# Therapeutic modulation of ROCK overcomes metabolic adaptation of cancer cells to OXPHOS inhibition and drives synergistic anti-tumor activity

**DOI:** 10.1101/2024.09.16.613317

**Authors:** Nicholas Blazanin, Xiaobing Liang, Iqbal Mahmud, Eiru Kim, Sara Martinez, Lin Tan, Waikin Chan, Nazanin Esmaeili Anvar, Min Jin Ha, Md Qudratullah, Rosalba Minelli, Michael Peoples, Philip Lorenzi, Traver Hart, Yonathan Lissanu

## Abstract

Genomic studies have identified frequent mutations in subunits of the SWI/SNF chromatin remodeling complex including *SMARCA4* and *ARID1A* in non-small cell lung cancer. Previously, we and others have identified that *SMARCA4*-mutant lung cancers are highly dependent on oxidative phosphorylation (OXPHOS). Despite initial excitements, therapeutics targeting metabolic pathways such as OXPHOS have largely been disappointing due to rapid adaptation of cancer cells to inhibition of single metabolic enzymes or pathways, suggesting novel combination strategies to overcome adaptive responses are urgently needed. Here, we performed a functional genomics screen using CRISPR-Cas9 library targeting genes with available FDA approved therapeutics and identified ROCK1/2 as a top hit that sensitizes cancer cells to OXPHOS inhibition. We validate these results by orthogonal genetic and pharmacologic approaches by demonstrating that KD025 (Belumosudil), an FDA approved ROCK inhibitor, has highly synergistic anti-cancer activity in vitro and in vivo in combination with OXPHOS inhibition. Mechanistically, we showed that this combination induced a rapid, profound energetic stress and cell cycle arrest that was in part due to ROCK inhibition-mediated suppression of the adaptive increase in glycolysis normally seen by OXPHOS inhibition. Furthermore, we applied global phosphoproteomics and kinase-motif enrichment analysis to uncover a dynamic regulatory kinome upon combination of OXPHOS and ROCK inhibition. Importantly, we found converging phosphorylation-dependent regulatory cross-talk by AMPK and ROCK kinases on key RHO GTPase signaling/ROCK-dependent substrates such as PPP1R12A, NUMA1 and PKMYT1 that are known regulators of cell cycle progression. Taken together, our study identified ROCK kinases as critical mediators of metabolic adaptation of cancer cells to OXPHOS inhibition and provides a strong rationale for pursuing ROCK inhibitors as novel combination partners to OXPHOS inhibitors in cancer treatment.

## Introduction

Lung cancer is a devastating disease that remains the top cause of cancer mortality accounting for a quarter of all cancer related deaths which is more than mortality due to colon, breast and pancreas cancers combined^1,2^. However, many patients with lung cancer still lack effective treatment options, underscoring the dire need for novel therapeutic approaches.

Genomic sequencing studies across numerous solid tumor types have demonstrated a high frequency of genetic alterations in multiple subunits of the SWI/SNF chromatin remodeling complex including *SMARCA4* (also known as *BRG1*) and *ARID1A* ranging from 16% in early stage disease to 33% in advanced lung cancer^3-7^. Furthermore, a meta-analysis of 44 genomic studies has shown that 20% of all solid tumors have mutations in subunits of the SWI/SNF complex making it one of the most frequently mutated complexes in cancer^6^. The SWI/SNF (SWItch/Sucrose Non-Fermenting) complex is a large multi-protein assembly that uses the energy derived from ATP hydrolysis to remodel nucleosomes and facilitate major chromatin dependent cellular processes such as DNA replication, repair and transcription^8,9^. There is intense effort to identify synthetic lethal or other vulnerabilities in SWI/SNF-mutant lung cancer. Most mutations in the SWI/SNF complex are inactivating and cannot be directly targeted therapeutically. In this regard, several vulnerabilities have been reported that enhanced sensitivity of *SMARCA4*-mutant lung cancers such as inhibition of Aurora kinase A^10^, CDK4/6^11^, EZH2^12^, ATR^13^, and KDM6 methyltransferase ^14^. However, none of these have progressed into advanced clinical studies.

Several reports have shown that SWI/SNF-mutant lung cancers have a blunted response to energy stress and rely on oxidative phosphorylation (OXPHOS) for their bioenergetic and biosynthetic needs, suggesting OXPHOS can be an attractive therapeutic target ^15,16^. However, it has so far been challenging to develop highly efficacious OXPHOS inhibitors with favorable safety profile as single agent cancer therapeutics. Routine use of metformin and related class of compounds have become standard care for the treatment of metabolic disorders (i.e diabetes) as they demonstrate robust safety with limited toxicity suggesting OXPHOS inhibition could be a beneficial approach in the clinic. However, metformin and other OXPHOS inhibitors display several limitations. These include inadequate potency ^17,18^, “off target” toxicity issues^19^, and lack of suitable in vivo pharmacokinetics that restrict their clinical use as therapeutic agents. The recent discovery of IACS-10759, a highly selective and potent small-molecule inhibitor of complex I of the mitochondrial electron transport chain with favorable in vivo pharmacokinetics^20^, has shown promise in several genetically defined preclinical cancer models including SWI/SNF-mutant lung cancer^15,21,22^. However, phase I clinical trials revealed dose-restricting toxicities (i.e lactic acidosis; peripheral neuropathy) with modest anti-tumor activity at tolerable doses putting into question the further development of OXPHOS inhibitors for clinical use^23^.

Due to current limitations of OXPHOS inhibitors as single agents, further clinical optimization through discovery of potent and efficacious combination agents will be required. While several combination agents with OXPHOS inhibitors have been proposed, such as tyrosine kinase inhibitors (i. e BRAF, cKIT, BTK) in melanoma, gastrointestinal tumors and lymphoma, respectively^24-26^, currently no proposed combination agent exists for patients harboring SWI/SNF-mutant lung tumors sensitive to OXPHOS inhibition.

To address this question and nominate a combination agent that can be translated rapidly in the clinic, we performed CRISPR-Cas9 screens using a sgRNA library against genes whose gene products have targeted FDA approved therapeutics or agents in advanced clinical development (FDAome-library). By using low concentrations of IACS-10759 that correspond to clinically tolerated doses in three *SMARCA4*-mutant cell lines, we identified several gene targets that are sensitive to OXPHOS inhibition. Validation of these targets through drug screening identified ROCK inhibitor KD025 (Belumosudil) as a novel, clinically available therapeutic that potently synergizes with IACS-10759. We demonstrate that KD025 and IACS-10759 have a profound impact on cell growth and survival by causing a severe energetic stress due to impingement on energy producing pathways. Mechanistically, we showed that ROCK inhibition abolished the adaptive increase in glycolysis upon OXPHOS inhibition with significant rewiring of cellular metabolic pathways. Global phosphoproteomics and kinase-motif enrichment analysis uncovered a dynamic regulatory kinome upon combination of OXPHOS and ROCK inhibition. Among several prominent signaling pathways and kinases observed in the combination treatment, an emergent phosphorylation-dependent regulatory cross-talk between AMPK and RHO GTPase signaling that converge on key RHO/ROCK-dependent substrates such as PPP1R12A, NUMA1, and PKMYT1 is proposed to drive cell cycle inhibition. In conclusion, we identified ROCK kinases as critical mediators of metabolic adaptation of cancer cells to OXPHOS inhibition and provide the preclinical basis for future clinical investigation of the combination of OXPHOS and ROCK inhibitors as cancer therapeutics.

## Results

### Integrative functional chemo-genomic screens and pharmacologic studies identifies ROCK inhibition as a novel combination strategy that synergizes with OXPHOS inhibition

To discover clinically relevant combination agents that could synergize with OXPHOS inhibition, we performed chemo-genomic CRISPR-Cas9 screens using a focused “FDA-ome” library that contains 1607 sgRNAs targeting 200 protein coding genes whose gene products have FDA approved therapeutics or therapeutics in clinical testing (Supplementary Data 1). We carried out CRISPR screens in three cell lines with damaging *SMARCA4* mutations as these are more sensitive to IACS-10759 compared to *SMARCA4* wild-type cells due to increased dependance on OXPHOS^15^. We determined the IC_20_ (inhibitory dose corresponding to 20% growth inhibition) for IACS-10759 treatment as this dose is well tolerated in patients ^23^. Each cell line was transduced with the lentiviral-based FDA-ome CRISPR library at a low multiplicity of infection (MOI) (<0.3), followed by selection with puromycin. Cells were treated with either dimethyl sulfoxide (DMSO) as a control or with IACS-10759 for the duration of the experiment. Cells collected after puromycin selection and seeding were referenced as time = 0 (T0), and cells were cultured for at least 10 population doublings and collected at both early and late time points. Genomic DNA was extracted, barcode labeled, and final PCR products were submitted for deep-sequencing and analyses (Fig. 1A). Processing of sequencing reads for each screen assured an adequate depth and distribution (>1000-fold of gRNA numbers) for subsequent bioinformatic analysis (Supplementary Fig. 1A). Next, we performed analysis using the BAGEL algorithm to calculate a Bayes factor for each gene^27^ and Pearson’s correlation coefficients were calculated based on the Bayes factor distributions. Notably, H1299, A549, and H2023 screens had a high correlation (correlation index >0.8) among the different groups indicated that the results of these 3 screens were of comparable quality (Supplementary Fig. 1B). Precision-recall curves were also used to evaluate each screens performance and we routinely observed a value >0.8, a metric commonly used to determine a quality screen (Supplementary Fig. 1C)^27^. We also compared changes in 50 essential and 50 nonessential control genes included in the FDA-ome library from the starting reference time point (T0) to the early and late time points ^27^. Overall, sgRNAs distributions targeting essential genes were reduced while those targeting nonessential genes did not change regardless of time point analyzed, indicating that our CRISPR screens worked well in each cell line (Supplementary Fig. 1D). Taken together, these results demonstrated that all three screens were of high quality and performance allowing for reliable identification of candidate co-essential genes to OXPHOS inhibition.

**Figure 1:**
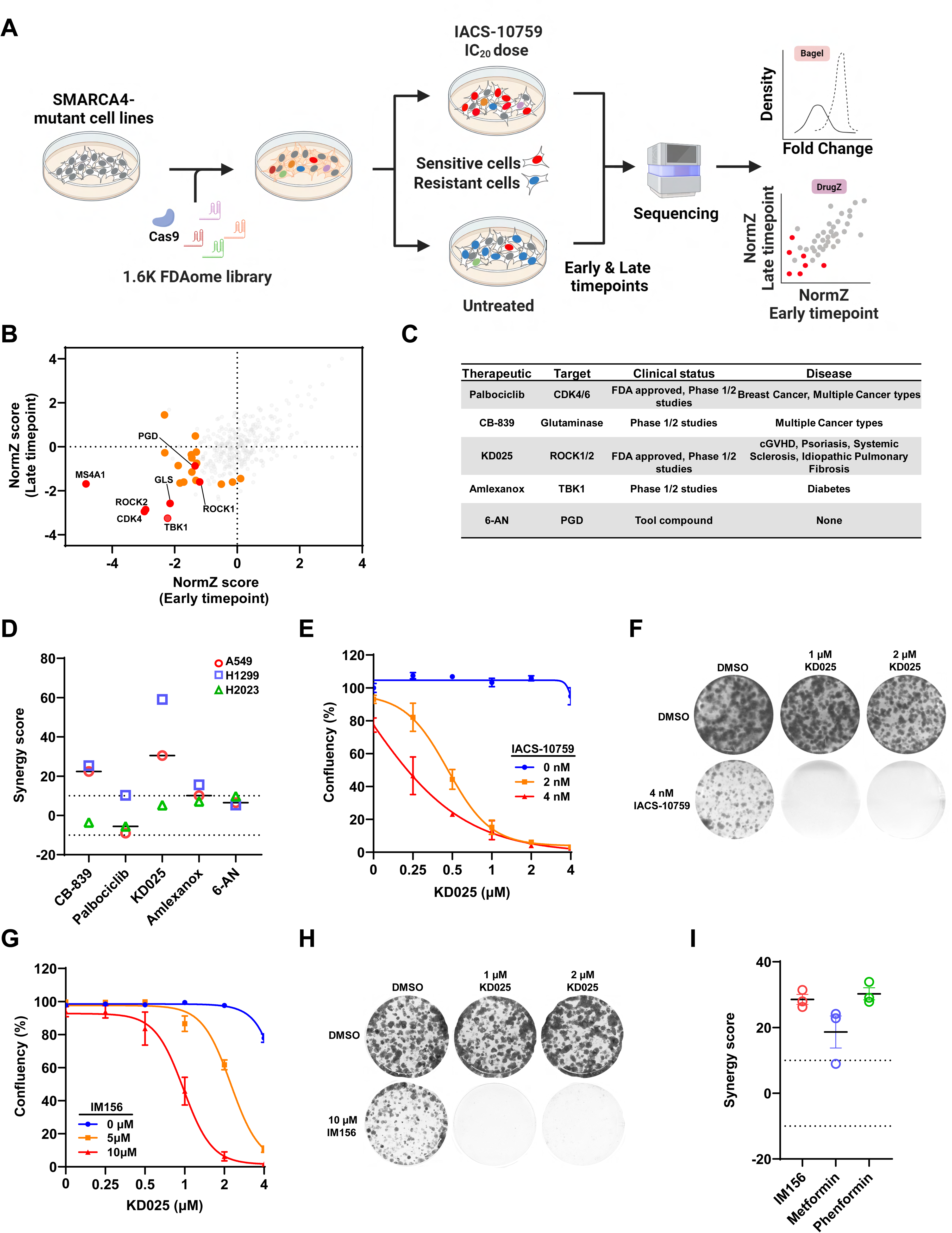
CRISPR and clinical drug screening identifies ROCK1/2 as a synergistic combination agent with OXPHOS inhibition. **(A)** Schematic representation of the workflow for CRISPR screens performed in A549, H1299, and H2023 SMARCA4-mutant lung cancer cells. Image was created using BioRender.com. (**B**) NormZ scores of candidate genes common at both early and late timepoints from FDAome library CRISPR screening results in H1299 cells cultured in presence or absence of 4 nM IACS-10759. The NormZ score was used to define a possible synthetic lethal interaction with IACS-10759. All genes targeted by the FDAome library were scored according to the fold change of levels of their respective sgRNAs. Genes whose loss of function led to IACS-10759 sensitivity appear on the bottom left quadrant, and genes whose loss of function led to IACS-10759 resistance appear on the top right quadrant. High-confidence candidate genes are shaded in orange and those selected for further analysis are indicated in red. **(C**) List of selected therapeutics and respective gene targets that are FDA approved or in clinical development tested in combination with IACS-10759 for synergistic effects on cell growth. (**D**) Overall synergy scores determined in A549, H1299, and H2023 exposed to therapeutics against selected CRISPR hits in combination with IACS-10759. An overall synergy score >10: the interaction between two drugs is likely to be synergistic. (**E**) Dose-response curves of H1299 cells exposed to increasing concentrations of KD025 in the presence or absence of IACS-10759. Cell growth (measured as % Confluency) was assayed 5 days after drug exposure. (**F**) Representative clonogenic growth assays of H1299 cells cultured in the presence of KD025 and IACS-10759 alone or in combination for 12 days. Surviving cells after the treatment were fixed and visualized by crystal violet staining. (**G**) Dose-response curves of H1299 cells exposed to increasing concentrations of KD025 in the presence or absence of IM156. Cell growth (measured as % Confluency) was assayed 5 days after drug exposure. (**H**) Representative clonogenic growth assays of H1299 cells cultured in the presence of KD025 and IM156 alone or in combination for 12 days. Surviving cells after the treatment were fixed and visualized by crystal violet staining. Representative images of three independent experiments are shown. (**I**)Overall synergy scores determined in H1299 cells exposed to indicated OXPHOS inhibitors in combination with KD025. An overall synergy score >10: the interaction between two drugs is likely to be synergistic. For panels (**D**, **E**, **G**, **I)** are presented as mean +/-SEM of three biological replicates (n=3).

To identify genes whose depletion caused fitness defects and reduced viability with IACS-10759, a normalized sgRNA depletion score was calculated for each gene using DrugZ by comparing IACS-10759 and the DMSO-treated groups in each cell line at both early and late time points^28^. Genes from each cell line were ranked by their NormZ scores (Supplementary Data 2). We determined high-confidence hits at each timepoint by including genes with a p-value ≤ 0.05 for each cell line or genes in at least two cell lines with a p-value threshold of ≤ 0.10. This analysis identified a total of 59 unique genes across all three cell lines (Supplementary Data3). Out of this list, we prioritized genes that were depleted in both early and late timepoints in at least one cell line that also had multiple sgRNAs (n≥3) depleted in IACS-10759-treated group compared to DMSO (Fig. 1B, Supplementary Fig. 2A-E). Upon passing these stringent criteria, we identified 5 genes as potential hits in IACS-10759-treated *SMARCA4*-mutant lung cancer cells. Notably, our top hit from this analysis was PGD, a metabolic enzyme essential in generating NADPH for redox homeostasis, included in the FDA-ome library for quality control, as previous CRISPR screens demonstrated PGD as a synthetic lethal target to OXPHOS deficiency ^29^. Other lead candidate co-essential genes included TBK1, CDK4, GLS, and ROCK1/ROCK2. TBK1 is a non-conical IKK serine/threonine kinase that regulates NFKB involved in innate immunity with additional diverse roles in autophagy, proliferation, apoptosis and glucose metabolism^30,31^. CDK4 is a serine/threonine kinase involved in G1 cell cycle phase progression. CDK4/6 inhibition is synthetic lethal in SMARCA4-mutant lung cancer ^11^. GLS is mitochondrial enzyme that promotes the breakdown of glutamine to glutamate to fuel the tricarboxylic (TCA) cycle via anaplerosis or fatty acid synthesis via reductive carboxylation ^32^. ROCK1 and ROCK2 are related serine/threonine kinases that regulate the actin cytoskeleton and are involved in multiple cellular processes including cell cycle, DNA damage, migration, invasion and glucose homeostasis^33^. We prioritized these targets for further analysis by identifying therapeutics that are FDA approved or in advanced clinical development (Fig. 1C).

To determine whether therapeutics targeting candidate co-essential genes sensitize OXPHOS inhibition, we employed a combinatorial drug screening strategy with IACS-10759 in A549, H1299, and H2023 *SMARCA4*-mutant cell lines by performing 5-day growth assays with a matrix titration of several doses of each compound alone or in combination. We determined the statistical significance of growth inhibition by the combination of IACS-10759 and each therapeutic by determining an overallsynergy score (>10 indicates synergy) by Bliss estimation using SynergyFinder^34^ (Fig. 1D). This approach revealed that the combination of IACS-10759 with 6-aminonicotinamide (6-AN), a cell-permeable compound that suppresses PGD activity by competing with nicotinamide for NAD^+^/NADP^+29,35^, was additive/weakly synergistic to suppress cell growth, further demonstrating predictability of our screens and in agreement with published results (Supplementary Fig. 3A)^29^. Overall, combinatorial drug screening revealed that pharmacological inhibition with KD025, a ROCK inhibitor, was our most promising therapeutic with IACS-10759 was highly synergistic in all three *SMARCA4*-mutant lung cancer cell lines (Fig. 1D, Supplementary Fig. 3B-E). Further, this combination completely suppressed cell growth in all three cell lines, while individual drug exposure had little effect on cell growth at the doses examined. (Figure 3E, Supplementary Fig. 4A-B). We corroborated this response by performing 12 to 14-day clonogenic assays which similarly showed robust and highly significant synergy in the KD025 and IACS-10759 combination (Fig. 1F, Supplementary Fig. 4C,D). While we designed these studies using sub-therapeutic doses of IACS-10759 as single agent that is expected to be clinically well-tolerated, we sought to expand the repertoire of OXPHOS inhibitors to maximize the potential of KD025 as a suitable combination agent in the clinic.Importantly, synergistic inhibition on cell growth with KD025 was also observed with other clinically available small-molecule agents targeting OXPHOS (biguanide class of OXPHOS inhibitors) including metformin, phenformin, and IM156^36^ (Fig. G-I, Supplementary Fig. 4E-J). In line with these pharmacologic results, genetic perturbation of both ROCK1 and ROCK2 by shRNA knockdown in H1299 cells treated with IACS-10759 similarly caused a synergistic anti-proliferative response (Supplementary Fig. 5A, B). Taken together, these results demonstrate that KD025 has synergistic anti-tumor activity in combination with IACS-10759 and with other clinically relevant OXPHOS inhibitors. The demonstration of a synergistic combination effect of KD025 with OXPHOS inhibitors in OXPHOS-sensitive *SMARCA4*-mutant lung cancer cells is highly promising and could enable expansion of the therapeutic indications for inhibitors of ROCK and OXPHOS.

### ROCK blockade combined with OXPHOS inhibition causes cell cycle growth arrest, apoptosis, and severe energy deficiency

To further extend these observations and gain additional insight into the strong synergistic inhibitory action on cell growth between ROCK and OXPHOS blockade, we used a cell cycle reporter (FUCCI)^37^ to monitor different phases of the cell cycle in real-time in H1299 cells (Fig. 2A). Treatment of KD025 or IACS-10759 individually did not significantly alter cell cycle phase distribution in line with the minimal effect on cell growth at the concentrations used (Fig. 2A). However, the combination of both compounds rapidly increased the percentage of cells in M-G1 and G1 cell cycle phases with a concomitant decrease in the percentage of cells in the G1/S cell cycle phase (Fig. 2A), suggesting these cells go into an early G1/G1 phase cell cycle arrest. Furthermore, combination of KD025 and IACS-10759 induced a significant cell death as determined by Annexin V and Cytotox Green, while individual treatment of each compound had no effect. (Fig. 2B, C). Notably, significant cell death occurred much later and at higher doses compared to cell cycle G1 arrest, suggesting growth inhibition as the primary response to combination treatment. Taken together, these data suggest that the potent, synergistic combination of KD025 and IACS-10759 displays features of synthetic lethality that is detrimental to *SMARCA4*-mutant lung cancer cell survival.

**Figure 2:**
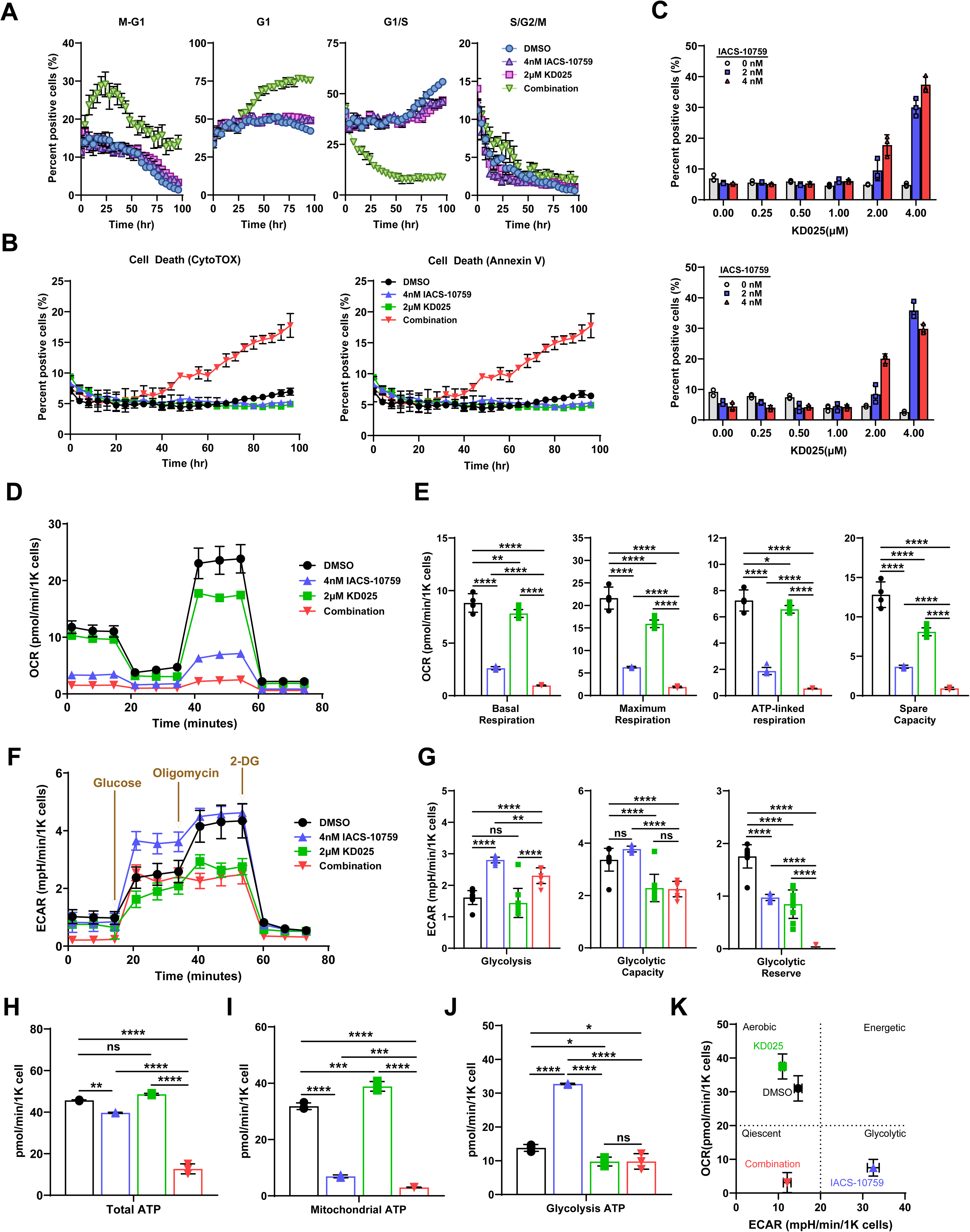
Combination of KD025 and IACS-10759 induces cell cycle growth arrest, cell death, and severe energy stress. **(A)** FUCCI cell cycle analysis showing percentage of H1299 cells in M-G1, G1, G1/S or S/G2/M following treatment with DMSO, IACS-10759, KD025, or the combination over a 96 hr timecourse. Percentage of cells at each cell cycle phase were quantitatively assessed by cell by cell analysis software from Incucyte S3. (**B**-**C**) Quantitation of apoptosis using either Cytotox green or Annexin V showing the percentage of cells over a 96 hr time course. (**B**) or at increasing doses of KD025 after 96 hr. (**C**) as assessed by cell-by-cell incucyte analysis following treatment with DMSO, IACS-10759, KD025 or the combination in H1299 cells. (**D**) Seahorse mitochondrial stress test assay showing a representative trace measuring mitochondrial oxygen consumption rate (OCR) in H1299 cells cultured with DMSO, KD025, IACS-10759, or the combination for 6 hr from which (**E**) basal respiration, maximum respiratory capacity, ATP-linked respiration, and spare capacity were calculated. Data indicate mean +/-S.E.M from DMSO (n=4), IACS-10759 (n=7), KD025 (n=11), and combination (n=7) independent experiments. (**F**) Seahorse glycolysis stress test assay showing a representative trace measuring extracellular acidification rate (ECAR) in H1299 cells cultured with DMSO, KD025, IACS-10759, or the combination for 6 hours from which (**G**), glycolysis, glycolytic capacity, and glycolytic reserve were calculated. Data indicate mean ± SEM from DMSO (n=10), IACS-10759(n=11), KD025(n=10), combination (n=9) independent experiments. (**H-J)** Quantitation of ATP production by Seahorse XF real-time ATP rate assay following treatment with DMSO, IACS-10759, KD025 or the combination for 6 hours in H1299 cells from which (**H**), total ATP production, (**I**) mitochondrial ATP production, and (**J**) glycolytic ATP production rates were calculated. Data shown are mean ± SEM from DMSO (n=10), IACS-10759(n=11), KD025 (n=10), combination (n=9) independent experiments. (**K**) Bioenergetic profile map in H1299 cells by plotting basal OCR and ECAR of the indicated treatment groups. For panels (**E**-**K)** One-way ANOVA was used corrected for multiple comparisons. ****indicates P-values <0.0001; ***<0.001; **<0.01; *<0.05; ns, not significant.

Given that *SMARCA4*-mutant lung cancer cells depend on elevated OXPHOS activity for survival^15^, we next profiled the IACS-10759 and KD025 combination for their effect on mitochondrial respiration and glycolytic capacities by oxygen consumption rate (OCR) and extracellular acidification rate (ECAR) assays by Seahorse metabolic analyzer. Consistent with previous results^15,20^, we found that even with low dose IACS-10759, H1299 cells had lower basal, ATP-linked, maximum respiration, and spare capacity rates that differed significantly from control cells (Fig. 2 D, E). KD025 decreased in mitochondrial respiration, in particular decreased maximum respiration, and spare capacity rates, although the effect was more modest when compared to IACS-10759. Importantly, combination of both compounds led to a profound decreases in mitochondrial respiration rates compared to each compound alone (Fig. 2D, E). By contrast, IACS-10759 caused a significant increase in glycolysis with glycolytic reserves being decreased (Fig. 2F, G), consistent with an adaptive shift to glycolysis due to suppression of OXPHOS^38,39^. On the other hand, glycolysis remained unchanged by KD025 but glycolytic capacity and glycolytic reserves were significantly decreased when compared to control cells (Fig. 2 F, G). Importantly, the adaptive increase in glycolysis stimulated by IACS-10759 is largely blocked in the combination with glycolytic reserves being depleted (Fig. 2G). Taken together, these results strongly suggest that addition of KD025 to IACS-10759 resulted in blockade of the normally observed physiological response of an increase in the glycolytic pathway.

Consistent with these data, total cellular ATP production was dramatically decreased in the combination while treatment of each compound individually had little to no effect on ATP production rates (Fig. 2H). Further analysis by Seahorse revealed that the while the proportion of ATP produced by mitochondria was suppressed by IACS-10759, there was a compensatory increase in glycolytic ATP production rates consistent with our ECAR results (Fig. 2I). By contrast, no significant effect of KD025 on mitochondrial and glycolytic ATP production rates were observed. However, the compensatory increase in glycolytic ATP production by IACS-10759 was significantly blocked in the combination, likely due to reduced glycolytic capacity and glycolytic reserves due to KD025 treatment (Fig. 2J). Taken together, IACS-10759 treatment derived almost all of their ATP from glycolysis with a glycolytic index (GI) consistent with the Warburg effect, that in turn was suppressed in the combination leading to a metabolic phenotype reminiscent of the quiescent state (Fig. 2K).

### Metabolic rewiring of glycolysis by IACS-10759 is impaired upon ROCK inhibition

Cancer cells use several carbon sources for survival, major players being glucose and glutamine, to fuel glycolysis and the tricarboxylic acid (TCA) cycle to continually sustain energy production (i.e ATP) and biosynthetic needs for survival ^40,41^. To understand how the combined treatment of IACS-10759 and KD025 impacts cancer cell metabolism, we profiled steady-state abundances of over 300 metabolites in H1299 cells treated for 24- and 48-hours using gas chromatography/mass spectrometry (GC/MS) (Supplementary Data 4). Combined treatment with both compounds caused a profound rewiring of the metabolic landscape when compared to each single agent individually (Supplementary Fig. 6A, B). Notably, while KD025 had little effect on metabolite abundance globally, IACS-10759 and the combination drastically altered the metabolic landscape although global metabolome abundance was completely distinct between the two groups. Enrichment analysis of significantly different metabolites showed key pathways involved in energy metabolism, including the TCA cycle, glycolysis/gluconeogenesis, pentose phosphate pathway, and pyruvate metabolism, as significantly changed by IACS-10759 or the combination (Supplementary Fig. 6C, D). Prompted by this observation and our ECAR/OCR results (Fig. 2D-K), we focused on metabolites involved in glycolysis, pentose phosphate pathway (PPP) and the TCA cycle. Overall, we noticed a dramatic rewiring of all three pathways in cells treated with the combination (Fig. 3A). Several glycolytic and TCA cycle intermediate metabolites were down-regulated in the combination compared to single agent and DMSO-treated cells including glucose-6-phosphate (G6P), pyruvate, citrate, isocitrate, and oxaloacetate (OAA) (Fig. 3A, B, Supplementary Fig. 6D). Unexpectedly, in contrast to reduced G6P levels, we observed up-regulated glucose levels in the combination (Fig.3A, B, Supplementary Fig. 6D), an effect also observed in KD025-treated cells, suggesting a blockade at the glucose to G6P rate-limiting step. By contrast, lactate levels were up-regulated by IACS-10759, further confirmation that OXPHOS inhibition induces the Warburg effect in these cells. However, the increase in lactate levels were blocked by the combination (Fig. 3A-C, Supplementary Fig. 6D). To confirm these results, we measured extracellular lactate levels in the media and observed increased lactate secretion following IACS-10759 treatment that was similarly blocked by the combination (Fig. 3C). Importantly, lactate secretion leading to lactic acidosis is one of the key adverse effects of OXPHOS inhibition and the observation that combined treatment of KD025 and IACS-10759, while being more efficacious, lacks elevated lactate levels is highly promising for improving tolerability and clinical translational. In addition, KD025 treatment alone and in combination with IACS-10759 led to decreases in glucose uptake, suggesting that altogether KD025 likely impinges on glycolysis through multiple rate-limiting steps (Fig. 3D).

**Figure 3:**
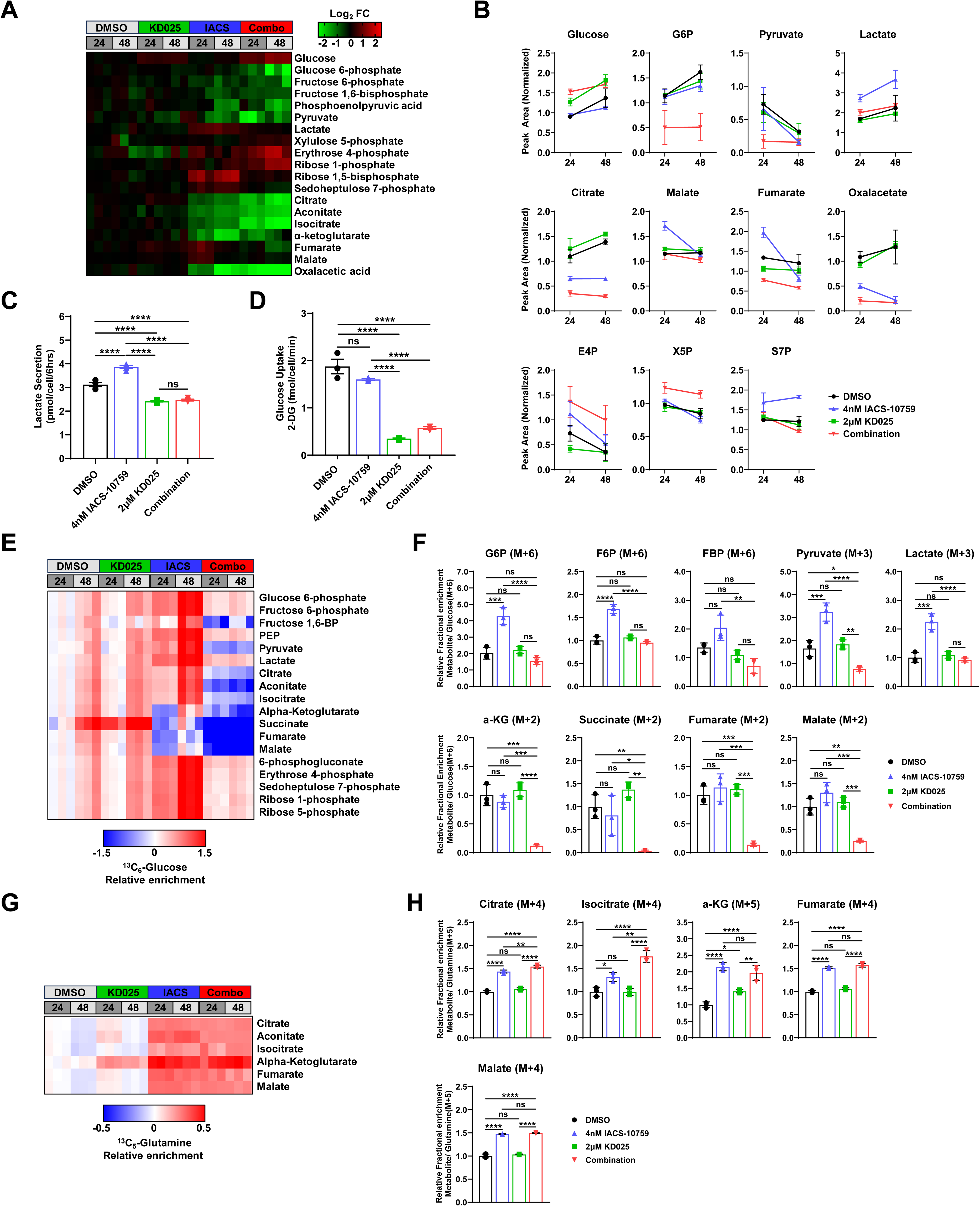
KD025 promotes metabolic reprogramming by suppressing adaptive increase in glycolysis due to OXPHOS inhibition. (A) Heatmap of significantly different metabolite abundances involved in glycolysis, pentose phosphate pathway, and TCA cycle following treatment with DMSO, KD025, IACS-10759, or the combination for 24- and 48- hr in H1299 cells. Log_2_ FC > 0 (red) represent an increase of metabolite abundance and Log_2_ FC < 0 (green) represent a decrease of metabolite abundance. (B) Relative abundance of select metabolites from glycolysis, pentose phosphate pathway, and TCA cycle pathways following treatment with DMSO, KD025, IACS-10759, or the combination for 24- and 48 hours. Metabolite abundance expressed as relative peak intensity. (**C**) Glucose uptake in H1299 cells following 6 hours treatment as measured by uptake of 2-deoxyglucose and normalized to cell number. (**D**) Lactate secretion into extraceullar media from H1299 cells were treated for 6 hours, and lactate levels accumulated in the media were measured and normalized to the endpoint cell number. (**E**) Heatmap of significantly labeled metabolites after isotope incorporation. Data reflect the relative sum of abundance of all ^13^C isotopologues. (**F**) Fractional isotopic incorporation of ^13^C_6_-glucose into glycolytic and TCA cycle metabolite intermediates as measured by GC/MS in H1299 cells following treatment with DMSO, KD025, IACS-10759 or the combination, and cultured in ^13^C_6_-glucose containing medium for 24 hours. m, number of labeled carbons. Data indicate mean ± SEM of three independent experiments. All data indicate mean ± SEM of three independent experiments (n=3). For panels (**C, D, F, H)** One-way ANOVA was used corrected for multiple comparisons. ****indicates P-values <0.0001; ***<0.001; **<0.01; *<0.05; ns, not significant.

To further support that ROCK inhibition causes a metabolic shift from consuming glucose in IACS-10759 treated cells, we performed stable isotope tracer analysis using ^13^C_6_-glucose (Fig. 3E, F). Importantly, ^13^C_6_-glucose labeling was significantly decreased at a number of rate-limiting glycolytic metabolites in cells treated with combination including fructose 1,6-bisphosphate (FBP) and pyruvate (Fig. 3E). This translated to a near complete shut down in ^13^C_6_-glucose labeling of TCA cycle intermediates. Notably, while IACS-10759 increased ^13^C_6_-glucose labeling in glycolysis, owing to the metabolic dependency on this pathway, labeling of several TCA cycle metabolites were decreased. This is consistent with the role of IACS-10759 as a complex I inhibitor^20^. Importantly, the overall lower metabolic activity of glycolysis and the TCA cycle in the combination was also evidenced by significant decreases in M+6 and M+3 isotopologs of glycolytic metabolites and M+2 isotopologs of TCA cycle metabolites (Fig. 3F, Supplementary Fig. 7A). We also performed isotope tracing using ^13^C_5-_glutamine and showed that IACS-10759 increased glutamine carbon enrichment in TCA cycle metabolites (Fig. 3G, H, Supplementary Fig. 7B), an effect that was not blocked in the combination. Thus, despite lack of effect on KD025 on glutamine utilization induced by IACS-10759, glutamine may still serve as an alternative compensatory fuel source via glutaminolysis in the combination. Overall, these observations were also confirmed through biochemical analysis of ATP/AMP, NADH/NAD+, and NADPH/NADP+ cellular pools as all were significantly depleted in the combination (Supplementary Fig. 7C-E). Taken together, these data support that ROCK inhibition induces a metabolic reprogramming that completely suppresses glucose utilization, thus depriving cancer cells an essential adaptive fuel source for survival during OXPHOS inhibition.

### Global phosphoproteomics analysis links energy stress and cell cycle arrest to RHO GTPase signaling

One of the most well-known and rapid cell signaling changes induced by OXPHOS inhibition is the activation of the master metabolic regulator AMPK^20,42,43^, which monitors cellular energy status by sensing increases in cellular AMP/ATP and ADP/ATP ratios^44^ Activated AMPK then phosphorylates a diverse array of substrates that shut down ATP-consuming anabolic pathways while simultaneously activating ATP-producing catabolic pathways to restore energy homeostasis^44^. On the other hand, KD025, by targeting ROCK1/2, can inhibit phosphorylation of various substrates, such as myosin light chain (MLC), MYPT (a regulatory subunit of MLC phosphatase, also known as PPP1R12A), and LIM-kinase-1 (LIMK1) to impact actin filament assembly and contraction, which in turn can regulate numerous cellular pathways related to metabolism, cell growth, migration, and survival ^33,45^. ROCK1/2 can also directly interact and phosphorylate substrates independent of cytoskeletal regulation^46,47^. Therefore, we hypothesized that investigation of the state of cellular phosphoproteome could elucidate the biochemical basis for the cancer cell growth inhibition effect of the combination of IACS-10759 and KD025. To this end, we evaluated global phosphorylation and protein changes by TMT mass spectrometry in H1299 *SMARCA4*-mutant cells treated acutely for 6 hours with the IACS-10759/KD025 combination, or each compound individually (Fig. 4A).

**Figure 4:**
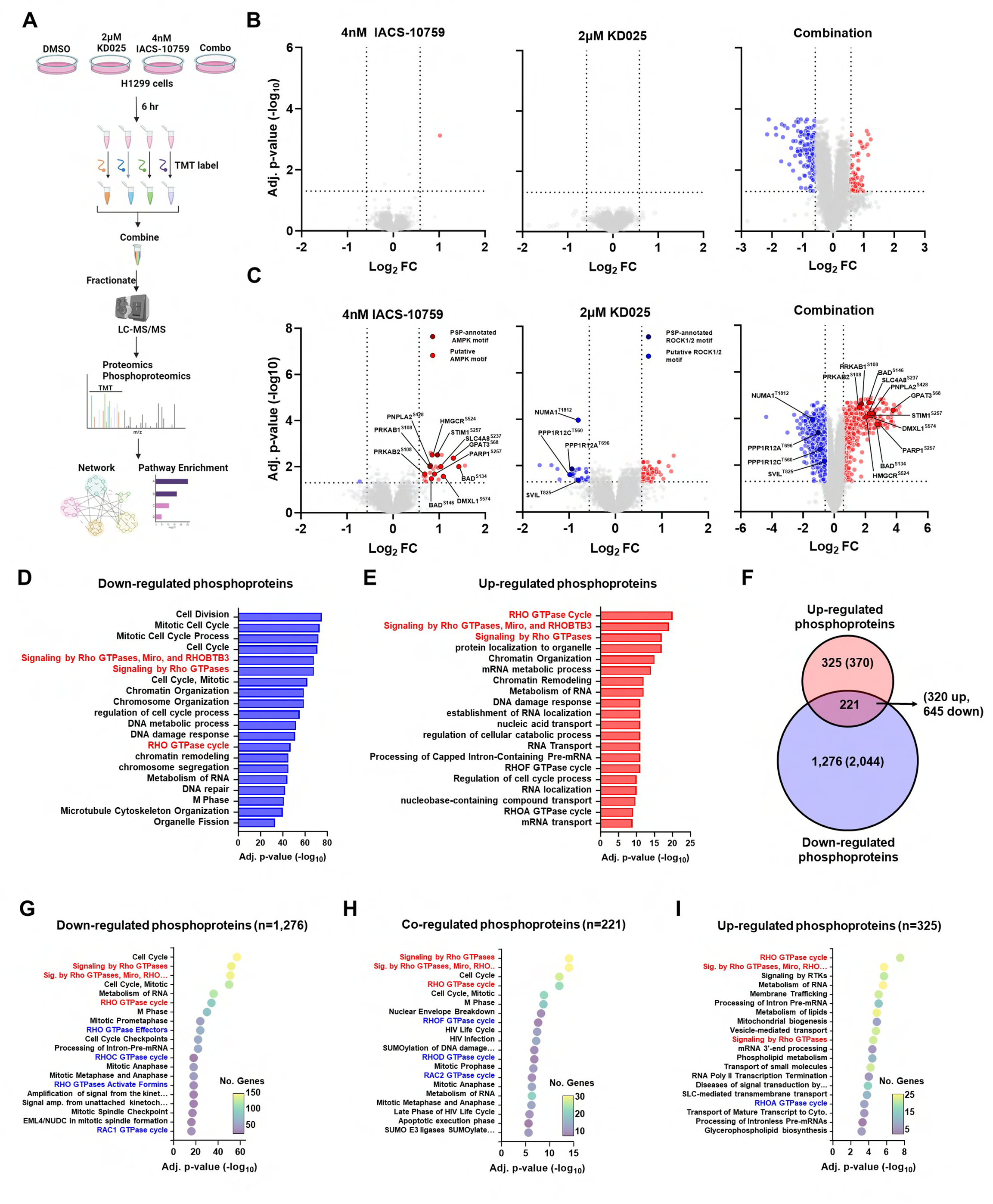
Phosphoproteomics uncovers RHO GTPase signaling as a top regulated pathway in the KD025 and IACS-10759 combination. (**A**) Experimental workflow for proteomic and phosphoproteomic analyses. Four replicates were analyzed for each treatment group. Image was created using BioRender.com. (**B**) Volcano plots of global proteome showing differentially expressed proteins (FDR < 0.05, 1.5 log_2_ FC) following treatment with KD025, IACS-10759, or the combination for 6 hours in H1299 cells. Volcano plots of global phosphoproteome showing differentially expressed phosphosites (FDR < 0.05, 1.5 log_2_ FC) following treatment with KD025, IACS-10759, or the combination for 6 hours in H1299 cells. PSP annotated and putative AMPK and ROCK regulated phosphosites are indicated. (**D**) Top Reactome, KEGG, GOBP gene sets significantly enriched (P > 0.05) in down-regulated phosphoproteins in the combination. RHO GTPase signaling-related pathways is highlighted in red (**E**) Top Reactome, KEGG, GOBP gene sets significantly enriched (P< 0.05) in up -regulated phosphoproteins in the combination. RHO GTPase signaling-related pathways are highlighted in red. (**F**) Venn diagram comparing overlap of proteins with down-regulated phosphosites, or in proteins with up-regulated phosphosites in the combination. The total number of phosphosites in each group are indicated. (**G-I**) Top enriched terms of Reactome gene sets significantly enriched (P < 0.05) in phosphoproteins with only down-regulated phosphosites (G), in phosphoproteins with up- and down-regulated phosphosites (H), or in phosphoproteins with only up-regulated phosphosites (I). Terms are sized and colored by significance of enrichment and number of associated genes. Less specific RHO GTPase signaling pathways are highlighted in red. More specific pathways related to RHO GTPase signaling are highlighted in blue.

Our global analysis quantified 9,050 proteins across all four treatment groups (Supplementary Fig. 8A, Supplementary Data 5), as well as 55,376 phosphopeptides corresponding to 17,539 phosphorylation sites on 4,474 proteins (Supplementary Fig. 8B, Supplementary Data 6). Our proteomic and phosphoproteomic data were of high quality as shown by tight clustering of treatment groups using principal component analysis and Pearson correlation (Supplementary Fig. 8C-D). Analysis of the global proteome revealed that 2.5% of proteins quantified were significantly changed in the combination (Fig. 4B). This included 57 differentially expressed up-regulated proteins, and 172 down-regulated proteins. By contrast, only 1 protein was significantly altered in response to IACS-10759, and none were significantly altered in response to the KD025 (Fig. 4B). Next, we performed pathway enrichment analysis using gene sets in the Reactome, Gene Ontology Biological Process (GOBP), and Kyoto Encyclopedia of Genes and Genomes (KEGG) databases. Importantly, pathways related to cell division and cell cycle regulation were the most significantly down-regulated pathways in the combination treatment (Supplementary Fig. 9A). By contrast, gene sets modestly up-regulated included MAPK and TGFbeta pathways, as well as mitochondrial oxidative phosphorylation and respiratory electron transport (Supplementary Fig. 9B). We further annotated functionally associated proteins with differential expression in the combination treatment by assessing networks comprised of protein-protein interaction (PPI) data using STRING^48^. Similarly, analysis revealed down-regulated PPI protein networks involved in cell cycle regulation, while up-regulated PPI networks involved oxidative phosphorylation and electron transport (Supplementary Fig. 9C-D). These results are consistent to the cell cycle arrest and metabolic dysregulation observed in H1299 cells, implicating these protein networks as sensitive to the combined treatment of IACS-10759 and KD025.

Next, we performed in depth analysis of the phosphoproteomics data. Here, we detected 27 phosphosites that were differentially regulated following IACS-10759 treatment with nearly all (26 phosphosites) up-regulated (Fig. 4C). KD025 treatment differentially regulated 94 phosphosites with 65 and 29 phosphosites up-regulated and down-regulated, respectively (Fig. 4C). However, combined treatment with IACS-10759 and KD025 revealed profound changes with 2,981 down-regulated phosphosites and 751 up-regulated phosphosites detected constituting 21.7% of the phosphoproteome (Fig. 4C). Next, we sought to enrich these data for phosphosites likely to be early, direct substrates of ROCK1/2 and AMPK. We selected only those that were down-regulated in response to KD025 and also contained the motif of ROCK1/2 with a βXpS/T or βXXpS/T consensus sequence (β = R, K, X = any amino acid, pS/T = phosphorylation site)(Supplementary Data 6)^33,45^. Similarly, to enrich for phosphosites more likely to be direct substrates of AMPK, we selected those that were up-regulated in response to IACS-10759 and contained the AMPK motif with a ϕββXXpS/TXXXϕ consensus sequence (hydrophobic, φ = M, L, I, F, or V; basic, β = R, K, or H, X = any amino acid, pS/T = phosphorylation site)(Supplementary Data 6)^49,50^. In addition, AMPK targets often contain slight variants of this consensus sequence^51^. We also functionally assigned known kinase-phosphosite interactions of AMPK and ROCK1/2 by using the PhosphositePlus (PSP) database ^52^. This analysis identified several known and suspected targets of AMPK and ROCK1/2 from treatment of IACS-10759 or KD025, respectively(Fig. 4C). IACS-10759 up-regulated AMPK targets PRKAB1(S108) and STIM1(S257) and AMPK putative targets PRKAB2 (S108), HMGCR (S524), GPAT(S68), PARP1 (S257), DMXL1(S574), PNPLA2 (S428), SLC4A8 (S237), and BAD (S134, S146) whereas KD025 down-regulated ROCK1/2 target PPP1R12A (T696) along with putative ROCK1/2 targets PPP1R12C (T560), NUMA1 (T1812) and SVIL (T825).

Similar to our proteomic analysis, we performed pathway enrichment using gene sets in the GOBP, KEGG, and Reactome databases and analysed PPI networks using STRING. After IACS-10759 treatment, we found enrichment of pathways related to membrane and vesicle trafficking, pH and ion regulation, and insulin/AMPK signaling in the up-regulated phosphoproteome (Supplementary Fig. 10A, D), all representing known adaptive responses to OXPHOS inhibition. No gene sets were enriched among the down-regulated phosphoproteome. Next, we found a different set of enriched pathways among up- and down-regulated phosphorylations differentially expressed following KD025 treatment. Down-regulated phosphoproteins were enriched in components related to smooth muscle contraction and focal adhesion, and nucleocytoplasmic transport (Supplementary Fig. 10B, D). On the other hand, DNA damage and repair, and cell cycle comprised the top enriched gene sets in KD025 up-regulated phosphoproteins (Supplementary Fig. 10C, D). By contrast, pathway enrichment of significant gene sets in the combination showed that down- and up-regulated phosphoproteins strongly shared a number of gene sets related to RHO GTPase signaling, RNA metabolism, chromatin remodeling and organization, and cell cycle-related processes (Fig. 4D, E). Importantly, analysis of PPI networks further supported these observations with RHO GTPase signaling as a top enriched pathway in both up- and down-regulated phosphoprotein networks in the combination (Supplementary Fig. 11A-D).

RHO GTPase signaling is comprised of a complex network of small guanine nucleotide binding proteins that regulate cell behavior by activating downstream effectors, many of which overlap with numerous cellular processes, including changes in cell cycle, cytoskeleton, and adhesion ^53^. We sought to further characterize changes in the RHO GTPase network by performing pathway enrichment analyses using Reactome gene sets. While the majority of proteins in the combination contained either significantly down-regulated (70%) or up-regulated (17.8%) phosphosites that made up the phosphoproteome (Fig. 4F), some were co-regulated (12.1%) and contained both up- and down-regulated phosphorylations (Fig. 4F). This suggests potential cross-talk mechanisms that may add an additional layer of protein regulation to this subset of phospho-regulated proteins. RHO GTPase signaling and RNA metabolism were top enriched pathways common to all three phosphoprotein-regulated subsets in the Reactome gene sets (Fig. 4G-I), while cell cycle-related processes comprised down-regulated and co-regulated phosphoproteins, and adaptative processes such as mitochondrial biogenesis, lipid metabolism, and vesicle and membrane trafficking comprised up-regulated phosphoproteins. Furthermore, each phosphoprotein-regulated subset was associated with specific subtypes of RHO GTPase pathways. Down-regulated phosphoproteins were primarily associated with RAC1 and RHOC GTPase cycles, RHO GTPase activate Formins, and RHO GTPase Effectors gene sets (Fig. 4G), whereas up-regulated phosphoproteins were associated with the RHOA GTPase cycle (Fig. 4I). By contrast, co-regulated phosphoproteins were primarily associated with RAC2, RHOD, and RHOF GTPase cycle gene sets (Fig. 4G-I). Taken together, we found that OXPHOS and ROCK1/2 inhibition led to widespread proteome and phosphoprotoeome changes in the combination, while relatively few phosphosite changes were seen by individual IACS-10759 or KD025 treatment. Top phospho-regulated pathways in the combination that emerged include RHO GTPase signaling, with cell cycle-related processes dominating down-regulated phosphorylations and up-regulated phosphoproteins being largely related to metabolic adaptation. Co-regulated phosphoproteins may serve as a central regulatory node that links RHO GTPase signaling to up- and down-regulated phosphorylations.

### Kinase profiling reveals AMPK-mediated cross-talk at key substrates involved in Rho GTPase signaling

Next, we sought to gain a comprehensive understanding of kinase regulation following combined treatment of IACS-10759 and KD025. Recently, Cantley and colleagues utilized positional scanning peptide arrays (PSPAs) to determine the substrate sequence specificity of 303 human serine-threonine kinases^54^. By employing an annotated data base of 89,752 phosphosites^54^, we determined how combined treatment of IACS-10759 and KD025 may alter the relative frequency of each kinase motif associated with up- and down-regulated phosphosites (Fig. 5A). Two approaches were performed for this analysis: First, we plotted the frequency factor and significance for each kinase and compared IACS-10759 or KD025 treatment with the combination (Fig. 5B). Second, we assigned a relative kinase activity score to each kinase, using the frequency factor and significance to represent how frequently the kinase motif was differentially regulated. We further partitioned each kinase into their respective motif class ^54^ (Supplementary Fig. 12A-D). Overall, we found significant dysregulation of several major kinase classes that could be broadly divided into three groups based on their phylogenic relationship: The first consists of CAMK kinases that were upregulated in the combination. The second consists of AGC and CAMK kinases that were upregulated by KD025 but significantly repressed in the combination. The third consists of TKL/STE/Atypical/Other kinases that were down-regulated in the combination. Notably, the most up-regulated CAMK kinases in the combination was AMPK (AMPKA1, AMPKA2) or AMPK-related kinases (TSSK1, TSSK2, NAUK1). (Fig. 5B, Supplementary Fig. 12A). AMPK was the only motif class whose kinase activity was significantly up-regulated by IACS-10759, although to a much lesser extent compared to the combination, in agreement with our initial phosphosite motif analysis. Other CAMK kinases up-regulated in the combination include related AMPK-like kinases MARK1-4, SIK, QSK, QIK, BRSK1-2 that belong to the MARK/SIK motif class, along with MAPKAPK2 (MK2), MAPKAPK3 (MK3), and MAPKAPK5 (MK5) of the PKRD/MAPKAPK motif class, which are known effectors of p38 MAPK^55^. Interestingly, another member of the PKRD/MAPKAPK motif class, CHK1 was strongly up-regulated, suggesting DNA damage checkpoint control and replicative stress in the combination. By contrast, CAMK kinases of the MLCK/DAPK motif class (smMLCK, caMLCK, skMLCK, MYLK4, DAPK2) were up-regulated in response to ROCK1/2 inhibition (Fig. 5B, Supplementary Fig. 12A). MLCKs and DAPK directly phosphorylate MLC to promote smooth muscle contraction, acting in concert with ROCK1/2 in a calcium/calmodulin-dependent manner^56^, suggesting compensatory activity in response to KD025 However, the combined effect of IACS-10759 and KD025 suppressed MLCK kinase activity. KD025 also up-regulated several AGC family kinases of the Akt/ROCK, S6K/RSK, AurAK/PKA, and PAK(II) motif classes that were significantly repressed in the combination (Fig. 5B, Supplementary Fig. 12A). These include members of the AKT/mTOR/SGK axis such as AKT1-3, SGK1, SGK3, and several RSKs (RSK2-4, MSK1-2, P90RSK), and PKA and cGMP-PKG signaling pathways (PKACA, PKACB, PKG1, PKG2).

**Figure 5.**
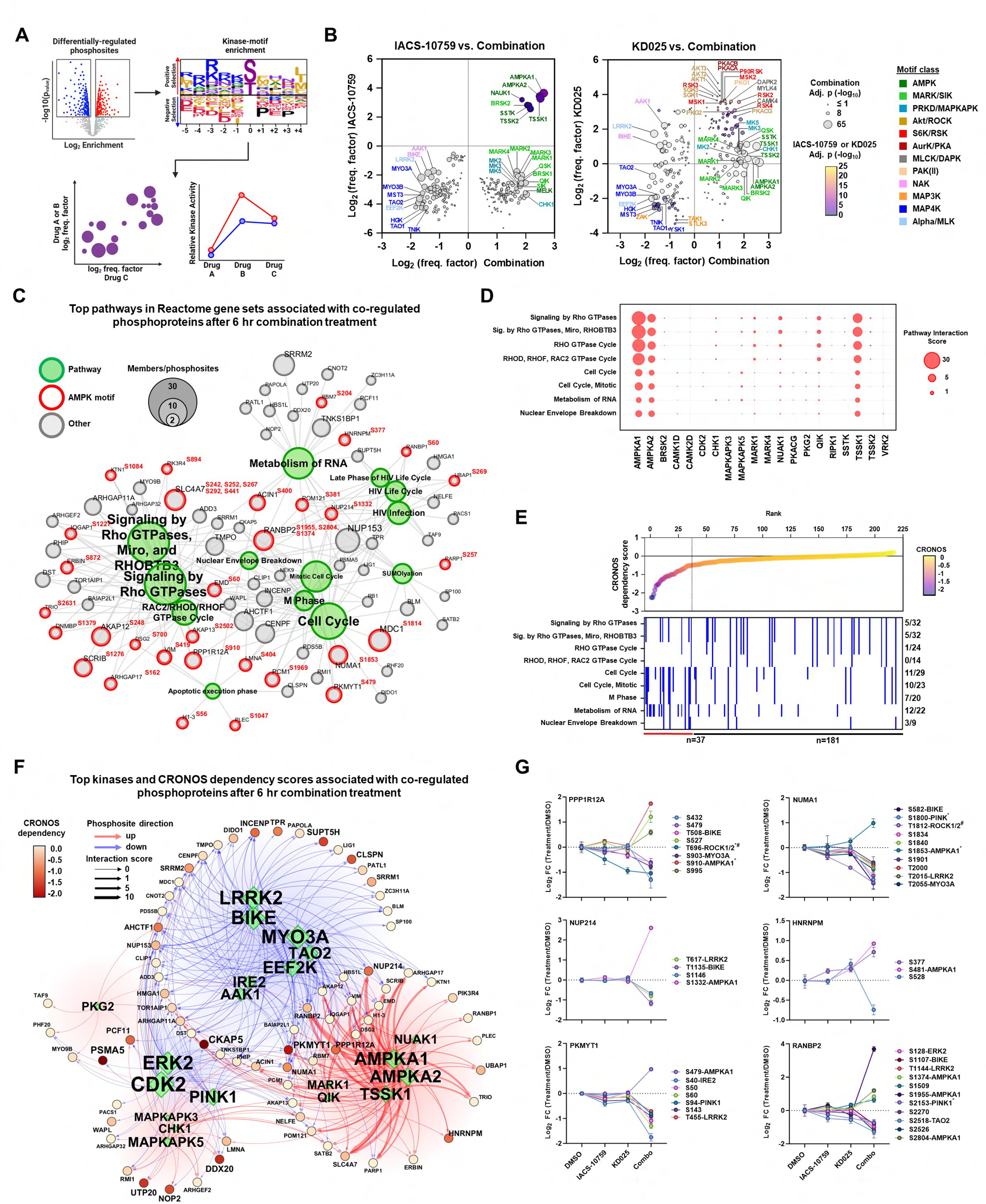
AMPK engages RHO GTPase signaling to regulate a dynamic phosphoproteome network at key substrates. (**A**) Schematic diagram of substrate scoring process and network analysis. Image was created using BioRender.com. (**B**) Kinase-motif enrichment [log2(frequency factor)] comparing IACS-10759 versus combination (left) and KD025 versus combination (right) phosphoproteomes derived from motifs of significant phosphosites shown in Fig. 4C. Fill color represents the significance of enrichment in either IACS-10759 or KD025 phosphoproteomes, and size represents the significance in the combination phosphoproteome. Select enriched kinases and kinase motif class are indicated^54^. The enrichments in these plots were determined using one-sided exact Fisher’s tests and corrected for multiple hypotheses using the Benjamini–Hochberg method. (**C**) Network of co-regulated phosphoprotein members in top enriched Reactome gene sets with up- and down-regulated phosphosites. Phosphoproteins are outlined in red if a differentially regulated phosphosite was identified with a AMPK motif. Phosphoprotein size is proportional to the total number of up- and down-regulated phosphosites, and pathways are sized by the number of phosphoprotein members. (**D**) Top kinase interaction scores for pathways in Reactome gene sets of co-regulated phosphoproteins. Kinase-pathway interactions were constructed using kinase-motif specificities ^54^, kinase-substrate specificities from PhosphoSitePlus ^52^, and kinase-substrate interactions from BioGRID^65^ for all phosphoproteins annotated for each pathway.(**E**) Annotated Chronos dependency rank plot ranked by DepMap dependency scores for co-regulated phosphoproteins. Chronos dependency scores for each phosphoprotein is represented as a point on the rank plot. Annotations indicate membership in Reactome gene sets. Phosphoproteins with a Chronos scores ≤–0.5 (dotted line) are considered essential, genetic dependent proteins. (**F**)Kinase-substrate networks and genetic dependency of co-regulated phosphoproteins in top Reactome gene sets after combined IACS-10759 and KD025 treatment. Kinase-substrate interaction scores were computed from kinase-pathway analysis in Figure 5E. Each phosphoprotein was annotated with genetic dependency scores from DepMap. The top 20 kinases are shown with nodes and text sized by their strength in the network. Edges are colored by direction of association with up- or down-regulated phosphosites. Edge width represents the magnitude of interaction score. Co-regulated phosphoprotein nodes and text are colored according to their CRONOS dependency score. (**G**) Phosphosites on essential, genetic dependent co-regulated proteins that interact with AMPK. Additional top kinases and their associated phosphosites computed by kinase-motif interaction analysis are also indicated. *indicates kinase-phosphosite interaction also annotated by PSP or by manual curation.

Lastly, combined treatment of IACS-10759 and KD025 down-regulated a diverse set of less characterized kinases (so-called “dark kinases”) predominantly of the Alpha/MLK, MAP4K, and NAK motif classes (Fig. 5B, Supplementary Fig 12D). Notable kinases of this group include EEF2K, LRRK2, MYO3A, MYO3B, BIKE, IRE2 and TAO1-2. Taken together, global motif analyses revealed important insights into the complexity of kinase regulation that occurs with OXPHOS and ROCK1/2 inhibition, many kinases of which have direct or indirect roles in actin cytoskeleton dynamics, cell cycle regulation and energy metabolism.

We next sought to validate the accuracy of our kinase-motif enrichment results by examining AMPK activation at T172 (Supplementary Fig. 13A). Indeed, immunoblot analysis demonstrated that AMPK activation was up-regulated by IACS-10759 after 6 hours treatment but subsided by 24 and 48 hours, consistent with the sharp, transient increases in AMPK activity that occur with OXPHOS inhibition^57^. By contrast, AMPK activation in the combination was even greater at 6 hours and activation was sustained over a 48-hour time period. In fact, AMPK total levels decreased with time in the combination, suggesting phosphorylated AMPK at T172 was a highly active form of AMPK. Encouraged, we further analyzed our kinase motif-enrichment results in our phosphoprotein subsets (Supplementary Fig. 13B). Unexpectedly, we found that co-regulated phosphoproteins were highly up-regulated with members of the AMPK motif class with AMPKA1 and AMPKA2 as top active kinases, at a level similar to up-regulated phosphoproteins.

Emerging evidence points to a role for AMPK not just as a master regulator for cellular metabolism, but also a key regulator of the actin cytoskeleton and cell cycle progression^58-63^. Thus, we reasoned that co-regulated phosphoproteins, where Rho GTPase signaling and cell cycle-related processes are top pathways, might be impacted by AMPK. Thus, we evaluated top Reactome gene sets for AMPK binding motifs. Using kinase-motif enrichment we scanned phosphosites for AMPKA1 and AMPKA2 kinase motifs^54^, and found 34.9% (37/106) of phosphoproteins contained the AMPK motif in the top Reactome datasets (Fig. 5C). When we investigated AMPK phosphorylations from our network, we found the AMPK motif to be highly enriched for hydrophobic residues (L, F, I, M, V) at +4 and -5 positions and basic residues (K and R) at the -3 position, as expected (Supplementary Fig. 13C). This is consistent with AMPK kinase-motif specificity ^54^, as well as published results^49,51,64^. Remarkably, AMPK-binding motifs were present in 50% (16/32) of the RHO GTPase-associated phosphoproteins. This decreased to 34.4% (10/29) and 27.3% (6/22) with AMPK-binding motifs, among cell cycle and RNA metabolism-associated phosphoproteins, respectively (Fig. 5C). Next, we used kinase-motif enrichment together with previously reported kinase-phosphosite and interaction data from PSP^52^ and BIOGRID^65^ to generate kinase-phosphosite interaction scores (see Materials and Methods and ^66^). Using this approach, we generated a pathway interaction score to elucidate which top kinases are likely to regulate both up- and down-regulated phosphorylations on co-regulated proteins in the top Reactome gene sets (Fig. 5D, Supplementary Fig. 13D). Indeed, our analysis suggests that AMPKA1 and to a lesser extent AMPKA2, TSSK1, and NUAK1, significantly impacted RHO GTPase signaling, RNA metabolism and cell cycle-related pathways on up-regulated phosphosites associated with co-regulated proteins. Top kinase interactions of down-regulated phosphosites revealed TAO2 and MYO3A preferentially interact with co-regulated proteins associated with RHO GTPase signaling, whereas LRRK2 and BIKE kinases preferentially interact with cell cycle and RNA metabolism pathways (Supplementary Fig. 13D). Although not well understood, evidence suggests that several of these kinases have unique roles in cytoskeletal control, mediating signaling between the actin and microtubule cytoskeleton network. For example, TAO kinases regulate microtubule plus end growth^67^ or interact with several substrates known to regulate the actin cytoskeleton ^68^.

Given that we found AMPK as a top interacting kinase in both RHO GTPase, cell cycle and RNA metabolism gene sets, we were interested in identifying which of its substrate phosphoproteins could impact cellular growth and viability and potentially explain the cellular phenotypic effects we observed. To this end, we evaluated the genetic dependencies of individual co-regulated phosphoproteins by taking advantage of the DepMap dataset (DepMap, https://depmap.org/portal/). Among the least dependent, non-essential genetic dependencies, we found significant enrichment in RHO GTPase signaling and RHO GTPase cycle and the more focused RHOF, RHOD, and RAC2 GTPase pathways (Fig. 5E, Supplementary Fig. 13F). More importantly, pathway enrichment of the more dependent, essential co-regulated phosphoproteins revealed RNA metabolism and cell cycle components as top enriched pathways, particularly mitotic, and M-phase gene sets (Fig. 5E, Supplementary Fig. 13F). We took advantage of our kinase-substrate and interaction network to associate top kinase interactions with genetic dependencies in the top Reactome gene sets (Fig. 5F). This revealed several genetic dependencies that were identified as AMPK and AMPK-related substrates; ERK2 and CDK2 substrates; and substrates of the less-well characterized kinases including LRRK2, MYO3A, TAO2, BIKE, and EEF2K. AMPK and AMPK-related kinases displayed the highest interaction scores with substrates compared to other top kinases, consistent with our pathway analysis, including those that have high genetic dependencies for cell survival according to DepMap. The co-regulated phosphoproteins with the highest genetic dependencies attributed to AMPK phosphorylation following combined treatment of KD025 and IACS-10759 after 6 hours were PPP1R12A, NUMA1, PKMYT1, NUP214, RANBP2, and HNRNPM (Fig. 5G). Remarkably, two are KD025-regulated phosphoproteins, ROCK substrate PPP1R12A (T696) and putative ROCK substrate NUMA1 (T1812) (Fig. 4C). Interestingly, both proteins are suggested to be AMPK substrates at S910 and S1853, respectively ^51,63^(Fig. 5G), further validating our network analysis. Taken together, these results highlight that AMPK can phosphorylate proteins involved in RHO GTPase signaling and cell cycle regulation and cross-talk with ROCK at key substrates critical for cell growth and survival.

### Anti-tumor activity of KD025 and IACS-10759 combination is synergistic in tumor xenograft models

Our study has revealed a potent vulnerability of *SMARCA4*-mutant cancer cells to dual inhibition of ROCK1/2 and OXPHOS. To validate these results in vivo, we first evaluated the anti-tumor effects of IACS-10759 and KD025 alone or in combination in mouse xenograft models of lung cancer. Thus, we decided to evaluate in vivo anti-tumor activity by treating mice bearing H1299 tumor xenografts with daily oral administration of 300 mg per kg body weight of KD025 and 1 mg kg body weight IACS-10759, both clinically relevant doses that have been shown to have minimal adverse effects ^23,69^. The dose chosen for KD025 was based on previous studies in other model systems that have shown KD025 to be effective and well tolerated in mice over multiple daily oral administrations. Individual daily treatments of both inhibitors did not significantly affect tumor growth in H1299 tumor xenografts (Fig. 6A, C). Importantly, co-administration of both compounds had a significant synergistic effect on suppressing tumor growth leading to stable disease in the H1299 xenograft model during 21 days of treatment, suggesting that low dose IACS-010759 and potentially other OXPHOS inhibitors in combination with KD025 could be viable treatment options for *SMARCA4*-mutant lung cancers. Similar results were evident in an additional lung cancer xenograft model, A549 where the combination of KD025 and IACS-10759 showed synergistic tumor growth inhibition activity (Fig. 6B, C). Importantly, both KD025 and IACS-10759 alone or as a combination regime was well tolerated in both lung cancer xenograft models, as both did not exhibit any significant weight loss or other overt toxicities over the course of the studies (Fig. 6D,E). Taken together, our data strongly suggests the inhibition of ROCK kinase helps to overcome the complex adaptive metabolic responses stimulated by OXPHOS inhibition, resulting in marked anti-tumor activity of the combination treatment regimen.

**Figure 6:**
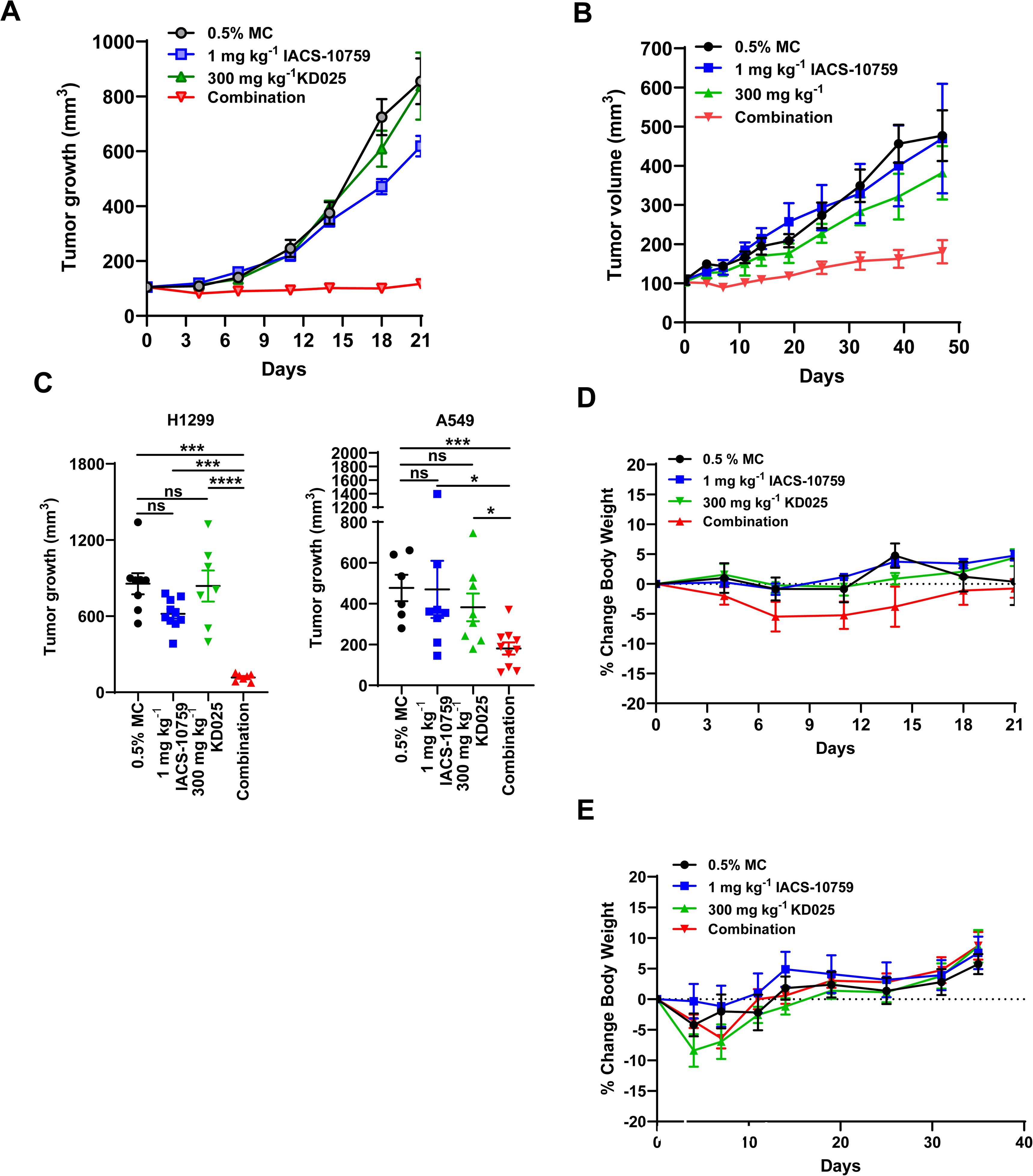
Combination of ROCK and OXPHOS inhibition displays synergistic anti-tumor activity in vivo. (**A**) In vivo xenograft in H1299 cells showing anti-tumor efficacy of IACS-10759 and KD025 alone or in combination after daily administration by oral gavage over 21 days. (**B)** In vivo xenograft in A549 cells showing anti-tumor efficacy of IACS-10759 and KD025 alone or in combination after daily administration by oral gavage over 47 days. (**C**) Comparison of final tumor volumes of H1299 (left) and A549 (right) tumor xenograft from (**A**, **B**). (**D-E**) Body weight changes of each treatment group over the during of the experiment in H1299 (**D**) and A549(**E**) tumor xenografts. For panel (**C**) Mann Whitney U-test was used. ****indicates P-values <0.0001; ***<0.001; **<0.01; *<0.05; ns, not significant.

## Discussion

In this report, we reveal the discovery of ROCK inhibition as synergistic combination partner to OXPHOS inhibition in *SMARCA4*-mutant lung cancer. Previously, we and others have shown that *SMARCA4*-mutant lung cancers are highly dependent on OXPHOS^15,16^. *SMARCA4*-mutant lung cancer is one of the worst prognosis subtypes of lung cancer with dismal outcome and poor response to chemotherapy, immunotherapy and even recently approved KRAS^G12C^ inhibitors ^70-74^. Unfortunately, initial efforts to utilize OXPHOS inhibitors in cancer have largely been disappointing due to limited efficacy and development of toxicity at doses required to induce tumor growth inhibition^17-19,75,76^ ^23^. This primarily stems from the rapid adaptation of cancer cells to inhibition of single metabolic enzymes or pathways and development of toxicities such as lactic acidosis upon higher doses of OXPHOS inhibitors. Taken together, it is unlikely that OXPHOS inhibitors alone can be efficaciously used as cancer therapeutics and requires the development of novel combination strategies that enhance therapeutic activity while avoiding toxicity. Here, we performed functional chemo-genomic screens with the main goal of identifying readily clinically translatable combination agents that would work with very low, and tolerable, doses of OXPHOS inhibitors. We identified ROCK1/2 as the most promising top hit and extensively validated our results using genetic and pharmacologic approaches in vitro and in vivo. Importantly, we have used several OXPHOS inhibitors including metformin that is extensively used in the clinic for decades, suggesting our main finding carries across various classes of OXPHOS inhibitors. Further, the FDA has recently approved the ROCK inhibitor KD025 (Belumosudil) for non-oncology indications. Thus, our finding lays the foundation for rapid clinical translation.

To decipher the mechanism of the profound degree of growth inhibition induced by the combination of OXPHOS and ROCK inhibition, we have performed comprehensive metabolic, metabolomics, and proteomic studies. In brief, we show that this combination induced a rapid, profound energetic stress and cell cycle arrest that was in part due to ROCK inhibition-mediated suppression of the adaptive increase in glycolysis upon OXPHOS inhibition. We also reveal a dynamic regulatory cross-talk by AMPK and ROCK kinases on key RHO/ROCK-dependent substrates such as PPP1R12A, NUMA1, PKMYT1 and RANBP2 that are all known regulators of cell cycle progression^77-80^.

The key question for future preclinical and clinical studies is: does the combination of OXPHOS and ROCK inhibitors lead to increased toxicity? So far, in our mouse models, we did not observe obvious signs of increased toxicity. One of the frequent dose-limiting toxicities of OXPHOS inhibitors is elevated blood lactate and lactic acidosis^23^. Excitingly, in the combination of OXPHOS and ROCK inhibitors, we observe lower lactate secretion than OXPHOS treatment alone due to the abolishment of the OXPHOS inhibition-induced adaptive increase in glycolysis by ROCK inhibitor. If this holds true in clinically settings, it will mark a major progress forward in the development of OXPHOS inhibitors as cancer therapeutics. It is important to note that another dose-limiting toxicity observed by OXPHOS inhibitors is peripheral neuropathy^23^, which we have not evaluated in the present study and needs to be carefully assessed in the future before trials in patients commence. On a positive note, in a recent clinical trial that carefully assessed plasma concentrations of lactate, IACS-10759 and peripheral neuropathy, elevated lactate was a stronger predictor of peripheral neuropathy^23^. As our combination regimen decreases lactate levels, it is likely that this adverse event will also occur less frequently in our combination than OXPHOS alone. However, this needs to be confirmed by future studies. In this study we have focused on *SMARCA4* mutant models as this genotype is associated with increased sensitivity to OXPHOS inhibition. Future work needs to be done to determine if the enhanced effect of our combination regimen is also observed in *SMARCA4* wild type lung cancers and other tumor types.

A key finding of our study is the demonstration that KD025 treatment leads to reduction in glucose uptake by cells. This is in line with previously demonstrated role of ROCK as an important regulator of cellular glucose uptake^81,82^. Interestingly, our metabolic flux analysis also revealed profound block at rate-limiting reactions in glycolysis by combination treatment. The exact biochemical basis for this phenomenon remains unknown and deserve detailed future investigations.

An additional key aspect of our study was the comprehensive proteomics and phosphoproteomics profiling studies to identify primary biochemical targets of the combination treatment. While the single agent treatments had minimal impact on the phosphoproteome, a remarkable rewiring of cellular signaling pathways was observed early after combination treatment. Detailed pathway analysis revealed the most prominent impact was the down-regulation of a vast array of cell cycle related pathways including key cell division regulators and executors upon combination treatment. This biochemical data is in line with our cellular phenotypic studies.

Further kinase motif profiling and kinase activity score determination utilizing a recently released database showed that the most profoundly elevated kinase family was AMPK and related kinases. AMPK is the master sensor and regulator of bioenergetic stress ^83^ and its phosphorylation signature is rapidly increased upon combination of KD025 and IACS-10759 in line with our metabolic studies. Importantly, we were able to perform phosphorylation site motif analysis across the altered phosphoproteome to identify major direct and indirect substrates of ROCK kinase signaling as well as AMPK that might serve as key nodes for signal propagation. While the majority of the phosphoproteome was down-regulated, a significant number was up-regulated. Interestingly, we also observed some phosphoproteins that showed both up- and down-regulated phosphosites suggesting they are dually regulated by different kinases. Detailed investigation of these unique proteins revealed that up-regulated sites were frequently phosphorylated by AMPK indicating these might constitute important regulators of downstream cellular behavior potentially accounting for the profound degree of synergy between OXPHOS and ROCK inhibitors. Some of these proteins that are heavily phospho-regulated are ROCK-dependent include PPP1R12A and NUMA1, which according to our DepMap analysis are critical mediators of cell growth and survival. PPP1R12A (MYPT1) is a regulatory subunit of protein phosphatase 1 (PP1), a key signaling molecule with roles in cell motility, morphology, and cell cycle including regulation of Rb ^79,80^. In our study, PPP1R12A showed highly interesting patterns of phosphorylation. While the ROCK motif phosphorylation site at T696 was significantly down-regulated, the AMPK phosphorylation site at S910 was highly up-regulated, suggesting a dual regulation by these two kinases which gets inactivated (ROCK) and activated (AMPK) upon combination of IACS-10759 and KD025. Indeed, previous studies have identified AMPK and AMPK-related member NUAK1 kinases as targets of PPP1R12A S910, where at least in the case of NUAK1 phosphorylation of S910 inactivates PPP1R12A preventing dephosphorylation of MLC by MLCP^51,84^. Another differentially regulated phosphoprotein is NUMA, a key cell cycle regulator^77,78^, with increased phosphorylation at AMPK S1853 and decreased phosphorylation at multiple other phosphosites including putative ROCK phosphosite T1812. NUMA coordinates mitotic spindle assembly by phase separation and is primarily regulated by Aurora A-mediated phosphorylation at S1969^77^. A number of the differentially regulated phosphosites discovered in our study reside in the regulatory C-terminus of NUMA and could regulate its activity or expression. Currently the impact of differentially regulated phosphorylation sites of PPP1R12A or NUMA on the enzymatic activity or protein-protein interactions of these proteins remains unknown and is a major future research direction. It has recently been proposed that a bidirectional relationship exists between metabolism and cell cycle regulation^85^. Our study further strengthens this concept by showing how early and profound changes in phosphorylation of cell cycle components occur during bioenergetic stress induced by combination treatment. It is important to note that in addition to the prominent phosphorylation signatures of AMPK kinases, many other kinase motifs were observed to be up- or down-regulated in our combination treatment including less characterized “dark kinases” of the Alpha/MLK, MAP4K, and NAK family such as EEF2K, LRRK2, MYO3A, MYO3B, BIKE, IRE2 and TAO1-2. Delineating the role of these kinases in the mechanism of action of drugs, energy metabolism, bioenergetic stress and response will be highly informative in the future.

In summary, our study identified ROCK kinases as critical mediators of metabolic adaptation of cancer cells to OXPHOS inhibition and provides a strong preclinical basis for pursuing ROCK inhibitors as novel combination partners to OXPHOS inhibitors in *SMARCA4*-mutant lung cancer treatment. This study is also expected to stimulate future mechanistic studies to understand the role of ROCK in energy stress sensing and metabolic adaptation in cancer cells.

## Acknowledgements

We thank the MD Anderson Cancer Center (MDACC) Core facilities, including the Advanced Technology Genomics Core (ATGC) and Metabolomics Facility Core, as well as the ThermoFisher Center for Multiplexed Proteomics at Harvard Medical School. We thank Shan Jiang for assistance with mouse colony maintenance. We also thank Dr Jack Roth and other members of the Department of Thoracic Surgery-Research section for valuable comments during the progress of this project. This study was in part supported by the generous philanthropic contributions to The University of Texas MD Anderson Lung Cancer Moon Shots Program (YL). This work is also supported by NIH grants R01CA272945 (YL) and R37CA251629 (YL).

## Author Contributions

Conceptualization: N.B. and Y.L.; Methodology: N.B., X.L., I.M., E.K., S.M., L.T., W.C., N.E.A., M.J.H., M.Q., R.M., M.P., P.L., T.H., Y.L. Software: I.M., E.K., N.E.A., M.J.H., M.Q., P.L., T.H.; Formal Analysis: N.B., I.M., E.K., L.T., W.C., N.E.A., M.J.H., M.Q.; Investigation: N.B., X.L., S.M., L.T., W.C.; Writing-Original Draft: N.B. and Y.L.; Supervision: N.B. and Y.L. Project Administration: N.B. and Y.L.; Funding Acquisition: Y.L.

## Material and Methods

### Cell culture

293 T cells were cultured with DMEM (Dulbecco’s modified Eagle medium, Hyclone, Cat# SH30243.02) containing 10% heat inactivated fetal bovine serum (Gibco, Cat# 1614007), 1% penicillin/streptomycin (Hyclone, Cat# SV30010), and 4 mM L-glutamine. All other cell lines were cultured in RPMI (Roswell Park Memorial Institute 1640 Medium, Hyclone, Cat# SH30027.01; no pyruvate) with 10% heat inactivated fetal bovine serum (Gibco, Cat# 1614007), 1% penicillin/streptomycin (Hyclone, Cat# SV30010), and 2 mM L-glutamine. Cell lines were cultured till 90% confluency and kept at low passage numbers and maintained at 37°C in a humidified 5% CO2-containing incubator. H1299 (Cat# CRL-5803), H2023 (Cat# CRL-5912) and A549 (Cat# CCL-185) were obtained from ATCC. Routine Mycoplasma testing was performed using Mycoplasma Detection Kit (ABM, Cat # G238).

### Compounds and Antibodies

KD025 (Cat# S7936), 6-aminonicatinimide (Cat# S9783), Palbociclib(Cat# S4482), CB-839(Cat# S7655), Amlexanox (Cat# S3648), IM156(Cat# S9604), Metformin(Cat# S1950), and Phenformin(Cat# S2542) were all purchased from Selleck Chemicals (Houston, TX). Antibodies against ROCK1 (Cat# 4035S), ROCK2 (Cat# 9029S), p-AMPK T172(Cat# 50081S), and AMPK (Cat# 2532S) were from Cell Signaling (Danvers, MA, USA). Antibody against β-actin (#A2228, 1:20,000) was from Millipore Sigma (Burlington, MA).

### Plasmids, Lentivirus production and infection

Individual shRNA vectors were obtained from the Gene Editing & Screening Core Facility (GES Core) at Memorial Sloan Kettering Cancer Center. All shRNAs were designed using the SplashRNA algorithm^86^ and cloned into LT3REPIR dox-inducible lentiviral vector. Sequence targeting both ROCK1 and ROCK2 was shROCK1/2 #1812 TAAGTTTACAAGATCTCCACCA. LT3REPIR was used as an empty vector control. Doxycycline working concentration of 1 µg/ml was used to induce shRNA mediated knockdown at various time points. Lentiviral transduction was performed with modifications using the protocol at https://portals.broadinstitute.org/gpp/public/resources/protocols. Briefly, 4 × 10^6^ 293 T cells were seeded in 10cm plates with 10 mL DMEM medium per dish. 24 hours later, cells were transfected with indicated lentiviral constructs, the packaging (psPAX2) and envelope (pMD2.G) plasmid by Lipofectamine 3000. Virus containing medium were collected (48-72 hours after transfection), filtered and stored at 4°C for no more than a week until infection. Infected cells were selected in medium containing puromycin for 48 hours and resistant cells subsequently grown for experiments.

### FDA-ome library construction and validation

All 1,607 gRNA sequences for the FDA-ome library were designed in a manner similar to the human TKOv3 library^87^ and cloned into the pLentiCRISPRv2 vector. The cloned plasmid pool yielded a 1000-fold representation of the library. A set of 250 control gRNAs were included in the library targeting 50 genes (5 gRNAs per gene) that exhibited non-essential or essential (according to Bayes Factor (BF) scores ^27,87,88^) fitness profiles and were expressed across all cell lines. Oligos were obtained from Eurofins.

### In vitro FDA-ome library CRISPR screens

A549, H1299, and H2023 cell lines were infected with the FDA-ome lentiviral library at a multiplicity of infection (MOI) of around 0.3. Twenty-four hours after infection, cells were selected with puromycin for 48–96 hours. Selected cells were divided into a control and an experimental group with three replicates each and maintained at around a 1000-fold library coverage throughout the screens. For each screen, the experimental groups were treated with IACS-10759 to achieve approximately 20% growth inhibition (IC_20_) throughout the course of the experiment, with control groups being treated with DMSO. Treatments were repeated 3 times throughout the course of each screen. At each passage, cell pellets were collected at around a 1000-fold library coverage for genomic DNA extraction, starting at day 0 after selection. Genomic DNA was extracted using the Wizard Genomic DNA Purification kit (Promega) according to the manufacturer’s instructions. gRNAs were amplified with TruSeq library prep kit (Illumina) according to manufacturer’s instructions. The resulting libraries were sequenced on an Illumina Nextseq500. For each screen, we sequenced both early and late time points defined as the population of cells following 1–2 and 2–3 treatment rounds with IACS-10759, respectively. At the end of the experiment, each cell line routinely went through a minimum of 10 population doublings in control cells.

### Data processing and DrugZ analysis of CRISPR screens

Raw sequencing read files of each sample were combined into one single fastq file. Adapter sequences were cut-off using cutadapt software (DOI:10.14806/ej.17.1.200) and only sgRNA sequence of each read was kept for the next steps. Data was mapped to gRNA library using bowtie ^89^ with allowing 3 mismatches for each read and removed sequences which are aligned to more than one reference sequence. Sequence mapping rates were ∼ 40% for all samples. We created a matrix of read count from mapping results to use for further analysis using an in-house script. Then, we processed each sample using the BAGEL2 pipeline ^90^. F-measures were calculated at Bayes Factor 5, which is a harmonic mean of precision and recall at BF 5 obtained through BAGEL ‘pr’ function, to determine the quality of screens. F-measures of all screens exceeded 0.9. To identify chemo-genetic interactions between drug and gene essentiality, we utilized DrugZ ^28^, which measures synthetic essentiality and buffering interaction between genes and a drug of interest. We ran DrugZ with read count information of a non-treated sample and the matched drug-treated sample with each cell line and time points. DrugZ results at P value ≤ 0.05 was considered significant for each time point.

### Non-targeted analysis of polar metabolites by HRLC/IC-MS

To determine the relative abundance of polar metabolites in H1299 cells, extracts were prepared and analyzed by ultra-high-resolution mass spectrometry (HRMS). H1299 cells were seeded in 10cm dishes in triplicate and approximately 80% confluent at the end of the experiment. H1299 cells were treated with freshly prepared 2µM KD025, 4nM IACS-10759 or the combination of both compounds for 24-48 hours. Equivalent volumes of DMSO were used for control groups. For ICMS analysis, metabolites were extracted using ice-cold 80/20 (v/v) methanol/water with 0.1% ammonium hydroxide. Extracts were centrifuged at 17,000 *g* for 5 min at 4°C, and supernatants were transferred to clean tubes, followed by evaporation to dryness under nitrogen. Dried extracts were reconstituted in deionized water, and 10 μL was injected for analysis by ion chromatography (IC)-MS. IC mobile phase A (MPA; weak) was water, and mobile phase B (MPB; strong) was water containing 100 mM KOH. A Thermo Scientific Dionex ICS-6000+ system included a Thermo IonPac AS11 column (4 µm particle size, 250 x 2 mm) with column compartment kept at 35°C. The autosampler tray was chilled to 4°C. The mobile phase flow rate was 360 µL/min, and the gradient elution program was: 0-2 min, 1% MPB; 2-25 min, 1-40% MPB; 25-39 min, 40-100% MPB; 39-50 min, 100% MPB; 50-50.5, 100-1% MPB. The total run time was 55 min. To assist the desolvation for better sensitivity, methanol was delivered by an external pump and combined with the eluent via a low dead volume mixing tee. Data were acquired using a Thermo Orbitrap IQ-X Tribrid Mass Spectrometer under ESI negative ionization mode. For HILIC analysis, the same samples were injected by liquid chromatography (LC)-MS. LC mobile phase A (MPA) was 95/5 (v/v) water/acetonitrile containing 20mM ammonium acetate and 20mM ammonium hydroxide (pH∼9), and mobile phase B (MPB) was acetonitrile. Thermo Vanquish LC system included a Xbridge BEH Amide column (3.5 µm particle size, 100 x 4.6 mm) with column compartment kept at 30°C. The autosampler tray was chilled to 4°C. The mobile phase flow rate was 350 µL/min, and the gradient elution program was: 0-1 min, 90% MPB; 1-20 min, 90-5% MPB; 20-25 min, 5% MPB; 25-26 min, 5-90% MPB. The total run time was 30 min. Data were acquired using a Thermo Orbitrap Fusion Tribrid Mass Spectrometer under ESI positive and negative ionization modes at a resolution of 240,000. All the raw data files were imported into Thermo Trace Finder 5.1 and Compound Discoverer 3.3 software for final analysis. The relative level of metabolites was normalized by total peak intensity.

### Isotope tracer analysis of polar metabolites by HRLC/IC-MS

To determine the incorporation of glucose and glutamine carbons into glycolysis, tricarboxylic acid (TCA) cycle, pentose phosphate pathway, nucleotides and amino acids, in H1299 cells, extracts were prepared and analyzed by high-resolution mass spectrometry (HRMS). H1299 cells were seeded in 10cm dishes in triplicate and approximately 80% confluent at the end of the experiment Before treatment, H1299 cells were washed with glutamine or glucose-free medium before incubated in fresh medium containing 11.1mM ^13^C_6_-glucose or 2mM ^13^C_5_-glutamine. Freshly prepared 2µM KD025, 4nM IACS-10759 or the combination of both compounds were added immediately and incubated for 24-48 hours. Equivalent volumes of DMSO were used for control groups. At the end of the experiment, H1299 cells were quickly washed with ice-cold deionized water to remove extra medium components. Metabolites were extracted using cold 80/20 (v/v) methanol/water with 0.1% ammonium hydroxide. Samples were centrifuged at 17,000 g for 5 min at 4°C, and supernatants were transferred to clean tubes, followed by evaporation to dryness under nitrogen. Samples were reconstituted in deionized water, and 10 μL was injected for analysis by ion chromatography (IC)-MS. IC mobile phase A (MPA; weak) was water, and mobile phase B (MPB; strong) was water containing 100 mM KOH. A Thermo Scientific Dionex ICS-6000+ system included a Thermo IonPac AS11 column (4 µm particle size, 250 x 2 mm) with column compartment kept at 35°C. The autosampler tray was chilled to 4°C. The mobile phase flow rate was 360 µL/min, and the gradient elution program was: 0-2 min, 1% MPB; 2-25 min, 1-40% MPB; 25-39 min, 40-100% MPB; 39-50 min, 100% MPB; 50-50.5, 100-1% MPB. The total run time was 55 min. To assist the desolvation for better sensitivity, methanol was delivered by an external pump and combined with the eluent via a low dead volume mixing tee. Data were acquired using a Thermo Orbitrap IQ-X Tribrid Mass Spectrometer under ESI negative ionization mode. For amino acids analysis, samples were diluted in 90/10 acetonitrile/water containing 1% formic acid, then 15 μL was injected into a Thermo Vanquish liquid chromatography (LC) system containing an Imtakt Intrada Amino Acid 2.1 x 150 mm column with 3 µm particle size. Mobile phase A (MPA) was acetonitrile containing 0.1% formic acid. Mobile phase B (MPB) was 50 mM ammonium formate. The flow rate was 300 µL/min (at 35°C), and the gradient conditions were: initial 15% MPB, increased to 30% MPB at 20 min, then increased to 95% MPB at 30 min, held at 95% MPB for 10 min, returned to initial conditions and equilibrated for 10 min. The total run time was 50 min. Data were acquired using a Thermo Orbitrap Fusion Tribrid mass spectrometer under ESI positive ionization mode at a resolution of 240,000. All the raw files were imported to Thermo Trace Finder 5.1 software for final analysis. The fractional abundance of each isotopologue is calculated by the peak area of the corresponding isotopplogy normalized by the sum of all isotopoloy areas^91^. The relative abundance of each metabolite was normalized by total peak intensity.

### Sample preparation for proteomic and phosphoproteomic analysis

H1299 cells were treated with freshly prepared 2µM KD025, 4nM IACS-10759, or the combination of both compounds for 6 hours. Equivalent volumes of DMSO were used for control groups. Samples for protein analysis were prepared essentially as previously described^92,93^. Proteomes were extracted using a buffer containing 200 mM EPPS pH 8.5, 8M urea, 0.1% SDS and protease inhibitors. Following lysis, 300 µg of each proteome was reduced with 5 mM TCEP. Cysteine residues were alkylated using 10 mM iodoacetimide for 20 minutes at RT in the dark. Excess iodoacetimide was quenched with 10 mM DTT. A buffer exchange was carried out using a modified SP3 protocol^94^. Briefly, ∼3000 µg of Cytiva SpeedBead Magnetic Carboxylate Modified Particles (65152105050250 and 4515210505250), mixed at a 1:1 ratio, were added to each sample. 100% ethanol was added to each sample to achieve a final ethanol concentration of at least 50%. Samples were incubated with gentle shaking for 15 minutes. Samples were washed three times with 80% ethanol. Protein was eluted from SP3 beads using 200 mM EPPS pH 8.5 containing Lys-C (Wako, 129-02541). Samples were digested overnight at room temperature with vigorous shaking. The next morning trypsin was added to each sample and further incubated for 6 hours at 37° C. Acetonitrile was added to each sample to achieve a final concentration of ∼33%. Each sample was labelled, in the presence of SP3 beads, with ∼750 µg of TMTPro reagents (ThermoFisher Scientific). Following confirmation of satisfactory labelling (>97%), excess TMT was quenched by addition of hydroxylamine to a final concentration of 0.3%. The full volume from each sample was pooled and acetonitrile was removed by vacuum centrifugation for 1 hour. The pooled sample was acidified and peptides were de-salted using a Sep-Pak 50mg tC18 cartridge (Waters). Peptides were eluted in 70% acetonitrile, 1% formic acid and dried by vacuum centrifugation.

### Phosphopeptide enrichment and reversed-phase separation

Phosphopeptide enrichment was performed using a High-Select Fe-NTA Phosphopeptide Enrichment Kit (ThermoFisher Scientific). Following enrichment, the phosphopeptides were fractionated using a High pH Reversed-Phase Peptide Fractionation Kit (Cat. No.: 84868; Pierce). The 8 resulting fractions were compressed into 4 fractions. Dried peptides were de-salted by Stage-Tip and re-dissolved in 5% formic acid/5% acetonitrile for LC-MS/MS. The flow through from the phosphopeptide enrichment was used for total proteome profiling. For reversed-phase separation TMT labeled peptides were solubilized in 5% acetonitrile/10 mM ammonium bicarbonate, pH 8.0 and ∼300 µg of TMT labeled peptides were separated by an Agilent 300 Extend C18 column (3.5 μm particles, 4.6 mm ID and 250 mm in length). An Agilent 1260 binary pump coupled with a photodiode array (PDA) detector (Thermo Scientific) was used to separate the peptides. A 45 minute linear gradient from 10% to 40% acetonitrile in 10 mM ammonium bicarbonate pH 8.0 (flow rate of 0.6 mL/min) separated the peptide mixtures into a total of 96 fractions (36 seconds). A total of 96 Fractions were consolidated into 24 samples in a checkerboard fashion and vacuum dried to completion. Each sample was desalted via Stage Tips and re-dissolved in 5% formic acid/ 5% acetonitrile for LC-MS3 analysis.

### Proteomic LC-MS3 analysis

Proteome data were collected on an Orbitrap Eclipse mass spectrometer (ThermoFisher Scientific) coupled to a Proxeon EASY-nLC 1000 LC pump (ThermoFisher Scientific). Fractionated peptides were separated using a 120 min gradient at 525 nL/min on a 35 cm column (i.d. 100 μm, Accucore, 2.6 μm, 150 Å) packed in-house. A FAIMS device was enabled using compensation voltages set to -40, -60, and -80. MS1 data were collected in the Orbitrap (120,000 resolution; maximum injection time 50 ms; AGC 4 × 10^5^). Charge states between 2 and 5 were required for MS2 analysis, and a 120 second dynamic exclusion window was used. Top 10 MS2 scans were performed in the ion trap with CID fragmentation (isolation window 0.5 Da; Turbo; NCE 35%; maximum injection time 35 ms; AGC 1 × 10^4^). Real-time search was used to trigger MS3 scans for quantification^95^. MS3 scans were collected in the Orbitrap using a resolution of 50,000, NCE of 55%, maximum injection time of 250 ms, and AGC of 1.25 × 10^5^. The close out was set at two peptides per protein per fraction^95^.

### Phosphoproteomic LC-MS/MS analysis

Phosphorylation data were collected on an Orbitrap Eclipse mass spectrometer (ThermoFisher Scientific) coupled to a Proxeon EASY-nLC 1000 LC pump (ThermoFisher Scientific). Fractionated peptides were separated using a 120 min gradient at 525 nL/min on a 35 cm column (i.d. 100 μm, Accucore, 2.6 μm, 150 Å) packed in-house. A FAIMS device enabled during data acquisition with compensation voltages set as −40, −60, and −80 V for the first shot and −45 and −65 V for the second shot^96^. MS1 data were collected in the Orbitrap (120,000 resolution; maximum injection time set to auto; AGC 4 × 10^5^). Charge states between 2 and 5 were required for MS2 analysis, and a 120 second dynamic exclusion window was used. Cycle time was set at 1 second. MS2 scans were performed in the Orbitrap with HCD fragmentation (isolation window 0.5 Da; 50,000 resolution; NCE 36%; maximum injection time 300 ms; AGC 2 × 10^5^).

### Proteomics M/S search and analysis

Raw files were converted to mzXML, and monoisotopic peaks were re-assigned using Monocle^97^. Searches were performed using the Comet search algorithm against a human database downloaded from Uniprot in May 2021. We used a 50 ppm precursor ion tolerance, 1.0005 fragment ion tolerance, and 0.4 fragment bin offset for MS2 scans collected in the ion trap, and 0.02 fragment ion tolerance; 0.00 fragment bin offset for MS2 scans collected in the Orbitrap. TMTpro on lysine residues and peptide N-termini (+304.2071 Da) and carbamidomethylation of cysteine residues (+57.0215 Da) were set as static modifications, while oxidation of methionine residues (+15.9949 Da) was set as a variable modification. For phosphorylated peptide analysis, +79.9663 Da was set as a variable modification on serine, threonine, and tyrosine residues.

Each run was filtered separately to 1% False Discovery Rate (FDR) on the peptide-spectrum match (PSM) level. Then proteins were filtered to the target 1% FDR level across the entire combined data set. Phosphorylation site localization was determined using the AScorePro algorithm^98^. For reporter ion quantification, a 0.003 Da window around the theoretical m/z of each reporter ion was scanned, and the most intense m/z was used. Reporter ion intensities were adjusted to correct for isotopic impurities of the different TMTpro reagents according to manufacturer specifications. Peptides were filtered to include only those with a summed signal-to-noise (SN) ≥ 160 across all TMT channels. An extra filter of an isolation specificity (“isolation purity”) of at least 0.5 in the MS1 isolation window was applied for the phosphorylated peptide analysis. For each protein or phosphorylation site, the filtered peptide TMTpro SN values were summed to generate protein or phosphorylation site quantification values. The signal-to-noise (S/N) measurements of peptides assigned to each protein were summed (for a given protein). These values were normalized so that the sum of the signal for all proteins in each channel was equivalent thereby accounting for equal protein loading. The resulting normalization factors were used to normalize the phosphorylation sites, again account for equal protein loading.

### Proteome and phosphoproteome enrichment analysis

Enrichment analysis using GOBP, KEGG, and Reactome Gene sets were performed using Metascape^99^. Overrepresentation was determined by one-tailed hypergeometric testing and P values adjusted by Benjamini-Hochberg with a minimum background set to three proteins or phosphoproteins. Enrichment was determined by the relative proportion of representation in the foreground to the proportion in the background. Protein-protein interaction (PPI) networks were generated using STRING v12^48^.

### Kinase library analysis and network construction

Differentially-regulated phosphosites (log_2_ fold change ≥ 1.5, Adj. P value ≤ 0.05) were analyzed for kinase-motif enrichment as reported previously ^54^ using the publically available software tools on PhosphositePlus (https://www.phosphosite.org/kinaseLibraryAction). Relative kinase activities and kinase-substrate interaction scores were determined as previously described ^66^, with slight modifications. Briefly, relative kinase activities were determined by the product min-max normalized adjusted P values and log_2_ enrichment factor for the most significant enrichment direction for each kinase. Individual phosphosites were scored for the 303 human kinases as described previously ^54^, and top kinases for each phosphosite was filtered for the top 25 scoring kinases by percentile.

As described ^66^, the first kinase-phosphosite link was assigned a score by taking the product of the kinase-phosphosite percentile and absolute value of the relative kinase activities for each cell line and comparison and defined as the kinase library score (KLS). Previously reported kinase-phosphosite evidence was added to the kinase library score by integrating the kinase-substrate dataset available from PhosphoSitePlus ^52^. A score of 0.5 was added to the KLS for each level of evidence, in vitro and in vivo, reported in the kinase-substrate set. The PSP score was added to the KLS. Next, kinase-substrate interactions were annotated from BioGRID^65^. Data from BioGRID (v4.4.213) was filtered for human interactions and the score was scaled by the number of references that reported the kinase-substrate interaction [score = n/(n+1), where n is the number of references reporting the interaction]. The BioGRID score for each kinase-substrate interaction was added to the KLS.

The final interaction score was determined by taking the product of the final KLS and the relative phosphosite differential regulation matched for each treatment comparison in H1299 cells. The relative phosphosite differential regulation was calculated by the product of the min-max normalized adjusted P values and log_2_ fold-change for each treatment comparison. Each differentially expressed phosphosite was filtered for the top three kinase interaction scores. This score represents the likelihood that a kinase is responsible for regulating that individual phosphosite. The final kinase interaction score for kinase-substrate interactions was defined as the sum of interaction scores for each phosphosite. Pathway interaction scores was defined as the product of the sum of all interaction scores for each phosphoprotein and the percentage of phosphoproteins with interaction scores in each pathway. All calculations were performed in Excel and networks were created in CytoScape (v3.10.2) using the final interaction score for each kinase-substrate as links and each kinase and substrate as nodes. Nodes were annotated by databases as described above.

### DepMap analysis

CRISPR Chronos dependency data was downloaded from the DepMap Public 24Q2 dataset (https://depmap.org/portal/). Foreground for overrepresentation analyses using Reactome gene sets of co-regulated phosphoproteins in the combination was determined for proteins with Chronos dependency scores ≤–0.5 for essential proteins and ≥ -0.5 for non-essential proteins in the H1299 cells. Overrepresentation was determined by one-tailed hypergeometric testing using the background of all detected phosphoproteins.

### Proliferation and colony formation assays

Single-cell suspensions of all cell lines were counted and plated into 6-well plates at a density of 0.5–4 × 10^3^ cells per well. Cells were cultured in a medium containing the indicated drugs for 10–14 days (refreshed every 4 days). At the endpoints, cells were fixed with 10% Formalin and stained with crystal violet (0.1%w/v in water) and photographed. All experiments were performed in triplicates. IncuCyte S3 (Essen Biosciences, Ann Arbor, MI, USA) was used to measure proliferation of cell lines. For IncuCyte S3 experiments, A549, H1299 and H2023cells were suspended in fresh growth media, plated in 96-well plates at 1-2.5 x 10^4^ cells/well, and allowed to settle overnight at 37 °C. On the following day, drugs were added, and confluency was assessed every 4 hours until the end of the experiment. Confluency was determined at each time point using the IncuCyte S3 analysis software.

### Mitochondrial respiration and glycolysis measurements

OCR and ECAR experiments were performed using the XF-96 Analyzer apparatus from Seahorse Bioscience. H1299 cells were plated (10,000 cells per well) in at least quadruplicate for each condition the day before the experiment. The next day, medium was replaced with reconstituted DMEM with 25 mM glucose and 2 mM glutamine (no sodium bicarbonate) adjusted to pH∼7.4 and incubated for 30 min at 37% in a CO2-free incubator. For the mitochondrial stress test (Seahorse 101706–100), oligomycin, FCCP, antimycin and rotenone were injected to a final concentration of 2 μM, 0.5 μM and 4 μM, respectively. For the glycolysis stress test (Seahorse 102194–100), glucose, oligomycin and 2-deoxyglucose were injected to a final concentration of 10 mM, 2 μM and 100 mM, respectively. OCR and ECAR were normalized to cell number as determined by CellTiter-Glo analysis at the end of the experiments. OCR and ECAR values were used to compute basal respiration, spare capacity, proton leak, ATP production, glycolysis and glycolytic capacity. Calculations were done by ExcelMacro Report Generator Version 3.0.3 provided by Seahorse Biosciences and two-sided Student’s t-test computed using GraphPad taking the mean and s.d values provided by Report Generator 3.0.3 and n=8 (4 values from 2 independent experiments). Formulas are described here. Basal Respiration= (Last rate measurement before first injection)-(Non-mitochondrial Respiration Rate), Spare Respiratory Capacity = (Maximal Respiration)-(Basal Respiration).Proton leak= (minimum rate measurement after Oligomycin injection)-(non-mitochondrial respiration). ATP production = (Last rate measurement before oligomycin injection)-(Minimum rate measurement after Oligomycin injection). For Glycolysis = (Maximum rate measurement before oligomycin injection)(Last rate measurement before glucose injection) and for glycolytic capacity = Maximum rate measurement after oligomycin injection)-(Last rate measurement before glucose injection).

### Immunoblotting

Cells were washed twice in ice-cold PBS, and proteins extracted in RIPA buffer with protease and phosphatase inhibitors for 30 min. Lysates were then collected and centrifuged at 14,000 rpm for 15 min at 4°C. Protein concentrations were measured using the BCA Protein Assay Kit (Thermo scientific). Lysates were mixed with 4X laemmli buffer, boiled at 95°C for 5 minutes and processed with Mini-Protean Gel Electrophoresis Systems (Bio-Rad) using Bis-Tris 4–20% gradient pre-cast gels (Biorad).

### Human lung xenografts anti-tumor efficacy studies

All animal experiments were performed according to standards outlined by following internationally recognized guidelines on animal welfare. All animal procedures were approved by the Institutional Animal Care Committee according to guidelines of the MDACC. To establish xenograft models, H1299 cells (2×10^6^ cells/mouse) and A549 cells (1 x10^6^ cells/mouse) were trypsinized and resuspended in 1X PBS, mixed with a 1:1 mix of Matrigel in a final volume of 200 μl, and injected subcutaneously into the flanks of nude female mice (Jackson Labs) at 6-8 weeks of age. Tumor volume was monitored visually and by palpation weekly. Mice were randomized to vehicle and treatment arms once the average tumor volume reached 100 mm^3^. KD025 (300 mg kg^-1^), and IACS-10759 (1 mg kg^-1^) was prepared in 0.5% methycellulose (MC). Briefly, drugs were dissolved in 0.5% MC for 30 minutes on ice followed by to five rounds of sonication; each pulsed for 15 seconds with a gap of 2 minutes, while on ice. Drugs were incubated at 4°C overnight while stirring. Experimental mice were treated with solutions of 0.5% MC, KD025, or IACS-10759, or both compounds by oral gavage once daily until the end of the experiment. Tumor volume and mouse body weight were measured two or three times weekly. Tumor diameter and volume were calculated based on caliper measurements of tumor length and height using the formula tumor volume = (length × width^2^)/2. The maximal tumor size permitted by our Institutional Animal Care and Use Committee is 2500 mm^3^, which was not exceeded in our experiments.

### Statistics and reproducibility

GraphPad Prism 8 software was used to generate graphs and statistical analyses. Statistical significance was determined by one-way ANOVA or unpaired Student’s t-test. Methods for statistical tests, the exact value of n, and definition of error bars were indicated in figure legends. All experiments have been reproduced in at least two independent experiments unless otherwise specified in the figure legends. All immunoblots and images shown are representatives of these independent experiments.

