## Supplementary Information for "Therapeutic modulation of ROCK overcomes metabolic adaptation of cancer cells to OXPHOS inhibition and drives synergistic anti-tumor activity"

### Supplementary Figure Legends

**Supplementary Figure 1. Analysis of CRISPR-Cas9 FDA-ome library screens in the presence of IACS-10759 in *SMARCA4*-mutant lung cancer cells. (A)** Box plots of $log_2$-transformed sgRNA normalized read counts from A549, H1299, and H2023 cell lines prior to drug treatment (baseline), at early timepoints, and at late timepoints with DMSO or 4nM IACS-10759 treatment. (**B**) Pearson correlation of all genes across all treatment groups in A549, H1299, and H2023 cell lines. Data indicate (n=2) independent experiements. (**C**) Precision-recall curves of BAGEL results for A549, H1299, and H2023 cell lines using the FDAome library. (**D**) Mean fold change distributions of core essential genes or non-essential genes included in FDA-ome library in A549, H1299, and H2023 cell lines at early and late time points, where fold change is calculated as $log_2$ (normalized read counts from early or late time points) $-log_2$ (normalized T0 read counts).

**Supplementary Figure 2. Chemo-genomic analysis of FDA-ome CRISPR hits** (**A**-**B**) Scatterplot of NormZ scores in A549 (**A**) and H2023 (**B**) cell lines similar to as shown in Figure.1B. (**C-E**) Rug plots showing NormZ score distributions of individual sgRNAs for all FDA-ome library genes in A549 (**C**) , H1299 (**D**) , and H2023 (**E**) cell lines at early and late time points following 4nM IACS-10759 treatment. In each plot, red lines show individual sgRNAs targeting the indicated genes and grey lines show all other genes in FDA-ome library.

**Supplementary Figure 3. Combinatorial drug screening of FDA-ome library targets with IACS-10759 using FDA approved and clinical grade therapeutics.** (**A**-**E**) Treatment of three *SMARCA4*-mutant cell lines with a 3 × 6 matrix titration using different concentrations of IACS-10759 and either 6-AN (**A**), KD025 (**B**), Palbociclib (**C**), Amlexanox (**D**) or CB-839 (**E**). Heatmaps of each cell line depict percent growth inhibition across all concentrations for indicated therapeutic. Synergy scores were determined using the Bliss independence model. A synergy score above 10 is considered a synergistic drug interaction. A score between -10 to 10 is considered additive. A score below -10 is considered antagonistic.

**Supplementary Figure 4. KD025 is highly synergistic with biguanide class of OXPHOS inhibitors.** (**A**-**B**) Dose-response curves of A549 (**A**) and H2023 (**B**) cell lines treated with increasing concentrations of KD025 in the presence or absence of IACS-10759. Cell growth (measured as % Confluency) was assayed 5 days after drug exposure. Data are presented as mean +/- SEM of three independent experiments. (**C-D**) Clonogenic growth assays of A549 (**C**) and H2023 (**D**) cell lines cultured in the presence of KD025 and IACS-10759 alone or in combination for 12 days. Surviving cells after the treatment were fixed and visualized by crystal violet staining. Representative images of three independent experiments are shown. (**E-F**) Dose-response curves of H1299 cells treated with increasing concentrations of KD025 in the presence or absence of Metformin (**E**) or presence or absence of Phenformin (**F**). Cell growth (measured as % Confluency) was assayed 5 days after drug exposure. Data are presented as mean +/- SEM of three independent experiments. (**G-H**) Clonogenic growth assays of H1299 cells cultured in the presence of KD025 and Metformin (**G**) or KD025 and Phenformin (**H**) alone or in combination for 12 days. Surviving cells after treatment were fixed and visualized by crystal violet staining. Representative images of three independent experiments are shown. (**I**-**K**) Treatment of H1299 cells with a 3 × 6 matrix titration using different concentrations of KD025 with either Metformin (**I**), Phenformin (**J**), or IM156 (**K**). Heatmaps of each combination depict percent growth inhibition across all concentrations for indicated therapeutic. Synergy scores were determined using the Bliss independence model. A synergy score above 10 is considered a synergistic drug interaction. A score between -10 to 10 is considered additive. A score below -10 is considered antagonistic.

**Supplementary Figure 5: Genetic validation of KD025 and IACS-10759 combination in H1299 cells. (A)** Immunoblot showing shRNA knockodown of both ROCK1 and ROCK2 in H1299 cells after dox addition (1μg/ml). **(B)** Clonogenic growth assay of H1299 cells stably transduced with ROCK1/2 shRNA and cultured in the presence or absence of doxycycline (1μg/ml). Cells were treated with IACS-10759 for 12 days. Surviving cells after treatment were fixed and visualized by crystal violet staining. Representative images of three independent experiments are shown.

**Supplementary Figure 6. Metabolomic analysis of KD025 and IACS-10759 combination in H1299 lung cancer cells. (A)** Heatmap and unsupervised clustering showing $log_2$ fold change in abundance changes in all quantitated metabolites following DMSO, 2μM KD025, 4nM IACS-10759 or the combined treatment for 24- and 48 hours. Yellow indicates up-regulated metabolites and blue indicates down-regulated metabolites. White indicates no change. **(B)** Principal component analysis (PCA) of all quantitated metabolites in H1299 cells following treatment with DMSO, IACS-10759, KD025, or the combination for 24 hours (top panel) and 48 hours (bottom panel) treatment. **(C)** Heatmap showing enriched metabolic pathways (FDR $<$ 0.05) from significantly different up- and down-regulated metabolite abundances ($log_2$ FC >1.5, P value < 0.05) by MSEA. The color is scaled according to significance (FDR), with blue representing non-significant. (**D**) Volcano plots of $log_2$ fold change metabolite abundance of KD025, IACS-10759, or the combination following 24- or 48-hours treatment. Metabolite abundance ($log_2$ FC (treatment/DMSO) > 1.5, P value < 0.05) were determined as significantly different as indicated by dotted lines. Red dots indicate glycolysis, TCA cycle, and PPP metabolites that were significantly up-regulated. Blue dots indicate glycolysis, TCA cycle, and PPP metabolites that were significantly down-regulated.

**Supplementary Figure 7. Depletion of ATP, NADH, and NADPH cellular pools in the IACS-10759 and KD025 combination. (A)** Isotope tracing diagram for $^{13}C_6$-glucose through glycolysis and into TCA cycle metabolic pathways. Solid red circle, $^{13}C$; open circle, $^{12}C$. (**B**) Isotope tracing diagram for $^{13}C_5$-glutamine through TCA cycle metabolic pathway. Solid red circle, $^{13}C$; open circle, $^{12}C$. (**C**) Relative ratios of NADPH/NADP+ (left) and NADH/NAD+ (right) following treatment with IACS-10759, KD025, or the combination for 48 hours. (**D**) Relative abundance of AMP (left) and ATP (right) following treatment with IACS-10759, KD025, or the combination for 48 hours. (E) Relative ratio of ATP/AMP following treatment with IACS-10759, KD025, or the combination for 48 hours.

**Supplementary Figure 8. Proteomic and phosphoproteomic analysis of KD025 and IACS-10759 combination in H1299 lung cancer cells. (A-B)** Quantitation of proteome (**A**) and phosphoproteome (**B**) across all treatment groups by TMT mass spectrometry. "Changed by" refers to $log_2$ fold change (FC) of proteins or phosphoproteins ($log_2$ FC $\geq$ 1.5, P value 0.05) compared to DMSO. (n =4) biological replicates (**C**) PCA plots of all quantitated proteins (left) or phosphoproteins (right) in H1299 cells following treatment with DMSO, 4nM IACS-10759, 2μM KD025, or the combination for 6 hours. Data indicate independent experiments(n=4) (**D**) Pearson correlation of all quantified proteins (left) or phosphoproteins (right) across all treatment groups. Data indicate four independent experiments (n=4).

**Supplementary Figure 9. Proteomic analysis of top enriched pathways and protein-protein interaction networks from combined treatment of IACS-10759 and KD025.** (**A**) Top Reactome, KEGG, GOBP pathway gene sets significantly enriched (P < 0.5) in down-regulated proteins. (**B**) Top Reactome, KEGG, GOBP pathway gene sets significantly enriched (P< 0.05) in up-regulated proteins. (**C**) PPI network of significantly down-regulated proteins from combined KD025 and IACS-10759 treatment. (**D**) PPI network of significantly up-regulated proteins from combined KD025 and IACS-10759 treatment. PPI, Protein-protein interaction; Top three enriched terms for each PPI cluster are indicated using GOBP, KEGG, and Reactome pathway gene sets. PPI, Protein-protein interaction.

**Supplementary Figure 10. Phosphoproteomic analysis of single agent IACS-10759 and KD025 treatment.** **(A)** Top Reactome, KEGG, GOBP pathway gene sets significantly enriched in up-regulated proteins of IACS-10759-treated cells. (**B**) Top Reactome, KEGG, GOBP pathway gene sets significantly enriched in down-regulated proteins of KD025-treated cells. (**C**) Top Reactome, KEGG, GOBP pathway gene sets significantly enriched in up-regulated proteins of KD025-treated cells. (**D**) PPI network with differentially expressed phosphoproteins either up-regulated by IACS-10769 (left), down-regulated by KD025 (middle), or up-regulated by KD025 (right). Top three enriched terms for each PPI cluster are indicated using GOBP, KEGG, and Reactome pathway gene sets. PPI, Protein-protein interaction.

**Supplementary Figure 11. Top protein-protein interaction networks from combined treatment of IACS-10759 and KD025.** (**A-B**) PPI network with differentially expressed phosphoproteins either up-regulated (**A**) or down-regulated (**B**) from combined treatment of KD025 and IACS-10759. Color scale represent the different MCODE clusters identified. PPI, Protein-protein interaction; MCODE, Molecular Complex Detection. (**C**) Top 3 down-regulated PPI clusters identified using MCODE with top three enriched terms for each cluster indicated from GOBP, KEGG, and Reactome gene sets. (**D**) Top 3 up-regulated PPI clusters identified using

MCODE with top three enriched terms for each cluster indicated from GOBP, KEGG, and Reactome gene sets.

**Supplementary Figure 12. Relative kinase activities and clustering of 303 Ser/Thr kinases on the basis of their motif class following IACS-10759, KD025, or combination treatment for 6 hours.** (**A**) Basophilic kinases subcategorized into 11 motif classes. (**B**) Proline-directed kinases subcategorized into 5 motif classes. (**C**) Acidophilic kinases subcategorized into 8 motif classes. (**D**) Unique kinases subcategorized into 13 motif classes.

**Supplementary Figure 13. Characterization of kinase regulation in top Reactome gene sets of co-regulated phosphoproteins (A)** Immunoblot analysis of p-AMPK T172 and AMPK in H1299 cells following a 6-48 hr timecourse with 4nM IACS-10759, 2μM KD025, or the combination. β-actin serves as a loading control. (**B**) Volcano plots showing kinase-motif enrichment of three phosphoprotein subsets in the combination, down-regulated phosphoproteins, co-regulated phosphoproteins, and up-regulated phosphoproteins. The enrichments in these plots were determined using one-sided exact Fisher's tests and corrected for multiple hypotheses using the Benjamini–Hochberg method. (**C**) Motif logo analysis of 37 up-regulated phosphosites with AMPK motif. (**D**) Top kinase interaction scores for pathways in Reactome gene sets co-regulated phosphoproteins. Kinase-pathway interactions were constructed using kinase-motif specificities [54], kinase-substrate specificities from PhosphoSitePlus [52], and kinase-substrate interactions from BioGRID[65] for all phosphoproteins annotated for each pathway. (**E**) Top significantly enriched Reactome gene sets in non-essential co-regulated phosphoproteins. (**F**) Top significantly enriched Reactome gene sets in essential co-regulated phosphoproteins.

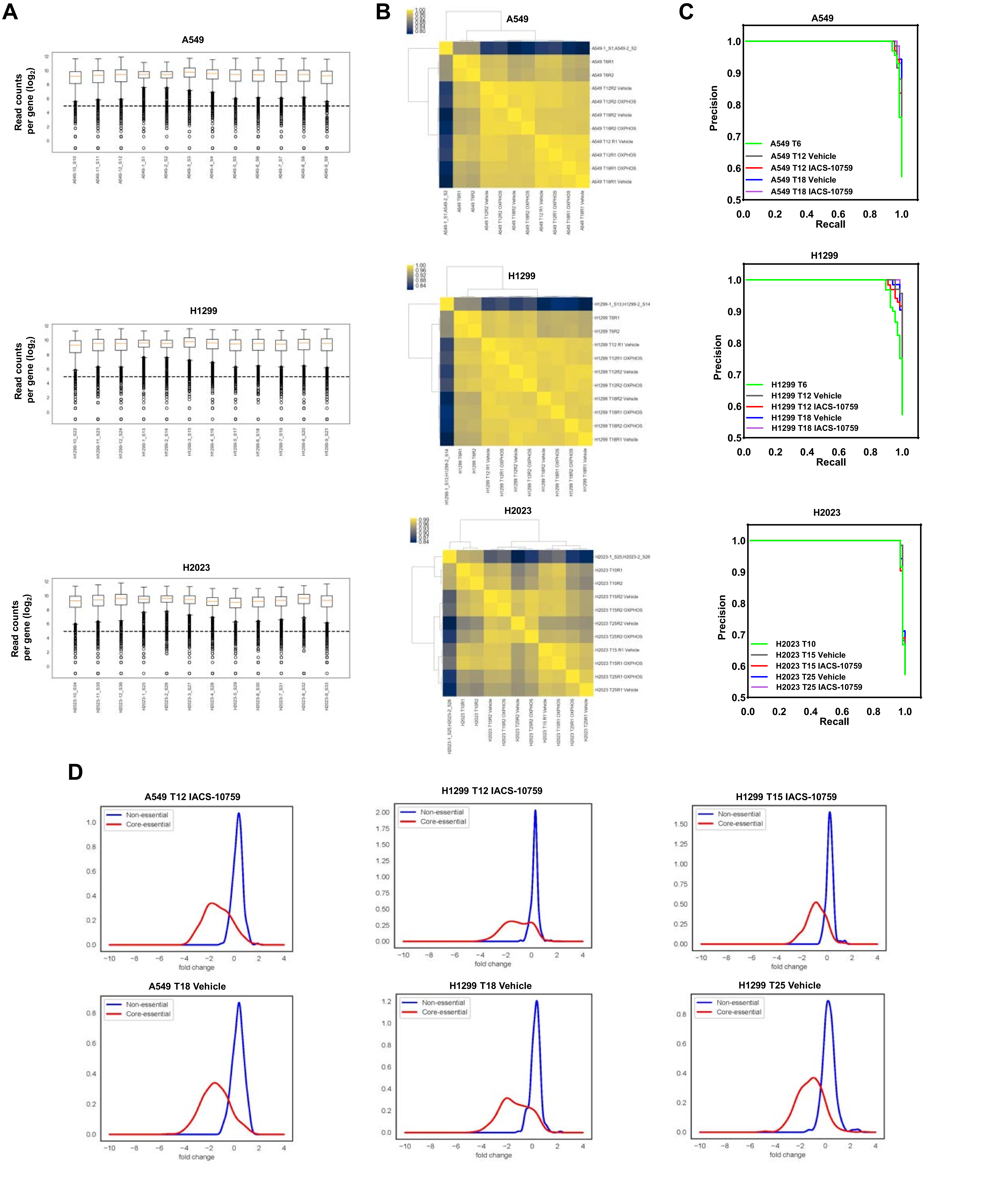

Supplementary Figure 1

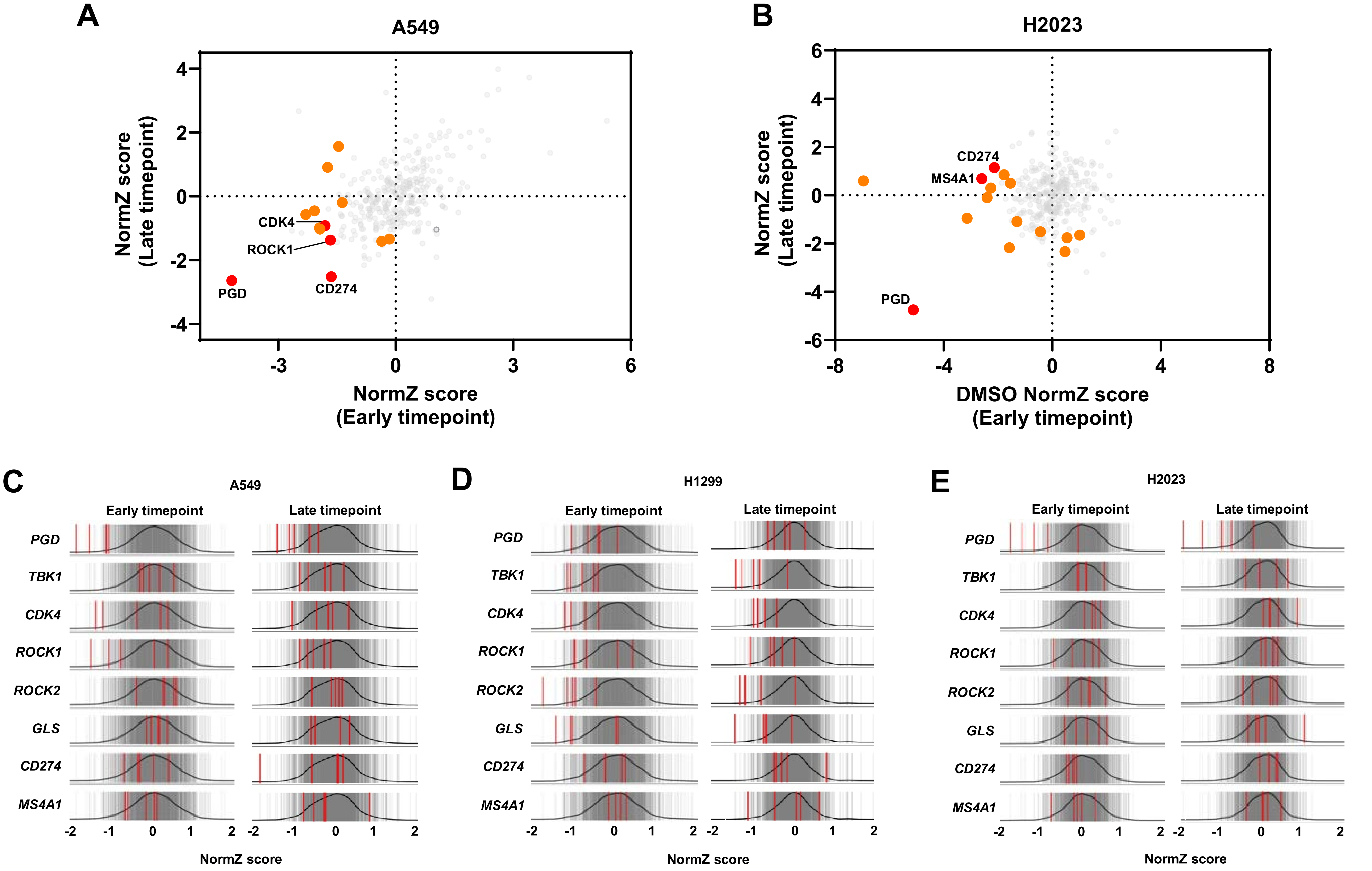

Supplementary Figure 2

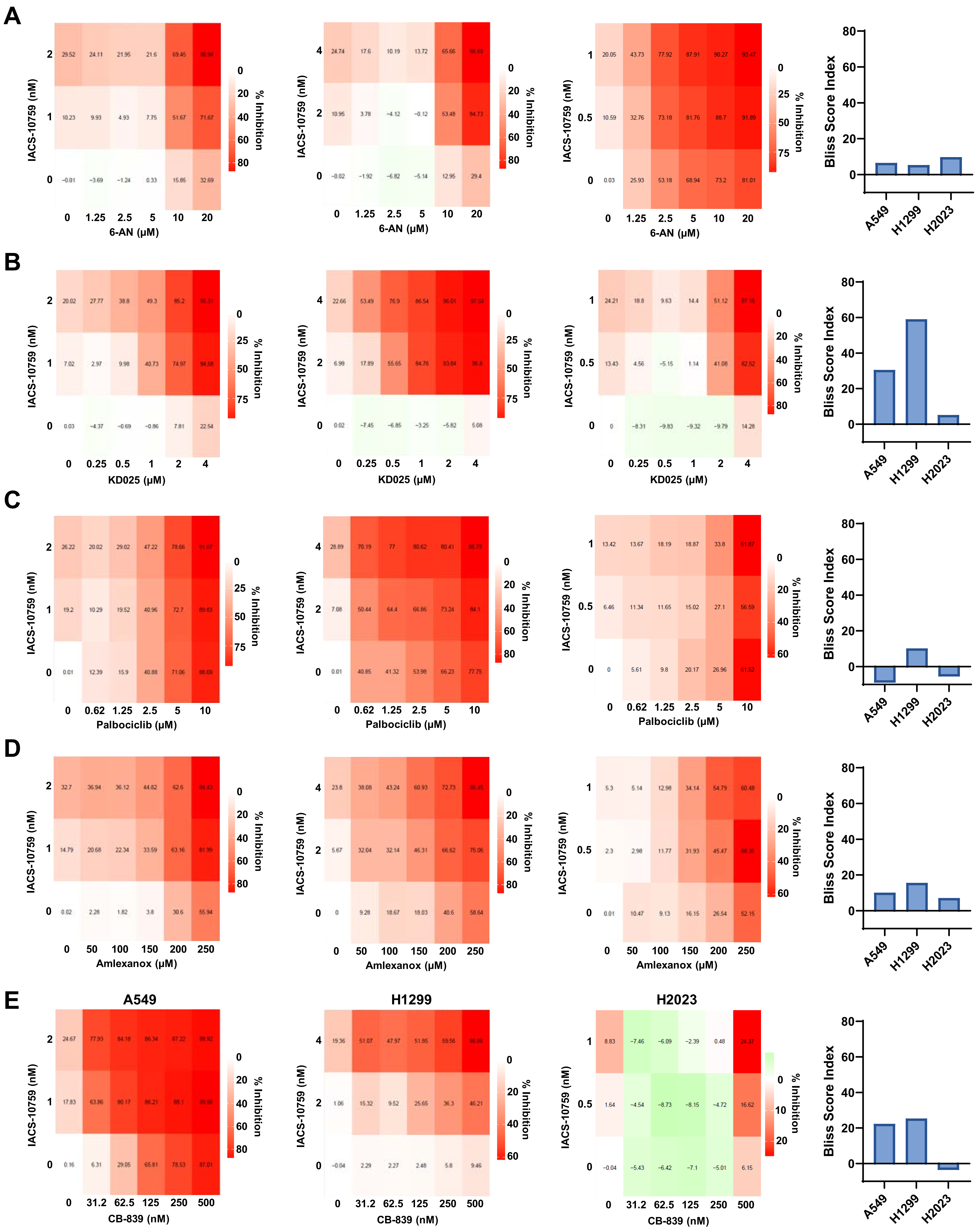

Supplementary Figure 3

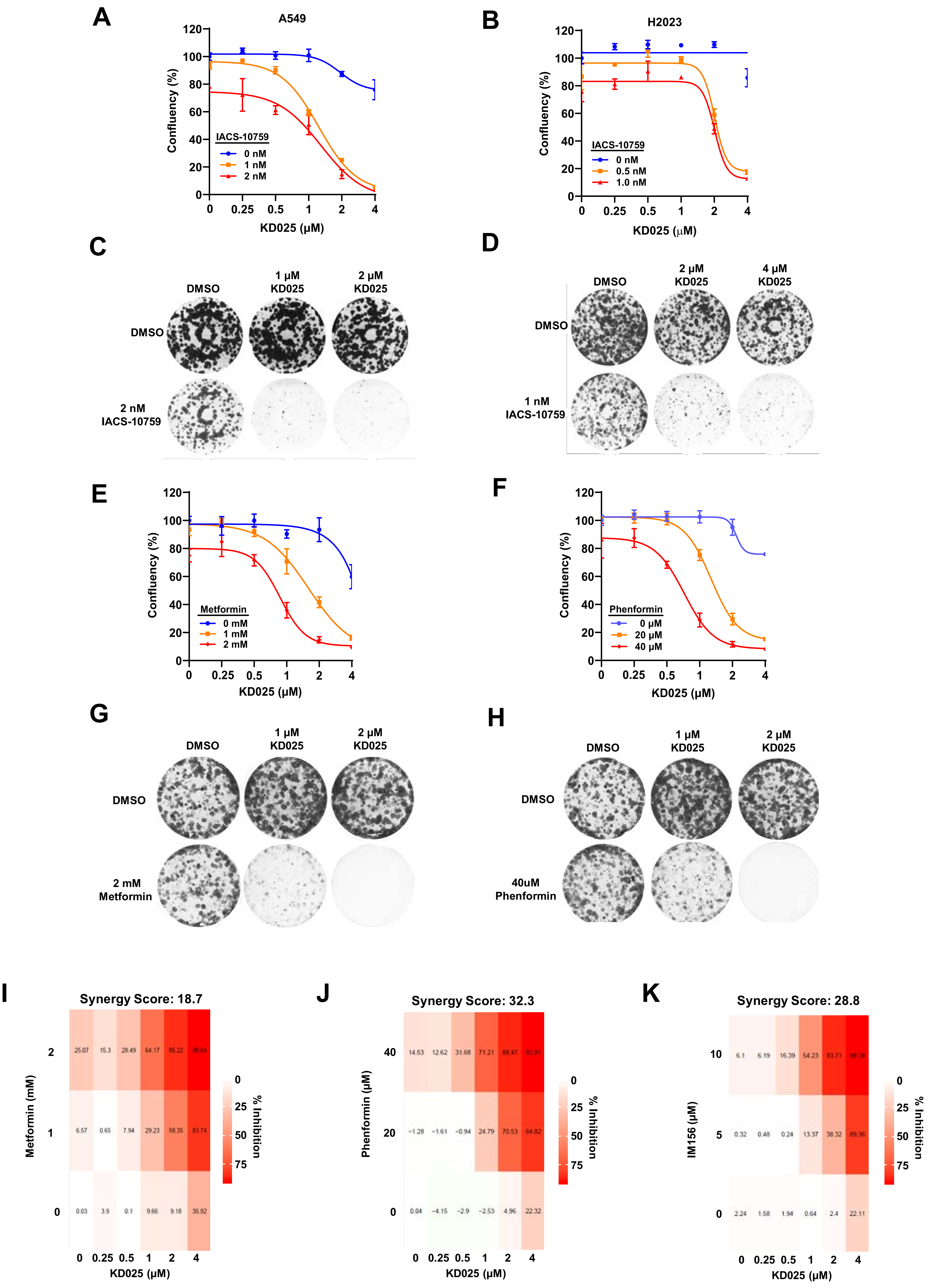

**Supplementary Figure 4**

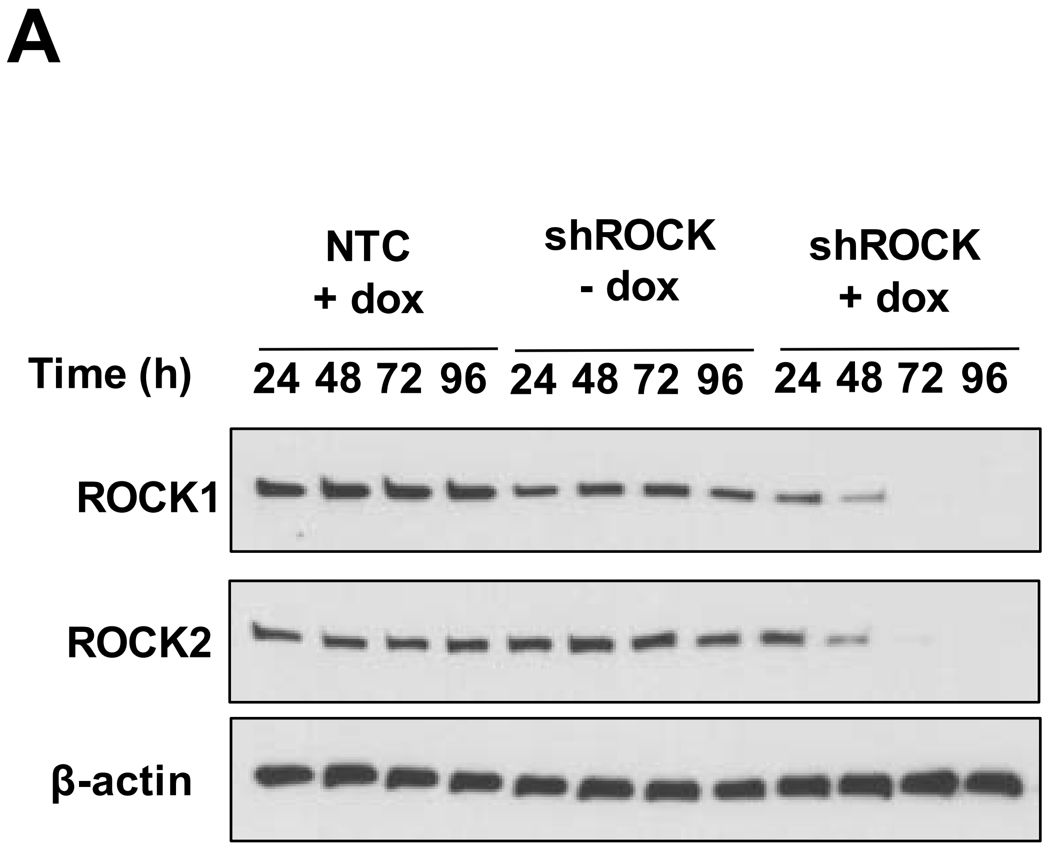

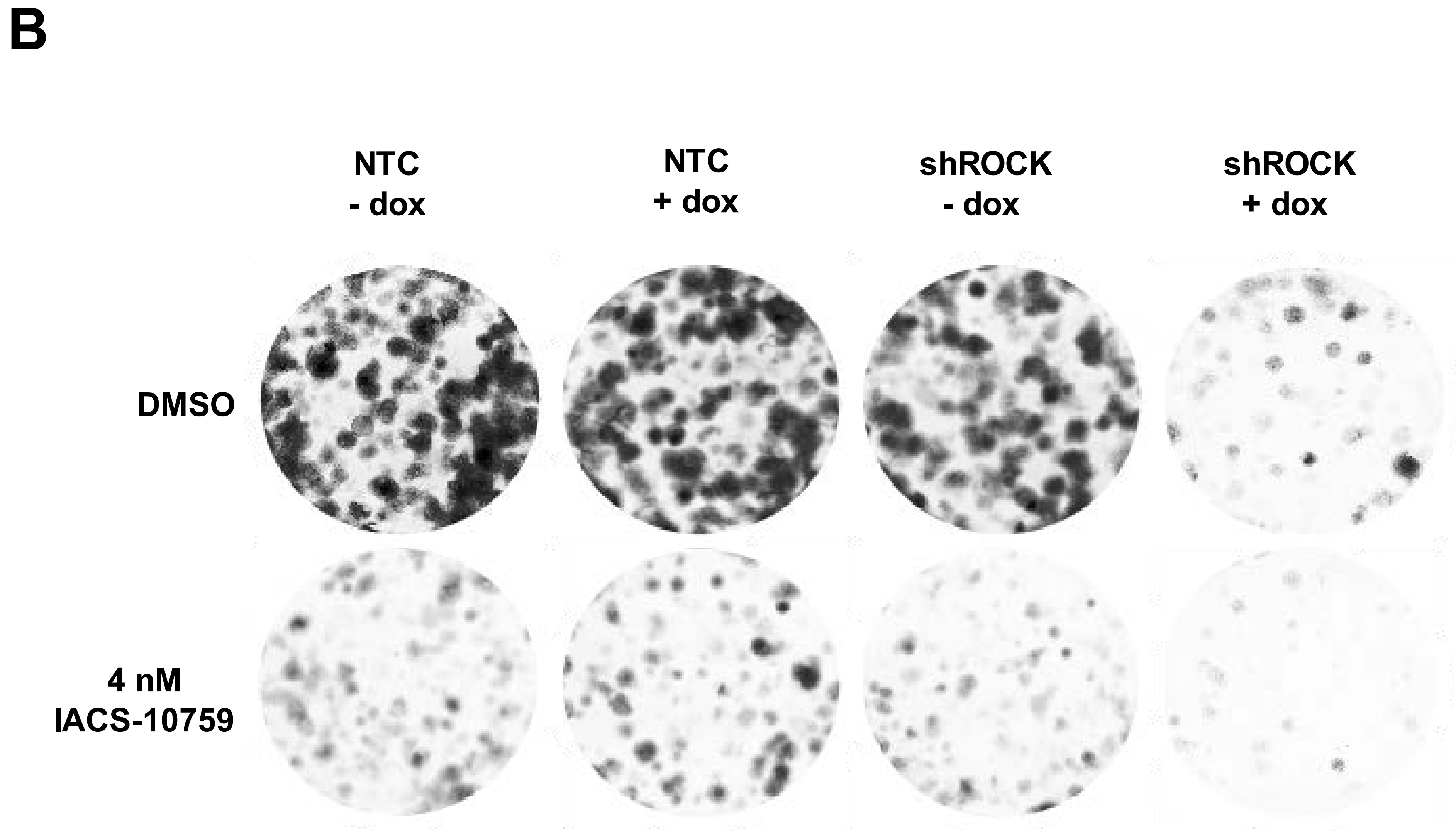

Supplementary Figure 5

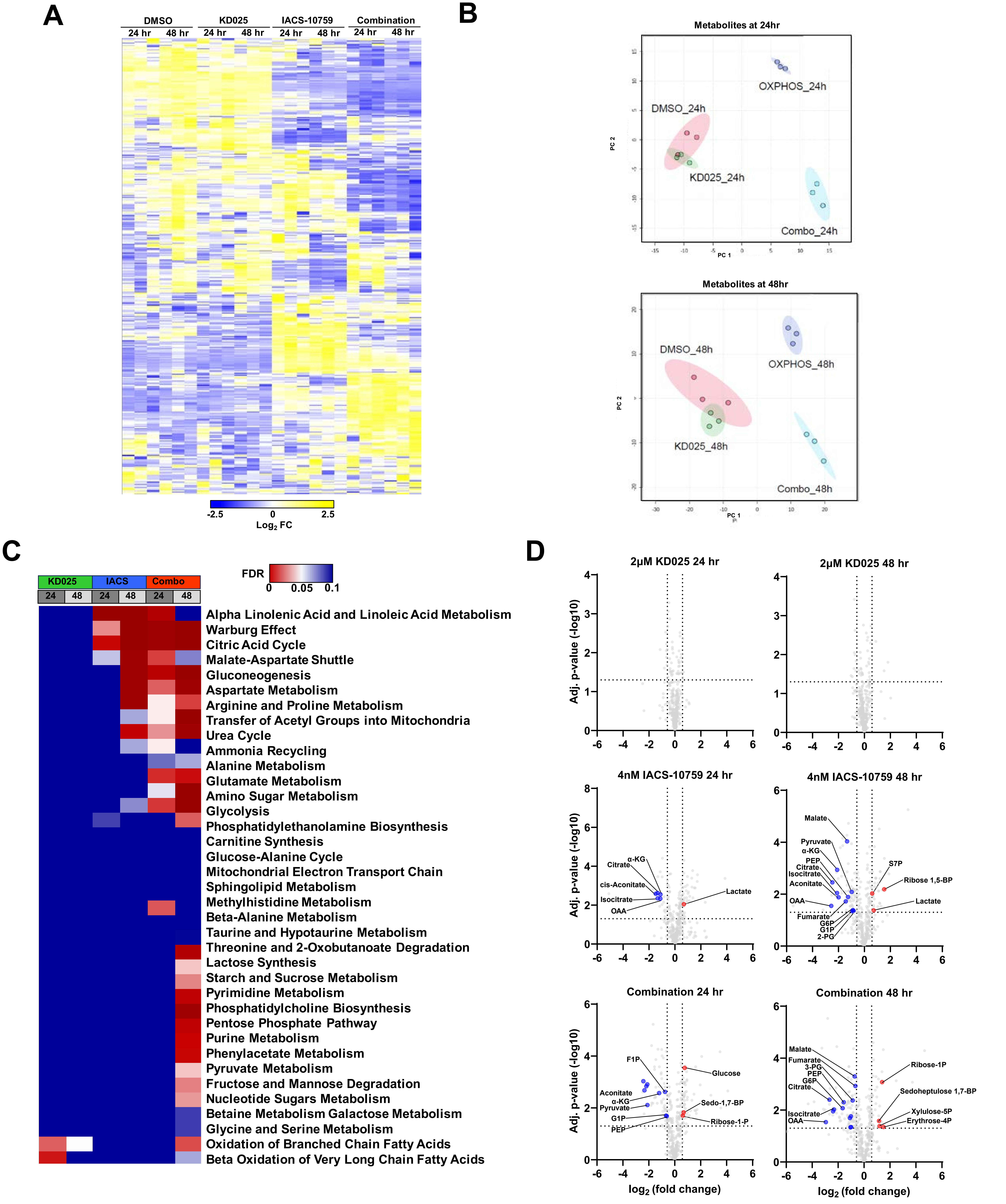

Supplementary Figure 6

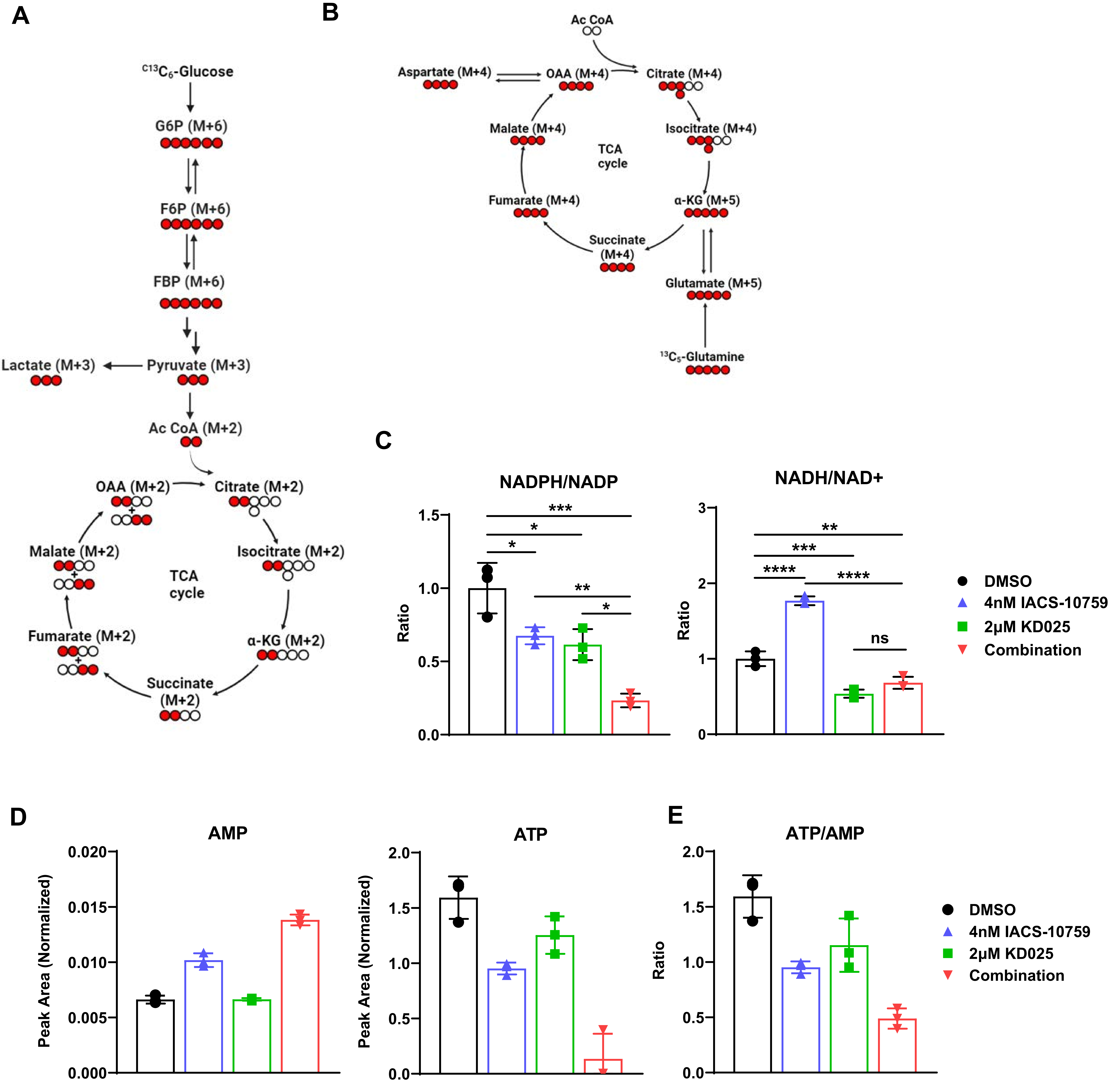

Supplementary Figure 7

**A**

| Proteome | |
|---|---|
| 9,050 | Total |
| 1 | Changed by IACS-10759 |
| 0 | Changed by KD025 |
| 229 | Changed by Combination |

**B**

Phosphoproteome

| Quantified | Changed by IACS-10759 | Changed by KD025 | Changed by Combination | |
|---|---|---|---|---|
| 55,376 | 87 | 296 | 9,927 | Phosphopeptides |
| 17,539 | 27 | 94 | 3,733 | Phosphosites |
| 4,474 | 22 | 87 | 1,822 | Phosphoproteins |

**C**

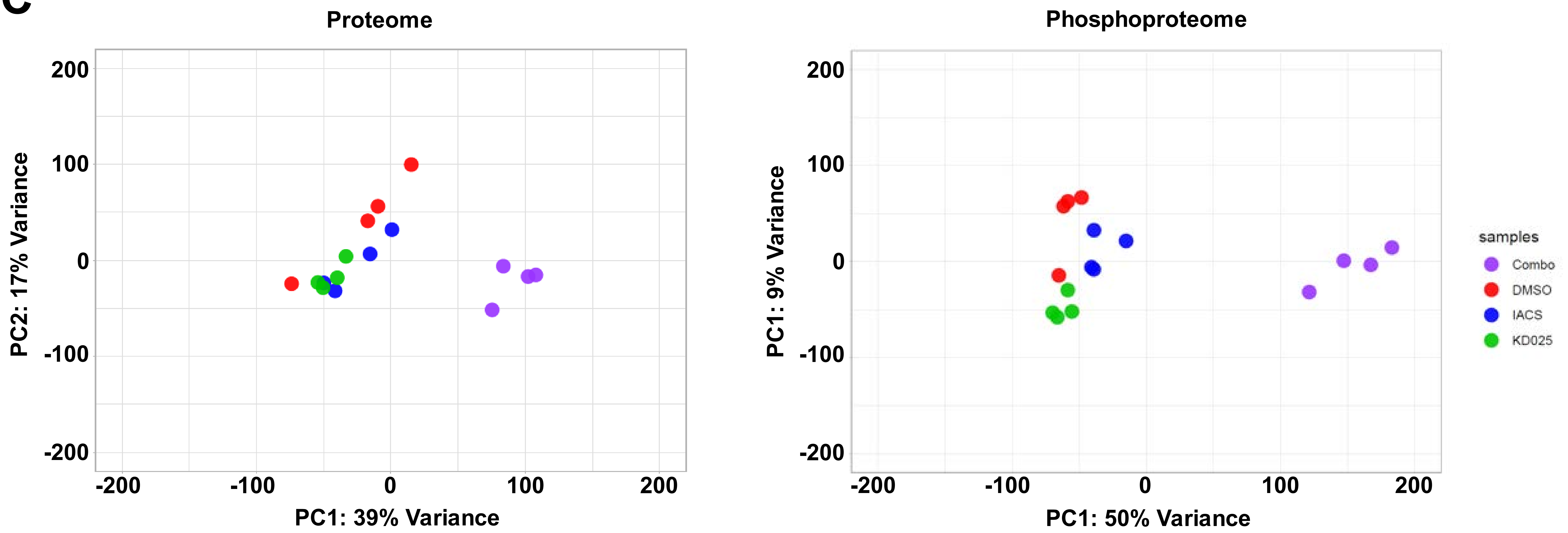

**D**

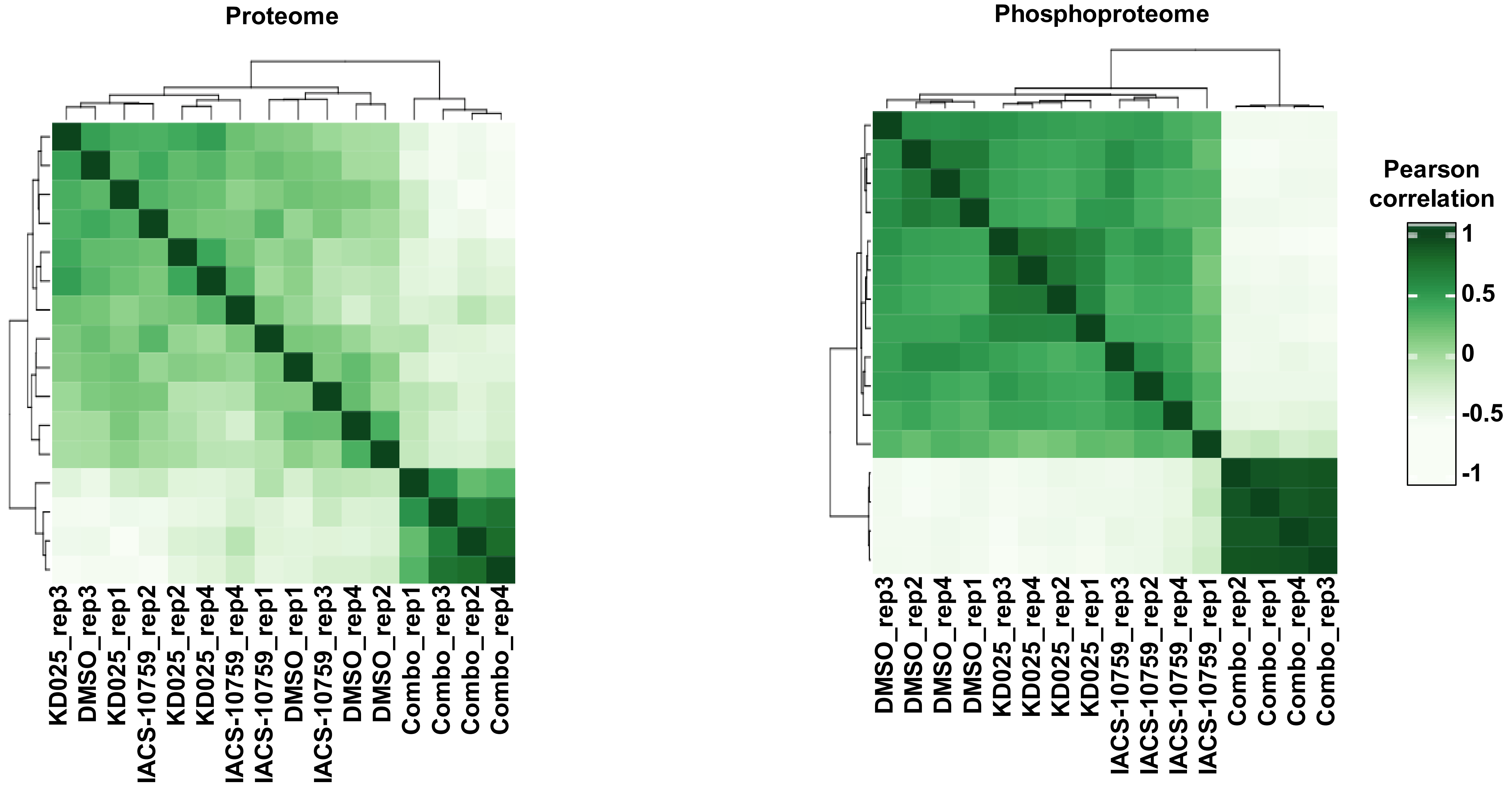

**Supplementary Figure 8**

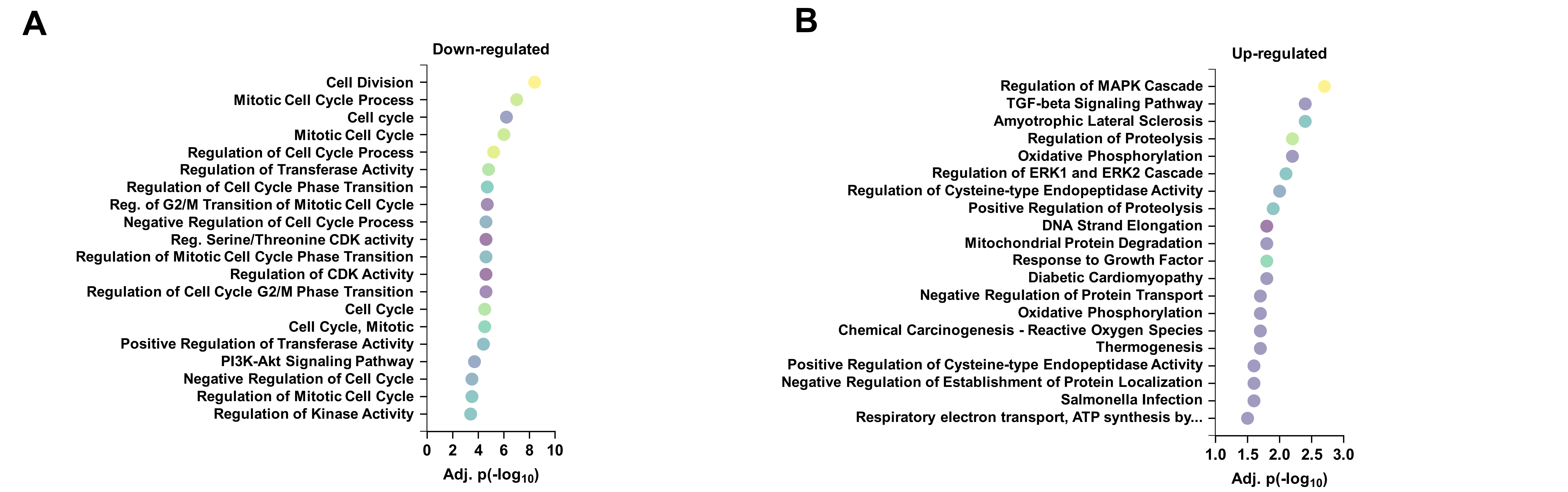

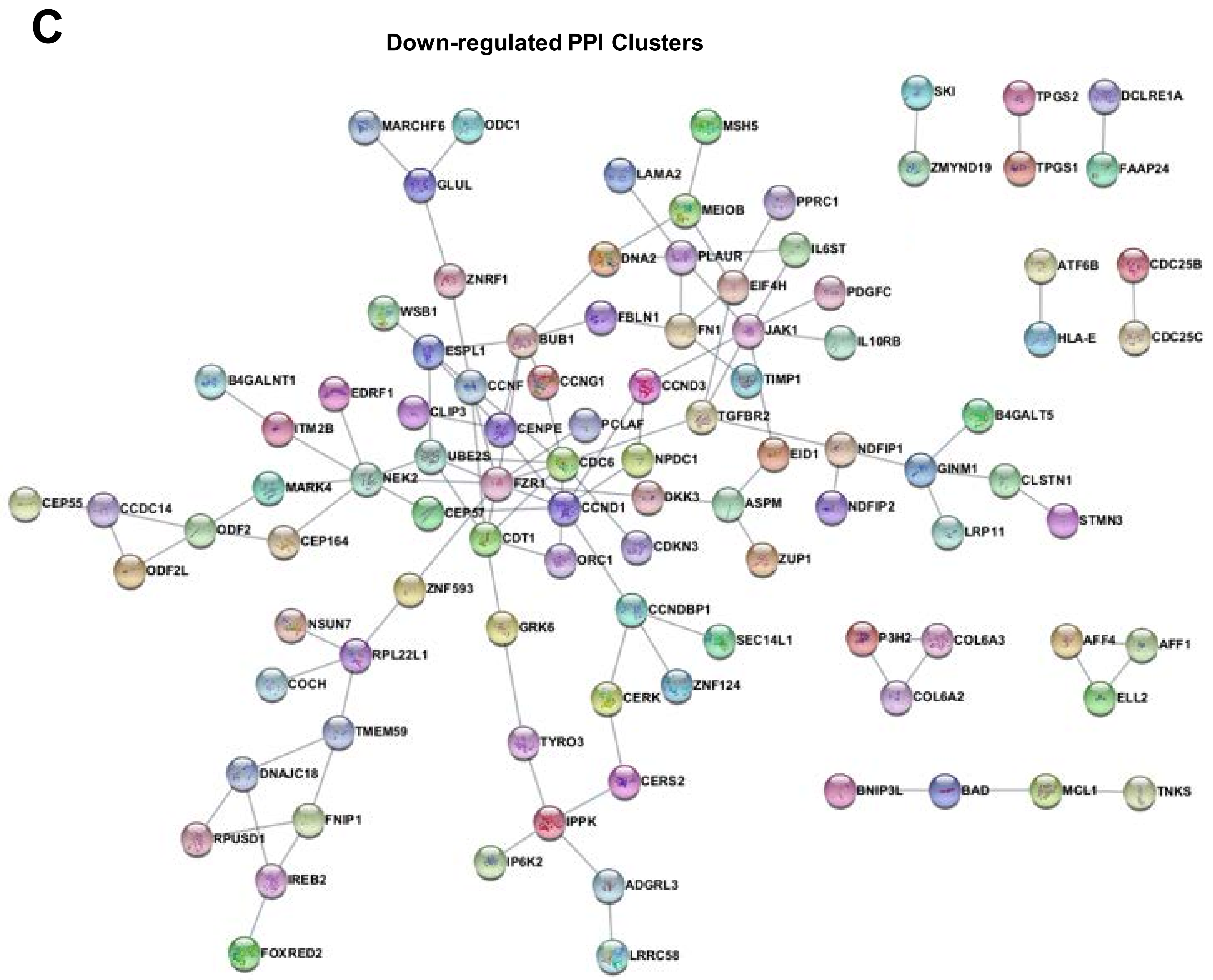

<u>Reactome</u>
1. Cell Cycle, Mitotic; 2. Cell Cycle; 3. S Phase

<u>KEGG</u>
1. Cell Cycle; 2. PI3K-Akt Signaling Pathway; 3. Focal Adhesion

<u>GOBP</u>
1. Cell Division; 2. Mitotic Cell Cycle Process; 3. Mitotic Cell Cycle

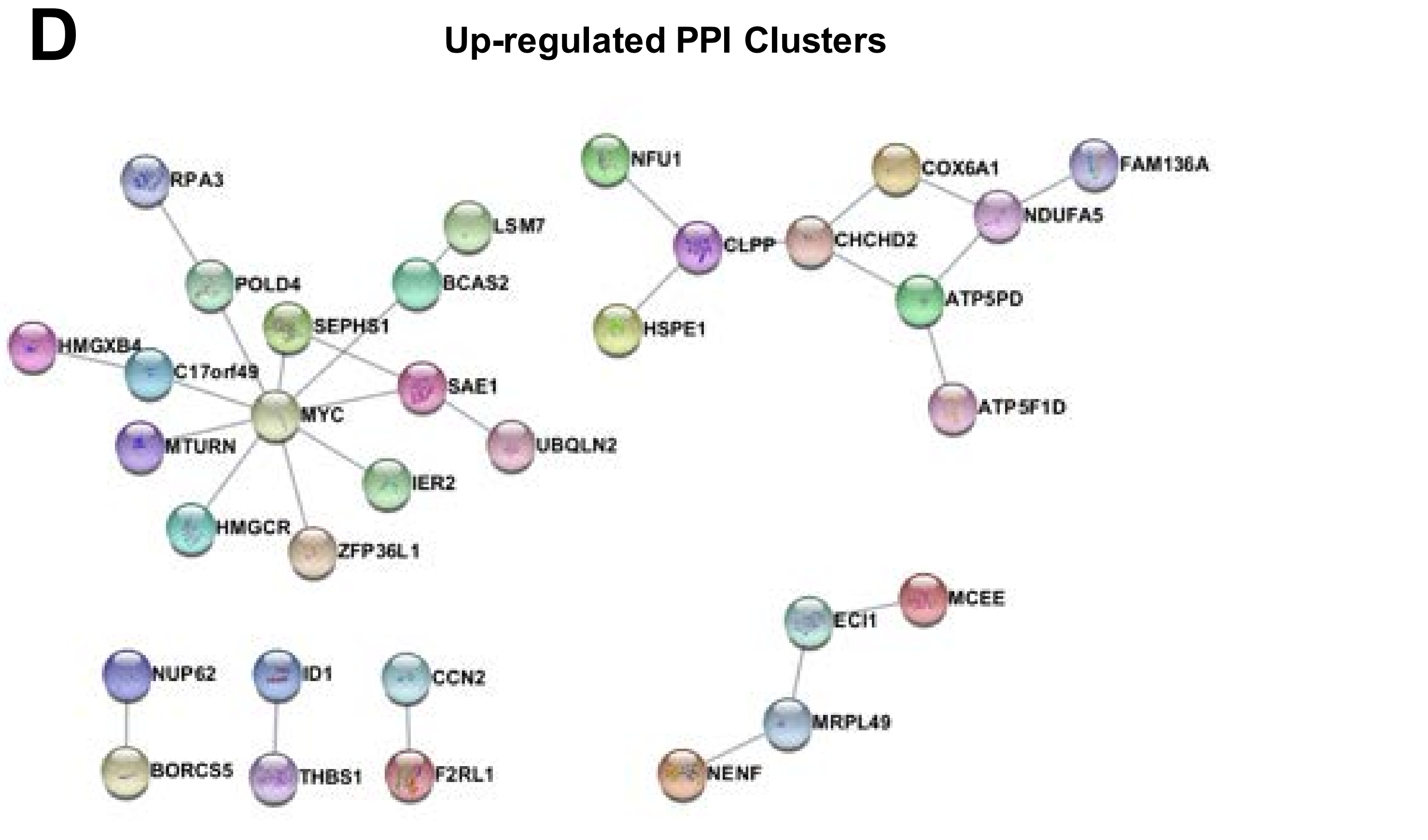

<u>Reactome</u>
1. Mitochondrial Protein Degradation; 2. Respiratory Electron Transport, ATP synthesis by Chemiosmotic Coupling, and Heat Production by Uncoupling Proteins; 3. Aerobic Respiration and Respiratory Electron Transport

<u>KEGG</u>
1. Amyotrophic lateral sclerosis; 2. Oxidative phosphorylation; 3. Diabetic cardiomyopathy

<u>GOBP</u>
1. Regulation of MAPK cascade; 2. Oxidative phosphorylation; 3. Response to growth factor

**Supplementary Figure 9**

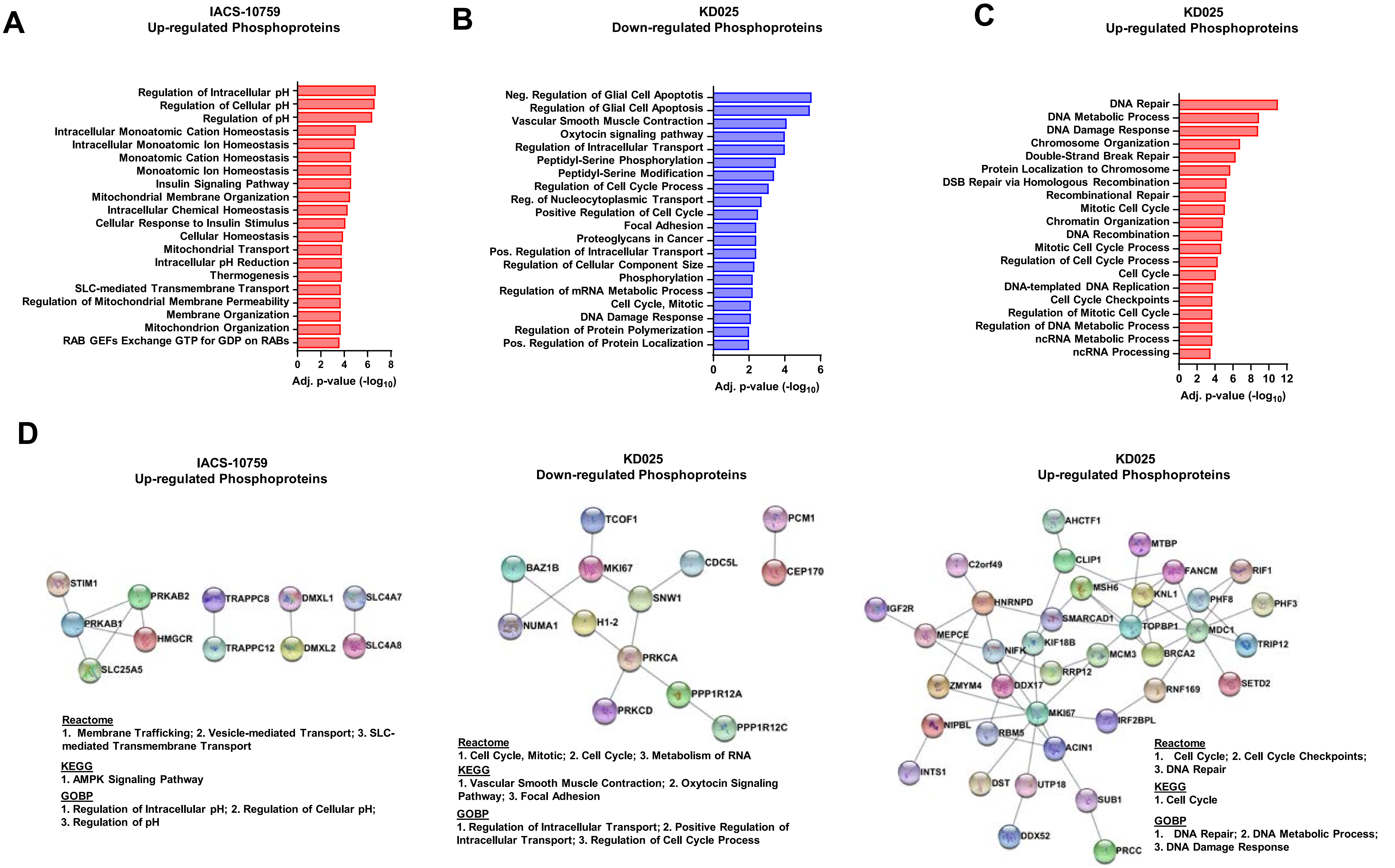

**Supplementary Figure 10**

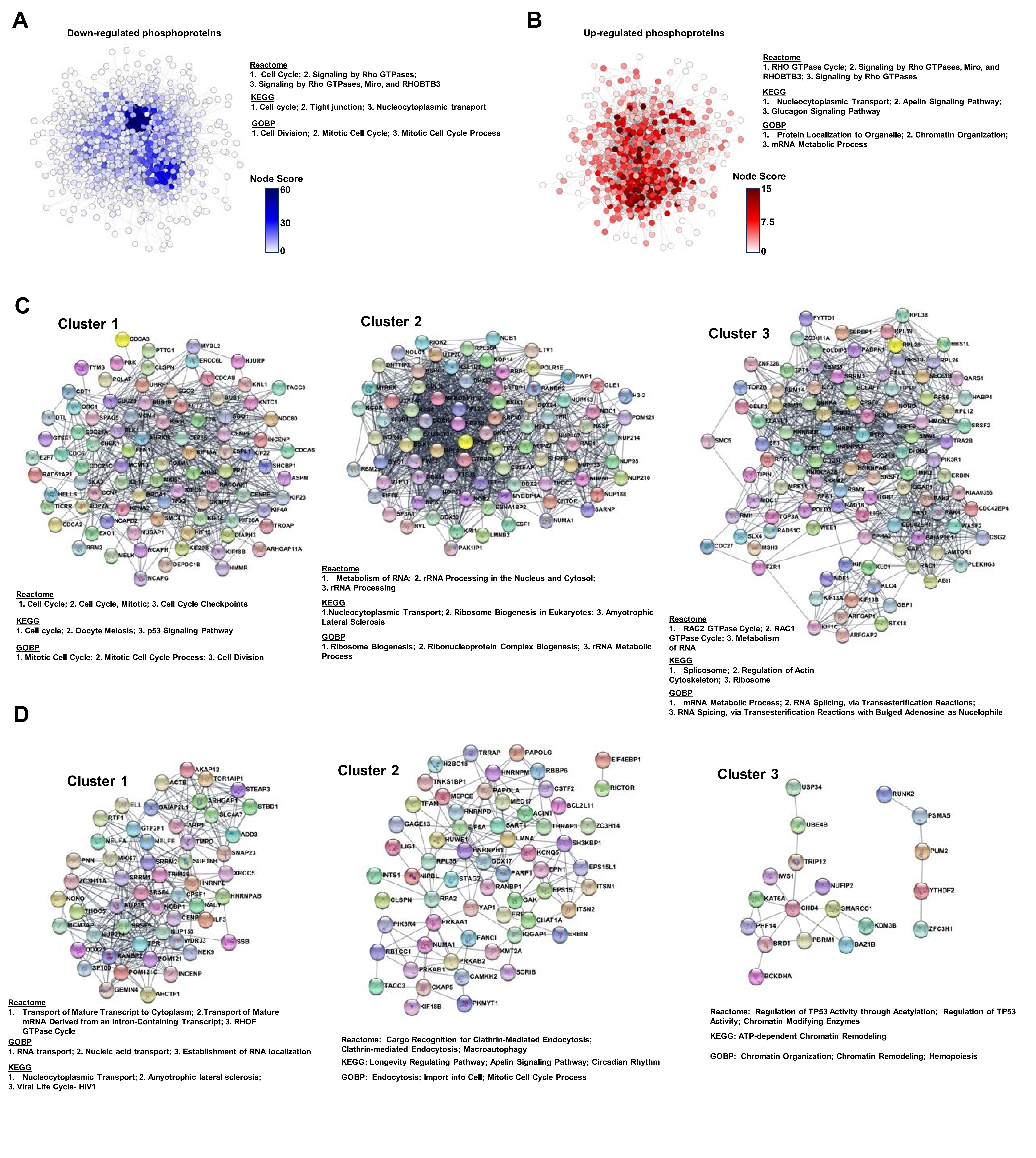

Supplementary Figure 11

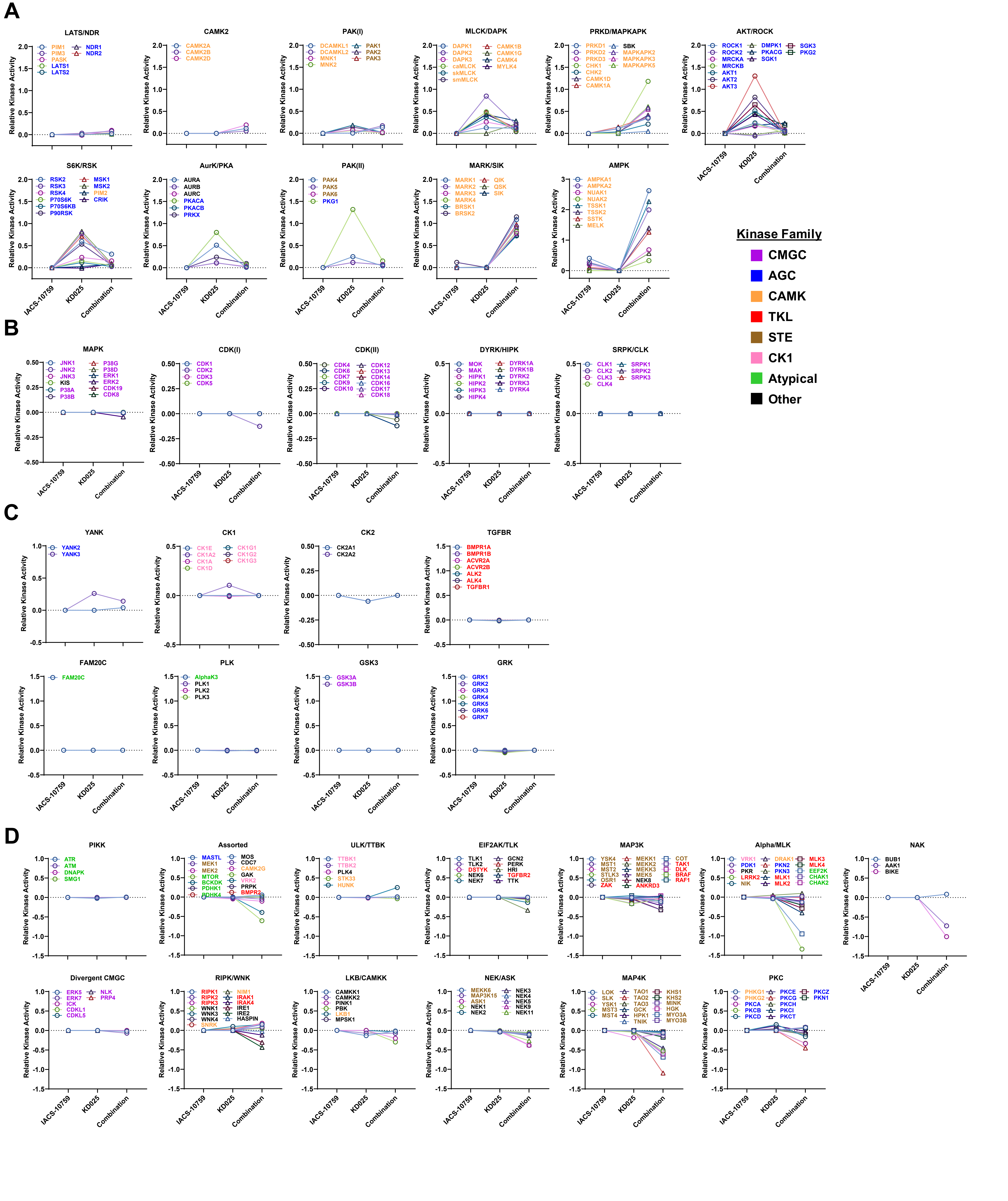

Supplementary Figure 12

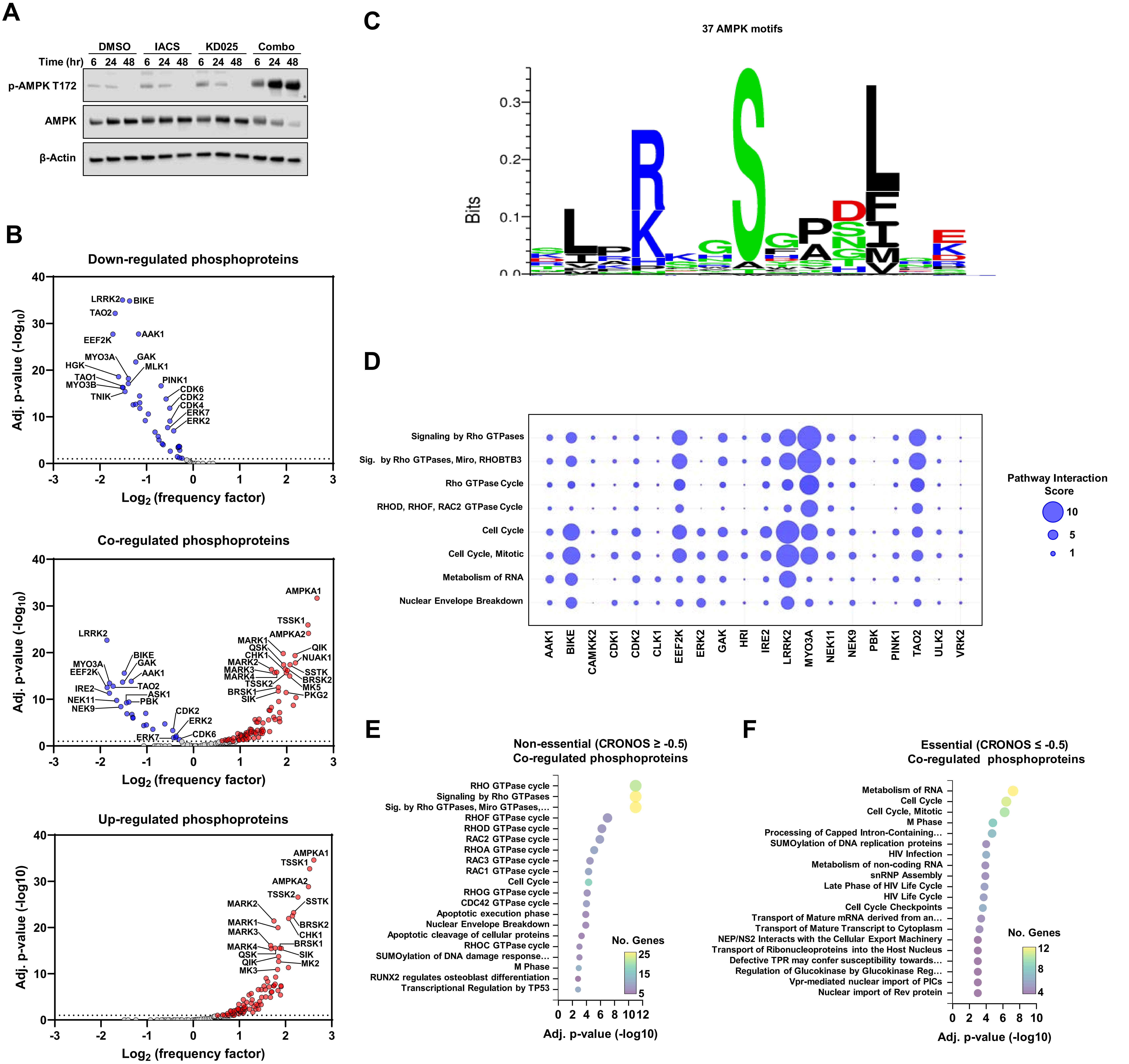

**Supplementary Figure 13**

| Gene Symbol | guide RNA | gRNA strand position | Target domain |
|---|---|---|---|
| AARS | AGAGCCCAGCATCTCGAAGA | chr16:70310893-70310912_AARS_+ | exon_03 |
| AARS | GTAGGAATTGGCATGTAAGA | chr16:70310519-70310538_AARS_- | exon_04 |
| AARS | GAAGGATAACTTCTGGGAGA | chr16:70305823-70305842_AARS_- | exon_05 |
| AARS | GGCCTGACAACACAGGGCGT | chr16:70303523-70303542_AARS_- | exon_07 |
| AARS | GGGCGTATCGGACAGCTCGG | chr16:70302239-70302258_AARS_+ | exon_08 |
| ABL1 | TGGGCTGCAAATCCAAGAAG | chr9:133710859-133710878_ABL1_+ | exon_01 |
| ABL1 | CATTGCGGGACACAGGCCCA | chr9:133730321-133730340_ABL1_- | exon_03 |
| ABL1 | GAAGAAATACAGCCTGACGG | chr9:133738383-133738402_ABL1_+ | exon_04 |
| ABL1 | CAAGAACTCTTCCACCTCCA | chr9:133747526-133747545_ABL1_- | exon_05 |
| ABL1 | TCAGTGATGATATAGAACGG | chr9:133748267-133748286_ABL1_- | exon_06 |
| ABL2 | GAAGCTTTGCATCGTCCCTA | chr1:179100598-179100617_ABL2_- | exon_03 |
| ABL2 | GCGAGATGGACAGCTGCCCA | chr1:179095568-179095587_ABL2_+ | exon_04 |
| ABL2 | TGTACACCATCACTCCACAG | chr1:179090939-179090958_ABL2_- | exon_05 |
| ABL2 | TTTCTAGGAAGATACCATGG | chr1:179089397-179089416_ABL2_- | exon_06 |
| ABL2 | TAATCCAGCAAATTCCCGTA | chr1:179087830-179087849_ABL2_+ | exon_07 |
| ADH7 | GAAGCAACCCTTCTCCATTG | chr4:100350736-100350755_ADH7_- | exon_02 |
| ADH7 | ATCTGTGCGACAGATTCCTG | chr4:100349758-100349777_ADH7_+ | exon_03 |
| ADH7 | AACAGCAGCGCCATATCCAG | chr4:100348942-100348961_ADH7_+ | exon_05 |
| ADH7 | GAAATGACAGGCAACAACGT | chr4:100341730-100341749_ADH7_- | exon_06 |
| ADH7 | TAGGTGAGCATCTTGGCTGA | chr4:100340196-100340215_ADH7_+ | exon_07 |
| ADORA2A | CTCCTCGGTGTACATCACGG | hr22:24829387-24829406_ADORA2A_ | exon_02 |
| ADORA2A | AGCACACCAGCACATTGCCC | hr22:24829438-24829457_ADORA2A_ | exon_02 |
| ADORA2A | AGGCAATGAAGAGGCAGCCG | hr22:24829597-24829616_ADORA2A_ | exon_02 |
| ADORA2A | GAGGGCAAGAACCACTCCCA | hr22:24836669-24836688_ADORA2A_ | exon_03 |
| ADORA2A | GGAGAGCCAGCCTCTGCCGG | hr22:24836851-24836870_ADORA2A_ | exon_03 |
| AKT1 | GACGTGGCTATTGTGAAGGA | chr14:105258955-105258974_AKT1_- | exon_01 |
| AKT1 | GCAGGATGTGGACCAACGTG | chr14:105246455-105246474_AKT1_- | exon_02 |
| AKT1 | GAAGGTGCGTTCGATGACAG | chr14:105243019-105243038_AKT1_+ | exon_03 |
| AKT1 | GGAGGAGATGGACTTCCGGT | chr14:105242060-105242079_AKT1_- | exon_04 |
| AKT1 | GGGCAAGGGCACTTTCGGCA | chr14:105241493-105241512_AKT1_- | exon_05 |
| AKT2 | TGGGTACAAGGAGAGGCCCG | chr19:40762881-40762900_AKT2_- | exon_03 |
| AKT2 | CATCGAGAGGACCTTCCACG | chr19:40761084-40761103_AKT2_- | exon_04 |
| AKT2 | GTGGATGCGGGCCATCCAGA | chr19:40748569-40748588_AKT2_- | exon_05 |
| AKT2 | TATCTCAAACTCCTTGGCAA | chr19:40747939-40747958_AKT2_- | exon_06 |
| AKT2 | CCGGGAGCGTGTCTTCACAG | chr19:40743962-40743981_AKT2_- | exon_09 |
| AKT3 | AACCCAACCTTCTTTCACAA | chr1:244006437-244006456_AKT3_+ | exon_02 |
| AKT3 | GAGAGAGCGGGTGTTCTCTG | chr1:243736314-243736333_AKT3_- | exon_09 |
| AKT3 | ATGGAACCGAAGCCTACCTC | chr1:243727005-243727024_AKT3_+ | exon_10 |
| AKT3 | CATCATCTGGTCCTCCACCA | chr1:243708877-243708896_AKT3_+ | exon_12 |
| AKT3 | ACTGCATGGACAATGAGAGG | chr1:243668599-243668618_AKT3_- | exon_14 |
| ALK | GATGCCCGAGAAGAAGGCGT | chr2:30142940-30142959_ALK_- | exon_01 |
| ALK | CATCTGGGAGGCCTCCTCGG | chr2:29917765-29917784_ALK_+ | exon_03 |
| ALK | TGGGCAATGTACCTTCCAGA | chr2:29754844-29754863_ALK_+ | exon_04 |
| ALK | AAACCGCAGCTTGTCTGCAG | chr2:29606631-29606650_ALK_- | exon_05 |
| ALK | AATGGGACAGTCCTCCAGCT | chr2:29551278-29551297_ALK_- | exon_06 |
| APOBEC1 | ATAGAAGACGTCAAACTCCC | chr12:7805395-7805414_APOBEC1_+ | exon_03 |
| APOBEC1 | TAGAAGACGTCAAACTCCCA | chr12:7805396-7805415_APOBEC1_+ | exon_03 |
| APOBEC1 | CAGGAGAAGAATCGAACCCT | chr12:7805415-7805434_APOBEC1_- | exon_03 |
| APOBEC1 | TCAGGAGAAGAATCGAACCC | chr12:7805416-7805435_APOBEC1_- | exon_03 |
| APOBEC1 | TACAACATCATCCACAGAGG | chr12:7803647-7803666_APOBEC1_+ | exon_04 |

| | | | |
|---|---|---|---|
| ARCN1 | GGTGTTCAGAGCCGTCAGAG | hr11:118454052-118454071_ARCN1_ | exon_03 |
| ARCN1 | AGTACGCTTGCCCATACTAG | :hr11:118455296-118455315_ARCN1_ | exon_05 |
| ARCN1 | CAGCCATTTCCACTCTCCGA | :hr11:118464319-118464338_ARCN1_ | exon_08 |
| ARCN1 | ACAGCCATTTCCACTCTCCG | :hr11:118464320-118464339_ARCN1_ | exon_08 |
| ARCN1 | GTCGACGAAATACCCTGGAG | hr11:118468478-118468497_ARCN1_ | exon_09 |
| AR | CAGCAGGAAGCAGTATCCGA | chrX:66765580-66765599_AR_+ | exon_01 |
| AR | TCTCTGCTAGACGACAGCGC | chrX:66765865-66765884_AR_+ | exon_01 |
| AR | AGAGAGACAGCTTGTACACG | chrX:66931485-66931504_AR_+ | exon_04 |
| AR | GGTGAAGGATCGCCAGCCCA | chrX:66937395-66937414_AR_- | exon_05 |
| AR | TCATGCACAGGAATTCCTGG | chrX:66941762-66941781_AR_- | exon_06 |
| ATM | GATGGCAGATATCTGTCACC | chr11:108117833-108117852_ATM_+ | exon_09 |
| ATM | GACACAATGCAACTTCCGTA | chr11:108121569-108121588_ATM_- | exon_11 |
| ATM | AAAGTCAAACAGCATACTGC | chr11:108122568-108122587_ATM_- | exon_12 |
| ATM | TTGCTTGAGCAACTGTACCA | chr11:108128312-108128331_ATM_+ | exon_16 |
| ATM | GGAGAGAGCCAAAGTACCAT | chr11:108138048-108138067_ATM_+ | exon_18 |
| ATP6V1A | TCTCTCCAACCAATTCGCTG | hr3:113499955-113499974_ATP6V1A_ | exon_03 |
| ATP6V1A | GGTTACACAGACATACCCGT | hr3:113503282-113503301_ATP6V1A_ | exon_04 |
| ATP6V1A | GAGGTATCATAATTCCCAGG | hr3:113503660-113503679_ATP6V1A_ | exon_05 |
| ATP6V1A | TGTAAAGGAGAAGTTCACCA | r3:113505108-113505127_ATP6V1A_ | exon_06 |
| ATP6V1A | AGACCTCTGGGAAGTCCCGG | hr3:113507707-113507726_ATP6V1A_ | exon_07 |
| ATR | GAAATCAAGCAACATCACGG | chr3:142285042-142285061_ATR_+ | exon_03 |
| ATR | CTTGTGTAACAAATGACAGG | chr3:142281881-142281900_ATR_+ | exon_04 |
| ATR | GTGATGGAATATCACCCAAA | chr3:142280144-142280163_ATR_- | exon_05 |
| ATR | TGTTCAGAAACTGGACCTGG | chr3:142278147-142278166_ATR_- | exon_07 |
| ATR | GCTGACTCTTACCTGACAGA | chr3:142272057-142272076_ATR_+ | exon_13 |
| AURKA | GCTAGTTTACCAGGTGCCGA | chr20:54961303-54961322_AURKA_+ | exon_03 |
| AURKA | TTACCAGGTGCCGATGGCAG | chr20:54961309-54961328_AURKA_+ | exon_03 |
| AURKA | TCTGAGCTGATGCTCCACTC | chr20:54958070-54958089_AURKA_+ | exon_05 |
| AURKA | CTTCGAATGACAGTAAGACA | chr20:54948568-54948587_AURKA_+ | exon_07 |
| AURKA | TTCAGGAGGACCACTCTCTG | chr20:54945701-54945720_AURKA_- | exon_08 |
| AURKB | GGGTGACAGGCTCTTTCCGG | chr17:8111101-8111120_AURKB_+ | exon_03 |
| AURKB | TGACGAGCAGCGAACAGCCA | chr17:8110070-8110089_AURKB_- | exon_06 |
| AURKB | AGGCCACCATACCTCAGGGA | chr17:8109797-8109816_AURKB_+ | exon_07 |
| AURKB | TGTGATGCACTCTCAAAGGG | chr17:8108562-8108581_AURKB_+ | exon_08 |
| AURKB | GGTCCGGGCCAACTCTCGGA | chr17:8108227-8108246_AURKB_- | exon_09 |
| AURKC | AGATGAACAGCGCACAGCCA | chr19:57744027-57744046_AURKC_+ | exon_04 |
| AURKC | TGGGAGGCGCTCTCAAAGGG | chr19:57746379-57746398_AURKC_- | exon_06 |
| AURKC | ACTGTGGGAGGCGCTCTCAA | chr19:57746383-57746402_AURKC_- | exon_06 |
| AURKC | TCGGGAGTGGGCCTGAACCC | chr19:57746730-57746749_AURKC_- | exon_07 |
| AURKC | GAAGCCATCTGAGCACAGGG | chr19:57746762-57746781_AURKC_- | exon_07 |
| AXL | AGCGCCAAGCACCAGGCCAG | chr19:41725334-41725353_AXL_- | exon_01 |
| AXL | GAACCTGGAGCTGACACCGA | chr19:41726617-41726636_AXL_- | exon_02 |
| AXL | CGTGTCGGAAAGCTGCAGGG | chr19:41727059-41727078_AXL_- | exon_03 |
| AXL | TGACCTGGAGCCGTGGCCAG | chr19:41727913-41727932_AXL_- | exon_04 |
| AXL | CTGCGAAGCCCATAACGCCA | chr19:41736897-41736916_AXL_+ | exon_05 |
| BCL2 | ACCTGGATCCAGGATAACGG | chr18:60985322-60985341_BCL2_- | exon_01 |
| BCL2 | GAAGAGCTCCTCCACCACCG | chr18:60985486-60985505_BCL2_+ | exon_01 |
| BCL2 | ACATCTCGGCGAAGTCGCGG | chr18:60985554-60985573_BCL2_+ | exon_01 |
| BCL2 | GTGGCCCAGATAGGCACCCA | chr18:60795864-60795883_BCL2_+ | exon_02 |
| BCL2 | GGTGATGCAAGCTCCCACCA | chr18:60795885-60795904_BCL2_+ | exon_02 |
| BCR | AGAGGAGAAGTCCTCCTCGC | chr22:23524169-23524188_BCR_- | exon_01 |

| | | | |
|---|---|---|---|
| BCR | GCAGCGCAGCCGTATCCAGG | chr22:23596006-23596025_BCR_- | exon_02 |
| BCR | GTGGCTCCACTGCTGCACGC | chr22:23603675-23603694_BCR_- | exon_04 |
| BCR | TCAGTTTGCAGAAATCTCCG | chr22:23610681-23610700_BCR_+ | exon_05 |
| BCR | GGTCGGTGAACAGGAAGACG | chr22:23626229-23626248_BCR_- | exon_09 |
| BIRC5 | GATGCGGTGGTCCTTGAGAA | chr17:76210435-76210454_BIRC5_- | exon_01 |
| BIRC5 | CAAGAACTGGCCCTTCTTGG | chr17:76210463-76210482_BIRC5_+ | exon_01 |
| BIRC5 | GCAGGCGCAGCCCTCCAAGA | chr17:76210477-76210496_BIRC5_- | exon_01 |
| BIRC5 | GGGCAGTCTCACCCGCTCCG | chr17:76210501-76210520_BIRC5_- | exon_01 |
| BIRC5 | GCAGTGGATGAAGCCAGCCT | chr17:76210768-76210787_BIRC5_- | exon_02 |
| BIRC8 | TCAGCCCTGAAGAGCCGCTA | chr19:53793088-53793107_BIRC8_- | exon_01 |
| BIRC8 | GAAGCCCAAGGAAGATCCTT | chr19:53793462-53793481_BIRC8_- | exon_01 |
| BIRC8 | GGAAGCCCAAGGAAGATCCT | chr19:53793463-53793482_BIRC8_- | exon_01 |
| BIRC8 | GCTAGCCAACTGGAAGCCCA | chr19:53793474-53793493_BIRC8_- | exon_01 |
| BIRC8 | GTGGAGGAGGGCTAGCCAAC | chr19:53793484-53793503_BIRC8_- | exon_01 |
| BLK | GAAGGTCAGCGCCCAAGACA | chr8:11400811-11400830_BLK_+ | exon_02 |
| BLK | TTTAGACAAGCATTTCGTGG | chr8:11405536-11405555_BLK_+ | exon_04 |
| BLK | GAAGGATGTCACCACCCAGG | chr8:11412271-11412290_BLK_+ | exon_07 |
| BLK | ACCAGCCTGAGAGACTGCCG | chr8:11412933-11412952_BLK_- | exon_08 |
| BLK | GCTGGTCCGACTCTACGCAG | chr8:11414279-11414298_BLK_+ | exon_09 |
| BMP15 | GCTTCCTTGGCTGTTCGCCA | chrX:50653942-50653961_BMP15_- | exon_01 |
| BMP15 | CATGCGAGTCAGCTGAACGC | chrX:50654008-50654027_BMP15_- | exon_01 |
| BMP15 | GAAGGCTAAATTTCTTCCCA | chrX:50659160-50659179_BMP15_+ | exon_02 |
| BMP15 | TGTCTCCGAGTACTACGCGA | chrX:50659398-50659417_BMP15_+ | exon_02 |
| BMP15 | TACGGGACACAGGAGGGCCG | chrX:50659488-50659507_BMP15_- | exon_02 |
| BMX | ATGAAGAGAGCCGAAGTCAG | chrX:15529600-15529619_BMX_+ | exon_04 |
| BMX | ACACATTACATGCTTCCCAG | chrX:15534344-15534363_BMX_- | exon_05 |
| BMX | GCTAGAGTGGTACTTGAAGA | chrX:15540514-15540533_BMX_- | exon_07 |
| BMX | GCTAGACTGGTACTTGAAGA | chrX:15540580-15540599_BMX_- | exon_07 |
| BMX | TGTATCCATACCATTTCCCA | chrX:15554542-15554561_BMX_- | exon_13 |
| BPIFB6 | GCACAGGTGGTGCCATGCTG | chr20:31619440-31619459_BPIFB6_+ | exon_01 |
| BPIFB6 | CCAGGGATGAAACCTATCAA | chr20:31620858-31620877_BPIFB6_+ | exon_02 |
| BPIFB6 | ACAGCCCTCACTCTTGAACA | chr20:31622658-31622677_BPIFB6_- | exon_04 |
| BPIFB6 | TCTGGGTCACTCACCAGCCC | chr20:31622945-31622964_BPIFB6_- | exon_05 |
| BPIFB6 | TAGGGCTACACTCACCACTG | chr20:31623474-31623493_BPIFB6_- | exon_06 |
| BRAF | AAGATGGCGGCGCTGAGCGG | chr7:140624487-140624506_BRAF_- | exon_01 |
| BRAF | GAGAGAAGAAACCAATTGGT | chr7:140507844-140507863_BRAF_- | exon_05 |
| BRAF | GAACTTCTGTACTACAACGC | chr7:140501243-140501262_BRAF_+ | exon_06 |
| BRAF | ATACCCAATAGAGTCCGAGG | chr7:140500158-140500177_BRAF_+ | exon_07 |
| BRAF | GATGACTTCCTTTCTCGCTG | chr7:140482849-140482868_BRAF_+ | exon_10 |
| BRD4 | TTAGGGTTGGAGGTCTCTGG | chr19:15383750-15383769_BRD4_+ | exon_02 |
| BRD4 | ACAGGAGGAGGATTCGGCTG | chr19:15376361-15376380_BRD4_+ | exon_05 |
| BRD4 | GGAGCCCAAGACCACCAAGC | chr19:15375475-15375494_BRD4_- | exon_06 |
| BRD4 | TTCGACTGATGACTCTGAGG | chr19:15367813-15367832_BRD4_- | exon_08 |
| BRD4 | CACGCGGCCCAGCTTCTCGC | chr19:15366256-15366275_BRD4_+ | exon_10 |
| BTK | GATGCTCTCCAGAATCACTG | chrX:100630246-100630265_BTK_+ | exon_02 |
| BTK | ACCGGAATCTGTCTTTCTGG | chrX:100629522-100629541_BTK_+ | exon_03 |
| BTK | CTGAAGAACTAAGGAAGCGG | chrX:100625008-100625027_BTK_- | exon_05 |
| BTK | ATGCAAATGATCTACAGCTG | chrX:100615627-100615646_BTK_- | exon_08 |
| BTK | GAATAGTAGCACTCACCCTG | chrX:100613589-100613608_BTK_+ | exon_11 |
| BTLA | AAAGGGATCTCCTGCTAAGA | chr3:112198543-112198562_BTLA_+ | exon_02 |
| BTLA | TGCCTGAGAAGGCACCAAGG | chr3:112190058-112190077_BTLA_- | exon_03 |

| | | | |
|---|---|---|---|
| BTLA | AAACAGGTAGTGATGAGTAG | chr3:112190091-112190110_BTLA_+ | exon_03 |
| BTLA | AACTCTCTGACACAGCAGGA | chr3:112188620-112188639_BTLA_- | exon_04 |
| BTLA | GCATCCTGAAACAAAGGTCA | chr3:112185116-112185135_BTLA_+ | exon_05 |
| BUD31 | AATCAACTCCCAGCCATCTG | chr7:99008750-99008769_BUD31_- | exon_03 |
| BUD31 | GGAGTTGATTGAGCCAACAC | chr7:99008760-99008779_BUD31_+ | exon_03 |
| BUD31 | GCTGAAACAGAACCGCATGA | chr7:99013760-99013779_BUD31_+ | exon_04 |
| BUD31 | AAAGAGGTCGAAGATGTAGC | chr7:99013839-99013858_BUD31_- | exon_04 |
| BUD31 | AAGCGGAAAGCCATCAGCAG | chr7:99013862-99013881_BUD31_+ | exon_04 |
| C3orf30 | TCAGCCAGAGCCTCTTGCGG | :hr3:118865049-118865068_C3orf30_ | exon_01 |
| C3orf30 | GAAGAACCAGGCCGAAAGGA | :hr3:118865141-118865160_C3orf30_ | exon_01 |
| C3orf30 | TCAGGTGTCTGGCCTGACGG | :hr3:118865432-118865451_C3orf30_ | exon_01 |
| C3orf30 | GGACCGCAGAATGTCTGGCG | :hr3:118865618-118865637_C3orf30_ | exon_01 |
| C3orf30 | TATGAGCTTGGTCAATAGCC | :hr3:118865999-118866018_C3orf30_ | exon_01 |
| C8B | ATGGGCTTGGAGGGCGCCGG | chr1:57431585-57431604_C8B_- | exon_01 |
| C8B | GGTGAAAGGCCACATTCCTT | chr1:57425829-57425848_C8B_- | exon_02 |
| C8B | CATGAAATGGACCAATACTG | chr1:57420383-57420402_C8B_- | exon_04 |
| C8B | ACCTGGCTGTCTCTTACCTG | chr1:57417704-57417723_C8B_+ | exon_05 |
| C8B | TGGGACCCACTACATCACAG | chr1:57411560-57411579_C8B_- | exon_07 |
| CA9 | GGATCAACAGAGGGAGCCAG | chr9:35673980-35673999_CA9_- | exon_01 |
| CA9 | AGGCGCAGTTCTGGGAGCGG | chr9:35675889-35675908_CA9_- | exon_03 |
| CA9 | AGAGCCCGGTACTCCCGCCC | chr9:35676105-35676124_CA9_- | exon_04 |
| CA9 | TCGCTTGGAAGAAATCGCTG | chr9:35677828-35677847_CA9_+ | exon_06 |
| CA9 | GACTACACCGCCCTGTGCCC | chr9:35679267-35679286_CA9_+ | exon_07 |
| CADM1 | GAGCGGATCCCAGTGTGCGG | hr11:115375073-115375092_CADM1_ | exon_01 |
| CADM1 | AAAGACGTGACAGTGATCGA | hr11:115111098-115111117_CADM1_ | exon_02 |
| CADM1 | ATGGTGGTGTAACTTTCCTG | hr11:115109231-115109250_CADM1_ | exon_03 |
| CADM1 | TATCCAGAAAGACACTGCGG | hr11:115102166-115102185_CADM1_ | exon_04 |
| CADM1 | AAGGTGCACAAGGAGGACGA | hr11:115099913-115099932_CADM1_ | exon_05 |
| CASP3 | AGTTTCTGAATGTTTCCCTG | :hr4:185553488-185553507_CASP3_ | exon_05 |
| CASP3 | GTTTGTGTGCTTCTGAGCCA | chr4:185553040-185553059_CASP3_- | exon_06 |
| CASP3 | CGTGGTACAGAACTGGACTG | chr4:185552286-185552305_CASP3_- | exon_07 |
| CASP3 | TCTAACACCCAGGCCTGCCG | chr4:185552304-185552323_CASP3_- | exon_07 |
| CASP3 | TCTTGGCGAAATTCAAAGGA | chr4:185550628-185550647_CASP3_- | exon_08 |
| CCR2 | GATAAACCGAGAACGAGATG | chr3:46399029-46399048_CCR2_- | exon_01 |
| CCR2 | TGGAAATTATTCCATCCTCG | chr3:46399604-46399623_CCR2_- | exon_01 |
| CCR2 | TCAACTGGACCAAGCCACGC | chr3:46399861-46399880_CCR2_+ | exon_01 |
| CCR2 | ACTCCTGGACCTCCACACAC | chr3:46401217-46401236_CCR2_- | exon_02 |
| CCR2 | ACTACACAAGGACTCCTCGA | chr3:46401262-46401281_CCR2_+ | exon_02 |
| CCT2 | AGACTTGGTAAAGAGCACCT | chr12:69980562-69980581_CCT2_+ | exon_03 |
| CCT2 | GGGTTGGAGAGAAGCCACGA | chr12:69981980-69981999_CCT2_+ | exon_06 |
| CCT2 | GAAAGGCTCTGGCAACCTGG | chr12:69983388-69983407_CCT2_+ | exon_07 |
| CCT2 | GAACGCCTAGCTCTTGTCAC | chr12:69987372-69987391_CCT2_+ | exon_10 |
| CCT2 | GACTCTAGAACAGTTTATGG | chr12:69991522-69991541_CCT2_+ | exon_12 |
| CCT7 | GAGGCTGTAAGAACTACCCT | chr2:73466864-73466883_CCT7_+ | exon_02 |
| CCT7 | GTACTGCCATACCAGCTGGG | chr2:73470250-73470269_CCT7_- | exon_04 |
| CCT7 | GCCACAGCAGGCTTACCTCG | chr2:73471840-73471859_CCT7_- | exon_06 |
| CCT7 | AAGAAGACTTTCTCTTACGC | chr2:73474920-73474939_CCT7_+ | exon_07 |
| CCT7 | CCTGTCAGCAAAGTACTGGG | chr2:73476234-73476253_CCT7_- | exon_08 |
| CD274 | GTTGAAGGACCAGCTCTCCC | chr9:5457287-5457306_CD274_+ | exon_03 |
| CD274 | TTGAAGGACCAGCTCTCCCT | chr9:5457288-5457307_CD274_+ | exon_03 |
| CD274 | TCTGAAGTGCAGCATTTCCC | chr9:5457308-5457327_CD274_- | exon_03 |

| | | | |
|---|---|---|---|
| CD274 | GAACTGACATGTCAGGCTGA | chr9:5462893-5462912_CD274_+ | exon_04 |
| CD274 | TGAATGTCAGTGCTACACCA | chr9:5465569-5465588_CD274_- | exon_05 |
| CD276 | GCTGACAGATACCAAACAGC | chr15:73994720-73994739_CD276_+ | exon_03 |
| CD276 | GGTGGCATCGGTGCCCACTA | chr15:73996042-73996061_CD276_- | exon_05 |
| CD276 | CACAAGGCAATGCATCCCTG | chr15:73996222-73996241_CD276_+ | exon_05 |
| CD276 | TGTGCACAGCGTCCTGCGGG | chr15:73996707-73996726_CD276_+ | exon_06 |
| CD276 | TCGCACCCAGCACCACCCGC | chr15:73996722-73996741_CD276_- | exon_06 |
| CD27 | TGGCACGGCCACATCCCTGG | chr12:6554263-6554282_CD27_+ | exon_01 |
| CD27 | AGAGAGGCACTACTGGGCTC | chr12:6554345-6554364_CD27_+ | exon_01 |
| CD27 | AGTGCCGACAGCTCTCACAG | chr12:6554693-6554712_CD27_- | exon_02 |
| CD27 | GACGACCGAGCGGTCAGCGA | chr12:6559449-6559468_CD27_- | exon_03 |
| CD27 | TGGGTAGAGAGAGTCCGGGC | chr12:6559769-6559788_CD27_- | exon_04 |
| CD28 | ATTGTCGTACGCTACAAGCA | chr2:204591386-204591405_CD28_- | exon_02 |
| CD28 | TGTAGGAATACTTGCAGCTA | chr2:204591417-204591436_CD28_- | exon_02 |
| CD28 | TTCCGGGCATCCCTTCACAA | chr2:204591454-204591473_CD28_+ | exon_02 |
| CD28 | GGGCTGGTAATGCTTGCGGG | chr2:204599577-204599596_CD28_- | exon_04 |
| CD28 | TAGGCTGCGAAGTCGCGTGG | chr2:204599606-204599625_CD28_- | exon_04 |
| CD38 | CCTCGTCGTGGTGCTCGCGG | chr4:15780139-15780158_CD38_+ | exon_01 |
| CD38 | CCACCGCGAGCACCACGACG | chr4:15780142-15780161_CD38_- | exon_01 |
| CD38 | CCAGCGGGACATGTTCACCC | chr4:15826554-15826573_CD38_+ | exon_03 |
| CD38 | ACTACTTGGTACTTACCCTG | chr4:15835922-15835941_CD38_- | exon_04 |
| CD38 | GATGTGCAAGATGAATCCTC | chr4:15850196-15850215_CD38_- | exon_08 |
| CD40LG | CATGCTGATGGGCAGTCCAG | hrX:135730451-135730470_CD40LG_ | exon_01 |
| CD40LG | GATTCTCTCGAATCTACCGG | hrX:135741381-135741400_CD40LG_ | exon_05 |
| CD40LG | GATCCAAGCCAAGTGAGCCA | hrX:135741515-135741534_CD40LG_ | exon_05 |
| CD40LG | AGCCAAGTGAGCCATGGCAC | hrX:135741521-135741540_CD40LG_ | exon_05 |
| CD40LG | AGGACGTGAAGCCAGTGCCA | hrX:135741535-135741554_CD40LG_ | exon_05 |
| CD40 | TCTGCTGGACCCGAAGCCCT | chr20:44751253-44751272_CD40_- | exon_04 |
| CD40 | GAGTGCTTATACCTTGTCCA | chr20:44751851-44751870_CD40_- | exon_05 |
| CD40 | TAGCTGTGAGACCAAAGACC | chr20:44755276-44755295_CD40_+ | exon_06 |
| CD40 | AGGGCTCTCAGCCGATCCTG | chr20:44756782-44756801_CD40_- | exon_07 |
| CD40 | AGAGAGTCGCATCTCAGTGC | chr20:44757646-44757665_CD40_+ | exon_09 |
| CD52 | GAAGAGGTGGATTATGGCAT | chr1:26646762-26646781_CD52_- | exon_02 |
| CD52 | GAAGCAGAAGAGGTGGATTA | chr1:26646768-26646787_CD52_- | exon_02 |
| CD52 | CTCAACTGAAGCAGAAGAGG | chr1:26646775-26646794_CD52_- | exon_02 |
| CD70 | GCTGAGCCTGTGCGAAGCGC | chr19:6590878-6590897_CD70_+ | exon_01 |
| CD70 | GTGCATCCAGCGCTTCGCAC | chr19:6590887-6590906_CD70_- | exon_01 |
| CD70 | ATGGGACCAAAGCAGCCCGC | chr19:6590938-6590957_CD70_+ | exon_01 |
| CD70 | GGCGGGAGAGCAGATTCCCA | chr19:6586205-6586224_CD70_+ | exon_03 |
| CD70 | AGAGCAGATTCCCACGGCCA | chr19:6586211-6586230_CD70_+ | exon_03 |
| CD80 | AAGCCATGGGCCACACACGG | chr3:119276561-119276580_CD80_- | exon_02 |
| CD80 | GCGCAGAGCCAGGATCACAA | chr3:119263494-119263513_CD80_+ | exon_03 |
| CD80 | GAGCCAGGATCACAATGGAG | chr3:119263499-119263518_CD80_+ | exon_03 |
| CD80 | TGAGATTAAGGTAATGGCCC | chr3:119248726-119248745_CD80_+ | exon_05 |
| CD80 | ATTAAGGTAATGGCCCAGGA | chr3:119248730-119248749_CD80_+ | exon_05 |
| CD86 | GTGGATGCGAATCATTCCTG | chr3:121822647-121822666_CD86_- | exon_03 |
| CD86 | ACAGTTCAGAATTCATCTGG | chr3:121822666-121822685_CD86_- | exon_03 |
| CD86 | GAATGAAACAGACAAGCTGA | chr3:121825246-121825265_CD86_- | exon_04 |
| CD86 | GGAAGTACAGCTGTAATCCA | chr3:121828150-121828169_CD86_- | exon_05 |
| CD86 | TCTTTCAGGAACCAACACAA | chr3:121836887-121836906_CD86_+ | exon_06 |
| CDC16 | AAAGTAGCTTCACTCTCTCG | hr13:115002153-115002172_CDC16_ | exon_02 |

| | | | |
|---|---|---|---|
| CDC16 | TTTGTAGCTGTAGGTAGCCA | :hr13:115007657-115007676_CDC16_ | exon_06 |
| CDC16 | TCAGGGATGACCGTTTCACT | :hr13:115009343-115009362_CDC16_ | exon_08 |
| CDC16 | GTAGAGCTGAATAAAGCCAA | hr13:115010423-115010442_CDC16_ | exon_09 |
| CDC16 | AGTGGGATGTTACTATCTCA | hr13:115012420-115012439_CDC16_ | exon_11 |
| CDH1 | GCTGCGGCTCCAAGGGCCCA | chr16:68771320-68771339_CDH1_- | exon_01 |
| CDH1 | AGAGGCCGCGTCCTGGGCAG | chr16:68772293-68772312_CDH1_+ | exon_02 |
| CDH1 | GCAATGCGTTCTCTATCCAG | chr16:68842722-68842741_CDH1_- | exon_05 |
| CDH1 | CAAGCCCGAATTCACCCAGG | chr16:68844186-68844205_CDH1_+ | exon_06 |
| CDH1 | GCTGAGGATGGTGTAAGCGA | chr16:68845653-68845672_CDH1_- | exon_07 |
| CDK1 | ACCCTTATACACAACTCCAT | chr10:62544469-62544488_CDK1_- | exon_03 |
| CDK1 | AATCCATGTACTGACCAGGA | chr10:62545512-62545531_CDK1_- | exon_04 |
| CDK1 | GAATCCATGTACTGACCAGG | chr10:62545513-62545532_CDK1_- | exon_04 |
| CDK1 | GATCTCCAGAAGTATTGCTG | chr10:62551667-62551686_CDK1_+ | exon_06 |
| CDK1 | GCACTCCCAATAATGAAGTG | chr10:62551914-62551933_CDK1_+ | exon_07 |
| CDK2 | AAGCAGAGAGATCTCTCGGA | chr12:56361670-56361689_CDK2_- | exon_02 |
| CDK2 | AATGGCAGAAAGCTAGGCCC | chr12:56362585-56362604_CDK2_- | exon_04 |
| CDK2 | TCTGAGGTTTAAGGTCTCGG | chr12:56362621-56362640_CDK2_- | exon_04 |
| CDK2 | TCAGGCATAGAAGTAACTCC | chr12:56364924-56364943_CDK2_- | exon_06 |
| CDK2 | TAACAAGCGGATTTCGGCCA | chr12:56365325-56365344_CDK2_+ | exon_07 |
| CDK4 | TTCGTGAGGTGGCTTTACTG | chr12:58145321-58145340_CDK4_- | exon_02 |
| CDK4 | GTTTCGGCTGGCAAGCCTGG | chr12:58144997-58145016_CDK4_+ | exon_03 |
| CDK4 | CCAGATGGCACTTACACCCG | chr12:58144708-58144727_CDK4_- | exon_04 |
| CDK4 | CAGAATCTACAGCTACCAGA | chr12:58144723-58144742_CDK4_- | exon_04 |
| CDK4 | GTACCACCGACTGCACTGGG | chr12:58143000-58143019_CDK4_+ | exon_07 |
| CDK6 | GGCCTTGAACACCTTCCCAT | chr7:92462548-92462567_CDK6_+ | exon_02 |
| CDK6 | ATGATGTTTCAGCTTCTCCG | chr7:92355085-92355104_CDK6_- | exon_04 |
| CDK6 | AACACTCCAGAGATCCACGG | chr7:92300772-92300791_CDK6_+ | exon_05 |
| CDK6 | TAGGCCAGTCTTCTTCTCCT | chr7:92247487-92247506_CDK6_+ | exon_07 |
| CDK6 | TCAGGCTGTATTCAGCTCCG | chr7:92244454-92244473_CDK6_+ | exon_08 |
| CDK7 | ATCTCTGGCCTTGTAAACGG | chr5:68531228-68531247_CDK7_- | exon_02 |
| CDK7 | AGCTCCAAATAGTAACTCGG | chr5:68558046-68558065_CDK7_- | exon_08 |
| CDK7 | TAGCTCCAAATAGTAACTCG | chr5:68558047-68558066_CDK7_- | exon_08 |
| CDK7 | TGTGATGCAAAGGTATTCCA | chr5:68568763-68568782_CDK7_- | exon_10 |
| CDK7 | TGACATCCAGGTGTTGGCCC | chr5:68572403-68572422_CDK7_- | exon_11 |
| CDK9 | GGCAAAGCAGTACGACTCGG | chr9:130548430-130548449_CDK9_+ | exon_01 |
| CDK9 | GCACCGCAAGACCGGCCAGA | chr9:130548998-130549017_CDK9_+ | exon_02 |
| CDK9 | AAGGATCTTGATCTCCCGCA | chr9:130549813-130549832_CDK9_- | exon_03 |
| CDK9 | GGTATATACTACCCTTGCAG | chr9:130550242-130550261_CDK9_- | exon_04 |
| CDK9 | TCACCGAGCAACAGCTCCGG | chr9:130550649-130550668_CDK9_- | exon_05 |
| CDX4 | AAAGAAGCAGGCATGTACCC | chrX:72667114-72667133_CDX4_+ | exon_01 |
| CDX4 | ACACGCTCCAGTCTTCTCGC | chrX:72667326-72667345_CDX4_- | exon_01 |
| CDX4 | GTCGGTCGAGCAGAAAGCGG | chrX:72667406-72667425_CDX4_- | exon_01 |
| CDX4 | TGGCGGGTCGCTTGTCCCGA | chrX:72667485-72667504_CDX4_+ | exon_01 |
| CDX4 | TCAGAGCTGGCAGTTAACCT | chrX:72673460-72673479_CDX4_+ | exon_02 |
| CHEK1 | TTGGTGCAAACCCTGGGAGA | hr11:125496694-125496713_CHEK1_ | exon_02 |
| CHEK1 | CTTCCATCAACTCATGGCAG | hr11:125499167-125499186_CHEK1_ | exon_04 |
| CHEK1 | GGAATGGTAATTCTTGCTGA | :hr11:125507376-125507395_CHEK1_ | exon_08 |
| CHEK1 | GGTGTGTCAGAGTCTCCCAG | hr11:125513716-125513735_CHEK1_ | exon_09 |
| CHEK1 | CATAAGGAAAGACCTGTGCG | :hr11:125514026-125514045_CHEK1_ | exon_10 |
| CHEK2 | GATTGGCAAATCCATCCTGA | chr22:29130394-29130413_CHEK2_+ | exon_02 |
| CHEK2 | CTTAGACTGCAAACTGGCCG | chr22:29126407-29126426_CHEK2_+ | exon_03 |

| | | | |
|---|---|---|---|
| CHEK2 | GCATACATAGAAGATCACAG | chr22:29121066-29121085_CHEK2_- | exon_05 |
| CHEK2 | TGCTGGGTATAACCGTGCTG | chr22:29091735-29091754_CHEK2_- | exon_12 |
| CHEK2 | AAGAAGCCTTAAGACACCCG | chr22:29090029-29090048_CHEK2_- | exon_14 |
| CKAP5 | TGTCACTGATTCCAATGCAG | chr11:46839925-46839944_CKAP5_- | exon_03 |
| CKAP5 | TAAAGCTAAAGCCAAGGAGC | chr11:46837885-46837904_CKAP5_- | exon_04 |
| CKAP5 | TGAAGCCAAACTAATTGCTG | chr11:46832622-46832641_CKAP5_- | exon_05 |
| CKAP5 | TGGGAACGAAGAAATCGAGT | chr11:46831351-46831370_CKAP5_+ | exon_06 |
| CKAP5 | AGTGGTACAAGCCCTGCAGG | chr11:46819659-46819678_CKAP5_- | exon_10 |
| CLDN17 | GAGGCATTAGACATAACTGG | chr21:31538255-31538274_CLDN17_+ | exon_01 |
| CLDN17 | AGTGCACAGGCTCTAACGAG | chr21:31538600-31538619_CLDN17_- | exon_01 |
| CLDN17 | GAATTGCATCCGACAAGCCA | chr21:31538761-31538780_CLDN17_- | exon_01 |
| CLDN17 | AAAGCTGATACTCTCCACTG | chr21:31538832-31538851_CLDN17_+ | exon_01 |
| CLDN17 | ACCAGCCCAGCAATTTGCAA | chr21:31538901-31538920_CLDN17_+ | exon_01 |
| CLRN1 | ACTGCCTCACACCCTCTCCG | chr3:150690276-150690295_CLRN1_+ | exon_01 |
| CLRN1 | ACAGTGGCTTTGATCCACAA | chr3:150690383-150690402_CLRN1_+ | exon_01 |
| CLRN1 | CCAAGGCTGTCACAACTCCG | chr3:150690411-150690430_CLRN1_+ | exon_01 |
| CLRN1 | CAAGGCTGTCACAACTCCGA | chr3:150690412-150690431_CLRN1_+ | exon_01 |
| CLRN1 | GGACCATGCAGAGTTTCAAA | chr3:150659401-150659420_CLRN1_+ | exon_02 |
| CPSF3 | AATAGGAGATCAATCTCAGC | chr2:9570121-9570140_CPSF3_- | exon_03 |
| CPSF3 | GATCTCAATCATGAACATGG | chr2:9572786-9572805_CPSF3_- | exon_05 |
| CPSF3 | GTCCACGATATTGTAAACAG | chr2:9576406-9576425_CPSF3_+ | exon_07 |
| CPSF3 | TGTAGAAGGGACACTTGCCA | chr2:9582084-9582103_CPSF3_+ | exon_09 |
| CPSF3 | CATAGACTAACCACATGAGG | chr2:9583782-9583801_CPSF3_- | exon_10 |
| CREBBP | GCTCAGAGGGCTCTTGCCCA | chr16:3900724-3900743_CREBBP_+ | exon_02 |
| CREBBP | ACGAGAGCAAGCAAACGGAG | chr16:3843477-3843496_CREBBP_- | exon_04 |
| CREBBP | TTTGTCGTGAAGATGCACAA | chr16:3842069-3842088_CREBBP_+ | exon_05 |
| CREBBP | GAGTTCCTTCACCTACCCAG | chr16:3832669-3832688_CREBBP_+ | exon_06 |
| CREBBP | AATCCAATGAACATTCCAGC | chr16:3831283-3831302_CREBBP_- | exon_07 |
| CST9 | GTTCAAGGCAAACTCCACTG | chr20:23586334-23586353_CST9_+ | exon_01 |
| CST9 | GAGAAGCAGTGACAGTGCCC | chr20:23586448-23586467_CST9_+ | exon_01 |
| CST9 | AGAGGAGGAAGGCTATGCCC | chr20:23586469-23586488_CST9_- | exon_01 |
| CST9 | AGCTGACAAAGCCATTCCGA | chr20:23584164-23584183_CST9_- | exon_02 |
| CST9 | CAGCTGACAAAGCCATTCCG | chr20:23584165-23584184_CST9_- | exon_02 |
| CTLA4 | TATCCAAGGACTGAGGGCCA | chr2:204735544-204735563_CTLA4_+ | exon_02 |
| CTLA4 | ATGCCCAGGTAGTATGGCGG | chr2:204735608-204735627_CTLA4_- | exon_02 |
| CTLA4 | AGAGGAGGAAGTCAGAATCT | chr2:204736117-204736136_CTLA4_- | exon_03 |
| CTLA4 | TAACTGCTGCAAGGATCCAG | chr2:204736135-204736154_CTLA4_- | exon_03 |
| CTLA4 | AAGAAGCCCTCTTACAACAG | chr2:204737439-204737458_CTLA4_+ | exon_04 |
| CTNNB1 | GATGGAGTTGGACATGGCCA | chr3:41266024-41266043_CTNNB1_+ | exon_03 |
| CTNNB1 | AAGGTTATGCAAGGTCCCAG | chr3:41266973-41266992_CTNNB1_- | exon_05 |
| CTNNB1 | ATGGCAGTGCGTTTAGCTGG | chr3:41267227-41267246_CTNNB1_+ | exon_06 |
| CTNNB1 | GAAGCTGAACAAGAGTCCCA | chr3:41275040-41275059_CTNNB1_- | exon_09 |
| CTNNB1 | TTGCTGCTGTGTCCCACCCA | chr3:41275763-41275782_CTNNB1_- | exon_10 |
| CXCL1 | AGAGAGCAGCGCGGGCCATG | chr4:74735187-74735206_CXCL1_- | exon_01 |
| CXCL1 | GAGAGAGCAGCGCGGGCCAT | chr4:74735188-74735207_CXCL1_- | exon_01 |
| CXCL1 | GGAGAGAGCAGCGCGGGCCA | chr4:74735189-74735208_CXCL1_- | exon_01 |
| CXCL3 | GGGACCTTACATTCACACTT | chr4:74904038-74904057_CXCL3_+ | exon_02 |
| CXorf66 | CCTGGAATCTAACTTACGAA | chrX:139038193-139038212_CXorf66_ | exon_03 |
| CXorf66 | AAGGTCAGTGAGTCTAGGCA | chrX:139038384-139038403_CXorf66_ | exon_03 |
| CXorf66 | AAAGGTCAGTGAGTCTAGGC | chrX:139038385-139038404_CXorf66_ | exon_03 |
| CXorf66 | TAAAGTGGCTTGAATAGCTG | chrX:139038491-139038510_CXorf66_ | exon_03 |

| | | | |
|---|---|---|---|
| CXorf66 | AGTGGCTTGAATAGCTGAGG | :hrX:139038494-139038513_CXorf66_ | exon_03 |
| DDOST | TGTCCAGCAGCACTAAGGTG | chr1:20987738-20987757_DDOST_+ | exon_01 |
| DDOST | TGAATGTGAGCTCAAAGCCC | chr1:20987461-20987480_DDOST_+ | exon_02 |
| DDOST | TAGAGATGATTTCCCAACGA | chr1:20981959-20981978_DDOST_+ | exon_05 |
| DDOST | TTTGGTGCTGGACATCCTGA | chr1:20981157-20981176_DDOST_- | exon_06 |
| DDOST | GGAGCCGCTGAAGATGACGC | chr1:20980781-20980800_DDOST_+ | exon_07 |
| DDX20 | TGACGGCGCGGCTACCATGG | :hr1:112298531-112298550_DDX20_+ | exon_01 |
| DDX20 | TAAAGAGGCGTATACTGCCT | chr1:112303136-112303155_DDX20_- | exon_04 |
| DDX20 | ATTGGGAGACATACATGCAT | :hr1:112305360-112305379_DDX20_+ | exon_09 |
| DDX20 | ACAGTGACCTACTGTTGCCG | :hr1:112305542-112305561_DDX20_+ | exon_10 |
| DDX20 | GAAGACAAGACTTAAAGAGG | :hr1:112309203-112309222_DDX20_+ | exon_11 |
| DGAT2L6 | CAGTCGGGAGAAGAAAGCCA | :hrX:69397434-69397453_DGAT2L6_- | exon_01 |
| DGAT2L6 | GATTGGAACACCCACAGTCA | :hrX:69419210-69419229_DGAT2L6_- | exon_02 |
| DGAT2L6 | GGGTACGAAACTGGACCCTA | :hrX:69419680-69419699_DGAT2L6_- | exon_03 |
| DGAT2L6 | TCCACCCACCACAATAACCA | :hrX:69421806-69421825_DGAT2L6_- | exon_05 |
| DGAT2L6 | CAATCAGGAGACCTTCCCTG | :hrX:69424200-69424219_DGAT2L6_- | exon_06 |
| DHFRL1 | TCATTCCTGAGCGGCGGCCT | chr3:93780264-93780283_DHFRL1_+ | exon_02 |
| DHFR | CCCGGCAGATACCTGAGCGG | chr5:79950211-79950230_DHFR_+ | exon_01 |
| DHFR | AGATACCTGAGCGGTGGCCA | chr5:79950217-79950236_DHFR_+ | exon_01 |
| DHFR | AACCTTAGGGAACCTCCACA | chr5:79933817-79933836_DHFR_- | exon_04 |
| DHX15 | AGAGTGCGAGGATGTCCAAG | chr4:24586008-24586027_DHX15_- | exon_01 |
| DHX15 | GAATGGGAAGCTTTCAAAGG | chr4:24578128-24578147_DHX15_+ | exon_02 |
| DHX15 | ATGTACTCCACACACCACTG | chr4:24572445-24572464_DHX15_+ | exon_03 |
| DHX15 | TACACCATAACGCTCCAGGA | chr4:24557967-24557986_DHX15_+ | exon_04 |
| DHX15 | TCAACAGGATGTGTACGCCC | chr4:24556469-24556488_DHX15_+ | exon_05 |
| DHX8 | GTTTGGCAAGTTCTTCCGCG | chr17:41561459-41561478_DHX8_- | exon_01 |
| DHX8 | ACCGGGACAAATATGGAGAG | chr17:41570255-41570274_DHX8_+ | exon_06 |
| DHX8 | TGGCCACACGACCCTCCCGC | chr17:41570852-41570871_DHX8_- | exon_07 |
| DHX8 | TAGACGGCGAAATCTTGTCG | chr17:41571089-41571108_DHX8_+ | exon_08 |
| DHX8 | GAAGAGCCTCCATTCCTGAG | chr17:41576259-41576278_DHX8_+ | exon_10 |
| DOT1L | GCTGAGACTGAAGTCGCCCG | chr19:2164201-2164220_DOT1L_+ | exon_01 |
| DOT1L | CAGGCCGTGCACTCACCGGA | chr19:2180752-2180771_DOT1L_- | exon_02 |
| DOT1L | TCAGCTTCGAGAGCATGCAG | chr19:2189727-2189746_DOT1L_+ | exon_04 |
| DOT1L | GATATGGCGCAGGAGTCCAG | chr19:2191057-2191076_DOT1L_- | exon_05 |
| DOT1L | CATCCCGGCCAAGTATGCGG | chr19:2193761-2193780_DOT1L_+ | exon_06 |
| DPAGT1 | AGAGGCGCGCAGCAATGAAG | ır11:118972242-118972261_DPAGT1_ | exon_03 |
| DPAGT1 | AGCACCGCTGATCACTCCCT | ır11:118971818-118971837_DPAGT1_ | exon_04 |
| DPAGT1 | GATGCAGGCCAAGTATCGGG | ır11:118971349-118971368_DPAGT1_ | exon_05 |
| DPAGT1 | TGCTGGCATGACCTTTGCCG | ır11:118968692-118968711_DPAGT1_ | exon_08 |
| DPAGT1 | GGTGGCAGAGAGCCTCCAGC | ır11:118967989-118968008_DPAGT1_ | exon_10 |
| DRD2 | CCTGATCGTCAGCCTCGCAG | chr11:113295142-113295161_DRD2_- | exon_02 |
| DRD2 | TAGGTGAGTGGAAATTCAGC | chr11:113288835-113288854_DRD2_- | exon_03 |
| DRD2 | GAAGGACAGGACCCAGACGA | :hr11:113287625-113287644_DRD2_+ | exon_04 |
| DRD2 | GATGGAGGAGTAGACCACGA | :hr11:113286281-113286300_DRD2_+ | exon_05 |
| DRD2 | GAGATCCAGACCATGCCCAA | chr11:113283373-113283392_DRD2_- | exon_07 |
| DSPP | GGTTCCTCAAAGCAAACCAC | chr4:88533256-88533275_DSPP_+ | exon_02 |
| DSPP | AATACCCAAGATGGTCACAA | chr4:88533549-88533568_DSPP_+ | exon_03 |
| DSPP | CATGATCATCTTCTTTCCCA | chr4:88534193-88534212_DSPP_- | exon_03 |
| DSPP | GACAATAACAGCAGTAGCCG | chr4:88535258-88535277_DSPP_+ | exon_04 |
| DSPP | GACAATGACAAATCAGACAG | chr4:88535420-88535439_DSPP_+ | exon_04 |
| ECD | GCTGTGAATAGGCGCATCAG | chr10:74916036-74916055_ECD_- | exon_05 |

| | | | |
|---|---|---|---|
| ECD | GCAGCCTGTATCCACTCCGC | chr10:74912101-74912120_ECD_+ | exon_07 |
| ECD | AAAGGTTTGTGCCAGACCGG | chr10:74912120-74912139_ECD_- | exon_07 |
| ECD | AGAGTGGTGAGGCAGTCACA | chr10:74908078-74908097_ECD_+ | exon_08 |
| ECD | CTGGTGAGAGATCTAACCAC | chr10:74899227-74899246_ECD_+ | exon_11 |
| EGFR | AGCGATGCGACCCTCCGGGA | chr7:55086967-55086986_EGFR_+ | exon_01 |
| EGFR | GATGTTCAATAACTGTGAGG | chr7:55210049-55210068_EGFR_+ | exon_02 |
| EGFR | ATAACCAGCCACCTCCTGGA | chr7:55211002-55211021_EGFR_- | exon_03 |
| EGFR | TGTGATCCAAGCTGTCCCAA | chr7:55218995-55219014_EGFR_+ | exon_05 |
| EGFR | CCAGGCAGTCGCTCTCCCGG | chr7:55220339-55220358_EGFR_- | exon_06 |
| EHMT2 | GCTGCACTCACCTCTCTCGG | chr6:31864689-31864708_EHMT2_+ | exon_01 |
| EHMT2 | TAGTGAAGAAACCCTGCCCA | chr6:31864554-31864573_EHMT2_- | exon_02 |
| EHMT2 | TCCAGCCACCACGAGCCCAG | chr6:31864099-31864118_EHMT2_- | exon_03 |
| EHMT2 | TCACTGGGAAAGGTGACCTC | chr6:31860605-31860624_EHMT2_- | exon_04 |
| EHMT2 | GGAGAAGTGACCCTGACGAA | chr6:31860290-31860309_EHMT2_- | exon_06 |
| EIF2S1 | TGAGGTGGAAGATGTAGTGA | chr14:67831526-67831545_EIF2S1_+ | exon_02 |
| EIF2S1 | TGAGGTGTTAGAATACACCA | chr14:67843182-67843201_EIF2S1_+ | exon_04 |
| EIF2S1 | AGCTGGAAAGCCTATTCCAG | chr14:67843211-67843230_EIF2S1_+ | exon_04 |
| EIF2S1 | GTAGTCATTACATACCGAGG | chr14:67849182-67849201_EIF2S1_- | exon_07 |
| EIF2S1 | TAACAGCCATAGCTTGACTG | chr14:67849238-67849257_EIF2S1_- | exon_07 |
| EIF4E | AGAAAGCTTACCTGTTCTGT | chr4:99823016-99823035_EIF4E_+ | exon_03 |
| EIF4E | GAAAGCTTACCTGTTCTGTA | chr4:99823017-99823036_EIF4E_+ | exon_03 |
| EIF4E | TTCTCCTCTTCTGTAGTCGG | chr4:99823090-99823109_EIF4E_+ | exon_03 |
| EIF4E | CTAGGAGGGTATACAAGGAA | chr4:99802278-99802297_EIF4E_- | exon_08 |
| EIF6 | GGCGATAGACGCGTGCACCA | chr20:33872010-33872029_EIF6_+ | exon_03 |
| EIF6 | GGAGGAGCGGCTCTCAGCCT | chr20:33868516-33868535_EIF6_- | exon_04 |
| EIF6 | GTGTTGCAGCTCCTGGTCGG | chr20:33868577-33868596_EIF6_+ | exon_04 |
| EIF6 | TCTGGCAGATGTGCTCAAGG | chr20:33867888-33867907_EIF6_- | exon_05 |
| EIF6 | AATGAGGGAATCCCGCATGC | chr20:33867374-33867393_EIF6_+ | exon_06 |
| ENTPD1 | AAAGGAGAATGACACAGGCG | chr10:97599513-97599532_ENTPD1_- | exon_03 |
| ENTPD1 | CAAGAGACACCCGTTTACCT | chr10:97602202-97602221_ENTPD1_- | exon_04 |
| ENTPD1 | ACTCTGGCAGAAACTGGCCA | chr10:97605323-97605342_ENTPD1_- | exon_06 |
| ENTPD1 | TAGTCACTTACCTCCTCCCA | chr10:97620331-97620350_ENTPD1_- | exon_08 |
| ENTPD1 | AATGAAATGGATGTGCTCCC | chr10:97624593-97624612_ENTPD1_- | exon_09 |
| EPHA2 | CAGCAGGGCGAAGCAGGCGC | chr1:16482389-16482408_EPHA2_+ | exon_01 |
| EPHA2 | GAACGTGGAGGAGCGCTCCG | chr1:16475188-16475207_EPHA2_- | exon_03 |
| EPHA2 | GCTCAGGGCACTCCAAGCAG | chr1:16464863-16464882_EPHA2_+ | exon_04 |
| EPHA2 | CATGAACTACACCTTCACCG | chr1:16464426-16464445_EPHA2_- | exon_05 |
| EPHA2 | ACTCGGCTCTGCTGCGGCGG | chr1:16462181-16462200_EPHA2_+ | exon_06 |
| ERBB2 | TTGGGTGCTCGCGGCTCCGG | chr17:37856544-37856563_ERBB2_- | exon_01 |
| ERBB2 | GGTGGCGGAGCATGTCCAGG | chr17:37863295-37863314_ERBB2_- | exon_02 |
| ERBB2 | ACAATCCGCAGCCTCTGCAG | chr17:37864634-37864653_ERBB2_- | exon_03 |
| ERBB2 | ACGCGCACTGTCTGTGCCGG | chr17:37866341-37866360_ERBB2_+ | exon_06 |
| ERBB2 | TGAGTCCATGCCCAATCCCG | chr17:37866670-37866689_ERBB2_+ | exon_07 |
| ERCC2 | GCGGGAGCTCAAACGCACGC | chr19:45873402-45873421_ERCC2_- | exon_02 |
| ERCC2 | TTATCGGCAGGCATATCCGC | chr19:45872241-45872260_ERCC2_- | exon_04 |
| ERCC2 | AAGGAACAGGTGCTCACCTC | chr19:45871871-45871890_ERCC2_+ | exon_05 |
| ERCC2 | ATCCAGGTTGTAGATGCCAG | chr19:45868156-45868175_ERCC2_+ | exon_07 |
| ERCC2 | GTAGCTATAAACCACCACAT | chr19:45867773-45867792_ERCC2_+ | exon_08 |
| ESR1 | TACCTGGAGAACGAGCCCAG | chr6:152129438-152129457_ESR1_+ | exon_01 |
| ESR1 | AATTCAGATAATCGACGCCA | chr6:152163736-152163755_ESR1_+ | exon_02 |
| ESR1 | GAAGTGGGAATGATGAAAGG | chr6:152201885-152201904_ESR1_+ | exon_03 |

| | | | |
|---|---|---|---|
| ESR1 | TCATGAGCGGCTTGGCCAA | chr6:152265420-152265439_ESR1_- | exon_04 |
| ESR1 | GAAGGTGGACCTGATCATGG | chr6:152332810-152332829_ESR1_- | exon_05 |
| ESR2 | ACAGATGTGATAACTGGCGA | chr14:64749409-64749428_ESR2_+ | exon_02 |
| ESR2 | TACGCATCGGGATATCACTA | chr14:64746752-64746771_ESR2_- | exon_03 |
| ESR2 | AGGCCTGGCAGCTCTTGCGC | chr14:64735555-64735574_ESR2_+ | exon_04 |
| ESR2 | GGTCAGGGACATCATCATGG | chr14:64727222-64727241_ESR2_+ | exon_05 |
| ESR2 | GAGCAAAGATGAGCTTGCCG | chr14:64723963-64723982_ESR2_+ | exon_06 |
| EZH2 | ACTGGGAAGAAATCTGAGAA | chr7:148544362-148544381_EZH2_- | exon_02 |
| EZH2 | ACCAAGAATGGAAACAGCGA | chr7:148543619-148543638_EZH2_- | exon_03 |
| EZH2 | GATCTGGAGGATCACCGAGA | chr7:148525834-148525853_EZH2_- | exon_06 |
| EZH2 | TCAGAAGGAAATTTCCGAGG | chr7:148524322-148524341_EZH2_+ | exon_07 |
| EZH2 | ATTCAGGAGGAAGTGCGCCT | chr7:148523675-148523694_EZH2_+ | exon_08 |
| F13B | TCAGGAGAACTCTATGCAGA | chr1:197036292-197036311_F13B_- | exon_01 |
| F13B | AAGGCTGGTCTCCAGAGCCA | chr1:197031997-197032016_F13B_- | exon_02 |
| F13B | TAGAATGTCTCACATACGGA | chr1:197030054-197030073_F13B_- | exon_04 |
| F13B | CCAGAATCTCCTGTATGCGA | chr1:197029498-197029517_F13B_- | exon_05 |
| F13B | AACGTATTTCTGCTGACCCA | chr1:197026455-197026474_F13B_+ | exon_06 |
| FGF4 | GCGCCACCAGGCTCTCCCAG | chr11:69589696-69589715_FGF4_+ | exon_01 |
| FGF4 | GTTGGGTGCAGTGGGTGCGG | chr11:69589745-69589764_FGF4_+ | exon_01 |
| FGF4 | CGTGGTGAGCATCTTCGGCG | chr11:69588845-69588864_FGF4_- | exon_02 |
| FGF4 | CCTGCTGGAGCTCTCGCCCG | chr11:69588875-69588894_FGF4_- | exon_02 |
| FGF4 | TCAGGGCGATGAACATGCCG | chr11:69588154-69588173_FGF4_+ | exon_03 |
| FGFR1 | TAGGTGGTGTCACTGCCCGA | chr8:38287220-38287239_FGFR1_+ | exon_03 |
| FGFR1 | ATCATCATCATCCTCCGAGG | chr8:38285922-38285941_FGFR1_+ | exon_04 |
| FGFR1 | AAGTTCAAATGCCCTTCCAG | chr8:38285518-38285537_FGFR1_- | exon_05 |
| FGFR1 | GGTGTAGTTGCCCTTGTCAG | chr8:38283698-38283717_FGFR1_+ | exon_06 |
| FGFR1 | GAACTCCACGTTGCTACCCA | chr8:38282138-38282157_FGFR1_+ | exon_07 |
| FGFR2 | GATACCACATTAGAGCCAGA | chr10:123353225-123353244_FGFR2_ | exon_02 |
| FGFR2 | CTTAGTCCAACTGATCACGG | hr10:123325106-123325125_FGFR2_ | exon_03 |
| FGFR2 | AAACGGGAAGGAGTTTAAGC | chr10:123310830-123310849_FGFR2_ | exon_05 |
| FGFR2 | ACTCACCCACAACATCCAGG | hr10:123298100-123298119_FGFR2_ | exon_06 |
| FGFR2 | GCCGGCAAATGCCTCCACAG | chr10:123279627-123279646_FGFR2_ | exon_07 |
| FGFR3 | CACGGCCACGCAGAGCGCGA | chr4:1795684-1795703_FGFR3_- | exon_01 |
| FGFR3 | GAGCAGTTGGTCTTCGGCAG | chr4:1801010-1801029_FGFR3_+ | exon_02 |
| FGFR3 | CATCCGGCAGACGTACACGC | chr4:1803439-1803458_FGFR3_+ | exon_05 |
| FGFR3 | GCTGCCGGCCAACCAGACGG | chr4:1803596-1803615_FGFR3_+ | exon_06 |
| FGFR3 | GGGCGCTAACACCACCGACA | chr4:1805424-1805443_FGFR3_+ | exon_07 |
| FGFR4 | ACCAAGCTCCACTTCCTCAG | chr5:176516677-176516696_FGFR4_- | exon_02 |
| FGFR4 | GATTGCCAGCTTCCTACCTG | chr5:176517560-176517579_FGFR4_+ | exon_03 |
| FGFR4 | AGACCTACTGACCTTGCTGG | chr5:176517819-176517838_FGFR4_- | exon_04 |
| FGFR4 | GGATGGACAGGCCTTTCATG | chr5:176518060-176518079_FGFR4_+ | exon_05 |
| FGFR4 | CCCGGCCAACACCACAGCCG | chr5:176519359-176519378_FGFR4_+ | exon_07 |
| FGR | CTGCAAGAAATTGGAGCCGG | chr1:27950394-27950413_FGR_- | exon_03 |
| FGR | GATGTGGAACTTCTCGCCCT | chr1:27949564-27949583_FGR_+ | exon_04 |
| FGR | CGGGACTGGGATCAGACCAG | chr1:27943478-27943497_FGR_- | exon_07 |
| FGR | GAACGGCAGCACTAAGGTGG | chr1:27942098-27942117_FGR_- | exon_09 |
| FGR | ATTGGTGGACATGGCAGCCC | chr1:27941363-27941382_FGR_- | exon_10 |
| FLT1 | GGTGTCCCAGTAGCTGACCA | chr13:29068960-29068979_FLT1_+ | exon_01 |
| FLT1 | AGCATAACTAAATCTGCCTG | chr13:29041186-29041205_FLT1_- | exon_03 |
| FLT1 | TCAGGGATCAAAGTGTCAAG | chr13:29008332-29008351_FLT1_+ | exon_05 |
| FLT1 | TACCATACAACTTCCGGCGA | chr13:29004184-29004203_FLT1_+ | exon_08 |

| | | | |
|---|---|---|---|
| FLT1 | AGACCCGGCTCTCTACCCAC | chr13:29001396-29001415_FLT1_- | exon_10 |
| FLT3 | GAAGTCATCATCATATCCCA | chr13:28644630-28644649_FLT3_- | exon_02 |
| FLT3 | AGTGTACGAAGCTGCCGCTG | chr13:28636131-28636150_FLT3_- | exon_03 |
| FLT3 | AAAGTAGGTATTCTCCAGCT | chr13:28631538-28631557_FLT3_+ | exon_04 |
| FLT3 | GGTGCTTTGCGATTCACAGG | chr13:28626689-28626708_FLT3_- | exon_05 |
| FLT3 | AAAGCTGTTCATGTGAACCA | chr13:28623818-28623837_FLT3_- | exon_07 |
| FLT4 | GCAGCGGCCGGAGATGCAGC | chr5:180076539-180076558_FLT4_- | exon_01 |
| FLT4 | CCCGACCTTGAACATCACGG | chr5:180058731-180058750_FLT4_- | exon_02 |
| FLT4 | CAAGGACAGCGAGGACACGG | chr5:180057720-180057739_FLT4_- | exon_03 |
| FLT4 | AGGTGGTGTGGGATGACCGG | chr5:180057055-180057074_FLT4_- | exon_05 |
| FLT4 | GCGATTTCGGGAGAGCACCG | chr5:180056274-180056293_FLT4_- | exon_07 |
| FMR1NB | AGTGAGCCACACGCATGGCA | hrX:147062973-147062992_FMR1NB_ | exon_01 |
| FMR1NB | TGGCTGAGGCCTGTCAGCCA | hrX:147063068-147063087_FMR1NB_ | exon_01 |
| FMR1NB | GAGAGCATGAGCATCCCAAA | hrX:147063127-147063146_FMR1NB_ | exon_01 |
| FMR1NB | TCTGGAAGAAGATTCCGCAT | hrX:147084788-147084807_FMR1NB_ | exon_02 |
| FMR1NB | AACCGGCCGATGATTTACAA | hrX:147106387-147106406_FMR1NB_ | exon_05 |
| FRK | GAGGACTTGAGCTTCCGAGC | chr6:116381281-116381300_FRK_- | exon_01 |
| FRK | GGATCCAAATGACTTCCTGA | chr6:116277747-116277766_FRK_- | exon_05 |
| FRK | GCTGCCAGAAATGTCCTCGT | chr6:116265462-116265481_FRK_- | exon_06 |
| FRK | CTCCAGATAGGCCATTCCAG | chr6:116265509-116265528_FRK_+ | exon_06 |
| FRK | TAGACACGAAATAAAGCTGC | chr6:116264303-116264322_FRK_- | exon_07 |
| FYN | AACAACTTCCACGCAGCCGG | chr6:112041085-112041104_FYN_- | exon_04 |
| FYN | TGGATACTACATTACCACCC | chr6:112024133-112024152_FYN_- | exon_08 |
| FYN | TGTGACAGGGAACTACTAGG | chr6:112021427-112021446_FYN_+ | exon_09 |
| FYN | GCTGGTCCAGCTCTATGCAG | chr6:112017528-112017547_FYN_- | exon_10 |
| FYN | GATCAGCAAACATTCTAGTG | chr6:112015657-112015676_FYN_- | exon_12 |
| GABRA1 | AAGGACCTATGACAGTCCTC | hr5:161300181-161300200_GABRA1_ | exon_06 |
| GABRA1 | CCATTAGGTTATTTAACCGG | hr5:161300200-161300219_GABRA1_ | exon_06 |
| GABRA1 | GCATTTGGAGGACTTCCCTA | hr5:161302593-161302612_GABRA1_ | exon_07 |
| GABRA1 | TGCATAAGCCACCTTAGGGA | hr5:161322726-161322745_GABRA1_ | exon_10 |
| GABRA1 | GATTCCAAATAGCAGCGGGA | hr5:161324337-161324356_GABRA1_ | exon_11 |
| GFRAL | AGTGGCAGAGGCATGTGTAG | chr6:55216079-55216098_GFRAL_+ | exon_05 |
| GFRAL | GCACTTGAAGGACCGTCCCA | chr6:55223833-55223852_GFRAL_- | exon_06 |
| GFRAL | CATGCCTTTGACATTAGACA | chr6:55263990-55264009_GFRAL_- | exon_07 |
| GFRAL | AACCAACAGAAGGATTCCAC | chr6:55264206-55264225_GFRAL_- | exon_08 |
| GFRAL | GCTTGACCATAACCAACAGA | chr6:55264216-55264235_GFRAL_- | exon_08 |
| GLRA2 | TCTTGGACAAGTTAATGGGA | chrX:14550432-14550451_GLRA2_+ | exon_02 |
| GLRA2 | TTGCAAGTAACGTTTACTGG | chrX:14592460-14592479_GLRA2_- | exon_03 |
| GLRA2 | ACAGTGGAATGATTCACGGC | chrX:14599334-14599353_GLRA2_+ | exon_04 |
| GLRA2 | GGACTTGAAGAACTTTCCGA | chrX:14622464-14622483_GLRA2_+ | exon_05 |
| GLRA2 | GCCCAGTGCGACCCTGGCAG | chrX:14627248-14627267_GLRA2_- | exon_07 |
| GLS | GCGTGAGCGCGACTCTGCGG | chr2:191745869-191745888_GLS_+ | exon_01 |
| GLS | GATTGCGAACGTCTGATCCC | chr2:191760346-191760365_GLS_+ | exon_03 |
| GLS | ATTGCTCCAGCATTTACCAT | chr2:191775006-191775025_GLS_- | exon_07 |
| GLS | GCTGCGACACTGGCTAATGG | chr2:191792074-191792093_GLS_+ | exon_12 |
| GLS | TAGGTGGAATAACACTCCCA | chr2:191827553-191827572_GLS_+ | exon_18 |
| GPR50 | TTACAGCCAATACAGCCATA | chrX:150345224-150345243_GPR50_ | exon_01 |
| GPR50 | ATAAAGATGATTAGAGCCGG | chrX:150345266-150345285_GPR50_ | exon_01 |
| GPR50 | AGACCTAATCGGCAACTCCA | chrX:150345316-150345335_GPR50_- | exon_01 |
| GPR50 | GGTGCGAGGATCGTACTCGA | chrX:150348561-150348580_GPR50_ | exon_02 |
| GPR50 | GTCAGCCTTGAAATGGACAG | chrX:150349383-150349402_GPR50_ | exon_02 |

| | | | |
|---|---|---|---|
| GPX6 | GAAGGTCTTTACTTTCCTGA | chr6:28473482-28473501_GPX6_- | exon_04 |
| GPX6 | CAGTCGCTAGCCCTTCCTAG | chr6:28472053-28472072_GPX6_+ | exon_05 |
| GPX6 | AAAGCAGTTCAATACCCACT | chr6:28472071-28472090_GPX6_- | exon_05 |
| GPX6 | GGTGGAACCAATGCATGACA | chr6:28472134-28472153_GPX6_+ | exon_05 |
| GPX6 | TGAAGGTCCATGATATCCGC | chr6:28472195-28472214_GPX6_- | exon_05 |
| GSK3A | GGAAGCTAGTGCCTGCGCCG | chr19:42746359-42746378_GSK3A_+ | exon_01 |
| GSK3A | GTCGTAGCCACTCTAGGCCA | chr19:42744252-42744271_GSK3A_- | exon_02 |
| GSK3A | TCAGCCTCACAATATTGCAG | chr19:42741039-42741058_GSK3A_+ | exon_03 |
| GSK3A | GGTGGCCCGCCACTTCACCA | chr19:42740793-42740812_GSK3A_- | exon_04 |
| GSK3A | TTGGCCTACATCCACTCCCA | chr19:42738784-42738803_GSK3A_- | exon_05 |
| GSK3B | AGAGAGTGATCATGTCAGGG | chr3:119812273-119812292_GSK3B_ | exon_01 |
| GSK3B | CGGCAGCAAGGTGACAACAG | chr3:119721057-119721076_GSK3B_ | exon_02 |
| GSK3B | TCTGCGGTTTAATATCCCGA | hr3:119634943-119634962_GSK3B_ | exon_05 |
| GSK3B | AGGACAACCAATATTTCCAG | chr3:119624640-119624659_GSK3B_ | exon_07 |
| GSK3B | TAAGGCACATCCTTGGACTA | chr3:119595262-119595281_GSK3B_ | exon_08 |
| GSX2 | GGATGAAGAAATCCGGCCCG | chr4:54966592-54966611_GSX2_- | exon_01 |
| GSX2 | GATGCGCATGGTTCACCCGC | chr4:54966871-54966890_GSX2_- | exon_01 |
| GSX2 | TGAGGCAGTGGAATCTCCGC | chr4:54967060-54967079_GSX2_- | exon_01 |
| GSX2 | ACGCAGAGGAACAGTCACGC | chr4:54967964-54967983_GSX2_+ | exon_02 |
| GSX2 | GGTGCACTACGCGCGCTCCG | chr4:54968008-54968027_GSX2_+ | exon_02 |
| GTF2B | GGCGGGACATACTAACCGGC | chr1:89357150-89357169_GTF2B_+ | exon_01 |
| GTF2B | TTAGTGGAGGACTACAGAGC | chr1:89352982-89353001_GTF2B_- | exon_02 |
| GTF2B | AAATCTCCATCACTCAGAAG | chr1:89329683-89329702_GTF2B_+ | exon_03 |
| GTF2B | ATATGAACAGAAGAGCCTGA | chr1:89325646-89325665_GTF2B_- | exon_05 |
| GTF2B | GAAAGCGCTAGAAACCAGTG | chr1:89323090-89323109_GTF2B_- | exon_06 |
| HAVCR2 | AGAAGTGGAATACAGAGCGG | hr5:156533947-156533966_HAVCR2_ | exon_02 |
| HAVCR2 | TCCAAGGATGCTTACCACCA | hr5:156531692-156531711_HAVCR2_ | exon_03 |
| HAVCR2 | ATAGCAGAGACACAGACACT | hr5:156525897-156525916_HAVCR2_ | exon_04 |
| HAVCR2 | GATGTAGATGCCTATTCTGA | hr5:156522378-156522397_HAVCR2_ | exon_05 |
| HAVCR2 | ATTGCAAAGCGACAACCCAA | hr5:156514120-156514139_HAVCR2_ | exon_07 |
| HDAC1 | GAGCAAGATGGCGCAGACGC | chr1:32757764-32757783_HDAC1_+ | exon_01 |
| HDAC1 | CCCTCACAAAGCCAATGCTG | chr1:32782268-32782287_HDAC1_+ | exon_03 |
| HDAC1 | GCACCATGCAAAGAAGTCCG | chr1:32792601-32792620_HDAC1_+ | exon_05 |
| HDAC1 | TAAAGGGCGTTCTCACCCGT | chr1:32793275-32793294_HDAC1_- | exon_06 |
| HDAC1 | GATGTTCCAGCCTAGTGCGG | chr1:32796199-32796218_HDAC1_+ | exon_08 |
| HDAC2 | ATGGCGTACAGTCAAGGAGG | chr6:114292053-114292072_HDAC2_ | exon_01 |
| HDAC2 | AAACCGACAACAGACTGATA | chr6:114277280-114277299_HDAC2_ | exon_05 |
| HDAC2 | GATGTATCAACCTAGTGCTG | chr6:114270212-114270231_HDAC2_ | exon_08 |
| HDAC2 | GATAAATCCAAGGACAACAG | chr6:114262896-114262915_HDAC2_ | exon_13 |
| HDAC2 | GGTGAGACTGTCAAATTCAG | hr6:114262206-114262225_HDAC2_- | exon_14 |
| HDAC3 | GACGTGGGCAACTTCCACTA | chr5:141016305-141016324_HDAC3_ | exon_01 |
| HDAC3 | TATGGGTCAATGCCAGGCGA | hr5:141016153-141016172_HDAC3_- | exon_02 |
| HDAC3 | CATGTGCCGCTTCCACTCCG | chr5:141014469-141014488_HDAC3_ | exon_03 |
| HDAC3 | GAGCAGAACTCAAAGAGCCC | hr5:141009660-141009679_HDAC3_- | exon_04 |
| HDAC3 | GTTGACATAGCAGAAGCCAG | hr5:141009283-141009302_HDAC3_- | exon_06 |
| HDAC6 | GAAGTGACACTGGAGTCCTG | chrX:48661172-48661191_HDAC6_- | exon_02 |
| HDAC6 | AATGGAAGAAGACCTAATCG | chrX:48661370-48661389_HDAC6_+ | exon_03 |
| HDAC6 | AGCCTCACCTGTCATCCCAG | chrX:48661612-48661631_HDAC6_- | exon_04 |
| HDAC6 | TGAGGTGGTCTGGTCCGGAG | chrX:48681649-48681668_HDAC6_- | exon_25 |
| HDAC6 | TCTGGACCAGACCACCTCAG | chrX:48681692-48681711_HDAC6_+ | exon_25 |
| HDAC8 | GCGGAAGATGGAGGAGCCGG | chrX:71792599-71792618_HDAC8_- | exon_01 |

| | | | |
|---|---|---|---|
| HDAC8 | GAAGGTGGCCATCTCCTCCA | chrX:71788689-71788708_HDAC8_+ | exon_03 |
| HDAC8 | ACTATGCAGCAGCTATAGGA | chrX:71787828-71787847_HDAC8_- | exon_04 |
| HDAC8 | GTGGATTTGGATCTGCACCA | chrX:71715014-71715033_HDAC8_- | exon_05 |
| HDAC8 | GACGTGTCTGATGTTGGCCT | chrX:71708864-71708883_HDAC8_- | exon_07 |
| HLA-G | ACAGCGACTCGGCGTGTCCG | chr6:29795932-29795951_HLA-G_+ | exon_02 |
| HLA-G | AGAGGAGACACGGAACACCA | chr6:29796005-29796024_HLA-G_+ | exon_02 |
| HLA-G | GCAGCCAATCATCCACTGGA | chr6:29796332-29796351_HLA-G_- | exon_03 |
| HLA-G | ACAAAGGAGAGCCTACCTGG | chr6:29796510-29796529_HLA-G_+ | exon_03 |
| HLA-G | CATAGTCAAAGACAGGGTGG | chr6:29797220-29797239_HLA-G_- | exon_04 |
| HSP90AA1 | GGTGTCGTTCCAGATGCCTG | 14:102552709-102552728_HSP90AA | exon_02 |
| HSP90AA1 | GATCTGTCAAGCTTTCATAC | 14:102552425-102552444_HSP90AA | exon_03 |
| HSP90AA1 | TTTCCGAAGACGTTCCACAA | 14:102549587-102549606_HSP90AA | exon_09 |
| HSP90AA1 | TAAGAACGACAAGTCTGTGA | 14:102548559-102548578_HSP90AA | exon_10 |
| HSP90AA1 | AGATGACGACACATCACGCA | 14:102548066-102548085_HSP90AA | exon_11 |
| HTR5A | TTGGAGACCAACCACAGCCT | chr7:154862661-154862680_HTR5A_ | exon_01 |
| HTR5A | ACTCCGAAGACCGAGAGCAG | chr7:154862712-154862731_HTR5A_- | exon_01 |
| HTR5A | GGTGGAAGGTGCGTACACGG | chr7:154862807-154862826_HTR5A_- | exon_01 |
| HTR5A | AATGAGGATGCCCACCATGA | chr7:154875977-154875996_HTR5A_- | exon_02 |
| HTR5A | AGTTGGAGTAGCCAAGCCAC | chr7:154876095-154876114_HTR5A_- | exon_02 |
| ICOSLG | GGTGACGTTGAACAAGCGCA | chr21:45656844-45656863_ICOSLG_ | exon_03 |
| ICOSLG | TGGGATGTGGTAGGTCACCA | chr21:45656952-45656971_ICOSLG_ | exon_03 |
| ICOSLG | GCAGCAGCCAATGTTCACGC | chr21:45655204-45655223_ICOSLG_ | exon_04 |
| ICOSLG | GACAGCCAGGATGCTCCACG | chr21:45651239-45651258_ICOSLG_ | exon_05 |
| ICOSLG | TGTCTCCGGACTCACAGCCC | chr21:45649947-45649966_ICOSLG_ | exon_06 |
| ICOS | CAAACATGAAGTCAGGCCTC | chr2:204801533-204801552_ICOS_+ | exon_01 |
| ICOS | AAATGCAGTTGCTGAAAGGG | chr2:204820455-204820474_ICOS_+ | exon_02 |
| ICOS | TGGGTAACCAGAACTTCAGC | chr2:204821401-204821420_ICOS_- | exon_03 |
| ICOS | ACAAAGGCTGCACATCCTAT | chr2:204821420-204821439_ICOS_- | exon_03 |
| ICOS | ACCGTTAGGGTCGTGCACAC | chr2:204822535-204822554_ICOS_- | exon_04 |
| IDH1 | ATGTAGATCCAATTCCACGT | chr2:209116156-209116175_IDH1_+ | exon_03 |
| IDH1 | ACCCATCCACTCACAAGCCG | chr2:209113133-209113152_IDH1_+ | exon_04 |
| IDH1 | TAGGCTCATCGACGACATGG | chr2:209106805-209106824_IDH1_- | exon_07 |
| IDH1 | ATCTGGACAAACCAGCACGC | chr2:209104681-209104700_IDH1_+ | exon_08 |
| IDH1 | TGAGGCTGGCTTCATGACCA | chr2:209103829-209103848_IDH1_- | exon_09 |
| IDH2 | AGAGCGAGCGCACGACCCGC | chr15:90645589-90645608_IDH2_+ | exon_01 |
| IDH2 | GATCAAGGTGGCGAAGCCCG | chr15:90634844-90634863_IDH2_- | exon_02 |
| IDH2 | ACACCCTCGCACCTTCCACA | chr15:90633699-90633718_IDH2_+ | exon_03 |
| IDH2 | AACTATCCGGAACATCCTGG | chr15:90631923-90631942_IDH2_- | exon_04 |
| IDH2 | GGGCATGTACAACACCGACG | chr15:90631593-90631612_IDH2_- | exon_05 |
| IGF1R | GATCAGGGACCAGTCCACAG | chr15:99251157-99251176_IGF1R_- | exon_02 |
| IGF1R | GCCGTTGCGGATGAAGCCCG | chr15:99434839-99434858_IGF1R_- | exon_03 |
| IGF1R | TCGGGCAAGGACCTTCACAA | chr15:99440001-99440020_IGF1R_- | exon_04 |
| IGF1R | ACTAAAGGGCGCCAAAGCAA | chr15:99452068-99452087_IGF1R_+ | exon_06 |
| IGF1R | GATGATGCGATTCTTCGACG | chr15:99454580-99454599_IGF1R_- | exon_07 |
| IGSF11 | GGAGCAGCAAAGGCGCCAGA | chr3:118753390-118753409_IGSF11_- | exon_01 |
| IGSF11 | GGGAGTATCCAGGTGGCCCG | chr3:118649068-118649087_IGSF11_ | exon_02 |
| IGSF11 | GGGCCACCTGGATACTCCCA | chr3:118649069-118649088_IGSF11_- | exon_02 |
| IGSF11 | TTTGTGACAGACCAGGTCCA | chr3:118624556-118624575_IGSF11_ | exon_05 |
| IGSF11 | TTGGCAGAAGAACACTTGGG | chr3:118621773-118621792_IGSF11_- | exon_07 |
| IKBKE | GTTGCGGGCCTTGTACACAC | chr1:206646635-206646654_IKBKE_- | exon_03 |
| IKBKE | TCAACACTACCAGCTACCTG | chr1:206647705-206647724_IKBKE_+ | exon_04 |

| | | | |
|---|---|---|---|
| IKBKE | ATAGGGCGGAAGCCGGCAGA | chr1:206648204-206648223_IKBKE_+ | exon_05 |
| IKBKE | TGCATCGCGACATCAAGCCG | chr1:206649560-206649579_IKBKE_+ | exon_06 |
| IKBKE | GATGAAGGGCAGGCTGCCAG | chr1:206650127-206650146_IKBKE_- | exon_07 |
| IL1B | GAGCTCAGGTACTTCTGCCA | chr2:113593786-113593805_IL1B_+ | exon_02 |
| IL1B | GCTCAGGTCATTCTCCTGGA | chr2:113590979-113590998_IL1B_+ | exon_04 |
| IL1B | AGCTGGATGCCGCCATCCAG | chr2:113591103-113591122_IL1B_+ | exon_04 |
| IL1B | GCTGGATGCCGCCATCCAGA | chr2:113591104-113591123_IL1B_+ | exon_04 |
| IL1B | AGAGGTGCTGATGTACCAGT | chr2:113588025-113588044_IL1B_+ | exon_07 |
| IL6R | GGAGGAAGCATGCTGGCCGT | chr1:154378097-154378116_IL6R_+ | exon_01 |
| IL6R | TGGGACTGTGCACTTGCTGG | chr1:154401895-154401914_IL6R_+ | exon_02 |
| IL6R | ACTCACAAACAACATTGCTG | chr1:154403005-154403024_IL6R_- | exon_03 |
| IL6R | CTGCCAGTTAGCAGTCCCGG | chr1:154407061-154407080_IL6R_+ | exon_04 |
| IL6R | CAACATCACAGTCACTGCCG | chr1:154407485-154407504_IL6R_+ | exon_05 |
| IL6 | GGAGAAGGCAACTGGACCGA | chr7:22767069-22767088_IL6_- | exon_03 |
| IL6 | TCAGGGCTGAGATGCCGTCG | chr7:22767226-22767245_IL6_- | exon_03 |
| IL6 | CATGTGTGAAAGCAGCAAAG | chr7:22768329-22768348_IL6_+ | exon_04 |
| IL6 | AGCTGCAGGCACAGAACCAG | chr7:22771086-22771105_IL6_+ | exon_06 |
| IL6 | TTGCCGAAGAGCCCTCAGGC | chr7:22771167-22771186_IL6_- | exon_06 |
| IMP3 | GAGCAAGGGCACGTACGCGT | chr15:75932097-75932116_IMP3_- | exon_02 |
| IMP3 | GGAGCAAGGGCACGTACGCG | chr15:75932098-75932117_IMP3_- | exon_02 |
| IMP3 | GCGCAGCTTGAGGAGCACGG | chr15:75932153-75932172_IMP3_+ | exon_02 |
| IMP3 | GGAGCTCTGCGACTTCGTCA | chr15:75932206-75932225_IMP3_- | exon_02 |
| IMP3 | ATACAGCTTGTCCAGCAGCG | chr15:75932258-75932277_IMP3_+ | exon_02 |
| INSR | AACCTGATGAACATCACCCG | chr19:7267586-7267605_INSR_- | exon_02 |
| INSR | ACAAATGCAAGAACTCGCGG | chr19:7184410-7184429_INSR_- | exon_03 |
| INSR | GAGGTGGCACACCTTGGGAC | chr19:7174694-7174713_INSR_+ | exon_04 |
| INSR | GGAAAGTGACACCAGAGCGT | chr19:7172348-7172367_INSR_+ | exon_05 |
| INSR | GAAGAAGTTTCAGGAACCAA | chr19:7170604-7170623_INSR_- | exon_06 |
| IRAK4 | TATGTGCGCTGCCTCAATGT | chr12:44161942-44161961_IRAK4_+ | exon_02 |
| IRAK4 | TCTGGACTTGAGGAGTCAGG | chr12:44166117-44166136_IRAK4_- | exon_04 |
| IRAK4 | GATGAACGACCCATTTCTGT | chr12:44166765-44166784_IRAK4_+ | exon_05 |
| IRAK4 | CATGAGATGCAAGATTGCTC | chr12:44172022-44172041_IRAK4_+ | exon_08 |
| IRAK4 | ATGGCACCAGAAGCTTTGCG | chr12:44176231-44176250_IRAK4_+ | exon_09 |
| ITK | TGATGAGCTGTTCTTCCAGG | chr5:156608006-156608025_ITK_- | exon_01 |
| ITK | ACAGAAGAAGCGCACGCTGA | chr5:156635896-156635915_ITK_+ | exon_02 |
| ITK | TGGGTGCTGGCCCTTAAAGA | chr5:156638358-156638377_ITK_+ | exon_03 |
| ITK | CAACGAAGAGTACTGCCTGC | chr5:156649959-156649978_ITK_+ | exon_06 |
| ITK | TGAAGGCTCCTTCTTTGCCC | chr5:156665118-156665137_ITK_- | exon_09 |
| JAK1 | CAGTGGAGAGTACACAGCAG | chr1:65348993-65349012_JAK1_- | exon_03 |
| JAK1 | ATGACGAGAACACCAAGCTC | chr1:65344776-65344795_JAK1_- | exon_04 |
| JAK1 | AAACAGATACTCCAGTGAGC | chr1:65339059-65339078_JAK1_+ | exon_05 |
| JAK1 | GCAGTTGCCAGAACTGCCCA | chr1:65335004-65335023_JAK1_- | exon_06 |
| JAK1 | GAGAAGGTTCCTCTGTCTGA | chr1:65332834-65332853_JAK1_+ | exon_07 |
| JAK2 | ATCTGCCTCAGATTTCCCAA | chr9:5022127-5022146_JAK2_- | exon_03 |
| JAK2 | TATCGGCATGGAATATCTCG | chr9:5044446-5044465_JAK2_+ | exon_05 |
| JAK2 | ATAGAGTTATAGATGGCCAG | chr9:5050809-5050828_JAK2_- | exon_06 |
| JAK2 | GAGGTGCTACTTCTTTACAG | chr9:5064975-5064994_JAK2_- | exon_09 |
| JAK2 | AATGAAGAGTACAACCTCAG | chr9:5069070-5069089_JAK2_+ | exon_11 |
| JAK3 | ACTAGGGATGCACTCACCGC | chr19:17955026-17955045_JAK3_+ | exon_01 |
| JAK3 | CCCGAGCCACATCTTCTCCG | chr19:17954626-17954645_JAK3_- | exon_02 |
| JAK3 | CCTTGCGTAGCCCGAAGCGG | chr19:17954242-17954261_JAK3_+ | exon_03 |

| | | | |
|---|---|---|---|
| JAK3 | TCAGCAGCTCTCCCGGCCGC | chr19:17953846-17953865_JAK3_+ | exon_04 |
| JAK3 | TGACGCGGAGGCGTATTCGG | chr19:17953344-17953363_JAK3_- | exon_05 |
| KCNK10 | AGCTGGACTTCACTTACCCA | chr14:88792746-88792765_KCNK10_+ | exon_01 |
| KCNK10 | GATGGAAGGCACCTCCCAAG | chr14:88729752-88729771_KCNK10_- | exon_02 |
| KCNK10 | AGGCACTGCCGAGGTCCCAG | chr14:88707080-88707099_KCNK10_+ | exon_03 |
| KCNK10 | GCCGGCCAAGATGAACAGGA | chr14:88658692-88658711_KCNK10_+ | exon_05 |
| KCNK10 | GGAGAACGGAATGATACCCA | chr14:88651935-88651954_KCNK10_- | exon_07 |
| KDM1A | GAATAGCAGAGACTCCGGAG | chr1:23346388-23346407_KDM1A_+ | exon_01 |
| KDM1A | GGATACTACAACTCACCCGA | chr1:23357124-23357143_KDM1A_- | exon_02 |
| KDM1A | TTCAAGACGACAGTTCTGGA | chr1:23370935-23370954_KDM1A_+ | exon_03 |
| KDM1A | TGGACCACAACAGACCCAGA | chr1:23376965-23376984_KDM1A_+ | exon_04 |
| KDM1A | CGAGTTGCCACATTTCGCAA | chr1:23383992-23384011_KDM1A_+ | exon_08 |
| KDR | GGGCTCCTTACCCACAGAGG | chr4:55991383-55991402_KDR_+ | exon_01 |
| KDR | GAGGTGACTGAGTGCAGCGA | chr4:55984887-55984906_KDR_- | exon_03 |
| KDR | AATTTCCTGGGACAGCAAGA | chr4:55981152-55981171_KDR_- | exon_05 |
| KDR | CTGGGAATACCCTTCTTCGA | chr4:55980295-55980314_KDR_- | exon_06 |
| KDR | GAAGAACAGCACATTTGTCA | chr4:55979480-55979499_KDR_- | exon_07 |
| KIF11 | GAAGTTAGTGTACGAACTGG | chr10:94366089-94366108_KIF11_+ | exon_02 |
| KIF11 | TGAAGAGTATACCTGGGAAG | chr10:94366973-94366992_KIF11_+ | exon_04 |
| KIF11 | CGTGGAATTATACCAGCCAA | chr10:94368783-94368802_KIF11_- | exon_05 |
| KIF11 | AATCCCTGTTGACTTTGGGA | chr10:94373213-94373232_KIF11_+ | exon_08 |
| KIF11 | TCTTGTGTAGGAGTATACGG | chr10:94381132-94381151_KIF11_+ | exon_10 |
| KIT | TCAGACTTAATAGTCCGCGT | chr4:55561740-55561759_KIT_+ | exon_02 |
| KIT | GAAAGAAGACAACGACACGC | chr4:55564490-55564509_KIT_+ | exon_03 |
| KIT | GCAAGCTATCTTCTTAGGGA | chr4:55565831-55565850_KIT_+ | exon_04 |
| KIT | CCAAACCTGAACACCAGCAG | chr4:55573360-55573379_KIT_+ | exon_06 |
| KIT | ACCACTAGCTTTCCAAACGG | chr4:55592078-55592097_KIT_- | exon_09 |
| KLK12 | AGAACTCACCAAGAACACAC | chr19:51537832-51537851_KLK12_+ | exon_02 |
| KLK12 | GAGGGCACCAGCCTGCGCTG | chr19:51537293-51537312_KLK12_- | exon_03 |
| KLK12 | GACGTGGCACTCGGTGCCAG | chr19:51535163-51535182_KLK12_+ | exon_04 |
| KLK12 | ACCGAGCAGATCCGGCACAG | chr19:51535323-51535342_KLK12_- | exon_04 |
| KLK12 | TGTGTGCAGGCGGCGTCCCG | chr19:51534062-51534081_KLK12_- | exon_05 |
| LAG3 | GCAGAAGGCTGAGATCCTGG | chr12:6882449-6882468_LAG3_- | exon_02 |
| LAG3 | GCGGTCCCTGAGGTGCACCG | chr12:6883105-6883124_LAG3_- | exon_03 |
| LAG3 | GTTCCGGAACCAATGCACAG | chr12:6883845-6883864_LAG3_- | exon_04 |
| LAG3 | GCCAAGTGGACTCCTCCTGG | chr12:6884543-6884562_LAG3_+ | exon_05 |
| LAG3 | GCAATGCCAGCTGTACCAGG | chr12:6886602-6886621_LAG3_+ | exon_06 |
| LAP3 | CTCCACCGCAGACATGACGA | chr4:17579169-17579188_LAP3_+ | exon_01 |
| LAP3 | TCTGGACCACCTCTGAAGGC | chr4:17583384-17583403_LAP3_+ | exon_03 |
| LAP3 | GAGGCCAACTAGCACCACGC | chr4:17583921-17583940_LAP3_- | exon_04 |
| LAP3 | TCTGTGGAGGTGGATCCCTG | chr4:17585141-17585160_LAP3_+ | exon_05 |
| LAP3 | AATGGGATCATTCCTCAGTG | chr4:17590469-17590488_LAP3_+ | exon_07 |
| LAS1L | GCCAGGCGACCACGATGCCG | chrX:64754460-64754479_LAS1L_+ | exon_01 |
| LAS1L | GCAGTAGAAGCCACTGCCAG | chrX:64753577-64753596_LAS1L_+ | exon_02 |
| LAS1L | GAGGAAGACAAAGTTTGCCA | chrX:64752470-64752489_LAS1L_- | exon_03 |
| LAS1L | TGTGGGTCAACTCATGGCGA | chrX:64751270-64751289_LAS1L_+ | exon_04 |
| LAS1L | CCAGGACACATTCTACACGT | chrX:64748171-64748190_LAS1L_+ | exon_07 |
| LCK | TATCCCATAGTCCCACTGGA | chr1:32740003-32740022_LCK_+ | exon_02 |
| LCK | GAGCCTTCGTAGGTAACCAG | chr1:32740374-32740393_LCK_- | exon_03 |
| LCK | GAAGGCGCAGTCCCTGACCA | chr1:32740937-32740956_LCK_+ | exon_05 |
| LCK | CCGGGAGAGCGAGAGCACCG | chr1:32741251-32741270_LCK_+ | exon_06 |

| | | | |
|---|---|---|---|
| LCK | AAGATCCGTAATCTGGACAA | chr1:32741577-32741596_LCK_+ | exon_07 |
| LDHA | TACGCTGGACCAAATTAAGA | chr11:18422457-18422476_LDHA_- | exon_04 |
| LDHA | TACAGAGAGTCCAATAGCCC | chr11:18427034-18427053_LDHA_- | exon_07 |
| LDHA | TAGACCTACCTTAATCATGG | chr11:18427109-18427128_LDHA_- | exon_07 |
| LGALS3 | GAAACCCAAACCCTCAAGGA | chr14:55604788-55604807_LGALS3_+ | exon_03 |
| LGALS3 | CATGCGCCAGGCCATCCTTG | chr14:55604802-55604821_LGALS3_- | exon_03 |
| LGALS3 | GTCTACCCAGGGCCACCCAG | chr14:55604976-55604995_LGALS3_+ | exon_03 |
| LGALS3 | TCTCACCAGTGGCCCAGCAG | chr14:55605073-55605092_LGALS3_- | exon_03 |
| LGALS3 | GCTGATAACAATTCTGGGCA | chr14:55607040-55607059_LGALS3_+ | exon_04 |
| LGALS9B | GCGGTGGAGATGGCCTTCAG | chr17:20370773-20370792_LGALS9B_ | exon_01 |
| LGALS9 | GACACACATGCCTTTCCAGA | chr17:25967730-25967749_LGALS9_+ | exon_03 |
| LGALS9 | ACACACATGCCTTTCCAGAA | chr17:25967731-25967750_LGALS9_+ | exon_03 |
| LUC7L3 | AGAAGCGCAGCAACGTGCGG | chr17:48797158-48797177_LUC7L3_+ | exon_01 |
| LUC7L3 | GTATTTGTGAACAATTCCGC | chr17:48814354-48814373_LUC7L3_- | exon_02 |
| LUC7L3 | GTAGAACGTAGGATCAGACG | chr17:48818542-48818561_LUC7L3_+ | exon_04 |
| LUC7L3 | ACTGCTAAGGTCCACAACGT | chr17:48821150-48821169_LUC7L3_+ | exon_06 |
| LUC7L3 | AGACACAAACACTGAATCGA | chr17:48827926-48827945_LUC7L3_+ | exon_10 |
| LYN | CAACGTCCAATAAACAGCAA | chr8:56854525-56854544_LYN_+ | exon_02 |
| LYN | TGAAAGACAAGTCGTCCGGG | chr8:56860232-56860251_LYN_- | exon_04 |
| LYN | GTTTGGCCACATAGTTGCTG | chr8:56863075-56863094_LYN_- | exon_05 |
| LYN | TTGGCACCAGGAAATAGCGC | chr8:56863286-56863305_LYN_+ | exon_06 |
| LYN | ACCAACTTGATGGACTCCCG | chr8:56866483-56866502_LYN_- | exon_08 |
| MAP2K1 | AGTTAACGGGACCAGCTCTG | chr15:66679742-66679761_MAP2K1_- | exon_01 |
| MAP2K1 | TTGAGAAGATCAGTGAGCTG | chr15:66727487-66727506_MAP2K1_- | exon_02 |
| MAP2K1 | TATGGTGCGTTCTACAGCGA | chr15:66729180-66729199_MAP2K1_- | exon_03 |
| MAP2K1 | AACATCCTAGTCAACTCCCG | chr15:66774107-66774126_MAP2K1_- | exon_06 |
| MAP2K1 | AATGGACAGCCGACCTCCCA | chr15:66779573-66779592_MAP2K1_- | exon_08 |
| MAP2K2 | CATCAACCCTACCATCGCCG | chr19:4123815-4123834_MAP2K2_- | exon_01 |
| MAP2K2 | CAAAGTCCAGCACAGACCCT | chr19:4117440-4117459_MAP2K2_- | exon_02 |
| MAP2K2 | TGATCTGGTGCTTCTCTCGG | chr19:4101236-4101255_MAP2K2_+ | exon_05 |
| MAP2K2 | GTAGGAGCGCGTGCCCACGA | chr19:4101023-4101042_MAP2K2_+ | exon_06 |
| MAP2K2 | ACAGCATCTCGCCTCGGCCG | chr19:4099224-4099243_MAP2K2_- | exon_07 |
| MAP2 | TAAGGATCAAGGCGGAGCAG | chr2:210517987-210518006_MAP2_+ | exon_04 |
| MAP2 | AGTTCAAGTAGTCACTGCTG | chr2:210543315-210543334_MAP2_+ | exon_05 |
| MAP2 | AGAACAGACTGTCACAGTGG | chr2:210545526-210545545_MAP2_+ | exon_06 |
| MAP2 | AGTGTGCCCAAACAAAGCTA | chr2:210557562-210557581_MAP2_- | exon_07 |
| MAP2 | TCTCCTGACACACCTCGCCG | chr2:210570370-210570389_MAP2_- | exon_11 |
| MAP3K8 | TGCCAAGTATACCTTTCCAA | chr10:30736820-30736839_MAP3K8_- | exon_04 |
| MAP3K8 | GTGACAAAGCATGTTCTCAA | chr10:30739379-30739398_MAP3K8_- | exon_05 |
| MAP3K8 | TAATTACCTCTGTTCCTCGG | chr10:30740661-30740680_MAP3K8_- | exon_06 |
| MAP3K8 | AGGAGGGATAGGCTGAGCGA | chr10:30747135-30747154_MAP3K8_- | exon_07 |
| MAP3K8 | TTCGTGCACAGGAAGCACCG | chr10:30749639-30749658_MAP3K8_- | exon_09 |
| MAP4 | TCAGAAACTAGCCAGATTGA | chr3:48040130-48040149_MAP4_- | exon_02 |
| MAP4 | AAACCAACACTCCTAGCCAA | chr3:48019390-48019409_MAP4_- | exon_04 |
| MAP4 | AATCTGCCAGGTCATCATCA | chr3:47963330-47963349_MAP4_+ | exon_06 |
| MAP4 | AAACAAGATAGTCCCAGCCA | chr3:47958095-47958114_MAP4_- | exon_08 |
| MAP4 | AAAGTGCAGCTTGCCGGCCG | chr3:47956388-47956407_MAP4_- | exon_09 |
| MAPK11 | GCCGGTAGAAGCCGGCGCGA | chr22:50708691-50708710_MAPK11_- | exon_01 |
| MAPK11 | GCAGAACGTACCGGGAGCTG | chr22:50706279-50706298_MAPK11_- | exon_02 |
| MAPK11 | GCTTCTGGACGTCTTCACGC | chr22:50706029-50706048_MAPK11_- | exon_03 |
| MAPK11 | CAGGTCGGCGCCCATCAGGG | chr22:50705878-50705897_MAPK11_- | exon_04 |

| | | | |
|---|---|---|---|
| MAPK11 | CTGCGGTCGCACCTACCCGG | chr22:50705670-50705689_MAPK11_- | exon_05 |
| MAPK12 | GGAGGTGACCAAGACGGCCT | chr22:50699878-50699897_MAPK12_- | exon_01 |
| MAPK12 | CTTGATGGCCACCTTAGCGC | chr22:50699683-50699702_MAPK12_- | exon_02 |
| MAPK12 | ACAGATCCTGGACTTCGGCC | chr22:50694613-50694632_MAPK12_- | exon_07 |
| MAPK12 | ACAGGCAAGACGCTGTTCAA | chr22:50694250-50694269_MAPK12_- | exon_08 |
| MAPK12 | CGCTGCACAAACTCAGCCGG | chr22:50694060-50694079_MAPK12_- | exon_09 |
| MAPK13 | CAAGTCATAGAAGTTGCGCA | chr6:36100438-36100457_MAPK13_- | exon_03 |
| MAPK13 | ATAGTACATCCACTCTGCTG | chr6:36103792-36103811_MAPK13_+ | exon_05 |
| MAPK13 | GCAGAAGCTGAACGACAAAG | chr6:36106200-36106219_MAPK13_+ | exon_09 |
| MAPK13 | GCTGAGTGAAATCCTTCCTG | chr6:36106514-36106533_MAPK13_- | exon_10 |
| MAPK13 | AGAACACGAGAAACTCACAG | chr6:36106798-36106817_MAPK13_+ | exon_11 |
| MAPK14 | GCTGAACAAGACAATCTGGG | chr6:35995970-35995989_MAPK14_+ | exon_01 |
| MAPK14 | AAAGAACCTACAGAGAACTG | chr6:36020556-36020575_MAPK14_+ | exon_02 |
| MAPK14 | CCTGTTAATGAGATAAGCGG | chr6:36068010-36068029_MAPK14_- | exon_09 |
| MAPK14 | TATATGGTCAACTTACCCAG | chr6:36070423-36070442_MAPK14_- | exon_10 |
| MAPK14 | AGCAAAGTAGGCATGTGCAA | chr6:36075298-36075317_MAPK14_- | exon_11 |
| MAPK1 | CCCGGCAGCCAACATGGCGG | chr22:22221724-22221743_MAPK1_- | exon_01 |
| MAPK1 | CCTACTGCCAGAGAACCCTG | chr22:22162048-22162067_MAPK1_- | exon_02 |
| MAPK1 | GATCCAGACCATGATCACAC | chr22:22153368-22153387_MAPK1_- | exon_04 |
| MAPK1 | TTTGGGAACAGCCTGTTCCA | chr22:22142560-22142579_MAPK1_+ | exon_06 |
| MAPK1 | TGACATTCAACCCACACAAG | chr22:22127228-22127247_MAPK1_- | exon_07 |
| MAPK3 | GCAGTTGCAGTACATCGGCG | chr16:30134383-30134402_MAPK3_- | exon_01 |
| MAPK3 | AGTAGGTCTGATGTTCGAAG | chr16:30133254-30133273_MAPK3_+ | exon_02 |
| MAPK3 | TGGAGGGCTTTAGATCTCGG | chr16:30129702-30129721_MAPK3_+ | exon_03 |
| MAPK3 | TGACCACACCGGCTTCCTGA | chr16:30129424-30129443_MAPK3_- | exon_04 |
| MAPK3 | GATCCAGGTAGTGCTTGCCA | chr16:30129009-30129028_MAPK3_+ | exon_05 |
| MAPK7 | GAAGGAGCGGGCTTTAAGCA | chr17:19282329-19282348_MAPK7_- | exon_02 |
| MAPK7 | GAATTCGCCATAGGGCACGG | chr17:19283233-19283252_MAPK7_- | exon_03 |
| MAPK7 | AAAGGCAAAGTCAAAGGGCG | chr17:19284604-19284623_MAPK7_- | exon_04 |
| MAPK7 | GCGGGAGCGAAAGGAACGGG | chr17:19285227-19285246_MAPK7_+ | exon_05 |
| MAPK7 | TGTGTTCTCAGGCACACCAA | chr17:19286146-19286165_MAPK7_+ | exon_06 |
| MAPK8 | TCTTTGCCAAGTGATTCAGA | chr10:49618103-49618122_MAPK8_+ | exon_05 |
| MAPK8 | TCGCTACTACAGAGCACCCG | chr10:49628311-49628330_MAPK8_+ | exon_06 |
| MAPK8 | TCAACGTAAGTCCTTACTGT | chr10:49634005-49634024_MAPK8_- | exon_08 |
| MAPK8 | GGGAACACACAATAGAAGAG | chr10:49635160-49635179_MAPK8_+ | exon_10 |
| MAPK8 | CCAAGAATGGAGTTATACGG | chr10:49639275-49639294_MAPK8_+ | exon_11 |
| MAPK9 | GAAGGATACGGTCAGTGCCT | chr5:179674430-179674449_MAPK9_- | exon_07 |
| MAPK9 | TCACGGCTTCTGCTTCGGCG | chr5:179668027-179668046_MAPK9_- | exon_09 |
| MAPK9 | TAGTGATTGATCCTGACAAG | chr5:179668103-179668122_MAPK9_- | exon_09 |
| MAPK9 | TGGGCATCATAAATTTGAGG | chr5:179666962-179666981_MAPK9_- | exon_10 |
| MAPK9 | TGAGGCCAGCGTCTGCTCAG | chr5:179663438-179663457_MAPK9_- | exon_12 |
| MAPT | GAGATCCCAGAAGGAACCAC | chr17:44051816-44051835_MAPT_+ | exon_03 |
| MAPT | TGGAGAGGAAATCCACAGGG | chr17:44060971-44060990_MAPT_- | exon_05 |
| MAPT | CACCCGGGAGCCCAAGAAGG | chr17:44073865-44073884_MAPT_+ | exon_10 |
| MAPT | AATATCAAACACGTCCCGGG | chr17:44087739-44087758_MAPT_+ | exon_11 |
| MAPT | AATATCACCCACGTCCCTGG | chr17:44096061-44096080_MAPT_+ | exon_13 |
| MCL1 | AGGCGCTGGAGACCTTACGA | chr1:150551365-150551384_MCL1_- | exon_01 |
| MCL1 | TCTGGCGTGAGGGTGGACGG | chr1:150551792-150551811_MCL1_+ | exon_01 |
| MCL1 | GCTTCCGCCAATCACCGCGC | chr1:150551827-150551846_MCL1_+ | exon_01 |
| MCL1 | ATCGGACTCAACCTCTACTG | chr1:150551960-150551979_MCL1_- | exon_01 |
| MCL1 | CATGTAGAGGACCTAGAAGG | chr1:150549927-150549946_MCL1_- | exon_03 |

| | | | |
|---|---|---|---|
| MDM2 | TTGGTAGTAGTCAATCAGCA | chr12:69214133-69214152_MDM2_+ | exon_05 |
| MDM2 | AACAGGTGTCACCTTGAAGG | chr12:69218175-69218194_MDM2_+ | exon_06 |
| MDM2 | GAGAACATTACCGGATTCGA | chr12:69222703-69222722_MDM2_- | exon_08 |
| MDM2 | AGGAAGCCAATTCTCACGAA | chr12:69233127-69233146_MDM2_- | exon_11 |
| MDM2 | CAGGAAGCCAATTCTCACGA | chr12:69233128-69233147_MDM2_- | exon_11 |
| MEP1A | ATAGGGCTTGAAATCCACAC | chr6:46777214-46777233_MEP1A_- | exon_06 |
| MEP1A | GTGGGACAGAACATTTCCAT | chr6:46787294-46787313_MEP1A_+ | exon_07 |
| MEP1A | TTTGAGAAGGCAAACATCTG | chr6:46794123-46794142_MEP1A_+ | exon_09 |
| MEP1A | AAGAATCCGAGACTCCAGTA | chr6:46797150-46797169_MEP1A_- | exon_10 |
| MEP1A | CAGCCGACAGAAGCGGTCGG | chr6:46803146-46803165_MEP1A_+ | exon_13 |
| MET | TCTGTGAGAATACACTCCAG | chr7:116340024-116340043_MET_- | exon_02 |
| MET | GCACCAGCCACACTGAACAA | chr7:116380982-116381001_MET_- | exon_05 |
| MET | GCTGGGACTTTGGATTTCGG | chr7:116395461-116395480_MET_+ | exon_06 |
| MET | AGCAGGAAGGAACTTTACAG | chr7:116399523-116399542_MET_+ | exon_10 |
| MET | AAAGCCAGTGATGATCTCAA | chr7:116403274-116403293_MET_+ | exon_11 |
| MKNK1 | GGTGCCGTGAGCCTACAGAA | chr1:47046224-47046243_MKNK1_- | exon_04 |
| MKNK1 | TCGGAGTAGGGTGTTTCGAG | chr1:47042281-47042300_MKNK1_- | exon_05 |
| MKNK1 | TGAGCGAGAAGCCAGCCGAG | chr1:47037780-47037799_MKNK1_- | exon_07 |
| MKNK1 | ACACCCTGCAGACCTCGCCC | chr1:47028326-47028345_MKNK1_+ | exon_11 |
| MKNK1 | GTGGTCAGAGCATTTACCTC | chr1:47025889-47025908_MKNK1_+ | exon_13 |
| MS4A1 | CAGGAGGATGTCTTCACTGG | chr11:60229940-60229959_MS4A1_+ | exon_02 |
| MS4A1 | TCATGAAGAAGCTTTGCGTG | chr11:60229967-60229986_MS4A1_- | exon_02 |
| MS4A1 | CTCATGAAGAAGCTTTGCGT | chr11:60229968-60229987_MS4A1_- | exon_02 |
| MS4A1 | TGGGCTCTTCCACATTGCCC | chr11:60230492-60230511_MS4A1_+ | exon_03 |
| MS4A1 | GTACCACACAGTCACACAGA | chr11:60230554-60230573_MS4A1_- | exon_03 |
| MTOR | GAGCCGGAATGAGGAAACCA | chr1:11319358-11319377_MTOR_- | exon_02 |
| MTOR | AGAGACTCTGTCCTTACCTA | chr1:11318525-11318544_MTOR_+ | exon_03 |
| MTOR | TGGCCGTCTTGCCATGGCAG | chr1:11317085-11317104_MTOR_- | exon_04 |
| MTOR | TGAGAATCAGACAGGCACGA | chr1:11316096-11316115_MTOR_+ | exon_05 |
| MTOR | TGAGACCTTGGCCAAAGAGA | chr1:11313979-11313998_MTOR_- | exon_06 |
| MYC | GCTGCACCGAGTCGTAGTCG | chr8:128750550-128750569_MYC_- | exon_02 |
| MYC | AGGGCGAGCAGAGCCCGGAG | chr8:128750706-128750725_MYC_- | exon_02 |
| MYC | GAAGGGTGTGACCGCAACGT | chr8:128750729-128750748_MYC_- | exon_02 |
| MYC | GAAGGCGCTGGAGTCTTGCG | chr8:128751137-128751156_MYC_- | exon_02 |
| MYC | CGAGGACGGAGAGAAGGCGC | chr8:128751149-128751168_MYC_- | exon_02 |
| NAMPT | CCGGCCCGAGATGAATCCTG | chr7:105925321-105925340_NAMPT_ | exon_01 |
| NAMPT | AGGAAATGTTCTCTTCACGG | chr7:105913023-105913042_NAMPT_ | exon_04 |
| NAMPT | GTCCTGGTATCCAATCACAG | chr7:105909725-105909744_NAMPT_ | exon_05 |
| NAMPT | CAAACCTCCACCAGAACCGA | chr7:105894882-105894901_NAMPT_- | exon_09 |
| NAMPT | AGGGCCGATTATCTTTACAT | chr7:105893529-105893548_NAMPT_ | exon_10 |
| NDST4 | TGAGAAAGGCCCTCTTCCTG | chr4:115997544-115997563_NDST4_- | exon_02 |
| NDST4 | TCTCAACAAGGAATTTGCAC | chr4:115891588-115891607_NDST4_- | exon_04 |
| NDST4 | AATCAACATGGGCTATGCTG | chr4:115858626-115858645_NDST4_- | exon_05 |
| NDST4 | AACAACCCACAAGTCTGTCG | chr4:115856462-115856481_NDST4_+ | exon_06 |
| NDST4 | ATGAAGATCCAGCTGCTCTG | chr4:115760672-115760691_NDST4_- | exon_11 |
| NELFB | GACTGGTGGAACTCCAGCCG | chr9:140150428-140150447_NELFB_- | exon_02 |
| NELFB | GATGCCGTCCCTGCAGCCCG | chr9:140150829-140150848_NELFB_+ | exon_03 |
| NELFB | GCTGTATCGAGCCTGCGCCG | chr9:140151317-140151336_NELFB_+ | exon_04 |
| NELFB | CGTGGGTGAAATCTGCACCG | chr9:140157638-140157657_NELFB_+ | exon_05 |
| NELFB | GCGGTTCGTGGACAGCAAGA | chr9:140158729-140158748_NELFB_+ | exon_06 |
| NFKB1 | GAACAAGAAGTCTTACCCTC | chr4:103459086-103459105_NFKB1_+ | exon_05 |

| | | | |
|---|---|---|---|
| NFKB1 | ACTGGAAGCACGAATGACAG | chr4:103498084-103498103_NFKB1_+ | exon_07 |
| NFKB1 | CAAGGAGATGGACCTCAGCG | chr4:103500081-103500100_NFKB1_+ | exon_08 |
| NFKB1 | ACAGCTGGATGTGTGACTGG | chr4:103501736-103501755_NFKB1_+ | exon_09 |
| NFKB1 | CCGAAGCTGGACAAACACAG | chr4:103505897-103505916_NFKB1_- | exon_11 |
| NOL6 | GCAACAGTGAGGGTTTCAGG | chr9:33469509-33469528_NOL6_+ | exon_05 |
| NOL6 | GGCCTGGTGGAAGTCAGCCA | chr9:33468394-33468413_NOL6_+ | exon_10 |
| NOL6 | AGAGGCAGACCAGCCTGAGG | chr9:33467391-33467410_NOL6_- | exon_13 |
| NOL6 | ACACTCACAGTGCCAAGAGG | chr9:33467104-33467123_NOL6_+ | exon_14 |
| NOL6 | ACAGGATGGACCCTCAGCCC | chr9:33464037-33464056_NOL6_- | exon_22 |
| NOS2 | GAGGTAGAAGCTCATCTGGA | chr17:26114747-26114766_NOS2_+ | exon_05 |
| NOS2 | ATTGCGCCAGGCCTGCTTGG | chr17:26110012-26110031_NOS2_+ | exon_06 |
| NOS2 | TACCTGAGTGAATTCCACGT | chr17:26108059-26108078_NOS2_+ | exon_08 |
| NOS2 | GAAGCAGAATGTGACCATCA | chr17:26101459-26101478_NOS2_- | exon_12 |
| NOS2 | TATGCTGATGCGCAAGACAA | chr17:26099441-26099460_NOS2_- | exon_14 |
| NOTCH1 | TGAAGCGGCCAATGGCACGG | hr9:139438486-139438505_NOTCH1_ | exon_02 |
| NOTCH1 | GAACGCCGGGACATGCCACG | hr9:139418346-139418365_NOTCH1_ | exon_03 |
| NOTCH1 | GCACTGGCCACCGTTGGCGC | hr9:139417579-139417598_NOTCH1_ | exon_04 |
| NOTCH1 | TCCTGCCAGAACACCCACGG | hr9:139405179-139405198_NOTCH1_ | exon_17 |
| NOTCH1 | TCCTCGCCCTGCAAGAACGG | hr9:139402797-139402816_NOTCH1_ | exon_20 |
| NR2C2 | AATCCAGATAGTCACCGCAG | chr3:15055125-15055144_NR2C2_+ | exon_03 |
| NR2C2 | AAACAGGTCTTCTTGATCCA | chr3:15065624-15065643_NR2C2_+ | exon_07 |
| NR2C2 | TAATAGGCTGAAACAAGCCA | chr3:15070087-15070106_NR2C2_+ | exon_08 |
| NR2C2 | GCAGATGGGATAGACACCAG | chr3:15071843-15071862_NR2C2_+ | exon_09 |
| NR2C2 | GGTGGAGAGACTCATGACCT | chr3:15076251-15076270_NR2C2_- | exon_11 |
| NT5E | GAGCCACGCCACCCATGCAG | chr6:86160025-86160044_NT5E_- | exon_01 |
| NT5E | GACGCCGGCGACCAGTACCA | chr6:86160101-86160120_NT5E_+ | exon_01 |
| NT5E | GATCGAGCCACTCCTCAAAG | chr6:86176819-86176838_NT5E_+ | exon_02 |
| NT5E | CAAAGGGCAATACAGCAGCC | chr6:86200248-86200267_NT5E_- | exon_07 |
| NT5E | AGGTGACCAAGATATCAACG | chr6:86203557-86203576_NT5E_+ | exon_09 |
| NTRK1 | CCGGGTGCATCGCAGTCCCG | chr1:156830863-156830882_NTRK1_- | exon_01 |
| NTRK1 | ACGGAGACCACTCTTCACGA | chr1:156834525-156834544_NTRK1_- | exon_03 |
| NTRK1 | AGGGCACAAGAACAGTGCAG | chr1:156837915-156837934_NTRK1_- | exon_05 |
| NTRK1 | GGGCACCTGGACCTTCAGCG | chr1:156838306-156838325_NTRK1_- | exon_06 |
| NTRK1 | TGTGCAGCTGCACACGGCGG | chr1:156843435-156843454_NTRK1_+ | exon_08 |
| NUDT1 | ATGCCCAGGAGAACTCGCTG | chr7:2284255-2284274_NUDT1_- | exon_03 |
| NUDT1 | GCAAGAAGGAGAGACCATCG | chr7:2284326-2284345_NUDT1_+ | exon_03 |
| NUDT1 | GGAGGAGAGCGGTCTGACAG | chr7:2289501-2289520_NUDT1_+ | exon_04 |
| NUDT1 | AGTGGACGCCCTGCACAAGG | chr7:2289519-2289538_NUDT1_+ | exon_04 |
| NUDT1 | CCTGGACTACACACTCCGCG | chr7:2290600-2290619_NUDT1_+ | exon_05 |
| NUP133 | GACCGGCGAGAAGAGCACTG | chr1:229643841-229643860_NUP133_ | exon_01 |
| NUP133 | GAAGCCCTAACATTGGCTGA | chr1:229641789-229641808_NUP133_ | exon_02 |
| NUP133 | AGTAGAATGTGCTTCACCTG | chr1:229636506-229636525_NUP133_ | exon_04 |
| NUP133 | TGGTGAAGATACCTACACAG | chr1:229635487-229635506_NUP133_ | exon_05 |
| NUP133 | AATGATAGGAACACCACCAC | chr1:229622229-229622248_NUP133_ | exon_11 |
| OPN5 | CAGGCGCTCGTCCTGAGGCA | chr6:47749825-47749844_OPN5_- | exon_01 |
| OPN5 | ATCCAGCCCATCCATACCAG | chr6:47759608-47759627_OPN5_- | exon_03 |
| OPN5 | TTTGGAAGAGGACTTAACCT | chr6:47763217-47763236_OPN5_- | exon_04 |
| OPN5 | TTTGCCTGTTGCCAAACTGG | chr6:47776070-47776089_OPN5_+ | exon_05 |
| OPN5 | AGCAGAAGACTTCCTGACTG | chr6:47779427-47779446_OPN5_- | exon_06 |
| ORAOV1 | CCTCTCATCCGCCATCACGA | chr11:69489957-69489976_ORAOV1_- | exon_01 |
| ORAOV1 | CATATTCGATGCCATCGTGA | chr11:69489971-69489990_ORAOV1_ | exon_01 |

| | | | |
|---|---|---|---|
| ORAOV1 | TATCGGGAAGGCTATGAAGA | :hr11:69488054-69488073_ORAOV1_ | exon_02 |
| ORAOV1 | GCACAGTTGCACCACTGAGA | :hr11:69486524-69486543_ORAOV1_ | exon_03 |
| ORAOV1 | GAAGACTTAGACAAGATCAG | :hr11:69482679-69482698_ORAOV1_ | exon_04 |
| P4HB | GACGTGGTCCTCCTCCTCGG | chr17:79818270-79818289_P4HB_+ | exon_01 |
| P4HB | GTTGGCCAAGGTGGACGCCA | chr17:79817156-79817175_P4HB_- | exon_02 |
| P4HB | GCTGCCACCACCCTGCCTGA | chr17:79813390-79813409_P4HB_- | exon_03 |
| P4HB | AGGCCGGAACAACTTTGAAG | chr17:79805196-79805215_P4HB_- | exon_05 |
| P4HB | CAAGTACAAGCCCGAATCGG | chr17:79804367-79804386_P4HB_- | exon_07 |
| PAK1 | GATGCTGGAACCCTAAACCA | chr11:77103459-77103478_PAK1_- | exon_02 |
| PAK1 | ACACATCCAGAACAGCCTGC | chr11:77090343-77090362_PAK1_+ | exon_04 |
| PAK1 | GGGCGTGGAGCAATCACTGG | chr11:77069959-77069978_PAK1_+ | exon_06 |
| PAK1 | GGTTTGAGAAGATTGGACAA | chr11:77064583-77064602_PAK1_- | exon_08 |
| PAK1 | ACAGCAATGGATGTGGCCAC | chr11:77060294-77060313_PAK1_- | exon_09 |
| PARP1 | AAGAAGACAGCGGAAGCTGG | :hr1:226589926-226589945_PARP1_- | exon_02 |
| PARP1 | AGCTAGGCATGATTGACCGC | :hr1:226578260-226578279_PARP1_- | exon_04 |
| PARP1 | CAAGCAGCAAGTGCCTTCTG | :hr1:226574038-226574057_PARP1_- | exon_06 |
| PARP1 | GACACAGACACCCAACCGGA | :hr1:226573225-226573244_PARP1_- | exon_07 |
| PARP1 | AGAAACCAGCGCCTCCGTGG | :hr1:226570800-226570819_PARP1_- | exon_08 |
| PARP2 | TCAGCGTTCGAATTCCATGG | chr14:20811785-20811804_PARP2_+ | exon_01 |
| PARP2 | AGGACAGAAGACAAGCAAGA | chr14:20813225-20813244_PARP2_+ | exon_02 |
| PARP2 | GGACCCAGAGTGTACAGCCA | chr14:20813593-20813612_PARP2_+ | exon_03 |
| PARP2 | TATTAGAAGATGATGCCCAG | chr14:20818728-20818747_PARP2_+ | exon_05 |
| PARP2 | AGGCAATCTCAACAAGGCCA | chr14:20819243-20819262_PARP2_+ | exon_06 |
| PARP3 | AGGGCCCTCAGTCTGTACCC | chr3:51977394-51977413_PARP3_- | exon_02 |
| PARP3 | ACTTATCGAAGTACAGGCAG | chr3:51978546-51978565_PARP3_+ | exon_04 |
| PARP3 | GGAGATGTTCAAGAACACCA | chr3:51978889-51978908_PARP3_+ | exon_05 |
| PARP3 | GAGCAAGCAACAGATTGCAC | chr3:51979042-51979061_PARP3_+ | exon_06 |
| PARP3 | TGAGCAGGAGAAGACGGTGG | chr3:51979555-51979574_PARP3_+ | exon_07 |
| PDCD1LG2 | CCAGGCTCAACATTAGCAGG | chr9:5522558-5522577_PDCD1LG2_- | exon_02 |
| PDCD1LG2 | GTGGAACGAGGCCTTCCCTA | chr9:5534934-5534953_PDCD1LG2_- | exon_03 |
| PDCD1LG2 | TGTGGAACGAGGCCTTCCCT | chr9:5534935-5534954_PDCD1LG2_- | exon_03 |
| PDCD1LG2 | GGCCAGGATACTTCTGCCAG | chr9:5549421-5549440_PDCD1LG2_- | exon_04 |
| PDCD1LG2 | TGTTCTGGAATACTCACGTG | :hr9:5549551-5549570_PDCD1LG2_+ | exon_04 |
| PDCD1 | TCCAGGCATGCAGATCCCAC | :hr2:242800978-242800997_PDCD1_- | exon_01 |
| PDCD1 | TCACCCTGAGCTCTGCCCGC | :hr2:242794776-242794795_PDCD1_- | exon_02 |
| PDCD1 | CGTGTCACACAACTGCCCAA | :hr2:242794904-242794923_PDCD1_- | exon_02 |
| PDCD1 | AGCCGGCCAGTTCCAAACCC | :hr2:242794437-242794456_PDCD1_- | exon_03 |
| PDCD1 | TCAGTGGCTGGGCACTCCGA | :hr2:242793245-242793264_PDCD1_- | exon_05 |
| PDGFRA | AGGGTAATGAAAGCTGGCAG | chr4:55127278-55127297_PDGFRA_- | exon_03 |
| PDGFRA | GGATTATTTAGTCATCGTGG | :hr4:55129868-55129887_PDGFRA_+ | exon_04 |
| PDGFRA | ATCTCCACTGTCTTTCACCG | chr4:55133538-55133557_PDGFRA_- | exon_06 |
| PDGFRA | CGTGCTAAGGAAGAAGACAG | :hr4:55136825-55136844_PDGFRA_+ | exon_08 |
| PDGFRA | GACTTGGTCGATGATCACCA | :hr4:55138578-55138597_PDGFRA_+ | exon_09 |
| PDGFRB | TGCACCCACAATGACTCCCG | hr5:149515165-149515184_PDGFRB_ | exon_03 |
| PDGFRB | ATTAGGGAGGAAGCCCACGG | hr5:149514554-149514573_PDGFRB_ | exon_04 |
| PDGFRB | CCCACATTACTTCTTTGCGG | hr5:149513434-149513453_PDGFRB_ | exon_05 |
| PDGFRB | AAAGGCCATCAACATCACCG | hr5:149513155-149513174_PDGFRB_ | exon_06 |
| PDGFRB | GTTCAAAGACAACCGCACCC | hr5:149512371-149512390_PDGFRB_ | exon_07 |
| PDILT | CAAGCCTGTGCACATCCTGG | chr16:20410502-20410521_PDILT_- | exon_02 |
| PDILT | AGTGGACATTACCATAGAGA | chr16:20396057-20396076_PDILT_- | exon_03 |
| PDILT | GAAGAAGCCAACGATGACCA | chr16:20387393-20387412_PDILT_+ | exon_04 |

| | | | |
|---|---|---|---|
| PDILT | ATGACTCCAAACGTTAGCTC | chr16:20386208-20386227_PDILT_+ | exon_05 |
| PDILT | TGTGATCGAATACAACACTG | chr16:20384336-20384355_PDILT_- | exon_06 |
| PDK1 | TTGCCGCAGAAACATAAATG | chr2:173423463-173423482_PDK1_- | exon_02 |
| PDK1 | AAAGTACTGAACATTCTGGC | chr2:173429341-173429360_PDK1_- | exon_04 |
| PDK1 | ACCAGGACAGCCAATACAAG | chr2:173433476-173433495_PDK1_+ | exon_07 |
| PDK1 | ATGGAACACCATGCCAACAG | chr2:173435472-173435491_PDK1_+ | exon_08 |
| PDK1 | TCAAAGGAACGCCACCTCCT | chr2:173451017-173451036_PDK1_- | exon_09 |
| PGD | GCGAGTCACTCACTGGGCCA | chr1:10459176-10459195_PGD_- | exon_01 |
| PGD | CGCCGCTTACCACAAAGCCG | chr1:10459752-10459771_PGD_- | exon_02 |
| PGD | ACCAGGAGGATGATCCGCCG | chr1:10460570-10460589_PGD_- | exon_03 |
| PGD | GGTGACATCATCATTGACGG | chr1:10463146-10463165_PGD_+ | exon_04 |
| PGD | GTGCCGAGACCTCAAGGCCA | chr1:10464223-10464242_PGD_+ | exon_05 |
| PHB2 | TTCGCGCACACCGTAGGCCA | chr12:7079593-7079612_PHB2_+ | exon_01 |
| PHB2 | GCAGGACACTATCCTGGCCG | chr12:7079375-7079394_PHB2_- | exon_02 |
| PHB2 | CAAGTGCTCAGACCTACCTT | chr12:7078644-7078663_PHB2_+ | exon_03 |
| PHB2 | GGGTCGAGACAACACTCGCA | chr12:7077715-7077734_PHB2_+ | exon_04 |
| PHB2 | GAAGCACTGAGCAAGAACCC | chr12:7076386-7076405_PHB2_- | exon_07 |
| PIGF | GGTCCTAACAAACATAAGCA | chr2:46839406-46839425_PIGF_+ | exon_04 |
| PIGF | CATGGTCTTTCCCAATCCAG | chr2:46819618-46819637_PIGF_+ | exon_05 |
| PIGF | CACTGGATTGGGAAAGACCA | chr2:46819620-46819639_PIGF_- | exon_05 |
| PIGF | TGGAGCACTTCCTATTCCAC | chr2:46819637-46819656_PIGF_- | exon_05 |
| PIGF | CTGGAGACTATTCTCCCATA | chr2:46819692-46819711_PIGF_+ | exon_05 |
| PIK3CA | TTATTAATGTAGCCTCACGG | hr3:178916724-178916743_PIK3CA_ | exon_02 |
| PIK3CA | GTTCGAACAGGTATCTACCA | hr3:178922297-178922316_PIK3CA_ | exon_06 |
| PIK3CA | AGGAACACTGTCCATTGGCA | hr3:178927972-178927991_PIK3CA_ | exon_08 |
| PIK3CA | GAATAGGCAAGTCGAGGCAA | hr3:178938864-178938883_PIK3CA_ | exon_14 |
| PIK3CA | ACAGCCACACACTACATCAG | hr3:178947182-178947201_PIK3CA_ | exon_18 |
| PIK3CB | TAAGAAATGGTAGCTTCCCG | hr3:138478025-138478044_PIK3CB_ | exon_02 |
| PIK3CB | TGACAAGAAGTTGTGACCCA | hr3:138474639-138474658_PIK3CB_ | exon_03 |
| PIK3CB | TGTAGCGTGGGTAAATACGA | hr3:138431119-138431138_PIK3CB_ | exon_09 |
| PIK3CB | GAAGCTCTAGGGCCTCCCGG | hr3:138413692-138413711_PIK3CB_ | exon_13 |
| PIK3CB | AAAGAGCACTTGGTAATCGG | hr3:138409874-138409893_PIK3CB_ | exon_14 |
| PIK3CD | AGAGCGGCTCATACTGGGCG | chr1:9775613-9775632_PIK3CD_- | exon_04 |
| PIK3CD | GATGTGCCAATTCTGCGAGG | chr1:9775962-9775981_PIK3CD_+ | exon_05 |
| PIK3CD | GTCCACCAAGGACGTGCCGC | chr1:9776518-9776537_PIK3CD_+ | exon_06 |
| PIK3CD | ATCTGCAGCTGCCTGCACAG | chr1:9777020-9777039_PIK3CD_+ | exon_07 |
| PIK3CD | GATGCGGAACGGCTGCTCCA | chr1:9777620-9777639_PIK3CD_- | exon_08 |
| PIK3CG | GAAGGCTTGGGACTCCTCGG | hr7:106508437-106508456_PIK3CG_ | exon_02 |
| PIK3CG | GCTTCAGCAGAAATCTGGCA | hr7:106513017-106513036_PIK3CG_ | exon_03 |
| PIK3CG | AGTCCAGACACTATCAGCAG | hr7:106513210-106513229_PIK3CG_ | exon_04 |
| PIK3CG | TGGACTGAAAGCAGGAGCGC | hr7:106515224-106515243_PIK3CG_ | exon_05 |
| PIK3CG | AAATGCAACCATATGGCAGG | hr7:106522615-106522634_PIK3CG_ | exon_07 |
| PIM1 | GTGGGCCCGCTACTGGGCAG | chr6:37138584-37138603_PIM1_+ | exon_02 |
| PIM1 | GAAGGACCGGATTTCCGACT | chr6:37138777-37138796_PIM1_+ | exon_03 |
| PIM1 | AAGTCGAAGAGATCTTGCAC | chr6:37139036-37139055_PIM1_- | exon_04 |
| PIM1 | GAGTGTATAGCCCTCCAGAG | chr6:37140778-37140797_PIM1_+ | exon_05 |
| PIM1 | TGGATCTCAGCAGTTTCCTG | chr6:37141814-37141833_PIM1_- | exon_06 |
| PIM2 | TGTTGACCAAGCCTCTACAG | chrX:48776091-48776110_PIM2_- | exon_01 |
| PIM2 | GCTCCGAAGGTACTCACCAA | chrX:48775041-48775060_PIM2_+ | exon_03 |
| PIM2 | ATCCTGATAGACCTACGCCG | chrX:48772368-48772387_PIM2_- | exon_04 |
| PIM2 | GATGCCCAGTGACCAGACAG | chrX:48771766-48771785_PIM2_+ | exon_05 |

| | | | |
|---|---|---|---|
| PIM2 | CTGGATGCAAACACCAGCCG | chrX:48771473-48771492_PIM2_- | exon_06 |
| PIM3 | GCTGCAGGATCTTCACCGGG | chr22:50354658-50354677_PIM3_- | exon_01 |
| PIM3 | GCGGGTAGCCGCATCGCCGA | chr22:50354853-50354872_PIM3_+ | exon_02 |
| PIM3 | GAAGGAGCGGGTGACCGAGT | chr22:50354986-50355005_PIM3_+ | exon_03 |
| PIM3 | GCTGTGGCAGTGGCGCACGG | chr22:50355313-50355332_PIM3_- | exon_04 |
| PIM3 | GAGGCCGCCTGCTCTTCCGG | chr22:50356478-50356497_PIM3_+ | exon_05 |
| PLA2G2F | AGAACCCGCAACATGGCAGA | chr1:20465909-20465928_PLA2G2F_+ | exon_01 |
| PLA2G2F | TATGAAGAAGTTCTTCACCG | chr1:20466671-20466690_PLA2G2F_+ | exon_02 |
| PLA2G2F | GTAGCAACCGTAGCCCACGA | chr1:20470020-20470039_PLA2G2F_- | exon_03 |
| PLA2G2F | TACCAGGAACTCTTTGACCA | chr1:20471098-20471117_PLA2G2F_+ | exon_04 |
| PLA2G2F | CATGAACCAGACGTACCGAG | chr1:20474750-20474769_PLA2G2F_+ | exon_05 |
| PLK1 | ACCGGCGAAAGAGATCCCGG | chr16:23690358-23690377_PLK1_+ | exon_01 |
| PLK1 | AGCCAAGCACAATTTGCCGT | chr16:23691467-23691486_PLK1_- | exon_02 |
| PLK1 | GAAAGGGCACAGTTTCGAGG | chr16:23692332-23692351_PLK1_+ | exon_03 |
| PLK1 | CCTGCCTGACCATTCCACCA | chr16:23695324-23695343_PLK1_+ | exon_05 |
| PLK1 | GCTCTGTGATAACAGCGTGG | chr16:23700566-23700585_PLK1_+ | exon_08 |
| PNPLA5 | TACCTGGGCGCCCACCACGT | chr22:44287687-44287706_PNPLA5_- | exon_01 |
| PNPLA5 | TCAGCGAAATGCCCAGCCGC | chr22:44287007-44287026_PNPLA5_+ | exon_02 |
| PNPLA5 | AAAGTATAAGGTGCAGACCA | chr22:44285721-44285740_PNPLA5_+ | exon_03 |
| PNPLA5 | ACAGGGCCAGACCTACCTCG | chr22:44285193-44285212_PNPLA5_+ | exon_04 |
| PNPLA5 | AAGGCTACCTGGATGCCCTG | chr22:44283490-44283509_PNPLA5_- | exon_05 |
| POLR1C | GATGGCGGCTTCTCAGGCGG | chr6:43484847-43484866_POLR1C_+ | exon_01 |
| POLR1C | GGACCAGGACCGCTTCGAGA | chr6:43485094-43485113_POLR1C_+ | exon_02 |
| POLR1C | AAACTCACTGGAGTTTGACA | chr6:43487103-43487122_POLR1C_+ | exon_03 |
| POLR1C | CCAGGCTGATCTCTTTCCAG | chr6:43488050-43488069_POLR1C_+ | exon_06 |
| POLR1C | ATTGAGGTGCAGGAAGTCCA | chr6:43488491-43488510_POLR1C_+ | exon_07 |
| POLR2B | GTTTGGCAATATGTACGACG | chr4:57845142-57845161_POLR2B_+ | exon_02 |
| POLR2B | GAGCACCATCTCTTTCCCAA | chr4:57860620-57860639_POLR2B_- | exon_05 |
| POLR2B | AAATAGCCACCAGGATCCAA | chr4:57860991-57861010_POLR2B_- | exon_06 |
| POLR2B | AATCACTGACACCAACATGA | chr4:57871558-57871577_POLR2B_- | exon_09 |
| POLR2B | TAGGAGAATTTAAACGACGC | chr4:57876568-57876587_POLR2B_- | exon_12 |
| POLR2E | AATCCGCAAGACCATCATGC | chr19:1095260-1095279_POLR2E_- | exon_01 |
| POLR2E | TTGGGCTTTGAACTCCTCCA | chr19:1094006-1094025_POLR2E_+ | exon_02 |
| POLR2E | GCAGGAGGAGAACATCACAC | chr19:1091838-1091857_POLR2E_- | exon_03 |
| POLR2E | GGAGCTGCTCATCAACATCA | chr19:1090918-1090937_POLR2E_- | exon_04 |
| POLR2E | TGAGCACGTCGTCATGACCA | chr19:1090117-1090136_POLR2E_- | exon_05 |
| POLR3C | AGAGTAATTGCCCATGACAC | chr1:145609209-145609228_POLR3C_ | exon_02 |
| POLR3C | TTGGTAGTATAGATGTACCG | hr1:145608520-145608539_POLR3C_ | exon_03 |
| POLR3C | TTATACAGATGGCAAGACCA | chr1:145608282-145608301_POLR3C_ | exon_04 |
| POLR3C | GAGCGCAGTTGCTAACAGGA | chr1:145601756-145601775_POLR3C_ | exon_06 |
| POLR3C | GGAAAGTCTGGCGACAGTGG | chr1:145598264-145598283_POLR3C_ | exon_09 |
| PORCN | GATGGGAGGAATGTCGGCAG | chrX:48369810-48369829_PORCN_- | exon_03 |
| PORCN | TATCCAGGGCCCGAAGACGA | chrX:48370834-48370853_PORCN_- | exon_05 |
| PORCN | GATCCCTACAGAGCTGCCGG | chrX:48370966-48370985_PORCN_+ | exon_06 |
| PORCN | TCTGTAGACTCACCTTACCA | chrX:48372523-48372542_PORCN_- | exon_07 |
| PORCN | GAAGGAGACAGCACTCTCGT | chrX:48372642-48372661_PORCN_- | exon_08 |
| PPM1D | GGAGGCGGCACAGTTTGCCC | chr17:58678105-58678124_PPM1D_+ | exon_01 |
| PPM1D | AAGGAAAGAGAACGAATCGA | chr17:58701076-58701095_PPM1D_+ | exon_02 |
| PPM1D | ATTGAGATGGCATCTTGTGG | chr17:58725390-58725409_PPM1D_- | exon_04 |
| PPM1D | GGAGCATACGCTGCCTCCAG | chr17:58734013-58734032_PPM1D_- | exon_05 |
| PPM1D | TGGAGGAGGATCCATGGCCA | chr17:58740360-58740379_PPM1D_+ | exon_06 |

| | | | |
|---|---|---|---|
| PPP3R2 | GATGGTGGGCAACAACCTGA | chr9:104356831-104356850_PPP3R2_ | exon_01 |
| PPP3R2 | GGAGCTGCGCCACAACCCGT | chr9:104357044-104357063_PPP3R2_ | exon_01 |
| PPP3R2 | GTTGTGGCGCAGCTCCGGCA | hr9:104357048-104357067_PPP3R2_ | exon_01 |
| PPP3R2 | AAGTGGGAGCACATCTCCGC | hr9:104357160-104357179_PPP3R2_ | exon_01 |
| PPP3R2 | AAACGAGGCCAGTTACCCGG | chr9:104357179-104357198_PPP3R2_ | exon_01 |
| PRKAA1 | GTGATGGAATATGTCTCAGG | chr5:40775549-40775568_PRKAA1_- | exon_03 |
| PRKAA1 | TTTGAAACATATGCTGCAGG | chr5:40767605-40767624_PRKAA1_- | exon_06 |
| PRKAA1 | GAGGTAGATATATGGAGCAG | chr5:40767757-40767776_PRKAA1_- | exon_06 |
| PRKAA1 | AGAATGGTACTCTTTCAGGA | chr5:40765030-40765049_PRKAA1_+ | exon_07 |
| PRKAA1 | GCAACTATCGATCTTGCCAA | chr5:40763048-40763067_PRKAA1_- | exon_09 |
| PRKCA | GAGGCAGAAGAACGTGCACG | chr17:64299047-64299066_PRKCA_+ | exon_01 |
| PRKCA | AAACAAGGCTTCCAGTGCCA | chr17:64302224-64302243_PRKCA_+ | exon_02 |
| PRKCA | GCTCCACACTAAATCCGCAG | chr17:64683346-64683365_PRKCA_+ | exon_06 |
| PRKCA | GCTCCGAAACTCCAAAGGAA | chr17:64684510-64684529_PRKCA_- | exon_07 |
| PRKCA | TACAACGTACCCATTCCGGA | chr17:64685103-64685122_PRKCA_+ | exon_08 |
| PRKCB | ATGCCCATTACTTACATCGG | chr16:23999907-23999926_PRKCB_- | exon_03 |
| PRKCB | GCTGTATGGACTCATCCACC | chr16:24043531-24043550_PRKCB_+ | exon_04 |
| PRKCB | ACGGACCACACGGAGCGCCG | chr16:24046796-24046815_PRKCB_+ | exon_05 |
| PRKCB | GTAGGGATCTGACAGGCCAT | chr16:24104148-24104167_PRKCB_- | exon_06 |
| PRKCB | CTGAAGTTCAGAAATCCCAA | chr16:24105579-24105598_PRKCB_- | exon_07 |
| PRKCD | GATGCGCAGGAACGGCGCCA | chr3:53212440-53212459_PRKCD_- | exon_02 |
| PRKCD | GGCCAGCACCGACACACCCA | chr3:53213734-53213753_PRKCD_- | exon_03 |
| PRKCD | CTGAACAGACATCAACACCT | chr3:53215245-53215264_PRKCD_- | exon_04 |
| PRKCD | TTCCCAACGATGAACCGCCG | chr3:53215498-53215517_PRKCD_+ | exon_05 |
| PRKCD | GCTGCCGCAGTGGTCACAGA | chr3:53217538-53217557_PRKCD_- | exon_08 |
| PRKCE | AAGGCCATTGAACACTACCA | chr2:45879241-45879260_PRKCE_- | exon_01 |
| PRKCE | ACCAGATGAAGTCTCTGCAA | chr2:46203710-46203729_PRKCE_- | exon_03 |
| PRKCE | AATGCCTCTTACCTTGACAC | chr2:46206142-46206161_PRKCE_- | exon_04 |
| PRKCE | ACCAACACTTACCTGGTCGG | chr2:46207513-46207532_PRKCE_- | exon_05 |
| PRKCE | AACTGTGGAGTGGATGCCAG | chr2:46228590-46228609_PRKCE_+ | exon_07 |
| PRKCG | CCTCTGAATCGCCTACGCCG | chr19:54385766-54385785_PRKCG_- | exon_01 |
| PRKCG | GGTGCACAAGCCCGTAGAGG | chr19:54392966-54392985_PRKCG_- | exon_04 |
| PRKCG | GTGGACCACACCGAGCGCCG | chr19:54393196-54393215_PRKCG_+ | exon_05 |
| PRKCG | GACGAAACAGAAGACCCGAA | chr19:54395019-54395038_PRKCG_+ | exon_06 |
| PRKCG | GGAGCTGCTCAAGGCGCCCG | chr19:54395868-54395887_PRKCG_+ | exon_07 |
| PRKCH | GGAAGACGGCCAACTCGAGG | chr14:61789062-61789081_PRKCH_- | exon_01 |
| PRKCH | GTAGGTGGATCTCGAGCCAG | chr14:61857939-61857958_PRKCH_+ | exon_02 |
| PRKCH | AGCGCCAAAGGGCTATGCGA | chr14:61909871-61909890_PRKCH_+ | exon_03 |
| PRKCH | AAACAGGGTTATCAGTGCCA | chr14:61912423-61912442_PRKCH_+ | exon_04 |
| PRKCH | GCAGGGATGGGTCTCCAACC | chr14:61920030-61920049_PRKCH_+ | exon_07 |
| PRKCI | GTGAGGAGATGCCGACCCAG | chr3:169940450-169940469_PRKCI_+ | exon_01 |
| PRKCI | TGTCTCGAACCTCATTGCAA | chr3:169953066-169953085_PRKCI_- | exon_02 |
| PRKCI | ACTGAGATGATACTGTACAC | chr3:169977764-169977783_PRKCI_- | exon_03 |
| PRKCI | ACCGTAGAGGTGCACGCCGC | chr3:169985712-169985731_PRKCI_+ | exon_05 |
| PRKCI | TCAACAGGCAATGAACACCA | chr3:169998008-169998027_PRKCI_+ | exon_09 |
| PRKCQ | AATCCGAAGAAATGGCGACA | chr10:6557077-6557096_PRKCQ_+ | exon_02 |
| PRKCQ | CAGCGAGTAGAGCTCCACGG | chr10:6552999-6553018_PRKCQ_+ | exon_03 |
| PRKCQ | TTTGCTTTGCATCAGCGCCG | chr10:6540463-6540482_PRKCQ_- | exon_05 |
| PRKCQ | AACACTGTGGGACCCTGCTG | chr10:6533679-6533698_PRKCQ_- | exon_08 |
| PRKCQ | GAAGCTAATGGCTGAAGCGC | chr10:6528023-6528042_PRKCQ_- | exon_09 |
| PRKCSH | GCTACCCATGTGCTGGGCCG | chr19:11546962-11546981_PRKCSH_ | exon_02 |

| | | | |
|---|---|---|---|
| PRKCSH | GGTGGCCGAACCGTCCAGGC | :hr19:11547246-11547265_PRKCSH_ | exon_03 |
| PRKCSH | TCTGCCACAGACTGCTGCGA | :hr19:11548878-11548897_PRKCSH_ | exon_05 |
| PRKCSH | GAAGTCTCTGGAAGACCAGG | :hr19:11553230-11553249_PRKCSH_ | exon_07 |
| PRKCSH | GACGCGGTCGTAGAAAGAGG | :hr19:11557923-11557942_PRKCSH_ | exon_10 |
| PRKCZ | CAGCATTAAAGACGACTCGG | chr1:2080342-2080361_PRKCZ_+ | exon_05 |
| PRKCZ | TAAGCCAGTTATCGATGGGA | chr1:2082234-2082253_PRKCZ_+ | exon_06 |
| PRKCZ | GGAGTGTAATCCGACCAGGA | chr1:2087492-2087511_PRKCZ_- | exon_07 |
| PRKCZ | CATTGAGTACGTCAACGGCG | chr1:2100969-2100988_PRKCZ_+ | exon_08 |
| PRKCZ | TGTGCCCGTCCGCATCCAGG | chr1:2103581-2103600_PRKCZ_- | exon_09 |
| PRPF19 | ACTGGAGCAGATTAGGGACA | :hr11:60673834-60673853_PRPF19_ | exon_01 |
| PRPF19 | CAACAACCAGCCTCTCTCCG | chr11:60671208-60671227_PRPF19_- | exon_02 |
| PRPF19 | GAAGTCACTGCTGCCCGAGA | chr11:60670214-60670233_PRPF19_- | exon_04 |
| PRPF19 | GGCACAATGAGGCCAGCCTG | :hr11:60670089-60670108_PRPF19_ | exon_05 |
| PRPF19 | GCGCTCCGTGGTTAGCACAG | :hr11:60668977-60668996_PRPF19_ | exon_07 |
| PRSS37 | TGATGAGGACGCCCACACAG | hr7:141539178-141539197_PRSS37_ | exon_02 |
| PRSS37 | ATGAGCATGAGGTCATCCTG | hr7:141537782-141537801_PRSS37_ | exon_03 |
| PRSS37 | TGAGCATGAGGTCATCCTGT | hr7:141537783-141537802_PRSS37_ | exon_03 |
| PRSS37 | TTCTCGATCAGACATCACGG | hr7:141536993-141537012_PRSS37_ | exon_04 |
| PRSS37 | TTTGCAGATGACAGTAGCAA | hr7:141536306-141536325_PRSS37_ | exon_05 |
| PSMA7 | CGTCAAGAAGGGCTCGACCG | chr20:60718266-60718285_PSMA7_- | exon_01 |
| PSMA7 | AGTGCGGAAGATCTGTGCTT | chr20:60715904-60715923_PSMA7_- | exon_02 |
| PSMA7 | TGTGGAGGACCCGGTCACTG | chr20:60714875-60714894_PSMA7_- | exon_03 |
| PSMA7 | CAGGCATGGTATGTGCCCGA | chr20:60714135-60714154_PSMA7_+ | exon_04 |
| PSMA7 | GAGGCGAGATCAATCCCTCA | chr20:60712422-60712441_PSMA7_- | exon_06 |
| PSMB1 | TCGCAGCTGCAAAGGGCCCG | :hr6:170862244-170862263_PSMB1_- | exon_01 |
| PSMB1 | GATGGAACCGCACAGAGCCG | :hr6:170862264-170862283_PSMB1_ | exon_01 |
| PSMB1 | TAATAAGGCCATGACTACGG | :hr6:170852782-170852801_PSMB1_ | exon_04 |
| PSMB1 | TGAGAGAGATGTGTACACTG | :hr6:170844382-170844401_PSMB1_ | exon_06 |
| PSMB1 | GCTGAGAGAGATGTGTACAC | :hr6:170844384-170844403_PSMB1_ | exon_06 |
| PSMD1 | GGGCGCAGCCATGATCACCT | :hr2:231921730-231921749_PSMD1_- | exon_01 |
| PSMD1 | AAAGGAAGACAACCTCCTGA | :hr2:231936959-231936978_PSMD1_- | exon_07 |
| PSMD1 | AATGGCAACATACCTTCCCA | :hr2:231944881-231944900_PSMD1_- | exon_12 |
| PSMD1 | ACATAACCAAACCTAGGGCC | :hr2:231948297-231948316_PSMD1_ | exon_14 |
| PSMD1 | ACCAGAGCCACAATAAGCCA | :hr2:231949773-231949792_PSMD1_ | exon_15 |
| PSMG3 | GTTTGGGAAGATGGGCACCC | chr7:1608843-1608862_PSMG3_- | exon_01 |
| PSMG3 | GAAGGCCGTACACACCACCT | chr7:1608892-1608911_PSMG3_+ | exon_01 |
| PSMG3 | ATCGAAGCAGAAGACGGAGG | chr7:1608933-1608952_PSMG3_- | exon_01 |
| PSMG3 | GATATCGAAGCAGAAGACGG | chr7:1608936-1608955_PSMG3_- | exon_01 |
| PSMG3 | CAGAGCAGTCCTCCTCGCCG | chr7:1607408-1607427_PSMG3_- | exon_02 |
| PTCH1 | GGAAATCTGCTCCAGAGCGA | chr9:98270446-98270465_PTCH1_+ | exon_01 |
| PTCH1 | GAACCTCGAGACCAACGTGG | chr9:98268707-98268726_PTCH1_- | exon_02 |
| PTCH1 | CATGATCACACACTTACAGG | chr9:98244214-98244233_PTCH1_+ | exon_05 |
| PTCH1 | GAGGTAAACCTCCTTTGCGG | chr9:98242852-98242871_PTCH1_- | exon_06 |
| PTCH1 | AAAGTATATGCACTGGCAGG | chr9:98242303-98242322_PTCH1_- | exon_07 |
| PTGS2 | TCAAGACAGATCATAAGCGA | chr1:186646814-186646833_PTGS2_- | exon_05 |
| PTGS2 | TTCAAGACAGATCATAAGCG | chr1:186646815-186646834_PTGS2_- | exon_05 |
| PTGS2 | TGAGCATCTACGGTTTGCTG | chr1:186645752-186645771_PTGS2_- | exon_07 |
| PTGS2 | GATGCTCAGGGACTTGAGGA | :hr1:186645764-186645783_PTGS2_ | exon_07 |
| PTGS2 | CCAAAGATGGCATCTGGCCG | :hr1:186643786-186643805_PTGS2_ | exon_10 |
| PTK2 | GCAATAGTGAGCCAACCACC | chr8:141900684-141900703_PTK2_- | exon_03 |
| PTK2 | GATTCCGCCTCAGTCACCTG | chr8:141889662-141889681_PTK2_- | exon_04 |

| | | | |
|---|---|---|---|
| PTK2 | AATAGATGGGTACTGCCGGC | chr8:141810603-141810622_PTK2_- | exon_12 |
| PTK2 | TCAGCTTCACAATATGAGGA | chr8:141756918-141756937_PTK2_+ | exon_18 |
| PTK2 | AGGGTAGGAGGACAATTTGG | chr8:141745428-141745447_PTK2_+ | exon_22 |
| PTPN11 | GGAGGAACATGACATCGCGG | hr12:112856908-112856927_PTPN11_ | exon_01 |
| PTPN11 | GTAGGATCTGCACAGTTCAG | hr12:112888288-112888307_PTPN11_ | exon_03 |
| PTPN11 | GTGCGCACTGGTGATGACAA | hr12:112891117-112891136_PTPN11_ | exon_04 |
| PTPN11 | CTGACAGCGAATCATAACAT | hr12:112891172-112891191_PTPN11_ | exon_04 |
| PTPN11 | ACGGCGTGCCCAGCGACCCT | hr12:112924331-112924350_PTPN11_ | exon_11 |
| PTPN6 | GATGCAGAGACCCTGCTCAA | chr12:7060800-7060819_PTPN6_+ | exon_02 |
| PTPN6 | GGTGTCCTGCAGGACCGCGA | chr12:7061264-7061283_PTPN6_+ | exon_03 |
| PTPN6 | AAAGCACGAAGTCTCCAGGC | chr12:7064064-7064083_PTPN6_- | exon_04 |
| PTPN6 | CCAGGGTGGACGCTACACAG | chr12:7064310-7064329_PTPN6_+ | exon_05 |
| PTPN6 | GGAGTCCGAGGATACAGCCA | chr12:7064603-7064622_PTPN6_+ | exon_06 |
| PVR | AACTGGCATGGCCCGAGCCA | chr19:45147390-45147409_PVR_+ | exon_01 |
| PVR | CAACCCGAACATCCTCAGCG | chr19:45150735-45150754_PVR_- | exon_02 |
| PVR | TTGAGGGCACCAATATCCAG | chr19:45153268-45153287_PVR_- | exon_03 |
| PVR | GGGTGGCCTCATTCTGGCCA | chr19:45157215-45157234_PVR_- | exon_04 |
| PVR | ACAGTGCCAAAGGACCTCAC | chr19:45162133-45162152_PVR_- | exon_06 |
| RABGGTB | TCCTGTATGAACTTTGACGG | hr1:76257194-76257213_RABGGTB_ | exon_06 |
| RABGGTB | TGTGAACGACAATTACCCTC | hr1:76257944-76257963_RABGGTB_ | exon_07 |
| RABGGTB | GAACGACAATTACCCTCAGG | hr1:76257947-76257966_RABGGTB_ | exon_07 |
| RABGGTB | ATGTCAAGATGAAGAAACGG | hr1:76259870-76259889_RABGGTB_ | exon_08 |
| RABGGTB | TGTCAAGATGAAGAAACGGG | hr1:76259871-76259890_RABGGTB_ | exon_08 |
| RAC1 | TCACATCTAGTGGTATCCTG | chr7:6441489-6441508_RAC1_+ | exon_05 |
| RAC1 | TCTAGTGGTATCCTGAGGTG | chr7:6441494-6441513_RAC1_+ | exon_05 |
| RAC1 | CTATCCGCAGGGTCTAGCCA | chr7:6441624-6441643_RAC1_+ | exon_05 |
| RAC1 | GCAGGGTCTAGCCATGGCTA | chr7:6441630-6441649_RAC1_+ | exon_05 |
| RAC1 | TTAGCCATGGCTAGACCCTG | chr7:6441631-6441650_RAC1_- | exon_05 |
| RAF1 | GCCGAACAAGCAAAGAACAG | chr3:12660016-12660035_RAF1_- | exon_02 |
| RAF1 | TATGAAAGCACTCAAGGTGA | chr3:12653504-12653523_RAF1_- | exon_03 |
| RAF1 | TGTCAGAAATTCCTGCTCAA | chr3:12650364-12650383_RAF1_- | exon_05 |
| RAF1 | ATACGACGCATAGTCAAAGA | chr3:12647730-12647749_RAF1_+ | exon_06 |
| RAF1 | GGAGAGGGAACCTTCAGATG | chr3:12645713-12645732_RAF1_+ | exon_07 |
| RARA | AGAGTGGTCAGAGCGCCTGG | chr17:38487582-38487601_RARA_- | exon_02 |
| RARA | GCAGTCGGCAGTACTGGCAG | chr17:38506125-38506144_RARA_- | exon_04 |
| RARA | TGAGAGCTACACGCTGACGC | chr17:38508217-38508236_RARA_+ | exon_05 |
| RARA | GAGGGTGATCTGGTCGGCGA | chr17:38508713-38508732_RARA_- | exon_06 |
| RARA | GGGCAGCAGCTGGTTGGCGA | chr17:38510681-38510700_RARA_- | exon_07 |
| RARB | TAGAAATCCAGGATTTGCCC | chr3:25470259-25470278_RARB_- | exon_01 |
| RARB | GCAGGGTTTGTACACTCGAG | chr3:25502750-25502769_RARB_- | exon_02 |
| RARB | TCTGCCAGGACAAATCATCA | chr3:25502774-25502793_RARB_+ | exon_02 |
| RARB | CAGGCAGGCGGCCTTCAGCA | chr3:25622185-25622204_RARB_- | exon_05 |
| RARB | TAGCATCTCACCCAGCTCAG | chr3:25638047-25638066_RARB_+ | exon_08 |
| RARG | AAAGAGTCGCTCCTTATTGG | chr12:53621303-53621322_RARG_+ | exon_03 |
| RARG | AAGCATGGCTTGTAGACCCG | chr12:53609491-53609510_RARG_+ | exon_04 |
| RARG | GCTACAGAAGTGCTTCGAAG | chr12:53609095-53609114_RARG_- | exon_05 |
| RARG | GTTAGAAGAGCTCATCACCA | chr12:53608292-53608311_RARG_- | exon_06 |
| RARG | CCAGATCCAGCTGCACGCGG | chr12:53607983-53608002_RARG_+ | exon_07 |
| RAX | GGCAAGCGAGAAGCTCCCGT | chr18:56940385-56940404_RAX_+ | exon_01 |
| RAX | TGCGCGCCAGCCATGGCCGA | chr18:56940404-56940423_RAX_- | exon_01 |
| RAX | CGAAGCGAAACTGTCAGAGG | chr18:56939751-56939770_RAX_- | exon_02 |

| | | | |
|---|---|---|---|
| RAX | TGACAGTTTCGCTTCGCCGG | chr18:56939755-56939774_RAX_+ | exon_02 |
| RAX | TTCAGGAAGGGCGGAGGCGG | chr18:56936423-56936442_RAX_+ | exon_03 |
| RCC1 | TTGGTGCTGACACTAGGCCA | chr1:28858347-28858366_RCC1_+ | exon_06 |
| RCC1 | CTTGCAGCTCCACTTTCCCA | chr1:28858757-28858776_RCC1_- | exon_07 |
| RCC1 | CCTGAGGCCACCTTTACCAC | chr1:28861640-28861659_RCC1_- | exon_08 |
| RCC1 | TGATGGTGACCTCTACACCT | chr1:28861798-28861817_RCC1_+ | exon_09 |
| RCC1 | AGAGGCCGAAGCCGTACACG | chr1:28862501-28862520_RCC1_- | exon_10 |
| RD3 | ATAGCAGGCCCACAATAGGA | chr1:211654468-211654487_RD3_+ | exon_02 |
| RD3 | GCTGGCTGGCCAGCACACCC | chr1:211654557-211654576_RD3_- | exon_02 |
| RD3 | GCGGTCAGAAAGGTCTGCAC | chr1:211654591-211654610_RD3_- | exon_02 |
| RD3 | GAAGGGCGAGATGCGCGCGC | chr1:211652480-211652499_RD3_+ | exon_03 |
| RD3 | GAAGGTGGCCAGGCTGCCGC | chr1:211652507-211652526_RD3_+ | exon_03 |
| RET | GAAGGCGACGTCCGGTGCCG | chr10:43572712-43572731_RET_+ | exon_01 |
| RET | ATTGGGCCTCTACTTCTCGA | chr10:43595911-43595930_RET_+ | exon_02 |
| RET | GAAGAAGGAGAAGTATACGC | chr10:43597883-43597902_RET_- | exon_03 |
| RET | CTTCCGCTTGAACTCCACCA | chr10:43600619-43600638_RET_- | exon_04 |
| RET | GGCCCAGCAGACCTTCCGGG | chr10:43601928-43601947_RET_+ | exon_05 |
| RNF17 | AGTGCACCAGGTGTGGAAGG | chr13:25338433-25338452_RNF17_+ | exon_01 |
| RNF17 | GTCCACATTGAAGTTCACAA | chr13:25341417-25341436_RNF17_- | exon_02 |
| RNF17 | TAGACAACGCTACTACCCAA | chr13:25348971-25348990_RNF17_+ | exon_03 |
| RNF17 | TCAGGGCTAGATCCCATCTG | chr13:25367427-25367446_RNF17_- | exon_10 |
| RNF17 | CTTGATCTTGAACTGGACAG | chr13:25399840-25399859_RNF17_- | exon_16 |
| ROCK1 | CAGCAGCAACATGTCGACTG | chr18:18690862-18690881_ROCK1_- | exon_01 |
| ROCK1 | AAGGCATAAATCCACCAGGA | chr18:18629169-18629188_ROCK1_- | exon_04 |
| ROCK1 | TTGGATGCAATCCATTCCAT | chr18:18625268-18625287_ROCK1_- | exon_05 |
| ROCK1 | GAAGCCTGATAACATGCTGC | chr18:18624122-18624141_ROCK1_- | exon_06 |
| ROCK1 | GGAGCGTTTCCCAAGCCCAC | chr18:18619439-18619458_ROCK1_+ | exon_09 |
| ROCK2 | GAGCCGCCAGAGGAAGCTGG | chr2:11484172-11484191_ROCK2_- | exon_01 |
| ROCK2 | TGCTTACCTGAACCACCCAG | chr2:11389780-11389799_ROCK2_+ | exon_04 |
| ROCK2 | TCTGGATGCAATACACTCCA | chr2:11375929-11375948_ROCK2_- | exon_05 |
| ROCK2 | GAGGTTCTGAAATCACAAGG | chr2:11367451-11367470_ROCK2_- | exon_06 |
| ROCK2 | GAGTTGATTCACTTTCTCAG | chr2:11355456-11355475_ROCK2_+ | exon_15 |
| RPA1 | GAACATCCGTCCCATTACTA | chr17:1747213-1747232_RPA1_+ | exon_03 |
| RPA1 | ACAGTTGAACCCTCTCGTGG | chr17:1747888-1747907_RPA1_+ | exon_04 |
| RPA1 | GTAAGGAGTGAGGCTGGCAA | chr17:1779060-1779079_RPA1_- | exon_07 |
| RPA1 | GTACCTGGAGCAACTCCCGA | chr17:1780547-1780566_RPA1_+ | exon_08 |
| RPA1 | GGACGACCATCATTTACCTA | chr17:1782619-1782638_RPA1_+ | exon_10 |
| RPL8 | TGAGCGGCACGGCTACATCA | chr8:146017391-146017410_RPL8_- | exon_02 |
| RPL8 | GTGCCGCTCAGCGAAATCCA | chr8:146017401-146017420_RPL8_+ | exon_02 |
| RPL8 | CCGGAAGACCACCTTGGCGA | chr8:146017246-146017265_RPL8_+ | exon_03 |
| RPL8 | GATAACCTTCTTGGAGCCGG | chr8:146016687-146016706_RPL8_+ | exon_04 |
| RPL8 | CGCCCGGCCAGCCTTCAAGA | chr8:146015800-146015819_RPL8_+ | exon_05 |
| RPS5 | CCTGTCCCAGGATGACCGAG | chr19:58899494-58899513_RPS5_+ | exon_01 |
| RPS5 | GCCCTACCTCGCCTGTGAGC | chr19:58904540-58904559_RPS5_- | exon_02 |
| RPS5 | ACCTGGTTCACACGGCGCAG | chr19:58904837-58904856_RPS5_- | exon_03 |
| RPS5 | TCACCTGGTTCACACGGCGC | chr19:58904839-58904858_RPS5_- | exon_03 |
| RPS5 | CATTAAGAAGAAGGACGAGC | chr19:58906069-58906088_RPS5_+ | exon_05 |
| RPS6KB1 | CATAGACCTGGACCAGCCAG | hr17:57970632-57970651_RPS6KB1_ | exon_01 |
| RPS6KB1 | AATGAAAGCATGGACCATGG | hr17:57987932-57987951_RPS6KB1_ | exon_02 |
| RPS6KB1 | TCAGAAACTAGTGTGAACAG | hr17:57990076-57990095_RPS6KB1_ | exon_03 |
| RPS6KB1 | AGTAAAGCATCCCTTCATCG | hr17:58003861-58003880_RPS6KB1_ | exon_05 |

| | | | |
|---|---|---|---|
| RPS6KB1 | TTACTTGGCAGAAATCTCCA | hr17:58008986-58009005_RPS6KB1_ | exon_07 |
| RRM1 | GTAATCCAAGGCTTGTACAG | chr11:4127294-4127313_RRM1_+ | exon_03 |
| RRM1 | TAATGGCAAACACTCTCCCA | chr11:4128702-4128721_RRM1_+ | exon_04 |
| RRM1 | TACTGGCAGCTACATTGCTG | chr11:4139670-4139689_RRM1_+ | exon_08 |
| RRM1 | AAGAGCAACCAGCAGAACCT | chr11:4144534-4144553_RRM1_+ | exon_12 |
| RRM1 | AAACTACTATCCTGTACCAG | chr11:4148042-4148061_RRM1_+ | exon_13 |
| RS1 | ATTGGGATTATCGTCTACCG | chrX:18675762-18675781_RS1_- | exon_02 |
| RS1 | GCATGCAAGTGCGATTGCCA | chrX:18674829-18674848_RS1_- | exon_03 |
| RS1 | TTGGCTGCAGGATGAAGGCG | chrX:18674869-18674888_RS1_- | exon_03 |
| RS1 | ATTTCAGGGATCCTCACCCA | chrX:18662656-18662675_RS1_- | exon_05 |
| RS1 | ATGCGGACGTGCCAGCCCAG | chrX:18660170-18660189_RS1_+ | exon_06 |
| RXFP3 | GGTGCGGATTCTCATCAGCG | chr5:33937082-33937101_RXFP3_+ | exon_01 |
| RXFP3 | CTTACACATGGCCTTGCCGA | chr5:33937294-33937313_RXFP3_- | exon_01 |
| RXFP3 | GATCCACACACACAGCGCCT | chr5:33937489-33937508_RXFP3_- | exon_01 |
| RXFP3 | GAAGAAGGACAGGACAACGA | chr5:33937834-33937853_RXFP3_- | exon_01 |
| RXFP3 | GGCGTCGTGGTCTACAGCGG | chr5:33938196-33938215_RXFP3_+ | exon_01 |
| RXRA | GGTGAGGGAGGAGTTCACCT | chr9:137293490-137293509_RXRA_- | exon_02 |
| RXRA | GAGGACGCCATTGAGGCCCA | chr9:137300047-137300066_RXRA_- | exon_03 |
| RXRA | TATGGAGTGTACAGCTGCGA | chr9:137300794-137300813_RXRA_+ | exon_04 |
| RXRA | CAAGACCGAGACCTACGTGG | chr9:137309125-137309144_RXRA_+ | exon_05 |
| RXRA | GATCCCGTCCTTCACGGCGA | chr9:137320996-137321015_RXRA_- | exon_07 |
| RXRB | AGGCGGAGAACAACAAACCC | chr6:33168073-33168092_RXRB_- | exon_01 |
| RXRB | ACTGGAAAGGGAGAGCCCAG | chr6:33167025-33167044_RXRB_+ | exon_02 |
| RXRB | GTGGCTTCACATCTTCAGGG | chr6:33166181-33166200_RXRB_+ | exon_03 |
| RXRB | GGACAACAAAGACTGCACAG | chr6:33165614-33165633_RXRB_- | exon_04 |
| RXRB | GTCCACAGGCATCTCCTCGG | chr6:33164298-33164317_RXRB_+ | exon_05 |
| SAMD4B | GAAACGGGTCACCCGTACCC | :hr19:39847623-39847642_SAMD4B_· | exon_05 |
| SAMD4B | TAGCCAACCGCTCTTCCAGG | :hr19:39860545-39860564_SAMD4B_ | exon_06 |
| SAMD4B | CAGCCGGAACACCTTCCAGG | :hr19:39866489-39866508_SAMD4B_· | exon_07 |
| SAMD4B | TTATCACAGAACGTCACCAA | :hr19:39867336-39867355_SAMD4B_· | exon_09 |
| SAMD4B | GGAGGCTCAGCACCTGGCAG | :hr19:39868278-39868297_SAMD4B_ | exon_10 |
| SERPINA7 | TCATTCAAGAACTTTGCCAG | rX:105280604-105280623_SERPINA7 | exon_01 |
| SERPINA7 | AGCTGCAGAAATGCTCACAG | rX:105280822-105280841_SERPINA7 | exon_01 |
| SERPINA7 | AGTGCCCATGATGCACCAGA | ırX:105279278-105279297_SERPINA7 | exon_02 |
| SERPINA7 | TCTGGACTCACAGAGGACAA | ırX:105278245-105278264_SERPINA7 | exon_03 |
| SERPINA7 | AGGGAAAGTTGTGAACCCAA | ırX:105277502-105277521_SERPINA7 | exon_04 |
| SERPINB12 | ACCCAAGCGTACCATACCAA | ır18:61223508-61223527_SERPINB12 | exon_01 |
| SERPINB12 | TGGGTTGAATGTCAATCCCA | r18:61228414-61228433_SERPINB12 | exon_04 |
| SERPINB12 | AATCAAGGAACTCTTCAGCA | r18:61231215-61231234_SERPINB12 | exon_05 |
| SERPINB12 | CCTGGAAATGAGGTACACCA | r18:61232758-61232777_SERPINB12 | exon_06 |
| SERPINB12 | GAGATCATAGCTGTCTTCCA | ır18:61233937-61233956_SERPINB12 | exon_07 |
| SF3B3 | ACTGGCCAGGAACGATGCGA | chr16:70563044-70563063_SF3B3_- | exon_03 |
| SF3B3 | GGACCACATGATTTAAACCA | chr16:70566456-70566475_SF3B3_- | exon_05 |
| SF3B3 | GGAGGGTCAGATGGTCCAAG | chr16:70569216-70569235_SF3B3_+ | exon_06 |
| SF3B3 | AATGTGTCTCCTTCTTCCAG | chr16:70575639-70575658_SF3B3_- | exon_09 |
| SF3B3 | TCTCTGAGAGTCCTAAGACA | chr16:70578407-70578426_SF3B3_+ | exon_10 |
| SGK3 | AATGCAAAGAGATCACACCA | chr8:67705971-67705990_SGK3_+ | exon_02 |
| SGK3 | GGATGAAAGAAGTTCTCAGA | chr8:67734673-67734692_SGK3_+ | exon_06 |
| SGK3 | TACCTCACAGAACATCAACC | chr8:67740897-67740916_SGK3_+ | exon_07 |
| SGK3 | ACATACTGAGGATTTCCAGA | chr8:67740925-67740944_SGK3_- | exon_07 |
| SGK3 | CAGGCTGTAAGACTCACTCC | chr8:67759500-67759519_SGK3_- | exon_15 |

| | | | |
|---|---|---|---|
| SLC13A1 | GTGGCTCAGCAACACCTCGA | hr7:122809328-122809347_SLC13A1_ | exon_04 |
| SLC13A1 | ACCAACCAGGAACAAATCCG | r7:122768915-122768934_SLC13A1_ | exon_10 |
| SLC13A1 | ACAAGAATGGCTATATCCCA | r7:122763214-122763233_SLC13A1_ | exon_12 |
| SLC13A1 | CATGCTGGTAATGAACCCAG | r7:122759235-122759254_SLC13A1_ | exon_13 |
| SLC13A1 | CTCATAGCAGGAGCCCACGA | r7:122755591-122755610_SLC13A1_ | exon_15 |
| SLC36A3 | TCAGAGAATGTCCATCCTGC | hr5:150682806-150682825_SLC36A3_ | exon_01 |
| SLC36A3 | GATGCCTCTTACCAACAAGC | r5:150678142-150678161_SLC36A3_ | exon_02 |
| SLC36A3 | AGAAGCAGGCCTCTGACCTC | r5:150675724-150675743_SLC36A3_ | exon_03 |
| SLC36A3 | CCTGCCCGAACACCTGGCTG | hr5:150672948-150672967_SLC36A3_ | exon_04 |
| SLC36A3 | GCAGATGTTGGAGGTCACGT | r5:150666990-150667009_SLC36A3_ | exon_06 |
| SMO | TGTAGCGCAGCGGCTCGCAG | chr7:128829199-128829218_SMO_- | exon_01 |
| SMO | ATGTATACGGCACACAGCAG | chr7:128843266-128843285_SMO_- | exon_02 |
| SMO | CCAGGGTGAAGAGCGTGCAG | chr7:128845235-128845254_SMO_- | exon_03 |
| SMO | CACGGCAGACGATCTCTCGG | chr7:128845573-128845592_SMO_- | exon_04 |
| SMO | GATGACAAAGATGATGACGC | chr7:128846011-128846030_SMO_- | exon_05 |
| SPC24 | GGTGCCGAGAAGCAGCTGCG | chr19:11266295-11266314_SPC24_- | exon_01 |
| SPC24 | ACATAGAGGAGGTGAGCCAG | chr19:11266417-11266436_SPC24_- | exon_01 |
| SPC24 | GGAGGAGGACACCCGTCTGA | chr19:11259783-11259802_SPC24_- | exon_02 |
| SPC24 | CCAGAGCCTTCTCAATGCGA | chr19:11259867-11259886_SPC24_- | exon_02 |
| SPC24 | TGGCAGAACCTACACGGCCG | chr19:11258676-11258695_SPC24_+ | exon_03 |
| SPEM1 | CGGCCGCTCAACCATGGCCA | chr17:7323705-7323724_SPEM1_- | exon_01 |
| SPEM1 | GGGCCTCGGCCGCTCAACCA | chr17:7323711-7323730_SPEM1_- | exon_01 |
| SPEM1 | TAGCTCTGGAGCCGATTCCG | chr17:7323943-7323962_SPEM1_+ | exon_02 |
| SPEM1 | GCTCCTAGCCACAAGAACGG | chr17:7324640-7324659_SPEM1_+ | exon_03 |
| SPEM1 | GCTTCGGGAACTGACCCGGG | chr17:7324804-7324823_SPEM1_+ | exon_03 |
| SRC | TAGCAACAAGAGCAAGCCCA | chr20:36012562-36012581_SRC_+ | exon_04 |
| SRC | GTCATAGAGGGCCACAAAGG | chr20:36014490-36014509_SRC_- | exon_05 |
| SRC | TCAATGCAGAGAACCCGAGA | chr20:36022624-36022643_SRC_+ | exon_07 |
| SRC | GTCTGACTTCGACAACGCCA | chr20:36024584-36024603_SRC_+ | exon_08 |
| SRC | TCCAGCCGCAGCGACTCCCG | chr20:36026197-36026216_SRC_- | exon_09 |
| SRSF2 | CCAAGAGGGAATCCAAATCC | chr17:74732303-74732322_SRSF2_- | exon_02 |
| SRSF2 | GGATTCCCTCTTGGACACTG | chr17:74732309-74732328_SRSF2_+ | exon_02 |
| SRSF2 | GATTCCCTCTTGGACACTGG | chr17:74732310-74732329_SRSF2_+ | exon_02 |
| SRSF2 | AGGACTCCTGGACCGAGACC | chr17:74732333-74732352_SRSF2_+ | exon_02 |
| SRSF2 | ACGAGATCTGGAGACCGACG | chr17:74732369-74732388_SRSF2_+ | exon_02 |
| STAT3 | CTACAGTGACAGCTTCCCAA | chr17:40500453-40500472_STAT3_- | exon_02 |
| STAT3 | GCAGGAAGCGGCTATACTGC | chr17:40498643-40498662_STAT3_+ | exon_03 |
| STAT3 | GAAGGCGTGATTCTTCCCAC | chr17:40497603-40497622_STAT3_+ | exon_04 |
| STAT3 | GGAACAGATGCTCACTGCGC | chr17:40489798-40489817_STAT3_- | exon_07 |
| STAT3 | ACAGCACCGGCCGATGCTGG | chr17:40485949-40485968_STAT3_- | exon_09 |
| SYK | TTTCGGCAACATCACCCGGG | chr9:93606228-93606247_SYK_+ | exon_02 |
| SYK | GTGCAGCAGGCACAGGGCGT | chr9:93624517-93624536_SYK_- | exon_04 |
| SYK | TGAGCCAGAACTTGCACCCT | chr9:93636539-93636558_SYK_+ | exon_08 |
| SYK | GGACCGAAAGCTGCTGACGC | chr9:93637045-93637064_SYK_+ | exon_09 |
| SYK | ATGGAAAGTGGCCTGTCAAG | chr9:93650041-93650060_SYK_+ | exon_12 |
| TBK1 | TTTGAACATCCACTGGACGA | chr12:64854019-64854038_TBK1_- | exon_03 |
| TBK1 | ACAGTGTATAAACTCCCACA | chr12:64858156-64858175_TBK1_- | exon_04 |
| TBK1 | AATGGACCAATTGACTGGAG | chr12:64873850-64873869_TBK1_+ | exon_07 |
| TBK1 | AACGTGGATGTACTTTAGGG | chr12:64879242-64879261_TBK1_- | exon_10 |
| TBK1 | GATGAAGATCAACCTGGAAG | chr12:64883833-64883852_TBK1_+ | exon_13 |
| TEC | AAACACTTACTTCACTGCGG | chr4:48158687-48158706_TEC_+ | exon_09 |

| | | | |
|---|---|---|---|
| TEC | GGAAGGTGCAATGTGCGAGG | chr4:48147456-48147475_TEC_- | exon_13 |
| TEC | AATGACCTTGTCTCTGTCGG | chr4:48147185-48147204_TEC_+ | exon_14 |
| TEC | TCAGGTGGACACCACTTCAC | chr4:48140971-48140990_TEC_+ | exon_16 |
| TEC | ATGTGGGAAGTATTCACGGA | chr4:48140812-48140831_TEC_- | exon_17 |
| TEK | AAAGCCAAACCTTACCAGAA | chr9:27109636-27109655_TEK_- | exon_01 |
| TEK | GAACCAGCACCAGGATCCGC | chr9:27157971-27157990_TEK_+ | exon_02 |
| TEK | AGGAGGAAACCTCTTCACCT | chr9:27169581-27169600_TEK_+ | exon_04 |
| TEK | TATGGGAAGGACGTGTGAGA | chr9:27172723-27172742_TEK_+ | exon_05 |
| TEK | TGTGGCACAGGAACACCCAT | chr9:27173313-27173332_TEK_- | exon_06 |
| TKTL2 | GTGGAGGATCACTACAGGGA | chr4:164393181-164393200_TKTL2_- | exon_01 |
| TKTL2 | AGGCGGCCGAGTTATCACAG | chr4:164393200-164393219_TKTL2_- | exon_01 |
| TKTL2 | TATCTAGCCGCCAATACCAA | chr4:164393481-164393500_TKTL2_- | exon_01 |
| TKTL2 | AGAAGATGGAGTCTCCCAGA | chr4:164393590-164393609_TKTL2_- | exon_01 |
| TKTL2 | GGAGACTCCATCTTCTCCAG | chr4:164393594-164393613_TKTL2_+ | exon_01 |
| TNFRSF14 | CAAGGAGGACGAGTACCCAG | chr1:2489221-2489240_TNFRSF14_+ | exon_02 |
| TNFRSF14 | ACTTGCTTAGGCCATTGAGG | chr1:2489861-2489880_TNFRSF14_- | exon_03 |
| TNFRSF14 | GGTGGCGTAAGCGCGGCACG | chr1:2491367-2491386_TNFRSF14_- | exon_04 |
| TNFRSF14 | CTGGTGCTGACATTCCTCCA | chr1:2492129-2492148_TNFRSF14_- | exon_05 |
| TNFRSF14 | CTGGGACCAGCAGCTCCCAC | chr1:2493141-2493160_TNFRSF14_+ | exon_06 |
| TNFRSF4 | GCCGGGCTTCTACAACGACG | chr1:1149088-1149107_TNFRSF4_- | exon_02 |
| TNFRSF4 | GCAGGCCTGGTTGTCGCCTG | chr1:1148032-1148051_TNFRSF4_+ | exon_04 |
| TNFRSF4 | ACAGTGATGGGCCTGGCCGG | chr1:1147390-1147409_TNFRSF4_+ | exon_05 |
| TNFRSF4 | GGCCATGAGGCACTCACCAG | chr1:1147067-1147086_TNFRSF4_+ | exon_06 |
| TNFRSF4 | CCCAGGTCAGATCTTGGCCA | chr1:1146929-1146948_TNFRSF4_+ | exon_07 |
| TNFRSF8 | GGCGCTACGAGCCTTCCCAC | hr1:12123697-12123716_TNFRSF8_- | exon_01 |
| TNFRSF8 | ACTATGACAAGGCTGTCAGG | hr1:12144564-12144583_TNFRSF8_- | exon_02 |
| TNFRSF8 | ACCTCGAGAACAAGTCACGC | hr1:12157257-12157276_TNFRSF8_ | exon_03 |
| TNFRSF8 | GCCCGACTACTACCTGGACG | hr1:12172013-12172032_TNFRSF8_- | exon_07 |
| TNFRSF8 | ACCTCGAGAACAGCTCACGC | hr1:12172054-12172073_TNFRSF8_ | exon_07 |
| TNFRSF9 | GCGCTGGAGAAACTATTTGG | chr1:7998822-7998841_TNFRSF9_+ | exon_03 |
| TNFRSF9 | TCCAGGGTTTCACTGCCTGG | chr1:7998313-7998332_TNFRSF9_- | exon_04 |
| TNFRSF9 | AACGTGGCATCTGTCGACCC | chr1:7997758-7997777_TNFRSF9_- | exon_05 |
| TNFRSF9 | AAGGAGAGGGACGTGGTCTG | chr1:7995144-7995163_TNFRSF9_- | exon_06 |
| TNFRSF9 | TGGGACGAAGGAGAGGGACG | chr1:7995151-7995170_TNFRSF9_- | exon_06 |
| TNFSF11 | GTTCTTCTATTTCAGAGCGC | hr13:43148637-43148656_TNFSF11_ | exon_01 |
| TNFSF11 | TGCAGATAGAGCTTACCAGA | hr13:43175114-43175133_TNFSF11_ | exon_04 |
| TNFSF11 | GAAGCACCAAGTATTGGTCA | hr13:43180876-43180895_TNFSF11_ | exon_05 |
| TNFSF11 | TGGAGAGGAAATCAGCATCG | hr13:43180955-43180974_TNFSF11_ | exon_05 |
| TNFSF11 | GATCCGGATCCAGTAAGGAG | hr13:43180988-43181007_TNFSF11_ | exon_05 |
| TNFSF13B | GCTGTCTTGCTGCCTCACGG | 13:108922411-108922430_TNFSF13 | exon_01 |
| TNFSF13B | AGAGCTGCAGGGCCACCACG | 13:108922486-108922505_TNFSF13 | exon_01 |
| TNFSF13B | GAAGCTCCAGCTGTCACCGC | 13:108922553-108922572_TNFSF13 | exon_01 |
| TNFSF13B | TACTGATAAGACCTACGCCA | 13:108955810-108955829_TNFSF13 | exon_05 |
| TNFSF13B | AATTAGATGTCCCATGGCGT | 13:108955824-108955843_TNFSF13 | exon_05 |
| TNFSF4 | GAGGAACAAGCTATTGCTGG | hr1:173176237-173176256_TNFSF4_ | exon_01 |
| TNFSF4 | GATGACTTCCATGTGAATGG | hr1:173155707-173155726_TNFSF4_ | exon_03 |
| TNFSF4 | CACATGGAAGTCATCCAGGG | hr1:173155712-173155731_TNFSF4_ | exon_03 |
| TNFSF4 | AGTCAGAGAGGCCACCATCA | hr1:173155775-173155794_TNFSF4_ | exon_03 |
| TNFSF4 | CCTGAAGGGCTACTTCTCCC | hr1:173155876-173155895_TNFSF4_ | exon_03 |
| TNFSF9 | GCCAGCCCGAGACTCCGCGA | chr19:6531243-6531262_TNFSF9_+ | exon_01 |
| TNFSF9 | GCAGGCGTGTCCCTGACGGG | chr19:6534658-6534677_TNFSF9_+ | exon_03 |

| | | | |
|---|---|---|---|
| TNFSF9 | GAGCTACAAAGAGGACACGA | chr19:6534684-6534703_TNFSF9_+ | exon_03 |
| TNFSF9 | GAAGATGGACGCCCAGGCGC | chr19:6534942-6534961_TNFSF9_- | exon_03 |
| TNFSF9 | CAGCCGGACTCCCTTCACCG | chr19:6535046-6535065_TNFSF9_+ | exon_03 |
| TNF | AGGCACTCACCTCTTCCCTC | chr6:31543695-31543714_TNF_- | exon_01 |
| TNF | GAGACACTTACTGACTGCCT | chr6:31544348-31544367_TNF_- | exon_02 |
| TNF | GCTGAGAGATAACCAGCTGG | chr6:31544966-31544985_TNF_+ | exon_04 |
| TNF | AGATAGATGGGCTCATACCA | chr6:31545180-31545199_TNF_- | exon_04 |
| TNF | TATGAGCCCATCTATCTGGG | chr6:31545183-31545202_TNF_+ | exon_04 |
| TOMM34 | GCGGAAACTCTCATTGCCGG | chr20:43588912-43588931_TOMM34_ | exon_01 |
| TOMM34 | GCAGCATGTCACTTGAAGGA | chr20:43585063-43585082_TOMM34_ | exon_02 |
| TOMM34 | GCAGATGCTCGCCGCAGCAG | chr20:43583808-43583827_TOMM34_ | exon_03 |
| TOMM34 | TGAGCTGAAACAGGCACCAA | chr20:43580560-43580579_TOMM34_ | exon_04 |
| TOMM34 | GCTGGATGGAAAGAACGTGA | chr20:43572135-43572154_TOMM34_ | exon_06 |
| TONSL | AAAGCCAAGGCGCAGAGGGC | chr8:145669625-145669644_TONSL_ | exon_02 |
| TONSL | GGCGCTCTCCGATCTTGCGG | chr8:145669300-145669319_TONSL_ | exon_03 |
| TONSL | AATAGCCAAGCTCTTCTCAA | chr8:145668540-145668559_TONSL_ | exon_04 |
| TONSL | CTTCCAGGGACACTGGCCCA | chr8:145668177-145668196_TONSL_ | exon_05 |
| TONSL | GCTCCCAGAAGCCTGTGCAG | chr8:145667451-145667470_TONSL_ | exon_07 |
| TOP1 | GTACCTGGGAATCGTTGTGG | chr20:39657725-39657744_TOP1_- | exon_01 |
| TOP1 | AACACAAAGATCGAGAACAC | chr20:39690055-39690074_TOP1_+ | exon_03 |
| TOP1 | GAGAAGACCAAACACAAAGA | chr20:39704848-39704867_TOP1_+ | exon_04 |
| TOP1 | GAAGAAGAGCGCTATCCTGA | chr20:39721116-39721135_TOP1_+ | exon_09 |
| TOP1 | AGAAATACGAGACTGCTCGG | chr20:39741438-39741457_TOP1_+ | exon_14 |
| TOP2A | ATGCTGCGGACAACAAACAA | chr17:38572298-38572317_TOP2A_- | exon_04 |
| TOP2A | TATGGGAAGAGCTGGTGAGA | chr17:38569190-38569209_TOP2A_- | exon_07 |
| TOP2A | TTAAACACACTTCCCACCTG | chr17:38567956-38567975_TOP2A_+ | exon_08 |
| TOP2A | TGTACGCTTATCCTGACTGA | chr17:38564337-38564356_TOP2A_- | exon_12 |
| TOP2A | TAATCAGCAAGCCTTTGATG | chr17:38563158-38563177_TOP2A_+ | exon_14 |
| TOP2B | GCAGCCACCCGACTTGGCCA | chr3:25705768-25705787_TOP2B_+ | exon_01 |
| TOP2B | GTCAGTGGAGCCATTGACGC | chr3:25686793-25686812_TOP2B_- | exon_02 |
| TOP2B | TGTAGGAATGAATTGCAGGG | chr3:25685233-25685252_TOP2B_- | exon_03 |
| TOP2B | ATGCTGCTGACAATAAACAG | chr3:25683854-25683873_TOP2B_- | exon_04 |
| TOP2B | AAAGTGATGGCCAATTGTGA | chr3:25670570-25670589_TOP2B_+ | exon_14 |
| TPH2 | AATGACGACAAAGGCAACAA | chr12:72335388-72335407_TPH2_+ | exon_02 |
| TPH2 | TCCAGAGAACATTTGGACAG | chr12:72338229-72338248_TPH2_+ | exon_03 |
| TPH2 | CTAACTCAGAGATCTTCCGA | chr12:72338391-72338410_TPH2_- | exon_04 |
| TPH2 | TAAATACTGTGGCTACAGAG | chr12:72366431-72366450_TPH2_+ | exon_06 |
| TPH2 | TGGCTTCACGGTGAGGCCGG | chr12:72372739-72372758_TPH2_+ | exon_07 |
| TRIM24 | CAGGACAATGGAGGTGGCGG | chr7:138145287-138145306_TRIM24_ | exon_01 |
| TRIM24 | GAGGACAACGCAGAAGCCAA | chr7:138199972-138199991_TRIM24_ | exon_03 |
| TRIM24 | TCTAAATGGGCAGTTTCCAG | chr7:138223492-138223511_TRIM24_ | exon_07 |
| TRIM24 | ATGCATCACACCTTGCACGA | chr7:138235834-138235853_TRIM24_ | exon_08 |
| TRIM24 | AGAGGCAACAGGTGCAACGG | chr7:138239620-138239639_TRIM24_ | exon_09 |
| TRPC7 | AGCGCCGGCACACAACGCTG | chr5:135693025-135693044_TRPC7_ | exon_02 |
| TRPC7 | TCCGGCTCAGACTTGGACGG | chr5:135651317-135651336_TRPC7_ | exon_03 |
| TRPC7 | GAAGTGGCATACTTCACCTA | chr5:135587339-135587358_TRPC7_ | exon_06 |
| TRPC7 | GCGAGAGAAGCTCAGCACGA | chr5:135583341-135583360_TRPC7_ | exon_07 |
| TRPC7 | GTTGAGCAACACTACCACCA | chr5:135567084-135567103_TRPC7_ | exon_08 |
| TRPV1 | CTCGGAGGAGGCTTTCCCGG | chr17:3495464-3495483_TRPV1_- | exon_02 |
| TRPV1 | CTGGCAGTTATTCTGAGCAA | chr17:3494548-3494567_TRPV1_+ | exon_03 |
| TRPV1 | GCGGCAAACGGACAGCCTGA | chr17:3494300-3494319_TRPV1_- | exon_04 |

| | | | |
|---|---|---|---|
| TRPV1 | GACGGCCGACATCAGCGCCA | chr17:3493304-3493323_TRPV1_- | exon_06 |
| TRPV1 | GCAGGACAGGTCGTACAGCG | chr17:3491545-3491564_TRPV1_+ | exon_07 |
| TSSK1B | TGTGGCCATCAACAAGGAGG | :hr5:112769675-112769694_TSSK1B_ | exon_01 |
| TSSK1B | GTGGCAGTACTTGATGGCCA | hr5:112770153-112770172_TSSK1B_ | exon_01 |
| TSSK1B | GCCAGAATCTCAATTTCCCG | hr5:112770343-112770362_TSSK1B_ | exon_01 |
| TSSK1B | TTTCTCCAAGAAGTCTGCGG | hr5:112770372-112770391_TSSK1B_ | exon_01 |
| TSSK1B | GAGGACAGCAGCGTCATCCA | hr5:112770516-112770535_TSSK1B_ | exon_01 |
| TUBB | ACCCAAAGTGTCTGACACCG | chr6:30691355-30691374_TUBB_+ | exon_04 |
| TUBB | TGAGCTGGCCAGGGAAACGG | chr6:30691559-30691578_TUBB_- | exon_04 |
| TUBB | TGAGGTATCGGCCGTGGCGG | chr6:30691754-30691773_TUBB_- | exon_04 |
| TUBB | ACCGTGGCTGCTGTCTTCCG | chr6:30691773-30691792_TUBB_+ | exon_04 |
| TUBB | GGAGGAGGATTTCGGTGAGG | chr6:30692135-30692154_TUBB_+ | exon_04 |
| TUBGCP2 | AGCCTGCTGCGTGTCCACGG | r10:135116384-135116403_TUBGCP2 | exon_01 |
| TUBGCP2 | TTTGACAACAGGTACACCAG | r10:135113508-135113527_TUBGCP2 | exon_02 |
| TUBGCP2 | CAGCATCAACGTCCCTGCCG | r10:135113014-135113033_TUBGCP2 | exon_03 |
| TUBGCP2 | GGTGCACAGGATCCTCCCAG | r10:135107094-135107113_TUBGCP2 | exon_05 |
| TUBGCP2 | AATCAGGTGCTCCTTCACCA | r10:135106652-135106671_TUBGCP2 | exon_06 |
| TXN | GGTGAAGCAGATCGAGAGCA | chr9:113018694-113018713_TXN_- | exon_01 |
| TYMS | ATCCAACACATCCTCCGCTG | chr18:657851-657870_TYMS_+ | exon_01 |
| TYMS | GTCAGGAAGGACGACCGCAC | chr18:657875-657894_TYMS_+ | exon_01 |
| TYMS | ACACGTTTGGTTGTCAGCAG | chr18:659652-659671_TYMS_- | exon_02 |
| TYMS | TCACCACATAGAACTGGCAG | chr18:670729-670748_TYMS_- | exon_05 |
| TYMS | TTTCGAAGAATCCTGAGCTT | chr18:672887-672906_TYMS_- | exon_07 |
| TYRO3 | AGACACTGTCAGCTTCACCG | chr15:41853346-41853365_TYRO3_- | exon_02 |
| TYRO3 | TCAGTGGAGCGCTCTGACGC | chr15:41853744-41853763_TYRO3_+ | exon_03 |
| TYRO3 | AGAGGAACTACGAAGATCGG | chr15:41854859-41854878_TYRO3_+ | exon_04 |
| TYRO3 | GCTGCTGGAAAGCTTTGTCA | chr15:41857254-41857273_TYRO3_- | exon_06 |
| TYRO3 | GGTGCGCTGTGCCAATGCCT | chr15:41859665-41859684_TYRO3_+ | exon_07 |
| T | ACAGAGCGCGAACTGCGCGT | chr6:166580949-166580968_T_- | exon_02 |
| T | TCACCAACAAGCTCAACGGA | chr6:166580089-166580108_T_- | exon_03 |
| T | TGGCTGGTGATCATGCGCTG | chr6:166579246-166579265_T_+ | exon_04 |
| T | TGATCACAAAGAGATGATGG | chr6:166578132-166578151_T_- | exon_06 |
| T | GAGAGGGCACCTCCAAACTG | chr6:166576033-166576052_T_+ | exon_07 |
| UBQLN3 | TCAGGCCCTCATTGCTACGG | chr11:5528865-5528884_UBQLN3_- | exon_02 |
| UBQLN3 | GGAGACCTAAGCTAAAGGCA | chr11:5530380-5530399_UBQLN3_+ | exon_02 |
| UBQLN3 | GAGACCTAAGCTAAAGGCAG | chr11:5530381-5530400_UBQLN3_+ | exon_02 |
| UBQLN3 | CAAGAGGCAGCACCGTGCCA | chr11:5530497-5530516_UBQLN3_- | exon_02 |
| UBQLN3 | AGGTGGAGAAGCCCTGCCAC | chr11:5530761-5530780_UBQLN3_- | exon_02 |
| UTS2R | GACCGAGCCCAGCTCCCTGG | chr17:80332317-80332336_UTS2R_+ | exon_01 |
| UTS2R | ACATGGAGGCCACCGCACGC | chr17:80332440-80332459_UTS2R_- | exon_01 |
| UTS2R | GACCACGTAGACGTACATGG | chr17:80332454-80332473_UTS2R_- | exon_01 |
| UTS2R | GAGCAGCGAGCGCTACGCTG | chr17:80332632-80332651_UTS2R_+ | exon_01 |
| UTS2R | ACAGCGCTGGAAGCGGGCGC | chr17:80333243-80333262_UTS2R_- | exon_01 |
| VCP | CAACAATTAACCGATTGGGA | chr9:35068292-35068311_VCP_+ | exon_02 |
| VCP | AATAACCTTCGTGTACGCCT | chr9:35067903-35067922_VCP_- | exon_03 |
| VCP | TTAGCTCTCACCTTTCCGGA | chr9:35066661-35066680_VCP_+ | exon_04 |
| VCP | GACACAGTGATCCACTGCGA | chr9:35065270-35065289_VCP_- | exon_05 |
| VCP | AAAGAGGGCAGGATGTCTCA | chr9:35064169-35064188_VCP_+ | exon_06 |
| VEGFA | GCTCTACCTCCACCATGCCA | chr6:43739028-43739047_VEGFA_+ | exon_01 |
| VEGFA | GGAGGGCAGAATCATCACGA | chr6:43742108-43742127_VEGFA_+ | exon_02 |
| VEGFA | GGTGATGTTGGACTCCTCAG | chr6:43745377-43745396_VEGFA_- | exon_03 |

| | | | |
|---|---|---|---|
| VEGFA | ATGCGGATCAAACCTCACCA | chr6:43746200-43746219_VEGFA_+ | exon_04 |
| VEGFA | TCAGATGTGACAAGCCGAGG | chr6:43752274-43752293_VEGFA_+ | exon_07 |
| VTCN1 | GAGCAGGATGAAATGTTCAG | chr1:117699348-117699367_VTCN1_- | exon_03 |
| VTCN1 | GTGATACAATGGCTGAAGGA | chr1:117699420-117699439_VTCN1_- | exon_03 |
| VTCN1 | TGGGCATCCCAAGTTGACCA | chr1:117695877-117695896_VTCN1_- | exon_04 |
| VTCN1 | TTGGGATGCCCAGACCACTG | chr1:117695885-117695904_VTCN1_+ | exon_04 |
| VTCN1 | ACACCGCAAGGTCTCTGAGC | chr1:117695933-117695952_VTCN1_+ | exon_04 |
| WDR3 | GATGGGCTACACTTAGCTGT | chr1:118477150-118477169_WDR3_+ | exon_03 |
| WDR3 | AAGGTCCCACCATTTCACCA | chr1:118481117-118481136_WDR3_- | exon_05 |
| WDR3 | AGAGGATGGTGCCTTTGAGA | chr1:118483524-118483543_WDR3_+ | exon_07 |
| WDR3 | TCTGCAAGATGAAATCCAGC | chr1:118485114-118485133_WDR3_+ | exon_10 |
| WDR3 | TCAGTGAATACAATTCCACC | chr1:118486085-118486104_WDR3_- | exon_11 |
| WDR43 | AGAGCAGCAATGGCGGCGGG | chr2:29117556-29117575_WDR43_+ | exon_01 |
| WDR43 | TTCGCTTTGGCCTCTACCGA | chr2:29117649-29117668_WDR43_+ | exon_01 |
| WDR43 | ACAGAGTCAACTGCATACAG | chr2:29129342-29129361_WDR43_+ | exon_03 |
| WDR43 | ATACAGTGGCATCAAGACAG | chr2:29129356-29129375_WDR43_+ | exon_03 |
| WDR43 | TTTCCAGTTCTTCTTACCCA | chr2:29152523-29152542_WDR43_+ | exon_11 |
| WEE1 | GTCGCCGGTCAAGTCGCCGG | chr11:9595846-9595865_WEE1_+ | exon_01 |
| WEE1 | TCTACGACGACACTGTCCTG | chr11:9597601-9597620_WEE1_- | exon_02 |
| WEE1 | TCATCAACAGAGCCCGCCAA | chr11:9598187-9598206_WEE1_- | exon_04 |
| WEE1 | AGTTCGATATTTCTCTGCGT | chr11:9598746-9598765_WEE1_+ | exon_05 |
| WEE1 | TCAGACAGGGTAGATTACCT | chr11:9608102-9608121_WEE1_+ | exon_09 |
| XPO1 | GTGGTGAATTGCTTATACCA | chr2:61760923-61760942_XPO1_- | exon_02 |
| XPO1 | CTTCCAAGGAACCAGTGCGA | chr2:61749748-61749767_XPO1_- | exon_04 |
| XPO1 | AACAAGAATGGCCCAAACAT | chr2:61727003-61727022_XPO1_- | exon_07 |
| XPO1 | TCTCAGGGAAACTCTTATGG | chr2:61722592-61722611_XPO1_- | exon_11 |
| XPO1 | ACTTCCAGAAAGCAACGGAG | chr2:61721083-61721102_XPO1_+ | exon_12 |

| GENE | A549 T12 sumZ | numOb | normZ | val svr | ank svr | dr svnt | val sur | ank sur | dr sup |
|---|---|---|---|---|---|---|---|---|---|
| AARS | -3.64 | 10 | -0.9 | 0.184 | 47 | 0.916 | 0.816 | 281 | 0.943 |
| ABL1 | 0.79 | 10 | 0.05 | 0.521 | 172 | 0.916 | 0.479 | 156 | 0.943 |
| ABL2 | -0.79 | 10 | -0.29 | 0.387 | 118 | 0.916 | 0.613 | 210 | 0.943 |
| ADH7 | -3.74 | 10 | -0.92 | 0.179 | 44 | 0.916 | 0.821 | 284 | 0.943 |
| ADORA2A | -4.15 | 10 | -1.01 | 0.157 | 42 | 0.916 | 0.843 | 286 | 0.962 |
| AKT1 | -3.18 | 10 | -0.8 | 0.212 | 59 | 0.916 | 0.788 | 269 | 0.943 |
| AKT2 | 4.62 | 10 | 0.88 | 0.809 | 282 | 0.938 | 0.191 | 46 | 0.943 |
| AKT3 | -0.45 | 10 | -0.21 | 0.415 | 131 | 0.916 | 0.585 | 197 | 0.943 |
| ALK | -0.32 | 10 | -0.19 | 0.426 | 139 | 0.916 | 0.574 | 189 | 0.943 |
| APOBEC1 | -8.26 | 10 | -1.89 | 0.029 | 10 | 0.916 | 0.971 | 318 | 0.993 |
| AR | -8.54 | 10 | -1.95 | 0.026 | 7 | 0.916 | 0.975 | 321 | 0.993 |
| ARCN1 | 5.84 | 10 | 1.14 | 0.872 | 298 | 0.955 | 0.128 | 30 | 0.943 |
| ATM | 2.08 | 10 | 0.33 | 0.629 | 213 | 0.916 | 0.371 | 115 | 0.943 |
| ATP6V1A | 12.74 | 10 | 2.62 | 0.996 | 324 | 1 | 0.004 | 4 | 0.295 |
| ATR | 2.44 | 10 | 0.41 | 0.658 | 229 | 0.916 | 0.342 | 99 | 0.943 |
| AURKA | 2.42 | 10 | 0.4 | 0.656 | 226 | 0.916 | 0.344 | 102 | 0.943 |
| AURKB | -1.78 | 10 | -0.5 | 0.309 | 95 | 0.916 | 0.691 | 233 | 0.943 |
| AURKC | 4.38 | 10 | 0.82 | 0.795 | 278 | 0.931 | 0.205 | 50 | 0.943 |
| AXL | 3.02 | 10 | 0.53 | 0.703 | 245 | 0.916 | 0.297 | 83 | 0.943 |
| BCL2 | 1.55 | 10 | 0.22 | 0.585 | 199 | 0.916 | 0.415 | 129 | 0.943 |
| BCR | -3.17 | 10 | -0.8 | 0.212 | 60 | 0.916 | 0.788 | 268 | 0.943 |
| BIRC5 | 6.13 | 10 | 1.2 | 0.885 | 302 | 0.957 | 0.115 | 26 | 0.943 |
| BIRC8 | -1.84 | 10 | -0.51 | 0.304 | 90 | 0.916 | 0.696 | 238 | 0.943 |
| BLK | 2.42 | 10 | 0.4 | 0.657 | 227 | 0.916 | 0.343 | 101 | 0.943 |
| BMP15 | -4.69 | 10 | -1.13 | 0.13 | 34 | 0.916 | 0.87 | 294 | 0.967 |
| BMX | 1.88 | 10 | 0.29 | 0.613 | 207 | 0.916 | 0.387 | 121 | 0.943 |
| BPIFB6 | 3.51 | 10 | 0.64 | 0.738 | 261 | 0.916 | 0.262 | 67 | 0.943 |
| BRAF | 3.84 | 10 | 0.71 | 0.761 | 271 | 0.916 | 0.239 | 57 | 0.943 |
| BRD4 | 1.8 | 10 | 0.27 | 0.606 | 205 | 0.916 | 0.394 | 123 | 0.943 |
| BTK | -2.32 | 10 | -0.62 | 0.269 | 80 | 0.916 | 0.731 | 248 | 0.943 |
| BTLA | -1.12 | 10 | -0.36 | 0.36 | 111 | 0.916 | 0.64 | 217 | 0.943 |
| BUD31 | 0.41 | 10 | -0.03 | 0.488 | 161 | 0.916 | 0.512 | 167 | 0.943 |
| C3orf30 | -0.7 | 10 | -0.27 | 0.395 | 121 | 0.916 | 0.605 | 207 | 0.943 |
| C8B | -2.27 | 10 | -0.6 | 0.273 | 82 | 0.916 | 0.727 | 246 | 0.943 |
| CA9 | -5.82 | 10 | -1.37 | 0.086 | 24 | 0.916 | 0.914 | 304 | 0.974 |
| CADM1 | -0.06 | 10 | -0.13 | 0.448 | 146 | 0.916 | 0.552 | 182 | 0.943 |
| CASP3 | -4.8 | 10 | -1.15 | 0.126 | 33 | 0.916 | 0.874 | 295 | 0.967 |
| CCR2 | -5.67 | 10 | -1.34 | 0.091 | 27 | 0.916 | 0.909 | 301 | 0.974 |
| CCT2 | 6.18 | 10 | 1.21 | 0.887 | 303 | 0.957 | 0.113 | 25 | 0.943 |
| CCT7 | -4.46 | 10 | -1.07 | 0.141 | 37 | 0.916 | 0.859 | 291 | 0.962 |
| CD27 | -1.54 | 10 | -0.45 | 0.327 | 100 | 0.916 | 0.673 | 228 | 0.943 |
| CD274 | -7.12 | 10 | -1.65 | 0.05 | 15 | 0.916 | 0.95 | 313 | 0.992 |
| CD276 | 0.09 | 10 | -0.1 | 0.461 | 151 | 0.916 | 0.539 | 177 | 0.943 |
| CD28 | -1.34 | 10 | -0.41 | 0.342 | 106 | 0.916 | 0.658 | 222 | 0.943 |
| CD38 | -5.24 | 10 | -1.24 | 0.107 | 29 | 0.916 | 0.893 | 299 | 0.974 |
| CD40 | 2.19 | 10 | 0.35 | 0.638 | 216 | 0.916 | 0.362 | 112 | 0.943 |
| CD40LG | -4.96 | 10 | -1.18 | 0.118 | 31 | 0.916 | 0.882 | 297 | 0.971 |
| CD52 | 2.15 | 6 | 0.48 | 0.684 | 237 | 0.916 | 0.316 | 91 | 0.943 |
| CD70 | -2.22 | 10 | -0.59 | 0.276 | 84 | 0.916 | 0.724 | 244 | 0.943 |
| CD80 | 2.42 | 10 | 0.4 | 0.657 | 228 | 0.916 | 0.343 | 100 | 0.943 |
| CD86 | 0.69 | 10 | 0.03 | 0.512 | 169 | 0.916 | 0.488 | 159 | 0.943 |
| CDC16 | 3.04 | 10 | 0.54 | 0.704 | 246 | 0.916 | 0.296 | 82 | 0.943 |
| CDH1 | 4.75 | 10 | 0.9 | 0.817 | 283 | 0.943 | 0.183 | 45 | 0.943 |
| CDK1 | -4.24 | 10 | -1.03 | 0.152 | 41 | 0.916 | 0.848 | 287 | 0.962 |
| CDK2 | 4.47 | 10 | 0.84 | 0.8 | 281 | 0.931 | 0.2 | 47 | 0.943 |
| CDK4 | -7.87 | 10 | -1.81 | 0.035 | 11 | 0.916 | 0.965 | 317 | 0.993 |
| CDK6 | 4.24 | 10 | 0.79 | 0.786 | 277 | 0.928 | 0.214 | 51 | 0.943 |
| CDK7 | -3.19 | 10 | -0.8 | 0.211 | 56 | 0.916 | 0.789 | 272 | 0.943 |
| CDK9 | 0.45 | 10 | -0.02 | 0.492 | 162 | 0.916 | 0.508 | 166 | 0.943 |
| CDX4 | -0.87 | 10 | -0.3 | 0.38 | 115 | 0.916 | 0.62 | 213 | 0.943 |
| CHEK1 | 1.76 | 10 | 0.26 | 0.603 | 204 | 0.916 | 0.397 | 124 | 0.943 |
| CHEK2 | -3.75 | 10 | -0.92 | 0.178 | 43 | 0.916 | 0.822 | 285 | 0.943 |
| CKAP5 | -3.53 | 10 | -0.87 | 0.191 | 49 | 0.916 | 0.809 | 279 | 0.943 |
| CLDN17 | -1.81 | 10 | -0.51 | 0.306 | 93 | 0.916 | 0.694 | 235 | 0.943 |
| CLRN1 | -0.59 | 10 | -0.24 | 0.404 | 130 | 0.916 | 0.596 | 198 | 0.943 |
| CPSF3 | 1.43 | 10 | 0.19 | 0.576 | 196 | 0.916 | 0.424 | 132 | 0.943 |
| CREBBP | 1.17 | 10 | 0.13 | 0.553 | 186 | 0.916 | 0.447 | 142 | 0.943 |
| CST9 | 3.87 | 10 | 0.72 | 0.763 | 272 | 0.916 | 0.237 | 56 | 0.943 |
| CTLA4 | 2.58 | 10 | 0.44 | 0.669 | 231 | 0.916 | 0.331 | 97 | 0.943 |
| CTNNB1 | 1.41 | 10 | 0.19 | 0.573 | 193 | 0.916 | 0.427 | 135 | 0.943 |
| CXCL1 | 16.22 | 4 | 5.39 | 1 | 327 | 1 | ##### | 1 | ##### |
| CXCL3 | 0.45 | 2 | 0.1 | 0.539 | 179 | 0.916 | 0.461 | 149 | 0.943 |
| CXorf66 | 1.59 | 10 | 0.22 | 0.589 | 200 | 0.916 | 0.411 | 128 | 0.943 |
| DDOST | -1.85 | 10 | -0.51 | 0.304 | 88 | 0.916 | 0.696 | 240 | 0.943 |
| DDX20 | -3.65 | 10 | -0.9 | 0.184 | 46 | 0.916 | 0.816 | 282 | 0.943 |
| DGAT2L6 | 0.69 | 10 | 0.03 | 0.513 | 170 | 0.916 | 0.487 | 158 | 0.943 |
| DHFR | 2.94 | 6 | 0.7 | 0.757 | 269 | 0.916 | 0.243 | 59 | 0.943 |
| DHFRL1 | -1.15 | 2 | -0.67 | 0.252 | 73 | 0.916 | 0.748 | 255 | 0.943 |
| DHX15 | 2.05 | 10 | 0.32 | 0.627 | 211 | 0.916 | 0.373 | 117 | 0.943 |
| DHX8 | 1.42 | 10 | 0.19 | 0.574 | 194 | 0.916 | 0.426 | 134 | 0.943 |
| DOT1L | -0.68 | 10 | -0.26 | 0.396 | 122 | 0.916 | 0.604 | 206 | 0.943 |
| DPAGT1 | -3.47 | 10 | -0.86 | 0.194 | 50 | 0.916 | 0.806 | 278 | 0.943 |
| DRD2 | 0.73 | 10 | 0.04 | 0.516 | 171 | 0.916 | 0.484 | 157 | 0.943 |
| DSPP | 3.26 | 10 | 0.58 | 0.72 | 256 | 0.916 | 0.28 | 72 | 0.943 |
| ECD | 1.76 | 10 | 0.26 | 0.603 | 203 | 0.916 | 0.397 | 125 | 0.943 |
| EGFR | 12.7 | 10 | 2.61 | 0.995 | 323 | 1 | 0.005 | 5 | 0.295 |
| EHMT2 | -3.22 | 10 | -0.81 | 0.209 | 55 | 0.916 | 0.791 | 273 | 0.943 |
| EIF2S1 | 3.11 | 10 | 0.55 | 0.709 | 249 | 0.916 | 0.291 | 79 | 0.943 |
| EIF4E | 1.42 | 8 | 0.22 | 0.589 | 201 | 0.916 | 0.411 | 127 | 0.943 |
| EIF6 | -0.79 | 10 | -0.29 | 0.387 | 119 | 0.916 | 0.613 | 209 | 0.943 |
| ENTPD1 | 0.07 | 10 | -0.1 | 0.46 | 150 | 0.916 | 0.54 | 178 | 0.943 |
| EPHA2 | -8.48 | 10 | -1.94 | 0.026 | 8 | 0.916 | 0.974 | 320 | 0.993 |
| ERBB2 | -4.37 | 10 | -1.06 | 0.146 | 39 | 0.916 | 0.854 | 289 | 0.962 |
| ERCC2 | 3.43 | 10 | 0.62 | 0.732 | 259 | 0.916 | 0.268 | 69 | 0.943 |
| ESR1 | -3.37 | 10 | -0.84 | 0.2 | 52 | 0.916 | 0.8 | 276 | 0.943 |
| ESR2 | 4.39 | 10 | 0.83 | 0.796 | 279 | 0.931 | 0.204 | 49 | 0.943 |
| EZH2 | 11.54 | 10 | 2.36 | 0.991 | 322 | 1 | 0.009 | 6 | 0.464 |
| F13B | 5.74 | 10 | 1.12 | 0.868 | 294 | 0.955 | 0.132 | 34 | 0.943 |
| FGF4 | -3.62 | 10 | -0.89 | 0.186 | 48 | 0.916 | 0.814 | 280 | 0.943 |
| FGFR1 | 3.9 | 10 | 0.72 | 0.765 | 273 | 0.916 | 0.235 | 55 | 0.943 |
| FGFR2 | 1.07 | 10 | 0.11 | 0.545 | 182 | 0.916 | 0.455 | 146 | 0.943 |
| FGFR3 | 5.61 | 10 | 1.09 | 0.862 | 293 | 0.955 | 0.138 | 35 | 0.943 |
| FGFR4 | 1.73 | 10 | 0.26 | 0.601 | 202 | 0.916 | 0.399 | 126 | 0.943 |
| FGR | -2.81 | 10 | -0.72 | 0.236 | 66 | 0.916 | 0.764 | 262 | 0.943 |
| FLT1 | -0.42 | 10 | -0.21 | 0.418 | 135 | 0.916 | 0.582 | 193 | 0.943 |
| FLT3 | 0.32 | 10 | -0.05 | 0.481 | 159 | 0.916 | 0.519 | 169 | 0.943 |
| FLT4 | 0.63 | 10 | 0.02 | 0.507 | 167 | 0.916 | 0.493 | 161 | 0.943 |
| FMR1NB | 5.41 | 10 | 1.05 | 0.852 | 291 | 0.955 | 0.148 | 37 | 0.943 |
| FRK | 0.89 | 10 | 0.07 | 0.53 | 174 | 0.916 | 0.47 | 154 | 0.943 |
| FYN | -6.27 | 10 | -1.46 | 0.072 | 17 | 0.916 | 0.928 | 311 | 0.976 |
| GABRA1 | -3.18 | 10 | -0.8 | 0.212 | 57 | 0.916 | 0.788 | 271 | 0.943 |
| GFRAL | -1.91 | 10 | -0.53 | 0.299 | 87 | 0.916 | 0.701 | 241 | 0.943 |
| GLRA2 | -1.59 | 10 | -0.46 | 0.323 | 99 | 0.916 | 0.677 | 229 | 0.943 |
| GLS | 1.17 | 10 | 0.13 | 0.553 | 187 | 0.916 | 0.447 | 141 | 0.943 |
| GPR50 | 0.18 | 10 | -0.08 | 0.469 | 152 | 0.916 | 0.531 | 176 | 0.943 |
| GPX6 | 3.72 | 10 | 0.68 | 0.752 | 264 | 0.916 | 0.248 | 64 | 0.943 |
| GSK3A | 8.27 | 10 | 1.66 | 0.951 | 314 | 0.991 | 0.049 | 14 | 0.943 |
| GSK3B | -3.3 | 10 | -0.83 | 0.205 | 53 | 0.916 | 0.795 | 275 | 0.943 |
| GSX2 | -4.83 | 10 | -1.15 | 0.124 | 32 | 0.916 | 0.876 | 296 | 0.967 |
| GTF2B | -2.55 | 10 | -0.67 | 0.253 | 75 | 0.916 | 0.747 | 253 | 0.943 |
| HAVCR2 | 6.09 | 10 | 1.19 | 0.883 | 300 | 0.957 | 0.117 | 28 | 0.943 |
| HDAC1 | -3.45 | 10 | -0.86 | 0.196 | 51 | 0.916 | 0.804 | 277 | 0.943 |
| HDAC2 | -0.24 | 10 | -0.17 | 0.433 | 141 | 0.916 | 0.567 | 187 | 0.943 |
| HDAC3 | 0.45 | 10 | -0.02 | 0.492 | 163 | 0.916 | 0.508 | 165 | 0.943 |
| HDAC6 | 1.24 | 10 | 0.15 | 0.559 | 188 | 0.916 | 0.441 | 140 | 0.943 |
| HDAC8 | 3.77 | 10 | 0.69 | 0.756 | 266 | 0.916 | 0.244 | 62 | 0.943 |
| HLA-G | -2.67 | 10 | -0.69 | 0.245 | 70 | 0.916 | 0.755 | 258 | 0.943 |
| HSP90AA1 | 3.01 | 10 | 0.53 | 0.702 | 243 | 0.916 | 0.298 | 85 | 0.943 |
| HTR5A | 0.24 | 10 | -0.06 | 0.474 | 156 | 0.916 | 0.526 | 172 | 0.943 |
| ICOS | -0.29 | 10 | -0.18 | 0.429 | 140 | 0.916 | 0.571 | 188 | 0.943 |
| ICOSLG | -1.44 | 10 | -0.43 | 0.335 | 103 | 0.916 | 0.665 | 225 | 0.943 |
| IDH1 | -2.64 | 10 | -0.68 | 0.247 | 72 | 0.916 | 0.753 | 256 | 0.943 |
| IDH2 | 2.99 | 10 | 0.53 | 0.7 | 241 | 0.916 | 0.3 | 87 | 0.943 |
| IGF1R | 3.02 | 10 | 0.53 | 0.703 | 244 | 0.916 | 0.297 | 84 | 0.943 |
| IGSF11 | -5.94 | 10 | -1.39 | 0.082 | 21 | 0.916 | 0.918 | 307 | 0.974 |
| IKBKE | -0.43 | 10 | -0.21 | 0.417 | 133 | 0.916 | 0.583 | 195 | 0.943 |
| IL1B | -2.67 | 10 | -0.69 | 0.245 | 71 | 0.916 | 0.755 | 257 | 0.943 |
| IL6 | 0.57 | 10 | 0 | 0.502 | 165 | 0.916 | 0.498 | 163 | 0.943 |
| IL6R | 0.23 | 10 | -0.07 | 0.473 | 154 | 0.916 | 0.527 | 174 | 0.943 |
| IMP3 | 4.55 | 9 | 0.91 | 0.819 | 284 | 0.943 | 0.181 | 44 | 0.943 |
| INSR | -3.18 | 10 | -0.8 | 0.212 | 58 | 0.916 | 0.788 | 270 | 0.943 |
| IRAK4 | 0.86 | 10 | 0.07 | 0.527 | 173 | 0.916 | 0.473 | 155 | 0.943 |
| ITK | 3.75 | 10 | 0.69 | 0.754 | 265 | 0.916 | 0.246 | 63 | 0.943 |
| JAK1 | 0.99 | 10 | 0.1 | 0.538 | 177 | 0.916 | 0.462 | 151 | 0.943 |
| JAK2 | -1.82 | 10 | -0.51 | 0.306 | 92 | 0.916 | 0.694 | 236 | 0.943 |
| JAK3 | -1.8 | 10 | -0.5 | 0.307 | 94 | 0.916 | 0.693 | 234 | 0.943 |
| KCNK10 | 3.25 | 10 | 0.58 | 0.72 | 254 | 0.916 | 0.28 | 74 | 0.943 |
| KDM1A | 9.39 | 10 | 1.9 | 0.971 | 318 | 0.999 | 0.029 | 10 | 0.891 |
| KDR | 0.3 | 10 | -0.05 | 0.479 | 157 | 0.916 | 0.521 | 171 | 0.943 |
| KIF11 | 0.99 | 10 | 0.1 | 0.538 | 178 | 0.916 | 0.462 | 150 | 0.943 |
| KIT | -2.56 | 10 | -0.67 | 0.253 | 74 | 0.916 | 0.747 | 254 | 0.943 |
| KLK12 | 1.45 | 10 | 0.2 | 0.577 | 197 | 0.916 | 0.423 | 131 | 0.943 |
| LAG3 | -1.17 | 10 | -0.37 | 0.356 | 109 | 0.916 | 0.644 | 219 | 0.943 |
| LAP3 | -0.34 | 10 | -0.19 | 0.425 | 138 | 0.916 | 0.575 | 190 | 0.943 |
| LAS1L | -3.07 | 10 | -0.78 | 0.219 | 62 | 0.916 | 0.781 | 266 | 0.943 |
| LCK | -0.2 | 10 | -0.16 | 0.436 | 142 | 0.916 | 0.564 | 186 | 0.943 |
| LDHA | -0.55 | 6 | -0.27 | 0.394 | 120 | 0.916 | 0.606 | 208 | 0.943 |
| LGALS3 | 0 | 10 | -0.12 | 0.454 | 148 | 0.916 | 0.546 | 180 | 0.943 |
| LGALS9 | 10.38 | 4 | 3.41 | 1 | 325 | 1 | 3E-04 | 3 | 0.036 |
| LGALS9B | 1.29 | 2 | 0.5 | 0.692 | 238 | 0.916 | 0.308 | 90 | 0.943 |
| LUC7L3 | 5.59 | 10 | 1.08 | 0.861 | 292 | 0.955 | 0.139 | 36 | 0.943 |
| LYN | 2.26 | 10 | 0.37 | 0.643 | 221 | 0.916 | 0.357 | 107 | 0.943 |
| MAP2 | 4.07 | 10 | 0.76 | 0.775 | 275 | 0.922 | 0.225 | 53 | 0.943 |
| MAP2K1 | 11.39 | 10 | 2.33 | 0.99 | 321 | 1 | 0.01 | 7 | 0.464 |
| MAP2K2 | -4.52 | 10 | -1.09 | 0.138 | 35 | 0.916 | 0.862 | 293 | 0.962 |
| MAP3K8 | -2.47 | 10 | -0.65 | 0.259 | 78 | 0.916 | 0.741 | 250 | 0.943 |
| MAP4 | -2.84 | 10 | -0.73 | 0.233 | 65 | 0.916 | 0.767 | 263 | 0.943 |
| MAPK1 | 6.1 | 10 | 1.19 | 0.884 | 301 | 0.957 | 0.116 | 27 | 0.943 |
| MAPK11 | 2.08 | 10 | 0.33 | 0.629 | 212 | 0.916 | 0.371 | 116 | 0.943 |
| MAPK12 | -2.29 | 10 | -0.61 | 0.271 | 81 | 0.916 | 0.729 | 247 | 0.943 |
| MAPK13 | -1.01 | 10 | -0.33 | 0.369 | 114 | 0.916 | 0.631 | 214 | 0.943 |
| MAPK14 | 5.04 | 10 | 0.97 | 0.833 | 286 | 0.951 | 0.167 | 42 | 0.943 |
| MAPK3 | -0.62 | 10 | -0.25 | 0.401 | 125 | 0.916 | 0.599 | 203 | 0.943 |
| MAPK7 | 9.3 | 10 | 1.88 | 0.97 | 317 | 0.999 | 0.03 | 11 | 0.891 |
| MAPK8 | -0.63 | 10 | -0.25 | 0.401 | 124 | 0.916 | 0.599 | 204 | 0.943 |
| MAPK9 | 0.31 | 10 | -0.05 | 0.48 | 158 | 0.916 | 0.52 | 170 | 0.943 |
| MAPT | 2.4 | 10 | 0.4 | 0.655 | 224 | 0.916 | 0.345 | 104 | 0.943 |
| MCL1 | -1.72 | 10 | -0.49 | 0.313 | 96 | 0.916 | 0.687 | 232 | 0.943 |
| MDM2 | 9.08 | 10 | 1.83 | 0.967 | 316 | 0.999 | 0.033 | 12 | 0.909 |
| MEP1A | -0.6 | 10 | -0.25 | 0.403 | 126 | 0.916 | 0.597 | 202 | 0.943 |
| MET | -1.84 | 10 | -0.51 | 0.304 | 89 | 0.916 | 0.696 | 239 | 0.943 |
| MKNK1 | -4.47 | 10 | -1.08 | 0.141 | 36 | 0.916 | 0.859 | 292 | 0.962 |
| MS4A1 | -2.76 | 10 | -0.71 | 0.239 | 67 | 0.916 | 0.761 | 261 | 0.943 |
| MTOR | 2.03 | 10 | 0.32 | 0.625 | 210 | 0.916 | 0.375 | 118 | 0.943 |
| MYC | 7.05 | 10 | 1.4 | 0.919 | 309 | 0.972 | 0.081 | 19 | 0.943 |
| NAMPT | 1.27 | 10 | 0.16 | 0.562 | 189 | 0.916 | 0.438 | 139 | 0.943 |
| NDST4 | -7.54 | 10 | -1.74 | 0.041 | 12 | 0.916 | 0.959 | 316 | 0.992 |
| NELFB | 0.55 | 10 | 0 | 0.501 | 164 | 0.916 | 0.499 | 164 | 0.943 |
| NFKB1 | -10.66 | 10 | -2.41 | 0.008 | 4 | 0.659 | 0.992 | 324 | 0.999 |
| NOL6 | 5.78 | 10 | 1.12 | 0.87 | 297 | 0.955 | 0.13 | 31 | 0.943 |
| NOS2 | 4.04 | 10 | 0.75 | 0.773 | 274 | 0.922 | 0.227 | 54 | 0.943 |
| NOTCH1 | 0 | 10 | -0.12 | 0.453 | 147 | 0.916 | 0.547 | 181 | 0.943 |
| NR2C2 | -0.86 | 10 | -0.3 | 0.381 | 116 | 0.916 | 0.619 | 212 | 0.943 |
| NT5E | -1.08 | 10 | -0.35 | 0.364 | 112 | 0.916 | 0.636 | 216 | 0.943 |
| NTRK1 | -1.63 | 10 | -0.47 | 0.32 | 98 | 0.916 | 0.68 | 230 | 0.943 |
| NUDT1 | 2.71 | 10 | 0.47 | 0.679 | 236 | 0.916 | 0.321 | 92 | 0.943 |
| NUP133 | 3.25 | 10 | 0.58 | 0.72 | 255 | 0.916 | 0.28 | 73 | 0.943 |
| OPN5 | -1.03 | 10 | -0.34 | 0.368 | 113 | 0.916 | 0.632 | 215 | 0.943 |
| ORAOV1 | -4.28 | 10 | -1.04 | 0.15 | 40 | 0.916 | 0.85 | 288 | 0.962 |
| P4HB | -0.81 | 10 | -0.29 | 0.385 | 117 | 0.916 | 0.615 | 211 | 0.943 |
| PAK1 | 1.32 | 10 | 0.17 | 0.566 | 190 | 0.916 | 0.434 | 138 | 0.943 |
| PARP1 | 6.55 | 10 | 1.29 | 0.901 | 306 | 0.963 | 0.099 | 22 | 0.943 |
| PARP2 | -2.53 | 10 | -0.66 | 0.255 | 76 | 0.916 | 0.745 | 252 | 0.943 |
| PARP3 | 0.58 | 10 | 0.01 | 0.503 | 166 | 0.916 | 0.497 | 162 | 0.943 |
| PDCD1 | 3.06 | 10 | 0.54 | 0.706 | 247 | 0.916 | 0.294 | 81 | 0.943 |
| PDCD1LG | -3.06 | 10 | -0.77 | 0.219 | 63 | 0.916 | 0.781 | 265 | 0.943 |
| PDGFRA | -1.5 | 10 | -0.44 | 0.33 | 102 | 0.916 | 0.67 | 226 | 0.943 |
| PDGFRB | -2.99 | 10 | -0.76 | 0.224 | 64 | 0.916 | 0.776 | 264 | 0.943 |
| PDILT | -1.66 | 10 | -0.47 | 0.318 | 97 | 0.916 | 0.682 | 231 | 0.943 |
| PDK1 | -0.13 | 10 | -0.15 | 0.442 | 144 | 0.916 | 0.558 | 184 | 0.943 |
| PGD | -18.98 | 10 | -4.19 | ##### | 1 | 0.004 | 1 | 327 | 1 |
| PHB2 | -6 | 10 | -1.41 | 0.08 | 19 | 0.916 | 0.92 | 309 | 0.974 |
| PIGF | 2.97 | 10 | 0.52 | 0.699 | 240 | 0.916 | 0.301 | 88 | 0.943 |
| PIK3CA | 1.11 | 10 | 0.12 | 0.549 | 183 | 0.916 | 0.451 | 145 | 0.943 |
| PIK3CB | -5.64 | 10 | -1.33 | 0.092 | 28 | 0.916 | 0.908 | 300 | 0.974 |
| PIK3CD | -1.83 | 10 | -0.51 | 0.305 | 91 | 0.916 | 0.695 | 237 | 0.943 |
| PIK3CG | -2.01 | 10 | -0.55 | 0.291 | 86 | 0.916 | 0.709 | 242 | 0.943 |
| PIM1 | -7.47 | 10 | -1.72 | 0.043 | 13 | 0.916 | 0.957 | 315 | 0.992 |
| PIM2 | -0.36 | 10 | -0.19 | 0.423 | 136 | 0.916 | 0.577 | 192 | 0.943 |
| PIM3 | -3.09 | 10 | -0.78 | 0.217 | 61 | 0.916 | 0.783 | 267 | 0.943 |
| PLA2G2F | -0.59 | 10 | -0.24 | 0.404 | 129 | 0.916 | 0.596 | 199 | 0.943 |
| PLK1 | 1.88 | 10 | 0.29 | 0.613 | 208 | 0.916 | 0.387 | 120 | 0.943 |
| PNPLA5 | -5.78 | 10 | -1.36 | 0.087 | 26 | 0.916 | 0.913 | 302 | 0.974 |
| POLR1C | -2.26 | 10 | -0.6 | 0.274 | 83 | 0.916 | 0.726 | 245 | 0.943 |
| POLR2B | 2.16 | 10 | 0.35 | 0.636 | 215 | 0.916 | 0.364 | 113 | 0.943 |

| | | | | | | | | | | | | | | | | | | | | | | | | | | | | | | | | | | | | | | | | | | | | | | | | | | | | | | |
|---|---|---|---|---|---|---|---|---|---|---|---|---|---|---|---|---|---|---|---|---|---|---|---|---|---|---|---|---|---|---|---|---|---|---|---|---|---|---|---|---|---|---|---|---|---|---|---|---|---|---|---|---|---|---|
| **POLR2E** | -1.29 | 10 | -0.4 | 0.346 | 107 | 0.916 | 0.654 | 221 | 0.943 | -6.07 | 10 | -1.32 | 0.0926 | 21 | 0.903 | 0.907 | 307 | 0.959 | 3.52 | 10 | 0.65 | 0.742 | 262 | 0.922 | 0.258 | 66 | 0.933 | 1.28 | 10 | 0.26 | 0.602 | 207 | 0.927 | 0.398 | 121 | 0.944 | 8.46 | 10 | 1.93 | 0.973 | 320 | 0.992 | 0.027 | 8 | 0.913 | -3.68 | 10 | -0.96 | 0.17 | 49 | 0.958 | 0.83 | 279 | 0.969 |
| **POLR3C** | 3.18 | 10 | 0.57 | 0.715 | 251 | 0.916 | 0.285 | 77 | 0.943 | -4.53 | 10 | -0.99 | 0.162 | 45 | 0.903 | 0.838 | 283 | 0.945 | 1.7 | 10 | 0.29 | 0.614 | 219 | 0.912 | 0.386 | 109 | 0.933 | -1.03 | 10 | -0.28 | 0.388 | 127 | 0.927 | 0.612 | 201 | 0.944 | 5.72 | 10 | 1.28 | 0.899 | 301 | 0.976 | 0.101 | 27 | 0.913 | 4.53 | 10 | 0.92 | 0.822 | 276 | 0.969 | 0.178 | 52 | 0.935 |
| **PORCN** | 2.38 | 10 | 0.39 | 0.653 | 223 | 0.916 | 0.347 | 105 | 0.943 | 3.67 | 10 | 0.8 | 0.788 | 272 | 0.948 | 0.212 | 56 | 0.945 | 0.19 | 10 | -0.01 | 0.497 | 170 | 0.912 | 0.503 | 158 | 0.933 | -2.38 | 10 | -0.6 | 0.274 | 83 | 0.927 | 0.726 | 245 | 0.944 | -1.28 | 10 | -0.38 | 0.351 | 97 | 0.923 | 0.649 | 231 | 0.913 | -3.66 | 10 | -0.95 | 0.171 | 50 | 0.958 | 0.829 | 278 | 0.969 |
| **PPM1D** | 6.53 | 10 | 1.28 | 0.901 | 305 | 0.963 | 0.099 | 23 | 0.943 | 8.61 | 10 | 1.88 | 0.97 | 313 | 1 | 0.03 | 15 | 0.619 | -1.74 | 10 | -0.39 | 0.348 | 107 | 0.912 | 0.652 | 221 | 0.933 | -2.53 | 10 | -0.64 | 0.262 | 75 | 0.927 | 0.738 | 253 | 0.944 | -2.92 | 10 | -0.77 | 0.22 | 52 | 0.923 | 0.78 | 276 | 0.924 | -0.3 | 10 | -0.18 | 0.428 | 133 | 0.958 | 0.572 | 195 | 0.948 |
| **PPP3R2** | -9.14 | 10 | -2.08 | 0.019 | 6 | 0.916 | 0.981 | 322 | 0.997 | -2.11 | 10 | -0.46 | 0.323 | 107 | 0.903 | 0.677 | 221 | 0.945 | -6.47 | 10 | -1.32 | 0.0927 | 22 | 0.912 | 0.907 | 306 | 0.965 | -6.32 | 10 | -1.53 | 0.0632 | 20 | 0.927 | 0.937 | 308 | 0.98 | -1.51 | 10 | -0.44 | 0.331 | 94 | 0.923 | 0.669 | 234 | 0.913 | -6.15 | 10 | -1.52 | 0.0641 | 22 | 0.94 | 0.936 | 306 | 0.993 |
| **PRKAA1** | 2.22 | 10 | 0.36 | 0.64 | 218 | 0.916 | 0.36 | 110 | 0.943 | 1.88 | 10 | 0.41 | 0.66 | 236 | 0.903 | 0.34 | 92 | 0.945 | 3.74 | 10 | 0.69 | 0.756 | 265 | 0.932 | 0.244 | 63 | 0.933 | 7.68 | 10 | 1.76 | 0.961 | 314 | 1 | 0.0389 | 14 | 0.894 | -3.88 | 10 | -1 | 0.159 | 37 | 0.923 | 0.841 | 291 | 0.945 | -0.29 | 10 | -0.18 | 0.429 | 134 | 0.958 | 0.571 | 194 | 0.948 |
| **PRKCA** | 2.68 | 10 | 0.46 | 0.677 | 235 | 0.916 | 0.323 | 93 | 0.943 | -3.85 | 10 | -0.84 | 0.2 | 62 | 0.903 | 0.8 | 266 | 0.945 | 3.54 | 10 | 0.66 | 0.744 | 263 | 0.922 | 0.256 | 65 | 0.933 | 5.35 | 10 | 1.22 | 0.888 | 291 | 0.986 | 0.112 | 37 | 0.944 | 2.7 | 10 | 0.56 | 0.713 | 251 | 0.923 | 0.287 | 77 | 0.913 | -7.72 | 10 | -1.88 | 0.03 | 10 | 0.85 | 0.97 | 318 | 0.997 |
| **PRKCB** | -0.18 | 10 | -0.16 | 0.438 | 143 | 0.916 | 0.562 | 185 | 0.943 | -0.62 | 10 | -0.13 | 0.446 | 157 | 0.903 | 0.554 | 171 | 0.945 | 0.63 | 10 | 0.08 | 0.532 | 184 | 0.912 | 0.468 | 144 | 0.933 | 1.5 | 10 | 0.31 | 0.622 | 218 | 0.927 | 0.378 | 110 | 0.944 | -4.13 | 10 | -1.06 | 0.145 | 34 | 0.923 | 0.855 | 294 | 0.951 | -2.31 | 10 | -0.64 | 0.26 | 81 | 0.958 | 0.74 | 247 | 0.948 |
| **PRKCD** | -3.24 | 10 | -0.81 | 0.208 | 54 | 0.916 | 0.792 | 274 | 0.943 | -8.52 | 10 | -1.86 | 0.0315 | 8 | 0.903 | 0.968 | 320 | 0.988 | 1.49 | 10 | 0.25 | 0.598 | 211 | 0.912 | 0.402 | 117 | 0.933 | 4.37 | 10 | 0.99 | 0.838 | 283 | 0.965 | 0.162 | 45 | 0.944 | 5.83 | 10 | 1.3 | 0.904 | 302 | 0.976 | 0.0964 | 26 | 0.913 | 2.18 | 10 | 0.39 | 0.65 | 208 | 0.958 | 0.35 | 120 | 0.935 |
| **PRKCE** | 0.36 | 10 | -0.04 | 0.484 | 160 | 0.916 | 0.516 | 168 | 0.943 | 0.61 | 10 | 0.13 | 0.553 | 195 | 0.903 | 0.447 | 133 | 0.945 | 0.97 | 10 | 0.15 | 0.559 | 197 | 0.912 | 0.441 | 131 | 0.933 | -2.68 | 10 | -0.67 | 0.25 | 70 | 0.927 | 0.75 | 258 | 0.944 | -0.48 | 10 | -0.19 | 0.423 | 128 | 0.923 | 0.577 | 200 | 0.913 | 5.37 | 10 | 1.12 | 0.868 | 290 | 0.973 | 0.132 | 38 | 0.935 |
| **PRKCG** | 3 | 10 | 0.53 | 0.701 | 242 | 0.916 | 0.299 | 86 | 0.943 | 2.08 | 10 | 0.45 | 0.675 | 240 | 0.903 | 0.325 | 88 | 0.945 | 2.02 | 10 | 0.35 | 0.638 | 228 | 0.914 | 0.362 | 100 | 0.933 | -1.76 | 10 | -0.46 | 0.324 | 105 | 0.927 | 0.676 | 223 | 0.944 | -1.02 | 10 | -0.32 | 0.374 | 107 | 0.923 | 0.626 | 221 | 0.913 | 2.9 | 10 | 0.55 | 0.709 | 237 | 0.958 | 0.291 | 91 | 0.935 |
| **PRKCH** | 4.17 | 10 | 0.78 | 0.782 | 276 | 0.926 | 0.218 | 52 | 0.943 | 1.34 | 10 | 0.29 | 0.616 | 221 | 0.903 | 0.384 | 107 | 0.945 | -4.31 | 10 | -0.9 | 0.185 | 46 | 0.912 | 0.815 | 282 | 0.945 | 0.12 | 10 | -0.01 | 0.494 | 166 | 0.927 | 0.506 | 162 | 0.944 | -1.15 | 10 | -0.35 | 0.363 | 101 | 0.923 | 0.637 | 227 | 0.913 | -0.75 | 10 | -0.28 | 0.388 | 121 | 0.958 | 0.612 | 207 | 0.948 |
| **PRKCI** | 0.94 | 10 | 0.08 | 0.534 | 175 | 0.916 | 0.466 | 153 | 0.943 | 2.78 | 10 | 0.61 | 0.728 | 262 | 0.908 | 0.272 | 66 | 0.945 | -0.97 | 10 | -0.24 | 0.407 | 132 | 0.912 | 0.593 | 196 | 0.933 | -2.43 | 10 | -0.61 | 0.27 | 81 | 0.927 | 0.73 | 247 | 0.944 | 0.93 | 10 | 0.14 | 0.556 | 184 | 0.923 | 0.444 | 144 | 0.913 | -6.27 | 10 | -1.55 | 0.0608 | 20 | 0.94 | 0.939 | 308 | 0.993 |
| **PRKCQ** | -0.43 | 10 | -0.21 | 0.417 | 134 | 0.916 | 0.583 | 194 | 0.943 | 2.36 | 10 | 0.52 | 0.697 | 250 | 0.903 | 0.303 | 78 | 0.945 | -2.9 | 10 | -0.62 | 0.268 | 73 | 0.912 | 0.732 | 255 | 0.933 | 0.68 | 10 | 0.12 | 0.547 | 182 | 0.927 | 0.453 | 146 | 0.944 | -5.31 | 10 | -1.34 | 0.0905 | 20 | 0.923 | 0.91 | 308 | 0.966 | -0.14 | 10 | -0.15 | 0.442 | 140 | 0.958 | 0.558 | 188 | 0.948 |
| **PRKCSH** | 7.35 | 10 | 1.46 | 0.928 | 310 | 0.979 | 0.072 | 18 | 0.943 | 5.8 | 10 | 1.27 | 0.897 | 295 | 0.992 | 0.103 | 33 | 0.945 | -2.34 | 10 | -0.51 | 0.306 | 88 | 0.912 | 0.694 | 240 | 0.933 | 3.97 | 10 | 0.89 | 0.813 | 278 | 0.955 | 0.187 | 50 | 0.944 | 1.79 | 10 | 0.35 | 0.635 | 217 | 0.923 | 0.365 | 111 | 0.913 | 4.47 | 10 | 0.91 | 0.819 | 275 | 0.969 | 0.181 | 53 | 0.935 |
| **PRKCZ** | 3.12 | 10 | 0.55 | 0.71 | 250 | 0.916 | 0.29 | 78 | 0.943 | 2.01 | 10 | 0.44 | 0.67 | 239 | 0.903 | 0.33 | 89 | 0.945 | -5.34 | 10 | -1.1 | 0.136 | 33 | 0.912 | 0.864 | 295 | 0.956 | -2.11 | 10 | -0.54 | 0.295 | 94 | 0.927 | 0.705 | 234 | 0.944 | -1.61 | 10 | -0.46 | 0.323 | 89 | 0.923 | 0.677 | 239 | 0.913 | 0.63 | 10 | 0.03 | 0.512 | 163 | 0.958 | 0.488 | 165 | 0.948 |
| **PRPF19** | 3.55 | 10 | 0.65 | 0.741 | 262 | 0.916 | 0.259 | 66 | 0.943 | 2.29 | 10 | 0.5 | 0.692 | 247 | 0.903 | 0.308 | 81 | 0.945 | -2.45 | 10 | -0.53 | 0.298 | 83 | 0.912 | 0.702 | 245 | 0.933 | -2.19 | 10 | -0.56 | 0.289 | 91 | 0.927 | 0.711 | 237 | 0.944 | 0.89 | 10 | 0.13 | 0.552 | 182 | 0.923 | 0.448 | 146 | 0.913 | -1.91 | 10 | -0.55 | 0.291 | 89 | 0.958 | 0.709 | 239 | 0.948 |
| **PRSS37** | 1.13 | 10 | 0.13 | 0.55 | 184 | 0.916 | 0.45 | 144 | 0.943 | -3.33 | 10 | -0.73 | 0.233 | 73 | 0.903 | 0.767 | 255 | 0.945 | -3.88 | 10 | -0.81 | 0.208 | 54 | 0.912 | 0.792 | 274 | 0.945 | -5.76 | 10 | -1.4 | 0.0813 | 23 | 0.927 | 0.919 | 305 | 0.98 | 8.28 | 10 | 1.88 | 0.97 | 319 | 0.992 | 0.0298 | 9 | 0.913 | 0.59 | 10 | 0.02 | 0.509 | 162 | 0.958 | 0.491 | 166 | 0.948 |
| **PSMA7** | 2 | 10 | 0.31 | 0.622 | 209 | 0.916 | 0.378 | 119 | 0.943 | -0.98 | 10 | -0.21 | 0.415 | 141 | 0.903 | 0.585 | 187 | 0.945 | 4.05 | 10 | 0.75 | 0.775 | 270 | 0.933 | 0.225 | 58 | 0.933 | 7.39 | 10 | 1.69 | 0.955 | 313 | 0.998 | 0.0451 | 15 | 0.894 | 5.95 | 10 | 1.33 | 0.909 | 303 | 0.976 | 0.0915 | 25 | 0.913 | -1.77 | 10 | -0.52 | 0.302 | 95 | 0.958 | 0.698 | 233 | 0.948 |
| **PSMB1** | -0.59 | 10 | -0.24 | 0.404 | 128 | 0.916 | 0.596 | 200 | 0.943 | 2.26 | 10 | 0.49 | 0.689 | 246 | 0.903 | 0.311 | 82 | 0.945 | -0.76 | 10 | -0.19 | 0.423 | 140 | 0.912 | 0.577 | 188 | 0.933 | -3.66 | 10 | -0.9 | 0.183 | 47 | 0.927 | 0.817 | 281 | 0.944 | 9.21 | 10 | 2.11 | 0.982 | 323 | 0.992 | 0.0176 | 5 | 0.913 | 4.04 | 10 | 0.81 | 0.792 | 268 | 0.966 | 0.208 | 60 | 0.935 |
| **PSMD1** | 2.52 | 10 | 0.42 | 0.664 | 230 | 0.916 | 0.336 | 98 | 0.943 | -0.18 | 10 | -0.04 | 0.484 | 170 | 0.903 | 0.516 | 158 | 0.945 | -0.95 | 10 | -0.23 | 0.408 | 133 | 0.912 | 0.592 | 195 | 0.933 | -0.25 | 10 | -0.1 | 0.46 | 154 | 0.927 | 0.54 | 174 | 0.944 | 1.47 | 10 | 0.27 | 0.607 | 210 | 0.923 | 0.393 | 118 | 0.913 | 0.43 | 10 | -0.01 | 0.494 | 155 | 0.958 | 0.506 | 173 | 0.948 |
| **PSMG3** | 6.41 | 10 | 1.26 | 0.896 | 304 | 0.963 | 0.104 | 24 | 0.943 | -1.64 | 10 | -0.36 | 0.36 | 126 | 0.903 | 0.64 | 202 | 0.945 | 2.23 | 10 | 0.4 | 0.654 | 231 | 0.914 | 0.346 | 97 | 0.933 | 1.32 | 10 | 0.27 | 0.606 | 209 | 0.927 | 0.394 | 119 | 0.944 | 6.52 | 10 | 1.47 | 0.929 | 309 | 0.978 | 0.0713 | 19 | 0.913 | 6.5 | 10 | 1.38 | 0.916 | 306 | 0.976 | 0.0843 | 22 | 0.935 |
| **PTCH1** | 1.46 | 10 | 0.2 | 0.578 | 198 | 0.916 | 0.422 | 130 | 0.943 | 1.75 | 10 | 0.38 | 0.649 | 229 | 0.903 | 0.351 | 99 | 0.945 | -2.38 | 10 | -0.52 | 0.303 | 86 | 0.912 | 0.697 | 242 | 0.933 | -1.31 | 10 | -0.35 | 0.363 | 119 | 0.927 | 0.637 | 209 | 0.944 | 0.35 | 10 | 0 | 0.502 | 163 | 0.923 | 0.498 | 165 | 0.913 | 2.48 | 10 | 0.45 | 0.675 | 222 | 0.958 | 0.325 | 106 | 0.935 |
| **PTGS2** | 10.99 | 10 | 2.24 | 0.988 | 320 | 1 | 0.012 | 8 | 0.507 | 5.42 | 9 | 1.25 | 0.894 | 294 | 0.992 | 0.106 | 34 | 0.945 | 1.33 | 10 | 0.22 | 0.586 | 206 | 0.912 | 0.414 | 122 | 0.933 | 5.62 | 9 | 1.35 | 0.911 | 298 | 0.986 | 0.0887 | 30 | 0.944 | 4.84 | 10 | 1.07 | 0.857 | 287 | 0.976 | 0.143 | 41 | 0.913 | 7 | 10 | 1.49 | 0.932 | 311 | 0.978 | 0.0681 | 17 | 0.935 |
| **PTK2** | 3.29 | 10 | 0.59 | 0.722 | 257 | 0.916 | 0.278 | 71 | 0.943 | 6.78 | 10 | 1.48 | 0.93 | 302 | 1 | 0.0695 | 26 | 0.874 | 5.93 | 10 | 1.13 | 0.87 | 293 | 0.971 | 0.13 | 35 | 0.933 | 2.45 | 10 | 0.53 | 0.703 | 245 | 0.935 | 0.297 | 83 | 0.944 | -4.81 | 10 | -1.22 | 0.111 | 25 | 0.923 | 0.889 | 303 | 0.958 | 9.89 | 10 | 2.15 | 0.984 | 321 | 0.996 | 0.0157 | 7 | 0.663 |
| **PTPN11** | 10.73 | 10 | 2.19 | 0.986 | 319 | 1 | 0.014 | 9 | 0.522 | 9.41 | 10 | 2.05 | 0.98 | 319 | 1 | 0.02 | 9 | 0.619 | -0.99 | 10 | -0.24 | 0.405 | 128 | 0.912 | 0.595 | 200 | 0.933 | -2.03 | 10 | -0.52 | 0.301 | 95 | 0.927 | 0.699 | 233 | 0.944 | 0.19 | 10 | -0.03 | 0.487 | 156 | 0.923 | 0.513 | 172 | 0.913 | 2.86 | 10 | 0.54 | 0.706 | 232 | 0.958 | 0.294 | 96 | 0.935 |
| **PTPN6** | 4.85 | 10 | 0.93 | 0.823 | 285 | 0.944 | 0.177 | 43 | 0.943 | 3.66 | 10 | 0.8 | 0.788 | 271 | 0.948 | 0.212 | 57 | 0.945 | -0.82 | 10 | -0.21 | 0.418 | 137 | 0.912 | 0.582 | 191 | 0.933 | 0.15 | 10 | -0.01 | 0.497 | 168 | 0.927 | 0.503 | 160 | 0.944 | 4.95 | 10 | 1.09 | 0.863 | 288 | 0.976 | 0.137 | 40 | 0.913 | 1.33 | 10 | 0.19 | 0.576 | 183 | 0.958 | 0.424 | 145 | 0.943 |
| **PVR** | 6.9 | 10 | 1.36 | 0.914 | 308 | 0.97 | 0.086 | 20 | 0.943 | 9.33 | 10 | 2.04 | 0.979 | 318 | 1 | 0.0208 | 10 | 0.619 | 2.83 | 10 | 0.51 | 0.696 | 249 | 0.914 | 0.304 | 79 | 0.933 | 4.44 | 10 | 1 | 0.841 | 285 | 0.965 | 0.159 | 43 | 0.944 | -0.29 | 10 | -0.15 | 0.442 | 135 | 0.923 | 0.558 | 193 | 0.913 | -1.16 | 10 | -0.38 | 0.352 | 112 | 0.958 | 0.648 | 216 | 0.948 |
| **RABGGTB** | 4.82 | 8 | 1.04 | 0.851 | 289 | 0.955 | 0.149 | 39 | 0.943 | -7.83 | 8 | -1.91 | 0.0281 | 7 | 0.903 | 0.972 | 321 | 0.988 | 9.9 | 10 | 1.91 | 0.972 | 315 | 1 | 0.0279 | 13 | 0.703 | 2.86 | 10 | 0.63 | 0.736 | 254 | 0.943 | 0.264 | 74 | 0.944 | 7.45 | 9 | 1.78 | 0.963 | 316 | 0.992 | 0.0373 | 12 | 0.913 | -2.3 | 10 | -0.64 | 0.261 | 82 | 0.958 | 0.739 | 246 | 0.948 |
| **RAC1** | -10.98 | 10 | -2.48 | 0.007 | 3 | 0.659 | 0.993 | 325 | 0.999 | 12.23 | 10 | 2.67 | 0.996 | 322 | 1 | 0.0038 | 6 | 0.207 | 12.84 | 10 | 2.49 | 0.994 | 322 | 1 | 0.0063 | 6 | 0.345 | 9.35 | 10 | 2.15 | 0.984 | 320 | 1 | 0.0156 | 8 | 0.632 | 6.86 | 10 | 1.55 | 0.939 | 314 | 0.978 | 0.0608 | 14 | 0.913 | 1.79 | 10 | 0.3 | 0.617 | 196 | 0.958 | 0.383 | 132 | 0.935 |
| **RAF1** | 8.48 | 10 | 1.7 | 0.956 | 315 | 0.992 | 0.044 | 13 | 0.943 | 1.91 | 10 | 0.42 | 0.662 | 237 | 0.903 | 0.338 | 91 | 0.945 | 1.27 | 10 | 0.21 | 0.581 | 205 | 0.912 | 0.419 | 123 | 0.933 | 4.95 | 10 | 1.12 | 0.869 | 288 | 0.986 | 0.131 | 40 | 0.944 | 8.88 | 10 | 2.03 | 0.979 | 321 | 0.992 | 0.0214 | 7 | 0.913 | 0.12 | 10 | -0.09 | 0.466 | 146 | 0.958 | 0.534 | 182 | 0.948 |
| **RARA** | 2.59 | 10 | 0.44 | 0.67 | 232 | 0.916 | 0.33 | 96 | 0.943 | 5.14 | 10 | 1.12 | 0.869 | 287 | 0.99 | 0.131 | 41 | 0.945 | 0.14 | 10 | -0.02 | 0.493 | 168 | 0.912 | 0.507 | 160 | 0.933 | -6.43 | 10 | -1.55 | 0.06 | 18 | 0.927 | 0.94 | 310 | 0.98 | -3.49 | 10 | -0.91 | 0.182 | 41 | 0.923 | 0.818 | 287 | 0.932 | 3.55 | 10 | 0.7 | 0.758 | 258 | 0.958 | 0.242 | 70 | 0.935 |
| **RARB** | 2.9 | 10 | 0.51 | 0.693 | 239 | 0.916 | 0.307 | 89 | 0.943 | 8.6 | 10 | 1.88 | 0.97 | 312 | 1 | 0.0303 | 16 | 0.619 | -2.43 | 10 | -0.53 | 0.3 | 85 | 0.912 | 0.7 | 243 | 0.933 | -1.14 | 10 | -0.31 | 0.378 | 122 | 0.927 | 0.622 | 206 | 0.944 | -2.31 | 10 | -0.63 | 0.266 | 68 | 0.923 | 0.734 | 260 | 0.92 | 1.56 | 10 | 0.24 | 0.597 | 187 | 0.958 | 0.403 | 141 | 0.935 |
| **RARG** | -2.04 | 10 | -0.56 | 0.289 | 85 | 0.916 | 0.711 | 243 | 0.943 | 2.53 | 10 | 0.55 | 0.71 | 257 | 0.903 | 0.29 | 71 | 0.945 | -6.52 | 10 | -1.33 | 0.0912 | 21 | 0.912 | 0.909 | 307 | 0.965 | -9.05 | 10 | -2.17 | 0.015 | 7 | 0.702 | 0.985 | 321 | 0.999 | -4.35 | 10 | -1.11 | 0.134 | 32 | 0.923 | 0.866 | 296 | 0.952 | 3.57 | 10 | 0.7 | 0.759 | 259 | 0.958 | 0.241 | 69 | 0.935 |
| **RAX** | 1.42 | 10 | 0.19 | 0.575 | 195 | 0.916 | 0.425 | 133 | 0.943 | 1.13 | 10 | 0.25 | 0.598 | 213 | 0.903 | 0.402 | 115 | 0.945 | -5.61 | 10 | -1.15 | 0.124 | 30 | 0.912 | 0.876 | 298 | 0.959 | 1.07 | 10 | 0.21 | 0.583 | 200 | 0.927 | 0.417 | 128 | 0.944 | -3.27 | 10 | -0.85 | 0.196 | 45 | 0.923 | 0.804 | 283 | 0.926 | -2.51 | 10 | -0.69 | 0.246 | 72 | 0.958 | 0.754 | 256 | 0.948 |
| **RCC1** | 2.62 | 10 | 0.45 | 0.672 | 233 | 0.916 | 0.328 | 95 | 0.943 | 1.71 | 10 | 0.37 | 0.646 | 228 | 0.903 | 0.354 | 100 | 0.945 | 0.13 | 10 | -0.02 | 0.492 | 167 | 0.912 | 0.508 | 161 | 0.933 | 0.87 | 10 | 0.16 | 0.565 | 194 | 0.927 | 0.435 | 134 | 0.944 | 2.55 | 10 | 0.53 | 0.7 | 244 | 0.923 | 0.3 | 84 | 0.913 | 3.84 | 10 | 0.77 | 0.778 | 264 | 0.964 | 0.222 | 64 | 0.935 |
| **RD3** | 0.23 | 10 | -0.07 | 0.473 | 155 | 0.916 | 0.527 | 173 | 0.943 | 1.76 | 10 | 0.38 | 0.649 | 231 | 0.903 | 0.351 | 97 | 0.945 | -3.94 | 10 | -0.82 | 0.205 | 53 | 0.912 | 0.795 | 275 | 0.945 | -4.59 | 10 | -1.12 | 0.131 | 35 | 0.927 | 0.869 | 293 | 0.97 | 2.37 | 10 | 0.48 | 0.685 | 235 | 0.923 | 0.315 | 93 | 0.913 | 3.27 | 10 | 0.64 | 0.738 | 249 | 0.958 | 0.262 | 79 | 0.935 |
| **RET** | -5.99 | 10 | -1.4 | 0.08 | 20 | 0.916 | 0.92 | 308 | 0.974 | -5.59 | 10 | -1.22 | 0.111 | 24 | 0.903 | 0.889 | 304 | 0.956 | -0.21 | 10 | -0.09 | 0.466 | 157 | 0.912 | 0.534 | 171 | 0.933 | -0.26 | 10 | -0.1 | 0.459 | 153 | 0.927 | 0.541 | 175 | 0.944 | -4.53 | 10 | -1.15 | 0.125 | 29 | 0.923 | 0.875 | 299 | 0.957 | -0.78 | 10 | -0.29 | 0.386 | 120 | 0.958 | 0.614 | 208 | 0.948 |
| **RNF17** | 5.85 | 10 | 1.14 | 0.873 | 299 | 0.955 | 0.127 | 29 | 0.943 | 0.27 | 10 | 0.06 | 0.523 | 184 | 0.903 | 0.477 | 144 | 0.945 | -1.41 | 10 | -0.32 | 0.373 | 114 | 0.912 | 0.627 | 214 | 0.933 | -1.45 | 10 | -0.38 | 0.351 | 115 | 0.927 | 0.649 | 213 | 0.944 | -1 | 10 | -0.32 | 0.376 | 109 | 0.923 | 0.624 | 219 | 0.913 | -7.29 | 10 | -1.78 | 0.0374 | 11 | 0.85 | 0.963 | 317 | 0.993 |
| **ROCK1** | -7.23 | 10 | -1.67 | 0.047 | 14 | 0.916 | 0.953 | 314 | 0.992 | -6.27 | 10 | -1.37 | 0.0856 | 17 | 0.903 | 0.914 | 311 | 0.959 | -5.82 | 10 | -1.2 | 0.116 | 27 | 0.912 | 0.884 | 301 | 0.961 | -6.64 | 10 | -1.6 | 0.0545 | 16 | 0.927 | 0.945 | 312 | 0.98 | -0.43 | 10 | -0.18 | 0.428 | 130 | 0.923 | 0.572 | 198 | 0.913 | 3.77 | 10 | 0.75 | 0.774 | 262 | 0.964 | 0.226 | 66 | 0.935 |
| **ROCK2** | 3.79 | 10 | 0.7 | 0.757 | 267 | 0.916 | 0.243 | 61 | 0.943 | -0.72 | 10 | -0.16 | 0.438 | 152 | 0.903 | 0.562 | 176 | 0.945 | -14.61 | 10 | -2.93 | 0.0017 | 3 | 0.182 | 0.998 | 325 | 1 | -11.99 | 10 | -2.86 | 0.0021 | 3 | 0.231 | 0.998 | 325 | 0.999 | 1.83 | 10 | 0.36 | 0.639 | 219 | 0.923 | 0.361 | 109 | 0.913 | 1.06 | 10 | 0.13 | 0.552 | 174 | 0.958 | 0.448 | 154 | 0.948 |
| **RPA1** | 3.24 | 10 | 0.58 | 0.719 | 253 | 0.916 | 0.281 | 75 | 0.943 | -2.5 | 10 | -0.54 | 0.293 | 94 | 0.903 | 0.707 | 234 | 0.945 | 2.76 | 10 | 0.5 | 0.692 | 245 | 0.914 | 0.308 | 83 | 0.933 | 4.09 | 10 | 0.92 | 0.821 | 281 | 0.955 | 0.179 | 47 | 0.944 | 3.91 | 10 | 0.85 | 0.802 | 274 | 0.954 | 0.198 | 54 | 0.913 | 4.09 | 10 | 0.82 | 0.795 | 269 | 0.966 | 0.205 | 59 | 0.935 |
| **RPL8** | -2.46 | 10 | -0.65 | 0.259 | 79 | 0.916 | 0.741 | 249 | 0.943 | -2.36 | 10 | -0.52 | 0.303 | 98 | 0.903 | 0.697 | 230 | 0.945 | 3.55 | 10 | 0.66 | 0.744 | 264 | 0.922 | 0.256 | 64 | 0.933 | 1.92 | 10 | 0.41 | 0.659 | 227 | 0.932 | 0.341 | 101 | 0.944 | 5.2 | 10 | 1.15 | 0.876 | 291 | 0.976 | 0.124 | 37 | 0.913 | -1.52 | 10 | -0.46 | 0.323 | 104 | 0.958 | 0.677 | 224 | 0.948 |
| **RPS5** | 7.98 | 10 | 1.6 | 0.945 | 311 | 0.991 | 0.055 | 17 | 0.943 | -2.33 | 10 | -0.51 | 0.306 | 99 | 0.903 | 0.694 | 229 | 0.945 | -1.91 | 10 | -0.42 | 0.336 | 103 | 0.912 | 0.664 | 225 | 0.933 | 2.73 | 10 | 0.6 | 0.726 | 249 | 0.943 | 0.274 | 79 | 0.944 | 0.43 | 10 | 0.02 | 0.509 | 167 | 0.923 | 0.491 | 161 | 0.913 | 3.16 | 10 | 0.61 | 0.729 | 248 | 0.958 | 0.271 | 80 | 0.935 |
| **RPS6KB1** | -8.37 | 10 | -1.92 | 0.028 | 9 | 0.916 | 0.972 | 319 | 0.993 | -4.56 | 10 | -0.99 | 0.16 | 44 | 0.903 | 0.84 | 284 | 0.945 | -3.02 | 10 | -0.64 | 0.26 | 70 | 0.912 | 0.74 | 258 | 0.933 | 5.5 | 10 | 1.25 | 0.894 | 293 | 0.986 | 0.106 | 35 | 0.944 | -0.51 | 10 | -0.2 | 0.42 | 127 | 0.923 | 0.58 | 201 | 0.913 | -1.04 | 10 | -0.35 | 0.363 | 117 | 0.958 | 0.637 | 211 | 0.948 |
| **RRM1** | 3.5 | 10 | 0.63 | 0.737 | 260 | 0.916 | 0.263 | 68 | 0.943 | 1.81 | 10 | 0.4 | 0.654 | 233 | 0.903 | 0.346 | 95 | 0.945 | 1.45 | 10 | 0.24 | 0.595 | 208 | 0.912 | 0.405 | 120 | 0.933 | 2.1 | 10 | 0.45 | 0.675 | 234 | 0.932 | 0.325 | 94 | 0.944 | 5.42 | 10 | 1.21 | 0.886 | 295 | 0.976 | 0.114 | 33 | 0.913 | -0.27 | 10 | -0.17 | 0.431 | 136 | 0.958 | 0.569 | 192 | 0.948 |
| **RS1** | 0.05 | 10 | -0.11 | 0.458 | 149 | 0.916 | 0.542 | 179 | 0.943 | 1.86 | 10 | 0.41 | 0.657 | 235 | 0.903 | 0.343 | 93 | 0.945 | -9.16 | 10 | -1.86 | 0.0317 | 11 | 0.912 | 0.968 | 317 | 0.998 | 0.22 | 10 | 0.01 | 0.504 | 169 | 0.927 | 0.496 | 159 | 0.944 | -2.53 | 10 | -0.68 | 0.249 | 64 | 0.923 | 0.751 | 264 | 0.924 | 4.91 | 10 | 1.01 | 0.844 | 285 | 0.969 | 0.156 | 43 | 0.935 |
| **RXFP3** | 1.05 | 10 | 0.11 | 0.543 | 181 | 0.916 | 0.457 | 147 | 0.943 | -0.33 | 10 | -0.07 | 0.471 | 166 | 0.903 | 0.529 | 162 | 0.945 | 2.5 | 10 | 0.45 | 0.674 | 238 | 0.914 | 0.326 | 90 | 0.933 | -1.07 | 10 | -0.29 | 0.384 | 124 | 0.927 | 0.616 | 204 | 0.944 | 2.76 | 10 | 0.57 | 0.717 | 254 | 0.923 | 0.283 | 74 | 0.913 | 3.06 | 10 | 0.59 | 0.722 | 245 | 0.958 | 0.278 | 83 | 0.935 |
| **RXRA** | 5.74 | 10 | 1.12 | 0.868 | 295 | 0.955 | 0.132 | 33 | 0.943 | 8.43 | 10 | 1.84 | 0.967 | 311 | 1 | 0.0329 | 17 | 0.633 | 1.48 | 10 | 0.25 | 0.597 | 210 | 0.912 | 0.403 | 118 | 0.933 | -3.23 | 10 | -0.8 | 0.211 | 60 | 0.927 | 0.789 | 268 | 0.944 | 0.35 | 10 | 0 | 0.501 | 161 | 0.923 | 0.499 | 167 | 0.913 | 1.46 | 10 | 0.22 | 0.588 | 185 | 0.958 | 0.412 | 143 | 0.943 |
| **RXRB** | -5.12 | 10 | -1.22 | 0.112 | 30 | 0.916 | 0.888 | 298 | 0.974 | 0.24 | 10 | 0.05 | 0.521 | 183 | 0.903 | 0.479 | 145 | 0.945 | 5.58 | 10 | 1.06 | 0.855 | 284 | 0.971 | 0.145 | 44 | 0.933 | 6 | 10 | 1.37 | 0.914 | 300 | 0.986 | 0.0856 | 28 | 0.944 | -3.84 | 10 | -0.99 | 0.161 | 38 | 0.923 | 0.839 | 290 | 0.945 | 4.84 | 10 | 0.99 | 0.84 | 282 | 0.969 | 0.16 | 46 | 0.935 |
| **SAMD4B** | 2.41 | 10 | 0.4 | 0.656 | 225 | 0.916 | 0.344 | 103 | 0.943 | -0.83 | 10 | -0.18 | 0.428 | 150 | 0.903 | 0.572 | 178 | 0.945 | 5.77 | 10 | 1.1 | 0.863 | 289 | 0.971 | 0.137 | 39 | 0.933 | 8.18 | 10 | 1.88 | 0.97 | 316 | 1 | 0.03 | 12 | 0.817 | 4.1 | 10 | 0.89 | 0.814 | 279 | 0.954 | 0.186 | 49 | 0.913 | 6.57 | 10 | 1.39 | 0.918 | 307 | 0.976 | 0.082 | 21 | 0.935 |
| **SERPINA7** | 2.22 | 10 | 0.36 | 0.641 | 219 | 0.916 | 0.359 | 109 | 0.943 | -1.93 | 10 | -0.42 | 0.337 | 113 | 0.903 | 0.663 | 215 | 0.945 | 4.16 | 10 | 0.78 | 0.781 | 274 | 0.933 | 0.219 | 54 | 0.933 | -2.51 | 10 | -0.63 | 0.264 | 77 | 0.927 | 0.736 | 251 | 0.944 | 5.35 | 10 | 1.19 | 0.883 | 293 | 0.976 | 0.117 | 35 | 0.913 | -1.28 | 10 | -0.41 | 0.342 | 109 | 0.958 | 0.658 | 219 | 0.948 |
| **SERPINB1** | -1.17 | 10 | -0.37 | 0.356 | 108 | 0.916 | 0.644 | 220 | 0.943 | -4.42 | 10 | -0.96 | 0.168 | 48 | 0.903 | 0.832 | 280 | 0.945 | 4.11 | 10 | 0.77 | 0.779 | 273 | 0.933 | 0.221 | 55 | 0.933 | 0.57 | 10 | 0.09 | 0.537 | 179 | 0.927 | 0.463 | 149 | 0.944 | -2.21 | 10 | -0.6 | 0.273 | 69 | 0.923 | 0.727 | 259 | 0.917 | -5.55 | 10 | -1.38 | 0.0833 | 28 | 0.94 | 0.917 | 300 | 0.993 |
| **SF3B3** | 8.19 | 10 | 1.64 | 0.95 | 313 | 0.991 | 0.05 | 15 | 0.943 | 1.03 | 10 | 0.23 | 0.589 | 208 | 0.903 | 0.411 | 120 | 0.945 | 4.38 | 10 | 0.82 | 0.794 | 275 | 0.943 | 0.206 | 53 | 0.933 | -3.53 | 10 | -0.87 | 0.192 | 50 | 0.927 | 0.808 | 278 | 0.944 | 0.59 | 10 | 0.06 | 0.524 | 171 | 0.923 | 0.476 | 157 | 0.913 | -6.93 | 10 | -1.7 | 0.0447 | 17 | 0.85 | 0.955 | 311 | 0.993 |
| **SGK3** | 3.08 | 10 | 0.54 | 0.707 | 248 | 0.916 | 0.293 | 80 | 0.943 | 0.83 | 10 | 0.18 | 0.572 | 202 | 0.903 | 0.428 | 126 | 0.945 | 2.75 | 10 | 0.5 | 0.691 | 244 | 0.914 | 0.309 | 84 | 0.933 | -0.21 | 10 | -0.09 | 0.463 | 156 | 0.927 | 0.537 | 172 | 0.944 | 6.01 | 10 | 1.34 | 0.911 | 305 | 0.976 | 0.0893 | 23 | 0.913 | 0.66 | 10 | 0.04 | 0.515 | 165 | 0.958 | 0.485 | 163 | 0.948 |
| **SLC13A1** | 1.01 | 10 | 0.1 | 0.54 | 180 | 0.916 | 0.46 | 148 | 0.943 | -0.94 | 10 | -0.2 | 0.419 | 144 | 0.903 | 0.581 | 184 | 0.945 | 6.07 | 10 | 1.15 | 0.876 | 294 | 0.971 | 0.124 | 34 | 0.933 | 2.83 | 10 | 0.62 | 0.734 | 253 | 0.943 | 0.266 | 75 | 0.944 | -1.01 | 10 | -0.32 | 0.375 | 108 | 0.923 | 0.625 | 220 | 0.913 | 2.87 | 10 | 0.55 | 0.707 | 235 | 0.958 | 0.293 | 93 | 0.935 |
| **SLC36A3** | 2.1 | 10 | 0.33 | 0.631 | 214 | 0.916 | 0.369 | 114 | 0.943 | 1.48 | 10 | 0.32 | 0.627 | 225 | 0.903 | 0.373 | 103 | 0.945 | -5.36 | 10 | -1.1 | 0.135 | 32 | 0.912 | 0.865 | 296 | 0.956 | -1.74 | 10 | -0.45 | 0.326 | 106 | 0.927 | 0.674 | 222 | 0.944 | -2.77 | 10 | -0.74 | 0.231 | 59 | 0.923 | 0.769 | 269 | 0.924 | 6.63 | 10 | 1.41 | 0.92 | 308 | 0.976 | 0.08 | 20 | 0.935 |
| **SMO** | -4.4 | 10 | -1.06 | 0.144 | 38 | 0.916 | 0.856 | 290 | 0.962 | 2.36 | 10 | 0.52 | 0.697 | 251 | 0.903 | 0.303 | 77 | 0.945 | -0.43 | 10 | -0.13 | 0.448 | 151 | 0.912 | 0.552 | 177 | 0.933 | 1.72 | 10 | 0.36 | 0.642 | 223 | 0.932 | 0.358 | 105 | 0.944 | -3.09 | 10 | -0.81 | 0.208 | 48 | 0.923 | 0.792 | 280 | 0.925 | -1.26 | 10 | -0.4 | 0.344 | 111 | 0.958 | 0.656 | 217 | 0.948 |
| **SPC24** | 0.97 | 10 | 0.09 | 0.537 | 176 | 0.916 | 0.463 | 152 | 0.943 | 3.26 | 10 | 0.71 | 0.762 | 265 | 0.94 | 0.238 | 63 | 0.945 | 1.72 | 10 | 0.29 | 0.616 | 220 | 0.912 | 0.384 | 108 | 0.933 | 1.52 | 10 | 0.31 | 0.623 | 219 | 0.927 | 0.377 | 109 | 0.944 | -3.79 | 10 | -0.98 | 0.164 | 40 | 0.923 | 0.836 | 288 | 0.945 | 3.82 | 10 | 0.76 | 0.777 | 263 | 0.964 | 0.223 | 65 | 0.935 |
| **SPEM1** | 5.07 | 10 | 0.97 | 0.834 | 287 | 0.951 | 0.166 | 41 | 0.943 | 8.04 | 10 | 1.75 | 0.96 | 309 | 1 | 0.0397 | 19 | 0.65 | -0.69 | 10 | -0.18 | 0.428 | 142 | 0.912 | 0.572 | 186 | 0.933 | -0.15 | 10 | -0.08 | 0.469 | 157 | 0.927 | 0.531 | 171 | 0.944 | -4.08 | 10 | -1.05 | 0.147 | 35 | 0.923 | 0.853 | 293 | 0.951 | -2.73 | 10 | -0.74 | 0.23 | 65 | 0.958 | 0.77 | 263 | 0.948 |
| **SRC** | 3.62 | 10 | 0.66 | 0.745 | 263 | 0.916 | 0.255 | 65 | 0.943 | 2.89 | 10 | 0.63 | 0.736 | 263 | 0.915 | 0.264 | 65 | 0.945 | 0.89 | 10 | 0.13 | 0.552 | 194 | 0.912 | 0.448 | 134 | 0.933 | 1.37 | 10 | 0.28 | 0.61 | 211 | 0.927 | 0.39 | 117 | 0.944 | -1.37 | 10 | -0.4 | 0.343 | 96 | 0.923 | 0.657 | 232 | 0.913 | -2.77 | 10 | -0.75 | 0.228 | 63 | 0.958 | 0.772 | 265 | 0.948 |
| **SRSF2** | -0.6 | 10 | -0.25 | 0.403 | 127 | 0.916 | 0.597 | 201 | 0.943 | 14.89 | 10 | 3.25 | 0.999 | 324 | 1 | 0.0006 | 4 | 0.0471 | 15.58 | 10 | 3.04 | 0.999 | 325 | 1 | 0.0012 | 3 | 0.131 | -1.84 | 10 | -0.48 | 0.317 | 101 | 0.927 | 0.683 | 227 | 0.944 | -0.28 | 10 | -0.15 | 0.442 | 136 | 0.923 | 0.558 | 192 | 0.913 | 0.57 | 10 | 0.02 | 0.507 | 160 | 0.958 | 0.493 | 168 | 0.948 |
| **STAT3** | 1.14 | 10 | 0.13 | 0.551 | 185 | 0.916 | 0.449 | 143 | 0.943 | -0.49 | 10 | -0.11 | 0.457 | 160 | 0.903 | 0.543 | 168 | 0.945 | -0.5 | 10 | -0.14 | 0.443 | 148 | 0.912 | 0.557 | 180 | 0.933 | -1.22 | 10 | -0.33 | 0.371 | 121 | 0.927 | 0.629 | 207 | 0.944 | -5.64 | 10 | -1.42 | 0.0784 | 18 | 0.923 | 0.922 | 310 | 0.972 | -2.09 | 10 | -0.59 | 0.277 | 87 | 0.958 | 0.723 | 241 | 0.948 |
| **SYK** | -6.21 | 10 | -1.45 | 0.074 | 18 | 0.916 | 0.927 | 310 | 0.976 | 0.29 | 10 | 0.06 | 0.526 | 185 | 0.903 | 0.474 | 143 | 0.945 | -4.18 | 10 | -0.87 | 0.192 | 49 | 0.912 | 0.808 | 279 | 0.945 | 1.68 | 10 | 0.35 | 0.638 | 222 | 0.932 | 0.362 | 106 | 0.944 | 2.1 | 10 | 0.42 | 0.662 | 225 | 0.923 | 0.338 | 103 | 0.913 | 0.35 | 10 | -0.03 | 0.487 | 153 | 0.958 | 0.513 | 175 | 0.948 |
| **T** | 3.3 | 10 | 0.59 | 0.723 | 258 | 0.916 | 0.277 | 70 | 0.943 | -0.68 | 10 | -0.15 | 0.442 | 153 | 0.903 | 0.558 | 175 | 0.945 | -3.45 | 10 | -0.73 | 0.233 | 61 | 0.912 | 0.767 | 267 | 0.933 | -0.14 | 10 | -0.07 | 0.47 | 158 | 0.927 | 0.53 | 170 | 0.944 | 0.39 | 10 | 0.01 | 0.505 | 165 | 0.923 | 0.495 | 163 | 0.913 | 0.88 | 10 | 0.09 | 0.536 | 170 | 0.958 | 0.464 | 158 | 0.948 |
| **TBK1** | -0.08 | 10 | -0.14 | 0.446 | 145 | 0.916 | 0.554 | 183 | 0.943 | -4.76 | 10 | -1.04 | 0.149 | 39 | 0.903 | 0.851 | 289 | 0.945 | -11.04 | 10 | -2.23 | 0.013 | 6 | 0.642 | 0.987 | 322 | 0.999 | -13.63 | 10 | -3.25 | 0.0006 | 1 | 0.191 | 0.999 | 327 | 0.999 | 2.18 | 10 | 0.44 | 0.669 | 229 | 0.923 | 0.331 | 99 | 0.913 | 1.89 | 10 | 0.32 | 0.625 | 198 | 0.958 | 0.375 | 130 | 0.935 |
| **TEC** | -10.15 | 10 | -2.3 | 0.011 | 5 | 0.706 | 0.989 | 323 | 0.999 | -2.6 | 10 | -0.57 | 0.285 | 88 | 0.903 | 0.715 | 240 | 0.945 | -7.08 | 10 | -1.45 | 0.0742 | 16 | 0.912 | 0.926 | 312 | 0.968 | -1.99 | 10 | -0.51 | 0.305 | 98 | 0.927 | 0.695 | 230 | 0.944 | -1.97 | 10 | -0.55 | 0.292 | 77 | 0.923 | 0.708 | 251 | 0.914 | -3.98 | 10 | -1.03 | 0.153 | 43 | 0.958 | 0.847 | 285 | 0.972 |
| **TEK** | 5.39 | 10 | 1.04 | 0.851 | 290 | 0.955 | 0.149 | 38 | 0.943 | -4.77 | 10 | -1.04 | 0.149 | 38 | 0.903 | 0.851 | 290 | 0.945 | 2.68 | 10 | 0.48 | 0.686 | 242 | 0.914 | 0.314 | 86 | 0.933 | -2.76 | 10 | -0.69 | 0.245 | 68 | 0.927 | 0.755 | 260 | 0.944 | 1.29 | 10 | 0.23 | 0.589 | 201 | 0.923 | 0.411 | 127 | 0.913 | 1.56 | 10 | 0.25 | 0.597 | 188 | 0.958 | 0.403 | 140 | 0.935 |
| **TKTL2** | 4.42 | 10 | 0.83 | 0.797 | 280 | 0.931 | 0.203 | 48 | 0.943 | -3.89 | 10 | -0.85 | 0.198 | 61 | 0.903 | 0.802 | 267 | 0.945 | -1.19 | 10 | -0.28 | 0.39 | 118 | 0.912 | 0.61 | 210 | 0.933 | 4.42 | 10 | 1 | 0.841 | 284 | 0.965 | 0.159 | 44 | 0.944 | 3.85 | 10 | 0.83 | 0.798 | 272 | 0.954 | 0.202 | 56 | 0.913 | 1.1 | 10 | 0.14 | 0.555 | 176 | 0.958 | 0.445 | 152 | 0.948 |
| **TNF** | -6.42 | 10 | -1.5 | 0.067 | 16 | 0.916 | 0.933 | 312 | 0.977 | -6.62 | 10 | -1.44 | 0.0744 | 13 | 0.903 | 0.926 | 315 | 0.961 | 0.96 | 10 | 0.14 | 0.557 | 196 | 0.912 | 0.443 | 132 | 0.933 | -3.05 | 10 | -0.76 | 0.224 | 64 | 0.927 | 0.776 | 264 | 0.944 | -1.61 | 10 | -0.46 | 0.322 | 88 | 0.923 | 0.678 | 240 | 0.913 | 2.46 | 10 | 0.45 | 0.674 | 217 | 0.958 | 0.326 | 111 | 0.935 |
| **TNFRSF14** | -2.51 | 10 | -0.66 | 0.256 | 77 | 0.916 | 0.744 | 251 | 0.943 | 0.21 | 10 | 0.05 | 0.518 | 182 | 0.903 | 0.482 | 146 | 0.945 | -3.16 | 10 | -0.67 | 0.251 | 67 | 0.912 | 0.749 | 261 | 0.933 | 0.83 | 10 | 0.15 | 0.561 | 190 | 0.927 | 0.439 | 138 | 0.944 | 1.7 | 10 | 0.32 | 0.627 | 216 | 0.923 | 0.373 | 112 | 0.913 | 4.65 | 10 | 0.95 | 0.83 | 280 | 0.969 | 0.17 | 48 | 0.935 |
| **TNFRSF4** | -2.69 | 10 | -0.69 | 0.244 | 68 | 0.916 | 0.756 | 260 | 0.943 | -2.65 | 10 | -0.58 | 0.282 | 87 | 0.903 | 0.718 | 241 | 0.945 | 0.47 | 10 | 0.05 | 0.519 | 177 | 0.912 | 0.481 | 151 | 0.933 | 3.69 | 10 | 0.82 | 0.795 | 275 | 0.943 | 0.205 | 53 | 0.944 | -3.47 | 10 | -0.9 | 0.183 | 42 | 0.923 | 0.817 | 286 | 0.932 | 0.36 | 10 | -0.03 | 0.487 | 154 | 0.958 | 0.513 | 174 | 0.948 |
| **TNFRSF8** | -11.8 | 10 | -2.65 | 0.004 | 2 | 0.655 | 0.996 | 326 | 0.999 | -4.94 | 10 | -1.08 | 0.14 | 32 | 0.903 | 0.86 | 296 | 0.945 | 3.03 | 10 | 0.55 | 0.71 | 254 | 0.914 | 0.29 | 74 | 0.933 | 1.32 | 10 | 0.27 | 0.605 | 208 | 0.927 | 0.395 | 120 | 0.944 | 0.59 | 10 | 0.06 | 0.525 | 172 | 0.923 | 0.475 | 156 | 0.913 | -2.44 | 10 | -0.67 | 0.251 | 75 | 0.958 | 0.749 | 253 | 0.948 |
| **TNFRSF9** | 0.2 | 10 | -0.07 | 0.47 | 153 | 0.916 | 0.53 | 175 | 0.943 | 1.17 | 10 | 0.26 | 0.601 | 214 | 0.903 | 0.399 | 114 | 0.945 | -1.23 | 10 | -0.29 | 0.386 | 117 | 0.912 | 0.614 | 211 | 0.933 | 1.17 | 10 | 0.23 | 0.592 | 204 | 0.927 | 0.408 | 124 | 0.944 | -5.1 | 10 | -1.29 | 0.0989 | 22 | 0.923 | 0.901 | 306 | 0.962 | -3.31 | 10 | -0.87 | 0.192 | 52 | 0.958 | 0.808 | 276 | 0.957 |
| **TNFSF11** | -0.44 | 10 | -0.21 | 0.416 | 132 | 0.916 | 0.584 | 196 | 0.943 | -1.25 | 10 | -0.27 | 0.393 | 133 | 0.903 | 0.607 | 195 | 0.945 | -1.95 | 10 | -0.43 | 0.333 | 101 | 0.912 | 0.667 | 227 | 0.933 | -2.88 | 10 | -0.72 | 0.236 | 67 | 0.927 | 0.764 | 261 | 0.944 | 1.92 | 10 | 0.38 | 0.646 | 221 | 0.923 | 0.354 | 107 | 0.913 | -0.48 | 10 | -0.22 | 0.412 | 127 | 0.958 | 0.588 | 201 | 0.948 |
| **TNFSF13B** | -1.14 | 10 | -0.36 | 0.358 | 110 | 0.916 | 0.642 | 218 | 0.943 | 4.79 | 10 | 1.05 | 0.852 | 283 | 0.983 | 0.148 | 45 | 0.945 | -0.97 | 10 | -0.24 | 0.406 | 130 | 0.912 | 0.594 | 198 | 0.933 | -3.4 | 10 | -0.84 | 0.2 | 54 | 0.927 | 0.8 | 274 | 0.944 | 0.07 | 10 | -0.06 | 0.475 | 151 | 0.923 | 0.525 | 177 | 0.913 | 3.91 | 10 | 0.78 | 0.783 | 265 | 0.966 | 0.217 | 63 | 0.935 |
| **TNFSF4** | 3.79 | 10 | 0.7 | 0.757 | 268 | 0.916 | 0.243 | 60 | 0.943 | 4.62 | 10 | 1.01 | 0.843 | 281 | 0.981 | 0.157 | 47 | 0.945 | -5.08 | 10 | -1.05 | 0.147 | 37 | 0.912 | 0.853 | 291 | 0.956 | 1.25 | 10 | 0.25 | 0.599 | 206 | 0.927 | 0.401 | 122 | 0.944 | 1.41 | 10 | 0.26 | 0.601 | 208 | 0.923 | 0.399 | 120 | 0.913 | 1.99 | 10 | 0.34 | 0.635 | 201 | 0.958 | 0.365 | 127 | 0.935 |
| **TNFSF9** | -3.66 | 10 | -0.9 | 0.183 | 45 | 0.916 | 0.817 | 283 | 0.943 | -4 | 10 | -0.87 | 0.191 | 56 | 0.903 | 0.809 | 272 | 0.945 | 3.18 | 10 | 0.58 | 0.72 | 255 | 0.918 | 0.28 | 73 | 0.933 | 4.51 | 10 | 1.02 | 0.846 | 286 | 0.967 | 0.154 | 42 | 0.944 | -2.14 | 10 | -0.59 | 0.279 | 70 | 0.923 | 0.721 | 258 | 0.914 | -0.41 | 10 | -0.21 | 0.418 | 130 | 0.958 | 0.582 | 198 | 0.948 |
| **TOMM34** | 2.2 | 10 | 0.36 | 0.639 | 217 | 0.916 | 0.361 | 111 | 0.943 | 0.06 | 10 | 0.01 | 0.505 | 179 | 0.903 | 0.495 | 149 | 0.945 | -2.94 | 10 | -0.63 | 0.266 | 71 | 0.912 | 0.734 | 257 | 0.933 | 2.21 | 10 | 0.48 | 0.683 | 237 | 0.932 | 0.317 | 91 | 0.944 | -0.95 | 10 | -0.31 | 0.38 | 112 | 0.923 | 0.62 | 216 | 0.913 | 0.99 | 10 | 0.11 | 0.545 | 173 | 0.958 | 0.455 | 155 | 0.948 |
| **TONSL** | 2.66 | 10 | 0.46 | 0.675 | 234 | 0.916 | 0.325 | 94 | 0.943 | -1.02 | 10 | -0.22 | 0.412 | 137 | 0.903 | 0.588 | 191 | 0.945 | 6.66 | 10 | 1.27 | 0.898 | 301 | 0.974 | 0.102 | 27 | 0.933 | -1.53 | 10 | -0.4 | 0.343 | 113 | 0.927 | 0.657 | 215 | 0.944 | -0.27 | 10 | -0.14 | 0.443 | 137 | 0.923 | 0.557 | 191 | 0.913 | -1.71 | 10 | -0.51 | 0.307 | 98 | 0.958 | 0.693 | 230 | 0.948 |
| **TOP1** | 5.77 | 10 | 1.12 | 0.869 | 296 | 0.955 | 0.131 | 32 | 0.943 | 9.12 | 10 | 1.99 | 0.977 | 316 | 1 | 0.0232 | 12 | 0.619 | -0.86 | 10 | -0.21 | 0.415 | 135 | 0.912 | 0.585 | 193 | 0.933 | 3.54 | 10 | 0.79 | 0.785 | 270 | 0.943 | 0.215 | 58 | 0.944 | 6.49 | 10 | 1.46 | 0.928 | 308 | 0.978 | 0.0723 | 20 | 0.913 | -6.22 | 10 | -1.54 | 0.0622 | 21 | 0.94 | 0.938 | 307 | 0.993 |
| **TOP2A** | 3.2 | 10 | 0.57 | 0.716 | 252 | 0.916 | 0.284 | 76 | 0.943 | -1.77 | 10 | -0.39 | 0.35 | 121 | 0.903 | 0.65 | 207 | 0.945 | 4 | 10 | 0.74 | 0.772 | 269 | 0.933 | 0.228 | 59 | 0.933 | 2.33 | 10 | 0.51 | 0.693 | 242 | 0.935 | 0.307 | 86 | 0.944 | 2.62 | 10 | 0.54 | 0.706 | 248 | 0.923 | 0.294 | 80 | 0.913 | -2.39 | 10 | -0.66 | 0.254 | 77 | 0.958 | 0.746 | 251 | 0.948 |
| **TOP2B** | 0.64 | 10 | 0.02 | 0.509 | 168 | 0.916 | 0.491 | 160 | 0.943 | 1.35 | 10 | 0.3 | 0.616 | 222 | 0.903 | 0.384 | 106 | 0.945 | 1.45 | 10 | 0.24 | 0.596 | 209 | 0.912 | 0.404 | 119 | 0.933 | -2.76 | 10 | -0.69 | 0.245 | 69 | 0.927 | 0.755 | 259 | 0.944 | -2.84 | 10 | -0.75 | 0.226 | 56 | 0.923 | 0.774 | 272 | 0.924 | 5.53 | 10 | 1.15 | 0.876 | 294 | 0.973 | 0.124 | 34 | 0.935 |
| **TPH2** | 2.29 | 10 | 0.37 | 0.646 | 222 | 0.916 | 0.354 | 106 | 0.943 | 1.4 | 10 | 0.31 | 0.62 | 224 | 0.903 | 0.38 | 104 | 0.945 | -0.51 | 10 | -0.15 | 0.442 | 147 | 0.912 | 0.558 | 181 | 0.933 | -6.67 | 10 | -1.61 | 0.0536 | 15 | 0.927 | 0.946 | 313 | 0.98 | 2.32 | 10 | 0.47 | 0.682 | 234 | 0.923 | 0.318 | 94 | 0.913 | -9.71 | 10 | -2.34 | 0.0098 | 5 | 0.579 | 0.99 | 323 | 1 |
| **TRIM24** | -5.9 | 10 | -1.38 | 0.083 | 22 | 0.916 | 0.917 | 306 | 0.974 | -4.86 | 10 | -1.06 | 0.145 | 35 | 0.903 | 0.855 | 293 | 0.945 | 1.59 | 10 | 0.27 | 0.606 | 215 | 0.912 | 0.394 | 113 | 0.933 | -1.71 | 10 | -0.44 | 0.328 | 109 | 0.927 | 0.672 | 219 | 0.944 | 0.69 | 10 | 0.08 | 0.533 | 175 | 0.923 | 0.467 | 153 | 0.913 | 4.01 | 10 | 0.81 | 0.79 | 266 | 0.966 | 0.21 | 62 | 0.935 |
| **TRPC7** | 1.4 | 10 | 0.18 | 0.573 | 192 | 0.916 | 0.427 | 136 | 0.943 | -3.69 | 10 | -0.81 | 0.21 | 68 | 0.903 | 0.79 | 260 | 0.945 | -1.78 | 10 | -0.4 | 0.345 | 106 | 0.912 | 0.655 | 222 | 0.933 | -4.1 | 10 | -1.01 | 0.157 | 41 | 0.927 | 0.843 | 287 | 0.954 | 3.27 | 10 | 0.7 | 0.757 | 263 | 0.937 | 0.243 | 65 | 0.913 | -0.06 | 10 | -0.13 | 0.449 | 142 | 0.958 | 0.551 | 186 | 0.948 |
| **TRPV1** | 2.23 | 10 | 0.36 | 0.641 | 220 | 0.916 | 0.359 | 108 | 0.943 | 2.36 | 10 | 0.52 | 0.697 | 249 | 0.903 | 0.303 | 79 | 0.945 | -2.06 | 10 | -0.45 | 0.325 | 99 | 0.912 | 0.675 | 229 | 0.933 | 0.29 | 10 | 0.03 | 0.51 | 173 | 0.927 | 0.49 | 155 | 0.944 | -0.73 | 10 | -0.25 | 0.4 | 116 | 0.923 | 0.6 | 212 | 0.913 | 0.84 | 10 | 0.08 | 0.531 | 168 | 0.958 | 0.469 | 160 | 0.948 |
| **TSSK1B** | -2.67 | 10 | -0.69 | 0.245 | 69 | 0.916 | 0.755 | 259 | 0.943 | -0.01 | 10 | 0 | 0.499 | 176 | 0.903 | 0.501 | 152 | 0.945 | -5.95 | 10 | -1.22 | 0.111 | 26 | 0.912 | 0.889 | 302 | 0.963 | -2.27 | 10 | -0.58 | 0.282 | 86 | 0.927 | 0.718 | 242 | 0.944 | 2.65 | 10 | 0.55 | 0.709 | 249 | 0.923 | 0.291 | 79 | 0.913 | 2.53 | 10 | 0.47 | 0.68 | 224 | 0.958 | 0.32 | 104 | 0.935 |
| **TUBB** | 3.83 | 10 | 0.71 | 0.76 | 270 | 0.916 | 0.24 | 58 | 0.943 | 3.58 | 10 | 0.78 | 0.783 | 268 | 0.948 | 0.217 | 60 | 0.945 | 6.69 | 10 | 1.28 | 0.899 | 302 | 0.974 | 0.101 | 26 | 0.933 | 5.01 | 10 | 1.14 | 0.872 | 289 | 0.986 | 0.128 | 39 | 0.944 | 1.26 | 10 | 0.22 | 0.586 | 198 | 0.923 | 0.414 | 130 | 0.913 | 6.19 | 10 | 1.31 | 0.904 | 303 | 0.973 | 0.0959 | 25 | 0.935 |
| **TUBGCP2** | -1.43 | 10 | -0.43 | 0.335 | 104 | 0.916 | 0.665 | 224 | 0.943 | 0.3 | 10 | 0.07 | 0.526 | 186 | 0.903 | 0.474 | 142 | 0.945 | -4.54 | 10 | -0.94 | 0.173 | 42 | 0.912 | 0.827 | 286 | 0.946 | 3.32 | 10 | 0.74 | 0.77 | 261 | 0.943 | 0.23 | 67 | 0.944 | 0.98 | 10 | 0.15 | 0.561 | 188 | 0.923 | 0.439 | 140 | 0.913 | -2.36 | 10 | -0.65 | 0.257 | 79 | 0.958 | 0.743 | 249 | 0.948 |
| **TXN** | 0.6 | 2 | 0.17 | 0.568 | 191 | 0.916 | 0.432 | 137 | 0.943 | 0.62 | 2 | 0.3 | 0.62 | 223 | 0.903 | 0.38 | 105 | 0.945 | 1.06 | 2 | 0.42 | 0.663 | 235 | 0.914 | 0.337 | 93 | 0.933 | 2.69 | 2 | 1.37 | 0.915 | 301 | 0.986 | 0.0851 | 27 | 0.944 | -0.03 | 2 | -0.09 | 0.463 | 144 | 0.923 | 0.537 | 184 | 0.913 | -5.39 | 2 | -2.87 | 0.002 | 3 | 0.22 | 0.998 | 325 | 1 |
| **TYMS** | 6.75 | 10 | 1.33 | 0.909 | 307 | 0.968 | 0.091 | 21 | 0.943 | 5.9 | 10 | 1.29 | 0.901 | 297 | 0.992 | 0.0989 | 31 | 0.945 | 14.03 | 10 | 2.73 | 0.997 | 323 | 1 | 0.0032 | 5 | 0.209 | 12.26 | 10 | 2.84 | 0.998 | 324 | 1 | 0.0023 | 4 | 0.185 | -2.66 | 10 | -0.71 | 0.239 | 62 | 0.923 | 0.761 | 266 | 0.924 | -5.5 | 10 | -1.37 | 0.085 | 29 | 0.94 | 0.915 | 299 | 0.993 |
| **TYRO3** | -1.53 | 10 | -0.44 | 0.328 | 101 | 0.916 | 0.672 | 227 | 0.943 | -0.45 | 10 | -0.1 | 0.461 | 162 | 0.903 | 0.539 | 166 | 0.945 | -6.97 | 10 | -1.42 | 0.0775 | 17 | 0.912 | 0.923 | 311 | 0.968 | -5.48 | 10 | -1.33 | 0.0916 | 25 | 0.927 | 0.908 | 303 | 0.979 | 2.2 | 10 | 0.44 | 0.671 | 230 | 0.923 | 0.329 | 98 | 0.913 | -0.41 | 10 | -0.21 | 0.418 | 131 | 0.958 | 0.582 | 197 | 0.948 |
| **UBQLN3** | 4.93 | 9 | 1 | 0.841 | 288 | 0.955 | 0.159 | 40 | 0.943 | -4.91 | 10 | -1.07 | 0.142 | 34 | 0.903 | 0.858 | 294 | 0.945 | -7.97 | 9 | -1.71 | 0.0441 | 13 | 0.912 | 0.956 | 315 | 0.992 | -6.34 | 9 | -1.61 | 0.0534 | 14 | 0.927 | 0.947 | 314 | 0.98 | -5.15 | 10 | -1.3 | 0.0968 | 21 | 0.923 | 0.903 | 307 | 0.962 | -4.04 | 9 | -1.09 | 0.138 | 40 | 0.958 | 0.862 | 288 | 0.976 |
| **UTS2R** | -0.34 | 10 | -0.19 | 0.424 | 137 | 0.916 | 0.576 | 191 | 0.943 | -0.99 | 10 | -0.22 | 0.414 | 139 | 0.903 | 0.586 | 189 | 0.945 | -4.93 | 10 | -1.02 | 0.154 | 40 | 0.912 | 0.846 | 288 | 0.956 | -1.07 | 10 | -0.29 | 0.384 | 123 | 0.927 | 0.616 | 205 | 0.944 | -0.57 | 10 | -0.21 | 0.415 | 121 | 0.923 | 0.585 | 207 | 0.913 | 1.07 | 10 | 0.13 | 0.552 | 175 | 0.958 | 0.448 | 153 | 0.948 |
| **VCP** | 8.03 | 10 | 1.61 | 0.946 | 312 | 0.991 | 0.054 | 16 | 0.943 | 0.33 | 10 | 0.07 | 0.529 | 187 | 0.903 | 0.471 | 141 | 0.945 | -2.33 | 10 | -0.51 | 0.306 | 89 | 0.912 | 0.694 | 239 | 0.933 | -7.07 | 10 | -1.7 | 0.0442 | 9 | 0.927 | 0.956 | 319 | 0.98 | 2.6 | 10 | 0.54 | 0.705 | 247 | 0.923 | 0.295 | 81 | 0.913 | -7.2 | 10 | -1.76 | 0.0391 | 14 | 0.85 | 0.961 | 314 | 0.993 |
| **VEGFA** | -5.82 | 10 | -1.37 | 0.086 | 23 | 0.916 | 0.914 | 305 | 0.974 | -2.84 | 10 | -0.62 | 0.268 | 85 | 0.903 | 0.732 | 243 | 0.945 | 5.06 | 10 | 0.95 | 0.83 | 280 | 0.967 | 0.17 | 48 | 0.933 | 3.7 | 10 | 0.83 | 0.796 | 276 | 0.943 | 0.204 | 52 | 0.944 | -0.16 | 10 | -0.12 | 0.454 | 140 | 0.923 | 0.546 | 188 | 0.913 | -2.05 | 10 | -0.58 | 0.28 | 88 | 0.958 | 0.72 | 240 | 0.948 |
| **VTCN1** | -5.8 | 10 | -1.36 | 0.086 | 25 | 0.916 | 0.914 | 303 | 0.974 | -0.24 | 10 | -0.05 | 0.48 | 168 | 0.903 | 0.52 | 160 | 0.945 | 0.57 | 10 | 0.07 | 0.527 | 182 | 0.912 | 0.473 | 146 | 0.933 | -1.49 | 10 | -0.39 | 0.347 | 114 | 0.927 | 0.653 | 214 | 0.944 | -0.3 | 10 | -0.15 | 0.44 | 134 | 0.923 | 0.56 | 194 | 0.913 | 0.9 | 10 | 0.09 | 0.537 | 171 | 0.958 | 0.463 | 157 | 0.948 |
| **WDR3** | -1.43 | 10 | -0.43 | 0.335 | 105 | 0.916 | 0.665 | 223 | 0.943 | -1.74 | 10 | -0.38 | 0.352 | 122 | 0.903 | 0.648 | 206 | 0.945 | 6.61 | 10 | 1.26 | 0.896 | 299 | 0.974 | 0.104 | 29 | 0.933 | 3.51 | 10 | 0.78 | 0.783 | 267 | 0.943 | 0.217 | 61 | 0.944 | 2.94 | 10 | 0.62 | 0.732 | 257 | 0.929 | 0.268 | 71 | 0.913 | 2.33 | 10 | 0.42 | 0.663 | 211 | 0.958 | 0.337 | 117 | 0.935 |
| **WDR43** | -0.67 | 10 | -0.26 | 0.397 | 123 | 0.916 | 0.603 | 205 | 0.943 | -2.86 | 10 | -0.62 | 0.267 | 83 | 0.903 | 0.733 | 245 | 0.945 | 11.25 | 10 | 2.18 | 0.985 | 318 | 1 | 0.0147 | 10 | 0.48 | 1.04 | 10 | 0.2 | 0.58 | 198 | 0.927 | 0.42 | 130 | 0.944 | 5.65 | 10 | 1.26 | 0.896 | 299 | 0.976 | 0.104 | 29 | 0.913 | -0.39 | 10 | -0.2 | 0.42 | 132 | 0.958 | 0.58 | 196 | 0.948 |
| **WEE1** | 1.84 | 10 | 0.28 | 0.61 | 206 | 0.916 | 0.39 | 122 | 0.943 | 1.34 | 10 | 0.29 | 0.616 | 219 | 0.903 | 0.384 | 109 | 0.945 | -6.24 | 10 | -1.28 | 0.101 | 25 | 0.912 | 0.899 | 303 | 0.965 | 3.64 | 10 | 0.81 | 0.792 | 274 | 0.943 | 0.208 | 54 | 0.944 | 0.02 | 10 | -0.07 | 0.471 | 149 | 0.923 | 0.529 | 179 | 0.913 | 9.83 | 10 | 2.14 | 0.984 | 320 | 0.996 | 0.0162 | 8 | 0.663 |
| **XPO1** | 18.9 | 10 | 3.94 | 1 | 326 | 1 | ##### | 2 | 0.007 | 6.22 | 10 | 1.36 | 0.913 | 299 | 0.997 | 0.0872 | 29 | 0.945 | 2.77 | 10 | 0.5 | 0.692 | 246 | 0.914 | 0.308 | 82 | 0.933 | 6.08 | 10 | 1.39 | 0.917 | 303 | 0.986 | 0.0828 | 25 | 0.944 | 3.83 | 10 | 0.83 | 0.797 | 270 | 0.954 | 0.203 | 58 | 0.913 | 6.44 | 10 | 1.36 | 0.913 | 305 | 0.976 | 0.0865 | 23 | 0.935 |

High confidence hits (P-value ≤ 0.05 in either early or late time points for one cell line or P-value ≤ 0.01 at either early or late timepoint for 2 cell lines)

| A549 | | H1299 | | H2023 | |
|---|---|---|---|---|---|
| Early (Day 12) | Late (Day 18) | Early (Day 12) | Late (Day 18) | Early (Day 15) | Late (Day 25) |
| AR | BTLA | AR | BRAF | BCL2 | FYN |
| CA9 | CD274 | BCL2 | CDK4 | BTLA | PGD |
| CD274 | LCK | BRAF | GLS | CA9 | PPP3R2 |
| CDK4 | PGD | CA9 | LCK | CD274 | VCP |
| EPHA2 | ROCK1 | CDK4 | MS4A1 | CXCL1 | |
| FYN | | CXCL1 | PPP3R2 | EPHA2 | |
| NDST4 | | EPHA2 | ROCK1 | FGF4 | |
| PGD | | FGF4 | ROCK2 | FYN | |
| PPP3R2 | | FYN | TBK1 | MS4A1 | |
| ROCK1 | | GLS | TPH2 | NDST4 | |
| TEC | | MS4A1 | VCP | PGD | |
| | | NDST4 | | TPH2 | |
| | | PGD | | UBQLN3 | |
| | | PPP3R2 | | | |
| | | ROCK2 | | | |
| | | TBK1 | | | |
| | | TEC | | | |
| | | UBQLN3 | | | |

| Metabolite | Method | Metabolite abundance | | | | | | | | | | | | | | | | | | | | | | | | | |
|---|---|---|---|---|---|---|---|---|---|---|---|---|---|---|---|---|---|---|---|---|---|---|---|---|---|---|---|
| | | DMSO 0h 1 | DMSO 0h 2 | DMSO 0h 3 | DMSO 24h 1 | DMSO 24h 2 | DMSO 24h 3 | DMSO 48h 1 | DMSO 48h 2 | DMSO 48h 3 | KD025 24h 1 | KD025 24h 2 | KD025 24h 3 | KD025 48h 1 | KD025 48h 2 | KD025 48h 3 | OXPHOS 24h | OXPHOS 24h | OXPHOS 24h | OXPHOS 48h | OXPHOS 48h | OXPHOS 48h | Combo | Combo | Combo | Combo | Combo | Combo 48h 3 |
| (2R)-2-Hydroxy-3-(phosphonatooxy)propanoate | HILIC | 4.388901e-02 | 4.795234e-02 | 5.217186e-02 | 4.512890e-02 | 4.437392e-02 | 3.800034e-02 | 5.485923e-02 | 5.747767e-02 | 3.781530e-02 | 3.812795e-02 | 4.780915e-02 | 4.314875e-02 | 4.482995e-02 | 4.483548e-02 | 4.654346e-02 | 3.795125e-02 | 4.853251e-02 | 3.018891e-02 | 2.129954e-02 | 2.848852e-02 | 2.963688e-02 | 3.70302 | 2.7608 | 2.6234 | 2.4601 | 2.2914 | 2.617495e-02 |
| 1-Pyrroline-2-carboxylic acid | HILIC | 9.072932e-05 | 8.875973e-05 | 9.422309e-05 | 7.138778e-05 | 7.116045e-05 | 6.721553e-05 | 6.397417e-05 | 6.532372e-05 | 5.352241e-05 | 7.787024e-05 | 8.523694e-05 | 7.102449e-05 | 6.175914e-05 | 6.059821e-05 | 6.329795e-05 | 5.379754e-05 | 4.576922e-05 | 4.843442e-05 | 3.071068e-05 | 3.201672e-05 | 3.795393e-05 | 5.51186 | 4.87814 | 5.41804 | 3.62271 | 3.2732 | 3.255019e-05 |
| 2'-Deoxyguanosine 5'-monophosphate | HILIC | 5.189971e-03 | 3.512378e-03 | 3.760765e-03 | 3.778769e-03 | 3.816289e-03 | 4.729207e-03 | 5.779016e-03 | 5.125279e-03 | 3.547276e-03 | 4.275076e-03 | 5.139398e-03 | 4.871686e-03 | 5.064854e-03 | 4.533909e-03 | 4.091831e-03 | 6.352102e-03 | 5.717030e-03 | 8.576276e-03 | 5.631893e-03 | 5.164410e-03 | 6.372419e-03 | 7.28733 | 8.13221 | 8.1469 | 7.49626 | 7.59631 | 5.822638e-03 |
| 2,3-Diphosphoglyceric acid | HILIC | 7.988024e-04 | 3.633074e-04 | 4.749338e-04 | 2.197258e-04 | 1.833555e-04 | 1.875686e-04 | 2.038711e-04 | 2.425939e-04 | 8.392951e-05 | 1.860328e-04 | 2.373046e-04 | 1.609644e-04 | 1.483740e-04 | 9.842493e-05 | 9.181434e-05 | 1.382121e-04 | 1.764010e-04 | 1.578359e-04 | 3.026533e-04 | 2.319919e-05 | 2.740614e-05 | 4.34656 | 6.85222 | 3.86148 | 2.69461 | 2.13735 | 3.228619e-05 |
| 2-Aminomuconic acid | HILIC | 4.867162e-04 | 5.707651e-04 | 5.933627e-04 | 4.723447e-04 | 4.611408e-04 | 4.339868e-04 | 4.247152e-04 | 4.203446e-04 | 3.233439e-04 | 4.891734e-04 | 5.320772e-04 | 4.657322e-04 | 4.119783e-04 | 4.175765e-04 | 4.414444e-04 | 3.216666e-04 | 3.028318e-04 | 3.372083e-04 | 1.720087e-04 | 1.940532e-04 | 2.324144e-04 | 3.75989 | 3.29449 | 3.56789 | 2.27343 | 2.00298 | 2.211721e-04 |
| 2-Aminomuconic acid semialdehyde | HILIC | 2.765234e-04 | 1.878943e-04 | 1.961622e-04 | 1.428525e-04 | 1.516501e-04 | 1.365150e-04 | 1.891404e-04 | 1.942816e-04 | 5.723063e-04 | 1.554593e-04 | 1.658610e-04 | 1.696782e-04 | 1.853331e-04 | 2.138409e-04 | 1.696809e-04 | 1.594242e-04 | 1.623756e-04 | 1.517245e-04 | 3.645573e-04 | 2.151803e-04 | 1.789071e-04 | 2.21869 | 1.63485 | 1.82604 | 1.60859 | 1.57892 | 5.051083e-04 |
| 2-Hydroxybutyric acid | HILIC | 1.885970e-04 | 1.009154e-04 | 9.153247e-05 | 8.280627e-05 | 7.848572e-05 | 8.140122e-05 | 8.472192e-05 | 7.765674e-05 | 2.574216e-04 | 9.825159e-05 | 7.025775e-05 | 7.927161e-05 | 7.815742e-05 | 9.392829e-05 | 7.488846e-05 | 9.699595e-05 | 9.870536e-05 | 8.866851e-05 | 1.792238e-04 | 1.093873e-04 | 1.004835e-04 | 9.14278 | 9.24188 | 8.83857 | 8.37424 | 9.10771 | 8.415856e-05 |
| 2-Hydroxyglutarate | HILIC | 4.505759e-04 | 4.819086e-04 | 4.368142e-04 | 4.351602e-04 | 4.318169e-04 | 4.418375e-04 | 3.965823e-04 | 3.850087e-04 | 3.616421e-04 | 4.131625e-04 | 4.321609e-04 | 4.017477e-04 | 3.022289e-04 | 2.855466e-04 | 3.272581e-04 | 8.678203e-04 | 5.480974e-04 | 5.213963e-04 | 1.752375e-04 | 2.866795e-04 | 1.704269e-04 | 4.41022 | 3.68949 | 3.92799 | 3.98207 | 3.48559 | 3.688676e-04 |
| 2-Methylglutaric acid | HILIC | 3.196605e-04 | 1.411083e-04 | 1.156979e-04 | 9.170391e-05 | 7.442574e-05 | 9.602925e-05 | 5.251738e-05 | 4.994881e-05 | 1.181125e-04 | 1.183759e-04 | 5.133396e-05 | 8.231920e-05 | 1.236437e-04 | 2.495600e-04 | 1.601621e-04 | 3.401966e-04 | 7.089633e-05 | 8.792176e-05 | 2.007873e-04 | 3.549729e-04 | 1.895525e-04 | 9.27315 | 1.31405 | 9.34286 | 2.07126 | 1.54983 | 1.867691e-04 |
| 2-Phospho-D-glyceric acid | HILIC | 4.291382e-02 | 4.699754e-02 | 5.100719e-02 | 4.413947e-02 | 4.340759e-02 | 3.716562e-02 | 5.368010e-02 | 5.627597e-02 | 3.699498e-02 | 3.730186e-02 | 4.676034e-02 | 4.219894e-02 | 4.364833e-02 | 4.372501e-02 | 4.533988e-02 | 3.711661e-02 | 4.745469e-02 | 2.939500e-02 | 2.072225e-02 | 2.772805e-02 | 2.882423e-02 | 3.62334 | 2.70308 | 2.57302 | 2.38874 | 2.23033 | 2.547671e-02 |
| 2-Propylglutaric acid | HILIC | 3.600791e-04 | 9.565309e-05 | 1.030781e-04 | 9.528766e-05 | 5.690587e-05 | 1.259730e-04 | 4.480437e-05 | 5.625760e-05 | 6.951606e-05 | 9.436217e-05 | 5.255073e-05 | 5.920315e-05 | 7.874416e-05 | 1.070301e-04 | 5.983383e-05 | 1.713700e-04 | 5.718130e-05 | 1.340003e-04 | 1.592429e-04 | 2.087879e-04 | 8.550006e-05 | 8.90509 | 1.05016 | 8.48615 | 1.04060 | 7.46994 | 1.688061e-04 |
| 3-Dehydro-L-gulonate | HILIC | 4.063601e-05 | 1.784237e-05 | 1.707558e-05 | 1.580822e-05 | 1.633376e-05 | 1.376571e-05 | 1.360885e-05 | 1.108772e-05 | 8.911554e-06 | 2.418332e-05 | 1.069345e-05 | 1.097084e-05 | 7.201987e-06 | 8.882873e-06 | 6.079198e-06 | 2.811667e-05 | 2.155730e-05 | 2.705918e-05 | 8.237167e-06 | 1.713293e-05 | 1.936854e-05 | 1.41789 | 1.56424 | 1.41892 | 1.29525 | 1.11143 | 9.059639e-06 |
| 3-Hydroxybutyric acid | HILIC | 2.580965e-04 | 5.957537e-05 | 5.491526e-05 | 4.510945e-05 | 4.303356e-05 | 4.772327e-05 | 3.690506e-05 | 3.462919e-05 | 1.422679e-04 | 5.238618e-05 | 3.693961e-05 | 4.388465e-05 | 4.861566e-05 | 5.974369e-05 | 4.515139e-05 | 4.386530e-05 | 3.710063e-05 | 4.252331e-05 | 1.065419e-04 | 5.285360e-05 | 4.634289e-05 | 4.38499 | 5.15322 | 4.75711 | 4.86119 | 4.89984 | 4.907649e-05 |
| 3-Hydroxymethylglutaric acid | HILIC | 2.314626e-04 | 1.408431e-04 | 1.259502e-04 | 1.173491e-04 | 1.010843e-04 | 1.327822e-04 | 1.132142e-04 | 1.043602e-04 | 1.270606e-04 | 1.322859e-04 | 1.137740e-04 | 1.154325e-04 | 1.166420e-04 | 1.203680e-04 | 1.155402e-04 | 3.427968e-04 | 1.039888e-04 | 1.178239e-04 | 1.265349e-04 | 1.749676e-04 | 1.161975e-04 | 1.09622 | 1.04063 | 9.99916 | 1.40636 | 9.74193 | 1.109167e-04 |
| 3-Sulfinoalanine | HILIC | 8.085827e-05 | 6.786547e-05 | 2.389328e-05 | 2.713962e-05 | 1.672097e-05 | 5.164981e-05 | 1.460104e-05 | 1.389511e-05 | 1.812345e-05 | 1.421214e-05 | 1.387997e-05 | 1.397191e-05 | 1.508643e-05 | 1.095698e-04 | 1.313564e-05 | 4.701443e-05 | 1.375126e-05 | 1.505194e-05 | 1.328301e-05 | 1.177185e-05 | 1.505969e-05 | 3.27719 | 1.90766 | 2.16686 | 1.86223 | 1.45708 | 1.893568e-05 |
| 5'-Phosphoribosyl-N-formylglycinamide | HILIC | 2.338743e-04 | 2.995581e-04 | 3.677421e-04 | 2.003045e-04 | 2.171475e-04 | 2.552104e-04 | 3.112693e-04 | 3.125458e-04 | 1.654785e-04 | 2.315223e-04 | 3.428101e-04 | 2.498088e-04 | 4.362883e-04 | 3.922903e-04 | 4.010761e-04 | 2.816298e-04 | 2.804925e-04 | 3.667840e-04 | 2.456045e-05 | 3.475744e-05 | 2.749404e-05 | 2.01075 | 1.87280 | 2.40049 | 1.50064 | 1.27305 | 1.159541e-04 |
| 5-L-Glutamyl-L-alanine | HILIC | 5.504365e-04 | 2.138572e-04 | 2.872328e-04 | 2.786345e-04 | 2.964878e-04 | 2.579526e-04 | 4.760534e-04 | 4.105190e-04 | 3.871919e-04 | 5.473599e-04 | 2.607292e-04 | 2.606319e-04 | 3.399399e-04 | 3.601654e-04 | 3.600063e-04 | 6.553541e-04 | 7.388059e-04 | 6.713982e-04 | 3.974193e-04 | 4.950994e-04 | 5.416019e-04 | 8.44470 | 8.07747 | 8.86872 | 9.77941 | 8.51538 | 9.146182e-04 |
| 5-L-Glutamyl-taurine | HILIC | 8.287553e-05 | 1.046780e-04 | 1.070568e-04 | 1.090780e-04 | 1.031366e-04 | 9.728846e-05 | 1.403874e-04 | 1.318436e-04 | 6.359764e-05 | 1.004805e-04 | 1.111491e-04 | 1.052452e-04 | 9.800239e-05 | 8.572810e-05 | 1.027079e-04 | 1.481573e-04 | 1.454786e-04 | 1.543304e-04 | 1.576518e-04 | 1.963325e-04 | 2.394013e-04 | 8.51829 | 7.80878 | 8.65277 | 7.32087 | 6.55087 | 6.366367e-05 |
| 5-Thymidylic acid | HILIC | 4.415260e-05 | 2.779573e-05 | 4.096958e-05 | 1.607285e-05 | 1.353098e-05 | 2.092817e-05 | 1.769021e-05 | 1.507953e-05 | 1.401834e-05 | 2.405319e-05 | 2.488790e-05 | 2.375021e-05 | 3.202265e-05 | 2.711222e-05 | 2.689748e-05 | 8.046695e-05 | 2.060419e-05 | 1.472415e-04 | 2.465752e-05 | 2.590019e-05 | 2.125872e-05 | 2.00099 | 1.71796 | 1.91834 | 2.11713 | 1.93326 | 8.600340e-06 |
| 6-Phosphogluconic acid | HILIC | 4.028600e-04 | 3.907573e-04 | 3.138706e-04 | 4.142344e-04 | 2.859926e-04 | 4.454839e-04 | 4.515015e-04 | 3.606219e-04 | 2.757341e-04 | 3.390889e-04 | 2.735149e-04 | 3.502729e-04 | 3.290122e-04 | 2.177706e-04 | 3.227378e-04 | 4.214455e-04 | 4.952643e-04 | 2.356349e-04 | 2.663269e-04 | 3.665628e-04 | 3.571512e-04 | 3.63329 | 4.09281 | 3.62935 | 3.47448 | 3.40642 | 1.760241e-04 |
| ADP | HILIC | 5.208569e-03 | 5.553968e-03 | 5.541395e-03 | 6.133494e-03 | 5.102160e-03 | 6.775318e-03 | 4.204409e-03 | 4.083614e-03 | 2.386372e-03 | 4.584610e-03 | 4.800204e-03 | 4.035706e-03 | 2.785132e-03 | 2.798157e-03 | 2.379560e-03 | 7.045640e-03 | 7.602841e-03 | 4.182674e-03 | 4.187345e-03 | 3.724459e-03 | 4.634204e-03 | 6.01604 | 7.32006 | 8.30494 | 5.47838 | 5.42072 | 3.940587e-03 |
| ADP-glucose | HILIC | 1.326130e-04 | 1.568892e-04 | 1.561270e-04 | 2.188166e-04 | 1.898865e-04 | 1.963663e-04 | 2.430099e-04 | 2.111575e-04 | 1.945459e-04 | 2.112328e-04 | 2.334341e-04 | 2.270816e-04 | 2.194095e-04 | 1.544196e-04 | 1.950741e-04 | 3.080719e-04 | 3.683541e-04 | 2.830468e-04 | 6.548074e-04 | 6.441828e-04 | 5.793762e-04 | 2.70099 | 2.82362 | 2.14571 | 2.52434 | 2.36007 | 2.119755e-04 |
| ADP-ribose 2'-phosphate | HILIC | 1.110708e-04 | 1.385158e-04 | 1.402246e-04 | 1.902082e-04 | 1.993521e-04 | 2.280114e-04 | 3.581013e-04 | 3.226251e-04 | 2.994362e-04 | 1.891240e-04 | 3.081718e-04 | 3.079043e-04 | 2.974601e-04 | 2.748353e-04 | 3.106690e-04 | 2.493365e-04 | 2.967796e-04 | 1.991501e-04 | 2.307019e-04 | 2.366255e-04 | 2.481821e-04 | 2.81377 | 3.07935 | 2.86799 | 2.89796 | 2.81829 | 2.495330e-04 |
| AICAR | HILIC | 3.604903e-05 | 3.558784e-05 | 3.890561e-05 | 8.130613e-05 | 4.977911e-05 | 6.258389e-05 | 1.066557e-03 | 4.855802e-04 | 5.349956e-04 | 5.404542e-05 | 5.251031e-05 | 4.738302e-05 | 2.997341e-03 | 1.881251e-03 | 2.468277e-03 | 5.404083e-05 | 5.929427e-05 | 4.978030e-05 | 9.569694e-07 | 3.504436e-06 | 3.593943e-06 | 2.36390 | 2.93491 | 3.09400 | 1.40373 | 1.46267 | 5.791407e-06 |
| Acetylcysteine | HILIC | 3.200989e-05 | 1.699295e-05 | 9.438754e-06 | 8.934218e-06 | 9.268201e-06 | 6.855980e-06 | 2.070371e-05 | 1.583032e-05 | 2.058814e-05 | 9.607447e-06 | 1.459942e-05 | 1.170205e-05 | 2.044694e-05 | 2.057717e-05 | 2.239780e-05 | 4.466183e-05 | 1.571217e-05 | 1.297466e-05 | 7.759426e-06 | 5.527107e-06 | 7.888444e-06 | 3.03173 | 2.31705 | 2.29829 | 2.08901 | 2.40955 | 1.569065e-05 |
| Adenosine diphosphate ribose | HILIC | 5.470972e-04 | 6.909173e-04 | 7.583005e-04 | 5.809633e-04 | 5.606499e-04 | 6.991276e-04 | 4.772255e-04 | 4.793631e-04 | 2.589805e-04 | 4.499708e-04 | 4.843939e-04 | 4.574391e-04 | 3.457757e-04 | 4.759209e-04 | 3.415028e-04 | 8.914127e-04 | 8.743647e-04 | 7.185751e-04 | 4.413552e-04 | 4.209225e-04 | 5.941103e-04 | 9.23249 | 1.04784 | 1.18894 | 8.76885 | 9.02453 | 1.310448e-03 |
| Adenosine monophosphate | HILIC | 5.000545e-03 | 3.434911e-03 | 3.665589e-03 | 3.677694e-03 | 3.692068e-03 | 4.579250e-03 | 5.642674e-03 | 4.964223e-03 | 3.397854e-03 | 4.188948e-03 | 5.068354e-03 | 4.773910e-03 | 4.876690e-03 | 4.241672e-03 | 3.926172e-03 | 6.173002e-03 | 5.535844e-03 | 8.343752e-03 | 5.320671e-03 | 4.967266e-03 | 6.134056e-03 | 7.12430 | 7.92065 | 7.92264 | 7.22339 | 7.31515 | 5.568947e-03 |
| Adenosine triphosphate | HILIC | 9.984208e-04 | 1.313801e-03 | 1.424643e-03 | 1.570730e-03 | 1.547194e-03 | 1.494855e-03 | 2.005983e-03 | 1.864004e-03 | 1.596228e-03 | 1.255393e-03 | 1.828590e-03 | 1.739526e-03 | 1.583939e-03 | 1.104830e-03 | 1.494862e-03 | 1.578934e-03 | 1.940520e-03 | 1.114985e-03 | 1.666299e-03 | 1.912585e-03 | 1.822849e-03 | 1.78375 | 1.74044 | 1.64915 | 1.52540 | 1.55615 | 1.223525e-03 |
| Alpha-D-Glucose 1,6-bisphosphate | HILIC | 3.246075e-03 | 3.354972e-03 | 3.463883e-03 | 4.255043e-03 | 4.018964e-03 | 3.967506e-03 | 4.445071e-03 | 4.005158e-03 | 2.607425e-03 | 4.070407e-03 | 4.134089e-03 | 4.349424e-03 | 3.250519e-03 | 2.461197e-03 | 3.343223e-03 | 4.027345e-03 | 4.524930e-03 | 3.008818e-03 | 2.033401e-03 | 2.830094e-03 | 3.236048e-03 | 4.01256 | 3.59664 | 3.60932 | 2.57528 | 2.48260 | 2.375896e-03 |
| Arabinonic acid | HILIC | 7.396273e-05 | 1.263726e-04 | 9.005385e-05 | 7.387108e-05 | 4.972675e-05 | 6.266740e-05 | 7.635511e-05 | 5.766853e-05 | 3.499759e-05 | 5.810469e-05 | 6.570734e-05 | 6.752718e-05 | 4.683670e-05 | 3.830745e-05 | 4.546036e-05 | 5.691823e-05 | 7.334108e-05 | 5.700640e-05 | 2.955215e-05 | 3.544648e-05 | 4.550885e-05 | 8.90551 | 6.00312 | 5.20629 | 3.95683 | 5.44628 | 3.414659e-05 |
| Beta-Citryl-L-glutamic acid | HILIC | 3.215418e-03 | 3.561429e-03 | 3.637430e-03 | 3.990623e-03 | 3.935193e-03 | 3.791767e-03 | 4.836013e-03 | 4.823901e-03 | 3.650903e-03 | 3.664443e-03 | 4.503827e-03 | 4.125252e-03 | 4.573439e-03 | 4.145815e-03 | 4.775717e-03 | 1.920967e-03 | 2.013385e-03 | 2.055574e-03 | 8.417285e-04 | 1.028981e-03 | 1.062166e-03 | 1.34229 | 1.14474 | 1.20573 | 5.98218 | 5.16407 | 5.514550e-04 |
| Beta-D-Glucose 6-phosphate | HILIC | 1.606916e-02 | 1.827098e-02 | 1.725322e-02 | 1.809241e-02 | 1.718094e-02 | 1.765045e-02 | 2.097642e-02 | 1.956809e-02 | 1.395383e-02 | 1.556320e-02 | 1.455936e-02 | 1.636727e-02 | 1.823486e-02 | 1.468471e-02 | 1.853283e-02 | 1.285416e-02 | 1.317984e-02 | 1.373283e-02 | 6.356418e-03 | 1.133603e-02 | 1.187116e-02 | 1.46866 | 1.14744 | 1.05410 | 1.04459 | 1.03454 | 3.166544e-03 |
| Butyric acid | HILIC | 2.664049e-05 | 2.560544e-05 | 2.309132e-05 | 2.074685e-05 | 2.244845e-05 | 2.018552e-05 | 2.444825e-05 | 2.360772e-05 | 5.280392e-05 | 2.333963e-05 | 2.234879e-05 | 2.227226e-05 | 3.570276e-05 | 4.150337e-05 | 3.184443e-05 | 1.928819e-05 | 1.853098e-05 | 2.394555e-05 | 3.506929e-05 | 2.965550e-05 | 2.662106e-05 | 1.55536 | 1.74223 | 1.49172 | 2.66421 | 2.51461 | 2.563922e-05 |
| CDP | HILIC | 1.075674e-03 | 1.488018e-03 | 2.304816e-03 | 6.217634e-04 | 7.215472e-04 | 4.986605e-04 | 5.718005e-04 | 8.472600e-04 | 3.297599e-04 | 5.331843e-04 | 1.166195e-03 | 7.710030e-04 | 4.360049e-04 | 8.217020e-04 | 4.071236e-04 | 3.845735e-04 | 2.883834e-04 | 4.072406e-04 | 2.719923e-04 | 3.090999e-04 | 2.906489e-04 | 8.93770 | 4.43306 | 6.61368 | 5.66290 | 4.51307 | 4.504389e-04 |
| CDP-Ethanolamine | HILIC | 6.503938e-05 | 9.653039e-05 | 8.957783e-05 | 6.781932e-05 | 5.938873e-05 | 6.526635e-05 | 4.333590e-05 | 4.567125e-05 | 1.671639e-05 | 4.962382e-05 | 6.949722e-05 | 6.360066e-05 | 4.062085e-05 | 3.168782e-05 | 4.539519e-05 | 1.314383e-04 | 1.421549e-04 | 1.234460e-04 | 1.162904e-04 | 1.345975e-04 | 1.390669e-04 | 1.10382 | 1.00616 | 1.17797 | 8.96774 | 8.92577 | 8.891625e-04 |
| Citric acid | HILIC | 3.505966e-02 | 2.034618e-02 | 2.207017e-02 | 2.888927e-02 | 2.606072e-02 | 2.980703e-02 | 3.217426e-02 | 2.986579e-02 | 2.653366e-02 | 2.414833e-02 | 2.639339e-02 | 2.611554e-02 | 2.754150e-02 | 2.535906e-02 | 2.778581e-02 | 1.124916e-02 | 1.167714e-02 | 1.062910e-02 | 4.708068e-03 | 5.398280e-03 | 5.845191e-03 | 7.16435 | 6.35207 | 5.59780 | 4.45575 | 5.02017 | 4.446548e-03 |
| CoA | HILIC | 6.726610e-04 | 4.000962e-04 | 3.684033e-04 | 3.892156e-04 | 3.490585e-04 | 3.161892e-04 | 2.549808e-04 | 2.725291e-04 | 2.309020e-04 | 1.838625e-04 | 2.726506e-04 | 2.677506e-04 | 2.007645e-04 | 1.870832e-04 | 1.963934e-04 | 2.893520e-04 | 2.434259e-04 | 1.358358e-04 | 1.758278e-04 | 1.801607e-04 | 1.668521e-04 | 2.25460 | 1.96775 | 1.86733 | 1.87635 | 1.58745 | 1.435228e-04 |
| Cyclic AMP | HILIC | 2.259671e-04 | 4.274586e-04 | 4.639713e-04 | 1.263376e-04 | 1.693848e-04 | 1.470001e-04 | 1.421772e-04 | 1.967160e-04 | 1.276746e-04 | 1.983035e-04 | 4.000922e-04 | 1.816308e-04 | 1.875112e-04 | 2.879722e-04 | 1.978687e-04 | 1.640991e-04 | 1.350876e-04 | 1.691203e-04 | 5.202896e-05 | 6.115327e-05 | 6.335980e-05 | 2.99817 | 2.26556 | 3.00013 | 2.33428 | 1.57106 | 1.835985e-04 |
| Cysteic acid | HILIC | 4.224219e-04 | 2.344859e-05 | 3.984647e-05 | 2.502176e-05 | 1.619167e-05 | 3.039157e-05 | 2.952172e-05 | 3.033589e-05 | 2.042339e-05 | 3.360710e-05 | 3.105759e-05 | 2.322886e-05 | 3.098582e-05 | 3.885041e-05 | 3.121194e-05 | 9.611031e-05 | 1.920211e-05 | 4.058714e-05 | 1.596275e-05 | 2.830628e-05 | 2.447971e-05 | 5.75627 | 3.20780 | 4.51921 | 3.51465 | 2.97497 | 3.304509e-05 |
| Cytidine monophosphate | HILIC | 1.267801e-03 | 1.396084e-03 | 2.184375e-03 | 1.001711e-03 | 1.168704e-03 | 1.138843e-03 | 1.139655e-03 | 1.279881e-03 | 7.729771e-04 | 8.572838e-04 | 1.319115e-03 | 1.088165e-03 | 1.179580e-03 | 1.644763e-03 | 1.130319e-03 | 1.094383e-03 | 1.017030e-03 | 1.399775e-03 | 8.733012e-04 | 9.133285e-04 | 1.125358e-03 | 1.59851 | 1.34652 | 1.61506 | 1.47163 | 1.50193 | 1.511635e-03 |
| Cytidine monophosphate N-acetylneuraminic acid | HILIC | 1.270431e-05 | 1.378724e-05 | 1.103318e-05 | 1.542531e-05 | 1.463084e-05 | 1.449275e-05 | 1.594968e-05 | 1.192775e-05 | 1.441946e-05 | 1.476353e-05 | 1.508648e-05 | 1.242858e-05 | 1.859431e-05 | 1.541404e-05 | 1.613311e-05 | 1.092358e-05 | 1.254507e-05 | 1.420857e-05 | 1.999516e-05 | 1.449766e-05 | 1.519667e-05 | 8.65993 | 7.51981 | 7.93711 | 9.29841 | 8.94734 | 9.449706e-06 |
| Cytidine triphosphate | HILIC | 1.719185e-04 | 3.678253e-04 | 5.981796e-04 | 1.645750e-04 | 1.877728e-04 | 1.207101e-04 | 3.695999e-04 | 5.199337e-04 | 2.967490e-04 | 1.398227e-04 | 4.573602e-04 | 4.206886e-04 | 2.688527e-04 | 3.365445e-04 | 2.920199e-04 | 1.023715e-04 | 1.070270e-04 | 1.347935e-04 | 1.373059e-04 | 2.346271e-04 | 1.573786e-04 | 3.69283 | 1.51841 | 1.48952 | 2.31707 | 2.27189 | 2.344620e-04 |
| D-Erythrose 4-phosphate | HILIC | 2.262227e-04 | 1.340772e-04 | 1.356573e-04 | 1.107186e-04 | 1.013387e-04 | 1.501295e-04 | 7.378232e-05 | 7.439603e-05 | 2.686336e-05 | 6.833521e-05 | 8.055238e-05 | 5.853981e-05 | 3.891301e-05 | 9.131119e-05 | 4.129264e-05 | 1.977130e-04 | 2.174657e-04 | 1.393377e-04 | 8.640442e-05 | 7.546411e-05 | 1.164149e-04 | 1.72924 | 2.28227 | 2.75227 | 1.86919 | 1.98662 | 1.079635e-04 |
| D-Glucaric acid | HILIC | 1.250568e-04 | 8.996170e-05 | 6.378104e-05 | 5.523466e-05 | 4.375440e-05 | 5.388131e-05 | 5.606307e-05 | 4.615710e-05 | 2.798773e-05 | 5.614621e-05 | 5.317578e-05 | 5.227375e-05 | 3.819301e-05 | 3.755571e-05 | 3.986162e-05 | 2.032425e-04 | 5.683015e-05 | 1.045372e-04 | 2.505173e-05 | 3.072430e-05 | 4.004029e-05 | 5.99374 | 5.20709 | 4.39997 | 3.80048 | 3.94697 | 3.331213e-05 |
| D-Glucose | HILIC | 1.660461e-02 | 1.123014e-02 | 1.212615e-02 | 1.156633e-02 | 1.278560e-02 | 1.174624e-02 | 1.556509e-02 | 1.755180e-02 | 2.160562e-02 | 1.833628e-02 | 1.674097e-02 | 1.573656e-02 | 2.274238e-02 | 2.628359e-02 | 2.352853e-02 | 1.207308e-02 | 1.287275e-02 | 1.285982e-02 | 1.469270e-02 | 1.540346e-02 | 1.472813e-02 | 1.98022 | 1.99575 | 2.14200 | 2.35306 | 2.15976 | 2.344194e-02 |
| D-Glucuronic acid 1-phosphate | HILIC | 3.459133e-04 | 3.402467e-04 | 3.092267e-04 | 4.024064e-04 | 3.802250e-04 | 4.053307e-04 | 4.475280e-04 | 3.965187e-04 | 3.087973e-04 | 3.000500e-04 | 2.785953e-04 | 2.779437e-04 | 3.579765e-04 | 2.919181e-04 | 4.194369e-04 | 3.235977e-04 | 3.842633e-04 | 1.565868e-04 | 1.527869e-04 | 2.237071e-04 | 2.154663e-04 | 1.42752 | 1.74054 | 1.66593 | 1.44574 | 1.29193 | 1.991266e-04 |
| D-Mannose 1-phosphate | HILIC | 1.989155e-04 | 3.153027e-04 | 3.073329e-04 | 2.858557e-04 | 2.759716e-04 | 2.801999e-04 | 2.581924e-04 | 2.705920e-04 | 1.663889e-04 | 2.905692e-04 | 2.662824e-04 | 2.826127e-04 | 2.494847e-04 | 2.345936e-04 | 2.597404e-04 | 2.662649e-04 | 2.934528e-04 | 2.518924e-04 | 1.346879e-04 | 1.562764e-04 | 1.443126e-04 | 2.99096 | 2.65790 | 2.65804 | 2.17812 | 2.02006 | 1.853811e-04 |

[illegible]

| | | | | | | | | | | | | | | | | | | | | | | | | | | | | |
|---|---|---|---|---|---|---|---|---|---|---|---|---|---|---|---|---|---|---|---|---|---|---|---|---|---|---|---|---|
| Benzylideneacetone | IC-MS | 1.255878e-03 | [illegible] | [illegible] | [illegible] | [illegible] | [illegible] | [illegible] | [illegible] | [illegible] | [illegible] | [illegible] | [illegible] | [illegible] | [illegible] | [illegible] | [illegible] | [illegible] | [illegible] | [illegible] | [illegible] | [illegible] | [illegible] | [illegible] | [illegible] | [illegible] | [illegible] | [illegible] |
| beta-Aspartylaspartic acid | IC-MS | 6.741826e-05 | [illegible] | [illegible] | [illegible] | [illegible] | [illegible] | [illegible] | [illegible] | [illegible] | [illegible] | [illegible] | [illegible] | [illegible] | [illegible] | [illegible] | [illegible] | [illegible] | [illegible] | [illegible] | [illegible] | [illegible] | [illegible] | [illegible] | [illegible] | [illegible] | [illegible] | [illegible] |
| beta-D-Mannopyranose | IC-MS | 9.166377e-04 | [illegible] | [illegible] | [illegible] | [illegible] | [illegible] | [illegible] | [illegible] | [illegible] | [illegible] | [illegible] | [illegible] | [illegible] | [illegible] | [illegible] | [illegible] | [illegible] | [illegible] | [illegible] | [illegible] | [illegible] | [illegible] | [illegible] | [illegible] | [illegible] | [illegible] | [illegible] |
| Betaine | IC-MS | 1.268832e-02 | [illegible] | [illegible] | [illegible] | [illegible] | [illegible] | [illegible] | [illegible] | [illegible] | [illegible] | [illegible] | [illegible] | [illegible] | [illegible] | [illegible] | [illegible] | [illegible] | [illegible] | [illegible] | [illegible] | [illegible] | [illegible] | [illegible] | [illegible] | [illegible] | [illegible] | [illegible] |
| Caprolactam | IC-MS | 1.396204e-03 | [illegible] | [illegible] | [illegible] | [illegible] | [illegible] | [illegible] | [illegible] | [illegible] | [illegible] | [illegible] | [illegible] | [illegible] | [illegible] | [illegible] | [illegible] | [illegible] | [illegible] | [illegible] | [illegible] | [illegible] | [illegible] | [illegible] | [illegible] | [illegible] | [illegible] | [illegible] |
| Caprylic acid | IC-MS | 2.202585e-04 | [illegible] | [illegible] | [illegible] | [illegible] | [illegible] | [illegible] | [illegible] | [illegible] | [illegible] | [illegible] | [illegible] | [illegible] | [illegible] | [illegible] | [illegible] | [illegible] | [illegible] | [illegible] | [illegible] | [illegible] | [illegible] | [illegible] | [illegible] | [illegible] | [illegible] | [illegible] |
| capuride | IC-MS | 2.125101e-04 | [illegible] | [illegible] | [illegible] | [illegible] | [illegible] | [illegible] | [illegible] | [illegible] | [illegible] | [illegible] | [illegible] | [illegible] | [illegible] | [illegible] | [illegible] | [illegible] | [illegible] | [illegible] | [illegible] | [illegible] | [illegible] | [illegible] | [illegible] | [illegible] | [illegible] | [illegible] |
| carglumic acid | IC-MS | 9.651610e-05 | [illegible] | [illegible] | [illegible] | [illegible] | [illegible] | [illegible] | [illegible] | [illegible] | [illegible] | [illegible] | [illegible] | [illegible] | [illegible] | [illegible] | [illegible] | [illegible] | [illegible] | [illegible] | [illegible] | [illegible] | [illegible] | [illegible] | [illegible] | [illegible] | [illegible] | [illegible] |
| CHLORIC ACID | IC-MS | 1.798429e-04 | [illegible] | [illegible] | [illegible] | [illegible] | [illegible] | [illegible] | [illegible] | [illegible] | [illegible] | [illegible] | [illegible] | [illegible] | [illegible] | [illegible] | [illegible] | [illegible] | [illegible] | [illegible] | [illegible] | [illegible] | [illegible] | [illegible] | [illegible] | [illegible] | [illegible] | [illegible] |
| Cholesterol sulfate | IC-MS | 1.141491e-03 | [illegible] | [illegible] | [illegible] | [illegible] | [illegible] | [illegible] | [illegible] | [illegible] | [illegible] | [illegible] | [illegible] | [illegible] | [illegible] | [illegible] | [illegible] | [illegible] | [illegible] | [illegible] | [illegible] | [illegible] | [illegible] | [illegible] | [illegible] | [illegible] | [illegible] | [illegible] |
| cis-4-Hydroxy-D-proline | IC-MS | 7.301190e-04 | [illegible] | [illegible] | [illegible] | [illegible] | [illegible] | [illegible] | [illegible] | [illegible] | [illegible] | [illegible] | [illegible] | [illegible] | [illegible] | [illegible] | [illegible] | [illegible] | [illegible] | [illegible] | [illegible] | [illegible] | [illegible] | [illegible] | [illegible] | [illegible] | [illegible] | [illegible] |
| Creatine | IC-MS | 2.312346e-02 | [illegible] | [illegible] | [illegible] | [illegible] | [illegible] | [illegible] | [illegible] | [illegible] | [illegible] | [illegible] | [illegible] | [illegible] | [illegible] | [illegible] | [illegible] | [illegible] | [illegible] | [illegible] | [illegible] | [illegible] | [illegible] | [illegible] | [illegible] | [illegible] | [illegible] | [illegible] |
| Creatinine | IC-MS | 4.343628e-04 | [illegible] | [illegible] | [illegible] | [illegible] | [illegible] | [illegible] | [illegible] | [illegible] | [illegible] | [illegible] | [illegible] | [illegible] | [illegible] | [illegible] | [illegible] | [illegible] | [illegible] | [illegible] | [illegible] | [illegible] | [illegible] | [illegible] | [illegible] | [illegible] | [illegible] | [illegible] |
| Crotonic acid | IC-MS | 2.653335e-04 | [illegible] | [illegible] | [illegible] | [illegible] | [illegible] | [illegible] | [illegible] | [illegible] | [illegible] | [illegible] | [illegible] | [illegible] | [illegible] | [illegible] | [illegible] | [illegible] | [illegible] | [illegible] | [illegible] | [illegible] | [illegible] | [illegible] | [illegible] | [illegible] | [illegible] | [illegible] |
| Cytarabine | IC-MS | 5.120688e-04 | [illegible] | [illegible] | [illegible] | [illegible] | [illegible] | [illegible] | [illegible] | [illegible] | [illegible] | [illegible] | [illegible] | [illegible] | [illegible] | [illegible] | [illegible] | [illegible] | [illegible] | [illegible] | [illegible] | [illegible] | [illegible] | [illegible] | [illegible] | [illegible] | [illegible] | [illegible] |
| Cytosine_RT1 | IC-MS | 1.612034e-03 | [illegible] | [illegible] | [illegible] | [illegible] | [illegible] | [illegible] | [illegible] | [illegible] | [illegible] | [illegible] | [illegible] | [illegible] | [illegible] | [illegible] | [illegible] | [illegible] | [illegible] | [illegible] | [illegible] | [illegible] | [illegible] | [illegible] | [illegible] | [illegible] | [illegible] | [illegible] |
| Cytosine_RT2 | IC-MS | 1.292602e-04 | [illegible] | [illegible] | [illegible] | [illegible] | [illegible] | [illegible] | [illegible] | [illegible] | [illegible] | [illegible] | [illegible] | [illegible] | [illegible] | [illegible] | [illegible] | [illegible] | [illegible] | [illegible] | [illegible] | [illegible] | [illegible] | [illegible] | [illegible] | [illegible] | [illegible] | [illegible] |
| D-(+)-Malic acid | IC-MS | 6.776436e-03 | [illegible] | [illegible] | [illegible] | [illegible] | [illegible] | [illegible] | [illegible] | [illegible] | [illegible] | [illegible] | [illegible] | [illegible] | [illegible] | [illegible] | [illegible] | [illegible] | [illegible] | [illegible] | [illegible] | [illegible] | [illegible] | [illegible] | [illegible] | [illegible] | [illegible] | [illegible] |
| D-(+)-Octopine | IC-MS | 1.472324e-05 | [illegible] | [illegible] | [illegible] | [illegible] | [illegible] | [illegible] | [illegible] | [illegible] | [illegible] | [illegible] | [illegible] | [illegible] | [illegible] | [illegible] | [illegible] | [illegible] | [illegible] | [illegible] | [illegible] | [illegible] | [illegible] | [illegible] | [illegible] | [illegible] | [illegible] | [illegible] |
| D-(+)-Proline | IC-MS | 1.994590e-02 | [illegible] | [illegible] | [illegible] | [illegible] | [illegible] | [illegible] | [illegible] | [illegible] | [illegible] | [illegible] | [illegible] | [illegible] | [illegible] | [illegible] | [illegible] | [illegible] | [illegible] | [illegible] | [illegible] | [illegible] | [illegible] | [illegible] | [illegible] | [illegible] | [illegible] | [illegible] |
| D-(+)-Tryptophan | IC-MS | 1.358514e-04 | [illegible] | [illegible] | [illegible] | [illegible] | [illegible] | [illegible] | [illegible] | [illegible] | [illegible] | [illegible] | [illegible] | [illegible] | [illegible] | [illegible] | [illegible] | [illegible] | [illegible] | [illegible] | [illegible] | [illegible] | [illegible] | [illegible] | [illegible] | [illegible] | [illegible] | [illegible] |
| D-(-)-3-Phosphoglyceric acid | IC-MS | 1.815613e-04 | [illegible] | [illegible] | [illegible] | [illegible] | [illegible] | [illegible] | [illegible] | [illegible] | [illegible] | [illegible] | [illegible] | [illegible] | [illegible] | [illegible] | [illegible] | [illegible] | [illegible] | [illegible] | [illegible] | [illegible] | [illegible] | [illegible] | [illegible] | [illegible] | [illegible] | [illegible] |
| D-benzylpenicilloic acid | IC-MS | 3.375304e-05 | [illegible] | [illegible] | [illegible] | [illegible] | [illegible] | [illegible] | [illegible] | [illegible] | [illegible] | [illegible] | [illegible] | [illegible] | [illegible] | [illegible] | [illegible] | [illegible] | [illegible] | [illegible] | [illegible] | [illegible] | [illegible] | [illegible] | [illegible] | [illegible] | [illegible] | [illegible] |
| D-Glucose 6-phosphate_RT1 | IC-MS | 4.267378e-04 | [illegible] | [illegible] | [illegible] | [illegible] | [illegible] | [illegible] | [illegible] | [illegible] | [illegible] | [illegible] | [illegible] | [illegible] | [illegible] | [illegible] | [illegible] | [illegible] | [illegible] | [illegible] | [illegible] | [illegible] | [illegible] | [illegible] | [illegible] | [illegible] | [illegible] | [illegible] |
| D-Glucose 6-phosphate_RT2 | IC-MS | 9.547487e-05 | [illegible] | [illegible] | [illegible] | [illegible] | [illegible] | [illegible] | [illegible] | [illegible] | [illegible] | [illegible] | [illegible] | [illegible] | [illegible] | [illegible] | [illegible] | [illegible] | [illegible] | [illegible] | [illegible] | [illegible] | [illegible] | [illegible] | [illegible] | [illegible] | [illegible] | [illegible] |
| D-PANTOTHENIC ACID | IC-MS | 3.064282e-03 | [illegible] | [illegible] | [illegible] | [illegible] | [illegible] | [illegible] | [illegible] | [illegible] | [illegible] | [illegible] | [illegible] | [illegible] | [illegible] | [illegible] | [illegible] | [illegible] | [illegible] | [illegible] | [illegible] | [illegible] | [illegible] | [illegible] | [illegible] | [illegible] | [illegible] | [illegible] |
| D-Sedoheptulose 7-phosphate | IC-MS | 3.481869e-04 | [illegible] | [illegible] | [illegible] | [illegible] | [illegible] | [illegible] | [illegible] | [illegible] | [illegible] | [illegible] | [illegible] | [illegible] | [illegible] | [illegible] | [illegible] | [illegible] | [illegible] | [illegible] | [illegible] | [illegible] | [illegible] | [illegible] | [illegible] | [illegible] | [illegible] | [illegible] |
| D-Xylulose 5-phosphate_RT1 | IC-MS | 1.586836e-04 | [illegible] | [illegible] | [illegible] | [illegible] | [illegible] | [illegible] | [illegible] | [illegible] | [illegible] | [illegible] | [illegible] | [illegible] | [illegible] | [illegible] | [illegible] | [illegible] | [illegible] | [illegible] | [illegible] | [illegible] | [illegible] | [illegible] | [illegible] | [illegible] | [illegible] | [illegible] |
| D-Xylulose 5-phosphate_RT2 | IC-MS | 5.738032e-04 | [illegible] | [illegible] | [illegible] | [illegible] | [illegible] | [illegible] | [illegible] | [illegible] | [illegible] | [illegible] | [illegible] | [illegible] | [illegible] | [illegible] | [illegible] | [illegible] | [illegible] | [illegible] | [illegible] | [illegible] | [illegible] | [illegible] | [illegible] | [illegible] | [illegible] | [illegible] |
| Decanoic acid | IC-MS | 5.419735e-04 | [illegible] | [illegible] | [illegible] | [illegible] | [illegible] | [illegible] | [illegible] | [illegible] | [illegible] | [illegible] | [illegible] | [illegible] | [illegible] | [illegible] | [illegible] | [illegible] | [illegible] | [illegible] | [illegible] | [illegible] | [illegible] | [illegible] | [illegible] | [illegible] | [illegible] | [illegible] |
| Dimethyl sulfoxide | IC-MS | 8.352481e-03 | [illegible] | [illegible] | [illegible] | [illegible] | [illegible] | [illegible] | [illegible] | [illegible] | [illegible] | [illegible] | [illegible] | [illegible] | [illegible] | [illegible] | [illegible] | [illegible] | [illegible] | [illegible] | [illegible] | [illegible] | [illegible] | [illegible] | [illegible] | [illegible] | [illegible] | [illegible] |
| DL-Alanine | IC-MS | 4.177048e-04 | [illegible] | [illegible] | [illegible] | [illegible] | [illegible] | [illegible] | [illegible] | [illegible] | [illegible] | [illegible] | [illegible] | [illegible] | [illegible] | [illegible] | [illegible] | [illegible] | [illegible] | [illegible] | [illegible] | [illegible] | [illegible] | [illegible] | [illegible] | [illegible] | [illegible] | [illegible] |
| DL-Glutamine | IC-MS | 1.023917e-02 | [illegible] | [illegible] | [illegible] | [illegible] | [illegible] | [illegible] | [illegible] | [illegible] | [illegible] | [illegible] | [illegible] | [illegible] | [illegible] | [illegible] | [illegible] | [illegible] | [illegible] | [illegible] | [illegible] | [illegible] | [illegible] | [illegible] | [illegible] | [illegible] | [illegible] | [illegible] |
| DL-Homoserine | IC-MS | 2.514874e-04 | [illegible] | [illegible] | [illegible] | [illegible] | [illegible] | [illegible] | [illegible] | [illegible] | [illegible] | [illegible] | [illegible] | [illegible] | [illegible] | [illegible] | [illegible] | [illegible] | [illegible] | [illegible] | [illegible] | [illegible] | [illegible] | [illegible] | [illegible] | [illegible] | [illegible] | [illegible] |
| DL-Beta-Leucine_RT1 | IC-MS | 2.870436e-03 | [illegible] | [illegible] | [illegible] | [illegible] | [illegible] | [illegible] | [illegible] | [illegible] | [illegible] | [illegible] | [illegible] | [illegible] | [illegible] | [illegible] | [illegible] | [illegible] | [illegible] | [illegible] | [illegible] | [illegible] | [illegible] | [illegible] | [illegible] | [illegible] | [illegible] | [illegible] |
| DL-Beta-Leucine_RT2 | IC-MS | 1.977874e-03 | [illegible] | [illegible] | [illegible] | [illegible] | [illegible] | [illegible] | [illegible] | [illegible] | [illegible] | [illegible] | [illegible] | [illegible] | [illegible] | [illegible] | [illegible] | [illegible] | [illegible] | [illegible] | [illegible] | [illegible] | [illegible] | [illegible] | [illegible] | [illegible] | [illegible] | [illegible] |
| Docosahexaenoic Acid | IC-MS | 1.195086e-03 | [illegible] | [illegible] | [illegible] | [illegible] | [illegible] | [illegible] | [illegible] | [illegible] | [illegible] | [illegible] | [illegible] | [illegible] | [illegible] | [illegible] | [illegible] | [illegible] | [illegible] | [illegible] | [illegible] | [illegible] | [illegible] | [illegible] | [illegible] | [illegible] | [illegible] | [illegible] |
| Docosanoic acid_RT1 | IC-MS | 1.330350e-03 | [illegible] | [illegible] | [illegible] | [illegible] | [illegible] | [illegible] | [illegible] | [illegible] | [illegible] | [illegible] | [illegible] | [illegible] | [illegible] | [illegible] | [illegible] | [illegible] | [illegible] | [illegible] | [illegible] | [illegible] | [illegible] | [illegible] | [illegible] | [illegible] | [illegible] | [illegible] |
| Docosanoic acid_RT2 | IC-MS | 1.349002e-04 | [illegible] | [illegible] | [illegible] | [illegible] | [illegible] | [illegible] | [illegible] | [illegible] | [illegible] | [illegible] | [illegible] | [illegible] | [illegible] | [illegible] | [illegible] | [illegible] | [illegible] | [illegible] | [illegible] | [illegible] | [illegible] | [illegible] | [illegible] | [illegible] | [illegible] | [illegible] |
| Docosatetraenoic acid | IC-MS | 4.252882e-04 | [illegible] | [illegible] | [illegible] | [illegible] | [illegible] | [illegible] | [illegible] | [illegible] | [illegible] | [illegible] | [illegible] | [illegible] | [illegible] | [illegible] | [illegible] | [illegible] | [illegible] | [illegible] | [illegible] | [illegible] | [illegible] | [illegible] | [illegible] | [illegible] | [illegible] | [illegible] |
| Drostanolone propionate | IC-MS | 7.122495e-05 | [illegible] | [illegible] | [illegible] | [illegible] | [illegible] | [illegible] | [illegible] | [illegible] | [illegible] | [illegible] | [illegible] | [illegible] | [illegible] | [illegible] | [illegible] | [illegible] | [illegible] | [illegible] | [illegible] | [illegible] | [illegible] | [illegible] | [illegible] | [illegible] | [illegible] | [illegible] |
| e,z-2,6-DODECADIEN-1-AL | IC-MS | 3.460672e-04 | [illegible] | [illegible] | [illegible] | [illegible] | [illegible] | [illegible] | [illegible] | [illegible] | [illegible] | [illegible] | [illegible] | [illegible] | [illegible] | [illegible] | [illegible] | [illegible] | [illegible] | [illegible] | [illegible] | [illegible] | [illegible] | [illegible] | [illegible] | [illegible] | [illegible] | [illegible] |
| Erucic acid | IC-MS | 3.431492e-04 | [illegible] | [illegible] | [illegible] | [illegible] | [illegible] | [illegible] | [illegible] | [illegible] | [illegible] | [illegible] | [illegible] | [illegible] | [illegible] | [illegible] | [illegible] | [illegible] | [illegible] | [illegible] | [illegible] | [illegible] | [illegible] | [illegible] | [illegible] | [illegible] | [illegible] | [illegible] |
| Fumaric acid | IC-MS | 2.563598e-04 | [illegible] | [illegible] | [illegible] | [illegible] | [illegible] | [illegible] | [illegible] | [illegible] | [illegible] | [illegible] | [illegible] | [illegible] | [illegible] | [illegible] | [illegible] | [illegible] | [illegible] | [illegible] | [illegible] | [illegible] | [illegible] | [illegible] | [illegible] | [illegible] | [illegible] | [illegible] |
| g-Guanidinobutyrate | IC-MS | 7.730754e-05 | [illegible] | [illegible] | [illegible] | [illegible] | [illegible] | [illegible] | [illegible] | [illegible] | [illegible] | [illegible] | [illegible] | [illegible] | [illegible] | [illegible] | [illegible] | [illegible] | [illegible] | [illegible] | [illegible] | [illegible] | [illegible] | [illegible] | [illegible] | [illegible] | [illegible] | [illegible] |
| gamma-Aminobutyric acid | IC-MS | 1.229312e-03 | [illegible] | [illegible] | [illegible] | [illegible] | [illegible] | [illegible] | [illegible] | [illegible] | [illegible] | [illegible] | [illegible] | [illegible] | [illegible] | [illegible] | [illegible] | [illegible] | [illegible] | [illegible] | [illegible] | [illegible] | [illegible] | [illegible] | [illegible] | [illegible] | [illegible] | [illegible] |
| glu-asp | IC-MS | 9.598090e-05 | [illegible] | [illegible] | [illegible] | [illegible] | [illegible] | [illegible] | [illegible] | [illegible] | [illegible] | [illegible] | [illegible] | [illegible] | [illegible] | [illegible] | [illegible] | [illegible] | [illegible] | [illegible] | [illegible] | [illegible] | [illegible] | [illegible] | [illegible] | [illegible] | [illegible] | [illegible] |
| Gluconic acid | IC-MS | 2.563394e-04 | [illegible] | [illegible] | [illegible] | [illegible] | [illegible] | [illegible] | [illegible] | [illegible] | [illegible] | [illegible] | [illegible] | [illegible] | [illegible] | [illegible] | [illegible] | [illegible] | [illegible] | [illegible] | [illegible] | [illegible] | [illegible] | [illegible] | [illegible] | [illegible] | [illegible] | [illegible] |
| glyceraldehyde 3-phosphate | IC-MS | 9.175883e-05 | [illegible] | [illegible] | [illegible] | [illegible] | [illegible] | [illegible] | [illegible] | [illegible] | [illegible] | [illegible] | [illegible] | [illegible] | [illegible] | [illegible] | [illegible] | [illegible] | [illegible] | [illegible] | [illegible] | [illegible] | [illegible] | [illegible] | [illegible] | [illegible] | [illegible] | [illegible] |
| glycerin | IC-MS | 3.786265e-04 | [illegible] | [illegible] | [illegible] | [illegible] | [illegible] | [illegible] | [illegible] | [illegible] | [illegible] | [illegible] | [illegible] | [illegible] | [illegible] | [illegible] | [illegible] | [illegible] | [illegible] | [illegible] | [illegible] | [illegible] | [illegible] | [illegible] | [illegible] | [illegible] | [illegible] | [illegible] |
| Glycerophosphoglycerol | IC-MS | 8.245351e-05 | [illegible] | [illegible] | [illegible] | [illegible] | [illegible] | [illegible] | [illegible] | [illegible] | [illegible] | [illegible] | [illegible] | [illegible] | [illegible] | [illegible] | [illegible] | [illegible] | [illegible] | [illegible] | [illegible] | [illegible] | [illegible] | [illegible] | [illegible] | [illegible] | [illegible] | [illegible] |
| glycidyl oleate_RT1 | IC-MS | 1.466120e-03 | [illegible] | [illegible] | [illegible] | [illegible] | [illegible] | [illegible] | [illegible] | [illegible] | [illegible] | [illegible] | [illegible] | [illegible] | [illegible] | [illegible] | [illegible] | [illegible] | [illegible] | [illegible] | [illegible] | [illegible] | [illegible] | [illegible] | [illegible] | [illegible] | [illegible] | [illegible] |
| glycidyl oleate_RT2 | IC-MS | 2.762528e-04 | [illegible] | [illegible] | [illegible] | [illegible] | [illegible] | [illegible] | [illegible] | [illegible] | [illegible] | [illegible] | [illegible] | [illegible] | [illegible] | [illegible] | [illegible] | [illegible] | [illegible] | [illegible] | [illegible] | [illegible] | [illegible] | [illegible] | [illegible] | [illegible] | [illegible] | [illegible] |
| glycidyl oleate_RT3 | IC-MS | 1.247524e-04 | [illegible] | [illegible] | [illegible] | [illegible] | [illegible] | [illegible] | [illegible] | [illegible] | [illegible] | [illegible] | [illegible] | [illegible] | [illegible] | [illegible] | [illegible] | [illegible] | [illegible] | [illegible] | [illegible] | [illegible] | [illegible] | [illegible] | [illegible] | [illegible] | [illegible] | [illegible] |
| Glycidyl Stearate | IC-MS | 3.447533e-04 | [illegible] | [illegible] | [illegible] | [illegible] | [illegible] | [illegible] | [illegible] | [illegible] | [illegible] | [illegible] | [illegible] | [illegible] | [illegible] | [illegible] | [illegible] | [illegible] | [illegible] | [illegible] | [illegible] | [illegible] | [illegible] | [illegible] | [illegible] | [illegible] | [illegible] | [illegible] |
| Guanine_RT1 | IC-MS | 8.564577e-04 | [illegible] | [illegible] | [illegible] | [illegible] | [illegible] | [illegible] | [illegible] | [illegible] | [illegible] | [illegible] | [illegible] | [illegible] | [illegible] | [illegible] | [illegible] | [illegible] | [illegible] | [illegible] | [illegible] | [illegible] | [illegible] | [illegible] | [illegible] | [illegible] | [illegible] | [illegible] |
| Guanine_RT2 | IC-MS | 4.635655e-04 | [illegible] | [illegible] | [illegible] | [illegible] | [illegible] | [illegible] | [illegible] | [illegible] | [illegible] | [illegible] | [illegible] | [illegible] | [illegible] | [illegible] | [illegible] | [illegible] | [illegible] | [illegible] | [illegible] | [illegible] | [illegible] | [illegible] | [illegible] | [illegible] | [illegible] | [illegible] |
| Guanosine | IC-MS | 3.759194e-04 | [illegible] | [illegible] | [illegible] | [illegible] | [illegible] | [illegible] | [illegible] | [illegible] | [illegible] | [illegible] | [illegible] | [illegible] | [illegible] | [illegible] | [illegible] | [illegible] | [illegible] | [illegible] | [illegible] | [illegible] | [illegible] | [illegible] | [illegible] | [illegible] | [illegible] | [illegible] |
| Hypoxanthine_RT1 | IC-MS | 2.270777e-02 | [illegible] | [illegible] | [illegible] | [illegible] | [illegible] | [illegible] | [illegible] | [illegible] | [illegible] | [illegible] | [illegible] | [illegible] | [illegible] | [illegible] | [illegible] | [illegible] | [illegible] | [illegible] | [illegible] | [illegible] | [illegible] | [illegible] | [illegible] | [illegible] | [illegible] | [illegible] |
| Hypoxanthine_RT2 | IC-MS | 5.130811e-03 | [illegible] | [illegible] | [illegible] | [illegible] | [illegible] | [illegible] | [illegible] | [illegible] | [illegible] | [illegible] | [illegible] | [illegible] | [illegible] | [illegible] | [illegible] | [illegible] | [illegible] | [illegible] | [illegible] | [illegible] | [illegible] | [illegible] | [illegible] | [illegible] | [illegible] | [illegible] |
| Imagabalin | IC-MS | 4.639983e-04 | [illegible] | [illegible] | [illegible] | [illegible] | [illegible] | [illegible] | [illegible] | [illegible] | [illegible] | [illegible] | [illegible] | [illegible] | [illegible] | [illegible] | [illegible] | [illegible] | [illegible] | [illegible] | [illegible] | [illegible] | [illegible] | [illegible] | [illegible] | [illegible] | [illegible] | [illegible] |
| L(-)-Carnitine | IC-MS | 6.792386e-03 | [illegible] | [illegible] | [illegible] | [illegible] | [illegible] | [illegible] | [illegible] | [illegible] | [illegible] | [illegible] | [illegible] | [illegible] | [illegible] | [illegible] | [illegible] | [illegible] | [illegible] | [illegible] | [illegible] | [illegible] | [illegible] | [illegible] | [illegible] | [illegible] | [illegible] | [illegible] |
| L-(+)-Arginine | IC-MS | 3.707236e-02 | [illegible] | [illegible] | [illegible] | [illegible] | [illegible] | [illegible] | [illegible] | [illegible] | [illegible] | [illegible] | [illegible] | [illegible] | [illegible] | [illegible] | [illegible] | [illegible] | [illegible] | [illegible] | [illegible] | [illegible] | [illegible] | [illegible] | [illegible] | [illegible] | [illegible] | [illegible] |
| L-(+)-Tartaric acid | IC-MS | 2.958422e-04 | [illegible] | [illegible] | [illegible] | [illegible] | [illegible] | [illegible] | [illegible] | [illegible] | [illegible] | [illegible] | [illegible] | [illegible] | [illegible] | [illegible] | [illegible] | [illegible] | [illegible] | [illegible] | [illegible] | [illegible] | [illegible] | [illegible] | [illegible] | [illegible] | [illegible] | [illegible] |
| L-(-)-methionine | IC-MS | 2.019881e-04 | [illegible] | [illegible] | [illegible] | [illegible] | [illegible] | [illegible] | [illegible] | [illegible] | [illegible] | [illegible] | [illegible] | [illegible] | [illegible] | [illegible] | [illegible] | [illegible] | [illegible] | [illegible] | [illegible] | [illegible] | [illegible] | [illegible] | [illegible] | [illegible] | [illegible] | [illegible] |
| L-alpha-Glycerylphosphorylcholine | IC-MS | 2.553385e-02 | [illegible] | [illegible] | [illegible] | [illegible] | [illegible] | [illegible] | [illegible] | [illegible] | [illegible] | [illegible] | [illegible] | [illegible] | [illegible] | [illegible] | [illegible] | [illegible] | [illegible] | [illegible] | [illegible] | [illegible] | [illegible] | [illegible] | [illegible] | [illegible] | [illegible] | [illegible] |
| L-Aspartic acid_RT1 | IC-MS | 1.598874e-03 | [illegible] | [illegible] | [illegible] | [illegible] | [illegible] | [illegible] | [illegible] | [illegible] | [illegible] | [illegible] | [illegible] | [illegible] | [illegible] | [illegible] | [illegible] | [illegible] | [illegible] | [illegible] | [illegible] | [illegible] | [illegible] | [illegible] | [illegible] | [illegible] | [illegible] | [illegible] |
| L-Aspartic acid_RT2 | IC-MS | 5.846423e-04 | [illegible] | [illegible] | [illegible] | [illegible] | [illegible] | [illegible] | [illegible] | [illegible] | [illegible] | [illegible] | [illegible] | [illegible] | [illegible] | [illegible] | [illegible] | [illegible] | [illegible] | [illegible] | [illegible] | [illegible] | [illegible] | [illegible] | [illegible] | [illegible] | [illegible] | [illegible] |
| L-Glutamic acid | IC-MS | 1.281686e-02 | [illegible] | [illegible] | [illegible] | [illegible] | [illegible] | [illegible] | [illegible] | [illegible] | [illegible] | [illegible] | [illegible] | [illegible] | [illegible] | [illegible] | [illegible] | [illegible] | [illegible] | [illegible] | [illegible] | [illegible] | [illegible] | [illegible] | [illegible] | [illegible] | [illegible] | [illegible] |
| L-Glutathione (reduced) | IC-MS | 6.072074e-03 | [illegible] | [illegible] | [illegible] | [illegible] | [illegible] | [illegible] | [illegible] | [illegible] | [illegible] | [illegible] | [illegible] | [illegible] | [illegible] | [illegible] | [illegible] | [illegible] | [illegible] | [illegible] | [illegible] | [illegible] | [illegible] | [illegible] | [illegible] | [illegible] | [illegible] | [illegible] |
| L-Hydroxyproline | IC-MS | 4.026357e-03 | [illegible] | [illegible] | [illegible] | [illegible] | [illegible] | [illegible] | [illegible] | [illegible] | [illegible] | [illegible] | [illegible] | [illegible] | [illegible] | [illegible] | [illegible] | [illegible] | [illegible] | [illegible] | [illegible] | [illegible] | [illegible] | [illegible] | [illegible] | [illegible] | [illegible] | [illegible] |
| L-Iditol | IC-MS | 4.461955e-04 | [illegible] | [illegible] | [illegible] | [illegible] | [illegible] | [illegible] | [illegible] | [illegible] | [illegible] | [illegible] | [illegible] | [illegible] | [illegible] | [illegible] | [illegible] | [illegible] | [illegible] | [illegible] | [illegible] | [illegible] | [illegible] | [illegible] | [illegible] | [illegible] | [illegible] | [illegible] |
| L-Phenylalanine | IC-MS | 8.358456e-04 | [illegible] | [illegible] | [illegible] | [illegible] | [illegible] | [illegible] | [illegible] | [illegible] | [illegible] | [illegible] | [illegible] | [illegible] | [illegible] | [illegible] | [illegible] | [illegible] | [illegible] | [illegible] | [illegible] | [illegible] | [illegible] | [illegible] | [illegible] | [illegible] | [illegible] | [illegible] |
| L-Serine | IC-MS | 3.011714e-04 | [illegible] | [illegible] | [illegible] | [illegible] | [illegible] | [illegible] | [illegible] | [illegible] | [illegible] | [illegible] | [illegible] | [illegible] | [illegible] | [illegible] | [illegible] | [illegible] | [illegible] | [illegible] | [illegible] | [illegible] | [illegible] | [illegible] | [illegible] | [illegible] | [illegible] | [illegible] |
| L-Threonic acid_RT1 | IC-MS | 1.024138e-03 | [illegible] | [illegible] | [illegible] | [illegible] | [illegible] | [illegible] | [illegible] | [illegible] | [illegible] | [illegible] | [illegible] | [illegible] | [illegible] | [illegible] | [illegible] | [illegible] | [illegible] | [illegible] | [illegible] | [illegible] | [illegible] | [illegible] | [illegible] | [illegible] | [illegible] | [illegible] |
| L-Threonic acid_RT2 | IC-MS | 2.374458e-03 | [illegible] | [illegible] | [illegible] | [illegible] | [illegible] | [illegible] | [illegible] | [illegible] | [illegible] | [illegible] | [illegible] | [illegible] | [illegible] | [illegible] | [illegible] | [illegible] | [illegible] | [illegible] | [illegible] | [illegible] | [illegible] | [illegible] | [illegible] | [illegible] | [illegible] | [illegible] |
| L-Tyrosine | IC-MS | 5.443707e-04 | [illegible] | [illegible] | [illegible] | [illegible] | [illegible] | [illegible] | [illegible] | [illegible] | [illegible] | [illegible] | [illegible] | [illegible] | [illegible] | [illegible] | [illegible] | [illegible] | [illegible] | [illegible] | [illegible] | [illegible] | [illegible] | [illegible] | [illegible] | [illegible] | [illegible] | [illegible] |
| Lauric acid | IC-MS | 3.041321e-04 | [illegible] | [illegible] | [illegible] | [illegible] | [illegible] | [illegible] | [illegible] | [illegible] | [illegible] | [illegible] | [illegible] | [illegible] | [illegible] | [illegible] | [illegible] | [illegible] | [illegible] | [illegible] | [illegible] | [illegible] | [illegible] | [illegible] | [illegible] | [illegible] | [illegible] | [illegible] |
| lentiginosine | IC-MS | 1.842214e-04 | [illegible] | [illegible] | [illegible] | [illegible] | [illegible] | [illegible] | [illegible] | [illegible] | [illegible] | [illegible] | [illegible] | [illegible] | [illegible] | [illegible] | [illegible] | [illegible] | [illegible] | [illegible] | [illegible] | [illegible] | [illegible] | [illegible] | [illegible] | [illegible] | [illegible] | [illegible] |
| Lignoceric acid | IC-MS | 2.165618e-04 | [illegible] | [illegible] | [illegible] | [illegible] | [illegible] | [illegible] | [illegible] | [illegible] | [illegible] | [illegible] | [illegible] | [illegible] | [illegible] | [illegible] | [illegible] | [illegible] | [illegible] | [illegible] | [illegible] | [illegible] | [illegible] | [illegible] | [illegible] | [illegible] | [illegible] | [illegible] |
| Linoleic acid | IC-MS | 2.911545e-03 | [illegible] | [illegible] | [illegible] | [illegible] | [illegible] | [illegible] | [illegible] | [illegible] | [illegible] | [illegible] | [illegible] | [illegible] | [illegible] | [illegible] | [illegible] | [illegible] | [illegible] | [illegible] | [illegible] | [illegible] | [illegible] | [illegible] | [illegible] | [illegible] | [illegible] | [illegible] |
| Lithocholic acid | IC-MS | 9.146350e-05 | [illegible] | [illegible] | [illegible] | [illegible] | [illegible] | [illegible] | [illegible] | [illegible] | [illegible] | [illegible] | [illegible] | [illegible] | [illegible] | [illegible] | [illegible] | [illegible] | [illegible] | [illegible] | [illegible] | [illegible] | [illegible] | [illegible] | [illegible] | [illegible] | [illegible] | [illegible] |
| Maleamic acid | IC-MS | 6.099988e-03 | [illegible] | [illegible] | [illegible] | [illegible] | [illegible] | [illegible] | [illegible] | [illegible] | [illegible] | [illegible] | [illegible] | [illegible] | [illegible] | [illegible] | [illegible] | [illegible] | [illegible] | [illegible] | [illegible] | [illegible] | [illegible] | [illegible] | [illegible] | [illegible] | [illegible] | [illegible] |
| Margaric acid_RT1 | IC-MS | 9.652727e-04 | [illegible] | [illegible] | [illegible] | [illegible] | [illegible] | [illegible] | [illegible] | [illegible] | [illegible] | [illegible] | [illegible] | [illegible] | [illegible] | [illegible] | [illegible] | [illegible] | [illegible] | [illegible] | [illegible] | [illegible] | [illegible] | [illegible] | [illegible] | [illegible] | [illegible] | [illegible] |
| Margaric acid_RT2 | IC-MS | 7.060382e-04 | [illegible] | [illegible] | [illegible] | [illegible] | [illegible] | [illegible] | [illegible] | [illegible] | [illegible] | [illegible] | [illegible] | [illegible] | [illegible] | [illegible] | [illegible] | [illegible] | [illegible] | [illegible] | [illegible] | [illegible] | [illegible] | [illegible] | [illegible] | [illegible] | [illegible] | [illegible] |
| Megastigma-5,8(Z)-dien-4-one | IC-MS | 4.844046e-04 | [illegible] | [illegible] | [illegible] | [illegible] | [illegible] | [illegible] | [illegible] | [illegible] | [illegible] | [illegible] | [illegible] | [illegible] | [illegible] | [illegible] | [illegible] | [illegible] | [illegible] | [illegible] | [illegible] | [illegible] | [illegible] | [illegible] | [illegible] | [illegible] | [illegible] | [illegible] |
| Mephedrone | IC-MS | 1.798013e-03 | [illegible] | [illegible] | [illegible] | [illegible] | [illegible] | [illegible] | [illegible] | [illegible] | [illegible] | [illegible] | [illegible] | [illegible] | [illegible] | [illegible] | [illegible] | [illegible] | [illegible] | [illegible] | [illegible] | [illegible] | [illegible] | [illegible] | [illegible] | [illegible] | [illegible] | [illegible] |
| Methyl (9E)-9-octadecenoate | IC-MS | 5.328803e-04 | [illegible] | [illegible] | [illegible] | [illegible] | [illegible] | [illegible] | [illegible] | [illegible] | [illegible] | [illegible] | [illegible] | [illegible] | [illegible] | [illegible] | [illegible] | [illegible] | [illegible] | [illegible] | [illegible] | [illegible] | [illegible] | [illegible] | [illegible] | [illegible] | [illegible] | [illegible] |
| Methyl acetoacetate | IC-MS | 2.778013e-04 | [illegible] | [illegible] | [illegible] | [illegible] | [illegible] | [illegible] | [illegible] | [illegible] | [illegible] | [illegible] | [illegible] | [illegible] | [illegible] | [illegible] | [illegible] | [illegible] | [illegible] | [illegible] | [illegible] | [illegible] | [illegible] | [illegible] | [illegible] | [illegible] | [illegible] | [illegible] |
| Methyl stearate | IC-MS | 9.447357e-05 | [illegible] | [illegible] | [illegible] | [illegible] | [illegible] | [illegible] | [illegible] | [illegible] | [illegible] | [illegible] | [illegible] | [illegible] | [illegible] | [illegible] | [illegible] | [illegible] | [illegible] | [illegible] | [illegible] | [illegible] | [illegible] | [illegible] | [illegible] | [illegible] | [illegible] | [illegible] |
| Methylmalonic acid | IC-MS | 3.736227e-04 | [illegible] | [illegible] | [illegible] | [illegible] | [illegible] | [illegible] | [illegible] | [illegible] | [illegible] | [illegible] | [illegible] | [illegible] | [illegible] | [illegible] | [illegible] | [illegible] | [illegible] | [illegible] | [illegible] | [illegible] | [illegible] | [illegible] | [illegible] | [illegible] | [illegible] | [illegible] |
| Myristic acid | IC-MS | 2.596089e-03 | [illegible] | [illegible] | [illegible] | [illegible] | [illegible] | [illegible] | [illegible] | [illegible] | [illegible] | [illegible] | [illegible] | [illegible] | [illegible] | [illegible] | [illegible] | [illegible] | [illegible] | [illegible] | [illegible] | [illegible] | [illegible] | [illegible] | [illegible] | [illegible] | [illegible] | [illegible] |
| Myristoleic acid | IC-MS | 8.207504e-05 | [illegible] | [illegible] | [illegible] | [illegible] | [illegible] | [illegible] | [illegible] | [illegible] | [illegible] | [illegible] | [illegible] | [illegible] | [illegible] | [illegible] | [illegible] | [illegible] | [illegible] | [illegible] | [illegible] | [illegible] | [illegible] | [illegible] | [illegible] | [illegible] | [illegible] | [illegible] |
| Myristyl sulfate | IC-MS | 3.122624e-04 | [illegible] | [illegible] | [illegible] | [illegible] | [illegible] | [illegible] | [illegible] | [illegible] | [illegible] | [illegible] | [illegible] | [illegible] | [illegible] | [illegible] | [illegible] | [illegible] | [illegible] | [illegible] | [illegible] | [illegible] | [illegible] | [illegible] | [illegible] | [illegible] | [illegible] | [illegible] |
| N'-nitrosoanatabine | IC-MS | 2.316384e-04 | [illegible] | [illegible] | [illegible] | [illegible] | [illegible] | [illegible] | [illegible] | [illegible] | [illegible] | [illegible] | [illegible] | [illegible] | [illegible] | [illegible] | [illegible] | [illegible] | [illegible] | [illegible] | [illegible] | [illegible] | [illegible] | [illegible] | [illegible] | [illegible] | [illegible] | [illegible] |
| N-(3-carboxypropanoyl)-N-hydroxycadaverine | IC-MS | 6.122425e-05 | [illegible] | [illegible] | [illegible] | [illegible] | [illegible] | [illegible] | [illegible] | [illegible] | [illegible] | [illegible] | [illegible] | [illegible] | [illegible] | [illegible] | [illegible] | [illegible] | [illegible] | [illegible] | [illegible] | [illegible] | [illegible] | [illegible] | [illegible] | [illegible] | [illegible] | [illegible] |
| N-(tert-Butoxycarbonyl)-L-leucine | IC-MS | 2.134628e-03 | [illegible] | [illegible] | [illegible] | [illegible] | [illegible] | [illegible] | [illegible] | [illegible] | [illegible] | [illegible] | [illegible] | [illegible] | [illegible] | [illegible] | [illegible] | [illegible] | [illegible] | [illegible] | [illegible] | [illegible] | [illegible] | [illegible] | [illegible] | [illegible] | [illegible] | [illegible] |
| N-[3-Carboxy-2-(carboxymethyl)-2-hydroxypropanoyl]glut | IC-MS | 1.698686e-03 | [illegible] | [illegible] | [illegible] | [illegible] | [illegible] | [illegible] | [illegible] | [illegible] | [illegible] | [illegible] | [illegible] | [illegible] | [illegible] | [illegible] | [illegible] | [illegible] | [illegible] | [illegible] | [illegible] | [illegible] | [illegible] | [illegible] | [illegible] | [illegible] | [illegible] | [illegible] |
| N-Acetyl-1-aspartylglutamic acid | IC-MS | 5.071543e-05 | [illegible] | [illegible] | [illegible] | [illegible] | [illegible] | [illegible] | [illegible] | [illegible] | [illegible] | [illegible] | [illegible] | [illegible] | [illegible] | [illegible] | [illegible] | [illegible] | [illegible] | [illegible] | [illegible] | [illegible] | [illegible] | [illegible] | [illegible] | [illegible] | [illegible] | [illegible] |
| N-acetyl-L-2-aminoadipic acid 6-phosphate | IC-MS | 3.710731e-04 | [illegible] | [illegible] | [illegible] | [illegible] | [illegible] | [illegible] | [illegible] | [illegible] | [illegible] | [illegible] | [illegible] | [illegible] | [illegible] | [illegible] | [illegible] | [illegible] | [illegible] | [illegible] | [illegible] | [illegible] | [illegible] | [illegible] | [illegible] | [illegible] | [illegible] | [illegible] |
| N-Acetyl-L-methionine | IC-MS | 1.841799e-04 | [illegible] | [illegible] | [illegible] | [illegible] | [illegible] | [illegible] | [illegible] | [illegible] | [illegible] | [illegible] | [illegible] | [illegible] | [illegible] | [illegible] | [illegible] | [illegible] | [illegible] | [illegible] | [illegible] | [illegible] | [illegible] | [illegible] | [illegible] | [illegible] | [illegible] | [illegible] |
| N-Acetylalanine | IC-MS | 6.859127e-04 | [illegible] | [illegible] | [illegible] | [illegible] | [illegible] | [illegible] | [illegible] | [illegible] | [illegible] | [illegible] | [illegible] | [illegible] | [illegible] | [illegible] | [illegible] | [illegible] | [illegible] | [illegible] | [illegible] | [illegible] | [illegible] | [illegible] | [illegible] | [illegible] | [illegible] | [illegible] |
| N-Acetylneuraminic acid | IC-MS | 1.839598e-04 | [illegible] | [illegible] | [illegible] | [illegible] | [illegible] | [illegible] | [illegible] | [illegible] | [illegible] | [illegible] | [illegible] | [illegible] | [illegible] | [illegible] | [illegible] | [illegible] | [illegible] | [illegible] | [illegible] | [illegible] | [illegible] | [illegible] | [illegible] | [illegible] | [illegible] | [illegible] |
| N-Acetylornithine | IC-MS | 7.361266e-05 | [illegible] | [illegible] | [illegible] | [illegible] | [illegible] | [illegible] | [illegible] | [illegible] | [illegible] | [illegible] | [illegible] | [illegible] | [illegible] | [illegible] | [illegible] | [illegible] | [illegible] | [illegible] | [illegible] | [illegible] | [illegible] | [illegible] | [illegible] | [illegible] | [illegible] | [illegible] |
| N-Methylisopelletierine | IC-MS | 2.663529e-04 | [illegible] | [illegible] | [illegible] | [illegible] | [illegible] | [illegible] | [illegible] | [illegible] | [illegible] | [illegible] | [illegible] | [illegible] | [illegible] | [illegible] | [illegible] | [illegible] | [illegible] | [illegible] | [illegible] | [illegible] | [illegible] | [illegible] | [illegible] | [illegible] | [illegible] | [illegible] |
| N-Undecanoylglycine | IC-MS | 1.116487e-04 | [illegible] | [illegible] | [illegible] | [illegible] | [illegible] | [illegible] | [illegible] | [illegible] | [illegible] | [illegible] | [illegible] | [illegible] | [illegible] | [illegible] | [illegible] | [illegible] | [illegible] | [illegible] | [illegible] | [illegible] | [illegible] | [illegible] | [illegible] | [illegible] | [illegible] | [illegible] |
| N3,N4-Dimethyl-L-arginine | IC-MS | 4.595198e-04 | [illegible] | [illegible] | [illegible] | [illegible] | [illegible] | [illegible] | [illegible] | [illegible] | [illegible] | [illegible] | [illegible] | [illegible] | [illegible] | [illegible] | [illegible] | [illegible] | [illegible] | [illegible] | [illegible] | [illegible] | [illegible] | [illegible] | [illegible] | [illegible] | [illegible] | [illegible] |
| N3-Metyladenine | IC-MS | 8.883119e-03 | [illegible] | [illegible] | [illegible] | [illegible] | [illegible] | [illegible] | [illegible] | [illegible] | [illegible] | [illegible] | [illegible] | [illegible] | [illegible] | [illegible] | [illegible] | [illegible] | [illegible] | [illegible] | [illegible] | [illegible] | [illegible] | [illegible] | [illegible] | [illegible] | [illegible] | [illegible] |
| Nervonic acid | IC-MS | 5.405332e-04 | [illegible] | [illegible] | [illegible] | [illegible] | [illegible] | [illegible] | [illegible] | [illegible] | [illegible] | [illegible] | [illegible] | [illegible] | [illegible] | [illegible] | [illegible] | [illegible] | [illegible] | [illegible] | [illegible] | [illegible] | [illegible] | [illegible] | [illegible] | [illegible] | [illegible] | [illegible] |
| Nonanoic acid | IC-MS | 1.950656e-03 | [illegible] | [illegible] | [illegible] | [illegible] | [illegible] | [illegible] | [illegible] | [illegible] | [illegible] | [illegible] | [illegible] | [illegible] | [illegible] | [illegible] | [illegible] | [illegible] | [illegible] | [illegible] | [illegible] | [illegible] | [illegible] | [illegible] | [illegible] | [illegible] | [illegible] | [illegible] |
| NP-011548 | IC-MS | 3.428359e-03 | [illegible] | [illegible] | [illegible] | [illegible] | [illegible] | [illegible] | [illegible] | [illegible] | [illegible] | [illegible] | [illegible] | [illegible] | [illegible] | [illegible] | [illegible] | [illegible] | [illegible] | [illegible] | [illegible] | [illegible] | [illegible] | [illegible] | [illegible] | [illegible] | [illegible] | [illegible] |
| Octodrine | IC-MS | 2.962081e-03 | [illegible] | [illegible] | [illegible] | [illegible] | [illegible] | [illegible] | [illegible] | [illegible] | [illegible] | [illegible] | [illegible] | [illegible] | [illegible] | [illegible] | [illegible] | [illegible] | [illegible] | [illegible] | [illegible] | [illegible] | [illegible] | [illegible] | [illegible] | [illegible] | [illegible] | [illegible] |
| Oleic acid | IC-MS | 6.115786e-02 | [illegible] | [illegible] | [illegible] | [illegible] | [illegible] | [illegible] | [illegible] | [illegible] | [illegible] | [illegible] | [illegible] | [illegible] | [illegible] | [illegible] | [illegible] | [illegible] | [illegible] | [illegible] | [illegible] | [illegible] | [illegible] | [illegible] | [illegible] | [illegible] | [illegible] | [illegible] |
| Ornithine | IC-MS | 4.190196e-04 | [illegible] | [illegible] | [illegible] | [illegible] | [illegible] | [illegible] | [illegible] | [illegible] | [illegible] | [illegible] | [illegible] | [illegible] | [illegible] | [illegible] | [illegible] | [illegible] | [illegible] | [illegible] | [illegible] | [illegible] | [illegible] | [illegible] | [illegible] | [illegible] | [illegible] | [illegible] |
| Osbond acid | IC-MS | 6.559208e-04 | [illegible] | [illegible] | [illegible] | [illegible] | [illegible] | [illegible] | [illegible] | [illegible] | [illegible] | [illegible] | [illegible] | [illegible] | [illegible] | [illegible] | [illegible] | [illegible] | [illegible] | [illegible] | [illegible] | [illegible] | [illegible] | [illegible] | [illegible] | [illegible] | [illegible] | [illegible] |
| oxolamine | IC-MS | 2.689609e-04 | [illegible] | [illegible] | [illegible] | [illegible] | [illegible] | [illegible] | [illegible] | [illegible] | [illegible] | [illegible] | [illegible] | [illegible] | [illegible] | [illegible] | [illegible] | [illegible] | [illegible] | [illegible] | [illegible] | [illegible] | [illegible] | [illegible] | [illegible] | [illegible] | [illegible] | [illegible] |
| p-Cresylsulfate | IC-MS | 2.969697e-04 | [illegible] | [illegible] | [illegible] | [illegible] | [illegible] | [illegible] | [illegible] | [illegible] | [illegible] | [illegible] | [illegible] | [illegible] | [illegible] | [illegible] | [illegible] | [illegible] | [illegible] | [illegible] | [illegible] | [illegible] | [illegible] | [illegible] | [illegible] | [illegible] | [illegible] | [illegible] |
| P-DMEA_RT1 | IC-MS | 5.603179e-04 | [illegible] | [illegible] | [illegible] | [illegible] | [illegible] | [illegible] | [illegible] | [illegible] | [illegible] | [illegible] | [illegible] | [illegible] | [illegible] | [illegible] | [illegible] | [illegible] | [illegible] | [illegible] | [illegible] | [illegible] | [illegible] | [illegible] | [illegible] | [illegible] | [illegible] | [illegible] |
| P-DMEA_RT2 | IC-MS | 1.670377e-04 | [illegible] | [illegible] | [illegible] | [illegible] | [illegible] | [illegible] | [illegible] | [illegible] | [illegible] | [illegible] | [illegible] | [illegible] | [illegible] | [illegible] | [illegible] | [illegible] | [illegible] | [illegible] | [illegible] | [illegible] | [illegible] | [illegible] | [illegible] | [illegible] | [illegible] | [illegible] |
| Palmitic acid_RT1 | IC-MS | 6.344278e-02 | [illegible] | [illegible] | [illegible] | [illegible] | [illegible] | [illegible] | [illegible] | [illegible] | [illegible] | [illegible] | [illegible] | [illegible] | [illegible] | [illegible] | [illegible] | [illegible] | [illegible] | [illegible] | [illegible] | [illegible] | [illegible] | [illegible] | [illegible] | [illegible] | [illegible] | [illegible] |
| Palmitic acid_RT2 | IC-MS | 3.551299e-02 | [illegible] | [illegible] | [illegible] | [illegible] | [illegible] | [illegible] | [illegible] | [illegible] | [illegible] | [illegible] | [illegible] | [illegible] | [illegible] | [illegible] | [illegible] | [illegible] | [illegible] | [illegible] | [illegible] | [illegible] | [illegible] | [illegible] | [illegible] | [illegible] | [illegible] | [illegible] |
| Pentadecanoic acid | IC-MS | 4.937272e-04 | [illegible] | [illegible] | [illegible] | [illegible] | [illegible] | [illegible] | [illegible] | [illegible] | [illegible] | [illegible] | [illegible] | [illegible] | [illegible] | [illegible] | [illegible] | [illegible] | [illegible] | [illegible] | [illegible] | [illegible] | [illegible] | [illegible] | [illegible] | [illegible] | [illegible] | [illegible] |
| Pentonic acid | IC-MS | 1.637112e-03 | [illegible] | [illegible] | [illegible] | [illegible] | [illegible] | [illegible] | [illegible] | [illegible] | [illegible] | [illegible] | [illegible] | [illegible] | [illegible] | [illegible] | [illegible] | [illegible] | [illegible] | [illegible] | [illegible] | [illegible] | [illegible] | [illegible] | [illegible] | [illegible] | [illegible] | [illegible] |
| phendimetrazine | IC-MS | 2.654624e-04 | [illegible] | [illegible] | [illegible] | [illegible] | [illegible] | [illegible] | [illegible] | [illegible] | [illegible] | [illegible] | [illegible] | [illegible] | [illegible] | [illegible] | [illegible] | [illegible] | [illegible] | [illegible] | [illegible] | [illegible] | [illegible] | [illegible] | [illegible] | [illegible] | [illegible] | [illegible] |
| Phosphoenolpyruvic acid | IC-MS | 8.431383e-04 | [illegible] | [illegible] | [illegible] | [illegible] | [illegible] | [illegible] | [illegible] | [illegible] | [illegible] | [illegible] | [illegible] | [illegible] | [illegible] | [illegible] | [illegible] | [illegible] | [illegible] | [illegible] | [illegible] | [illegible] | [illegible] | [illegible] | [illegible] | [illegible] | [illegible] | [illegible] |
| Phosphoric acid_RT1 | IC-MS | 3.778771e-02 | [illegible] | [illegible] | [illegible] | [illegible] | [illegible] | [illegible] | [illegible] | [illegible] | [illegible] | [illegible] | [illegible] | [illegible] | [illegible] | [illegible] | [illegible] | [illegible] | [illegible] | [illegible] | [illegible] | [illegible] | [illegible] | [illegible] | [illegible] | [illegible] | [illegible] | [illegible] |
| Phosphoric acid_RT2 | IC-MS | 2.977162e-04 | [illegible] | [illegible] | [illegible] | [illegible] | [illegible] | [illegible] | [illegible] | [illegible] | [illegible] | [illegible] | [illegible] | [illegible] | [illegible] | [illegible] | [illegible] | [illegible] | [illegible] | [illegible] | [illegible] | [illegible] | [illegible] | [illegible] | [illegible] | [illegible] | [illegible] | [illegible] |
| Platelet-activating factor | IC-MS | 3.779009e-04 | [illegible] | [illegible] | [illegible] | [illegible] | [illegible] | [illegible] | [illegible] | [illegible] | [illegible] | [illegible] | [illegible] | [illegible] | [illegible] | [illegible] | [illegible] | [illegible] | [illegible] | [illegible] | [illegible] | [illegible] | [illegible] | [illegible] | [illegible] | [illegible] | [illegible] | [illegible] |
| Propionylcarnitine | IC-MS | 2.652843e-04 | [illegible] | [illegible] | [illegible] | [illegible] | [illegible] | [illegible] | [illegible] | [illegible] | [illegible] | [illegible] | [illegible] | [illegible] | [illegible] | [illegible] | [illegible] | [illegible] | [illegible] | [illegible] | [illegible] | [illegible] | [illegible] | [illegible] | [illegible] | [illegible] | [illegible] | [illegible] |
| Pyrazolidine | IC-MS | 2.971578e-04 | [illegible] | [illegible] | [illegible] | [illegible] | [illegible] | [illegible] | [illegible] | [illegible] | [illegible] | [illegible] | [illegible] | [illegible] | [illegible] | [illegible] | [illegible] | [illegible] | [illegible] | [illegible] | [illegible] | [illegible] | [illegible] | [illegible] | [illegible] | [illegible] | [illegible] | [illegible] |
| Pyridoxine | IC-MS | 8.492463e-04 | [illegible] | [illegible] | [illegible] | [illegible] | [illegible] | [illegible] | [illegible] | [illegible] | [illegible] | [illegible] | [illegible] | [illegible] | [illegible] | [illegible] | [illegible] | [illegible] | [illegible] | [illegible] | [illegible] | [illegible] | [illegible] | [illegible] | [illegible] | [illegible] | [illegible] | [illegible] |
| Pyrogallol | IC-MS | 6.355518e-05 | [illegible] | [illegible] | [illegible] | [illegible] | [illegible] | [illegible] | [illegible] | [illegible] | [illegible] | [illegible] | [illegible] | [illegible] | [illegible] | [illegible] | [illegible] | [illegible] | [illegible] | [illegible] | [illegible] | [illegible] | [illegible] | [illegible] | [illegible] | [illegible] | [illegible] | [illegible] |
| Pyrophosphoric Acid | IC-MS | 4.352471e-03 | [illegible] | [illegible] | [illegible] | [illegible] | [illegible] | [illegible] | [illegible] | [illegible] | [illegible] | [illegible] | [illegible] | [illegible] | [illegible] | [illegible] | [illegible] | [illegible] | [illegible] | [illegible] | [illegible] | [illegible] | [illegible] | [illegible] | [illegible] | [illegible] | [illegible] | [illegible] |
| Pyruvic acid | IC-MS | 3.897336e-03 | [illegible] | [illegible] | [illegible] | [illegible] | [illegible] | [illegible] | [illegible] | [illegible] | [illegible] | [illegible] | [illegible] | [illegible] | [illegible] | [illegible] | [illegible] | [illegible] | [illegible] | [illegible] | [illegible] | [illegible] | [illegible] | [illegible] | [illegible] | [illegible] | [illegible] | [illegible] |
| S-Adenosylhomocysteine | IC-MS | 1.737445e-04 | [illegible] | [illegible] | [illegible] | [illegible] | [illegible] | [illegible] | [illegible] | [illegible] | [illegible] | [illegible] | [illegible] | [illegible] | [illegible] | [illegible] | [illegible] | [illegible] | [illegible] | [illegible] | [illegible] | [illegible] | [illegible] | [illegible] | [illegible] | [illegible] | [illegible] | [illegible] |
| Seryl-4-hydroxyproline | IC-MS | 1.634384e-05 | [illegible] | [illegible] | [illegible] | [illegible] | [illegible] | [illegible] | [illegible] | [illegible] | [illegible] | [illegible] | [illegible] | [illegible] | [illegible] | [illegible] | [illegible] | [illegible] | [illegible] | [illegible] | [illegible] | [illegible] | [illegible] | [illegible] | [illegible] | [illegible] | [illegible] | [illegible] |
| Stearic acid_RT1 | IC-MS | 3.442226e-01 | [illegible] | [illegible] | [illegible] | [illegible] | [illegible] | [illegible] | [illegible] | [illegible] | [illegible] | [illegible] | [illegible] | [illegible] | [illegible] | [illegible] | [illegible] | [illegible] | [illegible] | [illegible] | [illegible] | [illegible] | [illegible] | [illegible] | [illegible] | [illegible] | [illegible] | [illegible] |
| Stearic acid_RT2 | IC-MS | 5.137285e-02 | [illegible] | [illegible] | [illegible] | [illegible] | [illegible] | [illegible] | [illegible] | [illegible] | [illegible] | [illegible] | [illegible] | [illegible] | [illegible] | [illegible] | [illegible] | [illegible] | [illegible] | [illegible] | [illegible] | [illegible] | [illegible] | [illegible] | [illegible] | [illegible] | [illegible] | [illegible] |
| Styrene | IC-MS | 4.803443e-04 | [illegible] | [illegible] | [illegible] | [illegible] | [illegible] | [illegible] | [illegible] | [illegible] | [illegible] | [illegible] | [illegible] | [illegible] | [illegible] | [illegible] | [illegible] | [illegible] | [illegible] | [illegible] | [illegible] | [illegible] | [illegible] | [illegible] | [illegible] | [illegible] | [illegible] | [illegible] |
| Sulfuric acid_RT1 | IC-MS | 9.541475e-03 | [illegible] | [illegible] | [illegible] | [illegible] | [illegible] | [illegible] | [illegible] | [illegible] | [illegible] | [illegible] | [illegible] | [illegible] | [illegible] | [illegible] | [illegible] | [illegible] | [illegible] | [illegible] | [illegible] | [illegible] | [illegible] | [illegible] | [illegible] | [illegible] | [illegible] | [illegible] |
| Sulfuric acid_RT2 | IC-MS | 2.974037e-03 | [illegible] | [illegible] | [illegible] | [illegible] | [illegible] | [illegible] | [illegible] | [illegible] | [illegible] | [illegible] | [illegible] | [illegible] | [illegible] | [illegible] | [illegible] | [illegible] | [illegible] | [illegible] | [illegible] | [illegible] | [illegible] | [illegible] | [illegible] | [illegible] | [illegible] | [illegible] |
| Taurine | IC-MS | 1.417300e-03 | [illegible] | [illegible] | [illegible] | [illegible] | [illegible] | [illegible] | [illegible] | [illegible] | [illegible] | [illegible] | [illegible] | [illegible] | [illegible] | [illegible] | [illegible] | [illegible] | [illegible] | [illegible] | [illegible] | [illegible] | [illegible] | [illegible] | [illegible] | [illegible] | [illegible] | [illegible] |
| Thiosulfic acid | IC-MS | 2.829008e-04 | [illegible] | [illegible] | [illegible] | [illegible] | [illegible] | [illegible] | [illegible] | [illegible] | [illegible] | [illegible] | [illegible] | [illegible] | [illegible] | [illegible] | [illegible] | [illegible] | [illegible] | [illegible] | [illegible] | [illegible] | [illegible] | [illegible] | [illegible] | [illegible] | [illegible] | [illegible] |
| trans-10-Heptadecenoic acid | IC-MS | 8.620837e-04 | [illegible] | [illegible] | [illegible] | [illegible] | [illegible] | [illegible] | [illegible] | [illegible] | [illegible] | [illegible] | [illegible] | [illegible] | [illegible] | [illegible] | [illegible] | [illegible] | [illegible] | [illegible] | [illegible] | [illegible] | [illegible] | [illegible] | [illegible] | [illegible] | [illegible] | [illegible] |
| trans-3-Indoleacrylic acid | IC-MS | 1.444009e-04 | [illegible] | [illegible] | [illegible] | [illegible] | [illegible] | [illegible] | [illegible] | [illegible] | [illegible] | [illegible] | [illegible] | [illegible] | [illegible] | [illegible] | [illegible] | [illegible] | [illegible] | [illegible] | [illegible] | [illegible] | [illegible] | [illegible] | [illegible] | [illegible] | [illegible] | [illegible] |
| trans-Aconitic acid | IC-MS | 9.471110e-04 | [illegible] | [illegible] | [illegible] | [illegible] | [illegible] | [illegible] | [illegible] | [illegible] | [illegible] | [illegible] | [illegible] | [illegible] | [illegible] | [illegible] | [illegible] | [illegible] | [illegible] | [illegible] | [illegible] | [illegible] | [illegible] | [illegible] | [illegible] | [illegible] | [illegible] | [illegible] |
| UDP-N-acetylglucosamine | IC-MS | 1.614997e-04 | [illegible] | [illegible] | [illegible] | [illegible] | [illegible] | [illegible] | [illegible] | [illegible] | [illegible] | [illegible] | [illegible] | [illegible] | [illegible] | [illegible] | [illegible] | [illegible] | [illegible] | [illegible] | [illegible] | [illegible] | [illegible] | [illegible] | [illegible] | [illegible] | [illegible] | [illegible] |
| Uracil | IC-MS | 6.970307e-03 | [illegible] | [illegible] | [illegible] | [illegible] | [illegible] | [illegible] | [illegible] | [illegible] | [illegible] | [illegible] | [illegible] | [illegible] | [illegible] | [illegible] | [illegible] | [illegible] | [illegible] | [illegible] | [illegible] | [illegible] | [illegible] | [illegible] | [illegible] | [illegible] | [illegible] | [illegible] |
| Uric acid | IC-MS | 1.821198e-04 | [illegible] | [illegible] | [illegible] | [illegible] | [illegible] | [illegible] | [illegible] | [illegible] | [illegible] | [illegible] | [illegible] | [illegible] | [illegible] | [illegible] | [illegible] | [illegible] | [illegible] | [illegible] | [illegible] | [illegible] | [illegible] | [illegible] | [illegible] | [illegible] | [illegible] | [illegible] |
| vinylglycine | IC-MS | 7.573395e-04 | [illegible] | [illegible] | [illegible] | [illegible] | [illegible] | [illegible] | [illegible] | [illegible] | [illegible] | [illegible] | [illegible] | [illegible] | [illegible] | [illegible] | [illegible] | [illegible] | [illegible] | [illegible] | [illegible] | [illegible] | [illegible] | [illegible] | [illegible] | [illegible] | [illegible] | [illegible] |
| Xanthine | IC-MS | 5.316736e-04 | [illegible] | [illegible] | [illegible] | [illegible] | [illegible] | [illegible] | [illegible] | [illegible] | [illegible] | [illegible] | [illegible] | [illegible] | [illegible] | [illegible] | [illegible] | [illegible] | [illegible] | [illegible] | [illegible] | [illegible] | [illegible] | [illegible] | [illegible] | [illegible] | [illegible] | [illegible] |
| Ximenic acid | IC-MS | 5.549265e-04 | [illegible] | [illegible] | [illegible] | [illegible] | [illegible] | [illegible] | [illegible] | [illegible] | [illegible] | [illegible] | [illegible] | [illegible] | [illegible] | [illegible] | [illegible] | [illegible] | [illegible] | [illegible] | [illegible] | [illegible] | [illegible] | [illegible] | [illegible] | [illegible] | [illegible] | [illegible] |
| Y-Aminobutyric acid (GABA) | IC-MS | 6.709867e-04 | [illegible] | [illegible] | [illegible] | [illegible] | [illegible] | [illegible] | [illegible] | [illegible] | [illegible] | [illegible] | [illegible] | [illegible] | [illegible] | [illegible] | [illegible] | [illegible] | [illegible] | [illegible] | [illegible] | [illegible] | [illegible] | [illegible] | [illegible] | [illegible] | [illegible] | [illegible] |
| Beta-Alanine | IC-MS | 6.991005e-04 | [illegible] | [illegible] | [illegible] | [illegible] | [illegible] | [illegible] | [illegible] | [illegible] | [illegible] | [illegible] | [illegible] | [illegible] | [illegible] | [illegible] | [illegible] | [illegible] | [illegible] | [illegible] | [illegible] | [illegible] | [illegible] | [illegible] | [illegible] | [illegible] | [illegible] | [illegible] |

| Protein Identification | | TMT Quantitation | | | | | | | | | | | | | | | |
|---|---|---|---|---|---|---|---|---|---|---|---|---|---|---|---|---|---|
| Gene | Uniprot | DMSO_1 | DMSO_2 | DMSO_3 | DMSO_4 | IACS_1 | IACS_2 | IACS_3 | IACS_4 | KD025_1 | KD025_2 | KD025_3 | KD025_4 | Combo_1 | Combo_2 | Combo_3 | Combo_4 |
| ACTN4 | O43707 | 6.20954 | 6.12299 | 5.7296 | 5.84045 | 6.29036 | 6.07775 | 6.12107 | 5.99345 | 6.43828 | 5.90566 | 6.00305 | 6.03694 | 6.80369 | 6.76372 | 6.80971 | 6.85375 |
| PLEC | Q15149 | 6.2188 | 6.25297 | 5.90772 | 6.08305 | 6.13341 | 6.14663 | 6.12378 | 5.99908 | 6.10885 | 6.11178 | 6.14589 | 6.07038 | 6.63471 | 6.71241 | 6.70668 | 6.64386 |
| MACF1 | Q9UPN3 | 6.17538 | 6.28099 | 6.28471 | 6.08344 | 6.12238 | 6.27919 | 6.18053 | 6.3001 | 6.07643 | 6.18493 | 6.30074 | 6.14924 | 6.32211 | 6.4765 | 6.49728 | 6.28604 |
| HSP90AA1 | P07900-2 | 6.11965 | 6.20256 | 6.50696 | 5.89883 | 6.24661 | 6.58719 | 6.37366 | 6.56783 | 6.34564 | 6.09355 | 6.27979 | 6.32379 | 5.94805 | 6.30857 | 6.07036 | 6.12697 |
| FLNA | P21333-2 | 6.17369 | 6.24872 | 6.49961 | 6.06141 | 6.30643 | 6.40518 | 6.01369 | 6.4621 | 6.53137 | 6.41853 | 6.46469 | 6.48008 | 6.03823 | 6.20556 | 5.88237 | 5.80834 |
| VIM | P08670 | 6.33314 | 6.41839 | 5.75826 | 6.59792 | 6.1633 | 5.83302 | 6.69287 | 6.03125 | 6.37424 | 5.86739 | 5.91376 | 6.11599 | 6.42262 | 6.41094 | 6.36226 | 6.70465 |
| HSPA8 | P11142 | 5.9562 | 6.0328 | 6.37764 | 6.17614 | 6.167 | 6.3156 | 6.4116 | 6.23498 | 6.33773 | 5.9529 | 6.17352 | 6.28644 | 6.20974 | 6.50954 | 6.2594 | 6.59878 |
| LMNA | P02545 | 6.21947 | 6.18716 | 5.60743 | 6.23531 | 6.21461 | 6.0534 | 6.3744 | 5.98673 | 6.01254 | 5.89633 | 5.77349 | 5.71739 | 7.11893 | 6.87725 | 6.90563 | 6.81992 |
| HSPD1 | P10809 | 6.32505 | 6.25851 | 5.85364 | 6.53207 | 6.49876 | 6.08476 | 6.4324 | 6.22563 | 6.30549 | 5.95614 | 6.06367 | 6.04336 | 6.78167 | 6.0953 | 6.22076 | 6.32279 |
| DST | Q03001 | 6.22037 | 6.30533 | 6.44412 | 6.31057 | 6.12753 | 6.31235 | 6.24074 | 6.21168 | 6.29036 | 6.35965 | 6.35961 | 6.30604 | 6.07198 | 6.12975 | 6.22436 | 6.08556 |
| HNRNPM | P52272 | 6.24061 | 6.21939 | 5.84563 | 6.51415 | 6.24734 | 5.92159 | 6.40968 | 5.81254 | 6.18692 | 5.93917 | 5.91006 | 5.91179 | 6.84647 | 6.52614 | 6.80541 | 6.66309 |
| VCP | P55072 | 6.04258 | 6.1083 | 6.1312 | 6.12729 | 6.28201 | 6.25469 | 6.17282 | 6.19865 | 6.24551 | 6.12918 | 6.40372 | 6.39951 | 6.53967 | 6.39778 | 6.29597 | 6.27112 |
| TUBA1A | Q71U36 | 6.61156 | 6.50179 | 6.99077 | 6.00877 | 6.40165 | 6.91044 | 6.2018 | 6.84102 | 6.41004 | 6.40131 | 6.58853 | 6.61562 | 5.47846 | 5.54754 | 5.10871 | 5.382 |
| DYNC1H1 | Q14204 | 6.31724 | 6.14461 | 6.80585 | 6.10313 | 6.30618 | 6.55402 | 6.30372 | 6.47651 | 6.38904 | 6.33232 | 6.57838 | 6.56938 | 5.86157 | 6.00472 | 5.65903 | 5.65716 |
| CLTC | Q00610 | 6.18353 | 6.29504 | 6.38594 | 5.94547 | 6.27031 | 6.40247 | 6.27267 | 6.37706 | 6.30245 | 6.31379 | 6.30102 | 6.22676 | 6.16199 | 6.12048 | 6.3851 | 6.05593 |
| MYOF | Q9NZM1 | 6.32826 | 6.58598 | 6.21638 | 6.53752 | 6.07103 | 6.3279 | 6.4273 | 6.16124 | 6.09378 | 6.35291 | 6.20781 | 6.24582 | 5.90537 | 6.22031 | 6.13812 | 6.18026 |
| MYH9 | P35579 | 6.26727 | 6.26921 | 6.68484 | 6.02289 | 6.2643 | 6.48239 | 6.32898 | 6.4809 | 6.53296 | 6.28726 | 6.45947 | 6.42288 | 5.84309 | 6.12569 | 5.67829 | 5.84959 |
| LRPPRC | P42704 | 6.4307 | 6.51437 | 5.80903 | 6.61693 | 6.41209 | 6.019 | 6.53584 | 6.17031 | 6.12338 | 5.92415 | 5.93299 | 6.00084 | 6.44823 | 6.20768 | 6.52048 | 6.33399 |
| HSPA9 | P38646 | 6.41308 | 6.21456 | 5.87665 | 6.53073 | 6.48346 | 6.07186 | 6.26969 | 6.18383 | 6.30681 | 5.91941 | 6.13891 | 6.16615 | 6.94007 | 6.02086 | 6.23462 | 6.22932 |
| SYNE2 | Q8WXH0-2 | 6.44737 | 6.37679 | 5.7586 | 6.60682 | 6.19282 | 6.06826 | 6.33142 | 6.07515 | 6.0828 | 6.21246 | 5.93829 | 5.87029 | 6.75198 | 6.29385 | 6.64188 | 6.35124 |
| SMCHD1 | A6NHR9 | 6.38569 | 6.45516 | 5.84585 | 6.57573 | 6.25939 | 6.03438 | 6.41941 | 6.06467 | 6.11609 | 5.98847 | 5.7351 | 5.89042 | 6.60486 | 6.47192 | 6.54982 | 6.60303 |
| FLNB | O75369-8 | 6.14571 | 6.10591 | 6.19369 | 5.94173 | 6.26412 | 6.27956 | 6.17217 | 6.16987 | 6.34132 | 6.22541 | 6.41564 | 6.33956 | 6.37272 | 6.50979 | 6.28592 | 6.23687 |
| HSP90B1 | P14625 | 6.27412 | 6.3257 | 5.90873 | 6.24906 | 6.362 | 6.12817 | 6.40036 | 6.00048 | 6.07544 | 5.93089 | 5.91722 | 5.84546 | 6.76797 | 6.4695 | 6.76165 | 6.58325 |
| NUMA1 | Q14980 | 6.38732 | 6.50171 | 5.85302 | 6.60798 | 6.17749 | 6.0382 | 6.29515 | 5.97889 | 5.99805 | 6.0898 | 5.91592 | 5.90511 | 6.51787 | 6.55989 | 6.66378 | 6.50982 |
| UBR4 | Q5T4S7 | 6.25603 | 6.18603 | 6.83758 | 6.10445 | 6.32624 | 6.65503 | 6.30558 | 6.52373 | 6.36844 | 6.41252 | 6.50863 | 6.45662 | 5.71376 | 5.90966 | 5.71737 | 5.71833 |
| EEF1A2 | Q05639 | 6.43611 | 6.46934 | 7.00241 | 6.15156 | 6.39203 | 6.68663 | 6.32049 | 6.43371 | 6.55342 | 6.458 | 6.51393 | 6.51168 | 5.54358 | 5.60592 | 5.4795 | 5.44168 |
| NCL | P19338 | 5.89424 | 6.22999 | 5.71512 | 6.15033 | 6.17008 | 5.98707 | 6.29328 | 6.02137 | 6.10985 | 6.01151 | 5.92522 | 6.01738 | 6.76313 | 6.77981 | 6.92038 | 7.01124 |
| ACTN1 | P12814 | 6.20282 | 6.19519 | 5.91952 | 5.96778 | 6.26727 | 6.15313 | 6.22572 | 6.05891 | 6.62723 | 5.92474 | 6.03738 | 6.11816 | 6.44405 | 6.4259 | 6.62699 | 6.7052 |
| EEF1A1 | P68104 | 6.42774 | 6.08791 | 6.9708 | 6.18912 | 6.16802 | 6.45361 | 6.70888 | 6.17328 | 6.84351 | 6.07862 | 6.34429 | 6.4975 | 5.47085 | 5.96641 | 5.50527 | 6.11419 |
| PRPF8 | Q6P2Q9 | 6.26485 | 6.38642 | 5.93255 | 6.71294 | 6.18781 | 6.01972 | 6.34416 | 5.99366 | 6.15385 | 6.0917 | 5.93442 | 5.9163 | 6.48448 | 6.38985 | 6.52819 | 6.65911 |
| IQGAP1 | P46940 | 5.97683 | 6.1344 | 6.34675 | 5.69599 | 6.38617 | 6.38203 | 6.04477 | 6.12511 | 6.44031 | 6.36161 | 6.34075 | 6.29334 | 6.29189 | 6.6444 | 6.31394 | 6.22172 |
| PKM | P14618-2 | 6.1787 | 6.17552 | 6.78832 | 5.92029 | 6.32487 | 6.90998 | 6.11691 | 6.72921 | 6.38117 | 6.64952 | 6.44447 | 6.4672 | 5.71352 | 5.86955 | 5.63843 | 5.69234 |
| ILF3 | Q12906-7 | 6.24068 | 6.35258 | 5.9189 | 6.53628 | 6.28853 | 6.08123 | 6.33345 | 6.02061 | 6.31011 | 5.94472 | 5.95981 | 5.8996 | 6.61928 | 6.33699 | 6.64264 | 6.5146 |
| UTRN | P46939 | 6.29109 | 6.53936 | 6.08639 | 6.52732 | 6.21652 | 6.08531 | 6.19026 | 6.15245 | 6.18207 | 6.21486 | 6.15905 | 6.28755 | 6.43819 | 6.21006 | 6.19194 | 6.22759 |
| SNRNP200 | O75643 | 6.30938 | 6.2697 | 5.91061 | 6.57974 | 6.31337 | 6.07699 | 6.40565 | 5.90131 | 6.24088 | 5.95535 | 6.0613 | 5.91487 | 6.53336 | 6.28557 | 6.5489 | 6.69303 |
| HNRNPU | Q00839 | 6.09597 | 6.11917 | 5.61314 | 6.66032 | 5.97862 | 5.73861 | 6.31231 | 5.8813 | 6.25633 | 5.88863 | 5.87706 | 6.17253 | 6.58617 | 6.92924 | 6.92719 | 6.9634 |
| MYH10 | P35580-3 | 6.13104 | 6.13187 | 6.58198 | 5.83477 | 6.36709 | 6.41115 | 6.18153 | 6.32933 | 6.41716 | 6.33539 | 6.44276 | 6.1804 | 6.3074 | 6.29546 | 6.05578 | 5.99688 |
| XPO1 | O14980 | 5.9383 | 6.10879 | 5.9557 | 6.13238 | 6.02715 | 6.17751 | 6.19966 | 6.09393 | 6.18868 | 5.93304 | 6.01835 | 6.06864 | 6.56527 | 6.83752 | 6.76049 | 6.9946 |
| RANBP2 | P49792 | 6.30857 | 6.36417 | 5.87915 | 6.5392 | 6.28212 | 6.05435 | 6.30676 | 6.00679 | 6.15536 | 6.1581 | 5.97114 | 5.98196 | 6.56172 | 6.3405 | 6.45169 | 6.63843 |
| GOLGB1 | Q14789-2 | 6.4944 | 6.45761 | 5.99058 | 6.47167 | 6.24999 | 6.10722 | 6.38721 | 6.04533 | 6.12091 | 6.05245 | 6.13295 | 6.13247 | 6.56471 | 6.16543 | 6.34127 | 6.28579 |
| MAP4 | P27816 | 6.07998 | 6.0679 | 6.36621 | 5.77684 | 6.07057 | 6.37074 | 5.9525 | 6.57468 | 6.33351 | 6.46033 | 6.49288 | 6.51449 | 5.92422 | 6.76351 | 5.97285 | 6.27879 |
| HSPA4 | P34932 | 5.86059 | 5.95537 | 6.55848 | 5.81209 | 6.11212 | 6.42418 | 6.08531 | 6.13344 | 6.19472 | 6.13938 | 6.33286 | 6.2019 | 6.64017 | 6.53641 | 6.61375 | 6.39922 |
| ATP5F1A | P25705 | 6.38255 | 6.43005 | 5.91465 | 6.51749 | 6.3133 | 5.96446 | 6.49709 | 6.04154 | 6.23611 | 5.91013 | 5.98192 | 5.8888 | 6.7055 | 6.21903 | 6.62695 | 6.37044 |
| COPA | P53621-2 | 6.28005 | 6.42129 | 6.29135 | 6.062 | 6.31183 | 6.48627 | 6.29088 | 6.45846 | 6.26209 | 6.47275 | 6.38001 | 6.33689 | 5.93185 | 6.16847 | 5.92772 | 5.91808 |
| PRKDC | P78527 | 5.96878 | 6.0925 | 5.89515 | 6.27506 | 6.08437 | 6.01378 | 6.30435 | 5.95742 | 6.14481 | 6.22452 | 6.13841 | 6.0741 | 6.69595 | 6.59932 | 6.72965 | 6.80184 |
| SPTBN1 | Q01082 | 6.20388 | 6.31515 | 6.00783 | 6.01992 | 6.18402 | 6.27914 | 6.03671 | 6.17948 | 6.04903 | 6.04535 | 6.02818 | 5.9157 | 6.78552 | 6.54828 | 6.79394 | 6.60788 |
| ENO1 | P06733 | 6.12532 | 6.0871 | 6.58122 | 5.74589 | 6.26268 | 6.8741 | 6.39568 | 6.88699 | 6.42443 | 6.27271 | 6.332 | 6.39505 | 5.72409 | 6.21199 | 5.57596 | 5.84481 |
| NPM1 | P06748 | 6.36791 | 6.23968 | 5.77974 | 6.68855 | 6.43249 | 5.85273 | 6.46219 | 6.11125 | 6.81139 | 5.6477 | 6.06599 | 6.35392 | 6.33616 | 6.14428 | 6.14563 | 6.56041 |
| MDN1 | Q9NU22 | 6.47864 | 6.47899 | 6.16102 | 6.94684 | 6.10012 | 6.02277 | 6.34306 | 5.98467 | 6.18654 | 6.03465 | 6.24183 | 6.26041 | 6.11314 | 6.03849 | 6.26634 | 6.34249 |
| MDC1 | Q14676 | 6.25436 | 6.21915 | 5.75436 | 6.74747 | 6.05376 | 5.80151 | 6.31644 | 6.20061 | 5.91387 | 5.89731 | 5.85268 | 6.05359 | 6.32214 | 7.00472 | 6.70748 | 7.099 |
| GAPDH | P04406 | 6.14789 | 6.08262 | 6.71838 | 5.73778 | 6.4885 | 6.62712 | 6.4455 | 6.68402 | 6.47329 | 6.45351 | 6.62906 | 6.45728 | 5.61648 | 6.06204 | 5.74896 | 5.6276 |
| EIF4G1 | Q04637 | 6.34231 | 6.4363 | 6.63316 | 6.16174 | 6.2659 | 6.55596 | 6.33729 | 6.41661 | 6.64574 | 6.32514 | 6.5822 | 6.65883 | 5.57082 | 5.79876 | 5.46938 | 5.79984 |
| ATP5F1B | P06576 | 6.21185 | 6.18226 | 5.73367 | 6.50544 | 6.40837 | 5.9517 | 6.56065 | 5.98023 | 6.47181 | 5.8654 | 5.95769 | 6.04166 | 6.56022 | 6.26324 | 6.56041 | 6.7454 |
| FLNC | Q14315 | 6.25186 | 6.2243 | 6.57415 | 5.96981 | 6.21597 | 6.3338 | 6.05348 | 6.42181 | 6.55139 | 6.44522 | 6.45011 | 6.6446 | 5.98485 | 6.05282 | 5.87459 | 5.95123 |
| SYNE1 | Q8NF91-4 | 6.3779 | 6.29548 | 5.98126 | 6.50555 | 6.15667 | 6.02321 | 6.32264 | 5.9335 | 6.21362 | 6.14578 | 6.00819 | 6.05591 | 6.62979 | 6.24046 | 6.57006 | 6.54 |
| ATP2A2 | P16615 | 6.26378 | 6.40876 | 5.94914 | 6.76343 | 6.14294 | 5.94979 | 6.41444 | 5.83282 | 6.14424 | 5.94465 | 5.66742 | 5.81156 | 6.45129 | 6.44999 | 6.90276 | 6.90299 |
| LARS1 | Q9P2J5 | 6.26975 | 6.2471 | 6.53178 | 5.92925 | 6.26065 | 6.56669 | 6.28883 | 6.35991 | 6.28323 | 6.39964 | 6.48155 | 6.37084 | 6.10519 | 6.18748 | 5.93731 | 5.78079 |
| AHNAK2 | Q8IVF2 | 6.24534 | 6.37414 | 6.24756 | 6.0163 | 6.17411 | 6.34112 | 6.24064 | 6.30607 | 6.22358 | 6.2495 | 6.48908 | 6.265 | 6.21329 | 6.27461 | 6.23623 | 6.10345 |
| IGF2R | P11717 | 6.40579 | 6.54887 | 6.25271 | 6.40886 | 6.23505 | 6.28588 | 6.22955 | 6.0073 | 6.25682 | 6.49897 | 6.3641 | 6.28547 | 6.02068 | 6.03278 | 6.14487 | 6.0223 |
| HNRNPR | O43390-2 | 6.32559 | 6.43554 | 6.05505 | 6.52134 | 6.21367 | 6.11184 | 6.34803 | 6.08651 | 6.06288 | 6.08106 | 5.99616 | 6.09615 | 6.43875 | 6.27058 | 6.50594 | 6.45091 |
| EZR | P15311 | 6.14211 | 6.16203 | 6.65612 | 5.88709 | 6.29856 | 6.4704 | 6.38655 | 6.42132 | 6.52967 | 6.33947 | 6.37756 | 6.36625 | 5.96646 | 6.11576 | 6.00794 | 5.87271 |
| KTN1 | Q86UP2 | 6.31836 | 6.52302 | 6.06406 | 6.60885 | 6.03416 | 5.91581 | 6.27198 | 5.79197 | 6.10041 | 6.04509 | 5.95842 | 5.95001 | 6.48395 | 6.64559 | 6.77339 | 6.51493 |
| HNRNPA1 | P09651 | 6.49725 | 6.8803 | 5.68399 | 7.01016 | 6.42965 | 6.33435 | 6.19466 | 7.12705 | 6.14904 | 5.85536 | 5.78905 | 5.84831 | 5.99629 | 6.06078 | 6.19763 | 5.94613 |
| TPR | P12270 | 6.57829 | 6.47745 | 5.7038 | 6.53053 | 6.41666 | 6.18717 | 6.10433 | 6.27765 | 6.11368 | 6.04656 | 5.98474 | 6.15457 | 6.82227 | 6.0781 | 6.22499 | 6.2992 |
| UBA1 | P22314 | 5.87732 | 5.86134 | 6.44562 | 5.35324 | 6.10779 | 6.60261 | 5.86019 | 6.41014 | 6.05465 | 6.19647 | 6.25208 | 6.18469 | 6.60731 | 7.02194 | 6.60268 | 6.56193 |
| AKAP9 | Q99996-6 | 6.49591 | 6.42804 | 6.50619 | 6.2511 | 6.1206 | 6.5235 | 6.27973 | 6.29462 | 6.27843 | 6.6034 | 6.3686 | 6.41077 | 5.92951 | 5.9089 | 5.8831 | 5.71763 |
| DNMT1 | P26358-2 | 5.77748 | 6.32042 | 6.28895 | 6.27612 | 5.96809 | 6.26549 | 6.17415 | 6.07149 | 6.20703 | 6.31648 | 6.48959 | 6.44179 | 6.05662 | 6.51278 | 6.47028 | 6.36325 |
| LMNB1 | P20700 | 6.34229 | 6.422 | 5.70619 | 6.62731 | 6.32631 | 5.93855 | 6.39773 | 5.90162 | 6.09249 | 5.95332 | 5.91362 | 5.91746 | 6.74945 | 6.4633 | 6.64474 | 6.60359 |
| CAND1 | Q86VP6 | 5.70527 | 5.7551 | 6.1672 | 5.31771 | 6.06176 | 6.42893 | 5.94997 | 6.28149 | 6.0562 | 6.12535 | 6.22399 | 5.9902 | 6.98245 | 7.18266 | 7.02273 | 6.74899 |
| EPRS1 | P07814 | 6.10044 | 6.27696 | 6.2524 | 5.94041 | 6.16586 | 6.57428 | 6.00035 | 6.44002 | 6.1808 | 6.475 | 6.12689 | 6.32103 | 6.3351 | 6.62242 | 6.08599 | 6.10208 |
| PARP1 | P09874 | 5.56965 | 5.98002 | 5.85788 | 5.74407 | 5.88741 | 6.06305 | 5.97042 | 5.82535 | 6.0722 | 6.20351 | 6.26907 | 6.14028 | 6.93443 | 7.12782 | 7.3174 | 7.03744 |
| HNRNPK | P61978-2 | 6.20124 | 6.15518 | 5.86553 | 6.13801 | 6.31465 | 6.08839 | 6.09209 | 6.26053 | 6.26615 | 6.04568 | 6.16889 | 6.1396 | 7.04149 | 6.32417 | 6.55354 | 6.56184 |
| SPTAN1 | Q13813-3 | 6.27679 | 6.32807 | 6.11756 | 6.01776 | 6.29159 | 6.19734 | 5.91124 | 6.27278 | 6.13592 | 6.11921 | 6.16802 | 6.11943 | 6.9063 | 6.28867 | 6.54618 | 6.30315 |
| AHNAK | Q09666 | 6.34386 | 6.664 | 6.05637 | 6.43245 | 6.18487 | 6.05941 | 6.40316 | 6.21578 | 6.29126 | 5.9625 | 6.06551 | 6.14175 | 6.1757 | 6.39138 | 6.25038 | 6.36163 |
| AARS1 | P49588 | 6.39336 | 6.21437 | 6.75765 | 5.81434 | 6.46801 | 6.9176 | 6.22673 | 6.7709 | 6.45173 | 6.47417 | 6.48064 | 6.45681 | 5.83469 | 5.79912 | 5.45602 | 5.48388 |
| CNOT1 | A5YKK6 | 6.21146 | 6.27497 | 6.48617 | 6.0916 | 6.22307 | 6.44242 | 6.25878 | 6.43123 | 6.31192 | 6.42032 | 6.47472 | 6.43306 | 5.99206 | 6.05325 | 5.99661 | 5.89834 |
| SON | P18583-5 | 6.09288 | 6.06403 | 5.76597 | 6.57731 | 5.8587 | 5.75295 | 6.22874 | 5.74758 | 5.91245 | 5.84398 | 5.80491 | 5.81512 | 6.6231 | 7.2038 | 7.19794 | 7.51055 |
| KPNB1 | Q14974 | 6.16909 | 6.00157 | 6.61534 | 6.34223 | 6.32129 | 6.45529 | 6.35665 | 6.11734 | 6.477 | 6.01685 | 6.37704 | 6.53282 | 6.105 | 5.94195 | 5.91566 | 6.2549 |
| SF3B3 | Q15393 | 6.17741 | 6.33268 | 5.93173 | 6.45835 | 6.25073 | 6.09002 | 6.35262 | 5.96807 | 6.20313 | 6.07968 | 5.97207 | 5.99406 | 6.60369 | 6.28473 | 6.72551 | 6.57552 |
| DDB1 | Q16531 | 5.97011 | 6.19121 | 6.15003 | 6.02735 | 6.20772 | 6.1465 | 6.04448 | 6.11405 | 6.04134 | 6.08874 | 5.98492 | 6.02165 | 6.84047 | 6.63351 | 6.90969 | 6.62823 |
| ADAR | P55265-4 | 6.36069 | 6.40517 | 5.76652 | 6.57591 | 6.25538 | 6.03522 | 6.26769 | 6.04786 | 6.09284 | 6.00207 | 5.93181 | 5.90669 | 6.69865 | 6.51181 | 6.58871 | 6.55299 |
| DDX39B | Q13838-2 | 6.41546 | 6.38623 | 5.89485 | 6.53151 | 6.37642 | 6.11134 | 6.40237 | 5.92814 | 6.1982 | 6.06158 | 6.01415 | 5.93978 | 6.6039 | 6.22184 | 6.57441 | 6.33982 |
| H2BC18 | Q5QNW6-2 | 6.43911 | 6.55731 | 5.46733 | 6.61519 | 6.15506 | 5.8351 | 6.58715 | 6.04204 | 5.97032 | 5.58519 | 5.3698 | 5.60217 | 6.37236 | 7.22069 | 7.0635 | 7.11768 |
| VCL | P18206 | 6.25818 | 6.20523 | 6.83491 | 5.90297 | 6.34842 | 6.65648 | 6.49908 | 6.59785 | 6.52008 | 6.26738 | 6.37669 | 6.46034 | 5.84613 | 5.96148 | 5.57063 | 5.69416 |
| AFDN | P55196-5 | 6.42337 | 6.48978 | 6.16188 | 6.38491 | 6.30986 | 6.15259 | 6.34686 | 6.11527 | 6.33474 | 6.20561 | 5.90141 | 6.01664 | 6.36001 | 6.3644 | 6.28339 | 6.14928 |
| CAD | P27708 | 6.35709 | 6.28797 | 6.79284 | 5.99036 | 6.4698 | 6.89309 | 6.23743 | 6.6566 | 6.33022 | 6.32921 | 6.59636 | 6.68978 | 5.55766 | 5.87441 | 5.47628 | 5.4609 |
| CANX | P27824-2 | 6.20825 | 6.16771 | 5.96464 | 6.33579 | 6.1071 | 5.85895 | 6.33517 | 5.70211 | 6.08137 | 6.14081 | 6.03209 | 5.91335 | 6.98693 | 6.50842 | 6.70923 | 6.94807 |
| GANAB | Q14697-2 | 6.28532 | 6.27201 | 5.81458 | 6.47345 | 6.45624 | 6.28835 | 6.21781 | 6.13436 | 5.91779 | 5.74017 | 6.00706 | 5.8717 | 6.78984 | 6.49327 | 6.67357 | 6.56447 |
| ACTC1 | P68032 | 6.4941 | 6.16774 | 6.15382 | 5.76807 | 6.2146 | 5.84585 | 6.32566 | 6.06909 | 6.32491 | 5.73083 | 6.11286 | 6.32388 | 6.59424 | 6.52048 | 6.60349 | 6.75038 |
| EPB41L2 | O43491 | 6.12972 | 6.4802 | 5.79524 | 6.28287 | 6.38625 | 6.33486 | 6.27034 | 6.24218 | 6.10328 | 6.03374 | 5.87994 | 5.91696 | 6.49741 | 6.67612 | 6.47317 | 6.4977 |
| HNRNPA2B1 | P22626 | 6.35831 | 6.26763 | 5.77974 | 6.36392 | 6.35342 | 5.99173 | 6.15607 | 5.91596 | 6.17939 | 6.12656 | 6.18353 | 6.21934 | 6.93016 | 6.18098 | 6.48342 | 6.50984 |
| HDLBP | A0A024R4E5 | 6.6783 | 6.57619 | 6.53366 | 6.51079 | 6.44263 | 6.43718 | 6.33873 | 6.29481 | 6.64224 | 6.53348 | 6.49882 | 6.31503 | 5.71913 | 5.53699 | 5.55761 | 5.38444 |
| TP53BP1 | Q12888-2 | 6.28274 | 6.45853 | 5.86541 | 6.34863 | 6.28838 | 6.09891 | 6.26004 | 6.01418 | 5.85896 | 5.93737 | 5.8177 | 5.84563 | 6.92805 | 6.70134 | 6.66542 | 6.62872 |
| PRRC2C | Q9Y520-7 | 6.33489 | 6.48262 | 6.6486 | 6.43608 | 6.3731 | 6.47823 | 6.24164 | 6.63486 | 6.33315 | 6.33738 | 6.6057 | 6.48628 | 5.61277 | 5.81705 | 5.55084 | 5.62681 |
| CHD4 | A0A2R8Y521 | 6.30175 | 6.52786 | 6.02634 | 6.60416 | 6.12307 | 5.971 | 6.5348 | 5.94471 | 6.0136 | 6.08408 | 6.21542 | 6.06533 | 6.11325 | 6.59026 | 6.51608 | 6.36828 |
| HUWE1 | Q7Z6Z7 | 6.06616 | 6.28835 | 6.35273 | 6.01687 | 6.22154 | 6.4224 | 6.18573 | 6.28251 | 6.14302 | 6.18001 | 6.15396 | 6.18964 | 6.36017 | 6.36728 | 6.37328 | 6.39633 |
| EFTUD2 | Q15029 | 6.29193 | 6.22586 | 5.8321 | 6.53035 | 6.32265 | 6.06492 | 6.68105 | 6.17296 | 6.24928 | 5.7411 | 5.92957 | 6.01129 | 6.41352 | 6.31158 | 6.54278 | 6.67907 |
| EEF2 | P13639 | 6.25149 | 6.11903 | 6.9851 | 6.14204 | 6.4208 | 6.60889 | 6.33067 | 6.5212 | 6.78383 | 6.22147 | 6.62474 | 6.73064 | 5.49769 | 5.75791 | 5.38641 | 5.61808 |
| DDX5 | P17844 | 6.38852 | 6.3446 | 6.09898 | 6.67175 | 6.50345 | 6.2203 | 6.88423 | 6.10137 | 6.36489 | 5.95863 | 5.91103 | 5.9981 | 6.12757 | 5.95254 | 6.22318 | 6.25086 |
| RAD50 | Q92878-2 | 6.16131 | 6.12394 | 5.92387 | 6.44546 | 6.31381 | 6.0044 | 6.24729 | 5.84197 | 6.22273 | 6.02511 | 5.99222 | 5.85482 | 6.871 | 6.48661 | 6.79928 | 6.68619 |
| VDAC1 | P21796 | 5.6892 | 6.10586 | 5.49944 | 6.31466 | 6.39652 | 6.05662 | 6.54289 | 6.47139 | 6.41257 | 6.33378 | 6.24424 | 6.34623 | 6.1398 | 6.19505 | 6.71372 | 6.53803 |
| TCOF1 | Q13428-4 | 6.11104 | 6.2281 | 5.74037 | 6.54392 | 5.92987 | 5.77981 | 6.33604 | 5.7917 | 5.96982 | 5.84275 | 5.73827 | 5.79282 | 6.47592 | 7.27139 | 7.10053 | 7.34764 |
| SF3B2 | Q13435 | 6.07521 | 6.36198 | 5.81198 | 6.52073 | 6.13431 | 5.90476 | 6.40056 | 6.02738 | 6.07965 | 6.0619 | 5.89556 | 6.01027 | 6.64902 | 6.62842 | 6.68046 | 6.75784 |

| | | | | | | | | | | | | | | | | | |
|---|---|---|---|---|---|---|---|---|---|---|---|---|---|---|---|---|---|
| HSP90AB1 | P08238 | 6.18338 | 6.15397 | 6.66879 | 6.07753 | 6.27416 | 6.72271 | 6.32446 | 6.63182 | 6.36247 | 6.18142 | 6.65663 | 6.51974 | 5.85541 | 5.85951 | 5.70668 | 5.82133 |
| GARS1 | P41250 | 6.52497 | 6.28252 | 6.59278 | 6.0488 | 6.51765 | 6.67456 | 6.34181 | 6.60897 | 6.28719 | 6.0882 | 6.3759 | 6.37744 | 5.88732 | 5.96973 | 5.72395 | 5.6982 |
| CCT8 | P50990 | 6.26826 | 6.1219 | 6.53374 | 5.95618 | 6.47338 | 6.53484 | 6.32298 | 6.5014 | 6.39484 | 6.24045 | 6.41946 | 6.3235 | 6.05206 | 5.96631 | 6.02896 | 5.86174 |
| ANLN | Q9NQW6 | 5.66745 | 5.99098 | 6.11585 | 5.76672 | 6.08495 | 6.38464 | 6.37675 | 6.17831 | 6.19768 | 5.93798 | 6.3139 | 6.19654 | 6.55851 | 6.61509 | 6.79474 | 6.81991 |
| TOP2B | Q02880 | 6.38477 | 6.31623 | 5.85097 | 6.64695 | 6.12447 | 5.90496 | 6.38697 | 5.95638 | 6.06787 | 5.85887 | 5.68293 | 5.62473 | 6.46228 | 6.81342 | 7.04296 | 6.87523 |
| MATR3 | A8MXP9 | 6.42266 | 6.43349 | 6.02314 | 6.82375 | 6.26907 | 6.07509 | 6.58862 | 6.07656 | 6.17386 | 5.8646 | 6.01371 | 5.93368 | 6.42614 | 6.27603 | 6.29109 | 6.30851 |
| GPI | P06744 | 6.40971 | 6.15614 | 6.91734 | 5.79989 | 6.64567 | 6.90911 | 6.19566 | 6.27695 | 6.19655 | 6.37058 | 6.37179 | 6.38519 | 5.7187 | 6.34877 | 5.70255 | 5.59541 |
| YWHAE | P62258 | 6.08127 | 5.79234 | 6.61232 | 5.60878 | 6.29977 | 6.48443 | 6.26999 | 6.31781 | 6.45506 | 6.2341 | 6.38096 | 6.30792 | 6.29069 | 6.2455 | 6.39759 | 6.22147 |
| HNRNPL | P14866 | 6.28059 | 6.2333 | 5.92353 | 6.65777 | 6.37034 | 6.01414 | 6.17908 | 6.06432 | 6.13063 | 5.88315 | 6.09187 | 6.09422 | 6.91195 | 6.17456 | 6.48284 | 6.50771 |
| VARS1 | P26640 | 6.19788 | 6.24595 | 6.67233 | 5.9312 | 6.35452 | 6.33939 | 6.17237 | 6.363 | 6.59643 | 6.34359 | 6.37717 | 6.36379 | 6.01328 | 6.24311 | 5.89746 | 5.88852 |
| HNRNPC | P07910 | 6.12945 | 6.40646 | 5.96233 | 6.79846 | 6.32424 | 6.31823 | 6.62228 | 5.90832 | 6.15825 | 6.00324 | 5.87248 | 5.78221 | 6.52375 | 6.34668 | 6.50321 | 6.34042 |
| DDX17 | Q92841 | 6.25625 | 6.30714 | 5.84301 | 6.49278 | 6.29787 | 6.09177 | 6.44873 | 6.01374 | 6.07472 | 6.07718 | 6.08631 | 5.93661 | 6.83832 | 6.3055 | 6.5106 | 6.41946 |
| RRBP1 | Q9P2E9 | 6.27832 | 6.56018 | 5.78596 | 6.39435 | 6.10008 | 5.92003 | 6.13417 | 6.00505 | 6.02249 | 6.14502 | 5.85752 | 5.79835 | 6.21528 | 7.09073 | 6.75188 | 6.94058 |
| P4HB | P07237 | 6.58323 | 6.29759 | 5.91687 | 6.36192 | 6.5305 | 6.15299 | 6.10215 | 6.32794 | 6.20023 | 6.02394 | 6.09541 | 6.09624 | 6.8215 | 6.14402 | 6.17371 | 6.17175 |
| TPI1 | P60174-3 | 6.29374 | 6.04284 | 6.76179 | 5.65067 | 6.55627 | 6.83281 | 5.94707 | 6.6256 | 6.24728 | 6.4471 | 6.31651 | 6.27788 | 6.07497 | 6.26559 | 5.84478 | 5.81513 |
| KHSRP | Q92945 | 6.36252 | 5.70835 | 6.03451 | 5.47928 | 6.74836 | 5.70926 | 5.8348 | 6.49638 | 6.08242 | 6.11456 | 6.35973 | 6.6045 | 7.2255 | 6.31122 | 6.36388 | 6.56474 |
| HSPA5 | P11021 | 6.21656 | 6.2095 | 5.83182 | 6.15796 | 6.5017 | 6.0645 | 6.36636 | 6.1501 | 6.26606 | 5.96959 | 5.78915 | 5.82202 | 6.97448 | 6.51639 | 6.57186 | 6.59197 |
| RIF1 | Q5UIP0 | 6.51858 | 6.50507 | 6.05682 | 7.05956 | 6.14483 | 5.87639 | 6.25151 | 6.00221 | 6.17439 | 5.97172 | 5.78502 | 6.13553 | 6.30736 | 6.18752 | 6.43549 | 6.58801 |
| HADHA | P40939 | 6.33117 | 6.28678 | 5.78547 | 6.55548 | 6.26862 | 6.05315 | 6.41616 | 5.96861 | 6.23002 | 6.02674 | 5.95833 | 5.8698 | 6.64221 | 6.37173 | 6.73707 | 6.49865 |
| CCT7 | Q99832 | 6.19241 | 5.96446 | 6.68049 | 5.94705 | 6.4695 | 6.53425 | 6.54401 | 6.3393 | 6.58796 | 6.11867 | 6.37915 | 6.26967 | 5.98852 | 6.02459 | 5.95686 | 6.00312 |
| DHX9 | Q08211 | 6.31144 | 6.45659 | 5.94575 | 6.67174 | 6.16383 | 6.08628 | 6.30623 | 5.81378 | 6.14643 | 6.0479 | 6.07046 | 6.01893 | 6.57042 | 6.23753 | 6.63367 | 6.51903 |
| FASN | P49327 | 6.33486 | 6.16153 | 6.90536 | 6.01021 | 6.48296 | 6.7806 | 6.42596 | 6.69713 | 6.69296 | 6.38643 | 6.72634 | 6.58657 | 5.45612 | 5.56107 | 5.39969 | 5.39221 |
| GOLGA4 | Q13439-5 | 6.66455 | 6.66324 | 6.24684 | 6.47926 | 6.46996 | 6.17397 | 6.12549 | 6.0542 | 6.63532 | 6.36726 | 6.3752 | 6.41384 | 5.35643 | 5.66335 | 5.6157 | 5.69539 |
| SF3B1 | O75533 | 6.23436 | 6.3462 | 5.8083 | 6.47234 | 6.21987 | 5.9503 | 6.52149 | 5.93855 | 6.20791 | 5.83023 | 5.78736 | 5.80421 | 6.84338 | 6.53731 | 6.73199 | 6.76618 |
| NONO | Q15233 | 6.32943 | 6.36831 | 5.91968 | 6.56984 | 6.30407 | 6.04381 | 6.51203 | 6.03565 | 6.13564 | 5.98495 | 5.82235 | 5.98404 | 6.46527 | 6.35619 | 6.49532 | 6.67342 |
| PHB2 | Q99623 | 6.40964 | 6.44492 | 5.81288 | 6.54975 | 6.2785 | 5.98794 | 6.4682 | 5.94904 | 6.08035 | 6.05589 | 5.83636 | 5.90158 | 6.58276 | 6.41806 | 6.67955 | 6.54458 |
| MKI67 | P46013 | 6.47028 | 6.46798 | 5.95445 | 7.06607 | 6.03194 | 5.88346 | 6.35712 | 6.20258 | 6.10162 | 6.07876 | 6.11886 | 6.32835 | 5.76838 | 6.40057 | 6.27136 | 6.49823 |
| DDX3X | O00571 | 6.37132 | 6.51441 | 6.58926 | 6.5289 | 6.41754 | 6.55255 | 6.40727 | 6.4178 | 6.44717 | 6.53304 | 6.40843 | 6.42307 | 6.58604 | 5.55882 | 5.4976 | 5.6468 |
| XRCC5 | P13010 | 5.69337 | 6.00282 | 5.98646 | 5.90958 | 6.09845 | 6.02993 | 6.12732 | 5.82554 | 5.98305 | 5.85667 | 6.2452 | 6.02776 | 6.96252 | 7.02638 | 7.23422 | 6.99073 |
| YLPM1 | P49750 | 6.42562 | 6.29612 | 6.0078 | 6.65997 | 6.16515 | 5.96606 | 6.43531 | 6.01275 | 6.21541 | 6.01557 | 6.13036 | 6.07746 | 6.59341 | 6.27171 | 6.36229 | 6.36499 |
| HSPA1B | P0DMV9 | 5.8831 | 6.08182 | 6.25048 | 5.82786 | 6.08132 | 6.43115 | 6.13304 | 6.21 | 6.3104 | 6.03809 | 6.32234 | 6.22595 | 6.60789 | 6.58822 | 6.58187 | 6.42647 |
| TUBB3 | Q13509 | 6.42613 | 6.27977 | 6.88428 | 6.15749 | 6.48399 | 6.97718 | 6.29216 | 7.13267 | 6.42524 | 6.43752 | 6.66213 | 6.74705 | 5.33559 | 5.56152 | 4.92408 | 5.2732 |
| TOP1 | P11387 | 6.11403 | 5.98126 | 5.69491 | 6.31159 | 5.91692 | 5.67656 | 6.36608 | 5.79505 | 5.92505 | 5.81071 | 5.86033 | 5.69371 | 6.52919 | 7.42155 | 7.29224 | 7.61081 |
| RPS2 | P15880 | 6.42167 | 6.21279 | 6.50332 | 6.32979 | 6.32444 | 6.46136 | 6.19763 | 6.23081 | 6.41424 | 6.49044 | 6.58093 | 6.4291 | 5.82233 | 5.92067 | 5.76769 | 5.89278 |
| ACO2 | Q99798 | 6.42695 | 6.42846 | 5.93596 | 6.57423 | 6.36824 | 6.13697 | 6.26941 | 6.19895 | 6.09708 | 6.03166 | 6.17368 | 5.90182 | 6.71691 | 6.17984 | 6.3944 | 6.16544 |
| GLUD1 | P00367 | 6.27506 | 6.26459 | 5.64356 | 6.39993 | 6.47648 | 5.98376 | 6.43902 | 6.09188 | 6.35771 | 5.94222 | 5.91962 | 5.84472 | 6.84793 | 6.20832 | 6.83283 | 6.47236 |
| MYBBP1A | Q9BQG0-2 | 6.04968 | 5.94836 | 5.59511 | 6.65163 | 6.065 | 5.78725 | 6.45941 | 6.04465 | 6.25494 | 5.87816 | 6.01013 | 6.09192 | 6.59611 | 6.72657 | 6.74095 | 7.10014 |
| PABPC4 | Q13310-3 | 6.10153 | 6.52678 | 6.5851 | 6.21388 | 6.51958 | 6.62369 | 6.33877 | 6.59072 | 6.40451 | 6.3575 | 6.57617 | 6.55895 | 5.69173 | 5.92721 | 5.56268 | 5.62121 |
| RPS3 | P23396 | 6.58309 | 6.58382 | 6.56906 | 6.0829 | 6.57359 | 6.59313 | 6.33884 | 6.41494 | 6.56337 | 6.54325 | 6.3823 | 6.21331 | 5.80721 | 5.55202 | 5.75047 | 5.44868 |
| TJP1 | Q07157 | 6.43153 | 6.53185 | 5.99203 | 6.45708 | 6.30093 | 6.33383 | 6.01814 | 6.39122 | 6.23491 | 6.09261 | 6.09528 | 6.17824 | 6.27944 | 6.24991 | 6.27421 | 6.1388 |
| MCM4 | P33991 | 5.59995 | 6.26954 | 5.82562 | 5.82057 | 6.06159 | 6.41488 | 5.85426 | 6.27103 | 5.83965 | 6.1122 | 6.05481 | 5.99537 | 6.91496 | 7.14472 | 6.90036 | 6.9205 |
| EIF4A2 | Q14240-2 | 6.46133 | 6.59352 | 6.87609 | 6.39295 | 6.51091 | 6.81027 | 6.60711 | 6.55558 | 6.69324 | 6.57196 | 6.61611 | 6.64364 | 5.11737 | 5.37465 | 5.03422 | 5.14105 |
| TCP1 | P17987 | 6.16519 | 6.28188 | 6.44607 | 5.92529 | 6.528 | 6.81538 | 6.30023 | 6.65519 | 6.48313 | 6.12519 | 6.31642 | 6.19979 | 5.87934 | 6.08546 | 5.93001 | 5.86343 |
| SACS | Q9NZJ4 | 6.34984 | 6.21039 | 6.80778 | 6.31998 | 6.16912 | 6.47787 | 6.05942 | 6.2835 | 6.60944 | 6.52701 | 6.61948 | 6.69255 | 5.56957 | 5.92899 | 5.69759 | 5.68263 |
| USP7 | Q93009 | 5.98169 | 6.10331 | 5.85158 | 6.09941 | 6.1772 | 6.13393 | 6.21195 | 5.98222 | 6.07129 | 6.06058 | 5.95886 | 5.88268 | 6.91373 | 6.68432 | 7.04072 | 6.84654 |
| ACTB | A0A2R8Y793 | 6.42265 | 6.23267 | 6.2054 | 6.28686 | 6.74806 | 6.37883 | 6.11828 | 6.38174 | 6.36313 | 5.82006 | 5.85692 | 6.1758 | 6.13085 | 6.26776 | 6.34491 | 6.26608 |
| NT5E | P21589 | 6.12554 | 6.27622 | 5.75635 | 6.20065 | 6.24067 | 6.26216 | 6.17911 | 6.27011 | 6.02217 | 6.18498 | 6.05712 | 5.94161 | 6.64603 | 6.81541 | 6.61353 | 6.40834 |
| IMMT | Q16891-4 | 6.49096 | 6.29383 | 5.91393 | 6.51274 | 6.4352 | 6.01092 | 6.30322 | 6.15221 | 6.13674 | 5.96063 | 6.02135 | 5.93015 | 6.79123 | 6.2771 | 6.40354 | 6.36626 |
| NUP214 | P35658 | 6.3865 | 6.11066 | 6.08315 | 6.23887 | 6.24834 | 6.05504 | 6.10716 | 6.09827 | 6.35133 | 6.12119 | 6.0907 | 6.18746 | 6.63779 | 6.24548 | 6.39333 | 6.64472 |
| PDS5B | Q9NTI5 | 5.98357 | 6.06425 | 5.78059 | 6.27319 | 6.00564 | 5.91018 | 6.19122 | 5.84578 | 5.92779 | 5.95892 | 5.92459 | 5.76499 | 6.8806 | 7.29086 | 7.12395 | 7.07386 |
| CCT3 | P49368 | 6.22893 | 6.10318 | 6.66952 | 5.95083 | 6.46903 | 6.70685 | 6.32369 | 6.5747 | 6.27376 | 6.28646 | 6.37605 | 6.36302 | 6.0025 | 6.06177 | 5.81077 | 5.79896 |
| RNF213 | Q63HN8-4 | 6.1029 | 6.34143 | 6.54199 | 6.05884 | 6.14587 | 6.44089 | 6.16221 | 6.21036 | 6.30516 | 6.48183 | 6.37434 | 6.28343 | 6.02361 | 6.42673 | 6.0128 | 6.08762 |
| HNRNPUL1 | Q9BUJ2 | 6.08446 | 6.24201 | 5.56248 | 6.4303 | 6.17226 | 5.87977 | 6.68272 | 5.67836 | 6.16202 | 5.72976 | 5.59894 | 5.66435 | 6.64467 | 7.04323 | 6.99685 | 7.42782 |
| HSPH1 | Q92598 | 6.05358 | 6.15365 | 6.64731 | 5.86357 | 6.28517 | 6.56307 | 6.0881 | 6.27395 | 6.14707 | 6.29937 | 6.47733 | 6.34149 | 6.39419 | 6.1514 | 6.2205 | 6.04024 |
| ECPAS | Q5VYK3 | 6.2467 | 6.06277 | 6.47153 | 5.88672 | 6.28959 | 6.56627 | 6.27869 | 6.3942 | 6.23199 | 6.40584 | 6.52586 | 6.38871 | 6.22663 | 6.13147 | 5.98662 | 5.90641 |
| VDAC2 | P45880 | 5.9647 | 5.95198 | 5.67035 | 6.47999 | 6.39661 | 6.12061 | 6.40769 | 6.10455 | 6.356 | 6.18359 | 6.36332 | 6.19924 | 6.30186 | 6.09495 | 6.71128 | 6.69327 |
| ALDOA | P04075-2 | 6.34354 | 6.11585 | 6.68789 | 5.68631 | 6.65503 | 6.94558 | 6.00311 | 6.69792 | 6.50943 | 6.36666 | 6.52044 | 6.43577 | 5.98919 | 5.98196 | 5.48623 | 5.57509 |
| TRIM28 | Q13263 | 5.97744 | 6.29025 | 5.83318 | 6.33182 | 6.26384 | 6.11507 | 6.15563 | 6.08327 | 5.99462 | 6.11748 | 6.04347 | 6.02932 | 6.7964 | 6.65673 | 6.683 | 6.70948 |
| SMC1A | Q14683 | 6.256 | 6.51107 | 5.86368 | 6.38287 | 6.19711 | 6.16827 | 6.3989 | 6.17136 | 6.02887 | 5.9503 | 5.82701 | 5.77874 | 6.60701 | 6.54913 | 6.65546 | 6.65422 |
| DARS1 | P14868 | 6.19156 | 6.24329 | 6.53663 | 5.81297 | 6.47326 | 6.49024 | 6.18041 | 6.49207 | 6.22708 | 6.48071 | 6.3719 | 6.25147 | 6.13526 | 6.28364 | 6.06417 | 5.76534 |
| GCN1 | Q92616 | 6.42609 | 6.42833 | 6.66937 | 6.10737 | 6.33612 | 6.68799 | 6.23035 | 6.8245 | 6.47259 | 6.41136 | 6.62791 | 6.55109 | 5.54411 | 5.84341 | 5.43431 | 5.40511 |
| EIF4G2 | P78344 | 6.33816 | 6.49819 | 6.53734 | 6.30048 | 6.50356 | 6.66745 | 6.21403 | 6.53568 | 6.33965 | 6.36897 | 6.46501 | 6.38296 | 5.7272 | 5.84444 | 5.64295 | 5.63392 |
| TMPO | P42166 | 6.01026 | 6.21454 | 5.92929 | 6.69837 | 6.04496 | 6.17424 | 6.08615 | 6.07295 | 5.98049 | 5.96167 | 6.24829 | 6.1096 | 6.55795 | 6.46319 | 6.79428 | 6.45256 |
| CALR | P27797 | 6.30393 | 6.20611 | 6.07774 | 6.46048 | 6.35774 | 5.90356 | 6.08646 | 5.79917 | 5.9473 | 6.09372 | 6.05468 | 5.95647 | 7.0143 | 6.28631 | 6.57349 | 6.87854 |
| STIP1 | P31948-2 | 6.24698 | 6.26122 | 6.45844 | 5.71368 | 6.50571 | 6.66443 | 5.97828 | 6.67163 | 6.25335 | 6.35792 | 6.54778 | 6.39436 | 6.4731 | 5.99588 | 5.73819 | 5.73905 |
| CCAR2 | Q8N163 | 6.36523 | 6.16882 | 6.07026 | 6.47431 | 6.37022 | 5.96259 | 6.21407 | 6.04384 | 6.31032 | 5.90944 | 5.99118 | 6.00002 | 6.98339 | 6.25805 | 6.52127 | 6.35698 |
| SFPQ | P23246 | 6.30772 | 6.1187 | 5.97722 | 6.69852 | 6.17809 | 5.66411 | 6.67935 | 5.91311 | 6.48489 | 5.64992 | 5.95667 | 6.03405 | 6.50802 | 6.33594 | 6.78877 | 6.70493 |
| NASP | P49321 | 5.53318 | 5.79266 | 6.01147 | 5.25444 | 6.0239 | 6.32671 | 5.76409 | 6.1417 | 5.84248 | 6.03325 | 6.28165 | 6.19084 | 7.72086 | 7.05929 | 7.15816 | 6.86533 |
| TRAP1 | Q12931 | 6.32707 | 6.335 | 5.99241 | 6.69187 | 6.32444 | 5.99017 | 6.4564 | 5.96098 | 6.07573 | 5.89285 | 6.0212 | 5.95822 | 6.54171 | 6.30322 | 6.62882 | 6.49992 |
| SYNCRIP | O60506 | 6.18744 | 6.22612 | 6.10264 | 6.38459 | 6.27092 | 6.21933 | 6.31518 | 6.22201 | 6.38668 | 6.11084 | 6.19113 | 6.10533 | 6.16533 | 6.35816 | 6.39457 | 6.35971 |
| NUP98 | P52948 | 6.43466 | 6.29582 | 5.81803 | 6.54252 | 6.42615 | 6.14172 | 6.23357 | 5.9047 | 6.1316 | 6.1827 | 6.05985 | 5.98828 | 6.77535 | 6.30817 | 6.35397 | 6.40731 |
| XRCC6 | P12956 | 5.8498 | 6.11975 | 5.88678 | 5.84701 | 5.95124 | 6.08007 | 6.07199 | 5.86791 | 5.90432 | 6.07441 | 6.00937 | 5.92605 | 7.03846 | 7.07273 | 7.39379 | 6.90633 |
| PSMD2 | Q13200 | 6.17691 | 6.26667 | 6.37902 | 6.05094 | 6.30141 | 6.58775 | 6.27993 | 6.46165 | 6.23458 | 6.33825 | 6.34933 | 6.20888 | 6.14326 | 6.08423 | 6.04865 | 6.08853 |
| TLN1 | Q9Y490 | 6.37951 | 6.18675 | 6.71094 | 5.74296 | 6.4181 | 6.65622 | 6.25143 | 6.54978 | 6.36361 | 6.50036 | 6.44995 | 6.09824 | 5.92451 | 5.7026 | 5.60319 |
| NSF | P46459 | 6.24962 | 6.15183 | 5.91148 | 6.08895 | 6.16672 | 6.27233 | 6.22714 | 6.27105 | 6.22264 | 6.07904 | 6.14746 | 5.93173 | 6.58256 | 6.34957 | 6.53982 | 6.80807 |
| PDIA3 | P30101 | 6.45354 | 6.28599 | 5.88489 | 6.18987 | 6.48053 | 6.04859 | 6.29881 | 6.02458 | 6.07897 | 5.95104 | 6.08059 | 5.82318 | 7.10002 | 6.45143 | 6.5697 | 6.27827 |
| NUP93 | Q8N1F7 | 6.27759 | 6.22486 | 6.03115 | 6.49482 | 6.13055 | 6.08868 | 6.32433 | 6.16202 | 6.29248 | 6.13758 | 6.17294 | 6.05737 | 6.54762 | 6.09913 | 6.59973 | 6.35913 |
| MCM7 | P33993 | 5.63031 | 6.06056 | 6.10866 | 5.95755 | 6.02244 | 6.20761 | 6.12846 | 6.14175 | 6.09105 | 6.13372 | 6.22922 | 6.0501 | 6.61469 | 6.77665 | 6.90215 | 6.94509 |
| ACADVL | P49748-3 | 6.28327 | 6.38507 | 5.9201 | 6.52999 | 6.31945 | 6.08096 | 6.51846 | 6.1078 | 6.05834 | 5.99667 | 5.93892 | 5.95319 | 6.62867 | 6.35181 | 6.59612 | 6.33119 |
| RPN1 | P04843 | 6.34961 | 6.38615 | 5.85391 | 6.48916 | 6.27333 | 6.14335 | 6.29939 | 6.0361 | 6.16265 | 6.05671 | 5.89746 | 5.82006 | 6.60592 | 6.42365 | 6.70118 | 6.50139 |
| GLG1 | Q92896-2 | 6.58541 | 6.33875 | 5.92198 | 6.53602 | 6.54519 | 6.19056 | 6.46841 | 5.94067 | 6.37272 | 5.96392 | 5.88738 | 5.72652 | 6.49088 | 6.29041 | 6.37242 | 6.36876 |
| LDHB | P07195 | 6.38274 | 5.99268 | 7.00318 | 5.75525 | 6.49252 | 6.6228 | 6.41784 | 6.51088 | 6.48716 | 6.16229 | 6.4874 | 6.34193 | 5.81532 | 5.92316 | 5.80609 | 5.79877 |
| ZNF638 | Q14966 | 6.14793 | 6.33774 | 5.98226 | 6.91987 | 6.06798 | 5.99944 | 6.38773 | 6.0811 | 6.01572 | 6.00865 | 5.85526 | 5.82767 | 6.39692 | 6.51531 | 6.87798 | 6.57843 |
| LAP3 | P28838 | 6.45888 | 6.49596 | 6.17352 | 6.51408 | 6.66808 | 6.28613 | 6.36708 | 6.39412 | 6.29918 | 6.10415 | 6.23009 | 6.06263 | 6.2654 | 5.78673 | 6.04552 | 5.84847 |
| PA2G4 | Q9UQ80 | 5.99924 | 6.04847 | 6.55362 | 5.97814 | 6.28739 | 6.5904 | 6.38118 | 6.33357 | 6.62968 | 6.3381 | 6.67357 | 6.66119 | 5.67783 | 6.09321 | 5.85588 | 5.8985 |
| AP2B1 | P63010-2 | 6.28456 | 6.30494 | 6.37335 | 6.26222 | 6.34536 | 6.40276 | 6.36698 | 6.30388 | 6.30571 | 6.13346 | 6.28157 | 6.20007 | 6.117 | 6.18253 | 6.08621 | 6.04941 |
| AKAP12 | Q02952 | 6.63714 | 6.51003 | 6.49312 | 6.3354 | 6.32547 | 6.09123 | 5.97865 | 6.2165 | 6.2925 | 6.02322 | 6.38028 | 6.49041 | 6.55703 | 5.82606 | 5.92771 | 5.91525 |
| POLR2A | P24928 | 6.68142 | 6.7811 | 6.28837 | 7.02665 | 6.32471 | 6.23885 | 6.57284 | 6.3168 | 6.52804 | 6.3259 | 6.28436 | 6.14581 | 5.62689 | 5.58553 | 5.60721 | 5.66552 |
| IARS1 | P41252 | 6.30621 | 6.08379 | 6.42407 | 5.78105 | 6.5711 | 6.39154 | 6.17077 | 6.1494 | 6.39243 | 6.49228 | 6.47494 | 6.28028 | 6.30705 | 6.26372 | 5.95374 | 5.95764 |
| SMARCA1 | P28370 | 6.54216 | 6.59888 | 5.99223 | 6.77866 | 6.1863 | 6.08232 | 6.46072 | 6.00593 | 6.12648 | 6.03203 | 5.89734 | 5.99941 | 6.44518 | 6.1487 | 6.42901 | 6.27465 |
| MSN | P26038 | 6.05128 | 6.01075 | 6.42559 | 5.83094 | 6.33072 | 6.48526 | 6.21392 | 6.22585 | 6.59981 | 6.49942 | 6.45341 | 6.50872 | 6.00529 | 6.15865 | 6.13456 | 6.06584 |
| DDX46 | Q7L014 | 5.916 | 6.14878 | 5.92201 | 6.01247 | 6.15945 | 5.99105 | 6.19676 | 5.82661 | 6.18408 | 6.08101 | 5.92525 | 5.83243 | 6.76361 | 6.87229 | 7.15669 | 7.01151 |
| SRRM2 | Q9UQ35 | 6.266 | 6.22791 | 5.97974 | 6.5739 | 6.18304 | 5.99114 | 6.47694 | 5.80086 | 6.34433 | 5.82364 | 6.06063 | 5.99105 | 6.53698 | 6.5468 | 6.40522 | 6.79182 |
| MAP1B | P46821 | 5.93507 | 5.99692 | 6.37863 | 5.70503 | 6.07857 | 6.428 | 5.96109 | 6.29628 | 6.07904 | 6.49288 | 6.422 | 6.4251 | 5.95993 | 7.003 | 6.28212 | 6.55635 |
| DIDO1 | Q9BTC0 | 6.23828 | 6.32513 | 5.91241 | 6.74275 | 6.00324 | 6.0242 | 6.32103 | 5.79348 | 6.06627 | 6.1849 | 5.90478 | 6.02351 | 6.62463 | 6.53922 | 6.58571 | 6.71045 |
| PSMD1 | Q99460 | 6.17675 | 6.27608 | 6.32726 | 6.22276 | 6.32528 | 6.37493 | 6.12151 | 6.3451 | 6.11178 | 6.20342 | 6.26308 | 6.20491 | 6.37171 | 6.18898 | 6.23295 | 6.2535 |
| TRRAP | Q9Y4A5 | 6.52814 | 6.69869 | 6.10521 | 7.06899 | 6.19771 | 6.04261 | 6.48689 | 6.09378 | 6.55167 | 6.48965 | 6.30528 | 6.24976 | 5.88304 | 5.54571 | 5.89353 | 5.85933 |
| RACK1 | P63244 | 6.36441 | 6.5008 | 6.56349 | 6.18189 | 6.51787 | 6.44733 | 6.17539 | 6.54652 | 6.52813 | 6.60972 | 6.61928 | 6.57431 | 5.62852 | 5.59183 | 5.60678 | 5.54373 |
| NPEPPS | P55786 | 6.3442 | 6.08706 | 6.75799 | 5.72576 | 6.34531 | 6.60676 | 6.2385 | 6.56759 | 6.30629 | 6.52205 | 6.41278 | 6.44209 | 5.89173 | 6.1453 | 5.78935 | 5.81724 |
| SMC4 | Q9NTJ3 | 6.50376 | 6.41433 | 6.93341 | 6.43748 | 6.37401 | 6.52162 | 6.34445 | 6.22926 | 6.48418 | 6.56067 | 6.53309 | 6.47035 | 5.62987 | 5.54274 | 5.58362 | 5.43717 |
| H4C1 | P62805 | 6.21044 | 6.24154 | 5.36345 | 6.48863 | 6.16784 | 6.02917 | 6.52394 | 6.35497 | 5.96914 | 5.9009 | 5.57114 | 5.91772 | 6.78353 | 6.88812 | 6.48105 | 7.10844 |
| HERC2 | O95714 | 6.21732 | 6.24482 | 6.57823 | 6.34865 | 6.09751 | 6.3414 | 6.40886 | 6.32704 | 6.45563 | 6.60857 | 6.64086 | 6.67542 | 5.69983 | 5.88261 | 5.66524 | 5.808 |
| OPA1 | O60313-2 | 6.46724 | 6.36222 | 5.87346 | 6.62756 | 6.39179 | 5.99986 | 6.49354 | 6.06649 | 6.20581 | 5.95819 | 6.02417 | 5.87314 | 6.62866 | 6.145 | 6.44092 | 6.44195 |
| CCT2 | P78371 | 6.15133 | 6.1335 | 6.61085 | 6.05346 | 6.5033 | 6.81305 | 6.40209 | 6.64417 | 6.50352 | 6.0188 | 6.45551 | 6.43786 | 5.79554 | 5.96431 | 5.72312 | 5.7896 |
| ACIN1 | Q9UKV3-5 | 6.26468 | 6.19668 | 5.85673 | 6.62225 | 6.14976 | 5.87947 | 6.26853 | 6.10791 | 6.12003 | 6.01174 | 5.95704 | 6.06088 | 6.81947 | 6.485 | 6.47024 | 6.72958 |

| | | | | | | | | | | | | | | | | | |
|---|---|---|---|---|---|---|---|---|---|---|---|---|---|---|---|---|---|
| HEATR1 | Q9H583 | 6.29583 | 6.15205 | 5.76517 | 6.62592 | 6.34888 | 5.96482 | 6.52113 | 5.86647 | 6.25454 | 5.84871 | 5.94135 | 5.84334 | 6.62968 | 6.42046 | 6.63022 | 6.89143 |
| CKAP5 | Q14008 | 6.19109 | 6.32374 | 6.57248 | 6.20712 | 6.26589 | 6.50851 | 6.27521 | 6.3924 | 6.43335 | 6.65506 | 6.53755 | 6.50856 | 5.80695 | 5.88548 | 5.62767 | 5.80894 |
| OGDH | Q02218 | 6.3243 | 5.97638 | 5.89163 | 6.51623 | 6.37503 | 5.7708 | 6.88903 | 5.73875 | 6.46023 | 5.67106 | 5.71216 | 5.70261 | 6.53665 | 6.55299 | 6.77957 | 7.10258 |
| MDH2 | P40926 | 6.40195 | 6.37021 | 5.77348 | 6.39007 | 6.41853 | 6.08388 | 6.33843 | 6.20701 | 6.215 | 6.11292 | 6.0961 | 6.03776 | 6.77745 | 6.194 | 6.32877 | 6.25445 |
| XPO5 | Q9HAV4 | 5.93525 | 5.88917 | 6.07919 | 5.59856 | 5.99784 | 6.16697 | 5.96861 | 5.99437 | 6.41475 | 6.26983 | 6.27941 | 6.14847 | 6.75089 | 6.93091 | 6.82475 | 6.75105 |
| CCT6A | P40227 | 6.17705 | 6.26864 | 6.63006 | 6.02898 | 6.40091 | 6.61194 | 6.28146 | 6.48115 | 6.30538 | 6.25538 | 6.6154 | 6.48535 | 5.96533 | 5.87113 | 5.72804 | 5.89381 |
| TRIP11 | Q15643 | 6.58426 | 6.36395 | 6.22329 | 6.63573 | 6.29804 | 6.16822 | 6.27294 | 6.14511 | 6.42772 | 6.07312 | 6.49114 | 6.38498 | 6.65144 | 5.72816 | 5.85808 | 5.69381 |
| GTF2I | P78347 | 6.10058 | 6.27674 | 5.7745 | 6.602 | 6.05638 | 5.76023 | 6.70439 | 6.12928 | 6.07608 | 5.7655 | 5.77854 | 6.27846 | 6.33806 | 6.75935 | 6.51802 | 7.08189 |
| NSUN2 | Q08J23 | 5.31617 | 5.76386 | 6.03762 | 5.27412 | 5.7483 | 6.17683 | 5.75253 | 5.92532 | 5.91443 | 5.91687 | 6.12909 | 6.11236 | 7.2116 | 7.54411 | 7.69921 | 7.47758 |
| RPL4 | P36578 | 6.18396 | 6.17832 | 5.68209 | 5.83963 | 6.02772 | 6.21627 | 6.29121 | 6.63464 | 6.52649 | 6.13692 | 6.69112 | 6.8106 | 5.97156 | 6.92957 | 5.76441 | 6.11548 |
| CSE1L | P55060 | 5.64407 | 5.94423 | 6.15879 | 5.7728 | 6.04494 | 6.21648 | 5.93381 | 6.22504 | 6.34126 | 6.05147 | 6.18322 | 6.19749 | 6.36082 | 7.0504 | 6.77256 | 7.10262 |
| UBR5 | O95071 | 6.29847 | 6.35528 | 6.15399 | 6.38528 | 6.42083 | 6.23765 | 6.39543 | 6.14636 | 6.17628 | 6.03874 | 5.91891 | 6.04973 | 6.50788 | 6.23023 | 6.29942 | 6.38552 |
| BIRC6 | Q9NR09 | 6.34111 | 6.31856 | 6.97018 | 6.08195 | 6.31284 | 6.74678 | 6.17408 | 6.56299 | 6.389 | 6.65548 | 6.35589 | 6.43088 | 5.68098 | 5.78993 | 5.64753 | 5.74183 |
| PCNT | O95613 | 6.28481 | 6.39325 | 6.40441 | 6.3765 | 6.10409 | 6.29477 | 6.12477 | 6.15977 | 6.27619 | 6.38337 | 6.38012 | 6.27211 | 6.2007 | 6.18033 | 6.10591 | 6.0589 |
| HYOU1 | Q9Y4L1 | 6.22265 | 6.46196 | 5.97077 | 6.44991 | 6.38088 | 6.26926 | 6.38413 | 6.16411 | 6.01868 | 5.93309 | 6.03117 | 5.92428 | 6.61264 | 6.36702 | 6.48423 | 6.32521 |
| NOMO3 | P69849 | 6.45285 | 6.45659 | 5.96985 | 6.69099 | 6.26354 | 6.16619 | 6.47607 | 6.05847 | 6.15936 | 5.98701 | 5.93852 | 5.9202 | 6.33705 | 6.11515 | 6.58888 | 6.41928 |
| ACACA | Q13085-4 | 6.47266 | 6.4891 | 7.01843 | 6.26138 | 6.23193 | 6.4689 | 6.23689 | 6.58497 | 6.58694 | 6.57571 | 6.56394 | 6.60114 | 5.43834 | 5.58432 | 5.39424 | 5.49111 |
| PUF60 | Q9UHX1-2 | 6.27342 | 6.18319 | 6.06079 | 6.34306 | 6.38108 | 6.01174 | 6.39749 | 6.05553 | 6.35505 | 5.70133 | 5.82043 | 6.05014 | 6.45604 | 6.54567 | 6.59801 | 6.76704 |
| DDX1 | Q92499 | 5.74196 | 6.08052 | 6.11815 | 5.72102 | 5.93805 | 6.23097 | 6.08237 | 6.20477 | 6.08597 | 6.21988 | 6.41212 | 6.08844 | 6.80471 | 6.78119 | 6.7795 | 6.71037 |
| BPTF | Q12830 | 6.39615 | 6.47196 | 6.04353 | 6.89843 | 6.00839 | 5.99185 | 6.39152 | 5.79365 | 6.09974 | 6.20973 | 6.11708 | 6.1354 | 6.26473 | 6.53376 | 6.31685 | 6.32723 |
| EIF3I | Q13347 | 6.00858 | 5.96888 | 6.5438 | 6.01519 | 6.56177 | 6.74928 | 6.25046 | 6.89348 | 6.62533 | 6.86069 | 7.00849 | 7.07544 | 5.21037 | 5.4033 | 5.25041 | 5.57453 |
| CPSF1 | Q10570 | 6.4673 | 6.58878 | 6.23543 | 6.85744 | 6.14308 | 6.06335 | 6.26824 | 5.89434 | 5.99951 | 6.17936 | 6.08176 | 6.126 | 6.36297 | 6.07541 | 6.37147 | 6.28557 |
| LMNB2 | Q03252 | 6.40969 | 6.3653 | 5.92646 | 6.72154 | 6.18193 | 6.07322 | 6.33695 | 5.96907 | 6.5428 | 5.76837 | 5.76975 | 5.89288 | 6.6179 | 6.3394 | 6.53731 | 6.54744 |
| CHD3 | Q12873-3 | 6.3454 | 6.48486 | 5.98742 | 6.6992 | 6.31239 | 5.94033 | 6.39096 | 6.10734 | 6.12248 | 6.06579 | 5.91971 | 5.85505 | 6.24238 | 6.72634 | 6.4383 | 6.36205 |
| PHF3 | Q92576 | 6.29052 | 6.52419 | 5.96294 | 6.93713 | 6.04338 | 5.99059 | 6.40314 | 6.00742 | 6.15987 | 6.34408 | 5.98649 | 6.16685 | 6.0818 | 6.37685 | 6.36501 | 6.35973 |
| MTHFD1 | P11586 | 6.22979 | 6.0157 | 6.75182 | 5.70782 | 6.4807 | 6.74025 | 6.14891 | 6.75473 | 6.43535 | 6.4457 | 6.70577 | 6.70141 | 5.92433 | 5.84859 | 5.58067 | 5.52847 |
| PGK1 | P00558 | 6.0583 | 5.74905 | 6.49343 | 5.68305 | 6.47617 | 6.65952 | 6.46426 | 6.4963 | 6.57035 | 6.10501 | 6.31496 | 6.46855 | 5.97817 | 6.52303 | 5.7965 | 6.16333 |
| EIF3A | Q14152 | 6.35806 | 6.5886 | 6.73116 | 6.20223 | 6.31449 | 6.51702 | 6.3406 | 6.28166 | 6.42794 | 6.56995 | 6.54182 | 6.57325 | 5.55789 | 5.80696 | 5.54038 | 5.64798 |
| ACLY | P53396 | 5.93594 | 6.19691 | 6.69301 | 5.69569 | 6.1201 | 6.68168 | 6.00249 | 6.42034 | 6.29555 | 6.27525 | 6.48344 | 6.35145 | 6.0929 | 6.43648 | 6.19405 | 6.12472 |
| ASCC3 | Q8N3C0 | 6.32585 | 6.22349 | 6.50544 | 6.0623 | 6.39506 | 6.58709 | 6.16377 | 6.4203 | 6.23657 | 6.40524 | 6.31993 | 6.32315 | 6.01139 | 6.08381 | 5.89527 | 6.04134 |
| CCT5 | P48643 | 6.25453 | 6.09748 | 6.3921 | 6.0799 | 6.57424 | 6.68549 | 6.49829 | 6.47716 | 6.5754 | 6.92385 | 6.20261 | 6.26554 | 5.81803 | 6.13734 | 5.94221 | 6.07584 |
| CUL4B | Q13620 | 5.84939 | 6.24973 | 6.07509 | 6.04703 | 6.1149 | 6.45586 | 6.12767 | 6.06255 | 6.08799 | 6.30032 | 6.22198 | 6.11526 | 6.50317 | 6.59259 | 6.68809 | 6.50838 |
| MCM5 | P33992 | 5.6355 | 6.08704 | 5.82565 | 5.85587 | 5.90709 | 6.1221 | 6.10462 | 6.02396 | 5.96514 | 6.02744 | 5.96353 | 5.76655 | 7.04044 | 7.13086 | 7.51401 | 7.0302 |
| KMT2A | Q03164-3 | 6.40868 | 6.47257 | 5.98259 | 6.91383 | 6.18287 | 6.04987 | 6.49917 | 6.08018 | 6.27393 | 6.21183 | 6.07173 | 6.04167 | 6.90962 | 6.37921 | 6.3102 | 6.21205 |
| GPD2 | P43304 | 6.33903 | 6.17483 | 5.79105 | 6.44013 | 6.30015 | 5.98309 | 6.39994 | 6.20578 | 6.16311 | 5.97405 | 5.88072 | 5.69032 | 6.73409 | 6.40099 | 6.70213 | 6.82058 |
| PPFIBP1 | Q86W92 | 6.02049 | 6.01254 | 6.15893 | 5.89385 | 6.26924 | 6.14166 | 6.10198 | 6.00125 | 6.55945 | 6.18734 | 6.1484 | 6.01787 | 7.03354 | 6.69973 | 6.52402 | 6.22971 |
| KARS1 | Q15046-2 | 6.36462 | 6.6104 | 6.49892 | 6.28773 | 6.52055 | 6.60676 | 6.27885 | 6.64107 | 6.26539 | 6.20081 | 5.99783 | 6.2347 | 5.75987 | 6.12394 | 5.70226 | 5.90632 |
| PPIA | P62937 | 5.50948 | 5.70902 | 6.1675 | 5.56254 | 6.63245 | 6.40513 | 6.38985 | 7.03285 | 6.78502 | 6.39424 | 6.54463 | 6.87911 | 5.54157 | 6.19782 | 5.78553 | 6.46326 |
| POR | P16435 | 6.3529 | 6.51097 | 5.98463 | 6.65719 | 6.00463 | 6.06102 | 6.25162 | 5.87642 | 6.08214 | 5.99435 | 6.11304 | 5.97984 | 6.56505 | 6.36877 | 6.63527 | 6.56217 |
| MSH6 | P52701 | 5.3231 | 5.68566 | 5.84529 | 5.25587 | 5.73863 | 6.28866 | 5.91443 | 5.82585 | 5.97317 | 6.03205 | 6.07278 | 5.86171 | 7.32392 | 7.5739 | 7.79871 | 7.48628 |
| SND1 | Q7KZF4 | 6.37115 | 6.4208 | 6.15501 | 6.37744 | 6.34768 | 6.24652 | 6.27889 | 6.4553 | 6.27928 | 6.21885 | 6.31898 | 6.2102 | 6.04009 | 6.21403 | 5.99674 | 6.06904 |
| MARS1 | P56192 | 6.08612 | 6.07275 | 6.55376 | 5.84937 | 6.33322 | 6.41078 | 6.28734 | 6.25151 | 6.47235 | 6.42238 | 6.40532 | 6.27861 | 6.23813 | 6.21346 | 6.19038 | 5.93452 |
| NCAPD2 | Q15021 | 6.49385 | 6.66188 | 6.96051 | 6.65732 | 6.30282 | 6.54515 | 6.38691 | 6.73252 | 6.65227 | 6.70405 | 6.86598 | 6.91715 | 4.99754 | 5.21649 | 4.70089 | 5.20466 |
| SSB | P05455 | 5.20363 | 5.70096 | 5.79282 | 5.1978 | 5.52916 | 6.0567 | 5.81033 | 5.9036 | 5.73251 | 6.04689 | 5.92393 | 5.7546 | 7.46084 | 7.93432 | 8.24927 | 7.70416 |
| CLIP1 | P30622 | 6.38804 | 6.40517 | 7.00471 | 6.28658 | 6.62685 | 6.69015 | 6.11138 | 6.86168 | 6.53313 | 6.38597 | 6.51503 | 6.8229 | 5.65437 | 5.31708 | 5.16887 | 5.2281 |
| GBF1 | Q92538 | 6.0883 | 6.03086 | 6.27114 | 5.92644 | 6.24379 | 6.35282 | 6.07156 | 6.36519 | 6.26359 | 6.29878 | 6.25388 | 6.28888 | 6.24639 | 6.4361 | 6.36442 | 6.49786 |
| NBAS | A2RRP1 | 6.36877 | 6.41161 | 6.02598 | 6.4852 | 6.23933 | 6.06673 | 6.4644 | 5.88643 | 6.02767 | 6.13524 | 6.08991 | 5.90857 | 6.47517 | 6.2788 | 6.62873 | 6.50748 |
| GART | P22102 | 6.30733 | 6.30215 | 6.94813 | 6.19894 | 6.464 | 6.85516 | 6.27307 | 6.74442 | 6.34545 | 6.27007 | 6.42426 | 6.57507 | 5.56233 | 5.80096 | 5.3661 | 5.56254 |
| PDS5A | Q29RF7 | 6.061 | 6.00845 | 5.9662 | 6.34115 | 6.2721 | 6.15149 | 6.28241 | 5.91638 | 5.92514 | 6.12208 | 6.14796 | 6.04792 | 6.6531 | 6.73269 | 6.63393 | 6.738 |
| HCFC1 | P51610-4 | 6.26834 | 6.26009 | 5.79104 | 6.36659 | 6.48395 | 6.32807 | 6.09595 | 6.13034 | 5.99061 | 6.10818 | 5.87528 | 5.96686 | 6.95417 | 6.46139 | 6.4822 | 6.43692 |
| TUFM | P49411 | 6.28057 | 6.25924 | 5.72379 | 6.52521 | 6.38039 | 6.25014 | 6.29604 | 6.30098 | 6.11046 | 5.97608 | 5.87942 | 6.03615 | 6.57769 | 6.5279 | 6.42652 | 6.44941 |
| SEPTIN9 | Q9UHD8 | 6.13313 | 6.131 | 6.37881 | 5.93843 | 6.41429 | 6.53621 | 6.18829 | 6.38673 | 6.29521 | 6.47887 | 6.27889 | 6.21837 | 6.40118 | 6.25094 | 6.02098 | 5.94869 |
| TDRD12 | A0A2R8Y872 | 6.56314 | 6.47623 | 7.42659 | 6.59851 | 6.44184 | 6.81409 | 6.37931 | 6.61561 | 6.7155 | 6.8306 | 6.73415 | 6.99958 | 4.84884 | 4.96503 | 4.73237 | 4.85862 |
| NAT10 | Q9H0A0 | 6.3625 | 6.34055 | 5.90377 | 6.76924 | 6.07234 | 5.79424 | 6.48232 | 5.94867 | 6.13305 | 5.76219 | 5.97316 | 6.00209 | 6.44893 | 6.38756 | 6.91889 | 6.70051 |
| FRYL | O94915 | 6.14706 | 6.07549 | 6.63087 | 5.86585 | 6.23907 | 6.58901 | 6.21686 | 6.57893 | 6.40027 | 6.48277 | 6.44672 | 6.35181 | 5.99418 | 6.10395 | 5.90669 | 5.97046 |
| NAA15 | Q9BXJ9 | 6.10923 | 6.29873 | 6.83883 | 6.05207 | 6.30797 | 6.6263 | 6.20618 | 6.40054 | 6.52275 | 6.40812 | 6.68712 | 6.52594 | 5.7425 | 5.88569 | 5.62738 | 5.76064 |
| MTREX | P42285 | 6.30862 | 6.35351 | 5.93646 | 6.47125 | 6.42746 | 6.31428 | 6.37997 | 6.03043 | 6.10872 | 6.02047 | 6.00601 | 5.8109 | 6.59829 | 6.26583 | 6.48773 | 6.48007 |
| GCC2 | Q8IWJ2 | 6.48683 | 6.41453 | 6.2432 | 6.51479 | 6.53572 | 6.29218 | 6.14856 | 6.2262 | 6.52065 | 6.29358 | 6.36255 | 6.53974 | 6.31617 | 5.72985 | 5.71596 | 5.65948 |
| EIF3B | P55884-2 | 6.32682 | 6.32938 | 6.92977 | 6.25535 | 6.17786 | 6.38225 | 6.57646 | 6.41312 | 6.59159 | 6.20763 | 6.624 | 6.49388 | 5.52224 | 5.76089 | 5.63897 | 5.7698 |
| USP9X | Q93008-3 | 6.27436 | 6.23323 | 6.6738 | 5.94585 | 6.19085 | 6.72249 | 6.21372 | 6.40145 | 6.14977 | 6.41555 | 6.4306 | 6.49336 | 6.02571 | 6.09658 | 5.83157 | 5.90111 |
| PRDX1 | Q06830 | 5.83836 | 5.27326 | 6.45161 | 4.91062 | 6.42971 | 6.42119 | 6.04787 | 6.86438 | 6.04928 | 6.63291 | 6.46003 | 6.90147 | 6.4201 | 6.72258 | 6.19809 | 6.37854 |
| EIF4A3 | P38919 | 6.24693 | 6.35554 | 6.12422 | 6.4876 | 6.42066 | 6.16173 | 6.42457 | 6.12773 | 6.28346 | 6.22743 | 5.99665 | 6.00533 | 6.26738 | 6.36815 | 6.31344 | 6.18919 |
| ANXA6 | P08133 | 6.43124 | 6.1295 | 6.83259 | 5.82533 | 6.4444 | 6.79661 | 6.35049 | 6.83582 | 6.29819 | 6.45753 | 6.5242 | 6.41076 | 5.6899 | 5.85934 | 5.56288 | 5.55121 |
| RPS27A | P62979 | 6.57987 | 6.74269 | 6.31559 | 6.75575 | 6.29625 | 6.23094 | 6.29939 | 6.46421 | 6.04326 | 6.18238 | 6.13383 | 6.33264 | 5.39019 | 6.31038 | 5.705 | 6.21764 |
| H3C1 | P68431 | 6.31942 | 6.24515 | 5.55567 | 6.38899 | 6.09061 | 5.86646 | 6.55582 | 6.30163 | 5.86696 | 5.90398 | 5.71077 | 5.93886 | 6.62966 | 6.84603 | 6.80691 | 6.97308 |
| SVIL | O95425 | 6.13459 | 6.40126 | 5.79627 | 6.10464 | 6.35377 | 6.22628 | 6.28945 | 6.03643 | 5.9652 | 6.12581 | 5.6258 | 5.76204 | 6.85846 | 6.78787 | 6.87677 | 6.65535 |
| QARS1 | P47897 | 6.34218 | 6.16816 | 6.49092 | 5.92121 | 6.56879 | 6.41975 | 5.9875 | 6.36742 | 6.50891 | 6.43026 | 6.46359 | 6.42025 | 6.16623 | 6.1367 | 5.95227 | 5.65587 |
| TPP2 | P29144 | 6.38478 | 6.47219 | 6.25466 | 6.58327 | 6.42681 | 6.39905 | 6.37713 | 6.23151 | 6.18044 | 6.02817 | 5.97024 | 6.13392 | 6.21995 | 6.12592 | 6.15686 | 6.05511 |
| RPS3A | P61247 | 6.20127 | 6.55457 | 6.67109 | 6.33171 | 6.26404 | 6.31067 | 6.37724 | 6.1475 | 6.40242 | 6.58963 | 6.54592 | 6.37302 | 5.68132 | 5.96166 | 5.78129 | 5.80664 |
| TARS1 | P26639 | 6.30425 | 6.10036 | 6.9388 | 5.95119 | 6.3611 | 6.66544 | 6.45316 | 6.54422 | 6.47819 | 6.36565 | 6.49058 | 6.50119 | 5.76791 | 5.74626 | 5.68562 | 5.64607 |
| THOC2 | Q8NI27 | 6.13507 | 6.14878 | 5.83839 | 6.52205 | 6.06549 | 5.89633 | 6.33616 | 5.72048 | 6.09964 | 5.98034 | 5.87601 | 5.77445 | 6.58262 | 7.04615 | 6.87578 | 7.10225 |
| CCT4 | P50991 | 6.29756 | 6.13464 | 6.48253 | 5.97499 | 6.42996 | 6.69656 | 6.14039 | 6.60308 | 6.38295 | 6.4012 | 6.2578 | 6.40727 | 6.09902 | 6.025 | 5.71351 | 5.95355 |
| GTF3C1 | Q12789 | 6.41815 | 6.56731 | 6.12408 | 6.91681 | 6.25398 | 6.07421 | 6.40826 | 6.09856 | 6.20501 | 6.13719 | 6.18764 | 6.02271 | 6.06848 | 6.11386 | 6.26394 | 6.13982 |
| AHCTF1 | Q8WYP5-2 | 6.49735 | 6.45269 | 5.72701 | 6.71253 | 6.26633 | 5.79198 | 6.46307 | 6.06084 | 6.02297 | 5.91266 | 5.75349 | 5.82683 | 6.46583 | 6.66156 | 6.87239 | 6.53827 |
| DICER1 | Q9UPY3 | 6.41502 | 6.36906 | 6.72105 | 6.03262 | 6.31089 | 6.61308 | 6.08371 | 6.40783 | 6.61136 | 6.51579 | 6.61358 | 6.42131 | 5.76425 | 5.82716 | 5.57636 | 5.71693 |
| BOD1L1 | Q8NFC6 | 6.24868 | 6.35286 | 6.07283 | 6.58302 | 6.03309 | 5.96542 | 6.11663 | 5.8636 | 6.11014 | 6.14888 | 6.06642 | 6.01794 | 6.12704 | 6.94757 | 6.54656 | 6.79933 |
| CS | O75390 | 6.41267 | 6.08978 | 6.01189 | 6.4031 | 6.34361 | 6.01407 | 6.4522 | 6.18472 | 6.11357 | 6.03507 | 6.12247 | 5.96903 | 6.66833 | 6.24307 | 6.44627 | 6.49015 |
| ATM | Q13315 | 5.79535 | 6.14537 | 6.0859 | 5.78124 | 6.05432 | 6.25379 | 6.02511 | 5.86607 | 6.10554 | 6.43011 | 6.24469 | 6.03351 | 6.87768 | 6.74296 | 6.9356 | 6.62275 |
| DDX21 | Q9NR30 | 6.39056 | 6.37914 | 6.02718 | 6.98749 | 6.22163 | 5.94163 | 6.41697 | 6.04537 | 6.19785 | 5.78852 | 5.9604 | 5.89777 | 6.0664 | 6.35738 | 6.70634 | 6.61538 |
| SLC25A5 | P05141 | 6.57924 | 6.58296 | 5.86051 | 6.69746 | 6.29232 | 5.89222 | 6.34334 | 6.00275 | 6.19503 | 6.01426 | 5.93031 | 5.96552 | 6.49608 | 6.25744 | 6.51892 | 6.37166 |
| PI4KA | P42356 | 6.38398 | 6.64378 | 6.46109 | 6.48636 | 6.17571 | 6.32038 | 6.32047 | 6.00223 | 6.16149 | 6.35016 | 6.25871 | 6.32857 | 6.03084 | 5.93002 | 6.1546 | 5.9916 |
| IPO7 | O95373 | 6.27396 | 5.98132 | 6.21513 | 6.41409 | 6.17413 | 6.44535 | 6.16359 | 5.92772 | 6.54874 | 6.29018 | 6.38406 | 6.45403 | 5.99196 | 6.14622 | 6.03787 | 6.55165 |
| UTP20 | O75691 | 6.30867 | 6.31625 | 5.81142 | 6.60185 | 6.05843 | 5.82695 | 6.46753 | 6.1134 | 6.17131 | 5.96143 | 5.98884 | 6.04054 | 6.4041 | 6.42204 | 6.54417 | 6.96306 |
| MTHFD1L | Q6UB35 | 6.42668 | 6.37202 | 6.05167 | 6.6462 | 6.31369 | 5.91296 | 6.42951 | 6.05339 | 6.23205 | 6.01877 | 6.06824 | 6.14534 | 6.57573 | 6.06399 | 6.30721 | 6.38256 |
| DCTN1 | Q14203-4 | 6.30663 | 6.23591 | 6.74792 | 6.02851 | 6.46367 | 6.64376 | 6.10459 | 6.67904 | 6.46228 | 6.5619 | 6.48504 | 6.5933 | 5.80061 | 5.72825 | 5.55765 | 5.60094 |
| CENPF | P49454 | 6.24943 | 6.06309 | 5.86915 | 6.19672 | 6.27736 | 6.07603 | 6.43832 | 6.05501 | 6.17342 | 5.93831 | 6.12882 | 6.0272 | 6.93157 | 6.28643 | 6.62132 | 6.66781 |
| BAZ1B | Q9UIG0 | 6.4892 | 6.44625 | 5.85912 | 6.99454 | 6.10866 | 5.80716 | 6.35757 | 6.13113 | 6.09394 | 5.97501 | 6.04554 | 5.96167 | 6.44193 | 6.25991 | 6.59771 | 6.43063 |
| RPL3 | P39023 | 6.43029 | 6.33315 | 6.34625 | 6.24105 | 6.18092 | 6.29369 | 6.26668 | 6.50915 | 6.63143 | 6.34654 | 6.48091 | 6.24966 | 5.79376 | 6.18612 | 5.78503 | 5.92537 |
| TOP2A | P11388-4 | 6.48421 | 6.65377 | 6.01187 | 7.00697 | 6.11432 | 5.99923 | 6.45386 | 6.06981 | 6.34594 | 6.34244 | 6.24537 | 6.37087 | 5.7373 | 5.98095 | 6.00987 | 6.17321 |
| SMC2 | O95347 | 6.22324 | 6.36516 | 6.52372 | 6.13433 | 6.51061 | 6.57599 | 6.19471 | 6.51799 | 6.38458 | 6.36383 | 6.31219 | 6.19147 | 5.90778 | 6.01216 | 5.81941 | 5.96282 |
| MVP | Q14764 | 6.11208 | 6.13685 | 6.49125 | 5.647 | 6.33624 | 6.69113 | 6.16888 | 6.75177 | 6.36906 | 6.57314 | 6.58942 | 6.4926 | 5.98476 | 6.32846 | 5.73537 | 5.59199 |
| NUP205 | Q92621 | 6.31877 | 6.16896 | 6.0307 | 6.64022 | 6.18292 | 5.99146 | 6.48089 | 6.06391 | 6.13582 | 5.87567 | 6.00503 | 5.90052 | 6.56325 | 6.29114 | 6.67944 | 6.6713 |
| SF3A1 | Q15459 | 6.38422 | 6.3204 | 5.77982 | 6.53278 | 6.4366 | 6.0915 | 6.43188 | 6.14924 | 6.1796 | 5.78751 | 5.87409 | 5.99627 | 6.8556 | 6.29859 | 6.38951 | 6.49239 |
| ERBIN | Q96RT1-9 | 6.4965 | 6.41386 | 6.30439 | 6.64582 | 6.31367 | 6.00695 | 6.39632 | 6.06805 | 6.16714 | 6.20136 | 6.32301 | 6.23329 | 6.27121 | 5.84055 | 5.99038 | 6.3275 |
| PRMT1 | Q99873 | 5.66557 | 6.0918 | 5.94152 | 5.88985 | 6.12583 | 6.28652 | 6.1076 | 6.05198 | 5.79947 | 6.24592 | 5.99982 | 5.81097 | 7.02069 | 6.81034 | 7.36751 | 6.7846 |
| RAN | P62826 | 6.02364 | 6.08309 | 6.37834 | 6.17145 | 6.04547 | 6.42492 | 6.00498 | 6.085 | 6.24135 | 6.27666 | 6.37103 | 6.30678 | 6.41201 | 6.34154 | 6.5264 | 6.30733 |
| AP2A1 | O95782 | 6.11566 | 6.27721 | 6.11913 | 6.44447 | 6.08385 | 6.17069 | 6.24772 | 6.06562 | 6.24851 | 6.32033 | 6.25678 | 6.29957 | 6.34488 | 6.30019 | 6.35817 | 6.34722 |
| VPS13C | Q709C8 | 6.5306 | 6.24059 | 6.79142 | 6.29964 | 6.232 | 6.39292 | 6.31394 | 6.40433 | 6.57127 | 6.52079 | 6.33704 | 6.37144 | 5.69159 | 5.845 | 5.55762 | 5.89981 |
| H1-5 | P16401 | 6.18661 | 6.16349 | 5.34936 | 6.26325 | 6.03959 | 5.8409 | 6.27933 | 6.05932 | 5.78217 | 5.87606 | 5.64058 | 5.74239 | 6.67983 | 7.60328 | 7.07974 | 7.41408 |
| NIPBL | Q6KC79 | 6.16662 | 6.42327 | 5.85788 | 6.82312 | 5.9746 | 6.04486 | 6.20834 | 5.92231 | 5.99866 | 5.98429 | 5.90842 | 5.92572 | 6.4088 | 6.65253 | 6.86495 | 6.83562 |
| DDX23 | Q9BUQ8 | 6.19688 | 6.3283 | 5.81496 | 6.45951 | 6.06557 | 5.95703 | 6.21953 | 5.93088 | 5.91451 | 6.14832 | 5.8722 | 5.95869 | 6.59012 | 6.7435 | 7.00722 | 6.79277 |
| SMC3 | Q9UQE7 | 6.16235 | 6.40479 | 5.67452 | 6.47133 | 6.20178 | 6.0653 | 6.45109 | 6.1196 | 6.25746 | 5.87748 | 5.57053 | 5.88086 | 6.50834 | 6.631 | 6.74541 | 6.97816 |
| AP3D1 | O14617-5 | 6.12685 | 6.23001 | 6.6977 | 6.12914 | 5.92571 | 6.34289 | 6.02265 | 6.36366 | 6.19509 | 6.44053 | 6.13128 | 6.35434 | 5.70226 | 6.67895 | 6.28456 | 6.37439 |
| IPO5 | O00410-3 | 6.35006 | 5.95582 | 6.85196 | 6.04596 | 6.2162 | 6.50611 | 6.27645 | 6.41829 | 6.55841 | 6.20668 | 6.5584 | 6.5827 | 5.73376 | 5.96276 | 5.77319 | 6.00325 |
| MCM3 | P25205-2 | 5.58071 | 6.26489 | 5.99259 | 6.10577 | 6.18249 | 6.47234 | 6.18302 | 5.97859 | 5.7778 | 5.82294 | 5.99414 | 5.76531 | 7.05494 | 7.21492 | 6.92867 | 6.68088 |

| | | | | | | | | | | | | | | | | | |
|---|---|---|---|---|---|---|---|---|---|---|---|---|---|---|---|---|---|
| KNTC1 | P50748 | 6.48831 | 6.21996 | 6.87211 | 6.06035 | 6.31469 | 6.64778 | 6.41135 | 6.59749 | 6.48336 | 6.53046 | 6.43034 | 6.42343 | 5.5641 | 5.77861 | 5.54944 | 5.62822 |
| SUPT6H | Q7KZ85 | 6.2369 | 6.28475 | 6.15786 | 6.85163 | 6.34884 | 6.11083 | 6.43385 | 6.10635 | 6.27462 | 6.18698 | 6.25013 | 6.11075 | 6.25639 | 5.87169 | 6.21579 | 6.30263 |
| MIA3 | Q5JRA6 | 6.46185 | 6.35383 | 6.11902 | 6.43484 | 6.18822 | 5.91864 | 6.46604 | 6.0521 | 6.13156 | 5.86331 | 6.21269 | 6.11149 | 6.77824 | 6.15927 | 6.31499 | 6.43392 |
| MGA | Q8IWI9-3 | 6.28729 | 6.71531 | 6.15171 | 6.86764 | 6.2164 | 6.18512 | 6.43966 | 6.15474 | 6.14173 | 6.08587 | 5.9999 | 6.19389 | 5.89656 | 6.13037 | 6.08471 | 6.4491 |
| ILF2 | Q12905 | 6.35335 | 6.23778 | 5.92941 | 6.42919 | 6.27342 | 5.98082 | 6.20117 | 5.97736 | 6.2524 | 6.08997 | 6.00996 | 6.06546 | 6.73425 | 6.2827 | 6.60683 | 6.57593 |
| LARP1 | Q6PKG0 | 6.4524 | 6.5282 | 6.79496 | 6.21687 | 6.49264 | 6.57304 | 6.26108 | 6.33327 | 6.43225 | 6.34636 | 6.357 | 6.14867 | 5.73536 | 6.02453 | 5.65358 | 6.54979 |
| ITGB1 | P05556 | 6.37721 | 6.53253 | 5.93379 | 6.24551 | 6.41916 | 6.16012 | 6.42331 | 5.96968 | 6.18689 | 5.97502 | 5.79659 | 5.88637 | 6.66643 | 6.22408 | 6.54443 | 6.65889 |
| SMARCC2 | Q8TAQ2 | 6.09249 | 6.23102 | 6.01613 | 6.3357 | 6.37567 | 6.16244 | 6.32453 | 5.92898 | 6.16325 | 6.0422 | 6.07779 | 5.84523 | 6.77107 | 6.562 | 6.58661 | 6.4849 |
| USP34 | Q70CQ2 | 5.87166 | 6.16703 | 6.25875 | 6.32265 | 6.14444 | 6.16951 | 6.20911 | 5.91109 | 6.00917 | 6.19468 | 6.14804 | 6.12511 | 6.55954 | 6.57173 | 6.73051 | 6.60697 |
| RBM14 | Q96PK6 | 6.27098 | 6.42631 | 5.98521 | 6.81017 | 6.30599 | 5.84349 | 6.31698 | 5.95102 | 6.13292 | 5.90068 | 5.9875 | 5.99869 | 6.72841 | 6.16398 | 6.65231 | 6.52537 |
| SYMPK | Q92797 | 6.31632 | 6.51046 | 5.80586 | 6.61631 | 6.2702 | 6.12976 | 6.20493 | 6.21738 | 5.93459 | 6.09813 | 5.89678 | 5.94643 | 6.71776 | 6.36906 | 6.44198 | 6.52405 |
| ANXA5 | P08758 | 6.26274 | 6.04475 | 6.93857 | 5.54167 | 6.55254 | 6.93298 | 6.23401 | 6.53192 | 6.1625 | 6.37969 | 6.63278 | 6.44761 | 5.95669 | 6.28595 | 5.5614 | 5.53418 |
| HSD17B4 | P51659 | 6.30562 | 6.53794 | 5.89451 | 6.4982 | 6.29448 | 6.14051 | 6.34886 | 6.00989 | 6.04284 | 6.05855 | 5.89588 | 5.83782 | 6.67607 | 6.29701 | 6.71153 | 6.4503 |
| ASNS | P08243 | 6.28803 | 6.24639 | 6.76905 | 5.82634 | 6.50163 | 6.63754 | 6.41857 | 6.86764 | 6.66159 | 6.23166 | 6.41551 | 6.47768 | 5.70056 | 5.57245 | 5.74885 | 5.63651 |
| CIT | O14578-4 | 6.57232 | 6.74815 | 6.62011 | 6.77236 | 6.35005 | 6.3192 | 6.41324 | 6.43416 | 6.56837 | 6.50568 | 6.33912 | 6.44153 | 5.52932 | 5.32156 | 5.50519 | 5.55964 |
| YARS1 | P54577 | 6.24966 | 5.96444 | 6.88119 | 5.70065 | 6.62199 | 6.88563 | 6.09658 | 6.4893 | 6.35243 | 6.33334 | 6.48984 | 6.30366 | 6.39327 | 5.96695 | 5.67175 | 5.59932 |
| CUL3 | Q13618 | 5.72841 | 6.03037 | 6.28526 | 5.68758 | 5.96398 | 6.25067 | 6.12352 | 6.2167 | 6.13967 | 6.29346 | 6.28874 | 6.18055 | 6.58496 | 6.74864 | 6.85977 | 6.61772 |
| PFAS | O15067 | 6.09102 | 6.01461 | 6.93763 | 5.96955 | 6.2491 | 6.74507 | 6.26743 | 6.59192 | 6.44409 | 6.22091 | 6.36923 | 6.49086 | 5.72461 | 6.17136 | 5.83017 | 5.88245 |
| PSIP1 | O75475 | 6.24762 | 6.20139 | 5.57371 | 6.58159 | 6.07671 | 5.7448 | 6.50773 | 5.87128 | 6.11091 | 5.77535 | 5.63084 | 5.97336 | 6.52879 | 7.02667 | 6.96614 | 7.1791 |
| POTEF | A5A3E0 | 6.55703 | 6.30772 | 6.21218 | 5.9634 | 6.41265 | 6.30293 | 6.09892 | 6.03243 | 5.93255 | 6.17581 | 6.18376 | 6.42375 | 6.62963 | 6.26447 | 6.26471 | 6.23807 |
| TNPO1 | Q92973 | 6.05909 | 5.9427 | 6.37702 | 5.88737 | 6.23933 | 6.15999 | 6.10285 | 6.17966 | 6.30272 | 6.15815 | 6.40837 | 6.26055 | 6.317 | 6.32227 | 6.59888 | 6.68407 |
| NSD2 | O96028 | 6.35413 | 6.53428 | 6.03344 | 6.97393 | 6.23994 | 6.03944 | 6.42782 | 6.07099 | 6.30053 | 6.14028 | 6.23075 | 6.13939 | 5.90279 | 6.21575 | 6.29393 | 6.10262 |
| PRKCSH | P14314-2 | 6.63502 | 6.31305 | 5.92211 | 6.25926 | 6.86998 | 6.12235 | 6.22741 | 6.30602 | 6.1935 | 6.01613 | 6.01112 | 5.74846 | 7.14181 | 6.06371 | 6.16417 | 6.0059 |
| COPG1 | Q9Y678 | 6.30226 | 6.32875 | 6.43173 | 5.99135 | 6.3928 | 6.53405 | 6.3242 | 6.37867 | 6.43658 | 6.47828 | 6.50354 | 6.43692 | 6.0021 | 5.90245 | 5.78976 | 5.76656 |
| LRBA | P50851 | 6.26588 | 6.6329 | 6.87628 | 6.58114 | 6.44207 | 6.77119 | 6.1848 | 6.42019 | 6.14023 | 6.34017 | 6.38439 | 6.12082 | 5.76849 | 5.92117 | 5.70113 | 5.44916 |
| NUP155 | O75694 | 6.35444 | 6.3073 | 6.04331 | 6.61503 | 6.31482 | 6.05042 | 6.37929 | 5.95339 | 6.18862 | 5.98286 | 6.17823 | 6.058 | 6.63149 | 6.12094 | 6.44406 | 6.3778 |
| MYCBP2 | O75592 | 6.51433 | 6.6198 | 6.64224 | 6.5144 | 6.33199 | 6.42306 | 6.34424 | 6.41895 | 6.27702 | 6.58636 | 6.34948 | 6.25843 | 5.70025 | 5.81151 | 5.59221 | 5.61573 |
| PDIA4 | P13667 | 6.24765 | 6.52215 | 5.84236 | 6.39339 | 6.49826 | 6.16192 | 6.23928 | 6.06795 | 5.97466 | 6.00641 | 5.88699 | 5.82252 | 6.80288 | 6.42743 | 6.68006 | 6.4261 |
| GTPBP4 | Q9BZE4 | 6.29107 | 6.17557 | 6.01085 | 6.64838 | 6.30456 | 5.86104 | 6.54322 | 5.93674 | 6.33156 | 5.89858 | 6.01381 | 5.84667 | 6.55454 | 6.28808 | 6.80529 | 6.49003 |
| RARS1 | P54136 | 6.43216 | 6.17789 | 6.50583 | 5.83949 | 6.46823 | 6.52845 | 6.2662 | 6.48806 | 6.34456 | 6.19445 | 6.3085 | 6.40029 | 6.07105 | 6.27524 | 5.84623 | 5.85337 |
| PABPC1 | P11940 | 6.47046 | 6.66473 | 6.701 | 6.23648 | 6.42734 | 6.59602 | 6.32111 | 6.56671 | 6.3015 | 6.49924 | 6.49576 | 6.49417 | 5.54949 | 5.66264 | 5.56562 | 5.44773 |
| PSMC3 | P17980 | 6.42435 | 6.29224 | 6.38961 | 6.04575 | 6.66864 | 6.53819 | 6.05508 | 6.65056 | 6.14048 | 6.25257 | 6.49889 | 6.31664 | 6.43977 | 5.84801 | 5.72089 | 5.71834 |
| KNL1 | Q8NG31 | 6.34892 | 6.44801 | 6.2634 | 6.66176 | 6.2698 | 6.22673 | 6.36705 | 6.19021 | 6.39422 | 6.25615 | 6.29634 | 6.44163 | 6.06498 | 5.7153 | 5.93408 | 6.1214 |
| FLII | Q13045 | 6.39389 | 6.25627 | 6.95908 | 6.17181 | 6.36804 | 6.49637 | 6.32263 | 6.46166 | 6.60865 | 6.58954 | 6.61252 | 6.64522 | 5.56161 | 5.62788 | 5.36243 | 5.5624 |
| SRRT | Q9BXP5 | 6.01257 | 6.37355 | 5.69298 | 6.50606 | 6.15034 | 6.09327 | 6.13643 | 5.99405 | 5.95691 | 6.03434 | 5.94491 | 6.08865 | 6.84943 | 6.65524 | 6.62461 | 6.88664 |
| NUP153 | P49790 | 6.38843 | 6.22921 | 6.02984 | 6.70264 | 6.18103 | 5.86985 | 6.38986 | 5.89491 | 6.02857 | 6.1221 | 5.97764 | 5.85336 | 6.85376 | 6.43135 | 6.50635 | 6.5411 |
| NME2 | P22392-2 | 6.84443 | 6.4338 | 7.06815 | 6.06672 | 6.9567 | 6.96288 | 6.4934 | 7.04685 | 6.76626 | 6.39797 | 6.14588 | 5.98387 | 5.2005 | 5.19618 | 5.3145 | 5.12191 |
| PLIN3 | O60664 | 6.33478 | 5.62982 | 6.46585 | 5.29912 | 6.38289 | 6.54707 | 6.23406 | 6.83101 | 6.3457 | 6.24546 | 6.4393 | 6.73468 | 6.45566 | 6.191 | 5.89779 | 5.96582 |
| NNT | Q13423 | 6.4746 | 6.44683 | 5.90038 | 6.44096 | 6.22449 | 5.9879 | 6.2872 | 5.97232 | 5.94308 | 6.21221 | 5.72091 | 5.72363 | 6.68128 | 6.65189 | 6.75783 | 6.57449 |
| PHB | P35232 | 6.3351 | 5.26928 | 5.44521 | 5.28855 | 6.36871 | 5.49736 | 6.07948 | 6.38956 | 5.44512 | 6.01867 | 5.65158 | 6.38763 | 6.7358 | 7.65575 | 7.85277 | 7.57943 |
| COPB2 | P35606 | 6.20073 | 6.34489 | 6.64142 | 6.21503 | 6.24443 | 6.51905 | 6.37634 | 6.14972 | 6.36934 | 6.33597 | 6.41857 | 6.4617 | 5.8627 | 6.08656 | 5.92994 | 5.84361 |
| PTPRF | P10586 | 6.25904 | 8.02474 | 6.12484 | 6.75363 | 6.07656 | 5.93045 | 6.05776 | 6.7356 | 6.44445 | 6.56438 | 6.29731 | 6.53135 | 5.3715 | 5.55778 | 5.75768 | 5.51293 |
| RPSA | P08865 | 6.48622 | 5.376 | 6.23394 | 4.87826 | 6.22217 | 5.91738 | 6.07901 | 6.62508 | 5.75665 | 6.70409 | 6.27051 | 7.07072 | 5.78106 | 7.04776 | 6.66922 | 6.88193 |
| SLC3A2 | P08195-4 | 6.40482 | 6.49935 | 5.95217 | 6.34443 | 6.58379 | 6.18269 | 6.30852 | 6.10045 | 6.14821 | 5.78905 | 5.97914 | 6.20947 | 6.72433 | 6.13197 | 6.28868 | 6.35295 |
| GOT2 | P00505 | 6.28967 | 6.34956 | 5.87936 | 6.37833 | 6.58242 | 5.93982 | 6.66954 | 5.85431 | 5.7803 | 6.70499 | 6.08543 | 5.84005 | 6.48687 | 6.37357 | 6.58756 | 6.19823 |
| PCBP2 | Q15366-3 | 6.35916 | 6.15184 | 6.47168 | 6.05986 | 6.31144 | 6.54826 | 5.93886 | 6.30898 | 6.53136 | 6.63774 | 6.53802 | 6.36252 | 6.12489 | 5.98963 | 5.8078 | 5.85794 |
| KIF20B | Q96Q89-3 | 6.66703 | 6.66405 | 6.47478 | 7.11528 | 6.54445 | 5.9657 | 6.68481 | 6.11016 | 6.49462 | 6.23302 | 6.45238 | 6.41391 | 5.48675 | 5.27961 | 5.62234 | 5.79109 |
| DYNC2H1 | Q8NCM8-2 | 6.6328 | 6.71509 | 7.27779 | 6.51568 | 6.34637 | 6.37663 | 6.42848 | 6.5133 | 6.79782 | 6.86097 | 7.12257 | 7.05224 | 4.79245 | 5.0325 | 4.69494 | 4.84037 |
| TPX2 | Q9ULW0 | 6.34344 | 6.15258 | 5.76606 | 6.40103 | 6.07096 | 5.94175 | 6.42705 | 5.83552 | 6.08356 | 5.8469 | 5.9515 | 5.91062 | 6.96711 | 6.53128 | 6.85455 | 6.92209 |
| OGA | O60502 | 6.17498 | 6.26031 | 6.90905 | 5.95798 | 6.42741 | 6.68567 | 6.39314 | 6.47847 | 6.20742 | 6.28608 | 6.43023 | 6.23798 | 5.96403 | 6.08031 | 5.74301 | 5.76391 |
| SUPT5H | O00267 | 5.55166 | 6.02641 | 5.99971 | 5.74048 | 5.95551 | 6.11289 | 6.19904 | 5.79785 | 6.22928 | 5.90629 | 6.24291 | 5.98316 | 6.93039 | 6.93177 | 7.33161 | 7.06105 |
| HELLS | Q9NRZ9 | 6.15605 | 6.44043 | 6.43326 | 6.61097 | 6.26988 | 6.08506 | 6.22106 | 6.15565 | 6.43701 | 6.36534 | 6.3694 | 6.43013 | 5.9669 | 5.91177 | 6.0162 | 6.1309 |
| RPL5 | P46777 | 6.06707 | 6.11998 | 6.30374 | 6.27702 | 6.11434 | 6.14689 | 6.49291 | 6.15587 | 6.70545 | 5.95132 | 6.22773 | 6.298 | 5.73307 | 6.5805 | 6.23415 | 6.59195 |
| GDI2 | P50395 | 6.25453 | 5.97259 | 7.02512 | 5.88173 | 6.30394 | 6.73469 | 6.34042 | 6.62098 | 6.57966 | 6.53873 | 6.46535 | 6.67744 | 5.72925 | 5.68901 | 5.52265 | 5.6639 |
| PITRM1 | Q5JRX3-2 | 6.25378 | 6.44388 | 5.65216 | 6.67311 | 6.26488 | 6.23307 | 6.34812 | 6.22622 | 6.13825 | 6.1415 | 5.96993 | 5.92916 | 6.54184 | 6.25699 | 6.52194 | 6.40519 |
| RPN2 | P04844 | 6.32483 | 6.5067 | 5.88196 | 6.52676 | 6.23341 | 6.08491 | 6.44937 | 6.00705 | 5.91433 | 6.1086 | 5.90914 | 5.86159 | 6.61587 | 6.35013 | 6.76233 | 6.46301 |
| EXOSC10 | Q01780 | 6.33802 | 6.37143 | 5.95034 | 6.68353 | 6.28103 | 6.0156 | 6.45008 | 5.97904 | 6.11186 | 5.97949 | 5.98488 | 5.8977 | 6.52123 | 6.44396 | 6.58687 | 6.40494 |
| SMARCC1 | Q92922 | 5.99508 | 5.95582 | 5.9331 | 6.64546 | 6.18673 | 5.68467 | 6.84485 | 5.73426 | 6.60641 | 5.4716 | 5.8271 | 6.00198 | 6.3172 | 6.7601 | 6.73 | 7.30565 |
| PLXNA1 | Q9UIW2 | 6.4374 | 6.44948 | 5.92727 | 6.38154 | 6.14758 | 6.32268 | 6.18275 | 5.98429 | 6.04828 | 6.12706 | 6.17081 | 5.95392 | 6.41772 | 6.49343 | 6.47976 | 6.47605 |
| RFC1 | P35251 | 5.95441 | 6.22986 | 5.95233 | 6.26905 | 5.92177 | 5.98279 | 6.10757 | 6.03078 | 5.87581 | 6.06984 | 6.00651 | 6.00308 | 6.18039 | 7.29887 | 6.87083 | 7.24614 |
| CLASP1 | Q7Z460 | 6.30605 | 6.23393 | 6.4528 | 6.17346 | 6.25142 | 6.43008 | 6.17861 | 6.26396 | 6.33082 | 6.28537 | 6.31182 | 6.24499 | 6.28887 | 6.13092 | 6.12927 | 5.98763 |
| PRRC2A | P48634 | 6.75805 | 6.87373 | 6.7819 | 6.71342 | 6.45098 | 6.24719 | 6.25452 | 6.28425 | 6.37117 | 6.50303 | 6.48804 | 6.60285 | 5.38676 | 5.54663 | 5.32344 | 5.41405 |
| SHMT2 | P34897 | 6.26922 | 6.40272 | 5.62898 | 6.38737 | 6.47483 | 6.12061 | 6.35955 | 6.48501 | 6.09759 | 5.94637 | 6.13517 | 5.93742 | 6.81802 | 6.45026 | 6.24285 | 6.24403 |
| ABCF1 | Q8NE71 | 6.48936 | 6.41608 | 6.61244 | 6.37577 | 6.36199 | 6.3789 | 6.38745 | 6.56094 | 6.45479 | 6.26867 | 6.33221 | 6.24623 | 5.3218 | 6.24777 | 5.62762 | 5.91795 |
| CUX1 | P39880-3 | 6.45391 | 6.10252 | 6.13259 | 6.4774 | 6.25823 | 5.96295 | 6.16416 | 5.9684 | 6.24887 | 6.19689 | 6.30264 | 6.22933 | 6.90172 | 6.07573 | 6.31237 | 6.2123 |
| ANXA1 | P04083 | 5.99104 | 6.23201 | 6.70551 | 5.82813 | 6.34738 | 7.05537 | 6.10475 | 7.01396 | 6.11244 | 6.42292 | 6.07883 | 6.60321 | 5.94479 | 6.24133 | 5.45848 | 5.85985 |
| TASOR | Q9UK61 | 6.4964 | 6.64153 | 5.92963 | 6.77144 | 6.10463 | 6.06975 | 6.29842 | 6.11087 | 6.36483 | 6.14147 | 5.96065 | 5.96697 | 6.46072 | 6.13558 | 6.1759 | 6.3712 |
| MICAL3 | Q7RTP6 | 6.10397 | 6.3574 | 6.46133 | 6.14676 | 6.11239 | 6.33152 | 6.21585 | 6.19198 | 6.36033 | 6.53311 | 6.44862 | 6.45683 | 6.00807 | 6.15904 | 6.07126 | 6.04154 |
| ELAC2 | Q9BQ52 | 6.04716 | 6.04358 | 6.17243 | 6.11906 | 6.00801 | 6.32011 | 6.098 | 6.18292 | 6.11203 | 6.13048 | 6.07573 | 6.04775 | 6.39583 | 6.56411 | 6.95235 | 6.73046 |
| KIF23 | Q02241 | 6.38917 | 6.5866 | 6.38129 | 7.01373 | 6.09885 | 6.34299 | 6.68283 | 6.17217 | 6.06723 | 6.22527 | 6.19732 | 6.28216 | 5.57825 | 5.58752 | 6.10498 | 6.18963 |
| DNM1L | O00429-6 | 6.42323 | 6.13484 | 6.85704 | 5.99678 | 6.42666 | 6.81601 | 6.13683 | 6.77345 | 6.30915 | 6.45443 | 6.49168 | 6.34431 | 5.85971 | 5.95369 | 5.59875 | 5.42344 |
| CLUH | A0A494C0R8 | 6.38976 | 6.26583 | 7.192 | 6.41774 | 6.3947 | 6.75717 | 6.30377 | 6.74431 | 6.68736 | 6.49716 | 6.91912 | 6.77304 | 5.28675 | 5.18754 | 5.00852 | 5.17523 |
| NOL11 | Q9H8H0 | 6.4114 | 6.27202 | 6.10208 | 6.78242 | 6.38533 | 6.0346 | 6.27098 | 5.97371 | 6.46525 | 5.95012 | 6.04507 | 5.90551 | 6.64821 | 5.8899 | 6.62789 | 6.23551 |
| RTN4 | Q9NQC3 | 6.21455 | 6.35517 | 5.82616 | 6.23298 | 5.84702 | 6.06793 | 6.03383 | 6.28307 | 5.95184 | 5.93493 | 5.9281 | 6.11354 | 6.42594 | 6.68123 | 6.87242 | 7.23128 |
| ATIC | P31939 | 6.23153 | 6.05356 | 6.78828 | 5.8048 | 6.42496 | 6.61914 | 6.34959 | 6.83671 | 6.50902 | 6.35449 | 6.56778 | 6.45909 | 5.81523 | 5.84999 | 5.77851 | 5.55732 |
| GSN | P06396-2 | 6.39799 | 6.02515 | 6.70542 | 5.69764 | 6.3 | 6.85972 | 6.15764 | 6.62574 | 6.27804 | 6.49062 | 6.51518 | 6.4528 | 6.06931 | 6.24869 | 5.65573 | 5.52034 |
| ERAP1 | Q9NZ08-2 | 6.39924 | 6.50358 | 5.95331 | 6.3531 | 6.30452 | 6.10509 | 6.22192 | 6.00787 | 6.04312 | 6.15377 | 6.02306 | 5.90274 | 6.65773 | 6.19148 | 6.81699 | 6.36249 |
| NUP160 | Q12769 | 6.64979 | 6.47992 | 6.27778 | 6.91999 | 6.35298 | 6.18302 | 6.56604 | 6.03253 | 6.43758 | 6.08129 | 5.79937 | 5.7238 | 6.29763 | 5.80029 | 6.2415 | 6.1565 |
| FKBP4 | Q02790 | 6.33821 | 6.08424 | 6.56899 | 5.92929 | 6.48634 | 6.63919 | 6.22179 | 6.32233 | 6.26176 | 6.43941 | 6.53802 | 6.36725 | 6.18073 | 5.9205 | 5.90622 | 5.79575 |
| EPHA2 | P29317 | 6.14214 | 6.44656 | 5.95458 | 6.56863 | 6.04719 | 6.07016 | 6.30673 | 6.12051 | 5.98724 | 5.9553 | 5.99947 | 5.84446 | 6.47191 | 6.49681 | 6.8315 | 6.75681 |
| USP5 | P45974 | 5.85585 | 5.93956 | 6.60297 | 5.69856 | 5.93724 | 6.47451 | 5.94592 | 6.46979 | 6.04431 | 6.30923 | 6.59002 | 6.40018 | 6.37446 | 6.61603 | 6.49736 | 6.244 |
| EIF3D | O15371 | 6.2489 | 6.31414 | 6.42114 | 6.28478 | 6.26972 | 6.32596 | 6.71693 | 6.02859 | 6.65136 | 6.17531 | 6.21206 | 6.37712 | 5.61632 | 6.15796 | 5.87319 | 6.32652 |
| PFN1 | P07737 | 6.01286 | 5.7093 | 6.75563 | 5.41421 | 6.66006 | 6.50492 | 6.13602 | 6.91289 | 6.43274 | 6.67791 | 6.79537 | 6.87271 | 5.62583 | 5.95128 | 5.67046 | 5.86782 |
| HNRNPDL | O14979 | 6.3488 | 6.35066 | 5.90738 | 6.49205 | 6.51457 | 6.16351 | 6.51124 | 6.0558 | 6.30422 | 5.94406 | 6.07865 | 6.09092 | 6.53421 | 6.1339 | 6.31676 | 6.25327 |
| NOP56 | O00567 | 6.18683 | 5.94967 | 5.47108 | 6.68311 | 6.07522 | 5.51801 | 6.81593 | 5.68746 | 6.2099 | 5.34787 | 5.59932 | 5.83817 | 6.30359 | 7.32893 | 7.29227 | 7.69264 |
| EP400 | Q96L91 | 6.78585 | 6.72427 | 6.32637 | 7.16683 | 6.2509 | 5.9222 | 6.52115 | 5.92255 | 6.50538 | 6.47526 | 6.27388 | 6.31143 | 5.73678 | 5.55224 | 5.72114 | 5.80375 |
| TFRC | P02786 | 6.0002 | 6.32781 | 6.05883 | 6.43116 | 6.31746 | 6.11354 | 6.72041 | 6.39423 | 6.11449 | 5.89262 | 6.00812 | 6.05765 | 6.21177 | 6.16173 | 6.67117 | 6.5188 |
| EEF1G | P26641-2 | 6.27862 | 6.25025 | 6.71319 | 5.96326 | 6.27087 | 6.37549 | 5.99786 | 6.43081 | 6.55027 | 6.3746 | 6.4505 | 6.56884 | 6.01502 | 6.04954 | 5.91802 | 5.79287 |
| NAP1L1 | P55209 | 6.44466 | 6.18618 | 6.95249 | 6.19924 | 6.60631 | 6.83419 | 6.11942 | 6.75418 | 6.67293 | 6.52188 | 6.84958 | 6.93839 | 5.63675 | 5.2742 | 4.89194 | 5.11767 |
| NUP188 | Q5SRE5 | 6.40089 | 6.46956 | 6.09112 | 6.52698 | 6.37192 | 6.25183 | 6.24504 | 6.2704 | 6.25697 | 6.1785 | 6.11268 | 6.13232 | 6.00187 | 6.17163 | 6.10124 | 6.41705 |
| MTOR | P42345 | 6.43886 | 6.42558 | 6.55204 | 6.42804 | 6.13541 | 6.61253 | 6.09211 | 6.20111 | 6.44845 | 6.58262 | 6.63279 | 6.68886 | 5.67419 | 5.91698 | 5.51154 | 5.65889 |
| TUBB2B | Q9BVA1 | 6.57158 | 6.18654 | 7.02544 | 6.03738 | 6.59454 | 6.86406 | 6.0131 | 6.83351 | 6.43333 | 6.38833 | 6.69108 | 6.714 | 5.75775 | 5.4327 | 5.14461 | 5.31204 |
| RECQL | P46063 | 5.15161 | 5.44385 | 5.72676 | 5.05883 | 5.57042 | 6.15742 | 5.73907 | 5.77523 | 5.75173 | 6.01565 | 6.07169 | 5.80473 | 7.61508 | 8.24409 | 8.24568 | 7.62816 |
| FTSJ3 | Q8IY81 | 5.88908 | 6.24208 | 5.53778 | 6.72759 | 5.82655 | 5.97748 | 6.34904 | 6.10895 | 6.30609 | 5.98878 | 6.02141 | 6.08059 | 6.34563 | 6.8504 | 6.73276 | 7.01579 |
| CAST | P20810-6 | 6.25315 | 6.28945 | 6.61343 | 5.72395 | 6.17231 | 6.51386 | 5.98215 | 6.54195 | 6.36735 | 6.40296 | 6.30526 | 6.43936 | 6.06448 | 6.39758 | 5.86218 | 6.0706 |
| TNKS1BP1 | Q9C0C2 | 6.14233 | 6.28884 | 6.33487 | 5.84478 | 6.13723 | 6.27982 | 5.97238 | 6.30578 | 6.28626 | 6.5743 | 6.4303 | 6.39818 | 6.51893 | 6.25637 | 6.145 | 6.08462 |
| SLC25A13 | Q9UJS0-2 | 6.42596 | 6.09284 | 5.89493 | 6.51075 | 6.36145 | 6.00324 | 6.57789 | 6.07111 | 6.10582 | 6.03928 | 6.02109 | 5.89213 | 6.74519 | 6.3767 | 6.53673 | 6.34489 |
| CHD7 | Q9P2D1 | 6.43975 | 6.41461 | 6.29986 | 7.0583 | 5.97927 | 6.04095 | 6.47501 | 6.07334 | 6.164 | 6.05647 | 5.96332 | 6.0899 | 5.85374 | 6.302 | 6.26398 | 6.52549 |
| ITPR2 | Q14571 | 6.41964 | 6.41315 | 5.84607 | 6.5978 | 6.21398 | 5.84511 | 6.3244 | 5.92291 | 5.96179 | 6.11103 | 5.8228 | 5.76691 | 6.66876 | 6.59211 | 6.86914 | 6.6244 |
| ARFGEF1 | Q9Y6D6 | 6.26007 | 6.17459 | 6.85579 | 6.27074 | 6.1671 | 6.54668 | 6.20991 | 6.36645 | 6.30601 | 6.5148 | 6.50603 | 6.44105 | 5.90743 | 5.88705 | 5.64811 | 5.93816 |
| EMC1 | Q8N766 | 6.39208 | 6.55028 | 6.13757 | 6.85427 | 6.20719 | 5.9092 | 6.41995 | 6.10324 | 6.21418 | 6.0865 | 5.81234 | 5.89851 | 6.35624 | 6.04552 | 6.49103 | 6.5219 |
| ABCE1 | P61221 | 6.30686 | 6.26815 | 6.9307 | 6.02967 | 6.36642 | 6.81927 | 6.37891 | 6.5907 | 6.64812 | 6.38546 | 6.31121 | 6.22986 | 5.64676 | 5.71259 | 5.77522 | 5.6001 |
| UGGT1 | Q9NYU2 | 6.27751 | 6.44142 | 5.75755 | 6.51082 | 6.26937 | 6.16395 | 6.28664 | 6.08733 | 6.06687 | 5.96048 | 5.75628 | 5.60624 | 6.70118 | 6.64774 | 6.76931 | 6.69734 |
| ITGA6 | P23229 | 6.2613 | 6.35722 | 6.11737 | 6.57463 | 6.27318 | 6.26785 | 6.48815 | 6.2457 | 6.14305 | 5.98497 | 6.36477 | 6.34176 | 6.22794 | 5.93138 | 6.08002 | 6.34072 |
| ATAD2 | Q6PL18 | 6.45398 | 6.43341 | 5.99812 | 6.81021 | 6.23053 | 5.82295 | 6.26784 | 5.85097 | 6.51797 | 6.25828 | 6.26162 | 6.18769 | 6.39413 | 5.91415 | 6.31641 | 6.28175 |

| | | | | | | | | | | | | | | | | | |
|---|---|---|---|---|---|---|---|---|---|---|---|---|---|---|---|---|---|
| RPLP0 | P05388 | 6.3874 | 6.41087 | 6.45532 | 6.19441 | 6.29864 | 6.52682 | 6.07994 | 6.39788 | 6.4883 | 6.68371 | 6.45582 | 6.68168 | 5.7642 | 5.87012 | 5.66038 | 5.64453 |
| MCM2 | P49736 | 5.64013 | 6.18711 | 5.9454 | 5.7906 | 6.03589 | 6.12982 | 5.99965 | 6.16955 | 6.01357 | 6.01822 | 6.00565 | 6.09177 | 6.89597 | 6.9822 | 7.07023 | 7.02425 |
| RPL7 | P18124 | 5.99524 | 6.44866 | 5.75314 | 5.80967 | 6.13953 | 6.33854 | 6.52053 | 6.6362 | 7.03493 | 6.50832 | 6.43614 | 6.41606 | 5.96933 | 6.24142 | 5.62749 | 6.12479 |
| ESYT1 | Q9BSJ8 | 6.17909 | 6.5275 | 6.06057 | 6.44053 | 6.33292 | 6.14313 | 6.33359 | 6.15083 | 6.07621 | 6.18267 | 6.16603 | 5.96514 | 6.45562 | 6.32488 | 6.46406 | 6.19721 |
| PPP1CC | P36873-2 | 6.20955 | 6.13981 | 6.28134 | 6.51567 | 6.15622 | 6.24107 | 6.1733 | 6.04589 | 6.27725 | 6.29417 | 6.3597 | 6.46885 | 6.25544 | 5.95189 | 6.39498 | 6.23486 |
| CARS1 | P49589-3 | 6.11092 | 6.31411 | 6.5906 | 6.03609 | 6.23387 | 6.71921 | 6.14038 | 6.6483 | 6.04659 | 6.4672 | 6.36807 | 6.36847 | 5.84323 | 6.31996 | 5.97865 | 5.81434 |
| NDUFS1 | P28331-2 | 6.38954 | 6.27953 | 5.75626 | 6.55187 | 6.32582 | 5.72783 | 6.63805 | 5.88274 | 6.46493 | 5.60971 | 5.77515 | 5.8522 | 6.52683 | 6.63684 | 6.57386 | 7.00883 |
| PNPT1 | Q8TCS8 | 6.37219 | 6.30743 | 5.99398 | 6.83238 | 6.37585 | 5.95785 | 6.43699 | 5.94718 | 6.35243 | 6.14361 | 6.06263 | 6.01692 | 6.59128 | 5.96164 | 6.38661 | 6.26103 |
| SLC25A3 | Q00325 | 6.36049 | 6.39121 | 5.79058 | 6.68638 | 5.96107 | 5.76069 | 6.40334 | 5.97761 | 6.10452 | 5.91486 | 5.84866 | 5.87803 | 6.50299 | 6.82134 | 6.72547 | 6.87274 |
| UFL1 | O94874 | 6.32155 | 6.55137 | 5.89361 | 6.43256 | 6.37999 | 6.29204 | 6.44078 | 6.01922 | 5.96432 | 6.06991 | 5.90061 | 5.92025 | 6.60267 | 6.26569 | 6.57166 | 6.37376 |
| FSCN1 | Q16658 | 6.32618 | 6.44243 | 6.80735 | 5.87003 | 6.35021 | 6.82875 | 6.29847 | 6.57562 | 6.39818 | 6.65474 | 6.31935 | 6.3749 | 5.7606 | 5.74223 | 5.76237 | 5.48859 |
| TOMM40 | O96008 | 6.03747 | 5.99341 | 5.65477 | 6.89413 | 6.47722 | 5.69578 | 7.23785 | 5.79634 | 6.68109 | 5.22806 | 5.58053 | 5.97751 | 6.26511 | 6.44862 | 6.82884 | 7.20326 |
| MYO6 | A0A0A0MRM8 | 6.22791 | 6.41964 | 6.63995 | 6.01135 | 6.30577 | 6.65911 | 6.21083 | 6.46654 | 6.37542 | 6.5788 | 6.52656 | 6.21063 | 5.9356 | 6.02284 | 5.7045 | 5.70455 |
| WDR1 | O75083 | 6.16326 | 6.04385 | 6.56161 | 5.67001 | 6.68881 | 6.61554 | 6.35413 | 6.60088 | 6.26189 | 6.2687 | 6.18827 | 6.07489 | 6.12239 | 6.22402 | 6.17231 | 5.98943 |
| LDHA | P00338-3 | 6.17809 | 6.0851 | 6.82332 | 5.74032 | 6.45486 | 6.8578 | 6.27322 | 6.56361 | 6.29653 | 6.33334 | 6.37375 | 6.4259 | 5.77132 | 6.33036 | 5.73416 | 5.75832 |
| NOP58 | Q9Y2X3 | 6.12213 | 6.22407 | 5.58507 | 6.67806 | 6.11357 | 5.87838 | 6.28429 | 5.79098 | 6.03152 | 5.74049 | 5.69135 | 5.68557 | 6.55246 | 7.21163 | 7.2155 | 7.19493 |
| KIF5B | P33176 | 6.27021 | 6.1999 | 6.92895 | 5.9668 | 6.3729 | 6.52638 | 6.40924 | 6.65259 | 6.54918 | 6.45087 | 6.47361 | 6.42815 | 5.78267 | 5.72271 | 5.73843 | 5.52739 |
| AQR | O60306 | 6.28626 | 6.3159 | 6.25961 | 6.55879 | 6.18666 | 6.11036 | 6.41669 | 5.86203 | 6.2284 | 6.1413 | 6.15666 | 6.10481 | 6.45195 | 6.24912 | 6.33701 | 6.33446 |
| ITGA3 | P26006 | 6.47307 | 6.50067 | 5.88382 | 6.24456 | 6.34075 | 6.07078 | 6.33927 | 5.96534 | 6.2115 | 6.06631 | 5.83525 | 5.97433 | 6.62676 | 6.66433 | 6.37857 | 6.42468 |
| RUVBL2 | Q9Y230 | 6.32461 | 6.52773 | 6.44556 | 6.3872 | 6.443 | 6.24087 | 6.38793 | 6.40904 | 6.42842 | 6.09356 | 6.22357 | 6.29296 | 5.9656 | 5.88625 | 5.98512 | 5.95857 |
| MYO1C | O00159 | 6.06696 | 6.13964 | 5.87165 | 5.92423 | 6.22505 | 6.05124 | 6.38026 | 5.89997 | 6.52489 | 6.2753 | 6.13319 | 6.09176 | 6.72084 | 6.54524 | 6.66554 | 6.48423 |
| XRN2 | Q9H0D6 | 6.24147 | 5.98619 | 6.00766 | 6.38736 | 6.43833 | 5.96387 | 6.35535 | 5.87339 | 6.37651 | 6.12715 | 6.22798 | 6.00117 | 6.65268 | 6.29274 | 6.47698 | 6.59117 |
| WDR11 | Q9BZH6 | 6.27259 | 6.27007 | 6.48168 | 6.09926 | 6.30939 | 6.3423 | 6.33465 | 6.17969 | 6.38873 | 6.59083 | 6.42511 | 6.35446 | 5.97265 | 5.99048 | 6.0462 | 5.94194 |
| DDX27 | Q96GQ7 | 6.15368 | 6.22653 | 5.71018 | 6.65537 | 6.27999 | 5.93761 | 6.2965 | 6.07752 | 6.06644 | 6.0558 | 5.88698 | 5.8578 | 6.43084 | 6.57984 | 6.99625 | 6.78868 |
| DDOST | P39656 | 6.27427 | 6.2284 | 6.04129 | 6.53371 | 6.47783 | 5.88002 | 6.44275 | 5.83909 | 6.14905 | 6.41677 | 6.06288 | 5.80666 | 6.57231 | 6.01778 | 6.79772 | 6.45946 |
| POLA1 | P09884 | 5.63745 | 6.04499 | 6.13919 | 5.8966 | 5.80521 | 6.25673 | 6.08382 | 5.98911 | 6.09541 | 6.06502 | 6.15691 | 6.11911 | 6.80406 | 6.954 | 7.03537 | 6.91704 |
| PDCD6IP | Q8WUM4 | 6.67706 | 6.40398 | 6.86323 | 5.99993 | 6.51792 | 6.78302 | 6.19238 | 6.84389 | 6.32871 | 6.16693 | 6.32751 | 6.63645 | 5.7116 | 5.68205 | 5.41891 | 5.44644 |
| ATRX | P46100 | 6.35975 | 6.35003 | 5.97655 | 6.54073 | 6.15134 | 5.92234 | 6.29448 | 5.71391 | 6.1477 | 6.24808 | 6.09235 | 5.93956 | 6.64231 | 6.58548 | 6.60674 | 6.42865 |
| LRP1 | Q07954 | 6.27954 | 6.81394 | 5.94044 | 6.18163 | 6.3204 | 6.30501 | 6.07492 | 6.09606 | 5.86177 | 6.3567 | 6.07854 | 5.97353 | 6.38754 | 6.49355 | 6.53898 | 6.29746 |
| PSMA1 | P25786-2 | 6.08299 | 6.15873 | 6.32808 | 5.97805 | 6.39448 | 6.44654 | 6.0799 | 6.3866 | 6.13828 | 6.24468 | 6.36839 | 6.28588 | 6.64977 | 6.19741 | 6.21666 | 6.04358 |
| SIPA1L3 | O60292 | 6.05481 | 6.35315 | 5.91587 | 6.42788 | 6.03818 | 5.97171 | 6.3103 | 5.89303 | 5.9154 | 5.99537 | 5.99479 | 6.02429 | 6.86289 | 6.5631 | 6.80455 | 6.87468 |
| PSMC5 | P62195 | 6.21878 | 6.03154 | 6.25147 | 6.07156 | 6.49789 | 6.24204 | 6.63309 | 6.29058 | 6.42995 | 5.78365 | 6.33076 | 6.27538 | 6.0914 | 6.14561 | 6.29148 | 6.41483 |
| ARID2 | Q68CP9 | 6.48199 | 6.42942 | 6.10667 | 6.84305 | 6.39238 | 6.25708 | 6.3717 | 6.18351 | 6.05798 | 6.34831 | 6.23037 | 6.12693 | 6.10751 | 5.90265 | 6.19956 | 5.9609 |
| DLST | P36957 | 6.42564 | 6.42197 | 5.99411 | 6.70345 | 6.75443 | 6.293 | 6.48201 | 6.20849 | 6.08827 | 6.0743 | 5.87548 | 5.76077 | 6.77813 | 5.97219 | 6.23921 | 5.92855 |
| ACSL4 | O60488 | 6.30589 | 6.35203 | 6.231 | 6.80381 | 6.50167 | 6.35951 | 6.67473 | 6.00418 | 6.27928 | 6.04944 | 6.40312 | 6.3394 | 5.88065 | 5.75352 | 6.04112 | 6.02062 |
| ACSL3 | O95573 | 6.31869 | 6.39902 | 5.99718 | 6.62666 | 6.19379 | 6.08626 | 6.40968 | 5.9066 | 6.02606 | 5.85609 | 5.90304 | 5.91429 | 6.6261 | 6.39838 | 6.6683 | 6.66986 |
| CCAR1 | Q8IX12 | 6.3156 | 6.2718 | 5.874 | 6.42117 | 6.21494 | 6.04555 | 6.36822 | 6.06992 | 6.0685 | 6.04922 | 5.89699 | 5.67288 | 6.80811 | 6.49015 | 6.76565 | 6.66731 |
| ITGAV | P06756 | 6.40403 | 6.37394 | 6.0158 | 6.39461 | 6.21186 | 6.24879 | 6.23619 | 6.13422 | 6.26921 | 6.24955 | 6.22023 | 6.16735 | 6.3989 | 6.23981 | 6.22012 | 6.21539 |
| DHX15 | O43143 | 6.00319 | 6.25286 | 6.10334 | 6.41241 | 6.02356 | 6.01894 | 6.24999 | 5.96577 | 6.18762 | 6.09253 | 6.05708 | 6.10295 | 6.44149 | 6.39982 | 6.90284 | 6.78563 |
| PSAT1 | Q9Y617 | 6.37641 | 6.15709 | 6.68653 | 5.68939 | 6.64789 | 6.92666 | 6.20023 | 6.69056 | 6.36938 | 6.36646 | 6.39488 | 6.45723 | 5.96083 | 5.92573 | 5.69962 | 5.4511 |
| PGAM1 | P18669 | 6.17989 | 6.14592 | 6.85078 | 5.91271 | 6.50803 | 6.81015 | 6.09967 | 6.98877 | 6.24753 | 6.53732 | 6.55921 | 6.61447 | 5.69689 | 5.90753 | 5.5001 | 5.44103 |
| DNMBP | Q6XZF7 | 6.48371 | 6.13274 | 6.72468 | 6.0196 | 6.4416 | 6.74861 | 6.44488 | 6.6464 | 6.45316 | 6.2501 | 6.36258 | 6.30575 | 5.92246 | 5.80332 | 5.69653 | 5.56388 |
| MED12 | Q93074-2 | 6.44042 | 6.61197 | 6.20985 | 6.54793 | 6.3496 | 6.11488 | 6.50441 | 5.93983 | 6.36442 | 6.41008 | 6.06262 | 6.11015 | 6.13277 | 5.81212 | 6.18429 | 6.20464 |
| SERPINH1 | P50454 | 6.31824 | 6.49567 | 6.19157 | 6.59132 | 6.35079 | 6.27786 | 6.31323 | 5.88245 | 6.03106 | 6.04369 | 6.02258 | 5.88768 | 6.57777 | 6.32969 | 6.46417 | 6.22202 |
| ROCK2 | O75116 | 6.41628 | 6.364 | 6.97007 | 6.22715 | 6.35475 | 6.5848 | 6.39464 | 6.45796 | 6.34027 | 6.21064 | 6.42026 | 6.20029 | 5.82723 | 5.97651 | 5.55371 | 5.70143 |
| HNRNPAB | Q99729-3 | 6.23206 | 6.42587 | 5.85163 | 6.80462 | 6.12591 | 6.00601 | 6.35928 | 6.12938 | 6.14448 | 6.13629 | 5.99121 | 6.17858 | 6.53843 | 6.22707 | 6.27955 | 6.56964 |
| CAPRIN1 | Q14444 | 6.49873 | 6.49591 | 6.7901 | 6.27895 | 6.56074 | 6.67119 | 6.11428 | 6.65381 | 6.60574 | 6.55725 | 6.65096 | 6.69867 | 5.6161 | 5.32099 | 5.21208 | 5.27449 |
| EIF3E | P60228 | 6.52314 | 6.40652 | 6.56535 | 6.20351 | 6.44248 | 6.63129 | 6.40965 | 6.5934 | 6.45264 | 6.51325 | 6.35135 | 6.42765 | 5.59103 | 5.79246 | 5.45891 | 5.63739 |
| POLR1A | O95602 | 6.51243 | 6.39246 | 6.11966 | 6.97412 | 6.25092 | 6.02845 | 6.46529 | 6.09867 | 6.2338 | 5.9503 | 6.14092 | 6.06818 | 6.30054 | 5.86338 | 6.28096 | 6.3199 |
| COPB1 | P53618 | 6.2942 | 6.15693 | 6.30076 | 5.98213 | 6.46059 | 6.47204 | 6.14335 | 6.40939 | 6.51534 | 6.4707 | 6.64901 | 6.68369 | 5.92128 | 5.9063 | 5.79134 | 5.84296 |
| PCM1 | Q15154-5 | 6.52034 | 6.56356 | 6.577 | 6.46697 | 6.38746 | 6.65917 | 6.29851 | 6.5154 | 6.32233 | 6.46453 | 6.54365 | 6.66544 | 5.48442 | 5.6704 | 5.35506 | 5.50576 |
| EHBP1L1 | Q8N3D4 | 6.47557 | 6.50455 | 6.36357 | 6.42002 | 6.24232 | 6.11982 | 6.38115 | 6.33976 | 6.53302 | 6.42812 | 6.50775 | 6.4124 | 5.9602 | 5.83576 | 5.69145 | 5.78453 |
| RRP12 | Q5JTH9 | 5.97366 | 5.94545 | 5.41439 | 6.5605 | 5.96669 | 6.03378 | 6.42037 | 6.05046 | 6.71312 | 5.77704 | 5.73239 | 5.88177 | 6.57165 | 6.93769 | 6.82396 | 7.1971 |
| H1-3 | P16402 | 6.02101 | 5.98787 | 5.48934 | 6.36264 | 6.01373 | 5.7504 | 6.35923 | 5.94241 | 5.89802 | 5.69651 | 5.55062 | 5.65572 | 6.72899 | 7.65766 | 7.31914 | 7.5667 |
| RNF40 | O75150 | 5.85808 | 6.33032 | 6.28237 | 6.39341 | 6.29726 | 6.11478 | 6.58671 | 6.01724 | 6.31819 | 6.18589 | 6.21115 | 6.26472 | 6.19329 | 6.23431 | 6.29184 | 6.42043 |
| PSMD11 | O00231 | 6.31264 | 6.39141 | 6.16301 | 6.10514 | 6.67022 | 6.53908 | 6.22389 | 6.69393 | 6.17266 | 6.20508 | 6.04752 | 5.80655 | 6.20891 | 6.20189 | 6.1823 | 6.07577 |
| TUBB6 | Q9BUF5 | 6.3524 | 6.07545 | 6.92653 | 6.0841 | 6.47461 | 6.55072 | 6.44727 | 7.1492 | 6.56046 | 6.25271 | 6.72169 | 6.84329 | 5.54775 | 5.49577 | 5.0593 | 5.45874 |
| PYGL | P06737 | 6.42262 | 6.43331 | 6.94731 | 6.13328 | 6.17123 | 6.88991 | 6.05223 | 6.42909 | 6.51368 | 6.53069 | 6.72375 | 6.53975 | 5.4982 | 5.83341 | 5.40599 | 5.47556 |
| PDCD11 | Q14690 | 6.22528 | 6.576 | 5.79912 | 6.92809 | 6.14861 | 6.0283 | 6.39438 | 6.20291 | 6.0906 | 6.04156 | 5.653 | 6.0162 | 6.45056 | 6.32917 | 6.56072 | 6.55549 |
| SMG1 | Q96Q15 | 6.23171 | 6.35958 | 6.73984 | 6.24262 | 6.22708 | 6.4871 | 6.30743 | 6.27215 | 6.56793 | 6.52335 | 6.57617 | 6.71895 | 5.68319 | 5.52897 | 5.67871 | 5.85521 |
| DDX24 | Q9GZR7 | 6.10921 | 6.0142 | 5.94289 | 6.71155 | 6.18681 | 5.9483 | 6.54119 | 5.96405 | 6.09106 | 5.92282 | 5.9647 | 5.91524 | 6.52878 | 6.72121 | 6.57838 | 6.85962 |
| MCM6 | Q14566 | 5.7313 | 6.01683 | 6.10641 | 5.85374 | 5.83481 | 6.01199 | 6.00783 | 5.85248 | 6.03398 | 5.96209 | 6.45631 | 6.09768 | 7.08365 | 6.80355 | 7.26071 | 6.88664 |
| FAM120A | Q9NZB2-6 | 6.52179 | 6.42379 | 6.60582 | 6.41389 | 6.38718 | 6.50321 | 6.19332 | 6.49204 | 6.55577 | 6.42759 | 6.4017 | 6.39761 | 5.80258 | 5.77956 | 5.45288 | 5.64128 |
| SYNM | O15061 | 6.25232 | 6.24194 | 5.94463 | 6.47271 | 6.16917 | 6.11529 | 6.16801 | 6.14011 | 6.11404 | 6.02017 | 6.00128 | 5.94847 | 7.00995 | 6.4496 | 6.44682 | 6.5055 |
| ZNF318 | Q5VUA4 | 6.31629 | 6.50029 | 6.3692 | 6.96951 | 6.09666 | 6.08054 | 6.37488 | 5.96162 | 6.18109 | 6.4436 | 6.34576 | 6.15913 | 6.2782 | 5.8715 | 5.98512 | 6.06663 |
| VCPIP1 | Q96JH7 | 6.1144 | 6.35127 | 6.3752 | 6.15476 | 6.28075 | 6.51682 | 6.10552 | 6.4301 | 6.20387 | 6.26094 | 6.39983 | 6.42055 | 6.20071 | 6.13405 | 6.09114 | 5.96006 |
| KDM3B | Q7LBC6 | 5.76785 | 5.99535 | 5.92323 | 5.93162 | 5.91645 | 6.06589 | 6.07378 | 6.00642 | 5.96455 | 6.3336 | 6.2493 | 6.10018 | 6.70172 | 6.97758 | 7.06129 | 6.93119 |
| PSMA6 | P60900 | 6.01957 | 6.00191 | 6.13958 | 6.06093 | 6.49691 | 6.33036 | 6.40419 | 6.14915 | 6.38447 | 5.94218 | 6.16907 | 5.91082 | 6.33515 | 6.38487 | 6.62948 | 6.64138 |
| RPL6 | Q02878 | 6.1128 | 6.5517 | 6.07819 | 6.27804 | 6.40624 | 6.79672 | 6.2772 | 6.64633 | 6.40956 | 6.17932 | 6.4529 | 6.22258 | 5.55301 | 6.6648 | 5.78821 | 5.58242 |
| PSMD3 | O43242 | 6.26758 | 6.33158 | 6.12352 | 6.25024 | 6.43886 | 6.50866 | 6.14002 | 6.25803 | 6.06784 | 6.22053 | 6.13889 | 6.08137 | 6.31871 | 6.37559 | 6.17261 | 6.30598 |
| ZC3H14 | Q6PJT7-2 | 6.30096 | 6.44373 | 6.06379 | 6.68476 | 6.4459 | 6.35036 | 6.12133 | 5.98367 | 6.15195 | 6.10927 | 6.13481 | 6.19661 | 6.75896 | 5.997 | 6.12829 | 6.12859 |
| LETM1 | O95202 | 6.43226 | 6.42049 | 5.72281 | 6.51884 | 6.23215 | 6.04552 | 6.35431 | 6.14868 | 6.14391 | 5.88138 | 5.88706 | 5.76445 | 6.72328 | 6.42862 | 6.65422 | 6.64201 |
| SART3 | A0A499FI31 | 5.86354 | 5.89278 | 5.86887 | 6.22498 | 6.1529 | 5.99075 | 6.38654 | 5.92997 | 6.0953 | 5.87425 | 5.82907 | 5.67893 | 7.03996 | 7.0025 | 7.05644 | 7.11325 |
| HNRNPF | P52597 | 5.99204 | 6.14204 | 5.59989 | 6.312 | 6.37122 | 6.05736 | 6.76297 | 6.15557 | 6.61545 | 5.62954 | 5.87766 | 6.05522 | 6.36436 | 6.4693 | 6.53064 | 7.06475 |
| RUVBL1 | Q9Y265 | 6.21416 | 6.15384 | 6.45474 | 6.30187 | 6.43962 | 6.43705 | 6.26677 | 6.31035 | 6.31897 | 6.3507 | 6.3526 | 6.23178 | 6.21417 | 5.99891 | 5.92022 | 6.03424 |
| AP3B1 | O00203 | 6.4107 | 6.31192 | 6.69467 | 6.15814 | 6.19108 | 6.46047 | 6.38027 | 6.66792 | 6.23051 | 6.51482 | 6.42062 | 6.20165 | 5.86923 | 6.02044 | 5.86389 | 5.60367 |
| PNPLA6 | Q8IY17 | 6.07836 | 6.40443 | 5.88143 | 6.38652 | 5.84578 | 5.78231 | 6.12882 | 5.89495 | 6.30664 | 6.36423 | 6.29547 | 5.94608 | 6.60907 | 6.70397 | 6.70609 | 6.66595 |
| RPA1 | P27694 | 5.54805 | 5.63567 | 5.92346 | 5.31173 | 5.7852 | 6.27399 | 5.72385 | 5.72953 | 6.09268 | 6.20598 | 6.06224 | 5.86365 | 7.30736 | 7.40779 | 7.81068 | 7.31814 |
| CSDE1 | O75534-4 | 6.59877 | 6.64852 | 7.02174 | 6.483 | 6.41931 | 6.60554 | 6.39535 | 6.67702 | 6.84197 | 6.75701 | 6.86986 | 6.77792 | 5.0298 | 5.17709 | 4.81526 | 4.88183 |
| TKT | P29401-2 | 5.75858 | 5.73718 | 6.21775 | 5.38304 | 6.0799 | 6.54091 | 5.91355 | 6.43185 | 6.12223 | 6.86156 | 6.04862 | 5.84224 | 6.39045 | 7.61198 | 7.18319 | 6.87696 |
| PMPCA | Q10713 | 6.21154 | 6.21564 | 5.7881 | 6.62087 | 6.40415 | 5.98231 | 6.5672 | 6.01357 | 6.21858 | 5.78243 | 5.87672 | 5.8901 | 6.47468 | 6.39263 | 6.73315 | 6.82833 |
| UCHL1 | P09936 | 6.14792 | 4.94795 | 6.13231 | 4.34652 | 6.26906 | 5.86099 | 5.73231 | 7.16003 | 5.69528 | 6.72983 | 5.91411 | 6.95688 | 6.23283 | 7.54947 | 7.16972 | 7.1548 |
| FUBP1 | Q96AE4-2 | 6.43045 | 5.83036 | 6.13743 | 5.6789 | 6.74483 | 5.88464 | 6.00449 | 6.23629 | 6.26411 | 6.1539 | 6.39912 | 6.32068 | 7.19671 | 6.086 | 6.40697 | 6.22512 |
| SDHA | P31040 | 6.28556 | 6.33503 | 5.56987 | 6.43172 | 6.34551 | 6.11157 | 6.17788 | 6.08205 | 6.104 | 6.12413 | 5.89755 | 5.96841 | 6.77203 | 6.4246 | 6.6423 | 6.7278 |
| SPEN | Q96T58 | 6.4537 | 6.66689 | 6.02921 | 7.05264 | 6.0528 | 5.83558 | 6.21872 | 6.00192 | 6.18124 | 5.91933 | 6.1275 | 6.12854 | 5.84196 | 6.30775 | 6.6679 | 6.51432 |
| MYO1B | O43795 | 6.16083 | 6.3899 | 6.03044 | 6.25944 | 6.1802 | 6.10631 | 6.12215 | 5.83805 | 6.1971 | 6.16682 | 6.1556 | 6.14931 | 6.5622 | 6.45088 | 6.78018 | 6.45057 |
| AP1G1 | O43747-2 | 6.23989 | 6.43411 | 6.7884 | 6.63755 | 6.0547 | 6.64291 | 6.24162 | 6.28598 | 6.07871 | 6.3938 | 6.3041 | 6.25429 | 5.92458 | 5.97487 | 5.79701 | 5.94747 |
| DDX54 | Q8TDD1-2 | 6.42008 | 6.52356 | 5.87824 | 6.72495 | 5.95657 | 5.88987 | 6.4629 | 6.13048 | 6.24925 | 6.15877 | 6.11792 | 6.38029 | 5.92132 | 6.37666 | 6.27154 | 6.5376 |
| CEP170 | Q5SW79 | 6.18051 | 5.92533 | 6.55174 | 6.02298 | 6.1617 | 6.17905 | 6.16749 | 6.04967 | 6.65187 | 6.53846 | 6.67838 | 6.49023 | 6.04787 | 6.09692 | 6.01067 | 6.24713 |
| POLD1 | P28340 | 5.66651 | 6.02933 | 6.08894 | 5.61229 | 5.86901 | 6.2864 | 6.01089 | 5.87136 | 6.02412 | 6.13403 | 6.32287 | 6.05537 | 7.03222 | 7.04033 | 7.15303 | 6.8033 |
| POLE | Q07864 | 5.97468 | 6.38134 | 6.33241 | 6.41885 | 5.98948 | 6.24248 | 6.08696 | 6.12622 | 6.29223 | 6.31259 | 6.35507 | 6.54667 | 6.04198 | 6.22395 | 6.20017 | 6.47491 |
| ACAT1 | P24752 | 6.27682 | 6.02236 | 5.81825 | 6.27798 | 6.47557 | 6.01108 | 6.50771 | 6.30719 | 6.19542 | 5.98673 | 6.25473 | 6.2531 | 6.54759 | 6.22973 | 6.46759 | 6.36815 |
| UBE4B | O95155-4 | 6.0563 | 6.3572 | 6.64797 | 6.14131 | 6.1787 | 6.5755 | 6.24919 | 6.47512 | 6.25955 | 6.26187 | 6.22515 | 6.15726 | 6.05351 | 6.11672 | 6.10371 | 6.14094 |
| NCOR2 | C9J0Q5 | 6.43168 | 6.19281 | 6.37723 | 6.53378 | 6.37593 | 6.08557 | 6.16924 | 5.91814 | 6.28686 | 6.41513 | 6.4438 | 6.38184 | 6.38465 | 6.15128 | 5.96163 | 5.89044 |
| UBA2 | Q9UBT2 | 5.70204 | 5.73911 | 6.26488 | 5.16661 | 5.62935 | 6.26318 | 5.67297 | 6.1651 | 5.96743 | 5.99564 | 6.34916 | 5.92732 | 6.85443 | 7.64241 | 7.71102 | 6.94936 |
| EIF5B | O60841 | 6.52721 | 6.35266 | 6.52797 | 5.90023 | 6.30244 | 6.38765 | 6.10724 | 6.3358 | 6.34426 | 6.35211 | 6.28481 | 6.26996 | 5.53422 | 6.59928 | 5.8715 | 6.30265 |
| PSMC1 | P62191 | 5.82982 | 6.29293 | 6.26595 | 6.20537 | 6.06537 | 6.65068 | 6.71573 | 6.40089 | 6.30349 | 6.14235 | 6.5233 | 6.2669 | 5.86187 | 6.37818 | 6.0453 | 6.05187 |
| HSPA4L | O95757 | 6.14499 | 6.16654 | 6.59734 | 5.99279 | 6.3041 | 6.46512 | 6.19117 | 6.26501 | 6.24838 | 6.43564 | 6.47329 | 6.21512 | 6.21756 | 6.11437 | 6.12706 | 6.04151 |
| NRDC | O43847-2 | 6.21562 | 6.32789 | 7.11386 | 6.10948 | 6.37403 | 6.7161 | 6.25301 | 6.60231 | 6.31821 | 6.45994 | 6.73246 | 6.55741 | 5.54112 | 5.73692 | 5.47573 | 5.46591 |
| ANKLE2 | Q86XL3 | 6.37957 | 6.49301 | 6.32395 | 6.56811 | 6.10746 | 6.07972 | 6.26155 | 5.9815 | 6.21552 | 6.05965 | 6.07848 | 6.12496 | 6.51461 | 6.17579 | 6.38159 | 6.25452 |
| LONP1 | P36776 | 6.31909 | 6.35311 | 5.90007 | 6.47621 | 6.39225 | 6.11313 | 6.25379 | 6.11512 | 6.04321 | 5.99055 | 5.93495 | 5.70203 | 6.88994 | 6.36254 | 6.78271 | 6.37132 |
| RAB2B | Q8WUD1 | 6.21091 | 6.25852 | 5.99538 | 6.43165 | 6.24268 | 6.15638 | 6.50966 | 5.94095 | 6.17136 | 6.25806 | 6.0337 | 5.89669 | 6.44756 | 6.12381 | 6.86962 | 6.45306 |
| HLTF | Q14527 | 5.31542 | 5.82714 | 6.13464 | 5.55769 | 5.66554 | 6.1904 | 5.95347 | 5.92004 | 6.3086 | 6.34442 | 6.36407 | 6.30362 | 7.0341 | 7.16499 | 6.8684 | 7.04747 |
| DNAJC13 | O75165 | 6.00159 | 6.49243 | 6.06216 | 6.35623 | 5.88322 | 6.48355 | 6.26067 | 6.10773 | 5.93227 | 6.36174 | 6.58971 | 6.64642 | 5.93596 | 6.20212 | 6.32999 | 6.35421 |
| GMPS | P49915 | 5.91495 | 5.90034 | 6.3899 | 5.61354 | 5.92414 | 6.40161 | 5.76655 | 6.34263 | 6.04591 | 6.17862 | 6.29948 | 6.2895 | 6.33793 | 7.11499 | 6.95356 | 6.52635 |

| | | | | | | | | | | | | | | | | | |
|---|---|---|---|---|---|---|---|---|---|---|---|---|---|---|---|---|---|
| PLS3 | P13797 | 6.2182 | 6.00679 | 6.47953 | 5.79273 | 6.35216 | 6.53408 | 5.97793 | 6.36217 | 6.2097 | 6.22634 | 6.15957 | 6.28542 | 6.81457 | 6.34267 | 6.25675 | 5.98138 |
| ME2 | P23368 | 6.3804 | 6.22011 | 6.09879 | 6.66621 | 6.1655 | 5.946 | 6.45815 | 5.93429 | 6.21631 | 6.13301 | 6.04447 | 6.00638 | 6.47379 | 6.20515 | 6.4872 | 6.56424 |
| NCBP1 | Q09161 | 6.03331 | 6.24756 | 5.82039 | 6.21621 | 6.15419 | 6.2781 | 6.04078 | 5.98552 | 6.07304 | 6.05817 | 6.05692 | 5.92973 | 6.95158 | 6.49382 | 7.00907 | 6.6516 |
| CD2AP | Q9Y5K6 | 6.36445 | 6.25265 | 7.05638 | 6.15045 | 6.46166 | 6.71255 | 6.22393 | 6.28744 | 6.41706 | 6.20203 | 6.47499 | 6.42364 | 5.90294 | 5.8961 | 5.48679 | 5.68694 |
| DIS3 | Q9Y2L1 | 5.55409 | 5.77467 | 5.96321 | 5.45477 | 6.05732 | 5.85844 | 5.94295 | 5.92314 | 6.17194 | 5.97662 | 6.1514 | 5.91964 | 7.2272 | 7.25225 | 7.85009 | 6.92229 |
| NUP54 | Q7Z3B4 | 6.29186 | 6.23439 | 6.04959 | 6.26204 | 6.50866 | 6.04197 | 6.14854 | 6.19182 | 6.2401 | 6.03195 | 6.2959 | 6.29533 | 6.82829 | 5.84967 | 6.5434 | 6.18648 |
| USP10 | Q14694-2 | 6.28309 | 6.24613 | 6.53804 | 6.09136 | 6.36101 | 6.48586 | 6.19925 | 6.37996 | 6.36598 | 6.31103 | 6.40577 | 6.47255 | 5.83709 | 6.03536 | 5.97137 | 6.01614 |
| THOC1 | Q96FV9 | 6.23722 | 6.41537 | 6.08296 | 6.58266 | 6.33253 | 6.04587 | 6.16154 | 5.79804 | 6.33395 | 6.09163 | 6.15284 | 6.04383 | 6.45915 | 6.1764 | 6.63652 | 6.44951 |
| DOCK7 | Q96N67 | 6.39016 | 6.42327 | 6.4955 | 6.17552 | 6.21244 | 6.5399 | 6.10551 | 6.3455 | 6.20021 | 6.33702 | 6.1788 | 6.15054 | 6.06562 | 6.12917 | 6.1561 | 6.09474 |
| CTNNA1 | P35221 | 6.62108 | 6.45777 | 6.2628 | 6.40677 | 6.10716 | 6.28008 | 6.64671 | 6.12434 | 6.52269 | 6.14305 | 6.21939 | 6.13047 | 5.89258 | 5.97748 | 6.00285 | 6.20479 |
| LIG3 | P49916 | 5.84011 | 6.05559 | 5.83228 | 6.01071 | 5.96194 | 6.18767 | 6.15423 | 5.93581 | 5.85373 | 6.04419 | 6.28159 | 6.14386 | 6.86074 | 7.01374 | 7.04749 | 6.77632 |
| SUN1 | O94901-9 | 6.3744 | 6.29867 | 5.85285 | 6.55944 | 6.11142 | 6.01554 | 6.34043 | 5.8523 | 6.10625 | 6.96209 | 5.81357 | 5.73127 | 7.00923 | 6.50208 | 6.945 | 6.52544 |
| KIAA1671 | Q9BY89 | 5.88406 | 5.99361 | 5.81123 | 6.05864 | 6.08265 | 6.18012 | 6.19779 | 5.81749 | 5.92695 | 5.92256 | 5.73026 | 5.97653 | 7.36792 | 6.85229 | 7.12644 | 7.07146 |
| LUZP1 | Q86V48 | 6.04498 | 5.989 | 5.93743 | 6.28419 | 6.04188 | 5.94278 | 6.09201 | 5.93333 | 6.53389 | 6.22741 | 6.38112 | 6.20672 | 6.71674 | 6.54639 | 6.63859 | 6.48354 |
| PFKP | Q01813 | 5.94921 | 6.22206 | 6.8926 | 6.08303 | 6.40286 | 6.97752 | 6.52597 | 6.4375 | 6.60515 | 6.26019 | 6.47764 | 6.42738 | 5.63758 | 5.78182 | 5.5767 | 5.74279 |
| EHMT2 | Q96KQ7-2 | 6.31813 | 6.5777 | 6.04564 | 6.74301 | 6.35398 | 6.1051 | 6.25796 | 6.08778 | 6.29996 | 6.16001 | 6.13698 | 6.03517 | 6.3762 | 6.06205 | 6.30457 | 6.13577 |
| PWP2 | Q15269 | 6.3699 | 6.27339 | 5.934 | 6.78251 | 6.40928 | 5.80965 | 6.48255 | 5.90961 | 6.19186 | 5.93528 | 6.01154 | 5.98307 | 6.64391 | 6.14904 | 6.62773 | 6.48668 |
| UQCRC1 | P31930 | 5.90214 | 5.59198 | 5.40414 | 5.8054 | 6.42781 | 5.97983 | 6.32141 | 6.76093 | 6.09772 | 6.3128 | 6.12948 | 6.47867 | 6.49421 | 6.55812 | 6.77281 | 6.96255 |
| PKN2 | Q16513-2 | 6.47193 | 6.33059 | 6.54432 | 6.37012 | 6.30052 | 6.47735 | 6.3607 | 6.23898 | 6.31814 | 6.59004 | 6.3867 | 6.48324 | 5.94712 | 5.80628 | 5.76332 | 5.61063 |
| INF2 | Q27J81 | 6.22213 | 6.40258 | 6.34149 | 6.22267 | 5.98698 | 6.45486 | 6.10678 | 6.33587 | 6.50425 | 6.42267 | 6.58543 | 6.34166 | 5.86401 | 6.07625 | 6.20714 | 5.92525 |
| AXL | P30530 | 6.63049 | 6.51848 | 6.61017 | 7.15739 | 6.31268 | 6.50542 | 6.58841 | 6.31315 | 6.72001 | 6.68201 | 6.66381 | 7.04441 | 4.91143 | 4.97123 | 4.96907 | 5.40185 |
| LAS1L | Q9Y4W2 | 6.46584 | 6.04005 | 6.24333 | 6.56736 | 6.19089 | 5.67044 | 6.58086 | 5.85887 | 6.4889 | 5.95043 | 6.42258 | 6.05563 | 6.58461 | 5.86524 | 6.56375 | 6.45122 |
| SPAG9 | O60271-4 | 6.40889 | 6.35721 | 7.01904 | 6.19515 | 6.52437 | 6.67024 | 6.41161 | 6.7507 | 6.46037 | 6.36026 | 6.77836 | 6.55751 | 5.56977 | 5.47369 | 5.25588 | 5.20695 |
| AHCY | P23526 | 6.17573 | 6.09119 | 6.85285 | 5.93904 | 6.4776 | 6.77236 | 6.38338 | 6.65641 | 6.49122 | 6.4104 | 6.27973 | 6.22766 | 5.71386 | 5.82757 | 5.90799 | 5.79302 |
| THADA | Q6YHU6 | 6.41507 | 6.32587 | 6.93841 | 6.20593 | 6.27006 | 6.8067 | 6.3911 | 6.8087 | 6.61791 | 6.63025 | 6.54381 | 6.64404 | 5.42759 | 5.47101 | 5.16914 | 5.33444 |
| DHX16 | O60231 | 5.90466 | 6.42041 | 6.09447 | 6.11034 | 6.1151 | 6.18748 | 6.15567 | 5.9256 | 6.18159 | 6.11939 | 6.0392 | 6.11786 | 6.56627 | 6.55887 | 6.96007 | 6.54302 |
| ZFR | Q96KR1 | 6.49943 | 6.55507 | 6.23112 | 6.71788 | 6.50675 | 6.12708 | 6.62043 | 6.00989 | 6.10684 | 5.97645 | 5.88073 | 5.78635 | 6.28676 | 6.15052 | 6.31199 | 6.2327 |
| TXNDC5 | Q8NBS9 | 6.46889 | 6.29381 | 5.89118 | 6.61903 | 6.38838 | 6.1512 | 6.14951 | 6.28954 | 6.28356 | 6.0175 | 5.98684 | 6.47178 | 6.67903 | 5.87483 | 6.10582 | 6.32911 |
| PDIA6 | Q15084-2 | 6.44717 | 6.49307 | 6.02447 | 6.10849 | 6.42372 | 6.20822 | 6.18175 | 6.13312 | 6.04013 | 6.04134 | 5.91576 | 5.95216 | 7.05115 | 6.27893 | 6.28238 | 6.41817 |
| MSH2 | P43246 | 5.37133 | 5.9684 | 5.82501 | 5.48154 | 5.84094 | 6.18124 | 5.83525 | 5.83352 | 5.87386 | 6.15375 | 6.3443 | 5.94151 | 7.33109 | 7.48829 | 7.42215 | 7.10782 |
| EIF5 | P55010 | 6.07221 | 6.0087 | 6.82958 | 6.039 | 6.45888 | 6.6369 | 6.45195 | 6.48507 | 6.51204 | 6.18357 | 6.51014 | 6.42689 | 5.91327 | 5.90918 | 5.72517 | 5.83745 |
| RBM25 | P49756 | 6.24636 | 6.23566 | 5.92469 | 6.55964 | 6.13464 | 5.92641 | 6.36902 | 5.85583 | 6.19451 | 5.9368 | 6.01794 | 6.03225 | 6.78698 | 6.34977 | 6.7576 | 6.67189 |
| THRAP3 | Q9Y2W1 | 6.14139 | 6.42303 | 5.96274 | 6.54629 | 5.91874 | 5.87965 | 6.32498 | 5.8649 | 6.16796 | 6.16632 | 6.26667 | 6.28504 | 6.6294 | 6.30582 | 6.45979 | 6.65728 |
| AP2M1 | Q96CW1 | 6.07413 | 6.33089 | 6.45529 | 6.57281 | 6.10118 | 6.29969 | 6.32605 | 5.91192 | 6.30286 | 6.41801 | 6.17366 | 6.406 | 6.1844 | 6.012 | 6.44954 | 5.98155 |
| ERO1A | Q96HE7 | 6.34129 | 6.49367 | 5.89929 | 6.53526 | 6.38804 | 6.17224 | 6.26842 | 6.17986 | 5.95447 | 6.12446 | 5.91071 | 5.93796 | 6.41628 | 6.49253 | 6.51176 | 6.37377 |
| DHX38 | Q92620 | 5.75321 | 6.19827 | 5.90134 | 5.9839 | 6.12496 | 6.15503 | 6.11633 | 5.85288 | 5.95948 | 6.13972 | 5.97076 | 6.00384 | 6.7613 | 6.91531 | 7.24832 | 6.91538 |
| PHIP | Q8WWQ0 | 6.17434 | 6.31841 | 5.70704 | 6.97831 | 5.77335 | 5.98899 | 6.30204 | 6.08793 | 5.98213 | 6.11929 | 5.84535 | 6.1286 | 6.32903 | 6.78938 | 6.67328 | 6.80253 |
| CPSF2 | Q9P2I0 | 6.38027 | 6.18957 | 6.14211 | 6.54073 | 6.27675 | 6.09629 | 6.28089 | 5.98372 | 6.25144 | 6.04624 | 6.087 | 6.16263 | 6.6531 | 5.86596 | 6.48716 | 6.55613 |
| G3BP1 | Q13283 | 6.43043 | 6.50842 | 6.68859 | 6.33924 | 6.46027 | 6.4708 | 6.24274 | 6.51501 | 6.57523 | 6.54307 | 6.74658 | 6.77922 | 5.72265 | 5.40362 | 5.18437 | 5.38976 |
| PPIB | P23284 | 6.17905 | 6.44451 | 5.65986 | 6.369 | 6.32208 | 6.07872 | 6.23814 | 6.18372 | 6.19998 | 6.02492 | 6.04358 | 6.09923 | 6.67956 | 6.32461 | 6.75583 | 6.39721 |
| PLAA | Q9Y263 | 6.20067 | 6.17499 | 6.48204 | 6.35844 | 6.38846 | 6.28359 | 6.24677 | 6.22518 | 6.30001 | 6.42397 | 6.62382 | 6.33625 | 6.06904 | 5.93875 | 6.11212 | 5.83592 |
| LMAN2 | Q12907 | 5.95548 | 6.39452 | 5.89136 | 6.35572 | 6.07273 | 6.05944 | 6.10516 | 6.10501 | 5.99301 | 5.96773 | 6.11864 | 6.24185 | 6.75088 | 6.71097 | 6.71956 | 6.55793 |
| STAT1 | P42224 | 6.08022 | 6.45786 | 6.64291 | 6.05042 | 6.23633 | 6.92598 | 6.26357 | 6.84752 | 6.10316 | 6.57143 | 6.2728 | 6.34019 | 5.60785 | 6.02522 | 5.6713 | 5.90322 |
| PRPH | P41219-2 | 6.53641 | 6.32349 | 6.3733 | 6.59401 | 6.66638 | 5.7909 | 6.41997 | 6.11695 | 6.45316 | 5.75891 | 5.92551 | 5.98416 | 6.39454 | 6.07205 | 6.30849 | 6.28177 |
| EDC4 | Q6P2E9 | 5.97789 | 5.96711 | 6.67584 | 5.91313 | 6.17159 | 6.46344 | 6.11839 | 6.0792 | 6.56948 | 6.36801 | 6.58984 | 6.7143 | 6.21092 | 6.14179 | 6.08237 | 5.95671 |
| DBN1 | Q16643 | 6.44183 | 6.34007 | 6.03477 | 6.17319 | 6.41494 | 6.11041 | 6.13167 | 6.21499 | 6.21365 | 6.17193 | 6.32564 | 6.10551 | 7.01643 | 6.04401 | 6.12527 | 6.13568 |
| YWHAZ | P63104 | 6.18917 | 5.86502 | 6.62909 | 5.65494 | 6.36051 | 6.51138 | 6.05386 | 6.22701 | 6.48919 | 6.51307 | 6.55089 | 6.2918 | 6.4185 | 6.172 | 6.14345 | 5.93012 |
| CRYBG1 | Q9Y4K1 | 6.00187 | 6.27164 | 6.10409 | 6.07261 | 5.96329 | 6.25192 | 6.01915 | 6.18787 | 6.24684 | 6.17042 | 6.32585 | 6.23347 | 6.37236 | 6.56422 | 6.66827 | 6.54613 |
| ARID1A | O14497 | 6.39346 | 6.32537 | 6.10755 | 6.40196 | 6.23527 | 6.09961 | 6.37022 | 6.10689 | 6.08534 | 6.16 | 6.23928 | 6.06533 | 6.75723 | 6.05815 | 6.37545 | 6.21889 |
| HADHB | P55084 | 6.12576 | 6.2694 | 5.75332 | 6.43527 | 6.42759 | 6.10263 | 6.59601 | 6.07767 | 6.14944 | 5.99395 | 5.80618 | 5.67773 | 6.75873 | 6.65184 | 6.60088 | 6.57359 |
| EEA1 | Q15075 | 6.87522 | 6.61024 | 6.20894 | 6.32375 | 6.43737 | 6.06273 | 6.16162 | 6.08051 | 6.35115 | 6.23371 | 6.5722 | 6.41054 | 6.56472 | 5.64899 | 5.824 | 5.6343 |
| HK1 | P19367-3 | 6.36799 | 6.38454 | 6.09545 | 6.48717 | 6.31186 | 6.11328 | 6.17027 | 6.01135 | 6.23414 | 6.13853 | 5.82029 | 5.87349 | 6.55316 | 6.27481 | 6.67615 | 6.4875 |
| DLD | P09622 | 6.30637 | 6.56115 | 5.74857 | 6.50484 | 6.32629 | 6.05639 | 6.13077 | 6.12866 | 6.00404 | 6.06575 | 5.97278 | 5.98339 | 6.86512 | 6.40781 | 6.50734 | 6.43072 |
| SLC25A12 | O75746 | 6.41058 | 6.37058 | 5.98197 | 6.64599 | 6.20632 | 5.93342 | 6.41424 | 5.92572 | 6.11309 | 6.12302 | 6.08157 | 5.94052 | 6.6402 | 6.24365 | 6.48518 | 6.48395 |
| ETFA | P13804 | 6.45692 | 6.41076 | 5.74065 | 6.41341 | 6.70093 | 6.36365 | 6.1943 | 6.44016 | 6.39811 | 6.22624 | 6.17851 | 5.88621 | 6.55741 | 5.90056 | 6.09064 | 6.04154 |
| PPL | O60437 | 6.33649 | 6.40277 | 5.75915 | 5.97973 | 6.26219 | 5.99395 | 5.98091 | 6.15881 | 5.99446 | 6.29426 | 5.99006 | 5.84966 | 7.35402 | 6.84098 | 6.5245 | 6.27807 |
| EHMT1 | Q9H9B1 | 6.2672 | 6.51875 | 6.02999 | 7.03722 | 6.05106 | 6.11214 | 6.57747 | 6.14669 | 6.22538 | 6.13674 | 6.21069 | 6.13913 | 6.1549 | 6.09577 | 6.18727 | 6.1096 |
| IARS2 | Q9NSE4 | 6.396 | 6.46871 | 5.79316 | 6.66102 | 6.27972 | 5.98992 | 6.29781 | 6.15415 | 6.09702 | 6.07823 | 5.89101 | 5.83675 | 6.70176 | 6.36986 | 6.52452 | 6.46038 |
| FERMT2 | Q96AC1-3 | 6.12795 | 5.98652 | 6.36665 | 5.91385 | 6.19463 | 6.35942 | 6.30564 | 6.10179 | 6.47358 | 6.64866 | 6.28811 | 6.22581 | 6.49439 | 6.19639 | 6.50168 | 5.81495 |
| UPF1 | Q92900 | 6.42364 | 6.26891 | 6.94666 | 6.38379 | 6.17056 | 6.43315 | 6.28275 | 6.14123 | 6.47828 | 6.67117 | 6.60886 | 6.623 | 5.72501 | 5.77525 | 5.52643 | 5.54131 |
| WDR62 | O43379-4 | 6.21707 | 6.08184 | 7.02664 | 6.04881 | 6.18699 | 6.40705 | 6.12804 | 6.34339 | 6.54781 | 6.69826 | 7.01581 | 7.01028 | 5.47443 | 5.76638 | 5.50052 | 5.54666 |
| UTP14A | Q9BVJ6 | 6.22934 | 6.31995 | 6.05182 | 6.88392 | 6.15917 | 5.9403 | 6.50033 | 6.08219 | 6.16459 | 5.96971 | 6.02854 | 6.20997 | 6.35965 | 6.16238 | 6.49448 | 6.44367 |
| IGF2BP2 | Q9Y6M1 | 6.23867 | 6.35531 | 6.43814 | 6.06198 | 6.4612 | 6.61412 | 6.15108 | 6.47692 | 6.40685 | 6.4751 | 6.40193 | 6.45415 | 5.92542 | 5.9761 | 5.84811 | 5.71492 |
| UBTF | P17480 | 6.2994 | 6.40631 | 5.92178 | 6.62432 | 6.07408 | 6.07125 | 6.16731 | 6.06745 | 6.02177 | 6.25832 | 5.88955 | 5.97503 | 6.53926 | 6.559 | 6.5217 | 6.60347 |
| SRPRA | P08240 | 6.31321 | 6.10157 | 5.64129 | 6.49162 | 6.07494 | 5.78892 | 6.82902 | 5.88741 | 6.22992 | 5.4018 | 5.58392 | 5.84036 | 6.52832 | 6.95514 | 7.00508 | 7.32748 |
| POLR2B | P30876 | 6.60194 | 6.67453 | 6.33799 | 6.73119 | 6.34082 | 6.19398 | 6.56967 | 6.48019 | 6.30284 | 6.19163 | 6.34136 | 6.19328 | 5.79221 | 5.71818 | 5.68698 | 5.84321 |
| CORO1C | Q9ULV4-3 | 6.19428 | 6.5332 | 6.06144 | 6.38662 | 6.43402 | 6.51493 | 6.15782 | 6.2274 | 5.99534 | 5.94618 | 6.01596 | 5.91166 | 6.49776 | 6.48823 | 6.44922 | 6.18594 |
| RPS7 | P62081 | 6.31373 | 6.50724 | 6.87634 | 6.26605 | 6.33018 | 6.31727 | 6.27122 | 6.39522 | 6.43109 | 6.61323 | 6.68397 | 6.55688 | 5.62319 | 5.7131 | 5.62548 | 5.47582 |
| HECTD1 | A0A087X2H1 | 6.44376 | 6.32421 | 7.3641 | 6.32022 | 6.44858 | 6.81 | 6.2794 | 6.46007 | 6.69255 | 6.5947 | 6.76014 | 7.0256 | 5.25319 | 4.97946 | 4.97578 | 5.26825 |
| CEBPZ | Q03701 | 6.29437 | 6.26742 | 5.85327 | 6.66901 | 6.19337 | 5.76756 | 6.39176 | 5.89251 | 6.00559 | 5.81625 | 5.95372 | 5.8629 | 6.55721 | 6.59066 | 6.95059 | 6.93382 |
| HEXB | P07686 | 6.57861 | 5.9921 | 5.89727 | 5.79861 | 6.63043 | 5.84688 | 6.0811 | 6.47396 | 5.85511 | 6.47256 | 6.12338 | 6.42436 | 6.65516 | 6.41778 | 6.37077 | 6.38194 |
| AK2 | P54819 | 5.85244 | 5.11164 | 5.62219 | 5.2992 | 6.75678 | 5.16571 | 5.9897 | 6.69423 | 5.7174 | 6.46249 | 6.11129 | 6.63998 | 6.98287 | 6.84913 | 7.41082 | 7.3341 |
| SEC31A | D6REX3 | 6.25737 | 6.01066 | 6.88883 | 5.85517 | 6.07482 | 6.42803 | 6.08978 | 6.38264 | 6.14923 | 6.40732 | 6.50839 | 6.06394 | 6.50026 | 6.26232 | 6.21515 | 5.9061 |
| ZC3HAV1 | Q7Z2W4 | 6.32962 | 6.4296 | 6.36894 | 6.44385 | 6.0818 | 6.1941 | 6.2408 | 6.00211 | 6.23679 | 6.48136 | 6.46613 | 6.30079 | 6.16722 | 6.11509 | 6.12645 | 6.01534 |
| USO1 | O60763-2 | 6.28708 | 6.15947 | 6.3945 | 6.0351 | 6.09331 | 6.01864 | 6.07179 | 6.3957 | 6.34771 | 6.36204 | 6.19866 | 6.29871 | 6.416 | 6.27847 | 6.43295 | 6.20987 |
| EIF2A | Q9BY44 | 6.14637 | 6.38403 | 6.58127 | 6.36764 | 6.22411 | 6.48664 | 6.33484 | 6.33351 | 6.56026 | 6.32173 | 6.4931 | 6.36062 | 5.52207 | 5.88656 | 5.80646 | 6.19078 |
| KIF11 | P52732 | 6.55368 | 6.37806 | 6.83827 | 6.20325 | 6.38621 | 6.60211 | 6.28288 | 6.6281 | 6.90983 | 6.4756 | 6.76309 | 6.69708 | 5.42745 | 5.37523 | 5.17046 | 5.3087 |
| MOGS | Q13724 | 6.07617 | 6.08542 | 5.88218 | 6.73334 | 6.1666 | 5.97072 | 6.13189 | 5.88917 | 6.08519 | 6.15943 | 6.09288 | 6.20696 | 6.69689 | 6.61966 | 6.54808 | 6.65542 |
| UMPS | P11172 | 6.24959 | 6.1104 | 6.68379 | 5.87421 | 6.293 | 6.61522 | 6.18779 | 6.71082 | 6.49676 | 6.40268 | 6.61139 | 6.50185 | 5.78911 | 5.80899 | 5.86063 | 5.80375 |
| U2SURP | O15042 | 6.22525 | 6.32989 | 5.85747 | 6.75096 | 6.08413 | 5.95848 | 6.29084 | 5.97684 | 6.04436 | 6.14994 | 6.05335 | 6.01412 | 6.30569 | 6.55641 | 6.62915 | 6.77311 |
| ARF3 | P61204 | 6.33887 | 6.48491 | 6.79464 | 6.54182 | 6.28953 | 6.44079 | 6.60993 | 6.24058 | 6.33523 | 6.30008 | 6.31741 | 6.21124 | 5.73022 | 5.95619 | 5.80302 | 5.60557 |
| SARS1 | P49591 | 6.33848 | 6.07914 | 6.87846 | 5.9936 | 6.55071 | 6.73631 | 6.44449 | 6.59486 | 6.45095 | 6.31621 | 6.62943 | 6.37399 | 5.84881 | 5.84152 | 5.59092 | 5.33212 |
| EIF2S3 | P41091 | 6.37788 | 6.25284 | 6.62913 | 6.16461 | 6.35068 | 6.39091 | 6.14697 | 6.29324 | 6.3642 | 6.43443 | 6.41819 | 6.3154 | 6.0847 | 6.07788 | 5.97507 | 5.72388 |
| SNRNP70 | P08621 | 6.13463 | 6.34023 | 5.63627 | 6.5058 | 6.2215 | 5.85799 | 6.40413 | 6.08063 | 6.01117 | 5.83195 | 5.86807 | 5.68838 | 6.61015 | 6.81734 | 7.09766 | 6.89409 |
| RCC1 | P18754-2 | 6.23544 | 6.4231 | 5.78591 | 6.51576 | 6.0947 | 5.99656 | 6.39204 | 6.19858 | 6.00687 | 6.11363 | 5.84178 | 6.0228 | 6.50095 | 6.42724 | 6.82078 | 6.62386 |
| HTT | P42858 | 6.11581 | 6.20528 | 6.56754 | 5.8297 | 6.23476 | 6.67441 | 6.17066 | 6.50059 | 6.53372 | 6.4788 | 6.3548 | 6.41252 | 5.82791 | 6.10604 | 6.01948 | 5.96796 |
| SETX | Q7Z333-4 | 6.29363 | 6.49023 | 6.19375 | 6.6535 | 6.15874 | 6.2604 | 6.32592 | 6.23564 | 6.19453 | 6.37573 | 6.27799 | 6.2655 | 5.92429 | 6.11493 | 6.07687 | 6.15833 |
| GOLGA2 | Q08379 | 6.63129 | 6.54424 | 5.99658 | 6.83024 | 6.54603 | 6.30206 | 6.33776 | 6.22927 | 6.24568 | 6.27316 | 6.26149 | 6.12177 | 6.34189 | 5.67676 | 5.88503 | 5.77675 |
| RSL1D1 | O76021 | 6.16042 | 6.23907 | 5.67934 | 6.52177 | 6.07032 | 5.66286 | 6.35732 | 5.73042 | 5.97812 | 5.78477 | 5.57133 | 5.68847 | 6.58874 | 7.34411 | 7.17312 | 7.44982 |
| AARS2 | Q5JTZ9 | 6.3422 | 6.26508 | 5.92388 | 6.60371 | 6.26952 | 5.98819 | 6.20967 | 6.04341 | 6.15927 | 6.16576 | 5.95484 | 5.96224 | 6.74524 | 6.40881 | 6.47114 | 6.48702 |
| MYO9B | Q13459 | 6.26369 | 6.21359 | 6.82582 | 6.0094 | 6.26751 | 6.76466 | 6.26756 | 6.57953 | 6.27728 | 6.59497 | 6.50705 | 6.4285 | 5.69651 | 6.00006 | 5.74624 | 5.55763 |
| PARP14 | Q460N5 | 6.18124 | 6.26069 | 6.85346 | 6.28405 | 5.96999 | 6.40777 | 6.21007 | 6.29018 | 6.71853 | 6.91871 | 6.87814 | 6.78753 | 5.53716 | 5.67676 | 5.50672 | 5.51901 |
| PRRC2B | Q5JSZ5 | 6.68719 | 6.46006 | 6.69749 | 6.46607 | 6.04602 | 6.29303 | 6.38173 | 6.08666 | 6.48618 | 6.67579 | 6.7326 | 6.61769 | 5.67783 | 5.76877 | 5.47023 | 5.45265 |
| IPO4 | Q8TEX9-2 | 6.30204 | 6.08209 | 6.84436 | 6.17895 | 6.29962 | 6.90143 | 6.36884 | 6.75367 | 6.54913 | 6.1662 | 6.64194 | 6.79916 | 5.56586 | 5.7197 | 5.19691 | 5.63011 |
| ASPM | Q8IZT6 | 6.78842 | 6.83616 | 7.0749 | 7.06801 | 6.23409 | 6.54921 | 6.6601 | 6.84948 | 6.92558 | 6.80226 | 7.0461 | 7.20637 | 4.21245 | 4.78559 | 4.14361 | 4.81769 |
| MYO5A | Q9Y4I1-3 | 6.15269 | 6.36256 | 6.38579 | 5.94005 | 6.18903 | 6.23781 | 6.19332 | 6.16417 | 6.44954 | 6.62505 | 6.55515 | 6.49341 | 5.8166 | 6.15168 | 6.06677 | 6.21637 |
| NOLC1 | Q14978-2 | 6.17613 | 6.45189 | 5.76181 | 6.88334 | 5.99593 | 5.94418 | 6.26033 | 6.09178 | 6.13716 | 5.95897 | 5.80685 | 6.1226 | 6.42251 | 6.34372 | 6.63568 | 7.00712 |
| PYGB | P11216 | 6.63749 | 6.42381 | 7.0227 | 6.31539 | 6.24105 | 6.69288 | 6.0616 | 6.34678 | 6.46236 | 6.68694 | 6.69106 | 6.61907 | 5.52194 | 5.62656 | 5.41467 | 5.2357 |
| WASHC2A | Q641Q2 | 5.9671 | 6.0323 | 6.41301 | 5.78539 | 6.01653 | 6.3675 | 6.02796 | 6.23222 | 6.35894 | 6.53932 | 6.41971 | 6.46644 | 6.31845 | 6.47148 | 6.26899 | 6.31466 |
| LTN1 | O94822-3 | 6.04755 | 5.87847 | 6.41325 | 5.83089 | 6.14829 | 6.41594 | 6.02926 | 6.3609 | 6.55312 | 6.54694 | 6.44862 | 6.65646 | 6.09289 | 6.36308 | 6.02551 | 6.18882 |
| PSMC6 | P62333 | 6.15722 | 6.26555 | 6.35119 | 6.04686 | 6.34034 | 6.3115 | 6.18018 | 6.21776 | 6.20727 | 6.24128 | 6.49182 | 6.51275 | 6.64194 | 5.98328 | 6.05069 | 6.00037 |
| WAPL | Q7Z5K2-3 | 5.94028 | 6.47598 | 6.03202 | 6.44362 | 6.07105 | 6.18194 | 6.20621 | 6.02451 | 6.06847 | 6.08104 | 5.98636 | 6.11099 | 6.56026 | 6.62684 | 6.61312 | 6.5773 |
| PSME4 | Q14997 | 6.02116 | 6.08627 | 6.25771 | 6.35278 | 6.09353 | 6.20756 | 6.26963 | 6.31999 | 6.28475 | 6.18187 | 6.37309 | 6.38327 | 6.23328 | 6.14658 | 6.2182 | 6.57031 |
| PSMC4 | P43686 | 6.26503 | 6.20045 | 6.54874 | 6.12723 | 6.70443 | 6.76225 | 6.19863 | 6.40128 | 6.16125 | 6.28793 | 6.42149 | 6.35815 | 6.28064 | 5.71784 | 5.70117 | 5.8635 |
| SEPTIN7 | Q16181-2 | 6.15707 | 6.17787 | 6.57936 | 6.11263 | 6.23652 | 6.4518 | 6.20076 | 6.28594 | 6.2666 | 6.20384 | 6.32627 | 6.08786 | 6.19919 | 6.37897 | 6.24144 | 6.09388 |

| | | | | | | | | | | | | | | | | | |
|---|---|---|---|---|---|---|---|---|---|---|---|---|---|---|---|---|---|
| IDE | P14735 | 6.04658 | 6.35454 | 6.62693 | 5.88624 | 6.442 | 6.6553 | 6.1641 | 6.67595 | 6.25237 | 6.54915 | 6.32244 | 6.40388 | 5.84085 | 6.11738 | 5.82786 | 5.83441 |
| SLK | Q9H2G2-2 | 6.27782 | 6.51212 | 6.68842 | 6.12734 | 6.27331 | 6.72809 | 6.28687 | 6.62822 | 6.29648 | 6.70819 | 6.44804 | 6.44087 | 5.60382 | 6.07269 | 5.48905 | 5.41866 |
| SRCAP | Q6ZRS2 | 6.30278 | 6.15241 | 6.04374 | 6.81719 | 6.009 | 5.9226 | 6.31625 | 5.89449 | 6.15491 | 6.25702 | 6.03927 | 5.95573 | 6.44344 | 6.49711 | 6.60232 | 6.59175 |
| UACA | Q9BZF9 | 6.42953 | 6.57099 | 6.35957 | 6.35681 | 6.63498 | 6.51097 | 6.35661 | 6.58796 | 6.50042 | 6.44185 | 6.56585 | 6.40389 | 5.83633 | 5.5719 | 5.40627 | 5.46607 |
| PREP | P48147 | 6.33863 | 6.04524 | 6.73017 | 5.67339 | 6.32775 | 6.84898 | 6.21894 | 6.86931 | 6.32979 | 6.36927 | 6.19557 | 6.12868 | 5.91939 | 6.20099 | 5.87458 | 5.92932 |
| SCRIB | Q14160-3 | 6.53532 | 6.35091 | 6.0809 | 6.38077 | 6.32506 | 5.90907 | 6.53108 | 6.03794 | 6.62037 | 6.13297 | 6.35562 | 6.16415 | 6.48892 | 5.87469 | 6.1165 | 6.09574 |
| DPP3 | Q9NY33 | 6.17637 | 6.20899 | 6.72582 | 5.71632 | 6.44463 | 6.74608 | 6.28867 | 6.66231 | 6.41801 | 6.27751 | 6.62415 | 6.47023 | 5.72878 | 6.05043 | 5.72833 | 5.73338 |
| H2AX | P16104 | 6.35704 | 6.22234 | 5.63626 | 6.40349 | 6.06512 | 5.6206 | 6.62765 | 5.85111 | 6.06256 | 5.94614 | 5.85506 | 5.83998 | 6.96794 | 6.62098 | 6.83025 | 7.09347 |
| MON2 | Q7Z3U7-5 | 6.38711 | 6.13349 | 6.87285 | 5.96942 | 6.17694 | 6.49383 | 6.22058 | 6.32531 | 6.51711 | 6.47955 | 6.44428 | 6.66586 | 6.05831 | 5.92955 | 5.80805 | 5.51775 |
| PGD | P52209 | 6.23439 | 6.255 | 6.86887 | 5.86433 | 6.37587 | 6.74101 | 6.19336 | 6.69848 | 6.26973 | 6.35163 | 6.44333 | 6.43328 | 5.66223 | 6.14942 | 5.74641 | 5.71267 |
| KMT2B | Q9UMN6 | 6.27751 | 6.72642 | 5.72352 | 6.96628 | 6.05045 | 5.87063 | 6.54092 | 6.25903 | 6.44509 | 6.38131 | 6.38746 | 6.2493 | 5.94577 | 6.2011 | 5.87574 | 6.09947 |
| FXR1 | P51114 | 6.48239 | 6.8325 | 6.61208 | 6.48063 | 6.46356 | 6.66513 | 6.09504 | 6.56166 | 6.29985 | 6.59388 | 6.46577 | 6.34599 | 5.55885 | 5.56287 | 5.59159 | 5.38819 |
| TRIO | O75962 | 6.37475 | 6.36551 | 6.87925 | 6.14341 | 6.32459 | 6.50939 | 6.3271 | 6.4899 | 6.58416 | 6.33491 | 6.60739 | 6.51335 | 5.6717 | 5.82138 | 5.58056 | 5.47263 |
| OLA1 | Q9NTK5 | 6.19831 | 6.14993 | 6.95541 | 6.08832 | 6.4776 | 6.74766 | 6.39957 | 6.5934 | 6.50244 | 6.50676 | 6.49812 | 6.50807 | 5.53953 | 5.76547 | 5.38055 | 5.68887 |
| PSMA7 | O14818 | 6.23304 | 6.33021 | 6.23063 | 5.88212 | 6.49502 | 6.45636 | 6.27076 | 6.40503 | 6.11241 | 6.14972 | 6.20459 | 6.07905 | 6.5225 | 6.31842 | 6.26214 | 6.04801 |
| SERBP1 | Q8NC51 | 6.22436 | 6.51874 | 6.49147 | 5.92818 | 6.33773 | 6.62123 | 6.18209 | 6.65896 | 6.64757 | 6.29462 | 6.52318 | 6.31889 | 5.71729 | 5.87285 | 5.76223 | 5.90064 |
| VIRMA | Q69YN4 | 6.41305 | 6.41622 | 6.01311 | 6.7204 | 6.15633 | 6.13852 | 6.36382 | 6.11732 | 6.0868 | 6.05661 | 6.04866 | 6.01657 | 6.32241 | 6.20742 | 6.32578 | 6.59698 |
| NCAPG | Q9BPX3 | 6.39232 | 6.4122 | 6.92228 | 6.32571 | 6.22174 | 6.60402 | 6.46136 | 6.46214 | 6.59597 | 6.59789 | 6.47375 | 6.67486 | 5.45462 | 5.60908 | 5.28352 | 5.50851 |
| IMPDH2 | P12268 | 6.27432 | 6.18689 | 6.82335 | 5.97018 | 6.38708 | 6.53192 | 6.05799 | 6.73127 | 7.00362 | 6.31479 | 6.69026 | 6.58472 | 5.73733 | 5.80189 | 5.46467 | 5.43973 |
| FAT1 | Q14517 | 6.51341 | 6.40958 | 6.31989 | 6.81567 | 6.24131 | 6.32417 | 6.59384 | 6.22456 | 6.4163 | 5.9511 | 6.19569 | 6.05009 | 6.30876 | 5.78555 | 5.92739 | 5.92269 |
| UGP2 | Q16851 | 6.33947 | 6.2452 | 6.56839 | 5.91708 | 6.44167 | 6.76784 | 6.20773 | 6.61407 | 6.26823 | 6.38195 | 6.35598 | 6.25934 | 5.99562 | 6.07384 | 5.77874 | 5.78485 |
| OSBPL8 | Q9BZF1 | 6.21637 | 6.37383 | 5.80446 | 6.30941 | 5.97268 | 5.92402 | 6.27723 | 5.83703 | 5.95816 | 6.07108 | 6.06435 | 6.2451 | 6.5959 | 5.65645 | 6.8751 | 6.91883 |
| NDUFS2 | O75306 | 6.20498 | 6.11271 | 5.81642 | 6.34235 | 6.35858 | 6.0634 | 6.2653 | 6.20711 | 6.06289 | 6.16881 | 5.91875 | 5.87361 | 6.74039 | 6.37097 | 6.70665 | 6.7871 |
| ATP2B1 | P20020-1 | 6.18383 | 6.45774 | 5.9549 | 6.39578 | 6.06119 | 6.07904 | 6.25154 | 5.92174 | 6.02309 | 6.08936 | 5.98746 | 5.80432 | 6.65502 | 6.52572 | 6.79657 | 6.8127 |
| UQCRC2 | P22695 | 6.30098 | 6.33065 | 5.99275 | 6.73186 | 6.2822 | 6.09721 | 6.56959 | 6.04347 | 6.26562 | 5.95049 | 6.00316 | 6.03946 | 6.48985 | 6.10688 | 6.5123 | 6.28353 |
| MPDZ | O75970 | 6.25193 | 6.29203 | 6.29195 | 6.29648 | 6.12046 | 6.41862 | 5.97558 | 6.51593 | 6.17477 | 6.22437 | 6.35401 | 6.35163 | 6.25528 | 6.00172 | 6.21874 | 6.25651 |
| CPD | O75976 | 6.79084 | 6.55566 | 6.31107 | 6.84062 | 6.24116 | 6.09105 | 6.22495 | 6.13645 | 6.03584 | 6.1411 | 6.33165 | 6.23746 | 6.19417 | 6.06477 | 5.97823 | 5.82497 |
| GSPT1 | P15170-3 | 6.42864 | 6.28339 | 6.63363 | 6.10277 | 6.49602 | 6.83774 | 6.27642 | 6.58403 | 6.4838 | 6.36583 | 6.52574 | 6.50821 | 5.74768 | 5.71575 | 5.44509 | 5.56526 |
| ZFYVE16 | Q7Z3T8 | 6.35303 | 6.58539 | 6.8849 | 6.39884 | 6.29557 | 6.47105 | 6.05527 | 6.18523 | 6.53841 | 6.65185 | 6.89815 | 7.00671 | 5.51255 | 5.38473 | 5.33905 | 5.43927 |
| HNRNPD | Q14103-2 | 6.17109 | 6.35975 | 5.68876 | 6.42314 | 6.18154 | 6.17294 | 6.17559 | 6.18868 | 6.11726 | 6.0211 | 5.78379 | 6.06584 | 6.66704 | 6.51415 | 6.75331 | 6.71602 |
| PNN | Q9H307 | 6.2633 | 6.2968 | 5.94252 | 6.6899 | 6.03855 | 5.92201 | 6.2698 | 5.96001 | 6.18101 | 6.00256 | 6.11336 | 6.00059 | 6.6205 | 6.32657 | 6.58453 | 6.78798 |
| GDI1 | P31150 | 6.29654 | 6.12195 | 6.88203 | 5.97434 | 6.15598 | 6.75152 | 6.23264 | 6.5587 | 6.41831 | 6.47483 | 6.69425 | 6.74502 | 5.79659 | 5.71193 | 5.53932 | 5.64602 |
| FKBP10 | Q96AY3 | 6.66913 | 6.51231 | 6.26413 | 6.66589 | 6.61958 | 6.0182 | 6.1126 | 6.00414 | 6.13744 | 6.02539 | 6.23664 | 6.1342 | 6.66212 | 5.96889 | 5.90083 | 6.06851 |
| ATP5F1C | P36542 | 6.36593 | 6.44902 | 5.71219 | 6.62881 | 6.38714 | 6.00065 | 6.64892 | 5.93164 | 6.21569 | 5.91058 | 6.00542 | 6.00606 | 6.49308 | 6.08812 | 6.65908 | 6.49767 |
| CLPX | O76031 | 6.51988 | 6.36017 | 5.95207 | 6.75523 | 6.42933 | 6.09812 | 6.29626 | 6.18523 | 6.29339 | 6.07368 | 6.07846 | 5.78857 | 6.43325 | 6.03503 | 6.37044 | 6.33088 |
| INTS1 | Q8N201 | 6.43415 | 6.48576 | 5.79376 | 6.82759 | 6.25592 | 6.22775 | 6.37697 | 6.34278 | 6.14752 | 6.15802 | 6.14703 | 6.21314 | 6.11567 | 6.19585 | 6.03104 | 6.24706 |
| HNRNPUL2 | Q1KMD3 | 6.18255 | 6.55426 | 5.8151 | 6.63669 | 6.09999 | 5.98812 | 6.38136 | 5.92593 | 5.93339 | 6.21227 | 5.82508 | 5.84792 | 6.70174 | 6.58543 | 6.62947 | 6.68069 |
| PRPF6 | O94906 | 6.26394 | 6.18555 | 6.0388 | 6.5916 | 6.16674 | 5.85415 | 6.51191 | 5.85305 | 6.31096 | 5.92498 | 6.19684 | 5.98091 | 6.55732 | 6.29964 | 6.47788 | 6.78572 |
| WDR3 | Q9UNX4 | 6.32923 | 6.30525 | 5.94327 | 6.7925 | 6.0226 | 5.91412 | 6.44906 | 5.88155 | 6.10612 | 5.85022 | 6.05295 | 5.94283 | 6.64546 | 6.40356 | 6.74647 | 6.61481 |
| TBL3 | Q12788 | 6.48753 | 6.39998 | 6.07218 | 6.91837 | 6.26163 | 6.14808 | 6.5989 | 6.23904 | 6.36958 | 5.83412 | 5.80653 | 5.83651 | 6.44016 | 6.01699 | 6.34566 | 6.22475 |
| DIAPH1 | O60610 | 6.22721 | 6.23269 | 6.65555 | 6.02394 | 6.28251 | 6.64478 | 6.25005 | 6.63014 | 6.45482 | 6.49666 | 6.51947 | 6.40458 | 5.8144 | 6.00258 | 5.5498 | 5.81082 |
| MACROH2A1 | O75367 | 6.24328 | 6.28387 | 5.67306 | 6.69383 | 6.15979 | 5.79609 | 6.36024 | 5.8276 | 6.2252 | 5.91676 | 5.83306 | 5.74058 | 6.64191 | 6.70348 | 7.05427 | 6.847 |
| CHD1 | O14646 | 6.15463 | 6.42774 | 6.05932 | 6.64297 | 6.34512 | 6.16962 | 6.33787 | 5.91668 | 6.12325 | 6.04251 | 6.10788 | 5.90242 | 6.37567 | 6.29063 | 6.53002 | 6.57365 |
| ITPR1 | Q14643 | 6.2693 | 6.30373 | 5.98709 | 6.7392 | 6.08342 | 5.95219 | 6.2308 | 5.87769 | 6.29714 | 6.08906 | 5.94948 | 5.94921 | 6.57312 | 6.48545 | 6.61155 | 6.60607 |
| NOL6 | Q9H6R4 | 6.23668 | 6.18208 | 6.03664 | 6.85301 | 6.12627 | 6.00606 | 6.19556 | 6.10058 | 6.23491 | 5.96652 | 6.0291 | 6.13223 | 6.60141 | 6.12218 | 6.59935 | 6.57743 |
| SAMHD1 | Q9Y3Z3 | 5.16298 | 5.44898 | 5.77879 | 5.09301 | 5.60909 | 6.14534 | 5.81969 | 5.75776 | 5.97633 | 6.07806 | 6.1293 | 5.81755 | 7.47536 | 7.82149 | 8.19862 | 7.68765 |
| DYNC1LI1 | Q9Y6G9 | 6.33571 | 6.06678 | 6.87997 | 5.99119 | 6.50599 | 6.70994 | 5.96508 | 6.68864 | 6.22062 | 6.33664 | 6.71009 | 6.73706 | 5.94634 | 5.80302 | 5.50829 | 5.59464 |
| CDK5RAP2 | Q96SN8 | 6.63753 | 6.76641 | 6.85747 | 6.75946 | 6.44445 | 6.36441 | 6.29817 | 6.4769 | 6.57971 | 6.7051 | 6.92636 | 6.64838 | 5.26684 | 4.98922 | 5.1307 | 5.1489 |
| ALDH18A1 | P54886 | 6.20781 | 6.36617 | 5.96275 | 6.60478 | 6.15528 | 5.94302 | 6.20727 | 6.14474 | 6.18702 | 5.99735 | 6.2205 | 6.27147 | 6.53698 | 6.24889 | 6.57219 | 6.37533 |
| CAP1 | Q01518 | 5.9937 | 5.95863 | 6.62472 | 5.70857 | 6.36403 | 6.74984 | 5.94543 | 6.27459 | 6.06489 | 6.24888 | 6.56523 | 6.40065 | 6.46889 | 6.38995 | 6.21888 | 6.02312 |
| RANGAP1 | P46060 | 6.39551 | 6.23362 | 6.32564 | 6.4358 | 6.33016 | 6.32261 | 6.41918 | 6.23447 | 6.56711 | 6.11849 | 6.33644 | 6.48397 | 6.02996 | 5.80465 | 5.89157 | 6.07083 |
| ANAPC1 | Q9H1A4 | 6.49617 | 6.10706 | 6.45467 | 6.3382 | 6.31839 | 6.20762 | 6.28024 | 6.18357 | 6.16966 | 6.22108 | 6.34786 | 6.31789 | 6.20429 | 6.11165 | 5.93964 | 6.30201 |
| UBR1 | Q8IWV7 | 5.85123 | 6.16549 | 6.54495 | 5.72156 | 6.0097 | 6.51509 | 6.08722 | 6.14612 | 6.1312 | 6.16181 | 6.24168 | 6.20056 | 6.37596 | 6.64387 | 6.67687 | 6.52671 |
| RDX | P35241 | 6.13249 | 5.8726 | 6.79292 | 5.86308 | 6.39038 | 6.45048 | 6.33411 | 6.21206 | 6.75332 | 6.23702 | 6.8671 | 6.36106 | 6.21283 | 6.14106 | 5.88618 | 5.49331 |
| PES1 | O00541 | 6.2197 | 6.21381 | 5.78645 | 7.12166 | 6.0067 | 5.98069 | 6.39411 | 5.94714 | 6.32746 | 6.02908 | 5.88002 | 6.10945 | 6.34599 | 6.23998 | 6.4319 | 6.96585 |
| MTA2 | O94776 | 6.22785 | 6.06933 | 6.01756 | 6.3728 | 6.09756 | 5.9434 | 6.50547 | 5.77235 | 6.27901 | 6.09939 | 6.11493 | 5.98123 | 6.89871 | 6.47199 | 6.37162 | 6.77679 |
| TXNRD1 | Q16881 | 6.07632 | 6.06046 | 6.5845 | 5.71573 | 6.17766 | 6.8936 | 6.31455 | 6.49909 | 6.10385 | 6.24266 | 6.36311 | 6.23534 | 6.23118 | 6.33151 | 6.23525 | 5.93519 |
| MOV10 | Q9HCE1 | 6.36509 | 6.13493 | 6.4789 | 6.03803 | 6.15835 | 6.38716 | 6.37218 | 6.32783 | 6.40704 | 6.6126 | 6.57178 | 6.35069 | 5.85606 | 6.13271 | 5.99889 | 5.80774 |
| NUP133 | Q8WUM0 | 6.38476 | 6.18356 | 5.8101 | 6.47086 | 6.32177 | 5.76828 | 6.50045 | 5.93312 | 6.17436 | 5.87521 | 5.98999 | 6.02136 | 6.87625 | 6.38699 | 6.7249 | 6.57805 |
| DARS2 | Q6PI48 | 6.19903 | 6.17691 | 5.67583 | 6.43575 | 6.26913 | 6.02949 | 6.22123 | 6.03198 | 6.17043 | 5.97324 | 5.88392 | 5.73299 | 6.93459 | 6.55784 | 6.90983 | 6.7978 |
| RSF1 | Q96T23 | 6.50398 | 6.35464 | 5.99295 | 6.61934 | 6.3263 | 6.0303 | 6.31296 | 6.13853 | 6.23519 | 6.06017 | 6.20364 | 5.96876 | 6.45086 | 6.07844 | 6.38952 | 6.33444 |
| DHX30 | Q7L2E3-2 | 6.28335 | 6.34444 | 6.00904 | 6.68095 | 6.36843 | 6.22203 | 6.31422 | 6.23188 | 6.29443 | 6.24835 | 6.33552 | 6.27425 | 6.09255 | 5.9703 | 6.09782 | 6.23244 |
| TSR1 | Q2NL82 | 5.58406 | 5.73532 | 5.8257 | 5.83027 | 6.01021 | 6.20908 | 5.95251 | 6.08888 | 6.03658 | 5.94283 | 6.20959 | 6.18494 | 6.82644 | 7.18701 | 7.1728 | 7.20379 |
| RPS19 | P39019 | 6.53443 | 6.49539 | 6.49821 | 6.1495 | 6.32913 | 6.38974 | 6.30707 | 6.47272 | 6.51043 | 6.48279 | 6.68892 | 6.68603 | 5.58679 | 5.82872 | 5.46438 | 5.47574 |
| KIF4A | O95239 | 5.59067 | 6.27447 | 6.05611 | 6.1116 | 5.85839 | 6.22985 | 6.15915 | 6.2246 | 6.02688 | 6.26498 | 6.10143 | 6.20734 | 6.56588 | 6.85692 | 6.74994 | 6.7218 |
| SRP72 | O76094 | 6.00101 | 6.22136 | 5.89706 | 5.95933 | 6.05516 | 6.13448 | 6.04555 | 5.97576 | 6.03862 | 6.10182 | 5.93307 | 5.8771 | 6.47499 | 7.33297 | 7.00703 | 6.9447 |
| TRIM25 | Q14258 | 6.29897 | 6.44438 | 6.43842 | 6.0807 | 6.45611 | 6.83332 | 6.11967 | 6.87781 | 6.34194 | 6.3934 | 6.45049 | 6.04303 | 5.6627 | 6.17969 | 5.8935 | 5.48589 |
| SRP68 | Q9UHB9 | 6.13158 | 6.22075 | 6.40928 | 6.06477 | 6.18997 | 6.39947 | 6.2687 | 5.94819 | 6.45364 | 6.0967 | 6.10667 | 6.14067 | 6.51735 | 6.3862 | 6.36748 | 6.29859 |
| SLC4A7 | Q9Y6M7-13 | 6.1364 | 6.44582 | 6.04376 | 6.55299 | 6.02832 | 6.11984 | 6.33721 | 6.13845 | 6.18937 | 6.12128 | 5.93573 | 6.19774 | 6.39928 | 6.36547 | 6.42526 | 6.56307 |
| HNRNPH2 | P55795 | 6.25635 | 6.21539 | 5.78063 | 6.48917 | 6.2894 | 6.1641 | 6.14318 | 6.13833 | 6.27876 | 6.06976 | 6.28635 | 6.40131 | 6.62688 | 6.16264 | 6.40147 | 6.29627 |
| SPG11 | Q96JI7 | 6.85976 | 6.81864 | 7.05841 | 6.64898 | 6.39637 | 6.38339 | 6.10211 | 6.34468 | 6.69362 | 6.85985 | 6.98581 | 6.80323 | 4.87276 | 5.09381 | 4.97718 | 5.10139 |
| KIF2C | Q99661 | 5.87501 | 6.28488 | 6.6453 | 6.22966 | 5.93928 | 6.46225 | 6.37946 | 6.08383 | 6.43588 | 6.62116 | 6.67399 | 6.61017 | 5.85592 | 5.81431 | 5.91469 | 6.17421 |
| EPS15L1 | Q9UBC2-2 | 6.17979 | 5.78962 | 6.20314 | 5.26326 | 6.61098 | 6.13109 | 6.10035 | 6.50289 | 6.33783 | 6.42378 | 6.62533 | 6.49563 | 6.76494 | 6.22482 | 6.29202 | 6.05454 |
| TOPBP1 | Q92547 | 6.20413 | 6.7935 | 6.32083 | 6.92284 | 6.15228 | 6.15866 | 6.2456 | 5.99065 | 6.19998 | 6.3694 | 6.19695 | 6.29965 | 5.97574 | 5.72794 | 6.25851 | 6.17577 |
| RBBP6 | Q7Z6E9 | 7.43746 | 6.04179 | 5.95277 | 6.08518 | 5.39212 | 5.33874 | 8.32682 | 6.95082 | 6.54313 | 5.63657 | 5.36906 | 5.39974 | 5.6776 | 7.03195 | 6.23296 | 6.58329 |
| RPS9 | P46781 | 6.3978 | 6.49125 | 6.0978 | 6.0472 | 6.30099 | 6.24296 | 6.24514 | 6.66663 | 6.84084 | 6.33923 | 5.92853 | 6.11058 | 6.17169 | 6.31712 | 5.68712 | 6.11514 |
| CSTF3 | Q12996 | 6.27516 | 6.26956 | 5.86011 | 6.55805 | 6.28782 | 6.07206 | 6.73525 | 5.93472 | 6.10069 | 6.0522 | 5.98467 | 5.9647 | 6.69911 | 6.17096 | 6.70272 | 6.33221 |
| BRD4 | O60885 | 6.32251 | 6.49595 | 6.16932 | 6.79971 | 6.20481 | 6.1059 | 6.18659 | 5.93288 | 6.14818 | 6.26853 | 6.18781 | 6.27982 | 6.17352 | 6.06038 | 6.25622 | 6.40487 |
| ITPRID2 | P28290 | 6.44573 | 6.50903 | 6.00997 | 6.48327 | 6.11617 | 6.27673 | 6.17975 | 6.16367 | 6.2188 | 6.43438 | 6.39304 | 6.29799 | 6.20371 | 6.06659 | 6.03134 | 6.16985 |
| DIP2B | Q9P265 | 5.97858 | 6.11373 | 6.38761 | 6.03021 | 6.00671 | 6.37264 | 5.95086 | 6.37965 | 6.27692 | 6.38558 | 6.60801 | 6.57282 | 6.28395 | 6.32627 | 6.13539 | 6.19107 |
| TPM3 | P06753 | 6.33783 | 6.07204 | 6.52819 | 6.38734 | 6.71053 | 6.36182 | 6.63036 | 6.5314 | 6.63277 | 6.50253 | 6.1958 | 6.29662 | 6.10465 | 6.21162 | 5.37742 | 6.11908 |
| TPM1 | P09493-3 | 6.71613 | 5.97099 | 7.04739 | 5.98807 | 6.52884 | 6.1023 | 5.95653 | 6.81581 | 6.46693 | 6.04433 | 6.71911 | 6.54578 | 6.49965 | 6.54035 | 5.42437 | 5.63343 |
| PRPF3 | O43395 | 6.24465 | 6.55061 | 6.03169 | 6.61742 | 6.14063 | 6.13396 | 6.36995 | 6.13575 | 6.02875 | 6.00123 | 5.87129 | 5.90252 | 6.41344 | 6.50033 | 6.4327 | 6.62508 |
| DPYSL2 | A0A1C7CYX9 | 5.9696 | 5.99596 | 6.51797 | 5.80489 | 6.31083 | 6.70084 | 6.42944 | 6.73637 | 6.37566 | 6.43823 | 6.45811 | 6.43564 | 5.78061 | 6.29928 | 5.68945 | 6.05713 |
| DPYSL3 | Q14195-2 | 6.30149 | 6.1951 | 6.70012 | 5.94801 | 6.38892 | 6.7369 | 6.14698 | 6.67764 | 6.42697 | 6.54534 | 6.69321 | 6.37123 | 5.76928 | 5.83556 | 5.58776 | 5.67548 |
| LIMA1 | Q9UHB6-4 | 6.39994 | 6.57467 | 5.96559 | 6.41909 | 6.4374 | 6.22903 | 6.10188 | 6.25258 | 6.20348 | 5.89309 | 6.00504 | 5.98016 | 6.60935 | 6.17576 | 6.43354 | 6.3194 |
| RBM28 | Q9NW13 | 6.40917 | 6.21225 | 6.00568 | 6.55064 | 6.15581 | 5.66716 | 6.39982 | 5.99218 | 6.26156 | 6.02465 | 5.9898 | 5.96083 | 6.55248 | 6.38486 | 6.64791 | 6.78519 |
| WDHD1 | O75717 | 5.91695 | 6.1146 | 6.14366 | 5.99341 | 5.8217 | 6.05922 | 6.10264 | 6.60424 | 6.09045 | 6.10554 | 6.35592 | 6.29172 | 6.66073 | 6.65511 | 6.72616 | 6.95795 |
| CDC5L | Q99459 | 6.51389 | 6.57973 | 5.91322 | 6.67152 | 6.12166 | 6.15074 | 6.51691 | 5.85932 | 6.04443 | 6.05167 | 5.7969 | 5.97505 | 6.31194 | 6.48481 | 6.25427 | 6.75395 |
| ERC1 | X6RLX0 | 6.27037 | 6.39485 | 6.59018 | 6.05664 | 6.29606 | 6.59994 | 5.89448 | 6.53829 | 6.35975 | 6.36853 | 6.5001 | 6.36387 | 6.27594 | 5.96769 | 5.78357 | 5.73976 |
| HARS1 | P12081 | 6.43119 | 6.16561 | 6.82991 | 6.00169 | 6.23236 | 6.85959 | 6.26151 | 6.67421 | 6.31754 | 6.61919 | 6.70217 | 6.54641 | 5.71114 | 5.70649 | 5.36076 | 5.58021 |
| SUPT16H | Q9Y5B9 | 5.97523 | 6.14444 | 5.9016 | 6.10716 | 6.16178 | 6.09361 | 6.19045 | 5.84289 | 6.20837 | 6.30158 | 6.01341 | 5.85694 | 6.78308 | 6.59396 | 7.00795 | 6.81754 |
| TALDO1 | P37837 | 5.98657 | 6.0018 | 6.25421 | 5.22291 | 6.05095 | 6.49431 | 5.77632 | 6.53148 | 5.81079 | 6.14303 | 5.97604 | 6.04458 | 6.33902 | 7.60303 | 7.09441 | 6.67055 |
| NCLN | Q969V3-2 | 6.48772 | 6.72558 | 5.95934 | 7.00241 | 6.24281 | 6.01911 | 6.40674 | 6.15498 | 6.08449 | 5.88675 | 6.084 | 5.90135 | 6.24207 | 6.1433 | 6.40742 | 6.25193 |
| LAMB1 | P07942 | 6.44935 | 6.20096 | 6.39567 | 6.6965 | 6.23163 | 6.2402 | 6.37089 | 5.96396 | 6.26063 | 6.05732 | 6.40619 | 6.32695 | 6.22309 | 5.84451 | 6.04758 | 6.28458 |
| USP36 | Q9P275 | 6.21823 | 6.2182 | 5.67362 | 6.97737 | 6.0704 | 5.80704 | 6.35001 | 5.79099 | 5.93924 | 5.9361 | 5.94813 | 5.86581 | 6.22625 | 6.77441 | 7.08283 | 7.12136 |
| STRAP | Q9Y3F4-2 | 6.24579 | 6.21396 | 6.74284 | 6.02043 | 6.65439 | 6.65922 | 6.11143 | 6.82017 | 6.40266 | 6.45216 | 6.69857 | 6.57464 | 5.54092 | 5.71575 | 5.44688 | 5.70017 |
| FYCO1 | Q9BQS8-4 | 6.64472 | 6.60275 | 6.23837 | 6.53153 | 6.11021 | 6.03148 | 5.98881 | 6.09373 | 6.6211 | 6.54228 | 6.57463 | 6.5353 | 6.09936 | 5.95936 | 5.69756 | 5.72881 |
| NUP85 | Q9BW27 | 6.37852 | 6.18179 | 6.06861 | 6.56074 | 6.15035 | 6.08746 | 6.55936 | 6.18425 | 6.20928 | 6.01862 | 6.13796 | 6.18427 | 6.33554 | 6.27809 | 6.25937 | 6.40577 |
| NCOR1 | O75376 | 6.27796 | 6.32426 | 6.30653 | 6.60382 | 6.12922 | 6.16586 | 6.27301 | 6.21157 | 6.30576 | 6.19787 | 6.05356 | 6.19257 | 6.49892 | 6.09168 | 6.15437 | 6.21305 |
| EIF2S1 | P05198 | 6.37167 | 6.42402 | 6.52755 | 6.10664 | 6.31504 | 6.47361 | 6.39882 | 6.59712 | 6.29592 | 6.37515 | 6.3462 | 6.30957 | 5.70066 | 6.0902 | 5.81795 | 5.84985 |
| ANKHD1 | Q8IWZ3-6 | 6.27927 | 6.25691 | 6.6741 | 6.04579 | 6.2036 | 6.38663 | 6.25138 | 6.40185 | 6.69217 | 6.55689 | 6.65306 | 6.65521 | 5.7499 | 5.81283 | 5.69728 | 5.68124 |
| SSRP1 | Q08945 | 5.78688 | 6.13629 | 5.78824 | 6.41024 | 5.96128 | 6.0725 | 6.24537 | 5.74983 | 6.09776 | 6.11458 | 6.08443 | 6.13695 | 6.69437 | 6.70885 | 7.01378 | 6.99866 |
| BCLAF1 | Q9NYF8 | 6.34065 | 6.38456 | 5.74376 | 6.80095 | 6.27779 | 5.98148 | 6.28254 | 5.90892 | 6.15249 | 6.07802 | 6.1149 | 6.13352 | 6.60391 | 6.33041 | 6.47028 | 6.39579 |
| ATP13A1 | Q9HD20 | 6.23454 | 6.4929 | 6.03154 | 6.6601 | 6.14109 | 5.86883 | 6.32891 | 5.91847 | 6.04056 | 6.07585 | 6.04641 | 5.96773 | 6.60121 | 6.52284 | 6.55941 | 6.50962 |

| | | | | | | | | | | | | | | | | | |
|---|---|---|---|---|---|---|---|---|---|---|---|---|---|---|---|---|---|
| GSTP1 | P09211 | 5.97854 | 5.9226 | 6.92443 | 5.41878 | 6.38407 | 6.75302 | 5.97813 | 6.75638 | 6.20447 | 6.54783 | 6.64399 | 6.70824 | 6.07136 | 6.02737 | 5.93069 | 5.75011 |
| PLD3 | Q8IV08 | 6.74645 | 6.88356 | 6.20312 | 6.53952 | 6.29481 | 6.34977 | 6.36837 | 6.20082 | 6.25923 | 6.42335 | 6.25976 | 6.46048 | 5.77451 | 5.84589 | 5.71555 | 5.67481 |
| KIF21A | Q7Z4S6-4 | 5.84038 | 5.98013 | 6.62355 | 5.75804 | 5.88662 | 6.36336 | 5.95695 | 6.20457 | 6.25317 | 6.29733 | 6.67001 | 6.86672 | 6.30274 | 6.58694 | 6.11047 | 6.29903 |
| DKC1 | O60832 | 6.01871 | 6.079 | 5.57839 | 6.3567 | 6.05042 | 5.64463 | 6.33486 | 5.88496 | 6.08515 | 5.97258 | 5.73861 | 5.73431 | 6.6716 | 7.16648 | 7.31454 | 7.36907 |
| RAD23B | P54727 | 6.54188 | 6.10528 | 6.31479 | 5.25756 | 6.44871 | 6.26328 | 6.00568 | 6.67934 | 6.15881 | 6.17794 | 6.24973 | 6.55788 | 6.54963 | 6.36205 | 6.0427 | 6.28475 |
| MFN2 | O95140 | 6.88053 | 6.93975 | 6.54027 | 7.22883 | 6.45385 | 6.09268 | 6.63805 | 6.0317 | 6.68358 | 6.41926 | 6.43969 | 6.50771 | 5.44034 | 5.19408 | 5.20505 | 5.30464 |
| ALMS1 | Q8TCU4 | 6.69665 | 6.44155 | 6.91418 | 6.46797 | 6.3197 | 6.48761 | 6.23535 | 6.29763 | 6.58773 | 6.30062 | 6.78376 | 6.77313 | 5.45711 | 5.49709 | 5.32485 | 5.41507 |
| GAK | O14976 | 6.13771 | 6.13735 | 6.64959 | 5.96521 | 6.01616 | 6.36816 | 6.0618 | 6.16244 | 6.32357 | 6.48124 | 6.5218 | 6.61227 | 6.07014 | 6.31734 | 6.01411 | 6.16112 |
| NUP107 | P57740 | 6.50209 | 6.34901 | 6.08132 | 6.63025 | 6.30318 | 6.07456 | 6.43507 | 6.16105 | 6.27151 | 6.00089 | 5.88887 | 6.10777 | 6.4831 | 5.9934 | 6.50629 | 6.21165 |
| RPL10 | P27635 | 6.39082 | 6.06348 | 6.56954 | 6.38768 | 6.26932 | 6.30656 | 6.54847 | 6.01167 | 6.90601 | 6.28359 | 6.59982 | 6.19889 | 5.81596 | 5.93403 | 5.77496 | 5.93918 |
| WASHC5 | Q12768 | 6.05663 | 6.2411 | 6.25299 | 6.03676 | 6.23982 | 6.28368 | 6.12113 | 6.45345 | 6.21739 | 6.42083 | 6.32765 | 6.3284 | 6.16082 | 6.52932 | 6.16896 | 6.16107 |
| SUCLG2 | Q96I99 | 6.32382 | 6.37405 | 5.68701 | 6.11232 | 6.61257 | 6.14515 | 6.02066 | 6.30745 | 6.07728 | 6.03507 | 5.96872 | 6.16173 | 7.01428 | 6.43359 | 6.32331 | 6.40298 |
| TCERG1 | O14776 | 6.12102 | 6.39279 | 5.85027 | 6.32532 | 6.33679 | 6.14307 | 6.37015 | 6.20518 | 6.09339 | 6.02711 | 5.85535 | 5.76639 | 6.8956 | 6.58937 | 6.51817 | 6.51003 |
| PSMD13 | Q9UNM6 | 6.15932 | 5.50409 | 6.18603 | 5.54261 | 6.34216 | 6.39214 | 6.41872 | 6.19078 | 5.91333 | 6.04542 | 6.01611 | 6.42078 | 6.31534 | 6.79255 | 6.71507 | 7.04555 |
| SCFD1 | Q8WVM8 | 6.4638 | 6.32299 | 6.0443 | 6.32885 | 6.21812 | 6.21428 | 6.52371 | 5.80903 | 6.05373 | 5.92225 | 6.04755 | 5.91754 | 6.59047 | 6.30042 | 6.7256 | 6.51735 |
| SEC16A | O15027-5 | 6.18815 | 6.05393 | 6.94444 | 6.1147 | 6.04187 | 6.43607 | 6.02316 | 6.22769 | 6.26347 | 6.55705 | 6.70455 | 6.63796 | 6.17708 | 6.14514 | 5.74963 | 5.7351 |
| SH3KBP1 | Q96B97 | 6.14922 | 6.09177 | 6.68785 | 5.76079 | 6.22449 | 6.53804 | 6.07122 | 6.74268 | 6.56863 | 6.46462 | 6.49007 | 6.45664 | 6.02214 | 6.11178 | 5.78385 | 5.83621 |
| SEC63 | Q9UGP8 | 6.21096 | 6.39222 | 5.85207 | 6.35384 | 6.40748 | 6.03547 | 6.47849 | 6.2704 | 5.9695 | 5.98317 | 5.70061 | 5.56963 | 6.76495 | 6.63048 | 6.87664 | 6.5041 |
| PTBP1 | P26599-3 | 6.4833 | 6.77532 | 6.2631 | 6.87712 | 6.84179 | 6.43329 | 6.41415 | 6.46302 | 6.3426 | 5.74025 | 5.59589 | 5.65148 | 6.24098 | 5.9793 | 5.96086 | 5.93755 |
| CDK11B | P21127 | 5.9912 | 6.39259 | 6.06571 | 6.49317 | 6.09831 | 6.00505 | 6.09042 | 6.0211 | 5.99779 | 6.01757 | 6.25049 | 6.1167 | 6.4596 | 6.51353 | 6.64538 | 6.84138 |
| RPL8 | P62917 | 6.0174 | 6.4589 | 6.35579 | 5.96213 | 5.9796 | 7.09588 | 6.23276 | 6.53594 | 7.50751 | 6.19463 | 6.10703 | 5.89693 | 6.00175 | 6.19306 | 5.63632 | 5.82438 |
| API5 | Q9BZZ5 | 6.00674 | 6.01884 | 5.83962 | 5.8864 | 5.99741 | 5.92084 | 6.08248 | 5.87493 | 6.11699 | 6.01648 | 6.01569 | 5.92317 | 7.13426 | 7.03872 | 7.22688 | 6.90052 |
| NF1 | P21359 | 6.26064 | 6.27183 | 6.31936 | 6.33486 | 6.02923 | 6.40269 | 6.27244 | 6.2492 | 6.32911 | 6.28856 | 6.13291 | 6.26167 | 6.27624 | 6.19643 | 6.29059 | 6.08426 |
| ACAD9 | Q9H845 | 6.35103 | 6.41515 | 6.10492 | 6.78484 | 6.4474 | 5.95986 | 6.39413 | 5.93626 | 6.18089 | 6.0315 | 5.92705 | 5.91397 | 6.48311 | 6.13584 | 6.54583 | 6.38821 |
| CFL2 | Q9Y281 | 5.9822 | 6.00355 | 6.44819 | 5.95783 | 6.30353 | 6.71426 | 6.09458 | 6.51899 | 6.18302 | 6.46517 | 6.62589 | 6.58393 | 6.1745 | 6.21163 | 5.82148 | 5.91123 |
| PLOD3 | O60568 | 6.32918 | 6.29783 | 6.11305 | 6.56796 | 6.20487 | 6.03067 | 6.31608 | 6.00774 | 6.0744 | 6.03484 | 6.24029 | 6.14984 | 6.38319 | 6.18241 | 6.72941 | 6.33826 |
| ZMYM3 | Q14202-2 | 6.23173 | 6.5109 | 6.35598 | 6.80822 | 6.17985 | 6.14111 | 6.5281 | 5.8015 | 6.32339 | 6.4275 | 6.44691 | 6.06067 | 6.08166 | 5.98491 | 6.14285 | 5.97473 |
| RABGAP1 | Q9Y3P9 | 6.09582 | 6.19522 | 6.81959 | 6.1929 | 6.29577 | 6.55199 | 6.21367 | 6.21973 | 6.26246 | 6.50962 | 6.61228 | 6.35998 | 6.14463 | 6.05446 | 5.72437 | 5.7475 |
| USP24 | Q9UPU5 | 6.23837 | 6.23254 | 6.79787 | 6.11918 | 6.11509 | 6.53885 | 6.16428 | 6.46432 | 6.26359 | 6.39109 | 6.54268 | 6.5715 | 5.96912 | 5.80608 | 5.91949 | 5.86595 |
| INTS3 | Q68E01 | 6.3766 | 6.3685 | 6.23047 | 6.42531 | 6.26473 | 6.06096 | 6.30102 | 6.04197 | 6.1327 | 6.02643 | 6.06302 | 5.91872 | 6.44227 | 6.37399 | 6.52378 | 6.44953 |
| SRSF1 | Q07955-2 | 6.5718 | 6.11189 | 5.98868 | 6.47007 | 6.64345 | 5.82889 | 6.17992 | 5.94564 | 6.40523 | 5.81948 | 6.21752 | 5.83612 | 7.23794 | 6.05569 | 6.31533 | 6.37236 |
| PPP1R12A | O14974 | 6.41011 | 6.37874 | 6.57108 | 6.12509 | 6.36173 | 6.67495 | 6.16438 | 6.55492 | 6.44646 | 6.29858 | 6.45965 | 6.33351 | 5.93574 | 5.98947 | 5.6713 | 5.6243 |
| NOP2 | P46087-4 | 6.05643 | 6.14555 | 5.86059 | 6.63741 | 5.85636 | 5.77223 | 6.42811 | 5.83615 | 6.10209 | 5.75916 | 5.84952 | 5.9198 | 6.39079 | 6.9646 | 7.17023 | 7.25098 |
| NUDC | Q9Y266 | 6.52882 | 6.1645 | 6.93844 | 5.84493 | 6.48342 | 6.70877 | 6.27494 | 6.53738 | 6.69463 | 6.3425 | 6.47838 | 6.45158 | 5.76242 | 5.55882 | 5.63737 | 5.59311 |
| TTF2 | Q9UNY4 | 6.49962 | 6.27711 | 6.53237 | 6.54722 | 6.14154 | 6.27584 | 6.50368 | 6.48474 | 6.66772 | 6.3535 | 6.60703 | 6.66577 | 5.87045 | 5.43618 | 5.64528 | 5.49195 |
| HSD17B10 | Q99714 | 6.34001 | 6.10718 | 5.71175 | 6.51574 | 6.41629 | 5.91171 | 6.40845 | 5.96752 | 6.36824 | 6.15412 | 6.24991 | 6.15684 | 6.69798 | 6.03459 | 6.67366 | 6.28602 |
| NCOA2 | Q15596 | 5.75985 | 5.98422 | 6.09386 | 5.68342 | 6.20173 | 6.28354 | 6.2933 | 6.13206 | 5.99239 | 6.0325 | 6.14629 | 5.72798 | 7.03766 | 6.96218 | 6.99492 | 6.6741 |
| FH | P07954 | 6.38989 | 6.19663 | 5.91512 | 6.10634 | 6.46808 | 6.0782 | 6.36581 | 6.06769 | 6.43539 | 5.93822 | 6.16671 | 5.9272 | 7.33099 | 6.04681 | 6.48536 | 6.08156 |
| TOMM70 | O94826 | 6.39132 | 6.49596 | 5.72746 | 6.6262 | 6.39504 | 6.20754 | 6.47489 | 6.17793 | 6.05272 | 6.03702 | 5.58655 | 5.56541 | 6.67266 | 6.54442 | 6.46514 | 6.57973 |
| CTNND1 | O60716-3 | 6.28833 | 6.32423 | 6.05176 | 6.39182 | 6.16627 | 6.2269 | 6.36434 | 5.97328 | 6.31798 | 6.2698 | 6.29928 | 6.2393 | 6.45223 | 6.35926 | 6.17178 | 6.10342 |
| BTAF1 | O14981 | 5.38486 | 5.69609 | 5.90191 | 5.26467 | 5.79333 | 6.11268 | 5.9305 | 5.80747 | 5.88847 | 6.31602 | 6.15886 | 5.96427 | 7.4897 | 7.45324 | 7.56268 | 7.27525 |
| SIN3A | Q96ST3 | 6.44601 | 6.79182 | 5.91787 | 6.85462 | 6.3453 | 6.21649 | 6.24623 | 6.22352 | 5.94708 | 6.00489 | 5.93689 | 5.97572 | 6.1323 | 6.28225 | 6.37158 | 6.30744 |
| NCKAP1 | Q9Y2A7-2 | 6.20051 | 6.09973 | 6.5491 | 5.90401 | 6.07996 | 6.50216 | 6.33749 | 6.31594 | 6.53976 | 6.47415 | 6.48964 | 6.51841 | 5.96216 | 6.14446 | 6.04119 | 5.84134 |
| SMC6 | Q96SB8 | 6.38997 | 6.29974 | 6.12562 | 6.42293 | 6.15253 | 5.86907 | 6.20029 | 5.92769 | 6.18349 | 5.9267 | 6.07643 | 6.14095 | 6.71512 | 6.20956 | 6.85788 | 6.50894 |
| FARSB | Q9NSD9 | 6.21763 | 6.19361 | 6.51294 | 5.85369 | 6.53125 | 6.58235 | 6.03794 | 6.5552 | 6.47199 | 6.47975 | 6.47859 | 6.52725 | 6.0186 | 5.99124 | 5.68627 | 5.86172 |
| CDK1 | P06493 | 6.32859 | 6.17057 | 6.55445 | 5.97942 | 6.39692 | 6.882 | 6.37682 | 6.97668 | 6.26844 | 6.55315 | 6.59545 | 6.60969 | 5.68983 | 5.65603 | 5.27646 | 5.68551 |
| ATXN2 | A0A2R8Y5A6 | 6.29767 | 6.13678 | 6.67749 | 5.77416 | 6.56376 | 6.37313 | 6.19459 | 6.36519 | 6.28253 | 6.43245 | 6.37256 | 6.4588 | 6.41006 | 5.9758 | 5.93614 | 5.74889 |
| NIBAN2 | Q96TA1 | 6.65361 | 6.05613 | 6.9176 | 5.64811 | 6.49068 | 6.60972 | 6.25327 | 6.50828 | 6.5301 | 6.14648 | 6.43858 | 6.25187 | 6.14334 | 6.07987 | 5.77906 | 5.4933 |
| RARS2 | Q5T160 | 6.48507 | 6.44344 | 5.71822 | 6.61389 | 6.39295 | 6.06171 | 6.50926 | 6.06477 | 6.2413 | 5.95135 | 5.91013 | 5.99016 | 6.63124 | 6.15483 | 6.45456 | 6.37711 |
| CHAMP1 | Q96JM3 | 6.35053 | 6.2999 | 5.91933 | 6.40304 | 6.19974 | 5.97452 | 6.14602 | 6.01261 | 6.11439 | 6.25652 | 6.0375 | 6.08092 | 6.95796 | 6.25505 | 6.51026 | 6.48172 |
| CHD8 | Q9HCK8 | 6.0466 | 6.35004 | 6.06054 | 6.53806 | 5.98984 | 6.20899 | 6.30322 | 6.08025 | 6.06107 | 6.24703 | 6.54955 | 6.23639 | 6.10754 | 6.46144 | 6.47086 | 6.28858 |
| ATP1A3 | P13637-3 | 6.47519 | 6.60243 | 6.42049 | 6.76622 | 6.07079 | 6.10948 | 6.32716 | 5.92921 | 6.49701 | 6.34921 | 6.52505 | 6.31312 | 5.90627 | 5.84602 | 5.83256 | 6.02977 |
| ATP1A1 | P05023-4 | 6.66194 | 6.6966 | 6.172 | 6.74989 | 6.39872 | 6.25419 | 6.37651 | 6.06177 | 6.24647 | 6.28524 | 5.98929 | 5.96436 | 5.96845 | 6.13223 | 6.03113 | 6.0112 |
| UBA6 | A0AVT1 | 6.1181 | 6.02313 | 6.84314 | 5.91114 | 6.30497 | 6.58555 | 6.1529 | 6.47315 | 6.20477 | 6.43098 | 6.64368 | 6.62723 | 6.01578 | 6.09465 | 5.85751 | 5.71331 |
| DNAAF5 | Q86Y56 | 6.01381 | 6.10716 | 6.66325 | 5.80277 | 6.36536 | 6.67366 | 6.30107 | 6.72169 | 6.41348 | 6.3473 | 6.68397 | 6.50321 | 5.87882 | 5.85639 | 5.77402 | 5.89404 |
| SCP2 | P22307 | 5.93063 | 5.85319 | 5.68882 | 6.09741 | 6.65287 | 5.88594 | 6.42597 | 6.30209 | 6.11299 | 5.88047 | 5.98479 | 6.07736 | 6.72308 | 6.4934 | 6.74445 | 7.14655 |
| ARHGEF1 | Q92888-3 | 6.41266 | 6.4483 | 6.60431 | 6.3144 | 6.27333 | 6.7397 | 6.27164 | 6.70083 | 6.37724 | 6.41659 | 6.47211 | 6.51792 | 5.73728 | 5.79549 | 5.42303 | 5.49517 |
| EHD1 | Q9H4M9 | 5.78976 | 5.96385 | 6.03469 | 5.79129 | 6.30738 | 6.18222 | 6.52908 | 6.29883 | 6.72903 | 5.79388 | 6.1968 | 6.3265 | 6.22487 | 6.68766 | 6.42471 | 6.71945 |
| EML4 | Q9HC35 | 6.26034 | 6.09007 | 6.7625 | 5.76424 | 6.11634 | 6.32665 | 6.14502 | 6.25498 | 6.15796 | 6.51098 | 6.44269 | 6.33162 | 6.1819 | 6.39885 | 6.17864 | 6.07721 |
| NUP50 | Q9UKX7 | 6.39157 | 6.08578 | 6.22767 | 6.86375 | 6.35496 | 6.07495 | 6.12478 | 6.12319 | 6.20746 | 6.16206 | 6.25597 | 6.29833 | 6.82014 | 5.93315 | 5.87114 | 6.20512 |
| ATAD3B | Q5T9A4 | 6.40878 | 6.26952 | 6.11256 | 6.7432 | 6.29162 | 6.0894 | 6.5947 | 5.84797 | 6.01844 | 5.9899 | 6.05128 | 5.99193 | 6.39075 | 6.27647 | 6.42763 | 6.49586 |
| ZCCHC8 | Q6NZY4 | 6.43177 | 6.16295 | 5.86844 | 6.55135 | 6.60078 | 6.37543 | 6.08045 | 6.14899 | 6.3941 | 5.96564 | 6.35263 | 6.24352 | 6.81477 | 5.87056 | 6.02251 | 6.1161 |
| ESF1 | Q9H501 | 6.05268 | 6.02303 | 6.05159 | 6.57564 | 6.03251 | 5.7613 | 6.45842 | 5.91303 | 6.26177 | 5.78086 | 5.95436 | 5.84794 | 6.84069 | 6.57011 | 6.92394 | 6.95162 |
| FHOD1 | Q9Y613 | 6.16775 | 6.16236 | 6.47099 | 6.03003 | 6.19469 | 6.76157 | 6.12308 | 6.33836 | 6.29535 | 6.57465 | 6.44975 | 6.28254 | 5.98251 | 6.32773 | 5.94455 | 5.89408 |
| SQOR | Q9Y6N5 | 6.1291 | 6.39416 | 5.72927 | 6.69555 | 5.85666 | 6.17886 | 6.15997 | 6.01002 | 5.82595 | 6.33301 | 5.7692 | 6.10856 | 6.68908 | 6.70215 | 6.54878 | 6.86967 |
| SMARCA5 | O60264 | 6.52552 | 6.49522 | 6.07189 | 6.83719 | 6.26751 | 6.03608 | 6.38827 | 5.93974 | 6.09547 | 6.05291 | 5.89775 | 5.9937 | 6.51795 | 6.20988 | 6.49037 | 6.18056 |
| BZW1 | Q7L1Q6-3 | 6.41443 | 6.44307 | 6.51775 | 6.00585 | 6.4739 | 6.64482 | 6.54393 | 6.46877 | 6.47266 | 6.27847 | 6.2127 | 6.2242 | 5.60199 | 5.98349 | 5.75694 | 5.95703 |
| EIF4A1 | P60842 | 6.76038 | 6.73231 | 6.93408 | 6.42775 | 6.50927 | 6.71433 | 6.72766 | 6.72323 | 6.89976 | 6.12968 | 6.38996 | 6.36694 | 5.04739 | 5.34968 | 5.00096 | 5.28663 |
| LMO7 | Q8WWI1 | 6.41539 | 6.39926 | 6.07612 | 6.61575 | 6.13489 | 5.89853 | 6.45528 | 6.00081 | 6.14588 | 6.17092 | 6.09568 | 6.07747 | 6.48881 | 6.29554 | 6.49147 | 6.23819 |
| DNM2 | P50570 | 6.28415 | 6.15897 | 6.69785 | 5.89451 | 6.23696 | 6.44434 | 6.15988 | 6.40903 | 6.3806 | 6.37991 | 6.33102 | 6.32167 | 6.21922 | 6.16485 | 6.08426 | 5.83278 |
| NFKB2 | Q00653 | 6.66934 | 6.36056 | 6.88824 | 6.25876 | 6.58459 | 6.85298 | 6.42365 | 6.73574 | 6.2622 | 6.07877 | 6.01877 | 6.10225 | 5.80608 | 5.75542 | 5.54108 | 5.66157 |
| PTCD3 | Q96EY7 | 6.40246 | 6.25669 | 5.88119 | 6.81525 | 6.33826 | 6.12335 | 6.41606 | 6.17305 | 6.00943 | 6.01744 | 5.99733 | 6.01428 | 6.42388 | 6.21798 | 6.43431 | 6.47904 |
| MTA1 | Q13330 | 6.34943 | 6.35726 | 5.96582 | 6.49893 | 6.31671 | 5.81779 | 6.5182 | 5.90374 | 6.37739 | 6.07741 | 6.18791 | 5.74536 | 6.55846 | 6.28393 | 6.43592 | 6.60574 |
| VWA8 | A3KMH1 | 6.45663 | 6.47811 | 5.87284 | 6.65747 | 6.28948 | 6.14466 | 6.27498 | 5.97689 | 6.18601 | 5.93551 | 5.86434 | 5.91423 | 6.49907 | 6.38603 | 6.49875 | 6.55599 |
| MED14 | O60244 | 6.3322 | 6.33716 | 6.03637 | 6.80133 | 6.43871 | 6.23842 | 6.28927 | 6.11778 | 6.12633 | 6.03604 | 5.92666 | 5.951 | 6.44025 | 6.20504 | 6.43182 | 6.29162 |
| PAFAH1B1 | P43034 | 6.32486 | 6.2345 | 6.84394 | 5.9767 | 6.3737 | 6.74079 | 6.3676 | 6.39774 | 6.33163 | 6.46756 | 6.58326 | 6.3491 | 5.88966 | 5.61745 | 5.95636 | 5.54514 |
| RPAP3 | Q9H6T3 | 6.11808 | 6.34446 | 6.76733 | 6.09782 | 6.23971 | 6.63554 | 6.1117 | 6.5061 | 6.18344 | 6.49683 | 6.44162 | 6.48639 | 5.85866 | 5.99927 | 5.81211 | 5.90094 |
| PUS7 | Q96PZ0-2 | 5.39699 | 5.79761 | 6.21931 | 5.31009 | 5.81608 | 6.38376 | 5.53182 | 6.14047 | 5.82244 | 6.14938 | 6.30288 | 6.29245 | 6.77746 | 7.52945 | 7.29115 | 7.23867 |
| WDR33 | Q9C0J8 | 6.35591 | 6.40814 | 6.14176 | 6.89001 | 6.13612 | 5.98693 | 6.47508 | 6.0756 | 6.01634 | 6.04801 | 6.02345 | 6.18394 | 6.31112 | 6.18005 | 6.41338 | 6.35418 |
| UAP1 | Q16222 | 6.29281 | 6.033 | 7.1035 | 6.20464 | 6.39611 | 6.61953 | 6.25407 | 6.82268 | 6.48447 | 6.06647 | 6.68351 | 6.99395 | 5.39571 | 5.31003 | 5.74462 | 5.59493 |
| SUGP2 | Q8IX01 | 6.29593 | 6.51188 | 5.86247 | 6.91346 | 6.00033 | 5.9913 | 6.44403 | 5.94433 | 6.09495 | 6.14368 | 5.87744 | 5.98142 | 6.34378 | 6.51412 | 6.54197 | 6.53892 |
| HSDL2 | Q6YN16 | 6.27864 | 6.3771 | 5.95118 | 6.67376 | 5.91867 | 5.7314 | 6.1073 | 6.11487 | 6.17136 | 6.22972 | 6.04653 | 6.1532 | 6.63925 | 6.42425 | 6.58085 | 6.60191 |
| DYNC1I2 | Q13409-2 | 6.19199 | 6.00637 | 6.73145 | 6.30009 | 6.2324 | 6.47416 | 6.24981 | 6.41853 | 6.26851 | 6.42187 | 6.48939 | 6.71465 | 5.72598 | 6.22788 | 5.5681 | 5.97881 |
| ELP1 | O95163 | 6.30954 | 6.09903 | 6.7144 | 6.13979 | 6.38405 | 6.64583 | 6.30678 | 6.49234 | 6.50921 | 6.56515 | 6.68197 | 6.55362 | 5.68608 | 5.65985 | 5.67474 | 5.57761 |
| ZMYND8 | Q9ULU4-7 | 6.34518 | 6.4612 | 6.04774 | 6.80367 | 6.17325 | 6.1828 | 6.14333 | 5.99884 | 5.96793 | 6.13039 | 6.14871 | 6.20684 | 6.45766 | 6.2693 | 6.19484 | 6.46834 |
| UBR2 | Q8IWV8 | 6.14424 | 6.40752 | 6.80111 | 6.28504 | 6.21865 | 6.71798 | 6.06044 | 6.48398 | 6.30929 | 6.44878 | 6.44384 | 6.4481 | 5.76412 | 6.05709 | 5.71141 | 5.69842 |
| EIF3H | O15372 | 6.22849 | 6.50172 | 6.88594 | 6.19323 | 6.37251 | 6.54636 | 6.36659 | 6.55128 | 6.55153 | 6.48077 | 6.71643 | 6.69815 | 5.35177 | 5.60365 | 5.52223 | 5.42934 |
| HNRNPH1 | G8JLB6 | 6.26018 | 6.33396 | 5.66237 | 6.81082 | 6.26469 | 6.50086 | 5.98109 | 6.5261 | 6.16646 | 6.1142 | 6.16304 | 6.2791 | 6.32114 | 6.1878 | 5.99277 | 6.43542 |
| RBM12B | Q8IXT5 | 6.29451 | 6.52311 | 5.83263 | 6.71593 | 6.24598 | 6.08183 | 6.26772 | 6.06792 | 6.14321 | 6.0849 | 6.01152 | 6.02213 | 6.55286 | 6.23884 | 6.52875 | 6.38817 |
| BAIAP2L1 | Q9UHR4 | 6.16346 | 6.28967 | 6.15882 | 5.86115 | 6.18361 | 6.41468 | 6.15568 | 6.295 | 6.74035 | 6.42493 | 6.32003 | 6.68222 | 6.12795 | 6.21387 | 5.93439 | 6.03422 |
| ATP5PO | P48047 | 6.28732 | 6.27474 | 5.88204 | 6.66581 | 6.30675 | 5.9225 | 6.62629 | 6.14053 | 6.35092 | 5.73767 | 5.79856 | 5.99332 | 6.68922 | 6.17656 | 6.64096 | 6.50681 |
| DHX57 | Q6P158 | 6.19439 | 6.33587 | 6.69976 | 6.12063 | 6.08911 | 6.589 | 6.05982 | 6.25199 | 6.60681 | 6.50428 | 6.66099 | 6.43696 | 5.64478 | 6.01789 | 5.95494 | 5.83277 |
| NXF1 | Q9UBU9 | 6.17471 | 6.25452 | 6.00086 | 6.61514 | 6.01663 | 5.76689 | 6.53366 | 5.81421 | 6.24977 | 6.11294 | 5.89976 | 6.0382 | 6.64783 | 6.29822 | 6.92085 | 6.6558 |
| PTK2 | Q05397 | 6.11802 | 6.06769 | 6.75243 | 5.99268 | 6.19737 | 6.40899 | 6.35056 | 6.34193 | 6.44662 | 6.44517 | 6.76023 | 6.39048 | 5.96345 | 6.08429 | 5.97009 | 5.70999 |
| TRIP12 | Q14669-3 | 6.29685 | 6.46032 | 6.09261 | 6.87255 | 6.11872 | 6.00969 | 6.30478 | 6.16475 | 6.29497 | 6.05033 | 5.9693 | 6.0265 | 5.87919 | 6.55995 | 6.3108 | 6.58869 |
| SCAF11 | Q99590 | 6.23527 | 6.38777 | 5.92921 | 6.81607 | 6.034 | 5.80751 | 6.23431 | 5.95751 | 6.24643 | 6.29373 | 6.1186 | 6.20701 | 6.36072 | 6.39122 | 6.51843 | 6.46219 |
| RPL23A | P62750 | 5.86706 | 6.43827 | 6.3222 | 6.27925 | 6.19878 | 6.87304 | 6.06 | 6.46944 | 6.60212 | 6.17324 | 6.05207 | 6.24923 | 5.67761 | 6.59286 | 6.02969 | 6.11516 |
| UGDH | O60701 | 5.45793 | 5.75675 | 6.144 | 5.2298 | 5.88739 | 6.56484 | 5.83031 | 6.07689 | 5.79633 | 6.23088 | 6.21565 | 6.15623 | 7.15121 | 7.10481 | 7.33076 | 7.0662 |
| PPP2R1A | P30153 | 6.183 | 6.02307 | 6.35399 | 5.60757 | 6.33222 | 6.4372 | 6.01684 | 6.6176 | 6.16793 | 6.2442 | 6.48766 | 6.54475 | 6.30058 | 6.34925 | 6.03303 | 6.3011 |
| BOP1 | Q14137 | 6.34179 | 6.52073 | 6.05002 | 6.9339 | 6.22963 | 6.0245 | 6.33089 | 5.98738 | 6.04833 | 6.19781 | 6.08373 | 6.25202 | 6.21237 | 6.10322 | 6.31014 | 6.37353 |
| EIF3C | Q99613 | 6.40314 | 6.61181 | 6.62287 | 6.33896 | 6.38098 | 6.65167 | 6.33664 | 6.39106 | 6.40336 | 6.47201 | 6.53341 | 6.42978 | 5.42654 | 5.73193 | 5.6353 | 5.63054 |
| ZNF462 | Q96JM2-3 | 6.40577 | 6.743 | 6.18357 | 6.87669 | 6.12453 | 5.9799 | 6.32451 | 6.04897 | 6.19423 | 6.18293 | 6.33824 | 6.1035 | 6.16759 | 5.87247 | 6.16989 | 6.28419 |
| CREBBP | Q92793 | 6.08134 | 6.3258 | 6.18105 | 6.29845 | 6.21981 | 6.28588 | 6.06905 | 6.14055 | 6.16712 | 6.31013 | 6.32449 | 6.03911 | 6.48874 | 6.34423 | 6.4014 | 6.32286 |
| SF3A3 | Q12874 | 6.46276 | 6.36621 | 5.80771 | 6.63286 | 6.47939 | 6.36393 | 5.98987 | 6.2101 | 6.13803 | 6.04881 | 6.18654 | 6.16957 | 6.79537 | 6.09363 | 6.11553 | 6.13968 |
| GTF3C4 | Q9UKN8 | 6.10649 | 6.34154 | 6.15206 | 6.25762 | 6.23279 | 6.29262 | 6.01604 | 5.95972 | 6.08602 | 6.29852 | 6.10711 | 6.1317 | 6.66119 | 6.1455 | 6.87998 | 6.3311 |

| | | | | | | | | | | | | | | | | | |
|---|---|---|---|---|---|---|---|---|---|---|---|---|---|---|---|---|---|
| GEMIN5 | Q8TEQ6 | 6.3316 | 6.70144 | 6.74477 | 6.18246 | 6.33736 | 6.49698 | 6.25781 | 6.66006 | 6.39824 | 6.35422 | 6.46982 | 6.41467 | 5.66412 | 5.66612 | 5.79221 | 5.52813 |
| IDH1 | O75874 | 6.28751 | 6.18752 | 6.68533 | 5.54142 | 6.3782 | 6.75086 | 6.11417 | 6.52823 | 6.2946 | 6.46913 | 6.72435 | 6.26295 | 6.16266 | 6.03733 | 5.91848 | 5.65725 |
| PSMD7 | P51665 | 6.37366 | 6.32812 | 6.43144 | 6.25337 | 6.34807 | 6.5343 | 6.03844 | 6.47164 | 6.15659 | 6.18569 | 6.45554 | 6.37016 | 6.11742 | 6.01558 | 5.89298 | 6.02699 |
| WDR43 | Q15061 | 6.348 | 6.29845 | 5.78049 | 6.82905 | 6.40199 | 6.16338 | 6.37499 | 5.97247 | 6.14156 | 5.96732 | 5.8785 | 6.04711 | 6.8085 | 6.12405 | 6.30673 | 6.5574 |
| EIF3L | Q9Y262 | 6.32213 | 6.46892 | 6.64089 | 6.23315 | 6.39851 | 6.55502 | 6.30021 | 6.48066 | 6.46246 | 6.38122 | 6.44525 | 6.4112 | 5.6429 | 5.96165 | 5.67529 | 5.62055 |
| NDUFA9 | Q16795 | 5.97639 | 6.10501 | 5.23876 | 6.292 | 6.39657 | 6.30616 | 6.79261 | 6.5772 | 6.25595 | 5.82467 | 5.75929 | 5.74552 | 6.98121 | 6.57188 | 6.5007 | 6.67609 |
| RTCB | Q9Y3I0 | 5.65335 | 5.84603 | 5.92317 | 5.64758 | 6.26639 | 6.22132 | 6.45329 | 6.10934 | 6.26012 | 5.98253 | 6.28266 | 5.97296 | 6.53647 | 6.97075 | 6.79713 | 7.07691 |
| USP14 | P54578 | 6.02562 | 6.1078 | 6.49049 | 6.03781 | 6.20903 | 6.47127 | 6.36623 | 6.52743 | 6.36895 | 6.07761 | 6.52923 | 6.56843 | 5.80033 | 6.29474 | 5.80273 | 6.3223 |
| PDHA1 | P08559-4 | 6.35976 | 6.35826 | 5.87596 | 6.6828 | 6.16371 | 5.89204 | 6.55541 | 5.88389 | 6.25296 | 5.8408 | 5.78794 | 5.91723 | 6.85974 | 6.22683 | 6.70868 | 6.634 |
| XAB2 | Q9HCS7 | 6.31985 | 6.33301 | 6.08258 | 6.84417 | 6.41828 | 6.15474 | 6.37868 | 5.99326 | 6.19544 | 6.19491 | 6.23864 | 6.20743 | 6.23036 | 5.89706 | 6.31061 | 6.20097 |
| RAI1 | Q7Z5J4 | 6.55075 | 6.2732 | 6.03991 | 6.82984 | 6.06229 | 6.14319 | 6.34423 | 5.78196 | 6.12724 | 6.10725 | 6.27098 | 6.00577 | 6.1879 | 6.37823 | 6.47726 | 6.42 |
| DDX42 | Q86XP3 | 6.18665 | 6.33933 | 6.073 | 6.31145 | 6.27834 | 5.97875 | 6.36013 | 5.89722 | 6.22019 | 6.07185 | 6.06407 | 5.73442 | 6.82445 | 6.3218 | 6.75928 | 6.57907 |
| RNF20 | Q5VTR2 | 5.99748 | 6.52017 | 6.20524 | 6.11921 | 6.47893 | 6.38091 | 6.44931 | 6.1701 | 6.29174 | 6.56793 | 6.1893 | 5.88847 | 6.33775 | 6.17001 | 6.14727 | 6.08616 |
| RAB11B | Q15907 | 6.46083 | 6.33136 | 6.3833 | 6.24558 | 6.26549 | 6.30582 | 6.1752 | 5.99244 | 6.34327 | 6.29058 | 6.30687 | 6.38292 | 5.96166 | 5.87449 | 6.36956 | 6.31064 |
| SMU1 | Q2TAY7 | 6.2504 | 6.28487 | 5.79933 | 6.40635 | 6.39884 | 5.96758 | 6.43553 | 5.95043 | 6.02737 | 6.01459 | 6.00696 | 5.7814 | 6.66789 | 6.44171 | 6.88851 | 6.67825 |
| ATXN2L | Q8WWM7-3 | 6.34089 | 6.39644 | 6.56466 | 6.25395 | 6.47861 | 6.63973 | 5.86425 | 6.44327 | 6.46632 | 6.60445 | 6.49023 | 6.70823 | 6.0706 | 5.77568 | 5.37141 | 5.66229 |
| ACAA2 | P42765 | 6.2869 | 6.26246 | 5.68579 | 6.41299 | 6.49308 | 6.27296 | 6.13576 | 6.29524 | 6.16285 | 5.93504 | 5.93631 | 6.10297 | 6.89411 | 6.19798 | 6.51201 | 6.41354 |
| AIFM1 | O95831 | 6.27945 | 6.29505 | 5.71893 | 6.51642 | 6.20966 | 6.03161 | 6.53771 | 6.05052 | 6.17379 | 5.97578 | 5.95022 | 5.9151 | 6.66392 | 6.6149 | 6.53167 | 6.53527 |
| DNTTIP2 | Q5QJE6 | 6.33401 | 6.37995 | 5.91477 | 6.81776 | 6.22367 | 5.91124 | 6.45668 | 6.17439 | 6.03265 | 6.03743 | 5.78927 | 5.97735 | 6.59421 | 6.1828 | 6.53778 | 6.63604 |
| BLM | P54132 | 6.29264 | 6.13728 | 5.85429 | 6.5075 | 6.07225 | 6.02906 | 6.37012 | 5.96471 | 6.13147 | 6.01334 | 5.80923 | 5.84256 | 6.52968 | 6.63105 | 6.97775 | 6.8371 |
| PPA2 | Q9H2U2 | 6.43653 | 6.10206 | 5.85767 | 6.06814 | 6.54012 | 5.93747 | 6.13291 | 6.32437 | 6.02147 | 6.11193 | 5.97426 | 6.01562 | 7.04142 | 6.52586 | 6.6173 | 6.29286 |
| PSPC1 | Q8WXF1 | 6.18977 | 5.92587 | 5.80342 | 6.17261 | 6.40688 | 5.87952 | 6.16567 | 6.28369 | 6.19545 | 5.9208 | 6.22305 | 6.02545 | 7.23614 | 6.39538 | 6.52255 | 6.65377 |
| AGPS | O00116 | 6.25827 | 6.38722 | 6.19197 | 6.73618 | 6.19539 | 6.09028 | 6.41005 | 6.08846 | 6.14719 | 6.01574 | 6.06133 | 5.97855 | 6.244 | 6.2113 | 6.47836 | 6.50572 |
| ITSN2 | Q9NZM3 | 6.19089 | 6.21746 | 6.02515 | 5.92123 | 6.29165 | 6.23612 | 6.29649 | 6.14151 | 6.46619 | 6.26103 | 6.22648 | 6.24521 | 6.51504 | 6.32986 | 6.45465 | 6.18103 |
| ACSF2 | Q96CM8-2 | 6.29748 | 6.19192 | 5.90652 | 6.28464 | 6.20345 | 6.01951 | 6.23104 | 5.94994 | 6.07939 | 5.97388 | 6.04326 | 5.82997 | 7.06812 | 6.60545 | 6.82577 | 6.48965 |
| EIF4B | P23588 | 6.53469 | 6.58913 | 6.7433 | 6.25603 | 6.23917 | 6.53733 | 6.306 | 6.26452 | 6.65195 | 6.60406 | 6.77953 | 6.77028 | 5.52137 | 5.56843 | 5.21996 | 5.41421 |
| GOLGA3 | Q08378 | 6.33994 | 6.35109 | 6.43604 | 6.47565 | 6.31395 | 6.2767 | 6.14036 | 6.69707 | 6.55173 | 6.39184 | 6.51568 | 6.32678 | 6.11504 | 5.76322 | 5.64492 | 5.66628 |
| FBL | P22087 | 6.40702 | 6.20158 | 5.90569 | 6.4562 | 5.92141 | 5.80731 | 6.4315 | 5.79687 | 6.27769 | 6.10466 | 6.22614 | 6.18847 | 6.68175 | 6.26997 | 6.56571 | 6.75803 |
| DDX41 | Q9UJV9 | 5.96586 | 6.19592 | 5.94137 | 6.2294 | 6.07435 | 5.95877 | 6.23048 | 5.87298 | 6.20535 | 6.24005 | 6.17998 | 5.99404 | 6.90077 | 6.61924 | 6.72748 | 6.66398 |
| ECT2 | Q9H8V3 | 5.9254 | 6.18278 | 6.20357 | 6.07824 | 5.99467 | 6.11558 | 6.35481 | 6.33949 | 6.30735 | 5.98599 | 6.17951 | 6.18747 | 6.63045 | 6.32879 | 6.62553 | 6.56039 |
| AHSA1 | O95433 | 6.38785 | 6.15507 | 6.9243 | 5.90761 | 6.4782 | 6.69364 | 6.53548 | 6.48471 | 6.43906 | 6.18609 | 6.65754 | 6.19536 | 5.96276 | 5.68832 | 5.68817 | 5.61583 |
| PRDX4 | Q13162 | 6.34897 | 6.02915 | 5.97063 | 5.83053 | 6.46918 | 6.1228 | 5.90168 | 6.47802 | 6.00478 | 6.15 | 6.00555 | 6.31111 | 6.83237 | 6.47728 | 6.31926 | 6.74869 |
| EHD4 | Q9H223 | 6.06113 | 5.98619 | 6.35195 | 5.86814 | 6.06679 | 6.31637 | 6.01374 | 6.40153 | 6.67059 | 6.4212 | 6.60882 | 6.62529 | 6.47473 | 6.0287 | 6.03934 | 6.06551 |
| ZC3H18 | E7ERS3 | 6.24761 | 6.46745 | 5.93128 | 6.7794 | 6.17793 | 5.98249 | 6.32368 | 6.01356 | 6.21552 | 6.09501 | 5.95233 | 5.78643 | 6.5823 | 6.42374 | 6.48795 | 6.5333 |
| RPS6KA1 | Q15418-2 | 6.04419 | 6.19905 | 6.75947 | 5.8326 | 6.44872 | 6.74052 | 6.15915 | 6.45847 | 6.38563 | 6.42843 | 6.59392 | 6.46717 | 5.93471 | 5.79354 | 6.01997 | 5.73446 |
| SMPD4 | Q9NXE4-4 | 6.42907 | 6.45016 | 6.11973 | 6.77563 | 6.33715 | 6.05901 | 6.38609 | 5.95465 | 6.30679 | 6.20946 | 6.30778 | 6.24849 | 6.26654 | 5.91502 | 6.11774 | 6.11669 |
| PSMC2 | P35998 | 6.21835 | 5.88662 | 6.80087 | 6.07114 | 6.36972 | 6.44945 | 6.40308 | 6.16263 | 6.04811 | 6.07916 | 6.11147 | 6.24625 | 6.30368 | 6.17123 | 6.16286 | 6.51538 |
| PAPSS1 | O43252 | 5.56371 | 5.73719 | 6.13801 | 5.42374 | 5.89555 | 6.21524 | 5.88138 | 6.29134 | 6.00868 | 6.0958 | 6.30099 | 6.24554 | 6.85886 | 7.15144 | 7.4467 | 6.74582 |
| XRN1 | Q8IZH2 | 6.09231 | 6.23684 | 6.50506 | 5.92785 | 6.34141 | 6.84362 | 6.24889 | 6.44602 | 6.32869 | 6.45509 | 6.86383 | 6.42763 | 5.92607 | 6.25112 | 5.63435 | 5.47122 |
| PCNA | P12004 | 5.98282 | 6.24958 | 6.14527 | 5.64891 | 5.7235 | 5.97412 | 5.93431 | 6.03219 | 6.3444 | 6.11335 | 5.7953 | 6.08485 | 6.76018 | 6.90392 | 7.11731 | 7.18999 |
| PBRM1 | Q86U86 | 6.40871 | 6.40604 | 6.11271 | 6.66742 | 5.90892 | 5.90223 | 6.40256 | 5.92768 | 6.22923 | 6.22765 | 5.87965 | 5.91 | 6.51797 | 6.36834 | 6.58863 | 6.54618 |
| CACYBP | Q9HB71 | 6.36859 | 6.03644 | 6.51961 | 5.86716 | 6.82755 | 6.73256 | 6.41841 | 6.44166 | 6.35425 | 6.4632 | 6.32364 | 6.27926 | 6.09261 | 5.92298 | 5.70636 | 5.64574 |
| ACSL1 | P33121 | 6.33132 | 6.3945 | 5.90171 | 6.70215 | 6.15722 | 5.92833 | 6.3706 | 5.76685 | 6.08353 | 6.10717 | 5.95334 | 5.99676 | 6.57634 | 6.43476 | 6.58417 | 6.71125 |
| ZNF292 | O60281 | 6.25213 | 6.34029 | 5.96342 | 6.8768 | 5.97675 | 6.09223 | 6.48913 | 5.78812 | 6.09778 | 6.03926 | 5.9701 | 6.16977 | 6.34374 | 6.48274 | 6.64148 | 6.38625 |
| ARHGEF2 | Q92974 | 6.44156 | 6.49991 | 6.317 | 6.19783 | 6.44814 | 6.59306 | 6.21811 | 6.324 | 6.15577 | 6.41948 | 6.46702 | 6.54394 | 5.96785 | 5.9282 | 5.6762 | 5.80192 |
| USP39 | Q53GS9 | 6.06811 | 6.39005 | 5.9567 | 6.58838 | 6.00525 | 5.97224 | 6.03739 | 5.90368 | 5.90664 | 5.98181 | 5.90212 | 5.91314 | 6.67882 | 6.80564 | 6.84601 | 7.04404 |
| MME | P08473 | 6.38312 | 6.5323 | 5.95675 | 6.06087 | 6.32687 | 6.13535 | 6.36686 | 6.09883 | 6.21077 | 6.22448 | 5.88629 | 5.78068 | 6.60516 | 6.48035 | 6.58809 | 6.36324 |
| STAG2 | Q8N3U4-2 | 6.33554 | 6.54641 | 5.93036 | 6.54975 | 6.15378 | 6.17179 | 6.20731 | 6.18116 | 6.02252 | 6.24756 | 5.80997 | 5.78812 | 6.6991 | 6.30106 | 6.52042 | 6.53515 |
| NOTCH2 | Q04721 | 6.32494 | 6.23063 | 6.05711 | 6.4255 | 6.22908 | 6.05117 | 6.32852 | 6.08143 | 6.15363 | 5.90395 | 6.10717 | 6.20829 | 6.71399 | 6.29256 | 6.43019 | 6.46183 |
| PCK2 | Q16822 | 6.34376 | 6.04996 | 5.8864 | 6.29084 | 6.3932 | 6.00613 | 6.29738 | 5.83244 | 6.12184 | 5.9832 | 5.97041 | 5.82527 | 7.25216 | 6.52303 | 6.74996 | 6.47403 |
| UBAP2L | Q14157-5 | 6.38878 | 6.32992 | 6.74836 | 6.12099 | 6.35882 | 6.45704 | 6.08414 | 6.43197 | 6.50736 | 6.64468 | 6.87646 | 6.60786 | 5.67971 | 5.89861 | 5.44455 | 5.42075 |
| CACNA2D1 | P54289 | 6.21881 | 6.14943 | 6.07073 | 6.25362 | 6.2536 | 5.96267 | 6.17014 | 5.94294 | 6.209 | 6.3003 | 6.13183 | 5.86707 | 6.87531 | 6.41162 | 6.78341 | 6.39951 |
| UHRF1 | Q96T88-2 | 6.26518 | 6.23217 | 6.20858 | 6.66237 | 6.0535 | 5.8946 | 6.26523 | 5.79429 | 6.32814 | 6.21719 | 6.54509 | 6.40269 | 6.27806 | 6.09257 | 6.34221 | 6.41812 |
| ATL3 | Q6DD88 | 6.37946 | 6.22934 | 5.90596 | 6.36054 | 6.26007 | 6.19107 | 6.32817 | 6.02903 | 6.08081 | 6.12596 | 5.81025 | 5.98259 | 6.61352 | 6.48831 | 6.55365 | 6.66127 |
| TIMM44 | O43615 | 6.34433 | 6.69461 | 5.90878 | 6.9006 | 6.21685 | 6.2833 | 6.21751 | 6.20828 | 5.929 | 5.91237 | 6.095 | 5.97135 | 6.29851 | 6.46246 | 6.39181 | 6.16525 |
| PLCB4 | Q15147-4 | 6.18352 | 6.77806 | 6.28391 | 6.67819 | 6.25706 | 6.20793 | 6.46741 | 6.44722 | 6.59227 | 6.25952 | 6.52378 | 6.85495 | 5.59458 | 5.47874 | 5.51754 | 5.87533 |
| GRIPAP1 | Q4V328 | 6.48121 | 6.08884 | 6.79909 | 5.90715 | 6.57616 | 6.59579 | 5.98515 | 6.6873 | 6.39911 | 6.45475 | 6.49961 | 6.51061 | 6.37801 | 5.72252 | 5.57265 | 5.34205 |
| HBS1L | Q9Y450 | 6.40698 | 6.27998 | 6.65568 | 6.17195 | 6.41475 | 6.82474 | 6.18573 | 6.40155 | 6.31677 | 6.42575 | 6.32172 | 6.38658 | 5.82068 | 5.82081 | 5.93165 | 5.63468 |
| DDX18 | Q9NVP1 | 6.28674 | 6.24936 | 5.64184 | 6.85993 | 6.0799 | 5.8755 | 6.36754 | 5.96216 | 6.07428 | 5.78077 | 5.91109 | 5.85161 | 6.35403 | 6.71988 | 6.86289 | 7.12247 |
| RAB7A | P51149 | 6.147 | 6.19326 | 6.01894 | 6.23875 | 6.25054 | 6.04635 | 6.4582 | 5.85999 | 6.15309 | 6.11856 | 6.18496 | 5.92185 | 6.61853 | 6.43831 | 6.79402 | 6.55765 |
| RPRD2 | Q5VT52 | 6.47145 | 6.37844 | 6.16139 | 6.48209 | 6.32843 | 6.1906 | 6.29552 | 5.85045 | 6.32374 | 6.15306 | 6.21168 | 5.96764 | 6.54828 | 6.14938 | 6.22013 | 6.26774 |
| KPNA2 | P52292 | 6.4489 | 6.37947 | 6.78295 | 6.64278 | 6.14777 | 6.3997 | 6.65709 | 6.47909 | 6.51892 | 6.45295 | 6.66161 | 6.86392 | 5.29936 | 5.31288 | 5.19813 | 5.7545 |
| NBN | O60934 | 6.02899 | 6.4434 | 6.00609 | 6.46358 | 5.97977 | 6.15645 | 6.14995 | 5.936 | 6.05856 | 6.16177 | 5.88142 | 5.96122 | 6.7479 | 6.36888 | 6.69583 | 6.96017 |
| AKAP13 | Q12802-2 | 5.86919 | 5.9223 | 6.55301 | 6.00059 | 6.00625 | 6.37443 | 6.07152 | 6.30674 | 6.50986 | 6.69932 | 6.64566 | 6.78021 | 5.93144 | 6.22508 | 6.07248 | 6.03193 |
| RPL28 | P46779-3 | 6.30261 | 6.24949 | 6.24781 | 6.27224 | 6.17598 | 6.46916 | 6.1921 | 6.24121 | 6.7865 | 6.46427 | 6.51256 | 6.40495 | 5.86376 | 6.20408 | 5.80536 | 5.80792 |
| ZYX | Q15942 | 6.29996 | 5.32488 | 6.70285 | 5.10656 | 6.70938 | 6.37355 | 5.78551 | 6.95464 | 6.50052 | 6.58373 | 6.87351 | 6.88033 | 6.75926 | 5.89701 | 5.71638 | 5.73732 |
| POLR1B | Q9H9Y6-3 | 6.47796 | 6.21014 | 6.22973 | 6.82892 | 6.0969 | 5.8483 | 6.48276 | 5.78685 | 6.5009 | 5.97229 | 6.30618 | 6.2509 | 6.38623 | 5.79485 | 6.36335 | 6.46375 |
| ARCN1 | P48444 | 6.40062 | 6.39237 | 6.7 | 6.27668 | 6.41993 | 6.44472 | 6.29214 | 6.41043 | 6.40397 | 6.51218 | 6.76257 | 6.55191 | 5.77775 | 5.68739 | 5.53313 | 5.43421 |
| NOC2L | Q9Y3T9 | 6.39119 | 6.28658 | 6.01447 | 6.68503 | 6.282 | 5.80558 | 6.59515 | 5.99273 | 6.42351 | 5.93351 | 6.02507 | 5.84037 | 6.61538 | 6.0701 | 6.45348 | 6.58584 |
| RBM15 | Q96T37 | 6.32403 | 6.4361 | 5.957 | 6.76983 | 6.0275 | 5.84047 | 6.39322 | 5.76351 | 6.14375 | 6.11511 | 5.99608 | 6.20361 | 6.4398 | 6.20894 | 6.76816 | 6.61288 |
| INA | Q16352 | 6.44553 | 6.36612 | 5.81748 | 6.42756 | 6.43918 | 6.19569 | 6.23639 | 5.95001 | 6.23264 | 5.98978 | 6.06226 | 6.08107 | 7.05976 | 6.00215 | 6.41832 | 6.27607 |
| ETFB | P38117 | 6.3529 | 5.96304 | 5.80033 | 6.05237 | 6.40142 | 6.11256 | 6.40358 | 6.35439 | 6.29334 | 5.91103 | 5.66779 | 5.96458 | 7.10127 | 6.45232 | 6.5851 | 6.58397 |
| DEK | H0Y993 | 6.04394 | 6.44261 | 6.09203 | 6.50735 | 6.17093 | 6.18479 | 6.25245 | 6.10539 | 5.96749 | 6.2763 | 5.98449 | 6.02284 | 6.52671 | 6.56123 | 6.4233 | 6.43815 |
| FUBP3 | Q96I24 | 6.58596 | 6.41303 | 6.39231 | 6.51545 | 6.68507 | 6.42402 | 6.06333 | 6.13067 | 6.2674 | 6.06573 | 6.62186 | 6.35252 | 6.56083 | 5.64819 | 5.68747 | 5.58617 |
| RBM39 | Q14498 | 6.08271 | 6.40694 | 5.7667 | 6.54164 | 6.18296 | 5.99583 | 6.27455 | 5.89813 | 5.91286 | 5.88084 | 5.82098 | 5.76643 | 6.80426 | 6.71721 | 6.9801 | 6.96787 |
| UBE3A | Q05086 | 5.84438 | 5.98542 | 6.20459 | 5.90711 | 6.0411 | 6.33167 | 6.16964 | 6.13901 | 6.14496 | 6.16182 | 6.23293 | 6.27035 | 6.67664 | 6.70166 | 6.67334 | 6.51538 |
| YBX3 | P16989 | 6.48571 | 6.36202 | 6.45245 | 6.51746 | 6.30176 | 6.16689 | 6.59338 | 6.36876 | 6.76262 | 6.03733 | 6.35557 | 6.31232 | 5.28145 | 6.21979 | 5.67418 | 6.1083 |
| PRC1 | O43663 | 6.06476 | 6.05325 | 5.88593 | 6.20296 | 5.99745 | 6.38976 | 6.2772 | 6.1929 | 6.12725 | 6.12877 | 6.09906 | 6.17014 | 6.71293 | 6.40588 | 6.57811 | 6.71365 |
| XPO7 | Q9UIA9 | 5.55269 | 5.66348 | 6.168 | 5.43954 | 5.92348 | 6.23418 | 5.79063 | 6.06913 | 6.07139 | 6.11708 | 6.28535 | 6.05474 | 7.15741 | 7.33963 | 7.19249 | 6.94077 |
| PDLIM7 | Q9NR12 | 5.98331 | 6.05436 | 5.76242 | 5.65644 | 6.28423 | 6.31339 | 6.13298 | 6.12382 | 6.38178 | 6.38848 | 6.13237 | 6.18033 | 6.90843 | 6.63896 | 6.61674 | 6.44196 |
| TUBGCP2 | Q9BSJ2-4 | 6.28843 | 6.24522 | 6.89635 | 6.23894 | 6.22278 | 6.69026 | 6.21357 | 6.56453 | 6.3895 | 6.4203 | 6.86493 | 6.53607 | 5.56617 | 5.8 | 5.53011 | 5.53282 |
| ILVBL | A1L0T0 | 6.27781 | 6.28886 | 5.85852 | 6.67547 | 6.2451 | 5.97762 | 6.42171 | 5.93229 | 6.14979 | 6.00106 | 5.9374 | 5.89828 | 6.47529 | 6.52894 | 6.69651 | 6.63536 |
| CHD2 | O14647 | 6.28319 | 6.07423 | 6.0505 | 6.52992 | 6.31148 | 5.75116 | 6.37088 | 5.70179 | 6.08679 | 6.20542 | 5.86477 | 5.64086 | 6.6674 | 6.69154 | 7.10565 | 6.66441 |
| WNK1 | Q9H4A3-7 | 6.35414 | 6.3189 | 6.7108 | 5.93258 | 6.29623 | 6.57325 | 6.12018 | 6.55188 | 6.4962 | 6.65005 | 6.58284 | 6.42417 | 5.78414 | 5.89026 | 5.60192 | 5.71246 |
| SIPA1L1 | O43166-2 | 6.42908 | 6.49497 | 6.03879 | 6.57961 | 6.19955 | 5.94199 | 6.46817 | 5.97984 | 6.17217 | 5.97446 | 5.9728 | 5.83172 | 6.524 | 6.29349 | 6.56473 | 6.53462 |
| SEPTIN2 | Q15019-2 | 5.89843 | 5.83956 | 6.61098 | 5.71322 | 6.42205 | 6.6788 | 6.25896 | 6.4487 | 6.39007 | 6.46728 | 6.28161 | 6.22432 | 6.41109 | 6.11891 | 6.17954 | 6.05647 |
| KIF22 | Q14807 | 6.27189 | 6.17068 | 5.67697 | 6.24026 | 6.25152 | 6.03284 | 6.40841 | 6.20178 | 6.14102 | 5.92557 | 5.73164 | 5.84277 | 6.70926 | 6.666 | 6.89114 | 6.83824 |
| IGF2BP1 | Q9NZI8 | 6.24768 | 6.34013 | 6.42318 | 6.05625 | 6.60763 | 6.87326 | 6.32416 | 6.52556 | 6.49919 | 6.32376 | 6.73873 | 6.16419 | 5.92266 | 5.92363 | 5.57524 | 5.45476 |
| HTATSF1 | O43719 | 6.16373 | 6.3443 | 6.11688 | 6.59134 | 6.03276 | 6.05247 | 6.16938 | 5.87138 | 6.05977 | 6.33915 | 6.31847 | 6.26202 | 6.55666 | 6.33551 | 6.39405 | 6.39212 |
| VPS35 | Q96QK1 | 6.03472 | 6.14558 | 5.88906 | 5.83876 | 6.15866 | 6.49433 | 5.91066 | 6.45728 | 6.2128 | 6.29292 | 6.02223 | 6.21565 | 6.30571 | 6.92023 | 6.53662 | 6.5648 |
| PRPF40A | O75400 | 6.22845 | 6.41479 | 5.8483 | 6.6742 | 6.27387 | 5.93941 | 6.48266 | 5.88775 | 6.06211 | 5.8704 | 5.92263 | 5.93959 | 6.83583 | 6.55329 | 6.46832 | 6.59842 |
| RRP1B | Q14684 | 6.18423 | 6.17321 | 5.83124 | 6.76974 | 6.16393 | 5.841 | 6.50155 | 5.74491 | 5.97318 | 5.85866 | 6.11088 | 5.97612 | 6.65943 | 6.55014 | 6.91423 | 6.74757 |
| TRAPPC8 | Q9Y2L5 | 6.14154 | 6.1889 | 6.46733 | 6.00831 | 6.22667 | 6.69885 | 6.03881 | 6.4429 | 6.37312 | 6.47851 | 6.42909 | 6.50337 | 5.98938 | 6.1675 | 5.88902 | 5.95671 |
| CRYBG3 | Q68DQ2 | 6.18724 | 6.1773 | 6.69176 | 6.06663 | 6.10605 | 6.36169 | 6.08095 | 6.37544 | 6.50957 | 6.5903 | 6.89863 | 6.74872 | 5.96503 | 5.76858 | 5.78007 | 5.69204 |
| THOC5 | Q13769 | 6.35322 | 6.49308 | 5.93361 | 6.68894 | 6.16029 | 6.0208 | 6.43835 | 6.05681 | 6.11241 | 6.01687 | 6.02999 | 5.91226 | 6.6396 | 6.26232 | 6.5176 | 6.36386 |
| CLCC1 | Q96S66 | 6.29637 | 6.34186 | 6.09645 | 6.60459 | 6.19848 | 6.03575 | 6.11548 | 6.09974 | 6.03637 | 6.03239 | 6.0105 | 5.89614 | 6.59552 | 6.20581 | 6.62693 | 6.80763 |
| HMOX2 | P30519 | 6.36964 | 6.07826 | 5.93673 | 5.93132 | 6.62334 | 5.86392 | 6.29736 | 6.37879 | 6.021 | 6.01581 | 6.12025 | 6.10721 | 6.82222 | 6.47014 | 6.52528 | 6.43874 |
| RPL10A | P62906 | 6.27509 | 6.52747 | 6.29521 | 6.27503 | 6.17594 | 6.59954 | 6.37872 | 6.32071 | 6.45422 | 6.40123 | 6.3251 | 6.42473 | 5.85231 | 5.84447 | 5.83052 | 6.01972 |
| MAPRE1 | Q15691 | 6.08293 | 5.74132 | 6.8034 | 5.53488 | 6.39366 | 6.56575 | 6.02409 | 6.65593 | 6.17648 | 6.40513 | 6.72661 | 6.886 | 6.13029 | 5.98554 | 5.84196 | 6.04602 |
| CNN3 | Q15417 | 6.1417 | 6.09153 | 7.09516 | 5.98603 | 6.07903 | 6.74051 | 5.9393 | 6.72817 | 6.64474 | 6.75019 | 6.98799 | 7.01711 | 5.88603 | 5.63718 | 5.09392 | 5.18139 |
| PELP1 | Q8IZL8 | 6.48202 | 6.31606 | 5.97262 | 6.82822 | 6.19863 | 6.03542 | 6.27478 | 5.94099 | 6.02875 | 5.91941 | 6.20919 | 6.18015 | 6.71262 | 6.37892 | 6.32906 | 6.19316 |
| DECR1 | Q16698 | 6.33262 | 6.20117 | 5.80459 | 6.39706 | 6.18537 | 5.78714 | 6.35623 | 6.14641 | 6.21101 | 5.92215 | 5.78733 | 5.86432 | 6.80055 | 6.61773 | 6.82908 | 6.75722 |
| RPL7A | P62424 | 6.28422 | 6.14391 | 6.15461 | 5.98219 | 6.33273 | 6.24494 | 6.38669 | 6.40287 | 6.63082 | 6.51035 | 6.3042 | 6.24711 | 5.87293 | 6.28753 | 6.08566 | 6.12923 |
| PUM3 | Q15397 | 6.20413 | 6.23899 | 5.85061 | 6.9513 | 6.13732 | 5.63918 | 6.41974 | 5.87798 | 6.01585 | 5.86491 | 5.88089 | 6.20555 | 6.3787 | 6.44179 | 6.87907 | 7.01399 |
| RPL18A | Q02543 | 6.32525 | 6.273 | 6.34314 | 6.39368 | 6.20951 | 6.34678 | 6.39298 | 6.32887 | 6.61408 | 6.59503 | 6.4411 | 6.51333 | 5.74676 | 5.83415 | 5.69062 | 5.95174 |

| | | | | | | | | | | | | | | | | | |
|---|---|---|---|---|---|---|---|---|---|---|---|---|---|---|---|---|---|
| DHX8 | Q14562 | 6.47167 | 6.78319 | 5.94208 | 6.85533 | 6.18794 | 6.07828 | 6.39497 | 6.18766 | 6.3112 | 6.15295 | 6.23179 | 6.10036 | 5.89568 | 6.14284 | 6.17873 | 6.08533 |
| CDC42BPB | Q9Y5S2 | 6.4401 | 6.3793 | 7.03167 | 6.0422 | 6.34271 | 6.66294 | 6.27671 | 6.29924 | 6.49267 | 6.56107 | 6.65054 | 6.54451 | 5.83879 | 5.56183 | 5.6003 | 5.27544 |
| ATR | Q13535 | 6.02243 | 6.35039 | 6.22997 | 6.33351 | 6.319 | 6.28914 | 6.24878 | 6.10832 | 6.25894 | 6.3752 | 6.2702 | 6.20311 | 6.16907 | 6.25399 | 6.20366 | 6.3643 |
| GFM1 | Q96RP9 | 6.38738 | 6.21426 | 5.83925 | 6.49656 | 6.46322 | 6.0457 | 6.3284 | 6.15595 | 6.1367 | 5.83433 | 5.8497 | 5.81196 | 6.77373 | 6.4376 | 6.80566 | 6.41959 |
| RFC3 | P40938 | 6.04669 | 6.41837 | 6.31189 | 6.21914 | 6.22718 | 6.31757 | 6.1713 | 6.14148 | 5.97331 | 6.20992 | 6.2699 | 5.98892 | 6.54844 | 6.27774 | 6.5536 | 6.32454 |
| KLC1 | Q07866-6 | 6.37976 | 6.40603 | 6.80315 | 5.84015 | 6.47507 | 6.7808 | 6.27737 | 6.757 | 6.58503 | 6.45097 | 6.59529 | 6.54963 | 5.85281 | 5.5627 | 5.30185 | 5.38237 |
| CSNK2A1 | P68400 | 6.16139 | 6.43859 | 6.41063 | 6.498 | 6.38829 | 6.57838 | 6.20513 | 6.44051 | 6.06064 | 6.22002 | 6.33253 | 6.09239 | 6.06862 | 6.02959 | 6.19809 | 5.87719 |
| KIF1B | O60333-2 | 6.31249 | 6.35823 | 7.01666 | 6.16867 | 6.23461 | 6.72574 | 6.3632 | 6.34763 | 6.4124 | 6.48757 | 6.48101 | 6.55175 | 5.6411 | 5.87578 | 5.4078 | 5.61534 |
| MED23 | Q9ULK4 | 6.03548 | 6.25532 | 5.97817 | 6.34078 | 6.01061 | 6.07408 | 6.06007 | 5.93619 | 6.26518 | 6.2503 | 6.23495 | 6.2275 | 6.58915 | 6.45226 | 6.69253 | 6.59745 |
| EMSY | Q7Z589-7 | 6.56299 | 6.44399 | 6.16434 | 6.74803 | 6.18837 | 6.13974 | 6.12359 | 5.95938 | 6.17866 | 6.36473 | 6.09533 | 6.11072 | 6.4502 | 5.96898 | 6.29353 | 6.20741 |
| NARS1 | O43776 | 6.21478 | 6.14811 | 6.5972 | 5.83883 | 6.05827 | 6.52691 | 6.23517 | 6.48315 | 6.46003 | 6.38672 | 6.62992 | 6.51508 | 5.89581 | 6.21718 | 5.77809 | 6.01476 |
| PRPF19 | Q9UMS4 | 5.8888 | 6.34528 | 5.33933 | 6.37259 | 6.45974 | 6.41319 | 6.08769 | 6.64665 | 6.14622 | 6.38204 | 5.88362 | 6.05005 | 6.22427 | 6.34228 | 6.62844 | 6.78982 |
| NAA25 | Q14CX7 | 6.14612 | 6.38814 | 6.80056 | 5.97771 | 6.30585 | 6.71293 | 6.17895 | 6.78696 | 6.31472 | 6.61419 | 6.6354 | 6.40849 | 5.66814 | 5.80556 | 5.62842 | 5.62786 |
| CD44 | P16070 | 6.43233 | 6.47085 | 5.9028 | 6.54887 | 6.18629 | 6.10726 | 6.17604 | 6.49173 | 6.21375 | 5.91393 | 5.87981 | 5.93222 | 6.46515 | 6.54075 | 6.17282 | 5.6554 |
| TBCD | Q9BTW9 | 6.49534 | 6.13617 | 7.00457 | 6.24417 | 6.39176 | 6.89812 | 6.28652 | 6.63322 | 6.44796 | 6.33285 | 6.65276 | 6.68663 | 5.50721 | 5.56687 | 5.29029 | 5.42558 |
| INTS4 | Q96HW7 | 6.22917 | 6.32017 | 5.98897 | 6.53765 | 6.24619 | 6.06099 | 6.21364 | 5.95856 | 6.13813 | 6.24219 | 6.17711 | 6.0045 | 6.51987 | 6.2548 | 6.5765 | 6.53156 |
| CTTN | Q14247 | 6.19886 | 6.29882 | 6.41169 | 6.29873 | 6.32826 | 6.63797 | 6.03967 | 6.75085 | 6.01509 | 5.93438 | 6.44291 | 6.14453 | 6.25666 | 6.11647 | 6.18897 | 5.93616 |
| BAZ1A | Q9NRL2 | 6.45799 | 6.6076 | 6.15845 | 7.06223 | 6.12091 | 5.88945 | 6.65406 | 6.18397 | 6.23967 | 6.08767 | 6.16679 | 6.19253 | 5.99018 | 5.88645 | 6.07607 | 6.22596 |
| NCAPH | Q15003 | 6.41304 | 6.32106 | 6.73135 | 6.27407 | 6.34148 | 6.63018 | 6.04387 | 6.77183 | 6.40277 | 6.50306 | 6.67938 | 6.85531 | 5.65257 | 5.6371 | 5.38895 | 5.35398 |
| TRIP10 | Q15642-2 | 6.17935 | 6.39225 | 6.92996 | 6.01851 | 6.42989 | 6.88639 | 6.26104 | 6.46042 | 6.4511 | 6.61991 | 6.59162 | 6.7724 | 5.6945 | 5.67866 | 5.26752 | 5.36648 |
| SACM1L | Q9NTJ5 | 6.32931 | 6.28851 | 6.12371 | 6.52018 | 6.2153 | 6.08819 | 6.39315 | 6.00402 | 6.12151 | 6.00907 | 6.04814 | 5.85681 | 6.6188 | 6.33316 | 6.64224 | 6.4079 |
| ARPC2 | O15144 | 6.28448 | 6.40505 | 6.48422 | 6.10289 | 6.27009 | 6.3893 | 6.40416 | 6.55638 | 6.15788 | 6.38643 | 6.21098 | 6.11266 | 5.92284 | 6.02297 | 6.33294 | 5.95674 |
| GNL2 | Q13823 | 6.36236 | 6.24568 | 6.10183 | 6.77265 | 6.18572 | 5.8971 | 6.37033 | 5.91728 | 6.22535 | 6.03152 | 6.2022 | 6.15919 | 5.98577 | 6.35283 | 6.50866 | 6.68153 |
| PLRG1 | O43660 | 6.31591 | 6.41516 | 5.81329 | 6.7768 | 6.04362 | 5.84837 | 6.56142 | 6.0488 | 6.2828 | 6.42004 | 5.88716 | 5.98411 | 6.08361 | 6.47686 | 6.55814 | 6.48391 |
| CKAP4 | Q07065 | 6.38481 | 6.51838 | 6.1805 | 6.43844 | 6.37448 | 5.93155 | 6.2758 | 6.09381 | 6.1909 | 5.9541 | 5.96987 | 5.77601 | 6.71789 | 6.25797 | 6.57304 | 6.36244 |
| GFPT1 | Q06210 | 6.36397 | 6.36234 | 6.77434 | 6.00069 | 6.69079 | 7.02573 | 6.3369 | 6.71157 | 6.49119 | 6.08233 | 6.32038 | 6.36655 | 5.47925 | 6.03216 | 5.48327 | 5.47854 |
| MSH3 | P20585 | 5.65405 | 6.05214 | 5.87361 | 5.8399 | 6.0146 | 6.23968 | 6.18314 | 5.86567 | 5.99652 | 6.10872 | 6.26847 | 5.96585 | 6.91143 | 6.99452 | 7.1405 | 6.89119 |
| TWF1 | Q12792 | 6.28597 | 6.50134 | 6.83451 | 6.45683 | 6.35898 | 6.87616 | 6.35362 | 6.74597 | 6.46177 | 6.29606 | 6.24595 | 6.37035 | 5.45216 | 5.63236 | 5.63809 | 5.48989 |
| NOL10 | Q9BSC4 | 6.17467 | 6.1351 | 5.86186 | 6.72188 | 6.14199 | 5.89628 | 6.50571 | 5.78492 | 6.19709 | 6.05547 | 5.92931 | 6.07469 | 6.62515 | 6.34388 | 6.82854 | 6.72344 |
| ZNF644 | Q9H582 | 6.49098 | 6.6607 | 6.08145 | 6.87738 | 6.11497 | 6.14571 | 6.29051 | 5.99775 | 6.252 | 6.29661 | 6.06097 | 6.29132 | 6.09311 | 6.14843 | 5.98846 | 6.20965 |
| WDR44 | Q5JSH3 | 6.21781 | 5.94672 | 6.74138 | 5.70722 | 6.36771 | 6.51815 | 6.45631 | 6.53201 | 6.3907 | 6.34354 | 6.45257 | 6.28097 | 6.08374 | 6.15761 | 5.97215 | 5.8314 |
| RPL12 | P30050 | 6.37786 | 6.40444 | 6.36815 | 6.14172 | 6.44868 | 6.59764 | 6.22901 | 6.26481 | 6.50051 | 6.40099 | 6.53112 | 6.51108 | 5.86127 | 5.88067 | 5.73445 | 5.74758 |
| VAPA | Q9P0L0 | 6.42763 | 6.1219 | 5.90858 | 6.19713 | 6.32731 | 6.12082 | 6.24046 | 6.44377 | 6.06379 | 6.2642 | 5.86101 | 6.00727 | 6.97523 | 6.4243 | 6.33455 | 6.28206 |
| RPS4X | P62701 | 6.74143 | 6.3447 | 6.63161 | 6.71623 | 6.7562 | 6.46905 | 6.48255 | 6.23953 | 6.51326 | 6.0028 | 6.01418 | 6.13984 | 5.71082 | 5.87305 | 5.54807 | 5.81667 |
| EIF2S2 | P20042 | 6.02796 | 6.30071 | 6.53697 | 6.16382 | 6.11732 | 6.44383 | 6.16718 | 6.39771 | 6.24 | 6.59586 | 6.24395 | 6.52171 | 5.80666 | 6.21415 | 6.06163 | 6.16056 |
| PRDX6 | P30041 | 6.14061 | 5.27879 | 6.74754 | 4.99758 | 6.54945 | 6.25921 | 5.83016 | 7.22997 | 5.8051 | 6.94941 | 6.33829 | 7.15478 | 5.94278 | 6.62718 | 6.05172 | 6.09743 |
| ASH2L | Q9UBL3 | 5.99945 | 6.38179 | 5.67903 | 6.43811 | 6.18092 | 6.18396 | 6.25984 | 6.17537 | 6.19617 | 6.10303 | 5.92653 | 6.08248 | 6.66488 | 6.45834 | 6.65219 | 6.61791 |
| SNW1 | Q13573 | 5.87035 | 6.04295 | 5.73888 | 6.43703 | 6.10459 | 5.76926 | 6.86704 | 5.83402 | 6.50944 | 5.57447 | 5.79579 | 5.8319 | 6.30651 | 6.86315 | 6.95355 | 7.50106 |
| MDH1 | P40925-3 | 6.32641 | 6.2663 | 6.98747 | 5.86665 | 6.36167 | 6.77664 | 6.25546 | 6.33292 | 6.31738 | 6.56645 | 6.65146 | 6.31822 | 5.83502 | 5.95008 | 5.70996 | 5.47791 |
| CKB | P12277 | 6.22577 | 6.22216 | 6.95624 | 6.03393 | 6.59558 | 6.8348 | 6.13785 | 6.46488 | 6.27887 | 6.40234 | 6.31841 | 6.46215 | 5.85337 | 5.99226 | 5.59322 | 5.62817 |
| SRP54 | P61011 | 6.29619 | 6.30435 | 6.26963 | 6.20369 | 6.28766 | 6.23501 | 6.26335 | 6.12962 | 6.29844 | 6.25794 | 6.19548 | 6.09973 | 6.2753 | 6.23629 | 6.41214 | 6.23838 |
| FARSA | Q9Y285 | 6.07273 | 6.22441 | 6.73426 | 6.0786 | 6.25277 | 6.63103 | 6.08625 | 6.45496 | 6.64489 | 6.50004 | 6.79528 | 6.72567 | 5.81664 | 5.70137 | 5.65483 | 5.62627 |
| CLPTM1 | O96005 | 6.23748 | 6.56754 | 6.03589 | 6.82873 | 6.09979 | 6.15151 | 6.29942 | 5.99534 | 6.30088 | 6.24906 | 6.15391 | 6.30485 | 6.07518 | 6.17157 | 6.10397 | 6.42486 |
| VDAC3 | Q9Y277 | 6.21267 | 6.31087 | 5.82742 | 6.63041 | 6.47722 | 5.98113 | 6.44573 | 6.05327 | 6.24252 | 6.1 | 5.75092 | 5.89367 | 6.68663 | 6.14245 | 6.94398 | 6.33111 |
| BRD2 | P25440-2 | 6.32536 | 6.57302 | 6.09412 | 6.64974 | 6.09533 | 6.15 | 6.35873 | 5.9093 | 6.38172 | 6.20222 | 6.17683 | 6.23292 | 6.06309 | 6.05479 | 6.20623 | 6.5266 |
| TPM3 | P06753-2 | 6.60656 | 6.14509 | 6.3796 | 6.18463 | 6.85581 | 6.18508 | 6.0093 | 6.74078 | 6.37341 | 5.96173 | 6.66906 | 6.54473 | 6.49212 | 5.84808 | 5.30319 | 5.70082 |
| DGLUCY | Q7Z3D6-2 | 6.33921 | 6.21634 | 5.67441 | 6.01649 | 6.33256 | 6.1521 | 6.26013 | 5.88853 | 6.02463 | 6.28621 | 6.08175 | 6.13582 | 6.69827 | 6.57847 | 6.85642 | 6.45867 |
| RCN1 | Q15293 | 6.5309 | 6.25956 | 6.04772 | 6.23247 | 6.54529 | 6.01427 | 6.10441 | 6.11703 | 6.1731 | 6.01503 | 6.4019 | 6.15254 | 7.1063 | 5.95909 | 6.1844 | 6.156 |
| TTC37 | Q6PGP7 | 6.46573 | 6.35773 | 6.61491 | 6.03863 | 6.30653 | 6.65829 | 6.15169 | 6.67768 | 6.42629 | 6.55487 | 6.27226 | 6.2896 | 5.85252 | 6.08755 | 5.64635 | 5.59936 |
| NEMF | O60524 | 6.03154 | 5.92346 | 6.30602 | 5.9504 | 5.95108 | 6.32345 | 6.07574 | 6.22158 | 6.03126 | 6.44971 | 6.53737 | 6.43697 | 6.17783 | 6.75929 | 6.65856 | 6.16574 |
| SUPV3L1 | Q8IYB8 | 6.14285 | 6.28219 | 6.05435 | 6.60423 | 6.1453 | 6.03561 | 6.39957 | 5.8021 | 5.98093 | 6.04093 | 5.96897 | 6.01232 | 6.77692 | 6.32041 | 6.66423 | 6.76908 |
| EIF4G3 | A0A0U1RQK7 | 6.15439 | 6.50805 | 6.63431 | 6.06849 | 6.02739 | 6.45602 | 6.11112 | 6.31001 | 6.36742 | 6.587 | 6.38197 | 6.52154 | 6.01172 | 6.28933 | 5.83537 | 5.93586 |
| ECH1 | Q13011 | 6.15947 | 6.10831 | 5.94267 | 6.32647 | 6.26403 | 6.03219 | 6.60232 | 6.02115 | 6.42385 | 5.9583 | 6.08423 | 5.95875 | 6.67461 | 6.29427 | 6.53785 | 6.61154 |
| RAI14 | Q9P0K7 | 6.15123 | 6.23962 | 6.15587 | 6.02474 | 6.43355 | 6.28155 | 5.85044 | 6.20347 | 6.22282 | 6.43081 | 6.34137 | 6.20319 | 6.7212 | 6.27018 | 6.246 | 6.22396 |
| PSMA2 | P25787 | 6.08175 | 6.21794 | 6.20664 | 6.0744 | 6.40225 | 6.69871 | 6.24118 | 6.6111 | 6.3267 | 6.07686 | 6.2912 | 6.04854 | 6.43389 | 6.14236 | 6.15659 | 5.98988 |
| ITSN1 | Q15811-2 | 6.04387 | 6.13737 | 6.33972 | 5.95572 | 6.23947 | 6.38209 | 6.04594 | 6.23645 | 6.36782 | 6.41762 | 6.55944 | 6.26447 | 6.61352 | 6.14996 | 6.24028 | 6.00626 |
| SEC23A | Q15436 | 6.07415 | 6.27901 | 6.31523 | 5.98746 | 6.16629 | 6.21242 | 6.26981 | 6.14701 | 6.0222 | 6.22621 | 6.14067 | 6.09403 | 6.33767 | 6.63168 | 6.68123 | 6.41492 |
| RALY | Q9UKM9 | 6.52447 | 6.48413 | 5.86161 | 6.7116 | 6.44891 | 5.92857 | 6.46923 | 6.00812 | 6.12596 | 5.95722 | 5.74377 | 5.76623 | 6.65747 | 6.15973 | 6.6587 | 6.49429 |
| PARP4 | Q9UKK3 | 6.4589 | 6.34802 | 6.72164 | 6.09146 | 6.2352 | 6.63813 | 6.23087 | 6.5332 | 6.56991 | 6.49152 | 6.49229 | 6.41857 | 6.59842 | 5.8996 | 5.58621 | 5.58605 |
| FKBP15 | Q5T1M5 | 5.97743 | 6.05892 | 6.19055 | 5.78393 | 6.17326 | 6.20387 | 5.83471 | 6.02443 | 6.38434 | 6.62369 | 6.69952 | 6.56032 | 6.53029 | 6.28025 | 6.34103 | 6.33346 |
| DDX10 | Q13206 | 6.23877 | 5.95764 | 5.69783 | 6.69347 | 6.09239 | 5.86746 | 6.44292 | 6.07437 | 6.06423 | 5.52828 | 5.67976 | 5.66432 | 6.94832 | 6.68998 | 7.10969 | 7.25057 |
| PLOD2 | O00469-2 | 6.52405 | 6.27943 | 5.82368 | 6.42103 | 6.65688 | 6.06134 | 6.40033 | 6.00394 | 6.02897 | 6.03572 | 6.05794 | 5.84519 | 6.51303 | 6.66243 | 6.55487 | 6.13115 |
| MMS19 | Q96T76-8 | 6.1241 | 6.15707 | 6.56852 | 5.91647 | 6.20356 | 6.61409 | 6.08098 | 6.46011 | 6.31976 | 6.41433 | 6.54956 | 6.36062 | 5.88949 | 6.28176 | 5.85644 | 6.20314 |
| CTR9 | Q6PD62 | 6.17717 | 6.31935 | 5.93344 | 6.9416 | 6.2575 | 6.2499 | 6.38443 | 5.95913 | 6.1258 | 5.94916 | 5.98384 | 5.97906 | 6.21885 | 6.3419 | 6.68333 | 6.49553 |
| ISYNA1 | Q9NPH2 | 5.75786 | 5.62034 | 6.60057 | 5.37381 | 6.51667 | 6.91084 | 6.56863 | 6.6898 | 6.06018 | 6.47973 | 6.5434 | 6.24732 | 6.47052 | 6.21043 | 6.00056 | 5.94935 |
| SART1 | O43290 | 6.43622 | 6.4064 | 5.93904 | 6.54357 | 5.97332 | 5.84331 | 6.40057 | 5.82586 | 6.13515 | 6.09696 | 6.29563 | 5.81737 | 6.26825 | 6.8403 | 6.58 | 6.59806 |
| PITPNB | P48739-3 | 6.16208 | 6.12234 | 6.86974 | 6.1186 | 6.36935 | 6.81821 | 6.27958 | 6.6582 | 6.3454 | 6.38311 | 6.53753 | 6.61238 | 5.81658 | 5.80003 | 5.57817 | 5.5287 |
| EEF1D | P29692-2 | 6.59633 | 6.27463 | 6.68119 | 5.76119 | 6.2859 | 6.42264 | 5.90211 | 6.64324 | 6.52326 | 6.45768 | 6.44087 | 6.50044 | 6.11407 | 5.9181 | 5.53849 | 5.93986 |
| DCAF1 | Q9Y4B6 | 5.89388 | 6.18903 | 6.04099 | 6.25165 | 6.20207 | 5.93211 | 6.12598 | 5.95157 | 5.9425 | 6.00041 | 5.92371 | 5.99061 | 6.97944 | 6.53594 | 7.16827 | 6.87184 |
| LUC7L2 | Q9Y383 | 5.63983 | 6.08198 | 5.95816 | 5.40853 | 5.97797 | 6.22523 | 5.80319 | 5.89523 | 6.07363 | 6.20095 | 6.09358 | 5.91911 | 6.89391 | 7.44863 | 7.30176 | 7.07833 |
| PKM | P14618 | 6.32369 | 6.19415 | 6.98249 | 5.89277 | 6.59377 | 7.10077 | 6.46396 | 6.14082 | 6.5407 | 6.70503 | 6.38423 | 6.30415 | 5.89733 | 5.41469 | 5.78401 | 5.27742 |
| ASPH | Q12797 | 6.31191 | 6.35176 | 5.90119 | 6.34446 | 6.17529 | 5.83564 | 6.27689 | 6.01258 | 6.10329 | 6.16332 | 5.93153 | 5.71099 | 6.76165 | 6.60189 | 6.90722 | 6.61037 |
| SNRPB | P14678-3 | 6.21388 | 6.4144 | 5.95264 | 6.36159 | 6.46649 | 6.11893 | 6.35989 | 6.12802 | 6.23328 | 6.29563 | 6.06738 | 5.96941 | 6.77059 | 5.8855 | 6.4422 | 6.32018 |
| NCAPD3 | P42695 | 5.89819 | 6.55189 | 6.28221 | 6.69928 | 6.09248 | 6.44705 | 6.3245 | 6.2683 | 6.02115 | 6.48976 | 6.16146 | 6.48651 | 5.96659 | 6.12863 | 5.95044 | 6.23157 |
| CCDC88C | Q9P219 | 6.23533 | 6.62851 | 6.60927 | 6.52088 | 6.30863 | 6.32961 | 6.21961 | 6.60995 | 6.37674 | 6.50816 | 6.65202 | 6.59168 | 5.585 | 5.69734 | 5.45334 | 5.67392 |
| MCCC2 | Q9HCC0 | 6.35099 | 6.46914 | 5.97948 | 6.73805 | 6.51049 | 5.89042 | 6.22433 | 6.04486 | 6.08062 | 6.11269 | 6.14204 | 6.11126 | 6.62942 | 6.12066 | 6.43088 | 6.16468 |
| SDHB | P21912 | 6.2211 | 5.99442 | 5.64673 | 6.61056 | 6.45036 | 5.80312 | 6.84442 | 5.6937 | 6.68653 | 5.71849 | 5.59867 | 6.00138 | 6.67537 | 6.29749 | 6.50113 | 6.86654 |
| HDGFL2 | Q7Z4V5-2 | 5.97574 | 6.19203 | 5.97745 | 5.75823 | 6.08341 | 5.99908 | 6.07808 | 5.77385 | 5.92275 | 6.10803 | 6.33252 | 6.16493 | 6.91331 | 7.0706 | 6.86409 | 6.7859 |
| OXCT1 | P55809 | 5.99484 | 6.14449 | 5.64293 | 6.20081 | 6.33148 | 5.9683 | 6.50846 | 6.40295 | 6.58121 | 6.27772 | 6.37382 | 5.96553 | 6.647 | 6.14836 | 6.60487 | 6.20724 |
| SETDB1 | Q15047 | 6.10956 | 6.55367 | 6.12271 | 6.70144 | 6.09135 | 6.19912 | 6.37851 | 6.2097 | 6.01833 | 6.49215 | 6.38029 | 6.35698 | 6.18189 | 5.86638 | 6.18449 | 6.15345 |
| RAD21 | O60216 | 6.35376 | 6.53551 | 6.02898 | 6.50113 | 6.23752 | 6.24381 | 6.34651 | 6.28469 | 6.10704 | 5.97435 | 6.02339 | 5.75999 | 6.42199 | 6.34579 | 6.39892 | 6.43661 |
| ELAVL1 | Q15717-2 | 6.36999 | 6.45819 | 5.78361 | 6.51776 | 6.36448 | 6.20297 | 6.25851 | 6.3272 | 5.8164 | 6.11801 | 5.94891 | 6.07427 | 6.75052 | 6.45334 | 6.32169 | 6.23416 |
| POLRMT | O00411 | 6.30686 | 6.33802 | 6.21747 | 7.08014 | 6.25951 | 6.29461 | 6.43319 | 6.07606 | 6.37156 | 6.17304 | 6.18041 | 6.26462 | 6.30075 | 5.48238 | 6.11759 | 6.10379 |
| GEMIN4 | P57678 | 6.30239 | 6.59922 | 6.66045 | 6.26682 | 6.23437 | 6.44675 | 6.17554 | 6.38383 | 6.39885 | 6.33483 | 6.49458 | 6.3902 | 5.71233 | 5.9142 | 5.97574 | 5.7099 |
| KIF4B | Q2VIQ3 | 5.66208 | 6.12947 | 6.22301 | 5.95739 | 6.15292 | 6.36205 | 6.2014 | 5.80782 | 5.97898 | 6.60685 | 6.49643 | 6.21703 | 6.55369 | 6.51276 | 6.6537 | 6.4844 |
| XPOT | O43592 | 5.69478 | 5.91614 | 6.31171 | 5.61057 | 5.7501 | 6.27412 | 5.97026 | 6.39474 | 6.28315 | 6.55593 | 6.28542 | 6.2694 | 6.20447 | 6.60752 | 7.13132 | 6.74037 |
| METAP2 | P50579 | 6.0235 | 6.02966 | 6.79366 | 6.08021 | 6.01239 | 6.17647 | 6.28932 | 6.1366 | 6.41228 | 6.31672 | 6.87514 | 6.71385 | 5.84858 | 6.31944 | 5.79798 | 6.17419 |
| GLYR1 | Q49A26 | 6.11081 | 6.2016 | 5.86123 | 6.5469 | 6.05361 | 5.99302 | 6.57228 | 6.06823 | 6.0052 | 6.06676 | 5.87663 | 5.794 | 6.64649 | 6.58094 | 6.77111 | 6.8512 |
| RBMXL1 | Q96E39 | 6.39751 | 6.50636 | 5.79647 | 6.84099 | 5.98526 | 5.87296 | 6.32439 | 6.01928 | 6.04952 | 5.98419 | 5.92154 | 5.9505 | 6.59623 | 6.47941 | 6.55762 | 6.71778 |
| FKBP5 | Q13451 | 5.47387 | 5.98316 | 6.10168 | 5.52001 | 5.94374 | 5.98024 | 5.91473 | 5.82055 | 5.82902 | 6.04078 | 6.06349 | 5.83681 | 7.59233 | 7.04808 | 7.6058 | 7.24572 |
| TMF1 | P82094-2 | 6.3193 | 6.08255 | 5.94374 | 6.21879 | 6.40086 | 6.21747 | 6.25776 | 6.1914 | 6.2996 | 6.06607 | 6.56459 | 6.34143 | 6.69889 | 5.96436 | 6.19423 | 6.23896 |
| RFC5 | P40937 | 6.01733 | 6.16152 | 6.1998 | 6.03017 | 6.34205 | 6.38373 | 6.1596 | 5.9529 | 6.2198 | 6.20625 | 6.10243 | 6.20454 | 6.68644 | 6.43666 | 6.54311 | 6.35366 |
| KDM2A | Q9Y2K7 | 6.63385 | 6.65413 | 6.14942 | 6.9472 | 6.11653 | 5.99114 | 6.36518 | 6.05934 | 6.32501 | 6.24744 | 6.16117 | 6.17029 | 5.93345 | 5.85148 | 6.15154 | 6.24282 |
| RANBP3 | Q9H6Z4 | 5.39915 | 5.51342 | 5.9622 | 5.38187 | 5.52358 | 5.94607 | 5.89619 | 5.74174 | 5.98159 | 6.13698 | 6.24477 | 6.19172 | 7.43547 | 7.42892 | 7.72703 | 7.48931 |
| PAK2 | Q13177 | 6.1775 | 6.02382 | 6.71713 | 6.00276 | 6.6248 | 6.67356 | 6.33926 | 6.69834 | 6.54095 | 6.41122 | 6.49162 | 6.60597 | 5.82922 | 5.7192 | 5.5168 | 5.62786 |
| DPP9 | Q86TI2-2 | 6.03142 | 6.21728 | 6.86469 | 5.77422 | 6.38601 | 6.83141 | 6.19183 | 6.35348 | 6.36982 | 6.55922 | 6.70655 | 6.41554 | 5.80183 | 5.96096 | 5.95728 | 5.57844 |
| RPL15 | P61313 | 6.25178 | 6.24067 | 6.20636 | 6.11142 | 6.29848 | 6.33312 | 6.4308 | 6.22046 | 6.67456 | 6.60338 | 6.56261 | 6.44733 | 5.89535 | 6.06709 | 5.78869 | 5.86789 |
| SAE1 | Q9UBE0 | 5.42164 | 5.47478 | 5.96069 | 4.97033 | 5.87385 | 6.28032 | 5.56351 | 6.21376 | 5.88238 | 5.90767 | 6.00541 | 5.89477 | 7.20503 | 7.96005 | 7.95182 | 7.43397 |
| GRSF1 | Q12849 | 6.5822 | 6.40415 | 6.165 | 6.47644 | 6.75662 | 6.24474 | 6.51564 | 6.43756 | 6.21175 | 6.01573 | 6.33618 | 6.34796 | 6.30434 | 5.46621 | 5.81981 | 5.91566 |
| LRRC59 | Q96AG4 | 6.5131 | 6.41514 | 6.00406 | 6.67389 | 6.14962 | 5.97976 | 6.52087 | 5.97148 | 6.11857 | 6.0475 | 6.09767 | 5.94693 | 6.11511 | 6.2889 | 6.72376 | 6.43365 |
| ADNP | Q9H2P0 | 6.26275 | 6.42523 | 5.96726 | 6.94922 | 6.08755 | 6.02835 | 6.17134 | 5.76067 | 6.14684 | 6.16158 | 6.10953 | 6.2399 | 6.43153 | 6.4459 | 6.41884 | 6.39349 |
| SNRPA | P09012 | 6.44419 | 6.18869 | 5.99962 | 6.37027 | 6.32212 | 5.61033 | 6.43377 | 6.16192 | 6.30895 | 5.95288 | 5.81345 | 6.0268 | 6.69805 | 6.47238 | 6.54068 | 6.65588 |
| CUL5 | Q93034 | 5.73799 | 5.93831 | 6.11099 | 5.66823 | 5.93109 | 6.18717 | 5.98686 | 5.99925 | 6.14743 | 6.15416 | 6.14497 | 6.09075 | 6.90247 | 7.16725 | 7.03017 | 6.80292 |
| SF1 | Q15637-6 | 5.97177 | 6.36571 | 5.8843 | 6.02968 | 6.28475 | 6.22825 | 6.005 | 6.09135 | 6.1983 | 6.36365 | 6.31712 | 6.13091 | 6.97589 | 6.3142 | 6.50143 | 6.3377 |
| WASHC4 | Q2M389 | 5.99199 | 6.48311 | 6.19554 | 5.93913 | 6.0726 | 6.38266 | 5.94961 | 6.20514 | 6.18704 | 6.43192 | 6.40423 | 6.44604 | 6.28048 | 6.55468 | 6.37355 | 6.10229 |

| | | | | | | | | | | | | | | | | | |
|---|---|---|---|---|---|---|---|---|---|---|---|---|---|---|---|---|---|
| INTS11 | Q5TA45 | 6.24925 | 6.45479 | 6.17455 | 6.72522 | 6.28576 | 6.33135 | 6.21149 | 6.10629 | 6.0107 | 6.23689 | 6.45784 | 6.44591 | 6.11868 | 5.90647 | 6.27461 | 6.0102 |
| PTPN1 | P18031 | 6.36544 | 6.68203 | 5.85363 | 6.84086 | 6.06732 | 6.11469 | 6.27393 | 6.08481 | 6.00426 | 6.0279 | 5.70865 | 5.94803 | 6.4914 | 6.33748 | 6.59018 | 6.6094 |
| NFXL1 | Q6ZNB6 | 6.20511 | 6.03551 | 6.02381 | 6.73094 | 6.18471 | 5.77689 | 6.20064 | 5.70898 | 6.11806 | 6.18679 | 6.08465 | 5.87206 | 6.67647 | 6.67171 | 6.80228 | 6.72139 |
| USP15 | Q9Y4E8 | 6.18794 | 6.13577 | 6.59159 | 6.10825 | 6.22868 | 6.4048 | 6.18228 | 6.36005 | 6.20519 | 6.28732 | 6.41675 | 6.39367 | 6.22457 | 5.95216 | 6.04059 | 6.28038 |
| G6PD | P11413-2 | 6.38355 | 6.03843 | 6.81781 | 5.93962 | 6.50678 | 6.69244 | 6.26469 | 6.56289 | 6.45402 | 6.39646 | 6.50038 | 6.3835 | 5.91761 | 5.88236 | 5.66287 | 5.59659 |
| TARDBP | Q13148 | 6.05302 | 6.01079 | 5.89127 | 6.22144 | 6.27884 | 5.8709 | 6.2541 | 5.86853 | 6.33882 | 6.11607 | 6.21474 | 6.15681 | 7.0244 | 6.208 | 6.88672 | 6.60556 |
| APMAP | Q9HDC9 | 6.26896 | 6.41125 | 5.93539 | 6.38117 | 6.2332 | 6.02323 | 6.25097 | 5.95103 | 6.25542 | 6.03267 | 5.80011 | 5.77588 | 6.85869 | 6.45494 | 6.85445 | 6.51264 |
| NUDCD1 | Q96RS6 | 5.49541 | 6.30539 | 6.48383 | 5.61975 | 5.97617 | 6.157 | 5.92587 | 6.40611 | 5.99647 | 6.04843 | 6.30064 | 6.02631 | 6.5597 | 6.87083 | 7.14118 | 6.6869 |
| ALDH3A2 | P51648-2 | 6.49496 | 6.62297 | 5.80096 | 6.57513 | 6.52972 | 6.40169 | 6.29921 | 6.39363 | 5.74603 | 5.93816 | 6.06919 | 5.77799 | 6.37357 | 6.43132 | 6.5299 | 6.01556 |
| GLDC | P23378 | 6.15649 | 6.18001 | 5.80507 | 6.55856 | 6.37612 | 6.1313 | 6.46083 | 5.98702 | 5.89927 | 5.84647 | 6.03576 | 5.77727 | 6.79609 | 6.48334 | 6.76354 | 6.74285 |
| ALDH7A1 | P49419-2 | 6.30904 | 6.34031 | 5.81497 | 6.17974 | 6.33617 | 6.3807 | 6.42732 | 6.14211 | 6.12517 | 5.92197 | 6.13404 | 5.86609 | 6.60026 | 6.36654 | 6.58286 | 6.47269 |
| DDX6 | P26196 | 6.35517 | 6.38443 | 6.54925 | 6.03612 | 6.31054 | 6.45338 | 6.20743 | 6.2714 | 6.39073 | 6.50922 | 6.49617 | 6.44587 | 5.77634 | 6.1158 | 5.80589 | 5.89226 |
| ABCF2 | Q9UG63-2 | 6.86632 | 6.58901 | 6.95301 | 6.75738 | 6.57047 | 6.68213 | 6.52557 | 6.56088 | 6.79097 | 6.53405 | 6.55737 | 6.53915 | 5.02091 | 5.1274 | 5.04532 | 4.88006 |
| KIF20A | O95235 | 6.73901 | 7.03242 | 6.59673 | 7.79443 | 5.99311 | 6.13663 | 6.59561 | 6.11691 | 6.4153 | 6.45877 | 6.46404 | 6.6242 | 5.0837 | 5.09183 | 5.20993 | 5.64738 |
| SOGA1 | O94964-2 | 6.42515 | 6.57925 | 6.69473 | 6.56783 | 6.2827 | 6.70536 | 6.37321 | 6.68151 | 6.53436 | 6.75282 | 6.69214 | 6.70927 | 5.16092 | 5.35072 | 5.11089 | 5.37913 |
| ERCC6L | Q2NKX8 | 6.41374 | 6.2267 | 6.85455 | 6.32727 | 6.1615 | 6.7548 | 6.21073 | 6.63371 | 6.46682 | 6.45594 | 6.75287 | 6.809 | 5.47575 | 5.61965 | 5.31982 | 5.51715 |
| GALNT2 | Q10471 | 6.4772 | 6.38278 | 6.05826 | 6.73341 | 6.04021 | 5.72461 | 6.54847 | 5.99963 | 6.27452 | 5.87565 | 6.27051 | 6.02342 | 6.40972 | 6.17885 | 6.5501 | 6.45266 |
| CPT1A | P50416 | 6.35664 | 6.31586 | 5.92039 | 6.78935 | 6.25562 | 5.93054 | 6.60947 | 5.63291 | 6.33071 | 5.86778 | 5.8703 | 5.89002 | 6.33672 | 6.40203 | 6.81241 | 6.67926 |
| CUL7 | Q14999 | 6.57484 | 6.65809 | 6.96118 | 6.38569 | 6.36795 | 6.8193 | 6.31991 | 6.59908 | 6.66332 | 6.89321 | 6.93383 | 6.79881 | 4.99404 | 5.28355 | 4.79461 | 4.9526 |
| IQGAP3 | Q86VI3 | 6.29663 | 6.43308 | 6.78904 | 6.4175 | 6.10913 | 6.42864 | 6.4138 | 6.69897 | 6.58707 | 6.46572 | 6.63023 | 6.69026 | 5.49244 | 5.63591 | 5.3625 | 5.54907 |
| SMARCD2 | Q92925 | 6.10764 | 6.41512 | 6.0805 | 6.22099 | 6.1078 | 6.10888 | 6.25185 | 6.16267 | 6.05756 | 6.34571 | 5.97917 | 5.99573 | 6.52925 | 6.35363 | 6.71593 | 6.56756 |
| ARID1B | Q8NFD5-3 | 6.1771 | 6.49495 | 6.26309 | 6.25636 | 6.09374 | 6.20926 | 6.27034 | 5.90847 | 6.25884 | 6.19088 | 6.10864 | 5.97549 | 6.75213 | 6.28265 | 6.40274 | 6.35531 |
| RPL11 | P62913 | 6.31588 | 6.35034 | 6.67317 | 6.35355 | 6.46131 | 6.74897 | 6.36062 | 5.95669 | 6.27939 | 6.43047 | 6.59129 | 6.47 | 5.88541 | 5.82513 | 5.63632 | 5.66146 |
| ANPEP | P15144 | 6.15114 | 6.42009 | 5.87433 | 6.22119 | 6.27376 | 6.07578 | 5.84754 | 5.65825 | 6.23692 | 6.1769 | 6.09257 | 6.0368 | 6.91406 | 6.82894 | 6.64749 | 6.54423 |
| ZNF598 | Q86UK7 | 6.09789 | 6.3964 | 6.71459 | 6.2348 | 6.0573 | 6.47805 | 6.23071 | 6.31696 | 6.64157 | 6.52322 | 6.65109 | 6.70669 | 5.60772 | 5.9777 | 5.67853 | 5.68677 |
| ADH5 | P11766 | 6.24734 | 5.89896 | 6.74596 | 5.8197 | 6.33036 | 6.69163 | 6.16419 | 6.38033 | 6.27249 | 6.45293 | 6.54144 | 6.45873 | 6.07781 | 6.16822 | 5.92854 | 5.82137 |
| BAZ2A | Q9UIF9 | 6.84866 | 6.90095 | 6.30528 | 7.21048 | 6.34728 | 6.11777 | 6.50274 | 5.86283 | 6.43359 | 6.13524 | 6.18576 | 6.14566 | 5.3256 | 5.89127 | 5.6986 | 6.0883 |
| UBE2O | Q9C0C9 | 6.27739 | 6.18284 | 7.10603 | 6.08355 | 6.26503 | 6.68145 | 6.28294 | 6.58045 | 6.53834 | 6.54264 | 6.57128 | 6.64083 | 5.7923 | 5.473 | 5.41818 | 5.56374 |
| IWS1 | Q96ST2 | 6.05841 | 6.32737 | 5.97073 | 6.61735 | 6.11545 | 6.35358 | 6.2014 | 5.96202 | 6.04969 | 6.10987 | 6.05119 | 6.23681 | 6.62988 | 6.54366 | 6.24841 | 6.52418 |
| CTNNB1 | P35222 | 6.23206 | 6.41103 | 6.15239 | 6.48504 | 6.05055 | 6.26221 | 6.12605 | 5.99263 | 6.31875 | 5.99136 | 6.12104 | 6.44047 | 6.29255 | 6.27301 | 6.38915 | 6.46171 |
| GATAD2A | Q86YP4-3 | 6.36097 | 6.57931 | 6.00562 | 6.66086 | 6.34039 | 6.07141 | 6.219 | 6.16568 | 6.08976 | 6.16424 | 6.15835 | 6.42438 | 6.30453 | 5.96709 | 6.17529 | 6.31313 |
| CMTR1 | Q8N1G2 | 5.78462 | 6.02234 | 5.95626 | 5.33464 | 5.86663 | 6.03787 | 5.49328 | 5.84372 | 6.28771 | 6.34665 | 6.45262 | 6.16508 | 6.84153 | 7.11721 | 7.38131 | 7.06851 |
| HEATR5B | Q9P2D3 | 6.47437 | 6.25345 | 6.98723 | 6.4107 | 6.18515 | 6.64633 | 6.32761 | 6.52724 | 6.46607 | 6.49397 | 6.48542 | 6.35992 | 5.55424 | 5.83064 | 5.38399 | 5.61367 |
| PPP6R3 | Q5H9R7-5 | 6.35598 | 6.04898 | 6.59599 | 6.17577 | 6.28038 | 6.70061 | 6.22699 | 6.38487 | 6.36717 | 6.35968 | 6.33953 | 6.46646 | 6.10984 | 5.8772 | 5.82663 | 5.88394 |
| ANXA4 | P09525 | 6.28648 | 6.14087 | 6.64942 | 5.75904 | 6.39407 | 6.80092 | 6.09716 | 6.62983 | 6.33546 | 6.48792 | 6.48732 | 6.3762 | 5.92247 | 6.41958 | 5.67712 | 5.53615 |
| OGT | O15294 | 5.42216 | 5.22432 | 5.41132 | 5.15554 | 6.79957 | 6.78026 | 6.56355 | 6.73308 | 6.38079 | 6.07081 | 6.46237 | 6.17568 | 7.08513 | 6.58041 | 6.69553 | 6.45947 |
| SARS2 | Q9NP81-2 | 6.52915 | 6.55955 | 5.74307 | 6.49499 | 6.45948 | 6.14192 | 6.14015 | 6.09008 | 6.17367 | 6.09643 | 6.11243 | 6.15697 | 6.43942 | 6.26105 | 6.4742 | 6.12745 |
| PFKM | P08237-3 | 6.40528 | 6.21668 | 6.8429 | 5.82222 | 6.39175 | 6.91556 | 6.39752 | 6.57396 | 6.27135 | 6.61686 | 6.49779 | 6.44886 | 5.55799 | 5.94735 | 5.48858 | 5.60534 |
| RPS10 | P46783 | 6.39795 | 6.56236 | 6.61746 | 6.3417 | 6.37652 | 6.60611 | 6.2003 | 6.64139 | 6.55338 | 6.75473 | 6.54227 | 6.5836 | 5.5334 | 5.63445 | 5.26313 | 5.39124 |
| GNL3 | Q9BVP2 | 6.27354 | 6.3862 | 5.73281 | 6.87326 | 5.91733 | 5.94321 | 6.47307 | 5.82616 | 6.28755 | 5.6104 | 5.59068 | 5.84727 | 6.261 | 6.75318 | 6.93795 | 7.28639 |
| EGFR | P00533 | 6.43501 | 6.78518 | 5.98522 | 6.63489 | 6.35062 | 6.2986 | 6.3587 | 6.37164 | 6.19487 | 6.05179 | 6.10836 | 6.04757 | 6.18684 | 6.86914 | 6.03566 | 6.28591 |
| PAICS | P22234-2 | 6.27627 | 6.1963 | 7.11184 | 6.03678 | 6.38742 | 6.87072 | 6.26758 | 6.86805 | 6.40664 | 6.29302 | 6.4713 | 6.55418 | 5.71319 | 5.56595 | 5.60255 | 5.3782 |
| CIC | I3L2J0 | 6.46712 | 6.6413 | 6.2604 | 6.47338 | 6.23414 | 6.1665 | 6.10361 | 5.94993 | 6.36957 | 6.45717 | 6.62151 | 6.28153 | 6.37956 | 5.8694 | 5.86062 | 5.86426 |
| CTPS1 | P17812 | 6.49998 | 6.32462 | 6.81032 | 6.17425 | 6.61836 | 6.95065 | 6.15484 | 6.91158 | 6.52704 | 6.2254 | 6.48159 | 6.43408 | 5.70671 | 5.3819 | 5.30021 | 5.49848 |
| NCAPG2 | Q86XI2 | 6.25593 | 6.27609 | 6.27734 | 6.52332 | 6.25576 | 6.21517 | 6.27901 | 6.17696 | 6.37901 | 6.48924 | 6.33964 | 6.2709 | 5.94749 | 6.07415 | 6.08315 | 6.15684 |
| APLP2 | Q06481 | 6.77207 | 6.75624 | 6.20743 | 6.64032 | 6.31581 | 6.09099 | 6.33495 | 6.13857 | 6.23651 | 6.47879 | 6.50701 | 6.18063 | 5.75992 | 5.86776 | 5.78515 | 5.92784 |
| ANKFY1 | Q9P2R3-4 | 6.58093 | 6.69184 | 6.51043 | 6.47726 | 6.42885 | 6.38733 | 6.41013 | 6.19627 | 6.32606 | 6.3929 | 6.55741 | 6.13075 | 5.90287 | 5.57023 | 5.78915 | 5.6476 |
| NUP210 | Q8TEM1 | 6.50182 | 6.39046 | 5.76272 | 6.6561 | 6.25549 | 5.90915 | 6.24811 | 6.06541 | 6.073 | 5.92938 | 5.92876 | 5.97181 | 6.70448 | 6.49362 | 6.49652 | 6.61317 |
| DNAJC7 | Q99615 | 6.40455 | 6.16179 | 6.57416 | 6.14363 | 6.22285 | 6.53429 | 6.30544 | 6.24154 | 6.42325 | 6.56642 | 6.52877 | 6.40724 | 5.93794 | 5.91915 | 5.92406 | 5.70492 |
| ANXA2 | P07355-2 | 6.03514 | 6.39405 | 6.39005 | 6.14178 | 6.26987 | 6.42261 | 6.2639 | 6.0805 | 6.21087 | 6.34339 | 6.36875 | 6.36755 | 6.21564 | 6.21283 | 6.14056 | 6.14251 |
| PRPF4B | Q13523 | 5.92962 | 6.311 | 5.52589 | 6.53135 | 5.93098 | 5.75379 | 6.17034 | 5.89854 | 5.81661 | 5.71691 | 5.44256 | 5.71246 | 6.48776 | 7.67166 | 7.21054 | 7.88999 |
| GPATCH1 | Q9BRR8 | 5.88768 | 6.24443 | 6.06245 | 6.2543 | 5.95209 | 6.12216 | 6.09203 | 5.95213 | 6.00796 | 6.06365 | 6.12024 | 6.07479 | 6.40684 | 6.98305 | 6.84185 | 6.93436 |
| NOC4L | Q9BVI4 | 6.14403 | 6.25598 | 5.81974 | 6.79812 | 6.21567 | 5.98408 | 6.29331 | 6.24437 | 6.24589 | 5.98611 | 5.89474 | 6.19416 | 6.39241 | 6.23106 | 6.6315 | 6.66883 |
| BCAP31 | P51572-2 | 6.14155 | 6.14882 | 5.75676 | 6.56078 | 6.04743 | 5.67777 | 6.55867 | 5.93613 | 6.21316 | 5.69726 | 5.81096 | 5.974 | 6.56109 | 6.88713 | 6.97212 | 7.05639 |
| ACTR3 | P61158 | 6.06898 | 5.98835 | 6.77756 | 6.24268 | 6.25521 | 6.59759 | 6.36164 | 6.23828 | 6.24769 | 6.4083 | 6.4731 | 6.36616 | 6.00395 | 5.944 | 5.97155 | 6.05497 |
| ZFC3H1 | O60293 | 6.20113 | 6.19122 | 6.04975 | 6.84205 | 6.0143 | 6.00449 | 6.52762 | 6.04488 | 6.18024 | 6.206 | 6.09113 | 5.97111 | 6.34715 | 6.4426 | 6.31957 | 6.56676 |
| RAP1B | P61224 | 6.02532 | 6.33421 | 5.84847 | 6.48838 | 6.37878 | 6.39414 | 6.39181 | 6.11468 | 6.20278 | 5.98376 | 5.92244 | 5.98059 | 6.45546 | 6.50862 | 6.50953 | 6.46102 |
| UTP18 | Q9Y5J1 | 6.13267 | 6.23352 | 5.95797 | 7.00891 | 6.4672 | 5.88043 | 6.66617 | 6.24041 | 6.19972 | 5.99261 | 6.03049 | 6.02053 | 6.28208 | 6.1442 | 6.38217 | 6.36093 |
| CROCC | Q5TZA2 | 6.19798 | 5.99734 | 5.96774 | 7.77477 | 6.26214 | 6.21001 | 6.03773 | 6.40042 | 6.09726 | 6.11339 | 6.21064 | 6.09098 | 7.07114 | 6.49743 | 6.64191 | 6.42912 |
| HDGF | P51858 | 5.87176 | 5.85571 | 6.20138 | 5.13435 | 5.94014 | 6.16358 | 5.54898 | 6.23882 | 6.19611 | 6.35087 | 6.43522 | 5.91913 | 7.06542 | 7.37483 | 7.12587 | 6.57783 |
| BAG6 | P46379-3 | 6.39436 | 6.11882 | 6.83686 | 5.69818 | 6.53462 | 6.41253 | 6.12903 | 6.55025 | 6.14557 | 6.75543 | 6.67546 | 6.83793 | 5.84138 | 5.58599 | 5.57404 | 5.90954 |
| BYSL | Q13895 | 5.99345 | 5.99408 | 6.29253 | 6.19183 | 6.10034 | 5.91522 | 6.23565 | 6.11338 | 6.30554 | 6.00503 | 6.22448 | 5.87063 | 6.88209 | 6.19095 | 7.00904 | 6.67575 |
| NRDE2 | Q9H7Z3 | 5.90431 | 6.3659 | 6.032 | 6.38741 | 5.82658 | 5.79924 | 6.06636 | 5.57636 | 6.09881 | 6.00761 | 6.02373 | 6.13896 | 6.85743 | 6.96072 | 6.98423 | 6.97037 |
| PLXNB2 | O15031 | 6.45909 | 6.5452 | 6.02375 | 6.76389 | 5.99504 | 6.08634 | 6.13566 | 6.03358 | 6.23784 | 6.34624 | 6.30683 | 6.20272 | 6.31364 | 6.23834 | 6.23974 | 6.07211 |
| ATP6V1A | P38606 | 6.64862 | 6.4819 | 6.75018 | 6.36216 | 6.28075 | 6.84725 | 5.94877 | 6.35992 | 6.61867 | 6.70891 | 6.78623 | 6.64871 | 5.39329 | 5.45083 | 5.43218 | 5.28162 |
| CAPN2 | P17655 | 6.08006 | 6.39847 | 6.70338 | 6.00452 | 6.6065 | 6.98524 | 6.37173 | 6.67823 | 6.36104 | 6.47219 | 6.65969 | 6.575 | 5.81837 | 5.91776 | 5.23926 | 5.12856 |
| UTP6 | Q9NYH9 | 6.24046 | 6.27409 | 5.9119 | 6.77363 | 6.45098 | 5.87329 | 6.40087 | 5.89827 | 6.2051 | 6.0487 | 5.8216 | 5.73571 | 6.82308 | 6.15645 | 6.81096 | 6.57489 |
| NOL9 | Q5SY16 | 6.21731 | 6.25746 | 5.86034 | 6.69342 | 6.09783 | 5.91811 | 6.44713 | 6.02253 | 6.27694 | 5.81175 | 6.10975 | 5.89692 | 6.68631 | 6.36027 | 6.68881 | 6.65511 |
| RPS16 | P62249 | 6.28955 | 6.60288 | 6.6223 | 6.34376 | 6.35758 | 6.60672 | 6.63145 | 6.33258 | 6.69221 | 6.51429 | 6.56842 | 6.409 | 5.59744 | 5.48655 | 5.62968 | 5.3156 |
| ZW10 | O43264 | 6.51615 | 6.56582 | 6.23487 | 6.55943 | 6.46724 | 6.44683 | 6.20716 | 6.50139 | 5.94032 | 6.28973 | 6.06462 | 6.23094 | 6.01765 | 6.07756 | 5.81545 | 6.06485 |
| NIBAN1 | Q9BZQ8 | 6.09549 | 6.31869 | 6.25917 | 5.87547 | 6.27406 | 6.26687 | 5.98458 | 6.48597 | 6.26666 | 6.21217 | 6.28873 | 6.33961 | 6.47172 | 6.30401 | 6.35609 | 6.20071 |
| NUCB1 | Q02818 | 7.79992 | 6.05475 | 6.13837 | 5.61574 | 6.99824 | 4.70372 | 5.63137 | 7.26687 | 5.43847 | 7.335 | 6.12563 | 7.63512 | 5.56413 | 6.05833 | 5.55124 | 6.0831 |
| LTA4H | P09960 | 6.15519 | 5.99185 | 6.76969 | 5.59701 | 6.46155 | 6.93346 | 6.15936 | 6.67466 | 6.21591 | 6.43849 | 6.51707 | 6.48351 | 5.90112 | 6.24988 | 5.84353 | 5.60773 |
| PLCG1 | P19174-2 | 6.50538 | 6.06728 | 7.24192 | 6.06437 | 6.26366 | 6.84892 | 6.46664 | 6.36496 | 6.43123 | 6.70729 | 6.91495 | 6.44219 | 5.43181 | 5.45825 | 5.41511 | 5.37605 |
| SH3GL1 | Q99961 | 6.54887 | 6.25555 | 6.89025 | 6.13868 | 6.51375 | 6.76724 | 6.2024 | 6.47333 | 6.50612 | 6.25122 | 6.68515 | 6.6015 | 5.97825 | 5.65471 | 5.34519 | 5.18779 |
| OSBP | P22059 | 6.15396 | 6.54959 | 6.70445 | 6.55474 | 6.15658 | 6.67417 | 6.41624 | 6.36531 | 6.65751 | 7.03822 | 7.06226 | 6.80413 | 5.13053 | 5.45985 | 4.95663 | 5.31583 |
| CCDC88A | Q3V6T2 | 6.2361 | 6.28128 | 6.64181 | 5.98273 | 6.25642 | 6.52971 | 6.27819 | 6.55839 | 6.56432 | 6.45417 | 6.48218 | 6.59829 | 6.06079 | 5.83111 | 5.65444 | 5.59008 |
| RASA1 | P20936 | 6.14967 | 6.30981 | 6.67221 | 5.88029 | 6.27525 | 6.84238 | 6.19586 | 6.74943 | 6.30468 | 6.26812 | 6.29295 | 6.31677 | 5.9166 | 6.20931 | 5.84332 | 5.77336 |
| UTP4 | Q969X6 | 6.46978 | 6.41174 | 5.95957 | 6.78637 | 6.26415 | 5.89783 | 6.486 | 5.99451 | 6.35794 | 5.98291 | 5.99988 | 5.93851 | 6.42127 | 6.01117 | 6.52949 | 6.48889 |
| ATF7IP | Q6VMQ6-4 | 6.76766 | 6.56821 | 6.25199 | 6.68143 | 6.47074 | 6.14233 | 6.03452 | 6.27969 | 6.15022 | 6.34436 | 6.22884 | 6.3479 | 6.21732 | 5.58547 | 5.79267 | 6.13664 |
| CFL1 | P23528 | 6.19113 | 6.08265 | 6.54395 | 5.65258 | 6.19986 | 6.53624 | 6.04778 | 6.67209 | 6.16933 | 6.40311 | 6.19146 | 6.40475 | 6.25559 | 6.22402 | 6.01742 | 6.40803 |
| POGZ | Q7Z3K3 | 6.47041 | 6.73682 | 6.00513 | 6.80945 | 6.27472 | 6.04969 | 6.33405 | 5.80483 | 5.93509 | 6.24264 | 6.17905 | 6.18033 | 6.34538 | 6.13327 | 6.26898 | 6.23015 |
| DAP3 | P51398 | 6.57244 | 6.27908 | 5.95056 | 6.82612 | 6.6219 | 6.29362 | 6.65029 | 6.07446 | 6.25644 | 5.9335 | 5.99907 | 5.8811 | 6.42504 | 5.83066 | 6.25532 | 6.1504 |
| AGL | P35573 | 6.04799 | 6.0302 | 6.57781 | 5.65687 | 6.30465 | 6.34263 | 6.11721 | 6.26374 | 6.17638 | 6.27669 | 6.55437 | 6.42604 | 6.38451 | 6.34415 | 6.27665 | 6.22012 |
| CIP2A | Q8TCG1 | 6.10544 | 6.21639 | 6.89722 | 5.87068 | 6.15803 | 6.62691 | 6.3607 | 6.7624 | 6.53809 | 6.47095 | 6.73007 | 6.86347 | 5.5509 | 5.67168 | 5.39228 | 5.7848 |
| DNAJA1 | P31689 | 6.38124 | 6.26243 | 6.71155 | 6.60367 | 6.04927 | 6.56743 | 6.48462 | 6.58157 | 6.60603 | 6.32608 | 6.32283 | 6.57325 | 5.59854 | 5.8631 | 5.34359 | 5.7248 |
| SBDS | Q9Y3A5 | 6.17838 | 6.2545 | 6.8349 | 5.91093 | 6.03764 | 6.56349 | 6.22806 | 6.31349 | 6.41642 | 6.39486 | 6.55831 | 6.39323 | 5.91946 | 6.04889 | 6.04693 | 5.9005 |
| PDHB | P11177 | 6.08112 | 6.18185 | 5.52541 | 6.36926 | 6.63876 | 6.05618 | 6.54159 | 6.16158 | 6.2734 | 6.28324 | 6.39028 | 6.14497 | 6.61038 | 6.2294 | 6.354 | 6.29404 |
| LYAR | Q9NX58 | 6.04642 | 6.21395 | 5.9894 | 6.51537 | 5.94204 | 5.59035 | 6.22631 | 5.69278 | 5.97859 | 5.80183 | 5.7959 | 5.88018 | 6.6259 | 7.242 | 7.12224 | 7.33675 |
| TAGLN2 | P37802-2 | 5.74767 | 4.70039 | 6.35903 | 4.64351 | 6.62184 | 5.62336 | 5.95423 | 7.33661 | 5.9142 | 6.74523 | 6.30915 | 7.53995 | 5.5126 | 7.33888 | 6.37195 | 7.28141 |
| TRMT10C | Q7L0Y3 | 6.48393 | 6.38488 | 5.9242 | 6.50208 | 6.26229 | 6.04756 | 6.40257 | 6.00252 | 6.03429 | 5.95275 | 5.82296 | 5.72015 | 6.82317 | 6.40625 | 6.62747 | 6.60292 |
| RPL9 | P32969 | 6.45178 | 6.17252 | 6.59409 | 6.42894 | 6.21103 | 5.96604 | 6.40942 | 6.07618 | 6.96836 | 6.49436 | 6.31224 | 6.39681 | 5.71005 | 5.93723 | 5.69344 | 6.17749 |
| LNPEP | Q9UIQ6 | 6.29131 | 6.54563 | 6.30342 | 6.60782 | 6.43506 | 6.14785 | 6.21362 | 6.09045 | 6.15452 | 6.42908 | 6.07656 | 5.91896 | 6.12857 | 6.09514 | 6.3805 | 6.18151 |
| CALU | O43852-4 | 6.21213 | 6.15602 | 5.74719 | 6.42354 | 6.76227 | 6.25849 | 6.20541 | 6.36465 | 6.31815 | 5.85697 | 5.90177 | 6.02673 | 7.12292 | 6.22685 | 6.21619 | 6.20071 |
| TAF2 | Q6P1X5 | 6.21496 | 6.4014 | 6.19639 | 6.38213 | 6.09542 | 6.11278 | 6.37819 | 6.02721 | 6.08733 | 6.00861 | 6.11604 | 5.99902 | 6.01277 | 6.57182 | 6.69066 | 6.70526 |
| RNH1 | P13489 | 6.2216 | 5.86061 | 6.85058 | 5.66835 | 6.30525 | 6.42777 | 6.15266 | 6.54265 | 6.57612 | 6.50596 | 6.76496 | 6.70346 | 5.71739 | 5.83335 | 5.79923 | 6.07008 |
| AFG3L2 | Q9Y4W6 | 6.38371 | 6.58712 | 5.72728 | 6.84294 | 6.2755 | 5.85466 | 6.75159 | 6.05361 | 6.13369 | 5.75114 | 5.59435 | 5.86364 | 6.58482 | 6.36709 | 6.52273 | 6.70614 |
| SGO2 | Q562F6 | 6.38523 | 6.593 | 5.93809 | 6.66398 | 6.01298 | 5.91465 | 6.39476 | 6.05639 | 6.22074 | 6.07925 | 6.03104 | 5.90405 | 5.9262 | 6.70727 | 6.50948 | 6.6629 |
| PSMD12 | O00232 | 6.14883 | 6.18185 | 6.23127 | 6.23486 | 6.34946 | 6.53 | 6.36939 | 6.51851 | 5.98653 | 6.03609 | 6.31845 | 6.22943 | 6.17467 | 6.33978 | 6.26096 | 6.0899 |
| CMAS | Q8NFW8 | 5.86016 | 6.11199 | 5.78537 | 6.45353 | 6.12776 | 5.91205 | 6.12312 | 5.89453 | 5.86023 | 5.99199 | 5.90165 | 5.93263 | 7.04479 | 6.85203 | 7.10719 | 7.04098 |
| SWAP70 | Q9UH65 | 6.23129 | 6.4087 | 6.82882 | 5.99278 | 6.05124 | 6.74965 | 6.07705 | 6.44283 | 6.43694 | 6.61936 | 6.28782 | 6.53884 | 5.83698 | 5.84645 | 5.82729 | 5.82397 |
| HMGA1 | P17096 | 5.92 | 5.88486 | 5.56409 | 5.78524 | 5.85226 | 5.72215 | 6.01509 | 5.78884 | 5.99924 | 5.89213 | 5.44696 | 5.37866 | 7.08513 | 7.95136 | 7.87042 | 7.84356 |
| EIF5A | P63241-2 | 6.2882 | 5.94457 | 6.54023 | 5.46058 | 6.66819 | 6.70312 | 5.93892 | 7.27485 | 6.3904 | 6.4537 | 6.47623 | 6.66082 | 5.98153 | 5.91857 | 5.57045 | 5.72966 |
| RPS24 | P62847-4 | 6.00779 | 6.30511 | 6.03738 | 6.20151 | 5.97639 | 6.03457 | 6.85343 | 5.99033 | 7.10132 | 5.82626 | 6.29319 | 6.44377 | 5.49879 | 6.81594 | 6.11617 | 6.49804 |
| ATAD5 | Q96QE3 | 6.75751 | 6.84758 | 6.1278 | 7.18252 | 6.1531 | 6.01092 | 6.51387 | 5.93421 | 6.28531 | 6.14402 | 6.10194 | 6.35203 | 5.68451 | 5.86729 | 5.8439 | 6.19348 |

| | | | | | | | | | | | | | | | | | |
|---|---|---|---|---|---|---|---|---|---|---|---|---|---|---|---|---|---|
| TIMELESS | Q9UNS1 | 5.81661 | 6.21285 | 6.30013 | 5.6625 | 5.93852 | 6.15221 | 6.13028 | 6.2144 | 6.33829 | 6.21582 | 6.55966 | 6.11774 | 6.5695 | 6.47639 | 6.59079 | 6.70432 |
| COPG2 | Q9UBF2 | 6.4206 | 6.5913 | 6.54599 | 6.12523 | 6.42663 | 6.53236 | 6.1826 | 6.3259 | 6.18838 | 6.48081 | 6.63905 | 6.20131 | 5.98207 | 6.02852 | 5.72025 | 5.60902 |
| POP1 | Q99575 | 6.10295 | 6.42534 | 6.04766 | 6.56047 | 6.16571 | 6.18873 | 6.34081 | 6.01641 | 5.90527 | 6.02478 | 6.05483 | 5.93627 | 6.40169 | 6.46896 | 6.74812 | 6.61201 |
| WIZ | O95785 | 6.31541 | 6.03838 | 5.97489 | 6.75391 | 6.25427 | 6.02049 | 6.31252 | 6.09149 | 6.27496 | 6.00706 | 6.13432 | 6.16416 | 6.58447 | 6.37494 | 6.37268 | 6.32604 |
| CLASP2 | O75122-3 | 6.29971 | 6.0594 | 6.44111 | 6.14451 | 6.24555 | 6.3418 | 6.05771 | 6.5065 | 6.4685 | 6.34317 | 6.42561 | 6.36953 | 6.20212 | 6.19992 | 5.89281 | 6.00205 |
| SMARCD1 | Q96GM5 | 6.45692 | 6.6773 | 6.24821 | 6.58837 | 6.22034 | 5.93439 | 6.20937 | 6.11999 | 6.06704 | 6.03686 | 6.28851 | 6.10546 | 6.44614 | 6.3792 | 6.24507 | 5.97682 |
| RBM17 | Q96I25 | 6.36185 | 6.23967 | 6.01509 | 6.74471 | 6.23997 | 6.08933 | 6.23819 | 6.06674 | 6.22808 | 6.08119 | 6.19449 | 6.14827 | 6.66632 | 5.9606 | 6.47916 | 6.24636 |
| VPS35L | Q7Z3J2 | 6.01057 | 6.10943 | 6.3154 | 5.77748 | 6.15718 | 6.30148 | 6.03156 | 6.01193 | 6.08208 | 6.54431 | 6.20985 | 6.2264 | 6.51657 | 6.66893 | 6.66252 | 6.37429 |
| TUBB1 | Q9H4B7 | 6.54252 | 6.23606 | 7.10035 | 6.0706 | 6.41148 | 6.87995 | 6.27678 | 6.54865 | 6.47536 | 6.41032 | 6.82922 | 6.98374 | 5.69765 | 5.34965 | 4.94024 | 5.24743 |
| UPF2 | Q9HAU5 | 6.04558 | 6.57456 | 6.72561 | 6.29523 | 5.87093 | 6.51391 | 6.13838 | 6.26594 | 6.39938 | 6.57889 | 6.28121 | 6.66014 | 5.92902 | 6.02668 | 5.94547 | 5.74907 |
| BAG3 | O95817 | 6.62702 | 5.74913 | 6.86484 | 5.38635 | 6.4582 | 6.61939 | 5.82609 | 6.94566 | 6.18952 | 6.5452 | 6.98542 | 6.69167 | 6.33626 | 5.93825 | 5.39831 | 5.43869 |
| GPRIN1 | Q7Z2K8 | 6.50656 | 6.43775 | 6.12672 | 6.34572 | 6.39919 | 5.9759 | 6.23092 | 6.2369 | 6.53255 | 6.09488 | 6.1569 | 6.23184 | 6.32566 | 6.02279 | 6.07753 | 6.29821 |
| RPL32 | P62910 | 5.86115 | 6.38811 | 5.70591 | 5.983 | 6.12907 | 6.26749 | 6.63301 | 6.43066 | 7.12531 | 6.10129 | 6.23179 | 6.14199 | 6.03321 | 6.46701 | 5.92318 | 6.57782 |
| RB1CC1 | Q8TDY2 | 6.18179 | 6.07422 | 6.35095 | 6.00875 | 6.3681 | 6.53693 | 6.168 | 6.05648 | 6.23112 | 6.19657 | 6.42473 | 6.1974 | 6.43592 | 6.27782 | 6.37603 | 6.11519 |
| DBT | P11182 | 6.09604 | 6.16158 | 5.86526 | 6.32978 | 6.3207 | 6.08039 | 6.43553 | 5.86561 | 6.26248 | 6.09405 | 6.08637 | 6.06345 | 6.80754 | 6.38588 | 6.64468 | 6.50066 |
| PPWD1 | Q96BP3 | 5.85382 | 6.085 | 6.21744 | 5.84238 | 5.84293 | 5.95894 | 5.94153 | 5.83219 | 6.38711 | 6.1208 | 6.32216 | 5.94228 | 6.86973 | 6.66198 | 7.26332 | 6.85839 |
| STXBP3 | O00186 | 6.38992 | 6.54496 | 5.9548 | 6.39298 | 6.2383 | 6.229 | 6.42193 | 6.07536 | 6.20676 | 6.26429 | 6.10847 | 6.00765 | 6.41592 | 6.2348 | 6.30974 | 6.20509 |
| ITPR3 | Q14573 | 6.46435 | 6.40188 | 5.84513 | 6.65753 | 6.12531 | 5.92761 | 6.35988 | 5.82869 | 6.13445 | 6.21687 | 5.90512 | 6.01237 | 6.63415 | 6.35695 | 6.69453 | 6.43517 |
| KIFC1 | Q9BW19 | 5.95106 | 6.22095 | 5.94594 | 6.10487 | 6.13882 | 6.19088 | 6.23151 | 5.92166 | 6.1011 | 6.20643 | 6.14229 | 5.8972 | 6.69053 | 6.65655 | 6.79993 | 6.80029 |
| RPS8 | P62241 | 6.10922 | 6.22036 | 6.57057 | 5.87328 | 6.49034 | 6.41537 | 6.02414 | 6.40079 | 6.88147 | 6.45055 | 6.39384 | 6.32809 | 5.64064 | 6.28999 | 6.12997 | 5.78139 |
| PMPCB | O75439 | 6.38624 | 6.3483 | 5.75317 | 6.33285 | 6.59349 | 6.14024 | 6.30069 | 6.17664 | 6.14561 | 6.08857 | 6.03292 | 5.9296 | 6.74743 | 6.23944 | 6.45661 | 6.32819 |
| APP | P05067 | 6.64862 | 6.80168 | 6.25446 | 6.83871 | 6.54538 | 6.46924 | 6.45732 | 6.13754 | 6.26792 | 6.40328 | 6.33529 | 6.31416 | 5.64939 | 5.5516 | 5.62028 | 5.70511 |
| EHD2 | Q9NZN4 | 5.99478 | 6.1747 | 6.34136 | 6.29025 | 6.18318 | 6.6645 | 6.18458 | 6.54069 | 6.4061 | 6.43919 | 6.84668 | 6.79499 | 6.16736 | 6.02529 | 5.4686 | 5.47776 |
| TASOR2 | Q5VWN6 | 6.57806 | 6.38535 | 6.14668 | 6.97072 | 6.32051 | 6.19473 | 6.41327 | 6.14959 | 6.12069 | 5.82395 | 5.87963 | 6.03259 | 6.38767 | 5.85107 | 6.31234 | 6.43315 |
| SPATA5 | Q8NB90 | 6.40703 | 6.24799 | 6.6148 | 6.08185 | 6.44222 | 6.51508 | 6.06384 | 6.32593 | 6.48796 | 6.29241 | 6.5083 | 6.31747 | 6.10877 | 5.8756 | 5.99583 | 5.71492 |
| TARS2 | Q9BW92 | 6.30081 | 6.9104 | 5.68335 | 6.77087 | 6.34429 | 6.26936 | 6.24538 | 6.58973 | 5.9415 | 5.97172 | 6.06738 | 6.16716 | 6.12732 | 6.1466 | 6.26002 | 6.2041 |
| MED24 | O75448 | 6.25317 | 6.52897 | 5.8965 | 6.45008 | 6.28879 | 6.10337 | 6.21533 | 6.33539 | 6.06958 | 6.19658 | 5.90254 | 5.90371 | 6.54481 | 6.44791 | 6.42782 | 6.43546 |
| SEC24B | O95487-3 | 6.14649 | 6.18738 | 6.47386 | 6.35276 | 6.28115 | 6.50105 | 6.14801 | 6.40982 | 6.26016 | 6.25587 | 6.58429 | 6.22282 | 5.96907 | 6.12045 | 5.99455 | 6.09226 |
| HIP1R | O75146 | 5.95167 | 6.42455 | 6.42861 | 6.42702 | 6.0573 | 6.25381 | 6.35102 | 6.41347 | 6.1207 | 6.61294 | 6.45794 | 6.49294 | 5.82178 | 6.15924 | 6.07079 | 5.95621 |
| LEMD2 | Q8NC56 | 6.35188 | 6.27197 | 5.83704 | 6.3707 | 6.16545 | 6.21671 | 6.06914 | 6.2211 | 6.24689 | 6.31541 | 6.01628 | 6.36493 | 6.95625 | 6.33319 | 6.15815 | 6.10494 |
| NPLOC4 | Q8TAT6-2 | 6.22233 | 6.49424 | 6.12061 | 6.04464 | 6.48324 | 6.34191 | 6.03863 | 6.24378 | 6.17505 | 6.23212 | 6.16963 | 5.94988 | 6.63374 | 6.23882 | 6.32604 | 6.28535 |
| TBL2 | Q9Y4P3 | 6.31667 | 6.52485 | 6.07789 | 6.57647 | 6.07982 | 6.02921 | 6.47622 | 5.78685 | 6.02217 | 5.98192 | 5.97266 | 6.06771 | 6.40248 | 6.35839 | 6.70771 | 6.61899 |
| GNS | P15586 | 6.45058 | 6.54285 | 5.86331 | 6.46074 | 6.36722 | 6.4681 | 6.24804 | 6.29867 | 6.13769 | 6.19121 | 6.31618 | 6.09163 | 6.49178 | 6.10044 | 6.14879 | 5.82277 |
| CPT2 | P23786 | 6.37904 | 6.35839 | 5.84336 | 6.70024 | 6.28838 | 5.96411 | 6.12073 | 6.00375 | 6.04757 | 6.14549 | 5.95024 | 6.24424 | 6.75309 | 6.43229 | 6.45273 | 6.31635 |
| PAF1 | Q8N7H5 | 6.10494 | 6.39834 | 5.9319 | 6.48685 | 6.11925 | 6.29478 | 6.2939 | 6.07342 | 6.37103 | 6.30611 | 6.11824 | 5.99858 | 6.47215 | 6.2014 | 6.31962 | 6.50949 |
| KANK2 | Q63ZY3 | 6.24548 | 5.90536 | 5.96682 | 5.45768 | 6.31824 | 5.99196 | 6.04451 | 6.04547 | 6.7169 | 6.32804 | 6.27327 | 6.16706 | 7.20012 | 6.54885 | 6.57879 | 6.21144 |
| SLTM | Q9NWH9 | 6.17706 | 6.27867 | 6.03141 | 6.63817 | 6.0074 | 5.73069 | 6.33762 | 5.87985 | 5.97171 | 6.03085 | 5.92153 | 5.77772 | 6.45711 | 6.77696 | 7.09654 | 6.8867 |
| SKIV2L | Q15477 | 6.22765 | 6.37339 | 6.70611 | 6.09082 | 6.38212 | 6.84861 | 6.25599 | 6.82592 | 6.34949 | 6.36862 | 6.57607 | 6.46043 | 5.69384 | 5.83521 | 5.57499 | 5.43074 |
| AP1M1 | Q9BXS5-2 | 6.21754 | 6.56154 | 6.85625 | 6.5358 | 6.08025 | 6.52726 | 6.33624 | 6.6222 | 6.10993 | 6.45791 | 6.23868 | 6.36049 | 5.64519 | 5.99929 | 5.64108 | 5.81035 |
| ZZEF1 | O43149 | 6.12054 | 6.32792 | 6.54105 | 6.08755 | 6.05694 | 6.62063 | 6.19383 | 6.4584 | 6.36653 | 6.4697 | 6.66267 | 6.45774 | 5.9893 | 5.91483 | 5.72863 | 6.00374 |
| GSS | P48637 | 6.15462 | 6.0009 | 6.90483 | 5.53773 | 6.36263 | 6.84148 | 6.20914 | 6.86019 | 6.20807 | 6.53569 | 6.57979 | 6.33334 | 6.01277 | 6.00573 | 5.99327 | 5.45981 |
| MLH1 | P40692 | 5.60681 | 5.87908 | 6.13841 | 5.55502 | 5.98119 | 6.27504 | 6.20543 | 5.92747 | 6.01992 | 6.38886 | 6.37998 | 6.23387 | 6.8032 | 6.84791 | 7.05063 | 6.70718 |
| KHDRBS1 | Q07666 | 6.33383 | 6.44066 | 5.9205 | 6.60497 | 6.39627 | 6.14997 | 6.24055 | 6.16597 | 6.25742 | 6.02878 | 6.3146 | 6.11209 | 6.60795 | 6.04177 | 6.38609 | 5.99858 |
| UBE4A | Q14139-2 | 6.21365 | 6.3151 | 6.16027 | 6.51026 | 6.18955 | 6.19465 | 6.03118 | 6.00493 | 6.01533 | 6.30281 | 5.98071 | 6.15045 | 6.63068 | 6.2744 | 6.58676 | 6.36327 |
| MCM3AP | O60318 | 6.33145 | 5.9164 | 5.85846 | 6.77761 | 6.17261 | 6.05531 | 6.49403 | 5.70782 | 6.10129 | 6.17668 | 6.18364 | 6.23419 | 6.61978 | 6.27652 | 6.41021 | 6.68399 |
| TSFM | P43897 | 6.31842 | 6.27186 | 6.03325 | 6.58055 | 6.38659 | 6.03853 | 6.28635 | 5.88606 | 6.38074 | 6.00936 | 5.71597 | 5.66203 | 6.75136 | 6.11096 | 6.89503 | 6.67293 |
| HP1BP3 | Q5SSJ5 | 6.25443 | 6.45697 | 5.37286 | 6.11938 | 6.15632 | 5.74391 | 6.32057 | 5.97362 | 5.71018 | 5.93616 | 5.6455 | 5.53888 | 6.52494 | 7.61275 | 7.24325 | 7.39029 |
| FANCI | Q9NVI1 | 5.69425 | 6.12038 | 5.8688 | 5.86282 | 6.16851 | 6.22147 | 6.30606 | 6.38134 | 6.20239 | 6.21409 | 6.15872 | 6.24908 | 6.50792 | 6.50653 | 6.71277 | 6.82486 |
| HNRNPH3 | P31942 | 6.10369 | 6.14255 | 5.89586 | 6.28617 | 6.34051 | 6.09692 | 6.17215 | 6.2643 | 6.30915 | 6.43246 | 6.43031 | 6.38666 | 6.48486 | 6.03686 | 6.14768 | 6.46987 |
| RBM22 | Q9NW64 | 6.40226 | 6.2839 | 6.08733 | 6.56278 | 6.10565 | 5.91177 | 6.17843 | 5.92194 | 6.2932 | 6.24147 | 6.17615 | 6.00559 | 6.34962 | 6.35904 | 6.62642 | 6.49447 |
| MRPL13 | Q9BYD1 | 6.34777 | 6.51824 | 5.92234 | 6.69379 | 6.65343 | 6.16908 | 6.44161 | 6.21914 | 6.13313 | 6.12793 | 6.03555 | 5.95321 | 6.61508 | 5.95176 | 6.22144 | 5.9965 |
| DLAT | P10515 | 6.37172 | 6.32315 | 5.88561 | 6.45784 | 6.43604 | 6.11014 | 6.33558 | 6.08498 | 6.15251 | 5.92108 | 5.79969 | 6.05186 | 6.64937 | 6.41218 | 6.63728 | 6.37095 |
| NOL8 | Q76FK4 | 6.11696 | 6.4251 | 6.00732 | 6.81348 | 5.95944 | 5.91859 | 6.39851 | 5.85054 | 6.08269 | 6.04931 | 5.78348 | 6.09744 | 6.48976 | 6.41009 | 6.75267 | 6.84461 |
| ACOX3 | O15254 | 6.25868 | 6.28342 | 6.13613 | 6.66237 | 6.36903 | 5.91802 | 6.45281 | 5.88463 | 6.04141 | 6.25265 | 6.07912 | 5.91791 | 6.65484 | 6.03823 | 6.67276 | 6.37801 |
| PAPOLA | P51003 | 5.3237 | 5.78153 | 6.16204 | 5.31855 | 5.44857 | 5.95691 | 5.83533 | 5.87274 | 5.96916 | 6.13782 | 6.29794 | 6.18095 | 7.08823 | 7.38499 | 7.90553 | 7.33599 |
| PSMA3 | P25788 | 5.96407 | 6.3696 | 6.36523 | 6.03511 | 6.44259 | 6.14104 | 6.41426 | 6.25221 | 6.17674 | 5.97334 | 6.03733 | 5.92401 | 6.6578 | 6.35577 | 6.46979 | 6.42114 |
| CPNE8 | Q86YQ8 | 6.32218 | 6.4784 | 6.06009 | 6.54082 | 6.52041 | 6.40863 | 6.46519 | 6.38843 | 6.36061 | 5.98641 | 6.24059 | 6.19445 | 5.91173 | 6.1741 | 5.69038 | 6.25759 |
| EPG5 | Q9HCE0 | 6.33239 | 6.22206 | 6.96779 | 6.39459 | 6.23762 | 6.50121 | 6.30786 | 6.38328 | 6.53736 | 6.33045 | 6.4835 | 6.51694 | 5.58849 | 5.78404 | 5.73128 | 5.68117 |
| NAMPT | P43490 | 6.2842 | 6.19002 | 6.88659 | 6.10598 | 6.29571 | 6.67151 | 6.06271 | 6.85452 | 6.20483 | 6.18286 | 6.16914 | 6.38121 | 5.57562 | 6.27097 | 6.02297 | 5.74118 |
| ADAM9 | Q13443 | 6.18506 | 6.20969 | 6.16898 | 6.30781 | 6.16989 | 5.98118 | 6.54054 | 6.12499 | 6.01538 | 6.04667 | 6.0569 | 6.11061 | 6.46324 | 6.53178 | 6.53723 | 6.55464 |
| HNRNPA3 | P51991 | 6.55091 | 6.44036 | 5.84654 | 6.93954 | 6.33003 | 6.16233 | 6.18354 | 6.09577 | 6.0987 | 5.83488 | 6.00492 | 6.02999 | 6.53274 | 6.11801 | 6.35604 | 6.47571 |
| PPP2CB | P62714 | 5.86771 | 5.89856 | 6.74966 | 5.69194 | 6.20703 | 6.55969 | 6.5029 | 6.40774 | 6.39593 | 6.46369 | 6.74514 | 6.67773 | 5.84684 | 5.90731 | 6.12841 | 5.94973 |
| SORL1 | Q92673 | 6.49018 | 6.4026 | 6.62431 | 6.6576 | 6.43029 | 6.36612 | 6.63405 | 6.16646 | 6.62381 | 6.78445 | 6.70299 | 6.27616 | 5.39365 | 5.57146 | 5.3466 | 5.52928 |
| RPS11 | P62280 | 6.43001 | 6.42094 | 6.47222 | 6.25314 | 6.56343 | 6.44011 | 6.33979 | 6.28973 | 6.55344 | 6.54221 | 6.50428 | 6.22019 | 5.80658 | 5.90874 | 5.68243 | 5.57276 |
| CBX1 | P83916 | 6.17789 | 6.46571 | 5.82034 | 6.66569 | 6.11113 | 6.43514 | 5.82118 | 6.1915 | 6.00351 | 6.04163 | 5.9413 | 6.33633 | 6.54861 | 6.55287 | 6.15534 | 6.73183 |
| SRSF4 | Q08170 | 6.26605 | 6.36687 | 5.93692 | 6.49455 | 6.21125 | 5.98131 | 6.29111 | 5.92891 | 6.23419 | 6.25517 | 5.90795 | 5.9168 | 6.69054 | 6.26026 | 6.52022 | 6.73791 |
| EIF3G | O75821 | 6.29427 | 6.24177 | 6.77773 | 6.26845 | 6.38074 | 6.65421 | 6.33699 | 6.61447 | 6.58199 | 6.40007 | 6.70679 | 6.40924 | 5.5699 | 5.70078 | 5.44736 | 5.61525 |
| SYNJ2 | O15056 | 6.1996 | 6.10643 | 6.63311 | 6.07297 | 6.23246 | 6.58782 | 6.10289 | 6.61505 | 6.56413 | 6.72036 | 6.56796 | 6.57436 | 5.66603 | 6.30593 | 5.58073 | 5.47018 |
| AKAP8L | Q9ULX6 | 6.02137 | 6.15988 | 6.07604 | 6.5518 | 6.23398 | 6.24369 | 6.12271 | 5.83756 | 6.21862 | 6.20296 | 6.09439 | 5.98831 | 6.78374 | 6.37644 | 6.57723 | 6.5113 |
| TBC1D5 | Q92609-2 | 6.3505 | 6.33463 | 6.659 | 6.20852 | 6.41234 | 6.52435 | 6.27663 | 6.40579 | 6.34381 | 6.45102 | 6.71 | 6.59229 | 5.83428 | 5.77248 | 5.45275 | 5.67162 |
| STAT3 | P40763-3 | 6.18181 | 6.11703 | 6.6561 | 5.84696 | 5.99567 | 6.52752 | 6.12078 | 6.42513 | 6.35109 | 6.57374 | 6.54807 | 6.54842 | 5.92469 | 6.16089 | 6.04645 | 5.97563 |
| SLC25A24 | Q6NUK1 | 6.47644 | 6.57518 | 5.58477 | 6.53225 | 6.59265 | 6.16356 | 6.42779 | 6.25535 | 5.86183 | 5.93075 | 5.54671 | 5.77719 | 6.50287 | 6.59365 | 6.42372 | 6.75529 |
| ETF1 | P62495 | 6.31117 | 6.08517 | 6.94124 | 6.09044 | 6.57427 | 6.49234 | 6.19765 | 6.48769 | 6.70714 | 6.36001 | 6.49695 | 6.35936 | 5.77835 | 5.58308 | 5.79203 | 5.74312 |
| UNC45A | Q9H3U1 | 6.16295 | 6.25645 | 6.92913 | 6.22363 | 6.25613 | 6.53478 | 6.34275 | 6.53644 | 6.73766 | 6.52868 | 6.51139 | 6.63665 | 5.55725 | 5.76356 | 5.36638 | 5.65616 |
| MAGED2 | Q9UNF1 | 5.99379 | 6.04706 | 6.36793 | 5.60264 | 6.08793 | 6.49443 | 6.39684 | 6.37562 | 6.26079 | 6.32628 | 6.41933 | 6.03303 | 6.4395 | 6.42265 | 6.46866 | 6.26352 |
| RUFY1 | Q96T51 | 6.25186 | 6.20842 | 6.82933 | 5.8839 | 6.49856 | 6.41202 | 6.26542 | 6.34115 | 6.70104 | 6.08841 | 6.60026 | 6.43029 | 6.16839 | 5.87547 | 5.74008 | 5.70541 |
| RAB3GAP1 | Q15042 | 6.05192 | 6.14103 | 6.60555 | 6.01225 | 5.98463 | 6.30115 | 6.20746 | 6.33847 | 6.22043 | 6.46405 | 6.57258 | 6.57458 | 5.99591 | 6.26288 | 6.15411 | 6.113 |
| AP2A2 | O94973-2 | 6.06312 | 6.23925 | 6.0185 | 6.27207 | 5.95868 | 6.3151 | 6.07012 | 5.98402 | 6.33995 | 6.49637 | 6.19021 | 6.12686 | 6.33398 | 6.3039 | 6.80821 | 6.47966 |
| BMS1 | Q14692 | 6.21048 | 6.36717 | 5.65787 | 6.70324 | 6.26477 | 5.92873 | 6.46889 | 5.88429 | 6.23045 | 5.92381 | 5.70704 | 5.76232 | 6.62683 | 6.6575 | 6.62559 | 6.98102 |
| HGS | O14964 | 6.52253 | 6.31885 | 6.47877 | 6.17719 | 6.55548 | 6.77243 | 6.25715 | 6.57057 | 6.26506 | 6.54515 | 6.62458 | 6.46013 | 6.01886 | 5.65714 | 5.35559 | 5.42052 |
| LAMB3 | Q13751 | 6.66694 | 6.44378 | 6.16648 | 6.61348 | 6.81442 | 6.20646 | 6.57881 | 6.34999 | 6.19204 | 5.91015 | 6.00234 | 6.27798 | 6.3533 | 5.6857 | 5.80535 | 5.93279 |
| SPATS2L | Q9NUQ6-3 | 5.8306 | 6.28055 | 6.05587 | 6.05329 | 5.89571 | 6.45719 | 6.10659 | 6.60885 | 6.31217 | 6.71407 | 6.48133 | 6.80607 | 6.02321 | 6.22887 | 5.99536 | 6.15027 |
| IK | Q13123 | 6.26282 | 6.41745 | 5.91897 | 6.62133 | 6.06113 | 5.96939 | 6.33782 | 5.83399 | 6.05145 | 6.07247 | 6.10773 | 6.00935 | 6.62639 | 6.36122 | 6.5762 | 6.77228 |
| RBM4 | Q9BWF3 | 6.40119 | 6.32181 | 6.18751 | 6.51871 | 6.45538 | 5.87423 | 6.34242 | 6.31505 | 6.23245 | 5.97359 | 6.48802 | 6.31952 | 6.47264 | 5.74017 | 6.20831 | 6.14899 |
| FARP1 | Q9Y4F1-2 | 6.53321 | 6.51337 | 6.07381 | 6.78791 | 6.04142 | 5.99019 | 6.29269 | 5.97345 | 6.18826 | 6.17326 | 6.32099 | 6.11637 | 6.22412 | 6.31077 | 6.2933 | 6.16688 |
| SMARCAL1 | Q9NZC9 | 5.27581 | 5.88452 | 6.09337 | 5.39512 | 5.94058 | 6.35231 | 5.91544 | 5.79754 | 5.81981 | 6.43356 | 6.26459 | 6.055 | 7.13024 | 7.43401 | 7.1319 | 7.07619 |
| RHPN2 | Q8IUC4 | 6.74399 | 6.55314 | 7.05941 | 6.36206 | 6.66083 | 6.67505 | 6.15853 | 6.7862 | 6.36931 | 6.57563 | 6.39973 | 6.51702 | 5.40618 | 5.3325 | 5.10032 | 5.30011 |
| PPA1 | Q15181 | 6.20455 | 6.1845 | 6.78422 | 5.91101 | 6.7716 | 6.91145 | 5.9031 | 7.37065 | 6.6662 | 6.5778 | 6.72696 | 6.72794 | 5.39367 | 5.39521 | 5.33282 | 5.13833 |
| ACOT7 | O00154 | 5.81656 | 5.76034 | 6.0734 | 6.04702 | 6.32054 | 6.4205 | 6.74648 | 6.25409 | 6.40883 | 5.8099 | 6.82955 | 6.47701 | 6.14818 | 6.51344 | 6.1243 | 6.24986 |
| DSG2 | Q14126 | 6.42964 | 6.52245 | 6.16093 | 6.66473 | 6.3762 | 6.2678 | 6.26769 | 6.38106 | 6.38535 | 6.18604 | 6.13717 | 6.17361 | 6.13353 | 5.77596 | 6.09586 | 6.04199 |
| H0YHG0 | H0YHG0 | 6.42384 | 5.86981 | 5.98637 | 6.3257 | 6.46276 | 6.17293 | 6.33164 | 5.69368 | 6.1781 | 6.24512 | 5.74966 | 5.93799 | 6.98336 | 6.44744 | 6.6001 | 6.5915 |
| PFKL | P17858-2 | 6.10209 | 6.32624 | 6.59005 | 6.02052 | 6.34803 | 6.79176 | 6.61546 | 6.73239 | 6.58545 | 6.21242 | 5.94694 | 6.134 | 5.49893 | 6.20539 | 5.78082 | 6.10953 |
| COTL1 | Q14019 | 6.30427 | 5.87856 | 6.8186 | 5.79008 | 6.38652 | 6.565 | 6.5074 | 6.48123 | 6.8727 | 6.17689 | 6.46451 | 6.51268 | 5.65095 | 5.9976 | 5.71117 | 5.88186 |
| SEC23IP | Q9Y6Y8 | 6.11978 | 6.32456 | 6.78202 | 5.81559 | 6.1857 | 6.76219 | 6.19604 | 6.59516 | 6.57607 | 6.43226 | 6.52063 | 6.50886 | 5.88358 | 5.86162 | 5.79054 | 5.64539 |
| SERPINB6 | P35237 | 6.17265 | 6.26461 | 6.69596 | 5.72066 | 6.36254 | 7.07968 | 6.15947 | 6.91209 | 6.04607 | 6.70854 | 6.13621 | 6.31344 | 5.76563 | 6.32604 | 5.74648 | 5.58992 |
| RPS18 | P62269 | 6.52197 | 6.36348 | 6.68561 | 6.44331 | 6.46124 | 6.34456 | 6.51752 | 6.39582 | 6.54317 | 6.24899 | 6.38872 | 6.29908 | 5.67486 | 5.87378 | 5.51346 | 5.72444 |
| KIF15 | Q9NS87 | 6.50483 | 6.46715 | 7.04651 | 6.48271 | 6.46593 | 6.67348 | 6.25478 | 6.75129 | 6.59753 | 6.73276 | 6.79571 | 6.84998 | 5.05697 | 5.31492 | 4.97329 | 5.03216 |
| SEC24C | P53992 | 6.07953 | 6.10464 | 6.42647 | 6.1176 | 6.24652 | 6.78025 | 6.19628 | 6.03153 | 6.04818 | 6.19854 | 6.21641 | 6.19493 | 6.4459 | 6.30197 | 6.37992 | 6.23134 |
| CAT | P04040 | 6.39823 | 6.12912 | 5.72182 | 6.13805 | 6.44537 | 6.39823 | 6.1428 | 6.24009 | 6.13521 | 6.08514 | 6.10258 | 6.09468 | 6.83102 | 6.22628 | 6.55024 | 6.36114 |
| CAMSAP1 | Q5T5Y3-3 | 6.5841 | 6.47584 | 6.98008 | 6.14095 | 6.08388 | 6.37173 | 6.28007 | 6.45671 | 6.30303 | 6.27653 | 6.31546 | 6.38644 | 5.76413 | 6.00235 | 5.83506 | 5.74364 |
| KDM5C | P41229 | 5.81743 | 6.33143 | 6.11505 | 6.23859 | 6.37091 | 6.38076 | 6.24153 | 6.42172 | 6.24025 | 6.27631 | 6.50072 | 6.46489 | 6.09816 | 6.07462 | 6.27194 | 6.15569 |
| RACGAP1 | Q9H0H5 | 6.66407 | 6.32035 | 6.44152 | 6.89512 | 5.96627 | 6.15372 | 6.332 | 6.01677 | 6.45433 | 6.40068 | 6.57242 | 6.3003 | 5.88015 | 5.74136 | 5.79034 | 6.0706 |
| ZFHX4 | Q86UP3-5 | 6.14337 | 6.51614 | 6.21144 | 6.80177 | 6.33054 | 6.12951 | 6.46117 | 6.10067 | 6.29831 | 6.21501 | 6.32938 | 6.3972 | 6.22552 | 5.75354 | 5.90546 | 6.18098 |
| IGF2BP3 | O00425 | 6.37197 | 6.25003 | 6.26136 | 6.16948 | 6.51122 | 6.35911 | 6.32835 | 6.20278 | 6.49484 | 6.52902 | 6.17164 | 6.14976 | 6.03171 | 6.09623 | 6.05368 | 6.01884 |
| BUB1B | O60566-3 | 6.61849 | 6.39845 | 7.39045 | 6.49885 | 6.41278 | 6.52931 | 6.59634 | 6.48673 | 7.09538 | 6.46638 | 6.79676 | 6.98352 | 5.03688 | 4.97121 | 4.71507 | 5.00341 |

| | | | | | | | | | | | | | | | | | |
|---|---|---|---|---|---|---|---|---|---|---|---|---|---|---|---|---|---|
| HIP1 | O00291 | 6.40581 | 6.34245 | 6.04011 | 6.08196 | 6.51673 | 6.19956 | 6.33447 | 6.17321 | 6.3012 | 6.37375 | 6.08763 | 6.05884 | 6.39838 | 6.28151 | 6.29808 | 6.10633 |
| PHF14 | A0A0U1RRH6 | 6.59788 | 6.69737 | 6.31946 | 6.87332 | 5.72417 | 5.99532 | 6.27936 | 6.00714 | 6.24015 | 6.44511 | 6.31359 | 6.39433 | 5.99276 | 6.05548 | 6.06258 | 6.00199 |
| COG1 | Q8WTW3 | 6.3722 | 6.32972 | 6.43986 | 6.53012 | 6.2312 | 6.42169 | 6.46492 | 6.34971 | 6.38094 | 6.36462 | 6.3054 | 6.17715 | 5.97493 | 6.19233 | 5.82412 | 5.64107 |
| SCPEP1 | Q9HB40 | 6.31903 | 5.84386 | 5.86003 | 5.76192 | 6.56083 | 5.81576 | 6.34017 | 6.34152 | 5.91936 | 6.27346 | 5.79931 | 6.17157 | 6.45747 | 6.90666 | 6.80941 | 6.81965 |
| GPATCH8 | Q9UKJ3 | 6.27387 | 6.39962 | 6.24822 | 6.2445 | 5.82145 | 5.99692 | 6.10565 | 5.73157 | 5.84114 | 5.98509 | 5.92518 | 5.6869 | 6.37535 | 7.12599 | 6.99541 | 7.24313 |
| SPECC1L | Q69YQ0 | 6.76723 | 6.45322 | 6.51325 | 6.26079 | 6.20157 | 6.22549 | 6.32858 | 5.85921 | 6.61879 | 6.22371 | 6.30596 | 6.5038 | 6.35203 | 6.10378 | 5.51538 | 5.76722 |
| DHX37 | Q8IY37 | 6.05204 | 6.43753 | 6.27465 | 6.90359 | 6.13617 | 5.86617 | 6.39108 | 5.94007 | 6.31179 | 6.31444 | 6.06748 | 6.27592 | 6.02743 | 6.04025 | 6.71969 | 6.24169 |
| ECHS1 | P30084 | 6.43755 | 6.29893 | 5.71776 | 6.28464 | 6.54543 | 6.00111 | 6.41978 | 6.11343 | 6.11186 | 6.01459 | 5.98176 | 5.96384 | 6.78444 | 6.5201 | 6.48726 | 6.31753 |
| CPVL | Q9H3G5 | 6.3662 | 6.32979 | 5.79364 | 6.09022 | 6.3831 | 6.27248 | 6.39176 | 6.35831 | 6.0644 | 6.16758 | 6.06385 | 5.93654 | 6.63116 | 6.43384 | 6.50931 | 6.20783 |
| PXDN | Q92626 | 7.33546 | 7.11652 | 5.93587 | 7.53517 | 6.37047 | 6.17254 | 6.62329 | 6.07128 | 6.5859 | 6.41904 | 6.33852 | 6.52216 | 5.15296 | 5.08728 | 5.15392 | 5.57963 |
| RAB1A | P62820 | 6.03031 | 6.03844 | 6.15103 | 6.17849 | 6.11079 | 6.09979 | 6.45471 | 6.21289 | 6.32255 | 6.03154 | 6.20549 | 6.01173 | 6.27647 | 6.66882 | 6.71777 | 6.48916 |
| YTHDC2 | Q9H6S0 | 6.32325 | 6.32841 | 6.55044 | 6.28844 | 6.15786 | 6.30273 | 6.20224 | 6.29068 | 6.45543 | 6.60265 | 6.56331 | 6.56631 | 5.89826 | 5.87544 | 5.82103 | 5.77351 |
| HADH | Q16836-2 | 5.87036 | 6.20252 | 5.75127 | 6.5048 | 6.20097 | 5.98393 | 6.28469 | 6.03152 | 6.17618 | 6.14126 | 6.12521 | 6.05038 | 6.59642 | 6.54402 | 6.71927 | 6.81717 |
| HSPG2 | P98160 | 6.84616 | 7.0479 | 6.38118 | 6.9241 | 6.34475 | 6.62967 | 6.65887 | 6.23121 | 6.71161 | 6.93574 | 6.88098 | 6.65337 | 4.95925 | 5.06397 | 4.81036 | 4.92088 |
| JMJD1C | Q15652 | 6.10171 | 6.61249 | 6.13766 | 6.7295 | 6.10539 | 6.30356 | 6.22925 | 6.22428 | 6.11825 | 6.17893 | 5.99634 | 5.9333 | 6.2537 | 6.34038 | 6.44749 | 6.28775 |
| WRNIP1 | Q96S55 | 5.99717 | 6.21405 | 6.09936 | 5.91606 | 6.1525 | 6.05097 | 6.00471 | 5.96422 | 6.07052 | 6.11875 | 6.09761 | 5.94903 | 7.01466 | 6.57721 | 7.139 | 6.63418 |
| URB1 | O60287 | 6.38876 | 6.40796 | 5.7861 | 7.01501 | 6.21675 | 5.8581 | 6.341 | 6.16237 | 6.38972 | 5.90359 | 6.09448 | 6.24176 | 6.24566 | 6.06105 | 6.39783 | 6.48987 |
| CDCA2 | Q69YH5 | 6.94232 | 7.01963 | 6.54968 | 7.49303 | 6.20732 | 5.98506 | 6.61356 | 6.11191 | 6.46767 | 6.54058 | 6.45578 | 6.58552 | 4.90356 | 5.30326 | 5.40188 | 5.41925 |
| PHGDH | O43175 | 6.33156 | 6.0032 | 6.69152 | 5.67624 | 6.50578 | 6.82396 | 5.94732 | 6.66268 | 6.42201 | 6.5102 | 6.55859 | 6.42109 | 6.248 | 6.01994 | 5.64904 | 5.52885 |
| FXR2 | P51116 | 6.27545 | 6.51916 | 6.49871 | 6.24983 | 6.26137 | 6.28226 | 6.20754 | 6.36797 | 6.25028 | 6.4473 | 6.71658 | 6.37033 | 5.80405 | 6.13384 | 5.83549 | 5.77983 |
| TNPO2 | O14787 | 5.82653 | 5.6797 | 6.2927 | 5.7621 | 5.9279 | 6.23505 | 5.86379 | 6.31997 | 6.42558 | 6.23771 | 6.3684 | 6.49879 | 6.52486 | 6.3646 | 6.80421 | 6.86811 |
| MRPL39 | Q9NYK5 | 6.40795 | 6.3489 | 5.80676 | 6.54373 | 6.56928 | 6.36498 | 6.40605 | 6.38917 | 6.05354 | 6.08109 | 5.90766 | 5.86331 | 6.47893 | 6.19252 | 6.29668 | 6.28945 |
| CNP | P09543 | 6.26598 | 6.46057 | 6.00735 | 6.26408 | 6.20059 | 5.94317 | 6.31311 | 5.94367 | 6.29361 | 6.42184 | 6.10686 | 6.1989 | 6.51256 | 6.35264 | 6.4713 | 6.24375 |
| VPS39 | Q96JC1 | 6.28396 | 6.67684 | 6.37027 | 6.03076 | 6.06215 | 6.53089 | 6.05454 | 6.26932 | 6.27721 | 6.53208 | 6.37364 | 6.22098 | 5.93917 | 6.39298 | 6.10744 | 5.87776 |
| ADAM10 | O14672 | 6.36089 | 6.50123 | 6.00492 | 6.5819 | 6.16695 | 6.25246 | 6.48989 | 5.9535 | 5.98787 | 6.1542 | 6.11639 | 5.95395 | 6.48315 | 6.09794 | 6.38684 | 6.50793 |
| FANCD2 | Q9BXW9-1 | 6.02198 | 6.15438 | 6.13258 | 6.05876 | 5.96844 | 6.03268 | 6.18435 | 5.9073 | 6.54093 | 6.38527 | 6.22087 | 6.44831 | 6.49753 | 6.03848 | 6.77287 | 6.63526 |
| LARP7 | Q4G0J3-3 | 5.77022 | 6.54567 | 5.87524 | 6.1469 | 6.14126 | 5.98485 | 6.05426 | 5.80638 | 5.78568 | 6.08448 | 6.03158 | 5.80169 | 6.58511 | 7.20983 | 7.25988 | 6.91697 |
| ARHGAP1 | Q07960 | 6.57525 | 6.16877 | 6.71846 | 5.68327 | 6.5544 | 6.4148 | 6.1819 | 6.44196 | 6.56004 | 6.51569 | 6.55031 | 6.28469 | 6.21691 | 5.71931 | 5.803 | 5.61123 |
| CPSF6 | Q16630-2 | 6.23975 | 6.47523 | 5.947 | 6.3192 | 6.30295 | 5.99087 | 5.99389 | 5.88998 | 6.22922 | 6.28725 | 6.02589 | 5.98926 | 6.76707 | 6.54012 | 6.53608 | 6.46625 |
| PTPN13 | Q12923-4 | 6.36585 | 6.11187 | 6.19073 | 6.60731 | 6.25797 | 6.18533 | 6.33121 | 5.85141 | 6.39584 | 6.36241 | 6.20674 | 6.25425 | 6.44927 | 6.09782 | 6.25786 | 6.07413 |
| COX4I1 | P13073 | 6.38878 | 6.33544 | 5.67703 | 6.59683 | 6.58824 | 6.07045 | 6.60127 | 6.10987 | 6.19515 | 5.97795 | 5.81728 | 5.95651 | 6.63253 | 6.13064 | 6.44363 | 6.4784 |
| TBRG4 | Q969Z0 | 6.30187 | 6.35561 | 5.92464 | 6.86172 | 6.25453 | 6.30972 | 6.25482 | 6.21301 | 6.0133 | 5.96518 | 6.12618 | 6.15053 | 6.25392 | 6.14458 | 6.42459 | 6.44582 |
| RPL24 | P83731 | 6.52434 | 6.32601 | 6.39675 | 5.94413 | 6.274 | 6.39128 | 6.3389 | 6.2732 | 6.39533 | 6.4854 | 6.37742 | 6.35905 | 5.87557 | 6.16873 | 5.91821 | 5.95166 |
| DDX39A | O00148 | 6.39281 | 6.37799 | 6.09147 | 6.95207 | 6.44049 | 5.96003 | 6.25596 | 5.87597 | 6.22071 | 6.12293 | 6.17012 | 6.04819 | 6.39908 | 6.06953 | 6.44769 | 6.17496 |
| ECE1 | P42892 | 6.4253 | 6.32404 | 6.11835 | 6.27047 | 6.12758 | 6.24336 | 6.10817 | 6.22528 | 5.99669 | 6.59202 | 6.42913 | 6.37952 | 6.27365 | 6.43341 | 6.18204 | 5.87099 |
| RPTOR | Q8N122 | 6.15993 | 6.51571 | 6.65378 | 6.35025 | 6.07899 | 6.23945 | 6.14199 | 6.39105 | 6.50355 | 6.47253 | 6.4333 | 6.69792 | 5.79519 | 5.65557 | 6.01906 | 5.89176 |
| NCOA5 | Q9HCD5 | 6.32183 | 6.25243 | 5.74655 | 6.75827 | 6.00062 | 5.984 | 6.66445 | 5.97089 | 6.33175 | 5.76776 | 5.72997 | 6.11654 | 6.30658 | 6.5178 | 6.5255 | 7.00508 |
| RBMX | P38159-2 | 6.31868 | 6.31941 | 5.88067 | 6.82217 | 5.99299 | 5.97926 | 6.37764 | 6.02944 | 6.24679 | 6.21699 | 6.09465 | 6.22173 | 6.55698 | 6.22615 | 6.3553 | 6.36765 |
| PRPF4 | O43172 | 5.65734 | 5.97997 | 6.02311 | 6.4985 | 6.42441 | 6.25151 | 6.33721 | 5.79637 | 6.20905 | 5.91671 | 6.49473 | 6.10879 | 6.56686 | 6.40047 | 6.70491 | 6.63006 |
| WDR36 | Q8NI36 | 6.17824 | 6.41774 | 5.79203 | 6.82595 | 6.2977 | 6.00489 | 6.34702 | 6.26214 | 5.99772 | 5.84452 | 5.87712 | 5.99214 | 6.49099 | 6.21513 | 6.72223 | 6.73444 |
| SUCLA2 | Q9P2R7 | 6.58547 | 6.32342 | 6.07243 | 6.43857 | 6.48994 | 5.8725 | 6.28224 | 5.98014 | 6.33553 | 5.96512 | 6.00837 | 5.83042 | 6.79376 | 6.08053 | 6.61346 | 6.3279 |
| MYO18A | Q92614 | 6.18058 | 6.18547 | 6.44612 | 6.14889 | 6.24063 | 6.19145 | 6.17323 | 5.92543 | 6.58034 | 6.58946 | 6.08588 | 5.90422 | 6.46221 | 6.30563 | 6.22727 | 6.35318 |
| CEP192 | Q8TEP8 | 6.55435 | 6.4745 | 6.90581 | 6.65 | 6.21577 | 6.55133 | 6.21681 | 6.55057 | 6.66857 | 6.62808 | 7.03308 | 7.02068 | 5.14013 | 5.28357 | 4.92421 | 5.18254 |
| AATF | Q9NY61 | 6.31178 | 6.27163 | 6.00986 | 6.89783 | 6.28026 | 5.8356 | 6.25942 | 6.02508 | 6.11747 | 5.99308 | 6.13696 | 6.20341 | 6.37148 | 5.94645 | 6.56493 | 6.77475 |
| TACC3 | Q9Y6A5 | 6.92546 | 6.18705 | 7.12564 | 5.69537 | 6.49132 | 6.6544 | 5.80362 | 6.7327 | 6.86402 | 6.52158 | 7.05068 | 7.15087 | 5.67145 | 5.23455 | 4.83364 | 5.05765 |
| HMMR | O75330-3 | 6.26527 | 6.24574 | 6.7614 | 5.93192 | 6.22553 | 6.37593 | 6.47102 | 6.37248 | 6.97748 | 6.69774 | 6.65619 | 6.67183 | 5.60733 | 5.66791 | 5.52077 | 5.55144 |
| BCL2L2 | Q92843-2 | 6.69869 | 6.82613 | 5.93984 | 6.58398 | 6.68456 | 6.27423 | 6.31981 | 6.21464 | 6.22504 | 6.1909 | 5.77919 | 5.62759 | 6.39526 | 5.90261 | 6.22579 | 6.11173 |
| RTF1 | Q92541 | 5.25632 | 5.86479 | 5.83128 | 5.15897 | 5.67757 | 6.02253 | 5.5837 | 6.00647 | 5.91209 | 6.22745 | 6.18164 | 5.86828 | 7.47836 | 7.87929 | 7.7097 | 7.34158 |
| ABCC1 | P33527 | 6.41139 | 6.60651 | 6.01699 | 6.72144 | 6.11996 | 5.9837 | 6.23238 | 5.94406 | 6.32022 | 6.17407 | 5.98636 | 6.08384 | 6.32733 | 6.1601 | 6.44332 | 6.46834 |
| GNAS | P63092-3 | 6.33051 | 6.58029 | 6.17383 | 6.49084 | 6.45468 | 6.30747 | 6.36659 | 6.29892 | 6.46354 | 6.24708 | 5.91328 | 6.15167 | 5.95 | 6.16589 | 6.00125 | 6.10417 |
| RICTOR | Q6R327-3 | 6.42388 | 6.18263 | 6.58682 | 6.05624 | 6.35671 | 6.82075 | 6.30133 | 6.6045 | 6.34834 | 6.63735 | 6.41545 | 6.33933 | 5.86227 | 5.82853 | 5.56599 | 5.66987 |
| VPS11 | A0A087WXL6 | 6.12429 | 6.34018 | 6.49579 | 5.9009 | 6.10092 | 6.40731 | 6.26633 | 6.30933 | 6.34237 | 6.40852 | 6.49052 | 6.10986 | 5.93366 | 6.12733 | 6.615 | 6.0277 |
| SLC4A1AP | Q9BWU0 | 5.478 | 5.91081 | 5.99801 | 5.74494 | 6.01106 | 6.15567 | 5.93503 | 5.89021 | 6.2771 | 6.12092 | 6.28483 | 6.39325 | 7.08406 | 6.6512 | 6.79484 | 7.27006 |
| ARHGEF10 | O15013 | 6.39826 | 6.26841 | 6.86095 | 6.57766 | 6.19359 | 6.70676 | 6.36242 | 6.42441 | 6.28277 | 6.32614 | 6.40713 | 6.6907 | 5.54101 | 5.81614 | 5.36067 | 5.78299 |
| IDH3A | P50213 | 6.18398 | 5.8046 | 5.43777 | 6.05449 | 6.44709 | 5.71011 | 6.68889 | 6.40484 | 6.19243 | 6.25903 | 5.62184 | 5.58978 | 6.47019 | 6.79859 | 7.00187 | 7.35909 |
| CAPN1 | P07384 | 6.39066 | 6.50364 | 6.68694 | 5.83905 | 6.36594 | 6.74218 | 6.2366 | 6.66643 | 6.41014 | 6.62426 | 6.30262 | 6.35527 | 5.68959 | 5.84134 | 5.67918 | 5.66616 |
| TBCB | Q99426 | 6.39157 | 6.05423 | 6.72373 | 5.97465 | 6.31822 | 6.41339 | 6.21695 | 6.78743 | 6.44482 | 6.28091 | 6.24247 | 6.73469 | 5.56239 | 6.10995 | 5.69077 | 6.05385 |
| DHX36 | Q9H2U1 | 6.33114 | 6.29822 | 6.65998 | 6.22603 | 6.24568 | 6.26671 | 6.32081 | 6.34473 | 6.48078 | 6.27999 | 6.55935 | 6.41584 | 6.02938 | 6.01813 | 5.85683 | 5.66642 |
| DHX29 | Q7Z478 | 6.40617 | 6.45704 | 6.73371 | 6.2637 | 6.37427 | 6.55408 | 6.31124 | 6.20067 | 6.53388 | 6.45513 | 6.50764 | 6.20049 | 5.72579 | 6.00441 | 5.72637 | 5.54541 |
| LIG1 | P18858 | 5.20412 | 5.41313 | 5.9281 | 5.15185 | 5.29954 | 5.76274 | 5.93086 | 5.8231 | 6.3761 | 5.95524 | 6.40487 | 6.04373 | 7.77415 | 7.3672 | 8.06057 | 7.5047 |
| MPRIP | Q6WCQ1-2 | 5.88034 | 6.26649 | 5.9083 | 5.81199 | 6.27411 | 5.92661 | 5.87267 | 5.99808 | 6.33082 | 6.32794 | 6.53852 | 6.21162 | 6.84132 | 6.6407 | 6.57587 | 6.59463 |
| PKP4 | Q99569 | 6.48745 | 6.97982 | 6.1853 | 6.98382 | 6.21788 | 6.22135 | 6.30129 | 6.52313 | 6.08148 | 5.84561 | 6.09473 | 5.98927 | 6.17022 | 6.10439 | 6.02612 | 5.78814 |
| KIF2A | O00139-4 | 5.79235 | 6.25325 | 6.15647 | 5.89902 | 6.23321 | 6.47107 | 6.17908 | 6.60582 | 6.40184 | 5.5646 | 6.23397 | 6.5271 | 6.31046 | 6.42218 | 6.2873 | 6.26229 |
| PRPF31 | Q8WWY3 | 6.17084 | 6.14744 | 5.87369 | 6.29392 | 6.08007 | 5.97347 | 6.36524 | 5.91425 | 6.12919 | 5.9164 | 5.776 | 5.99003 | 6.80398 | 6.81884 | 6.87328 | 6.87334 |
| SBNO1 | A3KN83 | 5.4635 | 6.12209 | 6.47424 | 5.56572 | 5.58871 | 6.23027 | 5.75713 | 6.04293 | 6.13337 | 6.39362 | 6.67097 | 6.30871 | 6.31558 | 6.75286 | 7.14367 | 7.03662 |
| RNMT | O43148-2 | 5.97708 | 6.22782 | 5.84158 | 5.89552 | 6.26409 | 5.9749 | 6.3404 | 6.05891 | 6.18909 | 6.00008 | 5.67425 | 5.68978 | 7.01376 | 6.90869 | 6.94842 | 6.99565 |
| HDAC2 | Q92769 | 6.45422 | 6.61352 | 6.0446 | 6.72875 | 6.39315 | 6.33999 | 6.2992 | 6.30261 | 6.30493 | 6.05866 | 5.9247 | 5.99492 | 6.42445 | 5.88794 | 6.24581 | 5.98255 |
| RBM12 | Q9NTZ6 | 5.57987 | 5.76186 | 5.96657 | 5.31398 | 5.97646 | 6.342 | 5.57622 | 6.10945 | 6.05586 | 6.24985 | 6.30432 | 6.12017 | 7.44796 | 7.17883 | 7.1095 | 6.90711 |
| FBN1 | P35555 | 6.18916 | 6.36218 | 5.63465 | 6.54836 | 6.42609 | 6.49966 | 6.76698 | 6.44609 | 5.9568 | 6.38505 | 6.31997 | 5.93897 | 6.38765 | 6.03927 | 5.93662 | 6.1618 |
| PDPR | Q8NCN5 | 6.39489 | 6.25909 | 5.8315 | 6.39512 | 6.38849 | 6.02564 | 6.50082 | 6.13343 | 6.10193 | 6.1691 | 6.09974 | 5.77044 | 6.6117 | 6.24478 | 6.64568 | 6.42767 |
| LPP | Q93052 | 6.4437 | 5.97137 | 6.92296 | 5.61949 | 6.48682 | 6.59691 | 5.83902 | 6.74523 | 6.55436 | 6.29929 | 6.61655 | 6.50745 | 6.56361 | 5.74209 | 5.52004 | 5.52612 |
| ATG7 | O95352 | 6.3096 | 6.03695 | 6.99382 | 6.02066 | 6.21766 | 6.64031 | 6.24193 | 6.51317 | 6.6045 | 6.43701 | 6.60027 | 6.53022 | 5.78675 | 5.75953 | 5.79114 | 5.51651 |
| UBR7 | Q8N806 | 5.59557 | 5.67601 | 6.43029 | 6.50125 | 5.85932 | 6.44756 | 5.73586 | 6.29188 | 6.1195 | 6.27404 | 6.71384 | 6.33282 | 6.55999 | 7.16065 | 6.66142 | 6.53999 |
| BRIX1 | Q8TDN6 | 6.21477 | 6.02862 | 5.87231 | 6.61521 | 6.21779 | 5.65908 | 6.23371 | 5.90139 | 6.17005 | 5.5255 | 5.76118 | 5.88419 | 6.77115 | 6.82644 | 7.09055 | 7.22807 |
| LEMD3 | Q9Y2U8 | 6.33693 | 6.36145 | 6.07402 | 6.75337 | 6.34944 | 6.0693 | 6.30283 | 5.7745 | 6.13274 | 6.13357 | 5.99052 | 6.03722 | 6.67968 | 6.19599 | 6.40614 | 6.40231 |
| CDK5RAP3 | Q96JB5-4 | 6.43135 | 6.37328 | 6.12457 | 6.28666 | 6.29644 | 6.12907 | 6.09746 | 6.21032 | 6.24516 | 6.17256 | 5.93541 | 5.86185 | 6.78363 | 6.11962 | 6.4859 | 6.44674 |
| TMX3 | Q96JJ7 | 6.39533 | 6.27435 | 5.92374 | 6.4659 | 6.25539 | 5.87242 | 6.40557 | 5.91821 | 6.23942 | 5.94299 | 5.87178 | 5.84016 | 6.81877 | 6.13198 | 6.95527 | 6.68872 |
| PLEKHA5 | Q9HAU0-6 | 6.3299 | 6.44416 | 6.23778 | 6.02755 | 6.13252 | 6.14126 | 6.29831 | 6.39714 | 6.31119 | 6.45024 | 6.14542 | 6.16971 | 6.4579 | 6.24092 | 6.18559 | 6.03041 |
| ERP29 | P30040 | 6.09975 | 5.78086 | 5.78402 | 5.51991 | 6.77879 | 5.8664 | 5.99078 | 6.59983 | 5.97277 | 6.47743 | 6.30096 | 6.3981 | 6.98057 | 6.4206 | 6.65457 | 6.37465 |
| CYB5R3 | P00387-3 | 6.27636 | 6.44956 | 5.97844 | 6.62667 | 6.48496 | 6.25334 | 6.20648 | 6.14968 | 6.02129 | 5.90593 | 6.1474 | 6.03209 | 6.41104 | 6.49853 | 6.44432 | 6.11363 |
| CAPZA2 | P47755 | 6.21899 | 6.17273 | 6.6489 | 6.29116 | 6.40324 | 6.60777 | 6.09548 | 6.17396 | 6.55127 | 6.29499 | 6.46587 | 6.09308 | 6.17818 | 5.86791 | 6.08613 | 5.85035 |
| TMED10 | P49755 | 6.47939 | 6.79989 | 5.64747 | 6.43585 | 6.78617 | 6.50736 | 6.11565 | 6.32685 | 6.03689 | 5.83024 | 5.79704 | 5.75182 | 6.62489 | 6.4489 | 6.38247 | 6.0291 |
| PACS1 | Q6VY07 | 6.50633 | 6.30595 | 6.47143 | 6.39065 | 6.22569 | 6.66613 | 6.02298 | 6.34441 | 6.20651 | 6.4728 | 6.57776 | 6.484 | 5.65318 | 6.32145 | 5.84012 | 5.51062 |
| RBM19 | Q9Y4C8 | 6.18287 | 6.42721 | 5.99378 | 6.94437 | 6.16633 | 6.0255 | 6.44212 | 5.87751 | 6.22065 | 5.97423 | 5.8014 | 5.952 | 6.35924 | 6.26663 | 6.7074 | 6.65877 |
| TRMT1 | Q9NXH9 | 5.24525 | 5.55798 | 6.17624 | 5.68025 | 5.81664 | 6.16664 | 6.12178 | 5.95465 | 5.96847 | 6.1091 | 6.24346 | 5.93744 | 7.04293 | 7.06608 | 7.83189 | 7.0812 |
| SNRPD2 | P62316 | 6.11567 | 6.21872 | 5.92624 | 6.71637 | 6.09707 | 5.94737 | 6.93789 | 5.59841 | 6.27324 | 5.61107 | 5.68006 | 5.80221 | 6.42326 | 6.65474 | 6.79382 | 7.20386 |
| POFUT1 | Q9H488 | 6.40364 | 6.16093 | 5.95054 | 6.57037 | 6.46383 | 5.92624 | 6.34126 | 6.19888 | 6.13786 | 5.97684 | 5.89864 | 5.90618 | 6.79057 | 6.28341 | 6.53046 | 6.46033 |
| METAP1 | P53582 | 6.54777 | 6.32296 | 6.55828 | 6.20491 | 6.62028 | 6.54941 | 6.3708 | 6.76043 | 6.51496 | 6.18079 | 6.44253 | 6.41483 | 5.892 | 5.39029 | 5.55581 | 5.67394 |
| PSME1 | Q06323 | 6.12452 | 5.23124 | 6.46754 | 4.68463 | 6.48454 | 6.06446 | 5.48929 | 7.06715 | 5.6711 | 6.67701 | 6.72159 | 6.98417 | 6.80896 | 6.71916 | 6.36454 | 6.44011 |
| RPS13 | P62277 | 6.26868 | 6.47172 | 6.35421 | 6.24145 | 6.1708 | 6.38739 | 6.06433 | 6.48875 | 6.41924 | 6.46975 | 6.47975 | 6.54016 | 5.77735 | 6.16588 | 5.76184 | 5.9387 |
| WDR82 | Q6UXN9 | 6.31751 | 6.59482 | 6.24396 | 6.68691 | 6.25841 | 6.44748 | 6.24439 | 6.11948 | 6.40604 | 6.2886 | 6.19488 | 6.11875 | 6.2685 | 5.74554 | 6.07754 | 5.98719 |
| POLR3A | O14802 | 6.17142 | 6.42772 | 6.47402 | 6.24521 | 6.02243 | 6.15648 | 6.13243 | 6.15487 | 6.3491 | 6.27787 | 6.36778 | 6.59144 | 6.1869 | 6.16998 | 6.05048 | 6.22187 |
| RPL17 | P18621-3 | 6.27294 | 6.47345 | 6.60711 | 6.11328 | 6.23775 | 6.24638 | 6.42033 | 6.10094 | 6.60873 | 6.15616 | 6.50178 | 6.56335 | 5.84337 | 6.19768 | 5.73696 | 5.91978 |
| WARS1 | P23381 | 6.27833 | 6.09663 | 6.75083 | 5.75511 | 6.66122 | 6.64549 | 6.3453 | 6.40455 | 6.62631 | 6.32406 | 6.63665 | 6.414 | 6.02546 | 5.63607 | 5.71458 | 5.68542 |
| SPTLC1 | O15269 | 6.313 | 6.35595 | 5.98719 | 6.63373 | 6.33963 | 6.08377 | 6.37769 | 6.0632 | 5.87691 | 5.93778 | 5.92719 | 6.01699 | 6.47323 | 6.46922 | 6.59 | 6.55451 |
| NSD1 | Q96L73 | 6.58246 | 6.84483 | 6.28956 | 7.4756 | 6.04666 | 5.79947 | 6.28325 | 6.238 | 6.38418 | 6.03908 | 6.58536 | 6.42726 | 5.32539 | 5.82605 | 5.94389 | 5.90896 |
| QRSL1 | Q9H0R6 | 6.30188 | 6.37832 | 5.9934 | 6.5545 | 6.52595 | 6.29716 | 6.42555 | 6.1498 | 6.41306 | 6.04889 | 6.09907 | 6.04407 | 6.61989 | 5.6888 | 6.3243 | 6.13536 |
| LTBP4 | Q8N2S1 | 7.09947 | 6.36106 | 4.8848 | 7.07842 | 5.47824 | 5.92525 | 7.29811 | 6.22697 | 7.0706 | 5.9278 | 5.69811 | 6.0908 | 6.74257 | 5.7371 | 6.2172 | 6.16349 |
| MAN2A1 | Q16706 | 6.35814 | 6.29275 | 6.22725 | 6.8006 | 6.16428 | 6.0536 | 6.51972 | 5.82108 | 6.10011 | 6.14863 | 6.30624 | 6.18544 | 6.47077 | 6.0566 | 6.32127 | 6.17351 |
| POLR1E | Q9GZS1 | 6.33187 | 6.45027 | 5.85268 | 7.01257 | 5.97101 | 5.88576 | 6.70791 | 5.80558 | 6.28442 | 5.63547 | 6.01423 | 6.04638 | 6.42349 | 6.07019 | 6.56526 | 6.94292 |
| SCAF8 | Q9UPN6-2 | 6.53781 | 6.51982 | 5.90803 | 6.40692 | 6.3404 | 6.09802 | 6.36224 | 5.92589 | 6.09345 | 6.00469 | 5.84093 | 5.97607 | 6.78267 | 6.28502 | 6.56352 | 6.35451 |
| RRP1 | P56182 | 6.48745 | 6.55288 | 5.37755 | 6.66375 | 6.23552 | 5.65194 | 6.4721 | 6.12903 | 6.45852 | 5.60377 | 5.83683 | 5.99834 | 6.66466 | 6.79656 | 6.43594 | 7.53516 |
| TACC1 | O75410-2 | 6.59031 | 6.38876 | 6.18823 | 6.6491 | 6.38158 | 6.15911 | 6.206 | 6.17998 | 6.13817 | 5.94021 | 6.34258 | 6.13221 | 6.63425 | 5.87345 | 6.07602 | 6.12004 |
| SYNE3 | Q6ZMZ3 | 6.44969 | 6.37988 | 5.69655 | 6.38163 | 6.07453 | 5.91476 | 6.28319 | 5.93651 | 5.94225 | 6.11597 | 5.75865 | 5.55641 | 6.97665 | 6.85379 | 6.93425 | 6.74529 |
| NAPA | P54920 | 6.3849 | 6.03375 | 6.1434 | 5.87076 | 6.20051 | 6.1272 | 6.23655 | 6.37254 | 5.92815 | 6.19376 | 6.26832 | 6.28494 | 6.32572 | 6.1321 | 6.85916 | 6.63823 |

| | | | | | | | | | | | | | | | | | |
|---|---|---|---|---|---|---|---|---|---|---|---|---|---|---|---|---|---|
| DLGAP5 | Q15398 | 6.18709 | 6.11781 | 6.94245 | 6.20651 | 6.34902 | 6.42685 | 6.61748 | 6.92965 | 6.4847 | 6.3499 | 6.30291 | 6.78477 | 5.65493 | 5.59154 | 5.37343 | 5.68093 |
| MAP7D3 | Q8IWC1 | 6.48286 | 6.43956 | 6.42992 | 6.63157 | 6.13481 | 6.35612 | 6.16716 | 6.45808 | 6.26876 | 6.19532 | 6.56009 | 6.62456 | 5.69693 | 5.82694 | 5.70704 | 6.02027 |
| TRA2B | P62995 | 6.4419 | 6.22759 | 5.78744 | 6.4476 | 6.51103 | 5.91326 | 6.45954 | 6.09738 | 6.40292 | 5.96246 | 5.9477 | 5.70015 | 6.99282 | 6.17934 | 6.49405 | 6.43483 |
| CEP350 | Q5VT06 | 6.27481 | 6.16464 | 6.38663 | 6.22873 | 6.10465 | 6.38918 | 6.24658 | 6.27918 | 6.42231 | 6.70445 | 6.35563 | 6.44431 | 5.88346 | 6.04561 | 6.07747 | 5.99235 |
| SGPL1 | O95470 | 6.28554 | 6.50542 | 5.99621 | 6.62144 | 6.17985 | 6.00025 | 6.29726 | 5.82389 | 6.05157 | 6.15361 | 5.79751 | 5.64794 | 6.72399 | 6.44999 | 6.9154 | 6.55012 |
| ARPC1B | O15143 | 6.01609 | 6.09826 | 6.5118 | 6.05194 | 6.4149 | 6.32538 | 6.31321 | 6.37475 | 6.29934 | 6.4126 | 6.26098 | 6.35096 | 6.16498 | 6.14677 | 6.21199 | 6.04605 |
| SEC23B | Q15437 | 6.14942 | 6.17078 | 6.6398 | 6.15388 | 6.20534 | 6.57617 | 6.15451 | 6.36114 | 6.39091 | 6.20156 | 6.40445 | 6.37935 | 5.95273 | 6.21842 | 6.34434 | 5.69718 |
| EWSR1 | Q01844-5 | 6.22805 | 6.34421 | 5.97045 | 6.36017 | 6.37241 | 6.17078 | 6.03123 | 6.30614 | 6.16728 | 6.07072 | 6.00699 | 5.81692 | 6.69894 | 6.55785 | 6.35651 | 6.54132 |
| ENO2 | P09104 | 6.12966 | 6.00237 | 6.91628 | 5.70645 | 6.63029 | 6.98425 | 6.06892 | 6.88899 | 6.29435 | 6.71291 | 6.69479 | 6.52182 | 5.6037 | 5.72605 | 5.64081 | 5.47833 |
| WBP11 | Q9Y2W2 | 6.28099 | 6.56552 | 6.24465 | 6.09041 | 6.08114 | 6.23486 | 6.38832 | 6.11384 | 6.11712 | 6.15864 | 5.82894 | 5.75139 | 6.67755 | 6.46266 | 6.76003 | 6.24393 |
| SHPRH | Q149N8 | 6.22774 | 6.57842 | 6.01182 | 6.80522 | 6.32926 | 6.22633 | 6.43679 | 5.9871 | 6.40113 | 6.08106 | 6.09886 | 6.15705 | 6.16178 | 6.03312 | 6.19232 | 6.27201 |
| SAFB2 | Q14151 | 6.40175 | 6.25649 | 5.90001 | 6.65959 | 6.24594 | 5.85938 | 6.4622 | 5.95507 | 6.28183 | 5.98874 | 6.05632 | 5.86235 | 6.63468 | 6.26082 | 6.75846 | 6.41635 |
| FKBP8 | Q14318 | 6.44447 | 6.42898 | 6.12601 | 6.73933 | 6.39778 | 6.05352 | 6.45606 | 6.00161 | 6.1265 | 6.13736 | 6.14621 | 6.01654 | 6.26766 | 5.9792 | 6.53107 | 6.14769 |
| EEF1B2 | P24534 | 6.80112 | 6.18922 | 6.7644 | 5.47876 | 6.70659 | 6.21148 | 5.83476 | 6.76318 | 6.3308 | 6.48528 | 6.73391 | 6.60984 | 6.43561 | 5.76113 | 5.40504 | 5.4889 |
| KDM5A | P29375 | 6.43676 | 6.69515 | 6.32842 | 7.10382 | 6.32156 | 6.08632 | 6.73008 | 5.77358 | 6.10489 | 6.18043 | 5.98687 | 6.22112 | 5.98508 | 5.86058 | 6.14229 | 6.04305 |
| MTIF2 | P46199 | 6.38513 | 6.40993 | 6.08924 | 6.6947 | 6.25862 | 6.04445 | 6.3821 | 6.04627 | 6.07499 | 6.00179 | 6.10593 | 5.82196 | 6.66238 | 6.16441 | 6.55866 | 6.29945 |
| MRPS9 | P82933 | 6.5443 | 6.37613 | 5.92441 | 6.5903 | 6.24989 | 6.01891 | 6.33869 | 6.07069 | 6.19845 | 6.04078 | 6.1335 | 6.07411 | 6.43162 | 6.12709 | 6.49806 | 6.38308 |
| FIP1L1 | Q6UN15 | 6.18485 | 6.26839 | 5.81057 | 6.76021 | 6.23094 | 5.96898 | 6.30762 | 5.86304 | 6.19298 | 5.79378 | 5.59649 | 5.84789 | 6.60249 | 6.63259 | 6.7592 | 7.17998 |
| ABCF3 | Q9NUQ8 | 6.50243 | 6.48823 | 6.90301 | 6.3403 | 6.18673 | 6.71438 | 6.35109 | 6.71679 | 6.63982 | 6.58348 | 6.89402 | 6.86986 | 5.15335 | 5.39976 | 4.8902 | 5.36655 |
| STRBP | Q96SI9 | 6.49723 | 6.49454 | 6.04124 | 7.08926 | 6.29062 | 6.00579 | 6.5238 | 5.78605 | 6.2656 | 6.18473 | 6.19959 | 5.96595 | 6.19395 | 6.05459 | 6.13503 | 6.27204 |
| PPP4R3B | Q5MIZ7 | 5.84066 | 6.11122 | 6.19041 | 5.9942 | 6.31232 | 6.14692 | 5.99463 | 6.01314 | 6.27549 | 6.2243 | 6.10279 | 6.03441 | 6.7008 | 6.56985 | 6.68485 | 6.804 |
| PPP4R3A | Q6IN85 | 5.8491 | 6.08033 | 5.95473 | 5.83552 | 6.41864 | 6.28531 | 6.21524 | 5.91756 | 5.96105 | 6.02854 | 5.84298 | 5.58782 | 7.07134 | 6.7172 | 7.38621 | 6.84842 |
| PML | P29590 | 6.35185 | 6.37227 | 5.98841 | 6.16571 | 6.24589 | 5.82517 | 6.02069 | 6.0025 | 6.24764 | 6.06302 | 6.43521 | 6.20815 | 6.95587 | 6.33322 | 6.58269 | 6.20172 |
| MYEF2 | Q9P2K5 | 6.61316 | 6.55776 | 5.93201 | 7.00372 | 6.47005 | 5.93868 | 6.25063 | 6.01397 | 6.21367 | 5.95186 | 6.27492 | 5.80068 | 6.66282 | 5.88664 | 6.29511 | 6.13431 |
| CCDC86 | Q9H6F5 | 5.8085 | 5.98312 | 5.66431 | 6.58738 | 5.82835 | 5.60713 | 6.24248 | 5.8335 | 5.86096 | 5.69012 | 5.81488 | 5.94279 | 7.08081 | 7.02386 | 7.56602 | 7.46578 |
| QSOX2 | Q6ZRP7 | 6.39173 | 6.28796 | 5.95745 | 6.75606 | 6.3674 | 6.06352 | 6.38128 | 6.25298 | 6.36099 | 6.02666 | 6.00328 | 6.12561 | 6.38169 | 6.01237 | 6.36743 | 6.26358 |
| NSDHL | Q15738 | 6.31227 | 6.39332 | 5.80743 | 6.4294 | 6.43992 | 6.0661 | 6.43575 | 6.15308 | 5.95595 | 5.88074 | 5.97389 | 5.83208 | 6.66972 | 6.52714 | 6.56487 | 6.55835 |
| BAIAP2 | I3L4C2 | 6.4331 | 6.50137 | 6.53006 | 6.14599 | 6.65396 | 6.25035 | 6.57716 | 6.71093 | 6.62662 | 6.57253 | 6.35954 | 6.17056 | 5.99518 | 5.55174 | 5.58131 | 5.33963 |
| PIKFYVE | Q9Y2I7 | 6.25073 | 6.29743 | 6.6412 | 6.46579 | 6.15468 | 6.37362 | 6.05235 | 6.29122 | 6.34686 | 6.27343 | 6.50903 | 6.16772 | 6.06965 | 6.05386 | 6.11216 | 5.94027 |
| BCAM | P50895 | 6.5022 | 6.44498 | 6.27132 | 6.62197 | 6.10107 | 6.18703 | 6.34796 | 6.04893 | 6.24379 | 5.96318 | 6.0237 | 6.0873 | 6.54492 | 6.2382 | 6.21273 | 6.1607 |
| DHX40 | Q8IX18 | 5.81894 | 6.30158 | 6.13819 | 6.29901 | 6.03229 | 6.22932 | 6.27494 | 5.91496 | 6.24511 | 6.40935 | 6.2475 | 6.29294 | 6.38011 | 6.25389 | 6.61564 | 6.54625 |
| NEK9 | Q8TD19 | 6.39067 | 6.21893 | 6.72885 | 5.94942 | 6.19762 | 6.80119 | 6.13344 | 6.61163 | 6.28079 | 6.44245 | 6.38546 | 6.72331 | 5.78408 | 5.89246 | 5.73544 | 5.72426 |
| VAC14 | Q08AM6 | 6.15209 | 5.93022 | 6.44435 | 6.20036 | 6.10837 | 6.34025 | 6.11569 | 6.29992 | 5.98117 | 6.55549 | 6.09813 | 6.07948 | 6.5061 | 6.46438 | 6.58368 | 6.14033 |
| MRC2 | Q9UBG0 | 6.10203 | 6.46456 | 6.15545 | 6.28704 | 6.21215 | 6.47506 | 6.00623 | 6.0212 | 5.87002 | 6.35443 | 6.39407 | 6.20672 | 6.15669 | 6.49392 | 6.39341 | 6.40702 |
| CLIC1 | O00299 | 6.28788 | 5.93564 | 7.08552 | 5.57544 | 6.50533 | 6.75133 | 6.0932 | 6.53855 | 6.31503 | 6.57075 | 6.64107 | 6.6673 | 6.04313 | 5.76735 | 5.57326 | 5.64922 |
| ACTL6A | O96019 | 6.06072 | 6.1552 | 6.07801 | 6.56135 | 6.30007 | 6.20402 | 6.39592 | 6.29574 | 6.38746 | 6.13054 | 6.22177 | 6.33755 | 6.08321 | 6.13625 | 6.17737 | 6.47482 |
| EBNA1BP2 | Q99848 | 6.11531 | 6.31758 | 5.66597 | 6.58416 | 6.03159 | 5.82046 | 6.36757 | 5.92241 | 5.80353 | 5.83608 | 5.78283 | 5.91652 | 6.66938 | 7.02272 | 7.09028 | 7.05362 |
| DNAJC10 | Q8IXB1 | 6.14917 | 6.31165 | 5.92329 | 6.32667 | 6.08188 | 6.06686 | 6.11702 | 6.01157 | 5.98355 | 6.00591 | 5.99501 | 6.12183 | 6.87977 | 6.66811 | 6.82866 | 6.52904 |
| ERP44 | Q9BS26 | 6.30501 | 6.04269 | 6.01139 | 6.17596 | 6.21452 | 6.00982 | 6.18818 | 6.19758 | 6.25677 | 5.91796 | 6.06785 | 6.127 | 6.7736 | 6.2872 | 6.57962 | 6.54484 |
| AGO2 | Q9UKV8 | 6.41597 | 6.24534 | 6.64273 | 6.38361 | 6.47407 | 6.46497 | 6.00788 | 6.15927 | 6.41845 | 6.35704 | 6.53446 | 6.33251 | 6.03081 | 5.94201 | 5.95546 | 5.63542 |
| NMT1 | P30419 | 6.23263 | 6.51949 | 6.59198 | 5.91629 | 6.28978 | 6.6489 | 6.20338 | 6.29829 | 6.39243 | 6.56472 | 6.47362 | 6.52908 | 5.61398 | 6.29661 | 5.72514 | 5.70369 |
| CORO1B | Q9BR76 | 6.27735 | 6.50202 | 6.25698 | 5.9045 | 6.64225 | 6.46146 | 6.10211 | 6.29287 | 6.44845 | 6.229 | 6.21124 | 5.99362 | 6.41591 | 6.24512 | 6.04817 | 5.96894 |
| DOCK5 | Q9H7D0 | 6.09267 | 6.15459 | 6.77397 | 6.03067 | 6.32651 | 6.6118 | 6.15152 | 6.47206 | 6.42752 | 6.50331 | 6.78343 | 6.54478 | 5.73927 | 5.92152 | 5.66714 | 5.79924 |
| PLBD1 | Q6P4A8 | 6.39313 | 6.45344 | 6.07378 | 6.42913 | 6.11008 | 5.97735 | 6.46752 | 6.01753 | 6.29196 | 6.35566 | 6.19801 | 6.27556 | 6.11209 | 6.28495 | 6.3075 | 6.25353 |
| TCF25 | Q9BQ70 | 6.28062 | 6.32222 | 6.52218 | 6.43905 | 6.18824 | 6.43502 | 6.24312 | 6.2389 | 6.34472 | 6.11609 | 6.39205 | 6.22019 | 5.89746 | 6.06282 | 6.20015 | 6.09717 |
| DOCK1 | Q14185 | 6.23504 | 6.23302 | 6.90766 | 6.09667 | 6.32589 | 6.64575 | 6.15125 | 6.5007 | 6.39539 | 6.49645 | 6.40655 | 6.25552 | 6.00342 | 6.01594 | 5.79423 | 5.53653 |
| PPIE | Q9UNP9 | 6.46583 | 6.31963 | 5.96561 | 6.33807 | 6.47161 | 6.34968 | 6.27463 | 6.55522 | 6.34954 | 6.06273 | 6.4966 | 6.16285 | 6.60638 | 5.83175 | 5.8272 | 5.92267 |
| DDX47 | Q9H0S4 | 6.22551 | 6.24693 | 5.85473 | 6.65615 | 6.42671 | 5.99906 | 6.58734 | 6.04928 | 6.18977 | 6.10355 | 5.8577 | 5.84646 | 6.61886 | 6.15269 | 6.64357 | 6.5417 |
| PSMB5 | P28074 | 6.22319 | 6.76446 | 6.29669 | 6.13398 | 6.38062 | 6.5306 | 6.14915 | 6.41488 | 6.10669 | 6.27227 | 6.07961 | 5.83682 | 6.2747 | 5.94852 | 6.66236 | 5.92545 |
| SMC5 | Q8IY18 | 6.36596 | 6.54806 | 5.92502 | 6.53435 | 6.11216 | 5.94901 | 6.39032 | 6.02586 | 6.20053 | 5.91018 | 5.93396 | 6.13957 | 6.51199 | 6.39256 | 6.49728 | 6.56319 |
| PICALM | Q13492 | 5.96145 | 6.12698 | 6.57056 | 6.36227 | 5.85378 | 6.0853 | 6.4188 | 6.17059 | 6.1127 | 6.18269 | 6.42395 | 6.57193 | 6.11393 | 6.23691 | 6.35781 | 6.45035 |
| CPSF3 | Q9UKF6 | 6.34923 | 6.26292 | 5.8232 | 6.67257 | 6.23184 | 6.12743 | 6.63408 | 6.0862 | 6.34502 | 5.90425 | 5.78068 | 6.07589 | 6.26777 | 6.31382 | 6.44194 | 6.68318 |
| FNDC3B | Q53EP0 | 6.0843 | 6.28831 | 6.06988 | 6.46768 | 6.34992 | 6.11004 | 6.6059 | 6.12748 | 6.04615 | 5.8365 | 6.0871 | 5.93343 | 6.72358 | 6.20033 | 6.57307 | 6.49631 |
| SIPA1 | Q96FS4 | 6.21862 | 6.59866 | 5.90326 | 6.33617 | 6.10802 | 6.32267 | 6.1261 | 6.26808 | 6.21958 | 6.28919 | 5.96001 | 6.35639 | 6.05673 | 6.44843 | 6.388 | 6.40011 |
| HACL1 | Q9UJ83 | 6.26163 | 5.95641 | 5.91432 | 6.36586 | 6.69336 | 6.2407 | 6.37423 | 6.23125 | 6.07349 | 6.09982 | 6.01447 | 6.03437 | 6.77075 | 6.22455 | 6.31956 | 6.42522 |
| SRI | P30626 | 5.73289 | 4.95812 | 5.73704 | 4.77178 | 5.33697 | 5.09759 | 5.56321 | 5.51642 | 4.93469 | 5.43565 | 5.02082 | 5.59835 | 4.93186 | 11.2322 | 8.43947 | 10.693 |
| CUL2 | Q13617-2 | 5.95697 | 6.07916 | 6.46634 | 5.91685 | 6.13773 | 6.5695 | 6.10742 | 5.71994 | 6.21966 | 6.62523 | 6.34986 | 6.01973 | 6.31309 | 6.39247 | 6.54591 | 6.58017 |
| RBM27 | Q9P2N5 | 6.00769 | 6.29155 | 5.82749 | 6.22452 | 6.31327 | 6.13824 | 6.0704 | 6.15333 | 6.06041 | 6.11513 | 5.84859 | 5.91288 | 6.84265 | 6.62096 | 6.78865 | 6.78424 |
| DIAPH3 | Q9NSV4 | 6.53678 | 6.24577 | 7.14062 | 6.29788 | 6.00254 | 6.41629 | 6.52951 | 6.23702 | 6.68952 | 6.30734 | 6.86164 | 7.1486 | 5.48237 | 5.37773 | 5.19852 | 5.52787 |
| U2AF2 | P26368 | 5.93099 | 6.16856 | 5.75838 | 6.48896 | 6.30887 | 6.07098 | 6.56513 | 6.0995 | 6.08632 | 5.91445 | 5.82376 | 5.73016 | 6.63595 | 6.554 | 6.93362 | 6.92657 |
| PSMD6 | Q15008 | 6.12075 | 6.08758 | 6.37382 | 6.09331 | 6.39223 | 6.36672 | 6.33044 | 5.91659 | 6.10827 | 5.87618 | 6.16578 | 6.10936 | 6.52659 | 6.58657 | 6.41315 | 6.53265 |
| NOP14 | P78316 | 6.07294 | 6.26931 | 5.53445 | 6.68694 | 6.31218 | 5.54759 | 6.52816 | 6.35994 | 6.51604 | 5.73129 | 5.84405 | 5.77033 | 6.71779 | 6.54416 | 6.56818 | 6.99664 |
| RBM6 | P78332 | 6.52205 | 6.5495 | 6.34041 | 6.93659 | 6.03398 | 5.85229 | 6.03492 | 5.89456 | 6.47455 | 5.96763 | 6.06924 | 6.32236 | 5.99202 | 6.1811 | 6.2711 | 6.5577 |
| LRWD1 | Q9UFC0 | 5.92678 | 6.04325 | 6.34806 | 6.40542 | 6.16114 | 5.83141 | 6.05967 | 6.13528 | 6.27533 | 6.19768 | 6.28137 | 6.46059 | 6.24687 | 6.16741 | 6.76712 | 6.69262 |
| DRG2 | P55039 | 6.30909 | 6.22463 | 6.89885 | 6.28235 | 6.35279 | 6.87871 | 6.43025 | 6.52095 | 6.21296 | 6.6699 | 6.75069 | 6.51056 | 5.62048 | 5.5351 | 5.42637 | 5.37633 |
| TBC1D4 | O60343 | 6.44855 | 6.37026 | 7.039 | 6.13091 | 6.23914 | 6.58132 | 6.10797 | 6.56477 | 6.38142 | 6.19578 | 6.62256 | 6.52733 | 5.91936 | 5.62702 | 5.51828 | 5.72635 |
| AIMP1 | Q12904-2 | 6.15947 | 6.07226 | 6.63958 | 5.91588 | 6.35117 | 6.48744 | 6.08089 | 6.37684 | 6.27135 | 6.40058 | 6.75879 | 6.71014 | 6.44102 | 5.94204 | 5.86948 | 5.52311 |
| NDC80 | O14777 | 6.67137 | 6.92095 | 6.90341 | 6.68042 | 6.48744 | 6.96714 | 6.53551 | 6.85268 | 6.62829 | 6.80062 | 6.68966 | 6.86104 | 4.76387 | 4.96054 | 4.45009 | 4.82698 |
| RBM10 | P98175-5 | 6.37127 | 6.34204 | 5.96846 | 6.44428 | 6.40821 | 6.1146 | 6.47909 | 5.95476 | 6.12007 | 6.12423 | 5.89593 | 5.77228 | 6.78162 | 6.18354 | 6.69561 | 6.34401 |
| DCTN2 | Q13561 | 6.63479 | 5.08733 | 6.2374 | 4.60018 | 6.35739 | 5.40702 | 5.91587 | 7.415 | 5.57392 | 6.53284 | 5.88089 | 7.45591 | 6.08086 | 7.15949 | 6.6028 | 7.05832 |
| ECI2 | O75521 | 6.307 | 6.25052 | 5.75389 | 6.48897 | 6.30351 | 6.30662 | 6.2482 | 5.9669 | 6.14946 | 6.2081 | 6.05108 | 5.95296 | 7.04894 | 6.13164 | 6.44806 | 6.38414 |
| MRPL3 | P09001 | 6.16982 | 6.50514 | 5.98865 | 7.22844 | 6.55083 | 6.29003 | 6.31675 | 6.19889 | 5.94351 | 6.1699 | 5.80566 | 6.02134 | 5.83149 | 6.13771 | 6.17046 | 6.67138 |
| YME1L1 | Q96TA2 | 6.56934 | 6.65534 | 6.05798 | 7.00404 | 6.08478 | 5.85998 | 6.2536 | 5.76311 | 6.23061 | 6.18799 | 6.33255 | 6.21554 | 6.10815 | 5.89086 | 6.46994 | 6.31619 |
| MCMBP | Q9BTE3-2 | 5.74147 | 5.83628 | 6.22285 | 5.39753 | 6.25479 | 6.41032 | 5.81348 | 6.31833 | 6.05329 | 6.02353 | 6.26366 | 6.28512 | 6.92771 | 6.8637 | 6.7219 | 6.86602 |
| EIF2AK4 | Q9P2K8 | 6.41624 | 6.42733 | 6.81706 | 5.97332 | 6.21119 | 6.55557 | 6.23767 | 6.32711 | 6.43386 | 6.55439 | 6.67948 | 6.50569 | 5.83002 | 5.81018 | 5.6748 | 5.54609 |
| TUT7 | Q5VYS8 | 5.99995 | 6.12611 | 6.68204 | 6.04507 | 6.14771 | 6.49127 | 6.45421 | 6.21604 | 6.32298 | 6.60007 | 6.56825 | 6.56432 | 5.86852 | 6.24464 | 5.82553 | 5.84329 |
| PCF11 | O94913 | 6.16326 | 6.46685 | 5.90682 | 6.67279 | 5.98297 | 6.10217 | 6.26479 | 6.21475 | 6.08594 | 6.13339 | 6.15935 | 5.95191 | 6.14208 | 6.4881 | 6.68883 | 6.57602 |
| EIF4H | Q15056 | 6.62805 | 6.78221 | 7.21761 | 6.16129 | 6.52865 | 7.08832 | 5.9607 | 6.6435 | 6.63839 | 6.49516 | 7.01073 | 6.96863 | 5.32209 | 5.14712 | 4.6304 | 4.77715 |
| KDM1A | O60341 | 6.07607 | 6.36658 | 6.2456 | 6.11158 | 6.28315 | 5.96237 | 6.57239 | 5.9772 | 6.34673 | 5.99231 | 6.25409 | 5.88647 | 6.45943 | 6.20869 | 6.75858 | 6.49875 |
| CCN1 | O00622 | 6.14664 | 4.42339 | 3.23571 | 5.87571 | 4.16359 | 5.02784 | 8.58902 | 4.93768 | 5.74614 | 4.648 | 4.05643 | 4.87041 | 10.9011 | 6.83742 | 10.2553 | 10.2856 |
| COBLL1 | Q53SF7-2 | 5.76918 | 5.79215 | 5.97662 | 5.79969 | 6.08945 | 6.24707 | 5.97084 | 5.52856 | 6.11709 | 6.32278 | 6.35734 | 6.49227 | 7.26714 | 6.71514 | 6.75855 | 6.79612 |
| RPL13A | P40429 | 6.06736 | 6.38651 | 5.98644 | 5.95242 | 5.97272 | 6.28454 | 6.47832 | 6.45823 | 6.83263 | 6.50651 | 6.40105 | 6.33138 | 5.8678 | 6.25712 | 5.92467 | 6.2923 |
| CUL1 | Q13616 | 5.6828 | 5.84104 | 6.03948 | 5.67259 | 6.11483 | 6.30673 | 6.17564 | 5.94957 | 6.11394 | 6.32349 | 6.15769 | 5.96658 | 6.88005 | 6.74771 | 7.24418 | 6.78367 |
| ACTR2 | P61160 | 6.17013 | 5.9871 | 6.66779 | 6.16391 | 6.23933 | 6.45757 | 6.44246 | 6.5035 | 6.35832 | 6.05449 | 6.31853 | 6.51749 | 5.8658 | 6.14612 | 5.93027 | 6.17719 |
| USP4 | Q13107 | 6.05368 | 6.05247 | 6.77724 | 6.02948 | 6.03158 | 6.59141 | 6.13848 | 6.53538 | 6.26265 | 6.51707 | 6.5731 | 6.67825 | 5.84556 | 6.15151 | 5.81703 | 5.94512 |
| ARMC8 | Q8IUR7 | 6.3288 | 6.52899 | 6.3554 | 6.14425 | 6.30961 | 6.14849 | 6.03276 | 6.15435 | 6.12606 | 6.21284 | 6.17662 | 5.97992 | 6.61106 | 6.353 | 6.5985 | 6.53935 |
| VPS13A | Q96RL7 | 6.26069 | 6.49261 | 6.17133 | 6.26522 | 6.01236 | 6.06726 | 6.1773 | 5.80183 | 6.39906 | 6.46112 | 6.11662 | 6.16644 | 6.36296 | 6.43313 | 6.462 | 6.35006 |
| CBX5 | P45973 | 5.92196 | 6.14414 | 5.75686 | 6.45774 | 6.27716 | 5.92495 | 6.42538 | 5.8578 | 6.20092 | 5.92559 | 5.99029 | 5.91191 | 6.82431 | 6.67982 | 6.71309 | 6.98808 |
| PCYOX1 | Q9UHG3 | 6.25491 | 6.26291 | 5.70922 | 6.39809 | 6.29345 | 5.71812 | 6.5311 | 5.98923 | 6.12266 | 5.96847 | 5.81552 | 5.82542 | 6.77852 | 6.45799 | 7.2247 | 6.64969 |
| NFS1 | Q9Y697 | 6.30253 | 6.37054 | 6.01348 | 6.64442 | 6.40797 | 5.92387 | 6.48041 | 5.95063 | 6.19028 | 6.20719 | 6.15367 | 6.03785 | 6.55525 | 6.08549 | 6.39224 | 6.28418 |
| UBE3C | Q15386 | 6.14009 | 6.36103 | 6.36206 | 6.32939 | 6.30762 | 6.44809 | 6.25898 | 6.36882 | 6.13421 | 6.43204 | 6.05482 | 6.16662 | 6.104 | 6.09796 | 6.2034 | 6.23088 |
| CHERP | Q8IWX8 | 6.47141 | 6.46291 | 6.03553 | 6.72439 | 6.325 | 6.16032 | 6.16096 | 5.92849 | 6.18277 | 5.97724 | 6.12712 | 6.12938 | 6.62408 | 5.94797 | 6.17417 | 6.56826 |
| ZBTB10 | Q96DT7 | 6.21642 | 6.65794 | 6.44039 | 6.78031 | 6.22266 | 6.18664 | 6.36022 | 5.99703 | 6.10997 | 6.31263 | 6.13436 | 6.11838 | 6.22032 | 6.00879 | 6.081 | 6.15294 |
| NOC3L | Q8WTT2 | 6.18434 | 6.25261 | 5.44946 | 6.70617 | 6.03283 | 5.79362 | 6.54558 | 6.14177 | 6.25255 | 5.94736 | 5.68289 | 5.75192 | 6.76199 | 6.56991 | 6.67293 | 7.25407 |
| COL12A1 | Q99715 | 6.28238 | 6.19027 | 6.02232 | 6.82 | 6.41129 | 6.17931 | 6.41973 | 6.21748 | 6.36295 | 6.07583 | 6.18468 | 6.35901 | 6.14163 | 5.78019 | 6.27846 | 6.27447 |
| VPS33A | Q96AX1 | 6.59796 | 6.59647 | 6.56377 | 6.42083 | 6.25726 | 6.30199 | 6.41629 | 6.35097 | 6.49742 | 6.655 | 6.55402 | 6.31011 | 5.68382 | 5.75012 | 5.48351 | 5.56047 |
| PSAP | P07602-3 | 6.82371 | 6.00332 | 5.84643 | 5.86922 | 6.12975 | 5.51345 | 5.70845 | 6.08368 | 5.59221 | 5.81303 | 5.68567 | 5.81652 | 6.22966 | 8.10155 | 6.69148 | 8.09188 |
| EPHX1 | P07099 | 6.38584 | 6.42451 | 5.92666 | 6.36987 | 6.11369 | 5.72465 | 6.35313 | 5.78704 | 6.08428 | 6.01976 | 6.18816 | 5.83878 | 6.67599 | 6.59042 | 6.87957 | 6.63766 |
| COMT | P21964 | 6.04029 | 6.22114 | 5.92214 | 6.26675 | 6.28441 | 6.24062 | 6.15098 | 6.13015 | 5.86729 | 6.20183 | 5.88577 | 6.15753 | 6.75822 | 6.70795 | 6.59512 | 6.5698 |
| FAM3C | Q92520 | 6.55032 | 6.33753 | 5.71372 | 6.26436 | 6.46692 | 6.27047 | 6.44406 | 6.22088 | 6.07007 | 5.93783 | 5.82234 | 5.82581 | 6.94689 | 6.23602 | 6.36586 | 6.52691 |
| COL6A1 | P12109 | 6.66074 | 6.71948 | 5.92028 | 6.57457 | 6.64002 | 6.1976 | 6.42414 | 6.18263 | 6.04838 | 6.36486 | 6.17503 | 6.16549 | 5.98518 | 5.87793 | 5.88306 | 6.18061 |
| CNDP2 | Q96KP4 | 6.11019 | 5.93389 | 6.93968 | 5.82472 | 6.49596 | 6.84779 | 6.06541 | 6.70337 | 6.29726 | 6.46031 | 6.74172 | 6.63544 | 5.91969 | 5.92674 | 5.63301 | 5.46483 |
| GGH | Q92820 | 6.54741 | 6.3933 | 6.18978 | 6.51301 | 6.48252 | 6.11303 | 6.37797 | 6.29188 | 6.16789 | 6.01889 | 6.06686 | 5.87252 | 6.41784 | 6.07474 | 6.27324 | 6.19911 |
| LARS2 | Q15031 | 6.33733 | 6.42262 | 5.94488 | 6.68884 | 6.35943 | 6.06525 | 6.4312 | 6.14716 | 5.9201 | 6.18462 | 5.91603 | 5.81235 | 6.6611 | 6.21211 | 6.5558 | 6.34119 |
| ABCB1 | P08183 | 6.11105 | 6.53635 | 6.04737 | 6.37647 | 5.79332 | 5.71209 | 6.11128 | 5.79136 | 6.59694 | 6.63114 | 6.38953 | 6.38258 | 6.30613 | 6.23487 | 6.53948 | 6.44004 |

| | | | | | | | | | | | | | | | | | |
|---|---|---|---|---|---|---|---|---|---|---|---|---|---|---|---|---|---|
| IMPA1 | P29218-3 | 6.21611 | 6.23767 | 6.8676 | 5.65616 | 6.49167 | 6.96007 | 6.30926 | 6.93423 | 6.37664 | 6.41893 | 6.39312 | 6.2524 | 5.85977 | 5.87564 | 5.67302 | 5.47771 |
| NELFE | P18615 | 5.60154 | 5.93035 | 6.07756 | 5.73982 | 6.0058 | 6.1419 | 6.3926 | 6.11961 | 5.96484 | 6.17845 | 6.20432 | 5.99423 | 6.89875 | 7.00584 | 6.91296 | 6.83143 |
| INCENP | Q9NQS7 | 6.1475 | 6.49582 | 5.73292 | 6.87903 | 6.32432 | 5.88629 | 6.41859 | 5.92537 | 6.09662 | 6.11299 | 6.07391 | 5.93373 | 6.28311 | 6.31146 | 6.42465 | 6.95369 |
| RBM42 | Q9BTD8 | 6.03402 | 6.26339 | 5.97765 | 6.45728 | 6.05244 | 5.92033 | 6.2574 | 6.18772 | 6.32336 | 6.25398 | 6.23397 | 6.12761 | 6.49629 | 6.3994 | 6.48623 | 6.52894 |
| PSMD4 | P55036 | 6.26657 | 6.29616 | 6.45023 | 6.15943 | 6.36924 | 6.44551 | 6.00661 | 6.29749 | 6.37317 | 6.40241 | 6.45412 | 6.18136 | 6.33924 | 5.92382 | 5.94067 | 6.09397 |
| MRPL11 | Q9Y3B7 | 6.57671 | 6.54892 | 5.93697 | 6.51598 | 6.39612 | 5.77606 | 6.4917 | 6.11834 | 6.21798 | 6.01004 | 5.85356 | 5.94064 | 6.57266 | 6.2066 | 6.41601 | 6.4217 |
| IRF2BPL | Q9H1B7 | 6.26917 | 6.00361 | 6.1149 | 5.74707 | 6.50845 | 6.25195 | 5.92205 | 6.1514 | 6.0655 | 5.97526 | 5.84623 | 5.89913 | 7.52263 | 6.48498 | 6.6094 | 6.62826 |
| ADK | P55263 | 6.12783 | 6.07525 | 6.83134 | 5.75944 | 6.39443 | 6.84462 | 6.1554 | 6.5294 | 6.26659 | 6.53588 | 6.53635 | 6.4099 | 5.83245 | 6.06881 | 6.05313 | 5.5792 |
| WDR70 | Q9NW82 | 5.63551 | 5.9635 | 6.19538 | 5.77226 | 5.91387 | 6.36948 | 5.64547 | 5.93638 | 6.22664 | 6.26054 | 6.08408 | 6.12568 | 6.49447 | 7.20704 | 7.17345 | 6.99624 |
| GNPDA1 | P46926 | 6.1831 | 6.12362 | 6.93013 | 6.01019 | 6.21094 | 6.7534 | 6.24512 | 6.29249 | 6.4155 | 6.50439 | 6.57635 | 6.40003 | 6.11621 | 5.79104 | 5.94745 | 5.50004 |
| ASPSCR1 | Q9BZE9-2 | 5.77213 | 5.8006 | 6.76078 | 5.56626 | 5.93003 | 6.2767 | 5.76708 | 6.04801 | 6.20292 | 6.19227 | 6.4119 | 6.14778 | 7.11407 | 6.45958 | 6.8308 | 6.71909 |
| TAF15 | Q92804 | 6.37074 | 6.35454 | 5.69462 | 6.38877 | 6.3968 | 5.94136 | 6.34446 | 6.0779 | 6.24703 | 5.98811 | 5.82322 | 5.96362 | 6.88045 | 6.51951 | 6.35788 | 6.65101 |
| CEP250 | Q9BV73 | 6.08641 | 6.30644 | 6.59113 | 6.0107 | 6.02697 | 6.47053 | 6.08149 | 6.24474 | 6.44348 | 6.44097 | 6.56573 | 6.65297 | 5.9817 | 6.10441 | 5.99433 | 5.99802 |
| PTPN23 | Q9H3S7 | 6.22915 | 6.64484 | 6.49576 | 6.07913 | 6.27518 | 6.59597 | 5.87967 | 6.67558 | 6.44405 | 6.34624 | 6.49558 | 6.26042 | 5.82096 | 5.96007 | 6.00158 | 5.79584 |
| PPP2R5D | Q14738 | 6.27994 | 6.39366 | 6.65625 | 6.09635 | 6.41486 | 6.70101 | 6.18837 | 6.56664 | 6.17649 | 6.28346 | 6.48093 | 6.33357 | 5.93814 | 5.92747 | 5.70557 | 5.85731 |
| TUBB | P07437 | 6.33135 | 6.19952 | 6.69036 | 5.78066 | 6.37014 | 6.79715 | 6.15613 | 7.26198 | 6.58739 | 6.41964 | 6.59905 | 6.98252 | 5.40995 | 5.77765 | 5.14305 | 5.49346 |
| WFS1 | O76024 | 6.47896 | 6.42619 | 5.94568 | 6.63581 | 6.28027 | 5.88633 | 6.51701 | 6.05676 | 6.0443 | 6.10545 | 5.90416 | 5.86752 | 6.49336 | 6.5213 | 6.49783 | 6.33908 |
| NDUFS3 | O75489 | 6.22016 | 6.31996 | 5.91 | 5.90556 | 6.47162 | 6.09474 | 6.31943 | 5.97211 | 5.68367 | 6.13119 | 6.29449 | 6.21059 | 6.73343 | 6.39146 | 6.68711 | 6.6545 |
| STAU1 | O95793 | 6.38077 | 6.61908 | 6.60928 | 6.21697 | 6.2175 | 6.59486 | 6.29028 | 6.34826 | 6.51747 | 6.61341 | 6.7642 | 6.70218 | 5.47944 | 5.73328 | 5.38511 | 5.52791 |
| STAU2 | Q9NUL3 | 6.19118 | 6.2878 | 6.6199 | 5.92923 | 6.13446 | 6.44092 | 6.40339 | 6.44543 | 6.17869 | 6.57515 | 6.38722 | 6.3651 | 6.58955 | 6.35749 | 5.91077 | 5.78374 |
| AKAP2 | Q9Y2D5-6 | 6.58779 | 6.27863 | 6.58728 | 6.21158 | 6.36541 | 6.10731 | 5.7112 | 5.64813 | 6.14965 | 5.7761 | 6.08992 | 6.18366 | 7.35093 | 6.32321 | 6.42001 | 6.20919 |
| MACROH2A2 | Q9P0M6 | 6.16831 | 6.37303 | 5.56458 | 6.48875 | 6.1193 | 5.93584 | 6.41935 | 6.14043 | 6.08502 | 6.02788 | 5.81002 | 5.61963 | 6.47421 | 6.79914 | 7.26894 | 6.70561 |
| CHD6 | Q8TD26 | 6.47791 | 6.47398 | 6.24592 | 6.51196 | 6.11576 | 6.04873 | 6.2507 | 5.97853 | 6.23788 | 6.46839 | 6.0834 | 6.44696 | 6.54715 | 6.47636 | 6.07198 | 6.46438 |
| FLOT1 | O75955 | 6.49298 | 7.06131 | 6.20931 | 6.89008 | 6.3373 | 6.16395 | 6.3858 | 6.12238 | 6.21937 | 6.6779 | 6.37235 | 6.33779 | 5.71124 | 5.58127 | 5.86109 | 5.57588 |
| ALDH5A1 | P51649-2 | 6.46819 | 6.34557 | 5.75143 | 6.46315 | 6.30645 | 5.98166 | 6.4723 | 6.12486 | 6.05745 | 6.00659 | 5.95359 | 5.84561 | 6.78118 | 6.35565 | 6.69665 | 6.38966 |
| LSS | P48449 | 6.53458 | 6.49491 | 6.1032 | 6.61042 | 6.26145 | 6.14008 | 6.38629 | 6.09568 | 6.05291 | 6.37393 | 6.03701 | 5.94174 | 6.14383 | 6.38529 | 6.34079 | 6.0979 |
| OAS3 | Q9Y6K5 | 5.94325 | 6.56453 | 6.26555 | 6.00438 | 6.70064 | 6.65578 | 5.9583 | 6.53183 | 6.36211 | 6.46414 | 6.54791 | 6.12174 | 6.18255 | 6.15563 | 5.84324 | 5.69842 |
| GNL3L | Q9NVN8 | 6.70916 | 6.64165 | 6.55524 | 6.57488 | 6.51849 | 6.50868 | 6.72029 | 6.21936 | 6.43678 | 6.40182 | 6.51442 | 6.3586 | 5.42083 | 5.4956 | 5.40531 | 5.51887 |
| URB2 | Q14146 | 6.28032 | 6.36519 | 5.85687 | 6.59609 | 6.1234 | 5.90301 | 6.56736 | 5.91039 | 6.2325 | 5.88166 | 5.9597 | 5.91591 | 6.65196 | 6.25377 | 6.64732 | 6.85454 |
| ACTR1B | P42025 | 6.31626 | 6.21754 | 6.63189 | 6.15186 | 6.28348 | 6.71378 | 6.09683 | 6.61804 | 6.4387 | 6.32579 | 6.4612 | 6.58988 | 5.77031 | 6.13887 | 5.62653 | 5.61906 |
| MAD1L1 | Q9Y6D9 | 6.26514 | 6.18773 | 6.24667 | 6.40479 | 6.43498 | 6.09755 | 6.24204 | 6.33743 | 6.24756 | 5.99479 | 6.23149 | 6.16911 | 6.74272 | 5.84482 | 6.30249 | 6.2507 |
| HSPE1 | P61604 | 5.7199 | 4.8911 | 5.22544 | 4.87633 | 6.09257 | 5.2344 | 5.85368 | 6.71335 | 5.72977 | 6.21663 | 5.67113 | 6.27932 | 6.74003 | 8.31309 | 7.99346 | 8.4498 |
| XPO4 | Q9C0E2 | 6.23657 | 6.18409 | 6.35286 | 5.96504 | 6.11486 | 6.16204 | 6.16006 | 6.08485 | 6.30681 | 6.21374 | 6.06071 | 5.95762 | 6.55706 | 6.37185 | 6.70112 | 6.57073 |
| PGM1 | P36871 | 6.21939 | 6.06359 | 6.64766 | 5.70111 | 6.42586 | 6.94 | 6.07714 | 6.799 | 6.36208 | 6.68602 | 6.45298 | 6.35253 | 5.77888 | 6.28945 | 5.70969 | 5.49462 |
| ALDH1L2 | Q3SY69 | 6.58605 | 6.35508 | 5.72168 | 6.3319 | 6.32492 | 5.98595 | 6.2926 | 6.02732 | 6.03256 | 5.99635 | 5.74382 | 5.85191 | 7.06206 | 6.49404 | 6.57192 | 6.62183 |
| SNTB2 | Q13425 | 5.92318 | 6.88213 | 5.78901 | 5.60687 | 5.74081 | 6.26972 | 6.37388 | 5.9861 | 6.29038 | 6.18965 | 6.13527 | 6.10257 | 6.02399 | 6.27771 | 7.68278 | 6.72595 |
| SRGAP2 | O75044 | 6.05058 | 6.12555 | 6.65203 | 6.05185 | 6.17287 | 6.77262 | 6.28624 | 6.27589 | 6.30252 | 6.22653 | 6.3827 | 6.3327 | 6.03439 | 6.25944 | 5.93295 | 6.14114 |
| ZNF148 | Q9UQR1 | 6.63246 | 6.46659 | 6.04158 | 7.08761 | 6.59269 | 5.97871 | 6.08467 | 6.0553 | 6.12587 | 6.47497 | 5.98015 | 6.17053 | 6.18872 | 6.13916 | 6.16596 | 6.45502 |
| MORC3 | Q14149 | 5.799 | 6.19846 | 5.99218 | 6.19039 | 5.90908 | 6.12403 | 5.94775 | 5.82611 | 6.18825 | 6.22598 | 6.19463 | 5.99118 | 7.08135 | 6.70086 | 6.79723 | 6.83353 |
| DYNC1LI2 | O43237 | 6.28987 | 6.07298 | 6.91276 | 5.83315 | 6.32119 | 6.67423 | 6.06425 | 6.66825 | 6.45101 | 6.40229 | 6.52123 | 6.39755 | 6.02808 | 5.97773 | 5.61131 | 5.77414 |
| LCOR | Q96JN0-3 | 6.40814 | 6.3784 | 6.17006 | 6.83914 | 6.24074 | 5.88307 | 6.04056 | 5.98274 | 5.99835 | 5.9753 | 5.8119 | 5.82241 | 6.01272 | 6.71201 | 6.69558 | 7.02887 |
| CSNK1A1 | P48729 | 6.65799 | 6.66444 | 6.64312 | 6.59906 | 6.49548 | 6.37028 | 6.79203 | 6.46753 | 6.78725 | 6.33755 | 6.25291 | 6.30642 | 5.1936 | 5.40005 | 5.35444 | 5.67785 |
| EPS15 | P42566 | 6.2642 | 5.99389 | 6.53082 | 5.69751 | 6.56083 | 6.36512 | 6.05005 | 6.537 | 6.64327 | 6.29121 | 6.60458 | 6.51056 | 6.52332 | 5.93672 | 5.67529 | 5.81563 |
| MAGEA4 | P43358 | 6.24562 | 6.02717 | 6.74729 | 5.94574 | 6.41633 | 6.68976 | 6.09844 | 6.46243 | 6.35354 | 6.40108 | 6.51279 | 6.43156 | 6.04508 | 6.07113 | 5.76003 | 5.79202 |
| LRRFIP1 | Q32MZ4 | 6.27079 | 6.11466 | 6.68901 | 5.77928 | 6.5736 | 6.65727 | 5.94005 | 6.62888 | 6.53595 | 6.28903 | 6.79416 | 6.50797 | 6.21672 | 5.85457 | 5.66136 | 5.48671 |
| GPKOW | Q92917 | 5.69592 | 6.16552 | 5.84462 | 5.86516 | 6.3979 | 6.49774 | 6.06279 | 6.18518 | 6.15266 | 6.0335 | 6.3969 | 6.21944 | 7.04434 | 6.76139 | 6.42206 | 6.25487 |
| FNDC3A | Q9Y2H6 | 6.23251 | 6.25556 | 6.07752 | 6.54396 | 6.20179 | 5.82544 | 6.41166 | 5.8417 | 6.29763 | 6.00328 | 6.02117 | 6.11065 | 6.724 | 6.41907 | 6.50837 | 6.52568 |
| APPL1 | Q9UKG1 | 6.48298 | 6.51666 | 6.88546 | 6.00776 | 6.5767 | 6.74712 | 6.33586 | 6.70691 | 6.51635 | 6.64216 | 6.50723 | 6.53964 | 5.61417 | 5.2678 | 5.37748 | 5.27572 |
| CUL4A | Q13619 | 6.20905 | 6.16612 | 6.60285 | 6.10421 | 6.44389 | 6.17334 | 6.40133 | 5.98329 | 6.23081 | 6.29008 | 5.9956 | 5.91972 | 6.50794 | 6.11691 | 6.51093 | 6.34393 |
| IVD | P26440 | 6.17493 | 5.92977 | 6.17205 | 6.58215 | 6.28958 | 6.09539 | 6.36461 | 6.05114 | 6.14381 | 6.09728 | 6.10384 | 6.1683 | 6.80957 | 6.09576 | 6.40597 | 6.51584 |
| NSFL1C | Q9UNZ2-5 | 5.91811 | 6.104 | 6.40371 | 5.88595 | 6.15021 | 6.59327 | 5.76105 | 6.75273 | 6.35867 | 6.28097 | 6.50614 | 6.84053 | 6.59626 | 6.06677 | 5.69793 | 6.08369 |
| XPNPEP1 | Q9NQW7-3 | 6.43888 | 6.26489 | 6.98529 | 6.03723 | 6.3617 | 6.81896 | 5.99824 | 6.46125 | 6.18463 | 6.52148 | 6.69144 | 6.57211 | 5.88036 | 5.82347 | 5.44104 | 5.51901 |
| SLC12A2 | P55011 | 6.49123 | 6.42588 | 6.33429 | 6.74295 | 6.0961 | 6.1321 | 6.25864 | 5.80661 | 6.29925 | 6.29992 | 6.33901 | 6.44712 | 6.12609 | 5.84138 | 6.18537 | 6.17409 |
| TXNL1 | O43396 | 6.13611 | 6.30852 | 7.18925 | 6.39187 | 6.15163 | 6.9127 | 6.01255 | 6.34197 | 5.88798 | 6.59322 | 6.27345 | 6.70893 | 5.82574 | 5.78762 | 5.5077 | 5.97074 |
| AASS | Q9UDR5 | 6.46034 | 6.40118 | 5.67603 | 6.49395 | 6.44263 | 6.24369 | 6.23653 | 5.9972 | 5.90019 | 6.00202 | 6.01692 | 5.99284 | 6.59261 | 6.44977 | 6.49206 | 6.60206 |
| TCF20 | Q9UGU0 | 6.34631 | 6.37631 | 6.2028 | 6.865 | 6.15177 | 6.15519 | 6.33369 | 5.99567 | 5.89703 | 6.29895 | 6.03713 | 5.97406 | 6.33814 | 6.33642 | 6.5195 | 6.17204 |
| SFXN1 | Q9H9B4 | 6.36269 | 5.88023 | 5.81065 | 6.1691 | 6.23841 | 5.78729 | 6.49897 | 5.95617 | 6.1527 | 6.09783 | 5.83437 | 6.00307 | 6.73051 | 6.51721 | 6.86494 | 7.09587 |
| STUB1 | Q9UNE7 | 6.42382 | 6.35829 | 6.58686 | 6.1561 | 6.20844 | 6.49139 | 5.94846 | 6.5306 | 6.42184 | 6.76998 | 6.80156 | 6.53453 | 6.10763 | 5.58575 | 5.48396 | 5.5908 |
| SETD2 | Q9BYW2 | 6.5909 | 6.72357 | 6.45115 | 7.21571 | 6.18149 | 5.94038 | 6.42766 | 6.04792 | 6.47104 | 6.06678 | 6.22333 | 6.22747 | 5.70318 | 5.73485 | 6.15683 | 5.83774 |
| LARP4B | Q92615 | 6.342 | 6.42281 | 6.47452 | 5.93402 | 6.54022 | 6.25615 | 6.20335 | 6.39347 | 6.30578 | 6.26628 | 6.43833 | 6.5308 | 5.97858 | 6.04631 | 5.94135 | 5.92603 |
| IPO9 | Q96P70 | 6.15056 | 6.04261 | 6.78721 | 5.97845 | 6.34353 | 6.81342 | 6.2928 | 6.42731 | 6.43863 | 6.6088 | 6.47636 | 6.2428 | 6.01145 | 5.76947 | 5.91493 | 5.70168 |
| TMED2 | Q15363 | 6.35045 | 6.30179 | 5.92035 | 6.44169 | 6.05865 | 5.88874 | 6.01907 | 6.08788 | 6.07353 | 5.90208 | 6.12796 | 6.10267 | 6.65402 | 6.57475 | 6.91061 | 6.58577 |
| DDX56 | Q9NY93 | 6.48099 | 6.25614 | 5.93172 | 6.85901 | 6.00325 | 5.79563 | 6.12213 | 5.96334 | 6.23685 | 5.95086 | 6.06691 | 6.1049 | 6.38312 | 6.44569 | 6.5894 | 6.81008 |
| AKT2 | P31751 | 6.17453 | 6.39107 | 6.72676 | 5.9363 | 6.29894 | 6.9042 | 6.23972 | 6.58194 | 6.26109 | 6.52637 | 6.49938 | 6.35546 | 5.60287 | 6.15274 | 5.73018 | 5.61844 |
| VPS41 | P49754 | 6.28667 | 6.22939 | 6.3455 | 5.96834 | 6.41363 | 6.40695 | 6.1702 | 6.16671 | 6.32963 | 6.55368 | 6.23733 | 6.23385 | 6.32907 | 6.30346 | 6.21243 | 5.81317 |
| EHHADH | Q08426 | 6.3077 | 6.16814 | 6.02683 | 6.38314 | 6.16779 | 5.86283 | 6.37887 | 6.00344 | 6.34532 | 6.16433 | 6.2213 | 6.03637 | 6.49988 | 6.30724 | 6.70102 | 6.42581 |
| SLC25A1 | P53007 | 6.44054 | 6.41229 | 5.91003 | 6.69937 | 6.14044 | 5.96245 | 6.42723 | 5.82406 | 6.07053 | 6.00737 | 6.0635 | 5.89922 | 6.64971 | 6.28719 | 6.72663 | 6.47943 |
| THOP1 | P52888 | 6.23604 | 6.0848 | 6.90963 | 5.76918 | 6.43292 | 6.76696 | 6.27849 | 6.49143 | 6.54658 | 6.63721 | 6.60908 | 6.50495 | 5.6067 | 5.95807 | 5.68202 | 5.48596 |
| WDR46 | O15213 | 6.11494 | 6.15956 | 5.58654 | 6.44694 | 6.1009 | 5.91583 | 6.20502 | 6.16285 | 5.93841 | 5.86173 | 5.60606 | 5.7462 | 6.78074 | 7.06005 | 7.28583 | 7.0284 |
| EIF2B3 | Q9NR50 | 6.03029 | 5.69685 | 6.79342 | 6.02233 | 6.47034 | 6.57981 | 6.08845 | 6.45097 | 6.60696 | 6.71781 | 6.68935 | 6.82945 | 6.05907 | 5.68352 | 5.66014 | 5.62123 |
| TMEM214 | Q6NUQ4 | 6.4169 | 6.41564 | 5.74531 | 6.76039 | 6.30646 | 6.16386 | 6.44565 | 6.05138 | 6.04491 | 6.21852 | 5.9475 | 5.97282 | 6.38946 | 6.37869 | 6.38824 | 6.35426 |
| SUGT1 | Q9Y2Z0 | 6.1116 | 6.07906 | 6.52585 | 6.07554 | 6.31222 | 6.68748 | 6.08144 | 6.39771 | 6.5134 | 6.43727 | 6.38831 | 6.4483 | 6.28171 | 6.12804 | 5.83818 | 5.69389 |
| NUDT21 | O43809 | 4.85552 | 5.03939 | 5.76124 | 5.51071 | 6.44209 | 5.53933 | 6.02158 | 6.58912 | 6.11636 | 6.79942 | 6.33359 | 6.93329 | 7.13593 | 6.53687 | 7.36176 | 7.02383 |
| KIFBP | Q96EK5 | 6.26473 | 6.3895 | 6.61221 | 6.32417 | 6.28636 | 6.43643 | 6.49097 | 6.53948 | 6.52926 | 6.2501 | 6.34475 | 6.23189 | 5.8098 | 5.8714 | 5.82048 | 5.69847 |
| TELO2 | Q9Y4R8 | 6.56567 | 6.6271 | 6.74074 | 6.34095 | 6.45461 | 6.61469 | 5.97846 | 6.64845 | 6.81727 | 6.70566 | 6.59634 | 6.8552 | 5.29393 | 5.50815 | 5.16413 | 5.08866 |
| TXLNA | P40222 | 6.58957 | 6.29568 | 6.86633 | 6.18164 | 6.47837 | 6.56731 | 6.23562 | 6.46504 | 6.99938 | 6.70647 | 6.63858 | 6.75688 | 5.32045 | 5.39514 | 5.08996 | 5.41359 |
| RFC2 | P35250 | 6.00235 | 6.16814 | 6.13112 | 6.20456 | 6.47842 | 6.30722 | 6.09027 | 6.36744 | 6.13395 | 6.17772 | 6.01705 | 6.32046 | 6.52714 | 6.19894 | 6.35448 | 6.54144 |
| CYFIP2 | Q96F07 | 6.47136 | 6.08749 | 6.64007 | 5.95663 | 6.32045 | 6.42613 | 6.215 | 6.34663 | 6.42199 | 6.47308 | 6.62076 | 6.47468 | 5.95163 | 5.91667 | 5.92605 | 5.75138 |
| NOP9 | Q86U38 | 5.989 | 6.23729 | 5.62867 | 6.39667 | 6.34841 | 6.09742 | 6.31954 | 6.20951 | 6.11092 | 6.18228 | 6.02841 | 5.89233 | 6.74767 | 6.44822 | 6.5025 | 6.86115 |
| PRDX5 | P30044 | 5.91078 | 5.47735 | 5.98415 | 5.31233 | 6.41141 | 6.11295 | 6.02541 | 6.69253 | 6.31649 | 6.62039 | 6.37562 | 6.5654 | 6.36685 | 6.49896 | 6.76677 | 6.56261 |
| RIOX1 | Q9H6W3 | 5.89275 | 6.30144 | 5.84811 | 6.89258 | 5.99763 | 5.65969 | 6.40886 | 5.94012 | 6.02179 | 6.02258 | 6.06345 | 6.10833 | 6.78905 | 6.39534 | 6.71561 | 6.94267 |
| COLGALT1 | Q8NBJ5 | 6.26249 | 6.3052 | 5.83615 | 6.91883 | 5.97408 | 5.87035 | 6.48479 | 5.93228 | 6.19752 | 5.58495 | 5.84903 | 5.97194 | 6.54239 | 6.47923 | 6.81422 | 6.97657 |
| DDX31 | Q9H8H2 | 6.16954 | 5.90159 | 5.81278 | 6.61251 | 6.04446 | 5.7249 | 6.39569 | 5.87535 | 6.30142 | 6.10469 | 6.14925 | 6.0241 | 6.7663 | 6.66031 | 6.64604 | 6.81109 |
| MORC2 | Q9Y6X9 | 5.80545 | 5.98125 | 6.14882 | 6.3684 | 5.99065 | 6.02025 | 6.30052 | 5.80462 | 6.05068 | 6.12784 | 6.10205 | 5.94032 | 6.95613 | 6.48496 | 7.04906 | 6.86899 |
| AKAP8 | O43823 | 6.56176 | 6.35602 | 5.9455 | 6.69277 | 6.0583 | 6.06037 | 6.19733 | 6.0544 | 6.31754 | 6.13531 | 6.04896 | 6.25324 | 6.69368 | 6.09881 | 6.17107 | 6.35494 |
| SPTLC2 | O15270 | 6.4214 | 6.68242 | 6.34769 | 6.82829 | 6.26986 | 6.00816 | 6.17955 | 6.05877 | 6.15957 | 6.16695 | 6.23519 | 6.08617 | 6.30089 | 5.84603 | 6.44801 | 5.96105 |
| KIDINS220 | Q9ULH0 | 6.65797 | 6.53925 | 6.31527 | 6.84466 | 6.11779 | 6.05917 | 6.22277 | 5.92288 | 6.26883 | 6.1518 | 6.0434 | 6.33353 | 6.14705 | 6.09188 | 6.14162 | 6.14211 |
| FBXL18 | Q96ME1-4 | 6.05386 | 5.88094 | 6.70787 | 5.77476 | 6.00258 | 6.54758 | 6.01622 | 6.25347 | 6.2868 | 6.61735 | 6.53277 | 6.4135 | 6.15986 | 6.43058 | 6.35857 | 5.96327 |
| TIMM50 | Q3ZCQ8-2 | 6.51228 | 6.34163 | 5.94053 | 6.70461 | 6.56091 | 6.0818 | 6.21805 | 6.24822 | 6.17791 | 5.98565 | 6.27599 | 6.42376 | 6.54385 | 5.98198 | 6.01909 | 5.98373 |
| WDR5 | P61964 | 6.0919 | 6.45802 | 5.99895 | 6.30402 | 6.50899 | 6.31957 | 6.31704 | 6.16963 | 6.00578 | 6.04929 | 5.93442 | 5.74548 | 6.82967 | 6.45157 | 6.61892 | 6.19676 |
| CC2D1A | Q6P1N0 | 6.169 | 6.26698 | 6.85336 | 6.26071 | 6.18614 | 6.59604 | 6.31596 | 6.27526 | 6.51043 | 6.77147 | 6.67934 | 6.71103 | 5.61855 | 5.85504 | 5.18416 | 5.74653 |
| ANAPC4 | Q9UJX5-3 | 6.2275 | 6.17278 | 6.40615 | 6.54349 | 6.252 | 6.12111 | 6.54001 | 6.15576 | 6.41794 | 6.06033 | 6.10706 | 6.38334 | 5.93226 | 6.07907 | 6.18114 | 6.42006 |
| TNPO3 | Q9Y5L0-1 | 5.91373 | 6.01341 | 6.15583 | 5.66788 | 6.11215 | 6.47895 | 6.07827 | 6.37146 | 6.37118 | 6.32951 | 6.19175 | 6.26126 | 6.45692 | 6.60489 | 6.39887 | 6.59394 |
| NDC1 | Q9BTX1 | 6.27418 | 6.2245 | 5.69313 | 6.80237 | 6.08005 | 5.87301 | 6.50695 | 6.05489 | 6.21641 | 6.01001 | 5.87651 | 5.90784 | 6.59363 | 6.31379 | 6.68918 | 6.88354 |
| ETHE1 | O95571 | 6.38317 | 6.16864 | 6.10864 | 6.36026 | 6.41538 | 5.91738 | 6.06601 | 6.27136 | 6.19737 | 6.06218 | 6.17259 | 6.22733 | 6.72953 | 5.98617 | 6.49128 | 6.44273 |
| MED1 | Q15648 | 6.28298 | 6.52174 | 5.95067 | 6.74028 | 6.10466 | 5.98314 | 6.26337 | 5.83945 | 5.84287 | 6.04209 | 6.09299 | 5.97553 | 6.16016 | 6.73206 | 6.67053 | 6.79748 |
| POLR1G | O15446-2 | 5.82766 | 5.81173 | 5.77107 | 6.70272 | 5.80598 | 5.84859 | 6.43756 | 5.6719 | 6.00764 | 5.71593 | 5.66775 | 5.92488 | 6.33249 | 7.4753 | 7.21999 | 7.77881 |
| P3H1 | Q32P28 | 6.48758 | 6.59674 | 5.93721 | 6.54478 | 6.24195 | 6.29053 | 6.02245 | 6.09697 | 5.81522 | 6.15995 | 6.16718 | 6.15926 | 6.59417 | 6.27859 | 6.31284 | 6.29458 |
| NVL | O15381 | 6.14589 | 6.56103 | 6.27159 | 6.82628 | 6.14295 | 5.92216 | 6.20339 | 5.86917 | 6.41704 | 6.08654 | 6.15033 | 6.27114 | 6.23826 | 6.06373 | 6.3864 | 6.4441 |
| U2AF1 | Q01081-2 | 5.96157 | 6.1841 | 5.93224 | 6.32593 | 5.992 | 5.76626 | 6.43301 | 5.79639 | 6.24251 | 6.23761 | 6.13828 | 5.89746 | 6.56738 | 6.72144 | 6.85112 | 6.9527 |
| CDC42 | P60953 | 6.33954 | 5.97095 | 6.16077 | 5.86088 | 6.37641 | 6.16421 | 6.17368 | 6.53072 | 6.28237 | 6.43335 | 6.27538 | 6.39328 | 6.25909 | 6.39907 | 6.18287 | 6.19741 |
| TFIP11 | Q9UBB9 | 6.09714 | 6.25302 | 6.09584 | 6.24289 | 6.14061 | 6.06789 | 6.4414 | 5.9859 | 6.06151 | 6.23881 | 6.05807 | 6.13126 | 6.61988 | 6.44955 | 6.731 | 6.38523 |
| CYRIB | Q9NUQ9 | 6.3464 | 6.32603 | 6.49711 | 6.15516 | 6.17449 | 6.54522 | 6.09664 | 6.61911 | 6.15668 | 6.45325 | 6.46052 | 6.22926 | 5.93431 | 6.27207 | 5.72079 | 6.01295 |
| ERCC3 | P19447 | 5.93798 | 6.15582 | 6.1313 | 6.42657 | 6.08779 | 6.24745 | 6.47799 | 5.95511 | 6.19431 | 6.32915 | 6.21182 | 6.24619 | 6.38948 | 6.18363 | 6.59856 | 6.42685 |
| CORO7 | P57737-3 | 6.33013 | 6.04681 | 6.26994 | 5.64202 | 6.67415 | 6.59114 | 6.48724 | 6.64347 | 6.28058 | 6.10357 | 6.04304 | 6.10495 | 6.42895 | 6.17041 | 6.1301 | 6.0535 |

| | | | | | | | | | | | | | | | | | |
|---|---|---|---|---|---|---|---|---|---|---|---|---|---|---|---|---|---|
| ZC3H13 | Q5T200 | 6.19417 | 6.37459 | 6.03172 | 6.70128 | 5.86515 | 5.98054 | 5.83496 | 5.39872 | 6.07351 | 5.87051 | 6.19792 | 6.234 | 6.34535 | 6.95057 | 6.84979 | 7.09721 |
| EXOC4 | Q96A65 | 6.34688 | 6.4268 | 6.57814 | 5.97918 | 6.43056 | 6.55693 | 6.29817 | 6.45959 | 6.50554 | 6.56588 | 6.4132 | 6.30643 | 5.82558 | 5.89237 | 5.6337 | 5.78106 |
| AKAP1 | Q92667 | 6.41609 | 6.22777 | 6.18132 | 6.59944 | 6.16552 | 5.74882 | 6.29777 | 5.88306 | 6.20369 | 5.8136 | 6.08795 | 5.92706 | 6.73434 | 6.2949 | 6.71017 | 6.70851 |
| NCEH1 | Q6PIU2 | 6.22468 | 5.92602 | 6.08313 | 6.64278 | 6.38755 | 5.84492 | 6.42757 | 6.143 | 5.81535 | 5.60241 | 5.92492 | 5.77038 | 6.76286 | 6.58725 | 6.78243 | 7.07475 |
| DOCK10 | Q96BY6 | 5.80036 | 5.89636 | 6.06973 | 5.85696 | 5.78984 | 6.27005 | 6.2232 | 6.20051 | 6.47268 | 6.29974 | 6.40208 | 6.39749 | 6.60815 | 6.80481 | 6.4363 | 6.47175 |
| POGLUT3 | Q7Z4H8 | 6.45661 | 6.35893 | 6.10501 | 6.64615 | 6.36303 | 6.2536 | 6.44168 | 6.06441 | 5.99559 | 5.9421 | 6.39961 | 6.28102 | 6.25145 | 5.9929 | 6.25572 | 6.1922 |
| EIF3M | Q7L2H7 | 6.39955 | 6.38593 | 6.74605 | 6.17051 | 6.51872 | 6.68067 | 6.20948 | 6.49151 | 6.38102 | 6.44763 | 6.5735 | 6.37611 | 5.52984 | 5.71826 | 5.69285 | 5.67837 |
| ATP6V1C1 | P21283 | 6.31485 | 6.26246 | 6.56445 | 6.10545 | 6.36959 | 6.45982 | 6.058 | 6.80829 | 6.28843 | 6.65748 | 6.55545 | 6.66494 | 5.78907 | 5.89612 | 5.52074 | 5.68486 |
| CALM1 | P0DP23 | 7.16554 | 5.8561 | 6.0953 | 5.07229 | 5.62569 | 5.4494 | 5.61694 | 6.81146 | 5.8568 | 7.10726 | 6.5433 | 7.20436 | 5.77507 | 6.99588 | 6.35294 | 6.47167 |
| LGALS1 | P09382 | 5.29127 | 4.68042 | 4.29317 | 4.28719 | 4.98886 | 4.10075 | 5.42894 | 6.83702 | 4.27862 | 5.39643 | 4.30094 | 6.12296 | 5.43127 | 12.2436 | 9.51697 | 12.8016 |
| CDKAL1 | Q5VV42 | 6.36887 | 6.31685 | 5.96467 | 6.73422 | 5.99857 | 6.16585 | 6.47362 | 6.14867 | 6.07596 | 6.2157 | 6.20312 | 6.23439 | 6.21663 | 6.30369 | 6.11586 | 6.46332 |
| PTPN12 | Q05209 | 6.23615 | 6.16666 | 6.7203 | 6.14545 | 6.35032 | 6.67461 | 6.0443 | 6.75683 | 6.44376 | 6.35299 | 6.32877 | 6.69853 | 5.70876 | 5.96362 | 5.62187 | 5.78706 |
| MRPS22 | P82650 | 6.39381 | 6.34974 | 5.56586 | 6.52327 | 6.53975 | 6.24148 | 6.31396 | 6.40707 | 6.32732 | 6.18142 | 6.18606 | 6.30805 | 6.31844 | 5.94127 | 6.16041 | 6.24207 |
| ZMPSTE24 | O75844 | 6.15923 | 6.07008 | 5.72438 | 6.47038 | 6.35413 | 6.21301 | 6.35685 | 6.11242 | 6.1775 | 6.01633 | 5.82505 | 5.83316 | 6.96736 | 6.61249 | 6.59656 | 6.51105 |
| CHAF1A | Q13111 | 6.50996 | 6.8825 | 6.19888 | 7.33648 | 6.17241 | 6.14613 | 6.28636 | 6.02446 | 6.04994 | 6.44123 | 6.23229 | 6.42699 | 5.81762 | 5.80312 | 5.57985 | 6.09179 |
| RAP1GDS1 | P52306-5 | 6.42355 | 5.72432 | 6.95508 | 5.37355 | 6.4068 | 6.49053 | 6.3513 | 6.57292 | 6.28823 | 6.60009 | 6.35351 | 6.94218 | 5.70629 | 6.09638 | 5.72955 | 5.98572 |
| SF3A2 | Q15428 | 6.35863 | 6.43547 | 5.79803 | 6.58804 | 6.50774 | 6.21729 | 6.07581 | 6.00716 | 6.06968 | 6.38575 | 5.92528 | 5.91386 | 6.52104 | 6.14784 | 6.60916 | 6.43922 |
| MAT2A | P31153 | 5.88487 | 6.14506 | 6.71559 | 5.57881 | 6.09257 | 6.30313 | 6.14651 | 6.42076 | 6.10678 | 6.13669 | 6.4233 | 6.29741 | 6.23654 | 6.59973 | 6.64238 | 6.26987 |
| CRKL | P46109 | 6.10115 | 5.92191 | 6.52555 | 5.62417 | 6.46643 | 6.32757 | 5.80083 | 6.83326 | 6.46404 | 6.57634 | 6.72702 | 6.92544 | 5.79521 | 5.9775 | 5.9198 | 6.01379 |
| CDC37 | Q16543 | 6.01884 | 5.84269 | 6.67352 | 5.77566 | 6.13102 | 6.4724 | 6.06508 | 6.43349 | 6.24285 | 6.11188 | 6.43117 | 6.36777 | 6.51964 | 6.38152 | 6.31364 | 6.21882 |
| MTDH | Q86UE4 | 6.10968 | 6.58296 | 5.91983 | 6.49285 | 5.94633 | 5.92486 | 6.33993 | 6.01117 | 6.03212 | 6.1451 | 5.83717 | 5.74312 | 6.19873 | 7.13325 | 6.8286 | 6.7543 |
| NIFK | Q9BYG3 | 6.24992 | 6.44435 | 5.83457 | 6.94145 | 6.18705 | 5.833 | 6.34234 | 6.04819 | 5.83442 | 5.67519 | 5.80633 | 5.93105 | 6.40586 | 6.4919 | 7.00857 | 6.96583 |
| TBCE | Q15813 | 6.2258 | 6.16761 | 6.86276 | 5.91349 | 6.56858 | 6.84124 | 6.39503 | 6.87187 | 6.37792 | 6.28353 | 6.58476 | 6.47092 | 5.87433 | 5.61458 | 5.40927 | 5.53833 |
| PRKAR1A | P10644 | 5.68414 | 5.93387 | 6.22638 | 6.09443 | 5.98129 | 6.21648 | 5.96897 | 5.94645 | 6.50984 | 6.78862 | 6.85461 | 6.7105 | 6.3428 | 6.12487 | 6.43511 | 6.18164 |
| GLS | O94925-3 | 6.41416 | 6.06417 | 6.02609 | 6.86184 | 6.37059 | 5.87434 | 6.46899 | 5.8959 | 6.23347 | 5.91309 | 5.83321 | 6.03344 | 6.76881 | 6.07956 | 6.53072 | 6.63161 |
| SNRPA1 | P09661 | 6.0592 | 6.3973 | 5.87458 | 6.52775 | 6.2079 | 6.13765 | 6.36613 | 6.1793 | 5.86947 | 5.92741 | 6.07691 | 6.31396 | 6.79045 | 6.31837 | 6.47393 | 6.47967 |
| INPPL1 | O15357 | 6.27899 | 6.18285 | 6.48919 | 5.75761 | 6.39697 | 6.57154 | 6.16379 | 6.45953 | 6.19975 | 6.52137 | 6.3849 | 6.43908 | 6.00976 | 6.38038 | 5.83184 | 5.93245 |
| BCAR1 | P56945-6 | 6.44085 | 6.34081 | 7.37456 | 5.89898 | 6.33273 | 6.73395 | 6.38918 | 6.92818 | 6.38104 | 6.26237 | 6.91558 | 6.78866 | 5.44084 | 5.44866 | 5.22914 | 5.09446 |
| RAB3GAP2 | Q9H2M9 | 5.71381 | 6.07721 | 6.41089 | 5.74252 | 6.0175 | 6.2202 | 5.80594 | 6.26206 | 6.2335 | 6.2432 | 6.14816 | 6.4035 | 6.52794 | 6.82309 | 6.71385 | 6.65665 |
| FMNL3 | Q8IVF7 | 6.16513 | 6.50029 | 6.24588 | 6.12502 | 6.43608 | 6.26905 | 6.34205 | 5.90461 | 6.61694 | 6.3795 | 6.19765 | 6.24271 | 5.98328 | 6.0212 | 6.33811 | 6.23248 |
| AGK | Q53H12 | 6.23597 | 6.52191 | 5.88894 | 6.63217 | 6.29703 | 5.99272 | 6.20213 | 5.8354 | 6.22675 | 5.91309 | 6.00127 | 5.80068 | 6.89957 | 6.30417 | 6.7056 | 6.54257 |
| ATP2C1 | P98194-7 | 6.60853 | 6.46188 | 6.20741 | 6.77593 | 6.27705 | 6.11164 | 6.11385 | 6.00449 | 6.44091 | 6.15336 | 6.00714 | 6.07963 | 6.10358 | 6.09942 | 6.4658 | 6.08938 |
| ZNF512B | Q96KM6 | 6.54723 | 6.49198 | 5.81296 | 6.89793 | 5.78577 | 5.65522 | 6.26358 | 6.16284 | 5.94255 | 6.35982 | 5.59566 | 5.89429 | 6.30182 | 6.71775 | 6.78555 | 6.78507 |
| YWHAG | P61981 | 6.23329 | 6.21152 | 6.24131 | 5.61738 | 6.59925 | 6.58558 | 6.58125 | 6.52026 | 6.61955 | 6.52932 | 6.59096 | 6.11765 | 6.27215 | 6.30542 | 6.19675 | 6.07836 |
| LMAN1 | P49257 | 6.15958 | 6.28116 | 5.80108 | 6.28923 | 5.9961 | 5.84332 | 6.20851 | 5.94938 | 6.20479 | 6.0985 | 5.93189 | 6.06186 | 6.67066 | 6.61819 | 7.05437 | 6.83137 |
| SAMD4B | Q5PRF9 | 6.3057 | 6.36046 | 6.94652 | 5.96575 | 6.24278 | 6.70786 | 6.22383 | 6.53842 | 6.44505 | 6.49417 | 6.82258 | 6.69336 | 5.78809 | 5.78587 | 5.37312 | 5.30643 |
| COPS2 | P61201-2 | 6.10489 | 6.24547 | 6.34123 | 6.04272 | 6.42195 | 6.52339 | 6.16272 | 6.33624 | 6.14974 | 6.16547 | 6.26065 | 6.09473 | 6.5281 | 6.26749 | 6.32857 | 6.02664 |
| MAP7D1 | Q3KQU3 | 6.14904 | 6.31601 | 6.614 | 5.92632 | 6.08193 | 6.41455 | 6.0488 | 6.51436 | 6.30826 | 6.58617 | 6.40787 | 6.48441 | 5.93612 | 6.25728 | 6.02125 | 5.93364 |
| ESYT2 | H7BXI1 | 6.38321 | 6.43409 | 5.78861 | 6.47066 | 6.30449 | 6.1842 | 6.25918 | 5.8002 | 6.12826 | 6.18506 | 5.78806 | 5.68684 | 6.84611 | 6.41601 | 6.77738 | 6.54767 |
| HK2 | P52789 | 5.86327 | 5.92871 | 6.06131 | 5.82775 | 6.98049 | 7.07316 | 7.23695 | 6.59686 | 5.5513 | 5.48114 | 5.68063 | 5.59225 | 6.77246 | 6.27389 | 6.63732 | 6.44251 |
| VPS53 | Q5VIR6 | 6.34027 | 6.33935 | 6.4987 | 5.852 | 6.45795 | 6.66093 | 6.09612 | 6.45675 | 6.31094 | 6.51153 | 6.31399 | 6.41735 | 6.05814 | 6.09382 | 5.80608 | 5.78607 |
| TOR1AIP1 | Q5JTV8 | 6.34357 | 6.43466 | 5.85984 | 6.88131 | 6.32667 | 5.91967 | 6.2323 | 5.99808 | 6.20196 | 5.79773 | 6.05712 | 5.91216 | 6.72608 | 6.34656 | 6.49238 | 6.46992 |
| GNL1 | P36915 | 6.08723 | 6.08395 | 7.00161 | 6.20255 | 5.91579 | 6.40594 | 6.4053 | 6.64624 | 6.27981 | 6.4593 | 6.62139 | 6.37135 | 5.5905 | 6.22696 | 5.85226 | 5.84984 |
| NXF2 | Q9GZY0 | 6.4062 | 6.5309 | 6.23989 | 6.83924 | 6.32795 | 6.23841 | 6.4338 | 5.87607 | 6.23861 | 6.5144 | 6.30699 | 6.42738 | 6.07162 | 5.52251 | 6.07591 | 5.95013 |
| ERLIN1 | O75477 | 6.43763 | 6.5047 | 6.12119 | 6.48394 | 6.06612 | 6.25799 | 6.25355 | 6.01798 | 6.23162 | 5.86836 | 6.03011 | 6.21449 | 6.41196 | 6.12478 | 6.474 | 6.50158 |
| CALCOCO2 | Q13137-4 | 6.8335 | 6.87803 | 7.50191 | 6.37989 | 6.60613 | 6.94273 | 6.31375 | 6.79423 | 6.54423 | 6.51679 | 6.77829 | 6.82841 | 4.75153 | 4.87177 | 4.51245 | 4.94637 |
| OPTN | Q96CV9 | 6.44938 | 6.24862 | 6.92981 | 6.21656 | 6.54298 | 6.90095 | 6.07151 | 6.6055 | 6.26314 | 6.45189 | 6.74718 | 6.90265 | 5.95558 | 5.49332 | 5.08603 | 5.13491 |
| SEPTIN10 | Q9P0V9-2 | 6.14914 | 6.25325 | 6.45753 | 6.04651 | 6.37047 | 6.56278 | 6.17999 | 6.57619 | 6.38123 | 6.46103 | 6.27304 | 6.34141 | 5.9864 | 6.14928 | 6.09039 | 5.72136 |
| DDX19B | Q9UMR2 | 6.35279 | 6.02459 | 6.7177 | 6.03805 | 6.54759 | 6.65151 | 6.31018 | 6.40245 | 6.40641 | 6.43648 | 6.59554 | 6.65215 | 5.88391 | 5.80773 | 5.65883 | 5.71409 |
| SETD3 | Q86TU7 | 6.52869 | 6.34703 | 6.98533 | 6.36647 | 6.09244 | 6.33095 | 6.27105 | 6.37756 | 6.64447 | 6.46819 | 6.80608 | 6.86183 | 5.49217 | 5.66964 | 5.41165 | 5.34646 |
| RNPEP | Q9H4A4 | 5.98998 | 6.17983 | 6.89345 | 5.79497 | 6.18747 | 6.81316 | 6.26789 | 6.69214 | 6.3125 | 6.37638 | 6.45449 | 6.35346 | 5.82533 | 6.23996 | 5.88725 | 5.73174 |
| NDUFV3 | P56181-2 | 6.40525 | 6.43999 | 5.96525 | 6.79906 | 6.01582 | 5.88954 | 6.151 | 5.8528 | 6.21541 | 6.14159 | 6.06155 | 6.09202 | 6.80384 | 6.25204 | 6.53124 | 6.38362 |
| ARMT1 | Q9H993 | 5.94161 | 5.91473 | 6.8824 | 5.41376 | 6.35013 | 7.07418 | 6.16402 | 6.67794 | 6.15899 | 6.66335 | 6.27069 | 6.39382 | 5.90099 | 6.51824 | 5.74503 | 5.93013 |
| ALDH6A1 | Q02252 | 6.19614 | 6.45279 | 5.81368 | 6.28158 | 6.38293 | 6.14706 | 5.99525 | 6.07201 | 6.14695 | 6.11267 | 5.79962 | 5.96762 | 7.16053 | 6.50601 | 6.52523 | 6.43994 |
| LSG1 | Q9H089 | 6.6598 | 6.47537 | 6.27485 | 6.37845 | 5.99774 | 6.05343 | 6.31338 | 6.08144 | 5.91802 | 5.94605 | 5.72707 | 5.82693 | 6.62945 | 6.50405 | 6.57288 | 6.64107 |
| MARK3 | P27448 | 6.18696 | 6.18462 | 6.39651 | 6.13973 | 5.95865 | 6.19977 | 6.15593 | 6.05571 | 6.22881 | 6.34076 | 6.58286 | 6.60182 | 6.32157 | 6.06906 | 6.14266 | 6.43458 |
| CHORDC1 | Q9UHD1 | 5.95606 | 6.14063 | 6.69562 | 5.95679 | 6.36597 | 6.85622 | 6.25732 | 6.4142 | 6.33629 | 6.45332 | 6.75942 | 6.49217 | 5.92536 | 5.84621 | 5.68632 | 5.8581 |
| CKAP2L | Q8IYA6 | 6.25209 | 5.70578 | 6.16209 | 5.87452 | 5.94665 | 6.08114 | 6.15727 | 5.89017 | 6.46096 | 6.36023 | 6.51167 | 6.54221 | 6.20242 | 6.65761 | 6.37569 | 6.8195 |
| PCCA | P05165 | 6.40473 | 6.29724 | 6.0365 | 6.57789 | 6.03367 | 5.95573 | 6.3076 | 5.87754 | 6.27232 | 6.22975 | 5.9951 | 5.84212 | 6.94019 | 6.23107 | 6.85624 | 6.14231 |
| PGM2 | Q96G03 | 6.08816 | 6.19976 | 6.70807 | 5.71837 | 6.12766 | 6.58295 | 6.08175 | 6.46586 | 6.25671 | 6.39417 | 6.5409 | 6.48393 | 6.03837 | 6.10849 | 6.44898 | 5.75588 |
| CBR1 | P16152 | 6.23589 | 6.00424 | 6.8161 | 5.83828 | 6.58773 | 6.88733 | 6.04518 | 6.58507 | 6.24698 | 6.56638 | 6.73776 | 6.56116 | 6.04712 | 5.93541 | 5.58872 | 5.31664 |
| TYK2 | P29597 | 6.42143 | 6.64857 | 6.4645 | 6.58042 | 6.34077 | 6.40453 | 6.1338 | 6.03008 | 6.55032 | 6.57888 | 6.62931 | 6.67025 | 5.70129 | 5.62354 | 5.47917 | 5.74314 |
| MPP6 | Q9NZW5 | 6.26334 | 6.22153 | 5.93189 | 6.57769 | 6.14638 | 6.07795 | 6.19688 | 6.11371 | 6.28436 | 5.98591 | 6.01598 | 6.17296 | 6.63509 | 6.13486 | 6.60319 | 6.63828 |
| PRPSAP1 | Q14558-2 | 6.18055 | 6.21848 | 6.91728 | 6.04627 | 6.27175 | 6.76608 | 6.0955 | 6.71323 | 6.17446 | 6.43252 | 6.41149 | 6.44058 | 5.82677 | 6.09437 | 5.66256 | 5.74814 |
| CNN2 | Q99439 | 6.40216 | 6.09515 | 6.78027 | 5.97597 | 6.6226 | 6.58717 | 6.27218 | 6.92822 | 6.38756 | 6.12956 | 6.71033 | 6.64348 | 6.0269 | 5.73365 | 5.22489 | 5.47991 |
| ANAPC7 | Q9UJX3 | 6.39083 | 6.71939 | 6.25095 | 6.68506 | 6.51684 | 6.50532 | 6.48127 | 6.45012 | 6.40849 | 6.1302 | 6.45139 | 6.05558 | 5.59167 | 5.83553 | 5.79155 | 5.73581 |
| ZFYVE26 | Q68DK2 | 6.77514 | 6.79797 | 6.76269 | 6.76845 | 6.19809 | 6.41559 | 6.19597 | 6.28031 | 6.49606 | 6.65594 | 6.89213 | 6.68946 | 5.17448 | 5.45326 | 5.24545 | 5.19001 |
| CLIC4 | Q9Y696 | 6.14273 | 5.92662 | 6.70406 | 5.38559 | 6.54305 | 6.33151 | 5.74011 | 6.50202 | 6.26637 | 6.30312 | 6.10792 | 6.441 | 6.55347 | 6.63066 | 6.27101 | 6.15077 |
| KMT2D | O14686-3 | 6.40601 | 6.65641 | 6.08666 | 6.67603 | 5.95515 | 6.08258 | 6.24406 | 6.08891 | 6.29776 | 6.41195 | 6.25823 | 6.47829 | 6.19031 | 6.03885 | 6.10487 | 6.07501 |
| NSUN5 | Q96P11-2 | 5.9404 | 6.1232 | 5.94929 | 6.05411 | 6.2473 | 6.04436 | 6.31183 | 6.02915 | 6.29512 | 6.40835 | 6.10905 | 6.21776 | 6.62023 | 6.39983 | 6.6952 | 6.55483 |
| MFN1 | Q8IWA4 | 6.64589 | 6.29072 | 6.21883 | 7.04951 | 6.59727 | 6.53866 | 6.38392 | 6.14407 | 6.25829 | 6.98939 | 6.0183 | 6.51543 | 6.16609 | 5.93606 | 5.69881 | 6.14877 |
| LDLR | P01130 | 5.67915 | 5.89289 | 5.99894 | 6.40203 | 5.41774 | 5.48121 | 5.79907 | 5.60332 | 6.05944 | 6.30318 | 6.45732 | 6.12376 | 7.27288 | 6.85173 | 7.46883 | 7.1885 |
| TAP2 | X5CMH5 | 6.20122 | 6.33618 | 5.84245 | 6.59214 | 6.12399 | 6.07964 | 6.23584 | 6.01881 | 5.90846 | 6.0491 | 6.16369 | 6.01872 | 6.30429 | 6.58606 | 6.88202 | 6.6574 |
| LAMC1 | P11047 | 6.54388 | 6.59363 | 6.17282 | 6.83674 | 6.48189 | 6.46567 | 6.23828 | 6.31673 | 6.20751 | 6.17045 | 6.16033 | 6.36862 | 6.16984 | 5.72017 | 5.65323 | 5.90019 |
| FECH | P22830-2 | 6.49537 | 6.61659 | 5.83865 | 6.39434 | 6.30387 | 5.81293 | 6.3688 | 5.98716 | 6.15363 | 5.99709 | 6.00291 | 5.96329 | 6.92255 | 6.49923 | 6.41904 | 6.22456 |
| CRNKL1 | Q9BZJ0 | 6.33478 | 6.4545 | 6.04396 | 6.77609 | 6.32587 | 5.93319 | 6.37153 | 5.86615 | 6.219 | 6.09645 | 5.94188 | 5.90692 | 6.29481 | 6.39534 | 6.54748 | 6.49205 |
| SMARCA2 | P51531 | 6.32134 | 6.58806 | 5.69172 | 6.51743 | 6.26107 | 5.85284 | 6.20891 | 6.04542 | 5.92128 | 6.20335 | 5.71612 | 5.93438 | 6.5896 | 6.66938 | 6.82229 | 6.65682 |
| CKAP2 | Q8WWK9 | 6.13102 | 5.95717 | 6.22693 | 6.09731 | 5.80364 | 5.92748 | 6.13657 | 5.99994 | 6.74936 | 6.63979 | 6.89714 | 6.86476 | 6.00192 | 6.19245 | 6.01568 | 6.35885 |
| TTK | P33981 | 6.36713 | 6.66458 | 6.92825 | 6.29984 | 6.08332 | 6.75306 | 6.4733 | 6.71561 | 6.74769 | 6.67948 | 6.64565 | 6.65961 | 5.06429 | 5.33632 | 5.0659 | 5.51598 |
| GALNT7 | Q86SF2 | 6.33453 | 6.42372 | 5.87649 | 6.49905 | 6.23113 | 5.92862 | 6.57969 | 5.90646 | 6.17264 | 6.03773 | 6.02016 | 5.80501 | 6.56554 | 6.35837 | 6.81835 | 6.44443 |
| RHOT2 | Q8IXI1 | 6.60334 | 6.5494 | 5.98548 | 6.42887 | 6.54588 | 6.30486 | 6.66387 | 6.20553 | 6.28299 | 5.99615 | 5.80688 | 5.83557 | 6.27925 | 5.96842 | 6.36835 | 6.17516 |
| DCAF8 | Q5TAQ9 | 6.0343 | 6.15314 | 6.47576 | 6.11725 | 6.2704 | 6.43635 | 6.18164 | 6.09008 | 6.2637 | 6.16154 | 6.52092 | 6.31229 | 6.58403 | 5.92911 | 6.30935 | 6.16014 |
| PBXIP1 | Q96AQ6 | 6.69321 | 6.97465 | 6.5039 | 6.62251 | 6.12626 | 6.18858 | 6.29019 | 6.20793 | 6.09818 | 6.47241 | 6.14703 | 6.20636 | 5.87045 | 5.96004 | 5.91733 | 5.72097 |
| DLG1 | Q12959-7 | 6.65276 | 6.52215 | 6.23251 | 6.76854 | 5.92036 | 6.35261 | 6.27081 | 6.08954 | 6.19604 | 6.08323 | 6.13752 | 5.75064 | 6.17686 | 6.14586 | 6.17125 | 6.52932 |
| NT5DC2 | Q9H857-2 | 6.64931 | 6.72204 | 5.89228 | 7.054 | 6.15491 | 6.0703 | 6.15733 | 6.19297 | 6.47053 | 6.4956 | 6.16412 | 6.3727 | 6.03117 | 5.64886 | 6.05006 | 5.87383 |
| HERC1 | Q15751 | 6.24195 | 6.4885 | 6.9766 | 6.174 | 5.95747 | 6.39954 | 5.95981 | 6.42655 | 6.65487 | 6.39263 | 6.43614 | 6.8768 | 5.66362 | 5.9593 | 5.66892 | 5.7233 |
| GLOD4 | Q9HC38-2 | 6.44233 | 5.9297 | 6.90522 | 5.85962 | 6.5287 | 6.6375 | 6.01502 | 6.5744 | 6.41408 | 6.37171 | 6.63175 | 6.61559 | 6.0946 | 5.84261 | 5.49723 | 5.63995 |
| NACA | E9PAV3 | 6.75553 | 6.5873 | 7.11467 | 6.10877 | 6.3614 | 6.71516 | 6.25775 | 6.4902 | 6.85192 | 6.67507 | 7.06753 | 7.11073 | 5.16297 | 5.23578 | 4.79384 | 4.9114 |
| CYP51A1 | Q16850 | 6.86967 | 6.74389 | 6.43162 | 6.73886 | 6.54831 | 6.11811 | 6.46017 | 5.83852 | 6.39337 | 6.14922 | 6.5018 | 6.31418 | 5.70449 | 5.56694 | 5.91086 | 5.70999 |
| CYFIP1 | Q7L576 | 6.08364 | 6.19908 | 6.57743 | 5.85693 | 6.06543 | 6.57985 | 6.25932 | 6.37676 | 6.56056 | 6.55071 | 6.5427 | 6.36413 | 6.00452 | 6.12949 | 6.06675 | 5.78268 |
| COG2 | Q14746 | 6.49384 | 6.31713 | 6.44718 | 6.6892 | 6.25798 | 6.28891 | 6.31132 | 6.29069 | 6.58781 | 6.43886 | 6.13074 | 6.14783 | 6.14214 | 5.82437 | 5.95363 | 5.67837 |
| RIPOR1 | Q6ZS17-4 | 6.16695 | 6.03544 | 6.82168 | 6.15888 | 6.34673 | 6.72503 | 6.25057 | 6.68537 | 6.45236 | 6.3834 | 6.48706 | 6.48486 | 5.96213 | 5.69629 | 5.6583 | 5.68496 |
| PARK7 | Q99497 | 6.47925 | 5.77257 | 6.81831 | 5.00985 | 7.07842 | 6.37983 | 5.96953 | 7.03505 | 6.2866 | 6.53381 | 6.33466 | 6.64243 | 6.188 | 6.19837 | 5.69136 | 5.58197 |
| SUGP1 | Q8IWZ8 | 6.29316 | 6.46604 | 6.23057 | 6.31361 | 6.11296 | 5.94925 | 6.31155 | 6.04332 | 6.35369 | 6.51433 | 6.23815 | 6.40478 | 6.48878 | 6.01166 | 6.20307 | 6.06507 |
| CDCP1 | Q9H5V8 | 6.02358 | 6.27559 | 6.00888 | 6.58225 | 6.09882 | 5.97559 | 6.37725 | 5.93293 | 6.11081 | 6.14435 | 5.9913 | 5.93167 | 6.63856 | 6.32511 | 6.79506 | 6.78824 |
| PIK3R4 | Q99570 | 6.29485 | 6.28146 | 6.8034 | 6.21165 | 6.44989 | 6.50064 | 6.36907 | 6.56109 | 6.54713 | 6.42143 | 6.76265 | 6.47834 | 5.75078 | 5.66247 | 5.57295 | 5.33221 |
| CDC16 | Q13042 | 6.27888 | 6.38952 | 6.14767 | 6.20633 | 6.19785 | 6.16347 | 6.34542 | 6.2974 | 6.21102 | 5.91561 | 6.08376 | 6.11621 | 6.35242 | 6.26914 | 6.40997 | 6.61533 |
| DDX28 | Q9NUL7 | 6.43895 | 6.14548 | 5.96712 | 7.09649 | 6.11262 | 5.84855 | 6.51833 | 5.83644 | 6.07205 | 6.24325 | 6.17667 | 5.96391 | 6.53148 | 6.31336 | 6.35647 | 6.37883 |
| POM121 | Q96HA1 | 6.34592 | 6.20421 | 6.0353 | 6.78418 | 6.09558 | 5.73844 | 6.46427 | 5.97317 | 6.17508 | 6.11906 | 6.17788 | 6.13193 | 6.48954 | 6.28066 | 6.48428 | 6.50052 |
| CGN | Q9P2M7 | 6.45597 | 6.54496 | 6.40792 | 6.79544 | 6.28064 | 6.30509 | 6.00982 | 6.32675 | 6.34663 | 6.31438 | 6.06642 | 6.41071 | 6.28191 | 5.80162 | 5.85212 | 5.7996 |
| MELTF | P08582 | 6.43912 | 6.54278 | 5.91775 | 6.81448 | 6.45869 | 6.53234 | 6.05935 | 6.26742 | 6.2938 | 6.07725 | 6.2949 | 5.69176 | 6.29335 | 6.06091 | 6.24051 | 6.01558 |
| KPNA4 | O00629 | 6.02321 | 6.07116 | 6.52588 | 6.18655 | 6.24989 | 6.4924 | 5.88759 | 5.90919 | 6.32655 | 6.07936 | 6.23626 | 6.25238 | 6.48144 | 6.14659 | 6.48797 | 6.64358 |
| GUF1 | Q8N442 | 6.39578 | 6.48718 | 5.79423 | 6.79056 | 6.36521 | 6.21275 | 6.22103 | 6.12138 | 6.22317 | 6.02677 | 5.89051 | 5.93707 | 6.53965 | 6.38382 | 6.41137 | 6.19952 |
| THUMPD1 | Q9NXG2 | 5.23013 | 5.52699 | 6.07753 | 5.3599 | 5.49946 | 6.19813 | 5.45077 | 5.65755 | 5.73094 | 6.24688 | 6.22715 | 6.08258 | 7.2898 | 7.71065 | 7.99256 | 7.71897 |
| ZC3H11A | O75152 | 6.20293 | 6.02211 | 6.39328 | 6.91638 | 6.19327 | 6.01444 | 6.30589 | 6.00376 | 5.97676 | 6.00057 | 5.88295 | 6.03622 | 6.30185 | 6.3965 | 6.5506 | 6.80248 |

| | | | | | | | | | | | | | | | | | |
|---|---|---|---|---|---|---|---|---|---|---|---|---|---|---|---|---|---|
| PSMA4 | P25789 | 5.97376 | 6.21139 | 6.58041 | 6.09247 | 6.36904 | 6.22562 | 6.26176 | 6.21871 | 6.3824 | 5.94312 | 6.3967 | 6.09495 | 6.08393 | 6.14444 | 6.52631 | 6.49499 |
| ACO1 | P21399 | 6.32701 | 6.09287 | 6.78761 | 5.87284 | 6.2868 | 6.85035 | 6.22674 | 6.44945 | 6.33926 | 6.60146 | 6.5544 | 6.21038 | 5.74639 | 6.08272 | 5.8149 | 5.65683 |
| FBN2 | P35556 | 6.57455 | 6.77112 | 6.04708 | 6.02672 | 6.56673 | 6.55039 | 6.6481 | 6.35782 | 6.01456 | 5.85391 | 6.59808 | 6.42434 | 6.48691 | 5.88329 | 5.74441 | 6.06199 |
| CPNE3 | O75131 | 6.25216 | 6.17865 | 6.90905 | 5.92309 | 6.43295 | 6.64641 | 6.34122 | 6.26489 | 6.38952 | 6.48119 | 6.5668 | 6.37962 | 5.86341 | 5.9546 | 5.81778 | 5.59865 |
| PPP2R1B | P30154-2 | 6.28424 | 6.06072 | 6.79521 | 5.70102 | 6.45745 | 6.5706 | 6.1345 | 6.45823 | 6.11109 | 6.35369 | 6.39645 | 6.47586 | 6.02174 | 6.00694 | 6.12174 | 6.05051 |
| UFD1 | Q92890-1 | 5.73013 | 6.38825 | 5.79042 | 6.1428 | 6.33765 | 6.41891 | 6.16941 | 6.69016 | 6.37822 | 6.44774 | 6.02501 | 6.22977 | 6.03779 | 6.46633 | 6.1747 | 6.57271 |
| USP19 | O94966-5 | 6.4158 | 6.36698 | 6.58136 | 6.25079 | 6.35478 | 6.37676 | 6.54934 | 6.24808 | 6.41139 | 6.55262 | 6.37189 | 6.23622 | 5.72923 | 5.84816 | 5.94743 | 5.75917 |
| ZMYM2 | Q9UBW7 | 6.41621 | 6.7375 | 6.23051 | 6.55008 | 6.23727 | 6.33136 | 6.31858 | 5.92479 | 6.23066 | 6.36478 | 6.18537 | 6.25365 | 6.17558 | 6.10238 | 5.91776 | 6.02351 |
| ERCC2 | P18074 | 6.35031 | 6.30391 | 6.64647 | 6.42868 | 6.18404 | 6.31932 | 6.40576 | 6.43104 | 6.68977 | 6.62343 | 6.68418 | 6.43682 | 5.47503 | 5.6153 | 5.6127 | 5.79324 |
| AKR1A1 | P14550 | 6.04961 | 5.88538 | 6.46112 | 5.14856 | 6.58757 | 6.60155 | 5.99431 | 7.16989 | 6.26976 | 6.66554 | 6.56446 | 6.69882 | 5.97717 | 6.16385 | 5.97315 | 5.78927 |
| BRCA1 | P38398-7 | 6.37392 | 6.46851 | 6.45353 | 6.88219 | 5.8242 | 5.80332 | 6.43745 | 5.74134 | 6.30952 | 6.47647 | 6.31294 | 6.385 | 6.25846 | 6.10646 | 6.13683 | 6.02985 |
| FLOT2 | Q14254 | 6.37923 | 6.51521 | 6.01154 | 6.42517 | 6.3482 | 6.14954 | 6.30553 | 6.1345 | 6.36442 | 6.00958 | 6.03664 | 6.06096 | 6.29988 | 6.42093 | 6.33847 | 6.2002 |
| TES | Q9UGI8 | 6.47066 | 6.19998 | 6.70883 | 5.52198 | 6.53326 | 6.70235 | 6.11454 | 6.69343 | 6.41174 | 6.47061 | 6.42441 | 6.35408 | 6.3031 | 5.93615 | 5.61094 | 5.54394 |
| SLMAP | Q14BN4 | 6.25098 | 6.20881 | 6.02847 | 6.29368 | 6.31793 | 5.96935 | 6.32849 | 6.14906 | 6.13244 | 5.99858 | 6.03839 | 6.00601 | 6.80603 | 6.20377 | 6.7061 | 6.5619 |
| RELCH | Q9P260-2 | 6.17966 | 6.3068 | 7.00835 | 6.01236 | 6.35389 | 6.55084 | 5.98698 | 6.28337 | 6.10825 | 6.37252 | 6.5081 | 6.2781 | 6.15294 | 6.38876 | 5.7108 | 5.7983 |
| MAGOHB | Q96A72 | 6.4161 | 6.0602 | 5.97038 | 5.92178 | 6.3529 | 5.92262 | 6.01774 | 6.06098 | 6.31397 | 6.19899 | 6.12646 | 6.31635 | 6.85172 | 6.05398 | 7.02526 | 6.39056 |
| INTS6 | Q9UL03 | 6.27408 | 6.39906 | 5.8418 | 6.72325 | 6.2317 | 5.84735 | 6.4568 | 6.07333 | 6.18618 | 6.26818 | 5.8961 | 5.90586 | 6.65728 | 6.27259 | 6.46438 | 6.50208 |
| PDE12 | Q6L8Q7 | 6.48224 | 6.22916 | 6.21866 | 6.55125 | 6.36448 | 6.25258 | 6.35555 | 6.41432 | 6.50205 | 6.0218 | 6.11498 | 6.0758 | 6.01457 | 6.06219 | 6.19142 | 6.14893 |
| ADGRE5 | P48960 | 6.68626 | 6.42974 | 6.06106 | 6.51126 | 6.41969 | 6.34996 | 6.27229 | 6.32314 | 6.43984 | 6.11445 | 6.30789 | 6.31181 | 6.30215 | 5.84268 | 5.68636 | 5.94141 |
| CTCF | P49711 | 6.20879 | 6.40992 | 5.90848 | 6.95847 | 5.99047 | 6.06165 | 6.24103 | 6.15196 | 5.84432 | 6.13822 | 5.95315 | 6.03075 | 6.2941 | 6.51604 | 6.79951 | 6.49314 |
| PRKAA1 | Q13131-2 | 5.48444 | 5.63747 | 5.94364 | 5.49226 | 5.84211 | 6.19937 | 5.89657 | 6.15835 | 6.0224 | 6.22277 | 6.33105 | 6.33766 | 7.02594 | 7.26668 | 6.99919 | 7.14009 |
| ITGB5 | P18084 | 6.44503 | 6.64221 | 6.42527 | 6.37214 | 6.21011 | 6.1555 | 6.20129 | 6.02474 | 6.36788 | 6.47791 | 6.30338 | 6.10004 | 6.24552 | 5.83067 | 6.19928 | 5.99903 |
| ZC3H7A | Q8IWR0 | 5.45666 | 6.755 | 6.95498 | 6.26755 | 6.39585 | 6.40254 | 6.15788 | 6.15802 | 6.39657 | 6.7649 | 6.53167 | 6.5698 | 5.44979 | 5.71635 | 5.49667 | 5.52576 |
| CASK | O14936 | 6.38099 | 6.73289 | 6.11609 | 6.66795 | 6.22347 | 6.09696 | 6.11197 | 6.16267 | 6.24728 | 6.1822 | 5.99012 | 6.21155 | 6.23596 | 6.11368 | 6.23918 | 6.28705 |
| RBBP7 | Q16576 | 5.89579 | 6.00614 | 6.0356 | 5.88837 | 6.56775 | 6.09612 | 6.15167 | 6.23932 | 6.1227 | 5.91368 | 6.33341 | 6.14336 | 6.95561 | 6.63831 | 6.63062 | 6.38156 |
| G3BP2 | Q9UN86 | 6.28499 | 6.42527 | 6.62589 | 6.12205 | 6.55654 | 6.37942 | 6.36886 | 6.38443 | 6.7114 | 6.57329 | 6.56618 | 6.5707 | 5.69783 | 5.46282 | 5.56766 | 5.70267 |
| REPS1 | Q96D71 | 6.44483 | 6.25795 | 6.12776 | 5.882 | 6.40791 | 6.22211 | 6.11731 | 6.20807 | 6.81855 | 6.56068 | 6.28991 | 6.70694 | 6.07994 | 5.84166 | 6.05913 | 5.97521 |
| CDC40 | O60508 | 6.50071 | 6.53012 | 5.89164 | 6.70755 | 6.16623 | 6.12837 | 6.39083 | 6.10339 | 6.14253 | 6.192 | 6.11561 | 5.98537 | 6.40402 | 5.99705 | 6.48296 | 6.26161 |
| MRE11 | P49959-3 | 6.272 | 6.16867 | 6.03813 | 6.34323 | 6.3336 | 6.1734 | 6.14154 | 6.08201 | 6.03782 | 6.2272 | 6.16039 | 5.95348 | 6.73326 | 6.44853 | 6.4367 | 6.45004 |
| ARHGEF12 | Q9NZN5 | 6.51185 | 6.46653 | 6.82641 | 6.00706 | 6.36101 | 6.64567 | 6.03679 | 6.32032 | 6.17606 | 6.22432 | 6.32082 | 6.26666 | 6.06034 | 6.03085 | 5.79528 | 5.95005 |
| EXOSC2 | Q13868 | 6.27604 | 6.33655 | 5.96419 | 6.67691 | 6.27818 | 6.19585 | 6.07647 | 6.10229 | 6.13122 | 6.15043 | 6.19838 | 6.21004 | 6.28461 | 6.07956 | 6.73807 | 6.30121 |
| RPS6 | P62753 | 6.1123 | 6.36357 | 6.0523 | 5.96632 | 6.05794 | 6.30594 | 6.39749 | 6.69724 | 6.74438 | 6.56869 | 6.31443 | 6.47173 | 5.7835 | 6.18323 | 5.87057 | 6.11035 |
| VPS26A | O75436 | 6.14577 | 6.5051 | 6.03748 | 6.15677 | 6.157 | 6.6236 | 6.26401 | 6.58968 | 6.21673 | 5.94779 | 6.28662 | 6.11053 | 5.72154 | 6.46168 | 6.58801 | 6.1877 |
| WDR18 | Q9BV38 | 6.32077 | 6.51723 | 5.87059 | 6.58098 | 6.40625 | 6.0232 | 6.5698 | 6.00081 | 6.16578 | 5.88206 | 5.86863 | 6.12808 | 6.52023 | 6.06262 | 6.64026 | 6.44273 |
| TRIP4 | Q15650 | 6.40011 | 6.39684 | 6.64699 | 6.03821 | 6.36808 | 6.51065 | 6.18836 | 6.37871 | 6.42557 | 6.48069 | 6.35841 | 6.2804 | 5.92292 | 6.08535 | 5.7121 | 5.80661 |
| RB1 | P06400 | 5.31977 | 5.71381 | 6.20876 | 5.48437 | 5.31399 | 6.10826 | 5.62244 | 5.93167 | 5.8942 | 6.23972 | 6.27271 | 6.06664 | 6.77409 | 7.64679 | 8.00476 | 7.39803 |
| TNRC6B | Q9UPQ9 | 6.17919 | 6.30658 | 6.8799 | 6.1577 | 6.22329 | 6.36485 | 5.8981 | 6.25702 | 6.41459 | 6.61601 | 6.86143 | 6.72463 | 6.20134 | 5.67731 | 5.7049 | 5.53315 |
| CSPP1 | Q1MSJ5 | 6.31695 | 6.22299 | 6.56334 | 6.28871 | 5.96372 | 6.73515 | 6.51184 | 6.564 | 6.49899 | 6.50144 | 6.4871 | 6.53843 | 5.67948 | 5.82791 | 5.5457 | 5.75424 |
| PPFIA1 | Q13136 | 6.09748 | 5.97059 | 5.89731 | 5.99992 | 6.25544 | 6.10842 | 6.09674 | 5.8394 | 6.48152 | 6.20956 | 6.12048 | 6.10839 | 6.8637 | 6.62822 | 6.76376 | 6.5591 |
| CDH2 | P19022 | 6.64695 | 6.77202 | 6.08026 | 6.70811 | 6.1676 | 5.97818 | 6.00051 | 6.11997 | 6.37 | 6.32967 | 6.05924 | 6.32636 | 6.31289 | 6.04912 | 5.91146 | 6.16767 |
| PRPS2 | P11908-2 | 6.49255 | 6.07278 | 6.99659 | 6.10512 | 6.69371 | 6.71676 | 6.42201 | 6.5649 | 6.23642 | 6.16214 | 6.52883 | 6.2768 | 5.90329 | 5.73118 | 5.59909 | 5.49785 |
| CALD1 | Q05682-4 | 5.82712 | 5.93046 | 6.34592 | 6.08063 | 5.88299 | 5.92866 | 6.04615 | 6.12717 | 6.11129 | 5.73641 | 6.16664 | 6.17064 | 7.02348 | 6.81809 | 6.99936 | 6.80499 |
| FEN1 | P39748 | 5.46195 | 5.88677 | 6.04133 | 5.65489 | 5.74814 | 6.46996 | 5.5168 | 6.13759 | 5.82899 | 6.23571 | 6.43728 | 6.23927 | 6.37442 | 7.70998 | 7.48678 | 6.77011 |
| ACBD5 | Q5T8D3-3 | 6.36813 | 6.11673 | 6.01809 | 6.58906 | 6.39806 | 5.9395 | 6.40456 | 5.98914 | 6.37597 | 6.10375 | 6.28284 | 6.14926 | 6.71153 | 5.90139 | 6.38699 | 6.26501 |
| GSTO1 | P78417 | 6.38145 | 6.06705 | 6.88366 | 5.87713 | 6.41548 | 6.71079 | 5.96217 | 6.74828 | 6.18658 | 6.52052 | 6.32966 | 6.62819 | 5.86729 | 5.96128 | 5.59092 | 5.86956 |
| PSMA5 | P28066 | 6.01746 | 5.11649 | 6.06523 | 5.00666 | 6.1184 | 6.06835 | 6.22708 | 6.18408 | 5.9763 | 6.29015 | 6.34388 | 6.6889 | 6.56511 | 7.16652 | 7.17885 | 6.98652 |
| NEFH | P12036 | 6.7052 | 6.21176 | 5.75621 | 6.11729 | 6.39434 | 5.77833 | 6.54937 | 6.04758 | 6.34657 | 5.85742 | 6.22462 | 6.15676 | 6.99709 | 6.26155 | 6.42803 | 6.1679 |
| HERC4 | Q5GLZ8 | 6.65094 | 6.59676 | 7.00962 | 6.29936 | 6.45567 | 6.71586 | 6.26467 | 6.96161 | 6.79665 | 6.74496 | 6.902 | 7.01198 | 4.71905 | 4.98772 | 4.80199 | 5.08115 |
| MRPS35 | P82673 | 6.24236 | 6.50407 | 5.80176 | 6.78043 | 6.25633 | 6.05412 | 6.54208 | 6.22382 | 5.8891 | 6.07146 | 5.93626 | 6.08512 | 6.51842 | 6.19412 | 6.53323 | 6.3673 |
| NUSAP1 | Q9BXS6 | 5.97862 | 6.24735 | 5.54266 | 6.09102 | 6.22968 | 6.39591 | 6.58783 | 6.01709 | 6.10261 | 5.88618 | 5.99233 | 5.94133 | 6.51257 | 6.84646 | 6.5694 | 7.05895 |
| PTPN11 | Q06124-1 | 6.26409 | 6.41781 | 6.64528 | 6.06107 | 6.10649 | 6.45243 | 6.18323 | 6.45589 | 6.11053 | 6.52867 | 6.86886 | 6.64205 | 5.75447 | 5.9642 | 5.83708 | 5.70784 |
| CLGN | O14967 | 6.30999 | 6.28633 | 5.93765 | 6.62338 | 6.06322 | 5.99365 | 6.21486 | 6.1129 | 6.11308 | 6.22127 | 6.13674 | 5.89781 | 6.59519 | 6.479 | 6.57563 | 6.43933 |
| QKI | Q96PU8 | 5.87937 | 5.45736 | 6.07601 | 5.78806 | 6.48521 | 6.09669 | 6.56548 | 6.26697 | 6.2879 | 6.46214 | 6.4976 | 6.7249 | 6.99418 | 6.33105 | 6.49332 | 6.49376 |
| RPL30 | P62888 | 6.3834 | 6.40006 | 6.60545 | 6.46187 | 6.30687 | 6.46611 | 6.46682 | 6.18938 | 6.513 | 6.41545 | 6.74103 | 6.29022 | 5.73899 | 5.72838 | 5.87984 | 5.41315 |
| TLN2 | Q9Y4G6 | 6.17871 | 6.24467 | 6.45935 | 5.97804 | 6.00442 | 6.57409 | 6.15653 | 6.34153 | 6.21327 | 6.48461 | 6.30172 | 6.40602 | 6.27725 | 6.25606 | 5.99889 | 6.12486 |
| RRM2 | P31350-2 | 6.75065 | 6.24782 | 7.11219 | 5.91755 | 6.71901 | 6.69659 | 6.1189 | 6.64065 | 6.31761 | 6.20616 | 6.54068 | 7.08545 | 5.59582 | 5.56971 | 5.28659 | 5.19462 |
| FMNL2 | Q96PY5-3 | 6.34161 | 6.48097 | 6.3251 | 6.24644 | 6.02788 | 6.18391 | 6.36161 | 5.97763 | 6.50891 | 6.24736 | 6.33988 | 6.22821 | 6.23469 | 6.12086 | 6.42025 | 5.95468 |
| MTHFD2 | P13995 | 6.51654 | 6.2684 | 5.84387 | 6.5261 | 6.84967 | 6.17179 | 6.58473 | 6.58394 | 6.50837 | 5.77878 | 6.18242 | 5.96458 | 6.31051 | 5.71145 | 6.02098 | 6.17785 |
| CDC73 | Q6P1J9 | 6.24298 | 6.25687 | 6.10751 | 6.71528 | 6.09768 | 5.64659 | 6.39241 | 5.77854 | 6.41246 | 5.92754 | 6.35206 | 6.01502 | 6.68642 | 6.16115 | 6.70764 | 6.49984 |
| BCAT2 | O15382 | 6.51641 | 6.08763 | 5.89673 | 6.08625 | 6.71413 | 6.08229 | 6.3815 | 6.09493 | 5.95837 | 6.12533 | 5.92057 | 5.75805 | 6.66978 | 6.50798 | 6.60898 | 6.59107 |
| EPS8 | Q12929 | 6.19764 | 6.28267 | 6.41526 | 6.21599 | 6.07049 | 6.19037 | 6.03112 | 6.2132 | 6.32737 | 6.75221 | 6.50357 | 6.28523 | 6.07618 | 6.37271 | 6.1114 | 5.95461 |
| SEL1L | Q9UBV2 | 6.69798 | 6.64656 | 5.76607 | 6.79351 | 6.48224 | 6.26396 | 5.90364 | 6.30318 | 6.14056 | 6.06471 | 5.90343 | 5.74437 | 6.29574 | 6.07516 | 6.77697 | 6.14192 |
| VPS51 | Q9UID3 | 6.43406 | 6.62296 | 6.7627 | 6.00344 | 6.48392 | 6.49433 | 6.25248 | 6.57709 | 6.37205 | 6.36724 | 6.63496 | 6.44655 | 5.79039 | 5.87885 | 5.53876 | 5.34023 |
| CCDC124 | Q96CT7 | 6.50857 | 6.36332 | 6.92185 | 6.20137 | 6.41749 | 6.17436 | 6.63239 | 6.43706 | 6.36752 | 6.34675 | 6.6328 | 6.34685 | 5.61315 | 5.84883 | 5.80773 | 5.37995 |
| MRTO4 | Q9UKD2 | 6.39871 | 6.42993 | 6.65436 | 6.47722 | 6.09946 | 6.15613 | 6.09249 | 6.14624 | 6.02701 | 6.2309 | 6.30144 | 6.17353 | 5.99391 | 6.41613 | 6.2476 | 6.15496 |
| STT3B | Q8TCJ2 | 5.98934 | 6.28204 | 5.79866 | 6.57863 | 5.83335 | 5.8898 | 6.42871 | 6.01695 | 6.18017 | 6.07627 | 5.62912 | 5.92742 | 6.57136 | 7.01404 | 6.92844 | 6.85571 |
| STT3A | P46977 | 6.47624 | 6.34856 | 6.01042 | 6.5109 | 6.20453 | 6.04491 | 6.41586 | 5.75204 | 5.93747 | 6.11293 | 6.00553 | 5.71476 | 6.59419 | 6.46042 | 6.91361 | 6.49764 |
| YES1 | P07947 | 6.36535 | 6.0908 | 6.2043 | 6.51151 | 6.27169 | 6.41711 | 6.46268 | 6.02463 | 6.162 | 6.25093 | 6.11263 | 6.01126 | 6.35138 | 6.16057 | 6.16092 | 6.44225 |
| PSMB1 | P20618 | 6.41697 | 6.6748 | 6.02377 | 6.27075 | 6.73224 | 6.88552 | 6.21832 | 7.04772 | 5.99664 | 5.60799 | 6.35233 | 6.00034 | 5.78925 | 6.07075 | 5.87155 | 6.04106 |
| TACC2 | O95359-3 | 6.23712 | 5.99228 | 6.59929 | 5.84526 | 6.67347 | 6.49439 | 6.15064 | 6.28633 | 6.54119 | 6.32165 | 6.70124 | 6.14658 | 6.35704 | 6.22267 | 5.85191 | 5.57894 |
| AP1B1 | Q10567-3 | 6.24799 | 6.26386 | 6.68464 | 6.22903 | 6.02634 | 6.73147 | 6.34248 | 6.63863 | 6.21173 | 6.40741 | 6.47048 | 6.40471 | 5.80176 | 5.75768 | 5.84499 | 5.93681 |
| WRN | Q14191 | 6.24943 | 6.43426 | 6.20065 | 6.71446 | 6.07993 | 6.10488 | 6.1712 | 6.16765 | 6.12083 | 6.21802 | 6.19273 | 6.43135 | 6.37204 | 5.99722 | 6.25616 | 6.28921 |
| UBAP2 | Q5T6F2 | 6.15052 | 6.1744 | 6.93692 | 6.02973 | 6.27919 | 6.69695 | 6.09019 | 6.60796 | 6.59296 | 6.69532 | 7.13235 | 7.09463 | 5.61399 | 5.55416 | 4.95751 | 5.39322 |
| DIEXF | Q68CQ4 | 5.94822 | 5.91469 | 5.90101 | 6.28703 | 6.1584 | 5.89713 | 6.28507 | 5.9498 | 5.963 | 5.84609 | 6.09803 | 6.13174 | 7.02947 | 6.61725 | 7.05014 | 6.92293 |
| CENPE | Q02224 | 7.09975 | 6.86545 | 7.68018 | 7.05871 | 5.99604 | 6.38081 | 6.59915 | 6.46746 | 6.97575 | 7.10202 | 7.118 | 7.23981 | 4.33983 | 4.41357 | 4.21773 | 4.44575 |
| SPDL1 | Q96EA4 | 5.60986 | 5.58137 | 6.12447 | 5.45493 | 6.09918 | 6.52178 | 6.08162 | 6.23895 | 6.15155 | 6.01743 | 6.21131 | 6.09663 | 7.60012 | 6.67248 | 6.96484 | 6.5735 |
| DTX3L | Q8TDB6 | 6.28633 | 6.29249 | 6.68111 | 6.25343 | 6.10643 | 6.26495 | 6.21264 | 6.44729 | 6.57309 | 6.83389 | 6.75235 | 6.74444 | 5.39051 | 5.74732 | 5.64758 | 5.76613 |
| GOLGA5 | Q8TBA6 | 6.56736 | 6.56789 | 5.96688 | 6.47867 | 6.10622 | 6.02328 | 6.11013 | 6.00002 | 6.16949 | 5.96665 | 6.18088 | 6.01133 | 6.72121 | 6.36158 | 6.51652 | 6.25189 |
| FBLL1 | A6NHQ2 | 6.23272 | 6.32324 | 6.0155 | 6.91508 | 5.811 | 5.79689 | 6.4219 | 5.89701 | 6.4014 | 6.15746 | 6.13191 | 6.23853 | 6.43493 | 6.08989 | 6.49152 | 6.64103 |
| DDX50 | Q9BQ39 | 6.25505 | 6.40851 | 5.8208 | 6.80947 | 6.11258 | 5.62911 | 6.28284 | 5.93484 | 6.17223 | 6.06337 | 6.27257 | 6.05413 | 6.33227 | 6.55059 | 6.92893 | 6.37271 |
| ACADM | P11310-2 | 6.40289 | 6.29554 | 5.81186 | 6.22101 | 6.47444 | 6.13628 | 6.19891 | 6.14409 | 6.28439 | 6.01533 | 5.94529 | 5.99297 | 7.09529 | 6.20923 | 6.48173 | 6.29076 |
| NELFB | Q8WX92-2 | 5.73194 | 5.96624 | 5.86519 | 5.79185 | 6.02623 | 6.12601 | 6.16207 | 6.20587 | 6.21978 | 5.94611 | 5.85043 | 6.03855 | 7.14037 | 6.93665 | 7.11142 | 6.88329 |
| ARFGEF2 | Q9Y6D5 | 6.80799 | 6.60333 | 6.74248 | 6.67234 | 6.13712 | 6.35624 | 6.10691 | 6.57595 | 5.96452 | 6.41966 | 6.46244 | 6.34073 | 5.66604 | 5.92541 | 5.78359 | 5.43525 |
| EED | A0A5F9ZI63 | 6.16731 | 6.3626 | 6.25431 | 6.97039 | 5.72923 | 5.98223 | 6.30878 | 5.92248 | 6.36807 | 6.55261 | 6.45732 | 6.54479 | 5.90442 | 5.91214 | 6.2872 | 6.27611 |
| IPO8 | O15397 | 5.65074 | 5.76712 | 6.2029 | 5.63431 | 5.76747 | 6.18668 | 5.73048 | 6.2721 | 6.10246 | 6.26636 | 6.20648 | 6.35809 | 6.77337 | 7.02163 | 7.07841 | 6.98138 |
| MICAL2 | O94851 | 6.49091 | 6.38244 | 5.95676 | 6.63534 | 6.2551 | 5.99094 | 6.49684 | 5.92813 | 6.05739 | 5.7386 | 5.78722 | 5.92466 | 6.61824 | 6.32573 | 6.72494 | 6.68674 |
| RSBN1L | Q6PCB5 | 6.08495 | 6.47918 | 5.82972 | 6.80993 | 6.10352 | 6.15688 | 6.37249 | 5.75793 | 5.93393 | 6.15641 | 5.95318 | 5.9348 | 6.02549 | 6.8742 | 6.84475 | 6.68265 |
| SAFB | Q15424-3 | 6.44434 | 6.57748 | 6.06749 | 6.59211 | 6.19207 | 5.9363 | 6.4466 | 5.91218 | 6.51487 | 6.19486 | 6.0935 | 5.9277 | 6.72752 | 5.95077 | 6.42627 | 6.22936 |
| NELFA | Q9H3P2 | 5.85 | 5.98918 | 6.20981 | 5.81121 | 6.07897 | 6.09554 | 6.05238 | 5.92854 | 6.46995 | 6.14662 | 6.38347 | 6.05034 | 7.04202 | 6.50826 | 7.04521 | 6.33849 |
| PTK7 | Q13308-6 | 6.51116 | 6.55591 | 6.20151 | 6.66505 | 5.67589 | 5.81328 | 6.22663 | 5.90008 | 6.29376 | 6.2768 | 6.04427 | 6.46936 | 6.24627 | 6.24674 | 6.39116 | 6.4821 |
| CYCS | P99999 | 6.8199 | 6.59767 | 6.11913 | 6.4273 | 6.37909 | 5.70231 | 6.52118 | 5.86811 | 6.16967 | 5.78868 | 5.90697 | 5.67963 | 6.83436 | 6.09067 | 6.53529 | 6.56003 |
| HECTD4 | J3KPF0 | 6.56041 | 6.46374 | 6.93535 | 6.33356 | 5.97756 | 6.51348 | 6.09967 | 6.48476 | 6.63255 | 6.74591 | 6.77844 | 6.71718 | 5.39569 | 5.77301 | 5.26535 | 5.32336 |
| PIEZO1 | Q92508 | 6.41707 | 6.41591 | 6.01985 | 6.81738 | 6.0175 | 5.90999 | 6.03268 | 6.09092 | 6.32966 | 5.99579 | 6.03146 | 6.15719 | 6.22558 | 6.35631 | 6.49191 | 6.6908 |
| VPS4B | O75351 | 6.29772 | 6.31288 | 6.77552 | 6.15105 | 6.39718 | 6.9762 | 5.91548 | 6.86654 | 6.24128 | 6.60565 | 6.46745 | 6.78743 | 5.84617 | 5.83242 | 5.12627 | 5.40075 |
| ANKRD17 | O75179 | 6.32366 | 6.35043 | 6.79381 | 5.9821 | 6.20156 | 6.48751 | 6.24699 | 6.50556 | 6.58593 | 6.77515 | 6.68299 | 6.65132 | 5.47474 | 5.69408 | 5.47432 | 5.76986 |
| OTUD4 | Q01804 | 6.10631 | 6.27805 | 6.40522 | 6.19686 | 6.26417 | 6.50736 | 6.43192 | 6.27503 | 6.55115 | 6.86442 | 6.48152 | 6.5506 | 5.61931 | 5.98424 | 5.68455 | 5.7993 |
| NCSTN | Q92542 | 6.3516 | 6.50942 | 6.1105 | 6.47334 | 6.18755 | 6.14192 | 6.26092 | 6.04587 | 6.36559 | 6.24765 | 6.27611 | 6.08471 | 6.36185 | 6.26276 | 6.35134 | 5.96888 |
| RBM26 | A0A087X0H9 | 5.77634 | 6.29937 | 6.14843 | 6.27881 | 5.99295 | 5.91734 | 6.0944 | 5.95681 | 5.92875 | 6.05616 | 5.75908 | 6.06079 | 6.98934 | 6.61771 | 7.1881 | 6.93564 |
| CC2D1B | Q5T0F9 | 5.93093 | 6.07315 | 6.37383 | 5.69497 | 6.02541 | 6.55111 | 6.07853 | 6.41666 | 6.15424 | 6.51264 | 6.43259 | 6.44009 | 6.3728 | 6.69178 | 6.11157 | 6.1397 |
| ARHGDIA | P52565 | 6.00555 | 5.9016 | 6.92084 | 6.05312 | 6.40591 | 6.89565 | 6.35616 | 6.82238 | 6.3636 | 6.33455 | 6.61843 | 6.59848 | 5.73536 | 5.71068 | 5.77852 | 5.49919 |
| TM9SF2 | Q99805 | 6.68462 | 6.68372 | 6.1551 | 6.65455 | 6.61879 | 6.12825 | 6.35134 | 5.86361 | 6.20371 | 5.82726 | 6.07496 | 5.87109 | 6.41966 | 6.19496 | 6.30665 | 5.96174 |
| SPTBN2 | O15020 | 6.44492 | 6.19331 | 6.19053 | 6.24196 | 6.00839 | 5.81977 | 6.14273 | 5.93343 | 6.25723 | 6.02287 | 6.20498 | 6.2894 | 6.75452 | 6.31526 | 6.65927 | 6.52145 |
| NLN | Q9BYT8 | 6.5203 | 6.45977 | 6.3059 | 6.444 | 6.36745 | 6.23676 | 6.3042 | 6.39563 | 6.39866 | 6.2262 | 6.19908 | 6.17133 | 6.11928 | 5.82707 | 6.04809 | 5.97629 |
| POFUT2 | Q9Y2G5 | 6.39904 | 6.53415 | 6.11299 | 6.62009 | 6.14398 | 6.06737 | 6.25691 | 6.15564 | 5.91614 | 5.93451 | 6.02138 | 5.93149 | 6.40815 | 6.29242 | 6.66979 | 6.53596 |
| MTAP | Q13126 | 6.24514 | 5.89231 | 6.87121 | 6.01895 | 6.28813 | 6.74928 | 5.93956 | 6.62606 | 6.23049 | 6.29727 | 6.72683 | 6.48703 | 6.09048 | 5.77925 | 5.98208 | 5.77595 |

| | | | | | | | | | | | | | | | | | |
|---|---|---|---|---|---|---|---|---|---|---|---|---|---|---|---|---|---|
| DMAP1 | Q9NPF5 | 6.43574 | 6.65372 | 6.02184 | 6.70597 | 6.14848 | 5.83155 | 6.46057 | 6.03332 | 6.28584 | 6.28773 | 6.18243 | 6.133 | 6.14368 | 6.40724 | 6.20408 | 6.06481 |
| ACAP2 | Q15057 | 6.66558 | 6.21089 | 6.89971 | 5.94607 | 6.41701 | 6.91956 | 6.09998 | 6.49685 | 6.34822 | 6.4771 | 6.32342 | 6.2719 | 5.74867 | 5.67271 | 5.74145 | 5.76085 |
| PPP1R13L | Q8WUF5 | 6.57482 | 6.08418 | 6.7354 | 6.08925 | 6.4542 | 6.72968 | 6.06502 | 6.70067 | 6.76935 | 6.3041 | 6.81442 | 6.7008 | 5.53132 | 5.78958 | 5.38936 | 5.26788 |
| DNAJC8 | O75937 | 5.55719 | 5.77165 | 6.11927 | 5.6109 | 6.23825 | 6.37747 | 6.44461 | 5.74675 | 6.04151 | 6.07042 | 6.44719 | 6.25921 | 6.66963 | 7.19193 | 6.96188 | 6.49214 |
| PCCB | P05166-2 | 5.98932 | 6.05583 | 5.61953 | 6.41286 | 6.21846 | 6.0193 | 6.48006 | 5.95142 | 5.79458 | 5.99231 | 5.59849 | 5.78533 | 7.41003 | 6.65026 | 7.05373 | 6.96848 |
| PTGES2 | Q9H7Z7 | 6.63845 | 6.13081 | 5.91552 | 6.62326 | 6.19033 | 6.03071 | 6.20516 | 5.97624 | 5.96152 | 6.09615 | 6.22949 | 6.13009 | 6.8541 | 6.33059 | 6.44823 | 6.23936 |
| RPL34 | P49207 | 6.3915 | 6.30622 | 5.9512 | 6.00409 | 6.30638 | 6.17771 | 6.70195 | 6.44541 | 7.1003 | 5.97309 | 6.32793 | 6.2515 | 5.77039 | 6.25046 | 5.91943 | 6.12246 |
| RAB10 | P61026 | 6.21497 | 6.58961 | 6.29571 | 6.59782 | 6.08943 | 5.94802 | 6.34302 | 5.97525 | 6.14634 | 6.09565 | 5.80548 | 6.08704 | 6.40553 | 6.18603 | 6.73545 | 6.48464 |
| RAB14 | P61106 | 6.19722 | 6.54138 | 5.98682 | 6.25826 | 6.37371 | 6.34799 | 6.28124 | 6.25725 | 6.23266 | 6.15665 | 6.1746 | 5.88261 | 6.11078 | 6.37554 | 6.47116 | 6.35214 |
| KDM1B | Q8NB78 | 6.23807 | 6.71436 | 6.3694 | 6.4488 | 6.36096 | 6.27998 | 6.19771 | 6.05312 | 6.30646 | 6.14613 | 6.09438 | 6.17107 | 5.99712 | 6.29 | 6.25319 | 6.07925 |
| PEPD | P12955 | 6.52005 | 5.83114 | 6.8318 | 5.37763 | 6.64406 | 6.56272 | 6.06951 | 6.2932 | 6.39381 | 7.11755 | 6.88874 | 6.46405 | 6.26306 | 5.89536 | 5.51434 | 5.33299 |
| YY1 | P25490 | 5.69303 | 6.01836 | 6.35022 | 6.15933 | 6.19556 | 6.05127 | 5.90939 | 6.19113 | 6.36082 | 6.50988 | 6.65782 | 6.4103 | 6.33983 | 6.47969 | 6.49466 | 6.17872 |
| PSME2 | Q9UL46 | 6.19415 | 5.97816 | 6.62036 | 5.51681 | 6.07132 | 6.47213 | 5.71182 | 6.34691 | 6.03932 | 6.49029 | 5.97944 | 6.35561 | 6.75446 | 6.44793 | 6.6351 | 6.38621 |
| ARHGAP12 | Q8IWW6 | 6.00797 | 6.24968 | 6.49004 | 5.99074 | 6.46322 | 6.20011 | 6.03087 | 6.59617 | 6.51831 | 6.21336 | 6.25579 | 6.42709 | 6.24984 | 6.41581 | 5.89383 | 5.99719 |
| RFTN1 | Q14699 | 6.45331 | 6.44574 | 6.31711 | 6.47829 | 6.41632 | 6.1717 | 6.33748 | 6.0292 | 6.13957 | 5.9442 | 6.3038 | 6.15685 | 6.20541 | 6.14052 | 6.34831 | 6.11218 |
| NHLRC2 | Q8NBF2 | 6.12475 | 6.26021 | 6.90198 | 5.98739 | 6.14582 | 6.47554 | 6.23951 | 6.64603 | 6.36567 | 6.27186 | 6.27242 | 6.59996 | 5.97734 | 6.03959 | 5.7917 | 5.90023 |
| RCC2 | Q9P258 | 5.70075 | 5.94005 | 6.32709 | 5.62916 | 5.77428 | 6.27194 | 5.99522 | 6.10651 | 6.43618 | 6.16566 | 6.11709 | 6.01156 | 6.39757 | 7.36724 | 7.13107 | 6.62864 |
| SIRT1 | Q96EB6 | 5.40868 | 5.90841 | 6.01583 | 5.66285 | 5.61716 | 6.07879 | 5.77911 | 5.95974 | 5.98348 | 6.20734 | 6.63619 | 6.35431 | 7.17792 | 6.95981 | 7.05909 | 7.19129 |
| RPS15A | P62244 | 6.36354 | 6.27406 | 6.61386 | 6.263 | 6.45531 | 6.26017 | 6.73717 | 6.14101 | 6.7969 | 5.90726 | 6.1689 | 6.21716 | 5.46775 | 6.14969 | 5.96688 | 6.21733 |
| H2AC6 | Q93077 | 6.02542 | 5.95773 | 5.62159 | 6.71759 | 5.73397 | 5.78394 | 6.68583 | 6.20811 | 6.06405 | 6.15836 | 6.10903 | 6.03761 | 6.38404 | 6.54716 | 6.69729 | 7.26829 |
| GSTK1 | Q9Y2Q3 | 5.91031 | 4.72577 | 5.4476 | 5.26674 | 6.13441 | 5.28491 | 5.99787 | 6.36006 | 5.3388 | 6.09826 | 5.39435 | 6.50711 | 6.59755 | 8.30077 | 8.10064 | 8.53485 |
| RPL22 | P35268 | 6.30434 | 6.46415 | 6.37069 | 6.30289 | 5.75086 | 6.44195 | 6.42097 | 6.30854 | 6.60516 | 6.54361 | 6.63182 | 6.29641 | 5.77965 | 5.70376 | 6.2155 | 5.8597 |
| AIP | O00170 | 5.84077 | 5.6952 | 6.6877 | 5.46067 | 6.28046 | 6.57661 | 5.92783 | 6.58589 | 6.32352 | 6.37363 | 6.52881 | 6.4643 | 6.72128 | 6.29664 | 6.203 | 6.03367 |
| MICU1 | Q9BPX6-3 | 6.37581 | 6.55018 | 5.79213 | 6.72017 | 6.13619 | 6.25393 | 6.33086 | 6.04246 | 5.94855 | 6.13003 | 5.93508 | 6.10957 | 6.40311 | 6.25734 | 6.4853 | 6.52929 |
| RCC1L | Q96I51 | 6.46066 | 6.56116 | 6.0771 | 6.75669 | 6.23205 | 6.26496 | 6.55761 | 6.19355 | 6.15185 | 6.21665 | 6.20018 | 5.61142 | 6.1525 | 6.04295 | 6.40019 | 6.12048 |
| ALDH4A1 | P30038 | 6.4871 | 6.32473 | 5.94117 | 6.559 | 6.39489 | 6.11284 | 6.13731 | 6.09794 | 6.08014 | 6.35506 | 6.29243 | 5.96106 | 6.92225 | 5.89434 | 6.3134 | 6.12633 |
| PMS1 | P54277 | 5.50949 | 6.05836 | 6.6925 | 5.84705 | 5.5594 | 6.35031 | 6.00517 | 6.00991 | 6.09985 | 6.27112 | 6.81013 | 6.72949 | 6.37095 | 6.4448 | 6.54791 | 6.69356 |
| PGAM5 | Q96HS1 | 6.41194 | 6.47225 | 5.86389 | 6.86735 | 6.38423 | 6.12745 | 5.97912 | 6.24109 | 5.95812 | 5.91352 | 5.88329 | 6.1025 | 6.40812 | 6.31682 | 6.48247 | 6.58781 |
| USP47 | Q96K76 | 6.34507 | 6.27344 | 7.09007 | 6.02806 | 6.55329 | 6.89282 | 6.28726 | 6.71499 | 6.18657 | 6.42634 | 6.60429 | 6.55149 | 5.60005 | 5.59875 | 5.41511 | 5.43241 |
| RAVER1 | E9PAU2 | 5.87771 | 5.98668 | 5.96352 | 6.16864 | 6.14364 | 6.07075 | 6.00894 | 6.28489 | 6.15845 | 6.08837 | 6.32318 | 6.24636 | 6.97611 | 6.61804 | 6.62594 | 6.45878 |
| XRCC1 | P18887 | 5.70827 | 5.94026 | 5.73339 | 5.67203 | 5.49187 | 5.97677 | 5.90044 | 5.878 | 6.03305 | 6.43641 | 6.47279 | 6.24249 | 6.91521 | 7.3179 | 7.27393 | 7.00718 |
| RANBP6 | O60518 | 5.93293 | 6.21966 | 6.18182 | 6.182 | 6.07351 | 6.14823 | 6.10171 | 6.0718 | 6.21942 | 6.21225 | 6.28288 | 6.07097 | 6.61115 | 6.40826 | 6.78981 | 6.49362 |
| FDFT1 | P37268 | 6.57005 | 7.2434 | 6.58371 | 7.28241 | 6.31253 | 6.30869 | 6.62131 | 6.4774 | 6.32828 | 6.5412 | 6.29866 | 6.17833 | 5.17134 | 5.36971 | 5.35512 | 5.35785 |
| CDC42BPA | Q5VT25-6 | 6.29903 | 6.35979 | 6.66339 | 6.19332 | 6.39971 | 6.53316 | 6.32369 | 6.42501 | 6.27904 | 6.49041 | 6.66425 | 6.43369 | 5.84399 | 5.92058 | 5.67379 | 5.49715 |
| BRD8 | Q9H0E9-2 | 6.72489 | 6.69943 | 6.50504 | 7.35362 | 6.04162 | 6.14916 | 6.32954 | 6.13749 | 6.24297 | 6.50822 | 6.3957 | 6.74849 | 5.61503 | 5.4929 | 5.23227 | 5.82363 |
| LRSAM1 | Q6UWE0 | 6.42766 | 6.21107 | 6.93591 | 6.18904 | 6.11807 | 6.66664 | 6.45273 | 6.87202 | 6.52196 | 6.35674 | 6.72239 | 6.4353 | 5.75843 | 5.68781 | 5.46532 | 5.17891 |
| DNMT3A | Q9Y6K1 | 6.90992 | 6.59727 | 5.99818 | 6.65031 | 6.28634 | 6.19356 | 6.65016 | 6.05527 | 6.28603 | 6.32419 | 6.09161 | 6.00735 | 6.1637 | 5.79308 | 6.06132 | 5.93172 |
| PRIM2 | P49643 | 5.62363 | 6.1889 | 6.31095 | 6.07348 | 6.31604 | 6.48645 | 6.01976 | 5.87004 | 6.16124 | 6.24564 | 6.41678 | 6.37979 | 6.54099 | 6.33511 | 6.45724 | 6.57398 |
| SCFD2 | Q8WU76 | 6.35924 | 6.51422 | 5.87789 | 6.27403 | 6.25207 | 6.18603 | 6.36447 | 5.7867 | 6.04336 | 6.11715 | 5.94889 | 6.17396 | 6.65698 | 6.51287 | 6.47561 | 6.45653 |
| CSNK2A2 | P19784 | 6.04036 | 6.28864 | 6.60881 | 6.45093 | 6.06588 | 6.3676 | 6.17093 | 6.19291 | 5.80883 | 6.04228 | 6.33743 | 6.25931 | 6.304 | 6.45814 | 6.38544 | 6.21854 |
| CTDP1 | Q9Y5B0 | 5.86966 | 6.27667 | 6.17713 | 6.03308 | 5.70575 | 6.04983 | 5.81301 | 6.27695 | 6.13777 | 6.16693 | 5.9808 | 6.21943 | 6.7111 | 7.14783 | 6.73865 | 6.69541 |
| PUM1 | Q14671-3 | 6.58857 | 6.57731 | 6.59066 | 5.95528 | 6.20458 | 6.52586 | 6.42654 | 6.40182 | 6.30207 | 6.12266 | 6.48098 | 6.34418 | 5.78085 | 5.81743 | 6.03137 | 5.84985 |
| ABCD3 | P28288 | 6.26468 | 6.46493 | 5.92148 | 6.83851 | 5.97843 | 5.76236 | 6.24625 | 5.89734 | 6.05274 | 5.87847 | 6.43872 | 6.28286 | 6.3831 | 6.20492 | 6.77738 | 6.60784 |
| SNX6 | Q9UNH7 | 6.10218 | 6.30411 | 6.56613 | 5.97125 | 6.50491 | 6.69046 | 6.10427 | 6.61294 | 5.96623 | 6.5195 | 6.44981 | 6.36139 | 6.22458 | 6.05541 | 5.79216 | 5.768 |
| PDP1 | Q9P0J1-2 | 6.55961 | 6.1553 | 5.87932 | 6.3692 | 6.44296 | 6.12321 | 6.25631 | 6.19985 | 6.12231 | 5.7683 | 5.873 | 5.89329 | 6.88365 | 6.24774 | 6.76707 | 6.45887 |
| SNX1 | Q13596-3 | 6.46152 | 6.23638 | 6.81948 | 6.0203 | 6.56557 | 6.58801 | 6.1311 | 6.61525 | 6.4727 | 6.27471 | 6.45728 | 6.36548 | 5.95253 | 5.74723 | 5.78596 | 5.5065 |
| SNX2 | O60749 | 6.14774 | 6.68401 | 6.69013 | 6.54524 | 6.21718 | 6.66814 | 6.07486 | 6.61235 | 6.04662 | 6.1856 | 6.49771 | 6.64351 | 5.92555 | 5.83527 | 5.90563 | 5.32047 |
| SPATA5L1 | Q9BVQ7 | 6.23413 | 6.09478 | 6.85795 | 6.15628 | 6.41934 | 6.53104 | 6.30416 | 6.38115 | 6.60326 | 6.24344 | 6.63434 | 6.82313 | 5.80279 | 5.76679 | 5.68855 | 5.45887 |
| TCEA1 | P23193 | 6.03586 | 5.70121 | 6.44054 | 5.54035 | 5.93453 | 5.96249 | 6.17637 | 6.09486 | 6.36864 | 6.1862 | 6.50771 | 5.95323 | 6.81344 | 6.97094 | 6.85597 | 6.45767 |
| DUS3L | Q96G46 | 5.48965 | 5.51696 | 6.01751 | 5.15352 | 5.62405 | 6.26082 | 5.93564 | 5.96512 | 6.0193 | 6.07898 | 6.33902 | 6.06329 | 7.17635 | 7.68895 | 7.65904 | 7.01179 |
| INTS2 | Q9H0H0 | 6.46898 | 6.06247 | 5.98883 | 6.77358 | 6.33642 | 5.96464 | 6.13438 | 5.50319 | 5.8424 | 6.30871 | 6.13202 | 6.15292 | 6.37379 | 6.41792 | 6.73689 | 6.80287 |
| RECQL5 | O94762 | 6.07281 | 6.29015 | 6.18648 | 6.36047 | 6.10511 | 6.26111 | 6.16734 | 6.00364 | 6.46298 | 6.53237 | 6.37512 | 6.30866 | 6.31755 | 6.15723 | 6.31692 | 6.08208 |
| PSMB2 | P49721 | 5.67492 | 5.21988 | 5.96926 | 5.33338 | 6.13864 | 5.99106 | 6.52827 | 6.4507 | 5.80483 | 5.65475 | 5.70563 | 6.45257 | 6.04311 | 7.76817 | 7.2111 | 8.05372 |
| EMD | P50402 | 6.32231 | 6.47945 | 5.48046 | 6.48955 | 6.34545 | 6.03231 | 6.18372 | 6.42822 | 5.91625 | 6.19988 | 5.59291 | 5.9609 | 6.91053 | 6.59009 | 6.41009 | 6.65789 |
| PDCD4 | Q53EL6 | 6.11535 | 6.57939 | 5.93457 | 5.32251 | 6.25615 | 6.75508 | 5.76627 | 6.22373 | 5.89126 | 7.12713 | 6.05422 | 5.92795 | 6.37422 | 6.68944 | 6.88079 | 6.10194 |
| CGAS | Q8N884 | 6.43863 | 6.37958 | 6.13231 | 6.82436 | 6.26934 | 6.25497 | 6.41353 | 6.10653 | 6.44036 | 6.21849 | 6.24368 | 6.31738 | 5.89559 | 5.94401 | 5.9673 | 6.15394 |
| DDX20 | Q9UHI6 | 6.29143 | 6.28104 | 6.60702 | 6.10372 | 6.28287 | 6.57524 | 6.20183 | 6.72005 | 6.26917 | 6.27225 | 6.49413 | 6.64665 | 5.83881 | 5.99689 | 5.74558 | 5.67331 |
| SMARCB1 | Q12824 | 6.13707 | 6.23664 | 6.25103 | 6.36555 | 6.37006 | 5.8308 | 6.44565 | 6.03203 | 6.31973 | 6.03161 | 6.02756 | 6.03297 | 6.61226 | 6.28057 | 6.58424 | 6.44222 |
| QSER1 | Q2KHR3 | 6.04798 | 6.36141 | 6.48557 | 6.67336 | 6.12742 | 5.99389 | 6.35859 | 5.97451 | 5.98509 | 6.38485 | 6.20084 | 6.45068 | 6.20536 | 6.03454 | 6.16307 | 6.55282 |
| PLAU | P00749 | 5.50604 | 5.3988 | 5.37347 | 5.6985 | 6.01007 | 6.09446 | 6.65464 | 6.20077 | 5.81497 | 5.64317 | 5.90528 | 5.68253 | 8.02477 | 7.29188 | 8.01174 | 6.68888 |
| DAG1 | Q14118 | 6.64725 | 6.43728 | 6.16988 | 6.67676 | 5.96387 | 5.9958 | 6.19204 | 6.09373 | 5.9992 | 6.48345 | 6.39711 | 6.50549 | 6.22054 | 6.08622 | 6.08923 | 6.04216 |
| PANK4 | Q9NVE7 | 6.36053 | 6.07513 | 6.95185 | 5.99785 | 6.4042 | 6.86022 | 6.30249 | 6.8824 | 6.5882 | 6.58474 | 6.4262 | 6.3883 | 5.57088 | 5.72595 | 5.40642 | 5.47463 |
| COG5 | Q9UP83-2 | 6.27942 | 6.00606 | 6.35547 | 6.37596 | 6.23241 | 6.4976 | 6.30391 | 6.2463 | 6.46955 | 6.44945 | 6.38035 | 6.38492 | 6.39554 | 5.94396 | 5.99178 | 5.68732 |
| EEF1AKNMT | Q8N6R0 | 6.42615 | 6.29962 | 6.69789 | 6.21549 | 6.21518 | 6.53555 | 6.43261 | 6.55681 | 6.55312 | 6.42546 | 6.77222 | 6.56176 | 5.61434 | 5.7311 | 5.46055 | 5.50215 |
| TUT4 | Q5TAX3 | 6.08189 | 6.22214 | 6.95503 | 6.03372 | 5.93983 | 6.44755 | 6.21189 | 6.35478 | 6.3163 | 6.85145 | 6.63137 | 6.5053 | 5.59995 | 5.99344 | 5.89666 | 5.95871 |
| ZMYM4 | Q5VZL5 | 6.19083 | 6.48542 | 6.20975 | 6.91272 | 6.21196 | 6.14116 | 6.30334 | 5.79353 | 6.10938 | 6.47594 | 6.28199 | 5.91887 | 6.38991 | 6.06622 | 6.35051 | 6.15849 |
| NFRKB | Q6P4R8-2 | 6.48205 | 6.65206 | 6.26621 | 6.66218 | 6.2107 | 5.92078 | 6.29952 | 6.14784 | 6.18556 | 6.28688 | 6.20086 | 6.15114 | 6.28145 | 6.04299 | 6.10696 | 6.10281 |
| MIS18BP1 | Q6P0N0 | 6.67933 | 6.56334 | 6.22969 | 6.97959 | 6.42556 | 6.00367 | 6.54404 | 6.25789 | 6.61422 | 6.49736 | 6.14732 | 6.38339 | 5.57706 | 5.49903 | 5.6989 | 5.89961 |
| ATP6V1B2 | P21281 | 6.27147 | 6.8164 | 6.43163 | 6.29988 | 6.1935 | 6.63611 | 6.34937 | 6.30134 | 6.6018 | 6.94164 | 6.87595 | 6.78494 | 5.13035 | 5.29213 | 5.2364 | 5.83708 |
| VPS8 | Q8N3P4 | 6.07042 | 6.24685 | 6.42385 | 6.15168 | 6.08499 | 6.27297 | 6.09923 | 6.22279 | 6.23961 | 6.65797 | 6.36232 | 6.26563 | 6.1956 | 6.32043 | 6.16398 | 6.22168 |
| TEX10 | Q9NXF1 | 5.96006 | 6.11398 | 5.59772 | 6.72058 | 5.7699 | 5.91251 | 6.34354 | 6.02595 | 6.10686 | 6.05781 | 6.13098 | 6.1782 | 6.55193 | 6.90377 | 6.58293 | 7.04327 |
| SLU7 | O95391 | 6.24024 | 6.34657 | 6.23107 | 6.5995 | 6.03828 | 5.76999 | 6.1583 | 6.08613 | 6.19254 | 6.1899 | 6.21607 | 6.16512 | 6.10589 | 6.51501 | 6.59071 | 6.55467 |
| PRIM1 | P49642 | 5.91555 | 6.10777 | 6.36452 | 5.76712 | 6.14139 | 6.46464 | 6.079 | 6.32324 | 6.23564 | 6.33499 | 6.43337 | 6.37271 | 6.31628 | 6.34494 | 6.55414 | 6.2447 |
| TGM2 | P21980 | 6.31611 | 5.72443 | 7.21956 | 5.6985 | 6.80043 | 6.56292 | 6.19355 | 6.49109 | 6.47395 | 6.43255 | 6.70365 | 6.32855 | 6.08398 | 5.64151 | 5.80428 | 5.52494 |
| PPAT | Q06203 | 6.01859 | 6.24101 | 6.92571 | 5.91024 | 6.44125 | 6.67969 | 6.20577 | 6.58258 | 6.24003 | 6.3789 | 6.63432 | 6.20916 | 5.88968 | 5.91766 | 5.86976 | 5.85565 |
| TM9SF4 | Q92544 | 6.14099 | 6.52953 | 5.97183 | 6.65381 | 6.11777 | 5.81519 | 6.30619 | 5.99183 | 6.15824 | 6.16628 | 6.2303 | 6.05637 | 6.2871 | 6.3679 | 6.68936 | 6.5173 |
| NDUFA10 | O95299 | 5.89699 | 6.12251 | 5.73826 | 6.56608 | 6.47354 | 6.00349 | 7.00678 | 6.12371 | 6.28772 | 5.81189 | 5.73829 | 5.8034 | 6.43919 | 6.45101 | 6.631 | 6.90613 |
| PLCB3 | Q01970 | 6.16217 | 6.75768 | 6.3987 | 6.01482 | 6.1659 | 6.3115 | 6.27735 | 6.34617 | 6.12462 | 6.18412 | 6.51101 | 6.71127 | 6.03133 | 6.3692 | 5.87045 | 5.76371 |
| TRIM24 | O15164 | 6.42176 | 6.6389 | 5.87675 | 6.82972 | 6.22208 | 5.97581 | 6.2417 | 6.13438 | 6.13159 | 6.3413 | 6.2943 | 6.39384 | 6.1172 | 5.9959 | 6.16671 | 6.21805 |
| POLDIP3 | Q9BY77 | 6.27793 | 6.10478 | 6.18353 | 6.6955 | 6.20662 | 5.85766 | 6.8168 | 5.7826 | 6.3888 | 5.76766 | 5.95432 | 5.93745 | 6.27628 | 6.50611 | 6.59127 | 6.65267 |
| PPAN | Q9NQ55-3 | 6.30591 | 6.23559 | 5.82853 | 6.7328 | 6.00452 | 5.78422 | 6.53412 | 6.2813 | 6.12619 | 6.01984 | 5.92058 | 6.09265 | 6.21227 | 6.64853 | 6.70207 | 6.57088 |
| WDR19 | Q8NEZ3 | 6.1361 | 6.22634 | 6.4831 | 5.90098 | 6.29228 | 6.52439 | 6.32186 | 6.20948 | 6.20282 | 6.49026 | 6.6159 | 6.17197 | 6.35331 | 6.30213 | 6.05228 | 5.71681 |
| PARN | O95453 | 6.20514 | 6.35094 | 5.96658 | 6.63339 | 6.25198 | 6.10987 | 6.24612 | 6.10966 | 6.15102 | 6.04247 | 6.01624 | 5.96649 | 6.54287 | 6.34613 | 6.49556 | 6.56553 |
| FHL2 | Q14192 | 6.25346 | 6.03108 | 6.65113 | 6.15171 | 6.50468 | 6.6921 | 6.12659 | 6.20407 | 6.38587 | 6.06513 | 6.87892 | 6.39255 | 6.18747 | 5.7546 | 5.86094 | 5.8597 |
| TM9SF3 | Q9HD45 | 6.40309 | 6.91836 | 6.32383 | 6.70827 | 6.35327 | 6.28255 | 6.477 | 5.64566 | 5.79293 | 5.97271 | 5.91762 | 5.70388 | 6.67083 | 6.19311 | 6.37091 | 6.26601 |
| NT5C2 | P49902 | 6.35255 | 6.24348 | 6.82949 | 6.09186 | 6.32503 | 6.66878 | 5.92989 | 6.96888 | 6.01032 | 6.46719 | 6.40195 | 6.57771 | 5.54041 | 5.97561 | 5.70552 | 5.91133 |
| RRP9 | O43818 | 6.17565 | 5.58045 | 6.01806 | 6.61855 | 6.10124 | 5.99576 | 6.33706 | 5.5915 | 5.87891 | 6.10875 | 6.20092 | 6.15372 | 6.90961 | 6.79222 | 6.55979 | 6.97783 |
| GNE | Q9Y223-2 | 6.12179 | 6.19751 | 6.70742 | 6.05324 | 6.26842 | 6.79603 | 6.16673 | 6.75723 | 6.00238 | 6.33636 | 6.65745 | 6.28726 | 5.94633 | 6.14584 | 5.82237 | 5.73364 |
| RBBP4 | Q09028 | 6.16135 | 5.86676 | 6.03537 | 6.33591 | 6.11697 | 6.02902 | 5.90753 | 6.34385 | 6.36961 | 6.02948 | 6.41217 | 6.50416 | 6.99599 | 6.16006 | 6.1477 | 6.58406 |
| CWC22 | Q9HCG8 | 6.37461 | 6.21222 | 6.2855 | 6.56666 | 6.18937 | 5.8609 | 6.14858 | 5.79426 | 6.24634 | 6.16641 | 6.15461 | 6.01098 | 6.63166 | 6.20207 | 6.76081 | 6.39502 |
| PAK1IP1 | Q9NWT1 | 6.05443 | 6.14284 | 5.75322 | 6.67799 | 6.11609 | 5.90151 | 6.2345 | 5.7656 | 6.03935 | 5.99473 | 5.84647 | 5.93343 | 6.64388 | 6.49135 | 7.40303 | 7.00159 |
| LACTB | P83111 | 6.21506 | 6.59831 | 5.78148 | 6.7798 | 6.12314 | 6.02357 | 6.1682 | 5.99757 | 6.04947 | 6.1037 | 5.9289 | 6.32212 | 6.51858 | 6.26585 | 6.47636 | 6.6479 |
| ESS2 | Q96DF8 | 5.96867 | 5.9651 | 6.37913 | 5.98273 | 5.76461 | 6.05126 | 6.23229 | 6.14268 | 6.22554 | 6.25183 | 6.33454 | 6.50656 | 6.706 | 6.61262 | 6.39714 | 6.47931 |
| MRPL44 | Q9H9J2 | 6.39053 | 6.45866 | 6.01429 | 6.78373 | 6.1937 | 5.98488 | 6.48908 | 6.05763 | 5.98613 | 5.96096 | 6.00618 | 6.11323 | 6.52218 | 6.25557 | 6.33125 | 6.45201 |
| ILK | Q13418 | 6.44498 | 6.25461 | 6.81309 | 5.96005 | 6.54657 | 6.49231 | 6.27878 | 6.53762 | 6.53416 | 6.63758 | 6.78978 | 6.5953 | 5.82113 | 5.47722 | 5.40891 | 5.4079 |
| RPL23 | P62829 | 6.09035 | 6.08678 | 6.27739 | 6.22681 | 6.12659 | 6.13491 | 6.30054 | 6.32647 | 6.23378 | 6.31476 | 6.65673 | 6.46964 | 6.04817 | 6.54469 | 5.98262 | 6.17976 |
| DCAF13 | Q9NV06 | 5.99835 | 6.00081 | 6.15485 | 6.81201 | 6.16554 | 6.21359 | 6.5268 | 6.05248 | 6.18559 | 6.0453 | 6.24359 | 5.90636 | 6.37277 | 5.91485 | 6.77506 | 6.63203 |
| RHOC | P08134 | 6.11781 | 6.23969 | 6.20372 | 6.29629 | 6.18058 | 6.21974 | 6.34442 | 6.25943 | 6.1824 | 6.09269 | 6.26432 | 6.238 | 6.36859 | 6.45214 | 6.26283 | 6.27735 |
| DRG1 | Q9Y295 | 6.40318 | 6.20678 | 6.60582 | 5.98667 | 6.21837 | 6.36648 | 6.22179 | 6.29137 | 6.61331 | 6.34023 | 6.4843 | 6.39642 | 5.90096 | 5.9616 | 6.08459 | 5.91813 |
| CAPZA1 | P52907 | 6.3902 | 6.58326 | 6.54752 | 6.44073 | 6.44009 | 6.52921 | 5.92385 | 6.32924 | 6.1033 | 6.12329 | 6.34327 | 6.36423 | 6.13109 | 5.89336 | 6.08457 | 5.77278 |
| FASTKD2 | Q9NYY8 | 6.55621 | 6.42929 | 6.30953 | 6.94792 | 6.47565 | 5.97535 | 6.60299 | 5.9405 | 6.48896 | 5.93635 | 5.9924 | 6.01074 | 6.23937 | 5.58393 | 6.27436 | 6.23644 |
| TFB2M | Q9H5Q4 | 6.47461 | 6.39458 | 5.99445 | 6.97438 | 6.47148 | 6.20223 | 6.57663 | 6.22901 | 6.07981 | 5.73687 | 5.92213 | 6.13373 | 6.27596 | 5.9128 | 6.29294 | 6.3284 |
| CENPC | Q03188 | 6.07089 | 6.02964 | 5.80698 | 6.54506 | 6.13765 | 5.79046 | 6.29424 | 5.74182 | 6.16525 | 6.14948 | 5.83367 | 5.85084 | 6.55295 | 6.99242 | 7.11012 | 6.92851 |
| DNPEP | Q9ULA0 | 6.30118 | 6.2597 | 6.47217 | 5.9539 | 6.24094 | 6.7739 | 6.01129 | 6.41041 | 6.31996 | 6.32403 | 6.42643 | 6.52085 | 6.01657 | 6.1603 | 5.95535 | 5.85303 |
| TMEM43 | Q9BTV4 | 6.30716 | 6.55313 | 5.75019 | 6.50967 | 6.09609 | 5.94956 | 6.22081 | 6.04002 | 6.22857 | 6.02373 | 5.89242 | 5.87239 | 6.6425 | 6.45851 | 6.91242 | 6.54282 |

| | | | | | | | | | | | | | | | | | |
|---|---|---|---|---|---|---|---|---|---|---|---|---|---|---|---|---|---|
| MRPL37 | Q9BZE1 | 6.50028 | 6.24937 | 5.86742 | 6.73459 | 6.58202 | 6.13345 | 6.30282 | 6.24427 | 6.21247 | 6.2092 | 6.00929 | 5.87513 | 6.56901 | 6.04165 | 6.25955 | 6.20948 |
| ZC3H15 | Q8WU90 | 5.95215 | 6.14056 | 6.57904 | 5.96718 | 6.19841 | 6.69692 | 6.13107 | 6.20624 | 6.40953 | 6.71032 | 6.57453 | 6.57562 | 6.02748 | 6.24119 | 5.71903 | 5.87071 |
| DPM1 | O60762 | 6.34756 | 6.41359 | 5.93345 | 6.5965 | 6.34703 | 6.2339 | 6.33525 | 6.07541 | 6.00483 | 5.84268 | 5.84638 | 5.85925 | 6.63324 | 6.47842 | 6.55817 | 6.49434 |
| SSBP1 | Q04837 | 6.30497 | 6.24116 | 5.95426 | 6.54869 | 6.20272 | 5.98188 | 6.29481 | 6.0666 | 5.99301 | 6.01169 | 6.16288 | 5.99364 | 7.04788 | 6.34924 | 6.31104 | 6.53553 |
| ZNF326 | Q5BKZ1 | 6.18997 | 6.74135 | 5.94693 | 6.76517 | 6.05554 | 6.1988 | 6.20498 | 6.02545 | 5.89365 | 6.12974 | 6.07093 | 6.18559 | 6.61499 | 6.25778 | 6.37343 | 6.34569 |
| TRMT1L | Q7Z2T5 | 6.06976 | 6.36209 | 6.2201 | 6.30581 | 6.04496 | 6.02581 | 5.91897 | 5.70798 | 6.10381 | 6.36826 | 6.30744 | 6.04713 | 6.9036 | 6.40014 | 6.5985 | 6.61566 |
| BSG | P35613 | 6.26145 | 6.66322 | 5.68866 | 6.78334 | 6.07113 | 5.91783 | 6.36199 | 6.30486 | 6.23098 | 6.00896 | 5.49654 | 5.94103 | 6.08354 | 6.77611 | 6.51109 | 6.89927 |
| SAR1A | Q9NR31 | 6.38128 | 6.21759 | 6.64271 | 6.28773 | 6.43541 | 6.31775 | 6.20317 | 6.51945 | 6.34372 | 6.42618 | 6.39055 | 6.58941 | 6.05031 | 6.03242 | 5.54742 | 5.61493 |
| NPAT | Q14207 | 6.58624 | 6.65019 | 5.96689 | 7.02628 | 6.32138 | 6.04164 | 6.39606 | 6.29157 | 6.23251 | 5.87845 | 6.13109 | 6.11632 | 6.17961 | 6.01989 | 6.02913 | 6.13275 |
| DNAJA2 | O60884 | 6.32186 | 6.0782 | 6.44755 | 6.17956 | 6.07581 | 6.43874 | 6.30623 | 6.15144 | 6.4912 | 6.48378 | 6.47757 | 6.37913 | 6.14009 | 5.85189 | 6.07109 | 6.10585 |
| EXOC5 | O00471 | 6.2915 | 6.28322 | 6.47738 | 5.86126 | 6.49073 | 6.46955 | 6.05302 | 6.55525 | 6.55041 | 6.5756 | 6.31026 | 6.3231 | 6.053 | 6.16932 | 5.81755 | 5.71885 |
| TRIP13 | Q15645 | 6.40313 | 6.44733 | 6.82324 | 6.23976 | 6.49797 | 6.77894 | 6.28086 | 6.73861 | 6.64797 | 6.19713 | 6.40668 | 6.80181 | 5.46957 | 5.36472 | 5.33127 | 5.571 |
| RBM34 | P42696 | 6.22161 | 6.40483 | 6.03084 | 6.80154 | 6.07704 | 5.65488 | 6.35638 | 6.01308 | 6.04325 | 5.83139 | 5.91543 | 5.94488 | 6.2568 | 6.80326 | 6.84852 | 6.79626 |
| PDIA5 | Q14554 | 6.46396 | 6.76795 | 5.60844 | 6.2928 | 6.36761 | 6.14653 | 6.23193 | 6.26765 | 5.86595 | 5.75873 | 5.95422 | 5.76126 | 6.95594 | 6.40504 | 6.87225 | 6.27975 |
| ALYREF | Q86V81 | 6.11874 | 5.80874 | 5.52831 | 6.61036 | 5.95693 | 5.65401 | 7.01763 | 6.02039 | 6.33341 | 5.55727 | 5.8393 | 5.93806 | 6.52475 | 6.85387 | 6.61659 | 7.62163 |
| GET3 | O43681 | 6.19626 | 6.18591 | 6.48396 | 5.94021 | 6.23033 | 6.69814 | 5.9061 | 6.36483 | 6.12909 | 6.46253 | 6.27314 | 6.37098 | 6.57646 | 6.04469 | 6.19006 | 5.9473 |
| SRP14 | P37108 | 5.76582 | 6.14638 | 5.6598 | 5.76616 | 5.99764 | 6.06865 | 5.95628 | 6.19873 | 5.89681 | 6.51859 | 5.68043 | 6.06233 | 6.80194 | 7.20438 | 7.21456 | 7.0615 |
| STRN | O43815 | 6.0152 | 6.40146 | 6.49916 | 6.02072 | 6.38535 | 6.54612 | 6.19075 | 6.10001 | 6.31924 | 6.26019 | 6.11276 | 6.21787 | 6.23775 | 6.34599 | 6.19337 | 6.15406 |
| ABR | Q12979 | 6.30799 | 6.29198 | 6.7511 | 5.69268 | 6.38649 | 6.63513 | 6.31559 | 6.54462 | 6.44455 | 6.31991 | 6.35583 | 6.18242 | 6.10294 | 6.2828 | 5.76339 | 5.62258 |
| IDH3B | O43837 | 6.4839 | 6.71596 | 6.00593 | 6.97506 | 6.19884 | 5.94569 | 6.32441 | 6.05073 | 6.06444 | 6.1865 | 6.31196 | 6.18815 | 6.07306 | 6.2478 | 6.04283 | 6.18474 |
| GOLGA1 | Q92805 | 6.73063 | 6.51599 | 6.05293 | 6.51014 | 6.71408 | 6.20372 | 6.23727 | 6.33542 | 6.57406 | 6.16217 | 6.13738 | 6.51113 | 6.42408 | 5.53559 | 5.56697 | 5.78844 |
| TONSL | Q96HA7 | 6.21896 | 6.13646 | 6.03976 | 6.34954 | 5.93171 | 6.02132 | 6.16552 | 6.05625 | 6.17904 | 6.09745 | 6.23124 | 6.31049 | 6.60048 | 6.36371 | 6.79361 | 6.50446 |
| INTS10 | Q9NVR2 | 5.98475 | 6.28799 | 6.30794 | 6.74186 | 6.26582 | 5.83989 | 6.03963 | 5.58492 | 6.2781 | 6.12768 | 6.26624 | 6.05237 | 6.67941 | 6.32495 | 6.75955 | 6.4589 |
| GTF3C3 | Q9Y5Q9 | 6.26555 | 6.68888 | 6.23468 | 6.5187 | 6.20947 | 6.21155 | 6.45238 | 6.65403 | 6.23007 | 6.25301 | 6.24109 | 6.16309 | 5.97883 | 6.00862 | 6.22382 | 6.26624 |
| TIAL1 | Q01085-2 | 6.4503 | 6.28487 | 5.88705 | 6.49768 | 6.45411 | 6.30921 | 6.09433 | 6.31221 | 6.08639 | 6.0768 | 6.22425 | 6.2098 | 6.51429 | 6.11265 | 6.23008 | 6.25595 |
| YWHAH | Q04917 | 6.20066 | 6.22888 | 6.70982 | 5.90093 | 6.14206 | 6.39945 | 6.05382 | 6.45597 | 6.46932 | 6.27369 | 6.66212 | 6.42471 | 5.97399 | 6.09968 | 6.0125 | 5.99241 |
| SCAF4 | O95104 | 6.1088 | 6.59792 | 5.9573 | 6.66626 | 6.15391 | 5.86167 | 6.31021 | 6.22054 | 6.08742 | 6.03818 | 5.98956 | 5.99966 | 6.42354 | 6.35886 | 6.53282 | 6.69335 |
| ACAD10 | Q6JQN1-5 | 6.54926 | 6.34574 | 5.95127 | 6.48709 | 6.12379 | 5.92504 | 6.40678 | 6.11874 | 5.81304 | 5.90952 | 5.82868 | 6.02414 | 6.92861 | 6.63041 | 6.45571 | 6.50219 |
| DDX58 | O95786 | 6.28376 | 6.19763 | 6.75563 | 6.11463 | 6.14803 | 6.66271 | 6.3036 | 6.29856 | 6.12799 | 6.53257 | 6.53235 | 6.58865 | 5.91344 | 5.98361 | 6.03924 | 5.51758 |
| ADSL | P30566 | 6.10253 | 6.12538 | 6.85839 | 5.95966 | 6.40788 | 6.99007 | 5.86961 | 6.68256 | 6.58068 | 6.6147 | 6.60787 | 6.53979 | 5.72598 | 5.92698 | 5.5428 | 5.46511 |
| CDC23 | Q9UJX2 | 6.32727 | 6.29837 | 6.58162 | 6.18585 | 6.2072 | 6.39688 | 6.27709 | 5.96467 | 6.1469 | 6.46384 | 6.04503 | 6.0591 | 6.34231 | 5.84389 | 6.4616 | 6.3984 |
| BCCIP | Q9P287-2 | 5.92704 | 6.08838 | 6.42143 | 5.73895 | 6.19709 | 6.39883 | 5.98066 | 6.05936 | 6.18823 | 6.03233 | 6.17653 | 6.1147 | 6.65477 | 6.69632 | 6.90137 | 6.42403 |
| PRKAR2A | P13861 | 6.29669 | 6.33896 | 6.05419 | 6.11652 | 6.57062 | 6.31305 | 6.10305 | 6.35419 | 6.30741 | 6.20476 | 6.3069 | 6.15101 | 6.34959 | 6.06714 | 6.50318 | 5.96275 |
| FOCAD | Q5VW36 | 6.27914 | 6.70711 | 6.7056 | 6.25246 | 5.86106 | 6.23168 | 5.98426 | 6.67785 | 6.33977 | 6.39565 | 6.58132 | 6.73051 | 5.5602 | 5.73987 | 5.9296 | 6.02393 |
| MECP2 | P51608-2 | 6.22995 | 6.25166 | 5.73794 | 6.54971 | 5.98064 | 5.97067 | 6.27248 | 5.90837 | 6.04564 | 6.00499 | 5.61829 | 5.66423 | 6.68837 | 7.015 | 7.19763 | 6.86445 |
| BCL2L13 | Q9BXK5 | 6.43278 | 5.8886 | 5.726 | 6.31853 | 6.01914 | 5.71544 | 6.12357 | 5.79533 | 5.91081 | 5.93782 | 6.02078 | 6.07082 | 7.05151 | 6.94375 | 6.92967 | 7.11547 |
| TGFBI | Q15582 | 7.86193 | 6.41186 | 3.73245 | 7.80152 | 4.84786 | 5.65602 | 8.83628 | 5.79504 | 7.33048 | 5.98733 | 5.09951 | 5.8852 | 7.03701 | 4.45234 | 6.33783 | 6.92733 |
| AIMP2 | Q13155 | 6.21377 | 5.52368 | 6.59675 | 5.30451 | 6.49622 | 6.07911 | 5.82545 | 6.64292 | 6.07636 | 6.59078 | 6.60747 | 6.98158 | 6.47318 | 6.3341 | 6.18216 | 6.07196 |
| GOLIM4 | O00461 | 6.25674 | 6.45459 | 5.73121 | 6.27837 | 6.36586 | 6.37662 | 6.441 | 6.18114 | 6.30862 | 5.98289 | 5.77964 | 5.8291 | 6.73685 | 6.32651 | 6.54424 | 6.40661 |
| ADD1 | P35611-3 | 6.10336 | 6.19053 | 6.3473 | 5.7732 | 5.9346 | 6.38436 | 6.05074 | 6.29275 | 6.07375 | 6.41042 | 6.16233 | 6.08268 | 6.76763 | 6.67602 | 6.42511 | 6.32522 |
| PC | P11498 | 6.1712 | 6.37676 | 6.00208 | 6.4409 | 6.14367 | 6.00685 | 6.14564 | 5.70161 | 6.22575 | 6.10731 | 6.28737 | 6.08578 | 6.8938 | 6.30859 | 6.59872 | 6.50398 |
| BICD2 | Q8TD16-2 | 6.27082 | 6.11864 | 6.66446 | 6.05093 | 6.47921 | 6.64581 | 6.1963 | 6.45123 | 6.82159 | 6.49207 | 6.81754 | 6.64588 | 5.95429 | 5.4896 | 5.47856 | 5.42307 |
| ITGA2 | P17301 | 6.13852 | 6.49514 | 5.84309 | 6.23747 | 6.49701 | 6.23794 | 6.30142 | 6.06993 | 5.96246 | 6.05986 | 5.90815 | 5.72735 | 6.85876 | 6.52165 | 6.76934 | 6.37193 |
| GIGYF2 | Q6Y7W6 | 6.58172 | 6.65684 | 6.82347 | 6.34412 | 6.50947 | 6.66143 | 6.08065 | 6.27215 | 6.5622 | 6.4507 | 6.60445 | 6.72928 | 5.66994 | 5.37545 | 5.32886 | 5.34927 |
| TBC1D9B | Q66K14 | 6.37749 | 6.2276 | 7.01166 | 6.10549 | 6.13826 | 6.5554 | 6.18541 | 6.33926 | 6.53831 | 6.29166 | 6.65478 | 6.78754 | 5.74193 | 5.79841 | 5.64568 | 5.60112 |
| NUFIP2 | Q7Z417 | 6.33064 | 6.30669 | 6.3206 | 6.2094 | 6.47635 | 6.34127 | 6.2787 | 5.93049 | 6.64429 | 6.26416 | 6.28484 | 6.45758 | 6.0967 | 5.96268 | 5.80765 | 6.28794 |
| MAP2 | P11137 | 5.96565 | 6.22925 | 6.68476 | 5.58497 | 5.95547 | 6.67487 | 6.02948 | 6.55282 | 6.07782 | 6.6835 | 6.41308 | 6.23928 | 6.37146 | 6.84063 | 6.05087 | 5.64608 |
| FNBP1 | Q96RU3 | 6.75676 | 6.54924 | 7.28022 | 6.42034 | 6.65911 | 6.6325 | 6.38133 | 6.67401 | 6.37102 | 6.59817 | 6.56784 | 6.42473 | 5.3706 | 5.23374 | 4.98001 | 5.10039 |
| ABCC4 | O15439 | 6.42094 | 6.86849 | 6.11652 | 6.84249 | 6.0456 | 6.02628 | 6.35933 | 6.06962 | 6.33722 | 6.1564 | 5.74231 | 6.0201 | 6.15534 | 6.2421 | 6.18954 | 6.40772 |
| PAPSS2 | O95340-2 | 5.51709 | 5.75803 | 6.19783 | 5.34716 | 5.6484 | 6.41408 | 5.97597 | 6.17149 | 5.91231 | 6.32737 | 6.31204 | 6.35706 | 6.55436 | 7.30018 | 7.22299 | 6.98363 |
| TBC1D15 | Q8TC07-3 | 5.71457 | 5.92249 | 5.99307 | 5.68218 | 5.93213 | 6.32811 | 5.93078 | 6.24562 | 6.27996 | 6.26514 | 6.21622 | 6.43644 | 6.93364 | 6.56186 | 6.7831 | 6.77469 |
| RALB | P11234-2 | 6.55818 | 6.434 | 5.9499 | 6.49875 | 6.28365 | 6.19287 | 6.48113 | 6.23277 | 6.22893 | 6.01137 | 5.93915 | 5.91955 | 6.22592 | 6.48982 | 6.39044 | 6.16356 |
| SEC24A | O95486 | 6.17551 | 6.35753 | 6.67488 | 6.11445 | 6.28976 | 6.44895 | 6.18299 | 6.38298 | 6.2655 | 6.42198 | 6.54001 | 6.54155 | 5.91001 | 6.14021 | 5.79056 | 5.76313 |
| ATXN10 | Q9UBB4 | 6.39609 | 5.88785 | 6.94994 | 5.68269 | 6.3764 | 6.53385 | 6.32425 | 6.8389 | 6.27023 | 6.63009 | 6.56849 | 6.74448 | 5.75812 | 5.84431 | 5.47622 | 5.71809 |
| TOP3B | O95985 | 6.37009 | 6.23417 | 6.60817 | 6.43088 | 6.17951 | 6.12999 | 6.30042 | 6.00915 | 6.7514 | 6.48755 | 6.52703 | 6.52506 | 5.74215 | 5.75787 | 6.07057 | 5.87601 |
| RRS1 | Q15050 | 6.10737 | 6.28691 | 5.86697 | 6.69482 | 5.87868 | 5.8203 | 6.32475 | 6.0202 | 6.00953 | 5.58522 | 5.9479 | 5.77508 | 6.58311 | 6.97752 | 7.14993 | 6.87172 |
| MRPL38 | Q96DV4 | 6.18258 | 6.59598 | 5.96266 | 6.83613 | 6.2986 | 6.13244 | 6.57422 | 6.3589 | 6.43618 | 6.23651 | 5.82376 | 6.00889 | 6.40084 | 5.82571 | 6.23881 | 6.08778 |
| GPS1 | Q13098-7 | 6.22116 | 6.1825 | 6.61885 | 5.70594 | 6.30719 | 6.43359 | 6.22361 | 5.84138 | 6.22046 | 6.35783 | 6.51822 | 6.27151 | 6.58061 | 6.10279 | 6.30217 | 6.1122 |
| TTLL12 | Q14166 | 5.76285 | 6.07479 | 6.66177 | 5.84609 | 6.23267 | 7.0697 | 5.80985 | 6.63265 | 6.05302 | 6.28397 | 6.49247 | 6.6549 | 6.1898 | 6.36967 | 6.06489 | 5.80092 |
| OTUB1 | Q96FW1 | 6.28575 | 5.50708 | 6.61438 | 5.18352 | 6.49866 | 6.63118 | 6.12487 | 6.76014 | 6.41676 | 6.71405 | 6.37094 | 6.45669 | 6.0959 | 6.3507 | 6.07372 | 5.91566 |
| ITCH | Q96J02 | 6.32684 | 6.44355 | 6.80913 | 6.5297 | 6.48157 | 6.72226 | 6.23925 | 6.12901 | 6.4585 | 6.4044 | 6.56587 | 6.3212 | 5.61404 | 5.68201 | 5.78039 | 5.49229 |
| HSD17B11 | Q8NBQ5 | 6.30467 | 6.57865 | 6.36749 | 6.59217 | 6.20508 | 6.19074 | 6.39106 | 5.72387 | 6.09569 | 6.50387 | 6.0965 | 5.88976 | 6.23949 | 6.16496 | 6.61679 | 6.03922 |
| BRAT1 | Q6PJG6 | 5.88242 | 5.5389 | 6.52289 | 5.51309 | 5.93979 | 6.14486 | 5.93546 | 6.08029 | 6.22554 | 6.39125 | 6.75057 | 6.36988 | 6.3804 | 6.62182 | 6.86065 | 6.84219 |
| CCDC22 | O60826 | 6.04306 | 6.17769 | 6.58872 | 5.85015 | 6.43338 | 6.35035 | 5.90642 | 6.46593 | 6.2767 | 6.55668 | 6.40157 | 6.51704 | 6.07472 | 6.19174 | 6.07095 | 6.09491 |
| C1QBP | Q07021 | 6.36734 | 6.70064 | 5.54759 | 6.72103 | 6.25771 | 5.89926 | 6.06377 | 6.52109 | 5.93724 | 6.05613 | 5.71547 | 6.22561 | 6.72956 | 6.50506 | 5.81898 | 6.93352 |
| PGM3 | O95394-4 | 6.16145 | 6.2039 | 6.88097 | 5.86074 | 6.35596 | 6.85509 | 6.29514 | 6.76078 | 6.33761 | 6.54393 | 6.39716 | 6.43068 | 5.81795 | 5.67123 | 5.81715 | 5.61026 |
| ITGA5 | P08648 | 6.00197 | 6.38655 | 6.08511 | 6.53248 | 6.05881 | 5.9697 | 6.11757 | 6.11734 | 6.37395 | 6.19546 | 6.08312 | 6.05777 | 6.64864 | 6.23122 | 6.62274 | 6.51755 |
| GLB1 | P16278 | 6.60853 | 6.48512 | 6.03999 | 6.55853 | 6.37797 | 6.11874 | 6.17028 | 6.10557 | 6.05197 | 6.07273 | 6.04757 | 5.8779 | 6.64751 | 6.12642 | 6.59788 | 6.11329 |
| JAK1 | P23458 | 6.92506 | 7.25657 | 6.8737 | 7.23338 | 6.56364 | 6.40881 | 6.76519 | 6.2628 | 6.71794 | 6.89845 | 6.84856 | 6.74353 | 4.67784 | 4.57853 | 4.48481 | 4.76118 |
| DNAJC11 | Q9NVH1 | 6.70051 | 6.44649 | 5.97562 | 6.53763 | 6.45192 | 6.15585 | 6.31369 | 6.04715 | 6.17888 | 5.78876 | 5.90144 | 5.76322 | 6.49286 | 6.1996 | 6.7333 | 6.31309 |
| TAOK1 | Q7L7X3 | 6.1057 | 6.26265 | 6.26288 | 5.92909 | 5.94552 | 5.94522 | 6.15668 | 5.85754 | 6.45245 | 6.60039 | 6.64884 | 6.62975 | 6.24532 | 6.36006 | 6.29117 | 6.30675 |
| RBBP5 | Q15291 | 6.42436 | 6.31851 | 6.07048 | 6.80115 | 6.14859 | 6.2211 | 6.33939 | 5.78433 | 6.10368 | 5.85346 | 5.78226 | 5.73381 | 6.39609 | 6.48878 | 6.7626 | 6.77142 |
| RETSAT | Q6NUM9 | 6.44837 | 6.6629 | 6.038 | 6.7058 | 6.24532 | 6.15237 | 6.52155 | 6.09003 | 6.24897 | 6.30826 | 6.25034 | 6.31644 | 6.07046 | 6.0041 | 5.97999 | 5.95709 |
| RRAS | P10301 | 6.56403 | 6.40803 | 6.20628 | 6.51813 | 6.16182 | 6.09883 | 6.6025 | 6.0405 | 6.48201 | 5.9352 | 5.93148 | 6.04835 | 6.13585 | 6.1711 | 6.38278 | 6.31311 |
| ALDH9A1 | P49189-3 | 6.24021 | 6.11513 | 5.83875 | 6.11114 | 6.36125 | 6.35317 | 6.53677 | 6.40106 | 6.45477 | 6.04962 | 5.92041 | 6.31214 | 5.86627 | 6.51345 | 6.33779 | 6.58807 |
| CXXC1 | Q9P0U4-2 | 6.28765 | 6.39042 | 6.14988 | 6.61875 | 6.28259 | 6.09669 | 6.29677 | 6.03282 | 6.13331 | 6.15088 | 5.98192 | 6.147 | 6.21525 | 6.30593 | 6.56717 | 6.34297 |
| H2AZ1 | P0C0S5 | 6.18897 | 6.31121 | 5.83644 | 6.95884 | 6.21869 | 5.83313 | 6.74025 | 5.79547 | 6.21499 | 5.65216 | 5.62234 | 5.79957 | 6.41693 | 6.71942 | 6.67971 | 7.01188 |
| PRPSAP2 | O60256 | 6.18654 | 6.20389 | 7.08297 | 5.87697 | 6.39795 | 6.76742 | 6.4233 | 6.44609 | 6.21894 | 6.32979 | 6.52547 | 6.33944 | 5.88923 | 5.8665 | 5.79719 | 5.64829 |
| ATP5MF-PTCD1 | G3V325 | 6.16598 | 6.43583 | 5.6911 | 6.78643 | 6.14126 | 6.20367 | 6.4714 | 5.89548 | 6.41464 | 6.07426 | 6.1232 | 6.30624 | 6.5879 | 6.18205 | 6.16038 | 6.36018 |
| AAK1 | Q2M2I8 | 6.16423 | 6.04187 | 6.5815 | 5.86682 | 6.45757 | 6.50728 | 5.88994 | 6.25047 | 6.40072 | 6.56231 | 6.77424 | 6.63357 | 6.0197 | 6.20432 | 5.74722 | 5.89823 |
| ARHGAP35 | Q9NRY4 | 5.92452 | 6.0346 | 6.4012 | 5.92169 | 5.95505 | 6.42594 | 6.08119 | 6.51968 | 6.35817 | 6.33883 | 6.25941 | 6.49702 | 6.34697 | 6.50639 | 6.27037 | 6.15896 |
| HSPA14 | Q0VDF9 | 6.38933 | 6.23164 | 6.71131 | 5.90479 | 6.48688 | 6.40715 | 6.1326 | 6.79878 | 6.56606 | 6.33089 | 6.69571 | 6.47053 | 5.85355 | 5.76948 | 5.65481 | 5.5965 |
| FMR1 | Q06787 | 6.45784 | 6.78745 | 6.30265 | 6.25486 | 6.28896 | 6.32768 | 6.43038 | 6.24469 | 6.15423 | 6.399 | 6.2056 | 6.2865 | 6.10294 | 5.92813 | 5.90485 | 5.92424 |
| MPP5 | Q8N3R9 | 6.32839 | 6.19091 | 6.43797 | 6.25888 | 6.3839 | 6.36519 | 6.48076 | 6.2057 | 6.16493 | 6.14864 | 6.48063 | 6.10676 | 6.30828 | 6.18532 | 6.02209 | 5.93166 |
| ZPR1 | O75312 | 6.67916 | 6.07236 | 6.7953 | 5.77241 | 6.52613 | 6.77971 | 6.0093 | 6.98551 | 6.35323 | 6.3736 | 6.66174 | 6.6701 | 6.0434 | 5.53246 | 5.21064 | 5.43498 |
| ANXA11 | P50995 | 6.06279 | 6.24616 | 6.67655 | 5.77197 | 6.43343 | 6.71379 | 6.00994 | 6.7757 | 6.32567 | 6.35831 | 6.32687 | 6.36619 | 6.04189 | 6.29976 | 5.8732 | 5.71777 |
| FAF2 | Q96CS3 | 6.31643 | 5.96633 | 5.94788 | 6.32632 | 5.89371 | 5.93955 | 6.22938 | 6.05397 | 6.30485 | 6.11292 | 6.12485 | 6.20167 | 6.63447 | 6.56501 | 6.74629 | 6.63636 |
| PHACTR4 | Q8IZ21-2 | 6.18412 | 6.30026 | 6.28919 | 6.06025 | 6.18994 | 6.02822 | 6.49513 | 6.19674 | 6.72044 | 6.30619 | 6.16478 | 6.27714 | 6.16258 | 6.33194 | 6.10277 | 6.1903 |
| ZNF574 | Q6ZN55-2 | 5.39461 | 5.58405 | 5.72597 | 6.41402 | 5.87791 | 5.85659 | 6.87566 | 5.47976 | 6.45586 | 5.04382 | 5.64412 | 5.89441 | 6.22647 | 7.76464 | 7.58468 | 8.17743 |
| STAG1 | Q8WVM7 | 6.24843 | 6.55703 | 6.04051 | 6.81857 | 6.04509 | 5.80934 | 6.18846 | 5.66615 | 5.99084 | 6.02395 | 6.11258 | 6.15071 | 6.59158 | 6.35367 | 6.56921 | 6.83388 |
| ST13 | P50502 | 6.68966 | 6.30126 | 6.67157 | 5.96571 | 6.88467 | 6.82385 | 6.4188 | 6.91872 | 6.8437 | 6.13681 | 6.34235 | 6.19723 | 5.59789 | 5.49314 | 5.2009 | 5.51372 |
| KIF14 | Q15058 | 6.42564 | 6.49725 | 6.21544 | 6.75946 | 6.20273 | 6.4826 | 6.77929 | 6.18491 | 6.45768 | 6.4286 | 6.23455 | 6.46406 | 5.65874 | 5.79695 | 5.74675 | 5.66536 |
| SGTA | O43765 | 6.34958 | 5.25011 | 6.41204 | 4.86222 | 6.21544 | 5.96381 | 5.63371 | 6.82776 | 5.78241 | 6.72679 | 6.36149 | 7.00826 | 6.57763 | 6.85129 | 6.66446 | 6.513 |
| PPM1G | O15355 | 5.4302 | 5.64354 | 5.95221 | 5.04141 | 5.52362 | 6.297 | 5.59676 | 6.23358 | 5.8325 | 6.06264 | 6.40264 | 6.26827 | 6.80439 | 7.87039 | 7.68726 | 7.35356 |
| GTF3C2 | Q8WUA4 | 6.19863 | 6.5584 | 5.98958 | 6.5761 | 6.05075 | 6.10889 | 6.11154 | 6.13751 | 5.92961 | 6.32205 | 6.05359 | 5.82569 | 6.35289 | 6.64152 | 6.70755 | 6.4357 |
| GNPAT | O15228 | 6.23867 | 6.42367 | 6.23225 | 6.71193 | 6.44612 | 6.21189 | 6.26761 | 5.82239 | 6.26406 | 6.29347 | 6.2384 | 5.80268 | 6.3179 | 6.12271 | 6.3375 | 6.26874 |
| EML3 | Q32P44 | 6.45072 | 6.35674 | 6.85299 | 5.92631 | 6.46653 | 6.52482 | 6.13053 | 6.45684 | 5.94249 | 6.31894 | 6.38022 | 6.5487 | 6.11541 | 6.06924 | 5.70246 | 5.75707 |
| RFC4 | P35249 | 5.97399 | 6.06743 | 6.04552 | 5.95368 | 6.24878 | 6.17579 | 6.18696 | 6.11925 | 6.06868 | 6.29732 | 6.24601 | 6.19347 | 6.64637 | 6.54643 | 6.67629 | 6.55403 |
| SRPRB | Q9Y5M8 | 6.54882 | 6.27985 | 5.79284 | 6.41812 | 6.50277 | 5.98089 | 6.53884 | 5.99435 | 6.43385 | 6.04621 | 5.74372 | 5.71424 | 6.62571 | 6.27694 | 6.61326 | 6.48961 |
| NR2C2 | P49116 | 6.32728 | 6.47248 | 6.05176 | 6.68445 | 6.13252 | 6.10078 | 6.22546 | 5.90591 | 6.11938 | 6.18452 | 6.11922 | 6.03675 | 6.46966 | 6.23504 | 6.54987 | 6.38491 |
| KLC4 | Q9NSK0-3 | 6.58017 | 6.08993 | 7.03822 | 6.28387 | 6.58926 | 6.56304 | 6.04882 | 6.3607 | 6.67666 | 6.47432 | 6.224 | 6.44527 | 6.08769 | 5.67241 | 5.43852 | 5.42711 |
| ARHGEF40 | Q8TER5 | 6.34786 | 6.39104 | 6.29438 | 6.10865 | 5.99819 | 6.58659 | 6.3003 | 6.40227 | 6.46016 | 6.6401 | 6.50521 | 6.23431 | 5.97311 | 6.24697 | 5.99339 | 5.51747 |
| TDRD7 | Q8NHU6 | 6.0575 | 6.26498 | 6.69106 | 6.00142 | 5.87247 | 6.42595 | 6.10678 | 6.28188 | 6.23147 | 6.42182 | 6.6157 | 6.27736 | 6.07031 | 6.13895 | 6.64857 | 5.8938 |
| VPS50 | Q96JG6 | 6.30202 | 6.1688 | 6.74018 | 6.0325 | 6.26557 | 6.36992 | 6.22886 | 6.40731 | 6.44326 | 6.45573 | 6.6239 | 6.57443 | 5.92017 | 5.81691 | 5.88502 | 5.76543 |
| MAP2K3 | P46734-3 | 6.19344 | 5.89591 | 7.22328 | 6.09341 | 6.21111 | 6.68794 | 6.21397 | 6.3992 | 6.54549 | 6.03128 | 6.36474 | 6.31411 | 5.95114 | 6.02314 | 6.05901 | 5.79282 |

| | | | | | | | | | | | | | | | | | |
|---|---|---|---|---|---|---|---|---|---|---|---|---|---|---|---|---|---|
| TMOD3 | Q9NYL9 | 6.57501 | 5.93917 | 6.39875 | 5.67368 | 6.70574 | 6.21283 | 5.78594 | 6.80586 | 6.25108 | 6.45245 | 6.55992 | 6.96256 | 6.41841 | 5.963 | 5.55433 | 5.74127 |
| DHX35 | Q9H5Z1 | 6.44515 | 6.34794 | 5.63788 | 6.23081 | 6.42003 | 6.30167 | 6.28366 | 5.72193 | 6.05171 | 5.94905 | 5.97118 | 6.06579 | 6.67282 | 6.3308 | 6.99311 | 6.57646 |
| PPP1R18 | Q6NYC8 | 6.22922 | 6.43318 | 5.99939 | 6.2516 | 6.30526 | 6.28432 | 5.78189 | 5.90545 | 6.22406 | 6.2681 | 6.03508 | 6.29253 | 6.97893 | 6.47378 | 6.27368 | 6.26352 |
| NOM1 | Q5C9Z4 | 6.24673 | 6.10991 | 5.41974 | 6.56336 | 6.04536 | 5.79165 | 6.61861 | 6.14161 | 6.6311 | 5.99077 | 6.01691 | 6.08539 | 6.29799 | 6.46087 | 6.50114 | 7.07885 |
| IPO11 | Q9UI26-2 | 5.79379 | 5.55082 | 6.29213 | 5.39215 | 5.81039 | 6.22046 | 5.95123 | 6.11476 | 6.1051 | 6.1694 | 6.43614 | 5.97436 | 6.88374 | 7.09295 | 7.15477 | 7.05781 |
| CCNB1 | P14635 | 6.50305 | 5.8036 | 6.7975 | 6.26284 | 6.12448 | 6.25288 | 6.45658 | 6.52411 | 6.79921 | 6.21729 | 6.67952 | 6.47234 | 5.96622 | 5.67897 | 5.64517 | 5.81624 |
| FAM193A | A0A1B0GVL4 | 6.24122 | 6.57192 | 6.55063 | 6.32042 | 6.15083 | 6.5181 | 6.1526 | 6.59681 | 6.10571 | 6.77913 | 6.25344 | 6.60205 | 5.39152 | 6.17223 | 5.61475 | 5.97864 |
| TSC2 | P49815 | 6.57181 | 6.78808 | 7.2157 | 6.67077 | 6.21292 | 6.75034 | 6.13494 | 6.32487 | 6.67803 | 6.82616 | 6.62626 | 6.91716 | 5.01463 | 5.29078 | 4.83129 | 5.14628 |
| MRPS27 | Q92552-2 | 6.55113 | 6.44253 | 5.98611 | 6.71548 | 6.3208 | 5.99506 | 6.44649 | 6.19017 | 6.09249 | 5.71532 | 6.15785 | 6.1102 | 6.50435 | 6.03955 | 6.49549 | 6.23699 |
| NOSIP | Q9Y314 | 6.29037 | 6.00349 | 7.00512 | 6.38114 | 6.0917 | 6.47518 | 5.97017 | 6.39046 | 6.53716 | 6.79458 | 6.92636 | 7.02803 | 5.3468 | 5.82252 | 5.52931 | 5.40759 |
| HERC5 | Q9UII4 | 6.30171 | 6.62178 | 6.84865 | 6.25542 | 6.35178 | 6.34415 | 6.32765 | 6.19077 | 6.55936 | 6.75422 | 6.55228 | 6.50991 | 5.40374 | 5.77304 | 5.61673 | 5.58879 |
| PLK1 | P53350 | 6.9299 | 7.12738 | 6.75244 | 7.30596 | 6.19482 | 6.54369 | 6.68198 | 6.34451 | 6.68401 | 6.74597 | 6.54784 | 6.82301 | 4.53335 | 5.01945 | 4.65502 | 5.11066 |
| WDR75 | Q8IWA0 | 6.2237 | 6.1438 | 6.08983 | 6.89611 | 6.10341 | 5.85212 | 6.37738 | 5.94673 | 6.28864 | 6.01087 | 6.01751 | 6.02466 | 6.36136 | 6.09779 | 6.81177 | 6.75434 |
| ANKRD27 | Q96NW4 | 6.16529 | 6.17498 | 6.60757 | 6.47505 | 5.98438 | 6.39437 | 6.10742 | 6.92659 | 6.41859 | 6.54549 | 6.4858 | 6.59095 | 5.87572 | 5.90463 | 6.24247 | 6.1007 |
| COPS3 | Q9UNS2 | 5.97188 | 5.8758 | 6.4523 | 5.68651 | 6.41651 | 6.53111 | 6.27035 | 6.13006 | 6.16304 | 6.30019 | 6.50192 | 6.231 | 6.33223 | 6.48565 | 6.51909 | 6.13234 |
| OS9 | Q13438 | 6.32081 | 6.17863 | 6.01281 | 6.47775 | 6.55646 | 6.10128 | 6.27604 | 6.38766 | 6.23147 | 6.25024 | 6.2245 | 5.93931 | 6.595 | 5.95401 | 6.15003 | 6.34442 |
| EFL1 | Q7Z2Z2 | 6.18315 | 5.85726 | 6.50993 | 5.64027 | 6.2856 | 6.82075 | 5.86197 | 6.69903 | 6.4682 | 6.34135 | 6.26918 | 6.28949 | 6.32298 | 6.41622 | 5.96054 | 6.07407 |
| PNP | P00491 | 6.07919 | 6.293 | 6.64281 | 5.92736 | 6.54678 | 6.71729 | 6.15619 | 6.7312 | 6.25068 | 6.28597 | 6.6327 | 6.73182 | 5.87813 | 5.92973 | 5.6872 | 5.50993 |
| CLPB | Q9H078-2 | 6.3829 | 6.44203 | 5.76905 | 6.7282 | 6.35715 | 5.96933 | 6.64971 | 6.04508 | 6.18797 | 5.9052 | 6.09298 | 5.94205 | 6.67132 | 5.97888 | 6.54604 | 6.33209 |
| ZNF609 | O15014 | 5.9828 | 6.3088 | 5.95316 | 6.20523 | 5.97468 | 5.97499 | 6.2018 | 5.86811 | 6.12785 | 6.5248 | 6.09191 | 6.29192 | 6.20311 | 6.83037 | 6.59076 | 6.86969 |
| PTBP3 | O95758-4 | 6.44497 | 6.2555 | 6.18278 | 6.4491 | 6.45435 | 6.21285 | 6.27131 | 6.32732 | 6.14534 | 6.17924 | 6.00059 | 6.10289 | 6.46795 | 5.90921 | 6.37341 | 6.22321 |
| UBQLN2 | Q9UHD9 | 6.03958 | 4.70227 | 5.27132 | 4.4159 | 5.51605 | 4.55836 | 5.45812 | 7.03945 | 4.84991 | 5.8677 | 5.274 | 7.13777 | 6.24713 | 9.74171 | 8.52773 | 9.35299 |
| MLEC | Q14165 | 6.45779 | 6.45566 | 5.78088 | 6.43217 | 6.34808 | 6.06418 | 6.26532 | 6.04364 | 6.12797 | 6.1169 | 5.80588 | 5.754 | 6.68702 | 6.22748 | 6.86167 | 6.57134 |
| USP28 | Q96RU2-2 | 5.96133 | 6.40973 | 5.88887 | 6.41581 | 6.09651 | 6.20114 | 6.03487 | 6.24441 | 6.06835 | 6.26004 | 5.90556 | 6.15458 | 6.64846 | 6.55335 | 6.65858 | 6.4984 |
| GLRX3 | O76003 | 6.10881 | 6.03453 | 6.84399 | 5.67722 | 6.53735 | 7.05172 | 6.13216 | 6.77203 | 6.28833 | 6.55281 | 6.71268 | 6.66427 | 5.73937 | 5.78309 | 5.39155 | 5.71011 |
| PSMD5 | Q16401 | 6.33455 | 6.08624 | 6.72534 | 5.822 | 6.27844 | 6.80997 | 6.00977 | 6.56426 | 6.38735 | 6.52825 | 6.32278 | 6.24893 | 5.8877 | 6.21269 | 5.87592 | 5.90582 |
| DBR1 | Q9UK59 | 5.52177 | 5.82342 | 6.31176 | 5.41876 | 5.77216 | 6.01913 | 5.62579 | 5.94999 | 6.039 | 6.42851 | 6.19471 | 6.15667 | 7.0663 | 7.41147 | 7.25936 | 6.94119 |
| CAVIN1 | Q6NZI2 | 6.31181 | 6.32401 | 5.76093 | 6.64492 | 6.48046 | 6.08632 | 6.76237 | 6.15285 | 6.41713 | 5.72463 | 5.6971 | 5.84403 | 6.32129 | 6.4104 | 6.427 | 6.63475 |
| MRPL4 | Q9BYD3 | 6.3381 | 6.46001 | 5.8663 | 6.91835 | 6.46667 | 6.24503 | 6.55916 | 6.18377 | 6.28761 | 6.20422 | 5.92933 | 6.0825 | 6.21505 | 6.04488 | 6.22642 | 5.97261 |
| FCHO2 | Q0JRZ9 | 6.34725 | 6.13036 | 6.70684 | 5.99053 | 6.36747 | 6.51008 | 6.1762 | 6.26194 | 6.65614 | 6.52531 | 6.50969 | 6.67004 | 6.00534 | 5.87973 | 5.56227 | 5.7008 |
| USP13 | Q92995 | 5.86072 | 6.04059 | 6.51225 | 5.89972 | 5.83627 | 6.56806 | 6.23621 | 6.50867 | 6.35791 | 6.58893 | 6.6609 | 6.69084 | 6.10471 | 5.95485 | 6.05463 | 6.12473 |
| CLINT1 | Q14677-3 | 6.17804 | 6.13403 | 6.6551 | 6.4789 | 6.33387 | 6.54707 | 6.45367 | 6.18843 | 6.30167 | 6.08495 | 6.2432 | 6.13737 | 6.09147 | 6.17571 | 5.97787 | 6.01863 |
| SHTN1 | A0MZ66-3 | 6.48242 | 6.05494 | 6.81101 | 5.95459 | 6.27949 | 6.41183 | 6.25134 | 6.55802 | 6.5651 | 6.29893 | 6.70114 | 6.5306 | 6.11848 | 5.62106 | 5.71365 | 5.64742 |
| MYL12B | O14950 | 5.87758 | 6.38171 | 6.5682 | 5.8285 | 6.50184 | 7.00817 | 6.27222 | 7.03617 | 6.51389 | 6.40037 | 6.50582 | 6.75065 | 5.68127 | 6.04625 | 5.10093 | 5.52644 |
| TWF2 | Q6IBS0 | 6.04514 | 6.13491 | 6.43912 | 5.9421 | 6.09588 | 6.64988 | 6.7126 | 6.18449 | 6.32248 | 5.84152 | 6.14417 | 6.37814 | 5.83186 | 6.64463 | 6.01108 | 6.62198 |
| ZC3HC1 | Q86WB0 | 6.03591 | 6.20064 | 5.90051 | 6.41779 | 6.08496 | 6.1583 | 6.35546 | 5.99175 | 5.934 | 6.04193 | 6.11364 | 5.96753 | 6.85451 | 6.38627 | 6.82859 | 6.72822 |
| IKBIP | Q70UQ0-4 | 6.43619 | 6.54802 | 5.57062 | 6.70316 | 6.21114 | 6.00138 | 6.4983 | 6.20252 | 5.99091 | 6.17695 | 5.89161 | 5.92988 | 6.34039 | 6.31588 | 6.67933 | 6.50374 |
| GOPC | Q9HD26 | 6.35326 | 6.49777 | 6.40299 | 6.83515 | 6.69435 | 6.7075 | 6.22524 | 6.42468 | 6.45528 | 6.62116 | 6.38046 | 6.39775 | 5.45052 | 5.77206 | 5.32293 | 5.4599 |
| SLC25A11 | Q02978 | 6.25728 | 6.28263 | 5.7985 | 6.58754 | 6.19395 | 5.82441 | 6.62156 | 5.86226 | 6.5289 | 5.96369 | 5.99598 | 5.74445 | 6.64814 | 6.29839 | 6.83843 | 6.5539 |
| SHCBP1 | Q8NEM2 | 6.1649 | 6.73311 | 5.8951 | 6.41914 | 6.4207 | 6.42749 | 6.06438 | 6.45245 | 6.09123 | 6.2759 | 6.14399 | 6.41974 | 6.14243 | 5.96243 | 6.05644 | 6.33059 |
| DNAJC9 | Q8WXX5 | 5.94905 | 5.78027 | 6.30101 | 5.84621 | 5.7732 | 6.27404 | 5.99963 | 6.4859 | 6.39822 | 6.32502 | 6.40504 | 6.40321 | 6.11787 | 6.86223 | 6.71255 | 6.36653 |
| SQSTM1 | Q13501 | 5.62182 | 5.26946 | 5.79804 | 6.016 | 6.44696 | 6.35605 | 6.47089 | 6.28426 | 6.34251 | 5.76096 | 6.01523 | 6.08771 | 7.52996 | 6.52102 | 6.66168 | 6.81743 |
| EIF3F | O00303 | 6.28436 | 5.46279 | 6.11002 | 5.09415 | 6.42896 | 6.05107 | 6.01909 | 6.75651 | 5.80497 | 6.58017 | 6.17806 | 6.6618 | 5.69083 | 7.19304 | 6.8209 | 6.86329 |
| ITGB4 | P16144 | 6.23971 | 6.57834 | 6.04674 | 6.13203 | 6.02755 | 5.90182 | 6.09671 | 5.95104 | 6.36344 | 6.34294 | 6.05369 | 6.18543 | 6.58324 | 6.6453 | 6.51215 | 6.33987 |
| HNRNPA0 | Q13151 | 6.4446 | 6.53887 | 5.97403 | 7.01824 | 6.25289 | 6.2108 | 6.23474 | 6.06473 | 6.16336 | 6.16503 | 6.04793 | 6.18746 | 6.3887 | 5.92907 | 6.20699 | 6.17255 |
| INTS14 | Q96SY0 | 6.21165 | 6.26968 | 5.91671 | 6.41113 | 6.24994 | 6.10687 | 6.07196 | 6.0428 | 6.12326 | 6.13875 | 6.10938 | 5.93044 | 6.7563 | 6.56143 | 6.56351 | 6.53618 |
| KEAP1 | Q14145 | 6.75331 | 6.48257 | 6.73028 | 6.57068 | 6.50296 | 6.28671 | 6.47991 | 6.43977 | 6.36307 | 6.56249 | 6.62227 | 6.48677 | 5.36605 | 5.42837 | 5.33485 | 5.58996 |
| PPP1R7 | Q15435 | 5.87171 | 6.01648 | 6.3299 | 5.66366 | 6.45321 | 6.67632 | 6.1018 | 7.04498 | 6.8726 | 6.83218 | 6.53499 | 6.27913 | 5.86272 | 5.96927 | 5.87138 | 5.61967 |
| PXN | P49023-3 | 6.427 | 6.03709 | 6.92578 | 5.94504 | 6.69018 | 6.74408 | 6.24118 | 6.74865 | 6.51945 | 6.17291 | 6.89377 | 6.58945 | 6.11556 | 5.58909 | 5.11602 | 5.24477 |
| LRCH4 | O75427 | 6.21795 | 6.39463 | 6.08624 | 6.37375 | 6.31885 | 6.35542 | 6.38882 | 5.92591 | 6.13146 | 6.34565 | 6.33167 | 6.30152 | 6.27557 | 5.92236 | 6.35695 | 6.27327 |
| SDAD1 | Q9NVU7 | 6.15213 | 6.35291 | 6.05554 | 6.89699 | 6.28004 | 5.96251 | 6.51366 | 6.00405 | 6.35026 | 6.07374 | 5.93005 | 6.0516 | 6.19437 | 6.29303 | 6.48081 | 6.40832 |
| ANP32B | Q92688 | 5.57043 | 5.80537 | 6.15929 | 5.15594 | 5.80184 | 6.33378 | 5.73937 | 6.27005 | 6.07881 | 6.27998 | 6.47338 | 6.0971 | 6.67639 | 7.30744 | 7.43925 | 6.81158 |
| HAUS6 | Q7Z4H7 | 6.4509 | 6.35 | 6.86668 | 6.34681 | 6.14682 | 6.81369 | 6.17267 | 6.48736 | 6.42865 | 6.3383 | 6.73842 | 6.54076 | 5.94801 | 5.54895 | 5.30621 | 5.51579 |
| NKRF | O15226-2 | 6.4298 | 6.52708 | 6.03296 | 6.85097 | 6.04703 | 5.78705 | 6.48249 | 5.80013 | 6.16004 | 6.03348 | 6.19376 | 6.09667 | 6.4734 | 5.99418 | 6.59969 | 6.49127 |
| METTL16 | Q86W50 | 5.41075 | 5.74469 | 5.77047 | 5.34989 | 5.49734 | 6.02208 | 5.93189 | 6.07134 | 5.91486 | 6.08723 | 6.36974 | 6.00904 | 7.19568 | 7.33437 | 7.75878 | 7.53186 |
| RAPGEF6 | Q8TEU7-4 | 6.20739 | 5.80264 | 6.86967 | 6.24969 | 6.18489 | 6.56548 | 6.2955 | 6.48559 | 6.43136 | 6.6309 | 6.6198 | 6.47794 | 5.8395 | 5.83714 | 5.77529 | 5.72721 |
| PSMD8 | P48556 | 5.93663 | 5.65058 | 6.23513 | 5.5204 | 6.45695 | 6.18593 | 6.06113 | 6.80286 | 5.75899 | 6.56869 | 6.25401 | 6.65785 | 6.36329 | 6.48456 | 6.64497 | 6.41802 |
| ATP13A3 | Q9H7F0 | 6.63813 | 6.99802 | 6.58599 | 7.00314 | 6.28354 | 6.16368 | 6.23396 | 6.09508 | 6.30871 | 6.6111 | 6.33618 | 6.42074 | 5.38941 | 5.40343 | 5.53215 | 5.99672 |
| MRPL19 | P49406 | 6.45812 | 6.51667 | 6.00416 | 6.57002 | 6.61365 | 5.8152 | 6.57529 | 6.06803 | 6.09129 | 5.99146 | 5.76803 | 5.77132 | 6.64298 | 6.23218 | 6.54414 | 6.33746 |
| SENP6 | Q9GZR1 | 5.78945 | 6.19206 | 6.3728 | 6.26139 | 5.9142 | 6.22592 | 6.07988 | 6.00577 | 6.30318 | 6.38569 | 6.23337 | 6.32981 | 6.2429 | 6.37046 | 6.60357 | 6.68955 |
| ZNHIT6 | Q9NWK9 | 6.4727 | 6.5025 | 6.65436 | 6.76741 | 6.43739 | 6.47341 | 6.55137 | 6.53801 | 6.5022 | 6.08713 | 6.52812 | 6.36112 | 5.72326 | 5.2491 | 5.66879 | 5.48312 |
| NKTR | P30414 | 6.05711 | 6.18383 | 6.11693 | 6.68729 | 5.73495 | 5.8056 | 6.23954 | 5.77798 | 6.02198 | 5.88911 | 6.04447 | 5.77893 | 6.63347 | 7.07456 | 6.77343 | 7.18084 |
| EIF2B1 | Q14232 | 6.16943 | 6.00583 | 6.75595 | 6.018 | 6.46939 | 6.86159 | 6.18669 | 6.47787 | 6.44715 | 6.72605 | 6.32615 | 6.49883 | 5.77925 | 5.98306 | 5.56539 | 5.72935 |
| TPT1 | P13693 | 6.30346 | 5.08266 | 6.8311 | 5.32065 | 6.36837 | 6.23951 | 5.99086 | 6.79887 | 6.34473 | 6.85896 | 6.96466 | 7.40599 | 5.98908 | 5.98232 | 5.70287 | 5.81592 |
| UNC13B | O14795-2 | 6.31024 | 5.81625 | 6.58123 | 5.85947 | 6.35086 | 6.5078 | 6.3302 | 6.21434 | 6.46295 | 6.68709 | 6.7675 | 6.5335 | 5.91909 | 6.27129 | 5.80089 | 5.5873 |
| CAMSAP2 | Q08AD1 | 6.26488 | 6.2244 | 6.80656 | 6.06858 | 6.11808 | 6.64679 | 6.25151 | 6.36538 | 6.31455 | 6.32731 | 6.52216 | 6.60278 | 5.89487 | 5.87678 | 5.81806 | 5.89732 |
| MIA2 | Q96PC5 | 6.41087 | 6.28143 | 6.15784 | 6.48411 | 6.26754 | 5.92759 | 6.31504 | 5.84199 | 6.11234 | 6.07479 | 6.15192 | 6.04823 | 6.7645 | 6.14742 | 6.58727 | 6.42712 |
| MRPS31 | Q92665 | 6.52626 | 6.6063 | 6.08469 | 6.78607 | 6.11879 | 5.86084 | 6.51886 | 5.98362 | 6.05879 | 6.17107 | 6.04948 | 5.95458 | 6.49543 | 6.04805 | 6.46642 | 6.27076 |
| RPS15 | P62841 | 6.42497 | 6.26423 | 6.3231 | 6.36304 | 6.01768 | 5.87884 | 6.71241 | 6.09402 | 6.79522 | 6.18244 | 5.56147 | 6.2589 | 5.63246 | 6.36837 | 5.99947 | 7.12337 |
| WDR74 | Q6RFH5 | 6.29902 | 6.39971 | 6.41332 | 6.81497 | 6.26998 | 6.01519 | 6.10079 | 6.35007 | 6.41459 | 5.95394 | 6.30108 | 6.1639 | 6.3221 | 5.82932 | 6.11185 | 6.24016 |
| MRPS5 | P82675 | 6.37241 | 6.07237 | 5.52618 | 6.62928 | 6.28785 | 6.03224 | 6.46906 | 6.13988 | 6.09386 | 6.15219 | 5.83657 | 5.8846 | 6.47981 | 6.7731 | 6.60204 | 6.64856 |
| ATP1B3 | P54709 | 6.49144 | 6.30852 | 6.2344 | 6.40191 | 6.21787 | 5.95543 | 6.14573 | 6.28051 | 6.64831 | 6.59533 | 6.4444 | 6.70948 | 6.01626 | 5.65065 | 5.85125 | 6.04851 |
| NFKB1 | P19838-2 | 6.26043 | 5.8547 | 6.7639 | 5.94369 | 6.27989 | 6.78342 | 6.2784 | 6.62721 | 6.41253 | 6.33762 | 6.60239 | 6.58319 | 6.20161 | 5.69027 | 5.72872 | 5.65203 |
| FKBP9 | O95302-3 | 6.47494 | 6.20257 | 6.09384 | 6.62467 | 6.62114 | 6.07599 | 6.41935 | 6.01996 | 6.51428 | 5.74121 | 6.30084 | 6.04416 | 6.54354 | 5.89675 | 6.04384 | 6.38293 |
| DTYMK | P23919 | 6.2556 | 6.02465 | 6.84398 | 5.75856 | 6.23357 | 6.74746 | 6.04528 | 6.51387 | 6.33976 | 6.80335 | 6.76277 | 6.18755 | 6.08918 | 5.67468 | 5.88921 | 5.83054 |
| RPL13 | P26373 | 6.18444 | 5.92071 | 5.97031 | 5.94101 | 5.92927 | 6.0888 | 6.39877 | 6.41061 | 6.91255 | 6.39521 | 6.48042 | 6.37113 | 6.3943 | 6.44809 | 5.65279 | 6.50159 |
| KPNA1 | P52294 | 6.04447 | 5.76719 | 6.39266 | 6.23764 | 6.39637 | 6.4185 | 6.19959 | 6.21263 | 6.09592 | 6.14041 | 6.17637 | 6.02776 | 6.63323 | 6.41095 | 6.46431 | 6.38198 |
| CTNNBL1 | Q8WYA6 | 6.08912 | 6.27342 | 5.71843 | 6.11829 | 6.18337 | 6.01679 | 6.33338 | 5.94318 | 6.12145 | 6.09879 | 5.71418 | 6.04521 | 7.11785 | 6.53003 | 6.81654 | 6.87999 |
| LRPAP1 | P30533 | 6.67523 | 6.48111 | 6.26001 | 6.37963 | 6.49438 | 6.2682 | 6.31117 | 6.2741 | 6.46593 | 6.08897 | 6.12236 | 5.95529 | 6.53369 | 5.6934 | 6.12951 | 5.86703 |
| MAP2K2 | P36507 | 6.2163 | 5.95887 | 6.79731 | 5.9702 | 6.54474 | 6.96282 | 6.25849 | 7.00855 | 6.26455 | 6.46433 | 6.63844 | 6.55422 | 5.96292 | 5.98074 | 5.21578 | 5.20174 |
| DHODH | Q02127 | 6.52309 | 6.33888 | 5.70651 | 6.63181 | 6.48435 | 6.35996 | 6.37026 | 6.36286 | 5.96769 | 5.85519 | 6.01653 | 5.97875 | 6.69809 | 6.01224 | 6.37244 | 6.32133 |
| FABP5 | Q01469 | 6.0864 | 5.77856 | 6.72247 | 5.84839 | 6.99404 | 6.77474 | 6.71591 | 6.27004 | 7.06399 | 5.75721 | 6.37355 | 6.60132 | 5.4494 | 6.11428 | 5.38676 | 6.06295 |
| CCDC47 | Q96A33 | 6.50119 | 6.59257 | 6.24719 | 6.60422 | 6.05302 | 6.06388 | 6.4476 | 5.93427 | 5.8767 | 6.17169 | 5.95771 | 5.7194 | 6.53192 | 6.13746 | 6.56601 | 6.59518 |
| WDR61 | Q9GZS3 | 6.00769 | 6.23779 | 6.10281 | 5.98342 | 6.48491 | 6.42913 | 5.70813 | 6.45789 | 6.33281 | 6.37152 | 6.46588 | 6.44364 | 6.38248 | 6.06094 | 6.04455 | 6.48639 |
| PDXDC1 | Q6P996 | 6.22425 | 6.23879 | 6.08639 | 6.55766 | 6.08333 | 6.28704 | 6.14226 | 6.46269 | 6.40888 | 6.68486 | 6.57558 | 6.66137 | 6.17481 | 6.04598 | 5.78413 | 5.58198 |
| SUCLG1 | P53597 | 6.30953 | 6.32544 | 5.9464 | 6.66477 | 6.39928 | 6.14399 | 6.27259 | 6.17339 | 6.15066 | 6.25967 | 6.27637 | 6.11203 | 6.51715 | 6.053 | 6.21995 | 6.17579 |
| COPS5 | Q92905 | 6.12442 | 5.92761 | 6.59425 | 5.78229 | 6.29311 | 6.48036 | 6.28889 | 6.32014 | 5.96176 | 6.5132 | 6.39897 | 6.12557 | 6.37147 | 6.32026 | 6.34563 | 6.15206 |
| CAPNS1 | P04632 | 6.08965 | 6.41315 | 6.01919 | 4.81125 | 6.00075 | 5.77232 | 6.66275 | 6.64623 | 6.03346 | 6.16395 | 5.78618 | 7.10599 | 5.33781 | 7.94547 | 6.82882 | 8.18304 |
| STAG3 | Q9UJ98 | 6.38944 | 6.61414 | 6.23333 | 6.30603 | 6.36837 | 6.27437 | 6.30766 | 6.33229 | 6.2546 | 6.43234 | 6.44098 | 6.42806 | 5.7847 | 5.85062 | 5.72706 | 6.25601 |
| CDC27 | P30260-2 | 6.54817 | 6.58167 | 6.54768 | 6.67401 | 6.38198 | 6.36025 | 6.54577 | 6.11033 | 6.56352 | 6.37273 | 6.31491 | 6.29363 | 5.70806 | 5.64569 | 5.68038 | 5.67123 |
| STRIP1 | Q5VSL9 | 5.95496 | 6.18634 | 6.4562 | 5.94806 | 5.98201 | 6.43558 | 6.14766 | 6.06906 | 6.32502 | 6.46163 | 6.46179 | 6.42251 | 6.40781 | 6.35075 | 6.25937 | 6.13125 |
| CHCHD3 | Q9NX63 | 6.68047 | 6.16699 | 5.91805 | 6.52682 | 6.55505 | 6.08834 | 6.25668 | 6.20003 | 6.25252 | 6.04773 | 6.15144 | 5.94925 | 7.10014 | 5.85808 | 6.15925 | 6.08916 |
| TAF6 | P49848-3 | 6.11597 | 6.54501 | 6.22055 | 6.60253 | 6.32387 | 6.36344 | 6.1686 | 6.36647 | 6.34723 | 6.59825 | 6.43031 | 6.3385 | 6.25236 | 5.64495 | 5.71458 | 5.96738 |
| TEP1 | Q99973 | 6.14821 | 6.4924 | 6.74197 | 6.07572 | 6.22224 | 6.67327 | 6.09816 | 6.57442 | 6.33476 | 6.53482 | 6.65405 | 6.36787 | 5.64034 | 6.23262 | 5.73529 | 5.47386 |
| NUF2 | Q9BZD4 | 6.47144 | 6.63901 | 6.97441 | 6.64508 | 6.25539 | 6.79467 | 6.44142 | 6.66787 | 6.80648 | 6.54644 | 6.52297 | 6.38799 | 5.08465 | 5.20516 | 5.11349 | 5.44352 |
| PCID2 | Q5JVF3 | 6.60673 | 6.48337 | 6.35459 | 6.54617 | 6.4249 | 6.21411 | 6.35683 | 6.15954 | 6.40731 | 6.17813 | 6.28891 | 5.95826 | 6.12669 | 5.70828 | 6.20193 | 5.98425 |
| ALDH1B1 | P30837 | 6.45903 | 6.34195 | 5.69883 | 6.42495 | 6.70282 | 6.14075 | 6.42586 | 6.43724 | 6.01741 | 5.82226 | 5.86997 | 6.09397 | 6.96692 | 5.98286 | 6.30174 | 6.31341 |
| NMD3 | Q96D46 | 6.15169 | 6.19056 | 6.33649 | 6.00676 | 5.92725 | 5.97386 | 5.98627 | 6.17396 | 6.28835 | 5.81656 | 6.17368 | 5.90493 | 6.96709 | 6.60543 | 6.8624 | 6.63473 |
| CDK2 | P24941 | 6.16935 | 5.56552 | 6.48794 | 5.65251 | 6.09377 | 6.3093 | 6.25385 | 6.4505 | 6.3864 | 6.55085 | 6.83363 | 6.99072 | 5.99984 | 5.83822 | 6.17751 | 6.24009 |
| CDK12 | Q9NYV4 | 5.96707 | 6.02234 | 5.93895 | 6.72037 | 5.87547 | 5.83877 | 6.13589 | 5.68942 | 5.89388 | 5.79382 | 5.89298 | 6.15107 | 6.33412 | 7.24097 | 7.22311 | 7.28178 |
| MYO9A | B2RTY4-4 | 6.26645 | 6.16839 | 6.58807 | 6.38309 | 6.0121 | 6.41548 | 6.18317 | 6.33193 | 6.36169 | 6.46699 | 6.28812 | 6.53489 | 5.94937 | 6.20959 | 5.87587 | 5.96482 |
| CCDC91 | Q7Z6B0 | 6.46703 | 5.69176 | 6.84487 | 5.1248 | 6.49279 | 5.84084 | 5.54728 | 7.15467 | 6.09035 | 6.97878 | 6.97079 | 7.31859 | 6.3347 | 5.95833 | 5.63218 | 5.55224 |
| PARD3 | Q8TEW0 | 6.59036 | 6.13626 | 6.3384 | 6.12005 | 6.18798 | 6.13328 | 6.26118 | 6.2058 | 6.20289 | 6.08224 | 6.18767 | 6.22556 | 6.37997 | 6.33663 | 6.40761 | 6.20413 |
| POLG | P54098 | 6.5676 | 6.54355 | 6.10326 | 6.79683 | 6.15118 | 5.70475 | 6.26411 | 5.94799 | 6.33683 | 6.51435 | 6.30986 | 6.37534 | 6.32088 | 5.8628 | 6.18247 | 6.01819 |
| TUBB4B | P68371 | 6.25093 | 6.08631 | 6.78045 | 5.86504 | 6.69884 | 7.07622 | 6.26959 | 6.92184 | 6.08927 | 6.12878 | 6.6415 | 6.83441 | 5.69737 | 5.8149 | 5.16844 | 5.67611 |

| | | | | | | | | | | | | | | | | |
|---|---|---|---|---|---|---|---|---|---|---|---|---|---|---|---|---|
| TRMT2A | Q8IZ69 | 5.51567 | 6.12558 | 5.9024 | 5.39776 | 6.06601 | 6.19195 | 5.92544 | 6.08562 | 6.12364 | 6.26702 | 6.05396 | 5.91931 | 7.16124 | 7.17294 | 7.17585 | 6.91562 |
| ANK2 | Q01484 | 6.04962 | 6.30292 | 6.68806 | 5.74272 | 6.35204 | 6.63171 | 5.88967 | 6.23698 | 6.1661 | 6.16942 | 6.39493 | 6.49955 | 6.19963 | 6.25654 | 6.2739 | 6.1462 |
| HEATR5A | Q86XA9 | 6.22798 | 6.21234 | 6.64007 | 6.25237 | 5.99661 | 6.72155 | 6.15772 | 6.17318 | 6.31886 | 6.62486 | 6.62123 | 6.72525 | 6.07487 | 5.96604 | 5.58382 | 5.70324 |
| DNAJC2 | Q99543 | 5.88819 | 5.92592 | 6.2782 | 5.86899 | 5.85106 | 6.35492 | 5.83538 | 6.3656 | 6.1056 | 6.50353 | 6.4988 | 6.43144 | 6.4483 | 6.52844 | 6.49695 | 6.61867 |
| OSBPL6 | Q9BZF3-5 | 5.59242 | 5.95233 | 6.0351 | 6.23741 | 6.08706 | 6.0761 | 6.0067 | 6.03803 | 6.40045 | 6.38022 | 6.78107 | 6.66906 | 6.22929 | 6.24166 | 6.55732 | 6.71577 |
| TJP2 | Q9UDY2-7 | 6.33937 | 6.16058 | 5.84139 | 6.16103 | 6.17603 | 5.99642 | 6.18637 | 6.14676 | 6.29682 | 6.17607 | 6.05815 | 6.02375 | 7.02051 | 6.61094 | 6.56516 | 6.24064 |
| MAP1S | Q66K74 | 6.21208 | 6.28363 | 6.48291 | 5.98809 | 6.19422 | 6.54601 | 6.32832 | 6.63999 | 6.4691 | 6.62803 | 6.69028 | 6.43079 | 5.61694 | 6.06491 | 5.74171 | 5.68297 |
| MMUT | P22033 | 6.28409 | 6.43208 | 5.66803 | 6.48212 | 6.1486 | 5.99025 | 6.46917 | 6.07436 | 6.00347 | 6.0189 | 5.99736 | 5.77998 | 6.69809 | 6.53188 | 6.67944 | 6.74218 |
| PTPRK | Q15262-2 | 6.56603 | 6.39583 | 6.27743 | 6.42663 | 6.33435 | 6.09599 | 6.25461 | 5.96679 | 6.32562 | 5.95961 | 6.15199 | 6.03418 | 6.1443 | 6.32835 | 6.21967 | 6.5186 |
| FTO | Q9C0B1 | 5.81397 | 5.68969 | 6.58608 | 5.39476 | 6.17054 | 6.33283 | 5.81852 | 6.285 | 5.913 | 6.1803 | 6.48788 | 6.48872 | 6.55681 | 6.81484 | 7.01786 | 6.44921 |
| GNB1 | P62873 | 6.53142 | 6.48478 | 5.8258 | 6.18505 | 6.63665 | 6.35645 | 6.60046 | 6.25726 | 6.41675 | 6.26953 | 6.07123 | 5.83904 | 6.21667 | 5.85137 | 6.38366 | 6.07387 |
| HEATR6 | Q6AI08 | 6.41013 | 6.59471 | 6.62893 | 6.00562 | 6.31198 | 6.58153 | 6.03903 | 6.6787 | 6.37431 | 6.33897 | 6.18762 | 6.4982 | 5.95862 | 6.00511 | 5.56421 | 5.82233 |
| PPIL4 | Q8WUA2 | 5.9557 | 6.22372 | 6.15117 | 6.06315 | 6.27213 | 6.36661 | 6.17674 | 5.8581 | 6.02233 | 5.9528 | 5.77572 | 5.82003 | 6.35853 | 6.86478 | 7.148 | 6.99049 |
| MMS22L | Q6ZRQ5 | 5.95121 | 6.12053 | 6.10948 | 6.08918 | 6.0292 | 6.14138 | 6.23844 | 6.00438 | 6.20562 | 6.1369 | 5.99726 | 6.00252 | 6.86091 | 6.65969 | 6.73246 | 6.72084 |
| ELOB | Q15370 | 5.81846 | 4.90944 | 6.37656 | 4.23989 | 5.5 | 5.6566 | 5.53398 | 6.22936 | 5.69354 | 6.05765 | 5.61731 | 6.62811 | 5.99355 | 8.98166 | 7.85181 | 8.91209 |
| OCIAD1 | Q9NX40 | 6.48604 | 6.24817 | 6.09159 | 6.59581 | 6.61894 | 6.29006 | 6.38711 | 6.0565 | 6.04481 | 6.36022 | 6.4464 | 6.12026 | 6.65573 | 5.88723 | 5.85262 | 5.85852 |
| SMS | P52788 | 6.33816 | 5.91294 | 7.17626 | 5.93727 | 6.64317 | 6.63455 | 6.10011 | 6.65884 | 6.87621 | 6.46044 | 6.63817 | 6.86953 | 5.51965 | 5.39752 | 5.33522 | 5.50198 |
| INTS13 | Q9NVM9 | 5.99077 | 6.44184 | 5.95462 | 6.75624 | 6.24111 | 6.30421 | 6.14665 | 6.35091 | 6.07432 | 6.45622 | 6.00447 | 6.07823 | 6.3194 | 6.35608 | 6.11089 | 6.41403 |
| PACSIN2 | Q9UNF0 | 6.18883 | 6.27557 | 6.64863 | 5.62485 | 6.22749 | 6.31146 | 5.78585 | 6.52022 | 6.41146 | 6.69531 | 6.70211 | 6.78754 | 6.16684 | 5.97419 | 6.01875 | 5.6609 |
| SEPTIN6 | Q14141 | 5.98526 | 5.86537 | 6.48462 | 5.73549 | 6.35845 | 6.33336 | 6.3125 | 6.28127 | 6.29549 | 6.15291 | 6.85804 | 6.06911 | 6.71589 | 6.1998 | 6.36749 | 5.98496 |
| CLSPN | Q9HAW4 | 6.2393 | 6.11909 | 6.46001 | 6.20264 | 6.0443 | 5.97817 | 6.21736 | 5.77884 | 6.71836 | 6.30982 | 6.5737 | 6.18557 | 6.64234 | 6.13608 | 6.05343 | 6.34098 |
| ASH1L | Q9NR48-2 | 6.65387 | 6.91938 | 6.71143 | 7.4205 | 5.941 | 5.82898 | 6.2813 | 6.18867 | 7.30379 | 6.97968 | 6.77205 | 6.91062 | 4.87134 | 5.1017 | 5.08754 | 5.02814 |
| GTF2F2 | P13984 | 5.51153 | 5.66885 | 6.32643 | 5.57888 | 5.73715 | 6.15683 | 5.87882 | 5.96821 | 6.19894 | 6.05728 | 6.59679 | 6.17251 | 6.81808 | 7.093 | 7.32666 | 6.91004 |
| APC | P25054 | 6.19963 | 6.42936 | 6.54083 | 6.69468 | 6.07294 | 6.18887 | 6.1122 | 6.28741 | 6.28212 | 6.44154 | 6.27328 | 6.28048 | 5.82876 | 6.08424 | 6.05859 | 6.22506 |
| RCN2 | Q14257-2 | 6.59433 | 6.04603 | 6.56888 | 5.86654 | 6.84842 | 6.37748 | 5.63932 | 6.0216 | 6.32389 | 6.5748 | 6.3512 | 5.63727 | 6.83008 | 6.25677 | 5.88735 | 6.17604 |
| ZNF622 | Q969S3 | 5.79265 | 5.81359 | 5.78762 | 5.67942 | 5.96279 | 5.94864 | 5.68499 | 6.00826 | 6.09439 | 6.32357 | 6.07053 | 6.12767 | 7.18584 | 7.15391 | 7.27089 | 7.09525 |
| COG4 | Q9H9E3 | 6.53061 | 6.23885 | 6.59807 | 6.4219 | 6.25081 | 6.45881 | 6.24182 | 6.49622 | 6.66972 | 6.65163 | 6.75432 | 6.48082 | 5.50057 | 5.90215 | 5.49444 | 5.30926 |
| PHF6 | Q8IWS0 | 5.7922 | 6.01199 | 5.91195 | 5.9536 | 6.12077 | 6.49928 | 5.9557 | 5.86482 | 6.18656 | 6.26084 | 6.40056 | 5.91829 | 6.71704 | 6.61572 | 7.22014 | 6.57052 |
| ATAD3A | Q9NVI7-2 | 6.20311 | 6.31516 | 6.11676 | 6.72319 | 5.90752 | 5.87254 | 6.21357 | 5.84664 | 6.26294 | 6.17104 | 6.29447 | 6.23497 | 6.56644 | 6.16967 | 6.51576 | 6.58621 |
| IFT122 | Q9HBG6-6 | 6.45296 | 6.62914 | 6.96792 | 6.33031 | 6.36706 | 6.7708 | 6.27751 | 6.36214 | 6.47408 | 6.83816 | 6.59089 | 6.63019 | 5.21606 | 5.23034 | 5.37033 | 5.49213 |
| STK10 | O94804 | 6.00345 | 6.11259 | 6.92196 | 6.07281 | 6.55467 | 6.7757 | 6.36118 | 6.17836 | 6.03133 | 6.28256 | 6.71236 | 6.39411 | 6.21172 | 5.8859 | 5.9019 | 5.59941 |
| ASL | P04424 | 6.28836 | 6.21646 | 6.6381 | 5.73667 | 6.55195 | 6.9144 | 5.98917 | 6.71616 | 6.34652 | 6.65293 | 6.30422 | 6.20454 | 5.89756 | 5.96714 | 5.69012 | 5.8857 |
| RIPK1 | Q13546 | 6.52861 | 6.13481 | 7.06487 | 6.1816 | 6.2975 | 6.51571 | 6.28733 | 6.51681 | 6.93436 | 6.58771 | 6.7404 | 6.44462 | 5.65743 | 5.32066 | 5.52045 | 5.26712 |
| RMDN1 | Q96DB5 | 6.47463 | 5.90573 | 5.81562 | 6.51059 | 6.38983 | 5.85802 | 6.45882 | 6.00577 | 6.08386 | 5.89005 | 6.00333 | 5.94746 | 6.70519 | 6.46303 | 6.7044 | 6.78367 |
| AASDHPPT | Q9NRN7 | 6.26653 | 6.21945 | 6.70543 | 5.82033 | 6.23407 | 6.49937 | 6.00071 | 6.81157 | 6.4087 | 6.54593 | 6.68105 | 6.63158 | 5.79386 | 5.9782 | 5.71371 | 5.6895 |
| ORC1 | Q13415 | 7.38141 | 7.2988 | 6.87223 | 7.81971 | 5.96398 | 5.94071 | 6.39425 | 6.10055 | 6.81225 | 6.67563 | 6.81113 | 6.60804 | 4.71371 | 4.77601 | 4.83173 | 4.99986 |
| CARMIL1 | Q5VZK9 | 6.23499 | 6.17303 | 6.49624 | 5.83274 | 6.16243 | 6.29208 | 6.38921 | 6.21725 | 6.41759 | 6.63302 | 6.57526 | 6.31872 | 6.05578 | 6.44213 | 5.97766 | 5.78187 |
| ARHGAP5 | Q13017 | 5.94079 | 6.33743 | 6.80648 | 6.11429 | 5.77943 | 6.225 | 6.26525 | 6.08925 | 6.37662 | 6.72087 | 6.64124 | 6.96676 | 5.58278 | 6.08633 | 6.0395 | 6.02798 |
| SRSF7 | Q16629 | 6.49808 | 6.05193 | 6.08883 | 6.37257 | 6.35324 | 5.85186 | 6.5788 | 5.58725 | 6.47598 | 6.09534 | 6.2249 | 5.74964 | 7.0768 | 6.06867 | 6.28431 | 6.6418 |
| DIS3L2 | Q8IYB7 | 6.29436 | 6.45049 | 6.95172 | 6.12528 | 6.33588 | 6.50594 | 6.16241 | 6.68578 | 6.23108 | 6.49198 | 6.522 | 6.66199 | 5.73282 | 5.76906 | 5.51176 | 5.56746 |
| NUMB | P49757 | 5.89826 | 6.37527 | 5.9831 | 6.00348 | 6.53713 | 6.30929 | 6.268 | 6.41504 | 5.93379 | 6.27395 | 6.23042 | 6.07446 | 6.41829 | 6.35879 | 6.47134 | 6.44939 |
| OCRL | Q01968 | 6.16713 | 6.18118 | 6.78769 | 5.85142 | 6.25306 | 6.36974 | 6.07101 | 6.25219 | 6.47098 | 6.45191 | 6.68273 | 6.42391 | 5.99762 | 5.954 | 6.04986 | 6.03556 |
| POLR3E | Q9NVU0 | 6.29996 | 6.32626 | 6.14487 | 6.11111 | 6.17893 | 6.36378 | 6.18581 | 6.0827 | 6.25466 | 6.29282 | 6.27421 | 6.0311 | 6.56887 | 6.18428 | 6.48094 | 6.2197 |
| GBE1 | Q04446 | 6.43542 | 6.37922 | 6.71345 | 6.05975 | 6.54628 | 6.97128 | 5.97082 | 7.06795 | 6.12966 | 6.3406 | 6.32975 | 6.3256 | 5.32377 | 6.32694 | 5.56213 | 5.5174 |
| SMG9 | Q9H0W8 | 6.72035 | 6.60902 | 7.05897 | 6.16699 | 6.28035 | 6.85561 | 6.29615 | 7.0159 | 6.82223 | 6.68001 | 6.76167 | 7.13994 | 4.89665 | 5.18561 | 4.57146 | 4.93909 |
| THBS1 | P07996 | 4.92678 | 4.59662 | 4.74479 | 5.16973 | 4.85647 | 4.9527 | 5.50638 | 4.92043 | 4.82763 | 4.26645 | 4.52326 | 4.54167 | 11.2203 | 9.54003 | 11.5155 | 9.89116 |
| PSMD14 | O00487 | 5.75341 | 5.13684 | 6.03091 | 5.22669 | 6.50718 | 6.13204 | 6.31059 | 6.52298 | 6.17095 | 6.27621 | 6.22633 | 6.7294 | 6.28215 | 6.63568 | 7.20265 | 6.85599 |
| AGO3 | Q9H9G7 | 6.20622 | 6.2593 | 6.5996 | 6.18518 | 6.51701 | 6.65653 | 6.0801 | 6.07998 | 6.35036 | 6.32124 | 6.39707 | 6.20143 | 6.12858 | 6.08127 | 5.92472 | 6.01141 |
| TNIK | Q9UKE5 | 6.09034 | 6.15422 | 5.89191 | 6.17843 | 6.44461 | 6.6907 | 6.08339 | 6.154 | 6.12333 | 6.74921 | 6.09268 | 5.97279 | 6.47933 | 6.49902 | 6.09497 | 6.30106 |
| SNX8 | Q9Y5X2 | 6.4941 | 6.74946 | 6.59236 | 6.29382 | 6.46372 | 6.7812 | 6.20998 | 6.44798 | 6.18837 | 6.59385 | 6.70277 | 6.63185 | 5.52652 | 5.58879 | 5.62199 | 5.11323 |
| FGD6 | Q6ZV73 | 6.19301 | 6.39237 | 6.53338 | 6.47655 | 6.20075 | 6.48896 | 6.22167 | 6.55593 | 6.1816 | 6.17448 | 6.28127 | 6.53838 | 6.04131 | 6.07951 | 5.78283 | 5.858 |
| KAT7 | O95251 | 5.83867 | 6.4135 | 6.04469 | 6.77755 | 6.29342 | 6.48293 | 6.50202 | 5.88674 | 6.16676 | 6.32015 | 6.69498 | 6.15798 | 6.39795 | 5.84899 | 6.34142 | 5.83224 |
| DPF2 | Q92785 | 6.07195 | 6.13396 | 5.80132 | 6.33614 | 6.19564 | 5.94849 | 6.43673 | 5.81431 | 6.20539 | 6.38343 | 6.37772 | 6.11383 | 6.54175 | 6.61693 | 6.40154 | 6.62085 |
| HOOK3 | Q86VS8 | 6.3534 | 6.1379 | 6.71304 | 6.16084 | 6.64811 | 6.64869 | 5.92194 | 6.60783 | 6.40626 | 6.30061 | 6.4375 | 6.68764 | 6.16375 | 5.90718 | 5.3531 | 5.5522 |
| SCRN1 | Q12765-2 | 6.66408 | 6.12141 | 7.24104 | 6.01149 | 6.65547 | 6.71345 | 6.10811 | 6.8396 | 6.47952 | 6.52445 | 6.82891 | 6.97063 | 5.70308 | 5.33502 | 4.74578 | 5.05795 |
| CCNB2 | O95067 | 6.84584 | 6.68192 | 6.90677 | 6.55926 | 6.52061 | 6.41928 | 6.37799 | 6.1176 | 6.57322 | 6.31565 | 6.49856 | 6.159 | 5.50995 | 5.43447 | 5.41177 | 5.66812 |
| DOP1B | Q9Y3R5 | 6.24295 | 5.91017 | 6.83633 | 6.1515 | 6.22552 | 6.71229 | 6.06018 | 6.51099 | 6.51207 | 6.6007 | 6.63847 | 6.77351 | 5.72124 | 5.92174 | 5.33233 | 5.85002 |
| YTHDF3 | Q7Z739 | 6.27792 | 6.14809 | 6.45535 | 6.17581 | 6.0974 | 6.37583 | 5.95296 | 6.51547 | 6.24029 | 6.34151 | 6.45309 | 6.60021 | 6.18089 | 5.99981 | 6.13925 | 6.04611 |
| KIF18A | Q8NI77 | 6.27149 | 6.52544 | 6.01508 | 6.58398 | 6.07474 | 5.99205 | 6.24232 | 6.00968 | 6.20969 | 6.26648 | 6.13273 | 6.25765 | 6.21434 | 6.3082 | 6.43245 | 6.46369 |
| COPS4 | Q9BT78 | 5.771 | 4.93874 | 5.74463 | 4.83784 | 6.1023 | 5.85907 | 6.28159 | 6.35927 | 5.54117 | 6.23095 | 5.77913 | 6.59797 | 6.48 | 7.90797 | 7.69751 | 7.87086 |
| CPSF7 | Q8N684-3 | 6.28556 | 6.61703 | 5.63502 | 6.62658 | 6.28965 | 6.119 | 5.86392 | 6.28965 | 6.19432 | 6.3463 | 5.79361 | 5.97407 | 6.45829 | 6.50547 | 6.62458 | 6.37695 |
| FHL1 | Q13642-4 | 6.13315 | 6.14485 | 6.68612 | 5.50795 | 6.26833 | 6.55248 | 6.13818 | 6.26152 | 6.29191 | 6.71613 | 6.40014 | 6.13977 | 5.948 | 6.35706 | 6.21069 | 6.24371 |
| TGFB1 | P01137 | 6.46797 | 7.2633 | 5.87828 | 6.94431 | 6.35419 | 6.39685 | 5.9848 | 6.62251 | 5.86059 | 6.40428 | 6.18152 | 6.33889 | 5.86025 | 5.78269 | 5.83834 | 5.82124 |
| USP48 | Q86UV5-8 | 5.82915 | 5.73671 | 6.40729 | 5.89156 | 5.88978 | 6.02285 | 6.01446 | 5.60065 | 6.19595 | 6.41803 | 6.60303 | 6.03259 | 6.78807 | 6.6203 | 7.45814 | 6.49143 |
| KIF13B | Q9NQT8 | 6.44853 | 6.30399 | 6.65493 | 6.1459 | 6.22432 | 6.34887 | 6.35912 | 6.64771 | 6.22143 | 6.6682 | 6.23424 | 6.57902 | 5.71516 | 5.99404 | 5.65428 | 5.80026 |
| STXBP4 | Q6ZWJ1 | 6.36154 | 6.16757 | 6.85956 | 6.10425 | 6.42894 | 6.52675 | 6.31843 | 6.87681 | 6.57877 | 6.53764 | 6.78865 | 6.72539 | 5.60997 | 5.43977 | 5.34097 | 5.33498 |
| TTC13 | Q8NBP0 | 6.22317 | 6.33819 | 5.94922 | 6.4426 | 5.91195 | 6.17695 | 6.24696 | 5.88262 | 6.20066 | 6.19685 | 6.03827 | 6.12074 | 6.75333 | 6.46669 | 6.40858 | 6.64322 |
| GAPVD1 | Q14C86-6 | 6.48781 | 6.43273 | 6.85835 | 6.11533 | 6.28469 | 6.73928 | 6.39579 | 6.59364 | 6.60399 | 6.39556 | 6.5948 | 6.71951 | 5.5782 | 5.50317 | 5.44588 | 5.25126 |
| ANAPC5 | Q9UJX4 | 6.37032 | 6.36844 | 6.54434 | 6.63276 | 6.32896 | 6.45898 | 6.11743 | 6.39998 | 6.27767 | 6.42805 | 6.50465 | 6.63527 | 5.80542 | 5.67726 | 5.79979 | 5.65068 |
| ME3 | Q16798 | 6.49401 | 6.33729 | 5.80287 | 6.64447 | 6.08388 | 5.8359 | 6.20649 | 6.17376 | 6.15555 | 6.01303 | 6.13504 | 6.06773 | 6.84794 | 6.23664 | 6.51696 | 6.44845 |
| GNAI2 | P04899-4 | 6.64906 | 6.78469 | 6.18611 | 6.60607 | 6.47801 | 6.34563 | 6.25416 | 6.17736 | 6.36157 | 6.14749 | 5.97753 | 5.76791 | 5.99578 | 5.98996 | 6.42027 | 5.85837 |
| ARID4B | Q4LE39 | 6.10891 | 6.42421 | 5.94085 | 6.81532 | 6.19533 | 5.98666 | 6.28569 | 6.20209 | 5.94061 | 6.03838 | 6.15758 | 5.89541 | 6.59895 | 6.37268 | 6.68347 | 6.35388 |
| CISD1 | Q9NZ45 | 6.20061 | 6.36013 | 5.81505 | 6.52617 | 6.37637 | 5.81918 | 6.81092 | 6.04819 | 6.25987 | 5.62785 | 5.74655 | 5.94613 | 6.43025 | 6.68209 | 6.63595 | 6.71468 |
| MCCC1 | Q96RQ3 | 6.53014 | 6.26853 | 5.8203 | 6.45155 | 6.41021 | 5.88419 | 6.30486 | 5.86626 | 6.12196 | 6.0334 | 6.05977 | 5.80251 | 6.85704 | 6.29102 | 7.03461 | 6.26364 |
| BTF3 | P20290 | 6.34643 | 6.96557 | 7.03722 | 6.2263 | 6.55474 | 6.94892 | 6.45829 | 6.30318 | 6.41014 | 6.68847 | 7.07526 | 6.96936 | 4.77422 | 5.30822 | 4.88829 | 5.04539 |
| MED16 | Q9Y2X0-3 | 6.3936 | 6.44125 | 6.11403 | 6.72575 | 6.41478 | 6.05444 | 6.30163 | 5.86629 | 6.30921 | 6.341 | 6.01377 | 6.06546 | 6.32064 | 6.08675 | 6.29729 | 6.25412 |
| SPIN1 | Q9Y657 | 5.78729 | 6.45711 | 5.62496 | 6.14617 | 6.09134 | 6.01472 | 6.20079 | 6.20156 | 6.14396 | 6.34146 | 5.96859 | 5.9885 | 6.64711 | 6.48033 | 7.1182 | 6.7879 |
| EML2 | O95834-3 | 6.01955 | 5.99942 | 6.53748 | 5.54914 | 6.51247 | 6.80199 | 5.97879 | 6.28912 | 6.65637 | 6.61606 | 6.45896 | 6.11263 | 6.1256 | 6.46089 | 5.8089 | 6.07262 |
| PATL1 | Q86TB9 | 6.28794 | 6.28511 | 6.88035 | 5.93702 | 6.18155 | 6.72326 | 6.23638 | 6.58774 | 6.58293 | 6.50981 | 6.50184 | 6.45984 | 5.86551 | 5.56304 | 5.70291 | 5.69479 |
| SMARCAD1 | Q9H4L7 | 5.85881 | 6.48901 | 6.03814 | 6.247 | 6.11344 | 6.13354 | 6.17718 | 5.98621 | 5.93294 | 6.57554 | 6.1652 | 6.32147 | 6.60819 | 6.32086 | 6.58536 | 6.44712 |
| COPS6 | Q7L5N1 | 5.49593 | 4.84224 | 5.77245 | 5.01081 | 5.99905 | 6.0964 | 6.30473 | 6.35709 | 6.11763 | 6.44284 | 6.09999 | 6.64105 | 6.67688 | 7.15061 | 7.35255 | 7.63976 |
| RFX1 | P22670 | 6.10625 | 6.33259 | 6.15379 | 6.55994 | 6.13941 | 5.98833 | 5.89183 | 5.88968 | 6.20876 | 6.31025 | 6.05185 | 5.98538 | 6.42855 | 6.75742 | 6.64837 | 6.54759 |
| ZNF280C | Q8ND82 | 6.32906 | 6.2683 | 5.92895 | 6.72372 | 6.15255 | 6.07921 | 6.51705 | 5.9706 | 6.36497 | 5.83801 | 6.13962 | 5.92652 | 6.59274 | 6.36146 | 6.4285 | 6.37876 |
| SRBD1 | Q8N5C6 | 6.38347 | 6.11722 | 5.68912 | 6.73904 | 6.04177 | 5.90108 | 6.31908 | 5.98793 | 6.23348 | 6.21453 | 5.95902 | 6.09957 | 6.53917 | 6.47719 | 6.55264 | 6.74568 |
| MAN2B1 | O00754 | 6.64845 | 6.566 | 6.33768 | 6.6507 | 6.23108 | 6.15147 | 5.97615 | 6.09674 | 6.09364 | 6.44602 | 6.22741 | 6.62291 | 5.9872 | 6.18638 | 5.91415 | 5.86403 |
| NCOA6 | Q14686 | 6.58718 | 6.71543 | 5.98636 | 6.62147 | 6.33744 | 6.08686 | 6.31667 | 6.17942 | 6.4105 | 6.33948 | 6.08536 | 6.11953 | 6.27533 | 6.15127 | 5.80297 | 5.98472 |
| VBP1 | P61758 | 6.21207 | 5.92515 | 6.84909 | 5.4974 | 6.73573 | 6.55957 | 6.29115 | 6.59925 | 6.51959 | 6.46522 | 6.86206 | 6.49738 | 6.18243 | 5.58587 | 5.66814 | 5.54989 |
| ELMO2 | Q96JJ3 | 6.23532 | 6.31629 | 6.88814 | 5.80367 | 6.32365 | 6.67977 | 6.02229 | 6.79406 | 6.12347 | 6.495 | 6.59735 | 6.27825 | 6.03818 | 6.05346 | 5.76554 | 5.58557 |
| MAVS | Q7Z434 | 6.39093 | 6.08351 | 5.83033 | 5.86235 | 6.44315 | 5.7162 | 6.56572 | 6.44461 | 6.03993 | 5.88447 | 5.85052 | 5.99114 | 6.94897 | 6.56263 | 6.7721 | 6.61344 |
| VASP | P50552 | 6.24558 | 5.99587 | 6.48315 | 5.80135 | 6.80481 | 7.04829 | 6.2954 | 6.61332 | 6.55885 | 6.13489 | 6.89521 | 6.34317 | 6.28829 | 6.07508 | 5.246 | 5.17074 |
| NEDD4 | P46934-4 | 6.44758 | 6.311 | 6.73999 | 6.05501 | 6.31892 | 6.68378 | 6.25981 | 6.50424 | 6.45098 | 6.44894 | 6.58809 | 6.43531 | 5.63719 | 5.81707 | 5.60741 | 5.69467 |
| BCKDHA | P12694 | 6.28214 | 6.13998 | 5.84392 | 6.44579 | 6.45601 | 6.31799 | 6.54367 | 6.21283 | 6.24383 | 6.18284 | 6.35812 | 6.40702 | 6.33806 | 5.9466 | 6.22334 | 6.05786 |
| BRPF3 | Q9ULD4 | 6.74944 | 6.94749 | 6.20508 | 7.13378 | 5.8545 | 5.75499 | 6.29549 | 6.01673 | 6.61077 | 6.59754 | 6.26024 | 6.40261 | 5.80714 | 5.73973 | 5.76938 | 5.8551 |
| RAB5C | P51148-2 | 6.56897 | 6.75846 | 6.24665 | 6.41225 | 6.55734 | 6.33414 | 6.33742 | 6.3524 | 6.17907 | 6.23825 | 5.90355 | 5.74592 | 5.95098 | 6.03303 | 6.27859 | 6.10299 |
| EXOC7 | Q9UPT5 | 6.48083 | 6.53229 | 6.80005 | 5.91465 | 6.7478 | 6.85722 | 6.29074 | 6.42968 | 6.56624 | 6.85814 | 6.43692 | 6.41449 | 5.54222 | 5.665 | 5.16973 | 5.29399 |
| GTPBP1 | O00178 | 6.36866 | 6.1997 | 7.00348 | 6.07175 | 6.04163 | 6.52151 | 6.25948 | 6.32603 | 6.12772 | 6.56111 | 6.79078 | 6.66256 | 5.7836 | 5.91976 | 5.64659 | 5.71564 |
| AK3 | Q9UIJ7 | 6.59886 | 6.14104 | 5.81794 | 6.09311 | 6.55447 | 5.85998 | 6.164 | 6.36361 | 6.07921 | 6.09739 | 5.86706 | 5.9107 | 7.11128 | 6.36044 | 6.50077 | 6.48015 |
| FAM120B | Q96EK7 | 6.20437 | 6.21033 | 6.71549 | 6.3787 | 6.026 | 6.29636 | 6.36906 | 6.21422 | 6.54003 | 6.45262 | 6.43678 | 6.39766 | 6.06897 | 5.73977 | 5.91179 | 6.03784 |
| SOS1 | Q07889 | 6.23766 | 6.11274 | 6.83365 | 6.01743 | 6.1461 | 6.86821 | 6.4453 | 6.27356 | 6.40473 | 6.447 | 6.41417 | 6.5237 | 5.75675 | 6.18575 | 5.67009 | 5.66316 |
| TRIP6 | Q15654 | 6.38201 | 6.05481 | 6.78107 | 6.2592 | 6.51515 | 6.43626 | 6.07733 | 6.67138 | 6.5502 | 6.44232 | 6.41838 | 6.67349 | 5.82714 | 5.47208 | 5.60982 | 5.82937 |
| ACOT9 | Q9Y305-4 | 6.35027 | 6.45664 | 5.67744 | 6.72305 | 6.17391 | 6.06235 | 6.49812 | 6.03876 | 6.16382 | 5.96047 | 5.80899 | 5.92458 | 6.56735 | 6.41753 | 6.43326 | 6.74348 |
| VPS45 | Q9NRW7 | 6.43444 | 6.76353 | 6.13309 | 6.56318 | 6.25219 | 6.47887 | 6.4042 | 6.16876 | 6.17986 | 6.34146 | 6.59175 | 6.5021 | 5.62054 | 5.89736 | 5.56532 | 6.10336 |
| ROCK1 | Q13464 | 6.1781 | 6.18998 | 6.52761 | 5.63198 | 6.49621 | 6.34197 | 6.07497 | 6.33604 | 6.55102 | 6.56407 | 6.54835 | 6.55599 | 5.91304 | 6.19476 | 5.89351 | 6.0024 |
| PEX1 | O43933 | 6.4129 | 6.27621 | 6.49035 | 6.62289 | 6.16338 | 6.42982 | 6.47279 | 5.99684 | 6.52981 | 6.42339 | 6.64531 | 6.43894 | 5.87279 | 5.61804 | 5.77241 | 5.83414 |
| FAM91A1 | Q658Y4 | 6.31085 | 6.38843 | 6.42658 | 6.1802 | 6.19783 | 6.27408 | 6.22041 | 6.43853 | 6.37158 | 6.42645 | 6.50065 | 6.55862 | 5.90689 | 5.98901 | 5.96921 | 5.84065 |

| | | | | | | | | | | | | | | | | | |
|---|---|---|---|---|---|---|---|---|---|---|---|---|---|---|---|---|---|
| TRIM33 | Q9UPN9 | 6.56517 | 6.48006 | 6.52811 | 6.46602 | 6.21121 | 6.09431 | 6.19214 | 6.07162 | 6.46928 | 6.6198 | 6.64418 | 6.43207 | 5.95842 | 5.83137 | 5.64159 | 5.79462 |
| RALGAPB | Q86X10-3 | 6.32448 | 6.08136 | 6.77992 | 6.24317 | 6.34276 | 6.69408 | 6.29538 | 6.59251 | 6.42705 | 6.31045 | 6.33367 | 6.50758 | 6.04008 | 5.74207 | 5.75986 | 5.52558 |
| MCTS1 | Q9ULC4-3 | 6.09161 | 6.0267 | 7.13126 | 5.91411 | 6.32076 | 6.63573 | 6.20214 | 6.52892 | 6.13657 | 6.61593 | 6.80894 | 7.03695 | 6.11089 | 5.52725 | 5.46749 | 5.44475 |
| PI4K2A | Q9BTU6 | 6.3825 | 6.38838 | 6.47531 | 6.62189 | 5.90191 | 5.95245 | 6.41973 | 6.0244 | 6.55188 | 6.48098 | 6.48093 | 6.67196 | 6.03831 | 5.79211 | 5.85127 | 5.96599 |
| NR3C1 | P04150-3 | 6.00005 | 5.87128 | 6.30377 | 5.79405 | 6.10738 | 6.31994 | 6.09044 | 5.91648 | 5.9208 | 6.29023 | 6.24253 | 5.88434 | 6.88379 | 6.94882 | 6.76381 | 6.6623 |
| MTCH1 | Q9NZJ7 | 6.1654 | 6.58949 | 5.92426 | 6.90333 | 6.07839 | 6.05918 | 6.33956 | 6.19536 | 6.19453 | 6.15709 | 6.05514 | 6.05932 | 6.43825 | 6.07999 | 6.3611 | 6.39962 |
| NT5C3A | Q9H0P0 | 6.64262 | 6.50477 | 5.87239 | 6.72049 | 6.52251 | 6.01282 | 6.474 | 6.11817 | 6.38974 | 5.91464 | 5.76776 | 6.11035 | 6.35023 | 5.85748 | 6.40887 | 6.33317 |
| IFT140 | Q96RY7 | 6.20311 | 6.12726 | 6.47256 | 5.97365 | 6.21645 | 6.35361 | 6.21403 | 6.19773 | 6.33753 | 6.25335 | 6.32513 | 6.05162 | 6.39344 | 6.57951 | 6.4113 | 5.88971 |
| SNRNP40 | Q96DI7 | 5.76263 | 6.43617 | 5.72076 | 6.66238 | 6.45575 | 6.14953 | 6.86136 | 6.16341 | 6.67635 | 5.57664 | 6.09772 | 5.8423 | 6.01602 | 5.98954 | 6.84795 | 6.74149 |
| GYG1 | P46976 | 6.39444 | 6.51197 | 7.23192 | 6.31253 | 6.65204 | 7.1471 | 6.27671 | 6.89574 | 6.57435 | 6.54846 | 6.95443 | 6.51196 | 5.35305 | 4.94699 | 4.93066 | 4.75765 |
| PGRMC2 | O15173-2 | 6.45081 | 5.87427 | 5.70969 | 6.00536 | 6.07229 | 5.60957 | 5.9961 | 6.15477 | 5.40188 | 6.2365 | 5.71721 | 6.34703 | 6.76262 | 7.26887 | 7.03325 | 7.35979 |
| TAF4 | O00268 | 6.57209 | 6.4287 | 6.16338 | 6.56984 | 6.24807 | 6.13353 | 6.23579 | 6.11989 | 6.18506 | 6.56948 | 6.15567 | 6.15949 | 6.66216 | 5.9877 | 6.39723 | 6.01194 |
| SUB1 | P53999 | 5.84534 | 5.73054 | 6.29504 | 5.3614 | 5.81044 | 6.30554 | 6.17985 | 5.78651 | 6.16246 | 5.98012 | 6.75231 | 6.0754 | 5.91556 | 7.24377 | 7.27916 | 7.27656 |
| TMEM209 | Q96SK2 | 6.15445 | 6.12571 | 5.6875 | 6.72705 | 5.97077 | 5.82184 | 6.541 | 6.00657 | 6.05162 | 5.43458 | 5.80834 | 6.12951 | 6.36968 | 7.01905 | 6.94848 | 7.20386 |
| XPC | Q01831 | 6.13022 | 6.03827 | 5.66581 | 6.63816 | 5.94383 | 5.77611 | 6.10262 | 6.16045 | 6.15463 | 6.21029 | 5.75398 | 5.97926 | 6.24656 | 7.20426 | 6.92467 | 7.07089 |
| CWF19L1 | Q69YN2 | 5.45972 | 5.92773 | 6.10017 | 5.56824 | 5.80047 | 6.09973 | 5.61307 | 6.08777 | 5.93514 | 6.34832 | 6.33183 | 6.11697 | 6.84078 | 7.26208 | 7.46877 | 7.03921 |
| PDLIM5 | Q96HC4 | 5.90505 | 5.98831 | 6.66419 | 5.85219 | 6.38549 | 6.33876 | 6.23278 | 6.19624 | 6.3998 | 6.23859 | 6.5323 | 6.48116 | 6.35223 | 6.27125 | 6.20878 | 5.95286 |
| RBPJ | Q06330 | 5.53842 | 6.19764 | 5.9049 | 5.85136 | 5.88444 | 6.14323 | 5.86136 | 5.88915 | 6.02949 | 6.5231 | 6.25497 | 6.18238 | 6.56525 | 7.10235 | 7.1691 | 6.90284 |
| VAV2 | P52735-2 | 6.15711 | 6.21942 | 7.25232 | 6.02533 | 6.08454 | 6.44287 | 6.23977 | 6.37387 | 6.44375 | 6.5796 | 6.72666 | 6.43031 | 5.75 | 5.83246 | 5.65175 | 5.79026 |
| SCML2 | Q9UQR0 | 6.22098 | 6.6538 | 6.13176 | 7.05623 | 6.3229 | 5.98705 | 6.35219 | 5.94203 | 6.11349 | 6.26188 | 6.0225 | 6.07196 | 6.19327 | 5.89607 | 6.33826 | 6.43562 |
| SDF4 | Q9BRK5 | 7.44335 | 7.52781 | 6.87507 | 7.38452 | 7.00794 | 6.54111 | 6.42329 | 6.4065 | 7.21035 | 6.79674 | 7.04056 | 6.84601 | 4.25598 | 4.24395 | 3.78405 | 4.21277 |
| MOSPD2 | Q8NHP6 | 6.15511 | 6.2726 | 5.94795 | 6.47167 | 5.92925 | 6.34748 | 6.21035 | 5.97378 | 5.92721 | 6.40331 | 5.97608 | 5.88528 | 6.47198 | 6.40776 | 6.61828 | 7.00192 |
| HM13 | Q8TCT9-2 | 6.58 | 6.14548 | 6.07091 | 6.46572 | 6.17635 | 5.77028 | 6.59049 | 5.89581 | 6.251 | 5.97073 | 6.09704 | 5.7983 | 6.45277 | 6.39423 | 6.67758 | 6.6633 |
| RPS12 | P25398 | 6.40527 | 5.05497 | 6.40147 | 5.26126 | 6.68128 | 5.73993 | 5.69546 | 6.63848 | 5.73607 | 6.80263 | 6.38738 | 7.00294 | 5.98223 | 6.84063 | 6.69202 | 6.67797 |
| PTGES3 | Q15185 | 6.14615 | 6.2902 | 6.24135 | 6.07708 | 6.67076 | 6.753 | 6.1939 | 6.46613 | 6.14355 | 6.19415 | 6.52551 | 6.17079 | 6.33181 | 5.89988 | 5.67263 | 6.22311 |
| BAG2 | O95816 | 6.39512 | 6.06213 | 6.73079 | 5.89978 | 6.41409 | 6.14005 | 6.19741 | 6.17427 | 6.12136 | 6.34646 | 6.75903 | 6.6253 | 6.55345 | 5.63523 | 6.11365 | 5.83188 |
| UCHL5 | Q9Y5K5-3 | 6.19076 | 6.14922 | 6.37973 | 5.9517 | 6.69266 | 6.30985 | 5.99988 | 6.16738 | 6.09519 | 6.18559 | 6.28355 | 6.31211 | 6.59428 | 6.23987 | 6.24952 | 6.1927 |
| SET | Q01105 | 5.61291 | 5.85907 | 6.00941 | 5.50755 | 6.20138 | 6.29001 | 5.92354 | 5.84774 | 5.98662 | 6.10383 | 6.18696 | 5.84579 | 7.6473 | 7.05793 | 7.15828 | 6.7617 |
| RDH11 | Q8TC12 | 6.56091 | 7.0045 | 6.31874 | 6.70523 | 6.41171 | 6.00499 | 6.19075 | 6.03158 | 6.15714 | 5.89705 | 5.98439 | 6.06326 | 6.14203 | 6.22538 | 6.39976 | 5.90259 |
| CABIN1 | Q9Y6J0 | 6.29199 | 6.24865 | 5.93877 | 6.7747 | 6.3881 | 6.08842 | 6.41889 | 5.9564 | 6.30003 | 6.09729 | 6.0521 | 6.20074 | 6.50311 | 6.02494 | 6.39164 | 6.32426 |
| PON2 | A0A0J9YXF2 | 6.82484 | 6.53128 | 5.8949 | 6.31529 | 6.40859 | 6.03883 | 6.19125 | 6.24293 | 6.0944 | 6.10304 | 5.65127 | 5.77465 | 6.69461 | 6.27763 | 6.59886 | 6.35763 |
| TXNL4A | P83876 | 6.01909 | 5.99549 | 5.9921 | 6.01304 | 6.50519 | 6.23162 | 6.11952 | 6.26055 | 6.48013 | 6.29827 | 6.16281 | 6.07779 | 6.75765 | 6.05108 | 6.72737 | 6.30829 |
| RPF2 | Q9H7B2 | 6.11672 | 6.50188 | 5.57415 | 6.93328 | 6.05509 | 5.83586 | 6.26294 | 6.30804 | 5.77234 | 5.87164 | 5.6837 | 6.05064 | 6.48045 | 6.80156 | 6.82794 | 6.92377 |
| RPS17 | P08708 | 6.273 | 5.59622 | 6.40816 | 5.43882 | 6.08267 | 6.56827 | 6.35823 | 6.33001 | 6.70087 | 6.46977 | 6.38807 | 6.37707 | 5.74008 | 6.0928 | 5.53095 | 6.54501 |
| LUC7L3 | O95232 | 5.56697 | 5.98005 | 5.77631 | 6.45264 | 5.73406 | 5.76704 | 6.83718 | 5.46965 | 6.41493 | 5.50726 | 5.62472 | 5.97647 | 6.30928 | 7.42903 | 7.18813 | 7.96627 |
| PPP1R8 | Q12972 | 6.07583 | 5.96823 | 5.95598 | 6.07024 | 6.07062 | 6.09176 | 6.15336 | 6.24785 | 6.22926 | 6.09618 | 6.82507 | 6.11102 | 6.82195 | 6.59276 | 7.0496 | 6.64028 |
| CTNNA2 | P26232 | 6.46094 | 6.32364 | 6.34277 | 6.60098 | 6.54137 | 6.12524 | 6.06392 | 6.04895 | 6.71181 | 6.4219 | 6.18765 | 6.43852 | 6.0711 | 5.69255 | 5.7244 | 6.24428 |
| EFHD2 | Q96C19 | 6.876 | 6.43915 | 6.58054 | 6.17226 | 6.65613 | 6.51191 | 6.01477 | 6.50479 | 6.58191 | 6.14961 | 5.98574 | 6.33544 | 6.18832 | 5.81388 | 5.52879 | 5.66076 |
| IRGQ | Q8WZA9 | 6.27621 | 5.07169 | 6.96816 | 4.78895 | 6.69559 | 6.09298 | 5.84427 | 7.29677 | 5.61923 | 6.99257 | 5.99197 | 7.38408 | 6.29262 | 6.44472 | 6.1131 | 6.12709 |
| NAP1L4 | Q99733-2 | 6.41358 | 6.26145 | 6.73916 | 5.90267 | 6.40678 | 6.86782 | 6.07217 | 6.89089 | 6.55774 | 6.42015 | 6.94484 | 6.91695 | 5.45275 | 5.49039 | 5.23621 | 5.42645 |
| LRRC8A | Q8IWT6 | 6.3851 | 6.45839 | 5.98229 | 6.58354 | 6.31794 | 5.75814 | 6.44942 | 6.02971 | 6.40713 | 6.13893 | 6.37764 | 6.04811 | 6.38159 | 5.96836 | 6.48955 | 6.22416 |
| NDUFV2 | P19404 | 6.37184 | 5.18924 | 5.54607 | 4.96428 | 5.68349 | 4.41824 | 5.48073 | 6.55587 | 4.87746 | 6.13679 | 5.56333 | 6.90059 | 6.66984 | 8.60903 | 8.29365 | 8.73954 |
| ZC3H7B | Q9UGR2 | 6.10976 | 6.36486 | 6.61648 | 6.2006 | 6.28696 | 6.59664 | 6.33226 | 6.29472 | 6.42441 | 6.6578 | 6.57946 | 6.61779 | 5.67912 | 5.91651 | 5.6994 | 5.62322 |
| TPM4 | P67936 | 6.13975 | 6.22129 | 6.2891 | 6.33977 | 6.80457 | 6.90531 | 5.92836 | 6.85449 | 6.1116 | 6.3196 | 6.5904 | 6.63506 | 6.21628 | 6.27485 | 4.99108 | 5.37848 |
| SRPK1 | Q96SB4-3 | 6.43267 | 6.13992 | 6.85756 | 6.51634 | 6.02077 | 6.21452 | 6.27644 | 5.80922 | 6.59136 | 6.72163 | 6.6126 | 6.80352 | 5.50324 | 5.65527 | 5.99777 | 5.84717 |
| VPS18 | Q9P253 | 6.23007 | 6.27207 | 6.65616 | 6.11519 | 6.4298 | 6.16714 | 6.40076 | 6.4257 | 6.21411 | 6.4813 | 6.49946 | 6.32907 | 5.95855 | 5.95774 | 6.01178 | 5.85112 |
| KIF5A | Q12840 | 6.32112 | 6.27107 | 6.58435 | 6.10647 | 6.34246 | 6.75097 | 6.4047 | 6.6774 | 6.41653 | 6.65125 | 6.34209 | 6.40258 | 5.61439 | 5.83683 | 5.66642 | 5.61136 |
| AHDC1 | Q5TGY3 | 5.96743 | 6.47361 | 5.77862 | 6.69293 | 5.68126 | 5.8631 | 6.36536 | 5.98539 | 6.58303 | 6.38794 | 5.97589 | 6.05086 | 6.37769 | 6.58882 | 6.38558 | 6.84249 |
| YWHAQ | P27348 | 6.28595 | 6.05208 | 6.82383 | 5.77569 | 6.07511 | 6.65156 | 6.03848 | 6.49543 | 6.3341 | 6.5604 | 6.64402 | 6.41963 | 6.17434 | 6.09219 | 5.91086 | 5.66635 |
| STXBP1 | P61764-2 | 6.25476 | 6.35092 | 6.45806 | 6.13426 | 6.25465 | 6.53908 | 6.10539 | 6.38797 | 6.33327 | 6.33786 | 6.35997 | 6.34532 | 5.99222 | 5.96232 | 6.22832 | 5.95564 |
| TDRKH | Q9Y2W6 | 6.35104 | 6.23472 | 5.98473 | 6.6582 | 6.36008 | 6.27607 | 6.35924 | 6.2027 | 6.1904 | 6.24375 | 6.22099 | 5.74281 | 6.42855 | 6.17793 | 6.3665 | 6.20229 |
| CELF1 | Q92879-4 | 6.27686 | 6.60534 | 6.16588 | 6.36333 | 6.24541 | 6.07667 | 6.35035 | 6.05915 | 6.08842 | 6.25031 | 6.40651 | 6.19994 | 6.472 | 5.92832 | 6.31045 | 6.20104 |
| EXOSC7 | Q15024 | 6.13807 | 6.01657 | 6.13923 | 6.20291 | 6.15385 | 6.18761 | 6.22714 | 5.84758 | 6.14636 | 6.0425 | 6.27004 | 6.1231 | 6.83046 | 6.37759 | 6.7625 | 6.53448 |
| FDXR | P22570-3 | 6.44102 | 6.28248 | 5.99439 | 6.3789 | 6.44156 | 6.07343 | 6.13343 | 6.23666 | 6.21657 | 6.15603 | 5.86386 | 5.88695 | 6.8898 | 6.30729 | 6.26055 | 6.43707 |
| SREBF1 | P36956-4 | 6.82757 | 6.75814 | 6.15502 | 6.55509 | 6.31354 | 6.2174 | 6.08392 | 6.00661 | 6.95833 | 7.35268 | 6.95512 | 6.76837 | 5.19017 | 5.26227 | 5.23347 | 5.3623 |
| RBM33 | Q96EV2 | 5.98951 | 6.08175 | 6.05304 | 6.02453 | 5.89314 | 6.10193 | 6.14861 | 6.20923 | 6.1608 | 6.37964 | 6.1931 | 6.01126 | 6.99527 | 6.39047 | 6.66826 | 6.69946 |
| DIMT1 | Q9UNQ2 | 6.27257 | 6.38701 | 5.9916 | 6.6012 | 6.38444 | 6.34194 | 6.50892 | 6.05124 | 6.14419 | 5.95946 | 5.98466 | 6.09007 | 6.23191 | 5.91118 | 6.66335 | 6.47627 |
| ROBO1 | Q9Y6N7-2 | 6.36275 | 6.29186 | 6.23054 | 6.452 | 6.23079 | 6.01417 | 6.29554 | 6.15376 | 6.32558 | 6.30564 | 6.19886 | 5.99617 | 6.21751 | 6.50165 | 6.13279 | 6.30839 |
| CYC1 | P08574 | 6.26028 | 6.17242 | 6.11827 | 6.48296 | 6.61929 | 5.76537 | 6.60881 | 5.99252 | 6.27816 | 6.54604 | 6.28203 | 6.39151 | 6.42746 | 6.2545 | 6.44086 | 6.25951 |
| SCAF1 | Q9H7N4 | 6.003 | 6.3582 | 5.64054 | 6.69634 | 5.71237 | 5.73858 | 6.04851 | 6.00309 | 5.84641 | 6.03305 | 5.81225 | 6.18217 | 6.16613 | 7.19117 | 7.00023 | 7.56795 |
| PRKAG2 | Q9UGJ0 | 6.05415 | 5.99827 | 6.06188 | 5.47075 | 6.21719 | 6.18147 | 5.91567 | 6.37986 | 6.16517 | 6.22316 | 6.21667 | 6.26829 | 6.91615 | 6.48702 | 6.74707 | 6.69722 |
| MEPCE | Q7L2J0 | 6.16403 | 6.17621 | 6.05904 | 6.40065 | 5.96015 | 5.81664 | 6.04135 | 5.86313 | 6.05344 | 6.07839 | 6.10832 | 6.10329 | 6.71006 | 6.49935 | 7.15973 | 6.80622 |
| POLR3B | Q9NW08 | 5.95334 | 6.52133 | 6.18668 | 6.29209 | 6.23781 | 6.31819 | 6.12605 | 6.08572 | 6.26111 | 6.40476 | 6.30768 | 6.31334 | 6.24061 | 6.08098 | 6.356 | 6.31429 |
| OSBPL3 | Q9H4L5 | 5.66477 | 5.77858 | 6.04069 | 5.46679 | 5.82757 | 5.80163 | 5.81258 | 5.67263 | 6.36545 | 6.43578 | 6.34466 | 6.19686 | 7.23389 | 6.70272 | 7.45525 | 7.20015 |
| PPP1R21 | Q6ZMI0 | 6.22306 | 6.24529 | 6.30331 | 5.73173 | 6.17971 | 6.66003 | 6.07708 | 6.08345 | 6.35506 | 6.58692 | 6.77982 | 6.36568 | 6.54798 | 6.45223 | 5.84038 | 5.56827 |
| TRMT6 | Q9UJA5 | 5.348 | 6.20037 | 5.82644 | 5.42861 | 5.99574 | 6.3442 | 5.97024 | 6.15403 | 6.00667 | 6.19634 | 6.07339 | 6.13139 | 7.14402 | 6.92276 | 7.38263 | 6.87517 |
| WDR35 | Q9P2L0 | 6.22313 | 6.34628 | 6.63986 | 6.06475 | 6.5036 | 6.54302 | 6.31229 | 6.42662 | 6.63925 | 6.43954 | 6.30194 | 6.42595 | 5.95606 | 5.72146 | 5.71276 | 5.74348 |
| NAV1 | Q8NEY1 | 5.8399 | 6.08739 | 6.82653 | 6.14405 | 6.06277 | 6.43348 | 6.15426 | 6.25272 | 6.39413 | 6.5262 | 6.77782 | 6.52724 | 5.86804 | 5.99328 | 6.00904 | 6.10314 |
| LASP1 | Q14847 | 6.47727 | 5.9727 | 6.69346 | 5.82869 | 6.68476 | 6.62705 | 6.00172 | 6.94844 | 6.59966 | 6.35828 | 6.80541 | 6.44955 | 6.2912 | 5.85437 | 5.28739 | 5.12008 |
| SCCPDH | Q8NBX0 | 6.45417 | 6.3534 | 5.80362 | 6.68694 | 6.64826 | 6.20039 | 6.48258 | 5.97086 | 6.14248 | 6.51906 | 6.06646 | 5.62296 | 6.65669 | 6.52012 | 6.36443 | 6.11359 |
| NELFCD | Q8IXH7 | 5.91254 | 5.9839 | 6.13809 | 5.7497 | 6.25626 | 6.19666 | 6.17891 | 5.92966 | 5.90088 | 5.93819 | 6.21177 | 5.99857 | 7.20414 | 6.71624 | 6.94178 | 6.74271 |
| CHID1 | Q9BWS9-2 | 6.18716 | 6.09033 | 5.98948 | 6.71159 | 6.33272 | 6.05347 | 6.41211 | 5.82197 | 6.26761 | 6.15302 | 6.05659 | 5.73771 | 6.66834 | 6.27748 | 6.68134 | 6.55907 |
| HMGB1 | P09429 | 5.93437 | 5.93783 | 6.30932 | 5.69413 | 6.17725 | 7.03066 | 5.97253 | 6.50098 | 5.64774 | 5.9791 | 6.34538 | 6.06257 | 6.17036 | 7.75545 | 6.47923 | 6.00311 |
| LACTB2 | Q53H82 | 6.29262 | 6.22382 | 6.36681 | 6.08697 | 6.39452 | 6.21459 | 6.47981 | 6.04436 | 6.29128 | 6.20059 | 6.5511 | 6.23139 | 6.40254 | 6.07048 | 5.98702 | 6.1621 |
| ATP6V0A1 | Q93050 | 6.1496 | 6.61781 | 6.44893 | 6.58577 | 6.00951 | 6.16366 | 6.28781 | 6.50702 | 6.13718 | 6.33047 | 6.49618 | 6.18132 | 5.88311 | 6.08471 | 6.13303 | 5.98391 |
| GATD3B | A0A0B4J2D5 | 6.30335 | 5.91321 | 5.65575 | 6.07713 | 6.26934 | 5.97199 | 6.02163 | 6.16324 | 5.79316 | 6.30143 | 6.31242 | 6.51135 | 6.91197 | 6.50784 | 6.6398 | 6.64639 |
| SEC24D | O94855-2 | 5.9737 | 6.11195 | 6.51372 | 5.77599 | 6.17025 | 6.53803 | 6.25691 | 6.25223 | 5.98747 | 6.22752 | 6.30952 | 6.05995 | 6.62786 | 6.59766 | 6.33456 | 6.26689 |
| ANKRD52 | Q8NB46 | 6.6878 | 6.49567 | 7.2703 | 6.39152 | 6.41346 | 6.78101 | 6.25562 | 6.52593 | 6.85371 | 6.61341 | 7.03542 | 6.85105 | 4.90435 | 4.86753 | 5.03832 | 5.01488 |
| TXLNG | Q9NUQ3 | 6.38865 | 6.45822 | 6.70873 | 6.27793 | 6.32033 | 6.70465 | 6.07247 | 6.46196 | 6.43922 | 6.43761 | 6.66211 | 6.74284 | 5.35066 | 5.80528 | 5.37683 | 5.79249 |
| EXOC2 | Q96KP1 | 6.42222 | 6.3399 | 6.54105 | 5.80856 | 6.43588 | 6.3797 | 6.24187 | 6.38293 | 6.96744 | 6.75745 | 6.43925 | 6.31103 | 5.94975 | 5.84061 | 5.69522 | 5.48714 |
| ACOT2 | P49753 | 6.24495 | 6.32491 | 5.98429 | 6.42235 | 6.44626 | 6.12477 | 6.07377 | 6.20895 | 6.2559 | 6.12951 | 5.85514 | 5.93043 | 6.7134 | 6.3732 | 6.54544 | 6.36673 |
| PKN1 | Q16512-2 | 6.80887 | 6.4115 | 6.93749 | 6.28859 | 5.96594 | 6.23692 | 6.19156 | 6.54641 | 6.70337 | 6.52635 | 7.11466 | 6.92071 | 5.58083 | 5.1846 | 5.27037 | 5.31183 |
| PPP1R10 | Q96QC0 | 6.37713 | 6.30618 | 6.1386 | 6.82716 | 5.76915 | 5.82498 | 6.19423 | 5.78291 | 6.35115 | 6.50913 | 6.16325 | 6.5473 | 6.1265 | 6.12917 | 6.44069 | 6.51247 |
| DBNL | Q9UJU6-3 | 6.39849 | 6.21444 | 6.6493 | 5.76039 | 6.55048 | 6.74603 | 6.05785 | 6.81732 | 6.22112 | 6.51193 | 6.477 | 6.70549 | 6.26812 | 5.6397 | 5.46594 | 5.5164 |
| ARPC4 | P59998-3 | 5.95404 | 5.68915 | 6.47232 | 5.8363 | 6.07899 | 6.63883 | 6.07521 | 6.45112 | 6.1116 | 6.33241 | 6.51525 | 6.3667 | 5.93312 | 6.76454 | 6.3616 | 6.41883 |
| TBK1 | Q9UHD2 | 6.47244 | 6.29774 | 6.70762 | 5.98627 | 6.31083 | 6.87031 | 6.30077 | 6.6643 | 6.32825 | 6.37506 | 6.16232 | 6.48364 | 5.63102 | 5.94675 | 5.49088 | 5.97181 |
| DOCK9 | Q9BZ29-4 | 6.31029 | 6.3183 | 7.055 | 6.28182 | 6.23196 | 6.20995 | 6.09352 | 6.12296 | 6.62217 | 6.446 | 6.73782 | 6.56848 | 5.6793 | 5.73792 | 5.982 | 5.60251 |
| TOR4A | Q9NXH8 | 6.87353 | 6.8946 | 6.27272 | 6.9598 | 6.15726 | 5.79962 | 6.54254 | 6.14853 | 7.20982 | 6.6979 | 6.67797 | 6.53288 | 5.56039 | 5.24647 | 5.2019 | 5.22409 |
| MYL6 | P60660 | 6.37866 | 5.26245 | 5.68164 | 4.44093 | 5.91256 | 5.40737 | 5.4292 | 7.27964 | 5.18185 | 6.39721 | 5.6446 | 7.03687 | 5.9419 | 8.78871 | 7.05764 | 8.15879 |
| CDK5RAP1 | Q96SZ6-3 | 6.58062 | 6.53633 | 6.43038 | 6.73668 | 6.37478 | 6.13026 | 6.63367 | 6.25339 | 6.13488 | 6.03341 | 6.37591 | 6.25608 | 6.01482 | 5.45958 | 5.9504 | 6.09879 |
| TMTC3 | Q6ZXV5 | 6.22554 | 6.3945 | 5.77203 | 6.5396 | 6.13069 | 5.86746 | 6.20765 | 5.87895 | 6.0567 | 5.89437 | 5.98878 | 5.99781 | 6.66966 | 6.68627 | 6.95323 | 6.73675 |
| APEH | P13798 | 6.6728 | 6.15823 | 6.58556 | 5.89114 | 6.68113 | 6.91103 | 6.05509 | 6.55732 | 6.38169 | 6.42907 | 6.5182 | 6.56609 | 5.75205 | 5.99123 | 5.4022 | 5.44717 |
| UBA3 | Q8TBC4 | 5.81054 | 5.77763 | 6.08428 | 5.21105 | 5.91031 | 6.41871 | 5.74074 | 6.52372 | 5.7511 | 6.13591 | 6.19968 | 6.16086 | 6.822 | 7.71308 | 7.25272 | 6.48767 |
| CACTIN | Q8WUQ7-2 | 5.92335 | 6.22345 | 6.1497 | 6.50272 | 5.68167 | 6.13769 | 6.1153 | 5.82403 | 6.2041 | 6.15817 | 6.41268 | 6.36746 | 6.49211 | 6.37717 | 6.79296 | 6.63745 |
| PRDM2 | Q13029 | 6.55438 | 6.44648 | 6.08911 | 7.2125 | 5.80562 | 6.0033 | 6.48238 | 6.12358 | 5.81802 | 6.17868 | 6.26104 | 6.29366 | 6.07336 | 6.14394 | 6.25871 | 6.25526 |
| ERMP1 | Q7Z2K6 | 6.15409 | 6.42892 | 5.65762 | 6.18367 | 6.11249 | 5.97622 | 6.40211 | 6.08361 | 6.03318 | 6.09764 | 5.86769 | 5.90365 | 6.64105 | 6.63043 | 6.76838 | 7.05925 |
| TSPYL4 | Q9UJ04 | 6.46616 | 6.27737 | 6.98164 | 5.96304 | 6.08281 | 6.86077 | 6.36207 | 6.45166 | 6.51852 | 6.47942 | 6.56684 | 6.42189 | 5.67998 | 5.70783 | 5.50022 | 5.67976 |
| INTS5 | Q6P9B9 | 6.43133 | 6.33292 | 5.87883 | 6.41806 | 6.27373 | 5.90709 | 6.22546 | 6.13735 | 6.21762 | 5.85087 | 6.18998 | 6.06521 | 6.65074 | 6.32262 | 6.51914 | 6.57904 |
| RAB3B | P20337 | 6.41794 | 6.62188 | 6.03572 | 6.60158 | 6.21165 | 6.46229 | 6.18111 | 6.15938 | 6.21105 | 6.22497 | 5.71102 | 5.97788 | 6.24378 | 6.2009 | 6.35959 | 6.37924 |
| COIL | P38432 | 6.1931 | 6.40544 | 6.01779 | 6.90781 | 5.9752 | 5.82763 | 6.35771 | 5.9339 | 6.15225 | 5.96761 | 6.04762 | 6.06626 | 6.55085 | 6.37147 | 6.60111 | 6.62423 |
| COQ8B | Q96D53 | 6.57664 | 6.70981 | 5.94732 | 6.4661 | 6.37274 | 6.02669 | 6.24769 | 6.01943 | 6.08973 | 6.06638 | 6.01986 | 5.85025 | 6.67911 | 6.24955 | 6.50051 | 6.17821 |
| ELOA | Q14241 | 6.06791 | 6.23057 | 5.9456 | 6.46106 | 6.23207 | 5.90422 | 6.21648 | 6.05477 | 6.00602 | 5.84612 | 6.04998 | 5.86087 | 6.46076 | 7.02613 | 6.85529 | 6.78214 |
| ACOX1 | Q15067-2 | 6.25647 | 5.75018 | 5.93804 | 5.998 | 6.05683 | 6.29421 | 6.3738 | 5.83909 | 6.17469 | 6.23759 | 6.1852 | 6.10417 | 6.84058 | 6.62262 | 6.49196 | 6.83656 |
| TUBA1C | Q9BQE3 | 6.41916 | 6.00015 | 7.54484 | 6.49693 | 6.4599 | 6.92437 | 6.42661 | 6.12474 | 6.36299 | 6.31912 | 6.76969 | 6.67186 | 5.82229 | 5.26131 | 5.21415 | 5.18189 |
| IFT172 | Q9UG01 | 6.28763 | 6.68117 | 7.10516 | 6.2182 | 6.15181 | 6.63527 | 6.02242 | 6.48086 | 6.35747 | 6.98822 | 6.85426 | 6.77579 | 5.06321 | 5.78403 | 5.33544 | 5.25904 |

| | | | | | | | | | | | | | | | | | |
|---|---|---|---|---|---|---|---|---|---|---|---|---|---|---|---|---|---|
| ZNF217 | O75362 | 6.14599 | 6.22059 | 6.379 | 6.77967 | 6.15059 | 6.15091 | 6.70331 | 6.1622 | 6.47121 | 6.46955 | 6.26588 | 6.56634 | 5.87945 | 5.67229 | 5.82991 | 6.15312 |
| CRYZ | Q08257 | 6.28283 | 5.87729 | 6.24679 | 6.33499 | 6.26705 | 6.36625 | 6.0927 | 6.34807 | 6.14437 | 6.48544 | 6.32685 | 6.18305 | 6.25117 | 6.2269 | 6.29028 | 6.27597 |
| OGFR | Q9NZT2 | 5.4323 | 5.72083 | 6.16567 | 5.46668 | 6.03818 | 6.69644 | 5.92805 | 6.44312 | 5.95608 | 6.19671 | 6.29477 | 6.05652 | 7.03409 | 7.09448 | 6.84294 | 6.63313 |
| KDM3A | Q9Y4C1 | 5.75497 | 5.9943 | 6.07913 | 5.5773 | 5.74694 | 6.35968 | 5.79571 | 5.81859 | 5.99795 | 6.44159 | 6.3708 | 6.16038 | 6.97145 | 7.35948 | 6.80357 | 6.76819 |
| SLC9A3R1 | O14745 | 6.56009 | 6.09113 | 6.59793 | 5.51857 | 6.68053 | 6.45595 | 5.91391 | 6.78707 | 6.11781 | 6.50741 | 6.67182 | 6.77194 | 6.67994 | 5.78295 | 5.31504 | 5.54791 |
| VPS33B | Q9H267 | 6.17573 | 6.5226 | 6.68542 | 5.79425 | 6.28147 | 6.81286 | 6.1415 | 6.63065 | 6.38827 | 6.72318 | 6.42048 | 6.3165 | 5.85272 | 6.00751 | 5.57247 | 5.67439 |
| ATP5PD | O75947 | 6.03307 | 5.07291 | 4.95181 | 4.98628 | 5.84716 | 4.98642 | 5.39428 | 6.98517 | 4.94192 | 6.28052 | 5.29433 | 6.36359 | 6.43463 | 9.05762 | 8.15632 | 9.21398 |
| CARM1 | Q86X55 | 5.76511 | 5.87145 | 6.66659 | 5.72013 | 5.89654 | 6.37901 | 6.05353 | 6.09633 | 6.54094 | 6.49071 | 6.52578 | 6.24476 | 6.33716 | 6.50863 | 6.55103 | 6.35228 |
| PLEKHG2 | Q9H7P9 | 6.1965 | 6.13151 | 6.67827 | 5.85832 | 6.38013 | 6.33855 | 6.34207 | 6.56399 | 6.15019 | 6.46575 | 6.65178 | 6.49217 | 6.07338 | 6.00056 | 5.93453 | 5.74227 |
| NIT2 | Q9NQR4 | 6.43755 | 6.10956 | 6.64885 | 5.76907 | 6.32081 | 6.55059 | 6.05506 | 6.68236 | 6.59333 | 6.25935 | 6.33086 | 6.58444 | 6.03405 | 6.1834 | 5.81018 | 5.63055 |
| NAGK | Q9UJ70-2 | 6.27297 | 6.12826 | 6.81364 | 5.98701 | 6.34184 | 6.82975 | 6.00603 | 6.68981 | 6.2929 | 6.22016 | 6.4014 | 6.39055 | 5.82162 | 6.16371 | 5.81868 | 5.82167 |
| FAH | P16930 | 6.50957 | 5.92172 | 6.90569 | 5.46458 | 6.50195 | 6.64101 | 5.96222 | 7.06992 | 6.20616 | 6.57408 | 6.41232 | 6.60103 | 6.08061 | 5.86598 | 5.61893 | 6.56423 |
| HELZ2 | Q9BYK8 | 6.49014 | 6.39706 | 6.48233 | 6.2811 | 6.13953 | 6.34701 | 6.54068 | 6.94558 | 6.28911 | 6.72242 | 6.45368 | 6.40723 | 5.96385 | 5.80912 | 6.13843 | 5.59275 |
| GON4L | Q3T8J9-3 | 6.44607 | 6.86927 | 6.50926 | 6.95089 | 6.14848 | 6.35467 | 6.38812 | 6.35644 | 6.22954 | 6.25356 | 6.3318 | 6.15794 | 5.58604 | 5.79729 | 5.46038 | 6.16024 |
| RO60 | P10155 | 5.6354 | 6.25114 | 6.06291 | 5.55509 | 6.17554 | 6.42888 | 6.00869 | 6.31477 | 5.7407 | 6.41898 | 6.269 | 6.12176 | 6.86776 | 6.97782 | 6.75205 | 6.41951 |
| INTS8 | Q75QN2 | 6.51419 | 6.42461 | 6.03985 | 6.4577 | 6.23342 | 5.89057 | 6.45373 | 5.79944 | 6.33894 | 6.14261 | 6.19073 | 5.98786 | 6.66584 | 6.01333 | 6.65197 | 6.19522 |
| RGPD8 | O14715 | 6.53974 | 6.38912 | 6.04579 | 6.81291 | 6.18031 | 5.92862 | 6.47842 | 6.45726 | 6.29151 | 6.08515 | 5.90253 | 6.04151 | 6.25344 | 6.15653 | 6.1114 | 6.32576 |
| NBEAL1 | Q6ZS30 | 6.05748 | 6.02561 | 6.57755 | 6.36018 | 6.42079 | 6.25508 | 6.21177 | 6.06696 | 6.41306 | 6.24055 | 6.3149 | 6.49269 | 6.08162 | 6.31919 | 6.14457 | 6.01799 |
| WDR81 | Q562E7 | 6.52619 | 6.45578 | 6.69069 | 5.88426 | 6.56996 | 6.45741 | 6.28728 | 6.76691 | 6.73138 | 6.72441 | 6.79912 | 6.78544 | 5.34893 | 5.77268 | 5.02109 | 5.17848 |
| SFSWAP | Q12872-2 | 6.28146 | 6.35215 | 6.01609 | 6.69178 | 6.13622 | 5.81436 | 6.36312 | 5.86302 | 5.93867 | 6.0387 | 6.03514 | 5.96554 | 6.38846 | 6.61543 | 6.69698 | 6.80287 |
| BRCA2 | P51587 | 7.11102 | 7.08504 | 6.63513 | 6.91753 | 6.16108 | 6.36574 | 6.34179 | 6.31852 | 6.72848 | 6.59237 | 6.70627 | 6.70331 | 5.11707 | 4.82354 | 5.07058 | 5.32253 |
| SUZ12 | Q15022 | 6.47323 | 6.51344 | 6.29855 | 6.92405 | 6.20584 | 5.99016 | 6.43975 | 5.83282 | 6.41042 | 6.33317 | 6.47142 | 6.35621 | 5.87951 | 5.75765 | 6.00983 | 6.10396 |
| VPS16 | Q9H269 | 6.46335 | 6.51142 | 6.52304 | 6.05497 | 6.35017 | 6.6568 | 6.37195 | 6.39742 | 6.49609 | 6.62488 | 6.38438 | 6.22011 | 5.87436 | 5.75236 | 5.78182 | 5.53689 |
| SNX4 | O95219 | 6.25485 | 6.36304 | 6.40137 | 6.01293 | 6.05188 | 6.34522 | 6.1357 | 6.56364 | 5.94834 | 6.70729 | 6.19419 | 6.62549 | 6.06468 | 6.36152 | 5.93129 | 6.03856 |
| KRR1 | Q13601 | 6.06036 | 6.25229 | 5.81218 | 6.62479 | 6.00721 | 5.88285 | 6.45805 | 5.95859 | 5.9906 | 5.85819 | 5.85571 | 5.69667 | 6.73918 | 6.6588 | 7.04131 | 7.10322 |
| CTSA | P10619 | 6.47393 | 6.23331 | 6.21101 | 5.99315 | 6.40521 | 6.06975 | 6.02059 | 6.19023 | 6.01808 | 6.43909 | 6.35335 | 6.17413 | 6.67551 | 6.08887 | 6.84264 | 5.81114 |
| ERLEC1 | Q96DZ1 | 6.40234 | 6.52079 | 5.86319 | 6.49206 | 6.49315 | 6.20835 | 6.19321 | 6.1469 | 6.00852 | 6.19753 | 6.2684 | 5.8495 | 6.73038 | 6.21751 | 6.33683 | 6.07133 |
| CWF19L2 | Q2TBE0 | 5.45916 | 6.08349 | 5.85851 | 5.90646 | 5.67543 | 5.91022 | 5.87996 | 5.94483 | 5.86726 | 6.10193 | 6.06934 | 5.86715 | 6.62621 | 7.58075 | 7.43468 | 7.73464 |
| DENND4C | Q5VZ89-7 | 6.35046 | 6.30436 | 6.74564 | 5.95968 | 6.17604 | 6.3818 | 6.34521 | 6.36974 | 6.79248 | 6.78031 | 6.42289 | 6.65004 | 5.68164 | 5.73678 | 5.56641 | 5.73652 |
| DDX51 | Q8N8A6 | 6.58174 | 6.1361 | 6.0449 | 6.95173 | 6.09391 | 5.74066 | 6.42865 | 5.57134 | 6.2968 | 6.03989 | 6.11959 | 6.20332 | 6.57032 | 6.21045 | 6.51761 | 6.49298 |
| SNX9 | Q9Y5X1 | 6.23328 | 5.84593 | 6.83012 | 6.24197 | 6.11822 | 6.5384 | 5.81006 | 6.22732 | 6.26016 | 6.3721 | 6.39076 | 6.56181 | 6.44748 | 6.1433 | 6.10889 | 5.87021 |
| CRAT | P43155 | 6.3242 | 6.30147 | 5.9353 | 6.36361 | 6.34851 | 5.75704 | 6.63701 | 5.98823 | 6.24358 | 5.90758 | 5.77329 | 5.62526 | 6.74058 | 6.53657 | 6.86054 | 6.65724 |
| H2BC20P | Q6DN03 | 6.0885 | 5.93998 | 5.67874 | 6.27108 | 5.94528 | 5.70139 | 6.35294 | 5.86627 | 5.84057 | 5.66313 | 5.97593 | 5.96869 | 7.00253 | 7.14882 | 7.39659 | 7.15956 |
| BRD7 | Q9NPI1-2 | 6.05686 | 6.67408 | 5.98358 | 6.54257 | 5.97534 | 6.04783 | 6.13037 | 5.96584 | 5.79272 | 5.99359 | 6.19122 | 6.1483 | 6.33425 | 6.75241 | 6.5616 | 6.84945 |
| HSPA13 | P48723 | 6.54827 | 6.59586 | 6.05966 | 6.46342 | 6.25991 | 6.05237 | 6.05877 | 6.23023 | 6.1273 | 5.96809 | 5.90707 | 6.11895 | 6.65096 | 6.22053 | 6.51324 | 6.22535 |
| RPL26 | P61254 | 5.73996 | 6.28015 | 6.23394 | 6.24049 | 5.85995 | 6.43886 | 6.89644 | 6.48628 | 6.75005 | 5.85361 | 5.92289 | 6.43107 | 5.36366 | 6.69431 | 5.9529 | 6.85545 |
| RREB1 | Q92766-2 | 6.52701 | 6.30416 | 5.93076 | 6.85763 | 6.31509 | 6.00396 | 6.41596 | 6.24173 | 6.06647 | 6.1587 | 5.95255 | 5.92506 | 6.33247 | 6.23492 | 6.37387 | 6.35965 |
| ADAM17 | P78536 | 6.65666 | 6.7986 | 6.20111 | 6.89771 | 6.51871 | 6.2424 | 6.56921 | 6.23015 | 6.29908 | 6.36951 | 6.02527 | 6.19972 | 5.90785 | 5.64209 | 5.68493 | 5.75699 |
| RPL18 | Q07020 | 6.15487 | 6.41318 | 5.65039 | 5.8155 | 6.13904 | 6.1549 | 6.60858 | 6.51003 | 6.73169 | 6.30924 | 6.52694 | 6.12414 | 6.11761 | 6.60979 | 6.07439 | 6.05971 |
| P4HA2 | O15460 | 6.33819 | 6.7925 | 6.27454 | 6.72923 | 6.37986 | 6.312 | 6.23056 | 6.02225 | 6.07094 | 6.70987 | 6.21107 | 6.35616 | 6.05285 | 5.77083 | 5.88351 | 5.86564 |
| LLGL1 | Q15334 | 6.13397 | 6.37458 | 6.11089 | 6.10978 | 6.27228 | 6.25815 | 5.89698 | 6.09364 | 6.06647 | 6.22764 | 6.04962 | 6.32542 | 6.97115 | 6.11819 | 6.58947 | 6.42608 |
| HLA-A | P04439-2 | 6.49395 | 6.46062 | 6.17319 | 6.56416 | 6.45807 | 6.09886 | 6.16649 | 6.22857 | 6.16832 | 6.1037 | 6.11572 | 6.06241 | 6.4969 | 6.11947 | 6.23268 | 6.05689 |
| RPS5 | P46782 | 6.38382 | 6.39163 | 6.2688 | 5.92224 | 6.47262 | 6.36341 | 6.27823 | 6.50341 | 6.58032 | 6.45153 | 6.50981 | 6.41306 | 6.0781 | 5.99829 | 5.69649 | 5.68823 |
| NHSL1 | Q5SYE7-2 | 6.1864 | 6.14229 | 6.87169 | 5.97122 | 6.10721 | 6.38743 | 6.42064 | 6.3906 | 6.23574 | 6.59434 | 6.42707 | 6.59628 | 6.02995 | 6.03885 | 5.84851 | 5.83278 |
| PLOD1 | Q02809-2 | 6.62185 | 6.49904 | 5.69689 | 6.28559 | 6.25191 | 6.09069 | 6.28171 | 6.12095 | 6.14277 | 6.17331 | 5.92583 | 6.00966 | 6.40594 | 6.57563 | 6.55027 | 6.36797 |
| ATAD1 | Q8NBU5 | 6.32442 | 6.30238 | 6.09902 | 6.50767 | 6.26497 | 5.99451 | 6.25638 | 6.152 | 5.97293 | 5.93161 | 6.14454 | 5.88276 | 6.6174 | 6.35973 | 6.73117 | 6.45852 |
| HSPB1 | P04792 | 5.63139 | 5.05297 | 5.93423 | 4.40473 | 6.36883 | 6.10732 | 5.74837 | 7.57879 | 6.13017 | 7.44956 | 6.82678 | 7.27163 | 4.94679 | 7.13399 | 6.40355 | 7.01091 |
| CRIM1 | Q9NZV1 | 7.07842 | 6.38704 | 6.68773 | 7.56386 | 7.02382 | 6.42158 | 6.60596 | 6.2926 | 6.34502 | 6.16701 | 6.65305 | 6.38084 | 6.56856 | 4.58553 | 5.05175 | 5.08723 |
| CLYBL | Q8N0X4 | 6.29668 | 6.21369 | 5.73837 | 6.44226 | 6.19519 | 5.91372 | 6.14227 | 6.31666 | 6.14223 | 5.95663 | 6.0263 | 5.87967 | 6.86554 | 6.50001 | 6.76904 | 6.60174 |
| CPM | P14384 | 6.03444 | 5.99038 | 5.8047 | 6.3216 | 6.33998 | 5.86634 | 6.75097 | 5.85776 | 6.56662 | 5.98277 | 6.08524 | 6.03745 | 6.61581 | 6.365 | 6.90201 | 6.47893 |
| NAPG | Q99747 | 6.3792 | 5.41179 | 6.18241 | 5.01644 | 6.76147 | 6.02407 | 5.9659 | 6.83284 | 6.07302 | 6.5945 | 6.30307 | 6.62668 | 6.86918 | 6.34697 | 6.25992 | 6.35253 |
| MVD | P53602 | 6.34594 | 6.10847 | 6.99577 | 5.69339 | 6.12906 | 6.65836 | 6.16487 | 6.48401 | 6.45102 | 6.6559 | 6.63312 | 6.63843 | 5.79147 | 5.82784 | 5.82182 | 5.60353 |
| HDAC1 | Q13547 | 6.37376 | 6.12311 | 6.25696 | 6.22896 | 6.12509 | 5.72093 | 6.39348 | 5.98642 | 6.03398 | 5.98212 | 6.32878 | 5.98682 | 6.68389 | 6.27551 | 7.11643 | 6.38376 |
| C9orf78 | Q9NZ63 | 5.8376 | 6.0018 | 6.0611 | 5.4786 | 6.17198 | 6.25608 | 5.72168 | 6.24495 | 6.37405 | 6.40224 | 6.54822 | 6.479 | 6.65599 | 6.68788 | 6.59859 | 6.48022 |
| SNX14 | Q9Y5W7 | 6.65603 | 7.17731 | 6.61084 | 7.06179 | 6.80843 | 6.47898 | 6.47428 | 6.3317 | 6.49469 | 6.75735 | 6.47039 | 6.36778 | 5.13902 | 5.15351 | 4.93414 | 5.08374 |
| PRKACB | P22694 | 6.21789 | 6.35825 | 6.43318 | 6.27074 | 6.36739 | 6.49329 | 6.12585 | 6.55292 | 6.00269 | 6.21805 | 6.38882 | 5.90947 | 6.18709 | 6.08595 | 6.37695 | 6.01148 |
| RRM1 | P23921 | 6.39156 | 6.14332 | 6.93737 | 6.18552 | 6.46194 | 6.97019 | 6.4127 | 6.71061 | 6.79925 | 6.72751 | 6.75318 | 6.59307 | 5.34553 | 5.22985 | 5.33732 | 5.00108 |
| VTA1 | Q9NP79 | 6.15687 | 5.96071 | 6.75152 | 5.35067 | 6.26623 | 6.21439 | 6.28522 | 6.42974 | 6.11698 | 6.43286 | 6.52827 | 6.49421 | 6.08015 | 6.12519 | 6.3453 | 6.4617 |
| AACS | Q86V21 | 6.27074 | 6.25124 | 7.25718 | 6.35275 | 6.20722 | 6.91055 | 5.96333 | 6.64115 | 6.51703 | 6.88076 | 6.8032 | 6.79675 | 5.25196 | 5.71607 | 4.96291 | 5.21716 |
| UTP15 | Q8TED0 | 6.15939 | 6.37607 | 5.80658 | 6.71054 | 6.44363 | 6.11408 | 6.451 | 6.16969 | 6.12358 | 5.95585 | 5.94935 | 5.78809 | 6.4922 | 6.1877 | 6.58506 | 6.68718 |
| PIK3C3 | Q8NEB9 | 6.45605 | 6.19477 | 6.68271 | 5.8943 | 6.34757 | 6.55797 | 6.29934 | 6.51448 | 6.31806 | 6.44504 | 6.5767 | 6.44548 | 5.96993 | 5.88074 | 5.68908 | 5.72781 |
| MAT2B | Q9NZL9 | 5.2817 | 5.83519 | 6.24305 | 5.27778 | 6.08653 | 6.28153 | 6.23522 | 6.1494 | 6.52809 | 5.9465 | 6.28542 | 6.37311 | 6.5807 | 6.98874 | 6.9468 | 6.96024 |
| TUBG1 | P23258 | 6.37635 | 6.35742 | 6.8168 | 6.27185 | 6.3108 | 6.66824 | 6.19345 | 6.85238 | 6.45598 | 6.58327 | 6.80222 | 6.81951 | 5.23826 | 5.4206 | 5.28203 | 5.55083 |
| NUP88 | Q99567 | 6.40617 | 6.24497 | 6.18417 | 6.46392 | 6.32296 | 6.36985 | 6.31185 | 6.13309 | 6.37665 | 5.7951 | 6.02016 | 6.11556 | 6.53417 | 6.08795 | 6.13745 | 6.49598 |
| MTMR6 | Q9Y217 | 6.08027 | 6.05129 | 6.661 | 6.36141 | 6.16163 | 6.36317 | 6.24011 | 6.33626 | 6.19337 | 6.47437 | 6.59828 | 6.72905 | 5.75521 | 6.16458 | 5.76698 | 6.06304 |
| SYNRG | Q9UMZ2 | 6.57261 | 6.44126 | 6.96588 | 6.45886 | 6.32935 | 6.83862 | 6.26945 | 6.23437 | 6.14499 | 6.4332 | 6.86345 | 6.74513 | 5.78498 | 5.53784 | 5.22055 | 5.15947 |
| TXN | P10599 | 6.27909 | 5.59887 | 6.53425 | 5.11798 | 6.43985 | 6.81188 | 5.76303 | 7.13361 | 6.39531 | 6.85121 | 6.50621 | 6.97486 | 6.00801 | 6.08134 | 5.61201 | 5.89251 |
| AAAS | Q9NRG9 | 6.54728 | 6.34922 | 6.25096 | 6.8924 | 6.44895 | 5.8571 | 6.33049 | 6.10718 | 6.11087 | 5.97974 | 6.06093 | 6.11292 | 6.25646 | 6.08624 | 6.27554 | 6.33373 |
| NMI | Q13287 | 6.35889 | 6.1905 | 6.62098 | 5.80512 | 6.29758 | 6.92086 | 6.18575 | 6.57163 | 6.43502 | 6.84999 | 6.53623 | 6.20398 | 5.66141 | 5.95011 | 5.75225 | 5.65967 |
| DROSHA | Q9NRR4 | 6.25768 | 6.18663 | 5.63658 | 6.69382 | 6.15031 | 5.82921 | 6.32405 | 5.84411 | 6.66287 | 6.67919 | 6.04961 | 6.19819 | 6.35785 | 6.17831 | 6.4578 | 6.49379 |
| RMND1 | Q9NWS8 | 6.47568 | 6.54005 | 5.87205 | 7.15422 | 6.28022 | 5.85918 | 6.65614 | 5.72312 | 6.0657 | 5.92367 | 5.49583 | 5.45196 | 6.52594 | 6.42741 | 6.66129 | 6.88754 |
| PEX5 | P50542-2 | 6.77164 | 6.04407 | 6.95211 | 6.03936 | 6.24954 | 6.39581 | 5.94433 | 7.04577 | 6.42888 | 6.51204 | 6.92848 | 7.30377 | 5.52743 | 5.21505 | 5.08186 | 5.55985 |
| NCOA7 | Q8NI08-2 | 5.40918 | 6.24745 | 6.2331 | 6.0538 | 5.68418 | 6.30645 | 5.79601 | 6.31594 | 6.21631 | 6.57409 | 6.27726 | 6.7374 | 6.36915 | 6.91339 | 6.25603 | 6.61025 |
| SCD | O00767 | 7.02868 | 7.01062 | 6.5331 | 6.81544 | 5.93815 | 6.07194 | 6.74583 | 5.85506 | 6.98002 | 7.19448 | 6.99999 | 6.72246 | 5.03314 | 5.19559 | 4.85396 | 5.02155 |
| FAM50A | Q14320 | 5.9231 | 5.93858 | 6.51265 | 5.60769 | 5.96762 | 6.45858 | 5.76766 | 6.40546 | 6.12431 | 6.50231 | 6.5393 | 6.56523 | 5.94476 | 6.73926 | 6.97798 | 6.0255 |
| TERF2 | Q15554 | 6.24899 | 6.13927 | 6.1893 | 6.88241 | 6.32391 | 5.85403 | 6.33207 | 5.84404 | 6.50916 | 6.17005 | 6.33734 | 6.17097 | 6.28754 | 6.19532 | 6.3008 | 6.2148 |
| CAPZB | P47756-2 | 6.33083 | 6.29352 | 6.36296 | 6.34581 | 6.52242 | 6.29711 | 6.1047 | 6.30464 | 6.26525 | 6.26052 | 6.19943 | 6.19631 | 6.03604 | 6.30958 | 6.35047 | 5.82043 |
| RPL38 | P63173 | 6.30636 | 6.47363 | 6.56626 | 6.29952 | 6.16603 | 6.50158 | 6.28059 | 6.08704 | 6.46967 | 6.50215 | 6.53665 | 7.27548 | 5.51308 | 5.63991 | 5.77576 | 5.60631 |
| YTHDC1 | Q96MU7 | 6.24323 | 6.46807 | 5.86044 | 6.80422 | 6.10611 | 5.99524 | 6.42135 | 6.019 | 6.08183 | 5.85303 | 6.00896 | 5.97863 | 6.42844 | 6.66081 | 6.70352 | 6.36714 |
| DNAJB11 | Q9UBS4 | 6.07847 | 6.15092 | 5.91381 | 6.25929 | 6.31931 | 6.11703 | 6.48462 | 6.00892 | 6.12366 | 6.02709 | 6.08797 | 6.01635 | 6.61306 | 6.68285 | 6.59539 | 6.52124 |
| LPCAT1 | Q8NF37 | 6.40711 | 6.11945 | 6.0388 | 6.45294 | 6.02386 | 5.79184 | 6.17769 | 5.66823 | 6.35144 | 6.15924 | 6.06149 | 5.79527 | 6.78748 | 6.36908 | 6.96786 | 6.82823 |
| VPS13D | Q5THJ4 | 6.05441 | 6.10457 | 6.39982 | 6.2258 | 5.77795 | 6.44039 | 5.94435 | 5.92342 | 6.14697 | 6.27955 | 6.56496 | 6.63282 | 6.34883 | 6.54159 | 6.31303 | 6.30155 |
| MAP3K4 | Q9Y6R4 | 6.28096 | 5.95665 | 6.82126 | 6.1549 | 6.24762 | 6.75441 | 6.17426 | 6.29127 | 6.44893 | 6.54605 | 6.57466 | 6.67777 | 5.7977 | 6.00024 | 5.50544 | 5.76787 |
| CTTNBP2NL | Q9P2B4 | 5.96968 | 5.98845 | 6.23532 | 5.71898 | 6.0559 | 6.02661 | 5.7135 | 6.41923 | 6.26441 | 6.21819 | 6.61548 | 6.51009 | 7.10118 | 6.6949 | 6.35511 | 6.11298 |
| DAZAP1 | Q96EP5 | 6.34404 | 6.50124 | 6.30989 | 6.61385 | 6.85378 | 6.51357 | 6.57483 | 6.66764 | 6.66963 | 6.13891 | 6.56435 | 6.48879 | 5.7696 | 5.41538 | 5.26233 | 5.31217 |
| UBE2K | P61086 | 6.28538 | 5.6663 | 6.96236 | 5.3282 | 6.57639 | 6.84938 | 6.17265 | 6.72814 | 6.15837 | 6.54684 | 6.64897 | 6.60094 | 6.11267 | 6.05291 | 5.7591 | 5.5514 |
| QPCTL | Q9NXS2 | 6.04605 | 6.34631 | 5.88488 | 6.36917 | 5.9883 | 6.10084 | 6.13096 | 6.30779 | 5.67104 | 6.1084 | 6.18393 | 5.99956 | 6.95195 | 6.80917 | 6.42879 | 6.67286 |
| CAB39 | Q9Y376 | 6.54846 | 6.46911 | 6.90795 | 5.92313 | 6.51634 | 6.54875 | 6.14303 | 6.41816 | 6.57349 | 6.50522 | 6.75912 | 6.50699 | 5.63824 | 5.73465 | 5.383 | 5.42433 |
| RPL27 | P61353 | 6.27864 | 6.54258 | 6.21669 | 6.10221 | 6.22405 | 6.37575 | 6.44661 | 6.1899 | 6.64082 | 6.25381 | 6.60195 | 6.50815 | 5.9066 | 6.21709 | 5.8005 | 5.69465 |
| MRPL15 | Q9P015 | 5.98065 | 5.9172 | 5.46966 | 6.60669 | 6.1953 | 5.82309 | 6.67974 | 6.22721 | 6.19435 | 5.79276 | 5.99122 | 5.92136 | 7.08443 | 6.61352 | 6.4631 | 7.03975 |
| APEX1 | P27695 | 5.60802 | 5.5418 | 6.22615 | 5.67067 | 5.90091 | 6.39331 | 5.89555 | 5.51619 | 6.0001 | 6.08787 | 6.48621 | 6.12713 | 6.867 | 7.71642 | 7.17018 | 6.79248 |
| CDV3 | Q9UKY7 | 6.18119 | 6.28796 | 6.71392 | 5.85742 | 6.59191 | 6.83615 | 5.97218 | 6.72145 | 6.47097 | 6.39886 | 6.64142 | 6.80292 | 5.89831 | 5.82933 | 5.33671 | 5.45931 |
| TRMT5 | Q32P41 | 5.91508 | 5.93659 | 5.85272 | 5.89912 | 5.8055 | 6.20175 | 5.98984 | 6.16039 | 6.0266 | 6.45762 | 6.17325 | 6.20011 | 6.502 | 7.10295 | 6.89566 | 6.88081 |
| DES | P17661 | 6.20812 | 6.04546 | 6.00034 | 6.52427 | 6.39182 | 6.19652 | 6.06455 | 6.65761 | 6.40241 | 6.03264 | 6.52158 | 6.45738 | 6.76762 | 5.73033 | 5.95973 | 6.03962 |
| IST1 | P53990-5 | 6.52356 | 5.94122 | 7.0496 | 5.49268 | 6.48243 | 6.80201 | 6.19241 | 6.79253 | 6.30117 | 6.59577 | 6.54454 | 6.77356 | 6.05691 | 5.80641 | 5.36441 | 5.2808 |
| CDYL | Q9Y232 | 6.32649 | 6.22188 | 6.05294 | 6.74683 | 6.16973 | 5.85275 | 6.34471 | 5.90259 | 6.09378 | 6.28873 | 5.89699 | 6.06012 | 6.61919 | 6.16395 | 6.51982 | 6.73949 |
| VRK1 | Q99986 | 5.84312 | 6.0168 | 5.89311 | 6.09508 | 5.7027 | 5.66123 | 6.16794 | 6.07593 | 6.32507 | 6.1196 | 6.43616 | 6.20616 | 6.89921 | 6.76541 | 6.9531 | 6.83937 |
| RPL31 | P62899 | 6.48533 | 6.34758 | 6.31311 | 6.27846 | 6.1652 | 6.19052 | 6.40808 | 6.46342 | 6.79357 | 6.2245 | 6.51438 | 6.34152 | 5.76059 | 6.01553 | 5.82247 | 5.87576 |
| REXO1 | Q8N1G1 | 6.2561 | 6.27273 | 6.1577 | 6.93286 | 5.93675 | 5.86769 | 6.20091 | 5.90981 | 6.27387 | 6.3957 | 6.20899 | 6.52669 | 5.91681 | 6.27208 | 6.3842 | 6.48711 |
| RHEB | Q15382 | 6.5285 | 6.34764 | 6.6147 | 6.04932 | 6.22313 | 6.57379 | 6.4528 | 6.63843 | 6.5048 | 6.20649 | 6.23336 | 6.34657 | 5.74533 | 6.04178 | 5.56868 | 5.92468 |
| RPL21 | P46778 | 6.51392 | 6.17988 | 6.28966 | 6.28161 | 6.66016 | 6.19088 | 6.50429 | 6.14262 | 6.65507 | 6.29648 | 6.6635 | 6.04081 | 5.89685 | 5.98693 | 5.95547 | 5.74186 |
| HECTD3 | Q5T447 | 6.02397 | 6.21725 | 7.04942 | 6.2041 | 6.0526 | 6.54264 | 6.35595 | 6.40233 | 6.37658 | 6.59104 | 6.90691 | 6.80882 | 5.59501 | 5.75457 | 5.24884 | 5.86996 |
| NECTIN2 | Q92692 | 6.36893 | 6.5544 | 6.31866 | 6.66861 | 6.11802 | 6.18196 | 6.27107 | 6.19833 | 6.30431 | 6.16832 | 6.30571 | 6.07567 | 6.26011 | 5.97492 | 6.01192 | 6.21905 |
| MYDGF | Q969H8 | 6.13068 | 6.41153 | 5.92637 | 6.32712 | 6.40998 | 6.34171 | 6.42865 | 5.97437 | 5.99813 | 5.88122 | 5.88443 | 5.95515 | 6.46414 | 6.42276 | 6.88882 | 6.55493 |
| SNRPD3 | P62318 | 6.44911 | 6.3601 | 5.77835 | 6.68834 | 6.26771 | 5.99456 | 6.55489 | 6.01767 | 6.32858 | 5.66664 | 5.88065 | 5.86779 | 6.5357 | 6.22414 | 6.64784 | 6.73794 |

| | | | | | | | | | | | | | | | | | |
|---|---|---|---|---|---|---|---|---|---|---|---|---|---|---|---|---|---|
| CTSC | P53634 | 5.99264 | 5.91147 | 5.72698 | 5.93083 | 6.44617 | 6.20621 | 6.43834 | 6.86432 | 6.2469 | 6.52367 | 6.73424 | 6.34241 | 6.09085 | 6.23795 | 6.21788 | 6.08914 |
| INTS7 | Q9NVH2 | 6.14366 | 6.3722 | 6.11914 | 6.74635 | 5.97704 | 6.02866 | 6.23316 | 6.00362 | 6.0714 | 6.23378 | 6.26932 | 6.15202 | 6.5695 | 6.16553 | 6.55304 | 6.36157 |
| TFG | Q92734 | 6.28068 | 5.53442 | 6.75068 | 5.41416 | 6.06543 | 6.53967 | 6.27612 | 6.41104 | 5.8903 | 6.55359 | 6.91271 | 6.90941 | 6.3818 | 6.28956 | 5.85989 | 5.93052 |
| TRIM56 | Q9BRZ2 | 6.40074 | 6.09223 | 6.53618 | 5.97045 | 6.0139 | 6.53696 | 6.27526 | 6.46207 | 6.214 | 6.29872 | 6.40262 | 6.40694 | 6.14769 | 5.98842 | 5.98053 | 6.27328 |
| STRN3 | Q13033 | 6.08349 | 5.9227 | 6.25583 | 5.82921 | 6.25747 | 6.25507 | 5.95074 | 6.41773 | 6.27873 | 6.04779 | 6.55512 | 6.4617 | 6.65131 | 6.57037 | 6.37329 | 6.08945 |
| RPF1 | Q9H9Y2 | 6.65433 | 6.11347 | 5.96929 | 7.15317 | 6.03418 | 5.91608 | 6.37133 | 6.21861 | 6.43772 | 5.85601 | 6.23464 | 6.15551 | 6.24615 | 5.89785 | 6.47484 | 6.26682 |
| SH3BP4 | Q9P0V3 | 6.5353 | 6.28633 | 6.3074 | 6.69235 | 6.41303 | 6.2293 | 6.48261 | 5.74392 | 6.56327 | 6.31066 | 6.17428 | 6.24858 | 6.15317 | 5.77979 | 6.29362 | 5.78639 |
| UBXN4 | Q92575 | 6.59887 | 6.44639 | 5.93376 | 6.68618 | 6.21749 | 5.96913 | 6.5873 | 5.72371 | 6.09369 | 6.1926 | 6.04262 | 6.20527 | 6.25446 | 6.26153 | 6.6277 | 6.15929 |
| RAD18 | Q9NS91 | 6.25057 | 6.34572 | 6.11528 | 6.57398 | 6.39366 | 6.12416 | 6.46869 | 6.26798 | 6.13893 | 5.9466 | 6.33644 | 6.41524 | 6.31167 | 5.84554 | 6.19501 | 6.27053 |
| LIPE | Q05469 | 6.65424 | 6.4826 | 6.95533 | 6.26852 | 6.3306 | 6.45533 | 6.40554 | 6.62121 | 6.71778 | 6.61607 | 6.59095 | 6.48808 | 5.47976 | 5.26167 | 5.19142 | 5.4809 |
| ANO6 | Q4KMQ2-2 | 6.2552 | 6.41794 | 6.12931 | 6.42234 | 6.22207 | 6.10723 | 6.3045 | 5.97111 | 6.0371 | 6.2608 | 6.04198 | 5.98943 | 6.46209 | 6.35938 | 6.59421 | 6.4253 |
| KIAA2013 | Q8IYS2-2 | 6.51134 | 6.72424 | 5.99356 | 6.63197 | 6.08796 | 6.00083 | 6.35809 | 5.74748 | 6.18704 | 6.39927 | 6.22287 | 6.2675 | 6.32125 | 6.18816 | 6.19617 | 6.16226 |
| LYN | P07948 | 6.47573 | 6.39307 | 6.19085 | 6.37525 | 6.30172 | 6.20142 | 6.14291 | 5.98771 | 6.371 | 6.25878 | 6.10284 | 5.82781 | 6.43097 | 6.29494 | 6.40299 | 6.24202 |
| MET | P08581-2 | 6.43972 | 6.42791 | 6.2753 | 6.46783 | 6.27596 | 6.0523 | 6.47727 | 6.17273 | 6.24907 | 6.13972 | 6.20304 | 6.12496 | 6.26364 | 6.02292 | 6.27654 | 6.13111 |
| CDK13 | Q14004 | 5.99049 | 5.81464 | 5.78354 | 6.41169 | 5.92086 | 6.0121 | 6.2194 | 5.70169 | 6.06919 | 6.00826 | 5.80964 | 6.21465 | 6.51048 | 7.1173 | 6.93386 | 7.48221 |
| CDK5 | Q00535 | 6.42002 | 6.27348 | 6.65859 | 6.06006 | 6.45621 | 6.93689 | 6.59264 | 6.41485 | 6.32502 | 6.49812 | 6.39452 | 6.63572 | 5.59232 | 5.97318 | 5.62463 | 5.80965 |
| RPLP2 | P05387 | 6.84232 | 5.78124 | 5.43808 | 5.22279 | 5.55427 | 4.95997 | 4.97257 | 6.57202 | 4.78527 | 6.27186 | 5.35232 | 6.5175 | 5.61024 | 9.88542 | 7.18871 | 9.04541 |
| MYO1E | Q12965 | 6.49787 | 6.14746 | 7.01538 | 5.95803 | 6.23843 | 6.5978 | 6.54111 | 6.23398 | 6.45439 | 6.94411 | 6.43375 | 6.3685 | 5.78234 | 5.88812 | 5.46714 | 5.43159 |
| EDRF1 | Q3B7T1 | 6.86777 | 7.12609 | 7.82286 | 7.41827 | 6.80575 | 6.92786 | 6.63417 | 6.74757 | 7.15406 | 7.31541 | 7.62635 | 7.16504 | 3.43921 | 3.82301 | 3.34894 | 3.77763 |
| ANAPC2 | Q9UJX6 | 6.33523 | 6.23914 | 6.53405 | 6.11888 | 6.34376 | 6.23952 | 6.10187 | 6.02108 | 6.47452 | 6.34679 | 6.28315 | 6.20493 | 6.16502 | 6.12406 | 6.20531 | 6.26268 |
| PIK3C2A | O00443 | 6.30255 | 6.22136 | 6.73674 | 6.68873 | 5.92886 | 6.56555 | 6.50238 | 6.55703 | 6.14929 | 6.50077 | 6.53849 | 6.66625 | 5.57545 | 5.72906 | 5.69521 | 5.64228 |
| PAK4 | O96013 | 6.45205 | 6.12837 | 6.40645 | 6.55922 | 6.07274 | 6.52886 | 6.32609 | 6.61682 | 6.26168 | 6.35749 | 6.1151 | 6.11886 | 5.75732 | 6.45714 | 6.0478 | 5.79402 |
| CHUK | O15111 | 6.36844 | 6.29112 | 7.01339 | 6.10359 | 6.21299 | 6.41127 | 6.25484 | 6.16238 | 6.43204 | 6.63794 | 6.52693 | 6.39606 | 6.05932 | 5.81271 | 5.57411 | 5.74286 |
| UROD | P06132 | 6.36417 | 5.40364 | 6.778 | 5.00693 | 6.40161 | 6.58755 | 5.8109 | 6.87796 | 5.89018 | 6.65917 | 6.62113 | 6.93133 | 5.89373 | 6.63353 | 6.17533 | 5.96484 |
| CIAPIN1 | Q6FI81 | 6.34634 | 5.95404 | 6.91429 | 5.81359 | 6.42341 | 6.67842 | 6.05053 | 6.92604 | 6.28598 | 6.55753 | 6.72541 | 6.91653 | 5.73606 | 5.83014 | 5.36305 | 5.47865 |
| NUDT19 | A8MXV4 | 6.38677 | 6.57585 | 6.23195 | 7.00338 | 6.36397 | 6.24419 | 6.13231 | 6.31481 | 5.92888 | 5.93998 | 6.19029 | 6.02898 | 6.41255 | 5.88448 | 6.43842 | 5.92318 |
| YEATS2 | Q9ULM3 | 6.21699 | 6.09285 | 6.08027 | 6.93738 | 6.34966 | 6.38227 | 6.62834 | 5.9778 | 6.37752 | 6.53138 | 6.45306 | 6.35634 | 6.07575 | 5.97643 | 5.73758 | 5.82638 |
| MACF1 | A0A0A6YYJ5 | 6.27405 | 6.51764 | 6.07462 | 6.69873 | 6.15917 | 6.39249 | 6.56401 | 6.00733 | 6.1271 | 6.02518 | 6.10187 | 6.20396 | 6.01016 | 6.12126 | 6.42344 | 6.29898 |
| GRWD1 | Q9BQ67 | 5.88982 | 6.06543 | 6.36185 | 6.68629 | 6.07214 | 5.94889 | 5.86231 | 6.50318 | 7.01007 | 6.55987 | 6.92794 | 6.69517 | 5.79384 | 5.6328 | 5.89991 | 6.09048 |
| FBXO38 | Q6PIJ6 | 6.16803 | 6.53284 | 6.13819 | 6.96941 | 6.46497 | 6.02631 | 6.42639 | 5.8905 | 6.47836 | 6.30885 | 6.17804 | 6.15616 | 5.94179 | 5.93158 | 6.39581 | 5.99277 |
| ERLIN2 | O94905 | 6.33476 | 6.36946 | 5.84562 | 6.34453 | 6.35801 | 5.99555 | 6.65507 | 6.16496 | 6.05861 | 5.90003 | 5.61453 | 5.83236 | 6.68843 | 6.43218 | 6.72155 | 6.68435 |
| OBSL1 | O75147 | 6.3975 | 6.15492 | 6.53632 | 6.28701 | 6.11349 | 6.16989 | 6.34751 | 5.98859 | 6.32485 | 6.39349 | 6.45895 | 6.31446 | 6.16076 | 6.08369 | 6.33888 | 5.9297 |
| DHRS7 | Q9Y394 | 6.39067 | 6.24375 | 6.04824 | 6.61501 | 6.24993 | 6.01593 | 6.69126 | 5.72802 | 6.18033 | 5.70729 | 6.00616 | 5.76243 | 6.72094 | 6.32371 | 6.74486 | 6.57147 |
| TANC1 | Q9C0D5 | 6.05919 | 6.27064 | 6.61129 | 6.27473 | 6.13558 | 6.4865 | 6.24189 | 6.38301 | 6.28069 | 6.45081 | 6.80836 | 6.67614 | 5.91436 | 5.83126 | 5.77861 | 5.79695 |
| RPS21 | P63220 | 6.6956 | 6.75047 | 6.03871 | 5.60005 | 7.13751 | 6.85832 | 5.43866 | 7.81907 | 5.85936 | 6.516 | 6.89755 | 6.68591 | 4.95347 | 6.5877 | 5.83016 | 5.23145 |
| NUCB2 | P80303 | 7.40074 | 5.41171 | 5.66299 | 5.52993 | 6.26117 | 4.6613 | 4.69824 | 7.03931 | 4.66582 | 7.19327 | 5.85997 | 7.55319 | 6.46489 | 7.5862 | 6.6454 | 7.36587 |
| LENG8 | Q96PV6 | 6.34391 | 6.66407 | 6.22263 | 7.17034 | 6.28416 | 6.01727 | 6.48162 | 5.90479 | 6.08589 | 6.40106 | 6.28849 | 6.20776 | 6.21802 | 5.66692 | 6.06373 | 5.97935 |
| SPECC1 | Q5M775 | 6.40931 | 6.67309 | 6.3622 | 6.85334 | 6.79007 | 5.95067 | 6.28341 | 6.42334 | 6.39068 | 6.25286 | 6.31481 | 6.09306 | 6.10559 | 5.73914 | 5.66382 | 5.69407 |
| MORF4L1 | Q9UBU8-2 | 6.53187 | 6.95777 | 6.92829 | 6.83378 | 5.96859 | 6.60016 | 6.71323 | 6.47967 | 7.05762 | 6.78537 | 6.67143 | 6.58451 | 4.73628 | 5.09862 | 4.98279 | 5.07003 |
| MED13 | Q9UHV7 | 6.37129 | 6.5595 | 6.30234 | 6.87541 | 6.08813 | 6.27161 | 6.20728 | 5.92838 | 6.17241 | 6.43751 | 6.20575 | 6.197 | 6.45912 | 5.80924 | 6.02979 | 6.08524 |
| NCOA3 | Q9Y6Q9 | 5.54605 | 6.21284 | 6.20164 | 5.69637 | 5.9055 | 6.07097 | 5.69711 | 6.16697 | 6.21735 | 6.32046 | 6.50105 | 6.46242 | 6.95446 | 6.34154 | 6.86587 | 6.83941 |
| ME1 | P48163 | 6.03601 | 6.23586 | 6.73406 | 6.09682 | 6.37502 | 7.11168 | 6.05904 | 6.52528 | 6.44643 | 6.76008 | 6.6643 | 6.62834 | 5.5602 | 5.6527 | 5.62813 | 5.48603 |
| TBCA | O75347 | 6.41675 | 5.86916 | 7.03499 | 5.33756 | 6.79777 | 6.86966 | 6.12782 | 6.92924 | 6.51785 | 6.60569 | 6.62053 | 6.77459 | 5.89865 | 5.57242 | 5.26952 | 5.35779 |
| SPART | Q8N0X7 | 6.16767 | 6.37883 | 6.57955 | 6.18375 | 5.89145 | 6.75514 | 6.22063 | 6.71041 | 6.37417 | 6.66344 | 6.48688 | 6.93804 | 5.4877 | 5.92081 | 5.25768 | 5.98387 |
| PDE4DIP | Q5VU43-13 | 5.93761 | 5.80898 | 6.35924 | 5.87272 | 6.29815 | 6.46463 | 5.87779 | 6.64852 | 6.52193 | 6.50789 | 6.51489 | 6.66497 | 6.3254 | 6.17609 | 6.23573 | 5.78546 |
| PRKCA | P17252 | 6.80323 | 6.94662 | 7.73894 | 6.79217 | 5.9065 | 6.82434 | 6.53293 | 6.58137 | 6.0256 | 6.59477 | 6.57341 | 6.19384 | 4.71864 | 5.35151 | 5.39985 | 5.0163 |
| RABGGTA | Q92696 | 6.32377 | 6.23451 | 7.08943 | 6.03822 | 6.60402 | 6.79466 | 6.36031 | 6.75474 | 6.50599 | 6.45718 | 6.70819 | 6.46518 | 5.29646 | 5.65366 | 5.41943 | 5.29423 |
| SMG6 | Q86US8 | 6.13134 | 6.48817 | 6.81945 | 6.46472 | 6.11285 | 6.56375 | 6.01439 | 6.37857 | 6.63626 | 6.859 | 6.56595 | 6.77011 | 5.47005 | 5.69804 | 5.55532 | 5.47204 |
| MAGEA10 | P43363 | 5.46171 | 5.70835 | 5.83188 | 5.42632 | 5.70892 | 6.39407 | 5.70893 | 6.25373 | 5.83082 | 6.12454 | 6.32253 | 6.14238 | 7.20071 | 7.56627 | 7.14097 | 7.17787 |
| CCZ1B | P86790 | 6.36445 | 6.43195 | 6.88661 | 6.3625 | 6.48574 | 6.61708 | 6.1874 | 6.57582 | 6.76493 | 6.7168 | 6.82374 | 6.72388 | 5.2028 | 5.36362 | 5.29542 | 5.19725 |
| WDR26 | Q9H7D7 | 6.02065 | 6.46931 | 6.36816 | 6.66959 | 6.05828 | 6.5762 | 5.86877 | 6.38803 | 5.98369 | 6.21594 | 5.99914 | 6.09704 | 6.47961 | 6.15931 | 6.09841 | 6.54788 |
| SEH1L | Q96EE3-1 | 6.39518 | 6.30956 | 6.31043 | 6.59246 | 6.31678 | 6.24782 | 6.38011 | 5.80183 | 6.26924 | 6.17514 | 6.21801 | 6.11775 | 6.29951 | 5.96502 | 6.3078 | 6.29338 |
| GHITM | Q9H3K2 | 5.88702 | 5.70164 | 5.51386 | 6.23171 | 6.44245 | 6.63089 | 7.02679 | 6.54644 | 5.91182 | 5.68653 | 5.64578 | 5.81817 | 6.5299 | 6.82087 | 6.62654 | 6.97957 |
| CEP97 | Q8IW35-2 | 6.39419 | 6.11021 | 7.05123 | 6.31407 | 6.34233 | 6.58364 | 6.11115 | 6.71912 | 6.47409 | 6.63477 | 6.93398 | 7.07414 | 5.3677 | 5.48579 | 5.06575 | 5.33785 |
| TFAM | Q00059 | 6.04198 | 6.19816 | 5.55673 | 6.97356 | 5.96577 | 6.10312 | 6.89796 | 5.97603 | 6.15868 | 5.53752 | 5.73184 | 5.68694 | 6.64552 | 6.89133 | 6.6642 | 6.97068 |
| TTC28 | Q96AY4 | 6.50427 | 6.3989 | 6.77022 | 6.03577 | 6.18678 | 6.48829 | 6.18605 | 6.42096 | 6.70755 | 6.14426 | 6.49928 | 6.4408 | 5.98598 | 5.87267 | 5.67073 | 5.68749 |
| ANTXR2 | P58335 | 6.09501 | 6.05356 | 5.94503 | 6.59536 | 6.30313 | 6.2575 | 6.519 | 5.94056 | 6.42667 | 6.43365 | 6.60657 | 6.1697 | 6.17854 | 6.1507 | 6.03188 | 6.29314 |
| L1RE1 | Q9UN81 | 6.60698 | 6.54312 | 6.77892 | 6.28824 | 6.17922 | 6.41573 | 6.16918 | 6.11215 | 6.40847 | 6.55762 | 6.23048 | 6.39025 | 5.94521 | 5.79779 | 5.75649 | 5.82015 |
| RAB1B | Q9H0U4 | 5.89829 | 6.42076 | 6.11359 | 6.3816 | 5.73347 | 6.09443 | 6.43838 | 5.75326 | 6.23491 | 6.18177 | 6.17661 | 6.32737 | 6.40424 | 6.43605 | 6.55101 | 6.85426 |
| STEAP3 | Q658P3-2 | 6.428 | 6.75126 | 6.29621 | 6.7722 | 6.03216 | 6.08088 | 6.22205 | 6.1589 | 6.93155 | 6.48543 | 6.35323 | 6.68335 | 5.54861 | 5.72604 | 5.86492 | 5.6652 |
| ESD | P10768 | 5.84448 | 6.13371 | 6.5175 | 5.66544 | 6.67223 | 7.09562 | 6.18639 | 7.1523 | 6.54449 | 6.64979 | 6.51221 | 6.66026 | 5.66542 | 5.81405 | 5.47978 | 5.40632 |
| ERCC4 | Q92889 | 5.25758 | 5.72493 | 5.99656 | 5.79309 | 5.49348 | 6.22954 | 5.8621 | 5.81345 | 6.02392 | 6.41667 | 6.28239 | 6.21856 | 6.61692 | 7.33553 | 7.59512 | 7.34016 |
| TGFBRAP1 | Q8WUH2 | 6.30522 | 6.30043 | 6.59034 | 6.20219 | 6.26054 | 6.49569 | 5.99571 | 6.29665 | 6.20211 | 6.50063 | 6.65999 | 6.74478 | 5.77267 | 5.82731 | 6.13706 | 5.70868 |
| ZNF384 | Q8TF68 | 6.26453 | 6.32429 | 5.70307 | 7.02791 | 6.29785 | 6.02449 | 6.38468 | 6.14701 | 5.87984 | 6.36829 | 6.11487 | 6.0536 | 6.291 | 6.194 | 6.59597 | 6.32863 |
| CRTAP | O75718 | 6.69481 | 6.90856 | 6.37474 | 6.68376 | 6.62815 | 6.39909 | 6.37233 | 6.05545 | 6.04138 | 6.08856 | 6.05263 | 6.10258 | 5.98547 | 6.03348 | 5.66092 | 5.91808 |
| OSBPL1A | Q9BXW6 | 6.6852 | 6.20771 | 6.58631 | 6.13245 | 6.36156 | 6.42602 | 6.23991 | 6.45207 | 6.67179 | 6.65501 | 6.87186 | 6.41147 | 5.76974 | 5.73905 | 5.28421 | 5.50565 |
| LAMA5 | O15230 | 6.80794 | 7.04268 | 6.52602 | 6.79313 | 5.89351 | 6.2436 | 6.34302 | 5.85663 | 6.82013 | 7.21257 | 6.85008 | 6.42589 | 5.16906 | 5.78622 | 5.10862 | 5.12088 |
| CAMK2G | Q13555-6 | 6.73487 | 6.54825 | 7.02225 | 6.36299 | 6.08279 | 6.40343 | 6.51029 | 6.54794 | 5.86675 | 6.57214 | 6.36401 | 6.36089 | 5.38146 | 6.01991 | 5.67322 | 5.54882 |
| CAMK2D | Q13557-11 | 6.36404 | 6.09631 | 6.89552 | 5.81805 | 5.86882 | 6.48827 | 6.2747 | 6.33095 | 6.03143 | 6.67412 | 6.55628 | 6.39229 | 6.47221 | 5.92291 | 6.13918 | 5.67491 |
| TAOK2 | Q9UL54-2 | 6.29901 | 6.35498 | 6.22048 | 6.37635 | 6.23747 | 6.58577 | 5.91312 | 6.07991 | 6.2394 | 6.32784 | 6.39807 | 6.44862 | 5.90195 | 6.41992 | 6.08994 | 6.10718 |
| UBN1 | Q9NPG3 | 6.317 | 6.51721 | 5.97812 | 6.7264 | 6.04637 | 5.95381 | 6.40339 | 5.84999 | 6.14424 | 6.3357 | 6.0415 | 6.25061 | 6.25463 | 6.23217 | 6.59947 | 6.34938 |
| YARS2 | Q9Y2Z4 | 6.37187 | 6.60553 | 5.93352 | 6.71333 | 5.97615 | 6.08119 | 6.38838 | 6.28024 | 6.28296 | 6.04633 | 6.01454 | 6.14077 | 6.42157 | 6.10095 | 6.45757 | 6.18509 |
| HIVEP2 | P31629 | 6.2257 | 6.053 | 6.45696 | 6.08495 | 6.37932 | 6.43654 | 6.20139 | 6.31376 | 6.35438 | 6.27238 | 6.22059 | 6.25125 | 6.44569 | 6.1328 | 5.96346 | 6.20784 |
| EEFSEC | P57772 | 5.8956 | 5.59787 | 6.48649 | 5.79167 | 6.08755 | 6.40928 | 6.06895 | 6.36489 | 6.22066 | 6.44587 | 6.40373 | 6.13337 | 6.30635 | 6.63933 | 6.69248 | 6.4559 |
| MPHOSPH8 | Q99549 | 5.90507 | 6.33507 | 5.9205 | 6.74903 | 6.08091 | 6.09555 | 6.17392 | 5.79371 | 5.93213 | 5.97926 | 6.10307 | 5.96101 | 6.45307 | 7.16926 | 6.54032 | 6.80811 |
| GATB | O75879 | 6.34878 | 6.28206 | 5.89368 | 6.49699 | 6.04496 | 6.23315 | 6.51508 | 6.04694 | 6.26127 | 6.27654 | 6.09781 | 5.96541 | 6.76621 | 6.18563 | 6.34424 | 6.24126 |
| APPL2 | Q8NEU8 | 6.21252 | 6.44554 | 6.87195 | 6.12709 | 6.28944 | 6.59175 | 6.37997 | 6.45307 | 6.2933 | 6.78163 | 6.35645 | 6.6202 | 5.78428 | 5.6761 | 5.58028 | 5.53643 |
| CAPN7 | Q9Y6W3 | 6.35619 | 6.34623 | 6.9961 | 6.16831 | 6.272 | 6.56294 | 6.52584 | 6.35288 | 6.42081 | 6.76741 | 6.85139 | 6.72941 | 5.34174 | 5.64226 | 5.22984 | 5.43664 |
| RPL35A | P18077 | 6.27666 | 6.26564 | 6.47963 | 6.29402 | 6.1299 | 6.22816 | 6.47404 | 6.17024 | 6.75804 | 6.47674 | 6.50565 | 6.47106 | 5.86687 | 5.79593 | 5.88619 | 5.92125 |
| VLDLR | P98155 | 6.18554 | 6.45294 | 5.97227 | 6.12152 | 6.31215 | 6.47977 | 6.22674 | 6.23547 | 5.89846 | 6.15513 | 6.29206 | 6.42994 | 6.3352 | 6.22929 | 6.20183 | 6.47167 |
| ADSS2 | P30520 | 6.28865 | 6.24081 | 6.96381 | 5.8816 | 6.42427 | 7.04352 | 6.14476 | 6.783 | 6.46859 | 6.59687 | 6.78727 | 6.646 | 5.53077 | 5.56966 | 5.36253 | 5.26788 |
| RPS23 | P62266 | 6.44198 | 6.29265 | 6.35999 | 6.10448 | 6.44524 | 6.3133 | 6.35078 | 6.48189 | 6.73732 | 6.51938 | 6.3287 | 6.19197 | 5.82673 | 6.01811 | 5.8723 | 5.71517 |
| EIF2B4 | Q9UI10 | 6.34032 | 6.45361 | 6.77111 | 6.34413 | 6.32718 | 6.58765 | 6.14495 | 6.79233 | 6.2508 | 6.46807 | 6.67122 | 6.31037 | 5.4742 | 5.82499 | 5.80953 | 5.42955 |
| GLMN | Q92990 | 6.15723 | 6.05125 | 6.77949 | 6.14461 | 6.19572 | 6.58351 | 6.18377 | 6.48808 | 6.46599 | 6.42837 | 6.86973 | 6.68206 | 5.63882 | 5.85793 | 5.61221 | 5.86124 |
| LTV1 | Q96GA3 | 6.22891 | 6.47549 | 6.94287 | 6.28285 | 6.38048 | 6.70652 | 6.42069 | 6.81171 | 6.56227 | 6.62103 | 6.79816 | 6.64304 | 4.82897 | 5.29886 | 5.36023 | 5.63792 |
| C2CD5 | Q86YS7-3 | 6.44555 | 6.25367 | 6.38108 | 6.01283 | 6.2343 | 6.60415 | 6.59281 | 6.20058 | 6.28591 | 6.55648 | 6.21326 | 6.08682 | 6.08614 | 6.15887 | 5.95775 | 5.9298 |
| HOOK1 | Q9UJC3 | 6.5599 | 6.29678 | 6.87961 | 6.08844 | 6.31601 | 6.85037 | 5.96023 | 6.34094 | 6.26486 | 6.61281 | 6.46274 | 7.01964 | 6.13659 | 5.63113 | 5.14601 | 5.43393 |
| ASAP1 | Q9ULH1-2 | 6.04562 | 5.97665 | 6.78954 | 6.01731 | 6.39333 | 6.61388 | 6.36231 | 6.3109 | 6.46964 | 6.66056 | 6.32885 | 6.36787 | 6.00014 | 6.02461 | 5.79288 | 5.84592 |
| ASAP2 | O43150 | 6.36836 | 6.3736 | 6.99257 | 5.74967 | 6.11703 | 6.7219 | 6.07603 | 6.32132 | 6.73155 | 6.65052 | 6.63256 | 6.47804 | 5.73153 | 5.93361 | 5.6863 | 5.43541 |
| AP3S1 | Q92572 | 6.19298 | 5.93684 | 6.77023 | 5.77234 | 6.36242 | 6.54853 | 6.03458 | 6.52595 | 6.19549 | 6.47079 | 6.43005 | 6.44914 | 5.76321 | 6.51772 | 5.93934 | 6.0904 |
| LARP4 | Q71RC2-4 | 6.17416 | 6.28948 | 6.62337 | 6.37361 | 6.24058 | 6.60856 | 6.31625 | 6.47413 | 6.46871 | 6.03086 | 6.43312 | 6.50896 | 6.0894 | 5.82365 | 5.71052 | 5.83463 |
| PPIF | P30405 | 6.1998 | 5.98426 | 6.1061 | 7.20571 | 6.56039 | 5.86447 | 5.90875 | 6.20323 | 6.31477 | 5.99199 | 5.96009 | 6.05863 | 6.88709 | 6.01051 | 6.20469 | 6.63951 |
| KIF18B | Q86Y91 | 6.90068 | 6.77895 | 6.16759 | 7.31524 | 6.24072 | 5.97497 | 6.60626 | 6.00482 | 6.20784 | 5.68915 | 5.91987 | 5.7477 | 5.96518 | 5.88386 | 6.15696 | 6.44021 |
| TTC1 | Q99614 | 6.79275 | 6.26134 | 6.55122 | 6.15003 | 6.74126 | 6.56868 | 5.95007 | 6.80822 | 6.57125 | 6.4742 | 6.70141 | 6.62389 | 6.02952 | 5.26801 | 5.15742 | 5.35074 |
| TRAF2 | Q12933-2 | 6.14912 | 6.25952 | 6.97047 | 6.1128 | 6.04697 | 6.62516 | 6.05474 | 6.64006 | 6.18894 | 6.40806 | 6.55943 | 6.97908 | 5.56743 | 5.72978 | 6.08969 | 5.61876 |
| URGCP | Q8TCY9 | 6.44433 | 6.32557 | 6.81015 | 6.3921 | 6.28385 | 6.44162 | 6.32923 | 6.54616 | 6.69705 | 6.67317 | 6.80288 | 6.70671 | 5.33309 | 5.57388 | 5.11429 | 5.52593 |
| ABCA3 | Q99758 | 6.62385 | 6.89289 | 6.16751 | 6.75757 | 6.0054 | 6.03025 | 6.46053 | 6.2544 | 6.3144 | 6.48907 | 6.19624 | 6.04041 | 5.85728 | 6.16028 | 6.02277 | 5.72715 |
| CTBP2 | P56545 | 5.78044 | 5.34213 | 5.9503 | 6.17486 | 6.27517 | 5.47912 | 7.02925 | 5.90648 | 6.13826 | 5.57792 | 5.89237 | 5.51435 | 6.63349 | 7.07123 | 7.29414 | 7.94051 |
| SETD1A | O15047 | 6.2136 | 6.43692 | 6.16083 | 6.65478 | 6.30248 | 5.58953 | 6.37237 | 5.70183 | 5.96319 | 6.08455 | 6.22271 | 5.92261 | 6.52304 | 6.56735 | 6.77106 | 6.51316 |
| NGLY1 | Q96IV0 | 6.0585 | 6.37047 | 6.91727 | 6.36682 | 6.19324 | 6.50608 | 6.12079 | 5.99676 | 6.09734 | 6.53764 | 6.52391 | 6.62077 | 5.56939 | 6.10381 | 6.06136 | 5.95585 |
| APBB2 | Q92870-4 | 5.98082 | 5.95947 | 6.02067 | 6.01733 | 6.06336 | 6.15892 | 6.24556 | 5.85855 | 6.06367 | 6.42853 | 6.29641 | 6.16101 | 6.57302 | 6.6373 | 6.76497 | 6.7704 |
| REXO4 | Q9GZR2 | 6.08709 | 6.35487 | 5.81105 | 6.93064 | 6.28735 | 5.96715 | 6.32003 | 5.88298 | 6.14403 | 5.76914 | 5.60843 | 5.89252 | 6.37658 | 6.83202 | 6.69841 | 7.0377 |
| REXO2 | Q9Y3B8 | 6.51028 | 6.06964 | 6.75265 | 6.01022 | 6.75474 | 6.60832 | 6.12633 | 6.64078 | 6.87004 | 6.19819 | 6.44801 | 6.10437 | 6.00566 | 5.64549 | 5.67654 | 5.57872 |
| H6PD | O95479-2 | 6.34766 | 6.46301 | 5.77475 | 6.33878 | 6.2533 | 6.14154 | 6.17096 | 5.86899 | 6.02843 | 5.98669 | 5.95715 | 5.63402 | 6.89598 | 6.25765 | 6.97543 | 6.90567 |
| PAXBP1 | Q9Y5B6 | 6.38182 | 6.51503 | 6.04104 | 6.72519 | 6.24355 | 5.92879 | 6.13883 | 5.82701 | 5.92513 | 5.97364 | 6.10398 | 5.93369 | 6.75026 | 6.08416 | 6.83818 | 6.5897 |

| | | | | | | | | | | | | | | | | | |
|---|---|---|---|---|---|---|---|---|---|---|---|---|---|---|---|---|---|
| TMPO | P42167 | 6.31215 | 6.60198 | 5.7986 | 6.75518 | 6.0747 | 6.15524 | 6.14554 | 6.85224 | 5.78892 | 5.79651 | 5.89138 | 5.77937 | 6.11864 | 6.55529 | 6.79372 | 6.58054 |
| PLS1 | Q14651 | 6.16183 | 6.00009 | 6.63694 | 5.75064 | 6.1127 | 6.49776 | 6.11761 | 6.49504 | 6.55456 | 6.35336 | 6.50078 | 6.41049 | 6.47816 | 5.93006 | 6.06614 | 5.93383 |
| DDX55 | Q8NHQ9 | 6.1102 | 6.14138 | 5.86041 | 6.36839 | 6.02311 | 5.94206 | 6.1435 | 5.99036 | 6.12332 | 5.95321 | 6.29916 | 6.0706 | 6.13407 | 7.18082 | 6.96261 | 6.6968 |
| PPM1B | O75688 | 6.47666 | 6.26831 | 6.75518 | 6.07688 | 6.46196 | 5.75832 | 6.18582 | 6.72306 | 6.40472 | 6.59045 | 6.43014 | 6.4812 | 5.62778 | 5.69208 | 5.47057 | 5.59685 |
| PIGT | Q969N2 | 6.56306 | 6.76622 | 6.11662 | 6.43289 | 6.51643 | 6.14026 | 6.76543 | 5.9249 | 5.83967 | 6.07179 | 6.00104 | 5.80005 | 6.53376 | 6.08797 | 6.55054 | 5.88939 |
| RRP36 | Q96EU6 | 6.57853 | 6.25923 | 6.01016 | 6.89671 | 5.95878 | 5.9001 | 6.40954 | 6.10899 | 6.26075 | 5.8547 | 6.3703 | 6.14179 | 6.25436 | 6.0228 | 6.36349 | 6.60978 |
| POLR2E | P19388 | 5.84436 | 6.13851 | 6.05268 | 6.19265 | 6.68755 | 6.18355 | 6.79329 | 6.47183 | 7.17019 | 6.14175 | 6.18267 | 6.31473 | 5.8907 | 5.71464 | 5.98896 | 6.23193 |
| TMEM33 | P57088 | 6.24526 | 6.3579 | 6.00553 | 6.58526 | 6.18139 | 5.91122 | 6.35299 | 5.93689 | 6.21095 | 6.10279 | 5.98375 | 5.65721 | 6.76544 | 6.09615 | 6.90894 | 6.69833 |
| DDAH1 | O94760 | 6.22739 | 5.77242 | 6.28277 | 5.35416 | 6.45142 | 6.57052 | 5.8648 | 6.99723 | 6.33756 | 6.49766 | 6.31045 | 6.36219 | 6.30857 | 6.43883 | 6.0984 | 6.12564 |
| SMG8 | Q8ND04-2 | 6.14871 | 6.16059 | 7.18829 | 6.34078 | 6.16858 | 6.43671 | 6.30247 | 6.16876 | 6.41604 | 6.41093 | 6.66257 | 6.92245 | 5.77329 | 5.7365 | 5.39047 | 5.77286 |
| EXOC1 | Q9NV70 | 6.75385 | 6.51238 | 6.48164 | 5.79936 | 6.73103 | 6.61954 | 6.22085 | 6.46776 | 6.73708 | 6.62936 | 6.58272 | 6.34793 | 5.76741 | 5.48035 | 5.62871 | 5.24004 |
| GBA | P04062 | 6.16234 | 6.60202 | 6.0353 | 6.4166 | 6.31874 | 6.32457 | 6.51563 | 6.31477 | 6.21854 | 6.29816 | 6.04519 | 6.1033 | 6.17757 | 6.22447 | 6.12088 | 6.12193 |
| PHF8 | Q9UPP1 | 5.997 | 6.37668 | 6.22726 | 6.83125 | 6.09778 | 6.2703 | 6.2837 | 6.15217 | 6.27066 | 6.26502 | 6.37091 | 6.14128 | 5.93343 | 6.00397 | 6.4337 | 6.34491 |
| PHLDB1 | Q86UU1-2 | 6.52438 | 5.59754 | 6.4881 | 5.94906 | 6.22433 | 6.72417 | 6.11963 | 6.41322 | 6.60922 | 6.40798 | 6.52616 | 6.83043 | 5.73114 | 6.01405 | 5.73856 | 6.10204 |
| EIF3K | Q9UBQ5 | 6.2664 | 5.04402 | 6.37012 | 4.99813 | 6.28938 | 6.12211 | 6.19033 | 6.93186 | 6.08695 | 6.40658 | 6.30307 | 6.85774 | 5.23745 | 7.08628 | 6.77687 | 7.0327 |
| RPL14 | P50914 | 6.09668 | 6.1322 | 6.34374 | 6.71793 | 6.45866 | 6.30769 | 6.37075 | 6.13764 | 7.23991 | 5.9669 | 6.1197 | 6.35804 | 5.06246 | 6.2161 | 6.26162 | 6.20997 |
| MYG1 | Q9HB07 | 6.03802 | 6.17828 | 6.59641 | 5.56124 | 6.59237 | 6.88787 | 5.88502 | 7.01842 | 6.2817 | 6.38061 | 6.4679 | 6.6301 | 6.01304 | 6.06215 | 5.71542 | 5.69145 |
| GNAQ | P50148 | 6.51572 | 6.6389 | 6.27211 | 6.60872 | 6.10784 | 6.20853 | 6.36153 | 5.87717 | 6.46901 | 6.14164 | 6.25094 | 6.04106 | 6.21127 | 5.84566 | 6.35284 | 6.09706 |
| ARID4A | P29374 | 6.34406 | 6.43458 | 5.8338 | 6.78768 | 5.93735 | 5.85133 | 6.62533 | 6.0082 | 6.12007 | 6.27133 | 5.8848 | 6.02688 | 6.48209 | 6.15906 | 6.47914 | 6.75431 |
| RALGAPA1 | Q6GYQ0-6 | 6.37889 | 6.27842 | 6.88518 | 6.0565 | 6.17697 | 6.45035 | 6.28366 | 6.60784 | 6.41675 | 6.50926 | 6.63142 | 6.48542 | 5.87131 | 5.7874 | 5.65081 | 5.52982 |
| PAK1 | Q13153 | 6.38858 | 6.10942 | 6.89535 | 5.76662 | 6.53509 | 6.89372 | 6.22456 | 6.6107 | 6.49987 | 6.63743 | 6.4732 | 6.39964 | 5.89344 | 5.71301 | 5.53373 | 5.42565 |
| HJURP | Q8NCD3 | 6.11232 | 6.33476 | 6.52809 | 6.46401 | 6.40369 | 6.29443 | 6.59207 | 6.23103 | 6.15509 | 6.45901 | 6.131 | 6.47157 | 5.88888 | 5.87505 | 5.84867 | 6.21033 |
| NUP35 | Q8NFH5 | 5.99229 | 5.75863 | 5.97876 | 6.35677 | 6.65694 | 5.81995 | 5.97989 | 6.36239 | 6.35634 | 5.95975 | 6.13233 | 6.28536 | 6.96792 | 6.17256 | 6.55945 | 6.66067 |
| ETFDH | Q16134 | 6.51526 | 6.44182 | 5.90875 | 6.61934 | 6.48815 | 6.21106 | 6.4563 | 5.94507 | 6.10439 | 6.13997 | 5.78764 | 5.78114 | 6.85293 | 5.88046 | 6.61671 | 6.251 |
| EXOSC3 | Q9NQT5 | 6.24251 | 6.3577 | 5.96095 | 6.63027 | 6.24105 | 6.25966 | 6.35479 | 6.06592 | 6.14124 | 6.04527 | 6.2218 | 6.11665 | 6.60991 | 6.10208 | 6.46183 | 6.18836 |
| BUB1 | O43683 | 7.04342 | 6.87111 | 7.61794 | 7.00442 | 6.60948 | 6.62612 | 6.77779 | 6.57952 | 7.34179 | 7.0309 | 7.2376 | 7.02164 | 3.99459 | 4.23645 | 3.80907 | 4.19816 |
| PPP1R9A | Q9ULJ8-3 | 6.4306 | 6.09772 | 6.14619 | 6.66865 | 6.12122 | 5.96569 | 6.50925 | 6.0204 | 6.1189 | 5.88226 | 6.10397 | 6.02104 | 6.72774 | 6.16815 | 6.62596 | 6.39225 |
| MAPK1 | P28482 | 6.28899 | 6.31724 | 6.9372 | 5.64256 | 6.15105 | 6.57282 | 6.16817 | 6.41483 | 6.24224 | 6.59438 | 6.90661 | 6.89664 | 6.03234 | 5.74868 | 5.49133 | 5.5949 |
| EPB41 | P11171 | 6.27457 | 6.53601 | 6.07258 | 6.19111 | 6.14274 | 6.11208 | 6.31116 | 5.88708 | 6.03924 | 6.26248 | 6.26705 | 6.31736 | 6.54327 | 6.49254 | 6.22224 | 6.32846 |
| WDR12 | Q9GZL7 | 6.11193 | 6.07738 | 5.99487 | 6.45408 | 6.13763 | 6.07213 | 6.40255 | 6.04251 | 6.09427 | 5.94036 | 6.15073 | 6.04922 | 6.49647 | 6.29563 | 6.9141 | 6.76614 |
| RAE1 | P78406 | 6.08837 | 6.50736 | 5.82576 | 6.56542 | 6.47505 | 6.24199 | 6.20168 | 6.39447 | 5.92361 | 6.21485 | 6.26509 | 6.40772 | 6.31659 | 6.14954 | 6.08407 | 6.33843 |
| DECR2 | Q9NUI1 | 6.21332 | 6.21442 | 5.81934 | 6.32671 | 6.39393 | 5.83809 | 6.33709 | 6.11498 | 5.89654 | 6.17437 | 6.39234 | 6.02222 | 6.61125 | 6.1533 | 6.75364 | 6.73847 |
| MRM3 | Q9HC36 | 5.96961 | 6.11114 | 5.72323 | 6.98214 | 6.21012 | 6.25947 | 6.48797 | 6.07161 | 6.17523 | 6.14632 | 6.09872 | 6.20561 | 6.59699 | 6.18235 | 6.33893 | 6.44056 |
| PEBP1 | P30086 | 6.00668 | 5.92162 | 6.73285 | 5.47038 | 6.71123 | 6.64242 | 6.07389 | 7.01757 | 6.5618 | 6.71823 | 6.93517 | 6.59941 | 6.01995 | 5.66857 | 5.4907 | 5.42955 |
| LLGL2 | Q6P1M3 | 6.36487 | 6.31189 | 6.39499 | 6.46322 | 6.04541 | 6.11606 | 6.29131 | 5.99883 | 6.37132 | 6.21247 | 6.14529 | 6.11992 | 6.34396 | 6.18407 | 6.51364 | 6.12276 |
| ATP5PB | P24539 | 6.29895 | 5.72584 | 5.97478 | 5.94986 | 6.30543 | 5.72013 | 6.44388 | 6.02289 | 5.73844 | 6.21942 | 6.00695 | 6.15608 | 6.93064 | 6.55225 | 7.22975 | 6.72469 |
| KRI1 | Q8N9T8 | 6.03127 | 5.77878 | 6.16535 | 6.86673 | 6.2971 | 5.59945 | 6.26561 | 5.80516 | 6.48223 | 5.91442 | 5.92211 | 5.90042 | 6.64477 | 6.67535 | 6.57565 | 7.07559 |
| TRIOBP | Q9H2D6 | 6.43371 | 6.31734 | 6.15199 | 5.97404 | 6.25281 | 6.2679 | 5.91532 | 6.41287 | 6.4126 | 6.51099 | 6.57426 | 6.14389 | 6.29304 | 6.31689 | 6.26178 | 5.76058 |
| PPP4R1 | Q8TF05 | 6.33875 | 6.32102 | 7.27023 | 5.72737 | 6.58763 | 6.57657 | 6.4706 | 6.87932 | 6.54967 | 6.27866 | 6.66119 | 6.5354 | 6.56466 | 5.69176 | 5.23971 | 5.20747 |
| LOXL2 | Q9Y4K0 | 6.3613 | 6.54799 | 6.71732 | 7.06878 | 6.70216 | 6.42345 | 6.98501 | 6.32481 | 6.55535 | 5.8999 | 6.12468 | 6.16508 | 5.75437 | 5.17608 | 5.61156 | 5.58216 |
| PHLDB2 | Q86SQ0 | 6.59284 | 6.56265 | 6.34382 | 6.51407 | 6.39779 | 6.32429 | 6.32624 | 6.38944 | 6.21051 | 6.24898 | 6.09204 | 6.20249 | 5.98436 | 5.91419 | 6.0298 | 5.86564 |
| FAM111A | Q96PZ2 | 6.55831 | 6.90943 | 6.74189 | 6.72241 | 5.94864 | 6.27877 | 6.35467 | 6.14641 | 6.81489 | 6.54298 | 6.50346 | 6.52541 | 5.21286 | 5.4249 | 5.67284 | 5.64214 |
| GALNT1 | Q10472 | 6.29064 | 6.64607 | 6.2811 | 7.06457 | 6.05335 | 6.19591 | 6.4258 | 5.98486 | 6.24573 | 6.32917 | 6.17173 | 6.26572 | 6.04608 | 6.01634 | 5.9897 | 5.99323 |
| PSMB7 | Q99436 | 6.23921 | 6.23368 | 6.39476 | 5.9053 | 6.61569 | 6.4562 | 6.24468 | 6.78948 | 6.27913 | 6.44386 | 6.29804 | 6.19612 | 6.23301 | 5.89305 | 5.79763 | 5.98014 |
| MPHOSPH10 | O00566 | 6.12463 | 6.3973 | 5.74804 | 6.76936 | 5.99047 | 5.88716 | 6.33908 | 5.99665 | 6.01907 | 5.91014 | 5.63587 | 6.14235 | 6.65889 | 6.77966 | 6.94906 | 6.65226 |
| P4HA1 | P13674-2 | 6.47764 | 6.55988 | 5.81969 | 6.5978 | 6.9014 | 6.10269 | 6.31345 | 6.20121 | 5.83117 | 6.08146 | 5.81455 | 5.71034 | 6.69228 | 6.39157 | 6.2793 | 6.22558 |
| WASH2P | Q6VEQ5 | 6.23246 | 6.08927 | 6.43038 | 6.14941 | 6.20557 | 6.43791 | 6.15942 | 6.16502 | 6.42002 | 6.95608 | 6.8602 | 6.80269 | 5.75937 | 5.9662 | 5.64481 | 5.72118 |
| FYTTD1 | Q96QD9 | 6.60234 | 6.23335 | 6.23827 | 7.10658 | 6.3877 | 5.63814 | 6.71884 | 6.02352 | 6.70952 | 5.99721 | 6.18143 | 5.96997 | 6.2627 | 5.79545 | 6.19173 | 5.94326 |
| TUBGCP3 | Q96CW5 | 6.25238 | 6.46833 | 6.79378 | 6.15777 | 6.39561 | 6.78188 | 6.12393 | 6.62248 | 6.15684 | 6.49654 | 6.43501 | 6.449 | 5.87135 | 5.81752 | 5.75316 | 5.42442 |
| CPOX | P36551 | 6.14822 | 6.24805 | 5.96644 | 6.50174 | 6.63199 | 6.13627 | 6.47241 | 5.92218 | 6.10192 | 5.85146 | 6.01301 | 5.90242 | 6.97674 | 6.1581 | 6.50136 | 6.46768 |
| MRPS30 | Q9NP92 | 6.24981 | 6.60063 | 5.90779 | 6.98651 | 6.12256 | 5.99661 | 6.30466 | 6.1321 | 6.09566 | 6.15309 | 5.78866 | 6.08548 | 6.45385 | 6.13545 | 6.70303 | 6.2841 |
| CSTF2T | Q9H0L4 | 6.55226 | 5.41893 | 5.70155 | 5.49218 | 6.38822 | 5.25422 | 5.63711 | 6.90733 | 5.47467 | 6.39358 | 6.16969 | 7.00241 | 7.03851 | 6.86574 | 6.94651 | 6.7621 |
| NRBP1 | Q9UHY1 | 6.53154 | 6.14375 | 6.78185 | 6.07278 | 6.41291 | 6.72031 | 6.23947 | 6.68087 | 6.33139 | 6.7185 | 6.67406 | 6.64732 | 5.72461 | 5.49171 | 5.46976 | 5.35915 |
| ZRANB2 | O95218 | 5.49609 | 6.23755 | 5.71599 | 5.58672 | 6.16173 | 6.32628 | 5.55229 | 5.99454 | 5.81576 | 6.38257 | 6.07252 | 6.23454 | 7.10239 | 7.37622 | 6.90442 | 7.04039 |
| MEAK7 | Q6P9B6 | 6.1962 | 6.18387 | 5.89157 | 6.13706 | 6.18477 | 6.5029 | 6.08865 | 6.13487 | 6.02851 | 5.75925 | 6.07314 | 6.0816 | 6.33927 | 6.77597 | 6.68247 | 6.93991 |
| FAHD1 | Q6P587 | 6.1725 | 5.77253 | 5.9322 | 6.18179 | 6.44652 | 5.95043 | 5.93182 | 6.09608 | 6.15333 | 6.1519 | 6.24532 | 6.40991 | 6.85525 | 6.33382 | 6.70456 | 6.66204 |
| ORC3 | Q9UBD5-2 | 5.97529 | 6.29229 | 6.15866 | 6.17692 | 5.9509 | 6.05724 | 6.22496 | 5.96061 | 6.11706 | 6.14863 | 6.24666 | 6.14901 | 6.33968 | 6.64615 | 6.91259 | 6.64335 |
| LIN54 | Q6MZP7 | 5.98421 | 6.15779 | 6.16762 | 6.58517 | 6.32761 | 6.11688 | 6.46666 | 5.98751 | 5.99284 | 6.33188 | 6.1748 | 6.17532 | 6.76119 | 6.13915 | 6.20148 | 6.41153 |
| FOXK1 | P85037 | 5.75124 | 5.95508 | 5.94807 | 5.53726 | 6.25046 | 6.28831 | 6.01069 | 6.39302 | 5.99214 | 6.08594 | 6.16309 | 5.9034 | 7.1784 | 6.82358 | 6.95813 | 6.76119 |
| ZNFX1 | Q9P2E3 | 6.43293 | 6.18441 | 6.98394 | 6.25341 | 6.2483 | 6.87058 | 6.27155 | 6.25969 | 6.47835 | 6.64804 | 6.56025 | 6.43032 | 5.55373 | 5.56824 | 5.40079 | 5.44558 |
| TPP1 | O14773 | 6.15835 | 6.36784 | 6.01848 | 6.20176 | 6.66981 | 6.42156 | 6.24866 | 6.31908 | 5.97188 | 6.43044 | 6.06176 | 6.18369 | 6.58503 | 6.05444 | 5.87576 | 6.43145 |
| MFF | Q9GZY8 | 6.65439 | 5.55656 | 6.88073 | 5.93642 | 6.93549 | 5.49389 | 6.06393 | 6.02805 | 5.87534 | 6.73367 | 6.22565 | 6.82361 | 5.86853 | 6.34694 | 6.12101 | 6.45579 |
| RAB31 | Q13636 | 6.07007 | 6.06482 | 6.11358 | 6.09085 | 6.14813 | 6.11347 | 6.37427 | 5.90577 | 6.22592 | 6.26752 | 6.02317 | 5.9263 | 6.59933 | 6.47243 | 6.92434 | 6.68002 |
| GALC | P54803 | 6.41458 | 6.59476 | 5.65203 | 6.38325 | 6.28765 | 6.16596 | 6.25851 | 5.88329 | 6.08836 | 6.07929 | 5.90405 | 6.13497 | 6.43771 | 6.60714 | 6.5285 | 6.57994 |
| SLC30A1 | Q9Y6M5 | 6.21756 | 6.47654 | 5.91421 | 6.77827 | 6.03868 | 5.62409 | 6.03957 | 5.65799 | 5.98637 | 6.21271 | 5.87823 | 6.04029 | 6.84468 | 6.6151 | 6.96144 | 6.71426 |
| BLMH | Q13867 | 6.16528 | 6.30651 | 6.87403 | 6.05902 | 6.34297 | 6.98412 | 6.07375 | 6.34628 | 6.35142 | 6.65402 | 6.71496 | 6.77924 | 5.91254 | 5.40532 | 5.41136 | 5.61919 |
| ARHGAP18 | Q8N392 | 6.40728 | 6.40169 | 7.03708 | 5.98661 | 6.06769 | 6.48749 | 6.12475 | 6.6895 | 6.17671 | 6.43747 | 6.33385 | 6.47882 | 5.69697 | 5.87092 | 5.74001 | 6.06316 |
| SH3D19 | Q5HYK7 | 5.58349 | 6.15889 | 6.15691 | 5.64183 | 6.1598 | 6.42342 | 6.3222 | 6.22475 | 6.67846 | 6.68882 | 6.56108 | 6.60665 | 6.57454 | 6.35985 | 5.8141 | 6.04523 |
| CSNK2B | P67870 | 5.37001 | 5.60456 | 5.69782 | 5.43637 | 6.76497 | 6.07627 | 6.03422 | 7.36738 | 6.07381 | 6.72838 | 6.36035 | 6.40849 | 6.15376 | 6.45793 | 6.62455 | 6.84112 |
| PLAUR | Q03405 | 6.05629 | 6.17985 | 6.02155 | 6.42258 | 6.46838 | 6.21701 | 6.36632 | 6.1582 | 5.54592 | 5.58918 | 6.07341 | 5.45125 | 6.96034 | 6.47888 | 7.14226 | 6.86856 |
| UBE2Z | Q9H832 | 6.50155 | 5.87557 | 6.83302 | 5.99918 | 6.31423 | 6.65904 | 6.01206 | 6.45979 | 6.06873 | 6.21712 | 6.68382 | 6.52355 | 6.08625 | 6.22046 | 5.76236 | 5.78325 |
| TIGAR | Q9NQ88 | 6.13843 | 6.13989 | 6.81815 | 6.2362 | 6.17265 | 6.82501 | 6.05066 | 6.38991 | 6.57428 | 6.68793 | 6.77586 | 6.53406 | 5.72853 | 5.64693 | 5.67559 | 5.60593 |
| BUB3 | O43684 | 5.76873 | 6.19993 | 5.89452 | 5.87236 | 6.23455 | 6.17688 | 6.15522 | 6.33819 | 6.46118 | 6.59896 | 6.35933 | 5.77399 | 6.6704 | 6.17802 | 6.58188 | 6.73585 |
| COG7 | P83436 | 6.3692 | 5.94716 | 6.31774 | 5.96183 | 6.36717 | 6.34365 | 6.41962 | 6.46968 | 6.2994 | 6.48247 | 6.53589 | 6.28426 | 6.34911 | 6.10633 | 6.13289 | 5.6136 |
| MEIOC | A2RUB1 | 6.71499 | 6.51108 | 7.08885 | 6.47041 | 6.25218 | 6.72906 | 6.16743 | 6.61274 | 6.78575 | 6.72271 | 6.88122 | 7.10323 | 5.24012 | 4.91573 | 4.72397 | 5.08053 |
| SRSF10 | O75494 | 6.19904 | 6.51691 | 5.82031 | 6.61102 | 6.57315 | 6.228 | 6.28141 | 6.08845 | 5.80298 | 6.2368 | 6.46699 | 6.39926 | 6.10563 | 6.15754 | 6.14512 | 6.36739 |
| PPP6R1 | Q9UPN7 | 6.5161 | 6.16338 | 6.8751 | 6.00673 | 6.45341 | 6.72409 | 5.83829 | 6.86787 | 6.43257 | 6.14252 | 6.34909 | 6.60569 | 6.41244 | 5.64739 | 5.41871 | 5.54662 |
| GTPBP10 | A4D1E9 | 6.51394 | 6.42142 | 6.07165 | 6.74865 | 6.25565 | 6.02331 | 6.58567 | 6.15493 | 6.20112 | 6.19451 | 6.03535 | 6.04919 | 6.43982 | 5.92516 | 6.38569 | 5.99395 |
| RELA | Q04206 | 6.58513 | 6.43235 | 6.77666 | 5.89602 | 6.347 | 6.91545 | 6.07242 | 6.31879 | 6.32734 | 6.37852 | 6.60215 | 6.82662 | 5.83324 | 5.72504 | 5.69419 | 5.26908 |
| EIF2AK2 | P19525 | 6.41917 | 6.48601 | 6.44188 | 6.155 | 6.3897 | 6.48795 | 6.28599 | 6.60646 | 6.5306 | 6.78228 | 6.3676 | 6.36633 | 5.72241 | 5.86228 | 5.53059 | 5.56575 |
| USP11 | P51784 | 5.87463 | 6.32968 | 6.4786 | 5.96772 | 6.11133 | 6.565 | 5.90137 | 6.49886 | 6.07553 | 6.67149 | 6.71253 | 6.88231 | 5.98289 | 6.13022 | 5.69606 | 6.12179 |
| ELP2 | Q6IA86-6 | 6.35146 | 6.62107 | 7.01772 | 6.03114 | 6.30297 | 6.98859 | 6.22953 | 6.62858 | 6.51966 | 6.51421 | 6.5957 | 6.68155 | 5.39458 | 5.43328 | 5.35827 | 5.3317 |
| CYB5R1 | Q9UHQ9 | 6.3761 | 6.57341 | 5.80304 | 6.47491 | 6.04607 | 5.96834 | 6.27924 | 6.11491 | 6.09119 | 6.13709 | 5.74006 | 5.90987 | 6.77067 | 6.54635 | 6.61777 | 6.55096 |
| CTSB | P07858 | 6.53738 | 6.49807 | 6.01759 | 6.24094 | 6.77966 | 6.16085 | 6.42525 | 6.62399 | 6.19196 | 6.3542 | 6.38169 | 6.02738 | 5.85247 | 5.97714 | 5.92468 | 6.00674 |
| RINT1 | Q6NUQ1 | 6.5501 | 6.44929 | 6.25137 | 6.28619 | 6.44105 | 6.15294 | 6.53587 | 6.03517 | 5.97132 | 6.25643 | 5.91184 | 5.84197 | 6.33792 | 6.06639 | 6.57342 | 6.33873 |
| PIGG | Q5H8A4 | 6.32052 | 6.40793 | 5.98669 | 6.53691 | 6.37064 | 6.09731 | 6.40491 | 5.97362 | 6.06762 | 6.11508 | 5.94891 | 5.67939 | 6.66872 | 6.40818 | 6.54519 | 6.46837 |
| RESF1 | Q9HCM1 | 6.90967 | 6.7709 | 6.5515 | 7.50118 | 6.16715 | 6.03879 | 6.48209 | 6.35061 | 6.46182 | 6.60402 | 6.89726 | 6.55609 | 5.05386 | 5.03329 | 5.09383 | 5.52795 |
| BEND3 | Q5T5X7 | 6.50414 | 6.53702 | 5.88721 | 7.0206 | 6.06351 | 6.01739 | 6.37229 | 5.75171 | 6.208 | 5.61293 | 5.97139 | 5.94053 | 6.78408 | 6.26733 | 6.45452 | 6.60736 |
| ODF2 | Q5BJF6 | 6.77391 | 7.00284 | 7.11644 | 7.48476 | 6.20095 | 6.38687 | 6.58828 | 6.80604 | 6.95232 | 6.88894 | 6.60889 | 7.09037 | 4.7003 | 4.30978 | 4.5244 | 4.56492 |
| RCOR1 | Q9UKL0 | 6.28285 | 6.59116 | 6.25386 | 6.61796 | 6.10801 | 6.33489 | 6.13883 | 6.11551 | 6.11041 | 6.35752 | 6.12348 | 6.03961 | 6.14816 | 6.19511 | 6.3459 | 6.23674 |
| S100A10 | P60903 | 5.62013 | 4.93414 | 5.60865 | 4.76233 | 6.11551 | 5.59898 | 5.85632 | 6.36629 | 5.86876 | 6.53326 | 6.2205 | 6.58026 | 5.88418 | 7.92214 | 8.28143 | 7.84713 |
| RAB21 | Q9UL25 | 6.23839 | 6.36336 | 6.12099 | 6.51085 | 6.32062 | 6.24009 | 6.34302 | 6.09247 | 6.07648 | 6.38606 | 5.94719 | 5.72975 | 6.40067 | 6.13547 | 6.67017 | 6.42441 |
| SEC13 | P55735-3 | 6.17429 | 6.26517 | 6.30902 | 6.29294 | 6.17522 | 6.61578 | 6.05448 | 6.51413 | 6.13839 | 6.082 | 6.04968 | 6.14517 | 6.33622 | 6.53616 | 6.1098 | 6.20156 |
| PNPO | Q9NVS9 | 5.88172 | 5.12392 | 6.34083 | 5.19488 | 6.93875 | 6.10102 | 6.10312 | 6.9111 | 6.20944 | 7.01398 | 7.31727 | 6.81245 | 6.0394 | 5.78064 | 6.19815 | 6.03331 |
| PPP4R2 | Q9NY27 | 5.79884 | 5.94407 | 5.88869 | 5.76949 | 6.40498 | 6.20658 | 6.16759 | 5.81768 | 5.97172 | 5.75626 | 6.06412 | 6.20257 | 7.34399 | 6.70261 | 6.86513 | 7.09568 |
| CSK | P41240 | 6.07531 | 6.24551 | 6.84334 | 5.75671 | 6.48609 | 6.7325 | 6.19518 | 6.63633 | 6.46063 | 6.4064 | 6.48549 | 6.34919 | 5.95473 | 5.96065 | 5.8086 | 5.60333 |
| RIOK2 | Q9BVS4 | 6.39143 | 6.28187 | 7.22879 | 6.25101 | 6.40697 | 6.37305 | 6.28093 | 6.57312 | 6.46593 | 6.65831 | 6.6999 | 6.90652 | 5.39513 | 5.59606 | 5.22833 | 5.26265 |
| ASXL1 | Q8IXJ9 | 6.70897 | 6.3661 | 6.62977 | 6.6303 | 6.17674 | 6.1663 | 6.47928 | 6.61182 | 6.57016 | 6.28744 | 6.35732 | 6.67085 | 5.65439 | 5.68449 | 5.39079 | 5.61526 |
| AMPD2 | Q01433 | 6.18002 | 6.34316 | 6.68571 | 5.99031 | 6.48294 | 6.81463 | 6.39196 | 6.73168 | 6.59032 | 6.5164 | 6.14862 | 6.24271 | 5.72957 | 5.97306 | 5.46358 | 5.71535 |
| NUDT5 | Q9UKK9 | 6.14738 | 5.6672 | 6.61247 | 5.51765 | 6.59573 | 7.02791 | 6.0995 | 6.70898 | 6.3283 | 6.52118 | 6.62803 | 6.6173 | 6.15305 | 6.10106 | 5.55516 | 5.7191 |
| HDAC7 | Q8WUI4-5 | 6.05962 | 6.21937 | 6.81669 | 5.76185 | 6.12004 | 6.74047 | 6.11486 | 6.7103 | 6.47491 | 6.63925 | 6.64559 | 6.70883 | 5.62235 | 5.9321 | 5.69301 | 5.74077 |
| COG3 | Q96JB2 | 6.2431 | 6.22126 | 6.23878 | 6.70175 | 6.33141 | 6.23959 | 6.21492 | 6.08975 | 6.16302 | 6.46841 | 6.05705 | 6.42911 | 6.22596 | 6.03602 | 6.25781 | 6.08208 |
| A0A494C100 | A0A494C100 | 5.96147 | 6.42146 | 6.33093 | 6.2911 | 6.27348 | 6.53872 | 6.4587 | 6.34564 | 6.27355 | 6.30696 | 6.72335 | 6.56451 | 5.82377 | 5.88306 | 5.87616 | 5.92714 |
| MAGT1 | Q9H0U3 | 6.39924 | 6.15184 | 5.72244 | 6.67358 | 6.39078 | 5.68304 | 6.68922 | 5.88549 | 6.22931 | 5.60874 | 5.58382 | 5.74774 | 6.16894 | 7.04832 | 6.90835 | 7.10914 |
| MTG1 | Q9BT17 | 6.77523 | 6.76356 | 6.19361 | 7.05361 | 6.37523 | 6.3905 | 6.46528 | 5.93237 | 6.54081 | 6.36917 | 6.1873 | 6.41873 | 5.76904 | 5.524 | 5.55011 | 5.69146 |

| | | | | | | | | | | | | | | | | | |
|---|---|---|---|---|---|---|---|---|---|---|---|---|---|---|---|---|---|
| TPD52L2 | O43399-7 | 6.41904 | 5.55258 | 6.72043 | 5.62045 | 6.31319 | 6.24706 | 5.76164 | 6.48253 | 6.14282 | 6.46016 | 7.02879 | 7.04197 | 6.68451 | 6.1876 | 5.65089 | 5.68634 |
| NDUFA13 | Q9P0J0-2 | 6.00728 | 5.28134 | 5.34757 | 5.92836 | 6.25274 | 5.78035 | 7.10317 | 5.98878 | 6.18447 | 5.15606 | 5.55152 | 5.99825 | 6.22335 | 7.5341 | 7.38516 | 8.2775 |
| NUDCD3 | Q8IVD9 | 5.9593 | 5.89319 | 6.73634 | 5.71648 | 6.19281 | 6.57353 | 6.09948 | 6.40503 | 6.37812 | 6.1304 | 6.81175 | 6.43165 | 6.18385 | 5.39435 | 6.08109 | 6.01263 |
| NDUFAF7 | Q7L592 | 6.37391 | 6.2781 | 5.88744 | 6.33001 | 6.37994 | 6.05291 | 6.44568 | 6.14182 | 6.17703 | 6.0277 | 6.1664 | 5.88586 | 6.76281 | 6.26886 | 6.49394 | 6.32761 |
| IFI16 | Q16666 | 5.63315 | 6.0188 | 5.76517 | 5.86433 | 5.93195 | 5.99517 | 5.93512 | 5.94766 | 6.11025 | 6.51941 | 6.178 | 6.09996 | 6.93275 | 6.97078 | 7.29011 | 6.80739 |
| ATP6V1H | Q9UI12 | 6.44451 | 6.86443 | 6.45547 | 6.3202 | 6.40215 | 6.53525 | 6.40826 | 6.69656 | 6.57662 | 6.63372 | 6.55453 | 6.45793 | 5.32862 | 5.54704 | 5.38643 | 5.38827 |
| MRPL1 | Q9BYD6 | 6.39027 | 6.53157 | 6.01751 | 6.8784 | 6.40238 | 5.80608 | 6.32072 | 5.86423 | 6.16839 | 6.22783 | 5.74697 | 5.88149 | 6.65162 | 5.96752 | 6.67315 | 6.47187 |
| GFM2 | Q969S9 | 6.39855 | 5.93537 | 5.99667 | 6.69251 | 6.28566 | 6.16438 | 6.41113 | 5.77082 | 6.18828 | 5.87929 | 6.16828 | 6.14869 | 6.62868 | 6.49756 | 6.27906 | 6.55507 |
| SCYL2 | Q6P3W7 | 6.24647 | 6.49306 | 6.56979 | 6.35675 | 6.08869 | 6.48741 | 6.04915 | 6.51192 | 6.36128 | 6.46595 | 6.16995 | 6.5221 | 6.00063 | 5.79205 | 5.85431 | 6.03048 |
| SENP3 | Q9H4L4 | 6.25992 | 6.18732 | 5.8456 | 6.72124 | 6.24398 | 5.92894 | 6.47639 | 5.9509 | 6.13075 | 6.00943 | 5.93333 | 5.92163 | 6.55688 | 6.36125 | 6.76835 | 6.70408 |
| BLVRB | P30043 | 5.85383 | 5.44109 | 6.38008 | 5.40257 | 6.50666 | 6.72039 | 6.28726 | 6.97753 | 6.31807 | 6.87209 | 6.50693 | 6.95799 | 5.83897 | 6.2151 | 5.76565 | 5.95577 |
| NLE1 | Q9NVX2 | 6.36669 | 6.35651 | 6.27728 | 6.39243 | 5.79453 | 6.02723 | 6.11588 | 6.20855 | 6.26315 | 6.42533 | 6.39088 | 6.39134 | 6.12575 | 6.0498 | 6.55967 | 6.25497 |
| TRAPPC11 | Q7Z392 | 6.06996 | 6.20599 | 6.60203 | 5.85059 | 6.33795 | 6.58683 | 6.04251 | 6.6466 | 6.60648 | 6.6146 | 6.26241 | 6.33639 | 6.00614 | 6.10379 | 5.93848 | 5.78924 |
| RPAP1 | Q9BWH6 | 5.91254 | 6.18618 | 7.02001 | 5.85553 | 6.01961 | 6.37158 | 6.29658 | 6.58792 | 6.43837 | 6.47352 | 6.6653 | 6.68909 | 5.84191 | 5.89845 | 5.64768 | 6.09573 |
| MED17 | Q9NVC6 | 6.16057 | 6.15543 | 6.17016 | 6.36604 | 6.42587 | 6.11607 | 6.04109 | 5.86824 | 6.32034 | 6.20722 | 6.33171 | 6.06042 | 6.5147 | 6.09975 | 6.7126 | 6.44977 |
| EXOC8 | Q8IYI6 | 6.32286 | 6.55183 | 6.72928 | 6.14332 | 6.18607 | 6.52466 | 6.2443 | 6.29415 | 6.40639 | 6.39373 | 6.71386 | 6.40189 | 5.95413 | 5.85931 | 5.72065 | 5.55358 |
| DIP2A | Q14689 | 6.7864 | 6.58117 | 6.36643 | 6.41199 | 6.38637 | 6.38187 | 6.41535 | 6.10761 | 6.23136 | 6.18671 | 6.08307 | 6.09043 | 6.0203 | 6.10138 | 5.9181 | 5.93146 |
| SIN3B | O75182 | 6.01839 | 6.76424 | 6.20311 | 6.98671 | 6.24074 | 6.47239 | 5.97577 | 6.33454 | 6.04407 | 6.1951 | 6.74845 | 6.22414 | 5.73276 | 5.91684 | 6.36927 | 5.7735 |
| MTCL1 | Q9Y4B5 | 6.38406 | 6.15166 | 6.49034 | 6.30363 | 5.96489 | 6.54363 | 6.14433 | 6.70465 | 6.58804 | 6.68489 | 6.55857 | 6.924 | 5.63088 | 5.69553 | 5.60032 | 5.63059 |
| AP3M1 | Q9Y2T2 | 6.47119 | 6.24089 | 6.92518 | 6.19215 | 6.24332 | 6.42297 | 6.32631 | 6.54144 | 6.47571 | 6.36379 | 6.29896 | 6.00245 | 5.86648 | 5.95262 | 6.01314 | 5.6634 |
| ALDH16A1 | Q8IZ83 | 6.4331 | 6.54561 | 6.67042 | 5.95499 | 6.50994 | 7.12119 | 5.84387 | 6.54999 | 6.52988 | 6.34894 | 6.87576 | 6.90347 | 5.57147 | 5.62186 | 5.12368 | 5.39582 |
| MARCKS | P29966 | 6.02299 | 5.63901 | 5.8633 | 5.26109 | 5.7968 | 5.07858 | 5.62802 | 6.27108 | 5.46254 | 6.19049 | 5.92892 | 6.54843 | 8.23039 | 7.31628 | 7.51673 | 7.24537 |
| TOM1L2 | Q6ZVM7 | 6.31294 | 6.0795 | 6.74801 | 5.9135 | 6.51311 | 6.49091 | 6.08834 | 6.6446 | 6.52644 | 6.62807 | 6.67809 | 6.72807 | 6.05479 | 5.72522 | 5.37011 | 5.4983 |
| NECAP2 | Q9NVZ3 | 6.2788 | 5.94995 | 6.85593 | 5.93332 | 6.06544 | 6.50415 | 6.28326 | 6.7042 | 6.44944 | 6.44582 | 6.85574 | 6.74614 | 5.92844 | 5.77869 | 5.63993 | 5.58076 |
| YWHAB | P31946 | 6.02814 | 5.89519 | 6.67447 | 5.71443 | 6.29246 | 6.66518 | 6.19343 | 6.39973 | 6.38529 | 6.26805 | 6.34657 | 6.29369 | 6.67318 | 6.07005 | 6.16584 | 5.93429 |
| GOLPH3 | Q9H4A6 | 6.32902 | 6.26369 | 6.05099 | 6.27473 | 6.32514 | 6.2701 | 6.29268 | 6.14765 | 6.60452 | 6.26497 | 6.47408 | 6.16637 | 6.27268 | 6.27141 | 6.05805 | 5.93393 |
| TOMM34 | Q15785 | 6.33447 | 5.55984 | 6.74978 | 5.88746 | 6.45401 | 6.57527 | 6.13889 | 6.41126 | 6.41536 | 6.33083 | 6.65193 | 6.50572 | 6.04109 | 5.96342 | 5.84164 | 5.75304 |
| CHMP4B | Q9H444 | 6.12339 | 6.07818 | 6.75748 | 5.70998 | 6.39831 | 6.20807 | 6.05572 | 6.97783 | 6.70263 | 6.4705 | 6.48906 | 6.32913 | 5.95198 | 5.95869 | 5.81492 | 5.97411 |
| CEP131 | Q9UPN4 | 6.16246 | 6.43941 | 6.77841 | 6.54731 | 6.19946 | 6.32267 | 6.55423 | 6.43297 | 6.49431 | 6.31087 | 6.78646 | 6.93165 | 5.40864 | 5.77112 | 5.39246 | 5.46759 |
| SUN2 | Q9UH99 | 6.41775 | 6.49694 | 6.0433 | 6.78242 | 6.19448 | 6.2759 | 6.35321 | 5.94508 | 6.30372 | 5.91865 | 5.91604 | 6.11862 | 6.3527 | 6.14667 | 6.50936 | 6.22516 |
| PDLIM1 | O00151 | 6.50503 | 6.02985 | 6.43227 | 5.2926 | 6.76638 | 6.35205 | 5.94764 | 6.05959 | 6.38055 | 6.20246 | 6.70975 | 6.61376 | 6.8627 | 6.02328 | 5.9147 | 5.9074 |
| ARHGAP17 | Q68EM7 | 6.31881 | 6.33917 | 7.01497 | 6.21159 | 6.3887 | 6.87245 | 6.12299 | 6.40758 | 6.21874 | 6.68943 | 6.62665 | 6.6455 | 5.84478 | 5.57707 | 5.39605 | 5.32552 |
| PKP2 | Q99959 | 6.40819 | 6.55664 | 6.44521 | 6.56746 | 6.06522 | 6.04774 | 6.12549 | 6.211 | 6.38888 | 6.222 | 6.28117 | 6.36655 | 6.04495 | 5.92838 | 6.10986 | 6.23126 |
| MAN2B2 | Q9Y2E5 | 6.43549 | 6.35291 | 6.04117 | 6.44711 | 6.27058 | 6.199 | 6.30346 | 6.37549 | 6.05517 | 6.50513 | 6.25704 | 6.32795 | 5.78227 | 6.13569 | 6.40553 | 6.10599 |
| GATAD2B | Q8WXI9 | 6.46311 | 6.52463 | 6.00889 | 6.40802 | 6.64483 | 6.28624 | 6.19266 | 6.34402 | 6.20003 | 6.30011 | 6.15543 | 6.27602 | 6.48251 | 5.9216 | 5.88367 | 5.90822 |
| PRKAG1 | P54619 | 5.66979 | 5.56326 | 6.10995 | 5.32848 | 5.99869 | 6.1381 | 5.81755 | 5.90905 | 6.09291 | 6.16648 | 6.41832 | 6.33865 | 7.10993 | 6.71091 | 7.47876 | 7.14917 |
| CBS | P35520-2 | 6.09154 | 6.09916 | 7.07543 | 5.27398 | 6.53822 | 6.54007 | 6.40128 | 6.45771 | 6.58054 | 6.38942 | 6.72563 | 6.41226 | 6.06963 | 5.95981 | 5.59023 | 5.79508 |
| SPAG1 | Q07617 | 6.26995 | 6.35305 | 6.65893 | 6.35682 | 6.33573 | 6.6022 | 6.07146 | 6.35242 | 6.18523 | 6.32842 | 6.17668 | 6.62971 | 6.22537 | 5.90411 | 5.80877 | 5.74115 |
| SMARCE1 | Q969G3 | 6.51118 | 6.79577 | 5.94741 | 6.38272 | 6.60657 | 6.04606 | 5.90466 | 6.00183 | 6.06381 | 6.04731 | 6.30672 | 5.75592 | 6.96435 | 5.93687 | 6.10148 | 6.62734 |
| DDRGK1 | Q96HY6 | 6.31625 | 6.8248 | 5.77998 | 6.75249 | 6.00492 | 6.27592 | 6.1741 | 6.11385 | 5.93331 | 6.05638 | 5.88869 | 6.22089 | 6.20056 | 6.46744 | 6.22188 | 6.76853 |
| TNRC18 | O15417 | 6.49679 | 7.16926 | 6.50548 | 6.79239 | 6.1606 | 6.08665 | 6.35581 | 6.06009 | 6.33056 | 7.01351 | 6.58216 | 6.4022 | 5.16261 | 5.84098 | 5.39013 | 5.64778 |
| TRUB2 | O95900 | 6.28196 | 6.70241 | 5.99181 | 6.78643 | 6.68151 | 6.30677 | 6.61421 | 6.62194 | 6.1362 | 6.27442 | 5.95695 | 6.05286 | 6.01523 | 5.59845 | 6.05472 | 5.92412 |
| NAA35 | Q5VZE5 | 6.02089 | 6.39224 | 6.41089 | 5.93842 | 6.1651 | 6.35914 | 6.14907 | 6.26687 | 6.12805 | 6.64264 | 6.67283 | 7.01466 | 5.86414 | 6.00286 | 5.83674 | 6.13545 |
| TOR1AIP2 | Q8NFQ8 | 6.12294 | 6.17511 | 5.93395 | 6.24376 | 6.33881 | 6.23388 | 6.43944 | 6.36378 | 5.98165 | 5.79569 | 6.38466 | 6.06823 | 6.5777 | 6.33841 | 6.67073 | 6.33125 |
| VAT1 | Q99536 | 6.38907 | 5.97788 | 6.12667 | 5.69531 | 6.36377 | 6.36085 | 6.04299 | 6.16096 | 6.40437 | 6.51629 | 6.34928 | 6.21164 | 6.54578 | 6.53584 | 6.19285 | 6.12647 |
| MICAL1 | Q8TDZ2-4 | 6.17675 | 6.30389 | 6.97834 | 6.1903 | 6.21795 | 6.82166 | 6.45534 | 6.47905 | 6.3543 | 6.44967 | 6.67085 | 6.3753 | 5.66479 | 5.6953 | 5.60559 | 5.56092 |
| ATP6V0D1 | P61421 | 6.06869 | 5.69119 | 5.75668 | 5.63772 | 6.05091 | 5.98222 | 6.58093 | 6.61671 | 5.51553 | 6.0044 | 5.9312 | 6.31471 | 5.71601 | 7.90018 | 6.44475 | 7.78817 |
| GSR | P00390 | 5.85049 | 6.00711 | 8.0887 | 5.51853 | 6.27787 | 6.074 | 9.2076 | 6.13638 | 6.02806 | 6.0104 | 6.70707 | 5.46008 | 4.99547 | 5.23797 | 7.31684 | 5.08345 |
| GUSB | P08236-2 | 6.65995 | 6.62155 | 5.639 | 6.41629 | 6.33166 | 6.03789 | 6.39941 | 6.03389 | 6.21853 | 6.3394 | 6.0586 | 6.2135 | 6.15276 | 6.15527 | 6.36774 | 6.35457 |
| MRPL24 | Q96A35 | 5.94819 | 6.12166 | 5.5797 | 6.6191 | 6.68293 | 5.92073 | 6.69549 | 6.46826 | 6.16909 | 5.95367 | 6.16029 | 5.96912 | 6.12933 | 6.24852 | 6.72724 | 6.6067 |
| SP140L | Q9H930 | 6.29776 | 6.63304 | 6.33481 | 6.63835 | 5.95132 | 6.15023 | 6.52916 | 6.33082 | 6.22848 | 6.21327 | 6.47553 | 6.04904 | 5.6171 | 6.21777 | 6.04356 | 6.28977 |
| SP100 | P23497-4 | 6.4146 | 6.51716 | 5.69022 | 6.48746 | 6.1351 | 6.18167 | 5.97774 | 5.75645 | 6.19171 | 6.48059 | 6.07304 | 6.08584 | 6.61082 | 6.53054 | 6.3054 | 6.56166 |
| LRRC47 | Q8N1G4 | 6.15632 | 6.57402 | 6.90765 | 6.00667 | 6.23469 | 6.49238 | 6.02841 | 6.2834 | 6.21342 | 6.26898 | 7.0848 | 6.4343 | 5.93337 | 5.89532 | 5.71281 | 5.77347 |
| MRPL46 | Q9H2W6 | 6.34816 | 6.41874 | 5.57893 | 6.25127 | 6.54985 | 5.78735 | 6.06483 | 6.65567 | 6.0306 | 6.42787 | 5.93258 | 6.28113 | 6.796 | 6.15891 | 6.23881 | 6.47929 |
| YJU2 | Q9BW85 | 5.71318 | 5.73993 | 6.07938 | 5.45017 | 5.88569 | 5.74261 | 5.74858 | 5.54437 | 6.15804 | 6.38447 | 6.54549 | 6.32073 | 7.01416 | 7.18974 | 7.37262 | 7.11084 |
| NHP2 | Q9NX24 | 6.54529 | 6.53814 | 6.27428 | 6.87953 | 6.3355 | 5.68743 | 6.60236 | 5.62085 | 5.781 | 6.01454 | 6.1108 | 5.74133 | 6.55478 | 6.42024 | 6.39331 | 6.5006 |
| VPS4A | Q9UN37 | 6.60322 | 6.62999 | 6.96002 | 6.4758 | 6.44273 | 6.97126 | 6.00753 | 6.83725 | 6.54729 | 6.58586 | 6.75421 | 6.3524 | 5.40943 | 5.36099 | 5.08963 | 4.97239 |
| RPS14 | P62263 | 6.44565 | 6.59403 | 6.53467 | 6.41853 | 6.00523 | 6.31465 | 6.38142 | 6.17667 | 6.66061 | 6.41973 | 6.59748 | 6.44901 | 5.79098 | 5.72937 | 5.74896 | 5.73303 |
| BCORL1 | Q5H9F3-3 | 6.5323 | 6.62957 | 6.2912 | 7.01233 | 6.25655 | 6.25033 | 6.60496 | 5.84513 | 6.44385 | 6.47603 | 6.40735 | 6.19395 | 5.59108 | 5.72992 | 5.73429 | 6.00117 |
| MAN1B1 | Q9UKM7 | 6.52502 | 6.50953 | 6.16568 | 6.7353 | 6.45367 | 6.08952 | 6.29228 | 6.16903 | 5.92914 | 6.25278 | 6.17997 | 5.92736 | 6.37141 | 6.28499 | 6.33657 | 5.83981 |
| TGOLN2 | O43493-7 | 6.28969 | 6.2312 | 6.029 | 6.31384 | 6.32089 | 5.96872 | 6.11765 | 5.89483 | 6.41073 | 6.35687 | 6.49923 | 6.46474 | 6.55915 | 6.13125 | 6.17634 | 6.23588 |
| SPATS2 | Q86XZ4 | 5.77283 | 6.06397 | 6.25962 | 5.83021 | 5.97018 | 6.60443 | 6.27782 | 6.43717 | 6.59848 | 6.859 | 6.81614 | 6.63518 | 5.77062 | 6.16559 | 5.91453 | 6.02422 |
| ZFHX3 | Q15911 | 6.35077 | 6.63706 | 6.11131 | 6.94206 | 6.36081 | 6.11066 | 6.41408 | 5.94883 | 6.00759 | 6.38731 | 5.88195 | 5.86969 | 6.33712 | 6.32463 | 6.35234 | 5.96379 |
| PNO1 | Q9NRX1 | 6.00093 | 6.22918 | 5.91266 | 6.32017 | 6.44349 | 6.34004 | 6.18639 | 6.09355 | 6.1451 | 6.16132 | 6.02142 | 5.93403 | 6.78029 | 6.26017 | 6.5486 | 6.62265 |
| BASP1 | P80723 | 6.45095 | 4.82135 | 5.90312 | 5.42334 | 6.63339 | 5.0825 | 5.50636 | 6.29237 | 5.70369 | 6.67924 | 6.62361 | 6.93631 | 7.07918 | 6.87023 | 7.02933 | 6.96503 |
| PPP2R2A | P63151-2 | 5.72033 | 5.89028 | 6.66592 | 5.74831 | 5.90691 | 6.65772 | 6.24668 | 6.33162 | 6.20943 | 6.36403 | 6.58943 | 6.49049 | 6.19395 | 6.21935 | 6.45284 | 6.31272 |
| TOP3A | Q13472 | 6.24493 | 6.41193 | 6.22352 | 6.8639 | 6.08027 | 5.82587 | 6.51775 | 5.72934 | 6.7696 | 6.22064 | 6.27362 | 6.15411 | 6.1888 | 5.7621 | 6.47444 | 6.25918 |
| VPS52 | Q8N1B4 | 6.17132 | 6.38406 | 6.53039 | 6.13164 | 6.12477 | 6.27492 | 6.40415 | 6.52376 | 6.3979 | 6.53078 | 6.37708 | 6.67097 | 5.74978 | 6.27252 | 5.7435 | 5.71244 |
| CCDC93 | Q567U6 | 5.88959 | 6.24786 | 6.54771 | 5.71495 | 6.20605 | 6.26146 | 6.09228 | 6.30626 | 6.51126 | 6.16297 | 6.39245 | 6.05752 | 6.59672 | 6.54688 | 6.45594 | 6.0101 |
| TRIM27 | P14373 | 6.35869 | 6.20719 | 5.72906 | 6.30806 | 6.74794 | 6.36133 | 6.7116 | 6.41493 | 6.36195 | 5.97831 | 6.03513 | 6.15495 | 6.38693 | 5.80981 | 6.33046 | 6.10367 |
| RAB11FIP1 | Q6WKZ4 | 6.43755 | 7.12702 | 6.68375 | 6.65061 | 5.87955 | 6.81177 | 6.65702 | 6.15434 | 5.39088 | 6.03422 | 6.15331 | 6.13551 | 5.50389 | 5.79124 | 6.37791 | 6.21143 |
| CEP290 | O15078 | 6.24613 | 6.37496 | 6.89595 | 6.14294 | 6.17167 | 6.48823 | 6.11524 | 6.62132 | 6.55145 | 6.21703 | 6.53831 | 6.95719 | 5.5692 | 5.68106 | 5.66812 | 5.7612 |
| ZC3H4 | Q9UPT8 | 6.24028 | 6.23684 | 5.787 | 6.62348 | 6.09787 | 5.86049 | 6.16755 | 5.84918 | 6.0505 | 6.00644 | 6.17172 | 5.8362 | 6.72936 | 6.68724 | 6.80697 | 6.84886 |
| MARK2 | Q7KZI7-16 | 5.98448 | 6.1501 | 6.28547 | 6.31029 | 6.0057 | 6.06212 | 6.22861 | 6.1474 | 6.36073 | 6.18943 | 6.32828 | 6.40731 | 6.43812 | 5.96995 | 6.57121 | 6.5608 |
| HEXIM1 | O94992 | 6.2461 | 6.60353 | 6.07367 | 6.11922 | 6.15215 | 6.80064 | 5.80017 | 6.23678 | 6.03885 | 6.37495 | 6.12712 | 6.05951 | 6.4168 | 6.11763 | 6.4236 | 6.40929 |
| STOML2 | Q9UJZ1 | 6.0804 | 5.43001 | 5.86871 | 5.36413 | 6.16602 | 5.50769 | 6.11909 | 6.46523 | 5.68286 | 6.37668 | 5.7041 | 6.64945 | 6.95041 | 6.94722 | 7.49384 | 7.19417 |
| ZEB1 | P37275-2 | 6.18722 | 6.15965 | 5.99682 | 6.53581 | 6.48639 | 6.30694 | 6.4384 | 6.10141 | 6.10778 | 5.82604 | 5.8722 | 6.16025 | 6.90732 | 6.11177 | 6.26987 | 6.53215 |
| CANT1 | Q8WVQ1 | 6.52092 | 6.42163 | 6.11518 | 6.99414 | 6.29784 | 6.0671 | 6.33741 | 6.26579 | 6.25915 | 6.65287 | 6.32357 | 6.682 | 5.91619 | 5.51086 | 5.80175 | 5.83361 |
| MRPS23 | Q9Y3D9 | 6.16979 | 6.22544 | 5.80912 | 6.97554 | 6.46491 | 5.89883 | 6.63313 | 6.20496 | 6.28145 | 6.10181 | 6.0459 | 6.16664 | 6.59601 | 5.80412 | 6.33445 | 6.28791 |
| SNAPC4 | Q5SXM2 | 6.03179 | 5.91719 | 6.32267 | 6.17598 | 5.96218 | 6.03389 | 6.17446 | 6.00098 | 6.43641 | 5.87384 | 6.52181 | 6.20436 | 6.58742 | 6.19092 | 6.84437 | 6.72172 |
| USP16 | Q9Y5T5 | 6.04246 | 5.82469 | 6.5034 | 5.88627 | 6.11054 | 6.47649 | 5.95994 | 6.11811 | 6.40021 | 6.89501 | 6.76241 | 6.59042 | 5.90034 | 6.60018 | 5.77026 | 6.15926 |
| MBD3 | O95983 | 6.15041 | 6.4638 | 6.47053 | 6.55207 | 5.9866 | 5.81528 | 6.23109 | 6.07474 | 6.36874 | 6.27659 | 6.28483 | 6.75294 | 6.52844 | 5.67477 | 6.1305 | 6.23866 |
| MAIP1 | Q8WWC4 | 6.37804 | 6.69098 | 5.87321 | 6.76598 | 6.44394 | 6.24816 | 6.14935 | 6.24092 | 6.06048 | 5.9324 | 6.59118 | 5.7625 | 6.47756 | 6.14868 | 6.52573 | 6.34088 |
| CAP2 | P40123 | 6.13625 | 6.10338 | 6.88769 | 5.96251 | 6.27542 | 6.48717 | 6.25426 | 6.68864 | 6.41598 | 6.26928 | 6.61584 | 6.46597 | 5.79313 | 5.94226 | 5.8784 | 5.82382 |
| TSPYL1 | Q9H0U9 | 6.24982 | 6.02839 | 6.29889 | 6.52706 | 6.33492 | 6.06167 | 6.40861 | 6.06255 | 6.41393 | 6.35049 | 6.50668 | 6.00573 | 6.38639 | 6.01381 | 6.25788 | 6.09316 |
| PRDX3 | P30048 | 6.17603 | 4.96299 | 5.30141 | 4.97728 | 5.93424 | 5.15472 | 5.77566 | 6.23683 | 5.14187 | 6.26907 | 5.65072 | 6.6175 | 7.04147 | 8.16412 | 8.26162 | 8.33448 |
| CLMN | Q96JQ2 | 6.17225 | 6.46692 | 6.17808 | 6.56571 | 6.00459 | 6.16994 | 6.12171 | 5.91084 | 5.94917 | 6.22726 | 6.104 | 6.36263 | 6.48949 | 6.0106 | 6.65833 | 6.60847 |
| NUMBL | Q9Y6R0 | 6.44723 | 6.41309 | 6.55894 | 6.15436 | 6.23503 | 6.65157 | 6.15876 | 6.57841 | 6.18394 | 6.4768 | 6.5081 | 6.21695 | 5.88411 | 6.03553 | 5.82041 | 5.87936 |
| NPC1 | O15118 | 6.41914 | 6.66423 | 6.35123 | 7.00623 | 6.14808 | 5.96005 | 6.3083 | 6.05216 | 6.20102 | 6.28684 | 6.36168 | 6.34285 | 5.85088 | 5.85221 | 6.00688 | 6.18822 |
| ABCB7 | O75027-2 | 6.43387 | 6.69269 | 6.17511 | 6.83922 | 6.3717 | 5.88752 | 6.2151 | 5.99558 | 6.15476 | 6.08219 | 5.838 | 5.78867 | 6.60273 | 6.07338 | 6.60825 | 6.24125 |
| KDM4A | O75164 | 6.04119 | 6.45631 | 6.5685 | 6.33592 | 5.49762 | 5.98407 | 6.20976 | 6.04645 | 6.47761 | 6.36764 | 6.62179 | 7.0627 | 5.9533 | 6.28506 | 6.21855 | 5.87351 |
| AHCYL2 | Q96HN2 | 5.97652 | 5.97681 | 6.58591 | 5.8979 | 6.42656 | 6.84762 | 6.45473 | 6.85714 | 5.96991 | 6.06871 | 6.30664 | 6.01089 | 6.33738 | 6.17719 | 6.1406 | 5.96549 |
| COQ5 | Q5HYK3 | 6.55054 | 6.30509 | 5.77541 | 6.60184 | 6.66242 | 6.07929 | 6.3187 | 5.90236 | 6.35493 | 6.37529 | 5.95996 | 5.60936 | 6.61974 | 6.1304 | 6.5716 | 6.18308 |
| PIK3CB | P42338 | 6.32413 | 6.46157 | 6.94719 | 6.00157 | 6.21393 | 6.79257 | 5.90607 | 6.58305 | 6.0997 | 6.57637 | 6.71981 | 6.68669 | 6.6501 | 5.77617 | 5.5939 | 5.66716 |
| EPM2AIP1 | Q7L775 | 6.08637 | 6.23007 | 5.98227 | 6.58736 | 5.95315 | 6.18109 | 5.8836 | 6.24517 | 6.24032 | 6.24314 | 6.04476 | 5.94398 | 6.89445 | 6.93687 | 6.85479 | 6.5926 |
| TCIRG1 | Q13488 | 6.29458 | 6.65329 | 6.16439 | 6.52499 | 6.39295 | 6.36267 | 6.33279 | 6.32082 | 6.1145 | 6.20871 | 5.81248 | 6.05777 | 6.15308 | 6.11216 | 6.2915 | 6.20334 |
| TAX1BP1 | Q86VP1 | 6.4515 | 6.37146 | 6.39725 | 6.35195 | 6.73011 | 6.7791 | 6.26471 | 6.97827 | 6.48044 | 6.53421 | 6.41961 | 6.51615 | 5.57562 | 5.32558 | 5.33906 | 5.48498 |
| MRPL45 | Q9BRJ2 | 6.1663 | 6.28427 | 5.78099 | 6.48722 | 6.1978 | 5.9271 | 6.56924 | 6.10333 | 6.17339 | 5.99225 | 5.79815 | 6.07943 | 6.58815 | 6.46858 | 6.67977 | 6.70403 |
| ZNF687 | Q8N1G0 | 6.30101 | 6.65057 | 5.8723 | 6.64146 | 5.9767 | 6.14957 | 6.43011 | 6.04731 | 6.23954 | 6.44362 | 5.7357 | 5.99532 | 6.38334 | 6.13442 | 6.56858 | 6.43047 |
| SLC27A4 | Q6P1M0 | 6.06522 | 6.58053 | 6.5895 | 6.51902 | 6.09461 | 6.42197 | 6.25494 | 6.14347 | 6.16074 | 6.51725 | 5.76331 | 6.0689 | 6.30195 | 6.42644 | 6.32019 | 6.67197 |
| STK3 | Q13188-2 | 6.61903 | 6.10233 | 6.76762 | 5.97183 | 6.5829 | 6.37933 | 6.20587 | 6.45537 | 6.5993 | 6.17496 | 6.67195 | 6.32383 | 6.00638 | 5.63144 | 5.79247 | 5.71538 |
| TBC1D24 | Q9ULP9 | 5.6987 | 6.04178 | 6.34467 | 5.72041 | 6.12513 | 6.17034 | 6.28313 | 6.04872 | 6.40878 | 6.34758 | 6.47179 | 6.33806 | 6.34147 | 6.38874 | 6.81145 | 6.45925 |
| NCBP3 | Q53F19 | 6.52602 | 6.53821 | 6.04336 | 6.72364 | 6.20831 | 6.29479 | 6.28492 | 6.1008 | 5.99903 | 6.23171 | 6.12037 | 6.16065 | 6.4487 | 6.1295 | 6.03064 | 6.15936 |
| MED15 | Q96RN5 | 6.08784 | 5.73398 | 6.19811 | 5.2229 | 6.77341 | 6.3318 | 6.02162 | 6.84161 | 6.19988 | 6.47735 | 6.51061 | 6.54019 | 6.74433 | 6.10033 | 6.00771 | 6.20833 |
| TXNRD2 | Q9NNW7 | 6.44157 | 6.22036 | 5.8076 | 6.74772 | 6.12847 | 5.77048 | 6.74006 | 5.76982 | 6.16325 | 5.9924 | 5.95413 | 5.90884 | 6.67044 | 6.45664 | 6.46801 | 6.7602 |

| | | | | | | | | | | | | | | | | | |
|---|---|---|---|---|---|---|---|---|---|---|---|---|---|---|---|---|---|
| TRPV2 | Q9Y5S1 | 6.4059 | 6.81853 | 6.08059 | 6.72221 | 6.14198 | 6.14648 | 6.3406 | 5.92514 | 6.17007 | 6.1805 | 5.72055 | 5.76758 | 6.52735 | 6.24316 | 6.34514 | 6.46421 |
| CEP128 | Q6ZU80 | 6.30175 | 6.45442 | 5.96975 | 6.48661 | 6.05058 | 6.17396 | 6.125 | 6.29607 | 6.17807 | 6.25424 | 6.27435 | 6.21683 | 6.89002 | 6.10925 | 6.11383 | 6.10525 |
| TRAF1 | Q13077 | 6.25654 | 5.95761 | 6.38757 | 5.6967 | 6.5919 | 6.44505 | 6.18123 | 6.48025 | 6.44933 | 6.11548 | 6.20126 | 5.89814 | 6.5226 | 6.0844 | 6.43482 | 6.29713 |
| BPNT1 | O95861 | 6.13983 | 5.91935 | 6.75344 | 5.58956 | 6.07016 | 6.81082 | 6.31544 | 6.39469 | 6.52888 | 6.65755 | 6.67626 | 6.44393 | 5.89192 | 6.12385 | 5.93116 | 5.75317 |
| FAM111B | Q6SJ93 | 5.60157 | 6.29835 | 6.50903 | 5.87553 | 6.10243 | 6.4737 | 5.98877 | 6.41833 | 6.26281 | 6.56285 | 6.28594 | 6.26309 | 6.04953 | 6.56463 | 6.4352 | 6.30824 |
| RABEP1 | Q15276 | 6.34727 | 5.96773 | 7.06783 | 5.77986 | 6.53038 | 6.3712 | 6.04949 | 6.82627 | 6.60542 | 6.29411 | 6.71141 | 6.88726 | 6.26235 | 5.438 | 5.41405 | 5.4474 |
| DOT1L | Q8TEK3-1 | 6.61282 | 6.56348 | 6.33515 | 7.14609 | 6.34491 | 6.04999 | 6.50753 | 6.38514 | 6.44653 | 5.80917 | 6.40352 | 6.20122 | 5.69828 | 5.77398 | 5.80239 | 5.9198 |
| NMT2 | O60551 | 6.3312 | 6.39782 | 6.70519 | 6.13601 | 5.86552 | 6.17046 | 6.39568 | 6.50391 | 6.41644 | 6.4912 | 6.6456 | 6.94686 | 5.65382 | 5.90209 | 5.53247 | 5.90571 |
| STAM2 | O75886 | 6.44858 | 6.26934 | 6.74546 | 6.33904 | 6.52627 | 6.59048 | 6.09219 | 6.38815 | 6.36889 | 6.45967 | 6.6202 | 6.58437 | 6.15831 | 5.5247 | 5.399 | 5.48536 |
| DNAJB12 | Q9NXW2-2 | 6.49536 | 6.75897 | 6.25691 | 6.98906 | 6.54943 | 6.37106 | 6.33634 | 6.21455 | 6.35564 | 5.9703 | 6.10274 | 6.13553 | 6.02233 | 5.57266 | 5.90246 | 5.96667 |
| PSMB4 | P28070 | 6.15197 | 6.08923 | 6.18059 | 5.98865 | 6.47902 | 6.33973 | 6.32843 | 6.427 | 6.33934 | 6.36896 | 6.22622 | 5.99393 | 6.51236 | 6.11156 | 6.46582 | 5.99719 |
| ACTR8 | Q9H981 | 6.66006 | 6.44124 | 6.71682 | 6.56703 | 6.06378 | 6.39421 | 5.85297 | 6.15772 | 6.53284 | 6.21421 | 6.71449 | 6.37467 | 5.6893 | 5.80251 | 5.95466 | 5.86349 |
| POLA2 | Q14181 | 5.69242 | 6.07219 | 6.08622 | 5.83167 | 5.84617 | 6.27216 | 5.83018 | 6.18337 | 6.27643 | 6.01692 | 6.27427 | 6.15393 | 6.79586 | 6.75211 | 7.02986 | 6.88623 |
| TMEM70 | Q9BUB7 | 6.24742 | 6.72167 | 5.92063 | 7.16531 | 6.21695 | 5.86721 | 6.13071 | 6.33327 | 6.15152 | 5.75189 | 5.91382 | 6.01526 | 6.20804 | 6.21695 | 6.7577 | 6.38166 |
| LAMB2 | P55268 | 6.90233 | 7.4405 | 6.41309 | 6.80843 | 6.42689 | 6.19409 | 6.26245 | 5.85594 | 6.57918 | 6.74023 | 6.303 | 6.46296 | 5.57666 | 5.40345 | 5.30698 | 5.32383 |
| ACSF3 | Q4G176 | 6.51076 | 6.61062 | 6.08217 | 6.66537 | 6.32201 | 6.20183 | 6.48943 | 5.73865 | 5.86775 | 6.04167 | 6.20503 | 6.10478 | 6.45344 | 6.32053 | 6.26141 | 6.12455 |
| ABHD10 | Q9NUJ1 | 6.08265 | 6.40953 | 5.66451 | 6.14647 | 6.09094 | 6.20141 | 6.4057 | 6.27306 | 6.33404 | 6.26772 | 5.9497 | 6.08909 | 6.56467 | 6.06967 | 6.90704 | 6.54379 |
| UHRF2 | Q96PU4 | 5.53424 | 6.32824 | 6.37149 | 6.24564 | 5.67321 | 6.1927 | 5.79033 | 5.86625 | 6.12474 | 6.09623 | 6.25404 | 6.18661 | 6.719 | 6.64187 | 7.23199 | 6.74341 |
| NANS | Q9NR45 | 5.73254 | 5.74651 | 6.30279 | 5.53682 | 6.50572 | 6.6047 | 5.90633 | 6.58631 | 6.30784 | 6.72441 | 6.66305 | 6.29402 | 6.25173 | 6.45143 | 6.48058 | 5.9052 |
| XPO6 | Q96QU8 | 5.55615 | 5.73521 | 6.13353 | 5.48264 | 5.5066 | 6.2359 | 5.68288 | 6.38332 | 6.39664 | 6.31101 | 6.44349 | 6.70423 | 6.24247 | 7.00274 | 6.97099 | 7.2122 |
| PLEKHG4 | Q58EX7 | 6.50315 | 6.54084 | 6.60935 | 6.32585 | 6.21261 | 6.52877 | 6.29968 | 6.53344 | 6.59196 | 6.47648 | 6.5716 | 6.46404 | 5.65771 | 5.60359 | 5.41516 | 5.66576 |
| VARS2 | A0A140TA36 | 6.35237 | 6.19077 | 5.94943 | 6.78576 | 6.48556 | 5.85382 | 6.51675 | 5.91315 | 6.36589 | 6.28873 | 6.05279 | 6.12088 | 6.59289 | 5.92136 | 6.30931 | 6.30054 |
| PPM1A | P35813-3 | 6.42601 | 6.0505 | 7.04389 | 5.58012 | 6.31055 | 6.42443 | 6.12452 | 6.56587 | 6.74317 | 6.56087 | 6.72599 | 6.59188 | 6.05799 | 5.95979 | 5.37293 | 5.36151 |
| GTF2E2 | P29084 | 5.41608 | 5.9777 | 6.21824 | 5.47903 | 5.45456 | 6.16247 | 5.75709 | 6.16674 | 5.81588 | 6.4335 | 6.16709 | 6.20157 | 6.48056 | 7.50439 | 7.55381 | 7.21128 |
| ARAP3 | Q8WWN8 | 6.47257 | 6.00763 | 6.67931 | 6.06522 | 6.11631 | 6.76845 | 6.18548 | 6.56762 | 6.15588 | 6.57329 | 6.58579 | 6.41809 | 5.94641 | 5.95359 | 5.76844 | 5.73593 |
| NSMCE4A | Q9NXX6 | 6.06302 | 6.50809 | 5.78747 | 6.50087 | 6.11559 | 6.16449 | 6.14897 | 6.3085 | 6.04843 | 6.04553 | 6.08142 | 6.38491 | 6.73755 | 6.19989 | 6.34804 | 6.55724 |
| SCAP | Q12770 | 6.33667 | 6.55471 | 6.21913 | 6.49413 | 6.03367 | 6.11468 | 6.30679 | 6.09235 | 6.06506 | 5.84891 | 5.98765 | 6.16795 | 6.43153 | 6.29944 | 6.38779 | 6.65955 |
| PARG | Q86W56 | 5.61695 | 6.12567 | 6.80662 | 5.34323 | 6.1823 | 6.67595 | 5.97084 | 6.25599 | 6.11802 | 5.96472 | 6.50959 | 6.4664 | 6.57863 | 6.69039 | 6.11687 | 6.57785 |
| SYNJ1 | O43426 | 6.15428 | 6.44601 | 6.59424 | 6.25447 | 5.95583 | 6.3477 | 6.09363 | 6.22914 | 6.50797 | 6.64525 | 6.48709 | 6.23811 | 6.06606 | 6.12252 | 5.94116 | 5.91654 |
| TMEM245 | Q9H330 | 6.33961 | 6.40573 | 6.01872 | 6.51841 | 6.01202 | 5.89152 | 6.19577 | 6.0266 | 5.8539 | 5.93215 | 5.95072 | 5.90181 | 6.69434 | 6.7358 | 6.82288 | 6.7 |
| TTC19 | Q6DKK2 | 6.57733 | 5.81395 | 5.75951 | 5.82606 | 5.94613 | 5.58763 | 6.58163 | 6.11497 | 5.56971 | 6.23065 | 5.75168 | 6.42535 | 6.97209 | 7.37967 | 6.92176 | 7.44186 |
| THG1L | Q9NWX6 | 6.53836 | 6.15889 | 6.16069 | 6.44643 | 6.21091 | 6.31998 | 6.12491 | 6.37523 | 6.24856 | 6.13282 | 6.40525 | 6.15906 | 6.13979 | 6.56561 | 6.03074 | 5.98278 |
| SLC4A2 | P04920 | 6.59265 | 6.7445 | 6.22523 | 7.02037 | 5.9166 | 6.00938 | 6.3799 | 6.01498 | 6.27904 | 6.25694 | 6.32805 | 6.23041 | 5.8762 | 6.0994 | 6.13922 | 5.88713 |
| PFN2 | P35080-2 | 5.97667 | 6.10131 | 6.75946 | 5.64084 | 6.68037 | 6.5187 | 5.95414 | 6.80022 | 6.63937 | 6.72149 | 6.56809 | 6.59297 | 5.71226 | 5.82815 | 5.8014 | 5.70456 |
| RSU1 | Q15404 | 5.35855 | 5.8502 | 6.41916 | 5.71312 | 6.30032 | 6.6183 | 6.24725 | 6.83576 | 6.63072 | 7.01612 | 6.97248 | 6.92778 | 5.76179 | 5.83711 | 5.88132 | 5.63001 |
| STOM | P27105 | 6.40219 | 6.46502 | 5.78861 | 6.33965 | 6.14947 | 5.97899 | 6.58487 | 6.29993 | 6.36182 | 6.30802 | 5.90596 | 6.12048 | 6.51814 | 6.31891 | 6.35927 | 6.09866 |
| RFX5 | P48382 | 5.90012 | 6.16296 | 5.85774 | 6.16211 | 6.06515 | 5.98299 | 5.99655 | 5.97553 | 6.28089 | 6.5746 | 6.04815 | 5.96776 | 6.78596 | 6.57771 | 6.84289 | 6.81888 |
| GPT2 | Q8TD30 | 6.35856 | 6.47601 | 6.18229 | 6.68289 | 6.47811 | 6.36692 | 6.25433 | 5.98735 | 6.51307 | 6.38413 | 6.36564 | 6.42753 | 6.03374 | 5.63469 | 5.93204 | 5.9547 |
| IL6ST | P40189 | 7.19927 | 7.24556 | 6.96493 | 7.14391 | 6.45669 | 6.26429 | 6.45917 | 6.54194 | 6.76162 | 6.91009 | 6.76155 | 6.67616 | 4.6767 | 4.78942 | 4.44921 | 4.69949 |
| MTERF3 | Q96E29 | 6.11256 | 6.25148 | 6.11179 | 6.67208 | 6.54097 | 6.28368 | 6.68805 | 6.3927 | 6.08405 | 6.14627 | 6.46941 | 6.2068 | 6.06241 | 5.56612 | 6.16317 | 6.24847 |
| PUS1 | Q9Y606 | 5.94408 | 6.20795 | 5.82818 | 6.45112 | 5.74628 | 6.00759 | 6.10973 | 5.91815 | 6.07311 | 6.22026 | 6.18628 | 6.13577 | 7.04502 | 6.51358 | 6.82477 | 6.78812 |
| BTF3L4 | Q96K17 | 6.24205 | 6.1395 | 6.87387 | 5.79695 | 6.24557 | 6.80013 | 6.3315 | 6.59782 | 6.58509 | 6.83281 | 6.71426 | 6.44069 | 5.55007 | 5.81205 | 5.6011 | 5.43652 |
| AFF4 | Q9UHB7 | 6.94989 | 7.30454 | 7.10496 | 7.70723 | 6.20096 | 6.31813 | 6.76899 | 5.7946 | 6.829 | 7.21866 | 7.07565 | 6.59191 | 4.38977 | 4.88089 | 4.18498 | 4.67985 |
| FAM98A | Q8NCA5 | 6.43353 | 6.62682 | 6.32381 | 6.42451 | 6.48949 | 6.29982 | 6.13011 | 6.62255 | 6.42693 | 6.11997 | 6.25023 | 6.26842 | 6.16088 | 6.0779 | 5.84965 | 5.49539 |
| DNAJA3 | Q96EY1 | 6.43611 | 6.58089 | 6.0058 | 6.96024 | 6.40484 | 5.90012 | 6.31972 | 6.31744 | 5.98629 | 5.90602 | 5.91281 | 5.85386 | 6.34797 | 6.39014 | 6.27069 | 6.40709 |
| BCAP29 | Q9UHQ4-2 | 6.28532 | 6.45875 | 5.64464 | 6.77599 | 6.11275 | 5.77939 | 6.54372 | 5.75811 | 6.14581 | 5.76078 | 5.94797 | 5.87274 | 6.49374 | 6.577 | 6.65232 | 7.19098 |
| ATP2B4 | P23634 | 6.47193 | 6.18944 | 6.18262 | 6.32038 | 6.36489 | 5.97411 | 6.26661 | 6.00588 | 6.47106 | 5.98618 | 5.73952 | 5.65806 | 6.80062 | 6.20645 | 6.91974 | 6.44252 |
| IMP3 | Q9NV31 | 6.35513 | 6.78002 | 6.19505 | 6.81315 | 6.17859 | 6.02349 | 6.44694 | 5.8045 | 5.80725 | 6.31041 | 6.05143 | 5.99843 | 6.67316 | 6.18909 | 6.14625 | 6.22712 |
| DENND5B | Q6ZUT9-2 | 6.31406 | 6.41541 | 6.97711 | 6.07935 | 6.15005 | 6.72897 | 6.26829 | 6.69414 | 6.42764 | 6.68701 | 6.56147 | 6.53855 | 5.71463 | 5.96114 | 5.23441 | 5.24778 |
| CTDSPL2 | Q05D32 | 6.54651 | 6.44215 | 5.76589 | 6.79265 | 6.17854 | 5.68898 | 6.52758 | 6.11509 | 6.3819 | 5.9687 | 5.76105 | 5.87101 | 6.63887 | 6.18354 | 6.3036 | 6.83394 |
| HMGCS1 | Q01581 | 6.72795 | 6.52168 | 7.6112 | 6.06833 | 6.37314 | 6.35647 | 6.30441 | 6.9411 | 6.60225 | 6.5334 | 6.87049 | 7.01088 | 4.99385 | 5.15319 | 4.99805 | 4.93361 |
| ACADSB | P45954 | 6.24965 | 6.53166 | 5.9444 | 6.20061 | 6.43106 | 5.87443 | 6.39341 | 5.86244 | 6.16701 | 5.86137 | 5.97759 | 6.25739 | 7.01313 | 6.44765 | 6.82922 | 6.65897 |
| RANBP9 | Q96S59 | 6.36077 | 6.43982 | 6.53869 | 6.47228 | 6.60181 | 6.40533 | 6.30586 | 6.20426 | 6.4439 | 6.28174 | 6.61299 | 6.53473 | 5.92972 | 5.50953 | 5.6358 | 5.72276 |
| ARFGEF3 | Q5TH69 | 6.65278 | 6.33583 | 6.8181 | 6.41148 | 6.20162 | 6.59979 | 5.95844 | 6.17596 | 6.44909 | 6.60225 | 6.70443 | 6.97455 | 5.5697 | 5.58583 | 5.51767 | 5.44247 |
| ACAT2 | Q9BWD1 | 6.04819 | 6.12403 | 7.04359 | 5.98684 | 6.30241 | 7.09879 | 5.94084 | 6.61432 | 6.17905 | 6.2882 | 6.60649 | 6.58661 | 5.84668 | 5.87267 | 5.48142 | 5.97989 |
| CDC20 | Q12834 | 6.14007 | 5.90637 | 6.70084 | 6.46481 | 6.49352 | 6.38384 | 6.41625 | 6.41357 | 6.77498 | 6.487 | 6.60984 | 6.74541 | 5.30978 | 5.35056 | 5.88713 | 5.91605 |
| BRPF1 | P55201-2 | 6.13931 | 6.50506 | 5.77767 | 6.96053 | 6.14437 | 6.129 | 6.31215 | 6.1498 | 6.16552 | 5.97143 | 6.02043 | 6.07696 | 6.43361 | 6.05468 | 6.49355 | 6.66591 |
| BRD1 | O95696-2 | 6.28394 | 6.43795 | 5.89416 | 6.65502 | 6.10102 | 5.92282 | 6.32975 | 5.97387 | 6.24325 | 6.10255 | 6.11126 | 6.24614 | 6.39689 | 6.36755 | 6.43549 | 6.49834 |
| KAT2A | Q92830 | 5.64668 | 5.97874 | 6.09725 | 6.23226 | 6.20496 | 6.24887 | 5.9917 | 5.95259 | 6.18497 | 6.31433 | 6.59732 | 6.20723 | 6.14693 | 6.30782 | 6.83911 | 7.04923 |
| PAPOLG | Q9BWT3 | 6.0179 | 6.23323 | 6.45983 | 6.16872 | 6.04782 | 6.00335 | 6.17332 | 5.9411 | 6.18224 | 6.5081 | 6.57086 | 6.25019 | 6.27729 | 6.36356 | 6.4307 | 6.3718 |
| STX5 | Q13190 | 6.53369 | 6.66985 | 6.29881 | 6.61017 | 6.02415 | 6.11059 | 6.52669 | 5.91954 | 6.16207 | 6.23304 | 6.19075 | 6.02948 | 6.08033 | 5.93063 | 6.4018 | 6.27839 |
| DSTN | P60981 | 5.79333 | 5.9395 | 6.5157 | 5.65963 | 6.48776 | 6.89022 | 6.1192 | 6.51126 | 6.41143 | 6.64958 | 6.44209 | 6.51463 | 6.0231 | 5.94985 | 5.90559 | 6.18714 |
| ACAD8 | Q9UKU7 | 6.70773 | 6.53876 | 6.04182 | 6.72307 | 6.42931 | 5.85421 | 6.2007 | 5.71489 | 6.0957 | 6.2474 | 6.10508 | 6.05276 | 6.55464 | 6.21016 | 6.39637 | 6.1274 |
| MRPS10 | P82664 | 6.15017 | 6.09365 | 5.92636 | 6.16434 | 6.3544 | 6.13401 | 6.44072 | 6.0436 | 6.28942 | 6.05512 | 6.17297 | 5.97327 | 6.59623 | 6.47951 | 6.5384 | 6.58783 |
| FBXO11 | Q86XK2-5 | 6.40772 | 6.9048 | 6.86817 | 7.16846 | 6.40174 | 6.25401 | 6.73285 | 6.07997 | 6.78627 | 6.45959 | 6.92373 | 6.76854 | 5.14109 | 4.73352 | 5.02434 | 5.34521 |
| ARPC3 | O15145 | 6.1 | 5.57094 | 6.75567 | 5.57688 | 6.60373 | 6.13335 | 6.15205 | 7.10627 | 5.78646 | 6.48444 | 6.38946 | 6.61565 | 5.84069 | 6.5011 | 6.18653 | 6.20228 |
| ERGIC3 | Q9Y282-3 | 6.69176 | 6.84642 | 5.88118 | 6.43571 | 6.44501 | 6.02042 | 6.25811 | 5.69054 | 5.97366 | 6.06563 | 6.3385 | 5.80288 | 6.8996 | 6.34373 | 6.32337 | 5.98349 |
| GLO1 | Q04760 | 6.41729 | 4.89316 | 6.58844 | 4.60231 | 6.21137 | 5.85367 | 5.58557 | 7.27138 | 5.21391 | 6.95419 | 6.24537 | 6.9636 | 5.96313 | 7.5408 | 6.76299 | 6.93281 |
| RTN3 | O95197 | 6.2314 | 6.43177 | 5.96153 | 6.20262 | 5.99974 | 6.10424 | 6.35 | 6.14677 | 5.8609 | 6.13861 | 6.04633 | 5.97906 | 6.48593 | 6.60875 | 6.66369 | 6.78866 |
| RNGTT | O60942 | 5.19792 | 5.62134 | 6.08066 | 5.63972 | 5.61324 | 5.99966 | 5.87774 | 5.88984 | 6.02943 | 6.03868 | 6.28331 | 5.95594 | 7.45404 | 7.49145 | 7.71943 | 7.10758 |
| ACTR10 | Q9NZ32 | 6.2769 | 6.01273 | 7.05708 | 6.23944 | 6.12583 | 6.52093 | 6.34642 | 6.20032 | 6.4341 | 6.17096 | 6.5188 | 6.40851 | 5.81406 | 6.06679 | 5.88567 | 5.92334 |
| GSK3A | P49840 | 6.70376 | 6.27241 | 6.94089 | 6.13972 | 6.24201 | 6.42204 | 6.20068 | 6.29855 | 6.65197 | 6.75088 | 6.5196 | 6.38673 | 5.64964 | 5.63654 | 5.59964 | 5.58494 |
| PYCR2 | Q96C36 | 6.53817 | 6.23739 | 5.78404 | 6.49395 | 6.30111 | 6.0915 | 6.39601 | 6.3468 | 6.14569 | 5.71219 | 6.16833 | 5.8384 | 6.64321 | 6.24257 | 6.59421 | 6.46644 |
| ZNF800 | Q2TB10 | 5.75758 | 5.62321 | 5.66798 | 6.30384 | 6.02669 | 5.6361 | 6.14451 | 5.52383 | 5.93283 | 5.45088 | 5.64892 | 5.87444 | 7.57638 | 6.94703 | 7.99229 | 7.89349 |
| POLR3C | Q9BUI4 | 6.19965 | 5.88539 | 6.22537 | 5.74991 | 6.30164 | 6.23099 | 5.85255 | 5.94137 | 6.24578 | 6.18809 | 6.12537 | 5.92492 | 6.66909 | 6.68923 | 7.02389 | 6.74674 |
| RPP38 | P78345 | 6.12458 | 6.32382 | 5.98004 | 6.61793 | 6.273 | 6.20278 | 6.32435 | 5.75003 | 6.1143 | 6.19478 | 5.68184 | 5.63399 | 6.67408 | 6.51766 | 7.01483 | 6.57199 |
| LIN9 | Q5TKA1-2 | 6.26884 | 6.32984 | 5.98197 | 6.34907 | 6.08099 | 6.12472 | 6.42429 | 6.10121 | 6.1536 | 6.06805 | 6.28183 | 6.02907 | 6.41627 | 6.38785 | 6.56446 | 6.43792 |
| GAR1 | Q9NY12 | 6.37361 | 6.55354 | 5.69299 | 6.67841 | 6.24711 | 6.06693 | 6.35973 | 6.26861 | 6.18136 | 5.64037 | 5.70148 | 5.89161 | 6.62709 | 6.34676 | 6.69969 | 6.67071 |
| MIOS | Q9NXC5 | 6.55774 | 6.74832 | 6.39461 | 5.90625 | 6.31919 | 6.65265 | 5.91447 | 6.55984 | 5.92597 | 6.43258 | 6.63222 | 6.33378 | 5.69397 | 6.11614 | 6.04407 | 5.76821 |
| NAE1 | Q13564-4 | 5.81807 | 5.81265 | 6.60987 | 5.70251 | 5.88976 | 6.42622 | 5.87807 | 6.36835 | 5.94014 | 6.33885 | 6.40931 | 6.39694 | 6.01366 | 6.95343 | 6.79786 | 6.64431 |
| IDH2 | P48735 | 6.59225 | 6.49191 | 5.83181 | 6.71887 | 6.2121 | 6.26989 | 6.08712 | 6.1756 | 5.95438 | 5.97976 | 5.73123 | 5.95822 | 6.67462 | 6.34565 | 6.28094 | 6.69565 |
| GFUS | Q13630 | 6.53416 | 6.23077 | 6.95436 | 5.85885 | 6.53675 | 6.89753 | 5.97159 | 6.64395 | 6.26196 | 6.27137 | 6.70643 | 6.2019 | 5.8085 | 5.94579 | 5.62309 | 5.553 |
| BRWD1 | Q9NSI6-2 | 6.37303 | 6.74669 | 6.24 | 7.25352 | 6.19301 | 5.77237 | 6.38034 | 5.73868 | 6.24597 | 6.26898 | 5.88145 | 6.20991 | 5.85353 | 6.29236 | 6.12815 | 6.42203 |
| SRSF9 | Q13242 | 6.41403 | 6.73815 | 5.76362 | 6.14824 | 7.44357 | 5.97717 | 5.69295 | 6.67027 | 7.86183 | 5.98767 | 5.93974 | 6.09064 | 6.43127 | 5.53889 | 5.74021 | 5.56174 |
| TRIM13 | O60858-3 | 6.25687 | 6.83085 | 6.55465 | 7.06849 | 6.0391 | 5.92502 | 6.59559 | 5.86777 | 6.00504 | 6.20369 | 6.63321 | 6.37875 | 5.69597 | 5.65236 | 6.17595 | 6.11671 |
| DDX49 | Q9Y6V7 | 6.00213 | 6.09696 | 5.93364 | 6.43919 | 6.39229 | 6.39468 | 6.11934 | 6.04333 | 6.26162 | 6.08498 | 6.12555 | 6.15871 | 6.621 | 6.46103 | 6.33644 | 6.52911 |
| LRRC40 | Q9H9A6 | 6.19518 | 6.35074 | 6.65617 | 6.08761 | 6.20781 | 6.52944 | 6.31019 | 7.0166 | 5.91067 | 6.50953 | 6.34071 | 6.27695 | 5.91634 | 6.30373 | 5.65342 | 5.73492 |
| TBC1D23 | Q9NUY8 | 6.30681 | 6.12023 | 6.34547 | 6.13088 | 6.2251 | 6.25937 | 6.54631 | 6.15393 | 6.84581 | 6.35939 | 6.42507 | 5.5687 | 6.07463 | 5.92608 | 5.89745 | 5.81478 |
| EIF3J | O75822 | 6.36455 | 6.37809 | 6.83024 | 5.80661 | 6.50521 | 6.67947 | 6.27898 | 6.64601 | 6.43614 | 6.38211 | 6.21686 | 6.17431 | 6.14055 | 5.71321 | 5.78979 | 5.65788 |
| CEP152 | O94986 | 6.79132 | 6.57407 | 6.40427 | 6.53762 | 6.35232 | 6.43942 | 6.07844 | 6.1911 | 6.46941 | 6.16362 | 6.74274 | 6.61344 | 6.08807 | 5.23996 | 5.77544 | 5.53874 |
| YAP1 | P46937-9 | 6.80564 | 6.49384 | 6.36268 | 4.62162 | 6.49398 | 4.66252 | 5.25113 | 7.45241 | 4.95471 | 6.5278 | 6.07049 | 7.44256 | 6.60583 | 7.6054 | 6.93152 | 7.27885 |
| CEP135 | Q66GS9 | 6.58851 | 6.52446 | 6.71586 | 6.7042 | 6.32362 | 6.72135 | 6.01266 | 6.63294 | 6.53024 | 6.08076 | 6.49902 | 6.45981 | 5.82426 | 5.41191 | 5.35353 | 5.6169 |
| TAF5 | Q15542 | 6.02212 | 6.46285 | 6.28967 | 6.92431 | 6.11171 | 6.18246 | 6.29075 | 6.00653 | 6.24254 | 5.95834 | 6.32753 | 6.14755 | 6.37593 | 5.90891 | 6.34242 | 6.40638 |
| LMAN2L | Q9H0V9-2 | 6.30623 | 6.17224 | 6.15475 | 6.39242 | 6.25151 | 6.02635 | 6.21103 | 5.71729 | 6.41667 | 5.90282 | 5.98073 | 5.92431 | 6.44783 | 6.34523 | 7.08007 | 6.67052 |
| MRI1 | Q9BV20 | 6.2418 | 6.2432 | 6.80229 | 5.7282 | 6.37621 | 6.67398 | 6.27495 | 6.77629 | 6.00943 | 6.49498 | 6.44328 | 6.53408 | 5.78539 | 6.13009 | 5.8079 | 5.67795 |
| GTF3C5 | Q9Y5Q8-3 | 6.15139 | 6.36489 | 6.01856 | 6.41831 | 6.2523 | 6.31991 | 6.01757 | 6.20033 | 6.06731 | 6.51724 | 6.08817 | 6.17521 | 6.21132 | 6.40115 | 6.45323 | 6.34312 |
| GRB2 | P62993 | 6.31638 | 5.7296 | 6.31526 | 5.52321 | 6.65525 | 6.12341 | 6.05858 | 6.76887 | 6.36368 | 6.57963 | 6.69325 | 6.56944 | 6.12613 | 6.20043 | 6.10623 | 5.87063 |
| FIG4 | Q92562 | 6.38745 | 6.3801 | 6.71165 | 6.36419 | 6.03026 | 6.50508 | 6.27232 | 6.42839 | 6.21159 | 6.36389 | 6.44865 | 5.99103 | 6.0195 | 6.00353 | 6.21315 | 5.66923 |
| TSNAX | Q99598 | 5.78038 | 5.80757 | 6.11137 | 5.61205 | 6.21233 | 6.58452 | 6.21686 | 6.46616 | 6.23206 | 6.10912 | 6.20653 | 6.32266 | 6.34052 | 6.59911 | 6.87589 | 6.52286 |
| BCS1L | Q9Y276 | 6.24927 | 6.25359 | 5.93951 | 6.44903 | 6.17569 | 5.92139 | 6.68714 | 6.00222 | 6.35581 | 5.97116 | 6.13337 | 5.9143 | 6.54208 | 6.1779 | 6.84916 | 6.37839 |
| EMC3 | Q9P0I2 | 6.55464 | 6.38223 | 5.97992 | 6.73944 | 6.02438 | 6.1338 | 6.07455 | 6.04089 | 5.92683 | 6.44363 | 6.02537 | 6.11025 | 6.25811 | 6.32553 | 6.53042 | 6.45 |
| MELK | Q14680 | 6.78892 | 6.61101 | 7.26942 | 6.62531 | 6.21904 | 6.68787 | 6.34411 | 6.53174 | 7.01602 | 6.99744 | 6.87794 | 6.74993 | 4.90689 | 4.97483 | 4.70857 | 4.69097 |
| PPIL2 | Q13356 | 5.9083 | 6.17202 | 6.54296 | 6.1811 | 6.1403 | 6.21963 | 6.16781 | 5.72395 | 6.51758 | 6.33084 | 6.20168 | 5.98921 | 6.63112 | 6.23772 | 6.68201 | 6.35376 |
| PGM2L1 | Q6PCE3 | 6.61746 | 6.13685 | 6.79519 | 5.79325 | 6.40428 | 6.82734 | 6.19421 | 6.31408 | 6.44579 | 6.31661 | 6.57342 | 6.07973 | 5.97615 | 5.85679 | 5.71839 | 5.95047 |
| BTN2A1 | Q7KYR7 | 6.27665 | 6.48472 | 5.96732 | 6.37362 | 6.35364 | 6.06718 | 6.20479 | 6.48559 | 6.23653 | 6.07047 | 6.04414 | 5.91812 | 6.6815 | 6.29243 | 6.34942 | 6.19387 |

| | | | | | | | | | | | | | | | | | |
|---|---|---|---|---|---|---|---|---|---|---|---|---|---|---|---|---|---|
| CD2BP2 | O95400 | 6.37658 | 6.21978 | 6.39536 | 5.99602 | 6.51928 | 6.31715 | 6.00467 | 6.44144 | 6.10984 | 5.96206 | 6.61475 | 6.11446 | 6.96036 | 5.9094 | 5.99041 | 6.06843 |
| PWP1 | Q13610 | 6.81781 | 5.97078 | 6.91159 | 6.97946 | 6.33847 | 6.94307 | 6.5214 | 6.32884 | 6.23935 | 6.30035 | 6.53807 | 6.69418 | 5.37645 | 5.41473 | 5.35128 | 5.27418 |
| PPHLN1 | Q8NEY8-3 | 6.225 | 6.17188 | 5.82161 | 6.821 | 6.09966 | 5.64074 | 6.92767 | 5.64071 | 6.23124 | 5.70391 | 5.70737 | 6.06869 | 6.45678 | 6.41131 | 6.92969 | 7.14274 |
| CNOT9 | Q92600 | 5.94164 | 5.33654 | 6.37169 | 5.27144 | 6.52359 | 5.85908 | 5.80431 | 6.6301 | 5.85563 | 6.84986 | 6.3885 | 6.85177 | 6.0799 | 6.75087 | 6.83138 | 6.65369 |
| WASF2 | Q9Y6W5 | 5.97903 | 6.47958 | 6.7929 | 6.29693 | 6.70195 | 6.83728 | 6.15171 | 6.45478 | 6.36832 | 6.38614 | 6.81424 | 6.3454 | 5.76113 | 5.85352 | 5.79141 | 4.98569 |
| SAMM50 | Q9Y512 | 6.30129 | 6.37882 | 5.92564 | 6.48155 | 6.58865 | 5.87955 | 6.10865 | 6.26457 | 6.06517 | 6.05562 | 5.83864 | 5.83538 | 6.66421 | 6.24964 | 6.74074 | 6.62187 |
| WDR45B | Q5MNZ6 | 6.71388 | 6.93124 | 6.42214 | 6.49494 | 6.09082 | 6.34927 | 6.37707 | 6.23358 | 6.49978 | 6.62638 | 6.54562 | 6.42273 | 5.59073 | 5.40922 | 5.72243 | 5.57017 |
| MANBA | O00462 | 6.18089 | 6.73679 | 6.17809 | 6.57769 | 6.43174 | 6.2014 | 6.33198 | 6.13926 | 6.10097 | 6.4399 | 6.1244 | 6.31954 | 6.13907 | 5.95427 | 6.15944 | 5.98457 |
| RELB | D6R992 | 6.18005 | 6.36574 | 7.07109 | 6.00265 | 6.53119 | 6.38756 | 6.26613 | 6.46564 | 6.84813 | 6.36322 | 6.96547 | 6.70723 | 5.53554 | 5.38105 | 5.57942 | 5.34988 |
| DHFR | P00374 | 6.15094 | 6.35416 | 6.78441 | 5.88225 | 6.36837 | 6.51443 | 6.43436 | 7.34657 | 7.40723 | 6.89706 | 7.58213 | 7.57801 | 4.48753 | 4.72279 | 4.66681 | 4.82294 |
| TSN | Q15631 | 6.35289 | 6.31674 | 6.21569 | 6.09105 | 6.47326 | 7.06472 | 5.91045 | 7.33142 | 6.13907 | 6.26755 | 6.31341 | 6.2791 | 5.31053 | 6.21011 | 6.19841 | 5.52559 |
| NSMAF | Q92636 | 6.35423 | 6.13125 | 7.28941 | 6.33151 | 6.57584 | 6.64445 | 6.02875 | 6.59387 | 6.4955 | 6.65276 | 6.55256 | 6.50432 | 5.39236 | 5.97768 | 5.4713 | 5.00422 |
| NDUFV1 | P49821 | 6.34879 | 6.02328 | 6.00542 | 6.63971 | 6.09207 | 5.81222 | 6.41476 | 5.4962 | 6.33242 | 6.01141 | 6.25297 | 6.00092 | 7.02919 | 6.30898 | 6.57501 | 6.65666 |
| MTX1 | Q13505 | 5.95012 | 5.74397 | 5.78794 | 5.8568 | 6.03975 | 6.01446 | 6.28387 | 6.46504 | 5.76934 | 6.38241 | 6.24174 | 6.77717 | 6.34338 | 6.57916 | 6.8891 | 6.87575 |
| CPSF4 | O95639 | 6.73341 | 6.71306 | 5.78424 | 6.9852 | 6.42624 | 6.2949 | 6.20751 | 5.95499 | 5.90712 | 6.26593 | 5.83766 | 6.18912 | 6.49241 | 5.9547 | 6.04507 | 6.20842 |
| STX4 | Q12846 | 6.20078 | 6.71996 | 5.93951 | 6.76078 | 5.88961 | 6.11284 | 6.38983 | 5.96291 | 6.2304 | 6.4412 | 6.33974 | 6.29415 | 6.05106 | 6.13822 | 6.34791 | 6.18109 |
| MTCH2 | Q9Y6C9 | 6.44871 | 6.29836 | 5.97552 | 6.95377 | 6.255 | 6.23296 | 6.17154 | 6.24843 | 6.23441 | 6.01091 | 6.42614 | 5.96769 | 6.18205 | 5.89779 | 6.37858 | 6.31814 |
| RPL36 | Q9Y3U8 | 5.88772 | 6.21468 | 5.89984 | 5.89805 | 6.22166 | 6.29283 | 6.78458 | 6.3888 | 7.09256 | 5.55842 | 6.10623 | 6.27153 | 6.2288 | 6.86123 | 5.75463 | 6.53841 |
| OPHN1 | O60890 | 6.0568 | 6.10789 | 6.45244 | 5.85489 | 6.10036 | 6.1952 | 6.29051 | 6.44933 | 6.16408 | 6.26015 | 6.39909 | 6.42511 | 6.19385 | 6.27072 | 6.52081 | 6.31275 |
| TENM3 | Q9P273 | 5.92459 | 6.12658 | 5.9024 | 6.59231 | 6.0149 | 6.39298 | 6.38621 | 6.06753 | 6.33285 | 5.97701 | 6.17673 | 5.90612 | 6.72793 | 6.51973 | 6.39586 | 6.55629 |
| GPX8 | Q8TED1 | 6.57636 | 6.48571 | 6.06924 | 6.64657 | 6.35464 | 6.14882 | 6.11799 | 6.38531 | 6.0563 | 5.96536 | 5.85968 | 6.06237 | 6.26871 | 6.42751 | 6.33683 | 6.23861 |
| FAM114A2 | Q9NRY5 | 6.10093 | 5.92345 | 6.4821 | 5.60861 | 6.32393 | 6.52904 | 6.04068 | 6.15699 | 6.53015 | 6.58846 | 6.82538 | 6.48781 | 6.33958 | 6.07576 | 5.93989 | 6.04724 |
| TAP1 | Q03518 | 6.17827 | 6.35407 | 6.22103 | 6.67161 | 6.07243 | 5.87036 | 6.29714 | 5.96311 | 6.13947 | 6.18663 | 5.86337 | 5.74449 | 6.64715 | 6.13941 | 6.74711 | 6.90435 |
| ERI3 | O43414 | 6.03352 | 5.18525 | 6.44249 | 5.10301 | 6.22824 | 6.17553 | 5.96006 | 6.49837 | 5.71577 | 6.48637 | 6.26963 | 6.95339 | 6.26249 | 6.89907 | 6.9549 | 6.83192 |
| NLRP2 | Q9NX02 | 6.49842 | 5.96788 | 7.31036 | 6.23096 | 6.39796 | 6.57802 | 6.46138 | 6.3315 | 6.56907 | 6.41987 | 6.93687 | 6.53893 | 5.50633 | 5.30665 | 5.64455 | 5.30126 |
| AUP1 | Q9Y679 | 6.31942 | 6.3058 | 6.15883 | 6.60189 | 6.12032 | 6.16998 | 6.32283 | 5.81855 | 6.2984 | 6.09682 | 6.4333 | 6.2582 | 6.3975 | 6.10818 | 6.38106 | 6.20892 |
| NT5DC1 | Q5TFE4 | 6.05896 | 6.17242 | 6.54166 | 5.92856 | 6.17719 | 6.25121 | 5.63716 | 6.81501 | 6.12087 | 6.66289 | 6.6991 | 6.65571 | 5.54537 | 6.35312 | 6.16492 | 6.21584 |
| SLC25A22 | Q9H936 | 6.57366 | 6.10441 | 5.97343 | 6.37219 | 6.28515 | 6.09116 | 6.21231 | 5.74653 | 6.10651 | 5.91991 | 6.05466 | 5.80233 | 6.77215 | 6.00879 | 7.09528 | 6.88153 |
| JAK2 | O60674 | 6.40921 | 6.78748 | 6.55385 | 6.91644 | 6.38336 | 6.22512 | 6.43387 | 6.16484 | 6.8136 | 6.74294 | 6.62219 | 6.52795 | 5.24582 | 5.52018 | 5.27083 | 5.38233 |
| GGCT | O75223 | 6.41179 | 6.39714 | 6.94124 | 6.04727 | 6.42914 | 6.90504 | 6.00628 | 6.44091 | 6.6233 | 6.51085 | 6.61737 | 6.53724 | 5.6236 | 5.55786 | 5.5998 | 5.35118 |
| PSME3 | P61289-2 | 5.24229 | 5.53301 | 5.84694 | 5.081 | 5.95209 | 6.08139 | 5.7453 | 5.95468 | 5.76382 | 5.98252 | 6.20656 | 6.10798 | 7.47965 | 7.66385 | 7.81085 | 7.54808 |
| KIN | O60870 | 6.00474 | 6.29022 | 6.12708 | 6.31543 | 6.14522 | 6.38508 | 6.3471 | 5.71284 | 6.47057 | 6.20811 | 6.41478 | 6.13391 | 6.43811 | 6.09566 | 6.61231 | 6.29886 |
| MGME1 | Q9BQP7 | 6.42006 | 6.391 | 6.09018 | 7.1588 | 6.34392 | 6.24921 | 6.64417 | 6.03869 | 6.29704 | 6.01065 | 6.16181 | 6.20248 | 6.00625 | 5.78895 | 5.98857 | 6.20823 |
| EIF2B2 | P49770 | 6.4986 | 6.72856 | 6.84403 | 5.69174 | 6.36797 | 6.7879 | 6.06278 | 7.05166 | 6.46053 | 6.75761 | 6.54113 | 6.6622 | 5.43267 | 5.50965 | 5.29635 | 5.30662 |
| ALG5 | Q9Y673 | 6.31704 | 6.1843 | 5.98093 | 6.3986 | 6.11596 | 5.65652 | 6.3132 | 5.95905 | 6.21056 | 5.9013 | 6.07463 | 5.75855 | 6.74964 | 6.45981 | 6.92207 | 6.99785 |
| CYP20A1 | Q6UW02 | 6.26827 | 6.34737 | 5.81681 | 6.42816 | 6.11784 | 5.99098 | 6.32259 | 5.8551 | 6.16698 | 6.18919 | 5.86158 | 5.81639 | 6.77052 | 6.45249 | 7.02134 | 6.57439 |
| LSM4 | Q9Y4Z0 | 6.11095 | 6.07357 | 6.04765 | 6.06817 | 6.45449 | 6.01886 | 6.29021 | 6.24194 | 6.28356 | 6.08155 | 6.29971 | 6.2705 | 6.54579 | 6.32543 | 6.53122 | 6.35641 |
| TAF1 | P21675-4 | 6.23175 | 6.50626 | 6.10154 | 6.8412 | 5.74835 | 5.81887 | 6.22018 | 6.27275 | 6.18923 | 6.03115 | 6.12782 | 6.36672 | 6.23123 | 6.17693 | 6.61678 | 6.51923 |
| MRPL47 | Q9HD33 | 6.3463 | 6.01786 | 5.85101 | 6.46945 | 6.53497 | 5.90141 | 6.75132 | 5.9341 | 6.27595 | 5.90044 | 6.00385 | 5.94815 | 6.57631 | 6.34241 | 6.65999 | 6.48648 |
| SRP9 | P49458 | 5.62557 | 5.20723 | 5.71913 | 5.145 | 6.34708 | 5.81628 | 6.02617 | 6.11919 | 6.37371 | 6.35443 | 6.23903 | 6.40662 | 6.70242 | 7.08239 | 7.29523 | 7.54052 |
| GTF2H1 | P32780 | 6.11963 | 6.57277 | 6.12724 | 6.91536 | 6.35797 | 6.16103 | 6.43271 | 6.06978 | 6.14601 | 6.00356 | 5.98242 | 6.04144 | 6.50798 | 5.8133 | 6.64712 | 6.10168 |
| SNRPE | P62304 | 6.16471 | 6.37693 | 5.73524 | 6.50682 | 6.08551 | 5.92941 | 6.28919 | 6.07418 | 6.12247 | 5.84795 | 5.90329 | 6.08318 | 6.92843 | 6.15426 | 7.01041 | 6.78802 |
| ATP5MG | O75964 | 6.39173 | 5.67204 | 5.86229 | 5.73386 | 6.45533 | 6.06027 | 6.2463 | 6.35487 | 5.67463 | 6.01466 | 5.92001 | 5.68929 | 6.79849 | 6.77542 | 7.40621 | 6.94461 |
| RPL19 | P84098 | 5.87668 | 6.61384 | 5.72628 | 6.05293 | 6.12267 | 6.36828 | 6.61273 | 6.48376 | 6.76482 | 6.06511 | 6.15254 | 6.43645 | 5.78206 | 6.68257 | 5.77659 | 6.4827 |
| CISD2 | Q8N5K1 | 6.53138 | 6.30742 | 5.99012 | 6.25651 | 6.15991 | 5.93385 | 6.10698 | 6.04904 | 6.26354 | 6.46077 | 6.33088 | 6.28089 | 6.64186 | 6.22193 | 6.20666 | 6.25825 |
| RPRD1B | Q9NQG5 | 5.95083 | 6.11704 | 5.8985 | 5.91768 | 6.20941 | 6.19279 | 5.87754 | 6.09426 | 6.37543 | 6.07142 | 6.27751 | 6.11699 | 6.79705 | 6.68321 | 7.05828 | 6.36206 |
| MT-CO2 | P00403 | 5.90398 | 5.76311 | 5.1972 | 5.63737 | 5.72396 | 5.9069 | 6.16442 | 6.53275 | 5.86928 | 5.36295 | 5.78091 | 5.77739 | 6.99606 | 7.94233 | 7.17418 | 8.26721 |
| SH3PXD2B | A1X283 | 5.72706 | 5.62208 | 6.54449 | 6.02039 | 5.90213 | 6.55982 | 6.24967 | 6.2827 | 6.34711 | 6.52769 | 6.48397 | 6.55968 | 6.12647 | 6.57283 | 6.01695 | 6.45695 |
| HSPA2 | P54652 | 6.0174 | 6.0092 | 6.34448 | 5.59116 | 5.9982 | 6.05602 | 6.07281 | 5.8956 | 6.19332 | 6.29159 | 6.37535 | 6.55372 | 6.84171 | 6.68595 | 6.57182 | 6.50168 |
| GADD45GIP1 | Q8TAE8 | 6.26572 | 6.24798 | 5.73053 | 6.38532 | 6.3481 | 6.44452 | 6.50589 | 6.43974 | 5.97945 | 6.18177 | 5.92725 | 6.03995 | 6.412 | 6.29852 | 6.27571 | 6.51755 |
| SYDE1 | Q6ZW31 | 6.65505 | 6.16 | 7.36488 | 6.6837 | 6.07148 | 6.26588 | 6.30947 | 6.35425 | 6.63515 | 6.70711 | 6.84586 | 7.09527 | 5.33823 | 5.25512 | 5.10385 | 5.15468 |
| TTC4 | O95801 | 6.25806 | 6.23925 | 6.88039 | 6.08685 | 6.03937 | 6.64422 | 6.18504 | 6.14988 | 6.47157 | 6.83006 | 6.2579 | 6.52632 | 6.06541 | 5.92135 | 5.58955 | 5.85477 |
| IRF2BP2 | Q7Z5L9 | 5.91876 | 6.39135 | 6.01342 | 5.78451 | 6.11742 | 6.38129 | 5.94268 | 6.18112 | 5.9248 | 6.06908 | 6.32346 | 6.09028 | 7.34234 | 6.4508 | 6.65961 | 6.40906 |
| ARAP1 | Q96P48 | 6.26485 | 6.19696 | 7.00424 | 6.06675 | 6.09205 | 6.4457 | 6.20299 | 6.23045 | 6.3242 | 6.65401 | 6.54468 | 6.37058 | 5.89766 | 6.16614 | 5.95117 | 5.58756 |
| LRRC8C | Q8TDW0 | 6.29611 | 6.83631 | 6.27492 | 6.92766 | 6.21563 | 6.04997 | 6.30665 | 5.82221 | 6.23642 | 6.04878 | 6.24581 | 6.32021 | 6.08811 | 5.92328 | 6.17495 | 6.23298 |
| FMNL1 | O95466-3 | 6.46467 | 6.49103 | 6.14714 | 6.28344 | 6.57699 | 6.46663 | 6.26742 | 6.29589 | 6.49007 | 5.98372 | 6.06714 | 5.8566 | 6.1398 | 6.11433 | 6.07069 | 6.28444 |
| PDHX | O00330 | 6.71824 | 6.36077 | 5.76957 | 6.43766 | 6.76587 | 6.14002 | 6.06981 | 6.348 | 6.31849 | 6.0849 | 6.15974 | 6.15353 | 6.82201 | 5.67659 | 6.15783 | 6.01697 |
| PGLS | O95336 | 5.8292 | 5.26559 | 5.81322 | 4.55184 | 5.80067 | 5.94664 | 5.51844 | 7.33908 | 5.076 | 6.19039 | 5.20441 | 6.19387 | 5.32105 | 10.1874 | 6.77327 | 8.98895 |
| GRHPR | Q9UBQ7 | 6.46657 | 6.31563 | 6.42968 | 5.83871 | 6.51861 | 6.30041 | 6.27216 | 6.48584 | 6.15226 | 6.29838 | 6.25486 | 6.18421 | 6.33161 | 6.11583 | 6.1 | 5.93522 |
| CAND2 | O75155 | 6.19379 | 5.80959 | 6.27241 | 5.44387 | 5.85055 | 6.2137 | 5.98486 | 6.48319 | 6.0491 | 6.47952 | 6.45405 | 6.21664 | 6.39954 | 6.94945 | 6.83848 | 6.36127 |
| YBX1 | P67809 | 6.35168 | 6.20778 | 7.11107 | 6.40971 | 6.21669 | 6.18786 | 6.41253 | 6.01059 | 6.81995 | 6.49853 | 6.91941 | 6.75123 | 5.38767 | 5.60338 | 5.17079 | 5.94114 |
| RAD54B | Q9Y620 | 5.58002 | 5.77505 | 6.2967 | 5.82419 | 6.27324 | 6.47312 | 6.20601 | 6.06192 | 6.40619 | 6.41228 | 6.59944 | 6.15018 | 6.2243 | 6.50328 | 6.68451 | 6.52958 |
| SRM | P19623 | 6.01409 | 6.40338 | 6.84205 | 5.90127 | 6.70631 | 7.05294 | 6.3552 | 7.05414 | 6.60993 | 6.21009 | 6.76378 | 6.45747 | 5.40713 | 5.48461 | 5.35303 | 5.3846 |
| UBXN6 | Q9BZV1 | 6.24078 | 6.13652 | 6.59016 | 5.98012 | 6.24568 | 6.49738 | 5.83972 | 6.63841 | 6.17771 | 6.63593 | 6.50695 | 6.40217 | 6.32505 | 5.86687 | 6.11803 | 5.79851 |
| CDK6 | Q00534 | 6.32467 | 6.11527 | 6.58172 | 5.74634 | 6.13175 | 6.54335 | 6.4277 | 6.6768 | 6.53471 | 6.27581 | 6.52153 | 6.30035 | 5.89684 | 6.00885 | 5.95402 | 5.96029 |
| FUS | P35637 | 6.41742 | 6.47759 | 6.67202 | 6.89585 | 6.22164 | 6.40431 | 5.80875 | 5.91059 | 6.22076 | 6.41392 | 6.4365 | 6.11652 | 6.38713 | 5.64994 | 5.96437 | 6.00269 |
| HARS2 | P49590 | 6.40858 | 5.99577 | 5.96757 | 6.53507 | 6.11374 | 5.84145 | 6.82663 | 6.00221 | 6.17417 | 5.8822 | 5.90274 | 5.94589 | 6.44054 | 6.71166 | 6.57915 | 6.67262 |
| DNAJC3 | Q13217 | 6.23736 | 6.54804 | 5.81663 | 6.49778 | 6.28104 | 6.11082 | 6.22731 | 6.11675 | 6.14451 | 6.05748 | 6.24791 | 6.0756 | 6.55082 | 6.12953 | 6.82971 | 6.12871 |
| RPS25 | P62851 | 6.41469 | 6.41143 | 6.5142 | 6.19943 | 6.47577 | 6.51106 | 6.21921 | 6.66079 | 6.60031 | 6.42508 | 6.03351 | 6.04422 | 5.61003 | 6.18367 | 5.7588 | 5.93779 |
| PPP2R5C | Q13362-4 | 6.227 | 6.33108 | 6.62622 | 6.23216 | 6.31248 | 6.4994 | 6.29992 | 6.08158 | 6.24975 | 6.36957 | 6.49913 | 6.40141 | 6.03075 | 6.0151 | 5.93168 | 5.89278 |
| L3MBTL2 | Q969R5 | 6.2355 | 6.34088 | 5.88633 | 6.87218 | 5.8215 | 6.36733 | 6.4358 | 5.90844 | 6.13807 | 5.94167 | 6.01224 | 5.9584 | 6.26987 | 6.52106 | 6.58559 | 6.70514 |
| CLIP2 | Q9UDT6 | 6.34849 | 6.44979 | 6.90712 | 5.80995 | 6.6399 | 6.80641 | 5.8387 | 6.77235 | 6.2332 | 6.65441 | 6.67741 | 6.55701 | 6.08686 | 5.66811 | 5.28591 | 5.26438 |
| GALK2 | Q01415 | 6.65595 | 5.93081 | 6.76765 | 6.01243 | 6.14526 | 6.46516 | 6.1982 | 6.38284 | 6.36122 | 6.07028 | 6.36185 | 6.49421 | 6.06491 | 6.13928 | 6.20785 | 5.74211 |
| FBXO17 | Q96EF6 | 6.2587 | 6.2323 | 6.28251 | 6.42134 | 6.1106 | 6.01514 | 6.32955 | 6.59704 | 6.49551 | 6.43754 | 6.09367 | 6.3615 | 6.08214 | 6.35784 | 6.37294 | 6.21798 |
| ADD2 | P35612 | 5.92449 | 5.99101 | 6.15518 | 5.5976 | 6.03853 | 6.02535 | 5.86792 | 6.34868 | 6.24092 | 6.2298 | 6.01021 | 6.01542 | 6.85681 | 6.90794 | 7.01464 | 6.7755 |
| YKT6 | O15498 | 6.15914 | 6.24932 | 6.68239 | 5.86272 | 6.11115 | 6.74024 | 6.36824 | 6.87793 | 6.346 | 6.52717 | 6.21062 | 6.35475 | 5.90892 | 6.12613 | 5.62271 | 5.85258 |
| COA7 | Q96BR5 | 6.71058 | 6.31502 | 6.1289 | 6.30514 | 7.03793 | 6.31616 | 5.77124 | 6.06228 | 5.80196 | 5.71227 | 5.87144 | 5.91625 | 7.37278 | 6.03191 | 6.34532 | 6.30081 |
| ARFGAP2 | Q8N6H7 | 6.69802 | 6.58327 | 6.63715 | 6.09977 | 6.33145 | 6.92263 | 6.65084 | 6.63256 | 6.28887 | 6.64687 | 6.51995 | 6.3329 | 5.54657 | 5.45955 | 5.31107 | 5.33853 |
| FAM160B1 | Q5W0V3 | 6.54678 | 6.37013 | 6.7459 | 6.01976 | 6.28343 | 6.61743 | 6.17814 | 6.68358 | 6.3248 | 6.59838 | 6.29528 | 6.1852 | 5.88661 | 5.94326 | 5.71578 | 5.60552 |
| PNPLA8 | Q9NP80 | 6.03928 | 6.43219 | 5.71415 | 6.68165 | 6.06341 | 5.982 | 6.24309 | 6.04153 | 5.85413 | 5.9536 | 6.04191 | 6.31809 | 6.72612 | 6.56189 | 6.74362 | 6.60332 |
| COL18A1 | P39060 | 7.23055 | 7.3415 | 6.32316 | 7.08993 | 6.72262 | 6.65567 | 6.69066 | 6.1798 | 6.23673 | 6.49544 | 6.40829 | 6.06961 | 5.06139 | 5.26945 | 5.10389 | 5.1213 |
| NISCH | Q9Y2I1 | 6.49451 | 6.37277 | 6.21811 | 6.12204 | 6.51481 | 6.38059 | 6.24199 | 6.43182 | 6.34272 | 6.1846 | 6.73009 | 6.41858 | 6.0107 | 5.69957 | 5.94345 | 5.89364 |
| SFMBT1 | Q9UHJ3 | 6.00718 | 6.17397 | 6.17612 | 6.805 | 6.20124 | 5.94976 | 6.44727 | 6.16215 | 6.31552 | 6.15856 | 6.36471 | 6.14157 | 6.23326 | 6.18502 | 6.28428 | 6.39438 |
| TBC1D10B | Q4KMP7 | 5.91583 | 6.31373 | 5.71573 | 6.01444 | 6.30324 | 5.89916 | 6.23546 | 6.01063 | 6.18367 | 6.51068 | 5.95733 | 6.3342 | 6.59915 | 6.7322 | 6.44395 | 6.83058 |
| PRMT5 | O14744 | 5.91554 | 6.00659 | 6.66193 | 5.97356 | 6.13907 | 6.64849 | 6.27978 | 6.00046 | 6.24832 | 6.2139 | 6.45023 | 6.43471 | 6.30577 | 6.20274 | 6.18753 | 6.33138 |
| MLYCD | O95822 | 6.33764 | 6.67798 | 5.87232 | 6.70256 | 5.96223 | 5.82836 | 6.22707 | 6.27663 | 5.83496 | 6.1337 | 6.23536 | 6.1664 | 6.47256 | 6.39136 | 6.53587 | 6.34501 |
| ECSIT | Q9BQ95 | 6.47018 | 6.38949 | 6.29883 | 6.3657 | 6.62002 | 6.21353 | 6.34578 | 6.36348 | 6.18978 | 6.0354 | 6.4655 | 6.1204 | 6.64266 | 5.83748 | 5.90576 | 5.73601 |
| EHD3 | Q9NZN3 | 6.24209 | 6.3502 | 6.42667 | 5.83717 | 6.42125 | 6.51228 | 6.19758 | 6.48559 | 6.50377 | 6.41912 | 6.24669 | 6.17112 | 6.41039 | 5.79575 | 6.15935 | 5.82097 |
| UGGT2 | Q9NYU1 | 6.37623 | 6.28898 | 5.96803 | 6.86031 | 6.32121 | 6.38135 | 6.28548 | 6.09031 | 6.34685 | 5.94702 | 5.8853 | 6.09332 | 6.46602 | 6.10062 | 6.18475 | 6.40421 |
| ITGA4 | P13612 | 6.41817 | 6.18622 | 6.45199 | 6.44201 | 6.58939 | 6.24874 | 6.37979 | 5.76032 | 6.61703 | 6.0949 | 6.64443 | 6.60718 | 6.05238 | 5.92186 | 5.79596 | 5.78963 |
| GRPEL1 | Q9HAV7 | 6.24055 | 5.1399 | 5.57426 | 5.32528 | 6.41221 | 5.14672 | 5.64226 | 6.25124 | 4.92409 | 6.22064 | 5.51806 | 6.79829 | 6.96535 | 7.71877 | 7.86291 | 8.25944 |
| RPS6KA3 | P51812 | 6.14483 | 6.07942 | 6.57552 | 5.74588 | 6.23856 | 6.89718 | 6.17589 | 6.91471 | 6.22854 | 6.39662 | 6.66198 | 6.33272 | 6.18889 | 6.18612 | 5.70229 | 5.53087 |
| SRGAP1 | Q7Z6B7 | 6.16208 | 5.98274 | 6.56635 | 6.21923 | 6.09753 | 6.81499 | 6.16177 | 6.49865 | 6.47059 | 6.52752 | 6.67115 | 6.66366 | 5.73627 | 5.81224 | 5.85653 | 5.7587 |
| TMEM201 | Q5SNT2 | 6.41707 | 6.74374 | 5.95381 | 7.44044 | 6.30823 | 6.04724 | 6.18608 | 5.78118 | 6.14057 | 6.28353 | 6.40278 | 6.23088 | 5.94594 | 5.93256 | 5.91387 | 6.27211 |
| MBD1 | Q9UIS9-9 | 6.09697 | 6.29351 | 5.90453 | 6.9044 | 5.89492 | 5.61851 | 6.5569 | 5.55588 | 6.00433 | 6.14924 | 6.11411 | 6.25236 | 6.70991 | 6.57311 | 6.66873 | 6.70258 |
| SEPTIN11 | Q9NVA2 | 5.89924 | 6.43737 | 6.32104 | 6.04955 | 6.3341 | 6.64271 | 6.00235 | 6.52984 | 6.30551 | 6.51825 | 6.33323 | 6.27331 | 6.24307 | 6.34722 | 6.02853 | 5.73467 |
| GPHN | Q9NQX3-2 | 6.53461 | 6.21008 | 6.63127 | 6.07542 | 6.65266 | 6.66883 | 5.98295 | 6.54247 | 6.5716 | 6.18739 | 6.72777 | 6.85184 | 6.04571 | 5.35258 | 5.46488 | 5.49993 |
| MED13L | Q71F56 | 6.34548 | 6.47754 | 6.43184 | 6.62456 | 6.21506 | 6.08533 | 6.15397 | 6.14769 | 6.30753 | 6.1531 | 6.15838 | 6.25943 | 6.30034 | 6.06311 | 6.02557 | 6.25108 |
| TSC1 | Q92574 | 6.54585 | 6.2983 | 6.57721 | 6.10155 | 6.26086 | 6.65798 | 6.23706 | 6.42202 | 6.60072 | 6.45002 | 6.82331 | 6.89791 | 5.65592 | 5.9104 | 5.30205 | 5.25886 |
| PKN3 | Q6P5Z2 | 6.5625 | 6.36446 | 6.67724 | 6.92864 | 6.52506 | 6.42638 | 6.72112 | 6.28161 | 6.58492 | 6.85775 | 6.11232 | 6.4325 | 5.29127 | 5.36593 | 5.25902 | 5.60928 |
| ATAD2B | Q9ULI0 | 6.2905 | 6.2446 | 6.09076 | 7.08514 | 5.96692 | 5.74046 | 6.22609 | 5.66263 | 6.36527 | 6.08816 | 6.28527 | 6.16257 | 6.5809 | 6.08083 | 6.38164 | 6.74828 |
| ARHGAP45 | Q92619 | 6.62058 | 6.28752 | 6.62204 | 5.80373 | 5.84258 | 6.59714 | 5.75056 | 6.11995 | 6.63111 | 6.48711 | 6.33535 | 6.68389 | 6.21975 | 6.0671 | 5.81511 | 6.11648 |
| ZC2HC1A | Q96GY0 | 6.27607 | 6.22763 | 6.90588 | 5.97952 | 6.12719 | 6.55335 | 6.16074 | 6.43858 | 6.27904 | 6.72088 | 6.56525 | 6.56261 | 5.75619 | 6.0729 | 5.74135 | 5.63283 |
| LATS1 | O95835 | 6.26814 | 6.4062 | 7.04919 | 6.3958 | 6.30068 | 6.82831 | 6.16532 | 6.59871 | 6.30777 | 5.98969 | 6.43659 | 6.98463 | 5.80868 | 5.525 | 5.43379 | 5.50151 |
| CCDC6 | Q16204 | 6.29473 | 6.22122 | 6.71422 | 5.79656 | 6.41216 | 6.83654 | 5.8166 | 6.34246 | 6.23156 | 6.50662 | 6.54025 | 6.55925 | 6.45551 | 5.93691 | 5.51256 | 5.82286 |

| | | | | | | | | | | | | | | | | | |
|---|---|---|---|---|---|---|---|---|---|---|---|---|---|---|---|---|---|
| TLE3 | Q04726-4 | 5.79974 | 6.11051 | 5.85157 | 5.48683 | 6.10014 | 6.5802 | 5.80936 | 6.32689 | 5.88783 | 5.95808 | 6.37899 | 5.98324 | 7.23232 | 6.82367 | 6.95575 | 6.71487 |
| IGF1R | P08069 | 6.22525 | 6.31939 | 6.00634 | 6.20198 | 6.01986 | 6.12575 | 6.17728 | 6.27441 | 6.07253 | 6.1247 | 6.12508 | 6.01439 | 6.78843 | 6.04236 | 6.83272 | 6.64952 |
| CCDC186 | Q7Z3E2 | 6.38555 | 6.27675 | 6.60478 | 6.00078 | 6.17329 | 6.6176 | 6.30882 | 6.67606 | 6.42049 | 6.8377 | 6.89309 | 7.04291 | 5.72072 | 5.74284 | 5.44123 | 4.85739 |
| EAPP | Q56P03 | 6.03127 | 6.65303 | 5.97094 | 6.13067 | 6.1524 | 6.60979 | 6.00104 | 6.18786 | 6.06855 | 6.14869 | 6.25239 | 6.16 | 6.68909 | 6.39652 | 6.08371 | 6.46404 |
| SLC12A9 | Q9BXP2 | 6.56122 | 6.6038 | 6.20931 | 6.63222 | 6.09744 | 5.88598 | 6.23675 | 6.39148 | 6.14605 | 6.46395 | 6.50555 | 6.42181 | 5.84684 | 6.01499 | 5.95921 | 6.02339 |
| PCBP1 | Q15365 | 6.08389 | 5.68864 | 6.18552 | 5.96253 | 6.43115 | 6.91533 | 5.54657 | 5.97298 | 6.17969 | 6.49857 | 6.49299 | 6.41359 | 6.50617 | 7.03625 | 5.86825 | 6.21787 |
| NFIC | P08651 | 6.29549 | 5.99669 | 6.18744 | 6.24464 | 6.13016 | 5.88646 | 6.50668 | 5.90924 | 6.32525 | 6.13915 | 6.19298 | 6.08626 | 6.61004 | 6.27581 | 6.61347 | 6.60024 |
| HIBCH | Q6NVY1 | 6.39009 | 6.87147 | 5.59323 | 6.85337 | 6.11241 | 6.0447 | 6.01313 | 6.46506 | 5.89046 | 6.2445 | 5.95348 | 5.83532 | 6.33671 | 6.48474 | 6.20096 | 6.71039 |
| GIPC1 | O14908 | 6.4614 | 6.40672 | 7.12957 | 6.57508 | 6.49259 | 6.38129 | 6.43637 | 6.24548 | 6.71741 | 6.55975 | 6.75328 | 6.68783 | 5.52861 | 5.30304 | 5.09406 | 5.22752 |
| ARHGEF11 | O15085-2 | 6.62634 | 5.77044 | 6.70594 | 6.08514 | 5.91436 | 6.44642 | 5.85499 | 6.23371 | 6.03949 | 6.6324 | 6.46266 | 6.47744 | 6.21467 | 6.24718 | 6.07601 | 6.2128 |
| MED27 | Q6P2C8 | 5.96343 | 6.4528 | 6.07059 | 6.26695 | 6.00427 | 6.30861 | 6.16339 | 6.14454 | 6.00218 | 6.15744 | 6.46878 | 6.28489 | 6.27271 | 6.25465 | 6.71535 | 6.46944 |
| DNM3 | Q9UQ16-4 | 5.97774 | 6.23598 | 6.64306 | 6.0533 | 6.21492 | 6.54543 | 6.0092 | 6.46196 | 6.32376 | 6.26074 | 6.44983 | 6.51606 | 6.05983 | 6.27249 | 6.1097 | 5.866 |
| RAB8A | P61006 | 6.22325 | 6.41618 | 6.09371 | 6.55414 | 5.93821 | 6.19908 | 6.30165 | 6.18832 | 5.98694 | 6.18747 | 6.29416 | 6.27621 | 6.01366 | 6.31243 | 6.5758 | 6.43879 |
| RAB35 | Q15286 | 6.20589 | 6.60308 | 6.18525 | 6.54282 | 6.13635 | 6.31398 | 6.3213 | 5.99406 | 5.99569 | 6.15197 | 5.62539 | 5.79184 | 6.41915 | 6.55146 | 6.52448 | 6.63731 |
| SEC61A1 | P61619 | 6.19838 | 5.95036 | 5.84229 | 6.59581 | 6.17421 | 6.09227 | 6.23432 | 5.47969 | 5.88512 | 6.16597 | 5.59617 | 5.84069 | 6.89284 | 6.76847 | 6.91315 | 7.37024 |
| CIZ1 | Q9ULV3 | 6.91127 | 7.04134 | 6.73744 | 7.50657 | 6.05518 | 6.25842 | 6.65643 | 6.04735 | 6.54842 | 6.49832 | 6.69486 | 6.58785 | 5.13844 | 4.99705 | 5.1326 | 5.18847 |
| TATDN1 | Q6P1N9 | 5.65862 | 5.77814 | 6.52601 | 5.76906 | 6.28364 | 6.54343 | 6.1122 | 6.86063 | 6.61543 | 6.87943 | 7.28224 | 7.2185 | 5.85835 | 5.48486 | 5.44758 | 5.68188 |
| DCBLD2 | Q96PD2-2 | 6.27981 | 6.69816 | 6.28883 | 6.88777 | 6.31929 | 5.7126 | 6.58728 | 5.85113 | 6.28808 | 6.38055 | 6.42824 | 6.22699 | 6.18422 | 5.8073 | 6.22272 | 5.83702 |
| MAPRE2 | Q15555 | 6.13026 | 6.25413 | 6.80519 | 5.9235 | 5.99427 | 6.56709 | 5.7647 | 6.75346 | 6.36419 | 6.52844 | 6.54878 | 6.66852 | 5.7799 | 6.01752 | 6.06735 | 5.8327 |
| ARPC5L | Q9BPX5 | 6.10952 | 5.8385 | 6.56472 | 5.77546 | 6.26126 | 6.30503 | 5.88338 | 6.65209 | 6.05611 | 6.47643 | 6.45245 | 6.5222 | 6.42564 | 6.2378 | 5.95479 | 6.48464 |
| RTRAF | Q9Y224 | 5.83699 | 4.357 | 5.72526 | 4.43188 | 5.77826 | 4.19884 | 4.56884 | 6.62868 | 4.28776 | 6.35048 | 5.47775 | 6.67355 | 6.61865 | 10.4958 | 9.07162 | 9.49865 |
| SFXN3 | Q9BWM7 | 6.71273 | 6.33065 | 5.92017 | 6.43936 | 6.29371 | 5.72503 | 6.27433 | 5.849 | 6.3064 | 6.03481 | 6.0477 | 6.20043 | 6.6895 | 6.40171 | 6.49645 | 6.27801 |
| GRAMD1A | Q96CP6 | 6.81866 | 6.86956 | 6.31344 | 7.50858 | 6.61458 | 6.12851 | 6.82223 | 6.58393 | 6.52953 | 6.33704 | 5.89818 | 6.34085 | 5.05996 | 5.24538 | 5.38703 | 5.54255 |
| OSBPL9 | Q96SU4-2 | 6.4075 | 6.47922 | 6.34238 | 6.15568 | 6.09521 | 6.55302 | 6.07777 | 6.09096 | 6.39689 | 6.61723 | 6.40351 | 6.31722 | 6.08173 | 6.12181 | 5.76879 | 6.09106 |
| GTF2H4 | Q92759 | 6.16494 | 6.40561 | 5.97849 | 6.47687 | 6.443 | 6.14849 | 6.58892 | 6.32927 | 6.12383 | 6.30412 | 6.10412 | 5.94377 | 6.31332 | 6.28797 | 6.16919 | 6.21809 |
| ARSB | P15848 | 6.51215 | 6.17458 | 6.01184 | 6.48821 | 6.28111 | 6.10305 | 6.30304 | 5.85365 | 6.21568 | 6.35667 | 6.29569 | 6.13711 | 6.18475 | 6.28078 | 6.31439 | 6.48729 |
| KAT6B | Q8WYB5 | 6.37398 | 6.15013 | 6.0861 | 6.3045 | 6.2903 | 5.92067 | 6.24446 | 5.50238 | 6.49311 | 6.41391 | 5.93145 | 6.16173 | 6.80752 | 6.35895 | 6.32174 | 6.63907 |
| STAM | Q92783 | 6.55381 | 6.08653 | 6.50456 | 5.90162 | 6.75274 | 6.31999 | 6.00165 | 6.46129 | 6.26004 | 6.32497 | 6.51181 | 6.67087 | 6.68315 | 5.68445 | 5.68885 | 5.59368 |
| IFIH1 | Q9BYX4 | 6.72588 | 6.48957 | 6.95523 | 6.08313 | 6.22534 | 7.04574 | 6.25041 | 6.47566 | 6.58975 | 6.49161 | 6.73625 | 6.61747 | 5.42021 | 5.43964 | 5.1383 | 5.3158 |
| KYAT3 | Q6YP21 | 6.887 | 6.93093 | 6.79021 | 6.8689 | 6.90305 | 7.07167 | 6.52375 | 6.64107 | 6.35382 | 6.88544 | 5.69267 | 6.64387 | 5.46123 | 5.45592 | 5.69636 | 5.19413 |
| FUCA2 | Q9BTY2 | 6.97408 | 7.0155 | 6.36222 | 6.97305 | 6.73547 | 6.42088 | 6.38967 | 6.42466 | 6.55925 | 6.23458 | 6.55179 | 6.36197 | 5.31191 | 5.09878 | 5.27632 | 5.30987 |
| RABGEF1 | Q9UJ41-3 | 6.41332 | 6.34851 | 6.74365 | 6.07473 | 6.4883 | 6.53821 | 6.31651 | 6.47756 | 6.38032 | 6.61764 | 6.2432 | 6.25462 | 5.86838 | 5.62539 | 5.66074 | 5.94892 |
| DDX52 | Q9Y2R4 | 6.04494 | 6.15121 | 5.89566 | 6.68351 | 6.19689 | 5.85679 | 6.63631 | 5.85368 | 5.80225 | 5.9212 | 6.03654 | 5.81769 | 6.8603 | 6.59731 | 6.72157 | 6.92416 |
| CHAF1B | Q13112 | 6.43714 | 6.35282 | 6.15711 | 6.67047 | 6.05884 | 5.89908 | 6.13332 | 5.80038 | 6.28191 | 6.08538 | 6.32579 | 6.31317 | 6.22741 | 6.06525 | 6.48896 | 6.70298 |
| SOD2 | P04179 | 6.16681 | 4.98801 | 5.68686 | 5.17261 | 6.2185 | 5.20422 | 5.80261 | 6.50849 | 5.42655 | 6.33392 | 5.87539 | 6.71119 | 7.05762 | 7.38619 | 7.85786 | 7.60318 |
| ATL2 | Q8NHH9 | 6.37234 | 6.43829 | 5.99123 | 6.54417 | 6.06816 | 6.33447 | 6.49925 | 5.92868 | 6.03405 | 6.12325 | 6.02188 | 6.06683 | 6.24117 | 6.16793 | 6.42214 | 6.74616 |
| KIF16B | Q96L93 | 5.98231 | 6.42666 | 6.46129 | 6.10007 | 6.24449 | 6.27696 | 6.19418 | 6.14249 | 6.3095 | 6.38784 | 6.01196 | 5.8778 | 6.52031 | 6.36909 | 6.41015 | 6.2849 |
| FNBP4 | Q8N3X1-2 | 6.20006 | 6.55267 | 5.99394 | 6.49676 | 6.12152 | 6.13646 | 5.89917 | 6.89804 | 6.09689 | 6.30697 | 6.03744 | 6.23644 | 6.67407 | 6.17312 | 6.47659 | 6.50896 |
| PITPNM1 | O00562 | 6.13854 | 6.43841 | 6.19226 | 5.93965 | 6.25716 | 6.27741 | 5.8878 | 6.01654 | 6.16755 | 6.16196 | 5.63005 | 6.0049 | 6.96419 | 6.4668 | 7.00142 | 6.45536 |
| NQO1 | P15559 | 6.21177 | 6.08615 | 6.64743 | 5.59394 | 6.51546 | 6.85597 | 6.64718 | 6.95672 | 6.45799 | 6.50828 | 6.19489 | 5.9143 | 5.9312 | 6.20237 | 5.80009 | 5.47626 |
| NAXE | Q8NCW5 | 6.21428 | 4.81612 | 5.73189 | 4.94785 | 5.9084 | 5.28948 | 5.84728 | 6.76345 | 5.11101 | 6.41521 | 5.62891 | 6.79002 | 6.44063 | 8.02806 | 7.88094 | 8.18647 |
| RNF216 | Q9NWF9-1 | 5.83722 | 6.06466 | 6.85361 | 5.94142 | 5.90962 | 6.44459 | 6.30626 | 6.48325 | 6.31406 | 6.41846 | 6.92965 | 6.67271 | 5.81051 | 5.98266 | 5.72596 | 6.30537 |
| DHRS7B | Q6IAN0 | 6.27157 | 6.22889 | 5.78895 | 6.82742 | 6.33183 | 5.93102 | 6.5485 | 5.83912 | 6.12764 | 5.94622 | 5.92949 | 5.84782 | 6.65954 | 6.31418 | 6.80956 | 6.59825 |
| HPRT1 | P00492 | 6.25972 | 6.09292 | 6.95225 | 5.74741 | 6.56207 | 6.64081 | 6.26326 | 6.866 | 6.48177 | 6.42731 | 6.64635 | 6.61273 | 5.82419 | 5.65916 | 5.40814 | 5.55589 |
| NADK2 | Q4G0N4 | 6.49438 | 6.51044 | 5.76966 | 6.70473 | 6.25064 | 6.25368 | 6.1508 | 6.07183 | 6.24491 | 6.10956 | 5.76075 | 5.77691 | 6.42035 | 6.50136 | 6.53259 | 6.44741 |
| VPS13B | Q7Z7G8 | 6.70077 | 6.73939 | 6.76902 | 6.90419 | 6.17374 | 6.17824 | 6.60464 | 6.36858 | 6.78627 | 6.47464 | 6.44921 | 6.6596 | 5.24414 | 5.27226 | 5.1431 | 5.53223 |
| DNAJC1 | Q96KC8 | 6.31522 | 6.64934 | 6.15236 | 6.98546 | 6.24541 | 6.18566 | 6.45817 | 5.98717 | 6.03364 | 6.23725 | 6.12663 | 6.06315 | 5.99309 | 6.08536 | 6.289 | 6.19308 |
| PRR11 | Q96HE9 | 7.00199 | 7.08239 | 6.48214 | 7.33597 | 6.45116 | 6.14275 | 6.78739 | 6.40868 | 6.71591 | 6.58853 | 6.28282 | 6.535 | 4.76291 | 5.01203 | 4.85467 | 5.55567 |
| OAT | P04181 | 5.64441 | 6.54534 | 5.66325 | 6.30341 | 6.60039 | 6.31315 | 6.6866 | 6.21454 | 5.96945 | 6.02157 | 5.91272 | 6.00671 | 6.71342 | 6.43867 | 6.84681 | 7.01954 |
| ZYG11B | Q9C0D3 | 6.002 | 6.25376 | 6.41314 | 6.11939 | 6.01568 | 6.04413 | 6.15383 | 6.2811 | 6.09108 | 6.15036 | 6.16289 | 6.44925 | 6.19532 | 6.57313 | 6.53945 | 6.55551 |
| PDK2 | Q15119 | 6.67782 | 6.71222 | 5.95372 | 6.49942 | 6.44184 | 6.23958 | 6.3804 | 6.07288 | 6.26504 | 6.4831 | 6.48453 | 6.07121 | 6.16713 | 5.68614 | 5.92196 | 5.94303 |
| COG6 | Q9Y2V7 | 6.18994 | 6.02532 | 6.30434 | 6.51656 | 6.11631 | 6.42718 | 6.07386 | 6.33932 | 6.31676 | 6.61677 | 6.31722 | 6.17686 | 6.3033 | 6.4502 | 6.03432 | 5.79171 |
| CHMP1A | F5H875 | 5.98727 | 6.13143 | 6.87968 | 6.23544 | 6.09214 | 6.57018 | 6.2286 | 6.59428 | 6.61347 | 6.52241 | 6.57269 | 6.73328 | 5.42645 | 6.2037 | 5.39967 | 5.80931 |
| GNA13 | Q14344 | 6.63879 | 6.81047 | 6.01606 | 6.71033 | 6.35816 | 6.21682 | 6.5214 | 6.33336 | 6.33542 | 6.28508 | 5.91486 | 5.78755 | 5.889 | 6.02628 | 5.96781 | 6.18862 |
| GLE1 | Q53GS7 | 6.40967 | 6.22902 | 6.20097 | 6.51368 | 6.20975 | 6.25303 | 6.1563 | 6.28088 | 6.30646 | 6.35652 | 6.33706 | 6.16947 | 6.51279 | 6.03606 | 6.27917 | 6.14913 |
| JPT2 | Q9H910-3 | 6.45849 | 5.4717 | 7.20175 | 5.29821 | 6.79781 | 6.07308 | 5.44491 | 7.04351 | 6.54104 | 6.57982 | 7.00824 | 7.27282 | 6.11712 | 5.76658 | 5.25473 | 5.67018 |
| PRCP | P42785 | 6.40712 | 6.32706 | 6.1097 | 6.50625 | 6.81711 | 6.42646 | 6.02102 | 6.1711 | 6.39666 | 6.48206 | 6.20971 | 5.95519 | 6.26017 | 5.91444 | 5.89077 | 6.10519 |
| GTF2B | Q00403 | 5.83309 | 5.79955 | 5.94901 | 5.55169 | 5.96997 | 6.14912 | 5.80869 | 6.0079 | 6.14504 | 6.54865 | 6.31995 | 6.17788 | 6.64654 | 7.20575 | 7.03271 | 6.85446 |
| CMSS1 | Q9BQ75 | 6.08532 | 6.16313 | 5.95107 | 6.40377 | 6.53654 | 6.31241 | 6.14947 | 6.26212 | 6.1519 | 5.89737 | 5.70093 | 5.73303 | 6.72706 | 6.54148 | 6.78606 | 6.59836 |
| DCTN4 | Q9UJW0 | 6.02994 | 6.02532 | 6.9283 | 6.35332 | 6.01221 | 6.61279 | 6.30048 | 6.49736 | 6.42293 | 6.54127 | 6.73201 | 6.57718 | 5.58357 | 6.08483 | 5.79245 | 5.50604 |
| TBC1D1 | Q86TI0 | 6.58893 | 6.20748 | 6.98309 | 6.1984 | 6.30093 | 6.66631 | 6.45098 | 6.39456 | 6.64943 | 6.63729 | 6.60253 | 6.40304 | 5.60083 | 5.16666 | 5.53678 | 5.61276 |
| SAP18 | O00422-2 | 6.19762 | 6.27359 | 5.93674 | 6.39767 | 6.25743 | 6.13977 | 6.19634 | 6.20924 | 6.34946 | 6.00129 | 6.38551 | 6.51075 | 6.72003 | 5.91741 | 6.27088 | 6.23627 |
| GTF2F1 | P35269 | 5.4719 | 5.78669 | 6.25739 | 5.39826 | 5.84359 | 6.35029 | 5.89722 | 6.01496 | 5.94957 | 6.2774 | 6.70467 | 6.28492 | 6.84395 | 7.17013 | 7.02934 | 6.71971 |
| SPAST | Q9UBP0 | 6.26702 | 6.22843 | 6.88451 | 6.13548 | 5.8476 | 6.50346 | 6.24958 | 6.40523 | 6.3987 | 6.52952 | 6.49592 | 6.4157 | 5.68189 | 6.22149 | 5.84842 | 5.88703 |
| MYO19 | Q96H55 | 6.43749 | 6.58321 | 6.54831 | 6.90472 | 6.19861 | 6.18923 | 6.2855 | 6.2385 | 6.22747 | 6.05653 | 5.98401 | 5.79562 | 6.35107 | 5.99908 | 6.10026 | 6.10039 |
| CPNE1 | Q99829 | 6.31646 | 6.05778 | 6.71691 | 5.74418 | 6.20851 | 6.75368 | 6.06349 | 6.72351 | 6.3268 | 6.39923 | 6.63013 | 6.44655 | 5.95751 | 6.07027 | 5.96167 | 5.62333 |
| EARS2 | Q5JPH6-2 | 6.60959 | 6.48525 | 5.935 | 6.77146 | 6.62759 | 6.03607 | 6.30681 | 6.36791 | 6.20318 | 5.94936 | 5.86435 | 6.06135 | 6.69704 | 5.69342 | 6.14308 | 6.24854 |
| ORC2 | Q13416 | 5.71175 | 5.96191 | 6.2349 | 6.04712 | 6.52883 | 6.48708 | 6.49861 | 6.22411 | 6.22965 | 5.97759 | 6.55009 | 5.95608 | 6.53231 | 6.40908 | 6.48284 | 6.16805 |
| SPAG5 | Q96R06 | 6.81843 | 6.4376 | 6.81508 | 6.21806 | 6.42172 | 6.44494 | 6.31264 | 6.68099 | 6.72395 | 6.7508 | 7.16207 | 7.02453 | 4.95765 | 4.9946 | 4.96108 | 5.27586 |
| KPNA3 | O00505 | 5.84963 | 6.01928 | 6.139 | 5.87322 | 6.12785 | 6.25169 | 5.97998 | 6.05198 | 6.1475 | 6.25821 | 6.16964 | 6.4934 | 6.36369 | 6.49929 | 6.78907 | 6.98658 |
| RPUSD3 | Q6P087 | 6.44495 | 6.39513 | 5.90369 | 6.75277 | 6.03168 | 6.30795 | 6.35589 | 6.2381 | 6.15431 | 6.39262 | 5.88747 | 6.48046 | 6.11118 | 6.13049 | 6.07501 | 6.33832 |
| PLD1 | Q13393 | 6.42028 | 6.65669 | 6.32047 | 6.58988 | 5.97003 | 6.24856 | 6.22132 | 6.03485 | 6.32787 | 6.60543 | 6.43471 | 6.30913 | 5.98795 | 6.02254 | 6.04301 | 5.8073 |
| CDKN2AIP | Q9NXV6 | 6.11771 | 6.0627 | 5.79682 | 6.35569 | 6.29139 | 5.95077 | 6.2737 | 5.898 | 6.29015 | 6.06239 | 6.05184 | 5.80093 | 7.1401 | 6.48052 | 6.72242 | 6.70487 |
| PCDHGA3 | Q9Y5H0 | 6.34509 | 6.41289 | 5.93327 | 6.47121 | 6.3921 | 6.30674 | 6.38082 | 6.09962 | 6.14114 | 6.37218 | 6.18756 | 6.2974 | 6.23822 | 6.20503 | 6.06829 | 6.14845 |
| USP33 | Q8TEY7 | 6.96327 | 6.99362 | 6.65145 | 7.21726 | 6.23834 | 6.17552 | 6.68907 | 6.23663 | 6.76427 | 7.13334 | 6.68405 | 6.76981 | 4.57743 | 4.55583 | 4.96017 | 5.38994 |
| EIF6 | P56537 | 6.0531 | 6.15613 | 6.23943 | 5.83928 | 6.28183 | 6.48135 | 6.12124 | 6.58028 | 6.54837 | 6.34051 | 6.45475 | 6.54082 | 6.31531 | 6.08575 | 5.95221 | 6.00964 |
| WDR48 | Q8TAF3 | 5.97366 | 6.04157 | 5.99462 | 6.2896 | 6.25952 | 6.26972 | 6.00496 | 6.13726 | 6.30515 | 6.22469 | 6.17409 | 6.29785 | 6.41106 | 6.48838 | 6.52248 | 6.6054 |
| FAM207A | Q9NSI2 | 6.27904 | 6.28944 | 6.18623 | 6.74583 | 6.32447 | 6.14546 | 6.86633 | 6.24992 | 6.40068 | 6.19063 | 6.21712 | 6.1253 | 6.07239 | 5.75581 | 6.09932 | 6.05205 |
| CHP1 | Q99653 | 6.40764 | 5.48219 | 5.7366 | 5.53894 | 6.40947 | 5.83556 | 6.22978 | 6.47502 | 5.76566 | 6.68111 | 6.06366 | 6.51511 | 6.44453 | 6.93882 | 6.86279 | 6.61311 |
| ICAM1 | P05362 | 6.01272 | 5.94322 | 5.68128 | 6.34386 | 6.26597 | 6.1778 | 6.19536 | 5.558 | 5.64844 | 5.94345 | 5.88106 | 5.87022 | 7.37442 | 6.75267 | 6.94785 | 7.40367 |
| QTRT1 | Q9BXR0 | 6.40679 | 6.26792 | 6.41557 | 6.0085 | 6.23115 | 6.48727 | 6.20745 | 6.50572 | 6.53527 | 6.41982 | 6.3499 | 6.53161 | 5.94029 | 5.96336 | 5.92854 | 5.80085 |
| CEMIP2 | Q9UHN6 | 6.26169 | 6.5214 | 6.34766 | 6.44933 | 6.25451 | 5.875 | 6.47432 | 6.15609 | 6.26558 | 6.48057 | 5.91564 | 6.05555 | 6.31107 | 6.12163 | 6.42544 | 6.08452 |
| FASTKD5 | Q7L8L6 | 6.61147 | 6.53671 | 5.71846 | 6.88654 | 6.51577 | 6.06009 | 6.44991 | 5.90991 | 6.16241 | 6.1318 | 5.8426 | 5.94021 | 6.41787 | 6.1392 | 6.42409 | 6.25297 |
| ELMOD2 | Q8IZ81 | 6.2944 | 6.57919 | 5.79507 | 6.6182 | 5.93987 | 6.07288 | 6.47247 | 5.94003 | 5.86027 | 6.03404 | 6.05257 | 5.99076 | 6.56265 | 6.33754 | 6.82797 | 6.6221 |
| RAB34 | Q9BZG1 | 6.28656 | 6.55426 | 5.58677 | 6.3403 | 6.12314 | 6.36775 | 6.1172 | 6.51068 | 6.1436 | 6.28781 | 5.73809 | 6.08161 | 6.36323 | 6.53566 | 6.41005 | 6.55328 |
| AFAP1 | Q8N556-2 | 6.11938 | 6.33814 | 5.91008 | 6.54601 | 5.95666 | 6.13492 | 6.22388 | 5.89344 | 6.19738 | 6.13701 | 5.96399 | 6.10761 | 6.65237 | 6.17238 | 6.7039 | 6.94287 |
| CERT1 | Q9Y5P4-3 | 6.26748 | 6.40713 | 6.47349 | 6.11661 | 6.14545 | 6.83642 | 6.19782 | 6.98236 | 6.21627 | 6.49506 | 6.3227 | 6.54406 | 5.92961 | 6.16793 | 5.40896 | 5.48866 |
| KDSR | Q06136 | 6.24683 | 6.58954 | 6.05211 | 6.37648 | 6.18458 | 6.12473 | 6.7149 | 5.34407 | 6.13453 | 5.81397 | 5.67642 | 5.6474 | 6.80305 | 6.20427 | 7.33695 | 6.75018 |
| GK | P32189 | 6.40017 | 6.21457 | 6.19161 | 6.42294 | 6.21036 | 5.82974 | 6.42625 | 5.94911 | 6.25524 | 5.94391 | 6.39066 | 6.10992 | 6.54969 | 6.25624 | 6.65798 | 6.19161 |
| MTO1 | Q9Y2Z2-6 | 6.87193 | 7.08608 | 6.56803 | 7.48313 | 6.15921 | 6.17612 | 6.08009 | 5.7549 | 6.98639 | 6.56836 | 6.62302 | 6.55958 | 5.18092 | 5.12637 | 5.29828 | 5.47759 |
| AKR1B1 | P15121 | 5.40934 | 5.87888 | 6.34453 | 5.68835 | 6.17994 | 6.81057 | 6.40357 | 6.91538 | 6.41354 | 6.67802 | 6.6152 | 6.68902 | 5.91045 | 5.96904 | 6.22412 | 5.87006 |
| USP3 | Q9Y6I4 | 5.84548 | 6.38212 | 6.54003 | 6.13873 | 6.20459 | 6.05126 | 6.25365 | 5.67934 | 6.21468 | 5.90286 | 6.49404 | 6.4116 | 6.2938 | 6.30198 | 6.53708 | 6.74876 |
| SUMF2 | Q8NBJ7 | 6.10274 | 5.52962 | 5.95788 | 6.0544 | 6.77515 | 5.95722 | 6.07232 | 6.65929 | 6.10578 | 6.53357 | 6.10869 | 6.52412 | 6.59633 | 6.06861 | 6.4527 | 6.50157 |
| LIMS1 | P48059-3 | 6.0608 | 6.2221 | 6.68384 | 5.94973 | 5.98781 | 6.72686 | 6.07722 | 6.17415 | 6.26976 | 6.69766 | 6.13876 | 6.44438 | 5.98404 | 6.28893 | 6.15363 | 6.14032 |
| RTN1 | Q16799 | 6.34679 | 6.6505 | 6.02671 | 6.46934 | 6.41269 | 6.48133 | 6.58477 | 6.10336 | 6.45038 | 6.38099 | 6.03972 | 5.6983 | 6.08048 | 5.96018 | 6.42805 | 5.88641 |
| TRIM32 | Q13049 | 6.27246 | 6.11226 | 6.62519 | 6.24915 | 6.10338 | 6.25141 | 6.36108 | 6.22403 | 6.34865 | 6.37765 | 6.50819 | 6.41497 | 5.83609 | 6.23081 | 6.28336 | 5.80132 |
| TP53BP2 | Q13625-3 | 6.04707 | 6.579 | 6.47495 | 6.45394 | 6.4131 | 6.37743 | 6.15423 | 6.27177 | 6.30596 | 6.26351 | 6.49004 | 6.53594 | 6.06248 | 5.96229 | 5.81931 | 5.78898 |
| RLF | Q13129 | 6.28281 | 6.72938 | 6.0453 | 7.18672 | 6.06952 | 6.14843 | 6.55924 | 6.25294 | 6.15675 | 6.4303 | 6.03421 | 5.87415 | 6.12674 | 5.95447 | 6.21164 | 5.93739 |
| TMED9 | Q9BVK6 | 6.25043 | 5.79864 | 5.96641 | 6.16234 | 5.87221 | 6.10283 | 6.39237 | 6.40145 | 5.72182 | 6.01998 | 6.09749 | 6.25125 | 6.77663 | 6.60226 | 6.63734 | 6.94657 |
| CCNH | P51946 | 5.92588 | 6.17352 | 6.35962 | 6.19421 | 6.17983 | 6.28393 | 6.07207 | 6.06575 | 6.06952 | 6.55332 | 6.54812 | 6.39044 | 6.32926 | 6.2732 | 6.38466 | 6.19667 |
| NEDD1 | Q8NHV4-3 | 6.08819 | 6.24262 | 6.55784 | 5.92036 | 6.34558 | 6.57197 | 6.0901 | 6.40912 | 6.4313 | 6.44322 | 6.42236 | 6.31503 | 5.98499 | 6.24044 | 6.18384 | 5.75305 |
| PYCR1 | P32322-3 | 6.49744 | 6.34126 | 5.78216 | 6.51106 | 6.63789 | 6.33647 | 6.53985 | 5.93345 | 5.83009 | 6.05351 | 6.19964 | 5.92975 | 6.57749 | 6.06734 | 6.4608 | 6.3018 |
| HAT1 | O14929 | 5.54964 | 5.74606 | 6.24906 | 5.50924 | 5.81398 | 6.19488 | 6.18658 | 5.73007 | 5.68854 | 6.20341 | 6.27788 | 5.92183 | 7.17904 | 7.16916 | 7.6687 | 6.91194 |
| SCYL1 | Q96KG9 | 5.90746 | 6.41208 | 6.62911 | 5.91053 | 5.93312 | 6.49469 | 6.17502 | 6.46881 | 6.22815 | 6.40523 | 6.71384 | 6.88141 | 5.85985 | 6.05834 | 6.0137 | 5.90867 |
| SEMA4B | Q9NPR2 | 6.14298 | 6.22934 | 5.83185 | 6.27961 | 6.07409 | 6.3498 | 6.25503 | 6.47587 | 5.99847 | 6.63312 | 6.28445 | 6.32546 | 6.0951 | 6.36834 | 6.13174 | 6.52475 |

| | | | | | | | | | | | | | | | | | |
|---|---|---|---|---|---|---|---|---|---|---|---|---|---|---|---|---|---|
| CBX3 | Q13185 | 6.41702 | 5.87388 | 6.27606 | 6.04935 | 6.36988 | 5.91129 | 6.28051 | 5.80082 | 6.55482 | 6.02094 | 6.56374 | 5.83548 | 7.00686 | 5.7345 | 6.78876 | 6.51609 |
| TGS1 | Q96RS0 | 6.24875 | 6.20412 | 6.25126 | 6.0569 | 6.11297 | 6.35487 | 6.11786 | 6.16423 | 6.35287 | 6.09229 | 6.10668 | 6.24013 | 6.2045 | 6.28371 | 6.54606 | 6.66279 |
| WLS | Q5T9L3-2 | 6.80583 | 7.07524 | 6.68283 | 6.99208 | 6.19236 | 5.93662 | 6.69926 | 6.08878 | 6.42311 | 6.35259 | 6.08983 | 6.19914 | 5.39259 | 5.60962 | 5.76536 | 5.69476 |
| VAPB | O95292 | 6.41764 | 5.2223 | 6.03793 | 5.63417 | 6.23787 | 5.18325 | 5.77607 | 5.97809 | 5.74965 | 6.39508 | 5.90209 | 6.71552 | 6.98054 | 6.8751 | 7.22714 | 7.66758 |
| TIPRL | O75663 | 6.08731 | 6.03423 | 6.89976 | 6.16592 | 6.13901 | 7.01411 | 6.18414 | 6.85406 | 6.23246 | 6.63649 | 6.56753 | 6.9189 | 5.32245 | 5.7444 | 5.44255 | 5.75666 |
| NT5DC3 | Q86UY8 | 6.64012 | 6.63014 | 5.89981 | 6.44374 | 6.25053 | 6.09366 | 6.38436 | 5.93207 | 6.1843 | 6.04137 | 5.81353 | 5.7571 | 6.69033 | 5.9007 | 6.61434 | 6.7239 |
| GLA | P06280 | 6.65004 | 6.30853 | 6.16495 | 6.34449 | 6.72026 | 6.34496 | 6.43695 | 6.16838 | 6.36084 | 6.13916 | 6.17387 | 5.93755 | 6.61587 | 5.74572 | 6.05746 | 5.83096 |
| NIPSNAP1 | Q9BPW8 | 6.22405 | 6.80848 | 5.1434 | 6.84008 | 6.80132 | 6.9344 | 6.00165 | 6.96875 | 5.85207 | 5.80082 | 5.90623 | 5.66886 | 6.16197 | 6.43916 | 6.59392 | 5.85482 |
| BARD1 | Q99728 | 6.44107 | 6.63278 | 5.8988 | 6.82818 | 6.42793 | 6.0263 | 6.39836 | 5.96796 | 6.39233 | 6.1378 | 5.90489 | 5.8705 | 6.33153 | 6.2085 | 6.3654 | 6.16769 |
| ARF4 | P18085 | 6.11408 | 6.16385 | 6.55668 | 6.21709 | 5.90868 | 6.32714 | 6.48794 | 6.16374 | 6.4355 | 6.58556 | 6.29902 | 6.61561 | 5.87255 | 6.07645 | 6.15023 | 6.02589 |
| NUDT9 | Q9BW91 | 6.35695 | 6.57437 | 6.01335 | 6.40201 | 6.25994 | 6.45058 | 5.90252 | 6.65615 | 6.11963 | 6.2781 | 6.19845 | 6.36434 | 6.50748 | 6.05215 | 5.776 | 6.08796 |
| ACBD3 | Q9H3P7 | 6.29977 | 6.31065 | 5.81294 | 6.32478 | 6.33741 | 6.29536 | 5.79603 | 6.02611 | 5.84452 | 6.459 | 6.59186 | 6.23609 | 6.91579 | 6.64692 | 6.22809 | 5.87467 |
| PREPL | Q4J6C6 | 6.39999 | 6.40466 | 6.77021 | 6.42148 | 6.52846 | 6.71942 | 6.53817 | 6.61572 | 6.71849 | 6.70944 | 6.68585 | 6.36737 | 5.45718 | 5.09682 | 5.40267 | 5.16407 |
| BRIP1 | Q9BX63 | 6.72978 | 7.03676 | 6.77028 | 6.72382 | 6.27473 | 5.923 | 6.04776 | 6.28636 | 6.66632 | 6.74511 | 6.38078 | 6.81806 | 5.69535 | 5.18936 | 5.3526 | 5.35992 |
| LSM14A | Q8ND56-2 | 6.32933 | 5.97166 | 6.55175 | 6.06561 | 6.30432 | 6.34438 | 6.22334 | 6.33775 | 6.87591 | 6.6671 | 6.74444 | 6.51528 | 5.69341 | 5.9664 | 5.57075 | 5.83856 |
| TMX1 | Q9H3N1 | 6.47099 | 6.44936 | 5.89553 | 6.3379 | 6.44451 | 5.84563 | 6.58014 | 5.93367 | 6.14304 | 5.91498 | 6.05576 | 5.54157 | 6.72203 | 6.43035 | 6.52304 | 6.7115 |
| PPIH | O43447 | 5.77083 | 5.45577 | 6.11453 | 5.56559 | 6.52235 | 6.24031 | 5.86487 | 6.66401 | 6.07812 | 6.44612 | 6.49474 | 6.90924 | 6.30779 | 6.54545 | 6.46416 | 6.55613 |
| MFAP1 | P55081 | 6.50904 | 6.52998 | 6.1108 | 6.81962 | 6.23842 | 5.99801 | 6.31137 | 5.81923 | 6.09362 | 5.9541 | 5.80487 | 6.13787 | 6.55623 | 6.25516 | 6.49609 | 6.36557 |
| SHC1 | P29353-6 | 6.59908 | 6.28253 | 6.73038 | 6.32931 | 6.19771 | 6.74195 | 6.08899 | 6.33147 | 6.3561 | 6.22199 | 6.54782 | 6.73506 | 5.76957 | 5.66822 | 5.81871 | 5.5811 |
| NSMCE3 | Q96MG7 | 5.99371 | 6.02537 | 5.9869 | 6.49958 | 6.29791 | 5.96964 | 6.15033 | 5.97191 | 6.45182 | 6.14802 | 6.29933 | 6.36501 | 6.69407 | 6.16924 | 6.61807 | 6.35911 |
| KIRREL1 | Q96J84-2 | 6.48289 | 6.27053 | 6.07038 | 6.52431 | 6.21309 | 6.09726 | 6.5864 | 6.01998 | 6.32253 | 5.99043 | 6.03863 | 6.08343 | 6.61494 | 6.26562 | 6.17622 | 6.24337 |
| CCNT1 | O60563 | 6.03958 | 6.37311 | 5.93728 | 6.33723 | 5.83936 | 6.17385 | 6.58957 | 5.7599 | 6.0759 | 6.12572 | 5.87514 | 5.99087 | 6.29315 | 6.80665 | 6.92096 | 6.86173 |
| STK38 | Q15208 | 6.16535 | 6.43142 | 6.95834 | 6.24273 | 6.13089 | 6.42903 | 6.22319 | 6.74028 | 6.4217 | 6.46486 | 6.79385 | 6.65541 | 5.3496 | 5.74938 | 5.71772 | 5.52625 |
| CHEK1 | O14757 | 6.59852 | 6.18228 | 7.31837 | 6.36383 | 6.18794 | 6.59333 | 6.60942 | 6.66059 | 6.59238 | 6.60468 | 6.67501 | 6.8607 | 5.10778 | 5.35987 | 5.05212 | 5.23318 |
| MINPP1 | Q9UNW1 | 6.1129 | 6.23768 | 6.11111 | 6.59077 | 6.04573 | 6.05473 | 6.20243 | 5.50559 | 6.12185 | 6.2436 | 6.21994 | 6.09345 | 6.71958 | 6.34482 | 6.5464 | 6.84943 |
| VPS26B | Q4G0F5 | 5.89722 | 5.97572 | 6.34407 | 6.20645 | 6.2394 | 6.51864 | 5.86323 | 5.85628 | 6.43483 | 6.42973 | 6.79113 | 6.35347 | 6.50285 | 6.36218 | 6.15549 | 6.0693 |
| RTCA | O00442-2 | 5.99629 | 6.07979 | 6.55793 | 5.8499 | 5.96786 | 6.48215 | 6.20547 | 6.4906 | 6.17054 | 6.29393 | 6.24952 | 6.64089 | 5.93956 | 6.75821 | 6.29063 | 6.0267 |
| SSU72 | Q9NP77 | 5.82972 | 5.84548 | 6.41617 | 5.92555 | 6.06115 | 6.26657 | 5.84194 | 6.19804 | 6.16683 | 6.44456 | 6.51127 | 6.43597 | 6.49006 | 6.39135 | 6.32091 | 6.85444 |
| PRPF39 | Q86UA1 | 5.39642 | 5.93438 | 6.25249 | 5.45148 | 6.0614 | 6.06081 | 5.71758 | 5.90279 | 5.92294 | 6.09552 | 6.02194 | 5.58825 | 7.76285 | 6.85508 | 7.85585 | 7.1202 |
| GOLM1 | Q8NBJ4 | 6.74399 | 6.33873 | 6.2762 | 6.42885 | 6.83049 | 6.30976 | 6.62836 | 6.31246 | 6.33256 | 6.37931 | 6.60001 | 6.35084 | 5.85607 | 5.59001 | 5.42561 | 5.59675 |
| STXBP2 | Q15833-3 | 6.38549 | 6.6695 | 6.34881 | 6.10926 | 6.0811 | 6.62572 | 5.89005 | 6.23947 | 6.17516 | 6.52675 | 6.29539 | 6.04036 | 6.18853 | 6.0324 | 6.44481 | 5.94718 |
| PPME1 | Q9Y570-4 | 6.11888 | 5.91296 | 7.06197 | 6.0406 | 6.34872 | 6.45954 | 6.15413 | 6.57295 | 6.15845 | 6.19527 | 6.69508 | 6.41987 | 5.91767 | 6.0081 | 6.16504 | 5.77077 |
| CDK7 | P50613 | 5.77395 | 6.1604 | 6.36841 | 6.01874 | 5.9609 | 6.40643 | 6.15933 | 6.00705 | 6.11202 | 6.31352 | 6.20861 | 6.22605 | 6.52279 | 6.36187 | 6.63027 | 6.76965 |
| STXBP5 | Q5T5C0 | 6.06061 | 6.23245 | 6.44468 | 5.84659 | 6.08127 | 6.87171 | 6.38581 | 6.42847 | 6.36326 | 6.39574 | 6.65112 | 6.29406 | 6.15825 | 6.22537 | 5.97473 | 5.58587 |
| UQCRFS1 | P47985 | 6.94815 | 6.28032 | 6.05233 | 6.21887 | 6.25329 | 5.78168 | 6.1881 | 5.77432 | 5.65968 | 5.90203 | 5.87482 | 5.66523 | 6.12902 | 7.19285 | 6.82836 | 7.25096 |
| BLVRA | P53004 | 6.32474 | 6.11614 | 6.70145 | 5.78392 | 6.40708 | 6.85675 | 6.42362 | 6.75514 | 6.23804 | 6.38079 | 6.44322 | 6.42377 | 5.76284 | 6.06027 | 5.70517 | 5.61705 |
| MANF | P55145 | 5.97849 | 5.45994 | 6.14911 | 6.29548 | 6.57618 | 6.55164 | 6.2407 | 6.27392 | 6.60344 | 5.9006 | 6.39299 | 6.19398 | 7.18782 | 6.38057 | 5.7508 | 6.06436 |
| NUP43 | Q8NFH3 | 5.83918 | 5.98074 | 5.8784 | 5.98652 | 6.73537 | 6.25161 | 6.37983 | 7.15735 | 6.63529 | 6.08083 | 6.46047 | 6.85422 | 5.82182 | 5.8598 | 6.04398 | 6.03459 |
| HPF1 | Q9NWY4 | 6.08042 | 6.22545 | 6.86326 | 5.67394 | 6.03503 | 6.32477 | 5.95899 | 6.20267 | 6.02133 | 6.30414 | 6.45242 | 6.32071 | 6.10052 | 6.79639 | 6.50601 | 6.13395 |
| LMTK2 | Q8IWU2 | 6.17373 | 6.04572 | 6.13204 | 6.18224 | 6.04165 | 6.01134 | 5.93263 | 5.74222 | 6.37024 | 6.4501 | 6.37355 | 6.44217 | 6.65968 | 6.29908 | 6.67888 | 6.46474 |
| NOA1 | Q8NC60 | 6.45891 | 6.5237 | 6.42238 | 7.08469 | 6.26438 | 5.60605 | 6.78763 | 6.00636 | 6.61494 | 5.78915 | 6.43882 | 6.36134 | 6.07934 | 5.63975 | 5.93631 | 5.98625 |
| KBTBD6 | Q86V97 | 5.55165 | 6.01457 | 6.03338 | 5.73318 | 5.94349 | 6.14159 | 5.97615 | 6.12295 | 6.08291 | 6.51555 | 6.4225 | 5.96859 | 6.82501 | 6.97227 | 7.11939 | 6.57684 |
| RTTN | Q86VV8 | 6.25589 | 6.03336 | 6.41972 | 6.36517 | 6.17935 | 6.79632 | 6.51345 | 6.5128 | 6.68058 | 6.25297 | 6.18957 | 6.44067 | 6.24103 | 6.02051 | 5.49148 | 5.60715 |
| FIS1 | Q9Y3D6 | 6.13837 | 5.72888 | 5.89276 | 5.7487 | 6.33337 | 5.53119 | 5.83103 | 6.3078 | 5.57909 | 6.33621 | 5.94305 | 6.51688 | 6.74554 | 6.9975 | 7.12926 | 7.24036 |
| SRSF11 | Q05519 | 6.15799 | 6.46162 | 6.02993 | 6.51091 | 6.17508 | 5.95459 | 6.19922 | 5.7878 | 5.98075 | 6.2757 | 6.30954 | 5.91474 | 6.31984 | 6.35198 | 6.77082 | 6.79949 |
| TRIM4 | Q9C037 | 6.55617 | 6.70566 | 6.43571 | 6.77972 | 6.20537 | 6.09619 | 6.44095 | 6.07436 | 6.24712 | 5.89237 | 6.00168 | 5.87333 | 6.34438 | 5.96966 | 6.3801 | 5.99723 |
| NDUFAF2 | Q8N183 | 6.617 | 6.32004 | 6.08587 | 6.69695 | 6.26363 | 5.97626 | 6.19259 | 6.30259 | 6.4002 | 6.15339 | 6.05207 | 6.11023 | 6.78627 | 5.93048 | 5.99707 | 6.11535 |
| QRICH1 | Q2TAL8 | 5.51926 | 5.7699 | 5.87883 | 5.14927 | 5.71032 | 6.08781 | 5.34531 | 6.18378 | 5.9696 | 6.38933 | 6.14675 | 6.21507 | 7.40079 | 7.18116 | 7.68097 | 7.37186 |
| MRPL28 | Q13084 | 6.45316 | 6.45118 | 5.9983 | 6.76321 | 6.54626 | 6.33535 | 6.31087 | 5.9643 | 6.12859 | 6.44377 | 6.0652 | 6.35156 | 6.14904 | 5.98833 | 5.99501 | 6.05586 |
| SEC11A | P67812-3 | 6.41566 | 6.37615 | 5.69876 | 6.29899 | 6.25774 | 6.10629 | 6.47445 | 5.8911 | 6.03839 | 5.99258 | 6.00442 | 6.10005 | 6.82845 | 6.42121 | 6.50572 | 6.59004 |
| ATP5ME | P56385 | 6.13947 | 5.62633 | 5.76538 | 5.7405 | 6.66079 | 5.79451 | 6.53671 | 6.40756 | 5.63637 | 5.97209 | 5.91504 | 6.11084 | 6.55379 | 6.88612 | 6.82676 | 7.42775 |
| UBLCP1 | Q8WVY7 | 6.16333 | 6.05374 | 6.33386 | 5.52641 | 6.112 | 6.69978 | 5.99626 | 6.32146 | 6.05455 | 6.45272 | 5.98391 | 6.24266 | 6.30005 | 6.94861 | 6.61853 | 6.19214 |
| LIAS | O43766 | 6.55476 | 6.22494 | 6.10291 | 7.1699 | 6.17955 | 6.08486 | 6.47035 | 6.60351 | 6.19347 | 5.81709 | 6.16483 | 6.37129 | 6.17489 | 5.7548 | 6.14946 | 5.98338 |
| BABAM2 | Q9NXR7 | 6.14639 | 5.95996 | 6.7466 | 6.22245 | 6.49634 | 6.6676 | 6.02747 | 6.32299 | 6.23958 | 6.31908 | 6.61445 | 6.27487 | 6.09728 | 5.91024 | 5.81697 | 6.13771 |
| MRPS2 | Q9Y399 | 6.54479 | 6.33498 | 6.06445 | 6.49578 | 6.18322 | 6.14899 | 6.45026 | 6.16295 | 6.31222 | 5.75397 | 6.08024 | 6.06496 | 6.48525 | 6.12967 | 6.55869 | 6.2296 |
| SPCS2 | Q15005 | 6.44039 | 6.79015 | 5.6432 | 6.66186 | 6.51843 | 6.40476 | 6.36267 | 6.38746 | 6.06339 | 5.95934 | 5.70653 | 5.92292 | 6.46212 | 6.21516 | 6.27811 | 6.18349 |
| BCAS3 | Q9H6U6-8 | 6.12223 | 6.01614 | 6.69654 | 6.00837 | 6.36009 | 6.71167 | 6.34755 | 6.55676 | 6.35053 | 6.54667 | 6.55294 | 6.18364 | 6.00662 | 5.99739 | 6.10853 | 5.43431 |
| IRAK4 | Q9NWZ3 | 6.00559 | 6.15063 | 6.60908 | 5.90672 | 6.20897 | 6.89266 | 5.99029 | 6.5247 | 6.38125 | 6.78389 | 6.63419 | 6.55751 | 5.90863 | 5.86575 | 5.89181 | 5.68832 |
| ATPAF2 | Q8N5M1 | 6.81301 | 6.56626 | 5.64092 | 6.82029 | 6.2449 | 6.15273 | 6.34082 | 6.07472 | 5.93831 | 6.18553 | 5.67398 | 5.73169 | 6.56706 | 6.37528 | 6.57906 | 6.29545 |
| GOT1 | P17174 | 6.18722 | 5.88638 | 6.80652 | 5.75634 | 6.34325 | 6.93287 | 6.06318 | 6.68022 | 6.57343 | 6.27186 | 6.48465 | 6.36095 | 5.92395 | 6.13633 | 5.75638 | 5.83648 |
| ACOT13 | Q9NPJ3 | 6.05878 | 6.46024 | 5.85089 | 6.63824 | 6.22563 | 6.16717 | 6.29029 | 5.97701 | 5.88552 | 6.4528 | 6.246 | 5.93213 | 6.68208 | 6.00804 | 6.6771 | 6.44809 |
| DIS3L | Q8TF46 | 6.44218 | 6.31052 | 7.05973 | 6.05576 | 6.19355 | 6.52795 | 5.96482 | 6.43798 | 6.65379 | 6.61071 | 6.53596 | 6.66096 | 5.64757 | 5.62904 | 5.5748 | 5.69468 |
| SNRNP27 | Q8WVK2 | 6.33984 | 6.3699 | 5.87864 | 6.1917 | 5.92025 | 6.0529 | 6.39165 | 6.24509 | 5.86921 | 5.74128 | 5.72726 | 5.66954 | 6.83948 | 6.73217 | 7.01763 | 7.01347 |
| RPP30 | P78346-2 | 5.96639 | 6.16627 | 6.02661 | 6.30911 | 6.16034 | 6.02423 | 6.13344 | 5.89943 | 6.33111 | 6.45684 | 6.09622 | 5.95368 | 6.86298 | 6.18244 | 6.72789 | 6.70302 |
| USP32 | Q8NFA0 | 5.88352 | 5.93187 | 6.58706 | 5.60871 | 5.8409 | 6.47274 | 6.13449 | 6.30348 | 6.41836 | 6.59721 | 6.69547 | 6.47019 | 6.36107 | 6.37579 | 6.11597 | 6.20317 |
| NDUFA12 | Q9UI09 | 6.1072 | 6.57467 | 5.77417 | 6.52631 | 6.22131 | 5.90053 | 6.52828 | 6.17163 | 6.12832 | 5.94518 | 5.90935 | 6.05343 | 6.96892 | 6.48753 | 6.36715 | 6.33602 |
| PRRC1 | Q96M27-2 | 6.07263 | 5.83928 | 5.91479 | 5.44934 | 6.13149 | 6.43641 | 5.90819 | 6.09022 | 6.17541 | 6.29882 | 6.45493 | 6.38422 | 7.02763 | 6.50528 | 6.81092 | 6.50045 |
| RPL7L1 | Q6DKI1 | 6.41334 | 6.33744 | 5.58468 | 6.74876 | 5.83879 | 5.83975 | 6.49326 | 6.23075 | 6.25967 | 6.10009 | 5.93724 | 6.2131 | 6.54321 | 6.26248 | 6.37392 | 6.82352 |
| MARCHF5 | Q9NX47 | 6.39949 | 6.51322 | 6.11704 | 7.10882 | 6.28125 | 6.1402 | 6.41578 | 6.09092 | 6.38983 | 6.26808 | 6.0838 | 5.99346 | 6.14684 | 5.84031 | 6.12681 | 6.08414 |
| NIP7 | Q9Y221 | 6.3883 | 6.53529 | 6.14314 | 6.94287 | 6.35776 | 6.04987 | 6.26935 | 6.01564 | 5.59183 | 6.00323 | 6.39103 | 6.22895 | 6.45447 | 6.05092 | 6.4257 | 6.15166 |
| RPL27A | P46776 | 6.10981 | 6.1409 | 6.57444 | 5.7278 | 5.7355 | 5.99188 | 6.55301 | 6.0308 | 6.6456 | 6.29531 | 6.79594 | 6.35661 | 6.27939 | 6.10698 | 6.0623 | 6.59374 |
| RBM7 | Q9Y580 | 6.06772 | 6.47666 | 5.80574 | 6.3538 | 6.63374 | 6.01153 | 6.10269 | 6.31331 | 6.37718 | 6.22014 | 6.01981 | 5.9466 | 6.6998 | 6.27392 | 6.5216 | 6.17575 |
| MRPS15 | P82914 | 6.27757 | 6.40901 | 5.67606 | 6.88144 | 6.10095 | 5.95708 | 6.24954 | 6.07566 | 6.04596 | 6.12985 | 5.90337 | 5.99046 | 6.57399 | 6.41825 | 6.64419 | 6.66662 |
| ANKRD11 | Q6UB99 | 6.2339 | 6.08067 | 5.9771 | 6.81018 | 6.09804 | 5.82185 | 6.52728 | 6.1007 | 6.11595 | 6.35712 | 6.09016 | 6.09785 | 5.87618 | 6.65316 | 6.42563 | 6.73424 |
| NSD3 | Q9BZ95 | 6.26205 | 6.61218 | 6.10188 | 6.86319 | 5.98191 | 6.01064 | 6.38153 | 6.10796 | 6.17388 | 6.2413 | 5.7876 | 6.28851 | 5.99466 | 6.72975 | 6.1355 | 6.32747 |
| WBP2 | Q969T9 | 6.39619 | 5.84544 | 7.14191 | 6.17731 | 6.42729 | 6.48038 | 6.02918 | 6.75401 | 6.94243 | 6.39899 | 7.06194 | 6.91166 | 5.61362 | 5.56556 | 5.24368 | 5.01039 |
| EHBP1 | Q8NDI1 | 5.7267 | 6.09401 | 6.42865 | 5.84838 | 6.04163 | 6.27428 | 6.23383 | 6.34329 | 6.39823 | 6.55193 | 6.48644 | 6.22683 | 6.45861 | 6.27552 | 6.28036 | 6.33133 |
| DFFA | O00273 | 6.17183 | 5.86483 | 6.88621 | 5.67733 | 6.16722 | 6.3622 | 5.92792 | 6.80986 | 6.39583 | 6.32129 | 6.55479 | 6.71408 | 6.3975 | 6.03354 | 5.97979 | 5.73579 |
| PCGF2 | P35227 | 6.48517 | 6.77651 | 6.06321 | 6.39137 | 6.20734 | 6.13081 | 6.0991 | 5.8943 | 6.48158 | 6.34073 | 6.26866 | 6.04689 | 6.43274 | 6.0129 | 6.13611 | 6.23256 |
| TSPYL5 | Q86VY4 | 6.08054 | 6.36862 | 6.34402 | 5.81084 | 6.06407 | 6.14179 | 6.09916 | 6.25714 | 6.33466 | 6.51876 | 6.58746 | 6.58792 | 5.96629 | 6.3109 | 6.25422 | 6.27359 |
| FGFR1 | P11362-21 | 6.40638 | 6.58774 | 6.55378 | 6.67908 | 6.5469 | 6.13767 | 6.15542 | 6.34639 | 6.43671 | 6.38977 | 6.52721 | 6.29093 | 5.83294 | 5.78725 | 5.65455 | 5.66727 |
| DUT | P33316 | 6.73764 | 6.1202 | 7.95637 | 6.31849 | 6.57989 | 6.42123 | 5.77076 | 6.27776 | 6.32261 | 6.3057 | 6.66327 | 6.76831 | 5.76417 | 5.45117 | 5.11419 | 5.42821 |
| BUD13 | Q9BRD0 | 6.34709 | 6.42237 | 5.86338 | 6.82432 | 6.09654 | 5.84462 | 6.15925 | 6.13966 | 6.23397 | 6.03886 | 6.17684 | 5.9756 | 6.55969 | 6.24818 | 6.40531 | 6.66432 |
| CNOT3 | O75175 | 5.95552 | 6.12898 | 6.33049 | 5.94539 | 5.86973 | 6.2326 | 6.22908 | 6.50833 | 5.73785 | 6.10275 | 6.36944 | 6.2253 | 6.4228 | 6.69403 | 6.77108 | 6.47664 |
| VWA5A | O00534 | 5.68513 | 6.32057 | 6.14721 | 5.66935 | 6.30322 | 6.70725 | 6.02664 | 6.82548 | 6.01836 | 6.68696 | 6.42566 | 6.30262 | 6.02185 | 6.8075 | 5.65095 | 6.40125 |
| CARS2 | Q9HA77 | 6.06671 | 6.35328 | 5.95701 | 6.57539 | 6.15585 | 6.30133 | 6.04694 | 6.46738 | 6.35813 | 6.11408 | 6.06614 | 5.7865 | 6.45984 | 6.15866 | 6.47676 | 6.65599 |
| PHKA2 | P46019 | 6.15095 | 6.46424 | 6.81509 | 6.21916 | 6.25366 | 6.72597 | 6.34487 | 6.84856 | 6.40401 | 6.58208 | 6.81258 | 7.05702 | 5.31292 | 5.5202 | 5.21429 | 5.27439 |
| HPS3 | Q969F9 | 6.42075 | 6.35775 | 6.98183 | 6.2511 | 6.13284 | 6.66014 | 6.05404 | 6.35006 | 6.3904 | 6.86987 | 6.63018 | 6.59505 | 5.68783 | 5.64677 | 5.39747 | 5.57392 |
| ADGRL2 | O95490-6 | 5.99422 | 6.25197 | 5.88035 | 6.40361 | 5.89608 | 6.09219 | 6.17947 | 6.02666 | 6.13401 | 6.21281 | 6.02881 | 6.10909 | 6.64398 | 6.47309 | 6.8853 | 6.78836 |
| FANCA | O15360 | 6.63379 | 6.58684 | 6.66187 | 6.66672 | 6.45015 | 6.30581 | 6.54046 | 6.91118 | 6.46662 | 6.89236 | 6.66328 | 6.77393 | 5.11876 | 5.30788 | 4.94468 | 5.07569 |
| MYLK | Q15746 | 6.43502 | 6.41584 | 6.43227 | 6.09409 | 5.65 | 5.90042 | 5.6264 | 5.63559 | 6.26317 | 6.39265 | 5.79685 | 6.16116 | 6.48133 | 6.11887 | 7.43062 | 7.16571 |
| STK24 | Q9Y6E0 | 6.23903 | 6.27941 | 6.75513 | 5.79865 | 6.3705 | 6.64579 | 6.09335 | 6.34594 | 6.36973 | 6.43338 | 6.43919 | 6.41692 | 6.31015 | 5.93679 | 5.90866 | 5.65739 |
| METTL1 | Q9UBP6 | 5.11675 | 5.58329 | 5.76459 | 5.08866 | 5.6124 | 6.38885 | 5.4476 | 5.70323 | 5.63756 | 5.98612 | 6.32471 | 6.04642 | 7.12775 | 8.24815 | 8.28668 | 7.63723 |
| MAP7 | Q14244-7 | 5.98533 | 6.13376 | 6.1109 | 5.6474 | 6.49515 | 6.4013 | 6.03587 | 6.48579 | 7.11697 | 6.55567 | 6.54357 | 5.94561 | 6.34401 | 6.29992 | 6.02924 | 5.8695 |
| HSPBP1 | Q9NZL4-3 | 6.39467 | 6.18089 | 6.90637 | 5.76967 | 6.52682 | 6.68673 | 6.08986 | 6.80471 | 6.11807 | 6.46093 | 7.16462 | 6.1869 | 5.53719 | 6.06215 | 5.5614 | 5.54902 |
| HMGCL | P35914 | 6.35251 | 6.3284 | 5.73727 | 6.16635 | 6.67369 | 5.97576 | 6.29261 | 5.93164 | 6.08316 | 5.95828 | 6.11234 | 6.19197 | 7.06812 | 6.19784 | 6.63924 | 6.29085 |
| PRKAR2B | P31323 | 6.27451 | 6.55877 | 6.18287 | 6.4785 | 6.42789 | 6.62682 | 6.07203 | 6.68668 | 6.28641 | 6.02591 | 6.32551 | 6.11577 | 6.07717 | 5.86157 | 6.20095 | 5.79861 |
| PPP6R2 | O75170 | 6.45576 | 6.37683 | 7.14369 | 6.04888 | 6.61388 | 6.76692 | 6.05604 | 6.43902 | 6.7105 | 5.88713 | 6.41356 | 6.40313 | 5.84243 | 5.53566 | 5.65742 | 5.64914 |
| ZNF532 | Q9HCE3 | 6.46208 | 6.49379 | 6.05488 | 6.67901 | 6.2938 | 6.12488 | 6.45192 | 5.80399 | 6.40012 | 6.46389 | 6.08149 | 5.85401 | 6.11175 | 5.8822 | 6.51139 | 6.33079 |
| DPP7 | Q9UHL4 | 6.74841 | 6.45709 | 6.12463 | 6.37271 | 6.49801 | 6.18677 | 6.26849 | 6.38833 | 6.24214 | 6.21137 | 6.23014 | 6.16558 | 6.3599 | 5.74055 | 6.11878 | 5.8871 |
| PPP5C | P53041 | 6.35359 | 6.3411 | 6.88549 | 6.05599 | 6.49864 | 6.88674 | 6.33552 | 6.52107 | 6.33899 | 6.55117 | 6.10037 | 6.37446 | 5.71227 | 5.92628 | 5.50562 | 5.61271 |
| PHF21A | Q96BD5-3 | 6.57565 | 6.52794 | 5.91696 | 6.49611 | 6.29701 | 6.17156 | 6.29072 | 6.12056 | 6.09006 | 6.46132 | 6.03382 | 6.16947 | 6.44381 | 6.22868 | 6.11042 | 6.06589 |
| CHD9 | Q3L8U1 | 6.25826 | 6.27498 | 6.48566 | 6.49673 | 6.21334 | 6.29717 | 5.92666 | 6.2162 | 6.14276 | 6.25367 | 6.21696 | 6.20348 | 6.08423 | 6.24636 | 6.25051 | 6.43303 |
| CDAN1 | Q8IWY9 | 6.12331 | 6.53115 | 7.30098 | 6.17851 | 6.32504 | 6.40256 | 6.14548 | 6.29927 | 6.35224 | 6.45643 | 6.59893 | 6.55649 | 5.67186 | 5.25282 | 5.57496 | 6.22998 |
| L2HGDH | Q9H9P8 | 6.66556 | 6.68089 | 5.9483 | 6.70715 | 6.49005 | 5.94787 | 6.26983 | 6.34282 | 6.15655 | 5.95364 | 5.79995 | 5.75829 | 6.3205 | 6.19627 | 6.65392 | 6.10839 |

| | | | | | | | | | | | | | | | | | |
|---|---|---|---|---|---|---|---|---|---|---|---|---|---|---|---|---|---|
| ZFYVE1 | Q9HBF4 | 6.61979 | 6.22333 | 6.63667 | 6.04687 | 6.34053 | 6.69211 | 6.13903 | 6.30436 | 6.4928 | 6.37188 | 6.16329 | 5.86239 | 6.17312 | 6.17595 | 6.05552 | 5.70233 |
| SNU13 | P55769 | 5.99878 | 5.92585 | 5.70245 | 5.95926 | 6.48462 | 5.89518 | 6.20944 | 6.60483 | 6.08522 | 6.27898 | 6.43366 | 6.60269 | 6.53042 | 6.44961 | 6.44086 | 6.39816 |
| EXT2 | Q93063-3 | 6.64235 | 6.64145 | 5.88626 | 7.03046 | 6.47641 | 6.17098 | 6.93474 | 6.50513 | 6.23904 | 6.21927 | 6.01643 | 6.13249 | 5.93125 | 5.68047 | 5.54908 | 5.9442 |
| MIPEP | Q99797 | 6.21399 | 6.23699 | 5.74181 | 6.69622 | 6.83851 | 6.52416 | 6.55169 | 6.3775 | 6.25091 | 5.94269 | 5.68236 | 5.96598 | 6.56226 | 6.06131 | 6.21262 | 6.14098 |
| GORASP2 | Q9H8Y8 | 5.79114 | 5.6793 | 5.37838 | 5.59753 | 6.12422 | 6.15856 | 6.16192 | 6.85118 | 6.23707 | 6.73938 | 6.25178 | 6.46045 | 6.01701 | 6.62972 | 6.89651 | 7.02585 |
| TBC1D2 | Q9BYX2 | 6.51732 | 5.9106 | 7.27133 | 6.25452 | 6.6188 | 6.52765 | 6.41858 | 6.47236 | 6.68433 | 6.31553 | 6.8552 | 6.82396 | 6.52475 | 5.05949 | 5.5009 | 5.24469 |
| DCPS | Q96C86 | 5.63903 | 5.48249 | 5.97092 | 5.02018 | 5.88282 | 6.25644 | 5.35706 | 6.34826 | 5.65113 | 6.2541 | 6.30643 | 6.34267 | 6.94818 | 7.943 | 7.61769 | 6.97961 |
| BICD1 | Q96G01 | 6.26319 | 6.2384 | 6.46764 | 5.94154 | 6.17128 | 6.37373 | 6.17475 | 6.15973 | 6.36683 | 6.34376 | 6.49504 | 6.26461 | 6.48474 | 6.09166 | 6.03648 | 6.12661 |
| LRCH3 | Q96II8-2 | 6.65155 | 5.94348 | 6.79615 | 6.4085 | 6.20867 | 6.29668 | 6.06654 | 6.58909 | 6.41581 | 6.16577 | 6.73805 | 6.80697 | 6.12448 | 5.65032 | 5.62058 | 5.51738 |
| SP110 | Q9HB58-6 | 6.51659 | 6.62158 | 5.92142 | 6.88784 | 6.06614 | 6.03609 | 6.31651 | 6.14888 | 6.28133 | 6.2082 | 6.03068 | 6.05216 | 5.75953 | 6.18771 | 6.72785 | 6.23749 |
| IL13RA2 | Q14627 | 6.64168 | 6.68101 | 6.0141 | 6.8676 | 6.57339 | 6.13245 | 6.83735 | 6.05932 | 6.12618 | 6.40603 | 6.41063 | 5.77896 | 5.84626 | 5.36666 | 6.11278 | 6.1456 |
| UBXN7 | O94888 | 5.78519 | 6.09053 | 5.90418 | 6.52843 | 6.33352 | 6.49437 | 5.50432 | 6.68948 | 5.95758 | 6.29032 | 6.55777 | 6.47841 | 6.71075 | 6.88887 | 6.17016 | 6.51612 |
| CNTNAP1 | P78357 | 6.34229 | 6.70236 | 6.39229 | 6.48173 | 6.05939 | 5.71064 | 6.20372 | 6.00092 | 6.65552 | 6.28679 | 6.15186 | 5.80926 | 6.54404 | 6.13086 | 6.84902 | 6.67931 |
| MCM10 | Q7L590 | 6.37181 | 6.60152 | 6.07998 | 6.54505 | 6.09518 | 6.05119 | 6.21716 | 5.93535 | 6.0806 | 6.19809 | 6.23075 | 6.31121 | 6.38263 | 6.08324 | 6.44375 | 6.37248 |
| BPNT2 | Q9NX62 | 6.26601 | 6.34869 | 5.81802 | 6.58276 | 6.22075 | 5.99181 | 6.12036 | 6.11524 | 6.25139 | 6.07027 | 5.94531 | 6.08105 | 6.93399 | 6.20371 | 6.68255 | 6.36809 |
| TARS3 | A2RTX5 | 6.23763 | 6.3956 | 6.95916 | 6.33071 | 6.07131 | 6.36094 | 6.5013 | 6.5352 | 6.06125 | 6.72589 | 6.62678 | 6.66025 | 5.60482 | 5.82252 | 5.41316 | 5.69348 |
| MASTL | Q96GX5 | 5.3281 | 5.87755 | 6.52482 | 5.14974 | 5.37578 | 6.34811 | 5.51089 | 5.94825 | 6.06815 | 6.25342 | 6.52125 | 6.20234 | 6.89702 | 7.01929 | 7.60748 | 7.36781 |
| SEPTIN8 | Q92599 | 5.98189 | 6.2124 | 6.33524 | 5.98033 | 6.2667 | 6.62198 | 6.12303 | 6.7523 | 6.23601 | 6.43576 | 6.33925 | 6.46703 | 6.17888 | 6.21127 | 6.09801 | 5.75992 |
| ARMC1 | Q9NVT9 | 6.88976 | 6.48658 | 6.47511 | 6.30683 | 6.9087 | 6.96183 | 6.96668 | 6.71118 | 7.23818 | 6.96688 | 6.7261 | 6.65899 | 4.70156 | 4.55159 | 4.71519 | 4.73484 |
| GABPA | Q06546 | 5.84909 | 5.76985 | 5.99 | 5.65368 | 5.96318 | 6.14934 | 5.59554 | 5.81652 | 6.21195 | 6.13343 | 6.38923 | 6.22764 | 7.12205 | 6.94906 | 6.94775 | 7.2317 |
| ANO10 | Q9NW15 | 6.73629 | 6.45345 | 6.0593 | 6.5429 | 6.11531 | 5.87257 | 6.05969 | 5.8493 | 6.36024 | 6.02672 | 6.23524 | 6.02489 | 6.51526 | 6.02448 | 6.67487 | 6.44951 |
| ATP6V1E1 | P36543 | 6.81518 | 6.60669 | 6.61766 | 6.214 | 6.5333 | 6.72258 | 6.43852 | 6.48734 | 6.49112 | 6.63632 | 6.74408 | 6.5111 | 5.59295 | 5.38331 | 5.11179 | 5.09405 |
| MRPS26 | Q9BYN8 | 6.66191 | 6.05576 | 5.82158 | 6.65542 | 6.35069 | 5.72494 | 6.60429 | 5.8833 | 6.32338 | 5.89983 | 6.24426 | 6.15109 | 6.50676 | 6.16872 | 6.43938 | 6.50871 |
| COX6B1 | P14854 | 6.74527 | 5.04637 | 5.55656 | 4.91613 | 6.00312 | 4.83931 | 5.04488 | 6.25242 | 4.73583 | 6.04938 | 4.8915 | 5.52817 | 5.2482 | 10.3345 | 8.57571 | 10.2327 |
| HAUS5 | O94927 | 6.88549 | 6.9171 | 7.50823 | 6.77329 | 6.37342 | 6.81582 | 6.23584 | 6.29527 | 6.67488 | 6.61666 | 7.13725 | 6.8176 | 4.64438 | 4.8961 | 4.63913 | 4.76953 |
| GTPBP6 | O43824 | 6.45915 | 6.55223 | 6.12534 | 7.02926 | 6.52032 | 6.56714 | 6.72103 | 6.29116 | 6.03089 | 6.15774 | 6.0675 | 5.98434 | 5.88823 | 5.70168 | 5.84297 | 6.06101 |
| TRAM1 | Q15629 | 6.0994 | 6.38062 | 5.85695 | 6.54851 | 5.8643 | 5.74262 | 6.33211 | 5.57086 | 6.00739 | 6.1154 | 5.72802 | 5.7577 | 6.87538 | 6.60492 | 7.41073 | 7.10508 |
| LRRFIP2 | Q9Y608 | 6.21373 | 6.1888 | 6.5268 | 5.58973 | 6.62953 | 6.59909 | 5.94992 | 6.15526 | 6.6504 | 6.34168 | 6.74017 | 6.30187 | 6.56809 | 6.0515 | 5.76599 | 5.72744 |
| CNOT10 | Q9H9A5-6 | 6.40159 | 6.57949 | 6.54661 | 6.20362 | 6.23531 | 6.52203 | 6.11727 | 6.30457 | 6.3996 | 6.37738 | 6.59917 | 6.42853 | 5.89554 | 5.75245 | 5.93263 | 5.70423 |
| COG8 | Q96MW5 | 6.23208 | 5.95867 | 6.31932 | 6.21198 | 6.60829 | 6.33238 | 6.62692 | 6.48138 | 6.51492 | 6.54401 | 6.23547 | 6.5383 | 5.99814 | 5.9917 | 5.90261 | 6.02381 |
| ESPL1 | Q14674 | 6.80231 | 6.6292 | 7.49424 | 7.02232 | 6.21018 | 6.88164 | 6.59714 | 6.43526 | 7.22393 | 7.19171 | 7.06555 | 7.37568 | 4.21487 | 4.21369 | 4.0276 | 4.61466 |
| SRSF3 | P84103-2 | 6.14277 | 6.063 | 5.26362 | 5.58339 | 6.22264 | 5.5589 | 5.78428 | 5.27692 | 5.77033 | 6.17362 | 7.18799 | 6.75586 | 8.04146 | 6.81677 | 6.76825 | 6.5902 |
| AGTPBP1 | Q9UPW5-3 | 6.57644 | 6.34118 | 6.75206 | 6.31011 | 6.40339 | 6.60386 | 6.57388 | 6.58346 | 6.40911 | 6.57436 | 6.55645 | 6.6552 | 5.43423 | 5.40655 | 5.22153 | 5.5982 |
| UBE2V2 | Q15819 | 6.21407 | 5.49969 | 6.16424 | 5.00655 | 6.59978 | 6.63424 | 5.78422 | 7.53783 | 5.61866 | 6.83452 | 6.09178 | 6.66604 | 6.18783 | 7.05595 | 5.79556 | 6.30903 |
| ARMC6 | Q6NXE6 | 6.34803 | 6.25135 | 6.86105 | 6.0261 | 6.70535 | 6.59885 | 5.93266 | 7.18409 | 6.37514 | 6.20838 | 6.77771 | 6.97675 | 5.48565 | 5.52292 | 5.17999 | 5.46596 |
| ARHGEF17 | Q96PE2 | 6.45916 | 6.61758 | 6.45548 | 6.47224 | 6.31207 | 6.46979 | 6.17342 | 6.20929 | 6.10097 | 6.2591 | 6.57551 | 6.41027 | 5.89212 | 5.40151 | 6.04521 | 6.1463 |
| SLC9A1 | P19634 | 6.35009 | 6.54654 | 6.0257 | 6.65782 | 6.09503 | 6.25049 | 6.37319 | 6.28819 | 6.13057 | 6.20004 | 6.20895 | 5.91487 | 6.28296 | 6.25287 | 6.28984 | 6.13285 |
| PHAX | Q9H814 | 5.70393 | 5.65159 | 6.0139 | 5.50296 | 5.83991 | 6.30813 | 5.82101 | 6.04092 | 6.27049 | 6.2798 | 6.29641 | 6.17859 | 7.09367 | 6.71995 | 7.13831 | 7.14043 |
| ARHGAP11A | Q6P4F7 | 6.76192 | 6.87628 | 6.29897 | 7.42589 | 6.45327 | 5.9748 | 6.44447 | 6.23743 | 6.06591 | 6.07022 | 5.99464 | 6.2335 | 5.48665 | 5.56056 | 6.06724 | 6.04824 |
| LAMP1 | P11279 | 6.29604 | 6.74898 | 5.9497 | 6.51406 | 6.08615 | 5.93068 | 6.16644 | 6.00719 | 6.32673 | 6.38329 | 6.24759 | 6.13107 | 6.25894 | 5.99408 | 6.73672 | 6.22233 |
| FZD6 | O60353 | 6.39308 | 6.42377 | 6.09439 | 6.55584 | 5.99791 | 6.56471 | 6.59956 | 6.04179 | 5.98902 | 6.42335 | 6.04492 | 5.9843 | 6.05345 | 6.31991 | 6.30384 | 6.21016 |
| ODR4 | Q5SWX8 | 6.3601 | 6.18292 | 6.1966 | 6.7643 | 5.92989 | 5.83758 | 6.53095 | 5.87502 | 6.00568 | 5.86095 | 6.03544 | 5.98761 | 6.79507 | 6.44851 | 6.70725 | 6.48214 |
| PRR12 | Q9ULL5 | 6.62616 | 6.30583 | 6.10447 | 6.76242 | 6.05286 | 5.86562 | 6.35411 | 5.74899 | 6.33323 | 6.44848 | 5.94453 | 6.16235 | 6.34285 | 6.13054 | 6.47818 | 6.33938 |
| TMX2 | Q9Y320 | 6.29234 | 6.57701 | 6.21717 | 6.77473 | 6.4194 | 5.97739 | 6.5958 | 6.10536 | 6.33239 | 5.94537 | 6.05428 | 5.94842 | 6.16303 | 5.99575 | 6.37231 | 6.22924 |
| E9PSI1 | E9PSI1 | 6.35542 | 6.64769 | 6.02355 | 6.5455 | 6.3144 | 5.99751 | 6.23587 | 6.4657 | 5.96825 | 6.17039 | 6.11521 | 6.01991 | 6.22763 | 6.16736 | 6.3225 | 6.42311 |
| INO80 | Q9ULG1 | 6.48348 | 6.54725 | 5.68412 | 6.55752 | 6.1756 | 5.77783 | 6.46121 | 6.06541 | 6.22053 | 6.20964 | 5.79571 | 5.91656 | 6.30655 | 6.50273 | 6.84528 | 6.45059 |
| DTL | Q9NZJ0 | 5.59287 | 6.02953 | 6.37004 | 6.01971 | 5.52365 | 5.91522 | 6.02131 | 5.63417 | 6.27616 | 6.28675 | 6.77287 | 6.6673 | 6.74818 | 6.67638 | 6.80198 | 6.66388 |
| MPP2 | Q14168-4 | 6.22709 | 6.61321 | 6.13789 | 6.69942 | 6.01784 | 6.1017 | 6.31117 | 6.16394 | 6.13202 | 6.16194 | 6.02736 | 6.08656 | 6.46151 | 6.16443 | 6.4079 | 6.28603 |
| ACY1 | Q03154 | 6.36424 | 4.99785 | 6.6245 | 4.6385 | 6.3726 | 5.9862 | 6.5274 | 7.14052 | 5.6482 | 7.24862 | 6.47658 | 6.89031 | 6.01687 | 7.19924 | 6.58394 | 6.18444 |
| BIRC2 | Q13490 | 6.21386 | 5.84031 | 6.50462 | 5.70452 | 6.47371 | 6.58055 | 6.05663 | 6.62656 | 6.25759 | 6.32631 | 6.53947 | 6.76037 | 6.47125 | 6.10685 | 5.80661 | 5.7308 |
| SLC38A10 | Q9HBR0 | 6.37518 | 6.42205 | 5.79187 | 6.62131 | 6.18515 | 5.95535 | 6.36509 | 6.13346 | 5.76202 | 6.22873 | 6.20514 | 5.92092 | 6.58772 | 6.44406 | 6.13206 | 6.86988 |
| SATB2 | Q9UPW6 | 6.10425 | 6.28884 | 5.91742 | 6.81873 | 6.05962 | 6.2611 | 6.24187 | 6.16493 | 6.22589 | 6.39582 | 6.21157 | 6.21852 | 6.96128 | 5.74255 | 5.99711 | 6.3905 |
| STK4 | Q13043 | 6.49353 | 5.42035 | 6.90372 | 6.12985 | 6.54075 | 6.3868 | 6.5368 | 6.84522 | 6.51633 | 6.11531 | 6.99396 | 6.67049 | 5.76233 | 5.59494 | 5.52202 | 5.5676 |
| BMP2K | Q9NSY1 | 5.99519 | 6.32909 | 6.11094 | 6.53656 | 6.32276 | 6.36912 | 6.23218 | 6.0774 | 6.71612 | 6.43492 | 6.61347 | 6.37041 | 5.91055 | 6.22764 | 5.93871 | 5.81495 |
| RXRB | P28702-3 | 5.84288 | 6.6129 | 6.21953 | 6.14752 | 5.80145 | 6.15113 | 6.09956 | 6.26235 | 6.53744 | 6.47625 | 6.30548 | 6.38084 | 5.97158 | 6.58404 | 6.44727 | 6.15979 |
| GAA | P10253 | 6.63738 | 6.88177 | 6.1481 | 6.18967 | 6.58787 | 6.35676 | 6.47639 | 6.44014 | 5.98033 | 6.02806 | 5.82895 | 5.76676 | 6.12492 | 5.98427 | 6.2584 | 6.31023 |
| PREB | Q9HCU5 | 6.14535 | 6.27788 | 6.00121 | 6.57184 | 6.16252 | 5.92203 | 6.45839 | 5.80343 | 5.92028 | 6.17334 | 6.10677 | 5.92653 | 6.42548 | 6.49057 | 6.91681 | 6.69758 |
| FBXO28 | Q9NVF7 | 6.71795 | 6.81321 | 6.66595 | 7.38812 | 6.20616 | 5.92742 | 6.71109 | 5.95921 | 6.44957 | 6.39759 | 6.51394 | 6.5677 | 5.3265 | 5.25212 | 5.48423 | 5.60925 |
| OSBPL10 | Q9BXB5 | 6.42477 | 6.58316 | 6.47948 | 6.48135 | 6.37929 | 6.75921 | 6.2272 | 6.4477 | 6.2038 | 6.47959 | 6.40604 | 6.43586 | 5.53962 | 5.73168 | 5.75846 | 5.66279 |
| SNAP23 | O00161 | 6.12451 | 6.35408 | 5.77772 | 6.20659 | 5.82672 | 5.52414 | 6.05416 | 6.2214 | 6.20456 | 6.15425 | 5.88722 | 6.0893 | 6.8044 | 6.74893 | 6.99724 | 7.02478 |
| PPP4C | P60510 | 5.6731 | 6.0806 | 6.33294 | 5.65988 | 6.0216 | 6.11474 | 6.27889 | 6.02986 | 5.81137 | 6.4346 | 5.96025 | 6.22917 | 6.60534 | 6.80103 | 6.91414 | 7.05251 |
| PPP1CB | P62140 | 6.20054 | 6.05239 | 6.31073 | 6.13483 | 6.15987 | 6.40902 | 6.06223 | 6.27803 | 6.28005 | 6.378 | 6.30603 | 6.23751 | 6.37783 | 6.37532 | 6.18751 | 6.2501 |
| UIMC1 | Q96RL1 | 6.18429 | 6.11063 | 6.16018 | 6.84187 | 5.60207 | 5.81239 | 6.12163 | 5.87521 | 6.08068 | 5.97508 | 6.10121 | 6.08225 | 6.18101 | 6.90955 | 7.00081 | 6.96114 |
| HELZ | P42694 | 6.36148 | 6.18966 | 6.71557 | 6.26471 | 6.25648 | 6.62134 | 6.20038 | 5.9131 | 6.43707 | 6.39047 | 6.49928 | 6.38873 | 5.7612 | 6.12989 | 6.02523 | 5.84541 |
| ATP7A | Q04656-3 | 6.29789 | 6.34434 | 5.95095 | 6.13125 | 6.43734 | 6.13816 | 6.47985 | 5.81783 | 6.15251 | 6.31843 | 6.02933 | 5.91668 | 6.75666 | 6.28726 | 6.65661 | 6.28492 |
| SMARCA4 | P51532 | 6.17996 | 6.73592 | 5.9198 | 6.64972 | 6.13299 | 6.20502 | 6.38716 | 6.02331 | 6.04629 | 6.21465 | 5.87698 | 5.81257 | 6.41558 | 6.33881 | 6.55526 | 6.50599 |
| CENPP | Q6IPU0 | 6.26012 | 6.13071 | 6.36859 | 6.86708 | 5.7209 | 6.00232 | 6.3389 | 5.68582 | 6.60212 | 6.17332 | 6.19604 | 6.46594 | 6.2172 | 6.06924 | 6.23248 | 6.66922 |
| HTATIP2 | Q9BUP3-3 | 6.41375 | 6.28708 | 6.18404 | 6.39375 | 6.11136 | 6.15281 | 6.31425 | 6.33131 | 6.39254 | 6.52364 | 6.65844 | 6.49874 | 6.15914 | 5.78892 | 5.95352 | 5.83671 |
| ZER1 | Q7Z7L7 | 6.3623 | 6.57308 | 6.72745 | 6.13363 | 6.39071 | 6.36357 | 5.91993 | 6.38604 | 6.44102 | 6.78653 | 6.59495 | 6.40389 | 5.61191 | 5.72005 | 5.5462 | 6.03874 |
| EDF1 | O60869 | 6.60965 | 6.61619 | 6.502 | 6.04075 | 6.6803 | 6.80699 | 6.45038 | 6.65513 | 6.50255 | 6.67581 | 6.54218 | 6.40148 | 5.69538 | 5.46257 | 5.16888 | 5.28975 |
| PPID | Q08752 | 5.85762 | 6.12018 | 6.74042 | 5.65204 | 6.3593 | 6.58036 | 5.99771 | 6.61122 | 6.34981 | 6.38363 | 6.78821 | 6.47169 | 6.24629 | 6.02501 | 5.89816 | 5.91835 |
| NDUFAF1 | Q9Y375 | 6.56617 | 6.28344 | 6.11781 | 6.54841 | 6.43658 | 5.79344 | 6.50748 | 5.80503 | 6.06331 | 5.91051 | 6.16829 | 6.12408 | 6.92704 | 6.24453 | 6.27896 | 6.22491 |
| PWWP2A | Q96N64 | 6.49519 | 6.58193 | 6.31096 | 7.18299 | 5.9052 | 5.86695 | 6.58415 | 5.46745 | 6.01727 | 6.21703 | 6.25303 | 6.20302 | 6.34053 | 6.16616 | 6.32854 | 6.0796 |
| PDAP1 | Q13442 | 6.2268 | 6.35717 | 6.34619 | 5.35787 | 6.47796 | 6.57878 | 6.22278 | 6.60664 | 6.49348 | 6.13355 | 5.85934 | 6.01473 | 6.33511 | 6.88009 | 6.08939 | 6.0201 |
| RRP7A | Q9Y3A4 | 6.27198 | 6.34636 | 6.07613 | 6.67794 | 6.46536 | 5.90086 | 6.68515 | 6.02898 | 6.25097 | 5.94903 | 5.82433 | 5.64158 | 6.49747 | 6.10802 | 6.62011 | 6.65574 |
| SREK1 | Q8WXA9-2 | 5.42545 | 6.16461 | 5.43651 | 6.5134 | 5.97635 | 5.96663 | 6.31513 | 6.27705 | 5.99509 | 6.42098 | 6.24502 | 6.01057 | 5.86951 | 6.82313 | 7.14106 | 7.4195 |
| UQCRB | P14927 | 6.08125 | 5.35537 | 5.93297 | 5.48926 | 6.72934 | 5.19757 | 5.62808 | 6.78083 | 5.37101 | 6.33246 | 5.74907 | 5.65908 | 6.75384 | 7.44494 | 7.32691 | 7.228 |
| PCMT1 | P22061-2 | 5.64926 | 4.54081 | 5.92496 | 4.86949 | 6.06535 | 5.44235 | 5.72575 | 7.15261 | 5.49642 | 7.15485 | 6.59812 | 7.49607 | 5.99253 | 7.3907 | 6.88456 | 7.61617 |
| SORD | Q00796 | 6.37819 | 6.08581 | 6.85101 | 6.30115 | 6.00273 | 6.64273 | 5.99705 | 6.24332 | 6.2327 | 6.27555 | 6.50618 | 6.60295 | 5.99216 | 5.9902 | 5.6852 | 6.21307 |
| STMN1 | P16949-2 | 6.6257 | 5.38867 | 6.83343 | 4.87938 | 7.19385 | 5.93168 | 5.30786 | 7.50246 | 5.84706 | 6.62806 | 6.46116 | 7.11022 | 6.39711 | 6.39959 | 5.63812 | 5.85565 |
| IFT80 | Q9P2H3-3 | 6.40155 | 6.47446 | 6.35713 | 6.61879 | 6.21176 | 6.64791 | 6.38307 | 6.17688 | 6.12042 | 6.27548 | 6.1243 | 6.41473 | 5.90469 | 6.03107 | 5.80743 | 6.05034 |
| CRK | P46108 | 5.54464 | 5.66887 | 6.35628 | 5.56609 | 5.91896 | 6.54078 | 5.86769 | 6.83436 | 5.99423 | 5.9683 | 6.93822 | 7.15247 | 6.1456 | 6.31858 | 5.78971 | 6.39523 |
| ZNF207 | O43670-4 | 5.58006 | 5.77006 | 5.95011 | 6.15675 | 6.16769 | 6.18888 | 6.28557 | 5.99893 | 6.17587 | 5.92998 | 6.00224 | 6.03567 | 6.95686 | 6.93363 | 6.96097 | 6.90672 |
| RAB11FIP5 | A0A1B0GTL5 | 6.26339 | 7.08293 | 6.11831 | 6.38671 | 6.36197 | 6.45676 | 6.48629 | 6.5607 | 6.28046 | 6.48777 | 6.29539 | 6.33931 | 5.2369 | 5.60364 | 5.95465 | 6.08483 |
| RIC8A | Q9NPQ8-3 | 6.18225 | 6.05362 | 6.76819 | 6.27269 | 6.24813 | 6.69749 | 6.0946 | 6.536 | 6.47462 | 6.32906 | 6.62822 | 6.87451 | 5.55987 | 5.87705 | 5.39944 | 6.00428 |
| HEATR3 | Q7Z4Q2 | 5.33457 | 5.45789 | 5.98351 | 5.45753 | 5.43149 | 5.9018 | 5.78294 | 5.80439 | 5.79961 | 5.89759 | 6.2498 | 6.02458 | 7.36029 | 7.50142 | 8.04183 | 7.97075 |
| HEXA | P06865 | 6.49091 | 6.2313 | 5.63788 | 5.76401 | 6.56743 | 6.0635 | 6.34113 | 6.44415 | 6.0094 | 6.27345 | 5.73447 | 5.84923 | 6.84699 | 6.60866 | 6.62337 | 6.51411 |
| DHX32 | Q7L7V1 | 6.40521 | 6.28387 | 6.70034 | 6.12932 | 6.63411 | 6.41635 | 6.38539 | 6.47748 | 6.57605 | 6.62576 | 6.60854 | 6.33414 | 5.58798 | 5.67773 | 5.45667 | 5.70106 |
| NUB1 | Q9Y5A7 | 6.1337 | 6.35523 | 6.60754 | 6.02171 | 5.95817 | 7.00925 | 6.05597 | 6.38449 | 6.09682 | 6.50773 | 6.31645 | 6.42162 | 6.52414 | 6.26845 | 6.14153 | 6.0972 |
| DGKA | P23743 | 5.7037 | 6.1758 | 6.52535 | 5.78862 | 6.08721 | 6.62733 | 6.21595 | 6.56077 | 6.17726 | 6.70423 | 6.81723 | 6.5438 | 5.84692 | 6.01756 | 6.22911 | 5.97917 |
| QTRT2 | Q9H974-4 | 6.342 | 6.31683 | 6.31182 | 6.30613 | 6.1272 | 6.13787 | 6.27065 | 6.13548 | 6.56433 | 6.2956 | 6.22932 | 6.41622 | 6.23616 | 5.92099 | 6.31276 | 6.07665 |
| PHF2 | O75151 | 6.38972 | 5.91179 | 6.03056 | 6.93541 | 6.29536 | 5.80321 | 6.16976 | 5.76397 | 5.94054 | 6.12939 | 5.74061 | 6.33413 | 6.26765 | 6.97821 | 6.63343 | 6.67628 |
| EPHB2 | P29323 | 6.19165 | 6.62022 | 5.88312 | 6.61001 | 6.03393 | 5.94926 | 6.11784 | 6.0757 | 6.27507 | 6.22685 | 5.81078 | 6.29049 | 6.5902 | 6.32978 | 6.63797 | 6.35712 |
| EPHB4 | P54760 | 6.40386 | 6.39821 | 6.19477 | 6.62779 | 6.38613 | 6.38888 | 6.38085 | 5.91604 | 6.58915 | 6.48172 | 6.24837 | 6.3168 | 6.63846 | 6.07341 | 5.95392 | 6.00163 |
| VPS28 | Q9UK41 | 6.04583 | 5.32417 | 6.29746 | 4.92036 | 5.96491 | 6.03978 | 5.99059 | 6.84246 | 5.59352 | 6.94244 | 6.14996 | 6.8131 | 5.65481 | 7.6985 | 6.66197 | 7.06012 |
| ANXA7 | P20073 | 6.52625 | 5.74924 | 6.48957 | 5.05085 | 6.67077 | 6.74766 | 5.87043 | 7.01216 | 6.00883 | 6.76004 | 6.46817 | 6.79652 | 5.93998 | 6.29389 | 5.82863 | 5.78701 |
| MTX2 | O75431 | 5.90356 | 5.14324 | 5.44366 | 5.47155 | 6.09906 | 5.49323 | 6.14851 | 6.4618 | 5.38202 | 6.34328 | 5.71392 | 6.67407 | 6.64192 | 7.51931 | 7.6531 | 7.90777 |
| NCAPH2 | Q6IBW4-4 | 5.98095 | 6.22359 | 6.61305 | 6.27372 | 6.20525 | 6.29395 | 6.13405 | 6.1934 | 6.45863 | 6.0974 | 6.65239 | 6.62036 | 6.29524 | 5.97263 | 5.94772 | 6.03768 |
| SEC22B | O75396 | 6.1534 | 6.26269 | 5.53128 | 6.49298 | 6.0483 | 5.96869 | 6.31773 | 6.13802 | 5.9854 | 5.90754 | 5.72745 | 6.00074 | 6.52486 | 6.8949 | 7.0502 | 6.99582 |
| CDCA8 | Q53HL2 | 6.72464 | 6.80475 | 6.4644 | 7.34021 | 6.19189 | 5.95433 | 6.69651 | 6.16976 | 6.63667 | 6.47353 | 6.29531 | 6.47364 | 5.27667 | 5.37032 | 5.33107 | 5.7963 |
| SDCBP | O00560 | 6.26443 | 6.60069 | 7.03296 | 6.03153 | 6.41292 | 6.57107 | 6.24298 | 6.58018 | 6.50411 | 6.44157 | 6.46341 | 6.45774 | 5.72703 | 5.8478 | 5.48089 | 5.34069 |
| HOXB4 | P17483 | 6.30527 | 5.93772 | 6.34084 | 6.6001 | 6.18786 | 6.02784 | 6.5584 | 5.933 | 6.105 | 6.35538 | 6.16478 | 6.1456 | 6.15414 | 6.22662 | 6.2944 | 6.66304 |
| DNAJB6 | O75190 | 5.91762 | 6.08374 | 6.61556 | 6.16851 | 5.98228 | 6.22349 | 6.18997 | 6.19195 | 6.36797 | 5.9666 | 6.58601 | 6.30678 | 5.78716 | 6.5808 | 6.59301 | 6.43855 |
| USP54 | Q70EL1 | 6.19597 | 6.37076 | 6.43043 | 6.70976 | 6.08993 | 6.15234 | 6.24175 | 6.07002 | 6.12495 | 6.41586 | 6.36546 | 6.53027 | 5.75703 | 6.16908 | 6.06559 | 6.31081 |
| DPP8 | Q6V1X1 | 6.22609 | 6.09652 | 6.97502 | 6.10226 | 6.16846 | 6.43214 | 6.1768 | 6.45056 | 6.34946 | 6.48982 | 6.55029 | 6.56888 | 5.75556 | 6.01437 | 6.04476 | 5.59903 |

| | | | | | | | | | | | | | | | | | |
|---|---|---|---|---|---|---|---|---|---|---|---|---|---|---|---|---|---|
| EXOSC9 | Q06265-2 | 6.1675 | 6.35049 | 6.08478 | 6.52808 | 6.19202 | 5.97404 | 6.3098 | 5.36649 | 6.26239 | 5.94541 | 5.72812 | 5.45581 | 6.40064 | 6.58616 | 7.53267 | 7.11559 |
| KTI12 | Q96EK9 | 5.96628 | 5.54863 | 7.1007 | 5.76006 | 6.31094 | 6.70245 | 6.37646 | 6.42008 | 6.27736 | 6.55143 | 6.88975 | 6.87476 | 6.23954 | 5.87529 | 5.44974 | 5.65652 |
| THNSL1 | Q8IYQ7 | 6.67436 | 6.3305 | 5.99682 | 6.76244 | 6.21236 | 5.94941 | 6.01627 | 5.91271 | 6.2003 | 6.16244 | 6.05267 | 5.97102 | 6.41914 | 6.32251 | 6.53493 | 6.48211 |
| ANKRD26 | Q9UPS8 | 6.36823 | 6.16927 | 6.57302 | 6.25242 | 6.24466 | 6.3988 | 6.14465 | 6.41528 | 6.32361 | 6.53368 | 6.22578 | 6.42536 | 6.29929 | 6.02348 | 5.82051 | 5.78198 |
| HINT1 | P49773 | 5.61419 | 4.65698 | 5.76461 | 4.2265 | 5.49106 | 5.0269 | 5.31185 | 7.46819 | 5.09914 | 6.29134 | 5.24555 | 6.70967 | 5.12772 | 10.1095 | 8.35669 | 9.50007 |
| ATP5MJ | P56378 | 6.02208 | 5.70255 | 5.40132 | 5.99147 | 6.44199 | 6.03099 | 6.88642 | 6.18002 | 5.79953 | 5.96529 | 6.18357 | 6.08202 | 6.77352 | 6.70569 | 6.86375 | 6.9698 |
| RDH13 | Q8NBN7 | 6.72305 | 5.87671 | 5.96883 | 6.46065 | 6.66003 | 5.86683 | 6.15172 | 5.85014 | 6.14985 | 5.87163 | 6.25903 | 6.14531 | 7.00276 | 6.18301 | 6.47626 | 6.3542 |
| DPH2 | Q9BQC3 | 6.01955 | 6.1415 | 7.13161 | 5.79871 | 6.15672 | 6.83954 | 6.27666 | 6.5178 | 6.20463 | 6.46467 | 6.81062 | 6.72744 | 5.94091 | 5.68318 | 5.58354 | 5.70292 |
| ATG13 | O75143-5 | 6.0915 | 6.13873 | 6.79889 | 6.28044 | 6.061 | 6.16337 | 5.87945 | 6.51273 | 6.15469 | 6.35969 | 6.20101 | 6.58947 | 6.20563 | 6.0899 | 6.33524 | 6.13827 |
| CRIP2 | P52943 | 6.6425 | 6.25197 | 6.69829 | 5.65251 | 6.43887 | 6.36268 | 5.93365 | 6.69634 | 6.30602 | 6.48313 | 6.29711 | 6.21128 | 6.31091 | 6.20698 | 6.06023 | 5.44754 |
| RCL1 | Q9Y2P8 | 6.53635 | 6.15303 | 6.03471 | 6.70922 | 6.74893 | 6.26876 | 6.29057 | 6.09947 | 6.21912 | 5.99712 | 6.00654 | 5.97513 | 6.85202 | 5.82847 | 6.30171 | 5.97886 |
| BBX | Q8WY36 | 5.76494 | 5.9732 | 5.98478 | 5.93753 | 5.89392 | 5.75278 | 6.46147 | 5.56475 | 6.01826 | 6.09409 | 6.19276 | 5.79479 | 6.50526 | 7.30915 | 7.33276 | 7.41956 |
| ASCC2 | Q9H1I8 | 6.61312 | 6.62786 | 6.41106 | 6.20648 | 6.13184 | 6.28388 | 5.79548 | 6.22391 | 6.44744 | 6.96489 | 6.23119 | 6.45005 | 5.81001 | 5.94957 | 5.65141 | 6.20181 |
| EDEM3 | Q9BZQ6 | 6.40094 | 6.55049 | 5.9336 | 6.72927 | 6.39862 | 6.0831 | 6.19529 | 5.89329 | 6.41208 | 6.32148 | 6.11156 | 6.20408 | 6.39182 | 5.98693 | 6.14797 | 6.23948 |
| BCAT1 | P54687-5 | 6.16555 | 5.49734 | 6.81495 | 5.87938 | 6.51582 | 6.57493 | 6.60353 | 6.65368 | 6.32631 | 6.16284 | 7.14261 | 6.5696 | 5.81352 | 5.64946 | 5.77547 | 5.855 |
| ALG2 | Q9H553 | 6.38688 | 6.66538 | 6.11759 | 6.48851 | 6.19612 | 5.97906 | 6.13436 | 6.01708 | 6.22321 | 6.20077 | 6.02237 | 5.6081 | 6.495 | 6.17701 | 6.78797 | 6.50059 |
| PRMT9 | Q6P2P2 | 5.72321 | 6.14973 | 6.33487 | 5.92876 | 6.05895 | 6.31267 | 5.88313 | 6.27765 | 6.1516 | 6.47439 | 6.5951 | 6.52892 | 6.45966 | 6.51133 | 6.3052 | 6.30483 |
| TTF1 | Q15361 | 6.74617 | 6.58878 | 6.05879 | 6.66429 | 6.13202 | 6.03694 | 6.36147 | 5.96796 | 6.17835 | 6.0012 | 5.75336 | 6.21998 | 6.06702 | 6.32277 | 6.16084 | 6.74007 |
| VPS36 | Q86VN1 | 5.85202 | 6.16541 | 6.69605 | 5.88775 | 6.24487 | 6.40522 | 6.08386 | 6.26913 | 6.60159 | 6.43053 | 6.58277 | 6.78986 | 5.70035 | 6.1526 | 6.0574 | 6.0806 |
| RPL37A | P61513 | 6.60077 | 6.85911 | 6.44897 | 6.47975 | 6.72774 | 6.69641 | 6.16447 | 6.58919 | 6.60635 | 6.50336 | 6.22036 | 6.1376 | 6.50863 | 5.40372 | 5.86698 | 5.1866 |
| GTSE1 | Q9NYZ3 | 5.85987 | 5.72711 | 6.37266 | 6.26745 | 5.95017 | 6.31052 | 6.03519 | 6.1324 | 6.46449 | 6.16174 | 6.50853 | 7.00119 | 6.15259 | 6.53079 | 6.02559 | 6.49972 |
| AK4 | P27144 | 6.29449 | 5.25516 | 5.79332 | 5.34294 | 6.54922 | 5.31812 | 5.90176 | 6.59573 | 5.58888 | 6.20627 | 5.91418 | 6.65226 | 7.22531 | 7.1265 | 7.24782 | 6.98806 |
| NACC1 | Q96RE7 | 5.48375 | 6.26411 | 5.8171 | 5.73943 | 6.12684 | 6.19881 | 5.63809 | 5.87625 | 5.90423 | 6.41873 | 6.36761 | 5.99197 | 6.95555 | 7.14878 | 6.90363 | 7.16512 |
| PPP3CA | Q08209-2 | 6.19732 | 6.18964 | 6.59377 | 5.72993 | 6.51942 | 6.44996 | 6.02384 | 6.5325 | 6.2216 | 6.07908 | 6.43398 | 6.3408 | 6.1944 | 6.23756 | 6.08619 | 6.17001 |
| IBTK | Q9P2D0 | 5.85635 | 6.54185 | 6.04371 | 6.32377 | 6.51056 | 6.34385 | 6.24234 | 6.60935 | 6.13507 | 6.41406 | 6.3242 | 5.97143 | 6.22274 | 5.95789 | 6.20077 | 6.30207 |
| AAMP | Q13685 | 6.33842 | 6.41344 | 6.80814 | 5.85576 | 6.58283 | 7.03714 | 6.09734 | 6.99148 | 6.53942 | 6.30505 | 6.65735 | 6.54099 | 5.53571 | 5.60942 | 5.28941 | 5.3981 |
| NPC2 | P61916 | 6.68237 | 6.17228 | 6.16295 | 5.85868 | 6.95895 | 5.92628 | 6.26034 | 6.28233 | 6.15058 | 6.14391 | 6.15401 | 6.61636 | 6.25949 | 6.11714 | 6.07269 | 5.82166 |
| PTPN2 | P17706 | 5.62294 | 6.25288 | 6.07394 | 6.54537 | 5.70323 | 6.41406 | 5.93229 | 6.0279 | 5.97406 | 6.18292 | 6.07721 | 6.30879 | 6.44466 | 6.98821 | 6.83935 | 6.61219 |
| EMC7 | Q9NPA0 | 6.48489 | 6.54949 | 6.17405 | 6.64003 | 6.0345 | 5.94762 | 6.4438 | 5.96837 | 6.05339 | 6.11945 | 6.02108 | 6.06053 | 6.34405 | 6.10944 | 6.57206 | 6.47726 |
| CCDC43 | Q96MW1 | 6.19849 | 6.45263 | 7.00795 | 6.26939 | 6.29886 | 6.7746 | 6.47374 | 6.551 | 6.04221 | 6.74725 | 6.72984 | 6.46732 | 5.43409 | 5.8819 | 5.3291 | 5.34162 |
| PBDC1 | Q9BVG4 | 6.79801 | 6.39754 | 6.74809 | 5.85645 | 6.62386 | 6.69696 | 5.95683 | 6.60399 | 6.5957 | 6.51833 | 6.53935 | 6.76723 | 5.9864 | 5.55345 | 5.14809 | 5.20974 |
| RTN4IP1 | Q8WWV3 | 6.75197 | 6.39987 | 5.84013 | 6.6612 | 6.89502 | 6.2839 | 6.06856 | 5.77928 | 5.94783 | 5.8817 | 5.81246 | 5.95552 | 6.92997 | 6.28789 | 6.40858 | 6.09611 |
| CDCA3 | Q99618 | 6.24318 | 5.3187 | 5.95203 | 5.38859 | 6.77008 | 5.38096 | 5.44163 | 6.72085 | 5.89758 | 6.25695 | 6.00375 | 6.71578 | 7.25825 | 6.96716 | 6.58506 | 7.09941 |
| FKBP3 | Q00688 | 6.33826 | 6.22536 | 6.32409 | 6.56459 | 6.60312 | 6.65903 | 5.88662 | 6.60054 | 6.32343 | 6.43836 | 6.70803 | 6.3543 | 6.34138 | 6.06792 | 6.06273 | 5.50225 |
| RANBP1 | P43487 | 6.78849 | 6.31622 | 6.94439 | 6.29351 | 6.69557 | 6.93291 | 6.09978 | 6.24741 | 6.69795 | 6.16676 | 6.68974 | 6.57216 | 5.4156 | 5.66934 | 5.05585 | 5.41432 |
| HCCS | P53701 | 5.97211 | 5.90496 | 5.87963 | 6.39669 | 6.62805 | 6.04522 | 6.26603 | 6.53074 | 6.22735 | 6.0014 | 6.18435 | 6.24257 | 6.93482 | 6.21371 | 6.24936 | 6.323 |
| GSPT2 | Q8IYD1 | 6.16805 | 6.30802 | 6.86968 | 5.87576 | 5.78131 | 6.79767 | 5.94573 | 6.78118 | 6.08633 | 6.58322 | 6.73426 | 6.82516 | 5.50953 | 6.01212 | 5.71297 | 6.00901 |
| PNKP | Q96T60 | 5.36032 | 5.76315 | 5.8608 | 5.78065 | 5.83947 | 5.87495 | 5.75818 | 6.13723 | 6.12406 | 6.08743 | 6.56385 | 6.24354 | 7.22729 | 7.17618 | 7.27634 | 6.92656 |
| POLR1C | O15160 | 6.30944 | 6.37677 | 6.49818 | 6.21395 | 6.11391 | 6.59543 | 5.8448 | 6.73598 | 6.34538 | 5.97407 | 6.49808 | 6.50418 | 6.43443 | 5.59799 | 5.95028 | 6.00715 |
| CNBP | P62633-6 | 6.44111 | 6.11805 | 6.29718 | 5.80007 | 6.4897 | 6.58292 | 5.92792 | 6.11755 | 6.66502 | 6.89977 | 6.64655 | 6.36349 | 6.26011 | 5.83816 | 5.74479 | 5.80761 |
| ITGA1 | P56199 | 6.03593 | 6.23267 | 5.79365 | 6.29514 | 6.05698 | 6.11163 | 6.19972 | 6.12688 | 5.90051 | 6.38938 | 6.02193 | 6.1986 | 6.57144 | 6.66369 | 6.93437 | 6.46749 |
| RFWD3 | Q6PCD5 | 6.59538 | 6.57989 | 7.45519 | 7.17984 | 6.61986 | 6.59942 | 6.42226 | 6.248 | 7.07244 | 6.52954 | 6.51392 | 6.65834 | 4.83683 | 4.8833 | 4.8609 | 4.94489 |
| PRR14L | Q5THK1 | 6.31417 | 6.30117 | 6.1305 | 6.65224 | 6.29921 | 6.1531 | 6.5372 | 6.25719 | 6.39686 | 6.41544 | 6.58393 | 6.58486 | 5.87701 | 5.57475 | 5.89 | 6.03238 |
| MICALL1 | Q8N3F8 | 6.37669 | 6.63105 | 6.74456 | 6.08531 | 6.09243 | 6.38289 | 6.23392 | 6.29382 | 6.20594 | 6.37478 | 6.75148 | 6.69144 | 5.8319 | 5.71218 | 5.79728 | 5.79434 |
| TYW1 | Q9NV66 | 6.45468 | 6.7512 | 6.39111 | 6.54687 | 6.1365 | 5.96765 | 6.21835 | 6.5048 | 5.99836 | 6.41851 | 6.38339 | 6.18547 | 6.07447 | 6.10384 | 5.78853 | 6.07626 |
| FBXO30 | Q8TB52 | 6.339 | 6.17692 | 6.84633 | 5.95081 | 6.53017 | 6.63409 | 6.12898 | 6.64477 | 6.4809 | 6.40949 | 6.55735 | 6.61514 | 6.01965 | 5.65639 | 5.55578 | 5.45423 |
| FRG1 | Q14331 | 5.70547 | 6.01455 | 6.08614 | 5.8696 | 5.75647 | 5.96885 | 5.99645 | 5.81433 | 6.0442 | 6.3629 | 6.05655 | 6.07118 | 6.62587 | 7.28787 | 7.28248 | 7.05708 |
| FASTKD1 | Q53R41 | 6.53461 | 6.08134 | 6.04695 | 7.02819 | 6.15637 | 6.04136 | 6.5616 | 6.03373 | 6.47533 | 6.54543 | 6.13418 | 6.47267 | 6.19694 | 5.77912 | 6.14917 | 6.463 |
| CSRP1 | P21291 | 6.43564 | 6.08146 | 6.69063 | 6.01891 | 6.57501 | 6.55012 | 6.18627 | 6.27408 | 6.65308 | 6.32324 | 6.39435 | 6.5084 | 5.96498 | 6.0643 | 5.79813 | 5.59732 |
| WDR6 | Q9NNW5 | 6.5702 | 6.60743 | 7.08564 | 6.47523 | 6.31978 | 6.59527 | 6.70893 | 6.21051 | 6.75475 | 6.73583 | 6.88932 | 6.68288 | 5.13851 | 5.05019 | 5.08978 | 5.08575 |
| CACUL1 | Q86Y37 | 5.73204 | 5.05921 | 5.65311 | 4.61646 | 5.57152 | 5.72913 | 5.49253 | 6.24145 | 5.74478 | 6.52947 | 6.33435 | 6.78444 | 7.18629 | 8.02126 | 7.91287 | 7.39109 |
| CHD1L | Q86WJ1 | 5.763 | 5.96983 | 6.39589 | 5.336 | 5.57593 | 6.38429 | 5.88884 | 5.93108 | 6.12454 | 6.09168 | 6.69359 | 6.32979 | 6.51382 | 7.04069 | 7.1 | 6.86103 |
| PLEKHO2 | Q8TD55 | 6.48258 | 5.90211 | 6.52847 | 6.03455 | 6.66583 | 6.05036 | 5.83504 | 6.26419 | 6.34839 | 6.69971 | 6.58971 | 6.24007 | 6.65184 | 6.2515 | 5.9356 | 5.52004 |
| PRPF18 | Q99633 | 6.23634 | 6.34019 | 6.12021 | 5.86968 | 6.1109 | 6.35155 | 6.17295 | 6.57456 | 6.27491 | 6.34172 | 6.14226 | 6.11426 | 6.06054 | 6.6389 | 6.42219 | 6.22884 |
| TPMT | P51580 | 6.17674 | 6.29418 | 6.74932 | 5.93866 | 6.28001 | 6.88007 | 6.17701 | 6.55078 | 6.25711 | 6.57655 | 6.46196 | 6.58515 | 5.8883 | 5.87565 | 5.38811 | 5.92038 |
| APTX | Q7Z2E3 | 5.72517 | 6.27979 | 5.92329 | 5.7111 | 5.98801 | 6.13057 | 5.92686 | 6.00611 | 6.06852 | 6.32068 | 6.27783 | 6.06062 | 7.07022 | 6.76253 | 7.08746 | 6.66124 |
| NIPSNAP2 | O75323 | 6.15946 | 6.00797 | 6.10633 | 6.39327 | 6.43477 | 5.89311 | 6.73643 | 5.92621 | 6.42956 | 5.8168 | 6.39381 | 5.61868 | 6.56602 | 6.00252 | 6.93243 | 6.58264 |
| MCU | Q8NE86 | 6.45706 | 6.11692 | 6.00248 | 6.38438 | 6.50337 | 5.78526 | 6.48687 | 5.87152 | 6.29097 | 6.00112 | 6.03746 | 5.88822 | 6.95119 | 6.15003 | 6.84785 | 6.2223 |
| TUFT1 | Q9NNX1 | 6.70889 | 5.90232 | 6.84451 | 6.52856 | 6.41865 | 6.72666 | 6.34229 | 7.16652 | 6.27822 | 6.05397 | 6.76496 | 6.70262 | 5.61877 | 5.37941 | 5.09319 | 5.47047 |
| SLAIN2 | Q9P270 | 6.2398 | 6.22931 | 6.41826 | 5.64844 | 6.66526 | 6.60068 | 5.76654 | 6.88194 | 6.53375 | 6.35949 | 6.50327 | 6.38226 | 6.49877 | 5.91612 | 5.62685 | 5.72925 |
| UFSP2 | Q9NUQ7 | 6.19989 | 6.39563 | 6.15307 | 6.61092 | 6.11827 | 5.75934 | 6.30045 | 6.10475 | 6.31827 | 5.87126 | 6.23958 | 5.84237 | 6.53191 | 6.25316 | 6.66571 | 6.63541 |
| AGRN | O00468 | 7.28021 | 7.13625 | 6.72003 | 7.25347 | 6.29625 | 5.67068 | 6.11659 | 5.89703 | 6.79892 | 7.0435 | 7.06055 | 6.61841 | 4.97821 | 4.91355 | 5.08046 | 5.1359 |
| RRP15 | Q9Y3B9 | 6.28684 | 6.40797 | 6.19982 | 7.14311 | 6.23455 | 5.84471 | 6.61332 | 6.12809 | 5.90586 | 6.20475 | 6.16358 | 5.88352 | 6.63903 | 6.18896 | 6.2616 | 5.89429 |
| DHX33 | Q9H6R0 | 6.18552 | 6.44017 | 6.14066 | 7.11599 | 6.18509 | 5.8647 | 6.31439 | 6.05764 | 6.18421 | 5.86294 | 6.1144 | 5.96746 | 6.3742 | 6.12198 | 6.40548 | 6.66517 |
| SERPINE1 | P05121 | 6.30032 | 5.22906 | 4.96587 | 6.59592 | 6.12132 | 6.24836 | 7.9408 | 6.55545 | 5.49641 | 4.36721 | 4.81467 | 5.00909 | 8.61529 | 5.71527 | 8.04102 | 7.98393 |
| SYAP1 | Q96A49 | 6.15197 | 5.9805 | 6.6337 | 5.8575 | 6.40389 | 6.68409 | 5.97737 | 7.00804 | 6.30412 | 6.64634 | 6.72624 | 6.92373 | 6.04185 | 5.77251 | 5.3415 | 5.54665 |
| COPS8 | Q99627 | 5.48182 | 4.8572 | 5.42877 | 4.40864 | 5.73753 | 5.31427 | 5.52207 | 6.87501 | 5.35885 | 6.3071 | 5.57877 | 6.68248 | 6.30288 | 9.07283 | 8.33945 | 8.73234 |
| HDHD3 | Q9BSH5 | 5.86371 | 5.20195 | 5.56037 | 5.43786 | 6.00676 | 5.66745 | 5.84263 | 6.40295 | 5.64203 | 6.42231 | 5.64121 | 6.584 | 6.46355 | 7.48908 | 7.80543 | 7.96871 |
| TRAPPC9 | Q96Q05-2 | 6.21747 | 6.49727 | 6.59805 | 6.37011 | 6.29827 | 6.60116 | 6.33684 | 6.14764 | 5.98985 | 5.9408 | 6.36009 | 5.87864 | 5.89017 | 6.31406 | 6.45364 | 6.10596 |
| PAM | P19021-5 | 6.76264 | 6.71188 | 6.37458 | 6.79925 | 6.54066 | 6.55419 | 6.7587 | 6.4826 | 6.0638 | 6.34909 | 6.29653 | 6.55913 | 5.39249 | 5.55257 | 5.22043 | 5.58146 |
| TFB1M | Q8WVM0 | 6.44174 | 6.41126 | 6.09845 | 7.06101 | 6.162 | 6.05157 | 6.70635 | 6.01458 | 6.04063 | 6.21146 | 5.95706 | 6.17733 | 6.31377 | 5.72215 | 6.30937 | 6.32128 |
| PSMB6 | P28072 | 6.07012 | 6.23748 | 6.10554 | 5.71227 | 6.61894 | 6.72998 | 6.49534 | 6.46796 | 5.99251 | 6.00788 | 6.59465 | 6.29934 | 5.903 | 6.39617 | 5.81137 | 6.55744 |
| DYNLL1 | P63167 | 6.049 | 5.5811 | 6.05967 | 5.32938 | 6.49371 | 6.25592 | 5.85687 | 6.55854 | 5.45656 | 6.02959 | 5.75316 | 6.0509 | 5.93145 | 8.10932 | 7.14389 | 7.34094 |
| THAP12 | O43422 | 5.92765 | 6.52943 | 6.18432 | 6.37892 | 6.02 | 6.01193 | 6.06144 | 5.88042 | 6.19535 | 6.32362 | 6.23334 | 6.14811 | 6.613 | 6.53166 | 6.41562 | 6.5452 |
| ELP3 | Q9H9T3 | 6.00755 | 6.1962 | 6.62893 | 6.11931 | 6.40488 | 7.30889 | 6.48519 | 6.43401 | 6.06294 | 6.11574 | 6.74064 | 6.53724 | 5.98111 | 6.03144 | 5.53713 | 5.4088 |
| MIGA1 | Q8NAN2 | 6.55424 | 6.27724 | 5.76288 | 6.55155 | 6.54624 | 5.45743 | 6.39586 | 5.6202 | 6.39013 | 6.34622 | 6.18273 | 6.07248 | 6.49689 | 6.39718 | 6.15367 | 6.79507 |
| HAUS3 | Q68CZ6 | 6.675 | 6.3895 | 7.08069 | 6.34076 | 6.3027 | 6.58677 | 6.20164 | 6.63309 | 6.42055 | 7.0274 | 6.81513 | 6.94881 | 5.137 | 5.37218 | 4.94768 | 5.12111 |
| WTAP | Q15007 | 6.59587 | 6.27026 | 6.28618 | 6.64919 | 6.55363 | 6.36668 | 5.95367 | 6.35587 | 6.01504 | 6.2106 | 6.34106 | 6.60915 | 6.70093 | 5.53965 | 5.77356 | 5.77866 |
| CCNK | O75909-4 | 6.50887 | 6.15016 | 5.98452 | 6.30865 | 6.36608 | 6.14088 | 6.25654 | 6.11319 | 6.14779 | 6.03022 | 5.86836 | 6.06294 | 6.62825 | 6.17466 | 6.53488 | 6.72403 |
| FIBP | O43427 | 6.28906 | 6.39041 | 6.65289 | 6.2677 | 6.24585 | 6.44866 | 6.24259 | 6.44653 | 6.39558 | 6.57862 | 6.50249 | 6.38453 | 5.72201 | 5.78481 | 5.70483 | 5.94344 |
| TAF1C | Q15572 | 6.28165 | 6.26228 | 6.43252 | 7.05704 | 6.18941 | 5.97696 | 6.45411 | 5.81809 | 6.47778 | 6.2579 | 6.26761 | 6.21278 | 6.19311 | 5.70397 | 6.17387 | 6.24091 |
| SMNDC1 | O75940 | 5.91423 | 6.35586 | 5.83038 | 6.1138 | 5.99201 | 6.13741 | 6.00987 | 5.83982 | 5.91786 | 5.91627 | 6.1905 | 6.03133 | 6.83378 | 6.88952 | 7.18362 | 6.84375 |
| CWC15 | Q9P013 | 6.23991 | 6.48204 | 5.75583 | 6.4999 | 6.14141 | 6.1379 | 6.15812 | 6.26551 | 6.23231 | 6.11519 | 6.11687 | 6.34467 | 6.84851 | 6.26269 | 6.01788 | 6.38124 |
| SAP30BP | Q9UHR5 | 6.13965 | 5.8817 | 5.97438 | 6.09153 | 5.98389 | 6.05878 | 6.01767 | 6.31882 | 5.87987 | 6.06378 | 6.33314 | 6.10301 | 6.91196 | 6.61863 | 6.74163 | 6.88157 |
| N4BP2 | Q86UW6 | 6.14371 | 6.13999 | 7.07775 | 5.84872 | 5.74337 | 6.79765 | 6.30604 | 6.43533 | 6.51146 | 6.26246 | 7.21953 | 6.72332 | 5.3049 | 5.87455 | 5.74552 | 5.8657 |
| LRIF1 | Q5T3J3 | 6.84569 | 6.9698 | 6.3647 | 7.49367 | 6.29023 | 5.95399 | 6.10785 | 6.09618 | 6.21598 | 6.31885 | 6.28637 | 6.31099 | 5.59251 | 5.7283 | 5.46329 | 5.96159 |
| TDP1 | Q9NUW8 | 5.71602 | 5.89674 | 5.85802 | 5.31961 | 5.76264 | 6.40182 | 5.59626 | 6.32957 | 6.00532 | 6.1789 | 6.25498 | 6.25085 | 6.51018 | 7.41949 | 7.37981 | 7.11977 |
| FADS1 | A0A0A0MR51 | 6.96979 | 6.95711 | 6.44469 | 6.79972 | 6.34076 | 6.30458 | 6.5776 | 6.07673 | 6.40821 | 6.58146 | 6.4052 | 6.33351 | 5.51159 | 5.56157 | 5.24311 | 5.48436 |
| STAMBP | O95630 | 6.28575 | 6.2701 | 6.73641 | 5.66009 | 6.62679 | 6.67327 | 6.08935 | 6.50253 | 6.63466 | 6.29726 | 6.362 | 6.31783 | 6.01229 | 5.88024 | 5.99653 | 6.5549 |
| SLC12A4 | Q9UP95 | 6.05822 | 6.71893 | 6.49514 | 6.38441 | 6.05591 | 6.44719 | 6.27537 | 6.33874 | 6.4875 | 6.29993 | 6.51211 | 6.4766 | 5.79737 | 6.09826 | 5.78499 | 5.76935 |
| EXOC6 | Q8TAG9 | 6.35088 | 6.28561 | 6.48534 | 6.00883 | 6.65291 | 6.58694 | 6.269 | 6.55753 | 6.71567 | 6.56717 | 6.63573 | 6.4135 | 5.76414 | 5.61024 | 5.58401 | 5.5125 |
| LMF2 | Q9BU23 | 6.27901 | 6.55109 | 5.84008 | 6.89195 | 6.17072 | 6.07415 | 6.33082 | 6.01969 | 6.196 | 6.1397 | 5.83114 | 6.01299 | 6.92657 | 5.94434 | 6.64817 | 6.14359 |
| POLD3 | Q15054 | 5.1225 | 6.05249 | 6.10469 | 5.68129 | 5.67759 | 6.14606 | 5.84245 | 5.67194 | 5.97741 | 6.15437 | 6.37851 | 6.14558 | 6.78766 | 7.30076 | 7.62941 | 7.32729 |
| FLCN | Q8NFG4 | 6.82732 | 6.50031 | 7.31008 | 6.35697 | 6.08031 | 6.67033 | 6.26453 | 6.55963 | 6.77002 | 6.99924 | 6.91013 | 6.99351 | 4.75326 | 5.00141 | 4.75463 | 5.24831 |
| TBC1D13 | Q9NVG8 | 5.88045 | 5.27372 | 6.72264 | 5.20892 | 5.91939 | 5.89874 | 5.83483 | 6.45337 | 6.02802 | 6.51821 | 6.44264 | 6.81791 | 6.22906 | 7.19704 | 7.00564 | 6.56943 |
| NOP53 | Q9NZM5 | 6.52599 | 6.79248 | 6.04651 | 6.89084 | 6.25534 | 5.92376 | 6.54842 | 6.2187 | 6.90622 | 6.00305 | 6.19273 | 6.04116 | 6.22579 | 5.79805 | 5.70328 | 5.92766 |
| NAA50 | Q9GZZ1 | 6.43019 | 6.10065 | 7.0285 | 5.83726 | 6.32839 | 6.6964 | 6.24959 | 6.8202 | 6.53307 | 6.33804 | 6.96239 | 6.58711 | 5.77097 | 5.58175 | 5.51993 | 5.21556 |
| ZNF281 | Q9Y2X9 | 6.11842 | 6.42588 | 6.35347 | 6.80415 | 6.04376 | 6.25236 | 6.68856 | 5.88481 | 6.25279 | 6.13976 | 6.44786 | 6.36113 | 6.10275 | 5.75435 | 6.17363 | 6.19631 |
| NMNAT1 | Q9HAN9 | 5.76375 | 5.7566 | 5.97779 | 5.56689 | 5.91149 | 5.6522 | 6.06614 | 5.89358 | 6.17165 | 6.15226 | 6.20204 | 6.0175 | 7.08028 | 7.18181 | 7.58674 | 7.01929 |
| KIAA0319L | Q8IZA0 | 6.54173 | 6.68861 | 5.98503 | 6.66248 | 6.41014 | 6.12872 | 6.42255 | 6.18656 | 6.33906 | 6.27708 | 6.09776 | 6.07942 | 6.14052 | 5.9656 | 6.04515 | 6.02958 |
| MEN1 | O00255 | 6.51424 | 6.44775 | 6.10999 | 6.97122 | 5.83979 | 6.00827 | 6.43187 | 5.84596 | 6.34108 | 6.12994 | 6.06711 | 6.06056 | 6.22163 | 6.20229 | 6.54339 | 6.26493 |
| KPNA6 | O60684 | 6.14283 | 6.0743 | 6.2076 | 5.81434 | 6.17625 | 6.36448 | 6.22435 | 6.10354 | 6.2703 | 6.06649 | 6.1178 | 6.12519 | 6.48096 | 6.3278 | 6.91456 | 6.5892 |
| DYNC2I2 | Q96EX3 | 6.72777 | 6.29813 | 6.97809 | 6.275 | 6.42312 | 7.23679 | 6.20522 | 6.86798 | 6.62999 | 6.47924 | 6.58767 | 6.89784 | 5.19441 | 5.41925 | 4.79501 | 4.98449 |
| SCO1 | O75880 | 6.63802 | 6.39862 | 5.90073 | 6.80547 | 6.30223 | 5.767 | 6.09658 | 6.10013 | 5.9988 | 6.23848 | 6.10437 | 5.76451 | 6.66823 | 6.2667 | 6.61451 | 6.33563 |
| RNPC3 | Q96LT9 | 6.32665 | 6.22475 | 6.01318 | 6.88448 | 6.11964 | 6.22456 | 6.17867 | 6.01697 | 5.85373 | 6.10294 | 6.05455 | 6.05168 | 6.86252 | 6.19764 | 6.38079 | 6.50726 |

| | | | | | | | | | | | | | | | | | |
|---|---|---|---|---|---|---|---|---|---|---|---|---|---|---|---|---|---|
| RASSF8 | Q8NHQ8 | 6.43654 | 6.22382 | 6.52471 | 6.59535 | 6.20134 | 6.08605 | 6.24059 | 6.11254 | 6.48788 | 6.23886 | 6.39722 | 6.13706 | 6.078 | 5.89682 | 6.206 | 6.13721 |
| PCYT1A | P49585 | 5.54147 | 5.35704 | 6.32621 | 5.41114 | 5.70112 | 6.53938 | 5.47722 | 6.31591 | 6.21527 | 6.37923 | 6.46187 | 6.18293 | 6.59757 | 7.28389 | 7.14026 | 7.0695 |
| LNPK | Q9C0E8 | 6.54113 | 6.26908 | 5.82149 | 6.31797 | 6.37627 | 6.03207 | 6.29053 | 6.14142 | 5.99964 | 6.07128 | 6.04812 | 6.0807 | 6.31697 | 6.5121 | 6.66695 | 6.51426 |
| CTIF | O43310-2 | 6.59003 | 6.29469 | 6.99806 | 6.73158 | 6.12403 | 6.60491 | 6.35996 | 6.95676 | 6.61142 | 6.43903 | 6.82371 | 6.64185 | 5.36822 | 5.3789 | 4.9306 | 5.14626 |
| HMGN5 | P82970 | 5.85842 | 6.34028 | 5.70116 | 6.08845 | 5.9212 | 5.99933 | 6.33623 | 5.76383 | 6.06278 | 6.16974 | 6.05265 | 5.76869 | 7.00664 | 6.94111 | 6.70786 | 7.28164 |
| GLT8D1 | Q68CQ7 | 6.40889 | 6.45355 | 5.66754 | 6.54235 | 6.50461 | 6.25175 | 6.4346 | 6.42315 | 6.256 | 6.38888 | 5.83381 | 6.0236 | 6.24189 | 6.02902 | 6.34005 | 6.2003 |
| PIGK | Q92643 | 6.29886 | 6.33081 | 6.04442 | 6.6182 | 6.14649 | 5.7649 | 6.25576 | 5.83635 | 6.25652 | 6.38724 | 5.8662 | 6.096 | 6.34212 | 6.23886 | 6.98601 | 6.53125 |
| SSR1 | P43307 | 6.5329 | 5.76047 | 6.01797 | 6.6857 | 6.61588 | 5.73139 | 6.60567 | 5.69887 | 6.53474 | 5.86543 | 5.86717 | 6.02459 | 6.90946 | 6.053 | 6.74134 | 6.35542 |
| SNX27 | Q96L92 | 6.35437 | 6.32906 | 6.20129 | 6.1436 | 6.25946 | 6.42007 | 6.44584 | 6.21257 | 6.19502 | 6.3275 | 6.50134 | 6.36758 | 6.20696 | 5.6597 | 6.31282 | 6.06282 |
| ABRAXAS2 | Q15018 | 6.31516 | 6.09546 | 6.8656 | 5.88234 | 6.51412 | 6.3594 | 6.26508 | 6.36487 | 6.42411 | 6.42555 | 6.61009 | 6.36938 | 5.80034 | 6.11857 | 5.77134 | 5.8186 |
| TAB2 | Q9NYJ8 | 6.09978 | 6.36523 | 6.62164 | 5.92358 | 6.65479 | 6.87031 | 6.40879 | 6.86939 | 6.2142 | 6.24031 | 6.26027 | 6.4603 | 6.48103 | 5.53923 | 5.36444 | 5.6267 |
| SKA1 | Q96BD8 | 6.64462 | 6.60149 | 7.19027 | 6.261 | 6.3173 | 6.81049 | 6.45822 | 6.48005 | 6.96491 | 6.58047 | 7.06119 | 6.90687 | 5.06737 | 4.92215 | 4.76045 | 4.97313 |
| WDR91 | A4D1P6 | 6.49627 | 6.52779 | 6.51015 | 5.95998 | 6.26333 | 6.07227 | 6.00379 | 5.99668 | 6.43204 | 6.60187 | 6.31436 | 6.29288 | 6.46189 | 6.20401 | 6.02993 | 5.83276 |
| CEP89 | Q96ST8 | 6.4507 | 6.44658 | 7.0247 | 6.6672 | 6.48035 | 6.67472 | 6.6812 | 6.62995 | 7.00685 | 6.72844 | 7.00969 | 7.20071 | 4.56547 | 4.88471 | 4.60239 | 4.94636 |
| CWC25 | Q9NXE8 | 6.14709 | 6.19812 | 6.01013 | 7.14647 | 5.54296 | 5.82631 | 6.57469 | 6.34014 | 5.92754 | 5.86759 | 6.18922 | 5.88631 | 6.34596 | 6.80873 | 6.53879 | 6.64996 |
| AURKA | O14965 | 6.26479 | 5.9271 | 6.65572 | 6.92865 | 6.13104 | 6.33105 | 6.42877 | 6.05044 | 6.42623 | 6.56353 | 7.09708 | 7.04788 | 5.30396 | 5.39051 | 5.20109 | 6.25214 |
| ZNF830 | Q96NB3 | 6.67765 | 6.4895 | 7.04227 | 6.98956 | 6.44398 | 6.085 | 5.87245 | 5.97477 | 5.93019 | 5.93861 | 5.6053 | 6.11249 | 6.33495 | 6.04659 | 6.29543 | 6.16126 |
| HTRA2 | O43464 | 6.36492 | 6.18051 | 5.746 | 6.53186 | 6.36074 | 6.12381 | 6.15307 | 6.46411 | 6.07602 | 6.15989 | 5.98818 | 6.09844 | 6.77847 | 5.95691 | 6.43748 | 6.57959 |
| SNRPD1 | P62314 | 6.28934 | 6.80335 | 5.85908 | 6.97503 | 6.26416 | 6.51081 | 6.38956 | 5.9721 | 6.13453 | 5.98116 | 5.79392 | 5.97323 | 6.11325 | 5.88798 | 6.87007 | 6.18244 |
| VPS29 | Q9UBQ0-2 | 5.97278 | 5.93547 | 6.22244 | 5.56787 | 6.18793 | 6.43642 | 5.91432 | 6.57034 | 6.34612 | 6.43903 | 6.33191 | 6.3232 | 6.18835 | 6.62683 | 6.64867 | 6.28831 |
| MAP4K5 | Q9Y4K4 | 6.17598 | 6.53171 | 6.81224 | 6.06003 | 6.32195 | 6.40889 | 6.26708 | 6.57212 | 6.45917 | 6.61979 | 6.20627 | 6.54857 | 5.6451 | 5.67276 | 5.91729 | 5.78105 |
| SLC39A10 | Q9ULF5 | 5.86609 | 6.29319 | 6.04919 | 6.16695 | 6.15579 | 5.98852 | 6.60282 | 6.2421 | 6.10788 | 6.10942 | 5.55237 | 5.96715 | 6.41519 | 6.61523 | 6.86379 | 7.00434 |
| BCKDHB | P21953 | 5.83899 | 6.00635 | 5.62634 | 6.30152 | 6.44923 | 6.11552 | 6.54397 | 5.72009 | 6.32891 | 6.26158 | 6.15619 | 5.81714 | 6.80915 | 6.05505 | 7.04467 | 6.92529 |
| LBR | Q14739 | 6.45584 | 6.21634 | 6.35887 | 6.90112 | 5.90999 | 5.77818 | 6.35894 | 5.89224 | 6.02584 | 5.58266 | 6.30457 | 5.89199 | 6.32066 | 6.70946 | 6.37517 | 6.91811 |
| GPR89A | B7ZAQ6 | 6.12554 | 6.15664 | 6.02376 | 6.64112 | 5.93481 | 5.9227 | 5.99393 | 5.87396 | 6.04647 | 6.18944 | 5.96957 | 6.1709 | 6.70014 | 6.45972 | 6.72601 | 7.06528 |
| WDR59 | Q6PJI9 | 6.05495 | 6.16568 | 6.75425 | 6.46605 | 6.08002 | 6.65825 | 6.44583 | 6.20279 | 6.3925 | 6.80913 | 6.76252 | 6.52565 | 5.65814 | 5.76291 | 5.81475 | 5.44658 |
| UBFD1 | O14562 | 6.48793 | 6.24746 | 6.81581 | 6.02842 | 6.41024 | 6.58799 | 6.13641 | 6.92133 | 6.09494 | 6.47751 | 6.72103 | 6.59497 | 5.84591 | 5.61809 | 5.53041 | 5.48152 |
| SAMD9 | Q5K651 | 6.17777 | 6.35516 | 6.86411 | 5.90699 | 6.0622 | 6.22273 | 6.32555 | 6.42794 | 6.36616 | 6.52731 | 6.44832 | 6.77523 | 5.74503 | 6.00652 | 5.85813 | 5.93085 |
| MRPS7 | Q9Y2R9 | 6.53441 | 6.58341 | 6.21231 | 6.43624 | 6.59016 | 6.36002 | 6.54067 | 6.18349 | 6.1452 | 6.05428 | 6.17889 | 5.73732 | 6.59876 | 6.01931 | 5.98431 | 5.84123 |
| ARMH3 | Q5T2E6 | 6.22366 | 5.93183 | 6.34268 | 6.17128 | 6.06207 | 6.32306 | 6.50808 | 6.09081 | 6.61657 | 6.439 | 6.55373 | 6.36399 | 5.99165 | 6.17844 | 6.0902 | 6.11284 |
| HSD17B12 | Q53GQ0 | 6.57137 | 6.13957 | 6.01914 | 6.46416 | 6.32682 | 5.96586 | 6.244 | 5.79252 | 6.28161 | 6.10075 | 5.9693 | 5.80436 | 6.77653 | 6.02315 | 7.20312 | 6.31772 |
| RAB18 | Q9NP72 | 5.92173 | 6.08016 | 6.20664 | 6.38869 | 6.41866 | 6.07075 | 6.45882 | 5.796 | 6.18186 | 5.72313 | 6.01444 | 5.914 | 6.65592 | 6.51627 | 6.86848 | 6.78444 |
| MRPL17 | Q9NRX2 | 6.70822 | 6.34945 | 6.0175 | 6.13251 | 6.75466 | 5.68767 | 6.31208 | 6.33718 | 6.34625 | 6.14121 | 6.41177 | 6.17596 | 6.83051 | 5.72479 | 6.15233 | 5.9179 |
| SF3B6 | Q9Y3B4 | 6.56588 | 6.37855 | 5.93986 | 6.37557 | 6.52115 | 6.48774 | 5.97322 | 6.64862 | 6.22259 | 5.94604 | 6.33599 | 6.01155 | 6.36927 | 5.84663 | 6.42058 | 5.95677 |
| PQBP1 | O60828 | 6.22441 | 6.01439 | 6.16098 | 6.06237 | 6.29072 | 6.36139 | 6.11744 | 6.18371 | 6.37075 | 6.21116 | 6.01532 | 5.96498 | 6.59856 | 6.52443 | 6.38344 | 6.51592 |
| RIN1 | Q13671 | 6.55073 | 6.25517 | 6.66935 | 5.85516 | 6.51904 | 6.71971 | 6.38842 | 6.66355 | 6.26809 | 6.53657 | 6.53671 | 6.50106 | 5.94742 | 5.7092 | 5.41822 | 5.4616 |
| ZFPL1 | O95159 | 6.30248 | 6.86815 | 5.75512 | 6.78157 | 6.4785 | 6.5529 | 6.56148 | 6.12983 | 6.0612 | 6.23673 | 6.15048 | 5.99278 | 6.40154 | 5.82327 | 5.9237 | 5.98028 |
| PM20D2 | Q8IYS1 | 6.09943 | 5.65727 | 6.50317 | 5.32885 | 6.55782 | 6.6363 | 5.95311 | 7.14203 | 6.29175 | 6.54686 | 6.89925 | 7.03809 | 5.85433 | 5.86882 | 5.74062 | 5.88229 |
| NSRP1 | Q9H0G5 | 6.09929 | 6.47097 | 5.85527 | 6.5777 | 6.2275 | 6.2705 | 6.20004 | 6.0182 | 6.10079 | 6.19401 | 6.00202 | 5.84332 | 6.38268 | 6.55405 | 6.67164 | 6.53201 |
| NDST1 | P52848 | 6.46467 | 6.32026 | 6.06227 | 6.65715 | 6.17831 | 6.06067 | 6.44024 | 6.35067 | 6.21047 | 6.01963 | 6.03254 | 6.19925 | 6.03451 | 6.28455 | 6.04758 | 6.63722 |
| BRD3 | Q15059 | 6.13785 | 6.53078 | 6.17813 | 6.60441 | 6.03491 | 6.12634 | 6.3918 | 6.15367 | 6.24456 | 6.23737 | 6.12206 | 6.39629 | 6.18698 | 6.23716 | 6.22144 | 6.19625 |
| IRF2BP1 | Q8IU81 | 5.98353 | 6.18059 | 5.89138 | 6.00985 | 6.2591 | 6.36079 | 5.96652 | 6.25542 | 6.30539 | 6.5505 | 6.36135 | 6.07256 | 5.95715 | 6.30645 | 6.35542 | 6.18399 |
| LRRC8E | Q6NSJ5 | 6.76445 | 6.59809 | 6.3095 | 6.98188 | 5.82693 | 6.15445 | 6.19422 | 6.02661 | 5.93968 | 6.68068 | 6.02362 | 6.58286 | 6.04964 | 5.52325 | 6.05108 | 6.29305 |
| PURA | Q00577 | 6.23321 | 6.20401 | 6.68813 | 6.0182 | 6.47533 | 6.57838 | 6.43765 | 6.23694 | 6.62602 | 6.42915 | 6.53737 | 6.26967 | 6.13228 | 5.90451 | 5.76759 | 5.46157 |
| TPM2 | A7XZE4 | 6.28209 | 6.29633 | 6.82162 | 6.48117 | 6.5436 | 6.58451 | 5.52117 | 7.13313 | 6.16417 | 6.33689 | 6.61745 | 6.85298 | 6.26235 | 5.60622 | 4.94529 | 5.55104 |
| EIF2B5 | Q13144 | 6.06422 | 6.05821 | 6.87404 | 5.7869 | 6.26327 | 6.71051 | 6.08101 | 6.34012 | 6.40076 | 6.55579 | 6.89744 | 6.51824 | 5.89336 | 5.92652 | 5.93256 | 5.69704 |
| SRPK2 | P78362-2 | 6.37693 | 6.5163 | 6.46082 | 6.1484 | 6.23902 | 6.06877 | 6.10878 | 6.09318 | 6.67788 | 6.37193 | 6.54636 | 6.47843 | 5.86134 | 6.05948 | 6.10138 | 5.89101 |
| KNOP1 | Q1ED39 | 5.94739 | 6.08041 | 5.76443 | 6.67276 | 5.62886 | 5.47509 | 6.07542 | 5.60541 | 6.30194 | 6.04806 | 6.10463 | 6.2807 | 6.26063 | 7.13167 | 7.26338 | 7.35922 |
| SCLY | Q96I15 | 5.66561 | 5.73967 | 6.689 | 5.66524 | 5.94607 | 6.60359 | 5.68184 | 5.82618 | 6.05299 | 6.66265 | 6.26308 | 6.44968 | 6.66767 | 6.76538 | 6.98914 | 6.3322 |
| IGBP1 | P78318 | 6.38585 | 5.94872 | 6.90762 | 5.95327 | 6.38781 | 6.37208 | 5.80399 | 6.9576 | 6.38181 | 6.53197 | 7.06896 | 7.18976 | 5.79611 | 5.66257 | 5.40119 | 5.25068 |
| SLC16A1 | P53985 | 6.37136 | 6.44755 | 5.72367 | 6.26954 | 6.07102 | 5.92833 | 6.17346 | 6.29094 | 6.1166 | 5.95452 | 5.83609 | 6.13175 | 6.75036 | 6.53709 | 6.78426 | 6.61345 |
| NDUFAF4 | Q9P032 | 6.4363 | 6.04104 | 6.05648 | 6.00719 | 6.71041 | 5.73277 | 5.95695 | 6.17772 | 6.1688 | 5.75211 | 6.2068 | 6.26016 | 7.16736 | 6.33852 | 6.63984 | 6.34755 |
| RC3H1 | Q5TC82 | 6.23362 | 6.15418 | 6.18277 | 6.25338 | 6.19602 | 5.97504 | 6.21211 | 6.29465 | 6.22439 | 6.4518 | 6.41267 | 6.84851 | 6.14263 | 5.94445 | 6.06113 | 6.41263 |
| PALLD | Q8WX93 | 6.30928 | 5.99302 | 6.64496 | 6.15743 | 6.4179 | 6.20472 | 6.01691 | 6.24008 | 6.54195 | 6.7267 | 6.57076 | 6.74016 | 6.43244 | 5.86129 | 5.48941 | 5.65298 |
| OXSR1 | O95747 | 6.38224 | 6.28575 | 6.62314 | 5.72439 | 6.47211 | 6.65132 | 6.10468 | 6.50266 | 6.47496 | 6.43638 | 6.19663 | 6.30738 | 6.06479 | 6.16376 | 5.83404 | 5.77578 |
| CDK17 | Q00537-2 | 5.59146 | 6.32593 | 6.4526 | 6.25421 | 5.98104 | 6.80156 | 6.65504 | 6.71496 | 5.93246 | 6.26934 | 6.36679 | 6.44136 | 6.0498 | 6.09336 | 5.93251 | 6.13758 |
| PRKD2 | Q9BZL6-3 | 6.34236 | 6.45492 | 7.37075 | 6.32502 | 6.35422 | 6.82413 | 6.1117 | 6.66298 | 6.21465 | 6.45533 | 6.59855 | 6.69117 | 5.4562 | 5.63222 | 5.20241 | 5.30339 |
| MRPL2 | Q5T653 | 6.27585 | 6.35481 | 5.77195 | 6.3376 | 6.51773 | 5.91149 | 6.80679 | 6.30572 | 6.20013 | 6.04761 | 5.76481 | 5.75867 | 6.5569 | 6.41577 | 6.43652 | 6.53765 |
| PTPRS | Q13332 | 7.12423 | 6.9229 | 6.68351 | 6.52762 | 6.14869 | 5.98769 | 6.36858 | 5.88931 | 6.88717 | 6.72507 | 6.36852 | 6.67879 | 5.54476 | 6.07641 | 5.18078 | 4.88597 |
| GPR107 | Q5VW38-2 | 6.13058 | 6.19275 | 5.93177 | 6.29734 | 6.41027 | 6.13431 | 6.12597 | 5.79131 | 6.30084 | 6.44728 | 6.22212 | 6.08939 | 6.65454 | 6.52927 | 6.41371 | 6.32856 |
| SHMT1 | P34896 | 6.22666 | 6.26483 | 6.90347 | 5.60342 | 6.52274 | 6.86256 | 6.20471 | 6.75368 | 6.6765 | 6.56313 | 6.50458 | 6.24626 | 5.77145 | 5.67556 | 5.8168 | 5.40365 |
| MTR | Q99707 | 6.7586 | 6.37442 | 7.43783 | 6.24595 | 6.6554 | 6.88749 | 6.37181 | 6.70335 | 6.84677 | 6.83145 | 6.82352 | 7.12556 | 4.64816 | 4.77541 | 4.60298 | 4.91131 |
| NAA10 | P41227 | 6.29995 | 6.44229 | 6.55032 | 5.84571 | 6.41355 | 6.89401 | 5.90295 | 7.2517 | 6.55048 | 6.74464 | 6.62028 | 6.96055 | 5.3472 | 5.70373 | 5.01624 | 5.4564 |
| THUMPD3 | Q9BV44 | 6.66452 | 6.59963 | 6.70973 | 6.1629 | 6.42185 | 6.7495 | 6.36043 | 6.84279 | 6.42614 | 6.30311 | 5.99402 | 6.04824 | 5.8474 | 5.79223 | 5.48434 | 5.59318 |
| GGCX | P38435 | 6.1943 | 6.31556 | 6.26681 | 6.56846 | 5.88113 | 6.14618 | 6.38673 | 5.85757 | 5.86019 | 6.34322 | 6.24356 | 6.12736 | 6.13346 | 6.57188 | 6.58243 | 6.52117 |
| TRA2A | Q13595 | 6.27726 | 6.55802 | 6.11113 | 6.7148 | 6.35671 | 5.76096 | 6.20835 | 5.72136 | 6.16423 | 6.02588 | 6.39079 | 6.30842 | 6.46636 | 5.78379 | 6.49613 | 6.65582 |
| NFYC | Q13952 | 5.80787 | 5.26481 | 5.95306 | 5.17885 | 5.70735 | 5.55078 | 5.92328 | 5.4973 | 5.71601 | 6.58069 | 6.52727 | 6.97995 | 7.63068 | 6.79815 | 7.79934 | 7.08461 |
| MBNL1 | Q9NR56 | 6.46559 | 6.59917 | 6.34025 | 5.59995 | 6.4648 | 6.72519 | 5.7531 | 5.94756 | 6.11026 | 6.15512 | 6.13512 | 6.21007 | 6.62506 | 6.28498 | 6.34138 | 6.24241 |
| PHKB | Q93100-3 | 6.3093 | 6.1973 | 6.7412 | 5.65039 | 6.31559 | 6.63126 | 6.13802 | 6.76677 | 6.56679 | 6.3609 | 6.36215 | 6.29321 | 6.00135 | 6.1623 | 5.68477 | 5.81869 |
| STBD1 | O95210 | 6.624 | 6.64062 | 6.09711 | 6.61654 | 6.07358 | 6.3639 | 6.33743 | 6.1009 | 5.95589 | 6.29447 | 5.86745 | 6.01689 | 6.27096 | 6.19274 | 6.24254 | 6.30499 |
| EPB41L5 | Q9HCM4 | 6.17441 | 6.32441 | 6.27374 | 6.48782 | 5.68652 | 5.99551 | 6.41277 | 5.72004 | 6.46826 | 6.27724 | 6.1604 | 6.1901 | 6.33558 | 6.27762 | 6.60379 | 6.6118 |
| JADE2 | Q9NQC1 | 5.97079 | 6.54791 | 6.30205 | 6.76977 | 5.82028 | 6.14284 | 6.11116 | 5.99116 | 6.4566 | 6.46157 | 6.47169 | 6.63775 | 5.79847 | 6.25375 | 5.93412 | 6.3301 |
| TBCC | Q15814 | 6.0872 | 6.18819 | 7.17256 | 5.84819 | 5.83435 | 6.44668 | 6.04112 | 6.72488 | 6.23245 | 6.39378 | 6.52873 | 6.87812 | 5.93625 | 5.91213 | 5.82259 | 5.9528 |
| ISG20L2 | Q9H9L3 | 6.0183 | 6.41259 | 5.82133 | 7.12231 | 5.8452 | 6.6356 | 6.43871 | 5.90713 | 6.05011 | 6.05278 | 6.13592 | 6.39871 | 6.44158 | 6.34028 | 6.60842 | 6.68968 |
| FGD4 | F8VWL3 | 6.36691 | 6.3801 | 6.40129 | 6.28018 | 5.93079 | 6.37043 | 6.00016 | 5.94917 | 6.14954 | 6.04704 | 6.40375 | 6.33657 | 6.19973 | 6.41885 | 6.25089 | 6.51461 |
| PRORP | O15091 | 6.16237 | 6.68109 | 5.80871 | 6.65412 | 6.35466 | 6.19649 | 6.67303 | 6.18757 | 5.84219 | 6.01739 | 5.93506 | 6.08412 | 6.4923 | 5.9883 | 6.32473 | 6.59787 |
| DNAJB14 | Q8TBM8 | 6.58918 | 6.78386 | 6.28453 | 6.93191 | 6.15287 | 6.19689 | 6.40307 | 6.27728 | 6.07152 | 6.50079 | 6.23454 | 6.26016 | 5.87091 | 5.59866 | 5.80503 | 6.0388 |
| NUTF2 | P61970 | 4.9443 | 4.92999 | 5.07702 | 4.04621 | 6.10281 | 6.09631 | 5.15272 | 8.94892 | 6.315 | 7.12695 | 5.90386 | 7.08291 | 4.49361 | 8.46151 | 6.48073 | 7.83715 |
| UBE2L3 | P68036-3 | 6.12314 | 5.60388 | 6.82997 | 5.41611 | 6.50637 | 6.3833 | 6.07585 | 6.69518 | 6.73349 | 6.98607 | 7.55237 | 6.80141 | 5.96924 | 5.45639 | 5.62476 | 5.24247 |
| SERPINB1 | P30740 | 6.0911 | 6.02201 | 6.9281 | 5.88213 | 5.9548 | 6.80345 | 6.04262 | 6.84868 | 6.18408 | 6.538 | 6.58422 | 6.45737 | 5.73757 | 6.24241 | 5.82312 | 5.86034 |
| ERCC6 | Q03468 | 5.99568 | 6.10527 | 6.43069 | 6.51636 | 5.83087 | 6.26452 | 6.24621 | 5.94277 | 6.53391 | 6.28561 | 6.05443 | 6.61215 | 6.35486 | 6.16004 | 6.35216 | 6.31445 |
| CHTF18 | Q8WVB6 | 5.82587 | 6.08191 | 5.97856 | 6.21274 | 6.16274 | 6.5633 | 6.05688 | 6.21432 | 6.03605 | 5.85517 | 6.15271 | 5.79172 | 6.77174 | 6.65311 | 6.7598 | 6.9434 |
| FIGNL1 | Q6PIW4 | 5.84889 | 6.46843 | 6.39696 | 5.91767 | 6.03106 | 6.18669 | 6.32792 | 6.41997 | 6.18356 | 6.2274 | 6.55861 | 6.2883 | 6.39734 | 6.20414 | 6.31669 | 6.22637 |
| MYSM1 | Q5VVJ2 | 6.00664 | 5.95643 | 6.49975 | 6.37935 | 6.06062 | 5.96132 | 6.23589 | 5.6222 | 6.44765 | 6.29099 | 6.18093 | 6.14071 | 6.48785 | 6.15501 | 6.46121 | 7.11346 |
| AP4E1 | Q9UPM8 | 6.2604 | 6.47469 | 6.74955 | 6.42921 | 6.14407 | 6.43028 | 6.52247 | 6.55469 | 6.28001 | 6.39591 | 6.43392 | 6.12791 | 5.99073 | 5.8101 | 5.65532 | 5.74073 |
| CHML | P26374 | 6.11603 | 6.00159 | 6.73503 | 5.90832 | 6.2109 | 6.45466 | 5.97226 | 6.64648 | 6.15415 | 6.53663 | 6.50864 | 6.84088 | 5.68925 | 5.74011 | 6.2211 | 6.26398 |
| CHM | P24386 | 6.02716 | 6.16906 | 6.91288 | 5.95199 | 6.53537 | 6.81961 | 6.41071 | 6.20447 | 6.30012 | 6.53352 | 6.37321 | 6.60751 | 5.82581 | 5.60729 | 5.97571 | 5.74557 |
| LAMC2 | Q13753 | 6.73427 | 6.36057 | 5.8889 | 6.49629 | 6.94451 | 6.31309 | 6.6654 | 6.48457 | 5.92967 | 6.14865 | 6.02153 | 6.09763 | 6.17661 | 5.50775 | 6.17914 | 6.05142 |
| TYMS | P04818 | 6.31 | 6.1452 | 7.50009 | 6.31921 | 6.53067 | 6.88884 | 6.48669 | 7.28314 | 7.01293 | 6.20028 | 7.3771 | 7.2272 | 4.5466 | 4.62953 | 4.34831 | 5.1942 |
| SURF6 | O75683 | 6.13792 | 6.30975 | 6.02084 | 6.64073 | 5.89616 | 5.43456 | 6.54595 | 5.74532 | 6.20502 | 5.44834 | 6.05898 | 5.90792 | 6.64054 | 6.71193 | 7.26524 | 7.03079 |
| SYCE1 | Q8N0S2 | 6.72711 | 5.19812 | 7.32638 | 5.03428 | 6.97934 | 6.0511 | 5.85113 | 6.90577 | 6.39557 | 6.86937 | 7.54619 | 7.09362 | 6.17351 | 5.31392 | 5.24367 | 5.2909 |
| SMG5 | Q9UPR3 | 6.70137 | 6.10969 | 6.72643 | 6.23295 | 6.00349 | 6.65612 | 6.34168 | 6.36535 | 6.04345 | 6.66942 | 6.60385 | 6.80544 | 5.92464 | 5.68458 | 5.51222 | 5.61933 |
| PRKCE | Q02156 | 6.81267 | 6.6725 | 7.19262 | 7.25303 | 6.10864 | 6.08989 | 7.62016 | 6.08374 | 6.56235 | 5.67778 | 5.7308 | 6.37093 | 4.83444 | 5.92658 | 5.4425 | 5.62137 |
| TKFC | Q3LXA3 | 6.12611 | 5.90278 | 6.54592 | 5.80802 | 6.03204 | 6.64124 | 5.9469 | 6.97923 | 6.29566 | 6.21166 | 6.21645 | 6.53922 | 6.01325 | 6.58927 | 6.26353 | 5.88871 |
| RALBP1 | Q15311 | 6.23547 | 6.0885 | 6.3434 | 5.91821 | 6.0315 | 6.25424 | 6.30748 | 5.95867 | 6.88874 | 6.28986 | 6.33455 | 6.33376 | 6.18751 | 6.0847 | 6.19572 | 6.5477 |
| RNF214 | Q8ND24 | 6.18814 | 6.12732 | 6.32116 | 5.65662 | 6.55806 | 6.67803 | 5.86538 | 6.7522 | 6.51732 | 6.63435 | 6.68056 | 6.82363 | 6.59638 | 5.61408 | 5.47969 | 5.50707 |
| ZNF507 | Q8TCN5 | 5.85396 | 6.21267 | 6.41669 | 6.28555 | 5.5587 | 6.37286 | 5.81714 | 6.71291 | 6.4252 | 6.27068 | 6.87972 | 6.5137 | 6.04621 | 6.032 | 6.23946 | 6.36257 |
| SALL3 | Q9BXA9 | 7.32139 | 6.84328 | 6.80304 | 7.84493 | 6.72182 | 6.50559 | 7.15615 | 5.9605 | 6.2857 | 5.84826 | 6.63772 | 6.55091 | 5.26125 | 4.31149 | 4.81196 | 5.13601 |
| EPN1 | Q9Y6I3 | 6.05941 | 6.28004 | 6.62134 | 5.76097 | 6.27372 | 6.2981 | 6.00304 | 6.97016 | 5.93055 | 6.71441 | 6.35648 | 7.05552 | 5.90013 | 5.96799 | 5.86023 | 5.9479 |
| SLC9A6 | Q92581-2 | 6.02897 | 6.70004 | 6.32624 | 6.22004 | 6.31625 | 6.27807 | 6.25539 | 6.13951 | 6.21057 | 6.36822 | 6.27122 | 6.22724 | 5.946 | 6.37051 | 6.33845 | 6.00327 |
| WDR55 | Q9H6Y2 | 5.65408 | 6.00981 | 6.03421 | 5.62055 | 6.22941 | 6.296 | 5.6497 | 5.99288 | 6.50561 | 6.36816 | 6.43325 | 6.46749 | 6.87703 | 6.55361 | 6.56858 | 6.73962 |
| ARHGEF28 | Q8N1W1-6 | 6.62026 | 6.39164 | 7.13438 | 6.20487 | 6.15098 | 6.77053 | 6.46199 | 6.30912 | 6.73741 | 6.571 | 6.78676 | 6.65178 | 5.38788 | 5.11872 | 5.4732 | 5.22948 |
| GET4 | Q7L5D6 | 6.02041 | 6.24204 | 6.56309 | 5.84746 | 5.80234 | 6.57721 | 6.59528 | 6.17556 | 6.04829 | 6.53966 | 6.36106 | 6.56605 | 6.20261 | 6.16057 | 6.08301 | 6.21536 |
| BZW2 | Q9Y6E2 | 6.53399 | 6.06551 | 6.80181 | 6.03859 | 6.32291 | 6.31518 | 6.37979 | 6.21152 | 6.38036 | 6.35166 | 6.69985 | 6.54643 | 5.96316 | 6.03957 | 5.72972 | 5.61993 |
| FNTA | P49354 | 5.99494 | 5.89401 | 6.73545 | 5.47269 | 6.52932 | 6.87738 | 5.99658 | 6.43805 | 6.28558 | 6.17118 | 6.48537 | 6.50765 | 6.34746 | 6.06427 | 6.07961 | 6.12046 |

| | | | | | | | | | | | | | | | | | |
|---|---|---|---|---|---|---|---|---|---|---|---|---|---|---|---|---|---|
| RDH14 | Q9HBH5 | 6.4054 | 6.227 | 5.81 | 6.44576 | 6.69357 | 6.15836 | 6.38159 | 6.25713 | 6.17817 | 6.17683 | 5.64567 | 5.91047 | 6.88507 | 6.3989 | 6.22619 | 6.19991 |
| NDUFB10 | O96000 | 6.19747 | 6.36086 | 5.87186 | 6.72943 | 6.68227 | 6.18822 | 6.57573 | 5.78217 | 5.8107 | 6.12281 | 6.50472 | 5.93762 | 6.49613 | 6.19948 | 6.49297 | 6.04756 |
| ANTXR1 | Q9H6X2 | 6.50264 | 6.35976 | 6.21711 | 6.6056 | 6.25637 | 6.17383 | 6.21667 | 5.91999 | 6.03799 | 6.13307 | 6.57731 | 6.39455 | 6.54601 | 6.31482 | 5.69044 | 6.05382 |
| CBR3 | O75828 | 6.24754 | 5.85953 | 6.85064 | 5.56639 | 6.88886 | 6.83902 | 5.90647 | 6.53262 | 6.4564 | 6.79129 | 6.70562 | 6.46495 | 6.04668 | 5.87635 | 5.5488 | 5.41884 |
| hCG_1984214 | I3L0E3 | 6.54404 | 6.52925 | 6.0124 | 6.80757 | 5.86711 | 6.32725 | 6.58922 | 6.23389 | 6.06146 | 6.29883 | 5.52489 | 5.99992 | 6.16828 | 6.35455 | 6.03263 | 6.6487 |
| WWP1 | Q9H0M0 | 6.28019 | 6.57204 | 6.69941 | 6.47104 | 6.19756 | 6.1943 | 6.48513 | 6.1043 | 6.47472 | 6.29625 | 6.49474 | 6.65837 | 5.97992 | 5.74225 | 5.80178 | 5.54801 |
| RAB12 | A0A3B3ITT1 | 6.32405 | 6.53249 | 6.13543 | 6.45162 | 6.19364 | 6.02541 | 6.03656 | 6.19796 | 6.31332 | 6.32082 | 6.21097 | 6.4592 | 6.01314 | 6.29367 | 6.00648 | 6.48525 |
| RAB6B | Q9NRW1 | 6.14364 | 6.27636 | 6.11037 | 6.53751 | 6.39234 | 5.99952 | 6.52404 | 5.81965 | 6.20682 | 6.38866 | 6.31255 | 6.12027 | 6.15956 | 6.05768 | 6.50373 | 6.44731 |
| RAB4B | P61018-2 | 6.28237 | 6.76173 | 6.20818 | 6.63047 | 6.409 | 6.67894 | 6.40593 | 6.19017 | 5.96825 | 6.52282 | 5.87432 | 5.84434 | 5.9677 | 6.01553 | 6.14386 | 6.09639 |
| SPR | P35270 | 6.23148 | 5.31706 | 6.48968 | 5.2556 | 6.19565 | 5.85026 | 6.0439 | 6.78282 | 5.7602 | 6.72762 | 6.44684 | 6.8746 | 6.27494 | 6.60757 | 6.50082 | 6.64095 |
| UBXN1 | Q04323 | 6.47351 | 6.24904 | 6.80527 | 5.8523 | 6.38946 | 6.44132 | 6.00438 | 6.60991 | 6.48762 | 6.56271 | 6.98314 | 6.84457 | 6.12787 | 5.50499 | 5.2927 | 5.37121 |
| TOLLIP | Q9H0E2 | 5.71644 | 6.01208 | 6.53805 | 5.31748 | 6.35732 | 5.81067 | 5.91444 | 6.67421 | 5.5896 | 7.15272 | 6.2765 | 6.82488 | 6.21292 | 6.76546 | 6.35816 | 6.47906 |
| NAA20 | P61599 | 6.15749 | 5.23433 | 6.37695 | 4.7821 | 5.94447 | 6.59707 | 6.11801 | 6.63797 | 6.20058 | 6.31154 | 6.61011 | 7.23982 | 5.74904 | 6.80613 | 6.39819 | 6.8362 |
| PBK | Q96KB5-2 | 6.22366 | 6.32183 | 7.55529 | 6.07317 | 6.48487 | 6.29348 | 6.36133 | 6.8107 | 6.85771 | 6.75556 | 6.9018 | 6.7697 | 5.53536 | 4.98292 | 5.01382 | 5.05882 |
| CROT | Q9UKG9-3 | 6.35898 | 5.93911 | 6.35316 | 5.84393 | 6.26758 | 5.85637 | 6.3235 | 6.1793 | 6.25935 | 6.24416 | 6.14543 | 6.00252 | 6.27769 | 6.44781 | 6.85864 | 6.64249 |
| MGAT1 | P26572 | 6.40999 | 6.75236 | 5.87184 | 7.16216 | 5.97445 | 6.21397 | 6.64408 | 5.19404 | 6.09928 | 6.31481 | 6.0511 | 6.07967 | 6.16122 | 5.92184 | 6.24472 | 6.18439 |
| CUSTOS | Q96C57 | 5.90145 | 6.01177 | 5.8033 | 6.28358 | 5.88613 | 5.84194 | 6.20966 | 5.90176 | 6.24994 | 6.02935 | 6.35254 | 6.11154 | 6.75217 | 6.54595 | 7.08676 | 7.03217 |
| TPBG | Q13641 | 6.22526 | 6.36524 | 6.53823 | 6.07303 | 6.30094 | 6.27239 | 5.99649 | 6.21198 | 6.08568 | 6.49698 | 6.61999 | 6.41564 | 6.71654 | 5.93535 | 6.05081 | 5.69547 |
| GARRE1 | O15063 | 6.33245 | 6.1953 | 6.63378 | 5.95168 | 6.63088 | 6.71576 | 6.09324 | 6.40807 | 6.20007 | 6.38928 | 6.36563 | 6.72013 | 6.34567 | 5.94735 | 5.63711 | 5.43361 |
| RGS12 | O14924 | 6.08424 | 6.31143 | 6.8752 | 6.14814 | 5.96742 | 5.92282 | 6.62058 | 5.84469 | 6.67542 | 6.48932 | 6.51493 | 6.78174 | 6.06402 | 5.73401 | 5.86839 | 6.09766 |
| TAOK3 | Q9H2K8 | 6.23328 | 6.16228 | 6.36152 | 5.76822 | 6.38505 | 6.34269 | 5.92166 | 6.12952 | 6.73622 | 6.64721 | 6.66676 | 6.10573 | 6.25523 | 6.19638 | 6.52949 | 5.55874 |
| PARP9 | Q8IXQ6-2 | 6.32281 | 6.56673 | 6.23774 | 6.22535 | 6.01698 | 6.45631 | 6.15201 | 6.30851 | 6.66691 | 6.82916 | 6.26971 | 6.28487 | 5.84446 | 5.85068 | 6.0211 | 5.94665 |
| PRPF38B | Q5VTL8 | 6.2177 | 6.3993 | 5.59743 | 6.61048 | 6.31167 | 5.99708 | 6.16871 | 5.79277 | 6.10559 | 5.65118 | 5.99738 | 5.76531 | 6.62142 | 6.69737 | 7.19011 | 6.8765 |
| EMG1 | Q92979 | 6.03009 | 6.38107 | 5.70657 | 6.59112 | 6.54739 | 6.30258 | 6.15708 | 6.98344 | 6.03096 | 6.47264 | 5.97421 | 5.81107 | 5.96225 | 5.77886 | 7.1555 | 6.11516 |
| SLC27A1 | Q6PCB7 | 6.41624 | 6.40627 | 5.95633 | 6.55131 | 5.87758 | 6.06988 | 6.55842 | 6.24019 | 6.15477 | 6.16438 | 5.87848 | 5.83987 | 6.51897 | 6.45746 | 6.62323 | 6.28663 |
| MTA3 | Q9BTC8 | 6.37639 | 6.71365 | 5.97588 | 6.78862 | 6.14567 | 5.96146 | 6.12679 | 6.09117 | 6.48746 | 6.27161 | 5.87279 | 6.05634 | 6.40963 | 5.90465 | 6.35739 | 6.46049 |
| UTP3 | Q9NQZ2 | 6.15532 | 6.39386 | 5.81737 | 6.80988 | 5.85652 | 5.73245 | 6.51476 | 5.85976 | 6.09405 | 5.95578 | 5.87568 | 5.88973 | 6.66467 | 6.81959 | 6.59952 | 6.93107 |
| CENPB | P07199 | 6.78063 | 6.49712 | 6.35609 | 7.18216 | 6.23768 | 5.82025 | 6.27678 | 6.04569 | 6.40397 | 6.30651 | 5.73626 | 6.14158 | 6.05529 | 5.93304 | 5.91956 | 6.30738 |
| RAD23A | P54725 | 6.44994 | 6.27396 | 6.63682 | 5.35189 | 6.16068 | 6.3066 | 5.343 | 6.82671 | 5.84166 | 6.54266 | 6.84115 | 7.03638 | 5.84071 | 6.38576 | 6.00461 | 6.15747 |
| FAR1 | Q8WVX9 | 6.6658 | 6.84744 | 6.5624 | 7.06652 | 6.19108 | 5.89924 | 6.58525 | 6.17273 | 6.59166 | 6.56687 | 6.25777 | 5.95149 | 5.48803 | 5.56666 | 5.62365 | 5.96339 |
| WARS2 | Q9UGM6 | 6.62048 | 6.41106 | 6.32754 | 6.63569 | 6.67585 | 5.9627 | 6.6001 | 5.96171 | 6.33243 | 5.79328 | 5.97483 | 5.68849 | 6.44475 | 5.72193 | 6.68735 | 6.16181 |
| CSTF1 | Q05048 | 6.2834 | 6.59537 | 6.08474 | 6.35478 | 6.42995 | 6.13781 | 6.50017 | 6.22498 | 6.16288 | 5.99991 | 5.77744 | 5.50426 | 6.82357 | 5.93981 | 6.90198 | 6.27895 |
| OSBPL11 | Q9BXB4 | 6.45016 | 6.31838 | 6.58988 | 6.60552 | 6.17899 | 6.57336 | 6.08331 | 6.31832 | 6.48157 | 6.75147 | 6.75465 | 6.30094 | 5.77456 | 5.66009 | 5.43933 | 5.71947 |
| ABCB10 | Q9NRK6 | 6.3525 | 6.22884 | 6.12472 | 6.6215 | 6.05275 | 5.83994 | 6.43492 | 5.74245 | 6.13488 | 6.23536 | 6.14561 | 5.97272 | 6.65025 | 6.15833 | 6.88104 | 6.42418 |
| POLD2 | P49005 | 5.12695 | 5.24556 | 5.98504 | 5.07105 | 6.32404 | 6.2537 | 5.88601 | 6.63477 | 6.41815 | 6.89103 | 6.55969 | 7.01957 | 6.38705 | 6.21561 | 7.04502 | 6.93676 |
| MINK1 | Q8N4C8 | 6.02926 | 5.96533 | 6.66668 | 6.20707 | 5.90166 | 5.86989 | 6.14496 | 5.85141 | 6.79756 | 6.88438 | 6.53354 | 6.67784 | 6.15746 | 6.19402 | 6.23366 | 5.88528 |
| TPST1 | O60507 | 6.46856 | 6.62786 | 6.09687 | 6.77114 | 6.25139 | 6.0111 | 6.41217 | 6.21952 | 6.18436 | 6.12491 | 6.20191 | 5.9563 | 6.26833 | 6.0371 | 6.08536 | 6.28314 |
| WDR76 | Q9H967 | 6.5066 | 6.8613 | 5.70451 | 6.91544 | 6.73606 | 6.06573 | 6.38265 | 6.0123 | 5.97903 | 5.97017 | 5.93806 | 5.90285 | 6.33704 | 6.13176 | 6.47586 | 6.08064 |
| DNAJB1 | P25685 | 5.69924 | 5.8244 | 6.68457 | 5.68302 | 5.52874 | 6.3587 | 5.77261 | 6.33071 | 6.46016 | 6.57781 | 6.80085 | 6.70818 | 6.42645 | 6.3057 | 6.41378 | 6.42508 |
| USP40 | A0A7I2YQ75 | 6.51579 | 6.11159 | 6.79377 | 6.0745 | 6.10114 | 6.73183 | 6.24456 | 6.70782 | 6.09022 | 6.51936 | 6.92208 | 6.39548 | 5.62489 | 5.8592 | 5.65154 | 5.65624 |
| AKT1 | P31749 | 6.36737 | 6.18337 | 7.03211 | 5.9699 | 6.03819 | 6.68568 | 6.079 | 6.28217 | 6.38587 | 6.4561 | 6.5328 | 6.60234 | 5.69908 | 6.03652 | 5.90255 | 5.74695 |
| POMGNT1 | Q8WZA1-2 | 6.27421 | 6.63191 | 5.9523 | 6.89982 | 6.34354 | 6.0395 | 6.34601 | 6.31606 | 6.56624 | 6.47659 | 5.78025 | 6.21441 | 5.98747 | 5.59926 | 6.25859 | 6.31384 |
| GCDH | Q92947 | 5.97295 | 6.40868 | 6.04926 | 6.67065 | 6.03438 | 6.15665 | 6.25188 | 6.02457 | 5.57197 | 6.11009 | 6.23514 | 6.17024 | 6.90087 | 6.33016 | 6.57148 | 6.54102 |
| EFR3A | Q14156 | 6.21687 | 6.5136 | 6.14329 | 6.58885 | 6.28746 | 6.2055 | 6.24874 | 6.4898 | 6.14242 | 6.09063 | 6.09129 | 6.13635 | 6.13743 | 6.03901 | 6.45259 | 6.20717 |
| PAXIP1 | Q6ZW49 | 6.34523 | 6.56835 | 6.29162 | 6.95912 | 6.34407 | 5.98305 | 6.25764 | 5.90521 | 6.24102 | 5.83585 | 5.7258 | 5.99776 | 6.57465 | 6.26903 | 6.34419 | 6.35741 |
| CSTB | P04080 | 5.52202 | 5.19501 | 6.35759 | 4.39332 | 6.43549 | 6.41938 | 5.44548 | 7.18861 | 5.53734 | 7.00778 | 6.20843 | 6.68386 | 5.69999 | 8.00296 | 6.56454 | 7.3382 |
| CCDC18 | E9PFB9 | 6.26856 | 6.35365 | 6.82259 | 6.6354 | 6.56091 | 6.47717 | 6.19936 | 6.8553 | 6.77857 | 6.70148 | 7.03792 | 7.0812 | 4.9368 | 5.16782 | 4.89939 | 5.22388 |
| EGLN1 | Q9GZT9 | 6.15953 | 6.22536 | 6.72078 | 6.24409 | 6.28152 | 6.33552 | 5.87198 | 6.35709 | 6.34446 | 6.68625 | 6.69475 | 6.80767 | 6.06447 | 5.84652 | 5.78665 | 5.57335 |
| PPIP5K2 | O43314 | 5.94423 | 5.73929 | 6.73867 | 5.89501 | 6.20939 | 6.60583 | 6.05118 | 6.4145 | 6.26779 | 6.67294 | 6.39932 | 6.6681 | 6.02711 | 6.32817 | 6.04093 | 5.99756 |
| CEP63 | Q96MT8 | 6.43522 | 6.45435 | 6.46853 | 6.7274 | 6.46133 | 6.3048 | 6.369 | 6.31693 | 6.2542 | 6.17185 | 6.39577 | 6.44211 | 6.06083 | 5.696 | 5.74502 | 5.69665 |
| GCC1 | Q96CN9 | 7.09834 | 6.18618 | 6.00563 | 6.43455 | 7.03703 | 6.11075 | 6.31446 | 6.38975 | 6.53707 | 6.12898 | 6.27414 | 6.08551 | 6.64685 | 5.85614 | 5.51298 | 5.38163 |
| C9orf64 | Q5T6V5 | 6.18367 | 6.00274 | 6.72761 | 5.62886 | 6.42189 | 7.29152 | 6.00993 | 6.60789 | 6.56338 | 6.6604 | 6.38346 | 6.44622 | 5.48523 | 6.08986 | 5.76014 | 5.73719 |
| IMPACT | Q9P2X3 | 6.54747 | 6.65578 | 7.00969 | 5.87641 | 6.3525 | 6.83258 | 6.32332 | 6.45169 | 6.20214 | 6.45922 | 6.23523 | 6.50492 | 5.42691 | 5.92 | 5.71012 | 5.49203 |
| MAK16 | Q9BXY0 | 6.43054 | 6.31543 | 5.86018 | 6.61692 | 6.32124 | 5.84417 | 6.31863 | 5.82244 | 6.22113 | 6.08405 | 5.93777 | 5.88417 | 7.07692 | 5.9472 | 6.57757 | 6.74164 |
| FLAD1 | Q8NFF5 | 6.40359 | 6.07686 | 6.42083 | 6.39729 | 6.06289 | 5.84573 | 6.0049 | 6.31686 | 6.225 | 6.13345 | 6.24409 | 6.19855 | 6.56653 | 6.21531 | 6.58518 | 6.30294 |
| GPD1L | Q8N335 | 5.97188 | 6.1535 | 6.81877 | 5.84183 | 6.23057 | 6.4148 | 6.5136 | 6.53074 | 6.15806 | 6.41508 | 6.23675 | 6.7738 | 5.98788 | 6.30979 | 5.68504 | 5.9579 |
| KIF13A | Q9H1H9-4 | 6.6456 | 6.41965 | 6.84174 | 6.2989 | 6.29276 | 6.84707 | 6.06111 | 6.71853 | 6.54021 | 6.53943 | 6.53688 | 6.95305 | 5.34094 | 5.14438 | 5.35324 | 5.46649 |
| KIF1C | O43896 | 6.65831 | 6.46226 | 6.78925 | 6.15314 | 6.36178 | 6.45034 | 6.18874 | 6.62153 | 6.31534 | 6.56404 | 6.40277 | 6.37462 | 5.69815 | 5.7364 | 5.56485 | 5.65848 |
| IPO13 | O94829 | 5.75391 | 5.98218 | 6.16253 | 5.17427 | 5.92828 | 6.41602 | 5.82036 | 6.07197 | 6.58044 | 6.44832 | 6.24828 | 6.24416 | 6.5595 | 6.88156 | 7.01849 | 6.70973 |
| LYPLA1 | O75608 | 5.66194 | 4.70677 | 6.03266 | 4.71312 | 6.39554 | 5.75775 | 5.95281 | 6.9827 | 5.75894 | 6.74068 | 5.82337 | 6.50086 | 5.76815 | 7.7006 | 7.58771 | 7.9164 |
| SNX5 | Q9Y5X3 | 6.51764 | 6.56312 | 6.95802 | 5.97507 | 6.4094 | 7.03104 | 6.02165 | 6.68911 | 6.49383 | 6.34457 | 6.47731 | 6.78666 | 5.80923 | 5.30209 | 5.43206 | 5.18919 |
| MYBL1 | P10243 | 5.97408 | 5.88535 | 5.94076 | 6.30196 | 6.47741 | 6.22168 | 6.22191 | 5.96634 | 6.05456 | 6.01978 | 5.65942 | 5.68911 | 6.98165 | 6.56839 | 6.8885 | 7.14912 |
| ARMCX3 | Q9UH62 | 6.41304 | 7.15942 | 5.92806 | 6.51263 | 6.3702 | 6.48805 | 6.1394 | 6.23783 | 5.96824 | 6.30317 | 5.95433 | 5.75654 | 6.26529 | 6.1846 | 6.50207 | 5.81712 |
| INTS12 | Q96CB8 | 6.10081 | 6.10288 | 6.11586 | 6.50727 | 6.61968 | 6.10223 | 6.47884 | 5.99323 | 5.91319 | 6.32597 | 6.15056 | 5.97632 | 6.53652 | 6.13694 | 6.36749 | 6.57223 |
| ITPA | Q9BY32 | 6.70147 | 5.81171 | 6.67875 | 4.87796 | 6.92576 | 6.42647 | 6.09877 | 7.30548 | 6.21899 | 6.78325 | 6.05293 | 6.47998 | 6.22588 | 6.24598 | 5.67197 | 5.49463 |
| DMXL1 | Q9Y485 | 6.34945 | 6.25875 | 6.83969 | 6.55136 | 6.2903 | 6.40453 | 6.3453 | 6.70877 | 6.60188 | 6.3023 | 6.94513 | 6.95222 | 5.41046 | 5.48446 | 5.27492 | 5.28048 |
| UBR3 | Q6ZT12-4 | 5.84579 | 6.35341 | 6.62696 | 5.95151 | 5.96095 | 6.52749 | 6.30753 | 6.02239 | 6.33038 | 6.50544 | 6.53279 | 6.51503 | 6.02238 | 6.29506 | 6.29133 | 5.91157 |
| ABHD16A | O95870 | 6.57648 | 6.44041 | 5.72196 | 6.7585 | 6.09071 | 5.89829 | 6.09299 | 6.3339 | 5.76898 | 6.15138 | 6.14946 | 5.97914 | 6.26196 | 6.55358 | 6.64653 | 6.57572 |
| NIF3L1 | Q9GZT8 | 5.93223 | 6.07848 | 6.61981 | 5.76286 | 6.05668 | 6.40838 | 6.25333 | 6.33404 | 6.29744 | 6.13714 | 6.64867 | 6.25143 | 6.42724 | 6.18017 | 6.31055 | 6.30156 |
| GCFC2 | P16383 | 5.73831 | 6.11722 | 6.03568 | 6.09549 | 5.92814 | 6.26801 | 6.19662 | 6.34181 | 6.06512 | 6.25321 | 6.38481 | 6.16114 | 6.77097 | 6.31809 | 6.73187 | 6.59351 |
| CSNK1E | P49674 | 6.88286 | 6.94908 | 6.84777 | 7.14241 | 6.51589 | 6.34939 | 6.35075 | 6.48709 | 6.65163 | 6.72124 | 6.60188 | 6.66774 | 4.89917 | 4.85193 | 4.96772 | 5.11345 |
| SPATA20 | Q8TB22-2 | 6.64878 | 6.56927 | 5.88351 | 6.39943 | 6.15365 | 5.99135 | 6.43743 | 6.09569 | 5.888 | 5.94537 | 6.23653 | 5.90746 | 6.68445 | 6.38421 | 6.39937 | 6.37551 |
| GIGYF1 | O75420 | 6.74859 | 6.60746 | 6.93071 | 6.30605 | 6.47276 | 6.37891 | 5.90696 | 6.86954 | 6.65937 | 6.72935 | 6.8517 | 6.72536 | 5.52415 | 5.35246 | 4.87323 | 5.0634 |
| BIN3 | Q9NQY0 | 6.34738 | 5.90431 | 6.56432 | 5.70367 | 6.15606 | 6.72267 | 5.83382 | 7.10909 | 6.40812 | 6.85959 | 6.66088 | 6.82103 | 6.15632 | 5.85632 | 5.23027 | 5.66615 |
| ABCC3 | O15438-4 | 6.39046 | 6.26779 | 6.41558 | 6.27561 | 6.21185 | 5.86365 | 5.87166 | 5.63262 | 6.22328 | 6.10713 | 6.03845 | 6.08869 | 6.78626 | 6.43551 | 6.88538 | 6.50608 |
| ADPRS | Q9NX46 | 6.06056 | 5.74757 | 6.30375 | 5.92745 | 6.23061 | 6.0859 | 6.55845 | 6.07739 | 6.70839 | 5.6713 | 5.92206 | 6.59174 | 6.18031 | 6.68706 | 6.35594 | 6.89153 |
| PACSIN3 | Q9UKS6 | 6.36334 | 6.68379 | 6.29947 | 6.2755 | 6.58257 | 6.59068 | 5.85755 | 6.73227 | 6.02341 | 6.07425 | 6.10161 | 6.29495 | 6.20113 | 5.89654 | 5.91211 | 6.11085 |
| MRPL9 | Q9BYD2 | 6.38027 | 6.47354 | 6.00636 | 6.53932 | 6.61842 | 6.11127 | 6.95036 | 5.93171 | 6.25158 | 5.85439 | 5.5751 | 5.76749 | 6.40664 | 6.03248 | 6.61179 | 6.48926 |
| SDF2L1 | Q9HCN8 | 6.6236 | 6.66391 | 5.51371 | 6.24438 | 6.82691 | 6.23932 | 5.81402 | 6.54344 | 6.14535 | 6.11313 | 5.84046 | 5.96424 | 6.81334 | 6.17674 | 6.12931 | 6.34814 |
| TOR1A | O14656 | 6.71252 | 6.58508 | 6.18556 | 6.47118 | 6.14371 | 6.16521 | 6.41266 | 5.97196 | 5.99726 | 6.41632 | 6.12867 | 5.68274 | 6.55127 | 5.9333 | 6.47436 | 6.1682 |
| KBTBD2 | Q8IY47 | 5.83198 | 6.05467 | 6.55553 | 5.70482 | 6.03699 | 6.61681 | 6.23836 | 6.16695 | 6.44379 | 6.63494 | 6.4077 | 6.19005 | 5.96669 | 6.25471 | 6.73706 | 6.15896 |
| MRPL40 | Q9NQ50 | 6.05902 | 6.40743 | 6.00193 | 6.7209 | 6.48695 | 6.15376 | 6.77548 | 6.10587 | 6.21184 | 6.01881 | 6.13453 | 5.90903 | 6.3749 | 5.97008 | 6.06684 | 6.60263 |
| NSMCE2 | Q96MF7 | 6.27387 | 6.45809 | 6.59545 | 6.63701 | 6.31427 | 6.2351 | 6.43752 | 5.9918 | 6.07463 | 6.15903 | 6.15688 | 6.20528 | 6.26761 | 5.83161 | 6.11286 | 6.249 |
| RAD51 | Q06609 | 6.77994 | 6.09757 | 6.48875 | 6.76887 | 6.33148 | 6.1038 | 6.59959 | 6.14681 | 6.7139 | 6.22723 | 6.6569 | 6.75999 | 5.30363 | 5.51617 | 5.49178 | 6.01359 |
| PTER | Q96BW5 | 5.89871 | 5.65878 | 6.57522 | 5.13745 | 6.80005 | 6.64007 | 6.13294 | 6.68181 | 6.649 | 6.53844 | 6.73091 | 6.35795 | 6.14046 | 5.94009 | 6.14187 | 5.97625 |
| PTPN14 | Q15678 | 6.09431 | 6.40155 | 6.6067 | 6.2469 | 6.0179 | 6.59744 | 6.242 | 6.49823 | 6.25644 | 6.39649 | 6.68837 | 6.7424 | 5.78768 | 5.83425 | 5.95487 | 5.63448 |
| TCAF1 | Q9Y4C2 | 6.76538 | 6.71681 | 7.43198 | 6.40977 | 6.23359 | 6.43747 | 6.42784 | 6.51388 | 6.86947 | 7.11347 | 6.96589 | 7.23731 | 4.63638 | 4.99963 | 4.43454 | 4.80659 |
| TRPC4AP | Q8TEL6 | 6.29257 | 6.40692 | 7.0893 | 6.08226 | 6.4223 | 7.11996 | 6.83919 | 6.83839 | 6.83641 | 6.29474 | 6.63964 | 6.34072 | 4.94708 | 5.20476 | 4.89827 | 5.74748 |
| C11orf68 | Q9H3H3-3 | 6.22465 | 6.21274 | 6.69639 | 5.67623 | 6.59075 | 7.03775 | 5.96059 | 6.72137 | 6.45142 | 6.39703 | 6.58472 | 6.52191 | 5.85604 | 5.71568 | 5.82851 | 5.52422 |
| BRAP | Q7Z569 | 6.37371 | 6.26025 | 7.02636 | 6.27543 | 6.35375 | 6.81732 | 6.1687 | 6.68582 | 6.60442 | 6.32601 | 6.51285 | 6.75427 | 5.54052 | 5.57018 | 5.2781 | 5.45232 |
| ATP6AP2 | O75787 | 6.0341 | 5.59949 | 5.72622 | 5.91846 | 6.30064 | 5.65106 | 6.58739 | 6.3325 | 6.02323 | 6.35848 | 5.98794 | 6.56314 | 6.36007 | 6.59616 | 7.01827 | 6.94285 |
| CEP85 | Q6P2H3 | 7.25429 | 6.27132 | 7.24874 | 6.33488 | 6.57179 | 6.75152 | 6.16272 | 6.71468 | 6.44654 | 6.71895 | 6.96969 | 7.14464 | 5.13779 | 4.83495 | 4.39262 | 5.04487 |
| STAT2 | P52630 | 6.55659 | 6.59932 | 6.92657 | 6.07021 | 6.41117 | 7.20852 | 6.4137 | 6.65144 | 6.51328 | 6.84633 | 6.23743 | 6.49463 | 5.23299 | 5.51698 | 5.1683 | 5.15255 |
| MALT1 | Q9UDY8 | 6.28736 | 5.98985 | 6.92344 | 6.62986 | 6.31294 | 6.70935 | 6.23436 | 6.51669 | 6.63092 | 6.42815 | 6.9371 | 7.09707 | 5.41123 | 5.13543 | 5.25896 | 5.4973 |
| PPM1F | P49593 | 6.36717 | 5.77658 | 7.08372 | 5.86564 | 6.38514 | 6.72569 | 6.08936 | 6.5626 | 6.58501 | 6.54481 | 6.49018 | 6.2742 | 5.82944 | 5.82929 | 5.59931 | 5.99187 |
| SMTN | P53814-5 | 5.78985 | 6.22052 | 6.27453 | 5.89617 | 5.84512 | 6.18234 | 6.05401 | 5.94434 | 6.32637 | 6.16791 | 6.6936 | 6.69167 | 6.36198 | 6.23877 | 6.51425 | 6.79859 |
| RASA2 | Q15283 | 6.41482 | 6.68355 | 6.75032 | 6.45242 | 6.67833 | 6.64392 | 6.25777 | 6.1965 | 6.31846 | 6.25801 | 6.76713 | 6.2965 | 5.66157 | 5.74804 | 5.56002 | 5.31265 |
| MRPS18B | Q9Y676 | 6.26561 | 6.73895 | 5.9854 | 6.60677 | 6.26586 | 6.05669 | 6.57372 | 6.39032 | 6.08848 | 5.91195 | 6.28547 | 6.14178 | 6.52331 | 6.0972 | 6.11243 | 5.95607 |
| ZNF106 | Q9H2Y7 | 6.31868 | 6.5022 | 6.24032 | 6.96553 | 5.84306 | 5.90144 | 6.60058 | 6.17401 | 6.3989 | 6.195 | 6.25276 | 6.20876 | 5.77242 | 5.97774 | 6.25205 | 6.39656 |
| SUOX | P51687 | 6.45907 | 6.21798 | 5.99136 | 6.55152 | 6.16176 | 5.90405 | 6.24794 | 5.91874 | 6.13914 | 6.34472 | 6.33333 | 6.18403 | 6.42911 | 6.2913 | 6.40297 | 6.42297 |
| PPOX | P50336 | 6.13227 | 6.5756 | 6.24281 | 6.63074 | 6.37528 | 5.94027 | 6.05368 | 6.02612 | 5.99131 | 5.90548 | 5.66461 | 6.22644 | 6.47136 | 6.86536 | 6.50454 | 6.39412 |
| PHC2 | Q8IXK0-5 | 6.00518 | 6.03969 | 6.26989 | 6.99535 | 5.84118 | 6.30136 | 6.44419 | 5.92153 | 6.08781 | 6.33354 | 6.16228 | 6.50448 | 6.02666 | 5.94501 | 6.595 | 6.52684 |
| KIFAP3 | Q92845 | 6.12916 | 6.03986 | 7.10567 | 6.01991 | 6.5723 | 6.80297 | 6.27377 | 6.58357 | 6.65229 | 6.63431 | 6.58499 | 6.08036 | 5.77404 | 5.53095 | 5.54998 | 5.66588 |
| EML1 | O00423-3 | 6.53379 | 6.24171 | 6.70869 | 5.72679 | 6.38221 | 6.73222 | 6.01824 | 6.53617 | 6.46687 | 6.49026 | 6.38812 | 6.34336 | 6.00021 | 6.22556 | 5.67871 | 5.52709 |
| ZNF654 | A0A1B0GWA0 | 6.32971 | 6.52024 | 6.47831 | 6.83203 | 6.31799 | 6.21984 | 6.48942 | 5.92905 | 6.15589 | 6.24663 | 6.205 | 6.38875 | 6.12974 | 5.50411 | 6.18214 | 6.07116 |

| | | | | | | | | | | | | | | | | | |
|---|---|---|---|---|---|---|---|---|---|---|---|---|---|---|---|---|---|
| ZC3H3 | Q8IXZ2 | 6.5481 | 6.53974 | 6.28293 | 6.9069 | 6.16526 | 6.26364 | 6.56395 | 6.60948 | 6.66076 | 6.61056 | 6.55821 | 6.42283 | 5.23277 | 5.63633 | 5.28549 | 5.71305 |
| SCAMP1 | O15126 | 6.17019 | 6.28701 | 6.14106 | 6.17702 | 6.00767 | 5.96666 | 6.30503 | 6.3001 | 6.44898 | 6.48665 | 6.10236 | 5.90878 | 6.06729 | 6.17702 | 6.76235 | 6.69182 |
| AMOT | Q4VCS5 | 6.89576 | 6.57633 | 6.15724 | 6.18827 | 6.55165 | 6.44747 | 6.25328 | 5.99977 | 6.45385 | 6.64354 | 6.48204 | 6.48839 | 6.01372 | 5.73605 | 5.41784 | 5.69479 |
| FNBP1L | Q5T0N5 | 6.08237 | 6.31517 | 6.89409 | 6.23393 | 6.3351 | 6.72241 | 6.2514 | 6.57105 | 6.54429 | 6.61811 | 6.53899 | 6.51548 | 5.89458 | 5.80739 | 5.44552 | 5.23011 |
| METTL3 | Q86U44 | 5.53267 | 5.53317 | 5.99012 | 5.84791 | 6.35146 | 6.62643 | 6.03927 | 5.74791 | 6.28333 | 5.93499 | 6.44979 | 6.05689 | 6.88181 | 6.81587 | 7.26883 | 6.63953 |
| SRRM1 | Q8IYB3 | 5.70263 | 6.39131 | 5.59819 | 6.53444 | 5.61888 | 5.78324 | 5.85058 | 5.88084 | 5.8023 | 6.00609 | 5.91625 | 6.31831 | 6.59454 | 7.49464 | 6.96564 | 7.54212 |
| C7orf50 | Q9BRJ6 | 5.87521 | 6.32302 | 5.76461 | 5.92114 | 5.94266 | 5.75474 | 6.01651 | 6.02585 | 5.9784 | 6.49906 | 6.07575 | 5.87475 | 6.59321 | 7.1126 | 7.38455 | 6.85796 |
| RBM45 | Q8IUH3 | 6.67809 | 6.55161 | 5.68056 | 6.54611 | 6.45744 | 6.23432 | 6.25059 | 5.88447 | 6.33746 | 6.08286 | 6.31773 | 6.02881 | 6.68424 | 6.11634 | 6.09878 | 6.0506 |
| TRPS1 | Q9UHF7-2 | 6.3329 | 6.47005 | 5.99501 | 6.46958 | 6.08667 | 6.17214 | 6.28358 | 6.23381 | 6.26342 | 6.33599 | 5.99492 | 6.1567 | 6.37566 | 6.28342 | 6.20154 | 6.3446 |
| PRAME | P78395 | 6.0343 | 6.81626 | 7.23493 | 6.66378 | 6.36637 | 6.65725 | 6.56329 | 6.04722 | 6.48102 | 7.32794 | 7.34021 | 6.87247 | 4.45556 | 5.40381 | 4.71438 | 5.02121 |
| CHTOP | Q9Y3Y2-3 | 5.88476 | 6.56402 | 4.9889 | 6.13235 | 5.4171 | 5.71223 | 6.34379 | 6.34116 | 7.25499 | 5.30306 | 6.07315 | 5.73482 | 7.59304 | 6.73442 | 6.02526 | 7.89694 |
| RBM5 | P52756 | 6.30691 | 6.14506 | 6.14293 | 6.48383 | 6.31127 | 6.16085 | 6.74857 | 6.22872 | 6.57743 | 6.35426 | 6.28429 | 6.05943 | 6.26478 | 6.16416 | 6.00804 | 5.75947 |
| FAN1 | Q9Y2M0 | 6.20707 | 6.28923 | 6.4056 | 6.29364 | 6.47577 | 5.97246 | 6.30297 | 6.01788 | 6.38482 | 6.06704 | 6.29097 | 6.35356 | 6.09123 | 5.94605 | 6.32118 | 6.58053 |
| PTGR1 | Q14914 | 6.00106 | 6.07356 | 6.65331 | 5.93619 | 6.30297 | 6.86731 | 6.43962 | 6.5083 | 6.242 | 6.3759 | 6.50545 | 6.38761 | 5.63622 | 6.38213 | 5.95369 | 5.73469 |
| TWNK | Q96RR1 | 6.01203 | 6.32286 | 6.36976 | 6.97289 | 6.02306 | 5.80705 | 6.45073 | 6.01541 | 6.33931 | 6.13288 | 5.98457 | 6.36762 | 6.26128 | 5.89003 | 6.58639 | 6.46413 |
| ILKAP | Q9H0C8 | 5.21429 | 5.54758 | 6.34816 | 5.37342 | 5.60285 | 6.1783 | 5.72817 | 5.86534 | 5.88262 | 6.53688 | 6.15211 | 6.44181 | 7.02603 | 7.52104 | 7.12671 | 6.98564 |
| TEX2 | Q8IWB9-2 | 6.57324 | 6.51437 | 5.99842 | 6.67129 | 6.06572 | 6.20323 | 6.28796 | 6.1419 | 5.52325 | 5.92515 | 5.76089 | 5.92732 | 6.49032 | 6.41143 | 6.75906 | 6.74643 |
| HMBS | P08397 | 6.11185 | 5.81184 | 6.96844 | 5.76754 | 6.11877 | 6.43415 | 5.81553 | 6.21465 | 6.56642 | 6.41051 | 6.83587 | 6.44637 | 6.01261 | 6.37716 | 6.16282 | 5.94548 |
| L3MBTL3 | Q96JM7-2 | 6.26031 | 6.10178 | 5.7285 | 6.6032 | 6.25494 | 5.93443 | 6.51569 | 6.05042 | 6.15361 | 6.1855 | 5.97161 | 6.04278 | 6.72433 | 6.34136 | 6.539 | 6.59255 |
| AK1 | P00568 | 6.32607 | 5.93811 | 6.67297 | 5.41227 | 6.74943 | 6.88063 | 6.40098 | 6.93776 | 6.44028 | 6.43907 | 6.56404 | 6.39518 | 5.95381 | 6.06231 | 5.46353 | 5.36356 |
| WDR4 | P57081 | 5.34865 | 5.40384 | 5.86077 | 5.2989 | 5.51964 | 6.25534 | 5.53339 | 5.74351 | 5.90313 | 5.82353 | 5.8501 | 5.66639 | 7.43888 | 8.02092 | 8.82514 | 7.50787 |
| ANKS1A | Q92625 | 6.33824 | 6.20903 | 6.66607 | 6.11616 | 5.99898 | 6.65444 | 6.28006 | 6.56311 | 6.5815 | 6.5567 | 6.35389 | 6.60173 | 5.87879 | 6.52396 | 5.718 | 5.85934 |
| FDPS | P14324 | 6.15539 | 6.1899 | 6.3032 | 5.61099 | 6.43761 | 6.99182 | 5.72841 | 7.04404 | 6.03041 | 6.25483 | 6.17926 | 6.2768 | 5.77846 | 6.74039 | 6.3549 | 5.9236 |
| NEDD4L | Q96PU5 | 6.25423 | 6.11935 | 6.65985 | 6.14416 | 5.89415 | 6.8449 | 6.24438 | 6.56572 | 6.58297 | 6.52869 | 6.79018 | 6.61369 | 5.62726 | 5.66684 | 5.54455 | 5.91908 |
| RAB5B | P61020 | 6.73239 | 6.36775 | 6.14914 | 6.19419 | 6.07128 | 6.02799 | 6.75968 | 6.26027 | 6.33391 | 6.39872 | 6.00492 | 6.03482 | 6.17998 | 6.19026 | 6.10425 | 6.19045 |
| SCARB2 | Q14108 | 6.72511 | 6.93387 | 6.04842 | 6.40611 | 6.21237 | 6.03515 | 6.45579 | 6.15529 | 6.01518 | 6.45356 | 5.75358 | 5.98057 | 6.46077 | 5.85841 | 6.4129 | 6.09292 |
| CPNE2 | Q96FN4 | 5.23437 | 5.67283 | 5.86708 | 5.01906 | 5.88497 | 6.19354 | 5.57629 | 5.69132 | 6.68391 | 6.97473 | 6.69187 | 6.48092 | 6.1612 | 6.80865 | 7.48294 | 7.57632 |
| TMLHE | Q9NVH6-8 | 6.47694 | 6.14389 | 5.96223 | 6.5479 | 6.42217 | 6.02246 | 6.18509 | 5.98761 | 6.15341 | 6.22894 | 6.10988 | 5.83277 | 6.6109 | 6.14287 | 6.76922 | 6.40372 |
| RAC1 | P63000-2 | 6.60636 | 6.06532 | 6.56615 | 6.12237 | 6.56308 | 6.55421 | 6.35309 | 5.87149 | 6.24528 | 6.05982 | 6.51765 | 6.121 | 6.32611 | 5.9643 | 5.98568 | 6.07809 |
| ATG16L1 | Q676U5 | 5.97731 | 6.00545 | 6.65687 | 6.28438 | 6.18756 | 6.3358 | 6.36565 | 6.4027 | 6.29231 | 6.24465 | 6.92978 | 6.82154 | 5.74736 | 5.9008 | 6.01515 | 5.83269 |
| PTPA | Q15257 | 6.22326 | 6.14456 | 6.65396 | 5.59168 | 6.25209 | 6.63886 | 6.03938 | 6.89382 | 6.36089 | 6.48333 | 6.48669 | 6.64809 | 5.72447 | 6.23847 | 5.64499 | 5.97545 |
| DPYD | Q12882 | 6.29066 | 6.39547 | 7.31456 | 6.05191 | 6.65666 | 6.13918 | 6.37548 | 6.42456 | 6.55293 | 6.54712 | 6.17857 | 6.13369 | 5.59779 | 6.22716 | 5.55728 | 5.55701 |
| ECI1 | P42126 | 5.75891 | 4.89791 | 5.04349 | 5.0446 | 5.66725 | 4.96137 | 6.07792 | 6.57464 | 4.78443 | 5.44997 | 5.02763 | 6.22539 | 6.27377 | 9.22428 | 9.59385 | 9.39458 |
| CA2 | P00918 | 5.98982 | 6.24681 | 7.08806 | 5.57361 | 6.61503 | 7.17467 | 6.37612 | 6.82254 | 6.64806 | 6.2157 | 6.24931 | 6.5477 | 5.85797 | 5.78629 | 5.52534 | 5.28299 |
| IKBIP | Q70UQ0 | 6.14044 | 6.56563 | 6.06599 | 6.64423 | 6.00238 | 6.01148 | 6.28883 | 6.10562 | 6.16382 | 6.24354 | 6.19066 | 6.12185 | 6.41524 | 6.07823 | 6.4778 | 6.48426 |
| DLG5 | Q8TDM6 | 7.02956 | 6.53491 | 6.51356 | 6.88254 | 6.31048 | 6.36904 | 6.53949 | 6.3806 | 6.26472 | 6.1134 | 6.02156 | 6.38699 | 5.89921 | 5.66136 | 5.47042 | 5.62217 |
| SLC16A3 | O15427 | 5.8119 | 6.05388 | 6.04253 | 6.19388 | 6.01365 | 6.03296 | 6.44605 | 5.88608 | 6.2119 | 6.27323 | 6.07589 | 6.10519 | 6.53918 | 6.7365 | 6.63004 | 6.94714 |
| CENPI | Q92674 | 6.34252 | 6.66967 | 6.09956 | 7.16407 | 6.45645 | 5.99589 | 5.9978 | 6.02056 | 6.11452 | 6.18212 | 6.38381 | 6.3405 | 6.13208 | 5.81873 | 6.16841 | 6.1133 |
| M6PR | P20645 | 6.21826 | 6.45007 | 6.45056 | 6.49088 | 6.18423 | 6.01027 | 6.0073 | 5.73467 | 6.52371 | 6.66232 | 6.66148 | 6.58671 | 6.16705 | 5.6702 | 6.1583 | 6.024 |
| GXYLT1 | Q4G148 | 6.61308 | 6.7205 | 5.9189 | 6.7998 | 6.29929 | 6.03648 | 6.41552 | 6.16049 | 6.00426 | 6.18088 | 6.1229 | 6.03501 | 6.53008 | 5.95927 | 6.23677 | 5.96675 |
| ADA | P00813 | 6.19815 | 5.25204 | 7.1177 | 4.9774 | 6.76751 | 6.69992 | 6.069 | 6.43137 | 6.82848 | 6.43127 | 6.89848 | 6.77692 | 6.33904 | 6.14951 | 5.50373 | 5.55946 |
| USE1 | Q9NZ43 | 6.1009 | 6.60954 | 5.88884 | 6.51185 | 6.01811 | 5.92224 | 6.4587 | 5.87664 | 5.97137 | 6.30654 | 6.1397 | 6.209 | 6.44029 | 6.32016 | 6.75253 | 6.47358 |
| SDE2 | Q6IQ49 | 6.11166 | 6.2647 | 6.45453 | 6.3538 | 6.28613 | 6.11124 | 6.27958 | 6.13823 | 6.57588 | 6.25113 | 6.65581 | 6.3349 | 6.23928 | 5.85146 | 6.02925 | 6.06242 |
| CFAP36 | Q96G28 | 6.79146 | 6.47231 | 7.34685 | 5.79837 | 6.39503 | 6.54521 | 5.9393 | 7.13384 | 6.61693 | 6.78692 | 6.97186 | 6.70217 | 5.4792 | 4.93036 | 5.00723 | 5.08288 |
| ZBTB40 | Q9NUA8 | 6.25736 | 5.99461 | 6.2611 | 6.80049 | 5.83531 | 5.90511 | 6.63047 | 5.74426 | 6.20042 | 6.15305 | 6.00278 | 5.96068 | 6.76206 | 6.28052 | 6.58712 | 6.62464 |
| PPIG | Q13427 | 6.20014 | 6.28319 | 5.96276 | 6.55282 | 6.06055 | 5.87736 | 6.30609 | 5.98528 | 6.23445 | 5.86204 | 5.76525 | 5.84192 | 6.54501 | 6.66694 | 6.9733 | 6.88288 |
| LTBP3 | Q9NS15 | 5.99357 | 6.45844 | 5.92163 | 6.21623 | 6.19338 | 6.22394 | 6.51277 | 6.03717 | 6.2519 | 6.57056 | 6.34694 | 6.3268 | 6.5927 | 6.27646 | 5.98165 | 6.09587 |
| ATG4B | Q9Y4P1-2 | 6.00822 | 5.96146 | 7.17323 | 5.82241 | 6.12693 | 6.93795 | 6.45101 | 6.40992 | 6.47894 | 6.65982 | 6.99949 | 6.93964 | 5.51992 | 5.63847 | 5.38301 | 5.48958 |
| WDR92 | Q96MX6 | 5.95453 | 6.11188 | 6.6203 | 5.76172 | 6.40914 | 6.54838 | 6.16107 | 6.60206 | 6.52449 | 6.83382 | 6.4277 | 6.45164 | 6.17926 | 5.9297 | 5.81389 | 5.67042 |
| TIMM13 | Q9Y5L4 | 6.08821 | 5.20568 | 5.54765 | 5.21352 | 5.9793 | 4.37865 | 5.12711 | 6.66694 | 4.53286 | 5.67266 | 4.94555 | 5.68085 | 5.31273 | 10.5839 | 8.61635 | 10.448 |
| CIAO1 | O76071 | 5.88789 | 6.37574 | 6.3403 | 6.149 | 6.28155 | 6.92352 | 6.08877 | 6.5198 | 6.53098 | 6.74988 | 6.35651 | 6.44133 | 5.83124 | 5.87497 | 5.82872 | 5.8198 |
| FAM114A1 | Q8IWE2 | 6.63412 | 5.8923 | 6.93108 | 5.66578 | 5.95648 | 6.73713 | 6.02061 | 6.52829 | 6.61579 | 6.32268 | 6.94352 | 6.57038 | 6.3058 | 5.59302 | 5.42064 | 5.86236 |
| TTI1 | O43156 | 6.66487 | 5.90355 | 6.59233 | 6.06473 | 6.20064 | 6.74796 | 6.00886 | 6.19749 | 6.53199 | 6.6904 | 6.8726 | 6.40639 | 5.8328 | 6.16509 | 5.3372 | 5.7831 |
| SNX12 | Q9UMY4-1 | 5.94531 | 5.59263 | 6.46377 | 5.37392 | 6.42645 | 6.49481 | 5.92106 | 6.61905 | 6.43706 | 6.92691 | 6.70773 | 6.90154 | 6.25323 | 6.08163 | 5.85395 | 6.00097 |
| ALAS1 | P13196 | 6.50969 | 6.44384 | 6.19069 | 7.07218 | 5.80684 | 5.8244 | 6.1116 | 5.4471 | 7.54302 | 6.69187 | 6.63599 | 6.89553 | 5.99452 | 5.28204 | 5.65457 | 5.8961 |
| AKAP17A | Q02040 | 6.2829 | 6.22412 | 6.25539 | 6.25785 | 6.20729 | 5.92771 | 6.41663 | 5.32148 | 6.0114 | 5.83164 | 6.0552 | 5.59144 | 6.84987 | 6.97332 | 7.40509 | 6.38866 |
| ANP32A | P39687 | 5.78394 | 5.73265 | 6.03373 | 5.30774 | 6.12176 | 6.24234 | 5.46345 | 6.18026 | 6.02975 | 6.20784 | 6.44156 | 6.205 | 7.05438 | 7.46789 | 6.97172 | 6.756 |
| HEG1 | Q9ULI3 | 5.93443 | 6.76808 | 6.4437 | 6.99011 | 6.06765 | 6.40535 | 6.10544 | 6.2901 | 6.07585 | 6.50497 | 6.42788 | 6.37122 | 5.82181 | 5.84833 | 5.63567 | 6.3094 |
| ERAL1 | O75616 | 6.32315 | 6.385 | 6.0603 | 6.3746 | 6.49612 | 6.33189 | 6.60018 | 6.00326 | 6.06437 | 6.11966 | 6.42654 | 6.10397 | 6.30081 | 5.97558 | 6.1294 | 6.30517 |
| EXD2 | Q9NVH0 | 6.27825 | 6.30499 | 6.11985 | 6.6577 | 6.08767 | 5.89701 | 6.60058 | 5.92228 | 5.95321 | 6.25173 | 5.95541 | 5.81075 | 6.58309 | 6.45154 | 6.70853 | 6.41742 |
| TTC21B | Q7Z4L5 | 6.71154 | 6.64035 | 6.68478 | 6.12919 | 6.42276 | 6.40636 | 6.2302 | 6.52259 | 6.55558 | 6.5366 | 6.53287 | 6.37471 | 5.61562 | 5.62178 | 5.54659 | 5.46849 |
| PIN1 | Q13526 | 6.29298 | 6.09223 | 6.40217 | 6.09926 | 5.91462 | 6.87262 | 5.73457 | 6.26661 | 6.38356 | 6.37939 | 6.50022 | 6.86729 | 6.11538 | 6.1663 | 5.78649 | 6.12631 |
| CNST | Q6PJW8 | 5.91028 | 6.19635 | 6.29358 | 6.87428 | 6.04936 | 6.05871 | 6.3036 | 6.09076 | 6.25689 | 6.37406 | 6.41658 | 6.11553 | 6.224 | 5.94186 | 6.3073 | 6.58687 |
| HSDL1 | Q3SXM5 | 6.49851 | 6.20161 | 5.93821 | 6.74537 | 5.9875 | 6.21251 | 6.24006 | 5.80593 | 6.0097 | 5.98681 | 6.20234 | 5.96028 | 6.757 | 6.4845 | 6.36016 | 6.60951 |
| TRIM14 | Q14142 | 6.19182 | 6.60601 | 6.31866 | 6.45036 | 6.26384 | 6.30646 | 6.57925 | 5.91737 | 6.50567 | 6.11846 | 6.7928 | 6.6172 | 5.68775 | 5.25136 | 6.13475 | 6.25825 |
| SLC1A5 | Q15758 | 6.35096 | 6.15624 | 5.90395 | 6.42275 | 6.41728 | 6.13357 | 6.51115 | 6.14325 | 6.59239 | 5.5927 | 6.23593 | 6.13512 | 6.59303 | 6.05634 | 6.20533 | 6.55002 |
| TBL1XR1 | Q9BZK7 | 6.0781 | 6.05325 | 6.08066 | 6.34332 | 6.36762 | 6.4081 | 5.87251 | 5.99369 | 6.24137 | 6.56347 | 6.27505 | 6.44065 | 6.37917 | 6.05222 | 6.475 | 6.37581 |
| CFAP298 | P57076 | 6.54553 | 6.38594 | 7.05722 | 6.27715 | 6.76782 | 6.87809 | 6.72274 | 6.90203 | 6.77756 | 6.58909 | 6.8507 | 6.75174 | 5.05471 | 4.77379 | 4.76496 | 4.90094 |
| AEBP2 | Q6ZN18 | 6.54771 | 6.69153 | 6.69231 | 6.57887 | 6.16215 | 6.20676 | 6.33618 | 6.20902 | 6.66272 | 6.24867 | 6.36434 | 6.61082 | 5.68103 | 5.80755 | 5.56336 | 5.63698 |
| HMCES | Q96FZ2 | 6.27321 | 6.02906 | 6.53279 | 5.81275 | 5.91569 | 6.15414 | 5.80228 | 6.21141 | 6.4415 | 6.51212 | 6.86149 | 6.54257 | 5.97128 | 6.83542 | 6.10957 | 5.99471 |
| KDM6A | O15550 | 6.346 | 6.30819 | 6.01828 | 6.56019 | 6.1127 | 6.02557 | 6.09336 | 5.61444 | 6.35354 | 6.21503 | 6.11489 | 6.06788 | 6.77487 | 6.44493 | 6.58703 | 6.3631 |
| LIN28B | Q6ZN17 | 6.01479 | 6.3973 | 6.1704 | 6.39942 | 6.18327 | 6.2076 | 6.11689 | 6.49119 | 6.29017 | 6.14135 | 6.3302 | 6.06614 | 6.05423 | 6.52018 | 6.41122 | 6.20566 |
| SPAG7 | O75391 | 5.95775 | 5.82821 | 6.5812 | 5.75634 | 5.86006 | 6.54034 | 5.624 | 6.1682 | 6.31816 | 6.48771 | 6.56327 | 6.53567 | 6.5231 | 6.71372 | 6.14668 | 6.39559 |
| DCK | P27707 | 5.95615 | 5.34612 | 6.99249 | 5.59313 | 6.28145 | 6.05412 | 6.28308 | 6.36236 | 6.4697 | 6.51846 | 6.46074 | 6.18078 | 6.28461 | 6.49932 | 6.73055 | 5.98695 |
| AIDA | Q96BJ3 | 5.96434 | 6.24891 | 6.28303 | 6.13355 | 6.24835 | 6.11325 | 6.45379 | 6.31273 | 6.22806 | 6.35169 | 6.77304 | 6.51218 | 6.4814 | 5.89749 | 6.04713 | 5.95105 |
| PLIN2 | Q99541 | 6.21284 | 5.76994 | 6.26611 | 6.58184 | 6.17404 | 5.84648 | 6.20135 | 6.28148 | 6.05274 | 5.87675 | 5.83066 | 5.98598 | 6.75616 | 6.75604 | 7.05156 | 6.35605 |
| ASAH1 | Q13510-2 | 6.43572 | 6.54082 | 5.83376 | 6.24583 | 6.16977 | 6.02492 | 5.98621 | 6.22596 | 5.79434 | 6.43566 | 6.12158 | 5.95545 | 6.87251 | 6.46931 | 6.22542 | 6.66271 |
| PIK3CA | P42336 | 6.39931 | 6.19796 | 7.13639 | 6.0004 | 6.44942 | 6.56003 | 6.19754 | 6.2693 | 6.69577 | 6.6192 | 6.65585 | 6.68206 | 5.6841 | 5.51444 | 5.58428 | 5.35396 |
| EXOC3 | O60645 | 6.45816 | 6.29974 | 6.73781 | 5.91905 | 6.65831 | 6.42409 | 6.49464 | 6.36672 | 6.31819 | 6.49953 | 6.34535 | 6.59752 | 5.86742 | 5.97528 | 5.62739 | 5.41081 |
| PIGS | Q96S52 | 6.34032 | 6.36068 | 6.26221 | 6.33529 | 6.24632 | 5.9337 | 6.46578 | 5.9169 | 6.11943 | 5.93539 | 6.06427 | 5.88419 | 6.62558 | 6.02491 | 6.89421 | 6.59083 |
| TXNDC12 | O95881 | 6.22683 | 5.04455 | 5.39069 | 5.2308 | 6.00593 | 5.18965 | 5.68222 | 6.43725 | 5.44787 | 6.54972 | 5.62833 | 6.48019 | 6.87497 | 7.98057 | 7.91789 | 7.91253 |
| KLHDC4 | Q8TBB5 | 6.37956 | 6.33149 | 6.84994 | 6.29711 | 6.26244 | 6.28222 | 6.28141 | 6.5729 | 6.31943 | 6.47349 | 6.38735 | 6.54953 | 5.3326 | 6.02968 | 5.90301 | 5.74785 |
| NUP58 | Q9BVL2 | 6.56612 | 5.90897 | 6.18137 | 5.95261 | 6.52237 | 5.95129 | 6.18555 | 6.52301 | 5.99906 | 6.15726 | 6.18877 | 6.2291 | 7.14995 | 6.20866 | 6.21169 | 6.06422 |
| MIER1 | Q8N108-12 | 6.26337 | 6.27347 | 6.39109 | 6.03634 | 6.27091 | 6.09541 | 6.01783 | 6.17387 | 6.13622 | 5.89314 | 6.25528 | 6.21166 | 6.53469 | 6.48469 | 6.51085 | 6.45115 |
| RABL6 | Q3YEC7 | 6.10411 | 6.45418 | 6.85959 | 6.16595 | 6.10184 | 6.296 | 5.92621 | 6.41931 | 6.02788 | 6.41727 | 6.58873 | 6.45512 | 5.38844 | 6.56676 | 5.81919 | 6.40943 |
| DCAF6 | Q58WW2-3 | 5.69227 | 6.32887 | 6.08428 | 5.60838 | 6.44571 | 6.17658 | 5.93902 | 6.02764 | 6.12832 | 6.00342 | 5.72883 | 6.14762 | 6.82487 | 6.81021 | 7.14081 | 6.91319 |
| ARIH1 | Q9Y4X5 | 6.06197 | 6.17502 | 6.54619 | 5.71035 | 6.17726 | 6.43666 | 6.11255 | 6.58891 | 6.21236 | 6.27804 | 6.42737 | 6.15319 | 5.95861 | 6.4834 | 6.3083 | 6.36981 |
| HDAC3 | O15379 | 6.60854 | 6.50966 | 6.48101 | 6.59609 | 6.41373 | 6.25643 | 6.27801 | 6.12225 | 6.14628 | 6.14466 | 6.27767 | 6.25061 | 6.09671 | 5.75854 | 6.12228 | 5.93756 |
| MRPL22 | Q9NWU5 | 6.54769 | 6.53562 | 5.78078 | 7.08897 | 6.26576 | 5.90889 | 6.70126 | 5.86006 | 5.97573 | 6.07078 | 5.8104 | 5.63853 | 6.50355 | 6.15349 | 6.55511 | 6.60336 |
| USP25 | Q9UHP3 | 6.42528 | 6.17247 | 6.87894 | 6.2493 | 6.29046 | 6.39839 | 6.40066 | 6.53031 | 6.14503 | 6.3508 | 6.5837 | 6.634 | 5.64655 | 5.8579 | 5.67386 | 5.76236 |
| ENOPH1 | Q9UHY7 | 6.15741 | 6.05115 | 6.90604 | 5.60619 | 6.25843 | 6.95299 | 6.10531 | 6.68773 | 6.37201 | 6.58758 | 7.03389 | 6.79442 | 5.68289 | 5.77325 | 5.56417 | 5.46654 |
| CEP41 | Q9BYV8 | 6.34225 | 6.32356 | 6.3477 | 6.01566 | 5.9729 | 6.98047 | 6.1755 | 6.67808 | 6.35751 | 6.81145 | 6.57635 | 6.70921 | 6.06051 | 5.64702 | 5.37691 | 5.62492 |
| POGLUT2 | Q6UW63 | 6.46288 | 6.1622 | 6.00705 | 6.61056 | 6.38144 | 6.15676 | 6.29117 | 5.69112 | 6.2797 | 6.03248 | 6.17533 | 5.82474 | 6.49686 | 6.4125 | 6.6262 | 6.389 |
| SIRT6 | Q8N6T7 | 5.90358 | 6.44342 | 6.13814 | 6.0262 | 6.22219 | 6.14021 | 6.04063 | 6.03792 | 6.34694 | 6.14645 | 6.37467 | 6.06065 | 6.65232 | 6.42702 | 6.63601 | 6.40366 |
| TOR1B | O14657 | 6.468 | 6.63908 | 5.98677 | 6.62603 | 6.47465 | 6.20567 | 6.12216 | 6.17645 | 6.11408 | 6.2301 | 6.07414 | 6.15445 | 6.53665 | 6.00872 | 6.22326 | 5.95981 |
| STX7 | O15400 | 6.29019 | 6.11018 | 6.16551 | 5.89782 | 6.24338 | 6.02384 | 6.17458 | 6.26124 | 6.25255 | 6.5871 | 6.52905 | 6.13199 | 6.18958 | 6.31365 | 6.37394 | 6.45538 |
| ATG2B | Q96BY7 | 6.18829 | 6.18053 | 6.86648 | 5.85046 | 6.47965 | 6.56701 | 6.11354 | 6.55967 | 6.81277 | 5.97243 | 6.62423 | 6.80719 | 6.27518 | 5.70786 | 5.44437 | 5.55032 |
| ISY1 | Q9ULR0-1 | 6.64159 | 6.68088 | 6.09247 | 6.68871 | 6.17513 | 6.28055 | 5.96368 | 6.46043 | 6.00864 | 6.1228 | 6.51607 | 6.68329 | 6.27546 | 5.75068 | 5.82792 | 5.8317 |
| NCKIPSD | Q9NZQ3 | 6.24456 | 6.54153 | 6.98524 | 6.41392 | 6.26773 | 6.34713 | 6.30316 | 6.5001 | 6.17439 | 6.192 | 6.64295 | 6.57414 | 5.86373 | 5.95803 | 5.40693 | 5.58445 |
| LRRK2 | Q5S007 | 5.87642 | 6.38256 | 6.8092 | 6.07031 | 6.40225 | 6.1016 | 6.40599 | 6.17969 | 6.74257 | 6.94199 | 6.42888 | 5.92976 | 6.50472 | 6.38973 | 5.984 | 5.75034 |
| RCOR3 | Q9P2K3-3 | 6.36682 | 6.58841 | 5.85037 | 6.36263 | 6.30666 | 6.11965 | 6.21195 | 6.09919 | 6.18074 | 6.40475 | 5.881 | 6.05228 | 6.68885 | 6.46633 | 6.16088 | 6.25949 |
| INPP4B | O15327 | 5.90734 | 6.07745 | 6.76186 | 5.86495 | 6.00254 | 6.73813 | 6.16578 | 6.55707 | 6.30562 | 6.66836 | 6.53199 | 6.70264 | 5.74652 | 6.4013 | 5.75655 | 5.81188 |
| BAZ2B | Q9UIF8 | 6.41945 | 6.32794 | 5.80886 | 6.74811 | 6.01153 | 5.84435 | 6.42607 | 5.86228 | 6.0362 | 6.23361 | 6.03314 | 6.46332 | 6.32254 | 6.50984 | 6.51369 | 6.43907 |
| ZNF451 | Q9Y4E5 | 5.9915 | 6.27262 | 6.13951 | 6.34029 | 6.42829 | 6.28869 | 6.54667 | 5.92555 | 6.04498 | 6.30702 | 6.38688 | 5.98126 | 6.35549 | 6.14287 | 6.66883 | 6.17955 |
| TRMU | O75648 | 6.48238 | 6.42554 | 6.03824 | 6.75445 | 6.04702 | 5.83089 | 6.42215 | 6.05661 | 6.32889 | 5.88524 | 6.0676 | 6.04623 | 6.64596 | 6.16613 | 6.36498 | 6.43769 |

| | | | | | | | | | | | | | | | | | |
|---|---|---|---|---|---|---|---|---|---|---|---|---|---|---|---|---|---|
| EXOC6B | Q9Y2D4 | 6.28773 | 6.00627 | 6.56398 | 5.96511 | 6.60459 | 6.33211 | 6.24686 | 6.17141 | 6.35106 | 6.86637 | 6.61614 | 6.46169 | 6.10407 | 6.05379 | 5.88638 | 5.48245 |
| SF3B4 | Q15427 | 5.85138 | 4.75703 | 5.56142 | 5.03238 | 5.7577 | 5.01014 | 5.55669 | 6.52271 | 5.37955 | 6.61608 | 5.68648 | 7.52261 | 7.07757 | 7.77083 | 7.8214 | 8.07602 |
| COPE | O14579 | 5.9456 | 5.06751 | 5.59021 | 4.86135 | 5.97494 | 6.23932 | 5.89897 | 7.38001 | 5.22538 | 6.35436 | 5.93714 | 7.12493 | 5.16209 | 8.45072 | 7.07427 | 7.71318 |
| ACADS | P16219 | 6.38886 | 6.08888 | 5.68691 | 5.98074 | 6.55046 | 5.7541 | 6.57408 | 6.2356 | 6.17665 | 6.19481 | 5.79956 | 5.7037 | 6.95235 | 6.41478 | 6.87883 | 6.61968 |
| EOGT | Q5NDL2 | 6.01994 | 6.64747 | 6.30942 | 6.5419 | 6.20806 | 5.76244 | 6.18279 | 5.73235 | 6.07068 | 6.29746 | 5.93097 | 5.97208 | 6.71769 | 6.39342 | 6.57167 | 6.64166 |
| DOCK6 | Q96HP0 | 6.33465 | 6.54847 | 5.93348 | 6.06331 | 6.05646 | 6.37774 | 6.17012 | 6.09075 | 5.93312 | 6.07024 | 5.9782 | 5.9482 | 6.60492 | 6.58529 | 6.59509 | 6.70995 |
| BUD31 | P41223-2 | 6.13646 | 6.53015 | 6.21058 | 6.02 | 6.78122 | 6.4272 | 6.28001 | 5.86122 | 5.90598 | 6.32856 | 5.80285 | 5.96772 | 6.62628 | 6.13212 | 6.6908 | 6.29884 |
| POLDIP2 | Q9Y2S7 | 6.15768 | 6.11257 | 6.26389 | 6.67927 | 6.2421 | 5.89811 | 6.49154 | 6.05974 | 6.30164 | 5.93785 | 6.30872 | 6.10483 | 6.61404 | 6.0032 | 6.39459 | 6.43023 |
| TTC17 | Q96AE7 | 6.80716 | 6.72877 | 6.1528 | 7.2319 | 6.37221 | 5.98813 | 6.06252 | 6.45136 | 6.13669 | 6.51073 | 6.20428 | 6.30858 | 5.83771 | 5.76537 | 5.71078 | 5.73102 |
| PMS2 | P54278 | 5.43962 | 6.12614 | 5.84087 | 5.64252 | 5.7981 | 6.17453 | 5.57569 | 5.93362 | 5.81056 | 6.20561 | 6.052 | 6.07217 | 7.4188 | 7.49357 | 7.37763 | 7.0386 |
| NXN | Q6DKJ4 | 6.36882 | 6.20535 | 7.08332 | 5.73425 | 6.24513 | 7.09798 | 5.94246 | 6.47656 | 6.24215 | 6.17048 | 6.69192 | 6.55981 | 5.75341 | 5.68025 | 5.71662 | 6.03148 |
| BCAS2 | O75934 | 5.84615 | 4.78968 | 4.73769 | 5.01569 | 5.00666 | 3.66331 | 4.96424 | 6.01667 | 3.84559 | 5.49176 | 4.65648 | 5.68389 | 6.00734 | 12.0526 | 10.1809 | 12.0414 |
| PNISR | Q8TF01 | 6.42037 | 6.46806 | 5.98236 | 6.56325 | 6.26096 | 5.92273 | 6.04331 | 6.03758 | 6.15448 | 5.84101 | 6.30303 | 5.88073 | 6.5426 | 6.6543 | 6.36452 | 6.56071 |
| NCDN | Q9UBB6-3 | 6.21596 | 6.37999 | 7.30833 | 5.87712 | 6.52061 | 6.90573 | 6.52204 | 6.54622 | 6.58334 | 6.59076 | 6.63761 | 6.58398 | 5.30035 | 5.36126 | 5.25733 | 5.40936 |
| MED4 | Q9NPJ6 | 6.57704 | 6.19353 | 6.08302 | 6.23856 | 6.3387 | 6.08512 | 5.84083 | 6.20866 | 6.10502 | 6.48845 | 6.44386 | 6.54443 | 6.35839 | 6.08977 | 6.14477 | 6.25986 |
| OGFOD1 | Q8N543 | 5.24331 | 5.57722 | 6.20533 | 5.43538 | 5.61908 | 6.2296 | 5.70081 | 6.19416 | 6.04399 | 6.03072 | 6.24375 | 6.37956 | 6.80063 | 7.40973 | 7.5701 | 7.31662 |
| KHDC4 | Q7Z7F0 | 5.70608 | 5.58024 | 5.88116 | 5.91939 | 5.95287 | 6.24161 | 5.53149 | 6.62801 | 6.35038 | 6.21307 | 6.6547 | 6.63105 | 7.41148 | 6.31455 | 6.40676 | 6.57718 |
| HIRA | P54198 | 6.42628 | 6.40418 | 5.89872 | 6.96943 | 6.11031 | 5.80538 | 6.33234 | 5.97704 | 6.17093 | 5.94259 | 6.12898 | 6.2885 | 6.25186 | 6.46732 | 6.31636 | 6.5098 |
| SIX5 | Q8N196 | 6.26281 | 6.34906 | 6.66397 | 6.32308 | 6.04443 | 6.17485 | 6.04971 | 6.46443 | 6.22436 | 6.7123 | 6.50563 | 6.61758 | 6.02034 | 6.21937 | 5.62321 | 5.74486 |
| GAN | Q9H2C0 | 6.17987 | 6.26892 | 7.10424 | 6.10152 | 6.26735 | 6.20629 | 6.1788 | 6.3336 | 6.54429 | 6.50522 | 6.74331 | 6.39084 | 5.64944 | 5.73959 | 6.01313 | 5.77359 |
| GSTM3 | P21266 | 6.13861 | 5.92435 | 6.91884 | 5.39722 | 6.40769 | 6.58723 | 6.5092 | 6.44902 | 6.45114 | 6.65457 | 6.46527 | 6.48859 | 5.82842 | 5.80933 | 6.04238 | 5.92815 |
| JPT1 | Q9UK76 | 6.6497 | 4.93431 | 6.92048 | 4.9905 | 6.64285 | 5.3576 | 5.31951 | 6.8361 | 4.66083 | 7.03677 | 6.3995 | 7.80858 | 5.75676 | 7.50928 | 6.05569 | 7.12155 |
| AK6 | Q9Y3D8 | 5.42951 | 5.89097 | 6.64814 | 5.58498 | 6.42936 | 6.31425 | 6.03952 | 6.06985 | 6.16435 | 6.6142 | 6.98438 | 6.61429 | 6.06139 | 6.70484 | 6.18927 | 6.26072 |
| SAR1B | Q9Y6B6 | 6.23056 | 5.7809 | 6.44927 | 6.21373 | 5.69926 | 5.93487 | 5.9209 | 6.18752 | 6.04989 | 6.48235 | 6.83748 | 6.75216 | 6.26659 | 6.55898 | 6.25765 | 6.3779 |
| RING1 | Q06587 | 6.12154 | 5.9135 | 5.93666 | 6.34204 | 6.27448 | 6.11669 | 6.17215 | 6.33416 | 6.25304 | 6.07904 | 6.30209 | 6.02418 | 6.83832 | 6.34858 | 6.50116 | 6.44238 |
| TRAPPC12 | Q8WVT3 | 6.57412 | 6.21386 | 6.72424 | 5.84455 | 6.41529 | 6.73401 | 5.89548 | 6.40941 | 6.77658 | 6.31873 | 6.57497 | 6.45933 | 6.02233 | 5.9563 | 5.53194 | 5.54885 |
| MON1B | Q7L1V2 | 6.22809 | 6.432 | 6.78813 | 6.3303 | 6.1417 | 6.48817 | 6.26677 | 6.19 | 6.02205 | 6.33581 | 6.29636 | 6.54283 | 6.07795 | 5.91825 | 6.04143 | 5.90016 |
| CXADR | P78310 | 6.33458 | 6.32766 | 5.94736 | 6.67999 | 6.06735 | 6.0474 | 6.38046 | 6.08919 | 6.02069 | 6.33798 | 6.24019 | 6.14994 | 6.27396 | 6.30988 | 6.42754 | 6.36581 |
| FOXK2 | Q01167 | 6.74296 | 6.39981 | 5.95799 | 6.66279 | 6.34345 | 6.39818 | 6.5164 | 6.49985 | 6.4081 | 6.24862 | 5.88446 | 6.0287 | 6.33708 | 6.07602 | 6.02395 | 5.47164 |
| IFI35 | P80217-2 | 5.70989 | 4.92885 | 6.04991 | 4.62986 | 6.15652 | 6.31208 | 6.03578 | 7.00997 | 5.9719 | 7.11283 | 6.37984 | 6.91381 | 6.39741 | 7.09157 | 6.70709 | 6.59269 |
| COX5A | P20674 | 5.85478 | 5.67173 | 4.60632 | 5.7305 | 5.58875 | 4.02396 | 5.22253 | 5.38292 | 4.47522 | 5.24693 | 4.36476 | 4.62392 | 4.95218 | 12.4977 | 8.36175 | 13.395 |
| MAPKAP1 | Q9BPZ7 | 6.47275 | 6.59593 | 6.71088 | 6.43251 | 6.0809 | 6.62959 | 6.44066 | 6.4859 | 6.63338 | 6.70895 | 6.49059 | 6.42944 | 5.65553 | 5.52576 | 5.6511 | 5.05612 |
| COMMD4 | Q9H0A8 | 6.15667 | 5.81227 | 6.47843 | 5.39291 | 6.48021 | 6.64619 | 6.2295 | 6.60061 | 6.42396 | 6.7544 | 6.4501 | 6.24857 | 6.13122 | 6.12926 | 6.21337 | 5.85232 |
| KDM2B | Q8NHM5 | 7.0527 | 6.92118 | 6.71742 | 7.10614 | 6.49686 | 6.14787 | 6.86231 | 5.81978 | 6.29111 | 5.59879 | 5.75075 | 5.56052 | 5.44755 | 6.05498 | 5.99734 | 6.17469 |
| TAMM41 | Q96BW9-3 | 6.41327 | 6.35079 | 6.26275 | 6.57495 | 6.18013 | 6.08769 | 6.46041 | 5.90738 | 6.12457 | 6.30479 | 5.81105 | 5.98819 | 6.37748 | 6.14329 | 6.64144 | 6.37183 |
| CMPK1 | P30085 | 6.07263 | 6.11782 | 6.82404 | 5.57415 | 6.77871 | 6.9448 | 6.12396 | 7.04305 | 6.56929 | 6.76399 | 7.04873 | 6.68782 | 5.88785 | 5.39659 | 5.03676 | 5.12983 |
| SYNJ2BP | P57105 | 6.75783 | 5.54964 | 6.05603 | 5.77269 | 6.20477 | 5.90582 | 5.90296 | 6.39627 | 5.71581 | 6.03028 | 5.62546 | 6.10632 | 6.58799 | 7.3346 | 7.07144 | 6.98209 |
| SASS6 | Q6UVJ0 | 6.42372 | 5.84377 | 6.49255 | 5.64817 | 6.48764 | 6.64675 | 6.12172 | 6.84479 | 6.76999 | 6.41502 | 6.78235 | 6.43928 | 6.24722 | 5.5642 | 5.51399 | 5.75882 |
| SLFN5 | Q08AF3 | 5.40266 | 5.65575 | 6.00283 | 5.48988 | 5.67444 | 6.37302 | 5.60451 | 5.79735 | 6.38976 | 5.97768 | 6.11569 | 5.79194 | 7.33697 | 7.67796 | 7.43319 | 7.27636 |
| CTNNAL1 | Q9UBT7 | 6.92968 | 6.60092 | 6.92138 | 6.8379 | 6.16742 | 6.02525 | 6.82989 | 6.34849 | 6.82054 | 6.6313 | 7.27044 | 6.99469 | 4.69231 | 4.83788 | 4.97664 | 5.11526 |
| OXR1 | Q8N573 | 6.45603 | 6.30383 | 7.04097 | 6.04649 | 6.01581 | 6.70764 | 6.45574 | 6.59042 | 6.48081 | 6.7441 | 6.52721 | 6.33648 | 5.86568 | 5.56332 | 5.42354 | 5.44194 |
| THYN1 | Q9P016 | 5.63651 | 5.93647 | 6.2581 | 5.34134 | 5.64938 | 5.83722 | 5.7997 | 5.6248 | 6.33017 | 6.52288 | 6.43657 | 6.18823 | 6.89022 | 7.36482 | 7.49307 | 6.59053 |
| ZBTB33 | Q86T24 | 6.68686 | 6.78474 | 6.38687 | 6.9572 | 5.98472 | 5.91905 | 6.64812 | 5.86296 | 6.55341 | 6.10582 | 6.45643 | 6.27884 | 5.65645 | 5.72519 | 5.88308 | 6.11023 |
| PIR | O00625 | 5.82889 | 6.10398 | 6.22063 | 5.55515 | 6.94848 | 6.89547 | 6.25797 | 7.28126 | 6.82212 | 5.95271 | 6.29493 | 6.5586 | 5.38548 | 6.27057 | 5.64505 | 5.9787 |
| LAMP2 | P13473-2 | 6.74396 | 6.61009 | 6.20949 | 6.88527 | 6.58011 | 5.75346 | 5.83498 | 6.00958 | 6.09815 | 6.39788 | 6.3332 | 5.93199 | 6.56424 | 5.65736 | 6.63395 | 5.75627 |
| EXOSC1 | Q9Y3B2 | 6.09534 | 5.53994 | 6.20647 | 6.25502 | 6.40517 | 5.9682 | 6.27993 | 5.6562 | 6.52032 | 6.13417 | 6.15762 | 6.31127 | 6.77859 | 6.07516 | 6.91244 | 6.70415 |
| TIMM23 | O14925 | 6.36914 | 6.16076 | 6.13699 | 6.85474 | 6.19925 | 5.90481 | 6.40159 | 6.04178 | 6.56316 | 6.30332 | 6.26539 | 5.93185 | 6.59131 | 5.8873 | 6.46409 | 5.92454 |
| TEX264 | Q9Y6I9 | 6.47514 | 6.51565 | 6.5611 | 6.45324 | 6.48192 | 5.95963 | 6.14723 | 5.75085 | 6.1419 | 6.2772 | 5.83306 | 5.86313 | 6.86887 | 6.10995 | 7.07608 | 6.38506 |
| MRPS16 | Q9Y3D3 | 6.20102 | 6.51948 | 5.89828 | 6.51669 | 6.41847 | 6.07583 | 6.55775 | 6.02428 | 6.21121 | 6.18985 | 5.85901 | 5.89331 | 6.72869 | 6.20257 | 6.41812 | 6.28544 |
| MRPL43 | Q8N983-4 | 6.5944 | 6.07623 | 6.04985 | 6.14025 | 6.20602 | 6.05011 | 6.19815 | 6.82755 | 5.89207 | 6.11809 | 6.25301 | 6.32342 | 6.94435 | 6.09995 | 6.04897 | 6.1776 |
| CLP1 | Q92989 | 6.02402 | 6.04866 | 6.16257 | 6.498 | 6.19138 | 6.15191 | 6.23387 | 5.91254 | 6.32431 | 6.11595 | 6.30792 | 6.17351 | 6.84362 | 6.02006 | 6.68952 | 6.30216 |
| NDUFB4 | O95168 | 5.98416 | 5.76132 | 5.71129 | 6.49968 | 6.50921 | 6.00346 | 6.56672 | 6.51818 | 5.91146 | 6.42895 | 5.88335 | 6.45509 | 6.6482 | 6.04706 | 6.59744 | 6.47446 |
| FAM172A | Q8WUF8 | 6.27413 | 6.28022 | 6.26983 | 6.15982 | 6.27742 | 6.1359 | 6.65721 | 6.02312 | 6.09368 | 6.1076 | 6.20067 | 6.21976 | 6.48247 | 6.05243 | 6.47913 | 6.28658 |
| DNPH1 | O43598 | 6.28892 | 5.58365 | 6.77823 | 5.56898 | 6.40633 | 6.72032 | 6.09695 | 6.6667 | 6.11192 | 6.81139 | 6.78076 | 6.5827 | 6.5068 | 6.15285 | 5.58128 | 5.36221 |
| TDRD3 | Q9H7E2-3 | 6.11686 | 6.64794 | 6.55418 | 5.95188 | 6.22313 | 6.07114 | 6.56384 | 6.42614 | 6.07477 | 6.56238 | 6.4231 | 6.71823 | 6.00761 | 6.13476 | 5.92166 | 5.60295 |
| AP2S1 | P53680 | 6.42384 | 6.54235 | 6.40506 | 6.36786 | 6.04912 | 6.32336 | 6.54288 | 6.09139 | 6.37102 | 6.30261 | 6.2689 | 6.1906 | 6.00862 | 6.0273 | 6.03971 | 6.04537 |
| TOMM22 | Q9NS69 | 5.90539 | 6.14898 | 5.737 | 6.47282 | 5.91599 | 6.0278 | 6.21356 | 6.36961 | 5.55996 | 5.59837 | 6.01216 | 6.41793 | 6.47038 | 7.34007 | 6.7563 | 7.05367 |
| GOLT1B | Q9Y3E0 | 6.15547 | 6.14739 | 6.08446 | 6.24618 | 6.15677 | 6.21172 | 6.32695 | 5.99605 | 6.03821 | 6.23033 | 5.81916 | 5.91484 | 6.54966 | 6.50771 | 6.6249 | 6.99021 |
| FAM169A | Q9Y6X4 | 6.12153 | 5.93393 | 5.62214 | 6.43882 | 6.13111 | 6.03987 | 6.00877 | 5.94628 | 6.22244 | 6.10484 | 6.12677 | 6.03778 | 7.41038 | 6.4939 | 6.7811 | 6.58035 |
| FBXO22 | Q8NEZ5 | 5.74126 | 5.98163 | 6.34075 | 5.65769 | 5.82916 | 6.65575 | 5.78475 | 6.0175 | 5.86656 | 6.53586 | 6.52858 | 6.38577 | 6.18554 | 7.04808 | 6.75957 | 6.68156 |
| SENP2 | Q9HC62 | 6.051 | 6.38958 | 6.50499 | 6.62441 | 6.3517 | 6.09168 | 6.46505 | 5.643 | 6.13942 | 6.35705 | 6.16878 | 6.1046 | 6.75193 | 6.15483 | 5.66375 | 6.53823 |
| SURF1 | Q15526 | 6.25882 | 6.13228 | 5.84215 | 6.60583 | 6.18361 | 5.9847 | 6.40503 | 6.11311 | 6.20157 | 6.0747 | 5.95167 | 6.04462 | 6.84859 | 6.21479 | 6.76836 | 6.37016 |
| DENND3 | E9PF32 | 6.33154 | 6.16388 | 7.01405 | 6.25569 | 5.99579 | 6.47795 | 6.29444 | 6.1239 | 6.3746 | 6.95185 | 6.54943 | 6.36135 | 6.00551 | 5.81189 | 5.68861 | 5.59952 |
| PISD | Q9UG56 | 6.25045 | 6.46857 | 5.95901 | 6.67334 | 6.34884 | 6.23339 | 6.49732 | 6.18253 | 6.10119 | 5.93247 | 5.74121 | 5.88762 | 6.5703 | 5.84641 | 6.6173 | 6.69005 |
| UQCC1 | Q9NVA1 | 6.50706 | 6.62984 | 6.39124 | 6.57694 | 6.60569 | 5.90987 | 6.30478 | 5.89651 | 5.99465 | 6.09663 | 5.74061 | 5.80621 | 6.93065 | 5.95059 | 6.42572 | 6.233 |
| NDUFA5 | Q16718 | 5.76679 | 4.72081 | 5.14117 | 4.72476 | 5.866 | 5.29079 | 5.33141 | 6.50664 | 5.10351 | 5.99282 | 5.32907 | 6.17638 | 5.78514 | 9.4888 | 9.04084 | 9.73505 |
| SYF2 | O95926 | 6.82188 | 7.00902 | 5.84448 | 6.47661 | 6.1427 | 6.67439 | 6.82045 | 6.62012 | 6.83357 | 6.31135 | 6.35051 | 6.20854 | 5.86581 | 5.36096 | 5.41607 | 5.24355 |
| HMGB3 | O15347 | 5.94258 | 5.49841 | 6.37978 | 5.7133 | 6.09569 | 6.15248 | 5.96271 | 6.31128 | 5.87002 | 6.47511 | 6.39553 | 6.17765 | 6.20673 | 7.57174 | 6.86737 | 6.37963 |
| GPC1 | P35052 | 6.40661 | 6.62697 | 6.21502 | 6.89659 | 6.03079 | 5.81841 | 6.1594 | 5.86991 | 6.47907 | 6.24507 | 6.38617 | 6.46237 | 6.48524 | 6.20919 | 5.91555 | 5.79364 |
| PTRH2 | Q9Y3E5 | 6.38442 | 6.45673 | 6.08834 | 6.60094 | 6.38993 | 6.28031 | 6.77646 | 5.89334 | 6.18788 | 6.49076 | 6.15815 | 5.81718 | 6.37943 | 5.52822 | 6.56832 | 5.99958 |
| DENR | O43583 | 5.72531 | 6.71738 | 6.70549 | 6.2199 | 6.60646 | 7.65445 | 5.94027 | 6.67397 | 6.25707 | 6.44151 | 6.81683 | 6.67331 | 5.81465 | 5.78542 | 4.99573 | 4.97225 |
| RUSF1 | Q96GQ5 | 6.2992 | 6.19646 | 5.80607 | 6.30014 | 6.24092 | 5.87379 | 6.60412 | 5.94477 | 6.19452 | 5.85416 | 5.89277 | 5.93028 | 6.89247 | 6.35586 | 6.92649 | 6.68798 |
| MECOM | Q03112-3 | 6.4777 | 6.52723 | 6.44572 | 7.12041 | 6.52206 | 6.20453 | 6.77853 | 6.38674 | 6.28361 | 6.3536 | 6.38438 | 6.19501 | 5.80178 | 5.43687 | 5.36511 | 5.71671 |
| MIB1 | Q86YT6 | 6.48104 | 6.61863 | 6.9521 | 6.55893 | 6.11711 | 6.19299 | 6.36692 | 6.37547 | 6.45351 | 6.64837 | 6.99404 | 6.57345 | 5.40113 | 5.49265 | 5.51528 | 5.25839 |
| LRRC8D | Q7L1W4 | 6.22395 | 7.00418 | 6.00344 | 6.69939 | 6.483 | 5.86839 | 6.45346 | 5.80663 | 6.5053 | 5.83399 | 5.80664 | 6.19463 | 6.17754 | 6.09949 | 6.49938 | 6.3406 |
| ATP11A | P98196 | 6.02545 | 6.42229 | 6.27995 | 6.84109 | 6.08638 | 6.24737 | 6.16838 | 5.87321 | 6.00036 | 6.27153 | 6.40031 | 6.03761 | 6.15543 | 6.31147 | 6.5218 | 6.35736 |
| PRKRA | O75569 | 6.53049 | 6.46194 | 6.568 | 6.08175 | 7.00572 | 6.55989 | 5.83359 | 6.7476 | 6.5173 | 6.6166 | 6.67448 | 6.668 | 5.57404 | 5.56406 | 5.2798 | 5.31674 |
| SPG7 | Q9UQ90 | 6.396 | 6.18185 | 5.80609 | 6.80316 | 6.21314 | 5.67355 | 6.49763 | 6.10036 | 6.06802 | 5.99766 | 6.08484 | 6.00333 | 6.85109 | 6.16638 | 6.80732 | 6.34956 |
| MAN2A2 | P49641 | 6.30101 | 6.54631 | 5.94166 | 6.82568 | 6.19288 | 5.85212 | 6.28217 | 6.53008 | 6.30149 | 5.96093 | 6.48932 | 6.05318 | 6.08578 | 6.09843 | 6.25372 | 6.28523 |
| DNAJC17 | Q9NVM6 | 5.87672 | 5.48458 | 6.12095 | 5.68265 | 5.79355 | 6.1443 | 5.67548 | 5.97983 | 6.07747 | 6.47273 | 6.54234 | 6.54389 | 6.84525 | 7.08219 | 7.28405 | 6.39401 |
| CBX8 | Q9HC52 | 6.63743 | 6.45934 | 5.79173 | 6.48044 | 6.10098 | 5.8387 | 6.48956 | 6.32382 | 6.29619 | 5.91036 | 6.22737 | 6.52662 | 6.02413 | 6.1539 | 6.82607 | 5.91337 |
| H1-2 | P16403 | 6.17649 | 6.10128 | 5.45869 | 6.21874 | 6.1383 | 5.84687 | 6.09179 | 5.74945 | 5.37972 | 5.43645 | 5.40454 | 5.61251 | 7.06324 | 7.90954 | 7.68618 | 7.72621 |
| ATP6V1D | Q9Y5K8 | 6.74281 | 7.02025 | 7.27885 | 6.56408 | 6.23902 | 6.55204 | 6.31093 | 6.36458 | 6.84775 | 6.72036 | 6.97388 | 6.56339 | 4.91057 | 5.20974 | 4.7187 | 4.98306 |
| GNPDA2 | Q8TDQ7 | 6.06491 | 5.98006 | 6.71289 | 5.78635 | 6.26294 | 6.81715 | 6.46219 | 5.86609 | 6.15786 | 6.50206 | 6.8836 | 6.84531 | 6.07775 | 6.00802 | 6.08371 | 5.48911 |
| CYP2U1 | Q7Z449 | 6.36669 | 6.75232 | 6.05231 | 6.27323 | 6.23044 | 5.85189 | 6.3242 | 6.03644 | 6.1032 | 6.19921 | 5.97649 | 5.94959 | 6.23552 | 6.74979 | 6.71554 | 6.18312 |
| RPLP1 | P05386 | 5.9889 | 5.11507 | 5.04156 | 4.6047 | 5.10298 | 4.78937 | 5.73582 | 6.99895 | 5.33729 | 6.23073 | 5.60058 | 7.54638 | 5.6825 | 9.39155 | 7.43442 | 9.39921 |
| IQSEC2 | Q5JU85 | 6.01437 | 6.48578 | 6.70487 | 6.07519 | 5.99258 | 6.6974 | 6.07595 | 6.24056 | 6.16026 | 6.26233 | 7.00315 | 6.70287 | 5.83005 | 6.35201 | 5.66747 | 5.73517 |
| CENPU | Q71F23 | 6.76712 | 6.70346 | 5.86703 | 6.93924 | 6.19843 | 5.94529 | 6.22145 | 5.89672 | 6.2875 | 6.39647 | 6.39049 | 6.033 | 5.94171 | 6.17752 | 6.3107 | 5.92783 |
| GCLC | P48506 | 6.62066 | 6.10711 | 7.09175 | 5.97238 | 6.56713 | 6.61971 | 6.3046 | 6.60221 | 6.43721 | 6.51391 | 6.3613 | 6.3077 | 5.98943 | 5.54994 | 5.46276 | 5.4922 |
| CCNA2 | P20248 | 6.21578 | 5.91924 | 6.19628 | 5.40796 | 5.91586 | 6.02406 | 6.0362 | 6.00092 | 6.26286 | 6.20957 | 6.10293 | 5.99276 | 6.94765 | 6.52333 | 7.22549 | 7.01911 |
| MLKL | Q8NB16 | 6.1975 | 6.19258 | 6.9409 | 6.23547 | 6.08599 | 6.32896 | 6.38462 | 6.28336 | 6.43878 | 6.6928 | 6.90501 | 6.38647 | 5.50286 | 6.0207 | 5.86823 | 5.53577 |
| HNRNPLL | Q8WVV9 | 5.91129 | 5.87471 | 6.08853 | 6.28316 | 6.09545 | 5.98905 | 6.33073 | 5.89424 | 6.09803 | 6.00663 | 6.22329 | 5.77131 | 7.1369 | 6.50927 | 7.3269 | 6.4605 |
| DHX34 | Q14147 | 5.8535 | 6.1967 | 6.28349 | 6.06393 | 5.96338 | 6.48992 | 5.83498 | 5.77342 | 6.40832 | 6.35308 | 6.46763 | 6.22511 | 6.38491 | 6.53182 | 6.61245 | 6.55736 |
| NUDT2 | P50583 | 6.48226 | 6.07847 | 6.69038 | 5.59918 | 6.16056 | 6.85766 | 6.06266 | 6.84706 | 6.40313 | 6.65617 | 6.37474 | 6.66303 | 5.97297 | 5.79996 | 5.45251 | 5.89926 |
| PPP2R5E | Q16537 | 6.3361 | 5.96759 | 6.78326 | 6.1694 | 6.53338 | 6.68976 | 6.16409 | 6.04414 | 6.56509 | 6.40865 | 6.5015 | 6.36419 | 5.79084 | 5.89232 | 5.76305 | 6.02664 |
| SQLE | Q14534 | 6.55349 | 6.59132 | 6.19028 | 6.76229 | 6.06934 | 6.08565 | 6.59351 | 5.99614 | 6.13629 | 6.1114 | 5.9778 | 6.12581 | 6.39434 | 5.9966 | 6.12923 | 6.28654 |
| MAPK14 | Q16539-2 | 5.99815 | 5.8942 | 7.20795 | 6.03678 | 6.28059 | 6.84548 | 6.32539 | 6.48079 | 6.35347 | 6.5445 | 7.21703 | 6.59297 | 5.45147 | 5.89728 | 5.35699 | 5.51696 |
| ACIN1 | E7EQT4 | 6.14181 | 6.06282 | 5.80866 | 6.92427 | 6.09339 | 5.61362 | 6.42976 | 5.5466 | 6.16308 | 6.21443 | 5.95825 | 5.83431 | 7.16143 | 6.40934 | 7.04667 | 6.59157 |
| KATNA1 | O75449 | 6.0583 | 6.0969 | 6.86795 | 6.38774 | 6.45987 | 6.34858 | 6.39269 | 6.89569 | 6.75117 | 6.63185 | 6.9761 | 6.52502 | 5.44156 | 5.52892 | 5.35198 | 5.28569 |
| NCOA1 | Q15788 | 5.8355 | 6.0048 | 6.16334 | 5.21136 | 6.43239 | 6.41712 | 5.71034 | 6.03575 | 6.01673 | 6.14225 | 6.40814 | 6.53544 | 7.04306 | 6.70513 | 6.49073 | 6.84791 |
| TTI2 | Q6NXR4 | 6.33177 | 6.36647 | 6.63266 | 6.08165 | 6.44948 | 6.67337 | 6.02106 | 6.77187 | 6.60019 | 6.63938 | 6.73505 | 6.71077 | 5.76631 | 5.64892 | 5.34245 | 5.2286 |
| SBF2 | Q86WG5 | 6.46519 | 5.80926 | 6.69385 | 6.36542 | 6.1302 | 6.16898 | 6.37278 | 6.27397 | 6.38182 | 6.84208 | 6.59548 | 6.37827 | 5.67007 | 6.09041 | 5.82375 | 5.93849 |
| COL6A2 | P12110 | 7.56434 | 7.58472 | 6.96289 | 7.32643 | 6.77949 | 6.37449 | 6.87197 | 6.44632 | 6.84679 | 6.77476 | 6.70166 | 6.68534 | 4.14804 | 4.4316 | 4.18651 | 4.31466 |
| ENAH | Q8N8S7 | 6.27485 | 6.31735 | 6.40118 | 6.22534 | 6.21582 | 6.35566 | 5.5093 | 6.56051 | 6.66866 | 6.72094 | 7.12477 | 6.58739 | 6.02983 | 6.08936 | 5.56118 | 5.35787 |
| LRP6 | O75581 | 6.3796 | 6.46908 | 5.7108 | 6.54631 | 6.0442 | 6.10688 | 6.61424 | 6.3893 | 6.31999 | 5.98814 | 6.16526 | 6.40637 | 6.28618 | 6.19674 | 6.35818 | 6.01872 |

| | | | | | | | | | | | | | | | | | |
|---|---|---|---|---|---|---|---|---|---|---|---|---|---|---|---|---|---|
| SLC25A4 | P12235 | 6.22943 | 6.21651 | 5.90665 | 6.77906 | 6.19066 | 5.84679 | 6.26566 | 6.1322 | 6.43634 | 5.91811 | 6.1226 | 5.83489 | 6.75199 | 6.01451 | 6.72749 | 6.62711 |
| SLC25A6 | P12236 | 6.42042 | 6.50075 | 5.90916 | 6.80306 | 6.25625 | 6.03271 | 6.43262 | 6.10514 | 6.04716 | 5.82312 | 6.02698 | 6.00408 | 6.40104 | 6.19743 | 6.60359 | 6.43649 |
| SLC25A31 | Q9H0C2 | 5.9256 | 6.22304 | 6.15253 | 6.62337 | 6.17166 | 5.97707 | 6.31562 | 5.98665 | 5.85574 | 6.16249 | 6.19771 | 6.27498 | 6.67316 | 6.43925 | 6.34427 | 6.67686 |
| ATP9B | O43861 | 6.15864 | 6.65776 | 6.77258 | 6.82508 | 5.69465 | 5.93288 | 6.36785 | 5.97931 | 6.35851 | 6.46944 | 6.55853 | 6.43389 | 5.93396 | 5.93731 | 5.93865 | 5.98096 |
| ARMC9 | Q7Z3E5 | 6.33554 | 6.2673 | 6.93407 | 6.05277 | 6.20888 | 6.67679 | 6.34991 | 6.94322 | 6.4245 | 6.51623 | 6.60923 | 6.63877 | 5.55439 | 5.6508 | 5.25548 | 5.58213 |
| HCFC2 | Q9Y5Z7 | 6.48876 | 6.44571 | 6.16887 | 6.57816 | 6.04907 | 6.10888 | 6.07515 | 5.76303 | 6.10518 | 6.41251 | 6.22443 | 5.81205 | 6.2077 | 6.30978 | 6.79691 | 6.45381 |
| SEPHS1 | P49903 | 4.98436 | 4.37518 | 5.70795 | 4.03801 | 5.55243 | 5.36218 | 5.30267 | 6.43548 | 5.09079 | 6.35883 | 6.06727 | 6.72637 | 7.99195 | 8.64849 | 9.0139 | 8.34415 |
| GPAT3 | Q53EU6 | 6.18014 | 6.12135 | 5.73313 | 6.47466 | 6.12781 | 5.9844 | 6.46886 | 5.94993 | 5.91591 | 5.97314 | 6.1802 | 5.96195 | 6.68863 | 6.74186 | 6.94089 | 6.55716 |
| RPA2 | P15927-3 | 5.67805 | 5.51427 | 5.7677 | 5.33974 | 5.94576 | 6.15032 | 5.71561 | 6.33284 | 6.1077 | 6.20678 | 6.32316 | 6.33464 | 6.81447 | 7.39639 | 7.47526 | 6.89731 |
| PEAK1 | Q9H792 | 6.29313 | 6.38493 | 6.03907 | 6.39208 | 6.05394 | 6.11737 | 6.10205 | 6.43783 | 6.46397 | 6.08115 | 6.2296 | 6.22162 | 6.20088 | 6.0877 | 6.58779 | 6.3069 |
| DGUOK | Q16854 | 6.75958 | 7.20902 | 6.30643 | 7.22087 | 6.95153 | 6.65007 | 6.96514 | 6.60333 | 6.40584 | 5.965 | 5.45389 | 5.10636 | 5.78982 | 5.5467 | 5.67181 | 5.39462 |
| ACTR1A | P61163 | 6.4889 | 6.55108 | 6.58641 | 6.07497 | 6.10885 | 6.55376 | 6.23607 | 6.42831 | 6.76452 | 6.35417 | 6.59133 | 6.55368 | 5.59428 | 6.01658 | 5.54755 | 5.54958 |
| XPNPEP3 | Q9NQH7 | 6.16137 | 6.30667 | 5.60181 | 6.84246 | 6.21045 | 6.23874 | 6.3076 | 6.47479 | 6.05122 | 6.20209 | 6.08701 | 5.94023 | 6.7927 | 6.28154 | 6.10289 | 6.39844 |
| CWC27 | Q6UX04 | 5.58522 | 6.01521 | 5.85229 | 6.0987 | 5.90647 | 6.3248 | 6.2174 | 6.29234 | 5.4825 | 6.41294 | 6.13825 | 6.03501 | 6.6369 | 7.09257 | 6.90115 | 7.00828 |
| MCM9 | Q9NXL9 | 6.2924 | 6.31282 | 6.30079 | 6.95655 | 6.17686 | 6.01399 | 6.49363 | 6.20599 | 6.37574 | 6.27938 | 6.32169 | 6.01119 | 6.04475 | 5.99832 | 6.09315 | 6.12273 |
| IKBKG | Q9Y6K9-2 | 6.74197 | 5.39915 | 6.97895 | 4.91689 | 6.78437 | 5.80521 | 5.55478 | 7.41837 | 5.96047 | 6.89575 | 7.36451 | 7.111 | 6.34725 | 5.6982 | 5.11174 | 5.9114 |
| LRCH1 | Q9Y2L9-2 | 6.61449 | 5.7684 | 5.92612 | 6.32398 | 6.31349 | 6.67207 | 6.01185 | 6.13169 | 6.46829 | 5.79836 | 6.53645 | 6.46884 | 6.52344 | 5.92058 | 6.12705 | 6.3949 |
| HMGB2 | P26583 | 5.35346 | 6.14916 | 5.81452 | 5.39267 | 5.62356 | 6.73573 | 5.12294 | 7.10596 | 5.25135 | 6.41892 | 6.59563 | 6.13458 | 5.97497 | 8.44682 | 7.37008 | 6.50965 |
| TERF2IP | Q9NYB0 | 6.38455 | 6.07537 | 6.59903 | 6.18898 | 6.00323 | 6.33538 | 6.04383 | 6.31999 | 6.11291 | 6.162 | 6.25551 | 6.27388 | 7.01637 | 6.21885 | 5.87186 | 6.13825 |
| ACOT8 | O14734 | 5.69125 | 5.95938 | 5.99135 | 6.36098 | 6.40345 | 6.62334 | 6.58693 | 6.11657 | 6.11421 | 6.17191 | 6.45655 | 6.15874 | 6.75105 | 5.79819 | 6.35843 | 6.45766 |
| ATP6V0A2 | Q9Y487 | 6.26236 | 6.31944 | 6.38097 | 6.85467 | 5.68434 | 6.10523 | 6.62996 | 6.28502 | 6.05926 | 6.3768 | 6.23659 | 6.09944 | 6.16056 | 6.27595 | 6.24997 | 6.01943 |
| HAUS4 | Q9H6D7 | 6.42614 | 5.5443 | 6.75983 | 5.55198 | 5.87495 | 6.28651 | 6.19776 | 6.54548 | 5.43571 | 6.7656 | 6.35975 | 7.28023 | 5.3875 | 6.80798 | 6.26638 | 6.5099 |
| EIF2D | P41214 | 6.29326 | 6.56071 | 6.80155 | 6.11745 | 6.29018 | 6.55744 | 6.62598 | 6.48792 | 6.61515 | 6.62285 | 6.42596 | 6.70795 | 5.5647 | 5.37141 | 5.42724 | 5.53023 |
| ZCCHC3 | Q9NUD5 | 5.69619 | 5.78207 | 6.07986 | 6.32325 | 6.14592 | 5.69524 | 7.87483 | 5.82165 | 7.18109 | 4.71859 | 5.42104 | 6.02807 | 5.06556 | 7.58295 | 6.53186 | 8.05183 |
| COMMD10 | Q9Y6G5 | 5.68876 | 5.04369 | 6.15454 | 4.67695 | 5.47462 | 5.72499 | 5.46633 | 6.49537 | 5.70974 | 6.93543 | 6.84406 | 6.88699 | 7.09069 | 7.81344 | 7.49593 | 7.29846 |
| MCAT | Q8IVS2 | 6.52618 | 6.8131 | 5.46532 | 7.49054 | 6.74486 | 6.41087 | 6.02427 | 6.55547 | 5.95967 | 5.64988 | 6.03015 | 5.79242 | 6.26472 | 6.16914 | 6.21913 | 5.88428 |
| SGPP1 | Q9BX95 | 6.09276 | 6.43153 | 5.84778 | 6.37997 | 6.24032 | 6.22311 | 5.97588 | 5.95666 | 6.14224 | 5.95659 | 5.65044 | 5.85483 | 6.95264 | 6.29132 | 6.8367 | 7.12824 |
| ACAA1 | P09110 | 6.7305 | 6.23753 | 6.18887 | 6.59327 | 6.20108 | 5.77993 | 6.3458 | 5.87989 | 6.19656 | 6.16526 | 5.99493 | 6.27563 | 6.88358 | 6.00095 | 6.39332 | 6.1329 |
| TMEM87B | Q96K49 | 6.31305 | 5.63594 | 6.0997 | 6.44196 | 5.8995 | 6.00464 | 6.71971 | 5.34322 | 6.11309 | 5.90759 | 5.87049 | 5.85904 | 7.37025 | 6.35438 | 6.82524 | 7.24221 |
| EXOSC8 | Q96B26 | 6.13753 | 6.32446 | 6.12821 | 6.77239 | 6.4048 | 6.032 | 5.99549 | 5.88877 | 6.40961 | 6.17686 | 5.92905 | 6.33209 | 6.86188 | 5.83868 | 6.41734 | 6.35083 |
| IFT81 | Q8WYA0 | 6.07291 | 6.70347 | 6.55559 | 6.25131 | 6.23977 | 6.67701 | 6.16201 | 6.60429 | 6.43575 | 6.72164 | 6.75875 | 6.59018 | 5.5085 | 5.77664 | 5.53623 | 5.40592 |
| CLASRP | Q8N2M8 | 6.09729 | 6.31742 | 6.35254 | 6.86304 | 5.69107 | 5.91108 | 6.27736 | 5.79996 | 6.03566 | 5.91762 | 6.38147 | 6.45883 | 6.6451 | 6.43774 | 6.17225 | 6.64155 |
| KATNB1 | Q9BVA0 | 6.10618 | 6.23547 | 6.53488 | 6.05522 | 6.23099 | 6.64133 | 6.27581 | 6.30288 | 6.27908 | 6.36261 | 6.31344 | 6.48783 | 6.00859 | 6.33622 | 5.9814 | 5.84808 |
| EXOSC5 | Q9NQT4 | 5.95684 | 6.10719 | 6.21903 | 6.13029 | 6.11569 | 6.14589 | 6.22605 | 6.2264 | 6.18065 | 6.07346 | 5.98942 | 6.32315 | 6.56517 | 6.35848 | 6.83677 | 6.54553 |
| UBE2N | P61088 | 6.01212 | 5.02368 | 6.39858 | 4.66556 | 5.75006 | 4.8795 | 4.96516 | 7.02505 | 4.67353 | 5.84624 | 5.32103 | 6.73319 | 5.63054 | 9.71845 | 7.98309 | 9.37422 |
| RPUSD4 | Q96CM3 | 6.46145 | 6.4063 | 6.24123 | 7.00392 | 6.13176 | 6.07076 | 6.57372 | 5.97083 | 6.59109 | 6.12223 | 6.21304 | 6.25279 | 6.16747 | 5.70347 | 6.12947 | 5.96047 |
| PRCC | Q92733 | 6.08233 | 6.23804 | 5.85945 | 6.06776 | 6.55884 | 6.09758 | 6.03704 | 6.27894 | 5.91641 | 6.10745 | 6.09806 | 6.04044 | 7.20536 | 6.58461 | 6.40504 | 6.42266 |
| MERTK | Q12866 | 6.48212 | 6.49888 | 6.46384 | 6.98254 | 6.57969 | 6.48032 | 6.27499 | 6.20866 | 6.24152 | 5.91142 | 6.42968 | 6.32892 | 6.07796 | 5.80744 | 5.60822 | 5.6286 |
| DIABLO | Q9NR28 | 6.20387 | 5.33872 | 5.18904 | 5.29333 | 5.94715 | 5.43134 | 5.69765 | 6.30439 | 4.97967 | 5.97144 | 5.53487 | 6.43667 | 6.28395 | 8.30842 | 8.82737 | 8.25211 |
| RMI1 | Q9H9A7 | 6.60276 | 6.42734 | 6.21322 | 6.4141 | 5.81859 | 5.8641 | 6.51896 | 6.23559 | 6.4278 | 6.33866 | 6.30367 | 6.27638 | 6.32618 | 5.81563 | 6.18181 | 6.23522 |
| INTS9 | Q9NV88 | 5.62315 | 6.3363 | 5.9269 | 6.26699 | 6.15308 | 6.25964 | 6.33528 | 6.22222 | 6.0979 | 6.21222 | 5.99562 | 6.24704 | 6.39997 | 6.3517 | 6.8779 | 6.69411 |
| SNAPC1 | Q16533 | 6.46244 | 6.26727 | 6.71568 | 6.62736 | 6.42029 | 5.94885 | 6.65166 | 5.99586 | 6.22946 | 6.50514 | 6.09105 | 5.80336 | 6.24845 | 5.73399 | 6.88619 | 6.31295 |
| GMIP | Q9P107 | 6.5657 | 6.32201 | 7.40687 | 5.93521 | 6.47122 | 6.4324 | 6.17193 | 6.27167 | 6.5147 | 6.60661 | 6.90169 | 6.49218 | 5.46055 | 5.82334 | 5.39136 | 5.23255 |
| MKRN2 | Q9H000 | 6.69397 | 6.75398 | 6.41402 | 6.42961 | 6.46161 | 6.5116 | 6.47211 | 6.64302 | 6.49014 | 6.30725 | 6.20145 | 6.38416 | 5.65116 | 5.25539 | 5.80111 | 5.54563 |
| SETD1B | Q9UPS6 | 6.2121 | 6.69236 | 6.05696 | 6.54845 | 6.04493 | 6.28859 | 6.39367 | 5.98979 | 6.08487 | 6.43112 | 5.71281 | 5.9666 | 6.5067 | 6.51659 | 6.33326 | 6.22118 |
| ORC4 | O43929 | 6.30923 | 6.38709 | 6.20404 | 6.4938 | 6.13982 | 6.50141 | 6.2723 | 6.17872 | 6.27944 | 6.55742 | 6.04689 | 6.02154 | 6.21981 | 5.85938 | 6.53584 | 5.99327 |
| ZNF184 | Q99676 | 6.27672 | 6.89202 | 6.08322 | 7.15283 | 6.44861 | 6.11957 | 6.43032 | 6.15766 | 5.84092 | 6.39015 | 6.40578 | 6.08723 | 5.60763 | 6.09044 | 5.91648 | 6.10043 |
| ZNF629 | Q9UEG4 | 6.23272 | 6.22376 | 6.06308 | 6.44209 | 5.88279 | 6.07236 | 6.28235 | 6.03824 | 6.34639 | 6.29381 | 6.21163 | 5.93227 | 6.39895 | 6.44673 | 6.72385 | 6.40897 |
| MBD2 | Q9UBB5 | 6.56658 | 6.0551 | 5.785 | 6.80137 | 6.23079 | 6.02699 | 6.47855 | 6.1233 | 6.03995 | 5.93859 | 6.17406 | 6.08637 | 6.71257 | 6.05548 | 6.27324 | 6.65206 |
| ERF | P50548 | 6.27714 | 6.47376 | 7.08705 | 5.8806 | 6.07037 | 6.60441 | 6.05219 | 6.47017 | 6.18153 | 6.75537 | 6.61361 | 6.93016 | 5.77843 | 6.19302 | 5.32412 | 5.30805 |
| GALK1 | P51570 | 5.75347 | 5.97045 | 6.36694 | 5.59818 | 6.42552 | 7.24015 | 6.38458 | 6.85501 | 6.85277 | 6.60625 | 6.739 | 6.38599 | 6.05806 | 5.82678 | 5.42859 | 5.50826 |
| URI1 | O94763-4 | 6.53209 | 6.39044 | 6.60831 | 6.02582 | 6.81064 | 6.42078 | 6.08808 | 6.65258 | 6.60022 | 6.36158 | 6.60004 | 6.60746 | 5.83899 | 5.47651 | 5.51003 | 5.4764 |
| LANCL2 | Q9NS86 | 6.02492 | 6.19999 | 6.96234 | 5.8978 | 6.28879 | 6.32173 | 5.594 | 6.4116 | 6.3018 | 6.51392 | 6.60459 | 6.5738 | 6.15235 | 5.91308 | 6.29243 | 5.94687 |
| CHST14 | Q8NCH0 | 7.07813 | 6.45163 | 6.20455 | 7.50027 | 6.36719 | 6.27291 | 6.39112 | 6.11352 | 6.67888 | 6.27896 | 6.33433 | 6.74809 | 5.41732 | 5.37124 | 5.25643 | 5.53542 |
| KDM4B | O94953 | 6.24882 | 6.69126 | 6.21361 | 6.55172 | 6.2446 | 6.16313 | 6.3298 | 6.21409 | 6.46607 | 6.64759 | 6.304 | 6.13409 | 5.69805 | 6.2563 | 6.06952 | 5.76736 |
| CASC3 | O15234 | 6.36774 | 6.40653 | 6.97167 | 6.31036 | 6.01429 | 6.54487 | 6.10921 | 6.53105 | 6.16661 | 6.45816 | 6.76193 | 6.60478 | 5.99201 | 5.50327 | 5.5023 | 5.75522 |
| RAD54L | Q92698 | 5.44376 | 5.86726 | 6.00987 | 5.34296 | 5.80856 | 6.20097 | 6.21961 | 5.80836 | 6.52524 | 5.95436 | 6.45844 | 6.56194 | 6.88815 | 7.02228 | 7.09297 | 6.79527 |
| H1-10 | Q92522 | 5.84457 | 6.04373 | 4.91949 | 5.79177 | 5.78216 | 5.75192 | 5.81154 | 5.83702 | 5.34432 | 5.72018 | 5.26185 | 5.54838 | 7.26244 | 8.68869 | 8.25754 | 8.13442 |
| AHCYL1 | O43865 | 6.06086 | 6.35865 | 6.71971 | 6.08025 | 6.23261 | 6.67958 | 6.15219 | 6.37957 | 5.7869 | 6.3222 | 5.72076 | 5.95091 | 6.32823 | 6.69563 | 6.29636 | 6.2356 |
| DCP1A | Q9NPI6 | 6.01724 | 5.62086 | 6.69444 | 5.50842 | 6.46725 | 6.4106 | 5.77529 | 6.84593 | 6.08519 | 7.001 | 6.61235 | 6.82699 | 6.20299 | 6.24539 | 5.85201 | 5.83405 |
| SDC4 | P31431 | 6.99719 | 6.79308 | 6.01571 | 6.42369 | 6.20367 | 5.97929 | 6.18792 | 6.31724 | 6.60746 | 5.9841 | 5.8802 | 6.14166 | 6.44257 | 5.96729 | 5.76993 | 6.28898 |
| ARPC5 | O15511 | 6.11106 | 6.03271 | 6.98059 | 5.60671 | 6.63198 | 6.24642 | 5.91069 | 6.48874 | 6.02226 | 6.36031 | 6.56243 | 6.3096 | 6.55349 | 6.21028 | 6.28544 | 5.68727 |
| ICE2 | Q659A1 | 6.19824 | 6.47272 | 6.65508 | 6.50793 | 6.04626 | 6.2416 | 6.40407 | 5.77418 | 6.1425 | 6.32888 | 6.52105 | 6.16111 | 6.48587 | 6.26196 | 5.84164 | 5.95692 |
| WDSUB1 | Q8N9V3 | 6.33848 | 6.30513 | 6.9159 | 5.86854 | 6.39296 | 6.94088 | 6.33802 | 6.16596 | 6.22588 | 6.49932 | 6.74091 | 6.84167 | 5.90268 | 5.45786 | 6.10478 | 4.96105 |
| ELL | P55199 | 6.09427 | 6.46544 | 6.68158 | 6.6397 | 5.69372 | 6.22288 | 6.12289 | 5.96341 | 6.19465 | 6.19929 | 6.24374 | 6.29532 | 6.34252 | 6.15879 | 6.51097 | 6.17083 |
| UNK | Q9C0B0 | 6.12853 | 6.24248 | 6.33368 | 5.7903 | 6.48515 | 6.60263 | 5.90513 | 6.59342 | 6.2288 | 6.52989 | 6.44375 | 6.03503 | 6.44526 | 6.39606 | 5.82117 | 6.01873 |
| SNRPGP15 | A8MWD9 | 5.78173 | 5.85565 | 5.51678 | 5.90295 | 5.88488 | 5.69869 | 6.35833 | 6.0711 | 6.6075 | 6.06565 | 6.25753 | 6.48026 | 6.92677 | 6.52686 | 6.87724 | 7.18808 |
| MAP4K4 | O95819-3 | 6.11783 | 6.30587 | 6.27024 | 6.25316 | 6.07593 | 6.42294 | 5.88353 | 5.97659 | 6.27272 | 6.44937 | 6.56648 | 6.50185 | 6.38135 | 6.09864 | 6.10223 | 6.32128 |
| FLYWCH2 | Q96CP2 | 5.97593 | 5.70965 | 6.72943 | 5.20878 | 6.26273 | 6.32024 | 5.68725 | 6.4046 | 5.98006 | 6.87041 | 6.95192 | 6.38832 | 6.62489 | 6.63956 | 6.50097 | 5.74526 |
| KAT6A | Q92794 | 6.39149 | 6.84097 | 5.9112 | 6.80386 | 5.62602 | 5.62658 | 6.11646 | 6.25164 | 5.95253 | 6.19133 | 6.51617 | 6.27029 | 6.1996 | 6.11972 | 6.15903 | 7.02312 |
| PASK | Q96RG2-2 | 6.34288 | 6.53121 | 7.36872 | 6.15617 | 6.57419 | 6.49953 | 6.10044 | 6.80065 | 6.65197 | 6.72744 | 6.95552 | 6.93121 | 5.32341 | 4.92102 | 5.34053 | 5.13512 |
| PDCD10 | Q9BUL8 | 6.40481 | 6.31706 | 6.52188 | 5.72349 | 6.05398 | 6.20266 | 6.47306 | 6.55914 | 6.51131 | 6.33417 | 6.65504 | 6.27401 | 6.45982 | 5.74552 | 6.02328 | 5.74076 |
| DENND2B | P78524 | 6.47427 | 6.12414 | 6.23894 | 6.65581 | 6.02827 | 6.13207 | 6.61085 | 5.82176 | 6.20648 | 6.41187 | 6.21505 | 6.3894 | 6.22808 | 6.08075 | 6.08793 | 6.29432 |
| EZH2 | Q15910-2 | 6.40285 | 6.99868 | 5.9645 | 7.38511 | 6.36893 | 6.64183 | 6.4628 | 6.18322 | 6.15775 | 6.27069 | 6.21131 | 6.32499 | 5.41227 | 5.60914 | 5.94301 | 5.66288 |
| MAD2L1 | Q13257 | 5.98675 | 6.14589 | 6.37442 | 6.0895 | 6.32131 | 6.50942 | 6.15679 | 6.25736 | 6.08327 | 6.24531 | 5.86555 | 6.38952 | 6.231 | 6.21144 | 6.58192 | 6.55058 |
| C17orf75 | Q9HAS0 | 6.28557 | 6.36802 | 6.47781 | 5.94559 | 6.68287 | 6.24043 | 5.98396 | 6.0739 | 6.3716 | 6.41524 | 6.33736 | 6.46668 | 6.3995 | 5.78307 | 6.02344 | 6.14496 |
| EPC2 | Q52LR7 | 6.6641 | 7.13188 | 5.86062 | 6.52907 | 6.05771 | 5.96237 | 6.58566 | 5.71586 | 6.89656 | 6.03735 | 6.20276 | 6.29432 | 6.29031 | 6.03832 | 5.4578 | 6.2753 |
| NFKBIL1 | Q9UBC1 | 6.39033 | 6.42379 | 5.88319 | 6.70468 | 6.5867 | 6.4475 | 6.37125 | 6.05294 | 6.75444 | 6.11065 | 6.60401 | 6.6105 | 5.92104 | 5.66885 | 5.85598 | 5.61416 |
| PPIP5K1 | Q6PFW1 | 6.40409 | 6.68922 | 6.78029 | 5.832 | 6.45824 | 6.77887 | 6.18014 | 6.61479 | 6.1903 | 6.40196 | 6.42129 | 6.36637 | 5.79102 | 5.78865 | 5.52107 | 5.78169 |
| TRIM47 | Q96LD4 | 6.07877 | 6.18413 | 6.80019 | 5.507 | 6.45662 | 6.60078 | 5.85173 | 6.3594 | 6.42691 | 6.24306 | 6.56453 | 6.58771 | 6.22436 | 6.04948 | 6.22875 | 5.8366 |
| DMXL2 | Q8TDJ6-3 | 6.35686 | 6.65366 | 6.75076 | 6.24131 | 6.24405 | 6.43551 | 6.1276 | 6.39533 | 6.5967 | 6.81522 | 6.54306 | 6.65093 | 5.4841 | 5.71284 | 5.59533 | 5.39675 |
| UBE2S | Q16763 | 6.74825 | 7.15324 | 7.28717 | 7.05374 | 6.29187 | 6.7722 | 6.51166 | 6.57256 | 6.88195 | 6.94867 | 6.8061 | 6.89248 | 4.00508 | 5.0841 | 4.41148 | 4.57943 |
| KIF3A | Q9Y496 | 6.62091 | 6.42233 | 6.74802 | 5.94539 | 5.94576 | 6.56552 | 6.27274 | 6.72168 | 6.10446 | 6.7292 | 6.64245 | 6.57151 | 5.20325 | 5.96201 | 5.43311 | 6.11164 |
| STIM1 | Q13586 | 6.56762 | 6.55629 | 5.80992 | 6.52218 | 6.32405 | 6.00765 | 6.14832 | 6.18147 | 5.77991 | 6.26243 | 6.05 | 5.76212 | 6.42275 | 6.36011 | 6.61704 | 6.62814 |
| MARS2 | Q96GW9 | 5.95017 | 5.99122 | 5.95731 | 6.74691 | 6.3149 | 6.21037 | 6.46988 | 6.23506 | 5.90724 | 6.02386 | 6.12391 | 6.21204 | 6.79467 | 6.09965 | 6.42876 | 6.53406 |
| CEP78 | Q5JTW2-2 | 6.53001 | 6.59327 | 6.79816 | 6.38412 | 6.20579 | 6.40828 | 6.13794 | 6.32436 | 6.48351 | 6.54426 | 6.62374 | 6.89156 | 5.57961 | 5.43738 | 5.52366 | 5.53436 |
| PYM1 | Q9BRP8 | 5.80531 | 5.92598 | 6.36719 | 5.94309 | 6.24376 | 6.40329 | 6.03968 | 6.55644 | 6.30577 | 6.67788 | 6.59901 | 6.91862 | 5.80002 | 6.29795 | 6.13794 | 5.97807 |
| ATG9A | Q7Z3C6 | 6.20924 | 6.17535 | 6.4858 | 6.34744 | 6.15716 | 6.25908 | 6.14292 | 6.0131 | 6.38595 | 6.50434 | 6.65524 | 6.52548 | 5.99332 | 6.13058 | 5.99074 | 6.02425 |
| TRAPPC10 | P48553 | 6.47117 | 6.34446 | 6.53804 | 6.35853 | 6.42597 | 6.46652 | 5.97738 | 6.1584 | 6.43815 | 5.97652 | 6.4819 | 6.0829 | 6.00048 | 6.47459 | 6.12338 | 6.58163 |
| PUM2 | Q8TB72 | 6.25181 | 6.33124 | 6.58683 | 5.94544 | 6.44845 | 6.41402 | 5.39503 | 6.09477 | 5.97015 | 6.60912 | 6.87541 | 6.41181 | 6.15234 | 6.20638 | 6.26323 | 6.04398 |
| ZHX3 | Q9H4I2 | 5.72948 | 6.40857 | 6.25096 | 6.08576 | 6.32302 | 6.48121 | 5.94846 | 6.14602 | 6.0797 | 6.1194 | 6.66189 | 6.42399 | 6.75452 | 6.48339 | 6.24149 | 5.86212 |
| ALCAM | Q13740 | 6.43448 | 6.44931 | 5.95735 | 6.3181 | 6.14911 | 6.21058 | 6.32926 | 6.10734 | 6.07671 | 6.21026 | 5.95214 | 6.18091 | 6.37282 | 6.43801 | 6.44636 | 6.36725 |
| PIK3R1 | P27986 | 6.49606 | 6.21599 | 6.95737 | 6.07907 | 6.41838 | 6.57204 | 5.87866 | 6.52914 | 6.60012 | 6.63765 | 6.49347 | 6.59711 | 5.88516 | 5.63094 | 5.45905 | 5.54979 |
| STX18 | Q9P2W9 | 6.10955 | 6.21154 | 6.11168 | 6.60774 | 6.15608 | 5.80368 | 5.87425 | 6.37531 | 6.24838 | 6.22051 | 5.79112 | 6.16582 | 6.32632 | 6.53859 | 6.78625 | 6.67319 |
| BNIP1 | Q12981-1 | 6.58123 | 6.55515 | 5.84613 | 6.4632 | 6.16061 | 6.0753 | 6.22798 | 5.95668 | 5.72481 | 6.40186 | 5.94398 | 6.44269 | 6.10389 | 6.33568 | 6.51216 | 6.66866 |
| SLC25A19 | Q9HC21 | 6.00053 | 6.04085 | 5.98802 | 6.35545 | 6.07593 | 5.46536 | 6.6897 | 5.57012 | 6.32724 | 5.99549 | 6.2523 | 5.98676 | 6.68502 | 6.46085 | 7.06732 | 7.03907 |
| PPP1R37 | O75864 | 5.9963 | 6.13406 | 6.57338 | 5.93131 | 6.20354 | 6.57117 | 5.94808 | 6.16067 | 6.3826 | 6.3327 | 6.54003 | 6.43821 | 6.1501 | 6.32292 | 6.17386 | 6.14108 |
| VPS72 | Q15906 | 6.3877 | 6.88413 | 6.0314 | 7.05258 | 5.97075 | 6.19159 | 6.1576 | 6.03419 | 5.95966 | 6.34698 | 6.25726 | 6.32236 | 6.5775 | 6.24051 | 5.6097 | 5.97609 |
| MNAT1 | P51948 | 5.87496 | 5.7792 | 6.30274 | 6.22116 | 6.4715 | 6.47793 | 6.2506 | 6.15267 | 6.38916 | 6.30566 | 6.75052 | 6.63278 | 6.14539 | 5.8542 | 5.96838 | 6.42315 |
| DDX60 | Q8IY21 | 6.57549 | 6.20151 | 6.73775 | 5.81875 | 6.63578 | 6.90005 | 6.2958 | 6.51458 | 6.75092 | 6.26041 | 6.6137 | 6.32266 | 5.45085 | 5.85739 | 5.82627 | 5.23809 |
| RALGAPA2 | Q2PPJ7 | 5.82941 | 6.39801 | 6.86658 | 6.34938 | 6.21028 | 6.59649 | 6.39352 | 6.52875 | 6.35251 | 6.30406 | 6.75669 | 6.7856 | 5.70728 | 5.8298 | 5.65517 | 5.43648 |
| NOP16 | Q9Y3C1 | 6.65467 | 6.49408 | 6.08202 | 7.12717 | 6.41727 | 6.10596 | 6.74047 | 5.94685 | 5.68325 | 5.42569 | 5.50984 | 5.86294 | 6.36366 | 6.25373 | 6.72908 | 6.60332 |
| PCYT2 | Q99447-3 | 6.22254 | 6.19344 | 6.67838 | 5.81433 | 6.51383 | 6.65041 | 6.33124 | 6.65169 | 6.31549 | 6.60049 | 6.40702 | 6.27082 | 5.98737 | 5.64365 | 5.99363 | 5.72566 |
| IMPDH1 | P20839-6 | 6.02805 | 6.10053 | 6.87282 | 5.76966 | 6.3595 | 6.44538 | 6.25213 | 6.72534 | 6.80467 | 6.64088 | 6.62418 | 6.52335 | 5.72967 | 5.69005 | 5.83908 | 5.59472 |
| NDUFB9 | Q9Y6M9 | 6.24738 | 6.16814 | 5.82018 | 6.44712 | 6.73008 | 5.91535 | 6.47754 | 6.12961 | 5.88471 | 6.11318 | 6.07933 | 5.78411 | 6.92973 | 6.45963 | 6.57881 | 6.23511 |

| | | | | | | | | | | | | | | | | | |
|---|---|---|---|---|---|---|---|---|---|---|---|---|---|---|---|---|---|
| PELO | Q9BRX2 | 5.92408 | 6.13874 | 6.70759 | 6.24185 | 6.63578 | 6.55223 | 6.51938 | 6.39323 | 6.24933 | 6.176 | 6.17959 | 6.46844 | 5.82578 | 5.84035 | 5.99493 | 6.1527 |
| AKR7A2 | O43488 | 6.19771 | 6.23599 | 6.17887 | 6.10285 | 6.18439 | 6.01167 | 6.06845 | 6.86884 | 6.33241 | 6.70972 | 6.75559 | 6.59495 | 6.07904 | 6.1769 | 5.67906 | 5.82357 |
| DPY19L4 | Q7Z388 | 6.29412 | 6.32531 | 6.04145 | 6.62264 | 6.30882 | 5.7044 | 6.37448 | 5.89559 | 5.88081 | 6.1706 | 6.18853 | 5.99282 | 6.42523 | 6.44525 | 6.79034 | 6.53961 |
| EPS8L2 | Q9H6S3-3 | 6.43697 | 6.16544 | 7.01206 | 5.82003 | 6.1403 | 6.54562 | 6.05152 | 6.2806 | 6.5571 | 6.44475 | 6.13887 | 6.53808 | 6.37952 | 5.84949 | 5.74377 | 5.89588 |
| RP2 | O75695 | 6.18631 | 6.53116 | 5.95744 | 6.43564 | 6.40689 | 6.1008 | 6.56442 | 6.05296 | 6.19588 | 5.96109 | 5.92675 | 5.85326 | 6.55176 | 6.48621 | 6.50498 | 6.28445 |
| PPP1R9B | Q96SB3 | 6.19005 | 6.72458 | 6.10406 | 6.89848 | 6.4399 | 6.07598 | 6.06028 | 5.93293 | 5.96778 | 6.18963 | 6.58169 | 6.13242 | 6.5874 | 5.57048 | 6.25988 | 6.28446 |
| MAPK3 | P27361 | 6.48795 | 5.45275 | 6.96201 | 5.82385 | 6.14525 | 6.41469 | 5.97419 | 6.41387 | 6.40782 | 6.48527 | 7.0075 | 6.725 | 6.18883 | 5.87664 | 5.8337 | 5.80067 |
| PLCD3 | Q8N3E9 | 6.25712 | 6.28438 | 6.18177 | 6.6173 | 6.6011 | 6.13257 | 6.1208 | 6.0592 | 6.34136 | 5.96215 | 6.12293 | 6.37986 | 6.45595 | 6.21043 | 6.02402 | 6.24908 |
| TXNDC9 | O14530 | 6.07876 | 5.90829 | 6.88078 | 6.2217 | 6.68801 | 6.72129 | 6.47728 | 6.85189 | 6.63037 | 6.61561 | 6.98363 | 7.04713 | 5.34434 | 5.26461 | 5.24156 | 5.04475 |
| FARS2 | O95363 | 6.44028 | 6.49612 | 6.14243 | 6.52416 | 6.52551 | 6.12546 | 6.19038 | 6.01891 | 6.12278 | 5.97403 | 5.88782 | 5.87633 | 6.24444 | 6.08317 | 6.67698 | 6.6712 |
| PLPBP | O94903 | 6.52362 | 6.43759 | 6.58962 | 5.82463 | 7.27196 | 6.84445 | 6.05001 | 6.96457 | 6.11882 | 6.66388 | 6.44308 | 6.59399 | 5.76404 | 5.48628 | 5.18262 | 5.24083 |
| LGMN | Q99538 | 6.70653 | 7.28335 | 6.48327 | 7.02475 | 6.72703 | 6.15569 | 6.16201 | 6.52234 | 6.38914 | 6.66171 | 6.45201 | 6.18883 | 5.43972 | 5.30617 | 5.06438 | 5.43309 |
| CBLL1 | Q75N03 | 6.52536 | 6.54886 | 5.99367 | 6.67448 | 6.14448 | 6.24127 | 5.93822 | 6.00279 | 5.91741 | 6.36767 | 6.3335 | 6.21994 | 6.41301 | 6.51041 | 5.81476 | 6.35416 |
| C11orf98 | E9PRG8 | 5.97993 | 6.44038 | 5.56814 | 6.08367 | 6.16439 | 5.78475 | 6.71382 | 6.33012 | 6.54425 | 5.88588 | 6.22104 | 5.74925 | 6.5353 | 6.70259 | 6.40312 | 6.89337 |
| CSNK1G3 | Q9Y6M4-2 | 6.42338 | 6.66186 | 6.49844 | 6.59588 | 6.21625 | 6.42867 | 6.50766 | 5.97456 | 6.00047 | 6.30052 | 6.33207 | 6.45847 | 5.88325 | 6.12262 | 5.6968 | 5.8991 |
| AP5Z1 | O43299 | 6.58597 | 6.58469 | 6.86132 | 6.25395 | 6.1567 | 6.4166 | 6.3707 | 6.68624 | 6.30223 | 6.52434 | 6.5548 | 6.83907 | 5.46079 | 5.4896 | 5.33818 | 5.57483 |
| SPRYD4 | Q8WW59 | 5.83589 | 6.20783 | 5.67839 | 6.28743 | 6.40584 | 6.20013 | 6.45629 | 6.3019 | 6.36774 | 6.35558 | 5.76816 | 6.09848 | 6.4191 | 6.05434 | 6.91545 | 6.64746 |
| AGFG1 | P52594-4 | 6.02741 | 6.04162 | 6.34973 | 6.05772 | 5.99546 | 6.78102 | 6.15775 | 6.33635 | 6.60401 | 5.99143 | 6.57844 | 6.52001 | 6.64136 | 5.89822 | 6.25222 | 5.76726 |
| GPAM | Q9HCL2 | 6.99209 | 6.75937 | 6.31444 | 7.27988 | 5.88493 | 5.70014 | 6.20181 | 5.79361 | 6.08176 | 6.22311 | 6.41994 | 6.48785 | 6.21444 | 5.51717 | 6.20957 | 5.91989 |
| DDHD2 | O94830 | 6.65963 | 6.41376 | 6.68079 | 6.32948 | 6.37753 | 6.62118 | 6.34047 | 6.984 | 6.68834 | 6.38825 | 6.52999 | 6.63187 | 5.70832 | 5.28619 | 4.98956 | 5.37064 |
| DENND1A | Q8TEH3-6 | 6.52785 | 6.18245 | 6.39721 | 6.17246 | 6.19544 | 6.53457 | 6.11586 | 6.39694 | 6.50188 | 6.95396 | 6.20451 | 6.53701 | 5.93581 | 5.86703 | 5.65323 | 5.82377 |
| CCDC50 | Q8IVM0 | 6.61411 | 6.43347 | 6.78418 | 6.06797 | 6.62438 | 6.86368 | 5.9964 | 7.15838 | 6.38147 | 6.44688 | 6.36687 | 6.20164 | 5.71787 | 5.54648 | 5.22266 | 5.57355 |
| CCDC59 | Q9P031 | 6.13775 | 5.84902 | 5.812 | 6.63429 | 6.29759 | 6.12287 | 6.18116 | 5.9058 | 5.66591 | 5.41947 | 5.97826 | 5.96187 | 6.64226 | 6.97364 | 7.38018 | 7.03792 |
| CZIB | Q9NWV4 | 6.04124 | 5.6434 | 7.01993 | 5.47325 | 6.32887 | 6.53807 | 6.05583 | 6.48475 | 6.63832 | 6.32661 | 7.06599 | 6.89439 | 6.24913 | 5.69177 | 5.72269 | 5.82575 |
| C17orf80 | Q9BSJ5 | 6.2037 | 6.74079 | 6.0809 | 6.83902 | 6.13403 | 6.16955 | 6.23606 | 6.25016 | 6.23272 | 6.26943 | 6.18593 | 6.03239 | 6.04066 | 6.02016 | 6.3701 | 6.19441 |
| SBF1 | O95248-4 | 6.10358 | 6.13037 | 6.52125 | 5.75343 | 6.51121 | 6.79688 | 6.42344 | 5.97322 | 6.29368 | 6.50682 | 6.75846 | 6.43621 | 5.91311 | 6.28037 | 5.80002 | 5.79794 |
| ARHGAP21 | Q5T5U3 | 6.42082 | 6.41511 | 6.15158 | 6.63476 | 6.04139 | 5.91002 | 6.2301 | 6.46072 | 6.12979 | 6.25823 | 5.92031 | 5.74407 | 6.49517 | 6.44832 | 6.3143 | 6.42532 |
| SCAMP3 | O14828 | 6.15904 | 6.48016 | 6.55311 | 6.64658 | 6.25474 | 6.29649 | 6.02024 | 6.05377 | 6.24787 | 6.41323 | 6.44604 | 6.27074 | 5.89305 | 6.11831 | 6.05036 | 6.09628 |
| MKLN1 | Q9UL63 | 6.27322 | 6.46388 | 6.96328 | 6.78662 | 6.32963 | 6.25204 | 6.44329 | 6.34795 | 6.34789 | 6.41937 | 7.05069 | 6.32626 | 5.37835 | 5.64128 | 5.35072 | 5.62552 |
| ARMC10 | Q8N2F6 | 6.31573 | 6.23358 | 6.03273 | 6.54969 | 5.97631 | 6.20957 | 6.14738 | 5.81902 | 5.94444 | 5.99394 | 5.97039 | 6.52875 | 6.47608 | 6.55871 | 6.84 | 6.38258 |
| MAPKAPK2 | P49137 | 6.7268 | 6.34545 | 6.74959 | 6.14856 | 6.26377 | 6.86508 | 6.14028 | 6.70498 | 6.5433 | 7.01372 | 6.6189 | 6.87427 | 4.95605 | 5.10046 | 5.35668 | 5.59211 |
| CDC7 | O00311 | 6.08625 | 6.55327 | 6.29834 | 6.37084 | 5.90823 | 6.08846 | 5.99716 | 6.10871 | 6.75621 | 6.88287 | 7.06031 | 6.80295 | 6.17785 | 5.26523 | 5.9295 | 5.71384 |
| ANKMY2 | Q8IV38 | 6.13498 | 6.6543 | 6.80118 | 6.17559 | 6.1794 | 7.22666 | 5.99536 | 6.62359 | 6.1761 | 6.6823 | 6.1545 | 6.56014 | 5.54588 | 5.86805 | 5.7173 | 5.50465 |
| ARFGAP1 | Q8N6T3 | 6.05415 | 6.49699 | 6.50011 | 6.0853 | 6.18944 | 6.52752 | 6.168 | 6.45475 | 6.40905 | 6.16895 | 6.43583 | 6.17999 | 6.28526 | 6.26746 | 6.5587 | 6.11848 |
| EIF4ENIF1 | Q9NRA8-3 | 6.01386 | 6.3171 | 6.6945 | 6.33174 | 6.30921 | 6.48523 | 5.89673 | 6.36969 | 6.49993 | 6.68789 | 7.33672 | 6.81963 | 5.75112 | 5.57988 | 5.57107 | 5.33569 |
| RBMS2 | Q15434 | 6.61651 | 6.37147 | 7.00812 | 6.1545 | 6.56209 | 6.47487 | 6.08419 | 6.38706 | 6.38884 | 6.52241 | 6.79846 | 6.56308 | 5.8429 | 5.40604 | 5.38983 | 5.42963 |
| UBAC2 | Q8NBM4 | 6.28472 | 6.12009 | 5.8461 | 6.81398 | 6.0426 | 5.91269 | 6.48664 | 6.29309 | 6.19303 | 5.97556 | 6.20552 | 6.29316 | 6.02621 | 6.27414 | 6.30424 | 6.92823 |
| MTMR2 | Q13614 | 6.42699 | 6.03791 | 6.7005 | 5.87262 | 6.46428 | 6.56489 | 6.17343 | 6.39407 | 6.2937 | 6.04443 | 6.70635 | 6.37812 | 6.47353 | 5.74044 | 5.91416 | 5.81459 |
| STARD13 | Q9Y3M8 | 6.40215 | 6.22045 | 7.10099 | 6.37692 | 6.32084 | 6.65792 | 6.43139 | 6.83799 | 6.63012 | 6.12319 | 6.66462 | 7.03436 | 5.22449 | 5.19391 | 5.32867 | 5.45201 |
| DTD1 | Q8TEA8 | 6.25522 | 5.88594 | 6.08755 | 5.59609 | 6.01655 | 6.23199 | 6.13392 | 6.36117 | 6.75401 | 6.36483 | 6.50376 | 6.36111 | 6.71896 | 6.37578 | 6.11388 | 6.23921 |
| TRNT1 | Q96Q11 | 5.84331 | 5.02441 | 6.0178 | 6.29465 | 6.24214 | 6.72224 | 6.23356 | 5.50468 | 6.2597 | 6.49846 | 6.55625 | 6.68911 | 6.90133 | 6.56711 | 6.24106 | 6.30422 |
| TANGO6 | Q9C0B7 | 6.01559 | 6.16247 | 6.19899 | 5.67142 | 6.21732 | 6.51734 | 6.02232 | 6.64776 | 6.1654 | 6.44038 | 6.40134 | 6.11984 | 6.26166 | 6.42572 | 6.29701 | 6.43544 |
| RRAGB | Q5VZM2 | 6.71273 | 6.83617 | 6.69894 | 6.76219 | 5.95738 | 6.6741 | 6.3621 | 6.30834 | 6.44983 | 6.80406 | 6.34395 | 6.51356 | 5.17937 | 5.54514 | 5.50669 | 5.34546 |
| RIOX2 | Q8IUF8 | 5.57534 | 6.39201 | 5.49323 | 6.31509 | 5.80421 | 5.9112 | 6.27409 | 6.91677 | 7.78351 | 6.21752 | 6.45851 | 6.42741 | 6.37238 | 5.6983 | 5.85073 | 6.50971 |
| MIDEAS | A0A1C7CYX1 | 6.27869 | 6.38261 | 5.9553 | 6.78778 | 6.27179 | 6.10595 | 6.22876 | 6.03218 | 6.18059 | 6.43784 | 6.01961 | 5.94161 | 6.33704 | 6.28837 | 6.3228 | 6.42909 |
| KIAA0100 | Q14667 | 6.62235 | 6.44645 | 6.02908 | 6.89255 | 6.20611 | 5.70972 | 6.40531 | 5.83509 | 6.01126 | 6.14627 | 6.23751 | 5.98614 | 6.22553 | 6.19377 | 6.50717 | 6.54567 |
| RABEPK | Q7Z6M1 | 6.31563 | 5.79086 | 7.36119 | 6.18693 | 6.45039 | 6.50897 | 5.6811 | 6.52955 | 6.46568 | 6.63303 | 6.98537 | 7.31042 | 5.56276 | 5.45549 | 5.43109 | 5.33154 |
| RAB5A | P20339 | 6.24813 | 6.42057 | 6.24036 | 6.52358 | 5.99766 | 6.34261 | 6.35084 | 6.15338 | 6.38604 | 5.9847 | 6.4882 | 5.98508 | 5.97539 | 6.02971 | 6.3939 | 6.47986 |
| PDCD5 | O14737 | 6.43918 | 4.96394 | 6.34995 | 5.05436 | 6.22828 | 5.39907 | 5.22561 | 7.24577 | 5.26095 | 6.79163 | 6.27401 | 7.51621 | 5.82839 | 7.62323 | 6.53057 | 7.17482 |
| TXNDC17 | Q9BRA2 | 6.02812 | 5.48576 | 6.21983 | 4.89986 | 6.76135 | 6.7635 | 6.07885 | 7.26868 | 6.18042 | 6.93018 | 6.73215 | 6.72369 | 5.90159 | 6.48484 | 5.59987 | 5.94132 |
| MAEA | Q7L5Y9 | 6.63959 | 6.57756 | 6.91573 | 6.69033 | 6.47941 | 6.49535 | 6.4615 | 6.27894 | 6.43264 | 6.26317 | 6.49757 | 6.37115 | 5.53819 | 5.19113 | 5.60261 | 5.56512 |
| CPNE7 | Q9UBL6 | 6.14579 | 5.87594 | 6.32904 | 5.58427 | 6.12589 | 6.19868 | 5.89105 | 5.94352 | 6.55411 | 6.70361 | 6.48111 | 6.61346 | 6.00606 | 6.59866 | 6.65587 | 6.29294 |
| PCIF1 | Q9H4Z3 | 6.08982 | 6.10461 | 5.87149 | 5.34729 | 6.06227 | 6.57718 | 5.50571 | 6.39774 | 6.07563 | 6.35758 | 5.96988 | 6.20047 | 6.51528 | 6.81009 | 7.27853 | 6.83643 |
| MTMR14 | Q8NCE2 | 6.43207 | 6.29445 | 6.50326 | 6.29113 | 6.39289 | 6.55584 | 6.47895 | 6.28076 | 6.13526 | 6.43578 | 6.22821 | 6.26764 | 6.18752 | 5.66847 | 5.97258 | 5.87517 |
| PTDSS1 | P48651 | 6.13756 | 6.23911 | 5.98889 | 6.88525 | 5.44345 | 5.53285 | 6.76462 | 5.79167 | 5.3766 | 6.05703 | 6.2087 | 6.02491 | 6.64768 | 6.59338 | 7.49543 | 6.81287 |
| SMG7 | Q92540 | 6.39182 | 6.41921 | 6.97608 | 6.12632 | 6.42282 | 6.50386 | 6.30967 | 6.54905 | 6.39151 | 6.4483 | 6.57355 | 6.63231 | 5.6612 | 5.67507 | 5.43945 | 5.4798 |
| DHPS | P49366 | 5.85248 | 5.80473 | 6.36946 | 5.36366 | 6.11913 | 6.56773 | 6.12089 | 7.07323 | 6.35721 | 6.67596 | 6.44526 | 6.86926 | 6.10385 | 6.1348 | 5.976 | 6.16634 |
| NOVA1 | P51513 | 6.50218 | 5.97114 | 5.88031 | 6.5378 | 6.37433 | 6.15295 | 6.05629 | 5.98541 | 6.59469 | 6.23037 | 6.03939 | 6.26504 | 6.84179 | 5.93753 | 6.60073 | 6.03003 |
| ALG1 | Q9BT22 | 6.30085 | 6.45259 | 5.85457 | 6.718 | 6.16548 | 5.9123 | 6.15072 | 5.84074 | 6.09345 | 5.81239 | 6.1471 | 6.02842 | 6.50036 | 6.39042 | 6.96059 | 6.67203 |
| EIF4E2 | O60573 | 5.97774 | 5.93646 | 6.69367 | 6.30729 | 6.59946 | 6.80906 | 6.08053 | 6.87389 | 6.75196 | 6.39986 | 6.87908 | 6.74762 | 5.74165 | 5.35418 | 5.36506 | 5.4825 |
| ARL6IP5 | O75915 | 6.33921 | 6.38514 | 5.65715 | 6.41062 | 6.10197 | 5.99909 | 7.02237 | 5.54012 | 5.7837 | 5.85748 | 5.78106 | 6.12595 | 7.00969 | 6.93733 | 6.27208 | 6.77704 |
| DNAJC16 | Q9Y2G8 | 6.35949 | 6.3459 | 5.97465 | 6.75658 | 6.44738 | 5.86643 | 6.40296 | 5.67869 | 6.24978 | 6.2987 | 6.19004 | 5.78383 | 6.49682 | 5.99968 | 6.65091 | 6.49817 |
| ATP1B1 | P05026 | 6.47892 | 6.53242 | 6.4267 | 6.60942 | 6.32433 | 6.46228 | 6.35462 | 5.78604 | 6.38422 | 6.00881 | 5.80122 | 5.92133 | 6.08302 | 6.19237 | 6.30514 | 6.32916 |
| WBP4 | O75554 | 5.73873 | 6.19906 | 5.9763 | 5.71239 | 6.21508 | 6.46071 | 5.99368 | 5.81573 | 6.26508 | 6.22589 | 6.225 | 5.85432 | 7.1029 | 6.72813 | 6.97334 | 6.51367 |
| BANF1 | O75531 | 5.92014 | 5.75894 | 4.60091 | 5.91917 | 5.30287 | 4.34677 | 4.94938 | 6.06993 | 4.5518 | 5.10149 | 4.75218 | 5.39175 | 4.37879 | 12.2653 | 7.93953 | 12.7511 |
| MESD | Q14696 | 6.33333 | 6.43262 | 5.97125 | 6.50196 | 5.75733 | 6.06716 | 6.24553 | 6.02224 | 5.57487 | 6.49793 | 5.87895 | 6.28321 | 6.39999 | 6.51071 | 6.60935 | 6.91358 |
| CCN2 | P29279 | 5.35384 | 5.01031 | 4.85731 | 6.1708 | 5.35374 | 5.14016 | 6.73329 | 5.44711 | 4.82564 | 4.15835 | 4.77994 | 4.77907 | 10.1914 | 7.6456 | 10.249 | 9.30449 |
| DENND10 | Q8TCE6 | 5.77909 | 6.01724 | 6.24002 | 6.02915 | 5.89893 | 6.59131 | 5.6412 | 6.75985 | 5.8501 | 6.76217 | 6.58851 | 6.69957 | 6.09239 | 6.48404 | 6.03115 | 6.53525 |
| DCP1B | Q8IZD4 | 6.22315 | 5.73952 | 6.25903 | 5.43379 | 6.05712 | 6.00102 | 5.72172 | 6.55417 | 6.53472 | 7.03696 | 7.20245 | 7.07704 | 6.15183 | 6.13994 | 5.99852 | 5.86902 |
| SPRY4 | Q9C004 | 6.02002 | 6.15353 | 6.22285 | 6.15125 | 6.2249 | 6.16116 | 6.29829 | 5.98218 | 5.90699 | 6.26903 | 6.05153 | 5.96206 | 6.51256 | 6.70859 | 6.56642 | 6.80865 |
| R3HCC1 | Q9Y3T6 | 6.76982 | 6.09756 | 6.86959 | 6.3101 | 6.10342 | 6.19229 | 6.50097 | 6.40112 | 6.7186 | 6.28328 | 6.39642 | 6.26716 | 5.817 | 5.76215 | 5.63324 | 5.87726 |
| WDFY1 | Q8IWB7 | 6.22033 | 6.49517 | 6.50932 | 6.43626 | 5.74885 | 6.15571 | 6.27838 | 5.81671 | 6.31531 | 6.53056 | 6.06695 | 6.21377 | 6.03479 | 5.92622 | 6.81776 | 6.43391 |
| CLCN7 | P51798 | 6.47702 | 6.88228 | 6.3866 | 7.04292 | 5.84001 | 5.95675 | 6.26146 | 5.77947 | 6.32837 | 6.13832 | 6.31452 | 5.96469 | 5.87903 | 5.99024 | 6.42345 | 6.33486 |
| FBH1 | Q8NFZ0-2 | 5.58844 | 6.00061 | 6.49179 | 5.70918 | 6.0527 | 6.28311 | 5.75956 | 5.90006 | 6.27126 | 6.52 | 6.67127 | 6.13387 | 6.19783 | 6.75188 | 6.784 | 6.88444 |
| CASP6 | P55212 | 6.09101 | 6.37766 | 7.04149 | 5.63118 | 6.11598 | 6.91872 | 6.17376 | 6.56946 | 6.1517 | 6.87609 | 6.44793 | 6.44348 | 5.6195 | 6.20125 | 5.77232 | 5.56848 |
| S100A11 | P31949 | 5.43077 | 4.73126 | 5.98964 | 4.33894 | 5.49783 | 5.50246 | 5.32779 | 6.55134 | 4.97374 | 6.49316 | 5.4666 | 6.43891 | 5.34285 | 10.121 | 8.22723 | 9.56646 |
| CDC123 | O75794 | 6.29072 | 6.18528 | 7.32783 | 6.32985 | 6.59004 | 6.90868 | 6.31394 | 6.64245 | 6.76961 | 6.53501 | 7.08632 | 7.18747 | 5.09959 | 4.80742 | 4.75833 | 5.16746 |
| SLC1A4 | P43007 | 6.13469 | 6.80612 | 5.54624 | 6.54251 | 6.02148 | 6.36487 | 6.00852 | 5.82425 | 6.03921 | 6.58171 | 6.02841 | 6.53662 | 6.41797 | 6.19245 | 6.11756 | 6.8374 |
| GSK3B | P49841-2 | 6.40824 | 5.77386 | 7.01415 | 6.03991 | 6.37318 | 6.77161 | 6.29969 | 5.97509 | 6.42072 | 6.55247 | 6.68956 | 6.3399 | 6.08936 | 6.08929 | 5.4634 | 5.69957 |
| OXA1L | Q15070 | 6.39166 | 6.17841 | 5.83291 | 6.38713 | 6.51139 | 6.10272 | 6.53658 | 5.99582 | 6.18539 | 5.87682 | 5.81437 | 5.90688 | 6.69143 | 6.10959 | 6.786 | 6.69288 |
| SLC25A20 | O43772 | 6.38957 | 6.31969 | 5.63029 | 6.52972 | 6.21213 | 5.96478 | 6.17814 | 6.07371 | 5.98317 | 6.1606 | 5.85283 | 5.49889 | 6.83121 | 6.57765 | 7.15569 | 6.64191 |
| TTC12 | Q9H892 | 6.40112 | 6.25709 | 6.48727 | 6.44858 | 6.42434 | 6.704 | 6.5687 | 6.93356 | 7.20421 | 6.6105 | 6.61982 | 6.65069 | 5.35476 | 5.34544 | 4.78511 | 5.20483 |
| NFX1 | Q12986 | 5.80585 | 6.28313 | 6.47789 | 6.09838 | 6.24215 | 6.42967 | 6.31325 | 6.33889 | 6.19242 | 6.73706 | 6.56232 | 6.64451 | 5.72048 | 6.15023 | 5.83085 | 6.17292 |
| NAA30 | Q147X3 | 6.30862 | 6.25032 | 6.60139 | 5.56902 | 6.43774 | 6.32709 | 6.25009 | 6.31815 | 6.74485 | 6.48942 | 6.34611 | 6.37955 | 6.3651 | 6.04697 | 5.87647 | 5.6891 |
| ICE1 | Q9Y2F5 | 6.78152 | 6.62813 | 6.36238 | 7.18939 | 6.22472 | 6.2302 | 6.29776 | 5.86451 | 6.3825 | 5.89643 | 6.29612 | 6.2977 | 6.12585 | 5.35308 | 5.87276 | 6.19695 |
| MTMR1 | Q13613 | 5.9183 | 6.30942 | 6.96994 | 5.70713 | 6.17935 | 6.75724 | 5.80927 | 6.49144 | 6.49519 | 6.85829 | 6.62956 | 6.58507 | 5.76321 | 5.96835 | 5.66633 | 5.89192 |
| SRSF6 | Q13247 | 6.46235 | 6.51915 | 6.04379 | 6.33353 | 6.37283 | 5.8925 | 6.57133 | 5.9349 | 6.09334 | 6.01072 | 6.25629 | 5.90023 | 6.96101 | 6.39182 | 6.23768 | 6.01853 |
| SGO1 | Q5FBB7 | 6.18337 | 6.24128 | 5.9424 | 6.81745 | 5.77319 | 5.49297 | 6.59053 | 5.52207 | 6.19527 | 6.06412 | 5.99268 | 5.91698 | 6.54769 | 6.45183 | 7.23127 | 7.03691 |
| SPTY2D1 | Q68D10 | 6.37785 | 6.27964 | 5.91933 | 6.92154 | 6.2887 | 5.58232 | 6.35635 | 6.26484 | 6.22238 | 6.11402 | 6.11506 | 6.0443 | 5.79085 | 6.61902 | 6.51924 | 6.58457 |
| TBL1X | O60907 | 6.52429 | 6.10571 | 6.08476 | 6.36727 | 6.48407 | 5.77692 | 6.37068 | 5.96299 | 6.22657 | 6.4911 | 6.38166 | 6.16199 | 6.36756 | 5.91751 | 6.47631 | 6.30062 |
| NPR2 | P20594 | 6.56837 | 6.80661 | 5.84061 | 6.48918 | 6.36101 | 6.25939 | 6.25562 | 5.8745 | 6.2662 | 6.42941 | 6.63606 | 6.22377 | 6.02623 | 6.01366 | 5.97398 | 5.97541 |
| EMC4 | Q5J8M3 | 6.05433 | 5.61189 | 5.86545 | 5.87023 | 6.37922 | 5.74561 | 5.98537 | 5.88135 | 5.80383 | 6.14447 | 5.83697 | 6.10806 | 6.74322 | 6.9743 | 7.60268 | 7.39302 |
| HIBADH | P31937 | 6.49863 | 6.10723 | 6.09753 | 5.97312 | 6.61191 | 6.19426 | 6.27416 | 6.28905 | 6.31361 | 5.83905 | 6.40348 | 5.72943 | 7.19718 | 6.13914 | 6.26818 | 6.06408 |
| SRSF2 | Q01130 | 5.86562 | 6.36204 | 6.07327 | 5.8546 | 6.0922 | 6.14006 | 5.83274 | 5.68507 | 6.27989 | 6.35194 | 6.34324 | 6.15024 | 7.16703 | 6.32384 | 6.68974 | 6.78848 |
| PEX19 | P40855 | 6.34667 | 5.07042 | 4.36193 | 4.23196 | 5.08398 | 4.56852 | 5.40797 | 6.94696 | 4.61522 | 5.57699 | 4.61915 | 7.01169 | 5.79669 | 10.9267 | 8.68406 | 10.7511 |
| TJAP1 | Q5JTD0 | 6.45435 | 6.50335 | 6.81592 | 6.18361 | 6.47629 | 6.62524 | 6.26001 | 6.41888 | 6.58808 | 6.76597 | 7.08515 | 6.68111 | 5.77707 | 5.23462 | 5.14354 | 4.98681 |
| AGPAT3 | Q9NRZ7-3 | 6.0762 | 6.42261 | 5.72762 | 6.56277 | 5.8735 | 5.67598 | 6.33726 | 6.03071 | 5.89315 | 6.16077 | 6.03015 | 5.85239 | 6.60511 | 6.85796 | 6.90213 | 6.99167 |
| CIRBP | Q14011 | 6.5346 | 6.16985 | 6.00981 | 6.38009 | 6.44108 | 5.9581 | 6.14169 | 6.08952 | 6.64493 | 6.29791 | 6.39014 | 6.40134 | 6.37314 | 5.80927 | 6.32807 | 6.03046 |
| WDR24 | Q96S15 | 6.44771 | 6.27663 | 6.47847 | 5.97007 | 6.1596 | 6.35648 | 5.95033 | 6.10503 | 6.3934 | 6.29838 | 6.31104 | 6.57072 | 6.07031 | 6.39439 | 6.49694 | 5.7205 |
| ARF5 | P84085 | 6.49339 | 6.83339 | 6.61212 | 6.69947 | 6.34944 | 6.53176 | 5.98461 | 6.24208 | 6.53406 | 6.88567 | 6.52899 | 6.40037 | 6.58954 | 5.65803 | 5.4782 | 5.07888 |
| PSMB3 | P49720 | 6.06771 | 6.0288 | 5.95867 | 5.99863 | 6.027 | 6.07863 | 6.38876 | 6.45027 | 6.42774 | 6.46894 | 6.23854 | 6.12454 | 6.35877 | 6.40754 | 6.43117 | 6.54429 |
| RNF114 | Q9Y508 | 6.12885 | 6.07952 | 6.78258 | 6.29647 | 6.64408 | 7.26539 | 6.06364 | 7.0791 | 6.29535 | 6.45216 | 6.86244 | 6.74336 | 5.57516 | 5.51023 | 5.01948 | 5.20218 |
| SKA3 | Q8IX90 | 6.78798 | 6.06933 | 7.13462 | 5.7892 | 6.49669 | 6.3662 | 6.56953 | 6.9977 | 6.35886 | 6.55514 | 7.04276 | 7.26209 | 5.78031 | 5.1516 | 5.0797 | 5.45829 |
| GGT7 | Q9UJ14 | 6.69439 | 6.32384 | 5.59211 | 6.23476 | 6.40032 | 5.94208 | 6.18899 | 6.24824 | 6.07923 | 6.35987 | 6.19653 | 5.66022 | 6.66865 | 6.49807 | 6.44211 | 6.47059 |
| TECR | Q9NZ01 | 5.99044 | 6.71021 | 5.48377 | 6.31526 | 6.07272 | 6.28754 | 6.18022 | 6.49235 | 6.02714 | 6.53783 | 5.84677 | 6.08685 | 6.323 | 6.8746 | 6.27585 | 6.49546 |

| | | | | | | | | | | | | | | | | |
|---|---|---|---|---|---|---|---|---|---|---|---|---|---|---|---|---|
| PHF20L1 | A0A0A0MQS0 | 6.35226 | 6.60892 | 5.9359 | 6.83174 | 5.82671 | 5.93802 | 6.05072 | 6.10663 | 5.8754 | 6.1387 | 5.94436 | 6.3027 | 6.08243 | 6.71308 | 6.28975 | 7.00267 |
| AMFR | Q9UKV5 | 7.08162 | 7.31549 | 6.56869 | 7.23937 | 6.18807 | 5.97481 | 6.72388 | 6.03461 | 6.43582 | 6.64345 | 6.3586 | 6.22823 | 5.15203 | 5.30812 | 5.13261 | 5.61459 |
| PPP1R13B | Q96KQ4 | 6.22134 | 6.49714 | 6.48937 | 6.42918 | 6.31635 | 6.08449 | 6.18572 | 6.54448 | 6.1884 | 6.02542 | 6.17587 | 5.77236 | 6.63306 | 6.19439 | 6.12629 | 6.11613 |
| DDX11 | Q96FC9 | 6.83654 | 6.68338 | 6.90439 | 6.1782 | 6.51367 | 6.76151 | 5.94186 | 6.50183 | 6.51619 | 6.49292 | 6.78201 | 6.67919 | 5.19289 | 5.38451 | 5.34342 | 5.28749 |
| DGCR8 | Q8WYQ5 | 7.09893 | 7.01975 | 5.91897 | 6.8158 | 6.22376 | 6.18739 | 5.96921 | 6.07344 | 6.41794 | 6.53392 | 6.39771 | 6.3183 | 5.83406 | 5.71943 | 5.8207 | 5.65069 |
| SHKBP1 | Q8TBC3 | 5.80763 | 6.18462 | 6.03293 | 6.09374 | 5.94083 | 6.36678 | 6.68711 | 6.14257 | 6.0298 | 6.17063 | 6.30959 | 6.06859 | 6.49887 | 6.40435 | 6.69134 | 6.57063 |
| LSM7 | Q9UK45 | 5.31435 | 4.57555 | 5.26802 | 4.92838 | 6.00558 | 4.9689 | 5.53443 | 7.11838 | 5.05424 | 5.52146 | 5.30286 | 6.47792 | 6.01239 | 9.171 | 8.68816 | 10.0584 |
| BCAR3 | O75815 | 5.94124 | 6.09615 | 7.04829 | 6.18889 | 6.23177 | 6.78241 | 6.4675 | 6.55269 | 6.38257 | 6.1827 | 6.76283 | 6.70097 | 5.79434 | 5.38502 | 5.60515 | 5.87749 |
| EIF4E | P06730-2 | 5.95087 | 5.90959 | 6.7664 | 5.69241 | 6.08614 | 6.37741 | 6.01359 | 7.33759 | 5.8717 | 7.12589 | 7.65599 | 7.30137 | 5.49897 | 5.81424 | 5.37126 | 5.22658 |
| RNPS1 | Q15287 | 6.35614 | 6.2024 | 5.92127 | 6.78772 | 5.94401 | 5.71735 | 6.36156 | 5.77677 | 6.30654 | 5.84486 | 6.06551 | 5.96064 | 6.88961 | 6.19981 | 6.7815 | 6.88431 |
| KIF3B | O15066 | 6.37847 | 6.17732 | 6.60201 | 6.41778 | 5.92405 | 6.67783 | 6.23775 | 6.55602 | 6.5248 | 6.73592 | 6.22517 | 6.7866 | 5.47473 | 6.11886 | 5.45298 | 5.70971 |
| POLB | P06746 | 5.19479 | 5.82418 | 5.72679 | 5.00643 | 5.96225 | 6.36661 | 5.93125 | 6.21576 | 6.40497 | 6.20942 | 6.20516 | 6.29653 | 7.34798 | 6.89507 | 7.27201 | 7.1408 |
| NUP62 | P37198 | 6.27658 | 4.97205 | 4.66102 | 4.55073 | 5.01168 | 5.14274 | 5.68133 | 6.31436 | 4.70849 | 5.55164 | 4.81193 | 6.78404 | 6.89634 | 9.62033 | 9.21733 | 9.7994 |
| TMEM131 | Q92545 | 6.7536 | 6.83417 | 6.37307 | 6.80991 | 6.48085 | 5.75848 | 6.67988 | 5.79119 | 6.01457 | 6.57377 | 6.04328 | 6.22326 | 6.04827 | 5.65275 | 6.01893 | 5.94404 |
| YIPF3 | Q9GZM5 | 6.38835 | 6.05009 | 6.0294 | 6.90074 | 6.21614 | 5.71627 | 6.42496 | 5.8247 | 6.43174 | 6.11854 | 6.23862 | 6.41657 | 5.87779 | 6.22947 | 6.57811 | 6.55852 |
| PDCD2 | Q16342 | 6.42107 | 6.46772 | 7.06907 | 6.38123 | 6.17829 | 6.7714 | 5.93065 | 6.52887 | 6.79226 | 6.381 | 7.10531 | 7.41642 | 5.4117 | 5.00404 | 4.96843 | 5.17257 |
| DAAM1 | Q9Y4D1 | 6.35382 | 7.18548 | 6.3315 | 6.52654 | 5.84234 | 5.66855 | 6.2011 | 6.26604 | 6.44124 | 6.01184 | 6.49862 | 6.18401 | 5.98221 | 5.84565 | 6.40365 | 6.25743 |
| ABL1 | P00519-2 | 5.95294 | 6.12397 | 6.16633 | 5.73888 | 5.92768 | 6.75444 | 6.63092 | 6.53591 | 5.9354 | 6.29113 | 6.72109 | 6.31208 | 6.15751 | 6.63321 | 5.79156 | 6.32695 |
| ABI1 | Q8IZP0 | 6.42223 | 5.9023 | 6.4388 | 5.88315 | 6.30969 | 6.97512 | 6.09145 | 6.25699 | 6.23877 | 6.58892 | 6.74372 | 6.44932 | 6.52711 | 6.08723 | 5.54147 | 5.54375 |
| RHBDF2 | Q6PJF5 | 6.1838 | 6.3669 | 6.14777 | 6.63307 | 6.37542 | 5.8551 | 6.20054 | 6.26865 | 6.47656 | 6.15708 | 6.17783 | 6.15693 | 6.35701 | 6.06793 | 6.29727 | 6.27813 |
| UGT8 | Q16880 | 6.67135 | 6.55334 | 6.04065 | 6.99498 | 6.38634 | 5.94743 | 6.08352 | 5.78095 | 6.1838 | 5.92195 | 6.32406 | 6.05428 | 6.39339 | 5.89917 | 6.70015 | 6.06461 |
| PIGO | Q8TEQ8 | 6.16222 | 6.60869 | 6.11189 | 6.54018 | 6.23243 | 6.06067 | 6.32502 | 5.93738 | 5.99327 | 6.25456 | 6.15084 | 6.00917 | 6.4387 | 6.34478 | 6.44356 | 6.38663 |
| RPS6KA5 | O75582 | 5.56726 | 5.50435 | 5.81705 | 5.36155 | 5.78844 | 5.97898 | 5.88716 | 6.1745 | 5.93119 | 6.13382 | 6.38757 | 5.98142 | 6.74972 | 7.75726 | 7.68917 | 7.29056 |
| CAMK4 | Q16566 | 6.32273 | 6.29445 | 6.9066 | 5.68527 | 5.95448 | 6.68884 | 6.03822 | 6.81361 | 6.29092 | 6.57625 | 6.62898 | 6.31617 | 5.74304 | 6.22272 | 5.85466 | 5.66306 |
| ECHDC1 | Q9NTX5 | 6.09618 | 5.91875 | 6.45659 | 6.04614 | 6.42754 | 6.51674 | 6.3481 | 6.45439 | 6.1082 | 6.454 | 6.31383 | 6.26261 | 6.06638 | 6.33228 | 6.19835 | 5.9999 |
| POLR3D | P05423 | 6.05267 | 5.74965 | 6.47392 | 6.0178 | 6.16305 | 5.88391 | 6.41495 | 5.85485 | 6.36634 | 6.17255 | 6.62445 | 6.20804 | 6.49414 | 6.22689 | 6.50099 | 6.79579 |
| ADAM15 | Q13444 | 7.47664 | 6.7532 | 6.44542 | 6.90369 | 6.37046 | 6.40969 | 6.03408 | 5.97064 | 6.45718 | 6.72321 | 6.33853 | 6.24651 | 5.47138 | 5.46397 | 5.62592 | 5.31347 |
| PIP4K2C | Q8TBX8 | 6.29917 | 6.19983 | 6.49393 | 5.59658 | 6.36365 | 6.76244 | 6.01426 | 6.1727 | 6.21883 | 6.11902 | 6.28841 | 6.33184 | 6.68274 | 6.35685 | 6.15081 | 5.94894 |
| PEX3 | P56589 | 6.65568 | 6.28174 | 6.35302 | 6.70393 | 6.34245 | 5.73778 | 7.06719 | 6.09845 | 6.73483 | 6.23368 | 6.18207 | 6.05782 | 6.05932 | 5.44975 | 6.15409 | 5.88819 |
| NUDT1 | P36639 | 5.94951 | 5.72244 | 6.75363 | 5.71748 | 6.39488 | 6.37761 | 6.08011 | 7.19491 | 6.55525 | 6.90263 | 6.80955 | 7.20085 | 5.38674 | 5.74348 | 5.51873 | 5.69219 |
| CLK1 | P49759-3 | 6.49969 | 6.53734 | 6.07601 | 6.48841 | 5.85734 | 5.42768 | 5.7807 | 5.75469 | 5.82762 | 6.10143 | 5.71017 | 5.92073 | 6.74721 | 6.68715 | 7.19069 | 7.39312 |
| FAM135A | Q9P2D6 | 6.09043 | 5.88396 | 6.95848 | 6.57407 | 6.06043 | 6.08661 | 6.70753 | 6.30589 | 6.42659 | 6.4702 | 6.83919 | 6.67404 | 5.57163 | 5.81524 | 5.54288 | 5.99281 |
| TGFBR3 | Q03167 | 6.55323 | 6.28297 | 6.1032 | 6.55971 | 6.65652 | 6.43009 | 6.25341 | 5.90056 | 6.25157 | 6.17249 | 6.55951 | 6.35179 | 6.05195 | 5.6852 | 5.91731 | 6.27048 |
| TOM1L1 | O75674 | 6.32763 | 6.40717 | 6.73868 | 5.97393 | 6.14321 | 6.67538 | 6.10087 | 6.87615 | 6.31731 | 6.43573 | 6.7446 | 6.53981 | 5.70107 | 5.86931 | 5.53026 | 5.61891 |
| SLC12A6 | Q9UHW9 | 6.11617 | 6.31113 | 6.181 | 6.32819 | 5.99003 | 6.28534 | 6.23252 | 6.33506 | 6.05595 | 6.62755 | 6.29266 | 6.12789 | 6.21894 | 6.27324 | 6.01255 | 6.6118 |
| EVI5 | O60447-2 | 6.50745 | 6.59849 | 6.79748 | 6.44775 | 6.27381 | 6.67752 | 6.2735 | 6.65874 | 6.68582 | 6.57841 | 6.70127 | 6.73604 | 5.3661 | 5.35133 | 5.0542 | 5.29207 |
| TTC27 | Q6P3X3 | 6.53423 | 6.31364 | 6.8752 | 5.88486 | 6.3054 | 6.56517 | 6.06153 | 6.61297 | 6.74008 | 6.25266 | 6.5714 | 6.18646 | 5.66945 | 5.94133 | 5.81813 | 5.6675 |
| LPCAT3 | Q6P1A2 | 6.24208 | 6.18906 | 5.8648 | 6.38014 | 6.32986 | 5.83519 | 6.33027 | 5.93546 | 5.97068 | 6.15112 | 5.68463 | 5.54659 | 6.95866 | 6.61377 | 7.10013 | 6.86756 |
| HRAS | P01112 | 6.40655 | 6.48846 | 6.15827 | 6.70434 | 6.1501 | 5.85125 | 6.06286 | 6.0312 | 6.10912 | 5.92072 | 6.3889 | 6.37012 | 6.19676 | 6.18003 | 6.45962 | 6.5217 |
| NDUFB7 | P17568 | 6.54673 | 6.14134 | 5.9872 | 6.44424 | 6.4224 | 6.16291 | 6.68484 | 6.07711 | 6.29544 | 5.88179 | 6.1321 | 5.78828 | 6.57448 | 6.05987 | 6.48066 | 6.32061 |
| ARNT | P27540 | 5.80438 | 6.23 | 6.55388 | 5.52246 | 5.25654 | 6.01347 | 5.64147 | 6.25865 | 6.18583 | 6.08112 | 6.92124 | 6.2595 | 6.15311 | 7.09468 | 7.14514 | 6.75853 |
| SEC14L1 | Q92503-2 | 7.55934 | 8.09783 | 8.56859 | 7.97595 | 5.76262 | 6.9856 | 6.68868 | 6.98444 | 7.25105 | 7.43982 | 8.09113 | 7.92286 | 2.43916 | 2.99978 | 2.35665 | 2.87651 |
| RPL36A | P83881 | 5.83275 | 5.8756 | 6.23069 | 5.83002 | 6.23685 | 6.87559 | 6.51899 | 6.65618 | 6.80104 | 6.07674 | 6.85524 | 6.0648 | 5.82357 | 6.7981 | 5.93925 | 5.58458 |
| ENPP1 | P22413 | 6.34753 | 6.51734 | 5.91108 | 6.5583 | 6.04853 | 6.226 | 5.75411 | 6.16804 | 6.28357 | 6.25877 | 6.3767 | 6.24741 | 6.66935 | 6.33537 | 6.01817 | 6.28106 |
| ATP6AP1 | Q15904 | 6.28302 | 6.35354 | 6.16871 | 6.85365 | 6.19048 | 5.98935 | 6.45667 | 6.48903 | 6.31515 | 5.89823 | 6.48666 | 5.97868 | 6.29552 | 5.82759 | 6.26983 | 6.1439 |
| TOX4 | O94842 | 6.96641 | 6.67508 | 6.08791 | 6.84576 | 6.23399 | 5.80265 | 6.37783 | 6.21107 | 6.1792 | 6.20291 | 6.49032 | 6.06414 | 6.32243 | 5.70292 | 5.91928 | 5.91813 |
| DHTKD1 | Q96HY7 | 5.99192 | 5.82726 | 5.66389 | 6.44369 | 6.07222 | 6.38933 | 5.98295 | 6.05705 | 6.17752 | 5.99602 | 6.25476 | 6.35171 | 6.92226 | 6.65988 | 6.42123 | 6.78829 |
| SOAT1 | P35610 | 6.03388 | 6.19991 | 5.95006 | 6.59055 | 6.02283 | 5.75297 | 6.45186 | 5.91462 | 6.18963 | 6.23728 | 6.54454 | 6.08986 | 6.58571 | 6.52538 | 6.56053 | 6.35041 |
| TRAPPC3 | O43617 | 6.12941 | 5.34424 | 6.97772 | 4.92306 | 6.36036 | 6.04286 | 6.1016 | 6.31075 | 6.13751 | 6.47687 | 6.81452 | 6.68398 | 5.72069 | 6.60818 | 6.64551 | 6.72274 |
| DNAJC19 | Q96DA6 | 6.32892 | 6.2237 | 5.68327 | 6.52201 | 6.07663 | 5.91768 | 6.2321 | 6.02405 | 6.12125 | 6.54835 | 5.91303 | 6.25103 | 6.78685 | 6.44353 | 6.37295 | 6.55463 |
| SP1 | P08047 | 5.77896 | 5.98152 | 6.02036 | 6.20986 | 6.03031 | 6.1791 | 5.91879 | 6.03915 | 5.96645 | 6.31779 | 6.51485 | 6.29528 | 6.85003 | 6.74149 | 6.62918 | 6.52687 |
| TRIM16 | O95361 | 6.37043 | 6.39959 | 6.84389 | 5.70321 | 6.53648 | 6.81706 | 6.11885 | 6.47477 | 6.29829 | 6.4481 | 6.74717 | 6.23141 | 5.68369 | 5.98841 | 5.87251 | 5.46612 |
| BCL9L | Q86UU0 | 6.60733 | 6.24091 | 6.80511 | 5.8118 | 6.90154 | 6.43891 | 5.55752 | 6.59136 | 5.94823 | 6.88253 | 6.1089 | 6.25535 | 6.14715 | 6.11684 | 5.7839 | 5.8026 |
| SLC33A1 | O00400 | 6.01283 | 6.54135 | 6.20109 | 6.7176 | 6.1542 | 6.13516 | 6.19036 | 5.83268 | 5.8554 | 6.14499 | 5.93518 | 5.75833 | 6.48255 | 6.49403 | 6.85275 | 6.6915 |
| NDUFB11 | Q9NX14-2 | 6.3515 | 6.23971 | 5.5309 | 6.21514 | 6.37929 | 5.70223 | 6.4537 | 6.35787 | 5.72439 | 5.86868 | 5.83969 | 6.12814 | 6.42901 | 7.11461 | 6.56145 | 7.10368 |
| RBM8A | Q9Y5S9 | 6.25003 | 5.65289 | 5.99136 | 6.43748 | 6.02813 | 5.52426 | 6.54355 | 5.65341 | 6.24125 | 6.21637 | 6.18182 | 6.02009 | 6.72202 | 6.86268 | 6.83581 | 6.83885 |
| TNFRSF1A | P19438 | 6.04074 | 6.3389 | 5.99371 | 6.44233 | 5.85685 | 6.10481 | 6.49295 | 5.90104 | 6.26134 | 6.64896 | 6.41292 | 6.63181 | 6.50239 | 6.18598 | 5.94201 | 6.24326 |
| MRPS25 | P82663 | 6.36702 | 6.15536 | 6.07278 | 6.79576 | 6.34373 | 6.0871 | 6.32396 | 6.04934 | 6.30573 | 6.10878 | 6.09176 | 6.24788 | 6.88908 | 5.89648 | 6.19807 | 6.06716 |
| GPATCH4 | Q5T3I0-3 | 6.10179 | 6.36769 | 6.21367 | 7.00574 | 5.95047 | 5.83805 | 6.0465 | 5.97536 | 5.91852 | 5.96184 | 6.09645 | 6.20179 | 6.27151 | 6.6941 | 6.37437 | 6.98214 |
| ADD3 | Q9UEY8 | 5.88677 | 5.70656 | 6.125 | 5.8532 | 5.94652 | 6.35507 | 5.73995 | 5.79021 | 6.19335 | 6.33026 | 6.02763 | 5.99992 | 7.08737 | 7.13113 | 6.99608 | 7.32797 |
| MVK | Q03426 | 6.06218 | 5.8298 | 6.73769 | 5.81277 | 5.79551 | 6.46416 | 6.01921 | 6.30161 | 6.38173 | 6.60741 | 6.73259 | 6.87042 | 5.97429 | 6.32957 | 6.08842 | 5.99266 |
| SH2D4A | Q9H788 | 6.09646 | 6.13078 | 6.95065 | 6.18806 | 6.16734 | 6.50502 | 6.60771 | 6.32237 | 6.43256 | 6.15547 | 6.4549 | 6.53519 | 5.77812 | 5.95241 | 6.00931 | 5.71366 |
| SNRNP48 | Q6IEG0 | 6.22372 | 6.25939 | 6.20344 | 6.70985 | 5.8126 | 5.76761 | 6.41914 | 5.94811 | 6.37182 | 6.51535 | 6.3007 | 6.00972 | 5.94235 | 6.10091 | 6.4367 | 6.67858 |
| SP3 | Q02447 | 6.43005 | 6.19641 | 6.46442 | 6.69968 | 6.22159 | 6.51178 | 6.33293 | 5.94268 | 5.88677 | 6.54511 | 6.18667 | 5.96814 | 6.49992 | 6.13247 | 6.04692 | 5.93447 |
| PTPRA | P18433 | 6.88707 | 7.14158 | 6.22644 | 6.56016 | 6.42384 | 6.54174 | 6.74606 | 6.49032 | 5.86291 | 6.32708 | 5.75506 | 5.52382 | 5.97428 | 6.14011 | 5.79503 | 5.6045 |
| DHCR24 | Q15392 | 6.22392 | 6.20162 | 5.95591 | 6.74728 | 6.23356 | 5.97685 | 6.442 | 5.74697 | 6.16113 | 6.20597 | 5.95519 | 5.85397 | 6.4042 | 6.16253 | 6.89112 | 6.83778 |
| RNF14 | Q9UBS8 | 6.35679 | 6.05604 | 6.78338 | 5.96408 | 6.57937 | 6.69344 | 5.80353 | 6.66608 | 6.40357 | 6.37879 | 6.69934 | 6.82908 | 5.75059 | 6.11031 | 5.47516 | 5.45044 |
| SENP1 | Q9P0U3 | 6.23828 | 6.36704 | 5.69528 | 6.80656 | 6.29543 | 6.13032 | 6.57097 | 5.95823 | 5.6811 | 5.99193 | 6.13431 | 5.85039 | 6.83459 | 6.2094 | 6.75293 | 6.48325 |
| CDK19 | Q9BWU1 | 6.07774 | 6.26346 | 6.21022 | 6.12293 | 6.14926 | 5.99623 | 6.40358 | 5.8132 | 6.22679 | 6.0066 | 5.97046 | 5.99426 | 6.71778 | 6.31961 | 6.77556 | 6.95232 |
| PEX14 | O75381 | 6.8703 | 6.28096 | 5.83042 | 6.3796 | 6.91155 | 6.02686 | 6.11976 | 6.36653 | 6.3049 | 5.80098 | 6.03115 | 5.84396 | 6.94123 | 6.02249 | 6.20592 | 6.06336 |
| TNRC6A | Q8NDV7 | 6.34803 | 6.32547 | 7.1005 | 6.49077 | 6.21104 | 6.53904 | 6.66129 | 6.23537 | 6.46376 | 6.87197 | 6.48613 | 6.9109 | 5.25018 | 5.23679 | 5.28986 | 5.5789 |
| NIPSNAP3A | Q9UFN0 | 6.08126 | 6.35991 | 5.80726 | 6.15362 | 5.93017 | 5.63259 | 9.31044 | 6.04379 | 5.4298 | 6.08982 | 5.52061 | 5.32785 | 6.27837 | 6.06258 | 6.93169 | 7.04024 |
| ERCC5 | P28715 | 5.34953 | 6.457 | 5.83186 | 6.24343 | 6.0815 | 6.03438 | 6.38713 | 5.92071 | 6.23281 | 6.14836 | 6.44192 | 6.31227 | 6.53805 | 6.64087 | 6.52733 | 6.85285 |
| SLX4 | Q8IY92 | 6.38427 | 6.43284 | 6.27817 | 7.05828 | 6.22102 | 6.51816 | 6.37892 | 5.56072 | 6.63144 | 6.63271 | 5.53644 | 6.06229 | 5.95697 | 6.03047 | 6.03102 | 6.28629 |
| GRK2 | P25098 | 6.22899 | 6.45541 | 7.37836 | 6.22265 | 6.36699 | 6.36555 | 6.29219 | 6.34076 | 6.4643 | 6.37946 | 6.67461 | 6.7915 | 5.56471 | 5.68241 | 5.22804 | 5.56408 |
| CCDC85C | A6NKD9 | 6.38742 | 5.83827 | 6.46481 | 5.99976 | 6.433 | 6.28573 | 5.7332 | 6.69787 | 6.76089 | 6.27996 | 6.82829 | 7.27668 | 6.19967 | 5.97013 | 5.33412 | 5.51019 |
| ATXN1L | P0C7T5 | 6.28894 | 5.95254 | 6.35326 | 6.44947 | 6.48136 | 6.18851 | 6.10779 | 6.18221 | 6.2736 | 6.32716 | 6.37533 | 6.71801 | 6.35914 | 5.83241 | 5.80143 | 6.30883 |
| ERGIC2 | Q96RQ1 | 6.45745 | 6.20625 | 5.93492 | 6.31506 | 6.52454 | 5.95358 | 6.46802 | 5.9976 | 6.27139 | 5.8127 | 6.10256 | 5.60492 | 6.64078 | 6.35406 | 7.1034 | 6.25277 |
| GOLM2 | Q6P4E1 | 6.44447 | 6.22992 | 5.73574 | 6.28787 | 6.42691 | 6.0029 | 6.43765 | 6.17013 | 6.75853 | 6.24957 | 6.06138 | 6.2684 | 6.38058 | 6.06844 | 6.07576 | 6.40176 |
| POLG2 | Q9UHN1 | 6.43284 | 6.17978 | 5.86421 | 6.6805 | 6.28404 | 6.06232 | 5.97392 | 5.7357 | 6.15146 | 6.12073 | 6.10606 | 6.27816 | 6.76531 | 6.05968 | 6.76178 | 6.5435 |
| GDAP1 | Q8TB36 | 6.25186 | 6.65151 | 5.41566 | 6.36068 | 6.09149 | 5.97793 | 6.19832 | 6.00899 | 6.20058 | 6.08184 | 5.9593 | 6.05111 | 6.75489 | 6.74874 | 6.5554 | 6.6917 |
| PIBF1 | Q8WXW3 | 6.33955 | 6.46665 | 6.91742 | 6.33343 | 6.59393 | 6.42464 | 6.60506 | 6.43934 | 6.48222 | 6.49851 | 6.63548 | 6.60876 | 5.71116 | 5.03838 | 5.64178 | 5.26366 |
| CASP8 | Q14790-4 | 6.38316 | 6.12983 | 6.96263 | 6.07475 | 6.44211 | 6.78373 | 6.2471 | 6.7321 | 6.41577 | 6.58065 | 6.69156 | 6.43692 | 5.55883 | 5.86571 | 5.30867 | 5.38647 |
| GMDS | O60547 | 6.29686 | 6.40832 | 6.9747 | 5.83988 | 6.06528 | 6.39782 | 6.04878 | 6.48135 | 6.34735 | 6.71596 | 6.54523 | 6.5052 | 5.84333 | 5.55267 | 6.12595 | 5.85132 |
| SUPT4H1 | P63272 | 5.14399 | 5.91513 | 5.53618 | 5.27189 | 6.35991 | 5.87737 | 6.07722 | 6.842 | 6.30101 | 5.93972 | 6.79328 | 5.99186 | 7.31642 | 7.01738 | 6.78797 | 6.82867 |
| TK1 | P04183 | 6.26456 | 6.16043 | 6.77582 | 6.1734 | 5.96135 | 6.98548 | 5.93474 | 6.6426 | 6.55921 | 6.23389 | 6.61029 | 6.897 | 5.84413 | 5.57052 | 5.68316 | 5.70344 |
| GALNT10 | Q86SR1 | 6.35889 | 6.40795 | 6.29076 | 6.79068 | 6.26526 | 6.05467 | 6.50919 | 6.01297 | 6.45863 | 6.25981 | 6.53772 | 6.09679 | 6.23473 | 6.16316 | 6.14276 | 5.41604 |
| HPCAL1 | P37235 | 6.73594 | 5.2865 | 5.91973 | 4.866 | 5.54003 | 5.51358 | 5.21715 | 6.30063 | 5.40279 | 7.02522 | 5.78539 | 6.79867 | 5.79337 | 8.09365 | 7.45825 | 8.26309 |
| NDUFS8 | O00217 | 6.76666 | 4.91886 | 5.40934 | 4.83118 | 5.74222 | 4.61562 | 5.86375 | 6.68572 | 4.90836 | 6.15539 | 5.47986 | 6.37922 | 6.85888 | 8.39772 | 8.32677 | 8.66043 |
| MGAT5 | Q09328 | 6.25924 | 6.23988 | 5.7501 | 6.71517 | 6.41232 | 6.17249 | 6.23837 | 6.1702 | 6.02779 | 6.26605 | 5.86603 | 6.13858 | 6.58492 | 6.4313 | 6.4043 | 6.32324 |
| COQ6 | Q9Y2Z9 | 6.32314 | 6.40082 | 6.00662 | 6.52846 | 6.17298 | 6.09357 | 6.2592 | 5.89976 | 6.23181 | 6.23393 | 6.23387 | 6.20111 | 6.49738 | 6.42856 | 6.33541 | 6.15337 |
| GM2A | P17900 | 6.68823 | 5.69406 | 5.30033 | 5.06416 | 5.92933 | 5.43653 | 5.44665 | 6.55801 | 5.15405 | 6.39084 | 5.09157 | 6.25147 | 6.34579 | 8.82829 | 7.52989 | 8.29081 |
| DPH1 | Q9BZG8 | 6.40323 | 6.08681 | 6.71641 | 5.78109 | 6.22996 | 6.4482 | 6.33155 | 6.42534 | 6.65283 | 6.21351 | 6.60268 | 6.98434 | 5.77641 | 5.82129 | 5.82099 | 5.70535 |
| SNX7 | Q9UNH6-3 | 6.35694 | 6.03306 | 6.65761 | 6.35179 | 6.30689 | 6.73807 | 6.29054 | 6.34315 | 6.09973 | 6.29734 | 6.70432 | 6.34612 | 5.65707 | 6.26565 | 5.75304 | 5.79869 |
| SEMA4C | Q9C0C4 | 6.34123 | 6.72638 | 6.75636 | 6.80432 | 5.8986 | 5.99298 | 6.26272 | 6.05858 | 6.00865 | 6.3168 | 6.31298 | 6.40282 | 5.87305 | 5.95634 | 6.31626 | 5.97193 |
| PEX11B | O96011 | 6.4931 | 6.0984 | 5.44604 | 6.25041 | 6.16933 | 5.83472 | 6.40679 | 6.28238 | 5.974 | 6.47038 | 6.08891 | 5.89182 | 6.84765 | 6.49518 | 6.63531 | 6.61559 |
| NUP37 | Q8NFH4 | 5.93829 | 6.05435 | 5.89752 | 6.24563 | 6.44558 | 6.24281 | 6.36489 | 6.46597 | 6.71114 | 6.57847 | 6.45124 | 6.3609 | 6.21787 | 5.60945 | 6.25349 | 6.16242 |
| ERGIC1 | Q969X5 | 6.51025 | 6.38559 | 5.81934 | 6.21229 | 6.14729 | 5.92703 | 6.29906 | 6.03722 | 6.2847 | 6.14016 | 5.91521 | 5.67785 | 6.68723 | 6.48813 | 6.7992 | 6.66944 |
| PIN4 | Q9Y237-2 | 6.14275 | 5.97724 | 6.43883 | 6.17093 | 5.94133 | 6.67205 | 6.51307 | 6.52846 | 6.57495 | 6.39244 | 6.23491 | 6.84326 | 5.63283 | 6.13721 | 5.82726 | 5.97248 |
| PTAR1 | Q7Z6K3 | 5.96191 | 5.99869 | 6.43419 | 5.76351 | 6.26961 | 6.53537 | 5.94934 | 6.60492 | 6.24234 | 6.65331 | 6.38562 | 6.3989 | 6.06116 | 6.37263 | 6.32628 | 6.04223 |
| ZWILCH | Q9H900 | 6.47075 | 6.37455 | 6.89297 | 5.97892 | 6.47109 | 6.50173 | 6.24633 | 6.69499 | 6.43743 | 6.56675 | 6.75858 | 6.83768 | 5.41746 | 5.51538 | 5.45442 | 5.38099 |
| STAMBPL1 | Q96FJ0-2 | 5.68243 | 6.21077 | 6.71189 | 6.49969 | 5.86313 | 6.77739 | 6.15331 | 6.28132 | 6.0894 | 6.20756 | 6.57609 | 6.59776 | 5.83966 | 6.13117 | 5.98819 | 6.39024 |
| BECN1 | Q14457 | 6.57416 | 6.53803 | 6.61587 | 6.31474 | 6.35327 | 6.40597 | 6.27578 | 6.55488 | 6.38537 | 6.14881 | 6.45621 | 6.61966 | 5.86936 | 5.73813 | 5.60824 | 5.54153 |
| HACD3 | Q9P035 | 6.08701 | 6.34401 | 5.97219 | 6.52336 | 6.11391 | 5.97694 | 6.28217 | 6.15027 | 5.91399 | 6.18071 | 5.61009 | 5.75871 | 6.76961 | 6.45468 | 7.09291 | 6.76945 |
| MMP14 | P50281 | 5.69784 | 5.8746 | 5.3362 | 5.7719 | 5.72057 | 5.46602 | 5.88904 | 6.19017 | 5.37101 | 5.96001 | 5.71711 | 5.85454 | 6.5039 | 8.38132 | 7.72197 | 8.5438 |

| | | | | | | | | | | | | | | | | | |
|---|---|---|---|---|---|---|---|---|---|---|---|---|---|---|---|---|---|
| NDUFA4 | O00483 | 6.1616 | 5.34373 | 5.16154 | 5.63532 | 5.39043 | 4.72286 | 5.4824 | 6.51244 | 4.76485 | 5.64418 | 5.01601 | 5.82854 | 5.75401 | 10.0897 | 8.44278 | 10.0496 |
| SOGA3 | Q5TF21 | 6.05049 | 6.58813 | 6.31717 | 6.1611 | 6.24434 | 6.27097 | 5.86735 | 6.36262 | 6.11753 | 6.20993 | 6.35106 | 6.18841 | 6.05027 | 6.22898 | 6.41671 | 6.57494 |
| TUBGCP4 | Q9UGJ1 | 6.33249 | 6.18385 | 6.87324 | 6.19644 | 6.2177 | 6.3914 | 6.49069 | 6.6915 | 6.60673 | 6.51431 | 6.62028 | 6.78654 | 5.54324 | 5.40521 | 5.67257 | 5.47381 |
| TMEM38B | Q9NVV0 | 6.28155 | 6.05143 | 6.07643 | 6.64696 | 6.06861 | 5.57859 | 6.36361 | 5.77462 | 6.29 | 5.83503 | 6.2836 | 6.12419 | 6.44101 | 6.28564 | 6.88151 | 7.01722 |
| ADGRG6 | Q86SQ4 | 6.13027 | 6.18426 | 6.17464 | 6.89221 | 6.0668 | 6.31934 | 6.47822 | 6.29256 | 5.73606 | 5.98486 | 6.11293 | 6.48482 | 6.53243 | 5.84882 | 6.17525 | 6.58653 |
| ADI1 | Q9BV57 | 5.12706 | 5.63483 | 6.32811 | 4.62695 | 6.56189 | 7.06412 | 6.00968 | 7.90117 | 6.35666 | 7.33464 | 6.83341 | 6.93064 | 5.00954 | 6.10373 | 6.06644 | 6.11112 |
| USP1 | O94782 | 6.35301 | 6.79921 | 6.84101 | 6.44736 | 5.99844 | 6.27908 | 5.90061 | 6.46888 | 6.59567 | 7.14369 | 6.88216 | 6.92347 | 5.10754 | 5.50503 | 5.34676 | 5.40807 |
| POLR2G | P62487 | 6.42503 | 6.33548 | 6.38266 | 6.60732 | 6.63718 | 6.38021 | 6.35229 | 6.5809 | 6.59734 | 6.33486 | 6.5453 | 6.47992 | 5.87672 | 5.61959 | 5.18895 | 5.65626 |
| ADNP2 | Q6IQ32 | 6.60751 | 7.23652 | 6.1566 | 6.96144 | 6.01779 | 6.1198 | 6.14725 | 5.85542 | 6.2722 | 6.21382 | 6.25804 | 6.25823 | 6.13545 | 5.62032 | 5.90375 | 6.23588 |
| MRPL23 | Q16540 | 6.09207 | 6.2383 | 5.46651 | 6.58324 | 6.48714 | 5.96704 | 6.59362 | 6.48458 | 6.4161 | 6.45051 | 5.91608 | 6.10384 | 6.49203 | 5.96038 | 6.44926 | 6.2993 |
| DERL1 | Q9BUN8 | 6.12414 | 6.59446 | 6.08313 | 6.41885 | 5.88176 | 6.14044 | 6.01898 | 6.2019 | 6.04394 | 6.22768 | 6.08371 | 6.37317 | 6.5436 | 6.40051 | 6.41008 | 6.45365 |
| CNOT6 | Q9ULM6 | 6.2958 | 6.67104 | 6.62956 | 6.20092 | 6.34004 | 6.49492 | 6.40471 | 6.51481 | 6.46148 | 6.52336 | 6.73248 | 6.71913 | 5.46708 | 5.60439 | 5.60977 | 5.33048 |
| DCTN6 | O00399 | 6.21195 | 6.2067 | 6.72301 | 5.94133 | 6.70372 | 6.57937 | 6.02769 | 6.78676 | 6.18847 | 6.69642 | 6.5831 | 6.57004 | 5.84374 | 6.00413 | 5.0768 | 5.85678 |
| GSE1 | Q14687 | 6.60013 | 6.48335 | 6.33762 | 6.80688 | 6.44867 | 6.2799 | 6.37295 | 6.14437 | 6.28015 | 6.35216 | 6.51709 | 6.34186 | 5.99821 | 5.7286 | 5.62837 | 5.6797 |
| PMM2 | O15305 | 6.44676 | 6.07827 | 7.04694 | 5.64549 | 6.7689 | 6.91585 | 6.32462 | 6.53248 | 6.46882 | 6.34831 | 6.70072 | 6.43512 | 5.7507 | 5.76684 | 5.62408 | 5.14611 |
| TMEM63B | Q5T3F8 | 6.30882 | 6.72878 | 6.79545 | 6.51793 | 6.46535 | 5.99257 | 6.56427 | 6.05739 | 6.05394 | 6.22703 | 6.0776 | 5.90993 | 6.198 | 5.95528 | 6.01291 | 6.13474 |
| TBCK | Q8TEA7 | 6.1969 | 6.25429 | 6.73944 | 6.26857 | 5.97806 | 6.615 | 6.08913 | 5.99175 | 6.29124 | 6.46228 | 6.64703 | 6.94651 | 5.74247 | 6.181 | 5.90934 | 5.68701 |
| LGALS3 | P17931 | 5.44257 | 4.53638 | 5.44975 | 4.82484 | 5.13739 | 4.10987 | 5.01294 | 6.83295 | 4.45049 | 6.20608 | 4.99741 | 6.26417 | 5.43435 | 11.3028 | 9.21149 | 10.7865 |
| ARL3 | P36405 | 6.35415 | 6.05697 | 6.9131 | 5.78501 | 6.426 | 6.73182 | 5.94438 | 6.7568 | 6.30858 | 6.83435 | 6.58764 | 6.71018 | 6.12171 | 5.69313 | 5.30748 | 5.46872 |
| RPS20 | P60866-2 | 6.48302 | 6.68907 | 6.74053 | 6.05473 | 6.33729 | 6.64934 | 6.48612 | 6.15907 | 6.74133 | 6.64927 | 6.04477 | 6.30657 | 5.49262 | 5.99237 | 5.35804 | 5.81587 |
| NSUN4 | Q96CB9 | 6.48399 | 6.22002 | 6.28387 | 6.69078 | 6.37067 | 5.9532 | 6.50158 | 5.9366 | 6.51513 | 6.11521 | 6.07965 | 5.77763 | 6.12757 | 5.91593 | 6.79084 | 6.23733 |
| MSL1 | Q68DK7 | 6.19928 | 6.60633 | 6.23271 | 6.29885 | 6.90507 | 6.37763 | 5.50971 | 6.57365 | 6.18998 | 5.63788 | 6.98249 | 5.98022 | 7.62894 | 5.69742 | 5.4703 | 5.70954 |
| C11orf54 | Q9H0W9 | 5.83192 | 5.55655 | 6.67307 | 4.93556 | 7.11138 | 6.49915 | 5.92311 | 6.69896 | 6.54037 | 6.72731 | 7.17096 | 7.29213 | 6.06408 | 5.89056 | 5.88197 | 5.20293 |
| MAP1A | P78559-2 | 6.01975 | 5.98788 | 6.70276 | 5.49399 | 5.99015 | 6.42539 | 6.13826 | 6.34424 | 6.77267 | 6.04973 | 6.77303 | 6.61391 | 6.01793 | 6.38272 | 6.09197 | 6.19563 |
| PTMS | P20962 | 6.49887 | 6.86317 | 6.54627 | 5.68409 | 6.63221 | 6.35429 | 5.88074 | 6.80071 | 6.29987 | 6.09588 | 6.70511 | 6.01156 | 6.48907 | 6.50599 | 5.78212 | 5.85006 |
| USP8 | P40818 | 6.16982 | 6.13012 | 6.95882 | 5.85368 | 6.02271 | 6.70832 | 6.22275 | 6.49685 | 6.48342 | 6.38505 | 6.80705 | 6.46362 | 5.71201 | 6.16397 | 5.5711 | 5.85068 |
| LIG4 | P49917 | 5.7479 | 5.91473 | 5.93288 | 5.51331 | 6.1566 | 6.038 | 6.06813 | 5.89748 | 6.00732 | 6.01473 | 5.92337 | 5.88271 | 7.26311 | 7.06734 | 7.52516 | 7.04723 |
| MTPAP | Q9NVV4 | 6.22273 | 6.23881 | 5.60971 | 6.74289 | 6.45738 | 6.25804 | 6.37525 | 6.0314 | 6.11286 | 6.12732 | 5.92938 | 5.67554 | 6.64021 | 6.37413 | 6.55454 | 6.64982 |
| TIA1 | P31483 | 6.45968 | 6.41629 | 5.75702 | 6.44592 | 6.25127 | 6.1982 | 6.08212 | 6.09203 | 5.86822 | 6.43846 | 6.27685 | 6.32462 | 6.71734 | 6.16812 | 6.26763 | 6.23624 |
| POLE2 | P56282 | 5.78444 | 6.41603 | 6.32871 | 5.88818 | 5.92544 | 6.68655 | 6.20647 | 5.87755 | 6.43243 | 6.41282 | 6.38936 | 6.32234 | 6.51012 | 6.10116 | 6.48646 | 6.23194 |
| LGALS3BP | Q08380 | 6.72483 | 6.42019 | 5.87084 | 5.81685 | 6.28976 | 6.24649 | 6.63553 | 6.59513 | 6.25706 | 6.28134 | 6.25777 | 5.81337 | 6.20059 | 6.62068 | 6.10087 | 5.8687 |
| CDK9 | P50750-2 | 5.81316 | 6.2489 | 6.01123 | 6.48635 | 6.63172 | 5.52512 | 5.97498 | 6.23982 | 6.29942 | 5.64581 | 6.34033 | 6.24235 | 6.99262 | 6.37655 | 6.69144 | 6.48022 |
| EXTL3 | O43909 | 6.42776 | 6.62241 | 6.00294 | 6.88175 | 5.9727 | 6.10265 | 6.66013 | 6.30063 | 6.40492 | 6.21624 | 6.28661 | 6.4812 | 5.61687 | 6.05972 | 5.75798 | 6.2055 |
| RAB11A | P62491 | 6.32928 | 6.50236 | 6.26613 | 6.27885 | 5.72954 | 5.98948 | 5.93712 | 6.0682 | 6.02075 | 6.51447 | 6.13832 | 6.33607 | 5.91026 | 6.42933 | 6.34813 | 7.20168 |
| USP6NL | Q92738-2 | 6.57928 | 6.74219 | 6.16655 | 6.73177 | 5.68964 | 5.79379 | 6.04814 | 6.0423 | 6.03335 | 6.35275 | 5.75895 | 6.09638 | 6.34261 | 6.00896 | 6.59788 | 6.92545 |
| MYO5C | Q9NQX4 | 6.08785 | 6.14413 | 6.01631 | 6.10189 | 5.92146 | 5.95112 | 5.78603 | 6.36224 | 5.95096 | 6.37576 | 6.57411 | 6.11625 | 6.95523 | 6.50911 | 7.5313 | 6.81625 |
| FEM1B | Q9UK73 | 6.47316 | 6.62476 | 7.34477 | 6.5909 | 6.36101 | 6.24308 | 6.24856 | 5.94458 | 6.44099 | 6.2654 | 6.59625 | 6.76677 | 5.2951 | 5.46763 | 5.56335 | 5.77369 |
| PHKA1 | P46020 | 6.7721 | 6.78951 | 7.2995 | 6.6173 | 6.20859 | 6.69191 | 6.50586 | 6.74997 | 6.60442 | 6.68924 | 6.72837 | 7.04333 | 4.7824 | 5.01426 | 4.78502 | 4.71823 |
| HMOX1 | P09601 | 6.60141 | 6.62154 | 6.3783 | 6.76785 | 6.41326 | 6.16461 | 6.46044 | 6.43703 | 6.35854 | 6.31498 | 6.45809 | 6.45231 | 5.79581 | 5.45158 | 5.53297 | 5.79126 |
| WASL | O00401 | 6.08236 | 6.18789 | 6.92061 | 5.54982 | 6.34037 | 6.62427 | 6.47597 | 6.09746 | 6.39317 | 6.63199 | 6.58867 | 6.56435 | 5.86161 | 6.01025 | 5.98319 | 5.68803 |
| GALNS | P34059 | 5.79377 | 5.3655 | 5.96341 | 5.60709 | 6.29329 | 5.46634 | 6.04604 | 6.49388 | 5.86511 | 6.66073 | 6.10379 | 6.58856 | 6.60419 | 7.06348 | 7.19524 | 6.88956 |
| CNPY2 | Q9Y2B0 | 6.57682 | 7.04039 | 5.78398 | 6.35337 | 6.69949 | 6.47325 | 5.91899 | 6.9405 | 5.54173 | 5.73539 | 6.18674 | 6.18512 | 6.28958 | 6.06325 | 6.19585 | 6.01555 |
| PPP2R5A | Q15172 | 6.3668 | 6.31603 | 6.62928 | 6.13565 | 6.15497 | 6.70438 | 6.37116 | 6.68494 | 6.14247 | 6.69063 | 6.14295 | 6.23368 | 5.97167 | 5.87333 | 5.87069 | 5.71139 |
| RNF169 | Q8NCN4 | 5.90779 | 6.13769 | 5.45337 | 6.57444 | 5.45786 | 5.71363 | 6.10961 | 6.08303 | 6.30368 | 6.36802 | 6.52547 | 6.80822 | 6.48192 | 6.29978 | 6.50765 | 7.08999 |
| PHRF1 | Q9P1Y6 | 6.32109 | 6.23991 | 6.18559 | 6.76512 | 6.13272 | 6.00849 | 6.3736 | 5.97647 | 6.20219 | 5.90306 | 6.0523 | 6.02835 | 5.97939 | 6.59222 | 6.8129 | 6.42658 |
| ARL6IP4 | Q66PJ3 | 5.67622 | 5.79336 | 6.35015 | 5.4329 | 5.23384 | 5.9845 | 5.54091 | 5.81012 | 6.30545 | 6.61025 | 6.74304 | 6.68892 | 6.27727 | 7.39666 | 7.24241 | 6.91399 |
| SCLT1 | Q96NL6 | 6.63573 | 6.42365 | 7.02339 | 6.45992 | 6.21667 | 6.65929 | 6.50048 | 6.77034 | 6.49849 | 6.55438 | 6.86679 | 6.93021 | 5.18109 | 5.05635 | 4.93005 | 5.29318 |
| MBOAT7 | Q96N66 | 6.08983 | 6.45225 | 5.70407 | 6.48355 | 6.34632 | 5.88545 | 6.50726 | 6.05612 | 6.03099 | 6.50222 | 5.87859 | 5.73755 | 6.44971 | 6.72465 | 7.27183 | 6.77962 |
| ABT1 | Q9ULW3 | 6.31898 | 6.22373 | 5.56654 | 6.64197 | 6.55512 | 6.08304 | 6.14248 | 6.09166 | 6.42441 | 6.65678 | 6.2504 | 5.67193 | 6.34084 | 6.1329 | 7.07184 | 6.82737 |
| ARPC1A | Q92747 | 5.98696 | 5.91915 | 6.75983 | 6.29264 | 6.02487 | 6.60685 | 6.25031 | 5.86898 | 6.15875 | 6.37737 | 6.5481 | 6.66448 | 6.11005 | 6.20441 | 6.15242 | 6.07483 |
| ARFIP1 | P53367 | 6.19524 | 5.46153 | 6.35507 | 5.72001 | 6.63908 | 6.08632 | 6.4996 | 6.19256 | 5.67381 | 6.52479 | 6.16335 | 6.28858 | 6.85837 | 6.54348 | 6.37969 | 6.41852 |
| TSC22D1 | Q15714 | 6.28105 | 5.77339 | 7.19963 | 5.74746 | 6.50426 | 6.76234 | 5.85824 | 6.79693 | 6.29408 | 6.48971 | 6.82848 | 6.67594 | 6.16888 | 5.93445 | 5.20597 | 5.2992 |
| DCUN1D1 | Q96GG9 | 5.84283 | 6.24404 | 6.74857 | 5.63328 | 6.32965 | 6.63508 | 5.95391 | 6.11123 | 6.05733 | 6.15159 | 6.53607 | 5.94997 | 6.73677 | 6.4327 | 6.46933 | 6.16767 |
| EDC3 | Q96F86 | 6.05905 | 5.99507 | 6.49907 | 5.87984 | 6.09434 | 6.65953 | 5.92782 | 6.52948 | 6.75748 | 6.57391 | 6.69289 | 7.04068 | 6.2353 | 5.76459 | 5.69426 | 5.59669 |
| APIP | Q96GX9 | 5.90695 | 5.85663 | 7.20689 | 5.58955 | 5.9678 | 6.08514 | 6.52165 | 6.06495 | 5.84436 | 6.08326 | 6.55936 | 6.15696 | 5.95448 | 6.57986 | 7.52529 | 6.09687 |
| ATP4A | P20648 | 6.60128 | 6.74059 | 6.16211 | 6.70136 | 5.99034 | 6.26096 | 6.44929 | 5.96331 | 5.97842 | 6.39046 | 6.28854 | 6.2632 | 5.8031 | 5.94317 | 6.14053 | 6.32333 |
| GNB4 | Q9HAV0 | 6.48871 | 6.47409 | 5.87166 | 6.49609 | 6.58097 | 6.46848 | 6.22197 | 6.435 | 6.45855 | 6.12045 | 6.23569 | 5.96963 | 6.06774 | 5.96255 | 6.13903 | 6.00938 |
| CRYZL1 | O95825 | 5.89209 | 6.01905 | 6.78961 | 5.61746 | 6.61292 | 6.56845 | 6.4202 | 6.65666 | 6.31707 | 6.35006 | 6.75682 | 6.60022 | 5.98561 | 5.78985 | 5.85343 | 5.77048 |
| TRPM4 | Q8TD43 | 6.39437 | 6.2525 | 5.92771 | 6.93058 | 6.08699 | 5.75866 | 6.58931 | 5.87461 | 6.37768 | 6.11411 | 5.94914 | 6.22025 | 6.42403 | 6.23348 | 6.59143 | 6.27515 |
| PLBD2 | Q8NHP8 | 6.44622 | 6.5054 | 5.96466 | 6.14713 | 6.55757 | 6.47513 | 6.22861 | 6.1045 | 5.99741 | 6.18891 | 6.33656 | 6.16929 | 6.29226 | 6.27043 | 6.09361 | 6.2223 |
| GDPD1 | Q8N9F7 | 5.9906 | 6.38819 | 6.06538 | 6.53665 | 6.25111 | 5.99239 | 6.20597 | 6.08374 | 5.98691 | 6.15992 | 5.8401 | 5.62175 | 6.63313 | 6.68868 | 6.85411 | 6.70139 |
| ETNK1 | Q9HBU6 | 6.46904 | 6.46042 | 7.06628 | 6.82486 | 6.35876 | 6.69361 | 6.17347 | 6.45609 | 6.52917 | 6.76547 | 6.56754 | 6.88989 | 4.88895 | 5.25339 | 5.14081 | 5.46225 |
| CBL | P22681 | 5.84121 | 5.89506 | 6.96554 | 6.22639 | 6.26862 | 6.9005 | 6.14203 | 6.49639 | 6.51228 | 6.31413 | 7.19759 | 6.7304 | 5.96388 | 5.25201 | 5.34918 | 5.94478 |
| NCBP2 | P52298 | 5.63882 | 6.06569 | 5.83388 | 6.23706 | 6.04752 | 6.256 | 5.81203 | 6.09727 | 6.10542 | 6.44332 | 6.25212 | 6.39273 | 6.69783 | 6.74022 | 6.71741 | 6.66267 |
| PITPNA | Q00169 | 6.31425 | 6.72969 | 6.63836 | 6.30972 | 6.17421 | 6.77552 | 6.21779 | 6.94616 | 6.32795 | 6.70705 | 5.98912 | 6.21225 | 5.63964 | 5.83354 | 5.49949 | 5.68527 |
| UBAP1 | Q9NZ09-4 | 6.43958 | 6.57315 | 6.95557 | 5.58914 | 6.16486 | 6.5383 | 5.43283 | 6.88222 | 5.86561 | 7.10511 | 6.86827 | 7.76088 | 6.02904 | 5.75885 | 5.36059 | 5.57598 |
| ITPKA | P23677 | 6.29607 | 6.39268 | 6.29218 | 6.38813 | 5.89662 | 6.13431 | 5.99904 | 5.85457 | 6.19419 | 6.23028 | 6.15669 | 6.08476 | 6.66815 | 6.16869 | 6.5771 | 6.66653 |
| COX7A2L | O14548 | 6.16602 | 5.67995 | 5.69528 | 5.69844 | 6.53437 | 5.87084 | 6.37064 | 6.73951 | 5.32271 | 6.23298 | 5.41809 | 6.12036 | 5.9063 | 7.51336 | 7.27536 | 7.45579 |
| ZZZ3 | Q8IYH5 | 6.40027 | 6.43773 | 6.5054 | 6.87231 | 6.10217 | 5.80989 | 6.3425 | 5.91223 | 6.40188 | 6.32276 | 6.20389 | 5.78255 | 6.45058 | 5.95959 | 6.18505 | 6.31119 |
| DCAF4 | Q8WV16 | 6.43978 | 6.86217 | 6.26466 | 6.82655 | 6.7146 | 6.42069 | 6.68327 | 6.49108 | 6.4496 | 6.37656 | 6.20651 | 6.03031 | 5.3981 | 5.61437 | 5.57088 | 5.65087 |
| APAF1 | O14727 | 6.43992 | 6.34679 | 7.05292 | 6.11575 | 6.32753 | 6.83689 | 6.1999 | 6.71124 | 6.27933 | 6.46175 | 6.48881 | 6.17911 | 5.50494 | 6.05232 | 5.62628 | 5.37651 |
| AMIGO2 | Q86SJ2 | 6.50208 | 6.52397 | 6.6017 | 6.54168 | 6.02207 | 6.04422 | 6.40541 | 5.99661 | 6.13356 | 6.17194 | 5.92273 | 5.88461 | 6.65404 | 6.43165 | 6.24446 | 5.91927 |
| MORF4L2 | Q15014 | 6.9753 | 6.97233 | 6.80617 | 7.24775 | 6.29377 | 6.04654 | 6.34397 | 6.9272 | 7.34673 | 6.80366 | 7.23979 | 6.63055 | 4.07753 | 5.06094 | 4.34224 | 4.88553 |
| MRM2 | Q9UI43 | 6.29236 | 6.44745 | 6.05185 | 6.93572 | 6.04526 | 5.93286 | 5.87271 | 5.98657 | 6.2745 | 6.13517 | 6.36355 | 6.10679 | 6.69291 | 6.21134 | 6.28099 | 6.36998 |
| LUC7L | Q9NQ29 | 5.29158 | 6.23879 | 6.68953 | 5.82183 | 5.58712 | 6.30382 | 6.19694 | 5.86442 | 5.93542 | 6.67218 | 5.69248 | 6.2809 | 6.75237 | 6.91308 | 6.92356 | 6.836 |
| LIMD1 | Q9UGP4 | 6.24243 | 6.14896 | 6.57754 | 5.93293 | 6.37621 | 6.41538 | 6.14935 | 6.31502 | 6.70843 | 6.28003 | 7.00398 | 6.86278 | 6.51146 | 5.66087 | 5.41509 | 5.39951 |
| CHMP2A | O43633 | 6.23733 | 6.38204 | 6.50747 | 6.07128 | 5.94702 | 6.4618 | 6.26835 | 6.68321 | 6.05835 | 6.68244 | 6.15503 | 6.06821 | 5.64138 | 6.662 | 6.07425 | 6.09982 |
| SHPK | Q9UHJ6 | 6.52701 | 6.17402 | 6.72123 | 5.67381 | 6.11283 | 6.84695 | 6.00975 | 6.59505 | 6.62574 | 6.21566 | 6.42214 | 6.65824 | 5.66627 | 5.80305 | 5.98871 | 5.95954 |
| ZGPAT | Q8N5A5-2 | 6.57959 | 6.1485 | 6.14687 | 6.23252 | 6.19817 | 6.18439 | 6.17064 | 6.04395 | 6.39117 | 6.21155 | 6.63871 | 6.17495 | 6.30724 | 6.37021 | 5.97419 | 6.22734 |
| HMGA2 | P52926 | 5.81549 | 5.88008 | 5.57784 | 6.12332 | 5.83638 | 5.62591 | 6.33946 | 5.98736 | 6.08758 | 6.13483 | 5.68505 | 5.89076 | 7.05913 | 7.655 | 7.2778 | 7.02398 |
| UHRF1BP1L | A0JNW5 | 6.45111 | 6.2196 | 6.37175 | 5.96622 | 6.33265 | 6.49103 | 6.28719 | 6.51979 | 6.53634 | 6.4242 | 6.35088 | 6.37545 | 5.73969 | 6.22654 | 5.86065 | 5.84691 |
| CASP4 | P49662 | 6.24139 | 5.97673 | 7.16846 | 5.74783 | 6.34663 | 6.75756 | 6.01716 | 6.68215 | 6.71551 | 6.26759 | 6.75292 | 6.73601 | 5.87328 | 5.41805 | 5.60968 | 5.68903 |
| TMEM87A | Q8NBN3 | 6.33013 | 6.36763 | 5.94299 | 6.488 | 6.29573 | 5.90262 | 6.01399 | 6.08563 | 6.31219 | 6.39382 | 5.95941 | 6.01041 | 6.37323 | 6.27066 | 6.77066 | 6.48291 |
| RNF10 | Q8N5U6-2 | 6.6071 | 6.86731 | 7.20416 | 6.69603 | 6.3264 | 6.56517 | 6.32533 | 6.29684 | 6.91688 | 7.24196 | 7.38927 | 6.99703 | 4.13799 | 5.07415 | 4.49969 | 4.8547 |
| PRKCD | Q05655-2 | 7.07273 | 7.43584 | 7.10129 | 7.20576 | 6.28209 | 6.89516 | 6.27734 | 6.97704 | 6.31389 | 5.97178 | 5.82437 | 6.11212 | 5.16717 | 5.16632 | 5.08449 | 5.1126 |
| TUT1 | Q9H6E5 | 5.50966 | 5.85778 | 6.39456 | 6.31593 | 6.23105 | 6.26843 | 6.09476 | 5.82119 | 5.92802 | 6.01537 | 6.24868 | 6.3822 | 6.74116 | 6.46065 | 6.68618 | 7.0444 |
| PARP2 | Q9UGN5 | 6.0434 | 6.22152 | 6.18096 | 6.13745 | 6.08561 | 6.45825 | 6.32326 | 5.92083 | 5.84337 | 6.34794 | 6.49895 | 6.01996 | 6.38833 | 6.62689 | 6.48937 | 6.41392 |
| DVL2 | O14641 | 6.39381 | 6.55722 | 6.63706 | 5.94053 | 6.48495 | 6.64791 | 6.59346 | 6.90529 | 6.20382 | 6.2305 | 7.03507 | 6.28389 | 5.44711 | 5.84938 | 5.46642 | 5.32356 |
| MAN2C1 | Q9NTJ4-4 | 6.04046 | 5.96286 | 6.78415 | 5.89239 | 6.76557 | 6.86505 | 6.10843 | 6.47188 | 6.6539 | 6.08515 | 6.3881 | 6.46869 | 5.65389 | 6.13525 | 6.12562 | 5.59863 |
| TNS3 | Q68CZ2 | 6.0736 | 5.91255 | 7.00621 | 6.1016 | 5.9672 | 6.15294 | 6.42031 | 6.18095 | 6.32999 | 6.59288 | 6.754 | 6.17225 | 6.4934 | 6.09211 | 6.22558 | 5.52443 |
| SPTBN4 | Q9H254 | 6.01552 | 6.19049 | 5.95591 | 6.35932 | 6.1237 | 6.13346 | 6.41989 | 5.96725 | 6.3213 | 5.68837 | 6.23737 | 5.87117 | 6.59001 | 6.42959 | 6.7904 | 6.90625 |
| HEBP2 | Q9Y5Z4 | 6.85495 | 5.35622 | 6.55513 | 4.85338 | 6.56922 | 6.55192 | 5.79264 | 6.78301 | 5.99303 | 6.62971 | 6.51473 | 6.67889 | 6.28707 | 6.61829 | 5.81327 | 6.14855 |
| INSR | P06213 | 6.01598 | 6.24499 | 6.03565 | 6.20379 | 6.01854 | 6.00019 | 5.9637 | 5.89701 | 6.12276 | 6.35006 | 6.0275 | 6.27314 | 6.42204 | 6.60745 | 6.93137 | 6.88583 |
| LPCAT4 | Q643R3 | 6.22274 | 6.51577 | 6.31715 | 6.76437 | 6.28274 | 6.25892 | 6.35907 | 5.77834 | 5.59418 | 6.05269 | 6.33211 | 6.05652 | 6.64717 | 6.45502 | 6.39662 | 5.96658 |
| SP2 | Q02086 | 6.13226 | 6.33443 | 6.31386 | 6.47748 | 5.92471 | 6.20596 | 6.23964 | 5.9197 | 6.11265 | 6.0446 | 6.28508 | 5.97231 | 6.39274 | 6.43368 | 6.67369 | 6.53722 |
| SLC9A7 | Q96T83 | 6.20689 | 6.55613 | 6.30896 | 6.64493 | 5.89698 | 6.3962 | 6.42638 | 6.05698 | 6.08166 | 6.04145 | 6.86152 | 6.63895 | 5.8393 | 6.07271 | 5.89356 | 6.07739 |
| HELB | Q8NG08 | 6.33951 | 5.848 | 7.20555 | 6.27131 | 6.0945 | 6.67924 | 6.1761 | 6.02098 | 6.00639 | 6.62511 | 6.59931 | 6.38925 | 5.78055 | 6.01988 | 6.1437 | 5.80063 |
| UPF3B | Q9BZI7 | 6.11714 | 6.24223 | 6.53795 | 6.16478 | 6.22145 | 6.31708 | 6.27104 | 6.41782 | 6.5159 | 6.35999 | 6.27011 | 6.39147 | 5.80093 | 6.49396 | 6.01304 | 5.86512 |
| SLC30A9 | Q6PML9 | 6.24322 | 6.29442 | 5.97816 | 6.68004 | 5.96675 | 5.87544 | 6.31734 | 5.67706 | 6.26347 | 5.92294 | 6.15064 | 5.84414 | 6.77582 | 6.65216 | 6.71487 | 6.64354 |
| CISD3 | P0C7P0 | 6.44086 | 6.75274 | 5.92089 | 7.21188 | 6.40152 | 6.11168 | 6.1449 | 5.75083 | 6.08441 | 6.21561 | 6.25646 | 5.7671 | 6.68082 | 6.19396 | 6.30888 | 5.75744 |
| MAGI3 | Q5TCQ9-1 | 5.91813 | 6.19418 | 6.47751 | 6.01025 | 6.43384 | 6.50992 | 6.25018 | 6.1122 | 6.21326 | 6.2566 | 6.18297 | 6.28988 | 6.39785 | 6.34083 | 6.27931 | 6.13308 |
| PEX13 | Q92968 | 6.40521 | 6.4922 | 6.21719 | 6.7223 | 6.26957 | 6.04426 | 6.32941 | 6.00657 | 6.53125 | 6.31275 | 6.22241 | 5.94808 | 6.3473 | 5.80095 | 6.30765 | 6.0429 |
| CAMK1D | Q8IU85 | 6.11882 | 5.8876 | 7.11811 | 6.23324 | 5.99924 | 6.47995 | 6.28043 | 6.55852 | 6.41053 | 6.97681 | 7.04392 | 6.50701 | 5.65287 | 5.729 | 5.56009 | 5.44384 |
| SNX16 | P57768 | 5.76146 | 6.11359 | 6.51456 | 6.54212 | 6.38648 | 6.46334 | 5.82986 | 6.4067 | 6.49482 | 6.57654 | 6.73922 | 6.86495 | 6.01542 | 5.95008 | 5.52232 | 5.81854 |
| ZFYVE21 | Q9BQ24-2 | 6.04891 | 6.37112 | 6.84116 | 6.12892 | 6.02555 | 6.48908 | 6.16764 | 6.289 | 6.21038 | 6.64378 | 6.97184 | 6.8322 | 5.35586 | 6.10593 | 5.85114 | 5.66748 |

| | | | | | | | | | | | | | | | | | |
|---|---|---|---|---|---|---|---|---|---|---|---|---|---|---|---|---|---|
| WWP2 | O00308 | 6.67697 | 6.88446 | 6.13259 | 6.52376 | 6.53691 | 6.32376 | 6.61512 | 6.57685 | 6.51349 | 6.18105 | 6.33157 | 6.32338 | 5.65473 | 5.66131 | 5.66577 | 5.39828 |
| RAB15 | P59190 | 6.46669 | 6.41996 | 5.93463 | 6.21031 | 6.47531 | 6.25633 | 6.42181 | 6.29453 | 6.46949 | 6.33859 | 6.58816 | 5.7688 | 6.13102 | 6.13867 | 6.71525 | 6.27046 |
| RAB13 | P51153 | 5.89866 | 6.66213 | 6.23854 | 6.83322 | 6.70346 | 6.2633 | 6.20535 | 5.91788 | 6.65453 | 5.94752 | 5.97664 | 6.46495 | 5.80153 | 5.95673 | 6.47969 | 5.99587 |
| RAB4A | P20338 | 6.47598 | 6.8705 | 6.72132 | 6.54837 | 6.11219 | 6.63489 | 6.37875 | 6.48701 | 6.33121 | 6.42109 | 5.91966 | 6.26167 | 5.74056 | 5.69722 | 5.62576 | 5.77383 |
| VAMP7 | P51809 | 6.24883 | 6.68751 | 6.41654 | 6.48749 | 5.98692 | 6.39938 | 6.27076 | 6.02625 | 6.27073 | 6.5655 | 6.69578 | 6.7404 | 5.69477 | 5.85744 | 5.81204 | 5.83965 |
| LANCL1 | O43813 | 6.20384 | 5.62069 | 6.98031 | 5.9643 | 6.50068 | 6.95541 | 5.92693 | 6.35341 | 6.42812 | 6.39268 | 6.44638 | 6.08927 | 5.83484 | 6.18418 | 6.10159 | 6.01737 |
| SPC25 | Q9HBM1 | 6.44518 | 6.36618 | 6.80937 | 6.13289 | 6.71167 | 6.80909 | 6.30019 | 6.88602 | 6.69436 | 6.69887 | 6.4226 | 6.51692 | 5.70046 | 5.16509 | 5.06536 | 5.27576 |
| RBIS | Q8N0T1 | 7.2448 | 7.47711 | 7.44236 | 7.5416 | 6.6658 | 6.62242 | 6.92449 | 6.9168 | 7.08875 | 7.26118 | 7.27925 | 7.21803 | 3.33766 | 3.68903 | 3.49014 | 3.80057 |
| ZNF516 | Q92618 | 6.50777 | 6.33054 | 5.72902 | 6.52997 | 6.20502 | 6.41971 | 6.13854 | 6.38263 | 6.29843 | 6.27505 | 6.37173 | 6.15277 | 6.24384 | 6.1924 | 5.99369 | 6.22891 |
| XXYLT1 | Q8NBI6 | 6.79318 | 6.56524 | 5.92076 | 7.06102 | 5.8261 | 5.74974 | 6.10976 | 6.20691 | 6.20802 | 6.39622 | 6.2149 | 5.8295 | 6.98084 | 5.8282 | 6.02245 | 6.28717 |
| PSMB9 | P28065 | 6.06591 | 6.61122 | 6.05997 | 5.44823 | 6.6521 | 6.82917 | 5.7392 | 6.96502 | 6.13991 | 6.41023 | 6.05134 | 6.31786 | 6.23371 | 6.072 | 6.23084 | 6.17329 |
| DHRS1 | Q96LJ7 | 6.5727 | 6.29616 | 5.79467 | 6.09986 | 5.82056 | 5.64831 | 6.18234 | 6.19173 | 6.02973 | 6.48344 | 6.40014 | 6.49718 | 6.42866 | 6.36671 | 6.78987 | 6.39795 |
| LRRC1 | Q9BTT6 | 6.32336 | 6.6398 | 6.29958 | 6.61582 | 6.58044 | 6.14167 | 6.17488 | 6.0201 | 6.25869 | 6.10335 | 6.3224 | 6.31698 | 6.58085 | 5.95545 | 5.68138 | 5.98523 |
| XIAP | P98170 | 6.65964 | 6.11803 | 6.7564 | 5.9089 | 6.48174 | 6.92352 | 6.10578 | 6.79852 | 6.46912 | 6.41922 | 6.57203 | 6.46876 | 5.81911 | 5.61222 | 5.52917 | 5.35783 |
| RUSC2 | Q8N2Y8 | 5.87834 | 6.01815 | 6.48028 | 5.87465 | 6.39091 | 6.5762 | 6.28925 | 6.73267 | 5.7399 | 5.81656 | 6.147 | 5.89818 | 6.93963 | 6.46822 | 6.7192 | 6.03087 |
| CENPH | Q9H3R5 | 6.36835 | 5.33704 | 6.87939 | 5.43888 | 6.46837 | 6.54272 | 5.72729 | 6.77192 | 6.07384 | 6.60094 | 6.66829 | 7.28598 | 6.10845 | 6.43122 | 6.11623 | 6.08108 |
| MAN1A2 | O60476 | 6.59996 | 6.38917 | 6.11901 | 6.64951 | 5.97813 | 6.12633 | 6.33914 | 6.15316 | 6.16937 | 6.11905 | 5.87021 | 6.10097 | 6.3338 | 6.41947 | 6.31163 | 6.32111 |
| COX6C | P09669 | 6.04036 | 6.02528 | 5.42273 | 6.1972 | 6.83593 | 6.56456 | 6.68735 | 6.82652 | 5.53528 | 5.92588 | 5.27194 | 5.90948 | 7.12189 | 6.2843 | 6.74327 | 6.60803 |
| MBTPS1 | Q14703 | 6.24989 | 6.94061 | 6.2507 | 7.28243 | 6.09724 | 5.60188 | 6.34352 | 6.04962 | 6.17061 | 6.15755 | 5.79606 | 5.44819 | 6.50181 | 6.45818 | 6.6061 | 6.04561 |
| EME1 | Q96AY2-2 | 6.33861 | 6.70383 | 6.6472 | 6.38046 | 6.20679 | 6.51182 | 6.05109 | 6.33819 | 6.58068 | 6.29525 | 6.50829 | 6.61984 | 6.40261 | 5.89727 | 5.7738 | 5.74428 |
| GTPBP2 | Q9BX10 | 6.24849 | 6.69456 | 6.67717 | 6.64108 | 6.0138 | 6.26435 | 6.28498 | 6.17218 | 6.9537 | 7.20709 | 6.61034 | 6.55708 | 5.261 | 5.2752 | 5.57993 | 5.55905 |
| GIT2 | Q14161 | 6.42674 | 6.31179 | 7.10987 | 6.59683 | 6.13162 | 7.09151 | 6.13084 | 6.31307 | 6.22264 | 6.71916 | 6.85334 | 6.44512 | 5.37971 | 5.43403 | 5.53661 | 5.29711 |
| GIT1 | Q9Y2X7 | 6.2772 | 6.51303 | 6.51419 | 5.92005 | 6.44975 | 6.67816 | 6.06964 | 6.70665 | 6.28196 | 6.49778 | 6.4329 | 6.47659 | 6.09786 | 5.92117 | 5.74718 | 5.41589 |
| P3H3 | Q8IVL6 | 6.62503 | 6.33385 | 6.12996 | 6.75325 | 6.11717 | 6.34353 | 6.29324 | 5.95072 | 6.19214 | 6.0995 | 6.34206 | 6.23884 | 6.41366 | 6.13294 | 6.1033 | 5.9308 |
| PHACTR2 | O75167-4 | 6.04276 | 6.25977 | 7.17101 | 5.88736 | 6.06433 | 6.14511 | 6.26703 | 6.30514 | 6.48841 | 6.48924 | 6.59721 | 6.30721 | 5.88615 | 6.16086 | 5.80877 | 6.11966 |
| TSR2 | Q969E8 | 6.81914 | 6.29588 | 6.99696 | 5.73664 | 6.6741 | 6.64807 | 5.85741 | 6.49814 | 7.0107 | 6.50584 | 7.09648 | 7.08286 | 5.80817 | 5.02389 | 4.9328 | 5.01291 |
| KMT2C | Q8NEZ4-3 | 6.39658 | 6.40353 | 6.23562 | 6.90109 | 6.06308 | 6.19598 | 6.38029 | 6.10166 | 6.5537 | 6.43357 | 6.56461 | 6.05418 | 5.98065 | 6.36943 | 6.23536 | 6.03066 |
| PDCD6 | O75340 | 5.58723 | 4.87565 | 5.88386 | 4.60887 | 5.9231 | 5.85848 | 5.73215 | 6.66016 | 5.46446 | 6.14017 | 6.06146 | 6.326 | 6.34821 | 8.31451 | 8.39747 | 7.81823 |
| MFSD10 | Q14728 | 6.00845 | 6.32145 | 5.80722 | 6.46658 | 6.30002 | 6.07975 | 6.25886 | 6.25945 | 6.22996 | 6.2867 | 5.77374 | 5.6209 | 6.57154 | 6.45416 | 7.01046 | 6.55078 |
| ZNF512 | Q96ME7 | 6.25639 | 6.46615 | 5.64551 | 6.61744 | 6.21121 | 5.70959 | 6.10905 | 6.6153 | 5.70036 | 6.02866 | 5.65944 | 5.90024 | 6.47286 | 6.95414 | 6.93484 | 6.71883 |
| DPF1 | Q92782-2 | 5.96133 | 6.41303 | 6.1414 | 6.2867 | 6.44383 | 5.90524 | 6.61651 | 6.10726 | 5.7051 | 6.19893 | 6.33892 | 6.19866 | 6.21844 | 6.63324 | 6.44989 | 6.38153 |
| CEP295 | Q9C0D2-3 | 6.0974 | 6.87426 | 5.90643 | 6.09951 | 5.42698 | 6.35037 | 6.80896 | 6.32442 | 6.48772 | 6.59642 | 6.93288 | 7.18679 | 5.0109 | 5.71035 | 6.24806 | 5.93856 |
| PTPN9 | P43378 | 5.92801 | 6.17035 | 6.19821 | 6.07047 | 6.27488 | 6.52113 | 6.13635 | 6.53097 | 6.2997 | 6.51424 | 6.72067 | 6.39289 | 5.92447 | 5.99746 | 6.49413 | 5.82607 |
| AP1S1 | P61966 | 6.31334 | 6.47335 | 6.57546 | 6.42794 | 6.36049 | 6.46895 | 7.1057 | 6.15491 | 5.78043 | 6.2773 | 6.82376 | 6.23541 | 5.34208 | 5.92859 | 5.81873 | 5.91357 |
| SETMAR | Q53H47 | 5.66859 | 5.33963 | 5.96937 | 5.8138 | 5.72323 | 5.91754 | 5.31252 | 5.97425 | 5.95007 | 5.91431 | 6.41748 | 6.46233 | 6.88346 | 7.34922 | 7.60725 | 7.69696 |
| NEK1 | Q96PY6-3 | 6.24422 | 6.28733 | 6.5562 | 6.34286 | 6.03645 | 6.44542 | 6.15757 | 6.37492 | 6.18992 | 6.5287 | 6.43673 | 6.11841 | 6.25246 | 6.02643 | 5.92126 | 6.08114 |
| ABHD11 | Q8NFV4 | 5.76705 | 4.80472 | 5.54076 | 5.07114 | 5.85572 | 5.28052 | 5.33749 | 6.29723 | 5.24896 | 6.7059 | 5.71245 | 6.85641 | 6.99273 | 8.09917 | 8.12976 | 8.29998 |
| YTHDF2 | Q9Y5A9 | 6.90844 | 6.39404 | 6.66969 | 6.17593 | 6.30259 | 6.21266 | 6.08046 | 6.56294 | 6.53543 | 6.20925 | 6.22931 | 6.43473 | 5.87563 | 5.70258 | 5.88714 | 5.82278 |
| ATXN3 | P54252-1 | 6.58766 | 5.61203 | 6.58331 | 5.00918 | 6.58703 | 6.63093 | 5.81596 | 6.60852 | 6.1397 | 6.66875 | 6.14536 | 6.8765 | 5.85503 | 6.67576 | 6.10713 | 6.09715 |
| RPS6KB1 | P23443 | 6.32013 | 6.30607 | 6.5147 | 6.29229 | 5.8106 | 6.68814 | 6.04367 | 6.53702 | 6.29393 | 6.54347 | 6.37846 | 6.43843 | 5.83406 | 6.10653 | 5.69746 | 6.19504 |
| PUS7L | Q9H0K6 | 5.44775 | 5.96699 | 5.84176 | 5.47023 | 5.84018 | 6.65175 | 5.51451 | 5.8759 | 6.18195 | 6.43772 | 6.34831 | 6.66779 | 6.50492 | 7.35134 | 7.15189 | 6.74702 |
| EEF1E1 | O43324 | 6.18398 | 5.23517 | 6.49261 | 4.59712 | 6.45539 | 5.92962 | 6.07401 | 6.40155 | 5.80535 | 6.52457 | 6.45213 | 6.7018 | 6.95157 | 6.90591 | 6.68093 | 6.60831 |
| SAAL1 | Q96ER3 | 5.22153 | 5.38462 | 6.1056 | 5.2274 | 5.8038 | 6.25459 | 5.68771 | 5.57743 | 5.60038 | 6.0691 | 6.38639 | 5.67934 | 7.78582 | 7.65855 | 8.03504 | 7.52269 |
| RAP2A | P10114 | 6.26292 | 6.54858 | 5.95338 | 6.58623 | 6.22158 | 5.74177 | 6.42418 | 6.31995 | 5.88787 | 6.33247 | 6.45861 | 5.87716 | 6.23145 | 6.67059 | 6.14436 | 6.33889 |
| RHOT1 | Q8IXI2-3 | 6.37051 | 6.49372 | 5.95255 | 6.74263 | 6.4095 | 5.99186 | 6.38409 | 5.83969 | 6.49989 | 6.09512 | 5.86955 | 6.00726 | 6.31489 | 6.11013 | 6.49178 | 6.42682 |
| ZFP91 | Q96JP5 | 6.06448 | 5.88173 | 6.02123 | 6.17635 | 5.70739 | 5.96331 | 6.0528 | 5.87551 | 6.46528 | 6.20588 | 6.09221 | 6.00611 | 6.74873 | 6.90001 | 7.13687 | 6.70211 |
| MAP4K3 | Q8IVH8 | 6.6391 | 6.15952 | 6.57012 | 6.32995 | 6.29901 | 6.7675 | 6.28969 | 6.54586 | 6.5128 | 6.8915 | 6.74342 | 6.57977 | 5.85665 | 5.8639 | 5.2657 | 5.68552 |
| PIK3R2 | O00459 | 6.54831 | 6.22719 | 6.3665 | 5.70123 | 6.60707 | 6.90141 | 6.49964 | 6.58963 | 6.34417 | 6.24554 | 6.25681 | 6.42527 | 6.24944 | 6.11467 | 5.36212 | 5.56099 |
| FOSL1 | P15407 | 5.5727 | 5.06427 | 5.61389 | 5.57452 | 7.09122 | 6.07664 | 6.29303 | 7.03354 | 5.66238 | 5.73575 | 5.8774 | 5.69579 | 7.81408 | 6.52996 | 6.9337 | 7.43112 |
| SNAP91 | O60641-4 | 5.7134 | 5.95001 | 6.28599 | 5.98738 | 5.98908 | 6.51488 | 5.80355 | 6.02979 | 6.37349 | 6.45319 | 6.10559 | 6.49272 | 6.53387 | 6.45504 | 6.95133 | 6.36069 |
| INPP5A | Q14642 | 6.25711 | 6.48131 | 6.16419 | 6.47873 | 6.10165 | 6.23428 | 6.37486 | 5.95126 | 6.09192 | 6.13915 | 6.38322 | 6.05652 | 6.2418 | 6.27313 | 6.46094 | 6.30993 |
| UVRAG | Q9P2Y5 | 6.76252 | 6.77745 | 6.88821 | 6.48467 | 6.37994 | 6.68129 | 5.98049 | 6.46465 | 6.14612 | 6.21032 | 6.3841 | 6.52597 | 5.60843 | 5.71847 | 5.70822 | 5.27913 |
| TRIM65 | Q6PJ69 | 6.16689 | 6.19948 | 6.50114 | 5.76291 | 6.66109 | 6.17461 | 5.77138 | 6.90555 | 6.29986 | 6.12154 | 6.78484 | 6.94686 | 5.90681 | 6.05782 | 5.75313 | 5.9861 |
| ENTPD5 | O75356 | 6.31272 | 6.1358 | 6.15856 | 6.85821 | 5.91293 | 6.00733 | 6.1769 | 5.98882 | 5.80397 | 6.05845 | 6.01229 | 6.02406 | 6.74511 | 6.58117 | 6.55483 | 6.66886 |
| GTF2E1 | P29083 | 5.39466 | 5.87387 | 5.55194 | 5.17924 | 5.72662 | 5.3669 | 5.54355 | 5.86718 | 6.08605 | 6.23864 | 5.93275 | 6.49942 | 7.14253 | 7.78032 | 7.31818 | 7.49816 |
| TPRG1L | Q5T0D9 | 6.44753 | 6.4659 | 6.53242 | 6.024 | 6.46014 | 6.48922 | 6.17933 | 6.40434 | 6.27987 | 6.87002 | 6.51351 | 6.91269 | 5.31165 | 5.7938 | 5.873 | 5.44259 |
| KRIT1 | O00522 | 6.63907 | 6.39836 | 7.09795 | 6.82 | 6.62719 | 6.32072 | 6.37482 | 6.38409 | 7.05377 | 6.86237 | 6.63777 | 6.16817 | 4.88623 | 5.38017 | 5.1205 | 5.22881 |
| ZEB2 | O60315 | 6.3689 | 6.26422 | 6.03937 | 7.35854 | 6.28941 | 6.06016 | 6.38037 | 6.22082 | 6.4082 | 6.58196 | 6.43038 | 6.2098 | 5.75614 | 5.81771 | 5.7994 | 6.01463 |
| DDAH2 | O95865 | 5.72244 | 4.44628 | 5.24711 | 4.26626 | 5.73433 | 5.15582 | 5.22907 | 7.05503 | 4.9867 | 6.9158 | 5.45207 | 7.08481 | 6.62469 | 9.24213 | 8.30508 | 8.53238 |
| LZTFL1 | Q9NQ48 | 6.4949 | 5.90842 | 6.63753 | 5.81712 | 6.67129 | 6.64895 | 6.08979 | 6.78063 | 6.23129 | 6.71844 | 6.54048 | 6.65879 | 5.95892 | 5.99255 | 5.44339 | 5.4075 |
| AVEN | Q9NQS1 | 6.49884 | 6.54967 | 7.02223 | 6.36163 | 6.02569 | 6.40861 | 5.97785 | 6.8242 | 6.45872 | 6.80464 | 6.35541 | 6.92408 | 5.88298 | 5.3677 | 5.09635 | 5.44139 |
| DISP2 | A7MBM2 | 6.34457 | 6.41736 | 5.96468 | 6.58811 | 5.96308 | 5.60447 | 6.17317 | 5.92482 | 6.25398 | 6.13351 | 5.84854 | 6.41911 | 6.17726 | 6.59367 | 6.50399 | 7.0897 |
| DENND4A | Q7Z401-2 | 6.52818 | 6.43401 | 6.67358 | 6.33879 | 6.26766 | 7.03048 | 6.38188 | 6.61087 | 6.86404 | 6.07651 | 6.66982 | 6.27533 | 5.35925 | 5.69548 | 5.40348 | 5.39064 |
| MTF2 | Q9Y483 | 6.40719 | 6.84143 | 6.35304 | 7.12678 | 6.11656 | 6.04779 | 6.77646 | 6.17007 | 6.22906 | 6.20493 | 5.638 | 6.16536 | 5.74293 | 5.93695 | 5.9987 | 6.24475 |
| CTC1 | Q2NKJ3 | 6.05733 | 6.0661 | 6.2022 | 5.95897 | 5.98946 | 6.42682 | 6.30891 | 6.04554 | 6.00205 | 6.2401 | 6.64055 | 5.93102 | 6.46402 | 6.44209 | 6.56595 | 6.6589 |
| STIM2 | Q9P246-2 | 6.82666 | 6.77161 | 6.15823 | 7.15747 | 5.6158 | 5.88029 | 6.24657 | 6.06614 | 6.50706 | 6.62256 | 6.36642 | 6.34113 | 5.75279 | 5.5606 | 6.30438 | 5.8223 |
| PEX6 | Q13608 | 6.58124 | 6.31301 | 6.00522 | 6.65763 | 6.53077 | 5.99537 | 6.39069 | 5.52999 | 5.94164 | 5.88589 | 6.15629 | 6.1728 | 6.381 | 6.52362 | 6.49605 | 6.43879 |
| EIPR1 | Q53HC9 | 6.13592 | 6.08432 | 6.14698 | 5.68417 | 6.67638 | 6.71859 | 6.18285 | 6.56036 | 6.55394 | 6.65559 | 6.64545 | 6.62018 | 6.00243 | 6.02099 | 5.74708 | 5.56478 |
| IRF3 | Q14653 | 6.55834 | 6.27756 | 7.27248 | 6.10743 | 6.29625 | 6.54111 | 6.20212 | 6.78101 | 6.57524 | 6.54604 | 6.61825 | 6.42196 | 5.60485 | 5.37676 | 5.53024 | 5.29037 |
| GNAI3 | P08754 | 6.18335 | 6.27952 | 6.29597 | 6.85181 | 5.85408 | 6.03948 | 6.28477 | 6.16248 | 6.38026 | 6.68106 | 6.47829 | 6.57946 | 5.99559 | 5.56308 | 6.35193 | 6.01887 |
| GNA12 | Q03113 | 6.50723 | 6.39659 | 6.32134 | 6.83272 | 5.87704 | 5.982 | 6.39938 | 6.22824 | 6.46998 | 6.32401 | 5.99817 | 6.4199 | 6.04229 | 6.58402 | 6.09898 | 6.42711 |
| COQ8A | Q8NI60 | 6.35697 | 6.41348 | 6.02111 | 6.57658 | 6.01983 | 5.97196 | 6.11641 | 6.00478 | 6.3167 | 5.91144 | 6.12501 | 5.99919 | 6.74845 | 6.44979 | 6.37137 | 6.59691 |
| MRPS34 | P82930 | 6.00889 | 6.28515 | 5.49528 | 6.2769 | 5.99986 | 5.81848 | 6.4993 | 6.19511 | 6.46978 | 6.31434 | 5.94471 | 6.05813 | 6.54701 | 6.45721 | 6.44958 | 7.18028 |
| RSPRY1 | Q96DX4 | 6.03688 | 5.75464 | 6.45365 | 7.02895 | 6.056 | 5.99653 | 6.87309 | 5.40968 | 6.42929 | 5.79199 | 6.44672 | 6.57285 | 6.0141 | 6.00291 | 6.29295 | 6.83986 |
| ELOC | Q15369 | 5.99173 | 5.24508 | 5.64376 | 4.72807 | 6.08976 | 5.98396 | 6.56793 | 6.94884 | 5.68318 | 6.61025 | 6.38766 | 6.45929 | 5.83328 | 8.05465 | 7.42158 | 7.25099 |
| LSR | Q86X29 | 6.42877 | 6.54456 | 5.86205 | 6.61363 | 6.16325 | 6.22362 | 6.03689 | 6.01361 | 5.84756 | 6.30541 | 6.22034 | 6.14866 | 6.50425 | 6.29565 | 6.66835 | 6.1234 |
| ANKRD12 | Q6UB98 | 6.18487 | 6.66194 | 5.72616 | 6.70722 | 6.23335 | 5.94014 | 6.19575 | 6.19064 | 6.01148 | 6.25259 | 5.8844 | 5.96149 | 5.77457 | 7.04039 | 6.52897 | 6.70603 |
| PSMD9 | O00233 | 6.18807 | 5.56147 | 6.29839 | 5.68957 | 6.61142 | 6.27049 | 6.762 | 6.88103 | 6.78722 | 6.03189 | 6.62206 | 6.87921 | 5.40612 | 6.13557 | 5.40202 | 6.47347 |
| UNG | P13051 | 6.2013 | 5.80192 | 6.3967 | 5.84281 | 5.97211 | 6.31695 | 6.3247 | 5.69924 | 6.12342 | 6.29246 | 6.75874 | 6.22812 | 6.3177 | 6.80443 | 6.57561 | 6.34379 |
| ULK1 | O75385 | 6.35159 | 5.60219 | 6.67676 | 6.06996 | 6.26012 | 6.5185 | 5.9508 | 6.1049 | 6.36746 | 6.40151 | 6.28292 | 6.7934 | 6.17429 | 6.22335 | 5.79841 | 6.42386 |
| NCK1 | P16333 | 6.44736 | 6.34142 | 6.42335 | 6.10396 | 6.52971 | 6.64027 | 5.99279 | 7.11205 | 6.28366 | 6.09394 | 6.61603 | 6.57954 | 6.07862 | 5.89654 | 5.46653 | 5.39422 |
| LEO1 | Q8WVC0 | 6.3457 | 6.65275 | 5.69293 | 7.0558 | 6.32609 | 6.33079 | 6.31957 | 6.13835 | 6.42162 | 6.26976 | 5.41936 | 6.06789 | 6.23272 | 6.23786 | 6.1099 | 6.3789 |
| CERS2 | Q96G23 | 8.65083 | 8.39133 | 7.85232 | 8.6802 | 7.8099 | 7.23306 | 7.36298 | 5.79094 | 5.9558 | 4.91394 | 3.93421 | 4.14264 | 4.789 | 4.77093 | 4.61076 | 5.11116 |
| REV3L | O60673 | 6.53152 | 7.1618 | 6.74296 | 7.17725 | 5.91396 | 5.8306 | 6.17269 | 6.23723 | 6.17924 | 6.57798 | 5.75854 | 6.7946 | 5.14093 | 5.50066 | 6.07138 | 6.20865 |
| RFX7 | Q2KHR2 | 6.66746 | 6.76696 | 6.54533 | 6.5441 | 6.22888 | 6.18842 | 6.0739 | 6.17807 | 6.52042 | 6.81236 | 6.71229 | 7.02928 | 5.60867 | 5.27725 | 5.22311 | 5.62349 |
| RASA3 | Q14644 | 6.72069 | 6.57119 | 7.02059 | 6.5911 | 6.49732 | 6.36959 | 6.36 | 5.97901 | 6.22931 | 6.15073 | 6.2909 | 6.30315 | 5.74838 | 5.61657 | 5.74535 | 5.80612 |
| ASCC1 | Q8N9N2-2 | 6.16475 | 6.29331 | 6.35961 | 5.94236 | 6.52232 | 6.76332 | 6.03021 | 6.64001 | 6.30856 | 6.55593 | 6.3141 | 6.29459 | 6.12119 | 6.07959 | 5.63971 | 5.97044 |
| HLA-A | A0A140T8Y4 | 6.4408 | 6.59036 | 6.19707 | 6.50828 | 6.60204 | 6.47645 | 6.49594 | 5.92201 | 6.21034 | 6.29686 | 6.31094 | 6.3628 | 6.35367 | 5.80093 | 5.55442 | 5.87709 |
| RPL29 | P47914 | 6.07595 | 6.74445 | 6.173 | 6.1163 | 6.55452 | 6.51401 | 6.48734 | 6.80004 | 6.5922 | 6.34331 | 6.09045 | 5.80454 | 5.54762 | 6.48585 | 5.91686 | 5.75357 |
| TSG101 | Q99816 | 6.37094 | 5.96575 | 6.65218 | 5.53719 | 6.53003 | 6.62458 | 6.24753 | 6.82818 | 6.25338 | 6.52481 | 6.809 | 6.78981 | 6.15181 | 5.74096 | 5.49637 | 5.4775 |
| FAM98B | Q52LJ0 | 5.40985 | 5.57302 | 6.03261 | 5.56332 | 5.91683 | 6.28746 | 5.79043 | 5.911 | 5.99348 | 6.23384 | 6.3492 | 6.2515 | 7.4252 | 7.12103 | 7.20003 | 6.9412 |
| DUS2 | Q9NX74 | 6.13239 | 5.94328 | 6.5116 | 5.33848 | 6.06449 | 6.26929 | 5.77056 | 5.87948 | 6.00373 | 6.57309 | 6.0779 | 6.60732 | 6.22222 | 6.43338 | 6.42583 | 6.74698 |
| ARHGEF18 | Q6ZSZ5 | 6.40364 | 6.74783 | 6.77771 | 6.63338 | 6.40234 | 6.6794 | 6.31048 | 6.95141 | 6.3302 | 6.34855 | 6.23203 | 6.22849 | 5.39297 | 5.8441 | 5.26176 | 5.4557 |
| FAM193B | Q96PV7 | 6.40211 | 6.41091 | 6.87311 | 6.41306 | 5.87912 | 6.05368 | 6.13733 | 6.22747 | 6.71124 | 6.53109 | 6.48265 | 6.58031 | 5.42437 | 6.02682 | 5.93598 | 5.91074 |
| RRP8 | O43159 | 6.11137 | 6.07881 | 5.98724 | 7.03492 | 5.92715 | 5.81888 | 6.30181 | 5.86627 | 6.10514 | 5.92704 | 5.93699 | 6.02459 | 5.75717 | 6.41351 | 6.81724 | 6.89187 |
| UBQLN4 | Q9NRR5 | 6.21357 | 5.24386 | 5.40016 | 5.11678 | 5.68007 | 4.44206 | 5.21452 | 6.60883 | 4.65319 | 6.37745 | 5.46027 | 7.10337 | 6.41123 | 9.21569 | 8.23344 | 8.82549 |
| MRTFB | Q9ULH7-5 | 6.32199 | 6.45348 | 6.55628 | 5.43418 | 6.51448 | 6.77059 | 5.96214 | 6.11288 | 6.47909 | 6.59408 | 6.33277 | 6.61158 | 6.34054 | 6.00274 | 5.65043 | 5.86275 |
| CNTLN | Q9NXG0-2 | 6.43763 | 6.159 | 6.75579 | 6.66752 | 6.0957 | 6.56944 | 6.48135 | 6.53401 | 6.40564 | 6.72224 | 6.72034 | 6.86905 | 5.18648 | 5.49104 | 5.31577 | 5.58899 |
| THOC6 | Q86W42 | 5.71464 | 5.85501 | 5.65535 | 6.62856 | 6.1668 | 5.92623 | 6.18901 | 6.21244 | 6.49992 | 6.65669 | 6.43511 | 6.19449 | 6.58418 | 5.94642 | 6.71181 | 6.62334 |
| PARVA | Q9NVD7 | 5.72006 | 6.23872 | 6.1714 | 5.64105 | 6.20993 | 6.63931 | 5.88725 | 6.37439 | 6.3095 | 6.78793 | 7.03904 | 6.66219 | 6.17675 | 6.04211 | 5.9518 | 6.14857 |
| TPD52 | P55327-3 | 6.59912 | 5.37801 | 6.64576 | 5.20776 | 6.22341 | 6.06676 | 5.76408 | 6.65106 | 6.27926 | 6.76808 | 6.4097 | 6.76012 | 6.77643 | 6.39769 | 6.09089 | 5.98187 |
| RAB27A | P51159 | 6.4449 | 6.58502 | 6.22647 | 6.49925 | 6.13572 | 6.09631 | 6.39662 | 6.18395 | 5.92009 | 6.5002 | 6.04469 | 6.01992 | 6.1721 | 6.09114 | 6.48966 | 6.19397 |
| PRDM10 | Q9NQV6-6 | 6.42665 | 6.23655 | 5.3813 | 6.72981 | 6.18253 | 6.03797 | 6.19732 | 6.30502 | 5.86019 | 6.22548 | 6.38918 | 5.95374 | 6.52047 | 6.29628 | 6.44149 | 6.81602 |
| TAB1 | Q15750 | 6.68916 | 6.51536 | 6.73224 | 5.43085 | 6.71881 | 6.62591 | 5.95233 | 6.73011 | 6.67766 | 6.63302 | 6.49502 | 6.59144 | 5.81722 | 5.54954 | 5.40149 | 5.43981 |
| HDAC8 | Q9BY41 | 5.58621 | 5.50141 | 6.49366 | 5.45959 | 6.02435 | 6.90908 | 6.15556 | 5.843 | 5.9387 | 6.26745 | 6.401 | 6.16063 | 6.7034 | 6.90769 | 6.95328 | 6.69499 |

| | | | | | | | | | | | | | | | | | |
|---|---|---|---|---|---|---|---|---|---|---|---|---|---|---|---|---|---|
| CNPY3 | Q9BT09 | 6.29432 | 6.63333 | 5.85698 | 6.42453 | 6.45109 | 6.16564 | 6.42761 | 5.91134 | 5.9255 | 5.98318 | 5.46066 | 5.92378 | 6.67704 | 6.7867 | 6.51933 | 6.55896 |
| ARHGAP4 | P98171-2 | 5.91991 | 6.50005 | 6.65162 | 5.99806 | 6.56901 | 6.68741 | 5.67993 | 6.29182 | 6.46482 | 6.87627 | 6.67315 | 6.10341 | 6.20949 | 6.0873 | 5.69891 | 5.58886 |
| C19orf53 | Q9UNZ5 | 5.73295 | 6.01819 | 5.76018 | 5.81881 | 5.84213 | 5.85559 | 5.77841 | 5.96435 | 6.1353 | 6.35455 | 6.02864 | 6.2668 | 6.50959 | 7.44074 | 7.22634 | 7.26744 |
| OMA1 | Q96E52 | 6.33169 | 6.56702 | 6.10361 | 6.70055 | 6.33346 | 6.3812 | 6.6413 | 5.60847 | 6.21089 | 6.22971 | 5.81321 | 5.59307 | 6.35016 | 6.23283 | 6.38787 | 6.51495 |
| PSMD10 | O75832 | 5.9769 | 5.39887 | 6.76003 | 5.16782 | 6.23976 | 6.18259 | 6.26375 | 6.46268 | 6.09557 | 6.62195 | 6.60407 | 7.08367 | 5.94305 | 6.51101 | 6.49706 | 6.19123 |
| HDHD5 | Q9BXW7 | 6.64481 | 6.1442 | 6.25418 | 6.68231 | 6.22546 | 5.71987 | 5.96955 | 6.08873 | 5.84156 | 6.39235 | 6.59712 | 6.62913 | 6.64145 | 5.86648 | 6.17925 | 6.12355 |
| GTF2H2C | Q6P1K8 | 6.4031 | 6.10408 | 6.23335 | 6.56249 | 6.46517 | 6.29889 | 6.42613 | 6.2023 | 6.34873 | 6.59962 | 6.3028 | 6.18801 | 5.92535 | 5.90018 | 6.02048 | 6.01933 |
| RAB32 | Q13637 | 6.38153 | 6.51952 | 6.1464 | 6.35848 | 6.00774 | 6.0689 | 6.41863 | 6.19645 | 6.17635 | 6.43365 | 6.16797 | 6.04618 | 6.40728 | 6.19753 | 6.45926 | 6.01412 |
| RPA3 | P35244 | 4.44276 | 4.59777 | 5.05203 | 4.04431 | 5.71946 | 5.78332 | 4.89904 | 7.04515 | 5.27086 | 6.31039 | 5.86568 | 6.74483 | 6.12691 | 10.0773 | 8.60705 | 9.41314 |
| NDEL1 | Q9GZM8-2 | 6.51131 | 6.23514 | 6.69151 | 6.22282 | 6.4341 | 6.42711 | 5.56392 | 6.70588 | 6.42389 | 6.64927 | 6.90106 | 7.0164 | 6.21474 | 5.27663 | 5.24954 | 5.47667 |
| BORCS6 | Q96GS4 | 6.51733 | 5.9069 | 6.18835 | 5.47946 | 6.31962 | 6.11646 | 5.63601 | 6.44507 | 5.89589 | 6.42307 | 6.31177 | 6.80932 | 6.88543 | 6.55694 | 6.35654 | 6.15185 |
| ASXL2 | Q76L83 | 6.28406 | 6.88185 | 6.09495 | 6.84544 | 6.06944 | 6.46135 | 6.06523 | 6.13023 | 6.38475 | 6.15051 | 5.97615 | 6.42239 | 6.18324 | 5.88018 | 6.11527 | 6.05496 |
| CALCOCO1 | Q9P1Z2 | 6.35116 | 6.12091 | 6.6156 | 5.58567 | 6.6671 | 6.92748 | 6.12855 | 6.9537 | 6.25053 | 6.65277 | 7.17431 | 6.59083 | 6.26658 | 5.48067 | 5.0561 | 5.17804 |
| FSD1 | Q9BTV5 | 5.99201 | 6.19214 | 7.13611 | 6.13174 | 6.0098 | 6.90108 | 5.78075 | 6.51746 | 5.94705 | 6.92014 | 6.758 | 6.69016 | 5.81792 | 6.05976 | 5.31106 | 5.83483 |
| SCAPER | Q9BY12 | 5.97104 | 6.35631 | 6.75319 | 6.03567 | 6.09942 | 6.66269 | 6.26759 | 6.18191 | 6.36665 | 6.76599 | 6.46368 | 6.56552 | 5.65801 | 6.04743 | 6.02229 | 5.78263 |
| RNF113A | O15541 | 5.90323 | 5.89597 | 6.08296 | 5.80225 | 6.20524 | 6.39731 | 5.52961 | 6.48749 | 6.12336 | 6.0659 | 6.90983 | 6.36373 | 6.86209 | 6.54723 | 6.50707 | 6.31671 |
| ABHD17B | Q5VST6-2 | 6.58354 | 6.47994 | 6.44214 | 7.18368 | 6.08447 | 5.72243 | 6.34952 | 5.75508 | 6.13197 | 6.53772 | 6.13599 | 6.62701 | 5.40045 | 5.96799 | 6.01126 | 6.58682 |
| DUSP23 | Q9BVJ7 | 6.06841 | 4.99789 | 6.41692 | 5.2107 | 6.45462 | 5.88114 | 5.58757 | 6.58312 | 5.80394 | 6.85598 | 6.48452 | 6.91408 | 6.02838 | 7.06484 | 6.51349 | 7.13443 |
| ZNF518A | Q6AHZ1 | 7.11966 | 6.98405 | 6.06578 | 6.72136 | 5.93236 | 5.74674 | 6.05321 | 6.15021 | 6.23088 | 6.94626 | 6.39716 | 6.37219 | 5.61625 | 6.18842 | 5.68296 | 5.79249 |
| STX10 | O60499 | 6.66202 | 6.15506 | 5.97408 | 6.45985 | 6.16954 | 5.83199 | 6.2099 | 6.14262 | 6.31859 | 6.38229 | 6.02319 | 6.54864 | 6.54723 | 6.62989 | 6.51156 | 6.33355 |
| RMDN3 | Q96TC7 | 6.36778 | 6.36175 | 6.03726 | 6.89448 | 6.44583 | 6.23151 | 6.46686 | 5.98025 | 5.99041 | 5.86406 | 5.75065 | 5.92308 | 6.71866 | 5.78143 | 7.01264 | 6.17335 |
| SNRPB2 | P08579 | 6.1997 | 6.47078 | 5.83085 | 6.57491 | 6.15101 | 6.05639 | 6.16638 | 6.17106 | 5.84678 | 6.19479 | 5.98487 | 5.90042 | 6.76616 | 6.50228 | 6.64276 | 6.54087 |
| LPCAT2 | Q7L5N7 | 6.41344 | 6.17659 | 6.29371 | 6.35293 | 6.01623 | 5.96191 | 6.15737 | 5.59571 | 6.05777 | 6.24569 | 6.09011 | 5.83803 | 6.67621 | 6.53675 | 7.02818 | 6.55936 |
| TAF7 | Q15545 | 6.57635 | 6.55053 | 6.98142 | 6.5972 | 6.31479 | 6.50578 | 5.92461 | 6.86228 | 6.74459 | 6.41827 | 6.74408 | 6.83748 | 5.4695 | 5.31636 | 4.97906 | 5.17772 |
| CYHR1 | E9PJD7 | 5.7536 | 6.28952 | 6.01758 | 6.3097 | 5.8468 | 6.08448 | 6.07393 | 5.63199 | 5.98795 | 6.2764 | 6.33094 | 6.16859 | 7.06192 | 6.4283 | 7.0664 | 6.67189 |
| NDUFA7 | O95182 | 6.78512 | 6.49191 | 5.73911 | 6.83294 | 6.45048 | 5.87289 | 5.86269 | 5.94789 | 6.19502 | 6.1736 | 6.07185 | 6.12796 | 7.05546 | 6.21093 | 6.13776 | 6.04441 |
| EIF1AX | P47813 | 5.96569 | 6.09928 | 6.13678 | 6.10168 | 6.25346 | 6.28207 | 6.60314 | 6.1424 | 6.58135 | 6.27499 | 6.32421 | 6.55396 | 5.69726 | 6.45093 | 6.28526 | 6.24754 |
| NR2C1 | P13056 | 6.4302 | 6.74077 | 5.64521 | 6.83985 | 6.44925 | 6.41619 | 6.33343 | 6.4013 | 6.13083 | 6.09544 | 6.19697 | 6.3743 | 6.21504 | 5.75541 | 6.00561 | 5.97019 |
| PLEKHM2 | Q8IWE5 | 5.92298 | 6.5752 | 6.44155 | 5.91356 | 6.16276 | 5.98971 | 5.64853 | 6.36627 | 6.45137 | 6.79416 | 6.55973 | 6.65955 | 6.22036 | 5.87162 | 6.32405 | 6.09859 |
| MRPL16 | Q9NX20 | 6.51425 | 6.86279 | 5.67148 | 6.67221 | 6.20471 | 6.08954 | 6.21996 | 6.14289 | 5.98498 | 5.91784 | 5.73446 | 5.89356 | 6.45219 | 6.43293 | 6.3968 | 6.80941 |
| PROCR | Q9UNN8 | 6.32283 | 6.2895 | 5.91987 | 6.38545 | 6.4135 | 6.33739 | 6.45727 | 6.13752 | 5.80289 | 6.27171 | 6.41019 | 6.16349 | 6.22166 | 6.20718 | 6.27309 | 6.38645 |
| PLEKHF2 | Q9H8W4 | 6.33439 | 6.70776 | 6.96485 | 6.32281 | 6.22074 | 6.52871 | 5.79802 | 6.60985 | 6.53686 | 6.24262 | 6.18859 | 6.43946 | 5.86932 | 5.97554 | 5.59379 | 5.66669 |
| UGCG | Q16739 | 6.69507 | 6.50335 | 6.62399 | 6.81673 | 6.90618 | 6.49045 | 6.8904 | 6.91325 | 6.18754 | 5.58438 | 5.94375 | 6.11214 | 5.5534 | 5.27542 | 5.55353 | 5.95042 |
| ITGB3 | P05106 | 5.59961 | 6.64669 | 6.61347 | 6.76742 | 6.08894 | 6.18757 | 6.3656 | 5.38705 | 5.92964 | 6.26353 | 5.79288 | 6.06171 | 6.47482 | 6.0273 | 6.56049 | 6.23328 |
| PHF19 | Q5T6S3 | 6.35137 | 6.6823 | 6.63027 | 7.00233 | 6.29878 | 6.41597 | 6.53256 | 6.59778 | 6.46657 | 6.46442 | 6.68398 | 6.75777 | 5.12453 | 5.35251 | 5.20432 | 5.43455 |
| FBLIM1 | Q8WUP2 | 6.59874 | 6.09169 | 6.62642 | 5.66845 | 6.70983 | 6.77463 | 5.8949 | 6.82547 | 6.08116 | 6.20227 | 7.02891 | 7.1572 | 6.10602 | 5.41881 | 5.49555 | 5.31996 |
| SLIRP | Q9GZT3 | 5.93232 | 6.48908 | 5.58324 | 6.58283 | 6.71912 | 6.25742 | 6.09183 | 6.58706 | 6.18394 | 6.24374 | 6.02486 | 6.27629 | 6.22088 | 5.9413 | 6.51457 | 6.35153 |
| COPS7A | Q9UBW8 | 6.16789 | 6.15491 | 6.12118 | 5.83525 | 6.2778 | 6.59939 | 6.197 | 6.58523 | 6.03148 | 6.24979 | 5.90253 | 5.94935 | 6.58708 | 6.41778 | 6.48043 | 6.4429 |
| SNX3 | O60493 | 6.10605 | 6.53356 | 6.67635 | 5.69463 | 6.38679 | 6.33364 | 5.94496 | 6.57554 | 6.34594 | 6.81278 | 7.15223 | 6.93462 | 6.02795 | 5.89646 | 5.67349 | 5.80501 |
| MTHFS | P49914 | 5.82942 | 6.36811 | 6.63415 | 6.45284 | 6.00654 | 6.00631 | 6.16242 | 6.00191 | 6.4752 | 6.23625 | 6.40964 | 6.40442 | 6.26132 | 6.02857 | 6.33559 | 6.38731 |
| ESCO2 | Q56NI9 | 6.63746 | 6.15304 | 6.13208 | 6.16586 | 6.11784 | 5.6544 | 6.13907 | 5.43948 | 6.55941 | 6.12432 | 5.87361 | 5.59581 | 7.29001 | 6.27952 | 7.10174 | 6.73634 |
| WDFY3 | Q8IZQ1 | 6.3351 | 6.16672 | 6.74004 | 5.95506 | 6.24583 | 6.3996 | 6.43598 | 6.00928 | 6.29787 | 6.20138 | 6.41037 | 6.35584 | 6.04251 | 6.26772 | 6.04667 | 6.09004 |
| ECD | O95905-3 | 6.62705 | 6.50342 | 7.53363 | 6.85047 | 6.58175 | 6.7396 | 6.57332 | 6.69158 | 7.12518 | 6.8299 | 7.03199 | 7.33765 | 4.51864 | 4.16758 | 4.00864 | 4.8796 |
| RAD51AP1 | Q96B01 | 5.70472 | 5.84141 | 6.65676 | 5.80937 | 5.75926 | 6.0606 | 5.9371 | 6.06417 | 6.67653 | 6.56409 | 6.64165 | 6.744 | 5.98216 | 6.62069 | 6.28914 | 6.64836 |
| SLBP | Q14493 | 5.187 | 5.60643 | 5.91739 | 4.7783 | 5.62151 | 5.7392 | 5.16169 | 5.66325 | 6.19445 | 6.28023 | 6.1492 | 5.99829 | 7.97833 | 7.6947 | 8.21348 | 7.81654 |
| NDRG3 | Q9UGV2 | 6.77311 | 6.16989 | 6.93757 | 5.66086 | 6.54367 | 6.87366 | 6.21207 | 7.06998 | 6.27734 | 6.59409 | 6.32618 | 6.91941 | 6.163 | 5.56829 | 5.03365 | 4.87722 |
| BAP1 | Q92560 | 6.33563 | 6.25525 | 6.20931 | 6.40249 | 6.4657 | 6.62096 | 5.98315 | 5.91323 | 5.97966 | 6.30289 | 6.17671 | 5.91475 | 6.20139 | 6.44034 | 6.48621 | 6.21231 |
| MPDU1 | O75352 | 6.3533 | 6.18863 | 5.96234 | 6.65162 | 6.02135 | 5.78346 | 6.4751 | 6.13967 | 6.10894 | 6.12423 | 5.98346 | 5.82503 | 6.6129 | 6.29583 | 6.6308 | 6.84335 |
| ABCD1 | P33897 | 6.57594 | 6.43359 | 6.25705 | 6.34621 | 6.18476 | 5.99414 | 6.39349 | 5.84239 | 5.76614 | 6.19742 | 6.15013 | 6.05785 | 6.54924 | 6.30818 | 6.72048 | 6.21638 |
| TIMMDC1 | Q9NPL8 | 6.43495 | 6.69178 | 5.70943 | 6.69806 | 6.27442 | 6.36743 | 6.63573 | 6.04294 | 6.01778 | 5.92168 | 5.85754 | 5.80178 | 6.61758 | 6.41669 | 6.21029 | 6.30192 |
| DNAJC21 | Q5F1R6-3 | 6.67458 | 6.47571 | 7.11205 | 6.37096 | 6.16917 | 6.41284 | 6.32385 | 6.32769 | 6.61878 | 6.77658 | 6.7124 | 6.45216 | 4.965 | 5.77226 | 5.15772 | 5.67823 |
| BPHL | Q86WA6 | 5.68057 | 6.29313 | 5.68158 | 6.33269 | 6.31761 | 6.19882 | 6.17018 | 6.34296 | 6.24526 | 6.05256 | 5.73911 | 6.07311 | 7.2655 | 6.59311 | 6.23244 | 6.78137 |
| APOOL | Q6UXV4 | 6.60615 | 6.2718 | 5.84008 | 6.41782 | 6.43734 | 6.23636 | 6.7205 | 5.70433 | 6.29655 | 6.68513 | 6.12138 | 5.7472 | 6.62765 | 5.88502 | 6.2071 | 6.19559 |
| SERINC3 | Q13530 | 6.23059 | 6.46081 | 6.60552 | 6.73125 | 5.76283 | 5.83939 | 6.04626 | 6.19022 | 6.37935 | 6.37297 | 6.71012 | 6.66345 | 5.4434 | 6.1797 | 5.93459 | 6.44954 |
| ARFRP1 | Q13795 | 5.94411 | 5.97016 | 5.88626 | 6.6552 | 6.09527 | 6.41329 | 6.40317 | 5.98020 | 5.91694 | 6.26919 | 6.73258 | 6.10597 | 6.71256 | 6.2612 | 6.40511 | 6.42868 |
| PFKFB3 | Q16875-4 | 5.95077 | 5.608 | 7.06301 | 5.44056 | 5.56814 | 6.12427 | 5.83338 | 6.25975 | 6.19115 | 6.35978 | 6.73757 | 6.37333 | 6.52062 | 6.64376 | 6.91097 | 6.41492 |
| THEM4 | Q5T1C6 | 6.2871 | 6.21782 | 6.13129 | 6.75135 | 5.80795 | 5.67178 | 6.35286 | 5.94888 | 6.19223 | 5.91293 | 6.15075 | 5.91289 | 7.08803 | 6.20317 | 6.72217 | 6.6488 |
| TNFRSF10A | O00220 | 6.36638 | 6.82918 | 6.00143 | 6.37777 | 5.88588 | 6.11387 | 6.62148 | 6.53265 | 6.02452 | 5.93792 | 5.84911 | 6.20259 | 6.02476 | 6.27435 | 6.42287 | 6.53523 |
| UHRF1BP1 | Q6BDS2 | 6.67117 | 6.08932 | 7.18333 | 5.77956 | 6.35388 | 6.39321 | 6.29751 | 6.49771 | 6.66522 | 6.23036 | 6.54671 | 6.79726 | 5.85927 | 5.41859 | 5.50816 | 5.70872 |
| LCMT1 | Q9UIC8-2 | 6.37315 | 5.70441 | 6.84268 | 5.94191 | 6.53276 | 6.89679 | 6.62838 | 6.37382 | 6.31852 | 6.28931 | 7.00276 | 6.72318 | 5.76863 | 6.06965 | 5.34576 | 5.18828 |
| HAX1 | O00165 | 6.75731 | 6.33575 | 5.89255 | 6.35674 | 6.67043 | 5.88532 | 6.07537 | 6.19663 | 6.12399 | 6.26788 | 6.52139 | 5.98388 | 6.32838 | 6.01571 | 6.28277 | 6.3059 |
| SORBS3 | O60504 | 6.33997 | 6.09616 | 6.54304 | 5.84081 | 6.60256 | 6.80428 | 6.09379 | 6.76834 | 6.56189 | 6.60888 | 6.04457 | 6.25207 | 6.0138 | 5.96277 | 5.674 | 5.79305 |
| CASKIN2 | Q8WXE0 | 6.28778 | 5.71939 | 6.71195 | 5.71032 | 6.01726 | 6.50183 | 6.10172 | 6.35739 | 6.01194 | 6.75091 | 6.25289 | 6.9752 | 5.78308 | 6.85202 | 5.55655 | 6.40978 |
| KCTD10 | Q9H3F6 | 6.74596 | 6.72691 | 6.98696 | 6.72039 | 6.53216 | 6.6481 | 6.90258 | 6.47516 | 6.80744 | 6.35437 | 6.76046 | 6.37309 | 5.01431 | 5.08171 | 4.86056 | 5.00983 |
| CCDC97 | Q96F63 | 6.16655 | 6.38475 | 6.25051 | 6.18115 | 6.52755 | 6.49564 | 6.00993 | 6.44767 | 6.45031 | 6.34274 | 6.54244 | 6.65478 | 6.40389 | 5.5805 | 5.72611 | 5.8355 |
| CALU | O43852-3 | 6.14735 | 6.2788 | 5.40349 | 6.01432 | 6.55584 | 6.64251 | 6.015 | 6.40631 | 5.81265 | 6.05222 | 5.97313 | 6.17997 | 7.22962 | 6.84219 | 5.87555 | 6.57105 |
| HS2ST1 | Q7LGA3 | 5.88576 | 6.43631 | 5.69923 | 6.99635 | 6.12591 | 5.58606 | 5.52631 | 6.05994 | 6.05614 | 6.02078 | 6.65282 | 6.99058 | 7.46119 | 6.30537 | 6.30078 | 5.89647 |
| ANGEL1 | Q9UNK9 | 6.56697 | 6.71326 | 6.37847 | 6.84548 | 6.8139 | 6.29722 | 6.31534 | 6.2837 | 6.35353 | 6.43432 | 6.31944 | 5.91102 | 5.71255 | 5.65003 | 5.74035 | 5.66441 |
| ZRANB3 | Q5FWF4 | 6.01199 | 5.88828 | 6.25688 | 6.40196 | 6.47739 | 6.30848 | 6.69639 | 5.59347 | 6.34292 | 6.30826 | 6.45018 | 6.28499 | 5.89906 | 6.01289 | 6.62796 | 6.43891 |
| F11R | Q9Y624 | 6.23932 | 6.4882 | 6.62162 | 6.71037 | 6.20534 | 6.27816 | 6.11292 | 6.37478 | 5.9086 | 6.13899 | 6.30464 | 6.01086 | 5.88747 | 6.51077 | 6.43325 | 5.7747 |
| CDC45 | O75419-3 | 6.07309 | 5.80605 | 6.31569 | 5.86475 | 6.01043 | 6.23556 | 5.96644 | 6.19137 | 6.48279 | 6.3169 | 6.41821 | 6.49696 | 6.07535 | 6.57612 | 6.53643 | 6.63386 |
| PIGU | Q9H490 | 6.54211 | 5.96958 | 5.64636 | 6.63555 | 6.36891 | 6.03566 | 6.65881 | 5.97082 | 6.25354 | 6.15781 | 6.00751 | 6.12298 | 6.47645 | 6.21786 | 6.70835 | 6.22772 |
| UQCRH | P07919 | 6.81433 | 6.20069 | 5.44644 | 5.65284 | 5.21028 | 4.69114 | 5.47694 | 5.84689 | 5.13991 | 6.07666 | 5.42733 | 6.17295 | 6.6282 | 8.77817 | 7.46472 | 8.97249 |
| ZNF24 | P17028 | 6.03746 | 6.21687 | 6.39994 | 6.61738 | 5.70613 | 6.11981 | 6.17049 | 6.34863 | 5.85338 | 6.25608 | 6.63491 | 6.11569 | 6.50641 | 6.56302 | 6.24676 | 6.20704 |
| MECR | Q9BV79 | 6.40986 | 6.04047 | 5.70841 | 6.31494 | 6.51979 | 5.81349 | 5.75522 | 6.54953 | 6.11702 | 6.06917 | 6.35236 | 6.03781 | 7.31628 | 6.79072 | 6.20789 | 5.99703 |
| GUK1 | Q16774-2 | 6.11498 | 5.89813 | 5.9935 | 6.36424 | 6.21343 | 6.33755 | 5.8975 | 6.55254 | 6.29717 | 6.73464 | 6.45712 | 6.48303 | 6.36559 | 6.2834 | 5.73126 | 6.27593 |
| CADPS2 | Q86UW7 | 6.4803 | 6.11261 | 7.12322 | 6.2924 | 6.30855 | 6.5048 | 6.34373 | 6.52831 | 6.53554 | 6.39414 | 6.66321 | 6.86501 | 5.53012 | 5.37251 | 5.43829 | 5.50726 |
| GPX4 | P36969 | 6.5648 | 6.1143 | 6.43159 | 5.76938 | 6.74246 | 6.38944 | 6.24809 | 6.70322 | 6.47112 | 6.35654 | 6.23298 | 5.99219 | 6.36119 | 5.91592 | 6.09365 | 5.61314 |
| IRAK1 | P51617 | 6.69084 | 6.04992 | 7.04123 | 6.03886 | 6.0538 | 6.46258 | 5.80055 | 6.66588 | 6.12527 | 6.83168 | 6.96894 | 7.46143 | 5.41106 | 5.63843 | 5.31764 | 5.44191 |
| PRPF38A | Q8NAV1 | 5.89532 | 5.95625 | 6.24061 | 7.04247 | 5.79066 | 6.02266 | 6.07142 | 5.54033 | 6.27443 | 5.99568 | 6.23257 | 6.14111 | 6.92067 | 6.09708 | 7.02833 | 6.75044 |
| DDX59 | Q5T1V6 | 5.64262 | 5.60509 | 6.22693 | 5.40989 | 5.66326 | 6.12296 | 5.68085 | 6.18645 | 6.16385 | 5.94117 | 6.25723 | 6.49575 | 6.63022 | 7.51965 | 7.43361 | 7.02046 |
| ALDH3B1 | P43353 | 6.21553 | 6.68476 | 6.12641 | 6.56119 | 6.29954 | 5.95107 | 6.15201 | 6.16636 | 6.64613 | 6.06737 | 6.37357 | 6.43245 | 5.94165 | 6.26866 | 5.85873 | 6.25457 |
| RPS26 | P62854 | 6.52786 | 6.57261 | 6.59715 | 6.04216 | 6.47344 | 6.58404 | 6.27149 | 6.44995 | 6.44804 | 6.56634 | 6.17528 | 6.43048 | 5.2769 | 6.25637 | 5.63002 | 5.69788 |
| STX16 | O14662 | 6.6377 | 6.48987 | 5.94161 | 6.58575 | 6.44648 | 6.28309 | 6.45409 | 6.17506 | 5.96002 | 6.43442 | 6.31719 | 6.22981 | 6.00842 | 6.08929 | 5.9885 | 5.9587 |
| RPS6KA4 | O75676 | 6.02746 | 6.1038 | 5.95052 | 5.65708 | 5.99305 | 6.34831 | 5.95055 | 5.99076 | 6.13178 | 6.28786 | 6.73768 | 6.04791 | 6.41825 | 6.92153 | 6.72972 | 6.70372 |
| ARL8A | Q96BM9 | 6.18563 | 6.57293 | 6.27846 | 6.37495 | 5.92694 | 5.89571 | 6.11586 | 6.01059 | 6.18449 | 6.43295 | 6.01096 | 6.34165 | 6.20745 | 6.53433 | 6.36952 | 6.55757 |
| BABAM1 | Q9NWV8 | 6.42534 | 6.23583 | 6.47587 | 5.83198 | 6.57837 | 6.54123 | 6.06145 | 6.7831 | 6.56151 | 6.5649 | 6.50603 | 6.41141 | 6.4731 | 5.44164 | 5.60916 | 5.49908 |
| NAGA | P17050 | 6.59549 | 6.46857 | 6.46281 | 5.71549 | 6.30942 | 6.31328 | 6.285 | 6.71386 | 6.05272 | 6.1714 | 6.40131 | 5.98997 | 6.22292 | 5.82103 | 6.24996 | 6.22677 |
| POMT2 | Q9UKY4 | 6.25938 | 6.57181 | 5.644 | 6.45262 | 6.48499 | 6.00517 | 6.1319 | 5.78429 | 6.46542 | 6.09762 | 6.01256 | 5.80215 | 6.56484 | 6.47713 | 6.94852 | 6.2976 |
| ATRIP | Q8WXE1 | 6.13983 | 6.10849 | 6.7525 | 6.23 | 6.23978 | 5.94371 | 5.9259 | 5.73787 | 6.25505 | 6.4044 | 6.50058 | 6.13232 | 6.3456 | 6.44814 | 6.3402 | 6.49562 |
| MRPS18A | Q9NVS2-2 | 6.14807 | 6.12719 | 5.77883 | 6.58663 | 6.01697 | 5.84693 | 6.38969 | 6.13369 | 6.43311 | 6.1963 | 6.11738 | 6.05205 | 6.59025 | 6.4434 | 6.74533 | 6.39421 |
| SASH1 | O94885 | 6.15466 | 6.85406 | 6.80016 | 6.28666 | 6.51143 | 6.64246 | 6.37042 | 6.48009 | 6.43024 | 6.4329 | 6.60574 | 6.25494 | 5.54883 | 5.59988 | 5.41441 | 5.61314 |
| SMPDL3A | Q92484 | 6.44095 | 6.18507 | 5.77177 | 5.92656 | 6.95318 | 6.30684 | 6.17646 | 6.24212 | 6.15961 | 6.22332 | 6.1921 | 5.81077 | 6.89916 | 6.27208 | 6.48059 | 5.95943 |
| VEZT | Q9HBM0 | 5.85449 | 6.46776 | 5.8799 | 6.60859 | 5.96411 | 6.20783 | 6.2802 | 5.91053 | 5.7904 | 6.45147 | 6.48993 | 6.30852 | 6.48334 | 6.18136 | 6.70461 | 6.41693 |
| POMT1 | Q9Y6A1 | 6.64875 | 6.50548 | 6.77667 | 7.04013 | 6.35169 | 5.92587 | 6.3923 | 6.61775 | 6.61188 | 6.23059 | 6.28644 | 6.0979 | 5.59293 | 5.13303 | 6.17288 | 5.61571 |
| PCNP | Q8WW12 | 5.69055 | 6.28532 | 6.03994 | 5.763 | 5.74645 | 6.72318 | 5.42253 | 6.80848 | 6.05869 | 6.51974 | 6.40457 | 6.41226 | 6.11822 | 7.19685 | 6.239 | 6.57121 |
| WDR47 | O94967-4 | 6.32524 | 6.459 | 6.60979 | 5.82412 | 6.65347 | 6.74963 | 6.67272 | 6.33918 | 6.23615 | 6.06978 | 6.54931 | 6.14448 | 5.88595 | 5.76643 | 6.00003 | 5.71471 |
| ADAM22 | Q9P0K1 | 6.08884 | 7.05403 | 5.94506 | 6.31821 | 5.96761 | 6.17946 | 6.2418 | 5.73216 | 5.65545 | 6.95089 | 6.35852 | 6.02763 | 6.30777 | 6.49888 | 6.46156 | 6.21215 |
| LZTR1 | Q8N653 | 5.82404 | 6.51273 | 6.64412 | 6.54405 | 5.81332 | 6.81069 | 6.48505 | 6.33108 | 6.43485 | 6.78031 | 6.97884 | 6.67679 | 5.21373 | 5.95591 | 5.73888 | 5.25561 |
| SENP5 | Q96HI0 | 6.50813 | 6.26829 | 6.27154 | 6.89083 | 6.43481 | 5.92028 | 6.448 | 5.87463 | 5.80543 | 6.39632 | 6.13688 | 6.0022 | 6.29757 | 6.40835 | 6.14065 | 6.19608 |
| MRPL50 | Q8N5N7 | 5.29701 | 5.19443 | 4.48845 | 5.78175 | 5.93035 | 4.27473 | 5.2667 | 6.11507 | 4.26157 | 5.50655 | 4.65705 | 4.78333 | 4.60875 | 12.4156 | 9.24572 | 12.173 |
| USP22 | Q9UPT9 | 6.3965 | 6.38268 | 6.97107 | 6.47016 | 6.19437 | 6.32176 | 6.09978 | 6.49579 | 6.48018 | 7.21488 | 7.27321 | 7.06005 | 4.73484 | 5.45949 | 5.07606 | 5.3692 |
| TCP11L1 | Q9NUJ3 | 6.33841 | 5.32059 | 6.345 | 5.26123 | 6.01192 | 5.9586 | 6.10782 | 7.03175 | 5.685 | 6.28549 | 6.30883 | 7.35425 | 6.07029 | 6.81087 | 6.46736 | 6.64259 |
| SAMD4A | Q9UPU9 | 6.67481 | 6.58384 | 7.27001 | 6.20578 | 6.05274 | 6.65701 | 6.30042 | 5.99325 | 6.18121 | 6.29593 | 6.56895 | 6.69396 | 5.61948 | 5.7425 | 5.68947 | 5.47065 |
| PTGR2 | Q8N8N7 | 6.22214 | 5.9381 | 6.81807 | 5.49647 | 6.61043 | 6.88338 | 6.25317 | 6.84006 | 6.29866 | 6.52505 | 6.60332 | 6.40472 | 5.76818 | 5.93577 | 5.50357 | 5.89891 |

| | | | | | | | | | | | | | | | | | |
|---|---|---|---|---|---|---|---|---|---|---|---|---|---|---|---|---|---|
| DNAL1 | Q4LDG9 | 6.33661 | 5.72619 | 7.71569 | 5.4198 | 6.41663 | 6.12178 | 6.00904 | 6.91029 | 5.88773 | 6.65245 | 6.76293 | 7.14707 | 5.68774 | 5.89839 | 5.56918 | 5.73847 |
| MPZL1 | O95297 | 6.20739 | 6.43177 | 5.88714 | 6.67371 | 6.03541 | 6.04049 | 5.83534 | 5.88467 | 6.48882 | 5.95988 | 5.95727 | 5.94216 | 6.71935 | 6.3472 | 6.723 | 6.86639 |
| ALKBH5 | Q6P6C2-1 | 5.92303 | 6.38344 | 6.13398 | 5.94513 | 5.85335 | 6.43782 | 6.16549 | 5.58311 | 5.9877 | 6.40685 | 6.50119 | 6.19408 | 6.4909 | 6.76523 | 6.87491 | 6.35381 |
| KAT14 | Q9H8E8 | 6.26711 | 6.30218 | 5.95562 | 6.65067 | 6.04785 | 5.8455 | 5.94724 | 5.93207 | 6.46169 | 5.79903 | 6.43365 | 5.77303 | 6.89183 | 6.62014 | 6.6631 | 6.40931 |
| HAUS8 | Q9BT25 | 6.47265 | 6.72475 | 7.0368 | 6.76685 | 6.02416 | 6.77415 | 6.20523 | 6.58163 | 6.33813 | 6.65398 | 6.84083 | 6.9774 | 5.00562 | 5.35595 | 4.86064 | 5.38121 |
| TPRN | Q4KMQ1 | 6.56757 | 6.75647 | 6.98201 | 7.03431 | 5.90601 | 5.98661 | 6.20104 | 6.36772 | 6.31905 | 6.22963 | 6.84299 | 6.72952 | 5.53433 | 5.42874 | 5.47002 | 5.64399 |
| ATG5 | Q9H1Y0 | 6.5769 | 5.62069 | 6.74602 | 5.319 | 6.4455 | 6.16059 | 5.94027 | 6.91351 | 6.04184 | 6.77652 | 6.74124 | 7.08368 | 6.02598 | 6.12415 | 5.79349 | 5.69062 |
| MAPK7 | Q13164 | 6.27692 | 6.05538 | 7.08349 | 6.36544 | 6.65739 | 7.03826 | 6.37641 | 6.0703 | 6.09093 | 6.08385 | 6.6072 | 6.54764 | 5.9865 | 5.99979 | 5.43603 | 5.32448 |
| GGA2 | Q9UJY4 | 6.87276 | 6.57119 | 7.17139 | 6.57282 | 6.76916 | 6.67928 | 6.35984 | 6.58978 | 6.23965 | 6.62223 | 6.92123 | 6.32645 | 5.34504 | 5.04259 | 5.14906 | 4.76754 |
| GGA3 | Q9NZ52 | 6.58743 | 6.06136 | 7.01299 | 5.94114 | 6.41834 | 7.18258 | 6.15974 | 6.5876 | 6.31436 | 6.72771 | 6.597 | 6.59693 | 5.6001 | 5.85129 | 4.89139 | 5.47004 |
| GGA1 | Q9UJY5 | 6.77795 | 6.67986 | 6.49639 | 6.67294 | 6.83172 | 7.29311 | 5.9436 | 6.51385 | 5.98348 | 6.35815 | 7.08913 | 6.11128 | 5.32351 | 5.62911 | 5.2479 | 5.04802 |
| C2orf69 | Q8N8R5 | 6.52596 | 6.11521 | 6.78046 | 6.83086 | 6.34709 | 6.59629 | 6.24874 | 6.4633 | 6.38995 | 6.72562 | 6.34928 | 6.51325 | 5.68032 | 5.42063 | 5.40607 | 5.60697 |
| MTERF4 | Q7Z6M4 | 6.5762 | 5.99728 | 6.5077 | 6.45098 | 6.55951 | 6.38665 | 6.97538 | 5.95895 | 6.39815 | 6.09412 | 6.05964 | 5.51115 | 6.12047 | 6.14894 | 6.04343 | 6.21146 |
| INPP5K | Q9BT40 | 6.18638 | 6.33423 | 5.95967 | 6.78702 | 6.25802 | 6.06499 | 6.26662 | 6.06079 | 6.15617 | 6.17873 | 6.05031 | 5.8098 | 6.31963 | 6.05961 | 6.93227 | 6.57577 |
| COX5B | P10606 | 6.25945 | 4.99256 | 5.27074 | 4.78553 | 5.34834 | 4.20218 | 4.77271 | 6.18738 | 4.19357 | 5.27531 | 4.25802 | 5.49637 | 5.64034 | 11.6503 | 9.59802 | 12.0691 |
| HMG20B | Q9P0W2 | 6.63812 | 6.59417 | 6.23986 | 7.08093 | 6.21519 | 6.42709 | 6.41972 | 6.20327 | 5.90775 | 6.73658 | 6.32624 | 6.61417 | 5.7844 | 6.08003 | 5.20331 | 5.52916 |
| MTFMT | Q96DP5 | 6.76396 | 6.64545 | 5.78632 | 6.54841 | 6.19518 | 5.52965 | 6.12162 | 5.89792 | 6.64662 | 6.0609 | 5.93881 | 5.93684 | 6.50113 | 6.75998 | 6.14481 | 6.52241 |
| PAWR | Q96IZ0 | 6.31257 | 6.04418 | 6.84813 | 5.65017 | 6.18656 | 6.40459 | 5.993 | 6.33536 | 6.23773 | 6.68623 | 6.66991 | 6.56485 | 6.19109 | 5.88128 | 5.83558 | 6.15877 |
| ZBTB11 | O95625 | 6.09153 | 6.33088 | 6.40246 | 7.25444 | 6.02049 | 5.8656 | 6.54527 | 5.75623 | 6.27807 | 6.47795 | 6.46533 | 6.50331 | 5.86909 | 5.65271 | 6.13621 | 6.35043 |
| RBSN | Q9H1K0 | 6.74784 | 6.58468 | 6.58708 | 6.08774 | 6.41346 | 6.39788 | 6.1918 | 5.95858 | 6.28317 | 6.19516 | 6.19254 | 6.7973 | 6.87104 | 5.81106 | 5.83528 | 6.04537 |
| MSRA | Q9UJ68 | 6.30697 | 6.09698 | 6.29055 | 5.96143 | 6.65329 | 6.53115 | 5.99008 | 6.7443 | 6.21968 | 6.02916 | 6.59031 | 6.31164 | 6.31983 | 6.23946 | 5.9031 | 5.81205 |
| SLC25A32 | Q9H2D1 | 6.0783 | 6.1631 | 5.74956 | 6.81948 | 6.4393 | 6.17121 | 6.86543 | 6.00736 | 6.25879 | 6.05045 | 6.05429 | 6.051 | 6.39278 | 6.03688 | 6.25348 | 6.60857 |
| CAV1 | Q03135 | 6.3733 | 6.92291 | 5.91474 | 6.86155 | 5.9901 | 6.27299 | 6.236 | 6.25008 | 6.0088 | 5.95839 | 5.70105 | 6.09879 | 6.24359 | 6.62074 | 6.27435 | 6.27263 |
| TRAFD1 | O14545 | 6.22767 | 6.46292 | 6.48805 | 6.41828 | 6.48252 | 6.57374 | 6.5297 | 6.24114 | 7.24528 | 5.69772 | 6.41041 | 7.1806 | 5.00713 | 5.7081 | 5.22182 | 6.10492 |
| MYCBP | Q99417 | 6.11857 | 4.75403 | 5.86479 | 5.13369 | 4.92364 | 3.77311 | 4.6303 | 5.93437 | 4.0757 | 5.67745 | 4.81849 | 5.21984 | 4.75476 | 12.8275 | 9.69684 | 11.797 |
| OFD1 | O75665 | 6.69413 | 6.83767 | 6.90555 | 6.43244 | 6.57232 | 6.58226 | 6.40375 | 7.03435 | 6.47864 | 6.67585 | 6.56528 | 6.65868 | 5.29773 | 5.17791 | 4.7289 | 4.95453 |
| BAG5 | Q9UL15-2 | 6.15204 | 6.08877 | 6.54515 | 6.03682 | 6.37936 | 6.1673 | 6.18666 | 6.03954 | 6.50512 | 6.27054 | 6.66448 | 6.05719 | 6.41202 | 5.96064 | 6.51433 | 6.02006 |
| TRMT11 | Q7Z4G4 | 6.26665 | 6.34921 | 6.46917 | 6.32704 | 6.14467 | 6.40736 | 6.20008 | 6.49594 | 6.47704 | 6.32226 | 6.29493 | 6.58832 | 6.12689 | 5.69919 | 5.89907 | 5.93221 |
| ZXDC | Q2QGD7 | 5.71714 | 6.13033 | 6.57531 | 6.27377 | 5.90955 | 5.76328 | 6.38175 | 6.28521 | 6.10413 | 6.35936 | 6.34878 | 6.14249 | 6.42307 | 6.5078 | 6.48854 | 6.58948 |
| TIMM21 | Q9BVV7 | 6.36526 | 6.51274 | 5.86557 | 6.70314 | 5.92874 | 5.80952 | 6.20135 | 6.19342 | 6.14209 | 6.18681 | 5.94346 | 6.20072 | 6.69591 | 6.18813 | 6.65345 | 6.4097 |
| SKP1 | P63208 | 6.45551 | 5.25947 | 5.57652 | 4.86749 | 5.31566 | 5.10301 | 5.20028 | 6.33209 | 5.54328 | 6.30259 | 5.46492 | 6.57105 | 6.76813 | 8.81878 | 7.73135 | 8.68986 |
| NCALD | P61601 | 6.0815 | 5.08149 | 5.94212 | 4.91816 | 5.9343 | 5.24837 | 5.64393 | 6.36507 | 5.55285 | 6.61748 | 5.9659 | 6.98161 | 6.17371 | 7.98656 | 7.71077 | 7.79618 |
| HMGCR | P04035-3 | 5.08973 | 5.11989 | 5.11965 | 5.48366 | 7.10401 | 6.93701 | 7.68378 | 7.1615 | 4.38486 | 4.77727 | 4.81038 | 4.60464 | 8.2064 | 7.6229 | 7.71135 | 8.18296 |
| HMGA1 | P17096-2 | 5.66025 | 5.77583 | 5.25867 | 6.27383 | 6.90427 | 6.64315 | 6.24464 | 6.29885 | 6.07331 | 5.54198 | 6.56832 | 5.54743 | 6.88898 | 7.64845 | 6.72794 | 5.94409 |
| MYBL2 | P10244 | 6.11944 | 6.12485 | 6.33679 | 6.39158 | 6.34793 | 5.89919 | 6.02546 | 5.88427 | 6.1548 | 6.52622 | 6.55684 | 6.12201 | 6.8267 | 6.15333 | 6.21137 | 6.31921 |
| FUT8 | Q9BYC5 | 6.54503 | 6.40178 | 5.66646 | 6.91958 | 6.57787 | 6.17063 | 6.53914 | 6.13719 | 5.97297 | 6.10448 | 5.77042 | 5.81026 | 6.0888 | 6.27535 | 6.43805 | 6.582 |
| RBFOX2 | O43251-8 | 6.49287 | 6.35251 | 6.13336 | 6.33163 | 6.20396 | 5.91096 | 5.9953 | 6.28328 | 6.27881 | 5.90103 | 6.0012 | 6.68952 | 6.49653 | 6.07058 | 6.20223 | 6.65622 |
| NUBP1 | P53384 | 6.01176 | 6.29446 | 7.14912 | 5.37787 | 6.09082 | 6.40389 | 6.38027 | 6.54357 | 6.36039 | 6.40021 | 6.62871 | 6.79898 | 6.30446 | 5.80965 | 5.9573 | 5.48854 |
| ELP4 | Q96EB1-2 | 6.35613 | 6.05387 | 6.56577 | 6.45788 | 6.54049 | 6.99169 | 6.12989 | 6.89975 | 6.52995 | 6.58772 | 6.75056 | 6.5074 | 6.50151 | 5.97397 | 5.102 | 5.05143 |
| ABHD12 | Q8N2K0-2 | 6.57684 | 6.3795 | 6.03004 | 6.77939 | 6.30315 | 6.12653 | 6.0784 | 5.9386 | 5.90612 | 6.38576 | 6.07377 | 5.83267 | 6.52228 | 6.09931 | 6.57211 | 6.39553 |
| DZIP3 | Q86Y13 | 5.95858 | 5.7158 | 6.81053 | 5.7278 | 6.35513 | 6.93868 | 6.93441 | 6.27637 | 6.11649 | 6.32807 | 6.92763 | 6.79641 | 5.79787 | 5.80691 | 5.60714 | 5.80219 |
| MTX3 | Q5HYI7 | 6.19843 | 5.59943 | 5.89089 | 5.58398 | 6.21565 | 5.22561 | 6.18351 | 6.38592 | 5.72043 | 6.40114 | 6.15327 | 6.49816 | 6.77567 | 6.91298 | 7.16185 | 6.99307 |
| MSI2 | A0A6Q8PF05 | 6.24377 | 6.07544 | 5.88888 | 5.89782 | 6.42673 | 6.59258 | 6.22037 | 6.69804 | 6.32459 | 6.54374 | 6.45506 | 6.6836 | 6.23788 | 6.15819 | 5.79147 | 5.76184 |
| SNX17 | Q15036 | 6.72494 | 6.80523 | 6.88157 | 6.04204 | 6.47252 | 6.9864 | 6.2731 | 6.15005 | 7.07824 | 6.92367 | 7.1011 | 6.71198 | 4.82646 | 5.19654 | 5.01973 | 4.80642 |
| SH2B1 | Q9NRF2 | 6.41488 | 6.47193 | 6.77452 | 6.19514 | 6.27771 | 6.2184 | 6.0682 | 6.43127 | 6.56191 | 6.67825 | 6.27126 | 6.60452 | 6.11286 | 6.05568 | 5.25683 | 5.60666 |
| EXO1 | Q9UQ84 | 6.3887 | 6.30038 | 6.54526 | 6.07657 | 5.85014 | 5.90173 | 5.89061 | 5.60374 | 7.25183 | 6.07717 | 6.66117 | 6.45672 | 6.47792 | 6.11862 | 6.1604 | 6.23905 |
| WDR20 | Q8TBZ3-7 | 6.45419 | 6.70488 | 6.33279 | 6.37353 | 6.30601 | 6.49683 | 6.36706 | 6.82363 | 6.36614 | 6.43371 | 6.21734 | 6.93518 | 5.77133 | 5.79992 | 5.94743 | 5.67003 |
| TICAM2 | Q86XR7-2 | 6.53887 | 6.61645 | 6.07961 | 6.82506 | 6.59438 | 5.84888 | 6.12648 | 6.1762 | 6.35974 | 5.59402 | 6.06279 | 5.82108 | 6.32844 | 6.03668 | 6.77645 | 6.11489 |
| POU2F1 | P14859-6 | 5.55238 | 5.5944 | 6.12294 | 5.63047 | 5.56252 | 6.36471 | 5.73561 | 5.65905 | 6.0946 | 6.11429 | 7.05673 | 6.89162 | 7.33043 | 6.83292 | 6.89875 | 6.55859 |
| PIH1D1 | Q9NWS0 | 5.63816 | 4.95883 | 5.77838 | 4.72743 | 5.70215 | 4.92693 | 5.64337 | 7.56493 | 5.41416 | 6.83724 | 6.19998 | 7.59342 | 5.87831 | 8.23097 | 7.22399 | 7.68176 |
| RMC1 | Q96DM3 | 6.20194 | 6.69724 | 6.52182 | 6.62094 | 6.1941 | 6.45998 | 6.18924 | 6.38775 | 6.46233 | 6.59162 | 6.05892 | 6.35451 | 5.77453 | 5.64594 | 6.20071 | 5.63844 |
| CCDC137 | Q6PK04 | 6.28303 | 6.492 | 5.86707 | 6.93672 | 6.01953 | 5.81602 | 6.22336 | 6.11441 | 6.22682 | 6.05496 | 6.43427 | 6.14653 | 6.29856 | 6.31786 | 6.54072 | 6.22815 |
| XRCC3 | O43542 | 5.82963 | 5.8099 | 7.06102 | 5.92166 | 6.19572 | 6.22162 | 6.12497 | 5.9188 | 6.10322 | 6.31257 | 7.00525 | 6.92813 | 6.03453 | 6.52546 | 5.8887 | 6.11882 |
| HIF1AN | Q9NWT6 | 6.27917 | 6.32078 | 6.52588 | 6.11435 | 6.0661 | 6.42003 | 6.38538 | 6.72624 | 6.26327 | 6.57331 | 6.6988 | 6.91134 | 5.61431 | 5.97008 | 5.5653 | 5.56566 |
| MRPL58 | Q14197 | 6.32543 | 6.2186 | 5.44487 | 6.83627 | 6.60698 | 6.37846 | 6.14432 | 5.87096 | 6.31173 | 6.37629 | 5.94717 | 6.07893 | 6.60022 | 6.15977 | 6.34042 | 6.35958 |
| SRP19 | P09132 | 5.89632 | 6.07748 | 6.42298 | 6.12651 | 5.97838 | 5.95228 | 6.15807 | 5.96933 | 6.10626 | 5.8821 | 6.57905 | 6.35474 | 6.48537 | 6.76207 | 6.55609 | 6.69296 |
| METTL15 | A6NJ78 | 6.21426 | 6.45393 | 6.02913 | 6.80506 | 6.20913 | 6.0164 | 6.38779 | 6.24695 | 6.14384 | 6.30474 | 5.86717 | 6.31078 | 6.67242 | 5.98717 | 5.99046 | 6.36077 |
| SLC2A1 | P11166 | 6.05742 | 6.94313 | 6.09578 | 6.57186 | 6.40098 | 6.01497 | 6.65461 | 5.94952 | 5.90745 | 5.9336 | 6.01102 | 5.97803 | 6.6785 | 6.26733 | 5.92005 | 6.61576 |
| HPDL | Q96IR7 | 6.42128 | 5.20552 | 5.67427 | 5.7829 | 6.29707 | 5.30581 | 6.20639 | 6.23868 | 5.84822 | 6.17012 | 5.92315 | 6.95277 | 6.86733 | 7.1665 | 6.72392 | 7.21607 |
| FAM160A2 | Q8N612-2 | 6.59769 | 6.50576 | 6.92709 | 6.04958 | 6.28756 | 6.4118 | 5.59676 | 6.23399 | 6.52854 | 6.64029 | 7.03888 | 6.69471 | 5.49862 | 5.9936 | 5.52529 | 5.46983 |
| FAF1 | Q9UNN5 | 5.84966 | 6.11906 | 6.31001 | 5.75681 | 6.32937 | 6.5579 | 6.16451 | 6.36471 | 6.48285 | 6.14241 | 6.41702 | 6.41164 | 6.25591 | 6.17274 | 6.33697 | 6.32842 |
| LONP2 | Q86WA8 | 6.11234 | 6.39367 | 5.69089 | 6.37721 | 6.47493 | 6.23027 | 6.57642 | 6.22603 | 6.13671 | 6.00185 | 5.91245 | 5.98777 | 6.71334 | 6.10972 | 6.52689 | 6.52951 |
| INPP5F | Q9Y2H2 | 6.12108 | 6.23953 | 6.78716 | 6.07526 | 6.38524 | 6.52954 | 6.03204 | 6.16443 | 6.22964 | 6.55939 | 6.30673 | 6.32354 | 6.03846 | 5.8437 | 6.03944 | 6.32479 |
| FBRS | J3KNZ9 | 6.10729 | 6.89673 | 6.26011 | 6.73816 | 6.32585 | 6.75376 | 6.67498 | 6.27651 | 6.26951 | 6.46655 | 5.93171 | 6.33874 | 5.59429 | 6.22599 | 5.66877 | 5.47104 |
| WAC | Q9BTA9 | 6.35818 | 6.37309 | 6.32392 | 6.73949 | 6.15001 | 6.38073 | 6.02445 | 5.96012 | 6.21699 | 6.11198 | 6.53262 | 5.96013 | 6.62429 | 5.99207 | 6.01565 | 6.23627 |
| SLC25A10 | Q9UBX3-2 | 6.03199 | 6.40155 | 6.1938 | 6.60249 | 5.8873 | 6.09341 | 6.38971 | 5.63188 | 5.88121 | 6.09021 | 6.01321 | 6.20021 | 6.84672 | 6.41038 | 6.56719 | 6.75875 |
| NDUFS7 | O75251 | 6.4323 | 6.58966 | 5.45187 | 6.18266 | 6.4028 | 6.24589 | 6.3932 | 6.09148 | 6.19959 | 5.63576 | 6.16224 | 5.48177 | 7.11993 | 6.1735 | 7.04142 | 6.39594 |
| CLPTM1L | Q96KA5 | 6.60277 | 6.67818 | 6.43178 | 6.75765 | 6.13057 | 6.23144 | 6.08204 | 5.67827 | 5.96653 | 6.46009 | 6.03169 | 5.96701 | 6.3351 | 6.33058 | 6.256 | 6.0603 |
| LRRC57 | Q8N9N7 | 5.80007 | 5.82376 | 6.26226 | 5.68454 | 5.93624 | 6.056 | 6.196 | 6.50001 | 6.5332 | 6.31115 | 6.44951 | 6.86273 | 6.51785 | 6.39004 | 6.35562 | 6.32104 |
| ZCCHC9 | Q8N567 | 5.90779 | 6.33594 | 5.9769 | 6.4946 | 6.22575 | 6.10902 | 6.16675 | 5.78683 | 6.13448 | 6.45857 | 6.37232 | 6.31817 | 6.15649 | 6.82031 | 6.62234 | 6.11374 |
| TRIM41 | Q8WV44 | 6.23881 | 6.35766 | 6.57537 | 6.80023 | 6.41319 | 6.39375 | 6.0665 | 5.95656 | 6.56795 | 6.63996 | 6.7154 | 6.15416 | 5.79244 | 5.88284 | 5.71207 | 5.7331 |
| CKMT1A | P12532 | 6.25153 | 6.70188 | 5.82839 | 6.36378 | 6.53261 | 6.07264 | 6.11292 | 5.92651 | 6.19926 | 6.27141 | 6.23153 | 5.95904 | 6.38165 | 6.12591 | 6.52787 | 6.51309 |
| PPP6C | O00743-3 | 6.01319 | 5.74002 | 6.63741 | 5.74549 | 6.34972 | 6.761 | 6.34653 | 6.49259 | 6.48216 | 6.41267 | 6.37996 | 6.72746 | 5.99382 | 6.08976 | 5.98854 | 5.83968 |
| MRPL21 | Q7Z2W9 | 6.34395 | 6.35601 | 6.12443 | 6.58826 | 6.55757 | 6.00464 | 6.78266 | 6.218 | 6.26905 | 5.80835 | 5.8953 | 5.5594 | 6.38333 | 6.09229 | 6.61051 | 6.40626 |
| PARP12 | Q9H0J9 | 6.91714 | 6.45303 | 6.63732 | 6.84683 | 6.39193 | 6.1313 | 6.48561 | 6.18956 | 6.59487 | 6.45201 | 6.40993 | 6.34047 | 5.65231 | 5.44579 | 5.52994 | 5.52195 |
| COX11 | Q9Y6N1 | 6.61694 | 6.05864 | 6.19336 | 6.55095 | 6.47335 | 6.22197 | 6.39231 | 5.42052 | 6.01515 | 6.19508 | 5.75982 | 6.00984 | 6.63613 | 6.49066 | 6.56328 | 6.40199 |
| TERF1 | P54274 | 6.21091 | 6.2186 | 6.12194 | 6.69429 | 5.96654 | 5.91031 | 6.35807 | 5.94203 | 6.25281 | 6.3647 | 6.08512 | 6.10124 | 6.53451 | 6.34113 | 6.46224 | 6.43557 |
| SAP130 | Q9H0E3-3 | 6.33225 | 6.49624 | 6.35157 | 6.53558 | 6.21697 | 6.25856 | 6.09769 | 6.04522 | 6.21787 | 6.30276 | 6.25562 | 5.9634 | 6.36388 | 6.04225 | 6.2628 | 6.25733 |
| SERINC1 | Q9NRX5 | 6.33369 | 6.6342 | 6.2475 | 6.28329 | 5.83032 | 5.97343 | 6.08172 | 6.11764 | 6.33345 | 6.72693 | 6.15074 | 6.22726 | 6.12604 | 6.46405 | 6.40732 | 6.06239 |
| SLC44A1 | Q8WWI5 | 6.49623 | 6.6188 | 6.08757 | 6.46598 | 6.04354 | 6.26049 | 6.14393 | 6.00552 | 6.41484 | 6.58266 | 6.32044 | 5.87309 | 6.96548 | 6.30441 | 6.16875 | 6.24827 |
| AP4B1 | Q9Y6B7 | 6.61787 | 6.35267 | 7.30854 | 6.45903 | 6.12018 | 6.51841 | 6.1638 | 6.32002 | 6.64475 | 6.69647 | 6.68079 | 6.7135 | 5.44705 | 5.26014 | 5.35548 | 5.34131 |
| C1orf52 | Q8N6N3 | 6.11344 | 5.73029 | 6.55793 | 6.24337 | 6.35342 | 5.66873 | 5.64223 | 6.21103 | 6.05775 | 6.36312 | 7.3138 | 6.39694 | 6.63761 | 5.93994 | 6.44399 | 6.32639 |
| CEP55 | Q53EZ4 | 6.71596 | 6.83572 | 7.08386 | 7.01181 | 6.39704 | 7.14571 | 6.3495 | 6.9511 | 6.65875 | 7.18321 | 7.22179 | 7.15904 | 4.54621 | 4.51492 | 3.88179 | 4.3436 |
| SEMA3C | Q99985 | 6.96988 | 6.47196 | 4.33687 | 6.80294 | 5.78395 | 5.66948 | 7.4331 | 5.95955 | 6.56977 | 5.95727 | 5.27767 | 6.67069 | 7.30033 | 5.75558 | 6.44132 | 6.59964 |
| GCSH | P23434 | 7.2083 | 6.56597 | 5.53712 | 6.8352 | 6.18909 | 5.62983 | 5.74429 | 6.21937 | 5.98461 | 6.33982 | 5.81619 | 6.09942 | 6.66345 | 6.61976 | 6.06493 | 6.48265 |
| RRM2B | Q7LG56-6 | 6.52916 | 6.17486 | 6.48316 | 5.89813 | 6.48434 | 6.63797 | 5.93913 | 6.64252 | 6.2798 | 6.24909 | 6.33455 | 6.767 | 6.02318 | 6.06774 | 5.79389 | 5.69549 |
| C2CD2 | Q9Y426 | 6.07823 | 6.29257 | 6.1283 | 6.19529 | 6.1784 | 6.34548 | 6.80735 | 5.86617 | 5.86192 | 6.02652 | 6.21489 | 5.83891 | 6.59586 | 6.30554 | 6.77452 | 6.49006 |
| BCR | P11274 | 6.37241 | 6.17496 | 6.6668 | 6.00234 | 6.43241 | 6.52178 | 6.18192 | 6.17459 | 6.49731 | 6.42582 | 6.59993 | 6.48439 | 6.08001 | 6.02566 | 5.65037 | 5.70929 |
| C8orf33 | Q9H7E9-2 | 6.1451 | 6.26351 | 6.67947 | 6.40691 | 6.30422 | 6.68759 | 5.93244 | 6.49874 | 6.54904 | 6.14938 | 7.0494 | 6.63612 | 5.6939 | 5.6427 | 5.60803 | 5.75347 |
| PAXX | Q9BUH6 | 6.23742 | 5.96482 | 6.59311 | 5.44776 | 6.53073 | 6.5242 | 5.83024 | 6.71959 | 6.16382 | 6.85937 | 6.29011 | 6.56192 | 6.0593 | 6.11943 | 6.09032 | 6.00785 |
| NEK6 | Q9HC98-2 | 6.37147 | 5.96045 | 7.1287 | 6.50131 | 6.19913 | 7.09394 | 6.34594 | 6.26862 | 6.41362 | 6.83036 | 6.83487 | 6.64971 | 5.36108 | 5.61359 | 5.00093 | 5.42627 |
| GTF2H3 | Q13889 | 6.22808 | 6.35235 | 6.11003 | 6.20087 | 6.13883 | 6.19491 | 6.21564 | 6.15206 | 6.21311 | 6.09742 | 6.24736 | 6.16964 | 6.40203 | 6.10262 | 6.65545 | 6.51959 |
| PHF10 | Q8WUB8 | 6.39957 | 6.50316 | 6.02224 | 6.51316 | 5.85697 | 5.92511 | 6.33841 | 6.03379 | 6.30647 | 6.06428 | 5.9499 | 5.86373 | 6.66603 | 6.38374 | 6.50022 | 6.67323 |
| C9orf40 | Q8IXQ3 | 6.01654 | 5.46646 | 6.32017 | 4.94296 | 5.48254 | 5.14077 | 5.49442 | 6.84069 | 4.95912 | 6.87846 | 6.09582 | 7.35427 | 6.35186 | 7.89159 | 7.62026 | 7.14405 |
| CNOT2 | Q9NZN8 | 6.59925 | 6.14892 | 6.71956 | 6.04854 | 6.47555 | 6.16952 | 5.84385 | 6.72152 | 6.57922 | 6.58368 | 6.45366 | 6.94623 | 6.10758 | 5.67164 | 5.37398 | 5.55728 |
| RC3H2 | Q9HBD1 | 6.53313 | 6.88266 | 7.0526 | 6.26422 | 5.92949 | 6.29544 | 6.65213 | 6.15433 | 6.81401 | 6.28056 | 7.38203 | 6.32223 | 4.93374 | 5.47884 | 5.60711 | 5.41746 |
| TARBP2 | Q15633 | 6.58951 | 6.62664 | 6.89722 | 5.74884 | 6.60391 | 6.63599 | 6.36206 | 6.3653 | 6.75391 | 6.11673 | 6.60408 | 6.39228 | 6.06572 | 5.76328 | 5.17498 | 5.29956 |
| GALNT4 | Q8N4A0 | 6.65669 | 6.51754 | 6.5266 | 6.29848 | 6.53307 | 5.22679 | 6.45216 | 5.75054 | 6.73786 | 6.07752 | 6.04709 | 5.66975 | 6.3413 | 6.02456 | 6.72964 | 6.41042 |
| NBEA | Q8NFP9 | 6.27624 | 6.46694 | 6.71251 | 6.04357 | 5.98731 | 6.72649 | 5.95622 | 5.93091 | 6.28688 | 6.26237 | 6.58528 | 6.50221 | 6.13988 | 6.2306 | 6.05748 | 5.83512 |
| ALDH2 | P05091 | 6.50142 | 6.1142 | 5.64998 | 6.28032 | 6.63742 | 5.83691 | 6.36285 | 6.17254 | 6.06113 | 5.91314 | 5.93874 | 5.93664 | 6.94001 | 6.41532 | 6.70539 | 6.53399 |
| STK39 | Q9UEW8 | 6.26476 | 6.43482 | 7.02119 | 6.13407 | 6.42215 | 6.50278 | 5.89106 | 6.49263 | 6.1953 | 6.62744 | 6.47717 | 6.37057 | 5.64178 | 5.56342 | 5.92567 | 6.03519 |
| NAPRT | Q6XQN6 | 6.81646 | 6.01768 | 6.89487 | 5.13339 | 6.33836 | 6.68708 | 6.21791 | 6.81101 | 5.90704 | 6.1467 | 6.48014 | 6.23448 | 5.84003 | 6.82246 | 5.93922 | 5.71316 |
| CDK4 | P11802 | 6.45733 | 5.91598 | 7.02252 | 5.91424 | 6.60175 | 6.8321 | 6.37395 | 6.74366 | 6.45664 | 6.56806 | 6.78455 | 6.80353 | 5.49903 | 5.56606 | 5.19815 | 5.26244 |
| ZNF593 | O00488 | 7.24048 | 7.44668 | 7.17147 | 7.58367 | 6.6109 | 6.10578 | 6.34046 | 6.7046 | 6.96834 | 6.51019 | 6.23247 | 7.00751 | 4.05136 | 4.89828 | 4.6009 | 4.5269 |
| SAP30 | O75446 | 6.65432 | 6.86484 | 6.37946 | 7.09192 | 6.36823 | 6.29482 | 6.25766 | 6.09222 | 6.3361 | 6.20336 | 6.26004 | 6.15258 | 5.70844 | 5.62585 | 5.89106 | 5.8191 |

| | | | | | | | | | | | | | | | | | |
|---|---|---|---|---|---|---|---|---|---|---|---|---|---|---|---|---|---|
| CYB5R4 | Q7L1T6 | 6.60876 | 5.82589 | 7.14215 | 5.56979 | 6.01806 | 6.67615 | 6.40674 | 6.17598 | 6.27541 | 6.83691 | 6.56911 | 6.88542 | 5.63195 | 6.08501 | 5.5245 | 5.76818 |
| RAB24 | Q969Q5 | 6.59333 | 6.59959 | 6.22728 | 6.69012 | 6.04259 | 6.49361 | 6.88312 | 5.93424 | 6.35987 | 6.43544 | 6.28986 | 6.08908 | 5.75109 | 5.43779 | 6.31293 | 6.1259 |
| PRXL2A | Q9BRX8 | 6.36174 | 6.24698 | 5.89876 | 6.47645 | 6.25761 | 6.29963 | 6.33412 | 5.93929 | 6.31062 | 5.91527 | 6.01504 | 5.52045 | 7.01171 | 6.44689 | 6.62729 | 6.33816 |
| PAK5 | Q9P286 | 6.28702 | 6.52366 | 6.79669 | 6.33391 | 6.28494 | 6.55569 | 6.10059 | 6.09231 | 6.39421 | 6.63174 | 6.34381 | 6.57231 | 5.85759 | 5.98776 | 5.54923 | 5.68854 |
| RANBP17 | Q9H2T7 | 5.47472 | 5.06289 | 6.24473 | 5.54719 | 5.83151 | 6.36811 | 6.08629 | 5.9675 | 6.17123 | 6.22043 | 6.17556 | 6.25124 | 6.77762 | 7.26782 | 7.28924 | 7.26392 |
| ATF7 | P17544-1 | 6.41632 | 6.48674 | 6.05877 | 6.11241 | 6.34294 | 5.9652 | 6.30196 | 5.97998 | 6.67254 | 6.36482 | 6.33902 | 5.95548 | 6.73071 | 5.94908 | 6.09529 | 6.22874 |
| VANGL1 | Q8TAA9 | 6.90677 | 7.16911 | 6.49329 | 7.2785 | 5.96915 | 6.00198 | 6.38311 | 5.75009 | 6.41131 | 6.61301 | 6.56663 | 6.66537 | 5.4951 | 5.41494 | 5.52922 | 5.35242 |
| STK26 | Q9P289 | 6.36411 | 6.04903 | 6.76806 | 5.89651 | 6.47263 | 6.13695 | 6.2208 | 6.32108 | 6.22623 | 6.31657 | 6.53122 | 6.43622 | 6.44946 | 5.93385 | 5.83773 | 6.03956 |
| ANK3 | Q12955 | 5.92647 | 6.27696 | 6.39522 | 5.89433 | 6.11653 | 6.37837 | 6.09047 | 6.18686 | 6.06303 | 6.1836 | 6.394 | 6.40034 | 6.63999 | 6.28112 | 6.76905 | 6.00366 |
| ZNF592 | Q92610 | 6.45617 | 6.32851 | 6.122 | 6.91005 | 5.9383 | 5.93926 | 6.42815 | 5.84441 | 6.11886 | 6.24507 | 6.28586 | 6.04763 | 6.51699 | 6.17849 | 6.47068 | 6.16957 |
| NUS1 | Q96E22 | 6.66492 | 6.62705 | 6.21959 | 6.68172 | 6.45043 | 6.30219 | 5.89559 | 5.90811 | 6.19207 | 6.51973 | 6.44488 | 6.1018 | 5.55074 | 5.96423 | 6.24956 | 6.2274 |
| CAPN5 | O15484 | 6.45498 | 6.4124 | 6.55264 | 6.02327 | 6.21722 | 5.78189 | 6.5186 | 6.21739 | 5.79589 | 6.59106 | 6.24434 | 6.17783 | 6.28721 | 6.36381 | 6.06095 | 6.30052 |
| JADE3 | Q92613 | 6.07099 | 6.6149 | 6.39132 | 6.85295 | 6.46778 | 6.36887 | 6.33398 | 5.72874 | 6.20982 | 5.88768 | 6.08968 | 6.10877 | 6.37842 | 5.93534 | 6.40294 | 6.15782 |
| HNRNPC | B4DY08 | 6.08493 | 5.76176 | 6.00748 | 6.55282 | 6.36088 | 5.46808 | 7.22456 | 6.03602 | 6.48835 | 4.93284 | 5.88425 | 5.73825 | 6.42121 | 7.16198 | 6.81777 | 7.05882 |
| ZNF48 | Q96MX3 | 6.4767 | 6.44679 | 6.3275 | 6.73227 | 5.91966 | 6.09143 | 6.24242 | 5.70289 | 5.99076 | 6.15277 | 6.47495 | 6.29354 | 6.24052 | 6.28867 | 6.06572 | 6.55343 |
| ETS1 | P14921 | 5.66994 | 5.08662 | 6.68507 | 6.22962 | 6.01864 | 5.9659 | 5.70696 | 6.12045 | 6.17641 | 5.74637 | 6.05538 | 6.04527 | 6.90416 | 7.61394 | 7.47846 | 6.49683 |
| BTRC | Q9Y297 | 6.77385 | 6.49347 | 7.2768 | 6.95991 | 6.39314 | 6.29437 | 7.01905 | 6.08688 | 6.58538 | 6.59684 | 6.09515 | 5.96174 | 5.17682 | 5.24942 | 5.68465 | 5.35251 |
| NOL7 | Q9UMY1 | 6.52382 | 6.69565 | 5.95479 | 7.304 | 5.85154 | 5.67756 | 6.30906 | 5.98543 | 6.87352 | 5.98658 | 5.97585 | 6.33453 | 6.26966 | 5.87889 | 5.72922 | 6.64989 |
| DNAJC14 | Q6Y2X3 | 5.94984 | 7.06483 | 5.58927 | 7.02342 | 6.17086 | 5.68151 | 6.01478 | 5.79023 | 6.17682 | 6.39431 | 6.35271 | 6.14907 | 6.5142 | 6.16371 | 6.56943 | 6.395 |
| ARGLU1 | Q9NWB6 | 6.13586 | 6.01684 | 6.03301 | 6.29645 | 6.06138 | 5.54281 | 6.21637 | 5.71579 | 6.08554 | 6.04964 | 6.20473 | 6.05446 | 6.62697 | 6.75706 | 7.31469 | 6.8884 |
| MACF1 | H3BQK9 | 5.78537 | 6.46847 | 5.93375 | 6.52228 | 5.9788 | 6.72123 | 5.95968 | 6.43697 | 5.64397 | 6.24822 | 6.21099 | 6.45006 | 6.20345 | 6.80812 | 6.41002 | 6.21862 |
| SETD5 | Q9C0A6 | 6.86594 | 6.69151 | 6.09741 | 7.59007 | 5.9433 | 5.77056 | 6.49706 | 5.81206 | 6.43005 | 6.65235 | 6.52056 | 6.32131 | 5.39561 | 5.46902 | 5.89858 | 6.04461 |
| ANKIB1 | Q9P2G1 | 6.78553 | 6.32775 | 7.52282 | 6.27436 | 5.71936 | 6.34251 | 5.63188 | 5.63819 | 6.69905 | 6.90986 | 6.73011 | 6.92175 | 5.35434 | 5.96379 | 5.84468 | 5.33403 |
| MOB4 | Q9Y3A3 | 5.93703 | 6.46563 | 6.30928 | 5.97765 | 6.17721 | 6.49778 | 5.90672 | 6.2617 | 6.14606 | 6.42802 | 6.23764 | 6.1799 | 6.33262 | 6.44887 | 6.34171 | 6.35218 |
| PALD1 | Q9ULE6 | 6.3759 | 6.51538 | 6.35571 | 6.38401 | 6.62999 | 6.13842 | 6.30747 | 6.36914 | 6.5924 | 6.30799 | 6.08848 | 6.40422 | 6.35194 | 5.90487 | 5.59936 | 5.67475 |
| WNK3 | Q9BYP7 | 6.06469 | 6.1982 | 6.75097 | 5.94217 | 6.19201 | 6.84291 | 6.02211 | 6.5562 | 6.4618 | 6.55885 | 6.73055 | 6.60325 | 5.64326 | 6.13733 | 5.77239 | 5.52332 |
| BOLA2 | Q9H3K6 | 6.09945 | 5.70614 | 6.69117 | 5.26663 | 6.42682 | 6.86375 | 5.97208 | 6.71684 | 6.12526 | 6.69292 | 6.28195 | 6.70607 | 5.95842 | 6.50685 | 5.94138 | 6.04427 |
| SERPINB9 | P50453 | 6.21872 | 6.07472 | 6.73119 | 5.94129 | 5.95538 | 6.71398 | 6.34828 | 6.30138 | 6.17824 | 6.69506 | 6.80798 | 6.3912 | 5.87435 | 6.1263 | 5.72227 | 5.91965 |
| ITPRIP | Q8IWB1 | 6.60761 | 6.73798 | 6.09021 | 7.10192 | 6.46137 | 6.05769 | 6.2095 | 6.09497 | 6.12406 | 6.18799 | 6.04796 | 6.0572 | 6.12144 | 5.95428 | 6.09038 | 6.05543 |
| CD46 | P15529-2 | 6.30231 | 6.63058 | 5.85128 | 6.47843 | 6.018 | 5.96403 | 6.43705 | 5.85246 | 6.26906 | 5.99405 | 5.84412 | 6.15219 | 6.41473 | 6.58962 | 6.55035 | 6.65173 |
| FAM98C | Q17RN3 | 5.95925 | 5.72839 | 6.91385 | 6.38755 | 6.08899 | 6.57613 | 6.21246 | 6.60992 | 6.82434 | 6.80387 | 7.08221 | 6.3802 | 5.46648 | 5.70987 | 5.29443 | 5.96208 |
| TLK1 | Q9UKI8 | 5.55913 | 6.55962 | 6.28085 | 6.30485 | 5.68786 | 6.46927 | 5.74994 | 6.43877 | 5.80182 | 6.3918 | 5.8574 | 6.11484 | 6.48434 | 6.69814 | 6.72087 | 6.88051 |
| ERH | P84090 | 5.70669 | 4.80231 | 4.31955 | 4.81364 | 4.45134 | 3.77871 | 4.55823 | 4.96359 | 3.5941 | 4.74932 | 3.99464 | 4.63671 | 4.73752 | 16.087 | 9.39277 | 15.4139 |
| UNC119B | A6NIH7 | 5.87303 | 5.77343 | 6.53709 | 5.76562 | 6.47175 | 6.09791 | 5.94095 | 6.6739 | 6.68027 | 6.48747 | 6.74822 | 6.87578 | 6.00682 | 6.16291 | 5.88549 | 6.01936 |
| ASMTL | O95671 | 6.21604 | 5.76877 | 6.71001 | 5.71317 | 6.30334 | 6.5751 | 6.18714 | 6.4244 | 6.565 | 6.31474 | 6.66347 | 6.75387 | 6.17746 | 5.99578 | 5.81286 | 5.81885 |
| RAD51C | O43502 | 5.55526 | 5.92099 | 6.39824 | 5.66204 | 5.89766 | 6.03626 | 5.83948 | 6.41921 | 6.44357 | 6.37029 | 6.84925 | 7.02634 | 6.00782 | 6.3076 | 6.51264 | 6.75336 |
| TICRR | Q7Z2Z1 | 6.68057 | 6.99469 | 6.11895 | 7.25946 | 6.26956 | 5.92662 | 6.96345 | 6.45544 | 6.70774 | 6.25417 | 6.54749 | 6.31657 | 5.05243 | 5.34679 | 5.57497 | 5.53109 |
| CHPF2 | Q9P2E5 | 6.5063 | 6.89585 | 5.87834 | 6.63229 | 6.29747 | 5.84764 | 6.04253 | 6.0195 | 6.5532 | 6.72755 | 6.08618 | 5.87238 | 6.44295 | 6.39121 | 6.10151 | 5.7051 |
| NKAP | Q8N5F7 | 6.55073 | 5.84955 | 6.05595 | 6.33959 | 6.69055 | 5.97347 | 6.26272 | 6.03982 | 5.94433 | 6.06727 | 5.75909 | 5.68807 | 6.28335 | 6.87176 | 6.75578 | 6.86798 |
| HPS6 | Q86YV9 | 6.75439 | 6.53359 | 6.99714 | 6.42176 | 6.16448 | 6.63978 | 6.29701 | 6.705 | 6.41325 | 6.63364 | 6.6457 | 6.72961 | 5.33049 | 5.46237 | 4.92036 | 5.35143 |
| PKLR | P30613 | 6.57394 | 6.62694 | 6.62838 | 5.42124 | 6.46238 | 7.04591 | 6.25299 | 7.21384 | 5.86754 | 6.83644 | 5.72015 | 6.54451 | 5.60694 | 6.39727 | 5.32751 | 5.47403 |
| FAM120C | Q9NX05 | 6.38632 | 6.57805 | 6.68438 | 6.18379 | 6.19456 | 6.88316 | 5.83112 | 6.11308 | 6.51164 | 6.49632 | 6.79329 | 6.13737 | 5.84458 | 5.85041 | 5.85532 | 5.65658 |
| TBP | P20226 | 5.80901 | 5.95458 | 6.03726 | 6.2909 | 6.07355 | 6.18065 | 5.83493 | 5.95626 | 6.26665 | 6.31175 | 6.25608 | 6.16819 | 6.81458 | 6.17318 | 6.99622 | 6.8762 |
| IDI1 | Q13907-2 | 6.07328 | 5.75891 | 6.60985 | 5.57879 | 6.30842 | 6.56908 | 5.86448 | 6.5173 | 6.00266 | 6.86765 | 6.18138 | 6.95418 | 6.05521 | 6.16266 | 6.10042 | 6.39574 |
| PGAP1 | Q75T13 | 5.96878 | 6.47654 | 6.0697 | 6.24457 | 5.84764 | 6.37112 | 6.60438 | 6.21683 | 5.97838 | 6.18795 | 5.89901 | 5.92352 | 6.51365 | 6.40794 | 6.89632 | 6.39365 |
| AFF1 | P51825-2 | 7.87665 | 7.91383 | 7.3625 | 7.59533 | 6.07954 | 6.5296 | 6.71419 | 6.06994 | 7.1622 | 7.15807 | 7.50603 | 6.76501 | 3.17113 | 4.45993 | 3.58404 | 4.05203 |
| H2AB2 | P0C5Z0 | 5.90149 | 6.0811 | 5.60975 | 6.3913 | 6.1636 | 5.32795 | 6.72401 | 5.93572 | 6.13726 | 6.09983 | 6.22724 | 5.51259 | 6.63408 | 7.01772 | 7.0506 | 7.18574 |
| TLE1 | Q04724 | 5.85369 | 5.89929 | 6.40356 | 6.32713 | 5.84015 | 6.06438 | 6.00149 | 6.25043 | 5.87854 | 6.28666 | 6.06578 | 5.79322 | 7.17371 | 6.91339 | 6.88594 | 6.36265 |
| PKD2 | Q13563 | 6.53043 | 6.25056 | 6.37336 | 6.51479 | 6.25529 | 6.03457 | 6.39611 | 5.8781 | 6.0034 | 6.04078 | 5.94981 | 5.99342 | 6.33273 | 6.35377 | 6.88471 | 6.20816 |
| GATM | P50440-2 | 6.31027 | 6.69197 | 6.04923 | 6.58016 | 6.47091 | 6.30418 | 6.09795 | 6.02902 | 6.47082 | 5.6634 | 6.09846 | 6.29122 | 6.67883 | 5.83209 | 6.2955 | 6.13597 |
| PXK | Q7Z7A4 | 6.16877 | 6.15176 | 6.46182 | 5.73426 | 6.23793 | 6.72386 | 6.36512 | 6.45634 | 6.40068 | 6.5275 | 5.95082 | 6.08546 | 6.46795 | 6.52309 | 5.89543 | 5.84921 |
| NBR1 | Q14596 | 6.19637 | 5.85577 | 6.2152 | 6.37391 | 6.39747 | 6.06658 | 5.9448 | 6.64178 | 5.63491 | 6.48238 | 6.57876 | 6.49516 | 5.97501 | 6.32516 | 6.409 | 6.40776 |
| MYL6B | P14649 | 6.55637 | 5.84696 | 6.72857 | 5.36365 | 6.55676 | 6.18602 | 6.34781 | 7.01088 | 6.26788 | 6.66556 | 6.28214 | 6.75797 | 5.56028 | 5.92794 | 5.94679 | 5.8944 |
| DPY19L1 | A0A1B0GW05 | 6.20523 | 6.57684 | 6.19607 | 6.56839 | 5.90267 | 6.03337 | 6.05495 | 5.83157 | 6.07169 | 6.45889 | 5.95313 | 6.24854 | 6.48889 | 6.29293 | 6.54626 | 6.57057 |
| PCNX3 | Q9H6A9 | 6.55011 | 6.46884 | 6.37617 | 7.00603 | 5.99069 | 5.95206 | 6.39395 | 6.23273 | 6.59298 | 6.32096 | 6.076 | 6.41973 | 6.03781 | 5.69188 | 6.00273 | 5.88732 |
| TMX4 | Q9H1E5 | 6.1729 | 6.36063 | 6.03749 | 6.38116 | 6.03046 | 6.01354 | 6.63342 | 6.08979 | 6.17393 | 6.25458 | 6.10834 | 6.04014 | 6.76062 | 5.82031 | 6.49944 | 6.62325 |
| CCDC174 | Q6PII3 | 5.63148 | 6.17736 | 5.61663 | 5.69905 | 6.17971 | 6.02768 | 5.82591 | 5.67536 | 6.03224 | 6.15778 | 6.02339 | 6.14525 | 6.9982 | 7.41942 | 7.14674 | 7.24379 |
| CDS2 | O95674 | 6.03422 | 6.59375 | 5.85239 | 6.33816 | 6.06412 | 5.92297 | 6.07218 | 6.07293 | 5.90824 | 6.27758 | 5.80746 | 5.98891 | 6.40406 | 7.03956 | 6.87646 | 6.74701 |
| TFCP2 | Q12800 | 6.31301 | 6.55095 | 6.10509 | 5.79198 | 5.8826 | 6.65252 | 6.16246 | 6.2574 | 6.30207 | 6.29993 | 6.30298 | 5.77343 | 6.44591 | 5.5804 | 6.29619 | 6.2831 |
| ANKRD13A | Q8IZ07 | 6.43969 | 6.53052 | 6.41263 | 6.30254 | 6.72421 | 6.561 | 6.52805 | 6.28607 | 6.31976 | 6.27096 | 6.21024 | 5.87182 | 5.81903 | 6.03565 | 5.76666 | 5.92117 |
| NFIB | O00712-5 | 5.74786 | 6.16328 | 5.89655 | 6.3437 | 6.16068 | 6.15759 | 5.89413 | 5.91855 | 5.96893 | 6.25466 | 6.28363 | 5.89333 | 7.06297 | 6.82334 | 6.70884 | 6.72193 |
| ZKSCAN8 | Q15776 | 5.94108 | 6.53071 | 6.07162 | 6.61074 | 6.01974 | 6.6059 | 6.78214 | 5.93661 | 5.80403 | 6.53494 | 6.48182 | 6.22616 | 6.18259 | 5.81514 | 6.32324 | 6.13352 |
| ZNF268 | Q14587 | 5.90998 | 5.51674 | 6.21443 | 6.73681 | 6.17566 | 6.21062 | 6.27732 | 5.77166 | 6.30915 | 6.693 | 6.43496 | 6.37998 | 6.49261 | 6.19963 | 6.17121 | 6.50622 |
| PICK1 | Q9NRD5 | 6.90223 | 6.3658 | 6.74686 | 6.33709 | 6.16817 | 7.08928 | 6.19391 | 6.71124 | 6.54827 | 6.65129 | 6.70249 | 6.28136 | 5.38373 | 5.6977 | 4.87801 | 5.34255 |
| MPST | P25325-2 | 6.23613 | 5.55154 | 6.26233 | 5.44059 | 6.21547 | 5.87918 | 6.1956 | 6.7222 | 5.96566 | 7.14252 | 6.13343 | 6.80182 | 6.28688 | 6.4544 | 6.42099 | 6.29125 |
| NRF1 | Q16656-4 | 5.76988 | 5.89185 | 5.59233 | 6.08883 | 6.27363 | 5.90966 | 6.71226 | 5.86761 | 6.38524 | 5.98041 | 5.6229 | 6.1919 | 7.00716 | 6.61559 | 6.72422 | 7.36653 |
| TIAM1 | Q13009 | 6.06917 | 6.36575 | 6.42729 | 6.13639 | 6.08637 | 6.4907 | 6.43102 | 5.9339 | 6.14619 | 6.28229 | 6.29942 | 5.90735 | 6.5812 | 6.0878 | 6.27456 | 6.48062 |
| MTHFSD | Q2M296 | 6.21747 | 6.21649 | 6.46877 | 6.0923 | 6.31197 | 6.55014 | 6.43119 | 6.33364 | 6.38599 | 6.74234 | 6.00165 | 6.04145 | 6.28935 | 6.26769 | 5.85275 | 5.79681 |
| DYRK1A | Q13627 | 6.22189 | 6.75937 | 6.2888 | 5.96678 | 6.59075 | 6.26823 | 6.24805 | 6.39328 | 6.05412 | 6.2896 | 5.78687 | 6.12382 | 6.38167 | 6.35703 | 6.18551 | 6.08423 |
| RAB2A | P61019 | 6.29703 | 6.5456 | 5.81099 | 6.32855 | 6.42574 | 6.38481 | 6.34088 | 5.93682 | 6.08719 | 6.05734 | 5.97766 | 6.01357 | 6.68093 | 6.1918 | 6.73337 | 6.18772 |
| TSEN2 | Q8NCE0 | 5.58388 | 6.13411 | 6.19714 | 5.8745 | 5.56565 | 6.20082 | 5.96961 | 5.76325 | 6.22423 | 6.19378 | 5.62804 | 6.43481 | 7.0503 | 6.79323 | 7.02006 | 7.36659 |
| TUBA4A | P68366 | 6.46004 | 5.85309 | 7.11717 | 5.95882 | 6.52572 | 6.84784 | 6.00265 | 6.62008 | 6.62071 | 6.36572 | 6.9753 | 6.82445 | 5.41199 | 5.77171 | 5.06842 | 5.57631 |
| SUV39H1 | O43463-2 | 6.093 | 6.21233 | 5.65868 | 6.77272 | 5.64347 | 5.49099 | 6.31788 | 5.73494 | 6.19483 | 6.17254 | 5.68079 | 6.43925 | 6.86455 | 6.4533 | 7.26527 | 7.00548 |
| MED18 | Q9BUE0 | 5.70019 | 5.3292 | 5.8125 | 5.35086 | 5.9496 | 6.5872 | 5.89332 | 5.98549 | 5.50899 | 6.29661 | 6.12879 | 6.02978 | 6.81817 | 7.89552 | 7.13435 | 7.57944 |
| RAD54L2 | Q9Y4B4 | 5.93731 | 6.63563 | 5.75999 | 6.46809 | 5.45648 | 6.41044 | 5.88878 | 5.65319 | 5.861 | 6.74439 | 6.46944 | 6.23834 | 6.33092 | 6.13309 | 6.32376 | 6.68916 |
| ALAD | P13716-2 | 6.12874 | 5.50545 | 6.85 | 5.46297 | 6.44689 | 6.2654 | 6.0441 | 6.67084 | 6.19986 | 6.90962 | 6.55799 | 6.86973 | 5.81833 | 6.02588 | 6.32934 | 5.91487 |
| CHMP4A | Q9BY43-2 | 6.19547 | 6.1098 | 7.11286 | 6.1966 | 6.19138 | 7.11867 | 5.96065 | 6.53143 | 6.29089 | 6.2603 | 7.14401 | 6.96473 | 5.57235 | 5.96791 | 5.39681 | 4.98616 |
| RFLNA | Q6ZTI6 | 5.9922 | 6.02001 | 6.08557 | 5.97059 | 6.17778 | 6.01225 | 5.86232 | 6.61394 | 6.70451 | 6.26184 | 6.22528 | 6.63427 | 6.74787 | 6.24794 | 6.20396 | 6.23966 |
| TRAPPC4 | Q9Y296 | 6.55958 | 5.58943 | 6.68742 | 5.5988 | 6.5818 | 6.25281 | 6.38386 | 6.61535 | 6.39512 | 6.52449 | 6.41108 | 6.59242 | 6.11997 | 6.15359 | 5.72132 | 5.81295 |
| VDR | P11473-2 | 6.81797 | 6.76031 | 7.56364 | 6.74562 | 5.71891 | 6.60991 | 5.95053 | 6.09235 | 6.81634 | 7.04434 | 7.51878 | 6.92454 | 4.78838 | 5.44101 | 4.56202 | 4.64536 |
| RSRC1 | Q96IZ7 | 6.07886 | 6.45555 | 5.79524 | 6.47218 | 6.15423 | 6.09257 | 5.97822 | 5.75252 | 6.14599 | 5.98462 | 6.14594 | 5.91957 | 6.73566 | 6.50559 | 6.99007 | 6.79318 |
| OSGEPL1 | Q9H4B0 | 6.79932 | 6.87756 | 6.10968 | 7.00048 | 6.19391 | 5.94757 | 5.96796 | 5.91826 | 6.24581 | 5.89719 | 6.04899 | 6.09157 | 6.41828 | 6.06229 | 6.18348 | 6.23765 |
| CD59 | P13987-2 | 5.74012 | 5.32903 | 5.53032 | 5.1241 | 5.23683 | 5.44938 | 5.79465 | 6.54413 | 5.81182 | 6.19433 | 5.69714 | 6.38102 | 6.97547 | 8.22669 | 8.04759 | 7.91739 |
| HIVEP1 | P15822 | 6.70654 | 6.65388 | 6.41025 | 6.40977 | 6.46182 | 6.2394 | 6.08588 | 6.26607 | 6.23849 | 6.30577 | 5.92021 | 6.123 | 6.39334 | 5.88652 | 6.17361 | 5.72545 |
| NFAT5 | O94916-5 | 6.36433 | 5.79682 | 6.53267 | 6.20638 | 6.21722 | 5.94566 | 6.19265 | 6.07936 | 6.63541 | 6.46233 | 6.21434 | 6.43856 | 6.7022 | 5.75501 | 5.95241 | 6.50465 |
| HS1BP3 | Q53T59 | 6.25322 | 5.84387 | 6.41857 | 5.35989 | 6.2112 | 6.34612 | 5.67581 | 6.72098 | 6.50714 | 6.8512 | 6.76383 | 6.33747 | 6.34964 | 6.56884 | 5.82941 | 5.9628 |
| MMAB | Q96EY8 | 6.72696 | 6.19028 | 6.14074 | 6.55197 | 6.37881 | 6.33923 | 6.14672 | 5.98093 | 6.12456 | 5.86502 | 6.01852 | 5.86156 | 6.79424 | 6.17448 | 6.40416 | 6.30182 |
| AAK1 | A0A096LP25 | 6.07409 | 6.11585 | 6.54215 | 5.66935 | 6.28053 | 6.63994 | 6.0259 | 5.86495 | 6.23893 | 6.41081 | 6.28635 | 6.06613 | 6.86039 | 6.50329 | 6.20667 | 6.21468 |
| THBS3 | P49746 | 6.45377 | 6.63402 | 6.07612 | 6.46965 | 6.30518 | 6.19226 | 6.64585 | 6.23295 | 6.52942 | 6.65657 | 6.78797 | 6.01473 | 5.65365 | 5.89086 | 5.68325 | 5.77373 |
| RTEL1 | Q9NZ71-2 | 5.56391 | 5.90186 | 6.84069 | 6.49202 | 5.84447 | 6.18683 | 6.06008 | 6.12454 | 6.28246 | 6.39618 | 6.69345 | 6.79508 | 5.9856 | 6.37131 | 6.49515 | 5.96636 |
| HSD17B8 | Q92506 | 6.62482 | 6.3295 | 6.33171 | 6.51795 | 6.15149 | 5.89119 | 6.23067 | 5.79633 | 6.16085 | 6.50755 | 6.12456 | 5.99954 | 6.46636 | 6.03265 | 6.56731 | 6.26753 |
| GRAMD1B | Q3KR37-4 | 6.27198 | 6.1491 | 6.06111 | 6.65165 | 6.47873 | 6.38528 | 6.71889 | 6.22046 | 5.85223 | 6.07814 | 5.87291 | 5.77222 | 6.52986 | 6.39799 | 6.58225 | 6.57719 |
| UXT | Q9UBK9-2 | 5.94399 | 4.7304 | 6.10823 | 5.37315 | 5.39655 | 5.93645 | 5.62335 | 6.75393 | 5.35146 | 6.41451 | 5.89577 | 7.16444 | 5.68789 | 8.25342 | 7.29723 | 8.06924 |
| ADSS1 | Q8N142 | 6.32083 | 6.44423 | 6.38621 | 5.86048 | 6.46 | 6.69128 | 6.36441 | 6.67797 | 6.24975 | 6.53424 | 6.26265 | 5.843 | 5.89558 | 6.23936 | 5.94319 | 5.82683 |
| ABCB8 | Q9NUT2 | 6.23322 | 6.67754 | 6.10046 | 7.13772 | 6.29662 | 5.76101 | 6.78196 | 5.69067 | 6.64084 | 6.00166 | 5.99574 | 5.8045 | 6.27872 | 6.15556 | 6.69368 | 5.75011 |
| PPP1CA | P62136 | 6.24166 | 5.9855 | 6.04747 | 6.14446 | 6.34808 | 5.9989 | 6.22853 | 6.02392 | 6.05852 | 6.10813 | 6.16822 | 6.68961 | 6.71179 | 6.58045 | 6.58365 | 6.08111 |
| HOMER3 | Q9NSC5 | 6.02747 | 6.49487 | 6.78249 | 6.10708 | 6.19865 | 6.29592 | 5.58141 | 6.46637 | 6.47153 | 6.67562 | 6.93978 | 6.93179 | 6.00584 | 5.9875 | 5.49282 | 5.54087 |
| TTC7B | Q86TV6-2 | 6.47047 | 6.30659 | 6.24287 | 6.66623 | 6.31319 | 6.11925 | 6.24559 | 6.01479 | 6.05526 | 6.48207 | 5.98082 | 6.24544 | 6.32483 | 6.05061 | 6.4228 | 6.05919 |
| TTC14 | Q96N46 | 6.09694 | 7.03909 | 6.50339 | 6.99356 | 5.55905 | 6.60666 | 5.87093 | 6.12258 | 5.83734 | 6.16247 | 6.62735 | 6.09185 | 6.34439 | 5.94352 | 6.27831 | 5.92259 |
| OTUD6B | Q8N6M0 | 6.39885 | 6.49364 | 6.77654 | 6.40397 | 6.48821 | 6.30665 | 6.19595 | 6.90707 | 6.19321 | 6.35826 | 6.49733 | 6.10713 | 5.53285 | 5.93428 | 5.71809 | 5.68794 |
| NAA16 | Q6N069 | 6.2221 | 6.56861 | 6.35104 | 6.47166 | 5.97526 | 6.70854 | 5.92288 | 6.55537 | 6.17099 | 7.02028 | 6.39161 | 6.61239 | 5.82745 | 5.96656 | 5.5635 | 5.67179 |
| MOCS1 | Q9NZB8 | 6.3693 | 6.25633 | 6.1646 | 6.24516 | 6.40936 | 5.85701 | 6.35776 | 5.95262 | 6.34065 | 6.5805 | 6.53055 | 6.30048 | 6.12689 | 6.15611 | 6.37891 | 5.97378 |
| REV1 | Q9UBZ9 | 6.83114 | 6.36674 | 6.35225 | 6.52267 | 6.27991 | 6.14389 | 6.18164 | 5.95927 | 6.22612 | 5.90477 | 5.90253 | 6.22155 | 6.36794 | 5.7874 | 6.64746 | 6.30472 |
| TPST2 | O60704 | 6.4887 | 5.98569 | 6.10877 | 6.37463 | 6.17732 | 6.26885 | 6.44669 | 5.73771 | 6.55516 | 6.32282 | 6.0863 | 6.1671 | 6.52979 | 6.11109 | 5.95428 | 6.68511 |
| STON1 | Q9Y6Q2-3 | 5.789 | 5.86644 | 5.76136 | 5.38579 | 6.10406 | 6.2 | 5.86233 | 6.15622 | 6.20788 | 6.33115 | 6.02418 | 6.0949 | 6.74274 | 7.29264 | 7.34169 | 6.83962 |

| | | | | | | | | | | | | | | | | | |
|---|---|---|---|---|---|---|---|---|---|---|---|---|---|---|---|---|---|
| ACACB | O00763 | 6.54944 | 6.2232 | 6.88403 | 6.15051 | 6.69548 | 6.60492 | 6.34701 | 6.48426 | 6.67983 | 6.8302 | 6.60928 | 6.25828 | 5.39631 | 5.34817 | 5.47722 | 5.46186 |
| DNAJB4 | Q9UDY4 | 5.93497 | 6.31826 | 6.54987 | 6.09414 | 6.2317 | 6.29822 | 5.88746 | 6.31543 | 5.93729 | 6.07299 | 6.12086 | 6.35756 | 6.49008 | 6.2989 | 6.58581 | 6.50645 |
| CD9 | P21926 | 6.54529 | 6.54067 | 6.28297 | 5.96793 | 6.31451 | 6.38443 | 6.27254 | 5.75708 | 6.36644 | 6.19118 | 5.37511 | 6.07562 | 6.76171 | 6.30911 | 6.51241 | 6.34301 |
| TBC1D10A | Q9BXI6 | 6.20845 | 6.26562 | 5.92237 | 6.31231 | 6.01978 | 6.12842 | 6.24419 | 5.84092 | 6.44582 | 6.42806 | 6.20127 | 6.15058 | 6.26325 | 6.48894 | 6.36267 | 6.71733 |
| SLC25A25 | Q6KCM7-3 | 6.47245 | 6.20072 | 5.58948 | 6.35872 | 6.49409 | 6.10346 | 6.34559 | 6.14429 | 6.44615 | 5.99562 | 6.43993 | 5.83364 | 6.66109 | 6.03039 | 6.81651 | 6.06784 |
| DOP1A | Q5JWR5 | 6.66502 | 6.04017 | 7.12185 | 5.80508 | 6.1586 | 6.65476 | 6.37144 | 5.72248 | 6.58098 | 6.98094 | 6.47095 | 6.80377 | 5.83741 | 5.69383 | 5.57672 | 5.51599 |
| ZMYND11 | Q15326 | 6.15616 | 6.2971 | 5.95625 | 7.14822 | 6.0027 | 6.26937 | 6.65439 | 5.94548 | 6.15598 | 5.91344 | 6.14788 | 5.98645 | 6.19559 | 6.157 | 6.27899 | 6.735 |
| EPC1 | Q9H2F5 | 6.1547 | 6.60117 | 6.20564 | 6.88982 | 6.16177 | 5.85956 | 6.15544 | 5.98793 | 6.14207 | 6.37289 | 6.57814 | 6.60543 | 5.95711 | 6.30457 | 6.04003 | 5.98373 |
| PDCL3 | Q9H2J4 | 6.76287 | 6.50475 | 7.08329 | 5.77356 | 6.96961 | 6.779 | 6.0998 | 6.90626 | 6.60752 | 6.31366 | 6.1473 | 6.28128 | 6.05593 | 5.16213 | 5.22089 | 5.33216 |
| ZADH2 | Q8N4Q0 | 6.24668 | 6.76073 | 5.67894 | 6.5346 | 6.02764 | 6.15054 | 6.36204 | 6.05088 | 5.96684 | 6.25399 | 5.8005 | 6.08069 | 6.5519 | 6.11723 | 6.91153 | 6.50527 |
| EFR3B | Q9Y2G0 | 6.10634 | 6.58299 | 6.37393 | 6.99098 | 5.88991 | 6.11558 | 6.26907 | 5.84179 | 6.55957 | 6.99 | 6.50341 | 6.39839 | 5.85498 | 5.95163 | 5.60168 | 5.96976 |
| CTBP1 | Q13363 | 6.44334 | 5.72009 | 6.1651 | 5.43125 | 6.46031 | 5.95187 | 5.47272 | 6.13692 | 5.78129 | 6.16504 | 5.95362 | 6.83502 | 6.46527 | 6.81258 | 7.01301 | 7.19258 |
| TGFB2 | P61812-2 | 6.10108 | 6.35864 | 6.05388 | 6.02987 | 6.22506 | 6.28512 | 6.57142 | 6.33501 | 5.30597 | 5.56198 | 6.04047 | 5.61027 | 6.72179 | 7.14977 | 6.97153 | 6.67813 |
| FAM126A | Q9BYI3 | 6.33516 | 6.89849 | 6.56773 | 6.80716 | 5.97419 | 5.89954 | 6.36728 | 5.96667 | 6.25779 | 6.09218 | 6.23022 | 6.5736 | 6.18252 | 5.75256 | 5.90342 | 6.19148 |
| ABRAXAS1 | Q6UWZ7 | 6.61033 | 6.17834 | 5.9942 | 6.61858 | 6.1168 | 5.79194 | 6.39267 | 6.2119 | 6.1388 | 5.7988 | 6.41212 | 6.07407 | 6.62051 | 6.21474 | 6.42498 | 6.40122 |
| ZBTB21 | Q9ULJ3 | 5.85025 | 5.96738 | 5.99625 | 6.37747 | 6.06324 | 6.08311 | 6.33229 | 5.82827 | 5.849 | 6.00564 | 5.89629 | 6.23094 | 7.11974 | 6.34703 | 6.84967 | 7.20343 |
| TACO1 | Q9BSH4 | 6.43581 | 6.57069 | 6.38154 | 7.09326 | 6.08053 | 6.36286 | 6.66581 | 6.01775 | 6.23426 | 5.67961 | 5.54568 | 5.52605 | 6.04929 | 6.41157 | 6.35843 | 6.58686 |
| RRN3 | Q9NYV6 | 6.69211 | 6.61557 | 7.42732 | 7.55034 | 5.8864 | 6.17984 | 6.21011 | 6.28666 | 6.47492 | 6.40798 | 6.87465 | 6.85339 | 5.11378 | 4.65258 | 5.33333 | 5.44104 |
| EMC2 | Q15006 | 6.1256 | 6.88752 | 6.14475 | 7.12791 | 6.18011 | 6.05679 | 6.24763 | 5.66095 | 5.97454 | 6.55615 | 6.39299 | 6.27345 | 5.86957 | 6.07776 | 6.28579 | 6.1385 |
| MAU2 | Q9Y6X3 | 6.51187 | 5.97942 | 5.74462 | 6.77207 | 6.20523 | 6.25874 | 6.16338 | 5.91572 | 5.8752 | 6.10591 | 5.84669 | 6.63943 | 6.45133 | 6.553 | 6.60009 | 6.37731 |
| TRMT13 | Q9NUP7 | 5.56941 | 5.76601 | 6.58413 | 5.60854 | 5.78379 | 6.23113 | 5.55177 | 5.93257 | 6.08715 | 6.13748 | 6.25799 | 6.12487 | 6.69057 | 6.82641 | 7.62485 | 7.22334 |
| RNF149 | Q8NC42 | 6.82923 | 6.82193 | 6.72442 | 7.70939 | 6.18425 | 6.21097 | 6.14773 | 6.30346 | 6.6477 | 6.93827 | 6.97862 | 7.02416 | 4.77851 | 4.93808 | 4.80412 | 4.95916 |
| PSMF1 | Q92530 | 5.72275 | 5.73966 | 5.92225 | 4.37877 | 6.41766 | 6.92158 | 6.04958 | 6.71408 | 5.87768 | 7.12287 | 6.57126 | 7.70096 | 5.30467 | 6.48277 | 6.35986 | 6.71361 |
| NME3 | Q13232 | 6.51823 | 6.66631 | 6.14153 | 5.96506 | 6.06885 | 6.17658 | 6.16576 | 5.86319 | 6.45517 | 5.96289 | 5.95255 | 5.87107 | 6.61898 | 6.65147 | 6.4626 | 6.45977 |
| FANCB | Q8NB91 | 5.88521 | 6.15269 | 6.31956 | 6.37416 | 6.09867 | 6.17565 | 6.57075 | 6.85937 | 6.25889 | 6.19473 | 6.57849 | 6.28876 | 6.52929 | 6.10218 | 6.61441 | 6.65721 |
| HPS5 | Q9UPZ3 | 6.56095 | 6.37673 | 6.33958 | 6.4126 | 6.07564 | 6.06156 | 6.89631 | 6.65999 | 6.78882 | 6.31747 | 6.8101 | 6.027 | 4.98069 | 5.84251 | 6.03843 | 5.81163 |
| B4GALNT1 | Q00973 | 6.49799 | 6.4215 | 6.06328 | 6.57184 | 6.41898 | 5.74443 | 6.49412 | 5.95484 | 6.75319 | 6.7622 | 6.47608 | 6.37805 | 5.90891 | 5.74565 | 5.94292 | 5.86601 |
| MRPL14 | Q6P1L8 | 6.46188 | 6.50783 | 5.73701 | 6.54357 | 6.38543 | 5.91584 | 6.74369 | 5.71028 | 6.36107 | 6.17097 | 6.0977 | 6.28283 | 6.21835 | 5.8777 | 6.59318 | 6.39267 |
| SLC39A14 | Q15043 | 6.26566 | 6.58497 | 6.58659 | 6.1878 | 6.47339 | 6.6487 | 6.38669 | 6.24683 | 5.98823 | 6.19181 | 6.3608 | 6.3254 | 6.24509 | 5.76862 | 5.88925 | 5.85018 |
| TOE1 | Q96GM8 | 6.09165 | 6.0933 | 5.97727 | 6.2812 | 6.14275 | 5.92146 | 6.64699 | 5.76891 | 6.18001 | 6.50581 | 6.13347 | 6.16312 | 6.62743 | 6.34144 | 6.81522 | 6.30996 |
| APOE | P02649 | 6.39343 | 6.65276 | 6.45324 | 5.98108 | 6.65839 | 6.19984 | 6.90306 | 6.77887 | 6.21668 | 6.28726 | 6.16639 | 6.53585 | 5.7938 | 5.67913 | 5.51642 | 5.7838 |
| AJUBA | Q96IF1 | 6.24819 | 6.28705 | 7.29301 | 6.33647 | 6.23837 | 6.56179 | 6.48285 | 6.44096 | 6.26315 | 6.04497 | 6.62691 | 6.37256 | 5.86301 | 5.7437 | 5.63862 | 5.55837 |
| CSNK1D | P48730-2 | 6.96186 | 7.05502 | 6.97047 | 7.07365 | 6.40518 | 6.70973 | 5.87713 | 6.58783 | 6.47212 | 6.58371 | 6.64826 | 6.49617 | 5.29295 | 5.41315 | 4.82879 | 4.62398 |
| MCRS1 | Q96EZ8-2 | 6.27008 | 6.15172 | 6.10426 | 6.83672 | 5.7662 | 6.12463 | 6.35325 | 6.02646 | 6.38681 | 6.52026 | 6.40715 | 6.37437 | 6.30751 | 6.16204 | 6.22118 | 5.98736 |
| CHEK2 | O96017 | 5.3206 | 5.7525 | 5.59933 | 5.48473 | 5.45729 | 6.48007 | 5.53692 | 6.16718 | 6.04072 | 6.27512 | 6.77542 | 6.44322 | 6.8101 | 7.22531 | 7.32863 | 7.30285 |
| GSDME | O60443 | 6.58292 | 6.00779 | 7.77494 | 6.0458 | 6.10602 | 6.91721 | 6.37423 | 5.77359 | 6.57951 | 6.35229 | 6.02543 | 6.10767 | 5.52976 | 6.30083 | 5.83169 | 5.69033 |
| RASSF1 | Q9NS23-5 | 6.71245 | 6.71543 | 6.75771 | 6.61569 | 6.27691 | 6.28828 | 6.14248 | 6.28008 | 6.54578 | 6.17099 | 6.40681 | 6.53808 | 5.60083 | 5.42153 | 6.20899 | 5.31796 |
| CCM2 | Q9BSQ5-2 | 6.42513 | 6.39929 | 6.82809 | 6.28467 | 6.47941 | 6.5512 | 6.04599 | 6.35885 | 6.31269 | 6.7677 | 6.59352 | 6.63714 | 5.80764 | 5.55129 | 5.58703 | 5.34338 |
| AARSD1 | Q9BTE6-3 | 6.21473 | 6.16525 | 6.98129 | 6.31198 | 6.22302 | 6.72856 | 6.18538 | 6.59847 | 6.489 | 6.2831 | 6.5965 | 6.68904 | 5.79525 | 5.81066 | 5.43515 | 5.49262 |
| ZWINT | O95229 | 6.69838 | 5.21096 | 6.20348 | 4.95784 | 6.97881 | 5.63297 | 5.56922 | 7.96602 | 5.88704 | 6.93261 | 6.64313 | 7.45947 | 5.58322 | 6.13734 | 5.71832 | 6.42118 |
| MCUR1 | Q96AQ8 | 6.64461 | 6.20868 | 5.75947 | 6.47402 | 6.62292 | 5.98299 | 5.99394 | 6.42745 | 5.99524 | 5.99889 | 6.182 | 5.67442 | 6.64459 | 6.32314 | 6.73386 | 6.33377 |
| IL17RA | Q96F46 | 6.34016 | 6.73103 | 6.2928 | 6.58679 | 6.03893 | 5.92171 | 6.25421 | 5.98141 | 6.29776 | 6.45865 | 6.08486 | 6.21104 | 6.26733 | 6.17997 | 6.27771 | 6.07563 |
| MTRR | Q9UBK8-1 | 6.52191 | 6.10707 | 7.14157 | 5.99494 | 6.40933 | 6.69013 | 6.31899 | 6.79831 | 6.62754 | 6.32274 | 6.75752 | 6.73885 | 5.5055 | 5.34162 | 5.33002 | 5.39395 |
| RETREG3 | Q86VR2 | 6.36123 | 6.73971 | 6.30198 | 6.50665 | 5.73688 | 6.78701 | 6.07794 | 5.71466 | 5.9318 | 6.03969 | 6.03004 | 5.61891 | 6.37297 | 6.60579 | 6.87975 | 6.29501 |
| OTUD7B | Q6GQQ9 | 6.1248 | 6.23351 | 6.68043 | 6.14206 | 6.12562 | 6.07426 | 6.00025 | 6.40897 | 6.36633 | 6.41742 | 6.38868 | 6.45209 | 6.09102 | 6.30494 | 6.0593 | 6.13031 |
| PEG10 | Q86TG7 | 6.31289 | 6.44816 | 6.47676 | 6.21322 | 6.63451 | 6.49819 | 6.23433 | 6.40304 | 6.24083 | 6.32498 | 6.23985 | 6.50839 | 5.59199 | 5.88542 | 5.8485 | 6.13893 |
| RBM3 | P98179 | 7.34708 | 7.2697 | 6.42509 | 7.42004 | 7.00536 | 6.24381 | 6.13883 | 6.55187 | 6.52915 | 6.3303 | 6.57199 | 6.69368 | 5.42275 | 4.87164 | 4.22937 | 4.94936 |
| MRPS6 | P82932 | 5.84385 | 6.22761 | 6.02618 | 6.27253 | 6.43624 | 5.84287 | 6.05254 | 6.31818 | 6.09216 | 6.48687 | 6.53273 | 6.60248 | 6.31404 | 6.05272 | 6.40344 | 6.49558 |
| RBBP9 | O75884 | 5.94152 | 5.87865 | 6.74553 | 5.38289 | 6.08928 | 6.99805 | 6.25011 | 6.5561 | 6.14825 | 6.84998 | 6.26814 | 6.59821 | 5.82685 | 6.53228 | 5.87657 | 6.0576 |
| CENPQ | Q7L2Z9 | 6.53789 | 6.60698 | 5.8116 | 6.83318 | 6.11315 | 6.26061 | 6.45715 | 5.92792 | 6.22948 | 6.05052 | 6.00611 | 6.14567 | 5.85185 | 6.26597 | 6.44417 | 6.45776 |
| IMP4 | Q96G21 | 5.94829 | 5.95465 | 6.10724 | 7.20685 | 6.21925 | 5.9176 | 6.28695 | 6.6077 | 6.20322 | 5.94882 | 6.23713 | 5.89414 | 6.48701 | 6.07844 | 6.38365 | 6.51905 |
| BCLAF3 | A2AJT9 | 6.29559 | 6.31428 | 5.8845 | 6.82656 | 6.06333 | 6.20228 | 6.27902 | 5.78512 | 5.89892 | 5.50818 | 5.90479 | 5.91416 | 6.80649 | 6.74239 | 7.07519 | 6.49921 |
| LRRC42 | Q9Y546 | 5.94706 | 6.04282 | 6.65194 | 5.87588 | 5.85637 | 6.19392 | 6.10838 | 6.35489 | 6.38879 | 6.45462 | 6.80456 | 6.37098 | 5.87692 | 6.24366 | 6.60356 | 6.22563 |
| CHMP5 | Q9NZZ3 | 6.41094 | 6.25319 | 7.09215 | 5.54105 | 6.50012 | 6.86545 | 6.32291 | 6.78225 | 6.12804 | 6.72399 | 6.34114 | 6.58348 | 5.59503 | 6.13796 | 5.45213 | 5.27017 |
| PAAF1 | Q9BRP4 | 6.02209 | 6.36324 | 7.1813 | 5.73023 | 6.19046 | 6.65483 | 6.17891 | 6.24313 | 5.9926 | 6.5591 | 6.62677 | 6.45653 | 6.34444 | 6.16058 | 5.86679 | 5.42898 |
| HLA-C | A2BF26 | 6.48989 | 6.31699 | 6.12938 | 6.50668 | 6.4668 | 6.58743 | 6.34965 | 6.45704 | 6.40627 | 6.29923 | 6.21373 | 6.16231 | 5.80822 | 5.74207 | 5.89525 | 6.16906 |
| PTBP2 | Q9UKA9-2 | 6.10229 | 6.30051 | 6.32563 | 6.14997 | 6.34827 | 5.74168 | 6.18205 | 5.89249 | 6.20195 | 5.85789 | 5.89205 | 6.11002 | 6.7568 | 6.23597 | 6.91424 | 6.9882 |
| LAMTOR2 | Q9Y2Q5 | 5.87859 | 5.39191 | 5.56773 | 5.106 | 6.16248 | 6.04427 | 5.9678 | 6.41917 | 5.72419 | 6.35658 | 6.0539 | 6.16331 | 6.3095 | 7.47483 | 7.67301 | 7.70673 |
| CENPV | Q7Z7K6 | 5.98068 | 6.38151 | 6.08136 | 7.01408 | 5.8987 | 6.07554 | 6.23045 | 5.81643 | 5.99546 | 6.15747 | 6.14941 | 6.34545 | 6.25162 | 6.18726 | 6.73877 | 6.6958 |
| IFNAR1 | P17181 | 7.51068 | 7.16696 | 6.63351 | 6.70275 | 6.50936 | 6.00596 | 6.27518 | 6.07803 | 7.04999 | 6.57728 | 6.72425 | 6.80343 | 4.50245 | 4.92778 | 5.28503 | 5.24735 |
| SPNS1 | Q9H2V7 | 6.07742 | 6.5357 | 5.92982 | 6.90538 | 6.29802 | 5.88992 | 6.28818 | 5.79761 | 5.96891 | 6.31413 | 6.30474 | 6.04266 | 6.45498 | 6.52899 | 6.21529 | 6.44825 |
| JUN | P05412 | 5.60134 | 5.97032 | 5.58061 | 6.19107 | 6.54885 | 6.32788 | 6.6572 | 6.13349 | 5.47898 | 4.75801 | 5.40615 | 5.43169 | 7.85774 | 6.86485 | 7.86049 | 7.33132 |
| RPUSD2 | Q8IZ73 | 5.75726 | 5.32 | 6.31302 | 5.20433 | 5.76282 | 6.11884 | 5.57315 | 5.81369 | 6.03673 | 6.397 | 6.43437 | 6.10064 | 6.94469 | 7.48777 | 7.45518 | 7.28049 |
| TRIM26 | Q12899 | 7.24104 | 7.03425 | 6.77773 | 6.75506 | 6.44986 | 7.02314 | 5.82224 | 6.64837 | 7.17478 | 6.6367 | 7.25649 | 6.33256 | 4.64921 | 4.72204 | 4.66458 | 4.81195 |
| FRMD5 | Q7Z6J6 | 6.09566 | 6.08381 | 5.82484 | 6.57347 | 6.01799 | 5.81406 | 6.22255 | 5.84014 | 6.15976 | 6.28001 | 6.02309 | 5.89633 | 7.03209 | 6.51478 | 6.87511 | 6.74631 |
| BRF1 | Q92994 | 5.36278 | 5.59797 | 6.27217 | 5.93809 | 5.43324 | 6.14035 | 5.8135 | 5.81146 | 6.30391 | 6.48845 | 6.60618 | 6.43264 | 6.48274 | 7.0355 | 7.08036 | 6.80067 |
| CCDC102A | Q96A19 | 6.43726 | 6.24267 | 6.20394 | 6.46153 | 5.64324 | 5.85238 | 5.92211 | 5.97953 | 5.99565 | 6.50117 | 6.20302 | 6.61551 | 6.89966 | 6.31734 | 6.46593 | 6.25906 |
| ANKZF1 | Q9H8Y5 | 5.95287 | 6.6424 | 6.33744 | 6.41423 | 6.27094 | 6.40341 | 6.18598 | 6.66105 | 6.4398 | 6.62851 | 6.57679 | 6.8715 | 5.69243 | 5.41933 | 5.88972 | 5.61361 |
| NF2 | P35240-2 | 6.06159 | 6.28479 | 5.94635 | 6.38101 | 6.20219 | 6.25031 | 6.31936 | 6.20197 | 6.31979 | 5.90801 | 6.14568 | 6.23982 | 6.35196 | 6.30404 | 6.39347 | 6.68966 |
| PARVB | Q9HBI1-2 | 6.08149 | 6.25647 | 6.33277 | 6.1369 | 6.07866 | 6.69076 | 6.0519 | 6.70156 | 6.10327 | 6.5007 | 6.37706 | 6.58536 | 5.9304 | 6.54648 | 5.62927 | 5.99693 |
| MRPL48 | Q96GC5 | 6.06607 | 6.21168 | 5.51454 | 6.58124 | 6.4482 | 6.186 | 6.58672 | 6.30307 | 6.5101 | 6.21045 | 6.02022 | 6.112 | 6.1545 | 5.8605 | 6.90991 | 6.32482 |
| SERPINE2 | P07093-3 | 6.66387 | 6.72408 | 5.26248 | 7.21311 | 5.98816 | 6.04936 | 6.6158 | 6.20749 | 6.32005 | 6.69172 | 5.8928 | 6.24298 | 5.79883 | 5.98396 | 5.9978 | 6.34751 |
| GNG12 | Q9UBI6 | 5.84069 | 5.3573 | 6.1824 | 5.49091 | 5.62317 | 5.49779 | 5.96348 | 6.14939 | 6.11556 | 5.80151 | 5.92462 | 6.38494 | 6.3534 | 8.04117 | 7.67251 | 7.60115 |
| ISOC1 | Q96CN7 | 6.37435 | 6.92816 | 6.74282 | 5.98638 | 6.61804 | 6.51357 | 6.0752 | 6.62536 | 6.20121 | 6.19574 | 6.22013 | 6.49135 | 6.1452 | 6.1081 | 6.04232 | 6.16927 |
| SLC35B2 | Q8TB61 | 6.16612 | 6.12286 | 5.75956 | 6.15716 | 6.39505 | 6.10804 | 6.18381 | 6.1416 | 6.27394 | 6.35418 | 6.17283 | 6.10337 | 6.50027 | 6.46526 | 6.45677 | 6.63919 |
| CTAG1A | P78358 | 5.65332 | 5.93699 | 6.31993 | 5.25245 | 6.28801 | 6.10184 | 5.9092 | 7.0777 | 5.83517 | 6.81462 | 6.3872 | 6.6786 | 5.80608 | 6.70222 | 6.84475 | 6.3919 |
| RBMS1 | P29558-2 | 6.15561 | 6.77356 | 6.74307 | 6.35814 | 6.33249 | 6.42705 | 5.9925 | 6.09856 | 6.32223 | 6.72244 | 6.62711 | 6.80242 | 6.2491 | 5.46547 | 5.38186 | 5.54837 |
| MRPS11 | P82912 | 6.40127 | 6.29046 | 6.13143 | 6.83244 | 6.07296 | 5.73251 | 7.06576 | 5.64364 | 6.33217 | 6.13058 | 6.15862 | 5.94398 | 6.46248 | 6.55986 | 6.23781 | 6.00402 |
| TAPT1 | Q6NXT6 | 6.20107 | 5.85647 | 6.17492 | 6.93108 | 5.98916 | 6.17287 | 6.71025 | 5.94536 | 6.16436 | 6.22534 | 6.0611 | 6.03075 | 6.50767 | 6.317 | 6.54186 | 6.17073 |
| SURF2 | Q15527 | 5.54597 | 6.13315 | 5.9505 | 5.60419 | 5.91009 | 6.15519 | 5.9018 | 5.79437 | 6.14337 | 5.86473 | 6.90682 | 6.07007 | 7.10953 | 7.43469 | 6.82335 | 6.65219 |
| RGS10 | O43665 | 6.1193 | 5.69323 | 7.0215 | 5.6115 | 6.5501 | 6.11436 | 5.78304 | 6.57805 | 6.24786 | 6.70884 | 6.8675 | 7.06654 | 5.80753 | 6.35426 | 5.76854 | 5.70785 |
| PRKCI | P41743 | 5.7206 | 6.02769 | 6.62894 | 6.01511 | 5.99959 | 6.43824 | 6.03829 | 6.45386 | 6.68217 | 6.30415 | 6.38957 | 6.72969 | 5.96338 | 6.13234 | 6.1933 | 6.28307 |
| TENT4B | Q8NDF8-5 | 6.41807 | 6.45548 | 6.52853 | 7.18811 | 6.2261 | 5.90746 | 6.26995 | 5.78816 | 6.32663 | 5.94716 | 6.22363 | 6.0668 | 6.25199 | 5.8882 | 6.10515 | 6.40857 |
| PWWP3A | Q2TAK8-3 | 6.42402 | 6.67351 | 6.21855 | 6.82884 | 6.01995 | 6.67743 | 6.50475 | 6.46413 | 6.45255 | 6.29986 | 6.14067 | 6.39581 | 5.65274 | 5.65851 | 5.7276 | 5.86109 |
| P3H2 | Q8IVL5 | 7.64044 | 7.09579 | 6.91654 | 7.67664 | 6.5782 | 6.20467 | 6.43841 | 6.59041 | 6.91712 | 6.59439 | 6.67388 | 6.45858 | 4.94461 | 4.32141 | 4.94938 | 4.64854 |
| COPS7B | Q9H9Q2 | 6.35475 | 5.31281 | 6.40142 | 5.9343 | 6.35281 | 6.6832 | 5.9001 | 6.11631 | 6.33814 | 6.94626 | 6.38021 | 6.7231 | 6.87056 | 6.22189 | 6.43226 | 6.03188 |
| PRKACA | P17612 | 6.35726 | 6.24413 | 6.84513 | 6.20647 | 6.42929 | 6.66903 | 6.31091 | 6.37829 | 6.21813 | 6.12322 | 5.77447 | 6.30767 | 6.63879 | 5.88797 | 5.94932 | 5.65591 |
| RANBP10 | Q6VN20 | 6.34785 | 6.73589 | 6.4645 | 6.35441 | 6.28147 | 6.45601 | 6.14191 | 5.98371 | 6.28218 | 6.28643 | 6.54237 | 6.52773 | 6.2017 | 6.22901 | 5.9931 | 5.77172 |
| ENTPD6 | O75354 | 6.86013 | 6.44982 | 6.29325 | 6.6344 | 6.1792 | 5.95583 | 6.55029 | 5.96532 | 6.62024 | 6.56207 | 6.89838 | 6.59662 | 5.68774 | 5.50227 | 5.73779 | 5.50665 |
| NDUFA3 | O95167 | 6.50535 | 6.5457 | 5.82784 | 6.2125 | 5.97837 | 5.51069 | 6.28712 | 6.12989 | 5.78323 | 5.58932 | 6.26921 | 5.2359 | 7.08658 | 7.07464 | 6.82605 | 7.13762 |
| MSL2 | Q9HCI7 | 6.56855 | 6.57968 | 6.11046 | 7.145 | 6.09873 | 6.07564 | 6.3126 | 5.91196 | 6.05253 | 6.22899 | 6.43619 | 6.05334 | 6.17327 | 6.06989 | 6.33715 | 5.84603 |
| ZDHHC13 | Q8IUH4 | 6.23281 | 6.5488 | 6.04811 | 6.89074 | 6.62352 | 6.20323 | 5.98212 | 5.73785 | 6.39222 | 6.27109 | 6.10641 | 5.69607 | 6.58642 | 5.85297 | 6.30998 | 6.51767 |
| MCPH1 | Q8NEM0 | 6.07561 | 5.93809 | 5.82137 | 6.03488 | 6.37542 | 5.94371 | 6.33936 | 5.94091 | 6.1745 | 6.1325 | 6.13717 | 5.88139 | 6.48985 | 6.59984 | 7.03498 | 7.08039 |
| SH3GLB2 | Q9NR46 | 6.5765 | 5.91733 | 6.97401 | 5.34875 | 6.53853 | 6.63991 | 6.05648 | 6.65707 | 6.49265 | 7.05308 | 6.80905 | 7.11264 | 5.98369 | 5.21562 | 5.40005 | 5.22465 |
| TAF9 | Q16594 | 6.48641 | 5.92599 | 6.54258 | 5.98273 | 6.47967 | 6.27768 | 6.28314 | 6.60701 | 6.29977 | 6.5455 | 6.81039 | 6.60767 | 6.10655 | 5.60797 | 5.70298 | 5.73396 |
| CLSTN1 | O94985 | 7.20378 | 7.47023 | 6.87567 | 7.27805 | 6.50905 | 6.43912 | 6.48617 | 5.95302 | 6.99973 | 7.18091 | 6.72126 | 6.5572 | 4.4876 | 4.60587 | 4.67416 | 4.55819 |
| COMMD7 | Q86VX2 | 6.09995 | 5.80565 | 6.21511 | 5.51704 | 6.09144 | 6.15872 | 5.81627 | 6.50027 | 5.96351 | 7.02416 | 6.30472 | 6.66867 | 6.5386 | 6.34102 | 6.53379 | 6.42108 |
| PLA2G4C | Q9UP65-3 | 6.65734 | 6.55621 | 6.38048 | 6.94696 | 6.25246 | 6.38793 | 6.05393 | 5.78107 | 6.13792 | 6.52406 | 6.417 | 6.15073 | 5.58061 | 6.42702 | 5.5634 | 6.18289 |
| CSTF2 | P33240 | 6.33487 | 4.93882 | 4.88065 | 5.37693 | 6.06794 | 4.8739 | 5.23124 | 7.04155 | 4.58647 | 6.82818 | 5.51315 | 7.55103 | 6.66625 | 8.38888 | 7.25796 | 8.46217 |
| SEC62 | Q99442 | 6.37911 | 6.54413 | 6.1787 | 6.04206 | 6.32595 | 6.18669 | 6.11109 | 5.81226 | 6.43156 | 6.09135 | 5.78041 | 5.88801 | 6.48857 | 7.01672 | 6.35659 | 6.36679 |
| ACP6 | Q9NPH0 | 6.23981 | 6.00618 | 6.08902 | 6.7599 | 6.61963 | 5.73197 | 6.32202 | 5.53512 | 6.35272 | 6.69341 | 6.23636 | 6.57099 | 6.65819 | 5.67189 | 6.22397 | 6.28881 |
| CD47 | Q08722 | 6.37387 | 6.54104 | 5.95871 | 6.34095 | 6.31279 | 6.21673 | 6.33288 | 5.86295 | 6.38009 | 6.17569 | 5.72081 | 5.92302 | 6.7232 | 6.47317 | 6.28783 | 6.37627 |
| SMURF2 | Q9HAU4 | 6.27078 | 6.04044 | 6.51136 | 6.36552 | 6.05709 | 6.00027 | 6.55642 | 6.14269 | 6.49783 | 6.48228 | 6.44825 | 6.3737 | 6.27292 | 6.01256 | 6.06023 | 5.90767 |
| MAP2K4 | P45985-2 | 6.12268 | 6.03984 | 7.04522 | 5.60428 | 6.32256 | 6.4885 | 5.92808 | 6.68886 | 6.1762 | 6.33173 | 6.52673 | 6.10454 | 6.10493 | 6.28522 | 5.9739 | 6.25673 |
| GMEB1 | Q9Y692 | 6.36221 | 6.33189 | 5.98722 | 6.39201 | 6.29332 | 5.92124 | 5.9188 | 6.08589 | 6.24496 | 5.94016 | 6.19076 | 6.03633 | 6.9706 | 6.36182 | 6.17844 | 6.78433 |

| | | | | | | | | | | | | | | | | | |
|---|---|---|---|---|---|---|---|---|---|---|---|---|---|---|---|---|---|
| ABCC5 | O15440 | 6.29281 | 6.49112 | 6.39255 | 6.54147 | 6.39274 | 6.20206 | 6.25711 | 5.92072 | 6.12343 | 6.08096 | 5.84864 | 6.26725 | 6.08494 | 6.19183 | 6.30135 | 6.61102 |
| IDH3G | P51553 | 6.38842 | 6.33718 | 6.28628 | 6.57359 | 6.08265 | 6.01582 | 6.1358 | 6.03113 | 6.26322 | 6.17582 | 6.26275 | 5.88038 | 6.46399 | 5.99353 | 6.87934 | 6.23011 |
| RHOG | P84095 | 6.21218 | 6.00984 | 5.8821 | 6.07377 | 6.40338 | 6.20123 | 6.35593 | 5.89416 | 6.24629 | 5.81673 | 6.396 | 6.51324 | 6.2855 | 6.64702 | 6.42066 | 6.64197 |
| EXOG | Q9Y2C4 | 6.25876 | 6.58621 | 6.02542 | 7.02486 | 6.04075 | 6.05723 | 6.1695 | 5.92841 | 6.06501 | 5.89419 | 6.30871 | 5.71444 | 6.59254 | 6.66419 | 6.31807 | 6.35171 |
| PSMG3 | Q9BT73 | 6.00545 | 5.87465 | 6.45957 | 5.48018 | 6.3864 | 6.87256 | 5.90668 | 6.9458 | 6.25732 | 6.70688 | 5.9875 | 6.31274 | 5.59042 | 6.50107 | 6.17287 | 6.53992 |
| RNF123 | Q5XPI4-2 | 6.00411 | 6.70117 | 6.64404 | 5.77028 | 6.07713 | 6.34579 | 6.39018 | 6.82714 | 6.28401 | 6.33422 | 6.23313 | 6.42589 | 6.01131 | 5.92875 | 6.04203 | 5.98082 |
| TEFM | Q96QE5 | 6.52026 | 6.3574 | 6.18587 | 6.46933 | 6.38191 | 5.82367 | 6.72321 | 5.91515 | 6.26057 | 5.72695 | 5.98571 | 5.46732 | 6.77777 | 6.40441 | 6.53457 | 6.46592 |
| ZMYM1 | Q5SVZ6 | 6.0759 | 6.5564 | 6.47819 | 6.82034 | 6.20462 | 5.98244 | 6.41167 | 5.48236 | 6.20035 | 6.86369 | 6.4124 | 6.66214 | 5.5961 | 5.79706 | 6.31676 | 6.1396 |
| LARP1B | A0A3B3ISF0 | 5.7438 | 6.36849 | 6.53062 | 6.84415 | 6.13912 | 5.90664 | 6.20922 | 6.39294 | 6.70071 | 6.40695 | 6.90028 | 6.5374 | 5.86122 | 6.24211 | 5.69479 | 5.52156 |
| COASY | Q13057-2 | 5.90671 | 6.64692 | 6.54649 | 6.42787 | 6.56002 | 6.46329 | 5.91354 | 7.41217 | 5.9976 | 6.20853 | 6.53019 | 6.0316 | 5.85425 | 5.87096 | 6.04844 | 5.58142 |
| UQCRQ | O14949 | 6.25938 | 6.36819 | 5.79962 | 6.53369 | 6.94431 | 6.21482 | 6.28456 | 6.6725 | 5.47798 | 5.75576 | 5.70755 | 5.70605 | 6.52732 | 6.78002 | 6.49256 | 6.4757 |
| PHF20 | Q9BVI0 | 6.24147 | 6.40129 | 6.00726 | 6.95877 | 5.95591 | 6.17456 | 6.29266 | 5.87709 | 6.47847 | 6.29037 | 6.1962 | 6.3534 | 6.5438 | 6.349 | 6.34462 | 6.43513 |
| ANKRD28 | O15084-1 | 6.50234 | 6.39945 | 6.93341 | 6.23555 | 6.46539 | 6.11397 | 6.06129 | 6.0276 | 6.48231 | 6.45156 | 7.00693 | 6.78755 | 5.91177 | 5.67273 | 5.46378 | 5.48438 |
| MT1X | P80297 | 6.51711 | 5.76416 | 7.15433 | 5.4148 | 6.62106 | 6.78335 | 5.95996 | 6.26752 | 6.56429 | 6.87571 | 7.12937 | 6.72164 | 6.20966 | 5.513 | 5.23579 | 5.26822 |
| CSRP2 | Q16527 | 6.28509 | 6.0142 | 6.38506 | 6.34655 | 6.4038 | 6.38014 | 6.16976 | 6.38182 | 6.20035 | 6.49339 | 6.69146 | 6.46301 | 5.96049 | 6.0858 | 5.78029 | 5.9588 |
| RAB22A | Q9UL26 | 6.33136 | 6.55252 | 6.28258 | 6.53856 | 6.08105 | 6.20804 | 6.17553 | 6.3279 | 6.14419 | 6.52334 | 6.02558 | 6.12624 | 6.15595 | 6.11225 | 6.33782 | 6.07708 |
| SMAD4 | Q13485 | 6.51809 | 6.21257 | 6.623 | 6.03017 | 6.0711 | 6.43715 | 6.16323 | 6.61687 | 6.14022 | 6.33664 | 6.83299 | 6.51586 | 5.8796 | 5.98827 | 5.83539 | 5.79886 |
| CD55 | P08174-7 | 6.71027 | 6.3365 | 6.13614 | 6.54857 | 5.88629 | 6.05165 | 6.12723 | 5.91431 | 6.31319 | 6.2664 | 5.96917 | 5.50022 | 6.93604 | 6.34915 | 6.29194 | 6.66293 |
| ZNF142 | P52746 | 6.64934 | 6.48769 | 6.14032 | 6.7295 | 6.08347 | 5.67131 | 6.56587 | 6.29573 | 6.0498 | 6.13571 | 6.11742 | 6.26081 | 6.27681 | 5.82309 | 6.425 | 6.28812 |
| TMED4 | Q7Z7H5 | 6.41743 | 6.17094 | 5.7189 | 6.40667 | 6.3977 | 6.512 | 6.15474 | 6.28444 | 5.84784 | 6.12973 | 6.07401 | 5.96835 | 6.67236 | 6.40469 | 6.57586 | 6.26434 |
| FAM136A | Q96C01 | 6.09822 | 4.68369 | 5.0829 | 4.77504 | 5.68028 | 4.08426 | 4.84138 | 6.17253 | 3.95281 | 5.73488 | 4.76077 | 5.65849 | 6.11424 | 11.0973 | 10.026 | 11.2372 |
| PSMG2 | Q969U7 | 6.1426 | 6.05708 | 6.67916 | 5.77919 | 6.3064 | 6.51144 | 6.03542 | 6.80047 | 6.20483 | 6.56417 | 6.4724 | 6.21243 | 6.17473 | 5.99656 | 6.4073 | 6.5558 |
| GLIPR1 | P48060 | 6.45155 | 7.15522 | 6.17667 | 7.30119 | 6.01667 | 6.49557 | 6.33132 | 6.20721 | 6.08193 | 5.42478 | 6.36964 | 6.30845 | 5.93127 | 5.71332 | 6.14728 | 5.88791 |
| CDKN2A | Q8N726 | 6.03113 | 6.48691 | 5.73706 | 6.19968 | 6.21067 | 6.47396 | 6.45585 | 6.09636 | 6.23315 | 6.01251 | 5.80183 | 5.58747 | 6.62837 | 6.75213 | 6.51025 | 6.78268 |
| SNX30 | Q5VWJ9 | 5.89752 | 6.38367 | 6.37126 | 6.09773 | 6.10087 | 6.34017 | 6.11722 | 6.54176 | 6.00679 | 6.45398 | 6.52459 | 6.3957 | 6.49063 | 5.98337 | 6.40174 | 5.89302 |
| MRPL27 | Q9P0M9 | 6.17756 | 6.76906 | 5.51633 | 6.71383 | 6.36001 | 6.11845 | 6.47581 | 6.4198 | 5.97556 | 6.18564 | 5.99321 | 5.99323 | 6.09494 | 6.35719 | 6.38124 | 6.46813 |
| ARIH2 | O95376 | 6.15685 | 5.78794 | 6.7482 | 6.04096 | 6.27281 | 6.36783 | 6.20954 | 6.38313 | 6.53418 | 6.20735 | 6.60889 | 6.70021 | 5.91885 | 5.96783 | 6.20973 | 5.8857 |
| SNRPF | P62306 | 5.89039 | 5.3086 | 5.86639 | 6.18054 | 6.20441 | 6.02103 | 6.14198 | 5.32649 | 6.42787 | 6.62917 | 6.67027 | 6.65066 | 7.07462 | 5.99716 | 6.57865 | 7.03178 |
| SUV39H2 | Q9H5I1 | 6.43132 | 6.52916 | 5.71853 | 6.88879 | 5.76239 | 5.75152 | 6.32815 | 5.70321 | 6.03632 | 6.57279 | 6.31995 | 6.32266 | 6.67916 | 6.1978 | 6.6411 | 6.11714 |
| PPT1 | P50897 | 6.77856 | 6.02626 | 6.23518 | 6.16464 | 6.45704 | 5.86689 | 6.71382 | 7.01837 | 6.13486 | 6.01189 | 6.06428 | 5.89056 | 5.85688 | 5.5719 | 6.247 | 5.96187 |
| IRS1 | P35568 | 5.65847 | 5.9625 | 6.43622 | 6.09728 | 6.28066 | 6.32876 | 6.16377 | 6.57173 | 6.11368 | 6.6727 | 6.4573 | 6.50289 | 6.46211 | 6.32436 | 6.06281 | 5.90476 |
| TAF1B | Q53T94 | 6.54525 | 6.50484 | 6.3461 | 6.94078 | 5.85888 | 6.14132 | 6.17393 | 6.58805 | 6.62713 | 6.13989 | 6.54609 | 6.89437 | 5.73753 | 5.73046 | 5.48473 | 5.74064 |
| RELL1 | Q8IUW5 | 5.75531 | 6.60815 | 5.91687 | 6.5044 | 6.28316 | 6.70899 | 6.7366 | 6.55503 | 5.93696 | 5.99243 | 6.75406 | 6.28091 | 6.58518 | 6.11879 | 5.80825 | 5.45492 |
| YIPF4 | Q9BSR8 | 6.45891 | 6.13305 | 5.96855 | 6.81809 | 6.12468 | 6.06999 | 6.50417 | 6.12028 | 6.05158 | 6.4376 | 6.209 | 6.01207 | 6.22099 | 6.10402 | 6.47383 | 6.29318 |
| MUL1 | Q969V5 | 6.17716 | 6.50424 | 6.59743 | 6.49637 | 6.6513 | 5.40643 | 7.3535 | 5.88343 | 6.28761 | 6.47189 | 6.65966 | 5.59117 | 5.96396 | 5.51159 | 6.68785 | 5.75643 |
| BANP | Q8N9N5 | 6.37353 | 6.76783 | 5.83494 | 6.78712 | 6.40602 | 6.34075 | 6.40974 | 6.39345 | 5.98593 | 5.99413 | 5.90645 | 5.64614 | 6.51813 | 6.05928 | 6.34984 | 6.22672 |
| RMND5A | Q9H871 | 6.32245 | 5.52903 | 6.44786 | 5.50067 | 6.30013 | 5.93734 | 5.98 | 6.27705 | 5.63907 | 6.45854 | 6.30138 | 7.21986 | 6.65218 | 6.33215 | 6.50957 | 6.59271 |
| UFC1 | Q9Y3C8 | 6.1396 | 5.70581 | 6.47657 | 5.83818 | 6.50167 | 5.78541 | 5.67346 | 6.2745 | 5.78676 | 6.87404 | 5.98807 | 6.57195 | 6.58732 | 6.25494 | 7.03067 | 6.66255 |
| DGAT1 | O75907 | 6.2448 | 6.5253 | 5.86294 | 6.44315 | 6.01645 | 6.2384 | 6.18162 | 6.12596 | 6.18408 | 6.2521 | 5.356 | 5.73297 | 6.53091 | 6.71716 | 6.72101 | 6.86715 |
| GFER | P55789 | 6.66029 | 5.85182 | 6.39487 | 6.23038 | 6.33477 | 5.68287 | 5.91916 | 6.14917 | 6.04298 | 6.84438 | 6.68109 | 6.30051 | 6.99635 | 5.6467 | 6.05722 | 6.20746 |
| BACH1 | O14867 | 6.60717 | 6.34444 | 6.5947 | 6.28068 | 6.26391 | 6.21515 | 5.99469 | 5.99448 | 6.08133 | 6.5156 | 6.26956 | 6.80086 | 6.22657 | 5.85538 | 5.97325 | 6.05274 |
| GAS2L3 | Q86XJ1 | 6.37562 | 6.26287 | 6.54452 | 6.53095 | 6.05541 | 6.30133 | 6.19487 | 5.80942 | 6.58331 | 6.51228 | 6.69307 | 7.01236 | 5.0332 | 6.14893 | 5.91346 | 6.0284 |
| ARF6 | P62330 | 6.08818 | 6.2279 | 6.3989 | 6.19415 | 6.21225 | 6.38481 | 6.34617 | 5.99474 | 6.39017 | 6.1414 | 6.69137 | 5.80993 | 6.32538 | 6.54584 | 6.06578 | 6.18303 |
| PGRMC1 | O00264 | 6.56915 | 5.74703 | 5.73082 | 5.64569 | 5.71282 | 4.88647 | 5.54443 | 5.59448 | 5.2188 | 6.38841 | 5.97806 | 6.44606 | 6.95683 | 7.67007 | 7.47457 | 8.03632 |
| EMC8 | O43402 | 6.44882 | 6.295 | 5.50722 | 6.22951 | 6.23637 | 6.17726 | 6.11776 | 6.66343 | 5.96348 | 6.22791 | 6.07017 | 6.49835 | 6.32763 | 6.50321 | 5.96364 | 6.77026 |
| RAB9A | P51151 | 6.29425 | 6.0252 | 5.71126 | 6.4128 | 6.29094 | 6.2976 | 6.23467 | 6.51843 | 6.18868 | 5.95713 | 6.36462 | 6.65741 | 6.35892 | 6.26027 | 6.17408 | 6.25373 |
| PLEK2 | Q9NYT0 | 6.51602 | 6.33205 | 6.8569 | 6.20557 | 6.39882 | 6.32259 | 6.11317 | 6.62698 | 6.42333 | 6.82708 | 6.73066 | 6.79121 | 5.81279 | 5.60651 | 5.25152 | 5.18478 |
| POLQ | O75417 | 6.51103 | 6.87405 | 6.60012 | 7.78415 | 6.51249 | 6.12458 | 6.39403 | 5.64298 | 6.36403 | 6.87688 | 7.08849 | 6.99875 | 4.78608 | 4.90709 | 5.08353 | 5.4517 |
| CEP162 | Q5TB80 | 6.33089 | 5.99675 | 6.65785 | 6.15634 | 6.252 | 6.41918 | 6.46145 | 6.29576 | 6.58246 | 6.86037 | 6.60636 | 6.11462 | 6.16572 | 5.71584 | 5.86698 | 5.51744 |
| ZMIZ1 | Q9ULJ6 | 5.25608 | 5.4705 | 6.03971 | 5.49839 | 5.4776 | 5.81066 | 5.84384 | 6.19803 | 5.80488 | 6.09937 | 6.66582 | 6.41957 | 7.2202 | 7.28994 | 7.83127 | 7.07412 |
| EYA3 | Q99504 | 5.86159 | 5.94184 | 5.84048 | 5.51156 | 6.51367 | 5.9321 | 6.19245 | 6.32256 | 5.93907 | 5.92478 | 6.58592 | 6.52956 | 7.11265 | 6.47508 | 6.82838 | 6.48831 |
| MRPS33 | Q9Y291 | 6.09261 | 6.14909 | 5.59947 | 6.83156 | 5.76736 | 5.81737 | 6.36974 | 5.87823 | 6.16403 | 5.98992 | 6.02094 | 6.02684 | 6.84636 | 6.76881 | 6.94443 | 6.73324 |
| SCRN2 | Q96FV2 | 6.25217 | 5.86102 | 6.50099 | 4.98227 | 6.75484 | 6.14718 | 5.78212 | 7.09788 | 6.36013 | 6.89101 | 6.59873 | 6.8462 | 6.39284 | 5.76494 | 5.71461 | 6.05307 |
| MMADHC | Q9H3L0 | 5.59936 | 5.71506 | 5.93913 | 5.8275 | 6.75811 | 7.17817 | 7.44362 | 7.50449 | 6.46875 | 6.22322 | 6.6303 | 6.67614 | 5.286 | 5.14043 | 5.56817 | 6.04156 |
| TBC1D20 | Q96BZ9-3 | 6.38096 | 6.47501 | 6.0212 | 6.61558 | 6.3149 | 5.8373 | 6.37233 | 6.01601 | 6.02853 | 6.80859 | 6.22876 | 6.15618 | 6.22312 | 6.3613 | 6.09904 | 6.06119 |
| PPIC | P45877 | 6.63805 | 6.04155 | 6.455 | 6.82343 | 6.68655 | 5.9613 | 6.38951 | 5.8069 | 6.56925 | 6.41976 | 6.39161 | 6.25711 | 6.54961 | 5.47307 | 5.80987 | 5.72742 |
| PATZ1 | Q9HBE1 | 6.06108 | 5.92397 | 5.78554 | 6.58967 | 6.15157 | 6.24241 | 6.57285 | 5.90741 | 6.28599 | 6.43663 | 6.47671 | 6.55171 | 6.09778 | 6.08105 | 6.54492 | 6.29071 |
| KCTD15 | Q96SI1 | 5.86998 | 5.58406 | 6.43461 | 5.06947 | 6.0337 | 6.10154 | 5.49226 | 6.44227 | 6.61794 | 6.96567 | 6.96824 | 6.91516 | 6.54314 | 6.12025 | 6.39395 | 6.44775 |
| ZNF646 | O15015 | 6.28526 | 6.10124 | 5.74733 | 7.06546 | 5.9808 | 5.90292 | 6.35044 | 5.92203 | 5.77185 | 6.26942 | 6.34512 | 6.37115 | 6.64374 | 6.41782 | 6.55482 | 6.2706 |
| FBXW8 | Q8N3Y1 | 6.25572 | 6.18684 | 6.70882 | 6.29553 | 6.04951 | 6.85498 | 6.92942 | 6.32086 | 6.69025 | 6.95179 | 6.29701 | 6.58235 | 5.63248 | 5.42398 | 5.61238 | 5.20807 |
| SNX19 | Q92543 | 6.49648 | 6.84167 | 6.32209 | 6.50188 | 6.12726 | 6.43933 | 6.5532 | 5.9464 | 6.40273 | 6.28 | 6.2861 | 6.13229 | 6.09568 | 5.70205 | 5.77348 | 6.09939 |
| TRAPPC6B | Q86SZ2 | 6.05588 | 5.36065 | 6.41458 | 5.08911 | 6.81677 | 6.51823 | 5.76962 | 6.35432 | 5.88358 | 6.37527 | 6.45304 | 6.71297 | 6.5773 | 6.49422 | 6.67978 | 6.44469 |
| SNIP1 | Q8TAD8 | 5.61086 | 6.176 | 5.53609 | 6.65464 | 5.88698 | 5.72839 | 6.06899 | 6.02334 | 6.37523 | 6.08523 | 6.09314 | 6.03298 | 6.3119 | 6.72196 | 7.11746 | 7.57682 |
| SLIT3 | O75094-4 | 5.89031 | 6.21876 | 6.30542 | 6.20354 | 6.22237 | 6.63568 | 6.01144 | 6.07936 | 6.56612 | 6.52819 | 6.24172 | 6.14753 | 6.19243 | 6.09576 | 6.65021 | 6.01116 |
| RHOXF2 | Q9BQY4 | 5.43061 | 4.67997 | 5.48024 | 4.73375 | 4.73865 | 4.64891 | 4.96008 | 6.26194 | 4.59188 | 5.91663 | 5.6762 | 7.19664 | 5.23277 | 10.8817 | 8.33889 | 11.2311 |
| PRDM15 | P57071 | 6.15876 | 6.35312 | 6.103 | 6.89714 | 5.84112 | 6.13372 | 6.29861 | 5.77545 | 5.78973 | 6.11462 | 6.22158 | 6.3392 | 6.63907 | 6.20172 | 6.30437 | 6.82879 |
| C1orf174 | Q8IYL3 | 6.17721 | 6.80222 | 5.97734 | 6.8007 | 5.71838 | 5.92336 | 6.12602 | 6.13356 | 6.11435 | 6.7791 | 6.41149 | 6.31803 | 6.00353 | 6.49787 | 5.92187 | 6.29496 |
| HDAC5 | Q9UQL6-3 | 3.58863 | 4.24916 | 12.4537 | 10.3939 | 14.9459 | 8.69288 | 3.63284 | 3.54675 | 3.78782 | 9.50167 | 4.27645 | 4.16883 | 4.54812 | 4.37363 | 4.10423 | 3.73557 |
| MRPL41 | Q8IXM3 | 6.2412 | 6.23959 | 5.35944 | 6.82138 | 6.59042 | 5.9549 | 6.88575 | 6.27796 | 6.28301 | 5.97278 | 5.91098 | 5.92443 | 6.80021 | 6.0395 | 6.42933 | 6.26914 |
| MPHOSPH6 | Q99547 | 6.2058 | 6.403 | 6.137 | 6.47679 | 6.62872 | 6.05775 | 6.38693 | 6.37032 | 6.04811 | 6.28403 | 6.09309 | 6.10522 | 6.69076 | 5.87118 | 6.10716 | 6.13415 |
| STX2 | P32856 | 6.45798 | 6.3635 | 6.50758 | 6.42836 | 5.88096 | 6.10932 | 6.24691 | 6.0273 | 6.11644 | 6.49292 | 6.37569 | 6.17309 | 6.11628 | 6.44394 | 5.92261 | 6.33711 |
| CCNT2 | O60583 | 6.06845 | 6.84479 | 5.79697 | 6.33112 | 6.18017 | 6.63536 | 6.242 | 6.07356 | 5.86009 | 5.96953 | 5.59572 | 5.76894 | 6.72555 | 7.11903 | 6.43412 | 6.3546 |
| SH3BGRL | O75368 | 5.83688 | 5.35064 | 6.51565 | 4.38436 | 6.75406 | 6.40473 | 5.8746 | 6.84891 | 5.84091 | 6.70447 | 6.11717 | 6.989 | 6.07207 | 7.24543 | 6.42084 | 6.64027 |
| DYNLRB1 | Q9NP97 | 5.58156 | 4.46821 | 6.16451 | 4.33333 | 5.76938 | 5.44077 | 5.4529 | 7.47085 | 5.13932 | 6.28172 | 5.3894 | 6.5383 | 4.97265 | 9.69828 | 8.01469 | 9.28413 |
| GOSR1 | O95249 | 6.17537 | 6.39944 | 5.63891 | 6.49667 | 6.52394 | 6.00039 | 6.04586 | 6.13777 | 6.243 | 6.29263 | 5.99443 | 6.29043 | 6.43036 | 6.49941 | 6.34661 | 6.48477 |
| TMCC3 | Q9ULS5 | 6.52525 | 6.45561 | 5.67644 | 6.22392 | 5.97796 | 5.73836 | 6.37674 | 5.7941 | 6.16068 | 6.18379 | 6.34684 | 6.19824 | 6.74308 | 6.0173 | 6.91353 | 6.66818 |
| PPIL3 | Q9H2H8 | 5.84375 | 5.56973 | 5.79663 | 4.88867 | 6.61272 | 6.05312 | 5.98886 | 6.89365 | 6.33071 | 6.63213 | 6.61768 | 6.63235 | 6.28733 | 6.79222 | 6.63846 | 6.82199 |
| MAOA | P21397 | 6.41746 | 6.23377 | 5.63821 | 7.17277 | 6.44852 | 5.99512 | 6.68564 | 6.11319 | 5.58677 | 6.46483 | 5.68015 | 5.93654 | 6.22333 | 6.31017 | 6.46463 | 6.62888 |
| UCK2 | Q9BZX2 | 6.33309 | 5.9697 | 6.98587 | 6.38375 | 6.35317 | 6.21563 | 6.45973 | 6.59345 | 6.88604 | 6.42813 | 6.74442 | 6.53463 | 5.59298 | 5.70985 | 5.26713 | 5.54243 |
| CEP43 | O95684 | 6.48794 | 5.88489 | 6.67872 | 5.95548 | 6.48139 | 6.58539 | 6.05236 | 6.66958 | 6.38193 | 6.33786 | 6.92248 | 6.67533 | 6.24326 | 5.56047 | 5.28288 | 5.80004 |
| NPHP3 | Q7Z494 | 6.12222 | 6.3781 | 6.53501 | 6.11939 | 6.3153 | 6.71965 | 6.7542 | 6.83594 | 6.50407 | 6.29719 | 6.60324 | 6.59729 | 5.49125 | 5.46111 | 5.3803 | 5.88575 |
| PPP1R2 | P41236 | 6.38091 | 6.05411 | 6.17111 | 5.97991 | 6.67526 | 6.46978 | 6.08651 | 7.0641 | 6.8342 | 5.91841 | 5.7532 | 6.59611 | 5.78931 | 6.40816 | 5.38517 | 6.43374 |
| EEPD1 | Q7L9B9 | 7.05623 | 6.29132 | 6.37599 | 6.55086 | 6.20294 | 5.92821 | 6.08307 | 5.92563 | 6.4212 | 6.53519 | 6.5664 | 6.87749 | 5.84444 | 5.75863 | 5.66055 | 5.92185 |
| BLOC1S3 | Q6QNY0 | 6.52044 | 5.8889 | 6.78956 | 5.79658 | 6.4402 | 6.88453 | 5.91602 | 6.46228 | 6.47078 | 6.32607 | 6.39623 | 6.33855 | 6.25954 | 6.10387 | 5.74193 | 5.66452 |
| PPM1L | Q5SGD2 | 6.15599 | 6.31012 | 6.04167 | 6.85302 | 5.91293 | 5.81851 | 6.50465 | 5.8689 | 6.31658 | 6.3895 | 6.18197 | 6.45035 | 6.30057 | 6.13668 | 6.20007 | 6.55847 |
| MMGT1 | Q8N4V1-2 | 6.24662 | 6.26843 | 5.84847 | 6.21672 | 6.24083 | 6.14813 | 5.828 | 6.2044 | 5.9112 | 6.51405 | 5.7048 | 6.29334 | 6.37853 | 6.61565 | 6.72701 | 6.85382 |
| DCAF7 | P61962 | 5.31049 | 5.9785 | 6.03271 | 5.72129 | 6.54503 | 6.49863 | 6.51895 | 7.39321 | 6.71905 | 7.25349 | 6.53458 | 6.7435 | 5.467 | 5.48464 | 5.75808 | 6.04086 |
| NDUFB8 | O95169 | 6.54636 | 5.85787 | 5.39563 | 5.59835 | 6.33416 | 5.52254 | 5.9189 | 6.01819 | 5.33244 | 5.26951 | 5.6164 | 5.37838 | 6.74897 | 8.38196 | 7.39768 | 8.68267 |
| MVB12A | Q96EY5 | 5.90918 | 6.27191 | 6.53423 | 6.01839 | 6.12275 | 6.53299 | 6.7112 | 6.32912 | 6.40652 | 6.811 | 7.05811 | 6.81665 | 6.06879 | 5.51612 | 5.52337 | 5.36966 |
| ABHD6 | Q9BV23 | 6.28663 | 6.4706 | 5.91492 | 6.56616 | 5.99303 | 5.78563 | 6.51737 | 6.13635 | 6.27258 | 6.13461 | 6.07457 | 6.05136 | 6.56503 | 6.41385 | 6.32697 | 6.49035 |
| NTMT1 | Q9BV86 | 6.28783 | 5.84281 | 6.56022 | 5.60325 | 6.80965 | 6.73675 | 6.22491 | 6.99454 | 6.12574 | 6.11464 | 7.25397 | 6.82034 | 5.77762 | 5.4657 | 5.73834 | 5.64368 |
| SUCO | Q9UBS9 | 6.07414 | 6.20334 | 5.75296 | 6.3707 | 5.94705 | 6.27376 | 6.22035 | 6.11638 | 5.80768 | 6.31634 | 5.85443 | 5.96236 | 6.86468 | 6.8858 | 6.86424 | 6.48579 |
| CYLD | Q9NQC7 | 6.58755 | 5.87021 | 6.71272 | 6.31591 | 6.48628 | 6.52741 | 6.53993 | 6.63509 | 6.14594 | 6.76644 | 6.43695 | 6.35486 | 5.29235 | 5.97321 | 5.74059 | 5.61455 |
| DCTD | P32321-2 | 6.01287 | 5.84975 | 7.10166 | 5.70488 | 6.66633 | 6.87994 | 6.0248 | 6.65626 | 6.40741 | 6.7749 | 6.75599 | 7.03023 | 5.72237 | 5.50168 | 5.56324 | 5.34769 |
| PAN3 | Q58A45 | 6.25972 | 6.15952 | 7.11778 | 6.02804 | 6.36484 | 5.99589 | 6.43356 | 6.16985 | 6.51389 | 6.33177 | 6.35053 | 6.35766 | 6.25391 | 6.10728 | 5.84055 | 5.71521 |
| ACP2 | P11117 | 6.55948 | 6.56685 | 6.03923 | 6.35498 | 6.56555 | 6.19413 | 6.22512 | 6.44602 | 6.27313 | 6.05056 | 5.77101 | 5.75984 | 6.44731 | 6.13118 | 6.39497 | 6.22066 |
| LRCH2 | Q5VUJ6 | 6.61749 | 6.18886 | 6.2011 | 5.57808 | 6.33142 | 5.98273 | 6.31057 | 6.35018 | 6.33138 | 6.19691 | 6.32302 | 6.2139 | 6.42435 | 6.42808 | 6.45001 | 6.07193 |
| ENDOD1 | O94919 | 6.07032 | 6.66954 | 5.8557 | 6.41301 | 6.02143 | 6.1729 | 6.5559 | 5.91003 | 5.97896 | 6.13533 | 5.66196 | 5.90159 | 6.5863 | 6.10035 | 6.99503 | 6.97164 |
| PRPS1 | P60891 | 6.48534 | 6.0771 | 6.50929 | 6.08254 | 6.28221 | 7.18265 | 6.25776 | 6.47734 | 5.82975 | 6.06293 | 6.62838 | 6.32947 | 6.2825 | 5.72573 | 5.74509 | 6.04192 |
| ULBP3 | Q9BZM4 | 6.58687 | 6.90962 | 6.50744 | 7.33336 | 6.04815 | 5.97284 | 6.35794 | 6.02184 | 6.1608 | 6.43141 | 6.29137 | 6.46655 | 6.01075 | 5.37969 | 5.77345 | 5.74792 |
| ZMAT2 | Q96NC0 | 6.70665 | 6.24336 | 5.81569 | 6.26861 | 5.98832 | 6.15165 | 6.23291 | 6.10078 | 6.38276 | 6.17811 | 5.71996 | 5.42207 | 6.87981 | 6.72942 | 6.7909 | 6.389 |
| MTBP | Q96DY7 | 6.02548 | 6.33646 | 6.66762 | 6.95801 | 5.7804 | 6.31009 | 6.67133 | 6.04878 | 6.13994 | 6.33098 | 6.2988 | 6.71752 | 5.7152 | 5.9219 | 6.07885 | 5.99865 |
| SLC26A2 | P50443 | 6.62789 | 6.77937 | 6.14483 | 6.5719 | 6.45616 | 6.14116 | 6.38216 | 6.18796 | 6.21884 | 6.31438 | 6.12713 | 5.90038 | 6.15668 | 5.94661 | 5.9891 | 6.05545 |
| NFYA | P23511 | 5.80317 | 6.18594 | 6.17206 | 5.98934 | 6.13541 | 6.35631 | 6.04235 | 6.31656 | 5.77858 | 6.19534 | 6.31503 | 6.15828 | 6.66733 | 6.71924 | 6.6283 | 6.53676 |
| MPI | P34949 | 5.51256 | 4.79802 | 6.1827 | 4.80089 | 6.11507 | 6.45278 | 6.06823 | 7.15688 | 5.76335 | 7.06284 | 6.77807 | 7.41864 | 5.80612 | 7.10134 | 6.29717 | 6.68535 |

| | | | | | | | | | | | | | | | | | |
|---|---|---|---|---|---|---|---|---|---|---|---|---|---|---|---|---|---|
| GBP1 | P32455 | 6.18849 | 5.96 | 6.63719 | 5.60614 | 6.15628 | 6.34408 | 6.40977 | 6.26964 | 6.08692 | 6.57107 | 6.40466 | 6.70582 | 5.99834 | 6.59549 | 6.07275 | 5.99336 |
| PSME3IP1 | Q9GZU8 | 5.19728 | 5.93666 | 6.32489 | 5.06986 | 5.49579 | 6.69439 | 5.7027 | 6.82326 | 5.88038 | 6.2684 | 5.86139 | 6.2666 | 7.20895 | 7.33133 | 6.85739 | 7.0807 |
| LLPH | Q9BRT6 | 6.03364 | 6.15425 | 5.89913 | 6.52854 | 5.94716 | 5.65276 | 6.22635 | 6.02656 | 6.0515 | 5.91051 | 6.0672 | 6.01273 | 6.10064 | 7.16609 | 7.16192 | 7.06101 |
| WIPI2 | Q9Y4P8 | 6.55858 | 6.4982 | 6.75692 | 5.9619 | 6.11882 | 6.64838 | 6.68469 | 6.32876 | 6.51989 | 6.24543 | 6.19114 | 6.34651 | 5.79657 | 6.01677 | 5.77124 | 5.5562 |
| DPCD | Q9BVM2 | 5.9568 | 5.98427 | 6.49554 | 5.5704 | 6.06485 | 6.75148 | 6.36349 | 6.5063 | 6.503 | 6.54556 | 6.40553 | 6.25063 | 6.01792 | 6.52164 | 6.03067 | 6.03192 |
| CRIP1 | P50238 | 6.00038 | 6.05348 | 6.53546 | 5.63602 | 6.83119 | 7.65076 | 5.91916 | 7.08681 | 6.01125 | 6.92213 | 7.0769 | 6.63215 | 5.98849 | 5.89531 | 4.94239 | 4.81812 |
| CARNMT1 | Q8N4J0 | 5.55478 | 5.62443 | 6.23055 | 5.7944 | 5.85658 | 6.50846 | 5.67652 | 5.99263 | 5.66984 | 6.3537 | 6.56829 | 6.31108 | 6.39666 | 7.60036 | 6.85807 | 7.00365 |
| CHSY3 | Q70JA7 | 6.47592 | 6.62373 | 6.19606 | 7.2539 | 6.55499 | 6.51537 | 6.18609 | 6.13113 | 6.07782 | 6.46456 | 5.89133 | 6.44873 | 6.02002 | 5.89277 | 5.50024 | 5.76732 |
| GALE | Q14376 | 5.79368 | 5.70474 | 6.54526 | 5.5088 | 5.86442 | 6.32969 | 5.83005 | 6.38199 | 6.12979 | 6.96153 | 6.89199 | 6.82248 | 6.73602 | 6.69067 | 6.14698 | 6.66191 |
| DUSP11 | O75319 | 6.49435 | 5.90246 | 6.22886 | 6.89236 | 6.382 | 6.35317 | 6.68215 | 5.72153 | 6.24711 | 5.87615 | 6.12008 | 6.16655 | 6.30786 | 5.79984 | 6.32037 | 6.50516 |
| B3GLCT | Q6Y288 | 6.28909 | 6.64732 | 5.95676 | 6.5808 | 6.68828 | 6.41932 | 5.94895 | 6.72579 | 5.72444 | 5.85424 | 6.14965 | 5.60027 | 6.37809 | 6.37351 | 6.46201 | 6.20149 |
| UPP1 | Q16831 | 5.77052 | 6.10346 | 6.30747 | 5.55705 | 6.11865 | 6.37797 | 6.04764 | 6.17941 | 6.06268 | 5.92914 | 6.16878 | 6.16143 | 6.34204 | 7.42073 | 6.95365 | 6.49936 |
| HAGH | Q16775 | 6.63599 | 6.05307 | 6.3903 | 6.08397 | 6.54224 | 6.22527 | 6.23704 | 6.38254 | 6.40118 | 6.25889 | 6.63798 | 6.24767 | 6.45268 | 6.05451 | 5.91291 | 5.48375 |
| ERRFI1 | Q9UJM3 | 5.67941 | 5.47329 | 6.32992 | 5.17511 | 6.26121 | 6.71116 | 5.81767 | 6.55751 | 5.77196 | 5.79404 | 6.35428 | 6.20028 | 7.8247 | 6.33038 | 7.08127 | 6.63782 |
| TCEAL4 | Q96EI5-2 | 5.97903 | 6.08857 | 6.67747 | 5.79627 | 5.99285 | 6.79155 | 5.60734 | 6.5758 | 6.29419 | 6.96149 | 7.07816 | 7.30643 | 5.75102 | 5.95577 | 5.70451 | 5.43955 |
| ITM2C | Q9NQX7 | 7.29152 | 8.09227 | 6.64491 | 7.34986 | 6.47407 | 6.43198 | 6.74749 | 5.97999 | 6.86068 | 7.09833 | 6.61721 | 6.58033 | 4.3541 | 4.72029 | 4.24207 | 4.5149 |
| KLHL36 | Q8N4N3 | 5.51505 | 5.97353 | 6.36749 | 5.54426 | 6.01875 | 6.4809 | 6.07081 | 5.85445 | 6.25617 | 6.24141 | 6.31832 | 6.33307 | 6.47329 | 6.88824 | 7.02197 | 6.64228 |
| PDCD2L | Q9BRP1 | 6.78947 | 5.7636 | 7.07466 | 5.8652 | 6.05564 | 6.25756 | 6.14497 | 6.64927 | 6.30139 | 6.31808 | 6.70116 | 7.03069 | 5.89714 | 5.77251 | 5.60422 | 5.77443 |
| MANSC1 | Q9H8J5 | 6.75832 | 6.34595 | 6.40567 | 6.35528 | 6.49146 | 6.07903 | 5.53496 | 6.59916 | 6.2078 | 6.44544 | 6.58661 | 6.46827 | 5.99604 | 5.85058 | 5.87264 | 6.00278 |
| POLR2I | P36954 | 6.98647 | 5.4207 | 6.02796 | 5.23091 | 6.23836 | 5.76185 | 5.7021 | 7.09863 | 5.95332 | 6.73948 | 5.87265 | 6.80025 | 5.19055 | 7.0414 | 6.67345 | 7.26191 |
| TRMT61A | Q96FX7 | 5.48013 | 5.92144 | 5.59857 | 5.60305 | 6.33694 | 6.86724 | 5.59427 | 5.96099 | 5.97603 | 6.0174 | 6.30871 | 6.18112 | 6.88373 | 6.79741 | 7.47793 | 6.99504 |
| CFDP1 | Q9UEE9 | 5.52964 | 5.64498 | 6.14123 | 5.39726 | 5.38549 | 6.14731 | 5.50778 | 6.36477 | 6.02302 | 6.05926 | 6.74145 | 6.26495 | 6.72772 | 7.44492 | 7.40074 | 7.21949 |
| MGAT4B | Q9UQ53-3 | 6.49061 | 6.18482 | 5.89644 | 6.8264 | 6.16208 | 6.11762 | 6.55943 | 5.86296 | 6.40407 | 6.35664 | 6.1521 | 6.07485 | 6.48898 | 5.97968 | 6.37401 | 6.06932 |
| TNIP1 | Q15025 | 6.97512 | 5.6689 | 6.92187 | 5.99099 | 6.53786 | 6.55192 | 6.17426 | 6.65641 | 6.80809 | 6.71383 | 6.67778 | 6.77352 | 5.5935 | 5.23025 | 4.86596 | 4.95973 |
| GSTM2 | P28161 | 6.17615 | 5.46696 | 7.02968 | 5.11236 | 6.51418 | 6.60516 | 5.9604 | 6.70218 | 6.1799 | 6.83553 | 6.53922 | 6.30971 | 6.04413 | 6.49987 | 5.96269 | 6.06187 |
| HYPK | Q9NX55 | 6.29451 | 5.2081 | 5.50454 | 4.82475 | 5.07262 | 4.66773 | 5.15991 | 6.21512 | 4.34703 | 6.08505 | 5.40962 | 6.61861 | 5.31072 | 10.6512 | 8.00616 | 10.6243 |
| RAB23 | Q9ULC3 | 6.1162 | 6.58919 | 6.26714 | 6.66922 | 6.04206 | 6.08051 | 6.60133 | 5.9997 | 6.59866 | 6.5903 | 6.26999 | 6.1275 | 6.17262 | 5.85411 | 6.3255 | 6.29598 |
| PALM3 | A6NDB9 | 6.08046 | 6.56199 | 6.41358 | 5.89296 | 6.22777 | 6.21991 | 6.41017 | 5.80685 | 5.87441 | 6.45298 | 6.71616 | 6.60033 | 6.58751 | 6.16109 | 5.97721 | 6.01662 |
| NDUFC2 | O95298 | 6.65833 | 5.58937 | 6.12409 | 6.14323 | 6.36073 | 5.72548 | 6.67536 | 6.1029 | 6.22105 | 6.07753 | 6.32314 | 5.76544 | 6.40445 | 6.75034 | 6.69357 | 6.38499 |
| SLC25A17 | O43808 | 6.42806 | 6.36659 | 6.01919 | 6.79782 | 5.9718 | 6.00403 | 6.1664 | 6.18123 | 6.09126 | 5.65226 | 5.77009 | 5.74191 | 6.89109 | 6.48432 | 6.5977 | 6.83625 |
| PMF1 | Q6P1K2 | 5.76606 | 4.88609 | 5.6297 | 4.90663 | 5.92772 | 4.68247 | 5.16285 | 6.98426 | 4.62623 | 6.19363 | 5.37808 | 7.41807 | 5.92729 | 9.01367 | 8.2352 | 9.26206 |
| SIGMAR1 | Q99720 | 6.3347 | 6.39269 | 5.56149 | 6.07526 | 6.05273 | 6.01376 | 6.29284 | 6.03326 | 6.4887 | 6.12284 | 6.00059 | 6.13376 | 6.63131 | 6.41335 | 6.66472 | 6.788 |
| CNNM4 | Q6P4Q7 | 6.4848 | 6.38278 | 6.0843 | 6.97715 | 6.412 | 6.10472 | 6.49346 | 5.85822 | 6.20215 | 6.15655 | 5.88123 | 5.88818 | 6.20283 | 6.29936 | 6.32815 | 6.24412 |
| MADD | Q8WXG6-4 | 6.25009 | 6.50925 | 6.43723 | 6.50476 | 6.11547 | 6.15669 | 6.45173 | 6.79617 | 6.31416 | 6.67152 | 6.36853 | 6.38904 | 5.78413 | 6.89422 | 5.63948 | 5.61753 |
| PANX1 | Q96RD7 | 6.2531 | 6.56927 | 5.93046 | 6.70676 | 6.34922 | 6.04514 | 5.67478 | 6.19288 | 5.77182 | 6.14897 | 6.13792 | 6.46121 | 6.28781 | 6.39322 | 6.23601 | 6.84144 |
| MOXD1 | Q6UVY6 | 6.67466 | 6.9171 | 6.54243 | 6.63226 | 6.00368 | 5.99346 | 6.70641 | 6.5646 | 5.7821 | 6.24977 | 6.18694 | 6.0991 | 5.88327 | 5.77429 | 6.23784 | 5.75209 |
| HEBP1 | Q9NRV9 | 5.48078 | 5.5563 | 6.09285 | 5.45585 | 6.56513 | 6.06684 | 5.96401 | 6.37424 | 7.05456 | 7.002 | 6.51184 | 7.25627 | 6.19909 | 6.02419 | 6.00523 | 6.39081 |
| STING1 | Q86WV6 | 6.68202 | 6.46037 | 5.70867 | 5.87363 | 6.07592 | 5.98643 | 6.18753 | 5.90025 | 6.16766 | 5.90064 | 5.90386 | 5.57283 | 6.88466 | 6.81802 | 7.13065 | 6.74685 |
| MTMR9 | Q96QG7 | 6.14622 | 6.70752 | 6.8142 | 6.09513 | 6.39726 | 6.13141 | 6.16383 | 6.89329 | 6.35278 | 6.46138 | 6.7743 | 6.35158 | 5.78638 | 5.79043 | 5.643 | 5.4913 |
| NME7 | Q9Y5B8 | 6.17828 | 5.34926 | 6.94581 | 6.28666 | 6.33173 | 6.98585 | 6.35138 | 6.39996 | 6.16871 | 6.44948 | 7.03651 | 6.74806 | 5.7617 | 5.56985 | 5.46172 | 5.97504 |
| WDR37 | Q9Y2I8-2 | 5.97662 | 6.43271 | 7.14521 | 6.42643 | 6.55925 | 6.51016 | 6.34138 | 6.56457 | 6.83144 | 6.25793 | 6.15711 | 5.99316 | 5.90656 | 5.90184 | 5.48458 | 5.51105 |
| C12orf29 | Q8N999 | 6.19474 | 5.95542 | 6.92597 | 6.09978 | 6.29117 | 6.54616 | 6.1014 | 6.6312 | 6.40848 | 6.59187 | 6.67107 | 6.99741 | 5.78153 | 5.55939 | 5.49786 | 5.74656 |
| TTC3 | P53804 | 6.6544 | 6.62749 | 6.68236 | 6.52056 | 6.20005 | 6.72017 | 6.14963 | 6.37944 | 6.54755 | 6.80998 | 6.89955 | 6.81058 | 5.00777 | 5.59893 | 5.0961 | 5.29543 |
| GCLM | P48507 | 5.98168 | 6.33478 | 6.98651 | 6.42494 | 6.40874 | 6.76557 | 6.24629 | 6.59125 | 5.77447 | 6.73155 | 6.59454 | 6.90559 | 5.64489 | 5.66012 | 5.35101 | 5.79809 |
| FARP2 | O94887 | 6.68467 | 6.71788 | 6.54693 | 6.86488 | 6.07136 | 6.10977 | 6.73649 | 6.21152 | 6.51255 | 6.31544 | 6.14139 | 6.21347 | 6.58795 | 5.53715 | 5.64972 | 5.99884 |
| SEZ6L2 | Q6UXD5-5 | 6.6715 | 6.59691 | 6.49305 | 6.30429 | 6.28837 | 6.41453 | 6.39936 | 6.56918 | 6.41823 | 6.59283 | 6.61715 | 6.49751 | 5.4976 | 6.07131 | 5.26912 | 5.29906 |
| DHDDS | Q86SQ9-2 | 6.28639 | 7.31244 | 6.54517 | 6.69675 | 6.4539 | 6.57917 | 6.24689 | 6.10481 | 6.23619 | 6.52455 | 5.96522 | 6.22667 | 5.86952 | 5.54261 | 5.75808 | 5.65165 |
| TST | Q16762 | 6.37303 | 6.40553 | 6.03967 | 6.73207 | 5.85873 | 5.94978 | 6.22256 | 6.10581 | 6.09662 | 6.16673 | 5.6773 | 5.96887 | 6.84199 | 6.43402 | 6.42987 | 6.69741 |
| WIPF2 | Q8TF74 | 6.30659 | 6.33989 | 6.65943 | 6.02119 | 6.01683 | 6.87301 | 6.20776 | 6.68691 | 6.69529 | 6.34913 | 6.71155 | 7.03328 | 5.78681 | 5.84285 | 4.92151 | 5.54797 |
| GMPPB | Q9Y5P6-2 | 6.30198 | 5.83153 | 6.68915 | 5.79273 | 6.34883 | 6.63897 | 6.06275 | 6.52301 | 6.34556 | 6.50861 | 5.9657 | 5.84637 | 6.31981 | 6.67535 | 6.10463 | 6.04503 |
| VMP1 | Q96GC9 | 6.18122 | 6.59816 | 6.5731 | 6.86254 | 6.26058 | 5.73404 | 6.06377 | 5.44754 | 5.90872 | 6.38768 | 5.32185 | 5.73192 | 6.74279 | 6.3749 | 7.04402 | 6.76717 |
| OCLN | Q16625 | 6.56554 | 6.57106 | 5.96047 | 6.93154 | 5.96532 | 6.36818 | 5.85061 | 6.29965 | 5.93125 | 5.50709 | 6.17969 | 6.523 | 5.82415 | 6.43975 | 6.05181 | 7.03089 |
| NME6 | O75414 | 6.16798 | 5.09565 | 5.71694 | 5.66644 | 6.1343 | 5.36701 | 5.99781 | 6.30161 | 5.27417 | 6.28766 | 6.04672 | 6.57188 | 6.48689 | 7.43655 | 7.52743 | 7.92098 |
| ZHX1 | Q9UKY1 | 5.86379 | 5.82817 | 5.65123 | 6.15553 | 6.24364 | 6.10623 | 6.48188 | 5.75209 | 5.95859 | 6.07217 | 5.88243 | 5.85508 | 7.05087 | 6.99903 | 6.85153 | 7.24774 |
| REEP4 | Q9H6H4 | 6.19032 | 6.58327 | 5.57435 | 6.62965 | 6.06392 | 6.20873 | 6.31916 | 6.03518 | 6.17363 | 6.75693 | 6.60106 | 6.3644 | 6.04411 | 5.99436 | 6.39786 | 6.36308 |
| THOC7 | Q6I9Y2 | 6.57977 | 6.36508 | 5.69684 | 6.52401 | 6.1727 | 6.29672 | 6.00511 | 6.21929 | 6.38832 | 5.97639 | 6.08024 | 6.27882 | 6.75217 | 5.85377 | 6.37432 | 6.43643 |
| OBI1 | Q5W0B1 | 5.90221 | 5.83682 | 5.9434 | 6.7248 | 6.11783 | 6.25813 | 6.26181 | 6.08658 | 5.75404 | 5.67217 | 6.10885 | 6.34404 | 6.73295 | 6.35657 | 6.84937 | 7.05042 |
| CASP8AP2 | Q9UKL3 | 7.30153 | 6.92554 | 6.4806 | 7.34883 | 6.70688 | 6.09023 | 6.68429 | 6.31543 | 6.8219 | 6.18712 | 6.65873 | 6.0173 | 5.20365 | 5.09476 | 5.11342 | 5.0498 |
| SPC24 | Q8NBT2 | 6.19875 | 5.07519 | 5.6881 | 4.70306 | 5.5576 | 5.63231 | 5.29814 | 6.96232 | 5.30739 | 6.34752 | 5.34086 | 7.22918 | 4.9907 | 9.61801 | 7.32206 | 8.72882 |
| FCHSD2 | O94868 | 6.42231 | 6.21882 | 6.68536 | 6.29461 | 6.58273 | 6.73665 | 6.1849 | 6.59233 | 6.39081 | 6.78546 | 6.78317 | 6.55613 | 5.78432 | 5.67878 | 5.25019 | 5.71946 |
| ANKRD50 | Q9ULJ7 | 5.96383 | 6.15706 | 6.59646 | 6.13874 | 6.05848 | 6.12126 | 6.14269 | 5.52471 | 6.44844 | 6.33662 | 6.27595 | 6.05033 | 6.62671 | 6.51836 | 6.73966 | 6.30069 |
| GMNN | O75496 | 5.38752 | 6.00909 | 6.22065 | 5.14841 | 6.07251 | 6.27683 | 4.98573 | 6.81061 | 6.53334 | 6.47312 | 6.74935 | 6.83879 | 6.60708 | 6.73598 | 6.53346 | 6.61753 |
| DAXX | Q9UER7 | 5.68451 | 5.90091 | 6.25084 | 6.47952 | 6.25405 | 6.03236 | 6.37938 | 6.38614 | 5.6808 | 6.37007 | 6.51828 | 6.30964 | 6.55423 | 6.4808 | 6.18301 | 6.53544 |
| ATP5PF | P18859-2 | 6.46762 | 5.15637 | 4.23162 | 5.77867 | 5.31721 | 3.57846 | 4.47505 | 5.39703 | 3.71721 | 4.55445 | 4.65038 | 4.39567 | 4.6889 | 14.331 | 9.65135 | 13.609 |
| NCKAP5L | Q9HCH0 | 6.37194 | 6.80677 | 6.63132 | 5.96263 | 6.00615 | 6.30243 | 6.20777 | 5.96056 | 6.26583 | 6.88093 | 6.55922 | 6.68305 | 6.29169 | 5.86447 | 5.38599 | 5.81924 |
| ARHGEF7 | Q14155-5 | 6.37174 | 5.92112 | 7.02084 | 5.90757 | 5.94968 | 6.82652 | 6.3817 | 5.89513 | 6.63534 | 6.3711 | 6.90848 | 6.12232 | 6.41114 | 5.66134 | 5.92578 | 5.69022 |
| ARHGEF6 | Q15052 | 5.98586 | 6.19349 | 6.66407 | 5.81639 | 6.49724 | 6.69503 | 5.57427 | 6.45173 | 6.38236 | 6.75951 | 6.28249 | 6.4076 | 6.17321 | 6.46311 | 5.98657 | 5.66707 |
| PLEKHM1 | Q9Y4G2 | 6.67273 | 7.00484 | 6.26766 | 6.23141 | 6.09697 | 6.44082 | 6.15149 | 6.27408 | 6.7448 | 6.67005 | 6.22067 | 6.28647 | 5.56121 | 5.85637 | 5.64118 | 5.87924 |
| PDE6D | O43924 | 5.08898 | 5.90801 | 5.95839 | 5.37758 | 6.45994 | 6.51158 | 6.23152 | 7.6336 | 6.86999 | 7.37908 | 7.07191 | 6.45706 | 5.33211 | 5.74632 | 6.10708 | 5.86687 |
| WWC2 | Q6AWC2-6 | 6.83342 | 6.09565 | 6.97333 | 6.21141 | 6.27046 | 6.72837 | 6.53641 | 6.52048 | 6.638 | 6.09675 | 6.2708 | 6.36587 | 5.82244 | 5.4263 | 5.47044 | 5.73986 |
| RNASEH2B | Q5TBB1 | 5.42079 | 5.8563 | 6.59117 | 5.91568 | 5.50002 | 6.07845 | 5.86677 | 5.70791 | 5.82468 | 6.22515 | 6.493 | 6.43986 | 6.45823 | 7.26543 | 7.35835 | 6.9982 |
| TRUB1 | Q8WWH5 | 6.01453 | 5.90997 | 6.25984 | 5.95002 | 6.14558 | 6.22976 | 5.90192 | 6.20868 | 6.19494 | 6.3201 | 6.31292 | 6.30787 | 6.92694 | 6.62383 | 6.63913 | 6.05398 |
| KANSL3 | Q9P2N6-3 | 6.72868 | 6.53909 | 6.40722 | 6.71115 | 6.28553 | 5.79446 | 6.45184 | 6.11956 | 6.61188 | 6.40225 | 6.35417 | 6.44477 | 5.97271 | 5.84235 | 5.63246 | 5.70187 |
| FASTKD3 | Q14CZ7 | 6.25538 | 6.16536 | 6.218 | 7.08085 | 6.17392 | 5.94833 | 6.64688 | 5.53504 | 6.27312 | 5.89557 | 6.00131 | 6.52361 | 6.55094 | 5.79623 | 6.68172 | 6.25373 |
| RERE | Q9P2R6 | 6.13979 | 6.50482 | 6.33897 | 6.31696 | 6.52237 | 6.23726 | 6.38223 | 6.0317 | 5.84203 | 6.22889 | 6.31983 | 5.92736 | 6.21644 | 6.22631 | 6.22226 | 6.54279 |
| COX15 | Q7KZN9 | 6.05589 | 6.59785 | 5.67714 | 6.38567 | 6.20144 | 6.19839 | 6.28434 | 6.31603 | 6.51146 | 5.8497 | 5.95524 | 6.08017 | 6.35677 | 6.22989 | 6.77271 | 6.52732 |
| TMEM126A | Q9H061 | 6.43971 | 5.97887 | 6.17667 | 6.4328 | 6.39179 | 5.7652 | 6.24899 | 6.27794 | 5.79162 | 6.39741 | 5.8422 | 6.36673 | 6.38859 | 6.51634 | 6.67742 | 6.28774 |
| CLK3 | P49761 | 6.13876 | 6.18883 | 6.06883 | 6.45262 | 6.31983 | 6.02565 | 6.45342 | 5.887 | 6.60106 | 6.04004 | 6.39795 | 6.19597 | 6.14953 | 6.07928 | 6.53578 | 6.46547 |
| ASF1A | Q9Y294 | 5.99091 | 6.46532 | 6.1404 | 5.97176 | 5.95598 | 6.39969 | 5.7409 | 6.34024 | 6.39253 | 6.66692 | 6.70567 | 6.53666 | 6.19576 | 6.21355 | 5.86705 | 6.41666 |
| TRAPPC5 | Q8IUR0 | 6.07017 | 6.00691 | 6.59886 | 5.96207 | 6.20567 | 6.875 | 6.47577 | 6.65472 | 6.46578 | 6.67771 | 6.7623 | 6.71667 | 5.68828 | 5.82032 | 5.63302 | 5.38674 |
| ALG10 | Q5BKT4 | 6.27349 | 6.92753 | 5.97054 | 6.43009 | 5.47227 | 6.22994 | 6.4288 | 6.0922 | 6.07418 | 6.07294 | 5.36109 | 6.1084 | 6.37247 | 6.50462 | 6.859 | 6.82242 |
| LIN7C | Q9NUP9 | 5.99736 | 6.43823 | 6.49358 | 6.53168 | 6.32166 | 6.27385 | 5.97852 | 6.36333 | 6.33829 | 6.00372 | 6.92654 | 6.37665 | 6.43683 | 5.62282 | 5.89743 | 5.99953 |
| IFT74 | Q96LB3 | 6.36589 | 6.20612 | 6.61868 | 5.7557 | 6.46797 | 6.9782 | 6.08423 | 6.87431 | 6.77445 | 6.36068 | 6.56434 | 6.70541 | 6.03823 | 5.57697 | 5.48761 | 5.14118 |
| COMMD1 | Q8N668 | 6.17867 | 5.97425 | 6.78701 | 5.77567 | 6.24202 | 6.31641 | 6.69446 | 6.20868 | 6.1382 | 6.08283 | 6.90604 | 5.63729 | 6.52316 | 6.81361 | 5.87336 | 5.84834 |
| FRMD6 | Q96NE9 | 6.04728 | 5.87115 | 6.2778 | 6.30776 | 6.48163 | 6.34047 | 6.44375 | 5.97572 | 5.79518 | 6.01176 | 6.15192 | 5.6938 | 6.86854 | 6.44085 | 6.7757 | 6.51669 |
| YEATS4 | O95619 | 6.48157 | 6.4916 | 6.34479 | 6.71854 | 6.43231 | 6.23151 | 6.41861 | 6.1243 | 6.27727 | 6.75595 | 5.95566 | 6.03196 | 6.0153 | 5.80444 | 5.90074 | 6.01545 |
| COMMD9 | Q9P000 | 5.65006 | 4.8624 | 5.9343 | 4.33657 | 5.64854 | 4.9777 | 5.46124 | 6.10332 | 4.64801 | 6.25793 | 5.42273 | 6.32701 | 5.62069 | 10.8668 | 7.93773 | 9.94498 |
| UTP11 | Q9Y3A2 | 6.48538 | 6.23962 | 5.15525 | 6.29179 | 6.38974 | 6.13677 | 6.26124 | 6.37458 | 5.80494 | 6.0412 | 5.51981 | 5.63641 | 6.96548 | 6.75455 | 7.15507 | 6.78814 |
| RASAL2 | Q9UJF2-2 | 6.23334 | 6.04231 | 6.51622 | 6.33175 | 6.20444 | 6.33829 | 6.08153 | 6.58998 | 6.00063 | 6.10138 | 6.33994 | 6.242 | 6.10359 | 6.31417 | 6.34394 | 6.21649 |
| PPRC1 | Q5VV67 | 7.28172 | 6.6871 | 7.61828 | 7.76511 | 6.28766 | 5.84797 | 7.11264 | 6.31804 | 6.63874 | 6.20614 | 7.06612 | 6.89048 | 4.79018 | 4.04009 | 4.66745 | 4.78227 |
| NDUFS5 | O43920 | 6.4719 | 6.40875 | 6.19219 | 6.50399 | 6.27544 | 6.18371 | 6.39759 | 5.80069 | 6.224 | 6.28575 | 5.78519 | 5.60806 | 6.68759 | 6.43755 | 6.40403 | 6.33358 |
| MTMR10 | Q9NXD2 | 6.10672 | 6.24963 | 6.25489 | 6.20156 | 6.22988 | 6.13983 | 6.22804 | 6.22968 | 6.14709 | 6.22698 | 6.12748 | 6.16823 | 6.54916 | 6.36135 | 6.34753 | 6.43193 |
| TBC1D22B | Q9NU19 | 6.62726 | 7.29101 | 6.76928 | 6.46921 | 6.18633 | 6.25222 | 6.06097 | 7.00456 | 6.15242 | 6.69453 | 6.32077 | 6.1099 | 5.31498 | 5.80533 | 5.4027 | 5.53855 |
| FEZ2 | Q9UHY8-2 | 6.43231 | 6.57863 | 7.36997 | 6.36758 | 6.28298 | 6.96684 | 6.50313 | 6.52998 | 6.90571 | 7.09787 | 7.47601 | 6.98587 | 4.81946 | 4.71688 | 4.2641 | 4.70269 |
| EIF1 | P41567 | 6.18157 | 5.59062 | 7.34432 | 5.80571 | 6.43014 | 6.0307 | 6.05762 | 6.29764 | 6.37123 | 7.35868 | 7.4848 | 7.22207 | 5.1102 | 5.54526 | 5.6153 | 5.55415 |
| SPG21 | Q9NZD8 | 6.24545 | 5.94913 | 6.24102 | 5.87815 | 6.29661 | 6.09847 | 6.13584 | 6.14032 | 6.17587 | 6.70229 | 6.20565 | 6.67669 | 5.80696 | 6.40234 | 6.43369 | 6.61151 |
| AGAP3 | Q96P47-4 | 5.86907 | 6.17583 | 6.36458 | 5.73175 | 6.15693 | 6.37117 | 6.39694 | 6.25585 | 6.4314 | 6.33859 | 6.68193 | 6.41937 | 6.34955 | 6.21511 | 6.28867 | 5.95327 |
| PIAS1 | O75925-2 | 5.69109 | 5.80455 | 6.23273 | 6.30552 | 6.16671 | 6.54486 | 6.16378 | 6.0915 | 5.87416 | 6.50402 | 6.55907 | 6.42583 | 6.40884 | 6.48987 | 6.42239 | 6.31509 |
| PARS2 | Q7L3T8 | 6.10464 | 6.56097 | 5.55089 | 6.5229 | 6.18388 | 6.06044 | 6.35158 | 6.10614 | 6.11945 | 6.109 | 5.96484 | 6.11596 | 6.5625 | 6.7491 | 6.54523 | 6.39248 |
| PCYOX1L | Q8NBM8 | 6.38968 | 6.20997 | 5.3055 | 6.79991 | 6.32059 | 5.7722 | 6.59869 | 6.01989 | 6.40459 | 6.31938 | 5.66633 | 5.69348 | 7.00324 | 5.93635 | 6.84272 | 6.71749 |
| YIF1B | Q5BJH7 | 6.37585 | 6.6554 | 5.72072 | 6.48962 | 6.24103 | 6.22664 | 6.4376 | 6.56915 | 6.12085 | 6.38584 | 5.98173 | 6.00013 | 6.19711 | 6.37798 | 6.20965 | 6.01069 |
| NOB1 | Q9ULX3 | 6.12987 | 6.65883 | 6.93259 | 6.16658 | 6.32093 | 7.23529 | 6.33903 | 6.70524 | 6.48999 | 6.61657 | 6.9522 | 6.98067 | 4.89928 | 5.39996 | 4.9328 | 5.24017 |
| KNSTRN | Q9Y448 | 6.35654 | 6.20073 | 7.13334 | 6.34372 | 6.26089 | 6.89166 | 6.65762 | 7.04455 | 6.59339 | 6.58292 | 7.50421 | 7.28027 | 5.27071 | 4.82903 | 4.35817 | 4.69224 |
| DHCR7 | Q9UBM7 | 6.22766 | 6.35721 | 6.25364 | 6.35244 | 6.04672 | 5.95476 | 6.466 | 5.75512 | 6.24334 | 6.68907 | 5.89044 | 5.85053 | 6.44398 | 6.26329 | 6.72025 | 6.48554 |
| TMEM248 | Q9NWD8 | 7.40967 | 7.30786 | 7.28789 | 7.51043 | 6.45166 | 6.4473 | 6.60931 | 6.44675 | 6.81955 | 7.02233 | 7.14327 | 7.20323 | 3.91163 | 4.13793 | 3.96047 | 4.33073 |

| | | | | | | | | | | | | | | | | | |
|---|---|---|---|---|---|---|---|---|---|---|---|---|---|---|---|---|---|
| AAR2 | Q9Y312 | 6.44104 | 6.75603 | 6.55311 | 6.2546 | 6.9428 | 6.77733 | 6.30971 | 6.55472 | 6.25584 | 6.06353 | 5.97883 | 5.69933 | 6.09742 | 5.90498 | 5.82288 | 5.58786 |
| COQ7 | Q99807 | 6.46516 | 5.6907 | 5.5923 | 5.69716 | 5.75059 | 5.5622 | 5.99712 | 5.86925 | 6.13723 | 6.12202 | 5.63857 | 6.01953 | 7.03405 | 7.08847 | 7.45281 | 7.88283 |
| RHOF | Q9HBH0 | 6.89447 | 6.83141 | 6.28402 | 6.62168 | 6.21417 | 6.04221 | 6.49242 | 6.24354 | 6.55819 | 6.47011 | 6.21082 | 5.97611 | 5.80565 | 5.67383 | 5.76535 | 5.91601 |
| PHF12 | Q96QT6 | 6.61132 | 6.6938 | 6.1681 | 6.5193 | 6.09587 | 5.98566 | 6.56853 | 5.93273 | 6.35292 | 6.17853 | 6.02439 | 6.35371 | 6.10176 | 6.11917 | 6.24539 | 6.04883 |
| SNX33 | Q8WV41 | 6.52332 | 6.36963 | 6.87644 | 6.55706 | 6.42998 | 6.86256 | 6.5134 | 6.79306 | 6.40126 | 6.42981 | 6.32654 | 6.43638 | 5.5826 | 5.45252 | 5.42652 | 5.01892 |
| COMMD5 | Q9GZQ3 | 5.74052 | 4.97014 | 6.0344 | 4.36098 | 5.89341 | 5.37665 | 5.36622 | 6.84162 | 5.06134 | 6.50586 | 5.82784 | 6.57345 | 6.40031 | 8.8335 | 8.07081 | 8.14295 |
| NMRAL1 | Q9HBL8 | 6.44938 | 6.47503 | 7.05393 | 6.10536 | 5.99983 | 6.88317 | 6.36631 | 7.12123 | 6.27731 | 6.63675 | 6.76561 | 6.52948 | 5.45421 | 5.41445 | 5.03162 | 5.43635 |
| PSPH | P78330 | 6.59969 | 6.48933 | 6.31413 | 5.8592 | 6.83125 | 7.40852 | 5.9105 | 7.49531 | 5.98948 | 6.39308 | 6.09795 | 6.04619 | 5.78777 | 6.15035 | 5.42335 | 5.2039 |
| DAPK3 | O43293 | 5.87242 | 6.31803 | 5.95683 | 6.25422 | 6.1092 | 6.47027 | 6.28305 | 6.119 | 6.17032 | 6.62788 | 6.26442 | 6.25451 | 6.13749 | 6.58063 | 6.108 | 6.47374 |
| ADRM1 | Q16186 | 6.38609 | 6.19404 | 6.33916 | 6.14029 | 6.22143 | 6.49101 | 6.07206 | 6.35163 | 6.28573 | 6.24291 | 6.44388 | 6.49868 | 6.3837 | 5.72414 | 6.19185 | 6.0334 |
| SLC25A16 | P16260 | 6.48744 | 6.43775 | 5.90101 | 7.03673 | 6.33383 | 6.24236 | 6.59602 | 6.02862 | 6.30486 | 5.90176 | 5.89052 | 6.18141 | 6.33273 | 5.96182 | 6.59365 | 5.76951 |
| PHKG2 | P15735 | 7.07575 | 6.77349 | 7.34956 | 6.95655 | 6.6383 | 6.86183 | 6.15722 | 6.48036 | 7.24762 | 7.41804 | 6.82581 | 7.00134 | 3.98383 | 4.60447 | 4.06372 | 4.5621 |
| UBE2Q1 | Q7Z7E8 | 6.06249 | 6.14452 | 7.04273 | 6.25748 | 5.90617 | 6.30722 | 6.21997 | 6.60516 | 6.16931 | 6.52456 | 6.92499 | 6.89058 | 5.78736 | 5.99191 | 5.7661 | 5.39945 |
| EFNB1 | P98172 | 6.54138 | 6.01639 | 5.86446 | 6.55271 | 6.12744 | 5.98506 | 6.37751 | 5.83832 | 5.80844 | 5.8918 | 6.24888 | 6.56876 | 6.28682 | 6.36271 | 6.8209 | 6.70842 |
| NTPCR | Q9BSD7 | 6.07522 | 6.05231 | 6.13578 | 5.98678 | 6.44784 | 6.71812 | 6.16596 | 6.89015 | 6.59585 | 6.5266 | 6.27563 | 6.24367 | 6.40304 | 5.92059 | 5.80919 | 5.75328 |
| CLNS1A | P54105 | 6.58684 | 5.91101 | 6.69452 | 5.38311 | 6.84961 | 6.53697 | 6.42571 | 7.27564 | 6.99907 | 6.57769 | 6.71371 | 6.51021 | 5.74954 | 5.36743 | 5.33657 | 5.08238 |
| POLR1D | P0DPB6 | 6.43329 | 5.4734 | 5.91955 | 5.34057 | 6.17485 | 5.55427 | 6.5622 | 6.59134 | 5.5374 | 6.34924 | 6.07897 | 6.95852 | 6.20192 | 7.14572 | 7.22212 | 7.36663 |
| SLC25A15 | Q9Y619 | 6.55774 | 6.56457 | 5.82887 | 6.42557 | 6.47656 | 5.8041 | 6.34431 | 5.81612 | 6.06427 | 5.83786 | 6.00283 | 6.27214 | 6.4143 | 6.40968 | 6.60654 | 6.57454 |
| MT-ND1 | P03886 | 5.70983 | 6.70445 | 6.02912 | 6.30827 | 6.04237 | 5.84435 | 6.34524 | 6.26543 | 6.02387 | 5.9109 | 5.75812 | 6.09574 | 6.60327 | 6.67374 | 6.94948 | 6.73582 |
| TROAP | Q12815 | 6.80752 | 6.36355 | 6.43867 | 6.26574 | 6.44541 | 6.00781 | 6.30426 | 6.89661 | 6.6806 | 6.0291 | 6.56771 | 6.80776 | 5.71483 | 5.67699 | 5.39718 | 5.59626 |
| ATOX1 | O00244 | 6.60438 | 6.4452 | 7.3469 | 5.77647 | 6.70375 | 7.33249 | 5.20163 | 7.19274 | 5.41201 | 5.8592 | 5.61905 | 6.07834 | 4.62596 | 7.60825 | 6.31185 | 5.88178 |
| PPP4R4 | Q6NUP7 | 6.17305 | 6.03977 | 7.07004 | 6.13264 | 6.69496 | 6.34113 | 6.39854 | 6.23614 | 6.42047 | 6.49954 | 6.22463 | 6.65164 | 6.06759 | 5.84788 | 5.61488 | 5.5871 |
| GULP1 | Q9UBP9 | 6.30642 | 6.26621 | 6.17978 | 6.91744 | 6.01052 | 6.38146 | 6.11232 | 5.53915 | 6.16395 | 6.4555 | 6.3669 | 6.32497 | 5.94406 | 6.35732 | 6.1941 | 6.47989 |
| MEMO1 | Q9Y316-3 | 5.92218 | 6.00807 | 6.95495 | 5.93656 | 6.13284 | 6.88901 | 6.14565 | 6.53203 | 6.42939 | 6.57453 | 7.17991 | 7.01727 | 5.76256 | 5.47444 | 5.11277 | 5.92785 |
| NR2F2 | P24468 | 6.40252 | 6.44578 | 6.16792 | 6.73695 | 6.28417 | 6.21919 | 6.38779 | 6.49239 | 5.81372 | 6.10276 | 5.70891 | 6.08568 | 6.39036 | 6.58201 | 6.4452 | 6.33465 |
| SELENOT | P62341 | 6.40179 | 6.50563 | 5.98183 | 6.21096 | 6.4116 | 6.49605 | 6.03792 | 6.12673 | 5.89342 | 6.25726 | 5.78296 | 5.72821 | 6.48038 | 6.28016 | 7.01792 | 6.38719 |
| ACP1 | P24666-2 | 6.12536 | 5.40258 | 6.90263 | 5.44134 | 6.57733 | 6.16367 | 6.2336 | 6.90921 | 6.19863 | 6.76514 | 6.54811 | 6.70575 | 6.09718 | 6.22413 | 6.04859 | 5.65674 |
| TULP3 | O75386-2 | 6.44026 | 6.69011 | 6.51995 | 7.00024 | 5.98224 | 6.08016 | 6.33912 | 6.31505 | 7.4414 | 7.55813 | 6.29852 | 6.8546 | 5.14757 | 4.6376 | 5.61468 | 5.08036 |
| OSGEP | Q9NPF4 | 6.10584 | 5.56345 | 6.1301 | 5.50055 | 5.85074 | 5.94736 | 5.69237 | 5.77854 | 6.08491 | 5.95667 | 6.26808 | 6.06787 | 7.02717 | 7.34987 | 7.72774 | 6.94875 |
| ATG101 | Q9BSB4 | 6.20159 | 6.50418 | 6.77163 | 6.22486 | 6.06593 | 6.90721 | 6.18622 | 6.48003 | 6.38013 | 6.30196 | 6.65715 | 6.66559 | 5.81876 | 5.77676 | 5.62832 | 5.42967 |
| LRIG2 | O94898 | 5.99609 | 6.32321 | 6.35985 | 6.87021 | 6.46715 | 6.20209 | 6.15303 | 5.99609 | 6.25373 | 5.89572 | 6.44804 | 6.30436 | 6.40218 | 6.15352 | 6.21116 | 5.96356 |
| CHMP1B | Q7LBR1 | 6.26814 | 6.13614 | 6.4864 | 5.58208 | 6.26299 | 6.59905 | 6.12873 | 6.4917 | 6.34838 | 6.5053 | 6.56924 | 6.21119 | 6.34853 | 6.18656 | 6.11037 | 5.7652 |
| RFT1 | Q96AA3 | 6.44994 | 6.4076 | 5.84336 | 6.46466 | 6.03119 | 5.94037 | 6.30448 | 6.19287 | 5.85696 | 5.48048 | 5.93521 | 6.0028 | 6.62929 | 6.72588 | 6.9762 | 6.75871 |
| RPL35 | P42766 | 6.04281 | 6.39606 | 5.7183 | 5.85266 | 6.09055 | 6.20976 | 6.40836 | 6.75665 | 6.70886 | 6.33602 | 6.40205 | 6.58908 | 6.0929 | 6.54334 | 5.71497 | 6.13762 |
| GPAA1 | O43292 | 6.52252 | 6.3754 | 5.89079 | 6.37554 | 6.12661 | 6.06328 | 6.33143 | 6.18522 | 6.08447 | 6.16172 | 5.97584 | 5.86852 | 6.54932 | 6.54771 | 6.66678 | 6.27486 |
| UBA5 | Q9GZZ9 | 6.42292 | 6.28658 | 7.02045 | 6.21031 | 6.64073 | 7.12471 | 6.29898 | 6.86186 | 6.65605 | 6.54833 | 6.36732 | 6.41182 | 5.16628 | 5.74711 | 5.33482 | 4.90173 |
| MLF2 | Q15773 | 5.5505 | 6.05747 | 6.16545 | 5.62352 | 5.91945 | 6.41151 | 6.27297 | 6.64407 | 6.06394 | 6.54571 | 6.6081 | 6.24678 | 6.5244 | 6.28571 | 6.41478 | 6.66564 |
| SURF4 | O15260 | 5.98732 | 5.95498 | 5.5757 | 6.12606 | 6.03671 | 6.17204 | 6.03621 | 6.17245 | 5.98245 | 6.37148 | 5.93693 | 6.10243 | 6.62292 | 6.83299 | 6.94953 | 7.13978 |
| PSMG1 | O95456 | 6.08448 | 6.39793 | 7.0306 | 5.96383 | 6.12653 | 6.6958 | 6.2923 | 6.55479 | 6.40939 | 6.67323 | 6.20283 | 6.56588 | 5.45916 | 5.54118 | 6.16697 | 5.83508 |
| CD151 | P48509 | 6.40677 | 6.54956 | 6.0682 | 6.26323 | 6.28299 | 6.08825 | 6.45902 | 6.22776 | 6.2543 | 6.17902 | 5.78264 | 6.11102 | 6.43088 | 6.39392 | 6.15731 | 6.34515 |
| CMTR2 | Q8IYT2 | 6.25482 | 6.07917 | 6.73422 | 6.05644 | 6.44159 | 6.54871 | 5.76602 | 6.47922 | 6.11312 | 6.7927 | 6.46807 | 5.96528 | 6.05584 | 6.2412 | 6.18172 | 5.82186 |
| MLST8 | A0A0A0MR05 | 6.46202 | 6.08186 | 6.18517 | 6.12453 | 6.4992 | 7.15774 | 5.53352 | 7.12611 | 6.94412 | 7.06986 | 6.98108 | 7.09262 | 5.74564 | 5.32449 | 4.87995 | 4.79208 |
| OSTC | Q9NRP0 | 6.3662 | 6.15915 | 5.67007 | 6.63088 | 6.22057 | 5.96376 | 6.56969 | 5.95742 | 6.08295 | 6.33737 | 5.88578 | 5.78291 | 6.50336 | 6.48878 | 6.81073 | 6.57037 |
| MIB2 | Q96AX9-14 | 6.26125 | 6.50698 | 6.32498 | 6.28088 | 5.70708 | 6.49208 | 5.74316 | 6.04613 | 6.69165 | 6.78009 | 6.71807 | 6.59842 | 5.90206 | 6.13842 | 5.97845 | 5.733 |
| MPC2 | O95563 | 6.31707 | 6.38578 | 5.80688 | 6.57354 | 6.32585 | 6.02184 | 6.81723 | 6.23665 | 6.30154 | 6.0715 | 5.63834 | 5.7138 | 6.62226 | 5.79036 | 6.66523 | 6.71212 |
| NDUFS4 | O43181 | 6.2928 | 6.13409 | 5.54578 | 7.12827 | 6.10526 | 5.63581 | 6.47392 | 5.79658 | 5.72175 | 6.04503 | 6.03649 | 6.15874 | 6.49798 | 6.54768 | 6.49069 | 7.38912 |
| CD81 | P60033 | 5.9441 | 6.20381 | 6.29617 | 6.28526 | 5.80953 | 5.74665 | 6.4803 | 5.9779 | 6.0725 | 6.30694 | 6.27974 | 6.23463 | 6.43578 | 6.67881 | 6.2129 | 7.03497 |
| AKAP10 | O43572 | 6.32306 | 6.38436 | 7.10481 | 6.56303 | 6.22516 | 6.24969 | 5.97329 | 6.80031 | 6.0435 | 6.48827 | 5.79136 | 6.02994 | 5.97234 | 5.80369 | 6.21805 | 6.02916 |
| GAGE5 | Q13069 | 5.91263 | 5.32467 | 5.31666 | 5.197 | 4.04871 | 3.3324 | 4.02414 | 5.17178 | 3.28928 | 5.05684 | 4.38084 | 4.45133 | 3.76142 | 16.7337 | 9.70004 | 14.2986 |
| PGS1 | Q32NB8 | 6.28981 | 6.41809 | 5.60064 | 6.3994 | 6.44162 | 6.10433 | 6.59622 | 6.31143 | 6.11939 | 6.40338 | 5.79501 | 5.92584 | 6.5936 | 6.00568 | 6.62358 | 6.37195 |
| PHPT1 | Q9NRX4 | 6.36478 | 5.64255 | 6.88454 | 4.8778 | 6.6281 | 6.36958 | 5.64637 | 6.92112 | 6.20863 | 6.84342 | 6.46829 | 6.76446 | 6.24805 | 6.29147 | 5.6102 | 6.23065 |
| SUMO3 | P55854-2 | 5.91469 | 5.14167 | 6.31324 | 5.03161 | 6.19264 | 5.40302 | 5.4302 | 6.05321 | 5.93534 | 6.86963 | 6.58982 | 6.97406 | 6.71299 | 7.62783 | 6.95865 | 6.85141 |
| SLC15A4 | Q8N697 | 6.30141 | 6.56103 | 6.14136 | 6.92527 | 6.11746 | 6.04517 | 5.72415 | 5.95271 | 6.11327 | 6.57858 | 6.05595 | 6.76285 | 6.36874 | 6.23189 | 6.04714 | 6.073 |
| SELENOS | Q9BQE4 | 6.69621 | 6.85853 | 5.77391 | 7.04091 | 6.35652 | 7.20071 | 6.00929 | 6.86493 | 6.10471 | 6.3255 | 5.88671 | 5.61145 | 5.55678 | 5.5529 | 5.92422 | 6.23673 |
| MAGED1 | Q9Y5V3-2 | 6.83396 | 6.80794 | 6.59162 | 6.40182 | 6.31643 | 6.53213 | 6.43492 | 7.16732 | 6.60545 | 6.87315 | 6.83838 | 7.46258 | 4.8991 | 5.27693 | 4.45675 | 4.50151 |
| SINHCAF | Q9NP50 | 5.81548 | 5.71397 | 5.52727 | 6.54416 | 6.4791 | 5.8238 | 6.54142 | 6.24614 | 5.53543 | 5.79811 | 5.83279 | 5.69553 | 7.57444 | 6.62989 | 7.08457 | 7.1579 |
| B3GAT3 | O94766 | 6.35509 | 6.386 | 5.89263 | 6.23017 | 6.04665 | 5.97221 | 6.42513 | 6.07078 | 6.25004 | 5.98212 | 6.00495 | 6.33167 | 6.37358 | 6.35057 | 6.62699 | 6.70142 |
| RUFY2 | Q8WXA3-1 | 6.5262 | 6.26739 | 6.31036 | 6.59644 | 5.96441 | 6.72478 | 6.23019 | 6.37734 | 6.29409 | 6.63243 | 6.41113 | 7.16079 | 6.52291 | 6.09183 | 5.20807 | 5.58163 |
| TADA3 | O75528 | 5.92251 | 6.55663 | 6.21449 | 6.48452 | 6.56515 | 5.78073 | 5.97592 | 5.69139 | 6.43435 | 7.51302 | 6.19423 | 6.91911 | 5.67971 | 5.79027 | 6.35295 | 6.82502 |
| TPM2 | Q5TCU3 | 6.78822 | 5.87486 | 6.60325 | 6.11682 | 6.93098 | 6.24454 | 6.19553 | 7.42069 | 6.37881 | 6.021 | 6.78572 | 6.5941 | 6.37963 | 5.56987 | 4.83443 | 5.26156 |
| WIZ | A0A669KBH9 | 6.28932 | 6.32586 | 6.1492 | 6.83711 | 6.21331 | 6.01248 | 6.22675 | 6.07268 | 6.2056 | 5.74462 | 6.28055 | 6.4154 | 6.64134 | 6.32188 | 5.92449 | 6.33942 |
| YBX2 | Q9Y2T7 | 5.96431 | 6.34807 | 6.53007 | 6.33861 | 5.68291 | 6.01547 | 6.174 | 6.49459 | 6.41284 | 6.7437 | 6.12156 | 6.19111 | 6.03817 | 6.32087 | 6.1429 | 6.48081 |
| CLTA | P09496-2 | 7.28636 | 5.35152 | 5.48557 | 4.7612 | 5.65464 | 4.61278 | 5.42653 | 6.85838 | 4.92827 | 5.66132 | 5.13207 | 5.97894 | 6.77481 | 9.60855 | 7.95085 | 8.5282 |
| TBC1D8B | Q0IIM8 | 6.70684 | 5.33939 | 5.64141 | 5.37632 | 5.19433 | 4.93868 | 5.15297 | 6.67447 | 5.23828 | 6.1608 | 5.78928 | 6.54275 | 6.43581 | 8.92212 | 7.77123 | 8.11533 |
| PPP1R12C | Q9BZL4 | 6.19737 | 5.94615 | 6.27963 | 6.2307 | 6.2053 | 7.13068 | 6.52882 | 6.3986 | 6.2457 | 6.73877 | 6.48807 | 6.61367 | 5.85539 | 6.04997 | 5.5773 | 5.51387 |
| PTRH1 | Q86Y79 | 6.65818 | 6.43703 | 5.41067 | 6.55233 | 6.01884 | 5.58737 | 6.59756 | 6.34497 | 6.08348 | 5.95036 | 6.8738 | 6.47636 | 7.37888 | 5.73494 | 5.8088 | 6.08644 |
| KIF5C | O60282 | 6.35283 | 6.28256 | 6.74573 | 6.3824 | 6.51977 | 6.20972 | 5.78674 | 6.42605 | 6.22663 | 7.15375 | 6.48915 | 6.67826 | 5.39822 | 6.06161 | 5.60206 | 5.68452 |
| PCDHB11 | Q9Y5F2 | 5.95702 | 5.82789 | 5.37706 | 5.06879 | 6.91644 | 6.66155 | 6.65268 | 5.84067 | 5.57826 | 5.89465 | 5.55973 | 5.73542 | 7.47383 | 7.60654 | 6.6704 | 7.17907 |
| SFN | P31947 | 6.35696 | 6.70686 | 6.49413 | 6.14698 | 6.12641 | 6.7453 | 6.14126 | 6.62723 | 6.4875 | 6.00933 | 5.98941 | 6.10845 | 6.40768 | 5.96117 | 5.8291 | 5.86223 |
| CUL9 | Q8IWT3 | 6.48022 | 6.60578 | 6.55517 | 6.40287 | 5.93074 | 6.56461 | 6.61917 | 6.71138 | 6.68658 | 7.1113 | 6.72763 | 6.37737 | 5.39555 | 5.65899 | 5.19501 | 4.97764 |
| IQGAP2 | Q13576 | 5.95044 | 6.13409 | 5.6575 | 5.93371 | 5.85342 | 6.21068 | 6.37472 | 6.29757 | 6.65055 | 6.24871 | 5.32178 | 5.79657 | 6.98848 | 6.87755 | 6.87938 | 6.82484 |
| H1-4 | P10412 | 6.3791 | 6.58229 | 5.43037 | 6.03031 | 6.30023 | 6.21311 | 6.09255 | 6.28549 | 5.82364 | 6.26864 | 5.6317 | 5.1836 | 6.42138 | 7.24519 | 7.04083 | 7.07155 |
| DERA | Q9Y315 | 6.60716 | 6.71628 | 7.06077 | 6.22587 | 6.48542 | 6.72608 | 6.32347 | 6.4869 | 6.52172 | 6.61953 | 6.87916 | 6.3186 | 5.49117 | 5.26596 | 5.37433 | 4.89759 |
| CDK16 | Q00536-2 | 6.20401 | 6.51519 | 6.66891 | 6.3828 | 6.439 | 6.69394 | 6.43254 | 6.48894 | 6.33744 | 6.29618 | 7.07569 | 6.333 | 5.55553 | 5.33379 | 5.58723 | 5.65581 |
| PCBD1 | P61457 | 6.57442 | 5.04274 | 6.87985 | 4.68341 | 6.33258 | 5.75805 | 5.5191 | 7.10272 | 5.41921 | 6.87264 | 6.21912 | 7.50818 | 5.01978 | 7.40495 | 6.86882 | 6.79446 |
| IQSEC1 | Q6DN90-3 | 6.20446 | 6.17562 | 6.92922 | 6.62715 | 6.45895 | 6.3027 | 6.55513 | 6.03379 | 6.24864 | 6.24674 | 6.33764 | 6.50553 | 5.96421 | 6.0095 | 5.49295 | 5.90776 |
| DCBLD1 | Q8N8Z6 | 6.39549 | 6.2732 | 5.99922 | 6.49942 | 6.20975 | 6.213 | 6.78449 | 6.20004 | 6.72574 | 6.19933 | 6.40214 | 6.46058 | 5.87965 | 6.05392 | 6.0744 | 5.82913 |
| ISCU | Q9H1K1 | 6.4069 | 6.07003 | 6.2287 | 5.94006 | 6.4336 | 5.94515 | 6.05119 | 6.02745 | 6.53446 | 6.28347 | 6.36656 | 6.78115 | 6.65768 | 6.04446 | 6.20803 | 6.02109 |
| BCCIP | Q9P287 | 5.68222 | 5.84895 | 6.70972 | 6.06785 | 5.70091 | 5.67678 | 5.99917 | 5.64519 | 6.08959 | 6.20157 | 6.50218 | 6.6313 | 6.54109 | 6.54208 | 7.44767 | 6.71372 |
| SPICE1 | Q8N0Z3 | 6.14317 | 6.07461 | 6.36457 | 6.63471 | 5.88324 | 6.29979 | 6.57133 | 6.68603 | 6.16352 | 6.4215 | 6.52959 | 6.54196 | 5.97103 | 6.04275 | 5.73664 | 5.93557 |
| HNRNPL | A0A3B3ITJ4 | 6.07294 | 6.47073 | 6.10282 | 6.68076 | 6.35579 | 5.81483 | 6.16244 | 6.07456 | 6.19921 | 6.52036 | 6.49216 | 6.11925 | 6.66091 | 5.84174 | 6.2736 | 6.15791 |
| PIGA | P37287 | 6.20478 | 6.47042 | 6.3156 | 6.88478 | 5.92625 | 6.03328 | 6.61706 | 6.2766 | 5.88676 | 6.21251 | 5.82073 | 6.14985 | 6.38095 | 6.05338 | 6.42965 | 6.3374 |
| ATF2 | P15336 | 6.1953 | 6.10911 | 6.08344 | 6.16258 | 6.59203 | 6.08103 | 6.23395 | 6.04287 | 6.21716 | 6.6896 | 6.6105 | 6.43901 | 6.72878 | 6.06537 | 5.97328 | 6.42599 |
| IMMT | Q16891-2 | 6.66909 | 5.6743 | 5.47776 | 7.00112 | 6.16155 | 5.82971 | 5.66638 | 5.82002 | 6.31539 | 6.21723 | 5.7334 | 6.36198 | 6.96914 | 6.49087 | 6.34974 | 7.26231 |
| MAPK8 | P45983-4 | 6.44594 | 6.28466 | 7.01576 | 6.00071 | 6.23592 | 6.77829 | 5.98598 | 6.63491 | 6.35252 | 6.35028 | 6.0925 | 6.36192 | 5.96391 | 5.84187 | 5.98623 | 5.66861 |
| VRK2 | Q86Y07 | 6.90475 | 7.06958 | 6.46336 | 6.96504 | 6.51404 | 6.09795 | 6.23374 | 5.84279 | 6.72408 | 6.3384 | 6.23286 | 6.53786 | 5.44573 | 5.41352 | 5.44862 | 5.76767 |
| VGF | O15240 | 7.02626 | 6.80292 | 6.24628 | 6.3498 | 6.98985 | 6.1089 | 6.32913 | 6.77122 | 5.81036 | 5.95163 | 6.10651 | 6.67221 | 6.32492 | 5.59304 | 5.41359 | 5.50338 |
| NPTN | Q9Y639 | 6.30639 | 6.87712 | 6.23685 | 6.99888 | 6.23925 | 6.37598 | 5.88712 | 6.05269 | 5.95157 | 6.89081 | 6.04362 | 6.3876 | 5.82384 | 5.82273 | 5.93751 | 6.16802 |
| ITPRIPL2 | Q3MIP1 | 6.41951 | 6.46904 | 6.41423 | 7.2123 | 6.08252 | 6.18836 | 6.46663 | 6.00931 | 6.39479 | 6.22121 | 6.13103 | 5.92621 | 6.03869 | 5.80773 | 6.22159 | 5.99684 |
| PHF23 | Q9BUL5 | 5.91077 | 6.25167 | 5.73408 | 6.25704 | 5.92379 | 6.21663 | 6.30437 | 5.9652 | 5.9813 | 6.75728 | 6.4542 | 5.68352 | 6.56044 | 6.92355 | 6.38982 | 6.68633 |
| KATNAL1 | Q9BW62 | 6.32421 | 6.04846 | 7.1227 | 5.95461 | 6.57559 | 6.81265 | 6.55583 | 7.07119 | 6.45849 | 6.4171 | 6.56231 | 6.46748 | 5.5384 | 5.36907 | 5.25884 | 5.46307 |
| ADARB1 | P78563-5 | 6.64723 | 6.61267 | 5.43928 | 6.29879 | 6.16637 | 6.14784 | 6.50867 | 6.19215 | 6.38731 | 6.05431 | 6.13789 | 5.82736 | 6.69071 | 6.10323 | 6.58896 | 6.19724 |
| CLIP1 | A0A3B3ISJ8 | 6.52331 | 6.61002 | 6.55188 | 6.30616 | 6.50481 | 6.52422 | 6.082 | 6.30335 | 5.76205 | 6.82867 | 6.27394 | 6.3727 | 6.0514 | 5.64485 | 5.78983 | 5.87081 |
| DMAC2L | Q99766 | 6.33854 | 5.88046 | 5.55194 | 5.61851 | 6.32388 | 5.7734 | 6.05213 | 6.36366 | 5.66898 | 6.12144 | 5.17174 | 6.15027 | 7.09619 | 7.29493 | 7.46618 | 7.12775 |
| AMOTL2 | Q9Y2J4-4 | 6.78117 | 6.1652 | 6.61297 | 6.35116 | 6.25469 | 6.14748 | 5.89606 | 6.57525 | 6.50617 | 6.0892 | 6.95896 | 6.57164 | 6.63651 | 5.30678 | 5.68608 | 5.56068 |
| SCAMP2 | O15127 | 6.34916 | 6.70242 | 6.70183 | 6.97598 | 6.25699 | 6.30271 | 6.34967 | 6.45027 | 6.076 | 6.82563 | 6.90989 | 6.60227 | 5.2568 | 5.54817 | 5.21233 | 5.47987 |
| TTC26 | A0AVF1 | 6.74932 | 6.25049 | 7.11951 | 6.70806 | 6.37673 | 6.8819 | 5.31076 | 6.51298 | 6.4554 | 6.96891 | 6.01354 | 6.92691 | 5.27974 | 5.61392 | 5.09539 | 5.73643 |
| TMEM131L | A2VDJ0-5 | 7.19894 | 7.38721 | 6.64537 | 7.44786 | 6.25855 | 6.12012 | 6.62025 | 6.15685 | 6.37027 | 6.65742 | 6.54725 | 6.4326 | 4.89018 | 5.14779 | 5.07772 | 5.04163 |
| ATRN | O75882 | 6.30572 | 6.87228 | 6.34842 | 7.02638 | 6.48787 | 6.36144 | 6.5327 | 6.08493 | 6.21109 | 6.39969 | 6.40746 | 6.30646 | 5.6463 | 5.76485 | 5.49598 | 5.74842 |
| DAGLB | Q8NCG7 | 6.58532 | 6.52476 | 6.21519 | 6.02002 | 6.09837 | 6.18109 | 5.79671 | 6.18237 | 6.19048 | 6.41859 | 6.67405 | 6.47181 | 6.47058 | 6.40491 | 5.83976 | 5.92598 |
| GNB2 | P62879 | 6.25883 | 6.25667 | 6.27561 | 6.08204 | 6.49046 | 6.0942 | 6.44104 | 6.01868 | 6.66594 | 6.24805 | 6.04653 | 5.94254 | 6.27885 | 5.96693 | 6.47096 | 6.46264 |
| USF2 | Q15853 | 6.73532 | 6.55275 | 6.32718 | 6.35861 | 6.35294 | 5.77944 | 6.37553 | 5.74301 | 6.23608 | 6.65496 | 6.64558 | 6.5586 | 6.25603 | 5.61029 | 6.88417 | 5.92951 |
| NUDT3 | O95989 | 6.16492 | 6.14489 | 6.91803 | 5.76221 | 6.52828 | 6.6058 | 6.35003 | 6.50277 | 6.55734 | 6.69349 | 6.78345 | 6.44744 | 5.92879 | 5.70843 | 5.65043 | 5.25371 |
| ERBB2 | P04626 | 6.30631 | 6.20169 | 6.69944 | 6.29649 | 5.68434 | 6.64981 | 6.19037 | 6.5914 | 5.91703 | 6.46009 | 6.72675 | 6.6661 | 6.07359 | 6.0991 | 5.74842 | 5.68906 |
| GPAT4 | Q86UL3 | 6.25087 | 6.35358 | 5.75265 | 6.7414 | 5.83219 | 5.96805 | 6.51293 | 5.97677 | 6.06883 | 6.05449 | 6.2598 | 5.89365 | 6.64052 | 6.5035 | 6.8023 | 6.38847 |

| | | | | | | | | | | | | | | | | | |
|---|---|---|---|---|---|---|---|---|---|---|---|---|---|---|---|---|---|
| PCLAF | Q15004 | 7.4072 | 7.21781 | 7.90876 | 7.37349 | 6.67105 | 6.52302 | 6.4004 | 6.30551 | 7.92055 | 7.22796 | 8.24017 | 7.85722 | 3.23952 | 3.4572 | 3.03057 | 3.21956 |
| PI4KB | Q9UBF8 | 6.51987 | 6.5763 | 6.85153 | 6.12235 | 6.57338 | 6.5154 | 6.68371 | 6.56995 | 6.11578 | 6.4768 | 6.37199 | 6.24597 | 5.71104 | 5.89168 | 5.4104 | 5.36384 |
| RECQL4 | O94761 | 5.77542 | 6.00482 | 6.34354 | 6.28911 | 6.11659 | 6.10819 | 6.33086 | 5.95428 | 6.308 | 6.92814 | 6.72494 | 6.27174 | 6.10089 | 6.06816 | 6.29207 | 6.38324 |
| MFSD5 | Q6N075-2 | 6.10846 | 6.39024 | 5.78159 | 5.73551 | 5.80218 | 5.91917 | 6.03887 | 5.75461 | 6.22397 | 6.26181 | 6.61766 | 6.18289 | 6.52773 | 6.73966 | 6.63628 | 7.27936 |
| CBLB | Q13191 | 6.7473 | 6.18874 | 6.96206 | 6.37873 | 6.3393 | 6.90222 | 6.29957 | 6.22549 | 6.22664 | 6.83075 | 6.86549 | 6.32102 | 5.42157 | 6.05974 | 5.24367 | 4.98773 |
| B3GALT6 | Q96L58 | 6.45477 | 6.21885 | 5.83708 | 6.86155 | 6.02357 | 6.03702 | 6.70203 | 5.79347 | 6.42855 | 6.31927 | 6.15811 | 6.23375 | 6.19963 | 5.97409 | 6.61411 | 6.14415 |
| FAM76B | Q5HYJ3 | 6.42568 | 6.37907 | 5.79629 | 6.54681 | 6.35944 | 5.52335 | 5.88519 | 6.00365 | 6.59277 | 5.82864 | 5.8795 | 5.90551 | 7.12888 | 6.79647 | 6.47066 | 6.47808 |
| JUND | P17535 | 6.33307 | 6.27236 | 6.18895 | 6.75398 | 6.08262 | 6.28488 | 6.29947 | 6.28832 | 5.66013 | 5.87503 | 6.39172 | 6.31771 | 6.28258 | 6.16184 | 6.11708 | 6.69025 |
| SECISBP2 | Q96T21 | 6.17577 | 6.48202 | 6.71532 | 6.96887 | 5.93359 | 6.10132 | 5.88647 | 6.38629 | 6.75469 | 6.49719 | 6.61815 | 6.79967 | 5.73224 | 5.87832 | 5.54314 | 5.52695 |
| SECISBP2L | Q93073 | 6.68278 | 6.56188 | 6.57608 | 6.11644 | 6.11496 | 6.14346 | 6.59859 | 6.52622 | 6.77994 | 6.88316 | 7.15519 | 6.5439 | 5.39127 | 5.22098 | 5.60699 | 5.09817 |
| POP4 | O95707 | 6.40681 | 7.05054 | 5.68366 | 6.99356 | 6.47978 | 6.44235 | 6.62402 | 7.05925 | 6.05946 | 5.7876 | 6.1931 | 6.14797 | 5.34392 | 6.21133 | 5.94679 | 5.56985 |
| KCTD9 | Q7L273 | 6.2525 | 6.01905 | 6.74394 | 5.92231 | 6.53908 | 6.97192 | 5.75434 | 6.68952 | 6.36567 | 6.54956 | 6.55501 | 6.70223 | 5.8551 | 5.65782 | 5.72944 | 5.6925 |
| CDC42EP4 | Q9H3Q1 | 6.36053 | 6.28426 | 6.49223 | 6.41935 | 6.53474 | 6.3651 | 5.94907 | 6.49477 | 6.09056 | 6.09091 | 6.32664 | 6.09695 | 6.93063 | 6.34923 | 5.85622 | 5.3588 |
| COPRS | Q9NQ92 | 7.42024 | 6.15585 | 5.59981 | 4.99904 | 4.82636 | 4.85222 | 5.21885 | 5.78023 | 5.50951 | 6.25419 | 6.34149 | 6.84781 | 5.75283 | 8.67418 | 7.31504 | 8.45235 |
| FERMT1 | Q9BQL6 | 6.05708 | 6.43092 | 6.54741 | 6.0164 | 5.76509 | 6.74797 | 5.95277 | 6.64799 | 6.90643 | 6.85289 | 6.28264 | 7.06095 | 5.54709 | 5.86915 | 5.75177 | 5.56344 |
| LSM1 | O15116 | 5.49916 | 4.41023 | 5.55958 | 4.25352 | 5.56751 | 5.35562 | 5.02023 | 7.56847 | 5.80092 | 6.21971 | 5.85937 | 6.30695 | 5.05438 | 9.41261 | 8.71655 | 9.29519 |
| DDX19A | Q9NUU7 | 5.96327 | 6.48649 | 7.35091 | 6.51842 | 6.29165 | 6.51239 | 5.9582 | 6.7964 | 6.38309 | 5.98052 | 6.90455 | 6.84182 | 5.40864 | 5.71611 | 5.4832 | 5.40433 |
| ZFAND5 | O76080 | 6.23303 | 5.62331 | 6.80229 | 5.84327 | 6.6974 | 6.95032 | 7.06798 | 6.56748 | 6.15908 | 6.69048 | 6.94132 | 6.22813 | 5.99485 | 5.40036 | 5.32364 | 5.47706 |
| BAP18 | Q8IXM2-3 | 5.99395 | 4.83445 | 5.29999 | 4.90122 | 5.85452 | 4.38473 | 5.18348 | 6.49256 | 4.46164 | 6.23878 | 5.15004 | 6.17517 | 6.70641 | 9.78698 | 9.24829 | 9.28779 |
| PARL | Q9H300 | 6.54473 | 6.55045 | 5.75324 | 6.18336 | 6.39414 | 6.05938 | 6.11461 | 5.58172 | 5.48435 | 6.19515 | 5.96371 | 6.14361 | 6.58043 | 6.21118 | 7.00152 | 7.23842 |
| KDM5B | Q9UGL1-2 | 6.26034 | 6.63011 | 6.69886 | 6.84689 | 6.37747 | 6.47786 | 6.71963 | 6.33209 | 6.35871 | 6.28755 | 5.9946 | 6.40957 | 5.35932 | 5.75796 | 5.74038 | 5.74866 |
| TIMM29 | Q9BSF4 | 6.97758 | 6.501 | 6.38354 | 6.51244 | 5.96726 | 5.70089 | 6.32159 | 5.78288 | 6.19843 | 5.77902 | 6.09835 | 6.28414 | 6.53374 | 5.89509 | 6.52689 | 6.53715 |
| ADAT2 | Q7Z6V5 | 5.25572 | 5.49601 | 6.56138 | 5.65337 | 6.6462 | 6.51694 | 5.31967 | 7.01913 | 6.24822 | 6.70628 | 6.77752 | 6.58475 | 6.74659 | 6.15287 | 6.08607 | 6.22929 |
| RABEP2 | Q9H5N1 | 6.91605 | 5.01862 | 5.69787 | 3.91425 | 5.03954 | 4.83545 | 5.71943 | 7.10333 | 5.46423 | 6.13561 | 5.73155 | 7.41404 | 6.86717 | 8.50267 | 7.23522 | 8.40496 |
| RALA | P11233 | 6.31588 | 6.59675 | 6.03764 | 6.30718 | 6.06639 | 5.93093 | 6.31706 | 6.11528 | 6.36341 | 6.38484 | 6.27851 | 5.83342 | 6.27192 | 6.49822 | 6.31238 | 6.3702 |
| CGGBP1 | Q9UFW8 | 5.40135 | 5.59754 | 6.12696 | 4.67314 | 5.97575 | 6.7182 | 5.43267 | 5.34854 | 7.49788 | 6.72574 | 6.25834 | 6.0934 | 7.22933 | 7.23307 | 6.98713 | 6.70093 |
| HMGN3 | Q15651 | 5.84714 | 5.79555 | 5.71464 | 6.59785 | 6.18115 | 6.14456 | 5.84323 | 5.99486 | 5.71406 | 5.7277 | 5.52878 | 5.83919 | 7.02258 | 7.59104 | 6.84888 | 7.60878 |
| SMYD2 | Q9NRG4 | 7.08266 | 6.42481 | 7.33572 | 6.29948 | 6.56648 | 7.01278 | 5.89771 | 7.05012 | 6.62429 | 6.43487 | 6.59755 | 6.52378 | 5.2288 | 5.21981 | 4.93259 | 4.76856 |
| RNF31 | Q96EP0 | 6.49124 | 5.73199 | 6.62146 | 5.84119 | 6.15595 | 6.47113 | 6.39702 | 6.61389 | 6.63016 | 6.39457 | 6.80493 | 6.58158 | 6.05928 | 5.59363 | 5.75627 | 5.85572 |
| BAHCC1 | Q9P281 | 6.54071 | 6.5426 | 6.30162 | 5.56644 | 6.17835 | 6.40992 | 6.0779 | 5.68221 | 6.48479 | 6.68041 | 5.64394 | 6.31405 | 6.10307 | 6.09142 | 6.20859 | 7.17397 |
| UBE2D3 | P61077-3 | 6.00287 | 5.50285 | 6.99042 | 5.93368 | 5.86019 | 6.54921 | 5.59797 | 6.89774 | 5.94592 | 7.19755 | 6.65134 | 6.55891 | 4.95496 | 6.92848 | 6.50626 | 5.92164 |
| DOCK11 | Q5JSL3 | 6.0992 | 6.02626 | 6.38329 | 5.88721 | 6.00751 | 6.38032 | 5.97778 | 6.26091 | 6.37044 | 6.59238 | 6.36208 | 6.50372 | 6.38762 | 6.67944 | 6.03955 | 6.0423 |
| MSL3 | Q8N5Y2 | 6.49512 | 6.41942 | 6.43987 | 6.83052 | 6.39753 | 6.16819 | 6.33535 | 6.08084 | 5.91072 | 6.49128 | 6.2576 | 6.01833 | 5.9764 | 5.99215 | 6.1519 | 6.03479 |
| DSE | Q9UL01 | 5.8842 | 6.44074 | 6.59806 | 5.90327 | 6.16853 | 6.17688 | 6.28925 | 5.98633 | 6.44917 | 5.89267 | 6.267 | 6.32015 | 6.56076 | 6.66613 | 6.10896 | 6.28789 |
| ASB7 | Q9H672 | 5.89298 | 5.89483 | 6.28282 | 6.4006 | 6.38939 | 5.92137 | 6.3753 | 6.32903 | 6.72747 | 6.53707 | 6.14462 | 6.06552 | 6.56433 | 5.98902 | 6.50219 | 5.98346 |
| HACE1 | Q8IYU2 | 5.92052 | 5.90584 | 6.68613 | 5.88047 | 5.73515 | 6.46329 | 6.18387 | 5.84357 | 6.63602 | 6.88947 | 6.71536 | 6.83619 | 6.2679 | 5.9865 | 5.66291 | 6.38681 |
| PCBP2 | F8VRH0 | 5.82759 | 5.33587 | 6.27041 | 6.11259 | 6.20899 | 6.6468 | 5.77962 | 6.12837 | 6.17807 | 6.30061 | 6.81482 | 7.01545 | 6.48878 | 6.2724 | 6.0619 | 6.55773 |
| PUS3 | Q9BZE2 | 5.73093 | 5.64376 | 5.96971 | 5.67311 | 5.64692 | 6.72911 | 5.67394 | 6.13832 | 5.78985 | 5.77254 | 6.13395 | 6.17993 | 7.37073 | 6.82294 | 7.71423 | 6.96703 |
| FAM3A | P98173 | 6.78142 | 6.55919 | 6.11952 | 6.95937 | 6.65605 | 6.30831 | 6.44076 | 6.29228 | 6.17945 | 6.32024 | 6.31184 | 6.36686 | 6.18297 | 5.47882 | 5.54146 | 5.50147 |
| SLC2A3 | P11169 | 6.79978 | 6.22998 | 6.07812 | 6.34776 | 6.54421 | 6.3551 | 6.00532 | 5.91967 | 5.38157 | 6.05397 | 5.78227 | 5.7517 | 6.7567 | 6.81515 | 6.57093 | 6.60778 |
| FKTN | O75072 | 6.06954 | 5.95663 | 6.34828 | 6.98455 | 6.40946 | 6.20863 | 6.44236 | 5.9883 | 6.36873 | 6.30073 | 5.79593 | 6.04046 | 6.66259 | 5.8637 | 6.11806 | 6.44205 |
| ZNF740 | Q8NDX6 | 5.916 | 6.4841 | 6.05996 | 6.60799 | 6.01285 | 6.0988 | 6.59285 | 5.83842 | 5.97512 | 6.65934 | 5.81938 | 6.36031 | 6.5514 | 6.19282 | 6.43437 | 6.3963 |
| ZNF724 | A8MTY0 | 6.54193 | 7.37183 | 6.80851 | 7.50114 | 5.87738 | 6.25921 | 5.8118 | 5.92005 | 6.4789 | 6.23404 | 6.65356 | 6.92291 | 4.89289 | 5.55888 | 5.75953 | 5.40744 |
| ZKSCAN1 | P17029 | 5.99165 | 6.31325 | 6.39858 | 6.04861 | 6.10752 | 6.58415 | 6.00875 | 5.95589 | 6.38651 | 6.08052 | 6.47564 | 5.95307 | 6.23104 | 6.11307 | 6.38681 | 6.05916 |
| ZNF829 | Q3KNS6-3 | 6.36 | 6.3668 | 6.19634 | 6.75225 | 6.01558 | 6.2267 | 6.31332 | 6.21304 | 6.28551 | 6.00524 | 6.58324 | 6.6425 | 6.012 | 5.87564 | 6.12723 | 6.02462 |
| RBAK | Q9NYW8 | 6.53976 | 6.22843 | 5.84665 | 7.00471 | 6.06321 | 5.86121 | 5.95633 | 6.00172 | 6.43728 | 6.31552 | 6.07053 | 6.03249 | 6.44513 | 6.10734 | 6.70417 | 6.38551 |
| C1GALT1C1 | Q96EU7 | 6.53251 | 6.50217 | 6.3126 | 6.68564 | 6.52513 | 6.21181 | 6.36925 | 6.28482 | 6.42083 | 6.10021 | 6.28627 | 5.86386 | 6.14527 | 5.65921 | 6.03559 | 6.06483 |
| PPCS | Q9HAB8 | 6.57641 | 5.61548 | 7.04466 | 5.583 | 6.23413 | 6.51818 | 5.80685 | 6.70125 | 6.25245 | 6.31053 | 6.65084 | 6.73055 | 6.02636 | 5.8437 | 6.17127 | 5.93434 |
| LDAH | Q9H6V9-2 | 6.82821 | 6.0101 | 6.31853 | 6.16824 | 5.71059 | 5.69325 | 6.37814 | 6.23955 | 5.88023 | 6.12466 | 5.84245 | 6.2937 | 7.009 | 6.36389 | 6.87767 | 6.26178 |
| ZCCHC4 | Q9H5U6 | 5.36395 | 6.34062 | 6.17633 | 6.10316 | 5.63978 | 6.13274 | 6.05975 | 5.96868 | 6.42006 | 6.15771 | 6.29982 | 6.0762 | 6.54561 | 7.42941 | 6.81371 | 6.47247 |
| RAP1A | P62834 | 6.19195 | 6.26825 | 6.19865 | 6.55297 | 5.95426 | 5.56226 | 6.71761 | 5.85503 | 6.61618 | 5.75047 | 6.10953 | 6.14999 | 6.31442 | 6.36605 | 6.71935 | 6.67303 |
| RAB39A | Q14964 | 6.50397 | 6.19489 | 6.1898 | 6.73721 | 6.21262 | 6.18181 | 6.24351 | 5.84694 | 6.0409 | 6.67242 | 5.66337 | 6.28449 | 6.4818 | 5.36271 | 6.16566 | 7.2179 |
| RAB6A | P20340-2 | 6.21417 | 6.30988 | 6.20466 | 6.15887 | 6.23011 | 6.01204 | 6.18216 | 5.9944 | 6.27673 | 6.37577 | 6.27994 | 5.85956 | 6.60812 | 6.15291 | 6.84206 | 6.29861 |
| RAB33B | Q9H082 | 6.1333 | 6.56311 | 6.12346 | 6.72746 | 6.45335 | 6.03214 | 6.6635 | 6.20945 | 6.11763 | 6.21568 | 5.76304 | 5.84492 | 6.24706 | 6.26344 | 6.5249 | 6.11757 |
| DNASE2 | O00115 | 6.34582 | 5.69492 | 5.66887 | 5.98347 | 5.74126 | 5.85123 | 6.16642 | 5.5803 | 5.91302 | 6.10503 | 6.66883 | 6.23881 | 6.11499 | 7.51904 | 7.65426 | 6.65372 |
| ZSCAN21 | Q9Y5A6 | 5.65893 | 6.22124 | 6.31243 | 6.33702 | 6.28337 | 5.99065 | 6.29203 | 6.39957 | 6.1903 | 6.06478 | 6.0516 | 5.35568 | 6.72304 | 6.77524 | 6.58241 | 6.76171 |
| MXRA7 | P84157 | 6.48032 | 5.4948 | 4.52516 | 5.22712 | 4.22145 | 2.86547 | 4.79476 | 5.26058 | 4.21208 | 4.78158 | 5.15754 | 4.85856 | 5.52944 | 13.0756 | 10.7164 | 12.7991 |
| SNX15 | Q9NRS6 | 5.95464 | 6.00009 | 6.41997 | 5.7816 | 6.49808 | 6.62516 | 5.92594 | 7.29251 | 6.42141 | 6.29698 | 6.75389 | 7.08323 | 5.86005 | 5.40347 | 5.93753 | 5.74544 |
| BUD23 | O43709-3 | 6.32965 | 6.03153 | 6.01335 | 5.72896 | 5.80551 | 5.83605 | 6.10206 | 6.4556 | 6.58097 | 6.39629 | 6.15369 | 6.43372 | 6.39278 | 6.54279 | 6.6405 | 6.55655 |
| MED26 | O95402 | 6.51117 | 6.44084 | 6.41935 | 6.90037 | 5.90776 | 5.58254 | 6.15941 | 5.87156 | 6.38916 | 6.43449 | 6.32408 | 6.4541 | 6.18059 | 5.74423 | 6.50369 | 6.17666 |
| SYNGR2 | O43760-2 | 5.70413 | 6.50802 | 5.70206 | 5.7904 | 6.03439 | 6.39368 | 6.19812 | 6.24695 | 6.07735 | 6.41983 | 6.43814 | 6.39736 | 6.31601 | 6.51593 | 6.54751 | 6.71012 |
| C8orf82 | Q6P1X6 | 6.49324 | 6.02093 | 6.21852 | 6.01573 | 6.72552 | 5.85163 | 6.2486 | 5.84816 | 5.97379 | 6.08067 | 6.19715 | 6.10276 | 6.72395 | 6.17075 | 6.70697 | 6.62163 |
| DAP | P51397 | 5.02201 | 4.40979 | 5.29487 | 3.95668 | 5.4456 | 4.65882 | 4.06757 | 5.80549 | 4.27818 | 5.45857 | 5.20072 | 6.24313 | 10.154 | 10.3139 | 10.0121 | 10.0786 |
| MAN1A1 | P33908 | 6.45506 | 6.6707 | 6.38183 | 7.17096 | 6.21449 | 6.17412 | 6.25829 | 5.8096 | 6.57118 | 6.2429 | 6.39935 | 6.01876 | 5.73565 | 5.8382 | 5.82597 | 6.23293 |
| RNASEH1 | O60930 | 6.11197 | 6.20555 | 5.98991 | 7.08628 | 6.56553 | 6.20469 | 5.9322 | 6.41771 | 5.89232 | 6.22492 | 6.60057 | 6.24394 | 6.16648 | 5.86173 | 6.06574 | 6.43046 |
| LCORL | Q8N3X6 | 6.50471 | 6.15503 | 6.19016 | 6.71028 | 6.10973 | 6.59379 | 6.39176 | 6.29334 | 6.10741 | 6.90843 | 6.74845 | 6.7098 | 5.37785 | 5.51751 | 5.85401 | 5.82773 |
| TRHDE | Q9UKU6 | 6.49574 | 6.7155 | 6.47358 | 6.71706 | 6.03807 | 5.55624 | 6.15408 | 6.17136 | 6.12212 | 6.13481 | 6.55007 | 6.41549 | 5.93627 | 5.94968 | 6.25384 | 6.3161 |
| ALKBH6 | Q3KRA9-2 | 6.47812 | 6.2272 | 7.37419 | 6.39318 | 6.5264 | 6.4061 | 6.42334 | 6.56045 | 6.44419 | 6.43805 | 6.65265 | 6.80246 | 5.49416 | 5.19493 | 5.40042 | 5.18417 |
| SPOUT1 | Q5T280 | 6.0939 | 6.29929 | 5.69817 | 6.47118 | 6.01739 | 5.72882 | 6.49244 | 6.0391 | 6.28507 | 5.96035 | 6.22196 | 6.06498 | 6.46328 | 6.74244 | 6.67282 | 6.74882 |
| FKBPL | Q9UIM3 | 5.65686 | 5.33878 | 6.47764 | 5.09248 | 6.37617 | 6.04636 | 5.72705 | 7.32143 | 5.82765 | 7.08614 | 6.91391 | 8.06909 | 5.8137 | 6.55791 | 5.05136 | 5.74348 |
| DUSP12 | Q9UNI6 | 5.82489 | 5.81504 | 6.23655 | 5.52935 | 5.86195 | 6.09781 | 5.80897 | 5.7902 | 5.7749 | 6.23441 | 6.38801 | 6.28672 | 7.03938 | 6.96654 | 7.60663 | 6.73866 |
| MAPRE3 | Q9UPY8 | 6.39967 | 5.84066 | 6.46837 | 5.55316 | 6.41999 | 6.66127 | 6.5656 | 6.54137 | 6.38799 | 6.43754 | 6.39992 | 6.59367 | 6.10862 | 6.19546 | 5.76656 | 5.66016 |
| ABCC10 | Q5T3U5 | 6.56909 | 7.09866 | 6.29409 | 6.90104 | 6.23546 | 6.04704 | 6.03097 | 6.15278 | 6.53102 | 6.41441 | 6.50944 | 6.3541 | 5.64478 | 5.61534 | 5.76515 | 5.83663 |
| ALG9 | Q9H6U8-3 | 6.17557 | 6.35881 | 5.73501 | 6.43811 | 5.99195 | 6.19366 | 5.59281 | 6.29586 | 5.66607 | 5.72689 | 6.15461 | 5.56141 | 6.88135 | 6.43833 | 6.87112 | 6.91844 |
| CDC6 | Q99741 | 7.70104 | 6.85201 | 7.37645 | 7.41411 | 6.1591 | 6.6922 | 6.87485 | 6.39683 | 6.89952 | 6.6292 | 7.5965 | 7.27852 | 3.93084 | 4.08558 | 3.82022 | 4.29304 |
| IBA57 | Q5T440 | 6.60243 | 6.42875 | 6.02841 | 6.76675 | 6.3019 | 6.27713 | 6.11694 | 6.17135 | 6.01448 | 6.10102 | 6.17166 | 5.97468 | 6.19833 | 5.92957 | 6.50888 | 6.40773 |
| CLCN3 | P51790-2 | 6.34589 | 6.48721 | 6.41297 | 6.48786 | 6.4029 | 5.97099 | 6.15585 | 6.11097 | 6.06795 | 6.72355 | 6.37291 | 6.22014 | 6.21978 | 6.06949 | 6.08612 | 5.86544 |
| MAX | P61244 | 6.6152 | 6.33715 | 5.72615 | 6.18091 | 6.64718 | 5.89098 | 6.16374 | 6.78594 | 6.282 | 6.09369 | 6.47613 | 6.20188 | 6.5835 | 5.89093 | 5.89498 | 6.22965 |
| PPM1D | O15297 | 6.37952 | 6.61045 | 5.90644 | 6.76367 | 6.00946 | 6.36584 | 6.54877 | 6.1011 | 5.98714 | 5.43593 | 6.03754 | 6.30265 | 6.23064 | 6.37476 | 6.14634 | 6.79975 |
| UBE2G1 | P62253 | 5.84866 | 5.56433 | 6.62947 | 4.80582 | 6.48551 | 5.71446 | 5.83301 | 7.23806 | 6.03375 | 7.00318 | 6.44098 | 7.31601 | 5.77959 | 6.74678 | 6.339 | 6.22141 |
| KANSL1 | Q7Z3B3 | 6.59574 | 6.58967 | 6.11543 | 6.9778 | 6.53036 | 6.14855 | 6.95747 | 5.58272 | 6.29223 | 6.28973 | 6.3582 | 6.12694 | 5.80377 | 6.06415 | 5.7307 | 5.83654 |
| ZFAND1 | Q8TCF1 | 6.37278 | 5.52778 | 7.31174 | 5.95632 | 5.84808 | 6.55236 | 5.85114 | 6.38787 | 6.40239 | 6.62626 | 7.00725 | 6.83571 | 5.99713 | 5.9653 | 5.48101 | 5.87689 |
| KAT5 | Q92993-3 | 6.16605 | 6.35021 | 6.37174 | 6.97648 | 5.82822 | 6.16389 | 6.19667 | 6.23403 | 6.36151 | 6.15918 | 6.07608 | 6.33443 | 6.37429 | 5.75685 | 6.41656 | 6.23378 |
| ACOT11 | Q8WXI4 | 6.30321 | 6.50789 | 6.08477 | 6.32316 | 6.32132 | 6.73351 | 5.81257 | 7.07438 | 6.46843 | 6.54069 | 6.33953 | 6.93725 | 5.29109 | 6.11868 | 5.79752 | 5.346 |
| FBXL4 | Q9UKA2 | 6.04986 | 6.79284 | 6.37309 | 6.92788 | 6.46536 | 5.99965 | 6.32043 | 6.02415 | 6.63429 | 6.18642 | 7.08338 | 6.87442 | 5.29513 | 5.41815 | 5.7321 | 5.82285 |
| GYS1 | P13807 | 6.21663 | 6.3294 | 6.48131 | 5.86161 | 6.51442 | 6.68626 | 6.30486 | 6.65355 | 6.2372 | 6.93819 | 6.41309 | 6.41056 | 5.75458 | 6.06869 | 5.5624 | 5.56723 |
| GABARAPL2 | P60520 | 6.38202 | 5.71024 | 6.2749 | 5.78457 | 6.42618 | 6.84295 | 6.27821 | 7.13048 | 6.53358 | 6.36582 | 6.72944 | 6.66416 | 5.75703 | 6.02021 | 5.36531 | 5.73489 |
| LPIN1 | Q14693-7 | 5.97338 | 6.22595 | 6.76977 | 6.36594 | 5.58297 | 6.17847 | 5.77869 | 6.24811 | 6.33029 | 6.85912 | 6.75234 | 6.65755 | 6.06242 | 6.13126 | 6.16348 | 5.92026 |
| NSMCE1 | Q8WV22 | 6.0066 | 6.16039 | 6.21519 | 6.38811 | 6.0588 | 6.25957 | 5.97425 | 5.8253 | 6.53615 | 6.22884 | 6.66125 | 6.54729 | 6.29055 | 6.00486 | 6.61747 | 6.22537 |
| COPZ1 | P61923-4 | 6.65988 | 5.69174 | 5.92619 | 5.39614 | 6.44082 | 6.62408 | 6.46912 | 6.6356 | 5.43348 | 6.12677 | 6.65104 | 7.07726 | 5.4771 | 7.44391 | 5.82289 | 6.12399 |
| ARL1 | P40616 | 6.82669 | 6.42747 | 5.92247 | 7.01239 | 6.55054 | 6.44782 | 6.55664 | 6.14856 | 6.26777 | 6.42666 | 6.22082 | 6.06984 | 5.6824 | 5.99749 | 5.96056 | 5.48189 |
| SDR39U1 | Q9NRG7 | 6.56019 | 6.74783 | 5.92449 | 6.39103 | 6.60857 | 5.70923 | 6.11763 | 5.82021 | 5.93263 | 6.3508 | 6.32646 | 6.12122 | 7.06976 | 5.93941 | 6.11033 | 6.27022 |
| ICA1 | Q05084 | 6.57327 | 6.53316 | 6.96817 | 6.09444 | 6.15355 | 7.42473 | 5.977 | 6.70822 | 6.22088 | 6.41962 | 6.47116 | 6.58864 | 5.5891 | 5.50638 | 5.23765 | 5.53404 |
| TSPAN6 | O43657 | 6.12079 | 6.5059 | 5.84084 | 6.04228 | 6.10902 | 6.21447 | 6.01573 | 6.71244 | 6.61923 | 6.25775 | 6.20487 | 6.19896 | 5.96646 | 6.37447 | 6.37644 | 6.44035 |
| SMAD2 | Q15796 | 6.62018 | 6.56531 | 6.65727 | 5.79629 | 6.60363 | 7.12376 | 6.00336 | 6.76407 | 6.38113 | 5.98544 | 6.39759 | 5.90981 | 5.81772 | 6.22589 | 5.65172 | 5.49684 |
| XPR1 | Q9UBH6 | 6.78805 | 6.30177 | 6.0275 | 6.68816 | 6.11864 | 6.20044 | 6.60741 | 6.31656 | 6.69664 | 5.66847 | 5.8234 | 5.94947 | 5.72941 | 6.21167 | 6.60409 | 6.26832 |
| SLC25A33 | Q9BSK2 | 5.6484 | 6.33543 | 5.48306 | 6.77418 | 6.41243 | 6.83541 | 6.89618 | 6.17889 | 5.98322 | 5.64756 | 6.48648 | 5.79018 | 6.65425 | 5.79116 | 6.54668 | 6.53649 |
| FAHD2B | Q6P2I3 | 5.63115 | 5.76411 | 5.62921 | 5.57992 | 6.43747 | 5.94542 | 6.34061 | 6.79302 | 6.30149 | 6.67914 | 6.55373 | 6.79759 | 6.45239 | 6.10013 | 6.50211 | 6.4925 |
| SH3BGRL3 | Q9H299 | 6.05364 | 4.94692 | 6.00997 | 4.69924 | 6.20091 | 6.30658 | 5.43132 | 7.32112 | 6.20556 | 6.74826 | 6.01348 | 6.83086 | 5.9278 | 7.34751 | 6.47916 | 7.47767 |
| ESRRA | P11474 | 6.35756 | 6.24231 | 6.46541 | 6.87922 | 6.37861 | 6.35651 | 6.36635 | 6.03636 | 6.27933 | 5.84369 | 6.50555 | 6.23402 | 6.03236 | 5.72056 | 6.21772 | 6.08444 |
| SPINDOC | Q9BUA3 | 6.63142 | 6.8331 | 6.44487 | 6.68713 | 6.70087 | 6.80569 | 5.63405 | 6.05282 | 6.06555 | 6.19682 | 6.26107 | 6.30723 | 6.24663 | 5.71858 | 5.55456 | 5.85962 |
| SPIRE1 | Q08AE8-2 | 6.24382 | 6.24019 | 6.43366 | 6.74264 | 5.79765 | 6.73127 | 5.1292 | 5.98234 | 6.32534 | 6.20995 | 6.81646 | 6.28058 | 6.7785 | 6.13522 | 6.0297 | 6.12348 |
| RBMXL3 | Q8N7X1 | 6.35815 | 6.2905 | 6.09985 | 6.7849 | 5.86302 | 6.13405 | 6.36295 | 5.95452 | 6.10805 | 5.9869 | 6.26312 | 6.23961 | 6.26835 | 6.22183 | 6.49782 | 6.66639 |
| ATG3 | Q9NT62 | 6.5555 | 6.48679 | 7.03607 | 5.81476 | 6.69888 | 6.60646 | 6.11225 | 7.31666 | 6.76106 | 6.16088 | 6.87618 | 6.47029 | 6.1578 | 5.55246 | 4.58054 | 4.81342 |
| AREL1 | O15033 | 6.42657 | 6.49543 | 6.25891 | 6.72582 | 6.31558 | 6.09228 | 6.113 | 5.08583 | 6.19698 | 6.35529 | 6.49222 | 5.99948 | 6.26349 | 6.03351 | 6.71677 | 6.42886 |

| | | | | | | | | | | | | | | | | | |
|---|---|---|---|---|---|---|---|---|---|---|---|---|---|---|---|---|---|
| ACAD11 | Q709F0 | 6.28981 | 6.3148 | 6.07092 | 6.70352 | 6.22998 | 6.20239 | 6.14923 | 5.87408 | 6.05937 | 6.10493 | 5.96175 | 5.75062 | 6.84712 | 6.25666 | 6.54481 | 6.64 |
| CETN3 | O15182 | 5.93845 | 7.19161 | 5.88643 | 6.74053 | 6.28299 | 6.80709 | 5.53556 | 7.37745 | 5.88205 | 6.9484 | 5.47638 | 7.09024 | 5.42012 | 6.14584 | 4.87754 | 6.39932 |
| ATPAF1 | Q5TC12 | 6.52692 | 6.17932 | 6.11094 | 6.59326 | 6.73658 | 6.76327 | 6.71469 | 6.41988 | 6.59742 | 6.03257 | 6.03887 | 6.63651 | 5.52565 | 5.2762 | 6.04723 | 5.80068 |
| KIF1B | O60333-3 | 6.57991 | 6.50206 | 6.71549 | 6.16355 | 6.00596 | 6.33922 | 6.76459 | 6.79688 | 6.68142 | 6.44694 | 6.51097 | 6.33366 | 5.1712 | 5.51951 | 5.7828 | 5.68584 |
| TAPBP | O15533-3 | 6.35935 | 6.89784 | 7.22973 | 6.66469 | 6.904 | 6.3002 | 5.6777 | 5.65486 | 6.17772 | 6.40274 | 5.5716 | 5.45358 | 6.7625 | 5.64071 | 6.24576 | 6.057 |
| ACTR5 | Q9H9F9 | 6.21137 | 6.3998 | 6.41312 | 6.66491 | 6.3993 | 6.20813 | 6.09794 | 6.32401 | 5.91373 | 6.26567 | 6.34356 | 6.40102 | 6.31631 | 6.20065 | 5.97792 | 5.86255 |
| MTFR1 | Q15390 | 6.9434 | 7.16309 | 7.00494 | 7.56509 | 6.89326 | 5.62008 | 6.52186 | 6.34475 | 6.80368 | 6.24976 | 6.9218 | 6.73283 | 4.37873 | 4.86239 | 4.86849 | 5.12581 |
| KLHL15 | Q96M94 | 7.24201 | 6.91492 | 7.18123 | 7.67278 | 6.11302 | 6.06075 | 6.71645 | 5.76095 | 7.00595 | 6.10396 | 6.39874 | 6.27921 | 4.76686 | 5.08313 | 5.20874 | 5.4913 |
| DCTN3 | O75935 | 6.25879 | 5.48474 | 6.34561 | 4.81254 | 6.38966 | 6.22581 | 6.11841 | 6.7335 | 5.43131 | 6.80173 | 6.06147 | 7.49328 | 5.6138 | 6.91978 | 6.50271 | 6.80689 |
| ACBD6 | Q9BR61 | 6.63717 | 6.34275 | 6.9618 | 5.96002 | 6.67333 | 6.56769 | 6.14018 | 6.88934 | 6.49645 | 6.51212 | 6.6345 | 7.09436 | 5.89272 | 5.14772 | 4.86004 | 5.18982 |
| FAM20B | O75063 | 6.51852 | 6.40323 | 5.65808 | 7.26289 | 6.27801 | 6.15145 | 6.10816 | 6.16361 | 6.06066 | 6.54303 | 5.85345 | 5.66839 | 6.1479 | 5.98858 | 6.68462 | 6.50943 |
| CDC34 | P49427 | 6.745 | 6.16878 | 7.22988 | 6.09314 | 6.60128 | 6.48206 | 6.54336 | 6.85036 | 7.07306 | 6.50922 | 7.21981 | 6.76277 | 5.13024 | 5.00298 | 4.68275 | 4.90532 |
| CDIPT | O14735 | 6.20799 | 6.67265 | 5.99148 | 6.49276 | 6.13084 | 6.00875 | 6.01279 | 6.09088 | 5.67196 | 5.97185 | 6.00279 | 6.10502 | 6.38722 | 6.48045 | 7.14627 | 6.6263 |
| UVSSA | Q2YD98 | 6.15166 | 6.02631 | 6.77693 | 6.66653 | 5.63953 | 6.23897 | 6.10572 | 5.57785 | 5.72458 | 6.51275 | 6.80557 | 6.35377 | 6.47356 | 6.42978 | 6.56533 | 5.95116 |
| NT5C | Q8TCD5 | 5.0907 | 5.37899 | 5.82573 | 5.56088 | 6.2522 | 6.35322 | 6.64635 | 7.23947 | 6.86904 | 7.03221 | 6.60386 | 7.00885 | 6.03592 | 6.02838 | 5.83561 | 6.2386 |
| P3H4 | Q92791 | 6.73952 | 6.48982 | 5.98089 | 6.64916 | 6.83055 | 6.3174 | 5.73565 | 6.66954 | 5.77807 | 6.54505 | 6.24251 | 6.1627 | 6.04482 | 5.70717 | 5.94364 | 6.16352 |
| ATXN1 | P54253 | 6.16528 | 6.25794 | 5.96383 | 6.10778 | 5.97256 | 5.8794 | 6.26767 | 5.75968 | 6.21811 | 6.40085 | 6.07886 | 6.14043 | 7.00994 | 6.34266 | 6.59765 | 6.83735 |
| TIGD1 | Q96MW7 | 6.56397 | 6.64889 | 5.96048 | 6.68968 | 6.85139 | 6.04443 | 6.70285 | 6.46771 | 6.33159 | 6.00818 | 6.27218 | 6.44143 | 5.82109 | 5.46177 | 5.61603 | 6.11832 |
| IFFO2 | Q5TF58 | 5.8108 | 6.40375 | 6.79689 | 6.6903 | 5.98948 | 5.93788 | 6.36814 | 6.11011 | 7.93482 | 6.54325 | 7.36601 | 7.16125 | 6.41727 | 4.60015 | 4.95441 | 4.91548 |
| ENDOG | Q14249 | 5.98011 | 6.16226 | 6.09183 | 6.61578 | 6.6349 | 6.07841 | 6.44717 | 5.9446 | 6.2107 | 6.35659 | 6.07342 | 6.70721 | 6.42384 | 5.81376 | 6.38987 | 6.06757 |
| WRAP53 | Q9BUR4 | 5.92007 | 6.28893 | 6.55355 | 5.71683 | 6.37086 | 6.3257 | 6.47703 | 6.45531 | 6.27476 | 6.64957 | 6.88826 | 6.39341 | 5.91869 | 5.63154 | 5.77062 | 6.36486 |
| MICB | Q29980 | 6.0658 | 6.22587 | 6.26539 | 6.61319 | 6.12164 | 6.55202 | 6.07937 | 6.41622 | 5.73166 | 6.12935 | 6.30755 | 6.19514 | 6.36486 | 5.97163 | 6.22271 | 6.70761 |
| ORC5 | O43913 | 6.23971 | 6.55041 | 6.17718 | 6.36567 | 6.34218 | 6.19569 | 6.03847 | 5.93668 | 5.93586 | 5.98327 | 6.31356 | 5.9416 | 6.63696 | 6.43362 | 6.53351 | 6.37563 |
| ARVCF | O00192 | 6.51072 | 6.61556 | 6.40537 | 7.05692 | 6.21983 | 6.2706 | 6.26586 | 6.19708 | 6.1641 | 6.16875 | 6.46531 | 6.31117 | 5.85258 | 5.81437 | 5.79957 | 5.88219 |
| DUSP3 | P51452 | 6.24223 | 5.90196 | 6.9391 | 5.9526 | 7.02419 | 6.97444 | 6.14125 | 6.7431 | 6.39659 | 6.30984 | 6.75298 | 6.22305 | 5.33248 | 5.91054 | 5.70451 | 5.45115 |
| PHF1 | O43189 | 6.07494 | 6.73069 | 6.5734 | 6.87772 | 6.13512 | 6.39491 | 6.57601 | 5.6057 | 5.8428 | 7.03918 | 6.61836 | 5.76036 | 6.01166 | 6.00067 | 6.37372 | 5.38475 |
| MCAM | P43121 | 6.26441 | 6.85634 | 6.32079 | 6.76892 | 6.08402 | 6.13324 | 5.95465 | 5.76825 | 6.03692 | 5.88356 | 6.18678 | 6.34903 | 6.47954 | 6.34168 | 6.31091 | 6.26096 |
| BRMS1 | Q9HCU9 | 6.38296 | 6.32384 | 6.16989 | 6.59241 | 6.20018 | 6.06686 | 6.2427 | 6.04495 | 6.28485 | 6.11719 | 6.04718 | 6.06714 | 6.48548 | 6.27348 | 6.25706 | 6.44381 |
| NCK2 | O43639 | 6.60416 | 6.36213 | 6.63679 | 6.62571 | 6.13408 | 6.60117 | 6.53189 | 6.16435 | 6.41942 | 6.39622 | 6.54101 | 6.40865 | 5.97543 | 5.64135 | 5.50219 | 5.45546 |
| PAIP1 | Q9H074 | 6.26761 | 5.67445 | 6.74927 | 6.07649 | 6.44276 | 6.68295 | 6.24364 | 6.40077 | 6.41973 | 6.41315 | 6.71467 | 6.79693 | 6.10492 | 5.96891 | 5.41052 | 5.63322 |
| ADPGK | Q9BRR6 | 6.22821 | 6.26413 | 5.93162 | 6.53987 | 6.15428 | 6.11962 | 6.48457 | 5.99405 | 6.27108 | 5.65204 | 5.92366 | 5.87068 | 6.68944 | 6.34637 | 6.61632 | 6.91406 |
| SLC9A3R2 | Q15599 | 6.49943 | 6.53761 | 6.06867 | 6.32238 | 6.60865 | 5.94308 | 5.87149 | 6.27345 | 6.28768 | 5.90475 | 6.09509 | 6.475 | 7.07131 | 5.80777 | 6.12684 | 6.1068 |
| MEGF8 | Q7Z7M0 | 6.78411 | 6.4155 | 6.42593 | 6.57665 | 6.11466 | 6.14732 | 6.4542 | 5.90548 | 6.03422 | 6.93337 | 6.64005 | 6.95601 | 5.44602 | 5.71828 | 5.73669 | 5.71152 |
| ZSWIM8 | A7E2V4-4 | 6.94251 | 6.61017 | 6.92233 | 6.42092 | 6.46646 | 6.9082 | 5.31478 | 6.65935 | 6.81477 | 6.82921 | 6.37093 | 6.42503 | 5.1448 | 5.46288 | 5.52 | 5.18767 |
| MTG2 | Q9H4K7 | 7.24655 | 7.19937 | 6.41819 | 6.96262 | 6.37562 | 6.41567 | 6.47068 | 5.96555 | 6.27747 | 6.49862 | 6.29024 | 6.47299 | 5.73921 | 4.85523 | 5.43673 | 5.37525 |
| CTBS | Q01459 | 6.50445 | 6.6403 | 5.72668 | 6.4736 | 6.45828 | 6.44807 | 6.07277 | 6.29777 | 6.08859 | 6.18887 | 6.3632 | 6.23082 | 6.24593 | 6.07646 | 6.01601 | 6.16818 |
| PDXK | O00764 | 6.398 | 6.43796 | 7.18528 | 5.86171 | 6.2829 | 7.89009 | 6.01638 | 7.11274 | 6.47126 | 6.27519 | 6.65207 | 6.31414 | 5.2094 | 5.52896 | 5.80231 | 4.56161 |
| RIPK2 | O43353 | 6.42345 | 5.93783 | 6.77769 | 5.90367 | 6.47388 | 6.06795 | 6.56569 | 6.30088 | 6.32511 | 6.26918 | 6.68326 | 6.61466 | 5.19743 | 6.34632 | 5.84829 | 6.26471 |
| NSA2 | O95478 | 6.83551 | 5.98092 | 6.05828 | 7.32948 | 6.45229 | 6.84026 | 6.43447 | 6.15339 | 6.84595 | 5.88168 | 6.38565 | 6.20347 | 5.23562 | 6.02465 | 5.64864 | 6.28977 |
| MAP3K6 | O95382 | 6.19878 | 6.63375 | 6.93346 | 6.37756 | 6.06565 | 6.58652 | 6.0945 | 6.65669 | 6.21617 | 6.92086 | 6.77966 | 6.5333 | 5.56764 | 5.74941 | 5.40247 | 5.28358 |
| MAP3K5 | Q99683 | 6.28873 | 5.56452 | 6.9227 | 6.53181 | 6.03463 | 6.84724 | 6.56065 | 6.24319 | 7.09283 | 6.1684 | 6.78352 | 7.08147 | 5.70934 | 5.64102 | 4.91584 | 5.6141 |
| PDF | Q9HBH1 | 6.48961 | 6.26855 | 6.23728 | 6.69015 | 6.03306 | 5.76297 | 6.08024 | 6.14553 | 6.13691 | 6.21188 | 6.21509 | 6.28224 | 6.71695 | 5.93793 | 6.53071 | 6.2609 |
| IFIT5 | Q13325 | 6.44278 | 6.43022 | 7.17306 | 6.31124 | 6.34357 | 6.73832 | 6.22252 | 6.90586 | 6.15798 | 6.09155 | 6.56895 | 6.38019 | 5.56416 | 5.70811 | 5.33781 | 5.62369 |
| SIK2 | Q9H0K1 | 6.30153 | 6.58011 | 7.28554 | 5.92553 | 5.85954 | 7.32983 | 6.33643 | 6.78413 | 6.63935 | 6.78474 | 6.9329 | 6.52227 | 5.27882 | 5.54755 | 5.02583 | 5.26586 |
| PDLIM3 | Q53GG5-2 | 6.68052 | 6.15072 | 6.35833 | 6.23231 | 6.58815 | 6.31131 | 5.90716 | 6.28056 | 6.27551 | 6.25 | 6.60922 | 6.99719 | 5.91481 | 5.96168 | 5.46634 | 6.01617 |
| HINT2 | Q9BX68 | 5.13811 | 4.93828 | 4.34893 | 4.52743 | 5.58967 | 4.69285 | 5.54387 | 7.39315 | 5.14589 | 5.67443 | 4.80369 | 5.26352 | 4.85889 | 10.6859 | 9.27122 | 12.1242 |
| UBQLN1 | Q9UMX0 | 6.09893 | 4.46305 | 4.86201 | 4.26933 | 4.87342 | 3.7979 | 5.39863 | 6.63573 | 4.35711 | 5.24904 | 4.77465 | 6.9702 | 5.50572 | 11.1592 | 9.87682 | 11.7083 |
| STARD7 | Q9NQZ5 | 6.39005 | 5.8788 | 6.05695 | 6.48406 | 6.5756 | 6.5946 | 6.27543 | 6.1443 | 5.96476 | 6.27075 | 6.25603 | 6.75728 | 6.43282 | 6.47228 | 5.47837 | 5.96791 |
| TM7SF2 | O76062 | 5.85235 | 5.91032 | 6.13598 | 6.6277 | 5.83878 | 5.67642 | 6.15677 | 5.55032 | 6.09761 | 6.06306 | 5.79218 | 6.06693 | 7.01598 | 6.86796 | 7.10746 | 7.24018 |
| PRMT7 | Q9NVM4 | 6.61929 | 5.93723 | 7.41864 | 5.86799 | 6.28487 | 7.11643 | 6.15715 | 6.57328 | 6.60676 | 5.88634 | 6.92236 | 7.65172 | 5.43898 | 4.83973 | 5.41961 | 5.25964 |
| MRPS28 | Q9Y2Q9 | 6.34909 | 6.26762 | 6.25152 | 6.82671 | 6.27057 | 6.00579 | 6.29757 | 6.00607 | 6.04916 | 6.47878 | 6.10422 | 6.30025 | 6.55615 | 6.03382 | 6.13636 | 6.06633 |
| PINX1 | Q96BK5 | 6.52535 | 6.57709 | 6.0199 | 6.95929 | 6.31304 | 6.44508 | 6.44581 | 5.93516 | 5.9478 | 5.40826 | 6.187 | 6.56324 | 5.69591 | 6.41489 | 6.09774 | 6.46443 |
| GLIPR2 | Q9H4G4 | 6.15381 | 6.51016 | 6.2195 | 6.4898 | 6.30158 | 6.06707 | 6.31991 | 5.9483 | 5.91832 | 6.25873 | 6.07914 | 6.17911 | 6.7357 | 6.33568 | 6.58873 | 5.89445 |
| CASP7 | P55210-3 | 6.15631 | 6.43084 | 6.74318 | 5.87061 | 6.3213 | 6.22601 | 5.82069 | 6.7991 | 5.90567 | 6.48703 | 6.93953 | 6.7651 | 6.166 | 5.85921 | 5.61642 | 5.89301 |
| NEPRO | Q6NW34 | 6.50589 | 6.75058 | 6.38929 | 7.03161 | 6.01276 | 6.11798 | 6.51519 | 5.95666 | 6.25296 | 5.97927 | 6.19509 | 5.9548 | 5.59008 | 5.73797 | 6.43432 | 6.63544 |
| RUNX1 | Q01196-8 | 5.87718 | 6.52679 | 6.11707 | 6.36763 | 6.1531 | 5.95066 | 6.04648 | 6.05553 | 5.86581 | 6.07221 | 6.20002 | 6.23412 | 6.91661 | 6.39919 | 6.57328 | 6.64432 |
| PTDSS2 | Q9BVG9 | 6.33151 | 6.0562 | 6.5223 | 7.11 | 5.76535 | 6.56345 | 6.02478 | 6.09092 | 6.07869 | 6.27309 | 5.60688 | 6.16095 | 6.49803 | 6.38887 | 6.36702 | 6.16197 |
| ZNF280D | Q6N043 | 6.3682 | 6.86981 | 6.14305 | 7.22087 | 6.15623 | 6.1543 | 6.47551 | 5.65303 | 6.28902 | 6.30468 | 5.9461 | 5.964 | 6.38161 | 5.93937 | 5.96965 | 6.16459 |
| MEF2D | Q14814-4 | 5.79966 | 6.04825 | 6.08642 | 6.0325 | 5.963 | 6.29421 | 5.91011 | 5.73804 | 5.97875 | 6.34564 | 6.56836 | 6.32503 | 6.76086 | 6.80485 | 6.70031 | 6.64402 |
| R3HDM1 | Q15032-2 | 6.67794 | 6.34282 | 6.77105 | 6.42358 | 6.14573 | 6.82563 | 6.0445 | 5.93191 | 6.57066 | 6.62105 | 6.46127 | 7.14129 | 5.94913 | 5.30472 | 5.20865 | 5.57516 |
| MTM1 | Q13496 | 6.10727 | 6.54002 | 6.44116 | 6.04615 | 6.18555 | 6.55091 | 6.5234 | 6.19291 | 6.17234 | 6.43208 | 6.31228 | 6.23645 | 5.9528 | 6.21842 | 6.21412 | 5.87415 |
| TAF3 | Q5VWG9 | 6.44878 | 6.50038 | 6.12769 | 6.69494 | 5.64755 | 5.65659 | 6.00234 | 5.76559 | 5.99164 | 5.87808 | 6.2332 | 5.96943 | 6.1701 | 6.8865 | 6.93434 | 7.09283 |
| CEP44 | Q9C0F1 | 5.81521 | 6.03353 | 7.01503 | 5.97862 | 6.01906 | 6.58508 | 5.77292 | 6.15665 | 6.23481 | 6.79774 | 7.08828 | 7.10217 | 5.64067 | 6.08602 | 5.98703 | 5.68717 |
| ZFYVE19 | Q96K21 | 5.81731 | 6.11468 | 6.87745 | 5.88794 | 5.8482 | 7.05172 | 6.01351 | 6.21327 | 5.96628 | 7.10999 | 6.65295 | 6.82579 | 5.52496 | 6.13338 | 5.72575 | 6.23681 |
| TPD52L1 | Q16890 | 6.50156 | 6.09116 | 7.04427 | 5.73322 | 6.37562 | 6.82481 | 5.89913 | 6.70684 | 6.04571 | 6.31659 | 6.50972 | 6.65955 | 6.20562 | 5.71172 | 5.80962 | 5.56486 |
| SYTL4 | Q96C24 | 6.8371 | 7.02709 | 6.44643 | 6.69966 | 5.4552 | 6.40084 | 6.0563 | 5.56551 | 6.18758 | 6.33957 | 6.08181 | 6.62953 | 6.40954 | 6.4239 | 6.23181 | 5.20814 |
| NDE1 | Q9NXR1-1 | 6.73105 | 6.46411 | 6.41059 | 5.99584 | 6.70363 | 6.31636 | 6.0509 | 6.77694 | 6.1561 | 6.71851 | 6.54613 | 6.55985 | 5.96268 | 5.65293 | 5.33264 | 5.62174 |
| ARFGAP3 | Q9NP61 | 6.28525 | 6.03692 | 6.50386 | 6.1462 | 6.59055 | 6.20694 | 6.35615 | 6.5343 | 6.50986 | 6.60752 | 6.16531 | 6.59057 | 5.94797 | 5.96731 | 5.64452 | 5.90675 |
| DCAF16 | Q9NXF7 | 5.48021 | 5.93496 | 6.06779 | 5.67441 | 6.06752 | 6.09351 | 5.79147 | 6.59321 | 5.79886 | 6.10077 | 6.41556 | 6.39645 | 6.88863 | 7.35648 | 6.51435 | 6.82581 |
| GLCCI1 | Q86VQ1 | 6.63057 | 5.81432 | 7.33275 | 6.17093 | 6.10994 | 7.34127 | 6.06479 | 6.0275 | 6.57118 | 6.94449 | 7.00226 | 6.83616 | 5.8759 | 5.16285 | 4.75525 | 5.3598 |
| SIX4 | Q9UIU6 | 5.91462 | 6.35729 | 5.97716 | 5.81577 | 6.23476 | 6.14958 | 6.00154 | 6.4435 | 6.30785 | 6.1761 | 6.83527 | 6.55155 | 6.4837 | 6.41358 | 5.806 | 6.53174 |
| KIAA1549L | Q6ZVL6 | 6.14424 | 6.50713 | 6.3065 | 6.66699 | 5.97592 | 6.3783 | 6.25378 | 5.79894 | 6.07645 | 5.65699 | 6.48614 | 6.47628 | 6.46662 | 5.89314 | 6.19143 | 6.72115 |
| RETREG2 | Q8NC44 | 5.94163 | 5.88896 | 5.4908 | 5.76693 | 5.95225 | 5.5576 | 5.6544 | 5.94219 | 5.97543 | 6.07909 | 5.56259 | 6.03723 | 7.77735 | 7.21836 | 7.83844 | 7.31675 |
| POLH | Q9Y253 | 6.3467 | 6.73415 | 6.46461 | 6.70098 | 6.07485 | 6.44987 | 5.91323 | 5.97439 | 6.37006 | 6.57843 | 7.02391 | 6.86016 | 5.43659 | 5.5302 | 5.78963 | 5.75225 |
| REEP5 | Q00765 | 6.1456 | 6.19337 | 6.07762 | 5.89193 | 6.36578 | 6.26854 | 6.36212 | 6.14613 | 6.18453 | 6.34115 | 6.03457 | 6.27667 | 6.80926 | 6.29927 | 6.50992 | 6.09354 |
| TBPL1 | P62380 | 5.26 | 6.04917 | 6.23041 | 5.56943 | 5.26626 | 6.16578 | 5.64362 | 5.83093 | 5.78635 | 6.36703 | 6.11374 | 6.11727 | 6.63534 | 7.81989 | 7.77301 | 7.37176 |
| SETD7 | Q8WTS6 | 6.08253 | 5.69863 | 6.52927 | 5.80467 | 6.39225 | 6.36053 | 6.18398 | 6.70624 | 5.94862 | 6.62325 | 6.61793 | 6.74679 | 6.26966 | 6.08086 | 5.72286 | 6.23194 |
| DNTTIP1 | Q9H147 | 6.36048 | 6.26326 | 6.36966 | 6.77606 | 6.14497 | 6.05351 | 6.34355 | 6.12011 | 6.29089 | 6.30662 | 6.3657 | 6.44395 | 6.09978 | 5.91728 | 6.19713 | 5.94705 |
| SMURF1 | Q9HCE7 | 6.80879 | 5.90706 | 6.25561 | 6.22964 | 5.87051 | 6.66687 | 6.10621 | 6.65407 | 6.21134 | 6.63427 | 6.9707 | 6.73798 | 5.49197 | 5.80339 | 5.56336 | 6.08823 |
| ST7 | Q9NRC1 | 6.68354 | 6.52158 | 6.07452 | 6.71675 | 6.16139 | 5.88601 | 6.28 | 5.72726 | 6.29149 | 5.99117 | 6.48679 | 6.08598 | 6.29742 | 6.03912 | 6.50272 | 6.25426 |
| GALM | Q96C23 | 5.58695 | 5.58817 | 6.34096 | 5.77118 | 6.63465 | 6.86416 | 5.97491 | 6.57245 | 6.20595 | 6.72947 | 6.80119 | 6.14277 | 6.14097 | 6.47655 | 6.00646 | 6.06421 |
| DCP2 | Q8IU60 | 5.27008 | 5.87588 | 7.74035 | 6.23335 | 5.8522 | 6.11331 | 5.52171 | 6.23483 | 6.83697 | 6.47771 | 6.71515 | 6.56685 | 5.85797 | 5.92582 | 6.32127 | 6.45654 |
| SLC30A5 | Q8TAD4 | 6.34499 | 6.29867 | 5.92333 | 6.43791 | 6.2946 | 6.01122 | 6.37231 | 5.84189 | 6.44071 | 5.70188 | 5.84095 | 5.80417 | 6.74156 | 6.557 | 6.66133 | 6.72749 |
| GMPPA | Q96IJ6-2 | 6.224 | 6.00854 | 6.75851 | 6.01626 | 6.07821 | 6.59402 | 6.1075 | 6.55894 | 6.42901 | 6.5859 | 6.4696 | 6.51229 | 6.13581 | 5.98135 | 5.78756 | 5.75251 |
| FANCC | Q00597 | 6.40998 | 6.01821 | 6.7723 | 6.19644 | 6.51372 | 6.56949 | 6.32923 | 6.96922 | 6.65898 | 6.75894 | 6.88505 | 7.06466 | 5.16214 | 5.12008 | 5.04765 | 5.52391 |
| JRK | O75564-2 | 6.01238 | 6.07853 | 5.79896 | 6.30483 | 5.942 | 6.20522 | 6.36437 | 6.44046 | 6.17002 | 6.38327 | 5.98577 | 6.16914 | 6.46235 | 6.38045 | 6.52578 | 6.77647 |
| ERO1B | Q86YB8 | 6.36178 | 6.22536 | 6.03485 | 6.63776 | 6.16847 | 6.00577 | 6.49366 | 5.90847 | 5.83204 | 6.10687 | 6.24921 | 6.06157 | 6.62711 | 6.25296 | 6.63049 | 6.40362 |
| STX6 | O43752 | 6.37495 | 6.71177 | 5.96031 | 6.52464 | 6.02835 | 6.5907 | 6.36967 | 6.18403 | 6.23738 | 6.24173 | 6.51171 | 6.18788 | 5.72866 | 5.99295 | 6.24422 | 6.11106 |
| PPP3CB | P16298-4 | 6.20705 | 6.21054 | 6.79796 | 5.92779 | 6.23201 | 6.33225 | 5.90914 | 6.13174 | 6.16801 | 6.56423 | 6.52046 | 6.4189 | 6.27298 | 6.0567 | 5.93764 | 6.3126 |
| NOVA2 | Q9UNW9 | 6.14293 | 6.3407 | 6.24692 | 6.03762 | 6.4628 | 5.93851 | 6.065 | 6.20599 | 5.96446 | 6.06194 | 6.29554 | 6.05395 | 6.7818 | 6.32904 | 6.68012 | 6.39266 |
| ZBTB47 | Q9UFB7 | 6.16025 | 6.58339 | 6.36579 | 6.94401 | 5.972 | 6.22967 | 5.90646 | 5.66027 | 6.19333 | 6.63989 | 6.4288 | 6.26541 | 6.18489 | 6.07469 | 6.31402 | 6.07713 |
| WDR73 | Q6P4I2 | 5.91784 | 5.82642 | 6.39414 | 5.66687 | 6.54887 | 6.81049 | 5.77456 | 7.00554 | 6.98057 | 6.28859 | 6.47431 | 6.80741 | 5.40887 | 5.7822 | 6.36189 | 5.95141 |
| SRFBP1 | Q8NEF9 | 5.99751 | 5.95777 | 5.69551 | 6.59007 | 5.82042 | 5.62404 | 6.21795 | 5.7876 | 6.14772 | 5.80556 | 6.16988 | 5.82534 | 7.05888 | 6.62149 | 7.20467 | 7.47557 |
| AXIN1 | O15169 | 6.8216 | 6.99677 | 6.85247 | 6.66526 | 6.19091 | 6.31734 | 6.55639 | 6.42707 | 6.50034 | 6.35609 | 6.42454 | 6.60147 | 5.21178 | 5.22577 | 5.38445 | 5.46773 |
| SPPL2A | Q8TCT8 | 6.76269 | 6.83346 | 6.43473 | 6.83919 | 6.37811 | 5.90616 | 6.51512 | 6.16531 | 5.94058 | 6.23703 | 6.66095 | 6.36757 | 5.83976 | 5.79593 | 5.60288 | 5.72054 |
| GDAP2 | Q9NXN4 | 6.02405 | 6.17151 | 6.40011 | 6.26266 | 6.12449 | 6.02024 | 6.13249 | 6.16751 | 6.28052 | 6.39216 | 6.38414 | 6.22094 | 6.77163 | 6.10577 | 6.2807 | 6.26108 |
| DGKE | P52429 | 6.24716 | 6.2647 | 6.66225 | 6.87829 | 5.90671 | 5.86928 | 6.04246 | 5.76808 | 7.13444 | 5.97086 | 5.84876 | 6.22622 | 6.30222 | 6.00736 | 6.49986 | 6.37137 |
| CTU2 | Q2VPK5 | 6.64246 | 6.3428 | 7.02143 | 6.66972 | 6.79241 | 6.38036 | 6.15048 | 7.12203 | 6.23147 | 6.17567 | 6.41013 | 6.81195 | 5.43423 | 5.45914 | 4.9166 | 5.43913 |
| RPP40 | O75818 | 6.05927 | 6.39107 | 6.69836 | 6.46866 | 6.17619 | 6.36134 | 5.98348 | 6.01899 | 6.43378 | 6.19713 | 5.95984 | 6.19363 | 6.39155 | 6.1319 | 6.50178 | 6.03303 |
| BBS1 | Q8NFJ9-2 | 6.77881 | 6.09376 | 6.35223 | 5.85952 | 6.2626 | 6.75496 | 6.44693 | 6.84287 | 6.36742 | 6.43149 | 6.3072 | 6.38905 | 5.68061 | 6.25752 | 5.96871 | 5.20632 |
| PI4K2B | Q8TCG2 | 6.68299 | 6.90783 | 6.64278 | 6.55729 | 6.52369 | 5.9978 | 6.48345 | 6.23868 | 6.44647 | 6.68856 | 6.74875 | 6.17613 | 5.33387 | 5.62038 | 5.2658 | 5.68553 |
| NAXD | Q8IW45-2 | 5.83736 | 6.07209 | 6.54673 | 6.05594 | 6.65588 | 6.09493 | 6.19168 | 6.59321 | 6.34491 | 6.33269 | 6.30436 | 6.15052 | 6.259 | 6.06908 | 6.27257 | 6.21904 |
| USP37 | Q86T82 | 5.20544 | 6.02189 | 6.13713 | 5.71805 | 5.41122 | 5.85049 | 5.83261 | 5.76433 | 5.95311 | 6.35199 | 6.51673 | 6.51738 | 6.64246 | 7.44413 | 7.26749 | 7.36554 |
| NDUFA8 | P51970 | 6.70808 | 5.13557 | 6.10273 | 5.51903 | 7.26245 | 5.64958 | 5.96837 | 6.25617 | 5.92783 | 6.2305 | 6.29674 | 6.27398 | 7.31854 | 6.22738 | 6.34805 | 6.77499 |
| IRF2 | P14316 | 5.9423 | 6.1068 | 6.43295 | 6.85479 | 6.17898 | 6.2658 | 6.04528 | 5.62086 | 6.19598 | 6.59176 | 6.5036 | 6.46435 | 6.6813 | 6.12675 | 6.08719 | 5.90129 |

| | | | | | | | | | | | | | | | | | |
|---|---|---|---|---|---|---|---|---|---|---|---|---|---|---|---|---|---|
| TMEM67 | Q5HYA8 | 6.35705 | 6.69629 | 5.88032 | 6.58206 | 5.89109 | 6.26085 | 6.26676 | 6.37583 | 5.90545 | 5.92946 | 6.00675 | 6.15286 | 6.36413 | 6.54997 | 6.29702 | 6.48411 |
| SIRT5 | Q9NXA8 | 6.63055 | 6.59251 | 5.81618 | 6.3435 | 6.17639 | 6.41205 | 6.18534 | 6.15504 | 6.2822 | 6.25524 | 5.86636 | 6.20268 | 6.23467 | 6.30533 | 6.40787 | 6.13409 |
| ZNF335 | Q9H4Z2 | 6.26361 | 6.38093 | 6.39728 | 6.70438 | 6.08409 | 5.82102 | 6.258 | 6.04114 | 5.94016 | 6.42692 | 6.18732 | 6.06546 | 6.06436 | 6.00171 | 6.87139 | 6.49222 |
| FAS | P25445 | 6.0968 | 6.54941 | 5.97967 | 6.60391 | 6.34007 | 6.3588 | 6.2683 | 5.86763 | 6.08519 | 6.10863 | 6.15119 | 6.06301 | 6.67351 | 6.09369 | 6.39912 | 6.36108 |
| SDSL | Q96GA7 | 6.33189 | 6.06883 | 6.94485 | 5.92015 | 6.28892 | 6.72528 | 6.14123 | 6.23622 | 6.24999 | 6.56593 | 6.42523 | 6.36078 | 5.91874 | 6.00838 | 5.93629 | 5.8773 |
| EP300 | Q09472 | 5.55797 | 6.19107 | 6.31121 | 6.93692 | 6.05238 | 6.31339 | 6.36355 | 6.25338 | 5.90449 | 6.26826 | 6.69526 | 5.81438 | 6.49369 | 6.75209 | 6.13681 | 5.95514 |
| SLC25A40 | Q8TBP6 | 6.14928 | 6.25184 | 6.14361 | 6.77512 | 6.05211 | 6.01157 | 6.25966 | 5.84533 | 6.30009 | 6.07834 | 5.60743 | 5.86512 | 6.75976 | 6.67735 | 6.54602 | 6.67738 |
| ZBTB1 | Q9Y2K1 | 6.47736 | 6.92824 | 6.0828 | 6.54728 | 5.93978 | 6.12143 | 6.2286 | 6.0197 | 6.6914 | 6.6997 | 6.31918 | 6.47097 | 6.02266 | 5.22212 | 6.08094 | 6.14785 |
| LDB1 | Q86U70 | 6.35742 | 6.55491 | 6.30199 | 6.05682 | 5.8337 | 7.20722 | 6.10148 | 4.997 | 6.2508 | 6.71023 | 6.14347 | 6.29472 | 6.64523 | 6.59802 | 6.56623 | 5.38073 |
| WDR45 | Q9Y484-2 | 6.32925 | 6.59167 | 6.48373 | 6.48367 | 5.93104 | 6.10144 | 5.92221 | 6.33953 | 6.38359 | 6.35795 | 6.90543 | 6.94232 | 5.70888 | 5.75778 | 6.07703 | 5.68448 |
| TRIM5 | Q9C035 | 6.14798 | 6.03004 | 7.26923 | 6.35203 | 6.44203 | 6.81165 | 6.22843 | 6.24232 | 6.57861 | 6.25415 | 6.705 | 6.45933 | 5.83926 | 5.56007 | 5.31683 | 5.76305 |
| CRELD2 | Q6UXH1-5 | 5.73633 | 6.86698 | 6.04221 | 6.51687 | 6.06392 | 6.5517 | 6.11888 | 6.13847 | 5.84015 | 6.5684 | 5.72361 | 6.09728 | 6.62498 | 6.60187 | 6.11989 | 6.38848 |
| MYPN | Q86TC9 | 5.97092 | 5.69459 | 7.03582 | 5.82348 | 6.17626 | 6.50379 | 6.40704 | 6.07938 | 6.45984 | 6.52348 | 6.77253 | 6.41301 | 6.27121 | 5.85043 | 5.88541 | 6.13282 |
| THOC3 | Q96J01 | 6.13219 | 6.18066 | 6.11391 | 6.20159 | 6.18035 | 6.57813 | 6.24941 | 5.68501 | 6.13261 | 6.80849 | 6.39381 | 6.65109 | 6.1136 | 5.9259 | 6.33133 | 6.32193 |
| CRELD1 | Q96HD1-2 | 6.21933 | 6.38997 | 5.79229 | 6.11365 | 6.19861 | 6.28313 | 6.40564 | 5.74695 | 5.95274 | 6.54132 | 5.92588 | 5.27105 | 7.01475 | 6.65003 | 6.94986 | 6.54481 |
| TMEM9B | Q9NQ34 | 6.04377 | 7.1194 | 6.44712 | 6.35527 | 6.20669 | 6.41423 | 6.52021 | 6.2931 | 6.56388 | 6.16622 | 6.03717 | 6.22479 | 5.87737 | 5.91395 | 5.84136 | 5.97546 |
| FNIP1 | Q8TF40 | 7.61737 | 7.02369 | 7.50254 | 7.30288 | 6.55178 | 6.55622 | 6.42051 | 6.82159 | 7.34579 | 6.93132 | 7.19177 | 7.40703 | 4.02257 | 4.05316 | 3.26913 | 3.98267 |
| PDZD8 | Q8NEN9 | 6.45025 | 6.44125 | 5.79819 | 6.84253 | 5.98067 | 5.85244 | 6.45465 | 5.96966 | 5.777 | 6.41847 | 5.55727 | 6.45459 | 6.26282 | 6.55023 | 6.61119 | 6.57881 |
| FSTL1 | Q12841 | 6.81316 | 6.32413 | 5.67856 | 6.36614 | 7.1372 | 6.47858 | 6.76007 | 6.79075 | 5.81564 | 5.26621 | 6.13122 | 5.47897 | 7.05805 | 5.49814 | 5.83298 | 6.57019 |
| USP12 | O75317 | 6.21421 | 6.31981 | 6.08582 | 6.27048 | 5.99131 | 5.53703 | 6.30121 | 6.20576 | 6.23599 | 6.92191 | 6.29061 | 6.54753 | 6.36068 | 5.94835 | 6.51643 | 6.25286 |
| FHL3 | Q13643 | 6.56437 | 6.07468 | 6.54199 | 5.75791 | 5.97274 | 6.48874 | 5.87025 | 6.08803 | 6.31986 | 6.27911 | 6.43315 | 6.73763 | 6.16573 | 6.17624 | 6.28107 | 6.24849 |
| ESCO1 | Q5FWF5 | 6.05256 | 5.90211 | 6.03687 | 6.14103 | 5.82746 | 5.68626 | 6.28834 | 5.86479 | 5.63387 | 5.89639 | 5.95931 | 5.48121 | 6.71621 | 7.46665 | 7.64346 | 7.40347 |
| RECK | O95980 | 5.60451 | 6.40351 | 5.72633 | 6.88676 | 6.46246 | 6.1509 | 6.46697 | 5.93319 | 6.09902 | 6.60675 | 6.19569 | 6.48522 | 6.34861 | 5.92047 | 6.21119 | 6.49843 |
| NDRG2 | Q9UN36 | 6.4376 | 5.80978 | 6.6388 | 5.15925 | 6.67185 | 6.17442 | 5.39788 | 6.87996 | 5.5894 | 7.14551 | 6.10689 | 7.04087 | 6.88619 | 6.55701 | 5.51991 | 5.98467 |
| NUDT16L1 | Q9BRJ7 | 6.34585 | 5.99648 | 5.95781 | 6.00897 | 6.50319 | 6.10161 | 6.37308 | 6.21707 | 6.17019 | 6.52638 | 6.36654 | 6.16977 | 6.43302 | 6.05722 | 6.43035 | 6.34246 |
| MRPL20 | Q9BYC9 | 6.09122 | 6.174 | 5.91112 | 6.82449 | 6.14506 | 6.11059 | 6.75015 | 5.85812 | 6.00743 | 6.03545 | 6.34275 | 6.05769 | 6.99427 | 5.91355 | 6.02996 | 6.75415 |
| SMN1 | Q16637 | 5.84779 | 6.27687 | 6.7149 | 6.83207 | 6.31477 | 6.37333 | 5.96833 | 6.00386 | 6.07967 | 5.83547 | 6.71207 | 6.69993 | 6.65087 | 5.96054 | 5.84929 | 5.88023 |
| ARHGAP29 | Q52LW3 | 6.22255 | 6.41593 | 7.05874 | 6.37495 | 6.1063 | 6.92131 | 6.0338 | 6.46934 | 6.4423 | 6.18543 | 6.96191 | 7.26604 | 5.77818 | 5.32519 | 5.19847 | 5.23954 |
| SYNPO | Q8N3V7-2 | 6.31981 | 5.99981 | 5.73851 | 6.46589 | 6.45905 | 6.06321 | 6.20505 | 5.65598 | 6.20965 | 5.97311 | 5.66916 | 5.84943 | 7.59121 | 6.37248 | 6.69298 | 6.73468 |
| SRSF5 | Q13243 | 6.60876 | 6.47472 | 5.73684 | 5.84411 | 6.53893 | 6.16014 | 6.47851 | 6.28965 | 6.48786 | 6.34516 | 6.20323 | 5.91432 | 6.53317 | 6.10983 | 6.37558 | 5.89917 |
| TADA2B | Q86TJ2 | 6.0489 | 6.57136 | 6.01601 | 6.91913 | 6.02312 | 5.96712 | 6.30141 | 6.1466 | 6.15478 | 6.35398 | 6.30622 | 6.57449 | 5.99617 | 5.95915 | 6.17683 | 6.48472 |
| STIL | Q15468-2 | 5.92329 | 6.55312 | 6.57155 | 6.2424 | 6.19896 | 6.29928 | 6.56014 | 6.51795 | 6.92283 | 6.85832 | 6.9147 | 7.20335 | 5.1329 | 5.85746 | 4.8978 | 5.34595 |
| GINS1 | Q14691 | 6.12019 | 5.84976 | 6.43496 | 5.35927 | 5.89327 | 6.41397 | 5.93242 | 5.81859 | 6.21495 | 6.69036 | 6.42068 | 6.79072 | 6.00224 | 6.84014 | 6.58802 | 6.63046 |
| PHF5A | Q7RTV0 | 6.57732 | 6.42231 | 6.27562 | 6.55707 | 6.098 | 6.08664 | 6.04212 | 5.89756 | 5.98469 | 6.32288 | 6.13984 | 5.96527 | 6.81908 | 6.08754 | 6.29802 | 6.42604 |
| ZCCHC7 | Q8N3Z6 | 5.92894 | 6.26837 | 6.19445 | 6.96051 | 5.81331 | 5.50449 | 6.52664 | 6.05836 | 5.58303 | 5.96534 | 6.27175 | 6.01081 | 6.53879 | 6.56824 | 7.10512 | 6.70184 |
| ZC3H8 | Q8N5P1 | 6.53699 | 6.1871 | 6.02324 | 6.27951 | 5.72525 | 5.73854 | 5.94872 | 6.45236 | 6.26001 | 6.21 | 6.26942 | 6.28689 | 6.50447 | 6.93023 | 6.84876 | 6.78848 |
| NARF | Q9UHQ1-2 | 6.75836 | 6.93072 | 7.07908 | 6.16691 | 6.06992 | 6.41387 | 5.71278 | 5.95397 | 6.78479 | 7.29018 | 6.9631 | 7.02909 | 4.74863 | 5.89345 | 5.25517 | 4.94998 |
| DPH5 | Q9H2P9-5 | 6.16486 | 6.13086 | 7.39038 | 5.58682 | 6.16694 | 6.422 | 6.38832 | 7.06793 | 6.1865 | 6.46316 | 6.67847 | 7.16664 | 5.36641 | 5.60186 | 5.6077 | 5.61116 |
| CCDC134 | Q9H6E4 | 6.40119 | 6.17423 | 6.44122 | 6.32843 | 6.04321 | 6.41681 | 6.53077 | 5.92035 | 6.38579 | 6.32941 | 5.80776 | 5.83441 | 6.67165 | 5.94877 | 6.54081 | 6.2252 |
| PTOV1 | Q86YD1 | 6.40891 | 6.14831 | 6.4108 | 6.84612 | 6.12112 | 6.2854 | 6.09163 | 6.27334 | 6.00793 | 6.39873 | 6.76594 | 6.67148 | 6.25577 | 5.54748 | 5.88907 | 5.87797 |
| TRIM23 | P36406 | 6.23076 | 6.4181 | 6.3661 | 6.37096 | 6.12229 | 6.29791 | 6.45984 | 6.28199 | 6.60391 | 6.29802 | 6.46952 | 6.40953 | 5.82708 | 5.86431 | 6.07044 | 5.90926 |
| TMEM161A | Q9NX61 | 6.43942 | 6.1752 | 5.68916 | 6.78645 | 6.13607 | 5.9725 | 6.26917 | 5.83902 | 6.22329 | 6.04512 | 5.91441 | 5.82473 | 6.9955 | 6.21898 | 6.53281 | 6.93818 |
| USP42 | Q9H9J4 | 6.29367 | 6.41545 | 5.97388 | 6.81154 | 6.08621 | 6.12614 | 6.14187 | 5.95227 | 6.7346 | 6.46451 | 6.23156 | 6.44416 | 5.83627 | 6.16118 | 5.93477 | 6.39191 |
| NADK | O95544-2 | 6.41597 | 5.81112 | 7.38028 | 6.23224 | 5.83034 | 6.4443 | 6.47816 | 6.81296 | 6.62091 | 6.65171 | 6.7503 | 6.90882 | 5.49097 | 5.47824 | 5.25774 | 5.43594 |
| RPAP2 | Q8IXW5 | 6.4837 | 6.28779 | 6.78477 | 6.20808 | 6.07264 | 6.36147 | 6.80842 | 6.61464 | 6.6625 | 6.48119 | 6.64864 | 6.94152 | 5.444 | 5.27741 | 5.35599 | 5.56723 |
| DOK1 | Q99704 | 6.93597 | 5.61552 | 7.10915 | 5.79408 | 5.9205 | 5.86868 | 6.19019 | 6.4682 | 5.74804 | 6.25163 | 6.99323 | 6.72294 | 5.67073 | 6.86209 | 6.03756 | 5.81149 |
| RIOK1 | Q9BRS2 | 6.08643 | 5.92131 | 6.71019 | 6.56436 | 5.92383 | 6.5666 | 5.99248 | 6.40063 | 5.92042 | 6.24361 | 5.93263 | 6.06088 | 5.73943 | 6.51935 | 6.9124 | 6.50546 |
| NDOR1 | Q9UHB4-2 | 6.3324 | 5.81416 | 7.13627 | 6.39117 | 5.91765 | 6.63919 | 6.13951 | 6.37445 | 6.252 | 6.5295 | 7.2552 | 6.66125 | 5.66266 | 5.87441 | 5.66806 | 5.35214 |
| STX3 | Q13277 | 6.66093 | 6.70742 | 6.11 | 7.03131 | 6.17267 | 6.18414 | 6.26082 | 6.63795 | 6.01586 | 6.33513 | 6.51228 | 6.0824 | 5.59576 | 5.74473 | 6.20977 | 5.73882 |
| ZMIZ2 | Q8NF64 | 5.74093 | 5.53726 | 6.145 | 4.92551 | 5.62638 | 5.67718 | 5.71842 | 5.77161 | 6.38109 | 6.6351 | 5.99753 | 5.92636 | 7.16067 | 7.5392 | 7.6236 | 7.59416 |
| UBE2G2 | P60604 | 5.24974 | 5.28632 | 4.65551 | 6.54695 | 6.09291 | 5.01263 | 5.59442 | 6.75663 | 4.36992 | 5.30533 | 5.00421 | 5.24009 | 5.04413 | 11.1419 | 7.42542 | 11.2739 |
| TMEM30A | Q9NV96 | 6.71301 | 6.74252 | 6.21822 | 6.88004 | 6.14562 | 6.25254 | 6.21505 | 6.13205 | 6.59308 | 6.70193 | 6.42621 | 6.44524 | 5.74293 | 5.55556 | 5.48448 | 5.75153 |
| HSF1 | Q00613 | 6.08938 | 6.36004 | 6.08581 | 5.4608 | 6.15241 | 6.49042 | 5.5764 | 6.57648 | 6.01217 | 6.48375 | 6.36095 | 6.6098 | 6.20764 | 6.73308 | 6.44489 | 6.35599 |
| SIRT7 | Q9NRC8 | 6.13001 | 6.77976 | 5.71369 | 6.58755 | 5.46261 | 6.41 | 6.28789 | 5.82523 | 5.94813 | 6.23821 | 6.35956 | 6.46576 | 6.29787 | 6.21227 | 6.73448 | 6.54699 |
| TAF1A | Q15573 | 5.8699 | 6.25283 | 6.13314 | 6.47449 | 6.10855 | 6.21667 | 6.04779 | 5.657 | 6.01581 | 6.03371 | 6.08788 | 6.20887 | 6.68511 | 6.15591 | 7.03198 | 7.02035 |
| RABGGTB | P53611 | 6.42454 | 5.40009 | 6.86247 | 5.51123 | 6.16915 | 6.83019 | 5.90198 | 6.79817 | 5.93658 | 6.42567 | 6.56637 | 7.06836 | 5.11063 | 6.61732 | 5.96449 | 6.41275 |
| UBE2I | P63279 | 5.52515 | 4.66109 | 6.09172 | 5.00578 | 6.27851 | 5.40396 | 5.41256 | 6.84832 | 5.65013 | 6.75758 | 6.22447 | 6.98272 | 6.83165 | 7.72364 | 7.34359 | 7.25912 |
| FAM50B | Q9Y247 | 5.73806 | 6.70972 | 6.51602 | 6.07656 | 6.32042 | 6.27687 | 6.12696 | 5.98199 | 6.55267 | 6.76906 | 6.8019 | 6.39273 | 5.83317 | 6.19054 | 6.00185 | 5.81146 |
| FAM83G | A6ND36 | 5.72677 | 5.68628 | 6.42925 | 5.63519 | 6.04924 | 5.83441 | 6.19811 | 5.81928 | 6.51961 | 5.93925 | 6.35215 | 6.2662 | 7.26578 | 5.95387 | 7.36297 | 6.96163 |
| PPIL1 | Q9Y3C6 | 6.17816 | 5.66556 | 5.88073 | 5.37096 | 6.16991 | 6.20664 | 5.65745 | 6.80565 | 5.57417 | 6.47542 | 5.90246 | 6.58054 | 6.82634 | 7.23414 | 6.88512 | 6.58976 |
| ATG4C | Q96DT6 | 6.27837 | 6.02818 | 6.66133 | 5.59239 | 5.27208 | 6.82925 | 6.12706 | 5.68705 | 7.01017 | 7.18907 | 6.36386 | 6.59388 | 6.18568 | 6.26251 | 6.03204 | 5.88708 |
| APOB | P04114 | 6.27791 | 7.22909 | 6.55147 | 7.26506 | 5.78957 | 5.56498 | 6.28761 | 6.27759 | 6.97883 | 6.55235 | 6.08865 | 6.96015 | 5.25439 | 5.32552 | 5.98034 | 5.6165 |
| STX12 | Q86Y82 | 6.37014 | 6.23145 | 6.56771 | 6.23163 | 6.21504 | 6.57854 | 6.35127 | 6.02349 | 6.27491 | 6.22284 | 6.7235 | 6.50814 | 6.03854 | 5.97367 | 6.08919 | 5.59993 |
| ZBTB3 | Q9H5J0 | 6.70768 | 5.81718 | 5.26092 | 5.96996 | 6.7829 | 6.11297 | 5.54105 | 6.59384 | 6.51103 | 6.44287 | 6.51021 | 6.25739 | 7.00233 | 6.00302 | 6.49768 | 5.98898 |
| MYORG | Q6NSJ0 | 6.41663 | 6.60928 | 6.06604 | 6.88309 | 6.36322 | 5.80903 | 6.51732 | 5.8571 | 6.19818 | 6.99855 | 6.35657 | 5.94484 | 6.2063 | 6.0541 | 6.13617 | 5.5836 |
| HDHD2 | Q9H0R4 | 6.2605 | 5.83038 | 7.38584 | 5.76973 | 6.77776 | 7.75532 | 6.51028 | 6.29712 | 6.09588 | 6.3898 | 6.51569 | 6.41556 | 5.43302 | 6.02264 | 5.52745 | 5.01302 |
| CASP2 | P42575 | 6.51469 | 6.00714 | 6.90711 | 5.76802 | 5.41664 | 6.20403 | 6.04437 | 6.54345 | 6.33313 | 6.31684 | 6.8504 | 6.92379 | 5.30836 | 6.42859 | 6.74017 | 5.69328 |
| CCDC9 | A0A2R8Y4Z8 | 6.49159 | 6.26927 | 6.66817 | 6.14567 | 6.32942 | 6.19137 | 6.49168 | 6.69538 | 6.52946 | 5.92214 | 6.38265 | 6.47563 | 6.00669 | 5.93488 | 5.85834 | 5.60765 |
| COA1 | Q9GZY4 | 6.30454 | 6.15193 | 5.69957 | 6.45688 | 6.29286 | 5.78738 | 6.43763 | 5.96788 | 6.29713 | 5.98917 | 5.96593 | 6.09947 | 6.90309 | 6.31744 | 6.70511 | 6.61398 |
| MICU2 | Q8IYU8 | 6.14765 | 6.24417 | 6.21929 | 7.34158 | 6.45523 | 5.95274 | 6.5381 | 5.81028 | 6.12977 | 6.07147 | 5.7426 | 5.92673 | 6.44282 | 5.77178 | 6.70351 | 6.50229 |
| PRKAB2 | O43741 | 5.33801 | 5.51898 | 5.73434 | 5.32259 | 6.09627 | 6.56014 | 5.76429 | 6.15283 | 6.26448 | 6.27727 | 6.82013 | 6.75574 | 7.08677 | 6.92407 | 6.9682 | 6.41588 |
| RHOA | P61586 | 5.81564 | 6.38184 | 5.78064 | 6.32207 | 6.95334 | 7.67496 | 6.78054 | 6.57574 | 6.46581 | 5.71537 | 6.30304 | 5.87111 | 6.04151 | 6.34913 | 5.63404 | 5.33521 |
| RHOB | P62745 | 6.3676 | 6.37775 | 6.98672 | 6.77175 | 6.40995 | 6.34721 | 6.33126 | 5.95802 | 6.147 | 6.18057 | 6.03972 | 6.41828 | 5.98403 | 5.59323 | 5.87699 | 6.20991 |
| DLGAP4 | Q9Y2H0 | 6.1551 | 6.11494 | 6.57008 | 5.97275 | 6.87551 | 6.84962 | 6.37149 | 6.86292 | 6.77949 | 6.46407 | 6.61274 | 6.16425 | 5.63113 | 5.55126 | 5.31166 | 5.713 |
| FBXO2 | Q9UK22 | 6.30328 | 5.97619 | 6.69142 | 5.8243 | 6.49429 | 6.93081 | 6.08159 | 7.45816 | 6.41528 | 6.10506 | 7.00254 | 5.99117 | 5.99249 | 5.8874 | 5.48673 | 5.3593 |
| SNAP29 | O95721 | 6.2132 | 5.70058 | 6.79942 | 5.68413 | 6.43841 | 6.5095 | 5.91901 | 6.57609 | 6.57994 | 6.89993 | 6.93096 | 7.25486 | 6.18722 | 5.45613 | 5.36403 | 5.48658 |
| SCAI | Q8N9R8-2 | 6.33729 | 6.3584 | 5.94069 | 6.8384 | 6.58256 | 5.97229 | 6.61515 | 5.94178 | 6.20166 | 6.16118 | 6.48513 | 6.02477 | 6.21161 | 6.17936 | 6.109 | 6.04072 |
| SMYD4 | Q8IYR2 | 5.71262 | 6.50278 | 6.97893 | 6.41704 | 6.35962 | 7.04738 | 6.04523 | 6.3201 | 6.54303 | 6.59288 | 6.48745 | 6.3639 | 5.88359 | 5.77386 | 5.24884 | 5.72274 |
| BMPR2 | Q13873 | 6.70802 | 6.76965 | 6.59232 | 7.11025 | 5.79811 | 6.27735 | 6.47793 | 6.43696 | 6.24726 | 6.88661 | 6.75356 | 7.08302 | 5.38694 | 5.43805 | 5.02426 | 5.00969 |
| UBE2M | P61081 | 6.07631 | 5.60837 | 6.74648 | 5.44985 | 6.14887 | 6.35434 | 6.02101 | 6.23001 | 6.7407 | 6.4494 | 6.97817 | 6.43241 | 6.30348 | 5.92258 | 6.38542 | 6.1526 |
| ANXA3 | P12429 | 6.16644 | 6.12338 | 6.6631 | 5.95302 | 6.23517 | 6.22538 | 6.07775 | 6.67324 | 6.94841 | 7.07901 | 6.28701 | 6.23001 | 6.01509 | 5.69773 | 6.09017 | 5.53509 |
| DIXDC1 | Q155Q3 | 6.72622 | 6.39231 | 6.12181 | 6.91003 | 6.2964 | 6.23084 | 6.19802 | 6.52933 | 6.44465 | 6.06602 | 6.24708 | 6.78197 | 6.31721 | 5.80016 | 5.45175 | 6.3862 |
| ABI2 | Q9NYB9 | 6.65706 | 6.08528 | 6.74371 | 5.74224 | 6.89899 | 6.47539 | 6.13291 | 6.59172 | 6.31836 | 6.43861 | 6.23226 | 6.23386 | 6.04671 | 5.75923 | 5.77628 | 5.86738 |
| PAFAH1B3 | Q15102 | 5.60063 | 3.86664 | 6.71439 | 4.81175 | 5.07562 | 4.89796 | 5.31768 | 6.14933 | 5.2293 | 6.19426 | 5.76913 | 6.64197 | 5.82033 | 9.7077 | 8.75863 | 9.44467 |
| LYPLA2 | O95372 | 5.43236 | 5.31265 | 4.89681 | 4.57687 | 5.45973 | 5.48043 | 5.4243 | 7.04825 | 4.84787 | 5.9159 | 5.09476 | 5.49184 | 4.72329 | 11.8259 | 7.59436 | 10.8747 |
| GRK6 | P43250-2 | 7.17265 | 7.53129 | 7.16296 | 7.61383 | 6.45874 | 5.68826 | 6.03272 | 6.46168 | 6.57834 | 6.94089 | 7.10442 | 6.56944 | 4.3095 | 5.06604 | 4.76408 | 4.54516 |
| AIFM2 | Q9BRQ8 | 5.85871 | 5.75935 | 5.80764 | 6.37146 | 6.03712 | 5.87329 | 6.45713 | 6.26078 | 6.2671 | 6.5325 | 6.13527 | 6.22432 | 6.24749 | 6.70782 | 6.70946 | 6.75055 |
| POLR2C | P19387 | 6.72671 | 6.19017 | 6.4514 | 6.23008 | 6.9358 | 6.25998 | 6.73859 | 6.41892 | 6.66053 | 6.37745 | 6.73371 | 6.58208 | 5.68054 | 5.23897 | 5.51286 | 5.19001 |
| CLUAP1 | Q96AJ1 | 6.25967 | 5.94864 | 7.0405 | 6.16252 | 6.12457 | 6.55817 | 6.20002 | 6.94071 | 6.78681 | 6.21347 | 6.71357 | 6.62506 | 5.53833 | 5.47082 | 5.56396 | 5.85319 |
| MACO1 | Q8N5G2 | 6.16708 | 6.56775 | 6.38907 | 6.90723 | 5.94574 | 5.91965 | 6.36804 | 5.79419 | 6.70399 | 6.2496 | 6.43518 | 6.33994 | 5.87566 | 5.78181 | 6.22291 | 6.33217 |
| KIAA1191 | Q96A73 | 6.01307 | 5.82489 | 5.80383 | 6.0851 | 6.03829 | 6.01761 | 5.50985 | 6.6229 | 5.63484 | 5.95762 | 5.98505 | 6.2575 | 7.74075 | 6.85868 | 6.51694 | 7.13309 |
| CHMP7 | Q8WUX9 | 6.23613 | 6.53244 | 7.01567 | 6.34287 | 6.37878 | 7.03947 | 6.03048 | 6.39965 | 6.49007 | 6.67501 | 6.42716 | 6.81114 | 5.9798 | 4.90903 | 4.95811 | 5.77418 |
| CD276 | Q5ZPR3 | 6.36748 | 6.25749 | 6.06702 | 6.29356 | 6.15889 | 6.18812 | 6.06648 | 6.16164 | 6.02272 | 5.99904 | 6.18783 | 6.28376 | 6.93918 | 6.45233 | 5.86719 | 6.68727 |
| GALT | P07902 | 6.47635 | 5.94295 | 7.29481 | 4.9577 | 6.35523 | 6.42785 | 5.95278 | 6.84387 | 6.45585 | 6.65066 | 6.4992 | 6.90733 | 5.90154 | 6.02813 | 5.86981 | 5.43593 |
| MMTAG2 | Q9BU76 | 6.06525 | 6.04164 | 5.81995 | 6.3332 | 5.7842 | 5.88525 | 6.03355 | 5.583 | 5.96092 | 6.05037 | 6.01314 | 6.13903 | 6.57159 | 7.58221 | 6.883 | 7.25372 |
| DRAP1 | Q14919-2 | 5.77615 | 6.1256 | 6.22019 | 5.51417 | 5.78323 | 6.41821 | 5.77098 | 6.05228 | 6.14146 | 6.52761 | 6.24595 | 6.49342 | 6.4388 | 6.95501 | 6.92776 | 6.60919 |
| L3HYPDH | Q96EM0 | 6.13994 | 6.14434 | 7.02296 | 5.60038 | 6.84161 | 6.96818 | 6.24742 | 6.37121 | 6.92396 | 6.33593 | 6.21334 | 6.44987 | 6.3319 | 5.54804 | 5.61157 | 5.24933 |
| GTF3C6 | Q969F1 | 6.13212 | 6.70891 | 6.01576 | 6.66496 | 6.68345 | 6.75448 | 5.605 | 6.71806 | 5.91797 | 5.82375 | 5.65562 | 6.72894 | 6.36099 | 6.43567 | 5.9043 | 5.89001 |
| BBS2 | Q9BXC9 | 6.031 | 6.56334 | 7.43273 | 6.54204 | 6.27163 | 5.73409 | 6.21643 | 6.34114 | 6.54246 | 7.14552 | 6.82094 | 5.95181 | 5.54577 | 5.77812 | 5.41992 | 5.66306 |
| ANKRD13C | Q8N6S4 | 6.81596 | 6.7308 | 6.83376 | 6.48923 | 6.24187 | 6.07336 | 6.41148 | 6.58771 | 6.60863 | 6.59344 | 7.00971 | 6.90666 | 5.20833 | 5.27541 | 5.10397 | 5.10966 |
| CLK4 | Q9HAZ1 | 6.4824 | 6.00758 | 5.86905 | 6.86845 | 5.55187 | 5.16274 | 5.92988 | 4.71178 | 6.01642 | 6.233 | 5.99513 | 5.76262 | 7.22169 | 6.91301 | 7.5427 | 7.73168 |
| RNF139 | Q8WU17 | 6.89836 | 6.31114 | 5.80142 | 6.38592 | 6.28317 | 5.70299 | 6.55999 | 6.09111 | 6.06578 | 6.12907 | 5.89371 | 5.59883 | 6.63888 | 6.66551 | 6.81567 | 6.15844 |
| UBE2C | O00762 | 6.12922 | 4.64159 | 6.49119 | 5.21586 | 5.72254 | 4.80777 | 5.998 | 7.95065 | 4.62093 | 6.28187 | 5.61577 | 7.23231 | 3.97623 | 8.93617 | 7.33845 | 9.04143 |
| TNS2 | Q63HR2-4 | 6.36786 | 6.55912 | 6.47371 | 6.19822 | 6.08911 | 6.19099 | 6.18642 | 6.15374 | 5.87817 | 6.56879 | 6.52446 | 6.10623 | 6.31995 | 6.48605 | 5.97374 | 5.92345 |
| RAPGEF1 | Q13905-3 | 6.71156 | 6.34751 | 6.79283 | 6.31831 | 6.35869 | 6.61203 | 6.11913 | 6.37887 | 6.4967 | 6.3277 | 6.45322 | 6.28632 | 6.19487 | 5.58129 | 5.51733 | 5.50364 |

| | | | | | | | | | | | | | | | | | |
|---|---|---|---|---|---|---|---|---|---|---|---|---|---|---|---|---|---|
| DDA1 | Q9BW61 | 6.18346 | 5.52494 | 6.40743 | 5.98997 | 6.20643 | 5.98495 | 5.4942 | 6.6509 | 5.83699 | 6.58097 | 7.00189 | 6.60854 | 6.33459 | 6.06284 | 6.38059 | 6.75133 |
| RAF1 | P04049-2 | 6.77362 | 6.83957 | 6.40255 | 6.15259 | 6.30837 | 7.04326 | 6.20339 | 6.68948 | 6.19808 | 6.37069 | 6.79645 | 6.52972 | 5.33432 | 5.52596 | 5.37943 | 5.45252 |
| ARAF | P10398 | 6.10587 | 6.71149 | 6.9856 | 6.22443 | 6.60593 | 6.56684 | 6.32869 | 6.46039 | 6.34505 | 6.84923 | 6.52739 | 6.50284 | 5.50413 | 5.13347 | 5.64226 | 5.50642 |
| HAUS1 | Q96CS2 | 6.5698 | 4.78357 | 6.27293 | 4.85387 | 5.90697 | 4.44786 | 5.3233 | 7.98661 | 4.64066 | 6.71441 | 6.03127 | 7.486 | 4.90938 | 8.54836 | 7.38668 | 8.13832 |
| PLEKHG3 | A1L390 | 6.53497 | 6.62793 | 6.5221 | 6.66571 | 6.29795 | 6.31736 | 6.59458 | 5.86497 | 6.10823 | 6.17946 | 6.20888 | 6.49776 | 5.99524 | 5.37496 | 6.05553 | 6.15435 |
| KRAS | P01116-2 | 5.6928 | 6.39403 | 5.95212 | 6.33968 | 6.0486 | 6.36661 | 6.06971 | 6.17163 | 6.0484 | 6.00074 | 6.0377 | 6.01874 | 6.35757 | 6.96954 | 6.59067 | 6.94146 |
| RAD1 | O60671 | 5.97626 | 6.40338 | 6.7874 | 5.60093 | 6.04145 | 6.36732 | 5.88199 | 6.08335 | 6.64804 | 6.31576 | 6.45456 | 6.24933 | 6.40805 | 5.95734 | 7.01633 | 5.80851 |
| SLF2 | Q8IX21-2 | 6.26082 | 6.56446 | 5.94244 | 6.96386 | 5.88277 | 5.88344 | 6.1976 | 6.12757 | 5.93005 | 6.02164 | 6.15384 | 6.08401 | 6.34585 | 6.32374 | 6.63684 | 6.68107 |
| WDR89 | Q96FK6 | 5.90746 | 7.08035 | 5.69715 | 6.94821 | 6.38117 | 6.79488 | 6.70564 | 6.2864 | 5.66907 | 6.08869 | 5.84274 | 5.96223 | 6.41315 | 5.66959 | 6.29134 | 6.26193 |
| PURB | Q96QR8 | 6.4861 | 6.11695 | 6.48702 | 6.24916 | 6.5938 | 6.3959 | 6.06439 | 6.18234 | 6.23964 | 6.11318 | 6.65652 | 6.52759 | 6.07684 | 5.73989 | 6.06413 | 6.00657 |
| S100A13 | Q99584 | 6.02625 | 5.10567 | 6.31843 | 4.3861 | 7.13701 | 6.34865 | 5.7712 | 6.61243 | 5.76096 | 6.51927 | 6.19282 | 6.26275 | 6.12166 | 7.30016 | 7.25443 | 6.88222 |
| APOBEC3D | Q96AK3 | 6.53369 | 6.29595 | 6.48275 | 6.65238 | 6.24835 | 6.28281 | 6.31433 | 6.16573 | 6.36878 | 6.41364 | 6.10412 | 6.20928 | 6.07974 | 5.88563 | 5.90775 | 6.05508 |
| C5orf22 | Q49AR2 | 5.36347 | 5.69033 | 6.12432 | 5.43071 | 6.08547 | 6.41908 | 5.35517 | 6.60634 | 5.90112 | 6.00067 | 6.80775 | 6.44221 | 6.43825 | 7.07407 | 7.2353 | 7.02574 |
| FAM83D | Q9H4H8 | 6.52192 | 7.0005 | 6.97409 | 7.12726 | 6.36043 | 6.47794 | 6.28125 | 6.84336 | 7.28879 | 6.5462 | 7.12251 | 6.83259 | 4.43354 | 4.97646 | 4.428 | 4.78515 |
| CHMP3 | Q9Y3E7 | 6.88864 | 6.41505 | 6.97845 | 6.22444 | 6.56654 | 6.77414 | 6.20889 | 6.35868 | 6.44568 | 6.51606 | 6.59257 | 6.2911 | 5.49576 | 5.44775 | 5.28056 | 5.5157 |
| C1orf131 | Q8NDD1-5 | 6.36242 | 6.57897 | 6.27057 | 6.796 | 6.40454 | 5.96325 | 6.37085 | 6.11497 | 6.10928 | 6.1506 | 6.30442 | 6.18655 | 5.73877 | 5.99068 | 6.53751 | 6.12063 |
| FCGRT | P55899 | 6.5521 | 6.36029 | 6.15748 | 5.66658 | 6.12401 | 6.31968 | 6.16238 | 6.56681 | 6.29214 | 6.19607 | 6.08592 | 5.99691 | 6.38533 | 6.90881 | 5.86606 | 6.35942 |
| DCTPP1 | Q9H773 | 6.03657 | 5.19097 | 5.8543 | 4.94664 | 5.00416 | 4.60068 | 4.87032 | 6.35763 | 4.67953 | 5.7823 | 5.08565 | 6.1754 | 5.23826 | 11.1783 | 8.48899 | 10.5103 |
| CCPG1 | Q9ULG6-5 | 7.15426 | 6.7097 | 5.8085 | 6.67 | 6.29835 | 6.11038 | 6.18017 | 6.39901 | 6.98971 | 5.92498 | 6.27311 | 5.8899 | 6.20426 | 5.68892 | 5.81028 | 5.88849 |
| UBTD2 | Q8WUN7 | 6.39466 | 6.03623 | 6.39964 | 5.79454 | 6.10593 | 6.27717 | 5.88469 | 5.83751 | 6.2195 | 6.52722 | 7.1298 | 6.64244 | 6.19213 | 6.17607 | 6.76364 | 5.61884 |
| NDUFAF3 | Q9BU61 | 5.06248 | 4.85626 | 4.77473 | 4.75692 | 6.03502 | 5.49478 | 6.41349 | 7.01182 | 5.46849 | 6.78279 | 5.88164 | 6.667 | 5.27599 | 8.03059 | 8.3304 | 9.15762 |
| DYNLT1 | P63172 | 6.26924 | 6.19745 | 6.61332 | 5.69985 | 6.17734 | 7.4234 | 6.26954 | 6.41042 | 6.50518 | 6.92184 | 6.52748 | 6.26222 | 5.88919 | 5.84645 | 5.6193 | 5.36778 |
| TMEM68 | Q96MH6 | 6.76151 | 6.34559 | 5.78651 | 6.57288 | 6.16369 | 5.87972 | 6.3791 | 5.91458 | 6.33379 | 6.13132 | 5.81737 | 5.58597 | 6.62385 | 6.64016 | 6.62487 | 6.4391 |
| DENND6A | Q8IWF6 | 6.42739 | 6.91615 | 6.81038 | 6.38652 | 6.14726 | 6.62316 | 6.36983 | 6.48579 | 6.46732 | 6.42599 | 6.65185 | 6.9072 | 5.51255 | 5.23069 | 5.44775 | 5.19018 |
| ZNF668 | Q96K58-2 | 6.24941 | 5.85429 | 6.03951 | 6.21629 | 6.18412 | 6.03177 | 6.55032 | 5.8766 | 6.44411 | 6.17901 | 5.6664 | 6.19172 | 6.46818 | 6.71949 | 6.86027 | 6.46851 |
| NUBPL | Q8TB37 | 6.49885 | 6.41034 | 6.03848 | 6.40651 | 6.30234 | 5.90077 | 6.39448 | 6.06056 | 5.98515 | 6.13236 | 5.924 | 6.08075 | 6.94545 | 6.3133 | 6.53974 | 6.06692 |
| RBL1 | P28749 | 5.32147 | 5.55753 | 6.00693 | 4.92139 | 5.84883 | 6.1602 | 5.46449 | 5.57399 | 5.89268 | 6.16394 | 6.30744 | 5.98373 | 7.60199 | 7.7852 | 7.77154 | 7.63845 |
| PLCD1 | P51178-2 | 5.94875 | 6.39211 | 5.9044 | 5.89029 | 6.05964 | 5.93371 | 6.20638 | 5.64543 | 6.1693 | 6.29931 | 6.51461 | 6.31743 | 6.88536 | 6.85077 | 6.55688 | 6.42563 |
| TMED5 | Q9Y3A6 | 6.19108 | 6.33865 | 5.45373 | 6.17929 | 5.65518 | 6.07793 | 6.23802 | 5.82015 | 5.95909 | 5.96041 | 5.84055 | 6.39456 | 6.88356 | 6.91141 | 6.96419 | 7.13221 |
| KCNMA1 | Q12791-3 | 6.56567 | 6.95754 | 6.08889 | 6.91274 | 6.36588 | 6.00418 | 6.23095 | 5.98119 | 6.43214 | 5.98119 | 5.93458 | 5.92781 | 6.15799 | 5.94279 | 6.1122 | 6.40427 |
| GID8 | Q9NWU2 | 5.52307 | 4.98174 | 4.57707 | 5.11923 | 4.45515 | 3.43726 | 4.20799 | 5.31436 | 3.58043 | 4.90394 | 4.10065 | 4.38976 | 4.04793 | 15.6586 | 10.107 | 15.5958 |
| SLC8A1 | P32418-2 | 6.13711 | 6.44435 | 6.22453 | 6.51584 | 6.26251 | 5.92011 | 6.50787 | 5.95757 | 6.16299 | 6.02683 | 6.34442 | 6.05488 | 6.64102 | 6.29954 | 6.32505 | 6.17539 |
| FAM162A | Q96A26 | 6.06033 | 6.19693 | 5.41827 | 6.0856 | 5.74775 | 6.60841 | 5.55207 | 6.55083 | 5.12546 | 7.17206 | 5.86171 | 6.50418 | 6.28489 | 7.41272 | 6.33904 | 7.07976 |
| KIAA1143 | Q96AT1 | 6.51363 | 6.16142 | 6.79307 | 5.81398 | 5.96116 | 6.15045 | 6.12922 | 6.78978 | 5.91145 | 6.43209 | 6.91038 | 5.96465 | 5.99512 | 6.01956 | 6.20162 | 6.25245 |
| TP53I3 | Q53FA7 | 7.05069 | 5.83419 | 6.78978 | 5.5649 | 6.72276 | 6.27065 | 5.83705 | 6.93044 | 6.38216 | 6.12612 | 6.56656 | 6.47819 | 5.89891 | 5.85596 | 6.30529 | 5.38634 |
| NUDT15 | Q9NV35 | 6.25991 | 6.04809 | 6.84317 | 5.64608 | 6.42577 | 6.83571 | 6.14534 | 7.17297 | 6.49152 | 7.04896 | 6.89999 | 7.20167 | 4.82365 | 5.54844 | 5.16058 | 5.44816 |
| HMGN1 | P05114 | 6.02225 | 6.06363 | 5.88206 | 6.44056 | 5.68361 | 5.6192 | 6.22081 | 5.67723 | 5.99236 | 6.13855 | 5.94442 | 6.00276 | 6.65509 | 7.4383 | 7.04824 | 7.17092 |
| SH3BP1 | Q9Y3L3 | 6.25186 | 6.25289 | 7.00968 | 5.4785 | 6.42816 | 6.31234 | 6.05141 | 6.60404 | 6.69062 | 6.60813 | 7.0952 | 6.03827 | 6.30034 | 5.79003 | 5.84063 | 5.24185 |
| MRGBP | Q9NV56 | 6.30186 | 6.40904 | 6.4036 | 6.32526 | 5.93386 | 5.96338 | 6.43241 | 5.93959 | 6.6479 | 6.62142 | 6.45431 | 6.42708 | 6.22523 | 5.8274 | 5.91353 | 6.17414 |
| FER | P16591 | 6.19736 | 6.41779 | 6.63646 | 6.34885 | 5.93431 | 6.82831 | 6.14637 | 6.46935 | 6.07239 | 6.68478 | 6.72209 | 6.93573 | 5.90983 | 5.93051 | 5.01111 | 5.75477 |
| BNC2 | Q6ZN30 | 6.563 | 6.65906 | 6.03294 | 7.19178 | 5.92393 | 6.09845 | 6.42217 | 5.75934 | 6.19497 | 6.2107 | 6.33863 | 6.06631 | 6.23212 | 5.69907 | 6.22326 | 6.38427 |
| CHST3 | Q7LGC8 | 6.32706 | 6.42758 | 5.47555 | 6.92831 | 6.4091 | 6.51581 | 6.71338 | 5.44805 | 6.34643 | 6.24992 | 6.07185 | 5.9879 | 6.51104 | 5.60025 | 6.58787 | 6.39988 |
| GPATCH11 | Q8N954 | 5.80192 | 5.39147 | 6.43716 | 5.73119 | 5.82479 | 5.83153 | 5.88551 | 5.847 | 6.4475 | 6.6896 | 6.63882 | 6.38376 | 6.93417 | 6.50408 | 6.88249 | 6.769 |
| GTF3A | Q92664 | 5.36046 | 5.80076 | 6.22467 | 5.87428 | 5.74294 | 6.35352 | 5.67886 | 5.93299 | 6.00376 | 6.16987 | 6.62809 | 6.50953 | 6.7995 | 7.08215 | 7.15929 | 6.97903 |
| CNNM3 | Q8NE01 | 6.69938 | 6.55659 | 6.61665 | 6.63444 | 6.41764 | 6.11453 | 5.97541 | 5.53351 | 6.61146 | 6.53059 | 6.81776 | 6.29455 | 5.7107 | 5.71203 | 5.68676 | 6.08801 |
| LAMTOR1 | Q6IAA8 | 6.39738 | 5.96285 | 5.59585 | 5.89944 | 6.1569 | 6.20702 | 6.1875 | 6.19516 | 5.64249 | 6.23918 | 6.60517 | 6.2735 | 6.06299 | 7.33899 | 6.58118 | 6.6544 |
| PSMG4 | Q5JS54-2 | 5.70933 | 4.66939 | 6.20447 | 4.20182 | 6.30974 | 6.55416 | 6.03541 | 7.71986 | 5.5995 | 6.15889 | 5.84646 | 6.76577 | 5.72999 | 8.8798 | 6.27146 | 7.34394 |
| SEPHS2 | Q99611 | 5.15489 | 5.30011 | 5.97301 | 5.09039 | 6.14279 | 6.30866 | 5.64138 | 6.1004 | 6.21837 | 6.27944 | 6.20586 | 6.25499 | 7.34407 | 7.30072 | 7.43634 | 7.24859 |
| BTBD9 | Q96Q07 | 6.38051 | 6.64616 | 6.58607 | 6.42569 | 6.42029 | 6.64223 | 6.05442 | 6.76788 | 5.94672 | 6.36331 | 6.79817 | 6.86101 | 5.3527 | 5.52225 | 5.4 | 5.8326 |
| RNF2 | Q99496 | 6.34239 | 6.36116 | 6.25751 | 6.78849 | 6.185 | 6.02048 | 6.25702 | 5.68016 | 6.57982 | 6.18062 | 6.35507 | 6.51321 | 6.26582 | 5.74643 | 6.40649 | 6.06034 |
| PAN2 | Q504Q3 | 6.76278 | 5.9692 | 7.17771 | 5.79515 | 6.33147 | 6.23884 | 6.82468 | 6.04436 | 6.79661 | 6.41425 | 6.80448 | 5.96027 | 6.22422 | 5.27069 | 6.14456 | 5.24072 |
| NDUFB6 | O95139 | 6.11453 | 5.92046 | 6.00849 | 6.40928 | 6.26148 | 5.87996 | 6.68072 | 5.79412 | 6.09265 | 6.03087 | 5.99886 | 5.90975 | 7.13872 | 6.436 | 6.65975 | 6.66435 |
| FTSJ1 | Q9UET6 | 6.27969 | 6.14345 | 7.29918 | 6.41894 | 6.74669 | 6.82386 | 5.80246 | 6.46241 | 6.05487 | 6.65437 | 6.92249 | 6.73819 | 5.54632 | 5.70867 | 5.21562 | 5.1828 |
| BCL9 | O00512 | 6.00666 | 5.49548 | 6.38394 | 4.88181 | 6.72078 | 5.96673 | 5.63254 | 6.40257 | 5.6938 | 6.37793 | 6.55964 | 6.69771 | 7.76335 | 6.54311 | 6.4101 | 6.46385 |
| MRPL35 | Q9NZE8 | 6.51363 | 6.38528 | 6.09163 | 5.86813 | 6.50591 | 5.88424 | 6.79833 | 6.12239 | 6.22936 | 5.68381 | 5.3 | 5.59319 | 7.18615 | 6.58665 | 6.38352 | 6.86777 |
| ZSCAN18 | Q8TBC5-3 | 6.88404 | 6.39581 | 6.21424 | 5.75219 | 6.63885 | 6.35874 | 5.6431 | 6.85542 | 6.34917 | 6.16841 | 6.56129 | 6.34867 | 6.46017 | 5.88847 | 5.93192 | 5.5495 |
| BTN3A3 | O00478 | 6.56226 | 6.39167 | 6.2891 | 6.50141 | 5.98184 | 5.81736 | 5.89664 | 6.15366 | 6.33109 | 5.95948 | 5.80819 | 5.66095 | 6.36968 | 6.49681 | 6.86042 | 6.91945 |
| TANC2 | Q9HCD6-2 | 6.09501 | 5.56249 | 6.82295 | 5.74884 | 6.06751 | 6.25755 | 5.90998 | 6.34226 | 6.38027 | 6.63686 | 6.76463 | 6.56219 | 5.79581 | 7.00379 | 5.93724 | 6.10765 |
| ATG2A | Q2TAZ0-3 | 6.25814 | 6.31143 | 6.49094 | 5.95726 | 6.32501 | 6.73792 | 6.61483 | 6.28226 | 6.20314 | 6.75069 | 6.50619 | 7.36652 | 5.551 | 5.62241 | 5.49637 | 5.52588 |
| BDP1 | A6H8Y1 | 6.54184 | 6.19511 | 6.5441 | 6.98202 | 6.0863 | 5.93115 | 6.27602 | 5.90244 | 6.80045 | 6.2952 | 6.55096 | 6.47947 | 5.72327 | 5.59931 | 5.88807 | 6.20428 |
| NCR3LG1 | Q68D85 | 6.9118 | 6.11394 | 6.09944 | 6.33019 | 6.4702 | 5.99007 | 6.14554 | 6.22841 | 6.41793 | 5.81985 | 5.91454 | 6.14268 | 6.74983 | 6.06114 | 6.10648 | 6.49798 |
| EPHX4 | Q8IUS5 | 6.38094 | 6.77862 | 6.28108 | 6.49164 | 5.70271 | 5.76441 | 6.18458 | 6.13857 | 5.92015 | 6.13822 | 5.97814 | 6.22869 | 6.48507 | 6.22629 | 6.85371 | 6.4472 |
| TRAF6 | Q9Y4K3 | 6.54424 | 5.57569 | 6.6874 | 6.03723 | 6.39288 | 6.55711 | 5.79351 | 6.49435 | 6.19492 | 6.47789 | 7.02378 | 6.88152 | 6.02683 | 5.54648 | 5.47802 | 6.28814 |
| INPP4A | Q96PE3 | 5.88447 | 5.98637 | 7.08311 | 6.2112 | 5.90118 | 6.27984 | 6.06012 | 6.48441 | 6.19987 | 6.67495 | 6.60835 | 6.79036 | 5.97172 | 6.22925 | 5.99986 | 5.63496 |
| CLU | P10909-2 | 7.11578 | 7.10641 | 6.20935 | 6.92782 | 6.55681 | 6.24807 | 6.39238 | 6.26027 | 6.14026 | 7.06551 | 6.30566 | 6.058 | 5.39327 | 5.60309 | 5.3077 | 5.30962 |
| SHOC2 | Q9UQ13 | 6.04554 | 6.11278 | 6.48914 | 6.3443 | 5.92675 | 6.13745 | 6.18199 | 5.95237 | 6.55558 | 6.4573 | 6.30035 | 6.23255 | 6.33769 | 6.24162 | 6.35151 | 6.33309 |
| FBXL6 | Q8N531 | 6.198 | 6.86707 | 6.39335 | 6.63111 | 5.82147 | 6.80423 | 6.18449 | 6.66669 | 6.40337 | 5.97877 | 6.88401 | 6.50498 | 5.59171 | 5.48564 | 5.59895 | 5.98618 |
| MAP2K1 | Q02750 | 6.10818 | 6.03954 | 6.85634 | 5.99795 | 6.25793 | 7.00468 | 6.38791 | 6.6864 | 6.48883 | 6.41098 | 7.07531 | 6.84228 | 5.66093 | 5.62944 | 5.11809 | 5.43521 |
| UXS1 | Q8NBZ7-2 | 6.41464 | 6.40991 | 6.17864 | 6.86434 | 6.16689 | 5.7822 | 6.49354 | 6.00781 | 6.10148 | 6.12489 | 6.23298 | 6.28752 | 6.32369 | 5.95464 | 6.33741 | 6.31943 |
| TSEN54 | Q7Z6J9 | 5.76288 | 5.76104 | 6.7852 | 5.89722 | 6.35444 | 6.25181 | 5.63879 | 6.06554 | 6.21298 | 6.15533 | 6.79484 | 6.43727 | 6.63162 | 6.37089 | 6.69351 | 6.18664 |
| CNOT8 | Q9UFF9 | 6.1338 | 6.22474 | 6.26996 | 5.8849 | 6.24336 | 6.51274 | 6.47854 | 5.98475 | 6.35192 | 6.1653 | 6.06605 | 6.77975 | 6.3484 | 6.4028 | 5.65447 | 6.49851 |
| SOWAHC | Q53LP3 | 6.63268 | 6.87998 | 6.94171 | 6.25624 | 6.61406 | 6.26466 | 6.56754 | 6.72282 | 6.49798 | 6.14764 | 6.56349 | 6.10668 | 6.29444 | 5.26967 | 5.09918 | 5.14121 |
| NGEF | Q8N5V2 | 6.61841 | 6.33839 | 6.65031 | 5.93826 | 6.2681 | 6.97914 | 6.47618 | 6.38046 | 6.66492 | 6.57183 | 7.09182 | 6.64253 | 5.39199 | 5.5057 | 5.45769 | 5.02427 |
| NDUFB5 | O43674 | 6.37793 | 6.47874 | 5.82809 | 6.58585 | 6.36127 | 5.97591 | 6.46675 | 6.03362 | 5.92178 | 6.15821 | 5.83405 | 5.94417 | 6.9582 | 6.26904 | 6.73792 | 6.06848 |
| MAML1 | Q92585 | 5.65888 | 5.93368 | 6.56671 | 5.55242 | 5.56406 | 6.47586 | 5.83748 | 6.26827 | 6.25939 | 6.55515 | 7.37533 | 6.94308 | 6.50534 | 6.13462 | 6.19729 | 6.17243 |
| C5orf51 | A6NDU8 | 6.46997 | 5.35092 | 6.50508 | 5.28834 | 5.84561 | 5.89861 | 5.88997 | 6.58999 | 5.57441 | 6.55442 | 6.30092 | 7.15917 | 6.16137 | 7.28422 | 6.52266 | 6.60432 |
| ZNF639 | Q9UID6 | 6.54308 | 6.52603 | 5.98474 | 6.79343 | 6.41249 | 6.39204 | 6.05496 | 6.11376 | 6.32562 | 6.2075 | 6.07607 | 6.22175 | 6.21956 | 6.0465 | 6.06568 | 6.01681 |
| LIMK2 | P53671-3 | 6.34366 | 5.03599 | 6.84807 | 5.30912 | 6.32344 | 5.82316 | 5.44933 | 6.19161 | 6.05369 | 6.8205 | 6.50784 | 7.32183 | 6.17867 | 6.47755 | 6.27005 | 7.04549 |
| FPGT | O14772 | 6.16843 | 6.26229 | 6.93935 | 5.83805 | 6.15081 | 6.6284 | 6.44994 | 6.72609 | 6.25366 | 6.33993 | 6.51038 | 6.50645 | 5.69415 | 6.11246 | 5.70063 | 5.719 |
| TCF12 | Q99081-3 | 5.59447 | 5.99081 | 6.52873 | 5.23177 | 5.64973 | 6.24185 | 5.86222 | 6.29312 | 6.26376 | 7.14654 | 6.4963 | 6.08604 | 6.51117 | 6.72275 | 6.97171 | 6.4991 |
| JPH1 | Q9HDC5 | 6.24635 | 6.40375 | 5.92643 | 6.57121 | 5.87123 | 6.17991 | 6.38822 | 6.14089 | 6.39059 | 6.13869 | 6.107 | 5.69022 | 6.6862 | 6.62881 | 6.39549 | 6.23502 |
| CC2D2A | Q9P2K1 | 7.2228 | 6.95616 | 7.20698 | 6.69087 | 6.10841 | 5.78415 | 6.44372 | 6.06114 | 6.95486 | 7.01781 | 6.93752 | 6.5218 | 4.90993 | 4.97725 | 4.99565 | 5.21096 |
| AURKB | Q96GD4-5 | 6.82298 | 6.56927 | 6.28461 | 6.29564 | 6.53083 | 6.50655 | 6.52261 | 6.65269 | 6.26544 | 5.87827 | 6.24028 | 6.12948 | 5.71448 | 6.10045 | 5.74152 | 5.7449 |
| BROX | Q5VW32 | 6.47085 | 6.06664 | 6.69886 | 5.73269 | 5.70151 | 6.83787 | 6.29646 | 6.27796 | 6.38292 | 6.35299 | 6.56947 | 6.34073 | 5.9177 | 6.16684 | 6.20248 | 5.98405 |
| CHN1 | P15882 | 6.24933 | 6.16011 | 7.33413 | 5.47767 | 6.74725 | 6.80348 | 6.01896 | 5.96033 | 6.88856 | 6.69565 | 6.46379 | 6.6515 | 5.36588 | 5.77099 | 6.17883 | 5.23355 |
| ASF1B | Q9NVP2 | 6.06977 | 6.22529 | 6.34301 | 5.8588 | 6.53378 | 6.00065 | 5.5602 | 7.02327 | 6.71002 | 6.79409 | 6.22297 | 7.15145 | 5.5719 | 5.38406 | 5.68923 | 5.86151 |
| PJA2 | O43164 | 6.79209 | 6.17701 | 7.93064 | 6.76094 | 6.28534 | 7.15832 | 6.9256 | 7.38158 | 6.06052 | 6.83039 | 7.04203 | 7.70496 | 4.12904 | 4.79296 | 3.67229 | 4.3563 |
| CLPP | Q16740 | 5.97228 | 4.991 | 4.81085 | 5.15137 | 5.74662 | 4.65819 | 6.08632 | 6.69435 | 4.82163 | 5.36638 | 4.91633 | 6.21183 | 6.07373 | 9.7716 | 8.85902 | 9.86851 |
| R3HCC1L | A0A384DVK4 | 6.80019 | 6.25792 | 6.8525 | 6.29505 | 6.23419 | 6.22528 | 6.2419 | 6.68985 | 6.39636 | 6.60414 | 6.53947 | 6.85378 | 5.56982 | 5.70405 | 5.22191 | 5.51359 |
| TRMT61B | Q9BVS5 | 6.32872 | 6.37623 | 5.86581 | 6.81102 | 6.28621 | 6.19666 | 6.49187 | 6.54772 | 5.74606 | 5.90075 | 5.87409 | 5.757 | 6.54319 | 6.18166 | 6.69134 | 6.40166 |
| MRRF | Q96E11 | 6.17157 | 6.49251 | 5.66513 | 6.83214 | 5.9812 | 5.70147 | 6.54871 | 5.86788 | 6.04125 | 6.29753 | 6.38316 | 6.408 | 6.4488 | 6.01271 | 6.38263 | 6.76429 |
| SOD1 | P00441 | 6.88485 | 5.59252 | 6.6528 | 4.71031 | 5.94785 | 5.73241 | 5.4452 | 7.30699 | 5.73776 | 6.5426 | 6.83196 | 6.61925 | 6.0536 | 6.96517 | 6.00368 | 6.97304 |
| OXNAD1 | Q96HP4 | 6.42279 | 6.33648 | 5.29595 | 6.78682 | 6.72983 | 5.77791 | 6.35592 | 6.10847 | 5.97706 | 6.0158 | 5.92809 | 5.96627 | 6.66939 | 6.31765 | 6.62863 | 6.68295 |
| TDRD9 | Q8NDG6 | 8.17124 | 7.65851 | 8.24565 | 7.46901 | 6.8973 | 6.69687 | 6.28443 | 6.72556 | 7.71234 | 7.97219 | 7.37685 | 7.57567 | 2.5065 | 3.29171 | 2.51223 | 2.90396 |
| UBAC1 | Q9BSL1 | 6.95293 | 6.14877 | 6.94212 | 6.28456 | 6.05947 | 6.22024 | 5.94792 | 6.77048 | 6.76361 | 6.91315 | 6.88073 | 7.07332 | 5.63323 | 4.64326 | 5.40443 | 5.36177 |
| ARL2 | P36404 | 6.09166 | 6.42537 | 6.59597 | 6.11555 | 6.01879 | 6.51767 | 6.58987 | 7.09274 | 6.50855 | 6.70019 | 6.43889 | 6.691 | 5.8295 | 5.45785 | 5.4101 | 5.51629 |
| RBMX2 | Q9Y388 | 5.78061 | 6.68235 | 6.16917 | 6.71172 | 5.89669 | 6.00573 | 6.53791 | 6.08748 | 6.43305 | 5.71668 | 5.95978 | 6.16348 | 5.78903 | 6.73211 | 6.74866 | 6.58553 |
| NKIRAS2 | Q9NYR9 | 6.51737 | 6.3432 | 7.09179 | 6.97722 | 6.42814 | 6.60918 | 6.26727 | 6.59058 | 6.92392 | 7.2016 | 6.92062 | 7.13541 | 4.62967 | 5.04374 | 4.36351 | 4.95678 |
| FRA10AC1 | Q70Z53 | 5.99022 | 5.96418 | 6.87994 | 6.10578 | 6.26437 | 6.32114 | 5.80964 | 5.79483 | 6.42681 | 6.54939 | 6.44446 | 6.58338 | 5.98899 | 6.47072 | 6.53621 | 5.86994 |
| TBC1D22A | Q8WUA7 | 6.1503 | 5.90974 | 6.62765 | 6.20386 | 6.53942 | 6.76018 | 6.15597 | 6.81287 | 6.24403 | 6.34601 | 6.37919 | 6.06555 | 5.80135 | 6.43525 | 5.80396 | 5.76468 |
| TRAM2 | Q15035 | 6.61118 | 6.73873 | 6.36425 | 7.14559 | 6.15617 | 6.0322 | 6.21782 | 5.99732 | 6.02013 | 6.07404 | 6.17121 | 5.83139 | 6.18836 | 5.94359 | 6.45067 | 6.05734 |
| SNX29 | Q8TEQ0 | 6.51845 | 6.63588 | 6.08228 | 6.23021 | 6.26016 | 6.11205 | 5.92339 | 5.62634 | 5.95982 | 6.5538 | 6.50805 | 6.60879 | 6.09803 | 6.35541 | 6.35718 | 6.17016 |
| NSUN6 | Q8TEA1 | 5.58962 | 5.89377 | 6.4923 | 6.06955 | 6.55868 | 6.58479 | 6.17461 | 6.69336 | 6.1992 | 6.65254 | 6.65644 | 6.97841 | 6.15414 | 5.94347 | 5.61196 | 5.74718 |
| OARD1 | Q9Y530 | 6.48854 | 6.2135 | 6.14592 | 6.14698 | 6.21501 | 5.95233 | 6.46309 | 6.37005 | 6.09423 | 5.94786 | 5.73101 | 5.92114 | 6.74763 | 6.52404 | 6.70602 | 6.33265 |
| ASTE1 | Q2TB18 | 6.13601 | 6.18323 | 6.40024 | 6.22097 | 6.14737 | 6.20021 | 6.26976 | 5.88287 | 6.71917 | 6.48404 | 6.73399 | 6.87594 | 5.82309 | 5.86035 | 6.07074 | 5.99201 |

| | | | | | | | | | | | | | | | | | |
|---|---|---|---|---|---|---|---|---|---|---|---|---|---|---|---|---|---|
| UBN2 | Q6ZU65 | 6.23394 | 6.50055 | 5.96485 | 6.95434 | 6.21009 | 6.0992 | 6.23981 | 6.0573 | 6.14351 | 6.49055 | 5.93832 | 6.36357 | 5.9262 | 6.11428 | 6.32376 | 6.43974 |
| TUBGCP6 | Q96RT7 | 5.71467 | 6.22739 | 7.04534 | 6.38871 | 5.66079 | 6.45644 | 6.14052 | 6.41907 | 6.21373 | 6.89996 | 6.96398 | 6.44509 | 5.57662 | 6.07651 | 5.91444 | 5.85673 |
| HBA1 | P69905 | 6.841 | 5.93769 | 5.57511 | 5.92881 | 4.46862 | 5.64473 | 8.49192 | 4.26809 | 5.62311 | 6.17991 | 5.55702 | 6.38675 | 6.6213 | 5.70695 | 8.43547 | 8.33353 |
| COA3 | Q9Y2R0 | 6.40886 | 5.59446 | 5.88239 | 5.26385 | 6.5235 | 5.36718 | 5.85172 | 6.34996 | 5.73395 | 6.07687 | 5.85866 | 6.02883 | 6.74973 | 7.63887 | 7.47509 | 7.19609 |
| UCHL3 | P15374 | 6.07803 | 5.30026 | 6.34102 | 4.45899 | 6.05758 | 6.42142 | 6.75049 | 7.58293 | 5.16896 | 6.56911 | 5.77346 | 6.82509 | 6.19964 | 7.1402 | 6.38707 | 6.94577 |
| CTSZ | Q9UBR2 | 6.55693 | 6.51328 | 6.11876 | 6.01107 | 6.49223 | 6.30375 | 6.11557 | 6.25202 | 6.13622 | 5.93093 | 5.8547 | 5.50197 | 6.47365 | 6.85108 | 6.46119 | 6.42666 |
| PBX3 | P40426 | 6.49157 | 6.084 | 6.41951 | 6.43111 | 6.31569 | 6.25736 | 6.05833 | 5.69015 | 6.24326 | 6.0546 | 6.57296 | 6.3645 | 6.31886 | 6.17329 | 5.78796 | 6.73685 |
| FAM219A | Q8IW50 | 5.75676 | 5.77747 | 5.94512 | 6.22887 | 5.14119 | 5.71607 | 6.46658 | 4.90027 | 5.83753 | 5.8835 | 6.20245 | 6.50447 | 6.23072 | 7.85643 | 7.44675 | 8.10582 |
| BAK1 | Q16611 | 5.83123 | 4.77568 | 5.3543 | 5.26718 | 5.90287 | 5.77027 | 5.8486 | 6.80409 | 5.29021 | 5.98041 | 5.46894 | 5.8447 | 6.42477 | 8.65841 | 8.06853 | 8.70983 |
| RER1 | O15258 | 6.20998 | 6.58036 | 5.78702 | 6.2843 | 6.09295 | 6.10544 | 6.38386 | 6.25633 | 5.94583 | 6.1575 | 6.08771 | 6.12556 | 6.32815 | 6.5335 | 6.61528 | 6.50624 |
| NFATC2IP | Q8NCF5 | 5.99203 | 5.93666 | 6.1132 | 5.66708 | 5.70922 | 6.0442 | 5.69965 | 6.47643 | 5.86718 | 6.25971 | 6.35201 | 6.33062 | 6.46285 | 7.1589 | 6.85993 | 7.07033 |
| MRPL53 | Q96EL3 | 6.21966 | 5.38435 | 5.64322 | 5.72565 | 6.44242 | 5.89843 | 5.89587 | 6.70885 | 5.3875 | 6.15351 | 5.92563 | 6.86539 | 6.39121 | 7.19077 | 7.13532 | 7.03222 |
| STX8 | Q9UNK0 | 6.28302 | 5.76816 | 5.83768 | 6.68993 | 6.20343 | 5.88164 | 6.07786 | 6.25118 | 5.64501 | 6.49722 | 6.10361 | 6.81682 | 6.06007 | 6.74053 | 6.29933 | 6.84451 |
| SETD9 | Q8NE22 | 6.11166 | 6.47578 | 5.76615 | 6.65143 | 6.1766 | 6.37907 | 6.36139 | 5.98256 | 6.0734 | 6.47897 | 6.29053 | 5.94877 | 6.21448 | 6.16977 | 6.3468 | 6.57264 |
| FKBP2 | P26885 | 6.16465 | 6.62324 | 5.52194 | 6.27084 | 6.99796 | 6.72234 | 6.19241 | 6.906 | 5.58411 | 6.15192 | 6.10909 | 5.89498 | 6.58667 | 6.21082 | 6.09486 | 5.96817 |
| MRPL12 | P52815 | 6.076 | 4.82979 | 4.85707 | 5.20204 | 5.22888 | 3.65035 | 5.27583 | 5.47598 | 4.24322 | 4.72324 | 4.32032 | 4.82667 | 5.22902 | 12.7936 | 9.57485 | 13.6932 |
| MAGI1 | Q96QZ7 | 6.50772 | 6.71083 | 6.43002 | 5.77519 | 6.30093 | 6.56426 | 6.15155 | 5.81649 | 6.08096 | 6.22759 | 6.14204 | 6.06337 | 6.51711 | 6.43536 | 6.2116 | 6.06499 |
| TEC | P42680 | 6.53595 | 6.48582 | 7.40966 | 6.7894 | 5.79766 | 6.30267 | 6.01018 | 6.64919 | 6.11865 | 6.59488 | 6.37216 | 6.83882 | 5.42469 | 5.72351 | 5.31226 | 5.63451 |
| ATN1 | P54259 | 6.2275 | 5.73082 | 6.85769 | 6.26264 | 6.24493 | 6.40236 | 6.22477 | 6.25194 | 7.02326 | 6.12224 | 7.07836 | 6.92258 | 6.34539 | 4.8274 | 5.61869 | 5.85943 |
| SESTD1 | Q86VW0 | 6.17131 | 6.47643 | 6.83821 | 6.23692 | 6.13505 | 6.60382 | 6.23929 | 6.3254 | 6.88947 | 6.86333 | 6.41836 | 6.381 | 5.49841 | 5.75529 | 5.58494 | 5.58276 |
| ALG6 | Q9Y672 | 6.29795 | 6.35683 | 5.53474 | 6.67131 | 6.01567 | 5.89532 | 6.24148 | 6.01865 | 6.00605 | 6.28793 | 5.65951 | 5.74554 | 6.76584 | 6.54605 | 7.06375 | 6.89339 |
| GALNT11 | Q8NCW6 | 6.47113 | 6.7103 | 5.81079 | 6.52831 | 6.28146 | 6.27754 | 6.47428 | 5.81495 | 6.38046 | 5.96091 | 6.07982 | 5.74391 | 6.71651 | 6.33197 | 6.58243 | 5.83523 |
| NUBP2 | Q9Y5Y2 | 6.13095 | 5.66658 | 6.51278 | 5.61447 | 6.36734 | 6.29705 | 6.15671 | 6.63133 | 6.42379 | 6.08555 | 7.2113 | 7.10453 | 6.03696 | 6.00261 | 5.87917 | 5.87886 |
| BAX | Q07812-2 | 5.89136 | 5.65708 | 6.25492 | 5.84861 | 5.97452 | 6.7021 | 6.40638 | 6.19338 | 6.19716 | 6.47939 | 6.11946 | 6.23819 | 5.48675 | 7.10581 | 6.54518 | 6.89969 |
| CNOT11 | Q9UKZ1 | 6.60273 | 6.0748 | 6.40882 | 5.51987 | 6.44563 | 6.31252 | 6.20168 | 6.70707 | 6.27224 | 6.45927 | 6.54638 | 6.84101 | 5.75231 | 5.70898 | 5.90343 | 6.24326 |
| POT1 | Q9NUX5 | 5.89085 | 6.2079 | 6.09033 | 6.10935 | 5.92456 | 6.39455 | 6.42076 | 6.16489 | 5.97811 | 6.1649 | 6.18063 | 6.32651 | 6.64128 | 6.63934 | 6.52313 | 6.34291 |
| MTRF1 | O75570 | 5.69628 | 6.15545 | 5.97854 | 6.73224 | 6.46143 | 6.48568 | 6.0737 | 6.09058 | 6.48156 | 6.1028 | 6.5788 | 5.96163 | 6.2192 | 6.33538 | 6.14698 | 6.49976 |
| CXorf38 | Q8TB03 | 6.8368 | 6.08907 | 7.15616 | 5.93761 | 6.54203 | 6.74753 | 6.1892 | 6.63684 | 6.42007 | 6.52518 | 6.76953 | 6.59888 | 5.64513 | 5.29459 | 5.09825 | 5.51315 |
| RAVER2 | Q9HCJ3 | 5.47913 | 6.01631 | 6.35626 | 5.51438 | 5.99481 | 6.05897 | 6.01545 | 6.33257 | 6.11291 | 6.03361 | 6.576 | 6.24518 | 6.29629 | 6.87577 | 7.44625 | 6.64612 |
| FBXL12 | Q9NXK8 | 5.66673 | 5.67996 | 6.05022 | 5.18598 | 5.87044 | 6.15655 | 5.4845 | 6.32336 | 6.03711 | 6.38175 | 6.03329 | 6.08803 | 7.30212 | 7.2383 | 7.77749 | 6.72418 |
| MTIF3 | Q9H2K0 | 5.93874 | 7.07599 | 5.48331 | 6.58917 | 5.6764 | 6.46692 | 6.08653 | 5.81818 | 6.24406 | 6.36002 | 5.65093 | 5.99959 | 6.43506 | 6.84433 | 6.80552 | 6.52526 |
| LSM6 | P62312 | 5.29847 | 5.47287 | 5.56049 | 5.16648 | 6.71987 | 6.19048 | 6.32569 | 6.87724 | 6.26712 | 6.62077 | 6.37909 | 6.17883 | 6.49727 | 6.58287 | 7.12234 | 6.74012 |
| NAGPA | Q9UK23 | 6.4544 | 6.83395 | 6.21446 | 6.59507 | 6.57971 | 5.94638 | 6.22463 | 5.97095 | 6.42279 | 6.51021 | 6.5937 | 6.53022 | 6.33745 | 5.56198 | 5.55592 | 5.66817 |
| UCKL1 | Q9NWZ5 | 6.34979 | 6.22907 | 6.73324 | 5.81197 | 6.38216 | 6.7834 | 6.25811 | 6.47608 | 6.46074 | 6.58797 | 6.79791 | 6.352 | 5.63473 | 5.73705 | 5.7206 | 6.58519 |
| SLC5A3 | P53794 | 6.10428 | 6.32489 | 6.37124 | 6.45721 | 5.86359 | 6.2861 | 6.20696 | 6.54544 | 6.32977 | 6.44043 | 6.39696 | 6.09805 | 6.11988 | 6.4185 | 6.29357 | 6.64312 |
| PMM1 | Q92871 | 7.70455 | 7.26257 | 8.10169 | 7.48896 | 5.92686 | 6.56949 | 5.91966 | 6.24201 | 6.99254 | 8.04352 | 7.59255 | 7.49205 | 3.46418 | 4.08854 | 3.43015 | 3.68066 |
| IDUA | P35475 | 6.21402 | 6.41218 | 6.13353 | 6.3905 | 5.98537 | 5.99522 | 6.02464 | 6.16383 | 6.54159 | 6.19283 | 6.05857 | 6.16295 | 6.61357 | 6.35531 | 6.67128 | 6.08462 |
| APOO | Q9BUR5 | 6.44425 | 6.52082 | 5.91305 | 6.12024 | 6.51064 | 5.95735 | 5.79302 | 6.42721 | 6.03494 | 6.40672 | 6.12709 | 6.05783 | 6.76604 | 6.41601 | 6.29877 | 6.206 |
| PRIMPOL | Q96LW4 | 6.39963 | 6.92753 | 7.12256 | 6.55196 | 6.19953 | 6.57238 | 6.17964 | 5.95317 | 6.99052 | 6.86176 | 6.4254 | 6.56038 | 5.28989 | 5.21884 | 5.58936 | 5.15746 |
| RASA4 | O43374 | 6.63041 | 6.59389 | 6.65981 | 6.08407 | 6.20634 | 6.279 | 6.40022 | 6.61291 | 6.39827 | 6.761 | 6.44889 | 6.31646 | 5.53528 | 5.95529 | 5.45727 | 5.66088 |
| LCLAT1 | Q6UWP7 | 6.34248 | 6.20168 | 5.87918 | 6.42196 | 5.82788 | 5.97639 | 6.29558 | 6.038 | 6.28575 | 5.974 | 6.40759 | 6.03658 | 6.86072 | 6.39605 | 6.70022 | 6.35595 |
| DDI2 | Q5TDH0-3 | 6.08349 | 6.13316 | 6.60774 | 6.13294 | 6.09203 | 6.68074 | 6.36162 | 6.4061 | 6.27915 | 6.09688 | 6.59268 | 7.15813 | 5.8998 | 5.88546 | 5.80851 | 5.78156 |
| DCAKD | Q8WVC6 | 6.61091 | 6.51837 | 5.80104 | 6.30792 | 6.55163 | 5.7838 | 6.77014 | 6.38191 | 6.32927 | 5.89351 | 5.91505 | 5.8122 | 6.58469 | 6.03248 | 6.72792 | 5.97917 |
| GNB1L | Q9BYB4 | 6.16882 | 6.29742 | 6.72311 | 5.81115 | 6.19298 | 6.76449 | 6.23745 | 6.85398 | 6.32431 | 6.32929 | 6.62672 | 6.88711 | 5.6288 | 5.95129 | 5.50556 | 5.69753 |
| SUMO1 | P63165 | 6.41234 | 5.94724 | 5.97164 | 5.88904 | 6.19608 | 5.75391 | 5.58647 | 6.31051 | 5.51195 | 6.59965 | 6.26679 | 6.56717 | 6.73401 | 6.74808 | 6.77632 | 6.7288 |
| CNKSR2 | Q8WXI2 | 5.91216 | 6.01695 | 6.58305 | 6.85245 | 6.56584 | 6.03983 | 6.21934 | 6.50718 | 6.66085 | 6.39172 | 6.77064 | 6.40453 | 5.84434 | 5.63248 | 5.5524 | 6.04623 |
| MGST3 | O14880 | 6.23252 | 6.47475 | 5.56145 | 6.63424 | 6.4317 | 6.86365 | 6.30761 | 6.16943 | 5.25591 | 6.22237 | 6.20196 | 5.74156 | 6.69792 | 6.68788 | 6.36977 | 6.14727 |
| RPRD1A | Q96P16 | 5.91142 | 6.40393 | 5.94543 | 6.69397 | 6.03074 | 5.79614 | 5.53282 | 6.66093 | 6.03554 | 6.34956 | 6.26083 | 6.30645 | 6.93663 | 6.17202 | 6.39982 | 6.56377 |
| TMA16 | Q96EY4 | 6.35379 | 6.26367 | 6.34993 | 6.32187 | 6.25499 | 6.02422 | 5.97911 | 5.89489 | 5.96826 | 5.90902 | 6.35659 | 6.04947 | 6.31579 | 6.75153 | 6.68649 | 6.52542 |
| MIX23 | Q4VC31 | 6.27385 | 5.50533 | 5.95144 | 5.65617 | 6.9755 | 5.73869 | 5.49512 | 6.27815 | 5.61461 | 6.60689 | 6.37281 | 6.70676 | 6.98686 | 6.35213 | 6.7491 | 6.73657 |
| NDUFB3 | O43676 | 6.00876 | 6.23247 | 5.61234 | 6.417 | 6.57779 | 5.72882 | 7.25934 | 6.32919 | 5.89555 | 5.12695 | 5.79 | 5.84882 | 6.60217 | 6.82589 | 6.92603 | 6.81887 |
| PDPK1 | O15530 | 6.79261 | 6.19456 | 6.82615 | 6.40473 | 6.5206 | 6.78209 | 6.28174 | 6.49682 | 6.88233 | 6.49676 | 6.38242 | 6.39941 | 5.76724 | 5.24579 | 5.42446 | 5.10229 |
| SZRD1 | Q7Z422 | 6.93781 | 5.92532 | 7.27053 | 5.60248 | 6.48454 | 6.27135 | 6.23397 | 6.32694 | 6.65944 | 6.21831 | 7.29877 | 6.3897 | 5.93564 | 5.83017 | 5.28245 | 5.33259 |
| SLC35B1 | P78383-2 | 5.84523 | 6.30987 | 6.15562 | 6.75316 | 6.70058 | 6.63201 | 6.23023 | 5.88608 | 6.01797 | 6.36556 | 6.15631 | 5.96101 | 6.5676 | 6.2801 | 6.05311 | 6.08557 |
| CREB1 | P16220-1 | 6.46976 | 5.94265 | 6.31635 | 6.25614 | 6.45432 | 5.84964 | 6.23441 | 6.21441 | 6.50455 | 5.85524 | 6.28818 | 6.3872 | 6.32113 | 6.02278 | 6.53397 | 6.34927 |
| DSN1 | Q9H410 | 6.41482 | 6.2227 | 6.37155 | 6.08865 | 5.74979 | 6.02716 | 5.98893 | 6.09133 | 6.67284 | 6.97027 | 6.42056 | 6.07045 | 6.94594 | 6.40622 | 6.21641 | 6.34239 |
| CLN5 | O75503 | 6.77395 | 6.5637 | 6.08711 | 6.51998 | 6.19162 | 6.02436 | 6.42876 | 5.97902 | 6.21467 | 6.3466 | 6.17434 | 6.14013 | 6.24997 | 5.90813 | 6.28094 | 6.1167 |
| CDCA5 | Q96FF9 | 6.4275 | 6.16065 | 5.98882 | 6.70764 | 5.88067 | 5.97889 | 6.23995 | 5.64945 | 6.33227 | 6.03865 | 6.04554 | 6.05294 | 6.52066 | 6.28818 | 6.9295 | 6.75869 |
| FBXO21 | O94952 | 6.15082 | 6.18917 | 7.19014 | 6.13641 | 6.36636 | 6.59828 | 6.20288 | 6.66132 | 6.22595 | 6.00995 | 6.38429 | 6.51269 | 5.81659 | 5.80311 | 5.75242 | 5.99961 |
| ZNF346 | Q9UL40-2 | 6.54906 | 6.71115 | 6.44367 | 6.56306 | 6.20559 | 6.57057 | 6.5816 | 6.25298 | 6.02737 | 6.31393 | 5.96903 | 5.79033 | 5.89364 | 6.14774 | 6.38421 | 5.97177 |
| TMEM11 | P17152 | 6.39649 | 5.91489 | 6.09431 | 6.06463 | 6.5454 | 7.08141 | 6.54067 | 5.75252 | 5.93068 | 6.02883 | 6.24074 | 5.45805 | 6.69926 | 6.13425 | 6.72479 | 6.39309 |
| NUP42 | O15504 | 6.13938 | 6.66279 | 5.85802 | 6.75377 | 6.03504 | 6.35673 | 5.42488 | 6.57369 | 6.07039 | 6.69639 | 5.4438 | 6.57911 | 6.20631 | 6.20137 | 6.12943 | 6.86889 |
| IQCB1 | Q15051 | 7.01546 | 6.97399 | 8.00806 | 6.93597 | 7.07 | 6.53315 | 6.45642 | 6.05734 | 6.00012 | 5.45297 | 5.77024 | 5.82547 | 5.38597 | 5.49472 | 5.21476 | 5.80536 |
| MGAT2 | Q10469 | 6.28499 | 6.07157 | 6.10264 | 6.95197 | 5.9974 | 6.52058 | 6.26651 | 5.8368 | 6.33999 | 6.37866 | 5.85521 | 6.48677 | 6.19869 | 6.25863 | 6.51032 | 5.93926 |
| CINP | Q9BW66-3 | 6.0914 | 4.96607 | 5.7003 | 4.71433 | 6.5151 | 6.32104 | 6.03563 | 8.05752 | 6.32946 | 7.27381 | 7.21052 | 7.62961 | 4.87659 | 5.9539 | 6.28925 | 6.03546 |
| TARBP1 | Q13395 | 5.66712 | 6.57925 | 5.89402 | 6.3667 | 5.98702 | 6.86448 | 5.50422 | 6.56713 | 6.06262 | 5.94849 | 6.15903 | 6.22957 | 7.7563 | 6.23601 | 5.97175 | 6.20629 |
| SRA1 | Q9HD15 | 6.6653 | 6.00844 | 7.02732 | 5.42682 | 6.73584 | 6.86487 | 6.82595 | 7.20786 | 6.2719 | 6.47221 | 6.63425 | 7.0059 | 5.69347 | 5.70331 | 5.47466 | 4.9819 |
| GTSF1 | Q8WW33 | 6.51856 | 5.53448 | 7.13627 | 5.2568 | 6.57083 | 5.96532 | 5.5173 | 7.23847 | 5.7992 | 6.92292 | 6.66521 | 8.51652 | 5.3233 | 5.99787 | 5.2963 | 5.74065 |
| OSTM1 | Q86WC4 | 6.72147 | 6.48987 | 5.97347 | 6.92077 | 6.51876 | 6.01508 | 6.67109 | 5.63524 | 6.6472 | 5.91227 | 6.45652 | 6.32148 | 5.93226 | 5.98507 | 5.46992 | 6.32951 |
| CFAP20 | Q9Y6A4 | 6.20826 | 6.24417 | 6.47313 | 6.4779 | 6.13449 | 6.19371 | 6.41852 | 5.59933 | 6.33246 | 6.15293 | 6.22225 | 6.2048 | 6.85881 | 5.54138 | 6.69844 | 6.23942 |
| FOSL2 | P15408 | 5.82699 | 5.88306 | 6.16625 | 6.3659 | 5.9206 | 5.96944 | 6.21519 | 5.92865 | 5.54435 | 5.28695 | 5.3178 | 5.92964 | 8.20979 | 6.85728 | 7.27297 | 7.40485 |
| HMG20A | Q9NP66 | 6.20563 | 6.38116 | 6.00169 | 6.97673 | 5.84672 | 5.92313 | 6.43307 | 5.62513 | 6.16793 | 5.92671 | 6.30805 | 6.5569 | 6.67155 | 6.18555 | 6.51995 | 6.2701 |
| MIF | P14174 | 4.90263 | 4.93444 | 5.92188 | 4.77896 | 5.53475 | 5.12195 | 5.85182 | 7.0121 | 5.8324 | 6.1358 | 6.56257 | 6.02668 | 4.3977 | 9.89445 | 7.86251 | 9.22937 |
| PASD1 | Q8IV76 | 6.07065 | 6.40319 | 6.15387 | 6.44462 | 6.15424 | 6.10734 | 6.79101 | 6.04035 | 5.9095 | 6.33751 | 6.07491 | 6.10591 | 6.28788 | 6.06792 | 6.38838 | 6.66271 |
| MED10 | Q9BTT4 | 5.87207 | 5.32917 | 4.56892 | 5.3738 | 4.95297 | 3.45141 | 4.74916 | 5.21174 | 3.96249 | 5.0354 | 4.38969 | 4.4923 | 4.4782 | 14.0158 | 9.82129 | 14.2955 |
| MRPL30 | Q8TCC3-2 | 6.31752 | 6.51334 | 5.80575 | 6.58908 | 6.19997 | 6.2735 | 6.54751 | 6.00425 | 6.02108 | 6.20495 | 5.6827 | 5.7305 | 6.45968 | 6.35711 | 6.76231 | 6.53075 |
| XYLT2 | Q9H1B5 | 6.32678 | 6.46203 | 6.30755 | 6.8879 | 6.22173 | 6.05506 | 6.31191 | 5.74904 | 6.62249 | 6.38562 | 6.00336 | 6.45688 | 5.94515 | 5.99888 | 6.02156 | 6.24406 |
| H1-0 | P07305 | 6.31446 | 6.11284 | 5.3881 | 6.36308 | 6.02183 | 5.82366 | 6.21082 | 6.06063 | 5.81486 | 5.61871 | 5.49983 | 5.53816 | 7.03993 | 7.61699 | 7.40497 | 7.17114 |
| SPCS3 | P61009 | 6.46593 | 6.27894 | 6.01192 | 6.44169 | 6.4178 | 6.00143 | 6.4791 | 6.12764 | 6.16561 | 6.31108 | 6.02964 | 5.92541 | 6.42316 | 6.08909 | 6.60313 | 6.22845 |
| MRPL42 | Q9Y6G3 | 6.11035 | 5.19488 | 5.3074 | 6.71335 | 6.35663 | 5.12194 | 7.48055 | 5.86975 | 6.88893 | 4.78042 | 5.35725 | 5.95411 | 6.58422 | 6.85052 | 7.12912 | 8.30058 |
| SNAPC2 | Q13487 | 5.94281 | 6.30286 | 6.42456 | 6.13348 | 5.7763 | 6.24223 | 5.95991 | 6.07472 | 6.09634 | 6.2833 | 6.47522 | 6.64173 | 6.50264 | 6.39816 | 6.54126 | 6.20448 |
| DEF8 | Q6ZN54 | 6.29977 | 6.6565 | 6.65597 | 6.99658 | 6.11008 | 5.92578 | 6.40458 | 6.42557 | 7.44066 | 6.38122 | 6.01764 | 6.5515 | 6.06436 | 5.30802 | 5.69983 | 5.06194 |
| TMEM115 | Q12893 | 5.89685 | 6.37047 | 5.95146 | 6.98224 | 6.00633 | 5.87981 | 6.11372 | 6.26364 | 5.93656 | 6.28631 | 6.27043 | 6.3183 | 6.47504 | 6.49023 | 6.1951 | 6.56352 |
| CRABP2 | P29373 | 7.46952 | 5.97305 | 6.33995 | 5.4401 | 6.62198 | 7.20923 | 6.40023 | 5.99132 | 6.5152 | 7.0335 | 6.40342 | 6.56866 | 5.85179 | 5.47618 | 4.92019 | 5.78567 |
| STRN4 | Q9NRL3-3 | 5.92854 | 6.38358 | 6.31287 | 5.79584 | 6.27651 | 6.78771 | 5.73889 | 6.5608 | 6.43514 | 6.13312 | 6.05107 | 6.53994 | 6.69196 | 6.03218 | 6.01209 | 6.31976 |
| STRIP2 | Q9ULQ0 | 6.19016 | 6.56126 | 6.57203 | 6.17443 | 5.93265 | 6.44246 | 5.91367 | 6.06768 | 6.77352 | 6.67702 | 6.34876 | 6.10058 | 5.98416 | 6.11304 | 6.2041 | 5.94448 |
| BLOC1S4 | Q9NUP1 | 5.72899 | 5.28921 | 6.65481 | 5.15365 | 6.1572 | 6.21801 | 5.26029 | 6.58326 | 5.50377 | 6.675 | 6.57769 | 7.11107 | 6.47322 | 7.31656 | 6.57934 | 6.71793 |
| RBM23 | Q86U06 | 5.7038 | 6.36813 | 6.28213 | 6.68565 | 5.27549 | 6.39305 | 6.00506 | 5.9156 | 6.93892 | 7.11898 | 6.98356 | 6.70786 | 5.86334 | 5.86129 | 6.0469 | 5.85024 |
| NEK7 | Q8TDX7 | 5.95418 | 6.89568 | 6.83596 | 5.448 | 6.98109 | 7.08657 | 5.93892 | 6.20988 | 6.34506 | 6.14525 | 6.79725 | 6.10241 | 6.45809 | 6.28884 | 5.25479 | 5.25804 |
| BOD1 | Q96IK1 | 5.99423 | 5.27396 | 5.75944 | 5.46439 | 5.56586 | 4.16999 | 4.94877 | 6.65227 | 4.67677 | 5.63154 | 4.78756 | 5.91282 | 5.02754 | 11.4472 | 8.42862 | 10.259 |
| C5orf24 | Q7Z6I8 | 6.48685 | 6.17584 | 6.58679 | 6.48389 | 6.20526 | 5.93961 | 6.16551 | 5.62109 | 5.86088 | 6.44966 | 6.51028 | 6.01162 | 6.18388 | 6.47344 | 6.62847 | 6.21694 |
| GSTCD | Q8NEC7 | 6.14569 | 6.29127 | 6.88653 | 5.88839 | 6.48943 | 6.82868 | 6.46098 | 6.85024 | 6.4099 | 6.6156 | 6.22675 | 6.50053 | 6.58049 | 5.70308 | 5.36841 | 5.65403 |
| SYVN1 | Q86TM6 | 6.45489 | 6.3885 | 5.95427 | 6.43582 | 6.58641 | 6.07824 | 6.08909 | 6.27213 | 6.04784 | 6.56618 | 5.3161 | 5.83889 | 6.83527 | 6.41482 | 6.46985 | 6.25168 |
| COMMD3 | Q9UBI1 | 5.80286 | 4.87088 | 5.73645 | 5.03143 | 5.3755 | 5.49608 | 5.4941 | 6.74913 | 5.31454 | 6.04091 | 5.42422 | 5.50885 | 5.7467 | 9.40906 | 7.84909 | 10.1502 |
| FYN | P06241 | 5.99541 | 6.40308 | 5.97744 | 6.91455 | 6.30762 | 6.67621 | 5.99411 | 6.07947 | 6.25217 | 6.01411 | 5.95397 | 6.60781 | 6.83109 | 6.179 | 5.76218 | 6.05178 |
| PRKD1 | Q15139 | 6.15331 | 6.20448 | 6.76027 | 6.16711 | 6.15893 | 6.13449 | 6.21313 | 6.25574 | 6.49694 | 6.67458 | 6.71422 | 6.65805 | 5.75317 | 6.02392 | 5.644 | 5.98766 |
| STK36 | Q9NRP7 | 6.79451 | 6.44475 | 6.9755 | 6.77367 | 6.56291 | 6.69055 | 6.36196 | 6.73937 | 6.57272 | 6.59964 | 7.1293 | 6.36335 | 5.32692 | 5.09376 | 4.71631 | 4.85478 |
| MACROD1 | Q9BQ69 | 6.20668 | 6.46512 | 6.35032 | 6.32062 | 6.90012 | 6.23254 | 5.76465 | 5.68745 | 6.08212 | 5.52408 | 6.25414 | 5.96618 | 6.80667 | 6.71699 | 6.79297 | 5.92935 |
| SEPSECS | Q9HD40 | 6.43999 | 6.98103 | 6.47374 | 5.67067 | 6.75507 | 6.52136 | 6.6274 | 6.95594 | 6.39775 | 6.20963 | 6.80727 | 6.08013 | 5.68032 | 5.53915 | 5.57501 | 5.28553 |
| PCBD2 | Q9H0N5 | 6.01064 | 5.33377 | 6.28537 | 5.58123 | 6.58566 | 5.59388 | 5.80132 | 6.6937 | 5.4554 | 6.77004 | 6.44381 | 6.71786 | 6.30436 | 6.84396 | 6.72209 | 6.85689 |
| NEURL4 | Q96JN8 | 5.93499 | 6.20051 | 6.93052 | 6.11684 | 5.84049 | 5.93929 | 6.6049 | 6.27596 | 6.32083 | 6.47089 | 6.12507 | 6.87268 | 5.71622 | 6.33705 | 6.28297 | 6.03078 |
| MIEF1 | Q9NQG6 | 6.53866 | 5.78089 | 5.557 | 6.86481 | 6.85818 | 5.65959 | 6.3387 | 5.72703 | 5.53279 | 5.94741 | 5.622 | 6.32505 | 6.84392 | 6.52865 | 6.6234 | 7.25194 |
| KLHL24 | Q6TFL4 | 5.55383 | 6.09899 | 5.2838 | 5.01957 | 5.69835 | 5.60797 | 5.36528 | 5.92194 | 6.25159 | 7.59918 | 6.58189 | 6.07751 | 6.77922 | 7.14028 | 7.4699 | 7.5507 |
| GHDC | Q8N2G8 | 6.23268 | 6.3408 | 5.75085 | 6.1857 | 6.23337 | 5.90208 | 6.18695 | 6.19506 | 6.00764 | 5.92882 | 5.88583 | 5.89991 | 6.97801 | 6.57543 | 6.96983 | 6.72704 |
| VMA21 | Q3ZAQ7 | 6.18861 | 5.36527 | 5.92551 | 5.11606 | 6.22035 | 5.44094 | 6.55031 | 6.34368 | 5.63608 | 6.40349 | 5.6518 | 5.47641 | 6.12267 | 8.05711 | 7.49797 | 8.00373 |

| | | | | | | | | | | | | | | | | | |
|---|---|---|---|---|---|---|---|---|---|---|---|---|---|---|---|---|---|
| ADGRL1 | O94910 | 7.24255 | 6.72128 | 6.00456 | 6.46093 | 6.0702 | 6.23646 | 6.65668 | 6.25628 | 6.38137 | 6.00102 | 6.23897 | 5.94445 | 6.0638 | 5.60029 | 6.14559 | 5.97558 |
| PRDM11 | A0A087WWZ6 | 5.95747 | 6.27812 | 6.17359 | 6.10329 | 6.10206 | 6.24946 | 6.24262 | 5.65284 | 6.20286 | 6.37226 | 6.42026 | 6.24621 | 6.64598 | 6.78864 | 6.39292 | 6.17142 |
| NAA40 | Q86UY6 | 6.44781 | 7.04488 | 6.27921 | 6.7843 | 5.55829 | 6.70234 | 5.69478 | 6.48333 | 6.54892 | 7.36035 | 5.95418 | 6.78285 | 5.20517 | 5.97412 | 5.29973 | 5.87973 |
| PLD2 | O14939 | 6.72534 | 6.83526 | 6.37873 | 6.64903 | 6.13131 | 5.95315 | 6.1454 | 6.2795 | 6.24715 | 6.35236 | 6.45203 | 6.29877 | 5.75325 | 5.96412 | 5.96697 | 5.86762 |
| IKBKB | O14920 | 6.16429 | 6.68076 | 6.82007 | 5.43058 | 6.15374 | 7.08353 | 6.42907 | 6.59012 | 6.70732 | 6.62082 | 6.78663 | 6.13006 | 5.52498 | 5.78562 | 5.55595 | 5.53646 |
| MAPK9 | P45984 | 6.08801 | 6.53566 | 6.41608 | 5.91915 | 5.92773 | 6.33436 | 6.00391 | 6.61448 | 6.78991 | 6.62835 | 6.48724 | 6.39933 | 5.87326 | 6.36068 | 5.55232 | 6.06954 |
| CUX1 | Q13948 | 6.21736 | 6.55369 | 6.16484 | 6.07594 | 6.22765 | 6.39218 | 6.32345 | 5.99791 | 5.74405 | 5.77923 | 6.10907 | 5.78139 | 7.19934 | 6.57165 | 6.56824 | 6.29401 |
| STK25 | O00506 | 6.4991 | 5.82965 | 7.11242 | 5.66108 | 6.2475 | 6.44693 | 6.35403 | 6.95107 | 6.62469 | 6.62442 | 6.63852 | 6.18007 | 5.85093 | 5.93764 | 5.63877 | 5.4032 |
| ARID3A | Q99856 | 6.25714 | 5.83873 | 6.26473 | 6.57216 | 6.22097 | 6.09888 | 6.13588 | 6.6259 | 6.20793 | 6.09655 | 6.10266 | 6.2587 | 6.81438 | 5.97834 | 6.19747 | 6.3296 |
| NR0B1 | P51843 | 6.54673 | 5.5005 | 6.12835 | 7.30301 | 6.91309 | 6.08345 | 6.3735 | 6.00078 | 6.05711 | 5.79254 | 6.22988 | 5.42893 | 6.93929 | 5.85884 | 6.15542 | 6.68857 |
| ANK1 | P16157-14 | 6.21629 | 6.23422 | 7.01237 | 6.09128 | 6.40871 | 6.32386 | 5.62302 | 6.63016 | 6.63738 | 6.52615 | 6.49042 | 6.43958 | 6.17087 | 6.45379 | 5.62862 | 5.11329 |
| ALDOC | P09972 | 5.45819 | 5.97404 | 6.64769 | 5.50309 | 6.25928 | 6.25988 | 6.98567 | 6.63443 | 6.22647 | 6.12396 | 6.39653 | 6.95224 | 5.7567 | 6.84538 | 6.04082 | 5.93561 |
| PALM2 | Q8IXS6-2 | 6.26357 | 5.80268 | 6.20232 | 5.86202 | 5.9993 | 6.1187 | 5.90064 | 5.85364 | 6.30339 | 5.81462 | 6.20029 | 5.7263 | 7.56738 | 6.67261 | 7.29012 | 6.42243 |
| WASHC2C | Q9Y4E1-5 | 6.06815 | 5.77347 | 6.9202 | 5.43886 | 5.76841 | 6.40716 | 6.02876 | 5.7573 | 6.01079 | 6.46984 | 6.72974 | 6.72915 | 6.37223 | 6.52289 | 6.72776 | 6.27531 |
| MB21D2 | Q8IYB1 | 6.02058 | 6.01655 | 6.57899 | 6.54229 | 6.09617 | 6.55349 | 6.13579 | 6.5007 | 6.39272 | 6.82525 | 6.73513 | 6.60755 | 5.78818 | 5.76968 | 5.85275 | 5.58418 |
| ARFIP2 | P53365 | 6.24794 | 5.20993 | 6.32133 | 5.32867 | 6.24802 | 5.91395 | 6.26195 | 6.21492 | 5.65679 | 6.61829 | 6.23931 | 6.44585 | 6.92672 | 6.83581 | 7.16476 | 6.36578 |
| PYCR3 | Q53H96 | 6.30869 | 5.62186 | 6.40712 | 6.23732 | 6.07969 | 6.58339 | 6.41063 | 5.93015 | 6.22447 | 6.06914 | 6.68551 | 6.10552 | 6.55402 | 6.03731 | 6.13031 | 6.61488 |
| SARAF | Q96BY9 | 7.1938 | 7.92334 | 7.25391 | 7.50286 | 6.65318 | 6.26341 | 6.33617 | 6.54702 | 6.06915 | 7.20359 | 6.83147 | 7.21689 | 3.99089 | 4.52821 | 4.03589 | 4.45023 |
| TSC22D4 | Q9Y3Q8 | 6.22386 | 6.06194 | 7.20558 | 5.76825 | 6.62807 | 6.40147 | 5.49253 | 6.57048 | 6.2056 | 6.85793 | 7.16908 | 6.87449 | 6.26822 | 5.41202 | 5.31814 | 5.54233 |
| PDIA3 | H7BZJ3 | 6.31839 | 6.22999 | 6.07282 | 5.96425 | 5.99201 | 5.6704 | 6.51066 | 6.04586 | 6.05288 | 6.36818 | 5.98115 | 6.05765 | 6.55192 | 6.19824 | 6.93482 | 7.05079 |
| JADE1 | Q6IE81 | 6.26759 | 5.91996 | 6.68833 | 6.55995 | 6.42062 | 5.95855 | 6.43627 | 5.76638 | 6.7724 | 5.86668 | 6.5095 | 6.49321 | 6.2218 | 5.92842 | 6.13989 | 6.05045 |
| ASRGL1 | Q7L266 | 5.91745 | 6.0086 | 7.1245 | 4.99862 | 6.433 | 6.59339 | 6.60138 | 7.70736 | 6.57405 | 6.58191 | 6.41168 | 6.85776 | 5.6721 | 6.56025 | 5.71398 | 5.14397 |
| TTC38 | Q5R3I4 | 5.83607 | 5.87884 | 6.85547 | 4.99639 | 6.4531 | 6.72678 | 6.60655 | 6.82618 | 6.54551 | 7.3418 | 6.30791 | 6.54175 | 5.36246 | 6.42795 | 5.96254 | 5.33071 |
| M0R2C6 | M0R2C6 | 6.26489 | 6.33766 | 5.99589 | 6.9478 | 6.18225 | 5.87567 | 6.40092 | 6.35135 | 6.27375 | 6.12969 | 6.24079 | 6.03753 | 6.33002 | 6.00043 | 6.16226 | 6.4691 |
| LZTS2 | Q9BRK4 | 6.39855 | 6.33132 | 5.97965 | 6.18652 | 6.73034 | 6.18736 | 6.19732 | 6.08554 | 5.89962 | 6.00476 | 6.33103 | 6.00504 | 6.77417 | 6.1256 | 6.37956 | 6.38362 |
| RPS19BP1 | Q86WX3 | 6.26051 | 6.72396 | 5.95896 | 6.92634 | 5.83196 | 5.45086 | 6.55553 | 5.65029 | 6.51674 | 6.07526 | 5.99293 | 5.89164 | 6.24445 | 6.13532 | 6.89029 | 6.89497 |
| ASB3 | Q9Y575-3 | 6.55329 | 6.58598 | 6.84162 | 6.2885 | 6.13445 | 6.78629 | 6.32573 | 6.23862 | 6.32852 | 6.18769 | 6.50878 | 6.41069 | 5.44592 | 6.00503 | 5.65075 | 5.70813 |
| KMT2E | Q8IZD2 | 6.7912 | 6.49328 | 5.93456 | 6.87399 | 5.91965 | 6.08082 | 6.25964 | 5.92106 | 6.01088 | 6.48976 | 5.74745 | 6.26749 | 6.07926 | 6.45655 | 6.73544 | 5.93895 |
| FAM160B2 | Q86V87 | 6.51983 | 6.43786 | 6.38315 | 5.47501 | 6.11346 | 6.44941 | 6.06262 | 6.30271 | 6.17527 | 5.89019 | 5.89117 | 6.3335 | 6.58083 | 6.77805 | 6.22997 | 6.37697 |
| AFG1L | Q8WV93 | 6.6689 | 6.25563 | 5.93884 | 6.68513 | 6.50158 | 6.06531 | 6.0469 | 5.78695 | 6.24252 | 6.14978 | 5.87427 | 5.59941 | 6.62872 | 6.46835 | 6.39844 | 6.68926 |
| ARRB2 | P32121-4 | 6.0853 | 6.01268 | 6.9844 | 6.27061 | 6.21007 | 6.55822 | 6.55467 | 6.81546 | 6.7655 | 6.6189 | 6.41542 | 6.62753 | 5.52897 | 5.39575 | 5.4947 | 5.66182 |
| MAP3K11 | Q16584 | 5.9642 | 6.38026 | 5.93351 | 6.8461 | 6.14603 | 5.15778 | 5.92682 | 6.33192 | 6.19337 | 6.70805 | 5.96254 | 5.80606 | 7.12067 | 6.56062 | 6.83451 | 6.12757 |
| RSBN1 | Q5VWQ0 | 6.42235 | 6.40574 | 6.09912 | 6.63479 | 5.89298 | 6.02616 | 6.26691 | 6.04028 | 6.0101 | 5.91523 | 6.16726 | 5.97254 | 6.02118 | 6.81915 | 6.97448 | 6.33172 |
| INO80D | Q53TQ3 | 6.30988 | 6.72496 | 5.96305 | 6.90695 | 6.34466 | 6.31144 | 6.48573 | 6.09374 | 6.19649 | 7.17738 | 5.99468 | 6.31137 | 5.59619 | 5.88956 | 5.77669 | 5.91723 |
| UBE2T | Q9NPD8 | 6.30171 | 5.87518 | 6.87311 | 5.79915 | 6.28819 | 6.97305 | 6.29969 | 6.90489 | 6.4484 | 7.19755 | 7.15388 | 6.98551 | 5.19348 | 5.31759 | 5.11944 | 5.26919 |
| DAD1 | P61803 | 6.87482 | 6.43526 | 6.07861 | 6.16647 | 6.11647 | 6.07641 | 6.10932 | 5.97975 | 5.52915 | 5.64847 | 5.70731 | 5.94165 | 6.66538 | 7.66207 | 6.13089 | 6.87799 |
| SERPINB8 | P50452 | 7.92794 | 6.76821 | 7.14513 | 5.77427 | 4.77536 | 7.45715 | 6.77539 | 6.57534 | 7.47321 | 6.51924 | 6.54012 | 6.33864 | 5.57161 | 4.85457 | 4.27911 | 5.22472 |
| ITPKB | P27987 | 6.31617 | 6.54316 | 6.23135 | 5.63754 | 6.02207 | 6.47057 | 5.98181 | 5.81003 | 7.31347 | 6.88582 | 6.14715 | 6.77549 | 6.24053 | 5.77928 | 5.76967 | 6.0759 |
| SNTB1 | Q13884 | 6.59342 | 6.2868 | 5.64767 | 6.60849 | 6.58074 | 5.67374 | 5.6624 | 5.97286 | 5.98696 | 6.02608 | 5.94073 | 5.8764 | 6.85566 | 6.42705 | 6.28408 | 6.67692 |
| TLK2 | Q86UE8-2 | 5.2617 | 6.06436 | 6.2154 | 5.84523 | 5.68783 | 6.10242 | 5.97761 | 5.84443 | 6.10984 | 6.42138 | 6.217 | 6.17575 | 7.03808 | 7.10854 | 7.10783 | 6.82259 |
| SLC20A1 | Q8WUM9 | 5.40797 | 5.75683 | 5.42828 | 6.13435 | 5.47596 | 5.41254 | 6.06626 | 5.76845 | 6.07154 | 5.578 | 5.86237 | 5.47838 | 7.91867 | 7.62108 | 8.27081 | 7.75211 |
| OSBPL2 | Q9H1P3 | 6.07316 | 5.72833 | 6.5661 | 5.88016 | 6.34025 | 5.84746 | 6.38963 | 5.75607 | 6.68596 | 6.6131 | 7.2207 | 6.5901 | 6.38574 | 6.27327 | 5.88176 | 5.76819 |
| METTL25 | Q8N6Q8 | 6.60514 | 6.37973 | 7.31999 | 6.64575 | 6.20746 | 6.57488 | 6.20342 | 6.6052 | 6.53401 | 6.59212 | 6.66883 | 6.35071 | 5.43485 | 5.17193 | 5.24995 | 5.45602 |
| ERI1 | Q8IV48 | 5.73835 | 6.59475 | 6.69474 | 5.30471 | 6.4161 | 6.16559 | 6.52022 | 6.59931 | 6.73252 | 6.52744 | 6.44633 | 6.25205 | 6.10165 | 6.04338 | 6.13792 | 6.16494 |
| SSH1 | Q8WYL5 | 6.30729 | 6.56426 | 6.59965 | 5.98462 | 6.55832 | 5.77971 | 6.11812 | 5.90826 | 6.26858 | 5.91797 | 5.28192 | 6.27157 | 7.03721 | 6.49178 | 6.24318 | 6.66757 |
| FMC1 | Q96HJ9 | 6.14894 | 6.41387 | 5.89938 | 6.25562 | 6.39237 | 5.92151 | 6.48867 | 5.74806 | 6.10807 | 6.17527 | 6.26179 | 5.70981 | 6.61319 | 6.20232 | 6.83348 | 6.82767 |
| ITPK1 | Q13572 | 6.53952 | 6.17152 | 7.25125 | 6.15968 | 5.85328 | 6.74329 | 6.4388 | 6.33237 | 6.68383 | 6.57871 | 7.29757 | 6.30702 | 5.25453 | 5.3325 | 5.70602 | 5.35013 |
| RGS19 | P49795 | 5.88854 | 6.18734 | 6.31876 | 6.74243 | 6.1657 | 6.11931 | 6.29497 | 6.09885 | 5.84802 | 6.12436 | 6.3243 | 6.23114 | 6.16745 | 6.4539 | 6.59591 | 6.43901 |
| SH2B3 | Q9UQQ2 | 6.47951 | 5.86744 | 6.42366 | 5.71518 | 5.99806 | 6.79422 | 6.2856 | 6.71984 | 6.02128 | 6.01214 | 6.07102 | 6.44066 | 6.23075 | 6.37451 | 6.49805 | 6.06809 |
| RBL2 | Q08999 | 5.83671 | 5.81933 | 6.18759 | 5.52258 | 5.81946 | 6.31665 | 6.24814 | 5.91131 | 5.85769 | 6.26833 | 6.7094 | 6.38228 | 6.29767 | 7.314 | 7.02793 | 6.48092 |
| POLK | Q9UBT6 | 5.31364 | 6.30352 | 6.28795 | 5.38431 | 5.58997 | 6.41442 | 5.35444 | 6.28652 | 6.8706 | 5.95396 | 6.69537 | 6.50026 | 6.55173 | 6.30907 | 7.09499 | 7.08924 |
| POC5 | Q8NA72 | 6.58302 | 5.87272 | 7.30648 | 6.40802 | 6.92139 | 7.15586 | 6.7926 | 6.43606 | 7.14071 | 5.48164 | 6.46455 | 5.83241 | 5.44202 | 4.99795 | 5.23866 | 5.92591 |
| PRKCH | P24723 | 6.77774 | 7.12462 | 7.33017 | 7.79366 | 6.00222 | 6.22434 | 6.5329 | 5.99797 | 6.60605 | 6.8284 | 6.93238 | 6.77604 | 4.53951 | 5.00311 | 4.98042 | 4.65046 |
| ELP6 | Q0PNE2 | 6.22218 | 6.17295 | 6.8214 | 6.00843 | 6.44421 | 6.71004 | 6.26444 | 6.51099 | 6.88992 | 6.72442 | 6.37199 | 6.37133 | 5.52111 | 5.7551 | 5.55241 | 5.65907 |
| LATS2 | Q9NRM7 | 6.35518 | 5.94988 | 7.03156 | 6.94863 | 6.07922 | 6.58039 | 6.03477 | 6.36327 | 6.5872 | 6.50552 | 6.54078 | 6.89444 | 5.39497 | 5.85172 | 5.40108 | 5.4814 |
| VPS25 | Q9BRG1 | 6.79524 | 5.09812 | 6.42809 | 4.96978 | 5.79249 | 6.36069 | 6.51142 | 6.34535 | 6.37061 | 6.85527 | 6.39752 | 7.04587 | 5.93569 | 6.54443 | 5.97403 | 6.57538 |
| RCN3 | Q96D15 | 8.25357 | 7.24979 | 6.66508 | 7.07225 | 6.46946 | 6.17157 | 6.31575 | 7.2871 | 6.94203 | 7.29524 | 6.71528 | 7.39286 | 4.05877 | 4.15587 | 3.5781 | 4.37728 |
| FKBP7 | Q9Y680 | 6.72036 | 6.12633 | 5.94839 | 6.30614 | 6.46156 | 5.49927 | 6.13875 | 5.73538 | 6.44968 | 5.90089 | 6.49898 | 6.29684 | 6.93359 | 6.44081 | 6.47929 | 6.06374 |
| HIRIP3 | Q9BW71 | 5.10034 | 5.20525 | 6.18464 | 4.82207 | 5.65507 | 6.38424 | 5.63148 | 5.95247 | 6.53312 | 6.06644 | 6.32165 | 6.5742 | 7.58005 | 7.36913 | 7.38035 | 7.23949 |
| APRT | P07741 | 6.02547 | 5.47094 | 6.53448 | 4.77144 | 6.2562 | 6.37908 | 5.55512 | 7.05749 | 5.87474 | 6.76733 | 6.14544 | 7.40165 | 5.97769 | 6.85766 | 6.3235 | 6.60176 |
| SLC41A3 | Q96GZ6-9 | 6.26991 | 6.40361 | 6.03032 | 6.72842 | 6.051 | 5.55556 | 6.38538 | 6.29431 | 5.90805 | 6.17842 | 5.87541 | 5.81885 | 6.63901 | 6.38075 | 6.7347 | 7.11632 |
| ACSS2 | Q9NR19-2 | 6.14395 | 5.46087 | 6.97949 | 5.67519 | 6.42317 | 6.54811 | 5.62423 | 6.31165 | 6.2846 | 6.59348 | 6.43184 | 6.45288 | 5.64825 | 7.02039 | 6.64804 | 5.75386 |
| PRPF40B | Q6NWY9 | 6.31826 | 6.57585 | 5.99602 | 6.08636 | 6.45583 | 6.28108 | 6.24164 | 5.89863 | 6.37812 | 5.56699 | 6.06138 | 5.58295 | 7.05382 | 6.36679 | 6.81137 | 6.32492 |
| MAFG | O15525 | 6.58605 | 6.77994 | 5.56468 | 6.5722 | 6.36434 | 6.39312 | 6.57368 | 6.4244 | 5.95532 | 6.10336 | 5.46108 | 6.06539 | 6.232 | 6.30718 | 6.27203 | 6.34522 |
| JUNB | P17275 | 6.46029 | 6.2313 | 6.07133 | 6.72246 | 6.4407 | 5.93901 | 5.89841 | 6.49131 | 6.44203 | 5.25872 | 6.17323 | 5.29386 | 6.87863 | 6.49721 | 7.01264 | 6.18888 |
| ACTR3B | Q9P1U1 | 6.15619 | 6.21795 | 6.88238 | 6.01443 | 6.2459 | 6.15255 | 6.13592 | 6.67015 | 5.99318 | 6.24596 | 6.72542 | 6.78866 | 5.73005 | 6.22521 | 5.77757 | 6.03848 |
| LPGAT1 | Q92604 | 6.54586 | 6.56836 | 5.94819 | 6.78161 | 6.38842 | 5.73122 | 6.28701 | 6.03608 | 5.88874 | 6.08172 | 5.90157 | 5.98468 | 6.60643 | 6.20093 | 6.28612 | 6.76305 |
| EPN2 | O95208 | 6.46431 | 6.16753 | 6.24022 | 6.29676 | 6.46025 | 6.08295 | 6.48557 | 6.4003 | 6.06605 | 6.13268 | 6.45369 | 6.37598 | 6.73956 | 5.9872 | 6.0046 | 5.64235 |
| UBP1 | Q9NZI7 | 6.52237 | 5.7147 | 5.74537 | 5.82694 | 6.77069 | 6.81342 | 6.17179 | 6.28245 | 6.03605 | 5.72156 | 6.71299 | 6.20826 | 6.45045 | 6.08274 | 6.97372 | 5.96649 |
| MYEF2 | A0A0A0MQW0 | 6.16366 | 6.4131 | 5.88827 | 6.73438 | 6.04882 | 6.45635 | 6.25869 | 6.26746 | 6.09188 | 5.92638 | 5.93439 | 5.917 | 6.5741 | 6.17516 | 6.7111 | 6.43926 |
| FAAP100 | Q0VG06 | 5.80481 | 5.93915 | 6.09791 | 6.2618 | 5.91593 | 6.16097 | 5.96417 | 6.4116 | 6.02063 | 6.18948 | 6.45502 | 6.21884 | 6.63889 | 6.8832 | 6.26731 | 6.77029 |
| ZNF768 | Q9H5H4 | 6.21942 | 6.32414 | 5.66371 | 6.15866 | 6.08738 | 6.30418 | 6.59387 | 5.75299 | 6.21291 | 6.24489 | 6.25575 | 5.76251 | 6.79795 | 6.52035 | 6.64013 | 6.46115 |
| ZNF595 | Q8IYB9 | 6.40778 | 6.45661 | 6.63225 | 6.76919 | 6.31594 | 6.28687 | 6.7941 | 5.7508 | 6.20453 | 6.17508 | 6.46715 | 6.43549 | 5.75172 | 5.5593 | 5.91356 | 6.07963 |
| ZNF667 | Q5HYK9 | 5.87662 | 6.23445 | 5.2116 | 6.44465 | 5.92725 | 6.42746 | 6.06907 | 6.0354 | 5.61375 | 6.31334 | 5.41537 | 5.75737 | 7.18904 | 7.53926 | 7.23865 | 6.70675 |
| ZNF567 | Q8N184 | 6.43197 | 6.56487 | 5.69691 | 6.68306 | 6.04699 | 6.04739 | 6.18276 | 6.48467 | 5.57457 | 5.98444 | 5.74727 | 5.86742 | 6.47461 | 6.79799 | 7.06303 | 6.35205 |
| ZNF146 | Q15072 | 5.53398 | 5.45998 | 5.78329 | 5.87173 | 5.9008 | 6.12764 | 6.1711 | 5.50733 | 6.10535 | 6.00733 | 5.89412 | 5.83191 | 7.75719 | 7.18678 | 7.48217 | 7.37929 |
| ZNF569 | Q5MCW4 | 5.94031 | 6.44317 | 5.86662 | 7.24589 | 5.86824 | 6.35776 | 6.21883 | 6.62867 | 6.07797 | 6.89839 | 5.85235 | 6.40011 | 5.56968 | 6.52897 | 6.46555 | 6.37723 |
| ZNF316 | A6NFI3 | 5.47067 | 6.51382 | 6.60981 | 6.64692 | 6.81765 | 6.15617 | 5.89422 | 5.90956 | 6.81997 | 5.47924 | 6.56414 | 6.48328 | 6.03438 | 6.56797 | 6.69428 | 6.26101 |
| XYLB | O75191 | 6.43238 | 5.7705 | 6.8431 | 5.47024 | 6.78152 | 6.49127 | 5.99678 | 6.84189 | 6.21275 | 6.32049 | 6.99498 | 7.12975 | 5.74016 | 5.3513 | 6.236 | 5.38687 |
| RRAGD | Q9NQL2 | 6.15281 | 7.48041 | 6.11261 | 6.01075 | 6.30519 | 6.39852 | 5.85189 | 6.10337 | 6.33718 | 7.09278 | 6.91121 | 6.20579 | 6.48201 | 6.19726 | 5.69124 | 5.66698 |
| MPG | P29372 | 5.89851 | 6.49835 | 6.73033 | 6.38476 | 5.96494 | 5.89568 | 6.03603 | 6.06371 | 6.47006 | 6.5563 | 6.84683 | 6.70911 | 5.67792 | 6.27516 | 5.94251 | 6.04981 |
| CFAP97 | Q9P2B7 | 5.42337 | 6.29596 | 8.39558 | 6.66689 | 6.54246 | 6.47992 | 6.19142 | 7.16351 | 6.47298 | 6.85642 | 6.77022 | 7.09067 | 4.93141 | 4.98959 | 5.57199 | 5.23759 |
| DENND4B | O75064 | 6.12392 | 5.45632 | 6.20388 | 6.29096 | 5.67597 | 6.12026 | 6.4487 | 5.83138 | 5.96574 | 6.88481 | 6.22429 | 6.29297 | 6.45336 | 6.4032 | 6.93795 | 6.68629 |
| NDUFAF6 | Q330K2 | 6.64507 | 6.55252 | 6.31335 | 6.97446 | 6.13821 | 5.43856 | 6.28194 | 5.97963 | 6.48038 | 5.93975 | 6.02932 | 5.73524 | 6.56295 | 5.87862 | 6.79958 | 6.25041 |
| GOLGB1 | Q14789 | 5.79171 | 6.30046 | 6.42361 | 6.47548 | 5.94979 | 5.74506 | 6.3908 | 6.15858 | 6.22139 | 6.26603 | 5.43953 | 6.34768 | 6.70528 | 6.32821 | 6.78083 | 6.67558 |
| DOHH | Q9BU89 | 6.25793 | 4.25098 | 5.68506 | 4.98607 | 5.66919 | 4.83549 | 5.95584 | 6.80029 | 5.05694 | 5.54954 | 5.17638 | 7.34219 | 5.79088 | 9.37998 | 7.18823 | 10.075 |
| CDCA7 | Q9BWT1 | 6.41607 | 7.24964 | 7.16468 | 8.41757 | 6.3059 | 6.4391 | 7.27511 | 6.39602 | 7.07239 | 7.15178 | 6.93479 | 6.4894 | 4.39299 | 4.26486 | 3.72748 | 4.30221 |
| ZNF655 | Q8N720-3 | 6.17887 | 6.42433 | 5.95467 | 7.01076 | 6.36695 | 6.00975 | 6.41375 | 6.23219 | 6.29832 | 6.2 | 6.15673 | 6.29168 | 6.43859 | 5.84436 | 6.2092 | 5.96985 |
| RBBP8 | Q99708-2 | 5.82429 | 6.21639 | 6.11337 | 6.58464 | 6.23861 | 5.67667 | 6.30262 | 6.13976 | 6.13541 | 5.6763 | 6.76202 | 6.51072 | 6.45027 | 6.40381 | 6.46238 | 6.50275 |
| C2CD2L | O14523-2 | 6.30397 | 5.99728 | 5.84565 | 6.40526 | 5.92312 | 6.07883 | 6.36294 | 6.1095 | 5.92389 | 6.1542 | 6.36628 | 6.1319 | 7.14744 | 6.35104 | 6.54452 | 6.35419 |
| TCF3 | P15923-3 | 5.98342 | 5.95026 | 6.9348 | 5.72068 | 5.78098 | 6.22822 | 5.87113 | 6.47912 | 6.41476 | 6.66094 | 7.76278 | 7.35666 | 5.72583 | 5.8066 | 5.81295 | 5.51087 |
| OPLAH | O14841 | 5.92352 | 6.02652 | 6.29812 | 5.35558 | 5.91616 | 6.90707 | 6.53328 | 6.36201 | 5.95655 | 6.12669 | 6.60191 | 6.62499 | 6.08817 | 6.66105 | 6.46627 | 6.15211 |
| TCTN3 | Q6NUS6 | 7.1299 | 7.21413 | 6.42851 | 7.0806 | 7.02185 | 6.0613 | 6.61211 | 6.25781 | 6.14393 | 6.85737 | 6.50362 | 6.84388 | 4.8834 | 4.68829 | 4.92622 | 5.34708 |
| POLR2D | O15514 | 6.55698 | 5.60406 | 6.5262 | 5.95653 | 6.05848 | 5.83506 | 6.32972 | 6.22976 | 5.67827 | 6.76813 | 6.63346 | 7.00913 | 5.26025 | 6.47276 | 6.48702 | 6.59417 |
| TAF10 | Q12962 | 6.14815 | 5.29366 | 6.06715 | 5.11753 | 6.22173 | 4.78061 | 5.0495 | 6.7105 | 5.08829 | 6.86248 | 6.2679 | 7.06998 | 6.14707 | 7.93908 | 7.43743 | 7.79894 |
| NPM3 | O75607 | 6.43348 | 6.49552 | 6.78374 | 6.59023 | 6.3047 | 6.27163 | 5.94211 | 6.37586 | 6.20273 | 5.76129 | 7.06988 | 6.54468 | 6.2752 | 5.57184 | 5.46577 | 5.91135 |
| RRAS2 | P62070-4 | 6.3963 | 6.40716 | 6.51059 | 6.451 | 5.88912 | 5.86704 | 5.83937 | 6.16237 | 6.14432 | 6.41877 | 6.14689 | 6.33973 | 6.86053 | 5.77588 | 6.59181 | 6.19912 |
| METTL17 | Q9H7H0-3 | 6.60362 | 5.95522 | 6.49449 | 6.62904 | 6.54006 | 6.20007 | 6.3549 | 5.7453 | 6.28724 | 6.27095 | 6.24816 | 5.98302 | 6.05582 | 5.71508 | 6.68598 | 6.23104 |
| SATB1 | Q01826-2 | 6.41714 | 6.78105 | 6.18842 | 6.95092 | 6.12474 | 5.76502 | 6.57162 | 5.90003 | 6.48178 | 5.64729 | 5.45648 | 6.42233 | 7.06698 | 5.6268 | 5.8342 | 5.76519 |
| ARL6IP1 | Q15041 | 6.39321 | 6.29773 | 5.64532 | 6.25492 | 6.49959 | 6.59068 | 6.3839 | 5.66526 | 5.7921 | 6.45975 | 5.86384 | 5.42929 | 6.54955 | 6.98207 | 6.42557 | 6.7672 |
| BET1 | O15155 | 6.03733 | 5.84952 | 6.27962 | 6.58818 | 5.96001 | 5.86889 | 6.09262 | 5.66613 | 6.19973 | 5.85892 | 6.08712 | 6.27154 | 6.72605 | 6.7564 | 7.13279 | 6.62515 |
| TIMM9 | Q9Y5J7 | 6.54952 | 5.46421 | 5.77606 | 5.58361 | 4.62252 | 4.49238 | 5.72669 | 5.41243 | 4.82915 | 5.5817 | 4.8684 | 5.27464 | 4.67461 | 10.703 | 8.70906 | 11.7321 |
| EXOSC6 | Q5RKV6 | 6.54936 | 6.2034 | 6.25546 | 6.09328 | 6.57604 | 6.0701 | 6.39993 | 6.26276 | 6.03754 | 5.87019 | 6.22685 | 6.33768 | 6.97343 | 5.58179 | 6.20778 | 6.35439 |
| PDZD11 | Q5EBL8-2 | 5.68465 | 5.02301 | 6.45495 | 5.1453 | 6.20869 | 5.64423 | 5.996 | 7.38027 | 5.6049 | 6.90848 | 6.47708 | 7.04426 | 6.04961 | 7.03089 | 6.45984 | 6.88782 |
| FRMD8 | Q9BZ67 | 5.92335 | 6.81228 | 5.4158 | 6.51686 | 6.08578 | 6.34124 | 6.18831 | 5.65521 | 6.42592 | 6.54027 | 5.70367 | 6.30878 | 6.13602 | 6.5895 | 6.69742 | 6.65959 |

| | | | | | | | | | | | | | | | | | |
|---|---|---|---|---|---|---|---|---|---|---|---|---|---|---|---|---|---|
| NIPA2 | Q8N8Q9 | 6.91913 | 5.91734 | 6.70485 | 6.26136 | 6.21398 | 5.71487 | 5.89937 | 6.80409 | 6.339 | 5.83126 | 5.44267 | 5.98713 | 6.07268 | 5.64911 | 6.66111 | 7.58206 |
| BCOR | Q6W2J9 | 7.29764 | 7.00336 | 6.8688 | 7.14818 | 6.02778 | 6.37641 | 6.60058 | 5.87888 | 6.54827 | 6.08898 | 6.64963 | 6.26705 | 5.63827 | 4.92331 | 5.16359 | 5.51925 |
| RPP25L | Q8N5L8 | 5.9155 | 6.13702 | 6.38213 | 6.34043 | 5.98029 | 5.63499 | 6.05525 | 5.91472 | 6.78621 | 6.63152 | 6.88369 | 6.7939 | 6.68539 | 6.17904 | 5.51813 | 6.16179 |
| THAP11 | Q96EK4 | 6.71553 | 6.59372 | 6.00433 | 6.88537 | 6.46451 | 6.26236 | 6.26472 | 6.24159 | 6.35494 | 6.63234 | 5.90041 | 6.1559 | 5.99779 | 5.77451 | 5.85121 | 5.90077 |
| CLCN5 | P51795 | 6.28439 | 6.22586 | 7.50129 | 6.38774 | 5.90287 | 7.1247 | 6.68898 | 6.16614 | 6.33205 | 6.37196 | 6.79445 | 5.74051 | 5.90904 | 5.28062 | 5.73856 | 5.55085 |
| UBE2R2 | Q712K3 | 6.23062 | 6.54241 | 7.02422 | 5.90465 | 7.12796 | 6.41995 | 6.18134 | 7.2436 | 6.84577 | 6.5569 | 6.75044 | 6.57724 | 5.57356 | 5.13469 | 5.03373 | 4.85291 |
| CCDC12 | Q8WUD4 | 6.4826 | 6.22074 | 5.90652 | 6.28413 | 6.1713 | 6.19256 | 5.97968 | 6.19146 | 6.21044 | 6.19005 | 6.36292 | 5.97092 | 6.94869 | 6.24867 | 6.2595 | 6.37982 |
| AGO1 | Q9UL18 | 5.77537 | 6.38565 | 6.30342 | 6.0549 | 6.70992 | 5.94366 | 6.18409 | 5.73126 | 6.52913 | 6.28953 | 6.49701 | 6.64001 | 5.72303 | 6.13885 | 6.62604 | 6.46813 |
| SPRED1 | Q7Z699 | 5.90884 | 6.60526 | 6.34234 | 6.48835 | 5.59328 | 6.45726 | 6.75237 | 6.09948 | 6.19405 | 6.52226 | 5.7945 | 6.23642 | 6.50358 | 6.05949 | 5.83301 | 6.60949 |
| KAT8 | Q9H7Z6-2 | 6.46772 | 6.60515 | 5.91794 | 6.80741 | 6.41152 | 6.17524 | 6.23267 | 5.9911 | 6.29001 | 6.00986 | 6.10579 | 6.44205 | 6.24509 | 5.79815 | 6.38078 | 6.11952 |
| OTULIN | Q96BN8 | 5.96111 | 5.8741 | 7.1041 | 6.12812 | 6.1535 | 6.9747 | 6.37398 | 6.88734 | 6.41719 | 6.28267 | 6.18462 | 6.48993 | 5.95664 | 5.50045 | 6.06892 | 5.64263 |
| CORO1A | P31146 | 6.46773 | 6.61623 | 6.28771 | 6.14298 | 6.07971 | 6.23325 | 6.36642 | 6.36984 | 5.93043 | 6.72526 | 6.42091 | 6.57229 | 6.02902 | 6.36033 | 5.75752 | 5.64037 |
| APPBP2 | Q92624 | 6.26872 | 7.10831 | 6.86083 | 6.51198 | 6.24471 | 6.87828 | 6.44793 | 6.22559 | 6.30785 | 6.58605 | 6.87791 | 7.55971 | 5.38809 | 4.80003 | 4.97272 | 4.9613 |
| LPIN2 | Q92539 | 6.54175 | 6.73887 | 6.78604 | 6.28916 | 6.43973 | 6.88796 | 6.14579 | 6.64543 | 6.73857 | 6.72098 | 7.03676 | 7.20117 | 4.96406 | 5.28862 | 4.86551 | 4.70959 |
| AUH | Q13825 | 6.21476 | 6.33331 | 6.53651 | 6.11566 | 6.29354 | 6.076 | 6.56117 | 5.86329 | 5.57474 | 6.09803 | 6.42716 | 5.66898 | 7.13691 | 6.53644 | 6.60308 | 5.96041 |
| DNAJC18 | Q9H819 | 7.39333 | 7.14629 | 7.3454 | 8.47483 | 6.90899 | 6.85175 | 6.85389 | 6.64534 | 6.74456 | 7.22021 | 7.0433 | 6.93723 | 3.55127 | 3.661 | 3.27467 | 3.94795 |
| ATE1 | O95260-2 | 5.77698 | 6.00623 | 6.78195 | 5.72142 | 5.92906 | 6.44756 | 6.44847 | 6.1363 | 6.21365 | 6.3616 | 6.15661 | 6.3137 | 6.40433 | 6.46715 | 6.4976 | 6.33739 |
| AKT3 | Q9Y243 | 6.01966 | 6.31758 | 6.81528 | 6.43618 | 5.66561 | 6.72604 | 5.96607 | 6.64387 | 6.28505 | 6.40768 | 6.74054 | 6.54707 | 5.33493 | 6.10172 | 5.91853 | 6.07419 |
| SMAD3 | P84022 | 6.69203 | 6.22305 | 6.41859 | 5.83733 | 6.40999 | 6.44041 | 6.0515 | 6.47506 | 5.71593 | 6.49476 | 6.07136 | 6.3661 | 6.16507 | 6.99346 | 6.0199 | 5.62545 |
| OIP5 | O43482 | 6.6738 | 6.55188 | 6.82677 | 6.67975 | 6.34782 | 6.4103 | 6.30221 | 6.39335 | 6.68262 | 6.8214 | 6.65696 | 6.80846 | 5.11394 | 4.95367 | 5.37658 | 5.40047 |
| PDK1 | Q15118 | 6.476 | 6.72229 | 5.85012 | 6.39209 | 6.77749 | 6.00418 | 6.48932 | 6.39023 | 5.6518 | 5.50916 | 5.96662 | 5.72854 | 6.37848 | 6.72602 | 6.69329 | 6.24436 |
| DBI | P07108-5 | 5.60097 | 5.43027 | 5.87867 | 5.08938 | 5.95004 | 5.19839 | 4.86254 | 6.44054 | 4.40574 | 5.30356 | 4.97358 | 4.62607 | 4.36912 | 13.8858 | 7.34022 | 10.6451 |
| APBB1 | O00213 | 6.13871 | 6.1763 | 6.23643 | 5.98043 | 6.86649 | 5.89502 | 6.20775 | 6.1541 | 6.21965 | 6.70106 | 6.83684 | 6.05496 | 6.5567 | 5.71455 | 6.04526 | 6.21575 |
| RAP2B | P61225 | 5.91164 | 6.38508 | 5.98269 | 6.2816 | 5.81709 | 6.18036 | 6.11079 | 5.94644 | 6.15195 | 6.05162 | 6.07575 | 6.25616 | 6.5754 | 6.67041 | 6.77074 | 6.83227 |
| RAP2C | Q9Y3L5 | 6.37435 | 6.73184 | 6.66972 | 6.87498 | 6.56688 | 6.55524 | 6.17237 | 6.25236 | 5.1833 | 6.2222 | 6.04232 | 5.95169 | 6.23065 | 6.64781 | 5.86473 | 5.65957 |
| KLC2 | Q9H0B6 | 6.12927 | 6.32431 | 5.85265 | 5.82082 | 6.00591 | 6.55724 | 6.68114 | 6.59094 | 7.47915 | 6.27853 | 6.42379 | 6.58134 | 6.46663 | 5.57864 | 5.46006 | 5.7696 |
| SLC30A7 | Q8NEW0 | 5.7109 | 6.2219 | 5.83979 | 5.82958 | 6.63963 | 5.68272 | 6.00409 | 6.59908 | 5.90459 | 6.29559 | 6.0403 | 6.38913 | 6.90545 | 6.71546 | 7.34292 | 6.82008 |
| EXOSC4 | Q9NPD3 | 6.27503 | 6.17979 | 6.18155 | 6.5138 | 6.59825 | 6.01483 | 6.05408 | 5.94675 | 6.47343 | 6.42443 | 5.92676 | 6.09152 | 6.64725 | 5.89053 | 6.60069 | 6.18131 |
| HUS1 | O60921 | 5.51153 | 5.69236 | 6.19919 | 5.7068 | 5.59708 | 5.84278 | 5.53755 | 5.57296 | 5.95983 | 5.99773 | 6.53744 | 6.0985 | 7.12963 | 7.57134 | 7.73281 | 7.31247 |
| FCHSD1 | Q86WN1 | 6.26605 | 6.21057 | 6.98498 | 5.93061 | 6.08471 | 6.65783 | 6.02178 | 6.37095 | 6.63775 | 6.37865 | 6.45571 | 6.59449 | 5.93162 | 5.76859 | 5.80454 | 5.90117 |
| ING3 | Q9NXR8 | 6.21506 | 6.03891 | 6.18613 | 6.99945 | 6.4407 | 5.94646 | 6.44105 | 6.62021 | 6.07019 | 6.40463 | 6.66541 | 5.75915 | 6.1834 | 5.97879 | 6.05352 | 5.99694 |
| ZFAND2B | Q8WV99 | 6.39804 | 6.3961 | 6.5227 | 6.39049 | 6.04019 | 6.24452 | 5.93013 | 6.48961 | 6.81399 | 6.37254 | 6.21237 | 7.01731 | 6.22796 | 5.54499 | 5.33423 | 6.06482 |
| PNMA1 | Q8ND90 | 6.79105 | 5.62184 | 6.07794 | 5.35829 | 5.2668 | 5.37547 | 6.0561 | 6.98777 | 5.9007 | 7.3047 | 6.37207 | 8.18046 | 4.62117 | 6.96246 | 6.17593 | 6.94724 |
| ZHX2 | Q9Y6X8 | 6.40005 | 6.37423 | 5.9125 | 6.79223 | 5.64957 | 6.10539 | 6.49048 | 5.76178 | 5.95167 | 6.20348 | 5.80146 | 6.27255 | 6.70081 | 6.17532 | 6.44074 | 6.96775 |
| HSD17B7 | P56937 | 6.98367 | 6.6219 | 6.99015 | 6.48229 | 6.59483 | 6.17212 | 6.0019 | 5.18671 | 6.17141 | 5.84218 | 6.01253 | 5.68981 | 6.64323 | 6.59566 | 6.67591 | 5.96569 |
| MTMR12 | Q9C0I1 | 6.48819 | 6.21581 | 6.3866 | 6.23076 | 6.49307 | 6.28315 | 5.86889 | 6.404 | 6.73492 | 6.5682 | 6.07349 | 6.86788 | 6.15885 | 5.58175 | 5.54678 | 6.09765 |
| TMEM199 | Q8N511 | 6.81582 | 6.84406 | 5.36226 | 6.74988 | 6.48894 | 6.68304 | 5.8759 | 6.23733 | 6.46183 | 5.6889 | 6.13824 | 5.72913 | 6.47599 | 5.76341 | 5.7256 | 6.95967 |
| DTWD1 | Q8N5C7 | 5.83713 | 6.45415 | 7.27295 | 6.06279 | 5.86179 | 6.46528 | 6.14544 | 6.42958 | 6.49603 | 5.94781 | 6.79861 | 7.35984 | 5.3685 | 5.8679 | 5.81683 | 5.81537 |
| TRMT10A | Q8TBZ6 | 5.57094 | 5.76903 | 6.048 | 5.46258 | 5.35467 | 6.1876 | 5.71334 | 6.23657 | 5.74065 | 6.03232 | 6.51477 | 6.65797 | 6.50874 | 7.06224 | 7.67666 | 7.46392 |
| TBCEL | Q5QJ74 | 5.74595 | 5.96142 | 6.23271 | 5.76066 | 6.60722 | 6.42981 | 5.95241 | 7.12041 | 6.43259 | 6.68154 | 7.03038 | 7.09701 | 6.19703 | 5.82123 | 5.60961 | 5.32002 |
| TMEM132A | Q24JP5-2 | 6.56988 | 6.82981 | 6.32827 | 6.73908 | 6.27483 | 6.32471 | 6.60687 | 6.41274 | 6.8275 | 6.45519 | 6.51024 | 6.42845 | 5.46318 | 5.49543 | 5.44331 | 5.29052 |
| RTL8A | Q9BWD3 | 5.75298 | 4.56474 | 5.8609 | 4.37437 | 5.74147 | 6.26329 | 5.65982 | 6.88984 | 5.55987 | 5.53734 | 6.21036 | 7.19236 | 6.52566 | 8.53922 | 7.19307 | 8.13471 |
| CENPT | Q96BT3 | 6.46712 | 6.44846 | 5.6917 | 6.92511 | 6.08142 | 5.53769 | 6.44927 | 5.87295 | 6.19739 | 6.35299 | 6.2134 | 6.34715 | 6.81004 | 6.01921 | 6.23273 | 6.35335 |
| AP5M1 | Q9H0R1 | 6.9032 | 6.71031 | 7.13699 | 6.55222 | 6.2172 | 6.02211 | 5.59101 | 6.29718 | 6.34894 | 6.67832 | 6.82326 | 6.42165 | 5.73725 | 5.61818 | 5.43144 | 5.51075 |
| CCDC51 | Q96ER9 | 6.20412 | 5.8695 | 6.0642 | 6.7883 | 5.69551 | 6.44311 | 6.46324 | 5.71419 | 6.14545 | 6.27268 | 6.30462 | 6.37704 | 6.66164 | 5.86543 | 6.22027 | 6.91068 |
| TTC39C | Q8N584 | 6.31783 | 6.03072 | 7.19928 | 6.69123 | 6.6222 | 6.06818 | 6.35613 | 6.37077 | 6.58226 | 5.9301 | 6.83323 | 6.82509 | 5.70311 | 5.60609 | 5.10744 | 5.75635 |
| AMBRA1 | Q9C0C7-5 | 5.53149 | 5.69371 | 6.27223 | 5.92581 | 6.01611 | 6.52863 | 6.3434 | 6.19133 | 6.19757 | 6.42624 | 6.44456 | 6.43502 | 6.55066 | 6.18353 | 6.8094 | 6.45032 |
| PRTFDC1 | Q9NRG1 | 6.34691 | 6.23136 | 6.86692 | 6.56291 | 6.47864 | 6.68625 | 6.2342 | 6.95618 | 6.47325 | 6.1849 | 6.4317 | 6.20966 | 6.21912 | 5.68243 | 5.89351 | 5.54207 |
| PPT2 | Q9UMR5-3 | 6.55346 | 6.81949 | 6.11205 | 5.85343 | 6.8164 | 6.20953 | 6.16041 | 6.35487 | 5.85569 | 6.07786 | 6.16991 | 5.79241 | 6.79485 | 6.23864 | 6.1911 | 5.99991 |
| FANCG | O15287 | 6.73498 | 6.41115 | 7.04711 | 6.45156 | 6.15384 | 6.04398 | 6.19873 | 6.90919 | 6.27025 | 7.14242 | 6.86312 | 7.24303 | 5.21072 | 5.00141 | 4.85762 | 5.4609 |
| LIPT1 | Q9Y234 | 6.4648 | 6.56865 | 6.03045 | 7.43639 | 5.93195 | 5.60346 | 5.80079 | 5.5129 | 6.57528 | 6.51459 | 6.35294 | 6.14645 | 6.38233 | 5.91317 | 6.41429 | 6.35157 |
| SCRN3 | Q0VDG4 | 6.01551 | 6.08824 | 6.23027 | 5.96691 | 6.17782 | 6.7317 | 5.99246 | 6.64326 | 6.19924 | 6.95068 | 6.79282 | 7.16375 | 5.2671 | 6.089 | 5.83346 | 5.85776 |
| PROSER3 | Q2NL68 | 6.80184 | 6.85225 | 6.75099 | 5.16387 | 6.73157 | 6.67363 | 6.27458 | 6.68794 | 6.32637 | 6.35087 | 7.34446 | 7.33298 | 5.57459 | 5.10169 | 5.13397 | 4.89839 |
| ADO | Q96SZ5 | 6.68935 | 5.77996 | 6.70069 | 6.04689 | 6.07667 | 6.10087 | 5.75836 | 6.48868 | 6.62729 | 6.59944 | 6.17585 | 6.84579 | 6.23653 | 6.22786 | 5.91249 | 5.7333 |
| RHBDD1 | Q8TEB9 | 6.33931 | 5.44469 | 6.08545 | 6.28448 | 5.91879 | 6.19173 | 6.08734 | 5.73747 | 5.91354 | 6.64898 | 5.97497 | 5.642 | 6.91859 | 6.92623 | 6.37828 | 7.50815 |
| ASDURF | L0R819 | 6.42601 | 5.73423 | 6.62128 | 5.23585 | 6.46 | 6.15186 | 5.89581 | 6.79713 | 6.32382 | 6.9425 | 7.165 | 7.04195 | 5.94605 | 5.82467 | 5.5682 | 5.86564 |
| GTPBP3 | Q969Y2-2 | 6.35698 | 6.37907 | 6.11296 | 6.63962 | 6.2798 | 6.35915 | 5.91327 | 6.00054 | 6.35065 | 5.9208 | 6.09242 | 6.40152 | 6.72139 | 5.91354 | 6.40354 | 6.15474 |
| PRMT3 | O60678 | 6.26698 | 6.51578 | 6.83899 | 6.25506 | 6.23125 | 6.51967 | 6.32264 | 6.79023 | 6.34131 | 6.35497 | 6.69319 | 6.93172 | 5.55124 | 5.45316 | 5.48312 | 5.6507 |
| CEPT1 | Q9Y6K0 | 6.25128 | 6.33823 | 6.15166 | 6.5827 | 5.88732 | 6.15792 | 6.32937 | 5.72739 | 6.17209 | 6.26037 | 5.85722 | 6.58757 | 6.62174 | 6.34507 | 6.94682 | 6.68326 |
| HBS1L | Q9Y450-2 | 6.3182 | 6.13039 | 6.9598 | 6.14735 | 5.87751 | 6.31648 | 5.94574 | 6.13529 | 6.88888 | 5.88553 | 6.37938 | 6.67551 | 6.62083 | 5.86929 | 5.87228 | 5.97753 |
| CDT1 | Q9H211 | 6.84125 | 6.98126 | 6.93741 | 7.12871 | 6.33373 | 6.23614 | 6.74612 | 7.09492 | 7.09557 | 7.43444 | 7.96845 | 7.35712 | 4.17463 | 4.1673 | 3.44687 | 4.05609 |
| IFIT3 | O14879 | 6.97643 | 6.71752 | 6.26254 | 6.01572 | 7.05401 | 6.76777 | 6.15895 | 6.92714 | 6.50902 | 6.38654 | 6.45879 | 6.24948 | 5.58917 | 5.79395 | 5.08989 | 5.04307 |
| GCA | P28676 | 6.60267 | 5.55799 | 6.48862 | 4.9067 | 6.46308 | 6.31702 | 5.66169 | 6.81478 | 5.7827 | 6.85431 | 6.86357 | 6.9974 | 6.09025 | 6.38354 | 6.07871 | 6.13695 |
| TRNAU1AP | Q9NX07 | 5.48319 | 5.71206 | 5.84827 | 5.55264 | 6.24479 | 6.49686 | 5.90207 | 6.60234 | 6.25428 | 6.6953 | 6.67342 | 6.45317 | 6.22234 | 6.78061 | 6.34223 | 6.73644 |
| GNA11 | P29992 | 6.51609 | 6.90626 | 5.90429 | 6.44935 | 6.44318 | 6.39136 | 6.31618 | 6.442 | 6.38313 | 6.24334 | 6.32848 | 6.07553 | 5.52743 | 5.63217 | 6.3449 | 6.09631 |
| GATD1 | Q8NB37 | 5.98218 | 5.77118 | 6.71065 | 5.39843 | 6.38495 | 6.5394 | 5.88729 | 6.54129 | 6.48606 | 6.75268 | 6.89636 | 6.88876 | 5.99916 | 5.8958 | 5.92354 | 5.94226 |
| KDM4D | Q6B0I6 | 6.09562 | 7.17042 | 5.83907 | 6.98573 | 6.17214 | 5.99871 | 6.52307 | 6.20118 | 6.44114 | 6.60884 | 6.32474 | 5.83039 | 6.50337 | 6.04436 | 6.3038 | 5.85742 |
| ACYP1 | P07311 | 6.09053 | 5.8963 | 6.10371 | 5.03183 | 7.07982 | 6.58344 | 6.13439 | 7.68073 | 6.23587 | 6.79762 | 7.08609 | 6.28671 | 5.96071 | 5.78721 | 5.80549 | 5.43956 |
| SPRED2 | Q7Z698 | 6.74643 | 6.77412 | 5.41834 | 6.42524 | 6.70551 | 6.49274 | 5.93526 | 6.2277 | 6.11836 | 5.77057 | 7.10769 | 5.90074 | 6.26113 | 5.80677 | 6.6076 | 5.70179 |
| TSR3 | Q9UJK0 | 7.18593 | 6.79782 | 6.81856 | 6.71338 | 5.88008 | 6.38444 | 6.6396 | 6.39204 | 7.54018 | 7.00947 | 6.93365 | 6.73864 | 4.65011 | 4.97731 | 4.55199 | 4.78678 |
| DPY19L3 | Q6ZPD9 | 6.21984 | 6.57727 | 5.59995 | 6.90937 | 6.05343 | 5.91807 | 6.50539 | 5.65682 | 5.99882 | 6.23023 | 5.97629 | 5.78541 | 6.73093 | 6.14877 | 6.95518 | 6.73424 |
| MAPK6 | Q16659 | 5.33351 | 5.1986 | 5.8428 | 5.09139 | 6.51621 | 6.85893 | 6.50948 | 7.31063 | 5.28468 | 5.28895 | 5.87847 | 5.70859 | 7.65222 | 7.30833 | 6.77401 | 7.44321 |
| PCGF3 | Q3KNV8-2 | 6.38025 | 6.48016 | 6.47045 | 6.43337 | 5.93863 | 5.89468 | 6.24771 | 5.75027 | 5.96623 | 6.16474 | 6.64329 | 6.01563 | 6.54279 | 5.99489 | 6.7501 | 6.3268 |
| ZNF579 | Q8NAF0 | 5.87951 | 6.58469 | 5.82309 | 6.39393 | 6.34774 | 5.84891 | 5.89775 | 6.481 | 5.90711 | 6.7542 | 6.27616 | 5.86985 | 6.5541 | 6.39209 | 6.53562 | 6.45425 |
| PLA2G15 | Q8NCC3 | 6.66208 | 6.47265 | 5.90804 | 6.2502 | 6.50674 | 5.93391 | 6.30854 | 6.04493 | 6.40508 | 6.19254 | 5.99759 | 5.79382 | 6.75789 | 6.21816 | 6.32442 | 6.22341 |
| SAMD8 | A0A5F9ZGS6 | 6.6048 | 6.78308 | 6.55948 | 7.23944 | 6.62011 | 5.95623 | 6.41546 | 4.9417 | 6.13336 | 6.03934 | 5.86435 | 5.72537 | 6.26915 | 5.80969 | 6.78165 | 6.25676 |
| CGNL1 | Q0VF96 | 6.16628 | 6.33125 | 6.31981 | 6.27926 | 6.22194 | 6.24481 | 6.71746 | 6.40594 | 6.31982 | 6.45781 | 6.11621 | 6.2406 | 6.16797 | 5.74472 | 5.90323 | 6.36289 |
| MRPL32 | Q9BYC8 | 6.76681 | 6.42022 | 6.06048 | 6.60809 | 5.85586 | 5.93549 | 6.26128 | 5.07215 | 6.13648 | 6.35891 | 6.28041 | 6.43947 | 6.4988 | 6.2797 | 6.56572 | 6.46014 |
| SPIN2B | Q9BPZ2 | 6.13051 | 6.71253 | 5.66274 | 7.21885 | 6.12673 | 6.02844 | 6.17082 | 6.20219 | 6.78109 | 6.51371 | 6.11375 | 6.35819 | 5.64037 | 5.87385 | 6.3231 | 6.1431 |
| HLA-B | P01889 | 6.10198 | 6.73702 | 6.20636 | 5.93118 | 6.73724 | 6.24078 | 6.4721 | 6.58684 | 5.9627 | 5.94424 | 5.97332 | 5.95409 | 6.24288 | 6.59489 | 5.6962 | 6.61817 |
| HLA-C | A0A140T9M0 | 5.93108 | 6.7991 | 6.17022 | 6.30995 | 6.29294 | 6.94824 | 6.35147 | 6.39634 | 5.84942 | 6.33341 | 6.52285 | 6.90002 | 5.28219 | 6.29865 | 5.48548 | 6.12866 |
| GLCE | O94923 | 5.98072 | 6.30169 | 6.14991 | 6.05737 | 6.11058 | 6.05861 | 6.68978 | 5.85232 | 7.16986 | 5.94298 | 6.11892 | 5.75682 | 7.211 | 5.79567 | 6.36492 | 6.43887 |
| TAF6L | Q9Y6J9 | 6.14857 | 6.38446 | 6.36624 | 6.34408 | 5.82514 | 6.1631 | 6.0935 | 6.38274 | 6.20705 | 6.6542 | 5.87979 | 6.183 | 6.77347 | 6.03986 | 5.97545 | 6.57936 |
| MOB3A | Q96BX8 | 6.57419 | 6.85868 | 6.89652 | 6.22529 | 6.3026 | 6.94471 | 6.23709 | 6.92168 | 6.27138 | 6.62242 | 6.40337 | 6.62843 | 5.25042 | 5.30626 | 5.27354 | 5.28343 |
| NUDCD2 | Q8WVJ2 | 6.06023 | 5.5808 | 6.23203 | 4.95795 | 6.796 | 6.74098 | 5.66567 | 7.42241 | 6.18943 | 6.87468 | 6.56249 | 6.92357 | 5.72618 | 6.73283 | 5.35937 | 6.17537 |
| DONSON | Q9NYP3 | 6.12427 | 6.31422 | 6.67461 | 6.38739 | 5.8241 | 6.20522 | 5.88394 | 5.53322 | 6.74745 | 6.23478 | 6.92913 | 6.44772 | 6.14148 | 6.07151 | 6.09241 | 6.38857 |
| APOL2 | Q9BQE5 | 6.76959 | 6.51418 | 6.22043 | 7.20415 | 6.52564 | 5.91353 | 7.28354 | 5.68311 | 5.84688 | 5.94228 | 6.38114 | 7.36811 | 5.71684 | 6.04707 | 5.20896 | 5.37454 |
| ZNF787 | Q6DD87 | 5.74342 | 6.31898 | 5.96457 | 5.71423 | 5.86752 | 6.46877 | 6.22216 | 6.34497 | 5.90409 | 6.13188 | 6.69386 | 6.21406 | 6.70879 | 6.65017 | 6.49946 | 6.55306 |
| MRPS18C | Q9Y3D5 | 6.1605 | 6.13781 | 5.73118 | 6.31712 | 6.39884 | 5.80137 | 6.09485 | 6.43723 | 5.8275 | 6.84371 | 6.43862 | 6.63305 | 6.61758 | 6.2885 | 6.10271 | 6.16944 |
| BIN1 | O00499 | 6.85389 | 6.08165 | 7.00238 | 6.50124 | 5.82642 | 6.22556 | 6.2816 | 6.69158 | 5.63087 | 6.90769 | 6.56624 | 6.93004 | 6.00208 | 5.65455 | 5.30309 | 5.54114 |
| CBFB | Q13951 | 6.46516 | 6.00541 | 6.78449 | 5.67996 | 6.47132 | 6.85446 | 6.27584 | 6.59541 | 6.99161 | 6.44221 | 6.27866 | 6.27787 | 5.79411 | 5.72823 | 5.73338 | 5.62279 |
| AMOTL1 | Q8IY63 | 6.47007 | 6.34816 | 5.4818 | 6.0343 | 6.67296 | 6.10542 | 5.79328 | 6.40695 | 6.47998 | 6.44496 | 6.4778 | 6.21663 | 6.94971 | 5.9983 | 5.81696 | 6.30273 |
| DTNA | Q9Y4J8 | 6.32948 | 6.27228 | 6.57744 | 6.179 | 5.92893 | 6.15785 | 6.46525 | 5.98362 | 5.83812 | 6.11597 | 6.20027 | 6.26702 | 6.33482 | 6.94805 | 6.50434 | 5.89757 |
| DTNB | O60941 | 6.14375 | 6.42664 | 6.33769 | 6.63153 | 6.41873 | 5.70272 | 6.16955 | 6.26075 | 7.02447 | 5.85553 | 5.94274 | 5.93366 | 6.21813 | 6.07214 | 6.71979 | 6.14217 |
| MRTFA | Q969V6 | 6.26699 | 5.89487 | 6.17638 | 5.74961 | 5.99467 | 6.49019 | 5.62268 | 5.80204 | 6.53244 | 6.60767 | 6.93111 | 7.03382 | 6.17978 | 6.18926 | 6.04818 | 6.48031 |
| UBE2J1 | Q9Y385 | 6.38695 | 6.40959 | 5.92315 | 6.93348 | 6.57156 | 6.25409 | 6.67548 | 6.48975 | 6.25299 | 5.86048 | 5.90724 | 6.34047 | 6.23973 | 5.55919 | 5.96887 | 6.22698 |
| ATP11B | Q9Y2G3 | 6.04853 | 6.24897 | 6.36753 | 6.6163 | 5.84985 | 6.17685 | 6.45136 | 6.22142 | 6.10604 | 6.67168 | 6.7384 | 6.39856 | 5.97699 | 5.87372 | 6.37186 | 5.88194 |
| BET1L | Q9NYM9 | 6.1973 | 5.15526 | 5.28298 | 5.41906 | 5.98197 | 5.61834 | 5.56485 | 6.06579 | 5.33104 | 5.57862 | 5.61601 | 6.95963 | 6.31665 | 8.3809 | 7.86842 | 8.66318 |
| TRIP12 | Q14669-2 | 6.95234 | 6.25697 | 6.18426 | 7.24813 | 6.60655 | 5.34122 | 5.46784 | 7.06872 | 6.48698 | 5.49628 | 6.18718 | 6.4259 | 6.27193 | 5.83719 | 6.63734 | 5.53118 |
| SCAMP4 | Q969E2 | 6.05024 | 5.79006 | 6.61153 | 5.93825 | 5.58874 | 6.09754 | 6.40451 | 5.66581 | 6.70535 | 6.37331 | 6.26201 | 6.83081 | 5.87554 | 6.4602 | 6.53121 | 6.81489 |
| MLLT10 | P55197 | 6.10991 | 6.91755 | 6.08039 | 7.26993 | 5.7374 | 5.9724 | 5.85386 | 6.00694 | 5.88265 | 6.00523 | 5.80724 | 6.17625 | 5.49105 | 6.87136 | 6.6125 | 7.20535 |
| PSMB8 | P28062 | 6.08669 | 6.16413 | 6.29768 | 6.04641 | 6.52788 | 6.43688 | 6.14216 | 6.20034 | 6.06923 | 6.42127 | 6.38351 | 5.91553 | 6.70764 | 5.93863 | 6.54864 | 6.11337 |
| TBC1D31 | Q96DN5 | 6.89469 | 6.40295 | 6.6258 | 6.31807 | 6.01575 | 6.48839 | 6.30682 | 6.18498 | 7.39941 | 6.77152 | 5.91346 | 7.03001 | 5.14763 | 5.54464 | 5.13815 | 5.81772 |
| KANSL2 | Q9H9L4 | 5.81512 | 6.21147 | 5.86455 | 6.82 | 6.00303 | 6.20104 | 6.77337 | 6.27718 | 6.21877 | 5.96697 | 6.17974 | 6.00151 | 6.20674 | 6.62378 | 6.42024 | 6.41649 |
| CLTB | P09497-2 | 6.51367 | 5.12899 | 6.23755 | 5.37801 | 5.81109 | 4.48103 | 4.85932 | 6.5077 | 5.10274 | 6.64401 | 6.0817 | 7.23666 | 6.00525 | 8.03188 | 7.81478 | 8.16563 |

| | | | | | | | | | | | | | | | |
|---|---|---|---|---|---|---|---|---|---|---|---|---|---|---|---|
| TUBGCP5 | Q96RT8 | 6.40165 | 6.79337 | 7.24893 | 6.72652 | 6.30135 | 6.74327 | 6.03088 | 6.80866 | 6.61636 | 6.82037 | 6.71729 | 6.69825 | 5.0889 | 5.18463 | 4.61478 | 5.20479 |
| CEP112 | Q8N8E3 | 6.1471 | 6.31725 | 6.77597 | 5.67261 | 6.51551 | 6.74059 | 6.26367 | 6.42211 | 6.59291 | 6.5352 | 6.06963 | 6.44709 | 6.12371 | 5.66818 | 6.08773 | 5.82073 |
| CENPN | Q96H22-3 | 6.73866 | 6.83323 | 6.43776 | 6.46158 | 5.36967 | 6.51526 | 6.07986 | 5.85386 | 6.23315 | 6.14353 | 6.53068 | 6.76834 | 6.23858 | 5.84347 | 5.85347 | 6.0989 |
| IFT88 | Q13099-1 | 6.59203 | 6.6618 | 6.63522 | 6.05797 | 6.4524 | 6.48375 | 6.13436 | 6.79763 | 6.56301 | 6.60556 | 6.71547 | 6.48023 | 5.48342 | 5.55621 | 5.30469 | 5.47626 |
| SLC45A4 | Q5BKX6-2 | 5.95089 | 6.55154 | 6.2427 | 6.83458 | 6.01096 | 5.77906 | 5.97113 | 5.80617 | 5.91167 | 6.55453 | 6.34137 | 6.84171 | 6.07451 | 6.22257 | 6.2796 | 6.62702 |
| MIER3 | Q7Z3K6-2 | 5.89664 | 6.11752 | 6.3187 | 6.13745 | 6.17835 | 6.23131 | 5.82156 | 6.01468 | 6.15539 | 6.10314 | 6.7538 | 6.38066 | 6.47485 | 6.70683 | 6.30296 | 6.40616 |
| HMGXB4 | Q9UGU5 | 5.25272 | 5.57504 | 4.67663 | 4.98053 | 5.28915 | 5.56595 | 5.12008 | 5.2585 | 4.83546 | 5.41012 | 4.6183 | 4.88558 | 8.83836 | 10.1971 | 9.36416 | 10.1323 |
| PKMYT1 | Q99640 | 6.96285 | 7.28043 | 6.44014 | 7.08194 | 6.35748 | 6.59223 | 6.42059 | 6.56644 | 6.35002 | 5.98977 | 5.99217 | 6.07263 | 5.51172 | 5.30432 | 5.61175 | 5.4655 |
| TRIT1 | Q9H3H1 | 5.97351 | 6.6287 | 5.533 | 5.87533 | 7.53622 | 6.74103 | 5.67904 | 6.55811 | 5.96847 | 5.8304 | 5.76642 | 5.87229 | 6.25541 | 6.5395 | 6.16398 | 7.07858 |
| PODXL | O00592 | 6.38622 | 6.39186 | 6.18387 | 6.34724 | 6.39318 | 6.34594 | 6.19147 | 6.03976 | 6.26817 | 6.29815 | 6.22225 | 6.15882 | 6.3712 | 5.99946 | 6.12079 | 6.28159 |
| KIAA1109 | Q2LD37 | 7.00879 | 6.70244 | 6.65497 | 6.81459 | 5.49168 | 6.11389 | 6.2131 | 5.70681 | 6.01585 | 6.51171 | 6.45191 | 6.29115 | 6.13702 | 5.96827 | 5.65023 | 6.2676 |
| CYP27C1 | Q4G0S4 | 6.14303 | 6.3091 | 5.67286 | 5.91684 | 5.97557 | 5.88384 | 6.59967 | 5.85182 | 6.40317 | 6.24619 | 6.22524 | 6.07041 | 6.61663 | 6.77616 | 6.86414 | 6.44534 |
| PATJ | Q8NI35 | 6.38574 | 6.70879 | 5.94228 | 6.00797 | 6.65118 | 6.35812 | 6.07174 | 6.36796 | 6.24422 | 6.30427 | 5.89356 | 6.23926 | 6.44508 | 6.31936 | 5.93266 | 6.1278 |
| CHMP2B | Q9UQN3 | 6.77062 | 6.38825 | 6.33729 | 5.79807 | 6.33166 | 6.30413 | 6.46681 | 6.96477 | 6.55155 | 6.74807 | 6.52525 | 7.03362 | 5.1352 | 5.67712 | 5.30433 | 5.66325 |
| PANK1 | Q8TE04 | 6.37085 | 6.2454 | 6.75072 | 6.16101 | 6.01313 | 6.51533 | 5.96259 | 6.87126 | 6.56491 | 6.4048 | 6.69095 | 6.46182 | 5.55812 | 5.84554 | 6.09961 | 5.48396 |
| STAT5B | P51692 | 6.17153 | 6.2881 | 6.64864 | 5.92011 | 6.29609 | 6.36383 | 6.38762 | 6.83251 | 6.33003 | 5.9192 | 6.18231 | 6.09932 | 6.31254 | 6.11562 | 6.19502 | 5.93752 |
| FAM117B | Q6P1L5 | 6.26294 | 6.27583 | 6.86223 | 5.94884 | 6.20146 | 6.54374 | 6.21786 | 6.26503 | 6.44663 | 6.5973 | 6.56558 | 6.65838 | 5.85415 | 5.78073 | 5.58838 | 5.93092 |
| KIAA0586 | Q9BVV6-3 | 6.08029 | 6.79422 | 6.40259 | 6.37661 | 6.43586 | 6.78697 | 6.43772 | 7.34031 | 7.00517 | 6.69287 | 6.17255 | 7.07506 | 4.99893 | 5.15805 | 4.88183 | 5.36098 |
| CTH | P32929 | 5.94352 | 5.65496 | 5.8366 | 5.05885 | 6.09353 | 5.97374 | 6.72791 | 7.09067 | 6.80466 | 6.80186 | 6.61433 | 6.57603 | 6.38165 | 6.18231 | 6.84209 | 5.41731 |
| TFAP2A | P05549 | 6.30112 | 6.62925 | 6.32294 | 6.8137 | 6.35592 | 6.37632 | 6.19503 | 5.67029 | 6.64112 | 6.61151 | 6.42851 | 6.29956 | 5.62303 | 5.88194 | 5.60425 | 6.24551 |
| KAT2B | Q92831 | 5.75474 | 6.42441 | 5.55016 | 6.44958 | 6.81209 | 6.47612 | 6.04636 | 6.55095 | 6.47485 | 6.20692 | 6.67059 | 6.26437 | 6.34097 | 6.41658 | 6.30653 | 5.25477 |
| NTM | Q9P121-4 | 6.29812 | 6.27569 | 5.98149 | 6.05141 | 6.49616 | 6.29557 | 6.02758 | 6.24279 | 6.18283 | 6.00786 | 5.81987 | 5.88094 | 6.77237 | 6.48133 | 6.72243 | 6.46356 |
| PUSL1 | Q8N0Z8 | 6.61203 | 6.28157 | 6.39398 | 6.99392 | 6.06716 | 5.72553 | 6.22856 | 5.75415 | 6.37522 | 6.53062 | 6.30474 | 6.94176 | 6.45369 | 5.69251 | 6.12994 | 6.41462 |
| U2AF1 | Q01081 | 6.19773 | 6.41607 | 5.76794 | 6.8497 | 6.35148 | 5.79804 | 5.63482 | 5.59176 | 6.20556 | 5.80896 | 5.67934 | 5.73128 | 7.01561 | 6.72904 | 7.07256 | 7.1501 |
| LRRCC1 | Q9C099 | 6.43068 | 6.58108 | 6.50188 | 6.596 | 6.01989 | 6.45046 | 6.54867 | 6.52971 | 6.09391 | 6.66911 | 6.70478 | 7.51231 | 5.08582 | 5.53686 | 5.32797 | 5.41087 |
| KLHL11 | Q9NVR0 | 6.04944 | 6.27746 | 6.67936 | 6.77353 | 6.20611 | 6.44357 | 5.93582 | 6.33257 | 6.23103 | 6.66137 | 6.87039 | 6.74541 | 6.08713 | 5.5107 | 5.63796 | 5.55814 |
| MRS2 | Q9HD23 | 6.59234 | 6.30444 | 5.89771 | 6.41279 | 6.37346 | 6.05265 | 6.65333 | 5.33892 | 6.2121 | 6.26155 | 6.52398 | 5.6174 | 6.12399 | 6.11652 | 6.86134 | 6.6575 |
| S100A16 | Q96FQ6 | 6.09028 | 6.98978 | 5.34168 | 5.79355 | 6.04222 | 4.99648 | 5.32972 | 6.38913 | 5.06107 | 6.41703 | 5.73859 | 6.68993 | 6.02722 | 7.7681 | 7.70801 | 7.61722 |
| IREB2 | P48200 | 7.15267 | 7.1113 | 7.6061 | 6.95664 | 6.03095 | 7.11273 | 6.14092 | 7.17421 | 6.96422 | 7.41 | 7.74292 | 7.34497 | 3.38022 | 4.63477 | 3.97581 | 3.26157 |
| N4BP1 | O75113 | 6.01682 | 5.88987 | 6.6751 | 5.91272 | 5.7035 | 5.73129 | 6.07393 | 6.1585 | 6.49401 | 6.86142 | 6.82296 | 6.51136 | 5.9567 | 6.85651 | 6.37794 | 5.95737 |
| PPTC7 | Q8NI37 | 6.57001 | 6.872 | 6.29839 | 6.73161 | 6.29923 | 5.48563 | 6.3692 | 5.92793 | 6.32797 | 5.86942 | 5.61561 | 6.67267 | 6.62377 | 5.83809 | 6.1599 | 6.33856 |
| DNA2 | P51530 | 6.81449 | 6.84065 | 8.01261 | 6.80395 | 6.04952 | 7.04292 | 6.00247 | 6.59917 | 7.6074 | 7.80184 | 7.97877 | 7.40217 | 3.31919 | 3.92059 | 3.58608 | 4.21817 |
| POLR3F | Q9H1D9 | 5.8185 | 5.93753 | 6.049 | 5.9001 | 5.80165 | 5.7698 | 5.53704 | 6.09445 | 6.20789 | 6.48516 | 6.46904 | 6.4004 | 6.30164 | 6.72096 | 6.94512 | 5.5617 |
| DDR1 | Q08345-5 | 6.40979 | 6.87903 | 6.69286 | 6.5543 | 6.18218 | 6.40696 | 6.2805 | 6.63191 | 6.25925 | 6.65403 | 6.20105 | 6.46278 | 5.32872 | 5.83406 | 5.50977 | 5.71282 |
| NSL1 | Q96IY1 | 6.15315 | 6.30265 | 6.46346 | 6.43305 | 6.33782 | 5.69236 | 6.33042 | 5.88584 | 6.27456 | 5.98752 | 6.40879 | 6.26301 | 7.10164 | 6.0925 | 6.05895 | 6.21428 |
| ANKH | Q9HCJ1 | 6.30222 | 6.41368 | 6.13169 | 6.75486 | 6.45881 | 5.92113 | 6.04317 | 5.9328 | 6.43924 | 6.20781 | 5.9719 | 6.31911 | 6.03985 | 6.14206 | 6.76157 | 6.16009 |
| STARD8 | Q92502-2 | 6.9593 | 7.18503 | 6.27839 | 7.21141 | 6.91098 | 6.92814 | 6.69231 | 6.76145 | 7.06078 | 7.35198 | 6.99725 | 7.55113 | 3.58052 | 4.13597 | 3.90419 | 4.49117 |
| GALNT14 | Q96FL9-3 | 6.75024 | 6.40764 | 5.94122 | 6.98097 | 5.91225 | 5.63241 | 6.86417 | 6.15291 | 6.03723 | 5.91992 | 6.04844 | 6.38431 | 6.35751 | 5.88176 | 6.37266 | 5.35636 |
| TMUB1 | Q9BVT8 | 6.84546 | 6.45384 | 5.37573 | 6.37004 | 6.50404 | 5.5732 | 6.24417 | 6.21177 | 5.2933 | 4.98028 | 5.79588 | 5.48828 | 5.62651 | 8.32304 | 7.06409 | 7.85039 |
| PHLDB3 | Q6NSJ2 | 6.43585 | 6.0684 | 6.00557 | 5.97092 | 6.41537 | 6.35629 | 6.51622 | 6.33827 | 6.1018 | 6.62093 | 6.82472 | 6.45724 | 6.1472 | 5.61602 | 5.57225 | 6.55295 |
| PCMTD1 | Q96MG8 | 5.11561 | 5.95744 | 6.45821 | 5.67691 | 5.77569 | 6.23129 | 5.47 | 5.97594 | 6.08193 | 6.67656 | 6.65701 | 6.31372 | 6.88117 | 7.1252 | 6.86729 | 6.73603 |
| GEMIN6 | Q8WXD5 | 6.34654 | 6.28846 | 6.35365 | 5.48346 | 6.76645 | 5.99533 | 6.05796 | 6.84025 | 6.62835 | 6.47682 | 7.28761 | 6.99609 | 5.94261 | 6.23859 | 5.87143 | 5.42639 |
| NDUFA11 | Q86Y39 | 6.08904 | 6.42356 | 5.56098 | 6.03476 | 6.44426 | 6.47262 | 6.46697 | 6.42768 | 5.98442 | 6.02982 | 5.61629 | 5.70904 | 6.536 | 6.53642 | 6.79333 | 6.87482 |
| LRRC41 | Q15345 | 5.45858 | 5.90445 | 6.01751 | 5.685 | 5.76909 | 6.03857 | 5.92371 | 5.96924 | 6.41336 | 6.23806 | 6.74764 | 6.04893 | 7.02603 | 6.7773 | 7.09356 | 6.89001 |
| CMC4 | P56277 | 6.56481 | 5.91872 | 5.5477 | 5.78127 | 6.36284 | 5.40776 | 5.76894 | 5.75697 | 5.04743 | 5.87513 | 6.45288 | 5.97479 | 7.29287 | 7.20841 | 7.43843 | 7.60105 |
| IRAK2 | O43187 | 6.32463 | 6.20572 | 6.73032 | 6.14622 | 5.84028 | 6.46695 | 6.07008 | 6.35074 | 6.37886 | 5.71113 | 6.40882 | 6.60864 | 6.30633 | 6.34974 | 5.96707 | 6.13448 |
| DMAC2 | Q9NW81-4 | 6.71479 | 6.07091 | 5.95496 | 6.37655 | 6.79618 | 5.87454 | 6.33403 | 6.27177 | 6.52073 | 6.42365 | 6.02223 | 6.37745 | 6.73082 | 5.56185 | 6.29218 | 5.67735 |
| PHLDA1 | Q8WV24 | 5.76456 | 5.37431 | 5.96898 | 5.19238 | 6.25414 | 6.18808 | 6.33902 | 6.01976 | 5.95595 | 5.395 | 5.66545 | 5.79495 | 7.58103 | 6.82742 | 7.54378 | 8.13517 |
| ZFX | P17010-3 | 6.86579 | 6.55134 | 6.53922 | 7.12987 | 6.12222 | 5.86537 | 6.44274 | 4.98961 | 6.29137 | 6.04422 | 6.5474 | 6.55341 | 6.24224 | 5.96434 | 5.26626 | 6.58459 |
| MAZ | P56270-2 | 5.79609 | 5.90473 | 6.3772 | 6.75566 | 5.86545 | 5.95925 | 5.9836 | 5.66039 | 6.14162 | 6.45777 | 6.48249 | 6.83469 | 7.04231 | 6.32633 | 5.96758 | 6.44484 |
| VEZF1 | Q14119 | 6.63767 | 5.76285 | 6.11475 | 6.78892 | 6.36366 | 6.09058 | 6.0805 | 5.73263 | 6.1639 | 6.18721 | 6.10824 | 6.14583 | 6.86866 | 6.10463 | 6.39523 | 6.45475 |
| MAP3K20 | Q9NYL2 | 5.95893 | 6.28294 | 6.25172 | 5.90328 | 6.2718 | 6.85137 | 6.06289 | 6.79641 | 6.57468 | 6.97059 | 6.22258 | 6.83982 | 5.62449 | 5.7703 | 5.72855 | 5.88965 |
| ZFAND3 | Q9H8U3 | 6.25694 | 6.90926 | 7.77366 | 6.85064 | 6.64058 | 6.68809 | 6.42667 | 6.96488 | 5.91397 | 6.73788 | 6.42611 | 6.2352 | 4.94532 | 5.29605 | 4.93364 | 5.00112 |
| RABGAP1L | Q5R372-3 | 6.42517 | 5.94384 | 7.12257 | 6.08759 | 6.31155 | 6.62224 | 5.98483 | 6.63143 | 6.55412 | 6.28958 | 6.09398 | 6.18475 | 6.26289 | 6.41529 | 5.40087 | 5.66928 |
| STEEP1 | Q9H5V9 | 5.59905 | 9.14431 | 5.89236 | 6.13843 | 5.56894 | 5.57634 | 5.85612 | 7.22774 | 6.18358 | 6.06076 | 6.29668 | 5.55183 | 5.87862 | 5.94565 | 7.5318 | 5.54779 |
| LTF | P02788 | 5.29057 | 6.68759 | 5.42045 | 6.17004 | 5.39907 | 7.18298 | 6.47102 | 5.55273 | 6.504 | 5.57381 | 6.37341 | 5.83798 | 5.01067 | 8.89804 | 7.1828 | 6.44484 |
| PRAF2 | O60831 | 6.37552 | 6.30582 | 6.32587 | 6.15391 | 6.33898 | 5.92985 | 6.51779 | 6.45898 | 6.17675 | 5.67839 | 6.22753 | 5.66816 | 7.08262 | 6.29428 | 6.31075 | 6.15479 |
| IFT27 | Q9BW83 | 5.99357 | 5.80426 | 6.30944 | 5.04371 | 6.49462 | 6.932 | 5.64437 | 7.05197 | 6.96358 | 7.38619 | 7.12333 | 6.94761 | 5.5988 | 5.66852 | 5.48728 | 5.55076 |
| SLC49A4 | Q96SL1 | 6.27789 | 6.60339 | 6.02583 | 6.91465 | 6.10182 | 6.22706 | 6.19524 | 6.03328 | 6.26884 | 6.6104 | 6.5136 | 6.15708 | 5.92528 | 5.84583 | 5.95607 | 6.34374 |
| TRIM58 | Q8NG06 | 6.59359 | 5.61337 | 7.96852 | 6.72754 | 6.17828 | 6.1189 | 5.76898 | 6.97906 | 6.84868 | 6.01865 | 6.88825 | 6.22138 | 5.20615 | 5.88174 | 5.69499 | 5.29193 |
| LYPLAL1 | Q5VWZ2 | 6.63611 | 6.11742 | 6.79964 | 6.17537 | 6.24697 | 6.4486 | 5.85055 | 6.386 | 6.07273 | 6.48821 | 5.91851 | 6.37093 | 5.96461 | 6.40423 | 6.2088 | 5.91133 |
| DCUN1D5 | Q9BTE7 | 5.56309 | 5.69541 | 6.31633 | 5.76869 | 5.32416 | 6.26605 | 5.3017 | 6.23954 | 5.84135 | 6.12663 | 6.46286 | 6.23325 | 6.57148 | 7.8352 | 7.05506 | 7.39919 |
| ERG28 | Q9UKR5 | 5.64136 | 5.03527 | 5.74688 | 5.94734 | 6.65292 | 5.9333 | 6.10188 | 5.30076 | 5.67668 | 6.51899 | 5.43822 | 5.9544 | 6.61582 | 7.62032 | 7.52466 | 8.28489 |
| ARL13B | Q3SXY8 | 6.34524 | 6.20458 | 6.82893 | 6.26052 | 6.55204 | 6.61807 | 6.41109 | 5.93908 | 6.38891 | 6.12147 | 5.90266 | 6.18486 | 5.6818 | 6.40802 | 6.07427 | 6.07848 |
| CHCHD7 | Q9BUK0-4 | 6.43888 | 6.97113 | 6.11744 | 7.03688 | 6.65571 | 5.71512 | 7.42637 | 5.51806 | 5.60987 | 6.18688 | 6.3503 | 6.41885 | 6.2807 | 6.11846 | 6.21142 | 4.94394 |
| GZF1 | Q9H116 | 6.18341 | 6.38842 | 6.51507 | 7.41371 | 5.89423 | 5.84261 | 6.4685 | 5.73721 | 6.06743 | 6.36874 | 6.37656 | 6.4057 | 5.93012 | 5.92502 | 6.00191 | 6.48136 |
| MTMR8 | Q96EF0 | 6.27498 | 6.4414 | 6.50552 | 6.54591 | 5.6796 | 6.33902 | 6.41147 | 6.42442 | 6.69085 | 6.6144 | 7.05651 | 6.70645 | 5.29335 | 5.70589 | 5.56826 | 5.74196 |
| KCMF1 | Q9P0J7 | 6.98313 | 5.83717 | 7.33749 | 6.25042 | 5.94183 | 6.41463 | 5.67457 | 6.17357 | 6.02569 | 6.44626 | 7.23371 | 7.40434 | 6.03672 | 5.80213 | 5.00589 | 5.43247 |
| MICALL2 | Q8IY33 | 6.39372 | 6.08358 | 6.12767 | 5.44926 | 5.96301 | 5.74792 | 5.9856 | 6.34679 | 6.5953 | 6.03128 | 6.22801 | 6.32595 | 7.21846 | 6.53409 | 6.42666 | 6.54267 |
| BCKDK | O14874 | 6.32877 | 6.91661 | 5.7881 | 7.08858 | 6.5377 | 6.06145 | 5.73518 | 6.14812 | 5.85739 | 5.56772 | 6.16156 | 5.7333 | 6.40046 | 6.89823 | 6.09027 | 6.68655 |
| OXSM | Q9NWU1 | 6.41361 | 8.73857 | 5.54753 | 6.69585 | 5.41017 | 5.40376 | 6.2575 | 6.7784 | 7.02338 | 6.09167 | 5.54821 | 5.98437 | 6.13005 | 5.80807 | 5.95784 | 5.29338 |
| UBL5 | Q9BZL1 | 6.61605 | 5.93626 | 6.93406 | 6.4845 | 6.48823 | 6.72248 | 6.87961 | 6.06244 | 6.73714 | 7.04431 | 7.20568 | 6.92098 | 5.65854 | 4.80798 | 4.42336 | 5.07836 |
| GRN | P28799 | 7.26583 | 6.48041 | 6.47598 | 7.15941 | 6.27772 | 5.91235 | 5.80197 | 5.70858 | 6.34971 | 6.38013 | 6.8576 | 6.39215 | 5.56703 | 5.68555 | 5.62773 | 6.05786 |
| ZNF219 | Q9P2Y4 | 6.08996 | 6.41939 | 6.33346 | 7.02621 | 6.59071 | 6.84572 | 5.73215 | 5.93557 | 6.18958 | 7.02907 | 5.90581 | 6.1851 | 5.4782 | 5.61417 | 6.62356 | 6.00135 |
| ANP32E | Q9BTT0 | 5.42036 | 5.2943 | 5.99262 | 5.18853 | 5.92505 | 6.37165 | 5.36148 | 5.82796 | 5.76993 | 6.22181 | 6.48008 | 6.22988 | 7.25093 | 7.5513 | 7.47366 | 7.64045 |
| CMC1 | Q7Z7K0 | 7.39661 | 6.8242 | 6.9343 | 6.30003 | 6.3466 | 4.96041 | 5.32801 | 5.30819 | 6.2275 | 6.05131 | 6.52056 | 6.2313 | 7.23123 | 6.16277 | 6.07766 | 6.09933 |
| ARHGEF5 | Q12774 | 6.42002 | 6.40973 | 6.77601 | 6.08456 | 7.42525 | 6.16732 | 5.50407 | 6.52137 | 6.44408 | 6.86538 | 6.52881 | 5.99593 | 6.34328 | 6.06433 | 5.3873 | 5.60239 |
| NDUFAF5 | Q5TEU4 | 6.22228 | 5.77784 | 5.9058 | 6.51403 | 5.68609 | 5.88586 | 6.70445 | 5.73954 | 6.43967 | 6.26958 | 6.62042 | 6.43334 | 6.68795 | 6.30568 | 6.34906 | 6.45842 |
| CARD10 | Q9BWT7 | 6.64874 | 6.72981 | 7.20273 | 6.96059 | 6.10818 | 6.54168 | 6.12568 | 7.00313 | 7.07745 | 6.8177 | 7.36159 | 7.23358 | 4.71876 | 4.47735 | 4.41037 | 4.58265 |
| DBF4 | Q9UBU7 | 6.53077 | 5.99627 | 6.23945 | 6.56144 | 5.95792 | 6.03785 | 5.687 | 5.61081 | 6.59116 | 5.92371 | 6.5243 | 6.44708 | 6.15159 | 6.32036 | 6.6338 | 6.7865 |
| BCL10 | O95999 | 6.60123 | 5.89552 | 6.95184 | 5.57042 | 6.55281 | 6.46962 | 5.6248 | 7.10896 | 6.19567 | 6.7846 | 6.91438 | 7.37689 | 5.39086 | 5.40861 | 5.19595 | 5.95782 |
| IL13RA1 | P78552 | 7.22798 | 6.78132 | 6.48375 | 6.99779 | 5.834 | 6.38353 | 6.68748 | 5.81561 | 7.16466 | 7.48129 | 7.76904 | 6.59571 | 5.07143 | 4.46482 | 4.86516 | 4.37642 |
| RSAD1 | Q9HA92 | 5.89343 | 6.47575 | 6.30767 | 7.08558 | 6.35689 | 6.38495 | 6.2533 | 6.31971 | 6.16161 | 5.93186 | 6.42778 | 5.9936 | 5.99752 | 5.9071 | 6.25719 | 6.24608 |
| ARL14EP | Q8N8R7 | 6.5483 | 6.28018 | 6.61885 | 6.56763 | 6.14229 | 6.29061 | 6.70851 | 5.81659 | 6.31892 | 6.68884 | 6.05912 | 6.21626 | 6.36744 | 5.7198 | 6.0468 | 5.60987 |
| SORT1 | Q99523 | 6.41817 | 6.66206 | 6.99795 | 6.91864 | 5.92543 | 6.25199 | 6.61882 | 6.22701 | 6.3079 | 7.02633 | 6.69392 | 6.74355 | 5.28851 | 5.06942 | 5.38157 | 5.46872 |
| SRRD | Q9UH36 | 6.06527 | 6.05465 | 7.40393 | 6.78002 | 6.10392 | 6.7256 | 6.00689 | 6.18025 | 6.0966 | 6.24373 | 6.71942 | 7.32429 | 5.72095 | 5.06039 | 5.36718 | 6.14691 |
| DGKD | Q16760 | 5.86999 | 6.00642 | 6.31345 | 6.56842 | 6.66695 | 5.93801 | 5.93204 | 6.10273 | 6.65624 | 6.46389 | 6.42364 | 6.37073 | 5.99351 | 6.05514 | 6.25407 | 6.38476 |
| TRABD | Q9H4I3 | 6.46469 | 6.82332 | 6.41601 | 7.23456 | 6.10833 | 5.78699 | 6.15758 | 5.94609 | 6.57547 | 6.15477 | 6.67916 | 6.26232 | 5.73779 | 5.56925 | 6.09322 | 5.99045 |
| WDR77 | Q9BQA1 | 5.72422 | 5.91757 | 6.31639 | 5.56246 | 5.96714 | 6.45772 | 5.6276 | 6.61723 | 6.45943 | 7.04579 | 6.55299 | 6.54764 | 6.40224 | 6.26697 | 5.84376 | 6.69084 |
| SNRNP35 | Q16560-2 | 6.03283 | 6.4218 | 6.05374 | 6.84808 | 6.15016 | 6.84821 | 6.62863 | 6.17372 | 6.20803 | 6.41002 | 6.23914 | 6.03093 | 5.9768 | 5.90809 | 6.05774 | 6.01207 |
| SRSF8 | Q9BRL6 | 6.14967 | 6.32368 | 5.5987 | 6.2557 | 5.99575 | 5.93081 | 6.12543 | 6.43258 | 6.34193 | 6.34364 | 6.23504 | 5.56301 | 6.94264 | 6.55384 | 6.61163 | 6.59595 |
| STK19 | P49842 | 6.34239 | 5.90136 | 6.07799 | 5.9819 | 5.66149 | 5.86942 | 6.283 | 5.95965 | 6.96641 | 5.91796 | 6.6278 | 7.35543 | 6.40135 | 6.33033 | 5.89332 | 6.43021 |
| BLZF1 | Q9H2G9 | 6.47552 | 7.04759 | 6.22874 | 6.87836 | 6.7773 | 7.14133 | 6.82318 | 6.59056 | 6.27446 | 6.25014 | 5.78566 | 7.00633 | 5.12621 | 5.02293 | 5.49388 | 5.0778 |
| MED25 | M0QZQ2 | 6.03276 | 6.26905 | 6.38083 | 6.21701 | 6.22174 | 5.7393 | 6.52685 | 6.33054 | 6.49879 | 5.95901 | 6.48999 | 5.98101 | 6.34503 | 6.41971 | 5.40962 | 6.17875 |
| NARS2 | Q96I59 | 6.37599 | 6.01475 | 5.79091 | 6.71348 | 6.3238 | 6.09691 | 6.29925 | 6.05886 | 6.33079 | 5.95313 | 6.00999 | 5.99289 | 6.3328 | 6.52037 | 6.65475 | 6.53134 |
| SLC29A1 | Q99808-2 | 6.02034 | 6.36199 | 5.91479 | 6.79312 | 5.53136 | 5.73155 | 6.05418 | 6.29385 | 6.66841 | 6.84927 | 5.83073 | 6.95829 | 5.99121 | 6.567 | 6.18607 | 6.24782 |
| MARF1 | Q9Y4F3 | 6.11125 | 6.4405 | 6.96123 | 6.09297 | 5.82592 | 6.9045 | 5.85588 | 6.55867 | 6.03144 | 6.61469 | 6.58673 | 6.6084 | 5.14936 | 6.81891 | 5.86629 | 5.57326 |
| TRAF7 | Q6Q0C0 | 6.41192 | 6.91532 | 6.72617 | 6.80709 | 6.54783 | 6.11175 | 6.4302 | 6.30572 | 6.57204 | 7.1548 | 6.85918 | 6.47054 | 5.31608 | 4.84278 | 5.1505 | 5.37808 |
| HMGN4 | O00479 | 5.64561 | 6.04396 | 5.609 | 7.09682 | 6.04704 | 5.75483 | 6.19492 | 5.34183 | 5.59948 | 5.57566 | 6.2114 | 6.44654 | 6.1022 | 7.38206 | 7.2979 | 7.65077 |
| DTX3 | Q8N9I9-2 | 6.30653 | 6.13634 | 6.5304 | 5.55446 | 6.17671 | 6.87208 | 6.24876 | 6.57069 | 6.46457 | 6.79158 | 6.56083 | 6.53586 | 5.88112 | 6.00961 | 5.71406 | 5.64642 |
| FZR1 | Q9UM11 | 6.73917 | 6.89105 | 7.18387 | 6.86529 | 6.01506 | 6.88958 | 6.60847 | 6.62652 | 7.12966 | 7.25161 | 7.64437 | 6.92913 | 4.03794 | 4.70132 | 4.14078 | 4.34619 |
| RTF2 | Q9BY42 | 6.27694 | 6.30582 | 6.97383 | 5.83415 | 6.19306 | 6.57048 | 6.43597 | 6.46795 | 6.6248 | 6.26847 | 6.64465 | 6.35269 | 5.59131 | 6.22108 | 5.81586 | 5.42293 |
| GMEB2 | Q9UKD1 | 6.84835 | 6.73477 | 6.20505 | 6.61378 | 6.78167 | 6.45943 | 6.62221 | 6.34767 | 6.51284 | 5.83144 | 5.96478 | 5.88801 | 6.51812 | 5.56559 | 5.61414 | 5.49217 |
| SFXN4 | Q6P4A7 | 6.74758 | 6.84906 | 5.61738 | 6.46744 | 6.11318 | 6.20842 | 5.93717 | 6.80293 | 5.70743 | 6.27393 | 5.41028 | 6.42776 | 6.14807 | 6.32231 | 6.46406 | 6.50299 |
| NCS1 | P62166 | 6.18164 | 5.01285 | 5.70159 | 4.69765 | 5.91658 | 5.21648 | 5.45271 | 6.64045 | 5.20314 | 6.62089 | 5.51603 | 6.74138 | 6.36509 | 8.30964 | 8.11112 | 8.31275 |

| | | | | | | | | | | | | | | | | | |
|---|---|---|---|---|---|---|---|---|---|---|---|---|---|---|---|---|---|
| SNUPN | O95149 | 6.77152 | 6.46159 | 6.70824 | 5.95423 | 6.91052 | 6.81148 | 6.70409 | 6.4664 | 5.83073 | 6.62369 | 7.00819 | 7.06148 | 5.1819 | 5.68111 | 4.75464 | 5.07017 |
| MGRN1 | O60291-2 | 6.50946 | 6.08836 | 6.68002 | 7.47992 | 6.09839 | 5.83147 | 5.72485 | 6.34054 | 6.6585 | 6.8356 | 5.31752 | 6.71804 | 5.86007 | 6.62393 | 5.47716 | 5.75617 |
| SENP7 | Q9BQF6 | 6.26665 | 6.42041 | 5.59385 | 6.46132 | 5.97318 | 6.11425 | 6.44679 | 5.94524 | 5.95975 | 6.38055 | 6.15635 | 6.23173 | 6.74728 | 6.58898 | 6.45039 | 6.26328 |
| NEU1 | Q99519 | 6.74854 | 6.81158 | 6.1253 | 6.71978 | 6.84162 | 6.62861 | 6.21247 | 5.75252 | 6.35222 | 6.47377 | 6.1339 | 5.55139 | 5.58585 | 5.87654 | 6.23297 | 5.95293 |
| PRR3 | P79522 | 6.33353 | 6.84658 | 5.7429 | 6.79777 | 7.34427 | 5.50473 | 6.64997 | 6.07007 | 6.31289 | 6.28543 | 5.82973 | 5.58934 | 6.02466 | 6.22859 | 6.34654 | 6.09302 |
| MANEA | Q5SRI9 | 6.33465 | 6.98963 | 5.99666 | 6.2368 | 6.06493 | 6.3458 | 6.27827 | 5.88332 | 6.14475 | 6.14653 | 6.33065 | 6.49282 | 6.41574 | 6.07302 | 5.98688 | 6.27954 |
| JAM3 | Q9BX67 | 6.04824 | 6.60747 | 5.90774 | 6.2986 | 5.61892 | 5.9225 | 6.19816 | 6.16001 | 6.17123 | 6.17126 | 5.87202 | 6.14274 | 6.99802 | 6.597 | 6.30967 | 6.97641 |
| SCO2 | O43819 | 6.43873 | 6.64728 | 6.17165 | 6.81325 | 6.21318 | 5.93211 | 5.74729 | 5.93995 | 6.09874 | 6.31037 | 6.57108 | 6.75014 | 6.45936 | 5.93683 | 6.04616 | 5.92389 |
| GALNT6 | Q8NCL4 | 6.44318 | 6.02347 | 5.40748 | 7.01422 | 6.43981 | 5.93981 | 6.48135 | 5.87597 | 6.11588 | 5.81672 | 6.10204 | 5.63571 | 6.68753 | 6.34193 | 6.63658 | 7.03832 |
| PRSS23 | O95084 | 7.72006 | 6.40268 | 5.83759 | 7.23298 | 5.52582 | 5.6286 | 7.68971 | 6.14843 | 7.04309 | 6.77239 | 6.09547 | 6.41335 | 5.75384 | 4.41754 | 5.38204 | 5.93639 |
| TMEM222 | Q9H0R3 | 6.60707 | 6.51842 | 5.91456 | 6.25635 | 6.08824 | 6.33156 | 6.30068 | 6.44077 | 5.92926 | 5.46405 | 6.42464 | 6.18792 | 5.47835 | 6.74582 | 6.74868 | 6.56362 |
| CLEC11A | Q9Y240 | 6.63249 | 6.12527 | 5.4787 | 5.64764 | 6.77948 | 5.95255 | 5.82848 | 7.5786 | 6.23162 | 7.64537 | 6.81846 | 6.89932 | 5.95593 | 6.11955 | 5.00761 | 5.2989 |
| FAT4 | Q6V0I7 | 6.34848 | 6.25034 | 5.76099 | 6.00131 | 6.53692 | 7.17641 | 6.87204 | 6.31505 | 6.02999 | 6.55594 | 6.01753 | 6.12106 | 6.5382 | 5.88148 | 5.36936 | 6.22489 |
| KLHDC3 | Q9BQ90 | 6.2254 | 6.60129 | 7.01959 | 6.23494 | 6.54889 | 5.93682 | 6.29937 | 6.47071 | 6.2947 | 6.50686 | 6.72519 | 6.81521 | 5.88373 | 5.48042 | 5.577 | 5.37988 |
| D2HGDH | Q8N465 | 6.42709 | 6.56518 | 6.26796 | 6.86195 | 5.90501 | 5.98761 | 6.14026 | 6.1024 | 6.29281 | 6.20999 | 5.90634 | 6.12781 | 6.51681 | 6.28974 | 5.91699 | 6.48205 |
| GPR108 | Q9NPR9 | 6.26726 | 5.92769 | 5.98518 | 6.54681 | 6.05137 | 5.92468 | 6.3447 | 5.39556 | 6.11492 | 6.44062 | 6.58511 | 6.02317 | 6.79735 | 6.84971 | 6.4762 | 6.26967 |
| ARRB1 | P49407 | 5.65825 | 6.05739 | 7.00028 | 5.47779 | 5.58398 | 6.89772 | 6.20225 | 6.46161 | 6.09251 | 6.10696 | 6.22924 | 6.41805 | 6.25785 | 7.40622 | 6.27513 | 5.87478 |
| FKBP14 | Q9NWM8 | 6.52794 | 6.11692 | 6.04044 | 6.37078 | 6.80869 | 5.80546 | 6.38124 | 6.77228 | 6.17782 | 5.92913 | 6.19203 | 5.89214 | 6.85437 | 5.81322 | 6.02355 | 6.29399 |
| HDAC4 | P56524 | 6.20545 | 6.14262 | 6.40072 | 6.46583 | 6.55506 | 6.28495 | 6.84978 | 6.60914 | 6.81032 | 6.14257 | 6.58514 | 6.20127 | 5.83822 | 6.07393 | 5.62141 | 5.21359 |
| PSEN1 | P49768 | 6.65365 | 6.54149 | 5.77859 | 6.69912 | 6.11695 | 5.52742 | 5.9657 | 7.03564 | 5.70232 | 6.35169 | 6.51786 | 6.05828 | 5.78348 | 6.84434 | 6.24032 | 6.18316 |
| PTPMT1 | Q8WUK0 | 6.20524 | 5.56427 | 5.95476 | 6.72576 | 6.22003 | 5.76741 | 6.47654 | 5.62471 | 6.57126 | 6.28309 | 6.26855 | 6.50078 | 6.93355 | 6.15524 | 6.25459 | 6.49424 |
| PPP3R1 | P63098 | 5.71072 | 4.76116 | 6.15349 | 4.76513 | 6.35219 | 5.23556 | 5.94103 | 6.77409 | 5.90105 | 6.87359 | 6.20654 | 7.05407 | 6.11803 | 6.91449 | 7.84573 | 7.39312 |
| KIFC3 | Q9BVG8-6 | 6.60857 | 7.27494 | 6.79219 | 6.61316 | 6.71685 | 6.63455 | 6.31348 | 6.7683 | 6.65579 | 6.86495 | 6.576 | 7.73335 | 5.09347 | 5.23049 | 4.64519 | 4.47872 |
| DYNC2LI1 | Q8TCX1 | 6.71148 | 6.00392 | 7.63251 | 6.45906 | 5.91522 | 6.66172 | 5.98725 | 6.35541 | 6.88306 | 6.84787 | 6.74197 | 6.95478 | 5.51824 | 5.43075 | 4.947 | 4.94976 |
| ELOVL5 | Q9NYP7-2 | 6.00975 | 7.5337 | 6.03761 | 7.32849 | 6.17641 | 6.3045 | 6.02222 | 6.06991 | 5.78743 | 6.48988 | 6.2124 | 5.86071 | 5.31852 | 5.99337 | 6.37079 | 6.48428 |
| TSSC4 | Q9Y5U2 | 6.29395 | 6.2878 | 6.0416 | 5.77231 | 6.42882 | 6.44361 | 5.6193 | 6.74143 | 6.18093 | 5.98794 | 6.92386 | 7.13051 | 6.46503 | 5.41399 | 6.15686 | 6.11206 |
| E2F7 | Q96AV8 | 6.42268 | 6.44363 | 6.60742 | 6.28417 | 5.63174 | 5.60797 | 6.15162 | 6.31032 | 5.84785 | 5.74394 | 6.31932 | 6.48444 | 6.45893 | 6.22229 | 6.88658 | 6.5051 |
| PITPNC1 | Q9UKF7 | 6.27689 | 6.36109 | 6.7704 | 6.05647 | 6.20044 | 6.49393 | 6.29042 | 6.45864 | 6.61734 | 6.88093 | 6.50468 | 6.5747 | 6.81321 | 5.60061 | 5.56011 | 5.54015 |
| MTFR2 | Q6P444 | 7.30708 | 7.39729 | 6.79928 | 7.42504 | 6.57801 | 6.218 | 6.70568 | 6.40373 | 6.38506 | 6.63385 | 6.68073 | 6.94208 | 4.86008 | 4.46632 | 4.58368 | 4.61409 |
| TMCC1 | O94876 | 6.33386 | 6.60046 | 5.91743 | 6.10138 | 6.44875 | 5.9875 | 6.15282 | 6.35251 | 6.00309 | 6.03601 | 6.40437 | 6.03759 | 6.79564 | 6.23249 | 6.43004 | 6.16604 |
| ZRSR2 | Q15696 | 6.52451 | 6.52793 | 5.70928 | 6.49192 | 5.63653 | 5.70859 | 6.45817 | 5.70666 | 5.83081 | 5.79062 | 5.93061 | 6.05093 | 6.38198 | 6.49975 | 7.52743 | 7.22429 |
| CPEB2 | Q7Z5Q1-6 | 6.18994 | 6.292 | 6.56286 | 6.30305 | 6.45636 | 6.37795 | 6.28454 | 6.1634 | 6.28141 | 6.51147 | 7.10392 | 6.85309 | 5.79432 | 5.58135 | 5.83492 | 5.40941 |
| DVL3 | Q92997 | 6.25197 | 6.09974 | 7.62632 | 6.70079 | 6.31372 | 6.73667 | 5.70428 | 6.73243 | 6.75115 | 6.68601 | 6.954 | 6.67119 | 4.91576 | 5.36669 | 5.19372 | 5.29554 |
| USP20 | Q9Y2K6 | 6.71274 | 6.51954 | 6.07658 | 6.78477 | 6.1122 | 6.16228 | 6.75576 | 5.95215 | 5.93813 | 6.25918 | 6.1529 | 5.93825 | 6.07666 | 5.97986 | 6.1732 | 6.4058 |
| CNTRL | Q7Z7A1 | 5.89324 | 5.9022 | 7.13987 | 6.43279 | 6.76729 | 6.93293 | 5.57644 | 6.6736 | 6.41266 | 6.0596 | 6.36439 | 6.80277 | 6.25002 | 5.50507 | 5.54526 | 5.74189 |
| PZP | P20742 | 5.21593 | 6.72871 | 5.31973 | 5.00224 | 4.81789 | 6.6257 | 6.28891 | 5.19154 | 7.04723 | 5.5769 | 6.55239 | 6.31568 | 6.52136 | 6.41844 | 9.32248 | 7.05485 |
| FANCE | Q9HB96 | 6.19271 | 6.23608 | 6.02839 | 6.10628 | 6.63231 | 6.20638 | 5.51624 | 7.16684 | 6.55236 | 6.39225 | 6.69083 | 6.37259 | 5.61045 | 6.33502 | 6.44757 | 5.51371 |
| TRAPPC13 | A5PLN9-5 | 6.11802 | 6.22055 | 6.38104 | 5.81041 | 6.29752 | 6.57122 | 6.39474 | 6.72866 | 6.45112 | 6.54159 | 6.39762 | 6.39089 | 6.04234 | 5.87076 | 5.81027 | 5.97326 |
| FBXO4 | Q9UKT5 | 6.55199 | 6.02194 | 6.94887 | 5.44288 | 6.26276 | 7.05995 | 5.78546 | 6.83603 | 5.85251 | 6.45 | 6.59136 | 6.51699 | 6.2727 | 6.14621 | 5.75849 | 5.50184 |
| INO80B | Q9C086 | 6.16568 | 6.29739 | 6.66648 | 7.13505 | 6.28618 | 5.65034 | 6.24881 | 5.52572 | 6.57886 | 6.41109 | 6.518 | 6.02065 | 5.72761 | 6.24323 | 5.92303 | 6.60187 |
| PITHD1 | Q9GZP4 | 6.53806 | 5.81187 | 7.13633 | 6.03267 | 5.65057 | 6.1541 | 6.0786 | 5.94679 | 6.39309 | 7.08629 | 6.58795 | 6.94443 | 5.69909 | 5.97831 | 6.31385 | 5.64803 |
| GRPEL2 | Q8TAA5 | 5.54993 | 5.51141 | 5.49947 | 6.08084 | 6.26093 | 5.80175 | 6.38555 | 6.17374 | 5.19511 | 4.81845 | 5.55884 | 5.54748 | 8.64096 | 7.21038 | 8.28229 | 7.48287 |
| AVL9 | Q8NBF6 | 6.05043 | 6.37272 | 6.47292 | 6.13178 | 6.25335 | 6.5491 | 6.30314 | 6.44861 | 6.76741 | 6.98891 | 6.25998 | 6.29898 | 5.84726 | 5.7321 | 6.01532 | 5.50799 |
| TRAPPC6A | O75865-2 | 6.28019 | 4.91284 | 6.40129 | 4.67399 | 5.84526 | 6.47563 | 6.06363 | 6.79602 | 5.69123 | 6.73573 | 6.1189 | 6.70664 | 5.52918 | 8.48082 | 6.45927 | 6.72836 |
| MCC | P23508 | 5.98916 | 5.26082 | 6.8709 | 5.55341 | 6.25148 | 5.82835 | 6.15876 | 6.82616 | 6.43396 | 5.45664 | 6.50062 | 6.53531 | 6.24036 | 6.17259 | 6.36479 | 7.55667 |
| ZSCAN25 | Q6NSZ9 | 6.91142 | 6.71198 | 6.18498 | 6.29691 | 6.16192 | 5.81639 | 6.76549 | 5.81075 | 7.16141 | 6.04109 | 6.50672 | 7.11613 | 5.78379 | 5.19371 | 5.93622 | 5.60111 |
| MKRN1 | Q9UHC7 | 6.74199 | 6.4617 | 7.4056 | 6.21812 | 6.63029 | 6.25303 | 5.83748 | 6.28683 | 6.56329 | 6.61116 | 6.73446 | 6.35726 | 5.39745 | 5.93594 | 5.20474 | 5.36065 |
| PIP4K2B | P78356 | 5.88116 | 6.41966 | 6.59038 | 6.08637 | 6.1943 | 6.0649 | 5.67097 | 6.23842 | 6.4999 | 6.13404 | 6.76819 | 6.23329 | 6.14211 | 6.53744 | 6.44657 | 6.09232 |
| ANKRD44 | Q8N8A2-2 | 6.12186 | 6.37696 | 6.62924 | 6.04934 | 6.10633 | 6.83109 | 6.11863 | 6.55644 | 6.54817 | 6.98349 | 6.30339 | 7.00152 | 5.78806 | 5.53477 | 5.67907 | 5.37162 |
| VRK3 | Q8IV63 | 5.95784 | 6.53524 | 5.78772 | 6.43122 | 6.19154 | 6.40582 | 6.14987 | 6.25953 | 6.4391 | 6.45693 | 6.37464 | 6.06587 | 6.54017 | 5.99525 | 6.5483 | 6.76099 |
| ANGEL2 | Q5VTE6 | 5.55023 | 6.34079 | 5.93841 | 6.29994 | 5.84663 | 6.03318 | 6.37288 | 6.01776 | 6.24094 | 6.20586 | 5.94866 | 5.77684 | 6.80993 | 6.63842 | 7.26368 | 6.71586 |
| HM13 | Q8TCT9 | 6.25065 | 6.74789 | 6.0981 | 6.62284 | 6.00882 | 6.14796 | 6.24595 | 6.23517 | 5.75389 | 6.18641 | 5.7251 | 5.91491 | 6.68518 | 6.42468 | 6.50102 | 6.45143 |
| CENPL | Q8N0S6 | 6.57512 | 6.39502 | 6.19417 | 7.05773 | 6.07323 | 5.92125 | 6.50884 | 5.91372 | 6.24508 | 6.23656 | 5.89959 | 6.22831 | 6.43727 | 5.92303 | 6.15507 | 6.236 |
| TMEM97 | Q5BJF2 | 5.88138 | 6.2881 | 5.83259 | 7.02324 | 6.72307 | 6.54902 | 6.3034 | 6.4086 | 6.39802 | 6.02051 | 6.1356 | 5.79624 | 6.51351 | 6.30219 | 6.05398 | 5.77055 |
| LYRM4 | Q9HD34 | 6.67288 | 6.41556 | 5.87937 | 6.6295 | 6.01368 | 5.8572 | 6.45886 | 5.99441 | 6.14082 | 6.01686 | 6.0407 | 5.81143 | 6.49061 | 6.48351 | 6.64439 | 6.45022 |
| GATAD1 | Q8WUU5 | 6.4213 | 6.46831 | 7.27299 | 7.02151 | 6.12594 | 6.0883 | 6.41529 | 5.88137 | 6.73532 | 6.55527 | 6.9019 | 6.497 | 5.01657 | 5.17133 | 5.69097 | 5.73663 |
| FAM9C | Q8IZT9 | 5.61965 | 5.67857 | 5.87728 | 4.74809 | 5.75524 | 5.98693 | 5.82423 | 6.19051 | 6.03136 | 6.09903 | 6.12933 | 5.85063 | 7.60847 | 8.04773 | 7.47508 | 7.07787 |
| FAM83H | Q6ZRV2 | 6.11956 | 6.56712 | 6.2199 | 6.13669 | 6.17722 | 6.53791 | 6.53812 | 6.22603 | 6.2968 | 5.94952 | 5.91142 | 6.06535 | 6.24328 | 6.47532 | 6.50049 | 6.03528 |
| CMIP | Q8IY22 | 6.28024 | 6.66368 | 7.27653 | 6.53893 | 6.19466 | 6.67318 | 6.1411 | 7.48266 | 6.71508 | 6.71989 | 7.06471 | 7.70846 | 4.61899 | 4.5663 | 4.60285 | 4.75278 |
| RBM15B | Q8NDT2 | 6.37599 | 6.3083 | 5.61384 | 6.49509 | 6.2095 | 5.72823 | 6.3407 | 5.8962 | 6.46813 | 6.13078 | 5.84654 | 6.05024 | 6.60786 | 6.18126 | 6.91832 | 6.82902 |
| MINDY2 | Q8NBR6 | 6.07986 | 6.26026 | 6.91981 | 5.83737 | 5.97852 | 5.92597 | 6.62289 | 6.68693 | 5.73369 | 5.70719 | 6.70626 | 6.36319 | 6.44257 | 6.09086 | 6.56216 | 6.08246 |
| CRLF3 | Q8IUI8 | 6.37354 | 6.45633 | 6.85642 | 6.04242 | 5.78209 | 6.41815 | 6.77268 | 6.13673 | 5.96594 | 6.01296 | 6.57359 | 6.34981 | 6.41845 | 5.86139 | 5.88862 | 6.09088 |
| GLMP | Q8WWB7 | 6.82049 | 6.78421 | 5.07041 | 6.82517 | 7.88243 | 6.26304 | 5.88378 | 5.53713 | 5.39157 | 6.38172 | 7.49233 | 6.54279 | 6.43645 | 5.95744 | 4.97992 | 5.75112 |
| JMJD6 | Q6NYC1-3 | 5.38787 | 5.94169 | 6.18706 | 6.16003 | 5.54688 | 5.27568 | 5.90205 | 5.91492 | 5.59402 | 5.87567 | 5.85504 | 5.86122 | 6.65491 | 7.2267 | 8.7778 | 7.83845 |
| TMEM177 | Q53S58 | 6.39189 | 6.05558 | 5.93381 | 6.14636 | 6.27606 | 6.32626 | 6.18396 | 5.59627 | 6.34531 | 6.11908 | 6.29344 | 5.53076 | 6.70331 | 6.55619 | 6.38946 | 7.15225 |
| SLC27A6 | Q9Y2P4 | 6.54517 | 6.33127 | 5.96252 | 6.08921 | 6.35547 | 5.92146 | 6.26388 | 5.75686 | 6.37481 | 6.01371 | 5.71226 | 5.63738 | 6.46579 | 6.87288 | 6.94545 | 6.74288 |
| MICOS13 | Q5XKP0 | 6.59707 | 5.85673 | 5.67586 | 6.44213 | 6.34033 | 5.8202 | 6.27901 | 5.85787 | 6.20065 | 6.02355 | 6.19103 | 6.14481 | 7.0644 | 6.62458 | 6.2763 | 6.60546 |
| PEF1 | Q9UBV8 | 5.10072 | 4.32786 | 5.6346 | 4.24344 | 5.35796 | 5.63894 | 4.86986 | 5.85941 | 5.1566 | 6.06883 | 6.03638 | 6.69009 | 6.12237 | 10.0047 | 9.54776 | 9.34047 |
| PTPRJ | Q12913-3 | 6.54798 | 6.39188 | 6.10335 | 6.79556 | 6.01491 | 5.73234 | 6.37466 | 6.22699 | 6.00569 | 5.80196 | 6.31193 | 6.22874 | 6.84494 | 6.1213 | 6.02668 | 6.4711 |
| APOBEC3C | Q9NRW3 | 6.34749 | 6.09801 | 6.63785 | 6.39268 | 6.4954 | 6.25912 | 6.6315 | 5.98963 | 6.62171 | 6.76948 | 6.32151 | 6.51898 | 5.82129 | 5.83795 | 5.33306 | 5.92436 |
| DSC2 | Q02487 | 6.81954 | 6.9935 | 5.58136 | 6.56597 | 5.87251 | 5.60834 | 6.6973 | 6.4598 | 5.95857 | 6.46045 | 6.42119 | 6.27751 | 5.89061 | 6.24167 | 5.48459 | 6.6671 |
| TMEM104 | Q8NE00 | 6.13873 | 6.73878 | 6.61056 | 6.24275 | 5.94306 | 5.96547 | 6.30496 | 5.82664 | 6.19631 | 6.13947 | 6.78522 | 6.0201 | 6.37807 | 6.17916 | 6.16852 | 6.36221 |
| KATNBL1 | Q9H079 | 6.17856 | 6.43096 | 6.13296 | 6.07226 | 6.18689 | 7.1105 | 6.68031 | 6.06255 | 6.58775 | 6.12798 | 6.53057 | 6.92765 | 5.58423 | 5.69198 | 6.62984 | 5.06501 |
| KIAA1217 | Q5T5P2 | 6.31166 | 6.15918 | 6.18037 | 6.21691 | 6.04117 | 6.50474 | 5.98332 | 6.44846 | 6.28201 | 5.5623 | 5.98804 | 5.78996 | 5.6659 | 6.89134 | 5.9903 | 6.08435 |
| CCNL2 | Q96S94 | 5.72049 | 5.99627 | 5.85925 | 6.39997 | 5.84222 | 5.88187 | 6.26188 | 6.197 | 6.13057 | 5.9184 | 5.18487 | 5.60653 | 7.21042 | 6.99788 | 7.30329 | 7.48909 |
| GMPR2 | Q9P2T1-2 | 6.07865 | 6.51565 | 6.88645 | 6.05861 | 6.40963 | 6.58872 | 6.26982 | 6.33294 | 6.42908 | 6.55706 | 6.46494 | 6.34704 | 5.7278 | 5.49928 | 6.01003 | 5.82432 |
| AGPAT5 | Q9NUQ2 | 6.6908 | 6.31351 | 6.8237 | 7.38154 | 6.32752 | 5.12503 | 6.53049 | 6.24193 | 6.06803 | 6.21424 | 5.57446 | 5.84132 | 5.84039 | 6.26377 | 6.38966 | 6.37363 |
| MTFR1L | Q9H019 | 6.73227 | 6.40471 | 6.42771 | 6.16411 | 6.6891 | 6.44032 | 6.51653 | 6.96975 | 6.4724 | 6.70703 | 7.00746 | 6.31339 | 5.28166 | 5.26688 | 5.09888 | 5.5078 |
| TMEM168 | Q9H0V1 | 6.33832 | 6.40366 | 6.48279 | 6.61571 | 5.86042 | 5.67919 | 6.49673 | 5.92712 | 6.83647 | 6.23011 | 6.45529 | 5.61329 | 6.22246 | 6.51583 | 6.59812 | 5.72448 |
| GLRX5 | Q86SX6 | 6.15172 | 4.89834 | 4.65099 | 4.73854 | 5.5996 | 4.68766 | 5.54046 | 7.01091 | 4.93902 | 5.55379 | 4.80131 | 5.95165 | 5.38897 | 10.7044 | 8.88392 | 10.4987 |
| FCSK | Q8N0W3-2 | 6.05479 | 5.91443 | 6.24839 | 5.90216 | 6.05821 | 6.80552 | 6.25289 | 6.3514 | 6.17974 | 6.68507 | 6.58989 | 6.55612 | 6.00931 | 6.0612 | 6.10417 | 6.2267 |
| TMEM161B | Q8NDZ6 | 6.07528 | 6.43116 | 6.32863 | 6.78141 | 6.22245 | 5.74643 | 5.99638 | 5.69589 | 6.40828 | 6.05532 | 6.09773 | 5.45843 | 6.8008 | 6.52728 | 6.50842 | 6.8661 |
| TTC5 | Q8N0Z6 | 6.38624 | 5.72139 | 7.16168 | 6.01404 | 6.16913 | 7.03922 | 6.17339 | 6.34909 | 6.71953 | 6.32283 | 6.77111 | 6.61115 | 5.47482 | 5.53833 | 5.8876 | 5.66045 |
| FOXJ3 | Q9UPW0 | 5.71888 | 5.84369 | 6.15608 | 5.73584 | 6.17155 | 6.28481 | 5.7103 | 6.32214 | 6.53499 | 6.5022 | 6.39731 | 6.4656 | 7.01946 | 6.44236 | 6.40299 | 6.2918 |
| RUSC1 | Q9BVN2 | 6.95713 | 5.4045 | 6.21661 | 5.7148 | 6.36448 | 7.29101 | 6.51085 | 6.61611 | 5.62252 | 6.68489 | 6.65157 | 5.97583 | 6.2078 | 6.04896 | 6.07086 | 5.6621 |
| RWDD1 | Q9H446 | 6.39407 | 6.68652 | 6.49552 | 6.06989 | 7.18266 | 6.91557 | 6.92151 | 7.6223 | 6.13186 | 6.48881 | 6.44924 | 6.6387 | 5.82611 | 5.56634 | 4.62578 | 4.98511 |
| BCL2L1 | Q07817 | 7.08982 | 6.43839 | 6.25265 | 6.16729 | 6.50564 | 6.03835 | 6.14237 | 6.39838 | 6.48059 | 6.68899 | 6.30073 | 6.51832 | 5.3873 | 5.97492 | 5.75018 | 5.86609 |
| ATP5IF1 | Q9UII2 | 6.93376 | 6.47554 | 5.4807 | 5.91118 | 6.76653 | 5.55454 | 6.03095 | 6.79995 | 6.14136 | 5.93216 | 5.84799 | 6.32423 | 7.38091 | 6.17265 | 6.12434 | 6.14753 |
| C19orf47 | Q8N9M1 | 6.42575 | 6.52574 | 5.70625 | 6.66637 | 6.56254 | 5.77493 | 6.38577 | 5.99861 | 6.1911 | 5.81139 | 5.57039 | 5.75441 | 6.62124 | 6.58673 | 6.68618 | 6.7326 |
| XRCC4 | Q13426 | 5.47233 | 6.40966 | 5.7897 | 5.42603 | 5.80369 | 6.5552 | 5.83134 | 6.43311 | 5.73181 | 6.54064 | 6.21773 | 5.9229 | 6.94434 | 7.14753 | 7.02286 | 6.75113 |
| GBA2 | Q9HCG7 | 6.52735 | 6.51376 | 5.95222 | 6.26795 | 5.8832 | 5.93876 | 6.42017 | 6.39128 | 6.18966 | 6.38817 | 5.53626 | 6.34579 | 6.27252 | 6.24544 | 6.33261 | 6.79484 |
| C20orf27 | Q9GZN8-2 | 6.21935 | 6.56458 | 6.82337 | 6.14439 | 6.433 | 6.95423 | 6.17849 | 7.06411 | 6.13522 | 6.3863 | 6.77731 | 6.30869 | 5.48513 | 5.76384 | 5.41364 | 5.34834 |
| IGHMBP2 | P38935 | 5.99535 | 6.2956 | 6.80209 | 6.66448 | 6.19052 | 6.41169 | 6.55707 | 6.0872 | 6.34493 | 6.58005 | 6.3968 | 6.55646 | 5.60088 | 6.06716 | 5.56004 | 5.88967 |
| CPLANE1 | Q9H799 | 7.60937 | 7.05442 | 8.1101 | 7.31724 | 5.90212 | 6.2601 | 7.03348 | 6.97596 | 7.17635 | 6.97043 | 7.63092 | 7.78128 | 3.14582 | 3.88231 | 3.47859 | 3.67152 |
| ZFP82 | Q8N141 | 6.32633 | 6.44594 | 5.94147 | 7.0361 | 6.36558 | 5.99859 | 6.68911 | 5.64905 | 5.7241 | 5.67227 | 6.39091 | 6.56114 | 6.75649 | 5.99873 | 6.06667 | 6.37752 |
| NFYB | P25208 | 6.12801 | 5.62046 | 6.16391 | 5.46346 | 5.8171 | 5.3795 | 5.13031 | 5.95081 | 5.33914 | 5.4125 | 6.56934 | 7.48167 | 7.26111 | 7.63913 | 6.75732 | 7.88623 |
| TTL | Q8NG68 | 6.52479 | 6.73847 | 7.25809 | 6.13581 | 5.99969 | 7.05383 | 6.58033 | 6.90212 | 6.93646 | 6.80423 | 6.08775 | 6.30071 | 5.14807 | 5.29347 | 5.04245 | 5.19373 |
| TIMM17A | Q99595 | 6.48184 | 5.56707 | 6.53163 | 6.58811 | 5.99245 | 6.10053 | 6.66932 | 6.20071 | 6.502 | 6.37065 | 6.28769 | 6.10071 | 6.29054 | 5.72379 | 6.44003 | 6.15291 |
| UBASH3B | Q8TF42 | 5.76611 | 5.96296 | 7.37514 | 6.37879 | 5.80091 | 7.16961 | 5.44186 | 6.47018 | 6.35133 | 6.63315 | 6.93537 | 7.0862 | 5.63417 | 6.01152 | 5.35065 | 5.63206 |
| ACTR6 | Q9GZN1 | 6.39824 | 6.48086 | 5.69845 | 6.54151 | 6.51409 | 6.0689 | 6.57641 | 5.85427 | 6.4874 | 6.41393 | 5.92155 | 6.3739 | 6.18719 | 5.66511 | 6.37971 | 6.43848 |
| CNPY4 | Q8N129 | 6.43091 | 6.23603 | 6.3346 | 6.31043 | 6.82142 | 6.43774 | 6.29407 | 5.70563 | 6.22561 | 6.14727 | 6.46978 | 6.11438 | 6.7701 | 6.15843 | 5.85572 | 5.68789 |
| FBXO3 | Q9UK99 | 6.23128 | 6.47555 | 6.33311 | 5.56787 | 6.47815 | 6.69187 | 5.96825 | 6.76986 | 6.08362 | 6.22984 | 5.94851 | 7.19703 | 5.44042 | 5.99361 | 6.3552 | 6.23581 |
| CCDC77 | Q9BR77 | 6.09558 | 6.7665 | 6.62476 | 6.85105 | 5.99429 | 6.66739 | 6.5327 | 6.5342 | 6.27466 | 6.45223 | 5.95262 | 6.40082 | 5.77733 | 5.36972 | 5.73277 | 5.97338 |
| ARL6 | Q9H0F7-2 | 6.23059 | 5.32885 | 6.91759 | 6.46342 | 6.49191 | 6.02852 | 6.24877 | 6.12928 | 6.01879 | 6.34348 | 5.90551 | 6.76886 | 6.15055 | 6.5322 | 5.71826 | 6.72344 |
| CNNM1 | Q9NRU3 | 6.29792 | 6.42727 | 6.7127 | 6.72511 | 5.6703 | 6.29418 | 6.39327 | 6.48833 | 6.64374 | 5.63211 | 6.21971 | 6.58804 | 6.20715 | 5.28483 | 5.65198 | 6.76336 |

| | | | | | | | | | | | | | | | | | |
|---|---|---|---|---|---|---|---|---|---|---|---|---|---|---|---|---|---|
| TENT2 | Q6PIY7 | 6.8219 | 6.33575 | 5.83619 | 6.95045 | 6.61185 | 6.57188 | 6.26858 | 5.67242 | 6.61121 | 6.00574 | 5.7293 | 6.29209 | 6.21168 | 5.91904 | 6.24181 | 5.9201 |
| PTP4A1 | Q93096 | 5.99919 | 6.11957 | 6.30014 | 6.21717 | 5.92928 | 6.61525 | 6.37229 | 5.9884 | 6.54324 | 6.75233 | 6.47325 | 6.65888 | 5.71893 | 5.93949 | 6.33567 | 6.03694 |
| C6orf47 | O95873 | 5.97053 | 6.19969 | 5.79438 | 5.3574 | 6.492 | 5.68464 | 5.66836 | 6.81411 | 6.22993 | 5.97053 | 6.20227 | 6.15953 | 6.90405 | 6.61541 | 6.60155 | 7.3356 |
| PTPN21 | Q16825 | 6.9736 | 6.25136 | 7.35513 | 6.31751 | 6.05897 | 6.80723 | 6.61314 | 6.3766 | 6.65696 | 6.11845 | 6.41455 | 6.9501 | 5.30359 | 5.61401 | 5.1413 | 5.04749 |
| SIL1 | Q9H173 | 6.25086 | 6.23979 | 5.94221 | 6.25076 | 7.18572 | 6.14896 | 6.62378 | 6.09213 | 6.65528 | 5.59501 | 5.82621 | 6.13334 | 7.09153 | 5.90638 | 6.06166 | 5.99639 |
| NEMP1 | O14524 | 6.67622 | 6.41138 | 6.33729 | 6.50982 | 6.34149 | 5.66172 | 6.53027 | 5.70458 | 6.36849 | 5.70061 | 6.25465 | 5.47402 | 6.90773 | 6.27057 | 6.6757 | 6.17546 |
| GPRASP2 | Q96D09 | 6.07084 | 6.87958 | 7.01822 | 5.50676 | 6.4198 | 6.19406 | 6.21309 | 6.54952 | 6.77203 | 6.07609 | 6.60994 | 6.62566 | 6.17087 | 5.63022 | 5.88214 | 5.38118 |
| TRIM11 | Q96F44 | 6.05907 | 6.47303 | 5.68226 | 6.03007 | 6.2883 | 6.18001 | 6.03441 | 5.828 | 5.81376 | 6.71948 | 5.69982 | 6.33758 | 6.50762 | 6.80943 | 6.18216 | 7.35502 |
| SDC1 | P18827 | 6.60151 | 6.70457 | 5.71246 | 6.67022 | 6.55823 | 5.71497 | 6.18952 | 6.18023 | 6.38766 | 5.76001 | 6.3904 | 6.15444 | 6.13468 | 6.31002 | 5.85422 | 6.67685 |
| EIF1AD | Q8N9N8 | 6.2352 | 6.28768 | 6.99102 | 5.98256 | 6.34421 | 7.00877 | 5.77266 | 7.52541 | 6.21419 | 6.52226 | 6.54086 | 6.52016 | 5.95335 | 5.84941 | 4.91358 | 5.33868 |
| QDPR | P09417 | 6.21473 | 5.93645 | 6.89874 | 6.00396 | 6.39406 | 6.69843 | 6.01116 | 5.64759 | 6.82386 | 6.03848 | 6.57117 | 6.054 | 6.34647 | 6.09384 | 6.62583 | 5.64123 |
| SAMD1 | Q6SPF0 | 6.24721 | 6.48072 | 6.08343 | 7.21235 | 6.21026 | 6.78552 | 5.78477 | 5.69857 | 5.95964 | 6.27675 | 6.4656 | 6.58804 | 5.84549 | 6.3112 | 5.95266 | 6.09779 |
| ENSA | O43768-4 | 6.16975 | 5.50004 | 7.21322 | 5.23089 | 6.75585 | 6.1651 | 5.50577 | 7.12012 | 6.19201 | 6.66893 | 6.6439 | 7.22727 | 5.78892 | 6.50183 | 5.55891 | 5.75749 |
| NEDD8-MDP1 | E9PL57 | 6.32297 | 5.43367 | 6.72245 | 5.10838 | 5.44501 | 5.53318 | 6.11427 | 6.13965 | 5.97064 | 6.72919 | 6.28266 | 6.57609 | 4.87495 | 7.99084 | 7.14235 | 7.6137 |
| FOXRED1 | Q96CU9 | 6.61084 | 6.18798 | 6.16524 | 6.16683 | 6.10714 | 5.47506 | 5.93537 | 5.39199 | 6.5525 | 5.85486 | 6.5954 | 6.78283 | 6.67573 | 6.56095 | 6.59133 | 6.34596 |
| EEF1AKMT2 | A0A494BZY7 | 6.06784 | 6.24634 | 7.07649 | 6.3387 | 6.38057 | 6.75418 | 6.55055 | 7.33108 | 5.85296 | 6.3491 | 6.36887 | 6.67747 | 5.67209 | 5.91662 | 4.89993 | 5.5172 |
| LSM12 | Q3MHD2-2 | 6.49476 | 6.38833 | 6.55444 | 5.76829 | 6.38006 | 6.6198 | 6.01589 | 6.39668 | 6.35566 | 6.62566 | 6.49519 | 6.80068 | 6.15645 | 5.71186 | 5.64226 | 5.59398 |
| NIPSNAP3B | Q9BS92 | 6.28653 | 6.58459 | 5.84089 | 6.42708 | 5.83764 | 5.62965 | 7.84626 | 6.16131 | 5.89275 | 5.88798 | 5.65582 | 5.98141 | 6.18297 | 5.98836 | 7.22345 | 6.57333 |
| ZBTB2 | Q8N680 | 6.30182 | 6.54499 | 6.83503 | 6.93658 | 6.44149 | 5.29171 | 6.29111 | 6.8896 | 5.84901 | 5.60435 | 5.88924 | 5.67231 | 6.87011 | 5.57355 | 6.72954 | 6.27956 |
| MRPL10 | Q7Z7H8-2 | 6.38408 | 5.79211 | 5.81777 | 6.70308 | 5.85003 | 5.91462 | 6.52211 | 5.29165 | 6.44495 | 6.05748 | 6.76888 | 5.73129 | 7.19374 | 6.45812 | 6.81039 | 6.25971 |
| MED30 | Q96HR3 | 6.31132 | 5.19264 | 6.06689 | 5.35101 | 6.73186 | 5.21912 | 5.11099 | 7.04897 | 5.14161 | 6.07722 | 6.15176 | 6.62841 | 6.64233 | 7.50255 | 7.4091 | 7.41424 |
| REEP3 | Q6NUK4 | 6.32671 | 5.98049 | 5.91781 | 6.34449 | 6.29379 | 5.81661 | 6.43278 | 5.79789 | 6.1517 | 6.00849 | 6.08597 | 5.75846 | 6.3197 | 6.83485 | 6.85613 | 7.07413 |
| ATL1 | Q8WXF7 | 6.03245 | 7.11872 | 6.24618 | 6.42176 | 6.41461 | 6.31233 | 5.67861 | 5.49337 | 5.96257 | 6.4387 | 6.32456 | 5.97101 | 6.35843 | 6.31716 | 6.50539 | 6.40414 |
| ANAPC10 | Q9UM13 | 6.03365 | 6.45839 | 6.39983 | 6.30771 | 5.74421 | 6.48944 | 6.26536 | 6.15649 | 6.26758 | 6.16039 | 6.43565 | 6.35856 | 6.2521 | 6.27345 | 6.09609 | 6.30109 |
| BRK1 | Q8WUW1-2 | 5.68949 | 5.35277 | 5.68472 | 4.80922 | 5.54514 | 5.64721 | 5.41278 | 6.46972 | 5.30018 | 6.1834 | 6.084 | 6.37408 | 4.71122 | 10.2933 | 6.91517 | 9.52755 |
| ATXN7L1 | Q9ULK2 | 6.19846 | 5.36198 | 6.02934 | 6.75869 | 6.32796 | 5.98541 | 6.59906 | 5.79283 | 6.35578 | 6.4774 | 5.7379 | 6.45549 | 6.31695 | 6.95489 | 5.98548 | 6.66238 |
| PLEKHA3 | Q9HB20 | 6.84564 | 6.67016 | 6.50636 | 6.6292 | 5.96426 | 6.25108 | 6.59618 | 6.80137 | 7.43113 | 6.92773 | 7.44102 | 7.09597 | 4.81326 | 4.93943 | 4.48139 | 4.60582 |
| RYBP | Q8N488 | 5.53119 | 5.9042 | 5.99024 | 6.20816 | 6.39058 | 5.47715 | 5.63932 | 5.25756 | 6.11123 | 6.10954 | 6.25338 | 6.42715 | 7.58725 | 6.85379 | 6.96536 | 7.29389 |
| GNPTAB | Q3T906 | 6.90563 | 7.126 | 6.42568 | 6.85524 | 6.15615 | 6.6832 | 6.444 | 5.97013 | 6.69625 | 6.21594 | 6.45663 | 6.56304 | 5.2396 | 5.27303 | 5.69266 | 5.29683 |
| LAMTOR3 | Q9UHA4 | 5.50266 | 5.84841 | 5.67204 | 5.60058 | 6.18454 | 5.98854 | 6.61939 | 6.05896 | 6.55657 | 6.84632 | 6.31439 | 6.51968 | 5.75537 | 6.26633 | 6.90641 | 6.46792 |
| MOB1B | Q7L9L4-2 | 6.37258 | 5.87937 | 6.95789 | 6.06518 | 6.3147 | 6.628 | 6.48957 | 6.46721 | 6.2196 | 6.43119 | 6.80567 | 6.90902 | 5.46096 | 5.75897 | 5.94364 | 5.29645 |
| RNF168 | Q8IYW5 | 6.27204 | 6.63188 | 6.63894 | 6.99113 | 6.01312 | 6.25929 | 6.42741 | 5.99225 | 6.39271 | 6.13308 | 6.68777 | 6.43591 | 5.68853 | 5.73963 | 5.78879 | 5.90754 |
| NENF | Q9UMX5 | 6.29434 | 5.01035 | 4.97913 | 5.00692 | 5.34825 | 3.96239 | 5.18062 | 6.39837 | 4.27081 | 5.73782 | 4.81125 | 5.7731 | 6.09093 | 10.9541 | 9.46757 | 10.714 |
| FTH1 | P02794 | 6.4747 | 6.34884 | 7.7414 | 6.62051 | 5.11321 | 5.3764 | 5.09649 | 6.96909 | 7.03105 | 6.68104 | 6.88682 | 6.45477 | 4.43127 | 6.79159 | 5.61062 | 6.37221 |
| UBE2W | Q96B02-3 | 6.44957 | 6.40712 | 6.90876 | 6.00106 | 6.62575 | 6.38858 | 6.97534 | 6.51316 | 6.46989 | 7.04645 | 7.29729 | 6.54662 | 5.16306 | 5.22484 | 5.10382 | 4.87868 |
| DEPDC7 | Q96QD5 | 6.33438 | 7.00932 | 7.23724 | 6.05281 | 6.48149 | 6.3025 | 6.6612 | 6.26483 | 6.73839 | 6.40989 | 7.18163 | 7.26666 | 5.01626 | 4.9145 | 5.02153 | 5.10737 |
| PPP1R14B | Q96C90 | 6.5344 | 5.52698 | 6.65977 | 5.09835 | 6.49067 | 6.43181 | 5.41643 | 6.67202 | 5.95712 | 6.79166 | 6.62178 | 6.96453 | 5.49816 | 6.64279 | 5.77277 | 6.92076 |
| PFDN5 | Q99471 | 6.13032 | 5.42622 | 5.56159 | 4.54325 | 5.60263 | 4.60817 | 4.67059 | 7.10299 | 4.46061 | 5.92186 | 4.90365 | 5.16104 | 4.14333 | 12.3684 | 7.91154 | 11.4838 |
| DXO | O77932 | 5.67012 | 5.62859 | 7.26897 | 5.48847 | 5.70267 | 5.71767 | 5.72939 | 6.19431 | 6.37126 | 6.19976 | 6.6607 | 6.14438 | 6.65397 | 7.09078 | 6.6477 | 6.83126 |
| MED19 | A0JLT2 | 6.0424 | 5.90045 | 5.75106 | 6.77297 | 5.82855 | 5.99926 | 6.52745 | 5.9233 | 6.31284 | 5.76491 | 5.95049 | 6.23238 | 5.75838 | 7.4046 | 6.60276 | 7.22821 |
| CEP83 | Q9Y592 | 6.45046 | 6.30555 | 7.02013 | 6.20918 | 5.88881 | 6.59405 | 6.31393 | 6.35601 | 6.64412 | 6.44533 | 6.61928 | 6.45122 | 5.86666 | 5.30329 | 5.83288 | 5.69908 |
| RWDD4 | Q6NW29 | 6.54803 | 5.7957 | 7.04749 | 6.18834 | 6.67789 | 6.65998 | 5.825 | 6.53556 | 6.38811 | 6.64616 | 6.74975 | 7.09275 | 6.36875 | 5.33485 | 4.7604 | 5.38124 |
| DEPDC1 | Q5TB30 | 6.80303 | 6.7325 | 6.73137 | 6.61829 | 6.59444 | 6.48866 | 6.66649 | 6.13383 | 6.88529 | 6.79699 | 6.53807 | 6.72149 | 4.82345 | 4.97147 | 4.91976 | 5.57486 |
| CHST12 | Q9NRB3 | 6.67051 | 6.75927 | 6.29587 | 7.01642 | 6.10371 | 6.17254 | 6.48545 | 6.26244 | 6.4746 | 6.6789 | 6.52918 | 6.82305 | 5.33615 | 5.55393 | 5.22751 | 5.61049 |
| VPS9D1 | Q9Y2B5-2 | 6.3055 | 6.7614 | 6.39172 | 6.51649 | 5.86913 | 6.20932 | 6.34553 | 6.38345 | 6.28502 | 6.56989 | 6.73304 | 6.66331 | 6.01757 | 5.78779 | 4.97732 | 6.18352 |
| OGFRL1 | Q5TC84 | 6.44352 | 6.73344 | 6.79172 | 6.97587 | 5.90271 | 6.42679 | 5.90512 | 6.35291 | 6.63466 | 6.48528 | 6.54047 | 6.54573 | 5.6679 | 5.87654 | 5.21912 | 5.49821 |
| DR1 | Q01658 | 6.15488 | 5.36199 | 6.22736 | 5.47288 | 6.02518 | 6.9478 | 5.31973 | 5.92399 | 5.86434 | 6.42809 | 6.75973 | 6.27486 | 6.71654 | 6.68183 | 6.76908 | 7.07173 |
| MRNIP | Q6NTE8 | 5.4794 | 5.69919 | 6.0837 | 6.0115 | 5.53424 | 6.53267 | 6.22808 | 5.80549 | 5.66675 | 6.59272 | 6.97068 | 6.4189 | 6.61041 | 7.42489 | 6.12449 | 6.81691 |
| DMWD | Q09019 | 6.91646 | 6.26512 | 7.4333 | 6.30222 | 6.07173 | 7.31879 | 6.54049 | 6.23711 | 6.80004 | 6.83243 | 7.57973 | 7.37302 | 4.55673 | 4.7317 | 4.39625 | 4.6449 |
| TMA7 | Q9Y2S6 | 6.00918 | 5.97935 | 6.66465 | 5.68006 | 6.01289 | 6.29466 | 6.17491 | 6.38249 | 6.3855 | 6.57722 | 6.61417 | 6.57453 | 5.52453 | 6.97821 | 5.93749 | 6.21017 |
| BICC1 | Q9H694 | 6.02117 | 6.78261 | 6.49927 | 6.14004 | 6.41096 | 6.48477 | 5.96733 | 7.18092 | 6.69502 | 6.28789 | 6.17715 | 6.12156 | 6.24478 | 5.67245 | 5.85976 | 5.45431 |
| TUSC3 | Q13454-2 | 6.62151 | 6.14876 | 6.20689 | 7.05394 | 5.87702 | 5.89587 | 6.03773 | 6.11781 | 6.11167 | 5.73959 | 6.14579 | 6.31435 | 6.0241 | 6.63865 | 6.32618 | 6.74013 |
| SYNJ2BP-COX16 | A0A087WYV9 | 6.77207 | 6.1896 | 6.2317 | 6.6039 | 6.69616 | 5.96661 | 6.35997 | 5.84374 | 6.01573 | 5.72264 | 6.46187 | 5.98264 | 6.80015 | 5.67792 | 6.57747 | 6.09782 |
| CTSL | P07711 | 6.49435 | 6.40957 | 6.02221 | 6.40604 | 6.05471 | 5.0823 | 5.57353 | 6.28697 | 6.0779 | 6.67083 | 5.87107 | 5.45793 | 6.07843 | 7.56123 | 6.82623 | 7.12669 |
| SUDS3 | Q9H7L9 | 6.20087 | 6.52261 | 6.30033 | 6.8177 | 6.53693 | 6.11269 | 6.23511 | 5.96506 | 6.02298 | 6.04089 | 6.01929 | 6.51938 | 6.61559 | 6.03104 | 5.97174 | 6.68784 |
| PFDN2 | Q9UHV9 | 5.92735 | 5.10709 | 6.47848 | 4.9365 | 6.3724 | 6.27382 | 5.76776 | 6.90092 | 5.84534 | 6.89887 | 6.90676 | 7.23793 | 6.28025 | 6.40964 | 6.31201 | 6.34488 |
| TRAPPC2L | Q9UL33 | 5.41293 | 5.16022 | 6.25921 | 5.22293 | 6.70809 | 6.07397 | 6.62344 | 6.74387 | 7.71498 | 7.10057 | 6.65742 | 7.26415 | 5.22448 | 5.32 | 6.16183 | 6.3519 |
| CHCHD6 | Q9BRQ6 | 6.75499 | 6.40423 | 6.4261 | 6.36117 | 6.2733 | 6.00574 | 6.06688 | 6.05393 | 6.70429 | 6.40996 | 6.66782 | 6.52191 | 6.67809 | 5.28361 | 5.77431 | 5.61367 |
| REST | Q13127 | 6.2058 | 6.04486 | 5.98203 | 6.97238 | 6.26145 | 5.85913 | 6.20877 | 6.10594 | 6.11894 | 6.12192 | 5.97226 | 5.84367 | 6.16513 | 7.10246 | 6.30398 | 6.73126 |
| CSGALNACT2 | Q8N6G5 | 6.48434 | 6.63266 | 5.91603 | 7.07109 | 5.96909 | 6.01391 | 6.26469 | 5.65636 | 5.84741 | 6.18782 | 6.12655 | 6.36621 | 6.57017 | 6.0273 | 6.12003 | 6.74633 |
| SNAPIN | O95295 | 6.50619 | 4.89973 | 6.13967 | 5.55484 | 6.29205 | 5.55396 | 5.42971 | 6.14236 | 5.23074 | 6.42501 | 5.68451 | 6.7573 | 5.95929 | 8.24025 | 7.47607 | 7.70834 |
| POLR1F | Q3B726 | 6.40065 | 6.50362 | 6.26242 | 7.19889 | 5.44615 | 6.13774 | 6.41464 | 5.32522 | 6.57577 | 5.83224 | 5.7554 | 6.06764 | 6.52975 | 6.66194 | 6.81664 | 6.07128 |
| HDAC6 | Q9UBN7 | 6.66944 | 6.26599 | 6.9076 | 6.82015 | 6.56116 | 6.51212 | 6.85886 | 6.46707 | 6.74593 | 5.92773 | 6.12403 | 6.10539 | 6.10569 | 6.22772 | 5.583 | 5.11812 |
| PDSS2 | Q86YH6 | 6.19611 | 6.2647 | 5.98266 | 6.49381 | 6.44497 | 5.79898 | 6.32171 | 5.923 | 6.31789 | 6.60828 | 6.09071 | 6.39589 | 6.67151 | 6.08474 | 6.3316 | 6.07344 |
| SCARB1 | Q8WTV0 | 6.33338 | 6.55677 | 5.91687 | 6.26382 | 6.75705 | 6.20096 | 6.40984 | 6.33533 | 6.79662 | 5.84432 | 6.4042 | 6.06336 | 6.27054 | 5.74772 | 5.98845 | 6.11077 |
| OVCA2 | Q8WZ82 | 6.23809 | 6.57349 | 6.55357 | 5.68155 | 6.49906 | 5.95762 | 6.17384 | 6.26572 | 6.40802 | 7.23524 | 6.47929 | 7.07562 | 5.94137 | 5.74011 | 6.02735 | 6.05005 |
| SH3BGRL2 | Q9UJC5 | 5.86765 | 5.74486 | 6.29268 | 5.5047 | 5.9167 | 6.50495 | 5.78706 | 6.50264 | 6.21437 | 6.39208 | 6.36447 | 6.59074 | 6.52362 | 7.04028 | 6.47381 | 6.27939 |
| GPALPP1 | Q8IXQ4 | 5.52503 | 5.92235 | 5.81538 | 5.80717 | 5.62554 | 6.33026 | 5.51627 | 6.43424 | 6.83076 | 5.90053 | 6.16625 | 6.11068 | 7.10043 | 7.0723 | 6.92865 | 6.91417 |
| ING1 | Q9UK53 | 6.54059 | 6.47077 | 5.65711 | 6.90688 | 5.66852 | 5.65156 | 6.24439 | 5.89434 | 6.11463 | 5.68394 | 5.77214 | 5.57486 | 6.53541 | 6.72478 | 7.11211 | 7.44798 |
| SMCR8 | Q8TEV9 | 6.48653 | 6.90348 | 7.09172 | 6.19667 | 6.4416 | 6.91612 | 5.94338 | 5.74188 | 7.16435 | 6.71991 | 6.35185 | 7.5976 | 5.19556 | 5.61984 | 4.96426 | 4.66528 |
| C1GALT1 | Q9NS00 | 6.34337 | 6.26627 | 5.63253 | 6.59654 | 6.37921 | 5.9422 | 6.35474 | 6.1218 | 6.40403 | 6.01677 | 6.29559 | 6.183 | 6.77422 | 6.22951 | 6.31556 | 6.14466 |
| SKIL | P12757 | 6.77111 | 7.49763 | 5.94429 | 7.51752 | 6.12204 | 6.27658 | 6.67719 | 6.37311 | 6.35877 | 6.87885 | 6.63423 | 7.27492 | 4.45577 | 5.12082 | 5.10106 | 4.99611 |
| TMCO1 | Q9UM00 | 6.11124 | 6.26705 | 5.66059 | 6.73152 | 6.07677 | 6.11456 | 6.85688 | 5.92101 | 5.91087 | 5.99933 | 6.00496 | 5.97312 | 6.04037 | 6.99258 | 6.80219 | 6.53695 |
| LZIC | Q8WZA0-2 | 6.18293 | 5.23378 | 7.05134 | 5.07431 | 6.25536 | 5.95263 | 5.77576 | 6.96509 | 5.89983 | 7.01414 | 6.95297 | 7.77609 | 5.92708 | 6.41625 | 5.72261 | 5.79983 |
| GMFB | P60983 | 5.52861 | 5.46564 | 6.15195 | 5.14824 | 6.40245 | 6.20665 | 5.81565 | 7.61045 | 6.49083 | 6.7879 | 6.37951 | 7.4052 | 5.94592 | 6.27234 | 5.96419 | 6.42449 |
| ZC3H6 | P61129 | 6.20175 | 6.59231 | 6.03774 | 6.32317 | 5.47295 | 5.76062 | 6.29366 | 5.97251 | 5.7088 | 5.84358 | 6.06988 | 5.60134 | 6.5021 | 7.52311 | 7.29945 | 6.79702 |
| GNPNAT1 | Q96EK6 | 6.22537 | 6.40117 | 6.88604 | 5.91114 | 6.38899 | 6.71237 | 6.39401 | 6.71269 | 6.10957 | 6.25913 | 6.33201 | 6.86626 | 5.79876 | 5.66674 | 5.53187 | 5.80388 |
| STK11IP | Q8N1F8 | 6.86342 | 6.78809 | 6.43448 | 6.44298 | 6.59455 | 5.63945 | 6.16649 | 5.93846 | 6.9622 | 5.71159 | 6.41116 | 5.93954 | 6.01816 | 5.73455 | 6.2759 | 6.07897 |
| REX1BD | Q96EN9 | 6.47487 | 5.0918 | 6.85249 | 5.1433 | 6.3399 | 6.07455 | 5.91699 | 6.96773 | 5.33612 | 5.75136 | 6.14458 | 7.29886 | 4.44187 | 7.79709 | 7.06854 | 7.29993 |
| CCDC25 | Q86WR0 | 5.89128 | 6.15732 | 6.5527 | 5.91668 | 6.34021 | 6.46935 | 6.30959 | 6.92534 | 5.98408 | 7.13251 | 6.51457 | 6.96156 | 5.4951 | 5.85865 | 5.48725 | 6.00382 |
| MFGE8 | Q08431 | 7.10901 | 6.52047 | 6.24836 | 7.01147 | 5.37062 | 5.85994 | 7.06215 | 5.88744 | 6.51422 | 6.84521 | 6.48123 | 6.87797 | 5.75485 | 5.39446 | 5.49696 | 5.56563 |
| PHYH | O14832 | 5.56592 | 5.76681 | 5.73036 | 5.78125 | 6.53926 | 6.30323 | 6.48372 | 6.15073 | 5.90506 | 6.06051 | 6.04452 | 5.65087 | 7.53501 | 6.35032 | 6.9886 | 7.14383 |
| CCNE2 | O96020 | 6.34522 | 6.49049 | 6.5495 | 6.70122 | 6.228 | 6.02391 | 6.18181 | 5.75236 | 6.36143 | 6.4932 | 6.61182 | 6.06765 | 6.19228 | 5.75702 | 5.98761 | 6.25649 |
| NRP1 | O14786 | 6.01375 | 6.57824 | 5.91437 | 6.47319 | 6.15561 | 6.25294 | 6.00517 | 6.14897 | 5.99186 | 6.66441 | 5.99385 | 6.36041 | 6.02926 | 6.08959 | 6.67616 | 6.65224 |
| PDE8A | O60658 | 6.52468 | 6.77185 | 6.18494 | 6.64608 | 6.00626 | 5.93793 | 6.06202 | 6.27052 | 5.88864 | 6.2836 | 6.53407 | 6.62283 | 6.45581 | 6.05129 | 5.7732 | 5.98628 |
| SDHC | Q99643 | 6.02869 | 5.6631 | 5.50644 | 6.01203 | 6.14575 | 5.93273 | 6.10012 | 6.00849 | 5.85331 | 5.92631 | 5.7633 | 5.76811 | 7.22128 | 7.33865 | 7.27146 | 7.46023 |
| ITM2B | Q9Y287 | 7.1338 | 7.23784 | 6.8916 | 7.4855 | 6.04104 | 6.17352 | 6.69963 | 6.20037 | 6.62678 | 6.95391 | 6.9961 | 6.9756 | 4.28865 | 4.55154 | 4.72319 | 5.02093 |
| SEC14L2 | O76054 | 6.0857 | 6.19953 | 6.85636 | 6.05338 | 6.76806 | 6.76148 | 6.01182 | 6.40742 | 6.07098 | 6.67419 | 6.25752 | 6.36268 | 6.01184 | 6.22052 | 5.91457 | 5.34394 |
| SGTB | Q96EQ0 | 6.24255 | 5.63221 | 6.14194 | 5.17968 | 6.65281 | 6.16146 | 6.01018 | 7.23999 | 5.93844 | 6.53373 | 6.51837 | 6.72474 | 6.54919 | 6.28117 | 6.17887 | 6.01469 |
| RNF220 | Q5VTB9 | 5.87914 | 5.36905 | 6.12044 | 5.70076 | 6.39827 | 5.87089 | 5.80511 | 6.16529 | 6.51894 | 6.52877 | 6.69769 | 6.60998 | 6.83115 | 6.56272 | 6.25585 | 6.68595 |
| C3orf38 | Q5JPI3 | 6.11094 | 5.84778 | 6.25009 | 5.51722 | 6.0803 | 6.55845 | 5.72136 | 5.97722 | 6.76149 | 5.93637 | 6.66254 | 6.7137 | 6.40607 | 5.90093 | 6.94062 | 6.61491 |
| TMEM200A | Q86VY9 | 6.29695 | 6.32919 | 6.00257 | 6.31627 | 6.19406 | 6.3013 | 6.64437 | 6.42478 | 5.74266 | 6.2042 | 5.85769 | 5.56734 | 6.49609 | 6.55599 | 6.64016 | 6.42637 |
| CMC2 | Q9NRP2 | 6.48235 | 5.26459 | 5.60893 | 5.6302 | 5.87512 | 5.18648 | 5.64996 | 6.32705 | 5.53655 | 6.40395 | 6.23597 | 6.33041 | 6.69381 | 7.54169 | 7.48513 | 7.74781 |
| PIAS3 | Q9Y6X2 | 7.06744 | 6.86362 | 5.66517 | 6.54155 | 5.88518 | 6.09387 | 6.23854 | 6.62877 | 6.07543 | 7.28884 | 6.42406 | 6.01579 | 5.89331 | 5.7191 | 5.92084 | 5.67848 |
| TIFA | Q96CG3 | 6.11231 | 6.24064 | 7.14008 | 6.1352 | 6.41783 | 6.8879 | 6.0539 | 6.46875 | 6.74036 | 6.62041 | 7.13871 | 7.0377 | 5.11237 | 5.03147 | 5.71332 | 5.14906 |
| TXNDC11 | Q6PKC3 | 6.91073 | 5.58594 | 6.36847 | 6.49171 | 6.15546 | 6.40503 | 6.06143 | 6.18834 | 5.62634 | 6.3331 | 6.58782 | 6.00078 | 7.02266 | 6.34807 | 5.69594 | 6.21818 |
| GSTZ1 | O43708 | 6.52211 | 6.30617 | 6.25205 | 6.3284 | 6.81336 | 5.93491 | 6.15227 | 6.10753 | 6.36135 | 5.73733 | 5.9919 | 5.67401 | 6.39887 | 6.39134 | 6.45435 | 6.57405 |
| MSANTD2 | Q6P1R3 | 5.77824 | 5.26295 | 6.54255 | 6.31889 | 5.7803 | 6.10058 | 6.28063 | 5.70766 | 6.06342 | 6.18079 | 5.95216 | 5.99447 | 7.28 | 6.98063 | 6.97236 | 6.80436 |
| UBL4A | P11441 | 6.15622 | 4.67268 | 6.40899 | 4.8159 | 6.6066 | 5.22591 | 5.34174 | 7.16246 | 5.32873 | 6.89018 | 5.94006 | 6.74491 | 6.65508 | 7.70732 | 7.17407 | 7.16915 |
| RGS17 | Q9UGC6 | 6.13152 | 6.52494 | 6.12704 | 6.50632 | 6.20658 | 6.13723 | 6.25682 | 6.45951 | 6.30635 | 6.3459 | 5.94702 | 5.87159 | 6.05029 | 6.48653 | 6.39973 | 6.2426 |
| ALG3 | Q92685 | 6.0916 | 6.48147 | 5.80134 | 6.63779 | 6.10918 | 5.91699 | 6.08099 | 6.32452 | 6.28682 | 5.991 | 5.801 | 6.17015 | 6.25962 | 6.35325 | 6.85935 | 6.83492 |
| TBC1D17 | Q9HA65 | 6.01423 | 6.73268 | 6.75434 | 5.72509 | 6.20432 | 7.40899 | 5.93538 | 6.99966 | 6.33185 | 6.03364 | 6.76011 | 6.34773 | 5.78284 | 6.28186 | 5.64124 | 5.04604 |
| MAP1LC3B | Q9GZQ8 | 5.71379 | 6.42782 | 6.64546 | 5.95947 | 5.65581 | 6.51275 | 6.6218 | 6.18775 | 6.77398 | 5.9378 | 6.14929 | 6.28738 | 6.61472 | 6.7484 | 5.56379 | 6.19989 |
| ZBTB38 | Q8NAP3 | 7.0901 | 6.6121 | 5.36609 | 5.87689 | 6.07853 | 6.3677 | 5.86062 | 5.87309 | 5.80426 | 6.56885 | 6.2668 | 6.21659 | 6.40244 | 6.8339 | 6.61177 | 6.17027 |
| SCYL3 | Q8IZE3 | 6.53913 | 6.05566 | 6.41365 | 6.29056 | 6.29322 | 5.73323 | 5.95422 | 5.40792 | 7.10903 | 6.57062 | 6.55656 | 6.79605 | 6.25309 | 6.24799 | 5.76924 | 6.00983 |

| | | | | | | | | | | | | | | | |
|---|---|---|---|---|---|---|---|---|---|---|---|---|---|---|---|
| SELENOF | O60613 | 6.01012 | 5.35911 | 5.51902 | 5.30824 | 6.17035 | 5.7871 | 5.84821 | 6.31478 | 5.49521 | 6.30727 | 5.91224 | 6.5456 | 7.27666 | 7.22586 | 7.37776 | 7.54247 |
| MT-ND5 | P03915 | 6.08809 | 6.23392 | 5.25573 | 6.61416 | 5.73773 | 5.79915 | 6.38904 | 6.49494 | 6.65614 | 5.37504 | 6.54758 | 5.61154 | 6.82557 | 6.80165 | 6.8088 | 7.6609 |
| PBX2 | P40425 | 6.87227 | 6.96693 | 5.77125 | 7.24081 | 6.74937 | 6.87296 | 5.9453 | 7.08163 | 5.96969 | 6.68459 | 6.63454 | 5.64743 | 4.96802 | 5.63409 | 5.59457 | 5.36655 |
| NES | P48681 | 6.54338 | 6.2789 | 5.91184 | 6.64002 | 6.15631 | 5.88457 | 6.40806 | 6.01913 | 5.95981 | 6.28538 | 6.53868 | 6.3707 | 6.6556 | 6.03418 | 6.17155 | 6.14189 |
| CDC25B | P30305-3 | 6.9373 | 7.28598 | 7.8666 | 6.2943 | 5.92627 | 7.12351 | 6.49263 | 6.87355 | 7.73222 | 7.60199 | 8.4235 | 7.87026 | 3.1027 | 3.70747 | 3.08931 | 3.67239 |
| MBD4 | O95243 | 6.61044 | 6.86388 | 6.84763 | 7.14262 | 6.18797 | 6.17073 | 6.46641 | 6.32094 | 7.01852 | 6.85149 | 6.50133 | 6.71198 | 4.90883 | 4.99599 | 5.17812 | 5.22314 |
| CTSF | Q9UBX1 | 7.26112 | 7.03093 | 6.14887 | 6.13918 | 6.82835 | 6.48828 | 6.2941 | 6.36214 | 6.67051 | 6.1551 | 5.89363 | 5.5835 | 6.61554 | 5.46343 | 5.54114 | 5.52417 |
| GPATCH2L | Q9NWQ4 | 6.04003 | 6.92825 | 6.62662 | 6.91164 | 6.04817 | 6.39789 | 6.84766 | 6.59049 | 5.70395 | 5.66407 | 5.99649 | 6.10746 | 5.54226 | 6.26611 | 5.89959 | 6.42932 |
| SLC38A2 | Q96QD8 | 5.96232 | 7.07209 | 5.82756 | 6.32593 | 5.5075 | 6.5518 | 5.96599 | 6.47586 | 6.27415 | 6.52871 | 5.7098 | 6.34247 | 6.45689 | 7.00885 | 5.77917 | 6.21093 |
| VPS26C | O14972 | 5.71615 | 5.76606 | 6.09115 | 6.12971 | 6.07683 | 6.18756 | 5.69724 | 6.0868 | 6.20122 | 6.38172 | 6.67157 | 6.42909 | 6.39103 | 6.89562 | 6.89465 | 6.38361 |
| CERS5 | Q8N5B7 | 6.84974 | 6.53238 | 5.55696 | 6.71159 | 6.30904 | 6.24884 | 6.70954 | 5.90486 | 5.82957 | 5.44905 | 5.3528 | 5.31397 | 7.11015 | 6.60608 | 6.64306 | 6.87236 |
| NKAPD1 | Q6ZUT1-2 | 6.01299 | 6.43986 | 6.28229 | 6.13635 | 6.11545 | 5.44407 | 6.57947 | 5.96763 | 6.3845 | 6.88964 | 6.65929 | 5.39004 | 5.50891 | 6.45107 | 7.42406 | 7.31439 |
| ATPSCKMT | Q6P4H8 | 6.24523 | 6.62238 | 6.26264 | 6.42455 | 6.23083 | 6.38274 | 6.70246 | 5.77085 | 6.17501 | 6.07342 | 5.54625 | 5.9487 | 6.36417 | 6.34852 | 6.51344 | 6.38881 |
| SEC61B | P60468 | 6.16768 | 7.43124 | 5.5473 | 7.10698 | 6.09763 | 6.75494 | 6.12492 | 6.96533 | 5.27598 | 6.01375 | 4.81974 | 5.63208 | 6.43999 | 6.63035 | 6.32894 | 6.66315 |
| LMCD1 | Q9NZU5 | 6.15394 | 6.11726 | 7.1593 | 5.89581 | 5.86432 | 6.71272 | 5.70486 | 6.6441 | 5.89373 | 6.62474 | 7.07669 | 7.13575 | 5.88899 | 6.19734 | 5.34194 | 5.58852 |
| TOMM7 | Q9P0U1 | 6.53548 | 5.57677 | 6.21968 | 6.43458 | 6.28523 | 5.35377 | 5.83239 | 5.98059 | 4.83805 | 5.80293 | 5.67251 | 6.8802 | 6.46654 | 6.71861 | 7.52685 | 7.87581 |
| SRXN1 | Q9BYN0 | 6.43388 | 5.26067 | 6.68917 | 5.12175 | 5.97936 | 5.83052 | 5.70479 | 7.93069 | 5.58416 | 6.42247 | 6.24813 | 7.91918 | 4.14242 | 7.41076 | 6.12956 | 7.1925 |
| POP7 | O75817 | 6.70967 | 6.64156 | 6.29599 | 6.48115 | 6.22673 | 6.24759 | 6.26276 | 6.31763 | 6.73909 | 5.9746 | 6.49875 | 5.94375 | 6.25826 | 5.79863 | 5.92755 | 5.67628 |
| LIMK1 | P53667 | 6.89687 | 6.43317 | 6.88471 | 5.94436 | 6.29553 | 6.86661 | 5.90202 | 6.72856 | 6.54494 | 6.24234 | 6.4715 | 6.36882 | 5.84439 | 5.71303 | 5.76975 | 5.0934 |
| NR2C2AP | Q86WQ0-2 | 5.60187 | 5.53178 | 6.58379 | 5.21422 | 6.30964 | 6.60716 | 6.05034 | 7.54833 | 6.52238 | 6.79035 | 6.78588 | 7.09067 | 5.33175 | 6.27891 | 5.74822 | 6.00472 |
| TECPR1 | Q7Z6L1-4 | 6.68544 | 6.64743 | 7.62449 | 5.76697 | 6.01619 | 6.37006 | 6.49965 | 6.6394 | 6.08009 | 6.30123 | 6.36454 | 6.90798 | 5.56399 | 5.40975 | 5.81148 | 5.31132 |
| CARD8 | Q9Y2G2-5 | 6.13285 | 5.52424 | 6.65944 | 5.0356 | 6.08505 | 6.20322 | 6.18171 | 6.74952 | 5.728 | 6.51862 | 6.40602 | 7.33479 | 6.68184 | 6.71013 | 5.93509 | 6.11388 |
| MBLAC2 | Q68D91 | 6.12605 | 6.57316 | 5.76615 | 6.17813 | 6.61866 | 6.03099 | 6.36802 | 6.50581 | 6.00061 | 6.01607 | 6.57611 | 6.50907 | 6.27314 | 6.18858 | 6.49749 | 5.77194 |
| SLC25A29 | Q8N8R3 | 6.23165 | 6.23423 | 6.04926 | 6.32637 | 6.01586 | 6.53335 | 6.39388 | 6.00099 | 6.27654 | 6.86092 | 6.27224 | 5.96521 | 6.54338 | 6.16518 | 5.82119 | 6.30976 |
| STARD3 | Q14849-3 | 6.65288 | 6.98473 | 6.14468 | 7.33569 | 6.4751 | 6.13692 | 6.54729 | 6.52106 | 6.65794 | 6.71515 | 6.80635 | 6.18037 | 5.63864 | 5.57346 | 5.4829 | 5.44684 |
| OGFOD3 | Q6PK18-2 | 6.44188 | 6.65795 | 5.51256 | 7.82373 | 6.1669 | 6.08743 | 7.12172 | 6.05525 | 6.41156 | 5.9614 | 6.42198 | 5.63612 | 5.88015 | 5.80263 | 6.08131 | 5.93742 |
| CALB2 | P22676 | 6.38908 | 6.48519 | 5.99615 | 4.60217 | 6.09751 | 5.88875 | 5.56262 | 7.06659 | 5.78537 | 7.07805 | 6.26415 | 7.11858 | 6.61224 | 7.31286 | 6.51945 | 7.12125 |
| ZNF483 | Q8TF39 | 7.66285 | 7.42571 | 5.95484 | 6.48815 | 6.89711 | 6.52615 | 7.38025 | 6.32366 | 6.20586 | 6.16518 | 6.4205 | 6.48924 | 4.4138 | 5.59975 | 4.88093 | 5.16602 |
| NDST2 | P52849 | 6.23686 | 6.42162 | 5.95876 | 6.71246 | 6.48923 | 5.8843 | 6.13367 | 6.03239 | 6.11141 | 6.14364 | 5.70624 | 6.25685 | 6.56406 | 6.10481 | 6.46063 | 6.78308 |
| RNF5 | Q99942 | 5.84951 | 5.727 | 5.84282 | 5.66059 | 5.91335 | 5.84083 | 6.24737 | 6.41033 | 5.5556 | 6.03953 | 5.82309 | 6.7155 | 6.59005 | 6.94811 | 7.42446 | 7.41186 |
| CUTC | Q9NTM9 | 5.69889 | 5.68569 | 5.80047 | 5.5848 | 6.12577 | 6.84277 | 5.30432 | 5.46551 | 6.04596 | 6.30215 | 5.52645 | 6.61076 | 7.13478 | 7.84128 | 7.13042 | 6.89998 |
| NRP2 | O60462 | 5.97939 | 6.44072 | 5.8267 | 6.35538 | 5.85973 | 6.23225 | 6.19517 | 5.98075 | 6.18367 | 5.84802 | 6.51875 | 6.52126 | 6.60489 | 6.40144 | 6.55129 | 6.50059 |
| AGTRAP | Q6RW13 | 6.95037 | 6.90675 | 5.79014 | 5.79106 | 5.82808 | 6.80813 | 5.92943 | 6.73346 | 6.49432 | 6.76425 | 5.88657 | 5.87881 | 6.02537 | 6.41571 | 6.0082 | 5.78933 |
| AGPAT1 | Q99943 | 6.30753 | 6.60494 | 5.70408 | 6.38624 | 5.8108 | 5.97666 | 6.4738 | 6.29376 | 6.23588 | 6.52192 | 5.88767 | 6.38333 | 6.35406 | 6.47569 | 6.29445 | 6.2892 |
| HSPBAP1 | Q96EW2 | 5.50374 | 6.11973 | 6.45854 | 5.46873 | 5.9619 | 6.18433 | 5.77396 | 6.39695 | 6.06429 | 6.50782 | 6.53872 | 6.59597 | 6.54446 | 6.88282 | 6.5247 | 6.47335 |
| AP5B1 | Q2VPB7 | 6.41327 | 5.70271 | 6.89429 | 5.94985 | 6.29514 | 5.83636 | 6.21096 | 6.41457 | 6.32705 | 6.96545 | 6.88799 | 7.51226 | 5.23849 | 5.72912 | 5.62168 | 6.00082 |
| TMEM185B | Q9H7F4 | 6.60864 | 7.33789 | 6.34639 | 6.384 | 5.82988 | 4.97322 | 5.86876 | 6.47739 | 6.27377 | 6.17868 | 7.0729 | 5.98415 | 6.06664 | 5.82631 | 6.50836 | 6.26302 |
| GGPS1 | O95749 | 5.81672 | 5.32104 | 6.44927 | 4.75342 | 6.42818 | 6.24673 | 6.08939 | 7.09527 | 6.08984 | 6.41777 | 6.34629 | 6.90416 | 6.38794 | 6.57717 | 6.73598 | 6.34082 |
| SLC25A35 | Q3KQZ1 | 6.48861 | 6.28412 | 6.33413 | 6.49151 | 6.06986 | 5.99146 | 6.34363 | 5.44525 | 5.84216 | 6.42903 | 5.93129 | 5.75375 | 6.97196 | 6.20117 | 6.85135 | 6.57071 |
| LDLRAP1 | Q5SW96 | 6.21078 | 6.84204 | 6.88897 | 6.41135 | 5.79708 | 6.44125 | 6.02012 | 5.96539 | 6.28464 | 6.76747 | 6.33343 | 6.51037 | 6.02433 | 5.84324 | 5.97484 | 5.68471 |
| GINM1 | Q9NU53 | 7.36016 | 7.81586 | 7.23979 | 7.86418 | 6.61578 | 6.13667 | 6.60287 | 6.2303 | 6.71527 | 6.84923 | 6.92938 | 6.8913 | 4.0253 | 4.42274 | 4.09036 | 4.21081 |
| CHST10 | O43529 | 6.35608 | 6.64471 | 6.68401 | 6.69933 | 6.02921 | 6.45483 | 6.83336 | 5.86818 | 6.6231 | 6.54412 | 7.06129 | 6.87271 | 5.54396 | 5.50403 | 5.20424 | 5.07684 |
| GLB1L2 | Q8IW92 | 6.23216 | 6.69709 | 6.02749 | 6.44558 | 6.16298 | 5.98601 | 6.71452 | 5.8344 | 6.1401 | 6.24178 | 6.35063 | 6.14228 | 6.09262 | 5.83397 | 6.41758 | 6.68081 |
| TIMM17B | O60830-2 | 6.26119 | 5.90784 | 6.19583 | 6.28405 | 6.09805 | 6.2765 | 6.67258 | 5.71424 | 6.06825 | 6.12574 | 5.77862 | 6.35658 | 6.67097 | 6.35069 | 6.7305 | 6.48768 |
| BEND7 | Q8N7W2 | 6.34948 | 6.62969 | 6.50623 | 6.88827 | 6.02497 | 6.206 | 6.03619 | 6.01728 | 6.50096 | 6.46275 | 5.8298 | 6.09502 | 5.98673 | 6.18547 | 6.11827 | 6.16289 |
| TOM1 | O60784 | 6.79421 | 5.53298 | 6.15668 | 5.16663 | 5.84682 | 6.60451 | 6.22578 | 6.70323 | 5.72978 | 6.34576 | 7.25027 | 7.09648 | 5.9909 | 7.1352 | 6.24037 | 5.1804 |
| TMEM205 | Q6UW68 | 5.91176 | 6.87766 | 5.84509 | 6.29748 | 5.69774 | 6.00713 | 5.74894 | 6.27333 | 5.82108 | 6.43796 | 5.87534 | 6.17586 | 6.39147 | 6.59616 | 6.67851 | 7.3645 |
| BAD | Q92934 | 8.65729 | 8.91428 | 7.37965 | 8.76754 | 7.2788 | 6.74926 | 6.27303 | 7.47371 | 7.44389 | 8.33792 | 7.32585 | 7.85941 | 1.68224 | 2.42477 | 1.43207 | 2.0003 |
| WDR7 | Q9Y4E6 | 6.51084 | 6.36771 | 6.43724 | 5.96544 | 6.36668 | 6.27629 | 6.19068 | 6.39782 | 6.77198 | 6.86522 | 6.57467 | 6.40915 | 6.05286 | 5.61305 | 5.74471 | 5.45568 |
| LETMD1 | Q6P1Q0-7 | 6.63061 | 6.58333 | 5.74557 | 7.10621 | 6.20263 | 6.21115 | 6.40804 | 6.69497 | 5.46632 | 5.96607 | 6.11204 | 6.03876 | 5.98078 | 5.98829 | 6.23947 | 6.62576 |
| NGDN | Q8NEJ9 | 6.43503 | 6.39678 | 5.70037 | 6.51882 | 6.15019 | 5.76258 | 6.31323 | 5.81573 | 6.22018 | 6.07896 | 5.67421 | 5.83762 | 6.74159 | 6.37163 | 7.07257 | 6.91051 |
| IAH1 | Q2TAA2 | 6.52645 | 5.64621 | 6.37366 | 5.39529 | 6.25365 | 6.48064 | 6.68341 | 6.97026 | 6.38048 | 6.53575 | 6.53233 | 6.87004 | 6.08104 | 6.1324 | 5.6834 | 5.45501 |
| CNOT6L | Q96LI5 | 6.05089 | 6.0577 | 6.40002 | 6.25967 | 6.12761 | 6.86518 | 6.39974 | 6.34723 | 6.11558 | 6.64513 | 6.74004 | 6.83514 | 5.71667 | 5.6919 | 5.97811 | 5.76936 |
| TRMT112 | Q9UI30 | 4.31088 | 4.58365 | 4.62601 | 4.90122 | 5.48323 | 5.1994 | 5.62529 | 8.72779 | 5.52209 | 6.34461 | 5.45682 | 6.92227 | 4.29216 | 9.64248 | 8.43036 | 9.93174 |
| ARHGAP19 | Q14CB8-2 | 6.2653 | 6.028 | 6.23533 | 5.81809 | 5.9774 | 5.72653 | 6.49994 | 5.75284 | 6.69032 | 6.32831 | 6.46015 | 6.10945 | 6.45965 | 6.63216 | 6.64621 | 6.37032 |
| KLHL13 | Q9P2N7-5 | 6.42911 | 6.54639 | 6.9953 | 6.32922 | 6.29705 | 6.31899 | 6.42025 | 6.2188 | 6.53479 | 6.93395 | 7.11282 | 6.97352 | 5.12946 | 5.33167 | 5.38581 | 5.04288 |
| DCAF15 | Q66K64 | 5.15381 | 5.75173 | 6.2095 | 5.86813 | 5.69562 | 6.13379 | 6.02704 | 6.04784 | 5.96215 | 6.81904 | 7.29872 | 6.63579 | 6.24699 | 6.25898 | 6.80685 | 7.08402 |
| COX19 | Q49B96 | 6.72889 | 5.67881 | 5.98878 | 5.79177 | 6.89958 | 5.49949 | 5.56991 | 6.76677 | 5.27169 | 6.13303 | 5.52726 | 6.13205 | 6.88422 | 7.20277 | 6.90281 | 7.02216 |
| FN3KRP | Q9HA64 | 5.67546 | 5.35559 | 6.07357 | 5.32964 | 6.54781 | 6.44049 | 5.91495 | 7.24152 | 6.97003 | 6.43746 | 6.05266 | 6.77473 | 6.69049 | 5.85858 | 6.00485 | 6.63215 |
| BSDC1 | Q9NW68 | 6.65613 | 6.14495 | 6.0845 | 5.6248 | 6.13508 | 6.03183 | 5.81312 | 7.22411 | 6.04209 | 5.63132 | 6.54516 | 6.30052 | 7.33724 | 6.68693 | 6.00838 | 5.73384 |
| METTL5 | Q9NRN9 | 6.44876 | 6.23449 | 7.26076 | 6.09785 | 6.62275 | 6.58037 | 5.60882 | 6.89616 | 7.38765 | 6.52275 | 6.49193 | 7.17258 | 4.97 | 5.07916 | 5.2231 | 5.40288 |
| ORMDL3 | Q8N138 | 6.07394 | 5.6833 | 5.4024 | 6.1849 | 5.87446 | 5.70394 | 6.15963 | 5.92161 | 6.17262 | 6.07946 | 5.93501 | 6.01981 | 6.34854 | 7.53094 | 7.39359 | 7.51585 |
| FAM32A | Q9Y421 | 6.14072 | 6.45247 | 6.06369 | 6.14025 | 6.26462 | 6.60794 | 6.17375 | 6.61151 | 6.29695 | 6.58596 | 6.42731 | 6.21505 | 5.56249 | 6.59052 | 6.07306 | 5.7937 |
| CCDC138 | Q96M89 | 6.96211 | 6.64559 | 6.80953 | 6.86967 | 6.41579 | 6.43059 | 6.0799 | 6.11494 | 6.60615 | 6.48818 | 7.25815 | 7.54022 | 4.64409 | 5.37968 | 4.44833 | 5.30709 |
| ZBTB7A | O95365 | 6.23129 | 6.63436 | 6.25964 | 6.42721 | 6.56612 | 5.92974 | 6.57952 | 6.17832 | 6.65614 | 6.30595 | 6.41816 | 5.80091 | 6.40911 | 5.6091 | 6.18234 | 5.81209 |
| DCXR | Q7Z4W1 | 6.35529 | 6.04926 | 6.14067 | 6.62944 | 6.35798 | 6.01416 | 6.46652 | 5.88425 | 6.1266 | 6.07313 | 6.2578 | 6.33168 | 6.27763 | 6.07615 | 6.63576 | 6.3237 |
| RBP7 | Q96R05 | 6.53406 | 6.5387 | 6.52822 | 5.86983 | 7.00887 | 7.12611 | 6.57063 | 7.48116 | 6.18999 | 6.89191 | 6.74214 | 6.47381 | 4.98171 | 5.09497 | 4.76441 | 5.20347 |
| PCGF6 | Q9BYE7 | 6.22667 | 5.68077 | 6.39189 | 6.46478 | 6.73785 | 6.48749 | 6.1227 | 6.18042 | 6.34199 | 6.32393 | 6.52998 | 6.55899 | 5.95809 | 5.89487 | 5.99299 | 6.1066 |
| ZNHIT2 | Q9UHR6 | 6.68088 | 6.19046 | 6.71305 | 5.67934 | 6.16787 | 7.75807 | 5.82405 | 6.20907 | 6.53293 | 6.82557 | 6.8035 | 6.67479 | 6.12523 | 5.11235 | 5.29635 | 5.40648 |
| TRIM3 | O75382 | 6.19992 | 6.40576 | 6.67935 | 5.94938 | 5.98412 | 6.34621 | 6.22835 | 5.7754 | 6.48756 | 6.99147 | 6.88228 | 6.56191 | 6.23571 | 5.96547 | 5.53705 | 5.77006 |
| FAM92A | A1XBS5 | 6.58141 | 6.46686 | 7.39269 | 6.32083 | 6.44737 | 6.4706 | 6.30685 | 6.65756 | 6.55971 | 6.39762 | 6.7717 | 6.98405 | 5.31335 | 5.07331 | 5.15558 | 5.1005 |
| SNX24 | Q9Y343 | 6.29215 | 6.27417 | 7.10968 | 6.00198 | 6.42639 | 6.85827 | 6.53131 | 6.42333 | 6.25381 | 6.92065 | 6.65402 | 6.47645 | 5.26019 | 5.31426 | 5.47337 | 5.72996 |
| RFK | Q969G6 | 5.30334 | 6.20276 | 6.48079 | 6.04664 | 6.205 | 7.14915 | 6.49079 | 7.11826 | 6.96512 | 7.00041 | 7.4271 | 7.30207 | 5.02419 | 5.23766 | 5.12837 | 4.91834 |
| R3HDM2 | Q9Y2K5 | 6.15682 | 7.13847 | 6.1715 | 6.25724 | 5.84919 | 6.82454 | 5.91309 | 6.30845 | 6.36125 | 6.37536 | 6.89007 | 6.42137 | 5.81783 | 5.99623 | 5.55239 | 5.9662 |
| CCDC117 | Q8IWD4 | 4.96834 | 5.27829 | 6.52117 | 5.5917 | 6.03851 | 6.13376 | 6.50591 | 6.20656 | 6.24089 | 6.23087 | 7.08277 | 6.59674 | 7.02357 | 6.80815 | 6.0477 | 6.72506 |
| C18orf21 | Q32NC0 | 6.51263 | 6.19285 | 6.01824 | 6.77819 | 6.15237 | 5.97739 | 6.28448 | 6.14462 | 5.94153 | 5.94994 | 6.36098 | 5.84689 | 6.54747 | 6.25173 | 6.61707 | 6.42362 |
| KIAA1841 | Q6NSI8 | 6.19672 | 6.73847 | 6.49464 | 6.72596 | 5.96144 | 6.40426 | 6.48766 | 6.40087 | 6.3108 | 6.40738 | 6.61688 | 6.75734 | 5.22764 | 5.39665 | 5.74742 | 6.12588 |
| SYPL1 | Q16563 | 6.70172 | 6.4069 | 5.85896 | 6.16488 | 6.34631 | 5.7563 | 6.27911 | 5.98898 | 6.49396 | 7.11056 | 6.70727 | 6.6279 | 5.74312 | 5.82155 | 6.17527 | 5.81721 |
| BOLA3 | Q53S33 | 6.56103 | 6.31906 | 6.50442 | 6.60699 | 6.61089 | 5.99231 | 5.24852 | 5.94665 | 6.42565 | 6.43578 | 6.36333 | 6.24301 | 6.50772 | 5.96535 | 6.16935 | 6.09992 |
| ROR1 | Q01973 | 6.32346 | 6.42948 | 6.33453 | 6.99945 | 6.42487 | 6.06151 | 6.57616 | 6.31624 | 6.69254 | 6.56965 | 5.45742 | 5.76498 | 6.12669 | 6.15394 | 6.57111 | 6.09794 |
| RPE | Q96AT9 | 5.53787 | 4.55811 | 6.19025 | 4.09953 | 4.59238 | 4.1893 | 5.26553 | 7.46337 | 4.80949 | 5.70495 | 5.06304 | 6.55506 | 4.77563 | 10.8234 | 8.80284 | 11.5692 |
| NBEAL2 | Q6ZNJ1 | 6.55741 | 6.60291 | 6.46498 | 6.80327 | 6.35704 | 6.37847 | 6.901 | 6.63782 | 7.33012 | 6.33346 | 6.30337 | 6.74173 | 5.45863 | 5.24794 | 4.86016 | 5.02169 |
| POLL | Q9UGP5 | 5.34301 | 5.83385 | 6.53247 | 5.6198 | 5.75049 | 6.90958 | 5.66994 | 6.07041 | 6.25986 | 6.49879 | 6.83358 | 6.46638 | 6.51998 | 6.65853 | 6.8697 | 6.16363 |
| GOSR2 | O14653-2 | 6.7281 | 7.09531 | 6.69967 | 6.1608 | 6.57951 | 6.57201 | 6.48502 | 5.96834 | 5.65683 | 6.25309 | 5.46537 | 5.79738 | 6.44919 | 5.63285 | 6.26946 | 6.18709 |
| MAP2K7 | O14733-2 | 5.72822 | 6.73722 | 6.13968 | 6.13182 | 5.64707 | 6.9212 | 6.79554 | 6.90554 | 6.59133 | 6.38811 | 7.11841 | 6.96873 | 5.3218 | 5.75694 | 5.23291 | 5.61546 |
| COX20 | Q5RI15-2 | 6.68279 | 6.56557 | 6.21124 | 6.56254 | 6.2382 | 6.25254 | 5.97303 | 6.12983 | 5.95572 | 6.00884 | 5.83461 | 5.80142 | 6.69827 | 6.0877 | 6.40989 | 6.58781 |
| R3HDM4 | Q96D70 | 5.12344 | 5.85351 | 5.66024 | 5.33806 | 6.9009 | 7.38468 | 6.68215 | 6.12529 | 6.14761 | 5.89815 | 6.15567 | 6.13387 | 6.99585 | 6.51829 | 6.54172 | 6.54056 |
| GEMIN8 | Q9NWZ8 | 6.20674 | 5.87672 | 6.80459 | 6.17495 | 6.56944 | 6.133 | 5.86779 | 6.86757 | 6.30431 | 6.3639 | 7.36965 | 7.22326 | 5.92888 | 5.28311 | 5.8368 | 5.18929 |
| SLC38A1 | Q9H2H9 | 6.24184 | 6.9193 | 6.06615 | 6.21744 | 6.16939 | 6.41175 | 6.44458 | 6.74573 | 6.26817 | 5.78122 | 5.77657 | 6.26423 | 6.04446 | 6.03369 | 6.50714 | 6.10833 |
| BBS7 | Q8IWZ6 | 6.74847 | 6.48321 | 6.45917 | 6.49749 | 6.76824 | 6.48201 | 6.03568 | 7.08511 | 6.86739 | 6.20843 | 6.41941 | 6.50849 | 5.21119 | 5.78153 | 5.23954 | 5.20464 |
| FDX1 | P10109 | 6.34709 | 6.32199 | 5.73633 | 5.55747 | 6.77919 | 6.10735 | 6.73057 | 5.1913 | 6.75529 | 5.84325 | 6.17414 | 7.404 | 6.44107 | 5.36283 | 6.8385 | 6.40962 |
| EXT1 | Q16394 | 6.29613 | 6.68882 | 6.11642 | 6.40173 | 6.28281 | 6.17848 | 6.76403 | 6.31669 | 6.13995 | 5.98889 | 5.54363 | 6.05612 | 6.38574 | 6.33912 | 6.3409 | 6.16053 |
| MGST1 | P10620 | 5.78145 | 6.3163 | 5.71609 | 6.32526 | 6.97761 | 6.11431 | 6.36879 | 6.33184 | 6.17139 | 5.53609 | 6.53852 | 6.11166 | 6.44031 | 6.36404 | 7.06647 | 5.83988 |
| TNK2 | Q07912 | 6.2683 | 6.99181 | 7.37713 | 6.99633 | 5.86325 | 6.62663 | 6.69021 | 6.2628 | 6.3661 | 6.72255 | 6.92012 | 6.8904 | 4.9075 | 4.97669 | 4.84797 | 5.29221 |
| CIR1 | Q86X95 | 6.05399 | 6.5955 | 5.99232 | 6.40807 | 6.20987 | 6.03997 | 6.15792 | 5.92564 | 5.91748 | 6.258 | 6.25219 | 5.99546 | 6.32345 | 6.95524 | 6.22869 | 6.68622 |
| KLHL22 | Q53GT1 | 5.86549 | 6.57233 | 6.53646 | 6.49811 | 5.71488 | 6.29499 | 6.40461 | 6.50122 | 6.19689 | 6.24189 | 6.37836 | 6.31428 | 6.17438 | 6.2027 | 6.12188 | 5.98154 |
| DPAGT1 | Q9H3H5 | 6.11218 | 5.75621 | 5.93372 | 6.5584 | 6.02411 | 5.87653 | 6.10983 | 6.04025 | 6.31165 | 5.92846 | 5.79807 | 5.68449 | 7.10649 | 6.87717 | 6.83397 | 7.04848 |
| SSX2IP | Q9Y2D8 | 6.56477 | 6.59254 | 6.91886 | 6.524 | 6.20856 | 6.39573 | 6.08958 | 6.68706 | 6.37789 | 7.07541 | 6.80682 | 6.46992 | 5.54966 | 5.7105 | 4.93775 | 5.09095 |
| GPAT2 | Q6NUI2-4 | 7.19251 | 7.17679 | 7.1449 | 7.97992 | 6.71623 | 6.956 | 7.15651 | 6.2274 | 7.58853 | 6.48305 | 7.59463 | 7.51398 | 3.73294 | 3.67975 | 3.35142 | 3.50544 |
| AFTPH | Q6ULP2-9 | 6.35643 | 5.94973 | 6.95912 | 6.34771 | 6.34232 | 6.8481 | 6.24938 | 6.24126 | 6.44835 | 6.58252 | 7.21813 | 6.79524 | 5.72802 | 5.11672 | 5.3304 | 5.48659 |
| GLRX2 | Q9NS18-2 | 6.58524 | 7.09771 | 6.54183 | 7.26423 | 6.38406 | 5.91326 | 6.10107 | 6.72097 | 6.59719 | 6.39279 | 7.06977 | 6.45516 | 5.51586 | 5.08331 | 5.2736 | 5.00394 |
| SLC26A6 | Q9BXS9 | 5.97103 | 6.65048 | 6.1329 | 6.73537 | 5.82755 | 6.23867 | 5.90695 | 5.62429 | 6.58275 | 6.22158 | 5.74405 | 6.34077 | 6.63846 | 6.44336 | 6.71916 | 6.22264 |
| YIF1A | O95070 | 6.22607 | 6.50816 | 6.17733 | 6.68692 | 6.40342 | 5.80756 | 5.90011 | 5.71775 | 6.14899 | 5.97329 | 6.10171 | 5.89014 | 6.63954 | 6.35494 | 6.99711 | 6.46695 |
| COMMD6 | Q7Z4G1-2 | 5.95687 | 5.0446 | 6.30204 | 4.48889 | 5.89336 | 5.80078 | 5.31935 | 6.82063 | 5.81679 | 6.43206 | 6.09072 | 6.46296 | 5.79091 | 8.66627 | 7.44166 | 7.67211 |
| TRAPPC1 | Q9Y5R8 | 6.2696 | 5.84825 | 6.75291 | 5.68259 | 6.43513 | 6.44546 | 6.3539 | 6.63841 | 6.38848 | 7.10038 | 6.76793 | 6.72199 | 5.75863 | 5.41011 | 6.13995 | 5.28628 |

| | | | | | | | | | | | | | | | | | |
|---|---|---|---|---|---|---|---|---|---|---|---|---|---|---|---|---|---|
| NMES1 | Q9C002 | 6.24938 | 5.7126 | 6.63092 | 6.40051 | 5.52626 | 5.54235 | 6.06117 | 6.03861 | 6.04121 | 5.28974 | 5.33957 | 5.27439 | 6.32812 | 8.00399 | 7.32611 | 8.23507 |
| TFAP4 | Q01664 | 6.17356 | 6.65506 | 6.52834 | 6.54205 | 7.45513 | 6.20411 | 6.58905 | 6.25506 | 6.35944 | 6.06758 | 6.15512 | 6.28988 | 6.70296 | 4.81064 | 5.11367 | 5.75836 |
| TECPR2 | O15040 | 5.67478 | 6.18805 | 6.88452 | 6.15402 | 6.27375 | 6.23549 | 6.15964 | 5.78289 | 6.48014 | 6.33721 | 6.2215 | 6.64146 | 6.09993 | 6.56547 | 6.29074 | 6.01042 |
| ZNF280B | Q86YH2 | 6.28209 | 6.8003 | 6.21854 | 6.95342 | 6.67478 | 6.15495 | 5.75914 | 5.92892 | 6.71107 | 6.15704 | 6.57562 | 6.14987 | 6.24788 | 5.66648 | 6.03748 | 5.68242 |
| IGSF8 | Q969P0 | 6.75542 | 6.64085 | 6.06725 | 7.01194 | 6.15747 | 6.33035 | 6.52208 | 6.12473 | 6.20219 | 6.50066 | 6.20748 | 6.28191 | 5.95347 | 5.2418 | 6.05805 | 5.94435 |
| FAM118B | Q9BPY3 | 5.95862 | 5.76245 | 6.1212 | 4.99005 | 5.67608 | 5.76486 | 5.64229 | 5.91804 | 6.37563 | 6.72419 | 6.48567 | 5.90178 | 7.08404 | 7.31261 | 7.5767 | 6.7058 |
| UBE2B | P63146 | 6.02508 | 5.59962 | 6.64982 | 5.21094 | 6.19385 | 5.74428 | 5.83849 | 6.79622 | 5.43721 | 7.21763 | 6.21418 | 7.55665 | 5.15477 | 7.36785 | 6.22676 | 6.76664 |
| GPCPD1 | Q9NPB8 | 6.26932 | 6.47837 | 6.32926 | 5.85324 | 5.99595 | 7.06944 | 6.36497 | 6.58972 | 6.09493 | 5.98303 | 6.62516 | 6.50046 | 6.13206 | 5.98035 | 5.81856 | 5.91518 |
| MRPL51 | Q4U2R6 | 6.21409 | 6.45052 | 5.58501 | 6.56609 | 6.63851 | 6.16488 | 6.2718 | 6.21855 | 5.92612 | 6.22304 | 5.74585 | 5.44062 | 6.63863 | 6.58336 | 6.55193 | 6.781 |
| RAPH1 | Q70E73 | 5.91883 | 5.53613 | 6.94587 | 5.76062 | 6.60291 | 6.59259 | 6.93639 | 6.06992 | 6.77322 | 6.02907 | 6.97 | 6.59242 | 6.38911 | 5.91862 | 5.38669 | 5.57763 |
| MRPS36 | P82909 | 6.09897 | 5.80566 | 6.32639 | 6.77424 | 6.10525 | 5.55155 | 5.81688 | 6.25498 | 6.21073 | 6.103 | 6.63247 | 6.84571 | 7.00922 | 6.27616 | 5.97543 | 6.21337 |
| SMAP2 | Q8WU79 | 6.20432 | 6.07283 | 7.48379 | 6.12036 | 6.16923 | 7.00382 | 6.11127 | 6.52937 | 6.23259 | 6.56116 | 6.76623 | 6.43585 | 6.52611 | 5.90508 | 5.18988 | 5.58811 |
| RIC1 | Q4ADV7 | 6.06379 | 7.16476 | 6.57197 | 6.22557 | 6.32343 | 6.47899 | 6.1661 | 5.94388 | 6.49894 | 6.58863 | 6.83649 | 6.56123 | 5.29217 | 5.8961 | 5.59473 | 5.79323 |
| LYRM2 | Q9NU23 | 5.91171 | 5.82711 | 6.48395 | 6.31816 | 6.08858 | 5.87337 | 6.34197 | 6.49514 | 5.51431 | 6.64063 | 5.91341 | 6.02458 | 6.44538 | 6.6018 | 6.75656 | 6.76333 |
| ALG11 | Q2TAA5 | 5.98356 | 6.44619 | 5.63359 | 7.07578 | 5.84037 | 5.68642 | 6.05801 | 5.86603 | 5.84799 | 5.99055 | 6.50309 | 6.09679 | 6.88734 | 6.47188 | 7.14996 | 6.46248 |
| VPS54 | Q9P1Q0 | 5.8453 | 6.13624 | 6.78 | 6.14563 | 5.91641 | 6.19903 | 5.78869 | 6.57457 | 7.05973 | 6.62644 | 6.12392 | 6.35965 | 5.70303 | 5.89802 | 6.29326 | 6.54108 |
| SSR4 | P51571 | 6.60704 | 6.71368 | 5.76216 | 6.57987 | 7.17926 | 6.52258 | 6.01647 | 6.57817 | 6.04972 | 5.5946 | 5.93581 | 5.72905 | 6.37087 | 6.10471 | 6.57467 | 5.68134 |
| PNPLA2 | Q96AD5 | 6.0689 | 5.89115 | 6.751 | 5.76276 | 6.96538 | 7.26089 | 6.99432 | 6.70126 | 6.09801 | 6.11698 | 6.6798 | 6.57781 | 5.80253 | 5.40241 | 5.44619 | 5.48061 |
| UNC93B1 | Q9H1C4 | 6.33988 | 6.53686 | 6.26204 | 6.23918 | 5.99339 | 5.90059 | 5.90655 | 6.41447 | 6.32672 | 6.47285 | 6.19121 | 6.07406 | 6.21557 | 6.44853 | 6.20376 | 6.47436 |
| IFI30 | P13284 | 5.6651 | 6.2306 | 5.90878 | 5.76403 | 5.85441 | 6.09177 | 5.18444 | 6.31023 | 5.62722 | 5.7344 | 5.89639 | 6.37843 | 6.75483 | 7.63173 | 6.82971 | 7.23792 |
| PVR | P15151 | 5.60959 | 7.063 | 5.48684 | 6.37518 | 6.08726 | 6.00986 | 6.26603 | 6.55374 | 6.50335 | 5.52377 | 5.23006 | 6.39316 | 6.71069 | 6.02654 | 6.9282 | 7.23273 |
| UBE3D | Q7Z6J8 | 6.59247 | 6.61172 | 7.31498 | 6.46166 | 6.26522 | 7.06604 | 6.4288 | 6.48352 | 6.51444 | 6.15249 | 6.59881 | 6.58492 | 5.13874 | 5.42259 | 5.18785 | 5.16929 |
| FBXO10 | Q9UK96 | 6.27367 | 6.87448 | 7.06372 | 6.69054 | 5.9881 | 6.14068 | 5.74613 | 6.07052 | 7.23833 | 7.53473 | 6.63592 | 6.96553 | 4.84518 | 5.13185 | 5.58293 | 5.21768 |
| SYT1 | P21579 | 6.28522 | 6.34431 | 5.79288 | 6.30942 | 6.39498 | 5.85479 | 6.21204 | 5.89255 | 6.37214 | 6.23943 | 5.73714 | 5.78264 | 6.53564 | 6.68977 | 7.00323 | 6.55381 |
| SNRPC | P09234 | 6.04476 | 4.97374 | 5.52863 | 5.4466 | 6.66144 | 5.25123 | 5.68412 | 6.36614 | 5.1527 | 6.43298 | 6.13885 | 7.00903 | 6.5397 | 7.30349 | 7.56215 | 7.90444 |
| PRXL2B | Q8TBF2-3 | 6.21295 | 6.17145 | 7.06145 | 5.92832 | 6.25788 | 6.8031 | 6.41458 | 6.89809 | 6.67933 | 5.86373 | 6.38525 | 6.54131 | 5.78233 | 5.77598 | 5.18312 | 6.04115 |
| TMEM14C | Q9P0S9 | 6.50151 | 6.2712 | 5.97244 | 6.41825 | 5.97592 | 6.85092 | 4.9151 | 6.11304 | 5.53929 | 6.58657 | 5.54063 | 6.17083 | 6.98113 | 6.86037 | 7.22951 | 6.97429 |
| CNTN1 | Q12860 | 5.86189 | 6.26761 | 6.23417 | 6.54067 | 6.44431 | 6.07718 | 6.05705 | 6.73175 | 5.82174 | 6.43663 | 5.81452 | 5.87634 | 6.43506 | 6.29692 | 6.63582 | 6.46832 |
| PIP5K1A | Q99755 | 6.58844 | 6.16818 | 6.69868 | 6.9642 | 6.5764 | 5.6031 | 6.72004 | 6.12031 | 6.49712 | 6.20679 | 6.3495 | 6.08271 | 6.49386 | 5.16394 | 6.04477 | 5.72198 |
| PIP5K1C | O60331-3 | 6.21267 | 5.91611 | 6.96627 | 5.25221 | 6.39117 | 7.45152 | 6.39135 | 5.97264 | 6.4534 | 7.0616 | 6.39796 | 7.01231 | 5.98124 | 5.74169 | 5.58174 | 5.21551 |
| PDCD7 | Q8N8D1 | 6.23036 | 5.85133 | 7.13253 | 7.701 | 7.33657 | 6.26679 | 6.2622 | 5.19903 | 5.71981 | 6.37202 | 6.12057 | 6.17842 | 6.07471 | 5.53064 | 6.01353 | 6.01047 |
| PFDN4 | Q9NQP4 | 6.82202 | 5.5178 | 6.53544 | 4.37506 | 5.84214 | 5.289 | 5.72027 | 6.83482 | 5.53159 | 6.73391 | 6.04093 | 6.4128 | 5.77005 | 7.94145 | 7.28099 | 7.35173 |
| RUFY3 | Q7L099-3 | 5.43126 | 6.03281 | 6.6758 | 5.46053 | 6.74433 | 6.73869 | 6.1157 | 6.73778 | 6.07651 | 6.899 | 6.92081 | 6.68327 | 6.10897 | 5.86854 | 5.47394 | 6.03204 |
| HSPA1L | P34931 | 6.41855 | 6.3828 | 6.12478 | 6.70888 | 5.78101 | 5.97814 | 6.17832 | 6.18475 | 6.59724 | 6.11424 | 6.65639 | 6.87851 | 6.30276 | 5.77468 | 6.30956 | 6.50939 |
| SLC39A11 | Q8N1S5 | 5.95655 | 6.55301 | 5.65352 | 5.80966 | 6.28189 | 5.23366 | 6.86815 | 6.01592 | 5.82099 | 6.20997 | 6.25814 | 6.45148 | 7.01953 | 6.47758 | 7.1231 | 6.26685 |
| TPM1 | H0YK48 | 6.58332 | 6.44925 | 6.91461 | 5.67381 | 5.85623 | 5.62609 | 5.81672 | 6.27377 | 6.33266 | 6.15887 | 6.62966 | 6.48199 | 7.2507 | 6.36363 | 5.78374 | 5.80494 |
| ASAP3 | Q8TDY4 | 6.04235 | 6.02007 | 6.94132 | 5.88186 | 6.24566 | 6.54188 | 6.18581 | 6.54476 | 6.58993 | 6.9831 | 6.70815 | 6.35808 | 5.27894 | 6.14759 | 6.06216 | 5.46835 |
| YBX3 | P16989-2 | 6.72231 | 6.02338 | 6.76845 | 6.38551 | 6.14233 | 5.98566 | 6.76256 | 6.07469 | 6.67122 | 6.29254 | 6.70895 | 6.45211 | 6.11396 | 5.56523 | 5.7111 | 5.64793 |
| PTMA | P06454-2 | 6.36031 | 6.08491 | 6.88917 | 5.91781 | 5.88917 | 6.38566 | 5.5194 | 7.23808 | 6.2942 | 6.23455 | 6.56165 | 6.49347 | 5.9251 | 6.62218 | 5.67112 | 5.91325 |
| NXT2 | Q9NPJ8-3 | 5.84037 | 5.6418 | 5.68806 | 6.65041 | 6.08851 | 5.78455 | 6.42422 | 6.32717 | 5.96948 | 6.05201 | 6.22253 | 6.58628 | 6.50392 | 6.49086 | 6.68838 | 7.04144 |
| MARCHF6 | O60337 | 7.36499 | 7.77719 | 6.81956 | 7.86275 | 6.41517 | 6.07445 | 6.28364 | 6.31594 | 6.51381 | 6.66931 | 6.40382 | 6.56521 | 4.58685 | 4.79334 | 4.52648 | 5.0275 |
| ZNF131 | P52739 | 6.62764 | 6.32169 | 6.20166 | 6.56739 | 6.07472 | 5.56486 | 6.30621 | 6.24493 | 6.2807 | 6.73657 | 5.89199 | 7.10177 | 5.42631 | 6.33063 | 5.87961 | 6.44333 |
| MSTO1 | Q9BUK6 | 5.88147 | 5.43468 | 6.51876 | 5.43058 | 6.75835 | 5.96873 | 6.32966 | 6.63281 | 6.95298 | 6.72049 | 7.07795 | 6.59655 | 5.87533 | 5.77568 | 5.88042 | 6.16556 |
| PCDHB2 | Q9Y5E7 | 6.07852 | 6.4236 | 5.69702 | 5.8873 | 7.17042 | 6.55856 | 6.71479 | 6.69598 | 5.65428 | 5.93005 | 5.76362 | 6.38748 | 6.46822 | 6.2615 | 6.05679 | 6.25186 |
| PCDHB14 | Q9Y5E9 | 6.11376 | 6.07632 | 6.21594 | 6.81697 | 6.33703 | 6.42908 | 6.79344 | 6.5275 | 6.5793 | 6.77773 | 6.25541 | 6.50229 | 6.57162 | 5.67685 | 5.63461 | 5.59215 |
| PCDHB6 | Q9Y5E3 | 6.27395 | 6.13014 | 5.88025 | 6.49465 | 6.5828 | 6.1337 | 6.45917 | 6.7212 | 5.74446 | 6.14727 | 5.68855 | 6.08259 | 6.42215 | 6.01056 | 6.37575 | 6.85279 |
| TYSND1 | Q2T9J0 | 6.83795 | 5.88971 | 6.43495 | 6.85299 | 5.77008 | 6.77488 | 6.45305 | 5.81139 | 5.91456 | 5.84016 | 6.12843 | 6.53765 | 6.05083 | 5.95839 | 6.31618 | 6.42881 |
| SPATS2L | C9JGM8 | 6.21374 | 6.55666 | 6.5332 | 6.03421 | 5.24828 | 5.93572 | 5.97082 | 6.2587 | 6.73631 | 6.06875 | 6.11608 | 6.70226 | 6.01283 | 5.60844 | 7.19326 | 6.81073 |
| SMARCD3 | Q6STE5 | 5.56478 | 6.1344 | 6.02897 | 5.78007 | 5.96504 | 6.48371 | 6.31318 | 6.10658 | 5.58226 | 5.90399 | 6.37936 | 5.87202 | 6.91699 | 6.91149 | 6.70845 | 7.34872 |
| RIDA | P52758 | 6.12689 | 4.75321 | 5.23395 | 5.003 | 5.60578 | 5.33222 | 5.42833 | 6.47583 | 5.33439 | 6.63812 | 5.97885 | 6.9932 | 6.20446 | 8.75252 | 8.09046 | 8.0488 |
| SRC | P12931-2 | 6.37272 | 6.18556 | 6.16505 | 6.12079 | 5.71027 | 6.10044 | 6.43455 | 6.12005 | 6.26057 | 6.3879 | 6.22855 | 6.28284 | 6.14766 | 6.49093 | 6.52024 | 6.47188 |
| PARD3B | Q8TEW8 | 5.81373 | 6.6787 | 6.08049 | 7.16574 | 5.84414 | 5.87323 | 6.11416 | 6.17878 | 6.34823 | 6.41649 | 5.46236 | 6.35703 | 6.54868 | 6.04771 | 6.228 | 6.84252 |
| SIDT2 | Q8NBJ9-2 | 6.56095 | 6.66266 | 6.45837 | 6.86666 | 5.64739 | 6.18023 | 6.32606 | 6.0499 | 5.96531 | 6.85855 | 6.15357 | 6.58551 | 5.80124 | 6.37974 | 5.98916 | 5.51472 |
| TCOF1 | Q13428-7 | 5.18104 | 5.48816 | 5.67097 | 6.89977 | 5.76968 | 5.90601 | 5.31551 | 5.54868 | 5.99214 | 5.79081 | 5.18605 | 5.24832 | 7.46517 | 7.7041 | 8.3894 | 8.44417 |
| CELF2 | O95319-3 | 6.56279 | 6.50158 | 5.94297 | 6.07398 | 6.22553 | 6.27199 | 5.1845 | 6.1232 | 6.24239 | 6.17063 | 6.54687 | 6.25476 | 6.18089 | 5.76594 | 6.46168 | 6.49029 |
| ZNF706 | Q9Y5V0 | 6.53458 | 6.74059 | 6.92299 | 6.41232 | 6.16747 | 6.34337 | 5.91465 | 6.55309 | 6.57661 | 6.7557 | 6.26333 | 6.35773 | 4.95783 | 5.892 | 5.57765 | 6.03008 |
| FOXO3 | O43524 | 6.4328 | 6.29843 | 7.09864 | 6.30756 | 6.22543 | 6.55903 | 6.3734 | 7.04856 | 6.28926 | 6.15306 | 6.86551 | 6.80469 | 5.81151 | 5.70918 | 5.02599 | 4.99696 |
| PPP2R5B | Q15173 | 5.9587 | 6.05355 | 6.51109 | 6.28013 | 5.58354 | 6.68596 | 5.69951 | 6.04467 | 6.17377 | 6.3296 | 7.37679 | 6.83387 | 6.00182 | 6.29612 | 6.43935 | 5.73152 |
| TUBB4A | P04350 | 6.39339 | 6.1041 | 7.30238 | 6.58161 | 6.12366 | 6.87658 | 5.53973 | 6.7665 | 6.9331 | 6.90446 | 6.4984 | 7.07645 | 5.62733 | 5.30184 | 5.03969 | 4.93079 |
| TRIM44 | Q96DX7 | 6.54458 | 6.72286 | 7.06825 | 6.37089 | 6.84899 | 6.44954 | 6.23278 | 6.74974 | 6.53196 | 7.19982 | 7.04494 | 7.07177 | 4.73635 | 5.11172 | 4.73925 | 5.03618 |
| MCL1 | Q07820 | 8.1306 | 6.51753 | 6.68421 | 8.00613 | 6.43114 | 7.16452 | 7.9033 | 6.37429 | 5.89721 | 6.10078 | 6.05894 | 6.39482 | 4.30527 | 4.49241 | 4.60624 | 4.93261 |
| CHCHD2 | Q9Y6H1 | 3.95829 | 4.0799 | 3.99225 | 4.14772 | 8.49044 | 8.02642 | 7.9231 | 8.29987 | 4.60756 | 4.23197 | 4.07059 | 4.51126 | 8.96932 | 8.01646 | 7.47037 | 9.20449 |
| MBTD1 | Q05BQ5 | 6.98387 | 6.35384 | 5.94576 | 7.4473 | 6.91812 | 6.39094 | 6.60535 | 5.6087 | 6.13953 | 6.62975 | 5.9636 | 5.98646 | 6.10725 | 5.57931 | 5.7729 | 5.56733 |
| ZNRD2 | O60232 | 6.46866 | 4.80527 | 5.88105 | 4.08432 | 5.99135 | 5.38214 | 6.0242 | 6.4481 | 4.70408 | 5.7334 | 5.10142 | 6.07339 | 5.69024 | 9.82349 | 7.77135 | 10.0175 |
| ZNF827 | Q17R98 | 5.959 | 6.50509 | 5.94541 | 6.5612 | 6.12742 | 5.68172 | 5.86411 | 6.28505 | 6.57983 | 5.94263 | 6.19074 | 6.33632 | 6.48521 | 5.99422 | 6.7678 | 6.77427 |
| P2RX4 | Q99571-2 | 6.70221 | 7.11496 | 6.54509 | 6.80912 | 5.66002 | 6.5528 | 6.25666 | 5.53619 | 5.54365 | 6.56347 | 6.35131 | 6.03877 | 6.34969 | 6.39529 | 5.99075 | 5.59003 |
| DYSF | O75923-13 | 6.23995 | 6.22161 | 6.50926 | 6.37956 | 6.04114 | 5.71979 | 6.51368 | 5.95204 | 6.13723 | 6.46925 | 6.24235 | 5.93617 | 6.20525 | 6.54278 | 6.4234 | 6.46653 |
| KCNQ5 | Q9NR82-6 | 6.65062 | 6.20587 | 6.47021 | 7.1999 | 6.05282 | 5.34256 | 6.49859 | 5.91509 | 6.4599 | 6.24313 | 6.47951 | 6.29051 | 6.03742 | 5.92226 | 6.0982 | 6.13342 |
| CCDC71L | Q8N9Z2 | 6.36196 | 6.20309 | 5.36341 | 6.69644 | 6.16237 | 6.262 | 6.48871 | 5.2884 | 6.75139 | 5.30384 | 5.84583 | 5.46735 | 6.59559 | 7.86388 | 6.33351 | 7.01223 |
| TSHZ3 | Q63HK5 | 6.50106 | 6.15435 | 5.72469 | 6.42908 | 6.1503 | 6.16973 | 6.43667 | 5.4076 | 6.21416 | 6.60509 | 6.57414 | 6.19198 | 6.3897 | 6.30468 | 6.18351 | 6.56326 |
| ZNF518B | Q9C0D4 | 6.75779 | 6.90387 | 5.90101 | 6.74409 | 6.22372 | 6.44958 | 6.86154 | 5.96021 | 6.25877 | 6.56613 | 6.44636 | 6.6544 | 5.57985 | 5.43855 | 5.1132 | 6.14094 |
| AKAP11 | Q9UKA4 | 6.48806 | 6.05966 | 6.12053 | 6.38355 | 6.59527 | 5.85712 | 6.55951 | 6.46611 | 5.87142 | 6.02916 | 6.74655 | 6.41377 | 5.93633 | 6.58537 | 5.49024 | 6.39735 |
| TBC1D25 | Q3MII6 | 5.81807 | 5.47263 | 6.41226 | 5.64694 | 6.42126 | 6.58299 | 6.39568 | 6.83461 | 6.26863 | 6.10281 | 5.83936 | 6.63941 | 6.33742 | 6.49287 | 6.18177 | 6.55328 |
| LCMT2 | O60294 | 6.16355 | 7.07382 | 6.63629 | 6.41045 | 6.33269 | 5.63526 | 6.80526 | 6.88638 | 6.60053 | 7.20841 | 6.50353 | 6.68683 | 6.48217 | 5.0181 | 5.17111 | 4.38563 |
| DST | Q03001-11 | 6.20685 | 7.00831 | 5.59408 | 5.94027 | 6.27799 | 6.20064 | 6.80462 | 6.15115 | 6.72345 | 5.94522 | 5.89794 | 5.68297 | 6.70348 | 6.61774 | 6.48509 | 5.76019 |
| CYTH1 | Q15438 | 6.1187 | 5.91871 | 6.81253 | 5.87191 | 6.15607 | 6.58992 | 6.40183 | 5.98507 | 6.04137 | 6.31725 | 7.25046 | 6.32982 | 5.91885 | 6.29425 | 6.13503 | 5.85824 |
| CYTH3 | O43739 | 5.92509 | 6.07031 | 7.06492 | 6.7594 | 6.14176 | 5.89716 | 6.82 | 6.82975 | 6.59375 | 6.36183 | 6.36421 | 6.36283 | 5.89824 | 5.70176 | 5.38229 | 5.82671 |
| SDHAF3 | Q9NRP4 | 7.10955 | 7.08275 | 6.86304 | 6.84657 | 6.01997 | 5.62238 | 6.23383 | 5.91658 | 7.00338 | 7.01956 | 5.92174 | 7.03081 | 5.84873 | 5.57847 | 4.59126 | 5.31137 |
| DDHD1 | Q8NEL9 | 5.55367 | 7.02893 | 7.49379 | 5.37886 | 5.36409 | 6.46498 | 6.00294 | 5.93607 | 6.75902 | 6.61658 | 6.39204 | 6.94355 | 6.22583 | 5.53623 | 6.22251 | 6.08091 |
| GLUL | P15104 | 6.28717 | 7.81743 | 7.08993 | 7.61527 | 6.44899 | 6.3934 | 6.07314 | 6.74295 | 7.17242 | 6.67842 | 7.18537 | 6.43008 | 4.85207 | 4.45235 | 4.40082 | 4.3602 |
| RPS3 | E9PJN9 | 6.16443 | 6.24297 | 7.39461 | 6.0764 | 5.36314 | 6.10033 | 6.44901 | 6.67659 | 6.50596 | 6.62028 | 6.92935 | 6.66622 | 6.12185 | 5.34732 | 5.98742 | 5.35411 |
| ZFYVE9 | O95405 | 6.40207 | 6.37473 | 6.65296 | 6.60397 | 6.64284 | 5.92971 | 6.24038 | 6.3474 | 6.54119 | 6.54186 | 6.41372 | 6.65398 | 5.5404 | 5.87599 | 5.87937 | 5.35945 |
| VAMP2 | P63027 | 6.46221 | 6.65 | 5.80431 | 6.38078 | 6.31306 | 5.51721 | 6.29884 | 6.39528 | 5.65506 | 5.88404 | 5.87394 | 6.00149 | 6.17634 | 7.14923 | 6.8566 | 6.58161 |
| VAMP3 | Q15836 | 6.23143 | 7.39246 | 5.23551 | 6.37633 | 6.59753 | 5.8781 | 6.41447 | 5.82352 | 6.43717 | 7.08435 | 6.22433 | 6.52083 | 5.77215 | 5.68023 | 6.25945 | 6.07215 |
| RNASEH2A | O75792 | 6.01327 | 5.73056 | 6.74361 | 5.20019 | 6.24281 | 6.43726 | 6.16757 | 6.9245 | 5.9735 | 6.68173 | 6.70877 | 6.15788 | 5.93319 | 6.28006 | 6.66987 | 6.13523 |
| KLHDC2 | Q9Y2U9 | 7.07102 | 6.59242 | 6.60862 | 6.59721 | 5.76635 | 6.4686 | 5.97071 | 5.79377 | 6.80453 | 7.37005 | 7.02356 | 6.88132 | 5.15776 | 5.11476 | 5.65231 | 5.12701 |
| CCDC127 | Q96BQ5 | 6.53956 | 6.51445 | 5.96976 | 5.8817 | 6.53698 | 6.17251 | 6.5593 | 5.7939 | 6.17093 | 5.86332 | 6.25917 | 6.02814 | 6.47983 | 6.06293 | 6.85599 | 6.31155 |
| CTPS2 | Q9NRF8 | 6.38613 | 6.34946 | 7.39901 | 6.66649 | 5.83653 | 6.36097 | 6.5459 | 6.07125 | 7.06345 | 6.04247 | 6.40941 | 6.60977 | 5.8615 | 5.66215 | 5.56793 | 5.16756 |
| TMED1 | Q13445 | 6.16699 | 5.88753 | 5.5772 | 6.07042 | 6.39573 | 6.029 | 6.12385 | 5.83521 | 6.38055 | 5.93885 | 5.62926 | 6.0457 | 6.88416 | 7.09529 | 7.11432 | 6.82595 |
| SYNCRIP | O60506-3 | 6.45251 | 6.16293 | 6.3781 | 6.4843 | 6.55786 | 6.06011 | 6.07012 | 6.70177 | 6.84535 | 6.20798 | 6.07297 | 6.01205 | 6.15817 | 6.10878 | 5.67011 | 6.05689 |
| HNRNPR | A0A6Q8PHG0 | 6.64969 | 6.42501 | 5.71521 | 6.90499 | 6.28013 | 6.21001 | 6.03164 | 6.15039 | 6.13989 | 5.6264 | 6.04657 | 5.92507 | 6.68088 | 6.05937 | 6.71174 | 6.45801 |
| LHPP | Q9H008 | 6.58283 | 5.64917 | 6.96188 | 5.62889 | 6.32282 | 7.01295 | 5.77125 | 5.67397 | 7.22086 | 6.09596 | 6.09114 | 6.57485 | 6.46939 | 5.84064 | 6.12309 | 5.98031 |
| NRIP1 | P48552 | 6.33501 | 6.15274 | 6.48694 | 6.44814 | 6.42845 | 6.13353 | 7.04118 | 6.14785 | 6.26577 | 5.78673 | 6.65503 | 6.29341 | 6.11962 | 5.8235 | 5.85281 | 6.02928 |
| SGSH | P51688 | 4.71424 | 5.48767 | 9.58196 | 5.46793 | 4.9046 | 4.73435 | 17.118 | 4.72661 | 5.04218 | 5.10256 | 6.22366 | 4.74207 | 4.39032 | 4.03729 | 9.90494 | 3.82167 |
| SNTA1 | Q13424 | 6.42802 | 6.59352 | 6.04697 | 6.62918 | 6.40018 | 5.91285 | 6.00321 | 6.25958 | 6.72467 | 6.66983 | 6.21093 | 6.04603 | 6.35583 | 5.65569 | 6.12017 | 5.94334 |
| MAST2 | Q6P0Q8 | 6.39004 | 6.67846 | 6.74756 | 5.80649 | 6.23353 | 6.55639 | 5.85314 | 6.76168 | 6.57748 | 5.99049 | 6.54517 | 6.85186 | 5.62571 | 5.61407 | 5.84558 | 5.92235 |
| ZFAND6 | Q6FIF0 | 6.66402 | 6.2707 | 7.18788 | 6.1052 | 6.10162 | 6.1185 | 6.29274 | 6.78672 | 6.80769 | 6.28362 | 7.20174 | 6.66269 | 5.29882 | 5.69099 | 5.59869 | 4.92838 |
| CHPF | Q8IZ52 | 7.00112 | 6.4187 | 6.4084 | 7.55974 | 6.00967 | 6.18853 | 6.86089 | 6.18776 | 6.43937 | 6.18255 | 6.66754 | 6.35343 | 5.37611 | 5.65032 | 5.68074 | 5.01512 |
| SIRT3 | Q9NTG7 | 6.87937 | 5.55451 | 6.01221 | 4.99174 | 5.79723 | 4.8332 | 6.22993 | 5.77296 | 5.33795 | 6.49613 | 5.59052 | 6.35216 | 7.20854 | 7.76628 | 7.41597 | 7.76131 |
| ALKBH1 | Q13686 | 7.15873 | 6.88781 | 7.06044 | 7.50464 | 6.79374 | 5.898 | 6.78263 | 6.00394 | 6.2078 | 6.54874 | 7.2067 | 6.24337 | 5.75197 | 4.93197 | 4.94714 | 4.97237 |
| CMTM6 | Q9NX76 | 6.56787 | 6.77065 | 5.77113 | 6.08781 | 5.76792 | 6.08433 | 6.39423 | 6.05456 | 5.76352 | 6.16032 | 5.71525 | 6.01293 | 6.72185 | 6.93687 | 6.72578 | 6.46499 |
| ICAM5 | Q9UMF0 | 6.01552 | 6.58122 | 6.13961 | 5.49053 | 6.3239 | 6.22106 | 5.83927 | 6.7449 | 6.971 | 5.89687 | 5.92653 | 6.20622 | 6.75621 | 6.93647 | 6.50639 | 5.4443 |
| CHURC1-FNTB | B4DL54 | 6.28974 | 6.22366 | 6.84309 | 5.6422 | 6.76483 | 6.07076 | 6.05573 | 6.90803 | 6.69616 | 5.5705 | 6.89949 | 5.88525 | 6.07359 | 6.08882 | 6.06623 | 5.92191 |
| EEF2K | O00418 | 6.64237 | 6.65338 | 6.60122 | 6.27926 | 7.30034 | 7.92582 | 6.59462 | 7.29269 | 6.67129 | 6.62177 | 6.74879 | 6.661 | 4.61071 | 4.82473 | 4.12289 | 4.44911 |
| ZNF277 | Q9NRM2 | 6.0815 | 6.18357 | 5.99386 | 6.14847 | 7.03612 | 6.56621 | 6.67652 | 6.21858 | 6.58563 | 6.11474 | 5.44972 | 5.28333 | 6.70066 | 5.83924 | 6.57134 | 6.55048 |

| | | | | | | | | | | | | | | | | | |
|---|---|---|---|---|---|---|---|---|---|---|---|---|---|---|---|---|---|
| PAG1 | Q9NWQ8 | 6.88553 | 6.26958 | 6.14887 | 6.60484 | 5.77893 | 5.64765 | 6.22279 | 5.54037 | 5.9754 | 6.31929 | 6.33874 | 6.04497 | 7.26406 | 6.39973 | 6.61134 | 5.94792 |
| ARL6IP6 | Q8N6S5 | 6.29124 | 6.25007 | 5.52902 | 6.75411 | 5.77001 | 5.75221 | 6.28556 | 5.75485 | 6.18104 | 5.84928 | 6.03109 | 5.65451 | 6.84746 | 6.81645 | 7.31629 | 6.91682 |
| HMGN2 | P05204 | 6.45776 | 6.02944 | 5.11682 | 6.63558 | 5.74021 | 5.66159 | 6.3304 | 5.43797 | 5.80654 | 6.26307 | 6.31547 | 5.94839 | 6.1389 | 6.79462 | 7.81259 | 7.51066 |
| MALSU1 | Q96EH3 | 6.03067 | 6.07754 | 5.86353 | 6.2629 | 6.41003 | 6.24952 | 6.42241 | 6.23764 | 5.7132 | 6.40765 | 5.29299 | 6.29161 | 5.95727 | 6.97388 | 6.61496 | 7.19419 |
| ING5 | A0A1B0GW41 | 5.99726 | 5.92464 | 6.29614 | 6.76482 | 5.9655 | 5.53216 | 6.40135 | 5.67215 | 6.37765 | 6.43213 | 6.32326 | 6.105 | 6.20181 | 5.92479 | 7.17056 | 6.91079 |
| MTHFD1L | B7ZM99 | 6.19612 | 6.37246 | 5.8593 | 7.33227 | 6.76396 | 6.40187 | 6.21997 | 5.86617 | 6.09286 | 6.3883 | 6.28181 | 6.2418 | 6.7093 | 5.63016 | 5.83614 | 5.80751 |
| SVBP | Q8N300 | 6.27863 | 6.41529 | 6.91475 | 6.00029 | 6.03321 | 7.24349 | 6.48357 | 6.81704 | 6.63078 | 7.62814 | 6.78287 | 7.30892 | 4.87027 | 5.52807 | 4.41591 | 4.64877 |
| DLX6 | P56179-3 | 6.67018 | 6.71735 | 6.53004 | 6.80308 | 6.13696 | 6.21632 | 6.33222 | 5.9376 | 6.8359 | 6.29837 | 6.0689 | 6.45521 | 5.57959 | 5.34394 | 5.87301 | 6.20133 |
| PIP4P1 | Q86T03-2 | 6.25564 | 6.28876 | 6.28967 | 6.41206 | 6.31555 | 6.17804 | 6.45339 | 6.083 | 6.65044 | 6.41653 | 6.34446 | 5.64858 | 6.18151 | 5.99961 | 6.41643 | 6.06633 |
| LSM8 | O95777 | 6.67756 | 6.49769 | 5.48954 | 5.29471 | 6.82434 | 6.69084 | 5.94484 | 6.33627 | 5.55814 | 5.63902 | 6.34749 | 6.13908 | 6.37145 | 7.08651 | 6.58352 | 6.51901 |
| TMED3 | Q9Y3Q3 | 6.12458 | 5.9159 | 6.412 | 7.28016 | 6.26922 | 6.42191 | 6.67133 | 5.71919 | 6.78788 | 6.09173 | 6.07946 | 6.49014 | 5.59011 | 5.57203 | 6.30194 | 6.27243 |
| TMTC4 | Q5T4D3-3 | 5.86533 | 6.71613 | 6.80539 | 6.91318 | 5.8175 | 6.14112 | 5.92373 | 5.39395 | 5.85878 | 6.50096 | 6.14011 | 5.43394 | 5.75676 | 5.95409 | 6.84558 | 6.93346 |
| CDS1 | Q92903 | 5.82806 | 6.36587 | 6.10405 | 6.9747 | 5.46314 | 5.63335 | 6.14162 | 5.45598 | 6.57726 | 6.7125 | 6.34142 | 5.36177 | 6.66476 | 6.13522 | 6.84034 | 7.39997 |
| SMARCA1 | A0A0A0MRP6 | 5.88057 | 5.89976 | 5.59476 | 6.43141 | 5.94243 | 5.43416 | 6.14061 | 6.33903 | 6.97354 | 6.34555 | 6.19745 | 6.1701 | 6.53058 | 6.26842 | 6.83199 | 7.01964 |
| PML | P29590-13 | 6.48335 | 6.28827 | 6.10461 | 6.83278 | 5.77036 | 5.27097 | 6.44289 | 6.21999 | 6.10146 | 6.25964 | 6.66267 | 6.34776 | 6.48394 | 6.04443 | 6.39778 | 6.28909 |
| ASB6 | Q9NWX5 | 6.81031 | 6.56874 | 7.2023 | 5.90176 | 6.26329 | 7.18629 | 6.27434 | 6.85017 | 7.00232 | 6.66657 | 6.6736 | 6.89523 | 5.01648 | 4.95755 | 4.6089 | 5.12216 |
| HABP4 | Q5JVS0 | 5.54457 | 5.90887 | 6.61965 | 6.2135 | 7.32401 | 7.17538 | 6.30919 | 6.5739 | 6.696 | 6.65105 | 7.55036 | 6.93049 | 5.81673 | 5.71056 | 4.56566 | 4.41009 |
| PCBP4 | P57723 | 6.24565 | 6.39562 | 6.46613 | 6.18357 | 6.08939 | 6.08145 | 5.69594 | 6.64637 | 6.87731 | 6.39904 | 5.93183 | 6.56765 | 6.09425 | 6.46223 | 6.05551 | 5.80805 |
| NFIA | Q12857-4 | 5.63637 | 5.83426 | 6.34404 | 7.16776 | 6.07305 | 6.08449 | 5.30433 | 5.95195 | 5.81618 | 6.47939 | 6.20489 | 6.22511 | 6.57276 | 7.11328 | 7.14215 | 6.04998 |
| CST3 | P01034 | 6.93832 | 6.83096 | 6.20721 | 5.3068 | 7.22426 | 6.19437 | 6.53989 | 7.0971 | 6.75857 | 6.35059 | 6.43526 | 6.57797 | 5.46883 | 5.58281 | 5.1842 | 5.30286 |
| STEAP2 | Q8NFT2 | 6.24776 | 6.20934 | 6.34024 | 6.91039 | 5.55612 | 5.93419 | 5.8703 | 5.90755 | 6.43692 | 6.31572 | 5.66697 | 6.11849 | 6.39432 | 6.28572 | 6.98545 | 6.82052 |
| TBC1D7 | Q9P0N9 | 6.42904 | 6.47678 | 7.25954 | 6.44648 | 6.18694 | 7.12232 | 6.33369 | 6.72881 | 6.70836 | 6.9461 | 6.7139 | 6.88564 | 4.78799 | 4.62264 | 5.30333 | 5.04844 |
| ZNF93 | P35789 | 6.46037 | 6.03215 | 6.7618 | 6.7276 | 5.78589 | 5.91656 | 6.34613 | 6.44471 | 6.19556 | 5.89347 | 6.46538 | 5.93492 | 6.45739 | 6.2452 | 5.71335 | 6.61953 |
| DAB2 | P98082 | 6.35651 | 5.67383 | 6.86252 | 5.97568 | 6.15811 | 5.70635 | 5.81629 | 6.3624 | 6.84885 | 5.92154 | 6.96538 | 6.85853 | 6.26232 | 6.22562 | 5.78858 | 6.2175 |
| N4BP2L2 | Q92802-2 | 7.33971 | 6.51016 | 6.74628 | 7.48169 | 7.33776 | 6.44773 | 6.85944 | 6.4276 | 6.94937 | 6.96956 | 6.47903 | 7.10826 | 4.38794 | 4.61211 | 3.76571 | 4.57764 |
| RRAGC | Q9HB90 | 7.21275 | 7.26208 | 6.47578 | 6.40952 | 5.9995 | 6.93579 | 6.2568 | 5.71777 | 6.73105 | 7.29572 | 5.90838 | 5.69329 | 5.30432 | 5.62927 | 5.79982 | 5.36817 |
| ETV3 | P41162 | 6.11132 | 7.31807 | 7.48312 | 6.64743 | 6.02562 | 6.40391 | 5.53364 | 5.24982 | 5.79692 | 6.26968 | 6.41508 | 6.60369 | 5.94375 | 5.4021 | 6.57322 | 6.22263 |
| SGK3 | Q96BR1 | 6.5712 | 6.47529 | 6.7102 | 6.30876 | 6.27691 | 6.7608 | 6.2949 | 6.70824 | 6.76881 | 6.66427 | 6.51958 | 6.72022 | 5.53446 | 5.63509 | 5.11073 | 4.94054 |
| RAB3D | O95716 | 6.2362 | 6.38589 | 5.97472 | 6.6276 | 5.83741 | 5.99092 | 6.30776 | 6.26806 | 6.31759 | 6.04669 | 6.2198 | 6.317 | 6.40856 | 6.31055 | 6.37118 | 6.38007 |
| RAB8B | Q92930 | 6.10571 | 6.10919 | 6.2779 | 6.55548 | 5.75055 | 5.9977 | 5.83965 | 6.25882 | 6.36031 | 6.22176 | 6.28284 | 6.30541 | 6.50353 | 6.46723 | 6.43989 | 6.52403 |
| MKS1 | Q9NXB0 | 6.38313 | 6.46371 | 6.38017 | 6.60631 | 7.38954 | 6.54108 | 6.10638 | 6.04713 | 6.21111 | 6.08275 | 6.04834 | 6.32249 | 5.82757 | 6.00728 | 6.02919 | 5.55382 |
| ZNF100 | Q8IYN0 | 6.26191 | 6.52644 | 7.77147 | 6.91416 | 6.69512 | 6.96377 | 7.3155 | 6.25988 | 5.55669 | 6.13044 | 6.02557 | 6.07063 | 5.17348 | 5.70192 | 5.46308 | 5.16995 |
| MOCS3 | O95396 | 6.18659 | 5.67271 | 6.33274 | 6.19681 | 6.44108 | 6.42619 | 6.25738 | 6.6847 | 6.31752 | 6.53093 | 6.73424 | 6.27584 | 5.88686 | 5.83454 | 6.15304 | 6.06882 |
| CDCA7L | Q96GN5 | 7.28554 | 6.58165 | 6.90487 | 7.66446 | 6.63374 | 6.84757 | 7.01208 | 6.23301 | 7.0786 | 6.42732 | 6.14586 | 6.17171 | 4.90298 | 4.78225 | 4.7975 | 5.43085 |
| ZNF780A | O75290-3 | 7.15473 | 6.15022 | 6.35978 | 7.34127 | 5.91627 | 5.57339 | 6.79363 | 6.83136 | 6.18418 | 6.58126 | 6.735 | 6.60348 | 5.49808 | 5.22143 | 5.82371 | 5.2322 |
| PFDN1 | O60925 | 5.88841 | 5.84795 | 5.71195 | 5.411 | 5.07723 | 4.00537 | 4.98592 | 6.12249 | 4.44751 | 5.07366 | 4.43055 | 5.10171 | 3.99815 | 12.3714 | 8.72443 | 12.8023 |
| ZNF362 | Q5T0B9 | 6.72093 | 6.34619 | 5.37558 | 6.3101 | 5.76177 | 5.32007 | 6.20041 | 6.29237 | 6.02996 | 6.82132 | 6.35875 | 6.99047 | 6.08332 | 6.17957 | 6.85947 | 6.34973 |
| NFKBIE | O00221 | 6.53736 | 4.75353 | 6.12479 | 4.889 | 5.51556 | 5.33596 | 5.54509 | 7.12025 | 5.64346 | 6.47715 | 6.11663 | 7.61713 | 6.08974 | 7.45876 | 7.31424 | 7.46134 |
| GATA6 | Q92908 | 5.8688 | 6.2688 | 6.26227 | 6.351 | 6.36886 | 6.74287 | 5.96712 | 6.47318 | 6.16217 | 6.23046 | 6.19505 | 5.86241 | 6.65329 | 6.18433 | 6.98562 | 5.42378 |
| FBRSL1 | A0A3B3IRR3 | 6.37456 | 5.6569 | 6.98748 | 5.66943 | 6.57544 | 6.2488 | 5.92361 | 6.99271 | 6.65703 | 6.19375 | 6.42196 | 6.71735 | 6.14053 | 5.80425 | 5.48496 | 6.15126 |
| HDGFL3 | Q9Y3E1 | 5.35452 | 6.12046 | 5.26933 | 5.74602 | 6.28606 | 6.27342 | 5.66889 | 5.93361 | 5.97333 | 6.46792 | 5.50292 | 5.99838 | 6.57348 | 7.75705 | 7.39169 | 7.68293 |
| SMPDL3B | Q92485 | 6.01459 | 6.42411 | 6.41042 | 6.2535 | 5.95207 | 5.81869 | 5.90721 | 5.8893 | 6.0024 | 6.11535 | 6.06256 | 6.36756 | 6.91918 | 6.83639 | 6.48876 | 6.53792 |
| LYSMD2 | Q8IV50 | 6.26333 | 6.19322 | 7.24679 | 5.70338 | 6.27889 | 6.14176 | 6.00352 | 6.61168 | 6.26315 | 6.24076 | 7.45889 | 6.84508 | 6.16348 | 5.57893 | 5.78385 | 5.22329 |
| ANO8 | Q9HCE9 | 6.57174 | 5.98519 | 6.30497 | 6.69133 | 5.71201 | 5.32309 | 6.47823 | 6.96507 | 6.34476 | 6.33249 | 6.23844 | 5.40236 | 5.78832 | 5.95918 | 6.98155 | 6.92525 |
| SEC11C | Q9BY50 | 6.7483 | 6.18503 | 6.02228 | 6.15662 | 6.79312 | 5.8009 | 6.74573 | 5.86592 | 6.30637 | 5.77907 | 6.72373 | 6.37204 | 6.776 | 5.70932 | 5.77721 | 6.23837 |
| DERPC | P0CG12 | 4.79127 | 3.99871 | 5.24149 | 3.92783 | 6.1742 | 5.5777 | 5.12913 | 7.34847 | 4.76756 | 5.95078 | 5.67528 | 6.77726 | 8.30823 | 9.23749 | 8.31028 | 8.78431 |
| TMEM223 | A0PJW6 | 6.25645 | 6.4006 | 6.04784 | 6.73264 | 6.08854 | 6.27723 | 6.19527 | 6.25262 | 6.20553 | 6.47767 | 5.63338 | 5.34005 | 7.04017 | 6.50772 | 6.46695 | 6.07733 |
| DYNC2I1 | Q8WVS4 | 5.90943 | 6.56872 | 6.52608 | 5.31145 | 6.0564 | 6.38065 | 6.49159 | 6.65744 | 6.01823 | 5.90763 | 6.69406 | 6.5368 | 5.51919 | 6.68741 | 7.01057 | 5.72435 |
| MLLT1 | Q03111 | 6.78958 | 6.8796 | 6.45997 | 7.03865 | 6.31646 | 5.96591 | 6.43554 | 6.1503 | 5.6421 | 6.52446 | 6.96542 | 6.23814 | 5.91972 | 5.89001 | 5.23489 | 5.54925 |
| ARHGAP31 | Q2M1Z3 | 5.87385 | 6.27852 | 6.85468 | 6.2472 | 6.23495 | 6.48399 | 6.35525 | 6.92695 | 6.04312 | 6.30784 | 6.52423 | 6.88361 | 6.16712 | 6.25161 | 5.42138 | 5.1457 |
| C1orf122 | Q6ZSJ8 | 5.95204 | 5.65856 | 6.68986 | 6.11926 | 6.20907 | 6.25843 | 5.89582 | 6.07937 | 6.25091 | 6.44586 | 6.79856 | 6.90834 | 5.60006 | 6.54899 | 5.91554 | 6.66935 |
| H2AC11 | P0C0S8 | 4.70122 | 8.32503 | 7.95775 | 8.97304 | 5.15871 | 6.95432 | 4.40059 | 6.54327 | 5.54683 | 5.35302 | 3.72366 | 5.00475 | 6.18909 | 6.53213 | 8.27005 | 6.36654 |
| IL1RAP | Q9NPH3-5 | 6.57786 | 6.50709 | 5.9188 | 6.3122 | 5.53764 | 5.50167 | 6.42992 | 6.3955 | 6.71653 | 5.9178 | 5.53143 | 6.31561 | 6.45832 | 6.59792 | 7.16427 | 6.11744 |
| HOMER1 | Q86YM7 | 6.39185 | 6.25756 | 6.68018 | 5.94967 | 6.44147 | 6.25413 | 5.81942 | 6.68822 | 6.0327 | 6.34403 | 6.85966 | 6.64317 | 5.7352 | 6.14866 | 5.71106 | 6.04302 |
| POLR2M | P0CAP2 | 6.42144 | 6.37607 | 6.76959 | 5.76639 | 6.87718 | 6.7747 | 5.60166 | 6.25263 | 6.5364 | 6.47572 | 7.10538 | 6.35864 | 6.06682 | 5.64707 | 5.56045 | 5.40987 |
| GATC | O43716 | 5.31673 | 4.85895 | 4.7141 | 4.96051 | 5.30524 | 4.27617 | 5.83789 | 6.76306 | 4.275 | 5.08303 | 4.43265 | 5.71218 | 5.59588 | 11.6134 | 8.74818 | 12.5071 |
| NOTCH1 | P46531 | 7.10656 | 7.65429 | 5.60818 | 6.38867 | 5.40629 | 6.63846 | 6.99688 | 5.04539 | 6.63698 | 5.81543 | 6.59304 | 6.65919 | 5.51071 | 6.31113 | 5.10705 | 6.52174 |
| CAPN15 | O75808 | 5.93895 | 5.90427 | 6.9489 | 6.38599 | 6.17438 | 5.768 | 6.8098 | 6.43833 | 6.53167 | 6.9576 | 6.58788 | 6.79976 | 5.98137 | 5.31959 | 5.64211 | 5.81141 |
| DERL2 | Q9GZP9 | 6.61888 | 6.40406 | 6.01159 | 6.54849 | 6.18969 | 6.00293 | 6.00462 | 5.83849 | 5.43365 | 6.53414 | 6.2852 | 6.30477 | 6.11205 | 6.5967 | 6.63666 | 6.47808 |
| DTX2 | Q86UW9 | 5.6832 | 6.04179 | 6.47209 | 6.13448 | 5.237 | 6.65984 | 6.32054 | 5.81014 | 6.91211 | 6.41081 | 6.65496 | 5.9591 | 6.72023 | 6.08754 | 6.27811 | 6.61804 |
| SLC25A23 | Q9BV35 | 6.29707 | 6.36975 | 6.38226 | 6.76624 | 5.69373 | 5.58562 | 6.95822 | 5.43026 | 6.32416 | 5.94573 | 5.9201 | 6.3181 | 7.22289 | 5.973 | 6.0833 | 6.72958 |
| YTHDF1 | Q9BYJ9 | 6.64775 | 6.723 | 6.90415 | 6.11184 | 6.20253 | 6.23668 | 6.28025 | 6.394 | 6.11343 | 6.53079 | 6.65842 | 6.17689 | 5.63048 | 5.8011 | 6.00472 | 5.58397 |
| ADCK5 | Q3MIX3 | 6.61653 | 6.61737 | 5.48745 | 6.49882 | 6.16853 | 5.77854 | 6.25211 | 5.97243 | 6.30269 | 6.34395 | 6.12721 | 5.28292 | 7.01811 | 6.63967 | 6.10154 | 6.79211 |
| CNOT4 | O95628-9 | 6.50709 | 6.48673 | 6.62692 | 5.94144 | 6.133 | 6.21941 | 5.82412 | 6.52246 | 6.26683 | 6.39106 | 6.50315 | 6.80235 | 5.38999 | 5.76677 | 6.15607 | 6.4626 |
| ICA1 | E7ENI6 | 6.37331 | 5.94091 | 5.75817 | 6.43829 | 6.13488 | 5.72538 | 6.71057 | 5.33909 | 6.2667 | 6.03624 | 5.7988 | 6.12695 | 6.98313 | 6.50592 | 6.74046 | 7.12121 |
| WDTC1 | Q8N5D0 | 5.73961 | 6.12848 | 6.53031 | 5.8322 | 6.22057 | 6.66989 | 5.86641 | 6.41989 | 6.32031 | 6.77808 | 6.45054 | 6.02793 | 6.70928 | 6.18255 | 6.22635 | 5.8976 |
| ALS2 | Q96Q42 | 6.13077 | 6.65674 | 6.57174 | 6.05148 | 6.37938 | 6.57658 | 5.94869 | 6.72191 | 6.49354 | 6.22312 | 6.56388 | 6.15529 | 5.58376 | 6.34761 | 5.92216 | 5.67334 |
| SMAD5 | Q99717 | 5.97331 | 6.11058 | 6.74021 | 6.06204 | 6.12993 | 6.74774 | 6.43308 | 6.32631 | 6.35782 | 6.1271 | 7.0423 | 6.3607 | 6.44473 | 5.66419 | 6.00713 | 5.47282 |
| FAR2 | Q96K12 | 6.91018 | 6.89998 | 6.91617 | 7.68881 | 6.18599 | 5.78025 | 6.25324 | 5.94887 | 7.21894 | 7.12107 | 7.50073 | 7.62001 | 4.10892 | 4.48912 | 4.68723 | 4.67049 |
| DNMT3B | Q9UBC3 | 7.36657 | 6.69591 | 6.36581 | 7.93826 | 6.4736 | 6.35359 | 6.40569 | 5.74349 | 6.50305 | 6.69042 | 6.18069 | 6.61709 | 5.96892 | 5.56238 | 4.99377 | 4.77077 |
| SLC25A36 | Q96CQ1 | 6.58221 | 6.45808 | 5.56716 | 6.56845 | 6.13293 | 6.48528 | 6.61474 | 5.84799 | 5.3499 | 5.95626 | 5.97034 | 5.83069 | 6.8869 | 6.18941 | 7.05496 | 6.5047 |
| TTPAL | Q9BTX7 | 6.40605 | 6.31135 | 7.04002 | 6.0339 | 6.39902 | 6.73016 | 6.15309 | 6.48672 | 6.60431 | 6.61436 | 6.47696 | 6.82084 | 5.85498 | 5.3905 | 5.26762 | 5.41014 |
| RUNDC1 | Q96C34 | 6.54991 | 8.33132 | 7.56469 | 5.29227 | 6.76831 | 5.97395 | 6.87662 | 7.94943 | 6.77023 | 6.88071 | 6.68904 | 6.81742 | 4.98094 | 3.73161 | 3.89407 | 4.92948 |
| SERBP1 | Q8NC51-3 | 5.44778 | 5.85942 | 5.67079 | 5.45905 | 6.43614 | 6.23606 | 5.39613 | 6.57063 | 6.88775 | 6.64241 | 6.2817 | 7.40128 | 6.1007 | 7.306 | 6.23569 | 6.04298 |
| MPV17 | P39210 | 6.57265 | 6.31599 | 5.73109 | 5.25297 | 6.04147 | 6.12131 | 5.86205 | 6.04644 | 5.44079 | 6.2942 | 5.6379 | 6.37784 | 6.49025 | 7.03067 | 7.3197 | 7.46468 |
| SLC7A5 | Q01650 | 6.35113 | 6.46311 | 5.34946 | 5.92485 | 6.77234 | 6.13448 | 6.78412 | 6.25963 | 5.9773 | 5.784 | 5.3663 | 5.90965 | 6.67644 | 7.05336 | 6.76096 | 6.43287 |
| FAHD2A | Q96GK7 | 5.91042 | 6.34344 | 5.76976 | 5.94678 | 6.35148 | 6.37514 | 6.08094 | 6.47126 | 6.39966 | 6.67083 | 6.70849 | 6.98336 | 6.52348 | 5.88304 | 6.29285 | 6.16905 |
| PNKD | Q8N490 | 5.73552 | 5.84704 | 6.38002 | 6.49755 | 6.21553 | 6.08019 | 5.95934 | 5.90395 | 5.99203 | 6.90992 | 6.30042 | 5.93781 | 7.15201 | 6.27023 | 6.36636 | 6.45208 |
| SIAE | Q9HAT2 | 7.37115 | 6.03926 | 5.32842 | 6.76603 | 5.84451 | 6.10014 | 6.31352 | 5.60828 | 6.71165 | 6.59903 | 5.55115 | 5.577 | 6.89886 | 5.9044 | 6.49249 | 6.89411 |
| TWSG1 | Q9GZX9 | 7.45272 | 6.89956 | 5.85067 | 6.49074 | 7.03599 | 5.99548 | 6.20283 | 6.78594 | 5.86865 | 6.10649 | 6.56927 | 6.22121 | 6.30172 | 5.4456 | 5.34512 | 5.42802 |
| VKORC1L1 | Q8N0U8 | 6.15237 | 6.05812 | 5.62319 | 6.36548 | 6.06351 | 6.13142 | 6.8435 | 5.68513 | 6.15064 | 5.69762 | 5.61823 | 5.73175 | 6.8119 | 6.85031 | 6.80927 | 7.40758 |
| MDM1 | Q8TC05 | 6.20237 | 6.32955 | 6.14002 | 6.74448 | 6.16255 | 5.88642 | 6.2202 | 6.21692 | 6.21288 | 6.09018 | 6.47062 | 6.56885 | 5.57244 | 6.61383 | 6.61707 | 5.95162 |
| TRIM37 | O94972 | 6.48732 | 6.62949 | 6.79721 | 6.35734 | 6.25624 | 6.31249 | 6.80154 | 6.35283 | 6.30579 | 6.7269 | 6.60073 | 6.57933 | 5.53907 | 5.44413 | 5.42072 | 5.38886 |
| STK17A | Q9UEE5 | 6.39793 | 6.17561 | 6.23419 | 5.99199 | 6.41987 | 7.32926 | 5.2055 | 6.42877 | 6.35189 | 6.89514 | 7.56874 | 7.49305 | 5.18936 | 5.91123 | 4.93942 | 5.46806 |
| NDUFA2 | O43678 | 6.00576 | 5.18032 | 5.74011 | 5.22457 | 6.71663 | 5.20257 | 5.92495 | 6.33739 | 5.92576 | 6.67262 | 6.60251 | 6.40689 | 6.26578 | 6.96739 | 7.86927 | 6.95748 |
| IFIT2 | P09913 | 7.37902 | 5.81906 | 6.70985 | 5.99234 | 7.62133 | 6.69554 | 6.08339 | 6.13604 | 6.01612 | 5.81951 | 7.37485 | 5.93297 | 5.72887 | 6.01174 | 5.46579 | 5.21359 |
| GTPBP8 | Q8N3Z3 | 6.26103 | 6.43007 | 6.07534 | 6.53138 | 7.01063 | 6.09407 | 6.72428 | 6.01348 | 6.20659 | 6.24413 | 6.39244 | 6.66378 | 5.93779 | 5.02865 | 6.50983 | 5.8765 |
| RPGRIP1L | Q68CZ1 | 6.97618 | 6.39944 | 7.2206 | 6.77616 | 5.96136 | 6.64062 | 7.17168 | 6.66136 | 6.58485 | 6.81346 | 7.21542 | 6.85301 | 4.8411 | 4.52309 | 4.92918 | 4.43801 |
| SLC38A9 | Q8NBW4 | 6.70676 | 6.25851 | 6.50671 | 6.5515 | 6.1751 | 6.29601 | 6.0502 | 5.93925 | 5.89155 | 6.45219 | 6.32532 | 6.53925 | 6.34202 | 5.98007 | 6.13232 | 5.85324 |
| PLCL2 | Q9UPR0 | 6.08687 | 6.43939 | 6.40873 | 5.93585 | 6.2124 | 6.33573 | 6.3297 | 6.4813 | 6.52213 | 6.06012 | 6.36517 | 6.53851 | 5.74495 | 6.03625 | 6.49571 | 6.00719 |
| MAGEA6 | P43360 | 6.61296 | 6.74437 | 7.39146 | 6.23209 | 6.18103 | 6.5173 | 6.95239 | 7.1332 | 6.99221 | 6.85838 | 7.32836 | 6.90737 | 4.20129 | 4.53674 | 4.30786 | 5.10298 |
| NACA4P | Q9BZK3 | 6.81216 | 6.68107 | 6.9798 | 5.82649 | 6.74593 | 6.24039 | 6.7103 | 6.27191 | 7.22975 | 6.653 | 7.04038 | 6.42303 | 5.41426 | 5.00627 | 5.01564 | 4.94963 |
| SLC39A1 | Q9NY26 | 6.52774 | 7.3411 | 6.29885 | 8.02934 | 6.52298 | 6.5806 | 6.32194 | 6.08344 | 6.80484 | 6.53224 | 5.87739 | 6.3076 | 5.4612 | 5.3015 | 5.30741 | 4.70181 |
| ETV6 | P41212 | 6.8156 | 6.44802 | 5.90167 | 6.86873 | 6.86199 | 5.73233 | 6.30408 | 7.06582 | 6.2165 | 6.08006 | 6.37985 | 6.39633 | 5.80818 | 5.52082 | 6.19387 | 5.40615 |
| MUTYH | Q9UIF7 | 5.56602 | 5.9912 | 6.60564 | 6.24768 | 6.23978 | 6.19441 | 5.59512 | 5.90956 | 6.63426 | 6.43277 | 6.596 | 6.77666 | 6.0876 | 6.49116 | 6.33961 | 6.29253 |
| APEX2 | Q9UBZ4 | 6.09874 | 6.3004 | 7.20214 | 6.45312 | 5.57197 | 6.19984 | 6.2872 | 6.57921 | 7.09964 | 7.10056 | 7.56658 | 7.47448 | 4.66813 | 5.0338 | 5.10858 | 5.25561 |
| TTC31 | Q49AM3 | 6.30748 | 6.42626 | 6.70695 | 6.64687 | 6.97657 | 6.58278 | 6.65107 | 6.05545 | 6.93844 | 7.55958 | 6.88355 | 6.67923 | 5.3161 | 4.74653 | 4.97426 | 4.54886 |
| ZDHHC20 | Q5W0Z9 | 7.04138 | 6.44817 | 5.87902 | 7.09944 | 6.99283 | 5.58858 | 5.83884 | 5.69732 | 6.75077 | 6.65455 | 6.50861 | 6.66098 | 5.13536 | 5.6885 | 6.53124 | 5.4844 |
| CCDC130 | P13994 | 5.48587 | 6.59962 | 6.06721 | 5.85377 | 5.80153 | 6.64095 | 5.81837 | 6.21877 | 6.62057 | 6.4754 | 6.20653 | 6.84739 | 6.41207 | 6.09497 | 5.93094 | 6.92605 |
| ENOSF1 | Q7L5Y1-5 | 6.52191 | 6.38104 | 7.03293 | 5.52245 | 6.21555 | 7.04575 | 6.0402 | 6.70055 | 6.48026 | 6.54413 | 6.68769 | 6.72765 | 5.45421 | 5.48689 | 5.54451 | 5.61427 |
| TMCO3 | Q6UWJ1 | 6.84166 | 6.42441 | 6.20316 | 7.49625 | 6.07492 | 5.59353 | 6.52153 | 6.28214 | 6.32198 | 6.08805 | 5.94632 | 6.35978 | 6.17371 | 5.80458 | 5.98064 | 5.88733 |
| ASNSD1 | Q9NWL6 | 6.3091 | 6.46196 | 7.50966 | 6.29994 | 6.40045 | 6.15135 | 6.36566 | 6.35699 | 7.05418 | 6.97497 | 6.82725 | 7.26954 | 4.76357 | 4.76985 | 5.15675 | 5.32878 |
| ELOVL1 | Q9BW60 | 5.81296 | 6.78492 | 5.75004 | 6.50915 | 5.46156 | 6.12257 | 6.68235 | 6.59904 | 5.93445 | 5.73397 | 6.42948 | 6.11527 | 6.21364 | 6.31302 | 7.03299 | 6.50458 |
| TRADD | Q15628 | 6.60986 | 6.46901 | 7.04925 | 6.22856 | 6.41998 | 6.42819 | 5.75597 | 6.4482 | 6.02975 | 6.93479 | 6.05198 | 6.33173 | 6.05816 | 6.08061 | 5.11964 | 5.98434 |
| SLC4A8 | Q2Y0W8 | 6.02272 | 7.0767 | 6.16777 | 6.98897 | 5.9787 | 6.04322 | 6.35392 | 6.16438 | 6.35971 | 6.72631 | 6.73422 | 6.38317 | 5.59223 | 6.08809 | 5.79627 | 5.52362 |

| | | | | | | | | | | | | | | | | | |
|---|---|---|---|---|---|---|---|---|---|---|---|---|---|---|---|---|---|
| SMPD1 | P17405 | 6.59275 | 6.58443 | 6.35757 | 6.26937 | 6.23819 | 5.97923 | 6.12864 | 6.01106 | 6.16369 | 6.42566 | 6.18433 | 6.76548 | 5.78756 | 5.95649 | 6.27572 | 6.27983 |
| TBKBP1 | A7MCY6 | 6.70096 | 6.58825 | 6.65699 | 6.85133 | 6.75703 | 7.51064 | 5.80289 | 8.13024 | 6.37139 | 5.51008 | 6.67083 | 6.12505 | 5.43848 | 4.60599 | 5.6013 | 4.67857 |
| TCEANC2 | Q96MN5 | 6.1075 | 6.86319 | 6.22478 | 5.19264 | 6.17205 | 6.40683 | 6.16554 | 5.73431 | 5.88307 | 7.32433 | 6.85559 | 6.23387 | 6.0886 | 6.15841 | 6.12886 | 6.46043 |
| PFN2 | P35080 | 6.33558 | 6.12999 | 6.96676 | 5.64572 | 6.48584 | 6.96729 | 6.03687 | 6.7257 | 6.70938 | 6.49995 | 6.46698 | 6.66332 | 5.70198 | 5.55776 | 5.53853 | 5.56833 |
| ENY2 | Q9NPA8 | 6.31278 | 5.18974 | 5.74793 | 5.48464 | 6.59968 | 5.14357 | 4.98763 | 6.50661 | 4.93804 | 6.62516 | 5.65994 | 6.53429 | 6.03971 | 8.56147 | 7.63224 | 8.03656 |
| GNAI1 | P63096 | 6.28619 | 6.82267 | 5.64294 | 6.61294 | 6.73522 | 6.51291 | 5.05842 | 6.16208 | 5.38189 | 6.60206 | 5.72054 | 5.93119 | 5.91363 | 6.62399 | 6.34781 | 6.64553 |
| OTUD7A | Q8TE49-2 | 5.30049 | 4.15294 | 4.46822 | 5.04991 | 4.81249 | 5.27453 | 4.7562 | 5.57585 | 5.45955 | 7.51869 | 6.87959 | 8.17387 | 7.88183 | 8.93895 | 8.20322 | 7.55368 |
| SDF2 | Q99470 | 5.91064 | 6.04729 | 6.06392 | 6.20216 | 5.80411 | 6.68272 | 6.06721 | 6.01105 | 5.93695 | 6.82567 | 7.02379 | 6.29199 | 6.68134 | 6.00197 | 6.3687 | 6.08049 |
| TOR2A | Q5JU69 | 6.13568 | 6.63858 | 6.12364 | 6.35303 | 6.26844 | 5.55679 | 6.16462 | 6.1627 | 6.19728 | 6.08458 | 5.85647 | 6.14615 | 6.40065 | 6.51215 | 6.83951 | 6.55973 |
| INPP1 | P49441 | 5.91362 | 5.93506 | 6.46179 | 6.15007 | 6.64968 | 6.98913 | 5.81003 | 7.00445 | 6.56064 | 6.54575 | 6.45582 | 6.09784 | 5.59893 | 5.61373 | 6.20667 | 6.00682 |
| VTI1B | Q9UEU0 | 6.74921 | 6.96995 | 6.61842 | 6.52684 | 6.06957 | 6.22428 | 6.55023 | 6.04481 | 6.12345 | 6.65641 | 5.6757 | 6.3575 | 5.49816 | 5.49259 | 5.70335 | 5.83953 |
| TEAD3 | Q99594 | 6.41999 | 6.36324 | 6.65257 | 7.50047 | 6.97435 | 5.19909 | 5.69446 | 6.04351 | 6.54821 | 6.60573 | 6.44043 | 6.43583 | 4.89854 | 5.64472 | 6.26815 | 6.31071 |
| TEAD1 | P28347 | 6.36677 | 6.34693 | 5.92296 | 6.64584 | 6.82839 | 6.02763 | 6.27478 | 5.84157 | 5.87929 | 5.8301 | 6.19897 | 5.94314 | 6.54611 | 5.91642 | 6.96168 | 6.4694 |
| WWTR1 | Q9GZV5 | 6.81952 | 4.63326 | 6.5798 | 5.23388 | 7.01276 | 4.83019 | 5.47589 | 7.36935 | 4.77478 | 5.78095 | 5.97812 | 7.10001 | 6.20938 | 7.66902 | 7.37836 | 7.15472 |
| APBA2 | Q99767 | 6.73538 | 7.40102 | 7.05936 | 7.34011 | 6.29519 | 6.65222 | 6.14833 | 6.77656 | 6.30535 | 6.61402 | 6.49978 | 6.38168 | 4.76638 | 5.51059 | 4.5587 | 4.95531 |
| PITPNM2 | Q9BZ72 | 6.89625 | 6.38812 | 7.1194 | 6.0399 | 5.28411 | 6.61152 | 6.30698 | 4.78227 | 5.7935 | 6.32665 | 5.99386 | 6.40592 | 5.81976 | 6.80286 | 6.15077 | 7.27813 |
| CETN2 | P41208 | 6.37001 | 6.30188 | 6.44866 | 5.64507 | 6.58854 | 6.26416 | 5.99748 | 6.77904 | 6.13401 | 6.58581 | 6.31693 | 6.3712 | 6.24146 | 6.26706 | 5.97454 | 5.71415 |
| C2orf76 | Q3KRA6 | 6.11627 | 5.8917 | 6.55879 | 5.23805 | 6.32572 | 6.86225 | 6.45134 | 7.03556 | 6.74513 | 6.95765 | 7.06996 | 6.63877 | 5.69238 | 5.54778 | 5.38871 | 5.47993 |
| MEX3D | Q86XN8-3 | 6.70755 | 6.64115 | 7.68119 | 6.48235 | 6.69218 | 6.43565 | 5.87031 | 6.92676 | 6.3376 | 6.62787 | 6.78335 | 7.22967 | 5.27241 | 4.94029 | 4.6624 | 4.70929 |
| FBXO6 | Q9NRD1 | 5.96641 | 6.07145 | 6.82053 | 6.51963 | 6.2512 | 6.57853 | 5.93952 | 6.66578 | 6.16051 | 6.12841 | 6.44155 | 6.63481 | 6.58177 | 6.1222 | 5.79583 | 6.22187 |
| ARMCX1 | Q9P291 | 7.02896 | 7.30835 | 7.13631 | 7.10959 | 6.66533 | 6.54118 | 6.83184 | 6.6403 | 6.86992 | 7.7902 | 7.05883 | 7.65735 | 3.50057 | 4.07816 | 3.86696 | 3.91615 |
| APBA3 | O96018 | 6.33638 | 6.01974 | 6.1675 | 6.07777 | 6.44929 | 6.84088 | 6.46858 | 5.98302 | 6.42982 | 6.45742 | 7.50033 | 6.34607 | 5.61818 | 5.8478 | 5.72776 | 5.72945 |
| DNAJC5 | Q9H3Z4 | 6.36942 | 6.40312 | 6.02538 | 6.43036 | 5.80857 | 5.84841 | 6.23 | 6.1298 | 6.41206 | 6.2731 | 6.2107 | 6.69874 | 6.09203 | 6.20864 | 6.42692 | 6.43277 |
| POLR2M | Q6EEV4 | 6.36568 | 4.71338 | 5.39578 | 4.79779 | 5.23119 | 4.53274 | 4.78752 | 7.0741 | 3.9272 | 6.3935 | 5.58148 | 5.88577 | 4.65375 | 12.6309 | 7.38792 | 10.6413 |
| PXMP4 | Q9Y6I8 | 6.08019 | 6.20597 | 5.4735 | 6.39694 | 6.08499 | 5.43715 | 6.2488 | 6.581 | 6.08996 | 6.55305 | 6.2039 | 6.5164 | 6.72601 | 6.54685 | 6.53609 | 6.3192 |
| EPB41L2 | O43491-4 | 6.42915 | 7.56329 | 5.6491 | 7.11333 | 6.57694 | 6.85756 | 5.9759 | 7.24597 | 5.66069 | 5.36861 | 6.32112 | 6.07134 | 5.24552 | 6.50652 | 6.16937 | 5.2456 |
| ZMAT5 | Q9UDW3 | 5.61595 | 5.96397 | 6.51383 | 6.78292 | 6.16068 | 5.79639 | 6.48306 | 5.59623 | 6.60182 | 6.24475 | 6.9859 | 6.75649 | 5.91821 | 5.6753 | 6.22177 | 6.34234 |
| CEBPB | P17676 | 5.50883 | 6.43642 | 6.10198 | 7.06588 | 6.31785 | 6.08024 | 6.49917 | 6.44994 | 5.45169 | 6.43549 | 5.944 | 6.09937 | 6.09877 | 6.18684 | 6.83861 | 6.48491 |
| HNRNPU | Q00839-2 | 5.13308 | 5.68084 | 5.52857 | 6.76496 | 5.62028 | 5.89392 | 6.07917 | 6.38493 | 5.64087 | 5.96062 | 5.96681 | 7.09486 | 6.56042 | 6.68499 | 7.02535 | 7.98035 |
| PHC3 | Q8NDX5-7 | 6.11805 | 6.25715 | 5.95332 | 6.53623 | 6.18393 | 5.98476 | 6.41975 | 5.8712 | 6.36532 | 6.47643 | 6.43792 | 6.36558 | 6.4297 | 6.09992 | 6.29888 | 6.20188 |
| MAPKBP1 | O60336 | 5.83122 | 6.17342 | 6.93143 | 5.92125 | 6.13499 | 6.82227 | 6.51694 | 5.73364 | 6.74974 | 6.71975 | 7.07988 | 6.92988 | 5.92392 | 5.75011 | 5.06633 | 5.71524 |
| MPP1 | Q00013 | 6.50998 | 6.29053 | 6.51053 | 6.35052 | 6.20964 | 6.10408 | 6.56273 | 6.3654 | 6.15847 | 6.51781 | 6.48426 | 6.81494 | 5.62999 | 5.74569 | 5.87634 | 5.86907 |
| SH3BP5L | Q7L8J4 | 6.54399 | 6.49517 | 6.52477 | 5.80456 | 6.93494 | 6.67712 | 6.07765 | 6.92733 | 6.31598 | 6.22451 | 6.57983 | 6.29955 | 6.41469 | 5.87859 | 5.56199 | 4.73933 |
| TIMM8B | Q9Y5J9 | 6.20742 | 5.36778 | 4.44549 | 5.40993 | 5.85659 | 4.86415 | 5.64465 | 6.7611 | 4.9757 | 5.27375 | 4.74861 | 5.40552 | 5.40645 | 10.8221 | 8.08496 | 10.7258 |
| CDK11A | Q9UQ88 | 6.06216 | 6.5464 | 5.74839 | 6.4502 | 6.01625 | 5.82229 | 5.98199 | 5.91837 | 6.29234 | 6.19075 | 6.10113 | 5.8924 | 6.64841 | 6.84061 | 6.85142 | 6.6369 |
| SAP30L | Q9HAJ7 | 6.79992 | 6.87646 | 5.81654 | 6.70553 | 6.08652 | 6.14342 | 6.22165 | 6.05341 | 6.00143 | 6.16335 | 6.17385 | 6.19545 | 6.46805 | 6.19952 | 6.20767 | 5.88723 |
| TMEM260 | Q9NX78 | 6.05366 | 6.50526 | 6.41588 | 6.75744 | 5.82661 | 5.77565 | 6.08912 | 6.32264 | 6.30907 | 6.01753 | 5.68779 | 5.68544 | 6.43429 | 6.17636 | 7.25908 | 6.68417 |
| CCNY | Q8ND76 | 5.79712 | 5.82624 | 5.94926 | 6.56764 | 6.05784 | 6.36089 | 6.58215 | 6.23607 | 5.65494 | 6.40687 | 6.63912 | 6.41982 | 6.25102 | 6.48973 | 6.1106 | 6.65068 |
| ARHGAP23 | Q9P227 | 6.42873 | 6.29561 | 5.85485 | 6.45713 | 6.49818 | 6.38417 | 6.33857 | 6.40948 | 6.45573 | 5.8594 | 6.12012 | 6.33366 | 5.93649 | 6.42281 | 5.96592 | 6.23914 |
| FPGS | Q05932 | 6.3008 | 5.83042 | 5.85748 | 6.52852 | 7.07188 | 6.63608 | 6.95735 | 6.75833 | 6.35205 | 5.78677 | 6.12558 | 6.40634 | 5.95529 | 5.41423 | 5.73801 | 6.28087 |
| RNASEL | Q05823 | 6.21841 | 6.05125 | 6.28961 | 6.45034 | 6.371 | 6.37029 | 6.28951 | 6.33826 | 6.23766 | 6.16957 | 6.61174 | 6.30144 | 6.14043 | 6.28956 | 6.03819 | 5.83273 |
| MAPKAPK3 | Q16644 | 6.251 | 6.30319 | 6.71311 | 6.15043 | 6.55482 | 6.48448 | 6.21858 | 6.63059 | 6.90567 | 6.5922 | 6.47435 | 6.27919 | 5.69879 | 5.76022 | 5.51109 | 5.4723 |
| IFT43 | Q96FT9 | 6.61222 | 5.56722 | 6.42985 | 5.63946 | 6.57736 | 6.55343 | 6.53934 | 6.56071 | 6.64383 | 6.67591 | 6.72768 | 6.79767 | 6.53095 | 5.52357 | 5.19188 | 5.42895 |
| RFX3 | P48380 | 6.36921 | 6.45562 | 6.05779 | 5.92022 | 5.81799 | 5.80853 | 6.92984 | 6.6052 | 6.434 | 6.17999 | 5.8261 | 6.15804 | 6.43276 | 6.33412 | 6.08312 | 6.58745 |
| ABHD15 | Q6UXT9 | 6.41509 | 6.2656 | 6.22916 | 6.17401 | 6.34067 | 6.2058 | 5.98949 | 6.46165 | 6.37125 | 6.35649 | 6.1084 | 6.40324 | 5.90672 | 6.26165 | 6.31888 | 6.1889 |
| GPSM2 | P81274 | 6.07058 | 6.37084 | 6.89814 | 6.47628 | 5.95636 | 6.71381 | 6.53549 | 6.66856 | 6.62903 | 7.25101 | 7.41526 | 7.55336 | 4.33507 | 5.22962 | 4.68571 | 5.21085 |
| TMEM109 | Q9BVC6 | 6.04927 | 6.43839 | 5.77481 | 6.63099 | 6.06657 | 5.95026 | 6.21161 | 6.16038 | 5.87568 | 6.07612 | 5.74419 | 5.98355 | 6.57411 | 6.72826 | 6.77371 | 6.96211 |
| RAB27B | O00194 | 6.44288 | 6.61852 | 6.05012 | 6.44536 | 6.57454 | 6.37823 | 6.29818 | 5.99961 | 6.05368 | 6.14695 | 6.02883 | 5.70069 | 6.43215 | 6.55629 | 6.35241 | 5.92155 |
| SBNO2 | Q9Y2G9 | 5.79451 | 5.90242 | 6.70503 | 5.6961 | 5.58983 | 6.43425 | 6.12914 | 5.94109 | 6.71154 | 6.62886 | 6.87317 | 6.61357 | 5.98358 | 6.10371 | 6.68979 | 6.2034 |
| RAD17 | O75943 | 5.62777 | 5.99073 | 5.79868 | 5.40835 | 5.94417 | 6.02421 | 6.01165 | 6.40188 | 6.27657 | 6.32666 | 6.38441 | 6.15841 | 6.94442 | 6.82667 | 7.40986 | 6.46557 |
| CARHSP1 | Q9Y2V2 | 6.77575 | 6.04577 | 6.71207 | 5.3892 | 6.99784 | 6.74559 | 5.72551 | 6.69491 | 5.99695 | 6.75925 | 6.99598 | 6.7396 | 6.0692 | 5.88049 | 5.47732 | 4.99455 |
| MAP3K1 | Q13233 | 6.98341 | 6.6265 | 6.17348 | 6.48396 | 6.26102 | 6.33377 | 6.14519 | 6.42178 | 6.63475 | 6.88274 | 6.71731 | 6.47681 | 5.99528 | 5.63836 | 4.9284 | 5.29724 |
| TAF8 | Q7Z7C8-2 | 6.07371 | 5.94515 | 6.41161 | 6.48222 | 6.25283 | 6.00731 | 6.37956 | 5.85707 | 6.42479 | 6.29868 | 6.47506 | 6.65716 | 6.81775 | 6.07909 | 5.9121 | 5.9259 |
| SLC22A18 | Q96BI1 | 5.90607 | 5.93526 | 5.83145 | 6.12717 | 6.39875 | 6.09708 | 6.10521 | 5.88565 | 6.37076 | 6.10339 | 5.74806 | 5.65419 | 7.01705 | 6.55547 | 7.38401 | 6.88043 |
| LRP11 | Q86VZ4 | 7.66838 | 7.5303 | 7.04349 | 7.46342 | 6.70646 | 6.00004 | 6.68786 | 6.04141 | 6.89802 | 7.3006 | 7.54972 | 7.16214 | 3.92558 | 4.3256 | 3.64969 | 4.04729 |
| MYC | P01106-2 | 4.37851 | 5.06141 | 4.92982 | 6.16136 | 5.2986 | 5.51219 | 6.04151 | 5.8606 | 4.46191 | 5.4076 | 5.02671 | 5.52099 | 9.33064 | 7.50289 | 10.1854 | 9.31987 |
| APLF | Q8IW19 | 6.0517 | 6.01297 | 5.93063 | 5.79051 | 6.63276 | 6.12549 | 5.44062 | 6.48641 | 5.517 | 5.48615 | 6.50957 | 5.76321 | 7.06921 | 7.69417 | 6.16607 | 7.32353 |
| CHRAC1 | Q9NRG0 | 6.35139 | 6.41895 | 6.56236 | 6.1357 | 6.17323 | 6.5264 | 5.71654 | 6.2572 | 5.88291 | 6.68482 | 6.92144 | 6.80121 | 6.29421 | 6.31917 | 5.61828 | 5.33619 |
| EDEM2 | Q9BV94 | 6.67362 | 6.83071 | 6.18048 | 6.76232 | 6.63862 | 6.52469 | 6.37051 | 6.42292 | 6.1036 | 6.37783 | 6.1669 | 5.83861 | 5.9946 | 5.60529 | 5.72876 | 5.78053 |
| BAG1 | Q99933 | 6.30953 | 6.50706 | 6.15805 | 5.91703 | 6.3219 | 6.76689 | 5.86398 | 6.76652 | 6.09045 | 6.59659 | 5.89963 | 5.9918 | 5.99593 | 6.62067 | 5.59407 | 6.59991 |
| CORO2A | Q92828 | 6.23927 | 6.56027 | 6.45651 | 5.93611 | 6.47771 | 6.37664 | 6.4075 | 6.42033 | 6.53465 | 6.45041 | 5.90007 | 5.79794 | 6.28257 | 6.02604 | 6.02011 | 6.11387 |
| DAB2IP | Q5VWQ8 | 6.73126 | 6.36443 | 6.78828 | 6.61361 | 6.11256 | 6.04645 | 6.21295 | 5.83449 | 6.06135 | 6.44473 | 6.21779 | 6.11276 | 6.34172 | 6.23354 | 6.22526 | 5.65881 |
| ZDHHC17 | Q8IUH5 | 6.60195 | 6.25178 | 5.79759 | 6.82414 | 6.29415 | 6.26133 | 6.65817 | 5.86257 | 6.25148 | 5.80915 | 6.1276 | 5.8917 | 6.90432 | 6.0088 | 6.72216 | 5.73312 |
| DIPK2A | Q8NDZ4 | 6.79272 | 6.77927 | 6.11779 | 6.93151 | 5.90066 | 5.85746 | 6.54823 | 5.69075 | 6.32023 | 6.18783 | 6.13494 | 6.41038 | 6.03735 | 5.47538 | 6.48099 | 6.33451 |
| BTBD8 | A0A1B0GUL6 | 6.19285 | 5.73832 | 6.46829 | 6.19432 | 6.59024 | 6.96119 | 6.10056 | 6.00108 | 6.91593 | 6.80461 | 6.03649 | 6.64646 | 5.37528 | 6.46035 | 6.20897 | 5.30508 |
| NUCKS1 | Q9H1E3 | 5.55239 | 5.58385 | 5.59702 | 5.68883 | 5.72979 | 5.94635 | 5.38639 | 6.45312 | 5.47622 | 6.28578 | 5.33324 | 5.55038 | 7.01125 | 8.77847 | 7.98322 | 7.64369 |
| CUEDC2 | Q9H467 | 6.07929 | 5.83315 | 6.91869 | 5.41747 | 6.68616 | 6.30882 | 6.24711 | 6.60979 | 6.29333 | 6.47623 | 6.84312 | 7.01329 | 5.95069 | 5.91068 | 5.62161 | 5.79158 |
| CHRNB1 | P11230 | 8.68878 | 8.83069 | 7.96913 | 8.48416 | 6.30665 | 6.34694 | 6.2172 | 6.71969 | 7.20076 | 7.20399 | 7.21396 | 8.19195 | 2.42824 | 2.42122 | 2.53689 | 3.23975 |
| ABHD17A | Q96GS6-2 | 6.32334 | 6.82184 | 6.07311 | 6.46381 | 5.80613 | 6.21074 | 5.93398 | 6.29629 | 5.94904 | 6.33985 | 6.38555 | 6.70573 | 5.86379 | 6.32823 | 6.06595 | 6.43263 |
| BLOC1S6 | Q9UL45 | 6.14583 | 5.94326 | 6.67971 | 5.18878 | 6.40946 | 6.44924 | 5.83025 | 6.67142 | 5.98864 | 6.62266 | 6.72882 | 6.76602 | 6.58987 | 6.10768 | 5.83668 | 6.04168 |
| ZFP36L1 | Q07352 | 4.33688 | 4.28409 | 4.8717 | 4.34261 | 5.81616 | 6.0078 | 5.46032 | 5.64493 | 4.59376 | 4.7636 | 5.5977 | 4.72774 | 11.7657 | 9.01805 | 9.63273 | 9.13621 |
| ZFP36L2 | P47974 | 6.70582 | 6.29072 | 6.90331 | 5.37146 | 5.78721 | 6.14075 | 6.23457 | 7.88786 | 7.25647 | 6.19831 | 5.24185 | 6.27009 | 7.11906 | 5.90876 | 5.86546 | 4.81829 |
| DPEP3 | Q9H4B8 | 6.3433 | 6.32109 | 6.18716 | 6.57853 | 6.21895 | 5.9782 | 6.37745 | 5.8781 | 6.17331 | 7.00687 | 6.55599 | 6.37686 | 6.26958 | 6.06395 | 6.06198 | 5.60867 |
| NACC2 | Q96BF6 | 6.65049 | 5.85304 | 6.19427 | 5.84649 | 5.94316 | 6.2482 | 5.93171 | 5.71481 | 6.16979 | 7.0927 | 7.10114 | 6.41982 | 6.24929 | 6.30631 | 6.03718 | 6.24161 |
| NLRX1 | Q86UT6 | 6.69037 | 6.53045 | 6.08492 | 5.91412 | 6.53267 | 6.229 | 6.13914 | 5.60062 | 5.66079 | 6.70535 | 6.16578 | 5.74635 | 6.21496 | 6.649 | 6.44131 | 6.69518 |
| YBEY | P58557 | 5.97241 | 5.65956 | 5.14304 | 5.86806 | 6.26317 | 5.53842 | 6.45408 | 6.21011 | 6.58964 | 5.65232 | 5.80516 | 6.23195 | 7.26212 | 6.97764 | 7.00635 | 7.36599 |
| ATF6B | Q99941 | 6.54175 | 7.72616 | 6.92741 | 6.90558 | 6.26374 | 7.15949 | 7.58948 | 6.78528 | 6.17682 | 7.00422 | 6.77629 | 6.98125 | 4.05266 | 4.71174 | 4.1357 | 4.26242 |
| GINS4 | Q9BRT9 | 6.00536 | 5.62109 | 6.20063 | 5.65083 | 5.72773 | 5.8318 | 5.73679 | 7.58389 | 6.05666 | 5.9866 | 6.64068 | 6.41347 | 6.49992 | 6.60842 | 6.46588 | 6.97027 |
| PLEKHA1 | Q9HB21 | 5.99613 | 6.08246 | 6.89392 | 6.06001 | 5.88639 | 6.36148 | 6.36889 | 6.60613 | 6.26966 | 6.27599 | 6.70228 | 6.90003 | 5.84275 | 6.24146 | 5.9255 | 5.58693 |
| PDSS1 | Q5T2R2 | 6.56621 | 6.04035 | 6.12451 | 6.56189 | 7.14206 | 5.96351 | 5.90536 | 7.07948 | 6.13391 | 5.90535 | 6.80539 | 6.13381 | 5.90903 | 5.69689 | 6.27977 | 5.7525 |
| SH3RF1 | Q7Z6J0 | 6.62626 | 6.4613 | 6.4423 | 6.57465 | 6.20006 | 6.62589 | 5.88909 | 6.62442 | 5.8549 | 6.51896 | 6.92111 | 6.51306 | 5.7777 | 5.79244 | 5.35934 | 5.81852 |
| ZNF451 | Q9Y4E5-4 | 6.34415 | 6.48598 | 7.28857 | 6.38163 | 5.68849 | 6.5258 | 6.63605 | 6.35818 | 6.72537 | 6.53017 | 6.68945 | 6.84155 | 5.19262 | 5.68499 | 5.62689 | 5.00009 |
| GLTP | Q9NZD2 | 6.84763 | 5.66959 | 7.58256 | 5.64975 | 6.1031 | 5.82007 | 6.34126 | 6.79728 | 6.03514 | 6.5828 | 6.58865 | 6.44315 | 5.28469 | 6.11589 | 5.85344 | 6.28499 |
| FGD1 | P98174 | 6.23001 | 6.75074 | 6.67489 | 5.91415 | 5.88727 | 6.87497 | 6.36326 | 7.15982 | 6.32995 | 6.67387 | 6.2987 | 6.40489 | 5.62878 | 5.78445 | 5.51314 | 5.51112 |
| CRACD | Q6ZU35 | 6.26553 | 6.83961 | 6.35392 | 6.1778 | 5.82045 | 6.1997 | 6.54618 | 5.7483 | 6.30771 | 6.27062 | 5.88973 | 6.24415 | 6.54394 | 6.43902 | 5.90525 | 6.44808 |
| PSTPIP2 | Q9H939 | 6.35397 | 5.73675 | 6.43996 | 5.98269 | 6.25816 | 6.3123 | 5.9835 | 6.67517 | 5.92908 | 5.24909 | 6.50844 | 6.96638 | 5.95401 | 6.96368 | 6.0954 | 6.59142 |
| DVL1 | O14640 | 6.4334 | 6.27374 | 6.13888 | 6.55468 | 6.21837 | 5.88042 | 5.69925 | 6.05501 | 6.47642 | 6.69299 | 6.92291 | 7.36551 | 5.59416 | 6.36296 | 5.6306 | 5.70071 |
| PPP1R11 | O60927 | 6.23212 | 6.41437 | 6.88433 | 5.87389 | 5.82305 | 6.44689 | 5.42779 | 7.16921 | 7.01035 | 6.81942 | 7.51994 | 7.04548 | 5.91626 | 5.09082 | 4.79697 | 5.52911 |
| MEGF9 | Q9H1U4 | 7.07082 | 7.08464 | 5.99864 | 6.48088 | 6.7547 | 6.71 | 5.7729 | 5.33269 | 6.29671 | 6.82629 | 6.2972 | 6.12942 | 6.19099 | 6.16472 | 5.52862 | 6.36079 |
| METTL2B | Q6P1Q9 | 6.39009 | 6.24929 | 6.86479 | 5.84486 | 6.49368 | 6.8251 | 6.57349 | 7.00649 | 6.29824 | 6.49314 | 6.34902 | 6.68755 | 5.40038 | 5.56607 | 5.26666 | 5.69117 |
| FBXL3 | Q9UKT7 | 5.71155 | 6.43106 | 6.75392 | 5.97603 | 5.90647 | 6.12216 | 5.3492 | 5.97588 | 6.40548 | 6.59245 | 6.50517 | 6.17177 | 5.64875 | 6.80578 | 6.91355 | 6.73078 |
| RABGAP1L | Q5R372-5 | 6.40827 | 6.01632 | 6.87539 | 6.117 | 6.04829 | 6.37808 | 6.4178 | 6.1958 | 6.72893 | 6.27629 | 6.59713 | 6.64288 | 6.25577 | 5.73405 | 5.76112 | 5.54687 |
| RIN3 | Q8TB24 | 5.72823 | 5.61804 | 6.06851 | 5.64851 | 5.63423 | 5.98419 | 5.91351 | 6.28581 | 6.80451 | 6.77702 | 6.56917 | 7.56503 | 6.61088 | 6.70785 | 6.54499 | 5.53953 |
| CAV2 | P51636 | 6.34588 | 6.11077 | 6.69046 | 6.67201 | 6.53739 | 6.06505 | 6.38312 | 5.44678 | 6.49951 | 6.1451 | 6.16895 | 6.06134 | 6.18052 | 5.55079 | 6.45808 | 6.68424 |
| ANTKMT | Q9BQD7 | 6.41399 | 5.99328 | 7.17766 | 5.93092 | 7.32417 | 6.40371 | 6.40107 | 5.72874 | 7.11936 | 6.35168 | 6.60062 | 6.69912 | 5.87664 | 5.18619 | 5.50624 | 5.2866 |
| ATXN7L3B | Q96GX2 | 5.4476 | 6.01585 | 5.96895 | 4.80512 | 7.40863 | 6.75379 | 6.16462 | 6.74588 | 5.55182 | 6.90912 | 6.68797 | 6.15515 | 5.96982 | 7.00652 | 6.04649 | 6.36267 |
| CASP3 | P42574 | 6.29678 | 7.8369 | 7.16222 | 6.25374 | 6.95984 | 6.95926 | 5.49206 | 6.35644 | 6.08996 | 6.62891 | 6.38325 | 6.21576 | 5.30891 | 5.10174 | 5.58726 | 5.36697 |
| NADSYN1 | Q6IA69 | 7.34886 | 6.57508 | 6.81322 | 6.78156 | 6.81981 | 6.62436 | 5.5795 | 6.09775 | 6.54745 | 6.59097 | 6.02892 | 5.39668 | 6.69648 | 6.39988 | 5.09561 | 4.60389 |
| TMEM147 | Q9BVK8 | 6.38313 | 5.7954 | 6.9686 | 6.68615 | 6.73547 | 6.01508 | 6.2286 | 5.78736 | 6.26566 | 6.00363 | 5.85616 | 5.72594 | 6.31644 | 5.72884 | 6.95109 | 6.55245 |
| TMEM120A | Q9BXJ8 | 6.00297 | 6.03128 | 6.14656 | 6.71207 | 6.17809 | 5.76384 | 6.02921 | 5.72235 | 6.60588 | 6.38054 | 5.98094 | 5.92216 | 6.68939 | 6.26076 | 7.00856 | 6.56538 |
| COA6 | Q5JTJ3-2 | 6.42408 | 5.51503 | 5.92284 | 5.83613 | 6.41683 | 4.80412 | 6.12084 | 6.435 | 5.44857 | 5.86708 | 5.33189 | 6.00276 | 7.39612 | 7.49696 | 7.28778 | 7.69398 |
| LTO1 | Q8WV07 | 6.79562 | 6.0391 | 7.2991 | 6.06452 | 6.54463 | 6.52068 | 5.66968 | 6.72298 | 6.88 | 6.92561 | 7.10445 | 7.35006 | 4.92128 | 5.21762 | 4.70252 | 5.24214 |
| ID1 | P41134 | 5.00431 | 4.95152 | 5.81929 | 4.10038 | 5.59974 | 5.27774 | 5.24824 | 6.49936 | 5.21914 | 6.51097 | 6.3385 | 6.45901 | 6.8702 | 8.9632 | 8.04159 | 9.09681 |
| NR1D1 | P20393 | 5.84569 | 6.23763 | 5.60679 | 5.17815 | 6.15634 | 6.63236 | 5.8825 | 5.90034 | 6.51816 | 6.05464 | 6.3936 | 5.89804 | 6.80979 | 6.65509 | 7.40213 | 6.82875 |
| NR1D2 | Q14995 | 5.67294 | 5.77685 | 10.8745 | 5.91068 | 7.51835 | 6.6918 | 10.099 | 5.33044 | 6.06059 | 4.94405 | 6.09797 | 5.80442 | 4.95982 | 4.45803 | 5.00995 | 4.79063 |

| | | | | | | | | | | | | | | | | | |
|---|---|---|---|---|---|---|---|---|---|---|---|---|---|---|---|---|---|
| WDR41 | Q9HAD4 | 5.66103 | 6.47403 | 6.51388 | 6.18905 | 5.52434 | 6.29488 | 6.36062 | 5.86605 | 6.38656 | 6.74851 | 6.79248 | 7.16988 | 5.47881 | 5.45904 | 6.66728 | 6.41358 |
| INTS6L | Q5JSJ4 | 5.14298 | 6.8953 | 5.38558 | 6.95352 | 5.82967 | 6.5703 | 6.26191 | 5.91077 | 6.67241 | 5.75473 | 5.81362 | 7.61493 | 6.52901 | 5.87807 | 6.61516 | 6.17204 |
| PLAUR | Q03405-2 | 7.24839 | 7.72203 | 6.73499 | 7.69989 | 6.75172 | 6.69973 | 6.7959 | 6.69188 | 6.10063 | 6.349 | 6.36823 | 6.43609 | 4.67588 | 4.28556 | 4.52745 | 4.91263 |
| NAB2 | Q15742 | 5.72145 | 6.1106 | 6.20043 | 5.59484 | 6.07034 | 5.81728 | 5.93111 | 5.81607 | 6.17703 | 6.62006 | 5.92313 | 5.9294 | 6.86091 | 7.20579 | 7.22848 | 6.79306 |
| TMEM192 | Q8IY95 | 6.72286 | 6.50872 | 6.46906 | 5.80647 | 6.48497 | 5.57743 | 6.14265 | 5.62696 | 6.49686 | 5.53025 | 6.20739 | 6.4661 | 6.714 | 5.81765 | 6.72871 | 6.69991 |
| LTBP2 | Q14767 | 6.39795 | 5.09114 | 4.76724 | 7.27965 | 5.29982 | 5.38543 | 6.66727 | 5.59505 | 7.05962 | 5.84517 | 6.74636 | 6.06764 | 8.50873 | 6.0717 | 6.8934 | 6.32384 |
| NT5C3B | Q969T7 | 6.2001 | 6.10483 | 6.9712 | 5.95836 | 7.26906 | 7.2003 | 6.23453 | 6.48034 | 6.26317 | 6.82797 | 6.64286 | 6.36789 | 5.78518 | 5.58009 | 4.85043 | 5.26369 |
| JAG1 | P78504 | 6.19608 | 6.81729 | 6.72488 | 6.40227 | 5.96942 | 5.83502 | 5.66744 | 5.24998 | 6.14514 | 5.94729 | 6.61451 | 6.40578 | 7.06339 | 6.31332 | 6.4129 | 6.23529 |
| FRMD4A | Q9P2Q2 | 6.11623 | 6.25277 | 6.78152 | 6.26843 | 5.9607 | 6.68265 | 6.1061 | 6.02285 | 6.11555 | 6.10862 | 6.62798 | 6.78942 | 6.12843 | 6.07608 | 5.64632 | 6.31636 |
| E2F6 | O75461 | 6.52609 | 6.48155 | 6.21336 | 7.03404 | 6.19502 | 5.88008 | 6.47153 | 5.61223 | 6.33265 | 6.16492 | 6.56007 | 6.65789 | 6.36716 | 5.31785 | 5.84546 | 6.34011 |
| NIT1 | Q86X76-3 | 6.02265 | 5.07978 | 6.60735 | 5.89446 | 6.85991 | 6.48205 | 5.6111 | 6.24932 | 5.87006 | 6.9341 | 7.12206 | 6.45427 | 6.1743 | 6.20043 | 6.49649 | 5.94166 |
| FBXL20 | Q96IG2 | 5.61078 | 6.38545 | 5.91557 | 6.63527 | 6.75616 | 5.71236 | 6.18471 | 5.90532 | 6.03428 | 5.84761 | 6.12511 | 5.75806 | 7.08134 | 6.85766 | 6.89862 | 6.2917 |
| TRAF3 | Q13114 | 6.41851 | 6.11262 | 6.45978 | 5.96446 | 5.94443 | 6.80246 | 6.11951 | 6.63839 | 5.85421 | 6.45741 | 5.86479 | 6.2629 | 5.86839 | 6.87153 | 6.46179 | 5.89881 |
| MOB2 | Q70IA6-3 | 6.19559 | 6.28147 | 6.97612 | 5.31225 | 5.98664 | 6.82825 | 6.27174 | 5.94367 | 6.16186 | 6.1181 | 7.08137 | 6.27782 | 6.22399 | 6.28024 | 6.08931 | 5.97159 |
| DGKH | Q86XP1 | 6.80762 | 6.14987 | 6.69376 | 6.56196 | 6.42501 | 6.59258 | 6.15244 | 6.79686 | 6.2234 | 6.5941 | 6.3617 | 6.05941 | 5.59748 | 5.60749 | 5.60936 | 5.76696 |
| METTL14 | Q9HCE5 | 5.44684 | 5.20666 | 6.37483 | 6.39965 | 5.93724 | 5.918 | 5.8855 | 5.21224 | 6.00978 | 6.10354 | 6.26043 | 6.29323 | 7.24699 | 6.86593 | 7.40613 | 7.43302 |
| RMND5B | Q96G75 | 6.61379 | 5.8828 | 7.42059 | 6.50902 | 6.28651 | 6.11249 | 6.58133 | 6.40163 | 7.09751 | 6.84903 | 8.23686 | 6.33543 | 4.5374 | 4.66793 | 5.42402 | 5.04367 |
| RSL24D1 | Q9UHA3 | 5.93106 | 6.43999 | 6.11176 | 7.03828 | 6.50684 | 5.97555 | 6.54544 | 5.71535 | 6.23151 | 5.90244 | 5.74071 | 6.19753 | 5.92317 | 6.25173 | 6.75394 | 6.73469 |
| RMDN2 | Q96LZ7-2 | 6.8992 | 6.72586 | 6.63982 | 6.49108 | 6.45222 | 6.12272 | 6.35324 | 6.54505 | 6.45356 | 5.6984 | 5.77555 | 5.63714 | 6.3202 | 5.77323 | 6.53903 | 6.47372 |
| BAHD1 | Q8TBE0 | 6.56821 | 6.82406 | 6.3288 | 7.62336 | 6.16871 | 5.62663 | 7.20641 | 5.8478 | 6.66475 | 6.41775 | 6.88328 | 6.95882 | 5.16012 | 4.95466 | 4.94563 | 5.82103 |
| CACHD1 | Q5VU97 | 6.23782 | 6.00991 | 6.26016 | 6.36204 | 5.69681 | 6.54038 | 6.12852 | 5.68803 | 6.44968 | 7.0414 | 6.76697 | 6.06972 | 6.22037 | 5.8186 | 6.42708 | 6.28251 |
| CIAO3 | Q9H6Q4 | 6.57214 | 6.24134 | 6.94876 | 5.7953 | 6.43049 | 7.15994 | 5.76628 | 6.59272 | 5.96964 | 6.98815 | 6.6977 | 6.56813 | 5.65528 | 5.65251 | 5.28052 | 5.68111 |
| RMI2 | Q96E14 | 6.36591 | 5.31135 | 6.14974 | 5.57231 | 5.3249 | 4.68054 | 5.71471 | 7.4823 | 5.7591 | 6.84998 | 5.70982 | 6.95451 | 5.91702 | 7.19419 | 6.95251 | 8.06111 |
| STYX | Q8WUJ0 | 6.27125 | 5.64147 | 6.59638 | 5.6612 | 6.54765 | 7.02427 | 6.53221 | 6.46437 | 5.95853 | 6.51216 | 6.20837 | 5.69239 | 5.71182 | 6.49679 | 6.44168 | 6.23945 |
| NCBP2AS2 | Q69YL0 | 5.84482 | 5.9469 | 5.24028 | 6.34083 | 6.6056 | 6.22873 | 6.55471 | 6.83376 | 6.22785 | 6.16613 | 6.16187 | 5.83871 | 7.08222 | 6.2586 | 5.85998 | 6.80903 |
| TFDP1 | Q14186 | 6.00512 | 6.33642 | 6.25716 | 5.66318 | 6.21811 | 6.40839 | 5.9403 | 6.36467 | 6.40354 | 6.46047 | 6.13331 | 5.77011 | 6.91141 | 6.43571 | 6.38063 | 6.31148 |
| TAF4B | Q92750 | 6.7677 | 6.09015 | 6.17299 | 6.16583 | 5.9052 | 6.34086 | 6.13979 | 5.52966 | 5.59526 | 5.89288 | 6.80816 | 5.78293 | 6.65028 | 6.06955 | 6.90416 | 7.21686 |
| STN1 | Q9H668 | 5.61687 | 6.39015 | 5.88108 | 6.17497 | 6.30626 | 6.18841 | 5.96169 | 6.27511 | 5.78978 | 5.71078 | 6.25758 | 5.87612 | 7.13839 | 6.83473 | 6.98103 | 6.61706 |
| PRKCZ | Q05513 | 6.58675 | 6.48326 | 7.28065 | 6.32084 | 6.08243 | 5.83963 | 6.44442 | 6.02 | 6.44957 | 6.95088 | 6.65091 | 6.56914 | 5.49765 | 5.7025 | 5.24645 | 5.87493 |
| HSF2 | Q03933 | 6.40365 | 6.19084 | 6.14912 | 6.15343 | 6.73097 | 6.96967 | 6.35654 | 6.25526 | 6.4886 | 7.70038 | 7.67645 | 6.99973 | 5.3845 | 4.9445 | 4.47945 | 5.11693 |
| SUPT20H | Q8NEM7-3 | 6.11018 | 5.63989 | 6.24387 | 7.23124 | 6.536 | 6.50104 | 6.04128 | 5.29227 | 5.47378 | 5.97872 | 6.90984 | 6.84818 | 6.17873 | 6.47558 | 6.16303 | 6.37637 |
| SNX25 | Q9H3E2 | 6.50771 | 6.44455 | 5.76648 | 6.7376 | 5.54945 | 5.87768 | 6.14423 | 6.35272 | 6.76926 | 6.16303 | 5.95157 | 6.3848 | 5.44735 | 6.65841 | 6.17022 | 7.07494 |
| TMEM175 | Q9BSA9 | 6.76569 | 6.53734 | 5.79335 | 6.83587 | 6.0273 | 5.60448 | 6.45191 | 6.094 | 6.38703 | 6.12797 | 6.08435 | 6.6693 | 5.98876 | 6.16418 | 6.2175 | 6.25096 |
| NEU3 | Q9UQ49-2 | 6.01214 | 6.67453 | 6.87707 | 6.21037 | 6.16356 | 6.16349 | 6.76255 | 6.30011 | 6.51971 | 6.96838 | 6.20871 | 6.3169 | 5.69642 | 6.00588 | 6.20268 | 5.9175 |
| TXN2 | Q99757 | 5.38782 | 5.16178 | 4.3971 | 5.95307 | 5.33537 | 4.8584 | 5.5434 | 6.71513 | 5.10151 | 6.33901 | 5.15738 | 6.22835 | 6.16875 | 9.77099 | 7.35239 | 10.5295 |
| DSCC1 | Q9BVC3 | 5.60077 | 6.42636 | 5.91204 | 6.07767 | 5.67398 | 5.64811 | 6.87463 | 6.40214 | 6.34534 | 5.60948 | 6.58227 | 6.39381 | 5.94888 | 6.85175 | 6.4729 | 7.17986 |
| SLC25A46 | Q96AG3 | 5.97137 | 6.38659 | 6.23504 | 6.03638 | 6.03195 | 6.09868 | 6.20678 | 5.54745 | 6.09562 | 6.2314 | 6.19248 | 6.677 | 6.65945 | 6.47317 | 6.45927 | 6.69738 |
| LSM14B | Q9BX40 | 6.75292 | 6.90731 | 7.37498 | 6.16039 | 6.41644 | 7.3476 | 6.32099 | 5.89244 | 6.52548 | 6.48498 | 6.5988 | 7.42306 | 4.83342 | 5.05035 | 4.87016 | 5.09017 |
| PARP16 | Q8N5Y8-3 | 6.16179 | 6.50434 | 6.44312 | 6.83092 | 6.44789 | 6.1345 | 6.63682 | 5.78131 | 6.2607 | 6.2446 | 6.28885 | 5.68202 | 6.09365 | 6.02286 | 6.58939 | 5.87724 |
| SHB | Q15464 | 6.06877 | 6.73694 | 5.79378 | 6.21035 | 6.31057 | 6.04313 | 6.06119 | 6.65565 | 6.74348 | 5.77962 | 6.41142 | 5.92487 | 5.63653 | 6.62244 | 7.09172 | 5.90955 |
| ENGASE | Q8NFI3 | 5.70913 | 5.80607 | 6.0304 | 5.92604 | 5.62832 | 5.6975 | 5.78664 | 6.14034 | 6.12571 | 6.4784 | 6.36948 | 6.34369 | 6.81968 | 6.99709 | 7.14718 | 6.99431 |
| ARG2 | P78540 | 5.27784 | 4.67726 | 5.72083 | 5.25483 | 7.12966 | 6.5106 | 6.75905 | 6.89966 | 6.07259 | 6.04946 | 6.19902 | 6.17691 | 7.19209 | 6.15819 | 6.47721 | 7.44479 |
| QSOX1 | O00391 | 6.03705 | 7.09729 | 5.35421 | 6.56569 | 6.14692 | 7.2302 | 6.78523 | 6.15983 | 6.82483 | 6.06426 | 5.87034 | 6.31603 | 6.11698 | 4.89533 | 5.29642 | 7.23939 |
| MAD2L2 | Q9UI95 | 6.52063 | 6.46249 | 6.12346 | 6.01395 | 6.54405 | 5.95362 | 6.43665 | 6.10761 | 6.27972 | 6.54796 | 6.4688 | 6.61239 | 6.5537 | 5.90033 | 6.07827 | 6.29682 |
| OSMR | Q99650 | 7.13171 | 7.25161 | 6.69108 | 7.76858 | 6.32455 | 6.10866 | 5.87559 | 6.18653 | 6.78054 | 6.40889 | 7.03515 | 6.75646 | 4.82005 | 4.63262 | 5.04591 | 5.18206 |
| FBXW5 | Q969U6 | 6.25911 | 5.74397 | 6.6516 | 6.11986 | 6.28312 | 6.75288 | 6.46673 | 6.3721 | 6.14995 | 6.57257 | 7.48405 | 6.43297 | 5.8396 | 5.79527 | 5.84318 | 5.23303 |
| EMB | Q6PCB8 | 6.47566 | 6.7356 | 6.30878 | 6.223 | 6.23217 | 6.61425 | 6.33213 | 6.23472 | 6.64316 | 6.57308 | 6.03611 | 6.30112 | 5.98992 | 6.14863 | 5.13086 | 6.02082 |
| SEC22A | Q96IW7 | 6.6633 | 7.01599 | 5.85705 | 5.65327 | 4.64565 | 7.3248 | 6.06163 | 5.96064 | 6.65054 | 5.89272 | 6.27865 | 6.17336 | 6.27241 | 6.11592 | 7.86855 | 5.5655 |
| PALB2 | Q86YC2 | 6.69853 | 6.54899 | 6.32141 | 7.08006 | 5.93214 | 6.23671 | 6.29776 | 5.51031 | 6.39811 | 6.55279 | 6.26137 | 6.21604 | 6.07263 | 5.52606 | 5.85184 | 6.49525 |
| CEP120 | Q8N960 | 7.33452 | 6.20368 | 6.4139 | 7.22759 | 6.00803 | 6.72789 | 6.94642 | 5.4288 | 7.70701 | 5.83407 | 5.59186 | 6.44084 | 5.33899 | 5.45237 | 5.51303 | 5.83101 |
| COQ9 | O75208 | 6.24615 | 5.2129 | 5.06052 | 5.32446 | 6.30487 | 5.21057 | 6.49 | 6.34455 | 5.31449 | 6.70753 | 5.12674 | 6.12052 | 6.73851 | 8.13216 | 7.95472 | 7.71131 |
| ZSWIM9 | Q86XI8-2 | 6.89145 | 6.3151 | 6.41674 | 7.01906 | 6.45202 | 5.64139 | 7.25624 | 5.60189 | 6.41173 | 6.23814 | 6.25764 | 6.59935 | 5.86135 | 5.60873 | 5.08215 | 6.34732 |
| C18orf25 | Q96B23 | 5.59119 | 6.03217 | 5.83107 | 5.49214 | 6.04957 | 5.9262 | 5.93339 | 6.28764 | 5.62893 | 6.03181 | 6.25892 | 6.07451 | 7.38927 | 6.84851 | 7.59778 | 7.02692 |
| FGFR1OP2 | Q9NVK5 | 7.25145 | 5.31175 | 4.9777 | 5.34898 | 6.34389 | 4.87598 | 5.64767 | 7.31054 | 5.90548 | 6.5892 | 6.02131 | 7.6056 | 6.45534 | 6.48946 | 6.4657 | 7.39992 |
| RNF6 | Q9Y252 | 7.18789 | 6.87804 | 6.6971 | 7.40605 | 6.51563 | 6.87265 | 7.23634 | 5.75965 | 6.53259 | 6.73432 | 6.40105 | 6.71711 | 4.71458 | 4.59186 | 5.26499 | 4.49016 |
| KIF3C | O14782 | 6.60578 | 6.40384 | 6.61987 | 5.88575 | 6.34549 | 6.31295 | 6.3207 | 6.80671 | 6.42441 | 6.30856 | 6.44347 | 6.5358 | 5.41024 | 5.83552 | 5.8806 | 5.86031 |
| STARD9 | Q9P2P6 | 6.33938 | 6.33026 | 7.05251 | 6.69017 | 6.00496 | 6.33115 | 6.40925 | 6.1614 | 6.3406 | 6.68248 | 6.22132 | 6.4169 | 5.4545 | 5.87989 | 5.65268 | 6.03253 |
| TMEM263 | Q8WUH6 | 5.40608 | 5.22238 | 4.78974 | 4.77964 | 5.82967 | 5.04636 | 7.15764 | 7.66767 | 8.58476 | 6.26385 | 6.32113 | 5.63688 | 7.06011 | 7.61481 | 6.83321 | 5.78606 |
| NDUFB1 | O75438-2 | 5.86157 | 5.59642 | 5.6841 | 6.02171 | 5.75221 | 5.08731 | 5.10166 | 6.64446 | 5.21575 | 5.55501 | 4.77151 | 5.05567 | 5.99559 | 9.37511 | 7.5086 | 10.7733 |
| PPP2R3A | Q06190 | 6.20877 | 6.21831 | 6.65229 | 6.40895 | 5.91164 | 5.53537 | 6.30612 | 6.62958 | 6.80033 | 6.52847 | 6.37648 | 6.51553 | 6.50242 | 6.23779 | 5.35353 | 5.60116 |
| POLR2H | P52434 | 5.86287 | 6.05062 | 5.66086 | 6.06571 | 6.3396 | 6.84809 | 6.15217 | 6.85175 | 6.49779 | 6.82944 | 6.62497 | 7.05634 | 5.63013 | 5.62402 | 5.71945 | 6.18619 |
| LYSMD3 | Q7Z3D4 | 5.99981 | 5.6686 | 5.32811 | 6.09223 | 6.52104 | 5.76732 | 5.86852 | 6.20788 | 5.959 | 5.75274 | 6.18603 | 5.93867 | 7.37823 | 6.78545 | 7.40611 | 7.14025 |
| SLC6A6 | P31641 | 6.14489 | 5.83988 | 6.38212 | 6.82833 | 6.3793 | 6.9124 | 6.56778 | 5.76441 | 6.83181 | 5.69648 | 6.88574 | 6.45997 | 5.42146 | 6.40629 | 6.46242 | 5.01672 |
| MAML2 | Q8IZL2 | 5.83514 | 5.86683 | 6.63909 | 4.99948 | 6.11617 | 6.00881 | 5.54734 | 6.29251 | 6.78199 | 6.11871 | 6.41428 | 6.13261 | 7.4236 | 6.93346 | 6.61976 | 6.27023 |
| PLCG2 | P16885 | 5.92726 | 6.49998 | 7.21999 | 6.07068 | 6.29097 | 7.05027 | 6.10441 | 6.95153 | 6.93785 | 6.12228 | 6.78213 | 6.617 | 5.8697 | 5.03491 | 5.78621 | 4.73484 |
| ZNF302 | Q9NR11-2 | 5.75748 | 7.82219 | 6.71947 | 7.89718 | 6.35332 | 5.42374 | 6.61446 | 5.75715 | 6.57943 | 7.2065 | 6.79336 | 7.13501 | 4.85874 | 5.23878 | 4.66469 | 5.17851 |
| PAFAH1B2 | P68402 | 6.57544 | 5.09734 | 6.50791 | 4.25945 | 6.1242 | 5.79661 | 4.6483 | 6.92481 | 4.79462 | 6.21622 | 5.34922 | 7.21761 | 5.75687 | 8.72859 | 8.35779 | 7.64503 |
| CPEB4 | Q17RY0 | 6.37305 | 5.79638 | 6.47299 | 5.91494 | 6.77833 | 6.97963 | 5.78124 | 6.67659 | 6.89103 | 6.44203 | 6.32852 | 6.56591 | 6.27219 | 5.40163 | 5.62018 | 5.70537 |
| CRAMP1 | Q96RY5 | 6.15205 | 6.91937 | 6.4977 | 6.85557 | 6.23601 | 5.87696 | 6.89689 | 5.93575 | 7.03862 | 6.06108 | 7.26354 | 5.2481 | 5.62728 | 6.39537 | 5.47342 | 5.52229 |
| CCDC85B | Q15834 | 6.63687 | 5.87348 | 6.5799 | 5.3887 | 6.31561 | 6.53345 | 5.81441 | 6.46711 | 6.47471 | 6.99023 | 7.54865 | 7.00385 | 6.12651 | 5.22436 | 5.47225 | 5.5499 |
| B4GALT1 | P15291 | 6.61625 | 6.45293 | 6.03856 | 6.67061 | 6.12181 | 6.21635 | 6.41513 | 5.91946 | 6.96367 | 6.61577 | 6.51621 | 6.27703 | 5.96894 | 5.66387 | 5.64084 | 5.90256 |
| CENPS | Q8N2Z9 | 6.172 | 7.15871 | 7.4033 | 6.42757 | 6.09268 | 6.46671 | 5.28816 | 6.16688 | 6.74294 | 6.3841 | 6.07061 | 6.35606 | 5.73511 | 5.63957 | 5.42433 | 6.47127 |
| A2M | P01023 | 4.84701 | 7.0326 | 5.06876 | 4.43066 | 4.29987 | 6.85082 | 5.83431 | 5.19752 | 7.64031 | 5.75659 | 6.20769 | 6.47827 | 6.71778 | 6.28238 | 9.77102 | 7.58443 |
| FBXO7 | Q9Y3I1 | 5.82118 | 5.49778 | 6.66453 | 6.3356 | 5.94861 | 6.83756 | 5.81466 | 5.98244 | 6.3628 | 6.84957 | 6.12986 | 6.60813 | 6.67338 | 6.15637 | 5.61891 | 6.69861 |
| F8A1 | P23610 | 5.63564 | 5.59736 | 6.20297 | 5.19495 | 6.33521 | 6.53741 | 5.47451 | 6.92149 | 5.78406 | 7.01368 | 6.88502 | 7.27551 | 5.60902 | 6.46878 | 6.63921 | 6.42516 |
| MAP3K7 | O43318 | 6.63191 | 6.51298 | 6.3499 | 6.15743 | 6.3901 | 6.66909 | 6.01924 | 6.67685 | 6.27181 | 6.66506 | 6.47973 | 6.50041 | 5.7978 | 5.85524 | 5.53131 | 5.49113 |
| RPS27 | P42677 | 5.98009 | 6.06902 | 7.00779 | 6.47944 | 6.57588 | 6.33819 | 7.05467 | 6.15683 | 6.55405 | 5.43235 | 6.64115 | 6.63981 | 4.92643 | 6.42511 | 5.81993 | 5.89925 |
| WEE1 | P30291 | 5.33296 | 5.74564 | 6.21797 | 5.3078 | 6.3006 | 6.04981 | 6.05748 | 5.35971 | 6.22558 | 6.55545 | 6.90767 | 5.78013 | 7.17872 | 6.96841 | 7.22038 | 6.79168 |
| PSRC1 | Q6PGN9 | 6.29962 | 6.9742 | 7.71623 | 6.47107 | 6.15665 | 6.93716 | 6.20789 | 6.87916 | 6.53696 | 6.82857 | 7.57282 | 8.22781 | 4.60168 | 4.36095 | 3.49399 | 4.73526 |
| MCOLN2 | Q8IZK6 | 6.82717 | 6.35147 | 6.59613 | 6.71767 | 6.46668 | 5.8761 | 6.58959 | 6.28862 | 6.26141 | 6.32328 | 6.48999 | 6.03687 | 5.65007 | 5.56261 | 5.95509 | 6.00725 |
| CASTOR2 | A6NHX0 | 6.42384 | 6.0527 | 6.58685 | 5.60472 | 6.05724 | 7.02197 | 6.39643 | 6.57887 | 6.37751 | 6.86792 | 6.48704 | 6.63754 | 5.54871 | 6.23703 | 5.55606 | 5.56557 |
| ELL2 | O00472 | 7.42138 | 7.307 | 7.55652 | 8.26716 | 6.48174 | 6.79104 | 7.40568 | 6.95891 | 7.32951 | 7.2527 | 7.0273 | 6.4535 | 3.22586 | 3.43733 | 3.38353 | 3.70085 |
| SMIM8 | Q96KF7 | 6.26361 | 6.28123 | 5.70428 | 6.79023 | 6.16792 | 6.74977 | 6.25426 | 7.52327 | 5.2047 | 5.93495 | 6.0254 | 6.24134 | 5.79457 | 6.50858 | 6.26315 | 6.29274 |
| PIP4K2A | P48426 | 5.99815 | 6.16462 | 6.3103 | 6.07202 | 6.22137 | 6.75842 | 6.84937 | 5.54823 | 5.94547 | 6.14889 | 6.0973 | 5.79397 | 6.25813 | 6.4524 | 6.95481 | 6.42656 |
| PEA15 | Q15121-2 | 6.28675 | 4.88705 | 6.40581 | 4.24504 | 5.72159 | 5.35364 | 5.44527 | 7.39643 | 4.76937 | 6.4784 | 5.66562 | 7.50109 | 5.05988 | 8.95834 | 7.27979 | 8.54593 |
| PPP1R15B | Q5SWA1 | 6.29041 | 6.6938 | 6.66005 | 7.20025 | 5.76802 | 5.70916 | 6.66033 | 6.03897 | 6.49019 | 5.96637 | 6.59099 | 6.33568 | 5.74231 | 5.74614 | 5.92823 | 6.17909 |
| UBE2E3 | Q969T4 | 5.89848 | 5.37861 | 5.85367 | 5.73365 | 5.34016 | 6.09945 | 6.37105 | 7.74713 | 7.40775 | 7.05142 | 5.66724 | 6.7358 | 6.46415 | 6.45414 | 7.15176 | 4.64352 |
| GTDC1 | Q4AE62 | 6.39593 | 6.73832 | 7.03879 | 6.20244 | 5.53493 | 7.48562 | 5.61462 | 5.78499 | 5.75737 | 6.61602 | 6.55024 | 6.49371 | 5.76234 | 6.58504 | 6.00964 | 5.43 |
| RNASET2 | O00584 | 6.6643 | 5.35383 | 5.99075 | 5.20814 | 6.43724 | 4.98496 | 5.42883 | 6.60464 | 5.25738 | 6.22132 | 5.62319 | 6.17454 | 6.93722 | 7.84538 | 7.53137 | 7.7369 |
| MRPL49 | Q13405 | 5.14643 | 5.0229 | 5.00485 | 4.911 | 5.8017 | 4.82142 | 6.01575 | 7.18372 | 4.47923 | 6.69498 | 5.51216 | 7.16151 | 5.5612 | 8.51478 | 8.53286 | 9.63553 |
| BICRAL | Q6AI39 | 6.32679 | 6.52505 | 6.13354 | 6.19069 | 5.81938 | 6.10924 | 6.0388 | 6.07189 | 6.11498 | 6.4751 | 6.32264 | 6.84878 | 6.75437 | 6.14356 | 6.14717 | 5.97803 |
| SLC12A7 | Q9Y666 | 6.05078 | 6.20897 | 6.39437 | 6.06311 | 6.32867 | 6.26986 | 5.74578 | 5.38929 | 6.73086 | 6.98903 | 6.69612 | 6.50872 | 5.9756 | 6.09176 | 5.87591 | 6.68117 |
| SLC12A6 | Q9UHW9-2 | 6.76504 | 6.67374 | 6.60199 | 6.64807 | 6.75673 | 6.16345 | 6.33561 | 5.70085 | 6.02322 | 6.09397 | 6.60221 | 6.3212 | 5.70821 | 5.51703 | 6.14392 | 5.94276 |
| BRAF | A0A2R8Y8E0 | 6.51387 | 6.19232 | 7.07647 | 5.84153 | 6.40678 | 6.63234 | 6.40914 | 6.89679 | 6.51098 | 6.18439 | 6.17294 | 6.75292 | 5.76264 | 5.78285 | 5.48371 | 5.38033 |
| PER1 | O15534 | 6.16188 | 6.69036 | 6.56317 | 5.6678 | 6.02995 | 6.46588 | 6.26668 | 6.4757 | 6.68819 | 6.65812 | 6.48664 | 6.3507 | 5.88978 | 5.8585 | 5.66916 | 6.07749 |
| NRAS | P01111 | 5.90972 | 6.06744 | 6.1549 | 6.73918 | 5.88133 | 5.68679 | 6.54046 | 6.41935 | 6.06447 | 5.85142 | 6.01893 | 6.54662 | 7.10387 | 6.37646 | 6.39053 | 6.24852 |
| USPL1 | Q5W0Q7 | 6.35002 | 6.52117 | 6.52848 | 7.16132 | 5.81694 | 5.77336 | 6.46963 | 6.15724 | 7.0344 | 6.36838 | 6.57468 | 6.61753 | 5.63324 | 5.34511 | 6.05063 | 5.59786 |
| C2orf49 | Q9BVC5 | 5.83993 | 5.81379 | 5.43428 | 5.72233 | 6.41957 | 5.91026 | 6.25747 | 5.82647 | 5.58396 | 6.09204 | 5.83047 | 5.99665 | 7.75049 | 7.37085 | 7.62651 | 6.52494 |
| MCEE | Q96PE7 | 5.34329 | 4.86251 | 5.0161 | 4.84866 | 5.80121 | 5.20901 | 5.45319 | 7.01805 | 5.09205 | 6.88507 | 5.78302 | 6.96822 | 6.36065 | 8.51095 | 8.35799 | 8.49001 |
| C12orf45 | Q8N5I9 | 6.25271 | 6.33456 | 7.10824 | 6.46653 | 6.20562 | 6.87102 | 5.97291 | 5.99564 | 6.31058 | 6.24941 | 7.37839 | 7.12936 | 5.77787 | 5.35719 | 5.09864 | 5.49132 |
| SPIN4 | Q56A73 | 5.80353 | 6.42785 | 5.73915 | 6.38458 | 6.03045 | 6.54435 | 6.99174 | 6.14715 | 5.89127 | 5.82849 | 6.47234 | 5.55115 | 6.51982 | 5.90014 | 7.13147 | 6.63653 |
| USP38 | Q8NB14 | 5.28935 | 5.80043 | 6.188 | 5.74201 | 5.72883 | 6.15523 | 5.95673 | 6.65702 | 5.94138 | 6.03041 | 6.30278 | 6.16462 | 6.66741 | 7.35988 | 6.93131 | 7.08459 |
| STEAP1 | Q9UHE8 | 5.96113 | 7.24986 | 6.31845 | 7.29283 | 6.11738 | 6.19344 | 5.83346 | 6.71071 | 6.32248 | 6.15757 | 5.94128 | 6.54669 | 5.33286 | 5.93254 | 5.92669 | 6.16264 |
| PPHLN1 | Q8NEY8-2 | 5.9962 | 6.40914 | 5.45468 | 6.61927 | 6.21617 | 6.69536 | 6.41872 | 6.54953 | 5.91966 | 6.27365 | 5.95966 | 6.14155 | 6.5996 | 6.08781 | 6.05609 | 6.60292 |
| MKNK1 | Q9BUB5 | 6.14863 | 6.88777 | 7.81744 | 6.43134 | 6.2617 | 6.9127 | 6.59729 | 6.53268 | 6.73666 | 6.86989 | 7.48555 | 6.83074 | 4.90031 | 4.77197 | 4.34576 | 4.46957 |

| | | | | | | | | | | | | | | | | | |
|---|---|---|---|---|---|---|---|---|---|---|---|---|---|---|---|---|---|
| EMC10 | Q5UCC4 | 7.00638 | 6.73356 | 5.51983 | 5.89185 | 7.14365 | 5.88257 | 6.16001 | 6.35884 | 5.87357 | 6.62273 | 5.45588 | 7.02154 | 6.19496 | 5.70523 | 6.38282 | 6.04656 |
| ZNF8 | P17098 | 5.82081 | 5.72838 | 5.76138 | 6.37851 | 5.78873 | 5.57521 | 6.90578 | 5.98753 | 6.10712 | 5.80849 | 5.71834 | 6.5729 | 7.14009 | 6.64956 | 6.76623 | 7.29092 |
| CCNL1 | Q9UK58 | 5.89069 | 5.97479 | 6.24503 | 6.52149 | 6.17324 | 5.81278 | 6.30561 | 5.38906 | 6.10538 | 4.98654 | 6.01684 | 5.18448 | 7.27901 | 7.16414 | 7.69465 | 7.25626 |
| FAM177A1 | Q8N128-2 | 6.38861 | 6.01795 | 6.31232 | 7.03484 | 6.09148 | 6.17253 | 6.70956 | 6.1586 | 6.20324 | 6.84928 | 6.14385 | 5.94501 | 5.77007 | 5.55737 | 6.22617 | 6.41912 |
| RYK | P34925-2 | 6.06231 | 6.59771 | 7.21242 | 7.12253 | 6.17558 | 5.91712 | 6.30203 | 5.6141 | 6.40848 | 6.45735 | 6.6518 | 7.0772 | 5.62837 | 5.17524 | 5.77817 | 5.81959 |
| RILPL1 | Q5EBL4 | 6.59412 | 6.30663 | 7.39428 | 5.91308 | 6.37827 | 6.66894 | 6.4354 | 6.01745 | 7.20481 | 6.77395 | 8.17803 | 5.3923 | 5.16004 | 4.93604 | 4.80203 | 5.84463 |
| TMED8 | Q6PL24 | 5.87228 | 5.99306 | 6.92218 | 6.15037 | 5.95299 | 6.83677 | 6.31489 | 5.22606 | 6.17371 | 6.24964 | 6.30295 | 7.09345 | 6.49297 | 5.8557 | 6.24869 | 6.31428 |
| AMD1 | P17707 | 6.90776 | 6.24136 | 8.3935 | 6.25045 | 6.28345 | 5.84357 | 5.99329 | 7.13252 | 7.01962 | 7.51832 | 7.16319 | 8.65522 | 3.49268 | 4.66113 | 4.00584 | 4.43811 |
| LSM2 | Q9Y333 | 5.75652 | 5.30093 | 5.63371 | 5.53499 | 6.78855 | 5.92573 | 6.60227 | 6.48474 | 6.60359 | 6.28667 | 6.37781 | 5.82103 | 6.58914 | 6.33136 | 7.49926 | 6.4637 |
| VTN | P04004 | 7.15015 | 4.85787 | 3.23793 | 7.58909 | 4.15028 | 5.58133 | 9.07061 | 5.10655 | 6.64181 | 5.93235 | 3.95489 | 4.59312 | 9.04676 | 5.09792 | 8.03871 | 9.95063 |
| NAB1 | Q13506 | 5.51422 | 5.49061 | 6.15936 | 5.16082 | 5.88498 | 6.23958 | 5.34232 | 6.50908 | 6.29416 | 6.38492 | 6.73317 | 6.45727 | 6.8288 | 6.7887 | 7.51697 | 6.69503 |
| MBOAT2 | Q6ZWT7 | 6.15475 | 6.71045 | 6.04923 | 7.06139 | 6.26699 | 6.06805 | 5.8722 | 6.0863 | 6.09178 | 6.48834 | 5.66993 | 6.12576 | 5.58722 | 6.69893 | 6.70257 | 6.36612 |
| NECTIN3 | Q9NQS3 | 6.26732 | 7.20006 | 5.72421 | 6.537 | 6.02821 | 5.94157 | 6.4765 | 5.72429 | 6.14543 | 6.36557 | 6.10791 | 6.17869 | 6.91569 | 6.16863 | 5.96223 | 6.25667 |
| RFFL | Q8WZ73 | 6.86951 | 6.92878 | 6.35605 | 6.95144 | 6.59144 | 6.08435 | 6.93988 | 5.81594 | 5.80419 | 6.02557 | 7.02548 | 6.40739 | 5.59338 | 5.18394 | 6.35921 | 5.06345 |
| WDCP | Q9H6R7 | 5.96168 | 6.12323 | 6.66632 | 6.45861 | 6.2923 | 6.68263 | 5.79913 | 6.38052 | 6.28428 | 6.73259 | 6.20657 | 6.9195 | 6.50102 | 5.95224 | 5.5085 | 5.53088 |
| C12orf65 | Q9H3J6 | 6.29325 | 6.43967 | 5.83791 | 6.7792 | 5.96656 | 5.52162 | 6.60456 | 5.61097 | 6.21349 | 5.78512 | 5.63429 | 6.42017 | 6.66547 | 6.56691 | 7.06904 | 6.59177 |
| FOXJ2 | Q9P0K8 | 5.68933 | 5.41152 | 5.8965 | 5.04575 | 6.27528 | 6.64913 | 5.58269 | 5.19352 | 6.37194 | 7.17979 | 6.26357 | 6.172 | 7.53862 | 7.20755 | 6.74914 | 6.77367 |
| ELAPOR2 | A8MWY0 | 6.5052 | 7.25744 | 6.08763 | 6.24929 | 5.97866 | 6.35493 | 6.06675 | 5.89975 | 6.2471 | 6.1346 | 6.822 | 5.77817 | 5.80437 | 6.51667 | 6.1092 | 6.18824 |
| AKT1S1 | Q96B36-3 | 6.91998 | 6.27003 | 6.29423 | 6.03422 | 6.03744 | 5.79027 | 5.59033 | 7.76049 | 6.53709 | 6.75503 | 6.64398 | 6.87368 | 6.49415 | 5.2562 | 5.2325 | 5.51038 |
| MED6 | O75586-2 | 6.45473 | 5.90092 | 6.7156 | 6.19893 | 6.29684 | 5.82657 | 6.31316 | 6.38108 | 6.15273 | 6.54805 | 6.59588 | 6.23445 | 6.87549 | 6.39174 | 6.1075 | 5.90635 |
| CADM4 | Q8NFZ8 | 6.12197 | 6.99896 | 6.61252 | 6.34894 | 6.40392 | 6.15101 | 6.68982 | 5.8954 | 6.08298 | 5.42353 | 6.31976 | 5.48885 | 6.90649 | 6.40163 | 5.98474 | 6.16949 |
| DNAJC15 | Q9Y5T4 | 6.12943 | 6.18043 | 5.94104 | 6.69792 | 6.63546 | 6.7035 | 7.01448 | 6.48484 | 5.67993 | 6.767 | 6.3365 | 6.25464 | 5.65714 | 5.69164 | 5.89361 | 5.93245 |
| C16orf72 | Q14CZ0 | 7.42735 | 6.769 | 7.05453 | 6.52424 | 6.7917 | 6.86874 | 6.38852 | 6.14968 | 6.53643 | 6.34836 | 7.08719 | 6.59294 | 4.87991 | 4.28212 | 5.00067 | 5.29862 |
| ACD | Q96AP0 | 6.50322 | 5.95135 | 6.94795 | 6.58575 | 6.66132 | 6.23455 | 6.1386 | 6.72112 | 6.32091 | 5.63019 | 6.2345 | 7.1424 | 6.81651 | 5.04377 | 5.24719 | 5.82067 |
| POLE3 | Q9NRF9 | 6.44137 | 6.46603 | 6.85375 | 6.65338 | 6.2599 | 6.50238 | 6.09797 | 6.62506 | 6.21897 | 7.48576 | 6.46793 | 5.91111 | 6.07974 | 5.60776 | 5.59019 | 5.7387 |
| KLHL42 | Q9P2K6 | 6.04963 | 5.75208 | 7.03018 | 5.66209 | 6.59796 | 7.47855 | 6.02262 | 6.08177 | 5.90259 | 6.22583 | 6.82969 | 6.06608 | 6.38179 | 5.92453 | 5.81406 | 6.18054 |
| MRPL55 | Q7Z7F7-2 | 6.21359 | 6.52414 | 5.61423 | 6.80344 | 5.80133 | 5.93762 | 6.47652 | 6.34604 | 6.26037 | 6.1836 | 6.3261 | 6.23158 | 6.28763 | 6.40622 | 6.44349 | 6.3261 |
| RP9 | Q8TA86 | 5.84898 | 5.91364 | 6.55256 | 5.84119 | 6.24785 | 6.19933 | 6.39941 | 4.9765 | 6.23191 | 6.41364 | 5.49908 | 5.76825 | 6.52795 | 6.976 | 7.26749 | 7.33624 |
| LIN37 | Q96GY3 | 6.47143 | 5.97922 | 6.23649 | 5.92588 | 5.90462 | 6.36966 | 6.8788 | 5.78833 | 6.22124 | 6.96058 | 6.57065 | 5.91071 | 5.98795 | 6.34336 | 6.62751 | 5.82357 |
| DEGS1 | O15121 | 6.45765 | 7.37231 | 6.52846 | 7.31253 | 7.18478 | 6.3019 | 6.52125 | 5.67303 | 5.78176 | 6.11053 | 6.46033 | 6.25233 | 5.92861 | 5.75792 | 5.48076 | 4.87587 |
| RASSF3 | Q86WH2 | 6.12409 | 6.657 | 7.1102 | 6.85179 | 6.33189 | 5.01797 | 6.35014 | 5.72746 | 6.95101 | 6.55738 | 7.41227 | 6.6384 | 5.39779 | 5.3121 | 6.10956 | 5.45095 |
| PDXP | Q96GD0 | 5.18978 | 5.63333 | 6.14105 | 6.17745 | 5.95469 | 6.96388 | 6.48189 | 6.9411 | 6.24898 | 6.64496 | 7.0143 | 6.83024 | 5.41379 | 6.83842 | 5.81491 | 5.71123 |
| SNAP47 | Q5SQN1 | 5.98759 | 5.76775 | 6.60794 | 6.07098 | 6.44846 | 6.50109 | 6.02711 | 5.75075 | 6.88951 | 6.83101 | 6.62458 | 6.10007 | 6.30876 | 6.09128 | 6.16902 | 5.82411 |
| ZKSCAN4 | Q969J2 | 5.88961 | 6.42538 | 6.14575 | 6.01769 | 5.61445 | 5.96885 | 6.43785 | 5.85296 | 5.87103 | 6.58867 | 6.33143 | 6.14965 | 6.91131 | 6.83881 | 6.41598 | 6.54059 |
| C1orf198 | Q9H425 | 7.28589 | 6.04016 | 6.89235 | 5.92692 | 6.09657 | 6.75719 | 6.49837 | 6.50126 | 6.39041 | 5.94046 | 7.34997 | 6.6942 | 5.71762 | 5.78372 | 4.74206 | 5.38284 |
| CBWD2 | Q8IUF1 | 6.4415 | 6.47821 | 6.74436 | 6.01166 | 6.28824 | 6.63953 | 6.64542 | 7.19896 | 6.54023 | 6.5917 | 6.89272 | 6.56902 | 5.20108 | 4.99778 | 5.19441 | 5.56518 |
| KHNYN | O15037 | 6.34313 | 6.41928 | 6.32629 | 5.16048 | 6.17587 | 6.96815 | 6.30307 | 6.5481 | 5.98916 | 6.74666 | 6.56608 | 5.94864 | 6.38918 | 6.60732 | 6.02799 | 5.48059 |
| DNAJC25 | Q9H1X3 | 6.51247 | 6.34833 | 6.02895 | 6.59893 | 5.77986 | 5.79716 | 6.84768 | 5.0244 | 6.02053 | 6.35828 | 5.65402 | 6.0679 | 6.75652 | 6.73786 | 6.82548 | 6.64163 |
| ZBTB9 | Q96C00 | 7.08152 | 6.88147 | 6.29102 | 6.74178 | 5.95347 | 5.86191 | 7.055 | 6.23391 | 7.0599 | 6.49332 | 6.25128 | 6.73521 | 5.23334 | 4.82537 | 5.58015 | 5.72133 |
| TOR3A | Q9H497 | 6.18736 | 6.65992 | 6.32324 | 7.42288 | 6.51714 | 6.49958 | 6.34419 | 6.08377 | 5.52218 | 6.03086 | 5.84604 | 6.13922 | 5.86453 | 5.84796 | 6.43801 | 6.27312 |
| STX1A | Q16623 | 6.26968 | 6.29971 | 7.16139 | 6.98133 | 6.13562 | 6.50723 | 6.69888 | 7.05169 | 6.45743 | 6.63696 | 6.93029 | 6.71817 | 5.27463 | 4.87078 | 4.67703 | 5.32918 |
| CNNM2 | Q9H8M5 | 6.43662 | 6.86741 | 6.06478 | 6.81785 | 5.74041 | 6.31225 | 5.90797 | 6.10296 | 6.13599 | 6.22855 | 5.76327 | 6.78781 | 5.86927 | 5.95835 | 6.24886 | 5.75765 |
| RAB5IF | Q9BUV8-5 | 6.45329 | 6.0684 | 6.5079 | 6.61702 | 6.18255 | 5.42602 | 6.38274 | 5.50815 | 6.23115 | 6.09812 | 6.25834 | 6.02472 | 6.4153 | 6.14126 | 7.11889 | 6.56614 |
| FRS2 | Q8WU20 | 6.46743 | 6.79061 | 6.33169 | 6.83462 | 6.12368 | 6.34051 | 6.8844 | 6.35033 | 6.49017 | 5.7147 | 5.8118 | 5.78305 | 5.99675 | 6.01068 | 5.92738 | 6.14219 |
| TYW5 | A2RUC4 | 5.99635 | 5.9818 | 6.69276 | 5.78618 | 6.09522 | 6.60558 | 6.13399 | 6.53853 | 6.81339 | 6.78726 | 7.1288 | 6.87624 | 5.58139 | 5.75889 | 5.79475 | 5.42886 |
| TOPORS | Q9NS56 | 6.40471 | 6.90405 | 5.90033 | 7.35148 | 5.88983 | 5.75196 | 6.69658 | 6.24462 | 6.03614 | 6.13225 | 6.19337 | 6.21205 | 5.91252 | 6.77447 | 5.44061 | 6.15501 |
| GINS2 | Q9Y248 | 4.93201 | 4.40932 | 5.3682 | 4.41537 | 4.94899 | 5.15272 | 5.84273 | 7.0107 | 5.27572 | 6.24303 | 5.39492 | 6.86322 | 4.59721 | 9.87332 | 9.05471 | 10.6178 |
| GRAMD4 | Q6IC98 | 6.18084 | 6.91821 | 5.79893 | 6.36592 | 5.88016 | 6.01965 | 5.93234 | 6.00417 | 6.20831 | 6.60957 | 5.84518 | 5.53398 | 6.45426 | 6.8088 | 6.48094 | 6.95874 |
| YIPF6 | Q96EC8 | 5.8791 | 6.18619 | 5.8625 | 6.22451 | 6.41506 | 5.85437 | 5.76868 | 6.56472 | 6.59499 | 6.2069 | 6.28941 | 6.3245 | 6.03512 | 6.59723 | 6.61291 | 6.5838 |
| MTPN | P58546 | 5.83975 | 5.10235 | 5.32633 | 4.28325 | 5.82846 | 5.38132 | 4.8969 | 7.37693 | 5.05663 | 5.74344 | 4.98095 | 5.61836 | 4.2349 | 11.8883 | 7.46541 | 10.9767 |
| RPS28 | P62857 | 6.68653 | 6.11973 | 6.63386 | 5.67282 | 6.95078 | 6.24342 | 5.74388 | 6.29535 | 7.21255 | 6.13643 | 7.02602 | 6.96438 | 5.84102 | 5.39953 | 5.65097 | 5.42274 |
| MARCKSL1 | P49006 | 6.04054 | 5.12438 | 5.75212 | 5.36841 | 6.62506 | 4.89751 | 5.47773 | 5.92354 | 5.43797 | 6.2491 | 6.17903 | 6.48069 | 7.95617 | 7.31779 | 7.74804 | 7.42194 |
| PSD3 | Q9NYI0 | 4.87958 | 7.02581 | 6.08265 | 5.3841 | 6.17977 | 5.49243 | 5.77601 | 5.69501 | 6.61751 | 6.21479 | 7.35753 | 6.93029 | 7.46175 | 5.67286 | 6.2168 | 7.0131 |
| PNMA2 | Q9UL42 | 6.19324 | 6.37713 | 6.93786 | 5.72421 | 6.22862 | 6.88446 | 6.06442 | 6.85384 | 6.23513 | 6.79991 | 6.46133 | 6.99708 | 5.86535 | 5.72643 | 4.91175 | 5.73924 |
| RIC8B | Q9NVN3-1 | 5.49791 | 5.90083 | 7.42545 | 7.27533 | 6.39019 | 6.81613 | 6.49939 | 6.67675 | 6.30946 | 5.95148 | 6.123 | 7.04295 | 5.56436 | 5.98301 | 4.6066 | 5.93716 |
| EMC6 | Q9BV81 | 6.36471 | 6.55667 | 6.0851 | 6.66215 | 6.16067 | 6.04479 | 6.19609 | 5.66483 | 6.50965 | 5.48203 | 6.55947 | 6.3981 | 6.23954 | 6.04371 | 6.27791 | 6.7546 |
| MRPL54 | Q6P161 | 6.16994 | 6.53636 | 6.35625 | 6.7359 | 6.02085 | 5.86314 | 6.79153 | 6.00641 | 6.01006 | 5.53616 | 6.6316 | 5.72823 | 6.3256 | 6.54117 | 6.43021 | 6.3166 |
| TNRC6C | Q9HCJ0-2 | 6.54163 | 5.89929 | 6.61273 | 5.86824 | 6.41377 | 6.70749 | 6.12517 | 5.98348 | 6.51793 | 6.5486 | 6.71001 | 6.87142 | 6.28005 | 5.70071 | 5.30584 | 5.92036 |
| BTBD10 | Q9BSF8-2 | 6.19522 | 6.78079 | 7.21951 | 6.79795 | 6.20908 | 6.80283 | 6.34106 | 6.65652 | 6.40805 | 6.57566 | 7.26036 | 7.21716 | 4.70734 | 5.10155 | 4.79394 | 4.93298 |
| CA13 | Q8N1Q1 | 5.58807 | 6.08321 | 6.45631 | 5.88579 | 6.54265 | 6.71603 | 6.80916 | 5.98998 | 6.24187 | 5.98901 | 7.37688 | 6.77227 | 6.17681 | 6.13944 | 5.16551 | 6.06701 |
| REEP2 | Q9BRK0-2 | 6.18034 | 6.22794 | 6.01309 | 5.82374 | 5.98976 | 6.07364 | 7.02774 | 6.51666 | 5.62876 | 6.14821 | 5.67966 | 5.76217 | 6.93151 | 7.2992 | 6.19266 | 6.50491 |
| SSH3 | Q8TE77 | 5.83068 | 6.26184 | 5.49874 | 6.3692 | 5.86072 | 6.91593 | 6.24559 | 6.52395 | 6.27049 | 5.65243 | 6.46616 | 5.94697 | 6.69013 | 6.65834 | 5.9892 | 6.81962 |
| TBC1D2B | Q9UPU7 | 6.6476 | 5.56123 | 6.48462 | 5.86567 | 5.96998 | 6.72752 | 6.31217 | 5.92046 | 6.13328 | 6.89697 | 7.49556 | 7.02429 | 6.25588 | 5.24511 | 5.93089 | 5.52876 |
| HLCS | P50747 | 6.37999 | 6.11854 | 7.1239 | 5.95491 | 6.10779 | 6.67793 | 6.34361 | 6.36573 | 6.5067 | 6.63072 | 7.02369 | 6.25191 | 5.85245 | 6.02441 | 5.23304 | 5.40468 |
| CHCHD1 | Q96BP2 | 5.25907 | 5.46856 | 6.22743 | 6.11672 | 6.96153 | 7.75254 | 6.79843 | 5.93943 | 5.52145 | 5.58452 | 7.87389 | 6.39526 | 6.69814 | 6.23023 | 5.47709 | 5.69572 |
| ZNF174 | Q15697 | 6.39091 | 6.93316 | 6.01616 | 6.92724 | 6.10008 | 6.28459 | 6.16734 | 6.16355 | 6.16133 | 6.33923 | 6.2491 | 6.5872 | 6.05451 | 6.04876 | 5.70199 | 5.87485 |
| DPH6 | Q7L8W6 | 6.10438 | 5.81422 | 6.77774 | 5.59391 | 6.94109 | 6.80427 | 6.11787 | 7.73655 | 7.31889 | 6.06194 | 7.09839 | 6.72051 | 5.15703 | 5.79011 | 4.86968 | 5.0934 |
| AP1AR | Q63HQ0 | 7.0423 | 6.51612 | 6.47349 | 6.12074 | 5.88144 | 6.09106 | 6.89388 | 6.00726 | 6.53032 | 6.75928 | 6.54538 | 5.96768 | 5.75506 | 5.94478 | 6.01101 | 5.46021 |
| RHOBTB3 | O94955 | 6.69057 | 6.68827 | 7.01934 | 7.34925 | 6.3541 | 6.47416 | 6.83085 | 6.0238 | 6.75314 | 6.95678 | 6.64795 | 6.43978 | 4.68498 | 5.07701 | 4.77419 | 5.23584 |
| NEIL3 | Q8TAT5 | 6.94381 | 7.07151 | 6.89195 | 6.43141 | 6.57371 | 6.11351 | 6.49335 | 5.9336 | 7.06535 | 7.1013 | 7.58929 | 6.86547 | 4.58857 | 5.18217 | 4.6544 | 4.5006 |
| ALKBH7 | Q9BT30 | 7.60633 | 6.43604 | 6.05069 | 6.86216 | 6.00409 | 5.59379 | 5.68685 | 6.1515 | 6.54968 | 6.60119 | 6.57144 | 7.50128 | 5.60592 | 5.78094 | 5.36654 | 5.63156 |
| IFT46 | Q9NQC8-2 | 6.5113 | 6.54234 | 6.38721 | 5.60217 | 7.08547 | 6.17818 | 5.87869 | 6.74923 | 6.14418 | 6.41714 | 6.54556 | 6.65012 | 6.60468 | 5.75007 | 5.42655 | 5.52712 |
| TEDC1 | Q86SX3 | 6.13797 | 6.83127 | 7.08746 | 6.71863 | 6.51308 | 6.36742 | 6.68068 | 6.12393 | 6.73386 | 7.04667 | 6.88288 | 7.81368 | 4.83283 | 4.69092 | 4.39932 | 5.13941 |
| NSUN3 | Q9H649 | 6.29126 | 6.66874 | 6.05795 | 7.26845 | 5.95716 | 5.8975 | 6.15102 | 6.46881 | 5.96284 | 6.34529 | 6.13004 | 6.22603 | 6.76898 | 5.74608 | 6.11977 | 5.94009 |
| TSPO | P30536 | 6.49029 | 5.63172 | 5.84511 | 5.86589 | 6.38624 | 5.7164 | 6.24831 | 6.00359 | 6.05775 | 6.07127 | 5.97616 | 5.77531 | 6.56313 | 6.84643 | 7.42756 | 7.09485 |
| LRRC20 | Q8TCA0 | 6.02519 | 4.66002 | 6.42263 | 4.87824 | 6.17629 | 6.11789 | 6.0169 | 7.17604 | 5.45041 | 7.25492 | 6.50878 | 6.6493 | 5.20656 | 7.45052 | 6.82983 | 7.17648 |
| GPN3 | Q9UHW5-2 | 6.54401 | 6.22419 | 6.66383 | 6.47205 | 5.83358 | 6.98515 | 6.56279 | 7.02075 | 6.90724 | 6.06057 | 6.44022 | 6.96106 | 6.42841 | 5.33377 | 5.3662 | 6.19618 |
| DEPDC1B | Q8WUY9 | 6.6984 | 6.54091 | 6.26369 | 6.13278 | 6.26227 | 6.06399 | 6.71062 | 6.44625 | 6.05598 | 6.60663 | 7.03436 | 6.89582 | 5.6722 | 5.40548 | 5.68268 | 5.52796 |
| DPM3 | Q9P2X0-2 | 6.37771 | 5.68159 | 5.19311 | 5.64301 | 5.70855 | 6.28564 | 6.19604 | 6.23528 | 5.7354 | 5.48659 | 5.93015 | 6.08858 | 6.35676 | 7.87409 | 6.78454 | 8.42296 |
| RWDD2B | P57060 | 6.64647 | 6.43844 | 7.73352 | 6.2295 | 5.88448 | 6.95099 | 5.82732 | 5.99541 | 5.6552 | 5.95861 | 7.30222 | 6.38785 | 5.48615 | 5.53488 | 5.55011 | 5.50886 |
| MED31 | Q9Y3C7 | 6.3135 | 4.8528 | 6.23194 | 5.89329 | 6.05826 | 5.13084 | 5.00533 | 5.91349 | 5.08425 | 6.3747 | 5.97252 | 6.67013 | 6.00565 | 7.97007 | 8.32962 | 8.19361 |
| CYB5B | O43169 | 6.02962 | 5.48245 | 4.93355 | 4.96364 | 5.06881 | 5.11602 | 5.95883 | 6.51139 | 4.84249 | 6.19115 | 4.95635 | 6.16328 | 6.75957 | 9.33168 | 8.38425 | 9.30692 |
| PET100 | P0DJ07 | 6.2924 | 5.26971 | 5.86993 | 5.6052 | 6.78692 | 6.1125 | 7.11146 | 6.26909 | 6.44189 | 5.66787 | 6.06676 | 5.66787 | 6.83897 | 6.16528 | 7.23658 | 6.59757 |
| SIRT2 | Q8IXJ6 | 6.55688 | 6.25124 | 6.2623 | 5.60609 | 6.62606 | 6.75265 | 6.12774 | 6.74407 | 6.55866 | 6.61042 | 6.36194 | 6.38279 | 6.16897 | 6.02312 | 5.52562 | 5.44146 |
| CCNC | P24863 | 6.70388 | 6.42856 | 5.93221 | 6.16768 | 6.65154 | 5.70307 | 6.25349 | 6.11878 | 6.54033 | 6.48656 | 5.5959 | 5.99247 | 6.61628 | 6.29149 | 6.50558 | 6.11219 |
| SPOP | O43791 | 6.42491 | 5.8297 | 5.31013 | 6.21574 | 5.21169 | 5.55998 | 6.2405 | 5.03815 | 6.43505 | 5.09431 | 5.83384 | 6.98321 | 8.3307 | 6.14142 | 7.77386 | 7.57682 |
| UQCC3 | Q6UW78 | 5.37481 | 6.26993 | 5.67942 | 6.29361 | 5.61913 | 5.83638 | 6.10899 | 6.19466 | 6.11268 | 5.373 | 6.04158 | 6.11799 | 6.886 | 7.49768 | 6.72573 | 7.8684 |
| PRKAR1B | P31321 | 5.4382 | 6.35838 | 7.28412 | 6.25245 | 6.16179 | 6.23722 | 6.16354 | 6.4923 | 7.12135 | 6.88133 | 6.56665 | 6.58938 | 5.74819 | 6.02152 | 5.21911 | 5.46447 |
| CTSV | O60911 | 6.74862 | 8.16861 | 6.42947 | 7.03328 | 6.4068 | 5.91612 | 6.30277 | 6.46394 | 6.42387 | 6.5445 | 6.32586 | 5.6427 | 5.74361 | 5.13636 | 5.37347 | 5.34 |
| GEN1 | Q17RS7 | 6.25498 | 6.06194 | 7.24289 | 7.20368 | 6.59888 | 6.93333 | 6.17224 | 6.52358 | 6.90309 | 6.46375 | 7.1445 | 7.03449 | 4.94355 | 4.95167 | 4.25723 | 5.3102 |
| CCDC14 | Q49A88 | 7.03931 | 7.97433 | 7.04364 | 7.08316 | 6.63366 | 6.3576 | 5.52633 | 5.67582 | 7.01124 | 7.00106 | 7.04809 | 7.60111 | 4.40497 | 4.40764 | 4.60859 | 4.58345 |
| SMYD5 | Q6GMV2 | 6.83274 | 6.64796 | 7.30593 | 6.59648 | 6.01143 | 5.91396 | 6.24691 | 6.47558 | 5.77656 | 6.97022 | 5.67464 | 6.33619 | 5.91598 | 5.77001 | 6.20788 | 5.31751 |
| TIMP1 | P01033 | 8.59737 | 9.80922 | 5.8103 | 8.23361 | 7.05525 | 8.40153 | 5.74761 | 8.23392 | 5.95706 | 5.85215 | 7.4401 | 5.74599 | 3.08477 | 3.73933 | 3.18243 | 3.10936 |
| TMEM165 | Q9HC07 | 6.0112 | 5.74766 | 5.77473 | 6.80612 | 6.73314 | 5.6016 | 7.0776 | 5.34666 | 7.50609 | 4.12974 | 5.36299 | 5.9353 | 5.19379 | 7.57364 | 6.65167 | 8.54807 |
| TEPSIN | Q96N21-2 | 6.40366 | 6.12993 | 6.72271 | 6.2985 | 6.03751 | 6.38178 | 6.10103 | 6.5837 | 6.2441 | 6.37238 | 6.98666 | 6.33767 | 5.94616 | 6.04122 | 5.85656 | 5.55642 |
| ADGRA3 | Q8IWK6 | 6.98354 | 6.78761 | 6.73322 | 6.33557 | 6.51622 | 6.16159 | 5.92879 | 5.49104 | 6.67582 | 6.4994 | 6.67784 | 6.75744 | 5.53173 | 5.53182 | 5.52581 | 5.86257 |
| PRICKLE3 | O43900 | 6.13055 | 6.53914 | 6.38273 | 6.49651 | 6.0479 | 6.19226 | 6.48125 | 6.33123 | 5.91127 | 6.47466 | 6.38299 | 5.92781 | 6.21438 | 6.04469 | 6.17403 | 6.26859 |
| ASS1 | P00966 | 6.03805 | 5.6647 | 6.0187 | 5.65494 | 6.09901 | 5.47923 | 6.10936 | 6.25164 | 7.25424 | 7.14897 | 6.3595 | 6.40484 | 6.79061 | 6.51256 | 6.60137 | 5.61228 |
| MMAA | Q8IVH4 | 6.65043 | 6.10107 | 6.02953 | 6.56435 | 6.2628 | 5.5484 | 6.78757 | 5.75773 | 6.21814 | 5.92238 | 6.33073 | 5.442 | 7.10392 | 6.05376 | 6.75763 | 6.46956 |
| TMEM94 | Q12767-2 | 7.00193 | 6.60025 | 5.99159 | 5.75564 | 5.9191 | 5.95162 | 6.66629 | 5.12028 | 6.97394 | 6.19206 | 6.50827 | 5.65976 | 6.80675 | 6.2202 | 6.57139 | 6.06094 |
| MCUB | Q9NWR8 | 6.90454 | 6.24185 | 5.83879 | 6.39569 | 6.72515 | 6.47938 | 6.4803 | 6.35896 | 5.99358 | 5.95727 | 5.7602 | 6.10343 | 6.53476 | 5.79696 | 6.30102 | 6.12812 |
| LIN7A | O14910 | 6.42344 | 6.62844 | 6.21992 | 6.3991 | 6.66119 | 7.48022 | 5.90117 | 6.59763 | 5.73316 | 5.83657 | 6.56739 | 5.84091 | 6.1295 | 5.2413 | 5.99059 | 6.13568 |
| KCNN4 | O15554 | 6.63053 | 6.59282 | 6.26177 | 6.34041 | 6.49628 | 6.21732 | 6.23139 | 5.70788 | 6.88004 | 6.62393 | 5.97795 | 5.27135 | 6.55867 | 5.91502 | 6.477 | 5.81766 |
| ITFG1 | Q8TB96 | 6.8066 | 6.71275 | 6.86304 | 6.53074 | 6.57536 | 6.66115 | 6.81197 | 6.29053 | 6.3926 | 6.29121 | 6.03801 | 5.91664 | 5.5211 | 5.56706 | 5.6823 | 5.33896 |
| YPEL5 | P62699 | 5.78183 | 6.35207 | 6.06953 | 6.18735 | 5.65992 | 6.51058 | 6.77619 | 6.83337 | 6.39889 | 6.64673 | 6.38959 | 6.44056 | 5.20511 | 6.10865 | 6.24844 | 6.39119 |

| | | | | | | | | | | | | | | | | | |
|---|---|---|---|---|---|---|---|---|---|---|---|---|---|---|---|---|---|
| PGAP6 | Q9HCN3 | 6.63523 | 6.46441 | 6.84624 | 7.0841 | 6.47857 | 6.20511 | 6.37668 | 5.65574 | 6.28107 | 7.41814 | 6.53319 | 6.42843 | 5.60791 | 5.58962 | 5.01197 | 5.38359 |
| GPATCH2 | Q9NW75 | 6.06767 | 5.66027 | 6.61858 | 6.54417 | 6.78397 | 6.38701 | 6.20388 | 6.19318 | 6.18556 | 6.20472 | 6.30851 | 6.12152 | 5.61049 | 6.68677 | 6.40203 | 6.02168 |
| PLEKHG5 | O94827-7 | 6.40065 | 7.00434 | 6.67175 | 6.68574 | 6.25631 | 6.74864 | 5.5142 | 6.85429 | 6.09123 | 5.09436 | 7.04144 | 5.44386 | 6.2742 | 5.55341 | 6.18567 | 6.1799 |
| PRNP | P04156 | 7.08693 | 6.83781 | 6.02906 | 6.41009 | 6.43437 | 6.05842 | 6.52666 | 5.66524 | 6.23211 | 6.3297 | 5.87104 | 5.75708 | 6.34714 | 6.02433 | 6.11865 | 6.27137 |
| ARHGAP10 | A1A4S6 | 5.49344 | 6.4805 | 6.61869 | 5.90779 | 7.68075 | 6.22185 | 6.15294 | 7.20029 | 5.19998 | 5.98784 | 7.41917 | 6.84956 | 5.74149 | 5.75296 | 5.7277 | 5.56506 |
| NHEJ1 | Q9H9Q4 | 5.89468 | 5.56091 | 5.04966 | 5.62084 | 6.26819 | 5.69215 | 5.66623 | 6.7269 | 5.69935 | 6.03536 | 6.46858 | 6.57626 | 6.67894 | 8.67483 | 6.80528 | 6.58184 |
| FOXM1 | A0A0D9SFF0 | 6.07377 | 6.14585 | 7.32418 | 5.27537 | 5.61946 | 5.46278 | 6.06502 | 5.96201 | 7.01244 | 5.97367 | 6.89436 | 6.76002 | 6.0989 | 6.31237 | 6.72848 | 6.2913 |
| PTPRG | P23470 | 6.19976 | 6.0879 | 5.84949 | 6.38447 | 6.08733 | 6.17194 | 6.05211 | 5.03966 | 6.53359 | 6.08505 | 6.09732 | 6.37324 | 7.22608 | 6.29245 | 6.78975 | 6.72986 |
| MIS12 | Q9H081 | 6.22266 | 5.52347 | 5.60485 | 5.5412 | 5.63758 | 5.01616 | 5.39919 | 5.97292 | 4.89477 | 5.87849 | 5.03595 | 6.12044 | 6.00829 | 9.2704 | 8.71682 | 9.15682 |
| ZFYVE27 | Q5T4F4-3 | 6.77726 | 6.44992 | 6.04259 | 6.56752 | 6.0534 | 6.40696 | 6.33145 | 6.01207 | 6.35555 | 6.13858 | 6.26484 | 6.31249 | 6.00591 | 5.96899 | 6.32354 | 5.98893 |
| SUSD5 | O60279 | 6.48452 | 6.45961 | 6.20913 | 5.88719 | 6.18411 | 5.56164 | 6.28527 | 6.07261 | 6.52961 | 6.62774 | 6.04798 | 6.59373 | 6.5076 | 6.3434 | 6.22631 | 5.97954 |
| SGCE | O43556-4 | 7.04314 | 6.55386 | 6.09696 | 6.20564 | 7.16269 | 6.06966 | 6.96231 | 5.95468 | 6.18445 | 6.55327 | 6.12672 | 5.81391 | 5.94252 | 5.72704 | 5.96674 | 5.63642 |
| HDX | Q7Z353 | 5.71542 | 5.87604 | 6.54057 | 6.04875 | 6.09888 | 6.44019 | 6.35434 | 6.19054 | 6.23019 | 7.17316 | 6.45175 | 6.27616 | 6.4942 | 5.46523 | 6.08958 | 6.55501 |
| MTERF1 | Q99551 | 6.79519 | 6.81779 | 6.48809 | 7.58082 | 6.43723 | 6.13794 | 6.76189 | 5.95562 | 6.3901 | 6.28645 | 6.20726 | 5.57293 | 5.96982 | 5.20222 | 5.81858 | 5.57807 |
| AMN1 | Q8IY45 | 5.545 | 6.06258 | 7.42136 | 5.41122 | 6.021 | 6.53097 | 6.7233 | 6.42751 | 6.69035 | 6.64638 | 6.33314 | 6.17823 | 5.77082 | 5.55341 | 6.51174 | 6.17299 |
| MBIP | Q9NS73 | 6.42942 | 5.81111 | 5.5897 | 6.77563 | 7.1406 | 6.21618 | 5.96431 | 6.46748 | 5.88677 | 6.15821 | 6.55616 | 6.00622 | 6.29412 | 6.24557 | 6.13697 | 6.32156 |
| E2F3 | O00716 | 5.95334 | 6.10036 | 6.70727 | 6.1654 | 6.14505 | 5.94145 | 5.81966 | 5.54585 | 5.63421 | 6.62336 | 6.64481 | 6.68107 | 6.95033 | 6.14942 | 6.51028 | 6.42812 |
| PIGN | O95427 | 5.96529 | 6.79445 | 5.9647 | 6.29498 | 6.18915 | 6.04979 | 6.32249 | 5.98742 | 5.76386 | 6.1692 | 5.71891 | 6.00996 | 6.6851 | 6.50404 | 6.76352 | 6.81713 |
| ATF6 | P18850 | 7.35519 | 5.9782 | 6.33519 | 7.44814 | 7.05603 | 6.36465 | 6.84032 | 6.11372 | 6.33995 | 6.52314 | 6.69974 | 6.72527 | 5.47338 | 5.084 | 4.30346 | 5.35963 |
| RAMAC | Q9BTL3 | 6.05108 | 5.82089 | 5.62848 | 5.75762 | 6.83999 | 5.76649 | 4.90664 | 6.63289 | 5.1535 | 6.46678 | 6.02724 | 7.33036 | 6.75047 | 7.14466 | 6.98154 | 6.74138 |
| VPS37B | Q9H9H4 | 5.22879 | 4.80649 | 6.92416 | 4.81643 | 5.76234 | 5.30461 | 5.14698 | 7.22611 | 5.18443 | 6.84582 | 5.96949 | 6.96899 | 5.22477 | 9.13056 | 7.51401 | 7.94602 |
| PUS10 | Q3MIT2 | 5.32094 | 5.94115 | 6.14076 | 5.27299 | 5.75198 | 6.32772 | 5.71267 | 5.67847 | 5.71394 | 6.38605 | 6.55108 | 5.8226 | 7.15975 | 7.67184 | 7.47858 | 7.06948 |
| DCUN1D3 | Q8IWE4 | 6.53601 | 5.25662 | 6.68216 | 7.10479 | 5.76683 | 5.42114 | 6.20139 | 5.45192 | 6.16663 | 6.43361 | 6.32466 | 6.22078 | 6.47119 | 6.43826 | 6.56702 | 6.95697 |
| PLA2G6 | O60733 | 6.09622 | 6.1887 | 7.03351 | 6.16545 | 5.90255 | 6.34136 | 6.4695 | 5.86671 | 6.144 | 6.21643 | 6.93251 | 6.32229 | 6.52999 | 6.0646 | 5.40981 | 6.31637 |
| MTFP1 | Q9UDX5 | 7.10947 | 5.92471 | 6.79935 | 6.68387 | 6.34622 | 5.84217 | 6.6236 | 6.22285 | 5.70574 | 5.7834 | 5.24351 | 5.96365 | 5.76623 | 6.58905 | 6.70334 | 6.69282 |
| MLLT3 | P42568 | 6.29601 | 6.72722 | 5.83559 | 6.79721 | 6.30957 | 6.13136 | 6.93653 | 6.32937 | 6.08857 | 6.31173 | 6.42695 | 6.19879 | 5.36542 | 6.19361 | 6.02417 | 6.0279 |
| GANC | Q8TET4 | 5.68066 | 6.79254 | 7.04655 | 6.19837 | 6.36287 | 6.87392 | 6.07234 | 6.07111 | 7.12195 | 6.0905 | 6.16175 | 6.29653 | 5.30721 | 6.50773 | 5.70709 | 6.70888 |
| FBF1 | Q8TES7-6 | 5.84498 | 6.69955 | 7.30724 | 6.12971 | 6.06891 | 6.41296 | 5.386 | 6.13847 | 6.5473 | 6.43721 | 6.13567 | 6.78212 | 6.62857 | 6.02776 | 5.74398 | 5.70957 |
| ARHGEF26 | Q96DR7 | 6.34465 | 6.47729 | 6.41729 | 5.99768 | 6.50428 | 6.16002 | 5.76778 | 5.55443 | 6.88471 | 6.67415 | 6.52507 | 6.41196 | 6.02159 | 6.53457 | 5.65225 | 6.07226 |
| PIAS2 | O75928 | 5.8667 | 5.82459 | 6.08821 | 6.8472 | 5.90776 | 6.76193 | 6.01389 | 5.30834 | 6.32241 | 5.71601 | 5.80168 | 6.31412 | 6.09077 | 7.29176 | 6.9571 | 6.88755 |
| FZD2 | Q14332 | 6.67849 | 6.9948 | 6.21661 | 6.98693 | 6.20896 | 6.62913 | 5.98539 | 5.98573 | 6.86231 | 6.19156 | 6.3299 | 5.96582 | 5.4998 | 5.81899 | 5.92975 | 5.71582 |
| SERAC1 | Q96JX3 | 6.17709 | 6.15978 | 5.78326 | 7.15278 | 5.93175 | 5.96726 | 6.3244 | 6.02839 | 5.93208 | 6.03261 | 6.47071 | 6.47567 | 6.09879 | 6.42551 | 6.34904 | 6.69087 |
| TRPM7 | Q96QT4 | 6.61703 | 6.95063 | 6.18452 | 6.35956 | 6.42872 | 5.89124 | 6.00293 | 5.87178 | 5.12738 | 7.17757 | 6.75728 | 6.28945 | 5.80383 | 5.56265 | 6.57084 | 6.40459 |
| RABL3 | Q5HYI8 | 5.99437 | 6.17295 | 5.47487 | 6.16075 | 6.48361 | 6.0391 | 5.88771 | 6.42349 | 6.3958 | 6.29584 | 5.98212 | 6.15659 | 6.54708 | 6.34467 | 6.95267 | 6.68837 |
| SZT2 | Q5T011 | 6.66247 | 6.3667 | 6.76067 | 6.11516 | 6.18339 | 6.56995 | 6.14023 | 6.33163 | 6.27599 | 6.39643 | 7.01422 | 6.35846 | 6.76852 | 6.38441 | 5.5642 | 5.10758 |
| TMEM62 | Q0P6H9 | 6.74212 | 6.19598 | 5.87743 | 7.14674 | 5.56649 | 5.82468 | 6.78534 | 6.18669 | 5.74047 | 6.47418 | 6.41262 | 6.17809 | 6.10699 | 5.98077 | 6.14709 | 6.63232 |
| KSR1 | Q8IVT5 | 6.23508 | 5.32166 | 7.25355 | 5.83765 | 6.30084 | 7.11012 | 6.57164 | 6.01679 | 6.73453 | 6.63034 | 6.45095 | 5.85009 | 6.19533 | 6.37031 | 5.75795 | 5.36318 |
| FKRP | Q9H9S5 | 5.53722 | 6.86102 | 6.27313 | 6.80221 | 6.09161 | 5.89719 | 6.37637 | 6.36479 | 6.28687 | 6.29644 | 6.35021 | 6.49231 | 6.03194 | 6.58128 | 6.53906 | 7.01835 |
| TXNDC15 | Q96J42 | 7.07153 | 6.69397 | 6.64711 | 7.20571 | 6.99326 | 6.86596 | 6.21833 | 5.96983 | 6.42008 | 6.49346 | 7.14947 | 6.27533 | 5.17455 | 4.70933 | 4.79307 | 5.319 |
| TXNRD3 | Q86VQ6 | 6.01805 | 6.21493 | 6.5068 | 5.8898 | 5.81 | 6.64709 | 6.24455 | 6.37679 | 6.78983 | 6.32205 | 6.09098 | 6.47958 | 5.68315 | 6.43311 | 5.92428 | 6.569 |
| NFU1 | Q9UMS0 | 5.58335 | 5.11161 | 3.94069 | 5.29062 | 4.99178 | 4.84047 | 5.64313 | 7.17424 | 5.02862 | 6.54439 | 5.63805 | 7.1944 | 5.87442 | 8.8504 | 8.22165 | 10.0722 |
| GID4 | Q8IVV7 | 5.82719 | 6.09044 | 7.29162 | 5.98035 | 6.78214 | 6.21823 | 7.01798 | 6.54794 | 5.61797 | 6.36928 | 6.80644 | 6.59748 | 5.73364 | 5.95497 | 5.38031 | 5.784 |
| DOLPP1 | Q86YN1 | 6.43845 | 5.26446 | 5.58623 | 6.62608 | 5.97554 | 6.20763 | 5.77192 | 5.92497 | 5.97723 | 6.23709 | 6.30355 | 6.27582 | 7.10746 | 6.69261 | 6.63015 | 6.98081 |
| ALG8 | Q9BVK2 | 6.17418 | 5.97954 | 5.87439 | 6.41993 | 6.06478 | 5.89012 | 6.61058 | 6.25657 | 5.69241 | 6.46308 | 5.80056 | 5.84731 | 6.50569 | 6.12339 | 7.26024 | 7.03721 |
| ABCD4 | O14678 | 6.72658 | 6.83579 | 6.48885 | 6.87978 | 6.37575 | 5.92681 | 6.96368 | 6.0668 | 6.6283 | 6.57166 | 6.1556 | 6.29692 | 5.48804 | 5.74625 | 5.92744 | 5.92177 |
| MNT | Q99583 | 7.51485 | 6.11282 | 5.63269 | 5.86056 | 6.47846 | 5.29963 | 6.26428 | 6.58183 | 6.5125 | 6.45089 | 5.34766 | 6.65809 | 6.85653 | 6.35977 | 6.07672 | 5.99273 |
| UBL3 | O95164 | 6.57641 | 5.97594 | 6.03234 | 6.23506 | 5.63001 | 6.14443 | 6.25942 | 6.68041 | 5.58306 | 6.10168 | 5.69607 | 6.38096 | 6.04716 | 6.78802 | 6.02407 | 7.84496 |
| BBS9 | Q3SYG4-3 | 7.46813 | 6.25528 | 6.61104 | 5.57341 | 5.4787 | 6.20443 | 7.23987 | 6.29245 | 6.43183 | 6.50972 | 6.81201 | 6.68106 | 5.75475 | 5.91121 | 5.1368 | 5.6393 |
| HDDC2 | Q7Z4H3 | 6.54195 | 5.38796 | 7.00967 | 5.40723 | 6.04294 | 6.49198 | 5.81588 | 7.21388 | 6.25534 | 6.17939 | 6.45489 | 6.87488 | 5.97708 | 6.33518 | 5.68478 | 6.32698 |
| GOLGA7 | Q7Z5G4 | 6.75063 | 6.72782 | 7.04582 | 6.62234 | 6.33825 | 6.37451 | 6.24197 | 6.0423 | 6.73768 | 6.30255 | 5.60599 | 6.25171 | 5.39291 | 5.57815 | 6.05816 | 5.9292 |
| ARPIN | Q7Z6K5-2 | 6.51692 | 6.12371 | 6.48606 | 5.99949 | 6.48421 | 6.4995 | 6.27163 | 6.59811 | 6.26418 | 6.28455 | 6.85536 | 6.77156 | 6.17076 | 6.00724 | 5.37577 | 5.29095 |
| SELENON | Q9NZV5 | 7.3454 | 7.18144 | 6.778 | 7.82077 | 6.72917 | 5.82369 | 6.57193 | 5.86905 | 6.34834 | 5.98193 | 6.54696 | 5.98447 | 5.23765 | 5.36568 | 5.46425 | 5.85127 |
| PAFAH2 | Q99487 | 6.1866 | 6.31689 | 6.45487 | 6.2596 | 6.51673 | 6.26311 | 6.51285 | 6.28081 | 6.377 | 6.27713 | 5.96279 | 6.17828 | 5.93848 | 5.91265 | 6.17573 | 6.38646 |
| DTWD2 | Q8NBA8 | 5.71872 | 5.57121 | 7.87365 | 6.01826 | 6.56399 | 6.50069 | 6.37816 | 6.12955 | 6.23851 | 6.01022 | 7.22185 | 7.83668 | 5.48709 | 5.24845 | 5.2295 | 5.97348 |
| MROH1 | Q8NDA8 | 7.01434 | 5.2799 | 5.07879 | 6.08221 | 6.99404 | 6.27097 | 5.58173 | 6.53035 | 7.07838 | 7.21029 | 6.93427 | 6.65617 | 5.85415 | 6.35812 | 6.19613 | 4.88016 |
| ST3GAL2 | Q16842 | 6.49405 | 6.49051 | 5.86277 | 6.57435 | 6.87582 | 6.23537 | 6.46714 | 5.71835 | 6.56205 | 6.14493 | 6.03304 | 5.96342 | 6.31343 | 6.14651 | 5.88721 | 6.23104 |
| SLC37A4 | O43826-2 | 6.08479 | 6.23887 | 6.15239 | 7.0026 | 6.32162 | 5.65983 | 5.36797 | 6.39154 | 6.2918 | 5.94837 | 5.80838 | 5.975 | 7.15263 | 6.20312 | 6.54377 | 6.85733 |
| TM2D3 | Q9BRN9 | 6.32651 | 5.32086 | 5.95365 | 6.13005 | 6.12673 | 6.49633 | 6.96495 | 5.54551 | 6.21676 | 5.76882 | 5.94701 | 6.6884 | 6.7704 | 6.10913 | 6.58948 | 7.04541 |
| CERCAM | Q5T4B2 | 7.10638 | 8.65705 | 6.1819 | 6.82572 | 6.53435 | 6.30331 | 6.71858 | 7.30415 | 7.04579 | 6.7501 | 6.84459 | 6.86411 | 3.83195 | 4.54758 | 3.9889 | 4.49554 |
| NDUFA6 | P56556 | 6.1733 | 6.33254 | 5.3675 | 6.01209 | 7.2729 | 6.14827 | 6.46403 | 6.43657 | 5.66719 | 5.78537 | 5.52937 | 5.34967 | 7.11453 | 6.78447 | 6.81621 | 6.74597 |
| ARHGAP39 | Q9C0H5-2 | 6.51603 | 5.91996 | 6.88821 | 7.18572 | 6.96751 | 5.58593 | 6.48456 | 5.60781 | 5.94708 | 6.18352 | 7.08506 | 6.14026 | 5.67775 | 6.06301 | 6.17718 | 5.57038 |
| FBXO5 | Q9UKT4 | 6.71586 | 6.24159 | 7.07446 | 6.17979 | 6.11669 | 6.29807 | 6.26862 | 6.49818 | 6.80122 | 6.65292 | 7.21124 | 6.68405 | 5.11633 | 5.30717 | 5.00141 | 5.83241 |
| DAPK1 | P53355-3 | 6.76188 | 6.99169 | 6.44541 | 6.17486 | 6.279 | 6.56666 | 6.31463 | 6.14837 | 6.43041 | 6.18268 | 6.02974 | 6.10792 | 5.69444 | 5.85637 | 5.74958 | 6.26637 |
| TANK | Q92844 | 6.21996 | 4.75274 | 6.92048 | 5.07605 | 7.12472 | 5.62477 | 6.10019 | 6.88542 | 6.25655 | 6.52966 | 6.98653 | 7.19599 | 7.24108 | 5.94382 | 5.64185 | 5.50019 |
| ISCA2 | Q86U28 | 5.83335 | 5.13728 | 6.43103 | 5.16974 | 5.07323 | 5.61474 | 5.75308 | 6.43432 | 5.6517 | 6.62707 | 6.14437 | 7.36681 | 5.40441 | 7.95329 | 7.71851 | 7.68707 |
| C15orf40 | Q8WUR7 | 6.12659 | 5.92592 | 6.51293 | 6.03842 | 6.67575 | 6.34732 | 6.24004 | 6.58292 | 6.66911 | 6.36045 | 6.77996 | 6.45247 | 5.82304 | 5.59094 | 6.03302 | 5.8411 |
| NECAP1 | Q8NC96 | 6.49664 | 5.4429 | 7.15163 | 5.79053 | 6.35361 | 6.58369 | 5.58419 | 6.62806 | 7.06258 | 6.37544 | 6.88943 | 6.44399 | 5.88193 | 5.6584 | 5.88692 | 5.67007 |
| UPRT | Q96BW1 | 6.12588 | 5.87201 | 6.61269 | 6.09658 | 6.58456 | 6.51905 | 5.96878 | 6.31147 | 6.18163 | 5.8981 | 6.92292 | 6.37688 | 6.33538 | 6.19651 | 5.58939 | 6.40818 |
| ZBTB24 | O43167 | 6.4349 | 6.30711 | 5.73066 | 7.07379 | 6.15882 | 5.75548 | 5.73361 | 5.27623 | 6.1993 | 6.76701 | 5.86318 | 5.80759 | 6.65178 | 6.74667 | 7.15583 | 6.33804 |
| PLGRKT | Q9HBL7 | 5.8675 | 5.79004 | 6.5527 | 5.94782 | 6.00601 | 6.69291 | 6.298 | 6.39741 | 5.39412 | 5.98466 | 6.11619 | 6.43688 | 6.37959 | 7.09946 | 7.00465 | 6.93206 |
| ZC3HAV1L | Q96H79 | 5.45762 | 5.94271 | 6.32017 | 6.34035 | 5.79836 | 6.45325 | 5.84846 | 6.43648 | 5.97578 | 6.79264 | 6.62102 | 6.79367 | 6.19431 | 6.69017 | 6.0577 | 6.27731 |
| UBXN2B | Q14CS0 | 6.17372 | 6.51918 | 7.42558 | 6.18968 | 6.81048 | 6.36741 | 5.89861 | 6.39804 | 6.28878 | 6.7813 | 7.62392 | 7.06683 | 5.47037 | 5.30486 | 4.44109 | 5.24014 |
| TSEN34 | Q9BSV6 | 5.9035 | 6.26273 | 6.35414 | 5.63712 | 5.79871 | 5.88173 | 6.25961 | 6.49121 | 6.65196 | 6.37755 | 6.22022 | 6.12336 | 6.70487 | 6.3336 | 6.97273 | 6.02696 |
| TAF1D | Q9H5J8 | 6.37067 | 6.17003 | 6.22964 | 6.85759 | 5.6991 | 5.97692 | 5.84832 | 5.20528 | 5.85268 | 6.49874 | 6.62232 | 6.47385 | 6.14317 | 6.35217 | 6.79939 | 6.90013 |
| PDRG1 | Q9NUG6 | 6.13446 | 5.20876 | 6.51291 | 5.1807 | 6.83081 | 5.48544 | 5.58747 | 7.2497 | 5.46847 | 7.19581 | 6.23308 | 7.69447 | 5.58386 | 7.04031 | 6.01462 | 6.57912 |
| CNOT7 | Q9UIV1 | 5.99593 | 5.94832 | 7.263 | 6.08933 | 6.20815 | 6.21161 | 6.04489 | 5.85248 | 6.55021 | 6.51971 | 7.0962 | 6.81341 | 6.02812 | 5.83114 | 5.68279 | 5.8647 |
| PDLIM2 | Q96JY6-5 | 6.76838 | 5.49118 | 7.29016 | 5.69321 | 6.84792 | 7.00098 | 5.8967 | 6.88192 | 6.24184 | 6.53509 | 7.25705 | 7.24814 | 5.84002 | 5.38419 | 4.56766 | 5.05556 |
| COX7A2 | P14406 | 5.70303 | 5.37753 | 5.93226 | 4.71549 | 5.96194 | 5.82159 | 6.95466 | 5.40334 | 5.94233 | 5.49307 | 5.41663 | 6.12725 | 6.72194 | 7.58162 | 7.8267 | 9.02059 |
| ZFP62 | Q8NB50 | 5.96144 | 6.3578 | 6.06453 | 7.05994 | 5.49211 | 5.72155 | 6.354 | 6.18559 | 5.22601 | 6.35663 | 6.09744 | 6.53066 | 6.77138 | 6.40379 | 6.70356 | 6.71359 |
| SLC25A51 | Q9H1U9 | 6.13493 | 6.14368 | 5.6651 | 6.64663 | 6.19332 | 6.07583 | 6.70023 | 5.76644 | 6.58709 | 6.06148 | 5.93038 | 6.00457 | 6.60776 | 6.31339 | 6.69505 | 6.47409 |
| NAV2 | Q8IVL1 | 6.47344 | 5.70621 | 6.88307 | 6.43892 | 6.40074 | 6.37333 | 6.77253 | 5.87606 | 6.1228 | 6.48648 | 6.77058 | 6.61867 | 5.36617 | 6.08566 | 6.2677 | 5.35764 |
| NAV3 | Q8IVL0 | 6.2604 | 6.36981 | 6.60146 | 6.41658 | 6.53852 | 7.3905 | 6.29072 | 6.56928 | 5.83475 | 6.54479 | 6.78459 | 5.83043 | 6.256 | 5.63126 | 5.96825 | 5.61267 |
| SLC19A1 | P41440 | 7.30423 | 6.37594 | 7.00822 | 7.33236 | 5.7688 | 6.61602 | 6.79744 | 6.76561 | 5.73724 | 5.79648 | 6.26434 | 6.73582 | 5.68634 | 4.76685 | 5.43944 | 5.60486 |
| PBLD | P30039 | 6.93331 | 6.29691 | 6.19672 | 6.17821 | 6.21989 | 6.71462 | 6.42315 | 6.9874 | 6.39015 | 6.62392 | 6.01763 | 6.89729 | 5.65499 | 5.84538 | 5.29104 | 5.32939 |
| CD58 | P19256 | 5.7319 | 6.80432 | 6.07452 | 6.0219 | 6.38162 | 6.71341 | 5.59254 | 6.3435 | 6.79821 | 6.01683 | 6.16101 | 6.38893 | 7.02219 | 6.02427 | 6.85554 | 5.06931 |
| FXN | Q16595 | 5.81829 | 5.07262 | 4.79752 | 5.34653 | 5.57962 | 3.98427 | 5.7421 | 6.53157 | 4.71069 | 5.73599 | 5.1302 | 5.6527 | 5.11371 | 10.9247 | 8.68753 | 11.172 |
| CEP164 | Q9UPV0 | 6.69031 | 7.51977 | 8.42815 | 7.80969 | 6.24184 | 5.94492 | 6.11241 | 6.7344 | 7.40661 | 7.60502 | 6.76336 | 6.95894 | 3.85992 | 4.0208 | 3.69056 | 4.21331 |
| DNAJB5 | O75953-5 | 6.00504 | 6.36954 | 6.92269 | 5.97672 | 6.04047 | 6.01798 | 6.33499 | 6.67084 | 6.55038 | 7.2801 | 6.93208 | 6.73861 | 5.14807 | 5.74107 | 5.46433 | 5.8071 |
| RSRC2 | Q7L4I2 | 5.37948 | 5.72859 | 5.82563 | 5.45319 | 5.80134 | 6.16459 | 5.53094 | 5.51794 | 5.50909 | 6.3177 | 6.4263 | 5.84978 | 7.25574 | 6.92321 | 8.00021 | 8.31629 |
| LAMTOR4 | Q0VGL1 | 6.10549 | 6.45797 | 6.26409 | 6.59075 | 6.55328 | 6.31524 | 6.42691 | 6.23773 | 6.14804 | 6.70316 | 6.4515 | 6.41957 | 5.79992 | 5.71951 | 5.81767 | 5.98918 |
| LY75 | O60449-2 | 5.55151 | 7.04059 | 6.11439 | 6.5858 | 6.73851 | 6.75047 | 5.80574 | 6.5696 | 6.31119 | 6.84418 | 6.69537 | 6.43596 | 5.25239 | 6.41619 | 5.21606 | 5.67206 |
| MORC4 | Q8TE76 | 6.75929 | 7.06139 | 5.77683 | 6.53438 | 6.21765 | 6.61096 | 6.66402 | 6.86929 | 6.76019 | 6.40152 | 6.7021 | 5.77203 | 4.93275 | 5.64203 | 5.18767 | 6.1079 |
| FAM8A1 | Q9UBU6 | 6.75114 | 6.81315 | 6.3524 | 6.55683 | 6.10018 | 6.04284 | 6.1823 | 6.21299 | 5.73666 | 6.36596 | 6.23008 | 6.4934 | 5.82869 | 5.92195 | 6.24779 | 6.16364 |
| GINS3 | Q9BRX5-3 | 5.86068 | 4.29027 | 5.75444 | 4.66117 | 4.88704 | 5.27687 | 4.83022 | 6.11409 | 5.38836 | 6.77245 | 6.03488 | 7.86464 | 6.13559 | 8.5175 | 8.93443 | 8.67738 |
| CAMKK2 | Q96RR4 | 6.45751 | 6.18941 | 6.71354 | 5.48989 | 6.10883 | 6.21211 | 6.01429 | 6.97551 | 6.3419 | 6.68578 | 7.24411 | 7.05479 | 5.60456 | 6.05634 | 5.80333 | 5.0481 |
| INO80E | Q8NBZ0 | 6.27019 | 6.53285 | 6.71981 | 6.88447 | 5.64665 | 5.30701 | 6.23637 | 6.60564 | 6.76404 | 6.54182 | 6.66294 | 6.42704 | 6.29732 | 5.74671 | 6.13952 | 5.43462 |
| NAT14 | Q8WUY8 | 6.25551 | 5.87351 | 6.0384 | 6.54771 | 6.34669 | 6.14211 | 6.17515 | 5.60306 | 5.99187 | 6.28112 | 5.9648 | 5.89788 | 7.04726 | 6.77124 | 6.40944 | 6.65427 |
| NEK3 | P51956 | 6.92369 | 6.57206 | 6.73197 | 5.92975 | 6.64493 | 6.97385 | 6.71255 | 6.95897 | 6.60717 | 6.96557 | 6.46504 | 6.93566 | 4.83147 | 5.1076 | 4.83315 | 4.80659 |
| FAAP24 | Q9BTP7 | 6.631 | 6.88557 | 7.18585 | 7.1898 | 5.63126 | 6.36366 | 6.40725 | 6.57845 | 6.83686 | 7.2372 | 7.6477 | 7.23461 | 4.74269 | 4.81538 | 4.22103 | 4.39169 |
| GRIP1 | Q9Y3R0 | 6.60092 | 6.29845 | 6.72822 | 6.64346 | 5.78763 | 6.09417 | 6.36161 | 6.45835 | 5.88346 | 6.31134 | 6.25603 | 6.5235 | 6.01125 | 5.7839 | 6.20369 | 6.05403 |
| ZBTB48 | P10074 | 6.28304 | 6.4191 | 6.42538 | 6.96677 | 6.1816 | 6.61258 | 6.49896 | 5.76 | 6.12818 | 6.19684 | 6.76048 | 6.36795 | 6.06665 | 5.57989 | 5.64803 | 6.10453 |
| TCEAL5 | Q5H9L2 | 6.16385 | 5.73307 | 6.86349 | 5.65091 | 6.1031 | 6.2011 | 5.45458 | 6.69188 | 5.99596 | 6.54987 | 6.83791 | 6.39274 | 6.69245 | 6.77642 | 6.02023 | 5.87245 |
| MFSD11 | O43934 | 6.58015 | 6.86957 | 6.3593 | 6.59432 | 5.78691 | 6.10654 | 6.74674 | 6.25556 | 6.23286 | 6.30255 | 6.6691 | 6.00216 | 5.97465 | 5.85412 | 5.60364 | 6.06183 |
| RUNX2 | Q13950 | 6.4691 | 6.07046 | 6.19385 | 6.62241 | 6.94609 | 6.08401 | 5.73849 | 5.66353 | 6.45695 | 6.42201 | 6.26335 | 6.2016 | 6.9862 | 6.17313 | 5.952 | 5.75683 |
| MRPS12 | O15235 | 6.03038 | 6.76588 | 6.04246 | 6.5784 | 5.95015 | 6.3367 | 6.80123 | 6.04682 | 5.80306 | 6.07571 | 5.62841 | 6.10274 | 6.47265 | 6.44482 | 6.28655 | 6.63404 |

| | | | | | | | | | | | | | | | | | |
|---|---|---|---|---|---|---|---|---|---|---|---|---|---|---|---|---|---|
| MRPS24 | Q96EL2 | 6.24389 | 6.81559 | 5.70889 | 6.84817 | 6.03828 | 6.24543 | 6.33115 | 6.46921 | 6.00527 | 6.40925 | 5.54756 | 5.76237 | 6.35378 | 6.24049 | 6.6377 | 6.34296 |
| ASPHD1 | Q5U4P2 | 6.1212 | 6.43316 | 6.16202 | 7.19275 | 6.08538 | 6.56376 | 6.87067 | 5.673 | 6.57693 | 6.23233 | 6.32489 | 6.35605 | 5.67315 | 6.16403 | 5.95315 | 5.61754 |
| ITGB1BP1 | O14713 | 6.7022 | 6.77322 | 6.63745 | 6.76096 | 5.94882 | 6.37045 | 6.1276 | 6.58453 | 8.04335 | 6.91546 | 7.65413 | 7.26581 | 4.48114 | 4.85404 | 4.36336 | 4.51749 |
| HAUS2 | Q9NVX0 | 6.14452 | 5.53419 | 6.99089 | 5.47776 | 6.43976 | 6.32212 | 6.40903 | 6.42178 | 5.86687 | 6.43694 | 6.22781 | 6.93988 | 5.67682 | 6.59691 | 6.02015 | 6.49458 |
| UBE2H | P62256 | 5.90279 | 6.29711 | 5.37336 | 5.82681 | 6.67509 | 6.72375 | 6.42324 | 6.83238 | 6.09673 | 6.44814 | 7.27561 | 6.86409 | 6.10647 | 5.91052 | 5.17461 | 6.0693 |
| TINF2 | Q9BSI4 | 6.02478 | 6.271 | 5.23167 | 5.70954 | 6.39849 | 5.69369 | 6.48209 | 6.36088 | 6.03082 | 6.11039 | 5.51674 | 6.06136 | 6.56924 | 6.54499 | 7.67534 | 7.31895 |
| MCOLN1 | Q9GZU1 | 6.51233 | 6.9751 | 7.03215 | 6.35449 | 5.9153 | 6.16618 | 5.56804 | 6.54332 | 6.72158 | 6.92123 | 6.15946 | 6.12198 | 5.38062 | 5.2944 | 5.88551 | 6.44831 |
| PTPN3 | P26045 | 6.40602 | 6.58648 | 6.37023 | 6.32883 | 6.51114 | 5.92589 | 6.37213 | 6.51484 | 6.49035 | 6.71969 | 6.56044 | 6.3156 | 5.50001 | 5.62495 | 5.9245 | 5.84889 |
| GPBP1 | Q86WP2-2 | 7.06232 | 6.77728 | 7.60361 | 6.44536 | 6.32724 | 6.71913 | 6.62821 | 5.98899 | 6.58626 | 6.65333 | 6.71447 | 6.71182 | 5.17828 | 4.8653 | 4.68884 | 5.04954 |
| PLP2 | Q04941 | 6.76906 | 6.53936 | 6.38133 | 6.32311 | 5.90944 | 6.31087 | 6.81345 | 6.09199 | 6.19572 | 6.93161 | 7.13778 | 6.58638 | 5.34278 | 5.58377 | 5.51383 | 5.56951 |
| SLC7A1 | P30825 | 6.20027 | 6.34788 | 6.23004 | 6.84756 | 6.22576 | 6.30434 | 6.29216 | 5.66545 | 5.86818 | 6.07645 | 6.67299 | 6.36718 | 5.88087 | 5.91696 | 6.43755 | 6.66634 |
| FAM102B | Q5T8I3 | 5.5623 | 6.29297 | 6.48028 | 6.3037 | 6.10574 | 6.19283 | 6.15716 | 5.75531 | 6.82125 | 6.39993 | 6.26408 | 6.53513 | 6.2828 | 6.63339 | 6.09426 | 6.11885 |
| PRR14 | Q9BWN1 | 6.62746 | 6.20055 | 5.75278 | 6.97829 | 5.97606 | 6.03265 | 6.13601 | 6.00626 | 6.42124 | 5.78096 | 6.07205 | 6.1235 | 6.27039 | 6.09177 | 6.78926 | 6.74077 |
| EVI5L | Q96CN4-2 | 7.33093 | 5.77522 | 6.94232 | 6.64636 | 6.37253 | 6.85381 | 6.66753 | 6.83437 | 5.72339 | 6.23571 | 6.59154 | 6.49791 | 5.44385 | 5.69458 | 4.93979 | 5.45016 |
| KBTBD4 | Q9NVX7-2 | 5.69447 | 5.68472 | 6.69654 | 5.46085 | 5.51854 | 6.16684 | 5.93071 | 5.64595 | 5.89303 | 6.33768 | 6.13253 | 6.11853 | 6.78427 | 7.58061 | 7.47801 | 6.87674 |
| PDE3B | Q13370 | 6.91576 | 6.59661 | 6.33898 | 6.97583 | 6.44736 | 6.18519 | 6.34277 | 5.90813 | 6.09477 | 5.90519 | 6.06202 | 6.08175 | 6.31001 | 5.74138 | 6.24702 | 5.84724 |
| KMT5A | Q9NQR1 | 6.78687 | 6.26395 | 6.6179 | 6.79373 | 6.24753 | 6.35914 | 6.67792 | 6.77033 | 6.36057 | 6.84272 | 6.76424 | 6.44747 | 4.64414 | 5.49225 | 5.2204 | 5.71086 |
| GORAB | Q5T7V8 | 6.72056 | 6.31739 | 6.43151 | 6.34219 | 5.77413 | 5.71811 | 6.1864 | 6.72772 | 5.80927 | 6.26568 | 6.24552 | 6.64952 | 6.7692 | 6.15104 | 5.80698 | 6.08478 |
| TOMM5 | Q8N4H5-2 | 5.43399 | 5.37497 | 5.16403 | 5.25463 | 6.33434 | 4.758 | 5.52326 | 4.68859 | 4.69274 | 4.28141 | 4.17654 | 4.95581 | 5.55936 | 11.1652 | 9.12229 | 12.5148 |
| ACP1 | P24666 | 6.14505 | 5.51667 | 6.71549 | 5.06509 | 6.6064 | 6.28532 | 6.24697 | 7.28255 | 5.56899 | 6.63282 | 6.63361 | 6.97321 | 5.91217 | 6.04427 | 5.88257 | 6.48881 |
| RRAD | P55042 | 6.81354 | 6.50665 | 6.85584 | 5.50848 | 6.22571 | 6.1706 | 6.74838 | 6.6032 | 7.01132 | 6.60443 | 7.02268 | 6.12808 | 5.56651 | 5.50626 | 5.24078 | 5.48752 |
| GPR39 | O43194 | 6.31773 | 6.10279 | 5.82589 | 6.75852 | 6.26303 | 5.97531 | 5.9162 | 5.62671 | 6.44952 | 6.19276 | 6.19389 | 5.47023 | 7.14807 | 6.66816 | 6.90156 | 6.18962 |
| DENND11 | A4D1U4 | 6.78218 | 7.29563 | 6.67881 | 6.74547 | 6.05394 | 6.10279 | 6.00955 | 5.92733 | 6.42561 | 6.16039 | 6.71891 | 6.74195 | 5.36527 | 5.72144 | 5.70316 | 5.56756 |
| SMIM19 | Q96E16 | 6.42436 | 5.77797 | 6.37058 | 6.73672 | 6.0067 | 6.36533 | 6.81989 | 6.01711 | 6.38013 | 6.15877 | 6.45555 | 6.17889 | 5.81178 | 6.42637 | 6.23488 | 5.83499 |
| FICD | Q9BVA6 | 6.39234 | 6.68591 | 6.51331 | 6.43987 | 6.10582 | 6.03443 | 6.54041 | 5.58112 | 6.72268 | 6.95675 | 5.89857 | 6.08113 | 5.93233 | 5.49715 | 6.10747 | 6.51071 |
| PSMB10 | P40306 | 6.23042 | 6.31986 | 6.17873 | 5.78571 | 6.26352 | 6.39721 | 6.01848 | 6.74762 | 6.39272 | 6.20415 | 6.00284 | 6.24169 | 6.57775 | 6.20535 | 6.24221 | 6.19172 |
| CPE | P16870 | 6.38034 | 6.12129 | 6.2071 | 6.35236 | 6.6763 | 6.25036 | 6.06119 | 6.30246 | 6.31835 | 6.28493 | 7.05857 | 6.61568 | 5.83778 | 6.09474 | 5.46156 | 5.97699 |
| XPA | P23025 | 5.63502 | 5.47565 | 6.22513 | 5.76397 | 6.32212 | 6.55709 | 5.78665 | 5.89432 | 6.23515 | 6.26674 | 6.1249 | 6.23941 | 6.50179 | 7.29373 | 6.90853 | 6.7698 |
| PCMTD2 | Q9NV79 | 5.80784 | 6.18251 | 5.59662 | 5.04048 | 5.84511 | 5.86173 | 6.0872 | 6.23334 | 6.82569 | 6.32108 | 6.61001 | 6.30926 | 6.77339 | 7.16502 | 6.15948 | 7.18125 |
| C9orf72 | Q96LT7 | 6.24601 | 5.39268 | 6.25276 | 6.1238 | 6.0521 | 6.55164 | 6.81663 | 6.89281 | 6.47041 | 5.36744 | 6.34381 | 6.62441 | 5.98615 | 6.112 | 6.40369 | 6.36367 |
| CARD19 | Q96LW7-2 | 6.51048 | 5.4664 | 5.70501 | 6.09343 | 6.39258 | 5.61129 | 6.00847 | 6.09161 | 5.53855 | 6.4843 | 6.12506 | 6.50667 | 7.20972 | 6.79539 | 6.82822 | 6.6328 |
| AMZ2 | Q86W34 | 6.03066 | 6.10808 | 6.41298 | 5.99297 | 6.32123 | 6.47551 | 6.22709 | 6.0965 | 6.85031 | 6.25045 | 6.33537 | 6.53488 | 6.28648 | 6.2445 | 5.78676 | 6.04624 |
| ELF2 | Q15723 | 6.30235 | 6.42913 | 5.71702 | 6.64071 | 6.68109 | 5.89853 | 6.64433 | 5.54344 | 6.50336 | 6.0839 | 6.67596 | 5.99668 | 6.4812 | 5.90684 | 6.08569 | 6.40977 |
| ODF2L | Q9ULJ1-4 | 6.55483 | 7.29412 | 7.00193 | 7.33342 | 6.4934 | 6.70314 | 6.68774 | 6.70807 | 6.66919 | 6.55011 | 7.14087 | 7.56266 | 4.19884 | 4.41056 | 4.30791 | 4.38319 |
| POC1B | Q8TC44 | 6.36754 | 6.03602 | 6.3053 | 6.54018 | 5.9235 | 6.30125 | 5.9549 | 6.31133 | 6.63954 | 6.61858 | 6.77473 | 6.73957 | 5.76209 | 6.01379 | 6.02251 | 5.68915 |
| BRD9 | Q9H8M2 | 6.10941 | 6.44472 | 6.00585 | 6.93879 | 6.13288 | 6.32783 | 6.6748 | 5.9667 | 5.97775 | 6.2969 | 5.93305 | 6.48923 | 5.59922 | 6.27975 | 6.19563 | 6.62751 |
| WRAP73 | Q9P2S5 | 6.3081 | 6.86635 | 7.12308 | 6.72707 | 6.01987 | 6.56096 | 6.65287 | 6.9486 | 6.8073 | 6.52427 | 6.83906 | 7.68068 | 4.66852 | 4.42589 | 5.02954 | 4.81784 |
| MRPS14 | O60783 | 6.13592 | 6.11511 | 5.48891 | 6.07169 | 5.98005 | 5.87297 | 6.60218 | 6.59694 | 6.30885 | 5.87691 | 6.07593 | 5.88098 | 7.50555 | 6.7628 | 6.77328 | 6.85194 |
| DGKZ | Q13574-1 | 6.25785 | 6.02329 | 6.8406 | 6.56924 | 5.89916 | 6.28373 | 6.33281 | 6.63347 | 6.49332 | 6.36793 | 6.58587 | 6.72162 | 5.83048 | 5.67427 | 5.72765 | 5.75872 |
| REEP6 | Q96HR9 | 6.11761 | 5.1546 | 5.71503 | 5.21298 | 5.74895 | 5.83889 | 5.66111 | 5.80472 | 4.90833 | 6.05545 | 5.95007 | 6.6319 | 6.32469 | 8.2538 | 8.07052 | 8.55134 |
| SMAP | O00193 | 5.79702 | 5.77992 | 6.00112 | 5.45333 | 6.12425 | 6.33673 | 5.60424 | 6.68655 | 5.88211 | 6.31785 | 6.55394 | 6.13274 | 7.03275 | 6.77679 | 6.79606 | 6.72461 |
| SESN1 | Q9Y6P5-2 | 6.25614 | 6.40149 | 7.31722 | 5.97296 | 6.45029 | 6.91454 | 6.19083 | 6.49794 | 5.95067 | 5.98117 | 7.15644 | 7.01231 | 5.37419 | 5.93038 | 5.26895 | 5.32448 |
| ITIH2 | P19823 | 5.69126 | 7.08149 | 4.85036 | 5.26066 | 4.493 | 6.16833 | 6.79733 | 5.0273 | 6.9651 | 5.26416 | 6.02137 | 5.83506 | 6.58744 | 5.96826 | 10.1082 | 7.88067 |
| MDFIC | Q9P1T7-1 | 6.7552 | 6.6619 | 6.53678 | 6.52094 | 5.85443 | 6.31509 | 6.48716 | 6.00239 | 6.35002 | 6.24848 | 6.19258 | 6.20412 | 6.25055 | 5.8761 | 5.9188 | 5.82544 |
| ZBTB20 | Q9HC78 | 6.6913 | 6.55393 | 5.89878 | 6.56663 | 5.65301 | 5.83241 | 6.47114 | 5.84169 | 6.04042 | 6.58166 | 5.49948 | 5.86645 | 6.7989 | 6.30844 | 7.062 | 7.05375 |
| SLC35B3 | Q9H1N7 | 6.11811 | 6.84833 | 5.5972 | 6.55377 | 6.11777 | 5.94904 | 6.54807 | 5.9631 | 6.1421 | 6.35544 | 6.20529 | 6.16355 | 6.1204 | 6.23436 | 6.82921 | 6.25427 |
| TSPYL2 | Q9H2G4 | 7.07765 | 6.36355 | 5.85639 | 7.59458 | 6.62911 | 6.80597 | 6.9004 | 6.17306 | 6.43677 | 6.62495 | 6.41624 | 5.58508 | 6.20089 | 5.10884 | 4.88011 | 5.34642 |
| LYST | Q99698 | 6.44143 | 7.01155 | 6.46684 | 6.1992 | 5.94787 | 6.38152 | 6.00875 | 6.04134 | 6.70875 | 7.07643 | 7.04033 | 7.04177 | 5.52414 | 5.44022 | 5.2669 | 5.40295 |
| KDM4C | Q9H3R0 | 5.47775 | 6.21757 | 6.45015 | 6.05158 | 6.21883 | 6.71385 | 6.49295 | 5.98154 | 5.78163 | 6.13867 | 6.22149 | 6.65907 | 6.44974 | 6.18568 | 6.13565 | 6.82383 |
| JMY | Q8N9B5 | 7.44354 | 6.17401 | 7.15317 | 6.03819 | 5.82465 | 6.10931 | 5.80867 | 6.18097 | 6.13738 | 6.1062 | 5.50932 | 5.46054 | 6.76222 | 5.7576 | 7.04031 | 6.49393 |
| COQ10B | Q9H8M1 | 6.92995 | 6.29708 | 5.10293 | 6.82656 | 6.95307 | 5.94596 | 6.28547 | 6.41714 | 6.72275 | 6.57267 | 6.45016 | 5.53352 | 6.10368 | 5.57902 | 5.74274 | 6.53729 |
| C4A | P0C0L4 | 6.41198 | 6.54846 | 5.38039 | 6.60875 | 5.77459 | 5.96271 | 6.22475 | 5.7489 | 6.12218 | 5.98862 | 6.64777 | 6.64317 | 6.36398 | 5.67024 | 7.18267 | 6.72084 |
| MAF1 | Q9H063 | 4.57792 | 4.63498 | 4.96139 | 4.47134 | 5.41684 | 5.15481 | 5.86921 | 5.64719 | 4.85301 | 4.96492 | 4.8217 | 4.61258 | 10.2812 | 10.4491 | 10.1065 | 9.17723 |
| TNKS | O95271 | 7.05332 | 6.61447 | 7.34765 | 6.71616 | 6.72214 | 6.71503 | 6.81836 | 6.24782 | 6.87188 | 6.35177 | 7.17715 | 7.19699 | 4.78048 | 4.56455 | 4.27667 | 4.54555 |
| DCAF11 | Q8TEB1 | 5.42503 | 5.81331 | 6.5288 | 6.02624 | 6.95902 | 6.43374 | 6.10962 | 6.26533 | 6.48432 | 5.83928 | 6.16254 | 5.48025 | 6.96725 | 6.27216 | 6.93087 | 6.30223 |
| KIAA1586 | Q9HCI6 | 6.6866 | 6.9559 | 6.96644 | 6.7609 | 6.15708 | 6.39926 | 6.76044 | 6.62273 | 6.75699 | 7.4749 | 7.3088 | 6.71212 | 4.75282 | 4.70626 | 4.50431 | 4.47444 |
| RAB28 | P51157 | 6.32252 | 5.88386 | 6.74945 | 5.95885 | 6.50211 | 6.62418 | 6.35889 | 6.46382 | 6.66192 | 6.33151 | 6.62917 | 6.78917 | 5.48054 | 5.57397 | 5.74181 | 5.92821 |
| ZNF414 | Q96IQ9-2 | 5.62749 | 5.80006 | 5.60875 | 5.80504 | 6.47744 | 6.42831 | 5.99745 | 6.44496 | 6.05281 | 7.13161 | 6.43962 | 6.36544 | 6.69985 | 6.43742 | 6.41435 | 6.2694 |
| RPS29 | P62273-2 | 5.86323 | 6.68098 | 6.52772 | 5.41179 | 6.32665 | 7.55396 | 6.01911 | 6.22721 | 7.03766 | 7.09062 | 6.87456 | 6.50515 | 5.88789 | 5.65174 | 5.23426 | 5.10745 |
| ADCY3 | O60266 | 6.85853 | 6.33056 | 6.85105 | 6.44419 | 5.61088 | 7.01342 | 6.58473 | 6.3822 | 6.29576 | 5.87093 | 6.8562 | 6.073 | 5.55207 | 6.00125 | 6.24614 | 5.02908 |
| RTKN | Q9BST9-2 | 6.39231 | 6.38274 | 6.90815 | 6.19938 | 6.77428 | 6.90376 | 6.1887 | 6.15401 | 6.80384 | 6.69278 | 6.6246 | 6.555 | 5.46855 | 5.43645 | 5.34961 | 5.16584 |
| UMAD1 | C9J7I0 | 5.39265 | 4.53714 | 4.86284 | 4.724 | 5.37298 | 4.48543 | 5.47895 | 6.54039 | 4.23382 | 4.78002 | 5.28611 | 5.69827 | 5.92917 | 12.5685 | 7.68898 | 12.4207 |
| CUTA | O60888-2 | 6.81518 | 4.86134 | 5.55263 | 4.9753 | 6.40033 | 5.83617 | 5.25811 | 8.30251 | 5.12271 | 6.30548 | 5.71586 | 5.44829 | 4.5136 | 9.30104 | 7.18511 | 8.40633 |
| ARHGAP33 | O14559 | 7.65606 | 7.03057 | 6.67332 | 7.23135 | 6.5858 | 6.15642 | 6.64513 | 6.69361 | 6.69403 | 5.87531 | 6.90083 | 6.29436 | 4.89362 | 4.93579 | 4.85593 | 4.87787 |
| C3orf33 | Q6P1S2 | 5.694 | 6.23554 | 6.48967 | 6.75281 | 5.51437 | 6.19619 | 6.7771 | 6.35556 | 5.75839 | 7.42045 | 6.10613 | 5.66097 | 6.30926 | 5.31958 | 6.97477 | 6.43519 |
| HIKESHI | Q53FT3 | 6.68322 | 5.92088 | 6.68694 | 5.19376 | 6.31266 | 6.18919 | 6.61736 | 6.97404 | 6.00346 | 6.23479 | 6.52557 | 6.44224 | 5.52183 | 6.5492 | 6.25017 | 5.8947 |
| ZNF428 | Q96B54 | 6.56803 | 6.26385 | 6.92249 | 5.92299 | 6.52488 | 6.45941 | 5.97704 | 6.00923 | 6.01174 | 6.68437 | 7.35899 | 6.37808 | 6.39707 | 5.86386 | 5.37373 | 5.28422 |
| CDC26 | Q8NHZ8 | 5.93704 | 5.43537 | 5.79077 | 5.36121 | 5.75402 | 5.21209 | 4.54265 | 6.56893 | 4.0372 | 6.81288 | 5.38575 | 7.05677 | 5.73754 | 9.64775 | 7.24019 | 9.47985 |
| PIAS4 | Q8N2W9 | 5.37792 | 6.11716 | 6.97767 | 5.94549 | 5.98381 | 6.88531 | 6.79503 | 5.71223 | 6.45726 | 6.82492 | 6.88971 | 6.87982 | 5.62726 | 5.67863 | 5.83041 | 6.01736 |
| C1D | Q13901 | 6.25036 | 6.23413 | 6.08563 | 6.68288 | 5.88089 | 6.01399 | 6.74298 | 5.70467 | 6.5063 | 5.82331 | 6.77353 | 6.09215 | 6.73483 | 5.83604 | 6.47094 | 6.16739 |
| CCHCR1 | Q8TD31-2 | 6.37297 | 6.6113 | 6.54326 | 6.40154 | 6.9085 | 6.74667 | 6.35192 | 6.28493 | 6.51644 | 6.71771 | 7.25961 | 6.89074 | 5.11616 | 5.28068 | 4.84784 | 5.14974 |
| RWDD2A | Q9UIY3 | 6.26613 | 6.36096 | 7.11295 | 5.73507 | 6.6592 | 6.94975 | 6.37118 | 6.92899 | 7.16137 | 7.19046 | 6.55322 | 6.90672 | 4.68261 | 5.55346 | 4.70114 | 4.86677 |
| SELENOO | Q9BVL4 | 6.12599 | 6.75225 | 6.99073 | 5.92844 | 6.05577 | 7.05305 | 6.57623 | 5.48802 | 5.53021 | 6.33571 | 7.05196 | 6.26295 | 5.94359 | 6.57656 | 6.01075 | 5.31779 |
| IMMP2L | Q96T52 | 5.60494 | 6.76178 | 5.62099 | 6.77312 | 6.97213 | 6.75528 | 6.8628 | 6.11546 | 5.83881 | 5.48859 | 6.60102 | 6.49932 | 6.21161 | 4.76977 | 6.60647 | 6.51794 |
| CEP95 | Q96GE4 | 6.09926 | 7.02355 | 6.65149 | 6.90498 | 6.51243 | 6.71312 | 6.1729 | 6.57577 | 6.42862 | 6.75699 | 6.49139 | 6.52428 | 4.9127 | 5.41743 | 5.4057 | 5.4094 |
| TRIM36 | Q9NQ86 | 6.91617 | 6.44085 | 6.4316 | 6.12554 | 6.31372 | 6.37881 | 6.04757 | 6.49785 | 6.10701 | 6.41667 | 6.68098 | 6.82696 | 5.41057 | 5.72223 | 5.64256 | 6.0409 |
| CRTC3 | Q6UUV7 | 6.13698 | 5.14467 | 6.97901 | 5.03642 | 6.17682 | 5.92791 | 5.44871 | 6.40792 | 6.91579 | 5.76416 | 6.41446 | 6.61829 | 7.76559 | 6.68639 | 5.92129 | 6.6556 |
| TMEM126B | Q8IUX1 | 6.48769 | 6.63848 | 6.46024 | 6.8117 | 6.24598 | 5.80648 | 6.36224 | 5.8931 | 5.93095 | 6.19204 | 6.00172 | 5.94033 | 6.34542 | 6.19255 | 6.41903 | 6.27205 |
| ASB1 | Q9Y576 | 6.10526 | 6.00067 | 6.66758 | 6.4028 | 6.07644 | 6.61536 | 6.28927 | 6.88042 | 6.18596 | 5.92788 | 6.19358 | 6.47914 | 5.89018 | 5.70559 | 6.12104 | 6.45884 |
| NRM | Q8IXM6 | 6.56518 | 6.71798 | 5.35242 | 5.89945 | 6.05066 | 6.1631 | 5.93604 | 6.54043 | 6.05074 | 5.5449 | 5.72243 | 5.79358 | 6.29512 | 7.05172 | 7.00542 | 7.31084 |
| ARSK | Q6UWY0 | 6.00379 | 6.1451 | 5.52987 | 5.99161 | 6.47227 | 6.19271 | 6.20178 | 6.59065 | 6.23091 | 6.35456 | 6.53993 | 6.52738 | 6.41345 | 6.24341 | 6.26515 | 6.29743 |
| HBD | P02042 | 7.45788 | 5.7221 | 4.01406 | 5.48446 | 4.64208 | 5.5092 | 8.11355 | 5.93899 | 5.82515 | 5.47437 | 5.36408 | 5.88046 | 6.22036 | 5.93419 | 9.14493 | 9.27414 |
| TBC1D32 | Q96NH3-4 | 6.32223 | 6.96339 | 7.46063 | 6.71908 | 6.78959 | 7.29673 | 6.28115 | 6.48893 | 6.73809 | 6.89226 | 6.55614 | 6.81812 | 4.48315 | 5.4227 | 3.85343 | 4.91439 |
| NPRL2 | Q8WTW4 | 6.43919 | 6.30298 | 6.63077 | 6.36959 | 5.90881 | 6.03655 | 5.76559 | 6.46325 | 6.46554 | 6.6026 | 6.9774 | 7.13526 | 5.85069 | 5.84807 | 5.57822 | 5.6255 |
| TMTC2 | Q8N394 | 6.24198 | 7.02049 | 5.42301 | 6.40995 | 6.63767 | 6.34617 | 6.06405 | 6.04672 | 6.01817 | 6.31945 | 5.97157 | 6.01755 | 6.89811 | 6.20712 | 6.4931 | 5.8849 |
| TEX30 | Q5JUR7 | 6.3107 | 6.01109 | 6.73019 | 6.2141 | 6.50832 | 6.81944 | 6.28733 | 6.13144 | 6.62962 | 6.95582 | 6.8092 | 6.70647 | 5.42726 | 5.67757 | 5.08814 | 5.69334 |
| TRDMT1 | O14717 | 6.6117 | 6.43276 | 6.68526 | 5.9155 | 6.23503 | 6.69165 | 6.04115 | 7.28957 | 6.43467 | 6.59624 | 6.51575 | 6.89163 | 5.51646 | 5.35925 | 5.34725 | 5.43614 |
| NAGLU | P54802 | 5.97817 | 6.45247 | 5.87121 | 6.71092 | 6.76727 | 6.21465 | 6.79465 | 6.20804 | 5.91421 | 5.75333 | 6.31622 | 6.31482 | 5.88496 | 6.06036 | 6.4145 | 6.34423 |
| CDK10 | Q15131 | 5.76241 | 5.97793 | 5.58526 | 5.7493 | 6.11271 | 6.27664 | 6.33062 | 5.8435 | 6.01584 | 6.20801 | 5.77173 | 6.11306 | 7.27349 | 6.83736 | 7.15673 | 6.9854 |
| TLCD1 | Q96CP7 | 6.3507 | 5.98422 | 5.79182 | 6.76745 | 5.71996 | 6.02178 | 6.36558 | 6.04554 | 6.05309 | 6.09983 | 6.32384 | 5.92846 | 6.32415 | 6.55563 | 6.6932 | 6.97477 |
| B4GALT3 | O60512 | 6.56185 | 6.67583 | 6.4177 | 7.63141 | 6.16608 | 5.84144 | 6.12898 | 6.55938 | 6.0112 | 6.29584 | 6.56044 | 6.30742 | 6.2177 | 5.62128 | 5.32171 | 5.68174 |
| SLC5A6 | Q9Y289 | 6.16373 | 6.24237 | 6.57842 | 6.62427 | 5.94027 | 5.9028 | 6.09241 | 6.40862 | 6.3775 | 6.06701 | 6.53005 | 6.26261 | 5.79531 | 6.09666 | 6.14616 | 6.77181 |
| SMAP1 | Q8IYB5-2 | 6.12783 | 6.47591 | 6.94651 | 5.89313 | 5.71124 | 6.34144 | 6.60681 | 6.42194 | 6.57773 | 6.34534 | 6.29279 | 6.43534 | 6.18376 | 6.11535 | 5.52741 | 5.99746 |
| KCTD2 | Q14681 | 6.48936 | 5.51478 | 6.72613 | 6.46048 | 6.48121 | 6.7539 | 6.19905 | 6.14988 | 6.3573 | 6.56592 | 7.10083 | 6.74499 | 5.96009 | 5.3648 | 5.14823 | 5.98306 |
| TTC9C | Q8N5M4 | 4.97706 | 5.42194 | 6.29697 | 4.99089 | 6.60212 | 5.83788 | 5.87716 | 7.48241 | 6.11559 | 7.59117 | 6.49337 | 6.85718 | 6.10498 | 6.07253 | 6.59224 | 6.6865 |
| TAF5L | O75529 | 6.53085 | 6.62297 | 6.38571 | 5.72251 | 6.25761 | 6.08698 | 6.01977 | 6.44732 | 5.93585 | 5.57 | 6.48794 | 6.48153 | 6.52712 | 6.45792 | 6.41862 | 6.04731 |
| CREB3L1 | Q96BA8 | 7.38337 | 6.78969 | 6.2744 | 7.09709 | 6.57102 | 6.18547 | 6.47232 | 6.38944 | 6.25528 | 7.36278 | 7.01824 | 7.20618 | 4.10276 | 5.34086 | 4.65632 | 4.8948 |
| RXYLT1 | Q9Y2B1 | 6.50475 | 6.80344 | 6.55089 | 7.27293 | 5.78459 | 5.94255 | 6.50333 | 5.76514 | 5.87654 | 6.33554 | 6.05411 | 6.59093 | 5.86936 | 5.90651 | 5.93317 | 6.30624 |
| MSANTD4 | Q8NCY6 | 6.81664 | 7.20079 | 6.80505 | 7.01026 | 6.51727 | 6.34549 | 6.69814 | 6.49218 | 6.72272 | 6.46084 | 5.9691 | 5.71644 | 5.61373 | 4.97072 | 5.28349 | 5.37716 |
| AMACR | Q9UHK6-5 | 6.50855 | 5.27589 | 6.14891 | 6.48726 | 6.24385 | 5.98687 | 5.70041 | 5.8368 | 7.02639 | 6.77706 | 7.08805 | 6.91642 | 5.75339 | 5.91313 | 5.84346 | 6.49359 |
| ZDHHC5 | Q9C0B5 | 6.29984 | 6.33515 | 5.44437 | 6.41526 | 6.46784 | 6.10531 | 6.44134 | 6.04902 | 6.25577 | 6.06314 | 5.84855 | 5.61939 | 6.38878 | 6.62547 | 6.71257 | 6.92819 |
| TAF12 | Q16514 | 6.04436 | 6.48286 | 6.1772 | 6.31157 | 5.95234 | 6.31497 | 6.41986 | 6.46779 | 6.19876 | 6.26751 | 6.6097 | 7.06043 | 6.04267 | 5.75424 | 5.82247 | 6.07329 |
| CNEP1R1 | Q8N9A8-2 | 6.72532 | 5.80837 | 6.3714 | 7.22946 | 5.48145 | 6.67851 | 5.9177 | 5.32265 | 6.88122 | 6.60849 | 5.78075 | 5.9814 | 6.10125 | 5.45839 | 6.69231 | 6.96133 |
| PRXL2C | Q7RTV5 | 6.67971 | 6.65756 | 6.86702 | 7.07668 | 5.87415 | 6.62284 | 6.33693 | 6.78957 | 6.75387 | 6.4637 | 7.42407 | 7.34806 | 4.46637 | 5.0075 | 4.55871 | 5.07328 |
| MAD2L1BP | Q15013-3 | 6.01434 | 6.30494 | 6.52503 | 6.25138 | 6.0003 | 6.17371 | 5.94764 | 6.5735 | 6.29059 | 6.97277 | 6.48324 | 6.80442 | 6.25232 | 5.86166 | 5.70847 | 5.83567 |

| | | | | | | | | | | | | | | | | | |
|---|---|---|---|---|---|---|---|---|---|---|---|---|---|---|---|---|---|
| GET1 | O00258 | 6.22148 | 6.14868 | 6.58613 | 6.28922 | 5.53586 | 6.01164 | 5.72877 | 5.95699 | 6.43475 | 6.7344 | 5.99864 | 6.58643 | 6.56157 | 6.43686 | 5.93963 | 6.82893 |
| NOCT | Q9UK39 | 5.60443 | 6.41043 | 6.54231 | 5.09694 | 6.18874 | 6.22971 | 6.81749 | 6.0604 | 5.83516 | 6.01124 | 6.27168 | 7.35217 | 6.53659 | 6.02525 | 6.71027 | 6.30719 |
| CDR2 | Q01850 | 6.74657 | 6.27433 | 5.91472 | 6.35905 | 6.51354 | 6.23675 | 6.0746 | 6.2902 | 6.24484 | 5.97097 | 6.11194 | 7.15254 | 6.28742 | 5.71811 | 6.05646 | 6.04797 |
| MRFAP1 | Q9Y605 | 7.26074 | 6.14252 | 7.62474 | 5.63649 | 6.67429 | 5.60437 | 5.0544 | 7.61549 | 6.00046 | 8.05349 | 7.20012 | 7.45622 | 3.5876 | 6.25459 | 4.76582 | 5.06867 |
| ATP5MK | Q96IX5 | 6.17702 | 6.05107 | 6.48632 | 7.47333 | 6.01574 | 6.39101 | 6.31267 | 6.3895 | 6.58828 | 6.56718 | 6.07345 | 6.45021 | 5.31835 | 5.99034 | 5.95125 | 5.76426 |
| CRTC2 | Q53ET0 | 6.69026 | 6.00759 | 6.89571 | 5.84235 | 7.45346 | 6.63862 | 5.68021 | 6.02149 | 5.6238 | 6.40127 | 6.90164 | 6.88549 | 6.00883 | 5.97906 | 5.53612 | 5.43409 |
| FAM241A | Q8N8J7 | 6.63254 | 6.13068 | 5.76746 | 5.61368 | 6.19508 | 6.01558 | 6.5908 | 6.65581 | 5.44513 | 6.45165 | 5.05709 | 5.73198 | 7.56618 | 6.61323 | 7.29472 | 6.23837 |
| SF3B5 | Q9BWJ5 | 5.79756 | 4.43685 | 5.10446 | 5.22353 | 5.96077 | 4.94738 | 5.1963 | 5.61236 | 5.24522 | 5.73633 | 4.94364 | 5.06134 | 5.34759 | 10.427 | 9.16076 | 11.7989 |
| IL10RB | Q08334 | 6.59913 | 7.4795 | 7.42056 | 7.16807 | 6.24843 | 6.2438 | 6.76515 | 6.4748 | 6.6294 | 6.88627 | 7.23117 | 7.10401 | 4.19049 | 4.62441 | 4.27258 | 4.66224 |
| XAGE1A | Q9HD64 | 6.72905 | 6.7053 | 7.18435 | 6.89587 | 6.40855 | 6.13423 | 6.75117 | 7.16128 | 7.29176 | 6.62101 | 7.09109 | 6.50355 | 4.98831 | 4.78564 | 4.2283 | 4.52053 |
| PIGBOS1 | A0A0B4J2F0 | 6.36905 | 6.28389 | 6.03151 | 6.25411 | 5.73424 | 5.62337 | 6.05119 | 6.36646 | 4.88861 | 5.82623 | 5.63485 | 6.02121 | 5.65635 | 8.78443 | 6.84451 | 7.62998 |
| TMEM59 | Q9BXS4 | 7.12148 | 7.49678 | 6.69692 | 7.74319 | 6.14194 | 5.7514 | 6.35701 | 6.13756 | 6.79562 | 6.81213 | 7.19177 | 6.92846 | 4.42324 | 4.93235 | 4.46408 | 5.00605 |
| HECA | Q9UBI9 | 6.1945 | 6.43916 | 6.42217 | 5.81401 | 5.99114 | 6.25709 | 6.2553 | 6.20607 | 6.44 | 6.58831 | 6.75846 | 6.40207 | 5.90611 | 6.14483 | 5.90826 | 6.27251 |
| ZDHHC15 | Q96MV8 | 6.11169 | 6.29494 | 6.03627 | 6.66969 | 5.9097 | 6.34895 | 6.23915 | 5.90631 | 5.98212 | 6.60313 | 6.36698 | 6.50259 | 6.03564 | 6.30609 | 6.25784 | 6.42891 |
| TMEM167A | Q8TBQ9 | 5.70021 | 6.74085 | 5.81033 | 6.5557 | 5.80616 | 6.41927 | 5.86099 | 6.4645 | 6.52446 | 5.99268 | 6.16948 | 6.26087 | 6.40998 | 6.33136 | 6.64506 | 6.30808 |
| FUT10 | Q6P4F1-2 | 6.11574 | 7.08297 | 5.91036 | 6.98524 | 5.93248 | 5.94553 | 5.9982 | 5.86929 | 6.16857 | 6.3397 | 5.97479 | 5.82823 | 6.39689 | 5.89026 | 7.10199 | 6.45977 |
| SCMH1 | Q96GD3 | 6.21516 | 6.82318 | 6.29426 | 6.8201 | 6.35444 | 6.42473 | 6.74775 | 5.9017 | 5.94213 | 6.56045 | 6.04999 | 6.05956 | 6.15834 | 5.85982 | 6.33641 | 6.35197 |
| C12orf4 | Q9NQ89 | 5.69601 | 5.99113 | 6.83649 | 6.13672 | 6.7517 | 6.66315 | 6.64024 | 6.16266 | 6.37269 | 6.72526 | 6.5789 | 6.86802 | 5.51131 | 5.9034 | 5.94226 | 5.22004 |
| CD63 | P08962 | 6.41463 | 6.46873 | 6.72192 | 6.46393 | 5.40219 | 5.87494 | 6.53358 | 5.62435 | 6.48559 | 6.98711 | 6.92437 | 6.60378 | 5.76972 | 5.90132 | 6.30524 | 5.5186 |
| COL1A2 | P08123 | 5.99559 | 6.36555 | 5.28227 | 6.12822 | 4.16246 | 4.46436 | 4.74235 | 4.89516 | 6.01368 | 6.32849 | 6.78917 | 6.23049 | 8.38435 | 8.07834 | 8.90945 | 7.23008 |
| DCUN1D4 | Q92564-3 | 5.46995 | 5.71217 | 6.2799 | 5.33907 | 5.45909 | 6.47063 | 5.57315 | 6.03214 | 5.94395 | 6.34176 | 6.33598 | 6.14521 | 7.0615 | 7.73889 | 7.28268 | 6.81393 |
| TFDP2 | Q14188 | 5.98075 | 6.4514 | 6.42264 | 6.38631 | 6.05756 | 6.09096 | 5.91298 | 6.27206 | 6.31746 | 6.22498 | 6.84072 | 5.52121 | 7.01486 | 6.50251 | 6.16352 | 5.8401 |
| ZNF410 | Q86VK4-5 | 5.91191 | 5.79508 | 6.49167 | 6.0544 | 5.83132 | 6.14821 | 5.96738 | 5.96338 | 6.33307 | 6.51841 | 6.45744 | 6.56145 | 5.7074 | 6.49032 | 6.90378 | 6.86479 |
| MCRIP1 | C9JLW8 | 6.53909 | 4.75745 | 6.46305 | 5.03978 | 6.48346 | 5.30351 | 5.49792 | 6.9537 | 5.50939 | 6.83518 | 5.67889 | 7.26498 | 5.9659 | 7.79064 | 6.60139 | 7.84068 |
| ISG15 | P05161 | 5.5249 | 4.81256 | 5.87162 | 5.14938 | 6.16493 | 6.12295 | 6.12886 | 6.33858 | 7.13819 | 7.15506 | 6.90096 | 6.93908 | 5.80069 | 6.08717 | 6.5912 | 7.27388 |
| FUCA1 | P04066 | 6.12765 | 5.73902 | 5.77895 | 6.35002 | 6.1415 | 6.0893 | 5.92033 | 5.8696 | 6.37109 | 6.21763 | 5.57437 | 5.9382 | 6.77863 | 6.71988 | 7.04351 | 7.34032 |
| ZCCHC17 | Q9NP64 | 6.18042 | 5.97842 | 5.74741 | 6.36629 | 5.7106 | 5.81096 | 6.16186 | 5.66636 | 6.01711 | 5.994 | 5.81731 | 5.99345 | 6.46114 | 7.32141 | 7.14389 | 7.63213 |
| CAAP1 | Q9H8G2 | 6.39778 | 6.27638 | 5.75831 | 6.6998 | 6.22259 | 5.70105 | 5.71159 | 5.75603 | 6.42691 | 6.24458 | 6.08872 | 6.10709 | 6.58286 | 6.47895 | 6.52584 | 7.02152 |
| ATP5F1EP2 | Q5VTU8 | 6.60512 | 6.73084 | 5.48178 | 7.03429 | 5.9617 | 5.51368 | 6.40908 | 6.37813 | 6.79835 | 6.16023 | 6.4459 | 6.2159 | 6.41986 | 5.5345 | 6.18642 | 6.12421 |
| NID1 | P14543 | 6.37785 | 8.01696 | 6.71774 | 6.62876 | 6.42437 | 5.91189 | 6.43155 | 6.62802 | 6.25202 | 7.50284 | 7.19645 | 6.13042 | 4.41836 | 4.72614 | 5.06176 | 5.57484 |
| GAGE12F | P0CL80 | 5.2612 | 4.38359 | 4.09454 | 4.93913 | 3.81163 | 2.35217 | 4.92276 | 4.47657 | 3.64761 | 2.70507 | 3.37566 | 3.88248 | 2.98243 | 19.6807 | 10.4321 | 19.0524 |
| HINT3 | Q9NQE9 | 5.87807 | 5.45239 | 6.19716 | 5.16569 | 5.85595 | 5.66335 | 5.8548 | 7.24896 | 6.27538 | 6.44938 | 5.80191 | 6.55963 | 6.52915 | 7.11506 | 7.14389 | 6.80921 |
| FAM110A | Q9BQ89 | 5.94121 | 6.95534 | 7.74999 | 6.5142 | 6.3065 | 6.32336 | 6.42761 | 6.46297 | 6.34962 | 6.66335 | 7.16779 | 7.01714 | 5.05512 | 5.30219 | 4.9553 | 4.80832 |
| TAX1BP3 | O14907 | 6.04277 | 4.52032 | 6.2019 | 5.08213 | 6.63468 | 5.76825 | 5.81666 | 6.71044 | 5.62319 | 6.94628 | 6.23449 | 7.09175 | 5.41842 | 7.47425 | 6.60581 | 7.82866 |
| CKS1B | P61024 | 5.24206 | 5.12301 | 6.43414 | 5.49313 | 6.66797 | 5.99673 | 5.92782 | 7.96582 | 5.78022 | 7.61855 | 7.20022 | 7.44282 | 4.37848 | 6.25375 | 5.73942 | 6.73585 |
| POP5 | Q969H6 | 5.90277 | 6.44047 | 6.37895 | 6.40789 | 6.25137 | 6.42817 | 6.18718 | 5.76216 | 5.47682 | 6.60541 | 6.13495 | 5.95201 | 6.32579 | 6.30874 | 6.9696 | 6.46772 |
| TUBE1 | Q9UJT0 | 6.41487 | 6.01308 | 7.19487 | 6.15974 | 6.95493 | 6.52258 | 6.24391 | 6.29211 | 6.48634 | 6.8225 | 6.72444 | 6.5028 | 4.95506 | 5.56153 | 5.33485 | 5.81636 |
| ICMT | O60725 | 6.1706 | 6.03859 | 5.80009 | 6.97936 | 5.72511 | 6.05104 | 6.25994 | 6.1404 | 6.10782 | 6.17602 | 6.2884 | 6.25727 | 6.58285 | 6.32183 | 6.47133 | 6.62933 |
| ADCY9 | O60503 | 5.73187 | 6.8829 | 6.95006 | 7.23449 | 6.4394 | 6.55587 | 5.63637 | 6.14935 | 6.1612 | 5.8935 | 5.83882 | 6.64227 | 6.0449 | 5.54909 | 6.08055 | 6.20937 |
| IPP | Q9Y573 | 6.4465 | 6.23159 | 6.68971 | 5.8223 | 6.37688 | 6.79425 | 6.07694 | 6.66127 | 6.81381 | 7.03001 | 6.5513 | 6.38252 | 5.53426 | 5.85561 | 5.47136 | 5.2617 |
| BDH2 | Q9BUT1 | 6.12777 | 6.46561 | 6.99879 | 5.79755 | 6.17513 | 7.03698 | 6.4761 | 5.72913 | 6.00153 | 6.079 | 6.47457 | 6.03878 | 6.19003 | 6.26429 | 6.1899 | 5.95483 |
| ATP6V0C | P27449 | 5.88913 | 6.03351 | 5.49119 | 6.49604 | 6.28022 | 5.34879 | 7.97545 | 5.53506 | 7.33784 | 4.42078 | 5.2895 | 6.40235 | 5.72872 | 6.78043 | 6.45307 | 8.58792 |
| B2M | P61769 | 6.3291 | 5.24599 | 5.46028 | 5.62424 | 6.14301 | 5.16449 | 5.19513 | 7.84943 | 5.1378 | 5.78019 | 5.67948 | 6.46951 | 5.64719 | 8.45746 | 7.53232 | 8.28439 |
| LRP12 | Q9Y561 | 6.49103 | 6.84711 | 6.32856 | 6.4368 | 6.51461 | 6.65637 | 6.38312 | 6.43451 | 6.24238 | 6.41132 | 6.06012 | 6.5187 | 5.91545 | 5.24959 | 5.38124 | 6.22908 |
| CALHM5 | Q8N5C1 | 6.14545 | 6.36292 | 6.65305 | 7.00883 | 6.91454 | 6.4838 | 5.77336 | 5.8213 | 5.68428 | 6.0018 | 7.01808 | 6.5208 | 5.34305 | 6.25144 | 5.85374 | 6.16356 |
| NDUFS6 | O75380 | 6.84174 | 6.08813 | 5.91381 | 5.78438 | 6.106 | 5.96892 | 5.30906 | 6.29864 | 5.33151 | 6.41376 | 5.88103 | 6.74412 | 6.84261 | 6.6897 | 6.49328 | 7.2933 |
| TMEM230 | Q96A57-2 | 6.39787 | 6.24011 | 6.08123 | 6.36807 | 6.50934 | 6.17341 | 6.63152 | 6.11822 | 6.08304 | 6.09569 | 5.88072 | 5.32111 | 6.58806 | 6.36718 | 6.59851 | 6.54592 |
| FUBP1 | Q96AE4 | 6.89993 | 6.22279 | 5.06606 | 6.39607 | 6.83603 | 6.19727 | 5.8165 | 6.39287 | 5.8404 | 5.95184 | 6.85446 | 5.78696 | 7.32703 | 5.66826 | 6.47252 | 6.27102 |
| UNKL | Q9H9P5-6 | 6.22568 | 6.32062 | 6.70813 | 6.13415 | 5.23095 | 5.48639 | 5.71883 | 5.44752 | 7.19211 | 6.32455 | 5.37061 | 6.86474 | 6.53288 | 5.91174 | 6.82526 | 7.70585 |
| SH3PXD2A | Q5TCZ1 | 6.06138 | 6.37016 | 6.62541 | 5.99845 | 6.05956 | 6.46908 | 6.52001 | 6.1396 | 6.68576 | 6.54213 | 6.7938 | 6.8306 | 5.53649 | 5.90717 | 5.91741 | 5.54299 |
| TRO | Q12816 | 7.06992 | 6.23835 | 5.96095 | 5.31157 | 6.77835 | 6.86148 | 6.18833 | 6.35918 | 6.64054 | 7.25874 | 5.63822 | 6.05852 | 4.73321 | 7.57316 | 5.62088 | 5.7086 |
| TFR2 | Q9UP52 | 5.96689 | 5.7851 | 6.37393 | 6.63856 | 5.83498 | 6.37008 | 6.16311 | 6.49697 | 6.57393 | 6.30306 | 6.23127 | 5.79684 | 5.96077 | 5.98865 | 6.71587 | 6.79999 |
| PPP1R35 | Q8TAP8 | 6.5873 | 5.92205 | 7.3622 | 6.6306 | 5.71316 | 6.17854 | 5.84346 | 7.36328 | 7.38799 | 6.38087 | 7.87618 | 8.12136 | 4.89898 | 4.46669 | 3.86383 | 5.40351 |
| WDR87 | Q6ZQQ6-2 | 6.13412 | 6.33614 | 4.64305 | 6.64755 | 5.83625 | 6.60852 | 5.14176 | 7.10212 | 6.08031 | 6.31913 | 6.57921 | 6.62311 | 6.31517 | 6.24896 | 6.57289 | 6.81171 |
| SYDE2 | Q5VT97 | 6.50675 | 6.62467 | 7.22303 | 6.75741 | 5.81977 | 6.59166 | 7.01065 | 5.99678 | 6.59333 | 5.69464 | 6.56278 | 6.59094 | 5.81833 | 5.54856 | 5.39114 | 5.43155 |
| SMARCC2 | Q8TAQ2-3 | 6.529 | 6.35063 | 6.30778 | 6.39721 | 6.35036 | 6.03154 | 5.83018 | 6.16247 | 5.85643 | 6.24069 | 5.593 | 5.21947 | 6.85234 | 6.87895 | 6.43256 | 6.9674 |
| LRRC8B | Q6P9F7 | 5.26827 | 5.49046 | 5.86683 | 6.94236 | 7.17121 | 6.91184 | 6.61422 | 5.77685 | 7.64199 | 6.20825 | 6.13754 | 6.06352 | 6.57157 | 5.40695 | 6.07936 | 5.84879 |
| NRBF2 | Q96F24 | 7.03226 | 5.0769 | 7.8378 | 5.38037 | 6.28806 | 6.52446 | 5.29689 | 7.12161 | 6.58541 | 5.46341 | 7.10136 | 7.61055 | 6.64838 | 5.37482 | 5.01932 | 5.63841 |
| RARA | P10276 | 6.06361 | 6.19876 | 6.00757 | 5.79312 | 6.05504 | 6.23225 | 6.54542 | 6.02075 | 6.3138 | 6.28256 | 6.44681 | 6.63459 | 6.24618 | 5.83253 | 6.9196 | 6.40742 |
| MTMR4 | Q9NYA4 | 6.49443 | 5.81364 | 6.4004 | 6.89912 | 6.1624 | 6.31843 | 7.20947 | 7.07298 | 6.11554 | 6.26092 | 6.45823 | 7.29835 | 5.50859 | 5.23063 | 5.36058 | 5.3963 |
| TMEM231 | Q9H6L2-2 | 6.17568 | 6.90138 | 5.51933 | 5.92091 | 6.20738 | 6.05872 | 6.82597 | 6.07943 | 6.24845 | 6.00784 | 5.868 | 5.77358 | 6.63344 | 6.31568 | 6.25966 | 7.20454 |
| PLXNA3 | P51805 | 5.78463 | 6.64328 | 6.15937 | 7.03324 | 5.57762 | 6.25073 | 6.43682 | 6.04866 | 6.19798 | 6.48 | 6.91557 | 5.97807 | 5.94181 | 6.63567 | 5.69727 | 6.21927 |
| AP3M2 | P53677 | 6.2103 | 5.96545 | 6.76431 | 6.7248 | 6.40541 | 6.64343 | 5.26892 | 6.82731 | 6.75137 | 7.73787 | 6.56344 | 7.29686 | 5.20299 | 6.28458 | 5.1089 | 4.24406 |
| CCAR2 | G3V119 | 6.36032 | 6.09692 | 6.37113 | 6.15554 | 5.18701 | 5.26428 | 5.45631 | 5.71415 | 6.16114 | 4.67242 | 5.72171 | 5.70601 | 6.62904 | 6.59918 | 7.49774 | 9.40708 |
| NXT1 | Q9UKK6 | 5.79029 | 6.04955 | 6.02376 | 6.51083 | 5.7195 | 6.07193 | 5.77814 | 6.29613 | 6.11181 | 6.41051 | 6.92527 | 6.29335 | 6.06881 | 6.56523 | 6.88734 | 6.49756 |
| AFDN | P55196-1 | 5.499 | 6.87611 | 5.87142 | 7.11772 | 5.75305 | 6.6753 | 6.12917 | 6.49938 | 6.21308 | 6.3833 | 5.70177 | 6.22075 | 5.52396 | 6.46072 | 6.67696 | 6.39829 |
| PCDHB10 | Q9UN67 | 6.03599 | 6.42302 | 5.81765 | 6.22041 | 7.62251 | 6.48792 | 6.27078 | 6.3892 | 6.11055 | 5.7287 | 5.47671 | 5.66342 | 6.31517 | 5.89105 | 6.35276 | 7.19416 |
| CLTCL1 | P53675 | 6.23658 | 7.00907 | 7.21005 | 6.28449 | 6.1995 | 6.95603 | 6.4182 | 6.44773 | 6.43933 | 6.46155 | 6.56664 | 5.35755 | 5.63831 | 5.70285 | 5.23513 | 5.83701 |
| BOK | Q9UMX3 | 5.9017 | 6.13315 | 5.54421 | 5.36971 | 6.2369 | 5.95831 | 6.54326 | 6.12106 | 6.28696 | 7.55993 | 6.18945 | 4.87933 | 7.83901 | 6.04659 | 6.30084 | 7.0896 |
| ZNF696 | Q9H7X3 | 5.3718 | 7.26776 | 5.24343 | 6.65165 | 6.63867 | 6.20811 | 6.53341 | 7.16161 | 6.06049 | 6.12945 | 6.56783 | 6.09417 | 6.39066 | 5.08223 | 6.21176 | 6.38697 |
| ALDH1A2 | O94788 | 6.66092 | 6.09721 | 5.9914 | 6.69176 | 6.75267 | 5.39166 | 6.52274 | 6.27566 | 6.11408 | 5.73762 | 6.09283 | 6.04394 | 6.71672 | 5.92247 | 6.69128 | 6.29705 |
| COMMD3-BMI1 | R4GMX3 | 5.86752 | 6.483 | 5.80712 | 6.84956 | 6.10116 | 6.27012 | 6.22067 | 6.18803 | 6.49699 | 6.5587 | 5.3453 | 5.98995 | 6.41445 | 6.13068 | 6.37431 | 6.90243 |
| CEP104 | O60308 | 6.2234 | 5.91635 | 6.87253 | 6.13378 | 7.18233 | 6.27377 | 6.51611 | 5.60076 | 6.65201 | 5.87048 | 6.19077 | 6.22672 | 6.10665 | 5.68696 | 5.89915 | 6.64823 |
| GOLPH3L | Q9H4A5 | 6.10073 | 5.59174 | 6.16776 | 6.8333 | 6.30294 | 6.18302 | 6.61875 | 6.39029 | 5.7962 | 6.40859 | 6.56595 | 6.43246 | 6.81748 | 5.70251 | 5.58626 | 6.10202 |
| LYN | P07948-2 | 6.87019 | 6.57269 | 5.23768 | 6.15416 | 6.42336 | 5.86645 | 5.93383 | 6.16923 | 6.06999 | 5.99491 | 5.84893 | 6.47206 | 6.77471 | 6.26557 | 6.67409 | 6.67216 |
| FLNB | O75369 | 5.79715 | 6.07962 | 6.73498 | 6.61843 | 5.79887 | 5.75462 | 6.10804 | 6.81918 | 6.26692 | 6.26424 | 6.94433 | 6.69148 | 5.9934 | 5.71544 | 5.9256 | 6.48771 |
| SMU1 | Q2TAY7-2 | 5.99491 | 6.15745 | 4.89557 | 5.7878 | 6.33116 | 7.59504 | 4.91597 | 6.66791 | 7.50459 | 7.37839 | 7.02595 | 7.42245 | 8.20029 | 5.58477 | 4.79909 | 3.73868 |
| RBM4B | Q9BQ04 | 6.57093 | 6.24221 | 6.39807 | 7.00362 | 5.74197 | 6.47865 | 5.34195 | 5.55509 | 6.69837 | 6.86373 | 6.80648 | 6.15421 | 5.91974 | 5.83203 | 5.41531 | 6.97763 |
| NOL12 | Q9UGY1 | 6.27531 | 5.82089 | 5.92536 | 6.55111 | 7.68408 | 6.03071 | 6.33396 | 6.01652 | 6.95044 | 4.84619 | 7.48603 | 5.27497 | 8.20393 | 6.12301 | 5.65033 | 4.82715 |
| CBFA2T2 | O43439-4 | 5.81379 | 5.77594 | 6.63454 | 5.5228 | 5.48412 | 6.0379 | 6.31556 | 6.49386 | 7.24618 | 6.51426 | 6.43433 | 7.25913 | 6.48357 | 6.38604 | 5.21349 | 5.38449 |
| NPRL3 | Q12980 | 6.393 | 6.42904 | 6.96666 | 7.15611 | 5.79856 | 7.4216 | 6.5453 | 6.4643 | 6.8683 | 6.75292 | 6.66375 | 6.37367 | 5.07227 | 5.39939 | 4.95478 | 4.74036 |
| ATXN2 | F8W0B5 | 6.15477 | 6.25482 | 6.7538 | 6.54996 | 6.43791 | 6.26294 | 6.26639 | 6.37081 | 6.65326 | 5.89916 | 6.42122 | 6.69118 | 5.52588 | 6.15829 | 5.19303 | 6.10659 |
| ATXN2L | Q8WWM7 | 5.99269 | 5.50835 | 6.84782 | 5.36256 | 6.43511 | 6.24902 | 5.95988 | 6.75212 | 6.33198 | 5.9846 | 7.22647 | 7.74096 | 5.67708 | 6.18174 | 6.07731 | 5.67232 |
| STARD3NL | O95772 | 7.18836 | 6.37323 | 6.35481 | 7.07808 | 7.27389 | 6.008 | 6.44343 | 6.70432 | 6.4698 | 5.99833 | 7.91519 | 6.13923 | 4.8394 | 5.31878 | 5.07113 | 4.82402 |
| ISCA1 | Q9BUE6-2 | 6.68725 | 6.36359 | 5.89002 | 7.40999 | 5.58175 | 5.73275 | 5.12291 | 5.84163 | 5.91276 | 5.97164 | 6.05239 | 6.64306 | 6.79339 | 5.62366 | 7.034 | 7.33921 |
| CEP170B | Q9Y4F5 | 5.74858 | 6.4837 | 6.40514 | 6.11874 | 6.22709 | 6.45307 | 6.02836 | 6.64965 | 6.7814 | 6.53658 | 6.30347 | 6.79789 | 5.92944 | 6.16079 | 5.71619 | 5.65989 |
| MEAF6 | Q9HAF1-3 | 6.27086 | 6.48271 | 6.20192 | 6.79908 | 6.12934 | 6.10263 | 6.8819 | 5.84703 | 5.87217 | 6.17877 | 6.52294 | 6.12701 | 6.36948 | 6.19889 | 5.92755 | 6.08772 |
| GPANK1 | O95872 | 5.94041 | 6.87891 | 6.17766 | 6.90215 | 6.09001 | 6.64716 | 6.91872 | 7.15169 | 7.24963 | 6.44223 | 6.16956 | 6.07511 | 5.65149 | 4.5063 | 5.42637 | 5.7726 |
| MBNL1 | Q9NR56-5 | 6.17226 | 6.88983 | 6.36788 | 6.22367 | 6.7477 | 6.2736 | 5.84833 | 6.71796 | 6.62959 | 5.86047 | 6.93747 | 5.95822 | 6.78607 | 5.65829 | 6.01419 | 4.91447 |
| B3GNTL1 | Q67FW5 | 6.46449 | 6.12221 | 6.884 | 5.33746 | 6.06574 | 6.84749 | 6.43189 | 4.83411 | 8.10259 | 6.01101 | 7.08878 | 5.87295 | 6.63973 | 6.74161 | 5.06084 | 5.4951 |
| ARMC5 | Q96C12 | 5.67604 | 5.6205 | 5.74689 | 6.46893 | 5.92205 | 5.77736 | 6.15088 | 6.23312 | 6.73014 | 6.64519 | 6.55884 | 6.81441 | 6.83859 | 6.05229 | 6.37986 | 6.38491 |
| SDHD | O14521 | 5.81694 | 5.2427 | 6.41964 | 5.75974 | 5.26636 | 5.60655 | 6.83941 | 7.12952 | 5.63506 | 5.35935 | 5.80975 | 6.13083 | 6.82766 | 7.10544 | 6.21718 | 8.83389 |
| AAGAB | Q6PD74 | 6.1715 | 6.51722 | 6.75912 | 6.37855 | 6.05121 | 7.11959 | 5.78445 | 6.61691 | 7.21372 | 5.57134 | 7.07306 | 7.03844 | 4.8709 | 5.66696 | 4.89 | 6.27704 |
| EWSR1 | H7BY36 | 6.42994 | 6.11535 | 5.1983 | 6.69625 | 6.91727 | 6.38436 | 5.72623 | 5.72749 | 6.45126 | 5.99472 | 6.41853 | 6.29739 | 6.61103 | 5.36218 | 6.62218 | 7.04754 |
| SHROOM1 | Q2M3G4 | 6.98712 | 6.80089 | 7.03574 | 6.45202 | 6.29596 | 5.06295 | 5.63774 | 7.24588 | 7.6778 | 6.07295 | 6.4859 | 6.75536 | 5.39039 | 5.48347 | 5.25037 | 5.36547 |
| FBXW11 | Q9UKB1 | 6.48212 | 6.15838 | 7.02874 | 6.50986 | 6.70862 | 6.15251 | 6.52807 | 6.10535 | 7.31169 | 6.71522 | 6.62074 | 6.53531 | 4.99022 | 5.05775 | 5.79639 | 5.29901 |
| PGK2 | P07205 | 6.37804 | 6.19097 | 6.71202 | 5.80267 | 6.62002 | 6.82167 | 6.31842 | 6.06374 | 6.19773 | 7.15026 | 6.39861 | 5.96392 | 5.77687 | 6.47549 | 5.34783 | 5.78175 |
| PROSER2 | Q86WR7 | 6.52246 | 6.39092 | 7.15634 | 6.32427 | 6.15167 | 5.98466 | 5.59803 | 7.0605 | 6.32706 | 6.48125 | 6.4202 | 6.24574 | 6.07184 | 5.70863 | 5.59307 | 5.96336 |
| EBP | Q15125 | 6.28999 | 6.81769 | 5.81503 | 7.33165 | 6.03462 | 5.43588 | 6.95883 | 5.90333 | 5.66047 | 5.79046 | 5.58301 | 5.99435 | 6.78163 | 5.73184 | 7.2834 | 6.58782 |
| LRP5 | O75197 | 6.13242 | 6.17766 | 5.97076 | 6.83347 | 5.73034 | 6.29511 | 6.86874 | 5.99246 | 6.18677 | 6.24068 | 5.86801 | 6.17968 | 6.48538 | 6.70506 | 6.40815 | 5.92528 |
| EFCAB7 | A8K855 | 6.16573 | 6.10828 | 6.31302 | 5.70725 | 6.3842 | 6.40384 | 6.43057 | 5.98983 | 7.0055 | 7.19087 | 6.34591 | 6.46214 | 6.2976 | 6.17305 | 5.40985 | 5.61237 |
| DST | F6QMI7 | 6.00804 | 6.50723 | 5.89079 | 6.15244 | 6.09675 | 6.86385 | 5.78386 | 6.36096 | 5.71555 | 6.04406 | 5.9753 | 6.2483 | 6.30866 | 6.58638 | 6.54162 | 6.91621 |
| PLEC | Q15149-4 | 6.62786 | 6.55051 | 6.92656 | 6.76073 | 5.76174 | 5.07945 | 6.69981 | 6.02022 | 6.12131 | 6.12642 | 7.16358 | 5.94777 | 6.58455 | 6.37188 | 6.90931 | 5.34831 |
| PLEC | H0YDN1 | 6.40676 | 5.93032 | 6.16482 | 5.96052 | 5.50662 | 6.11856 | 5.48165 | 5.90277 | 7.20314 | 6.60428 | 5.36619 | 7.27824 | 6.059 | 6.66131 | 6.57658 | 6.77922 |
| PLEC | E9PKG0 | 5.98057 | 5.50037 | 5.90431 | 6.20563 | 6.53746 | 5.99482 | 5.89521 | 6.63291 | 6.82778 | 5.78867 | 6.40461 | 6.03028 | 6.35089 | 6.71772 | 6.61726 | 6.61151 |

| | | | | | | | | | | | | | | | | | |
|---|---|---|---|---|---|---|---|---|---|---|---|---|---|---|---|---|---|
| C3 | P01024 | 5.19533 | 7.45981 | 4.80204 | 5.45667 | 3.77159 | 5.99524 | 6.45334 | 5.50307 | 7.43078 | 5.6484 | 6.15845 | 5.73312 | 6.52785 | 5.95786 | 10.8108 | 7.09569 |
| CYTH2 | Q99418 | 5.25276 | 6.34195 | 6.38236 | 5.96375 | 5.71698 | 6.05317 | 6.29166 | 5.51278 | 6.51347 | 6.6311 | 6.60821 | 6.7289 | 6.80106 | 6.30093 | 6.49804 | 6.40288 |
| CXCL2 | P19875 | 7.59363 | 7.28536 | 7.35412 | 7.49515 | 6.84154 | 5.73835 | 7.09511 | 6.86003 | 7.12273 | 7.24483 | 8.04328 | 7.02691 | 4.07638 | 2.90113 | 3.24784 | 4.0736 |
| SGF29 | Q96ES7 | 6.27002 | 5.42143 | 5.32305 | 6.35125 | 6.26199 | 4.82198 | 6.27605 | 6.02662 | 7.14788 | 5.91401 | 6.714 | 6.3659 | 7.46243 | 5.51191 | 6.55486 | 7.57661 |
| HSP90AB3P | Q58FF7 | 5.5146 | 6.02085 | 6.49907 | 6.44939 | 5.76185 | 7.01266 | 6.39778 | 6.71355 | 5.84609 | 6.4292 | 6.43856 | 6.70788 | 5.82146 | 6.24635 | 5.88864 | 6.25208 |
| APH1A | Q96BI3 | 5.98249 | 5.27152 | 5.18475 | 5.73851 | 6.36408 | 6.32963 | 6.91513 | 6.46271 | 5.89736 | 5.68794 | 6.14397 | 5.6813 | 6.99608 | 6.74813 | 6.82446 | 7.77194 |
| ITPKC | Q96DU7 | 6.06147 | 6.62335 | 6.67549 | 6.06722 | 5.52224 | 6.84415 | 6.71251 | 6.63597 | 6.6417 | 6.68574 | 6.1824 | 6.34986 | 5.47752 | 5.69735 | 6.16104 | 5.66199 |
| CELSR1 | Q9NYQ6 | 7.39077 | 6.44345 | 5.89487 | 7.15684 | 5.84757 | 6.27486 | 6.78668 | 6.33974 | 6.31514 | 5.78048 | 6.41984 | 5.52557 | 6.35338 | 5.3195 | 5.55145 | 6.59985 |
| RGS20 | O76081 | 6.41393 | 6.25051 | 6.15469 | 6.42651 | 6.49015 | 6.0983 | 6.18769 | 5.63434 | 6.03305 | 4.89352 | 6.94534 | 6.17059 | 6.5154 | 5.95075 | 6.81701 | 7.01823 |
| PIEZO2 | Q9H5I5-4 | 6.31104 | 6.45693 | 6.10837 | 5.74362 | 6.13082 | 6.4966 | 6.36525 | 6.03196 | 6.55951 | 6.19724 | 5.9963 | 6.13078 | 6.84099 | 6.04106 | 6.43941 | 6.1501 |
| ERCC6 | P0DP91 | 5.73644 | 6.19275 | 6.45501 | 6.66666 | 6.0446 | 5.59232 | 6.35935 | 6.19257 | 6.50672 | 5.72682 | 6.38522 | 6.20086 | 6.32761 | 6.3443 | 6.38237 | 6.88641 |
| MATN2 | O00339 | 6.26655 | 7.13712 | 6.24391 | 7.11811 | 6.47569 | 5.4143 | 6.66035 | 6.2445 | 6.09891 | 6.18225 | 5.61233 | 6.56337 | 5.97819 | 4.66508 | 6.20279 | 7.13655 |
| ALG13 | Q9NP73 | 6.37448 | 6.3428 | 6.87898 | 5.93056 | 6.30133 | 6.29447 | 6.85127 | 6.04514 | 6.67203 | 6.22572 | 6.33721 | 6.19659 | 5.63471 | 5.68406 | 6.13984 | 6.09079 |
| CCDC112 | Q8NEF3-2 | 4.43919 | 3.73919 | 16.1869 | 5.95908 | 11.5285 | 9.08698 | 13.9234 | 4.46574 | 4.62182 | 3.16798 | 6.20182 | 3.83651 | 2.52498 | 2.88708 | 3.68424 | 3.54671 |
| CEND1 | Q8N111 | 5.60804 | 4.83503 | 5.92373 | 5.7246 | 6.36391 | 5.8194 | 6.05933 | 5.77954 | 5.40521 | 7.38867 | 6.49552 | 7.32163 | 7.40289 | 6.94632 | 6.20369 | 6.7225 |
| SNAP25 | P60880-2 | 5.89902 | 4.89367 | 6.46917 | 5.71235 | 5.5546 | 5.92624 | 6.35507 | 5.02697 | 5.67848 | 7.06298 | 6.10155 | 6.48834 | 7.27636 | 7.65119 | 6.12984 | 6.68415 |
| SEPTIN3 | Q9UH03 | 5.82162 | 6.38193 | 7.43475 | 6.53154 | 6.15716 | 6.11083 | 5.47265 | 6.09667 | 5.34877 | 5.78577 | 6.30454 | 6.92341 | 6.28537 | 6.59396 | 5.87671 | 6.87433 |
| SMPD2 | O60906 | 6.03356 | 6.53513 | 5.75946 | 6.54788 | 5.73904 | 6.43372 | 6.43052 | 6.30554 | 5.84644 | 5.74584 | 7.4484 | 6.04145 | 6.85328 | 5.72028 | 5.95366 | 6.60579 |
| SELENOH | Q8IZQ5 | 5.79182 | 5.99586 | 5.26589 | 5.27937 | 5.63388 | 5.59531 | 6.04566 | 5.66334 | 6.25323 | 6.12434 | 5.95988 | 5.85208 | 7.21998 | 7.92353 | 8.27766 | 7.11817 |
| TGIF2 | Q9GZN2 | 6.22564 | 6.39557 | 7.17385 | 6.53259 | 7.91685 | 6.64051 | 6.30305 | 6.75027 | 7.5971 | 8.524 | 7.74561 | 6.96257 | 4.89169 | 3.82665 | 3.03478 | 4.37827 |
| NOMO1 | Q15155 | 5.32291 | 6.53105 | 6.36535 | 5.58144 | 6.92095 | 5.15273 | 6.84828 | 5.55036 | 6.48461 | 6.38308 | 5.11772 | 7.37981 | 7.3986 | 5.79608 | 6.60651 | 6.5605 |
| CHURC1 | Q8WUH1-2 | 6.00006 | 4.88766 | 6.08555 | 6.33899 | 6.13695 | 6.94447 | 7.64662 | 6.8512 | 7.19463 | 6.09677 | 7.19956 | 7.74682 | 4.09216 | 6.05405 | 5.24924 | 5.47527 |
| KIF23 | Q02241-3 | 7.16709 | 7.08551 | 7.05588 | 6.758 | 6.33044 | 6.45711 | 6.76632 | 5.75579 | 5.63592 | 6.3005 | 6.30042 | 6.04734 | 5.50787 | 5.28506 | 5.31425 | 6.23251 |
| BTBD2 | Q9BX70 | 6.30192 | 6.39443 | 7.74006 | 7.32826 | 5.91881 | 6.23896 | 6.16633 | 6.14987 | 6.57428 | 7.06891 | 7.01227 | 6.98886 | 5.02773 | 5.32646 | 5.30211 | 4.46074 |
| RNF130 | Q86XS8 | 8.09081 | 8.25074 | 7.37783 | 7.80382 | 5.44084 | 6.64064 | 6.13414 | 5.69985 | 7.94988 | 8.38546 | 7.45815 | 7.76984 | 2.95321 | 3.21067 | 3.3385 | 3.49564 |
| CDC37L1 | Q7L3B6 | 6.57679 | 5.72636 | 6.38269 | 5.30225 | 5.94503 | 5.37908 | 5.40508 | 6.95499 | 5.66063 | 6.51814 | 6.51693 | 7.31507 | 5.9765 | 6.90761 | 6.78991 | 6.64292 |
| CDSN | Q15517 | 6.49241 | 5.9627 | 5.60398 | 7.72332 | 7.39575 | 7.81673 | 6.08057 | 5.26651 | 6.05551 | 5.42659 | 5.09541 | 5.87706 | 8.50732 | 5.84623 | 5.29453 | 5.55538 |
| CCND1 | P24385 | 6.86223 | 6.10594 | 8.13413 | 7.45061 | 7.34945 | 7.48192 | 7.5709 | 7.7706 | 6.49112 | 6.60135 | 7.7499 | 7.08258 | 3.01642 | 3.44997 | 3.22849 | 3.65442 |
| TOP1MT | Q969P6 | 6.30721 | 6.23078 | 5.96287 | 6.89542 | 6.20611 | 6.46785 | 7.51241 | 6.66695 | 5.72912 | 6.07143 | 6.76472 | 6.72169 | 5.57109 | 5.08218 | 5.60834 | 6.20184 |
| FUNDC2 | Q9BWH2 | 6.79321 | 6.86125 | 5.73009 | 6.8112 | 6.53905 | 6.06143 | 6.59719 | 5.91734 | 6.60766 | 6.28431 | 6.13108 | 6.28192 | 5.86307 | 5.25838 | 6.09349 | 6.16933 |
| FAM120A | A0A494C0Y9 | 6.29972 | 6.66094 | 6.80477 | 6.82205 | 6.01001 | 6.43793 | 6.13644 | 6.36984 | 6.59815 | 6.67423 | 6.56014 | 6.70318 | 5.27447 | 5.79251 | 5.35381 | 5.5018 |
| PPP1R26 | Q5T8A7 | 5.6037 | 7.63733 | 5.31106 | 4.91214 | 5.80438 | 5.68092 | 6.90557 | 6.33452 | 5.83192 | 5.47065 | 6.63868 | 6.37806 | 6.42197 | 6.49284 | 5.70425 | 8.87202 |
| ALDH18A1 | P54886-2 | 6.22772 | 6.25141 | 5.92717 | 6.5676 | 6.58596 | 5.17026 | 6.40731 | 6.11854 | 5.79313 | 6.47036 | 6.1438 | 6.78055 | 6.27256 | 6.45437 | 6.85532 | 5.97392 |
| STK17B | O94768 | 6.31511 | 6.32648 | 6.69749 | 5.818 | 6.02131 | 6.0122 | 5.87989 | 6.51673 | 6.12905 | 6.37407 | 7.25785 | 7.12117 | 5.26289 | 6.25595 | 6.10216 | 5.90967 |
| DLX1 | P56177 | 6.75386 | 6.68134 | 6.21955 | 7.15965 | 6.40369 | 6.43359 | 6.14452 | 6.06868 | 6.08761 | 6.26632 | 6.21199 | 6.54332 | 5.73954 | 5.67559 | 5.89961 | 5.71113 |
| TRAF3IP1 | Q8TDR0 | 5.7307 | 6.13966 | 6.90429 | 6.2024 | 5.51853 | 6.20339 | 6.37969 | 6.24121 | 6.55844 | 6.66996 | 6.87842 | 6.47662 | 5.28504 | 6.44566 | 6.25827 | 6.10773 |
| ACTN4 | F5GXS2 | 6.8805 | 6.69769 | 5.71491 | 6.12214 | 6.54507 | 6.44364 | 5.24468 | 6.00849 | 6.53713 | 5.78356 | 5.39816 | 6.60485 | 6.31716 | 6.12065 | 6.65308 | 6.92829 |
| PRPF40A | O75400-3 | 6.16526 | 5.9843 | 6.23543 | 6.24765 | 6.25356 | 5.56068 | 6.51525 | 5.36391 | 6.519 | 5.81427 | 6.78415 | 5.78854 | 7.04359 | 5.95596 | 7.30636 | 6.46207 |
| MAFF | Q9ULX9 | 6.41252 | 5.79023 | 5.77111 | 5.73027 | 6.6934 | 6.88316 | 6.67412 | 5.26475 | 6.01589 | 6.34984 | 6.15681 | 6.06835 | 6.27381 | 5.16561 | 7.1022 | 7.64794 |
| MAGEA1 | P43355 | 7.20543 | 6.58169 | 6.95974 | 6.49878 | 6.31762 | 6.61183 | 6.29932 | 6.6671 | 6.63801 | 7.05961 | 6.5435 | 6.52289 | 5.04063 | 4.87451 | 4.84461 | 5.33473 |
| PHC1 | P78364 | 7.09949 | 6.68634 | 6.05938 | 6.74348 | 6.25239 | 6.03597 | 6.61793 | 6.10249 | 6.13348 | 6.72318 | 6.74526 | 6.28494 | 6.03017 | 5.09066 | 5.82949 | 5.56534 |
| SAC3D1 | A6NKF1 | 7.86404 | 7.12761 | 7.08619 | 7.83325 | 6.07257 | 6.34829 | 6.51143 | 6.80637 | 5.90682 | 6.78723 | 6.64944 | 6.34419 | 4.30368 | 4.14059 | 4.99412 | 5.22418 |
| CHST7 | Q9NS84 | 6.74317 | 6.75373 | 5.8819 | 6.99799 | 6.29942 | 6.73331 | 6.36603 | 5.54035 | 6.49451 | 6.09819 | 6.484 | 6.43363 | 5.65335 | 5.60529 | 5.87818 | 6.03693 |
| ZNF850 | A8MQ14 | 6.2328 | 5.63983 | 7.68326 | 7.80351 | 6.13431 | 5.6142 | 5.96055 | 5.53806 | 5.83906 | 6.99721 | 5.98424 | 7.21462 | 5.44975 | 5.13668 | 6.63125 | 6.14067 |
| ZNF92 | Q03936 | 7.10241 | 7.06286 | 7.5624 | 7.20798 | 6.34176 | 5.71786 | 7.52272 | 6.57717 | 5.7778 | 6.61133 | 6.84127 | 6.32769 | 4.99588 | 4.86798 | 5.01553 | 4.46735 |
| ZNF749 | O43361 | 6.24763 | 7.26479 | 6.55393 | 6.27324 | 5.96487 | 6.08291 | 7.18321 | 6.00335 | 6.18275 | 5.77188 | 5.81678 | 5.85882 | 6.37968 | 6.51114 | 5.84727 | 6.05774 |
| ZNF260 | Q3ZCT1 | 6.17469 | 6.54381 | 5.83684 | 6.5526 | 6.08376 | 6.76319 | 6.15878 | 6.09315 | 5.83796 | 6.43341 | 6.06859 | 6.07237 | 6.02132 | 6.04235 | 7.08434 | 6.23284 |
| ZNF16 | P17020 | 6.47673 | 6.45476 | 6.10857 | 6.60201 | 6.28592 | 6.5742 | 6.25117 | 4.82741 | 6.24416 | 6.13598 | 6.63766 | 6.39052 | 6.89834 | 5.90755 | 5.86371 | 6.34133 |
| ZNF445 | P59923 | 6.5822 | 7.53977 | 6.40771 | 7.3179 | 6.36832 | 6.48004 | 6.31486 | 5.99667 | 5.7221 | 6.94278 | 6.80099 | 5.28526 | 5.21625 | 5.08007 | 5.68905 | 6.25603 |
| ZNF354A | O60765 | 6.19609 | 6.41178 | 6.2546 | 7.18623 | 6.03352 | 5.93442 | 6.94767 | 5.93438 | 7.18186 | 6.26421 | 6.73833 | 7.22485 | 5.20374 | 5.39593 | 5.47517 | 5.61724 |
| ZNF836 | Q6ZNA1 | 6.43056 | 6.27833 | 5.939 | 6.39126 | 6.03026 | 6.00301 | 6.20958 | 5.90448 | 6.2784 | 6.37586 | 6.36662 | 6.49186 | 6.42923 | 6.13613 | 6.58879 | 6.14662 |
| ZNF180 | Q9UJW8 | 6.43531 | 5.73391 | 5.91384 | 6.71006 | 6.50531 | 6.0412 | 6.24545 | 5.97099 | 6.36404 | 6.86986 | 6.03723 | 6.41156 | 6.53867 | 6.34429 | 5.73762 | 6.14065 |
| ZNF7 | P17097-2 | 5.95789 | 6.62323 | 5.69747 | 6.44707 | 6.80447 | 5.89288 | 6.64644 | 5.95284 | 5.77657 | 6.04428 | 5.64324 | 5.35587 | 6.5405 | 6.26401 | 6.79259 | 7.56065 |
| ZNF33B | Q06732 | 6.47701 | 5.95649 | 6.05765 | 6.22856 | 5.5022 | 6.05551 | 5.81338 | 5.56655 | 6.54409 | 6.36883 | 6.18495 | 6.37457 | 6.69557 | 6.08825 | 6.44364 | 7.64274 |
| FADS3 | Q9Y5Q0 | 6.40776 | 6.51875 | 5.96819 | 6.73703 | 6.08923 | 6.13192 | 6.1836 | 5.85226 | 6.67991 | 6.10616 | 6.68034 | 6.6121 | 5.96013 | 6.01662 | 6.34094 | 5.71505 |
| SNAPC3 | Q92966 | 5.16747 | 5.85179 | 6.01748 | 6.51388 | 5.67239 | 6.00045 | 6.6044 | 6.16684 | 6.5231 | 5.80885 | 7.16433 | 6.29785 | 6.54812 | 6.13627 | 6.62738 | 6.8994 |
| CCNDBP1 | O95273 | 7.55078 | 5.96234 | 7.72356 | 6.18002 | 6.09356 | 6.44927 | 5.90872 | 6.57776 | 7.01159 | 7.75025 | 7.57028 | 7.76341 | 4.00928 | 4.68217 | 4.12641 | 4.64059 |
| ZNF564 | Q8TBZ8 | 6.26735 | 7.25956 | 5.63044 | 6.05291 | 6.14288 | 6.53561 | 5.337 | 6.11015 | 5.37089 | 6.84578 | 6.62819 | 6.35102 | 5.91725 | 7.01722 | 6.39163 | 6.14212 |
| PEDS1 | A5PLL7 | 7.40041 | 6.87368 | 5.55369 | 6.50793 | 7.17606 | 5.40518 | 6.65271 | 5.80932 | 6.64378 | 6.52821 | 6.93138 | 6.47043 | 5.14141 | 6.48463 | 5.5708 | 5.85038 |
| PEDS1-UBE2V1 | I3L0A0 | 7.0808 | 5.37012 | 6.91188 | 5.50984 | 6.62097 | 6.40528 | 5.94458 | 6.753 | 5.65067 | 7.0348 | 6.47832 | 7.168 | 6.17479 | 5.80151 | 5.36595 | 5.72947 |
| DYRK1B | Q9Y463 | 6.45787 | 6.57869 | 5.93525 | 6.06323 | 6.02311 | 6.33936 | 5.95372 | 6.34064 | 6.02321 | 7.33435 | 6.24233 | 5.85479 | 6.3137 | 6.76371 | 6.26651 | 5.50953 |
| ZNF408 | Q9H9D4 | 5.74874 | 6.87911 | 6.22613 | 7.05031 | 6.02383 | 7.01824 | 5.95895 | 7.16987 | 5.6702 | 6.36236 | 7.04014 | 6.71103 | 6.24347 | 5.55025 | 4.97881 | 5.36856 |
| RAB30 | Q15771 | 6.64134 | 6.95919 | 6.455 | 6.57587 | 6.2503 | 5.59927 | 6.73659 | 6.06407 | 6.34702 | 6.33384 | 6.36995 | 6.26978 | 5.81448 | 5.61986 | 5.89619 | 6.06725 |
| RAB43 | Q86YS6 | 6.12867 | 6.32234 | 6.578 | 6.79789 | 6.15082 | 5.62107 | 6.59024 | 6.28649 | 6.27252 | 6.22627 | 6.63632 | 6.12936 | 5.61729 | 6.02488 | 6.79811 | 5.81975 |
| ZFP30 | Q9Y2G7 | 6.64944 | 7.1855 | 5.93813 | 7.34285 | 6.44785 | 6.37579 | 6.7533 | 6.46166 | 6.46353 | 5.75952 | 5.78599 | 6.17291 | 6.06276 | 5.32202 | 5.64875 | 5.62999 |
| FITM2 | Q8N6M3 | 6.71032 | 6.38969 | 5.60166 | 6.51211 | 6.57869 | 5.52235 | 6.12687 | 6.29041 | 6.20229 | 6.06104 | 6.30955 | 5.90584 | 6.66928 | 6.10523 | 6.56903 | 6.44565 |
| ZFP14 | Q9HCL3 | 6.76056 | 6.69417 | 6.26151 | 7.04416 | 6.66725 | 6.17559 | 7.0562 | 6.74806 | 6.88558 | 5.98554 | 5.77483 | 6.49363 | 5.67381 | 5.33812 | 4.8083 | 5.63267 |
| DCLRE1B | Q9H816 | 6.65577 | 6.77248 | 6.31837 | 6.39959 | 5.86177 | 6.02502 | 6.79115 | 6.32136 | 6.29107 | 6.64145 | 6.48942 | 6.35596 | 5.32645 | 6.23056 | 5.14653 | 6.37305 |
| FANCL | Q9NW38-2 | 6.71802 | 5.51118 | 5.9545 | 6.3792 | 6.27216 | 6.53105 | 6.09864 | 5.56213 | 6.35635 | 7.08103 | 6.43541 | 6.39445 | 6.91289 | 4.74752 | 6.72204 | 6.26452 |
| WTIP | A6NIX2 | 7.14403 | 6.64205 | 7.31026 | 7.12689 | 5.95181 | 6.38683 | 6.36939 | 6.25698 | 6.67423 | 5.89099 | 6.65223 | 6.53472 | 5.56648 | 4.91714 | 5.02225 | 5.45471 |
| PSIP1 | V9GYT7 | 5.82303 | 6.18966 | 6.71075 | 5.53639 | 6.23843 | 5.64231 | 6.7598 | 5.49454 | 6.77324 | 6.98543 | 5.83366 | 6.08966 | 7.21438 | 6.36326 | 5.82666 | 6.51881 |
| LYSMD1 | Q96S90 | 5.42609 | 4.88584 | 5.96959 | 4.47619 | 6.07194 | 5.42142 | 5.14686 | 5.97386 | 6.18522 | 6.65216 | 7.44473 | 6.86345 | 7.86023 | 7.32453 | 7.36738 | 6.93051 |
| HOXC8 | P31273 | 6.04178 | 6.14882 | 6.4487 | 5.7857 | 5.80023 | 6.48831 | 5.69678 | 6.06325 | 6.4899 | 6.77021 | 6.88434 | 7.04487 | 6.16114 | 5.95797 | 6.16118 | 6.05683 |
| JMJD8 | Q96S16 | 6.98787 | 7.0666 | 6.30421 | 6.999 | 5.89447 | 6.31871 | 6.27606 | 6.6459 | 6.93494 | 7.04145 | 7.41236 | 6.83316 | 5.10745 | 4.43013 | 4.56064 | 5.18705 |
| ACTBL2 | Q562R1 | 6.54899 | 7.39856 | 7.20573 | 6.66394 | 7.65436 | 7.65868 | 5.99229 | 5.34408 | 5.28247 | 5.96556 | 5.89178 | 5.55295 | 6.88661 | 5.69102 | 5.4295 | 4.83348 |
| BMP1 | P13497 | 7.32095 | 6.76649 | 5.44775 | 6.81743 | 5.7651 | 5.9968 | 7.66934 | 5.6228 | 6.73728 | 6.73398 | 5.89301 | 5.81466 | 6.56493 | 4.73546 | 6.02295 | 6.09105 |
| BIRC3 | Q13489 | 5.71669 | 6.09716 | 6.40981 | 5.32869 | 6.80678 | 6.75297 | 5.84732 | 6.86051 | 7.62205 | 6.05451 | 7.12908 | 6.84912 | 6.11672 | 5.51259 | 5.38335 | 5.51265 |
| CYBC1 | Q9BQA9 | 6.54665 | 6.55922 | 5.66973 | 6.46194 | 6.21574 | 6.08739 | 6.30177 | 5.33174 | 5.94408 | 5.9469 | 5.98649 | 5.62381 | 6.98235 | 6.68933 | 7.34718 | 6.30567 |
| ALDH9A1 | P49189 | 6.04939 | 6.29866 | 5.61916 | 6.25064 | 6.02617 | 6.01557 | 7.24422 | 5.17964 | 7.4505 | 5.03399 | 5.53943 | 5.93128 | 6.44866 | 6.6408 | 6.27066 | 8.00123 |
| C19orf54 | Q5BKX5 | 6.40693 | 6.23572 | 6.72288 | 6.52552 | 6.59709 | 6.16792 | 5.48349 | 7.01572 | 6.39559 | 6.50852 | 6.90456 | 6.77851 | 6.6895 | 5.51609 | 5.32153 | 5.63041 |
| NR1H2 | P55055 | 6.12409 | 6.12182 | 5.95085 | 6.44304 | 5.9717 | 6.03813 | 6.32034 | 5.93774 | 6.13815 | 6.41118 | 6.42765 | 6.31131 | 6.20006 | 6.06986 | 6.96912 | 6.56495 |
| MTA1 | Q13330-3 | 6.62994 | 7.17481 | 6.35386 | 6.68826 | 6.40353 | 5.16218 | 6.44294 | 6.03992 | 5.64152 | 6.46513 | 5.93499 | 5.88732 | 6.51415 | 5.79779 | 6.65631 | 6.15736 |
| MTMR11 | A4FU01 | 6.98287 | 6.12417 | 6.00699 | 6.29328 | 6.70722 | 6.68217 | 6.98785 | 5.91865 | 6.42166 | 5.41731 | 6.01727 | 5.77153 | 5.88815 | 5.86315 | 6.5429 | 6.37484 |
| AKAP5 | P24588 | 7.26921 | 6.2311 | 6.54763 | 6.04547 | 6.04606 | 6.48015 | 5.98484 | 5.77527 | 5.75807 | 5.39916 | 5.83596 | 7.17299 | 5.90832 | 6.27595 | 6.58117 | 6.68864 |
| HIC2 | Q96JB3 | 6.01767 | 5.17317 | 6.94405 | 7.28456 | 6.94277 | 7.09166 | 6.51934 | 6.27103 | 6.40374 | 5.89741 | 6.45189 | 6.77166 | 6.27567 | 4.96418 | 6.08269 | 5.90851 |
| YRDC | Q86U90 | 6.12469 | 5.91379 | 6.05352 | 6.45969 | 7.1863 | 5.36794 | 5.98736 | 7.18202 | 6.45208 | 6.83799 | 6.16543 | 7.42494 | 6.21212 | 5.71086 | 5.44187 | 5.47939 |
| POGK | Q9P215 | 5.4736 | 6.1667 | 6.8599 | 5.58932 | 6.18892 | 7.18263 | 6.77681 | 5.9143 | 5.7092 | 6.50082 | 6.11712 | 5.15436 | 5.85597 | 6.7158 | 6.67462 | 7.11993 |
| ERICH1 | Q86X53 | 5.25645 | 5.76243 | 5.71797 | 5.38247 | 5.57531 | 6.00872 | 5.93644 | 5.76265 | 5.85822 | 6.34114 | 6.51654 | 6.57838 | 6.6556 | 7.72122 | 7.83272 | 7.29373 |
| MT-ND4 | P03905 | 5.64488 | 6.47849 | 5.78622 | 5.8847 | 6.0943 | 6.58093 | 6.52979 | 6.80996 | 6.00831 | 6.55503 | 5.52939 | 5.51081 | 6.5613 | 6.78138 | 6.69817 | 6.54635 |
| POLR1D | P0DPB5 | 6.75773 | 6.54878 | 5.89772 | 6.98169 | 6.5994 | 5.99594 | 6.33612 | 6.76635 | 6.81014 | 5.85732 | 6.56642 | 6.21673 | 5.18564 | 5.42404 | 5.71013 | 6.34585 |
| AP1S2 | P56377 | 6.04397 | 6.94346 | 7.01357 | 6.60094 | 6.17746 | 6.78697 | 6.67783 | 5.92261 | 6.5391 | 6.25008 | 6.3489 | 5.98016 | 5.55315 | 5.7857 | 5.66208 | 5.71402 |
| GABARAPL1 | Q9H0R8-2 | 6.51946 | 6.39211 | 6.39907 | 5.8207 | 7.11333 | 7.32825 | 6.53233 | 6.84261 | 6.89053 | 6.3279 | 5.8464 | 6.13625 | 5.69356 | 5.45677 | 5.29165 | 5.40909 |
| CEP57 | Q86XR8 | 6.3452 | 7.94214 | 6.18921 | 6.59952 | 5.98712 | 6.83389 | 7.73076 | 7.71689 | 6.92988 | 7.44704 | 6.57286 | 6.23745 | 4.35759 | 5.3436 | 4.05731 | 3.70953 |
| PYROXD1 | Q8WU10 | 6.52498 | 5.9276 | 7.21137 | 5.86489 | 6.1609 | 6.45655 | 6.0295 | 6.35162 | 7.01593 | 6.16479 | 6.78111 | 7.21046 | 5.38288 | 5.10954 | 6.2891 | 5.51876 |
| XRCC1 | F5H8D7 | 5.78275 | 5.62202 | 5.89996 | 5.81036 | 5.95599 | 5.78298 | 6.08134 | 6.23565 | 6.25736 | 6.28908 | 6.60671 | 5.963 | 6.39012 | 7.24439 | 6.4224 | 7.65589 |
| KYAT1 | Q16773 | 6.03566 | 5.84171 | 6.65736 | 5.4901 | 6.47074 | 7.01509 | 6.26237 | 6.55138 | 6.23222 | 6.38554 | 6.88586 | 6.78082 | 5.87037 | 5.90198 | 5.97637 | 5.64246 |
| CPQ | Q9Y646 | 5.8957 | 6.28021 | 5.8653 | 6.31503 | 6.1919 | 6.89359 | 6.91524 | 6.28795 | 5.11531 | 6.40306 | 6.31152 | 6.16803 | 5.76138 | 6.03566 | 6.72107 | 6.83903 |
| FGF1 | P05230 | 6.06001 | 6.06119 | 7.16735 | 6.41604 | 6.69296 | 6.22183 | 7.64424 | 7.8501 | 6.159 | 5.3544 | 7.24826 | 7.84297 | 4.99423 | 4.70977 | 5.00598 | 4.57169 |
| FGF2 | P09038 | 5.99454 | 6.50029 | 6.07103 | 6.05165 | 6.69527 | 6.27955 | 6.14064 | 6.19906 | 6.36051 | 5.92097 | 6.20375 | 5.71454 | 6.82203 | 6.48278 | 6.62428 | 5.93911 |
| RGPD5 | Q99666 | 6.72382 | 5.86391 | 6.50801 | 7.26216 | 6.18904 | 6.00978 | 6.06888 | 6.08672 | 6.61381 | 6.04206 | 5.70128 | 5.59079 | 5.73157 | 6.73487 | 6.02546 | 6.84784 |
| ZC3H14 | Q6PJT7-9 | 5.97156 | 6.20198 | 5.81596 | 6.81941 | 7.3473 | 6.52336 | 6.18152 | 6.07387 | 5.94649 | 5.70882 | 6.63899 | 6.87295 | 6.49648 | 5.5603 | 5.62695 | 6.21406 |
| REL | Q04864 | 6.25948 | 6.90387 | 7.0456 | 5.87532 | 5.6667 | 6.38229 | 6.75495 | 7.15723 | 6.86307 | 6.12295 | 5.7783 | 6.86631 | 5.34432 | 6.09523 | 5.55962 | 5.32477 |
| SMAD1 | Q15797 | 6.31957 | 5.72205 | 6.77194 | 6.17189 | 5.78483 | 6.6348 | 5.69209 | 6.68416 | 7.63476 | 6.08367 | 5.99803 | 6.43459 | 5.65836 | 6.37698 | 5.75622 | 6.27607 |
| SMAD9 | O15198 | 6.63087 | 6.45878 | 6.54415 | 6.42928 | 6.35926 | 6.58455 | 5.98815 | 5.94992 | 6.05025 | 6.66764 | 6.28257 | 6.47911 | 5.93371 | 6.0168 | 6.08642 | 5.53852 |
| ULK3 | Q6PHR2 | 5.91867 | 5.91658 | 7.34654 | 6.27294 | 6.41633 | 6.8 | 6.19976 | 5.70989 | 6.13028 | 6.84775 | 6.92786 | 6.77847 | 5.29239 | 5.75109 | 6.37036 | 5.32108 |

| | | | | | | | | | | | | | | | | | |
|---|---|---|---|---|---|---|---|---|---|---|---|---|---|---|---|---|---|
| PDK3 | Q15120-2 | 6.69968 | 6.20773 | 5.4764 | 7.19445 | 6.21725 | 6.61444 | 6.25605 | 6.60335 | 6.05037 | 6.15776 | 6.78893 | 5.73208 | 6.28635 | 5.65296 | 6.68618 | 5.37603 |
| KHDRBS3 | O75525 | 7.01569 | 6.44668 | 5.85694 | 6.64845 | 6.15793 | 6.02177 | 6.63889 | 6.27027 | 6.42075 | 6.00437 | 5.94507 | 6.00999 | 6.45231 | 5.43692 | 6.14345 | 6.53052 |
| RPL10 | F8W7C6 | 5.9353 | 5.97332 | 5.78588 | 5.43095 | 5.15395 | 6.54556 | 7.07994 | 7.81358 | 7.16532 | 6.19313 | 7.44498 | 5.88579 | 6.27483 | 6.07035 | 4.95453 | 6.29258 |
| FAM133B | Q5BKY9 | 5.63401 | 5.18558 | 6.2451 | 5.03427 | 5.00482 | 5.32105 | 5.36149 | 5.41533 | 6.10635 | 5.65749 | 7.06226 | 6.48619 | 7.71129 | 8.0586 | 7.98318 | 7.733 |
| WDR46 | A0A0G2JJL1 | 5.67435 | 6.08369 | 5.93437 | 6.34078 | 5.67145 | 5.42659 | 6.18259 | 6.38265 | 6.53161 | 5.84084 | 6.07526 | 6.1813 | 6.50022 | 6.74222 | 7.02433 | 7.40774 |
| COA8 | Q96IL0 | 6.46017 | 5.62309 | 6.95151 | 7.36866 | 7.25942 | 6.45511 | 5.17526 | 6.55465 | 6.61643 | 5.95619 | 5.79619 | 6.9121 | 6.43544 | 5.14454 | 5.03202 | 6.2592 |
| DHRS4 | Q9BTZ2 | 6.34268 | 7.08716 | 6.02812 | 6.44945 | 6.50118 | 6.57259 | 5.9896 | 6.72256 | 5.75791 | 6.42172 | 5.73156 | 5.56486 | 6.3386 | 6.24763 | 6.43867 | 5.80571 |
| PXMP2 | Q9NR77 | 6.24925 | 6.13165 | 5.72164 | 6.70495 | 7.29762 | 6.54696 | 5.86132 | 6.43785 | 6.06123 | 6.24348 | 5.18373 | 5.94235 | 6.53403 | 6.13919 | 6.3396 | 6.60514 |
| PNKD | Q8N490-2 | 6.04889 | 5.40277 | 5.59191 | 6.49831 | 6.76099 | 6.0862 | 6.27696 | 5.91122 | 5.96463 | 6.10294 | 5.76864 | 6.03251 | 7.21916 | 6.9278 | 6.88732 | 6.51974 |
| VKORC1 | Q9BQB6 | 5.72554 | 6.07307 | 5.84178 | 6.35751 | 6.16656 | 6.29814 | 6.41291 | 5.98013 | 6.12261 | 6.06388 | 6.46317 | 5.90838 | 6.96483 | 6.71057 | 6.42818 | 6.48274 |
| SIVA1 | O15304 | 6.16498 | 6.46195 | 7.47523 | 5.73949 | 5.77762 | 6.5003 | 6.00222 | 6.25069 | 6.51706 | 7.03525 | 7.25333 | 7.23363 | 4.77276 | 5.4076 | 5.82999 | 5.57789 |
| FAM126A | Q9BYI3-3 | 6.00499 | 5.80624 | 5.00375 | 5.87311 | 6.56874 | 7.07049 | 6.21275 | 5.41861 | 6.09276 | 6.9012 | 7.81436 | 5.83825 | 6.8885 | 6.9151 | 6.1182 | 5.47295 |
| ZNF608 | Q9ULD9 | 6.49416 | 7.92746 | 6.04741 | 6.24191 | 6.36294 | 6.80493 | 4.89719 | 5.03498 | 6.38085 | 7.12613 | 5.87284 | 5.36914 | 6.2415 | 6.75136 | 6.50703 | 5.94016 |
| TMOD1 | P28289 | 6.49788 | 6.5457 | 6.08501 | 5.5232 | 6.48836 | 6.23839 | 6.1242 | 6.48532 | 5.89455 | 6.54467 | 6.65653 | 6.12672 | 6.57267 | 6.37642 | 5.72351 | 5.75687 |
| DLG3 | Q92796-2 | 6.49676 | 6.47545 | 5.62111 | 6.6633 | 6.18802 | 6.49473 | 5.94698 | 6.52698 | 5.77857 | 5.88267 | 6.28566 | 6.66727 | 6.008 | 6.02331 | 6.89008 | 6.0511 |
| MAP4K2 | Q12851 | 6.63947 | 6.27999 | 6.32008 | 6.27124 | 5.65488 | 6.25538 | 6.44943 | 6.60916 | 5.99509 | 6.56347 | 7.19745 | 6.42329 | 5.75521 | 6.20744 | 5.69691 | 5.6815 |
| ZNF207 | O43670-2 | 5.87306 | 4.65201 | 7.11101 | 6.54032 | 6.49696 | 5.04185 | 5.95822 | 6.0763 | 6.92976 | 5.4115 | 6.43048 | 6.22786 | 7.05552 | 7.05979 | 6.52295 | 6.61241 |
| FTL | P02792 | 8.30203 | 7.22846 | 6.62593 | 6.88623 | 5.47758 | 4.95109 | 4.91862 | 7.15309 | 6.6897 | 6.66962 | 6.31298 | 6.72325 | 3.84082 | 6.40366 | 6.22391 | 5.59303 |
| ANAPC16 | Q96DE5 | 6.34097 | 5.39837 | 6.20947 | 4.54243 | 5.55545 | 5.27414 | 5.75067 | 6.7791 | 5.19313 | 6.33811 | 5.68043 | 7.14629 | 5.18806 | 8.48919 | 7.85341 | 8.16076 |
| PIK3R3 | Q92569 | 6.34257 | 6.15222 | 6.88221 | 6.53518 | 5.83773 | 7.52639 | 6.01585 | 6.61321 | 6.4887 | 7.6087 | 8.17698 | 7.3792 | 4.55015 | 4.96753 | 4.11693 | 4.80646 |
| VGLL4 | Q14135-4 | 4.79752 | 4.87995 | 6.78178 | 4.66696 | 6.08882 | 5.15715 | 5.51902 | 6.98545 | 5.42833 | 7.72004 | 7.37927 | 6.82337 | 6.86473 | 8.24766 | 6.15356 | 6.50639 |
| SERPINI1 | Q99574 | 6.24176 | 6.30811 | 6.75755 | 7.02436 | 5.97776 | 5.54594 | 6.44488 | 6.27177 | 5.9844 | 6.75004 | 5.73121 | 6.47323 | 6.46824 | 5.53151 | 5.69199 | 6.79724 |
| NMNAT3 | Q96T66 | 6.30118 | 5.99758 | 5.98651 | 6.91679 | 5.93016 | 6.1111 | 5.79021 | 5.93537 | 5.82056 | 6.59358 | 6.66784 | 6.52265 | 6.70603 | 6.37579 | 6.06956 | 6.27509 |
| TMEM242 | Q9NWH2 | 6.66621 | 5.81755 | 5.33334 | 6.09223 | 6.44057 | 6.07481 | 7.11044 | 7.23055 | 6.75719 | 6.08022 | 6.30845 | 5.79182 | 6.51718 | 5.41965 | 6.51828 | 5.84151 |
| ING2 | Q9H160 | 5.88565 | 7.95937 | 5.11401 | 7.15777 | 4.87444 | 6.65131 | 5.80168 | 6.56191 | 5.59224 | 6.42015 | 5.10886 | 6.43055 | 5.54177 | 7.37486 | 6.17693 | 7.34848 |
| SRF | P11831 | 4.8758 | 5.08691 | 6.53222 | 5.72844 | 5.18136 | 6.53945 | 5.66959 | 5.47033 | 6.01829 | 5.59646 | 7.73907 | 5.84019 | 7.21001 | 7.64161 | 7.25394 | 7.61632 |
| EEF1AKMT4-ECE2 | P0DPD8 | 6.61876 | 6.38844 | 6.94473 | 6.10514 | 5.74115 | 6.74569 | 6.61541 | 6.94159 | 6.22034 | 6.92911 | 7.0794 | 6.90638 | 5.28556 | 5.01271 | 4.63676 | 5.82883 |
| METTL26 | Q96S19 | 5.6191 | 5.31317 | 5.61823 | 4.91506 | 5.07126 | 4.94511 | 5.50322 | 6.80758 | 4.68223 | 6.43852 | 5.32381 | 6.68089 | 4.69455 | 11.2569 | 7.48694 | 9.64341 |
| EPDR1 | Q9UM22 | 6.48318 | 6.08091 | 5.47497 | 6.38629 | 5.6574 | 6.05262 | 6.10686 | 6.34209 | 6.19822 | 6.11134 | 6.69179 | 6.03916 | 6.94691 | 6.60925 | 6.10066 | 6.71834 |
| FAM171A1 | Q5VUB5 | 6.42441 | 6.37784 | 5.98505 | 6.53101 | 5.93345 | 5.99932 | 6.3493 | 6.14863 | 6.48962 | 6.06426 | 5.9657 | 6.40321 | 6.48486 | 6.16007 | 6.33987 | 6.3434 |
| TPCN1 | Q9ULQ1-3 | 6.26091 | 6.91392 | 6.06594 | 6.19599 | 6.35878 | 6.22347 | 6.53696 | 6.25462 | 6.1295 | 5.4692 | 5.84614 | 5.98285 | 6.4116 | 5.89885 | 6.60614 | 6.84513 |
| PAPLN | O95428 | 6.83574 | 5.6894 | 5.86866 | 7.43354 | 5.91947 | 6.33031 | 5.88634 | 6.75169 | 6.27634 | 6.61114 | 7.38712 | 6.49296 | 5.72961 | 5.25842 | 6.00264 | 5.52663 |
| WIPF1 | O43516-3 | 6.53411 | 5.6206 | 7.79668 | 5.75772 | 5.83733 | 6.37926 | 6.42638 | 5.79826 | 6.43953 | 6.03445 | 7.01564 | 6.70726 | 5.51388 | 6.2642 | 6.1413 | 5.7334 |
| MAP4 | E7EVA0 | 6.57166 | 6.0138 | 6.38092 | 5.9839 | 5.83708 | 6.28768 | 6.08335 | 6.44085 | 6.65774 | 6.66982 | 6.45204 | 6.75198 | 5.33403 | 6.61445 | 5.8913 | 6.0294 |
| COLGALT2 | Q8IYK4 | 6.36507 | 6.79143 | 6.09912 | 6.45555 | 6.31555 | 6.33715 | 6.81623 | 5.68652 | 6.57019 | 5.55367 | 6.39153 | 6.31658 | 6.26468 | 5.48338 | 6.5263 | 6.02704 |
| B3GNT2 | Q9NY97 | 7.00432 | 7.23442 | 6.88229 | 6.8107 | 6.68115 | 6.62168 | 6.75303 | 6.03767 | 5.92242 | 6.61446 | 5.97528 | 6.16294 | 5.24573 | 5.6302 | 5.06683 | 5.35689 |
| NOSIP | A0A075B6F9 | 5.73122 | 6.54106 | 7.75825 | 6.00604 | 6.37069 | 6.604 | 6.38961 | 6.59216 | 6.79002 | 6.38864 | 7.20692 | 6.52183 | 5.16158 | 5.69882 | 5.28765 | 4.95149 |
| GTF2IRD1 | Q9UHL9-3 | 6.93955 | 7.21695 | 7.37538 | 7.569 | 6.96391 | 6.34318 | 6.16677 | 5.70106 | 6.43937 | 6.55266 | 6.19467 | 6.48409 | 5.33251 | 4.48734 | 5.02163 | 5.21193 |
| RNPEPL1 | Q9HAU8 | 5.90915 | 5.87652 | 6.94797 | 6.10534 | 6.13499 | 7.01125 | 6.25743 | 6.44775 | 6.29118 | 6.5024 | 6.01937 | 7.07248 | 5.90403 | 6.6654 | 4.90716 | 5.94758 |
| VASN | Q6EMK4 | 7.60389 | 6.53059 | 6.67344 | 6.49949 | 5.52391 | 5.44102 | 5.72776 | 5.64254 | 6.21061 | 7.13735 | 7.32438 | 6.87761 | 5.44409 | 5.98043 | 5.14153 | 6.24138 |
| ST3GAL4 | Q11206-4 | 6.2402 | 6.68171 | 6.21848 | 6.90681 | 6.34859 | 5.98483 | 6.65446 | 5.71401 | 6.23546 | 6.8906 | 6.55109 | 6.06853 | 6.12862 | 5.69567 | 5.52805 | 6.1529 |
| TRIM62 | Q9BVG3 | 6.23782 | 7.08144 | 6.33912 | 5.9852 | 6.22675 | 6.69893 | 5.70616 | 7.1825 | 6.72386 | 7.06221 | 6.89379 | 7.82953 | 4.62588 | 5.61709 | 4.54071 | 5.24901 |
| ZDHHC3 | Q9NYG2-2 | 5.98561 | 6.73128 | 6.64726 | 6.85474 | 6.22002 | 6.21316 | 5.84125 | 5.54504 | 5.95919 | 6.07265 | 6.19225 | 5.90006 | 6.49784 | 6.54565 | 6.69842 | 6.09558 |
| TEAD2 | Q15562-4 | 6.99637 | 6.52733 | 6.90621 | 6.55224 | 6.48788 | 5.47025 | 5.69984 | 6.25042 | 6.34354 | 6.01496 | 5.73392 | 6.2108 | 5.94359 | 5.47477 | 6.77928 | 6.6086 |
| TEAD4 | Q15561 | 6.42312 | 5.92991 | 5.42079 | 6.53027 | 6.10119 | 6.14278 | 6.43186 | 5.88678 | 5.83146 | 6.48323 | 6.46683 | 6.58478 | 6.43765 | 6.34185 | 6.41093 | 6.57657 |
| TCEA2 | Q15560 | 6.05716 | 5.77462 | 6.3738 | 5.58941 | 6.46703 | 6.221 | 6.23471 | 6.01992 | 6.29682 | 6.51617 | 6.29023 | 6.17735 | 6.74735 | 6.6351 | 6.43063 | 6.1687 |
| G2E3 | Q7L622 | 6.53537 | 7.86168 | 6.91267 | 8.57594 | 6.20386 | 6.64025 | 7.76697 | 6.88825 | 6.06296 | 6.11773 | 6.80424 | 7.6484 | 3.82692 | 4.5585 | 3.29292 | 4.30334 |
| FBXL14 | Q8N1E6 | 6.86686 | 6.64603 | 6.59675 | 6.74784 | 6.02806 | 6.32928 | 6.40943 | 6.13411 | 7.0368 | 7.05344 | 7.17554 | 6.70991 | 4.67143 | 5.04027 | 5.13376 | 5.42047 |
| TRPT1 | Q86TN4 | 6.45428 | 6.0913 | 7.10702 | 6.39877 | 6.54912 | 6.7066 | 5.86805 | 5.96592 | 6.53841 | 6.93911 | 6.25185 | 6.77595 | 5.68554 | 5.88738 | 5.61628 | 5.36362 |
| CDC42EP3 | Q9UKI2 | 6.06669 | 5.80567 | 7.1654 | 6.27098 | 6.59145 | 6.51591 | 6.63771 | 7.86114 | 5.91486 | 5.76749 | 6.55964 | 6.98408 | 5.88569 | 5.21266 | 5.87566 | 4.88497 |
| GJC1 | P36383 | 6.55419 | 7.04081 | 7.167 | 6.13248 | 6.17091 | 5.61712 | 6.26127 | 6.55356 | 6.4542 | 5.99901 | 5.5868 | 5.94526 | 5.68872 | 5.79655 | 6.50664 | 6.52549 |
| CHSY1 | Q86X52 | 6.4325 | 6.22554 | 5.78128 | 6.83243 | 6.38768 | 6.34739 | 6.87568 | 6.19445 | 5.9832 | 6.0999 | 6.07728 | 5.9393 | 6.12794 | 6.38999 | 6.2967 | 6.00876 |
| SDCBP | O00560-2 | 5.59617 | 6.12114 | 6.66422 | 6.23575 | 5.59251 | 7.018 | 6.29567 | 6.47418 | 7.11634 | 6.99366 | 7.05019 | 6.79597 | 5.00582 | 6.16089 | 5.20673 | 5.67276 |
| MEX3C | Q5U5Q3 | 7.40682 | 7.24029 | 7.39092 | 5.76453 | 5.86764 | 6.68942 | 6.44256 | 6.71357 | 6.24752 | 7.20194 | 6.75852 | 7.27597 | 3.91464 | 4.21987 | 5.80542 | 5.06037 |
| PMAIP1 | Q13794 | 6.66963 | 6.66482 | 5.82625 | 6.24652 | 7.08749 | 6.09407 | 6.51072 | 6.24184 | 6.1203 | 5.89255 | 5.89879 | 5.37317 | 6.06592 | 6.28023 | 6.49935 | 6.52837 |
| TSPAN4 | O14817 | 6.74139 | 7.00801 | 6.68456 | 7.07382 | 5.97521 | 6.4057 | 6.95134 | 6.59866 | 5.47425 | 6.46006 | 5.82101 | 6.06674 | 5.07491 | 5.98745 | 6.00279 | 5.67409 |
| IL27RA | Q6UWB1 | 6.45163 | 6.71564 | 6.68683 | 7.06565 | 7.07928 | 6.46994 | 6.58803 | 6.25371 | 7.47393 | 7.00129 | 6.51294 | 5.72586 | 4.45772 | 5.06998 | 5.70134 | 4.74624 |
| SCARF2 | Q96GP6 | 6.79347 | 6.70928 | 6.2708 | 7.30307 | 5.99768 | 5.43413 | 6.43507 | 6.36265 | 5.68785 | 6.3726 | 6.78071 | 6.2113 | 6.08997 | 6.10175 | 5.71779 | 5.73188 |
| EPB41L2 | H0Y5B0 | 7.48852 | 5.75263 | 6.40882 | 6.53449 | 6.20008 | 6.04361 | 6.96843 | 5.37 | 5.45401 | 6.30399 | 6.49324 | 6.51389 | 5.84714 | 6.34938 | 6.73005 | 6.54173 |
| ABCA6 | Q8N139 | 6.28494 | 6.19316 | 6.77261 | 6.18202 | 6.71275 | 6.61961 | 5.98046 | 6.4291 | 5.62475 | 6.95521 | 5.79172 | 7.06311 | 6.12735 | 5.22132 | 5.92512 | 6.11677 |
| KIAA1522 | Q9P206-2 | 6.26045 | 6.55058 | 5.96317 | 6.46636 | 6.11976 | 5.88311 | 6.00845 | 6.55876 | 6.17322 | 6.58878 | 5.95804 | 6.50378 | 6.51406 | 6.67573 | 5.51707 | 6.25868 |
| MGA | H3BU53 | 5.5459 | 6.48934 | 5.71804 | 7.77761 | 6.65356 | 6.08022 | 5.69396 | 5.95774 | 6.28134 | 5.90367 | 6.49397 | 6.49322 | 5.12689 | 6.00015 | 6.91036 | 6.87403 |
| ZNF576 | Q9H609 | 6.39448 | 6.17087 | 7.09738 | 5.73886 | 6.29368 | 6.48761 | 5.76191 | 6.9243 | 6.70794 | 6.07913 | 6.47136 | 6.51427 | 5.77167 | 5.73023 | 5.91959 | 5.93671 |
| CSNK1G1 | Q9HCP0 | 6.26267 | 6.01366 | 5.89035 | 6.82687 | 6.13392 | 6.20623 | 5.8908 | 6.62353 | 6.79575 | 6.35276 | 6.43851 | 6.75871 | 5.6994 | 5.99681 | 6.46007 | 5.64997 |
| PLEKHA2 | Q9HB19 | 6.0344 | 5.71623 | 7.13228 | 5.80742 | 5.83151 | 7.35451 | 7.41102 | 5.65609 | 5.89582 | 6.69567 | 5.91206 | 6.40988 | 5.50414 | 6.66401 | 5.64415 | 6.33079 |
| IFNGR1 | P15260 | 6.79275 | 6.89663 | 7.39654 | 7.782 | 5.67109 | 5.80984 | 6.23693 | 5.93979 | 6.7175 | 6.74486 | 7.16897 | 6.88412 | 4.48784 | 5.12494 | 4.81116 | 5.53504 |
| TMEM259 | Q4ZIN3 | 6.86521 | 6.05807 | 6.17887 | 7.92511 | 6.21954 | 6.57087 | 6.09172 | 5.42298 | 5.81488 | 6.76937 | 6.55402 | 5.59117 | 5.62523 | 5.49929 | 6.36493 | 6.44874 |
| ZFP64 | Q9NTW7-5 | 6.72948 | 7.29793 | 6.28195 | 6.95099 | 5.96769 | 6.41546 | 6.05319 | 5.21557 | 5.32829 | 6.45096 | 5.24634 | 6.66824 | 7.08384 | 5.71502 | 6.22761 | 6.36745 |
| HLA-C | A0A140T912 | 6.78069 | 6.50222 | 5.54711 | 6.75996 | 6.57011 | 6.3795 | 6.47801 | 6.57515 | 6.10585 | 6.65012 | 6.01112 | 6.34082 | 6.24581 | 5.48367 | 5.73155 | 5.83831 |
| HLA-E | P13747 | 7.88233 | 7.04728 | 7.95238 | 6.72715 | 7.49347 | 6.20343 | 6.56314 | 5.99267 | 7.38 | 6.75029 | 7.67343 | 7.89547 | 3.95183 | 3.47339 | 2.90352 | 4.11022 |
| HLA-F | P30511-3 | 5.84655 | 5.21871 | 5.6276 | 6.25844 | 5.89573 | 5.63764 | 6.25692 | 6.28795 | 6.83634 | 6.40384 | 6.68662 | 6.41485 | 6.68267 | 6.38918 | 6.58036 | 7.58155 |
| MAP3K15 | Q6ZN16 | 6.51297 | 6.29436 | 7.02708 | 5.94311 | 6.10996 | 6.76782 | 6.16817 | 6.62585 | 6.67712 | 6.80037 | 6.82077 | 6.77272 | 5.52192 | 5.65251 | 5.16018 | 5.14508 |
| ZDHHC18 | Q9NUE0 | 6.75439 | 7.62691 | 7.24597 | 6.34725 | 7.08252 | 5.25853 | 6.87436 | 6.39364 | 6.65529 | 5.32043 | 6.56897 | 6.13479 | 5.42567 | 5.13155 | 5.68165 | 5.49806 |
| MPRIP | A0A494BZV2 | 6.3582 | 5.28415 | 7.44868 | 6.90501 | 5.46537 | 5.24861 | 5.77703 | 6.46373 | 6.80684 | 6.23721 | 6.89006 | 6.11207 | 7.01423 | 5.81527 | 5.24875 | 6.92479 |
| MIS18A | Q9NYP9 | 6.13194 | 6.59462 | 5.92782 | 6.54434 | 6.152 | 6.57998 | 6.14237 | 6.79101 | 5.99293 | 6.91278 | 5.94558 | 6.92684 | 5.83444 | 5.38196 | 5.63419 | 6.50722 |
| NSD2 | O96028-3 | 6.65616 | 6.07959 | 6.06955 | 7.99085 | 6.26438 | 5.12187 | 6.30617 | 5.2125 | 6.72012 | 7.04176 | 6.28494 | 5.60928 | 6.04892 | 5.45789 | 5.75496 | 7.38107 |
| IFIT1 | P09914 | 7.32086 | 7.19468 | 6.08387 | 6.1456 | 6.06167 | 7.1197 | 6.57002 | 6.66754 | 7.39425 | 5.90041 | 6.07856 | 6.44697 | 5.10425 | 5.2524 | 4.95732 | 5.70192 |
| CCNYL1 | Q8N7R7 | 6.30056 | 5.94305 | 5.36211 | 6.75855 | 6.29486 | 6.45271 | 5.54941 | 5.87799 | 5.6155 | 6.99501 | 6.23664 | 6.15045 | 5.86497 | 5.96606 | 7.57132 | 7.06081 |
| COPA | A0A3B3ITX2 | 6.19436 | 6.94354 | 7.42073 | 5.61171 | 5.59174 | 7.35506 | 5.65568 | 6.59913 | 6.1781 | 6.51997 | 5.9815 | 5.98835 | 5.82183 | 6.30887 | 5.71245 | 6.11697 |
| ATP8B2 | P98198-3 | 6.37532 | 6.60489 | 6.41579 | 6.44188 | 6.05486 | 5.72182 | 6.50574 | 5.76713 | 6.62923 | 6.7124 | 5.90004 | 6.09969 | 6.37259 | 6.08689 | 6.18099 | 6.13074 |
| ATP8B1 | O43520 | 6.26096 | 5.89698 | 6.54204 | 6.70142 | 6.82279 | 5.80002 | 6.22019 | 6.81173 | 5.73967 | 6.59505 | 6.43232 | 6.40205 | 5.39649 | 6.03068 | 6.97385 | 5.37375 |
| MICA | A0A140T986 | 6.14661 | 7.02645 | 6.7603 | 7.93775 | 5.66657 | 6.26779 | 6.78096 | 5.78258 | 6.19463 | 6.29971 | 6.46577 | 6.706 | 5.25613 | 5.40059 | 5.30108 | 6.00709 |
| CIB1 | Q99828-2 | 5.95205 | 5.40799 | 5.85169 | 5.18655 | 5.24981 | 4.44179 | 6.08238 | 5.55597 | 5.18186 | 5.92872 | 5.8535 | 5.89213 | 5.65977 | 10.1954 | 7.28447 | 10.2759 |
| ADCK1 | Q86TW2 | 6.13953 | 6.68265 | 5.87985 | 6.52262 | 6.61076 | 6.02305 | 6.37124 | 5.4296 | 5.9666 | 6.60036 | 5.95317 | 5.9094 | 7.13497 | 5.83936 | 6.72129 | 6.21555 |
| PPCDC | Q96CD2 | 6.61107 | 5.11381 | 6.84005 | 4.71013 | 5.54591 | 5.57325 | 5.05489 | 7.57332 | 4.69554 | 6.63522 | 5.49607 | 7.30336 | 5.3703 | 9.07607 | 6.82524 | 7.57578 |
| IL31RA | Q8NI17-2 | 6.19251 | 5.86363 | 6.83698 | 6.78443 | 6.37123 | 6.44469 | 6.07558 | 5.85503 | 6.26193 | 5.3908 | 6.95127 | 6.13614 | 6.83763 | 5.90864 | 6.04854 | 6.04097 |
| NOL3 | O60936-3 | 6.98542 | 6.23825 | 6.97072 | 4.59737 | 5.30059 | 5.34974 | 5.76746 | 7.12654 | 4.81128 | 7.02231 | 6.60058 | 7.16672 | 5.71437 | 7.53693 | 6.21699 | 6.59471 |
| NME2P1 | O60361 | 7.19747 | 5.02213 | 5.93824 | 4.82872 | 6.31469 | 6.11088 | 7.03727 | 5.94725 | 7.88587 | 6.43248 | 7.05758 | 6.49548 | 5.77769 | 5.95214 | 6.00001 | 6.00659 |
| SLC2A13 | Q96QE2 | 6.10275 | 6.47592 | 6.08822 | 6.91092 | 5.99976 | 6.65087 | 5.5994 | 6.54297 | 6.22083 | 5.91931 | 6.52652 | 6.00908 | 5.51033 | 6.2961 | 6.72753 | 6.41948 |
| DDB2 | Q92466 | 6.15463 | 6.22846 | 5.98973 | 6.13393 | 6.22763 | 6.07057 | 5.6883 | 5.63206 | 6.27272 | 5.98788 | 6.83854 | 5.77355 | 7.24009 | 6.6131 | 6.64279 | 6.50603 |
| TRIM38 | O00635 | 5.2392 | 6.23388 | 6.52978 | 6.18706 | 6.6821 | 6.55857 | 6.00883 | 5.3272 | 6.16564 | 6.43128 | 6.35832 | 6.1771 | 6.34459 | 6.67126 | 6.78185 | 6.30335 |
| ELFN1 | P0C7U0 | 5.41378 | 5.85413 | 5.76228 | 6.42679 | 6.30871 | 6.16296 | 5.92382 | 5.98554 | 6.80652 | 6.31216 | 5.85232 | 5.53172 | 7.05434 | 6.77923 | 7.37621 | 6.44949 |
| ZDHHC21 | Q8IVQ6 | 6.38076 | 5.98723 | 6.19752 | 6.34582 | 5.92015 | 6.21284 | 6.79372 | 5.91979 | 5.06373 | 5.58045 | 6.92487 | 6.01027 | 6.56787 | 7.45266 | 6.37522 | 6.2671 |
| ZNRF1 | Q8ND25 | 6.84297 | 6.78179 | 7.57014 | 5.92341 | 6.02322 | 5.7158 | 6.02319 | 7.01984 | 6.63418 | 7.93712 | 7.25961 | 8.43195 | 3.99172 | 4.52426 | 4.61389 | 4.7069 |
| NRG1 | Q02297-2 | 5.86398 | 5.42009 | 6.34834 | 6.41243 | 6.42929 | 6.18294 | 6.5625 | 5.59679 | 6.83511 | 6.1807 | 6.88531 | 6.51273 | 6.33216 | 6.05399 | 5.75107 | 6.63258 |
| OTUD5 | Q96G74 | 6.65412 | 6.07778 | 7.66136 | 6.11538 | 6.03245 | 6.29728 | 6.19 | 6.11445 | 6.61898 | 5.63708 | 7.6043 | 6.53877 | 5.79384 | 5.11914 | 5.40196 | 6.1431 |
| TENT4A | Q5XG87 | 6.53337 | 6.64404 | 6.25311 | 6.97819 | 6.02964 | 5.99445 | 6.35296 | 6.00363 | 6.18516 | 6.29059 | 6.82795 | 6.58778 | 5.96861 | 5.58345 | 5.90313 | 5.86393 |
| ELMO1 | Q92556 | 6.17645 | 5.76424 | 6.58945 | 6.29355 | 5.83683 | 6.7692 | 5.63816 | 6.70026 | 6.62244 | 6.69632 | 6.37596 | 6.52257 | 5.95862 | 6.2686 | 5.82806 | 5.9593 |
| DHRS11 | Q6UWP2 | 5.93655 | 5.76609 | 6.82484 | 6.23497 | 7.28993 | 7.31524 | 6.23696 | 5.56142 | 6.36515 | 7.09178 | 6.8612 | 6.45794 | 5.61097 | 5.63891 | 5.1487 | 5.65936 |
| RRAGA | Q7L523 | 5.94499 | 6.12359 | 7.37896 | 6.9299 | 6.52864 | 6.13731 | 6.18693 | 6.56946 | 7.07884 | 6.26357 | 6.64292 | 6.40622 | 5.37042 | 5.86629 | 5.4 | 5.17196 |
| RAB29 | O14966 | 6.50221 | 6.60612 | 6.3251 | 6.64563 | 6.30258 | 6.10793 | 6.43706 | 6.25229 | 5.95706 | 6.11805 | 5.90214 | 5.99147 | 5.90521 | 6.24851 | 6.58442 | 6.11421 |
| PANK3 | Q9H999 | 6.64142 | 7.00297 | 8.21728 | 7.15892 | 6.34377 | 6.81783 | 6.68684 | 6.25931 | 7.67322 | 6.25009 | 6.78997 | 7.98866 | 3.47985 | 4.19475 | 3.94083 | 4.55428 |
| SLC35E1 | Q96K37 | 6.41143 | 6.52704 | 6.43476 | 7.24643 | 5.81925 | 5.63218 | 6.36288 | 5.51911 | 6.29735 | 6.12131 | 6.49029 | 6.51166 | 6.58069 | 5.65661 | 6.42722 | 5.96178 |
| TMEM44 | Q2T9K0 | 6.37689 | 6.11947 | 6.03193 | 8.15123 | 5.88554 | 6.82168 | 6.39696 | 6.04323 | 6.14845 | 6.24737 | 6.54656 | 6.69502 | 5.782 | 5.85723 | 4.89179 | 6.00466 |
| MYL9 | P24844 | 6.76281 | 6.31065 | 7.91669 | 6.10916 | 5.47758 | 6.53046 | 4.84063 | 7.05205 | 6.27528 | 6.94177 | 7.02439 | 7.61847 | 5.74658 | 6.20251 | 4.9015 | 4.28946 |

| | | | | | | | | | | | | | | | | | |
|---|---|---|---|---|---|---|---|---|---|---|---|---|---|---|---|---|---|
| SH3RF2 | Q8TEC5 | 6.12101 | 7.36793 | 6.62793 | 7.00712 | 5.34668 | 5.97522 | 5.91561 | 5.42739 | 6.86021 | 7.62127 | 6.25105 | 6.67786 | 5.47931 | 5.98946 | 5.31781 | 6.01414 |
| TP53RK | Q96S44 | 6.12726 | 5.44523 | 6.87899 | 5.59206 | 5.81604 | 6.16608 | 5.75227 | 5.56506 | 6.27527 | 6.4982 | 6.71446 | 6.26436 | 6.58465 | 6.52788 | 7.68201 | 6.11016 |
| CSDE1 | O75534-3 | 6.57775 | 6.76109 | 7.27617 | 5.9809 | 6.11837 | 7.17743 | 6.0653 | 6.42165 | 7.87416 | 6.68493 | 6.78418 | 6.68573 | 4.86057 | 4.88652 | 4.83591 | 5.00935 |
| TBRG1 | Q3YBR2 | 5.93989 | 6.51394 | 5.61794 | 7.59854 | 6.75381 | 6.02062 | 6.18939 | 6.05453 | 6.04788 | 6.79637 | 5.92716 | 4.86116 | 6.59739 | 6.36639 | 6.85009 | 5.86489 |
| TTC39B | Q5VTQ0 | 6.30994 | 5.72141 | 6.13246 | 5.8243 | 5.48246 | 6.15515 | 6.28783 | 6.76256 | 6.29042 | 6.57015 | 6.1626 | 7.02902 | 5.82639 | 6.45363 | 6.96517 | 6.02651 |
| ADCY6 | O43306 | 6.26483 | 6.42654 | 6.5556 | 7.35366 | 5.97369 | 5.85252 | 6.17134 | 5.59211 | 6.99099 | 6.20372 | 5.88146 | 6.93035 | 6.75841 | 5.83849 | 5.81957 | 5.38672 |
| CBR4 | Q8N4T8 | 6.61128 | 6.4581 | 5.95186 | 6.94441 | 6.01005 | 6.13178 | 5.48701 | 5.18736 | 6.42973 | 6.61194 | 5.81066 | 5.80141 | 7.03051 | 6.20528 | 6.93512 | 6.3935 |
| LMLN | Q96KR4-3 | 6.14677 | 6.60869 | 6.17404 | 6.69315 | 6.45358 | 6.56526 | 6.26145 | 5.6761 | 6.5966 | 6.6212 | 5.8083 | 5.68842 | 6.81121 | 6.01994 | 6.01444 | 5.86087 |
| ATP23 | Q9Y6H3 | 6.38609 | 6.4193 | 6.10064 | 6.81049 | 6.65083 | 6.02493 | 5.831 | 6.239 | 6.10903 | 6.50112 | 6.28707 | 6.36228 | 5.72002 | 6.23965 | 5.44511 | 6.87344 |
| AKIP1 | Q9NQ31 | 6.6836 | 6.50237 | 6.78888 | 6.038 | 6.2074 | 6.58388 | 7.28137 | 6.42479 | 5.67454 | 7.93121 | 7.13155 | 7.98903 | 4.5332 | 4.35776 | 4.34688 | 5.52555 |
| MBTPS2 | O43462 | 5.54917 | 6.53423 | 6.68378 | 6.95066 | 6.59791 | 6.74162 | 5.91893 | 6.76355 | 6.13955 | 6.30857 | 6.78812 | 5.41428 | 5.54748 | 6.01378 | 6.60509 | 5.44327 |
| CFLAR | O15519 | 6.59429 | 5.32672 | 6.58162 | 6.40503 | 6.01381 | 6.24171 | 6.43616 | 7.131 | 6.35812 | 5.92002 | 6.93354 | 6.899 | 6.08343 | 5.48778 | 5.54033 | 6.04744 |
| NANP | Q8TBE9 | 6.28698 | 6.02457 | 6.81584 | 6.06578 | 5.99394 | 6.68286 | 5.50763 | 7.20641 | 6.12833 | 6.85423 | 6.09423 | 7.25394 | 5.24542 | 5.80579 | 6.07384 | 5.9602 |
| ATG12 | O94817 | 6.89466 | 5.90338 | 6.45871 | 5.40876 | 6.79344 | 5.83556 | 5.89266 | 7.02797 | 6.10576 | 6.25945 | 7.13971 | 6.7028 | 6.21961 | 5.6594 | 5.98803 | 5.71011 |
| NFATC3 | Q12968 | 6.41251 | 6.43224 | 7.03684 | 5.58781 | 6.3517 | 5.84727 | 6.09579 | 6.38846 | 6.30082 | 6.76434 | 6.91826 | 7.40035 | 5.81856 | 6.14087 | 5.01432 | 5.48987 |
| ROGDI | Q9GZN7 | 6.28305 | 6.1669 | 6.4414 | 5.32907 | 6.625 | 5.78695 | 6.16241 | 6.7933 | 6.42147 | 6.40108 | 6.27313 | 6.41704 | 5.98882 | 6.82062 | 6.23437 | 5.8554 |
| JRKL | Q9Y4A0 | 6.31714 | 5.59189 | 6.20963 | 7.15368 | 5.79759 | 6.02464 | 6.3899 | 6.80253 | 5.79014 | 6.39811 | 5.45325 | 5.63255 | 7.48612 | 6.82061 | 5.75974 | 6.37247 |
| PPP3CC | P48454-3 | 7.10947 | 5.64593 | 6.78832 | 5.27791 | 6.25697 | 6.18406 | 6.04549 | 6.22282 | 6.45827 | 6.63552 | 6.6633 | 6.24709 | 6.05345 | 5.82415 | 6.61552 | 5.97172 |
| ZMYM5 | Q9UJ78 | 6.35146 | 6.63557 | 6.62203 | 6.24394 | 5.98855 | 6.0444 | 6.09013 | 6.00633 | 6.37064 | 6.55346 | 6.80261 | 6.55747 | 5.88066 | 6.36603 | 5.72101 | 5.76571 |
| PARD6B | Q9BYG5 | 5.60206 | 5.84936 | 6.87916 | 6.10272 | 6.27914 | 6.25376 | 6.01255 | 6.46878 | 6.34405 | 5.80641 | 6.62955 | 6.48983 | 6.66383 | 5.88219 | 6.61992 | 6.11669 |
| SLC31A1 | O15431 | 6.61273 | 6.65134 | 6.31561 | 6.76856 | 6.47639 | 5.85821 | 6.07845 | 5.61989 | 6.37924 | 5.96142 | 6.63212 | 6.41566 | 5.65943 | 5.87099 | 6.34431 | 6.35566 |
| SLC25A28 | Q96A46 | 5.87026 | 6.07548 | 6.53669 | 6.53438 | 6.91768 | 6.68731 | 6.55272 | 6.50123 | 6.27399 | 6.53211 | 5.85003 | 5.06613 | 6.42759 | 6.49914 | 6.01475 | 5.66052 |
| SMIM15 | Q7Z3B0 | 6.73123 | 6.59297 | 6.37947 | 6.63013 | 5.82496 | 5.78784 | 6.13627 | 5.18056 | 6.30708 | 6.07745 | 5.84279 | 6.35635 | 6.24346 | 6.11059 | 6.58445 | 7.21441 |
| VIPAS39 | Q9H9C1 | 6.79759 | 6.14112 | 5.95956 | 6.54582 | 5.91018 | 5.95324 | 6.71865 | 6.45927 | 6.38441 | 6.56576 | 6.83666 | 7.01915 | 5.92427 | 5.77873 | 5.37383 | 5.63174 |
| CRY2 | Q49AN0 | 6.48506 | 7.28092 | 7.24019 | 7.28369 | 6.23272 | 6.0278 | 6.19556 | 6.02955 | 6.58966 | 6.82374 | 6.50786 | 6.38156 | 4.99197 | 5.40069 | 5.40595 | 5.12308 |
| C8orf37 | Q96NL8 | 5.94168 | 6.45415 | 6.56121 | 5.9923 | 5.84186 | 7.11767 | 5.81456 | 7.09707 | 6.21751 | 7.12287 | 6.22464 | 7.37533 | 5.56354 | 5.97431 | 5.16771 | 5.53359 |
| MMACHC | Q9Y4U1 | 5.55327 | 6.17659 | 6.01315 | 6.73886 | 6.36462 | 6.04882 | 6.43145 | 6.65882 | 6.28922 | 7.24257 | 7.65606 | 7.75815 | 6.07526 | 5.08306 | 4.51216 | 5.39794 |
| MAP3K2 | Q9Y2U5 | 6.80528 | 6.4437 | 7.0073 | 6.41602 | 5.62143 | 7.36091 | 5.81153 | 6.64232 | 6.88439 | 6.47062 | 6.65708 | 7.18531 | 5.26718 | 4.98098 | 5.24161 | 5.20433 |
| MXRA8 | Q9BRK3-2 | 7.01782 | 6.55414 | 6.74119 | 6.80047 | 6.14419 | 6.49643 | 6.49157 | 6.60879 | 6.19082 | 6.52986 | 6.51649 | 5.51029 | 5.7963 | 5.43632 | 5.21767 | 5.94766 |
| FOXC1 | Q12948 | 6.15637 | 6.36413 | 7.27572 | 7.46847 | 6.73003 | 6.56522 | 6.32531 | 5.56652 | 6.26201 | 6.06036 | 6.52708 | 7.08374 | 6.14632 | 5.11444 | 4.6641 | 5.69017 |
| RPUSD1 | Q9UJJ7 | 7.12369 | 6.9439 | 8.08191 | 7.15521 | 5.8269 | 5.9848 | 6.94913 | 6.07226 | 7.30122 | 6.09623 | 7.39088 | 7.24654 | 4.53554 | 4.54549 | 3.78101 | 4.96528 |
| ALKBH8 | Q96BT7-4 | 5.9271 | 7.45124 | 7.43733 | 6.64281 | 5.96795 | 6.58876 | 6.10311 | 6.30518 | 5.9777 | 6.74947 | 7.04898 | 5.99792 | 6.13447 | 5.03878 | 5.09462 | 5.53458 |
| GSKIP | Q9P0R6 | 5.85411 | 5.23374 | 6.04278 | 4.57777 | 5.42456 | 5.33324 | 5.90687 | 8.16622 | 4.63166 | 5.87985 | 5.23874 | 7.00788 | 5.3088 | 9.72459 | 6.86265 | 8.80654 |
| TADA2A | O75478 | 6.98289 | 5.78478 | 6.96473 | 6.71564 | 6.25539 | 6.32309 | 5.79253 | 5.9203 | 5.84036 | 6.4288 | 5.96668 | 6.85774 | 6.31802 | 5.05921 | 5.91036 | 6.87948 |
| RAB3IP | Q96QF0 | 5.94462 | 5.9214 | 6.51958 | 5.89576 | 6.10444 | 6.68495 | 6.70331 | 6.61386 | 6.47429 | 6.55343 | 6.83505 | 6.58871 | 5.99816 | 5.78996 | 5.64914 | 5.72333 |
| SLAIN1 | Q8ND83 | 6.55209 | 5.41006 | 6.9292 | 5.35241 | 5.84258 | 6.77292 | 6.26285 | 6.1118 | 6.13857 | 6.54872 | 6.69636 | 8.51369 | 6.60453 | 5.99661 | 5.08158 | 5.18602 |
| DNAH5 | Q8TE73 | 6.99753 | 6.65245 | 5.00497 | 6.16206 | 5.98279 | 6.9451 | 5.17917 | 6.92735 | 6.81948 | 6.0575 | 5.66646 | 6.5667 | 5.53586 | 6.57022 | 5.84311 | 7.08924 |
| ZNF22 | P17026 | 6.58259 | 6.4624 | 5.31212 | 6.51316 | 5.71445 | 5.62664 | 6.1552 | 5.55695 | 5.85212 | 6.11626 | 5.43323 | 6.03223 | 6.9416 | 7.43192 | 6.89854 | 7.37058 |
| OTUB2 | Q96DC9 | 5.94904 | 5.88418 | 6.5567 | 6.05486 | 6.25998 | 7.08405 | 5.94535 | 6.18193 | 6.20032 | 6.92432 | 6.63362 | 6.53899 | 5.93775 | 6.26103 | 5.46791 | 6.11999 |
| DNASE1L1 | P49184 | 6.10225 | 6.55586 | 6.92809 | 5.14401 | 4.76242 | 6.09026 | 6.68085 | 7.08184 | 5.8119 | 5.72493 | 6.20391 | 5.72079 | 6.5157 | 7.36783 | 7.3768 | 5.93256 |
| DUS1L | Q6P1R4 | 5.44683 | 4.9009 | 6.98355 | 6.35735 | 5.88881 | 7.15253 | 5.90763 | 6.12469 | 6.4183 | 6.24135 | 6.51553 | 6.84713 | 5.64489 | 6.88856 | 6.34401 | 6.33794 |
| RTN3 | O95197-5 | 6.96424 | 7.46276 | 5.81831 | 6.67343 | 6.39151 | 6.09698 | 6.19554 | 7.54427 | 6.09683 | 5.863 | 6.27409 | 6.30178 | 4.77882 | 7.46333 | 4.72091 | 5.35423 |
| CRBN | Q96SW2 | 5.15264 | 5.64558 | 6.20785 | 4.55824 | 7.05717 | 5.25486 | 6.98568 | 5.70392 | 6.26134 | 6.87823 | 7.3298 | 6.29567 | 6.55504 | 6.98973 | 6.58501 | 6.53923 |
| E2F4 | Q16254 | 6.53275 | 8.0471 | 6.60305 | 6.96046 | 7.00652 | 6.15735 | 4.69514 | 7.01806 | 5.6328 | 7.35504 | 6.65565 | 6.63793 | 5.72862 | 5.52502 | 4.47177 | 4.97276 |
| HNRNPA1 | H0YH80 | 6.36016 | 6.34616 | 5.98534 | 7.38038 | 5.47487 | 5.65502 | 6.00284 | 5.93239 | 5.74845 | 7.40187 | 6.01583 | 5.63221 | 6.4919 | 5.21803 | 7.0988 | 7.25575 |
| EEF2KMT | Q96G04 | 6.53754 | 5.84479 | 7.48254 | 6.37967 | 6.06391 | 6.54056 | 6.46604 | 6.09135 | 7.16223 | 6.16943 | 7.02231 | 7.26385 | 5.10682 | 5.36865 | 4.97777 | 5.52254 |
| GNPTG | Q9UJJ9 | 6.36684 | 5.52906 | 5.2475 | 6.58094 | 5.97976 | 6.56033 | 6.37213 | 6.15078 | 7.32263 | 5.73704 | 6.08482 | 5.89481 | 6.16907 | 6.27772 | 6.8783 | 6.84828 |
| PLEKHF1 | Q96S99 | 6.24667 | 5.28962 | 7.0111 | 7.10168 | 5.94311 | 6.39281 | 6.31542 | 6.74718 | 6.19809 | 6.17302 | 6.1929 | 7.22671 | 5.12132 | 6.28591 | 5.91137 | 5.84309 |
| LRRC58 | Q96CX6 | 7.63148 | 6.94774 | 8.73483 | 8.39221 | 7.38147 | 7.48117 | 6.69266 | 7.40958 | 6.12172 | 5.16934 | 7.09254 | 6.73017 | 3.37528 | 2.97584 | 3.65683 | 4.20714 |
| DPYSL5 | Q9BPU6 | 6.05403 | 5.89162 | 7.04958 | 5.9732 | 6.07474 | 7.10371 | 6.22395 | 6.80752 | 6.66344 | 6.72969 | 6.89568 | 6.72822 | 6.22816 | 4.96429 | 5.01583 | 5.59632 |
| TMEM9 | Q9P0T7 | 6.12768 | 6.77262 | 6.83146 | 6.50828 | 6.26016 | 6.32262 | 5.55982 | 6.39581 | 6.02227 | 6.63855 | 7.55105 | 7.18168 | 6.07019 | 5.01546 | 5.73537 | 5.00697 |
| LDLRAD3 | Q86YD5 | 7.16601 | 6.68535 | 6.86852 | 6.46254 | 6.27859 | 6.06152 | 6.7023 | 7.63748 | 6.32925 | 6.59699 | 6.81219 | 6.46142 | 5.04808 | 4.66283 | 4.73409 | 5.49283 |
| SLC6A15 | Q9H2J7 | 6.16071 | 6.76504 | 6.64826 | 6.29899 | 5.47958 | 7.29675 | 6.49752 | 6.33144 | 6.04663 | 6.58313 | 6.29716 | 6.50595 | 5.95676 | 5.50821 | 5.67968 | 5.94419 |
| FBLN1 | P23142 | 7.17905 | 7.05961 | 6.3585 | 7.54305 | 6.84724 | 6.23043 | 6.58829 | 6.52814 | 7.05806 | 6.30346 | 7.31218 | 6.34295 | 4.95139 | 4.52807 | 4.36913 | 4.80046 |
| POLDIP3 | Q9BY77-2 | 6.72729 | 6.29895 | 6.43019 | 6.00308 | 6.16878 | 5.76534 | 6.64244 | 6.08604 | 6.24655 | 5.41673 | 6.38614 | 5.60542 | 7.17271 | 6.28971 | 6.6349 | 6.12573 |
| CENPK | Q9BS16 | 6.54635 | 5.25684 | 6.0196 | 5.05518 | 6.0244 | 5.16569 | 5.07137 | 5.91875 | 5.6623 | 6.07723 | 5.91817 | 6.9673 | 6.62192 | 7.47983 | 7.76909 | 8.44598 |
| CEP57L1 | Q8IYX8 | 6.2551 | 6.62515 | 7.31465 | 7.13482 | 6.34706 | 6.7746 | 6.74865 | 6.53181 | 6.63561 | 6.85658 | 6.62355 | 6.77598 | 4.68135 | 5.09887 | 4.63546 | 4.96077 |
| POLI | Q9UNA4 | 6.1517 | 5.35203 | 7.48659 | 5.82594 | 5.92504 | 5.99414 | 5.66168 | 6.45901 | 5.84038 | 6.13728 | 6.63061 | 6.28137 | 6.59145 | 6.31134 | 6.79469 | 6.55676 |
| METTL18 | O95568 | 6.47071 | 6.48269 | 7.12499 | 7.35381 | 5.80602 | 6.17187 | 6.2856 | 5.81557 | 6.13418 | 7.26359 | 7.15136 | 7.13568 | 4.90155 | 5.60969 | 5.6909 | 4.60179 |
| OLFML2A | Q68BL7 | 7.79898 | 7.35111 | 6.30322 | 6.92277 | 6.53781 | 6.6223 | 6.59098 | 6.1425 | 6.16139 | 7.58436 | 7.03055 | 6.65242 | 4.19382 | 5.19754 | 4.44429 | 4.46596 |
| ETV5 | P41161-2 | 4.21389 | 4.45753 | 5.80071 | 4.36691 | 6.75436 | 6.35531 | 6.1601 | 6.33999 | 6.36091 | 5.4579 | 5.36863 | 5.77644 | 8.26112 | 7.60957 | 8.23248 | 8.48415 |
| SMO | Q99835 | 5.82325 | 6.95704 | 7.72113 | 6.90163 | 6.25443 | 6.43752 | 5.96706 | 6.18354 | 5.82355 | 6.55017 | 7.71241 | 6.66071 | 4.8217 | 5.38676 | 5.82321 | 4.97588 |
| RNF167 | Q9H6Y7 | 7.1484 | 6.83767 | 6.10691 | 7.03611 | 6.55991 | 5.78749 | 6.12627 | 6.61981 | 6.16668 | 6.44406 | 6.53668 | 6.31773 | 5.50488 | 5.88477 | 5.41415 | 5.50848 |
| PEX12 | O00623 | 6.15967 | 6.42231 | 6.02551 | 6.20766 | 6.46052 | 6.21021 | 6.27474 | 5.87971 | 6.2341 | 6.16352 | 5.81571 | 6.04802 | 6.40771 | 6.30628 | 6.65483 | 6.72953 |
| PHLPP1 | O60346 | 6.41687 | 6.52029 | 7.31493 | 7.13676 | 6.31654 | 5.95992 | 6.53259 | 5.86242 | 7.37416 | 6.53075 | 7.56215 | 6.64834 | 5.4813 | 4.92642 | 4.62899 | 4.78756 |
| MSRB3 | Q8IXL7 | 7.31448 | 6.03154 | 5.70892 | 6.11829 | 6.42623 | 6.35077 | 5.35618 | 6.36913 | 6.46647 | 6.15191 | 6.51279 | 5.92379 | 6.99963 | 6.09385 | 5.9988 | 6.17723 |
| PROSER1 | Q86XN7 | 6.30539 | 5.20428 | 6.05561 | 5.55295 | 6.83681 | 6.45263 | 6.18414 | 6.52807 | 5.69682 | 6.98324 | 6.38476 | 6.2191 | 6.74012 | 6.74522 | 6.12482 | 5.98602 |
| MED7 | O43513 | 6.38959 | 5.80082 | 5.6835 | 5.60259 | 5.75536 | 5.52959 | 5.84847 | 6.11638 | 5.05946 | 6.86194 | 5.38694 | 6.40725 | 6.96894 | 7.38829 | 6.38199 | 8.81888 |
| KLF5 | Q13887 | 5.50944 | 5.8597 | 6.6257 | 5.94049 | 6.46687 | 5.58002 | 5.21567 | 6.28255 | 6.07422 | 6.79346 | 7.19297 | 6.48728 | 6.76707 | 6.45499 | 6.55155 | 6.19804 |
| BRWD3 | Q6RI45 | 6.32655 | 7.38676 | 6.00126 | 7.26947 | 5.6464 | 6.1969 | 6.59459 | 5.91913 | 6.34688 | 7.22539 | 6.57855 | 6.14891 | 5.3302 | 5.45695 | 5.68399 | 5.88807 |
| RABAC1 | Q9UI14 | 6.61074 | 6.57784 | 6.54353 | 5.83304 | 5.84559 | 6.18517 | 6.32793 | 5.31305 | 7.00631 | 7.02872 | 7.71455 | 6.81202 | 4.94728 | 6.47807 | 4.91144 | 5.86472 |
| AGPAT4 | Q9NRZ5 | 5.89251 | 6.12577 | 6.10719 | 7.20357 | 6.34753 | 5.95826 | 6.41458 | 5.50034 | 6.52102 | 6.16327 | 5.35178 | 5.83797 | 6.54602 | 6.38474 | 7.1793 | 6.46616 |
| NFKBIB | Q15653 | 7.15174 | 6.08486 | 7.54679 | 5.75726 | 7.2744 | 6.6346 | 5.73477 | 5.76195 | 6.45587 | 6.08372 | 7.01286 | 7.40865 | 5.99393 | 4.64798 | 5.08597 | 5.36465 |
| ABCD2 | Q9UBJ2 | 7.07136 | 5.31451 | 7.40865 | 6.74886 | 5.69606 | 6.41946 | 5.64633 | 5.52583 | 6.96431 | 5.27638 | 5.69504 | 6.74083 | 6.96113 | 6.06318 | 6.09755 | 6.37053 |
| NTHL1 | P78549 | 5.84072 | 5.72789 | 6.08849 | 6.56909 | 6.12649 | 6.19822 | 5.97551 | 5.76145 | 6.56734 | 6.23151 | 7.18622 | 6.67086 | 6.22238 | 6.00268 | 7.04516 | 5.786 |
| RNASEH2C | Q8TDP1 | 5.91467 | 5.3114 | 5.61499 | 4.41625 | 4.54925 | 5.69941 | 6.26232 | 6.43607 | 5.0233 | 6.90424 | 6.27114 | 7.52461 | 6.40429 | 7.65737 | 7.2227 | 8.78801 |
| TBX18 | O95935 | 5.63407 | 4.80812 | 6.31574 | 5.97283 | 6.16004 | 6.00795 | 7.41408 | 6.06327 | 6.83681 | 7.39065 | 6.42339 | 6.4069 | 6.44238 | 5.72897 | 6.07366 | 6.32116 |
| NPEPL1 | Q8NDH3 | 4.8928 | 5.32339 | 4.73181 | 6.14836 | 5.65709 | 6.41828 | 7.2234 | 6.41561 | 7.49257 | 4.77886 | 4.8328 | 5.20609 | 6.36297 | 8.76466 | 7.45156 | 8.29975 |
| PFKFB2 | O60825 | 5.9438 | 5.75288 | 6.57555 | 6.10631 | 6.36253 | 6.33334 | 5.89833 | 6.31773 | 5.58011 | 6.73714 | 6.80923 | 6.27654 | 6.65772 | 6.28334 | 6.65098 | 5.71447 |
| TTC7A | Q9ULT0-4 | 6.54911 | 6.97853 | 6.76821 | 7.11448 | 6.14928 | 5.97272 | 6.10781 | 6.11927 | 6.07526 | 6.09765 | 6.31668 | 6.4841 | 5.75362 | 5.49533 | 5.89899 | 6.11896 |
| NET1 | Q7Z628 | 6.49067 | 6.48858 | 6.41557 | 6.53144 | 6.75171 | 5.96182 | 6.40313 | 5.96764 | 6.92445 | 6.66844 | 6.0965 | 5.96061 | 5.66913 | 5.6957 | 5.87455 | 6.10007 |
| STX17 | P56962 | 6.90388 | 6.60849 | 6.03982 | 7.5055 | 6.98417 | 5.65401 | 6.63928 | 6.66107 | 6.46399 | 5.9203 | 5.91126 | 6.55233 | 6.27017 | 5.4328 | 5.26023 | 5.19272 |
| PPP1R12B | O60237-6 | 5.50618 | 6.43018 | 6.85109 | 6.18618 | 5.89422 | 7.30447 | 6.52047 | 5.45783 | 6.58812 | 6.38455 | 6.4391 | 7.35482 | 6.1441 | 5.59489 | 6.35099 | 4.9928 |
| PIK3CD | O00329 | 5.76037 | 5.83531 | 6.79346 | 6.11903 | 6.31826 | 5.61614 | 6.49753 | 6.65667 | 6.96689 | 6.03232 | 6.79551 | 6.8384 | 6.79774 | 5.50882 | 6.0211 | 5.44246 |
| EIF2AK3 | Q9NZJ5 | 5.50045 | 6.56769 | 4.79553 | 5.25794 | 5.8529 | 5.76134 | 5.83045 | 5.44171 | 5.92709 | 6.41608 | 5.43488 | 6.60575 | 6.94068 | 7.91308 | 7.90433 | 7.85013 |
| SLC27A2 | O14975 | 6.14461 | 6.16565 | 5.40797 | 6.32747 | 5.71715 | 5.262 | 5.95313 | 5.67516 | 5.92204 | 6.48856 | 6.43819 | 6.42936 | 6.76372 | 6.97706 | 6.87088 | 7.45703 |
| OPA3 | Q9H6K4-2 | 6.21347 | 6.17594 | 5.63543 | 6.32218 | 6.48884 | 5.81624 | 6.17326 | 6.61095 | 6.11196 | 5.74352 | 5.6745 | 5.51373 | 7.16781 | 6.95496 | 6.7339 | 6.66329 |
| ATP6V1G1 | O75348 | 6.20802 | 6.19101 | 7.57165 | 6.30147 | 6.51935 | 6.2447 | 6.54438 | 6.43712 | 6.80854 | 7.24343 | 6.70096 | 6.6105 | 5.50646 | 4.6809 | 5.51456 | 4.91693 |
| C12orf60 | Q5U649 | 6.48834 | 6.35547 | 6.06149 | 7.19233 | 6.20102 | 5.92253 | 7.27133 | 7.05424 | 5.85192 | 7.09983 | 6.95008 | 8.2569 | 4.15107 | 5.2888 | 4.90963 | 4.94502 |
| NUFIP1 | Q9UHK0 | 7.10852 | 5.75496 | 7.25034 | 7.64561 | 5.63521 | 5.13412 | 6.6349 | 6.03117 | 6.59272 | 4.73615 | 6.48038 | 6.18858 | 6.12942 | 6.07811 | 6.72421 | 5.87562 |
| TMEM65 | Q6PI78 | 6.47297 | 7.05361 | 5.89659 | 5.49735 | 6.65555 | 6.95399 | 6.87944 | 7.13875 | 5.29565 | 4.6304 | 6.60753 | 5.7605 | 5.08964 | 7.40299 | 6.66304 | 6.002 |
| CUEDC1 | Q9NWM3 | 6.93503 | 6.22934 | 6.32002 | 5.99956 | 6.04776 | 6.23763 | 6.07733 | 6.39236 | 6.45297 | 6.25083 | 6.81313 | 6.97981 | 5.54149 | 5.9834 | 5.57045 | 6.16888 |
| PIGL | Q9Y2B2 | 7.13789 | 6.17373 | 6.0104 | 6.87015 | 6.1212 | 7.39003 | 6.12299 | 4.89142 | 6.27296 | 5.93972 | 6.24149 | 6.45244 | 6.31461 | 5.06315 | 6.8741 | 6.12373 |
| IFRD1 | O00458 | 6.00595 | 6.16358 | 7.03659 | 6.88324 | 6.18215 | 6.6046 | 6.47604 | 5.80259 | 6.69211 | 6.89724 | 6.69279 | 6.69668 | 5.67962 | 5.0655 | 5.68961 | 5.4317 |
| INPP5B | P32019 | 5.8338 | 6.6724 | 6.48616 | 7.27353 | 6.69592 | 6.32268 | 5.94855 | 6.15183 | 6.55358 | 6.30792 | 6.91216 | 5.99372 | 6.07266 | 5.5355 | 5.8406 | 5.39898 |
| NAF1 | Q96HR8 | 5.8262 | 6.1399 | 6.93491 | 6.01014 | 5.72687 | 6.61997 | 6.30683 | 6.53456 | 5.6135 | 6.2364 | 6.44915 | 6.72345 | 5.79868 | 5.75671 | 6.87768 | 6.44507 |
| AIF1L | Q9BQI0-2 | 6.36074 | 5.28811 | 6.97825 | 6.53127 | 6.07116 | 5.65961 | 5.96113 | 6.08691 | 6.43488 | 6.0759 | 6.57839 | 7.4649 | 6.29746 | 5.67235 | 6.08059 | 6.45836 |
| SLC4A11 | Q8NBS3-4 | 6.9812 | 7.49525 | 5.62179 | 6.70844 | 5.93307 | 6.73359 | 5.90625 | 6.33948 | 5.99659 | 5.8614 | 6.08906 | 5.68062 | 6.21083 | 6.80583 | 5.9597 | 5.6769 |
| GDE1 | Q9NZC3 | 6.23654 | 6.77496 | 5.97012 | 6.21028 | 6.19948 | 6.19129 | 6.41118 | 6.2573 | 5.96543 | 6.46294 | 5.94048 | 5.68535 | 6.61976 | 6.0994 | 6.51543 | 6.46006 |
| ALKBH2 | Q6NS38 | 5.79652 | 4.63597 | 6.05324 | 6.144 | 5.33351 | 5.31184 | 6.0496 | 6.25126 | 6.20715 | 6.94402 | 7.56129 | 6.61843 | 6.36178 | 6.31241 | 7.0131 | 7.40589 |
| LAYN | Q6UX15 | 6.84808 | 5.6491 | 7.31313 | 6.29917 | 5.99283 | 5.8938 | 6.47365 | 5.28751 | 6.74984 | 6.16906 | 6.10144 | 6.43072 | 6.59189 | 6.50928 | 6.15329 | 5.53722 |
| TRERF1 | Q96PN7 | 6.44299 | 6.20203 | 6.04436 | 6.62363 | 6.00952 | 5.71777 | 5.82093 | 6.39841 | 6.31847 | 5.91986 | 6.45311 | 6.15368 | 6.53062 | 6.50235 | 6.49661 | 6.36565 |
| FBXW2 | Q9UKT8 | 6.05007 | 5.67244 | 6.78123 | 6.26298 | 6.03835 | 6.80457 | 6.53654 | 6.24066 | 6.09475 | 6.73591 | 6.74825 | 6.35906 | 6.0098 | 6.29727 | 5.93217 | 5.43596 |
| CCDC115 | Q96NT0 | 5.5716 | 5.33566 | 5.15017 | 5.19287 | 6.32346 | 5.52572 | 6.42839 | 6.95886 | 5.92329 | 7.21759 | 5.92827 | 7.61444 | 6.40489 | 6.83279 | 6.39897 | 7.19304 |
| RNF187 | Q5TA31 | 8.25441 | 8.22095 | 8.60242 | 6.82277 | 4.69931 | 6.01953 | 6.73031 | 7.15453 | 8.17546 | 8.58132 | 7.28649 | 7.10893 | 2.59416 | 3.25529 | 2.74487 | 3.74923 |

| | | | | | | | | | | | | | | | | | |
|---|---|---|---|---|---|---|---|---|---|---|---|---|---|---|---|---|---|
| NTAN1 | Q96AB6 | 6.64952 | 7.0235 | 6.36374 | 6.26769 | 6.35557 | 5.59491 | 5.97863 | 7.20473 | 5.91396 | 6.67238 | 7.10924 | 7.44368 | 5.26477 | 5.71375 | 5.02301 | 5.42093 |
| ADAM20 | O43506 | 5.92798 | 5.99 | 6.6 | 6.33877 | 6.04905 | 6.10375 | 6.20241 | 6.15748 | 6.37022 | 6.63609 | 6.6829 | 6.89401 | 5.85244 | 6.45668 | 5.65801 | 6.0802 |
| PRKAB1 | Q9Y478 | 5.93782 | 6.59314 | 6.15325 | 6.02565 | 6.56488 | 6.96437 | 5.29912 | 6.95565 | 6.54392 | 5.93379 | 6.51744 | 6.13954 | 5.7992 | 5.89944 | 6.80281 | 5.86997 |
| ZBED5 | Q49AG3 | 5.90393 | 6.44542 | 6.304 | 6.204 | 5.94409 | 6.43782 | 6.23044 | 6.20371 | 6.676 | 6.69209 | 6.43105 | 6.93513 | 5.76549 | 5.96077 | 5.99916 | 5.86693 |
| ATP5MC2 | Q06055-2 | 6.31642 | 6.3017 | 5.42731 | 6.03008 | 6.20946 | 6.05757 | 5.83977 | 6.25425 | 6.00068 | 6.16671 | 5.16716 | 5.63697 | 6.81782 | 7.33592 | 7.33376 | 7.10441 |
| FAM234B | A2RU67 | 6.47592 | 6.17989 | 6.75303 | 7.44858 | 5.67917 | 6.25147 | 5.49005 | 6.2143 | 6.79192 | 5.67618 | 6.94015 | 6.82925 | 6.20054 | 6.2574 | 5.83735 | 4.97481 |
| STK38L | Q9Y2H1 | 6.83109 | 7.89874 | 6.64496 | 6.23004 | 6.50621 | 6.27693 | 7.2948 | 5.3744 | 5.64258 | 7.06207 | 6.80898 | 6.5221 | 4.89117 | 5.37883 | 5.79704 | 4.84005 |
| DCLK2 | Q8N568-3 | 5.97721 | 6.04127 | 5.82966 | 5.75576 | 6.42533 | 6.65334 | 5.62309 | 6.07172 | 6.15035 | 6.74728 | 6.03255 | 7.05715 | 6.674 | 6.5526 | 6.42745 | 5.98123 |
| TADA1 | Q96BN2 | 6.66419 | 6.4604 | 5.9578 | 6.44919 | 5.96392 | 6.19839 | 6.21619 | 6.48797 | 5.95232 | 5.78387 | 5.78274 | 6.41092 | 6.67373 | 6.15179 | 6.2812 | 6.56538 |
| STX11 | O75558 | 6.17277 | 7.19315 | 6.66401 | 6.84322 | 5.60039 | 5.83258 | 6.11888 | 6.37504 | 6.41369 | 6.36465 | 6.11907 | 6.11968 | 5.98704 | 5.81926 | 6.07873 | 6.29783 |
| ABL2 | P42684 | 5.71183 | 6.00444 | 6.10224 | 5.83275 | 6.2174 | 6.35178 | 5.9633 | 6.55872 | 6.10041 | 6.05182 | 6.3589 | 6.04782 | 6.61908 | 6.60994 | 6.75041 | 6.71915 |
| NQO2 | P16083 | 6.98107 | 4.78573 | 6.6777 | 5.03237 | 6.48544 | 5.48328 | 6.63692 | 7.37294 | 5.64633 | 7.69203 | 5.62886 | 6.52372 | 5.12536 | 6.88771 | 6.47194 | 6.56859 |
| PIGH | Q14442 | 6.60595 | 6.39934 | 6.67988 | 7.00351 | 6.51573 | 5.65119 | 5.96846 | 5.54622 | 7.27951 | 6.57327 | 5.39996 | 6.39169 | 6.19307 | 5.04086 | 7.01003 | 5.74132 |
| ERCC6L2 | Q5T890 | 5.80098 | 6.36728 | 5.86177 | 5.59659 | 5.71195 | 6.94407 | 6.43351 | 5.93268 | 6.05644 | 6.43385 | 6.46261 | 6.33709 | 6.09256 | 7.13014 | 6.54353 | 6.29496 |
| NCOA4 | Q13772-4 | 6.29225 | 6.38924 | 6.70126 | 7.98781 | 6.47074 | 7.04254 | 6.30903 | 5.73754 | 6.44072 | 6.07596 | 7.01343 | 6.74209 | 4.32613 | 6.23003 | 4.88551 | 5.35572 |
| BID | P55957-2 | 6.33406 | 4.77801 | 5.67919 | 4.72233 | 5.25365 | 4.15607 | 4.96387 | 6.93916 | 4.03713 | 5.93629 | 5.22242 | 6.53024 | 4.91619 | 11.4881 | 8.66124 | 10.3821 |
| TATDN3 | Q17R31-3 | 5.81002 | 5.797 | 6.96844 | 5.74687 | 5.96825 | 6.55781 | 6.16082 | 6.44981 | 6.13248 | 7.05997 | 7.03625 | 6.99985 | 5.86542 | 5.94222 | 5.78378 | 5.721 |
| CCDC167 | Q9P0B6 | 7.28265 | 6.59194 | 5.96421 | 6.18135 | 5.67069 | 6.12982 | 6.65775 | 5.60515 | 5.79877 | 5.6658 | 7.47152 | 6.12633 | 6.17123 | 6.12452 | 5.76526 | 6.79301 |
| CD99L2 | Q8TCZ2-5 | 6.95698 | 5.7309 | 6.03492 | 5.38577 | 6.36977 | 5.87364 | 5.82787 | 6.44883 | 5.82174 | 6.45918 | 5.90376 | 5.93852 | 7.38513 | 7.10938 | 6.10818 | 6.64544 |
| GRK5 | P34947 | 5.34232 | 5.73641 | 5.48445 | 4.90499 | 5.28679 | 6.75262 | 5.09667 | 5.18224 | 6.77383 | 5.97249 | 6.89416 | 7.3628 | 7.50535 | 7.22178 | 6.76126 | 7.72183 |
| PAGR1 | Q9BTK6 | 6.57766 | 5.64542 | 6.20882 | 5.8949 | 5.22531 | 6.46324 | 5.60713 | 6.56621 | 5.86969 | 7.1562 | 6.51043 | 6.73178 | 5.68664 | 5.99235 | 6.63175 | 7.23247 |
| BMPR1A | P36894 | 6.91228 | 7.23427 | 6.91367 | 7.11365 | 5.7525 | 5.82622 | 6.26093 | 6.89552 | 6.68423 | 7.21115 | 6.20214 | 6.25978 | 4.97472 | 4.61414 | 5.41882 | 5.72596 |
| RPS27L | Q71UM5 | 6.39993 | 7.19452 | 5.48378 | 6.40017 | 7.30469 | 7.72734 | 5.75222 | 8.10377 | 5.62445 | 6.07759 | 5.66108 | 5.57883 | 5.05598 | 6.74717 | 5.51585 | 5.37265 |
| LYRM7 | Q5U5X0 | 8.11646 | 6.51102 | 6.39383 | 5.5803 | 6.63355 | 5.68588 | 5.86417 | 5.93766 | 6.24123 | 6.10787 | 6.6402 | 5.67406 | 6.74801 | 6.22354 | 6.70019 | 4.94205 |
| FBXO42 | Q6P3S6 | 6.18775 | 6.05701 | 6.65129 | 6.32243 | 5.50066 | 5.60306 | 6.47458 | 6.41871 | 6.38979 | 7.02605 | 7.4152 | 7.19493 | 5.52096 | 5.64985 | 5.04204 | 6.54568 |
| KLHL12 | Q53G59-2 | 6.37557 | 7.18055 | 6.58726 | 6.01137 | 5.5821 | 6.66921 | 5.99948 | 6.22077 | 6.52311 | 7.02579 | 6.47546 | 6.81777 | 6.21452 | 4.96072 | 5.49752 | 5.8588 |
| KLHL18 | O94889 | 6.0006 | 6.41649 | 7.59801 | 5.46395 | 5.73745 | 6.38426 | 5.92193 | 6.70124 | 6.44871 | 5.74622 | 4.60855 | 6.16762 | 6.548 | 5.66156 | 6.36599 | 6.42941 |
| HSPB11 | Q9Y547 | 6.28003 | 6.05355 | 7.14796 | 5.63811 | 6.64834 | 6.5888 | 6.38296 | 6.06557 | 6.80472 | 6.70201 | 6.89216 | 6.19447 | 5.56287 | 5.42756 | 6.06243 | 5.54847 |
| TET2 | Q6N021 | 6.84428 | 6.38565 | 6.54451 | 6.84108 | 5.87735 | 6.42346 | 5.91374 | 5.88069 | 6.28125 | 6.16655 | 6.57303 | 6.43861 | 6.18105 | 5.84559 | 5.85195 | 5.95121 |
| MOCS2 | O96007 | 6.71701 | 5.22431 | 6.54416 | 6.41302 | 5.86724 | 5.34965 | 6.40519 | 6.11809 | 6.1568 | 7.38159 | 6.69253 | 7.07492 | 5.91089 | 5.71471 | 6.34427 | 6.08563 |
| GPN1 | Q9HCN4-5 | 6.6435 | 6.16212 | 7.07291 | 6.18952 | 6.36268 | 6.5388 | 5.69798 | 6.96427 | 6.45449 | 6.38518 | 7.21374 | 6.42976 | 5.58124 | 5.53255 | 5.14789 | 5.62336 |
| CHST2 | Q9Y4C5 | 6.241 | 6.50113 | 5.45135 | 6.03594 | 5.80061 | 6.11216 | 6.04954 | 6.78413 | 6.26191 | 6.77689 | 6.1664 | 6.82425 | 5.95019 | 5.85373 | 6.59157 | 6.5992 |
| STRADA | Q7RTN6 | 6.3388 | 6.27867 | 6.67743 | 6.5266 | 6.94805 | 6.50964 | 6.14504 | 6.86844 | 6.03882 | 6.70266 | 7.39982 | 7.11854 | 4.85725 | 5.75491 | 4.46468 | 5.37065 |
| RBFA | Q8N0V3 | 5.67215 | 5.49977 | 6.18347 | 7.46697 | 5.09364 | 6.24304 | 6.46736 | 6.66482 | 6.62404 | 6.33341 | 5.1058 | 7.14332 | 6.67005 | 6.96406 | 5.73379 | 6.13431 |
| RLIM | Q9NVW2 | 5.87321 | 6.93619 | 6.08689 | 7.07219 | 6.78093 | 6.94479 | 5.96571 | 6.4936 | 6.77845 | 6.82969 | 6.00748 | 7.05881 | 5.01382 | 4.57499 | 5.4998 | 6.08345 |
| CLK2 | P49760 | 4.69323 | 6.32414 | 6.4421 | 5.62564 | 6.20688 | 6.75599 | 6.51073 | 5.57265 | 6.52458 | 6.94184 | 6.08238 | 5.97423 | 5.92159 | 6.5446 | 7.27163 | 6.60778 |
| PAQR3 | Q6TCH7 | 6.1942 | 6.57605 | 6.26657 | 6.96615 | 6.48251 | 5.35279 | 6.04992 | 5.90858 | 6.57818 | 5.66912 | 5.64757 | 5.21218 | 6.72013 | 6.78814 | 6.20397 | 7.38393 |
| ZNF511 | Q8NB15-1 | 5.61028 | 5.92428 | 6.74647 | 5.15893 | 5.90371 | 6.63735 | 6.20926 | 6.85917 | 6.44008 | 6.13827 | 7.30693 | 6.01709 | 6.01764 | 6.17005 | 6.55326 | 6.30724 |
| NFE2L1 | Q14494 | 6.21347 | 6.47329 | 7.48185 | 7.46184 | 6.13919 | 6.46241 | 7.00217 | 6.51218 | 6.06607 | 6.35099 | 5.91901 | 6.91858 | 5.10655 | 5.66432 | 4.67446 | 5.55363 |
| AMDHD2 | Q9Y303-3 | 6.16461 | 6.49563 | 6.23576 | 5.74642 | 6.7317 | 7.03505 | 6.21684 | 7.21164 | 6.41392 | 6.3021 | 6.42903 | 6.21962 | 5.9945 | 5.97567 | 5.40913 | 5.41836 |
| SLC25A21 | Q9BQT8 | 6.13872 | 5.53785 | 5.63139 | 6.03895 | 6.45345 | 5.78115 | 6.3836 | 6.13348 | 5.85362 | 6.49482 | 6.32497 | 5.58419 | 7.30174 | 5.90268 | 6.85179 | 7.5876 |
| ELP5 | Q8TE02-4 | 5.97675 | 5.88116 | 6.72354 | 6.05567 | 6.17188 | 6.45232 | 6.3762 | 7.19209 | 6.60862 | 6.65491 | 6.6426 | 7.13407 | 5.80208 | 5.9674 | 5.22697 | 5.13373 |
| NPHP1 | O15259-4 | 6.22516 | 6.57052 | 6.88332 | 6.63323 | 5.90809 | 5.69319 | 6.29055 | 6.44117 | 6.34094 | 6.7068 | 6.53565 | 7.34419 | 5.32818 | 5.23575 | 5.47176 | 6.39152 |
| SMIM12 | Q96EX1 | 6.60038 | 5.23171 | 5.61819 | 5.1938 | 5.86888 | 5.15065 | 5.5778 | 6.33101 | 5.71442 | 5.75311 | 5.40129 | 5.90911 | 7.04194 | 7.73557 | 8.08105 | 8.79107 |
| ITPRIPL1 | Q6GPH6-2 | 6.26723 | 6.19278 | 6.38154 | 6.95537 | 6.3854 | 6.26096 | 6.88343 | 5.41847 | 6.4613 | 6.82033 | 6.47037 | 6.54224 | 5.54802 | 5.18016 | 6.17808 | 6.05433 |
| ELF1 | P32519 | 6.2754 | 6.74967 | 6.69231 | 6.6662 | 5.76161 | 5.96 | 6.56446 | 6.64415 | 5.70707 | 6.78047 | 6.27086 | 7.00472 | 5.59804 | 5.70556 | 5.53103 | 6.08845 |
| CCDC15 | Q0P6D6 | 6.73432 | 5.47625 | 7.01852 | 7.07025 | 5.27005 | 6.53283 | 6.51396 | 6.63132 | 7.7319 | 6.73031 | 6.44764 | 7.31931 | 4.84856 | 5.77457 | 4.70822 | 5.19201 |
| GKAP1 | Q5VSY0 | 5.8691 | 6.30292 | 7.11019 | 5.86796 | 6.05167 | 6.22507 | 5.75679 | 6.34319 | 6.74556 | 6.94777 | 7.10818 | 6.73739 | 5.50907 | 5.96848 | 5.84513 | 5.61153 |
| ANKRD54 | Q6NXT1 | 5.62617 | 6.1512 | 6.00119 | 5.83234 | 6.0434 | 6.5491 | 5.69423 | 6.73441 | 6.16293 | 6.42785 | 6.17153 | 6.85711 | 6.42808 | 6.01354 | 6.24937 | 7.05757 |
| ZCRB1 | Q8TBF4 | 6.55037 | 6.4332 | 6.30278 | 7.06724 | 5.99547 | 6.02689 | 6.63521 | 5.94784 | 6.4461 | 6.2548 | 6.28145 | 6.07258 | 6.25443 | 6.06208 | 5.63879 | 6.03079 |
| KLF13 | Q9Y2Y9 | 6.91342 | 6.10478 | 5.8626 | 7.38742 | 5.04448 | 6.18145 | 5.9162 | 6.99032 | 4.77071 | 6.9179 | 8.41388 | 7.2975 | 6.14084 | 4.88347 | 5.05824 | 6.11679 |
| HOXB7 | P09629 | 6.31794 | 6.53789 | 6.07137 | 7.39207 | 6.37733 | 7.05491 | 5.76745 | 5.79012 | 7.14327 | 7.31988 | 7.08912 | 6.47402 | 4.95609 | 5.05848 | 4.77185 | 5.8782 |
| ANKRD16 | Q6P6B7 | 6.95698 | 6.03481 | 7.17524 | 5.79366 | 6.47847 | 6.43649 | 5.8987 | 7.11261 | 6.77593 | 6.09898 | 7.09744 | 6.38069 | 5.92927 | 5.81933 | 4.72821 | 5.2832 |
| KMT5B | Q4FZB7 | 6.43855 | 6.74567 | 6.18662 | 7.27258 | 5.55395 | 5.92535 | 6.9221 | 6.48746 | 6.3674 | 6.67158 | 6.60819 | 6.28645 | 5.48235 | 5.64711 | 5.39062 | 6.01402 |
| TMEM106C | Q9BVX2 | 6.33236 | 5.56143 | 6.49641 | 6.01676 | 7.28819 | 6.69555 | 6.94404 | 7.03443 | 6.27923 | 7.2621 | 6.81847 | 6.92182 | 4.87698 | 4.55629 | 5.37503 | 5.44089 |
| FLVCR1 | Q9Y5Y0 | 6.66826 | 6.51951 | 6.70513 | 6.55669 | 6.41859 | 6.59876 | 6.68013 | 5.9867 | 6.47114 | 5.49004 | 6.11411 | 6.17691 | 4.93977 | 6.29013 | 5.5321 | 6.84905 |
| KLHL7 | Q8IXQ5 | 6.45835 | 6.1745 | 6.38723 | 7.48076 | 6.27073 | 6.15531 | 6.62273 | 5.55342 | 6.66268 | 6.45767 | 6.49309 | 6.42872 | 5.49093 | 5.63202 | 6.03238 | 5.69947 |
| WDR53 | Q7Z5U6 | 6.29228 | 5.9734 | 7.76525 | 6.42262 | 6.28833 | 6.03945 | 6.26549 | 6.9568 | 6.3561 | 6.61269 | 7.07187 | 6.95622 | 5.21194 | 5.77262 | 5.00732 | 5.00761 |
| SPRYD3 | Q8NCJ5 | 6.00739 | 6.805 | 7.84232 | 7.47073 | 6.23076 | 7.06912 | 6.24863 | 6.39882 | 6.84082 | 6.89831 | 6.37062 | 6.70959 | 4.01758 | 5.27142 | 4.83319 | 4.9857 |
| RGP1 | Q92546 | 6.22664 | 6.41221 | 7.0494 | 7.44542 | 6.04489 | 6.2725 | 6.3921 | 5.70672 | 6.74174 | 6.90324 | 6.55651 | 7.05907 | 4.98062 | 4.81509 | 6.01488 | 5.37896 |
| APOBEC3F | Q8IUX4 | 6.06598 | 5.89843 | 6.56331 | 6.27337 | 6.81429 | 6.20371 | 5.61572 | 6.00049 | 5.84586 | 7.1584 | 6.22087 | 6.05812 | 6.18429 | 6.57321 | 6.39749 | 6.12646 |
| WDR83 | Q9BRX9 | 5.65726 | 6.57132 | 5.72737 | 5.88603 | 6.43704 | 6.34798 | 5.49532 | 6.40405 | 5.90159 | 6.3088 | 5.84493 | 6.26989 | 7.09839 | 6.97135 | 6.54089 | 6.5378 |
| NHLRC3 | Q5JS37 | 6.49637 | 6.82655 | 6.22987 | 6.96687 | 6.2181 | 5.98141 | 6.05312 | 6.07831 | 5.58924 | 5.39086 | 6.13678 | 6.20916 | 6.48495 | 6.53561 | 7.04955 | 5.75325 |
| SRPX | P78539 | 6.90411 | 7.76173 | 5.50754 | 7.25423 | 5.61234 | 6.53112 | 7.2727 | 5.69245 | 6.93705 | 7.0517 | 6.30056 | 6.95891 | 5.07655 | 4.20008 | 4.48706 | 6.45187 |
| CDC25C | P30307-3 | 6.65672 | 6.40515 | 7.6495 | 7.25021 | 6.17525 | 6.69047 | 6.78815 | 6.43455 | 6.51046 | 7.74027 | 7.11971 | 7.94042 | 3.94102 | 4.45797 | 3.6437 | 4.59646 |
| STMN3 | Q9NZ72 | 7.50723 | 6.92508 | 7.55703 | 7.30338 | 5.79294 | 5.81924 | 6.87757 | 6.80468 | 7.53217 | 8.92076 | 7.61136 | 8.06004 | 3.39039 | 3.41904 | 3.00144 | 3.47764 |
| RILPL2 | Q969X0 | 6.18188 | 7.05488 | 7.1035 | 6.60472 | 6.42813 | 6.63993 | 5.9175 | 6.77586 | 6.88129 | 6.60928 | 6.59026 | 7.65618 | 4.40896 | 4.99273 | 5.12301 | 5.03189 |
| MT-CYB | P00156 | 5.831 | 6.15318 | 5.33863 | 6.59191 | 6.94258 | 6.11018 | 6.74089 | 6.21099 | 5.51303 | 6.21822 | 6.64522 | 5.58264 | 6.36621 | 7.65061 | 6.24274 | 5.86196 |
| USP30 | Q70CQ3 | 6.20927 | 6.88498 | 6.63989 | 6.74897 | 6.00291 | 6.27718 | 6.49495 | 5.92107 | 5.79364 | 6.18995 | 5.66609 | 5.53106 | 6.86396 | 6.4016 | 6.29118 | 6.0833 |
| TGIF1 | Q15583 | 3.87283 | 4.78737 | 4.7154 | 4.83855 | 4.94754 | 5.65968 | 4.88901 | 4.82098 | 4.86027 | 4.41351 | 4.81731 | 4.71454 | 11.7028 | 10.0572 | 10.8961 | 10.0069 |
| CLCN6 | P51797 | 6.434 | 6.56592 | 6.62855 | 6.05876 | 5.47005 | 5.74328 | 5.8153 | 5.62979 | 6.54356 | 6.50237 | 6.70368 | 5.98634 | 5.87725 | 7.18391 | 6.97807 | 6.77917 |
| MBOAT1 | Q6ZNC8 | 5.66237 | 6.34093 | 6.32343 | 6.8889 | 6.07449 | 5.23963 | 5.80726 | 5.88519 | 6.30439 | 6.59012 | 5.8532 | 5.76724 | 6.9246 | 6.44451 | 6.98851 | 6.90522 |
| WWC1 | Q8IX03-2 | 6.2521 | 5.93466 | 7.10546 | 6.22431 | 6.56232 | 6.56268 | 6.23236 | 6.23327 | 6.3479 | 6.31088 | 7.12507 | 6.7694 | 5.88836 | 5.23416 | 5.62633 | 5.59073 |
| UBTD1 | Q9HAC8 | 5.95722 | 6.78964 | 5.74645 | 6.57334 | 5.85998 | 6.34743 | 6.07744 | 6.22661 | 6.75386 | 6.58541 | 6.67285 | 6.83836 | 5.61883 | 6.18573 | 5.8709 | 5.89594 |
| TAF11 | Q15544 | 6.66505 | 6.08451 | 6.14625 | 6.1485 | 6.93945 | 6.13721 | 6.23608 | 6.16793 | 6.87099 | 6.29827 | 5.99299 | 5.52141 | 6.30471 | 5.25846 | 6.2792 | 6.949 |
| FOXN3 | O00409 | 6.90621 | 6.98104 | 6.22799 | 6.86208 | 6.2938 | 6.29093 | 5.79204 | 6.67026 | 6.02954 | 6.49791 | 7.47127 | 6.44765 | 4.83348 | 5.78905 | 5.24442 | 5.66233 |
| FOXG1 | P55316 | 6.16036 | 6.21928 | 6.00605 | 6.8472 | 6.08186 | 6.11627 | 5.83934 | 6.87566 | 6.41205 | 6.30298 | 6.2271 | 6.62785 | 6.19666 | 5.59636 | 5.52945 | 6.59153 |
| ADAT1 | Q9BUB4 | 6.77375 | 7.25026 | 7.75525 | 7.31214 | 7.26207 | 5.71171 | 6.64289 | 4.93065 | 6.09009 | 5.59089 | 5.30072 | 5.8152 | 5.31968 | 6.51396 | 6.38672 | 5.34403 |
| TGFBR1 | P36897 | 6.79701 | 6.69616 | 6.28609 | 6.91497 | 6.64041 | 5.9993 | 6.48967 | 6.22944 | 6.86704 | 6.19088 | 6.41647 | 5.87144 | 5.812 | 5.58902 | 6.5587 | 5.5414 |
| C16orf72 | A0A087X1J2 | 6.60533 | 6.72191 | 7.2052 | 6.60882 | 5.86828 | 6.17216 | 6.76145 | 6.38359 | 6.61088 | 6.74821 | 7.21878 | 7.35996 | 5.01992 | 4.74028 | 5.11117 | 4.86407 |
| AURKAIP1 | Q9NWT8 | 6.05398 | 6.72752 | 5.06674 | 6.49833 | 6.5832 | 6.2 | 6.93801 | 6.3708 | 6.57873 | 5.99621 | 6.17326 | 5.95994 | 5.5622 | 6.34786 | 6.16959 | 6.77361 |
| CBX4 | O00257 | 6.66393 | 7.13719 | 5.8675 | 8.00708 | 6.28246 | 5.88892 | 5.77803 | 5.80925 | 6.06905 | 6.31522 | 6.34677 | 6.47286 | 5.94954 | 5.4816 | 5.93421 | 5.99638 |
| LAT2 | Q9GZY6 | 6.66982 | 6.19234 | 5.47528 | 6.43188 | 5.96316 | 6.17214 | 5.72594 | 6.00653 | 6.35851 | 6.51006 | 6.38855 | 6.10362 | 6.58311 | 7.4278 | 5.83423 | 6.15705 |
| SFXN5 | Q8TD22 | 6.18128 | 6.08048 | 6.22431 | 6.79593 | 5.76938 | 5.82347 | 5.92275 | 5.76117 | 6.94099 | 6.50251 | 5.51354 | 5.8546 | 6.44775 | 6.55329 | 6.35593 | 7.27261 |
| PLCH1 | Q4KWH8 | 6.61655 | 7.16307 | 5.1937 | 5.76894 | 6.1465 | 6.95271 | 6.04838 | 5.934 | 5.84427 | 6.14825 | 6.25296 | 5.97004 | 6.45586 | 6.88345 | 6.26118 | 6.36013 |
| CLMP | Q9H6B4 | 6.20848 | 6.3299 | 5.98269 | 6.78524 | 6.47597 | 6.15218 | 6.05453 | 6.5821 | 6.06208 | 5.97784 | 5.73764 | 5.96125 | 5.93093 | 6.61789 | 6.70417 | 6.43712 |
| CENPM | Q9NSP4 | 6.36365 | 6.75067 | 6.87459 | 7.94247 | 6.23303 | 5.90192 | 5.67717 | 5.66434 | 6.58004 | 5.5067 | 6.92244 | 6.22907 | 5.98901 | 5.47007 | 5.57784 | 6.31698 |
| MTMR3 | Q13615 | 5.97253 | 6.25967 | 6.65353 | 6.44669 | 6.16824 | 7.04838 | 5.57407 | 5.56732 | 6.42799 | 6.44837 | 6.15509 | 5.61014 | 5.87278 | 5.76812 | 6.49069 | 7.53638 |
| TXNDC16 | Q9P2K2 | 6.89053 | 6.36787 | 6.42663 | 6.47485 | 6.373 | 5.91831 | 7.00325 | 5.41533 | 6.21718 | 6.04106 | 5.38384 | 6.37577 | 5.99602 | 6.55478 | 6.74464 | 5.81696 |
| EFCAB14 | O75071 | 6.48392 | 6.92704 | 6.52191 | 6.66862 | 5.47864 | 6.67249 | 6.36934 | 6.34118 | 6.19483 | 6.44567 | 6.12089 | 6.64908 | 5.85032 | 5.77756 | 5.55235 | 5.94616 |
| RNF115 | Q9Y4L5 | 6.52031 | 6.03413 | 6.63448 | 6.09287 | 7.16429 | 7.13892 | 6.32723 | 6.63711 | 6.49759 | 6.0916 | 6.89019 | 7.58405 | 5.11773 | 4.31062 | 5.48568 | 5.4732 |
| CCDC9B | Q6ZUT6 | 5.16905 | 7.21055 | 6.94802 | 6.55631 | 6.40325 | 6.62255 | 6.31393 | 5.10937 | 5.9073 | 6.28653 | 5.66032 | 6.32586 | 5.7921 | 5.79321 | 6.50137 | 7.40027 |
| BRCC3 | P46736 | 5.6064 | 6.31435 | 5.8528 | 6.71514 | 6.74695 | 6.41272 | 5.93979 | 6.17651 | 6.98606 | 6.42225 | 6.47412 | 6.20237 | 6.37424 | 5.95756 | 6.27534 | 5.54341 |
| OSCP1 | Q8WVF1 | 6.53402 | 5.55585 | 7.12854 | 6.3573 | 6.52408 | 6.10857 | 5.59871 | 7.1032 | 6.4413 | 6.51048 | 7.17788 | 7.60198 | 6.23761 | 4.91673 | 4.79411 | 5.40964 |
| CHKA | P35790 | 6.1688 | 6.2002 | 7.85762 | 6.52093 | 6.0933 | 6.59464 | 6.14156 | 6.59457 | 6.61253 | 6.52658 | 7.52608 | 6.94874 | 5.5189 | 5.18064 | 5.08872 | 4.42619 |
| PTTG1IP | P53801 | 6.59938 | 6.37552 | 6.48059 | 7.08879 | 5.76574 | 5.96376 | 6.24249 | 5.72934 | 6.0997 | 7.11911 | 6.26176 | 6.2596 | 5.62142 | 6.33917 | 5.67478 | 6.37884 |
| FAM210A | Q96ND0 | 7.10148 | 6.14576 | 6.11911 | 6.59791 | 6.51591 | 6.05685 | 6.44586 | 5.8781 | 5.77807 | 6.56131 | 5.58559 | 5.71844 | 6.73646 | 5.96438 | 6.43021 | 6.36456 |
| BCL7B | Q9BQE9 | 6.56366 | 6.65589 | 6.20874 | 7.02882 | 6.30612 | 5.90301 | 5.93783 | 6.4708 | 6.02026 | 6.40864 | 6.74261 | 6.6038 | 5.84479 | 5.66622 | 5.79759 | 5.84121 |
| PDCL | Q13371 | 6.06646 | 6.14363 | 6.75079 | 5.7669 | 6.61596 | 6.88859 | 6.16192 | 6.31846 | 6.3825 | 6.37356 | 6.58194 | 6.40451 | 6.01056 | 6.30533 | 5.47907 | 5.74982 |
| ATXN7 | O15265-2 | 6.90711 | 6.28421 | 6.11127 | 6.83398 | 5.82324 | 6.06895 | 6.43288 | 5.71721 | 6.00145 | 5.80706 | 7.15173 | 6.10632 | 5.8758 | 6.04392 | 6.35923 | 6.47565 |
| BIRC5 | O15392 | 6.41849 | 5.62579 | 6.13093 | 5.63007 | 6.80749 | 5.04144 | 6.15845 | 6.82346 | 5.71863 | 6.98736 | 7.32007 | 7.78715 | 5.42783 | 6.18158 | 5.71624 | 6.22503 |
| RWDD3 | Q9Y3V2 | 4.92802 | 6.99976 | 8.81416 | 5.51895 | 5.55075 | 6.73568 | 6.8792 | 6.35688 | 5.78843 | 6.82917 | 7.31251 | 5.90681 | 6.15961 | 5.83056 | 5.32863 | 5.06088 |
| MSANTD3 | Q96H12 | 7.37289 | 7.26781 | 6.44138 | 7.01221 | 6.7599 | 6.08457 | 6.66818 | 6.54479 | 6.40271 | 6.02463 | 5.82869 | 5.93245 | 5.80238 | 5.20265 | 5.32979 | 5.32498 |
| CDK8 | P49336 | 6.49594 | 6.21619 | 6.34872 | 6.9409 | 5.94283 | 6.27151 | 6.28039 | 5.88163 | 6.24107 | 6.32936 | 5.95685 | 6.66149 | 6.22306 | 5.94514 | 6.16553 | 6.09939 |
| NIPA1 | Q7RTP0 | 6.11496 | 6.09712 | 6.2766 | 6.48267 | 5.93339 | 6.00483 | 6.10734 | 5.46807 | 5.79508 | 5.98625 | 5.27351 | 6.00996 | 6.94509 | 6.86266 | 7.57996 | 7.06252 |

| | | | | | | | | | | | | | | | | | |
|---|---|---|---|---|---|---|---|---|---|---|---|---|---|---|---|---|---|
| NAP1L5 | Q96NT1 | 6.89327 | 6.66245 | 7.38665 | 5.65015 | 7.09607 | 6.00804 | 5.27373 | 6.77518 | 6.4658 | 6.34235 | 7.37361 | 7.12205 | 5.85628 | 5.27361 | 4.86908 | 4.95171 |
| NLRP1 | Q9C000 | 6.17247 | 6.51189 | 6.43416 | 5.97178 | 6.24632 | 6.02081 | 5.71676 | 5.70827 | 7.15818 | 5.85954 | 6.05025 | 6.53777 | 6.32339 | 5.7929 | 6.87864 | 6.61687 |
| TMEM128 | Q5BJH2 | 6.48296 | 6.58598 | 5.98611 | 6.83268 | 5.74705 | 6.42786 | 6.44381 | 5.77272 | 5.80857 | 6.32669 | 6.25348 | 6.03938 | 6.23747 | 6.27753 | 6.24748 | 6.53025 |
| ABHD14B | Q96IU4 | 4.3107 | 5.4662 | 5.57069 | 5.16468 | 7.08122 | 5.3548 | 7.01174 | 7.5712 | 5.99826 | 7.07354 | 5.48204 | 7.0006 | 5.40813 | 7.33569 | 6.32479 | 7.84572 |
| COA5 | Q86WW8 | 6.84266 | 6.55996 | 6.99819 | 7.41536 | 6.61336 | 6.78533 | 6.16453 | 5.85519 | 6.34402 | 6.29932 | 6.64104 | 6.11547 | 5.76532 | 4.93576 | 5.38307 | 5.28143 |
| NKX2-6 | A6NCS4 | 6.8059 | 7.04432 | 6.47538 | 6.63125 | 6.53796 | 6.29144 | 6.76722 | 6.13481 | 6.35457 | 6.38498 | 6.57628 | 6.26383 | 5.30638 | 5.33464 | 5.79573 | 5.29531 |
| DCLK1 | O15075 | 6.46369 | 5.94794 | 7.00024 | 5.74904 | 5.44553 | 6.66682 | 6.38453 | 5.87638 | 5.86149 | 6.34635 | 6.20875 | 6.41363 | 6.39758 | 6.59463 | 6.49187 | 6.1515 |
| NEO1 | Q92859 | 6.22752 | 6.69371 | 6.47539 | 7.16882 | 5.96769 | 6.13354 | 5.77793 | 5.76966 | 6.70277 | 6.31689 | 6.36683 | 6.28035 | 6.73612 | 5.95124 | 5.83011 | 5.60142 |
| LEF1 | Q9UJU2 | 6.41242 | 5.82808 | 6.69737 | 6.19284 | 5.38392 | 5.47153 | 6.29561 | 5.52759 | 6.54928 | 6.67519 | 6.08693 | 6.60011 | 6.81877 | 5.92548 | 6.73811 | 6.79675 |
| H3-3A | P84243 | 6.64346 | 6.68043 | 6.1227 | 6.49438 | 6.14837 | 6.04491 | 6.25788 | 5.90952 | 6.03637 | 5.97621 | 6.22626 | 6.32623 | 6.16706 | 6.48416 | 6.20312 | 6.27896 |
| DOCK4 | Q8N1I0-3 | 5.85646 | 6.41407 | 6.70955 | 6.71479 | 6.50427 | 6.52578 | 6.17703 | 5.57186 | 5.5283 | 7.02777 | 6.35186 | 6.76799 | 6.18843 | 6.84852 | 5.5581 | 5.25521 |
| ABCG2 | Q9UNQ0 | 5.6626 | 6.44596 | 5.60836 | 6.60879 | 6.31226 | 5.75713 | 5.90097 | 5.68966 | 6.19237 | 6.45071 | 6.88987 | 6.69796 | 6.32631 | 5.82069 | 6.36693 | 7.26945 |
| MID1 | O15344 | 6.25598 | 6.30954 | 6.93118 | 5.98677 | 5.84935 | 6.61485 | 6.17187 | 6.34708 | 6.28941 | 6.59615 | 6.54039 | 6.60205 | 5.66117 | 5.96627 | 5.9899 | 5.88803 |
| SYS1 | Q8N2H4 | 6.04053 | 6.19825 | 5.54306 | 6.32513 | 5.99362 | 5.93505 | 6.59765 | 6.35065 | 5.19611 | 6.3027 | 5.42297 | 6.34712 | 6.30023 | 7.56161 | 6.76863 | 7.11668 |
| GCNT2 | Q8N0V5 | 6.54372 | 6.37674 | 6.11495 | 6.79427 | 6.98324 | 6.23472 | 5.99182 | 7.11717 | 6.57478 | 6.47246 | 5.50705 | 5.29029 | 5.69953 | 6.28157 | 5.70556 | 6.31215 |
| PTRHD1 | Q6GMV3 | 6.45693 | 5.34853 | 5.68115 | 4.68125 | 5.81217 | 4.68968 | 5.37244 | 6.78021 | 5.04871 | 6.20201 | 4.86154 | 5.8388 | 4.90166 | 10.6348 | 7.94878 | 9.74133 |
| HGH1 | Q9BTY7 | 5.28141 | 5.88666 | 5.36043 | 4.92877 | 4.84137 | 4.91054 | 6.39011 | 6.30601 | 4.6369 | 4.31326 | 5.19227 | 6.39306 | 5.60914 | 9.74373 | 8.14433 | 12.062 |
| BRMS1L | Q5PSV4 | 6.18707 | 6.33843 | 5.87404 | 6.81863 | 6.24884 | 6.20666 | 6.23581 | 6.02553 | 6.18278 | 6.29757 | 6.09194 | 5.90353 | 6.35768 | 6.22312 | 6.54794 | 6.46043 |
| RNF146 | Q9NTX7 | 5.71442 | 5.64737 | 6.67598 | 5.66774 | 5.8129 | 7.31792 | 7.623 | 7.01744 | 6.6496 | 6.52093 | 7.02968 | 6.91352 | 5.4476 | 5.41538 | 5.02248 | 5.52403 |
| C6orf89 | Q6UWU4-2 | 6.72952 | 6.48249 | 6.16663 | 6.48184 | 6.58326 | 6.09698 | 6.15089 | 6.3752 | 6.15908 | 6.11607 | 5.92178 | 5.72717 | 6.66707 | 5.99139 | 5.94962 | 6.401 |
| AKAP7 | F5GXD1 | 6.37523 | 6.41486 | 6.39902 | 4.5908 | 6.3574 | 5.38218 | 7.09748 | 5.7309 | 5.71956 | 6.51577 | 7.464 | 5.61229 | 7.1726 | 6.23952 | 6.15918 | 6.76922 |
| TMEM184B | Q9Y519 | 6.12281 | 6.06568 | 6.27804 | 5.60614 | 6.25962 | 5.3757 | 5.72244 | 6.41915 | 6.99645 | 6.20076 | 6.35732 | 6.19507 | 6.15607 | 6.84914 | 6.93099 | 6.46462 |
| ROMO1 | P60602 | 6.48032 | 6.1015 | 5.88465 | 6.22996 | 6.7944 | 6.34652 | 6.26929 | 6.78057 | 6.28292 | 5.92395 | 5.88745 | 6.01268 | 6.7724 | 6.16386 | 6.10937 | 5.96015 |
| CDC42EP2 | O14613 | 5.87955 | 4.95228 | 5.54558 | 6.45858 | 5.18149 | 5.16053 | 5.68762 | 7.30219 | 5.30837 | 6.33754 | 6.75585 | 6.41674 | 5.13027 | 8.14231 | 7.13481 | 8.6063 |
| SAYSD1 | Q9NPB0 | 6.93407 | 5.49615 | 6.0488 | 6.03021 | 5.94398 | 5.80803 | 6.14911 | 6.05129 | 6.01282 | 6.48776 | 6.34589 | 6.32052 | 6.1237 | 6.8635 | 6.58616 | 6.79801 |
| PIMREG | Q9BSJ6 | 5.94118 | 6.29631 | 6.34266 | 6.34253 | 6.28879 | 6.16416 | 6.39898 | 6.01695 | 6.37884 | 5.71344 | 6.07552 | 6.26807 | 6.43459 | 6.58489 | 6.55679 | 6.19629 |
| ALKBH4 | Q9NXW9 | 7.3996 | 5.57449 | 7.01381 | 4.32526 | 6.57836 | 6.66467 | 7.16948 | 6.36257 | 7.04802 | 6.46221 | 7.34308 | 5.33339 | 5.79773 | 5.76396 | 4.82971 | 6.33367 |
| TMEM41A | Q96HV5 | 6.4248 | 6.71614 | 7.2049 | 6.14508 | 5.7903 | 5.62259 | 6.28625 | 6.01131 | 6.52372 | 5.67041 | 5.98729 | 6.48497 | 5.58876 | 6.22091 | 6.39287 | 6.9297 |
| GMCL1 | Q96IK5 | 5.87391 | 7.18723 | 5.6557 | 7.08742 | 5.29132 | 5.63456 | 6.30988 | 5.98612 | 6.75525 | 6.17906 | 6.3287 | 6.88771 | 6.27556 | 5.96342 | 6.09705 | 6.48711 |
| HASPIN | Q8TF76 | 6.43417 | 6.27322 | 6.28876 | 7.31021 | 5.90319 | 5.76533 | 7.09727 | 6.4702 | 6.5086 | 6.87748 | 5.99546 | 6.14545 | 5.56716 | 5.59628 | 5.67672 | 6.09049 |
| BTG3 | Q14201-2 | 6.6927 | 6.14684 | 6.86057 | 6.22373 | 7.24364 | 6.91856 | 6.85006 | 6.97445 | 5.84705 | 6.3538 | 6.69596 | 6.92898 | 4.92702 | 5.2476 | 4.24934 | 5.8397 |
| CHFR | Q96EP1 | 5.25737 | 5.65855 | 6.26975 | 5.9457 | 5.64964 | 6.93194 | 5.81344 | 6.34937 | 5.76527 | 5.9992 | 6.41338 | 6.84063 | 6.30697 | 6.85602 | 6.74501 | 7.19778 |
| MOCOS | Q96EN8 | 6.49832 | 7.25536 | 6.72798 | 6.08216 | 6.42161 | 7.47303 | 5.8748 | 6.00926 | 6.60046 | 5.60269 | 6.93241 | 6.25735 | 5.60768 | 5.75138 | 4.9848 | 5.92073 |
| IGFBP3 | P17936-2 | 6.80338 | 5.72889 | 6.08294 | 7.61021 | 5.08722 | 5.90744 | 8.03108 | 5.79132 | 5.60302 | 5.02324 | 5.72952 | 6.22419 | 7.55281 | 4.80787 | 6.83803 | 7.17886 |
| NUSAP1 | Q9BXS6-7 | 4.91054 | 5.59531 | 6.1365 | 4.45593 | 5.52459 | 5.59767 | 6.27683 | 5.61325 | 6.6492 | 7.16728 | 7.08402 | 7.03757 | 6.47942 | 6.82053 | 7.2759 | 7.37546 |
| B4GAT1 | O43505 | 5.96485 | 6.44812 | 6.0656 | 6.77764 | 6.61611 | 6.73418 | 6.53703 | 6.23839 | 6.19717 | 5.966 | 6.20722 | 5.93583 | 5.77094 | 6.08622 | 6.24999 | 6.2047 |
| SLF1 | Q9BQI6 | 6.56487 | 6.79164 | 6.49683 | 7.94821 | 6.04412 | 5.5236 | 6.39124 | 6.05471 | 6.28319 | 7.11488 | 6.0731 | 6.94118 | 4.80187 | 5.44866 | 5.51621 | 6.0057 |
| TMC7 | Q7Z402 | 6.89705 | 6.11821 | 6.51726 | 6.63287 | 6.18967 | 5.42861 | 6.43923 | 5.67775 | 6.74729 | 5.91 | 5.70087 | 6.2452 | 6.90388 | 5.95363 | 6.447 | 6.1915 |
| CCDC90B | Q9GZT6 | 6.04889 | 4.81907 | 4.93499 | 5.67597 | 5.21155 | 4.41748 | 5.36766 | 6.38222 | 4.52328 | 5.6823 | 4.25345 | 5.80621 | 5.51487 | 10.3718 | 10.3559 | 10.6344 |
| POGLUT1 | Q8NBL1 | 6.2982 | 6.71668 | 5.92772 | 6.64414 | 6.48836 | 6.28272 | 6.42242 | 6.34002 | 5.76062 | 5.94429 | 5.64091 | 5.95549 | 5.9901 | 6.53387 | 6.6717 | 6.38275 |
| C5orf34 | Q96MH7 | 7.22378 | 6.87622 | 8.22004 | 7.10436 | 6.07459 | 7.35914 | 7.03055 | 6.94163 | 7.20216 | 7.06272 | 7.88264 | 8.41134 | 3.14303 | 3.21978 | 2.78695 | 3.46108 |
| CCDC61 | Q9Y6R9 | 5.79592 | 6.23376 | 6.43014 | 6.78877 | 6.73122 | 6.56688 | 6.62532 | 6.25418 | 6.39937 | 6.34663 | 5.91315 | 6.14099 | 5.96992 | 5.62962 | 5.81763 | 6.35649 |
| DEAF1 | O75398 | 5.68828 | 6.80158 | 5.71276 | 6.43225 | 5.92058 | 6.2131 | 5.77213 | 6.22621 | 6.20923 | 6.10373 | 6.46068 | 5.69161 | 6.84068 | 6.13614 | 6.88202 | 6.90901 |
| B4GALT2 | O60909-3 | 6.9326 | 7.21924 | 5.14771 | 6.88504 | 5.5995 | 5.63422 | 6.61107 | 6.41307 | 6.10045 | 6.15608 | 5.95433 | 6.2245 | 6.08515 | 6.34946 | 6.14435 | 5.51365 |
| PLD6 | Q8N2A8 | 6.67135 | 6.59851 | 5.47449 | 7.02871 | 6.31231 | 5.83739 | 7.43796 | 6.99152 | 6.00316 | 4.57473 | 4.87938 | 5.91431 | 6.18526 | 5.52045 | 6.69884 | 7.87162 |
| RDH10 | Q8IZV5 | 7.12644 | 6.16617 | 6.31639 | 7.34308 | 6.25458 | 6.22041 | 6.09471 | 5.7026 | 6.54765 | 6.01203 | 7.02608 | 6.13696 | 5.89716 | 5.69531 | 5.42927 | 6.03118 |
| SSH2 | Q76I76 | 5.80243 | 5.98491 | 6.05242 | 5.54716 | 6.34883 | 6.16847 | 6.61249 | 6.56093 | 6.34353 | 5.94522 | 6.27611 | 7.00679 | 6.54375 | 5.80788 | 6.4287 | 5.5704 |
| FOXD1 | Q16676 | 6.10593 | 4.71639 | 5.72026 | 4.88395 | 6.17462 | 6.01927 | 5.87163 | 6.12468 | 5.36492 | 5.69772 | 6.95491 | 6.35046 | 8.2388 | 6.82444 | 7.74313 | 7.20889 |
| RNF25 | Q96BH1 | 6.38997 | 6.03819 | 7.19065 | 6.00217 | 6.38397 | 6.44532 | 6.41319 | 6.28838 | 6.5291 | 6.45239 | 6.57063 | 6.34457 | 6.02464 | 5.47596 | 5.66732 | 5.78354 |
| BTBD1 | Q9H0C5 | 6.88236 | 6.17878 | 6.71246 | 6.49424 | 6.56069 | 6.58603 | 7.10218 | 6.43872 | 6.26449 | 6.54811 | 7.14108 | 6.07858 | 4.85996 | 5.03224 | 5.26044 | 5.85964 |
| BRF2 | Q9HAW0 | 5.79996 | 5.99554 | 6.69749 | 6.03549 | 6.42208 | 5.91311 | 5.90031 | 6.2198 | 6.97202 | 6.28634 | 8.41985 | 6.45368 | 4.82875 | 6.16691 | 5.95356 | 5.93511 |
| NDUFC1 | O43677 | 5.73611 | 6.25594 | 5.26892 | 6.18517 | 5.93323 | 5.63189 | 5.42372 | 5.89849 | 4.90172 | 5.1265 | 4.96262 | 5.31633 | 5.76564 | 10.3885 | 7.42241 | 9.78284 |
| VHL | P40337 | 6.77778 | 5.76073 | 6.26281 | 5.2053 | 6.8603 | 5.96129 | 6.64668 | 6.55371 | 6.46722 | 6.73897 | 5.99387 | 8.14063 | 5.26715 | 6.09719 | 5.51088 | 5.75548 |
| ZNF585A | Q6P3V2 | 5.68971 | 6.47493 | 6.13918 | 6.28851 | 6.73113 | 5.66101 | 6.63276 | 5.47485 | 6.76402 | 7.4792 | 6.774 | 6.79233 | 6.09925 | 5.99956 | 5.13833 | 5.86123 |
| AMMECR1 | Q9Y4X0 | 5.79409 | 6.26416 | 6.13931 | 6.2228 | 5.5878 | 6.7585 | 6.14141 | 7.26417 | 6.13575 | 6.43249 | 6.92743 | 6.46378 | 5.38697 | 6.91243 | 6.02085 | 5.54804 |
| PHTF1 | Q9UMS5 | 6.58227 | 6.59828 | 5.95588 | 7.13585 | 6.05001 | 6.06741 | 6.06661 | 5.81592 | 6.42764 | 5.46025 | 6.30638 | 7.06882 | 6.55692 | 6.19009 | 5.99955 | 5.71912 |
| PHTF2 | Q8N3S3-2 | 6.47366 | 7.27265 | 6.76371 | 6.99283 | 5.80175 | 6.83366 | 6.27661 | 5.44204 | 6.81192 | 6.1485 | 6.89882 | 7.47509 | 4.83044 | 5.27956 | 4.81361 | 5.88517 |
| PMVK | Q15126 | 5.27939 | 5.80807 | 6.94077 | 6.26844 | 6.21875 | 6.85557 | 6.04317 | 6.65608 | 6.98276 | 6.22733 | 6.77947 | 6.31877 | 5.90841 | 6.46187 | 5.71811 | 5.53305 |
| CCS | O14618 | 6.22362 | 5.339 | 6.35799 | 5.13872 | 5.81529 | 5.87556 | 5.65988 | 7.2903 | 4.90702 | 6.59281 | 6.70888 | 6.88681 | 6.8304 | 7.05006 | 6.07204 | 7.25162 |
| TCF7L2 | Q9NQB0 | 6.51432 | 6.6847 | 6.29076 | 7.01667 | 5.59419 | 4.66731 | 5.86139 | 7.44369 | 5.92197 | 6.10471 | 6.59947 | 6.27956 | 5.86405 | 6.42108 | 6.11197 | 6.62416 |
| CLSTN3 | Q9BQT9-2 | 7.87204 | 6.55748 | 5.70717 | 6.49504 | 5.82536 | 7.30607 | 6.52827 | 5.73478 | 7.06332 | 7.46414 | 4.44142 | 7.14337 | 5.3473 | 6.59543 | 5.42763 | 4.49121 |
| EMP3 | P54852 | 6.07729 | 6.55896 | 6.20589 | 6.81492 | 6.31877 | 6.11579 | 6.48874 | 5.70459 | 5.89226 | 6.34339 | 5.38047 | 5.37553 | 6.67171 | 6.26273 | 7.03992 | 6.74903 |
| ARHGAP26 | Q9UNA1 | 6.33998 | 6.38397 | 6.04768 | 6.47403 | 6.94382 | 5.76821 | 6.9391 | 6.53957 | 5.90541 | 6.2944 | 5.81543 | 6.16331 | 5.93714 | 5.27205 | 6.63665 | 6.53927 |
| ZSWIM6 | Q9HCJ5 | 6.50825 | 6.20688 | 6.81583 | 6.84501 | 5.64453 | 6.79657 | 5.93993 | 7.13988 | 6.29667 | 6.56585 | 6.48723 | 6.29627 | 5.66837 | 5.6976 | 5.19951 | 5.89162 |
| RIOK3 | O14730 | 7.5239 | 6.9047 | 6.33026 | 6.64447 | 6.80434 | 5.94254 | 6.23602 | 7.46984 | 6.69253 | 5.69928 | 6.83765 | 5.9649 | 4.99613 | 5.45213 | 5.29385 | 5.20747 |
| ACVR1 | Q04771 | 6.46679 | 6.91639 | 5.95472 | 6.6684 | 6.10642 | 6.55972 | 6.7599 | 6.11223 | 6.456 | 6.82609 | 5.60987 | 6.61419 | 6.10901 | 5.2753 | 5.74528 | 5.81969 |
| FBXL19 | Q6PCT2 | 4.96134 | 6.07187 | 6.42072 | 6.41204 | 5.34943 | 5.69831 | 6.13666 | 5.50287 | 6.40615 | 6.69467 | 5.91335 | 5.89818 | 7.12833 | 7.44309 | 7.0069 | 6.9561 |
| DNAAF2 | Q9NVR5 | 6.64423 | 6.11965 | 6.45033 | 5.60625 | 6.35354 | 7.69484 | 6.03342 | 6.60319 | 6.77134 | 6.82006 | 6.94555 | 6.40776 | 5.21385 | 5.72666 | 5.17095 | 5.43838 |
| MEIOB | Q8N635-2 | 6.73402 | 7.22907 | 7.1856 | 6.53128 | 6.49508 | 6.50443 | 5.76096 | 6.56136 | 6.58523 | 7.04813 | 7.35148 | 8.39495 | 4.24959 | 4.61399 | 4.04297 | 4.71185 |
| CCND3 | P30281 | 6.75081 | 6.96616 | 7.47901 | 6.61565 | 6.22566 | 7.77859 | 5.55656 | 6.39173 | 6.79309 | 8.39498 | 7.9913 | 8.3196 | 3.47561 | 4.2734 | 2.84748 | 4.14036 |
| AHI1 | Q8N157 | 6.06903 | 5.36686 | 6.40706 | 6.61761 | 5.65108 | 6.37228 | 6.34191 | 6.13508 | 6.44229 | 5.95562 | 7.12806 | 7.24618 | 5.61548 | 6.49119 | 5.96918 | 6.1911 |
| VCAN | P13611 | 6.29006 | 6.55155 | 5.43723 | 7.95826 | 5.60672 | 6.07543 | 6.90152 | 6.23445 | 6.86127 | 6.06533 | 6.27132 | 6.62313 | 6.33688 | 5.23289 | 5.37056 | 6.1834 |
| TXNL4B | Q9NX01 | 5.15314 | 4.60211 | 5.80856 | 4.50959 | 5.35259 | 5.13056 | 5.63979 | 8.07045 | 6.06131 | 7.12716 | 6.50364 | 7.62126 | 4.46994 | 7.96129 | 7.50549 | 8.48311 |
| PACC1 | Q9H813-2 | 5.68272 | 6.15837 | 7.05865 | 7.04709 | 6.3474 | 7.16854 | 6.47265 | 5.77001 | 6.23401 | 6.59633 | 6.70267 | 7.57409 | 5.31032 | 5.56019 | 4.91093 | 5.40602 |
| PCDH9 | Q9HC56 | 6.68561 | 6.60497 | 6.44227 | 6.39962 | 4.82583 | 6.42315 | 6.32945 | 6.26395 | 6.28294 | 7.21637 | 6.01129 | 5.27526 | 6.39337 | 6.23677 | 6.74906 | 5.86009 |
| SARM1 | Q6SZW1 | 5.95982 | 5.79816 | 6.872 | 6.92721 | 6.07411 | 5.57303 | 6.3993 | 6.78499 | 5.57908 | 6.7724 | 6.33059 | 5.98485 | 5.72709 | 6.26573 | 6.22785 | 6.7238 |
| CCDC120 | Q96HB5-4 | 6.50424 | 6.30713 | 4.80366 | 5.79838 | 6.2502 | 8.40816 | 5.3832 | 5.96785 | 5.80594 | 6.92821 | 7.17408 | 6.81273 | 7.09224 | 6.1497 | 5.76606 | 4.84821 |
| DBF4B | Q8NFT6 | 6.28241 | 7.33012 | 7.19179 | 8.10912 | 6.53093 | 7.60029 | 7.70804 | 5.38531 | 6.34761 | 6.93709 | 6.7713 | 4.99711 | 4.84613 | 4.47191 | 5.4465 | 4.04435 |
| FOXP4 | Q8IVH2 | 6.12467 | 6.33505 | 6.98664 | 5.69267 | 6.10697 | 6.57134 | 5.94793 | 5.54028 | 6.38373 | 6.97611 | 6.50297 | 6.78825 | 6.23097 | 5.9488 | 5.91898 | 5.94463 |
| ULBP2 | Q9BZM5 | 5.62034 | 6.6614 | 6.99858 | 6.52643 | 5.93657 | 6.43296 | 6.9838 | 5.86259 | 7.73591 | 4.76874 | 7.14906 | 6.71233 | 5.59214 | 5.97825 | 5.44998 | 5.59091 |
| NUDT4 | Q9NZJ9 | 6.39597 | 6.1165 | 6.59722 | 6.10276 | 6.36878 | 7.45075 | 6.08042 | 7.22223 | 6.57068 | 6.83455 | 6.61942 | 7.01539 | 5.06654 | 5.37306 | 4.78515 | 5.40058 |
| USP43 | Q70EL4 | 6.37577 | 6.06585 | 5.19891 | 7.46209 | 6.63806 | 6.08383 | 5.40459 | 5.77234 | 6.20678 | 6.46861 | 5.59123 | 6.04572 | 6.38432 | 5.96391 | 6.97024 | 7.36774 |
| UFM1 | P61960-2 | 6.15249 | 5.90374 | 5.88682 | 5.47468 | 5.43207 | 5.23644 | 5.57596 | 6.42193 | 5.41312 | 6.19756 | 6.03629 | 6.67197 | 5.25177 | 8.46002 | 7.24114 | 8.64402 |
| PPP1R3D | O95685 | 6.33711 | 6.5831 | 7.26277 | 5.86628 | 6.42334 | 7.1958 | 6.38282 | 5.82114 | 6.91269 | 6.2995 | 6.97007 | 6.62452 | 5.20217 | 5.60083 | 5.22835 | 5.28949 |
| GPR155 | Q7Z3F1 | 5.77891 | 7.00082 | 6.58732 | 5.96862 | 5.48964 | 5.69195 | 6.25253 | 5.87293 | 6.388 | 6.40646 | 5.77641 | 6.5288 | 6.05703 | 6.97377 | 6.11804 | 7.10876 |
| TM7SF3 | Q9NS93 | 6.74484 | 7.4054 | 6.39169 | 6.42077 | 6.86233 | 6.24964 | 6.40715 | 6.81586 | 6.54643 | 6.33061 | 6.34386 | 5.6163 | 5.25881 | 5.45237 | 5.4328 | 5.72115 |
| ZNF771 | Q7L3S4 | 5.73696 | 6.39418 | 5.98103 | 6.24341 | 6.25444 | 6.2407 | 6.44972 | 6.22243 | 6.05301 | 6.38791 | 6.3446 | 6.2687 | 6.06472 | 6.3071 | 6.69907 | 6.35202 |
| TMEM129 | A0AVI4 | 6.33204 | 7.48931 | 6.52788 | 6.64827 | 4.88519 | 5.84561 | 6.696 | 5.45854 | 6.45336 | 5.85751 | 6.89504 | 6.18885 | 5.70146 | 6.09507 | 6.64384 | 6.28202 |
| PEX2 | P28328 | 5.86509 | 6.20915 | 5.86639 | 6.20803 | 6.44618 | 5.85877 | 7.19962 | 6.06567 | 6.3136 | 6.00437 | 5.73566 | 5.73378 | 6.88934 | 5.85934 | 6.89348 | 6.85153 |
| ZNF689 | Q96CS4 | 6.19152 | 6.45249 | 6.1069 | 6.27517 | 5.82475 | 6.6639 | 5.98119 | 6.204 | 6.07536 | 6.2954 | 5.83496 | 5.97232 | 6.34508 | 6.52173 | 6.81892 | 6.43632 |
| GUCD1 | Q96NT3-3 | 7.34872 | 7.27642 | 7.20862 | 7.78889 | 5.63745 | 6.41672 | 6.66995 | 5.29731 | 5.93976 | 6.79494 | 7.25461 | 6.2731 | 5.38025 | 4.85051 | 5.19271 | 4.67003 |
| LMBR1 | Q8WVP7-3 | 5.92345 | 6.262 | 5.93768 | 6.46938 | 5.74336 | 5.91811 | 6.70665 | 5.85132 | 5.98725 | 6.29071 | 5.49386 | 5.73989 | 6.58033 | 6.51033 | 6.79759 | 7.7881 |
| ZDHHC6 | Q9H6R6 | 5.92419 | 6.58405 | 6.01782 | 7.12622 | 5.80916 | 5.49847 | 6.08124 | 5.90749 | 5.91577 | 6.30568 | 6.28641 | 6.43726 | 6.09386 | 6.47065 | 6.85047 | 6.69126 |
| RBKS | Q9H477 | 5.99514 | 6.10154 | 7.42589 | 5.744 | 6.52452 | 6.45417 | 6.82272 | 6.40076 | 5.7678 | 6.08493 | 6.36607 | 5.9113 | 6.12461 | 5.86176 | 6.81286 | 5.60192 |
| FZD1 | Q9UP38 | 6.77842 | 6.16139 | 6.63437 | 6.82147 | 6.7315 | 6.23943 | 6.94873 | 6.65792 | 6.0227 | 5.88459 | 6.19307 | 6.23841 | 5.30512 | 5.68413 | 5.71154 | 5.9872 |
| COCH | O43405 | 7.3066 | 7.58152 | 7.16278 | 7.15406 | 5.37308 | 6.73033 | 6.88655 | 5.27223 | 6.9194 | 6.98892 | 7.58704 | 7.20674 | 4.77388 | 4.24905 | 4.34836 | 4.45945 |
| SUMF1 | Q8NBK3 | 6.4469 | 6.53646 | 6.23192 | 6.43087 | 6.2926 | 6.44231 | 6.16367 | 5.99125 | 5.19879 | 6.00878 | 6.20406 | 6.53531 | 6.58509 | 6.03919 | 6.27335 | 6.61944 |
| PTCD2 | Q8WV60 | 6.52892 | 6.78885 | 6.07548 | 6.28297 | 6.22195 | 5.89961 | 6.78509 | 5.26639 | 6.06265 | 6.21222 | 5.59881 | 5.99027 | 6.78415 | 5.91926 | 6.75922 | 6.82414 |
| POMK | Q9H5K3 | 6.08621 | 6.42071 | 5.79393 | 6.67189 | 6.28737 | 6.3242 | 6.3005 | 6.36752 | 6.00707 | 6.52134 | 5.6339 | 6.26789 | 6.5608 | 6.40509 | 6.2 | 6.1516 |
| ARHGAP32 | A7KAX9 | 6.87043 | 6.90846 | 6.23985 | 6.46945 | 6.27003 | 6.04631 | 6.3653 | 6.64516 | 6.57452 | 6.14426 | 5.52575 | 6.01684 | 6.22656 | 6.44043 | 5.51198 | 5.74465 |
| MRPL57 | Q9BQC6 | 5.98155 | 5.28558 | 6.4216 | 7.04747 | 7.43148 | 6.10162 | 7.07488 | 6.66565 | 5.94818 | 6.0883 | 6.4299 | 5.90109 | 5.0846 | 5.86053 | 5.10486 | 7.57273 |
| NUDT7 | P0C024 | 6.79666 | 6.17592 | 6.10375 | 5.53076 | 6.47255 | 5.98441 | 5.94097 | 6.31921 | 5.99962 | 6.09534 | 6.2188 | 6.00483 | 6.47238 | 5.86667 | 7.05811 | 6.96001 |
| RPIA | P49247 | 4.82064 | 5.96626 | 9.0476 | 5.6115 | 6.58426 | 5.13751 | 7.837 | 5.99107 | 7.58021 | 6.46362 | 8.64103 | 5.91449 | 4.72245 | 3.96286 | 7.59166 | 4.12784 |
| CLOCK | O15516 | 6.79696 | 6.45363 | 6.50724 | 6.24972 | 5.94872 | 6.37664 | 5.73422 | 5.85078 | 6.17048 | 6.57367 | 5.61113 | 6.0245 | 5.48475 | 6.5911 | 6.36926 | 7.25721 |
| TMUB2 | Q71RG4 | 6.68596 | 7.62591 | 6.86987 | 6.47702 | 6.28811 | 6.68602 | 5.35415 | 6.18004 | 6.65448 | 6.80562 | 6.82797 | 6.54013 | 5.47706 | 4.7055 | 4.91861 | 5.90353 |

| | | | | | | | | | | | | | | | | | |
|---|---|---|---|---|---|---|---|---|---|---|---|---|---|---|---|---|---|
| CNTNAP3 | Q9BZ76 | 5.68625 | 5.09631 | 6.41663 | 6.94209 | 6.00408 | 5.40938 | 5.76296 | 7.27264 | 7.71819 | 6.62961 | 6.47367 | 6.49279 | 6.23995 | 7.2717 | 5.34842 | 5.23532 |
| DCTN5 | Q9BTE1 | 5.70289 | 6.42574 | 6.56468 | 5.88184 | 6.45028 | 7.1345 | 6.19733 | 6.8347 | 6.80871 | 6.75862 | 6.33067 | 6.23424 | 5.36159 | 5.79414 | 5.78531 | 5.73474 |
| C12orf66 | Q96MD2 | 6.24432 | 6.55884 | 6.30828 | 6.43723 | 6.51005 | 6.69121 | 6.1238 | 6.9523 | 5.75749 | 5.85946 | 5.73391 | 5.74408 | 6.46464 | 6.45774 | 5.84432 | 6.31231 |
| PTS | Q03393 | 6.43144 | 6.38894 | 6.58539 | 5.29121 | 6.70276 | 7.05969 | 5.98477 | 7.75305 | 6.45795 | 6.56768 | 6.4845 | 6.54741 | 5.58268 | 5.85642 | 4.9356 | 5.37052 |
| FANCM | Q8IYD8 | 7.64554 | 7.16799 | 7.39159 | 7.30074 | 6.68768 | 5.51454 | 5.93812 | 6.28842 | 6.9848 | 6.4363 | 5.18552 | 6.51514 | 5.78895 | 4.68776 | 5.31068 | 5.15623 |
| SLC43A2 | Q8N370-3 | 6.33287 | 6.64088 | 6.0884 | 6.61681 | 5.65132 | 5.77416 | 6.09931 | 6.15885 | 6.71542 | 6.83623 | 6.69116 | 7.30016 | 5.48187 | 5.50089 | 5.5347 | 6.57797 |
| LIX1L | Q8IVB5 | 5.70584 | 5.97026 | 6.20996 | 6.13954 | 7.0603 | 7.55212 | 6.06172 | 5.75651 | 5.45531 | 7.07178 | 7.59501 | 6.82322 | 5.40797 | 5.39686 | 5.69316 | 6.10044 |
| JMJD7 | P0C870 | 5.86041 | 5.10316 | 5.97163 | 5.02998 | 5.88973 | 6.09248 | 5.46732 | 7.53664 | 6.00007 | 6.69616 | 6.84226 | 7.36214 | 6.72111 | 7.37245 | 6.21392 | 5.84054 |
| ETNPPL | Q8TBG4 | 6.02284 | 5.99471 | 5.46367 | 6.52479 | 5.76731 | 6.32327 | 6.94021 | 7.02097 | 5.63242 | 6.88041 | 5.81164 | 5.55463 | 7.05907 | 5.98763 | 7.0336 | 5.98284 |
| AFAP1L2 | Q8N4X5 | 5.47977 | 5.53744 | 6.63784 | 5.44981 | 6.51583 | 6.92207 | 5.86835 | 6.01009 | 6.42838 | 6.26449 | 6.38849 | 7.32017 | 6.09375 | 6.56331 | 6.55632 | 5.96388 |
| SLITRK5 | O94991 | 6.36818 | 7.67409 | 5.51821 | 5.71512 | 5.48712 | 6.23295 | 5.0231 | 7.09325 | 7.2265 | 6.47279 | 6.89387 | 6.69763 | 6.47391 | 5.95509 | 5.37922 | 5.78898 |
| TTC8 | Q8TAM2 | 6.93162 | 5.99794 | 6.61914 | 6.11811 | 6.35675 | 6.58228 | 5.98692 | 7.88439 | 7.4009 | 7.49508 | 7.36844 | 6.58872 | 4.74657 | 5.45401 | 4.20503 | 4.2641 |
| ZNF580 | Q9UK33 | 6.90385 | 6.90161 | 6.64574 | 7.17218 | 6.8088 | 6.39548 | 6.48785 | 6.27495 | 7.19968 | 8.1336 | 7.73847 | 7.61678 | 3.62568 | 4.33279 | 3.71968 | 4.04285 |
| RIPK4 | P57078 | 6.4553 | 6.09975 | 6.56684 | 6.40365 | 6.60109 | 6.75233 | 5.80352 | 6.39581 | 5.78542 | 6.20815 | 5.96454 | 5.61982 | 6.41176 | 6.74559 | 6.60133 | 5.58511 |
| SAMD9L | Q8IVG5 | 5.96799 | 6.31926 | 5.83728 | 6.1382 | 5.72877 | 7.27229 | 6.47196 | 6.94815 | 6.71029 | 6.45161 | 6.36212 | 6.28867 | 5.5823 | 5.97234 | 5.9269 | 6.02188 |
| LRRC14 | Q15048 | 6.2323 | 6.10856 | 6.20902 | 5.39621 | 6.30151 | 6.79535 | 5.869 | 6.3401 | 5.84551 | 6.44038 | 6.95215 | 6.51351 | 6.42149 | 6.51884 | 6.58733 | 5.46874 |
| STOML1 | Q9UBI4 | 6.83954 | 6.94527 | 6.45416 | 6.45969 | 6.5263 | 5.97251 | 6.19402 | 5.67973 | 6.84234 | 6.03981 | 6.31685 | 6.29641 | 6.48962 | 5.88615 | 5.18019 | 5.87743 |
| SIK3 | Q9Y2K2 | 6.37442 | 6.62581 | 7.8614 | 6.59324 | 7.15121 | 7.50223 | 7.87051 | 7.94231 | 5.40515 | 5.22065 | 5.52204 | 5.60759 | 5.33905 | 4.46732 | 5.90658 | 4.61049 |
| CLN3 | Q13286 | 6.13839 | 7.0343 | 6.51524 | 6.50327 | 6.13104 | 6.76879 | 5.92047 | 6.26826 | 6.42161 | 6.81264 | 6.00822 | 6.67956 | 5.72527 | 5.59719 | 5.95747 | 5.51827 |
| CABLES1 | Q8TDN4 | 6.61588 | 6.08611 | 7.78828 | 6.2373 | 5.63896 | 7.08606 | 6.79477 | 7.27355 | 7.10092 | 4.73639 | 5.72207 | 7.0165 | 5.78622 | 4.76351 | 5.52801 | 5.82548 |
| C7orf25 | Q9BPX7-2 | 6.38478 | 6.32778 | 6.37898 | 6.04412 | 6.06439 | 6.71581 | 6.0492 | 6.71912 | 6.3795 | 6.53137 | 7.34225 | 6.4941 | 5.24482 | 5.82713 | 5.69304 | 5.80362 |
| TSPAN15 | O95858 | 5.62051 | 5.99356 | 6.09622 | 6.62883 | 5.74282 | 6.34826 | 6.26347 | 5.98249 | 6.53258 | 5.90856 | 6.18544 | 6.18772 | 6.55428 | 6.51231 | 6.68675 | 6.75621 |
| CDC14A | Q9UNH5-2 | 6.57231 | 6.08635 | 6.95032 | 6.39573 | 6.41829 | 6.66852 | 6.38963 | 6.69734 | 6.4536 | 6.72174 | 6.65894 | 6.60998 | 6.54675 | 5.3917 | 4.91033 | 5.42948 |
| ZNF143 | P52747 | 5.88825 | 6.40768 | 6.51929 | 5.99238 | 6.04 | 6.26087 | 5.90494 | 6.0899 | 6.60175 | 5.95996 | 6.17551 | 6.01329 | 7.15962 | 6.57368 | 6.30334 | 6.10952 |
| TNFRSF12A | Q9NP84 | 7.09673 | 7.13605 | 6.94075 | 6.59495 | 5.15217 | 5.02194 | 6.23721 | 6.54697 | 6.16353 | 7.58979 | 6.62804 | 8.24207 | 3.27346 | 6.44541 | 4.87456 | 6.05637 |
| FAU | P62861 | 6.80834 | 6.45377 | 6.37771 | 5.69442 | 6.77682 | 6.31149 | 6.8826 | 6.47286 | 6.46549 | 6.15458 | 5.70167 | 5.70602 | 5.80265 | 6.63115 | 5.72583 | 6.0346 |
| RNF166 | Q96A37 | 6.35797 | 7.06907 | 7.58323 | 6.88946 | 5.22163 | 6.09074 | 6.39043 | 6.61441 | 6.74587 | 6.65944 | 7.20548 | 6.40996 | 4.74572 | 5.81335 | 5.1783 | 5.02495 |
| LCA5 | Q86VQ0 | 6.94386 | 6.16351 | 6.08688 | 6.62406 | 5.91229 | 6.15378 | 6.19056 | 5.9998 | 5.88996 | 6.32365 | 6.56778 | 6.55197 | 5.96888 | 6.03592 | 5.95455 | 6.63254 |
| CASZ1 | Q86V15 | 5.83602 | 6.03749 | 6.53199 | 6.74538 | 6.12274 | 5.85953 | 6.19379 | 5.91301 | 5.98044 | 6.53059 | 6.35935 | 6.4697 | 6.3892 | 6.09968 | 6.58222 | 6.34889 |
| TTLL7 | Q6ZT98 | 6.43613 | 6.42868 | 7.68643 | 7.05859 | 6.38932 | 6.22674 | 7.49225 | 6.55709 | 6.60347 | 6.54778 | 6.72565 | 5.95083 | 3.75642 | 5.58372 | 5.51754 | 5.03935 |
| NAALAD2 | Q9Y3Q0 | 6.80906 | 6.28759 | 6.01245 | 6.81318 | 5.77806 | 6.25391 | 6.43944 | 6.64391 | 6.12536 | 6.24692 | 5.27097 | 6.68262 | 6.02506 | 6.20304 | 6.55971 | 5.84872 |
| TYW3 | Q6IPR3 | 6.00241 | 6.99104 | 7.16818 | 5.99651 | 6.46018 | 7.45313 | 6.30813 | 6.3283 | 6.24005 | 6.02011 | 6.58822 | 5.8678 | 6.03636 | 5.85399 | 5.54904 | 5.13656 |
| HFM1 | A2PYH4 | 5.99454 | 7.18182 | 5.29055 | 6.19277 | 6.05757 | 6.18802 | 5.86777 | 6.30108 | 6.04747 | 5.84066 | 4.65498 | 5.31268 | 6.10597 | 6.81103 | 8.36079 | 7.79231 |
| WASF1 | Q92558 | 7.36275 | 6.29415 | 7.05849 | 6.14487 | 6.49655 | 6.59436 | 6.72221 | 6.13186 | 6.63227 | 6.52395 | 6.5156 | 6.88695 | 6.24074 | 4.64386 | 5.26423 | 5.18717 |
| PECR | Q9BY49 | 6.28621 | 5.97609 | 6.02203 | 6.7758 | 6.18941 | 6.39894 | 6.64368 | 5.38657 | 5.72513 | 6.03741 | 6.14188 | 6.32307 | 7.22279 | 6.40402 | 6.10787 | 6.35912 |
| EAF1 | Q96JC9 | 6.11857 | 6.14874 | 6.49975 | 6.26689 | 5.98397 | 5.92417 | 5.68326 | 6.16415 | 6.26548 | 6.35411 | 6.51285 | 6.53256 | 6.17146 | 6.35436 | 6.27894 | 6.74075 |
| LSM3 | P62310 | 5.91132 | 4.74138 | 6.02083 | 5.60637 | 5.18327 | 4.59796 | 5.39772 | 7.19933 | 5.46866 | 6.716 | 6.20856 | 6.45417 | 6.62098 | 8.401 | 7.20271 | 8.26974 |
| IGFBP2 | P18065 | 7.30449 | 6.56042 | 6.69782 | 7.33712 | 5.93913 | 6.02479 | 6.10529 | 7.29789 | 6.65738 | 6.37744 | 7.12057 | 7.59119 | 3.76628 | 4.47942 | 6.13715 | 4.60361 |
| IFRD2 | Q12894-2 | 6.09938 | 5.98414 | 6.13141 | 6.51699 | 6.37347 | 6.53529 | 6.34654 | 6.39148 | 6.66996 | 6.62288 | 6.48097 | 7.13454 | 5.6178 | 5.83015 | 5.91182 | 5.3532 |
| KPTN | Q9Y664 | 6.1821 | 6.20211 | 6.55915 | 6.00932 | 5.96307 | 6.81023 | 6.16527 | 6.75415 | 6.29886 | 6.43937 | 6.56608 | 6.6948 | 5.81265 | 6.23985 | 5.82148 | 5.48151 |
| CLEC16A | Q2KHT3 | 6.56891 | 7.02817 | 5.62106 | 6.07234 | 7.12842 | 5.05261 | 6.77775 | 7.37077 | 6.03567 | 6.7107 | 6.19929 | 5.50856 | 6.69981 | 5.77655 | 5.18093 | 6.26845 |
| ZNF467 | Q7Z7K2 | 6.0634 | 6.08041 | 5.62335 | 6.39022 | 6.55851 | 6.30997 | 5.75834 | 5.93059 | 5.94193 | 6.98465 | 6.5373 | 6.32527 | 6.35375 | 6.71946 | 5.94803 | 6.47481 |
| TJP3 | O95049-3 | 7.00404 | 5.40943 | 5.76615 | 6.43491 | 5.76416 | 5.12007 | 6.9326 | 6.01456 | 5.41615 | 6.47966 | 6.2688 | 6.27143 | 6.84028 | 7.31384 | 6.91324 | 6.05067 |
| ZNF526 | Q8TF50 | 5.17041 | 7.00568 | 6.5231 | 5.9499 | 5.42881 | 6.02474 | 5.40789 | 6.71809 | 5.703 | 6.69013 | 6.03103 | 5.90323 | 6.54865 | 6.80775 | 7.17498 | 6.91261 |
| ZNF226 | Q9NYT6 | 6.45279 | 6.63124 | 6.32503 | 6.51636 | 6.11145 | 6.62033 | 6.83024 | 6.36183 | 6.47725 | 6.99948 | 6.1087 | 6.22811 | 5.29377 | 5.67435 | 5.48634 | 5.88274 |
| ZNF234 | Q14588 | 5.77136 | 6.266 | 5.60333 | 7.26625 | 6.51951 | 6.39023 | 7.5774 | 5.88643 | 6.38594 | 6.01662 | 5.31689 | 5.87623 | 6.41924 | 6.6146 | 6.15548 | 5.93448 |
| SLC35A4 | L0R6Q1 | 6.52862 | 5.25697 | 5.49594 | 4.93503 | 5.48163 | 4.83296 | 6.24563 | 6.75653 | 4.88285 | 5.54585 | 5.38147 | 5.74534 | 5.73818 | 9.33973 | 8.64806 | 9.18518 |
| PLEKHA8 | Q96JA3 | 6.8904 | 7.38286 | 7.08132 | 6.10188 | 6.12716 | 6.92098 | 6.71014 | 6.01723 | 5.56128 | 6.57728 | 6.50492 | 7.07818 | 5.28637 | 5.60183 | 4.70263 | 5.45556 |
| ZNF597 | Q96LX8 | 5.92473 | 5.63093 | 6.69205 | 6.30313 | 6.07786 | 5.25376 | 6.51294 | 6.2456 | 6.6532 | 7.06352 | 5.74608 | 6.84006 | 6.44196 | 5.43394 | 6.11226 | 7.06798 |
| CEP76 | Q8TAP6 | 6.60792 | 6.31356 | 6.72275 | 5.97143 | 6.28333 | 6.60655 | 6.29734 | 6.20625 | 6.3779 | 6.5141 | 6.99896 | 7.19817 | 5.59033 | 5.40888 | 5.61749 | 5.28502 |
| NOL4 | O94818 | 5.64529 | 6.23764 | 5.73639 | 6.42 | 6.57275 | 5.72579 | 6.49199 | 5.67734 | 5.609 | 5.84123 | 6.40493 | 6.72905 | 6.7939 | 6.28911 | 6.8423 | 6.9833 |
| SLC26A11 | Q86WA9 | 6.43577 | 6.19635 | 6.01365 | 6.72369 | 5.81215 | 6.14335 | 6.15781 | 6.55453 | 6.11983 | 6.61361 | 6.40087 | 6.45671 | 6.31126 | 6.24637 | 6.35352 | 6.36052 |
| PEX16 | Q9Y5Y5-2 | 6.27394 | 5.89481 | 5.94436 | 6.58448 | 5.64546 | 5.78033 | 6.51622 | 6.02735 | 6.37262 | 6.54751 | 6.51169 | 6.42802 | 6.36662 | 6.63843 | 6.54005 | 6.9281 |
| ACADL | P28330 | 5.89551 | 6.79369 | 6.48006 | 6.27457 | 6.94562 | 5.91955 | 5.95085 | 5.86194 | 5.66596 | 5.69739 | 6.06198 | 6.51355 | 6.5898 | 6.6851 | 6.31534 | 6.3491 |
| ZC3H10 | Q96K80 | 5.98519 | 6.23085 | 6.15517 | 5.98503 | 5.85958 | 6.35462 | 6.25591 | 5.99274 | 6.30949 | 6.22499 | 6.61252 | 6.45981 | 6.57147 | 6.03138 | 6.47924 | 6.49202 |
| GGNBP2 | Q9H3C7 | 6.64644 | 6.96337 | 7.22151 | 6.7331 | 6.30543 | 6.43985 | 6.25238 | 6.27022 | 6.91953 | 6.94931 | 6.95417 | 7.46317 | 4.58362 | 4.88639 | 4.46044 | 4.95109 |
| TK2 | O00142 | 6.45204 | 6.98562 | 6.06715 | 7.13 | 5.73953 | 5.7651 | 5.98858 | 5.94806 | 6.63328 | 6.23428 | 5.92564 | 6.27842 | 6.63678 | 5.42893 | 6.55801 | 6.22858 |
| ZC3H12A | Q5D1E8 | 5.75408 | 6.62507 | 8.14242 | 6.1342 | 5.23934 | 7.17915 | 6.36981 | 5.47166 | 6.76928 | 6.38538 | 7.47584 | 6.56486 | 5.22682 | 5.34212 | 5.68577 | 5.63419 |
| ZC3H12C | Q9C0D7-2 | 5.57184 | 6.71385 | 7.1903 | 5.24787 | 6.565 | 6.97619 | 5.22783 | 5.42278 | 5.90866 | 6.47108 | 8.0121 | 8.80398 | 6.44237 | 5.04259 | 5.00757 | 5.39599 |
| CRIPT | Q9P021 | 6.21719 | 6.11194 | 6.47776 | 5.96433 | 6.74873 | 6.8959 | 6.26239 | 6.26064 | 6.61739 | 6.8657 | 6.78709 | 6.08762 | 5.48755 | 5.95096 | 5.57052 | 5.29289 |
| RPS6KC1 | Q96S38 | 6.57807 | 6.81846 | 6.73632 | 6.39556 | 5.94461 | 6.59211 | 5.623 | 5.83759 | 6.03124 | 6.40309 | 6.90242 | 6.456 | 6.41904 | 5.83033 | 5.86779 | 5.56439 |
| CCP110 | O43303 | 6.33666 | 5.6453 | 5.83809 | 6.01143 | 6.03453 | 5.75469 | 5.60561 | 5.71229 | 6.46269 | 6.07052 | 6.21795 | 7.59056 | 7.26874 | 6.87335 | 6.6073 | 5.9703 |
| GPC2 | Q8N158 | 5.92078 | 6.75049 | 6.18836 | 6.78589 | 5.69324 | 6.76603 | 6.16439 | 6.07974 | 5.53259 | 6.36759 | 6.33696 | 6.73739 | 5.97667 | 6.19576 | 5.78992 | 6.71419 |
| NABP1 | Q96AH0 | 6.86585 | 6.18775 | 6.68104 | 7.06821 | 6.68307 | 6.19487 | 6.37809 | 7.10749 | 6.16454 | 5.40763 | 5.95207 | 6.37603 | 5.77277 | 5.87326 | 5.88328 | 5.40405 |
| BORCS5 | Q969J3 | 5.58308 | 4.99435 | 5.16849 | 5.36374 | 5.38516 | 4.66205 | 5.3083 | 6.35803 | 5.03496 | 5.98421 | 5.14879 | 6.59044 | 6.19856 | 9.48635 | 8.94479 | 9.78871 |
| LRRC28 | Q86X40 | 7.06411 | 5.63608 | 6.25761 | 6.27465 | 6.35073 | 7.49721 | 6.74506 | 5.86975 | 6.70233 | 6.17249 | 6.64498 | 6.77193 | 4.90943 | 5.95914 | 5.87196 | 5.27254 |
| HPSE | Q9Y251 | 6.33446 | 6.69447 | 6.32257 | 5.76846 | 6.03907 | 5.91933 | 6.3937 | 6.13394 | 6.562 | 6.74503 | 6.555 | 6.08826 | 6.26079 | 6.02612 | 5.6961 | 6.46069 |
| ENG | P17813 | 6.23983 | 6.37633 | 6.3858 | 6.90964 | 5.84015 | 6.20163 | 6.27922 | 5.70103 | 6.36816 | 5.98848 | 6.69444 | 6.54656 | 6.5334 | 6.10572 | 5.7718 | 6.0578 |
| CASD1 | Q96PB1 | 5.68813 | 6.39725 | 6.28661 | 6.56965 | 5.75901 | 6.14322 | 7.68181 | 5.86623 | 6.36843 | 6.52454 | 6.07698 | 5.67585 | 6.18972 | 6.13071 | 6.88682 | 6.65506 |
| PLPP2 | O43688-2 | 6.12926 | 7.33662 | 6.31184 | 6.46099 | 4.76709 | 6.27527 | 6.13908 | 5.8625 | 6.61242 | 6.99305 | 6.39438 | 5.81496 | 4.93643 | 6.81365 | 6.28692 | 6.86553 |
| NLGN2 | Q8NFZ4 | 7.25664 | 6.83447 | 6.81358 | 5.77044 | 6.51073 | 5.97914 | 6.16301 | 6.34687 | 6.33102 | 6.91106 | 6.26039 | 5.99859 | 5.38366 | 5.33782 | 5.87053 | 6.23205 |
| PPP1R14A | Q96A00 | 6.57351 | 5.99475 | 6.09984 | 5.77805 | 6.66968 | 6.64379 | 6.50187 | 6.10372 | 6.41556 | 6.97732 | 7.13583 | 6.12243 | 6.48972 | 5.05229 | 5.882 | 5.55962 |
| AGFG2 | Q95081 | 6.75746 | 6.29121 | 7.37214 | 5.25573 | 6.30918 | 6.31157 | 5.7145 | 6.47656 | 6.30266 | 6.48024 | 6.28974 | 6.8253 | 6.77035 | 5.68339 | 5.9161 | 5.24387 |
| PLEKHH3 | Q7Z736 | 6.86465 | 6.4457 | 6.63589 | 7.09184 | 5.7073 | 6.99834 | 6.71708 | 6.10828 | 6.82723 | 7.54839 | 7.23081 | 6.52983 | 4.56089 | 5.48598 | 4.60346 | 4.64433 |
| VSIG10 | Q8N0Z9 | 6.83385 | 6.27521 | 6.83644 | 5.72393 | 5.9659 | 6.51108 | 5.12768 | 5.80153 | 6.43228 | 6.9687 | 7.41502 | 6.3821 | 6.52705 | 5.69981 | 5.96193 | 5.5375 |
| NEIL2 | Q969S2 | 6.08132 | 5.26302 | 6.44442 | 5.44534 | 5.7441 | 6.41486 | 6.1972 | 5.29442 | 5.13795 | 7.417 | 6.06612 | 5.98456 | 7.02355 | 7.21961 | 7.5036 | 6.76292 |
| GAS2L1 | Q99501 | 6.10479 | 6.848 | 5.55555 | 4.85909 | 5.41078 | 6.05789 | 8.49828 | 6.03022 | 4.62796 | 6.58208 | 5.38757 | 5.18826 | 6.50059 | 7.3081 | 7.90457 | 7.13628 |
| SLC35F6 | Q8N357 | 6.94001 | 6.54934 | 6.6369 | 6.17595 | 5.65685 | 5.62703 | 6.66139 | 5.63514 | 5.49023 | 7.44873 | 6.8393 | 6.71453 | 4.85413 | 6.18789 | 5.68501 | 6.89758 |
| PHLPP2 | Q6ZVD8 | 7.18052 | 6.6795 | 6.73614 | 8.07886 | 6.70938 | 5.786 | 5.8637 | 7.28834 | 7.25679 | 5.81981 | 6.60872 | 6.86246 | 5.10381 | 4.95671 | 4.41773 | 4.65153 |
| BNIP2 | Q12982-2 | 6.77215 | 7.53835 | 7.02487 | 6.77218 | 4.76621 | 7.65922 | 7.25809 | 7.03751 | 5.359 | 4.55126 | 5.43291 | 5.65678 | 6.6126 | 5.29264 | 6.94181 | 6.32442 |
| RIMS2 | Q9UQ26 | 6.3806 | 5.62462 | 7.28832 | 6.72547 | 5.98238 | 7.15019 | 6.17465 | 6.30722 | 5.39404 | 6.5359 | 6.59722 | 7.58011 | 6.18551 | 5.72742 | 5.78206 | 6.46429 |
| BBS10 | Q8TAM1 | 6.44966 | 6.23629 | 7.15505 | 6.0839 | 6.43176 | 6.65437 | 6.94292 | 6.58471 | 6.38536 | 6.48129 | 7.36487 | 6.13358 | 5.51048 | 5.16983 | 5.15578 | 5.26016 |
| TMEM164 | Q5U3C3 | 5.5858 | 5.55447 | 5.48869 | 5.97722 | 5.45297 | 5.65339 | 6.08466 | 6.07639 | 6.52166 | 5.71598 | 5.82665 | 6.61358 | 7.24814 | 7.28676 | 7.60184 | 7.31179 |
| ORMDL1 | Q9P0S3 | 5.69874 | 6.78127 | 5.27392 | 5.43184 | 5.79527 | 6.38641 | 6.01864 | 5.56132 | 5.36173 | 6.34231 | 5.52213 | 6.62486 | 5.72879 | 7.53826 | 7.8196 | 8.11491 |
| SLC25A14 | O95258-3 | 7.11661 | 5.47081 | 5.91832 | 6.89489 | 6.37051 | 4.58378 | 6.36978 | 6.81128 | 6.21914 | 6.3981 | 6.58725 | 6.53686 | 7.36836 | 6.12365 | 6.82238 | 5.30829 |
| SNF8 | Q96H20 | 6.33455 | 6.46453 | 7.01944 | 5.92672 | 6.11331 | 6.6122 | 5.91112 | 6.8281 | 6.19731 | 6.78153 | 6.84602 | 7.24379 | 5.20317 | 5.78087 | 5.26821 | 5.46911 |
| FCF1 | Q9Y324 | 6.24063 | 6.61843 | 5.7342 | 7.02652 | 5.34358 | 6.19144 | 6.35448 | 6.69511 | 5.66235 | 5.96712 | 5.5081 | 5.70973 | 6.73743 | 6.79778 | 6.50615 | 6.90694 |
| ADAT3 | Q96EY9 | 6.00977 | 5.40756 | 8.20099 | 4.97966 | 6.8255 | 6.75245 | 6.643 | 5.83952 | 6.49827 | 6.52458 | 6.98152 | 7.06886 | 6.33721 | 6.04762 | 5.72696 | 5.05653 |
| DDX43 | Q9NXZ2 | 6.59479 | 6.31808 | 6.76341 | 6.0568 | 6.16564 | 6.64924 | 6.60897 | 6.42576 | 6.13723 | 6.02002 | 6.45932 | 6.67843 | 5.91086 | 5.45579 | 6.008 | 5.74767 |
| UST | Q9Y2C2 | 7.62993 | 5.62166 | 5.18121 | 6.72648 | 6.11934 | 6.4277 | 5.96592 | 5.41006 | 6.30502 | 6.49444 | 6.23363 | 6.28697 | 6.59207 | 5.77902 | 6.41762 | 6.80893 |
| SLC25A30 | Q5SVS4 | 6.30367 | 6.52743 | 5.73315 | 7.15637 | 6.35466 | 6.32361 | 5.82567 | 6.64355 | 7.20652 | 6.55235 | 6.26318 | 6.10572 | 5.7269 | 5.98937 | 5.62444 | 5.66341 |
| LACC1 | Q8IV20 | 6.13006 | 6.23082 | 7.06647 | 5.9598 | 6.15622 | 6.83465 | 6.45875 | 7.39501 | 7.22257 | 5.56545 | 6.92419 | 5.5953 | 5.90592 | 4.94541 | 5.86372 | 5.74567 |
| TMEM167B | Q9NRX6 | 5.44722 | 6.37865 | 6.4242 | 6.31247 | 6.51233 | 6.184 | 6.02394 | 6.27245 | 6.36549 | 6.04489 | 5.7843 | 6.19601 | 6.8817 | 6.14795 | 6.66666 | 6.35774 |
| PRRT3 | Q5FWE3 | 6.16685 | 5.5279 | 7.15148 | 6.60252 | 6.21212 | 7.05169 | 6.11167 | 6.30713 | 6.60034 | 6.17077 | 6.52412 | 6.23523 | 5.92657 | 5.36127 | 5.74088 | 6.30947 |
| FKBP1A | P62942 | 5.36908 | 5.70332 | 6.40589 | 4.80858 | 6.44616 | 6.5044 | 5.72531 | 7.7935 | 5.61471 | 7.25668 | 7.24185 | 6.78525 | 5.53726 | 6.37708 | 6.04455 | 6.38637 |
| WDFY2 | Q96P53 | 6.49546 | 7.16031 | 6.50566 | 5.91391 | 6.48033 | 5.98466 | 6.01023 | 5.90072 | 5.60308 | 6.19508 | 6.49761 | 5.92958 | 5.73636 | 6.34351 | 6.96168 | 6.28182 |
| DCAF12 | Q5T6F0 | 5.40815 | 5.94371 | 5.88235 | 6.09145 | 5.65415 | 6.0569 | 5.75892 | 5.77723 | 6.51196 | 5.75213 | 6.27021 | 7.44234 | 6.1317 | 6.78139 | 7.10691 | 7.4305 |
| NABP2 | Q9BQ15 | 6.77143 | 6.36339 | 6.66186 | 6.42218 | 6.35024 | 6.03183 | 5.71772 | 5.75371 | 6.37107 | 7.04134 | 5.90132 | 6.03911 | 6.69367 | 5.3448 | 6.46269 | 6.07364 |
| RPGR | Q92834 | 5.92454 | 6.90628 | 6.74663 | 6.90249 | 6.89021 | 6.55131 | 6.31281 | 6.47989 | 6.14519 | 5.74049 | 7.64496 | 6.86339 | 4.87953 | 5.54219 | 4.56248 | 5.9076 |
| CAMLG | P49069 | 5.90548 | 5.77728 | 6.56147 | 7.78456 | 6.14439 | 5.35442 | 6.6325 | 6.19675 | 6.30081 | 5.85527 | 6.63034 | 5.56669 | 6.25044 | 6.68634 | 6.0079 | 6.34537 |
| CNTROB | Q8N137-2 | 6.94308 | 6.46106 | 6.94688 | 7.38314 | 5.90091 | 5.87761 | 5.56563 | 6.17348 | 6.13704 | 6.0583 | 7.3389 | 6.93401 | 5.93324 | 4.60102 | 5.85201 | 5.89369 |
| MND1 | Q9BWT6 | 5.65605 | 6.48184 | 6.78332 | 5.80407 | 5.99007 | 6.45452 | 5.58702 | 5.89604 | 6.11758 | 6.53823 | 7.43454 | 7.11299 | 5.96678 | 6.46128 | 6.01643 | 5.69925 |
| CDCA4 | Q9BXL8 | 5.97046 | 5.21269 | 6.46786 | 5.41481 | 5.95257 | 5.79507 | 5.79892 | 6.92777 | 6.15543 | 6.90907 | 6.94545 | 7.97717 | 5.1558 | 6.64528 | 5.96158 | 6.71008 |
| ZNF770 | Q6IQ21 | 6.71015 | 6.18491 | 5.71336 | 7.23668 | 5.82615 | 5.741 | 6.31842 | 5.71591 | 6.39128 | 6.29137 | 6.89443 | 6.16524 | 6.3659 | 6.01805 | 6.50701 | 5.92013 |
| C9orf85 | Q96MD7-1 | 6.05578 | 6.22736 | 6.36949 | 5.95343 | 5.87393 | 5.98059 | 6.21018 | 5.98631 | 6.37128 | 6.07239 | 6.16702 | 6.16205 | 6.52402 | 6.59903 | 6.9895 | 6.45765 |

| | | | | | | | | | | | | | | | | | |
|---|---|---|---|---|---|---|---|---|---|---|---|---|---|---|---|---|---|
| GC | P02774-3 | 5.01351 | 7.86775 | 5.1255 | 3.71094 | 3.73459 | 7.02965 | 6.53983 | 5.72122 | 7.00709 | 5.51696 | 7.03074 | 7.08279 | 5.12796 | 5.69093 | 11.2709 | 6.52962 |
| ATP13A2 | Q9NQ11 | 6.02936 | 6.06976 | 6.67489 | 6.38134 | 6.03213 | 6.20229 | 6.61178 | 6.13354 | 6.24297 | 6.66394 | 6.40168 | 6.16608 | 5.84746 | 5.98089 | 6.35822 | 6.20368 |
| PRMT2 | P55345 | 5.31737 | 6.47183 | 6.12526 | 5.27822 | 7.38795 | 5.64885 | 5.89814 | 6.58381 | 6.64217 | 5.85503 | 6.16104 | 6.97397 | 6.87301 | 6.14815 | 6.16154 | 6.47366 |
| USP45 | Q70EL2 | 5.37988 | 6.10436 | 6.88392 | 6.46257 | 6.26035 | 5.88735 | 5.22732 | 5.41134 | 6.27611 | 6.09794 | 6.3145 | 7.38329 | 6.84831 | 7.01172 | 6.11854 | 6.33249 |
| SLC7A2 | P52569-3 | 6.95501 | 7.0286 | 6.43556 | 6.63998 | 6.55565 | 5.75944 | 6.32888 | 6.37693 | 6.58851 | 6.01317 | 5.84601 | 6.77576 | 5.63935 | 5.48775 | 5.96026 | 5.60914 |
| METAP1D | Q6UB28 | 5.34668 | 5.60104 | 5.65311 | 5.40222 | 6.45012 | 5.86921 | 5.94625 | 6.74056 | 6.14634 | 6.8217 | 5.93426 | 6.32866 | 6.89005 | 6.63994 | 7.24356 | 6.98629 |
| CRYL1 | Q9Y2S2 | 6.56305 | 6.08298 | 7.09561 | 6.25447 | 6.36228 | 6.52232 | 5.97194 | 6.61798 | 5.95027 | 6.70502 | 6.34258 | 6.5303 | 5.68115 | 6.16694 | 5.72826 | 5.42484 |
| PIGM | Q9H3S5 | 6.44577 | 6.35326 | 5.66427 | 6.83883 | 6.17042 | 5.52211 | 5.93975 | 5.97706 | 5.87423 | 6.28934 | 6.06904 | 5.84735 | 6.91328 | 6.54899 | 6.58053 | 6.96578 |
| MED28 | Q9H204 | 5.71138 | 5.11769 | 5.28162 | 4.57292 | 4.8334 | 4.67711 | 4.70958 | 5.907 | 4.78062 | 5.90493 | 4.54703 | 5.03418 | 5.21593 | 12.3272 | 9.72272 | 11.6567 |
| VTI1A | Q96AJ9 | 6.26538 | 5.6488 | 6.58845 | 5.98746 | 5.56587 | 5.46875 | 6.02636 | 5.46311 | 5.98935 | 6.41392 | 6.48131 | 6.95984 | 7.0362 | 7.0568 | 6.59286 | 6.45555 |
| SPRYD7 | Q5W111 | 6.46147 | 6.54116 | 6.42599 | 6.58196 | 6.07779 | 5.85265 | 6.44281 | 6.0387 | 6.44966 | 6.0558 | 6.45843 | 6.13832 | 6.03427 | 5.67463 | 6.56351 | 6.20285 |
| GPM6A | P51674 | 6.20684 | 6.52769 | 5.9591 | 6.1313 | 5.95602 | 6.01776 | 6.40084 | 6.18585 | 6.25017 | 5.88084 | 5.63775 | 5.6636 | 6.92175 | 6.68075 | 6.37672 | 7.20303 |
| CRTC1 | Q6UUV9-2 | 6.13639 | 6.15773 | 7.52923 | 5.61228 | 6.62408 | 6.4524 | 6.09769 | 6.65703 | 6.02875 | 7.12933 | 6.41333 | 6.87733 | 6.23738 | 5.52211 | 4.93825 | 5.58671 |
| SERGEF | Q9UGK8 | 6.53375 | 6.63025 | 6.83524 | 6.15524 | 6.31265 | 6.39313 | 5.76105 | 5.88434 | 6.27907 | 6.1756 | 7.08183 | 6.86522 | 5.92767 | 5.44152 | 5.91189 | 5.81157 |
| EBAG9 | O00559-2 | 6.80886 | 5.88125 | 5.35324 | 6.62293 | 6.20663 | 6.86829 | 6.57433 | 6.4769 | 6.36241 | 5.73196 | 5.64591 | 5.61295 | 6.44248 | 6.86341 | 6.06156 | 6.48689 |
| GAREM1 | Q9H706 | 6.87142 | 6.65716 | 6.90135 | 5.75791 | 6.17209 | 6.77775 | 6.6609 | 6.61328 | 6.20491 | 6.5933 | 6.07482 | 6.33725 | 5.38786 | 5.90649 | 5.38139 | 5.70213 |
| LRRC49 | Q8IUZ0-2 | 6.19487 | 7.00738 | 7.63639 | 6.23708 | 6.21292 | 6.96623 | 5.5827 | 5.81166 | 7.47837 | 7.62687 | 6.27185 | 6.09645 | 4.80811 | 5.01773 | 5.00255 | 6.04886 |
| ARHGEF16 | Q5VV41 | 6.10721 | 6.15088 | 7.23636 | 6.25174 | 6.25933 | 5.8975 | 6.0648 | 6.79736 | 5.97193 | 6.17901 | 6.20096 | 6.5409 | 6.30336 | 6.19675 | 5.90996 | 5.93193 |
| HOXD8 | P13378 | 6.27208 | 6.59079 | 5.80086 | 5.57996 | 5.7544 | 6.13581 | 5.98758 | 5.99593 | 6.35746 | 6.32009 | 6.37578 | 6.96433 | 6.32913 | 6.48235 | 6.94119 | 6.11224 |
| SLC44A2 | Q8IWA5-2 | 5.83353 | 5.68067 | 6.00326 | 6.68051 | 6.14989 | 6.06507 | 6.08185 | 5.57342 | 5.50318 | 6.62116 | 5.73041 | 6.20079 | 7.39493 | 6.56382 | 6.48436 | 7.43317 |
| MSMO1 | Q15800 | 5.9185 | 6.51789 | 6.50203 | 6.79768 | 5.49075 | 5.91956 | 7.0606 | 6.79747 | 5.7291 | 6.31431 | 6.23211 | 6.45679 | 5.46487 | 6.67663 | 6.10845 | 6.01329 |
| STAB1 | Q9NY15 | 6.22759 | 6.93935 | 6.30175 | 6.05158 | 4.86286 | 6.03351 | 5.86209 | 5.5589 | 5.92436 | 6.11834 | 5.55432 | 6.10997 | 6.34221 | 6.75067 | 7.36359 | 7.99889 |
| SYTL2 | Q9HCH5-8 | 6.09536 | 6.43387 | 5.58107 | 6.19911 | 5.52772 | 6.53974 | 7.63751 | 5.69952 | 6.87769 | 7.01704 | 5.05407 | 5.96471 | 5.93203 | 6.49986 | 6.11385 | 6.82687 |
| SUPT3H | O75486-4 | 6.09306 | 6.08286 | 6.74515 | 7.34129 | 6.8672 | 6.85037 | 5.92993 | 5.55674 | 7.68907 | 6.18932 | 6.66168 | 5.34871 | 5.93062 | 6.57398 | 6.52794 | 4.51209 |
| MAPT | P10636-9 | 6.64796 | 6.90956 | 7.13349 | 5.78143 | 6.7092 | 7.01169 | 6.27725 | 6.24767 | 6.10752 | 6.5076 | 5.99615 | 6.54454 | 5.60856 | 5.50776 | 5.5111 | 5.49853 |
| DDT | P30046 | 3.52158 | 3.87786 | 3.9674 | 3.5709 | 5.89583 | 6.15136 | 6.5212 | 9.31516 | 7.12851 | 7.95059 | 5.66879 | 7.62137 | 4.81634 | 7.62265 | 7.50095 | 8.86951 |
| PCGF1 | Q9BSM1 | 6.45992 | 6.32914 | 5.83936 | 6.77123 | 6.88323 | 6.50072 | 5.91694 | 6.63948 | 5.85359 | 6.11423 | 6.09385 | 5.75842 | 6.58573 | 6.35712 | 5.93557 | 5.96147 |
| SLC38A7 | Q9NVC3 | 6.39725 | 6.72043 | 5.86426 | 6.3572 | 6.53283 | 6.55851 | 6.34835 | 6.2827 | 5.87541 | 6.16278 | 5.66333 | 6.0561 | 6.3276 | 6.36409 | 6.37489 | 6.11426 |
| TWIST1 | Q15672 | 6.44796 | 6.87684 | 5.28452 | 7.11861 | 6.76223 | 7.42677 | 7.22803 | 5.18958 | 5.2942 | 6.45276 | 6.94276 | 8.09307 | 4.41814 | 5.32607 | 5.2571 | 5.88137 |
| ZMYM6 | O95789 | 6.50871 | 6.17095 | 6.90735 | 7.59204 | 6.3438 | 5.91118 | 6.74256 | 6.07128 | 6.67198 | 6.36371 | 6.57664 | 6.86236 | 4.7343 | 5.45612 | 5.75749 | 5.32952 |
| POC1A | Q8NBT0 | 6.0077 | 6.2487 | 6.54986 | 6.61719 | 5.68297 | 6.41631 | 6.34431 | 6.35958 | 6.61057 | 6.8948 | 6.906 | 7.13111 | 5.16454 | 5.86527 | 5.39942 | 5.80166 |
| ORC6 | Q9Y5N6 | 6.51741 | 6.2606 | 5.70671 | 5.73987 | 7.52464 | 6.02642 | 5.62314 | 5.26176 | 6.96985 | 5.24399 | 7.06744 | 7.63377 | 5.42429 | 6.78119 | 6.08711 | 6.13182 |
| UBIAD1 | Q9Y5Z9 | 6.20632 | 7.15316 | 5.98073 | 6.53674 | 5.97679 | 5.28832 | 6.34141 | 7.18349 | 6.9816 | 5.82686 | 5.92668 | 5.88203 | 6.1076 | 5.40007 | 6.31855 | 6.88963 |
| SLC66A2 | Q8N2U9 | 6.13261 | 5.9094 | 5.74122 | 5.72899 | 6.15895 | 6.08119 | 5.95374 | 6.23182 | 6.15181 | 5.58389 | 5.5589 | 6.06163 | 6.72901 | 6.63712 | 8.03853 | 7.20119 |
| ENTPD4 | Q9Y227 | 6.60063 | 6.58431 | 6.24039 | 6.94606 | 5.88559 | 6.4517 | 6.2409 | 6.35754 | 6.8335 | 6.62997 | 4.757 | 6.15498 | 6.50398 | 6.1237 | 6.20478 | 5.48496 |
| PKNOX1 | P55347 | 6.3919 | 6.79726 | 6.16706 | 7.1421 | 6.14839 | 6.45136 | 5.85365 | 6.20168 | 6.0374 | 5.55492 | 6.47292 | 6.9111 | 6.31098 | 5.82456 | 5.56757 | 5.96713 |
| POMGNT2 | Q8NAT1 | 5.68097 | 6.38628 | 6.40638 | 7.28704 | 6.25935 | 5.61168 | 6.88376 | 6.72808 | 8.14525 | 5.15132 | 5.17313 | 6.01473 | 6.15659 | 6.40061 | 5.34936 | 6.36549 |
| KXD1 | Q9BQD3 | 6.58272 | 5.58017 | 5.84033 | 5.12369 | 5.41462 | 5.17953 | 5.67264 | 6.21574 | 5.36469 | 6.17433 | 5.56966 | 7.31762 | 5.47228 | 8.9081 | 7.50844 | 8.07543 |
| MRPS21 | P82921 | 6.4525 | 5.76044 | 6.12159 | 6.69304 | 6.0677 | 5.52305 | 6.53407 | 5.7594 | 6.47289 | 6.04192 | 6.18812 | 6.24209 | 6.84884 | 5.86262 | 6.69612 | 6.73561 |
| TMBIM6 | P55061-2 | 5.69385 | 6.43787 | 5.52897 | 5.8754 | 6.6383 | 5.61084 | 7.15158 | 5.83935 | 5.72684 | 5.49208 | 4.72065 | 5.6238 | 7.21166 | 6.97942 | 9.35117 | 6.11822 |
| CHMP6 | Q96FZ7 | 6.13177 | 7.00552 | 5.9685 | 6.20759 | 6.31494 | 6.85319 | 6.3494 | 6.65595 | 6.28744 | 5.66232 | 5.85145 | 5.35661 | 6.70222 | 6.43134 | 6.35873 | 5.86302 |
| RAB11FIP2 | Q7L804-2 | 7.02037 | 6.50478 | 8.33799 | 7.33387 | 6.65587 | 6.3337 | 5.62286 | 4.72251 | 6.2899 | 6.82265 | 6.73585 | 6.33874 | 4.95736 | 5.77401 | 4.9428 | 6.50672 |
| SCD5 | Q86SK9 | 6.58291 | 6.75387 | 6.23346 | 6.76753 | 5.97422 | 5.80963 | 6.48535 | 5.88919 | 6.17735 | 6.47887 | 6.51396 | 6.07996 | 5.90318 | 6.02667 | 6.10942 | 6.21444 |
| CNRIP1 | Q96F85 | 5.74635 | 5.23539 | 5.90718 | 5.35743 | 5.88481 | 6.5254 | 5.73472 | 6.93506 | 7.27592 | 7.43719 | 7.1216 | 7.0446 | 5.63197 | 5.73837 | 5.87451 | 6.54951 |
| ZFYVE28 | Q9HCC9 | 6.15994 | 5.91817 | 6.32497 | 5.96265 | 5.91445 | 7.41729 | 5.96728 | 5.78003 | 5.83964 | 6.55817 | 6.399 | 6.25158 | 6.47741 | 5.83651 | 6.38566 | 6.80724 |
| FAM210B | Q96KR6 | 5.93289 | 5.83153 | 6.03685 | 6.602 | 5.7266 | 5.85367 | 6.38463 | 6.09954 | 5.93746 | 6.78488 | 6.74846 | 6.42552 | 6.02848 | 6.48013 | 6.57991 | 6.54744 |
| DNAJC30 | Q96LL9 | 6.75317 | 6.45867 | 6.09213 | 7.15591 | 6.38386 | 6.17609 | 6.175 | 6.1225 | 6.12278 | 5.93686 | 5.9924 | 5.63341 | 6.67829 | 5.9488 | 6.1172 | 6.25294 |
| COQ4 | Q9Y3A0 | 7.16988 | 6.05485 | 7.13692 | 6.33641 | 4.79116 | 4.64702 | 5.5378 | 8.08755 | 5.54006 | 7.43144 | 7.1505 | 8.44483 | 4.12459 | 5.62139 | 5.95466 | 5.97093 |
| DIPK1A | Q5T7M9 | 6.29156 | 7.26077 | 6.38787 | 5.9844 | 5.3851 | 6.04615 | 6.37449 | 5.01211 | 6.74343 | 6.8208 | 6.00395 | 5.95601 | 7.07947 | 5.59442 | 6.88959 | 6.16988 |
| UEVLD | Q8IX04 | 6.09475 | 6.01809 | 7.45305 | 6.0467 | 6.14425 | 6.92044 | 6.35893 | 6.60169 | 6.32748 | 6.00938 | 6.33777 | 6.54687 | 5.45017 | 5.99004 | 5.54424 | 6.15616 |
| AHR | P35869 | 7.11076 | 6.75452 | 7.74364 | 6.4765 | 6.00898 | 6.38179 | 6.22317 | 5.90026 | 7.03813 | 7.5979 | 7.76441 | 7.7378 | 4.15748 | 4.62253 | 4.20167 | 4.28044 |
| ZBTB43 | O43298 | 5.26193 | 6.66629 | 5.98768 | 6.3209 | 5.94662 | 6.27573 | 5.64079 | 5.92551 | 5.27939 | 6.27994 | 6.23239 | 6.22905 | 6.76397 | 6.67237 | 7.38859 | 7.12887 |
| PIK3AP1 | Q6ZUJ8 | 6.20898 | 6.13935 | 7.07849 | 6.30732 | 5.83191 | 6.34082 | 6.7687 | 6.73779 | 6.32404 | 6.05538 | 6.46556 | 7.03819 | 5.91867 | 5.3733 | 5.69501 | 5.71649 |
| TOMM20 | Q15388 | 7.12614 | 6.44985 | 6.15699 | 7.03234 | 5.95618 | 6.73571 | 6.80864 | 5.65248 | 5.58083 | 6.26055 | 5.94995 | 5.83116 | 6.15896 | 6.12871 | 6.34332 | 5.82818 |
| TIGD2 | Q4W5G0 | 6.43848 | 6.82403 | 6.42949 | 7.24853 | 5.44739 | 5.97645 | 6.27334 | 6.56357 | 6.09909 | 6.60133 | 5.61332 | 4.82502 | 6.18296 | 6.11008 | 6.65177 | 6.61517 |
| SLC39A6 | Q13433 | 6.15376 | 7.26852 | 5.42743 | 6.63942 | 6.0877 | 6.429 | 6.71662 | 6.62989 | 6.02795 | 5.76785 | 5.86049 | 5.89541 | 5.4201 | 5.45344 | 7.02739 | 7.19502 |
| NME4 | O00746 | 6.33942 | 6.48128 | 5.92462 | 7.57266 | 6.39923 | 4.94731 | 5.56693 | 6.40452 | 7.26143 | 6.69419 | 6.75817 | 7.03656 | 5.68278 | 6.05008 | 6.57264 | 4.30817 |
| SHQ1 | Q6PI26 | 7.78887 | 6.72574 | 6.38979 | 6.28705 | 6.5734 | 6.06283 | 7.0885 | 7.2072 | 7.60901 | 7.05221 | 7.36267 | 6.68328 | 4.05009 | 4.72244 | 4.04519 | 4.35172 |
| UBL7 | Q96S82 | 5.99179 | 5.26306 | 5.47538 | 4.99989 | 4.72554 | 4.82222 | 6.18389 | 6.63256 | 4.47388 | 6.51606 | 4.96465 | 6.62353 | 6.55016 | 9.43895 | 6.56257 | 10.7759 |
| MTRF1L | Q9UGC7 | 6.58548 | 6.49344 | 6.41184 | 6.61813 | 6.21038 | 5.82198 | 5.81977 | 5.87872 | 6.61303 | 6.33496 | 5.78073 | 6.02959 | 6.47851 | 6.38143 | 6.31057 | 6.23145 |
| RPP25 | Q9BUL9 | 5.72295 | 5.94106 | 5.80494 | 5.8382 | 5.99569 | 5.88242 | 6.04828 | 6.114 | 5.76591 | 5.93328 | 6.34985 | 5.68324 | 7.2828 | 7.37003 | 7.11125 | 7.1561 |
| NUDT8 | Q8WV74 | 6.526 | 5.09503 | 4.39041 | 7.33038 | 7.97762 | 6.0083 | 6.36196 | 5.75681 | 6.11926 | 7.68148 | 5.96654 | 6.90111 | 7.22682 | 4.85461 | 6.27375 | 5.52994 |
| PIGB | Q92521 | 5.55956 | 6.41184 | 4.75477 | 6.3232 | 6.23108 | 6.47338 | 6.79556 | 6.46224 | 5.68946 | 5.90919 | 5.53415 | 6.54642 | 6.7811 | 6.17057 | 7.12927 | 7.22822 |
| MIF4GD | A9UHW6-3 | 6.88261 | 5.17959 | 6.20565 | 5.1599 | 6.41677 | 6.02541 | 6.31505 | 6.5415 | 5.72489 | 6.75949 | 6.12447 | 6.45253 | 5.98365 | 6.43211 | 6.50725 | 7.28911 |
| LSM11 | P83369 | 6.80569 | 7.10427 | 5.72028 | 6.01516 | 5.90315 | 6.38026 | 6.28386 | 6.06276 | 6.05916 | 6.61603 | 6.27001 | 6.47056 | 5.93773 | 6.24382 | 5.91306 | 6.21418 |
| ING4 | Q9UNL4 | 6.56676 | 6.67454 | 5.86138 | 6.5264 | 6.50013 | 5.70414 | 6.97481 | 5.75723 | 6.0205 | 6.96681 | 6.55884 | 5.94891 | 6.55822 | 5.16695 | 6.10861 | 6.10577 |
| APH1B | Q8WW43 | 6.43514 | 6.97746 | 6.7569 | 6.67515 | 6.34471 | 6.03695 | 6.59661 | 6.33845 | 6.90914 | 7.04774 | 6.63511 | 6.56389 | 4.98852 | 5.33597 | 5.12557 | 5.23269 |
| DFFB | O76075 | 4.97452 | 5.29008 | 6.97097 | 5.61552 | 4.7483 | 5.68381 | 5.62667 | 5.43362 | 5.27313 | 6.45327 | 5.89521 | 6.14517 | 6.98927 | 8.79203 | 8.04407 | 8.06437 |
| NRIP3 | Q9NQ35 | 7.32758 | 6.26588 | 7.50664 | 7.45689 | 7.63494 | 5.94061 | 7.51752 | 6.54973 | 7.56453 | 5.99935 | 7.06519 | 7.38524 | 4.4498 | 3.45228 | 3.95233 | 3.93149 |
| PRUNE1 | Q86TP1 | 6.39028 | 6.17655 | 6.58672 | 5.83898 | 6.88099 | 6.41872 | 6.58192 | 7.08692 | 6.67966 | 5.85193 | 6.34321 | 6.63321 | 6.14449 | 5.55418 | 5.47861 | 5.35362 |
| NPDC1 | Q9NQX5 | 7.88478 | 6.7679 | 7.13668 | 7.04644 | 6.29586 | 6.63978 | 7.59081 | 6.66703 | 6.96599 | 6.12139 | 7.61345 | 7.27067 | 3.91047 | 4.26307 | 3.6418 | 4.18389 |
| DLC1 | Q96QB1 | 6.12086 | 5.61917 | 6.64979 | 6.09271 | 6.10889 | 6.47036 | 6.33723 | 6.56625 | 6.14546 | 5.77892 | 6.02591 | 5.99625 | 6.71627 | 6.46664 | 6.70604 | 6.19926 |
| SLC35F2 | Q8IXU6 | 6.14681 | 6.66813 | 6.15931 | 6.58647 | 6.26645 | 5.92788 | 5.92333 | 6.29828 | 6.08843 | 6.32297 | 6.31505 | 6.31784 | 6.10067 | 6.22094 | 6.59681 | 6.06061 |
| RFXAP | O00287 | 6.16801 | 5.97459 | 6.121 | 6.52177 | 5.25871 | 5.98493 | 6.38179 | 6.15132 | 6.52787 | 6.37706 | 6.81987 | 6.25563 | 6.86951 | 5.87731 | 6.04076 | 5.97587 |
| ISG20 | Q96AZ6 | 6.46113 | 6.43948 | 6.22794 | 6.33999 | 6.79987 | 5.91452 | 6.22026 | 6.68168 | 6.77996 | 6.60279 | 6.49873 | 6.43307 | 6.01307 | 5.56668 | 5.65134 | 5.36949 |
| JDP2 | Q8WYK2-2 | 6.8562 | 7.89147 | 6.96039 | 7.38747 | 6.21761 | 6.25983 | 6.15047 | 6.09305 | 7.21696 | 7.58487 | 6.85535 | 7.09569 | 4.27645 | 4.53714 | 3.99811 | 4.61894 |
| CHKB | Q9Y259 | 6.90192 | 7.25058 | 6.5437 | 6.39097 | 6.61037 | 6.88983 | 5.7201 | 5.40746 | 5.36104 | 5.97154 | 6.19225 | 6.95012 | 5.84974 | 6.85071 | 5.34651 | 5.76319 |
| TLCD3A | Q8TBR7 | 5.25015 | 6.73453 | 5.54238 | 6.66944 | 6.64211 | 5.97594 | 6.17849 | 5.31634 | 7.27538 | 6.05339 | 6.02168 | 6.39571 | 6.31956 | 6.13787 | 6.99965 | 6.48738 |
| LMO4 | P61968 | 6.28975 | 6.41188 | 7.02004 | 6.05946 | 6.7004 | 6.21783 | 6.87522 | 6.77626 | 6.32411 | 6.4451 | 6.86291 | 6.35967 | 6.04483 | 4.9018 | 5.5041 | 5.20666 |
| OSTF1 | Q92882 | 6.38312 | 5.93806 | 6.92989 | 5.34493 | 6.24823 | 6.46991 | 5.99428 | 7.01437 | 6.23349 | 6.54157 | 7.06326 | 7.27669 | 5.34387 | 6.32363 | 5.25286 | 5.64184 |
| PGP | A6NDG6 | 6.42285 | 6.07759 | 7.33086 | 5.5686 | 7.61688 | 7.29673 | 6.57065 | 6.60454 | 6.9348 | 6.77758 | 5.91042 | 5.51825 | 5.67397 | 5.63744 | 5.35249 | 4.70633 |
| ATG4D | Q86TL0 | 6.48886 | 6.16247 | 7.64099 | 7.07176 | 6.38199 | 5.95657 | 6.62794 | 6.30266 | 6.76302 | 7.49447 | 7.22454 | 7.28403 | 4.28667 | 5.20449 | 4.59745 | 4.5121 |
| EGLN2 | Q96KS0 | 6.2208 | 6.70414 | 6.23862 | 7.17203 | 5.23754 | 5.58741 | 5.9265 | 7.86739 | 7.62558 | 7.59665 | 7.30753 | 7.20523 | 4.82378 | 4.66544 | 5.06577 | 4.75561 |
| ARMC7 | Q9H6L4 | 5.85047 | 4.86563 | 6.41476 | 5.77123 | 5.62492 | 4.96481 | 5.17357 | 6.73575 | 6.0665 | 6.45817 | 6.12966 | 6.08365 | 4.49974 | 9.38472 | 7.39673 | 8.57971 |
| STK32C | Q86UX6 | 6.04845 | 5.69544 | 7.09394 | 6.36921 | 6.2532 | 6.64586 | 6.05994 | 6.11534 | 5.97185 | 6.10958 | 7.07017 | 5.96859 | 5.66287 | 5.79124 | 5.97508 | 7.16924 |
| NUDT22 | Q9BRQ3 | 6.54235 | 9.90707 | 6.00628 | 4.33985 | 5.13422 | 5.12047 | 6.64557 | 13.1086 | 7.85828 | 6.79756 | 6.38548 | 7.14693 | 2.71357 | 4.25061 | 3.59808 | 4.44503 |
| TCEAL9 | Q9UHQ7 | 6.20382 | 5.27353 | 5.62251 | 4.86414 | 6.42963 | 6.71533 | 5.9684 | 6.60588 | 5.8385 | 6.62613 | 6.88904 | 6.7817 | 6.49324 | 6.83037 | 5.9309 | 6.92687 |
| MT-ATP6 | P00846 | 5.91683 | 6.19675 | 5.57228 | 6.54874 | 5.59708 | 5.86247 | 6.16002 | 6.42188 | 6.05936 | 6.13929 | 5.53113 | 6.17122 | 6.34889 | 6.58631 | 7.55179 | 7.33596 |
| AP5S1 | Q9NUS5 | 6.65012 | 6.01361 | 6.4023 | 5.84402 | 4.8397 | 6.20074 | 5.51677 | 6.71485 | 6.60699 | 6.38642 | 5.94041 | 6.47067 | 4.88595 | 8.38317 | 5.91336 | 7.23092 |
| BMT2 | Q1RMZ1 | 6.40394 | 6.09196 | 7.53189 | 6.12438 | 6.74964 | 6.68873 | 5.72782 | 6.13965 | 6.01248 | 6.93629 | 6.43355 | 7.09797 | 5.6346 | 5.73396 | 5.54568 | 5.14744 |
| TIMM22 | Q9Y584 | 6.91554 | 6.45212 | 6.17278 | 5.86487 | 6.60592 | 6.46875 | 5.7261 | 6.45111 | 5.99914 | 6.44231 | 5.66731 | 5.60188 | 6.47338 | 6.2794 | 5.89205 | 5.98733 |
| UBE3B | Q7Z3V4 | 6.41516 | 5.49559 | 6.19788 | 6.33702 | 6.22508 | 7.71676 | 6.46624 | 6.66182 | 6.87742 | 5.88539 | 6.32322 | 5.6672 | 5.79085 | 6.49337 | 5.76559 | 5.68141 |
| ISOC2 | Q96AB3-2 | 5.991 | 4.84032 | 5.56059 | 6.12067 | 6.34093 | 6.04444 | 6.15681 | 6.36231 | 5.39797 | 5.86015 | 4.98686 | 5.71751 | 6.0217 | 8.17618 | 7.82065 | 8.60191 |
| ZNF276 | Q8N554 | 5.28617 | 6.13551 | 6.05323 | 5.32748 | 6.24321 | 5.6598 | 5.95113 | 6.32919 | 5.95659 | 6.21746 | 7.43469 | 6.7257 | 6.35946 | 7.00639 | 7.24276 | 6.07123 |
| PDP2 | Q9P2J9 | 5.57135 | 5.21067 | 4.86655 | 5.64114 | 6.80251 | 6.62655 | 7.07951 | 7.19737 | 6.271 | 6.20001 | 5.82065 | 6.17303 | 6.41961 | 6.6272 | 6.04298 | 7.44985 |
| CCNG1 | P51959 | 7.82491 | 7.85822 | 8.21363 | 8.44413 | 6.05803 | 5.91988 | 7.95982 | 6.56999 | 8.10977 | 8.41665 | 8.25733 | 8.87399 | 1.59653 | 1.91153 | 1.89061 | 2.09497 |
| C15orf61 | A6NNL5 | 6.44785 | 5.25839 | 6.08736 | 5.78442 | 6.12415 | 6.09657 | 6.57259 | 6.44109 | 5.78066 | 5.92142 | 7.06014 | 7.01549 | 6.31589 | 6.82555 | 5.7982 | 6.47023 |
| POLR2L | P62875 | 5.36911 | 5.25351 | 5.53201 | 5.70502 | 5.93092 | 6.61875 | 6.18669 | 7.20801 | 6.26634 | 6.71982 | 6.40756 | 6.29547 | 4.94617 | 7.92099 | 5.96667 | 7.67295 |
| GPR176 | Q14439 | 6.00132 | 6.39249 | 6.46198 | 6.10835 | 5.96627 | 5.78382 | 6.37568 | 6.0535 | 6.24473 | 6.65229 | 6.58528 | 6.62359 | 6.08867 | 6.09726 | 6.02959 | 6.53518 |
| TBC1D14 | Q9P2M4 | 6.29166 | 6.43122 | 7.70717 | 5.9098 | 5.87193 | 6.89002 | 6.39701 | 6.77892 | 7.01234 | 6.68836 | 7.11084 | 6.64096 | 5.08118 | 4.93146 | 5.03736 | 5.21978 |
| TIPIN | Q9BVW5 | 5.76732 | 5.8111 | 6.30779 | 6.05368 | 6.2617 | 5.93107 | 5.47258 | 6.76064 | 5.15151 | 7.22312 | 6.52247 | 7.50698 | 6.77042 | 6.27173 | 5.58953 | 6.59838 |
| TNFRSF10B | O14763 | 7.09961 | 5.88033 | 5.40994 | 5.51282 | 6.788 | 5.56836 | 6.86974 | 6.03932 | 6.20133 | 6.62697 | 5.61123 | 4.99872 | 7.12229 | 6.68188 | 6.9207 | 6.66876 |
| MPPE1 | Q53F39 | 6.56096 | 6.44054 | 5.53263 | 7.28068 | 5.75733 | 5.59584 | 6.4256 | 5.94313 | 6.34684 | 5.918 | 5.79506 | 5.95598 | 6.26618 | 5.98812 | 6.92852 | 7.26459 |
| POLR3H | Q9Y535 | 5.62006 | 6.83462 | 6.29314 | 5.52637 | 6.22826 | 5.84464 | 6.25335 | 6.36976 | 6.37489 | 6.31356 | 5.44739 | 6.01413 | 6.86821 | 6.53577 | 6.39728 | 7.07856 |
| POLR3G | O15318 | 5.82272 | 5.71984 | 6.05767 | 5.69202 | 5.80254 | 7.16002 | 5.04003 | 5.77901 | 5.39339 | 6.93 | 6.84114 | 6.38667 | 6.26235 | 7.63308 | 6.52861 | 6.95093 |

| | | | | | | | | | | | | | | | | | |
|---|---|---|---|---|---|---|---|---|---|---|---|---|---|---|---|---|---|
| ZNF330 | Q9Y3S2 | 5.7528 | 6.34514 | 6.8108 | 6.39441 | 6.17356 | 6.91407 | 5.76435 | 6.70785 | 6.38967 | 7.17008 | 6.72798 | 6.2845 | 5.52337 | 5.85951 | 5.85151 | 5.33039 |
| JAK3 | P52333 | 6.52464 | 6.75212 | 7.19214 | 6.18167 | 6.87139 | 7.19151 | 5.95721 | 6.4729 | 5.75298 | 6.24803 | 6.60043 | 6.92069 | 4.83316 | 5.92962 | 4.78318 | 5.78831 |
| CCDC28A | Q8IWP9 | 6.22875 | 6.03479 | 7.03938 | 6.84302 | 5.68014 | 6.28593 | 6.90987 | 6.85578 | 5.68181 | 6.41981 | 6.62226 | 7.02397 | 5.87062 | 5.44609 | 5.70269 | 5.35509 |
| ORAI1 | Q96D31 | 5.70131 | 6.23732 | 6.49684 | 6.59707 | 6.15927 | 5.76297 | 5.59392 | 5.91238 | 6.21234 | 6.46087 | 6.60389 | 6.80315 | 6.10243 | 5.80433 | 6.55118 | 7.00075 |
| ELK3 | P41970 | 5.75421 | 6.26125 | 5.51427 | 5.55731 | 6.99038 | 6.82614 | 5.87228 | 7.10243 | 7.38482 | 5.0788 | 6.30705 | 5.51703 | 6.21898 | 6.11419 | 7.58581 | 5.91505 |
| RNF34 | Q969K3-2 | 5.77062 | 5.7285 | 7.67449 | 6.45453 | 5.98769 | 5.47482 | 6.65892 | 5.98502 | 6.29072 | 6.72875 | 7.00628 | 6.9387 | 6.19663 | 5.80413 | 5.43308 | 5.86711 |
| SERPINC1 | P01008 | 5.43402 | 7.4436 | 4.64816 | 4.93373 | 4.53693 | 6.8278 | 7.12437 | 6.35921 | 6.21524 | 5.40066 | 5.71288 | 5.60178 | 5.86766 | 5.71987 | 10.371 | 7.80307 |
| TRAIP | Q9BWF2 | 6.89894 | 6.78181 | 6.04881 | 7.33234 | 6.41271 | 6.28031 | 6.67088 | 6.05928 | 6.48034 | 6.99954 | 7.55527 | 7.21389 | 4.35028 | 4.99788 | 4.45968 | 5.45805 |
| SSR3 | Q9UNL2-2 | 6.18808 | 6.52813 | 6.08599 | 6.50056 | 6.10486 | 5.89405 | 6.44246 | 5.8326 | 6.22016 | 5.84985 | 6.11502 | 5.84099 | 6.41639 | 6.46804 | 6.72126 | 6.79155 |
| PAQR7 | Q86WK9 | 6.31465 | 6.43213 | 6.49788 | 7.56936 | 6.77429 | 5.88619 | 6.42051 | 6.61577 | 6.25181 | 5.76405 | 6.37883 | 6.71012 | 5.42 | 5.44564 | 5.64102 | 5.87776 |
| DESI1 | Q6ICB0 | 5.60892 | 5.39852 | 4.86084 | 5.23837 | 6.10725 | 5.54577 | 6.4899 | 8.28571 | 5.25923 | 6.79382 | 7.23995 | 7.86855 | 4.37099 | 6.93392 | 6.37391 | 7.62436 |
| TGFBR2 | P37173-2 | 6.84653 | 7.2445 | 7.2467 | 7.56953 | 6.37332 | 6.31215 | 6.346 | 6.39324 | 6.99177 | 6.89196 | 6.67599 | 6.65056 | 4.40425 | 4.91722 | 4.49376 | 4.64251 |
| EID1 | Q9Y6B2 | 7.2969 | 6.75522 | 7.84101 | 6.44499 | 6.70481 | 7.11619 | 6.21527 | 7.0444 | 7.39666 | 7.29729 | 7.90536 | 7.40873 | 3.43945 | 3.74663 | 3.40086 | 3.98622 |
| TIMM10 | P62072 | 6.26957 | 4.77631 | 4.80732 | 5.66388 | 4.62698 | 2.73332 | 4.60767 | 4.99274 | 3.9208 | 4.96827 | 4.53325 | 4.56254 | 4.42855 | 14.6934 | 10.1172 | 14.2982 |
| CREBRF | Q8IUR6-3 | 5.64468 | 6.30517 | 6.20292 | 5.38217 | 6.06862 | 5.68367 | 5.80846 | 5.52995 | 4.96876 | 5.65808 | 5.27203 | 6.01187 | 8.70113 | 7.35657 | 7.69009 | 7.71581 |
| FAM161A | Q3B820-3 | 6.55217 | 5.46205 | 6.95789 | 5.45819 | 6.23038 | 6.26449 | 5.84173 | 6.49917 | 6.45316 | 6.2912 | 6.27503 | 6.49027 | 6.48679 | 6.06676 | 5.30232 | 6.36841 |
| LARP6 | Q9BRS8 | 5.54243 | 6.1395 | 6.48534 | 5.50023 | 6.34031 | 7.17755 | 5.83144 | 6.70533 | 6.1165 | 5.6795 | 6.85278 | 5.76727 | 6.467 | 6.6636 | 6.05402 | 5.77719 |
| CCDC82 | Q8N4S0 | 5.68523 | 6.61957 | 5.97162 | 4.69269 | 6.06874 | 6.15243 | 6.39708 | 6.04724 | 6.13532 | 6.92467 | 7.47647 | 6.70549 | 6.76219 | 5.90192 | 7.00491 | 5.45442 |
| IQCJ-SCHIP1 | B3KU38 | 6.61135 | 5.89023 | 5.93869 | 5.88676 | 5.87037 | 6.33038 | 6.65041 | 6.27594 | 6.50402 | 5.83936 | 7.22631 | 6.7935 | 6.45525 | 5.20186 | 6.80617 | 5.7194 |
| HORMAD1 | Q86X24 | 5.3304 | 6.51803 | 6.10124 | 6.82914 | 5.71403 | 5.45148 | 7.03676 | 6.50071 | 6.66094 | 4.73797 | 6.08795 | 6.28048 | 6.54805 | 7.14087 | 6.70543 | 7.25651 |
| NOP10 | Q9NPE3 | 5.30258 | 6.51728 | 6.68542 | 6.14436 | 6.52847 | 6.53453 | 6.40881 | 6.59432 | 6.09221 | 6.17238 | 5.75008 | 6.35095 | 6.10151 | 6.29482 | 6.34635 | 6.17593 |
| GAS6 | Q14393-1 | 5.96957 | 6.0985 | 5.85401 | 5.68079 | 6.51951 | 6.11434 | 6.20608 | 6.31961 | 5.5996 | 7.05022 | 6.71879 | 6.30084 | 5.91387 | 6.96056 | 6.29523 | 6.39846 |
| SLC16A2 | P36021 | 6.31005 | 5.9287 | 5.62374 | 5.29562 | 6.65734 | 6.1875 | 6.55636 | 5.5744 | 7.08764 | 6.80935 | 6.81897 | 6.85963 | 7.02404 | 5.09498 | 6.26947 | 5.90221 |
| MED29 | Q9NX70 | 5.80504 | 5.11942 | 4.89499 | 4.89865 | 5.11251 | 5.2674 | 5.70775 | 6.36892 | 5.04399 | 5.42517 | 4.86619 | 4.8787 | 5.35409 | 11.7421 | 7.69616 | 11.8189 |
| NTAQ1 | Q96HA8 | 6.01165 | 5.29992 | 7.10421 | 5.46587 | 6.31346 | 6.82421 | 5.55004 | 7.04577 | 6.34576 | 6.67582 | 7.29295 | 7.21084 | 6.0067 | 5.62943 | 5.84165 | 5.38173 |
| ELOF1 | P60002 | 5.27319 | 5.68352 | 4.97622 | 6.41978 | 5.81204 | 5.61306 | 7.64718 | 5.2652 | 6.48879 | 5.0898 | 5.48099 | 5.52473 | 5.70107 | 9.17393 | 7.16267 | 8.68783 |
| C1QL1 | O75973 | 7.88498 | 6.882 | 6.4608 | 6.94094 | 6.52309 | 6.15088 | 5.87973 | 6.59962 | 7.03797 | 7.40513 | 6.74127 | 6.35836 | 4.75073 | 5.40402 | 4.38866 | 4.59182 |
| WASHC3 | Q9Y3C0 | 6.13308 | 5.70884 | 6.14173 | 5.64887 | 6.15022 | 6.35067 | 6.04851 | 6.64734 | 5.81315 | 6.32715 | 7.26078 | 6.74769 | 6.49647 | 5.97 | 6.48666 | 6.06884 |
| FBXO9 | Q9UK97 | 6.8664 | 6.45804 | 7.39785 | 6.3996 | 6.1882 | 6.55572 | 6.40512 | 6.8205 | 6.95124 | 6.79972 | 6.71868 | 7.08322 | 4.89551 | 5.0986 | 4.66053 | 4.70107 |
| MFSD14B | Q5SR56 | 5.91526 | 6.49205 | 5.9597 | 6.8384 | 6.7328 | 5.8865 | 6.30898 | 5.94782 | 6.08134 | 6.2227 | 6.1677 | 6.52139 | 6.132 | 6.36117 | 6.37547 | 6.05672 |
| SNX11 | Q9Y5W9 | 6.26082 | 5.90131 | 6.6032 | 6.9082 | 5.92404 | 5.98043 | 5.92927 | 5.97987 | 6.57241 | 6.51669 | 7.02503 | 6.86672 | 6.26119 | 5.22996 | 5.94869 | 6.09215 |
| KIF24 | Q5T7B8 | 6.56392 | 6.32314 | 6.86992 | 5.88149 | 6.29961 | 6.84199 | 6.6063 | 5.90025 | 6.61533 | 6.79299 | 6.63303 | 6.84052 | 5.00136 | 5.7212 | 5.74913 | 5.35982 |
| MKKS | Q9HB66 | 5.84057 | 5.11092 | 5.93916 | 5.53333 | 5.31359 | 4.53142 | 5.61923 | 5.28816 | 5.82828 | 6.00837 | 5.56216 | 6.34791 | 6.36167 | 8.52476 | 8.35678 | 9.8337 |
| AGPAT2 | O15120 | 7.073 | 7.29918 | 6.09144 | 6.46222 | 6.0156 | 6.133 | 5.7992 | 5.88263 | 6.12701 | 5.08108 | 6.47664 | 5.19161 | 6.61725 | 6.10574 | 7.10188 | 6.54253 |
| AP4M1 | O00189 | 5.49328 | 5.74063 | 7.23319 | 6.16452 | 6.85293 | 6.12117 | 6.06171 | 6.32294 | 7.13151 | 5.6335 | 7.37823 | 7.86113 | 5.63067 | 5.82369 | 5.06817 | 5.48272 |
| TMEM106B | Q9NUM4 | 6.34767 | 6.05486 | 6.12184 | 5.89204 | 6.20755 | 6.05362 | 6.32579 | 6.2381 | 6.04291 | 5.6988 | 6.72696 | 7.83316 | 4.87677 | 6.56543 | 6.83963 | 6.17487 |
| ZDBF2 | Q9HCK1 | 7.74413 | 6.28259 | 5.37895 | 7.36205 | 5.31195 | 6.552 | 5.86045 | 5.01254 | 7.053 | 6.8755 | 6.8957 | 6.33874 | 5.8222 | 6.43616 | 5.30102 | 5.77302 |
| ZNF556 | Q9HAH1 | 6.20682 | 5.75561 | 6.52534 | 6.55867 | 5.28648 | 5.83076 | 6.54001 | 6.38177 | 6.14353 | 5.22131 | 6.8583 | 6.20474 | 6.82343 | 6.48445 | 6.26822 | 6.91058 |
| MIIP | Q5JXC2 | 6.48187 | 6.46631 | 7.38973 | 7.50012 | 6.0967 | 6.93243 | 6.18654 | 6.41132 | 7.33077 | 6.75168 | 8.20425 | 8.60207 | 3.87645 | 4.3107 | 3.70951 | 3.74954 |
| CDKN1B | P46527 | 5.89641 | 5.58451 | 6.59885 | 5.26383 | 5.8451 | 5.92243 | 5.48331 | 6.769 | 5.61403 | 7.48689 | 7.15612 | 7.54149 | 6.31251 | 7.22863 | 5.73984 | 5.55705 |
| SUPT7L | O94864 | 6.02457 | 5.30382 | 7.10646 | 5.70082 | 6.19385 | 5.80091 | 6.69186 | 5.36384 | 6.0199 | 6.8022 | 5.9931 | 6.46558 | 7.21672 | 6.30552 | 6.57773 | 6.43311 |
| TPRKB | Q9Y3C4-3 | 6.48701 | 6.17383 | 6.00449 | 5.15715 | 5.96991 | 6.6849 | 6.16013 | 6.03489 | 6.02701 | 6.484 | 7.19076 | 5.94079 | 5.86572 | 6.44617 | 6.89964 | 6.4736 |
| COX7C | P15954 | 6.13634 | 5.39863 | 5.57344 | 6.32253 | 5.75236 | 5.29722 | 5.6503 | 5.86881 | 5.97022 | 5.47345 | 5.55531 | 5.83364 | 5.88615 | 8.97577 | 7.14685 | 9.15899 |
| FN1 | P02751 | 7.48119 | 7.25857 | 7.06307 | 7.33353 | 6.71767 | 6.68163 | 6.22316 | 6.59466 | 6.04678 | 7.04283 | 6.48265 | 7.16573 | 4.42268 | 5.49898 | 4.77713 | 3.85795 |
| TOMM6 | Q96B49 | 4.58781 | 4.32807 | 4.4802 | 6.65201 | 5.76492 | 6.2113 | 6.38658 | 5.41979 | 6.14164 | 5.97953 | 6.23268 | 6.76032 | 5.35882 | 8.51124 | 6.83518 | 10.3499 |
| CDIN1 | Q9Y2V0 | 5.4578 | 5.80257 | 6.64296 | 5.60925 | 6.36179 | 6.87313 | 6.18687 | 6.83993 | 7.18299 | 6.91987 | 6.87886 | 6.74792 | 6.00418 | 6.21855 | 5.42084 | 4.85248 |
| CHAC2 | Q8WUX2 | 6.43963 | 6.11204 | 6.70494 | 6.04452 | 7.29086 | 5.83653 | 6.2352 | 7.21192 | 6.20528 | 6.51926 | 6.37447 | 6.52161 | 5.95177 | 5.6376 | 5.44566 | 5.4687 |
| BLOC1S5 | Q8TDH9 | 5.83764 | 5.2255 | 6.47695 | 4.70345 | 6.62526 | 5.50457 | 4.9095 | 8.26969 | 5.38419 | 7.1647 | 7.06009 | 7.36625 | 6.33729 | 6.63559 | 6.63133 | 5.868 |
| BORCS7 | Q96B45 | 6.65966 | 4.87677 | 6.39093 | 4.56636 | 6.93353 | 5.77544 | 5.52526 | 6.44281 | 4.75521 | 5.82368 | 5.97138 | 6.42991 | 5.56491 | 8.94314 | 7.93243 | 7.40858 |
| UBXN2A | P68543 | 7.17435 | 7.10546 | 7.9543 | 7.31165 | 6.3902 | 5.62892 | 7.30211 | 6.10622 | 7.0119 | 6.67774 | 7.41693 | 7.05116 | 4.3114 | 4.25601 | 4.0451 | 4.25656 |
| CERS6 | Q6ZMG9-2 | 6.19196 | 6.20166 | 5.92214 | 6.53521 | 6.06407 | 5.97788 | 6.07838 | 6.10083 | 5.41069 | 5.90135 | 6.18636 | 6.11493 | 6.76904 | 6.71832 | 6.80979 | 7.0174 |
| SLC17A5 | Q9NRA2 | 6.42333 | 6.69634 | 6.86008 | 6.01421 | 6.19389 | 5.52917 | 6.30805 | 6.45034 | 5.36857 | 6.80782 | 6.25076 | 5.42715 | 6.35037 | 6.52064 | 6.02504 | 6.77424 |
| ZBTB44 | Q8NCP5 | 6.03233 | 6.68801 | 5.70818 | 6.97569 | 5.95585 | 5.34232 | 6.19491 | 5.39531 | 6.02132 | 6.71399 | 6.08722 | 6.62124 | 6.90511 | 6.82195 | 5.95559 | 6.58099 |
| UNC119 | Q13432 | 5.96142 | 6.24358 | 7.29959 | 5.95156 | 6.0125 | 6.41448 | 5.84882 | 5.3632 | 6.09522 | 8.59943 | 7.03948 | 6.39505 | 4.69418 | 5.38334 | 6.03344 | 6.66472 |
| ARL5B | Q96KC2 | 5.89633 | 5.78163 | 6.34441 | 6.04567 | 5.97883 | 6.67296 | 6.73542 | 6.62928 | 6.9915 | 6.34209 | 5.60729 | 6.31568 | 6.91888 | 5.78961 | 6.07397 | 5.87644 |
| ILRUN | Q9H6K1 | 6.79847 | 5.92255 | 8.72328 | 4.87713 | 5.89396 | 6.62434 | 5.90536 | 6.40005 | 6.05972 | 7.21414 | 6.8109 | 5.93896 | 6.0118 | 5.77116 | 5.96755 | 5.08062 |
| TMEM51 | Q9NW97 | 6.79239 | 7.475 | 7.24338 | 6.90124 | 6.25449 | 6.28012 | 5.91584 | 5.779 | 7.04263 | 6.95996 | 5.98112 | 6.5442 | 5.45675 | 5.83351 | 4.43591 | 5.10445 |
| SGMS1 | Q86VZ5 | 6.31303 | 6.43814 | 6.63457 | 7.10699 | 6.11936 | 5.21652 | 6.68259 | 5.05102 | 6.09789 | 6.12538 | 6.32399 | 6.21641 | 5.97744 | 5.81005 | 6.71477 | 7.17183 |
| DMAC1 | Q96GE9 | 6.45104 | 4.98724 | 5.89507 | 6.50698 | 6.43581 | 6.56579 | 6.4251 | 6.75246 | 5.89205 | 5.48463 | 6.70842 | 6.24119 | 6.56898 | 6.49032 | 6.30975 | 6.28518 |
| ERVK3-1 | A0A0U1RQV1 | 6.56188 | 4.57906 | 4.84915 | 5.02773 | 5.21713 | 4.144 | 4.35468 | 5.19974 | 4.38133 | 6.63793 | 5.75571 | 6.13401 | 4.59161 | 12.5744 | 8.15658 | 11.8351 |
| ZIC2 | O95409 | 6.42352 | 5.49685 | 6.02857 | 7.42143 | 6.53372 | 6.37855 | 5.79336 | 5.23554 | 6.86263 | 7.0844 | 6.98537 | 7.66766 | 5.68688 | 5.31871 | 5.50434 | 5.57845 |
| COX6A1 | P12074 | 6.0708 | 4.3956 | 4.83327 | 5.7904 | 5.77345 | 6.08148 | 7.10886 | 6.65527 | 4.61178 | 4.86872 | 5.5829 | 4.78345 | 6.65609 | 9.64743 | 7.30874 | 9.83177 |
| CHTF8 | P0CG13 | 5.54618 | 5.88307 | 6.05383 | 5.51348 | 6.21648 | 5.91133 | 5.90314 | 6.37979 | 6.14336 | 6.1558 | 6.22549 | 6.58465 | 6.66808 | 7.4341 | 6.78481 | 6.59641 |
| GEMIN2 | O14893 | 6.72306 | 4.8969 | 6.59686 | 5.84005 | 6.69506 | 6.32668 | 6.16148 | 6.24328 | 5.12047 | 6.95412 | 6.98559 | 6.94263 | 7.31835 | 6.20671 | 5.28441 | 5.70434 |
| SLC25A38 | Q96DW6 | 5.56921 | 5.88608 | 5.9603 | 7.10193 | 6.77437 | 6.57754 | 7.03767 | 6.24892 | 6.41625 | 6.02069 | 5.94622 | 5.96501 | 6.22003 | 5.80862 | 6.20356 | 6.2636 |
| FBXW9 | Q5XUX1-2 | 5.94504 | 5.52974 | 6.24382 | 6.31597 | 5.98558 | 6.90998 | 5.75814 | 7.25871 | 7.13259 | 5.99299 | 6.51379 | 6.43246 | 6.17377 | 6.76274 | 5.43227 | 5.6124 |
| CIAO2B | Q9Y3D0 | 6.27094 | 5.41377 | 5.94506 | 5.40603 | 5.7924 | 6.50317 | 5.36518 | 6.41567 | 5.47964 | 5.92242 | 5.86689 | 6.44789 | 5.39636 | 9.01701 | 6.88772 | 7.86986 |
| APLP1 | P51693-2 | 7.75621 | 8.40556 | 7.47306 | 7.00272 | 6.06402 | 6.96593 | 6.02464 | 6.89107 | 6.83898 | 7.12973 | 7.15654 | 6.45901 | 4.29356 | 4.01597 | 3.60098 | 3.922 |
| TPPP3 | Q9BW30 | 6.3852 | 6.06351 | 7.81239 | 5.53052 | 6.32892 | 6.15543 | 5.1417 | 6.91728 | 6.40189 | 6.6558 | 6.31844 | 6.55547 | 6.56495 | 6.46125 | 4.9805 | 5.72673 |
| KCTD20 | Q7Z5Y7 | 6.94999 | 6.9875 | 7.12844 | 6.70714 | 6.13554 | 6.75786 | 6.21144 | 6.95178 | 7.06745 | 6.75146 | 6.73752 | 7.66765 | 4.315 | 4.83276 | 4.43965 | 4.35883 |
| WDR13 | Q9H1Z4 | 5.68555 | 6.13951 | 7.02092 | 7.47761 | 5.30158 | 6.93914 | 5.80374 | 6.07297 | 7.02354 | 6.08473 | 5.6385 | 6.99117 | 6.80342 | 4.96697 | 6.10748 | 5.94317 |
| STK35 | Q8TDR2 | 6.90197 | 6.51897 | 6.3011 | 6.52727 | 6.46348 | 6.0197 | 6.58538 | 5.8807 | 6.4436 | 6.43115 | 6.33583 | 6.55116 | 5.72613 | 5.60596 | 5.79974 | 5.90787 |
| CLN6 | Q9NWW5 | 7.35102 | 5.81445 | 6.12049 | 6.66983 | 5.84866 | 5.85926 | 6.75357 | 5.74372 | 5.51837 | 6.25535 | 5.87506 | 6.0292 | 5.69458 | 6.90698 | 7.04037 | 6.5191 |
| GABPB1 | Q06547 | 6.08772 | 5.20953 | 6.20922 | 5.48772 | 6.08945 | 5.91983 | 5.9206 | 4.71658 | 6.06337 | 6.10077 | 5.85194 | 6.53336 | 7.19355 | 7.76829 | 7.56946 | 7.27859 |
| RABIF | P47224 | 5.01686 | 5.77146 | 8.0932 | 4.51995 | 5.94572 | 7.47882 | 5.76686 | 7.63286 | 7.05018 | 6.37236 | 6.39904 | 7.3554 | 6.17825 | 6.84028 | 5.84926 | 5.34949 |
| HDAC9 | Q9UKV0-7 | 5.40781 | 5.79217 | 6.54385 | 5.66858 | 6.20425 | 6.10583 | 6.95152 | 6.10794 | 5.40064 | 5.02605 | 5.71198 | 6.30385 | 6.88634 | 6.57559 | 7.71557 | 7.59801 |
| MT-CO3 | P00414 | 6.07452 | 6.09711 | 5.08446 | 7.10868 | 6.76573 | 6.08216 | 5.97043 | 7.22281 | 6.04968 | 5.72444 | 5.97417 | 5.83583 | 6.25712 | 6.3583 | 6.75047 | 6.64412 |
| ARPP19 | P56211 | 5.27344 | 6.08184 | 6.90784 | 5.35616 | 7.67445 | 6.39944 | 5.68625 | 7.54875 | 5.83289 | 6.33583 | 5.96202 | 7.46121 | 4.8839 | 6.93309 | 5.11274 | 6.55015 |
| SNX21 | Q969T3 | 6.60147 | 6.68907 | 6.59288 | 6.22471 | 6.64634 | 7.47127 | 5.83263 | 6.62665 | 6.98859 | 6.32225 | 5.97317 | 6.31587 | 5.63832 | 5.24033 | 5.29129 | 5.54517 |
| ALG12 | Q9BV10 | 6.49559 | 6.10075 | 5.95706 | 6.31185 | 6.30182 | 6.11593 | 6.98002 | 6.02474 | 5.51673 | 6.30673 | 6.04738 | 6.07535 | 6.56782 | 6.48332 | 6.26424 | 6.45069 |
| F2RL1 | P55085 | 5.56669 | 5.42103 | 4.24985 | 5.70386 | 5.94566 | 5.86115 | 6.2042 | 5.60375 | 5.60801 | 5.29918 | 5.10656 | 4.99356 | 9.94159 | 7.32238 | 9.01684 | 8.15568 |
| RPL39P5 | Q59GN2 | 5.93288 | 6.38528 | 6.13066 | 5.72631 | 5.87649 | 6.13289 | 6.07038 | 6.72125 | 6.57171 | 6.22005 | 6.27593 | 6.31691 | 6.61989 | 6.8301 | 5.92621 | 6.26307 |
| SCML1 | Q9UN30 | 6.25274 | 6.64994 | 6.3525 | 7.30567 | 6.28171 | 6.58261 | 6.16048 | 6.03034 | 5.7852 | 5.85002 | 5.98536 | 6.45244 | 5.77058 | 6.06565 | 5.71417 | 6.7606 |
| BLOC1S1 | P78537 | 5.40965 | 5.25514 | 5.74068 | 5.26281 | 5.37283 | 4.81657 | 5.13123 | 6.82511 | 4.76803 | 6.29069 | 5.79724 | 6.04186 | 5.11179 | 10.765 | 7.93323 | 9.47818 |
| CTDSP1 | Q9GZU7 | 6.61765 | 6.59122 | 6.37363 | 5.97546 | 6.51011 | 6.72534 | 5.96648 | 5.98833 | 5.48333 | 6.69215 | 6.17241 | 6.1523 | 6.75857 | 6.38523 | 5.95413 | 5.65817 |
| PCDHGC3 | Q9UN70 | 7.58807 | 6.24351 | 5.98282 | 7.88872 | 6.3923 | 6.44626 | 6.6183 | 5.80573 | 6.80759 | 6.78343 | 5.88318 | 6.93754 | 4.93269 | 5.50829 | 4.79695 | 5.38461 |
| NXPE3 | Q969Y0 | 6.60082 | 7.61421 | 6.06856 | 6.87322 | 6.32782 | 6.15949 | 6.0273 | 5.96728 | 5.62893 | 6.12719 | 6.87971 | 6.30031 | 6.12889 | 5.9641 | 5.74359 | 5.58857 |
| PCTP | Q9UKL6 | 6.89793 | 6.04873 | 6.84669 | 6.11463 | 6.13563 | 7.66126 | 5.77402 | 6.00178 | 6.53257 | 6.62096 | 6.24598 | 6.82179 | 5.94179 | 5.79368 | 5.87679 | 4.68607 |
| YIPF5 | Q969M3-3 | 6.28966 | 6.50053 | 5.64375 | 6.33477 | 5.7769 | 5.89719 | 5.73444 | 6.18639 | 5.94548 | 6.14771 | 5.87929 | 6.60195 | 6.76868 | 6.71243 | 6.69684 | 6.884 |
| TTYH3 | Q9C0H2-4 | 7.09211 | 6.99635 | 6.43688 | 7.28572 | 6.12354 | 5.98571 | 5.7745 | 5.37606 | 6.9544 | 7.65016 | 6.9336 | 6.24468 | 4.85687 | 5.86378 | 5.2575 | 5.16814 |
| NTN4 | Q9HB63 | 7.05089 | 6.488 | 5.42432 | 7.56262 | 5.53717 | 6.8555 | 6.69811 | 6.09171 | 6.09309 | 6.11161 | 6.07486 | 6.19948 | 6.56395 | 4.93154 | 5.07843 | 7.23872 |
| COX7B | P24311 | 6.03067 | 5.67523 | 5.90155 | 5.91341 | 6.57183 | 6.13709 | 6.29862 | 6.45314 | 6.16774 | 5.92197 | 5.90489 | 6.2918 | 6.26801 | 6.92049 | 6.77316 | 6.7704 |
| BCL7C | Q8WUZ0-2 | 6.75394 | 6.39931 | 7.00565 | 6.9873 | 6.85661 | 6.16147 | 5.95268 | 6.4526 | 6.07886 | 7.17547 | 7.19748 | 5.76932 | 5.50907 | 5.4541 | 4.50183 | 5.74431 |
| SLC7A6OS | Q96CW6 | 6.24715 | 6.71706 | 6.9872 | 6.60135 | 6.14246 | 6.93241 | 6.1149 | 6.15251 | 6.58278 | 6.69029 | 7.15201 | 6.67197 | 5.50441 | 5.04025 | 4.9314 | 5.53186 |
| SSBP4 | Q9BWG4 | 6.09837 | 5.85648 | 5.44363 | 5.28197 | 5.84244 | 5.81349 | 5.43418 | 6.10143 | 5.54258 | 6.38335 | 6.18302 | 6.88362 | 7.00499 | 7.8127 | 7.74659 | 6.57116 |
| PHLDA2 | Q53GA4 | 6.53011 | 6.67319 | 7.41995 | 7.188 | 5.53607 | 6.40001 | 5.09898 | 6.87967 | 7.18062 | 6.39413 | 8.19788 | 6.7854 | 4.79119 | 5.16438 | 4.45057 | 5.30984 |
| CDH13 | P55290-4 | 5.39009 | 5.96773 | 5.16203 | 6.53338 | 6.95347 | 6.87679 | 6.28798 | 6.88513 | 6.11755 | 5.84824 | 5.71069 | 5.61261 | 7.68284 | 7.38556 | 5.75417 | 5.83174 |
| MYNN | Q9NPC7 | 6.03699 | 5.92953 | 6.2772 | 5.56023 | 5.56666 | 4.75743 | 6.33304 | 6.19342 | 6.51395 | 5.93402 | 6.63387 | 6.53924 | 6.60358 | 7.21959 | 7.08341 | 6.81783 |
| CEBPZOS | A8MTT3 | 6.18506 | 7.35357 | 6.00265 | 5.69321 | 7.28624 | 6.03778 | 6.60652 | 6.13374 | 6.3298 | 6.49259 | 5.64488 | 5.37458 | 6.85198 | 5.76235 | 6.93863 | 5.30642 |
| TVP23C | Q96ET8 | 5.90406 | 6.26762 | 6.08591 | 6.64402 | 6.03562 | 5.92295 | 6.43708 | 5.70393 | 6.02817 | 6.50531 | 6.28688 | 6.35849 | 6.50383 | 6.5189 | 6.41557 | 6.38167 |
| FAXC | Q5TGI0 | 7.33215 | 6.48037 | 5.41907 | 6.10358 | 6.35796 | 5.72033 | 5.79559 | 6.70319 | 6.63932 | 6.67861 | 5.4131 | 4.48192 | 5.76987 | 6.67536 | 6.78154 | 7.64806 |
| RAB36 | O95755 | 7.7709 | 6.81967 | 6.91189 | 5.71188 | 5.72431 | 6.88658 | 6.0487 | 5.87333 | 6.08056 | 6.06203 | 6.61752 | 6.35729 | 6.44526 | 6.27746 | 4.90318 | 5.50945 |
| PTPRZ1 | P23471 | 6.28376 | 6.40336 | 5.64199 | 6.94714 | 6.39212 | 6.06983 | 7.04347 | 6.05206 | 6.844 | 6.31543 | 5.99396 | 5.93556 | 6.58659 | 5.8979 | 6.19253 | 6.3003 |
| MT-ND3 | P03897 | 5.82528 | 5.87153 | 4.92055 | 6.7097 | 6.00582 | 5.76539 | 6.09596 | 5.51551 | 6.40034 | 5.78087 | 6.55297 | 5.19661 | 7.20462 | 8.09392 | 6.52058 | 7.54033 |
| ZDHHC7 | Q9NXF8 | 6.2152 | 6.22976 | 5.21929 | 6.75563 | 6.4882 | 6.52848 | 6.73218 | 6.49588 | 6.53759 | 6.32558 | 6.20851 | 6.20482 | 6.72916 | 6.0716 | 5.43151 | 5.82661 |

| | | | | | | | | | | | | | | | | | |
|---|---|---|---|---|---|---|---|---|---|---|---|---|---|---|---|---|---|
| DYNC1I1 | O14576 | 5.97817 | 5.71861 | 6.84394 | 6.04491 | 6.34761 | 6.57179 | 6.45367 | 7.26694 | 6.59509 | 6.34266 | 6.65227 | 6.52421 | 4.94813 | 6.12097 | 5.6765 | 5.91453 |
| PRADC1 | Q9BSG0 | 6.62118 | 6.00349 | 6.41818 | 6.47282 | 6.49654 | 6.20237 | 5.74902 | 6.70366 | 6.54132 | 7.02321 | 6.62651 | 6.87795 | 5.12985 | 5.70513 | 5.56951 | 5.85924 |
| FAM126B | Q8IXS8 | 6.35471 | 5.87019 | 6.50105 | 7.79286 | 6.22937 | 5.255 | 7.53697 | 6.02874 | 6.10397 | 5.47796 | 5.72297 | 6.8438 | 6.89665 | 5.65626 | 6.32486 | 5.40463 |
| CD40 | P25942 | 6.88074 | 6.47211 | 5.90625 | 6.49284 | 6.28078 | 6.30104 | 5.99092 | 5.69002 | 5.7513 | 6.06641 | 6.20089 | 6.04879 | 6.10616 | 7.40012 | 5.93432 | 6.47732 |
| CLDND1 | Q9NY35 | 6.13717 | 6.91725 | 6.03411 | 6.04962 | 6.30222 | 6.19288 | 5.97177 | 6.24969 | 6.33331 | 6.21644 | 5.86428 | 6.11685 | 6.67413 | 6.17869 | 6.17296 | 6.58862 |
| ARRDC1 | Q8N5I2 | 6.62352 | 6.54077 | 6.23272 | 5.77693 | 6.48817 | 6.28908 | 6.20455 | 6.52005 | 6.58386 | 6.4574 | 6.4416 | 6.30466 | 5.77186 | 5.87786 | 6.00731 | 5.87965 |
| MIPOL1 | Q8TD10 | 6.17251 | 5.97229 | 6.69795 | 6.54294 | 5.82782 | 5.86382 | 6.39399 | 6.84922 | 6.97701 | 6.34179 | 6.69121 | 6.60094 | 5.75971 | 5.38417 | 6.07847 | 5.84618 |
| HIPK1 | Q86Z02 | 5.79487 | 6.03511 | 6.27447 | 6.2691 | 6.3493 | 6.2439 | 6.54341 | 6.45117 | 5.97626 | 6.24929 | 6.72157 | 6.82659 | 5.98096 | 6.10684 | 6.12466 | 6.05253 |
| CDKN3 | Q16667 | 6.71924 | 6.6938 | 7.93979 | 6.64339 | 6.94409 | 6.90994 | 6.96301 | 6.75527 | 7.14292 | 6.90212 | 7.14497 | 6.69467 | 4.04565 | 3.99435 | 4.15996 | 4.34681 |
| HOXC9 | P31274 | 7.27045 | 6.35863 | 6.54395 | 7.62867 | 7.12229 | 6.21728 | 6.04162 | 5.8252 | 5.92522 | 6.86359 | 7.31016 | 7.16731 | 5.37035 | 4.93024 | 4.72879 | 4.69624 |
| RAB34 | P0DI83 | 6.56471 | 6.8691 | 6.75047 | 6.45217 | 6.80829 | 6.58311 | 6.01453 | 6.55468 | 6.74911 | 6.87557 | 6.31296 | 6.57018 | 6.42544 | 5.35717 | 4.73231 | 5.38019 |
| MTRES1 | Q9P0P8 | 6.26367 | 6.21436 | 6.10863 | 5.94747 | 5.81448 | 5.33393 | 7.4086 | 5.87831 | 6.27182 | 6.24953 | 6.4498 | 5.83443 | 6.75225 | 6.45401 | 7.03609 | 5.98266 |
| KCNAB2 | Q13303-3 | 6.27639 | 6.10138 | 6.40978 | 5.86766 | 6.22249 | 6.75726 | 6.29205 | 6.17131 | 6.4099 | 6.66281 | 5.93096 | 6.17338 | 6.51095 | 6.21704 | 6.06568 | 5.93096 |
| PLEKHB2 | Q96CS7-2 | 6.0828 | 6.17192 | 7.23881 | 6.19487 | 6.58452 | 6.61974 | 7.38185 | 6.65845 | 5.74218 | 6.03769 | 6.07298 | 6.97087 | 5.83457 | 5.21982 | 5.72497 | 5.46395 |
| ARSD | P51689 | 6.4934 | 6.48441 | 6.35196 | 6.92249 | 6.22748 | 5.76263 | 5.9752 | 5.76058 | 6.34842 | 6.22823 | 6.25571 | 6.27345 | 6.35352 | 5.72832 | 6.58025 | 6.25395 |
| SLC35C1 | Q96A29 | 7.94776 | 6.35403 | 6.97062 | 6.5778 | 6.58373 | 5.015 | 5.46334 | 4.21201 | 5.5945 | 5.63175 | 4.94076 | 5.69219 | 7.77813 | 7.98448 | 7.40871 | 6.74519 |
| FADD | Q13158 | 6.58141 | 5.67748 | 7.47097 | 5.40483 | 6.31746 | 6.13437 | 5.9321 | 6.64991 | 7.04766 | 7.36655 | 6.85279 | 6.96484 | 5.74045 | 5.62292 | 4.77901 | 5.45725 |
| TLCD4 | Q96MV1 | 6.41042 | 6.60713 | 5.94262 | 6.79333 | 5.76924 | 5.7761 | 6.11474 | 6.00244 | 5.79995 | 6.31895 | 5.86815 | 6.10449 | 6.51296 | 6.4115 | 6.70191 | 6.86608 |
| LEPROT | O15243 | 6.44467 | 5.8619 | 6.94236 | 6.73352 | 5.77314 | 6.99486 | 7.17363 | 6.10822 | 7.01304 | 5.82925 | 6.36924 | 5.87455 | 5.4021 | 5.91366 | 5.52036 | 6.04549 |
| MT-ND2 | P03891 | 6.0256 | 6.50477 | 5.29669 | 6.84999 | 6.2166 | 6.24634 | 6.46025 | 6.72126 | 5.65017 | 6.36578 | 5.40894 | 5.72663 | 6.40832 | 6.87247 | 6.53419 | 6.71201 |
| C16orf70 | Q9BSU1 | 6.65937 | 6.72557 | 6.78793 | 5.98111 | 6.94915 | 6.44728 | 6.72688 | 7.08774 | 6.05162 | 6.38575 | 6.4281 | 6.17086 | 5.27278 | 5.11195 | 5.05815 | 6.15577 |
| CRISPLD1 | Q9H336 | 7.04651 | 7.50223 | 6.40677 | 7.69585 | 6.15713 | 6.66683 | 6.79181 | 6.10142 | 6.80251 | 7.86758 | 6.21792 | 6.90881 | 4.33841 | 4.42164 | 4.38527 | 4.68931 |
| PPP2R3C | Q969Q6 | 5.77903 | 7.6645 | 6.17429 | 6.33543 | 5.6855 | 7.41816 | 4.71117 | 6.26719 | 6.89237 | 6.86399 | 6.78392 | 7.61078 | 5.49049 | 5.57365 | 5.26611 | 5.48342 |
| ARL2BP | Q9Y2Y0 | 5.73551 | 5.93603 | 5.88389 | 5.58289 | 5.98949 | 6.28748 | 6.06645 | 7.19247 | 6.49795 | 6.90792 | 6.83504 | 6.84092 | 5.10555 | 6.6645 | 6.01106 | 6.46284 |
| CNIH4 | Q9P003 | 5.54869 | 6.84034 | 5.53202 | 7.78976 | 6.73067 | 7.4092 | 5.66184 | 6.02876 | 4.08871 | 5.11451 | 7.30605 | 6.45239 | 6.46682 | 6.87761 | 6.43381 | 5.71881 |
| SELENOI | Q9C0D9 | 5.23662 | 7.19113 | 6.78615 | 6.12862 | 5.51771 | 5.59417 | 6.59314 | 6.3753 | 5.53355 | 6.47042 | 6.3264 | 6.44391 | 5.77289 | 6.20906 | 7.11379 | 6.70714 |
| PHOX2B | Q99453 | 6.53128 | 6.05472 | 6.14393 | 6.73625 | 5.11997 | 5.4965 | 5.88815 | 6.05163 | 6.08438 | 7.33818 | 5.72816 | 6.80901 | 6.52315 | 6.31881 | 6.54767 | 7.62822 |
| MUC5B | Q9HC84 | 6.01393 | 6.48157 | 6.60351 | 6.3595 | 6.51051 | 6.35339 | 6.1314 | 6.47084 | 7.04447 | 6.38684 | 6.44423 | 6.18852 | 6.36429 | 5.34564 | 5.8518 | 6.44957 |
| PSMD4 | Q5VWC4 | 6.29669 | 6.93626 | 5.17548 | 5.64042 | 4.52879 | 6.22436 | 6.26207 | 7.10534 | 6.8017 | 6.51729 | 6.32564 | 7.28888 | 6.5552 | 5.87601 | 5.10893 | 7.35694 |
| MAGED4 | Q96JG8-4 | 6.30333 | 6.29414 | 5.80379 | 5.8089 | 5.21673 | 7.06997 | 7.1675 | 5.90478 | 6.01741 | 6.82508 | 7.14249 | 6.88151 | 5.95574 | 6.1461 | 5.70293 | 5.7596 |
| PRR36 | Q9H6K5 | 6.92176 | 6.7573 | 6.50412 | 6.44631 | 5.7988 | 5.291 | 6.79807 | 6.25494 | 6.50677 | 5.90836 | 5.88447 | 6.36895 | 6.40015 | 7.2837 | 5.53448 | 5.34082 |
| HSPA8 | E9PN89 | 6.26611 | 5.84964 | 7.31773 | 5.94414 | 5.91769 | 5.84438 | 5.87217 | 6.49061 | 6.53543 | 6.35345 | 5.70238 | 5.5586 | 6.34389 | 6.11013 | 6.63314 | 7.2605 |
| HSPA8 | E9PQK7 | 6.71568 | 6.1037 | 6.25504 | 6.29422 | 4.90533 | 5.75726 | 5.19515 | 5.69971 | 6.24979 | 7.28617 | 5.53026 | 6.74309 | 6.59995 | 6.59756 | 6.33691 | 7.73017 |
| SFXN2 | Q96NB2 | 4.93696 | 4.28956 | 7.31097 | 6.8935 | 5.21382 | 7.8677 | 7.49533 | 5.35961 | 5.29854 | 6.87662 | 6.02899 | 6.09847 | 7.33265 | 5.38982 | 7.0563 | 6.55118 |
| NDFIP2 | Q9NV92 | 5.95737 | 6.85241 | 7.39915 | 7.05472 | 6.33609 | 5.96945 | 5.71163 | 6.08705 | 6.5122 | 7.90875 | 8.1752 | 9.40498 | 4.44887 | 4.4636 | 4.13901 | 3.5795 |
| HDAC10 | Q969S8 | 6.35324 | 4.99405 | 6.41632 | 5.71313 | 5.62971 | 4.99314 | 5.61145 | 6.7786 | 7.09259 | 6.2147 | 9.09383 | 6.7541 | 6.3399 | 4.93609 | 5.05736 | 7.12178 |
| CAMSAP3 | Q9P1Y5-2 | 6.09284 | 6.46091 | 5.46424 | 6.78193 | 6.90364 | 7.39705 | 6.67988 | 6.73758 | 7.12943 | 6.90794 | 4.67783 | 6.03632 | 5.46331 | 5.79229 | 5.8834 | 5.59139 |
| NAP1L1 | P55209-3 | 6.47573 | 6.52293 | 7.06005 | 5.80949 | 5.74645 | 6.03431 | 5.97134 | 7.77363 | 6.96793 | 7.27538 | 6.57193 | 6.59176 | 5.72826 | 4.65515 | 5.44983 | 5.36583 |
| NCL | H7BY16 | 6.4429 | 6.73121 | 5.41114 | 6.45477 | 6.45526 | 6.49695 | 6.37633 | 6.0561 | 6.2974 | 5.63687 | 6.35703 | 5.81111 | 6.42974 | 6.03872 | 6.44253 | 6.56193 |
| ODC1 | P11926 | 8.68024 | 9.07862 | 10.1775 | 8.98882 | 5.68524 | 5.68434 | 6.26482 | 6.64413 | 7.05111 | 6.43778 | 8.08726 | 7.14616 | 2.76537 | 2.17109 | 2.48924 | 2.6483 |
| ANKRD30B | Q9BXX2 | 4.87333 | 7.34143 | 7.29748 | 4.65793 | 4.99987 | 7.22406 | 4.45085 | 7.54246 | 6.27562 | 4.75734 | 5.70219 | 10.8193 | 7.81064 | 5.2452 | 5.37888 | 5.62338 |
| SETX | Q7Z333 | 7.21573 | 7.16237 | 5.85177 | 6.36112 | 5.15246 | 6.06637 | 7.26221 | 4.86415 | 6.67732 | 6.19125 | 7.34669 | 5.75183 | 5.38585 | 5.77485 | 7.68001 | 5.25602 |
| TPM1 | A0A087WTJ7 | 6.44329 | 6.11374 | 6.7196 | 4.41608 | 6.26029 | 5.34004 | 4.97347 | 7.55105 | 3.70455 | 6.53386 | 4.99155 | 6.9343 | 6.316 | 9.07834 | 7.69546 | 6.92836 |
| ZBTB41 | Q5SVQ8 | 5.81554 | 5.65316 | 5.74648 | 7.12653 | 7.08557 | 6.35663 | 6.3056 | 6.12829 | 5.91972 | 6.09414 | 6.16087 | 5.69915 | 6.71905 | 6.65216 | 6.05726 | 6.47983 |
| TRPM6 | Q9BX84 | 7.43078 | 7.68104 | 6.49542 | 6.63432 | 6.12855 | 7.0089 | 6.76981 | 6.62637 | 6.01146 | 5.70314 | 6.1045 | 6.2906 | 5.35895 | 5.12491 | 5.53807 | 5.09318 |
| EIF3B | C9JQN7 | 7.62523 | 7.71441 | 8.42459 | 5.76423 | 6.47523 | 8.28694 | 4.31686 | 6.35802 | 7.59436 | 4.49129 | 4.91551 | 4.78802 | 6.77187 | 6.0479 | 5.37949 | 5.04603 |
| DPYS | Q14117 | 6.62824 | 6.76751 | 5.94116 | 6.26462 | 6.43093 | 6.425 | 6.16482 | 6.54665 | 5.93946 | 6.7275 | 6.40044 | 5.4518 | 5.76205 | 6.83211 | 5.34908 | 6.36862 |
| POLE4 | Q9NR33 | 4.39813 | 3.94173 | 7.51129 | 4.27262 | 4.99229 | 3.43194 | 5.40252 | 7.55928 | 5.74117 | 4.36529 | 5.49648 | 6.98047 | 6.26419 | 12.7464 | 8.00999 | 8.88616 |
| PCDHB5 | Q9Y5E4 | 6.69827 | 5.5503 | 5.27713 | 5.64059 | 7.31994 | 6.79793 | 6.75953 | 5.88963 | 6.0146 | 5.82778 | 5.72695 | 5.02376 | 7.63102 | 6.2132 | 7.31388 | 6.3155 |
| HNRNPM | M0R019 | 5.15592 | 7.41343 | 5.54503 | 4.78957 | 5.83071 | 5.87098 | 3.53805 | 5.93714 | 7.98596 | 8.33437 | 5.18994 | 8.42674 | 7.52574 | 4.94174 | 6.71624 | 6.79842 |
| HNRNPM | M0QZM1 | 7.23021 | 5.97531 | 5.67552 | 4.5632 | 6.30415 | 5.28605 | 7.08781 | 4.2926 | 5.08818 | 5.63354 | 6.67108 | 8.82468 | 7.72184 | 6.29707 | 7.29643 | 6.05234 |
| YWHAZ | E7ESK7 | 6.51615 | 5.23036 | 6.43924 | 6.33447 | 6.85156 | 5.97998 | 5.90113 | 6.97416 | 5.58045 | 7.89017 | 6.11942 | 6.20855 | 6.10335 | 5.18799 | 5.4448 | 7.23824 |
| GOLGA4 | Q13439 | 6.76028 | 6.27289 | 5.66362 | 5.73452 | 6.40567 | 6.93173 | 6.66898 | 5.86839 | 7.7438 | 6.69602 | 6.85295 | 7.32733 | 6.30024 | 4.92232 | 5.14404 | 4.70722 |
| TMEM208 | Q9BTX3 | 6.4872 | 6.10126 | 6.68398 | 7.18187 | 4.60761 | 5.95449 | 6.77315 | 5.49249 | 6.02422 | 6.25907 | 6.36753 | 6.62766 | 6.4439 | 6.30818 | 6.34142 | 6.34597 |
| KCTD17 | Q8N5Z5 | 6.1788 | 5.3074 | 5.86011 | 6.57266 | 6.42469 | 5.92358 | 5.02288 | 7.41319 | 6.62082 | 6.36098 | 6.92335 | 6.44558 | 6.76246 | 5.85875 | 5.96064 | 6.36411 |
| PRKD3 | O94806 | 6.03699 | 6.93675 | 6.67443 | 5.50696 | 5.89215 | 7.07275 | 4.68513 | 5.40705 | 6.21274 | 5.07249 | 7.31831 | 8.4627 | 5.91491 | 5.65511 | 6.56415 | 6.58737 |
| PEX7 | O00628 | 7.26864 | 6.4307 | 6.65432 | 5.58849 | 6.80422 | 5.83708 | 6.47397 | 7.07186 | 6.22416 | 5.8612 | 6.66731 | 6.94743 | 5.37276 | 4.42227 | 6.43228 | 5.9433 |
| JAGN1 | Q8N5M9 | 5.70499 | 6.63502 | 6.91128 | 7.06162 | 5.64545 | 6.23663 | 5.24294 | 5.84047 | 6.355 | 6.32292 | 6.82311 | 5.95683 | 4.9528 | 6.67032 | 7.25871 | 6.38192 |
| MYO1B | C9K0I9 | 6.6567 | 6.79022 | 7.64389 | 5.78531 | 5.70966 | 5.97778 | 5.15048 | 6.20636 | 5.93002 | 5.25856 | 5.78403 | 6.20284 | 6.23492 | 6.23384 | 7.05806 | 7.37732 |
| RBM4 | Q9BWF3-3 | 8.40498 | 6.53489 | 6.90887 | 4.19109 | 6.82482 | 6.17382 | 5.70834 | 6.70116 | 6.10622 | 7.0778 | 7.8004 | 7.67186 | 3.96553 | 4.87162 | 5.06104 | 5.99757 |
| BBS4 | Q96RK4-2 | 6.75206 | 6.68415 | 7.45762 | 6.17501 | 6.40185 | 6.05599 | 6.16123 | 7.00964 | 7.3494 | 6.54508 | 7.29545 | 7.43955 | 4.96985 | 4.87907 | 4.43188 | 4.39218 |
| PRC1 | G3V3N7 | 5.96526 | 6.66388 | 6.09892 | 5.07516 | 6.25662 | 5.18473 | 7.73851 | 6.54777 | 6.34029 | 4.98137 | 6.63692 | 6.2019 | 5.45713 | 7.02189 | 6.88548 | 6.94417 |
| CHD3 | H7C0J3 | 7.54248 | 7.16087 | 5.96626 | 6.05697 | 5.17698 | 7.54591 | 6.91827 | 6.73653 | 4.85514 | 5.08314 | 6.44414 | 5.58953 | 5.71173 | 8.34639 | 5.62698 | 5.23868 |
| MYO5B | Q9ULV0 | 5.56875 | 5.06225 | 7.03581 | 5.28288 | 6.28954 | 5.92299 | 6.17411 | 7.10164 | 5.78163 | 7.32894 | 6.60941 | 7.34462 | 6.32885 | 6.34634 | 5.97555 | 5.84669 |
| KIAA1671 | Q9BY89-2 | 6.74341 | 4.60015 | 6.7404 | 6.48337 | 6.4684 | 5.33137 | 6.00285 | 6.47087 | 5.90757 | 7.01261 | 6.02081 | 6.3408 | 5.54565 | 7.83413 | 4.88604 | 7.61155 |
| ARRDC3 | Q96B67 | 5.86366 | 7.01204 | 5.37974 | 6.3817 | 6.19922 | 5.25143 | 4.90288 | 5.98218 | 5.66848 | 6.45973 | 6.10484 | 6.29714 | 7.40828 | 7.32233 | 7.98844 | 5.7779 |
| SYNE3 | Q6ZMZ3-2 | 5.68147 | 6.96202 | 6.99449 | 5.85674 | 4.71443 | 6.8106 | 6.10382 | 6.37725 | 5.36163 | 6.00211 | 7.23717 | 5.60771 | 6.70305 | 5.32762 | 7.21785 | 7.04204 |
| SHMT1 | J3KRZ5 | 7.4912 | 6.8412 | 7.31734 | 4.41098 | 4.38882 | 5.77091 | 6.22447 | 6.12571 | 6.35731 | 6.80443 | 5.38597 | 6.27495 | 7.7156 | 4.76602 | 5.93144 | 8.19366 |
| SREBF2 | Q12772 | 7.43885 | 5.24617 | 6.56426 | 8.05146 | 6.03766 | 7.83337 | 6.15289 | 5.82868 | 7.50634 | 6.67847 | 5.2883 | 5.94996 | 6.08701 | 4.59545 | 5.46459 | 5.27654 |
| ULK2 | Q8IYT8 | 7.70145 | 5.56345 | 5.74491 | 6.60589 | 5.76506 | 6.33044 | 5.48482 | 6.58595 | 5.74492 | 6.01674 | 6.60477 | 5.37125 | 7.17503 | 6.59243 | 6.16291 | 6.54998 |
| ADGRL2 | A0A6I8PLG0 | 5.84509 | 6.35394 | 5.98417 | 6.60276 | 5.66714 | 6.09504 | 6.20595 | 5.82982 | 5.99137 | 6.53689 | 6.32902 | 6.12934 | 5.7851 | 6.86146 | 6.11324 | 7.51129 |
| HOMEZ | Q8IX15-3 | 7.53068 | 5.9272 | 6.16046 | 5.91252 | 5.43802 | 6.99292 | 5.35332 | 6.09012 | 6.04165 | 5.84026 | 7.15667 | 5.0274 | 7.03306 | 5.72948 | 8.39011 | 5.37615 |
| PPP2R5C | Q13362 | 5.91885 | 4.03313 | 5.22397 | 7.37456 | 6.1863 | 5.97093 | 8.79768 | 6.33072 | 7.78877 | 3.89411 | 4.53876 | 5.13073 | 6.90267 | 6.68739 | 5.88717 | 9.33426 |
| NOLC1 | Q14978-3 | 6.43341 | 7.12818 | 5.49335 | 7.37909 | 6.27243 | 5.34716 | 5.18744 | 4.94519 | 6.41965 | 7.46956 | 5.45798 | 7.04852 | 6.01605 | 6.27763 | 6.85373 | 6.27064 |
| TMEM267 | Q0VDI3 | 5.35979 | 6.88691 | 6.69071 | 6.68101 | 6.5849 | 6.6333 | 5.84867 | 6.83595 | 5.8734 | 4.57287 | 6.39718 | 7.2202 | 5.52738 | 5.90252 | 5.72494 | 7.26028 |
| TUBB2A | Q13885 | 7.44168 | 8.54185 | 6.60621 | 6.3346 | 5.88242 | 7.3364 | 6.52171 | 8.23591 | 5.45398 | 5.08092 | 4.80665 | 6.4078 | 5.5092 | 5.86316 | 4.18113 | 5.7964 |
| MAPK11 | Q15759 | 5.72916 | 5.81574 | 9.96582 | 6.02355 | 6.36069 | 7.64453 | 5.23795 | 5.07416 | 5.95781 | 6.60011 | 6.25203 | 7.46059 | 5.99631 | 4.90776 | 5.49316 | 5.48064 |
| MAPK12 | P53778 | 6.79753 | 6.10262 | 7.1694 | 6.38221 | 5.95652 | 6.03628 | 6.22129 | 8.28767 | 5.80916 | 6.33744 | 5.46862 | 6.85642 | 5.95991 | 5.19719 | 5.3063 | 6.11141 |
| CEP170P1 | Q96L14 | 7.06073 | 5.94681 | 5.62598 | 5.46919 | 4.29846 | 7.43822 | 5.56901 | 5.49417 | 8.82523 | 6.43543 | 5.59239 | 6.27144 | 7.79909 | 6.09537 | 5.53187 | 6.54662 |
| MAP7D1 | Q3KQU3-3 | 5.52224 | 4.98405 | 7.70935 | 5.76837 | 6.45417 | 6.48239 | 6.6311 | 5.74006 | 6.07857 | 7.11825 | 6.69041 | 5.46543 | 5.66021 | 5.23065 | 7.9491 | 6.51906 |
| FUNDC1 | Q8IVP5 | 8.46374 | 6.14018 | 6.71638 | 6.83198 | 6.01637 | 6.85434 | 7.46855 | 5.72882 | 6.29935 | 5.30675 | 6.39406 | 5.86295 | 6.22282 | 5.57614 | 6.26968 | 3.84788 |
| CLCN2 | P51788 | 6.44712 | 6.34411 | 6.88178 | 6.98642 | 5.52237 | 6.30694 | 6.29447 | 6.4571 | 6.00238 | 5.65215 | 5.69892 | 6.60177 | 6.3495 | 6.7935 | 5.59628 | 6.06519 |
| STAU2 | Q9NUL3-7 | 6.90056 | 7.16714 | 6.25444 | 6.62719 | 6.07583 | 6.21302 | 7.53975 | 5.99774 | 6.21311 | 6.66584 | 6.06452 | 6.74394 | 5.17479 | 6.06755 | 5.07745 | 5.21714 |
| SON | P18583-6 | 6.93838 | 5.89194 | 4.74163 | 7.22362 | 5.33295 | 4.99552 | 7.54202 | 6.27613 | 5.20174 | 4.60501 | 7.31506 | 5.22794 | 6.34464 | 6.87702 | 8.17676 | 7.30963 |
| PHF23 | Q9BUL5-3 | 8.03909 | 6.2121 | 6.28892 | 5.93325 | 5.65045 | 5.70097 | 9.83068 | 7.04498 | 6.49856 | 4.98991 | 5.29704 | 5.68416 | 5.23755 | 6.0969 | 6.12972 | 5.36574 |
| SMARCE1 | Q969G3-2 | 6.71127 | 6.07766 | 6.15629 | 5.78004 | 6.66509 | 6.326 | 6.33486 | 5.65193 | 6.69502 | 5.53114 | 6.32623 | 6.40801 | 7.14982 | 5.72368 | 6.614 | 5.84897 |
| VARS1 | A0A0G2JJT9 | 6.20074 | 6.2277 | 6.71533 | 6.69194 | 5.93383 | 4.83406 | 6.43848 | 5.65015 | 7.62187 | 5.69556 | 6.72631 | 6.73859 | 6.31241 | 6.16193 | 5.82185 | 6.22926 |
| FOCAD | S4R400 | 6.7392 | 6.50901 | 6.21295 | 6.1716 | 6.43382 | 6.82446 | 5.83772 | 6.8392 | 6.3465 | 6.61439 | 6.4451 | 5.77734 | 5.91328 | 6.35304 | 5.08519 | 5.8972 |
| TNKS1BP1 | A0A2R8Y5C4 | 7.10542 | 6.00475 | 5.58961 | 5.2686 | 7.76269 | 6.62582 | 6.44968 | 6.27376 | 6.85361 | 7.94656 | 6.62776 | 5.03877 | 5.10126 | 6.96211 | 4.26245 | 6.12714 |
| TSC22D2 | O75157 | 6.49441 | 4.67648 | 6.1712 | 7.47981 | 6.80262 | 6.03201 | 5.15235 | 6.49727 | 7.14067 | 6.94614 | 7.20747 | 7.20099 | 4.75977 | 7.03123 | 4.22199 | 6.18561 |
| PDIA4 | A0A499FI48 | 5.72222 | 6.2421 | 5.7189 | 5.41692 | 6.15349 | 6.7582 | 7.11847 | 5.33507 | 6.08964 | 7.04245 | 5.92705 | 6.02903 | 5.69691 | 7.21337 | 6.4385 | 7.09769 |
| GOLIM4 | F8W785 | 4.77737 | 6.41572 | 6.49355 | 6.08604 | 7.58215 | 5.93787 | 6.34375 | 6.10784 | 6.61555 | 4.77056 | 6.29855 | 6.33733 | 7.39764 | 6.02244 | 6.58487 | 6.22877 |
| ILVBL | E9PL44 | 7.36814 | 7.34169 | 6.19993 | 6.35627 | 5.99259 | 5.75065 | 7.90849 | 4.35915 | 6.864 | 4.87107 | 4.51831 | 5.88941 | 6.54604 | 5.96194 | 6.57196 | 7.50036 |
| HAUS7 | Q99871-2 | 6.75816 | 5.3333 | 6.09651 | 5.77595 | 6.19133 | 6.06921 | 6.28539 | 7.95842 | 5.20538 | 5.42247 | 4.82494 | 7.15035 | 5.93883 | 7.63123 | 6.14292 | 7.3156 |
| CHD1 | A0A087WVF4 | 5.41143 | 7.77072 | 4.05499 | 6.80064 | 4.28979 | 7.61309 | 5.7824 | 5.21992 | 4.26112 | 7.30978 | 5.94696 | 6.27468 | 6.95697 | 8.64869 | 5.12326 | 8.53557 |
| PDS5B | A9IYQ1 | 6.13415 | 6.3226 | 6.36789 | 6.65828 | 4.82335 | 5.66534 | 5.84581 | 5.69401 | 6.25621 | 5.42178 | 5.61493 | 5.76504 | 6.74097 | 6.89913 | 7.3852 | 8.40532 |
| TRPC1 | P48995 | 7.26418 | 6.78036 | 5.37318 | 6.70764 | 5.22815 | 5.99034 | 5.77959 | 5.94138 | 6.79733 | 6.78564 | 6.61718 | 5.8549 | 7.22967 | 5.05074 | 6.9712 | 5.62853 |
| SF3B2 | E9PJ04 | 6.54958 | 6.33994 | 6.95336 | 5.5986 | 6.53918 | 6.1503 | 7.60009 | 5.33362 | 7.15224 | 6.77478 | 5.80268 | 6.06046 | 5.95367 | 6.2545 | 5.55266 | 5.38435 |
| ENO3 | P13929 | 5.73176 | 5.12952 | 7.30416 | 5.92468 | 7.08218 | 7.38789 | 6.9817 | 8.06523 | 6.19067 | 6.61012 | 6.17112 | 6.14677 | 5.92136 | 5.00539 | 5.42129 | 4.92615 |
| ENO1 | P06733-2 | 5.34089 | 5.60532 | 7.25262 | 5.5328 | 6.88145 | 7.78372 | 5.1894 | 7.14192 | 6.96758 | 6.77758 | 9.21724 | 5.50548 | 7.25424 | 5.75184 | 4.0603 | 3.73762 |
| ATRX | P46100-4 | 5.7225 | 6.12428 | 5.21376 | 7.59758 | 6.02473 | 6.8729 | 7.09601 | 5.95364 | 6.15118 | 5.63313 | 6.54 | 5.20478 | 6.19477 | 7.24926 | 6.16528 | 6.25619 |
| ATRX | A0A096LNL9 | 6.07955 | 6.06085 | 5.49721 | 6.70411 | 6.9949 | 6.81612 | 5.5341 | 6.24774 | 6.67348 | 5.4045 | 5.98416 | 6.3302 | 7.50758 | 6.58988 | 6.47807 | 5.09754 |
| ATRX | A0A096LNL6 | 6.18943 | 6.36056 | 5.46311 | 6.876 | 6.725 | 5.63722 | 7.94556 | 6.32565 | 5.542 | 5.01529 | 5.63437 | 7.23908 | 6.73044 | 6.2987 | 6.96644 | 5.05115 |
| PABPC1 | A0A7I2YQ88 | 6.9917 | 6.65795 | 7.12912 | 6.05792 | 6.17068 | 6.29952 | 6.28403 | 6.66454 | 7.11611 | 6.32073 | 6.85291 | 6.85435 | 5.19216 | 4.9443 | 5.25991 | 5.20407 |
| ATP10D | Q9P241 | 6.06437 | 6.06995 | 6.74883 | 6.53854 | 8.75547 | 7.55483 | 7.00288 | 6.53661 | 5.39614 | 5.94502 | 5.14266 | 5.82131 | 5.47708 | 6.30192 | 5.49242 | 5.15196 |

| | | | | | | | | | | | | | | | | | |
|---|---|---|---|---|---|---|---|---|---|---|---|---|---|---|---|---|---|
| HCFC1 | P51610 | 6.12618 | 6.0587 | 5.34199 | 5.59381 | 5.84364 | 5.51023 | 6.04472 | 6.49804 | 5.81936 | 5.77445 | 6.34514 | 6.57955 | 7.26936 | 6.70458 | 6.69646 | 7.79378 |
| USF1 | P22415 | 6.19612 | 6.77991 | 5.43993 | 5.80467 | 6.0277 | 7.64243 | 6.05484 | 7.25272 | 8.64898 | 5.01749 | 7.7231 | 4.92346 | 6.6353 | 4.69023 | 5.44352 | 5.71959 |
| DST | A0A0U1RQJ2 | 6.47387 | 6.25896 | 5.68376 | 6.46274 | 6.09606 | 6.68157 | 5.34276 | 5.28755 | 5.77874 | 5.76895 | 5.85015 | 6.04019 | 6.15859 | 7.45369 | 7.73798 | 6.92445 |
| MACF1 | E9PLY5 | 6.4081 | 6.37365 | 7.27359 | 6.24762 | 5.19698 | 6.71257 | 6.01596 | 6.1931 | 7.11562 | 5.94032 | 6.42101 | 5.32242 | 6.14299 | 6.17828 | 6.71072 | 5.74705 |
| USF2 | Q15853-3 | 7.20521 | 3.98108 | 7.02096 | 7.97522 | 7.37431 | 5.44373 | 4.89897 | 6.65124 | 6.79082 | 5.92363 | 7.43113 | 5.27957 | 6.525 | 6.21769 | 5.12071 | 6.16073 |
| CENPO | Q9BU64 | 5.82199 | 5.88941 | 6.54837 | 6.07172 | 5.77197 | 6.92128 | 6.35231 | 7.00779 | 6.65139 | 6.00456 | 6.12744 | 7.23881 | 5.26534 | 6.04705 | 6.17356 | 6.10703 |
| TPP2 | A0A494C0U1 | 6.72378 | 6.16896 | 6.30128 | 6.57068 | 6.59975 | 5.85292 | 6.0817 | 6.07016 | 6.19659 | 5.78357 | 5.7445 | 5.877 | 6.72508 | 5.89699 | 6.62102 | 6.78603 |
| NUP85 | J3QL91 | 6.97271 | 7.56538 | 5.63508 | 6.07572 | 5.21785 | 5.13026 | 6.85114 | 6.16729 | 5.5491 | 5.98079 | 5.2124 | 5.23783 | 7.32896 | 6.52476 | 7.23061 | 7.32012 |
| FBXO38 | Q6PIJ6-2 | 6.50215 | 6.41622 | 6.26515 | 6.47799 | 6.21965 | 6.19463 | 6.0821 | 6.0862 | 6.28709 | 6.23484 | 6.1093 | 6.23535 | 6.32969 | 6.33409 | 6.18014 | 6.04541 |
| HSP90AA2P | Q14568 | 5.42042 | 7.39842 | 6.50549 | 5.66802 | 5.67952 | 7.85921 | 5.6381 | 6.11568 | 5.39413 | 5.63295 | 6.51079 | 7.0934 | 5.82428 | 5.98248 | 6.36718 | 6.90992 |
| BCL3 | P20749 | 7.15905 | 5.96851 | 7.65928 | 5.88828 | 6.61021 | 5.79399 | 6.04121 | 6.52493 | 5.1419 | 7.28953 | 7.08587 | 5.57416 | 6.7989 | 6.22646 | 4.81956 | 5.41816 |
| WNK4 | Q96J92 | 6.42722 | 7.25998 | 5.93326 | 5.2648 | 5.01188 | 6.96808 | 6.78042 | 6.81296 | 7.70809 | 5.13101 | 7.03282 | 6.56385 | 5.97506 | 5.15625 | 5.95726 | 6.01704 |
| SLC48A1 | Q6P1K1 | 6.60768 | 6.80424 | 7.14135 | 7.32141 | 5.37116 | 6.4993 | 5.96672 | 6.80698 | 4.39687 | 5.7088 | 4.10641 | 5.93202 | 6.99806 | 8.09071 | 5.37215 | 6.87612 |
| OCIAD1 | D6RDK6 | 7.15101 | 5.1188 | 6.49857 | 6.5515 | 5.98504 | 4.59765 | 7.43156 | 5.20013 | 10.6193 | 3.91075 | 6.1861 | 5.48574 | 6.14376 | 4.52765 | 6.62602 | 7.96642 |
| CD46 | P15529 | 6.65258 | 6.69613 | 5.49394 | 6.56435 | 6.70329 | 5.78988 | 6.2427 | 6.307 | 6.65728 | 6.01186 | 6.65186 | 5.83698 | 6.31433 | 5.80329 | 6.3876 | 5.88694 |
| ZBTB46 | Q86UZ6 | 7.67555 | 6.38537 | 5.67001 | 6.59825 | 6.50773 | 6.3282 | 5.86585 | 4.63808 | 7.0609 | 7.09985 | 5.22827 | 5.78614 | 6.09633 | 6.92345 | 6.97393 | 5.16208 |
| ZNF664 | Q8N3J9 | 5.45563 | 6.16184 | 4.64912 | 5.32505 | 6.06278 | 5.95964 | 6.25148 | 6.71474 | 5.24843 | 5.50252 | 6.40523 | 5.99989 | 6.84694 | 6.9379 | 8.78441 | 7.6944 |
| RNF213 | Q63HN8 | 6.4827 | 6.98761 | 6.52362 | 6.10013 | 6.31455 | 6.44177 | 6.84152 | 6.39162 | 5.97609 | 6.43296 | 5.69007 | 6.15099 | 5.9524 | 6.10979 | 5.77764 | 5.82655 |
| FLII | Q13045-2 | 6.06578 | 5.99529 | 6.94383 | 6.00189 | 6.22814 | 6.40592 | 6.4381 | 6.50142 | 6.57558 | 6.89298 | 7.11635 | 6.55118 | 5.65477 | 5.7558 | 5.2522 | 5.62077 |
| ALG13 | Q9NP73-2 | 6.47884 | 4.35075 | 5.5105 | 4.9508 | 6.25933 | 4.95612 | 5.40871 | 6.14837 | 6.13407 | 5.47043 | 5.6824 | 6.60938 | 6.7549 | 8.4543 | 8.59382 | 8.23728 |
| SLC20A2 | Q08357 | 6.49041 | 6.27635 | 4.43967 | 6.44333 | 6.73598 | 5.39201 | 7.0314 | 6.13197 | 6.973 | 5.46083 | 6.79895 | 6.82133 | 5.84458 | 6.22154 | 6.40659 | 6.53204 |
| SRGAP2C | P0DJJ0 | 7.55885 | 7.8373 | 6.81948 | 6.81234 | 6.42396 | 7.94359 | 5.80865 | 7.31313 | 4.43316 | 4.69323 | 6.47644 | 6.83093 | 7.64725 | 4.75089 | 4.21157 | 4.43923 |
| OTUD4 | Q01804-5 | 6.15822 | 5.33979 | 7.54283 | 5.38038 | 6.51921 | 6.77233 | 5.85117 | 4.88233 | 5.99327 | 6.81179 | 7.27605 | 7.26486 | 5.6647 | 7.43276 | 5.36395 | 5.74637 |
| MKI67 | P46013-2 | 7.20466 | 6.53551 | 5.89958 | 6.92614 | 6.26416 | 6.05079 | 5.97323 | 5.43244 | 5.65328 | 6.34174 | 5.87894 | 6.33036 | 6.06223 | 5.76705 | 6.15416 | 7.52573 |
| MAP2 | P11137-4 | 5.86016 | 5.7502 | 5.60006 | 6.48401 | 6.17596 | 6.66128 | 5.89618 | 6.90396 | 5.20841 | 6.76072 | 6.69286 | 6.97866 | 7.44979 | 6.47349 | 5.7954 | 5.30886 |
| BAP18 | Q8IXM2-2 | 5.5296 | 4.15581 | 4.94154 | 4.9988 | 4.68233 | 4.99165 | 5.09236 | 6.97957 | 4.29703 | 5.41953 | 5.33517 | 5.60444 | 6.81282 | 10.6152 | 10.3656 | 10.1785 |
| PARL | C9JNP8 | 5.97859 | 6.01393 | 5.72446 | 6.18296 | 5.82319 | 5.93113 | 6.13337 | 6.32967 | 5.91268 | 6.46 | 6.28106 | 6.36538 | 6.73839 | 6.50689 | 6.78796 | 6.83036 |
| PSMC3IP | Q9P2W1 | 5.66275 | 5.96459 | 6.12219 | 5.40375 | 6.57959 | 6.46507 | 5.4266 | 5.86703 | 5.96493 | 6.37759 | 6.76074 | 6.47601 | 7.00525 | 6.62425 | 6.99288 | 6.30677 |
| CHMP2A | M0R1T5 | 7.74826 | 5.64071 | 7.72582 | 6.8078 | 5.94471 | 6.57865 | 6.88003 | 6.78564 | 5.09985 | 6.90342 | 4.85158 | 8.08485 | 2.81175 | 6.46343 | 5.41939 | 6.2541 |
| HYOU1 | A0A494C039 | 7.66145 | 6.61359 | 7.29655 | 6.43563 | 5.87157 | 6.78537 | 6.75145 | 7.15221 | 6.22543 | 6.10032 | 5.56582 | 5.54395 | 6.84095 | 4.83935 | 6.21068 | 4.10568 |
| HYOU1 | J3QL06 | 7.67446 | 6.44312 | 6.4199 | 8.14672 | 7.7154 | 7.11275 | 7.03296 | 3.87704 | 6.65658 | 7.10203 | 3.69435 | 4.07991 | 6.63606 | 4.42213 | 5.79656 | 7.19004 |
| MACIR | Q96GV9 | 6.02447 | 4.75276 | 6.84521 | 5.61709 | 6.74313 | 7.61348 | 6.30184 | 6.59152 | 6.37571 | 7.36942 | 6.12398 | 6.17406 | 6.86986 | 5.59988 | 5.51455 | 5.48304 |
| CCDC8 | Q9H0W5 | 6.26148 | 6.30228 | 7.90583 | 6.74723 | 6.73242 | 7.78377 | 6.49914 | 6.04981 | 6.20315 | 5.85667 | 7.46755 | 7.26257 | 5.1073 | 4.76548 | 5.41617 | 3.63915 |
| SCNM1 | Q9BWG6 | 5.9952 | 6.41764 | 6.05104 | 6.64339 | 6.01924 | 7.20134 | 6.07932 | 6.72163 | 5.60913 | 7.15874 | 6.59648 | 7.87591 | 5.71508 | 5.96062 | 4.21687 | 5.73839 |
| CBX6 | O95503 | 6.80609 | 6.77889 | 5.61529 | 5.76706 | 7.14839 | 6.7885 | 6.4704 | 7.16875 | 7.46589 | 6.91035 | 5.79704 | 6.36534 | 5.7317 | 6.90381 | 4.46606 | 4.81645 |
| THAP4 | Q8WY91 | 6.36093 | 6.95578 | 9.12612 | 9.05827 | 6.02568 | 5.25767 | 7.38883 | 7.46461 | 6.14526 | 6.13028 | 7.36048 | 6.13704 | 4.46474 | 3.54969 | 4.36817 | 4.20646 |
| HMGA2 | H0YFY4 | 5.3726 | 6.16321 | 5.08564 | 5.93265 | 5.08246 | 5.23699 | 4.72217 | 5.97654 | 6.15785 | 6.29068 | 6.31298 | 5.37334 | 8.0798 | 7.93237 | 6.75144 | 9.52928 |
| NDFIP1 | Q9BT67 | 6.96466 | 6.93858 | 6.84964 | 6.02896 | 6.63414 | 7.13177 | 7.15344 | 6.17015 | 6.59394 | 7.5835 | 8.31862 | 6.6513 | 3.96769 | 4.05086 | 4.52244 | 4.39531 |
| UBTF | E9PLT2 | 6.3549 | 5.89709 | 5.68791 | 7.2439 | 5.09308 | 5.10987 | 6.15 | 5.42418 | 6.28243 | 6.66087 | 5.18556 | 6.56289 | 6.41363 | 6.57844 | 7.10638 | 8.24887 |
| FAM204A | Q9H8W3 | 5.99568 | 6.32713 | 6.94942 | 6.43057 | 5.95064 | 7.03532 | 6.33922 | 5.83205 | 6.6981 | 6.75441 | 6.45943 | 5.91478 | 5.56426 | 5.86592 | 5.78511 | 6.09798 |
| SFT2D3 | Q587I9 | 6.34006 | 6.53493 | 5.55163 | 6.8481 | 6.24034 | 6.55295 | 5.97337 | 6.80801 | 6.91276 | 6.00548 | 5.32605 | 6.0767 | 6.41544 | 6.37141 | 6.03305 | 6.00974 |
| PCDHAC2 | Q9Y5I4 | 6.94708 | 5.71892 | 6.88454 | 6.33357 | 5.38234 | 6.4958 | 6.04594 | 7.86212 | 8.6513 | 5.64229 | 6.73449 | 6.47276 | 5.00971 | 5.9274 | 4.90099 | 4.99075 |
| BCLAF1 | Q9NYF8-3 | 5.96027 | 7.20376 | 5.88218 | 6.89199 | 6.26067 | 5.8455 | 6.10598 | 6.10649 | 5.63334 | 5.44117 | 6.1507 | 6.79808 | 6.11289 | 6.23411 | 6.5089 | 6.86396 |
| NDUFV3 | P56181 | 6.40448 | 5.85507 | 6.6214 | 4.79422 | 7.29148 | 5.91136 | 6.01536 | 5.07255 | 5.00597 | 5.37227 | 5.90601 | 7.33608 | 8.6552 | 5.64645 | 6.63836 | 7.47374 |
| PPP2R2D | Q66LE6 | 6.40466 | 5.24108 | 6.68457 | 5.50193 | 6.50833 | 6.4382 | 6.92348 | 6.39661 | 5.86231 | 7.23166 | 6.7456 | 6.78304 | 5.5968 | 5.9277 | 6.44769 | 5.30636 |
| TMPPE | Q6ZT21 | 6.39073 | 5.94422 | 5.9787 | 7.02527 | 5.93903 | 5.58008 | 6.53146 | 6.2165 | 5.93644 | 5.84521 | 5.80394 | 5.63398 | 7.06117 | 6.5259 | 6.92786 | 6.6595 |
| FOSB | P53539 | 6.83579 | 5.33645 | 5.50898 | 6.15341 | 6.85125 | 5.75931 | 6.16398 | 6.1007 | 6.99089 | 5.69599 | 6.09431 | 5.37854 | 8.17094 | 5.66672 | 6.60258 | 6.69018 |
| SPTBN1 | Q01082-3 | 4.9555 | 7.0532 | 5.27015 | 7.39193 | 5.93152 | 6.13238 | 5.49739 | 5.51311 | 7.38405 | 6.54594 | 6.13999 | 4.94237 | 7.1115 | 7.03027 | 7.48618 | 6.51454 |
| SPTB | P11277-2 | 9.37833 | 5.94972 | 7.42347 | 7.37955 | 5.52939 | 5.46713 | 5.28941 | 5.00889 | 5.33121 | 5.50508 | 4.68682 | 6.15204 | 6.50328 | 7.22781 | 5.5629 | 7.60496 |
| PIP4P2 | Q8N4L2 | 5.65599 | 6.95997 | 5.80956 | 5.60417 | 6.26165 | 6.39563 | 6.94734 | 6.19559 | 6.11831 | 7.5226 | 6.24202 | 6.02484 | 5.9227 | 5.9901 | 6.57904 | 5.77048 |
| MAFK | O60675 | 7.54779 | 5.65341 | 6.11963 | 6.80538 | 5.52925 | 6.59212 | 7.70383 | 5.82479 | 5.96538 | 6.30457 | 4.5016 | 5.96962 | 5.70942 | 5.7277 | 7.0811 | 6.96443 |
| UBE2D1 | P51668 | 6.44388 | 6.59575 | 6.86417 | 5.16216 | 6.02914 | 6.31258 | 6.02348 | 6.78336 | 5.64869 | 6.28138 | 6.15098 | 6.44808 | 5.39185 | 7.20651 | 6.67453 | 6.88347 |
| GPATCH3 | Q96I76 | 5.75093 | 5.93701 | 6.29491 | 6.5462 | 6.92273 | 5.87082 | 6.22603 | 6.23358 | 6.26862 | 6.55791 | 7.28332 | 7.50449 | 5.89887 | 6.13433 | 5.34615 | 5.2241 |
| SLC10A4 | Q96EP9 | 6.33345 | 6.77369 | 7.31902 | 6.37414 | 6.25477 | 6.64173 | 4.53251 | 6.38595 | 6.52544 | 7.52797 | 5.65545 | 6.22684 | 5.99767 | 7.3438 | 5.49769 | 4.60985 |
| ZNF865 | P0CJ78 | 6.36673 | 6.43486 | 5.94411 | 5.90655 | 7.43198 | 6.46809 | 7.06414 | 5.55152 | 3.83624 | 4.67098 | 6.95367 | 6.35892 | 7.08532 | 6.90092 | 7.03494 | 5.99103 |
| MARK4 | Q96L34 | 7.41622 | 7.07546 | 7.04939 | 7.17145 | 5.71813 | 5.99839 | 6.35127 | 7.52023 | 7.37735 | 6.54189 | 6.92894 | 6.58608 | 4.79135 | 4.81121 | 4.1599 | 4.50275 |
| MYL6 | G3V1V0 | 6.44553 | 4.66355 | 6.97161 | 5.68883 | 6.96287 | 4.19352 | 4.70988 | 5.71737 | 4.87811 | 4.91657 | 3.93671 | 6.82647 | 6.99833 | 8.88978 | 9.35906 | 8.84181 |
| RTL5 | Q5HYW3 | 6.87821 | 6.391 | 4.77187 | 6.90147 | 6.91183 | 6.77171 | 6.33897 | 5.44204 | 6.74147 | 6.17642 | 6.92001 | 6.17534 | 6.13461 | 6.42969 | 5.78713 | 5.22824 |
| RACK1 | D6RHH4 | 6.618 | 6.50617 | 6.06486 | 8.04582 | 5.84393 | 4.86076 | 7.33013 | 7.17374 | 5.59699 | 4.04158 | 7.58481 | 6.33202 | 6.97748 | 4.56639 | 6.75806 | 5.69924 |
| EPN1 | Q9Y6I3-3 | 6.08422 | 5.72533 | 5.836 | 5.99123 | 8.03586 | 6.01548 | 5.54114 | 5.17991 | 6.07955 | 6.4468 | 5.78698 | 6.98925 | 7.04764 | 5.84613 | 7.21952 | 6.17497 |
| RTN4 | Q9NQC3-5 | 6.15939 | 5.90939 | 6.03444 | 5.86719 | 5.4845 | 5.74995 | 6.76538 | 6.62602 | 6.86783 | 5.98113 | 5.0148 | 6.94233 | 6.84259 | 6.71341 | 6.89774 | 6.1439 |
| RTN4 | Q9NQC3-2 | 6.61188 | 7.26479 | 6.13648 | 6.59491 | 6.41557 | 4.7233 | 7.03687 | 6.5775 | 6.3803 | 6.92926 | 5.11935 | 5.55226 | 5.86588 | 6.40655 | 6.56585 | 5.81925 |
| CIT | O14578-3 | 6.24033 | 8.81516 | 7.25625 | 5.76256 | 7.22578 | 4.4718 | 5.24849 | 6.21483 | 6.27036 | 7.65792 | 7.44367 | 5.90627 | 4.82909 | 6.4171 | 5.13679 | 5.10359 |
| KRT18 | F8VZY9 | 6.43478 | 6.75191 | 5.63921 | 5.91397 | 5.77104 | 5.5071 | 6.68305 | 6.66898 | 6.14057 | 5.62361 | 5.40585 | 6.34223 | 6.76175 | 6.33953 | 7.31291 | 6.70352 |
| PML | P29590-5 | 7.38218 | 5.11212 | 5.54429 | 5.62791 | 6.79805 | 6.59486 | 5.34096 | 8.30896 | 5.31747 | 7.45784 | 4.69565 | 5.84973 | 6.33753 | 7.13141 | 5.45672 | 7.04433 |
| PCBP2 | F8VZX2 | 7.33848 | 5.59764 | 7.10888 | 5.17581 | 5.48854 | 7.78032 | 5.41191 | 5.59586 | 5.73586 | 6.62845 | 7.20215 | 6.2605 | 6.44843 | 5.45469 | 6.15763 | 6.61483 |
| PCBP2 | F8W0G4 | 6.40075 | 5.22424 | 7.38863 | 5.86192 | 5.79684 | 5.9019 | 5.54304 | 4.75424 | 7.22148 | 6.57189 | 6.03339 | 6.908 | 5.90617 | 6.14999 | 6.85012 | 7.48737 |
| NFIC | P08651-5 | 4.39762 | 5.57172 | 7.52179 | 6.19092 | 6.35646 | 6.93624 | 6.23992 | 5.74519 | 7.72398 | 5.36752 | 5.79916 | 5.85263 | 6.51895 | 6.5666 | 5.54382 | 7.66747 |
| NFIX | Q14938 | 7.05531 | 7.1936 | 7.56785 | 7.35714 | 6.37982 | 7.88058 | 6.14429 | 4.6775 | 5.09219 | 6.35915 | 7.06509 | 5.67099 | 6.09179 | 5.63879 | 5.36849 | 4.45743 |
| ZNF627 | Q7L945 | 4.26781 | 8.07135 | 7.19302 | 7.17246 | 6.34637 | 6.52631 | 6.19869 | 6.54606 | 4.52981 | 4.57524 | 5.60666 | 7.31064 | 7.34136 | 7.06016 | 6.38618 | 4.8679 |
| ZNF791 | Q3KP31 | 7.19054 | 8.04142 | 6.0823 | 6.45561 | 6.79126 | 5.78844 | 7.00605 | 6.28305 | 5.65817 | 6.31461 | 6.54086 | 6.68167 | 5.07079 | 5.05626 | 5.63501 | 5.40397 |
| LMO7 | E9PMS6 | 6.08928 | 5.9985 | 5.8825 | 6.96514 | 5.65876 | 5.9643 | 6.2149 | 5.33023 | 6.042 | 6.28658 | 5.88803 | 6.46881 | 6.86459 | 6.42036 | 6.95203 | 6.97399 |
| TMEM158 | Q8WZ71 | 6.23278 | 5.95278 | 5.66623 | 7.29276 | 7.22996 | 7.03899 | 7.42863 | 6.42356 | 5.81227 | 6.52737 | 7.54543 | 6.98079 | 5.30106 | 5.1795 | 4.53153 | 4.85638 |
| ZNF568 | Q3ZCX4 | 5.70484 | 5.70496 | 5.76296 | 6.01428 | 5.46394 | 5.2895 | 6.7164 | 6.29904 | 4.76138 | 5.562 | 5.84364 | 7.02859 | 7.42685 | 8.32442 | 6.98358 | 7.11363 |
| ZNF625 | Q96I27-2 | 6.16995 | 6.54312 | 6.26403 | 7.05069 | 5.96168 | 6.02808 | 6.65635 | 6.04552 | 5.66567 | 5.80788 | 5.94968 | 6.65943 | 5.99625 | 6.57963 | 6.40965 | 6.21239 |
| ZFP1 | Q6P2D0 | 5.55913 | 6.57261 | 6.62195 | 6.26325 | 5.83723 | 6.56992 | 6.66614 | 4.48841 | 5.86191 | 6.95452 | 5.92644 | 6.02274 | 7.40513 | 5.93509 | 6.04858 | 7.26694 |
| ZNF624 | Q9P2J8 | 6.63492 | 6.09624 | 5.80639 | 6.15736 | 6.66314 | 6.19794 | 5.8929 | 5.62924 | 5.50233 | 6.28632 | 7.08142 | 6.32318 | 6.33281 | 7.15074 | 5.48244 | 6.76264 |
| ZNF23 | P17027 | 4.77215 | 7.25401 | 5.86923 | 8.79012 | 7.56574 | 4.05513 | 6.98853 | 5.86596 | 5.61776 | 4.17505 | 4.66136 | 4.63314 | 6.94781 | 7.96279 | 8.50139 | 6.33985 |
| ZNF286B | P0CG31 | 6.9061 | 6.39464 | 5.86923 | 6.82795 | 3.84439 | 4.8115 | 6.28815 | 5.95183 | 6.62245 | 7.02658 | 7.06543 | 6.73726 | 5.35075 | 6.5172 | 7.05455 | 6.732 |
| ZNF90 | Q03938 | 6.76752 | 7.70984 | 6.79617 | 8.40644 | 5.27017 | 6.37945 | 6.86134 | 5.86414 | 6.29232 | 6.86471 | 6.46387 | 5.97132 | 4.35271 | 5.0496 | 5.63576 | 5.31464 |
| ZNF391 | Q9UJN7 | 6.25992 | 6.3602 | 6.42934 | 6.19221 | 6.5997 | 5.72415 | 5.35399 | 5.01848 | 4.98845 | 5.44577 | 5.86855 | 7.41249 | 6.53087 | 6.39697 | 6.98019 | 8.43871 |
| ZNF3 | P17036 | 6.12387 | 7.69048 | 6.5359 | 7.97341 | 5.73981 | 6.47753 | 7.51047 | 6.23793 | 4.31157 | 5.83125 | 7.09676 | 3.96372 | 5.94178 | 7.01828 | 5.5754 | 5.97183 |
| ZFP37 | Q9Y6Q3-2 | 5.42873 | 6.92658 | 6.28081 | 6.10349 | 7.14233 | 5.61576 | 5.15436 | 5.31202 | 5.74341 | 5.22748 | 5.47753 | 4.87502 | 7.44253 | 7.86712 | 8.00779 | 7.39503 |
| ZNF121 | P58317 | 5.79191 | 6.08593 | 5.62154 | 7.11297 | 6.51462 | 5.98602 | 5.91909 | 6.22524 | 5.7085 | 5.62503 | 6.15064 | 5.86323 | 6.46076 | 6.80424 | 7.54024 | 6.59004 |
| ZNF350 | Q9GZX5 | 5.65018 | 6.8514 | 8.89077 | 5.71263 | 9.44965 | 4.4558 | 4.78934 | 6.98066 | 8.25936 | 6.91974 | 5.50901 | 6.17191 | 4.67179 | 5.01805 | 4.85495 | 5.72477 |
| ZSCAN30 | Q86W11 | 7.63463 | 6.0721 | 7.66322 | 4.56119 | 8.31127 | 5.78825 | 4.20979 | 6.79975 | 5.79972 | 6.7806 | 4.93008 | 4.46152 | 5.80467 | 9.31815 | 7.01877 | 4.84629 |
| ZKSCAN7 | Q9P0L1 | 6.3943 | 7.27828 | 5.51078 | 5.56379 | 5.45448 | 6.31645 | 7.05837 | 7.35072 | 6.5475 | 4.74959 | 7.13741 | 6.80944 | 5.72291 | 6.60196 | 6.08236 | 5.42165 |
| ZNF14 | P17017 | 6.22559 | 5.99904 | 6.34659 | 6.81328 | 6.619 | 6.62183 | 6.43748 | 5.86576 | 5.43025 | 6.10218 | 6.46014 | 6.63182 | 6.00712 | 6.21666 | 6.40612 | 5.77422 |
| DNM1 | Q05193 | 6.915 | 6.11005 | 6.80725 | 6.4538 | 6.26252 | 6.78992 | 6.83757 | 7.92554 | 5.82059 | 6.56608 | 7.05308 | 5.27646 | 5.54754 | 5.94885 | 5.61846 | 4.06729 |
| MTHFSD | Q2M296-3 | 6.14396 | 6.98825 | 6.07496 | 6.05766 | 5.61086 | 6.34282 | 5.44533 | 6.27374 | 6.35981 | 7.00725 | 6.50151 | 6.833 | 5.17877 | 6.28869 | 6.3146 | 6.57878 |
| PLD3 | M0R1F7 | 6.06542 | 6.06451 | 6.60561 | 7.48661 | 4.94077 | 5.97994 | 5.76811 | 7.15683 | 8.33413 | 5.87486 | 6.81254 | 5.71757 | 5.65992 | 5.77733 | 6.01348 | 5.74238 |
| RAB39B | Q96DA2 | 4.86953 | 6.8476 | 6.27237 | 6.56215 | 6.6503 | 4.76407 | 6.65609 | 5.30929 | 4.9619 | 7.37201 | 8.01236 | 5.89095 | 6.68529 | 5.46954 | 7.10046 | 6.57609 |
| TRIM7 | Q9C029 | 6.96034 | 6.56032 | 4.97398 | 8.35647 | 6.13494 | 6.16718 | 7.06612 | 6.57765 | 6.4411 | 6.4425 | 6.84798 | 6.70702 | 5.38802 | 4.29486 | 5.80505 | 5.27647 |
| ZNF670 | Q9BS34 | 6.29436 | 6.56151 | 6.28617 | 7.24053 | 5.66269 | 6.24081 | 6.29066 | 5.83234 | 5.83383 | 6.33517 | 5.86663 | 6.06764 | 6.39349 | 5.9398 | 7.40192 | 5.75246 |
| KIF7 | Q2M1P5 | 5.52823 | 6.67798 | 6.945 | 6.37998 | 5.12742 | 5.88495 | 7.96523 | 4.384 | 5.83511 | 7.78418 | 7.11364 | 7.03812 | 4.64606 | 6.43235 | 6.56655 | 5.69119 |
| GOLGB1 | Q14789-4 | 7.28644 | 4.75973 | 7.27847 | 6.92996 | 6.4507 | 5.82764 | 5.19728 | 5.83537 | 5.25437 | 6.79596 | 5.87628 | 6.00216 | 6.13086 | 6.21955 | 6.95758 | 7.19764 |
| ZNF799 | Q96GE5-2 | 5.90403 | 7.23045 | 7.897 | 7.34415 | 6.28241 | 5.73813 | 5.78428 | 6.83617 | 3.80995 | 4.92817 | 6.27205 | 7.59005 | 6.06953 | 5.51598 | 6.1101 | 6.68755 |
| ZNF607 | Q96SK3 | 5.57616 | 6.76103 | 6.00075 | 7.79888 | 6.58084 | 6.80558 | 6.17003 | 6.7889 | 5.67647 | 6.26131 | 5.94464 | 6.95279 | 5.04373 | 5.99506 | 6.3902 | 5.25364 |
| ZNF317 | Q96PQ6 | 6.08177 | 6.5879 | 6.47967 | 6.59289 | 6.06745 | 6.23651 | 6.29685 | 6.19399 | 6.13952 | 5.9083 | 6.19508 | 6.10191 | 6.47489 | 6.35243 | 6.58006 | 5.71078 |
| ANKLE1 | Q8NAG6 | 6.15653 | 5.88536 | 7.07233 | 6.93597 | 6.20097 | 6.80693 | 5.37513 | 5.79107 | 7.13698 | 6.32382 | 5.65493 | 6.41598 | 7.02411 | 6.10933 | 5.82635 | 5.28422 |
| SEC61A2 | Q9H9S3 | 5.79875 | 7.49189 | 5.5496 | 5.84433 | 6.60992 | 6.31146 | 6.94088 | 6.2937 | 5.76799 | 6.05747 | 5.59118 | 5.492 | 7.14082 | 7.23326 | 6.26338 | 5.61338 |
| ZNF136 | P52737 | 6.82788 | 6.58906 | 5.18176 | 6.55582 | 6.01328 | 5.95087 | 6.38577 | 7.74148 | 6.68457 | 5.31407 | 6.48099 | 6.30004 | 5.11877 | 6.59974 | 7.10883 | 5.14707 |
| THUMPD2 | Q9BTF0 | 4.95593 | 5.74676 | 6.49283 | 5.34348 | 5.71886 | 6.55029 | 5.57707 | 5.83508 | 5.82646 | 6.26336 | 6.15197 | 7.03956 | 6.44213 | 6.8536 | 7.81666 | 7.38594 |
| ZNF367 | Q7RTV3 | 6.6234 | 6.36874 | 6.58419 | 6.67212 | 5.95655 | 6.34293 | 6.3603 | 6.20997 | 6.61293 | 6.66987 | 6.094 | 5.95789 | 6.36007 | 5.38506 | 6.08381 | 5.71817 |
| ZNF669 | Q96BR6 | 6.23951 | 7.01163 | 6.03178 | 6.45029 | 7.58517 | 6.68024 | 5.59236 | 5.45289 | 6.26201 | 7.64358 | 6.50014 | 7.37255 | 4.68766 | 4.67853 | 6.29527 | 5.5164 |

| | | | | | | | | | | | | | | | | | |
|---|---|---|---|---|---|---|---|---|---|---|---|---|---|---|---|---|---|
| ARHGAP12 | Q8IWW6-3 | 6.08999 | 6.14877 | 6.17616 | 5.7854 | 6.65926 | 6.4032 | 6.22208 | 6.78744 | 6.32343 | 6.35115 | 6.47861 | 6.09345 | 6.35976 | 6.04676 | 6.31054 | 5.76402 |
| ARHGEF2 | V9GYF0 | 6.91888 | 5.36557 | 7.87872 | 5.39956 | 6.25491 | 5.64716 | 6.17733 | 7.74722 | 6.01829 | 5.87117 | 6.4914 | 7.90396 | 4.63114 | 6.55389 | 5.21632 | 5.92448 |
| ATP5F1B | F8W079 | 9.08271 | 6.27301 | 7.02826 | 6.3258 | 5.14208 | 4.87894 | 5.46553 | 4.15762 | 4.11613 | 4.93786 | 5.24944 | 5.13625 | 7.5461 | 10.5795 | 6.47458 | 7.60621 |
| MXRA7 | P84157-3 | 5.5859 | 4.98351 | 3.7268 | 5.88349 | 3.94792 | 3.19957 | 4.95309 | 5.33259 | 3.88081 | 4.54176 | 4.94511 | 6.24216 | 6.05332 | 12.3814 | 10.2158 | 14.1267 |
| NFKB2 | Q00653-4 | 5.94871 | 6.107 | 6.58356 | 5.77828 | 6.83401 | 6.842 | 7.08281 | 5.44637 | 6.55688 | 6.10543 | 6.04666 | 5.58374 | 6.82915 | 6.28435 | 5.87453 | 6.09652 |
| TMEM160 | Q9NX00 | 6.023 | 4.33495 | 5.12179 | 7.56003 | 6.46385 | 5.24893 | 6.60886 | 6.63604 | 8.13788 | 4.74673 | 5.96303 | 7.52009 | 7.59225 | 6.57938 | 6.44764 | 5.01554 |
| CAMK2A | Q9UQM7-2 | 5.49095 | 6.99596 | 6.8977 | 7.58304 | 7.40174 | 4.83622 | 6.77158 | 7.07459 | 5.15071 | 6.87678 | 5.58541 | 4.95651 | 5.17417 | 6.6965 | 5.83069 | 6.67745 |
| CAMK2D | Q13557-12 | 5.7141 | 6.41341 | 7.13752 | 6.14987 | 5.91667 | 6.65299 | 6.20091 | 6.50954 | 6.05681 | 6.25192 | 6.8556 | 6.3934 | 5.98185 | 6.03193 | 6.19525 | 5.53824 |
| SDHB | A0A087WWT1 | 8.03635 | 6.24217 | 5.49251 | 6.60872 | 5.45208 | 6.07807 | 5.27839 | 6.59245 | 6.72748 | 5.98137 | 5.81705 | 5.79358 | 6.38854 | 4.86799 | 7.55247 | 7.09076 |
| C19orf25 | Q9UFG5 | 6.72654 | 5.2207 | 4.78059 | 5.43118 | 6.30828 | 2.95755 | 5.30931 | 4.9172 | 3.70587 | 6.4662 | 4.37347 | 5.67768 | 4.70981 | 13.2209 | 8.75826 | 11.4365 |
| FAM174A | Q8TBP5 | 6.72468 | 6.29721 | 5.95839 | 5.27648 | 5.97646 | 5.6602 | 6.03001 | 5.32162 | 6.16818 | 6.11305 | 6.05236 | 6.51001 | 6.81422 | 5.79768 | 7.73576 | 7.5637 |
| TOP2B | H7BZ82 | 6.61254 | 6.45124 | 5.52883 | 6.80997 | 5.11877 | 4.35749 | 6.82054 | 5.6529 | 6.86881 | 6.42269 | 5.44377 | 6.13466 | 6.34343 | 6.3701 | 7.89861 | 7.16565 |
| POTEKP | Q9BYX7 | 2.7958 | 1.76318 | 21.6659 | 4.64235 | 13.1508 | 11.0517 | 18.4916 | 3.64628 | 2.48746 | 1.66901 | 6.05085 | 2.24354 | 1.90833 | 2.19441 | 3.25519 | 2.98356 |
| ANGPTL4 | Q9BY76 | 6.86041 | 5.57114 | 5.39875 | 6.48477 | 4.15576 | 6.10771 | 9.12497 | 4.46075 | 8.17131 | 6.14142 | 5.21056 | 5.98347 | 6.89878 | 5.55062 | 7.42311 | 6.45649 |
| METTL9 | Q9H1A3 | 6.435 | 6.35414 | 5.72352 | 6.85933 | 6.10475 | 6.24605 | 6.32495 | 6.15536 | 5.99707 | 5.88533 | 5.57576 | 6.52123 | 6.34047 | 6.43542 | 6.30284 | 6.73878 |
| RBBP5 | Q15291-2 | 5.60154 | 7.60489 | 6.09042 | 6.85788 | 5.61932 | 6.26061 | 6.31517 | 5.69078 | 6.73045 | 6.74645 | 5.12739 | 5.83555 | 6.59415 | 6.56289 | 5.7655 | 6.597 |
| CACNA1C | Q13936-2 | 4.91858 | 6.0642 | 6.75418 | 6.26348 | 5.56659 | 7.15349 | 6.09411 | 6.47132 | 6.36427 | 5.63025 | 6.53784 | 5.02516 | 8.18066 | 7.04202 | 6.39949 | 5.53439 |
| EIF4EBP1 | Q13541 | 4.58847 | 5.76961 | 4.27003 | 4.08849 | 3.92253 | 3.37673 | 4.08843 | 4.21546 | 3.45786 | 4.99481 | 3.89085 | 3.87616 | 4.90732 | 16.5347 | 9.94082 | 18.0777 |
| PYGO2 | Q9BRQ0 | 5.15555 | 5.97155 | 7.73387 | 5.87973 | 5.88647 | 5.89935 | 4.50041 | 6.72997 | 7.88969 | 6.48383 | 5.8883 | 5.73495 | 6.08556 | 7.01361 | 7.27545 | 5.87171 |
| MBD6 | Q96DN6 | 6.33367 | 6.27106 | 5.0754 | 5.65779 | 5.3243 | 6.9799 | 8.1237 | 6.04769 | 7.04935 | 7.5064 | 6.58745 | 7.47853 | 4.93769 | 6.7581 | 5.39023 | 4.47875 |
| SMIM20 | Q8N5G0-2 | 5.98433 | 5.95203 | 5.46063 | 5.3322 | 5.99488 | 5.00334 | 5.46375 | 6.64715 | 5.91102 | 5.87072 | 5.84524 | 5.51213 | 6.85321 | 8.38222 | 7.57518 | 8.21198 |
| PRMT6 | Q96LA8 | 5.02187 | 5.39312 | 5.70962 | 5.3062 | 5.7983 | 6.39897 | 6.27638 | 4.77406 | 6.05897 | 6.181 | 6.53238 | 5.8119 | 7.50801 | 7.79198 | 7.92441 | 7.51283 |
| H2AC21 | Q8IUE6 | 6.14901 | 5.72402 | 5.44662 | 8.73283 | 6.4125 | 5.08977 | 7.39355 | 4.67568 | 5.1826 | 7.03646 | 5.92807 | 5.6039 | 6.4698 | 7.48676 | 5.8957 | 6.77273 |
| H2AZ2 | Q71UI9-2 | 7.67371 | 5.97344 | 5.41943 | 6.65462 | 6.20626 | 5.6033 | 8.95222 | 6.56641 | 7.53839 | 5.31509 | 4.73557 | 7.33295 | 4.95119 | 5.5395 | 6.34401 | 5.19391 |
| DUSP12 | V9GYV5 | 6.75668 | 6.08397 | 4.38595 | 5.39215 | 5.16628 | 7.32487 | 5.4009 | 6.87226 | 5.32208 | 6.15847 | 5.07505 | 5.82048 | 7.48262 | 7.49752 | 6.89821 | 8.36252 |
| F3 | P13726-2 | 5.67373 | 6.88621 | 5.80578 | 6.80637 | 6.03977 | 4.86007 | 6.90793 | 6.43419 | 6.12044 | 6.5196 | 5.80502 | 5.54628 | 6.74462 | 5.8718 | 6.56811 | 7.41008 |
| GAP43 | P17677-2 | 7.38986 | 6.82238 | 5.66376 | 6.06608 | 6.2734 | 4.96655 | 5.52861 | 6.92371 | 6.77472 | 6.33205 | 6.30092 | 6.68203 | 6.14445 | 6.07209 | 6.04511 | 6.01429 |
| BAG6 | F6WML8 | 5.08949 | 6.6624 | 6.90514 | 5.446 | 6.32065 | 6.53388 | 7.05019 | 6.33419 | 6.09871 | 6.35595 | 6.49023 | 8.50389 | 6.31724 | 4.64695 | 5.94228 | 5.30282 |
| NETO2 | Q8NC67 | 7.10535 | 7.39814 | 6.48774 | 6.98052 | 6.668 | 6.32843 | 6.78859 | 6.12325 | 7.00143 | 7.49957 | 6.13912 | 6.51932 | 4.32997 | 5.08889 | 4.90398 | 4.63771 |
| CLCN3 | P51790 | 6.91312 | 5.73114 | 5.6462 | 5.41293 | 6.30313 | 7.3065 | 6.0979 | 6.05481 | 6.63133 | 6.79277 | 6.14443 | 6.23802 | 6.2845 | 5.74669 | 5.84585 | 6.8507 |
| HOMER2 | Q9NSB8 | 6.78264 | 5.81616 | 5.22628 | 6.04756 | 6.93573 | 6.21268 | 6.401 | 6.8836 | 6.80985 | 5.95282 | 6.51927 | 6.30702 | 6.73013 | 6.43542 | 5.22332 | 5.71651 |
| FRYL | O94915-2 | 4.99103 | 5.62046 | 4.52277 | 6.34428 | 6.5194 | 6.66082 | 6.71379 | 6.22815 | 6.77712 | 7.42663 | 6.40572 | 6.61466 | 6.67233 | 7.55603 | 5.42876 | 5.51804 |
| RBM41 | Q96IZ5 | 5.7698 | 6.54887 | 6.1492 | 5.53147 | 6.88796 | 6.00256 | 6.73385 | 5.65111 | 7.42143 | 5.58346 | 6.12846 | 6.75414 | 5.18658 | 6.3224 | 6.56379 | 6.76493 |
| NOTCH3 | Q9UM47 | 6.41937 | 5.80969 | 6.65172 | 6.41028 | 6.48845 | 6.15212 | 6.16195 | 6.52135 | 6.14755 | 6.6345 | 7.21707 | 6.25441 | 5.88719 | 5.42506 | 6.13461 | 5.68467 |
| NBPF20 | P0DPF2 | 4.96038 | 6.68412 | 6.09689 | 5.98191 | 6.50358 | 7.25394 | 5.76567 | 5.00209 | 4.98527 | 6.08126 | 7.18201 | 6.00679 | 6.74087 | 7.11241 | 6.61881 | 7.024 |
| TNIK | Q9UKE5-4 | 6.74276 | 5.367 | 6.43426 | 6.45469 | 5.62579 | 5.99313 | 6.58739 | 6.54055 | 6.81095 | 5.72907 | 6.64052 | 6.66461 | 5.07932 | 6.4647 | 6.92 | 6.84526 |
| NAA15 | A0A0B4J1W3 | 6.61429 | 5.79234 | 8.01441 | 5.99839 | 5.71336 | 5.66832 | 5.81854 | 6.11458 | 7.06654 | 6.23274 | 6.42094 | 4.99627 | 6.70467 | 5.44408 | 5.7148 | 7.68573 |
| GABARAP | O95166 | 6.24632 | 5.37978 | 6.83421 | 6.83099 | 6.24146 | 6.04666 | 7.14261 | 6.38521 | 7.58926 | 6.41676 | 7.69858 | 6.20937 | 5.6192 | 5.3972 | 4.85178 | 5.11061 |
| DHX30 | Q7L2E3 | 5.70192 | 6.54926 | 7.92139 | 5.5341 | 5.51237 | 5.56117 | 6.54672 | 6.28307 | 6.01297 | 7.42844 | 6.39157 | 7.04444 | 6.21935 | 5.28421 | 6.6722 | 5.33681 |
| GTF2A1 | P52655 | 5.91894 | 5.96014 | 5.18802 | 4.10638 | 6.13994 | 7.48422 | 5.75737 | 5.54584 | 5.82524 | 6.16172 | 6.35471 | 6.7027 | 7.53064 | 6.81513 | 7.3701 | 7.1389 |
| DNAJB1 | M0R128 | 5.73086 | 5.71671 | 6.69389 | 5.77542 | 5.42323 | 6.38636 | 6.40723 | 5.82297 | 6.14799 | 6.59218 | 5.87913 | 6.51609 | 6.06263 | 5.93802 | 7.26355 | 7.64373 |
| RPS7 | A0A2R8Y623 | 3.90403 | 4.0586 | 4.19705 | 14.2481 | 5.48691 | 10.989 | 4.33642 | 4.5539 | 4.95001 | 10.6488 | 6.27952 | 9.27713 | 4.80001 | 4.81114 | 4.16438 | 3.295 |
| MCM3 | Q7Z6P5 | 4.0536 | 5.96054 | 6.0981 | 6.92767 | 4.64253 | 7.33903 | 5.40895 | 5.37784 | 6.00272 | 6.72142 | 7.46879 | 5.95943 | 6.60372 | 7.56871 | 6.88712 | 6.97983 |
| GK5 | Q6ZS86 | 6.50693 | 8.10728 | 6.84109 | 6.62946 | 6.16503 | 6.51054 | 7.07643 | 5.79744 | 5.71574 | 5.58004 | 7.27219 | 6.1705 | 5.48077 | 5.75369 | 5.07128 | 5.32161 |
| ANXA6 | E5RK69 | 6.85387 | 6.57096 | 6.54944 | 5.7416 | 6.46103 | 6.22003 | 6.53867 | 6.55789 | 6.27174 | 6.87083 | 6.56459 | 6.6317 | 5.88327 | 5.9041 | 5.29108 | 5.08921 |
| UBAP2L | F8W726 | 6.79017 | 6.70443 | 6.05746 | 6.21207 | 6.60244 | 6.53227 | 5.56613 | 6.58264 | 6.58094 | 6.32452 | 6.75145 | 7.20141 | 5.36418 | 5.82933 | 5.28828 | 5.61228 |
| NIN | Q8N4C6-7 | 6.09486 | 6.78288 | 7.80494 | 5.22432 | 7.15665 | 6.5306 | 6.97698 | 6.44994 | 5.53638 | 7.07931 | 5.16116 | 7.97842 | 5.12965 | 6.64791 | 4.63615 | 4.80986 |
| EZH1 | Q92800-2 | 6.40667 | 7.31399 | 6.78443 | 9.3871 | 3.9579 | 6.2911 | 6.85056 | 5.33481 | 5.09887 | 7.42534 | 5.14893 | 7.10761 | 4.73114 | 4.77008 | 5.91537 | 7.47608 |
| CNOT4 | O95628-6 | 7.2673 | 6.61434 | 6.71676 | 5.90979 | 5.64131 | 7.41547 | 5.58514 | 6.24112 | 6.88243 | 7.82988 | 5.92409 | 6.7039 | 5.83621 | 5.87949 | 4.97847 | 5.47432 |
| SHLD3 | Q6ZNX1 | 5.92819 | 5.49861 | 5.23253 | 5.26751 | 5.45589 | 4.5955 | 5.60229 | 6.64214 | 7.74699 | 6.55658 | 5.96938 | 6.81671 | 5.84141 | 6.92944 | 8.78972 | 7.12708 |
| RPS6KB2 | Q9UBS0 | 5.54784 | 7.23454 | 7.2967 | 4.35584 | 6.09406 | 7.63567 | 4.98007 | 5.3925 | 7.46609 | 7.69666 | 5.69877 | 6.77113 | 4.8866 | 7.14912 | 6.50176 | 5.29265 |
| TRAPPC14 | Q8WVR3 | 6.32152 | 6.72365 | 6.4967 | 5.45711 | 6.13904 | 7.66528 | 6.68293 | 6.20138 | 6.11553 | 6.09569 | 5.39794 | 7.01355 | 6.01944 | 5.38097 | 6.0761 | 6.21318 |
| MYH16 | Q9H6N6 | 5.85845 | 6.74705 | 5.65451 | 6.71566 | 6.74818 | 8.17235 | 7.23667 | 6.61522 | 8.48777 | 7.06973 | 6.57279 | 4.59873 | 5.31477 | 4.78502 | 4.70052 | 4.72259 |
| OXLD1 | Q5BKU9 | 5.65113 | 4.97909 | 4.71533 | 5.51339 | 4.59724 | 4.48986 | 4.95672 | 4.28482 | 4.28195 | 5.05003 | 4.49865 | 4.41768 | 5.19394 | 13.7797 | 8.54595 | 15.0445 |
| UROS | P10746 | 5.84934 | 5.81007 | 6.25536 | 4.32524 | 7.02574 | 6.48632 | 7.05963 | 7.55667 | 5.23124 | 7.66116 | 6.53144 | 6.51567 | 5.17308 | 6.28338 | 6.02508 | 6.21056 |
| APBB2 | Q92870-2 | 7.67641 | 8.51311 | 7.67174 | 6.52682 | 5.00583 | 6.91278 | 7.51484 | 5.18243 | 5.34739 | 4.48653 | 5.13303 | 4.66409 | 6.0684 | 6.11759 | 6.47307 | 6.70595 |
| PPIP5K2 | H0Y9S9 | 5.72461 | 6.15573 | 6.59195 | 5.39082 | 6.26061 | 7.03025 | 7.05279 | 7.40468 | 5.72701 | 6.45552 | 6.66073 | 6.12363 | 4.32148 | 6.785 | 6.38486 | 5.93035 |
| ESRRB | O95718-1 | 6.39331 | 6.45314 | 7.09444 | 6.96798 | 6.02834 | 6.10202 | 6.60576 | 6.3754 | 6.6833 | 6.50748 | 7.04366 | 6.09309 | 5.33958 | 5.49608 | 5.70001 | 5.11641 |
| DHRS4L1 | P0CG22 | 5.44587 | 6.47531 | 5.85989 | 6.74278 | 6.89239 | 5.4123 | 5.98505 | 5.6668 | 6.42129 | 6.48939 | 5.91301 | 5.975 | 6.92871 | 6.13116 | 7.34515 | 6.31591 |
| CTU1 | Q7Z7A3 | 6.09571 | 7.12956 | 6.76219 | 6.21125 | 7.54189 | 8.94292 | 5.12759 | 7.00398 | 4.35074 | 5.5243 | 8.07397 | 6.09531 | 4.38038 | 6.24301 | 5.47681 | 5.04039 |
| RNF138 | Q8WVD3 | 5.48267 | 6.05029 | 5.46131 | 4.34226 | 6.54179 | 6.88208 | 5.09772 | 5.51808 | 6.12886 | 6.62772 | 8.41301 | 6.68526 | 7.50232 | 6.66112 | 5.66365 | 6.84186 |
| S100A4 | P26447 | 6.21577 | 4.7631 | 7.04079 | 5.19254 | 5.5324 | 6.26074 | 6.15621 | 6.21468 | 5.69754 | 6.76051 | 5.78918 | 7.48683 | 5.62315 | 8.06302 | 6.6416 | 6.56194 |
| MED11 | Q9P086 | 5.15454 | 4.614 | 4.44408 | 4.01755 | 5.89829 | 5.30979 | 5.38771 | 6.38904 | 5.14358 | 5.9574 | 6.00042 | 5.98052 | 4.42358 | 11.5073 | 8.16461 | 11.6076 |
| RILP | Q96NA2 | 5.21163 | 4.53251 | 3.64719 | 4.49695 | 5.13793 | 5.15009 | 5.94608 | 5.65466 | 6.64982 | 5.82529 | 7.03221 | 8.99013 | 8.78156 | 7.36218 | 8.27653 | 7.30523 |
| PRG4 | Q92954 | 6.53435 | 6.06249 | 5.19373 | 6.64684 | 6.2972 | 5.42798 | 6.59497 | 5.97256 | 6.25661 | 6.20676 | 5.07187 | 6.38495 | 7.44825 | 6.81639 | 6.54962 | 6.53545 |
| ZNF653 | Q96CK0 | 6.0271 | 7.00349 | 6.18728 | 6.45931 | 6.39128 | 7.02211 | 6.02736 | 6.16187 | 5.92263 | 5.69126 | 6.60716 | 5.57846 | 6.22582 | 6.62505 | 6.24424 | 5.82557 |
| PUM2 | Q8TB72-3 | 6.84999 | 6.3118 | 6.91766 | 6.51016 | 6.29525 | 6.32087 | 7.2463 | 5.96504 | 5.44433 | 6.36213 | 5.80923 | 5.68151 | 6.22928 | 5.76475 | 6.11925 | 6.17246 |
| CIAO2A | Q9H5X1 | 5.22225 | 4.95041 | 6.29551 | 5.74909 | 7.37196 | 6.87972 | 4.27699 | 6.22164 | 7.75676 | 6.35399 | 6.26293 | 6.48293 | 8.22692 | 5.18006 | 5.91053 | 6.85831 |
| TEX9 | Q8N6V9 | 6.83786 | 6.89292 | 6.42454 | 6.95996 | 6.88439 | 6.41303 | 6.66489 | 6.73914 | 6.75203 | 7.28016 | 7.27997 | 6.71612 | 4.24926 | 4.84036 | 4.261 | 4.80436 |
| PPFIBP1 | Q86W92-2 | 6.89759 | 5.46742 | 6.35703 | 6.21078 | 5.6693 | 7.18001 | 5.31795 | 6.62766 | 6.49738 | 5.95052 | 6.00599 | 6.26466 | 6.79255 | 6.23628 | 6.80744 | 5.71743 |
| CRK | P46108-2 | 5.56817 | 5.1975 | 6.13067 | 5.2582 | 6.97217 | 5.31642 | 4.84674 | 8.05446 | 6.36622 | 6.84417 | 7.5255 | 6.82523 | 5.55855 | 6.39907 | 7.02902 | 6.1079 |
| LENG1 | Q96BZ8 | 6.5899 | 7.80247 | 7.06365 | 5.32264 | 5.92336 | 5.15322 | 5.74024 | 6.78897 | 5.45851 | 7.41422 | 7.37728 | 5.89222 | 5.31031 | 4.43781 | 6.97162 | 6.75358 |
| PICALM | H0YEH1 | 6.02525 | 4.74279 | 6.30196 | 6.32861 | 7.49017 | 7.06816 | 6.50741 | 7.03561 | 5.84792 | 6.37249 | 7.73772 | 6.72951 | 6.31904 | 5.30601 | 4.93502 | 5.25234 |
| LYPLA1 | A0A087X1K9 | 4.93505 | 4.28681 | 4.81339 | 5.16218 | 5.36455 | 5.2333 | 5.41986 | 6.09948 | 7.45952 | 9.34375 | 4.98595 | 6.51584 | 6.98477 | 7.33277 | 7.72104 | 8.34174 |
| C1orf43 | Q9BWL3 | 5.61692 | 6.61468 | 6.92772 | 7.092 | 5.44877 | 6.62431 | 7.6713 | 6.4398 | 7.30136 | 7.19803 | 5.97883 | 5.81825 | 5.06993 | 5.19986 | 5.22687 | 5.77138 |
| B4GALNT1 | Q00973-3 | 6.40112 | 7.05374 | 7.98406 | 7.32523 | 6.54524 | 6.85447 | 7.37775 | 6.57026 | 6.37361 | 6.96911 | 6.82119 | 6.26121 | 4.10497 | 4.40095 | 4.28742 | 4.66966 |
| MFSD1 | Q9H3U5-6 | 6.11391 | 5.81928 | 4.29129 | 6.94335 | 6.35167 | 5.91122 | 6.3522 | 7.84206 | 6.77802 | 6.56156 | 6.10316 | 7.45217 | 6.67693 | 6.3681 | 5.49368 | 5.84141 |
| RTL8C | A6ZKI3 | 5.89207 | 5.29804 | 6.58897 | 4.76707 | 6.06929 | 6.80211 | 6.25882 | 6.38709 | 5.30257 | 6.24909 | 6.59428 | 6.06503 | 6.52107 | 8.29867 | 5.99319 | 6.91262 |
| CNK3/IPCEF1 | G9CGD6-2 | 6.25742 | 6.42313 | 6.34826 | 6.40643 | 7.80865 | 6.33227 | 6.90455 | 6.04235 | 6.66672 | 5.95904 | 6.18218 | 6.53783 | 5.55901 | 5.55765 | 5.61143 | 5.40308 |
| CEP85L | Q5SZL2-4 | 6.84323 | 6.48553 | 6.6047 | 5.8442 | 5.34326 | 7.45343 | 6.43503 | 6.21652 | 6.22644 | 7.79607 | 7.95066 | 5.70788 | 4.55471 | 5.60067 | 5.69621 | 5.24148 |
| MPV17L2 | Q567V2 | 6.0653 | 6.1183 | 6.62413 | 6.99554 | 6.39597 | 5.96468 | 7.24609 | 5.35278 | 6.11227 | 5.99292 | 5.96657 | 4.9464 | 6.9915 | 6.06169 | 6.36019 | 6.80566 |
| ICOSLG | O75144-2 | 5.87411 | 6.40169 | 6.05226 | 6.40265 | 5.92212 | 6.48163 | 6.03904 | 5.94797 | 5.81118 | 6.49946 | 6.51263 | 6.69324 | 6.04572 | 6.63813 | 5.93294 | 6.74524 |
| PROSER3 | A0A2R8Y8D9 | 6.55976 | 5.00097 | 7.3839 | 5.42109 | 6.17311 | 8.37836 | 5.26954 | 8.1235 | 5.70146 | 5.33927 | 8.01883 | 7.67054 | 5.69599 | 4.84844 | 4.93979 | 5.47545 |
| PROSER3 | Q2NL68-2 | 5.97135 | 5.77164 | 4.95723 | 6.1072 | 7.86818 | 7.16996 | 6.55268 | 5.86827 | 7.76483 | 6.7215 | 7.01106 | 7.27899 | 4.14347 | 6.21987 | 4.90109 | 5.69269 |
| PGGT1B | P53609 | 6.05585 | 5.61125 | 6.04781 | 6.29015 | 5.74618 | 7.29164 | 6.33491 | 6.17809 | 6.62874 | 6.71866 | 6.8668 | 7.75968 | 5.45564 | 5.8566 | 5.90404 | 5.25396 |
| PPIA | P62937-2 | 5.37366 | 4.82512 | 6.13782 | 5.45964 | 5.46881 | 6.3125 | 4.95629 | 7.45234 | 7.23584 | 7.45737 | 6.7543 | 7.46914 | 5.62971 | 6.44163 | 6.58482 | 6.44102 |
| IMPA2 | O14732 | 6.0268 | 6.39852 | 6.33172 | 6.59791 | 5.97114 | 6.55492 | 6.56154 | 6.65461 | 6.65821 | 6.60432 | 6.29016 | 7.03621 | 5.70306 | 5.62034 | 5.26014 | 5.73038 |
| TTLL4 | Q14679 | 6.75876 | 7.04354 | 7.94807 | 6.74064 | 6.95785 | 6.37114 | 6.10922 | 6.53348 | 6.59278 | 6.97359 | 6.77407 | 7.81856 | 4.43072 | 4.37372 | 4.15609 | 4.38828 |
| KDELR2 | P33947 | 5.58651 | 6.07796 | 5.69293 | 6.01093 | 5.39823 | 5.89674 | 6.01461 | 6.24791 | 6.40011 | 6.67024 | 6.08996 | 6.31308 | 6.90926 | 6.40465 | 7.06121 | 7.22567 |
| DDX46 | A0A0C4DG89 | 5.43227 | 6.00594 | 5.14954 | 5.17083 | 6.54255 | 5.53543 | 5.46046 | 6.28177 | 7.0433 | 7.67196 | 7.95062 | 5.67392 | 5.97997 | 6.25318 | 6.54904 | 7.29923 |
| BRD7 | Q9NPI1 | 6.61768 | 6.39428 | 6.40983 | 6.31198 | 5.56618 | 6.15799 | 6.26881 | 5.73959 | 5.22216 | 5.85347 | 5.7284 | 6.67764 | 6.25057 | 7.06532 | 6.87318 | 6.86391 |
| GPBP1L1 | Q9HC44 | 4.86344 | 4.69917 | 5.79561 | 5.82479 | 6.3125 | 6.30125 | 6.33591 | 6.63475 | 6.15866 | 7.35975 | 6.89901 | 6.16758 | 7.23396 | 6.4836 | 7.20901 | 5.72103 |
| PEG10 | Q86TG7-2 | 6.23201 | 6.21711 | 5.96888 | 5.58881 | 6.41294 | 6.67304 | 6.81462 | 6.30736 | 6.58058 | 5.69005 | 6.44217 | 5.76326 | 5.7289 | 6.79728 | 6.84255 | 5.94045 |
| JPT2 | Q9H910-2 | 5.77294 | 6.36626 | 5.86012 | 5.07182 | 8.55054 | 6.54442 | 5.50472 | 6.56777 | 7.31367 | 6.61271 | 6.89546 | 6.63855 | 6.02435 | 5.59049 | 6.67014 | 4.01604 |
| RAD50 | A0A494C0Y7 | 6.76044 | 6.74396 | 5.58335 | 6.96487 | 6.5605 | 6.48676 | 6.34756 | 5.74646 | 5.29897 | 6.00406 | 5.4837 | 5.29074 | 6.93015 | 5.37404 | 6.93923 | 7.4852 |
| CRADD | P78560 | 5.31701 | 5.13976 | 7.61017 | 5.23973 | 6.10935 | 6.38391 | 6.38174 | 7.74005 | 5.39649 | 7.27696 | 6.81198 | 7.41175 | 4.96995 | 6.05251 | 5.6995 | 6.45913 |
| RADX | Q6NSI4 | 6.71641 | 5.6815 | 5.84242 | 5.44939 | 6.46306 | 6.5992 | 6.37742 | 7.30532 | 6.97326 | 5.61621 | 6.03061 | 6.06013 | 6.32405 | 6.08357 | 6.13309 | 6.34437 |
| RBM38 | Q9H0Z9 | 7.56552 | 5.78806 | 5.01311 | 5.79361 | 6.39236 | 5.08996 | 4.53703 | 7.12577 | 5.85864 | 6.0168 | 6.45668 | 7.88932 | 6.81423 | 6.76914 | 6.64032 | 6.24945 |
| PLCH2 | O75038-4 | 5.97809 | 3.79406 | 5.93062 | 5.23708 | 5.97926 | 3.89519 | 6.68308 | 5.9478 | 5.76516 | 5.27007 | 4.93295 | 6.56382 | 6.84225 | 7.89736 | 8.48935 | 10.7938 |
| UNG | P13051-2 | 6.62514 | 5.90895 | 5.32206 | 6.72494 | 6.166 | 5.98351 | 6.9226 | 6.18989 | 6.62584 | 5.30673 | 6.82744 | 6.5858 | 6.54877 | 6.16991 | 6.11489 | 5.97753 |
| LRIG1 | Q96JA1 | 6.70287 | 6.17202 | 5.82972 | 6.53403 | 6.43274 | 5.63778 | 6.62108 | 6.13319 | 6.16095 | 5.89614 | 6.10156 | 6.00942 | 6.92756 | 5.92073 | 6.69399 | 6.22621 |
| SKI | P12755 | 8.14105 | 6.90494 | 7.81643 | 9.92879 | 6.00591 | 7.09304 | 6.79816 | 6.13431 | 7.47788 | 6.42361 | 8.03065 | 7.09499 | 1.67168 | 3.33373 | 3.43707 | 3.70776 |
| KLF16 | Q9BXK1 | 6.34782 | 5.71151 | 6.3478 | 5.97996 | 6.14217 | 5.94564 | 6.08937 | 7.01866 | 6.3526 | 6.82775 | 6.28369 | 6.87827 | 6.99257 | 5.56568 | 5.42389 | 6.09261 |
| RHNO1 | Q9BSD3 | 6.08056 | 6.18323 | 6.36466 | 6.04954 | 5.75122 | 6.03864 | 7.13071 | 6.43695 | 5.86967 | 6.50776 | 6.87454 | 8.72782 | 5.62228 | 5.73546 | 4.51057 | 6.11639 |
| FOXS1 | O43638 | 5.55452 | 5.19574 | 5.58851 | 4.53458 | 5.37476 | 6.78583 | 6.21465 | 6.87649 | 6.25255 | 7.17592 | 5.37529 | 4.59956 | 7.20813 | 10.5276 | 7.49938 | 5.23647 |

| | | | | | | | | | | | | | | | | | |
|---|---|---|---|---|---|---|---|---|---|---|---|---|---|---|---|---|---|
| ATP8A1 | Q9Y2Q0 | 6.52454 | 6.31302 | 6.08739 | 6.44247 | 6.56894 | 5.92014 | 5.82936 | 5.93393 | 5.99583 | 6.61888 | 7.67567 | 7.07979 | 6.31089 | 5.28134 | 5.61482 | 5.80298 |
| POM121C | A8CG34 | 6.75479 | 5.22904 | 6.05844 | 7.04322 | 6.2793 | 6.14891 | 6.56651 | 5.43627 | 6.40224 | 5.56842 | 6.36649 | 6.08677 | 6.8997 | 6.65232 | 5.93879 | 6.56878 |
| CSNK1G3 | Q9Y6M4-4 | 6.21455 | 6.12464 | 6.58644 | 6.9018 | 5.18773 | 6.00159 | 5.30681 | 7.25003 | 6.3701 | 6.41369 | 6.11808 | 6.31678 | 6.1937 | 7.10099 | 5.72046 | 6.19263 |
| HLA-A | A0A140T9X5 | 6.8937 | 6.79447 | 5.59095 | 6.24684 | 6.5976 | 6.68195 | 6.92196 | 6.43191 | 6.60274 | 6.24721 | 6.47801 | 6.22645 | 6.20166 | 5.26092 | 5.53633 | 5.28731 |
| YLPM1 | P49750-3 | 6.02516 | 6.10551 | 6.73865 | 6.97576 | 5.68472 | 5.5376 | 6.08357 | 5.30682 | 5.69992 | 6.04305 | 6.25766 | 6.50653 | 6.88333 | 6.74915 | 6.41861 | 6.98396 |
| DENND1B | Q6P3S1 | 8.2126 | 5.78321 | 6.84069 | 5.42949 | 5.60498 | 7.09244 | 6.47655 | 6.59925 | 5.80153 | 5.81529 | 6.35521 | 6.80031 | 4.92567 | 5.52373 | 6.69115 | 6.04891 |
| ATP5F1D | P30049 | 5.48793 | 4.65516 | 4.7172 | 4.25024 | 7.23529 | 4.31529 | 5.28038 | 7.35268 | 4.27555 | 5.46792 | 6.51608 | 5.81756 | 6.07451 | 9.66579 | 8.37478 | 10.5136 |
| SHFL | Q9NUL5 | 5.73457 | 7.41726 | 7.38206 | 6.52926 | 6.01491 | 7.36832 | 6.22921 | 7.59108 | 5.962 | 6.5617 | 5.15125 | 4.27801 | 5.47251 | 6.68725 | 6.28123 | 5.3394 |
| DONSON | H7C006 | 6.08498 | 6.13084 | 7.08894 | 5.85164 | 6.55672 | 6.62294 | 6.47305 | 3.85863 | 5.91374 | 5.51241 | 6.23893 | 6.80032 | 6.06285 | 9.03647 | 5.83393 | 5.93362 |
| GLRX | P35754 | 5.15905 | 5.3859 | 5.71465 | 5.03901 | 5.47522 | 4.69988 | 5.53956 | 5.3651 | 4.328 | 6.00715 | 4.91325 | 5.8473 | 4.68968 | 12.5804 | 7.09576 | 12.16 |
| KIF2C | Q5JR91 | 6.20365 | 5.72483 | 5.72669 | 5.94557 | 6.00269 | 6.28412 | 6.28332 | 6.36988 | 7.01318 | 5.34993 | 6.03669 | 5.46792 | 7.40229 | 6.34627 | 6.76832 | 7.07465 |
| WDR5B | Q86VZ2 | 6.45744 | 6.74618 | 5.97576 | 6.51707 | 6.486 | 6.00299 | 7.27145 | 6.74106 | 6.59048 | 6.5776 | 6.27522 | 5.97408 | 5.58881 | 5.76054 | 5.58027 | 5.45505 |
| TTC33 | Q6PID6 | 6.1253 | 5.76025 | 6.15605 | 6.38731 | 5.8237 | 6.0087 | 6.33734 | 6.35216 | 5.70472 | 6.18839 | 6.36604 | 5.9152 | 6.92232 | 6.6137 | 6.70453 | 6.63429 |
| SAE1 | M0R054 | 5.59021 | 5.98377 | 3.68819 | 5.24812 | 5.12101 | 5.68139 | 6.40132 | 6.14264 | 5.37579 | 4.74073 | 5.81194 | 6.67467 | 7.74225 | 8.74461 | 9.86939 | 7.18398 |
| KIAA0753 | Q2KHM9 | 6.42184 | 6.22511 | 6.49543 | 7.04557 | 5.53012 | 5.63518 | 6.34667 | 6.76652 | 6.06201 | 5.81446 | 6.41703 | 6.35537 | 5.96954 | 6.27677 | 5.81831 | 6.82006 |
| HNRNPK | P61978-3 | 6.38584 | 6.37169 | 5.55852 | 6.381 | 7.54088 | 6.13567 | 6.42335 | 5.21441 | 6.63912 | 5.98146 | 6.40473 | 6.66818 | 6.8267 | 5.28192 | 6.24395 | 5.94258 |
| PRKCSH | K7ELL7 | 7.02742 | 6.11815 | 5.74316 | 5.45301 | 6.92619 | 5.76676 | 6.15867 | 5.49801 | 6.17549 | 5.9206 | 6.91695 | 6.61651 | 6.38232 | 6.07016 | 6.24254 | 6.98405 |
| PDGFD | Q9GZP0 | 6.15569 | 5.79201 | 6.59736 | 6.76777 | 6.79923 | 6.34992 | 6.33209 | 6.84005 | 6.94696 | 5.88272 | 6.1746 | 6.38802 | 6.60272 | 5.92597 | 5.29948 | 5.14541 |
| ARMC10 | Q8N2F6-2 | 9.51563 | 8.25103 | 8.03572 | 9.40909 | 6.02744 | 5.13106 | 5.01048 | 4.81151 | 9.66361 | 9.10757 | 8.01393 | 7.35267 | 2.02484 | 2.60379 | 2.77968 | 2.26194 |
| ZFPM2 | Q8WW38 | 6.87002 | 6.07101 | 5.81615 | 6.2194 | 6.60652 | 6.34585 | 6.37321 | 6.34649 | 7.3439 | 5.42242 | 5.50709 | 6.26863 | 6.16459 | 6.2251 | 6.71995 | 5.69965 |
| MED9 | Q9NWA0 | 6.56389 | 4.89272 | 4.32029 | 4.38215 | 4.57454 | 3.08766 | 4.18754 | 5.05627 | 4.48395 | 3.83175 | 3.22799 | 4.20056 | 4.11544 | 17.1179 | 9.48816 | 16.4692 |
| LNX2 | Q8N448 | 6.21147 | 6.19074 | 6.14921 | 6.63014 | 6.35793 | 6.60688 | 6.07739 | 6.37509 | 6.78663 | 6.17437 | 6.41467 | 6.86055 | 5.7616 | 5.64933 | 5.82427 | 5.92973 |
| TGFB1I1 | O43294 | 7.3891 | 6.08977 | 5.0914 | 4.82921 | 9.13396 | 5.90698 | 5.015 | 6.26559 | 8.41647 | 7.07177 | 6.41538 | 7.08373 | 7.42543 | 5.28239 | 3.90553 | 4.67827 |
| IQCG | Q9H095 | 6.5758 | 7.71388 | 7.28027 | 6.88004 | 5.22567 | 6.67184 | 7.47456 | 7.4373 | 6.84621 | 7.72648 | 8.5806 | 8.90302 | 3.36601 | 4.07198 | 2.46836 | 2.77796 |
| API5 | Q9BZZ5-2 | 6.6616 | 6.28254 | 4.32747 | 6.39881 | 5.08058 | 6.32323 | 4.43575 | 5.59418 | 6.32005 | 5.62969 | 7.54386 | 8.33594 | 7.31785 | 7.35415 | 6.15276 | 5.84154 |
| AP2M1 | C9JTK4 | 5.62715 | 7.14186 | 5.44699 | 5.51416 | 5.40999 | 6.48506 | 6.59575 | 6.19102 | 7.20164 | 6.78002 | 5.72983 | 5.98235 | 5.6063 | 6.29079 | 6.58086 | 7.41623 |
| GPSM1 | Q86YR5 | 6.71477 | 6.32771 | 6.58154 | 6.19481 | 5.99482 | 6.61871 | 5.8402 | 7.06895 | 6.13742 | 6.44913 | 7.19342 | 6.77011 | 5.81884 | 5.54545 | 5.5156 | 5.22853 |
| ZC3HAV1 | Q7Z2W4-3 | 7.07276 | 5.97019 | 5.74481 | 5.5193 | 7.43936 | 5.68562 | 4.77498 | 7.49488 | 6.49507 | 5.79982 | 7.8069 | 7.34645 | 6.05809 | 5.79198 | 5.85275 | 5.14704 |
| SUPT5H | O00267-2 | 5.73501 | 5.87657 | 5.91072 | 6.08389 | 4.18756 | 5.65101 | 5.5066 | 6.84433 | 10.0207 | 6.00633 | 6.52872 | 6.31599 | 5.43808 | 6.81434 | 6.16276 | 6.9174 |
| RPL23A | H7BY10 | 6.17176 | 4.50884 | 6.34571 | 4.50414 | 5.51423 | 6.65872 | 5.58125 | 5.51889 | 6.30175 | 6.42371 | 7.40519 | 7.63061 | 7.40117 | 6.70167 | 6.44547 | 6.88689 |
| MEF2A | Q02078 | 6.12209 | 5.86275 | 5.64316 | 6.51928 | 6.17663 | 6.2725 | 6.32312 | 6.12704 | 5.73732 | 6.62055 | 6.43216 | 5.92297 | 6.12904 | 7.30781 | 6.38147 | 6.42211 |
| MEF2C | Q06413-5 | 5.46849 | 5.89879 | 6.24619 | 5.41692 | 5.31174 | 5.9563 | 6.50929 | 6.67085 | 6.20919 | 6.23952 | 6.54808 | 6.67267 | 6.55114 | 7.66751 | 5.74241 | 6.8909 |
| TFPT | P0C1Z6 | 6.14698 | 5.76544 | 7.98616 | 5.82527 | 5.74285 | 8.17566 | 6.91643 | 5.70087 | 8.58857 | 6.59726 | 6.04235 | 5.45568 | 5.73725 | 4.9721 | 4.76999 | 5.57714 |
| CARHSP1 | H3BNU9 | 9.18088 | 6.83728 | 9.02783 | 6.17483 | 5.28841 | 6.08331 | 4.42864 | 5.35407 | 5.08845 | 4.40811 | 5.49454 | 5.5736 | 8.93939 | 7.93862 | 5.28293 | 4.89912 |
| CCNF | P41002 | 8.44924 | 7.25399 | 9.18267 | 6.56565 | 6.55648 | 5.03364 | 8.1151 | 7.30954 | 6.41434 | 5.94264 | 11.9973 | 7.48038 | 1.11439 | 3.44043 | 2.20389 | 2.93831 |
| DNAH9 | Q9NYC9 | 5.38757 | 6.40747 | 6.33769 | 5.10674 | 6.87095 | 6.39653 | 6.79665 | 6.81113 | 6.24311 | 6.63911 | 5.75143 | 6.41546 | 6.39957 | 6.2988 | 5.5798 | 6.55798 |
| ZFYVE19 | Q96K21-3 | 5.7497 | 5.65803 | 7.66229 | 6.23039 | 5.94959 | 7.06097 | 4.91109 | 5.6202 | 6.39923 | 7.49285 | 7.23431 | 6.99399 | 6.22492 | 5.61047 | 6.09994 | 5.10202 |
| SMIM26 | A0A096LP01 | 5.89096 | 6.44984 | 6.23736 | 7.47644 | 6.29477 | 6.5901 | 6.99305 | 6.03369 | 6.09114 | 6.5937 | 6.07599 | 6.75842 | 5.05188 | 5.92687 | 5.21416 | 6.32164 |
| RCL1 | Q5VZU2 | 6.49026 | 6.3503 | 6.74481 | 7.36026 | 5.7023 | 5.54964 | 6.86508 | 5.81637 | 7.52055 | 7.27526 | 4.68298 | 5.43115 | 6.32694 | 4.08967 | 7.30303 | 6.49141 |
| PLPP6 | Q8IY26 | 6.67447 | 6.3898 | 6.63701 | 7.28182 | 5.84598 | 5.54094 | 6.31073 | 5.96763 | 4.91807 | 6.50789 | 6.5716 | 6.27954 | 6.18599 | 6.35199 | 6.31483 | 6.22169 |
| C1orf159 | Q96HA4 | 6.55238 | 6.98508 | 6.31094 | 7.08909 | 6.1575 | 5.90933 | 6.58117 | 6.23781 | 6.64181 | 6.07363 | 6.02948 | 6.46974 | 5.6091 | 5.27356 | 6.08905 | 5.99033 |
| MPDZ | H0YGQ3 | 6.38154 | 6.049 | 6.39179 | 6.7255 | 6.48179 | 6.25973 | 6.24877 | 6.56999 | 6.47327 | 6.69123 | 5.71434 | 6.54321 | 5.53194 | 6.05625 | 5.69584 | 6.1858 |
| RIOX2 | Q8IUF8-4 | 6.01643 | 6.65454 | 6.33839 | 6.67032 | 6.80378 | 6.45109 | 5.55068 | 6.38133 | 6.6198 | 6.39065 | 4.65562 | 7.40943 | 6.49738 | 5.77555 | 6.2716 | 5.51339 |
| WDR25 | Q64LD2 | 7.34007 | 7.07289 | 6.44878 | 5.94979 | 6.76005 | 5.59182 | 6.48357 | 5.59619 | 6.0727 | 6.23559 | 4.9542 | 5.62848 | 6.8048 | 6.31551 | 6.18917 | 6.55638 |
| P4HA1 | P13674 | 5.83324 | 8.37048 | 6.03974 | 5.62901 | 6.32689 | 5.97951 | 6.58308 | 6.33624 | 5.62817 | 5.8329 | 6.53557 | 6.61817 | 6.41085 | 6.36452 | 5.37861 | 6.13303 |
| CBARP | Q8N350 | 6.33234 | 6.73264 | 6.56953 | 6.65777 | 6.07377 | 5.29866 | 6.77306 | 5.5715 | 6.08249 | 5.34117 | 5.73373 | 6.33218 | 7.08458 | 6.7882 | 6.2614 | 6.36698 |
| PUDP | Q08623-4 | 5.43053 | 4.39869 | 6.07845 | 4.31574 | 6.02177 | 6.11109 | 5.49246 | 7.31747 | 4.98698 | 7.01632 | 6.56798 | 6.6602 | 5.90033 | 7.80155 | 8.01601 | 7.88441 |
| CLINT1 | Q14677 | 5.95 | 5.87021 | 6.58695 | 6.86193 | 7.16682 | 6.187 | 4.97796 | 6.15161 | 6.42054 | 6.91708 | 7.03167 | 5.68995 | 5.53818 | 6.36475 | 6.64987 | 5.63548 |
| CBX2 | Q14781 | 6.88132 | 6.66154 | 6.72776 | 7.30127 | 6.60776 | 6.23126 | 6.56216 | 5.45528 | 6.19129 | 6.34721 | 6.08277 | 5.92668 | 5.56286 | 5.89283 | 5.59279 | 5.97523 |
| TTLL1 | O95922 | 7.02874 | 7.28587 | 6.28284 | 6.22953 | 5.78982 | 6.65903 | 5.64583 | 6.21847 | 5.80389 | 6.59514 | 6.36195 | 7.24216 | 5.46911 | 5.11061 | 5.95419 | 6.3228 |
| PNMA8A | Q86V59 | 6.33946 | 5.95778 | 6.4313 | 6.98577 | 5.76268 | 5.19268 | 6.99042 | 5.66397 | 7.3734 | 6.43571 | 6.17364 | 6.92949 | 5.22192 | 6.08433 | 6.37268 | 6.08476 |
| MLLT6 | P55198 | 7.51812 | 6.20255 | 6.53668 | 6.52458 | 5.54568 | 6.45911 | 5.79325 | 6.8188 | 7.19365 | 5.25797 | 6.0161 | 5.32213 | 6.0095 | 6.34564 | 6.60009 | 5.85615 |
| TKT | A0A0B4J1R6 | 5.50571 | 5.4305 | 6.51624 | 5.22134 | 5.36421 | 6.38429 | 4.71192 | 5.23345 | 7.0164 | 5.66489 | 6.0092 | 7.26831 | 6.84677 | 6.03424 | 8.96747 | 7.82505 |
| UTP23 | Q9BRU9 | 6.55131 | 7.26414 | 4.86406 | 7.36567 | 6.34819 | 7.04011 | 6.18144 | 7.46161 | 5.46891 | 5.46608 | 6.17816 | 5.74514 | 5.47897 | 5.90553 | 6.34495 | 6.33573 |
| FLYWCH1 | Q4VC44-3 | 7.27652 | 7.19743 | 6.84815 | 8.06969 | 5.83051 | 6.35755 | 6.33419 | 5.89453 | 6.53259 | 7.6406 | 6.60808 | 5.95695 | 4.62106 | 4.8914 | 4.55444 | 5.38631 |
| WSB1 | Q9Y6I7 | 9.37438 | 7.3053 | 8.41205 | 7.21533 | 6.08444 | 7.74692 | 6.93037 | 4.78975 | 9.03268 | 5.76501 | 6.69559 | 5.22863 | 3.42703 | 4.24193 | 4.60481 | 3.1458 |
| NFATC1 | O95644 | 5.44674 | 5.38664 | 6.78063 | 5.06583 | 6.50971 | 7.02203 | 7.04779 | 6.20036 | 5.49758 | 6.46324 | 6.94022 | 8.05864 | 6.30965 | 5.95451 | 6.09457 | 5.22186 |
| KLHL8 | Q9P2G9 | 6.53891 | 6.21623 | 7.09149 | 6.93499 | 6.02499 | 7.01053 | 5.97842 | 6.56853 | 6.65531 | 6.60226 | 6.90133 | 6.68901 | 4.64347 | 5.11841 | 5.48702 | 5.53909 |
| POMP | Q9Y244 | 5.05126 | 7.14868 | 6.46815 | 7.08025 | 5.96257 | 6.64703 | 8.40785 | 6.06515 | 4.48504 | 6.37676 | 6.23354 | 7.95559 | 5.00522 | 6.58947 | 5.71927 | 4.80417 |
| TMBIM1 | Q969X1 | 6.82327 | 6.46654 | 6.67125 | 6.10577 | 6.16304 | 5.5686 | 6.76346 | 6.06597 | 5.97299 | 6.21094 | 6.37151 | 6.56906 | 6.08701 | 5.85306 | 6.51784 | 5.7907 |
| ZNF652 | Q9Y2D9 | 5.65398 | 7.16154 | 5.3993 | 6.25385 | 6.7056 | 6.63085 | 6.56495 | 6.957 | 5.77065 | 5.88838 | 6.77769 | 5.01665 | 5.3818 | 7.05137 | 6.80755 | 5.97886 |
| CA8 | P35219 | 7.13493 | 4.64812 | 7.63321 | 5.63004 | 5.24951 | 7.05952 | 6.33134 | 6.91024 | 5.44629 | 7.33059 | 6.54369 | 5.71213 | 5.06037 | 7.78892 | 5.29195 | 6.22913 |
| LARP1 | H0YC73 | 4.90863 | 5.76646 | 6.91836 | 5.72011 | 6.12479 | 7.13756 | 6.47619 | 5.33854 | 7.04756 | 7.03691 | 5.51084 | 5.84849 | 5.38424 | 6.93282 | 7.94275 | 5.90574 |
| ZSWIM1 | Q9BR11 | 6.54543 | 6.34955 | 6.986 | 5.33841 | 5.94156 | 6.83778 | 6.27329 | 6.42075 | 6.54517 | 7.36781 | 7.06846 | 7.06136 | 5.00632 | 5.21097 | 5.71312 | 5.33403 |
| CPPED1 | Q9BRF8 | 6.12658 | 6.20814 | 10.1224 | 7.81296 | 5.16216 | 7.37055 | 6.54917 | 6.25899 | 8.62558 | 4.88499 | 4.55929 | 5.43051 | 4.5297 | 3.70231 | 6.02887 | 6.62778 |
| CRY1 | Q16526 | 6.62715 | 6.78712 | 7.2799 | 6.87647 | 6.76882 | 5.11199 | 6.20984 | 7.27461 | 5.6337 | 6.07518 | 7.61397 | 8.00704 | 5.25267 | 5.23005 | 4.48377 | 4.76771 |
| FN3K | Q9H479 | 6.4476 | 6.35955 | 7.01805 | 4.29075 | 6.48731 | 6.0209 | 6.34116 | 6.68306 | 6.60125 | 6.28321 | 7.90708 | 6.59904 | 6.18031 | 5.58161 | 5.40662 | 5.79249 |
| ELOVL2 | Q9NXB9 | 5.81137 | 5.7432 | 5.88873 | 7.11295 | 5.26805 | 5.33415 | 6.53322 | 6.37248 | 5.62814 | 7.1092 | 5.91375 | 6.89347 | 6.39563 | 6.38322 | 6.98959 | 6.62286 |
| MAP3K3 | Q99759-2 | 6.77384 | 7.35972 | 5.86091 | 8.73825 | 4.34837 | 8.10937 | 5.50157 | 4.67543 | 7.2364 | 6.48252 | 5.30854 | 5.59235 | 5.72186 | 7.47252 | 5.86278 | 4.95558 |
| MIGA2 | Q7L4E1 | 4.48465 | 7.43355 | 7.13858 | 6.74019 | 4.73739 | 5.66915 | 4.57033 | 5.67921 | 6.73035 | 6.55657 | 7.33893 | 7.01357 | 6.51427 | 5.49913 | 7.20859 | 6.68554 |
| SF1 | H7C561 | 6.52879 | 4.97504 | 6.31979 | 7.01933 | 6.71321 | 5.63187 | 5.77957 | 7.02212 | 5.88735 | 6.86203 | 6.6332 | 6.30651 | 5.30356 | 6.59319 | 6.85371 | 5.57072 |
| FAM221A | A4D161 | 6.93968 | 5.95219 | 7.56331 | 6.07623 | 6.47009 | 6.93072 | 5.96862 | 6.77788 | 5.98491 | 6.67607 | 6.58636 | 6.91844 | 5.10442 | 5.60027 | 5.64825 | 4.80257 |
| SLC52A2 | Q9HAB3 | 6.36585 | 8.26915 | 6.13623 | 6.96918 | 5.77392 | 5.29327 | 6.58752 | 4.28129 | 7.17083 | 7.42066 | 5.35657 | 5.41043 | 7.27912 | 5.9245 | 5.06201 | 6.69948 |
| EIF4H | Q15056-2 | 7.18689 | 6.78505 | 8.26378 | 7.38234 | 7.95921 | 9.24472 | 5.71098 | 6.72256 | 6.54509 | 7.32035 | 8.23178 | 6.52411 | 2.9599 | 3.51328 | 2.72416 | 2.82578 |
| BAG4 | O95429 | 5.69937 | 6.25718 | 6.19581 | 4.10345 | 7.93496 | 5.55907 | 4.54217 | 6.54243 | 6.58916 | 7.15302 | 7.03903 | 7.4959 | 6.88521 | 6.35162 | 5.17177 | 6.47985 |
| MT2A | P02795 | 6.83789 | 5.98892 | 7.25934 | 6.29672 | 7.16493 | 7.042 | 5.96231 | 6.92333 | 7.12664 | 7.99114 | 7.61144 | 7.0828 | 4.92224 | 3.94003 | 3.81712 | 4.03318 |
| TSPAN9 | O75954 | 5.22995 | 6.9582 | 6.294 | 7.05563 | 4.72345 | 6.00291 | 5.57935 | 5.31094 | 5.77326 | 6.03088 | 5.9711 | 8.02791 | 7.73854 | 3.74768 | 8.86273 | 6.69347 |
| SFT2D1 | Q8WV19 | 5.70145 | 6.85351 | 7.67983 | 7.02913 | 6.70713 | 7.36759 | 6.22136 | 5.04311 | 5.74655 | 6.67382 | 7.31153 | 7.45103 | 5.02378 | 5.35047 | 5.17452 | 4.6652 |
| KLF12 | Q9Y4X4 | 6.13249 | 5.81968 | 5.69712 | 6.54985 | 4.98379 | 6.69888 | 6.01424 | 4.72708 | 6.29914 | 6.67326 | 7.65376 | 6.46163 | 6.92419 | 7.02872 | 6.75299 | 5.58316 |
| RTN3 | O95197-7 | 5.969 | 7.22544 | 7.49414 | 5.70574 | 7.00453 | 6.30051 | 6.83593 | 6.41918 | 5.23443 | 5.83934 | 5.76725 | 5.59333 | 6.08996 | 5.96716 | 5.86236 | 6.69169 |
| HNRNPA1 | F8W646 | 5.74465 | 5.96548 | 6.1697 | 5.87554 | 8.80158 | 4.84866 | 6.15046 | 6.35335 | 7.24258 | 4.95726 | 5.74562 | 7.4324 | 5.73065 | 4.69833 | 7.45024 | 6.83349 |
| HNRNPA3 | A0A7I2V2R3 | 6.18295 | 7.66467 | 6.10342 | 7.50483 | 5.73742 | 8.55033 | 4.9255 | 4.62689 | 5.33163 | 5.90272 | 6.20008 | 7.06252 | 4.73083 | 6.29025 | 7.11422 | 6.07174 |
| TRIM33 | H0Y612 | 7.66738 | 6.36415 | 5.9362 | 5.09674 | 5.78507 | 5.96281 | 5.6975 | 5.95306 | 6.72218 | 7.12352 | 6.03101 | 7.79271 | 5.15068 | 5.49188 | 6.12348 | 7.10162 |
| LTBP1 | Q14766-4 | 7.74983 | 7.983 | 4.95432 | 5.50587 | 5.13244 | 6.29588 | 7.80644 | 5.73536 | 5.37098 | 6.03629 | 5.4272 | 7.2995 | 4.98404 | 7.2394 | 4.68128 | 7.79817 |
| ZNF737 | O75373 | 6.54562 | 7.10264 | 6.91788 | 7.69446 | 5.9341 | 6.1587 | 5.50563 | 5.71366 | 7.50721 | 6.17224 | 7.8187 | 5.95688 | 5.18099 | 6.77239 | 4.88944 | 4.13159 |
| SRD5A3 | Q9H8P0 | 6.04657 | 5.28019 | 4.98734 | 6.44974 | 6.74892 | 5.42133 | 7.1574 | 6.29578 | 6.16113 | 5.71777 | 5.01509 | 6.60565 | 7.55536 | 6.69036 | 7.07616 | 6.7912 |
| TDP2 | O95551-2 | 5.11601 | 5.96703 | 9.14978 | 7.32274 | 5.79037 | 6.50132 | 5.34246 | 5.57942 | 7.02407 | 5.99707 | 6.31544 | 5.64209 | 5.53834 | 5.98805 | 6.20987 | 6.51591 |
| PPP2CB | E5RHC1 | 9.44335 | 5.98583 | 8.47033 | 6.75035 | 5.62981 | 5.13421 | 5.41441 | 4.66035 | 4.93859 | 3.87229 | 5.43112 | 5.25149 | 8.14017 | 8.03286 | 6.46069 | 6.38417 |
| TSPAN5 | P62079 | 5.64211 | 6.80008 | 6.13993 | 6.49958 | 6.53583 | 6.40222 | 6.63078 | 6.72894 | 6.15693 | 6.80066 | 6.35033 | 6.14364 | 5.62821 | 5.85622 | 5.75775 | 5.9268 |
| ZFP36 | P26651 | 5.80805 | 5.98592 | 6.7938 | 5.48638 | 6.08965 | 7.46375 | 6.32895 | 6.92079 | 6.44495 | 6.04986 | 6.59527 | 6.89477 | 5.752 | 5.97466 | 5.96553 | 5.4457 |
| SSNA1 | O43805 | 6.59294 | 4.69441 | 5.93309 | 4.28842 | 5.14535 | 3.86958 | 4.53171 | 6.11897 | 3.85853 | 5.46475 | 5.54988 | 5.30584 | 4.53972 | 12.6169 | 9.5398 | 11.9501 |
| CALR | K7EJB9 | 10.3219 | 6.89518 | 6.03079 | 7.27787 | 7.0038 | 5.85192 | 3.9243 | 3.94085 | 4.79496 | 4.43822 | 4.81838 | 5.21174 | 10.6631 | 6.53666 | 6.55043 | 5.73981 |
| DNAAF1 | Q8NEP3 | 6.17693 | 6.53137 | 6.30027 | 5.56015 | 6.76497 | 5.24842 | 6.69022 | 5.8109 | 6.34331 | 6.01655 | 4.87111 | 7.84236 | 7.27841 | 6.48842 | 4.92982 | 7.14678 |
| CORO7 | P57737 | 4.93833 | 5.1028 | 5.61451 | 6.26432 | 6.5746 | 8.52354 | 4.79107 | 6.76984 | 6.29298 | 6.50816 | 7.50191 | 4.66602 | 4.71729 | 7.30715 | 5.68187 | 8.7456 |
| RPP21 | A0A096LP39 | 5.51678 | 6.34705 | 6.4155 | 6.50267 | 5.64781 | 6.19239 | 6.64993 | 5.62672 | 6.22558 | 7.14469 | 6.80528 | 5.75723 | 6.24156 | 6.74678 | 6.33092 | 5.84911 |
| ZNF319 | Q9P2F9 | 6.25962 | 6.74801 | 5.92523 | 7.38759 | 6.33595 | 6.1488 | 6.56165 | 5.6824 | 5.86773 | 6.99203 | 7.23967 | 6.56421 | 5.6432 | 4.98 | 5.14579 | 6.51812 |
| COP1 | Q8NHY2 | 5.13281 | 6.46875 | 5.19604 | 6.96165 | 7.16958 | 5.66685 | 7.23062 | 6.20376 | 6.17962 | 6.26674 | 5.91861 | 5.99721 | 7.13713 | 5.51663 | 6.45435 | 6.49964 |
| PRKAA2 | P54646 | 5.80662 | 5.07367 | 5.91473 | 5.81821 | 5.71408 | 6.7879 | 6.06619 | 5.64311 | 6.60899 | 5.38075 | 6.75347 | 6.26576 | 7.01166 | 7.21091 | 6.8391 | 7.10485 |
| APOD | P05090 | 1.14005 | 33.2493 | 0.734335 | 10.8843 | 1.3147 | 1.96855 | 2.90334 | 3.67176 | 4.35899 | 3.52948 | 2.27193 | 2.47371 | 8.45605 | 3.91017 | 3.55157 | 15.5818 |
| NUDT16 | Q96DE0-4 | 6.19866 | 5.46618 | 5.82472 | 4.94943 | 6.34188 | 7.89783 | 6.02819 | 6.84851 | 6.40387 | 6.46526 | 7.25046 | 6.9533 | 4.85214 | 6.76108 | 6.61778 | 5.14071 |
| TPRA1 | Q86W33 | 6.14147 | 7.08835 | 5.79278 | 5.20929 | 6.10371 | 6.39702 | 5.62375 | 5.6774 | 6.02376 | 5.73834 | 6.20484 | 6.2579 | 6.39273 | 7.29604 | 6.42485 | 7.62777 |
| FRMD4B | C9JAW0 | 5.32823 | 7.72124 | 6.33323 | 5.65831 | 6.23072 | 6.63005 | 6.99686 | 5.73181 | 4.26835 | 7.51025 | 7.9789 | 5.4255 | 4.58037 | 6.46009 | 5.95371 | 7.19238 |
| GLI4 | P10075 | 6.13755 | 7.34165 | 6.04251 | 6.85218 | 6.60459 | 5.29077 | 6.41385 | 4.80398 | 6.53933 | 7.42407 | 7.12387 | 7.22323 | 6.08029 | 4.55673 | 5.85615 | 5.70925 |
| KCTD5 | Q9NXV2 | 6.38146 | 6.29218 | 6.83773 | 7.01253 | 6.04862 | 6.70047 | 6.24041 | 6.42103 | 6.66913 | 6.40964 | 7.00029 | 6.86253 | 5.51712 | 5.3017 | 4.90972 | 5.39543 |
| FBLN2 | P98095-2 | 6.94877 | 6.12792 | 5.01979 | 6.53927 | 6.67434 | 6.1204 | 6.04429 | 6.49044 | 6.78159 | 6.9393 | 6.92355 | 7.86459 | 5.25256 | 5.25271 | 4.71304 | 6.30745 |

| | | | | | | | | | | | | | | | | | |
|---|---|---|---|---|---|---|---|---|---|---|---|---|---|---|---|---|---|
| TMEM50A | O95807 | 6.11654 | 6.92541 | 6.85044 | 6.52435 | 6.60004 | 6.67095 | 7.42569 | 6.56072 | 6.05513 | 5.90161 | 6.4962 | 6.2527 | 5.60468 | 5.2515 | 4.66389 | 6.10015 |
| DCAF10 | Q5QP82 | 5.64618 | 6.26627 | 6.41329 | 5.72475 | 6.08448 | 6.05297 | 6.38335 | 6.1512 | 6.46968 | 6.03966 | 7.4487 | 7.5706 | 5.13858 | 6.03291 | 5.63097 | 6.94641 |
| RAPGEF2 | Q9Y4G8 | 7.20715 | 7 | 6.47141 | 6.07983 | 5.66975 | 5.82767 | 6.82105 | 7.51638 | 6.73199 | 7.46714 | 6.41917 | 4.72466 | 5.22541 | 6.57712 | 6.01084 | 4.25042 |
| NAT9 | Q9BTE0 | 6.05848 | 6.48649 | 6.29293 | 5.50748 | 7.07808 | 5.36048 | 6.68822 | 7.59718 | 6.29223 | 6.20487 | 5.38433 | 6.795 | 5.98321 | 6.06055 | 5.96188 | 6.24861 |
| SLC30A4 | O14863 | 5.88898 | 7.41926 | 6.41161 | 5.47898 | 5.90683 | 5.36431 | 7.92445 | 5.2342 | 7.78298 | 6.18387 | 4.82859 | 7.80412 | 7.35936 | 6.01425 | 4.99109 | 5.4071 |
| RNF13 | O43567 | 6.18382 | 6.59197 | 6.17547 | 6.82078 | 6.28496 | 6.47691 | 6.03518 | 6.13477 | 6.29437 | 6.29852 | 6.13866 | 6.31857 | 5.66938 | 5.88552 | 6.02881 | 6.66232 |
| GCAT | O75600-2 | 7.24509 | 7.3568 | 7.70365 | 5.97177 | 6.79658 | 4.382 | 6.19737 | 5.68828 | 4.62957 | 5.78058 | 7.35718 | 5.84535 | 5.59633 | 6.52873 | 6.67813 | 6.24259 |
| ATG14 | Q6ZNE5 | 5.96606 | 7.62171 | 7.63362 | 5.8155 | 5.43236 | 5.03432 | 6.9936 | 7.07858 | 5.48173 | 6.30321 | 6.61227 | 5.62529 | 4.76247 | 8.01464 | 5.01865 | 6.606 |
| FAM72D | Q6L9T8 | 6.44769 | 5.61863 | 6.85785 | 5.61856 | 6.42096 | 6.70385 | 7.01695 | 8.40115 | 7.66758 | 7.11003 | 6.59366 | 7.13751 | 5.32537 | 4.66491 | 3.6636 | 4.75168 |
| EIF3G | K7ER90 | 6.15772 | 6.92322 | 5.23887 | 6.92872 | 6.771 | 5.84594 | 7.40848 | 6.46244 | 5.57767 | 4.49314 | 7.18411 | 6.15651 | 6.73652 | 5.58642 | 6.55336 | 5.97589 |
| HSCB | Q8IWL3 | 6.98056 | 5.46364 | 6.33075 | 5.32498 | 6.59533 | 5.46459 | 5.3945 | 6.20838 | 5.12466 | 5.90196 | 5.84169 | 6.52825 | 6.46785 | 7.78815 | 7.5586 | 7.02611 |
| HNRNPD | Q14103-3 | 7.15069 | 5.28386 | 7.38218 | 6.51495 | 7.35627 | 7.13699 | 5.6041 | 5.63105 | 5.66771 | 5.64622 | 7.35047 | 6.29809 | 5.68274 | 6.74935 | 4.5161 | 6.02922 |
| COX17 | Q14061 | 6.4676 | 6.35244 | 4.6804 | 4.86286 | 4.51098 | 3.5744 | 4.21196 | 5.41429 | 3.28023 | 5.27377 | 4.28975 | 4.38865 | 4.63666 | 13.2245 | 9.67548 | 15.156 |
| ZC3H13 | Q5T200-2 | 7.57451 | 6.87348 | 5.91935 | 5.80352 | 5.66968 | 5.41703 | 6.09786 | 6.02554 | 6.41456 | 6.87189 | 5.8618 | 7.03966 | 5.94623 | 5.37167 | 6.40213 | 6.71108 |
| SRSF12 | Q8WXF0 | 7.03102 | 8.45446 | 5.6696 | 5.64751 | 6.05618 | 3.5302 | 4.80196 | 6.86526 | 6.49764 | 8.54137 | 5.97089 | 7.2289 | 7.36556 | 5.41965 | 6.06063 | 4.85917 |
| SFXN4 | Q6P4A7-3 | 5.96662 | 5.50472 | 6.21641 | 6.42576 | 4.96618 | 6.45799 | 5.96301 | 7.63711 | 6.70446 | 6.39593 | 6.50562 | 7.08696 | 5.44481 | 5.61891 | 6.67158 | 6.43393 |
| DTNBP1 | Q96EV8 | 5.72652 | 5.5249 | 5.82579 | 6.53198 | 6.28525 | 7.18425 | 6.77407 | 7.52267 | 6.31574 | 7.04821 | 6.95923 | 7.51968 | 4.67541 | 5.87763 | 4.83615 | 5.39253 |
| RASSF4 | Q9H2L5-2 | 6.27794 | 5.9257 | 7.14048 | 6.16191 | 6.31678 | 6.73782 | 6.26591 | 6.15321 | 7.05477 | 6.64203 | 6.92266 | 6.42718 | 5.89701 | 5.05107 | 5.67341 | 5.35212 |
| RASSF2 | P50749 | 6.80936 | 6.46388 | 6.54717 | 6.13889 | 6.11118 | 7.08462 | 5.89308 | 6.81568 | 6.45332 | 6.11188 | 5.87601 | 6.29685 | 6.36866 | 5.63088 | 5.81641 | 5.58215 |
| MDK | P21741 | 7.34246 | 7.19379 | 7.13416 | 7.3725 | 6.01431 | 6.4033 | 7.71372 | 6.789 | 6.16063 | 5.85893 | 5.50004 | 5.48545 | 4.78591 | 5.75682 | 4.96596 | 5.52304 |
| ERCC8 | Q13216 | 5.49958 | 5.87592 | 5.75746 | 5.43716 | 6.31786 | 5.55898 | 5.50647 | 5.58666 | 7.47656 | 4.87518 | 5.35966 | 6.07032 | 7.40854 | 7.7519 | 8.15541 | 7.36232 |
| PFKFB4 | Q16877 | 7.11454 | 7.1123 | 5.78777 | 5.29295 | 7.05646 | 7.0183 | 4.95471 | 5.89368 | 5.90109 | 6.97134 | 5.70543 | 6.02311 | 5.44807 | 8.24225 | 6.18884 | 5.28917 |
| SPSB2 | Q99619-2 | 7.05136 | 6.0726 | 4.72906 | 5.01285 | 7.25274 | 7.12817 | 5.53326 | 5.39801 | 6.50265 | 6.69226 | 6.90052 | 6.30042 | 7.18495 | 6.95824 | 4.99756 | 6.28534 |
| BLOC1S2 | Q6QNY1 | 4.46229 | 5.30496 | 7.91496 | 5.94033 | 7.00543 | 5.11609 | 5.36455 | 6.13552 | 5.52999 | 6.46146 | 1.85794 | 5.46034 | 6.48601 | 9.58005 | 7.67224 | 9.70783 |
| EVC2 | Q86UK5 | 6.34771 | 7.29847 | 6.19193 | 7.46258 | 5.90059 | 6.48998 | 6.95303 | 7.06342 | 7.60137 | 7.16254 | 6.46609 | 5.72819 | 4.65611 | 5.51451 | 4.25593 | 4.90754 |
| ST14 | Q9Y5Y6 | 6.32203 | 5.96812 | 5.86452 | 5.66729 | 5.73326 | 6.8594 | 6.12045 | 7.30178 | 7.43557 | 6.42424 | 6.40278 | 7.57326 | 6.0555 | 6.55499 | 4.90892 | 4.80787 |
| ADGRL3 | Q9HAR2-4 | 7.40401 | 7.15654 | 7.16532 | 7.1891 | 5.82934 | 6.02577 | 7.05793 | 7.9576 | 8.59655 | 7.75353 | 6.683 | 8.14761 | 3.13884 | 3.64791 | 3.0866 | 3.16034 |
| MTHFR | P42898-2 | 6.05265 | 7.08582 | 6.7234 | 5.53461 | 5.39459 | 6.59901 | 6.43639 | 6.36249 | 6.48697 | 6.55919 | 6.19901 | 7.62943 | 5.62076 | 5.45135 | 6.14977 | 5.71457 |
| PHF20L1 | A8MXR8 | 7.2045 | 7.10323 | 5.94226 | 8.67814 | 5.75745 | 6.2266 | 6.23516 | 5.72219 | 5.85307 | 5.91555 | 6.34847 | 6.4624 | 5.2883 | 6.09527 | 5.47231 | 5.69508 |
| SLC9B2 | Q86UD5 | 5.22365 | 5.54441 | 7.24291 | 6.59988 | 5.25629 | 6.68704 | 7.65535 | 6.89684 | 5.49538 | 5.83757 | 5.62086 | 7.22501 | 5.38366 | 5.88564 | 5.76099 | 7.67553 |
| GALNT3 | Q14435 | 6.49137 | 5.99535 | 6.21336 | 6.39901 | 7.44043 | 4.6035 | 6.29708 | 5.26978 | 6.5402 | 7.47271 | 5.39205 | 6.45538 | 5.37556 | 5.7768 | 7.713 | 7.5644 |
| RNF4 | P78317 | 6.50238 | 7.36538 | 7.70536 | 7.95797 | 5.57123 | 6.4289 | 6.12154 | 8.40234 | 5.82654 | 5.51225 | 8.54517 | 4.35081 | 4.75314 | 5.13836 | 4.94711 | 4.87151 |
| RBBP7 | Q5JP01 | 6.39612 | 5.39918 | 3.58682 | 4.96969 | 6.52662 | 5.00842 | 6.75096 | 7.97925 | 6.20093 | 7.37398 | 6.23122 | 7.44135 | 6.02874 | 9.3391 | 4.82592 | 5.94168 |
| RGL2 | O15211 | 6.25096 | 5.69872 | 7.01299 | 5.47134 | 5.74283 | 6.67904 | 5.67594 | 6.58063 | 5.86803 | 6.99878 | 6.42924 | 6.79223 | 5.85038 | 6.71343 | 5.57566 | 6.65981 |
| NPM1 | E5RGW4 | 7.71846 | 7.30107 | 4.38738 | 7.54769 | 6.69421 | 5.16966 | 6.26177 | 5.25076 | 7.61008 | 3.9261 | 5.91004 | 6.5332 | 6.51679 | 6.64586 | 6.25154 | 6.27537 |
| KCTD3 | Q9Y597 | 5.79421 | 6.13102 | 6.30557 | 6.03986 | 6.22435 | 6.61949 | 6.04523 | 6.02891 | 5.8155 | 6.41565 | 5.87612 | 6.05238 | 7.40169 | 6.03754 | 6.67036 | 6.54211 |
| OPA3 | Q9H6K4 | 5.93111 | 6.10048 | 5.66422 | 6.74989 | 5.75453 | 6.27662 | 6.12465 | 5.94878 | 6.02967 | 5.91983 | 5.61553 | 6.4737 | 6.80416 | 6.55026 | 7.03371 | 7.02286 |
| C2orf42 | Q9NWW7 | 5.96565 | 7.74743 | 6.20546 | 6.13372 | 6.42663 | 6.4925 | 5.46524 | 6.12412 | 6.06491 | 5.55053 | 6.36496 | 5.9983 | 6.43256 | 6.23819 | 6.30497 | 6.48482 |
| CCDC80 | Q76M96-2 | 6.78884 | 6.34248 | 4.381 | 7.96745 | 4.71514 | 6.31615 | 8.00368 | 5.64409 | 7.38036 | 5.9679 | 4.87342 | 7.03765 | 6.43718 | 5.48158 | 5.94877 | 6.71431 |
| TCTN2 | Q96GX1 | 6.54833 | 6.74273 | 6.25663 | 6.63774 | 5.99706 | 6.15852 | 6.15205 | 6.21083 | 6.30798 | 6.65502 | 6.1795 | 6.70273 | 6.12175 | 5.83282 | 5.42349 | 6.0728 |
| OTULINL | Q9NUU6 | 6.72131 | 5.97133 | 6.0893 | 6.72398 | 6.09996 | 5.80056 | 6.29901 | 5.11058 | 5.70813 | 6.06318 | 6.52196 | 5.88655 | 6.39216 | 7.06461 | 6.64592 | 6.93149 |
| NPHP4 | O75161 | 6.29936 | 7.44642 | 7.37638 | 5.48481 | 5.9109 | 6.6653 | 5.97203 | 5.06206 | 7.28264 | 7.37334 | 6.19112 | 7.45681 | 6.14187 | 5.58895 | 4.40472 | 5.34331 |
| FIZ1 | Q96SL8 | 5.46413 | 5.8279 | 6.02738 | 6.41388 | 5.98742 | 5.77089 | 6.06575 | 6.28988 | 6.32654 | 5.97643 | 6.35636 | 6.08548 | 6.72334 | 6.82721 | 7.19407 | 6.66335 |
| OGFOD2 | Q6N063 | 6.86913 | 5.83994 | 6.71329 | 6.45279 | 7.02926 | 5.86049 | 6.38397 | 6.69733 | 6.4263 | 6.0857 | 6.4393 | 5.90753 | 6.56606 | 5.74765 | 5.57053 | 5.41074 |
| KCTD1 | Q719H9 | 6.27961 | 5.80623 | 7.40589 | 2.63674 | 8.45142 | 8.01283 | 5.65542 | 4.16756 | 5.88541 | 5.11344 | 7.72648 | 8.56796 | 5.92788 | 6.20922 | 5.24469 | 6.90922 |
| COL22A1 | Q8NFW1 | 6.26617 | 6.19507 | 6.36732 | 5.97209 | 6.56728 | 7.18953 | 6.46421 | 6.3606 | 5.70388 | 6.41509 | 6.94664 | 6.46268 | 5.60837 | 6.37681 | 5.7572 | 5.34706 |
| SHC1 | P29353-3 | 7.09647 | 6.80096 | 7.33907 | 6.65393 | 6.32472 | 5.28735 | 6.66104 | 6.71159 | 6.46903 | 6.089 | 6.41781 | 6.56196 | 4.52326 | 5.43244 | 5.66681 | 5.96455 |
| CLIP3 | Q96DZ5 | 6.29653 | 8.69043 | 6.81879 | 6.35679 | 7.4045 | 7.32674 | 6.15672 | 6.00196 | 7.83545 | 7.93876 | 6.8813 | 7.69037 | 3.19546 | 4.49678 | 2.92162 | 3.9878 |
| SLC39A7 | Q92504 | 6.49106 | 5.63139 | 5.51317 | 6.30322 | 6.01387 | 5.67746 | 7.78063 | 5.93121 | 6.33739 | 5.98083 | 6.05888 | 6.27751 | 6.70982 | 6.11314 | 6.36009 | 6.82034 |
| TRIM21 | P19474 | 6.05701 | 6.67226 | 7.73434 | 5.04966 | 5.65815 | 7.68563 | 6.63966 | 5.62607 | 7.11241 | 6.32771 | 4.98555 | 6.08391 | 6.29879 | 7.52109 | 3.68254 | 6.8652 |
| TPGS1 | Q6ZTW0 | 6.84651 | 6.36569 | 7.62796 | 6.4418 | 6.25394 | 6.66536 | 5.86315 | 7.61169 | 6.62352 | 7.47382 | 6.96618 | 7.84342 | 4.00481 | 4.69542 | 3.93417 | 4.78255 |
| NOP2 | P46087-2 | 5.44525 | 5.45558 | 5.77712 | 7.3163 | 5.18766 | 5.9694 | 7.21098 | 5.13226 | 6.06361 | 5.87995 | 6.58111 | 7.14461 | 6.64683 | 6.74977 | 5.99771 | 7.44186 |
| DYNLL2 | Q96FJ2 | 6.03803 | 5.68444 | 6.07874 | 4.94328 | 6.61826 | 6.42265 | 5.99825 | 6.53779 | 5.52857 | 6.11032 | 5.4541 | 5.36265 | 6.60033 | 7.91717 | 7.4453 | 7.26011 |
| COX18 | Q8N8Q8 | 6.6791 | 5.94174 | 7.60368 | 7.81617 | 5.44197 | 5.72438 | 6.00879 | 6.39825 | 5.4326 | 3.96467 | 5.46528 | 4.87856 | 8.71264 | 8.07096 | 7.10309 | 4.75813 |
| ZNF816 | Q0VGE8 | 6.11277 | 7.16439 | 6.29335 | 7.77107 | 5.82746 | 5.26015 | 6.0986 | 6.39188 | 6.90918 | 6.66802 | 6.53599 | 6.22054 | 5.59054 | 5.84603 | 5.9996 | 5.31042 |
| GSDMD | P57764 | 6.83927 | 6.23183 | 6.5108 | 5.7241 | 6.41467 | 6.93584 | 5.70606 | 6.11053 | 6.56197 | 6.07893 | 6.25822 | 6.39774 | 5.79135 | 6.84987 | 5.66301 | 5.9258 |
| HNF4G | Q14541-2 | 6.37889 | 5.91038 | 6.62031 | 6.37533 | 6.25332 | 5.62929 | 6.61182 | 6.17986 | 6.51675 | 6.34202 | 6.12975 | 6.42469 | 5.87583 | 5.18661 | 6.32965 | 7.23552 |
| TAF2 | H0YEX2 | 5.41527 | 6.34846 | 7.03198 | 6.91436 | 5.06704 | 5.20336 | 4.00933 | 7.38335 | 4.15017 | 5.57349 | 6.68545 | 5.85364 | 7.44838 | 9.84203 | 5.12491 | 7.94878 |
| ZBTB18 | Q99592-2 | 6.57086 | 6.97447 | 7.45167 | 7.24172 | 6.59516 | 6.43781 | 6.40759 | 6.10111 | 5.54269 | 5.97792 | 6.34575 | 6.61978 | 5.3603 | 5.22377 | 5.55504 | 5.59436 |
| ABLIM1 | O14639 | 6.82488 | 5.09201 | 6.44992 | 6.2933 | 5.32613 | 6.80536 | 6.13871 | 6.06246 | 7.02232 | 6.20238 | 5.37546 | 5.86391 | 6.70962 | 7.45508 | 5.50477 | 6.8737 |
| B4GALT5 | O43286 | 7.23858 | 7.43665 | 6.43607 | 7.90985 | 7.20281 | 5.58893 | 5.81436 | 5.51392 | 7.17411 | 8.07908 | 8.11462 | 6.84209 | 4.3447 | 3.90021 | 3.87733 | 4.52668 |
| B4GALT6 | Q9UBX8 | 7.6314 | 6.25123 | 5.73804 | 6.31595 | 4.98758 | 5.1994 | 6.33071 | 7.15561 | 6.41297 | 5.31345 | 7.49069 | 7.48784 | 7.19679 | 5.86558 | 6.27404 | 4.34873 |
| UCK1 | Q9HA47-4 | 6.02581 | 6.46488 | 7.88724 | 5.4766 | 6.16849 | 6.05427 | 6.06924 | 6.02946 | 6.69734 | 6.62288 | 6.85778 | 7.09039 | 6.36519 | 5.23912 | 6.29492 | 6.45638 |
| TCEAL1 | Q15170-2 | 6.49437 | 5.9483 | 5.97574 | 5.29334 | 6.97921 | 6.60555 | 5.84424 | 6.0896 | 5.61981 | 6.76062 | 7.01242 | 7.05705 | 5.71206 | 6.30695 | 5.80822 | 6.49255 |
| CEBPG | P53567 | 5.95378 | 6.79681 | 6.76549 | 7.04009 | 7.40348 | 6.3763 | 5.79015 | 8.10653 | 7.38957 | 4.85224 | 5.41848 | 4.66472 | 5.60163 | 4.28013 | 5.51692 | 8.0437 |
| POTEH | Q6S545 | 6.07002 | 5.29985 | 6.14269 | 5.95841 | 5.93978 | 6.25789 | 6.23968 | 6.15773 | 6.26884 | 6.83737 | 6.63923 | 6.5384 | 6.37615 | 6.55085 | 6.4496 | 6.27349 |
| FEM1A | Q9BSK4 | 6.16161 | 6.63785 | 7.31423 | 6.67227 | 6.11903 | 6.00717 | 6.54958 | 6.98269 | 7.44374 | 5.55065 | 6.60239 | 7.40398 | 4.93185 | 6.00534 | 5.26426 | 5.25335 |
| FEM1C | Q96JP0 | 6.70194 | 6.75268 | 8.04556 | 6.67212 | 6.0489 | 6.47385 | 6.03935 | 6.13614 | 7.02526 | 5.98794 | 6.44967 | 7.11872 | 4.7189 | 4.88599 | 5.87815 | 5.06484 |
| PACS2 | Q86VP3-2 | 6.31479 | 5.42354 | 5.47193 | 6.11949 | 6.72434 | 6.48478 | 7.33724 | 5.56087 | 6.74731 | 8.9845 | 6.47482 | 6.46953 | 5.78583 | 4.47635 | 5.19708 | 6.42758 |
| ARL8B | Q9NVJ2 | 4.07994 | 4.26972 | 5.61046 | 4.98793 | 2.58751 | 4.08341 | 4.43947 | 4.59052 | 5.23452 | 4.91879 | 8.49057 | 9.38427 | 9.40702 | 8.56667 | 9.83288 | 9.51632 |
| SYNGR1 | O43759-3 | 6.30239 | 5.98563 | 3.84925 | 5.72091 | 6.02746 | 5.62479 | 5.99188 | 6.11035 | 6.09389 | 5.36504 | 8.05376 | 7.06936 | 6.01146 | 6.30711 | 7.49847 | 7.98826 |
| SNX13 | Q9Y5W8 | 6.49492 | 6.36189 | 6.15303 | 6.59993 | 5.87972 | 6.19503 | 6.66241 | 6.3383 | 6.20997 | 5.85004 | 6.79188 | 5.40661 | 6.43963 | 6.29694 | 6.29451 | 6.02519 |
| KLHL20 | Q9Y2M5 | 5.87366 | 6.30372 | 6.54496 | 6.13538 | 5.86757 | 6.17886 | 6.01058 | 6.70609 | 6.55574 | 6.70088 | 6.48859 | 6.98575 | 5.8253 | 6.025 | 5.64792 | 6.15002 |
| SEMA3E | O15041 | 6.81234 | 5.89154 | 4.83021 | 7.31798 | 4.96354 | 4.95827 | 8.62052 | 5.56821 | 6.07594 | 4.96926 | 4.51071 | 7.14928 | 8.20838 | 6.66423 | 6.49572 | 6.86526 |
| TNRC6B | Q9UPQ9-2 | 6.29446 | 4.23944 | 6.55435 | 6.64779 | 7.31463 | 7.6011 | 4.74176 | 5.17362 | 6.74523 | 7.11672 | 6.2082 | 7.82454 | 6.13585 | 5.80373 | 5.34838 | 6.2502 |
| TLCD2 | A6NGC4 | 6.58299 | 6.57145 | 5.65851 | 5.88813 | 6.51858 | 6.01705 | 6.63564 | 6.96039 | 5.63327 | 7.11172 | 6.56849 | 5.78749 | 5.98801 | 6.9642 | 4.9684 | 6.14569 |
| RCC1 | M0R1Q8 | 7.066 | 5.78688 | 4.73789 | 5.40348 | 5.81589 | 7.177 | 4.28728 | 7.19337 | 9.73408 | 9.31422 | 7.39517 | 9.54333 | 6.41998 | 4.20521 | 3.45614 | 2.46408 |
| ANAPC15 | P60006-2 | 6.44214 | 6.29014 | 6.0055 | 6.58647 | 6.66757 | 6.53373 | 5.44103 | 7.11803 | 5.88969 | 6.53297 | 6.02018 | 6.26463 | 6.74378 | 5.99572 | 5.51914 | 5.94929 |
| ADAM23 | O75077 | 7.22234 | 6.76856 | 6.90726 | 6.62873 | 6.17385 | 6.58885 | 6.81696 | 5.91786 | 6.55762 | 7.24162 | 7.45444 | 5.39261 | 5.02208 | 5.50957 | 4.90488 | 4.89278 |
| POLR2J | P52435 | 5.70385 | 5.8814 | 4.8353 | 5.30736 | 6.04183 | 4.94884 | 5.36967 | 7.32404 | 5.16249 | 6.17136 | 4.7169 | 5.64025 | 4.58602 | 9.88126 | 7.37238 | 11.057 |
| CYP2J2 | P51589 | 5.57778 | 5.19418 | 6.34171 | 5.56477 | 5.66459 | 5.59409 | 6.83317 | 4.66155 | 4.74977 | 6.84108 | 7.44724 | 6.55163 | 7.85404 | 6.84965 | 7.37136 | 6.90338 |
| EVPL | Q92817 | 5.86495 | 6.77277 | 6.50088 | 6.56877 | 5.9558 | 6.61056 | 5.44726 | 6.17715 | 5.9911 | 5.47395 | 6.75608 | 6.20114 | 8.2935 | 6.01867 | 5.83957 | 5.52785 |
| CCDC74B | Q96LY2 | 7.0048 | 6.96164 | 7.30231 | 6.86785 | 6.25932 | 6.86597 | 7.04458 | 6.15826 | 7.79928 | 7.23637 | 7.73991 | 7.14734 | 3.67097 | 4.73429 | 3.39286 | 4.21425 |
| POSTN | Q15063 | 5.32927 | 6.90341 | 5.05375 | 8.27399 | 4.56265 | 6.42154 | 7.38762 | 6.33518 | 5.48188 | 5.33631 | 6.64636 | 5.84061 | 6.2959 | 5.28289 | 6.99071 | 7.85793 |
| TBC1D8 | O95759 | 7.09601 | 6.25647 | 6.19203 | 6.04364 | 6.01712 | 6.79252 | 7.05098 | 6.41589 | 6.29932 | 6.8849 | 5.48159 | 6.71424 | 5.78021 | 5.41257 | 6.06319 | 5.49933 |
| LMBRD2 | Q68DH5 | 5.56811 | 7.06771 | 5.37066 | 4.32376 | 3.53484 | 6.60691 | 7.59291 | 4.78579 | 7.11266 | 5.2304 | 6.90026 | 7.4663 | 4.43905 | 5.85425 | 10.6321 | 7.51428 |
| TMEM186 | Q96B77 | 5.49171 | 7.06217 | 5.345 | 6.83254 | 6.88154 | 6.64876 | 6.2418 | 6.51886 | 5.10327 | 5.78808 | 6.32597 | 6.00895 | 7.00809 | 6.66162 | 7.73489 | 4.34674 |
| MBP | P02686 | 6.04096 | 5.29423 | 6.91013 | 5.27442 | 7.01007 | 4.91382 | 4.74149 | 6.02783 | 5.54114 | 7.77201 | 5.80147 | 5.95061 | 7.95755 | 7.06613 | 6.10273 | 7.59541 |
| CMBL | Q96DG6 | 7.14279 | 5.88458 | 6.08149 | 5.42382 | 6.88042 | 6.2414 | 5.51019 | 6.65707 | 7.3892 | 6.20526 | 6.58216 | 6.45948 | 6.15507 | 5.39174 | 6.00678 | 5.98856 |
| SERF2 | P84101 | 5.81279 | 7.08882 | 7.11313 | 5.68325 | 6.84319 | 6.73393 | 6.21041 | 6.83056 | 5.76415 | 7.16348 | 7.04072 | 6.76567 | 5.96524 | 6.47052 | 4.35588 | 4.15827 |
| ARL15 | Q9NXU5 | 6.20178 | 6.07979 | 5.91972 | 6.32731 | 6.16493 | 5.67502 | 6.29508 | 6.28612 | 6.66426 | 5.84192 | 6.61044 | 5.70014 | 5.80994 | 6.86142 | 6.42929 | 7.13283 |
| G6PC3 | Q9BUM1 | 6.17214 | 6.06665 | 5.71071 | 6.78787 | 6.51121 | 5.87268 | 6.19111 | 6.03185 | 5.96257 | 6.32235 | 6.43547 | 6.01556 | 6.2329 | 6.59058 | 6.05671 | 7.03964 |
| HOOK2 | Q96ED9-2 | 5.39686 | 6.40623 | 6.63109 | 5.3111 | 6.64266 | 6.16079 | 5.6429 | 6.50368 | 7.15595 | 6.63955 | 7.18057 | 6.39379 | 6.29574 | 6.15749 | 5.74017 | 5.7414 |
| WDR47 | O94967 | 6.14397 | 5.99996 | 6.53713 | 5.79296 | 7.21435 | 6.23773 | 6.76179 | 5.76212 | 5.75456 | 6.1036 | 6.2909 | 6.04903 | 7.32065 | 6.60884 | 5.84078 | 5.58164 |
| DPH7 | Q9BTV6 | 5.79304 | 6.78442 | 6.01059 | 6.91585 | 5.87839 | 6.54543 | 6.57665 | 8.07218 | 6.36602 | 6.46051 | 6.62034 | 6.8996 | 4.88344 | 5.80069 | 5.64594 | 4.74693 |
| CAB39L | Q9H9S4 | 6.91048 | 6.6924 | 6.1435 | 5.89889 | 5.58423 | 7.08003 | 4.70125 | 6.3776 | 6.91222 | 7.288 | 5.02359 | 7.96068 | 6.27226 | 6.25804 | 5.44388 | 5.45294 |
| OSBPL8 | F8W060 | 6.05414 | 5.96496 | 6.50611 | 6.19037 | 5.88904 | 5.64166 | 6.48618 | 5.92515 | 6.34638 | 5.61222 | 5.33638 | 5.50613 | 7.30293 | 6.2812 | 7.74694 | 7.2102 |
| ZNF326 | Q5BKZ1-3 | 6.75346 | 8.1551 | 5.36039 | 6.83766 | 6.43034 | 5.42404 | 6.17668 | 7.33285 | 5.39943 | 5.72313 | 5.42403 | 6.35812 | 5.15305 | 6.02428 | 7.47899 | 5.96847 |
| ZNF224 | Q9NZL3 | 6.79111 | 6.93624 | 5.5495 | 7.11783 | 6.66237 | 5.40405 | 3.99957 | 4.13334 | 7.68229 | 6.20856 | 8.58178 | 7.32268 | 4.34917 | 5.13002 | 7.04586 | 7.08564 |
| ZNF222 | Q9UK12-2 | 6.53947 | 7.45902 | 4.39045 | 7.70045 | 6.29273 | 8.22565 | 6.28273 | 5.56412 | 7.16014 | 6.94861 | 8.03072 | 5.24156 | 4.86904 | 5.88872 | 4.79123 | 4.61535 |
| TTC17 | Q96AE7-2 | 6.04662 | 6.40698 | 5.69417 | 8.49685 | 6.63776 | 6.29404 | 6.65809 | 6.21681 | 6.50093 | 6.08355 | 6.24912 | 6.3095 | 6.34519 | 4.82525 | 5.87671 | 5.35844 |
| DMTF1 | Q9Y222 | 6.39092 | 6.38612 | 6.8617 | 6.29244 | 6.54959 | 6.62466 | 6.21489 | 6.69506 | 6.58208 | 6.51668 | 6.86578 | 6.7834 | 5.44329 | 5.21004 | 5.32642 | 5.25694 |
| AZGP1 | P25311 | 2.73644 | 11.9609 | 1.57969 | 19.5019 | 4.64369 | 7.30564 | 4.30315 | 2.07198 | 4.45907 | 4.52779 | 1.87617 | 4.33486 | 9.96102 | 6.07112 | 3.85721 | 10.8093 |
| hCG_2039718 | K7EN88 | 7.26367 | 6.6173 | 5.69227 | 5.72943 | 5.26644 | 7.83739 | 5.8672 | 6.23983 | 6.68249 | 7.09913 | 6.23424 | 6.31851 | 4.71119 | 6.40055 | 5.67556 | 6.3648 |
| EID2 | Q8N6I1 | 5.51778 | 5.20577 | 8.41544 | 5.7351 | 5.7899 | 7.35298 | 6.594 | 7.34503 | 7.49362 | 7.51163 | 8.46507 | 8.81388 | 3.41844 | 4.3209 | 3.88122 | 4.13924 |

| | | | | | | | | | | | | | | | | | |
|---|---|---|---|---|---|---|---|---|---|---|---|---|---|---|---|---|---|
| KPNA5 | O15131 | 5.95382 | 6.30791 | 6.86511 | 6.76094 | 6.2973 | 6.32626 | 6.02243 | 6.36963 | 6.73638 | 6.07457 | 7.12526 | 6.7123 | 5.70307 | 5.39087 | 5.77155 | 5.58258 |
| FAM13B | Q9NYF5 | 7.42844 | 7.32483 | 9.44755 | 6.00467 | 5.07782 | 5.26217 | 6.72192 | 6.20451 | 6.85872 | 7.83798 | 7.16004 | 6.76541 | 4.26436 | 3.90766 | 4.17608 | 5.55783 |
| ACVR1B | P36896-4 | 6.21949 | 7.01167 | 5.1956 | 7.26743 | 6.57623 | 6.28778 | 5.90648 | 6.56485 | 6.34015 | 6.34398 | 5.9503 | 6.23086 | 5.72965 | 6.24832 | 7.00042 | 6.02678 |
| PLSCR1 | O15162 | 6.36387 | 5.52938 | 5.98191 | 7.38656 | 7.85848 | 6.60707 | 5.62219 | 5.10074 | 5.68464 | 6.60604 | 5.60267 | 5.96314 | 6.42211 | 5.48725 | 6.83311 | 6.95083 |
| TMEM143 | Q96AN5 | 6.48061 | 6.07303 | 6.00808 | 6.79258 | 5.94118 | 6.0837 | 6.08546 | 6.13818 | 6.07395 | 6.65215 | 5.92519 | 5.51129 | 6.69435 | 6.95607 | 6.42629 | 6.1579 |
| ABCB4 | P21439 | 5.54357 | 6.61961 | 6.36107 | 6.87067 | 5.41885 | 5.69587 | 6.03748 | 5.43587 | 6.39318 | 7.17105 | 6.40479 | 6.74882 | 6.30066 | 5.89114 | 6.76545 | 6.34191 |
| STAT1 | P42224-2 | 6.58652 | 6.75688 | 5.6577 | 4.66644 | 5.98913 | 6.856 | 5.65991 | 6.52883 | 6.75912 | 5.37514 | 6.82662 | 6.9234 | 6.08388 | 7.28553 | 6.81078 | 5.2341 |
| NUDT14 | O95848 | 7.67746 | 6.28469 | 7.8443 | 6.49882 | 6.48312 | 7.45899 | 4.73221 | 6.68256 | 6.31445 | 6.65253 | 5.62583 | 5.41908 | 5.09349 | 5.93944 | 5.51364 | 5.77936 |
| GSTM4 | Q03013 | 6.29377 | 4.46164 | 6.488 | 4.08398 | 6.0423 | 6.07529 | 5.72643 | 5.83993 | 5.51975 | 7.13041 | 6.36191 | 7.08929 | 5.50469 | 7.66971 | 8.46883 | 7.24406 |
| ACBD4 | Q8NC06-3 | 5.76219 | 6.89779 | 5.37814 | 5.19655 | 5.47668 | 5.24433 | 7.38787 | 6.71839 | 5.7731 | 7.18569 | 5.69918 | 5.91434 | 6.60856 | 7.95482 | 6.87054 | 5.93181 |
| LAMA2 | P24043 | 6.81868 | 7.41766 | 5.96249 | 7.23836 | 6.96592 | 6.07632 | 6.50279 | 6.5907 | 6.27905 | 7.49382 | 7.67179 | 6.98733 | 3.96968 | 4.73418 | 4.1975 | 5.09374 |
| MTDH | E5RJU9 | 6.89317 | 5.75786 | 6.38792 | 6.0426 | 5.91263 | 4.4361 | 6.80656 | 4.45705 | 7.5674 | 7.4184 | 6.0689 | 7.79441 | 5.41841 | 6.16126 | 7.17752 | 5.6998 |
| MT-ND6 | P03923 | 5.81337 | 6.32912 | 6.39455 | 4.18071 | 6.86909 | 6.73387 | 6.72051 | 6.92577 | 5.15246 | 6.91038 | 6.0345 | 5.61824 | 7.05847 | 6.21087 | 5.98482 | 7.06326 |
| UBASH3A | P57075 | 5.46132 | 6.28938 | 6.34544 | 6.37463 | 6.29039 | 6.90198 | 5.37989 | 7.93887 | 6.72236 | 7.0232 | 5.75425 | 7.03561 | 5.31146 | 6.03869 | 5.36031 | 5.77223 |
| SORCS2 | Q96PQ0 | 5.4192 | 7.50425 | 5.8442 | 7.85808 | 5.79607 | 3.99822 | 6.47293 | 6.53043 | 5.19113 | 6.1821 | 5.8793 | 6.25428 | 7.14656 | 6.98643 | 7.19327 | 5.74354 |
| TMEM120B | A0PK00 | 5.863 | 6.52341 | 6.23774 | 6.77986 | 6.77841 | 5.67023 | 5.94621 | 5.28838 | 6.20511 | 5.61322 | 6.75125 | 5.32925 | 6.58291 | 6.72455 | 6.60203 | 7.10444 |
| YIPF1 | Q9Y548 | 5.73227 | 6.48472 | 6.30582 | 6.34763 | 5.69762 | 5.91978 | 6.34958 | 5.80087 | 5.91822 | 6.64282 | 5.9266 | 6.41983 | 6.59696 | 6.59191 | 6.25164 | 7.01374 |
| SLC39A13 | Q96H72 | 5.80249 | 5.85565 | 6.62331 | 6.90782 | 6.78731 | 5.72148 | 6.07773 | 4.90045 | 6.07357 | 5.0501 | 6.25518 | 6.88577 | 7.27179 | 5.56262 | 7.08812 | 7.1366 |
| SELENOM | Q8WWX9 | 4.83738 | 4.91143 | 4.47853 | 4.97172 | 5.75127 | 4.0874 | 5.91708 | 7.20369 | 4.1771 | 5.40342 | 4.90566 | 5.7545 | 4.80468 | 12.4551 | 10.1356 | 10.2054 |
| SHARPIN | Q9H0F6 | 3.65255 | 3.61466 | 6.28166 | 3.2272 | 6.17887 | 5.24842 | 5.09453 | 9.36453 | 6.31123 | 2.45977 | 5.54081 | 9.54797 | 4.40303 | 9.49234 | 8.92954 | 10.6529 |
| DYNLT2B | Q8WW35 | 5.61991 | 5.96891 | 6.48603 | 4.98185 | 5.67834 | 5.82898 | 6.73321 | 6.06578 | 4.83676 | 8.82013 | 7.59844 | 6.25515 | 5.80121 | 6.15906 | 7.00458 | 6.16166 |
| PALM | O75781 | 6.19194 | 6.00469 | 6.35506 | 4.46777 | 6.76721 | 7.27239 | 5.98449 | 5.63415 | 4.5978 | 6.4856 | 7.02354 | 6.62023 | 7.72562 | 7.34355 | 5.74468 | 5.78127 |
| PTP4A2 | Q12974 | 6.42258 | 5.92496 | 6.81188 | 5.73189 | 6.4074 | 5.90717 | 6.85981 | 6.51185 | 6.72323 | 6.68454 | 6.40358 | 6.75987 | 6.77811 | 5.47769 | 5.31222 | 5.28322 |
| PLEKHJ1 | Q9NW61 | 8.28697 | 5.80062 | 6.65808 | 4.5905 | 5.63902 | 7.73358 | 6.17869 | 6.69085 | 8.07073 | 7.87141 | 7.56395 | 5.32174 | 5.06667 | 6.23159 | 4.49885 | 4.44773 |
| TTBK2 | Q6IQ55 | 8.15849 | 5.12896 | 2.95544 | 7.6577 | 9.40544 | 5.14226 | 6.06001 | 6.60471 | 5.45311 | 5.65059 | 3.68344 | 6.73137 | 7.33743 | 5.47457 | 8.47231 | 6.08417 |
| PAIP2 | Q9BPZ3 | 7.55064 | 6.19898 | 6.19951 | 4.97216 | 5.68441 | 6.26316 | 5.33265 | 7.29858 | 6.21601 | 7.4794 | 6.96931 | 7.3471 | 5.09516 | 6.78244 | 4.89035 | 5.72017 |
| PRRG1 | O14668 | 6.13913 | 6.62248 | 4.47641 | 6.66064 | 7.02094 | 6.33472 | 6.39808 | 5.93789 | 5.91681 | 6.01955 | 6.36817 | 6.10681 | 5.77602 | 6.03419 | 7.31492 | 6.87325 |
| ABCA1 | O95477 | 6.2124 | 7.14966 | 5.67591 | 6.19482 | 6.5802 | 5.71712 | 6.0806 | 6.03851 | 6.00744 | 6.98021 | 5.93462 | 6.01157 | 6.04498 | 6.44726 | 6.32074 | 6.60396 |
| STOX2 | Q9P2F5 | 6.24 | 7.22974 | 7.53906 | 6.65033 | 5.24721 | 4.97917 | 6.62841 | 5.46121 | 6.48389 | 6.24063 | 7.9039 | 6.35396 | 6.33764 | 5.48202 | 6.51314 | 4.70971 |
| MON1A | Q86VX9 | 6.4992 | 6.1568 | 7.33476 | 5.78869 | 5.21096 | 6.43034 | 6.20836 | 6.33812 | 7.03434 | 6.76952 | 6.68559 | 6.65088 | 5.76641 | 6.22382 | 5.45958 | 5.44264 |
| PCNX1 | Q96RV3 | 7.43402 | 7.76256 | 4.17331 | 9.56313 | 5.49207 | 4.74167 | 7.0328 | 6.05758 | 6.47065 | 5.30919 | 5.91683 | 7.37793 | 5.03931 | 3.8142 | 5.16665 | 8.6481 |
| CDKN2A | P42771-4 | 6.92444 | 5.47091 | 6.11793 | 4.46083 | 5.47565 | 4.95263 | 4.77646 | 6.6682 | 4.78841 | 5.92949 | 5.66342 | 4.99364 | 4.35695 | 11.7194 | 8.0731 | 9.62856 |
| RHBDF1 | Q96CC6 | 6.10453 | 4.6802 | 5.14561 | 5.79073 | 6.3244 | 5.86172 | 5.73095 | 6.60584 | 8.07722 | 6.30923 | 6.276 | 6.67127 | 6.65378 | 6.84267 | 6.29406 | 6.6318 |
| TTC30B | Q8N4P2 | 6.39551 | 6.6264 | 7.9636 | 6.04124 | 6.17581 | 5.66039 | 6.25862 | 5.83631 | 6.88509 | 7.20503 | 7.03137 | 7.27921 | 4.76532 | 5.43581 | 5.71685 | 4.72345 |
| SH3BGR | P55822 | 6.01309 | 6.43637 | 5.76883 | 5.56941 | 7.24112 | 6.64041 | 5.84968 | 4.87132 | 6.62724 | 7.11929 | 6.25733 | 7.67614 | 6.2531 | 6.51537 | 6.42333 | 5.63796 |
| SORBS1 | Q9BX66-12 | 6.26166 | 5.4034 | 5.5725 | 6.22118 | 5.64108 | 5.49208 | 6.6957 | 5.90764 | 6.21227 | 6.94949 | 5.69742 | 6.25128 | 6.67578 | 6.63074 | 7.80155 | 7.58626 |
| ZNF764 | Q96H86 | 6.1597 | 6.51016 | 6.06078 | 6.44095 | 7.18899 | 6.33048 | 6.15452 | 5.58895 | 6.06891 | 6.08874 | 6.553 | 6.37116 | 6.48884 | 6.6968 | 5.81234 | 5.48569 |
| SNCA | P37840 | 6.53748 | 6.37925 | 6.79566 | 6.24582 | 5.04435 | 6.3375 | 5.82294 | 7.0128 | 6.32753 | 6.71244 | 7.37534 | 6.92587 | 6.29182 | 6.02924 | 5.81226 | 4.34972 |
| ALKBH3 | Q96Q83 | 6.23316 | 6.50741 | 7.28146 | 5.72099 | 6.3772 | 6.59139 | 6.27176 | 6.69813 | 6.60171 | 6.6873 | 7.02077 | 7.38706 | 5.38767 | 5.17989 | 4.92789 | 5.1262 |
| SNX18 | Q96RF0 | 6.76632 | 5.90861 | 7.03418 | 5.94575 | 6.64519 | 6.52534 | 6.19231 | 5.67223 | 7.05548 | 6.49914 | 7.17362 | 6.38288 | 6.13503 | 5.5352 | 5.66203 | 4.86668 |
| VDAC1 | C9JI87 | 5.66201 | 5.49855 | 4.44185 | 6.87354 | 6.12525 | 5.55158 | 6.1273 | 5.85058 | 7.45639 | 5.36728 | 5.28166 | 6.31036 | 6.83242 | 6.80592 | 8.7801 | 7.03521 |
| SEMA4G | Q9NTN9-2 | 6.31741 | 7.36066 | 5.15585 | 7.59959 | 6.73478 | 6.78295 | 5.28063 | 5.64515 | 7.4705 | 7.4439 | 6.43168 | 5.48062 | 5.77567 | 6.52851 | 5.26443 | 4.72769 |
| MAPKAP1 | H0Y866 | 5.56731 | 5.59679 | 5.58068 | 6.07534 | 6.75669 | 6.05973 | 6.21313 | 5.52162 | 7.8055 | 6.2149 | 4.89549 | 5.85041 | 6.45956 | 6.4473 | 7.89949 | 7.05608 |
| DNAAF3 | Q8N9W5-3 | 5.88261 | 5.85015 | 6.46811 | 5.10075 | 7.04681 | 6.31952 | 6.30876 | 6.11434 | 7.16177 | 7.40191 | 6.10191 | 6.47001 | 6.42268 | 5.95311 | 6.29011 | 5.10745 |
| C2CD3 | Q4AC94 | 7.5611 | 7.86571 | 4.81422 | 4.93004 | 5.83108 | 8.39438 | 6.34732 | 4.79366 | 5.44465 | 6.06107 | 5.93041 | 5.98666 | 7.42584 | 7.15254 | 6.46199 | 4.99933 |
| NEK2 | P51955 | 5.71514 | 6.57011 | 7.22862 | 7.4413 | 6.13682 | 6.76869 | 5.80367 | 5.62125 | 7.55303 | 8.40541 | 8.8882 | 8.12361 | 3.96476 | 3.88405 | 3.93082 | 3.96454 |
| TSPOAP1 | O95153-3 | 5.97616 | 6.05361 | 6.67206 | 5.50971 | 6.14024 | 4.68796 | 7.56321 | 6.23369 | 6.46129 | 6.59557 | 6.08671 | 7.34581 | 6.43628 | 6.08143 | 6.14139 | 6.0149 |
| TEDC2 | Q7L2K0 | 6.43478 | 6.13916 | 6.96142 | 6.21693 | 6.14205 | 6.12212 | 6.75941 | 5.82216 | 6.71366 | 6.73031 | 7.11057 | 7.01084 | 5.56083 | 5.45204 | 5.45882 | 5.36491 |
| SPIRE2 | Q8WWL2 | 6.97789 | 7.51015 | 6.27557 | 6.26324 | 5.77949 | 5.39316 | 5.38301 | 7.00768 | 7.35567 | 6.63854 | 5.65707 | 7.12841 | 6.40327 | 6.07541 | 5.7244 | 5.42703 |
| LRP8 | Q14114 | 6.79438 | 5.8316 | 7.0199 | 5.73585 | 7.63981 | 5.78565 | 5.47807 | 5.93166 | 6.55711 | 5.067 | 7.0329 | 6.6904 | 5.97329 | 5.04296 | 6.55877 | 6.86065 |
| ETFRF1 | Q6IPR1 | 6.39989 | 5.79774 | 5.74377 | 6.45587 | 6.66129 | 6.11715 | 6.6539 | 5.76745 | 5.47214 | 5.85216 | 6.22081 | 6.1004 | 8.07536 | 6.52298 | 5.73425 | 6.42482 |
| ATP5F1C | P36542-2 | 6.54279 | 6.41078 | 5.50812 | 7.37278 | 6.20116 | 5.71072 | 6.45498 | 6.30071 | 8.1925 | 6.21843 | 5.16851 | 4.57976 | 5.75061 | 6.05446 | 7.17377 | 6.35994 |
| ECM1 | Q16610-4 | 6.1957 | 6.91038 | 5.28892 | 6.26147 | 5.90917 | 6.57404 | 6.17805 | 6.4693 | 6.43294 | 6.61767 | 6.45729 | 6.92706 | 6.22099 | 5.7815 | 5.53085 | 6.24468 |
| PGAP2 | Q9UHJ9-2 | 6.12075 | 6.08758 | 5.90339 | 6.49306 | 6.67557 | 5.89299 | 6.0195 | 6.31546 | 7.03646 | 6.11944 | 5.63656 | 6.89021 | 6.4315 | 5.51097 | 5.98676 | 6.8798 |
| PLAAT3 | P53816 | 6.35395 | 5.58764 | 7.73682 | 5.66741 | 6.93954 | 7.14295 | 6.73323 | 6.93548 | 6.62286 | 6.83649 | 7.64397 | 7.36206 | 4.00957 | 5.71388 | 4.03159 | 4.58256 |
| ZNF692 | Q9BU19-5 | 6.5793 | 6.78833 | 6.27537 | 7.05036 | 5.04155 | 6.37572 | 6.52486 | 5.9584 | 6.51407 | 6.51866 | 5.28529 | 5.7278 | 5.92916 | 7.32362 | 5.87637 | 6.23114 |
| TSKU | Q8WUA8 | 5.79234 | 6.05369 | 5.12383 | 5.46345 | 6.85408 | 5.15878 | 6.14368 | 7.13701 | 5.49504 | 6.44231 | 6.59248 | 8.49924 | 6.27298 | 5.70152 | 6.70016 | 6.56943 |
| NGRN | Q9NPE2 | 6.28716 | 6.54574 | 6.55203 | 6.57176 | 6.76523 | 5.31606 | 7.31624 | 6.49684 | 6.09322 | 5.97423 | 6.8211 | 6.62705 | 5.9756 | 5.05777 | 6.05636 | 6.4436 |
| MED31 | I3L2J1 | 6.0066 | 4.46276 | 5.78454 | 5.65219 | 5.8544 | 4.19334 | 4.83211 | 6.78553 | 4.91067 | 6.60062 | 5.67512 | 7.75562 | 6.25048 | 7.56865 | 7.82862 | 9.83877 |
| ITFG2 | Q969R8 | 4.36763 | 5.08263 | 8.1251 | 6.40825 | 4.63349 | 7.6672 | 6.92012 | 5.68403 | 7.95387 | 7.86857 | 6.2328 | 4.81423 | 6.90647 | 6.19104 | 6.98088 | 4.16368 |
| LRRC9 | Q6ZRR7 | 6.24241 | 6.44014 | 5.95591 | 4.80426 | 6.84797 | 6.77718 | 5.94226 | 6.17654 | 7.55252 | 6.58984 | 7.24542 | 6.82735 | 5.59631 | 5.74622 | 5.71004 | 5.54566 |
| TMEM102 | Q8N9M5 | 7.81132 | 5.35381 | 7.80532 | 6.39117 | 6.42142 | 5.07221 | 5.61965 | 7.09542 | 5.22375 | 5.46975 | 5.71538 | 6.82398 | 4.80759 | 7.6105 | 7.55505 | 5.22367 |
| TK1 | K7ERV3 | 5.23638 | 7.10873 | 9.49484 | 4.97387 | 5.5467 | 6.00329 | 5.96894 | 5.41311 | 6.05895 | 6.01465 | 6.34629 | 6.03606 | 5.89813 | 7.11785 | 5.93623 | 6.84598 |
| COMTD1 | Q86VU5 | 7.0091 | 5.78122 | 6.44792 | 6.81262 | 5.88951 | 5.25381 | 7.70016 | 5.67079 | 5.95234 | 6.361 | 5.76878 | 5.05816 | 5.59412 | 6.27579 | 7.25674 | 7.16796 |
| CDPF1 | Q6NVV7 | 7.31836 | 5.1969 | 8.02235 | 4.7802 | 5.1318 | 7.00426 | 5.56259 | 4.46157 | 8.06087 | 5.07004 | 6.30904 | 7.36303 | 6.46224 | 5.08112 | 7.7847 | 6.39093 |
| MEIS2 | O14770 | 6.55306 | 6.21392 | 5.8555 | 4.93223 | 6.36387 | 5.50076 | 5.126 | 5.67828 | 6.77126 | 7.47844 | 5.52346 | 7.73704 | 5.89174 | 6.93848 | 6.05286 | 7.3831 |
| A0A1W2PSA9 | A0A1W2PSA9 | 6.48025 | 6.09019 | 5.84464 | 6.28863 | 5.52243 | 5.72315 | 6.24737 | 6.03217 | 6.15957 | 6.33166 | 5.15687 | 5.98676 | 6.94787 | 6.74656 | 7.16727 | 7.27461 |
| CYBRD1 | Q53TN4 | 6.05226 | 6.49721 | 5.37067 | 6.74952 | 6.30522 | 6.73804 | 6.70681 | 5.70589 | 6.29677 | 5.44314 | 6.21691 | 6.30098 | 6.2978 | 6.84371 | 6.85513 | 5.61995 |
| CYB5A | P00167 | 5.47974 | 5.89706 | 4.8675 | 4.77401 | 6.08569 | 5.20346 | 6.20708 | 7.12026 | 4.24469 | 4.49843 | 4.50517 | 5.43003 | 5.80202 | 11.0908 | 8.83211 | 9.9619 |
| AMMECR1L | Q6DCA0 | 4.66894 | 5.91245 | 6.02268 | 4.6267 | 5.62017 | 5.47232 | 4.46919 | 7.42412 | 9.90275 | 5.81413 | 4.99093 | 7.12628 | 5.53708 | 6.71866 | 8.3788 | 7.3148 |
| ETAA1 | Q9NY74 | 6.06633 | 6.40224 | 5.48737 | 6.17954 | 6.20393 | 7.32157 | 5.99081 | 5.64894 | 6.99496 | 6.40723 | 6.83384 | 6.36996 | 5.82654 | 5.43394 | 6.20291 | 6.83096 |
| SDCCAG8 | A0A0G2JQM2 | 6.32398 | 5.39506 | 6.46098 | 5.56909 | 5.86288 | 5.58403 | 6.80526 | 7.51775 | 6.90064 | 5.62413 | 6.25818 | 6.67437 | 6.63632 | 5.60941 | 6.27502 | 6.50289 |
| SDCCAG8 | Q86SQ7 | 6.35632 | 5.53137 | 6.23138 | 6.28631 | 5.84325 | 6.35243 | 6.18038 | 4.83553 | 5.49005 | 7.12254 | 6.60987 | 7.09814 | 8.06184 | 6.59176 | 5.55246 | 5.85638 |
| NAT1 | P18440 | 6.78789 | 5.65603 | 5.79566 | 6.35977 | 5.85905 | 7.54133 | 7.19207 | 6.34511 | 6.67926 | 7.28619 | 6.14272 | 6.83182 | 5.92887 | 5.71325 | 5.60248 | 4.3585 |
| SLC2A4 | P14672 | 7.10794 | 7.713 | 5.82301 | 7.25265 | 6.37564 | 5.63877 | 5.70412 | 6.65747 | 5.61538 | 8.03318 | 6.34875 | 6.13948 | 4.91361 | 5.10682 | 5.55789 | 6.01228 |
| CTHRC1 | Q96CG8 | 6.94462 | 6.33539 | 6.00934 | 6.894 | 5.74054 | 5.88112 | 6.75612 | 5.40264 | 5.90491 | 6.44393 | 5.94782 | 5.95405 | 6.49723 | 6.72423 | 6.14278 | 6.4213 |
| RNLS | Q5VYX0 | 5.29429 | 7.02511 | 6.23491 | 7.96577 | 6.75882 | 5.19335 | 6.99385 | 5.63246 | 5.99037 | 5.32348 | 7.05564 | 7.38835 | 5.0629 | 5.79778 | 6.22766 | 6.05524 |
| NAA38 | Q9BRA0-2 | 6.30107 | 6.26487 | 5.8648 | 4.89209 | 4.63668 | 5.46289 | 5.01698 | 8.23159 | 6.31047 | 7.87798 | 6.70403 | 6.7804 | 5.02003 | 6.98637 | 5.81064 | 7.8391 |
| NEMP2 | A6NFY4 | 5.9416 | 9.06006 | 6.03773 | 4.07273 | 7.41186 | 6.1112 | 8.46172 | 4.43244 | 7.78581 | 6.9401 | 4.3603 | 6.81953 | 3.56664 | 6.04851 | 6.59598 | 6.35379 |
| F2R | P25116 | 6.26982 | 7.28269 | 6.29482 | 7.4663 | 6.27559 | 6.56138 | 6.10678 | 6.31231 | 6.30191 | 6.2076 | 6.07877 | 5.84697 | 5.3947 | 5.68663 | 5.71752 | 6.19621 |
| KLHL26 | Q53HC5 | 6.50525 | 5.90868 | 6.85076 | 5.95592 | 6.03571 | 6.48264 | 6.00419 | 7.3216 | 6.32969 | 5.72822 | 6.47807 | 6.89492 | 5.88123 | 5.93185 | 6.00667 | 5.68459 |
| VPS29 | Q9UBQ0 | 5.78066 | 6.09028 | 6.1149 | 5.92117 | 6.09637 | 6.29111 | 5.6554 | 6.08205 | 6.55082 | 6.42546 | 6.25606 | 6.00857 | 6.38779 | 6.54401 | 7.13902 | 6.65633 |
| HENMT1 | Q5T8I9 | 4.95742 | 7.84345 | 5.62365 | 7.43992 | 5.06347 | 5.76979 | 7.2673 | 8.40655 | 6.14599 | 5.76284 | 6.87139 | 6.00652 | 6.59229 | 6.70559 | 5.35251 | 4.19131 |
| PRELID1 | Q9Y255 | 5.76253 | 5.78363 | 4.5158 | 6.50037 | 5.75097 | 7.27746 | 7.58297 | 7.67744 | 8.10615 | 9.7394 | 8.80327 | 8.99319 | 2.45175 | 4.42543 | 2.5585 | 4.07116 |
| LRRC23 | Q53EV4 | 6.5016 | 7.35518 | 6.16403 | 6.3697 | 8.14389 | 6.16366 | 5.97685 | 6.99655 | 5.4114 | 6.26719 | 7.07056 | 7.36926 | 5.28265 | 5.16209 | 3.61902 | 6.14637 |
| ZNF324 | O75467 | 6.10108 | 6.01911 | 6.44937 | 6.63293 | 6.95858 | 6.4273 | 6.53703 | 5.68514 | 6.11988 | 5.47719 | 5.9501 | 5.7363 | 6.93419 | 6.56569 | 7.30898 | 5.09711 |
| TMSB10 | P63313 | 5.34879 | 5.79394 | 4.46245 | 5.57105 | 4.79127 | 4.57486 | 5.0071 | 6.83166 | 3.84459 | 5.16072 | 4.30069 | 5.68271 | 3.37852 | 14.6365 | 7.00097 | 13.6142 |
| TNFRSF10D | Q9UBN6 | 8.10546 | 7.36715 | 7.19668 | 7.61629 | 6.79665 | 6.17273 | 7.1444 | 5.76961 | 7.17596 | 6.43797 | 6.83522 | 5.6827 | 4.6213 | 4.07867 | 3.98177 | 4.11744 |
| PFDN6 | O15212 | 4.88733 | 4.62319 | 5.21187 | 5.00597 | 4.97604 | 4.3183 | 3.97875 | 7.53212 | 3.98733 | 6.39957 | 4.58367 | 6.42351 | 4.80098 | 12.6633 | 8.85371 | 11.7543 |
| MED22 | Q15528 | 5.55331 | 4.62159 | 5.14789 | 4.05641 | 5.17856 | 4.53136 | 4.89463 | 6.22173 | 4.54432 | 5.802 | 4.78377 | 5.49714 | 5.41701 | 11.7674 | 10.3793 | 11.6036 |
| CERK | Q8TCT0 | 7.3806 | 7.22731 | 8.34249 | 7.20171 | 5.53264 | 5.83804 | 6.0188 | 6.23247 | 8.73251 | 6.39603 | 8.17526 | 6.17164 | 3.70407 | 4.32563 | 4.43247 | 4.28831 |
| PGBD4 | Q96DM1 | 5.99955 | 6.22454 | 5.58655 | 5.7383 | 5.9917 | 6.23898 | 6.29343 | 6.18818 | 5.213 | 6.38731 | 5.99537 | 6.45614 | 6.40174 | 6.73747 | 7.60805 | 6.93968 |
| TRAK2 | O60296 | 6.02327 | 6.98818 | 6.30338 | 6.98179 | 6.49386 | 5.81974 | 7.97394 | 8.97623 | 7.35496 | 4.37863 | 6.95013 | 7.42252 | 5.10035 | 5.28929 | 4.81003 | 3.1337 |
| CCDC66 | A2RUB6 | 6.23454 | 6.20037 | 6.30062 | 6.63623 | 7.05661 | 7.20044 | 5.95282 | 7.29128 | 6.95762 | 6.13143 | 7.04579 | 6.95965 | 4.54301 | 5.55172 | 4.68481 | 5.25304 |
| POLR3K | Q9Y2Y1 | 6.04826 | 5.18242 | 6.41347 | 5.51079 | 6.41946 | 5.54652 | 6.18316 | 7.24431 | 5.15375 | 7.32551 | 6.67436 | 7.61019 | 5.73078 | 6.33823 | 6.47411 | 6.14467 |
| DKK3 | Q9UBP4 | 8.60488 | 6.73544 | 7.84689 | 6.80131 | 7.61706 | 6.06807 | 8.25283 | 7.06096 | 7.47581 | 7.27825 | 7.29083 | 6.32668 | 3.61403 | 3.56239 | 2.37861 | 3.08595 |
| TRMT44 | Q8IYL2 | 6.47415 | 6.52491 | 7.27244 | 6.86019 | 7.32715 | 7.80101 | 6.35336 | 5.61145 | 6.59396 | 5.37177 | 6.67694 | 5.66512 | 5.40179 | 5.75963 | 4.07078 | 6.23536 |
| KBTBD8 | Q8NFY9 | 6.1138 | 6.21884 | 6.88179 | 6.40203 | 5.82589 | 6.97297 | 6.00472 | 7.05262 | 6.63514 | 6.4461 | 6.97878 | 6.85301 | 4.94584 | 5.35065 | 5.44213 | 5.87568 |
| CABYR | O75952 | 7.10291 | 7.32085 | 6.05182 | 6.78367 | 5.68782 | 6.1897 | 6.33061 | 6.81035 | 5.47998 | 5.57437 | 8.50836 | 8.22804 | 5.58472 | 3.96798 | 5.39642 | 4.98241 |
| CAPG | P40121 | 6.20122 | 4.87732 | 7.04086 | 6.54763 | 6.2448 | 6.27596 | 6.20797 | 6.25835 | 6.90509 | 6.70766 | 9.21958 | 8.54566 | 5.47645 | 3.77384 | 5.6341 | 4.08351 |
| DBF4B | Q8NFT6-3 | 5.51412 | 6.17156 | 6.30972 | 6.4854 | 6.37839 | 6.29689 | 4.4598 | 7.40516 | 5.86338 | 6.40754 | 7.52993 | 6.80712 | 5.99635 | 5.97329 | 6.31093 | 6.09042 |
| MOSPD1 | Q9UJG1-3 | 6.05789 | 6.08777 | 5.96794 | 6.12818 | 6.65163 | 5.84961 | 6.37241 | 6.29277 | 6.09487 | 5.79299 | 6.19205 | 5.8721 | 6.74582 | 6.72604 | 6.80569 | 6.36223 |
| GSTA4 | O15217 | 3.69343 | 7.47527 | 8.50167 | 5.4789 | 5.92591 | 7.9903 | 7.03775 | 7.32551 | 7.81727 | 5.7943 | 6.18886 | 5.37665 | 4.97664 | 5.5784 | 5.02801 | 5.81114 |
| TCEA3 | O75764 | 5.88182 | 6.35246 | 6.74071 | 6.40377 | 6.25851 | 6.16043 | 6.17971 | 5.5524 | 6.35677 | 6.91099 | 6.39043 | 5.91504 | 5.63249 | 7.92048 | 5.61598 | 5.72801 |
| NUDT11 | Q96G61 | 6.5987 | 7.27781 | 6.41481 | 5.67365 | 7.07286 | 6.57645 | 5.40951 | 7.56517 | 6.93027 | 7.31565 | 7.15332 | 6.13186 | 5.4323 | 5.17671 | 4.47719 | 4.79373 |

| | | | | | | | | | | | | | | | | | |
|---|---|---|---|---|---|---|---|---|---|---|---|---|---|---|---|---|---|
| APOC1 | P02654 | 7.26046 | 6.15744 | 4.89344 | 6.48297 | 5.98798 | 4.44296 | 5.65463 | 5.48727 | 4.70119 | 6.15937 | 5.67593 | 4.89152 | 3.39198 | 10.9958 | 6.68641 | 11.1306 |
| NOS1AP | O75052 | 6.41815 | 7.11624 | 6.7559 | 6.46278 | 5.72853 | 5.96031 | 7.49433 | 5.93318 | 5.90914 | 7.63201 | 6.08024 | 5.14323 | 5.84592 | 4.92115 | 7.06512 | 5.53378 |
| LAPTM4A | Q15012 | 6.85841 | 7.27304 | 5.5402 | 6.65562 | 6.48367 | 7.1572 | 7.98676 | 5.61175 | 5.87631 | 6.06824 | 6.87287 | 7.49369 | 4.53575 | 5.79862 | 5.01941 | 4.76845 |
| EIF1B | O60739 | 6.18236 | 5.47344 | 7.39038 | 6.30629 | 6.92824 | 6.97629 | 5.09185 | 5.8285 | 5.64685 | 6.66641 | 6.66643 | 7.01422 | 5.94897 | 5.78276 | 5.72682 | 6.37021 |
| SULT1A3 | P0DMM9-3 | 5.01124 | 4.83185 | 6.72981 | 4.57661 | 7.38046 | 6.20683 | 5.26592 | 8.91184 | 6.98547 | 7.4491 | 6.26992 | 6.48918 | 7.13173 | 5.5919 | 5.67035 | 5.4978 |
| SDHAF4 | Q5VUM1 | 6.1682 | 5.34211 | 5.78839 | 5.27026 | 7.76031 | 6.3188 | 5.72546 | 6.91078 | 5.87287 | 6.32189 | 5.37379 | 6.07808 | 6.53892 | 7.02161 | 6.35065 | 7.15789 |
| RAP1GAP2 | Q684P5 | 5.23542 | 7.97744 | 8.34268 | 6.73065 | 6.23771 | 5.93277 | 7.65294 | 7.36791 | 5.53926 | 5.50728 | 5.93856 | 6.46141 | 5.52655 | 4.70193 | 5.10898 | 5.73849 |
| AGAP1 | Q9UPQ3 | 6.59007 | 6.65645 | 6.9218 | 5.41616 | 4.6162 | 4.62989 | 7.19751 | 4.63754 | 6.81317 | 6.41276 | 6.61958 | 7.84426 | 6.59801 | 8.44645 | 5.55272 | 5.04742 |
| TFPI | P10646 | 6.40596 | 6.15747 | 5.75163 | 6.74057 | 6.0309 | 6.16324 | 6.27089 | 5.9317 | 6.17097 | 6.45971 | 6.56951 | 6.04473 | 6.52053 | 5.31492 | 7.33205 | 6.13523 |
| USB1 | Q9BQ65 | 5.89237 | 6.35511 | 6.64578 | 5.75346 | 6.16473 | 6.44468 | 6.00145 | 6.10775 | 6.51716 | 6.35851 | 6.28224 | 6.58303 | 5.74178 | 5.95862 | 7.09473 | 6.09859 |
| FZD7 | O75084 | 7.63996 | 7.37436 | 5.76975 | 6.51234 | 4.80858 | 5.38247 | 7.12799 | 6.05299 | 6.43681 | 5.55131 | 6.63852 | 6.04037 | 5.83637 | 6.24591 | 5.94612 | 6.63613 |
| PIGX | Q8TBF5-2 | 6.44175 | 6.92199 | 6.13225 | 7.37237 | 5.77851 | 5.88135 | 5.86709 | 5.29431 | 5.5627 | 7.43643 | 7.42773 | 5.70569 | 6.58598 | 5.89824 | 6.49256 | 5.20104 |
| USP31 | Q70CQ4 | 6.82533 | 6.71454 | 6.22625 | 6.65954 | 6.60476 | 6.03141 | 5.78681 | 5.74091 | 5.51373 | 5.99735 | 6.53722 | 6.03128 | 6.37579 | 6.39168 | 6.32677 | 6.23662 |
| SPRED3 | Q2MJR0 | 7.16764 | 5.12626 | 5.78031 | 7.61652 | 3.54831 | 6.59157 | 6.84396 | 5.20209 | 5.88915 | 7.78107 | 7.98449 | 6.46863 | 6.42625 | 4.6238 | 6.28479 | 6.66516 |
| MPLKIP | Q8TAP9 | 6.24552 | 4.88496 | 7.60225 | 3.69159 | 6.87738 | 6.42843 | 6.1906 | 5.25815 | 7.5126 | 6.95517 | 7.03964 | 6.38685 | 5.58641 | 4.98179 | 6.16891 | 8.18976 |
| TMEM17 | Q86X19 | 5.56721 | 5.96898 | 5.03562 | 6.54613 | 6.36179 | 6.12178 | 5.99496 | 6.17955 | 5.46316 | 5.09332 | 6.01287 | 6.03229 | 6.61186 | 7.79414 | 6.76637 | 8.44997 |
| DNAJC24 | Q6P3W2 | 6.4442 | 5.68172 | 7.05998 | 5.34591 | 6.0993 | 6.44867 | 5.57661 | 6.8734 | 5.60451 | 7.14044 | 7.15036 | 7.58427 | 4.79473 | 6.07856 | 5.60181 | 6.51555 |
| TRAPPC2 | P0DI81-3 | 4.90641 | 5.84042 | 6.63659 | 5.21052 | 6.4909 | 6.765 | 5.5551 | 5.90733 | 5.40331 | 6.52338 | 6.87582 | 6.45315 | 5.29892 | 7.82979 | 6.79887 | 7.5045 |
| TASP1 | Q9H6P5 | 7.22238 | 7.03706 | 5.87517 | 5.7926 | 6.17083 | 5.78147 | 6.07719 | 8.00524 | 6.14247 | 5.43668 | 5.64271 | 5.71996 | 6.9424 | 5.57973 | 7.22072 | 5.35339 |
| ARHGAP24 | Q8N264 | 5.89204 | 5.71626 | 6.34417 | 5.88324 | 6.10886 | 7.44935 | 5.50303 | 6.9249 | 6.09865 | 6.54339 | 6.74529 | 5.97681 | 6.68267 | 7.65311 | 5.42965 | 5.94857 |
| ZCCHC14 | Q8WYQ9-2 | 5.65021 | 5.76859 | 7.42156 | 6.34589 | 5.57461 | 8.02712 | 6.73052 | 6.74022 | 5.2326 | 6.87111 | 6.93829 | 8.50516 | 5.49033 | 4.89759 | 4.27848 | 5.52772 |
| TLNRD1 | Q9H1K6 | 6.04884 | 6.5422 | 5.28513 | 5.40833 | 6.92699 | 6.05002 | 6.24157 | 6.35841 | 6.54989 | 8.27013 | 6.24968 | 5.42468 | 5.85152 | 6.50787 | 6.19866 | 6.08609 |
| MAP1LC3A | Q9H492-2 | 5.35327 | 5.02829 | 6.53221 | 5.65816 | 6.94427 | 6.44174 | 6.27466 | 6.46433 | 7.14231 | 5.07291 | 5.8873 | 6.86478 | 5.8334 | 6.62781 | 6.97176 | 6.90281 |
| RRNAD1 | Q96FB5 | 5.76819 | 4.97302 | 8.32561 | 6.93071 | 5.35814 | 5.85193 | 5.94138 | 6.29949 | 7.19915 | 6.15161 | 7.6454 | 7.12559 | 5.80171 | 5.33685 | 5.73489 | 5.55633 |
| ZNF581 | Q9P0T4 | 5.90878 | 6.21791 | 5.66827 | 7.12819 | 5.84056 | 5.56638 | 6.37931 | 5.76133 | 6.64619 | 6.64941 | 7.04307 | 7.02585 | 6.67359 | 5.17453 | 6.27571 | 6.04093 |
| RIBC2 | Q9H4K1 | 6.95991 | 6.2129 | 6.4328 | 6.77853 | 6.69078 | 6.23353 | 6.25266 | 6.4248 | 6.84986 | 6.86238 | 7.5315 | 6.36164 | 5.13528 | 4.60748 | 5.23154 | 5.43443 |
| ZBED6 | P86452 | 5.35063 | 5.90501 | 6.10696 | 4.48132 | 6.27034 | 7.16097 | 5.45853 | 6.90405 | 6.28428 | 6.03825 | 5.69897 | 6.59172 | 8.04944 | 7.73408 | 6.49641 | 5.46905 |
| RPL22L1 | Q6P5R6 | 7.31378 | 7.58687 | 8.53219 | 8.6382 | 5.93027 | 5.87845 | 6.54853 | 4.80728 | 5.97692 | 7.09588 | 7.55815 | 7.34134 | 4.17264 | 4.20865 | 3.96779 | 4.44307 |
| ZNF407 | Q9C0G0 | 5.64636 | 6.21268 | 5.28898 | 5.33996 | 6.1946 | 6.15555 | 6.09792 | 6.57483 | 6.92835 | 6.88859 | 5.51734 | 5.71506 | 6.3918 | 6.4977 | 7.37536 | 7.17493 |
| TFPI2 | P48307 | 7.09558 | 6.36443 | 5.40664 | 8.096 | 5.21477 | 6.77228 | 7.15409 | 6.84536 | 7.03296 | 6.85354 | 5.6711 | 6.36106 | 5.65084 | 4.15167 | 5.91781 | 5.41185 |
| AP3S2 | P59780 | 6.94927 | 8.64615 | 7.18903 | 5.13612 | 5.07945 | 6.77962 | 8.29683 | 6.1223 | 7.65273 | 5.4954 | 6.81546 | 6.21712 | 4.33904 | 4.09829 | 6.57752 | 4.60568 |
| SLC35C2 | Q9NQQ7-3 | 6.15949 | 6.73546 | 6.20107 | 6.02758 | 5.76748 | 7.04927 | 7.12632 | 6.91232 | 7.12458 | 4.80269 | 5.32999 | 5.56543 | 6.22 | 6.53997 | 6.83009 | 5.60829 |
| CIPC | Q9C0C6 | 5.94665 | 6.52281 | 5.68142 | 6.28772 | 6.30087 | 6.21127 | 5.79404 | 5.63602 | 6.14574 | 7.16712 | 6.19112 | 6.50341 | 6.45009 | 6.55986 | 5.9547 | 6.64715 |
| IER2 | Q9BTL4 | 4.32769 | 5.18629 | 4.85765 | 5.42941 | 6.0782 | 5.0429 | 6.15901 | 6.26354 | 4.90213 | 4.89709 | 4.661 | 5.79503 | 9.05131 | 8.47717 | 9.57098 | 9.30059 |
| PLSCR3 | Q9NRY6 | 5.23463 | 5.9713 | 4.78443 | 6.2564 | 6.46069 | 4.78119 | 5.89231 | 5.54154 | 6.39586 | 6.36223 | 6.59385 | 6.69073 | 7.57236 | 7.74143 | 5.622 | 8.09907 |
| MITD1 | Q8WV92 | 5.79361 | 6.85667 | 7.18341 | 6.12191 | 6.8969 | 6.31236 | 6.11529 | 6.15165 | 5.64847 | 6.96667 | 6.56381 | 6.04706 | 5.77478 | 5.58123 | 6.02033 | 6.96585 |
| MICU3 | Q86XE3 | 7.40689 | 7.10052 | 5.40394 | 7.36011 | 7.06952 | 6.0285 | 5.4991 | 6.19328 | 5.8399 | 6.15425 | 5.05538 | 6.44993 | 6.31984 | 6.2985 | 6.13667 | 5.68366 |
| INO80C | Q6PI98-4 | 4.33336 | 5.83597 | 4.87671 | 7.79769 | 6.60199 | 6.34275 | 6.00609 | 7.05882 | 5.3411 | 6.48757 | 8.13017 | 6.86245 | 4.84169 | 7.04444 | 6.11643 | 6.32278 |
| SLC39A8 | Q9C0K1 | 6.19336 | 6.77572 | 6.83673 | 7.17297 | 5.72324 | 5.88598 | 6.88605 | 5.93372 | 5.92409 | 6.37334 | 6.45659 | 5.89743 | 5.89736 | 6.30762 | 6.26751 | 5.46829 |
| CES2 | O00748-4 | 7.20934 | 6.59514 | 6.18303 | 6.03589 | 5.34519 | 5.68687 | 6.7346 | 8.46475 | 5.42243 | 4.61207 | 5.38067 | 5.46775 | 7.11332 | 7.34814 | 5.98381 | 6.417 |
| AGGF1 | Q8N302 | 6.0529 | 6.33699 | 5.6013 | 6.44213 | 5.30271 | 6.19577 | 6.32555 | 5.9614 | 6.00161 | 6.52285 | 6.49545 | 6.6169 | 6.5795 | 6.43325 | 6.81851 | 6.31017 |
| HMBOX1 | Q6NT76-5 | 5.93565 | 6.26077 | 6.16298 | 6.36113 | 6.43988 | 6.63676 | 6.04532 | 6.1461 | 6.23162 | 6.75958 | 6.60517 | 5.78459 | 6.05788 | 6.09864 | 6.65524 | 5.8187 |
| EFNA1 | P20827 | 7.3151 | 8.37737 | 7.48628 | 8.56173 | 6.87951 | 5.60666 | 6.57944 | 6.52815 | 5.72803 | 7.7378 | 6.94972 | 7.15938 | 4.30135 | 4.07163 | 3.84042 | 2.87743 |
| B9D1 | Q9UPM9 | 6.215 | 5.68246 | 6.43629 | 5.92375 | 5.40223 | 5.52423 | 5.66187 | 7.08607 | 6.80337 | 6.32433 | 6.28519 | 6.64415 | 6.30177 | 6.75698 | 7.01889 | 5.93582 |
| COL3A1 | P02461 | 6.59204 | 6.29521 | 5.61264 | 5.9397 | 4.66899 | 4.94227 | 4.90441 | 4.65131 | 6.91356 | 5.70361 | 6.40664 | 5.51675 | 7.98707 | 7.43778 | 9.24871 | 7.1793 |
| PODXL2 | Q9NZ53 | 6.45897 | 5.68531 | 5.8535 | 7.65119 | 6.31717 | 6.15067 | 6.2738 | 5.67376 | 6.43264 | 6.29119 | 7.67058 | 7.73402 | 5.61634 | 5.20862 | 5.30007 | 5.68215 |
| ERCC1 | P07992-3 | 6.12156 | 5.73862 | 6.50413 | 5.02241 | 6.49244 | 6.82331 | 6.78592 | 6.42876 | 6.38451 | 6.23034 | 7.64947 | 6.73633 | 5.48954 | 5.95966 | 5.79547 | 5.83753 |
| ARNTL2 | Q8WYA1 | 5.89538 | 6.91746 | 5.90567 | 7.21765 | 6.84735 | 6.41769 | 6.53987 | 7.25857 | 7.41275 | 6.03283 | 6.13914 | 5.25315 | 4.99629 | 5.68728 | 5.46922 | 6.00969 |
| TM2D1 | Q9BX74 | 4.94744 | 4.88796 | 5.65059 | 7.23665 | 6.25682 | 6.99606 | 4.93714 | 6.94542 | 7.85085 | 7.37658 | 5.95614 | 6.55638 | 5.24018 | 6.79032 | 5.98139 | 6.39006 |
| SPA17 | Q15506 | 7.1816 | 6.52604 | 7.04044 | 5.44128 | 7.16694 | 6.79174 | 5.89524 | 7.45828 | 6.41917 | 6.47472 | 6.58967 | 7.66277 | 4.79693 | 5.38862 | 4.24667 | 4.91989 |
| UBXN8 | O00124 | 4.84256 | 5.6807 | 5.69671 | 7.65266 | 5.57429 | 6.15983 | 6.3824 | 6.74202 | 4.71889 | 7.12935 | 6.62316 | 6.30302 | 7.18653 | 5.84734 | 7.2616 | 6.19894 |
| C18orf32 | Q8TCD1 | 6.3555 | 5.78856 | 5.16713 | 5.13039 | 5.59189 | 4.45795 | 5.39998 | 5.62146 | 5.34204 | 5.78689 | 5.70701 | 5.87763 | 6.08554 | 9.02207 | 8.65016 | 10.0158 |
| ATAT1 | Q5SQI0 | 5.79725 | 6.60747 | 6.1821 | 5.36164 | 6.1178 | 7.21705 | 6.2739 | 6.4188 | 6.36761 | 6.83935 | 6.35154 | 6.59734 | 6.95706 | 4.95595 | 5.73903 | 6.2161 |
| MOSPD3 | O75425 | 5.85972 | 5.50048 | 6.35396 | 6.09762 | 5.71649 | 5.67238 | 5.09385 | 5.74688 | 6.71795 | 5.42864 | 6.15838 | 7.37878 | 7.45973 | 6.02799 | 6.45872 | 8.32843 |
| ABITRAM | Q9NX38 | 5.25977 | 5.75992 | 6.9459 | 5.42957 | 5.75921 | 6.50654 | 6.27504 | 5.15297 | 6.36549 | 6.19495 | 6.56857 | 6.59959 | 6.50795 | 6.47153 | 7.03187 | 7.17112 |
| ARNTL | O00327 | 5.59959 | 5.86581 | 5.95907 | 6.63046 | 6.10093 | 6.37619 | 5.99123 | 5.96738 | 5.8954 | 6.61251 | 6.03601 | 5.93579 | 6.77183 | 7.05206 | 6.31262 | 6.89311 |
| FOXP2 | O15409-4 | 5.90422 | 6.48237 | 6.41591 | 5.79285 | 6.37975 | 6.32857 | 5.91103 | 6.55655 | 5.87143 | 6.11817 | 6.16294 | 5.76146 | 6.7711 | 6.96745 | 6.86118 | 5.71502 |
| HOXC10 | Q9NYD6 | 5.49244 | 6.19575 | 4.99294 | 6.28614 | 6.52838 | 6.48157 | 7.87527 | 5.66185 | 7.69769 | 7.32986 | 5.35845 | 6.23939 | 6.05082 | 6.05939 | 6.27968 | 5.47037 |
| LRR1 | Q96L50 | 5.8552 | 6.48151 | 7.23925 | 6.19609 | 5.81502 | 6.4248 | 5.81611 | 5.76225 | 6.65203 | 7.0058 | 7.16291 | 6.41168 | 5.54956 | 5.19162 | 5.93681 | 6.49935 |
| B3GNT9 | Q6UX72 | 9.26034 | 5.52191 | 4.24871 | 7.08741 | 5.69457 | 5.56619 | 6.46652 | 5.15591 | 9.09312 | 7.46427 | 4.01494 | 7.15639 | 6.59297 | 6.03117 | 5.60888 | 5.08156 |
| ATRAID | Q6UW56-3 | 5.61897 | 6.7266 | 6.3608 | 5.726 | 6.70479 | 5.88471 | 6.15549 | 5.59619 | 6.45378 | 6.38343 | 6.66464 | 6.14361 | 5.89761 | 6.25596 | 7.48688 | 5.94054 |
| CFAP251 | Q8TBY9 | 9.06185 | 5.97626 | 5.82333 | 5.65436 | 7.91795 | 6.56076 | 7.28675 | 6.05109 | 5.57973 | 6.50545 | 6.36716 | 7.60019 | 5.44795 | 3.75913 | 4.38075 | 6.0273 |
| TNFAIP8L1 | Q8WVP5 | 5.29871 | 5.76075 | 5.70496 | 5.45624 | 5.93723 | 6.94526 | 7.35849 | 7.15464 | 6.40474 | 7.2513 | 7.93944 | 7.79393 | 5.95871 | 5.4943 | 3.96293 | 5.5784 |
| TF | P02787 | 5.73132 | 6.51263 | 6.27168 | 5.84821 | 5.26827 | 6.81432 | 5.81652 | 6.40872 | 5.57308 | 4.7876 | 6.14507 | 7.44999 | 5.21118 | 6.63138 | 9.1793 | 6.35073 |
| CNTN3 | Q9P232 | 5.82988 | 5.8412 | 6.84366 | 6.01765 | 5.66763 | 5.80209 | 6.52669 | 7.23297 | 6.43428 | 7.13489 | 6.62679 | 6.9052 | 5.61357 | 5.94123 | 5.47678 | 6.10548 |
| ERN1 | O75460 | 6.30467 | 6.48084 | 6.27544 | 7.19903 | 6.70305 | 6.15333 | 6.86702 | 6.04954 | 6.09894 | 6.44177 | 5.9199 | 5.88346 | 6.04614 | 5.56009 | 5.7899 | 6.22689 |
| CD82 | P27701 | 6.6783 | 6.96245 | 6.28081 | 5.81154 | 5.77827 | 5.59258 | 5.8757 | 6.20285 | 6.69038 | 5.80498 | 6.86518 | 5.89336 | 5.55316 | 6.75586 | 6.62112 | 6.63346 |
| SKAP2 | O75563 | 6.90902 | 7.89557 | 7.80125 | 6.83771 | 6.27555 | 8.14268 | 6.10169 | 5.9405 | 6.49304 | 6.75495 | 7.66361 | 7.16428 | 4.56627 | 3.60109 | 3.96024 | 3.89255 |
| IFT57 | Q9NWB7 | 6.10131 | 6.67407 | 6.43361 | 5.25682 | 4.97439 | 6.97857 | 6.06841 | 7.60546 | 5.5979 | 6.74232 | 5.67949 | 6.59141 | 7.39417 | 4.66603 | 7.09722 | 6.13881 |
| SS18 | Q15532 | 6.83802 | 3.41583 | 4.78684 | 3.70522 | 4.62219 | 3.63962 | 3.69785 | 4.94553 | 3.14638 | 6.18333 | 4.90879 | 5.60024 | 6.30186 | 12.9766 | 11.0164 | 14.2153 |
| NUAK1 | O60285 | 5.62323 | 6.10265 | 6.68651 | 4.80406 | 5.38083 | 7.28485 | 7.28778 | 4.8791 | 5.14003 | 7.14669 | 7.13873 | 6.34415 | 6.84742 | 4.88317 | 7.35354 | 7.09728 |
| ANKRD40 | Q6AI12 | 6.19787 | 5.99438 | 6.8927 | 4.16794 | 6.51262 | 6.45547 | 5.42756 | 6.58854 | 6.35285 | 6.50432 | 6.79579 | 7.15371 | 5.3476 | 6.12054 | 7.27751 | 6.21058 |
| KATNIP | O60303 | 7.01641 | 6.49142 | 5.87256 | 6.2481 | 5.85658 | 8.78079 | 5.06925 | 6.90131 | 6.31124 | 4.65014 | 8.69565 | 7.03064 | 5.70882 | 4.4082 | 5.09531 | 5.86357 |
| ZDHHC23 | Q8IYP9 | 5.66052 | 6.22443 | 6.49101 | 6.35799 | 6.51981 | 5.91297 | 7.92076 | 5.71338 | 7.72539 | 5.03405 | 4.87054 | 5.47647 | 4.60272 | 5.87182 | 6.32324 | 9.2949 |
| ZDHHC23 | A0A1W2PRJ8 | 5.83121 | 6.08141 | 6.21241 | 6.40756 | 5.49211 | 5.45042 | 6.1372 | 7.0494 | 6.05698 | 5.8708 | 6.11061 | 6.99703 | 6.34783 | 6.43965 | 6.87732 | 6.63808 |
| NKX2-4 | Q9H2Z4 | 6.26004 | 6.88557 | 6.12458 | 6.92623 | 6.74048 | 7.40728 | 6.22961 | 5.99786 | 6.17039 | 6.03694 | 6.39705 | 5.92814 | 6.37151 | 5.40299 | 6.11763 | 5.00371 |
| N4BP2L1 | Q5TBK1 | 5.84256 | 6.40462 | 6.25837 | 6.24313 | 5.00558 | 7.14226 | 6.61791 | 7.12213 | 5.99469 | 6.51819 | 5.83938 | 7.52526 | 5.69379 | 5.16195 | 5.81214 | 6.81805 |
| ZCCHC2 | Q9C0B9 | 6.67949 | 6.55399 | 8.25142 | 6.37387 | 5.58731 | 6.80756 | 6.35925 | 6.4554 | 7.71097 | 7.34802 | 7.52819 | 7.10748 | 3.97014 | 4.70727 | 4.18373 | 4.37592 |
| FUT4 | P22083 | 6.74048 | 5.69598 | 6.15509 | 6.71707 | 6.1192 | 5.72211 | 6.43599 | 6.18308 | 5.79705 | 6.07287 | 5.89906 | 6.78653 | 6.83268 | 6.01093 | 7.2795 | 5.55238 |
| CDC42SE2 | Q9NRR3 | 5.93834 | 6.09426 | 6.48743 | 5.80595 | 5.6128 | 6.07812 | 5.71487 | 6.28137 | 6.07783 | 5.55636 | 6.25632 | 6.28826 | 6.26425 | 6.94556 | 7.05022 | 7.54806 |
| ANKRD39 | Q53RE8 | 5.81211 | 4.61864 | 6.592 | 4.51081 | 6.58446 | 5.72729 | 5.35577 | 7.70116 | 5.33948 | 7.15185 | 6.57882 | 7.89554 | 4.29412 | 7.33301 | 6.69227 | 7.81267 |
| ZBED8 | Q8IZ13 | 6.13275 | 6.5698 | 6.32333 | 5.67996 | 6.56589 | 6.82222 | 6.14207 | 5.9447 | 6.58301 | 6.34357 | 6.34545 | 7.0395 | 5.76071 | 6.47059 | 6.26774 | 5.90871 |
| IFRD1 | A0A6Q8PF03 | 7.49382 | 7.17321 | 6.49006 | 5.5714 | 4.80227 | 5.04826 | 5.95877 | 6.08696 | 5.17767 | 5.01174 | 5.26482 | 5.39161 | 7.36657 | 5.97231 | 8.13001 | 9.06054 |
| SUFU | Q9UMX1 | 5.24386 | 6.36772 | 6.77573 | 6.27311 | 6.86523 | 7.4733 | 7.21502 | 5.57834 | 5.93863 | 7.70883 | 5.87028 | 8.66421 | 6.22084 | 5.08036 | 4.29271 | 4.43182 |
| MFHAS1 | Q9Y4C4 | 5.39803 | 7.04748 | 5.82905 | 6.07576 | 5.07885 | 6.85167 | 6.98441 | 6.46382 | 6.04519 | 5.67732 | 5.98615 | 5.81708 | 7.52079 | 6.49052 | 7.40627 | 5.3276 |
| SVIP | Q8NHG7 | 3.66829 | 6.83679 | 6.17068 | 5.47697 | 4.78715 | 5.65921 | 5.98892 | 5.87169 | 5.08105 | 6.37598 | 4.55646 | 4.6071 | 7.47205 | 9.05687 | 8.20493 | 10.1859 |
| FAM185A | Q8N0U4 | 7.08984 | 6.08518 | 6.39496 | 6.99306 | 6.09345 | 6.24221 | 6.44764 | 5.47571 | 5.07912 | 6.3611 | 4.87728 | 6.29155 | 6.3418 | 6.76106 | 6.84269 | 6.62335 |
| DCHS1 | Q96JQ0 | 6.64909 | 6.26294 | 4.27881 | 5.71579 | 6.58882 | 6.78168 | 6.01609 | 5.93974 | 7.55269 | 6.83935 | 4.88936 | 7.94679 | 8.1096 | 4.46921 | 4.59194 | 5.37041 |
| SEMA4D | Q92854 | 6.59311 | 6.73363 | 5.99198 | 7.52573 | 6.37659 | 6.33515 | 5.93876 | 5.17578 | 6.59101 | 5.88989 | 5.56874 | 7.12844 | 6.62125 | 5.85782 | 4.73593 | 6.93618 |
| CLPTM1L | G5E9Z2 | 6.32994 | 5.91096 | 5.22802 | 6.15491 | 5.75595 | 6.80174 | 6.4708 | 5.94311 | 6.56834 | 5.64964 | 6.41532 | 6.18511 | 6.19088 | 5.78595 | 7.43282 | 7.17652 |
| LIN52 | Q52LA3 | 4.96466 | 6.0373 | 5.49137 | 4.76839 | 5.40072 | 4.43423 | 4.71723 | 5.31389 | 4.46688 | 6.08721 | 5.71785 | 5.10842 | 5.8643 | 11.6553 | 7.48553 | 12.4868 |
| ITGAE | P38570 | 7.28994 | 5.68072 | 5.85911 | 6.76306 | 6.10004 | 7.65764 | 6.98248 | 6.48473 | 6.14527 | 5.55852 | 6.79317 | 6.68725 | 8.02457 | 5.28282 | 5.41489 | 5.27579 |
| SLC11A2 | P49281-4 | 5.70649 | 5.93267 | 6.38851 | 6.66713 | 5.98683 | 6.03595 | 6.13256 | 5.59806 | 5.91626 | 6.53296 | 6.38358 | 6.55806 | 6.20892 | 6.27402 | 6.62386 | 7.05412 |
| RNF111 | Q6ZNA4-3 | 6.90866 | 5.48451 | 6.53556 | 6.08783 | 6.75217 | 7.26509 | 7.19965 | 5.21232 | 6.19878 | 5.70468 | 5.98626 | 7.04235 | 5.73807 | 6.51216 | 5.11732 | 6.25461 |
| SLC25A44 | Q96H78 | 7.38348 | 7.16128 | 5.79851 | 7.46529 | 6.00267 | 4.66263 | 6.37373 | 6.16771 | 5.60433 | 6.05353 | 5.43418 | 6.6342 | 5.99825 | 7.23411 | 6.23305 | 5.79306 |
| LRRC45 | Q96CN5 | 7.69702 | 6.59609 | 5.70455 | 5.34485 | 5.25178 | 5.96065 | 6.16097 | 7.57255 | 6.34456 | 6.55132 | 6.68164 | 7.72554 | 6.27106 | 5.73095 | 4.51898 | 5.88746 |
| DYM | Q7RTS9 | 6.31659 | 5.73302 | 6.49567 | 5.77081 | 5.58871 | 6.5069 | 5.79972 | 5.8462 | 6.06039 | 6.09994 | 7.09326 | 7.14738 | 6.34542 | 6.9659 | 5.87287 | 6.35724 |
| UQCC2 | Q9BRT2 | 6.68577 | 5.93796 | 6.0988 | 6.42125 | 7.45525 | 6.71291 | 6.49195 | 6.58816 | 5.64901 | 6.14529 | 7.26066 | 6.30322 | 6.70822 | 5.81931 | 5.09561 | 4.62661 |
| SLC10A7 | Q0GE19-1 | 6.63825 | 5.48107 | 6.01638 | 7.23026 | 5.77403 | 7.09763 | 5.76915 | 5.86428 | 5.40927 | 6.96839 | 5.30861 | 6.03568 | 7.26075 | 5.77757 | 6.3247 | 7.04398 |
| TAPBPL | Q9BX59-7 | 7.09191 | 4.44978 | 5.00076 | 4.94933 | 7.26603 | 5.68736 | 9.20868 | 5.35687 | 6.82613 | 7.18162 | 5.89484 | 4.89435 | 8.35777 | 7.24318 | 4.48267 | 6.1087 |
| KLHL9 | Q9P2J3 | 6.03085 | 6.3737 | 7.75534 | 6.83883 | 6.12081 | 7.34056 | 7.51796 | 5.82451 | 6.31488 | 6.90343 | 7.23614 | 6.63667 | 4.70223 | 5.11076 | 4.72686 | 4.56647 |
| TYRO3 | Q06418 | 6.55095 | 8.76356 | 8.73481 | 7.23809 | 5.98094 | 5.21367 | 5.90605 | 7.72034 | 8.46886 | 7.22617 | 7.33115 | 6.25583 | 2.96793 | 2.14736 | 4.21275 | 5.28154 |
| HMCN2 | Q8NDA2 | 6.83158 | 5.95877 | 5.32317 | 7.48925 | 7.07844 | 6.04958 | 7.90386 | 5.2551 | 7.72358 | 5.3789 | 4.80502 | 5.75539 | 5.70333 | 5.5911 | 6.56129 | 6.59163 |
| S100PBP | Q96BU1 | 5.921 | 6.41838 | 6.63273 | 7.6364 | 6.34697 | 5.97077 | 6.58241 | 6.13245 | 5.80959 | 4.34401 | 7.02841 | 5.88732 | 7.00499 | 5.95076 | 5.93313 | 6.40068 |
| SMAGP | Q0VAQ4 | 4.9726 | 5.15324 | 3.5813 | 5.60316 | 5.0979 | 5.39604 | 6.0626 | 6.99105 | 3.60949 | 4.99121 | 3.3749 | 6.00947 | 5.30172 | 11.8026 | 7.52586 | 14.5269 |

| | | | | | | | | | | | | | | | | | |
|---|---|---|---|---|---|---|---|---|---|---|---|---|---|---|---|---|---|
| SLC30A6 | Q6NXT4-2 | 6.1881 | 6.29766 | 5.05445 | 6.21778 | 6.60934 | 5.27805 | 6.49485 | 6.21533 | 6.62622 | 5.54712 | 5.64504 | 5.85662 | 6.67341 | 6.7161 | 7.49182 | 7.08812 |
| HELQ | Q8TDG4 | 5.65246 | 7.33572 | 6.60234 | 5.34626 | 5.74423 | 6.0366 | 5.9133 | 6.98694 | 6.42709 | 6.05843 | 7.69894 | 6.96326 | 5.82158 | 5.49536 | 5.78535 | 6.13212 |
| ACYP2 | P14621 | 6.0353 | 5.53556 | 5.81942 | 5.55511 | 5.90385 | 7.24933 | 6.57395 | 6.52565 | 6.73671 | 7.53407 | 6.22854 | 6.9048 | 6.91935 | 6.53136 | 5.83423 | 4.11277 |
| ENKD1 | Q9H0I2 | 6.65808 | 6.88778 | 6.04589 | 6.57728 | 6.56696 | 6.82108 | 6.34827 | 6.36559 | 6.7276 | 6.72298 | 6.39458 | 7.12898 | 5.22031 | 5.20064 | 5.19768 | 5.13632 |
| PAQR4 | Q8N4S7 | 7.12736 | 6.61832 | 5.757 | 6.50917 | 6.43676 | 5.70896 | 8.01449 | 6.7693 | 5.38116 | 6.33953 | 7.90515 | 6.62027 | 5.77878 | 4.46753 | 5.52291 | 5.04331 |
| PLLP | Q9Y342 | 5.47457 | 6.30983 | 5.61108 | 6.2747 | 6.03012 | 5.58041 | 5.54489 | 6.43724 | 8.54309 | 4.74051 | 5.05563 | 6.62293 | 6.58417 | 8.33438 | 6.0916 | 6.76487 |
| CCDC171 | Q6TFL3-4 | 6.35263 | 6.89164 | 6.315 | 6.59629 | 6.93372 | 5.79662 | 5.65724 | 6.65549 | 6.06131 | 5.87082 | 5.851 | 6.31019 | 6.07209 | 5.65542 | 5.81473 | 7.16582 |
| IP6K2 | Q9UHH9 | 7.21422 | 7.17577 | 8.22053 | 7.76411 | 6.2909 | 6.66516 | 6.5552 | 6.23265 | 8.00449 | 8.40359 | 9.01877 | 8.71848 | 2.1281 | 3.04669 | 1.84573 | 2.71562 |
| RBM18 | Q96H35 | 6.38229 | 6.7231 | 6.39897 | 6.60914 | 6.54739 | 6.28243 | 5.91462 | 6.1759 | 5.93517 | 6.30406 | 6.12936 | 6.34781 | 5.92223 | 5.48729 | 6.1367 | 6.70355 |
| LRP10 | Q7Z4F1 | 6.24281 | 6.62105 | 6.71705 | 7.76386 | 6.26841 | 7.03228 | 6.90106 | 6.27985 | 7.95878 | 6.88701 | 5.76493 | 6.26416 | 4.49639 | 4.84413 | 4.81932 | 5.13893 |
| FBXO45 | P0C2W1 | 6.16083 | 6.2087 | 6.37641 | 7.48725 | 5.95939 | 6.50058 | 5.5358 | 6.10342 | 6.50583 | 7.82025 | 5.48281 | 6.22945 | 6.58975 | 5.9902 | 5.65725 | 5.39206 |
| LDHD | Q86WU2 | 6.64892 | 5.79969 | 5.68109 | 6.36254 | 6.60915 | 6.03922 | 6.06714 | 6.26893 | 5.78091 | 6.02143 | 6.28245 | 6.15162 | 6.89754 | 6.44067 | 6.51965 | 6.42904 |
| TLR3 | O15455 | 5.83753 | 5.65926 | 5.74976 | 7.53857 | 6.41087 | 6.47417 | 6.75878 | 6.10379 | 6.04933 | 6.40877 | 5.16053 | 6.51104 | 5.86801 | 6.29601 | 7.17204 | 6.00156 |
| EIF4EBP2 | Q13542 | 6.07649 | 4.86185 | 5.05232 | 4.65473 | 4.92374 | 3.81536 | 4.96298 | 5.53248 | 3.84327 | 4.62269 | 3.64672 | 5.55315 | 4.14505 | 15.3989 | 9.55125 | 13.359 |
| DYNLT3 | P51808 | 5.72424 | 5.83775 | 6.68207 | 5.248 | 6.2326 | 7.11747 | 6.89216 | 6.26196 | 6.73286 | 6.85873 | 6.60516 | 6.94456 | 6.10733 | 5.60224 | 4.86054 | 6.29235 |
| DCLRE1A | Q6PJP8 | 7.48667 | 6.98332 | 7.98537 | 7.22708 | 7.8415 | 6.25877 | 4.73287 | 6.03138 | 8.48782 | 6.54002 | 5.90307 | 5.34411 | 4.70516 | 5.99338 | 4.22822 | 4.25126 |
| NUDT12 | Q9BQG2 | 6.82982 | 6.55398 | 6.29434 | 6.44818 | 6.43614 | 6.76164 | 6.15228 | 6.50863 | 6.50907 | 6.50176 | 6.95138 | 5.7674 | 5.79276 | 5.92664 | 5.47463 | 5.09134 |
| SFR1 | Q86XK3-2 | 5.07539 | 5.80638 | 5.93573 | 5.92719 | 6.47641 | 6.69948 | 6.24085 | 6.03183 | 6.01188 | 6.65039 | 6.00506 | 7.18184 | 5.84851 | 6.90618 | 7.07421 | 6.12866 |
| NUDT18 | Q6ZVK8 | 6.52162 | 5.30442 | 7.48324 | 6.70671 | 6.83877 | 5.88427 | 6.18607 | 5.90697 | 7.20921 | 6.42592 | 7.57917 | 3.43457 | 5.76703 | 6.67588 | 5.86102 | 6.21513 |
| C14orf93 | Q9H972 | 6.78014 | 6.69727 | 6.71393 | 7.51799 | 6.43859 | 6.35159 | 5.87962 | 6.32848 | 6.22696 | 6.56697 | 6.81543 | 6.2033 | 5.34848 | 5.29151 | 5.4417 | 5.39808 |
| EPB41L4A | Q9HCS5 | 6.71731 | 6.92022 | 6.71283 | 5.797 | 5.28814 | 6.39396 | 5.2279 | 6.85602 | 6.56652 | 7.09297 | 5.94896 | 6.2782 | 7.08581 | 5.21122 | 6.0161 | 5.88684 |
| TMEM183B | Q1AE95 | 6.83399 | 6.60702 | 6.15313 | 7.2911 | 6.48603 | 7.20278 | 6.1594 | 6.66669 | 9.62304 | 6.78883 | 7.85489 | 6.57628 | 3.26235 | 4.53498 | 3.78802 | 4.17146 |
| UBXN11 | Q5T124 | 6.85731 | 5.60633 | 7.38636 | 6.39372 | 6.94132 | 7.00343 | 6.02748 | 7.43877 | 6.19234 | 6.8153 | 8.2331 | 7.58515 | 5.06723 | 4.51376 | 3.91338 | 4.02501 |
| EEF1AKMT1 | Q8WVE0 | 5.86089 | 6.19137 | 6.479 | 5.55331 | 5.81404 | 7.0506 | 6.81221 | 6.39784 | 6.45612 | 6.29846 | 6.74979 | 7.37313 | 5.66682 | 5.72138 | 5.4598 | 6.11525 |
| COMMD2 | Q86X83 | 5.77681 | 5.93192 | 6.34216 | 5.31714 | 6.77536 | 6.27332 | 5.59676 | 5.83248 | 7.51231 | 5.08923 | 7.94371 | 5.02424 | 6.74427 | 7.51918 | 5.53071 | 6.79039 |
| HMGXB3 | Q12766 | 5.74093 | 6.28118 | 6.73436 | 7.28988 | 5.9006 | 6.50464 | 6.32429 | 6.21197 | 6.37522 | 6.94113 | 6.75768 | 7.05111 | 5.39727 | 5.41464 | 5.52156 | 5.55352 |
| VPS37C | A5D8V6 | 6.91963 | 4.4206 | 5.84106 | 6.0481 | 7.23289 | 3.61705 | 3.37651 | 8.57262 | 5.00022 | 5.10773 | 4.55397 | 6.39736 | 5.06847 | 10.299 | 9.12802 | 8.41679 |
| RCAN3 | Q9UKA8 | 6.95374 | 7.00083 | 7.01407 | 6.31778 | 6.57448 | 7.23296 | 6.22792 | 6.59532 | 6.84786 | 6.65575 | 6.9972 | 6.96843 | 4.07016 | 5.11844 | 5.02623 | 4.39882 |
| PRDM4 | Q9UKN5 | 5.52286 | 7.04973 | 5.21681 | 6.3497 | 6.4329 | 5.62672 | 7.52981 | 5.09823 | 6.11119 | 6.87152 | 6.45604 | 5.55656 | 6.32855 | 5.62713 | 5.96345 | 8.25881 |
| KLHDC10 | Q6PID8 | 6.42011 | 6.66918 | 7.8142 | 6.19288 | 6.45506 | 6.66898 | 6.45331 | 5.9004 | 5.85001 | 6.8563 | 5.69454 | 6.6272 | 6.0064 | 6.10451 | 5.14777 | 5.13917 |
| B3GNT5 | Q9BYG0 | 7.10272 | 7.0462 | 6.50404 | 7.6773 | 6.62324 | 7.21463 | 6.76069 | 5.88248 | 5.62832 | 6.00087 | 5.84116 | 6.5054 | 5.11883 | 5.59114 | 4.95408 | 5.5489 |
| SH3GLB1 | Q9Y371-2 | 6.46702 | 6.57255 | 6.63643 | 6.01277 | 6.1725 | 6.83862 | 6.59116 | 6.76483 | 6.20798 | 6.53317 | 6.69439 | 6.84432 | 5.66735 | 5.34854 | 5.13701 | 5.51136 |
| ZNF562 | Q6V9R5 | 6.9183 | 7.01655 | 6.69099 | 7.2401 | 6.33127 | 5.57118 | 6.15232 | 5.97756 | 6.27482 | 6.20854 | 7.20923 | 6.33216 | 5.51012 | 5.37275 | 5.69495 | 5.49916 |
| CAMKK1 | Q8N5S9-2 | 6.76901 | 6.39077 | 6.38418 | 6.19306 | 6.39655 | 6.2784 | 6.2996 | 6.77967 | 6.26243 | 6.2837 | 6.36038 | 6.18047 | 5.74518 | 6.44551 | 5.86435 | 5.36675 |
| AZI2 | Q9H6S1 | 5.65828 | 6.87811 | 8.54155 | 5.14797 | 5.88131 | 5.37596 | 3.9971 | 6.05482 | 5.79635 | 6.21715 | 8.18604 | 7.6523 | 6.62855 | 6.78088 | 4.56207 | 5.64156 |
| SLC39A3 | Q9BRY0 | 6.54215 | 6.30522 | 7.29837 | 7.55171 | 3.81576 | 5.17829 | 5.13854 | 5.83805 | 6.52197 | 4.88848 | 8.13589 | 7.25832 | 5.04678 | 6.37327 | 6.99224 | 7.11497 |
| SLC35D2 | Q76EJ3 | 6.968 | 6.59816 | 6.228 | 6.62147 | 6.04581 | 5.65507 | 6.19991 | 5.57609 | 6.10538 | 6.43769 | 5.89143 | 6.27607 | 6.24776 | 5.82179 | 6.33372 | 6.99366 |
| ZBTB7B | O15156-2 | 7.17288 | 6.1905 | 5.94176 | 6.87559 | 6.05367 | 5.81332 | 5.86504 | 5.22745 | 6.69925 | 6.76862 | 6.44212 | 5.99439 | 5.62444 | 5.43704 | 6.50638 | 7.08755 |
| ZFP90 | Q8TF47 | 6.26505 | 7.69877 | 5.84247 | 5.90482 | 5.06615 | 5.51526 | 5.92795 | 6.28547 | 5.52879 | 6.41497 | 6.38572 | 6.86283 | 6.96534 | 7.05719 | 6.92218 | 5.35702 |
| AP4S1 | Q9Y587 | 6.34185 | 6.74184 | 6.92556 | 6.31508 | 6.194 | 6.43157 | 6.46447 | 6.6119 | 6.20391 | 6.7786 | 6.42959 | 6.22872 | 5.88362 | 5.82828 | 5.15421 | 5.46679 |
| TPGS2 | Q68CL5-5 | 7.34697 | 6.91827 | 7.02656 | 6.70865 | 5.86912 | 7.25789 | 6.49369 | 6.05228 | 6.72054 | 6.73724 | 7.37405 | 7.41725 | 4.59489 | 5.00736 | 3.93355 | 4.54169 |
| IFT22 | Q9H7X7 | 5.89707 | 4.16413 | 4.05391 | 3.95764 | 4.12129 | 5.08789 | 4.97515 | 7.93344 | 5.77023 | 7.32209 | 4.9702 | 6.87165 | 4.28693 | 10.0913 | 8.89421 | 11.6029 |
| IGSF9B | Q9UPX0-2 | 5.46969 | 6.10196 | 5.90283 | 6.3469 | 6.52757 | 5.40869 | 5.14863 | 5.73803 | 5.88331 | 6.61784 | 5.22105 | 5.66947 | 6.39029 | 7.44215 | 8.52378 | 7.60781 |
| SIMC1 | Q8NDZ2-5 | 6.54478 | 6.7525 | 6.61251 | 7.07585 | 6.15286 | 6.28016 | 6.63322 | 6.75241 | 6.81329 | 6.11012 | 6.10526 | 6.27634 | 5.65887 | 5.40459 | 5.24382 | 5.58342 |
| CREM | Q03060-1 | 6.29377 | 6.30355 | 6.21852 | 6.51827 | 7.12591 | 5.75275 | 6.10401 | 6.4165 | 5.59641 | 6.49681 | 7.54506 | 5.73359 | 6.24255 | 5.96378 | 6.1294 | 5.55915 |
| ATF1 | P18846 | 5.79071 | 6.06562 | 7.7926 | 6.81317 | 5.10956 | 5.29078 | 4.67507 | 3.47068 | 6.95816 | 6.46811 | 6.19022 | 10.0587 | 7.13788 | 6.18496 | 5.6323 | 6.36145 |
| HOXD9 | P28356 | 5.82993 | 6.32007 | 5.83129 | 5.85863 | 5.89704 | 5.91888 | 6.21288 | 6.35024 | 5.7795 | 6.95137 | 5.38328 | 5.61533 | 6.07303 | 6.64486 | 8.04599 | 7.28767 |
| PKP3 | Q9Y446-2 | 5.5213 | 8.36485 | 4.18839 | 5.45672 | 5.90709 | 5.75714 | 5.72629 | 7.26435 | 6.67223 | 6.60877 | 6.21904 | 5.79835 | 7.06676 | 6.95442 | 6.58529 | 5.909 |
| PGPEP1 | M0R3H3 | 6.14806 | 5.91004 | 5.30003 | 6.2701 | 5.91977 | 5.15695 | 5.23697 | 6.75518 | 6.63568 | 5.89931 | 6.06346 | 6.27603 | 6.85388 | 7.11741 | 6.6671 | 7.79004 |
| HACD2 | Q6Y1H2 | 6.04308 | 6.26389 | 5.5269 | 6.93449 | 6.25223 | 5.93614 | 6.22414 | 6.04374 | 5.77493 | 6.55046 | 6.24603 | 5.8747 | 6.42259 | 6.53979 | 6.7342 | 6.63269 |
| KIAA0930 | Q6ICG6-2 | 6.1345 | 6.72948 | 7.16113 | 6.40856 | 6.36656 | 7.75812 | 6.69207 | 5.3544 | 6.81195 | 5.83452 | 7.3884 | 7.84239 | 4.92843 | 4.24194 | 5.50209 | 4.84546 |
| DNAJB13 | P59910 | 7.12404 | 7.21592 | 5.58062 | 6.51118 | 5.97388 | 5.22828 | 6.08417 | 6.02269 | 7.40056 | 5.74914 | 5.69052 | 6.32783 | 6.05377 | 4.77437 | 8.11086 | 6.15219 |
| RNF170 | Q96K19 | 5.77733 | 7.48961 | 8.79778 | 4.4492 | 5.96106 | 6.06948 | 7.81179 | 7.35863 | 6.72405 | 3.786 | 6.24507 | 6.73456 | 6.28086 | 4.14561 | 7.86183 | 4.50715 |
| NELL2 | Q99435-3 | 6.74792 | 6.60766 | 5.9751 | 6.88429 | 6.38611 | 6.65848 | 5.65246 | 5.8169 | 6.93326 | 6.4206 | 6.25034 | 6.33441 | 5.88595 | 6.01432 | 5.8392 | 5.593 |
| OTUD3 | Q5T2D3 | 7.11775 | 7.10979 | 6.03384 | 6.20489 | 6.46773 | 6.20322 | 6.07243 | 7.59507 | 6.43813 | 7.74993 | 6.17549 | 6.35417 | 5.1406 | 5.2892 | 5.08769 | 4.96007 |
| CYP24A1 | Q07973 | 5.57536 | 6.78251 | 4.53111 | 6.6183 | 5.82497 | 5.4471 | 5.38091 | 6.07325 | 6.33726 | 6.28318 | 5.38496 | 6.83317 | 6.38136 | 6.3035 | 8.68169 | 7.56138 |
| GPR180 | Q86V85 | 6.95651 | 6.91657 | 5.88233 | 6.4643 | 5.96281 | 6.25368 | 6.48826 | 5.99016 | 6.06673 | 6.31504 | 5.25297 | 4.92136 | 6.92426 | 5.96796 | 7.27853 | 6.35855 |
| ALG14 | Q96F25 | 6.56508 | 5.63765 | 5.93361 | 7.34413 | 5.15325 | 5.29389 | 5.52392 | 5.73677 | 6.08982 | 6.87238 | 6.83515 | 6.61805 | 6.33507 | 6.32851 | 6.8115 | 6.92122 |
| C16orf87 | Q6PH81 | 6.02346 | 6.34598 | 6.08651 | 6.81963 | 5.87837 | 6.09347 | 6.62151 | 5.81222 | 6.36409 | 6.68889 | 6.33091 | 6.64265 | 6.17283 | 5.5692 | 6.21454 | 6.33576 |
| DPPA2 | Q7Z7J5 | 6.37653 | 7.14058 | 5.03791 | 7.12744 | 6.06338 | 6.64947 | 6.17314 | 6.07029 | 6.4598 | 7.29714 | 6.48324 | 6.34014 | 5.42299 | 6.32951 | 5.71513 | 5.31332 |
| CHMP6 | I3L4G8 | 7.2359 | 5.97038 | 5.39741 | 6.07356 | 5.21977 | 5.7863 | 6.56255 | 5.6762 | 6.53347 | 5.986 | 6.36899 | 6.38463 | 7.07576 | 5.34215 | 8.3623 | 6.02461 |
| STK11 | Q15831 | 5.42683 | 6.57679 | 6.51409 | 7.43891 | 6.42133 | 6.06269 | 5.68714 | 7.41016 | 6.90275 | 6.58495 | 5.85156 | 5.93689 | 5.17833 | 5.89776 | 6.10267 | 6.00715 |
| NHSL1 | H0YDF6 | 7.72172 | 5.71739 | 7.66542 | 7.23271 | 4.52923 | 7.22545 | 6.02345 | 4.81549 | 8.20737 | 6.79841 | 5.52507 | 4.54459 | 7.45248 | 4.25584 | 4.14054 | 7.24384 |
| NIPAL2 | Q9H841 | 6.76945 | 6.6153 | 6.75759 | 6.94169 | 4.66528 | 6.32321 | 6.67259 | 6.45862 | 6.05392 | 6.54116 | 7.81296 | 5.98857 | 5.86945 | 5.42628 | 6.11915 | 4.98477 |
| LGALSL | Q3ZCW2 | 5.07344 | 5.94701 | 6.31392 | 6.00597 | 6.21634 | 5.53086 | 6.26197 | 6.492 | 4.98905 | 6.377 | 5.20418 | 6.8979 | 5.22984 | 8.56923 | 6.31141 | 8.57986 |
| SYPL2 | Q5VXT5 | 5.24938 | 6.15408 | 5.65538 | 6.28832 | 4.70086 | 6.74776 | 7.46858 | 7.01501 | 5.58036 | 5.99456 | 6.30123 | 6.24945 | 6.57092 | 6.69996 | 6.98687 | 6.33729 |
| PPM1K | Q8N3J5 | 6.48966 | 6.9176 | 7.535 | 6.78787 | 4.71497 | 5.41431 | 5.49825 | 5.67065 | 7.33339 | 6.65523 | 5.78644 | 7.55612 | 5.16217 | 5.30314 | 6.74478 | 6.4304 |
| RP1 | P56715 | 6.82686 | 6.13567 | 5.50222 | 5.96435 | 6.11141 | 7.04351 | 7.19062 | 5.33149 | 5.16312 | 4.6563 | 6.3883 | 5.57049 | 7.81068 | 6.56612 | 6.45279 | 7.28607 |
| CRPPA | A4D126 | 6.02469 | 7.12547 | 7.54349 | 5.31612 | 5.37868 | 6.39983 | 5.81245 | 6.15861 | 5.91475 | 6.25129 | 5.50302 | 7.3885 | 6.73697 | 5.90331 | 5.85368 | 6.68912 |
| KRTCAP2 | Q8N6L1-2 | 5.41107 | 6.1044 | 6.23634 | 7.0984 | 6.20524 | 5.91401 | 6.39313 | 6.01981 | 5.9503 | 5.71647 | 5.26325 | 6.13556 | 6.59004 | 7.00498 | 7.49585 | 6.46112 |
| POLR3GL | Q9BT43 | 5.77026 | 6.51981 | 6.74568 | 5.43703 | 5.80732 | 6.95025 | 6.3311 | 5.57147 | 6.58683 | 6.79055 | 6.80932 | 5.72904 | 7.15431 | 7.20624 | 5.23495 | 5.35585 |
| RFESD | Q8TAC1-2 | 5.65509 | 6.71546 | 7.46434 | 5.40493 | 7.18799 | 6.56849 | 6.4909 | 8.20689 | 7.55456 | 4.95428 | 7.6774 | 5.89972 | 5.42793 | 4.19792 | 6.1745 | 4.41961 |
| GTF2A2 | P52657 | 5.67965 | 5.46929 | 5.58554 | 4.67707 | 6.5417 | 6.6859 | 5.86204 | 5.73981 | 5.9357 | 5.6769 | 5.65461 | 6.07461 | 7.18942 | 8.03879 | 7.68934 | 7.49964 |
| TNKS2 | Q9H2K2 | 6.39176 | 7.16968 | 6.53306 | 7.22203 | 7.20209 | 6.92633 | 7.27665 | 6.12908 | 5.75781 | 6.08815 | 6.54585 | 6.3182 | 5.20806 | 5.79202 | 4.61513 | 4.8241 |
| PSMA3 | H0YJ03 | 5.75121 | 6.04309 | 6.88782 | 5.9575 | 7.41408 | 8.75806 | 4.2097 | 7.70406 | 7.69244 | 6.16165 | 5.82631 | 5.87558 | 8.31997 | 4.17913 | 4.57 | 4.64941 |
| SP4 | Q02446 | 6.34532 | 5.47476 | 7.32265 | 7.32254 | 5.10586 | 6.09858 | 6.32613 | 5.53543 | 5.77317 | 5.92609 | 6.0452 | 5.79676 | 6.82845 | 6.1294 | 7.13379 | 6.83317 |
| CYBA | P13498 | 5.73744 | 6.9536 | 5.50371 | 6.61292 | 5.70146 | 6.22603 | 6.06825 | 6.75587 | 6.87595 | 7.72927 | 6.30615 | 7.13837 | 4.30099 | 5.74799 | 4.97129 | 7.37071 |
| B4GALT4 | O60513 | 6.70791 | 5.73864 | 6.6653 | 5.61071 | 6.34531 | 8.06193 | 6.99929 | 4.27598 | 5.8501 | 7.39214 | 5.96222 | 5.08772 | 6.63796 | 5.79622 | 5.99781 | 6.87074 |
| SERPINF1 | P36955 | 6.44609 | 6.33671 | 4.76241 | 6.27983 | 4.91177 | 6.99632 | 7.26613 | 4.52151 | 7.29313 | 5.40332 | 5.99871 | 5.20046 | 6.58858 | 5.73996 | 9.24011 | 7.01498 |
| LRFN3 | Q9BTN0 | 6.39644 | 7.38286 | 6.4884 | 7.23692 | 4.77014 | 6.78441 | 6.20366 | 4.85553 | 8.11485 | 6.31222 | 6.65169 | 5.43501 | 5.41385 | 5.14915 | 6.72119 | 6.0837 |
| TRMT2B | Q96GJ1 | 5.71832 | 5.94511 | 5.56471 | 7.39725 | 6.82295 | 6.86347 | 5.77378 | 5.7371 | 6.55251 | 5.85563 | 6.34307 | 4.91244 | 5.33598 | 6.83493 | 6.12471 | 6.21804 |
| C8orf76 | Q96K31 | 7.75968 | 7.28999 | 7.36188 | 4.81553 | 6.80823 | 6.87512 | 6.7141 | 7.06626 | 5.42179 | 6.68653 | 6.95929 | 8.16007 | 4.81249 | 4.56439 | 4.32702 | 4.37763 |
| CACFD1 | Q9UGQ2 | 6.87918 | 6.49161 | 6.32324 | 6.14517 | 6.76253 | 6.60788 | 6.20939 | 5.70317 | 5.73262 | 6.03163 | 5.11308 | 6.61297 | 6.66582 | 6.68997 | 6.30724 | 5.72449 |
| DSEL | Q8IZU8 | 6.2843 | 6.90493 | 7.42819 | 6.77511 | 7.30948 | 5.72328 | 6.48966 | 6.6247 | 4.88433 | 6.45857 | 6.87864 | 6.06875 | 6.1756 | 4.20898 | 5.13375 | 6.65173 |
| MZT2B | Q6NZ67 | 5.78041 | 5.20391 | 6.55202 | 5.19181 | 5.71829 | 6.85206 | 7.77997 | 7.86873 | 6.21775 | 8.28197 | 6.16438 | 6.84568 | 6.28237 | 4.78653 | 5.51431 | 4.96073 |
| MINDY1 | Q8N5J2-3 | 5.44614 | 6.14015 | 6.92746 | 5.92598 | 5.76598 | 6.2962 | 5.70148 | 5.88571 | 5.85704 | 6.81079 | 8.22527 | 7.18304 | 5.82044 | 5.9387 | 5.11543 | 6.96019 |
| CTNS | O60931-2 | 6.06917 | 6.82073 | 5.06357 | 5.52764 | 5.64495 | 5.47197 | 6.20941 | 6.95882 | 5.91781 | 6.87173 | 5.59579 | 7.14434 | 6.48805 | 6.10602 | 7.3775 | 6.73249 |
| TBCCD1 | Q9NVR7 | 5.58745 | 5.48759 | 6.88379 | 7.23029 | 6.42952 | 6.65365 | 6.70093 | 6.6513 | 6.72405 | 6.44916 | 6.78017 | 7.22628 | 5.16804 | 5.24067 | 5.25828 | 5.52883 |
| MFSD3 | Q96ES6 | 5.30083 | 6.00929 | 6.06151 | 6.25112 | 5.63383 | 5.73706 | 7.07897 | 6.96485 | 6.99256 | 6.86579 | 5.79491 | 5.15381 | 6.19083 | 6.83065 | 6.71277 | 6.42121 |
| DESI2 | Q9BSY9 | 6.17364 | 5.74666 | 5.03802 | 4.63232 | 6.61407 | 5.71269 | 5.77776 | 5.81599 | 7.21063 | 5.18716 | 6.07797 | 6.67586 | 5.85486 | 7.74797 | 7.16554 | 8.56887 |
| ENOX2 | Q16206 | 6.54316 | 5.5689 | 6.81667 | 5.64337 | 6.14978 | 6.20786 | 6.98847 | 5.84692 | 6.37552 | 6.25398 | 6.70238 | 5.72553 | 6.30003 | 5.91172 | 7.00568 | 5.96002 |
| TNFAIP8 | O95379-4 | 5.79159 | 4.93128 | 7.93149 | 5.62778 | 6.1849 | 7.55157 | 5.96187 | 5.53403 | 6.53709 | 6.49805 | 6.73279 | 5.78401 | 6.34225 | 6.57477 | 5.16934 | 6.8472 |
| CCDC154 | A6NI56 | 4.64194 | 5.83056 | 7.45492 | 5.81627 | 7.49796 | 8.47086 | 6.0984 | 5.84706 | 6.38782 | 6.58797 | 7.89682 | 5.95158 | 6.06039 | 4.66798 | 5.22128 | 5.5682 |
| ZNF229 | Q9UJW7 | 6.56667 | 7.31578 | 4.99416 | 6.194 | 5.72163 | 5.43615 | 5.02776 | 7.54552 | 7.51137 | 7.51196 | 5.79451 | 6.78726 | 5.63536 | 5.5892 | 6.13262 | 6.23605 |
| TCTN1 | Q2MV58-2 | 7.35007 | 7.07429 | 6.38105 | 7.03348 | 6.20582 | 6.16804 | 6.32411 | 5.94097 | 6.18524 | 7.38438 | 6.35094 | 6.64873 | 5.49064 | 5.22729 | 5.05389 | 5.18106 |
| KCT2 | Q8NC54 | 6.11903 | 6.83875 | 5.44318 | 6.05748 | 5.74959 | 6.33673 | 5.44668 | 6.96998 | 5.85516 | 7.17328 | 5.79663 | 7.48168 | 6.47328 | 6.1526 | 5.64542 | 6.46053 |
| RHBDD2 | Q6NTF9-2 | 6.34774 | 6.49515 | 5.93868 | 6.95027 | 5.82139 | 5.48364 | 6.89777 | 7.00969 | 6.69542 | 6.11233 | 5.56431 | 5.17162 | 6.19091 | 6.30219 | 5.97881 | 7.04008 |
| SOCS2 | O14508 | 6.48903 | 5.10687 | 6.97049 | 5.93264 | 7.14982 | 7.05227 | 5.72994 | 6.16571 | 6.98156 | 7.29619 | 7.08569 | 7.27376 | 5.51884 | 4.70336 | 5.55695 | 4.98689 |
| MED21 | Q13503 | 4.94926 | 6.03583 | 5.01398 | 6.24967 | 7.27753 | 4.70462 | 4.506 | 5.72429 | 6.18054 | 4.80059 | 5.528 | 3.65985 | 4.29842 | 10.3038 | 6.35175 | 14.4158 |
| TBC1D12 | O60347 | 7.06611 | 6.56354 | 6.56649 | 6.17415 | 5.61662 | 6.95467 | 6.44855 | 5.8443 | 6.54662 | 5.94038 | 6.09665 | 5.97971 | 6.76273 | 5.95341 | 5.09758 | 6.3885 |
| MGMT | P16455 | 6.54118 | 4.04332 | 5.05201 | 6.21106 | 5.53751 | 5.10211 | 6.40918 | 6.49453 | 5.33996 | 6.18774 | 6.27946 | 7.10473 | 5.43941 | 9.50874 | 8.8225 | 5.92656 |
| MED20 | Q9H944 | 6.16988 | 6.40507 | 7.93766 | 4.56172 | 6.49874 | 5.80844 | 5.82079 | 5.55988 | 5.66885 | 5.50285 | 7.17043 | 5.48791 | 7.58666 | 6.84989 | 6.01184 | 6.95939 |
| ZNF394 | Q53GI3 | 6.27331 | 6.85807 | 7.98443 | 5.77607 | 6.37705 | 5.68264 | 6.36527 | 5.6157 | 6.27145 | 5.62346 | 5.02997 | 5.47256 | 5.98789 | 6.21799 | 7.30291 | 7.16125 |
| ANKRD46 | Q86W74-1 | 7.47077 | 5.39139 | 6.91333 | 5.37607 | 6.83248 | 5.58392 | 7.56793 | 6.3919 | 6.36582 | 6.40289 | 6.56095 | 6.28346 | 5.26978 | 5.42266 | 5.85905 | 6.30761 |
| RNF19A | Q9NV58 | 7.51002 | 6.41123 | 7.11069 | 7.33187 | 4.95697 | 5.38004 | 6.27608 | 5.63493 | 6.13086 | 6.07982 | 6.60588 | 7.58524 | 6.79977 | 4.48687 | 6.70459 | 4.99513 |

| | | | | | | | | | | | | | | | | | |
|---|---|---|---|---|---|---|---|---|---|---|---|---|---|---|---|---|---|
| RALGPS2 | Q86X27 | 6.75541 | 6.97717 | 6.81079 | 5.7067 | 6.25108 | 8.21425 | 7.13083 | 5.47519 | 6.15697 | 7.28107 | 5.97933 | 5.18885 | 5.69024 | 5.73066 | 5.29488 | 5.35657 |
| FAM217B | Q9NTX9 | 6.00494 | 8.46703 | 7.63002 | 6.83119 | 6.21488 | 5.11268 | 5.23011 | 6.68928 | 6.97282 | 7.95754 | 7.56774 | 7.86517 | 3.89065 | 5.72081 | 3.36189 | 4.48326 |
| GXYLT2 | A0PJZ3 | 6.08028 | 6.28548 | 6.10538 | 6.85732 | 6.22567 | 6.03562 | 6.15101 | 6.02885 | 6.207 | 6.68026 | 6.37455 | 6.37264 | 6.3565 | 6.05753 | 6.13588 | 6.04605 |
| C19orf12 | Q9NSK7 | 6.02902 | 4.96991 | 6.49482 | 5.35862 | 6.12287 | 6.46581 | 5.98161 | 6.9828 | 6.08425 | 6.67421 | 6.12989 | 6.59259 | 5.76522 | 7.25423 | 6.52692 | 6.5672 |
| MUC15 | Q8N387 | 6.86475 | 5.79003 | 6.60927 | 5.57906 | 5.92991 | 6.38256 | 5.98832 | 6.15138 | 6.17357 | 6.39675 | 6.42017 | 7.0464 | 6.33023 | 6.21264 | 6.5859 | 5.53906 |
| MLXIP | Q9HAP2 | 5.99553 | 6.45501 | 7.67213 | 6.03863 | 6.03192 | 6.63924 | 6.27568 | 6.27895 | 7.264 | 6.58355 | 6.61665 | 6.7072 | 4.89523 | 5.16542 | 5.51913 | 5.86173 |
| ZNF736 | B4DX44 | 5.74864 | 7.21478 | 6.69052 | 5.88737 | 5.53629 | 7.24249 | 7.04153 | 5.27889 | 5.24627 | 7.62507 | 7.09324 | 7.01631 | 5.93174 | 5.27023 | 4.97206 | 6.20459 |
| FAM241B | Q96D05-2 | 5.88078 | 5.005 | 4.64234 | 4.54017 | 4.58968 | 3.93135 | 4.69277 | 6.35462 | 2.98672 | 5.00948 | 5.56653 | 4.66143 | 5.54952 | 12.9366 | 11.2112 | 12.4419 |
| ZG16B | Q96DA0 | 1.52909 | 5.383 | 0 | 20.1748 | 3.40063 | 8.36336 | 8.79426 | 2.41076 | 2.67717 | 2.36978 | 2.76281 | 2.46121 | 4.56238 | 26.6321 | 2.73229 | 5.74643 |
| TMEM138 | Q9NPI0 | 6.33743 | 6.96315 | 5.58938 | 6.79634 | 6.76973 | 6.81622 | 6.44243 | 6.3091 | 5.84631 | 6.9617 | 6.26669 | 5.88469 | 4.97751 | 6.36376 | 5.747 | 5.92859 |
| MIEF2 | Q96C03-3 | 7.61576 | 8.5187 | 7.77543 | 4.59194 | 8.75738 | 5.7319 | 7.72758 | 6.86877 | 6.79023 | 3.05983 | 7.88232 | 4.88644 | 5.64059 | 5.24793 | 4.64657 | 4.25865 |
| FBXO46 | Q6PJ61 | 4.93482 | 6.87091 | 6.1361 | 6.13445 | 5.83175 | 6.20064 | 6.54333 | 5.05773 | 6.86951 | 7.13031 | 5.63924 | 6.44484 | 6.57086 | 6.59379 | 6.95173 | 6.08998 |
| C1orf167 | Q5SNV9-2 | 6.12835 | 5.97116 | 5.44405 | 6.12695 | 6.06857 | 5.37807 | 6.92633 | 6.60074 | 6.40389 | 5.60363 | 5.25097 | 5.51009 | 6.73172 | 6.62908 | 7.62456 | 7.60185 |
| UBE2F | Q969M7-2 | 6.15509 | 4.48489 | 5.98945 | 7.76266 | 6.70044 | 5.72085 | 5.75969 | 6.09523 | 5.28854 | 7.02122 | 7.02263 | 5.64918 | 5.87551 | 6.92228 | 5.89667 | 7.65567 |
| FHOD3 | Q2V2M9-4 | 5.80455 | 6.3723 | 6.10407 | 5.7732 | 6.15822 | 6.5333 | 6.12473 | 6.2383 | 6.728 | 7.1326 | 6.87698 | 6.83153 | 6.28985 | 5.89416 | 5.44016 | 5.69803 |
| MREG | Q8N565-2 | 6.49256 | 6.41083 | 6.13528 | 6.3373 | 5.663 | 5.70677 | 6.50799 | 6.83293 | 6.19666 | 6.63542 | 6.32581 | 7.02405 | 5.75857 | 5.45636 | 6.39926 | 6.11721 |
| NFIL3 | Q16649 | 6.20268 | 6.17063 | 6.15133 | 6.81775 | 6.30746 | 5.80452 | 6.84457 | 6.20142 | 6.7598 | 5.71061 | 6.43822 | 6.47796 | 6.40277 | 5.63592 | 6.21757 | 5.85679 |
| YOD1 | Q5VVQ6 | 5.94299 | 6.68662 | 6.62757 | 7.35664 | 6.05091 | 6.96484 | 5.96694 | 6.20798 | 6.88769 | 6.06881 | 7.34683 | 6.87088 | 5.80679 | 5.72553 | 4.63552 | 4.85346 |
| RIN2 | Q8WYP3-2 | 5.78065 | 6.30278 | 6.22459 | 5.62151 | 6.12002 | 7.21293 | 5.94812 | 6.93346 | 6.82974 | 6.60346 | 6.22891 | 6.55886 | 6.16952 | 6.03948 | 5.71633 | 5.70964 |
| UQCR11 | O14957 | 5.40396 | 6.11822 | 5.35886 | 5.69375 | 6.20849 | 6.15127 | 7.09572 | 5.9291 | 5.43802 | 6.1813 | 6.15994 | 6.04898 | 6.29036 | 7.3841 | 7.53605 | 7.00188 |
| SCOC | Q9UIL1 | 6.82265 | 5.20498 | 3.44405 | 5.67975 | 5.31707 | 5.691 | 5.42485 | 6.66648 | 5.17328 | 6.29873 | 5.22835 | 5.10608 | 4.14553 | 10.5703 | 8.69139 | 10.5355 |
| CTRB2 | Q6GPI1 | 6.53921 | 5.28561 | 5.88409 | 6.04395 | 6.66006 | 5.64044 | 6.4258 | 6.07778 | 6.62141 | 4.97356 | 4.38983 | 5.29513 | 6.1913 | 7.23495 | 8.24219 | 8.49469 |
| PIGW | Q7Z7B1 | 5.76614 | 6.97095 | 5.67684 | 6.59221 | 5.17133 | 5.63556 | 6.40324 | 5.95997 | 7.22026 | 5.70721 | 6.12235 | 5.97633 | 6.79724 | 6.235 | 5.95343 | 7.81194 |
| POLD4 | Q9HCU8 | 5.45235 | 4.97462 | 5.2743 | 4.67603 | 5.96171 | 5.37285 | 5.83346 | 6.58185 | 4.93307 | 6.49761 | 7.33909 | 5.53322 | 6.92322 | 7.97816 | 8.83828 | 7.83016 |
| MTHFD2L | Q9H903 | 6.41898 | 6.64056 | 5.58122 | 6.4722 | 6.27087 | 6.44123 | 6.00842 | 5.20694 | 6.75814 | 6.29785 | 5.7479 | 6.57295 | 6.68652 | 6.26382 | 6.04505 | 6.58734 |
| GPS2 | Q13227 | 7.19707 | 6.17819 | 7.22561 | 5.34935 | 5.93431 | 5.86249 | 5.97207 | 6.25533 | 5.96199 | 6.81615 | 6.68956 | 6.29727 | 7.336 | 5.36154 | 5.31109 | 6.25199 |
| ADRB2 | P07550 | 4.69902 | 6.1099 | 5.86907 | 7.71927 | 5.14832 | 6.60561 | 6.70237 | 5.93204 | 6.60612 | 4.23068 | 5.58516 | 6.39264 | 7.25755 | 6.36561 | 8.11105 | 6.66559 |
| ZNF225 | Q9UK10 | 6.92136 | 5.91104 | 6.60636 | 7.1834 | 6.29592 | 6.12149 | 6.65599 | 6.60551 | 7.56624 | 6.71177 | 6.36314 | 6.17944 | 5.22481 | 5.50137 | 5.29365 | 4.8585 |
| ADGRV1 | Q8WXG9 | 6.19313 | 5.04843 | 6.46008 | 7.45727 | 5.94325 | 5.52949 | 8.102 | 6.25894 | 6.23709 | 6.27702 | 6.19952 | 4.10206 | 7.45018 | 5.61334 | 7.12033 | 6.00786 |
| CKLF | Q9UBR5 | 4.9263 | 6.24766 | 6.40279 | 7.46965 | 6.94393 | 6.77586 | 6.72455 | 6.68579 | 5.77046 | 6.08522 | 5.37286 | 5.30456 | 5.35493 | 6.69241 | 6.82372 | 6.41932 |
| PET117 | Q6UWS5 | 5.99937 | 5.65853 | 5.47808 | 6.30078 | 6.1244 | 5.12917 | 5.71181 | 6.41748 | 5.54582 | 6.38924 | 5.39726 | 6.56184 | 5.98027 | 7.60517 | 7.67997 | 8.02081 |
| PPM1M | Q96MI6 | 6.52964 | 6.4113 | 6.52573 | 5.71655 | 5.09856 | 6.32462 | 6.9144 | 8.07103 | 8.53059 | 6.33126 | 8.71605 | 8.64225 | 3.67873 | 5.29377 | 3.31576 | 3.89976 |
| ZMYND19 | Q96E35 | 6.36105 | 7.51916 | 7.47075 | 7.06963 | 6.41966 | 5.5103 | 7.05725 | 7.93185 | 6.11323 | 6.9652 | 7.03572 | 8.42786 | 4.51982 | 2.90219 | 4.01066 | 4.68566 |
| ZBTB45 | Q96K62 | 5.9334 | 5.91956 | 6.40999 | 6.92838 | 5.34134 | 5.88407 | 5.95545 | 6.27652 | 6.58021 | 6.43802 | 6.50632 | 5.97534 | 6.45415 | 6.52123 | 6.67606 | 6.19996 |
| ABTB2 | Q8N961 | 5.07222 | 6.77204 | 6.94618 | 5.77257 | 4.34702 | 7.53883 | 6.2731 | 4.25541 | 5.87143 | 6.89338 | 6.26381 | 6.11514 | 6.25451 | 6.66165 | 6.34715 | 8.61556 |
| ZCCHC10 | Q8TBK6 | 5.9086 | 6.5689 | 5.70587 | 7.13541 | 6.56055 | 6.5356 | 6.50669 | 6.36328 | 6.3338 | 5.62814 | 6.13649 | 6.19018 | 5.96364 | 6.36663 | 6.07434 | 6.02188 |
| NUDT6 | P53370 | 7.76893 | 6.52327 | 6.21184 | 7.84838 | 5.66985 | 5.3117 | 4.77353 | 5.42941 | 5.60646 | 7.99693 | 6.18526 | 6.36432 | 6.84605 | 5.74624 | 5.99463 | 5.7232 |
| PIGQ | Q9BRB3-2 | 6.93682 | 6.24302 | 6.80692 | 7.05582 | 6.86916 | 5.37608 | 5.47283 | 6.8182 | 4.98811 | 5.9398 | 5.75529 | 6.08856 | 5.9811 | 6.83704 | 7.09434 | 5.7369 |
| EXTL2 | Q9UBQ6 | 6.15048 | 6.28149 | 5.2689 | 6.33922 | 6.4818 | 6.0686 | 7.08748 | 6.62202 | 6.26079 | 5.76717 | 5.95024 | 6.22056 | 6.03931 | 6.41634 | 6.08196 | 6.96365 |
| PARP8 | Q8N3A8 | 5.42883 | 6.12626 | 6.77907 | 6.29219 | 5.54696 | 5.93541 | 5.86301 | 6.24056 | 6.55053 | 6.85496 | 6.88499 | 5.71899 | 7.18203 | 6.26313 | 5.3911 | 6.94198 |
| PRELID3B | Q9Y3B1 | 5.71527 | 6.78297 | 7.11709 | 6.99682 | 8.35906 | 7.05654 | 7.8337 | 9.27049 | 7.22073 | 6.70723 | 5.31772 | 6.1422 | 4.27703 | 4.48866 | 1.12261 | 5.59189 |
| SLC35D1 | Q9NTN3 | 7.95973 | 4.8901 | 5.31063 | 7.19301 | 6.99752 | 5.14477 | 5.12379 | 7.07241 | 4.66627 | 6.06814 | 6.07465 | 5.18559 | 6.42311 | 5.62691 | 9.40143 | 6.86194 |
| ZNF839 | A8K0R7-5 | 5.80535 | 6.30497 | 5.80905 | 6.29505 | 6.15814 | 6.42283 | 6.24258 | 6.77627 | 5.91476 | 6.07965 | 7.02038 | 6.58489 | 5.9705 | 5.90507 | 6.02939 | 6.68114 |
| B3GALNT2 | Q8NCR0 | 7.55634 | 6.91329 | 5.8885 | 6.06024 | 6.07469 | 6.64748 | 6.42108 | 5.67298 | 6.30706 | 6.34024 | 6.40658 | 6.06278 | 5.62351 | 6.4123 | 5.52994 | 6.08299 |
| COMMD8 | Q9NX08 | 6.24204 | 5.74639 | 5.25439 | 5.34892 | 5.22966 | 6.01127 | 5.54497 | 6.36012 | 5.33108 | 5.81213 | 5.55385 | 5.722 | 5.46769 | 9.88639 | 7.12842 | 9.36068 |
| SHF | Q7M4L6 | 4.26846 | 7.84522 | 6.47152 | 6.19442 | 5.02012 | 3.42164 | 5.40985 | 8.74063 | 5.12898 | 8.89977 | 7.04867 | 7.71383 | 6.80978 | 6.32635 | 5.27361 | 5.42716 |
| RAB40C | Q96S21 | 6.73584 | 6.1053 | 6.63762 | 6.64174 | 6.47036 | 5.33375 | 6.21254 | 6.50204 | 5.984 | 5.79475 | 6.05717 | 6.04924 | 5.3927 | 6.73609 | 6.53867 | 6.8082 |
| GDPGP1 | Q6ZNW5 | 6.34717 | 6.21369 | 8.0643 | 6.13324 | 6.05368 | 4.65343 | 7.11535 | 7.0472 | 6.0159 | 4.9969 | 4.81192 | 6.3793 | 8.27458 | 5.081 | 5.21364 | 7.59869 |
| DCAF5 | Q96JK2 | 6.35075 | 6.6129 | 7.37726 | 7.11288 | 6.13062 | 6.32167 | 6.10195 | 6.27835 | 7.69858 | 7.31071 | 6.83537 | 6.74103 | 4.24111 | 5.21171 | 4.59662 | 5.07849 |
| ZNHIT1 | O43257 | 6.56138 | 6.83343 | 6.83171 | 6.59588 | 6.38383 | 6.40211 | 6.58697 | 6.97378 | 6.52667 | 6.86188 | 7.18261 | 5.91897 | 5.20031 | 4.3456 | 5.83405 | 4.96081 |
| SLCO2A1 | Q92959 | 6.11723 | 6.23095 | 6.29523 | 6.97342 | 6.44514 | 5.46555 | 6.49867 | 5.74481 | 6.93716 | 5.80298 | 5.64253 | 7.12851 | 6.38564 | 5.98257 | 6.20546 | 6.14415 |
| RASSF7 | Q02833 | 5.76584 | 5.55611 | 6.69087 | 4.91099 | 6.65832 | 6.76535 | 6.23107 | 6.1305 | 5.6579 | 6.62724 | 6.96159 | 7.48074 | 5.56661 | 6.41457 | 6.45951 | 6.12277 |
| DACH1 | Q9UI36 | 6.4925 | 6.24581 | 5.80972 | 6.37993 | 6.50684 | 5.87297 | 6.48852 | 5.98589 | 6.874 | 6.75463 | 6.07656 | 5.7969 | 6.78691 | 6.13266 | 6.40735 | 5.38882 |
| SPRTN | Q9H040 | 6.01644 | 5.50659 | 6.55519 | 7.3855 | 5.33792 | 5.6823 | 8.78751 | 6.52055 | 5.43815 | 5.40598 | 6.98785 | 5.55965 | 6.61829 | 5.99464 | 5.8544 | 6.34903 |
| EPOP | A6NHQ4 | 6.49193 | 6.61253 | 5.95754 | 6.83772 | 6.36963 | 6.50582 | 6.47056 | 5.59823 | 6.99111 | 8.10734 | 5.49621 | 6.71811 | 6.42598 | 4.55856 | 5.67905 | 5.17969 |
| ENPP4 | Q9Y6X5 | 6.08288 | 7.42863 | 6.32416 | 5.79404 | 5.87123 | 7.441 | 5.33116 | 5.90561 | 4.97598 | 5.79231 | 7.28825 | 8.02777 | 5.55075 | 6.00157 | 5.42325 | 6.7614 |
| MRPL52 | Q86TS9 | 6.1048 | 5.40911 | 5.04812 | 6.00518 | 8.08495 | 5.56917 | 5.46377 | 7.61051 | 4.79816 | 8.07669 | 5.64749 | 8.11909 | 6.64647 | 6.06539 | 6.09035 | 5.26073 |
| DHRSX | Q8N5I4 | 7.12208 | 5.76446 | 6.0005 | 8.50912 | 7.59958 | 4.90447 | 6.57506 | 7.94275 | 2.88244 | 7.20034 | 7.32549 | 4.54828 | 6.80888 | 5.92752 | 5.42708 | 5.46196 |
| PPP1R16A | Q96I34 | 6.53232 | 7.77911 | 6.05798 | 5.60736 | 6.23977 | 6.2084 | 5.79607 | 8.47561 | 6.30961 | 5.33511 | 5.48818 | 5.42362 | 6.52682 | 5.95965 | 7.12348 | 5.13691 |
| MRPL18 | Q9H0U6 | 5.38271 | 5.99863 | 5.65116 | 5.97448 | 7.35429 | 8.15795 | 6.67714 | 5.61532 | 7.49502 | 5.94771 | 4.93137 | 6.22461 | 6.67251 | 5.54301 | 6.58114 | 5.79294 |
| SLC50A1 | Q9BRV3 | 4.9948 | 7.21459 | 5.8388 | 6.62795 | 6.63447 | 4.20525 | 4.23323 | 6.36068 | 7.2003 | 8.12065 | 5.1178 | 6.52697 | 6.90989 | 5.97413 | 8.46269 | 5.57779 |
| G0S2 | P27469 | 3.80017 | 5.64642 | 6.71819 | 6.2582 | 5.00773 | 6.16506 | 5.17925 | 6.59957 | 6.62751 | 5.07296 | 6.82773 | 8.08071 | 5.79514 | 7.81423 | 6.25911 | 8.14801 |
| SLC23A2 | Q9UGH3 | 6.70681 | 7.59656 | 6.00778 | 6.21391 | 6.20535 | 5.92293 | 6.4432 | 6.25988 | 6.42516 | 6.05588 | 6.11558 | 5.42877 | 6.47439 | 5.79156 | 6.29112 | 6.26112 |
| TMEM35B | Q8NCS4 | 5.56726 | 5.47523 | 5.01029 | 6.94702 | 4.74446 | 5.54563 | 6.66805 | 3.96274 | 4.83717 | 6.45318 | 4.4568 | 8.10543 | 8.86388 | 8.21964 | 7.46294 | 7.68029 |
| ZUP1 | Q96AP4 | 7.00715 | 7.24767 | 8.46658 | 6.75364 | 6.85502 | 7.16994 | 7.01745 | 7.27671 | 6.1733 | 6.47543 | 7.67712 | 7.89702 | 2.91578 | 3.57764 | 4.18575 | 3.3038 |
| LPAR2 | Q9HBW0 | 7.16914 | 5.35792 | 6.20821 | 5.20207 | 6.53799 | 7.58604 | 6.09388 | 6.96398 | 5.93024 | 6.54987 | 5.8856 | 5.89066 | 6.29206 | 5.62229 | 6.08078 | 6.62928 |
| SYTL1 | Q8IYJ3 | 6.54258 | 5.84022 | 7.29256 | 6.77651 | 4.87103 | 7.47627 | 5.73265 | 4.22055 | 4.99098 | 7.62986 | 4.88731 | 6.54054 | 7.3587 | 5.67855 | 6.95463 | 7.20704 |
| DOCK8-AS1 | Q5T8R8 | 7.04861 | 6.89849 | 6.62742 | 6.4048 | 6.51215 | 6.06344 | 6.26676 | 7.02132 | 5.84358 | 5.50806 | 6.6684 | 6.92487 | 4.93086 | 5.51653 | 6.00829 | 5.7564 |
| C8orf88 | P0DMB2 | 6.78863 | 7.6764 | 8.1938 | 6.61994 | 7.19155 | 6.68233 | 6.53212 | 6.9757 | 8.27698 | 6.69232 | 6.35838 | 8.17661 | 3.06032 | 3.31636 | 3.84425 | 3.61431 |
| BNIP3L | O60238 | 9.54536 | 9.24723 | 5.31404 | 7.99385 | 4.80712 | 3.49096 | 4.97072 | 5.30977 | 8.01303 | 8.5 | 7.57826 | 9.72605 | 2.73245 | 5.11162 | 3.24671 | 4.41282 |
| FBXL17 | Q9UF56 | 6.1247 | 6.62314 | 6.87199 | 6.97586 | 4.54644 | 5.48937 | 6.19871 | 6.50442 | 5.47411 | 6.30379 | 5.70971 | 6.34364 | 5.89666 | 7.94374 | 6.71932 | 6.49807 |
| CCDC34 | Q96HJ3 | 6.01565 | 7.07651 | 6.74918 | 7.00008 | 6.62548 | 6.171 | 5.61436 | 6.69617 | 7.65333 | 7.09951 | 6.70058 | 6.94594 | 5.29975 | 5.01912 | 4.75041 | 4.58294 |
| UBE2A | P49459 | 5.77081 | 6.06927 | 6.10541 | 4.91314 | 6.13838 | 5.38796 | 5.30556 | 7.8682 | 5.84146 | 6.98524 | 5.50313 | 6.22882 | 5.51914 | 7.82339 | 7.34768 | 7.19239 |
| AMIGO1 | Q86WK6 | 7.43845 | 7.52254 | 7.2374 | 6.46629 | 7.88208 | 6.76452 | 4.78521 | 7.61535 | 6.13114 | 3.47891 | 7.33281 | 5.9939 | 6.89293 | 4.90559 | 4.02139 | 5.53149 |
| ADGRD1 | Q6QNK2-4 | 7.11516 | 4.84413 | 6.09603 | 6.84436 | 7.02733 | 6.68556 | 5.45437 | 7.47921 | 5.76036 | 4.26658 | 5.94762 | 5.88348 | 6.05322 | 6.74132 | 7.55139 | 6.24989 |
| CLUAP1 | I3L2E1 | 4.20834 | 5.23446 | 7.01612 | 6.16077 | 7.09911 | 9.15975 | 5.96606 | 5.86314 | 7.13341 | 5.94485 | 6.8596 | 6.11119 | 7.42925 | 5.55029 | 5.68396 | 4.57969 |
| PPARD | Q03181 | 3.19345 | 5.58436 | 5.19805 | 5.71456 | 4.19254 | 6.16992 | 6.68824 | 7.08887 | 7.08239 | 6.9294 | 7.05729 | 6.79887 | 8.1757 | 4.99561 | 7.88041 | 7.25036 |
| LEPROTL1 | O95214-2 | 6.89166 | 7.0787 | 6.48065 | 6.24549 | 5.73943 | 6.2018 | 7.39738 | 5.72367 | 7.39233 | 6.87071 | 7.39281 | 6.03055 | 5.07145 | 5.3633 | 4.91655 | 5.20351 |
| SLC16A7 | O60669 | 7.17835 | 6.9195 | 6.97219 | 5.90989 | 6.03567 | 6.23248 | 6.26617 | 5.88134 | 7.5293 | 5.96754 | 6.41957 | 5.33656 | 6.16163 | 6.06518 | 4.4684 | 6.65622 |
| HDDC3 | Q8N4P3 | 6.06437 | 4.67151 | 7.33254 | 3.9884 | 7.01019 | 5.29045 | 5.17263 | 5.8217 | 5.92366 | 7.2233 | 7.06292 | 6.59351 | 6.75277 | 7.74548 | 7.42806 | 5.91853 |
| BCL2L12 | Q9HB09 | 4.85621 | 7.09388 | 6.63255 | 4.86189 | 4.48852 | 3.71567 | 4.54325 | 7.22668 | 4.79562 | 3.49832 | 6.32815 | 8.05209 | 5.71162 | 9.29808 | 10.9054 | 7.99203 |
| AKTIP | Q9H8T0-2 | 6.37216 | 6.61417 | 7.00555 | 5.83913 | 6.82943 | 7.32414 | 6.10045 | 6.02207 | 6.06236 | 6.26563 | 5.8993 | 6.18361 | 5.49435 | 6.41925 | 5.65397 | 5.91443 |
| C6orf136 | Q5SQH8-4 | 6.46471 | 6.65062 | 6.4663 | 6.54563 | 5.92597 | 5.91544 | 6.83013 | 5.60714 | 6.39272 | 5.97126 | 5.83742 | 6.33822 | 5.89863 | 6.07467 | 6.27113 | 6.81002 |
| ZNF239 | Q16600 | 4.57236 | 5.84922 | 5.69879 | 6.0363 | 5.34895 | 7.40767 | 5.12957 | 5.09356 | 6.70179 | 5.746 | 6.4486 | 6.44076 | 8.42263 | 7.86373 | 7.15518 | 6.08489 |
| ZYG11A | Q6WRX3 | 7.00099 | 7.53502 | 7.37266 | 6.36664 | 6.11888 | 6.52567 | 7.47463 | 7.03904 | 6.52823 | 6.67533 | 6.92717 | 6.75201 | 4.10781 | 4.4339 | 4.86786 | 4.27415 |
| XRCC2 | O43543 | 5.29889 | 5.89549 | 6.02034 | 6.11377 | 5.56688 | 5.68136 | 5.84531 | 6.25934 | 6.45749 | 6.16759 | 6.28367 | 6.53212 | 6.74224 | 7.19485 | 6.96519 | 6.97546 |
| POLR2K | P53803 | 7.14603 | 6.94986 | 7.31059 | 6.57833 | 6.93371 | 6.88688 | 6.57515 | 6.45151 | 6.46661 | 5.93719 | 5.92329 | 5.76212 | 6.26805 | 5.077 | 4.61442 | 5.11928 |
| RPP14 | O95059 | 5.74462 | 7.05728 | 6.41827 | 6.52438 | 6.17695 | 5.88311 | 5.7912 | 5.80687 | 5.21923 | 6.29015 | 6.01387 | 6.271 | 6.50235 | 6.16801 | 7.02182 | 7.1109 |
| NAT8L | Q8N9F0 | 6.77083 | 6.92607 | 6.94346 | 7.28494 | 5.85274 | 7.46108 | 6.98213 | 4.18364 | 7.03733 | 6.6748 | 6.9136 | 5.85419 | 6.68652 | 4.35952 | 4.07171 | 5.99743 |
| CD70 | P32970-2 | 7.16706 | 5.4435 | 7.37907 | 5.40601 | 6.19664 | 6.33389 | 6.33183 | 5.42468 | 5.97622 | 6.7727 | 6.65839 | 6.48419 | 5.77855 | 8.17744 | 5.14288 | 5.32695 |
| DISP1 | Q96F81 | 5.80962 | 7.18734 | 6.82538 | 7.03506 | 6.36943 | 6.2931 | 6.1706 | 6.21972 | 6.41559 | 6.18675 | 5.78702 | 6.09833 | 5.95107 | 6.23353 | 5.97225 | 5.44521 |
| ANAPC11 | Q9NYG5 | 7.9702 | 6.01755 | 6.17336 | 5.81067 | 4.12801 | 6.13488 | 8.31583 | 5.86792 | 6.40598 | 7.14142 | 7.13022 | 5.65303 | 5.33115 | 7.50943 | 5.15148 | 5.25888 |
| TMEM258 | P61165 | 6.71951 | 7.81538 | 5.6389 | 8.17155 | 5.62499 | 6.12686 | 6.54373 | 6.80903 | 6.21603 | 5.10836 | 6.90682 | 5.07462 | 4.83146 | 6.63832 | 5.48808 | 6.28635 |
| PNKP | A0A1B0GVJ3 | 6.06597 | 6.49054 | 6.73614 | 5.19458 | 5.21039 | 6.1809 | 6.45489 | 4.63757 | 6.48505 | 7.2418 | 6.64972 | 6.85228 | 7.21965 | 7.0183 | 6.52826 | 5.03397 |
| AKIRIN2 | Q53H80 | 6.48877 | 6.36568 | 6.31608 | 6.70514 | 7.30005 | 6.37381 | 6.16929 | 7.14884 | 5.76114 | 9.03742 | 7.73097 | 7.03648 | 3.40126 | 5.18874 | 4.4737 | 4.50263 |
| TIMP2 | P16035 | 5.02436 | 7.13114 | 6.37415 | 6.87247 | 6.4876 | 4.41517 | 5.74229 | 7.30184 | 5.05795 | 7.0646 | 6.11227 | 6.60251 | 7.86364 | 6.81609 | 6.70537 | 4.42854 |
| IP6K1 | Q92551 | 6.88908 | 7.13857 | 6.14598 | 7.68811 | 5.74993 | 6.76698 | 5.88245 | 6.60416 | 5.63055 | 6.2177 | 6.07406 | 6.12437 | 5.26495 | 5.91268 | 6.27546 | 5.63497 |
| SPCS1 | A0A5F9YFS9 | 6.55626 | 6.80012 | 5.56938 | 6.71523 | 6.16822 | 6.45929 | 8.60536 | 5.26845 | 5.92078 | 5.68887 | 4.91838 | 6.21754 | 5.12291 | 6.7935 | 7.653 | 5.54271 |
| FAM25A | B3EWG3 | 4.88381 | 7.47299 | 7.40875 | 5.56117 | 6.11221 | 5.49382 | 4.73942 | 6.69101 | 5.31297 | 3.88729 | 7.54039 | 6.41379 | 8.73013 | 8.60285 | 5.94402 | 5.20539 |
| MT-ATP8 | P03928 | 5.24253 | 4.55492 | 5.49217 | 5.72763 | 5.74207 | 4.7284 | 4.97882 | 5.80461 | 3.74259 | 6.24167 | 4.91269 | 5.82226 | 6.2248 | 11.9169 | 8.83148 | 10.0365 |
| RAD51B | O15315 | 6.16174 | 5.97543 | 7.19806 | 6.54832 | 5.31082 | 6.41183 | 5.77207 | 6.03752 | 4.91016 | 5.11826 | 6.07436 | 8.59244 | 6.01045 | 6.19376 | 7.81362 | 5.87116 |
| GAB3 | Q8WWW8-2 | 7.62004 | 6.27239 | 6.91015 | 6.83944 | 6.82349 | 6.62506 | 5.72191 | 5.65285 | 6.17234 | 6.51339 | 6.50339 | 5.78148 | 6.55053 | 6.09538 | 4.79461 | 5.12355 |
| MLH3 | Q9UHC1 | 5.48255 | 6.42113 | 6.33551 | 5.50888 | 6.06298 | 7.09483 | 6.84101 | 6.05053 | 6.78152 | 6.83924 | 5.36395 | 4.84469 | 6.28335 | 7.72792 | 7.08163 | 5.28026 |
| TMEM181 | Q9P2C4 | 6.14479 | 6.5185 | 6.46265 | 5.89488 | 6.67632 | 6.1203 | 5.55179 | 5.7949 | 7.10842 | 6.61693 | 6.66375 | 6.36809 | 5.61013 | 5.22322 | 6.49638 | 6.74893 |
| CD99 | P14209 | 5.67687 | 4.3262 | 4.7442 | 4.92828 | 4.7731 | 3.59745 | 5.71333 | 5.83076 | 3.77619 | 5.32262 | 4.36218 | 6.60362 | 5.88812 | 12.6138 | 9.32336 | 12.5199 |

| | | | | | | | | | | | | | | | | | |
|---|---|---|---|---|---|---|---|---|---|---|---|---|---|---|---|---|---|
| GATA2 | P23769 | 6.27498 | 6.19766 | 5.99517 | 8.14866 | 6.43809 | 5.86258 | 6.64618 | 5.98937 | 6.4598 | 6.82922 | 6.47281 | 6.34532 | 6.80976 | 3.57831 | 5.9083 | 6.04379 |
| CCSER2 | Q9H7U1-3 | 7.25798 | 6.36473 | 7.84659 | 6.84561 | 3.4649 | 5.46712 | 7.4024 | 7.4744 | 5.797 | 6.67244 | 5.95964 | 5.74412 | 6.52233 | 5.44874 | 4.81671 | 6.91529 |
| FBXO33 | Q7Z6M2 | 4.48944 | 4.01399 | 7.70941 | 4.33607 | 5.79896 | 8.25466 | 5.48636 | 4.38003 | 6.04007 | 6.05143 | 6.58141 | 6.82825 | 6.24395 | 7.97725 | 7.70783 | 9.50087 |
| FAM200B | P0CF97 | 6.87162 | 6.30111 | 5.73826 | 5.89114 | 6.72223 | 7.50216 | 5.2685 | 5.47472 | 4.98027 | 6.30502 | 6.67544 | 7.43562 | 6.2987 | 5.32262 | 7.15193 | 6.06067 |
| SMDT1 | Q9H4I9 | 6.74664 | 7.24443 | 5.87936 | 7.34899 | 6.46827 | 6.12068 | 6.10419 | 4.85984 | 5.95835 | 5.52116 | 5.7661 | 6.02061 | 6.53987 | 5.78345 | 6.0702 | 7.56788 |
| ZDHHC9 | Q9Y397 | 6.78039 | 7.11833 | 7.12752 | 7.28878 | 6.53179 | 6.0047 | 6.86884 | 6.1753 | 6.66655 | 6.54661 | 6.85251 | 7.04841 | 4.79641 | 4.61049 | 4.47595 | 5.1074 |
| ATXN7L3 | Q14CW9-2 | 7.15263 | 6.55392 | 6.79757 | 6.4037 | 5.86848 | 5.60352 | 5.56509 | 4.88826 | 7.33375 | 6.24145 | 5.65666 | 5.7233 | 6.25888 | 7.15625 | 5.86815 | 5.92841 |
| FAM104A | Q969W3-2 | 6.13227 | 6.40474 | 6.90023 | 7.0784 | 6.3915 | 5.64665 | 6.19354 | 5.86311 | 6.76886 | 7.31636 | 7.19213 | 7.34564 | 5.57665 | 5.43551 | 4.66114 | 5.09326 |
| PKDREJ | Q9NTG1 | 6.33549 | 6.617 | 5.58015 | 6.25576 | 7.17616 | 5.88125 | 6.41666 | 5.70175 | 5.89893 | 5.96371 | 6.33002 | 5.57078 | 6.57244 | 6.1204 | 7.48166 | 6.09782 |
| IER3IP1 | Q9Y5U9 | 5.86139 | 5.91147 | 5.9695 | 6.25241 | 6.05241 | 5.77057 | 6.8555 | 5.42281 | 6.91573 | 5.00882 | 6.01954 | 5.72847 | 6.51413 | 6.56307 | 7.50431 | 7.64986 |
| DUSP22 | Q9NRW4 | 6.40357 | 6.6701 | 6.96446 | 5.63473 | 6.74028 | 6.80806 | 6.66387 | 6.2643 | 7.12646 | 6.28127 | 7.30785 | 7.09657 | 4.84143 | 5.30388 | 4.48706 | 5.40612 |
| ZNF124 | Q15973 | 6.99102 | 6.75892 | 7.74799 | 7.70578 | 5.51165 | 5.33766 | 6.31752 | 6.84534 | 6.3848 | 7.68671 | 6.56927 | 6.87501 | 4.75534 | 5.01644 | 4.14781 | 5.34872 |
| ZNF57 | Q68EA5 | 6.09154 | 5.42283 | 6.0974 | 6.1353 | 5.95718 | 6.56768 | 6.95053 | 6.05789 | 6.21647 | 5.70377 | 6.16713 | 5.86131 | 6.14472 | 6.90794 | 6.7467 | 6.97161 |
| TUSC2 | O75896 | 6.75234 | 5.66426 | 6.46857 | 5.71339 | 6.84121 | 6.43735 | 5.12672 | 5.14027 | 6.21649 | 8.04509 | 6.69379 | 5.83573 | 8.10094 | 5.13865 | 6.63774 | 5.18745 |
| VPS37A | Q8NEZ2 | 6.41136 | 6.25018 | 6.49501 | 5.35421 | 5.6177 | 5.67578 | 6.29717 | 6.76128 | 6.45863 | 7.05404 | 6.84672 | 7.19309 | 5.68323 | 5.90277 | 5.89981 | 6.09902 |
| RABL2A | Q9UBK7-3 | 6.13358 | 8.13158 | 7.17842 | 6.25468 | 5.13448 | 7.18183 | 5.90425 | 6.57532 | 8.22672 | 6.01156 | 5.88887 | 7.02976 | 4.25111 | 6.1629 | 4.63047 | 5.30445 |
| MEST | Q5EB52 | 5.47158 | 5.04995 | 6.55524 | 7.73295 | 5.10094 | 5.87944 | 8.2506 | 6.90206 | 4.71089 | 5.61092 | 6.82587 | 6.75369 | 6.90397 | 5.49549 | 7.16655 | 5.58987 |
| MCFD2 | Q8NI22 | 5.39192 | 4.74671 | 4.07395 | 4.06731 | 5.13953 | 3.5822 | 5.80479 | 6.13758 | 3.80269 | 4.36155 | 4.14161 | 5.96642 | 6.77544 | 13.9061 | 8.25817 | 13.844 |
| EYA4 | O95677 | 6.2065 | 6.13586 | 4.79305 | 6.42372 | 7.20661 | 7.15248 | 6.21996 | 5.62906 | 6.86079 | 6.29583 | 6.69405 | 5.10524 | 6.45268 | 7.81099 | 6.41434 | 4.59884 |
| NEXMIF | Q5QGS0 | 6.87167 | 6.83868 | 6.45745 | 7.22511 | 5.1477 | 6.41942 | 7.0563 | 6.15201 | 6.25006 | 6.51454 | 6.36295 | 5.71849 | 5.17415 | 5.38683 | 6.30265 | 6.12199 |
| TUBD1 | Q9UJT1 | 6.36667 | 6.25837 | 6.28811 | 6.68366 | 6.09385 | 7.22563 | 6.07437 | 6.37636 | 6.07755 | 7.30497 | 6.70439 | 7.04424 | 5.49652 | 5.72159 | 5.00983 | 5.27389 |
| GPRC5A | Q8NFJ5 | 6.40016 | 6.18179 | 7.69283 | 6.76855 | 5.63822 | 5.34605 | 6.36579 | 5.58183 | 6.62845 | 5.90589 | 6.09536 | 5.9214 | 6.25059 | 5.73413 | 6.83812 | 6.65084 |
| SNRNP25 | Q9BV90 | 7.06765 | 7.68231 | 7.60308 | 6.58744 | 6.95396 | 6.30459 | 6.73141 | 5.9285 | 5.47924 | 6.05009 | 6.06553 | 4.64975 | 5.77706 | 5.01706 | 5.50309 | 6.59925 |
| MED8 | Q96G25-2 | 5.87921 | 4.33972 | 6.05816 | 4.81501 | 5.84623 | 3.83626 | 5.49665 | 6.49679 | 5.06502 | 6.95495 | 5.65611 | 7.94965 | 7.23426 | 8.17684 | 6.49526 | 8.69988 |
| DNAJA4 | Q8WW22-2 | 7.14342 | 6.15457 | 7.20541 | 5.47892 | 5.35141 | 5.56344 | 6.79254 | 5.4036 | 6.82829 | 6.78933 | 6.94767 | 7.27823 | 5.49083 | 5.78083 | 5.88528 | 5.90622 |
| FBXW4 | P57775 | 6.88588 | 5.31164 | 6.31475 | 6.28059 | 5.86713 | 4.97908 | 5.95435 | 6.56638 | 6.20869 | 6.41453 | 6.38127 | 6.92674 | 6.68535 | 6.02278 | 7.06189 | 6.13895 |
| CHST11 | Q9NPF2 | 7.56045 | 6.57419 | 6.53947 | 6.70221 | 5.46795 | 6.59313 | 6.7398 | 4.94454 | 6.87142 | 6.01184 | 6.71854 | 7.28752 | 4.85266 | 6.15138 | 6.26929 | 5.61562 |
| CADPS | Q9ULU8-4 | 6.46358 | 6.35483 | 6.56676 | 6.34464 | 6.66633 | 6.07907 | 5.92336 | 6.31701 | 5.89729 | 6.40404 | 6.46116 | 6.31298 | 6.31139 | 6.44536 | 6.0449 | 5.4073 |
| NTN1 | O95631 | 5.92394 | 5.55897 | 6.80106 | 4.78111 | 5.7024 | 6.85042 | 6.67157 | 6.70383 | 5.90112 | 6.35072 | 6.14654 | 7.05619 | 5.86843 | 6.23451 | 7.02418 | 6.42501 |
| ZNF444 | Q8N0Y2 | 5.95444 | 5.11755 | 7.02735 | 7.20481 | 6.39244 | 6.17963 | 6.66393 | 5.72144 | 6.51666 | 6.76162 | 7.26919 | 6.59692 | 5.70964 | 6.28228 | 5.76207 | 4.84003 |
| ZNF205 | O95201 | 6.16961 | 6.70113 | 6.47952 | 7.39115 | 6.47052 | 6.83332 | 6.04018 | 6.55548 | 6.33069 | 6.64808 | 6.49033 | 7.09795 | 5.25571 | 5.58417 | 4.63436 | 5.31779 |
| CMTM3 | Q96MX0 | 6.21698 | 4.81703 | 5.88866 | 6.73643 | 5.75215 | 5.86808 | 6.60147 | 5.70102 | 5.55308 | 6.67011 | 5.97824 | 6.3954 | 6.52326 | 7.07145 | 7.23034 | 6.99631 |
| IMUP | Q9GZP8 | 6.32476 | 5.45574 | 7.44202 | 6.23235 | 6.40701 | 6.89926 | 6.21407 | 6.30443 | 6.79896 | 6.87047 | 6.84685 | 7.55444 | 5.26662 | 5.19841 | 4.82762 | 5.357 |
| COX10 | Q12887 | 6.42539 | 4.77158 | 5.91044 | 6.36797 | 6.49979 | 6.18339 | 6.90196 | 6.36407 | 5.85555 | 5.83964 | 6.85914 | 5.88458 | 6.11039 | 6.36348 | 6.39703 | 7.26559 |
| SH2D5 | Q6ZV89 | 6.1637 | 4.96054 | 7.92228 | 5.54793 | 6.50685 | 5.17186 | 8.51477 | 5.81926 | 6.45677 | 5.88332 | 5.76723 | 5.7145 | 7.62538 | 6.82644 | 5.63705 | 5.4821 |
| DUSP16 | Q9BY84 | 6.23001 | 5.28096 | 7.28253 | 6.16414 | 7.13671 | 6.93886 | 6.95155 | 6.72032 | 6.11741 | 6.3262 | 6.42645 | 6.78003 | 5.98826 | 5.28808 | 4.75947 | 5.60903 |
| ROBO2 | Q9HCK4-3 | 6.56403 | 6.87929 | 5.3375 | 5.75946 | 5.71855 | 5.20669 | 7.09551 | 5.74052 | 5.2541 | 5.58209 | 5.3072 | 5.91229 | 7.00072 | 7.46283 | 7.40923 | 7.76999 |
| YAF2 | Q8IY57-5 | 5.45427 | 5.15276 | 8.58283 | 6.09387 | 5.78472 | 6.16735 | 4.52701 | 6.06777 | 5.60488 | 5.86718 | 6.34923 | 7.29594 | 6.72173 | 7.26529 | 6.03014 | 7.03501 |
| JCAD | Q9P266 | 6.43559 | 7.39404 | 6.6307 | 7.47699 | 5.84033 | 6.27467 | 6.5025 | 6.27591 | 5.8576 | 5.673 | 6.21511 | 7.16114 | 5.51875 | 5.80387 | 5.97402 | 5.96577 |
| CD164 | Q04900 | 7.39214 | 6.31158 | 7.31293 | 8.21022 | 7.10201 | 6.04126 | 6.71923 | 5.69183 | 6.8945 | 7.3185 | 6.31193 | 6.98836 | 4.61964 | 4.25902 | 3.9893 | 4.83754 |
| ACBD7 | Q8N6N7 | 4.89848 | 4.3163 | 4.71149 | 4.4782 | 4.69593 | 5.10333 | 5.70165 | 5.87777 | 4.52608 | 6.19659 | 4.9083 | 4.72709 | 4.80148 | 13.4597 | 9.06697 | 12.5307 |
| KIAA0513 | O60268 | 6.81759 | 6.00522 | 7.24735 | 6.20266 | 6.36502 | 6.10616 | 6.91088 | 6.12842 | 6.36283 | 7.15883 | 8.03491 | 7.14833 | 4.0431 | 4.81497 | 5.43677 | 5.21695 |
| HERPUD2 | Q9BSE4 | 8.35629 | 5.57862 | 7.14889 | 6.78537 | 6.19029 | 7.49962 | 6.7168 | 5.12766 | 6.30692 | 6.27524 | 6.36068 | 5.83842 | 5.05545 | 6.25399 | 4.51132 | 5.99445 |
| GAGE13 | Q4V321 | 4.70409 | 5.36749 | 5.57772 | 5.21037 | 3.81807 | 2.66388 | 3.84672 | 5.75886 | 3.15367 | 5.19043 | 4.05466 | 4.63258 | 4.76601 | 17.9557 | 8.98433 | 14.3155 |
| ENTR1 | Q96C92 | 6.04531 | 7.07722 | 6.3705 | 5.16666 | 7.12808 | 6.13297 | 5.50087 | 6.01573 | 7.68369 | 6.04707 | 5.82046 | 6.83227 | 6.56573 | 5.83842 | 6.00996 | 5.76508 |
| ZBED9 | Q6R2W3 | 6.41173 | 7.07866 | 6.11342 | 7.82887 | 5.44134 | 6.59857 | 6.15234 | 7.51545 | 6.14658 | 7.00593 | 6.47411 | 6.28909 | 4.72784 | 6.24574 | 4.78079 | 5.18954 |
| ACOT9 | F8WDI2 | 5.22922 | 4.84577 | 6.28487 | 6.52664 | 5.55878 | 6.47839 | 6.68031 | 5.87147 | 5.81159 | 6.19123 | 5.03749 | 6.28311 | 8.03217 | 7.28404 | 6.84502 | 7.03989 |
| SLC35E2B | P0CK96 | 5.50621 | 6.19721 | 6.03267 | 7.20743 | 6.23385 | 6.66056 | 6.31817 | 6.10062 | 5.90527 | 7.2038 | 6.89338 | 6.36194 | 5.73345 | 6.05418 | 5.75212 | 5.83914 |
| MAP2K5 | Q13163 | 5.58821 | 6.22422 | 7.03493 | 6.37634 | 7.22556 | 6.0247 | 5.93354 | 5.79332 | 6.48828 | 6.49463 | 7.15244 | 7.17595 | 5.9075 | 5.68076 | 5.55485 | 5.34477 |
| CKS2 | P33552 | 5.89286 | 5.3765 | 6.9413 | 5.63634 | 5.10586 | 5.96466 | 5.49547 | 7.4375 | 5.12823 | 7.00152 | 5.92559 | 6.69477 | 3.57549 | 8.69854 | 5.87355 | 9.25181 |
| INSIG2 | Q9Y5U4 | 6.41496 | 6.34617 | 6.72884 | 6.64418 | 5.94537 | 6.20812 | 6.32594 | 5.28283 | 6.40599 | 6.17359 | 6.03459 | 6.41252 | 6.37078 | 6.26361 | 6.0187 | 6.42381 |
| UBE2J2 | Q8N2K1-3 | 6.7699 | 6.70581 | 6.43868 | 6.83337 | 6.54481 | 5.67288 | 6.85942 | 5.90464 | 6.18515 | 5.70659 | 7.72332 | 6.54434 | 5.57657 | 5.31637 | 5.18657 | 6.03157 |
| BBX | H7C1U1 | 4.80677 | 5.26574 | 7.18379 | 6.36131 | 6.99249 | 5.05831 | 5.51311 | 6.8834 | 7.26981 | 5.61579 | 6.31588 | 5.47917 | 6.49412 | 7.0811 | 6.4783 | 7.2009 |
| CGRRF1 | Q99675 | 6.06407 | 8.04613 | 7.06277 | 5.91943 | 5.39327 | 4.52342 | 9.08182 | 4.12875 | 6.68812 | 6.90683 | 6.46577 | 7.24216 | 5.78348 | 5.36901 | 5.45586 | 5.86912 |
| CREB5 | Q02930 | 5.81899 | 5.71656 | 6.21311 | 6.20152 | 6.28178 | 5.54516 | 6.67605 | 6.1418 | 5.90796 | 6.3138 | 7.57765 | 7.06019 | 6.56433 | 5.73461 | 5.63453 | 6.61196 |
| CDK2AP1 | O14519 | 5.44015 | 4.79918 | 6.02533 | 5.55562 | 6.83671 | 4.57051 | 5.81397 | 5.77765 | 5.95267 | 6.82069 | 6.7893 | 8.04415 | 5.58687 | 7.15054 | 7.30753 | 7.52914 |
| RAD52 | P43351 | 6.14288 | 6.3747 | 6.31906 | 6.10707 | 6.50513 | 6.24816 | 6.74499 | 6.09052 | 6.72755 | 5.4188 | 6.02147 | 6.26872 | 6.69725 | 5.87776 | 6.34658 | 6.10937 |
| IGHMBP2 | A0A6Q8PGC0 | 5.80955 | 5.39192 | 5.66217 | 6.19124 | 5.33861 | 5.38622 | 6.03451 | 6.39365 | 6.94397 | 6.05836 | 5.98329 | 6.22585 | 7.22363 | 6.22818 | 7.45756 | 7.67129 |
| C1orf109 | Q9NX04 | 6.09922 | 4.61439 | 6.43527 | 4.78821 | 6.78501 | 5.32332 | 5.84908 | 6.36758 | 7.34228 | 6.87219 | 5.596 | 6.59483 | 5.37606 | 7.66388 | 7.19367 | 7.09903 |
| EMC9 | Q9Y3B6 | 5.52832 | 5.21973 | 6.75112 | 6.38798 | 6.18804 | 6.20473 | 6.69905 | 6.84604 | 5.40912 | 5.18721 | 6.9041 | 7.82461 | 5.0867 | 6.44713 | 6.49553 | 6.82058 |
| C4orf3 | Q8WVX3-2 | 5.86037 | 5.9539 | 5.18164 | 6.02323 | 6.95512 | 7.38327 | 6.64015 | 6.36813 | 6.829 | 5.24089 | 4.84474 | 5.61825 | 5.98607 | 7.50238 | 6.00609 | 7.60675 |
| ARID3B | Q8IVW6 | 7.13357 | 8.12247 | 6.12358 | 7.71425 | 5.57036 | 4.71087 | 6.32539 | 5.42952 | 6.54435 | 7.16916 | 8.26077 | 5.8587 | 5.07528 | 6.7893 | 4.79969 | 4.37274 |
| SPRR2G | Q9BYE4 | 0.907881 | 70.0743 | 0.893704 | 9.52228 | 0.237164 | 0 | 1.18567 | 0.948212 | 0.652258 | 2.28694 | 0.555697 | 1.07625 | 2.0006 | 1.91064 | 1.30149 | 6.44695 |
| PIGC | Q92535 | 6.06164 | 6.25274 | 6.48134 | 6.4912 | 6.32071 | 5.725 | 5.88951 | 6.09582 | 6.73352 | 5.97053 | 6.36596 | 6.01023 | 6.39661 | 6.22875 | 6.77497 | 6.20147 |
| NLRP3 | Q96P20 | 5.4205 | 6.40177 | 6.66649 | 7.35005 | 6.87542 | 4.96519 | 6.07314 | 6.18812 | 6.53321 | 4.88538 | 5.78508 | 7.10451 | 6.45972 | 6.68869 | 6.15637 | 6.44636 |
| TSNARE1 | A0A590UJA6 | 4.64351 | 6.61307 | 4.33399 | 5.74869 | 7.78413 | 5.21964 | 4.97742 | 7.11944 | 5.86202 | 4.62499 | 6.83008 | 5.37513 | 8.47042 | 8.2299 | 6.90992 | 7.25764 |
| SAV1 | Q9H4B6 | 7.18133 | 6.19542 | 5.77234 | 6.45273 | 6.55837 | 6.35911 | 5.93082 | 6.2202 | 5.81483 | 7.83134 | 6.30282 | 6.67221 | 5.79534 | 5.20292 | 6.05401 | 5.65621 |
| GRAMD2B | Q96HH9-3 | 6.63325 | 6.46087 | 6.56225 | 6.61317 | 5.93507 | 6.91751 | 6.46265 | 5.39061 | 6.26687 | 5.72249 | 6.34442 | 6.41892 | 5.966 | 6.1234 | 6.40151 | 5.781 |
| RNF175 | Q8N4F7 | 6.88492 | 6.14456 | 6.02139 | 7.41581 | 5.31314 | 5.80186 | 7.51237 | 3.76192 | 5.19746 | 8.20609 | 6.97082 | 6.53869 | 5.88331 | 6.66406 | 5.2353 | 6.44831 |
| LHFPL4 | Q7Z7J7 | 7.05546 | 6.25354 | 5.71544 | 5.13348 | 5.20575 | 6.20831 | 6.11847 | 5.55985 | 6.19155 | 6.39358 | 6.3474 | 6.70517 | 6.48617 | 6.71706 | 6.60462 | 7.30415 |
| RBX1 | P62877 | 6.57251 | 6.21148 | 6.72207 | 7.04417 | 4.94078 | 5.57409 | 6.177 | 6.57796 | 6.33595 | 8.23319 | 5.05102 | 5.12631 | 6.99261 | 5.94273 | 7.13162 | 5.36652 |
| C20orf194 | Q5TEA3 | 6.8196 | 6.37096 | 7.89288 | 5.87626 | 6.48939 | 6.75863 | 6.6018 | 6.35114 | 6.91832 | 5.52837 | 6.28581 | 6.7499 | 4.43722 | 5.59684 | 5.87268 | 5.4502 |
| NECAB3 | Q96P71 | 5.09947 | 6.44064 | 7.75116 | 6.57191 | 6.46127 | 6.3952 | 6.33016 | 6.40034 | 7.63793 | 7.68096 | 8.31679 | 5.87699 | 4.3064 | 5.36026 | 4.19 | 5.18049 |
| CCSMST1 | Q4G0I0 | 6.39187 | 6.30814 | 5.53518 | 5.08757 | 5.45993 | 4.92564 | 5.65967 | 5.75912 | 3.67924 | 5.62578 | 6.15098 | 6.56809 | 6.47704 | 9.70163 | 10.066 | 9.10753 |
| LYRM1 | O43325 | 5.44228 | 7.27236 | 6.82106 | 6.82128 | 5.58348 | 5.28356 | 6.27997 | 6.25537 | 6.80107 | 5.59975 | 4.75866 | 7.54047 | 8.0859 | 5.58928 | 5.46859 | 6.39692 |
| ST3GAL3 | Q11203-4 | 6.17553 | 5.77217 | 5.48285 | 6.83028 | 5.64398 | 5.68894 | 5.91739 | 6.17942 | 6.47667 | 6.33922 | 5.87168 | 6.40478 | 6.54573 | 6.23149 | 7.19043 | 7.24944 |
| TOMM40L | Q969M1 | 5.99146 | 6.5362 | 6.03596 | 6.65732 | 6.18351 | 6.73638 | 5.69551 | 6.16427 | 5.66753 | 5.79949 | 6.5561 | 6.09581 | 6.60114 | 6.26474 | 6.80033 | 6.21426 |
| ARSG | Q96EG1 | 6.43146 | 6.72323 | 6.19284 | 6.46058 | 7.71329 | 5.97846 | 6.52681 | 5.045 | 5.58777 | 6.67001 | 7.89854 | 6.48868 | 6.04301 | 6.13419 | 5.35114 | 5.75499 |
| SUMO2 | P61956 | 5.53117 | 6.07326 | 4.80931 | 5.52978 | 5.79184 | 6.22666 | 4.79376 | 7.15953 | 5.15093 | 7.36655 | 5.30798 | 7.30175 | 6.53889 | 8.56871 | 6.11104 | 7.73883 |
| HYLS1 | Q96M11 | 5.92656 | 7.51281 | 7.29491 | 6.54024 | 5.24649 | 6.49662 | 6.85245 | 5.83284 | 8.07165 | 6.55486 | 7.50048 | 7.07665 | 3.17834 | 6.59489 | 3.98126 | 5.33895 |
| PEX26 | Q7Z412 | 6.15427 | 6.66467 | 5.894 | 6.14849 | 5.85378 | 6.13164 | 6.19266 | 6.785 | 5.85063 | 5.61648 | 6.61403 | 6.51589 | 6.56694 | 6.28307 | 6.76247 | 5.96598 |
| FOXRED2 | Q8IWF2 | 6.28448 | 6.8741 | 7.05362 | 7.69172 | 5.33539 | 6.08125 | 6.53997 | 6.85011 | 7.11327 | 6.2594 | 8.84656 | 7.37384 | 4.60153 | 5.54023 | 4.31483 | 3.23971 |
| S1PR1 | P21453 | 6.2774 | 6.65478 | 6.03482 | 6.81247 | 5.02193 | 6.22879 | 7.46553 | 6.82004 | 6.32912 | 5.78548 | 7.36128 | 7.01198 | 5.15519 | 5.34221 | 6.1941 | 5.50487 |
| KNG1 | P01042 | 5.54205 | 8.17934 | 4.67889 | 6.01665 | 3.83777 | 5.5811 | 9.30239 | 4.00438 | 6.46498 | 5.62481 | 3.18655 | 5.4819 | 7.97034 | 6.33403 | 8.96761 | 8.82722 |
| FASTK | Q14296 | 9.25894 | 7.2865 | 5.0535 | 7.1284 | 4.77995 | 7.81698 | 6.10106 | 5.90423 | 6.81769 | 3.79379 | 3.41865 | 7.38871 | 7.19996 | 7.64059 | 5.72858 | 4.68246 |
| GABPB2 | Q5SZG2 | 4.44605 | 4.1622 | 6.78569 | 6.21589 | 5.10816 | 4.96959 | 5.90004 | 6.18066 | 7.38162 | 6.40231 | 6.84623 | 7.97252 | 7.90199 | 6.61683 | 6.22076 | 6.88945 |
| LPIN3 | Q9BQK8-2 | 5.63603 | 6.55808 | 7.77124 | 6.38443 | 6.56987 | 8.27617 | 6.10488 | 6.4907 | 5.63621 | 5.8211 | 5.48879 | 6.37143 | 4.92829 | 5.90753 | 6.25051 | 5.80474 |
| YIPF2 | Q9BWQ6 | 6.453 | 5.33868 | 6.71907 | 6.85715 | 6.93597 | 6.44958 | 5.70508 | 6.74639 | 6.38877 | 5.84702 | 7.02017 | 5.44402 | 6.81069 | 6.64117 | 5.13629 | 5.50696 |
| TTC16 | Q8NEE8 | 5.5455 | 6.1138 | 5.94978 | 6.0648 | 5.52112 | 6.14749 | 5.91403 | 5.59799 | 6.00911 | 5.84713 | 5.84301 | 6.2698 | 6.8878 | 7.65137 | 7.25524 | 7.38203 |
| KLHL5 | Q96PQ7-3 | 6.16266 | 5.59542 | 8.45485 | 5.93662 | 7.67088 | 4.93021 | 5.44689 | 6.36153 | 6.26896 | 6.46733 | 7.78104 | 7.11833 | 5.49577 | 6.04355 | 4.80602 | 5.45994 |
| RGS3 | P49796 | 7.09775 | 7.97248 | 6.44234 | 7.23719 | 6.8123 | 5.20331 | 6.07246 | 6.88121 | 6.69587 | 5.63745 | 6.57887 | 7.97705 | 4.75906 | 5.2523 | 5.46593 | 3.91443 |
| RAD9A | Q99638 | 3.92546 | 5.72793 | 5.86962 | 5.50523 | 6.4585 | 7.19785 | 6.08415 | 6.00647 | 5.28511 | 5.45084 | 7.57487 | 6.5011 | 6.92501 | 6.53804 | 7.29275 | 7.65708 |
| WSB2 | Q9NYS7-2 | 5.5119 | 6.4304 | 7.09218 | 5.63453 | 5.94706 | 5.62151 | 6.01053 | 6.45209 | 6.76928 | 7.31338 | 6.82554 | 6.31445 | 5.27198 | 6.01186 | 5.8282 | 6.96511 |
| CENPW | Q5EE01 | 6.14634 | 6.64362 | 5.77223 | 6.98713 | 6.34935 | 6.36814 | 6.85761 | 5.97269 | 6.3737 | 5.97889 | 6.03616 | 6.25521 | 6.3318 | 6.00549 | 5.96226 | 5.95938 |
| PDE4A | P27815 | 8.40745 | 5.70235 | 7.02365 | 6.41046 | 5.5158 | 6.47404 | 5.31401 | 6.04168 | 6.67685 | 6.28891 | 5.51645 | 6.34284 | 6.51711 | 6.3824 | 5.24623 | 6.13976 |
| MYADM | Q96S97 | 7.33185 | 7.10287 | 6.57859 | 6.18867 | 7.66125 | 5.73665 | 5.6232 | 5.41388 | 5.28828 | 5.58621 | 6.75413 | 5.12248 | 5.89227 | 7.08042 | 7.09183 | 5.54742 |
| PEMT | Q9UBM1-2 | 6.62983 | 6.05714 | 5.34824 | 5.86027 | 6.11135 | 6.8162 | 5.976 | 5.76696 | 6.45291 | 6.20463 | 5.79845 | 5.51193 | 7.23431 | 6.37871 | 6.73663 | 7.11645 |
| MANBAL | Q9NQG1 | 6.27629 | 6.53936 | 5.87517 | 6.64213 | 6.17757 | 6.42482 | 6.3674 | 6.14792 | 6.00282 | 6.48881 | 5.86861 | 6.15461 | 6.35539 | 6.3042 | 6.10598 | 6.26892 |
| CTDNEP1 | O95476 | 7.42697 | 6.85176 | 6.13637 | 6.99426 | 5.76463 | 6.56464 | 5.64427 | 5.81424 | 6.13104 | 6.46118 | 5.51956 | 6.09719 | 5.13622 | 7.0042 | 5.81556 | 6.63792 |
| NBDY | A0A0U1RRE5 | 5.89558 | 4.58127 | 7.85633 | 5.56969 | 5.95422 | 4.61786 | 4.95196 | 6.10059 | 5.36132 | 6.7893 | 7.51709 | 5.79754 | 4.68204 | 8.36304 | 7.16656 | 8.79562 |
| PLAGL1 | Q9UM63 | 6.09063 | 7.53099 | 6.39964 | 5.95023 | 6.31439 | 8.53708 | 6.44213 | 5.97204 | 6.81644 | 7.25529 | 6.42472 | 6.60238 | 5.15316 | 5.10369 | 3.65395 | 5.75324 |
| CMTM8 | Q8IZV2 | 7.13231 | 7.71502 | 7.25539 | 6.78939 | 7.08391 | 6.32389 | 5.99682 | 8.44105 | 5.3122 | 5.85933 | 7.00813 | 6.26762 | 5.58177 | 3.98305 | 5.36291 | 3.88721 |
| ZBTB8OS | Q8IWT0 | 6.67497 | 5.66016 | 6.38941 | 4.94373 | 6.75315 | 6.02506 | 6.6938 | 8.03911 | 8.19211 | 6.18365 | 6.49544 | 7.25778 | 4.58452 | 6.36282 | 4.63765 | 5.10663 |
| TMEM268 | Q5VZI3-3 | 7.598 | 7.21813 | 6.2152 | 4.7313 | 6.17014 | 5.91236 | 5.45748 | 7.08644 | 6.23473 | 5.32024 | 7.1554 | 7.49142 | 5.44692 | 6.88501 | 4.55898 | 6.51825 |
| PNRC1 | Q12796 | 7.41903 | 8.4509 | 7.37693 | 7.71722 | 5.83904 | 5.87091 | 6.05101 | 5.92524 | 6.33037 | 7.04586 | 7.72207 | 6.78597 | 4.33607 | 4.2787 | 4.04996 | 4.80072 |

| | | | | | | | | | | | | | | | | | |
|---|---|---|---|---|---|---|---|---|---|---|---|---|---|---|---|---|---|
| ZNHIT3 | Q15649 | 6.92733 | 6.43694 | 7.06331 | 6.95788 | 6.66329 | 5.97622 | 6.63703 | 6.69366 | 6.6347 | 5.39246 | 6.56901 | 6.77182 | 5.71587 | 5.00146 | 5.05794 | 5.50106 |
| IPPK | Q9H8X2 | 6.6966 | 6.96284 | 7.3513 | 7.77709 | 6.21238 | 6.20833 | 6.04354 | 5.99036 | 6.7602 | 7.46566 | 7.62458 | 5.7193 | 5.22678 | 4.86718 | 4.56882 | 4.52504 |
| SPPL3 | Q8TCT6 | 6.71171 | 6.37783 | 6.07432 | 7.06852 | 6.07561 | 6.16512 | 5.9523 | 5.25117 | 6.18512 | 7.2585 | 7.15119 | 5.5925 | 6.10356 | 5.83036 | 6.24837 | 5.95381 |
| BORCS8 | Q96FH0 | 6.2182 | 5.35473 | 4.83354 | 5.85628 | 5.83177 | 4.70354 | 5.59653 | 6.33355 | 4.90595 | 5.98034 | 4.60456 | 5.24142 | 4.83859 | 10.1409 | 8.7108 | 10.8493 |
| CPTP | Q5TA50 | 5.94774 | 6.07539 | 6.34012 | 6.14856 | 6.48141 | 6.98295 | 6.75559 | 7.48326 | 5.97901 | 7.10525 | 5.30122 | 7.10244 | 4.82412 | 5.79473 | 5.48317 | 6.19503 |
| TLR8 | Q9NR97-2 | 7.78113 | 5.72043 | 5.99178 | 5.27079 | 5.08607 | 4.32732 | 5.57533 | 6.78956 | 5.88464 | 6.37837 | 5.99113 | 5.72486 | 7.27898 | 6.27732 | 7.65726 | 8.26503 |
| CEP126 | Q9P2H0 | 5.9975 | 6.49254 | 6.24554 | 7.21312 | 7.4392 | 5.76988 | 4.86789 | 7.18668 | 7.44954 | 6.65953 | 6.93915 | 6.93441 | 5.24734 | 4.87287 | 4.39792 | 6.2869 |
| ZDHHC4 | Q9NPG8 | 7.05824 | 5.12826 | 6.07218 | 4.93749 | 6.06146 | 7.42251 | 6.53856 | 5.07751 | 6.37587 | 7.43489 | 5.74224 | 6.08898 | 6.77839 | 6.90333 | 6.28982 | 6.09026 |
| SLC6A8 | P48029 | 6.64896 | 7.04925 | 6.41447 | 6.12664 | 5.23848 | 6.4025 | 5.64325 | 7.21738 | 6.97995 | 5.77023 | 6.28735 | 5.12024 | 6.08355 | 5.83214 | 6.50822 | 6.77741 |
| KCNE2 | Q9Y6J6 | 6.69869 | 6.22919 | 5.46522 | 5.25095 | 6.09992 | 4.86861 | 5.13079 | 7.15291 | 5.12001 | 7.21094 | 6.29989 | 6.60595 | 6.25708 | 7.88408 | 7.26151 | 6.46427 |
| SLC18B1 | Q6NT16 | 6.5315 | 7.4988 | 7.73247 | 4.94331 | 4.87324 | 6.22508 | 5.38716 | 7.15056 | 7.00577 | 5.84776 | 6.96961 | 6.03625 | 6.28754 | 5.05279 | 6.37332 | 6.08483 |
| TSPAN3 | O60637 | 6.60552 | 7.21669 | 6.75227 | 7.04784 | 7.30123 | 6.66228 | 5.84737 | 5.70378 | 6.09344 | 7.91394 | 7.75734 | 5.92704 | 4.36986 | 5.59056 | 4.76856 | 4.44229 |
| IGHV4-28 | A0A0C4DH34 | 6.4954 | 6.25186 | 5.96212 | 6.25483 | 6.76811 | 6.66157 | 5.05724 | 7.03511 | 6.38207 | 5.98834 | 6.96344 | 5.8661 | 7.71183 | 5.96826 | 4.98012 | 5.6536 |
| C2orf16 | C9JG08 | 5.42241 | 6.62246 | 6.529 | 5.93038 | 5.9365 | 6.02819 | 5.69259 | 5.93875 | 6.82002 | 6.0621 | 6.88945 | 7.7305 | 6.13892 | 6.28495 | 6.00376 | 5.97002 |
| TMBIM4 | Q9HC24 | 5.60841 | 6.8858 | 5.98258 | 4.9523 | 6.49593 | 5.45106 | 7.15504 | 8.37978 | 5.5251 | 5.97069 | 4.68173 | 7.71359 | 3.78614 | 6.60575 | 6.53957 | 8.26653 |
| ZSCAN29 | Q8IWY8 | 5.591 | 7.14078 | 6.65344 | 7.39676 | 7.59075 | 5.38286 | 4.86609 | 6.38171 | 6.35142 | 6.70677 | 5.87914 | 5.09042 | 6.03599 | 6.20208 | 6.10068 | 6.6301 |
| SHROOM3 | Q8TF72 | 6.6786 | 7.33522 | 7.38381 | 6.53715 | 9.69723 | 6.4331 | 6.99107 | 4.56549 | 5.7909 | 4.52176 | 6.77231 | 3.24452 | 6.29798 | 6.97236 | 5.5058 | 5.27272 |
| ZNF160 | Q9HCG1 | 6.82429 | 7.37654 | 4.43282 | 7.47281 | 6.39116 | 6.47076 | 6.89142 | 5.42675 | 5.62651 | 6.43928 | 4.86481 | 6.16278 | 6.78004 | 7.154 | 4.81794 | 6.86809 |
| SLC66A1 | Q6ZP29 | 5.85549 | 7.15622 | 6.20646 | 5.5533 | 5.69723 | 5.10789 | 7.35502 | 5.06982 | 6.65208 | 5.71864 | 5.71556 | 6.14599 | 6.22356 | 6.98767 | 7.15772 | 7.39735 |
| ZBTB5 | O15062 | 6.17991 | 6.52445 | 6.17161 | 6.35637 | 6.277 | 6.28249 | 6.09638 | 5.85491 | 6.32403 | 5.83647 | 5.80868 | 5.67733 | 6.71273 | 6.61708 | 6.88414 | 6.39644 |
| NCOR1 | H0Y459 | 6.05154 | 6.0626 | 6.17279 | 6.90345 | 6.29694 | 5.7545 | 6.16202 | 5.9954 | 6.27881 | 6.03055 | 6.82346 | 6.71549 | 6.61951 | 5.73241 | 6.36598 | 6.03456 |
| SPATA2 | Q9UM82 | 6.61153 | 5.80107 | 6.86173 | 5.6355 | 6.1019 | 6.47383 | 7.22573 | 6.71483 | 7.34353 | 5.53233 | 6.10926 | 7.01069 | 5.61046 | 5.58697 | 5.31463 | 6.06601 |
| B4GALT7 | Q9UBV7 | 5.25416 | 6.78861 | 6.62085 | 6.72065 | 6.12689 | 6.27846 | 6.57116 | 4.3495 | 6.97952 | 7.3625 | 5.30296 | 5.95271 | 6.29918 | 5.85324 | 6.71471 | 6.82489 |
| DHRS13 | Q6UX07 | 6.63249 | 6.1351 | 5.82147 | 6.33071 | 6.02174 | 6.06916 | 6.56097 | 5.95182 | 6.09409 | 6.55238 | 6.12681 | 5.90425 | 6.50795 | 6.34079 | 6.43192 | 6.51832 |
| RNF128 | Q8TEB7 | 5.57809 | 7.28972 | 7.18105 | 7.26421 | 6.26843 | 6.09863 | 6.78629 | 6.0425 | 6.47645 | 8.26798 | 6.53355 | 6.30748 | 4.13221 | 5.91372 | 5.62573 | 4.23396 |
| ST8SIA5 | O15466-2 | 6.96959 | 4.94082 | 5.89374 | 6.74704 | 6.44201 | 7.26875 | 6.31002 | 6.40782 | 6.50345 | 4.45533 | 6.87935 | 8.17899 | 5.32948 | 7.24658 | 4.57796 | 5.84906 |
| TMCO6 | Q96DC7-2 | 4.76975 | 5.23827 | 5.91688 | 5.48471 | 5.83735 | 6.3175 | 5.48058 | 5.88714 | 6.22986 | 7.20092 | 7.6189 | 6.39982 | 7.03505 | 6.81266 | 6.9574 | 6.81322 |
| PYURF | Q96I23 | 6.44124 | 4.91503 | 5.79448 | 5.51552 | 6.47347 | 4.58684 | 5.18879 | 6.23984 | 4.47508 | 6.23688 | 5.50415 | 7.29657 | 5.57949 | 8.3247 | 8.40558 | 9.02235 |
| METTL8 | Q9H825 | 6.19778 | 6.37306 | 5.67867 | 6.76329 | 7.18984 | 6.61287 | 6.36527 | 6.56984 | 6.61027 | 5.74472 | 5.21509 | 5.38724 | 6.4209 | 6.01318 | 6.34259 | 6.51539 |
| SEMA3A | Q14563 | 7.6695 | 7.07748 | 5.32535 | 6.22427 | 4.24208 | 5.55416 | 7.98034 | 5.37947 | 6.48683 | 6.2113 | 5.9929 | 4.31135 | 7.12134 | 7.04816 | 5.71729 | 7.64419 |
| SLC25A26 | Q70HW3 | 6.15544 | 5.59195 | 5.94913 | 6.65201 | 6.1376 | 5.63123 | 7.6606 | 6.35267 | 6.63833 | 5.43728 | 6.20561 | 5.54007 | 7.10528 | 5.68457 | 6.84442 | 6.41382 |
| ZSCAN26 | Q16670 | 5.84823 | 5.86418 | 5.88447 | 5.96852 | 6.65169 | 6.62449 | 5.93368 | 6.12275 | 6.02729 | 6.61136 | 6.36183 | 6.72115 | 6.60995 | 6.95115 | 6.09576 | 5.72349 |
| RNFT1 | Q5M7Z0 | 5.72358 | 4.56551 | 6.36394 | 5.7358 | 5.73208 | 5.46094 | 6.54788 | 3.88136 | 6.14986 | 7.16586 | 8.11203 | 7.67117 | 4.56882 | 8.35294 | 7.67822 | 6.29002 |
| AUNIP | Q9H7T9-2 | 5.46788 | 6.76111 | 3.68323 | 4.61779 | 6.26471 | 4.37872 | 6.73622 | 8.05496 | 6.78342 | 7.92327 | 9.05331 | 6.75283 | 5.53932 | 5.96195 | 4.9979 | 7.02336 |
| EVA1B | Q9NVM1 | 6.57826 | 6.67324 | 5.90661 | 5.71686 | 4.88592 | 5.69543 | 7.26083 | 7.1617 | 5.08351 | 4.73761 | 6.31663 | 7.10083 | 3.86875 | 7.35422 | 6.79081 | 8.86878 |
| JAM2 | P57087-3 | 5.98937 | 6.48909 | 5.9937 | 5.91137 | 6.29854 | 6.46826 | 6.61977 | 6.06617 | 6.45359 | 6.39144 | 5.58811 | 5.73504 | 6.76998 | 6.83822 | 6.04583 | 6.24451 |
| TBC1D19 | Q8N5T2 | 6.50247 | 5.90503 | 6.22638 | 5.83317 | 6.75986 | 6.64938 | 6.93489 | 6.75397 | 6.01377 | 5.52548 | 5.91518 | 7.11195 | 5.66098 | 6.59253 | 5.95045 | 5.66452 |
| TMEM123 | Q8N131 | 6.67922 | 8.44898 | 6.09572 | 9.04463 | 5.42307 | 7.90125 | 5.56826 | 4.90281 | 5.81866 | 5.72585 | 3.97504 | 5.69673 | 6.51797 | 5.07692 | 5.62883 | 7.49606 |
| TNC | P24821-4 | 7.56972 | 7.44572 | 3.6281 | 8.4341 | 5.32811 | 4.32394 | 6.6196 | 6.07557 | 8.0771 | 6.61693 | 4.65011 | 4.66574 | 7.33979 | 4.56906 | 6.42342 | 8.233 |
| TMEM18 | Q96B42 | 6.26494 | 6.49309 | 6.63475 | 7.02417 | 6.53151 | 5.30714 | 5.98793 | 6.10693 | 6.60403 | 5.95629 | 5.23235 | 5.85651 | 6.0038 | 5.9352 | 6.94489 | 7.11647 |
| SLC16A6 | O15403 | 6.28215 | 6.12585 | 6.53227 | 7.49792 | 6.9771 | 8.1062 | 6.13442 | 7.17172 | 6.1233 | 4.83513 | 5.32082 | 5.67664 | 4.72811 | 4.40363 | 6.75031 | 7.33442 |
| KDELR1 | P24390 | 5.62911 | 6.32268 | 6.20622 | 5.66035 | 5.76134 | 5.95995 | 6.33398 | 6.38801 | 6.76354 | 5.83258 | 5.95022 | 6.00197 | 6.41921 | 6.17751 | 7.30671 | 7.28663 |
| MEA1 | Q16626 | 6.11895 | 4.22822 | 8.59872 | 6.27568 | 4.03277 | 4.16293 | 4.99792 | 8.22316 | 4.90696 | 7.05695 | 5.33586 | 6.33494 | 4.23711 | 9.28092 | 7.68033 | 8.52858 |
| CACNB3 | P54284 | 7.13973 | 5.57432 | 6.58645 | 6.98382 | 6.09221 | 7.3858 | 5.22955 | 7.76719 | 4.9038 | 5.41331 | 6.09111 | 7.55541 | 6.27887 | 6.27585 | 5.52385 | 5.19872 |
| PLG | P00747 | 8.14753 | 6.31514 | 4.80657 | 7.54193 | 5.0038 | 5.70762 | 7.83021 | 6.84373 | 5.69054 | 5.97433 | 4.72553 | 6.82716 | 6.84831 | 4.99625 | 6.60963 | 6.13172 |
| ZNF649 | Q9BS31 | 5.79364 | 5.58335 | 5.4455 | 6.45045 | 6.0511 | 3.96983 | 6.36805 | 6.05783 | 6.9836 | 5.86786 | 6.05187 | 5.50877 | 7.52099 | 7.16875 | 8.0833 | 7.09511 |
| SEL1L3 | Q68CR1 | 7.0928 | 7.81491 | 5.84935 | 8.1349 | 6.88953 | 7.36911 | 5.92758 | 6.03055 | 7.37194 | 6.66317 | 5.68408 | 7.49755 | 4.79399 | 4.10102 | 5.02137 | 3.75813 |
| TAS2R42 | Q7RTR8 | 7.38273 | 6.54008 | 6.06592 | 4.39606 | 8.0319 | 5.61601 | 8.35749 | 6.83591 | 6.90804 | 6.1364 | 4.31104 | 6.15933 | 6.3057 | 6.39858 | 4.69792 | 5.8569 |
| RBM48 | Q5RL73-2 | 6.06209 | 6.60542 | 5.23845 | 5.33787 | 6.40122 | 6.32177 | 5.27533 | 5.98352 | 6.15275 | 6.80793 | 5.96469 | 6.04536 | 7.17506 | 6.7705 | 7.34813 | 6.50989 |
| MINDY4 | Q4G0A6 | 5.54066 | 6.37454 | 6.83203 | 7.17685 | 6.81412 | 6.26105 | 6.30983 | 6.18896 | 6.09209 | 6.1996 | 7.6768 | 7.31187 | 4.74638 | 5.85666 | 4.99862 | 5.61994 |
| SEMA4F | O95754 | 6.83101 | 6.85721 | 7.53192 | 9.27019 | 6.20035 | 6.93087 | 6.28015 | 7.64039 | 4.32691 | 4.49742 | 7.1409 | 5.40096 | 6.49983 | 3.7626 | 5.14175 | 5.68756 |
| RGMB | Q6NW40 | 6.45104 | 5.98808 | 5.35136 | 8.14021 | 6.56749 | 7.99094 | 5.91633 | 4.36527 | 3.98549 | 6.61663 | 6.9327 | 6.5212 | 5.61737 | 8.6423 | 6.06739 | 4.84619 |
| CMTM7 | Q96FZ5 | 5.14499 | 6.56458 | 8.03751 | 7.34391 | 4.83245 | 7.35275 | 7.43962 | 5.95172 | 6.7693 | 7.64887 | 6.94304 | 6.89012 | 4.22234 | 6.56358 | 3.83201 | 4.4632 |
| JTB | O76095 | 6.94788 | 6.13911 | 6.09777 | 6.63185 | 6.38969 | 6.04388 | 6.43448 | 5.48047 | 6.65966 | 5.85428 | 6.62247 | 5.96928 | 6.17966 | 6.18871 | 6.58208 | 5.77872 |
| PWWP2B | Q6NUJ5 | 6.92997 | 6.75702 | 6.01924 | 6.39465 | 6.11473 | 6.58993 | 6.01638 | 5.67138 | 6.01338 | 6.45513 | 6.05358 | 6.30678 | 6.30238 | 5.83056 | 6.34984 | 6.19908 |
| PLAT | P00750 | 5.81589 | 6.45249 | 5.23075 | 9.92171 | 5.7993 | 4.97009 | 6.82344 | 6.67445 | 6.59851 | 5.00974 | 6.39902 | 6.94521 | 4.39615 | 5.60225 | 6.74276 | 6.61825 |
| OASL | Q15646 | 6.35151 | 6.36412 | 5.25407 | 6.01756 | 6.79439 | 5.02095 | 6.38956 | 6.78405 | 6.28593 | 7.09445 | 6.13052 | 5.4088 | 6.54113 | 6.47183 | 5.94012 | 7.15101 |
| EFNB2 | P52799 | 5.82553 | 8.69756 | 6.11956 | 5.5384 | 7.49293 | 6.68823 | 6.45925 | 5.37886 | 6.89022 | 3.00846 | 6.1654 | 6.08299 | 7.049 | 6.55403 | 6.55992 | 5.48966 |
| ARL4A | P40617 | 6.86743 | 7.13497 | 6.4558 | 7.48221 | 6.37044 | 6.43466 | 7.74483 | 6.60029 | 7.0345 | 5.69001 | 5.13595 | 6.29113 | 4.30147 | 5.94482 | 4.95994 | 5.55154 |
| RNF181 | Q9P0P0 | 7.75245 | 6.06683 | 6.29259 | 3.89125 | 4.80808 | 4.39371 | 5.32783 | 6.43435 | 5.38636 | 6.13836 | 4.81265 | 5.70832 | 4.49736 | 14.0045 | 5.6805 | 8.80485 |
| TMEM243 | Q9BU79 | 6.9309 | 6.15389 | 5.98652 | 5.9324 | 6.40252 | 6.14246 | 7.37797 | 6.73489 | 5.90154 | 5.54023 | 7.27868 | 5.33562 | 5.29861 | 5.97472 | 5.73866 | 6.37039 |
| SIAH1 | H3BU09 | 5.87402 | 6.88093 | 5.95586 | 5.56834 | 5.57864 | 5.33419 | 6.37212 | 6.28421 | 7.72138 | 6.28228 | 5.99251 | 7.08586 | 5.72272 | 5.44446 | 7.16417 | 6.7383 |
| DHRS2 | Q13268-2 | 5.80972 | 5.02808 | 6.14624 | 6.17881 | 7.10582 | 6.31969 | 5.88006 | 5.25636 | 6.23188 | 4.20494 | 7.74119 | 5.08947 | 6.89922 | 5.96788 | 7.68502 | 8.45563 |
| COL6A3 | P12111 | 8.37011 | 7.12307 | 8.03252 | 4.80798 | 4.40924 | 5.63128 | 6.86896 | 9.29093 | 6.70002 | 10.5161 | 6.15197 | 6.83773 | 4.5053 | 2.83406 | 3.66228 | 4.25841 |
| BPGM | P07738 | 5.90274 | 6.6177 | 6.48714 | 6.17299 | 6.66006 | 7.59754 | 6.60756 | 6.11814 | 4.91348 | 7.03191 | 5.89619 | 6.25381 | 6.23719 | 5.77321 | 5.48544 | 6.24492 |
| PLAG1 | Q6DJT9 | 5.84574 | 7.24472 | 6.47579 | 6.81996 | 4.45441 | 7.73244 | 5.79668 | 6.08447 | 6.92089 | 6.36192 | 6.45407 | 6.85589 | 5.73212 | 5.66058 | 6.55446 | 5.00586 |
| TOB1 | P50616 | 6.97402 | 6.43768 | 7.92791 | 5.91105 | 6.82526 | 6.11412 | 6.15245 | 7.26418 | 7.79738 | 8.10603 | 7.78462 | 7.57273 | 3.90587 | 4.10029 | 3.09263 | 4.03378 |
| DNLZ | Q5SXM8 | 6.99501 | 6.07039 | 5.6937 | 6.88118 | 6.32418 | 6.53615 | 6.39721 | 5.40425 | 6.07178 | 6.47862 | 6.65257 | 6.00368 | 6.31457 | 5.67925 | 6.0923 | 6.40517 |
| IPMK | Q8NFU5 | 6.79744 | 6.55326 | 6.8258 | 6.2223 | 5.74356 | 6.55901 | 5.69894 | 6.67381 | 6.57347 | 7.24404 | 7.24569 | 7.31603 | 5.27595 | 5.61855 | 4.97411 | 4.67804 |
| THEM6 | Q8WUY1 | 6.96649 | 5.84107 | 5.92493 | 7.3363 | 6.31961 | 4.35193 | 5.61364 | 8.32738 | 5.86231 | 5.10978 | 6.41793 | 6.65671 | 6.51795 | 7.17306 | 5.76891 | 5.81199 |
| PROS1 | P07225 | 5.50165 | 5.99929 | 4.88678 | 5.80707 | 5.30833 | 7.82772 | 7.97439 | 6.20483 | 4.6239 | 5.2096 | 5.57087 | 5.72155 | 7.04777 | 8.0043 | 7.42563 | 6.88632 |
| NHLRC1 | Q6VVB1 | 6.34323 | 5.9991 | 6.84331 | 5.76903 | 6.07899 | 6.39597 | 6.50326 | 6.71077 | 6.68403 | 6.79118 | 7.29723 | 6.79021 | 5.73212 | 5.00726 | 5.5636 | 5.4907 |
| METTL6 | Q8TCB7 | 7.25237 | 5.65437 | 6.52922 | 7.7003 | 6.057 | 5.72148 | 5.64606 | 7.73116 | 4.8012 | 6.33481 | 8.3492 | 7.19411 | 5.53112 | 3.79654 | 6.37475 | 5.32634 |
| FDX2 | Q6P4F2 | 6.07338 | 4.34024 | 4.28022 | 4.34778 | 7.56637 | 4.8405 | 5.54513 | 5.61404 | 4.41272 | 5.96864 | 6.14107 | 4.82377 | 5.70576 | 10.8965 | 7.5844 | 11.8595 |
| SMIM4 | Q8WVI0 | 6.8363 | 6.27184 | 6.07784 | 6.27392 | 6.71136 | 5.83858 | 7.16032 | 7.10781 | 5.42795 | 5.65552 | 6.36692 | 6.5023 | 5.56111 | 5.35622 | 6.91962 | 5.93239 |
| DLK2 | Q6UY11 | 7.14621 | 6.05321 | 7.6454 | 8.09388 | 5.46572 | 5.72257 | 6.59809 | 8.14077 | 6.77351 | 6.45184 | 7.5761 | 6.72712 | 4.28342 | 4.6471 | 4.50267 | 4.17239 |
| N6AMT1 | Q9Y5N5 | 4.97368 | 4.18729 | 5.96901 | 4.75196 | 5.19033 | 5.0028 | 5.72343 | 7.21707 | 5.39815 | 6.81384 | 5.49299 | 6.76963 | 5.25413 | 9.21049 | 8.28657 | 9.75863 |
| ZNF720 | Q7Z2F6-2 | 5.75223 | 6.93341 | 5.10613 | 6.44548 | 6.26091 | 6.4356 | 7.31079 | 6.48414 | 5.72861 | 6.97891 | 6.04497 | 6.50146 | 6.76468 | 7.4455 | 5.8117 | 3.9955 |
| MARCHF3 | Q86UD3 | 7.06767 | 5.57364 | 6.66126 | 7.38187 | 6.5337 | 5.41583 | 7.95478 | 6.57889 | 6.30231 | 5.81039 | 7.27527 | 6.57933 | 5.14716 | 5.0936 | 5.37214 | 5.25214 |
| NSUN7 | Q8NE18 | 7.66229 | 6.79724 | 7.72921 | 8.66614 | 7.17456 | 6.7386 | 6.72911 | 4.64021 | 7.76327 | 5.26623 | 4.51708 | 5.76673 | 5.09411 | 4.5902 | 4.96783 | 5.89718 |
| ADIPOR1 | Q96A54 | 7.05834 | 7.89171 | 7.34754 | 7.78215 | 6.31503 | 5.11421 | 5.89447 | 5.92768 | 8.26953 | 8.11663 | 7.6543 | 9.17315 | 3.48585 | 2.89417 | 2.72512 | 4.35011 |
| MSH5 | O43196-3 | 7.72731 | 6.25917 | 6.32385 | 5.55621 | 7.30279 | 7.26667 | 5.68728 | 7.74509 | 7.16523 | 8.5366 | 8.53463 | 6.54371 | 3.75411 | 3.21599 | 3.00876 | 5.37261 |
| RHOU | Q7L0Q8 | 5.5486 | 6.20352 | 7.04835 | 6.81667 | 5.98141 | 6.60749 | 5.02243 | 6.20218 | 6.76893 | 5.66843 | 6.87957 | 6.39176 | 8.09268 | 5.49781 | 6.17784 | 5.09235 |
| TIRAP | P58753-3 | 5.7311 | 5.57963 | 5.77684 | 7.31791 | 6.11198 | 5.57754 | 6.0569 | 6.49092 | 7.5746 | 5.60307 | 6.19169 | 6.43027 | 6.57365 | 5.41544 | 6.59338 | 6.9751 |
| MT-CO1 | P00395 | 6.40402 | 7.22732 | 5.1693 | 6.80064 | 7.02277 | 6.64066 | 5.62698 | 7.21359 | 5.60825 | 5.44612 | 5.32931 | 5.49309 | 6.54027 | 6.14648 | 6.89645 | 6.43474 |
| MGLL | Q99685 | 6.74127 | 7.02484 | 8.09055 | 5.73549 | 6.73449 | 6.55858 | 7.98001 | 7.2743 | 5.60755 | 5.481 | 5.07271 | 6.79844 | 5.27459 | 4.89161 | 4.97722 | 5.75737 |
| CSKMT | A8MUP2 | 6.5982 | 5.03168 | 7.48025 | 5.46716 | 6.97036 | 5.99148 | 6.18223 | 6.42306 | 5.62573 | 5.79999 | 4.65656 | 8.42663 | 4.46375 | 6.72044 | 7.26338 | 6.89908 |
| TSC22D1 | H7C4J9 | 6.011 | 6.05828 | 6.85165 | 6.85232 | 5.87606 | 7.45432 | 5.2549 | 6.15069 | 7.69544 | 5.23668 | 6.91255 | 4.58357 | 6.31757 | 6.65998 | 5.79928 | 5.74092 |
| AAMDC | Q9H7C9 | 5.81749 | 4.23006 | 5.58249 | 4.35789 | 5.64535 | 5.98739 | 5.38843 | 7.30053 | 4.81931 | 7.21102 | 5.28377 | 5.78864 | 6.05789 | 10.1962 | 8.01619 | 8.31732 |
| SFT2D2 | O95562 | 6.05817 | 5.78198 | 6.15157 | 6.55762 | 6.3905 | 6.5471 | 6.30874 | 6.10853 | 6.32326 | 6.60602 | 6.06331 | 6.75267 | 5.91289 | 6.08821 | 6.13086 | 6.21858 |
| ARMCX5 | Q6P1M9 | 6.61886 | 7.14768 | 6.85231 | 7.15342 | 6.00573 | 6.16869 | 6.13528 | 5.52825 | 5.72252 | 7.43704 | 6.60364 | 6.96962 | 5.36652 | 5.06796 | 5.70643 | 5.51604 |
| CYP1B1 | Q16678 | 5.79765 | 7.26245 | 5.41617 | 6.59853 | 6.67147 | 6.29769 | 4.26672 | 5.88635 | 7.84386 | 5.07901 | 6.94864 | 5.70541 | 6.93688 | 5.17207 | 5.65394 | 8.46315 |
| ZBTB14 | O43829 | 6.78003 | 7.25975 | 6.41534 | 5.41419 | 6.48508 | 6.72272 | 5.81975 | 6.86442 | 6.36247 | 5.93691 | 5.93222 | 6.67724 | 5.74067 | 5.34483 | 5.97218 | 6.27218 |
| ITIH4 | Q14624 | 7.00696 | 6.85556 | 5.13781 | 5.07275 | 4.26817 | 5.92446 | 6.36209 | 6.54605 | 6.20864 | 5.40889 | 6.12974 | 6.42253 | 7.04947 | 5.56759 | 9.29969 | 6.73961 |
| GON7 | Q9BXV9 | 5.09192 | 4.14476 | 3.84773 | 6.24807 | 5.9448 | 2.88323 | 4.88203 | 5.64415 | 4.16188 | 4.42504 | 4.48748 | 5.35875 | 4.84161 | 16.268 | 7.74823 | 14.0223 |
| TSPAN33 | Q86UF1 | 7.32232 | 6.02446 | 5.87001 | 6.32979 | 5.75949 | 5.15325 | 6.75223 | 6.23945 | 6.37939 | 6.50853 | 6.2485 | 5.72336 | 5.80077 | 6.19545 | 6.51005 | 7.18297 |
| IDS | P22304 | 6.40782 | 6.17126 | 5.88476 | 5.352 | 6.23658 | 7.39515 | 6.83941 | 5.82116 | 5.8336 | 7.06993 | 6.85321 | 6.20263 | 5.85681 | 6.44573 | 6.48985 | 5.14011 |
| TMEM107 | Q6UX40-2 | 6.20574 | 5.71638 | 4.83556 | 7.85752 | 5.23442 | 5.0734 | 3.55878 | 4.87309 | 6.41272 | 6.6792 | 6.51896 | 6.7757 | 7.49354 | 8.06782 | 6.74625 | 7.95091 |
| GNG5 | P63218 | 6.14797 | 5.52258 | 5.23196 | 5.18989 | 6.72816 | 5.71163 | 6.13663 | 6.44059 | 5.34797 | 5.59153 | 5.1565 | 5.55467 | 6.09583 | 8.81285 | 7.73138 | 8.59985 |
| PABIR1 | Q96E09 | 5.59469 | 4.72762 | 6.48197 | 3.42724 | 5.69384 | 5.831 | 6.33294 | 5.39394 | 4.74253 | 5.23842 | 6.37615 | 8.14052 | 9.232 | 7.51617 | 6.7412 | 8.52977 |
| E4F1 | Q66K89 | 6.30866 | 6.87877 | 6.86413 | 7.19549 | 6.83242 | 6.67339 | 5.80219 | 5.94136 | 6.46339 | 6.74892 | 6.61107 | 5.7237 | 5.47957 | 6.01517 | 4.86285 | 5.59891 |
| WDR90 | Q96KV7 | 6.41227 | 5.71915 | 6.87274 | 7.53598 | 7.37385 | 6.25605 | 6.00185 | 5.99419 | 7.33659 | 6.97551 | 7.20478 | 6.67534 | 4.43904 | 4.40903 | 6.62816 | 4.16547 |
| LSAMP | Q13449 | 6.34893 | 5.8914 | 5.81922 | 7.21411 | 6.07429 | 6.77406 | 5.42912 | 5.96772 | 5.17033 | 6.24084 | 6.63995 | 5.83821 | 6.47295 | 6.93655 | 7.13984 | 6.04247 |
| TMEM53 | Q6P2H8 | 6.31201 | 6.26982 | 5.70686 | 6.06647 | 5.25275 | 6.28642 | 6.20304 | 6.11901 | 6.02682 | 6.06823 | 5.5904 | 5.63896 | 7.07982 | 7.26333 | 7.22949 | 6.88656 |
| TSPAN14 | Q8NG11 | 6.25824 | 6.24989 | 5.60909 | 6.70972 | 6.3995 | 8.10416 | 6.45453 | 5.38607 | 5.2889 | 4.95826 | 4.95766 | 7.04169 | 7.09326 | 6.17339 | 7.23753 | 6.07812 |

| | | | | | | | | | | | | | | | | | |
|---|---|---|---|---|---|---|---|---|---|---|---|---|---|---|---|---|---|
| GFRA1 | P56159-2 | 7.14401 | 7.45983 | 6.06085 | 6.67399 | 7.19414 | 6.77982 | 6.31098 | 5.96358 | 5.99464 | 5.41499 | 6.54782 | 5.54851 | 5.0828 | 6.31716 | 5.49014 | 6.01673 |
| MTURN | Q8N3F0 | 4.78629 | 5.4086 | 5.2303 | 4.68514 | 4.21051 | 5.54837 | 7.88669 | 6.3698 | 5.80056 | 6.98018 | 4.22392 | 6.72556 | 5.83064 | 9.51519 | 8.05521 | 8.74302 |
| BEAN1 | J3KRN6 | 8.37358 | 8.78316 | 8.56522 | 8.8046 | 7.00773 | 7.89478 | 9.04564 | 3.33904 | 7.03057 | 4.14739 | 4.11796 | 3.54173 | 5.13263 | 4.30396 | 4.49759 | 5.41445 |
| ACAP3 | Q96P50 | 6.12241 | 6.12642 | 6.18475 | 6.36676 | 6.62256 | 7.15837 | 6.0684 | 7.0879 | 6.27356 | 6.27782 | 6.77469 | 5.81222 | 4.78776 | 6.1885 | 6.53541 | 5.61249 |
| KLHL29 | Q96CT2 | 4.75633 | 5.97602 | 6.01418 | 5.70681 | 5.79233 | 6.32832 | 7.02191 | 5.75444 | 5.54386 | 4.02646 | 4.20337 | 6.73512 | 7.74053 | 7.34094 | 8.77426 | 8.28512 |
| POU3F2 | P20265 | 8.02498 | 5.69793 | 6.6321 | 7.69944 | 5.66346 | 4.08855 | 5.12964 | 6.53391 | 5.74414 | 5.31667 | 7.04085 | 7.12569 | 6.1444 | 6.77312 | 7.10415 | 5.28096 |
| SUGCT | Q9HAC7-3 | 6.598 | 6.08761 | 5.77701 | 7.28302 | 5.79956 | 6.42187 | 5.73921 | 5.80411 | 6.07502 | 6.72354 | 6.65694 | 5.95296 | 6.0512 | 6.79669 | 6.4456 | 5.78768 |
| RNF145 | Q96MT1-5 | 7.42208 | 6.92263 | 6.2372 | 7.2354 | 7.09139 | 6.78668 | 6.65839 | 6.276 | 5.87141 | 6.45124 | 6.05134 | 5.75012 | 4.97798 | 5.99667 | 5.65394 | 4.61752 |
| USP53 | Q70EK8 | 6.09254 | 6.12572 | 6.52037 | 6.1805 | 5.94084 | 6.61719 | 6.8482 | 6.19504 | 6.20643 | 5.58168 | 6.14654 | 5.76915 | 6.44113 | 6.62435 | 7.10833 | 5.60197 |
| F2 | P00734 | 7.71392 | 6.5104 | 4.50318 | 5.47692 | 5.44537 | 7.59342 | 8.73857 | 5.03138 | 6.96877 | 6.16685 | 3.49115 | 8.25491 | 5.4451 | 2.82434 | 8.34678 | 7.48894 |
| ST6GAL1 | P15907 | 6.81303 | 6.19366 | 5.2328 | 6.5769 | 5.58121 | 5.92567 | 7.81041 | 5.41471 | 5.53199 | 5.86142 | 6.03575 | 6.29074 | 5.11161 | 6.73551 | 8.15321 | 6.73139 |
| CTDSPL | O15194 | 6.36392 | 6.32589 | 6.05272 | 6.39114 | 6.37591 | 6.2477 | 7.41248 | 6.67635 | 5.81585 | 5.4146 | 5.58673 | 5.92964 | 6.21719 | 5.51132 | 7.00088 | 6.67768 |
| ZNF32 | P17041 | 6.64149 | 7.32473 | 6.62549 | 6.97194 | 6.17494 | 6.04353 | 6.11437 | 5.81476 | 5.72588 | 5.92985 | 5.84914 | 6.96262 | 6.20826 | 6.18413 | 5.45122 | 5.97764 |
| LHFPL6 | Q9Y693 | 6.91679 | 8.09165 | 7.63626 | 5.75686 | 5.77079 | 4.11237 | 7.46234 | 5.40051 | 6.55452 | 5.91475 | 7.1683 | 6.28071 | 5.97016 | 5.41897 | 5.55719 | 5.98783 |
| RCBTB1 | Q8NDN9 | 7.25701 | 5.91822 | 8.2966 | 4.95693 | 5.25581 | 5.46508 | 7.4845 | 6.75746 | 6.27677 | 6.70493 | 6.54123 | 6.91153 | 6.96617 | 3.77028 | 6.35097 | 5.08651 |
| PDGFC | Q9NRA1 | 8.57338 | 6.64639 | 9.07972 | 8.41452 | 5.97743 | 6.24474 | 6.44489 | 6.01658 | 6.50292 | 6.2694 | 7.1308 | 5.85433 | 3.19701 | 4.46968 | 4.52621 | 4.65201 |
| ZNF529 | Q6P280 | 6.98279 | 6.98217 | 5.91504 | 7.39843 | 5.58288 | 5.83824 | 5.41503 | 5.85088 | 6.07589 | 5.79372 | 4.84724 | 5.2463 | 6.64387 | 7.41541 | 7.10809 | 6.90402 |
| TMEM250 | H0YL14 | 6.44824 | 5.91026 | 6.04387 | 7.03328 | 6.07719 | 7.76136 | 5.40077 | 4.78818 | 5.69301 | 5.66872 | 6.05899 | 7.79195 | 6.2864 | 7.91856 | 6.18906 | 4.93015 |
| CHCHD4 | Q8N4Q1-2 | 7.87466 | 5.22975 | 6.20687 | 4.81326 | 6.30708 | 5.25589 | 6.61304 | 5.61699 | 4.05014 | 6.28186 | 6.43723 | 6.874 | 7.29759 | 7.25323 | 6.37298 | 7.51545 |
| MIEF1 | L0R8F8 | 7.06256 | 5.42068 | 6.30725 | 7.61376 | 7.01018 | 5.45105 | 8.58645 | 6.45081 | 5.71671 | 5.44233 | 3.73833 | 6.10336 | 6.94589 | 5.92533 | 4.89543 | 7.32988 |
| TTLL5 | Q6EMB2 | 6.74104 | 6.18471 | 5.15176 | 6.0642 | 8.13131 | 6.36662 | 6.04845 | 5.7295 | 6.51879 | 6.29415 | 5.9618 | 6.74206 | 6.41745 | 5.26573 | 6.2623 | 6.12013 |
| BRAWNIN | Q69YU5 | 6.60449 | 6.20758 | 5.53823 | 5.7707 | 5.97638 | 5.57942 | 4.87082 | 6.58552 | 4.8044 | 5.99787 | 5.5285 | 5.28744 | 5.12848 | 8.28372 | 8.79797 | 9.03849 |

| Phosphopeptide identification | | | | | TMT Quant Data | | | | | | | | | | | | | | | |
|---|---|---|---|---|---|---|---|---|---|---|---|---|---|---|---|---|---|---|---|---|
| **Phosphopeptide** | **Gene** | **Uniprot** | **Site** | **Motif** | **DMSO_1** | **DMSO_2** | **DMSO_3** | **DMSO_4** | **IACS_1** | **IACS_2** | **IACS_3** | **IACS_4** | **KD025_1** | **KD025_2** | **KD025_3** | **KD025_4** | **Combo_1** | **Combo_2** | **Combo_3** | **Combo_4** |
| AAAS_S495 | AAAS | Q9NRG9 | 495 | AQFPRFSPVLGRA | 6.7966 | 6.7425 | 6.0810 | 6.5925 | 6.3399 | 5.9616 | 6.4219 | 6.0478 | 6.3369 | 6.2046 | 5.8592 | 6.0276 | 6.2327 | 6.1126 | 6.0997 | 6.1432 |
| AAGAB_S310;S311 | AAGAB | Q6PD74 | 310;311 | NA | 6.4578 | 6.2594 | 6.8777 | 6.2193 | 6.2984 | 6.4757 | 6.0010 | 6.8358 | 6.5829 | 6.7463 | 6.6736 | 6.8579 | 5.6903 | 5.8223 | 5.0095 | 5.1920 |
| AAK1_S1093 | AAK1 | A0A096LP25 | 637 | GHRRILSDVTHSA | 5.7476 | 5.7773 | 6.5809 | 5.7927 | 5.5097 | 5.9630 | 6.0672 | 5.8631 | 6.3859 | 5.6521 | 5.8307 | 6.1322 | 7.0411 | 7.5238 | 7.1190 | 7.0134 |
| AAK1_S21 | AAK1 | A0A096LP25 | 901 | FAPPEVSPEQAKT | 6.1565 | 6.5889 | 6.8427 | 6.8420 | 6.1847 | 6.9516 | 6.4525 | 6.8420 | 6.9271 | 6.8545 | 7.0018 | 7.2140 | 4.6673 | 5.0751 | 4.4190 | 4.9803 |
| AAK1_S618;S624 | AAK1 | A0A096LP25 | 606 | QPKVQTTPPPAVQ | 5.7509 | 6.4520 | 6.8475 | 6.3342 | 6.1435 | 6.7424 | 6.4719 | 6.7520 | 6.6289 | 6.6783 | 6.8964 | 7.1215 | 4.9871 | 5.8619 | 4.9614 | 5.3702 |
| AAK1_S637 | AAK1 | A0A096LP25 | 389 | PIQPALTPRKRAT | 6.2730 | 6.7749 | 6.8456 | 6.8560 | 6.0387 | 7.2891 | 6.8824 | 6.5049 | 7.2946 | 7.4117 | 6.4819 | 6.9476 | 4.5803 | 5.4603 | 4.0133 | 4.3456 |
| AAK1_S678 | AAK1 | A0A096LP25 | 731 | PEKLGGSAESLIP | 6.2099 | 5.8948 | 6.3851 | 6.3323 | 5.9396 | 5.9583 | 6.2876 | 6.3113 | 6.2010 | 6.1028 | 6.2526 | 6.6424 | 6.4504 | 6.4588 | 6.2471 | 6.3259 |
| AAK1_S682 | AAK1 | A0A096LP25 | 682 | SGSPRTSQQNVYN | 5.4006 | 5.7372 | 6.7789 | 5.3123 | 5.5742 | 5.9319 | 5.7207 | 6.1254 | 6.3310 | 5.6034 | 6.1555 | 6.1344 | 7.1506 | 7.5503 | 7.3387 | 7.1549 |
| AAK1_S731 | AAK1 | A0A096LP25 | 875 | RPLLMESEEEDES | 5.3070 | 5.5607 | 6.4673 | 6.1154 | 5.6648 | 6.0758 | 5.5834 | 6.0686 | 7.3180 | 6.2550 | 6.4823 | 6.0104 | 6.7451 | 7.0634 | 6.8088 | 6.4741 |
| AAK1_S875 | AAK1 | A0A096LP25 | 1093 | KPFRPQSLSKYSR | 7.6339 | 5.8628 | 6.9214 | 6.4657 | 5.7306 | 6.9235 | 6.4732 | 5.5317 | 8.3476 | 6.8064 | 7.2879 | 7.6232 | 4.3903 | 4.8514 | 3.9884 | 5.1620 |
| AAK1_S901 | AAK1 | A0A096LP25 | 678 | TTTPSGSPRTSQQ | 6.9039 | 6.8962 | 6.4732 | 6.4979 | 8.3140 | 5.9880 | 5.5533 | 6.7302 | 6.5059 | 6.2550 | 5.7653 | 5.9281 | 6.2219 | 5.9210 | 5.5326 | 4.5134 |
| AAK1_T389 | AAK1 | A0A096LP25 | 620 | QKVGSLTPPSSPK | 5.4995 | 5.5407 | 6.7017 | 6.3507 | 6.0973 | 6.0539 | 5.1831 | 6.0598 | 6.1027 | 5.6534 | 5.8231 | 6.5764 | 7.3187 | 6.9759 | 7.1273 | 6.9356 |
| AAK1_T606 | AAK1 | A0A096LP25 | 21 | GLGSGSSGGGGST | 6.5703 | 6.2389 | 7.3850 | 6.6629 | 6.5582 | 6.4776 | 7.3457 | 6.5743 | 6.0929 | 6.9358 | 6.7747 | 6.0053 | 4.7374 | 5.9650 | 5.3265 | 4.3496 |
| AAK1_T620 | AAK1 | A0A096LP25 | 620;624 | NA | 6.2659 | 5.8619 | 6.6102 | 5.9849 | 6.4747 | 6.0913 | 6.2860 | 6.1170 | 6.2762 | 6.5276 | 6.4169 | 6.4115 | 6.3005 | 6.5495 | 5.9655 | 5.8602 |
| AAK1_T620;S624 | AAK1 | A0A096LP25 | 618;624 | NA | 6.7492 | 6.5095 | 7.1754 | 6.7967 | 5.9969 | 6.1485 | 6.3577 | 6.3701 | 6.9229 | 6.3003 | 6.5560 | 7.0988 | 5.0064 | 5.5702 | 5.0052 | 5.4361 |
| AASDH_S724 | AASDH | Q4L235 | 724 | NLKGLNSPVLIGK | 7.1858 | 7.6116 | 6.9594 | 7.0593 | 5.5876 | 7.1256 | 7.7233 | 6.7615 | 6.9915 | 7.3434 | 7.8023 | 8.3725 | 3.0108 | 4.3117 | 2.8507 | 3.3030 |
| AATF_S153 | AATF | Q9NY61 | 203 | RAGDRNSEDDGVV | 5.2711 | 6.2842 | 6.1292 | 6.3922 | 6.9355 | 5.5470 | 5.9521 | 6.7532 | 5.5847 | 5.9697 | 5.7405 | 5.8976 | 6.8915 | 6.6843 | 6.6891 | 7.2783 |
| AATF_S203 | AATF | Q9NY61 | 153 | PGFSVQSISDFEK | 6.6281 | 6.7666 | 5.8337 | 7.5166 | 5.8085 | 5.6490 | 6.1866 | 6.0062 | 6.7580 | 5.5283 | 6.0690 | 6.1004 | 6.6416 | 6.5930 | 5.8110 | 6.1034 |
| AATF_S273 | AATF | Q9NY61 | 273 | GGPEFSSALKNSH | 7.1717 | 7.2258 | 7.4670 | 6.7163 | 5.7844 | 6.1480 | 6.8481 | 6.1151 | 6.2939 | 6.7269 | 6.5222 | 5.4514 | 5.2126 | 6.1198 | 5.1008 | 5.0961 |
| AATF_S316;S320;S321 | AATF | Q9NY61 | 316;320;321 | NA | 6.0509 | 6.3349 | 6.1250 | 6.7210 | 5.9682 | 6.3373 | 6.0924 | 5.9737 | 6.0948 | 6.0846 | 6.2106 | 6.6946 | 6.2933 | 6.2602 | 6.2742 | 6.4845 |
| AATF_S320;S321 | AATF | Q9NY61 | 320;321 | NA | 5.8681 | 6.0768 | 5.9120 | 6.2245 | 5.8574 | 5.4256 | 6.4193 | 5.4440 | 6.5653 | 7.4298 | 7.1807 | 6.9085 | 6.2189 | 6.1833 | 6.3316 | 5.9543 |
| ABCA3_S866 | ABCA3 | Q99758 | 866 | QHERRASDWAVDS | 5.8136 | 5.7880 | 5.7520 | 6.1146 | 6.6930 | 4.9398 | 5.6725 | 6.2557 | 6.4393 | 7.4286 | 6.5870 | 7.1560 | 6.2793 | 7.0017 | 6.4914 | 5.5873 |
| ABCC1_S929 | ABCC1 | P33527-9 | 940 | ISRHHNSTAELQK | 5.8932 | 6.0492 | 5.6175 | 6.0783 | 6.1876 | 5.7951 | 6.0723 | 5.8709 | 5.9762 | 5.6032 | 5.7125 | 5.6049 | 7.4325 | 7.5339 | 7.5160 | 7.0568 |
| ABCC1_S940 | ABCC1 | P33527-9 | 929 | QLSSSSSYSGDIS | 6.3514 | 6.9994 | 6.4900 | 6.9272 | 5.7787 | 6.5542 | 6.5675 | 6.6861 | 6.5011 | 6.8661 | 6.7231 | 6.5760 | 5.0229 | 5.6597 | 5.1379 | 5.1588 |
| ABCC3_S895 | ABCC3 | O15438-4 | 895 | LLQRNGSFANFLC | 5.3862 | 5.6280 | 5.8548 | 5.1286 | 4.9448 | 6.0883 | 6.0567 | 5.9535 | 5.8013 | 6.9192 | 5.3425 | 6.6375 | 8.2021 | 7.5897 | 8.1660 | 6.3008 |
| ABCC4_S404 | ABCC4 | O15439 | 646 | QPPVPGTPTLRNR | 6.3657 | 6.5752 | 6.4159 | 6.8507 | 6.8207 | 5.9783 | 6.3640 | 6.3342 | 6.1299 | 6.0517 | 6.0207 | 6.1956 | 6.0864 | 6.1643 | 5.8873 | 5.7592 |
| ABCC4_S629 | ABCC4 | O15439 | 404 | RNRQLPSDGKKMV | 6.5934 | 6.6174 | 6.1956 | 6.8266 | 6.0144 | 5.8116 | 6.4420 | 6.1341 | 5.9159 | 5.8128 | 5.2988 | 5.8406 | 6.3920 | 6.4925 | 6.6426 | 6.9698 |
| ABCC4_S655 | ABCC4 | O15439 | 655 | LRNRTFSESSVWS | 5.9524 | 5.8033 | 5.8224 | 6.1476 | 5.5774 | 5.7599 | 5.9109 | 6.5293 | 9.0711 | 6.5501 | 6.2734 | 6.7843 | 6.8433 | 5.9248 | 5.2357 | 5.8141 |
| ABCC4_S664;S668 | ABCC4 | O15439 | 629 | SGIDFGSLLKKDN | 3.4774 | 3.4859 | 3.6286 | 4.0593 | 4.0198 | 3.9771 | 4.0387 | 3.8126 | 9.0098 | 9.0209 | 8.3232 | 8.5845 | 8.3129 | 8.6632 | 9.1415 | 8.4446 |
| ABCC4_T646 | ABCC4 | O15439 | 664;668 | NA | 6.6526 | 6.9149 | 7.0787 | 7.3639 | 5.8765 | 6.2537 | 6.3478 | 5.9156 | 6.6093 | 6.6470 | 6.1477 | 6.8609 | 6.0684 | 6.1259 | 4.4878 | 4.6495 |
| ABCE1_T42 | ABCE1 | P61221 | 42 | KLCIEVTPQSKIA | 6.9045 | 6.4641 | 7.0351 | 6.4749 | 5.4927 | 6.5591 | 7.6168 | 6.9110 | 6.7879 | 6.9649 | 6.3065 | 5.5488 | 5.3254 | 5.9525 | 4.7318 | 4.9239 |
| ABCF1_S105;T108;S109 | ABCF1 | Q8NE71 | 140 | ALIQDQSEEEEEE | 5.9637 | 6.2644 | 6.5853 | 6.4559 | 6.1159 | 6.2175 | 6.2798 | 6.2668 | 6.3182 | 6.4478 | 6.4271 | 6.5242 | 5.7249 | 6.4800 | 5.5740 | 6.3546 |
| ABCF1_S108 | ABCF1 | Q8NE71 | 109 | KLSVPTSDEEDEV | 5.9301 | 6.3340 | 6.4192 | 6.4164 | 5.9121 | 5.7303 | 5.9631 | 6.3013 | 7.0036 | 7.1461 | 7.0078 | 7.2156 | 5.2786 | 6.1626 | 5.5140 | 5.6652 |
| ABCF1_S109 | ABCF1 | Q8NE71 | 22 | GDGESTSPSDKVV | 6.0563 | 6.3995 | 6.8784 | 6.9496 | 6.1039 | 6.5914 | 6.4496 | 6.5442 | 6.5168 | 6.6796 | 6.7290 | 6.8566 | 5.0382 | 5.7523 | 5.0110 | 5.4438 |
| ABCF1_S140 | ABCF1 | Q8NE71 | 228 | KKAEQGSEEEGEG | 6.2171 | 6.0371 | 6.3545 | 6.4492 | 5.9079 | 6.5334 | 5.8943 | 6.2315 | 6.4676 | 6.6600 | 6.6869 | 6.8038 | 5.2480 | 6.4719 | 5.9302 | 6.1067 |
| ABCF1_S22 | ABCF1 | Q8NE71 | 621 | PDPPPLSPPVLGL | 5.7667 | 6.4052 | 6.7487 | 6.3144 | 6.3345 | 6.6933 | 6.2052 | 6.3651 | 5.9964 | 5.9719 | 6.4670 | 6.2747 | 5.6450 | 6.8243 | 6.0161 | 5.9714 |
| ABCF1_S228 | ABCF1 | Q8NE71 | 108 | KKLSVPTSDEEDE | 5.4093 | 5.2029 | 6.9019 | 5.5754 | 5.8926 | 5.6157 | 6.7585 | 5.8699 | 6.6452 | 7.3475 | 6.2349 | 8.0069 | 6.5948 | 5.7516 | 7.9113 | 4.2816 |
| ABCF1_S621 | ABCF1 | Q8NE71 | 105;108;109 | NA | 5.1705 | 5.3544 | 5.1467 | 5.2092 | 5.9901 | 6.0059 | 5.4946 | 5.9025 | 5.8540 | 5.6021 | 5.5197 | 5.6023 | 7.6496 | 8.9230 | 8.2732 | 8.3022 |
| ABCF1_T108;S109 | ABCF1 | Q8NE71 | 108;109 | NA | 7.1539 | 7.7539 | 7.4134 | 7.1348 | 6.4915 | 6.6148 | 6.8970 | 7.0234 | 5.9423 | 6.5425 | 6.2082 | 6.6404 | 3.8432 | 5.0354 | 4.2124 | 5.0930 |
| ABCF2_T218 | ABCF2 | Q9UG63-2 | 218 | LHGLGFTPAMQRK | 6.8216 | 6.1449 | 7.2458 | 7.7065 | 6.5134 | 5.8367 | 5.9796 | 6.5849 | 6.6485 | 6.4753 | 7.1997 | 6.9630 | 4.2989 | 5.5968 | 5.2759 | 4.7087 |
| ABHD15_S434 | ABHD15 | Q6UXT9 | 434 | LSRHRASFLGGRR | 4.7174 | 4.4481 | 5.1880 | 5.3161 | 5.7050 | 5.0604 | 5.7069 | 5.4796 | 5.6607 | 5.4186 | 4.8945 | 5.0216 | 9.6721 | 9.8030 | 9.4925 | 8.4155 |
| ABI1_S183 | ABI1 | Q8IZP0 | 183 | PTQKPPSPPMSGR | 6.2046 | 5.9359 | 6.6089 | 6.0355 | 6.2213 | 6.2565 | 6.1297 | 6.1021 | 6.6482 | 6.7141 | 6.4836 | 6.6166 | 6.2905 | 6.2491 | 5.7321 | 5.7713 |
| ABI1_S216 | ABI1 | Q8IZP0 | 225 | RLGSQHSPGRTAS | 6.7457 | 7.3820 | 6.6218 | 6.5232 | 6.8370 | 7.6766 | 5.9318 | 7.7927 | 6.5921 | 6.0276 | 7.3698 | 6.8772 | 4.2701 | 4.6132 | 4.3042 | 4.4350 |
| ABI1_S225 | ABI1 | Q8IZP0 | 216 | PNDYMTSPARLGS | 6.6893 | 6.8511 | 6.9546 | 6.7902 | 6.5608 | 7.3050 | 6.8935 | 7.2743 | 7.5851 | 7.2432 | 7.4361 | 7.3165 | 3.5418 | 4.4887 | 3.2325 | 3.8373 |
| ABI1_S231 | ABI1 | Q8IZP0 | 213 | PTVPNDYMTSPAR | 5.4251 | 5.5070 | 6.7591 | 5.4381 | 6.2728 | 6.5905 | 6.0490 | 6.7558 | 5.9324 | 7.1853 | 7.1590 | 7.0053 | 6.2358 | 6.0539 | 5.6155 | 6.0156 |
| ABI1_Y213 | ABI1 | Q8IZP0 | 231 | SPGRTASLNQRPR | 7.6577 | 6.5923 | 7.5806 | 5.9906 | 5.5127 | 6.5578 | 5.7428 | 6.5010 | 6.9510 | 7.1704 | 7.1561 | 7.3881 | 5.6989 | 6.1322 | 3.6683 | 3.6994 |
| ABI2_S183 | ABI2 | Q9NYB9 | 227 | MAPSQQSPVRTAS | 6.8225 | 6.7270 | 7.0428 | 6.3282 | 6.1638 | 6.8499 | 6.3317 | 6.5635 | 6.8697 | 7.0539 | 7.2947 | 7.3555 | 4.6407 | 4.9588 | 4.2756 | 4.7217 |
| ABI2_S227 | ABI2 | Q9NYB9 | 183 | PTQKPPSPPMSGK | 6.7724 | 5.6261 | 6.8617 | 5.7122 | 6.4883 | 5.9137 | 5.9912 | 6.2547 | 6.4342 | 6.3190 | 6.4658 | 6.7010 | 6.6172 | 6.1625 | 5.6732 | 6.0069 |
| ABI2_S368 | ABI2 | Q9NYB9 | 368 | PPTIGGSLPYRRP | 6.6930 | 6.7024 | 6.6618 | 6.0484 | 6.0644 | 6.3357 | 5.7394 | 6.7899 | 6.4554 | 6.7099 | 6.5363 | 6.4589 | 5.7010 | 6.2525 | 5.7508 | 5.1002 |
| ABL1_T800 | ABL1 | P00519-2 | 833 | SSPPNLTPKPLRR | 5.3878 | 5.8831 | 5.9170 | 5.7543 | 5.5556 | 6.4060 | 6.6579 | 5.7927 | 6.2372 | 6.1090 | 6.1118 | 6.5986 | 6.7505 | 7.0035 | 6.8329 | 7.0023 |
| ABL1_T833 | ABL1 | P00519-2 | 800 | VTRGTVTPPPRLV | 5.6338 | 5.5177 | 5.8278 | 5.8892 | 5.7518 | 6.0712 | 7.6331 | 7.4428 | 9.8455 | 5.9204 | 6.5022 | 5.9167 | 5.1075 | 5.6301 | 5.5865 | 5.7237 |
| ABL1_T871 | ABL1 | P00519-2 | 871 | AAAEPVTPTSKAG | 6.1695 | 6.9369 | 5.9515 | 7.4987 | 5.0051 | 5.4091 | 6.6356 | 5.8841 | 7.5182 | 6.5806 | 7.3233 | 6.7579 | 6.1252 | 6.0862 | 4.2713 | 5.8469 |
| ABL2_S620 | ABL2 | P42684 | 655 | KGGFFSSFMKKRN | 10.7357 | 6.4003 | 8.0673 | 6.8597 | 6.4191 | 5.6016 | 6.3108 | 5.2159 | 6.8055 | 5.3157 | 6.3277 | 5.1879 | 5.5725 | 6.6379 | 4.3011 | 4.2414 |
| ABL2_S631 | ABL2 | P42684 | 936 | KVPVLISPTLKHT | 6.6122 | 6.7395 | 6.7033 | 6.2415 | 4.4620 | 7.2370 | 7.2134 | 7.6027 | 6.7349 | 6.6830 | 7.0762 | 6.7583 | 4.1987 | 4.8363 | 4.1025 | 4.7986 |
| ABL2_S655 | ABL2 | P42684 | 631 | RKQRDKSPSSLLE | 6.0547 | 5.8367 | 6.4554 | 6.0396 | 5.7858 | 6.3644 | 6.1591 | 6.3585 | 6.3225 | 6.4165 | 6.2431 | 6.3735 | 6.3114 | 6.6130 | 6.2603 | 6.4055 |
| ABL2_S72 | ABL2 | P42684 | 620 | AQASSGSPALPRK | 6.0950 | 6.0752 | 6.7046 | 6.1800 | 5.8051 | 6.5830 | 6.2524 | 6.2067 | 6.7278 | 6.6155 | 6.7311 | 6.7793 | 5.6662 | 6.1775 | 5.2341 | 6.1665 |
| ABL2_S915 | ABL2 | P42684 | 915 | EQPGWPSPAKAAP | 7.1203 | 6.6404 | 7.1936 | 6.0668 | 6.0262 | 6.8763 | 7.0349 | 7.7834 | 6.1380 | 7.4091 | 6.4980 | 6.2153 | 4.2852 | 4.8567 | 4.3869 | 5.4689 |
| ABL2_S936 | ABL2 | P42684 | 72 | DKTGGSSPEALHR | 6.4005 | 6.7167 | 6.1403 | 6.2837 | 6.4546 | 6.5675 | 5.9695 | 6.8866 | 6.2189 | 5.7259 | 5.8536 | 6.5704 | 5.4313 | 6.6451 | 6.1158 | 6.0195 |
| ABLIM1_S706 | ABLIM1 | O14639 | 706 | RMDRGVSMPNMLE | 6.5851 | 6.5095 | 6.6881 | 5.7422 | 6.8733 | 6.1023 | 6.2074 | 6.7848 | 6.6855 | 6.8475 | 7.0766 | 6.7706 | 5.2195 | 5.4633 | 5.3101 | 5.1341 |
| ABR_S33 | ABR | Q12979-2 | 33 | ELGTPTSPGSGSP | 5.8996 | 6.2778 | 6.5055 | 6.5848 | 5.9954 | 5.9171 | 6.9660 | 6.7476 | 6.3581 | 7.0423 | 6.8135 | 7.0453 | 5.9903 | 5.2433 | 5.6245 | 4.9888 |
| ABRAXAS1_S336 | ABRAXAS1 | Q6UWZ7 | 48 | KNSITDSQMDDVE | 5.9047 | 6.3349 | 5.4923 | 7.0545 | 5.8303 | 5.7117 | 5.7763 | 6.7350 | 6.8542 | 6.4369 | 6.1450 | 5.4893 | 7.4159 | 6.3804 | 6.2882 | 6.1505 |
| ABRAXAS1_S48 | ABRAXAS1 | Q6UWZ7 | 336 | TDIPEASPASTPQ | 6.6605 | 6.3246 | 6.3058 | 6.4875 | 6.0517 | 7.1258 | 6.0785 | 6.3930 | 6.1221 | 6.4909 | 6.1322 | 5.8747 | 6.6478 | 6.1704 | 5.5991 | 5.5354 |
| ABRAXAS2_S280 | ABRAXAS2 | Q15018 | 280 | SLDPAFSPRMPSS | 5.3675 | 4.8870 | 5.1104 | 6.1148 | 6.8612 | 6.4319 | 6.3913 | 6.0143 | 5.7088 | 6.6640 | 6.0240 | 5.3317 | 7.3929 | 7.2817 | 6.9482 | 7.4702 |
| ABTB2_S365 | ABTB2 | Q8N961 | 365 | PLCPGASPARQAR | 5.9437 | 6.6772 | 7.5040 | 6.4992 | 6.3715 | 6.7674 | 6.8863 | 6.9339 | 6.6251 | 6.9327 | 7.2517 | 6.4827 | 4.2900 | 5.1665 | 4.5779 | 5.0902 |
| ABTB2_S67 | ABTB2 | Q8N961 | 67 | MNSRHNSWDTVNT | 5.6435 | 6.4715 | 7.6954 | 6.7843 | 5.8369 | 6.0152 | 5.9608 | 7.5353 | 6.5909 | 5.9472 | 6.7224 | 5.9744 | 5.9004 | 5.1827 | 5.2356 | 6.5033 |
| ACACA_S115 | ACACA | Q13085-4 | 117 | HIRSSMSGLHLVK | 5.0268 | 4.9292 | 4.8007 | 4.6188 | 7.4863 | 6.3242 | 5.9453 | 6.5739 | 4.3704 | 3.9618 | 4.7043 | 4.5081 | 9.5283 | 9.8425 | 9.1398 | 8.2395 |
| ACACA_S117 | ACACA | Q13085-4 | 66 | SVSEDNSEDEISN | 5.7776 | 6.2404 | 6.9012 | 6.1282 | 6.1791 | 6.6124 | 6.1074 | 6.5127 | 6.0020 | 6.3150 | 5.9272 | 6.1992 | 6.0975 | 6.5789 | 6.1308 | 6.2905 |
| ACACA_S60 | ACACA | Q13085-4 | 60 | SRFIIGSVSEDNS | 6.2460 | 6.6165 | 6.4560 | 6.2675 | 6.5561 | 6.3664 | 5.8781 | 6.9314 | 6.1617 | 5.9447 | 6.1817 | 6.0140 | 6.2136 | 6.3457 | 6.1255 | 5.6950 |
| ACACA_S60;S66 | ACACA | Q13085-4 | 62 | FIIGSVSEDNSED | 6.4854 | 5.9976 | 6.8360 | 6.2178 | 6.1928 | 6.0629 | 6.0533 | 6.4079 | 6.1958 | 6.7328 | 6.4406 | 6.1154 | 5.9236 | 6.3757 | 5.9206 | 6.0416 |
| ACACA_S62 | ACACA | Q13085-4 | 115 | ALHIRSSMSGLHL | 9.0176 | 8.2806 | 7.2076 | 8.2614 | 6.5175 | 5.7757 | 6.0202 | 5.4995 | 8.6546 | 7.7813 | 6.8818 | 7.2889 | 2.8507 | 4.0270 | 3.1186 | 2.8169 |
| ACACA_S62;S66 | ACACA | Q13085-4 | 62;66 | NA | 6.6010 | 6.6582 | 7.6033 | 6.9127 | 5.9894 | 6.4924 | 6.1537 | 6.4225 | 6.5868 | 6.7838 | 6.6471 | 6.5961 | 5.4378 | 5.2634 | 4.5154 | 5.3365 |
| ACACA_S66 | ACACA | Q13085-4 | 60;66 | NA | 7.2590 | 7.4813 | 7.6100 | 7.2091 | 6.6399 | 5.8445 | 6.0645 | 6.5708 | 7.3712 | 6.6545 | 6.8967 | 7.1435 | 4.4163 | 4.6812 | 4.0274 | 4.1300 |
| ACAP2_S334 | ACAP2 | Q15057 | 521 | KYSISLSPPEQQK | 5.8696 | 6.2439 | 6.7696 | 6.0629 | 6.3374 | 6.7338 | 6.4105 | 6.5972 | 6.4497 | 6.6795 | 6.4835 | 6.6220 | 5.8177 | 5.8191 | 5.3944 | 5.7094 |
| ACAP2_S521 | ACAP2 | Q15057 | 334 | FCFEVVSPTKSCM | 5.3066 | 6.8229 | 7.2201 | 6.4595 | 6.2465 | 7.7084 | 7.2873 | 6.1344 | 7.0311 | 6.9815 | 6.0864 | 6.9879 | 4.1729 | 4.7221 | 5.6957 | 5.1368 |
| ACBD3_S43 | ACBD3 | Q9H3P7 | 43 | PPLPPPSPPGSGR | 6.7170 | 5.7079 | 5.3767 | 5.6484 | 7.0741 | 5.5475 | 4.5462 | 7.5994 | 6.0288 | 5.8088 | 5.9330 | 6.6523 | 7.4131 | 8.1246 | 6.7972 | 5.0248 |
| ACBD5_T172 | ACBD5 | Q5T8D3-3 | 172 | GNVLTSTPNAKTV | 6.2942 | 6.5940 | 6.3345 | 6.4339 | 6.8165 | 6.6215 | 6.5400 | 6.5075 | 7.0782 | 6.6972 | 6.6107 | 6.7504 | 5.0032 | 5.2005 | 5.0264 | 5.4913 |
| ACBD5_T181 | ACBD5 | Q5T8D3 | 181 | GNVLTSTPNAKTV | 6.4686 | 6.8605 | 7.0907 | 7.1486 | 6.6745 | 7.4175 | 7.7446 | 6.4324 | 7.0501 | 6.7653 | 6.3817 | 7.0335 | 3.9302 | 4.4452 | 3.5926 | 4.9642 |
| ACD_S25 | ACD | Q96AP0 | 25 | GSETPSSPRAGQL | 6.4412 | 6.7139 | 6.4924 | 6.5065 | 6.2860 | 6.1521 | 6.1347 | 6.6586 | 6.0634 | 6.7832 | 6.6464 | 6.1742 | 5.6240 | 5.8870 | 5.8463 | 5.5902 |
| ACD_S338 | ACD | Q96AP0 | 349 | SCTPSLSPRSHVP | 6.1949 | 6.4372 | 5.9956 | 6.2029 | 5.8382 | 6.0076 | 6.1754 | 6.8879 | 7.0938 | 6.8439 | 6.8235 | 6.1609 | 6.6871 | 6.0825 | 4.9723 | 5.5964 |
| ACD_S338;S349 | ACD | Q96AP0 | 338 | HASRTPSSPLQSC | 5.0746 | 5.3230 | 5.8407 | 5.8301 | 6.1874 | 5.4776 | 6.3719 | 5.9027 | 6.3168 | 6.1833 | 5.4899 | 6.1569 | 8.0363 | 7.2972 | 7.4969 | 7.0149 |
| ACD_S349 | ACD | Q96AP0 | 338;349 | NA | 6.2869 | 6.2650 | 6.3085 | 6.9039 | 6.4496 | 6.3251 | 6.5204 | 6.6804 | 6.3647 | 6.0682 | 7.0503 | 7.0400 | 5.4285 | 5.3229 | 5.5413 | 5.4444 |
| ACIN1_S1004 | ACIN1 | Q9UKV3 | 898 | ADDSRISEDETER | 6.0638 | 6.0291 | 6.5973 | 6.6870 | 6.4793 | 6.3975 | 6.4528 | 6.0466 | 6.1522 | 6.2784 | 6.5454 | 6.2699 | 6.3244 | 6.1558 | 5.7417 | 5.7788 |
| ACIN1_S1104 | ACIN1 | Q9UKV3 | 1004 | RTAQVPSPPRGKI | 6.5081 | 6.6643 | 6.1958 | 6.6851 | 6.0081 | 5.9067 | 6.2854 | 6.1039 | 6.2772 | 6.3304 | 6.1254 | 6.2740 | 6.2071 | 6.1555 | 6.0062 | 6.2669 |
| ACIN1_S1161 | ACIN1 | Q9UKV3 | 414 | TASEEETPPPLLT | 5.9196 | 6.0860 | 5.6910 | 6.1637 | 6.0643 | 5.8699 | 6.0053 | 5.8085 | 6.0062 | 6.1138 | 5.9489 | 5.9101 | 7.0767 | 7.1340 | 7.1737 | 7.0282 |
| ACIN1_S166 | ACIN1 | Q9UKV3-5 | 885 | ADDSRISEDETER | 6.3718 | 6.3923 | 6.0873 | 6.9665 | 6.2492 | 6.0145 | 6.3037 | 5.9288 | 6.2889 | 6.0723 | 6.0575 | 5.9018 | 6.4108 | 6.0542 | 6.5050 | 6.3956 |
| ACIN1_S207 | ACIN1 | Q9UKV3 | 682 | SGQEVATPPVPQL | 6.1936 | 6.4776 | 6.1129 | 6.3857 | 6.1741 | 6.0566 | 6.3058 | 6.0160 | 6.1531 | 6.0843 | 6.2220 | 6.0712 | 6.4550 | 6.3871 | 6.4118 | 6.4933 |
| ACIN1_S386;S388 | ACIN1 | Q9UKV3 | 490 | SDTKAESPAEKVP | 6.0541 | 6.2304 | 5.7743 | 6.3720 | 6.2890 | 5.9125 | 6.1443 | 6.0475 | 6.2017 | 6.2347 | 5.8610 | 5.8802 | 6.8451 | 6.5732 | 6.8742 | 6.7058 |
| ACIN1_S400 | ACIN1 | Q9UKV3 | 825 | SFKRKISVVSATK | 5.5017 | 4.5985 | 4.8805 | 5.3571 | 7.9258 | 4.9234 | 4.3385 | 5.0984 | 9.1500 | 9.0548 | 7.8346 | 8.5901 | 6.6031 | 7.2436 | 3.9379 | 4.9620 |
| ACIN1_S410 | ACIN1 | Q9UKV3 | 561 | KEGRRASHTLLPS | 4.6012 | 4.1959 | 4.4373 | 5.2186 | 8.0471 | 5.1774 | 4.8013 | 5.7814 | 7.4751 | 7.7978 | 6.9069 | 6.9752 | 7.9393 | 7.8942 | 6.4690 | 6.2823 |
| ACIN1_S410;T414 | ACIN1 | Q9UKV3 | 838 | GVPAGNSDTEGGQ | 7.0516 | 7.4709 | 7.0173 | 7.7953 | 7.1055 | 7.0813 | 7.0263 | 7.5756 | 5.3538 | 5.0957 | 5.0284 | 5.2073 | 5.0570 | 4.9932 | 5.4863 | 5.6546 |
| ACIN1_S414 | ACIN1 | Q9UKV3 | 410 | LPEQTASEEETPP | 6.5085 | 6.5421 | 6.4062 | 6.9351 | 6.5855 | 5.9280 | 6.4346 | 6.4568 | 6.0241 | 6.1390 | 5.8041 | 6.3753 | 5.9196 | 6.2029 | 5.8519 | 5.8861 |
| ACIN1_S490 | ACIN1 | Q9UKV3 | 657 | NSRKSLSPGVSRD | 6.2662 | 6.5676 | 5.8363 | 6.4141 | 6.0588 | 5.7278 | 5.8351 | 5.8677 | 6.3813 | 6.1431 | 5.9863 | 5.8841 | 6.8727 | 6.7724 | 6.7828 | 6.6037 |
| ACIN1_S561 | ACIN1 | Q9UKV3 | 729 | LQPERGSPKKCEA | 7.2753 | 6.5780 | 6.5573 | 6.3161 | 6.5400 | 6.3592 | 6.6927 | 5.2185 | 5.8987 | 6.0181 | 6.1342 | 6.2010 | 6.6537 | 6.1168 | 5.6018 | 5.8386 |
| ACIN1_S655;S657 | ACIN1 | Q9UKV3 | 710 | VQARRLSQPESAE | 4.6173 | 4.6490 | 4.4420 | 5.0452 | 6.7770 | 5.6425 | 5.9656 | 5.9569 | 5.0982 | 4.9752 | 4.5887 | 4.7748 | 10.0753 | 8.5993 | 9.6829 | 9.1102 |
| ACIN1_S655;S661 | ACIN1 | Q9UKV3 | 1332 | RSRSRSTPVRDRG | 7.2751 | 7.0820 | 6.6388 | 7.1700 | 6.4266 | 6.1297 | 5.9797 | 6.2088 | 6.4609 | 6.9722 | 6.8980 | 6.7418 | 4.9337 | 5.3453 | 4.8821 | 4.8552 |
| ACIN1_S657 | ACIN1 | Q9UKV3 | 676 | ETKDPSSGQEVAT | 6.6024 | 6.5077 | 6.4985 | 6.6033 | 6.4455 | 5.8892 | 6.4661 | 6.7813 | 6.0957 | 5.6534 | 6.0172 | 5.7989 | 6.1522 | 6.1728 | 6.4113 | 5.9046 |
| ACIN1_S667 | ACIN1 | Q9UKV3 | 408 | VALPEQTASEEET | 6.8755 | 6.5422 | 6.8320 | 6.6415 | 6.1379 | 5.6843 | 6.2705 | 6.7291 | 6.2980 | 6.5994 | 5.6488 | 6.0661 | 5.6633 | 6.2647 | 6.1016 | 5.6452 |
| ACIN1_S676 | ACIN1 | Q9UKV3 | 207 | HSPRKSSSISEEK | 6.1353 | 5.7095 | 6.8393 | 7.2453 | 6.0560 | 6.1505 | 6.2489 | 6.1930 | 6.2305 | 6.6181 | 6.7119 | 6.8978 | 5.3482 | 5.4362 | 5.7626 | 6.4168 |
| ACIN1_S710 | ACIN1 | Q9UKV3 | 400 | EDRKKASLVALPE | 5.7100 | 5.2039 | 5.0544 | 5.6672 | 5.8886 | 5.3105 | 5.5101 | 5.2063 | 5.8572 | 5.9053 | 6.2100 | 5.1887 | 8.2379 | 8.0575 | 8.6132 | 8.3794 |
| ACIN1_S729 | ACIN1 | Q9UKV3 | 895 | DLHADDSRISEDE | 6.2250 | 6.3344 | 10.4689 | 5.8398 | 10.8513 | 6.0053 | 5.3237 | 5.9315 | 6.7542 | 6.4868 | 6.7664 | 6.9543 | 3.6081 | 4.1326 | 4.2607 | 4.0568 |
| ACIN1_S815 | ACIN1 | Q9UKV3 | 1104 | LLVDRPSETKTEE | 7.3471 | 7.0329 | 7.2401 | 7.3830 | 6.4026 | 6.3361 | 6.8494 | 6.5826 | 7.1702 | 6.0983 | 6.7073 | 6.3952 | 4.6426 | 4.5845 | 4.5749 | 4.6532 |
| ACIN1_S825 | ACIN1 | Q9UKV3 | 1161 | RRERTRSEREWDR | 7.7645 | 7.5598 | 7.7377 | 8.4340 | 6.2550 | 6.5212 | 6.8645 | 5.0968 | 5.6389 | 6.4494 | 7.6445 | 6.4740 | 4.5805 | 4.7005 | 4.0743 | 4.2044 |
| ACIN1_S825 | ACIN1 | Q9UKV3 | 987 | SISQQKSGVSITI | 5.9946 | 6.8193 | 6.8457 | 6.7525 | 6.5894 | 5.3762 | 6.0072 | 5.9161 | 6.9043 | 6.4271 | 6.5223 | 6.3425 | 5.7428 | 6.1415 | 5.8347 | 5.7839 |
| ACIN1_S838 | ACIN1 | Q9UKV3 | 166 | AELEEASAESEDE | 5.9418 | 6.4922 | 6.4824 | 6.1570 | 6.6243 | 5.5308 | 6.0494 | 6.6659 | 7.2529 | 5.7877 | 6.1204 | 4.5988 | 6.7637 | 7.5416 | 6.4481 | 5.5431 |
| ACIN1_S885 | ACIN1 | Q9UKV3 | 667 | SRDSSTSYTETKD | 6.2494 | 6.6512 | 6.0355 | 6.3166 | 6.3490 | 6.4033 | 7.0096 | 6.3925 | 6.0907 | 6.3836 | 6.5548 | 6.3095 | 5.7634 | 5.6801 | 5.7558 | 6.0549 |
| ACIN1_S895 | ACIN1 | Q9UKV3 | 815 | EEKKESSLPKSFK | 5.7400 | 5.4577 | 4.9763 | 5.2174 | 5.7665 | 5.9866 | 5.8640 | 4.7926 | 4.9725 | 5.5349 | 5.2408 | 4.7517 | 10.2423 | 8.2315 | 8.6800 | 8.5452 |
| ACIN1_S895;S898 | ACIN1 | Q9UKV3-5 | 825 | SFKRKISVVSTKG | 5.4429 | 5.5337 | 5.2260 | 5.6513 | 7.9588 | 5.3869 | 5.2789 | 6.0070 | 8.4626 | 7.7368 | 7.0873 | 7.4465 | 6.3675 | 6.2435 | 4.4128 | 5.7575 |
| ACIN1_S898 | ACIN1 | Q9UKV3 | 655;657 | NA | 6.3431 | 6.2981 | 6.0324 | 6.6745 | 6.0093 | 5.9186 | 6.1610 | 6.1095 | 6.4309 | 6.3279 | 5.9150 | 6.2528 | 6.3695 | 6.2850 | 6.3925 | 6.4800 |
| ACIN1_S987 | ACIN1 | Q9UKV3 | 410;414 | NA | 6.3042 | 6.4954 | 6.3273 | 6.3282 | 6.5471 | 5.7841 | 6.2473 | 6.4013 | 6.0995 | 6.2420 | 5.9930 | 5.8654 | 6.4623 | 6.3134 | 6.4458 | 6.1435 |
| ACIN1_T1332 | ACIN1 | Q9UKV3 | 655;661 | NA | 6.4364 | 6.6544 | 5.8070 | 6.5561 | 6.1322 | 5.7761 | 6.2298 | 5.8808 | 6.4722 | 6.1914 | 6.0332 | 6.1972 | 6.3689 | 6.4219 | 6.3071 | 6.5352 |
| ACIN1_T408 | ACIN1 | Q9UKV3 | 386;388 | NA | 5.9754 | 6.2965 | 6.2348 | 6.5925 | 6.0922 | 6.0173 | 5.9041 | 5.4778 | 6.3149 | 6.1360 | 5.8926 | 6.4326 | 6.9321 | 6.5907 | 6.6004 | 6.5101 |
| ACIN1_T682 | ACIN1 | Q9UKV3 | 895;898 | NA | 6.7780 | 6.2165 | 6.4391 | 6.8426 | 6.6872 | 6.5901 | 6.5333 | 6.5707 | 6.3655 | 7.0855 | 6.5621 | 6.3843 | 4.4771 | 5.2776 | 5.0925 | 6.0978 |
| ACLY_S455 | ACLY | P53396 | 481 | PQDSVPSPRSLQG | 6.0529 | 6.0603 | 6.6931 | 6.1803 | 5.9522 | 6.2338 | 6.3367 | 5.9072 | 6.5166 | 6.6619 | 6.7349 | 6.6238 | 5.8395 | 6.4184 | 5.6648 | 6.1236 |
| ACLY_S481 | ACLY | P53396 | 455 | APSRTASFSESRA | 4.5625 | 4.5877 | 4.9599 | 4.2974 | 5.7602 | 5.9701 | 5.7606 | 6.1475 | 4.7001 | 4.5040 | 4.6626 | 4.7535 | 10.2940 | 9.7039 | 10.0840 | 9.2520 |
| ACLY_T447 | ACLY | P53396 | 447 | ASGSTSTPAPSRT | 4.6747 | 5.1347 | 5.6409 | 5.2540 | 6.0287 | 6.3741 | 6.6060 | 5.9931 | 5.0770 | 5.4436 | 5.3022 | 5.3763 | 8.5200 | 8.7327 | 7.6934 | 8.1486 |
| ACOX1_S26 | ACOX1 | Q15067-2 | 26 | THILDGSPEKTRR | 5.6346 | 6.6594 | 5.7742 | 7.2711 | 6.0839 | 5.8777 | 6.8961 | 6.4878 | 6.7445 | 6.8562 | 6.5827 | 6.3842 | 5.2983 | 5.6681 | 5.9445 | 5.8368 |
| ACSL3_S683 | ACSL3 | O95573 | 683 | PVKIRLSPEPWTP | 6.1141 | 6.2145 | 5.8788 | 6.4700 | 6.0788 | 5.9361 | 6.0422 | 6.2632 | 6.0851 | 6.2747 | 6.1291 | 6.2144 | 6.2758 | 6.7997 | 6.5620 | 6.6614 |
| ACSL3_T688 | ACSL3 | O95573 | 688 | LSPEPWTPETGLV | 5.4365 | 5.9300 | 5.7919 | 6.5995 | 5.6903 | 6.5509 | 6.0134 | 6.6744 | 6.5165 | 6.3724 | 6.3239 | 6.5627 | 6.0356 | 6.7643 | 6.3825 | 6.3550 |
| ACSL4_S674 | ACSL4 | O60488 | 674 | PIKVRLSPEPWTP | 5.1961 | 6.1375 | 6.6279 | 6.9834 | 5.9492 | 5.5895 | 6.6518 | 6.3117 | 5.9366 | 5.4504 | 6.1979 | 6.3525 | 7.0199 | 6.2977 | 7.2082 | 6.0897 |
| ACSS2_S267 | ACSS2 | Q9NR19-2 | 30 | GRARSWSPPPEVS | 6.3374 | 6.4258 | 6.6423 | 5.8711 | 6.1364 | 6.6087 | 6.0349 | 5.9818 | 5.1167 | 6.3192 | 6.2289 | 6.2341 | 5.9982 | 7.2153 | 6.6237 | 6.2255 |
| ACSS2_S30 | ACSS2 | Q9NR19-2 | 267 | GDSTSQSPPIKRS | 6.1318 | 6.3934 | 6.7243 | 6.3366 | 5.8350 | 6.6284 | 6.0463 | 6.4651 | 6.3469 | 6.7293 | 6.6932 | 6.7321 | 5.6586 | 6.1338 | 5.6721 | 5.4732 |
| ACTB_S233 | ACTB | P60709 | 233 | MATAASSSSLEKS | 6.2942 | 6.0918 | 6.5586 | 6.1972 | 6.5437 | 6.3114 | 6.1105 | 6.1077 | 6.3275 | 6.3616 | 6.3286 | 6.2800 | 6.4819 | 6.0554 | 6.0167 | 5.9332 |
| ACTB_S235 | ACTB | P60709 | 323 | ITALAPSTMKIKI | 4.5028 | 4.3083 | 4.2490 | 4.2788 | 5.9689 | 5.5537 | 5.9598 | 6.0346 | 4.5284 | 4.3199 | 4.2747 | 4.4371 | 11.2026 | 9.9595 | 11.2889 | 9.1329 |
| ACTB_S323 | ACTB | P60709 | 33 | PRAVFPSIVGRPR | 5.1574 | 4.9416 | 4.7546 | 4.8183 | 5.1762 | 5.5051 | 5.9873 | 7.0911 | 8.0388 | 6.3167 | 6.2903 | 7.0594 | 7.2981 | 7.3386 | 5.6177 | 8.6086 |
| ACTB_S33 | ACTB | P60709 | 235 | TAASSSSLEKSYE | 6.1163 | 5.4136 | 7.0092 | 5.7975 | 5.7589 | 6.2585 | 5.6830 | 6.1014 | 5.4373 | 6.3106 | 6.4399 | 6.5340 | 6.3844 | 7.1649 | 7.2076 | 6.3830 |
| ACTL6A_S233 | ACTL6A | O96019 | 233 | EAVREGSPANWKR | 6.1782 | 6.3270 | 6.6099 | 6.3441 | 6.2406 | 6.2489 | 6.7197 | 6.1203 | 6.4142 | 6.5973 | 6.3936 | 6.6960 | 5.6784 | 5.9110 | 5.5355 | 5.9852 |
| ACTL6A_S86 | ACTL6A | O96019 | 86 | ENMEAISPLKNGM | 6.2311 | 6.6165 | 6.7353 | 6.2174 | 6.0510 | 6.5908 | 7.1529 | 6.3983 | 6.5008 | 7.2134 | 5.8589 | 5.9318 | 5.4272 | 5.8032 | 5.2865 | 5.9850 |
| ACTN4_S159 | ACTN4 | O43707 | 423 | KFRQKASIHEAWT | 6.2032 | 6.1645 | 5.6379 | 5.5712 | 6.6585 | 6.1763 | 5.3773 | 5.8263 | 6.6722 | 6.1025 | 6.1190 | 6.0097 | 6.9986 | 6.8042 | 7.0411 | 6.6373 |
| ACTN4_S423 | ACTN4 | O43707 | 159 | FAIQDISVEETSA | 6.0670 | 6.6252 | 6.7243 | 6.9779 | 6.2376 | 6.0467 | 6.2417 | 6.1311 | 6.4918 | 5.9939 | 6.4230 | 6.6036 | 5.7682 | 5.8189 | 5.9806 | 5.8682 |
| ACTN4_S656 | ACTN4 | O43707 | 656 | LRRQFASQANVVG | 5.9175 | 5.7011 | 6.3137 | 6.5021 | 6.0175 | 5.6530 | 6.2253 | 6.2905 | 6.5129 | 6.5073 | 5.4015 | 5.1375 | 7.6264 | 7.1139 | 6.9250 | 6.1549 |
| ACTN4_S816 | ACTN4 | O43707 | 816 | EFNRIMSLVDPNH | 5.0055 | 5.5466 | 5.5118 | 6.4549 | 5.7492 | 5.5618 | 5.5236 | 5.4215 | 6.1690 | 6.3564 | 5.6967 | 5.6505 | 7.4075 | 7.9756 | 7.1905 | 8.7788 |
| ACTR10_S414 | ACTR10 | Q9NZ32 | 414 | LMKRAFSTEKxxx | 5.9325 | 5.3498 | 6.3664 | 5.5537 | 6.4812 | 6.0285 | 5.8436 | 5.7722 | 5.4785 | 6.3194 | 6.2219 | 5.9867 | 7.7943 | 7.5104 | 7.1214 | 6.2393 |
| ACTR2_T122 | ACTR2 | P61160-2 | 122 | EPPMNPTKNREKI | 8.1256 | 6.9459 | 7.3135 | 6.0779 | 7.4559 | 5.8565 | 6.0182 | 5.3638 | 6.3723 | 5.6333 | 6.2030 | 5.2849 | 5.2253 | 7.2230 | 5.7295 | 5.1714 |
| ACTR8_S132 | ACTR8 | Q9H981 | 132 | TRRIPVSPEQARS | 6.0892 | 6.3487 | 6.4088 | 6.6085 | 5.9354 | 6.0711 | 6.2076 | 5.8935 | 6.6640 | 6.1622 | 6.3311 | 6.4261 | 6.1711 | 6.2385 | 6.2383 | 6.2058 |
| ACVR1_T209 | ACVR1 | Q04771 | 209 | TVARQITLLECVG | 5.2844 | 5.0065 | 6.3033 | 6.6136 | 6.0115 | 5.0528 | 4.6128 | 5.7104 | 4.3581 | 4.5156 | 3.7468 | 3.4040 | 10.0315 | 10.7540 | 12.5371 | 6.0578 |
| ACVR2A_S184 | ACVR2A | P27037 | 184 | PGPPPPSPLLGLK | 6.7608 | 7.2417 | 7.0563 | 7.6151 | 6.2898 | 6.3420 | 5.9164 | 6.5520 | 7.1018 | 6.7204 | 7.0537 | 7.1855 | 4.5473 | 4.6647 | 4.3776 | 4.5749 |
| ADAM10_T719 | ADAM10 | O14672 | 719 | PKPLPGTLKRRRP | 7.7087 | 6.7982 | 9.1088 | 6.9474 | 5.2747 | 6.6312 | 6.8672 | 5.3395 | 6.5609 | 7.2948 | 7.4745 | 6.6617 | 3.9259 | 5.7016 | 3.4756 | 4.2294 |
| ADAM17_S791 | ADAM17 | P78536 | 735 | PFPAPQTPGRLQP | 6.8735 | 6.3459 | 7.3268 | 7.0909 | 5.7832 | 6.5737 | 6.4763 | 5.6201 | 6.0876 | 6.2307 | 6.5526 | 6.5867 | 5.6814 | 6.0669 | 4.8814 | 5.8225 |
| ADAM17_T735 | ADAM17 | P78536 | 791 | SSTAAKSFEDLTD | 6.4603 | 6.7734 | 8.0357 | 6.8844 | 6.0986 | 6.1409 | 5.9942 | 6.7030 | 6.0764 | 6.9898 | 6.8126 | 6.7713 | 4.0683 | 5.4092 | 5.3822 | 5.3995 |
| ADAM19_T779 | ADAM19 | Q9H013 | 779 | KRKVINTPEILRK | 6.8912 | 7.7257 | 7.1711 | 6.3398 | 5.5076 | 6.6862 | 6.9178 | 6.7603 | 6.4180 | 6.5410 | 6.8507 | 6.6139 | 4.3794 | 5.3849 | 4.5220 | 5.2904 |
| ADAM22_S834 | ADAM22 | Q9P0K1 | 857 | FRPRSNSTETLSP | 5.4183 | 5.6102 | 5.6213 | 5.5226 | 5.9755 | 6.0201 | 5.9080 | 5.8471 | 5.9745 | 6.3492 | 5.8667 | 6.0759 | 7.2629 | 7.5999 | 7.3415 | 7.6063 |
| ADAM22_S857 | ADAM22 | Q9P0K1 | 834 | GRPRSNSWQGNLG | 5.9015 | 5.8938 | 5.9253 | 5.3699 | 5.9675 | 6.1564 | 6.1886 | 6.2156 | 6.6524 | 6.7923 | 6.3531 | 6.5798 | 6.4529 | 7.0918 | 6.4199 | 6.0393 |
| ADAM9_S752 | ADAM9 | Q13443 | 758 | SVPRHVSPVTPPR | 6.2042 | 6.6695 | 6.4904 | 6.4424 | 6.1597 | 6.3367 | 6.2839 | 6.2235 | 6.4467 | 6.3352 | 6.1951 | 6.3021 | 5.6193 | 6.3860 | 5.8389 | 6.0665 |
| ADAM9_S758 | ADAM9 | Q13443 | 799 | PQPKVSSQGNLIP | 5.1840 | 5.0721 | 4.8109 | 4.9204 | 4.7902 | 5.1480 | 4.6863 | 5.0010 | 5.1808 | 5.5993 | 4.4802 | 5.1001 | 10.8128 | 9.7968 | 9.7153 | 9.7017 |
| ADAM9_S758;T761 | ADAM9 | Q13443 | 761 | RHVSPVTPPREVP | 6.3156 | 6.6529 | 6.7172 | 6.5008 | 6.3586 | 6.9011 | 6.5463 | 6.6936 | 6.6523 | 6.4582 | 6.6980 | 6.3731 | 4.8561 | 5.8014 | 5.0678 | 5.4071 |
| ADAM9_S761 | ADAM9 | Q13443 | 769 | PREVPIYANRFAV | 6.3093 | 6.5602 | 6.6601 | 5.9694 | 5.6076 | 6.9742 | 6.0366 | 7.0523 | 7.5659 | 7.1936 | 7.3459 | 7.7399 | 5.0642 | 5.2193 | 3.9932 | 4.7083 |
| ADAM9_S799 | ADAM9 | Q13443 | 752 | PSRQPGSVPRHVS | 6.2089 | 5.9795 | 7.8197 | 6.4063 | 5.1351 | 6.6951 | 5.9618 | 7.1587 | 6.3188 | 7.2068 | 7.3534 | 7.6612 | 4.9355 | 5.7563 | 4.6884 | 4.7144 |

| | | | | | | | | | | | | | | | | | | | | | |
|---|---|---|---|---|---|---|---|---|---|---|---|---|---|---|---|---|---|---|---|---|---|
| ADAM9_Y769 | ADAM9 | Q13443 | 758;761 | | NA | 6.8056 | 7.2478 | 6.6236 | 7.1784 | 6.5886 | 6.7492 | 6.9810 | 6.8222 | 6.7377 | 6.9291 | 7.0178 | 7.3090 | 3.8360 | 4.7608 | 4.0795 | 4.3338 |
| ADAR_S642 | ADAR | P55265-4 | | 868 | MLLLSRSPEAQPK | 6.7047 | 6.9331 | 6.1734 | 6.9350 | 6.2803 | 6.0797 | 6.5556 | 6.2391 | 6.2327 | 6.2753 | 6.0512 | 6.2456 | 5.8760 | 5.8518 | 5.7869 | 5.7798 |
| ADAR_S642;T646 | ADAR | P55265-4 | | 657 | SFFSGKSPVTTLL | 6.5402 | 7.4925 | 6.7421 | 7.4829 | 6.3965 | 6.4494 | 6.8976 | 6.3108 | 6.4628 | 6.7579 | 6.7693 | 6.7369 | 4.3418 | 5.0752 | 4.6769 | 4.8670 |
| ADAR_S657 | ADAR | P55265-4 | | 234 | LQKEAGTPPLWKI | 7.5708 | 8.0686 | 6.4388 | 7.0849 | 6.8175 | 7.1800 | 7.7456 | 7.2948 | 8.0858 | 6.4859 | 6.6547 | 6.5905 | 3.4216 | 4.0035 | 2.4936 | 4.0634 |
| ADAR_S868 | ADAR | P55265-4 | | 642 | SEKTAESQTPTPS | 5.6668 | 6.3414 | 6.4131 | 5.7711 | 6.1569 | 5.7152 | 6.8705 | 7.6231 | 5.8544 | 6.5543 | 5.9482 | 5.6984 | 6.3361 | 6.0062 | 6.2244 | 6.8199 |
| ADAR_T234 | ADAR | P55265-4 | | 644 | KTAESQTPTPSAT | 5.9775 | 5.7728 | 5.6823 | 5.9179 | 5.6189 | 5.6058 | 6.2883 | 6.4253 | 5.7813 | 5.9860 | 5.2307 | 6.4110 | 7.6758 | 7.4944 | 6.9446 | 7.1874 |
| ADAR_T644 | ADAR | P55265-4 | | 646 | AESQTPTPSATSF | 5.8517 | 5.4907 | 6.2582 | 5.7960 | 6.0797 | 5.8900 | 5.4112 | 6.3878 | 6.3695 | 7.3762 | 7.3612 | 6.8906 | 6.0380 | 6.0808 | 6.2549 | 6.4635 |
| ADAR_T646 | ADAR | P55265-4 | 642;646 | | NA | 6.9011 | 7.0647 | 5.9915 | 6.4699 | 6.3830 | 5.1312 | 5.5450 | 6.4636 | 6.1396 | 4.9368 | 6.5108 | 4.9886 | 7.7868 | 6.7810 | 6.9162 | 5.9903 |
| ADARB1_S75 | ADARB1 | P78563-5 | | 75 | RNLDNVSPKDGST | 6.0544 | 6.1478 | 6.1185 | 6.4160 | 5.7461 | 5.8818 | 6.3764 | 5.7210 | 6.4893 | 6.4028 | 6.3035 | 6.0707 | 6.5681 | 6.7106 | 6.5386 | 6.4544 |
| ADARB1_T789 | ADARB1 | P78563-5 | | 789 | QDQFSLTPxxxxx | 5.6064 | 5.9040 | 5.4529 | 7.2850 | 6.7998 | 6.7990 | 6.2398 | 6.0888 | 6.3300 | 5.9779 | 5.8679 | 5.9312 | 6.5976 | 6.8773 | 5.9950 | 6.2475 |
| ADAT1_S227 | ADAT1 | Q9BUB4 | | 227 | QKSGPISPGIHSC | 5.9011 | 6.7568 | 6.6998 | 6.6023 | 5.3260 | 6.3124 | 5.6112 | 6.8139 | 6.8323 | 6.3446 | 7.2363 | 7.4764 | 4.6993 | 6.2756 | 5.1539 | 5.9581 |
| ADCY3_S521 | ADCY3 | O60266 | | 525 | ASSKSSSPALIET | 6.7993 | 7.0097 | 6.3930 | 7.1380 | 5.9746 | 6.0422 | 5.5660 | 6.2388 | 6.8750 | 6.6758 | 6.6481 | 7.0169 | 4.7992 | 5.2714 | 5.0933 | 5.4586 |
| ADCY3_S525 | ADCY3 | O60266 | | 580 | DRVVDASEDEHEL | 6.0564 | 6.8881 | 6.2377 | 7.0511 | 6.5180 | 6.8020 | 7.1757 | 6.4393 | 5.5306 | 6.2726 | 5.6992 | 6.7342 | 5.9135 | 5.0653 | 5.7743 | 5.8421 |
| ADCY3_S580 | ADCY3 | O60266 | | 521 | NGAPASSKSSSPA | 7.2389 | 7.0216 | 6.8955 | 6.8639 | 5.7135 | 6.7807 | 7.0593 | 5.5935 | 7.0733 | 6.7027 | 7.5859 | 6.9314 | 4.7813 | 4.9924 | 3.4179 | 5.3481 |
| ADCY6_S54 | ADCY6 | O43306 | | 576 | QRTRANSMEGLMP | 6.2934 | 6.6757 | 6.1698 | 6.1565 | 5.5667 | 5.6794 | 6.4831 | 6.1115 | 6.6530 | 6.3641 | 6.3050 | 6.4409 | 6.0408 | 6.4299 | 6.3776 | 6.2525 |
| ADCY6_S576 | ADCY6 | O43306 | | 54 | RDAEPPSPTPAGP | 6.0661 | 6.9456 | 6.0242 | 6.6982 | 6.2513 | 6.1256 | 6.3885 | 6.4017 | 6.1741 | 5.8815 | 6.4475 | 5.9779 | 5.6534 | 5.9223 | 6.1163 | 6.9258 |
| ADCY9_S1257 | ADCY9 | O60503 | | 1257 | IPQHQLSISPDIR | 6.3281 | 6.7296 | 7.0864 | 7.2389 | 6.2465 | 6.2524 | 6.1538 | 6.1153 | 6.5482 | 6.7825 | 6.9234 | 6.6039 | 5.3024 | 5.7291 | 4.8469 | 5.1125 |
| ADCY9_S1307 | ADCY9 | O60503 | | 1307 | AKDAHLSPKRPWK | 6.3483 | 6.9350 | 6.5315 | 6.7011 | 6.4649 | 6.8011 | 7.3064 | 6.6091 | 6.8928 | 6.7866 | 7.1179 | 7.0568 | 4.2508 | 4.7789 | 4.5429 | 4.8758 |
| ADCY9_S365 | ADCY9 | O60503 | | 365 | KRHATSSPKNRKK | 7.4948 | 7.4584 | 6.9824 | 7.1150 | 6.8990 | 6.3759 | 6.6562 | 4.8079 | 6.4219 | 5.5352 | 5.2732 | 6.4230 | 5.4839 | 5.3887 | 5.8882 | 5.7962 |
| ADCY9_S610 | ADCY9 | O60503 | | 691 | LTNSQTSLCEILQ | 6.3391 | 6.1882 | 6.4861 | 6.3282 | 6.3602 | 6.3608 | 6.4638 | 6.1038 | 5.2172 | 5.6026 | 6.2445 | 5.5330 | 6.4286 | 6.7438 | 6.8403 | 6.7597 |
| ADCY9_S688;S691 | ADCY9 | O60503 | | 610 | RGQGTASSGNVSD | 4.6273 | 5.1879 | 5.4260 | 5.1649 | 5.3231 | 5.8831 | 4.6381 | 4.8262 | 5.4910 | 5.0030 | 4.7752 | 5.4765 | 9.4867 | 9.3202 | 9.6589 | 9.7118 |
| ADCY9_S691 | ADCY9 | O60503 | 688;691 | | NA | 5.7157 | 5.4607 | 5.6320 | 5.7134 | 6.5628 | 5.8854 | 6.5406 | 5.5986 | 6.4835 | 5.2987 | 4.9231 | 5.6459 | 7.6232 | 7.5299 | 7.9599 | 7.4266 |
| ADD1_S12 | ADD1 | P35611-3 | | 631 | REQKEKSPPDQPA | 5.8746 | 6.5825 | 6.7127 | 6.5813 | 5.6576 | 6.0839 | 6.3515 | 6.4351 | 6.1531 | 6.8230 | 6.4211 | 6.8789 | 5.8937 | 6.3208 | 5.5050 | 5.7252 |
| ADD1_S358 | ADD1 | P35611-3 | | 358 | KSRSPGSPVGEGT | 5.8678 | 6.4003 | 5.9397 | 5.8772 | 5.6918 | 5.9308 | 5.9487 | 6.2028 | 5.9193 | 5.9342 | 5.7940 | 6.3988 | 6.5803 | 7.4397 | 6.9256 | 7.1490 |
| ADD1_S358;S366 | ADD1 | P35611-3 | | 663 | PTTGDDSDAATFK | 5.3087 | 5.5912 | 6.5605 | 5.9734 | 5.9259 | 6.6551 | 5.9993 | 6.2065 | 5.3786 | 5.8538 | 5.7095 | 5.6902 | 7.3235 | 7.2274 | 7.4640 | 7.1325 |
| ADD1_S366 | ADD1 | P35611-3 | | 755 | KKKKFRTPSFLKK | 7.3728 | 7.9675 | 7.3045 | 7.4422 | 6.1218 | 7.6338 | 7.7563 | 7.1760 | 7.7146 | 7.3390 | 7.8639 | 7.8427 | 2.3008 | 3.2328 | 2.0202 | 2.9109 |
| ADD1_S512 | ADD1 | P35611-3 | | 12 | RAAVVTSPPPTTA | 6.2123 | 6.3803 | 6.5671 | 6.4477 | 6.6903 | 7.3369 | 6.8289 | 7.1821 | 6.7834 | 6.5871 | 6.4855 | 6.7031 | 4.5280 | 5.2776 | 4.8380 | 5.1518 |
| ADD1_S631 | ADD1 | P35611-3 | | 645 | PHPPPSTPIKLEE | 5.6232 | 5.8708 | 6.4002 | 5.8435 | 5.8172 | 5.9364 | 6.4406 | 6.6809 | 6.1057 | 6.0028 | 5.7264 | 6.3743 | 6.2556 | 6.8446 | 6.6886 | 7.3892 |
| ADD1_S663 | ADD1 | P35611-3 | | 757 | KKFRTPSFLKKSK | 7.1105 | 6.9468 | 7.7266 | 6.2724 | 7.3358 | 7.8837 | 6.6927 | 4.1532 | 4.4483 | 5.8122 | 6.4079 | 6.8413 | 6.5258 | 6.6435 | 4.2935 | 4.9060 |
| ADD1_S757 | ADD1 | P35611-3 | | 366 | VGEGTGSPPKWQI | 5.7563 | 6.5126 | 6.6849 | 6.5200 | 5.4664 | 6.5752 | 5.7248 | 6.1042 | 7.2299 | 7.1956 | 6.8838 | 7.6472 | 5.1302 | 5.4579 | 5.4788 | 5.6321 |
| ADD1_T645 | ADD1 | P35611-3 | | 512 | EDGHRTSTSAVPN | 5.6498 | 5.8434 | 6.3720 | 6.9985 | 5.1200 | 5.4831 | 5.8754 | 6.2122 | 6.0106 | 5.8096 | 6.1853 | 5.9614 | 7.2809 | 7.6995 | 6.5810 | 6.9172 |
| ADD1_T755 | ADD1 | P35611-3 | 358;366 | | NA | 5.6922 | 6.3357 | 6.9425 | 7.3929 | 5.0454 | 5.9800 | 6.2494 | 6.5988 | 6.0993 | 6.3443 | 6.4210 | 5.8660 | 6.1577 | 6.2633 | 6.0886 | 6.5229 |
| ADD2_S396 | ADD2 | P35612 | | 617 | TKSPLVSPSKSLE | 5.8371 | 5.8265 | 6.4938 | 6.5407 | 6.5413 | 6.5113 | 6.4697 | 6.8722 | 6.4088 | 6.1786 | 6.4001 | 6.4028 | 5.9028 | 6.0884 | 5.5860 | 5.9400 |
| ADD2_S530 | ADD2 | P35612 | | 613 | KEAETKSPLVSPS | 5.4526 | 5.7232 | 5.9331 | 5.2837 | 6.0696 | 5.8890 | 6.0071 | 5.8824 | 5.3827 | 5.5160 | 5.7811 | 5.5461 | 7.9374 | 7.6025 | 8.2759 | 7.7178 |
| ADD2_S592 | ADD2 | P35612 | | 396 | EKTKHKSEVEIPA | 6.4707 | 5.2868 | 5.8548 | 6.5555 | 5.5764 | 4.6838 | 6.1167 | 6.1406 | 5.2352 | 5.2481 | 5.1541 | 5.7036 | 8.1008 | 8.2318 | 6.9409 | 8.7001 |
| ADD2_S600 | ADD2 | P35612 | | 600 | AKSAPASPVQSPA | 4.7658 | 5.1284 | 5.6072 | 4.9733 | 4.9403 | 6.2706 | 5.3258 | 6.1565 | 5.9971 | 5.7645 | 6.0653 | 6.2125 | 7.7238 | 7.7920 | 8.8151 | 8.4618 |
| ADD2_S613 | ADD2 | P35612 | | 592 | APEEPGSPAKSAP | 5.9262 | 6.3863 | 6.2523 | 5.4896 | 6.3339 | 6.2550 | 6.0553 | 6.0869 | 6.7700 | 6.0667 | 5.9059 | 6.2145 | 6.3565 | 6.7935 | 6.7454 | 6.3620 |
| ADD2_S613;S617 | ADD2 | P35612 | | 530 | IAEKSRSPSTESQ | 5.8003 | 7.0696 | 6.5451 | 6.5839 | 6.0994 | 6.6948 | 6.6462 | 7.5668 | 7.1688 | 7.4808 | 7.0669 | 6.5426 | 4.6183 | 5.2401 | 4.2320 | 4.6445 |
| ADD2_S617 | ADD2 | P35612 | | 697 | PMSPEGSPSKSPS | 5.2224 | 4.7998 | 6.1712 | 5.2147 | 6.1975 | 5.4945 | 5.6095 | 5.4713 | 5.9450 | 5.1956 | 5.8878 | 5.5858 | 8.8708 | 8.2323 | 7.6092 | 8.4925 |
| ADD2_S693;S697 | ADD2 | P35612 | 613;617 | | NA | 6.5106 | 6.6111 | 6.9770 | 6.3518 | 6.3336 | 6.8556 | 6.9706 | 6.6028 | 6.8957 | 6.7537 | 7.2365 | 7.1652 | 4.3250 | 4.7742 | 4.5705 | 5.0662 |
| ADD2_S697 | ADD2 | P35612 | 693;697 | | NA | 5.6552 | 6.5433 | 6.2754 | 6.0744 | 6.1801 | 6.8184 | 6.4390 | 6.6125 | 6.4266 | 6.3643 | 6.8609 | 6.5380 | 5.6579 | 5.8276 | 5.7522 | 5.9741 |
| ADD3_S42 | ADD3 | Q9UEY8 | | 442 | KTRWLNSPNTYMK | 7.2723 | 7.5116 | 7.2585 | 7.7496 | 5.9028 | 7.4884 | 7.5165 | 7.0442 | 8.1466 | 7.5721 | 7.5929 | 7.5887 | 2.5153 | 3.4688 | 2.2510 | 3.1206 |
| ADD3_S423 | ADD3 | Q9UEY8 | | 677 | VLSPEGSPSKSPS | 4.9248 | 5.0417 | 4.9223 | 5.2338 | 5.2977 | 5.0329 | 5.6727 | 5.1902 | 5.3646 | 4.9851 | 4.3950 | 4.7311 | 10.1624 | 9.5701 | 9.7779 | 9.6978 |
| ADD3_S442 | ADD3 | Q9UEY8 | | 42 | IRERNMSPDLRQD | 6.8092 | 7.4582 | 7.2518 | 7.0660 | 5.9851 | 6.5903 | 6.4701 | 6.8195 | 7.0060 | 7.5796 | 7.4367 | 7.4265 | 3.9281 | 4.3314 | 3.7049 | 4.1368 |
| ADD3_S64 | ADD3 | Q9UEY8 | | 423 | DDTVPLSPLKYMA | 5.7915 | 6.3255 | 7.0168 | 5.9532 | 6.3727 | 6.7957 | 6.4610 | 6.2416 | 6.6489 | 6.6922 | 7.2196 | 6.5590 | 5.0095 | 5.7394 | 5.2865 | 5.8870 |
| ADD3_S650 | ADD3 | Q9UEY8 | | 64 | VTQILQSPAFRED | 6.9728 | 8.0094 | 6.8216 | 8.0973 | 5.8916 | 6.2680 | 6.0548 | 7.1027 | 7.3232 | 7.3039 | 6.5864 | 7.2299 | 3.3835 | 4.2055 | 3.6952 | 5.0542 |
| ADD3_S673;S677 | ADD3 | Q9UEY8 | | 481 | SKVSGGTPIKIED | 7.6309 | 8.1319 | 7.2714 | 7.1902 | 5.4150 | 7.3289 | 7.5922 | 6.9422 | 7.7433 | 8.1308 | 7.8837 | 7.8169 | 2.0935 | 3.3299 | 2.2537 | 3.2457 |
| ADD3_S677 | ADD3 | Q9UEY8 | | 650 | KRVSRLSTSTTIE | 5.7677 | 6.8428 | 6.4721 | 5.7222 | 5.4800 | 5.9741 | 5.9886 | 6.4143 | 6.3050 | 6.4822 | 5.2820 | 6.8226 | 6.3201 | 6.8664 | 6.3549 | 6.9051 |
| ADD3_T481 | ADD3 | Q9UEY8 | 673;677 | | NA | 5.8455 | 5.6355 | 6.1412 | 6.5826 | 5.6468 | 7.1096 | 6.0881 | 6.9602 | 6.5339 | 7.3402 | 7.4885 | 6.9791 | 4.8043 | 5.8233 | 5.3177 | 5.7034 |
| ADGB_S94 | ADGB | Q8N7X0 | | 94 | KIELPPSLKIYSW | 6.9458 | 7.9927 | 4.8309 | 7.6220 | 6.8500 | 7.6286 | 4.9027 | 9.2190 | 5.7101 | 5.5921 | 5.9066 | 5.5717 | 3.5659 | 6.3684 | 6.3976 | 4.8958 |
| ADGRB1_T1252 | ADGRB1 | O14514 | | 1252 | TATITGTLKRPSL | 6.8907 | 6.3536 | 8.0011 | 6.6495 | 5.7898 | 5.9730 | 7.3277 | 6.0905 | 5.4969 | 7.3762 | 5.6911 | 5.1383 | 4.5040 | 7.7254 | 5.3419 | 5.6504 |
| ADGRE5_S831 | ADGRE5 | P48960 | | 831 | TRALRASESGIxx | 6.9536 | 6.1046 | 7.1396 | 6.6716 | 6.3386 | 6.3310 | 6.2032 | 6.9233 | 6.3322 | 7.1785 | 7.3338 | 7.3895 | 4.3843 | 5.8243 | 4.6123 | 4.2797 |
| ADGRG6_S1163 | ADGRG6 | Q86SQ4 | | 1180 | TSKSKSSSTTYFK | 6.6823 | 6.5293 | 7.8906 | 6.4813 | 4.6226 | 6.3062 | 5.8435 | 6.6921 | 6.7674 | 7.6612 | 7.4172 | 8.2897 | 4.0207 | 5.8876 | 4.1933 | 4.7151 |
| ADGRG6_S1163;S1165;S | ADGRG6 | Q86SQ4 | | 516 | EGLRLHTVNVRQL | 7.2707 | 8.1715 | 4.3378 | 7.5750 | 6.8249 | 8.2070 | 4.6777 | 10.3281 | 6.0416 | 5.8912 | 5.4968 | 5.4539 | 3.6248 | 5.7963 | 6.3069 | 3.9957 |
| ADGRG6_S1180 | ADGRG6 | Q86SQ4 | | 1147 | DWSKTATNIIKKS | 8.4564 | 7.2458 | 8.4385 | 7.0831 | 4.7637 | 5.2080 | 6.2490 | 5.1708 | 5.7820 | 5.6461 | 7.5044 | 7.8514 | 4.4488 | 7.1618 | 4.4387 | 4.5516 |
| ADGRG6_S1189 | ADGRG6 | Q86SQ4 | | 1189 | TYFKRNSHTDNVS | 5.2124 | 5.6039 | 6.6130 | 6.7356 | 6.6970 | 6.6692 | 5.7792 | 6.9201 | 6.6824 | 6.8770 | 6.5979 | 6.9691 | 5.1266 | 6.4240 | 5.5142 | 5.5785 |
| ADGRG6_T1147 | ADGRG6 | Q86SQ4 | | 1163 | LGKSLSSSSIGSN | 6.4013 | 5.5825 | 6.9640 | 5.3589 | 5.5943 | 5.2700 | 6.4294 | 5.7527 | 6.3510 | 6.1348 | 4.7725 | 5.1175 | 7.8747 | 8.0065 | 7.4216 | 6.9684 |
| ADGRG6_T516 | ADGRG6 | Q86SQ4 | 1163;1165;11 | | NA | 4.4706 | 5.3232 | 4.5162 | 5.0427 | 4.3953 | 5.5316 | 5.1739 | 5.0741 | 4.8689 | 4.9799 | 4.8374 | 4.9222 | 9.1465 | 9.7112 | 10.6313 | 11.3750 |
| ADGRL1_S1447 | ADGRL1 | O94910 | | 1156 | RRMWNDTVRKQTE | 10.5367 | 7.3545 | 8.2509 | 8.5716 | 4.8499 | 5.1606 | 6.6894 | 4.0274 | 6.5301 | 5.6202 | 5.7677 | 5.5736 | 5.2445 | 6.2134 | 4.5580 | 5.0514 |
| ADGRL1_S75 | ADGRL1 | O94910 | | 1447 | YQVRRPSHEGYLA | 6.0857 | 5.4717 | 5.2340 | 5.7694 | 5.3415 | 6.7203 | 7.6732 | 6.9208 | 6.2097 | 7.2453 | 5.5420 | 7.2288 | 6.4267 | 7.8283 | 5.5048 | 4.7978 |
| ADGRL1_T1156 | ADGRL1 | K7EK47 | | 75 | QPRGGTSPYNTLI | 6.0928 | 5.8873 | 6.8317 | 6.5220 | 6.7796 | 5.6037 | 6.8103 | 6.5098 | 6.1617 | 6.5006 | 6.6248 | 6.6094 | 5.4734 | 5.7570 | 5.8076 | 6.0284 |
| ADGRL2_S1349 | ADGRL2 | O95490-6 | | 1445 | QISRGNSDGYIIP | 6.0361 | 5.8961 | 5.7993 | 5.7932 | 6.1458 | 5.9729 | 6.5062 | 5.9434 | 5.7890 | 5.9015 | 5.8877 | 5.5551 | 7.2786 | 7.4501 | 7.0670 | 6.9781 |
| ADGRL2_S1445 | ADGRL2 | O95490-6 | | 1349 | IPQRTHSLLYQPQ | 6.8154 | 6.9481 | 7.5000 | 7.1430 | 6.0837 | 6.6313 | 7.4333 | 6.5721 | 7.2829 | 6.8154 | 7.1590 | 6.8798 | 4.3354 | 5.0145 | 3.4151 | 3.9709 |
| ADNP_S608 | ADNP | Q9H2P0 | | 921 | VIPEDASESEEKL | 6.0867 | 6.2439 | 6.0582 | 6.7944 | 6.1154 | 5.9113 | 6.4530 | 6.0058 | 5.9319 | 5.8268 | 5.6278 | 6.0645 | 6.8934 | 6.7397 | 6.5084 | 6.7390 |
| ADNP_S709 | ADNP | Q9H2P0 | | 970 | EWKDGASPSESGP | 5.9769 | 6.0629 | 6.4804 | 6.6479 | 5.7451 | 5.8228 | 6.1697 | 5.8984 | 7.5326 | 7.4869 | 6.8790 | 7.5797 | 5.3676 | 5.1694 | 5.3720 | 5.8089 |
| ADNP_S736 | ADNP | Q9H2P0 | | 953 | KLMHNASDSEVDQ | 5.6535 | 6.0214 | 6.0030 | 6.5778 | 6.2560 | 5.5817 | 5.8209 | 5.8114 | 6.9822 | 6.7796 | 7.2155 | 6.9183 | 5.9555 | 6.0242 | 6.1819 | 6.2171 |
| ADNP_S740 | ADNP | Q9H2P0 | | 608 | DIPVKSSPQAAVP | 7.1919 | 8.0062 | 6.9706 | 7.5630 | 6.5092 | 6.7334 | 7.0138 | 6.4699 | 7.0950 | 7.0205 | 7.3103 | 7.4237 | 3.2959 | 4.0681 | 3.2668 | 4.0616 |
| ADNP_S921 | ADNP | Q9H2P0 | | 98 | HFRNVHSEDFENR | 7.8871 | 7.0099 | 6.7988 | 8.6842 | 5.6585 | 6.6226 | 6.8755 | 6.1631 | 7.0155 | 7.9921 | 6.8721 | 6.2173 | 4.1648 | 4.5408 | 3.4332 | 4.0644 |
| ADNP_S953 | ADNP | Q9H2P0 | | 709 | PSRLNQSPSLAPV | 7.2149 | 6.6667 | 6.5683 | 7.1297 | 7.2397 | 6.7513 | 6.7087 | 6.8759 | 7.1959 | 7.0008 | 6.4777 | 6.4359 | 4.5050 | 4.7729 | 3.8768 | 4.5795 |
| ADNP_S953;S955 | ADNP | Q9H2P0 | | 736 | RKLDDDSDSPSFF | 6.4535 | 7.8685 | 7.1563 | 7.4893 | 6.3163 | 6.3761 | 6.0852 | 6.3394 | 5.2615 | 6.2633 | 6.1368 | 6.2415 | 5.4015 | 5.2887 | 5.3988 | 5.9234 |
| ADNP_S970 | ADNP | Q9H2P0 | | 740 | DDSDSPSFFEEKP | 8.2846 | 6.4695 | 6.4420 | 9.6836 | 7.0867 | 5.4133 | 5.7928 | 6.9541 | 6.1814 | 6.0580 | 6.3826 | 6.0946 | 5.4270 | 5.2350 | 3.7811 | 4.7138 |
| ADNP_S98 | ADNP | Q9H2P0 | 953;955 | | NA | 6.1352 | 6.5820 | 6.2400 | 6.7403 | 6.0796 | 5.6597 | 6.0414 | 6.2130 | 5.7896 | 6.2557 | 5.3810 | 5.9683 | 6.7405 | 6.6795 | 7.0200 | 6.4742 |
| ADRA1B_S470 | ADRA1B | P35368 | | 470 | RRGRHDSGPLFTF | 5.3990 | 5.7421 | 5.5819 | 6.1402 | 4.9859 | 5.8760 | 6.1424 | 5.9758 | 6.4109 | 6.5444 | 6.6510 | 6.3278 | 6.5966 | 7.8415 | 6.8783 | 6.9063 |
| ADRA2C_S358 | ADRA2C | P18825 | | 358 | SVEFFLSRRRRAR | 6.5828 | 6.5828 | 8.0653 | 6.1071 | 5.5281 | 6.2657 | 7.4237 | 4.8643 | 6.0628 | 4.8495 | 6.0038 | 5.9227 | 5.1667 | 6.8631 | 5.7420 | 7.9695 |
| ADRB2_S246 | ADRB2 | P07550 | | 246 | FHVQNLSQVEQDG | 7.3019 | 6.9831 | 8.1118 | 8.0413 | 6.3705 | 5.9079 | 5.8645 | 6.0803 | 7.2092 | 6.8431 | 6.5817 | 6.9634 | 4.4206 | 4.8309 | 4.4729 | 4.0169 |
| ADRM1_T37 | ADRM1 | Q16186 | | 37 | MSLKGTTVTPDKR | 7.0942 | 7.3753 | 7.3019 | 7.4637 | 6.5144 | 7.0442 | 7.1127 | 7.0401 | 7.9105 | 7.2838 | 7.8789 | 7.5226 | 2.9162 | 3.4809 | 2.6804 | 3.3802 |
| AEBP2_S18;S24 | AEBP2 | Q6ZN18 | | 390 | KNKRRRSLPRPHD | 4.9259 | 5.5312 | 5.9252 | 6.0768 | 6.4107 | 6.1909 | 6.1740 | 6.3589 | 7.7035 | 8.0054 | 7.8777 | 7.4591 | 5.8882 | 6.2013 | 4.2657 | 5.0055 |
| AEBP2_S206 | AEBP2 | Q6ZN18 | | 24 | SPLPPGSPGSAAR | 6.5059 | 6.7624 | 6.8606 | 6.4214 | 6.4814 | 6.4547 | 6.2187 | 6.5099 | 6.5847 | 6.8443 | 6.8433 | 6.7481 | 5.2495 | 5.4207 | 4.7830 | 5.3115 |
| AEBP2_S24 | AEBP2 | Q6ZN18 | | 206 | SGGRRGSLEMSSD | 6.1708 | 6.0961 | 6.3432 | 6.5715 | 5.9633 | 6.2281 | 6.0588 | 5.9651 | 6.5896 | 6.7092 | 6.3655 | 7.2548 | 6.2569 | 6.0982 | 5.7162 | 5.6126 |
| AEBP2_S371 | AEBP2 | Q6ZN18 | | 371 | PKAKEESPSKAGM | 5.9032 | 6.6345 | 7.0635 | 6.8678 | 5.6295 | 6.0167 | 6.4059 | 6.2627 | 6.2787 | 6.3067 | 7.0926 | 7.6028 | 5.1090 | 6.1495 | 5.0981 | 5.5788 |
| AEBP2_S390 | AEBP2 | Q6ZN18 | 18;24 | | NA | 6.2600 | 7.4020 | 6.2677 | 6.0722 | 6.6484 | 5.2178 | 6.0406 | 6.7322 | 7.8916 | 6.8994 | 6.5141 | 5.1738 | 5.3607 | 7.0955 | 5.4189 | 5.0051 |
| AFAP1_S265 | AFAP1 | Q8N556-2 | | 752 | GTSSPQSPVFRHR | 5.9746 | 6.1362 | 5.7237 | 6.0012 | 6.0437 | 5.7133 | 6.2030 | 5.9209 | 6.0978 | 5.8387 | 5.8627 | 5.9161 | 7.1904 | 6.9541 | 7.2937 | 7.1299 |
| AFAP1_S277;S282;S283 | AFAP1 | Q8N556-2 | | 632 | VVKRTGSNAAQYK | 5.8732 | 5.3229 | 6.1796 | 5.9499 | 6.3322 | 5.9394 | 6.3154 | 5.4323 | 6.8520 | 6.1229 | 5.9713 | 6.2497 | 7.4438 | 7.0333 | 6.1374 | 6.8446 |
| AFAP1_S277;S283 | AFAP1 | Q8N556-2 | | 265 | CPPPPSSPVHKAE | 5.9399 | 6.2257 | 6.1853 | 6.7870 | 6.6562 | 6.5839 | 6.0898 | 6.1072 | 6.1158 | 5.7232 | 5.8924 | 6.0444 | 6.3162 | 6.0615 | 6.7116 | 6.5600 |
| AFAP1_S283 | AFAP1 | Q8N556-2 | | 630 | AAVVKRTGSNAAQ | 6.7196 | 5.8735 | 6.0971 | 5.9511 | 6.3634 | 5.4980 | 6.4261 | 5.3137 | 6.7715 | 5.8798 | 5.6220 | 6.0954 | 7.5525 | 7.0418 | 6.1285 | 6.6661 |
| AFAP1_S342;S343 | AFAP1 | Q8N556-2 | | 283 | SSERPSSDGEGVV | 6.0797 | 6.4367 | 6.5442 | 6.9073 | 5.8470 | 5.9721 | 6.2092 | 5.8552 | 6.3688 | 6.3298 | 6.3759 | 6.3834 | 5.9690 | 6.4747 | 5.9853 | 6.2616 |
| AFAP1_S632 | AFAP1 | Q8N556-2 | | 795 | QAAPGSSPCRGHV | 6.4512 | 6.5356 | 6.4921 | 6.8487 | 6.4826 | 6.7770 | 6.8851 | 6.2229 | 7.1574 | 6.9960 | 6.9490 | 6.8357 | 4.6897 | 5.3153 | 4.3880 | 4.9735 |
| AFAP1_S763 | AFAP1 | Q8N556-2 | | 763 | HRTLENSPISSCD | 4.8372 | 5.3715 | 5.0030 | 5.8276 | 5.3306 | 5.6400 | 6.3526 | 5.9400 | 6.6718 | 6.2263 | 6.6418 | 6.5217 | 7.2963 | 7.4998 | 7.3294 | 7.5104 |
| AFAP1_S795 | AFAP1 | Q8N556-2 | | 747 | IEPKSGTSSPQSP | 6.3524 | 7.4458 | 6.6293 | 6.5718 | 5.9823 | 5.6387 | 5.7347 | 5.9739 | 6.2136 | 5.6288 | 6.6646 | 6.1203 | 6.1021 | 6.5028 | 5.5280 | 6.9110 |
| AFAP1_T341;S343 | AFAP1 | Q8N556-2 | 342;343 | | NA | 5.6345 | 6.1344 | 6.3267 | 7.0573 | 6.1922 | 6.3108 | 6.0890 | 5.6552 | 6.5650 | 6.3199 | 6.5577 | 6.3809 | 6.4017 | 6.0161 | 5.9220 | 6.4367 |
| AFAP1_T630 | AFAP1 | Q8N556-2 | 747;752 | | NA | 6.4274 | 6.7107 | 6.6623 | 7.0943 | 6.2604 | 5.5861 | 6.9590 | 6.5358 | 6.7519 | 6.0540 | 6.7050 | 6.2942 | 5.4349 | 5.4495 | 5.3695 | 5.7050 |
| AFAP1_T747 | AFAP1 | Q8N556-2 | 341;343 | | NA | 6.0957 | 6.4734 | 6.0961 | 6.6649 | 5.5742 | 5.7192 | 6.0280 | 5.9564 | 6.7244 | 6.5230 | 6.4013 | 5.6701 | 6.3497 | 6.0413 | 6.0116 | 6.7707 |
| AFAP1_T747;S752 | AFAP1 | Q8N556-2 | 277;282;283 | | NA | 5.2738 | 5.4879 | 5.5088 | 6.3306 | 6.2525 | 5.9239 | 6.6413 | 5.8975 | 5.8686 | 5.5276 | 6.1756 | 5.5860 | 7.5878 | 6.7490 | 7.9213 | 7.2679 |
| AFAP1_T752 | AFAP1 | Q8N556-2 | 277;283 | | NA | 5.4135 | 5.8239 | 6.2607 | 6.6515 | 6.1525 | 5.6979 | 5.5189 | 5.4784 | 6.4172 | 7.0282 | 7.3330 | 7.8107 | 6.7128 | 6.1578 | 5.5497 | 5.9932 |
| AFAP1L1_S329 | AFAP1L1 | Q8TED9 | | 329 | GVEVPRSPVLLCK | 6.5902 | 5.9432 | 7.3263 | 7.1242 | 6.5984 | 6.6809 | 6.3790 | 5.7032 | 6.7333 | 6.5783 | 6.7462 | 7.0937 | 5.1777 | 4.9417 | 5.0967 | 5.2871 |
| AFAP1L2_S372 | AFAP1L2 | Q8N4X5-4 | | 569 | RVYLDLTPVKSFL | 6.6210 | 6.4828 | 6.6691 | 6.3432 | 6.3758 | 6.6399 | 6.7800 | 6.8790 | 6.9167 | 7.1375 | 7.2908 | 5.9895 | 4.5080 | 5.2678 | 4.9976 | 5.1014 |
| AFAP1L2_S668 | AFAP1L2 | Q8N4X5-4 | | 668 | ATPPGASPPVKDR | 5.6089 | 6.5051 | 7.1754 | 6.2346 | 6.4543 | 6.2502 | 5.4853 | 6.4872 | 6.7147 | 6.9720 | 6.7296 | 6.8485 | 5.7692 | 5.8029 | 5.5323 | 5.4298 |
| AFAP1L2_S820 | AFAP1L2 | Q8N4X5-4 | | 820 | LKNRPLSVVVTGK | 7.7983 | 7.4221 | 8.0637 | 8.0162 | 7.4260 | 6.2220 | 5.6121 | 4.2497 | 7.9092 | 4.4794 | 5.3151 | 5.2953 | 5.6438 | 6.2299 | 5.4972 | 4.8200 |
| AFAP1L2_T569 | AFAP1L2 | Q8N4X5-4 | | 372 | NLGRKKSTSLEPV | 5.9995 | 4.9151 | 6.4182 | 5.8033 | 6.6991 | 6.0613 | 5.2021 | 6.3449 | 7.0199 | 7.0527 | 6.8346 | 7.6386 | 6.4083 | 6.5422 | 5.4267 | 5.6333 |
| AFDN_S1082 | AFDN | P55196-5 | | 1106 | TLLNQPSPMMQRI | 6.2719 | 6.3866 | 6.4471 | 6.6170 | 6.4595 | 6.3890 | 6.1639 | 6.5012 | 6.2794 | 6.0972 | 6.0864 | 6.3420 | 5.7354 | 6.1820 | 5.9817 | 6.0597 |
| AFDN_S1106 | AFDN | P55196-5 | | 215 | SFTRTISNPEVVM | 5.7134 | 6.5280 | 7.7740 | 5.4433 | 4.5554 | 7.0777 | 6.3252 | 7.1559 | 7.3793 | 9.0292 | 8.9255 | 8.9499 | 3.5410 | 4.8238 | 2.8707 | 3.9075 |
| AFDN_S1139 | AFDN | P55196-5 | | 1210 | LCTEEQTPPPRPE | 5.8379 | 6.5057 | 6.6729 | 6.8179 | 5.8192 | 6.5666 | 6.2246 | 6.3178 | 6.3804 | 6.4370 | 6.2028 | 6.6550 | 5.8751 | 6.0613 | 5.7920 | 5.8338 |
| AFDN_S1139;S1142 | AFDN | P55196-5 | | 1454 | LNPAPFSPLTAQQ | 6.4033 | 6.9678 | 6.5766 | 7.1471 | 6.2113 | 7.0440 | 6.8576 | 6.9662 | 6.8340 | 6.7942 | 7.0440 | 6.7563 | 4.2604 | 5.1188 | 4.2453 | 4.7731 |
| AFDN_S1142 | AFDN | P55196-5 | | 1139 | NSTQNGSPESPQL | 6.3087 | 6.4789 | 6.4044 | 6.4489 | 6.5819 | 6.3783 | 6.2655 | 6.6940 | 6.5229 | 6.4421 | 6.4218 | 6.1844 | 5.6087 | 6.1725 | 5.5530 | 5.5340 |
| AFDN_S1171;S1181 | AFDN | P55196-5 | | 1082 | LMTRTSSVVTLEV | 6.5644 | 6.3016 | 6.2886 | 6.2033 | 6.3380 | 5.8294 | 6.2463 | 5.7992 | 6.2929 | 5.8553 | 5.6231 | 5.5368 | 7.1383 | 6.7592 | 7.0019 | 6.2215 |
| AFDN_S1172 | AFDN | P55196-5 | | 1329 | HSSKSVTPASTLT | 6.2644 | 7.6128 | 7.4221 | 6.2966 | 5.5431 | 7.4371 | 7.1677 | 7.0648 | 6.4791 | 7.4447 | 7.3674 | 7.0697 | 3.7867 | 4.8491 | 3.8951 | 4.2995 |
| AFDN_S1181 | AFDN | P55196-5 | | 1181 | VANQPPSPGGKSA | 6.2565 | 6.2488 | 6.0943 | 6.7014 | 6.5812 | 5.8806 | 5.9470 | 6.0955 | 6.5740 | 6.4316 | 6.0687 | 6.5487 | 6.1393 | 6.2774 | 6.3362 | 5.8187 |
| AFDN_S1274 | AFDN | P55196-5 | | 1809 | ERQRLFSQGQDVS | 6.0800 | 5.2891 | 5.6305 | 5.3927 | 6.2822 | 5.9414 | 6.5398 | 5.7114 | 6.6825 | 4.7184 | 4.1080 | 4.1907 | 9.4241 | 8.9872 | 7.8526 | 7.1693 |
| AFDN_S1454 | AFDN | P55196-5 | | 1231 | YTREYFTFPASKS | 8.4443 | 8.1161 | 7.2397 | 7.8496 | 7.0930 | 6.1273 | 6.5304 | 5.8272 | 6.5562 | 6.1048 | 5.8651 | 5.5831 | 4.8428 | 5.4293 | 4.3064 | 4.0848 |
| AFDN_S1500 | AFDN | P55196-5 | | 423 | LYRLQLSVTEVGT | 5.8911 | 6.3104 | 5.9496 | 6.5076 | 6.0950 | 5.8935 | 6.1431 | 6.2078 | 6.0214 | 5.7482 | 5.5924 | 5.7207 | 7.0886 | 6.9903 | 7.2690 | 6.5715 |
| AFDN_S1511 | AFDN | P55196-5 | | 1500 | LQYITVSKEELSS | 5.4036 | 6.6420 | 6.7130 | 6.1259 | 4.8280 | 6.4090 | 6.8742 | 6.9529 | 4.9659 | 6.2946 | 7.2661 | 6.7239 | 5.1934 | 5.9810 | 6.6810 | 6.9454 |
| AFDN_S1706 | AFDN | P55196-5 | | 1789 | KSQDADSPGSSGA | 6.8231 | 6.8164 | 6.4476 | 6.7123 | 6.1523 | 6.7440 | 6.6775 | 6.0302 | 6.3382 | 6.7509 | 6.2342 | 6.8042 | 5.3427 | 5.6716 | 5.2666 | 5.1884 |
| AFDN_S1731 | AFDN | P55196-5 | | 1731 | LKTQVLSPDSLFT | 6.7275 | 7.2108 | 6.6897 | 7.8698 | 6.3450 | 6.9715 | 6.8460 | 6.7326 | 7.3141 | 7.6492 | 7.3687 | 7.2845 | 3.4036 | 4.4674 | 3.1649 | 3.9547 |
| AFDN_S1789 | AFDN | P55196-5 | | 1511 | SSGDSLSPDPWKR | 5.5658 | 7.0599 | 6.9701 | 6.7981 | 5.0319 | 7.2806 | 7.0915 | 6.6436 | 5.6589 | 6.9153 | 7.7161 | 7.6887 | 4.2913 | 5.0819 | 4.7337 | 5.4727 |
| AFDN_S1789;S1792 | AFDN | P55196-5 | | 1274 | IQRVTRSQEELRE | 7.0046 | 7.9489 | 7.3162 | 6.6029 | 5.8139 | 7.9389 | 7.0498 | 4.5504 | 5.1585 | 6.2422 | 7.7366 | 7.1192 | 4.8066 | 5.5316 | 4.7171 | 4.4625 |
| AFDN_S1809 | AFDN | P55196-5 | | 1706 | RDYEPPSPSPAPG | 6.4579 | 5.4495 | 6.0871 | 6.2987 | 6.1567 | 6.6334 | 7.2126 | 5.6180 | 5.7637 | 6.2555 | 6.5103 | 6.3116 | 6.5438 | 6.6782 | 5.7432 | 6.2798 |
| AFDN_S215 | AFDN | P55196-5 | | 511 | HALAKRSVDGGLM | 6.1486 | 6.2414 | 6.0429 | 6.7706 | 6.2447 | 5.9215 | 6.5594 | 6.3318 | 6.6976 | 6.6570 | 6.1836 | 6.1203 | 5.8694 | 6.5141 | 5.6222 | 6.0749 |
| AFDN_S268 | AFDN | P55196-5 | | 1468 | KPEKPSTLQRPQE | 8.7436 | 7.0179 | 7.2252 | 6.8035 | 5.5266 | 5.0867 | 6.7619 | 4.8575 | 7.2637 | 6.0613 | 6.1655 | 6.5726 | 5.7071 | 6.0345 | 4.5738 | 5.5986 |
| AFDN_S423 | AFDN | P55196-5 | | 1142 | QNGSPESPQLPWA | 5.6543 | 6.1077 | 6.5771 | 6.3721 | 6.6660 | 5.7981 | 6.8862 | 6.5775 | 6.3569 | 6.2118 | 6.2113 | 6.3518 | 5.7787 | 6.9378 | 5.7188 | 5.7939 |
| AFDN_S511 | AFDN | P55196-5 | | 556 | STTRLDSDRVSSA | 6.9383 | 6.5878 | 5.6357 | 6.1063 | 6.6618 | 6.5218 | 6.1569 | 6.2825 | 6.5625 | 6.3990 | 5.3999 | 6.7811 | 6.5182 | 6.6058 | 5.3268 | 5.5156 |
| AFDN_S556 | AFDN | P55196-5 | | 1188 | PGGKSAYASGTTA | 6.4238 | 5.7529 | 5.9368 | 7.4669 | 6.1743 | 5.9061 | 6.6756 | 5.4366 | 6.8463 | 6.3844 | 6.9164 | 7.2425 | 5.4344 | 6.3943 | 4.9790 | 6.0298 |
| AFDN_T1135 | AFDN | P55196-5 | | 1172 | RADHRSSPNVANQ | 4.5845 | 7.0913 | 6.8329 | 6.4432 | 5.4865 | 5.1252 | 5.3551 | 5.7054 | 6.0141 | 6.1805 | 6.2527 | 5.5368 | 8.2078 | 7.5860 | 6.0294 | 7.5685 |
| AFDN_T1210 | AFDN | P55196-5 | | 1135 | ELYNNSTQNGSPE | 5.7811 | 6.5232 | 6.4495 | 7.2913 | 6.5232 | 6.3838 | 5.7449 | 6.6376 | 6.0451 | 6.7452 | 6.8392 | 6.8381 | 5.1173 | 5.8185 | 5.2634 | 5.9986 |
| AFDN_T1231 | AFDN | P55196-5 | | 268 | YKTILLSTTDPAD | 6.3942 | 6.3111 | 6.2548 | 5.9858 | 6.3551 | 5.8990 | 6.5262 | 7.0128 | 5.1973 | 5.7230 | 6.5458 | 6.2267 | 6.5751 | 6.1264 | 6.5372 | 6.3294 |
| AFDN_T1329 | AFDN | P55196-5 | 1139;1142 | | NA | 5.5598 | 6.4628 | 6.5401 | 7.2117 | 6.1460 | 6.4151 | 6.2804 | 6.4775 | 6.5752 | 6.9331 | 6.7220 | 6.2823 | 5.7099 | 5.8614 | 5.3351 | 5.4875 |
| AFDN_T1468 | AFDN | P55196-5 | 1171;1181 | | NA | 5.8592 | 6.5887 | 6.9085 | 6.3031 | 5.8230 | 6.5135 | 6.7403 | 5.7285 | 6.0762 | 6.4173 | 6.6065 | 7.0542 | 5.3852 | 6.1527 | 5.4894 | 6.3537 |
| AFDN_Y1188 | AFDN | P55196-5 | 1789;1792 | | NA | 7.0051 | 7.0735 | 5.9317 | 7.3447 | 6.0767 | 5.6917 | 6.7842 | 6.8073 | 6.7195 | 6.3150 | 6.3591 | 6.4399 | 5.2856 | 5.7443 | 5.3095 | 5.1122 |
| AFF1_S1099 | AFF1 | P51825-2 | | 1099 | KVAQAPSPCIARS | 6.2783 | 6.5439 | 6.0797 | 6.7775 | 6.2445 | 6.3742 | 6.4392 | 6.1813 | 6.6352 | 6.5448 | 5.7463 | 6.0672 | 5.8211 | 6.4503 | 5.3601 | 6.4564 |
| AFF1_S206;S213;S219 | AFF1 | P51825-2 | | 757 | RDTKLLSPLRDTP | 8.1953 | 7.6941 | 7.9879 | 8.1362 | 5.9231 | 6.5019 | 7.5732 | 6.2060 | 6.9375 | 6.9758 | 7.7948 | 6.8707 | 2.6158 | 4.5274 | 3.0557 | 3.0048 |
| AFF1_S213;S219 | AFF1 | P51825-2 | | 383 | PLTAIHTPSTAEP | 6.2380 | 6.5629 | 6.7122 | 7.1972 | 6.2340 | 6.6121 | 6.1053 | 6.1739 | 6.6653 | 7.2404 | 6.8050 | 6.4571 | 5.3426 | 5.6194 | 4.3245 | 5.7102 |
| AFF1_S757 | AFF1 | P51825-2 | 213;219 | | NA | 6.3620 | 6.8911 | 6.9661 | 8.5140 | 6.7892 | 7.2842 | 5.3066 | 6.9193 | 7.0610 | 7.1152 | 6.2715 | 6.4434 | 4.2688 | 5.3918 | 3.7900 | 4.6260 |
| AFF1_T383 | AFF1 | P51825-2 | 206;213;219 | | NA | 6.7825 | 6.4710 | 6.0237 | 7.6265 | 6.4912 | 6.0129 | 6.9145 | 5.9714 | 6.5880 | 6.2977 | 6.6236 | 5.1068 | 5.1591 | 6.2525 | 5.5424 | 6.1363 |
| AFF4_S1043 | AFF4 | Q9UHB7 | | 694 | FRQRMFSPMEEKE | 6.9091 | 7.3086 | 6.8687 | 7.3348 | 6.0829 | 6.0736 | 6.5722 | 6.1303 | 6.8020 | 7.0790 | 7.1720 | 6.9880 | 4.3554 | 5.0875 | 4.5851 | 4.6508 |
| AFF4_S1055 | AFF4 | Q9UHB7 | | 1058 | GMPSPVSPKLSPG | 7.0067 | 7.2903 | 6.9597 | 7.4034 | 6.1612 | 5.9951 | 6.3519 | 6.1604 | 7.2412 | 7.4631 | 6.7008 | 6.8691 | 4.3306 | 5.1269 | 4.3156 | 4.6240 |
| AFF4_S1055;S1058 | AFF4 | Q9UHB7 | | 703 | EEKELLSPLSEPD | 6.3677 | 6.6377 | 6.3324 | 7.0168 | 6.2585 | 6.3026 | 6.3786 | 6.0714 | 6.6381 | 6.8943 | 6.5769 | 6.5915 | 5.1541 | 6.1022 | 5.3332 | 5.3439 |
| AFF4_S1058 | AFF4 | Q9UHB7 | | 674 | YPESNRTPVKPSS | 6.5836 | 6.9340 | 6.7284 | 7.3496 | 6.2595 | 6.1936 | 6.4788 | 6.3703 | 6.7279 | 7.0336 | 6.6530 | 6.8099 | 4.5652 | 5.5686 | 4.7159 | 5.0282 |
| AFF4_S1062 | AFF4 | Q9UHB7 | | 814 | EKDLLPSPAGPVP | 6.5918 | 6.8832 | 6.3390 | 6.8863 | 6.1252 | 6.2636 | 6.3097 | 6.3386 | 6.6888 | 6.6517 | 6.5959 | 6.4380 | 4.8988 | 6.2439 | 5.1774 | 5.5681 |
| AFF4_S222 | AFF4 | Q9UHB7 | | 1055 | KAVGMPSPVSPKL | 6.9287 | 6.9746 | 6.9083 | 7.0912 | 6.4161 | 6.2298 | 6.3954 | 6.3562 | 7.5343 | 6.8249 | 6.2380 | 6.6477 | 4.3759 | 5.5365 | 4.5073 | 5.0353 |
| AFF4_S32 | AFF4 | Q9UHB7 | | 336 | PLTAIHTPCKTEP | 6.5545 | 6.5060 | 6.7430 | 7.2551 | 5.7792 | 6.7346 | 6.4410 | 6.3581 | 6.8693 | 6.6370 | 6.7422 | 6.7380 | 5.0834 | 5.7503 | 4.5380 | 5.2705 |
| AFF4_S487 | AFF4 | Q9UHB7 | | 620 | KKESKSSPRPTAE | 6.5222 | 6.8790 | 6.5585 | 7.6297 | 6.1518 | 6.0154 | 6.7129 | 6.0081 | 6.6648 | 6.6658 | 6.5969 | 6.9464 | 4.5244 | 5.7488 | 4.9393 | 5.4359 |
| AFF4_S487;S491 | AFF4 | Q9UHB7 | | 32 | DAFPPSSPLFAEP | 6.2439 | 6.4807 | 6.8380 | 7.1072 | 6.1690 | 5.7099 | 6.3291 | 6.6583 | 6.5247 | 6.9258 | 6.5778 | 6.3963 | 5.6114 | 5.7372 | 5.2300 | 5.4608 |
| AFF4_S620 | AFF4 | Q9UHB7 | | 1062 | PVSPKLSPGNSGN | 6.1793 | 6.7033 | 6.8228 | 6.7042 | 5.9799 | 6.3670 | 6.6215 | 6.9261 | 6.7287 | 6.7822 | 6.5207 | 6.5992 | 5.2465 | 5.5709 | 4.9483 | 5.2994 |
| AFF4_S694 | AFF4 | Q9UHB7 | | 528 | KETSSATPGRDSK | 6.0990 | 6.6279 | 7.1726 | 7.0576 | 6.4111 | 6.0149 | 6.6735 | 5.6785 | 6.6102 | 6.2530 | 7.4570 | 6.8030 | 5.1152 | 5.5189 | 5.0981 | 5.4094 |
| AFF4_S703 | AFF4 | Q9UHB7 | | 836 | SRKRTISQSSSLK | 7.1774 | 7.4466 | 8.3369 | 8.7222 | 6.8705 | 6.8959 | 7.6148 | 6.0972 | 6.6302 | 7.1327 | 8.6823 | 7.6251 | 2.4246 | 3.1935 | 2.5368 | 2.6134 |
| AFF4_S703;S706 | AFF4 | Q9UHB7 | | 1043 | NNSQAPSPGLGSK | 6.3170 | 6.6724 | 5.6067 | 6.9599 | 6.0369 | 5.9710 | 6.3740 | 6.0094 | 6.3208 | 6.2553 | 6.1341 | 6.3177 | 5.6677 | 7.1346 | 5.5032 | 6.7193 |
| AFF4_S814 | AFF4 | Q9UHB7 | | 487 | VNPHKVSPASSVD | 6.2632 | 6.4046 | 6.3922 | 7.0454 | 6.2301 | 6.8944 | 6.1815 | 6.5367 | 6.6811 | 6.4524 | 7.5567 | 7.3025 | 4.8987 | 5.7661 | 4.3380 | 5.0565 |
| AFF4_S836 | AFF4 | Q9UHB7 | | 222 | PDANWDSPSRVPF | 6.9702 | 7.3115 | 6.2139 | 7.5938 | 5.9134 | 5.8166 | 6.4400 | 6.8787 | 7.4424 | 6.8081 | 6.5790 | 6.6647 | 4.3754 | 4.9330 | 5.0781 | 4.9813 |
| AFF4_T336 | AFF4 | Q9UHB7 | 703;706 | | NA | 6.8190 | 7.2868 | 7.0301 | 7.6589 | 6.1875 | 6.1812 | 6.6020 | 6.1665 | 6.9925 | 6.7902 | 6.7583 | 6.9133 | 4.3345 | 5.0474 | 4.4460 | 4.7859 |
| AFF4_T528 | AFF4 | Q9UHB7 | 1055;1058 | | NA | 6.5266 | 6.2871 | 5.9299 | 6.4461 | 6.4539 | 6.3531 | 6.7207 | 6.3439 | 6.7001 | 6.1141 | 6.0091 | 6.0764 | 5.7379 | 6.6056 | 5.4211 | 6.2745 |

| | | | | | | | | | | | | | | | | | | | | |
|---|---|---|---|---|---|---|---|---|---|---|---|---|---|---|---|---|---|---|---|---|
| AFF4_T674 | AFF4 | Q9UHB7 | 487;491 | NA | 5.9986 | 6.1355 | 6.4213 | 6.9440 | 6.2685 | 6.1740 | 6.3939 | 6.3748 | 7.6878 | 7.6429 | 7.7006 | 7.3985 | 4.3878 | 5.6957 | 4.4391 | 4.3370 |
| AFTPH_S151 | AFTPH | Q6ULP2-9 | 411 | DLDDSLSVKNGDS | 6.4315 | 5.8556 | 6.8711 | 6.1420 | 6.5475 | 6.0918 | 5.9278 | 6.1314 | 6.2805 | 5.9006 | 6.2638 | 6.0290 | 7.1002 | 6.4172 | 5.8100 | 6.1998 |
| AFTPH_S411 | AFTPH | Q6ULP2-9 | 151 | GTLESFSPGDFRT | 6.3482 | 6.4887 | 6.7324 | 7.0045 | 6.0012 | 6.3878 | 6.3287 | 6.5704 | 6.7201 | 6.5088 | 7.0162 | 7.1703 | 4.9893 | 5.5869 | 5.0772 | 5.0693 |
| AGAP1_S269 | AGAP1 | E7EUN2 | 733 | KLDPPPSPHANRK | 5.9306 | 6.2807 | 7.2298 | 6.2601 | 6.1958 | 6.6731 | 6.7985 | 5.4546 | 5.5926 | 6.3881 | 6.3464 | 6.1188 | 5.9374 | 6.6966 | 6.0871 | 6.0099 |
| AGAP1_S687 | AGAP1 | E7EUN2 | 1048 | RFVLMATPNLSRR | 6.1747 | 7.2233 | 5.8060 | 5.7680 | 5.7517 | 6.5823 | 6.0953 | 5.7915 | 6.4632 | 6.7801 | 5.5775 | 6.0248 | 5.7963 | 7.0750 | 6.6812 | 6.4089 |
| AGAP1_S733 | AGAP1 | E7EUN2 | 687 | ACAPISSPKTNGL | 6.3181 | 7.0158 | 6.9619 | 7.1553 | 6.1873 | 6.8121 | 7.2234 | 6.8004 | 6.8446 | 7.1352 | 7.3657 | 7.1174 | 3.9578 | 4.5774 | 3.9427 | 4.5849 |
| AGAP1_T1048 | AGAP1 | E7EUN2 | 269 | RLLRFFSGIFARR | 8.1638 | 7.0325 | 6.6130 | 5.8842 | 5.7449 | 5.8268 | 6.0771 | 6.7990 | 7.1993 | 6.6581 | 6.7018 | 6.8212 | 5.5635 | 5.7755 | 4.2369 | 4.9021 |
| AGAP1_T210 | AGAP1 | E7EUN2 | 210 | ASEKRNTLDVGEV | 7.2111 | 6.8986 | 7.8338 | 6.5100 | 5.8565 | 5.4987 | 6.3168 | 6.4954 | 7.1445 | 7.7021 | 6.6161 | 7.4643 | 4.6930 | 4.9132 | 4.0073 | 4.8386 |
| AGAP2_S1181 | AGAP2 | Q99490 | 523 | DRISASSPRVVGD | 6.6164 | 6.7239 | 6.5334 | 7.3509 | 5.8841 | 6.2707 | 6.5895 | 6.5254 | 6.2269 | 6.0332 | 6.2566 | 6.2247 | 5.6395 | 5.5969 | 5.8720 | 5.6559 |
| AGAP2_S523 | AGAP2 | Q99490 | 1181 | RRSSAASVGRADA | 9.2385 | 7.1163 | 8.5094 | 7.7200 | 5.8931 | 5.6822 | 5.9509 | 5.3459 | 6.8234 | 7.0861 | 6.2871 | 6.8981 | 3.9890 | 6.1100 | 3.5623 | 3.7877 |
| AGAP3_S574 | AGAP3 | Q96P47-4 | 574 | KLDPPPSPHSNRK | 5.9919 | 6.4701 | 6.5894 | 6.2279 | 5.9514 | 6.4906 | 6.0663 | 6.4189 | 6.0637 | 6.2452 | 6.3608 | 6.9088 | 6.0075 | 6.2878 | 6.0283 | 5.8913 |
| AGFG1_T177;S181 | AGFG1 | P52594-4 | 181 | KGTPSQSPVVGRS | 6.1787 | 6.0851 | 6.1273 | 5.7527 | 6.7888 | 6.5852 | 5.8484 | 6.7246 | 5.9582 | 5.7896 | 6.3255 | 6.0261 | 6.6923 | 6.4687 | 6.3786 | 6.2700 |
| AGFG1_T181 | AGFG1 | P52594-4 | 177;181 | NA | 6.6696 | 6.8917 | 6.8913 | 6.3341 | 6.0651 | 6.7406 | 6.5012 | 6.9694 | 7.1083 | 6.7756 | 7.1732 | 7.1978 | 4.8038 | 5.1711 | 4.1670 | 4.5401 |
| AGGF1_S306 | AGGF1 | Q8N302 | 344 | SGNTIESPLHENI | 5.6957 | 6.4478 | 5.9082 | 6.2352 | 6.5828 | 6.1846 | 5.8751 | 6.6231 | 6.2217 | 6.2443 | 6.5735 | 5.8105 | 6.5109 | 6.4635 | 6.4757 | 6.1474 |
| AGGF1_S344 | AGGF1 | Q8N302 | 306 | FSVEHTSCNEEEN | 6.0854 | 7.2675 | 7.5838 | 7.2369 | 7.3508 | 6.8593 | 6.4001 | 7.3321 | 6.8168 | 5.6517 | 5.3473 | 6.1739 | 5.3881 | 4.3149 | 4.8030 | 5.3881 |
| AGK_S285 | AGK | Q53H12 | 285 | ILRRLASYWAQPQ | 6.5140 | 7.0712 | 5.9453 | 5.6934 | 6.8292 | 5.4667 | 6.0682 | 6.1430 | 5.5118 | 5.2667 | 5.9400 | 5.6095 | 6.5819 | 7.3551 | 6.6414 | 7.3627 |
| AGO2_S387 | AGO2 | Q9UKV8 | 387 | KLMRSASFNTDPY | 6.4897 | 6.1266 | 6.7661 | 5.5911 | 5.8173 | 6.0853 | 6.1395 | 6.6128 | 6.7187 | 6.4588 | 6.7071 | 6.4463 | 6.4359 | 6.4022 | 5.6268 | 5.5758 |
| AGPS_T155 | AGPS | O00116 | 155 | SLNPSDTPPSVVN | 4.9806 | 7.0855 | 5.9347 | 6.2138 | 5.6238 | 6.5779 | 6.2058 | 7.8642 | 6.5061 | 5.9454 | 6.2829 | 7.2503 | 6.2111 | 5.9640 | 6.0065 | 5.3474 |
| AGTPBP1_S1154 | AGTPBP1 | Q9UPW5-3 | 1154 | RLKRLTSPLEYNL | 5.8741 | 6.1534 | 6.6934 | 6.0773 | 6.2281 | 6.4600 | 6.3250 | 6.3882 | 5.9725 | 5.6694 | 6.7853 | 6.5850 | 6.0198 | 6.2410 | 6.6331 | 5.8942 |
| AHCTF1_S1030 | AHCTF1 | Q8WYP5-2 | 1318 | TSFFLNSPEKEHQ | 6.7137 | 6.5922 | 6.3563 | 7.0520 | 6.4629 | 5.9314 | 6.6712 | 6.3146 | 6.6985 | 6.5962 | 6.3642 | 6.4864 | 5.1200 | 5.6862 | 5.3573 | 5.5969 |
| AHCTF1_S1116 | AHCTF1 | Q8WYP5-2 | 1292 | KLEVFTTPKKCAV | 7.4305 | 6.9536 | 6.1729 | 6.6028 | 6.1821 | 5.9273 | 6.9960 | 6.6923 | 6.9926 | 6.3466 | 5.9193 | 6.2795 | 4.9194 | 5.6007 | 5.1382 | 5.8461 |
| AHCTF1_S1177 | AHCTF1 | Q8WYP5-2 | 1548 | KLPISDSPPDTQE | 6.1233 | 6.5004 | 6.0310 | 7.2398 | 6.0257 | 6.2028 | 6.2062 | 6.2159 | 6.3852 | 6.6005 | 6.3960 | 6.4416 | 5.4520 | 6.2820 | 5.8479 | 6.0498 |
| AHCTF1_S1185 | AHCTF1 | Q8WYP5-2 | 1267 | LKETRISFVEEDV | 4.5326 | 4.8284 | 4.5831 | 5.2486 | 5.3407 | 4.8668 | 5.4218 | 5.4933 | 5.1366 | 4.7153 | 4.6375 | 5.1866 | 9.0249 | 9.2795 | 10.8132 | 10.8910 |
| AHCTF1_S1185;S1188 | AHCTF1 | Q8WYP5-2 | 1195 | SSQLKGSPQAISR | 6.4182 | 7.0257 | 6.0537 | 6.9855 | 5.9327 | 5.9722 | 6.3917 | 6.2479 | 6.1527 | 6.2727 | 6.2059 | 6.4323 | 5.4830 | 6.3585 | 5.9059 | 6.1616 |
| AHCTF1_S1190 | AHCTF1 | Q8WYP5-2 | 1933 | KVSTVTSPSRMIR | 7.5230 | 7.8969 | 6.8138 | 7.7569 | 6.6205 | 6.8134 | 7.2212 | 6.8443 | 7.4576 | 7.1725 | 7.2924 | 7.3273 | 2.8380 | 3.8228 | 3.0412 | 3.5581 |
| AHCTF1_S1195 | AHCTF1 | Q8WYP5-2 | 563 | LVAGLLSPRFVDV | 6.4799 | 6.7242 | 6.0895 | 6.9937 | 6.1622 | 6.3635 | 6.7171 | 6.0829 | 6.0604 | 6.3400 | 6.3187 | 6.5458 | 5.4468 | 5.9882 | 5.6379 | 6.0494 |
| AHCTF1_S1203 | AHCTF1 | Q8WYP5-2 | 1257 | SPSPGRSPQRLKE | 6.7437 | 6.5347 | 6.0988 | 7.1518 | 6.6346 | 6.0490 | 6.8602 | 6.2556 | 6.2741 | 5.8962 | 5.9221 | 6.1561 | 5.8035 | 6.1024 | 5.9220 | 5.5951 |
| AHCTF1_S1249 | AHCTF1 | Q8WYP5-2 | 1285 | PGAADDSKLEVFT | 8.3098 | 7.0915 | 6.6870 | 8.1256 | 6.8374 | 5.1649 | 6.6190 | 4.9332 | 6.1406 | 4.9938 | 5.5086 | 5.4550 | 5.6244 | 6.4533 | 5.7774 | 6.2785 |
| AHCTF1_S1257 | AHCTF1 | Q8WYP5-2 | 1576 | EEARNLSFNELYP | 6.1952 | 6.6294 | 6.7025 | 7.1029 | 6.6097 | 6.6479 | 6.9389 | 6.7820 | 6.9410 | 6.6846 | 6.6981 | 6.8314 | 4.2235 | 5.4089 | 4.4723 | 5.1316 |
| AHCTF1_S1267 | AHCTF1 | Q8WYP5-2 | 2155 | NNEPLFSPASEVP | 7.0813 | 7.5814 | 7.0531 | 7.6872 | 5.8621 | 6.8496 | 7.1203 | 6.7580 | 7.8456 | 7.5826 | 6.7449 | 6.5480 | 3.4599 | 4.3899 | 3.2043 | 4.2319 |
| AHCTF1_S1285 | AHCTF1 | Q8WYP5-2 | 1177 | QQSSMKSPLYLVS | 6.7765 | 6.9916 | 6.2967 | 6.9354 | 6.2731 | 6.0418 | 6.7011 | 6.4903 | 6.4436 | 6.4340 | 6.3743 | 6.5630 | 5.2620 | 5.6459 | 5.2976 | 5.4732 |
| AHCTF1_S1313;S1318 | AHCTF1 | Q8WYP5-2 | 1116 | TTPFNSSKIEEPS | 6.3751 | 6.3477 | 6.5054 | 7.4732 | 6.3077 | 6.2736 | 6.1944 | 6.0081 | 6.4011 | 6.2707 | 6.6406 | 6.9559 | 5.7885 | 5.4808 | 5.2472 | 5.7300 |
| AHCTF1_S1318 | AHCTF1 | Q8WYP5-2 | 2257 | IEIRLISPLASPA | 7.3321 | 7.6316 | 6.3430 | 7.9451 | 6.9726 | 6.5473 | 6.3014 | 6.7655 | 7.1960 | 7.2534 | 7.0614 | 7.0943 | 3.5333 | 4.1174 | 3.8461 | 4.0596 |
| AHCTF1_S1332 | AHCTF1 | Q8WYP5-2 | 1203 | QAISRASELHLLE | 6.1808 | 5.8070 | 5.7452 | 6.7937 | 6.1374 | 5.5946 | 6.4229 | 5.5006 | 6.2410 | 5.9093 | 5.4502 | 5.2197 | 7.4718 | 7.6125 | 6.8482 | 7.0651 |
| AHCTF1_S1499 | AHCTF1 | Q8WYP5-2 | 2216 | LKDDAQSVETLGK | 5.9453 | 6.2581 | 5.8106 | 6.6070 | 6.2936 | 5.5303 | 5.8755 | 5.9615 | 6.6512 | 6.1831 | 5.8555 | 6.0767 | 6.3238 | 7.1128 | 7.1966 | 6.3188 |
| AHCTF1_S1548 | AHCTF1 | Q8WYP5-2 | 2247 | EKESAWSPPPIEI | 6.8910 | 7.4685 | 6.9299 | 8.0055 | 6.4831 | 6.6828 | 7.6854 | 7.1683 | 7.0495 | 7.4609 | 7.6598 | 7.6771 | 2.8388 | 3.6493 | 2.9739 | 3.3762 |
| AHCTF1_S1576 | AHCTF1 | Q8WYP5-2 | 2048 | RRSTRNTPAKSEN | 8.5845 | 8.1040 | 7.0283 | 7.4910 | 6.6638 | 7.1901 | 7.4176 | 6.0584 | 6.7279 | 6.5402 | 7.3210 | 6.9627 | 3.2976 | 4.0625 | 2.7338 | 3.8164 |
| AHCTF1_S1933 | AHCTF1 | Q8WYP5-2 | 1979 | VRGREVSPSDVRE | 6.6792 | 6.6560 | 6.1527 | 7.2577 | 6.0363 | 5.5604 | 6.1151 | 6.3148 | 6.3979 | 6.0260 | 5.7751 | 5.9316 | 5.9255 | 6.4313 | 6.5191 | 6.2212 |
| AHCTF1_S1979 | AHCTF1 | Q8WYP5-2 | 2124 | RLLATASFTKSSR | 7.4024 | 7.5911 | 6.9335 | 8.0938 | 6.7169 | 6.9017 | 7.8899 | 6.9865 | 7.1641 | 7.0634 | 7.1266 | 7.0547 | 2.9609 | 4.1690 | 2.5893 | 3.3561 |
| AHCTF1_S2124 | AHCTF1 | Q8WYP5-2 | 1140 | PEAFFGTPISKAS | 6.8864 | 6.0084 | 6.8955 | 7.0092 | 5.5972 | 6.3530 | 6.1799 | 6.1001 | 6.6432 | 6.1980 | 6.4718 | 6.7744 | 5.2691 | 6.1357 | 5.4897 | 5.9886 |
| AHCTF1_S2155 | AHCTF1 | Q8WYP5-2 | 1249 | RSTPLASPSPSPG | 6.2360 | 6.6298 | 5.9722 | 7.0949 | 6.3472 | 6.0470 | 6.9859 | 6.5345 | 6.2723 | 5.8178 | 6.2553 | 6.0970 | 5.7989 | 5.7609 | 6.1685 | 5.9819 |
| AHCTF1_S2189 | AHCTF1 | Q8WYP5-2 | 1190 | RSLPSSSQLKGSP | 7.7385 | 8.7127 | 7.4294 | 8.3856 | 5.9091 | 7.1629 | 7.8778 | 6.7190 | 7.2758 | 6.8057 | 7.4397 | 7.9797 | 1.9147 | 3.5887 | 1.9680 | 3.0928 |
| AHCTF1_S2216 | AHCTF1 | Q8WYP5-2 | 2189 | SSQFVISPPALRS | 6.6342 | 7.0408 | 7.0327 | 7.0367 | 5.8978 | 6.3580 | 7.2208 | 6.1628 | 7.5020 | 7.5526 | 6.8314 | 6.4127 | 3.9811 | 5.3961 | 3.8729 | 5.0674 |
| AHCTF1_S2247 | AHCTF1 | Q8WYP5-2 | 1030 | RSLARNSILDQYG | 5.2636 | 4.9078 | 4.7755 | 4.6265 | 7.7976 | 5.4008 | 4.6251 | 5.0213 | 7.6757 | 7.5434 | 7.6785 | 9.5795 | 6.8136 | 8.0260 | 4.6522 | 5.6132 |
| AHCTF1_S2257 | AHCTF1 | Q8WYP5-2 | 1185 | LYLVSRSLPSSSQ | 7.1612 | 7.5735 | 6.9514 | 7.3570 | 5.6193 | 7.0748 | 6.5842 | 6.5968 | 6.9366 | 7.4109 | 8.0840 | 6.4638 | 3.1902 | 5.0771 | 3.7250 | 4.1942 |
| AHCTF1_S2257;S2261 | AHCTF1 | Q8WYP5-2 | 1332 | MDEGSQSLEKLDV | 6.3063 | 6.2947 | 6.2304 | 7.0473 | 5.9159 | 6.2272 | 6.5269 | 6.4996 | 6.8792 | 6.8057 | 6.4425 | 7.2414 | 4.5592 | 5.8263 | 5.6027 | 5.5947 |
| AHCTF1_S563 | AHCTF1 | Q8WYP5-2 | 1843 | IPQNSVTPRRGRR | 6.2301 | 6.6934 | 6.7260 | 6.5595 | 6.2079 | 6.3057 | 6.8598 | 6.6258 | 7.0309 | 6.2580 | 6.5479 | 6.7027 | 4.6376 | 5.8785 | 5.2453 | 5.4909 |
| AHCTF1_T1140 | AHCTF1 | Q8WYP5-2 | 1404 | VFASEVTPSDLQK | 6.7428 | 6.8274 | 6.6099 | 7.5301 | 5.9198 | 6.5852 | 6.7466 | 6.5457 | 6.3188 | 6.7671 | 6.5917 | 6.5962 | 4.1315 | 5.3885 | 5.0793 | 5.6196 |
| AHCTF1_T1292 | AHCTF1 | Q8WYP5-2 | 1499 | GDGGNSSLTISEG | 5.9100 | 6.7009 | 6.4164 | 6.8393 | 6.2654 | 6.6771 | 6.3813 | 7.1625 | 6.1291 | 7.0908 | 6.8871 | 7.4919 | 4.8183 | 5.4259 | 5.0571 | 4.7467 |
| AHCTF1_T1404 | AHCTF1 | Q8WYP5-2 | 2122;2124 | NA | 8.0374 | 8.1814 | 8.3687 | 9.4680 | 6.5166 | 6.5473 | 7.9792 | 5.9826 | 6.2106 | 6.8731 | 7.4549 | 6.3474 | 2.6264 | 3.8415 | 2.4512 | 3.1139 |
| AHCTF1_T1843 | AHCTF1 | Q8WYP5-2 | 1185;1188 | NA | 6.5342 | 6.8455 | 7.6507 | 8.2253 | 5.0416 | 6.8095 | 6.0738 | 6.3719 | 6.5975 | 7.3502 | 7.3150 | 6.7722 | 2.8260 | 5.6358 | 4.7654 | 5.1854 |
| AHCTF1_T2048 | AHCTF1 | Q8WYP5-2 | 2257;2261 | NA | 7.3689 | 6.9674 | 7.4562 | 6.4866 | 6.6754 | 6.7765 | 6.8081 | 7.5665 | 7.4434 | 7.7410 | 7.9098 | 8.5819 | 2.2477 | 3.9450 | 2.6907 | 3.3350 |
| AHCTF1_T2122;S2124 | AHCTF1 | Q8WYP5-2 | 1313;1318 | NA | 7.5993 | 7.1875 | 7.1684 | 7.7145 | 6.0158 | 7.0408 | 7.0477 | 5.7872 | 7.5849 | 6.2097 | 6.7043 | 7.2597 | 3.4253 | 4.2662 | 3.5289 | 5.4597 |
| AHCYL2_S107 | AHCYL2 | Q96HN2 | 107 | GGEALVSPDGTVT | 7.5617 | 6.8046 | 6.5960 | 7.5957 | 5.4889 | 6.4564 | 7.5939 | 6.8010 | 5.9382 | 8.0754 | 6.9334 | 6.3401 | 3.9120 | 5.1990 | 4.3138 | 4.3899 |
| AHDC1_S1064 | AHDC1 | Q5TGY3 | 1476 | PGSAARSPPYEGK | 6.2654 | 6.8059 | 6.1542 | 6.7297 | 5.8398 | 6.0188 | 6.7495 | 6.7396 | 6.4731 | 6.5412 | 6.5520 | 6.5078 | 5.3697 | 5.8783 | 5.6131 | 5.7618 |
| AHDC1_S1399 | AHDC1 | Q5TGY3 | 46 | PTRPPASPPDKAF | 6.2880 | 5.5448 | 6.3664 | 7.2349 | 6.4478 | 5.9010 | 6.3509 | 6.0277 | 6.7723 | 6.7831 | 7.0154 | 6.7575 | 5.5369 | 5.9365 | 4.9962 | 6.0407 |
| AHDC1_S1399;S1403 | AHDC1 | Q5TGY3 | 1064 | SRASTVSPGGYMV | 6.0730 | 6.6779 | 5.7525 | 6.4718 | 6.1623 | 6.5264 | 6.1867 | 6.6937 | 6.4784 | 6.5430 | 6.3118 | 6.0885 | 5.9475 | 6.1685 | 5.9962 | 5.9219 |
| AHDC1_S1476 | AHDC1 | Q5TGY3 | 1549 | LSDLTLSPVPRDS | 5.6979 | 5.9311 | 5.5116 | 6.7115 | 5.8468 | 6.6272 | 6.4283 | 6.5353 | 6.7609 | 6.4359 | 5.7524 | 7.1425 | 6.7952 | 6.2445 | 5.5681 | 6.0108 |
| AHDC1_S1507 | AHDC1 | Q5TGY3 | 596 | RKQKLASPQPSYA | 6.6391 | 7.2832 | 6.2292 | 6.7993 | 6.1383 | 6.7123 | 7.2563 | 5.3628 | 7.7003 | 7.0046 | 6.3161 | 6.1618 | 5.1399 | 5.3805 | 4.5468 | 5.3295 |
| AHDC1_S1549 | AHDC1 | Q5TGY3 | 1399 | GLQKAYSPTCSPT | 5.8736 | 5.3927 | 4.6099 | 6.0843 | 6.1456 | 5.9028 | 5.8018 | 5.6330 | 5.6133 | 5.2501 | 5.4279 | 4.5972 | 8.8976 | 8.0257 | 8.0850 | 8.6596 |
| AHDC1_S46 | AHDC1 | Q5TGY3 | 1507 | SAPHLASPPATPK | 5.8845 | 6.8094 | 5.9600 | 6.7952 | 6.0525 | 6.1910 | 6.8185 | 6.5432 | 6.1555 | 6.5873 | 6.7833 | 6.5308 | 5.3296 | 6.1043 | 5.3598 | 6.0952 |
| AHDC1_S596 | AHDC1 | Q5TGY3 | 1399;1403 | NA | 6.2675 | 6.7557 | 6.4854 | 6.7475 | 6.7393 | 6.6182 | 6.2758 | 6.2230 | 7.6622 | 6.4993 | 6.9202 | 6.7344 | 4.9975 | 5.4355 | 4.7861 | 4.8525 |
| AHNAK_S1010 | AHNAK | Q09666 | 177 | KDIDISSPEFKIK | 6.8510 | 6.9303 | 6.6058 | 6.7243 | 6.4935 | 6.6269 | 6.8817 | 6.6033 | 6.8741 | 6.7036 | 6.8618 | 7.0116 | 4.4757 | 4.9981 | 4.4921 | 4.8662 |
| AHNAK_S1068 | AHNAK | Q09666 | 2708 | IKAPKISMPDIDL | 3.7086 | 3.5702 | 3.4202 | 3.3281 | 4.0187 | 3.8220 | 3.8841 | 3.7195 | 3.8777 | 3.4153 | 3.2533 | 3.3013 | 15.0103 | 14.2369 | 14.4612 | 12.9724 |
| AHNAK_S1286 | AHNAK | Q09666 | 2580 | IKAPKISMPDFDL | 1.4485 | 1.3795 | 1.3530 | 1.5306 | 1.9175 | 1.7741 | 1.8363 | 1.7957 | 1.5075 | 1.3667 | 1.0859 | 1.3157 | 21.5881 | 20.0596 | 20.9617 | 19.0797 |
| AHNAK_S1324 | AHNAK | Q09666 | 3412 | PFLSISSPKVSMP | 6.4048 | 6.4256 | 6.6767 | 6.7195 | 5.9228 | 6.5748 | 7.2106 | 6.7458 | 6.8579 | 6.5957 | 6.8113 | 6.3448 | 4.8255 | 5.6616 | 5.1179 | 5.1045 |
| AHNAK_S1445 | AHNAK | Q09666 | 570 | TGTCRISMSEVDL | 5.5255 | 5.1354 | 5.4860 | 5.3541 | 5.9419 | 5.3580 | 5.5829 | 5.3682 | 5.8751 | 4.9006 | 4.6874 | 4.7410 | 9.5342 | 9.1598 | 8.9294 | 8.4204 |
| AHNAK_S1452 | AHNAK | Q09666 | 93 | HRKGDRSPEPGQT | 5.9639 | 6.1750 | 6.0811 | 6.2392 | 6.0816 | 6.1309 | 6.0183 | 6.3350 | 6.1647 | 6.0948 | 6.1151 | 6.0504 | 6.5943 | 6.7094 | 6.7778 | 6.4683 |
| AHNAK_S1580 | AHNAK | Q09666 | 5110 | AEAPLPSPKLEGE | 5.9626 | 5.7166 | 5.7993 | 5.9160 | 6.8338 | 6.5201 | 7.0085 | 6.5005 | 6.1895 | 5.6752 | 5.7355 | 5.6960 | 6.7277 | 6.5629 | 6.6286 | 6.5271 |
| AHNAK_S176 | AHNAK | Q09666 | 5752 | SKASLGSLEGEAE | 5.9677 | 5.8473 | 6.0989 | 6.0064 | 6.1154 | 5.8955 | 5.7311 | 5.8668 | 6.1787 | 6.1088 | 5.8316 | 5.8808 | 7.3950 | 7.2347 | 7.1420 | 6.6995 |
| AHNAK_S177 | AHNAK | Q09666 | 1445 | FKMPEMSIKPQKI | 11.9336 | 7.3860 | 9.2535 | 7.5556 | 5.0842 | 4.4455 | 6.0343 | 4.5103 | 7.3045 | 6.1737 | 7.0248 | 6.0635 | 4.5844 | 5.6725 | 3.4669 | 3.5065 |
| AHNAK_S1782 | AHNAK | Q09666 | 5731 | KGGVTGSPEASIS | 6.4896 | 6.6412 | 6.5743 | 6.6817 | 6.3700 | 6.2663 | 6.1531 | 6.3691 | 6.4649 | 6.3703 | 6.6374 | 6.6142 | 5.6218 | 5.6843 | 5.4934 | 5.5683 |
| AHNAK_S18 | AHNAK | Q09666 | 5794 | EFSGPSTPTGTLE | 5.9726 | 6.1721 | 6.0493 | 6.3437 | 5.7853 | 6.1626 | 6.0246 | 6.1938 | 6.2962 | 6.3628 | 6.1838 | 6.0553 | 6.4515 | 6.8341 | 6.5623 | 6.5502 |
| AHNAK_S210 | AHNAK | Q09666 | 490 | KGTKVKTPEMIIQ | 6.6871 | 7.2163 | 6.8784 | 7.2919 | 5.9695 | 6.3651 | 6.6603 | 6.6953 | 7.0179 | 7.1458 | 6.7690 | 7.0316 | 4.2224 | 4.6644 | 4.3730 | 5.0122 |
| AHNAK_S210;S216 | AHNAK | Q09666 | 216 | SGSGAASPTGSAV | 6.2884 | 6.4022 | 6.0782 | 6.2609 | 6.1480 | 6.2248 | 6.1902 | 6.2631 | 6.2628 | 6.2634 | 6.1219 | 6.0556 | 6.1824 | 6.5016 | 6.4660 | 6.2904 |
| AHNAK_S2138 | AHNAK | Q09666 | 4960 | LDVHMDSPDINIE | 6.0604 | 6.7222 | 6.0730 | 6.9159 | 5.9368 | 6.6110 | 6.6246 | 6.6148 | 6.2224 | 6.5544 | 6.4030 | 6.5792 | 5.3851 | 6.1106 | 5.4773 | 5.7094 |
| AHNAK_S216 | AHNAK | Q09666 | 5448 | LKGPRISAPNVDF | 6.1445 | 5.9319 | 6.5210 | 6.4157 | 6.2353 | 5.8003 | 6.0075 | 6.1520 | 6.4862 | 5.4600 | 5.4183 | 5.9687 | 7.1404 | 6.6525 | 6.9855 | 6.6802 |
| AHNAK_S2185 | AHNAK | Q09666 | 1580 | IQTHKISMPDVGL | 5.5448 | 5.4430 | 5.5384 | 5.3399 | 5.3646 | 5.1782 | 5.5674 | 5.3987 | 5.7536 | 5.5351 | 5.3659 | 5.7040 | 9.0728 | 9.5600 | 7.6879 | 7.9458 |
| AHNAK_S2313 | AHNAK | Q09666 | 4516 | PDVHFKSPQISMS | 6.6624 | 6.4430 | 6.4503 | 7.0489 | 6.7370 | 7.2091 | 7.6796 | 7.2957 | 7.7961 | 6.9489 | 6.7692 | 6.5428 | 3.8123 | 4.8001 | 3.5227 | 4.2818 |
| AHNAK_S2387 | AHNAK | Q09666 | 819 | IKVPKISMPDVDL | 5.3354 | 4.8248 | 4.7073 | 4.8665 | 5.6087 | 4.8454 | 5.2000 | 5.3400 | 5.9124 | 4.8104 | 4.5894 | 4.4538 | 10.8901 | 10.4518 | 9.4757 | 8.6882 |
| AHNAK_S2397 | AHNAK | Q09666 | 4386 | IKAPKISMPDIDF | 4.4245 | 3.8646 | 3.9251 | 3.7856 | 4.8943 | 3.9930 | 4.1119 | 3.8937 | 4.3504 | 4.0316 | 3.7807 | 3.7629 | 13.9469 | 13.3690 | 12.5787 | 11.2872 |
| AHNAK_S256 | AHNAK | Q09666 | 1452 | IKPQKISIPDVGL | 3.4476 | 2.9605 | 3.0800 | 3.4110 | 4.6506 | 3.7004 | 3.6088 | 3.7516 | 4.6315 | 4.3821 | 4.1666 | 4.1042 | 14.1539 | 14.0895 | 14.0547 | 11.8070 |
| AHNAK_S2580 | AHNAK | Q09666 | 3672 | IKAPKISMPDFDL | 3.0197 | 2.9745 | 2.5560 | 2.5549 | 3.3902 | 3.3076 | 2.8380 | 3.0721 | 3.1661 | 2.2092 | 2.4031 | 2.5262 | 18.0095 | 16.7006 | 16.5488 | 14.7235 |
| AHNAK_S2708 | AHNAK | Q09666 | 5552 | PAFNMASPESDFG | 6.5262 | 7.0617 | 6.6866 | 7.4835 | 6.2716 | 6.7783 | 7.2990 | 6.7323 | 7.1525 | 7.3606 | 7.5071 | 6.8782 | 3.7024 | 4.7055 | 3.5500 | 4.3045 |
| AHNAK_S2775 | AHNAK | Q09666 | 4903 | AKLSGPSLKMPSL | 8.0015 | 7.2987 | 7.7620 | 7.4362 | 6.0494 | 6.2633 | 5.9079 | 6.1001 | 7.2874 | 6.6563 | 7.3451 | 7.0210 | 4.7546 | 5.0604 | 3.2824 | 3.7737 |
| AHNAK_S3054 | AHNAK | Q09666 | 1068 | FRAPKMSLPDVDL | 6.4850 | 5.0012 | 5.6811 | 5.5503 | 7.3398 | 5.4390 | 5.1109 | 6.1549 | 7.7232 | 7.1169 | 6.4793 | 6.2773 | 7.4120 | 7.3472 | 5.9684 | 4.9136 |
| AHNAK_S3112 | AHNAK | Q09666 | 511 | VDLSLGSPKLKGD | 6.7528 | 6.5456 | 6.4044 | 6.8435 | 6.8021 | 5.6622 | 7.1583 | 6.4483 | 6.9138 | 6.3717 | 6.5950 | 6.3086 | 5.0603 | 5.3570 | 4.7067 | 5.1697 |
| AHNAK_S3182 | AHNAK | Q09666 | 5400 | LEASEGSIKLPKM | 7.3716 | 6.5050 | 7.9360 | 7.6428 | 6.8171 | 5.8977 | 6.3090 | 6.3859 | 7.4415 | 7.3500 | 6.9583 | 6.5790 | 4.5300 | 5.1195 | 3.4750 | 3.6814 |
| AHNAK_S3294 | AHNAK | Q09666 | 4430 | PSLDIDTPDVNIE | 5.2374 | 5.9022 | 6.5761 | 6.5413 | 5.0722 | 6.0872 | 5.9711 | 6.3837 | 5.5602 | 5.9052 | 6.1573 | 6.2600 | 6.6811 | 7.3386 | 7.3241 | 7.0023 |
| AHNAK_S332 | AHNAK | Q09666 | 638 | KMPTFSTPGAKGE | 6.5734 | 6.8514 | 6.5370 | 6.7552 | 5.8330 | 7.2693 | 8.0481 | 6.9621 | 7.2961 | 6.6685 | 6.6294 | 7.1007 | 3.8490 | 5.0550 | 3.8197 | 4.7520 |
| AHNAK_S3326 | AHNAK | Q09666 | 559 | TGPRLGSPSGKTG | 6.5139 | 6.9416 | 6.9335 | 6.8942 | 5.9481 | 6.3047 | 6.7307 | 6.1242 | 6.5881 | 6.9407 | 7.0998 | 7.4526 | 4.6776 | 5.2089 | 4.5462 | 5.0950 |
| AHNAK_S3360 | AHNAK | Q09666 | 4130 | LKAPKISMPEVDL | 6.1923 | 5.4505 | 5.1482 | 5.5857 | 6.4288 | 5.4211 | 5.6452 | 5.2883 | 6.0002 | 5.5031 | 4.9797 | 4.8149 | 9.0410 | 8.7773 | 8.0112 | 7.7126 |
| AHNAK_S3362 | AHNAK | Q09666 | 2387 | FKAPKISMPDLDL | 4.3075 | 4.0199 | 3.9820 | 3.9681 | 4.3073 | 4.1614 | 4.7419 | 4.3231 | 4.5748 | 3.9913 | 3.7823 | 3.8440 | 12.5482 | 13.3451 | 12.0384 | 12.0647 |
| AHNAK_S3412 | AHNAK | Q09666 | 5830 | TFGGLGSKSKGHY | 9.3391 | 7.4961 | 8.6959 | 7.5695 | 5.7361 | 5.9656 | 5.9083 | 6.5718 | 7.4607 | 7.7929 | 7.5177 | 7.6187 | 2.8981 | 4.1070 | 2.7199 | 2.6024 |
| AHNAK_S3416 | AHNAK | Q09666 | 41 | QEVTQNSPAARTG | 6.5312 | 6.6475 | 6.4571 | 6.9332 | 6.1453 | 6.2450 | 6.5078 | 6.4496 | 7.3560 | 6.9587 | 7.1009 | 7.1421 | 4.7524 | 5.3670 | 4.6302 | 4.7761 |
| AHNAK_S3672 | AHNAK | Q09666 | 5782 | PRHRSNSFSDERE | 6.5975 | 6.6970 | 6.2332 | 6.1389 | 6.0455 | 5.8715 | 5.9058 | 6.0142 | 6.6317 | 6.5826 | 6.6611 | 6.6034 | 6.0880 | 6.2035 | 5.8538 | 5.8723 |
| AHNAK_S41 | AHNAK | Q09666 | 561 | PRLGSPSGKTGTC | 5.9307 | 5.9714 | 7.3317 | 5.5040 | 6.3164 | 7.2406 | 5.0697 | 6.7740 | 6.5403 | 5.8265 | 7.0987 | 6.6945 | 7.3967 | 6.0592 | 4.8926 | 5.3533 |
| AHNAK_S4130 | AHNAK | Q09666 | 5031 | MPKIHMSGPKIKA | 8.6139 | 6.6299 | 7.4063 | 6.0481 | 4.9223 | 6.2188 | 5.7359 | 5.9132 | 8.6470 | 8.0558 | 7.9844 | 9.1355 | 3.8775 | 5.0620 | 2.6615 | 3.0879 |
| AHNAK_S4220 | AHNAK | Q09666 | 698 | VKTPKISMPDVDL | 5.5036 | 4.9688 | 4.7964 | 5.1522 | 6.1134 | 5.2292 | 5.6608 | 5.3174 | 5.3327 | 5.5410 | 4.7494 | 5.2982 | 9.3136 | 9.6137 | 9.1562 | 8.2534 |
| AHNAK_S4386 | AHNAK | Q09666 | 5298 | LSMPKVSGPDLDL | 2.0371 | 1.7626 | 2.4071 | 2.1161 | 2.4623 | 2.4937 | 2.7574 | 2.4046 | 1.9772 | 2.0375 | 1.8423 | 2.0218 | 18.3021 | 17.7615 | 19.1171 | 18.5000 |
| AHNAK_S440 | AHNAK | Q09666 | 1782 | IKAPKVSMPDVDL | 6.2079 | 5.2973 | 5.4380 | 5.4199 | 7.0439 | 5.2256 | 5.9390 | 5.6071 | 7.0624 | 6.0742 | 5.3411 | 5.0707 | 8.4026 | 7.8133 | 7.4645 | 6.5922 |
| AHNAK_S4516 | AHNAK | Q09666 | 5620 | AGGLHFSGPKVEG | 6.6843 | 6.5529 | 6.7588 | 7.2848 | 6.0122 | 5.5052 | 5.9612 | 6.0048 | 5.9593 | 5.2358 | 5.4785 | 5.8977 | 6.8513 | 6.4565 | 6.8997 | 6.4570 |
| AHNAK_S4522 | AHNAK | Q09666 | 886 | MKMPKFSMPGFKA | 7.7496 | 5.6724 | 5.3377 | 5.3883 | 6.3873 | 5.4260 | 5.9228 | 5.1278 | 8.2165 | 6.1479 | 6.7148 | 6.2483 | 7.4444 | 7.3951 | 5.3431 | 5.4780 |
| AHNAK_S4850 | AHNAK | Q09666 | 2185 | FKTPKISMPDVNL | 5.6815 | 5.1349 | 5.5387 | 5.2615 | 5.5415 | 5.0534 | 5.3256 | 4.8878 | 6.0781 | 4.8833 | 5.1589 | 5.0267 | 9.3360 | 10.4287 | 8.6187 | 8.0448 |
| AHNAK_S4903 | AHNAK | Q09666 | 5824 | GKLKFGTFGGLGS | 10.8488 | 7.5547 | 8.2742 | 6.5159 | 6.4142 | 4.9553 | 6.6528 | 4.1350 | 6.8346 | 5.4986 | 5.9450 | 5.7724 | 6.0241 | 6.3424 | 4.0546 | 4.1773 |
| AHNAK_S4908 | AHNAK | Q09666 | 2845 | DIDLNLTGPKIKG | 6.3861 | 6.4310 | 6.4800 | 7.0737 | 6.5192 | 6.7012 | 7.2447 | 6.4196 | 7.2216 | 7.1285 | 6.7035 | 7.0116 | 4.4147 | 5.1773 | 4.1706 | 4.9168 |
| AHNAK_S4953 | AHNAK | Q09666 | 3362 | KFNFSGSKVQTPE | 8.4051 | 7.1663 | 7.8522 | 6.8219 | 4.2074 | 5.9568 | 6.1747 | 5.7854 | 7.5004 | 7.7437 | 8.6062 | 9.0742 | 3.2434 | 5.3231 | 2.9152 | 3.2241 |
| AHNAK_S4960 | AHNAK | Q09666 | 5739 | EASISGSKGDLKS | 5.7687 | 5.7670 | 5.5347 | 5.2543 | 5.7952 | 5.9176 | 5.5213 | 5.4078 | 6.0034 | 5.8967 | 5.0881 | 5.0664 | 9.1826 | 8.6287 | 7.6347 | 7.5328 |
| AHNAK_S4993 | AHNAK | Q09666 | 5321 | LDASVPSMKVHAP | 8.7012 | 7.2050 | 8.2911 | 7.5802 | 5.2963 | 5.3706 | 6.4415 | 5.1786 | 7.2405 | 6.4433 | 7.2684 | 6.1643 | 4.9479 | 5.8018 | 4.0244 | 4.0449 |
| AHNAK_S5031 | AHNAK | Q09666 | 4522 | SPQISMSDIDLNL | 4.0859 | 4.0648 | 3.9913 | 3.9781 | 4.4290 | 4.5568 | 4.6380 | 4.3254 | 3.9392 | 3.7453 | 3.5141 | 3.5667 | 13.3593 | 12.1754 | 13.5032 | 12.1274 |
| AHNAK_S5054 | AHNAK | Q09666 | 1010 | PKFSMPSLKGEGP | 7.6610 | 7.1599 | 8.1385 | 6.6007 | 5.5148 | 5.8071 | 6.3789 | 6.1169 | 7.6600 | 8.0493 | 7.3768 | 7.5060 | 3.8614 | 5.0009 | 3.4409 | 3.7270 |
| AHNAK_S511 | AHNAK | Q09666 | 5763 | AEAEASSPKGKFS | 6.4929 | 6.4987 | 6.8510 | 7.2159 | 6.3059 | 6.1322 | 6.9030 | 6.6906 | 6.7942 | 6.5903 | 6.6796 | 6.9163 | 4.9607 | 5.3954 | 4.6112 | 4.9622 |
| AHNAK_S5110 | AHNAK | Q09666 | 4993 | PKADIKSPSLDVT | 6.1145 | 6.9632 | 6.7562 | 7.5797 | 6.9230 | 7.7988 | 7.0019 | 6.9253 | 7.1817 | 7.1204 | 7.8853 | 7.4578 | 3.5567 | 4.1593 | 3.2322 | 3.3441 |
| AHNAK_S5125 | AHNAK | Q09666 | 4766 | PKVDINTPDVDVH | 3.9736 | 5.8480 | 6.9338 | 6.3905 | 4.5678 | 6.5898 | 6.1383 | 6.7336 | 4.6802 | 5.9751 | 7.3354 | 6.6262 | 6.1451 | 6.6649 | 7.4391 | 7.9587 |
| AHNAK_S5186 | AHNAK | Q09666 | 2313 | FKTPKISMPDVDF | 6.5175 | 5.4046 | 5.5583 | 6.1182 | 6.3899 | 5.1136 | 6.1532 | 5.5372 | 6.8774 | 5.6418 | 5.3040 | 5.7102 | 7.7762 | 8.2161 | 6.9151 | 6.7664 |
| AHNAK_S5195 | AHNAK | Q09666 | 3182 | KADLDVSGPKVDV | 6.1869 | 6.3639 | 6.0018 | 6.5850 | 7.1319 | 7.6514 | 7.2953 | 6.7281 | 6.5932 | 6.3471 | 7.8305 | 7.4569 | 4.6378 | 5.1142 | 3.8886 | 4.1873 |
| AHNAK_S5279 | AHNAK | Q09666 | 1286 | KADIDVSGPKVDV | 6.5675 | 7.1509 | 6.4063 | 7.0609 | 6.2246 | 6.8123 | 7.3514 | 6.6799 | 7.0506 | 6.8617 | 7.1137 | 7.0125 | 3.9892 | 4.7397 | 3.9526 | 5.0261 |
| AHNAK_S5293 | AHNAK | Q09666 | 4564 | PKVGIDTPDIDIH | 5.1892 | 6.4713 | 7.7041 | 6.6536 | 4.6640 | 6.7564 | 6.9620 | 7.0581 | 5.5054 | 6.7044 | 6.8215 | 6.7020 | 5.0972 | 5.8908 | 5.7381 | 6.0819 |
| AHNAK_S5298 | AHNAK | Q09666 | 3366 | SGSKVQTPEVDVK | 7.8397 | 7.9396 | 6.8411 | 7.0052 | 4.8912 | 7.1192 | 7.9819 | 7.0843 | 7.7752 | 7.4187 | 7.8334 | 7.9999 | 2.7842 | 3.7977 | 2.2859 | 3.4029 |
| AHNAK_S5321 | AHNAK | Q09666 | 5589 | LGEGHLSVKGSGG | 7.3497 | 6.8854 | 7.7161 | 7.4099 | 6.1791 | 6.0618 | 6.4369 | 6.3029 | 6.9290 | 6.9986 | 7.3008 | 6.4005 | 4.3816 | 5.4405 | 4.1256 | 4.0815 |
| AHNAK_S5332 | AHNAK | Q09666 | 5857 | SGVSLASKKSRLS | 9.1375 | 7.4829 | 8.2123 | 7.3554 | 5.4105 | 5.5482 | 5.9489 | 5.5837 | 7.4564 | 6.8967 | 6.8992 | 7.4734 | 4.1380 | 5.4073 | 3.5322 | 3.5175 |
| AHNAK_S539 | AHNAK | Q09666 | 158 | TVTRRVTAYTVDV | 10.0351 | 6.7505 | 6.7844 | 6.4981 | 7.1152 | 5.4010 | 6.2659 | 5.3363 | 7.2375 | 5.4144 | 5.8078 | 5.9656 | 6.4467 | 6.0247 | 4.1138 | 4.8031 |
| AHNAK_S5393 | AHNAK | Q09666 | 5293 | LPSVNLSMPKVSG | 5.3614 | 5.0747 | 5.0899 | 5.4210 | 5.7725 | 5.3388 | 5.3061 | 5.0902 | 5.2387 | 5.1620 | 5.0860 | 5.0131 | 9.7891 | 9.4748 | 9.2531 | 8.5287 |
| AHNAK_S5400 | AHNAK | Q09666 | 5870 | SSSSNDSGNKVGI | 5.9455 | 5.7840 | 5.4361 | 6.1754 | 6.3777 | 5.6947 | 6.0078 | 6.1528 | 6.5361 | 6.3518 | 5.9502 | 6.3334 | 6.8896 | 7.1452 | 6.6160 | 6.6038 |
| AHNAK_S5448 | AHNAK | Q09666 | 5841 | HYEVTGSDDETGK | 6.6081 | 6.8295 | 6.4233 | 6.4222 | 6.6304 | 6.4658 | 6.3684 | 6.5958 | 6.4054 | 5.9445 | 6.1021 | 5.9635 | 5.7230 | 6.2603 | 5.8248 | 5.4328 |
| AHNAK_S5464 | AHNAK | Q09666 | 176 | AKDIDISSPEFKI | 6.8554 | 6.7191 | 6.2868 | 6.5968 | 6.3483 | 7.0652 | 6.8288 | 6.7984 | 7.4456 | 6.1765 | 7.2878 | 7.1743 | 4.2683 | 4.5663 | 4.9809 | 4.6018 |
| AHNAK_S5500 | AHNAK | Q09666 | 3054 | KADLDVSGPKVDI | 7.0942 | 6.1772 | 6.8684 | 6.9462 | 6.6807 | 7.1543 | 7.8425 | 6.2287 | 6.5048 | 7.0991 | 8.4765 | 6.9931 | 3.9508 | 4.4060 | 3.5802 | 3.9974 |
| AHNAK_S5519 | AHNAK | Q09666 | 4220 | KADLDVSGPKVDI | 6.3073 | 6.9894 | 6.6345 | 6.6080 | 6.8954 | 7.7774 | 7.0645 | 6.3759 | 6.3617 | 7.0697 | 7.6314 | 6.8423 | 4.2515 | 5.0167 | 4.0481 | 4.1264 |
| AHNAK_S5530 | AHNAK | Q09666 | 4850 | IKAPKISIPDVDL | 5.4851 | 5.4586 | 5.4670 | 5.0967 | 5.3169 | 5.1152 | 5.3881 | 5.4687 | 6.7379 | 5.6275 | 5.6634 | 6.3610 | 8.5011 | 8.0287 | 8.3671 | 7.9168 |
| AHNAK_S5552 | AHNAK | Q09666 | 5593 | HLSVKGSGGEWKG | 3.8029 | 4.0837 | 3.9197 | 4.0506 | 3.8815 | 3.9560 | 4.1552 | 3.9196 | 3.9026 | 3.9816 | 3.9890 | 3.9670 | 13.1406 | 13.3666 | 13.5101 | 12.3732 |
| AHNAK_S5589 | AHNAK | Q09666 | 5279 | LEGPDVSLKGPGV | 6.0962 | 5.5393 | 6.6950 | 6.0861 | 6.7850 | 5.4471 | 6.4553 | 5.9853 | 6.3026 | 6.1409 | 6.0972 | 5.7754 | 7.2623 | 6.7834 | 5.8797 | 6.6693 |
| AHNAK_S559 | AHNAK | Q09666 | 947 | IKAPKISMPDVDL | 5.2277 | 4.9554 | 4.5280 | 4.5401 | 5.6894 | 4.8995 | 4.8722 | 4.4608 | 5.0679 | 5.0009 | 4.5004 | 4.3803 | 11.7014 | 10.5614 | 10.4524 | 9.1623 |
| AHNAK_S5593 | AHNAK | Q09666 | 332 | KAGLRVSAPEVSV | 6.0909 | 6.2670 | 5.8608 | 6.3401 | 6.5395 | 6.7954 | 5.2944 | 6.3591 | 6.9046 | 4.7280 | 5.5773 | 5.3711 | 6.4235 | 7.6851 | 7.0353 | 6.7279 |
| AHNAK_S561 | AHNAK | Q09666 | 5851 | TGKLQGSGVSLAS | 4.0720 | 5.0389 | 4.9470 | 3.8980 | 4.7568 | 5.8290 | 4.9861 | 6.2565 | 5.7574 | 6.7851 | 5.5386 | 5.4178 | 8.9527 | 8.4820 | 10.7393 | 8.5427 |
| AHNAK_S5620 | AHNAK | Q09666 | 210 | TVIRLPSGSGAAS | 4.4275 | 4.6215 | 4.4442 | 4.9690 | 6.5670 | 5.2941 | 5.3148 | 6.3316 | 5.5819 | 4.7718 | 4.3751 | 5.4181 | 9.5786 | 8.9775 | 9.9681 | 9.3591 |
| AHNAK_S570 | AHNAK | Q09666 | 3360 | LPKFNFSGSKVQT | 7.5667 | 7.3637 | 8.0916 | 7.7732 | 5.2716 | 6.2172 | 5.8670 | 6.4952 | 6.9577 | 8.3182 | 8.0582 | 8.4232 | 3.1174 | 4.4386 | 3.0583 | 2.9821 |
| AHNAK_S5731 | AHNAK | Q09666 | 1469 | PKMKGDYDVTVPK | 6.2742 | 7.3443 | 6.8159 | 7.2996 | 6.7986 | 6.8410 | 6.3965 | 6.5109 | 6.3039 | 7.3161 | 7.4627 | 7.3921 | 4.0712 | 4.7148 | 3.8258 | 4.6325 |
| AHNAK_S5731;S5739 | AHNAK | Q09666 | 5125 | APDLELSLPAIHV | 5.7398 | 5.7567 | 5.6103 | 5.6902 | 5.6200 | 5.9116 | 6.2092 | 5.5898 | 6.8686 | 6.5395 | 6.1596 | 5.8753 | 7.2461 | 7.1457 | 6.7501 | 7.2873 |
| AHNAK_S5739 | AHNAK | Q09666 | 651 | GPDVHMTLPKGDI | 9.1781 | 7.5212 | 8.7906 | 7.1643 | 6.3974 | 5.1273 | 5.4973 | 4.2701 | 6.9240 | 6.0843 | 5.7114 | 5.0086 | 6.4885 | 6.5533 | 4.5359 | 4.7478 |
| AHNAK_S5749;S5752 | AHNAK | Q09666 | 5054 | GGEIDASLKAPDV | 9.4579 | 6.6784 | 7.9260 | 7.2912 | 5.2999 | 4.9205 | 6.2727 | 5.7531 | 7.3350 | 6.0717 | 7.3926 | 6.3240 | 5.4476 | 6.0561 | 3.5126 | 4.2607 |
| AHNAK_S5749;S5752;S5 | AHNAK | Q09666 | 3416 | ISSPKVSMPDVEL | 6.5956 | 4.5171 | 4.7667 | 5.2419 | 7.4645 | 5.4483 | 4.7667 | 6.5860 | 6.2212 | 5.5540 | 6.0388 | 6.0100 | 6.1684 | 10.3111 | 7.2629 | 7.0469 |
| AHNAK_S5752 | AHNAK | Q09666 | 3326 | GDIKAPSLDIKGP | 6.2738 | 6.0802 | 6.1010 | 6.0269 | 5.7935 | 5.9225 | 6.0097 | 6.1016 | 6.1117 | 6.3865 | 5.7661 | 5.8987 | 7.2190 | 7.1256 | 6.5673 | 6.6159 |
| AHNAK_S5752;S5763 | AHNAK | Q09666 | 2138 | KGDVDVSLPKLEG | 5.1918 | 4.4203 | 4.9913 | 5.4292 | 6.0796 | 5.3916 | 5.8800 | 5.3295 | 5.2575 | 5.0898 | 5.0262 | 5.0553 | 9.6559 | 9.5048 | 8.9559 | 8.7412 |
| AHNAK_S5763 | AHNAK | Q09666 | 3112 | KGDMDVSLPKVEG | 5.7110 | 5.8268 | 5.5705 | 6.0232 | 6.1638 | 5.8778 | 6.2689 | 6.3274 | 6.0848 | 6.3124 | 6.1462 | 6.1364 | 6.8302 | 7.0733 | 6.9254 | 6.7218 |
| AHNAK_S5780 | AHNAK | Q09666 | 4100 | PKVDIDTPDIDIH | 4.7606 | 5.8422 | 6.7270 | 5.7906 | 5.2140 | 6.3072 | 5.9538 | 6.5302 | 5.6248 | 5.7450 | 5.7730 | 5.5396 | 7.0737 | 7.3717 | 8.6393 | 7.1075 |
| AHNAK_S5782 | AHNAK | Q09666 | 247 | HSKLQVTMPGIKV | 8.8345 | 6.9447 | 8.4745 | 7.2460 | 4.6715 | 7.8524 | 6.1074 | 5.8883 | 5.2310 | 7.4494 | 8.2280 | 5.2662 | 4.0768 | 5.0354 | 3.3178 | 5.3758 |
| AHNAK_S5807 | AHNAK | Q09666 | 5332 | APGLNLSGVGGKM | 5.2002 | 6.0489 | 5.3218 | 5.8877 | 6.5516 | 6.0571 | 7.2835 | 6.3990 | 6.7385 | 6.3372 | 5.5768 | 5.5788 | 6.7199 | 7.0907 | 6.4736 | 6.7347 |
| AHNAK_S5830 | AHNAK | Q09666 | 2775 | IKMPKISMPGFKG | 7.5687 | 6.1958 | 5.2808 | 6.4922 | 5.9560 | 5.7670 | 6.1937 | 5.5856 | 7.2844 | 6.2796 | 7.0613 | 6.8145 | 6.2654 | 6.0198 | 5.4586 | 5.7767 |
| AHNAK_S5841 | AHNAK | Q09666 | 256 | GIKVGGSGVNVNA | 5.1607 | 4.7913 | 5.1677 | 4.0080 | 5.0280 | 5.3331 | 4.4893 | 4.9300 | 5.6638 | 5.0894 | 5.2301 | 5.4984 | 9.2618 | 10.1392 | 10.0957 | 10.1135 |

| | | | | | | | | | | | | | | | | | | | | |
|---|---|---|---|---|---|---|---|---|---|---|---|---|---|---|---|---|---|---|---|---|
| AHNAK_S5851 | AHNAK | Q09666 | 3294 | VNMPKISMPEIDL | 4.8628 | 5.9256 | 4.5521 | 4.2546 | 6.1676 | 5.6640 | 3.9897 | 5.7262 | 7.5117 | 5.5496 | 5.4580 | 6.5339 | 9.6177 | 10.6544 | 7.0506 | 6.4816 |
| AHNAK_S5851;S5857 | AHNAK | Q09666 | 1324 | FKAPKISMPDVDL | 6.3533 | 5.5152 | 5.0749 | 5.6620 | 6.7519 | 5.9682 | 6.1842 | 5.4891 | 6.9821 | 5.4802 | 5.8452 | 6.2699 | 7.8157 | 7.4045 | 7.2116 | 5.9920 |
| AHNAK_S5854;S5857 | AHNAK | Q09666 | 5530 | KGGLKGSEVGFHG | 3.4498 | 3.7745 | 3.9555 | 4.1376 | 3.3696 | 3.3908 | 3.6487 | 3.4767 | 4.0578 | 4.2298 | 4.0750 | 3.9003 | 13.6442 | 14.4841 | 13.9008 | 12.5048 |
| AHNAK_S5857 | AHNAK | Q09666 | 2397 | LDLHLKSPKAKGE | 6.8788 | 7.7128 | 8.2246 | 7.7193 | 6.0808 | 5.5695 | 5.3952 | 7.8022 | 6.5930 | 6.8764 | 5.9240 | 7.8976 | 4.2027 | 4.2892 | 4.2730 | 4.5608 |
| AHNAK_S5863 | AHNAK | Q09666 | 3716 | PKVDIDTPDINIE | 4.8508 | 5.5264 | 6.8614 | 5.8636 | 5.1122 | 5.7711 | 6.1035 | 7.0126 | 6.3332 | 5.9610 | 5.8464 | 6.0664 | 6.6758 | 7.1228 | 7.7274 | 7.1653 |
| AHNAK_S5864 | AHNAK | Q09666 | 4908 | PSLKMPSLEISAP | 5.9743 | 6.4002 | 6.0130 | 6.5272 | 5.9758 | 6.0735 | 5.9761 | 5.7714 | 6.1155 | 5.7569 | 5.4731 | 6.4909 | 7.2009 | 6.7130 | 7.1480 | 6.3904 |
| AHNAK_S5864;S5870 | AHNAK | Q09666 | 5464 | GPKVKGSLGATGE | 5.5323 | 5.8831 | 6.6398 | 5.3467 | 6.3492 | 5.7720 | 6.6102 | 7.0222 | 7.0110 | 6.2390 | 5.8332 | 5.1795 | 6.8988 | 6.2513 | 7.1303 | 6.3012 |
| AHNAK_S5870 | AHNAK | Q09666 | 539 | QVALKGSRVDIET | 4.7657 | 4.4982 | 4.9595 | 4.9299 | 5.1175 | 4.5318 | 4.9895 | 5.4565 | 5.0566 | 4.6355 | 4.8473 | 4.3446 | 10.6079 | 10.6556 | 10.9350 | 9.6691 |
| AHNAK_S698 | AHNAK | Q09666 | 4953 | PKVEAPSLDVHMD | 6.0425 | 6.8787 | 6.4088 | 5.2503 | 6.4425 | 6.2820 | 5.5131 | 6.4671 | 7.0470 | 5.8742 | 5.8289 | 6.6185 | 6.0166 | 6.0930 | 6.3855 | 6.8515 |
| AHNAK_S819 | AHNAK | Q09666 | 5393 | VGAPDLSLEASEG | 6.3835 | 6.0875 | 5.6038 | 6.2932 | 6.4113 | 6.2793 | 7.6246 | 5.8609 | 7.1215 | 5.5593 | 6.8616 | 5.1132 | 6.2862 | 5.2925 | 6.7185 | 6.5030 |
| AHNAK_S886 | AHNAK | Q09666 | 5519 | LPTGQISGPEIKG | 2.7502 | 3.0679 | 3.0434 | 3.2625 | 3.8320 | 3.4396 | 3.8408 | 4.2043 | 2.5611 | 2.5300 | 2.5327 | 2.7998 | 15.0987 | 15.7320 | 15.5429 | 15.7620 |
| AHNAK_S93 | AHNAK | Q09666 | 5807 | FEGGEVSLEGGKV | 6.3334 | 6.0965 | 6.3848 | 5.7364 | 5.7604 | 6.1784 | 6.4749 | 6.2549 | 6.8488 | 5.4674 | 5.7320 | 5.6718 | 6.8681 | 6.9003 | 6.3584 | 6.9334 |
| AHNAK_S947 | AHNAK | Q09666 | 5780 | KKPRHRSNSFSDE | 5.6278 | 5.3271 | 6.3832 | 6.0475 | 6.2631 | 6.2493 | 5.9518 | 6.8772 | 4.8841 | 6.0570 | 5.1295 | 6.5597 | 8.2956 | 6.9484 | 7.4967 | 6.8018 |
| AHNAK_T158 | AHNAK | Q09666 | 440 | MKVPKFSVSGAKG | 7.4204 | 6.0718 | 6.4370 | 5.6013 | 5.5887 | 5.4666 | 6.0996 | 5.3668 | 7.2146 | 6.4542 | 7.0184 | 6.5083 | 6.7845 | 6.9599 | 5.3756 | 5.6321 |
| AHNAK_T247 | AHNAK | Q09666 | 5195 | ISAPQVSIPDVNV | 5.6137 | 5.7694 | 6.2044 | 5.6682 | 5.6560 | 5.4244 | 5.9976 | 5.7650 | 6.1255 | 6.7308 | 5.9051 | 5.6536 | 7.9248 | 8.0196 | 6.9784 | 6.5634 |
| AHNAK_T2845 | AHNAK | Q09666 | 18 | LPNWQGSGSHGLT | 6.0879 | 6.0879 | 5.8169 | 7.1519 | 6.2459 | 7.7333 | 5.1593 | 7.6515 | 5.0972 | 7.1068 | 7.2422 | 5.5121 | 4.4932 | 6.8894 | 6.1302 | 5.5940 |
| AHNAK_T3366 | AHNAK | Q09666 | 5863 | SKKSRLSSSSSND | 7.1805 | 7.4524 | 7.1726 | 7.1391 | 6.3260 | 6.3749 | 6.4435 | 6.7008 | 6.8066 | 7.6823 | 7.4260 | 7.4413 | 3.8472 | 4.8071 | 3.6434 | 3.5564 |
| AHNAK_T3716 | AHNAK | Q09666 | 5184 | LDAKVKTPSFGIS | 6.0217 | 6.3594 | 6.0857 | 6.6110 | 5.4380 | 4.9600 | 6.0620 | 5.8369 | 5.8960 | 5.8161 | 4.9294 | 5.9780 | 6.8591 | 7.5143 | 7.4031 | 8.2293 |
| AHNAK_T4100 | AHNAK | Q09666 | 5186 | AKVKTPSFGISAP | 3.3589 | 2.5995 | 2.7391 | 3.7152 | 3.9870 | 4.1385 | 4.3699 | 3.5958 | 3.3571 | 2.9127 | 2.9090 | 3.6647 | 12.7700 | 13.9490 | 15.1087 | 16.8249 |
| AHNAK_T4430 | AHNAK | Q09666 | 5864 | KKSRLSSSSSNDS | 6.7430 | 6.3034 | 7.1193 | 6.7440 | 5.7635 | 7.1625 | 6.0977 | 6.0689 | 7.1604 | 6.8365 | 6.9342 | 6.6730 | 5.2273 | 5.7769 | 4.2910 | 5.0982 |
| AHNAK_T4564 | AHNAK | Q09666 | 5500 | MPGIKSSGCDVNL | 4.5857 | 4.0991 | 4.3705 | 4.4887 | 5.3026 | 4.7578 | 4.8775 | 4.4311 | 4.8095 | 4.4536 | 4.2868 | 4.0397 | 11.3287 | 11.0458 | 12.0255 | 11.0975 |
| AHNAK_T4766 | AHNAK | Q09666 | 210;216 | NA | 5.3999 | 5.8217 | 5.5061 | 5.8991 | 6.0690 | 5.7074 | 6.0093 | 5.7861 | 6.0473 | 5.6025 | 5.7339 | 5.5099 | 7.7184 | 8.0605 | 7.7670 | 7.3620 |
| AHNAK_T490 | AHNAK | Q09666 | 5752;5763 | NA | 6.4386 | 5.8132 | 6.4495 | 6.5456 | 6.4535 | 5.8641 | 6.3766 | 6.2599 | 6.7123 | 6.0045 | 5.8929 | 6.5664 | 6.3654 | 6.2078 | 6.1254 | 5.9244 |
| AHNAK_T5184 | AHNAK | Q09666 | 5731;5739 | NA | 5.6796 | 5.8524 | 5.6878 | 5.2243 | 6.4105 | 6.0016 | 5.8568 | 5.7095 | 6.1525 | 5.6903 | 5.8214 | 5.6253 | 8.1550 | 7.4570 | 7.5117 | 7.1643 |
| AHNAK_T5794 | AHNAK | Q09666 | 5864;5870 | NA | 5.8875 | 6.2187 | 6.0770 | 6.3282 | 6.0822 | 6.6920 | 6.0946 | 6.6031 | 6.6343 | 6.5375 | 6.8275 | 6.5144 | 5.8822 | 6.0919 | 5.5704 | 5.9583 |
| AHNAK_T5794;T5796 | AHNAK | Q09666 | 5851;5857 | NA | 3.8598 | 4.2298 | 4.4007 | 4.1218 | 5.8744 | 5.8997 | 4.8485 | 5.7881 | 5.1323 | 6.0323 | 5.0527 | 5.6320 | 10.4144 | 9.3301 | 10.6851 | 8.6983 |
| AHNAK_T5824 | AHNAK | Q09666 | 5749;5752 | NA | 5.4781 | 5.7697 | 6.0925 | 6.4483 | 4.9367 | 6.1392 | 5.5799 | 5.7720 | 5.3771 | 7.0626 | 6.5352 | 6.1968 | 6.4317 | 7.5031 | 7.2736 | 7.4036 |
| AHNAK_T638 | AHNAK | Q09666 | 5749;5752;57 | NA | 6.1427 | 5.6730 | 7.4613 | 6.3989 | 5.6988 | 6.7531 | 6.2941 | 4.8324 | 6.8945 | 7.2593 | 7.0328 | 7.4362 | 5.4466 | 5.1778 | 5.9603 | 5.5382 |
| AHNAK_T651 | AHNAK | Q09666 | 5854;5857 | NA | 4.9175 | 4.4415 | 4.8427 | 5.6005 | 4.6826 | 4.7644 | 4.9403 | 4.4796 | 4.7782 | 4.5022 | 4.8406 | 4.0747 | 11.4528 | 10.6152 | 10.3048 | 10.7622 |
| AHNAK_Y1469 | AHNAK | Q09666 | 5794;5796 | NA | 6.4309 | 6.7600 | 6.8086 | 6.0405 | 6.2023 | 7.1830 | 6.3251 | 5.7411 | 6.2028 | 6.4286 | 6.7213 | 6.9894 | 5.9031 | 6.2888 | 4.7553 | 5.2192 |
| AHNAK2_S1033 | AHNAK2 | Q8IVF2 | 4477 | PSFGMLSPGKSIE | 6.6017 | 6.5577 | 6.2351 | 6.1784 | 6.7135 | 7.4347 | 7.4373 | 7.0363 | 7.0703 | 6.3373 | 6.6711 | 6.6118 | 4.2264 | 5.1309 | 4.5073 | 5.2500 |
| AHNAK2_S1172 | AHNAK2 | Q8IVF2 | 593 | IPSLGWSPSKHTK | 6.5577 | 6.2799 | 6.2721 | 6.0160 | 6.6398 | 6.5220 | 6.3518 | 6.3970 | 6.6228 | 6.5245 | 6.5356 | 6.2537 | 5.6532 | 5.9086 | 5.7577 | 5.7076 |
| AHNAK2_S1198 | AHNAK2 | Q8IVF2 | 5175 | EVLTVESPEEEAM | 4.3057 | 4.4333 | 4.8660 | 4.5653 | 5.4937 | 5.7845 | 5.6818 | 5.8681 | 4.8061 | 4.7031 | 5.2030 | 4.8427 | 9.9474 | 9.1687 | 10.3027 | 10.0280 |
| AHNAK2_S1418 | AHNAK2 | Q8IVF2 | 4897 | PVMSPLSPGERVQ | 5.7482 | 5.9779 | 6.0993 | 6.3036 | 6.3411 | 7.0571 | 6.5689 | 6.5788 | 6.0696 | 6.2226 | 6.3631 | 6.4500 | 5.7451 | 6.1457 | 6.0013 | 6.3276 |
| AHNAK2_S1583 | AHNAK2 | Q8IVF2 | 842 | PKFKMPSFGVSAP | 7.3780 | 7.0363 | 7.9179 | 7.6032 | 7.4994 | 5.4861 | 5.5871 | 6.2209 | 6.4916 | 6.6838 | 5.7090 | 5.4753 | 4.7845 | 6.1105 | 5.2968 | 4.7197 |
| AHNAK2_S1710 | AHNAK2 | Q8IVF2 | 4419 | PKVEVTSPNLDVS | 4.9213 | 5.1358 | 5.3080 | 5.5234 | 6.2741 | 6.5627 | 6.5708 | 6.7776 | 5.3167 | 5.2915 | 5.4926 | 5.5780 | 7.3729 | 7.6361 | 8.0113 | 8.2273 |
| AHNAK2_S1913 | AHNAK2 | Q8IVF2 | 1418 | PKLQMPSFKVPKV | 7.2081 | 6.2632 | 7.7232 | 6.1728 | 4.8894 | 6.4344 | 7.1058 | 6.2940 | 7.4735 | 7.3049 | 8.4156 | 8.6252 | 3.5473 | 5.3086 | 3.1903 | 4.0437 |
| AHNAK2_S1997 | AHNAK2 | Q8IVF2 | 5707 | KLGFSSSPTKKSK | 6.5336 | 6.7694 | 6.3004 | 6.0278 | 6.8076 | 6.7362 | 6.4567 | 6.4710 | 6.7074 | 6.5220 | 6.5083 | 6.5163 | 5.2743 | 5.4704 | 5.2685 | 5.6302 |
| AHNAK2_S2023;S2026 | AHNAK2 | Q8IVF2 | 2179 | EASVDVSPPKVEA | 6.0845 | 6.0861 | 6.5145 | 6.5486 | 6.0598 | 6.2298 | 6.4184 | 6.1772 | 6.3831 | 6.5700 | 6.5271 | 6.8553 | 5.7434 | 5.8904 | 5.7728 | 6.1389 |
| AHNAK2_S2179 | AHNAK2 | Q8IVF2 | 1710 | TDLSIQSPSADLE | 4.8783 | 5.0879 | 5.5014 | 5.2374 | 5.9455 | 6.3597 | 6.3447 | 6.4706 | 5.3572 | 5.1409 | 5.2635 | 5.5615 | 8.0054 | 7.5003 | 8.5128 | 8.8329 |
| AHNAK2_S2327 | AHNAK2 | Q8IVF2 | 4185 | TDLSIQSPSADLE | 5.1874 | 5.2912 | 5.7050 | 5.5572 | 6.0131 | 6.0770 | 6.5114 | 6.3575 | 5.4223 | 5.7927 | 5.9698 | 5.8422 | 7.3576 | 7.1886 | 7.9536 | 7.7734 |
| AHNAK2_S2336 | AHNAK2 | Q8IVF2 | 2327 | PKFKMLSFGVSAL | 8.1974 | 7.6747 | 8.9975 | 8.9440 | 7.7371 | 5.7975 | 5.9048 | 5.5698 | 6.8973 | 6.1334 | 6.8325 | 5.9133 | 3.6507 | 5.0317 | 3.1251 | 3.5930 |
| AHNAK2_S260 | AHNAK2 | Q8IVF2 | 1198 | KVEADVSLPSMQG | 5.9108 | 6.3372 | 6.8739 | 6.4059 | 6.1963 | 7.2926 | 7.0236 | 7.0514 | 7.0510 | 7.2630 | 7.9286 | 6.9368 | 4.2545 | 4.9866 | 3.9630 | 4.5246 |
| AHNAK2_S280 | AHNAK2 | Q8IVF2 | 294 | RDAHDVSPTSTDT | 5.9279 | 6.3700 | 6.2321 | 6.4867 | 6.1082 | 6.5610 | 6.3009 | 6.5182 | 6.7754 | 6.6090 | 6.1795 | 6.6811 | 5.3572 | 6.1222 | 6.0522 | 5.7186 |
| AHNAK2_S294 | AHNAK2 | Q8IVF2 | 510 | RERRLSTPQRGKR | 7.0873 | 7.4681 | 6.7699 | 6.8685 | 6.2173 | 7.2079 | 7.3207 | 6.5121 | 7.2210 | 7.3965 | 7.3249 | 7.2246 | 3.4896 | 4.4142 | 3.4083 | 4.0692 |
| AHNAK2_S294;S298 | AHNAK2 | Q8IVF2 | 1583 | PKVQMPSFKMPKV | 7.0051 | 6.4896 | 7.6243 | 6.4332 | 5.1418 | 6.8325 | 7.3076 | 6.4503 | 7.5048 | 7.0983 | 8.2526 | 8.5650 | 3.4991 | 4.8832 | 2.9608 | 3.9517 |
| AHNAK2_S2996 | AHNAK2 | Q8IVF2 | 4713 | PKVSFSSTKTPKD | 8.4080 | 7.0133 | 7.9183 | 6.5462 | 5.0237 | 6.4462 | 6.3983 | 6.6026 | 7.5541 | 7.3160 | 8.1710 | 8.4379 | 3.4440 | 4.8244 | 2.7565 | 3.1395 |
| AHNAK2_S3161 | AHNAK2 | Q8IVF2 | 260 | SQRERLSWPKFQS | 5.6254 | 5.8615 | 6.1043 | 5.6612 | 5.9701 | 6.1762 | 5.9206 | 5.6661 | 5.9769 | 5.7619 | 6.3554 | 6.1046 | 7.3418 | 7.0094 | 7.5663 | 6.8982 |
| AHNAK2_S3408 | AHNAK2 | Q8IVF2 | 4962 | PKIKLPSFRWSPK | 6.1910 | 6.4592 | 7.3333 | 6.0070 | 6.0247 | 7.9368 | 6.3112 | 7.2555 | 6.4002 | 7.1407 | 8.6951 | 7.4689 | 4.2917 | 5.8677 | 3.3828 | 3.2342 |
| AHNAK2_S3491 | AHNAK2 | Q8IVF2 | 3408 | PKVDLKSPQVDIK | 4.8496 | 4.7098 | 5.0121 | 4.7446 | 6.1286 | 6.3107 | 6.4220 | 6.1356 | 5.1735 | 4.5018 | 4.8097 | 4.7955 | 9.1770 | 8.3413 | 9.8943 | 8.9939 |
| AHNAK2_S365 | AHNAK2 | Q8IVF2 | 5428 | TALCKESPGLWGA | 6.0925 | 7.1680 | 6.5615 | 7.1856 | 6.5266 | 6.8656 | 6.8706 | 6.7314 | 6.6932 | 6.8269 | 7.1242 | 6.6936 | 4.5237 | 5.2902 | 4.1086 | 4.7377 |
| AHNAK2_S375 | AHNAK2 | Q8IVF2 | 1997 | PKFKMPSFGVSAP | 7.7836 | 7.2525 | 7.8250 | 7.6476 | 7.2214 | 5.6086 | 5.3618 | 6.0063 | 6.6239 | 6.6838 | 5.6109 | 5.4955 | 4.9068 | 6.1629 | 5.4847 | 4.3249 |
| AHNAK2_S4185 | AHNAK2 | Q8IVF2 | 4425 | SPNLDVSLPSMEV | 6.4817 | 7.2334 | 7.0712 | 7.1967 | 6.0719 | 6.9261 | 6.9788 | 7.2233 | 7.6880 | 7.3975 | 8.1492 | 7.4564 | 3.1023 | 4.1748 | 3.1385 | 3.7103 |
| AHNAK2_S4419 | AHNAK2 | Q8IVF2 | 4710 | FKVPKVSFSSTKT | 9.5798 | 6.9584 | 7.9953 | 6.5104 | 7.0587 | 6.7241 | 6.5817 | 7.3785 | 7.6686 | 6.2982 | 6.1740 | 6.1795 | 4.4846 | 4.7156 | 2.8493 | 2.8433 |
| AHNAK2_S4425 | AHNAK2 | Q8IVF2 | 280 | GPQRSHSSSEAYE | 5.6414 | 6.3087 | 5.3855 | 6.0770 | 7.0563 | 7.0386 | 5.9178 | 7.6647 | 6.0881 | 6.2333 | 5.6701 | 5.6471 | 5.5434 | 6.6086 | 6.6327 | 6.4866 |
| AHNAK2_S4472 | AHNAK2 | Q8IVF2 | 4785 | FESSILSPCEDVT | 5.4565 | 5.3047 | 5.0568 | 5.6045 | 6.2594 | 6.9828 | 6.7972 | 6.6855 | 5.5733 | 5.2555 | 5.4174 | 5.1734 | 7.5465 | 7.3431 | 7.5291 | 8.0142 |
| AHNAK2_S4472;S4477 | AHNAK2 | Q8IVF2 | 497 | TPKFAFSTEKEPE | 8.9259 | 7.7985 | 9.5607 | 8.7434 | 7.1005 | 5.2879 | 5.6248 | 6.5884 | 7.3423 | 7.9996 | 5.8853 | 6.2099 | 2.5113 | 4.8919 | 3.0725 | 2.4571 |
| AHNAK2_S4477 | AHNAK2 | Q8IVF2 | 4931 | LVASLQTSVVAPG | 5.7432 | 6.0801 | 5.7030 | 6.1567 | 6.7313 | 7.2221 | 6.4006 | 6.8860 | 6.7735 | 5.6819 | 6.6550 | 6.1274 | 6.2087 | 5.4560 | 6.1794 | 5.9949 |
| AHNAK2_S4642 | AHNAK2 | Q8IVF2 | 1913 | PKVDMPSFKMPKV | 6.2395 | 6.8699 | 8.0936 | 6.7110 | 5.2537 | 6.1946 | 6.6354 | 6.0731 | 7.1924 | 7.3814 | 8.1370 | 8.5998 | 4.0905 | 5.3388 | 3.2410 | 3.9483 |
| AHNAK2_S4646 | AHNAK2 | Q8IVF2 | 2996 | VSAPGKSIEVSVD | 7.5166 | 6.8705 | 8.3493 | 7.2692 | 6.7258 | 5.9235 | 5.7113 | 6.8544 | 6.1125 | 7.7455 | 6.2840 | 5.7969 | 4.2246 | 5.6639 | 4.6964 | 4.2557 |
| AHNAK2_S4710 | AHNAK2 | Q8IVF2 | 693 | ASAPGKSMEASVD | 6.2490 | 6.3608 | 7.6994 | 7.1727 | 6.4728 | 5.9011 | 6.0858 | 5.8555 | 6.1214 | 7.4105 | 6.1346 | 5.6367 | 5.3621 | 6.4331 | 5.8112 | 5.2935 |
| AHNAK2_S4713 | AHNAK2 | Q8IVF2 | 1172 | PKFKMPSFGASAP | 6.7134 | 6.1451 | 6.9604 | 6.5636 | 6.2206 | 5.9682 | 5.9214 | 6.1833 | 6.3399 | 6.6288 | 5.9609 | 5.8465 | 5.6913 | 6.5997 | 6.4352 | 5.8217 |
| AHNAK2_S4782 | AHNAK2 | Q8IVF2 | 228 | ADGCRETPTKTLE | 6.4425 | 6.4147 | 6.0915 | 6.6532 | 6.8275 | 7.9287 | 6.7430 | 6.8211 | 7.1749 | 7.2682 | 8.0170 | 7.5408 | 4.0121 | 4.9274 | 3.3747 | 3.7627 |
| AHNAK2_S4785 | AHNAK2 | Q8IVF2 | 375 | TGAATGSRREERA | 6.1516 | 5.8832 | 8.1482 | 6.2098 | 5.6467 | 6.8640 | 6.5867 | 6.6861 | 7.0931 | 7.9599 | 8.1228 | 7.3882 | 3.7396 | 5.3067 | 4.0335 | 4.1801 |
| AHNAK2_S4894 | AHNAK2 | Q8IVF2 | 4894 | VGAPVMSPLSPGE | 5.4322 | 6.5199 | 5.8415 | 5.9321 | 6.0645 | 6.7386 | 6.0038 | 7.0925 | 6.2413 | 6.9296 | 5.8469 | 6.2732 | 6.1575 | 6.2792 | 6.3982 | 6.2491 |
| AHNAK2_S4897 | AHNAK2 | Q8IVF2 | 4782 | GPHFESSILSPCE | 5.6121 | 5.4048 | 6.7076 | 5.6807 | 5.8560 | 7.0170 | 6.5699 | 6.0744 | 6.4718 | 5.4286 | 6.2757 | 5.1654 | 6.4017 | 6.7558 | 7.1430 | 7.4357 |
| AHNAK2_S4962 | AHNAK2 | Q8IVF2 | 724 | LKTTDLSVQTPSA | 5.2801 | 5.3589 | 5.7177 | 5.8087 | 5.5656 | 5.6154 | 5.7563 | 5.6845 | 5.9323 | 5.2192 | 5.2831 | 5.4633 | 8.4116 | 8.1712 | 8.8609 | 7.8712 |
| AHNAK2_S4962;S4966 | AHNAK2 | Q8IVF2 | 4472 | PKFKMPSFGMLSP | 4.5438 | 4.5167 | 4.5735 | 3.9439 | 4.4462 | 3.7776 | 3.7985 | 3.9773 | 3.5365 | 3.8465 | 2.8822 | 3.2599 | 12.2405 | 12.9568 | 14.9233 | 12.7768 |
| AHNAK2_S497 | AHNAK2 | Q8IVF2 | 408 | PRLCEGTPQEGGL | 7.0654 | 7.3092 | 6.5478 | 6.8403 | 5.7176 | 7.0415 | 7.2502 | 6.4465 | 6.8900 | 6.7842 | 7.6062 | 7.3818 | 3.7535 | 4.8347 | 3.8579 | 4.6731 |
| AHNAK2_S509 | AHNAK2 | Q8IVF2 | 765 | PKVQRPSLKMPKV | 7.3701 | 6.8284 | 7.6573 | 6.4206 | 7.2267 | 6.5713 | 6.3566 | 7.4099 | 6.7645 | 7.0110 | 6.6524 | 6.1295 | 4.6378 | 5.0612 | 3.8141 | 4.0887 |
| AHNAK2_S5175 | AHNAK2 | Q8IVF2 | 5199 | PKFRMPSLRRSFR | 6.3777 | 6.1657 | 6.7976 | 5.9358 | 5.6277 | 6.7739 | 7.3668 | 6.5646 | 5.7042 | 5.5278 | 6.9120 | 7.6164 | 4.3559 | 6.8081 | 4.6681 | 6.7976 |
| AHNAK2_S5199 | AHNAK2 | Q8IVF2 | 923 | PKVEMPSFKMPKV | 8.0941 | 6.9754 | 7.5238 | 7.2385 | 5.7539 | 6.8097 | 6.8989 | 6.0311 | 7.2222 | 6.7414 | 7.6360 | 7.5438 | 4.1420 | 5.1112 | 2.8961 | 3.3818 |
| AHNAK2_S5428 | AHNAK2 | Q8IVF2 | 4254 | PKADVMTPVVEVS | 5.3508 | 4.6222 | 6.6122 | 5.9178 | 6.0356 | 6.5478 | 6.2561 | 6.8683 | 5.3056 | 5.3357 | 5.5768 | 4.8482 | 7.7051 | 8.4665 | 7.6407 | 6.9107 |
| AHNAK2_S5707 | AHNAK2 | Q8IVF2 | 3161 | VSAPGKSIEVLVD | 5.1874 | 5.2365 | 9.3056 | 9.1256 | 7.2929 | 7.6856 | 7.2274 | 5.4983 | 6.0001 | 7.8438 | 7.2056 | 4.4892 | 3.3655 | 5.3019 | 4.5219 | 4.7128 |
| AHNAK2_S5741 | AHNAK2 | Q8IVF2 | 2336 | VSALGKSIEASAD | 8.0024 | 7.2066 | 9.5160 | 7.1677 | 6.4377 | 5.6520 | 5.7437 | 6.6722 | 8.0003 | 7.5056 | 5.5359 | 6.7074 | 3.6366 | 5.3433 | 3.4469 | 3.4259 |
| AHNAK2_S589;S593 | AHNAK2 | Q8IVF2 | 3491 | VSAPGRSIEASLD | 6.4897 | 7.9056 | 8.4429 | 6.8471 | 5.4700 | 5.1406 | 5.4540 | 7.2061 | 6.4289 | 7.7459 | 6.0496 | 7.1976 | 3.7982 | 5.5165 | 5.3839 | 4.9235 |
| AHNAK2_S593 | AHNAK2 | Q8IVF2 | 851 | VSAPGKSMEDSVD | 7.2451 | 7.1937 | 7.9575 | 7.0462 | 7.4213 | 5.9009 | 6.2783 | 6.5476 | 6.4837 | 6.9770 | 6.1422 | 5.8442 | 4.2737 | 5.6358 | 4.9383 | 4.1143 |
| AHNAK2_S693 | AHNAK2 | Q8IVF2 | 1033 | KVEADLSLPSMQG | 5.9229 | 7.0475 | 6.9679 | 7.3805 | 6.1189 | 8.0059 | 7.3683 | 6.6999 | 5.8762 | 6.6020 | 6.9595 | 7.0192 | 4.1630 | 4.8252 | 4.3383 | 4.7050 |
| AHNAK2_S710 | AHNAK2 | Q8IVF2 | 710 | KVEADVSLLSMQG | 7.7770 | 7.0771 | 6.8715 | 6.6361 | 6.4131 | 7.1787 | 7.5540 | 5.7244 | 7.5317 | 5.4395 | 8.8052 | 6.5581 | 2.9805 | 5.7244 | 3.7188 | 4.0099 |
| AHNAK2_S724 | AHNAK2 | Q8IVF2 | 4775 | SSRLDLTGPHFES | 6.9263 | 7.4700 | 6.8119 | 7.6394 | 6.1255 | 7.5755 | 6.9783 | 6.9174 | 7.4106 | 6.8802 | 7.9811 | 6.4419 | 3.4498 | 4.3709 | 3.2017 | 3.8197 |
| AHNAK2_S765 | AHNAK2 | Q8IVF2 | 4642 | SGFEWSSKKVSMS | 7.3578 | 6.4260 | 7.0167 | 7.0219 | 5.8198 | 5.9824 | 6.9758 | 6.1258 | 7.6638 | 6.6253 | 7.3206 | 8.0106 | 4.7725 | 5.5879 | 3.2179 | 4.0752 |
| AHNAK2_S842 | AHNAK2 | Q8IVF2 | 5741 | DARESFSPEEKEE | 5.4277 | 6.0302 | 6.2167 | 6.1264 | 5.9659 | 6.3514 | 6.6053 | 6.6969 | 5.8034 | 6.2532 | 6.5954 | 6.3952 | 5.7454 | 6.0656 | 6.5929 | 7.1283 |
| AHNAK2_S851 | AHNAK2 | Q8IVF2 | 4646 | WSSKKVSMSSSEI | 5.6418 | 6.3270 | 7.1610 | 6.0193 | 5.7358 | 6.4739 | 5.9514 | 7.4338 | 6.9679 | 9.4762 | 8.5889 | 7.8461 | 3.9588 | 4.8078 | 3.7791 | 3.8312 |
| AHNAK2_S923 | AHNAK2 | Q8IVF2 | 365 | LGPWGDSLEETGA | 7.6517 | 7.4791 | 7.9663 | 7.3340 | 6.7002 | 5.9630 | 5.8537 | 5.9802 | 7.3593 | 7.1555 | 5.6052 | 5.9616 | 4.5638 | 5.6618 | 4.7713 | 3.9932 |
| AHNAK2_T228 | AHNAK2 | Q8IVF2 | 5172 | PDAEVLTVESPEE | 4.8311 | 5.5491 | 6.0022 | 4.8893 | 5.4015 | 6.0503 | 6.0815 | 5.7727 | 4.8885 | 4.4557 | 5.2614 | 5.6909 | 8.5975 | 8.0887 | 9.2716 | 9.1678 |
| AHNAK2_T408 | AHNAK2 | Q8IVF2 | 509 | ERERRLSTPQRGK | 7.0081 | 7.6417 | 6.4542 | 7.2090 | 6.0582 | 7.0638 | 7.6338 | 6.5257 | 7.1552 | 7.4051 | 7.0353 | 7.2495 | 3.7051 | 4.5699 | 3.4859 | 3.7994 |
| AHNAK2_T4254 | AHNAK2 | Q8IVF2 | 294;298 | NA | 6.7837 | 6.7716 | 6.7108 | 6.7464 | 6.8211 | 7.1190 | 6.8710 | 6.9578 | 5.8554 | 5.0880 | 5.0512 | 5.2584 | 5.7806 | 6.4227 | 5.8922 | 5.8701 |
| AHNAK2_T4775 | AHNAK2 | Q8IVF2 | 4962;4966 | NA | 6.6645 | 6.1526 | 8.7626 | 6.1977 | 6.5770 | 8.4808 | 6.6486 | 7.0741 | 6.3478 | 7.4766 | 9.4726 | 7.8217 | 3.3085 | 4.2404 | 2.3576 | 2.4170 |
| AHNAK2_T4931 | AHNAK2 | Q8IVF2 | 4472;4477 | NA | 8.0027 | 6.6955 | 7.4342 | 6.6862 | 6.6108 | 5.5958 | 5.9353 | 6.4614 | 7.2269 | 6.3157 | 5.0394 | 5.4816 | 5.0166 | 5.9316 | 5.8268 | 5.7395 |
| AHNAK2_T510 | AHNAK2 | Q8IVF2 | 589;593 | NA | 6.5796 | 7.3081 | 7.0146 | 6.1213 | 5.8154 | 5.8735 | 6.1297 | 6.6201 | 5.0480 | 5.2322 | 8.4136 | 5.8919 | 5.3582 | 7.8311 | 5.9891 | 4.7738 |
| AHNAK2_T5172 | AHNAK2 | Q8IVF2 | 2023;2026 | NA | 6.0895 | 6.5559 | 7.1699 | 6.6441 | 6.0333 | 6.0022 | 5.7707 | 5.9829 | 7.0903 | 6.3925 | 6.4273 | 6.3994 | 6.0151 | 6.3260 | 5.5331 | 5.5680 |
| AIDA_S162 | AIDA | Q96BJ3 | 162 | LLPRLPSEPGMTL | 4.8019 | 5.6282 | 6.3865 | 6.6740 | 6.7914 | 5.3265 | 5.3054 | 5.4944 | 6.6212 | 6.3841 | 5.7620 | 5.8195 | 7.1541 | 7.0731 | 7.7163 | 7.0614 |
| AIFM1_S268 | AIFM1 | O95831 | 268 | GTPRSLSAIDRAG | 6.8724 | 6.6501 | 5.7276 | 6.3937 | 6.4438 | 5.8886 | 6.1882 | 6.0475 | 6.4279 | 5.9021 | 5.5070 | 5.4039 | 6.7746 | 6.5085 | 6.9031 | 6.3611 |
| AIFM1_S292 | AIFM1 | O95831 | 263 | LIATGGTPRSLSA | 6.4154 | 6.9192 | 6.2234 | 6.2975 | 6.0161 | 6.2764 | 6.0161 | 5.7673 | 6.2048 | 6.0052 | 5.1599 | 5.5705 | 7.2603 | 6.4775 | 6.6192 | 6.7712 |
| AIFM1_T105 | AIFM1 | O95831 | 292 | KIGDFRSLEKISR | 7.3004 | 7.4341 | 6.3407 | 6.7315 | 6.6660 | 6.4096 | 6.5432 | 5.0618 | 5.1232 | 6.1173 | 6.3806 | 6.3245 | 6.2199 | 5.5572 | 5.5173 | 6.2726 |
| AIFM1_T263 | AIFM1 | O95831 | 105 | ISGLGLTPEQKQK | 5.2018 | 6.4441 | 6.5159 | 8.1404 | 7.5573 | 5.9642 | 7.2694 | 7.3376 | 6.9877 | 6.2665 | 7.7071 | 7.0568 | 4.0097 | 4.8403 | 4.3658 | 4.3353 |
| AJM1_S109 | AJM1 | C9J069 | 856 | VALAAGSPARPPP | 6.4853 | 6.0643 | 6.3051 | 6.4645 | 5.5817 | 5.6346 | 6.7417 | 5.7715 | 6.5295 | 7.2191 | 5.9179 | 6.4091 | 6.8111 | 6.5243 | 5.4778 | 6.0625 |
| AJM1_S856 | AJM1 | C9J069 | 109 | LTPAPASPPVLPR | 5.7915 | 6.6628 | 6.2636 | 5.9653 | 6.0479 | 6.7122 | 5.6112 | 6.6886 | 6.3999 | 6.5308 | 6.5802 | 5.6681 | 6.2024 | 5.9674 | 6.7444 | 6.1638 |
| AJUBA_S119 | AJUBA | Q96IF1 | 119 | PTAPALSPRSSFA | 6.3347 | 6.1522 | 6.4068 | 6.3255 | 6.2291 | 6.3667 | 6.2040 | 6.5425 | 6.4694 | 6.6269 | 6.5309 | 6.6715 | 5.8817 | 5.9151 | 5.6655 | 5.6777 |
| AJUBA_S137 | AJUBA | Q96IF1 | 230 | CQESRHSYPPALG | 5.9309 | 6.2364 | 7.2508 | 6.1967 | 5.8988 | 6.4326 | 6.6423 | 6.2039 | 6.2003 | 6.2477 | 6.8132 | 6.6975 | 5.8197 | 5.7785 | 5.7225 | 5.9282 |
| AJUBA_S230 | AJUBA | Q96IF1 | 137 | DASKPSSPRGSLL | 5.5406 | 6.0366 | 6.5014 | 5.6219 | 6.4859 | 7.6329 | 6.4371 | 6.0903 | 6.4007 | 6.5505 | 7.1129 | 6.6858 | 6.0681 | 6.2212 | 5.3226 | 5.2914 |
| AJUBA_S237 | AJUBA | Q96IF1 | 79 | ERNQRGSFEAPRY | 5.3835 | 5.6778 | 6.5472 | 5.7669 | 6.1640 | 6.0970 | 5.8988 | 5.7722 | 7.5469 | 6.3066 | 6.5485 | 6.4902 | 6.4818 | 6.8069 | 6.2113 | 6.3003 |
| AJUBA_S263 | AJUBA | Q96IF1 | 39 | PGKGRLSGLGGPR | 7.0594 | 7.1696 | 7.1616 | 7.3154 | 5.5032 | 6.7321 | 6.7148 | 6.6999 | 9.2701 | 7.1191 | 6.8779 | 8.3157 | 3.9666 | 3.5831 | 2.4967 | 4.0149 |
| AJUBA_S39 | AJUBA | Q96IF1 | 237 | YPPALGSPGALAG | 6.3727 | 6.7004 | 7.2876 | 6.3567 | 6.4292 | 5.8723 | 6.1512 | 6.6869 | 5.6769 | 6.3390 | 5.5573 | 6.1815 | 4.7974 | 6.5075 | 6.0897 | 6.9936 |
| AJUBA_S79 | AJUBA | Q96IF1 | 263 | AQPGRHSVTGYGD | 6.5184 | 6.6726 | 7.4501 | 6.8350 | 6.6556 | 6.5418 | 5.8547 | 6.9506 | 6.7460 | 5.8443 | 7.2999 | 6.4255 | 5.4629 | 5.3797 | 4.6158 | 4.7469 |
| AKAP1_S107 | AKAP1 | Q92667 | 533 | PRDKAITPPLPES | 6.0719 | 6.1904 | 6.0650 | 6.9029 | 6.3796 | 5.8687 | 6.1774 | 6.1852 | 5.9583 | 6.1583 | 6.1110 | 6.3428 | 6.3813 | 6.5828 | 6.2664 | 6.3578 |
| AKAP1_S151 | AKAP1 | Q92667 | 151 | LECPLSSPKGVLF | 6.1676 | 5.9488 | 6.2023 | 6.7032 | 6.4382 | 6.6451 | 6.6959 | 6.4420 | 6.1008 | 5.9073 | 6.0308 | 5.9562 | 6.2571 | 6.0418 | 6.2052 | 6.2576 |
| AKAP1_S231 | AKAP1 | Q92667 | 445 | LKSLLSSPTKDSK | 6.6027 | 6.4714 | 5.8117 | 6.5189 | 6.6094 | 5.9192 | 6.4076 | 6.1850 | 6.3605 | 6.0267 | 5.9650 | 6.0779 | 6.1351 | 6.1448 | 6.3388 | 6.4255 |
| AKAP1_S445 | AKAP1 | Q92667 | 107 | PCRRSESSGILPN | 4.8348 | 5.1222 | 5.2594 | 5.4020 | 5.2047 | 4.6805 | 5.0437 | 5.0363 | 5.5509 | 5.3494 | 5.1888 | 5.3265 | 9.1999 | 9.2048 | 10.1098 | 9.4864 |
| AKAP1_S592 | AKAP1 | Q92667 | 231 | VLELENSKGPSLA | 6.2555 | 6.4027 | 5.3866 | 6.6438 | 5.5589 | 5.7021 | 7.2117 | 6.4116 | 6.1826 | 5.8744 | 5.8971 | 5.8356 | 5.9229 | 6.7789 | 7.0338 | 6.9019 |
| AKAP1_T533 | AKAP1 | Q92667 | 592 | SLKKTESFQNAQA | 0.9824 | 1.2623 | 1.0113 | 1.2633 | 1.9130 | 2.0190 | 1.8913 | 1.5737 | 1.3040 | 1.3673 | 1.3336 | 1.3551 | 21.4564 | 19.4791 | 22.6087 | 19.1793 |
| AKAP10_S187 | AKAP10 | O43572 | 281 | SRNSPASPLKELS | 6.3893 | 6.1877 | 7.0595 | 6.1409 | 6.1676 | 6.4990 | 6.5729 | 6.3609 | 7.0475 | 6.5554 | 6.9187 | 6.9777 | 5.4043 | 5.1582 | 5.1380 | 5.4223 |
| AKAP10_S281 | AKAP10 | O43572 | 7 | MRGAGPSPRQSPR | 7.2024 | 7.8235 | 7.1322 | 7.0979 | 5.6694 | 7.1364 | 7.4668 | 7.2278 | 7.1938 | 7.4219 | 7.8269 | 7.7822 | 2.5798 | 3.9712 | 2.8820 | 3.5857 |
| AKAP10_S7 | AKAP10 | O43572 | 187 | SLAEPVSPSKKHE | 6.6413 | 6.4102 | 6.4964 | 6.3141 | 6.6715 | 6.4432 | 6.4121 | 6.5499 | 6.6197 | 6.0202 | 6.6118 | 6.5977 | 5.4455 | 5.8638 | 5.6443 | 5.2582 |
| AKAP11_S1242 | AKAP11 | Q9UKA4 | 448 | EDSGLFSPIRSSA | 5.9660 | 6.0743 | 5.9245 | 6.0597 | 6.5019 | 6.3678 | 5.9832 | 6.3802 | 6.8927 | 6.6470 | 7.0553 | 7.0708 | 5.8341 | 6.0246 | 5.5306 | 5.6873 |
| AKAP11_S133 | AKAP11 | Q9UKA4 | 1580 | TYAEKLSPLTGQA | 6.4810 | 6.1700 | 6.1757 | 6.0379 | 6.1663 | 6.3379 | 6.1571 | 6.1660 | 6.6510 | 6.3794 | 6.3582 | 6.1357 | 6.3154 | 6.2920 | 6.1960 | 5.9804 |
| AKAP11_S1337 | AKAP11 | Q9UKA4 | 456 | IRSSAFSPLGGCT | 5.2600 | 5.8742 | 6.4668 | 5.9668 | 6.2591 | 6.2701 | 6.0648 | 6.2367 | 6.4865 | 6.7819 | 7.1296 | 6.6027 | 5.7436 | 7.0244 | 5.6692 | 6.1635 |
| AKAP11_S1337;Y1345 | AKAP11 | Q9UKA4 | 1242 | QSQRSVSPTFLNP | 5.7674 | 5.9377 | 6.0142 | 6.0445 | 6.2397 | 6.0771 | 5.9511 | 6.3005 | 6.7031 | 6.5063 | 6.5646 | 6.7466 | 6.2480 | 6.5180 | 6.1985 | 6.1828 |
| AKAP11_S1580 | AKAP11 | Q9UKA4 | 433 | LCDAPDSPRPVKA | 8.1811 | 6.0426 | 6.9508 | 6.2236 | 6.6578 | 6.3479 | 7.8114 | 6.4583 | 6.4200 | 6.6931 | 7.1011 | 6.4890 | 5.3001 | 4.5331 | 4.0039 | 4.7862 |
| AKAP11_S1611 | AKAP11 | Q9UKA4 | 1100 | RNVIPDTPPSTPL | 5.7163 | 6.3106 | 6.8487 | 5.5414 | 7.0166 | 6.3934 | 6.4600 | 6.7809 | 6.9182 | 6.5301 | 6.3071 | 6.0574 | 5.5263 | 6.5619 | 5.3850 | 5.6462 |
| AKAP11_S18 | AKAP11 | Q9UKA4 | 981 | GEESQLTPEKSPK | 6.2599 | 7.0301 | 7.0136 | 7.2063 | 5.8784 | 6.4526 | 6.3038 | 6.8139 | 7.1261 | 6.4164 | 7.3068 | 6.8972 | 4.3107 | 4.6598 | 4.9632 | 5.3613 |
| AKAP11_S422;S433 | AKAP11 | Q9UKA4 | 1337 | LMYKYPSCESVTD | 4.6394 | 5.1538 | 6.0150 | 5.4203 | 5.3225 | 6.9946 | 5.6198 | 6.7821 | 6.2237 | 7.6799 | 6.4315 | 7.2041 | 6.3895 | 6.2519 | 7.1065 | 6.7651 |
| AKAP11_S433 | AKAP11 | Q9UKA4 | 1611 | HFFRQGSLASSKP | 6.8608 | 6.4609 | 7.9301 | 6.4883 | 4.8261 | 5.9222 | 6.0706 | 4.7274 | 6.8001 | 6.5249 | 7.6282 | 6.6120 | 5.8610 | 7.2855 | 5.1489 | 4.8529 |
| AKAP11_S448 | AKAP11 | Q9UKA4 | 18 | KASVRKSFSEDVF | 6.6000 | 6.6218 | 6.7488 | 7.1448 | 6.4341 | 6.1895 | 6.1955 | 6.2878 | 6.8972 | 7.0212 | 7.4489 | 7.5280 | 4.0223 | 5.0922 | 4.4106 | 5.3573 |
| AKAP11_S456 | AKAP11 | Q9UKA4 | 1485 | GLPKSLTDSCLFE | 1.9359 | 2.2828 | 2.2568 | 2.3815 | 3.2036 | 3.2117 | 3.3536 | 3.0122 | 2.2051 | 2.0719 | 2.1471 | 2.3022 | 17.7861 | 15.6434 | 19.4001 | 16.8059 |
| AKAP11_T1100 | AKAP11 | Q9UKA4 | 133 | MRVSPTSPRLRID | 5.9037 | 5.6946 | 6.1574 | 5.8164 | 6.1557 | 6.0877 | 6.1902 | 6.2629 | 5.8590 | 5.5099 | 5.5438 | 5.9700 | 7.1420 | 6.9411 | 7.6278 | 7.1380 |
| AKAP11_T1100;T1104 | AKAP11 | Q9UKA4 | 1337;1345 | NA | 6.8413 | 5.4336 | 6.7592 | 7.0096 | 6.9612 | 5.3258 | 6.0722 | 5.9146 | 6.2554 | 5.1771 | 5.8645 | 5.7250 | 6.4865 | 7.5412 | 5.9114 | 6.7214 |
| AKAP11_T1485 | AKAP11 | Q9UKA4 | 1100;1104 | NA | 5.6096 | 5.6376 | 6.3281 | 5.9277 | 6.4651 | 6.5838 | 5.9757 | 6.4332 | 6.1966 | 5.5440 | 5.9345 | 5.7799 | 6.4500 | 7.0825 | 6.8645 | 7.1871 |
| AKAP11_T981 | AKAP11 | Q9UKA4 | 422;433 | NA | 6.2016 | 6.1416 | 6.5986 | 6.1437 | 6.2838 | 7.2003 | 6.3065 | 6.3024 | 6.5147 | 6.9322 | 7.2765 | 6.1660 | 4.8204 | 6.2462 | 4.4921 | 6.3734 |
| AKAP12_S1251 | AKAP12 | Q02952 | 612 | GVTPWASFKKMVT | 7.6585 | 6.9439 | 7.1480 | 6.7383 | 7.0716 | 6.0579 | 6.0930 | 6.0985 | 6.7291 | 5.5815 | 5.9474 | 6.0811 | 6.3129 | 6.1429 | 4.7041 | 4.6911 |
| AKAP12_S1259 | AKAP12 | Q02952 | 514 | RMKVQGSPLKKLF | 7.4900 | 6.9347 | 6.8353 | 6.2762 | 7.1820 | 6.9764 | 6.8923 | 7.2137 | 7.6985 | 6.8307 | 6.8625 | 6.7338 | 4.1015 | 4.1947 | 3.6701 | 4.1076 |
| AKAP12_S128 | AKAP12 | Q02952 | 371 | AHEPRLSAEYEKV | 6.5532 | 6.3862 | 6.4323 | 6.3303 | 6.4550 | 5.8014 | 5.9790 | 7.1559 | 7.1980 | 5.8246 | 6.1159 | 6.2767 | 6.0769 | 5.3758 | 5.9888 | 6.0500 |
| AKAP12_S1328 | AKAP12 | Q02952 | 1395 | VSSLEGSPPPCLG | 6.4246 | 6.3952 | 6.5656 | 6.8512 | 6.3827 | 6.2213 | 6.3086 | 6.5408 | 6.4708 | 6.3140 | 6.3834 | 6.5961 | 5.6581 | 5.7846 | 5.4890 | 5.6139 |
| AKAP12_S1331 | AKAP12 | Q02952 | 1331 | HAKSPPSPVEREM | 6.5361 | 6.4637 | 6.1862 | 6.2435 | 6.1782 | 6.1806 | 5.9549 | 6.3704 | 6.2147 | 5.9035 | 6.1053 | 6.1456 | 6.7212 | 6.3789 | 6.1966 | 6.2206 |
| AKAP12_S1395 | AKAP12 | Q02952 | 806 | KEESWVSIKKFIP | 9.8437 | 7.8271 | 9.4304 | 7.3999 | 4.9697 | 5.2705 | 6.1439 | 5.6370 | 6.6622 | 5.9126 | 6.4833 | 6.9294 | 4.2409 | 5.3581 | 3.7048 | 4.1867 |
| AKAP12_S1418 | AKAP12 | Q02952 | 347 | GKAEVASEKLTAS | 7.1943 | 6.1442 | 6.0291 | 6.1273 | 6.2960 | 5.6255 | 5.9966 | 6.3327 | 6.3401 | 5.5660 | 5.8375 | 5.9127 | 7.2506 | 6.2809 | 6.6457 | 6.4210 |
| AKAP12_S1498 | AKAP12 | Q02952 | 1116 | KVVGQTTPESFEK | 6.5860 | 6.7050 | 6.4475 | 6.4662 | 6.4429 | 6.3145 | 5.9426 | 6.3982 | 6.9121 | 6.5699 | 6.5045 | 6.7750 | 5.7316 | 5.7330 | 5.0764 | 5.3948 |
| AKAP12_S1674 | AKAP12 | Q02952 | 505 | EVEMLSSQERMKV | 6.2613 | 6.2642 | 6.4436 | 6.3377 | 6.3877 | 5.9573 | 5.8751 | 6.1557 | 6.2242 | 6.0634 | 6.2582 | 6.2375 | 6.6436 | 6.3499 | 6.1969 | 6.3438 |
| AKAP12_S1727 | AKAP12 | Q02952 | 75 | AEQDELSLQEGDL | 5.9014 | 5.9292 | 6.5079 | 6.3139 | 6.2174 | 5.7813 | 5.9391 | 6.1401 | 6.4896 | 5.9637 | 5.7538 | 5.7284 | 7.1979 | 6.8621 | 6.8173 | 6.4568 |
| AKAP12_S219 | AKAP12 | Q02952 | 248 | TLKREQSHAEISP | 3.5547 | 3.4281 | 3.3898 | 3.6519 | 5.3953 | 4.9420 | 4.7825 | 4.5780 | 3.5611 | 3.4267 | 3.5893 | 3.4713 | 13.7162 | 12.0449 | 13.4722 | 12.9962 |
| AKAP12_S248 | AKAP12 | Q02952 | 283 | PSKSAESPTSPVT | 6.2209 | 6.3449 | 6.5204 | 6.4624 | 6.4851 | 6.6759 | 6.3406 | 6.4632 | 6.8254 | 6.7092 | 6.8332 | 7.1599 | 5.1229 | 5.3651 | 5.1059 | 5.3649 |
| AKAP12_S283 | AKAP12 | Q02952 | 504 | SEVEMLSSQERMK | 6.1264 | 6.2228 | 6.4116 | 6.2204 | 6.1761 | 6.0285 | 5.6995 | 5.9930 | 6.4686 | 6.0928 | 6.2655 | 6.4799 | 6.5599 | 6.3466 | 6.5982 | 6.3101 |
| AKAP12_S294 | AKAP12 | Q02952 | 135 | AVVHDITDDGQEE | 4.9247 | 5.4766 | 5.1701 | 5.4276 | 5.6313 | 5.5556 | 4.4416 | 5.7729 | 5.6066 | 5.4029 | 5.4237 | 5.2559 | 9.1364 | 8.8318 | 9.2145 | 8.7280 |
| AKAP12_S347 | AKAP12 | Q02952 | 598 | AEEGATSDGEKKR | 6.6410 | 6.3836 | 6.6923 | 6.5241 | 6.4270 | 6.4438 | 6.5496 | 6.6043 | 6.1023 | 6.2708 | 6.2410 | 6.3771 | 5.7764 | 5.9075 | 5.3941 | 5.6650 |
| AKAP12_S371 | AKAP12 | Q02952 | 887 | SKELSESQVHMMA | 5.3677 | 6.0259 | 6.0305 | 5.9922 | 6.2883 | 5.8754 | 5.6917 | 5.5685 | 6.0698 | 5.2155 | 5.5614 | 5.4812 | 8.1469 | 7.4688 | 7.6949 | 7.5213 |
| AKAP12_S386 | AKAP12 | Q02952 | 1484 | DCQAKSTPVIVSA | 7.4914 | 7.8801 | 6.9645 | 7.0782 | 6.6568 | 7.0947 | 7.5301 | 7.1642 | 7.7427 | 7.0382 | 7.3264 | 6.9513 | 3.2389 | 3.7283 | 2.6982 | 3.4160 |
| AKAP12_S422 | AKAP12 | Q02952 | 483 | TQGADLSPDEKVL | 6.9832 | 6.8138 | 6.9413 | 7.2244 | 6.4089 | 6.4731 | 6.9869 | 6.7429 | 7.0352 | 6.4168 | 7.2626 | 7.1966 | 4.3385 | 4.4202 | 3.7990 | 4.9567 |
| AKAP12_S483 | AKAP12 | Q02952 | 219 | GDHKDPSLGAGEA | 6.2123 | 6.1200 | 6.2214 | 6.2681 | 5.9785 | 5.5777 | 5.6427 | 5.7877 | 5.8088 | 5.4259 | 5.5149 | 5.8794 | 7.6548 | 6.6744 | 7.2009 | 8.0325 |
| AKAP12_S490 | AKAP12 | Q02952 | 128 | KEMATKSAVVHDI | 6.3533 | 6.7155 | 6.9276 | 6.1524 | 5.9223 | 6.2563 | 5.8039 | 6.3310 | 6.9980 | 5.2820 | 6.1988 | 5.8271 | 6.4846 | 6.0803 | 5.9910 | 6.6761 |

| | | | | | | | | | | | | | | | | | | | | | |
|---|---|---|---|---|---|---|---|---|---|---|---|---|---|---|---|---|---|---|---|---|---|
| AKAP12_S504 | AKAP12 | Q02952 | | 1259 | ETVSILSKTEGTQ | 5.5802 | 5.9912 | 6.4571 | 6.2577 | 6.1471 | 6.0728 | 5.7838 | 6.1235 | 5.6616 | 5.7644 | 5.9583 | 6.0567 | 7.0127 | 6.9340 | 7.0801 | 7.1188 |
| AKAP12_S505 | AKAP12 | Q02952 | | 1727 | GGLTKESPDTNGP | 6.4060 | 6.2152 | 6.5356 | 6.5331 | 6.2284 | 6.3572 | 6.1568 | 6.0049 | 6.6693 | 6.3874 | 6.5712 | 6.8299 | 5.5992 | 6.0827 | 6.3005 | 5.1227 |
| AKAP12_S514 | AKAP12 | Q02952 | | 295 | TSETGSTFKKFFT | 9.2114 | 7.4725 | 10.9332 | 8.6327 | 5.5483 | 5.3896 | 5.7276 | 5.0664 | 6.5245 | 6.7776 | 7.5468 | 5.6793 | 4.3108 | 5.3454 | 2.8756 | 2.9582 |
| AKAP12_S522 | AKAP12 | Q02952 | | 568 | KGESSASSPEEPE | 6.5739 | 6.5958 | 6.6139 | 5.9111 | 6.4037 | 6.6002 | 6.0859 | 6.4312 | 5.4059 | 5.9127 | 6.4552 | 6.1103 | 6.9494 | 5.8416 | 6.2270 | 5.8821 |
| AKAP12_S568 | AKAP12 | Q02952 | | 645 | KSATLSSTESTAS | 5.2088 | 5.7844 | 6.1785 | 5.5923 | 5.8662 | 5.9034 | 5.5743 | 6.6381 | 7.0912 | 7.4423 | 7.1564 | 7.6461 | 5.5454 | 6.1381 | 6.0703 | 6.1639 |
| AKAP12_S598 | AKAP12 | Q02952 | | 627 | KRVRRPSESDKED | 5.4900 | 5.1609 | 6.0297 | 6.0552 | 5.9893 | 5.2457 | 5.5535 | 6.2206 | 7.2613 | 6.3273 | 6.3072 | 6.6897 | 6.8101 | 6.2064 | 6.8243 | 7.8288 |
| AKAP12_S627 | AKAP12 | Q02952 | | 1674 | GGAGTKSVPEDDG | 6.3467 | 6.8332 | 6.2821 | 6.4661 | 6.0956 | 6.5295 | 6.6267 | 6.5128 | 6.9033 | 6.1544 | 6.5199 | 6.7060 | 5.5136 | 5.6813 | 5.1506 | 5.6784 |
| AKAP12_S627;S629 | AKAP12 | Q02952 | | 1760 | ESDKAITPQAQEE | 5.7958 | 6.1006 | 6.3518 | 6.4067 | 5.7531 | 6.1689 | 5.7855 | 6.7971 | 6.1922 | 6.0002 | 5.8358 | 5.9427 | 7.0829 | 6.6477 | 6.6754 | 6.4635 |
| AKAP12_S645 | AKAP12 | Q02952 | | 490 | PDEKVLSKPPEGV | 5.9219 | 6.6915 | 6.6929 | 6.2747 | 6.2775 | 5.2662 | 7.4208 | 6.2442 | 6.6734 | 7.0693 | 6.2789 | 6.6109 | 5.2635 | 5.7135 | 5.6316 | 5.9691 |
| AKAP12_S696 | AKAP12 | Q02952 | | 1251 | HTDKEVSVETVSI | 6.4370 | 6.9508 | 5.9480 | 6.4658 | 6.4777 | 6.2603 | 5.8479 | 6.6503 | 6.5676 | 6.5087 | 6.6873 | 6.6151 | 5.8452 | 5.6575 | 5.4283 | 5.6525 |
| AKAP12_S75 | AKAP12 | Q02952 | | 386 | PSEEQVSGSQGPS | 6.2484 | 6.3481 | 5.6295 | 6.2174 | 5.6186 | 6.1546 | 6.4946 | 5.2133 | 7.0189 | 6.5298 | 6.4143 | 6.7317 | 6.9393 | 6.4193 | 5.8490 | 6.1731 |
| AKAP12_S806 | AKAP12 | Q02952 | | 696 | KRARRGSSSDEEG | 6.2235 | 5.8121 | 6.0383 | 6.0852 | 6.1418 | 5.6570 | 5.2756 | 5.9295 | 7.3746 | 6.3875 | 6.7069 | 6.9821 | 6.8748 | 5.9676 | 6.1545 | 6.3889 |
| AKAP12_S887 | AKAP12 | Q02952 | | 1328 | LQSHAKSPPSPVE | 6.9636 | 7.6210 | 6.3647 | 7.5110 | 6.2742 | 6.5793 | 6.3540 | 6.4299 | 6.7540 | 6.6415 | 6.9643 | 7.5627 | 4.3875 | 4.8887 | 4.3893 | 4.3144 |
| AKAP12_T1116 | AKAP12 | Q02952 | | 1498 | TKKGLSSDLEGEK | 7.0239 | 7.7947 | 7.6349 | 7.2843 | 6.7933 | 7.0548 | 7.0000 | 7.9569 | 5.5828 | 5.9345 | 5.2117 | 4.3142 | 5.1546 | 5.1820 | 5.1843 | 4.8931 |
| AKAP12_T135 | AKAP12 | Q02952 | | 793 | SGVEHSTPDTEPG | 6.3475 | 7.0094 | 6.3520 | 6.9038 | 7.2991 | 8.3970 | 6.8974 | 6.9637 | 6.8239 | 7.1272 | 8.6127 | 6.7441 | 2.7848 | 4.4300 | 3.5749 | 3.7326 |
| AKAP12_T1484 | AKAP12 | Q02952 | | 294 | VTSETGSTFKKFF | 7.8512 | 7.1436 | 8.5811 | 7.5287 | 7.0216 | 5.8710 | 6.6793 | 5.6709 | 6.8658 | 6.4325 | 7.1776 | 6.6200 | 4.9264 | 4.5142 | 3.0786 | 4.0375 |
| AKAP12_T1760 | AKAP12 | Q02952 | | 1418 | VQSSEASFTLTAA | 6.2814 | 6.6953 | 7.7397 | 6.2950 | 6.6084 | 6.5633 | 7.0266 | 6.0328 | 7.7761 | 6.6713 | 6.8736 | 6.1679 | 4.7930 | 5.2445 | 3.9472 | 5.2840 |
| AKAP12_T295 | AKAP12 | Q02952 | | 422 | VAEVHVSTVEERT | 6.7559 | 4.7775 | 8.4021 | 8.3875 | 6.2332 | 7.3323 | 7.3079 | 8.4705 | 6.3798 | 5.2513 | 7.0637 | 5.7740 | 3.8200 | 5.6421 | 4.4942 | 3.9080 |
| AKAP12_T608;S612 | AKAP12 | Q02952 | | 522 | LKKLFTSTGLKKL | 8.5607 | 6.2429 | 7.8996 | 6.8764 | 7.0821 | 5.7347 | 5.5926 | 6.4139 | 6.9057 | 6.3510 | 6.0535 | 6.1151 | 5.9584 | 6.0288 | 3.9804 | 4.2041 |
| AKAP12_T612 | AKAP12 | Q02952 | 627;629 | | NA | 6.4695 | 6.2547 | 6.7541 | 6.2799 | 6.2175 | 5.9704 | 6.3792 | 5.6282 | 6.3057 | 6.1231 | 6.2795 | 6.5038 | 6.7127 | 6.1410 | 5.6107 | 6.3699 |
| AKAP12_T793 | AKAP12 | Q02952 | 608;612 | | NA | 8.1978 | 7.6967 | 8.2894 | 6.5036 | 6.0320 | 6.6974 | 6.3004 | 5.8765 | 7.7937 | 6.1865 | 7.0482 | 6.9402 | 4.5903 | 4.6597 | 3.3928 | 3.7948 |
| AKAP13_S1411 | AKAP13 | Q12802-2 | | 2732 | DSELSVSPKRNSI | 6.1051 | 6.1672 | 6.4370 | 6.2711 | 6.1887 | 6.1998 | 6.1305 | 6.1280 | 6.5641 | 6.4964 | 6.9715 | 6.9963 | 5.7833 | 5.8829 | 5.6320 | 6.0459 |
| AKAP13_S1565 | AKAP13 | Q12802-2 | | 1606 | FNYRSFSLEGLTG | 6.1421 | 5.8526 | 6.8977 | 6.0538 | 5.7580 | 6.1491 | 6.1914 | 6.2614 | 6.6278 | 6.7816 | 6.3777 | 6.6784 | 5.6367 | 5.9778 | 6.2544 | 6.3596 |
| AKAP13_S1606 | AKAP13 | Q12802-2 | | 1910 | SLSKSVSIQNITG | 6.5821 | 6.2117 | 6.8794 | 5.9984 | 5.9320 | 6.4002 | 6.4841 | 6.0687 | 6.8382 | 6.1066 | 6.7545 | 6.5785 | 5.6316 | 6.4611 | 5.0762 | 5.9967 |
| AKAP13_S1649 | AKAP13 | Q12802-2 | | 2471 | LPRRAETFGGFDS | 5.7238 | 5.5866 | 5.9650 | 5.3746 | 6.0389 | 5.7000 | 5.7543 | 5.9332 | 5.4252 | 5.2763 | 5.5089 | 4.9156 | 8.3356 | 8.3416 | 8.9592 | 7.1612 |
| AKAP13_S1687 | AKAP13 | Q12802-2 | | 790 | TFSLANSPGSESV | 5.6363 | 6.0392 | 6.3948 | 6.3141 | 6.0576 | 6.6015 | 6.1071 | 6.2582 | 6.7693 | 6.5191 | 6.6164 | 6.3552 | 5.8454 | 6.3506 | 6.0250 | 6.1102 |
| AKAP13_S1880 | AKAP13 | Q12802-2 | | 2713 | SPEEPPSPSAPSI | 5.8391 | 5.9322 | 5.9509 | 5.9153 | 6.0638 | 6.8181 | 6.0110 | 6.2036 | 5.8547 | 6.2482 | 5.9927 | 6.2265 | 6.7849 | 6.9291 | 6.6083 | 6.6216 |
| AKAP13_S1904 | AKAP13 | Q12802-2 | | 1880 | PKERPRSAVLLVD | 6.4875 | 6.6655 | 7.1700 | 6.2079 | 6.5035 | 5.8220 | 6.4202 | 6.3005 | 6.5170 | 6.3036 | 6.3074 | 6.4417 | 5.9317 | 6.2700 | 5.2231 | 5.4282 |
| AKAP13_S1910 | AKAP13 | Q12802-2 | | 1149 | DPQGVGTPEMIPL | 6.6243 | 6.3158 | 7.0338 | 6.2496 | 5.3911 | 6.7754 | 7.0766 | 6.9240 | 6.2205 | 5.8199 | 5.9362 | 6.2480 | 5.5574 | 6.7019 | 5.7965 | 5.3289 |
| AKAP13_S1933 | AKAP13 | Q12802-2 | | 983 | ALQLSNSPGASSA | 5.4804 | 6.7822 | 6.8893 | 7.2483 | 5.9438 | 6.7825 | 6.3294 | 6.2066 | 6.4817 | 6.7846 | 6.6211 | 7.1533 | 5.0162 | 5.5934 | 5.2955 | 5.3915 |
| AKAP13_S1933;S1936 | AKAP13 | Q12802-2 | | 1933 | WKFLSHSTDSLNK | 5.7743 | 6.4216 | 6.3366 | 6.0244 | 5.6503 | 6.8286 | 6.2149 | 6.8562 | 6.3919 | 6.1731 | 6.8971 | 6.8080 | 5.7030 | 6.0170 | 6.0238 | 5.8793 |
| AKAP13_S1936 | AKAP13 | Q12802-2 | | 1565 | SPFRRHSWGPGKN | 5.8634 | 5.9744 | 6.7872 | 6.0822 | 6.3704 | 6.8093 | 5.8679 | 6.3172 | 6.4508 | 7.0089 | 6.6456 | 6.6178 | 5.4927 | 6.3300 | 5.7346 | 5.6476 |
| AKAP13_S2502 | AKAP13 | Q12802-2 | | 1936 | LSHSTDSLNKISK | 7.4390 | 7.4238 | 7.1588 | 7.0768 | 8.2779 | 5.1799 | 5.2199 | 5.8433 | 7.7994 | 5.6984 | 4.6747 | 5.6908 | 5.1284 | 6.5964 | 5.1818 | 5.6107 |
| AKAP13_S2703 | AKAP13 | Q12802-2 | | 1649 | RVDSLVSLSEEDL | 5.4125 | 5.4822 | 6.5846 | 7.0994 | 6.2285 | 6.0059 | 5.8442 | 5.0816 | 7.1128 | 7.1306 | 6.7211 | 7.0786 | 5.9214 | 5.9347 | 5.8576 | 6.5045 |
| AKAP13_S2713 | AKAP13 | Q12802-2 | | 2703 | KLMRIPSFFPSPE | 8.9720 | 8.8118 | 8.2928 | 6.6586 | 5.3169 | 5.7420 | 5.1623 | 6.4322 | 6.1396 | 7.0451 | 7.5033 | 5.6151 | 3.0643 | 4.3397 | 5.7531 | 5.1513 |
| AKAP13_S2724 | AKAP13 | Q12802-2 | | 2502 | DLRRTESDSGLKK | 4.7409 | 4.5680 | 5.1638 | 5.1073 | 5.9624 | 6.1539 | 5.7701 | 5.5372 | 5.4117 | 5.2879 | 5.8297 | 5.6965 | 8.6741 | 9.5023 | 8.8007 | 7.7932 |
| AKAP13_S2732 | AKAP13 | Q12802-2 | | 1904 | RSQQSVSLSKSVS | 7.2045 | 7.7063 | 7.7950 | 7.1516 | 5.2743 | 6.4829 | 6.2779 | 6.8508 | 7.1757 | 7.2474 | 7.7433 | 8.3151 | 3.2406 | 4.9611 | 3.1025 | 3.4709 |
| AKAP13_S790 | AKAP13 | Q12802-2 | | 953 | LQEEQRTPPPGQD | 5.2533 | 6.1629 | 6.5599 | 6.2024 | 6.1003 | 6.2750 | 5.9207 | 6.8203 | 5.9895 | 6.4071 | 6.8748 | 6.5348 | 6.4973 | 6.0809 | 5.9463 | 6.3745 |
| AKAP13_S983 | AKAP13 | Q12802-2 | | 2724 | SIAKSGSLDSELS | 5.2861 | 5.7883 | 8.8070 | 6.5929 | 5.9709 | 5.8358 | 4.9285 | 6.4331 | 7.7760 | 5.2747 | 6.8040 | 5.2499 | 6.3893 | 7.3100 | 6.1421 | 5.4116 |
| AKAP13_T1149 | AKAP13 | Q12802-2 | | 1411 | LLASKQSPECENF | 5.8177 | 6.9121 | 7.0829 | 5.6481 | 6.3148 | 7.0868 | 6.2082 | 5.5582 | 6.4394 | 7.1382 | 7.3360 | 6.9866 | 4.2685 | 5.9436 | 5.7278 | 5.5312 |
| AKAP13_T2471 | AKAP13 | Q12802-2 | | 1687 | CTSAISSPLTKSI | 6.6743 | 7.4090 | 6.5884 | 6.8436 | 6.1805 | 6.9288 | 6.3394 | 6.9724 | 7.1987 | 7.2563 | 7.4526 | 7.0705 | 4.3160 | 4.6901 | 3.7272 | 4.3522 |
| AKAP13_T953 | AKAP13 | Q12802-2 | 1933;1936 | | NA | 7.0080 | 6.6283 | 7.4315 | 6.0464 | 6.3575 | 6.6691 | 6.4557 | 6.7314 | 6.9592 | 6.5263 | 7.1969 | 5.8621 | 4.7278 | 5.1241 | 5.3020 | 4.9738 |
| AKAP17A_S537 | AKAP17A | Q02040 | | 537 | VVPEDGSPEKRCP | 6.2859 | 6.6128 | 6.2058 | 6.2705 | 5.9401 | 5.6783 | 6.2696 | 5.6681 | 6.1155 | 5.8020 | 5.6970 | 5.8715 | 6.5825 | 6.9837 | 6.9952 | 7.0215 |
| AKAP17A_S584 | AKAP17A | Q02040 | | 584 | KCNREPSKGRGRA | 6.2883 | 6.0840 | 6.0083 | 6.3339 | 6.1314 | 5.5042 | 6.4201 | 5.1895 | 5.8957 | 5.6318 | 5.8460 | 5.9120 | 7.2213 | 7.3996 | 7.1003 | 7.0337 |
| AKAP17A_S633 | AKAP17A | Q02040 | | 633 | HAYKDDSPRRRST | 6.0502 | 6.2928 | 6.4403 | 7.2931 | 6.2723 | 7.3281 | 6.4810 | 6.0261 | 6.5165 | 6.3178 | 6.7693 | 6.6008 | 5.7278 | 5.7491 | 4.8759 | 5.2590 |
| AKAP2_S1009 | AKAP2 | Q9Y2D5-4 | | 979 | VLQSTQSPRTKNA | 6.0262 | 5.8503 | 5.9133 | 6.0424 | 6.4838 | 6.0426 | 6.2865 | 6.5674 | 6.5283 | 6.0013 | 6.0015 | 6.0563 | 6.7935 | 6.4153 | 6.5272 | 6.4640 |
| AKAP2_S352 | AKAP2 | Q9Y2D5-4 | | 862 | EPPQVSSPVQEKR | 5.8395 | 5.3803 | 6.0784 | 6.1952 | 6.5528 | 6.2280 | 5.8804 | 5.7142 | 6.1473 | 5.8000 | 6.1763 | 6.3146 | 7.6529 | 6.6947 | 6.4899 | 6.8554 |
| AKAP2_S366 | AKAP2 | Q9Y2D5-4 | | 1009 | KVKPPPSPTTEGP | 5.6578 | 5.9383 | 6.0086 | 6.0525 | 6.1594 | 5.3795 | 5.9013 | 6.1984 | 6.1681 | 5.9016 | 5.8314 | 6.3244 | 7.2211 | 6.7781 | 7.4359 | 7.0437 |
| AKAP2_S383 | AKAP2 | Q9Y2D5-4 | | 383 | SLQVPVSPSSTTS | 6.3353 | 5.6760 | 5.9628 | 6.5827 | 5.9918 | 6.0204 | 6.0614 | 6.3174 | 6.4343 | 5.4890 | 6.3571 | 6.4251 | 6.6570 | 6.3809 | 6.8266 | 6.4822 |
| AKAP2_S624 | AKAP2 | Q9Y2D5-4 | | 624 | AAKGQKSPGALET | 6.1733 | 6.4288 | 6.2164 | 6.6581 | 6.2370 | 6.4399 | 6.3796 | 6.3995 | 6.7086 | 6.7737 | 6.6989 | 6.5879 | 5.8193 | 5.5993 | 5.2365 | 5.6432 |
| AKAP2_S696 | AKAP2 | Q9Y2D5-4 | | 951 | RKQRTLSMIEEEI | 5.7989 | 5.5483 | 6.6624 | 6.2864 | 6.0681 | 5.7442 | 5.7777 | 6.1312 | 6.6104 | 6.1638 | 6.4205 | 6.0927 | 7.0438 | 6.7758 | 6.5348 | 6.3410 |
| AKAP2_S862 | AKAP2 | Q9Y2D5-4 | | 696 | SRSVNVSLTQEEL | 6.5218 | 6.1185 | 6.3998 | 6.6561 | 6.5448 | 6.3214 | 6.3057 | 6.4250 | 6.1976 | 6.3019 | 6.6142 | 6.5670 | 5.4627 | 5.7158 | 5.7781 | 6.0696 |
| AKAP2_S951 | AKAP2 | Q9Y2D5-4 | | 352 | IELSNSSPDPMAE | 6.3838 | 6.4484 | 7.0814 | 6.5175 | 5.8894 | 5.8588 | 5.8871 | 6.1904 | 6.7631 | 6.7016 | 6.7990 | 7.1956 | 5.3715 | 6.1191 | 5.1780 | 5.6154 |
| AKAP2_S979 | AKAP2 | Q9Y2D5-4 | | 366 | ERTNGHSPSQPRD | 6.3604 | 6.8404 | 6.2622 | 6.7633 | 6.1747 | 5.9340 | 6.2144 | 6.0371 | 6.0503 | 6.1454 | 6.5194 | 6.8475 | 6.2124 | 5.8707 | 5.8975 | 5.8703 |
| AKAP2_Y304 | AKAP2 | Q9Y2D5-4 | | 322 | VLVGGLSPPVHEA | 5.6867 | 5.5617 | 6.1665 | 6.3331 | 6.3218 | 5.3951 | 5.7891 | 6.7470 | 6.4034 | 6.2845 | 6.5344 | 6.3548 | 6.6402 | 6.8450 | 6.6827 | 6.2541 |
| AKAP2_Y304;S322 | AKAP2 | Q9Y2D5-4 | | 304 | SAVDGTYNGTSSP | 7.0583 | 5.5806 | 7.8993 | 7.3617 | 5.4154 | 6.9051 | 5.8089 | 6.1513 | 5.0249 | 7.1124 | 5.9170 | 6.7790 | 6.6829 | 4.8537 | 5.8539 | 5.5956 |
| AKAP2_Y322 | AKAP2 | Q9Y2D5-4 | 304;322 | | NA | 6.5181 | 5.7137 | 6.4404 | 5.7173 | 6.6549 | 6.6315 | 6.1760 | 4.7177 | 5.9527 | 6.9686 | 7.4713 | 6.8910 | 5.7980 | 7.1075 | 6.1392 | 5.1020 |
| AKAP5_S178 | AKAP5 | P24588 | | 81 | TRGAWASLKRLVT | 6.6872 | 6.7592 | 7.7376 | 6.5056 | 5.4380 | 6.8403 | 6.7269 | 6.5322 | 6.7691 | 7.5026 | 7.4957 | 7.5437 | 4.0100 | 5.3835 | 3.5756 | 4.4928 |
| AKAP5_S81 | AKAP5 | P24588 | | 178 | QATKAKSTQDLSE | 5.8658 | 5.8099 | 6.0785 | 6.1294 | 6.3906 | 6.7340 | 6.7596 | 6.3666 | 6.0224 | 5.7423 | 7.1695 | 6.8124 | 6.5755 | 6.8124 | 5.2134 | 5.5176 |
| AKAP8_S182 | AKAP8 | O43823 | | 339 | AAGDFRSGDEEFK | 5.9885 | 6.7517 | 6.4626 | 7.0527 | 5.3627 | 4.7699 | 5.8068 | 5.8416 | 6.1187 | 6.1379 | 5.3044 | 6.1044 | 6.8116 | 6.1305 | 7.2071 | 8.1490 |
| AKAP8_S323;S328;S339 | AKAP8 | O43823 | | 182 | PARERGSLDGFMR | 6.3370 | 8.1670 | 8.3922 | 7.8598 | 6.6518 | 7.0357 | 7.2686 | 7.6525 | 6.3831 | 5.7944 | 6.0760 | 6.7747 | 4.8500 | 3.7572 | 3.3656 | 3.6343 |
| AKAP8_S328 | AKAP8 | O43823 | | 328 | DSEGDFSENDDAA | 6.0092 | 5.6705 | 5.8120 | 6.3072 | 5.6956 | 6.2154 | 6.0575 | 5.8535 | 6.9561 | 6.9657 | 7.1520 | 6.6636 | 5.2134 | 6.6541 | 6.2351 | 6.5391 |
| AKAP8_S339 | AKAP8 | O43823 | 323;328;339 | | NA | 5.9478 | 5.8890 | 6.7651 | 6.1630 | 6.2652 | 6.1498 | 6.4006 | 5.7579 | 6.3501 | 5.3764 | 6.5330 | 6.5036 | 7.3508 | 6.5109 | 6.0236 | 6.0131 |
| AKAP9_S3580 | AKAP9 | Q99996-6 | | 43 | AQSDGQSPSKKQK | 6.1600 | 6.4326 | 6.7184 | 6.4108 | 6.3534 | 6.2954 | 6.5658 | 6.3557 | 6.7969 | 6.4680 | 6.8051 | 6.5814 | 5.0005 | 5.8335 | 5.6365 | 5.5860 |
| AKAP9_S3846 | AKAP9 | Q99996-6 | | 3846 | DLDYIRSPLPFQN | 6.4193 | 6.3880 | 5.9004 | 6.5898 | 6.0192 | 6.7822 | 6.7605 | 6.8731 | 6.4606 | 6.4049 | 6.0879 | 6.2507 | 5.6645 | 6.2537 | 5.5133 | 5.6318 |
| AKAP9_S43 | AKAP9 | Q99996-6 | | 3580 | EEPSLVSPSTSCG | 8.7151 | 7.4612 | 6.4389 | 6.5010 | 5.8289 | 6.9811 | 5.7554 | 7.8396 | 5.9418 | 5.0664 | 8.8732 | 5.3657 | 5.6820 | 5.7046 | 3.6713 | 4.1740 |
| AKIRIN1_S22 | AKIRIN1 | Q9H9L7 | | 22 | ALLSPGSPKRRRC | 7.3347 | 5.6368 | 7.4417 | 5.9153 | 7.0985 | 4.9458 | 5.4241 | 7.2854 | 6.3477 | 7.7049 | 7.1197 | 8.8036 | 4.4923 | 5.7202 | 3.8413 | 4.8883 |
| AKIRIN2_S111 | AKIRIN2 | Q53H80 | | 21 | PLLSPASPKRRRC | 6.5686 | 5.8863 | 7.2434 | 5.8102 | 6.5724 | 5.4877 | 5.7353 | 7.1397 | 6.2689 | 7.3723 | 7.2201 | 8.4165 | 4.3216 | 5.3904 | 4.8354 | 5.7313 |
| AKIRIN2_S21 | AKIRIN2 | Q53H80 | | 111 | TDPCCTSDAQPHA | 6.2932 | 6.2093 | 5.8892 | 5.8841 | 7.1489 | 6.8889 | 6.5957 | 6.8060 | 5.3454 | 7.6731 | 7.6607 | 6.8796 | 5.0584 | 5.0926 | 4.9320 | 5.6427 |
| AKNA_S1102 | AKNA | Q7Z591 | | 534 | SARGDLSPSSLTS | 6.4778 | 6.8740 | 5.5430 | 6.2242 | 5.4909 | 4.9592 | 5.4978 | 7.5934 | 5.8766 | 6.3562 | 7.6907 | 5.0912 | 4.9974 | 6.5717 | 7.5517 | 7.2042 |
| AKNA_S52 | AKNA | Q7Z591 | | 52 | LFPNATSPELLED | 6.4692 | 6.1432 | 6.4141 | 6.2577 | 5.7907 | 5.5903 | 6.2313 | 8.2533 | 6.0771 | 6.8238 | 5.1828 | 6.2643 | 6.6410 | 6.5727 | 5.1211 | 6.1674 |
| AKNA_S534 | AKNA | Q7Z591 | | 1102 | HQPLQGSPTRPAS | 6.9412 | 7.4524 | 7.1798 | 4.4817 | 6.4470 | 6.1176 | 7.0491 | 7.9466 | 6.4811 | 7.2479 | 7.7535 | 7.9068 | 3.7262 | 3.9477 | 3.6126 | 5.7086 |
| AKT1_S124 | AKT1 | P31749 | | 450 | AQMITITPPDQDD | 5.6393 | 6.2093 | 6.6197 | 5.9418 | 6.4107 | 6.2520 | 6.0544 | 6.3721 | 6.6116 | 6.2262 | 6.5084 | 6.1601 | 6.0484 | 6.4531 | 6.4199 | 6.0730 |
| AKT1_T443 | AKT1 | P31749 | | 124 | MDFRSGSPSDNSG | 6.0551 | 6.2313 | 6.3400 | 6.1749 | 6.0052 | 6.2475 | 6.4932 | 6.1768 | 6.9300 | 6.4104 | 6.7881 | 6.5753 | 5.9570 | 6.3172 | 5.4182 | 5.8797 |
| AKT1_T450 | AKT1 | P31749 | | 443 | YFDEEFTAQMITI | 6.7673 | 5.6742 | 5.7337 | 6.0698 | 5.6316 | 6.5759 | 6.4143 | 7.0693 | 6.3866 | 6.4887 | 6.4696 | 5.2807 | 6.0995 | 7.1714 | 5.2680 | 6.8992 |
| AKT1S1_S108;S112 | AKT1S1 | Q96B36-3 | | 203 | TQQYAKSLPVSVP | 7.9109 | 7.2891 | 7.9238 | 6.8313 | 6.7117 | 6.7780 | 6.2605 | 7.3349 | 6.4580 | 6.2629 | 7.0134 | 7.0721 | 3.8683 | 4.4900 | 3.6374 | 4.1577 |
| AKT1S1_S203 | AKT1S1 | Q96B36-3 | | 266 | PRPRLNTSDFQKL | 6.2667 | 5.4513 | 7.1586 | 5.9414 | 6.4882 | 6.2537 | 5.9085 | 6.7792 | 6.2891 | 5.6853 | 5.9596 | 6.1648 | 7.2173 | 6.6299 | 5.8872 | 5.9193 |
| AKT1S1_T266 | AKT1S1 | Q96B36-3 | 108;112 | | NA | 6.6436 | 6.1926 | 6.8174 | 6.2140 | 6.4843 | 6.2162 | 5.7669 | 6.4590 | 6.5516 | 6.3670 | 6.8661 | 6.9621 | 5.5678 | 5.9165 | 5.2244 | 5.7505 |
| AKT2_S447 | AKT2 | P31751 | | 451 | AQSITITPPDRYD | 5.9493 | 6.2025 | 6.7413 | 6.0451 | 6.0881 | 6.5757 | 6.2826 | 6.5925 | 6.3382 | 6.5660 | 6.4681 | 6.2235 | 5.6831 | 6.2388 | 6.0916 | 5.9137 |
| AKT2_T451 | AKT2 | P31751 | | 447 | DEFTAQSITITPP | 6.2488 | 6.5698 | 6.9639 | 5.4051 | 6.1703 | 6.4964 | 6.2842 | 6.3767 | 6.1404 | 6.9853 | 6.7556 | 6.1431 | 5.6352 | 6.1163 | 6.1031 | 5.6056 |
| AKT3_T447 | AKT3 | Q9Y243 | | 447 | AQTITITPPEKYD | 6.0358 | 6.2798 | 7.0316 | 5.7425 | 5.7831 | 6.3427 | 5.9041 | 7.2641 | 6.5884 | 6.5794 | 5.9503 | 5.7772 | 6.0320 | 6.7959 | 5.7373 | 6.1557 |
| ALAD_S244 | ALAD | P13716-2 | | 244 | RDAAKSSPAFGDR | 6.1900 | 5.4839 | 6.6106 | 5.2001 | 6.1666 | 6.4984 | 5.7198 | 6.6288 | 5.5778 | 6.9884 | 6.1847 | 7.2183 | 6.0225 | 6.5839 | 6.6762 | 6.2499 |
| ALDH18A1_S431 | ALDH18A1 | P54886 | | 431 | RLSLSTSKLNSLA | 5.6078 | 6.7195 | 5.7465 | 6.6029 | 6.2873 | 5.6179 | 6.1757 | 6.6381 | 5.5284 | 5.5917 | 5.3002 | 5.4088 | 6.7457 | 7.6021 | 6.8482 | 7.5790 |
| ALDH3A2_S293 | ALDH3A2 | P51648-2 | | 293 | HFKRILSLLEGQK | 4.5562 | 4.2342 | 5.3924 | 4.7337 | 5.1012 | 4.9352 | 5.2706 | 4.7956 | 4.4229 | 4.5522 | 4.2046 | 4.6273 | 11.6460 | 9.7487 | 11.4328 | 10.3462 |
| ALDOA_S93 | ALDOA | P04075-2 | | 93 | ADESTGSIAKRLQ | 5.8122 | 5.8403 | 6.8698 | 6.0248 | 6.2585 | 6.4844 | 5.6677 | 6.0761 | 6.5730 | 6.5312 | 6.2798 | 6.1774 | 6.4833 | 6.4566 | 6.5647 | 5.9001 |
| ALDOA_Y258 | ALDOA | P04075-2 | | 258 | DLKRCQYVTEKVL | 6.5324 | 7.3027 | 6.3421 | 6.4540 | 6.3320 | 6.9713 | 6.9043 | 7.5044 | 6.7854 | 6.3908 | 5.7154 | 6.0050 | 4.8250 | 6.0771 | 5.4207 | 4.4374 |
| ALG3_S11 | ALG3 | Q92685 | | 11 | LRKRGRSGSAAQA | 5.9022 | 5.7353 | 5.4980 | 6.0537 | 5.9449 | 5.8581 | 5.4105 | 5.9695 | 5.3563 | 4.9872 | 5.7758 | 5.6463 | 8.0431 | 7.5323 | 8.2760 | 8.0109 |
| ALG5_S62 | ALG5 | Q9Y673 | | 62 | LPSIWDSPTKQLS | 6.4109 | 6.6163 | 6.0919 | 6.8006 | 6.8263 | 6.0072 | 6.0545 | 6.1794 | 6.0114 | 7.4064 | 6.9513 | 6.2275 | 5.1577 | 5.6877 | 5.7377 | 5.8332 |
| ALKBH5_S252 | ALKBH5 | Q6P6C2-1 | | 361 | THRRRGSFSSENY | 4.7479 | 5.1259 | 5.4127 | 5.1925 | 6.3606 | 5.8966 | 5.4123 | 6.0696 | 7.0944 | 7.2380 | 6.5748 | 6.5728 | 7.2770 | 7.5753 | 6.5060 | 6.9438 |
| ALKBH5_S361 | ALKBH5 | Q6P6C2-1 | | 96 | RQMRLFSQDECAK | 4.4590 | 4.9525 | 5.8161 | 4.7748 | 5.5072 | 5.1136 | 6.9852 | 4.7170 | 5.0661 | 5.0372 | 5.1890 | 4.5506 | 9.4037 | 8.6835 | 10.3731 | 9.3714 |
| ALKBH5_S64 | ALKBH5 | Q6P6C2-1 | | 64 | RKYQEDSDPERSD | 5.1428 | 5.0505 | 5.5390 | 5.1864 | 5.2720 | 5.5083 | 5.4242 | 5.7984 | 6.2664 | 7.4607 | 6.9563 | 7.0645 | 6.8666 | 7.8139 | 7.6232 | 7.0270 |
| ALKBH5_S64;Y71 | ALKBH5 | Q6P6C2-1 | | 252 | LPVRRGSVTVLSG | 5.2662 | 4.5504 | 5.5844 | 5.3081 | 6.6602 | 6.4635 | 5.2662 | 6.4551 | 7.4389 | 6.2207 | 5.7309 | 5.0276 | 7.1668 | 6.6561 | 9.4650 | 6.7398 |
| ALKBH5_S96 | ALKBH5 | Q6P6C2-1 | 64;71 | | NA | 6.0944 | 6.3075 | 6.0723 | 6.4239 | 5.8732 | 6.1956 | 5.9844 | 6.0349 | 6.3547 | 6.2643 | 6.2607 | 6.4826 | 6.4561 | 6.4928 | 5.9691 | 6.7336 |
| ALMS1_S1635 | ALMS1 | Q8TCU4 | | 1635 | RQALLDSPLNKEV | 7.5638 | 7.8306 | 6.7471 | 7.7709 | 6.7881 | 6.3658 | 6.5399 | 7.2901 | 8.5883 | 6.7415 | 6.9013 | 6.6295 | 2.9546 | 3.9393 | 3.5480 | 3.8012 |
| ALMS1_S2105 | ALMS1 | Q8TCU4 | | 92 | RILPPLSPPQHRY | 6.5182 | 6.4165 | 7.3319 | 5.9436 | 6.3357 | 7.0369 | 6.0702 | 5.9926 | 8.0082 | 7.1580 | 6.9577 | 6.3163 | 4.8403 | 4.9770 | 4.5981 | 5.4987 |
| ALMS1_S2770 | ALMS1 | Q8TCU4 | | 2105 | EKSNIFSPQELPG | 5.8337 | 7.0338 | 6.6777 | 6.7426 | 6.0074 | 6.9443 | 8.5252 | 5.7951 | 6.6426 | 6.7303 | 6.5180 | 6.3531 | 4.3038 | 4.9003 | 5.1688 | 5.8232 |
| ALMS1_S464 | ALMS1 | Q8TCU4 | | 464 | LRMLRMSPDTVPK | 6.9213 | 7.1792 | 7.3824 | 6.5358 | 6.3163 | 6.0052 | 6.5351 | 6.8512 | 6.8981 | 6.9852 | 7.1541 | 7.3821 | 4.5297 | 4.3314 | 4.2460 | 4.7469 |
| ALMS1_S92 | ALMS1 | Q8TCU4 | | 2770 | LKQKTSSPSSFKM | 7.4954 | 7.2802 | 6.6187 | 6.9987 | 6.6687 | 7.0626 | 7.4770 | 6.6874 | 7.3100 | 7.2366 | 7.0266 | 7.0917 | 3.2990 | 3.8885 | 3.5418 | 4.3173 |
| ALS2_S1464 | ALS2 | Q96Q42 | | 466 | QGKKSSSLVDIRE | 6.0229 | 6.0116 | 6.1518 | 5.7053 | 5.6751 | 5.7662 | 6.1197 | 6.7023 | 7.1876 | 6.3462 | 6.2093 | 6.9271 | 5.9172 | 6.6358 | 6.2539 | 6.3679 |
| ALS2_S466 | ALS2 | Q96Q42 | | 492 | GLLSQVSPRLLRK | 7.1216 | 8.4729 | 6.7708 | 7.0495 | 5.1596 | 7.6402 | 7.1157 | 6.9657 | 4.1130 | 4.2890 | 5.0391 | 4.4709 | 5.3805 | 5.6592 | 6.7649 | 7.9874 |
| ALS2_S483;S492 | ALS2 | Q96Q42 | | 510 | TRTVVLTPTYSGE | 5.7761 | 6.4823 | 6.7251 | 6.2490 | 6.4992 | 6.6487 | 6.3444 | 6.1981 | 5.7465 | 5.4772 | 6.2108 | 5.9362 | 6.3476 | 6.6031 | 5.5164 | 7.2393 |
| ALS2_S492 | ALS2 | Q96Q42 | | 1464 | SDSRSESPEPGYV | 5.4963 | 7.1467 | 6.4999 | 5.2825 | 5.7771 | 6.3981 | 6.4641 | 5.7894 | 5.9785 | 7.8382 | 6.5704 | 6.0143 | 5.4873 | 6.7640 | 6.2202 | 6.2728 |
| ALS2_T510 | ALS2 | Q96Q42 | 483;492 | | NA | 6.0865 | 6.3804 | 6.9861 | 6.0026 | 6.2342 | 6.4999 | 5.8932 | 6.6026 | 7.0373 | 6.8354 | 6.8249 | 6.5476 | 5.1589 | 5.7474 | 5.6924 | 5.4707 |
| ALYREF_S239 | ALYREF | Q86V81 | | 8 | ADKMDMSLDDIIK | 6.9139 | 7.0132 | 6.4853 | 6.9493 | 6.4701 | 6.9607 | 7.0386 | 6.8340 | 6.8405 | 6.8916 | 6.8017 | 6.7840 | 4.2202 | 4.8521 | 4.0631 | 4.8816 |
| ALYREF_S8 | ALYREF | Q86V81 | | 239 | NSKQQLSAEELDA | 6.5654 | 5.7793 | 6.1940 | 5.6206 | 7.0506 | 5.6844 | 6.5008 | 7.0818 | 6.6542 | 6.1553 | 5.9912 | 6.5570 | 5.9145 | 6.9815 | 5.3700 | 5.8993 |
| AMBRA1_S639 | AMBRA1 | Q9C0C7-3 | | 693 | DSLRRLSPAAYYA | 6.2904 | 6.8663 | 7.5776 | 6.1834 | 7.4747 | 7.2402 | 9.2315 | 7.1310 | 6.0910 | 5.7357 | 6.9618 | 6.5435 | 3.9647 | 4.3192 | 4.2421 | 4.1466 |
| AMBRA1_S693 | AMBRA1 | Q9C0C7-3 | | 639 | ELSSSASPQEERT | 5.9866 | 6.5179 | 6.6632 | 6.4317 | 6.1404 | 6.4555 | 6.9703 | 6.5074 | 6.2944 | 6.6173 | 6.8449 | 7.2228 | 5.6475 | 5.3202 | 5.0531 | 5.3268 |
| AMER1_S175 | AMER1 | Q5JTC6 | | 318 | LFGDVTSLKSFDS | 6.0983 | 5.9310 | 6.3212 | 6.8021 | 6.3764 | 5.8968 | 6.0133 | 5.7452 | 6.3944 | 6.1791 | 5.9124 | 6.0145 | 6.6583 | 6.5323 | 6.4158 | 6.7089 |
| AMER1_S318 | AMER1 | Q5JTC6 | | 175 | LKGFFSSIRRHRK | 8.6853 | 7.4046 | 8.2712 | 6.6633 | 5.1102 | 6.0010 | 7.4139 | 5.9714 | 7.4461 | 5.7060 | 6.9032 | 6.4345 | 4.8469 | 5.0242 | 3.1910 | 4.9272 |
| AMER1_S868 | AMER1 | Q5JTC6 | | 868 | LPWGVSSLPRYLG | 6.7372 | 6.2759 | 6.2609 | 6.5587 | 7.4127 | 5.7090 | 6.1331 | 5.5847 | 6.7194 | 5.9189 | 6.4630 | 6.8786 | 5.2134 | 5.6497 | 6.5851 | 5.8996 |
| AMFR_S542 | AMFR | Q9UKV5 | | 542 | RVPLDLSPRLEET | 7.3079 | 8.3591 | 7.3329 | 7.5380 | 6.3410 | 6.7590 | 6.9938 | 6.6443 | 6.9593 | 7.4709 | 7.3324 | 6.7965 | 3.1863 | 4.0143 | 3.1737 | 3.7906 |
| AMMECR1L_S74 | AMMECR1L | Q6DCA0 | | 74 | TLGPGNSPITRMN | 6.3478 | 5.9312 | 6.8494 | 6.3019 | 6.5457 | 6.4203 | 6.0768 | 7.2110 | 6.0462 | 6.4335 | 7.0828 | 6.6126 | 5.5934 | 6.0754 | 5.5989 | 4.8730 |
| AMOT_S1041 | AMOT | Q4VCS5 | | 200 | VTSAPLSPPQPND | 6.7580 | 6.6006 | 6.8887 | 7.3547 | 6.6784 | 6.2648 | 6.3947 | 6.9300 | 6.7819 | 6.7071 | 7.0758 | 7.0656 | 4.8751 | 4.5187 | 4.4582 | 4.6479 |
| AMOT_S200 | AMOT | Q4VCS5 | | 823 | DKSWKGSLGILLG | 5.8758 | 7.2980 | 7.2827 | 6.9691 | 4.3557 | 7.2065 | 5.8824 | 7.2130 | 4.7063 | 5.9085 | 8.1582 | 8.3651 | 3.8853 | 4.1553 | 5.4032 | 7.3350 |
| AMOT_S823 | AMOT | Q4VCS5 | | 1041 | PGPHRLSIPSLTC | 5.4100 | 6.5177 | 6.3793 | 5.9579 | 6.2763 | 5.8626 | 5.5772 | 6.2251 | 7.4266 | 7.4415 | 6.2500 | 6.7150 | 6.2241 | 6.6590 | 5.5752 | 5.5025 |
| AMOT_T1061 | AMOT | Q4VCS5 | | 1061 | PVFHSNTLERKTP | 6.6131 | 6.7526 | 6.5323 | 5.7346 | 6.7592 | 5.9961 | 6.3332 | 7.1029 | 6.4625 | 6.8121 | 6.1327 | 5.9146 | 6.0490 | 6.1092 | 5.0479 | 5.6480 |
| AMOTL1_S322 | AMOTL1 | Q8IY63 | | 828 | EKTWKGSIGLLLG | 6.0269 | 6.8729 | 6.1920 | 5.8467 | 5.3550 | 6.4947 | 6.3727 | 5.9655 | 5.4587 | 5.7975 | 6.3068 | 6.2381 | 6.3518 | 6.3872 | 6.9129 | 7.4206 |
| AMOTL1_S805 | AMOTL1 | Q8IY63 | | 322 | KTKQMMSPVSKTQ | 6.9911 | 6.9817 | 6.0600 | 6.0351 | 6.7765 | 6.9765 | 6.1737 | 6.6832 | 6.9381 | 6.4410 | 6.6777 | 6.8184 | 5.0393 | 5.5042 | 4.8837 | 5.0197 |
| AMOTL1_S828 | AMOTL1 | Q8IY63 | | 805 | AATGTHSRQTSLT | 5.6383 | 6.3675 | 5.6004 | 6.0985 | 6.2381 | 6.6443 | 6.0115 | 6.5832 | 6.1466 | 6.3297 | 5.3823 | 5.8711 | 7.9017 | 6.6793 | 5.9044 | 6.6032 |
| AMOTL1_S906 | AMOTL1 | Q8IY63 | | 906 | STTPAHSPVLKHP | 5.7831 | 5.1901 | 4.5804 | 5.9395 | 7.9771 | 5.8611 | 5.5119 | 6.3049 | 5.2261 | 4.3738 | 4.6065 | 4.7134 | 9.4178 | 8.6470 | 9.0817 | 6.7855 |
| AMOTL1_T902;S906 | AMOTL1 | Q8IY63 | 902;906 | | NA | 6.0818 | 5.7584 | 6.3025 | 6.5718 | 6.2475 | 7.2590 | 6.3480 | 7.1263 | 6.8467 | 6.7036 | 6.1558 | 5.3164 | 6.0306 | 5.9985 | 5.6453 | 5.6078 |
| AMOTL2_S817 | AMOTL2 | Q9Y2J4-4 | | 817 | SSQRAASLDSVAT | 6.0333 | 7.1922 | 5.9025 | 5.4336 | 6.7315 | 6.4301 | 5.9607 | 6.8351 | 6.9923 | 5.9139 | 6.4981 | 6.3069 | 6.1381 | 5.6931 | 5.9412 | 5.9974 |
| AMOTL2_T106 | AMOTL2 | Q9Y2J4-4 | | 106 | GAGTGGTGSPQAS | 5.3648 | 5.5970 | 5.5117 | 5.5039 | 6.1424 | 5.9116 | 6.5064 | 6.1192 | 6.6313 | 6.3326 | 6.2491 | 6.2564 | 6.6473 | 7.5275 | 6.5105 | 7.1882 |
| AMPD2_S100 | AMPD2 | Q01433 | | 168 | RLERQISQDVKLE | 6.0827 | 6.2780 | 7.4208 | 5.6108 | 6.5286 | 6.6955 | 6.3400 | 6.5358 | 6.5220 | 6.8021 | 6.7421 | 6.5581 | 4.8428 | 6.2877 | 5.2073 | 5.5458 |
| AMPD2_S168 | AMPD2 | Q01433 | | 100 | PLQSARSLPGPAP | 6.5959 | 6.6484 | 7.4694 | 6.2970 | 6.9516 | 6.8037 | 6.3562 | 6.7562 | 6.4408 | 6.8249 | 6.6838 | 5.8915 | 4.0312 | 6.4044 | 4.8974 | 4.9476 |
| AMPD2_S190 | AMPD2 | Q01433 | | 289 | YEQGPDTPVSADA | 5.0724 | 6.6631 | 7.2742 | 6.2791 | 5.8858 | 6.0221 | 5.8281 | 5.9859 | 6.8863 | 5.6965 | 6.1090 | 6.6647 | 6.0759 | 6.0305 | 7.2219 | 6.3045 |
| AMPD2_T289 | AMPD2 | Q01433 | | 190 | DFLKTDSDSDLQL | 4.5323 | 4.9976 | 5.3074 | 5.2043 | 5.7365 | 5.3340 | 5.9214 | 5.8668 | 4.9881 | 5.5953 | 5.9637 | 5.1955 | 9.1308 | 8.5829 | 8.9486 | 8.6948 |
| AMPD3_S94 | AMPD3 | Q01432-4 | | 94 | KMIRSQSLSLQMP | 4.9134 | 6.5703 | 8.3123 | 5.8095 | 4.9260 | 7.1314 | 6.6805 | 6.3930 | 7.6660 | 7.6520 | 6.6903 | 6.7838 | 4.2587 | 6.1264 | 4.0047 | 6.0817 |
| ANAPC1_S1058 | ANAPC1 | Q9H1A4 | | 688 | DFEGSLSPVIAPK | 5.9085 | 6.2427 | 6.6628 | 6.6447 | 6.2223 | 6.4159 | 6.4588 | 6.3563 | 6.4498 | 6.4365 | 6.5284 | 6.6030 | 5.7449 | 5.7610 | 5.7626 | 5.8019 |
| ANAPC1_S1347 | ANAPC1 | Q9H1A4 | | 1347 | HREKHKSPSYQIK | 6.2425 | 6.6380 | 6.3809 | 6.5556 | 6.1150 | 6.2752 | 6.3053 | 6.1366 | 6.6141 | 6.3808 | 6.5130 | 6.6183 | 5.5122 | 5.8619 | 5.7574 | 6.0932 |
| ANAPC1_S355 | ANAPC1 | Q9H1A4 | | 569 | SSKLHDSLYNEDC | 6.3110 | 6.5888 | 6.6252 | 6.4003 | 6.4031 | 6.5513 | 6.0929 | 6.3874 | 7.4568 | 6.3484 | 6.3839 | 6.7140 | 5.0736 | 5.6416 | 5.2856 | 5.7362 |
| ANAPC1_S355;S362 | ANAPC1 | Q9H1A4 | | 377 | ISSHNQSPKRHSI | 6.7758 | 7.7889 | 6.4854 | 7.6384 | 6.9536 | 7.0791 | 6.3750 | 7.3721 | 6.9158 | 6.5814 | 7.5369 | 7.4282 | 3.4956 | 4.1926 | 3.5922 | 3.7890 |
| ANAPC1_S377 | ANAPC1 | Q9H1A4 | | 731 | NRSLCLSPSEASQ | 6.0929 | 5.9200 | 6.4725 | 6.5302 | 5.6778 | 6.0571 | 5.8109 | 6.4572 | 6.3929 | 6.7286 | 6.6344 | 6.4992 | 5.9769 | 6.1338 | 6.1344 | 6.4812 |
| ANAPC1_S51;S60 | ANAPC1 | Q9H1A4 | | 355 | ALSRAHSPALGVH | 6.4543 | 8.0883 | 6.6925 | 7.6123 | 6.4824 | 7.3573 | 6.9126 | 7.8098 | 6.9761 | 7.0908 | 7.6050 | 7.5430 | 2.6773 | 3.6648 | 3.3270 | 3.7065 |
| ANAPC1_S547;S563 | ANAPC1 | Q9H1A4 | | 1058 | VQYPELSDHEFIE | 5.7196 | 6.0481 | 5.9008 | 6.3936 | 6.3650 | 6.4239 | 6.0126 | 6.0168 | 6.1760 | 5.8443 | 6.1980 | 5.7640 | 6.7659 | 6.8288 | 6.8434 | 6.6992 |
| ANAPC1_S555 | ANAPC1 | Q9H1A4 | | 727 | SHLLNRSLCLSPS | 5.7103 | 6.1780 | 6.6799 | 6.6803 | 6.6302 | 6.4732 | 6.4561 | 5.6932 | 6.6430 | 6.2423 | 6.5424 | 6.4664 | 6.0274 | 5.9127 | 5.5547 | 6.1098 |
| ANAPC1_S563 | ANAPC1 | Q9H1A4 | | 555 | DEVVLLSPVPELR | 6.5868 | 6.3586 | 6.3110 | 6.6784 | 7.4094 | 5.5434 | 6.5600 | 6.3220 | 6.6491 | 6.2268 | 6.4148 | 6.2830 | 6.9859 | 6.1414 | 4.3253 | 5.2041 |
| ANAPC1_S569 | ANAPC1 | Q9H1A4 | | 60 | AAGLVGSLQEVTI | 5.9001 | 6.5606 | 5.6722 | 6.2304 | 5.9466 | 5.4762 | 6.0322 | 6.4554 | 5.6004 | 6.1360 | 6.1991 | 7.0454 | 6.5301 | 7.2726 | 6.5875 | 6.3552 |
| ANAPC1_S60 | ANAPC1 | Q9H1A4 | | 563 | VPELRDSSKLHDS | 6.7160 | 6.1286 | 6.7143 | 6.0592 | 6.7219 | 5.8948 | 6.3966 | 6.2661 | 6.9279 | 6.2805 | 6.4425 | 6.5513 | 5.6296 | 5.8632 | 5.6476 | 5.7601 |
| ANAPC1_S686;S688 | ANAPC1 | Q9H1A4 | 686;688 | | NA | 6.5050 | 7.2528 | 6.6481 | 7.2487 | 6.5851 | 6.9838 | 6.8952 | 7.2029 | 7.0718 | 6.9141 | 7.2582 | 7.2160 | 3.9783 | 4.5991 | 3.3275 | 4.3134 |
| ANAPC1_S688 | ANAPC1 | Q9H1A4 | 51;60 | | NA | 6.3409 | 6.4911 | 6.4271 | 6.3010 | 6.4418 | 6.5212 | 6.2432 | 6.3453 | 6.7142 | 6.1006 | 6.3125 | 6.4715 | 5.9143 | 5.9073 | 5.7032 | 5.7649 |
| ANAPC1_S727 | ANAPC1 | Q9H1A4 | 547;563 | | NA | 6.8195 | 8.0565 | 8.1538 | 5.5502 | 6.4163 | 5.7209 | 5.8175 | 5.8132 | 7.3962 | 6.1536 | 7.1071 | 7.5714 | 3.5858 | 5.3222 | 4.7250 | 5.7908 |
| ANAPC1_S731 | ANAPC1 | Q9H1A4 | 355;362 | | NA | 7.8387 | 7.7409 | 6.5419 | 7.4975 | 6.9694 | 6.6431 | 7.0646 | 6.6846 | 7.2929 | 6.5745 | 6.8734 | 6.7212 | 3.6722 | 4.6255 | 3.5815 | 3.6780 |
| ANAPC2_S218 | ANAPC2 | Q9UJX6 | | 534 | LHQFSFSPEREIR | 5.9058 | 6.8819 | 6.6412 | 6.8512 | 6.4975 | 6.4620 | 6.1245 | 6.4206 | 6.4939 | 6.2021 | 6.4637 | 6.4011 | 5.7639 | 5.8286 | 5.4336 | 5.6282 |
| ANAPC2_S314 | ANAPC2 | Q9UJX6 | | 218 | YYRLLQSPLCAGC | 6.4485 | 6.4527 | 7.3495 | 6.4648 | 6.3323 | 6.7121 | 6.5015 | 6.3086 | 6.7061 | 6.7583 | 6.7660 | 6.2276 | 4.9680 | 5.8115 | 4.7445 | 5.4480 |
| ANAPC2_S534 | ANAPC2 | Q9UJX6 | | 314 | GPARPASPEAGNT | 6.2508 | 7.3897 | 6.6275 | 5.5281 | 5.7164 | 6.6713 | 6.5750 | 5.4142 | 5.3616 | 5.7471 | 5.5237 | 7.6044 | 6.8466 | 6.3822 | 6.3910 | 5.9705 |
| ANAPC4_S778 | ANAPC4 | Q9UJX5-3 | | 778 | IKEEVLSESEAEN | 6.0295 | 6.2389 | 6.2034 | 6.5154 | 6.4436 | 6.5520 | 6.0608 | 6.6352 | 6.4710 | 6.3086 | 6.6592 | 6.2850 | 5.8961 | 6.1633 | 5.7584 | 5.7797 |
| ANAPC5_S195 | ANAPC5 | Q9UJX4 | | 195 | KEELDVSVREEEV | 6.2865 | 6.4874 | 6.7349 | 6.8073 | 6.2124 | 6.7379 | 6.4244 | 6.7583 | 7.1485 | 6.1746 | 6.6280 | 6.5735 | 4.9903 | 5.3177 | 5.1052 | 5.6131 |
| ANAPC5_T232 | ANAPC5 | Q9UJX4 | | 232 | DETKALTPASLQK | 6.6263 | 6.7323 | 6.6145 | 7.2346 | 5.9664 | 6.4317 | 6.6231 | 6.1706 | 6.5302 | 6.2795 | 6.6747 | 6.3054 | 5.4609 | 5.6691 | 5.2531 | 5.4275 |

| | | | | | | | | | | | | | | | | | | | | |
|---|---|---|---|---|---|---|---|---|---|---|---|---|---|---|---|---|---|---|---|---|
| ANK2_S1831 | ANK2 | Q01484 | 3793 | AMIVPSSPSKTPE | 6.0290 | 6.0618 | 6.8202 | 5.8410 | 6.1185 | 6.1360 | 6.1004 | 6.6078 | 6.4237 | 6.2702 | 6.2823 | 6.3479 | 6.1373 | 6.3466 | 6.3911 | 6.0864 |
| ANK2_S1846 | ANK2 | Q01484 | 846 | TEVLDVSDEEGDD | 5.8625 | 5.8000 | 6.3114 | 5.9296 | 6.1081 | 6.1787 | 6.0782 | 6.2201 | 6.3510 | 6.4762 | 5.8946 | 6.8828 | 6.1666 | 6.6464 | 6.5946 | 6.4992 |
| ANK2_S1858 | ANK2 | Q01484 | 1905 | ERHPPVSPSGKTD | 6.5093 | 6.4826 | 6.2335 | 6.0810 | 6.5155 | 6.9282 | 6.1102 | 6.9492 | 6.5682 | 6.3739 | 6.2151 | 6.3026 | 5.5785 | 5.8674 | 5.4773 | 5.8076 |
| ANK2_S1882 | ANK2 | Q01484 | 3814 | EKLYLQTPTSSER | 6.2858 | 6.7137 | 6.4350 | 6.4292 | 6.2251 | 6.6597 | 6.4222 | 6.6707 | 6.9130 | 6.2496 | 6.4845 | 6.2908 | 5.7055 | 5.6444 | 5.2831 | 5.5876 |
| ANK2_S1905 | ANK2 | Q01484 | 3845 | SSPRKTSLVIVES | 5.8017 | 5.1661 | 5.7991 | 5.4023 | 6.1310 | 5.4569 | 5.3924 | 6.0745 | 6.6799 | 6.8527 | 6.8564 | 6.5431 | 7.2041 | 7.9060 | 6.4391 | 6.2946 |
| ANK2_S1964 | ANK2 | Q01484 | 1882 | EKHSPVSPSTKTE | 6.2888 | 7.1530 | 7.0821 | 6.4609 | 6.3464 | 6.6764 | 6.5796 | 6.2869 | 6.5024 | 5.7753 | 6.4590 | 6.9021 | 5.3277 | 5.6935 | 5.0975 | 5.3684 |
| ANK2_S31 | ANK2 | Q01484 | 1858 | EKHSPVSPSAKTE | 5.5497 | 6.3536 | 7.6880 | 6.4518 | 6.5304 | 6.8305 | 6.1043 | 6.0811 | 6.3009 | 7.2289 | 7.6943 | 6.7537 | 3.5622 | 6.1079 | 5.3576 | 5.4049 |
| ANK2_S3793 | ANK2 | Q01484 | 1846 | ERHPPVSPSSKTE | 6.3753 | 6.5111 | 6.9041 | 5.8123 | 6.5179 | 6.5355 | 6.9887 | 6.8575 | 6.1903 | 6.6314 | 6.6220 | 6.5431 | 5.3666 | 5.3185 | 4.8359 | 5.9898 |
| ANK2_S3845 | ANK2 | Q01484 | 1964 | EKHLPVSPSGKTE | 6.3759 | 6.4013 | 6.8482 | 6.0306 | 5.7191 | 6.4143 | 5.8025 | 6.2413 | 6.6312 | 8.0090 | 6.8626 | 6.7026 | 4.6835 | 5.8443 | 5.6608 | 5.7729 |
| ANK2_S3910 | ANK2 | Q01484 | 31 | RPKKSDSNASFLR | 5.2163 | 4.9682 | 5.8002 | 5.2017 | 5.7073 | 5.4024 | 5.8776 | 6.0765 | 5.1024 | 4.8568 | 5.6950 | 5.2289 | 8.9138 | 8.8751 | 9.1024 | 7.9756 |
| ANK2_S846 | ANK2 | Q01484 | 1831 | PKTERHSTLSSSA | 6.0469 | 5.5276 | 6.3303 | 5.8498 | 4.2907 | 5.7136 | 5.7413 | 6.3990 | 6.8109 | 6.2739 | 6.4145 | 7.4343 | 4.2985 | 8.0466 | 7.2073 | 7.6148 |
| ANK2_T3814 | ANK2 | Q01484 | 3910 | IRRYVSSEGTEKE | 4.1437 | 4.7519 | 4.6617 | 4.5257 | 5.0428 | 5.1598 | 4.6919 | 5.2682 | 4.4938 | 4.4025 | 4.7483 | 5.0478 | 10.6415 | 10.3005 | 11.5514 | 10.5686 |
| ANK3_S131 | ANK3 | A0A087WZ26 | 131 | DSRFLVSFMVDAR | 6.9949 | 8.2850 | 4.5825 | 7.2231 | 7.1857 | 7.8723 | 5.1008 | 9.5683 | 5.7536 | 5.5476 | 5.8051 | 5.2135 | 3.9758 | 5.9631 | 6.1490 | 4.7796 |
| ANKHD1_S1670 | ANKHD1 | Q8IWZ3-6 | 2281 | SQRVSTSPVGLPS | 5.8980 | 6.0961 | 6.2248 | 6.3795 | 6.1037 | 6.2002 | 6.3333 | 6.4585 | 6.0626 | 6.6148 | 7.0726 | 6.5226 | 5.8746 | 6.4009 | 5.7527 | 6.0052 |
| ANKHD1_S1679 | ANKHD1 | Q8IWZ3-6 | 1538 | RANVVTTPSTNRK | 6.5789 | 6.9485 | 7.5184 | 6.5888 | 5.8279 | 7.1706 | 6.7942 | 6.8178 | 7.1196 | 7.1199 | 7.5625 | 7.5419 | 3.8471 | 4.6113 | 3.6360 | 4.3167 |
| ANKHD1_S2281 | ANKHD1 | Q8IWZ3-6 | 1670 | VSLPLSSPNIKLN | 7.0119 | 6.5566 | 6.5249 | 6.4326 | 5.9658 | 7.2620 | 6.9772 | 7.6301 | 7.8330 | 6.5316 | 7.1686 | 7.3745 | 3.9044 | 4.4225 | 3.8172 | 4.5868 |
| ANKHD1_S2281;S2291 | ANKHD1 | Q8IWZ3-6 | 2539 | LKGEIPSPQLTRP | 6.6637 | 6.4864 | 6.7678 | 6.3769 | 6.4518 | 7.0444 | 6.6207 | 7.0896 | 7.1353 | 7.0358 | 7.1437 | 6.8952 | 4.4023 | 5.2108 | 4.1855 | 4.4900 |
| ANKHD1_S2291 | ANKHD1 | Q8IWZ3-6 | 1679 | IKLNLTSPKRGQK | 6.5327 | 6.4298 | 7.0630 | 5.8555 | 6.6651 | 6.5948 | 6.5161 | 6.3843 | 7.2193 | 6.5062 | 6.5887 | 6.7628 | 5.3853 | 5.5174 | 4.9257 | 5.0533 |
| ANKHD1_S2539 | ANKHD1 | Q8IWZ3-6 | 95 | GGGASGSDEDEVS | 4.9872 | 5.9105 | 6.9755 | 6.2527 | 5.6887 | 5.4978 | 6.6150 | 6.3630 | 7.3758 | 6.7259 | 6.7389 | 6.8540 | 6.0528 | 6.5112 | 5.6104 | 5.8405 |
| ANKHD1_S2556 | ANKHD1 | Q8IWZ3-6 | 2291 | LPSIDPSGSSPSS | 5.7817 | 5.7017 | 6.5002 | 6.7041 | 6.4373 | 6.3051 | 6.4480 | 6.6958 | 6.3869 | 6.6490 | 6.3490 | 6.5251 | 6.0899 | 6.9584 | 4.9867 | 5.4811 |
| ANKHD1_S95 | ANKHD1 | Q8IWZ3-6 | 2556 | GRPMVASPNQRHQ | 6.7405 | 7.5999 | 7.0929 | 6.7150 | 6.6446 | 6.9940 | 6.4210 | 6.2638 | 6.6455 | 6.5371 | 7.2471 | 6.8411 | 4.8016 | 5.1039 | 4.1650 | 4.1871 |
| ANKHD1_T1538 | ANKHD1 | Q8IWZ3-6 | 2323 | GPAPVGTPSFNRQ | 6.9565 | 7.0645 | 6.8717 | 6.9227 | 5.6758 | 6.5665 | 7.0248 | 5.7589 | 7.3897 | 6.7489 | 6.7161 | 6.7757 | 4.0442 | 5.3680 | 4.5379 | 5.5781 |
| ANKHD1_T2323 | ANKHD1 | Q8IWZ3-6 | 2281;2291 | NA | 5.9665 | 6.7541 | 6.6328 | 6.8016 | 6.2272 | 6.3419 | 6.5086 | 6.6404 | 7.0818 | 6.3841 | 6.7043 | 6.6102 | 5.1970 | 5.8272 | 5.1460 | 5.1761 |
| ANKIB1_S304 | ANKIB1 | Q9P2G1 | 737 | GVAPADSPEAPRR | 6.6710 | 6.7346 | 6.6209 | 6.4834 | 6.2716 | 6.0887 | 5.8267 | 6.0367 | 6.6283 | 6.5633 | 6.6202 | 6.6982 | 5.7366 | 5.9653 | 5.3703 | 5.6843 |
| ANKIB1_S737 | ANKIB1 | Q9P2G1 | 304 | AWDTLPSPRTPRT | 5.1631 | 7.8369 | 6.7472 | 6.3538 | 6.1136 | 8.2564 | 6.2424 | 6.1205 | 6.5644 | 6.4878 | 7.3756 | 7.2729 | 6.2772 | 4.1099 | 4.6931 | 4.3850 |
| ANKIB1_S911 | ANKIB1 | Q9P2G1 | 911 | SPRAALSSSELLE | 6.2823 | 6.4582 | 8.1607 | 7.4371 | 5.8992 | 6.6880 | 7.1165 | 4.7273 | 7.1477 | 6.6256 | 4.4890 | 7.3832 | 5.6382 | 5.1870 | 4.8039 | 5.9560 |
| ANKLE2_S259 | ANKLE2 | Q86XL3 | 528 | AFAGPLSPAKAED | 7.3484 | 7.2667 | 6.3017 | 7.1615 | 6.8477 | 6.8427 | 7.0799 | 6.7037 | 7.3459 | 6.7637 | 6.7929 | 6.7222 | 4.1943 | 4.5671 | 3.6227 | 4.4390 |
| ANKLE2_S268 | ANKLE2 | Q86XL3 | 662 | NAARNNSPPTVGA | 6.7620 | 6.7848 | 7.0365 | 7.0735 | 6.4204 | 6.0072 | 6.5576 | 5.8811 | 6.5708 | 6.6626 | 7.0988 | 6.7825 | 4.9906 | 5.4407 | 4.8074 | 5.1234 |
| ANKLE2_S512 | ANKLE2 | Q86XL3 | 896 | SGPHSYSPGRNSV | 6.6536 | 6.9750 | 6.6459 | 7.4639 | 6.8910 | 7.4050 | 7.2343 | 6.9337 | 7.2987 | 6.6982 | 7.1117 | 6.9442 | 3.8796 | 4.5757 | 3.4303 | 3.8593 |
| ANKLE2_S528 | ANKLE2 | Q86XL3 | 268 | KTSLPLSPVKTAP | 7.1133 | 6.9347 | 6.3998 | 7.2386 | 6.8041 | 6.8732 | 6.8521 | 6.6875 | 7.0574 | 6.7132 | 6.7321 | 6.8434 | 4.2511 | 4.7419 | 4.1176 | 4.6400 |
| ANKLE2_S662 | ANKLE2 | Q86XL3 | 914 | AKPGLGSPGRYSP | 7.0422 | 6.4047 | 6.8815 | 6.4554 | 7.0011 | 6.7970 | 7.3359 | 6.7684 | 7.3990 | 6.7631 | 7.0359 | 7.0232 | 4.6496 | 4.5034 | 3.8535 | 4.0860 |
| ANKLE2_S807 | ANKLE2 | Q86XL3 | 259 | ICDYFPSPSKTSL | 7.8603 | 7.4168 | 6.8049 | 7.6018 | 6.6372 | 7.3742 | 7.1983 | 7.1015 | 7.2971 | 7.2448 | 7.1705 | 7.2631 | 3.0683 | 3.6719 | 2.7852 | 3.5041 |
| ANKLE2_S866 | ANKLE2 | Q86XL3 | 866 | SAVLCYSPSDRQS | 7.5000 | 7.4246 | 6.9450 | 8.1190 | 6.3994 | 6.7357 | 7.4607 | 6.2973 | 6.8481 | 6.9900 | 7.2162 | 7.3989 | 3.3478 | 3.8701 | 3.4633 | 3.9839 |
| ANKLE2_S875 | ANKLE2 | Q86XL3 | 875 | DRQSWPSPAVKGR | 7.2123 | 6.8982 | 7.0839 | 7.7579 | 7.2976 | 7.4158 | 7.5431 | 7.2103 | 7.6650 | 7.4068 | 7.3224 | 7.1851 | 2.6526 | 3.9201 | 2.4145 | 3.0147 |
| ANKLE2_S896 | ANKLE2 | Q86XL3 | 512 | VSRYGGSPRDPVL | 7.6393 | 7.6838 | 7.2171 | 9.2070 | 6.6995 | 6.8692 | 6.9413 | 6.6910 | 8.3266 | 7.7178 | 7.8090 | 6.4534 | 2.3336 | 4.0392 | 2.0472 | 2.3251 |
| ANKLE2_S914 | ANKLE2 | Q86XL3 | 807 | MSLSPSSPRHEDQ | 6.7270 | 6.7842 | 6.6228 | 6.9015 | 6.2694 | 6.7336 | 7.0273 | 6.8425 | 6.6061 | 7.3709 | 7.0956 | 7.2909 | 4.1660 | 4.5731 | 4.0577 | 4.9314 |
| ANKRD11_S1354 | ANKRD11 | Q6UB99 | 1419 | EDELDKTIELFST | 6.7037 | 6.6295 | 5.9947 | 7.3249 | 6.1925 | 5.7985 | 6.2460 | 5.6208 | 6.3227 | 5.8442 | 6.0184 | 6.0084 | 5.4958 | 6.9693 | 6.0033 | 6.8274 |
| ANKRD11_S1509 | ANKRD11 | Q6UB99 | 505 | SGCLKGSPLVLKD | 6.8759 | 6.8854 | 5.6865 | 6.6875 | 6.2743 | 5.8428 | 6.2440 | 6.2026 | 6.0928 | 6.1145 | 5.7950 | 5.9047 | 5.5935 | 6.8248 | 6.3630 | 6.6127 |
| ANKRD11_S1792 | ANKRD11 | Q6UB99 | 1878 | VVTVTPSPEGVFS | 5.8182 | 6.2661 | 5.1940 | 6.1453 | 5.6854 | 5.7042 | 6.3123 | 5.8625 | 6.1665 | 6.1971 | 5.9371 | 6.2036 | 7.0312 | 7.6189 | 6.9923 | 6.8653 |
| ANKRD11_S1847 | ANKRD11 | Q6UB99 | 1792 | NLYRSVSVDIRRT | 5.6902 | 5.8294 | 4.9664 | 5.5721 | 5.9760 | 5.6680 | 6.0961 | 6.2148 | 5.1399 | 4.9165 | 5.0504 | 5.0328 | 7.9155 | 8.6333 | 8.5692 | 8.7293 |
| ANKRD11_S1847;S1852 | ANKRD11 | Q6UB99 | 1983 | SDLLLKSPQRFPE | 6.0487 | 6.0621 | 6.6026 | 6.7289 | 5.8734 | 6.7502 | 6.2430 | 5.8954 | 6.4344 | 6.1873 | 7.1057 | 7.0780 | 5.1161 | 6.2079 | 5.7455 | 5.9208 |
| ANKRD11_S1847;S1852;! | ANKRD11 | Q6UB99 | 1509 | ATRDKDSPPRVLK | 6.6842 | 6.5489 | 5.6170 | 7.2344 | 6.5950 | 5.8346 | 6.7851 | 6.2839 | 6.3059 | 6.3775 | 6.3898 | 6.5054 | 5.3189 | 6.1373 | 5.6131 | 5.7690 |
| ANKRD11_S1847;S1859 | ANKRD11 | Q6UB99 | 935 | SVPGYLSEKDKKR | 7.6246 | 8.0737 | 6.8366 | 7.4714 | 6.4791 | 6.3145 | 6.9558 | 7.0895 | 5.4146 | 6.2659 | 5.5225 | 5.7997 | 3.8313 | 5.8767 | 4.6080 | 5.8362 |
| ANKRD11_S1878 | ANKRD11 | Q6UB99 | 1798 | SVDIRRTPEEEFS | 6.4986 | 6.9792 | 6.6479 | 7.6690 | 6.0358 | 5.8072 | 6.3887 | 6.6524 | 6.6553 | 6.7762 | 6.4910 | 7.2521 | 4.5369 | 5.2044 | 5.0395 | 5.3659 |
| ANKRD11_S1983 | ANKRD11 | Q6UB99 | 609 | SLSEKKSPFLSSA | 6.2994 | 6.2446 | 5.5734 | 6.1416 | 6.1543 | 5.5320 | 6.2962 | 6.4137 | 6.8105 | 6.6036 | 6.7787 | 6.7068 | 5.5419 | 6.8890 | 5.6757 | 6.3386 |
| ANKRD11_S1983;S1990 | ANKRD11 | Q6UB99 | 1847 | EKFACLSPGYYSP | 6.6583 | 7.3735 | 6.3678 | 6.6509 | 6.3715 | 5.3788 | 7.8131 | 7.3009 | 6.2058 | 7.4890 | 7.3866 | 6.9284 | 3.1606 | 6.0046 | 4.2613 | 4.6487 |
| ANKRD11_S276 | ANKRD11 | Q6UB99 | 276 | PLKVANSPTMVNL | 6.1094 | 6.6007 | 6.5102 | 7.4273 | 5.7227 | 6.2795 | 6.7451 | 6.1589 | 6.4759 | 5.6719 | 6.7622 | 6.2599 | 5.3225 | 6.3658 | 5.4015 | 6.1865 |
| ANKRD11_S505 | ANKRD11 | Q6UB99 | 136 | NSPVDTTPKHPSQ | 7.1164 | 6.4106 | 6.2862 | 6.4785 | 5.8406 | 6.3292 | 7.1593 | 6.1618 | 5.9786 | 5.1145 | 6.9196 | 7.2182 | 5.7569 | 5.6822 | 5.3316 | 6.2161 |
| ANKRD11_S609 | ANKRD11 | Q6UB99 | 73 | NGEQKDSDTEKQG | 6.3662 | 6.3308 | 6.4287 | 6.5491 | 5.9769 | 5.6425 | 6.5084 | 6.1089 | 5.9093 | 5.9719 | 5.8555 | 5.8988 | 5.9840 | 6.9368 | 6.3345 | 7.1978 |
| ANKRD11_S73 | ANKRD11 | Q6UB99 | 1354 | ERHRHSSSSSKKS | 7.0052 | 7.9330 | 6.1171 | 7.0041 | 7.1114 | 7.0374 | 4.5736 | 7.2283 | 6.0152 | 6.3895 | 5.4488 | 5.9068 | 5.3480 | 6.1085 | 6.0227 | 4.7506 |
| ANKRD11_S935 | ANKRD11 | Q6UB99 | 1847;1852 | NA | 6.8252 | 7.0777 | 5.6451 | 6.9502 | 5.8020 | 5.6933 | 5.9549 | 6.8985 | 5.9598 | 6.3624 | 5.9547 | 5.6475 | 5.9017 | 6.8045 | 5.8752 | 6.6473 |
| ANKRD11_T136 | ANKRD11 | Q6UB99 | 1983;1990 | NA | 6.2909 | 6.1935 | 6.2405 | 5.9657 | 6.4905 | 6.0741 | 6.6536 | 5.5429 | 6.2654 | 6.3202 | 6.2221 | 5.9961 | 6.3205 | 6.5848 | 6.3423 | 6.4968 |
| ANKRD11_T1419 | ANKRD11 | Q6UB99 | 1847;1852;18! | NA | 5.0676 | 4.8105 | 4.6954 | 5.1542 | 5.7042 | 5.4939 | 5.9281 | 5.1325 | 5.0233 | 5.1852 | 5.0084 | 4.9421 | 8.6050 | 9.1504 | 9.5326 | 10.5666 |
| ANKRD11_T1798 | ANKRD11 | Q6UB99 | 1847;1859 | NA | 5.5601 | 7.1199 | 6.8757 | 4.7611 | 5.9305 | 6.7444 | 5.8840 | 6.0763 | 6.7978 | 4.5586 | 5.0572 | 6.2799 | 6.6150 | 7.0224 | 7.5647 | 7.1525 |
| ANKRD12_S1189 | ANKRD12 | Q6UB98 | 67 | KLPFTISPSRNEE | 6.1866 | 6.7645 | 6.2969 | 6.6356 | 6.6591 | 5.9891 | 6.2950 | 6.5643 | 6.3141 | 6.1110 | 5.8096 | 5.8187 | 5.5985 | 6.5294 | 6.2299 | 6.1980 |
| ANKRD12_S1311 | ANKRD12 | Q6UB98 | 1189 | DNSLNRSPRSENE | 6.5283 | 6.3839 | 5.8166 | 6.2313 | 6.5212 | 6.0053 | 6.1954 | 5.8022 | 6.0360 | 6.0930 | 5.7178 | 5.8822 | 6.2681 | 7.1071 | 6.6302 | 6.7815 |
| ANKRD12_S1345 | ANKRD12 | Q6UB98 | 1311 | IEVRRCSMPSVIC | 5.0862 | 4.0692 | 4.8509 | 4.1709 | 6.8636 | 4.4949 | 4.6948 | 5.6538 | 7.5046 | 9.2950 | 8.3016 | 8.6398 | 6.7655 | 8.7817 | 5.5096 | 5.3180 |
| ANKRD12_S1799 | ANKRD12 | Q6UB98 | 1799 | PQPVQVSPSLLQA | 6.0121 | 7.0681 | 6.8670 | 6.1643 | 5.5679 | 6.1579 | 6.5354 | 5.9869 | 6.9529 | 6.5912 | 6.0719 | 6.9431 | 5.4152 | 6.0363 | 5.7057 | 5.9242 |
| ANKRD12_S568 | ANKRD12 | Q6UB98 | 568 | HTKTCLSPGSSEM | 6.3663 | 5.9522 | 7.0861 | 5.7008 | 5.7717 | 5.8063 | 6.4026 | 6.3300 | 6.3802 | 6.3023 | 7.0595 | 7.2729 | 6.2175 | 5.7129 | 5.2873 | 6.3513 |
| ANKRD12_S67 | ANKRD12 | Q6UB98 | 1345 | PGDTSPSPKPEVF | 5.5483 | 6.0356 | 5.8441 | 6.5990 | 5.9968 | 5.7872 | 5.8595 | 5.3605 | 5.7867 | 5.1823 | 5.8133 | 5.9106 | 7.3933 | 7.5077 | 7.4737 | 7.9014 |
| ANKRD13D_S552;T556 | ANKRD13D | Q6ZTN6-3 | 552;556 | NA | 6.0787 | 5.9156 | 6.1526 | 5.9067 | 6.2066 | 6.6393 | 6.4100 | 6.1266 | 5.8580 | 6.4512 | 7.2532 | 6.4261 | 5.7712 | 6.4772 | 6.0021 | 6.3249 |
| ANKRD17_S1696 | ANKRD17 | O75179 | 1709 | GKLTVASPKRGQK | 6.4657 | 6.4621 | 7.2398 | 6.0059 | 6.5292 | 6.6074 | 6.4775 | 6.6722 | 7.1426 | 6.6237 | 6.5645 | 6.5693 | 5.0066 | 5.4490 | 4.9565 | 5.2280 |
| ANKRD17_S1709 | ANKRD17 | O75179 | 2067 | LDCGSASPNKVAS | 5.7116 | 5.8063 | 7.1192 | 6.0679 | 6.1390 | 6.1514 | 6.0741 | 6.5968 | 6.8343 | 6.9803 | 7.0507 | 7.5020 | 5.6607 | 5.3948 | 5.2603 | 5.6505 |
| ANKRD17_S19 | ANKRD17 | O75179 | 1940 | PARATNSPKPHMV | 6.7086 | 7.1104 | 7.3175 | 6.3606 | 6.2662 | 7.0676 | 6.9645 | 7.1587 | 7.5855 | 7.3696 | 8.0826 | 7.9638 | 3.5174 | 4.1923 | 2.8009 | 3.5339 |
| ANKRD17_S1940 | ANKRD17 | O75179 | 2401 | SGVRAPSPAPSSV | 5.4892 | 5.5668 | 5.6069 | 4.9284 | 5.4719 | 6.8997 | 6.5455 | 6.7201 | 6.2337 | 5.7985 | 5.8012 | 6.3246 | 6.5408 | 7.4617 | 6.8435 | 7.7676 |
| ANKRD17_S2045 | ANKRD17 | O75179 | 19 | AAEGEGSPPAVAA | 5.8323 | 7.0707 | 7.0752 | 7.0570 | 6.0258 | 5.0036 | 6.4947 | 7.2072 | 6.6040 | 5.7936 | 6.5744 | 7.4713 | 5.4498 | 5.2472 | 5.7549 | 5.3383 |
| ANKRD17_S2067 | ANKRD17 | O75179 | 1696 | CTKSGPSPLSSPN | 5.5201 | 6.3246 | 7.8142 | 6.8378 | 6.2126 | 5.9981 | 5.5580 | 6.7252 | 6.8204 | 6.9811 | 6.6474 | 7.1837 | 4.8494 | 5.8756 | 5.0668 | 5.5850 |
| ANKRD17_S2401 | ANKRD17 | O75179 | 2045 | PVSSPSSPSPPAQ | 5.0078 | 6.2668 | 6.2340 | 6.4548 | 7.6483 | 5.2176 | 5.4777 | 8.1291 | 5.7619 | 7.4078 | 6.4592 | 5.9805 | 7.0122 | 6.6952 | 5.0231 | 5.2242 |
| ANKRD26_S257 | ANKRD26 | Q9UPS8 | 631 | SKESVNSPVFGKA | 6.2389 | 6.6838 | 6.4801 | 6.0080 | 6.2907 | 6.4578 | 6.1899 | 6.5822 | 6.3010 | 6.3018 | 6.5874 | 6.4736 | 5.9326 | 6.2044 | 5.7133 | 5.5545 |
| ANKRD26_S261 | ANKRD26 | Q9UPS8 | 261 | DDSWPTSDDEDLN | 5.8526 | 5.9996 | 6.8875 | 6.2923 | 6.5209 | 6.7104 | 6.1352 | 6.3523 | 6.2477 | 6.0194 | 6.9546 | 6.5937 | 5.7390 | 6.0481 | 5.9109 | 5.7357 |
| ANKRD26_S312;S314 | ANKRD26 | Q9UPS8 | 489 | PVAHMESPERYLH | 6.4889 | 7.0265 | 6.7847 | 6.8608 | 6.6791 | 6.1891 | 5.9325 | 6.0267 | 7.0748 | 7.6255 | 7.3195 | 7.4081 | 4.6959 | 4.4427 | 4.4779 | 4.9673 |
| ANKRD26_S489 | ANKRD26 | Q9UPS8 | 654 | DDDSSLSEIDEDE | 5.4814 | 5.8868 | 5.3944 | 6.4195 | 6.0258 | 6.5156 | 5.8946 | 5.6373 | 5.6438 | 6.4169 | 6.2116 | 7.2822 | 7.7590 | 7.9487 | 6.1778 | 5.3047 |
| ANKRD26_S587 | ANKRD26 | Q9UPS8 | 587 | LIQKRKSGETDHQ | 5.7282 | 6.0073 | 6.9095 | 7.8436 | 6.7296 | 6.6413 | 6.7960 | 6.5858 | 6.2932 | 5.4280 | 8.0496 | 7.2172 | 5.8030 | 4.8286 | 4.0593 | 5.0800 |
| ANKRD26_S631 | ANKRD26 | Q9UPS8 | 257 | KPGVDDSWPTSDD | 5.6014 | 6.7921 | 6.9828 | 6.5620 | 6.0401 | 6.8672 | 6.4886 | 5.7811 | 5.9684 | 6.5898 | 6.6273 | 6.1967 | 5.6141 | 6.2458 | 5.7834 | 5.8592 |
| ANKRD26_S654 | ANKRD26 | Q9UPS8 | 312;314 | NA | 6.0335 | 6.5803 | 6.4983 | 5.9563 | 6.2156 | 7.0644 | 6.5705 | 6.2770 | 6.4666 | 6.6655 | 6.6576 | 6.0501 | 5.4738 | 6.0716 | 5.2480 | 6.1708 |
| ANKRD27_T1023 | ANKRD27 | Q96NW4 | 1023 | RMLRRHTVEDAVV | 6.8005 | 6.7384 | 7.1289 | 7.6962 | 6.1328 | 5.0713 | 4.6490 | 6.1885 | 6.5503 | 6.7782 | 5.2419 | 4.3494 | 6.7910 | 6.8197 | 6.4834 | 6.5806 |
| ANKRD28_S1044 | ANKRD28 | O15084-1 | 1044 | NTSKTVSFEALPI | 6.3988 | 6.0611 | 7.3380 | 6.2438 | 6.1117 | 6.6180 | 6.1871 | 6.5003 | 6.5064 | 6.8761 | 6.4905 | 7.5213 | 4.7204 | 5.9268 | 5.0968 | 5.4030 |
| ANKRD33B_S405 | ANKRD33B | A6NCL7 | 405 | PAPRKASLLPLQR | 6.3661 | 5.9319 | 6.9581 | 6.7476 | 6.0533 | 5.9024 | 5.8541 | 6.3481 | 8.2245 | 7.2836 | 7.8932 | 7.9149 | 4.8277 | 5.4120 | 4.0456 | 4.2369 |
| ANKRD33B_S416 | ANKRD33B | A6NCL7 | 416 | QRLRRRSVRPGVV | 7.3469 | 6.8581 | 7.4250 | 6.8863 | 5.9801 | 5.8119 | 6.1436 | 6.2200 | 7.3747 | 6.4717 | 7.3143 | 7.5203 | 4.7156 | 5.2871 | 3.8570 | 4.7874 |
| ANKRD34A_S461 | ANKRD34A | Q69YU3 | 461 | YAGAPGSPRTKRK | 6.3013 | 6.6721 | 6.4452 | 6.7366 | 6.1341 | 6.7413 | 6.3265 | 5.7418 | 6.1341 | 6.4536 | 6.0155 | 6.9654 | 6.4984 | 6.3069 | 5.3448 | 5.1823 |
| ANKRD40_T199 | ANKRD40 | Q6AI12 | 199 | PSAILRTPESTKP | 5.7452 | 6.7133 | 7.6590 | 6.8103 | 7.2188 | 7.3254 | 6.5812 | 8.0143 | 7.6736 | 7.3375 | 8.2484 | 7.2861 | 3.7837 | 3.4589 | 2.9689 | 3.1755 |
| ANKRD50_S1111 | ANKRD50 | Q9ULJ7 | 1183 | SSLSNNSLKSSKN | 5.6827 | 5.7413 | 6.4765 | 5.8762 | 5.4777 | 6.3255 | 5.9508 | 6.4956 | 6.1361 | 6.1128 | 6.4509 | 6.3098 | 6.6486 | 7.1487 | 6.7408 | 6.4262 |
| ANKRD50_S1162;S1167 | ANKRD50 | Q9ULJ7 | 1111 | LNGCSPSPVHTME | 5.8978 | 6.7394 | 6.2095 | 5.9342 | 7.3682 | 6.7206 | 6.0142 | 6.5848 | 5.7965 | 6.9947 | 7.0298 | 6.5041 | 6.4441 | 6.5502 | 5.2836 | 3.9284 |
| ANKRD50_S1165 | ANKRD50 | Q9ULJ7 | 1167 | FSSPSESPDSTVD | 6.9076 | 6.6680 | 6.2248 | 7.5052 | 5.3482 | 5.7783 | 5.2989 | 6.2083 | 6.8223 | 5.9949 | 6.1525 | 6.0869 | 6.1033 | 7.0685 | 6.3725 | 5.4598 |
| ANKRD50_S1167 | ANKRD50 | Q9ULJ7 | 1165 | HAFSSPSESPDST | 4.9108 | 5.4567 | 6.5089 | 5.8547 | 5.1400 | 7.4277 | 5.6713 | 6.3151 | 5.8297 | 6.2880 | 6.9902 | 6.1338 | 6.8214 | 6.6464 | 7.5756 | 6.4297 |
| ANKRD50_S1183 | ANKRD50 | Q9ULJ7 | 1162;1167 | NA | 5.7174 | 6.0407 | 7.3866 | 7.2083 | 6.0396 | 6.0237 | 5.5837 | 6.9152 | 6.8139 | 7.6161 | 6.8676 | 6.7162 | 4.3982 | 6.2205 | 5.2566 | 5.1957 |
| ANKRD52_S1028 | ANKRD52 | Q8NB46 | 1028 | PPKDAVSPFSFSL | 7.0031 | 7.6762 | 7.5699 | 7.3794 | 6.0226 | 7.0933 | 6.4769 | 7.5944 | 6.9953 | 6.5025 | 7.2898 | 7.6138 | 3.5800 | 3.7042 | 3.5567 | 3.9419 |
| ANKRD54_S63 | ANKRD54 | Q6NXT1 | 63 | ASGGAQSPLRYLH | 6.2682 | 6.4862 | 6.7151 | 5.0477 | 6.4565 | 6.4062 | 5.5186 | 6.6150 | 6.7246 | 6.7595 | 6.7873 | 6.9112 | 5.9398 | 5.9982 | 5.7851 | 5.5808 |
| ANKS1A_S647 | ANKS1A | Q92625 | 318 | TKEVDKTPPPQPP | 6.2421 | 6.6467 | 7.0444 | 6.9559 | 6.4062 | 6.6381 | 6.6798 | 6.6854 | 6.3953 | 6.8194 | 7.0610 | 6.9984 | 4.9061 | 5.1862 | 4.4796 | 4.8553 |
| ANKS1A_S661 | ANKS1A | Q92625 | 647 | MGSRSESLSNCSI | 6.6989 | 6.1618 | 7.1996 | 6.0648 | 6.4848 | 7.1543 | 6.1489 | 6.5963 | 6.7964 | 6.4336 | 6.5452 | 6.6559 | 5.3192 | 5.6717 | 4.9898 | 5.0787 |
| ANKS1A_S663 | ANKS1A | Q92625 | 661 | KKRLEKSPSFASE | 6.0744 | 5.7176 | 6.6423 | 6.2871 | 6.1295 | 6.0564 | 6.1911 | 6.4780 | 6.4822 | 6.5304 | 6.4035 | 7.0070 | 5.7171 | 6.3345 | 5.8687 | 6.0804 |
| ANKS1A_T318 | ANKS1A | Q92625 | 663 | RLEKSPSFASEWD | 5.5911 | 6.2642 | 6.0991 | 6.4268 | 6.6194 | 6.4843 | 6.0876 | 5.9673 | 6.5209 | 6.2527 | 6.2490 | 6.7233 | 5.6581 | 6.6498 | 6.0301 | 6.3759 |
| ANKZF1_S361 | ANKZF1 | Q9H8Y5 | 361 | EAVRLHSPQTHWK | 6.4841 | 6.8219 | 6.9194 | 6.4290 | 6.3903 | 6.4157 | 6.6148 | 6.6102 | 7.0856 | 7.1463 | 7.3484 | 7.1673 | 4.2755 | 5.0635 | 4.3997 | 4.8283 |
| ANKZF1_S675 | ANKZF1 | Q9H8Y5 | 675 | QLGAPTSPIPDSA | 6.1312 | 6.7054 | 7.0620 | 6.3219 | 5.9713 | 6.6386 | 6.7122 | 6.5959 | 6.8526 | 7.4313 | 6.5636 | 6.3099 | 4.8311 | 4.9932 | 5.0180 | 5.8617 |
| ANLN_S102 | ANLN | Q9NQW6 | 323 | LPKTPISPLKTGV | 5.5848 | 5.8579 | 5.9594 | 5.4878 | 5.8342 | 6.1357 | 6.0076 | 5.9669 | 6.1339 | 5.9995 | 6.2431 | 6.2977 | 6.7806 | 7.0819 | 7.3863 | 7.2428 |
| ANLN_S141 | ANLN | Q9NQW6 | 792 | QRKNKASPQSEFM | 5.1477 | 5.5242 | 5.9288 | 5.7981 | 5.3930 | 5.8842 | 5.8676 | 5.7824 | 5.8028 | 5.3537 | 5.7977 | 5.7660 | 7.5004 | 8.1019 | 8.1996 | 8.1521 |
| ANLN_S172 | ANLN | Q9NQW6 | 661 | RAESGDSLGSEDR | 6.0056 | 6.6408 | 6.8433 | 7.2596 | 6.1312 | 6.6438 | 7.2585 | 6.8777 | 6.0556 | 7.1576 | 7.3157 | 6.5859 | 4.5256 | 5.2900 | 4.3743 | 5.0347 |
| ANLN_S225 | ANLN | Q9NQW6 | 99 | AKSCSPSPVSPQV | 5.9425 | 6.1469 | 6.3822 | 6.0061 | 6.5244 | 6.6386 | 6.5085 | 6.7539 | 6.2926 | 6.4102 | 6.8117 | 6.3686 | 5.5646 | 6.2314 | 5.7530 | 5.6650 |
| ANLN_S245 | ANLN | Q9NQW6 | 449 | DKGNIWSAEKGGN | 7.5511 | 8.0171 | 7.0085 | 7.4640 | 5.6472 | 7.3984 | 8.0505 | 7.3960 | 7.9562 | 7.4221 | 8.0748 | 7.8766 | 2.0657 | 3.1870 | 1.9720 | 2.9127 |
| ANLN_S252 | ANLN | Q9NQW6 | 401 | HRTPIITPNTKAI | 7.0921 | 7.4203 | 7.1747 | 7.4803 | 5.9410 | 7.4847 | 7.6568 | 7.1312 | 7.7976 | 7.4290 | 7.9782 | 8.0141 | 2.3899 | 3.5386 | 2.2391 | 3.2326 |
| ANLN_S339 | ANLN | Q9NQW6 | 485 | GVSKTQSLPVTEK | 6.9367 | 5.0262 | 5.1378 | 6.1377 | 8.0953 | 5.2860 | 5.1830 | 6.2003 | 8.0390 | 5.8152 | 5.2124 | 6.0800 | 7.3963 | 7.3998 | 6.1490 | 5.9054 |
| ANLN_S356 | ANLN | Q9NQW6 | 252 | ARINSSSVKQEAT | 7.6582 | 6.6898 | 7.6174 | 6.9942 | 6.9156 | 6.4450 | 6.6165 | 6.7675 | 8.7638 | 7.7497 | 7.4052 | 7.1151 | 3.6295 | 4.0268 | 2.6159 | 2.9897 |
| ANLN_S417 | ANLN | Q9NQW6 | 913 | PKRLLTSITTKSN | 11.7145 | 6.5788 | 7.2439 | 7.8219 | 5.0170 | 5.5177 | 6.9966 | 5.9124 | 9.0624 | 6.6945 | 7.1988 | 7.2537 | 3.3745 | 4.1924 | 2.5248 | 2.8960 |
| ANLN_S449 | ANLN | Q9NQW6 | 483 | HQGVSKTQSLPVT | 6.7744 | 4.3622 | 5.2645 | 6.0672 | 8.3245 | 5.6371 | 5.1828 | 6.4385 | 8.4091 | 5.8450 | 5.0204 | 5.7785 | 7.3739 | 7.4996 | 6.1318 | 5.8904 |
| ANLN_S485 | ANLN | Q9NQW6 | 54 | SNQQPLSGGEEKS | 6.1045 | 6.9413 | 7.5751 | 6.9191 | 6.7935 | 7.2445 | 7.1431 | 7.8403 | 5.6609 | 5.5777 | 5.9361 | 6.0978 | 4.8884 | 5.1116 | 4.9364 | 5.2296 |
| ANLN_S54 | ANLN | Q9NQW6 | 194 | LSNASATPVGRRG | 7.9759 | 6.8510 | 7.4758 | 7.0726 | 4.7343 | 6.2717 | 7.8428 | 6.3830 | 7.6271 | 6.9459 | 7.4032 | 8.1481 | 3.3677 | 4.3501 | 3.1696 | 4.3812 |
| ANLN_S642 | ANLN | Q9NQW6 | 642 | TSRSDESPKPGKF | 6.2856 | 6.2984 | 7.7840 | 6.9290 | 6.6880 | 7.0488 | 7.4883 | 6.1942 | 7.0734 | 6.6063 | 6.9825 | 7.1364 | 4.1319 | 4.8924 | 3.9551 | 4.5058 |
| ANLN_S65 | ANLN | Q9NQW6 | 102 | CSPSPVSPQVQPQ | 5.7603 | 6.1479 | 6.0587 | 6.1598 | 6.3832 | 6.2533 | 6.3521 | 6.4084 | 6.5124 | 6.2056 | 6.6289 | 6.6533 | 5.8171 | 6.6257 | 5.9417 | 6.0915 |
| ANLN_S661 | ANLN | Q9NQW6 | 141 | EATAASSVKTRMQ | 9.8309 | 6.5859 | 7.4506 | 8.0458 | 7.0790 | 5.8345 | 6.4467 | 5.9998 | 10.0292 | 6.8949 | 6.9593 | 6.7714 | 3.5419 | 4.0506 | 2.1195 | 2.3601 |
| ANLN_S72 | ANLN | Q9NQW6 | 472 | ETHCQSTPLKKHQ | 7.4137 | 7.3157 | 6.8739 | 7.3239 | 5.3203 | 6.8218 | 7.5412 | 6.5730 | 7.5972 | 7.0387 | 7.2674 | 7.6520 | 3.4182 | 4.3560 | 3.2020 | 4.2850 |
| ANLN_S792 | ANLN | Q9NQW6 | 72 | PSKKRCSDNTEVE | 6.8538 | 6.8919 | 6.9608 | 8.0799 | 6.4433 | 7.3407 | 7.3268 | 7.0680 | 6.6672 | 7.3885 | 7.3644 | 7.0149 | 3.3180 | 4.1655 | 3.0973 | 4.0188 |
| ANLN_S913 | ANLN | Q9NQW6 | 245 | SSASGASARINSS | 8.3911 | 7.2307 | 8.2320 | 7.2895 | 5.1453 | 6.1310 | 6.7426 | 6.2796 | 8.2363 | 7.3288 | 7.9175 | 7.9969 | 3.2235 | 4.1058 | 2.5253 | 3.2241 |
| ANLN_S97;S102 | ANLN | Q9NQW6 | 172 | PESSLFSPMPSEE | 6.6441 | 6.7886 | 6.8500 | 6.8027 | 5.3800 | 7.3644 | 7.2439 | 6.6775 | 7.5342 | 6.8949 | 7.1838 | 7.7229 | 3.7716 | 4.4093 | 3.7173 | 5.0148 |
| ANLN_S99 | ANLN | Q9NQW6 | 356 | REICLQSQSKDKS | 6.5343 | 6.4998 | 6.7229 | 7.4908 | 5.9587 | 5.9458 | 7.4957 | 5.7930 | 7.1796 | 6.5048 | 7.1044 | 7.1944 | 4.4575 | 5.2605 | 4.4587 | 5.3992 |
| ANLN_T194 | ANLN | Q9NQW6 | 417 | LFKQDTSSSTTHL | 6.8395 | 7.7254 | 6.5559 | 7.6202 | 5.6509 | 7.0214 | 6.8132 | 7.6144 | 7.4609 | 6.1078 | 7.3533 | 7.5048 | 3.9150 | 4.4476 | 3.0991 | 4.2707 |
| ANLN_T300 | ANLN | Q9NQW6 | 363 | QSKDKSTTPGGTG | 8.1159 | 8.0325 | 7.1336 | 7.2580 | 5.9122 | 7.1592 | 8.4247 | 7.0165 | 7.8362 | 7.4733 | 7.9100 | 7.2820 | 2.0220 | 3.3513 | 2.1560 | 2.9166 |
| ANLN_T320;S323 | ANLN | Q9NQW6 | 225 | SFAKQNSVQEQPG | 5.8246 | 6.4576 | 8.4918 | 6.4925 | 5.2771 | 6.4519 | 6.7312 | 6.4760 | 7.7575 | 7.5657 | 7.8822 | 7.0293 | 4.1461 | 4.8207 | 4.0073 | 4.5884 |
| ANLN_T323 | ANLN | Q9NQW6 | 65 | KSCTKPSPSKKRC | 5.9279 | 5.8782 | 6.0696 | 5.4076 | 6.1939 | 6.3463 | 6.1712 | 6.0756 | 6.1182 | 6.1940 | 6.3777 | 6.3193 | 6.4784 | 7.0563 | 7.0120 | 6.3738 |
| ANLN_T363 | ANLN | Q9NQW6 | 397 | RSTPHRTPIITPN | 5.6976 | 6.7734 | 7.2459 | 7.2317 | 7.2193 | 6.0926 | 6.0926 | 7.4186 | 7.1022 | 6.4907 | 7.6291 | 6.0412 | 4.6437 | 5.1730 | 4.5047 | 4.6437 |
| ANLN_T397 | ANLN | Q9NQW6 | 339 | IVKSTLSQTVPSK | 6.5239 | 5.6063 | 5.7463 | 5.7923 | 7.4020 | 6.4662 | 6.1276 | 5.9536 | 7.4906 | 6.3081 | 5.8352 | 5.9319 | 6.4160 | 7.3904 | 5.4305 | 5.5792 |
| ANLN_T397;T401 | ANLN | Q9NQW6 | 300 | TSPVKSTTSITDA | 6.1513 | 5.8756 | 6.3486 | 5.5937 | 6.9832 | 5.5042 | 5.9042 | 6.2987 | 7.2511 | 7.0389 | 7.2188 | 6.3805 | 5.4598 | 6.6758 | 5.6479 | 5.6677 |
| ANLN_T401 | ANLN | Q9NQW6 | 397;401 | NA | 6.3434 | 7.6521 | 7.2586 | 8.3648 | 7.3541 | 7.3019 | 6.7694 | 7.1872 | 7.0046 | 6.9370 | 7.5581 | 7.6978 | 2.8330 | 3.7021 | 2.6907 | 3.3451 |
| ANLN_T472 | ANLN | Q9NQW6 | 97;102 | NA | 6.3491 | 6.8237 | 6.5972 | 6.7788 | 6.4718 | 6.2307 | 6.4613 | 6.3852 | 5.6272 | 7.1829 | 7.3934 | 7.0406 | 5.3274 | 6.0886 | 4.2399 | 5.0021 |
| ANLN_T483 | ANLN | Q9NQW6 | 320;323 | NA | 6.4667 | 6.3264 | 7.3025 | 7.4164 | 6.3500 | 8.6018 | 7.3522 | 6.3049 | 7.5757 | 7.2234 | 8.1786 | 7.6088 | 3.3126 | 3.9817 | 2.8626 | 3.1357 |
| ANO6_S177 | ANO6 | Q4KMQ2-2 | 177 | DLKNRSSAFGTLN | 5.7234 | 5.8847 | 6.4207 | 6.4055 | 6.1068 | 5.8918 | 5.6940 | 6.0373 | 6.1948 | 6.1211 | 5.8019 | 6.0255 | 6.5439 | 6.9614 | 7.2711 | 6.9163 |
| ANO8_S641 | ANO8 | Q9HCE9 | 641 | ALLEEGSPTMVEK | 6.4540 | 5.9963 | 6.5422 | 6.6871 | 6.3320 | 5.8212 | 5.7004 | 5.8188 | 6.5301 | 4.9818 | 5.1944 | 5.4504 | 7.2052 | 6.8272 | 7.0253 | 7.4335 |
| ANTXR1_S381 | ANTXR1 | Q9H6X2 | 563 | RPPPRPSVxxxxx | 6.2445 | 6.1433 | 5.4545 | 6.1207 | 5.8878 | 5.7365 | 6.1070 | 5.9699 | 7.2150 | 7.3887 | 6.9550 | 6.9840 | 5.6735 | 6.5125 | 5.6944 | 5.9127 |
| ANTXR1_S563 | ANTXR1 | Q9H6X2 | 381 | WPTVDASYYGGRG | 7.1151 | 6.1715 | 7.4892 | 6.1923 | 6.5233 | 5.4634 | 5.7272 | 5.6139 | 5.7720 | 6.3802 | 6.3399 | 6.6992 | 5.8226 | 7.3744 | 5.9195 | 5.3963 |
| ANXA1_S182 | ANXA1 | P04083 | 37 | GPGSAVSPYPTFN | 5.9233 | 6.3491 | 6.4151 | 6.2077 | 6.6779 | 7.0520 | 6.1282 | 6.7725 | 6.5944 | 6.2823 | 6.4448 | 6.6444 | 5.6512 | 6.1300 | 5.1188 | 5.6083 |
| ANXA1_S37 | ANXA1 | P04083 | 182 | FRNALLSLAKGDR | 6.0892 | 5.4328 | 7.3701 | 5.5235 | 6.5375 | 6.2013 | 6.7830 | 4.8511 | 6.1159 | 6.1639 | 6.8951 | 7.8664 | 5.1660 | 6.3294 | 5.6570 | 7.0178 |
| AP1G1_S767 | AP1G1 | O43747-2 | 767 | FQLQLLSPSSSIV | 6.1221 | 6.1800 | 7.4501 | 6.2698 | 5.9800 | 6.6522 | 6.8290 | 5.8818 | 6.4115 | 6.5297 | 6.3436 | 6.5001 | 5.4719 | 5.4368 | 5.7553 | 6.1860 |
| AP1M1_T152 | AP1M1 | Q9BXS5-2 | 152 | APRPPATVTNAVS | 10.8351 | 7.7003 | 6.9890 | 9.8344 | 5.8516 | 5.5623 | 5.9240 | 5.4819 | 5.5783 | 4.7022 | 3.6090 | 2.8575 | 5.3332 | 8.5082 | 5.5301 | 5.7029 |
| AP1M1_T152;T154 | AP1M1 | Q9BXS5-2 | 154 | RPPATVTNAVSWR | 8.7545 | 6.3613 | 8.1333 | 8.3437 | 7.0892 | 4.6944 | 6.3214 | 4.3889 | 5.2237 | 4.2874 | 5.5552 | 4.5292 | 7.0510 | 6.6509 | 6.0425 | 6.5733 |
| AP1M1_T154 | AP1M1 | Q9BXS5-2 | 152;154 | NA | 8.3628 | 6.5539 | 6.5642 | 8.3696 | 7.2074 | 4.7770 | 6.2109 | 5.0683 | 5.6691 | 4.8073 | 5.4082 | 5.3025 | 6.9325 | 6.7590 | 6.3597 | 5.6476 |
| AP1S1_S147 | AP1S1 | P61966 | 147 | LQEEDESPRSVLE | 5.9761 | 5.8380 | 6.3209 | 6.1275 | 5.9393 | 6.3266 | 6.5764 | 6.7761 | 6.5702 | 5.9761 | 6.2235 | 6.4651 | 6.3305 | 6.1203 | 6.3333 | 6.1002 |
| AP2A1_S655 | AP2A1 | O95782 | 655 | STVSTPSPSADLL | 5.7850 | 5.8777 | 5.9692 | 6.1897 | 5.8793 | 5.6151 | 5.9694 | 6.0145 | 5.9685 | 5.8012 | 5.8952 | 6.1828 | 7.0053 | 7.3380 | 7.1711 | 7.3381 |
| AP2M1_T156 | AP2M1 | Q96CW1 | 156 | QITSQVTGQIGWR | 6.4457 | 5.9857 | 6.2437 | 7.1143 | 6.2029 | 5.5232 | 6.0032 | 5.5096 | 6.8068 | 5.9449 | 6.6774 | 6.6223 | 6.0448 | 6.3259 | 5.4929 | 7.0568 |
| AP3B1_S276 | AP3B1 | O00203 | 276 | GKNFYESDDDQKE | 6.0376 | 6.1032 | 6.7215 | 5.8071 | 6.2389 | 6.0558 | 6.3516 | 6.2072 | 6.4073 | 6.3771 | 6.1926 | 6.4177 | 6.2081 | 6.5380 | 6.1465 | 6.1900 |
| AP3B1_S609 | AP3B1 | O00203 | 661 | ELAKEWTPAGKAK | 7.0702 | 7.2791 | 7.2204 | 6.7136 | 6.4761 | 7.1418 | 7.5273 | 6.6583 | 7.6753 | 6.7548 | 7.1364 | 7.2891 | 3.3959 | 4.4436 | 3.0960 | 4.1220 |
| AP3B1_S750 | AP3B1 | O00203 | 609 | PAPLLESPFKDRD | 7.3824 | 5.5608 | 7.2853 | 6.7379 | 6.1174 | 6.7003 | 7.7554 | 6.1607 | 7.1938 | 6.4620 | 6.9690 | 5.7706 | 4.8857 | 5.4154 | 4.7963 | 4.8070 |
| AP3B1_S750;S752 | AP3B1 | O00203 | 810 | KTKQDRTPLTKDV | 6.7377 | 7.6864 | 6.8577 | 8.2219 | 5.9804 | 7.0491 | 7.1982 | 6.7251 | 7.2928 | 7.3823 | 7.7622 | 7.8624 | 2.7140 | 3.9056 | 2.9297 | 3.6946 |
| AP3B1_T661 | AP3B1 | O00203 | 750 | SKAKGKSDSEDGE | 5.4527 | 6.5213 | 7.3080 | 6.4245 | 5.8511 | 5.9204 | 6.1808 | 6.5084 | 6.1464 | 6.6040 | 7.1868 | 7.6098 | 5.1887 | 6.1767 | 5.1559 | 5.7645 |
| AP3B1_T810 | AP3B1 | O00203 | 750;752 | NA | 6.0509 | 6.1622 | 6.8538 | 6.1426 | 5.8570 | 6.1498 | 5.9933 | 5.9032 | 6.0324 | 6.3838 | 6.3143 | 6.4632 | 6.4918 | 6.8485 | 6.2105 | 6.1428 |
| AP3D1_S597 | AP3D1 | O14617-2 | 786 | LDFWLSTTPPPAP | 6.1995 | 5.9872 | 6.8274 | 6.4396 | 6.0789 | 6.2200 | 6.1966 | 6.3482 | 6.6399 | 6.7347 | 6.6458 | 6.4095 | 5.1355 | 6.4435 | 5.6129 | 6.0806 |
| AP3D1_S738 | AP3D1 | O14617-2 | 597 | YIKSSPSPQKRYQ | 5.9091 | 5.7442 | 6.4080 | 5.8152 | 6.1411 | 6.2887 | 5.7562 | 6.4550 | 6.2289 | 6.0652 | 6.3296 | 6.6068 | 5.8483 | 7.2317 | 6.3828 | 6.7892 |
| AP3D1_T786 | AP3D1 | O14617-2 | 738 | NTETSKSPEKDVP | 6.1020 | 5.9248 | 6.8575 | 5.7777 | 6.2819 | 7.4720 | 6.4716 | 6.4869 | 6.6192 | 6.1973 | 8.0613 | 7.8376 | 5.1033 | 6.0049 | 4.4369 | 4.3649 |
| AP3D1_T786;S801 | AP3D1 | O14617-2 | 786;801 | NA | 6.0158 | 6.1021 | 7.0902 | 6.2832 | 6.7717 | 6.6937 | 6.1268 | 6.1442 | 5.7316 | 6.2585 | 6.1369 | 5.8437 | 5.9871 | 6.6119 | 5.9191 | 6.2836 |
| AP3S1_S183 | AP3S1 | Q92572 | 183 | INIGDISIKVPNL | 6.5040 | 5.5962 | 9.8465 | 6.4908 | 5.9406 | 6.0505 | 8.3130 | 5.3574 | 6.2021 | 5.5376 | 6.3670 | 5.6775 | 5.1551 | 6.5209 | 4.9962 | 5.4446 |
| APBA3_S9 | APBA3 | O96018 | 9 | FPTISRSPSGPPA | 6.4423 | 6.7565 | 7.0257 | 6.6422 | 6.2133 | 6.5786 | 6.5811 | 6.6090 | 6.3044 | 6.8855 | 7.0437 | 6.8005 | 5.2213 | 5.5253 | 4.7097 | 4.6609 |

| | | | | | | | | | | | | | | | | | | | | | |
|---|---|---|---|---|---|---|---|---|---|---|---|---|---|---|---|---|---|---|---|---|---|
| APBA3_T32 | APBA3 | O96018 | | 32 | VPSEDLTPDSQWD | 5.5404 | 6.2794 | 6.0773 | 7.2859 | 6.5435 | 5.7925 | 5.1433 | 7.4672 | 6.2276 | 5.8149 | 6.7231 | 6.6972 | 6.0014 | 6.7524 | 5.3090 | 6.3450 |
| APBB1_S205 | APBB1 | O00213 | | 205 | PREHSKSASLLFG | 4.2775 | 4.7218 | 3.4369 | 3.7542 | 4.6893 | 4.5391 | 4.5159 | 5.2218 | 3.9617 | 3.9214 | 4.1149 | 5.1692 | 12.0816 | 10.8029 | 12.0522 | 12.7396 |
| APBB2_S123 | APBB2 | Q92870-4 | | 331 | PADLQGSRKGSLS | 5.7162 | 5.3501 | 5.9361 | 5.9816 | 5.7390 | 5.7913 | 5.8006 | 5.7037 | 6.9840 | 7.1635 | 6.8132 | 6.7310 | 6.3630 | 7.0170 | 6.2012 | 6.7085 |
| APBB2_S160 | APBB2 | Q92870-4 | | 34 | NRNSPATPPNTLN | 5.8591 | 6.2650 | 6.2675 | 5.9577 | 6.2274 | 6.1920 | 6.8561 | 6.4273 | 6.5507 | 6.7056 | 6.6007 | 6.7607 | 5.6950 | 6.3433 | 5.5096 | 5.7824 |
| APBB2_S228 | APBB2 | Q92870-2 | | 334 | QGSRKGSLSSVTP | 5.2664 | 5.1134 | 5.8196 | 5.5022 | 6.3406 | 5.2612 | 5.3220 | 5.9383 | 6.5128 | 6.4592 | 6.6292 | 6.7410 | 7.2552 | 7.7782 | 7.4124 | 6.6485 |
| APBB2_S229 | APBB2 | Q92870-4 | | 160 | QPRRTKSFLNYYA | 7.0958 | 5.7330 | 6.5048 | 6.5551 | 5.9628 | 6.4046 | 6.0639 | 6.4258 | 6.6260 | 6.3906 | 6.6803 | 6.6476 | 5.7726 | 6.0174 | 5.3022 | 5.8176 |
| APBB2_S331 | APBB2 | Q92870-4 | | 229 | VATVSSSPETKKD | 5.6072 | 5.4102 | 6.4451 | 6.3204 | 5.9651 | 6.4218 | 6.3263 | 5.7760 | 6.5834 | 6.5753 | 6.4281 | 6.8250 | 6.1662 | 6.5195 | 5.8201 | 6.8104 |
| APBB2_S334 | APBB2 | Q92870-4 | | 335 | QGSRKGSLSSVTP | 5.4057 | 5.4971 | 5.7206 | 6.2546 | 7.1019 | 6.2150 | 6.5976 | 5.8104 | 7.0639 | 6.6989 | 6.3147 | 6.0394 | 6.7072 | 6.3180 | 6.5877 | 5.6672 |
| APBB2_S335 | APBB2 | Q92870-4 | | 123 | DPNKNLSPTAVIN | 6.8986 | 7.1045 | 7.0228 | 6.4915 | 6.1510 | 5.9335 | 7.2797 | 6.4926 | 6.7716 | 6.3975 | 6.1145 | 6.5935 | 5.1986 | 5.3523 | 4.9063 | 5.2914 |
| APBB2_S335;S342 | APBB2 | Q92870-4 | | 410 | DDDDSCSINSDPE | 5.6626 | 5.4974 | 6.0261 | 6.4654 | 5.4953 | 5.9705 | 6.0613 | 5.9112 | 6.6680 | 6.7703 | 5.9417 | 6.3619 | 7.1640 | 7.0860 | 6.3724 | 6.5458 |
| APBB2_S410 | APBB2 | Q92870-4 | | 228 | QVATVSSSPETKK | 5.5554 | 6.4028 | 7.1915 | 6.1378 | 6.6137 | 8.9061 | 6.3080 | 5.1144 | 6.5144 | 6.8929 | 8.9152 | 6.4383 | 6.3512 | 5.4993 | 3.7989 | 3.3604 |
| APBB2_S44 | APBB2 | Q92870-4 | | 44 | TLNLRSSHNELLN | 5.0938 | 5.8008 | 5.2083 | 4.8963 | 5.9051 | 6.5609 | 5.8197 | 5.7537 | 4.8892 | 5.5423 | 5.4085 | 5.1534 | 7.8917 | 8.5214 | 9.2497 | 8.3050 |
| APBB2_T34 | APBB2 | Q92870-4 | 335;342 | | NA | 4.7935 | 4.6973 | 5.8803 | 4.8258 | 6.1203 | 5.9400 | 6.0264 | 6.2908 | 6.0736 | 6.9362 | 7.2388 | 6.7584 | 6.6235 | 7.2593 | 7.2495 | 7.2862 |
| APC_S1100 | APC | P25054 | | 1438 | PPSRSKTPPPPPQ | 5.8909 | 6.1748 | 6.9477 | 6.1744 | 6.5490 | 6.3238 | 6.5889 | 6.3735 | 6.3015 | 6.3927 | 6.5013 | 6.6439 | 5.7915 | 6.0221 | 5.5928 | 5.7311 |
| APC_S1371 | APC | P25054 | | 1100 | GQQECVSPYRSRG | 5.8737 | 6.0682 | 6.4174 | 6.7733 | 6.0448 | 6.6336 | 7.0196 | 6.4840 | 6.7772 | 6.2032 | 6.4225 | 6.4918 | 5.3676 | 6.1013 | 5.4260 | 5.8958 |
| APC_S1567 | APC | P25054 | | 2449 | FIKEAPSPTLRRK | 6.5009 | 6.6741 | 6.4960 | 6.6375 | 6.4015 | 6.0797 | 6.0945 | 6.2160 | 6.5698 | 6.2178 | 6.3484 | 6.7083 | 5.5850 | 5.9372 | 5.5845 | 5.9489 |
| APC_S1774 | APC | P25054 | | 2323 | LSRPIQSPGRNSI | 6.7021 | 6.5999 | 6.8579 | 6.9890 | 6.2048 | 6.3286 | 6.3120 | 6.0681 | 6.4373 | 6.6740 | 6.9219 | 7.2454 | 4.8829 | 5.5539 | 4.8147 | 5.4074 |
| APC_S2146 | APC | P25054 | | 1371 | GAQTPKSPPEHYV | 5.8800 | 6.1170 | 6.1747 | 6.0346 | 6.5042 | 5.9916 | 6.4653 | 6.3160 | 7.0414 | 6.5698 | 6.0875 | 6.7285 | 6.0252 | 6.1048 | 5.8057 | 6.1538 |
| APC_S2283 | APC | P25054 | | 2789 | PFNYNPSPRKSSA | 6.5625 | 6.7909 | 6.4395 | 6.4814 | 6.0882 | 6.9848 | 6.9598 | 6.3851 | 6.9611 | 6.6495 | 6.8669 | 7.1076 | 4.7936 | 5.3348 | 4.6740 | 4.9203 |
| APC_S2323 | APC | P25054 | | 2768 | FSSSSSSKHSSPS | 6.3069 | 6.6590 | 6.8461 | 6.9647 | 6.0413 | 6.4573 | 6.7560 | 6.2601 | 6.7630 | 6.8959 | 6.7967 | 6.7184 | 4.8654 | 5.2143 | 5.0882 | 5.3667 |
| APC_S2368 | APC | P25054 | | 2368 | GKMSYTSPGRQMS | 6.7853 | 7.2223 | 6.7214 | 6.9774 | 6.1976 | 6.8957 | 6.4448 | 7.0253 | 6.9548 | 7.3324 | 7.3777 | 7.6434 | 3.6238 | 4.4472 | 3.8978 | 4.4532 |
| APC_S2374 | APC | P25054 | | 2724 | LENRLNSFIQVDA | 7.9403 | 6.1113 | 7.1794 | 6.9600 | 4.2969 | 4.8676 | 4.3457 | 5.3358 | 7.6769 | 5.9747 | 6.1454 | 5.5455 | 7.5550 | 6.0333 | 4.8920 | 9.1401 |
| APC_S2449 | APC | P25054 | | 2374 | SPGRQMSQQNLTK | 5.9158 | 5.7021 | 5.4020 | 6.8714 | 6.7874 | 5.1091 | 4.8786 | 6.0791 | 7.1643 | 6.5136 | 5.5533 | 6.4944 | 6.5929 | 6.9650 | 6.6001 | 7.3708 |
| APC_S2724 | APC | P25054 | | 1567 | KDLLDDSDDDDIE | 5.8398 | 7.0699 | 6.6330 | 6.5901 | 5.3912 | 6.3288 | 6.5537 | 6.3079 | 6.8021 | 6.5706 | 5.5342 | 6.1844 | 6.2949 | 6.1181 | 5.7982 | 5.9829 |
| APC_S2768 | APC | P25054 | | 2283 | SVKSELSPVARQT | 7.3868 | 6.5704 | 6.3113 | 7.5003 | 5.9906 | 5.9434 | 6.2031 | 6.1198 | 6.9212 | 7.0681 | 6.5992 | 7.0006 | 4.7007 | 4.9709 | 5.0155 | 5.6982 |
| APC_S2789 | APC | P25054 | | 2835 | QSPKRHSGSYLVT | 5.9376 | 5.7507 | 6.4025 | 5.9336 | 6.2289 | 6.2347 | 6.1170 | 6.9967 | 6.3565 | 5.8703 | 6.1665 | 6.4462 | 6.2744 | 6.0581 | 6.6831 | 6.5434 |
| APC_S2835 | APC | P25054 | | 2146 | SGISLGSPFHLTP | 6.3389 | 7.0284 | 7.7751 | 7.1639 | 6.4745 | 7.0401 | 6.1786 | 7.0179 | 6.9710 | 5.9858 | 7.4871 | 6.8563 | 4.1599 | 4.8272 | 4.9953 | 3.6999 |
| APC_T1438 | APC | P25054 | | 1774 | KKKKPTSPVKPIP | 4.7989 | 5.5948 | 6.1021 | 6.3411 | 5.7121 | 6.7086 | 5.5745 | 7.1933 | 6.8067 | 6.7007 | 6.6635 | 5.7380 | 5.9161 | 7.0378 | 6.8732 | 6.2385 |
| APH1A_S257 | APH1A | Q96BI3 | | 257 | SRVMVYSALRIPP | 5.8825 | 6.0173 | 5.2278 | 5.4978 | 6.0663 | 5.9745 | 6.1671 | 6.2241 | 5.7570 | 5.1887 | 5.2146 | 5.4260 | 7.6574 | 7.6321 | 8.0654 | 8.0013 |
| API5_S464 | API5 | Q9BZZ5 | | 464 | EDTTSGSPPKKSS | 5.6646 | 5.9696 | 5.9966 | 6.0146 | 6.0523 | 6.1766 | 6.1824 | 5.8456 | 6.4080 | 6.2612 | 6.3298 | 6.0590 | 6.7827 | 6.9022 | 6.7245 | 6.6303 |
| API5_S470 | API5 | Q9BZZ5 | | 470 | SPPKKSSAGPKRD | 6.3155 | 6.0760 | 6.7563 | 6.3500 | 5.5196 | 6.2814 | 5.8222 | 6.1349 | 6.1867 | 6.4290 | 7.2466 | 7.9225 | 5.5843 | 6.0774 | 5.5287 | 5.7688 |
| APIP_S87 | APIP | Q96GX9 | | 87 | KDISGPSPSKKLK | 6.2668 | 6.2717 | 6.5169 | 5.5821 | 6.2039 | 6.4094 | 5.9826 | 6.4786 | 6.3242 | 6.1756 | 6.4390 | 6.4909 | 5.7498 | 6.5807 | 6.5252 | 6.0026 |
| APLP2_S213 | APLP2 | Q06481 | | 213 | KIIGSVSKEEEEE | 7.2907 | 8.1660 | 6.8408 | 7.3033 | 5.7691 | 5.5892 | 5.6236 | 5.4122 | 6.9117 | 8.2846 | 6.9104 | 7.2196 | 4.1919 | 5.3048 | 4.5780 | 4.6039 |
| APPL1_S401 | APPL1 | Q9UKG1 | | 401 | LEAVTPSPSFQQR | 5.9129 | 6.4724 | 7.0172 | 5.3970 | 5.9634 | 6.8948 | 6.0609 | 7.1724 | 6.4773 | 6.2394 | 6.9325 | 7.2426 | 5.3333 | 5.6389 | 5.1989 | 6.0461 |
| APPL2_T352 | APPL2 | Q8NEU8-3 | | 352 | YCFQITTPNGKSG | 6.0256 | 6.7208 | 6.5196 | 6.9493 | 6.3083 | 6.2129 | 7.1741 | 6.6079 | 7.7971 | 6.5865 | 6.8087 | 5.6935 | 5.2690 | 5.4003 | 5.2275 | 4.6991 |
| APRT_S15 | APRT | P07741 | | 15 | VEQRIRSFPDFPT | 5.4555 | 6.2029 | 7.5932 | 4.6647 | 5.4451 | 6.3193 | 6.7607 | 6.9120 | 6.1733 | 5.8726 | 5.0714 | 5.8448 | 6.2932 | 8.3632 | 6.2515 | 6.7764 |
| ARAF_S186 | ARAF | P10398 | | 213 | LQRIRSTSTPNVH | 6.0218 | 6.4982 | 6.7952 | 6.6105 | 6.1064 | 6.2825 | 5.9945 | 6.3671 | 6.3070 | 6.3227 | 6.1832 | 5.9946 | 6.1001 | 6.2061 | 6.1694 | 6.0410 |
| ARAF_S257 | ARAF | P10398 | | 186 | LTPQGPSPRTQHC | 6.7066 | 5.5214 | 5.9307 | 7.5449 | 5.8699 | 7.5104 | 7.2441 | 6.2463 | 6.5455 | 6.5192 | 6.4699 | 5.4178 | 5.3061 | 5.8304 | 4.9987 | 6.3384 |
| ARAF_T213 | ARAF | P10398 | | 257 | DGTPRGSPSPASV | 6.1398 | 6.5558 | 7.1167 | 6.3091 | 6.1078 | 6.9186 | 6.1512 | 7.2461 | 6.3342 | 6.3168 | 6.9503 | 6.0266 | 5.0628 | 6.0441 | 5.1882 | 5.5319 |
| ARAP1_S207 | ARAP1 | Q96P48 | | 229 | FPEFDDSDYDEVP | 6.2212 | 5.5632 | 6.6087 | 6.3678 | 6.0894 | 6.4961 | 6.4847 | 5.7901 | 6.2971 | 6.7849 | 6.9560 | 6.4263 | 5.7980 | 6.4935 | 5.5693 | 6.0537 |
| ARAP1_S229 | ARAP1 | Q96P48 | | 207 | GPPQPPSPPPCPP | 5.7169 | 5.5511 | 6.8923 | 5.9628 | 5.9785 | 6.9225 | 6.1260 | 6.3213 | 5.9844 | 6.4315 | 6.4485 | 6.8045 | 6.4354 | 6.4498 | 5.9051 | 6.0696 |
| ARAP2_Y77 | ARAP2 | Q8WZ64 | | 77 | MQDIPIYANVHKT | 7.0726 | 6.4794 | 6.2371 | 5.7489 | 5.8899 | 4.4540 | 7.1630 | 6.4378 | 7.1214 | 6.0509 | 6.9207 | 5.7651 | 6.1847 | 5.6006 | 6.1015 | 6.7724 |
| ARAP3_S128 | ARAP3 | Q8WWN8 | | 128 | GPGVSRSPEPSPR | 5.6942 | 6.0337 | 7.4150 | 6.4729 | 6.5380 | 6.2125 | 6.6760 | 6.2724 | 6.2109 | 6.8326 | 6.3732 | 6.5308 | 5.3448 | 6.8781 | 5.0978 | 5.4172 |
| ARAP3_S1337 | ARAP3 | Q8WWN8 | | 1444 | PLTSQKSLDQPFL | 6.6294 | 7.1373 | 9.2119 | 6.3989 | 6.4986 | 6.3781 | 6.1593 | 6.7246 | 6.3929 | 9.0111 | 7.3132 | 6.0379 | 3.0632 | 5.6611 | 3.7460 | 3.6364 |
| ARAP3_S1380 | ARAP3 | Q8WWN8 | | 1337 | VVLRRHSSSDLAR | 7.0885 | 5.8428 | 4.9481 | 6.3371 | 5.5363 | 6.3223 | 4.9135 | 7.2912 | 7.0934 | 7.4740 | 7.5136 | 6.2679 | 5.8774 | 7.8349 | 4.2709 | 5.3880 |
| ARAP3_S1444 | ARAP3 | Q8WWN8 | | 1380 | HNRRTLSMFFPMK | 8.3863 | 5.7390 | 10.7842 | 6.9501 | 6.2654 | 5.1943 | 4.9691 | 4.7622 | 7.2087 | 5.9763 | 6.1224 | 5.6629 | 5.2430 | 8.0881 | 4.2540 | 4.3940 |
| ARAP3_S539 | ARAP3 | Q8WWN8 | | 539 | ALGSGISKVQSLK | 5.6673 | 6.3900 | 6.3053 | 6.7654 | 5.1045 | 4.4829 | 6.1467 | 4.6560 | 6.9200 | 5.9552 | 5.5246 | 5.7207 | 7.2943 | 6.3496 | 8.3879 | 8.3296 |
| ARF3_S147 | ARF3 | P61204 | | 147 | DKLGLHSLRHRNW | 6.6810 | 6.8441 | 6.7381 | 6.5319 | 6.1691 | 6.4409 | 6.0872 | 6.1779 | 6.7152 | 6.7693 | 6.2874 | 6.3840 | 5.1870 | 5.9628 | 5.5629 | 5.4611 |
| ARFGAP1_S369 | ARFGAP1 | Q8N6T3-2 | | 369 | STERRSSDSWEVW | 5.5949 | 5.6986 | 6.0186 | 6.0971 | 6.3465 | 5.7773 | 5.6533 | 6.7081 | 6.9735 | 6.9721 | 6.7953 | 6.6624 | 6.0850 | 6.7688 | 5.9434 | 5.9050 |
| ARFGAP1_T135 | ARFGAP1 | Q8N6T3-2 | | 135 | SPAQNWTPPQPRT | 6.3091 | 6.2892 | 6.3031 | 6.8867 | 6.2399 | 6.5861 | 7.0353 | 6.8459 | 6.3411 | 6.6128 | 7.0452 | 7.1657 | 4.8361 | 5.7395 | 4.5238 | 5.2406 |
| ARFGAP1_T189 | ARFGAP1 | Q8N6T3-2 | | 189 | YVGFGNTPPPQKK | 6.9922 | 7.2174 | 6.6848 | 7.0697 | 6.2293 | 6.7315 | 6.9655 | 6.7247 | 7.7419 | 7.0985 | 7.6310 | 7.4764 | 3.6802 | 4.1791 | 3.4238 | 4.1541 |
| ARFGAP2_S146 | ARFGAP2 | Q8N6H7 | | 146 | SAVPNHSPEKKDS | 6.1444 | 6.4713 | 6.4966 | 7.0305 | 6.1804 | 6.8869 | 6.0310 | 7.1788 | 6.4701 | 6.4166 | 6.9382 | 6.6575 | 5.1866 | 5.7516 | 5.0215 | 5.1381 |
| ARFGAP2_S315 | ARFGAP2 | Q8N6H7 | | 315 | GLVSRSSVSHSVL | 7.2051 | 5.4935 | 6.4334 | 5.5242 | 6.6707 | 5.9749 | 5.6139 | 6.3193 | 7.3340 | 7.0482 | 7.0543 | 6.6812 | 6.1670 | 7.0549 | 4.5273 | 4.8982 |
| ARFGAP2_S368 | ARFGAP2 | Q8N6H7 | | 432 | AGAKAISSDMFFG | 8.1943 | 7.3629 | 9.0343 | 7.1848 | 6.2963 | 5.6270 | 5.7315 | 6.5593 | 7.6338 | 7.5838 | 6.5883 | 6.3018 | 3.5709 | 5.3071 | 3.4945 | 3.5295 |
| ARFGAP2_S432 | ARFGAP2 | Q8N6H7 | | 368 | PFSLGESFGSRWD | 8.2987 | 7.6098 | 7.7374 | 7.2208 | 8.5655 | 5.6979 | 6.3060 | 6.0816 | 7.5351 | 6.5865 | 6.8647 | 6.2583 | 4.1756 | 4.2594 | 3.1695 | 3.6331 |
| ARFGAP3_S143 | ARFGAP3 | Q9NP61 | | 143 | CVVPPLSPPPKEE | 6.0722 | 6.4363 | 6.6191 | 6.2156 | 6.2506 | 7.0012 | 6.2575 | 7.2049 | 5.9491 | 5.8784 | 6.2320 | 6.4612 | 5.5747 | 6.1755 | 5.8252 | 5.8466 |
| ARFGAP3_Y396 | ARFGAP3 | Q9NP61 | | 396 | VLKTTGYSDRPTA | 4.7904 | 4.9543 | 7.2926 | 6.0041 | 6.8301 | 6.5915 | 6.4903 | 6.7693 | 7.0146 | 6.8749 | 6.5476 | 6.0028 | 5.4132 | 6.5450 | 6.1523 | 5.7270 |
| ARFGEF1_S1079 | ARFGEF1 | Q9Y6D6 | | 1566 | KPLDTISQKSVDI | 6.1906 | 5.6359 | 6.9956 | 6.4504 | 6.2750 | 6.3749 | 6.4301 | 5.8075 | 6.3832 | 6.1621 | 6.2519 | 6.3052 | 6.2850 | 6.3035 | 5.7920 | 6.3572 |
| ARFGEF1_S1566 | ARFGEF1 | Q9Y6D6 | | 1079 | VRGREGSLTGTKD | 5.4303 | 4.3129 | 5.4932 | 4.9887 | 6.3468 | 5.1268 | 5.1107 | 5.3508 | 7.1424 | 7.7456 | 7.0034 | 7.0832 | 8.0189 | 8.3224 | 6.6110 | 5.9129 |
| ARFGEF1_S1569 | ARFGEF1 | Q9Y6D6 | | 664 | TINRYGSLNSLES | 5.9401 | 6.2633 | 6.1151 | 6.1173 | 5.7462 | 6.1641 | 6.0719 | 6.0748 | 6.2215 | 6.4491 | 6.3486 | 6.3402 | 6.3900 | 6.9789 | 6.0639 | 6.7149 |
| ARFGEF1_S397 | ARFGEF1 | Q9Y6D6 | | 1569 | DTISQKSVDIHDS | 7.2257 | 6.8069 | 6.9535 | 6.8002 | 6.6813 | 6.3807 | 6.4781 | 6.7016 | 5.6144 | 6.1950 | 5.8999 | 6.7129 | 5.2061 | 5.2143 | 5.2483 | 5.8811 |
| ARFGEF1_S52 | ARFGEF1 | Q9Y6D6 | | 397 | DRLSVSSNDTQES | 6.0217 | 7.2023 | 7.2437 | 5.9807 | 7.5762 | 6.5775 | 5.7687 | 6.7880 | 6.8969 | 6.9560 | 7.1401 | 7.2128 | 4.6661 | 5.6067 | 4.9425 | 5.2400 |
| ARFGEF1_S664 | ARFGEF1 | Q9Y6D6 | | 52 | AETEKQSPPHGEA | 6.2455 | 6.8156 | 7.2938 | 6.7074 | 5.2804 | 6.1997 | 6.1971 | 6.2977 | 6.4205 | 6.5485 | 6.7191 | 6.1706 | 5.6653 | 5.7664 | 5.5381 | 6.1344 |
| ARFGEF2_S1511;S1514 | ARFGEF2 | Q9Y6D5 | | 227 | IQAAAVSPKFVRL | 6.2738 | 5.1925 | 6.9977 | 6.3969 | 6.2867 | 6.3111 | 6.1194 | 5.8364 | 6.3551 | 6.4667 | 6.3715 | 6.3012 | 6.4859 | 6.8410 | 5.7710 | 5.9930 |
| ARFGEF2_S1528 | ARFGEF2 | Q9Y6D5 | | 1528 | RGQSQLSNPTDDS | 5.7979 | 6.2338 | 7.0941 | 6.7299 | 6.5383 | 6.2760 | 6.1734 | 6.4882 | 6.4499 | 6.2124 | 6.1184 | 6.3186 | 5.8521 | 6.3705 | 5.6843 | 5.6622 |
| ARFGEF2_S218 | ARFGEF2 | Q9Y6D5 | | 218 | IQSKPQSPVIQAA | 5.9529 | 4.8242 | 6.9006 | 4.7715 | 5.4396 | 4.1562 | 6.3017 | 4.4178 | 4.9476 | 5.5695 | 5.3392 | 5.3984 | 8.9146 | 7.8089 | 9.9232 | 9.3341 |
| ARFGEF2_S227 | ARFGEF2 | Q9Y6D5 | | 276 | APRERGSSLSGTD | 5.4445 | 5.6309 | 6.4804 | 5.8552 | 5.9177 | 6.4644 | 5.9447 | 6.6252 | 6.0612 | 5.4405 | 5.8418 | 5.9821 | 6.9012 | 7.0743 | 7.6477 | 6.6883 |
| ARFGEF2_S276 | ARFGEF2 | Q9Y6D5 | 614;616 | | NA | 6.6128 | 7.1086 | 6.3225 | 5.3197 | 6.1127 | 6.3597 | 6.1050 | 6.7136 | 5.4246 | 5.2934 | 7.1553 | 6.3911 | 6.4894 | 6.4515 | 6.0975 | 6.0428 |
| ARFGEF2_S614;T616 | ARFGEF2 | Q9Y6D5 | 1511;1514 | | NA | 5.9863 | 6.1261 | 6.9692 | 6.3241 | 6.0861 | 7.1581 | 5.8502 | 6.3041 | 6.7910 | 6.2430 | 6.6626 | 6.2926 | 5.7718 | 6.0462 | 5.5183 | 5.8703 |
| ARFGEF3_S1066 | ARFGEF3 | Q5TH69 | | 2079 | QGQMRHSFSAGPE | 6.0793 | 5.4167 | 6.2119 | 5.1782 | 5.8388 | 6.0872 | 6.4984 | 5.7359 | 8.7736 | 7.1624 | 7.3302 | 7.1446 | 6.2079 | 6.9536 | 4.3618 | 5.0194 |
| ARFGEF3_S1741 | ARFGEF3 | Q5TH69 | | 1741 | EFVKGPSPGEEKT | 5.4113 | 5.4624 | 6.6412 | 6.3661 | 5.8577 | 6.1862 | 5.6914 | 6.3769 | 6.2850 | 6.5804 | 6.3919 | 6.5349 | 6.0818 | 6.6695 | 6.4810 | 6.9824 |
| ARFGEF3_S2079 | ARFGEF3 | Q5TH69 | | 1066 | GSPPEHSPEQGRS | 5.3955 | 5.8303 | 6.7325 | 7.2098 | 7.7896 | 5.3780 | 4.8282 | 7.3372 | 5.6004 | 8.0495 | 8.0770 | 6.7375 | 5.6279 | 6.3851 | 3.1063 | 5.9153 |
| ARFGEF3_S2097 | ARFGEF3 | Q5TH69 | | 2097 | KRPRSGSTGSSLS | 5.7973 | 5.4905 | 6.1487 | 5.5285 | 6.2679 | 6.4641 | 5.7710 | 6.7234 | 7.0191 | 6.6934 | 6.3831 | 6.7772 | 6.3717 | 6.5349 | 6.0940 | 5.9352 |
| ARFIP1_S132 | ARFIP1 | P53367 | | 79 | PLPSVMSPSRVAA | 5.7992 | 5.9602 | 6.6608 | 5.8538 | 6.3862 | 6.2915 | 6.5249 | 6.7991 | 6.5772 | 6.8571 | 6.7679 | 7.3773 | 5.2945 | 6.1665 | 5.2494 | 5.4344 |
| ARFIP1_S39 | ARFIP1 | P53367 | | 39 | LKHSLPSGLGLSE | 2.7637 | 2.8085 | 2.8328 | 2.5247 | 3.8811 | 3.8135 | 3.1580 | 3.2529 | 2.8168 | 2.8721 | 2.9308 | 2.7547 | 18.0078 | 16.3510 | 16.1544 | 13.0772 |
| ARFIP1_S69;S79 | ARFIP1 | P53367 | | 132 | ELVRKWSLNTYKC | 7.2957 | 6.6287 | 6.7106 | 4.8650 | 5.6905 | 5.8234 | 5.4112 | 6.7345 | 7.1281 | 7.5541 | 6.4564 | 7.6552 | 5.5697 | 5.8846 | 4.7209 | 5.8713 |
| ARFIP1_S79 | ARFIP1 | P53367 | 69;79 | | NA | 5.1660 | 7.3877 | 7.3372 | 5.6113 | 5.9086 | 5.0898 | 4.6889 | 6.7745 | 7.8828 | 6.4957 | 5.1589 | 7.3786 | 4.3802 | 6.7171 | 6.0295 | 7.9933 |
| ARGLU1_S60 | ARGLU1 | Q9NWB6 | | 77 | DRERASSPPDRID | 6.6855 | 7.8241 | 7.1670 | 6.9557 | 7.0023 | 8.0721 | 5.9873 | 4.6770 | 5.5655 | 5.8847 | 6.5578 | 5.7898 | 6.3036 | 5.9379 | 5.2065 | 4.3831 |
| ARGLU1_S77 | ARGLU1 | Q9NWB6 | | 60 | SRSRSRSTNTAVS | 6.1141 | 6.0849 | 4.8754 | 5.8534 | 6.0686 | 5.4767 | 5.4194 | 6.2735 | 6.3101 | 5.9957 | 5.5032 | 5.4534 | 7.3575 | 7.7621 | 7.9859 | 7.4661 |
| ARGLU1_S94 | ARGLU1 | Q9NWB6 | | 94 | TVSKRSSLDEKQK | 4.9232 | 5.4861 | 4.9722 | 5.6699 | 5.9028 | 5.8309 | 6.3614 | 5.5413 | 5.2990 | 5.4482 | 5.8945 | 5.5344 | 7.8506 | 8.2621 | 8.8652 | 8.1582 |
| ARHGAP1_S223 | ARHGAP1 | Q07960 | | 223 | LKSTQKSPATAPK | 7.1666 | 7.0397 | 6.4106 | 5.9999 | 7.0452 | 7.2020 | 6.6565 | 6.7811 | 6.5068 | 6.8163 | 6.4315 | 6.3243 | 4.8754 | 5.6237 | 4.4518 | 4.6685 |
| ARHGAP1_S27 | ARHGAP1 | Q07960 | | 50 | SDDSKSSSPELVT | 4.9416 | 5.1962 | 4.8929 | 4.7639 | 5.8375 | 5.1787 | 5.5222 | 5.0327 | 5.3084 | 4.6729 | 4.9299 | 4.7289 | 10.0416 | 9.5302 | 10.2773 | 9.1452 |
| ARHGAP1_S50 | ARHGAP1 | Q07960 | | 27 | NQLKLASIDEKNW | 6.8440 | 6.6521 | 6.5304 | 6.1623 | 6.9089 | 6.7079 | 7.1620 | 6.5799 | 7.0627 | 7.1568 | 6.1700 | 6.4727 | 4.6704 | 5.3652 | 4.3018 | 5.2529 |
| ARHGAP1_S50;S51 | ARHGAP1 | Q07960 | | 51 | DDSKSSSPELVTH | 6.4737 | 5.7322 | 5.9881 | 5.3492 | 6.7892 | 6.4504 | 6.4643 | 6.4711 | 6.7686 | 5.8354 | 6.3019 | 6.0054 | 6.7093 | 6.2455 | 6.2396 | 6.1762 |
| ARHGAP1_S51 | ARHGAP1 | Q07960 | 50;51 | | NA | 6.1026 | 5.8447 | 6.3135 | 5.4046 | 5.9615 | 6.1102 | 5.9130 | 5.8077 | 6.3568 | 5.9092 | 5.6612 | 6.1581 | 7.1850 | 7.1447 | 6.7674 | 7.3597 |
| ARHGAP10_S591 | ARHGAP10 | A1A4S6 | | 591 | PTCLSASPPNAPP | 6.2966 | 6.2208 | 6.5072 | 6.7319 | 7.0121 | 7.2729 | 6.6352 | 6.4247 | 6.6826 | 6.6219 | 6.2592 | 6.7902 | 5.6101 | 5.2824 | 4.4755 | 5.1767 |
| ARHGAP10_S645 | ARHGAP10 | A1A4S6 | | 645 | DSLSSPSPVTTAV | 5.5636 | 6.0859 | 6.5361 | 5.7001 | 5.8679 | 6.7124 | 6.0167 | 6.9370 | 6.4915 | 6.1114 | 6.6185 | 7.0953 | 6.2934 | 6.1209 | 5.8897 | 5.9598 |
| ARHGAP11A_S285 | ARHGAP11A | Q6P4F7 | | 422 | GKAGCFSPKISHK | 8.1379 | 7.7595 | 6.5875 | 7.1413 | 6.7651 | 6.6453 | 7.5180 | 6.5553 | 7.4932 | 7.0495 | 6.8052 | 6.5253 | 3.5259 | 4.0473 | 3.4572 | 3.9865 |
| ARHGAP11A_S318 | ARHGAP11A | Q6P4F7 | | 285 | KRKRRQSVGDFVS | 6.2059 | 6.1028 | 5.6317 | 5.6657 | 5.8901 | 6.3402 | 6.5134 | 6.4072 | 8.1048 | 6.8818 | 6.7599 | 6.7273 | 5.7040 | 6.7381 | 4.5831 | 5.7441 |
| ARHGAP11A_S318;T323 | ARHGAP11A | Q6P4F7 | | 868 | SLGDTASPLVKSV | 7.3975 | 8.0243 | 6.6215 | 8.0632 | 6.3416 | 7.0493 | 7.4199 | 6.9496 | 7.3851 | 7.7177 | 7.4322 | 7.4542 | 2.4831 | 3.8444 | 2.3531 | 3.4634 |
| ARHGAP11A_S323 | ARHGAP11A | Q6P4F7 | | 620 | MNQRQSSVTNVGK | 5.3485 | 5.4066 | 5.3701 | 5.4016 | 5.9248 | 5.5299 | 5.6529 | 5.6189 | 5.5002 | 5.6487 | 5.8226 | 5.6404 | 8.1777 | 8.1110 | 8.8131 | 8.0326 |
| ARHGAP11A_S422 | ARHGAP11A | Q6P4F7 | | 318 | IAQLSESPVILTP | 6.0316 | 6.9854 | 6.1135 | 7.8205 | 6.8839 | 5.8700 | 6.2577 | 6.6663 | 7.1961 | 6.6932 | 6.8600 | 6.7026 | 4.9790 | 5.2933 | 4.4560 | 5.1909 |
| ARHGAP11A_S469 | ARHGAP11A | Q6P4F7 | | 469 | RLANQQSLKNRIE | 8.5014 | 7.3987 | 7.6163 | 7.2074 | 5.7812 | 6.8322 | 6.8880 | 6.0128 | 8.5811 | 7.9998 | 8.4571 | 7.9751 | 2.4509 | 3.5249 | 1.7957 | 2.9772 |
| ARHGAP11A_S484 | ARHGAP11A | Q6P4F7 | | 323 | ESPVILTPNAKRT | 8.2436 | 7.2107 | 7.0156 | 7.6729 | 5.9897 | 6.8335 | 6.5029 | 5.8647 | 7.8437 | 6.8252 | 6.7438 | 6.8632 | 3.7977 | 4.4926 | 3.8333 | 4.2669 |
| ARHGAP11A_S620 | ARHGAP11A | Q6P4F7 | | 484 | KTGLLFSPDVDEK | 7.0118 | 7.3404 | 6.6802 | 7.5408 | 6.0998 | 6.1192 | 7.4135 | 6.6617 | 7.1179 | 7.6249 | 7.1210 | 7.6613 | 3.4477 | 4.1541 | 3.6912 | 4.3147 |
| ARHGAP11A_S847 | ARHGAP11A | Q6P4F7 | | 847 | SVRRINSLLEYSR | 7.5032 | 6.9531 | 7.0028 | 7.1862 | 6.5387 | 6.2720 | 6.4826 | 6.0482 | 7.3374 | 6.8033 | 6.8009 | 6.7395 | 4.6424 | 4.8544 | 4.2389 | 4.5964 |
| ARHGAP11A_S868 | ARHGAP11A | Q6P4F7 | | 508 | SEETLLTPERLVG | 6.1016 | 7.3008 | 7.5121 | 7.8648 | 6.4744 | 6.8257 | 7.3300 | 6.8584 | 8.0199 | 7.6739 | 7.6451 | 7.5487 | 2.6708 | 3.9470 | 2.6926 | 3.5342 |
| ARHGAP11A_T508 | ARHGAP11A | Q6P4F7 | 318;323 | | NA | 7.4287 | 7.4255 | 6.9572 | 7.3054 | 6.3735 | 7.0301 | 7.6896 | 6.7891 | 6.9371 | 7.4091 | 7.3217 | 7.2987 | 3.3987 | 4.0991 | 2.5539 | 3.9824 |
| ARHGAP12_S103 | ARHGAP12 | Q8IWW6 | | 592 | EEEIPDSPGIEKH | 5.9216 | 5.9909 | 6.4736 | 5.8459 | 6.2177 | 6.5578 | 6.4635 | 6.3238 | 6.4884 | 6.4647 | 6.4660 | 6.3481 | 5.8715 | 6.4141 | 5.9110 | 6.2412 |
| ARHGAP12_S115 | ARHGAP12 | Q8IWW6 | | 176 | SQNRTRSFGHFPG | 5.6978 | 5.8720 | 6.0553 | 5.3821 | 5.0989 | 5.4116 | 5.5038 | 5.8925 | 6.2862 | 6.3191 | 6.5649 | 6.6960 | 6.4533 | 7.4609 | 7.2842 | 8.0214 |
| ARHGAP12_S165 | ARHGAP12 | Q8IWW6 | | 262 | LPPLPGSPAIQIN | 6.2890 | 6.3501 | 6.7904 | 6.3843 | 6.3813 | 6.6294 | 6.4602 | 6.3370 | 6.6837 | 6.7626 | 6.8700 | 6.7911 | 4.9252 | 5.6063 | 5.1367 | 5.6026 |
| ARHGAP12_S176 | ARHGAP12 | Q8IWW6 | | 231 | EQIRATTPPNQGR | 5.6388 | 5.8819 | 5.8442 | 5.0764 | 6.0157 | 5.8745 | 5.4032 | 5.9604 | 6.5024 | 6.2406 | 6.2183 | 6.3200 | 7.4726 | 7.5829 | 6.9047 | 7.0634 |
| ARHGAP12_S240 | ARHGAP12 | Q8IWW6 | | 165 | FNNDSHSPKVSSQ | 6.2544 | 6.2576 | 6.2704 | 5.8903 | 6.4348 | 6.6371 | 6.6778 | 6.6459 | 6.6577 | 6.8839 | 6.2484 | 6.4271 | 5.5898 | 6.1197 | 5.3751 | 5.6301 |
| ARHGAP12_S262 | ARHGAP12 | Q8IWW6 | | 240 | NQGRPDSPVYANL | 5.2750 | 5.9611 | 6.1447 | 5.5815 | 6.1513 | 5.9012 | 5.5373 | 5.6888 | 6.2339 | 6.8627 | 6.3653 | 6.2361 | 7.0932 | 7.4120 | 7.1030 | 6.4529 |
| ARHGAP12_S592 | ARHGAP12 | Q8IWW6 | | 115 | KLPELSSFGKPSS | 5.6396 | 6.1508 | 6.4355 | 6.1983 | 4.8648 | 6.4302 | 5.8451 | 6.7359 | 5.1969 | 5.3550 | 8.6333 | 6.7306 | 5.4867 | 7.8796 | 6.8993 | 5.5184 |
| ARHGAP12_T231 | ARHGAP12 | Q8IWW6 | | 103 | SLHLQRSTENVNK | 5.4714 | 6.2248 | 7.1362 | 6.4377 | 4.9813 | 6.4823 | 6.0450 | 6.3115 | 6.2250 | 6.7742 | 6.7631 | 7.2500 | 5.3721 | 6.3721 | 5.9207 | 6.2323 |
| ARHGAP12_T231;S240 | ARHGAP12 | Q8IWW6 | 231;240 | | NA | 5.7649 | 6.2132 | 6.5172 | 5.9243 | 6.1622 | 6.2382 | 6.0000 | 6.3730 | 6.3447 | 6.6993 | 6.5811 | 6.3486 | 5.9191 | 6.4075 | 6.2375 | 6.2690 |
| ARHGAP17_S162 | ARHGAP17 | Q68EM7 | | 676 | PPTRSPSPPTQHT | 6.0388 | 6.1925 | 6.4158 | 5.7187 | 6.6227 | 6.7850 | 5.7821 | 6.6655 | 6.3208 | 6.3119 | 6.1409 | 6.2659 | 6.3076 | 6.2563 | 6.2581 | 5.9174 |
| ARHGAP17_S484 | ARHGAP17 | Q68EM7 | | 484 | ERKRPASMAVMEG | 8.3061 | 6.5547 | 7.6424 | 6.2113 | 6.6279 | 6.9690 | 6.3966 | 6.8818 | 8.0272 | 6.4816 | 5.9718 | 6.3307 | 4.9305 | 5.4148 | 3.5369 | 3.7167 |
| ARHGAP17_S497 | ARHGAP17 | Q68EM7 | | 497 | DLVKKESFGVKLM | 8.8328 | 7.1113 | 6.8472 | 6.0547 | 6.5878 | 6.3857 | 6.3790 | 6.3727 | 8.3838 | 6.7461 | 6.5350 | 6.2426 | 5.2186 | 5.3090 | 3.3998 | 3.5941 |
| ARHGAP17_S575 | ARHGAP17 | Q68EM7 | | 575 | PSPGDGSPPKPKD | 6.1434 | 5.9937 | 7.1730 | 7.4641 | 5.8696 | 6.0893 | 6.0371 | 6.6451 | 6.5502 | 6.9152 | 7.2030 | 7.1847 | 5.1072 | 5.3303 | 4.7162 | 5.5780 |
| ARHGAP17_S625 | ARHGAP17 | Q68EM7 | | 513 | AHRRGGTLNRKHI | 9.6715 | 7.4479 | 7.7255 | 6.2615 | 6.1123 | 6.8652 | 6.6514 | 5.6511 | 7.5100 | 6.4057 | 7.1344 | 6.3424 | 4.7611 | 4.6618 | 3.4569 | 3.3412 |
| ARHGAP17_S674 | ARHGAP17 | Q68EM7 | | 674 | PKPPTRSPSPPTQ | 5.4744 | 7.4995 | 6.5841 | 5.7481 | 6.1545 | 5.3417 | 6.1613 | 6.6717 | 6.3118 | 6.6881 | 6.7866 | 6.2010 | 6.4254 | 6.5102 | 6.3406 | 5.1009 |
| ARHGAP17_S674;S676 | ARHGAP17 | Q68EM7 | | 625 | HNAAGPSPHTLRR | 5.5826 | 6.8549 | 7.9844 | 5.5520 | 5.3206 | 5.5724 | 6.5589 | 8.2667 | 6.3922 | 5.8445 | 5.9670 | 7.8006 | 6.7426 | 5.5486 | 4.8035 | 5.2084 |
| ARHGAP17_S676 | ARHGAP17 | Q68EM7 | | 162 | NQAHKSSGTNFQG | 1.6600 | 1.5368 | 1.3766 | 1.6355 | 2.2458 | 2.3260 | 2.1488 | 2.4218 | 1.5211 | 1.8016 | 1.5985 | 1.5886 | 19.1571 | 18.4247 | 22.1882 | 18.3689 |
| ARHGAP17_S702;S704 | ARHGAP17 | Q68EM7 | 674;676 | | NA | 5.6173 | 6.5288 | 6.2284 | 5.9540 | 5.8017 | 5.5693 | 6.3080 | 6.6714 | 6.6291 | 6.3408 | 6.3097 | 6.7934 | 5.9585 | 6.2058 | 6.4600 | 6.6238 |
| ARHGAP17_S702;S706 | ARHGAP17 | Q68EM7 | 702;704 | | NA | 5.6167 | 6.6166 | 6.2141 | 6.6180 | 6.1191 | 6.1203 | 6.1980 | 7.4826 | 7.4739 | 6.3451 | 6.8724 | 6.6544 | 4.6364 | 5.5396 | 5.6366 | 5.8563 |
| ARHGAP17_T513 | ARHGAP17 | Q68EM7 | 702;706 | | NA | 5.5141 | 5.9678 | 6.6772 | 6.7271 | 6.3888 | 6.3402 | 6.9737 | 6.5679 | 7.0310 | 6.7512 | 7.6627 | 7.4273 | 4.9805 | 5.6763 | 4.4958 | 4.8184 |
| ARHGAP19_S422 | ARHGAP19 | Q14CB8-2 | | 438 | LGNQMMSEKKKKN | 7.8967 | 6.7874 | 8.0200 | 7.0554 | 5.8674 | 6.4930 | 6.4239 | 6.5557 | 8.1094 | 7.4419 | 7.0929 | 7.5236 | 3.7579 | 4.4804 | 2.9972 | 3.4973 |
| ARHGAP19_S438 | ARHGAP19 | Q14CB8-2 | | 422 | KRARSRSFSGLIK | 6.3092 | 5.8582 | 5.9188 | 5.1781 | 5.6398 | 6.2630 | 5.5883 | 6.2187 | 7.5856 | 7.0192 | 7.0875 | 7.4885 | 5.9237 | 6.8947 | 5.1632 | 5.8634 |
| ARHGAP19_T404 | ARHGAP19 | Q14CB8-2 | | 404 | KQSLTQTPGREPS | 8.3232 | 8.8936 | 6.9604 | 6.9135 | 5.1810 | 6.7832 | 9.1835 | 7.6389 | 7.9041 | 7.8255 | 7.2503 | 8.0004 | 2.0459 | 2.6304 | 1.8123 | 2.6539 |
| ARHGAP21_S1099 | ARHGAP21 | Q5T5U3 | | 1860 | ARERLRTSTSDLS | 6.2076 | 6.6667 | 6.1519 | 5.9478 | 5.8626 | 6.6038 | 6.4555 | 5.9601 | 5.9835 | 5.6525 | 5.6448 | 6.1151 | 7.1930 | 6.6658 | 6.5171 | 6.3723 |
| ARHGAP21_S1129 | ARHGAP21 | Q5T5U3 | | 253 | EIQVPPSPTDVAK | 5.9589 | 6.3201 | 6.1167 | 6.3846 | 6.6779 | 6.3581 | 5.9791 | 6.4140 | 5.7054 | 5.7146 | 5.9839 | 5.8222 | 7.1802 | 6.7546 | 6.2437 | 6.3860 |
| ARHGAP21_S1514 | ARHGAP21 | Q5T5U3 | | 1861 | RERLRTSTSDLSR | 6.3463 | 6.8500 | 6.0794 | 6.2858 | 5.8751 | 6.4060 | 6.8848 | 5.9975 | 5.7414 | 5.9045 | 6.0512 | 6.3357 | 6.3932 | 6.3818 | 6.2038 | 6.2635 |
| ARHGAP21_S1605 | ARHGAP21 | Q5T5U3 | | 319 | TVSRTTSPPLSIP | 6.2651 | 7.4013 | 7.2988 | 7.3299 | 5.5929 | 6.3031 | 6.5742 | 6.5028 | 6.1610 | 6.6259 | 6.4542 | 5.9529 | 5.3036 | 5.4319 | 5.3818 | 5.4205 |
| ARHGAP21_S1669 | ARHGAP21 | Q5T5U3 | | 1129 | WRKGIPSIMRKTF | 13.2340 | 8.2674 | 7.8786 | 7.1735 | 6.2244 | 6.1596 | 6.6189 | 4.9723 | 6.6951 | 5.7270 | 6.1310 | 5.9004 | 4.1661 | 5.0752 | 2.8187 | 2.9578 |
| ARHGAP21_S1861 | ARHGAP21 | Q5T5U3 | | 954 | GKRVGGSIRPWKQ | 6.5651 | 6.8022 | 7.0715 | 7.2827 | 6.2205 | 6.0181 | 6.0856 | 6.2467 | 5.1388 | 5.8956 | 4.6315 | 4.6221 | 7.1751 | 7.2217 | 7.3428 | 5.6799 |
| ARHGAP21_S253 | ARHGAP21 | Q5T5U3 | | 1605 | LDSSRLSPEVQSV | 7.1711 | 7.5659 | 6.8219 | 7.0391 | 5.6275 | 6.3050 | 7.0583 | 6.5832 | 6.9879 | 6.3968 | 6.7059 | 6.2236 | 4.4722 | 5.1040 | 4.6790 | 5.2586 |
| ARHGAP21_S319 | ARHGAP21 | Q5T5U3 | | 1099 | KTQSPHSPKEESE | 6.9789 | 6.8944 | 6.5877 | 7.1981 | 6.2520 | 6.5266 | 6.1196 | 6.4621 | 5.9365 | 5.9213 | 5.9202 | 6.1002 | 5.5904 | 6.0083 | 5.9078 | 5.5960 |
| ARHGAP21_S616 | ARHGAP21 | Q5T5U3 | | 857 | LIRRQLSHDHESV | 6.0471 | 7.3829 | 6.5094 | 6.7986 | 5.7173 | 7.1066 | 6.1766 | 7.4228 | 6.1081 | 5.5238 | 5.5690 | 4.9290 | 5.8205 | 6.7519 | 6.3295 | 5.8069 |
| ARHGAP21_S857 | ARHGAP21 | Q5T5U3 | | 1669 | KSSRRNSEGSELS | 5.7334 | 4.9035 | 7.0573 | 6.1689 | 6.6753 | 5.6650 | 5.6289 | 5.6998 | 7.5985 | 7.2651 | 7.4965 | 6.4339 | 6.5297 | 5.9375 | 5.7471 | 5.4597 |
| ARHGAP21_S954 | ARHGAP21 | Q5T5U3 | | 616 | GISQDRSPLVKVR | 5.9112 | 6.3580 | 5.6696 | 6.1333 | 5.5038 | 6.9867 | 7.7791 | 6.5102 | 5.5195 | 5.7185 | 5.6291 | 5.7757 | 6.2377 | 6.5543 | 6.7752 | 6.9382 |
| ARHGAP21_T1860 | ARHGAP21 | Q5T5U3 | | 1514 | NSKHNKSPTLSCR | 7.0181 | 7.2157 | 6.1742 | 6.6842 | 6.2067 | 6.3462 | 5.9683 | 6.1875 | 6.0790 | 6.2251 | 6.1118 | 5.8766 | 5.7197 | 5.9933 | 5.9185 | 6.2750 |
| ARHGAP23_S1393 | ARHGAP23 | Q9P227 | | 1393 | RLRRPLSPETRRR | 6.2844 | 6.4678 | 6.6257 | 6.3646 | 6.0907 | 6.2772 | 6.3799 | 5.8979 | 6.1600 | 6.4492 | 6.2279 | 6.6054 | 6.0084 | 6.1690 | 6.0256 | 5.9663 |
| ARHGAP23_S1412 | ARHGAP23 | Q9P227 | | 611 | KAGRRSSYLLAIT | 6.6217 | 5.9548 | 5.5538 | 6.0024 | 6.5969 | 5.4692 | 5.7960 | 5.3292 | 6.7215 | 6.0154 | 6.6486 | 6.3844 | 7.0010 | 7.3220 | 6.1366 | 6.4464 |
| ARHGAP23_S372 | ARHGAP23 | Q9P227 | | 372 | GPGALVSPRFERC | 5.7321 | 6.4451 | 5.7697 | 6.3614 | 5.7526 | 5.9702 | 6.3171 | 6.5290 | 6.3098 | 6.5419 | 6.1305 | 6.5445 | 5.8111 | 6.6429 | 6.7820 | 6.3602 |
| ARHGAP23_S423 | ARHGAP23 | Q9P227 | | 437 | GLLHALSFRDSPF | 9.0800 | 7.3449 | 6.4791 | 6.5083 | 6.0035 | 6.1283 | 5.6841 | 6.0349 | 7.7003 | 5.9843 | 6.7051 | 6.4679 | 5.1017 | 6.2565 | 4.1076 | 4.4135 |
| ARHGAP23_S437 | ARHGAP23 | Q9P227 | | 554 | FIDEPTSPSIDLQ | 7.0267 | 7.2027 | 6.7017 | 7.0800 | 6.9446 | 7.5952 | 7.0845 | 6.3665 | 5.8366 | 5.9071 | 5.0831 | 5.9938 | 5.1642 | 5.6591 | 5.2545 | 5.0998 |
| ARHGAP23_S554 | ARHGAP23 | Q9P227 | | 423 | IGYRSYSPSFQRR | 6.3375 | 6.5306 | 5.9883 | 6.1843 | 5.9294 | 6.0792 | 6.4489 | 6.3927 | 6.2661 | 6.2236 | 6.2693 | 6.3364 | 6.1847 | 6.3368 | 6.2734 | 6.2188 |
| ARHGAP23_S611 | ARHGAP23 | Q9P227 | | 1412 | HTVVVQSPLTDLN | 6.0406 | 6.3846 | 7.1249 | 8.6600 | 4.5079 | 7.0229 | 7.1439 | 6.8758 | 5.4522 | 5.3051 | 7.1818 | 7.1486 | 5.9718 | 4.5411 | 5.7512 | 4.8875 |
| ARHGAP24_S415 | ARHGAP24 | Q8N264 | | 415 | KLDVSRSPPLMVK | 6.1577 | 6.3208 | 6.5034 | 6.0436 | 6.2037 | 6.7741 | 5.9540 | 6.9262 | 6.3035 | 6.5306 | 6.5482 | 6.5764 | 5.7821 | 6.0335 | 5.6424 | 5.6996 |
| ARHGAP29_S1019 | ARHGAP29 | Q52LW3 | | 930 | KSLFFSSKEDIHT | 5.7653 | 5.8305 | 7.0748 | 5.9014 | 6.7236 | 6.3976 | 6.5435 | 6.3365 | 6.2436 | 6.4302 | 7.0854 | 6.4971 | 6.0440 | 5.0555 | 5.7172 | 6.3536 |
| ARHGAP29_S1029 | ARHGAP29 | Q52LW3 | | 1144 | ASERRSSDSYPLA | 5.5394 | 5.3435 | 6.2558 | 5.4149 | 6.2084 | 5.9727 | 5.7635 | 6.6372 | 7.2844 | 7.2787 | 7.9214 | 7.8896 | 5.9615 | 6.0701 | 5.0665 | 5.3922 |
| ARHGAP29_S1144 | ARHGAP29 | Q52LW3 | | 1019 | TKIRPVSLPVDRL | 7.7258 | 6.1437 | 8.3127 | 6.8987 | 7.8853 | 5.4896 | 5.3468 | 6.6474 | 8.0300 | 5.3326 | 5.8949 | 6.0580 | 5.4170 | 5.6651 | 4.8467 | 4.3060 |
| ARHGAP29_S1185 | ARHGAP29 | Q52LW3 | | 1029 | DRLLLASPPNERN | 6.4358 | 6.3447 | 7.1420 | 6.8474 | 6.2077 | 6.8108 | 6.9545 | 7.2807 | 6.8920 | 6.6379 | 7.2214 | 7.2899 | 4.3244 | 4.7251 | 4.3724 | 4.5133 |
| ARHGAP29_S930 | ARHGAP29 | Q52LW3 | | 1185 | NEEKPASPSAAVP | 5.8854 | 5.1530 | 6.3797 | 6.6119 | 6.3545 | 6.2802 | 7.1526 | 7.3769 | 6.3943 | 6.0161 | 6.6451 | 6.2789 | 6.7187 | 6.1727 | 5.5371 | 5.0429 |
| ARHGAP31_S1036 | ARHGAP31 | Q2M1Z3 | | 1374 | DSKVLLSPIRSPT | 6.3278 | 6.6499 | 7.4839 | 6.0763 | 6.0791 | 5.8935 | 6.5043 | 6.0484 | 6.7958 | 6.2554 | 6.5837 | 6.5125 | 5.9109 | 5.8707 | 5.3575 | 6.6504 |
| ARHGAP31_S1178 | ARHGAP31 | Q2M1Z3 | | 863 | KGCGFPSPTREVE | 6.1947 | 6.2688 | 6.6040 | 6.1737 | 6.1383 | 6.4289 | 6.4334 | 6.1817 | 6.4759 | 6.2482 | 6.5399 | 6.5073 | 5.8997 | 5.7996 | 6.0194 | 6.0866 |
| ARHGAP31_S1374 | ARHGAP31 | Q2M1Z3 | | 1178 | LTGRRNSAPVSVS | 6.1798 | 4.8869 | 5.9748 | 6.1718 | 7.0006 | 6.1191 | 5.5475 | 5.9574 | 7.8304 | 6.9061 | 7.5676 | 6.7674 | 6.1375 | 6.7500 | 4.7887 | 5.4144 |

| | | | | | | | | | | | | | | | | | | | | | |
|---|---|---|---|---|---|---|---|---|---|---|---|---|---|---|---|---|---|---|---|---|---|
| ARHGAP31_S346 | ARHGAP31 | Q2M1Z3 | | 346 | TIRPAKSMDSLCS | 6.2270 | 5.7816 | 6.7788 | 6.6422 | 5.4042 | 6.5965 | 6.3706 | 5.7508 | 7.1062 | 6.9340 | 7.0052 | 6.5832 | 5.6312 | 5.9411 | 4.9582 | 6.2892 |
| ARHGAP31_S863 | ARHGAP31 | Q2M1Z3 | | 1036 | PNPAETSPISLAE | 6.3058 | 6.8400 | 6.2723 | 7.0738 | 6.0488 | 6.3681 | 6.7810 | 6.3286 | 7.1052 | 6.3297 | 6.6674 | 7.1133 | 4.9540 | 5.6429 | 4.3322 | 5.8372 |
| ARHGAP32_S587 | ARHGAP32 | A7KAX9 | | 952 | RDATNRSPTQIVK | 6.4643 | 6.6412 | 6.5965 | 6.4303 | 6.4151 | 6.2392 | 6.0888 | 6.5503 | 6.4609 | 6.4276 | 6.1425 | 5.9841 | 5.8286 | 6.2350 | 5.9148 | 5.5808 |
| ARHGAP32_S592 | ARHGAP32 | A7KAX9 | | 892 | KVVYAFSPKIGRK | 6.0913 | 6.6072 | 6.5479 | 5.7210 | 5.9731 | 6.0247 | 6.7281 | 6.4132 | 6.3081 | 6.1672 | 6.5911 | 6.5799 | 6.1283 | 6.3870 | 6.0490 | 5.6829 |
| ARHGAP32_S856 | ARHGAP32 | A7KAX9 | | 592 | KSLLVSSPSTKLL | 6.5765 | 6.2817 | 7.0251 | 6.7789 | 6.3637 | 6.4135 | 7.3929 | 7.4379 | 6.9940 | 7.4120 | 7.1036 | 7.7897 | 3.7899 | 4.8732 | 3.4345 | 4.3329 |
| ARHGAP32_S892 | ARHGAP32 | A7KAX9 | | 587 | SLSRPKSLLVSSP | 6.7047 | 6.6528 | 6.3821 | 6.9868 | 6.5929 | 6.0936 | 6.5407 | 6.4327 | 6.6560 | 6.2805 | 6.8295 | 6.1337 | 5.4341 | 6.0265 | 5.1720 | 5.0814 |
| ARHGAP32_S952 | ARHGAP32 | A7KAX9 | | 856 | ASSEPVSPLQEKL | 5.3271 | 5.5985 | 6.4592 | 5.7269 | 5.6975 | 6.1633 | 5.7733 | 5.7587 | 5.8271 | 5.7684 | 6.6206 | 6.4996 | 7.5669 | 7.5572 | 6.7099 | 6.9458 |
| ARHGAP32_T1236 | ARHGAP32 | A7KAX9 | | 595 | LVSSPSTKLLTLE | 6.9605 | 4.7293 | 5.2856 | 4.9276 | 7.1709 | 5.6639 | 4.6323 | 4.7132 | 6.1858 | 4.4623 | 3.3761 | 4.8811 | 8.2025 | 9.7702 | 9.1654 | 9.8734 |
| ARHGAP32_T595 | ARHGAP32 | A7KAX9 | | 1236 | FADKSPTPPNLPS | 6.9495 | 6.5398 | 6.8672 | 5.7125 | 6.3411 | 6.7386 | 5.8546 | 6.6813 | 6.1685 | 6.2325 | 6.3478 | 5.8753 | 6.2069 | 5.9582 | 5.8930 | 5.6333 |
| ARHGAP33_S780 | ARHGAP33 | O14559 | | 780 | VTPQAISPRGPTS | 7.9771 | 7.9610 | 7.8472 | 7.7105 | 6.5555 | 7.1407 | 7.4060 | 7.0280 | 7.4923 | 7.1160 | 7.3091 | 6.6655 | 2.7955 | 3.1236 | 2.6597 | 3.2123 |
| ARHGAP33_T564 | ARHGAP33 | O14559 | | 564 | TPTEPTTPKAPAS | 7.7339 | 7.0750 | 7.0479 | 7.4492 | 5.8716 | 6.8896 | 6.9106 | 6.7618 | 6.8047 | 6.4424 | 6.4821 | 6.8777 | 4.4229 | 5.0822 | 3.5508 | 4.5974 |
| ARHGAP35_S1070 | ARHGAP35 | Q9NRY4 | | 1070 | RDGQRKSVSSSPW | 5.8824 | 5.5169 | 6.0924 | 5.6388 | 6.4958 | 6.4936 | 5.9209 | 6.4942 | 6.8431 | 7.1826 | 6.6785 | 6.8076 | 6.1946 | 6.7293 | 5.4364 | 5.5928 |
| ARHGAP35_S1150 | ARHGAP35 | Q9NRY4 | | 970 | TEEVFNSPRAGSP | 6.1272 | 6.5835 | 7.0446 | 6.3228 | 6.0600 | 6.8013 | 5.8457 | 6.4275 | 6.6196 | 6.2475 | 6.5386 | 6.7653 | 5.3054 | 6.2410 | 5.3776 | 5.6923 |
| ARHGAP35_S1179 | ARHGAP35 | Q9NRY4 | | 975 | NSPRAGSPLCNSN | 5.8719 | 6.4418 | 6.6800 | 5.8658 | 6.1467 | 5.8475 | 5.5585 | 5.7763 | 6.1426 | 6.4622 | 6.2973 | 6.5640 | 7.4086 | 6.9690 | 5.3692 | 6.5986 |
| ARHGAP35_S970 | ARHGAP35 | Q9NRY4 | | 1150 | ERGRKVSIVSKPV | 5.9738 | 5.9631 | 6.3883 | 5.7341 | 7.3624 | 6.4283 | 5.2882 | 5.7758 | 7.2966 | 6.8624 | 7.0821 | 7.0835 | 5.6299 | 6.5972 | 5.2454 | 5.2889 |
| ARHGAP35_S975 | ARHGAP35 | Q9NRY4 | | 1179 | TSFSVGSDDELGP | 6.6767 | 6.5232 | 7.0513 | 6.1251 | 6.2133 | 6.1830 | 6.4068 | 6.6296 | 6.7951 | 6.9280 | 6.5329 | 6.4799 | 5.4839 | 5.3965 | 5.1669 | 5.4079 |
| ARHGAP39_S173 | ARHGAP39 | Q9C0H5-2 | | 286 | ELPGSSSPLLAQP | 5.9945 | 5.9998 | 6.4215 | 6.8072 | 6.1237 | 5.9370 | 6.5041 | 6.1953 | 6.1183 | 6.4029 | 6.3586 | 6.3257 | 6.2329 | 6.1974 | 6.2403 | 6.1408 |
| ARHGAP39_S273 | ARHGAP39 | Q9C0H5-2 | | 366 | NKQGPPSPCQQLV | 5.7626 | 5.9987 | 6.1495 | 6.2505 | 5.8985 | 5.9015 | 6.1515 | 6.4128 | 6.1531 | 6.3258 | 6.2192 | 6.4318 | 6.1784 | 6.6229 | 6.6049 | 6.9383 |
| ARHGAP39_S286 | ARHGAP39 | Q9C0H5-2 | | 500 | RKSRKPSLCQATS | 5.3721 | 5.1385 | 6.0881 | 5.4012 | 5.9224 | 5.7086 | 4.9841 | 6.2338 | 7.8606 | 7.3094 | 7.3480 | 7.6277 | 6.7958 | 7.3823 | 5.0198 | 5.8077 |
| ARHGAP39_S299 | ARHGAP39 | Q9C0H5-2 | | 384 | CPERFLSLEYSPA | 6.5124 | 6.0130 | 7.7163 | 6.4104 | 5.3012 | 5.8993 | 6.6474 | 6.3891 | 5.9514 | 6.2776 | 6.5165 | 7.1285 | 5.0829 | 6.0453 | 5.5438 | 6.5649 |
| ARHGAP39_S344 | ARHGAP39 | Q9C0H5-2 | | 388 | FLSLEYSPAGKEY | 5.4808 | 6.6432 | 7.0201 | 7.2727 | 5.6093 | 5.8227 | 6.5010 | 7.3587 | 7.4798 | 6.7579 | 6.7823 | 7.3672 | 4.6167 | 4.5881 | 4.2653 | 6.4341 |
| ARHGAP39_S366 | ARHGAP39 | Q9C0H5-2 | | 690 | TLRKPSSETDIEN | 5.5296 | 5.7540 | 6.1782 | 6.3752 | 5.4728 | 5.3556 | 5.7171 | 6.2200 | 5.9813 | 6.2272 | 5.9199 | 5.5777 | 7.5750 | 7.5290 | 7.8275 | 6.7599 |
| ARHGAP39_S384 | ARHGAP39 | Q9C0H5-2 | | 173 | EDSGSSSPPGVFL | 6.5197 | 7.1744 | 6.8693 | 6.2920 | 6.0088 | 6.1147 | 6.2830 | 6.2046 | 7.0367 | 6.2830 | 6.7841 | 7.2727 | 4.7965 | 5.7795 | 5.5627 | 5.0182 |
| ARHGAP39_S388 | ARHGAP39 | Q9C0H5-2 | | 604 | PVVRAFSEDEALA | 6.3510 | 6.6821 | 6.5639 | 6.3312 | 5.3173 | 6.6699 | 7.3161 | 6.0650 | 6.6997 | 5.9360 | 6.4594 | 7.1277 | 4.9466 | 5.4325 | 5.8300 | 6.2717 |
| ARHGAP39_S407 | ARHGAP39 | Q9C0H5-2 | | 273 | FPERRPSPFLKRA | 5.4859 | 5.9546 | 5.6815 | 5.9263 | 5.1502 | 6.8210 | 7.0572 | 6.6974 | 5.6383 | 5.4742 | 5.6192 | 5.9339 | 6.3423 | 7.0209 | 7.5124 | 7.6846 |
| ARHGAP39_S488 | ARHGAP39 | Q9C0H5-2 | | 344 | GGYQAGSPQRSPG | 5.7183 | 6.9766 | 6.6494 | 6.7811 | 6.0841 | 6.4206 | 5.5201 | 7.3410 | 6.7704 | 5.5480 | 6.8130 | 6.4419 | 5.6065 | 6.4219 | 5.1556 | 5.7515 |
| ARHGAP39_S500 | ARHGAP39 | Q9C0H5-2 | | 407 | VEQAGSSPKLRAG | 6.0464 | 6.2528 | 6.4251 | 6.1999 | 5.8061 | 6.5912 | 6.9379 | 5.7872 | 6.4509 | 6.6655 | 6.0842 | 6.3956 | 6.1433 | 5.7469 | 5.8734 | 6.5937 |
| ARHGAP39_S604 | ARHGAP39 | Q9C0H5-2 | | 299 | RKPSGDSQPSSPR | 6.4125 | 6.4288 | 6.7336 | 7.1580 | 6.0760 | 5.8341 | 6.6328 | 6.7938 | 6.6360 | 6.3843 | 6.7434 | 6.5077 | 5.5894 | 5.4747 | 5.0135 | 5.5814 |
| ARHGAP39_S690 | ARHGAP39 | Q9C0H5-2 | | 488 | LSSTGYSPGTRKR | 5.4162 | 5.9229 | 6.7756 | 6.0650 | 5.8163 | 7.3364 | 6.9981 | 5.6201 | 5.7931 | 7.5295 | 7.2453 | 6.0496 | 5.3822 | 5.3189 | 6.1562 | 6.5748 |
| ARHGAP4_S946 | ARHGAP4 | P98171-2 | | 946 | PSPGPRSPKAPPS | 6.5232 | 5.9829 | 7.5091 | 6.5162 | 6.5753 | 7.3652 | 6.4485 | 6.0490 | 6.9968 | 6.3508 | 7.0854 | 6.8636 | 4.8880 | 5.9792 | 4.1302 | 4.7365 |
| ARHGAP45_S23 | ARHGAP45 | Q92619 | | 569 | VSANAWSPVMRAR | 6.8936 | 7.1929 | 5.9103 | 7.6578 | 9.0205 | 5.0019 | 3.9278 | 8.9189 | 5.0232 | 6.8722 | 6.9738 | 7.1127 | 4.5530 | 5.6485 | 5.6378 | 3.6552 |
| ARHGAP45_S23;S25 | ARHGAP45 | Q92619 | | 23 | KKNRAGSPSPQPS | 5.8291 | 6.3111 | 6.9518 | 6.5028 | 5.4203 | 5.9630 | 6.1490 | 6.4739 | 5.9313 | 6.3485 | 6.3999 | 6.9553 | 6.0377 | 5.9411 | 6.1392 | 6.6459 |
| ARHGAP45_S569 | ARHGAP45 | Q92619 | | 951 | EATVSLSSLVDYP | 6.0806 | 5.4655 | 6.6502 | 6.0203 | 6.5477 | 6.1547 | 5.4245 | 5.6865 | 6.9612 | 6.0510 | 6.3734 | 5.7298 | 7.7301 | 6.4224 | 6.1957 | 6.5067 |
| ARHGAP45_S578 | ARHGAP45 | Q92619 | | 578 | MRARKSSFNVSDV | 5.6128 | 5.8850 | 5.2270 | 5.4680 | 5.8798 | 5.4949 | 6.2135 | 6.3002 | 5.5469 | 5.8451 | 6.8645 | 6.2759 | 7.9333 | 7.1921 | 7.4175 | 6.8437 |
| ARHGAP45_S951 | ARHGAP45 | Q92619 | 23;25 | | NA | 5.8123 | 5.4546 | 6.2837 | 5.7380 | 6.3447 | 6.4867 | 6.2443 | 7.2935 | 6.0843 | 6.7091 | 6.3592 | 6.3854 | 6.2611 | 6.4545 | 6.0123 | 6.0763 |
| ARHGAP5_S1202 | ARHGAP5 | Q13017 | | 1202 | EEDPLLSPVETWK | 6.4250 | 6.1858 | 6.5129 | 7.3103 | 7.3808 | 6.3013 | 7.5444 | 7.2234 | 6.3984 | 6.5487 | 6.8308 | 7.3675 | 4.7526 | 4.6687 | 4.2813 | 4.2680 |
| ARHGEF1_S25 | ARHGEF1 | Q92888-3 | | 791 | RPKPRPSPSSTRE | 6.1447 | 5.5293 | 6.7035 | 6.5255 | 6.2946 | 6.8771 | 6.5717 | 6.4346 | 6.6843 | 6.7651 | 6.8916 | 7.0610 | 5.3569 | 5.7595 | 4.8771 | 5.5234 |
| ARHGEF1_S791 | ARHGEF1 | Q92888-3 | | 25 | FARGAASPGPSRP | 6.2595 | 6.6416 | 6.7782 | 6.3747 | 5.8042 | 6.6132 | 6.8556 | 7.7465 | 7.1058 | 7.0048 | 6.7747 | 6.7586 | 4.4919 | 5.2818 | 4.5876 | 4.9213 |
| ARHGEF1_S878 | ARHGEF1 | Q92888-3 | | 878 | PGGGPLSPARTQE | 6.1847 | 6.6720 | 6.9531 | 6.7614 | 5.9620 | 6.5079 | 6.6749 | 6.4289 | 6.2979 | 6.3509 | 6.8640 | 6.7096 | 5.2862 | 5.6725 | 5.2561 | 5.4180 |
| ARHGEF1_T710 | ARHGEF1 | Q92888-3 | | 710 | SHSRTLTPTPDGK | 6.2688 | 6.1944 | 7.1691 | 6.2869 | 5.9834 | 6.2218 | 6.3901 | 6.8594 | 5.9644 | 6.3458 | 6.5278 | 7.0376 | 5.4605 | 6.0753 | 5.4467 | 5.7680 |
| ARHGEF10_S379 | ARHGEF10 | O15013 | | 379 | SFIRTKSLIAQDH | 6.5282 | 6.1568 | 7.6778 | 6.5645 | 5.6662 | 6.2055 | 6.0539 | 6.1227 | 6.6358 | 6.4387 | 6.6238 | 6.5546 | 5.5478 | 6.5857 | 5.2576 | 5.3806 |
| ARHGEF10_S59 | ARHGEF10 | O15013 | | 59 | EQFDFDSGDEIPE | 6.6292 | 6.4518 | 6.3561 | 6.5531 | 6.5986 | 6.2125 | 6.0084 | 6.3004 | 6.4338 | 6.4165 | 6.7359 | 6.4856 | 4.9694 | 5.9590 | 5.5878 | 6.3019 |
| ARHGEF11_S1335 | ARHGEF11 | O15085-2 | | 1520 | WTDGSLSPPAKEP | 5.9865 | 6.2963 | 7.4448 | 6.2496 | 6.1915 | 6.6375 | 6.5326 | 6.3861 | 6.2193 | 6.1718 | 6.5396 | 6.2610 | 5.7151 | 6.1072 | 5.6976 | 5.5637 |
| ARHGEF11_S150 | ARHGEF11 | O15085-2 | | 1335 | PPRTRNSGIWESP | 6.0987 | 5.9468 | 6.8036 | 5.9695 | 6.1172 | 6.2575 | 6.1838 | 6.1446 | 6.2958 | 6.3756 | 6.1590 | 6.6276 | 6.1394 | 6.6512 | 6.1683 | 6.0615 |
| ARHGEF11_S1518;S1520 | ARHGEF11 | O15085-2 | | 703 | SSLSTRSLENPTP | 5.8265 | 6.4516 | 6.6126 | 6.6664 | 4.9983 | 5.0725 | 5.6866 | 5.6296 | 6.4863 | 6.8230 | 6.6596 | 5.4229 | 6.7296 | 7.0415 | 7.0681 | 6.8249 |
| ARHGEF11_S1520 | ARHGEF11 | O15085-2 | | 708 | RSLENPTPPFTPK | 6.1176 | 6.4748 | 6.1727 | 5.7715 | 5.9873 | 6.1008 | 5.6981 | 6.2067 | 6.6566 | 6.5379 | 6.5097 | 6.3988 | 6.1991 | 6.3975 | 6.2340 | 6.5369 |
| ARHGEF11_S1559 | ARHGEF11 | O15085-2 | | 1559 | TADAAASPGPxxx | 5.9945 | 5.4939 | 6.0744 | 6.5629 | 6.9037 | 6.5922 | 6.8065 | 5.4154 | 6.0451 | 7.0235 | 6.3699 | 6.7040 | 5.0906 | 5.5885 | 6.7160 | 6.6188 |
| ARHGEF11_S291;S295 | ARHGEF11 | O15085-2 | | 295 | LDSPRTSPVIMAR | 5.9385 | 6.4053 | 6.9204 | 6.5429 | 8.8228 | 5.6897 | 5.5697 | 5.6956 | 5.6209 | 5.1307 | 5.0282 | 5.4058 | 7.6068 | 6.9555 | 6.9336 | 5.7336 |
| ARHGEF11_S295 | ARHGEF11 | O15085-2 | | 150 | ITSVIPSPPPPPP | 5.9292 | 7.1586 | 6.1751 | 5.8677 | 5.7007 | 7.4579 | 6.1684 | 6.0829 | 5.8891 | 6.7898 | 6.7644 | 7.4045 | 4.6236 | 5.7007 | 6.4584 | 5.8290 |
| ARHGEF11_S629 | ARHGEF11 | O15085-2 | | 722 | GRRSIESPSLGFC | 6.4973 | 5.7580 | 7.4393 | 6.8284 | 6.2656 | 6.3168 | 5.3758 | 4.9304 | 6.0289 | 7.4694 | 6.8053 | 7.0962 | 5.9356 | 5.9286 | 5.8844 | 5.4400 |
| ARHGEF11_S703 | ARHGEF11 | O15085-2 | | 629 | PGTQRLSTGSFPE | 5.6149 | 5.6178 | 5.8780 | 6.9214 | 5.9019 | 5.8211 | 5.3069 | 6.5865 | 5.7853 | 7.2443 | 6.9812 | 7.1636 | 7.2234 | 6.7181 | 5.7315 | 5.5042 |
| ARHGEF11_S703;T708 | ARHGEF11 | O15085-2 | 703;708 | | NA | 5.4447 | 5.8829 | 6.1172 | 6.1662 | 5.3398 | 6.0372 | 6.0299 | 6.6918 | 6.4852 | 6.4554 | 6.3558 | 6.9356 | 6.2461 | 6.6623 | 6.8773 | 6.2727 |
| ARHGEF11_S722 | ARHGEF11 | O15085-2 | 291;295 | | NA | 5.9235 | 6.4266 | 6.5712 | 5.7635 | 6.0309 | 6.2110 | 6.6700 | 5.8797 | 6.2184 | 6.5150 | 6.4509 | 6.6741 | 6.6038 | 6.4829 | 6.0180 | 5.5604 |
| ARHGEF11_T708 | ARHGEF11 | O15085-2 | 1518;1520 | | NA | 6.5839 | 5.9934 | 5.2903 | 7.0304 | 7.2306 | 5.1292 | 5.2591 | 7.7213 | 5.8930 | 7.9109 | 6.1920 | 6.7865 | 5.5218 | 6.0553 | 5.4688 | 5.9335 |
| ARHGEF12_S1176 | ARHGEF12 | Q9NZN5 | | 736 | RVTEHGTPKPFRK | 6.5502 | 6.6238 | 6.4607 | 6.3503 | 5.9459 | 6.3700 | 6.2335 | 6.2364 | 6.2644 | 6.0981 | 6.3986 | 6.1641 | 6.1234 | 6.4611 | 5.7238 | 5.9959 |
| ARHGEF12_S1288 | ARHGEF12 | Q9NZN5 | | 1176 | SVTGLQSPDRDLG | 6.0527 | 6.6211 | 6.5385 | 6.7688 | 6.3600 | 6.4406 | 6.0994 | 6.6564 | 6.7401 | 6.5324 | 6.9362 | 6.6422 | 5.3809 | 5.6384 | 5.4732 | 5.1190 |
| ARHGEF12_S22 | ARHGEF12 | Q9NZN5 | | 22 | KPIRHGSILNRES | 7.9901 | 6.9270 | 6.6164 | 5.7113 | 5.7251 | 5.9907 | 5.9855 | 5.5546 | 7.3250 | 6.4224 | 6.7246 | 6.8503 | 6.5474 | 6.4572 | 4.0232 | 5.1492 |
| ARHGEF12_S41 | ARHGEF12 | Q9NZN5 | | 41 | KVERIASHDFDPT | 7.2014 | 6.2635 | 7.0129 | 6.7193 | 7.7836 | 6.0479 | 5.2074 | 6.7757 | 8.1265 | 7.1248 | 6.3044 | 6.7129 | 5.0327 | 5.5976 | 3.7999 | 4.2897 |
| ARHGEF12_S608 | ARHGEF12 | Q9NZN5 | | 608 | GPPRRPSRHDNSA | 8.1143 | 7.2774 | 6.7937 | 5.3330 | 6.0027 | 7.2956 | 5.4925 | 6.0319 | 6.4562 | 7.0176 | 7.2221 | 7.1286 | 5.4087 | 6.2876 | 3.7291 | 4.4090 |
| ARHGEF12_T736 | ARHGEF12 | Q9NZN5 | | 1288 | PRYRTASQGPQTD | 6.0303 | 5.9607 | 6.2916 | 5.8412 | 5.8408 | 6.6877 | 6.2471 | 5.7170 | 5.5525 | 5.3486 | 6.3647 | 6.2533 | 7.3416 | 6.7463 | 7.0330 | 6.7436 |
| ARHGEF16_S107 | ARHGEF16 | Q5VV41 | | 107 | TPARHQSFGAAVL | 8.4374 | 6.9690 | 6.6317 | 6.9556 | 6.6987 | 6.1176 | 6.6260 | 5.6481 | 6.9670 | 5.1267 | 5.5697 | 6.5012 | 6.0964 | 6.8987 | 3.8712 | 4.8849 |
| ARHGEF16_S577 | ARHGEF16 | Q5VV41 | | 577 | GGGNRSSSVPHPF | 7.1913 | 6.0862 | 6.8860 | 6.9821 | 6.4053 | 6.8410 | 6.7847 | 6.9536 | 6.2372 | 5.8161 | 5.8796 | 6.4093 | 5.5552 | 5.4347 | 5.5717 | 4.9659 |
| ARHGEF17_S1331 | ARHGEF17 | Q96PE2 | | 348 | REWGSGSPPCVPG | 6.2119 | 6.5755 | 6.8672 | 6.8449 | 6.1846 | 6.1966 | 6.3515 | 6.4404 | 6.5934 | 6.7307 | 6.7586 | 6.5640 | 5.1619 | 5.7036 | 5.3599 | 5.4553 |
| ARHGEF17_S142 | ARHGEF17 | Q96PE2 | | 420 | GVYRSPSFGAGEG | 6.7052 | 6.3278 | 6.3423 | 6.1419 | 5.9372 | 6.1288 | 6.0195 | 6.2097 | 6.4727 | 6.2011 | 6.0991 | 6.4392 | 6.6379 | 6.3572 | 5.9201 | 6.0604 |
| ARHGEF17_S320;S332 | ARHGEF17 | Q96PE2 | | 914 | KLADILSPRLIRR | 7.1336 | 8.0255 | 7.3494 | 7.1752 | 4.4803 | 7.1120 | 8.0558 | 6.6779 | 6.5507 | 7.4051 | 7.5233 | 7.8191 | 2.8137 | 4.1006 | 3.2333 | 4.5447 |
| ARHGEF17_S348 | ARHGEF17 | Q96PE2 | | 546 | TLRRAKSFTCSEK | 7.2601 | 7.0202 | 7.0560 | 6.1839 | 5.8640 | 6.6175 | 6.1571 | 5.9300 | 6.4846 | 6.1928 | 6.4795 | 6.7735 | 5.6239 | 5.8451 | 4.8755 | 5.6362 |
| ARHGEF17_S420 | ARHGEF17 | Q96PE2 | | 79 | RPLRSLSPSVRQL | 5.7717 | 6.2703 | 5.8527 | 6.2131 | 6.0017 | 6.2959 | 6.2860 | 6.2335 | 6.0122 | 5.4024 | 5.5261 | 5.9012 | 7.0190 | 7.1284 | 7.0220 | 7.0638 |
| ARHGEF17_S461 | ARHGEF17 | Q96PE2 | | 461 | KSRQRKSLSNPDI | 6.5017 | 6.3272 | 7.1468 | 6.5093 | 5.6656 | 6.0172 | 6.2458 | 6.4153 | 6.2939 | 5.7455 | 6.4120 | 6.0615 | 6.0506 | 6.7072 | 6.0850 | 5.8155 |
| ARHGEF17_S507;S527 | ARHGEF17 | Q96PE2 | | 764 | NLLGSLSPKTGLP | 6.0954 | 5.8680 | 6.0118 | 6.2784 | 6.3930 | 6.5928 | 6.7206 | 6.5788 | 6.7132 | 6.1805 | 6.6337 | 6.3621 | 6.2340 | 6.4134 | 5.3466 | 5.5776 |
| ARHGEF17_S527 | ARHGEF17 | Q96PE2 | | 527 | PIPAPASPGTRPT | 6.3131 | 5.7282 | 6.4105 | 5.9684 | 6.1930 | 6.0199 | 5.9100 | 5.8705 | 5.9790 | 6.1868 | 6.2094 | 6.4838 | 6.5542 | 6.8975 | 6.6019 | 6.6738 |
| ARHGEF17_S546 | ARHGEF17 | Q96PE2 | | 569 | ALKSSSSELLLTG | 5.9948 | 5.9113 | 6.2354 | 7.0856 | 6.1656 | 5.8399 | 6.2194 | 5.6093 | 5.9597 | 6.4023 | 5.8427 | 6.6627 | 6.3051 | 7.1768 | 6.5770 | 6.0124 |
| ARHGEF17_S569 | ARHGEF17 | Q96PE2 | | 619 | SDAPPGSPDWAGD | 6.1978 | 5.5977 | 5.5592 | 6.2593 | 6.6253 | 6.4033 | 6.5693 | 6.7550 | 6.9661 | 5.7702 | 7.4958 | 6.2088 | 5.7768 | 6.0626 | 5.5109 | 6.2418 |
| ARHGEF17_S619 | ARHGEF17 | Q96PE2 | | 142 | PGSRRPSADSESP | 5.7624 | 5.8423 | 6.5062 | 6.0078 | 6.1043 | 5.6260 | 5.4421 | 6.4139 | 7.3116 | 6.6443 | 6.7100 | 6.6185 | 5.5386 | 7.8815 | 6.2233 | 5.3672 |
| ARHGEF17_S735 | ARHGEF17 | Q96PE2 | | 735 | EAYRSLSDPIPQR | 5.7264 | 6.7603 | 6.6064 | 6.3614 | 6.2839 | 6.1142 | 6.0232 | 6.6043 | 5.9457 | 5.9428 | 6.2371 | 6.3464 | 6.0164 | 6.3014 | 6.2167 | 6.5135 |
| ARHGEF17_S764 | ARHGEF17 | Q96PE2 | | 1331 | GSLRRSSMSLYTA | 6.4358 | 5.1623 | 5.8894 | 6.0564 | 6.2119 | 6.3883 | 6.7108 | 6.3430 | 7.2018 | 6.4329 | 7.0924 | 6.1738 | 6.0262 | 6.0759 | 5.8786 | 5.9204 |
| ARHGEF17_S79 | ARHGEF17 | Q96PE2 | 507;527 | | NA | 6.6863 | 6.6336 | 6.7376 | 6.9412 | 6.0390 | 6.7482 | 6.5359 | 6.8093 | 6.4136 | 6.3064 | 6.7467 | 6.6291 | 5.4079 | 5.3970 | 4.9413 | 5.0271 |
| ARHGEF17_S914 | ARHGEF17 | Q96PE2 | 320;332 | | NA | 6.3735 | 5.8430 | 6.1277 | 7.0409 | 6.8714 | 6.4412 | 6.1195 | 5.7738 | 6.1166 | 5.8804 | 7.5540 | 7.0388 | 5.6512 | 6.7607 | 5.0402 | 5.3671 |
| ARHGEF18_S1261 | ARHGEF18 | Q6ZSZ5 | | 1291 | RNRRSLSPILPGR | 6.0555 | 5.9377 | 7.0317 | 5.6898 | 6.1481 | 6.5118 | 6.1240 | 6.6904 | 6.2691 | 6.2587 | 6.4725 | 6.4609 | 5.9485 | 6.3344 | 6.0887 | 5.9783 |
| ARHGEF18_S1264 | ARHGEF18 | Q6ZSZ5 | | 948 | WRGPPNSPDLKLS | 6.9526 | 7.0789 | 6.9034 | 6.5545 | 6.3793 | 6.7626 | 6.8294 | 6.8157 | 6.9518 | 7.0925 | 7.1166 | 7.2844 | 4.0761 | 4.5918 | 3.9475 | 4.6630 |
| ARHGEF18_S1291 | ARHGEF18 | Q6ZSZ5 | | 921 | GYDCTNSPTKNGS | 6.1019 | 6.2860 | 6.8087 | 6.8559 | 6.1770 | 6.3142 | 6.5346 | 7.0201 | 6.6183 | 6.6097 | 6.6574 | 6.7698 | 5.0587 | 5.6342 | 5.0376 | 5.5159 |
| ARHGEF18_S1312 | ARHGEF18 | Q6ZSZ5 | | 1348 | LPATPLSAKEDAS | 5.7790 | 5.5684 | 6.9858 | 6.4431 | 5.6089 | 6.6375 | 5.2282 | 6.7428 | 5.6210 | 4.8354 | 3.7096 | 6.2244 | 9.2253 | 8.1521 | 6.6051 | 6.6335 |
| ARHGEF18_S1348 | ARHGEF18 | Q6ZSZ5 | | 1345 | PSPLPATPLSAKE | 5.4377 | 6.6266 | 7.5305 | 6.6231 | 5.1317 | 6.9607 | 5.2794 | 7.5622 | 7.9842 | 6.7989 | 8.2445 | 7.4953 | 4.7132 | 5.4448 | 3.8655 | 4.3016 |
| ARHGEF18_S921 | ARHGEF18 | Q6ZSZ5 | | 1312 | PGFPAPSPPPADS | 5.1924 | 6.3981 | 7.3644 | 6.9217 | 5.8420 | 6.5542 | 5.5283 | 6.2751 | 6.7833 | 6.6713 | 6.0379 | 6.8547 | 6.2390 | 6.1491 | 5.8950 | 5.2933 |
| ARHGEF18_S948 | ARHGEF18 | Q6ZSZ5 | | 1264 | RWESSASFDLKQQ | 6.2961 | 6.0811 | 9.3504 | 7.9697 | 6.0242 | 5.6047 | 5.8050 | 6.1317 | 6.8673 | 7.4154 | 7.2109 | 8.2585 | 3.9290 | 4.6815 | 4.1419 | 4.2325 |
| ARHGEF18_T1345 | ARHGEF18 | Q6ZSZ5 | | 1261 | GSQRWESSASFDL | 6.9914 | 6.3737 | 8.1025 | 6.1598 | 5.8854 | 6.1692 | 5.6277 | 6.1494 | 7.8312 | 8.3498 | 7.0174 | 7.5255 | 3.7268 | 5.8573 | 3.8551 | 4.3778 |
| ARHGEF2_S137 | ARHGEF2 | Q92974 | | 174 | RRILSQSTDSLNM | 6.4387 | 6.3850 | 6.6972 | 5.7363 | 5.2156 | 6.6543 | 7.0531 | 6.2805 | 6.8330 | 6.4886 | 6.3491 | 6.4196 | 6.0726 | 5.9802 | 5.1116 | 6.2846 |
| ARHGEF2_S143 | ARHGEF2 | Q92974 | | 696 | DSGGNTSPGVTAN | 6.1680 | 6.1002 | 6.3485 | 6.1081 | 6.2815 | 6.1180 | 6.3735 | 6.3689 | 6.5754 | 6.5784 | 6.2944 | 6.1583 | 5.9160 | 6.3176 | 6.3185 | 5.9748 |
| ARHGEF2_S151 | ARHGEF2 | Q92974 | | 143 | SRRGRSSLSLAKS | 8.8058 | 6.5133 | 6.9022 | 5.5918 | 5.4598 | 6.2219 | 6.6955 | 6.0019 | 9.9989 | 7.7333 | 7.3882 | 7.1003 | 4.8440 | 4.8771 | 2.4903 | 3.3755 |
| ARHGEF2_S163 | ARHGEF2 | Q92974 | | 163 | GHFNDESPLGLRR | 6.3351 | 7.1490 | 6.3987 | 6.6291 | 5.6433 | 7.5825 | 7.1267 | 7.0397 | 7.0769 | 6.9957 | 6.9630 | 6.7474 | 4.4586 | 5.0782 | 4.2848 | 4.4914 |
| ARHGEF2_S174 | ARHGEF2 | Q92974 | | 137 | RQSLLGSRRGRSS | 9.0351 | 7.6784 | 8.4953 | 6.7774 | 4.7154 | 6.5812 | 7.0129 | 6.3293 | 8.3927 | 7.0092 | 7.5752 | 6.6700 | 3.6884 | 4.0928 | 2.3880 | 3.5589 |
| ARHGEF2_S645 | ARHGEF2 | Q92974 | | 886 | VDPRRRSLPAGDA | 5.9343 | 5.5282 | 6.6302 | 6.1154 | 6.4694 | 6.2697 | 5.7609 | 6.1974 | 7.2076 | 7.1634 | 6.6591 | 7.1475 | 6.1276 | 5.9899 | 5.0390 | 5.7605 |
| ARHGEF2_S696 | ARHGEF2 | Q92974 | | 151 | SLAKSVSTTNIAG | 7.0336 | 5.9276 | 6.9249 | 6.8680 | 4.7964 | 6.9477 | 7.2770 | 5.1044 | 6.7422 | 6.0913 | 7.0620 | 6.5382 | 5.7292 | 5.8498 | 4.6498 | 6.4580 |
| ARHGEF2_S886 | ARHGEF2 | Q92974 | | 932 | ERQELGSPEERLQ | 6.4738 | 6.6867 | 6.5888 | 6.0770 | 5.6839 | 6.3985 | 6.4972 | 6.6080 | 6.6398 | 6.5570 | 6.5958 | 6.5789 | 5.6472 | 5.9162 | 5.7311 | 5.3199 |
| ARHGEF2_S932 | ARHGEF2 | Q92974 | | 645 | GLFRSESLESPRG | 4.2412 | 4.1329 | 4.5342 | 4.1344 | 5.4321 | 5.8909 | 5.7011 | 4.8774 | 4.2325 | 4.2475 | 4.3118 | 4.7419 | 11.0402 | 10.2884 | 11.5212 | 10.6721 |
| ARHGEF2_S956;S960 | ARHGEF2 | Q92974 | 956;960 | | NA | 5.4336 | 5.8220 | 5.9807 | 5.9773 | 6.0642 | 5.8063 | 6.1095 | 6.6654 | 6.7870 | 6.6003 | 6.1798 | 6.2855 | 6.3334 | 6.6709 | 6.4894 | 6.7948 |
| ARHGEF26_S124 | ARHGEF26 | Q96DR7 | | 422 | STWSQLSAVKRKG | 7.7021 | 6.9956 | 7.7021 | 7.5391 | 6.4447 | 6.0317 | 6.2554 | 5.9120 | 6.9478 | 6.5437 | 6.5730 | 6.8026 | 4.5793 | 5.5921 | 4.0588 | 4.3199 |
| ARHGEF26_S222 | ARHGEF26 | Q96DR7 | | 124 | KAVPGGSPKSPAN | 6.5454 | 6.3656 | 6.6265 | 6.9485 | 6.2311 | 6.1828 | 6.8423 | 6.3024 | 7.6883 | 6.5932 | 6.8500 | 7.0597 | 5.0276 | 5.1068 | 4.4770 | 5.1527 |
| ARHGEF26_S333 | ARHGEF26 | Q96DR7 | | 80 | SRTVHRSPLLLGA | 6.8178 | 6.4232 | 6.7780 | 6.6174 | 6.5202 | 6.2926 | 6.3303 | 6.5619 | 7.0381 | 6.3757 | 6.3917 | 6.0949 | 5.4585 | 5.7809 | 5.1053 | 5.4135 |
| ARHGEF26_S419 | ARHGEF26 | Q96DR7 | | 222 | PLQRLPSQENELL | 5.6887 | 6.3852 | 5.7330 | 6.7915 | 6.6931 | 5.5162 | 6.5077 | 6.4988 | 6.8608 | 6.2207 | 6.2167 | 6.3602 | 6.1272 | 6.5859 | 5.6855 | 6.1288 |
| ARHGEF26_S422 | ARHGEF26 | Q96DR7 | | 333 | FDSPTSSKKKNRM | 8.7078 | 6.8718 | 9.9751 | 10.2557 | 7.1394 | 4.8129 | 6.3868 | 5.7383 | 6.2326 | 7.3921 | 7.3457 | 6.6916 | 3.1665 | 4.5137 | 2.5161 | 2.2541 |
| ARHGEF26_S80 | ARHGEF26 | Q96DR7 | | 419 | PLRSTWSQLSAVK | 6.6644 | 6.8942 | 6.4717 | 6.9053 | 5.9480 | 6.8374 | 6.1580 | 6.2037 | 7.4649 | 6.4606 | 7.3266 | 6.2963 | 5.2253 | 6.9350 | 3.6725 | 4.5360 |
| ARHGEF28_S1538 | ARHGEF28 | Q8N1W1-6 | | 535 | NISRAESLPLSSN | 5.6960 | 5.5995 | 5.8031 | 5.5977 | 5.8787 | 5.8925 | 5.8852 | 6.1445 | 5.2717 | 5.4647 | 5.6591 | 5.8582 | 8.1231 | 7.9335 | 8.0684 | 7.1242 |
| ARHGEF28_S535 | ARHGEF28 | Q8N1W1-6 | | 1538 | KHGRQRSLPAVLL | 6.2792 | 6.5759 | 6.4765 | 6.4190 | 6.1596 | 6.3879 | 6.1425 | 5.5617 | 7.3043 | 5.9593 | 6.1860 | 6.3522 | 5.8940 | 6.2777 | 5.5151 | 6.5091 |
| ARHGEF28_S624 | ARHGEF28 | Q8N1W1-6 | | 624 | KVSRTFSFLMNRM | 7.9523 | 7.0885 | 7.2955 | 6.0640 | 5.6751 | 6.5703 | 6.7570 | 5.5587 | 7.0408 | 6.3648 | 7.3817 | 6.5706 | 5.3174 | 5.3089 | 4.3044 | 4.7501 |
| ARHGEF3_S127 | ARHGEF3 | Q9NR81-2 | | 127 | RNAAPSSTKRRDS | 6.6959 | 6.6777 | 6.6082 | 5.9835 | 5.0795 | 6.2202 | 5.9895 | 5.7949 | 7.4165 | 7.9329 | 7.8809 | 8.9981 | 4.0968 | 6.0652 | 3.7004 | 4.8597 |
| ARHGEF3_S79 | ARHGEF3 | Q9NR81-2 | | 79 | PLSRVTSLANLIP | 7.4972 | 5.7584 | 5.1590 | 4.7195 | 5.5054 | 4.6893 | 5.2855 | 4.4744 | 6.2673 | 4.8985 | 5.3516 | 6.0259 | 8.4456 | 10.5424 | 7.7337 | 7.6461 |
| ARHGEF40_S1187 | ARHGEF40 | Q8TER5 | | 214 | GPPGLPSPPLPEE | 6.0020 | 6.3285 | 6.1466 | 6.3090 | 6.4101 | 6.3645 | 6.5615 | 6.4148 | 6.5100 | 6.5413 | 6.7789 | 6.6574 | 5.7616 | 5.9240 | 5.5650 | 5.7246 |
| ARHGEF40_S1504 | ARHGEF40 | Q8TER5 | | 1504 | RGILGLSRQSHAR | 7.5507 | 6.7484 | 7.5666 | 6.1657 | 5.0915 | 6.1140 | 5.7670 | 6.2005 | 7.4977 | 7.8187 | 8.3958 | 8.4114 | 3.7256 | 6.0200 | 3.3255 | 3.6009 |
| ARHGEF40_S1513 | ARHGEF40 | Q8TER5 | | 931 | LALDLGSPAALRE | 6.6066 | 7.6273 | 6.8584 | 7.1394 | 6.6072 | 6.8591 | 7.5323 | 6.9090 | 7.4938 | 7.6124 | 6.8783 | 7.0601 | 3.2924 | 4.4336 | 3.3941 | 3.6960 |
| ARHGEF40_S214 | ARHGEF40 | Q8TER5 | | 401 | GDSAPLSPGDKED | 6.1089 | 6.3239 | 7.2549 | 6.6753 | 6.5325 | 6.4536 | 6.9382 | 6.4061 | 6.2898 | 6.8606 | 6.9069 | 7.0303 | 4.4976 | 5.7995 | 5.0268 | 4.8951 |
| ARHGEF40_S226 | ARHGEF40 | Q8TER5 | | 1513 | SHARALSDPTTPL | 5.9546 | 7.0620 | 6.2816 | 6.1974 | 6.4856 | 6.1513 | 5.5030 | 6.0118 | 6.3295 | 5.6105 | 5.9903 | 6.5950 | 6.0438 | 6.6159 | 6.7855 | 6.3824 |
| ARHGEF40_S262 | ARHGEF40 | Q8TER5 | | 262 | PTDAEGSPGLSRV | 6.4303 | 6.7344 | 6.3678 | 6.0574 | 6.4218 | 6.7052 | 6.5242 | 6.6526 | 6.5826 | 7.0797 | 6.4784 | 6.5114 | 5.1579 | 5.9505 | 5.2659 | 5.0801 |
| ARHGEF40_S401 | ARHGEF40 | Q8TER5 | | 1074 | GPWGVGTPRMERK | 7.6857 | 7.9669 | 6.8468 | 7.0176 | 5.7107 | 7.0169 | 7.4503 | 6.3982 | 7.5219 | 6.5866 | 6.5859 | 6.3817 | 4.3245 | 4.7909 | 3.3951 | 4.3203 |
| ARHGEF40_S931 | ARHGEF40 | Q8TER5 | | 961 | HVGEEASPRGYRR | 7.1527 | 7.1321 | 6.5643 | 7.3551 | 5.8031 | 6.5058 | 6.7267 | 6.4250 | 6.8527 | 7.0800 | 7.5117 | 7.1545 | 3.8721 | 4.9476 | 4.4848 | 4.4319 |
| ARHGEF40_S961 | ARHGEF40 | Q8TER5 | | 1187 | NGLAALSPLSKGS | 6.3629 | 7.0174 | 6.9509 | 7.1202 | 5.9333 | 6.5676 | 6.4754 | 6.8166 | 7.0803 | 7.2724 | 6.9033 | 6.7655 | 3.9916 | 5.6377 | 4.2943 | 4.8105 |
| ARHGEF40_S986 | ARHGEF40 | Q8TER5 | | 1492 | PHPGSSTPTLASR | 6.2455 | 6.7847 | 6.6235 | 6.1130 | 5.7522 | 7.4424 | 6.1445 | 6.9205 | 6.7912 | 6.3764 | 6.5910 | 6.3145 | 5.2283 | 6.6124 | 5.0332 | 5.0267 |
| ARHGEF40_T1074 | ARHGEF40 | Q8TER5 | | 226 | EALGTRSPGDGHN | 6.1403 | 7.0597 | 7.0559 | 5.7994 | 6.3596 | 6.9931 | 6.8424 | 6.5141 | 6.5208 | 6.6850 | 6.3982 | 7.4122 | 4.6164 | 4.5246 | 4.8742 | 6.2041 |
| ARHGEF40_T1492 | ARHGEF40 | Q8TER5 | | 270 | GLSRVRTVPTRKG | 7.5998 | 7.2067 | 8.4472 | 6.1029 | 4.0150 | 6.1524 | 6.0985 | 5.7263 | 8.2458 | 7.2288 | 8.1201 | 7.6357 | 4.2962 | 5.9851 | 3.0615 | 4.0780 |
| ARHGEF40_T1517 | ARHGEF40 | Q8TER5 | | 1517 | ALSDPTTPLxxxx | 7.7539 | 7.5087 | 8.1601 | 6.9054 | 5.5138 | 7.9341 | 7.9197 | 6.6795 | 8.2274 | 6.7372 | 6.0498 | 6.9415 | 3.0020 | 4.2976 | 2.4420 | 3.9274 |
| ARHGEF40_T270 | ARHGEF40 | Q8TER5 | | 986 | WGPRSPSPSLSSL | 5.3683 | 5.7264 | 6.2896 | 5.7506 | 5.3135 | 5.4652 | 5.4894 | 6.1379 | 6.1188 | 5.0689 | 5.8385 | 5.8219 | 8.1436 | 7.8378 | 8.3870 | 7.2427 |
| ARHGEF5_S11 | ARHGEF5 | Q12774 | | 11 | EAQRGASPPISAI | 6.2354 | 6.4023 | 6.2965 | 6.5993 | 6.4112 | 6.0721 | 6.5129 | 6.4018 | 6.4412 | 5.6102 | 6.3768 | 6.2793 | 6.3071 | 6.2271 | 6.1199 | 5.7071 |
| ARHGEF5_S1126 | ARHGEF5 | Q12774 | | 474 | SFLTEESPDKEID | 6.8929 | 5.9393 | 8.0616 | 7.3001 | 5.4957 | 6.5466 | 7.2216 | 6.5196 | 6.6237 | 8.6432 | 8.1968 | 7.4340 | 3.3680 | 4.2039 | 3.2490 | 4.3040 |
| ARHGEF5_S474 | ARHGEF5 | Q12774 | | 897 | ASAGRTSWPPATA | 6.2387 | 6.7619 | 6.8544 | 6.3132 | 6.4316 | 6.3903 | 6.0148 | 6.5152 | 6.7977 | 5.8397 | 6.2041 | 6.3778 | 5.6785 | 6.1486 | 5.7011 | 5.7323 |
| ARHGEF5_S897 | ARHGEF5 | Q12774 | | 1126 | ETPGPSSPRQPRK | 6.0427 | 6.0963 | 6.6195 | 6.3371 | 6.7310 | 6.3215 | 6.5757 | 5.8718 | 6.8030 | 6.2594 | 6.8694 | 6.0363 | 5.7440 | 5.9226 | 5.7532 | 6.0165 |
| ARHGEF5_T658 | ARHGEF5 | Q12774 | | 658 | SGRDYSTVSASPT | 5.8239 | 6.9254 | 6.6297 | 7.2492 | 6.5261 | 6.7624 | 6.7498 | 6.5625 | 7.2865 | 7.1191 | 7.6493 | 7.1612 | 3.8667 | 5.4502 | 3.9061 | 4.3318 |
| ARHGEF6_S225 | ARHGEF6 | Q15052 | | 488 | SASPRMSGFIYQG | 6.2593 | 6.1176 | 6.6831 | 5.7995 | 6.4733 | 6.1895 | 6.5185 | 6.2731 | 6.6366 | 5.8487 | 5.8699 | 5.9089 | 6.5006 | 6.4572 | 6.4520 | 6.0123 |
| ARHGEF6_S488 | ARHGEF6 | Q15052 | | 225 | SSERPLSPKAVKG | 6.3201 | 6.5555 | 6.5867 | 5.7908 | 6.1207 | 6.7555 | 6.2634 | 6.1386 | 6.5737 | 7.2578 | 6.7608 | 6.4703 | 5.1872 | 6.4044 | 5.3019 | 5.5126 |
| ARHGEF6_S649 | ARHGEF6 | Q15052 | | 684 | SSTRKDSIPQVLL | 5.6472 | 5.0548 | 5.8652 | 5.1307 | 6.9783 | 5.5513 | 5.7124 | 6.0980 | 6.7373 | 7.7593 | 7.0909 | 6.5065 | 6.8423 | 7.9938 | 5.5931 | 5.4390 |
| ARHGEF6_S684 | ARHGEF6 | Q15052 | | 649 | EYVIRKSTAALEE | 6.3257 | 6.0262 | 6.2785 | 6.5798 | 6.6173 | 6.3678 | 6.2114 | 6.1614 | 6.6833 | 5.2687 | 5.5002 | 5.8997 | 7.4130 | 6.8233 | 6.2997 | 5.5439 |
| ARHGEF7_S601 | ARHGEF7 | Q14155-5 | | 645 | SSSRKESAPQVLL | 6.6291 | 5.4106 | 6.1931 | 5.5817 | 5.8647 | 5.3844 | 5.7564 | 5.9957 | 7.5892 | 7.0670 | 6.1162 | 6.7590 | 6.7159 | 6.7916 | 6.3695 | 5.7762 |
| ARHGEF7_S645 | ARHGEF7 | Q14155-5 | | 601 | KPERKPSDEEFAS | 5.8596 | 5.9670 | 6.1059 | 5.7846 | 6.4118 | 6.2421 | 6.0622 | 6.2883 | 5.9766 | 5.3840 | 5.7866 | 5.7629 | 7.2598 | 7.3609 | 7.1372 | 6.6104 |
| ARID1A_S1184 | ARID1A | O14497 | | 1755 | RFSKVSSPAPMEG | 5.9004 | 6.2149 | 6.1217 | 6.4124 | 6.1190 | 5.8906 | 5.9696 | 5.8944 | 6.1578 | 6.1236 | 5.9264 | 6.1285 | 6.8383 | 6.7767 | 6.6870 | 6.8387 |
| ARID1A_S1604 | ARID1A | O14497 | | 604 | PPPQELSQDSFGS | 4.5703 | 5.4094 | 7.0214 | 6.2934 | 4.9798 | 6.5484 | 6.3834 | 6.8919 | 5.9385 | 7.2593 | 7.5862 | 7.2833 | 5.7099 | 6.7327 | 5.9971 | 5.3949 |
| ARID1A_S1751 | ARID1A | O14497 | | 696 | PGIRGPSPSPVGS | 5.8735 | 6.0064 | 5.9577 | 6.1937 | 6.3561 | 6.2792 | 6.2990 | 6.5205 | 5.9445 | 5.8470 | 6.0936 | 5.8869 | 6.9416 | 6.7362 | 6.6092 | 6.4549 |
| ARID1A_S1755 | ARID1A | O14497 | | 382 | RTPQPSSPMDQMG | 6.1072 | 6.2679 | 6.1779 | 6.2231 | 6.3585 | 6.3686 | 6.4307 | 6.4655 | 6.2177 | 6.3179 | 6.1965 | 6.3772 | 6.3964 | 6.1402 | 6.0126 | 5.9421 |
| ARID1A_S1944 | ARID1A | O14497 | | 772 | QPGSALSPRQPSG | 6.0098 | 5.7943 | 6.6955 | 6.7127 | 6.5218 | 6.3441 | 6.8892 | 6.6436 | 6.4271 | 6.6042 | 6.3324 | 6.5908 | 5.5837 | 5.7214 | 5.4742 | 5.6551 |
| ARID1A_S233 | ARID1A | O14497 | | 1751 | LDPGRFSKVSSPA | 6.4022 | 6.3246 | 5.4983 | 6.1935 | 6.6222 | 5.4371 | 5.6233 | 5.2210 | 5.9941 | 5.2352 | 4.8617 | 4.8758 | 8.4050 | 7.8381 | 7.9971 | 7.4707 |
| ARID1A_S301 | ARID1A | O14497 | | 1604 | RTSPSKSPFLHSG | 6.3804 | 6.5807 | 6.3647 | 6.6706 | 6.4744 | 6.8503 | 6.9563 | 6.9487 | 6.7719 | 6.5660 | 6.5531 | 6.4544 | 4.7215 | 5.4622 | 4.7312 | 5.5137 |
| ARID1A_S382 | ARID1A | O14497 | | 1944 | KFPFGISPAQSHR | 5.9727 | 6.3374 | 6.1961 | 6.7705 | 5.7952 | 6.1438 | 6.3037 | 6.2006 | 6.3434 | 7.0479 | 5.8682 | 6.8179 | 5.6848 | 6.3455 | 5.7514 | 6.4209 |
| ARID1A_S604 | ARID1A | O14497 | | 233 | PAYALSSPRGGTP | 6.6144 | 6.0524 | 6.1173 | 6.7854 | 6.6275 | 6.6493 | 6.5589 | 6.8618 | 7.1501 | 6.9319 | 6.5439 | 6.9634 | 4.6930 | 5.1538 | 5.0428 | 5.2539 |
| ARID1A_S696 | ARID1A | O14497 | | 1184 | GMSRSNSVGIQDA | 5.1960 | 5.9044 | 6.8429 | 7.6735 | 5.2499 | 5.1852 | 5.4621 | 6.8465 | 6.2783 | 6.2424 | 5.4585 | 5.7569 | 7.1737 | 7.0910 | 6.0590 | 7.5800 |
| ARID1A_S772 | ARID1A | O14497 | | 301 | LNQLLTSPSSARG | 5.5316 | 5.7494 | 6.9104 | 7.4738 | 6.6108 | 6.7876 | 7.1348 | 6.0769 | 5.8657 | 6.3046 | 8.2107 | 6.3668 | 5.4760 | 4.9094 | 4.6474 | 5.9443 |
| ARID1B_S1595 | ARID1B | Q8NFD5-3 | | 516 | MGRSQGSPMDPMV | 6.2090 | 6.7526 | 7.2702 | 6.9382 | 6.2206 | 6.4849 | 6.4993 | 6.7656 | 6.9434 | 7.0701 | 7.1650 | 7.0400 | 4.4464 | 4.7135 | 4.2991 | 5.1820 |
| ARID1B_S1612 | ARID1B | Q8NFD5-3 | | 1612 | RMSPSKSPFLPSM | 6.7470 | 7.1378 | 5.9731 | 6.7828 | 6.0150 | 6.6363 | 6.8633 | 6.8539 | 6.6985 | 6.4946 | 7.0001 | 6.7567 | 4.7846 | 5.5938 | 4.5723 | 5.0903 |
| ARID1B_S516 | ARID1B | Q8NFD5-3 | | 1595 | HISRAPSPASFQR | 4.7832 | 5.2017 | 6.3459 | 5.8217 | 6.8150 | 6.9382 | 6.3171 | 6.4216 | 6.6526 | 7.0875 | 6.8114 | 6.3653 | 6.1329 | 6.4796 | 5.6698 | 6.1566 |
| ARID2_S1300 | ARID2 | Q68CP9 | | 1300 | LNGRKYSDSSLPP | 4.6878 | 4.6859 | 4.9989 | 5.0256 | 6.5949 | 5.6143 | 5.9089 | 5.5710 | 5.3151 | 5.9945 | 5.4406 | 5.6621 | 8.9598 | 8.6645 | 8.5883 | 8.2878 |
| ARID2_T1726 | ARID2 | Q68CP9 | | 1726 | VGGTSSTPRAQKA | 5.9620 | 6.4918 | 5.9043 | 6.7462 | 6.5294 | 6.6474 | 7.1729 | 6.3624 | 6.2339 | 7.2280 | 6.3502 | 7.2367 | 5.4330 | 5.0999 | 5.1541 | 5.4479 |
| ARID4A_S1140;S1145 | ARID4A | P29374 | | 864 | KILGQSSPEKKIR | 6.6945 | 6.7227 | 6.3665 | 6.8174 | 5.9846 | 5.7850 | 6.6414 | 5.9295 | 6.3637 | 6.3738 | 6.0288 | 6.3890 | 5.6726 | 6.1797 | 5.9181 | 6.1324 |
| ARID4A_S1164 | ARID4A | P29374 | | 1164 | EKHPNSSPRTYKW | 6.1874 | 6.9860 | 5.8022 | 7.3121 | 6.2045 | 5.4864 | 6.1754 | 7.0048 | 6.4125 | 5.6362 | 6.6137 | 5.0807 | 5.7406 | 6.4818 | 6.9449 | 5.9306 |

| | | | | | | | | | | | | | | | | | | | | | |
|---|---|---|---|---|---|---|---|---|---|---|---|---|---|---|---|---|---|---|---|---|---|
| ARID4A_S864 | ARID4A | P29374 | 1140;1145 | | NA | 6.9188 | 7.0699 | 5.9913 | 6.5177 | 6.8057 | 7.0083 | 6.8420 | 5.9565 | 6.0810 | 6.2294 | 6.1289 | 6.3244 | 5.5054 | 5.9869 | 4.9081 | 5.7259 |
| ARID4B_S1029 | ARID4B | Q4LE39 | | 133 | NPEHFGTPVIGKK | 7.1972 | 7.4623 | 6.9229 | 7.1413 | 6.3490 | 6.7258 | 6.7631 | 6.9516 | 7.1654 | 6.9015 | 7.2222 | 6.5536 | 3.9862 | 4.6642 | 3.6182 | 4.3754 |
| ARID4B_S1153 | ARID4B | Q4LE39 | | 675 | PFQTNPSPEMVSK | 5.6462 | 6.3151 | 6.5466 | 6.8007 | 5.7880 | 6.2395 | 6.0224 | 6.1297 | 6.4638 | 6.6602 | 6.3414 | 6.5400 | 6.1212 | 5.9766 | 5.9712 | 6.4374 |
| ARID4B_S675 | ARID4B | Q4LE39 | | 1029 | PPTTPESPSSVTV | 6.3108 | 5.6046 | 6.5246 | 6.2747 | 6.6275 | 6.5841 | 6.1337 | 6.0315 | 6.0215 | 5.9293 | 6.9202 | 6.0108 | 6.0616 | 6.0810 | 6.2306 | 6.6536 |
| ARID4B_S838 | ARID4B | Q4LE39 | | 742 | SKIDHLTNNRNDL | 6.5245 | 5.8207 | 5.6993 | 6.8653 | 6.1208 | 6.3860 | 5.7869 | 6.2047 | 5.7879 | 6.5230 | 5.3862 | 6.6386 | 6.4209 | 6.6298 | 6.7981 | 6.4073 |
| ARID4B_T133 | ARID4B | Q4LE39 | | 838 | ECLKTGSPGKKEE | 6.2334 | 6.7810 | 6.5416 | 6.6382 | 6.3916 | 6.7548 | 6.2243 | 5.7034 | 6.0203 | 6.6769 | 6.2305 | 6.3151 | 6.2124 | 6.1321 | 5.9405 | 5.2041 |
| ARID4B_T742 | ARID4B | Q4LE39 | | 1153 | GKGTNSSDSEELS | 5.4355 | 5.8178 | 6.3599 | 6.1117 | 6.1591 | 6.0751 | 6.1138 | 6.0566 | 6.0980 | 6.2611 | 6.0021 | 6.7787 | 6.9516 | 6.6983 | 6.6912 | 6.3895 |
| ARL13B_T385 | ARL13B | Q3SXY8 | | 385 | PPVGWGTPKVTRL | 6.5650 | 6.1701 | 6.5027 | 6.9317 | 5.8439 | 6.3422 | 6.8610 | 5.7907 | 6.0730 | 6.1373 | 6.3573 | 6.4406 | 5.6072 | 6.1565 | 5.7163 | 6.5044 |
| ARL2_S45 | ARL2 | P36404 | | 45 | EDIDTISPTLGFN | 5.5101 | 6.1651 | 6.5073 | 5.9996 | 6.1669 | 6.6282 | 6.2808 | 6.6819 | 7.1705 | 7.4128 | 7.0351 | 6.8962 | 5.1468 | 5.5647 | 5.4744 | 5.3597 |
| ARL6IP4_S332 | ARL6IP4 | Q66PJ3 | | 332 | LDQWHRSAGEEED | 6.0701 | 6.5309 | 7.3349 | 7.1152 | 5.8145 | 6.5866 | 6.1694 | 7.2170 | 6.2561 | 5.8761 | 6.0152 | 6.5678 | 5.1138 | 5.9364 | 5.5550 | 5.8409 |
| ARL6IP5_S184 | ARL6IP5 | O75915 | | 184 | RLTDYISKVKExx | 6.0611 | 5.7617 | 5.3796 | 4.5531 | 5.5065 | 4.6083 | 5.9080 | 5.2237 | 4.9229 | 4.8649 | 5.2540 | 5.3685 | 9.3490 | 9.5643 | 10.2707 | 7.4036 |
| ARL6IP6_S65 | ARL6IP6 | Q8N6S5 | | 19 | RRRGPGTPGPVAR | 7.2159 | 7.2253 | 6.2913 | 7.1166 | 6.2835 | 6.5739 | 7.0770 | 6.6886 | 6.7481 | 6.4440 | 6.5748 | 6.6530 | 4.6495 | 5.1802 | 4.2784 | 5.0000 |
| ARL6IP6_S80 | ARL6IP6 | Q8N6S5 | | 80 | PPDGNGSPVLPDK | 6.4576 | 6.9619 | 6.4367 | 7.4165 | 6.2926 | 6.4904 | 6.7706 | 6.2779 | 6.3876 | 6.6662 | 6.5786 | 6.6028 | 4.7736 | 5.3884 | 5.2427 | 5.2558 |
| ARL6IP6_T19 | ARL6IP6 | Q8N6S5 | | 65 | FSAGAWSEPRKRS | 7.2884 | 7.7005 | 6.8771 | 7.8755 | 6.5779 | 7.0328 | 7.4863 | 6.8986 | 7.1001 | 7.2377 | 7.3989 | 7.2123 | 2.8959 | 4.0897 | 2.7388 | 3.5895 |
| ARMC1_S246 | ARMC1 | Q9NVT9 | | 246 | YLPEDESPTKEQD | 6.1436 | 6.3227 | 6.5004 | 6.5933 | 6.4885 | 6.4227 | 6.4230 | 6.5361 | 6.7559 | 6.8016 | 6.7053 | 6.7926 | 5.0960 | 5.6741 | 5.2340 | 5.5102 |
| ARMC1_S260 | ARMC1 | Q9NVT9 | | 260 | AVSRVGSHPEGGA | 1.8505 | 2.2799 | 2.1415 | 2.9332 | 2.7872 | 3.1884 | 3.3420 | 3.3874 | 2.5427 | 2.4800 | 2.0885 | 2.3729 | 17.0627 | 17.1233 | 19.0268 | 15.3928 |
| ARMC1_S267 | ARMC1 | Q9NVT9 | | 267 | HPEGGASWLSTAA | 3.8157 | 4.1173 | 4.8481 | 3.8420 | 3.6356 | 4.7408 | 4.1740 | 5.2084 | 3.5181 | 5.0566 | 3.4169 | 4.1862 | 12.3176 | 11.8682 | 13.3297 | 11.9249 |
| ARMC10_S45 | ARMC10 | Q8N2F6-2 | | 45 | GIRSSKSAEDLTD | 4.5920 | 4.5627 | 4.8002 | 4.8712 | 6.6025 | 6.1604 | 6.1450 | 6.0347 | 5.2546 | 5.2026 | 5.1119 | 5.0094 | 9.0914 | 9.1291 | 8.6695 | 8.7628 |
| ARMC10_S45 | ARMC10 | Q8N2F6 | | 45 | GIRSSKSAGALEE | 3.3272 | 3.1672 | 3.7347 | 4.2274 | 5.1234 | 5.4913 | 6.1950 | 5.4269 | 3.8750 | 3.7555 | 3.8054 | 3.9010 | 11.8527 | 11.8839 | 12.5678 | 11.6656 |
| ARMCX3_S110 | ARMCX3 | Q9UH62 | | 72 | DSNESKSIVWYPP | 7.0587 | 6.7877 | 6.0346 | 7.0239 | 6.4316 | 5.7511 | 6.1028 | 5.9584 | 6.1413 | 5.8228 | 5.6524 | 5.4258 | 6.6853 | 6.8895 | 6.2486 | 5.9856 |
| ARMCX3_S61 | ARMCX3 | Q9UH62 | | 61 | ARYNDWSDDDDDS | 5.9986 | 6.2206 | 6.5195 | 6.8991 | 5.7860 | 5.7491 | 6.1257 | 5.2479 | 5.6660 | 6.2198 | 6.4928 | 6.9140 | 5.9743 | 7.3078 | 5.7327 | 7.1462 |
| ARMCX3_S72 | ARMCX3 | Q9UH62 | | 110 | AVQKRASPNSDDT | 6.0782 | 6.0958 | 6.9903 | 6.4663 | 6.3020 | 6.2657 | 6.2764 | 6.6887 | 6.3672 | 6.6231 | 7.0168 | 6.5947 | 5.2493 | 6.0833 | 5.1884 | 5.7138 |
| ARMT1_S204 | ARMT1 | Q9H993 | | 204 | LSGGESSSQNTNV | 6.6557 | 6.0015 | 6.4482 | 6.0132 | 6.2772 | 6.1175 | 6.6177 | 5.9436 | 6.6775 | 6.3610 | 6.7603 | 6.8832 | 5.4856 | 5.1378 | 6.3337 | 6.2863 |
| ARNTL2_S565 | ARNTL2 | Q8WYA1 | | 565 | KELFPPSPSEMGE | 6.2879 | 5.8005 | 7.4462 | 6.2364 | 4.6730 | 6.0631 | 6.5607 | 6.6997 | 6.0270 | 6.3376 | 7.5389 | 5.3217 | 5.7662 | 6.4526 | 6.4852 | 6.3033 |
| ARPC1B_T80 | ARPC1B | O15143 | | 80 | RNAYVWTLKGRTW | 6.0017 | 8.4642 | 8.5938 | 5.0097 | 7.6965 | 6.4603 | 7.2828 | 5.6727 | 6.1762 | 5.0646 | 5.5082 | 5.8023 | 6.2509 | 6.4653 | 6.0914 | 3.4594 |
| ARPC5L_S64 | ARPC5L | Q9BPX5 | | 64 | HAALRNSPVNTKN | 6.1153 | 5.4583 | 6.7147 | 5.6232 | 5.6939 | 5.4890 | 5.9503 | 6.4287 | 6.5838 | 7.0612 | 6.6840 | 7.1726 | 5.8971 | 6.3654 | 6.0493 | 6.7132 |
| ARRDC1_S292 | ARRDC1 | Q8N5I2 | | 292 | VNHAPVSPRPGLG | 6.3988 | 6.1284 | 6.4243 | 5.9284 | 5.9458 | 6.5314 | 6.3692 | 6.9191 | 6.2161 | 7.4649 | 6.7783 | 6.7069 | 5.2459 | 5.9213 | 5.1949 | 5.8264 |
| ARVCF_S267 | ARVCF | O00192 | | 642 | MDRNFDTLDLPKR | 7.3151 | 7.5133 | 6.4833 | 7.6719 | 6.6400 | 5.5488 | 6.0711 | 6.0991 | 6.3360 | 5.7299 | 5.3103 | 5.4189 | 5.7338 | 6.2945 | 6.0459 | 5.7881 |
| ARVCF_S335 | ARVCF | O00192 | | 267 | LEDDTRSLAADDE | 5.7217 | 6.1113 | 6.8222 | 6.5820 | 6.8175 | 5.8655 | 6.1980 | 5.8706 | 6.6105 | 6.1759 | 6.2982 | 6.1736 | 6.0650 | 6.3703 | 5.8547 | 6.4630 |
| ARVCF_S864 | ARVCF | O00192 | | 864 | GPKGALSPGGFDD | 6.0425 | 6.5456 | 6.3194 | 7.3247 | 6.4596 | 6.2393 | 6.2975 | 7.0851 | 7.0682 | 7.4033 | 7.2009 | 6.8454 | 4.4519 | 5.4717 | 4.1332 | 5.1117 |
| ARVCF_T642 | ARVCF | O00192 | | 335 | ERGSMGSLDRLVR | 6.9404 | 7.2914 | 5.8549 | 6.8563 | 5.8671 | 5.8940 | 5.6732 | 6.1487 | 6.7460 | 6.4903 | 5.8825 | 6.1493 | 6.3598 | 6.3727 | 5.7093 | 5.7641 |
| ARVCF_T872 | ARVCF | O00192 | | 872 | GGFDDSTLPLVDK | 8.6178 | 8.1451 | 7.6052 | 8.3988 | 6.3226 | 6.1947 | 6.7734 | 6.1869 | 5.9191 | 6.6286 | 6.3134 | 5.1382 | 4.0789 | 4.5983 | 5.0661 | 4.0132 |
| ASAP1_S1011 | ASAP1 | Q9ULH1-2 | | 1011 | SQTGDVSPKAQQP | 5.8049 | 6.3602 | 6.5145 | 6.1757 | 5.6300 | 6.2375 | 5.9765 | 6.0422 | 5.9559 | 6.1223 | 6.7983 | 6.9874 | 6.2783 | 6.6526 | 6.2614 | 6.2023 |
| ASAP1_S1030 | ASAP1 | Q9ULH1-2 | | 761 | ALPGFSTPRDKQR | 8.2076 | 7.8343 | 7.1890 | 7.2115 | 5.6177 | 7.9093 | 7.3462 | 6.7073 | 7.6771 | 6.9799 | 7.3077 | 8.1248 | 2.5502 | 3.6888 | 2.2482 | 3.4004 |
| ASAP1_S842;S846 | ASAP1 | Q9ULH1-2 | | 846 | TLSDPPSPLPHGP | 6.3561 | 6.5454 | 5.8957 | 6.4908 | 6.2432 | 6.1523 | 6.2293 | 6.6088 | 6.4108 | 6.1812 | 6.2634 | 6.4199 | 5.9849 | 6.1986 | 5.8785 | 6.1411 |
| ASAP1_S846 | ASAP1 | Q9ULH1-2 | | 1030 | SHPLDLSPNVQSR | 6.9541 | 7.5776 | 7.0232 | 6.7107 | 6.1411 | 7.0738 | 7.0015 | 6.9979 | 7.1787 | 6.9408 | 6.2631 | 6.9785 | 4.1778 | 4.4787 | 3.8857 | 4.6168 |
| ASAP1_T761 | ASAP1 | Q9ULH1-2 | 842;846 | | NA | 6.6681 | 5.6811 | 5.5003 | 6.7912 | 6.5746 | 6.1505 | 6.2860 | 6.0388 | 6.1094 | 6.6981 | 6.4303 | 6.3005 | 6.4214 | 6.2047 | 6.6004 | 5.5446 |
| ASAP2_S701 | ASAP2 | O43150 | | 712 | DEKLQPSPNRRED | 6.4219 | 7.0649 | 6.6584 | 7.3713 | 5.8426 | 6.9895 | 6.5561 | 6.7696 | 7.1816 | 7.0150 | 7.4070 | 6.9861 | 4.0016 | 4.7895 | 4.1629 | 4.7820 |
| ASAP2_S712 | ASAP2 | O43150 | | 701 | HEDLDESDDDMDE | 6.4168 | 5.8841 | 7.0363 | 6.3328 | 6.7067 | 6.2100 | 6.3537 | 6.3410 | 5.8923 | 6.7321 | 6.7198 | 6.1695 | 5.7489 | 5.9668 | 5.7420 | 5.7473 |
| ASAP2_S822 | ASAP2 | O43150 | | 900 | PGADKSTPLTNKG | 6.5837 | 6.9364 | 6.8490 | 6.6270 | 6.1921 | 6.5973 | 6.2847 | 5.9311 | 6.3060 | 6.1385 | 6.2241 | 7.0205 | 5.5544 | 5.8437 | 5.3441 | 5.5674 |
| ASAP2_T798 | ASAP2 | O43150 | | 822 | SRQRSSSDPPAVH | 5.4336 | 5.4852 | 5.9581 | 6.8989 | 5.9805 | 5.8023 | 6.1983 | 6.0442 | 6.6794 | 6.2011 | 6.7502 | 6.1994 | 5.9855 | 7.2637 | 6.4863 | 6.6333 |
| ASAP2_T900 | ASAP2 | O43150-2 | | 798 | VGKDPLTPTPPPP | 6.6318 | 6.4494 | 6.3412 | 6.4981 | 5.9494 | 6.3271 | 7.0345 | 7.4452 | 7.3489 | 7.2509 | 6.2854 | 7.5113 | 4.6781 | 4.7582 | 4.7528 | 4.7376 |
| ASB1_S15 | ASB1 | Q9Y576 | | 15 | GRAGPGSAGRNLK | 6.7046 | 6.6960 | 8.1323 | 6.3399 | 5.8427 | 6.6429 | 6.4315 | 6.8408 | 6.7642 | 7.4701 | 7.7130 | 7.7962 | 3.8184 | 5.2250 | 3.5650 | 4.0173 |
| ASCC2_T233 | ASCC2 | Q9H1I8 | | 233 | EERGRLTPSDMPL | 6.1624 | 6.1279 | 6.3560 | 6.3597 | 6.4038 | 6.0246 | 5.6088 | 6.2826 | 6.3881 | 6.8506 | 5.8112 | 6.3916 | 6.6040 | 6.0313 | 6.4449 | 6.1524 |
| ASCC3_S2195 | ASCC3 | Q8N3C0 | | 209 | HLQEACTPELKPV | 7.0267 | 7.7267 | 7.1066 | 7.7836 | 6.4620 | 7.9301 | 7.7786 | 6.9260 | 7.5864 | 7.4232 | 7.5296 | 7.2893 | 2.5701 | 3.6219 | 1.9929 | 3.2463 |
| ASCC3_T209 | ASCC3 | Q8N3C0 | | 2195 | NTKVSDSLTDLAL | 6.6588 | 6.1821 | 6.3684 | 5.5535 | 6.2709 | 6.4208 | 6.3938 | 6.1100 | 6.5099 | 5.5751 | 6.2771 | 5.4915 | 6.4850 | 6.6596 | 6.5162 | 6.5274 |
| ASF1A_S166 | ASF1A | Q9Y294 | | 166 | LEDAESSNPNLQS | 5.1615 | 5.3029 | 5.3013 | 5.4950 | 5.7913 | 5.9916 | 5.7279 | 6.0160 | 5.9560 | 5.5815 | 5.8669 | 5.6962 | 7.6565 | 8.1443 | 8.2659 | 8.0451 |
| ASF1A_S192 | ASF1A | Q9Y294 | | 157 | INWEDNTEKLEDA | 6.4829 | 6.9427 | 6.7810 | 5.9317 | 6.7369 | 6.4846 | 6.9571 | 6.1010 | 6.1459 | 5.8868 | 7.2603 | 6.4169 | 4.5734 | 5.8969 | 5.0925 | 6.3093 |
| ASF1A_T157 | ASF1A | Q9Y294 | | 192 | WSTSENSLNVMLE | 5.1563 | 6.0097 | 8.5810 | 6.0511 | 4.7034 | 5.7197 | 4.9602 | 5.4739 | 4.7945 | 5.3441 | 6.4737 | 6.7886 | 9.1195 | 7.6447 | 7.5840 | 5.5954 |
| ASF1B_S198 | ASF1B | Q9NVP2 | | 179 | GLPLNCTPIKGLG | 6.3822 | 6.5040 | 6.4239 | 6.0484 | 6.4157 | 6.8525 | 6.6693 | 7.0785 | 7.4499 | 7.0159 | 7.2289 | 7.0841 | 4.4244 | 5.0879 | 4.3460 | 4.9883 |
| ASF1B_T179 | ASF1B | Q9NVP2 | | 198 | GLLPENSMDCIxx | 5.4934 | 6.4191 | 6.8274 | 6.1391 | 5.9017 | 6.7344 | 6.1674 | 6.0040 | 6.3642 | 6.6392 | 7.0253 | 6.5331 | 5.7282 | 6.4059 | 5.6545 | 5.9630 |
| ASH1L_S1170 | ASH1L | Q9NR48 | | 884 | KQGLSVSPFPKKR | 7.5356 | 7.2275 | 6.5034 | 7.4854 | 6.0181 | 6.1356 | 6.6898 | 6.0936 | 7.9342 | 7.0646 | 6.6961 | 6.9024 | 4.1814 | 4.7306 | 4.2297 | 4.5722 |
| ASH1L_S1681 | ASH1L | Q9NR48 | | 1748 | SAPPSSSPGRSHS | 6.5984 | 6.0742 | 6.7481 | 6.7186 | 6.2149 | 6.1837 | 6.6296 | 6.4741 | 6.6138 | 6.1134 | 6.1173 | 6.7756 | 5.4770 | 5.7074 | 5.4549 | 6.0990 |
| ASH1L_S1748 | ASH1L | Q9NR48 | | 179 | PLSKKLSPVHSEM | 6.2995 | 6.4428 | 6.4102 | 6.7269 | 6.1566 | 5.4976 | 6.3036 | 6.9513 | 6.3834 | 6.6141 | 6.3458 | 6.6530 | 5.8870 | 5.5302 | 5.6443 | 6.1536 |
| ASH1L_S179 | ASH1L | Q9NR48 | | 22 | EGFSRKSPSAIST | 6.5803 | 6.7125 | 7.0836 | 7.4521 | 5.7835 | 5.6456 | 6.1311 | 5.8107 | 7.5796 | 6.9347 | 7.0737 | 7.1445 | 4.8303 | 5.1947 | 4.8945 | 5.1486 |
| ASH1L_S1791 | ASH1L | Q9NR48 | | 529 | PPVYCTSPDFKMG | 6.0292 | 6.5448 | 7.5115 | 8.0376 | 6.2264 | 6.5420 | 6.5190 | 5.7535 | 7.4745 | 6.9012 | 7.2854 | 7.7373 | 4.6457 | 3.8302 | 4.1256 | 4.8360 |
| ASH1L_S1943 | ASH1L | Q9NR48 | | 918 | PFVATESPSKLES | 6.6274 | 7.0406 | 6.5872 | 7.6297 | 6.0657 | 5.4492 | 6.0584 | 6.3681 | 7.2646 | 7.4415 | 6.6642 | 7.0950 | 5.1110 | 4.9370 | 4.5895 | 5.0708 |
| ASH1L_S1953 | ASH1L | Q9NR48 | | 1953 | APVEIPSPSETPA | 6.6919 | 6.2500 | 6.5399 | 7.0331 | 5.7153 | 5.9343 | 5.5339 | 6.2818 | 6.6841 | 6.9808 | 6.4162 | 6.4617 | 5.3526 | 5.6884 | 5.6664 | 6.7696 |
| ASH1L_S1955 | ASH1L | Q9NR48 | | 1681 | SESTNCSPTRKRS | 6.8674 | 7.0038 | 6.7032 | 7.6681 | 6.9330 | 6.4335 | 6.1288 | 6.3297 | 7.3705 | 7.7497 | 7.2182 | 6.1562 | 4.5585 | 4.8276 | 4.1075 | 3.9443 |
| ASH1L_S22 | ASH1L | Q9NR48 | | 2966 | LFIPENSFRKxxx | 8.7847 | 6.1149 | 8.4031 | 8.6589 | 6.9268 | 5.6887 | 8.4180 | 6.2259 | 6.8362 | 6.3829 | 5.8430 | 5.8199 | 4.7631 | 4.0811 | 2.7280 | 4.3247 |
| ASH1L_S2966 | ASH1L | Q9NR48 | | 1943 | EQENNKSFNEAPV | 6.1427 | 6.6514 | 6.1747 | 6.6312 | 6.9074 | 6.9616 | 6.1297 | 6.4157 | 6.5234 | 6.4699 | 7.0269 | 6.7011 | 5.2416 | 5.3239 | 5.6634 | 5.0353 |
| ASH1L_S529 | ASH1L | Q9NR48 | | 1170 | PTLLPNSPSHLSE | 7.7323 | 7.6694 | 5.7559 | 8.5342 | 6.6087 | 6.5767 | 5.5655 | 7.9062 | 7.1178 | 6.3539 | 6.8830 | 6.3229 | 3.7965 | 5.3977 | 4.0657 | 3.7135 |
| ASH1L_S884 | ASH1L | Q9NR48 | | 1791 | KLTSSCSPHHIKR | 7.1463 | 7.8151 | 6.5146 | 7.7961 | 6.9473 | 6.5575 | 6.1286 | 6.9157 | 6.7080 | 6.7695 | 6.8130 | 6.5974 | 3.9956 | 4.8168 | 4.3201 | 4.1583 |
| ASH1L_S918 | ASH1L | Q9NR48 | | 1955 | VEIPSPSETPAKP | 5.8640 | 6.1775 | 6.2859 | 6.7862 | 5.9913 | 6.4633 | 5.9084 | 6.3884 | 6.4982 | 6.4553 | 7.3011 | 6.9644 | 5.6524 | 6.6640 | 5.1797 | 5.4197 |
| ASH2L_S101 | ASH2L | Q9UBL3 | | 623 | EVDGRRSPPWEPx | 6.0531 | 6.0913 | 5.6187 | 6.4213 | 5.6216 | 5.7459 | 6.3790 | 6.0897 | 6.4764 | 6.1470 | 6.1403 | 6.2808 | 6.4172 | 6.4881 | 6.9532 | 7.0762 |
| ASH2L_S291 | ASH2L | Q9UBL3 | | 101 | MDTQAGSVDEENG | 5.5060 | 5.5862 | 5.7622 | 5.9659 | 6.2117 | 5.8663 | 5.8045 | 6.1433 | 6.2929 | 6.4162 | 6.5283 | 5.9559 | 6.9935 | 7.3135 | 6.7355 | 6.9181 |
| ASH2L_S623 | ASH2L | Q9UBL3 | | 291 | GGIAAGSSGKGRG | 6.2124 | 7.2242 | 6.2242 | 6.9448 | 6.1689 | 6.7289 | 6.7784 | 7.2429 | 7.6657 | 6.5051 | 7.0759 | 6.4994 | 4.5435 | 4.8585 | 4.3395 | 4.9878 |
| ASMTL_S21 | ASMTL | O95671 | | 21 | VVLASASPRRQEI | 6.3982 | 6.2551 | 6.6490 | 5.8915 | 6.3658 | 6.7783 | 5.9288 | 6.9359 | 6.5998 | 6.3973 | 6.3938 | 6.5087 | 5.7789 | 5.9757 | 5.5379 | 5.6054 |
| ASPHD1_S158 | ASPHD1 | Q5U4P2 | | 158 | AYARRYSWAGMGR | 5.9319 | 5.1973 | 5.9716 | 5.1474 | 6.6076 | 5.4648 | 4.5509 | 5.9533 | 8.2468 | 9.2202 | 8.0365 | 7.3110 | 5.8251 | 6.3507 | 5.4819 | 4.7031 |
| ASPM_S170 | ASPM | Q8IZT6 | | 367 | IYQKILSPDSFIK | 6.9897 | 7.4831 | 7.0505 | 7.7516 | 6.5304 | 6.4625 | 7.0111 | 6.8405 | 7.7230 | 7.4209 | 7.5175 | 7.1960 | 2.9689 | 4.1520 | 3.0212 | 3.8811 |
| ASPM_S190 | ASPM | Q8IZT6 | | 392 | SVNPILSPNQFLK | 6.7555 | 6.9212 | 6.9208 | 7.0673 | 6.3918 | 6.6886 | 7.2631 | 6.9501 | 7.3970 | 7.2721 | 7.1621 | 7.3699 | 3.4206 | 4.6848 | 3.5583 | 4.1766 |
| ASPM_S222 | ASPM | Q8IZT6 | | 425 | NSQVPQSPEDWRK | 6.5543 | 7.0726 | 6.9090 | 7.6418 | 7.0264 | 6.7631 | 6.2881 | 6.9086 | 6.5912 | 6.4254 | 7.3344 | 7.5699 | 4.0865 | 4.7713 | 3.6737 | 4.3838 |
| ASPM_S222;S225 | ASPM | Q8IZT6 | | 622 | KTKNVTTPISKRI | 7.6325 | 8.1243 | 7.5815 | 7.3752 | 5.7427 | 7.3445 | 8.0632 | 6.9848 | 7.3774 | 7.1733 | 7.4557 | 7.5597 | 2.2681 | 3.7749 | 2.3447 | 3.1973 |
| ASPM_S280 | ASPM | Q8IZT6 | | 647 | DLSIFRTPISKTN | 8.4048 | 8.4153 | 8.3909 | 7.7183 | 4.2103 | 6.4376 | 8.3902 | 6.0547 | 7.5107 | 7.2042 | 7.6769 | 7.4291 | 2.7981 | 3.6659 | 2.4120 | 3.2812 |
| ASPM_S3235 | ASPM | Q8IZT6 | | 170 | SHNRRVSNIQNVN | 6.8745 | 6.2175 | 6.5194 | 6.7288 | 5.1481 | 6.5730 | 6.3228 | 5.4539 | 9.2605 | 7.9711 | 8.6294 | 8.2257 | 3.9110 | 5.6510 | 2.8863 | 3.6270 |
| ASPM_S367 | ASPM | Q8IZT6 | | 3235 | IKAIRLSLQVVNR | 7.4835 | 7.5603 | 6.0056 | 8.1821 | 5.8817 | 6.1768 | 7.5059 | 7.3596 | 7.9515 | 6.4619 | 6.6544 | 6.6422 | 3.0783 | 4.6004 | 3.5279 | 4.9281 |
| ASPM_S367;S370 | ASPM | Q8IZT6 | | 190 | KVDRVRSPLQACE | 6.7045 | 6.2045 | 7.8935 | 7.5736 | 6.5114 | 7.2536 | 7.8120 | 6.3268 | 6.1699 | 7.0621 | 6.4853 | 7.5313 | 3.6495 | 4.7693 | 3.9233 | 4.1294 |
| ASPM_S392 | ASPM | Q8IZT6 | | 222 | ENKIPISPISPAF | 6.3711 | 6.4083 | 6.7548 | 6.1607 | 6.4813 | 7.0228 | 6.0029 | 6.7464 | 6.6573 | 6.2652 | 6.1930 | 6.6980 | 5.4438 | 5.5568 | 5.5841 | 5.6536 |
| ASPM_S425 | ASPM | Q8IZT6 | | 605 | IKRIHFSPSEPKT | 7.3229 | 7.7116 | 6.7097 | 7.5781 | 6.5048 | 6.7694 | 7.5715 | 7.3942 | 7.6752 | 7.2328 | 6.9362 | 7.0655 | 2.8225 | 3.9067 | 3.2859 | 3.5129 |
| ASPM_S605 | ASPM | Q8IZT6 | | 280 | KAVTETSFNSVNV | 6.5959 | 6.1383 | 7.3580 | 7.3094 | 7.1553 | 7.2085 | 6.7563 | 7.7472 | 6.4833 | 6.7473 | 6.1086 | 6.7698 | 4.3501 | 4.5510 | 3.9177 | 4.8033 |
| ASPM_T622 | ASPM | Q8IZT6 | 222;225 | | NA | 6.2008 | 6.6794 | 6.9007 | 7.4757 | 6.7281 | 6.3348 | 7.2076 | 7.0645 | 6.7828 | 7.0060 | 7.1032 | 7.0614 | 4.1460 | 4.9870 | 3.6122 | 4.7098 |
| ASPM_T647 | ASPM | Q8IZT6 | 367;370 | | NA | 7.2354 | 7.8701 | 7.2856 | 8.5668 | 5.9256 | 6.9557 | 7.6074 | 6.8061 | 6.7560 | 7.0945 | 7.9308 | 7.8593 | 2.2408 | 3.8340 | 2.0398 | 3.9921 |
| ASPSCR1_S275 | ASPSCR1 | Q9BZE9-2 | | 594 | MSRAAGSPSPLPA | 5.4395 | 5.6727 | 5.7564 | 5.7498 | 5.6741 | 6.0153 | 6.0698 | 6.4125 | 5.7234 | 6.0236 | 6.0716 | 6.2529 | 7.2706 | 7.3267 | 7.2622 | 7.2788 |
| ASPSCR1_S594 | ASPSCR1 | Q9BZE9-2 | | 275 | LPKSLSSPGGPSK | 6.5912 | 6.8116 | 6.6453 | 6.5004 | 6.3835 | 6.9337 | 6.5773 | 6.8213 | 7.0678 | 6.9924 | 7.1683 | 7.3261 | 4.2581 | 4.8536 | 4.2834 | 4.7860 |
| ASXL1_S503 | ASXL1 | Q8IXJ9 | | 526 | KDQKRKSFEQAAS | 6.0877 | 5.8488 | 7.1135 | 7.5149 | 6.6484 | 6.5433 | 5.9508 | 6.5337 | 6.8013 | 6.0304 | 7.7124 | 7.5818 | 4.7211 | 4.5905 | 6.3776 | 3.9438 |
| ASXL1_S526 | ASXL1 | Q8IXJ9 | | 503 | LARASASPDRIPS | 6.5788 | 7.3241 | 6.6429 | 7.2809 | 6.5528 | 6.0072 | 6.5053 | 6.4384 | 6.3630 | 6.9057 | 7.1282 | 7.0589 | 4.4367 | 5.0799 | 4.8043 | 4.8930 |
| ASXL2_S1154 | ASXL2 | Q76L83 | | 395 | SKKLTASPSDPKV | 6.1280 | 6.4100 | 8.0906 | 7.0109 | 6.9390 | 6.7478 | 7.6164 | 5.9014 | 6.1530 | 6.0438 | 6.4626 | 6.3101 | 4.9085 | 5.2198 | 4.8016 | 5.2564 |
| ASXL2_S1319 | ASXL2 | Q76L83 | | 562 | ATLVDQSPESLKR | 6.5999 | 7.0375 | 6.7397 | 7.3462 | 6.5854 | 6.6723 | 6.6163 | 6.5769 | 6.6709 | 6.9102 | 6.8100 | 7.1299 | 4.4403 | 4.9084 | 4.2828 | 4.6734 |
| ASXL2_S139 | ASXL2 | Q76L83 | | 524 | QESLVTSPSKPKS | 6.6589 | 6.6519 | 6.6040 | 7.3613 | 6.1904 | 5.7255 | 6.4088 | 5.9765 | 6.7679 | 6.5441 | 6.9081 | 6.7951 | 5.4969 | 5.2530 | 5.0400 | 5.6175 |
| ASXL2_S146 | ASXL2 | Q76L83 | | 440 | EALQMSSPGRKEE | 6.3908 | 6.8464 | 6.5087 | 7.1438 | 6.2733 | 6.3296 | 6.1253 | 5.9379 | 6.5498 | 6.5794 | 6.7958 | 6.5011 | 5.2402 | 5.6640 | 5.4927 | 5.6210 |
| ASXL2_S395 | ASXL2 | Q76L83 | | 1319 | TPKLYGSPTQIGP | 5.7041 | 6.8547 | 6.4274 | 6.7423 | 6.4567 | 6.9207 | 6.8185 | 6.2298 | 6.8997 | 6.8009 | 6.2132 | 6.5002 | 5.1643 | 5.6821 | 5.2953 | 5.2899 |
| ASXL2_S440 | ASXL2 | Q76L83 | | 146 | PQSGCPSPTIPAG | 5.7960 | 6.4915 | 6.7843 | 7.0210 | 6.7199 | 5.6663 | 5.6261 | 5.8036 | 6.5879 | 6.8966 | 6.1950 | 6.3550 | 5.6503 | 6.6076 | 6.2670 | 5.5320 |
| ASXL2_S524 | ASXL2 | Q76L83 | | 648 | FPVSITSPNRTGA | 7.4434 | 7.3415 | 6.5694 | 7.1373 | 6.1988 | 7.2518 | 6.8451 | 7.2520 | 7.8936 | 7.4105 | 7.4684 | 7.7324 | 3.1719 | 3.7914 | 2.8971 | 3.5955 |
| ASXL2_S562 | ASXL2 | Q76L83 | | 139 | RKVSSSSPQSGCP | 6.9095 | 6.4123 | 5.5055 | 7.2320 | 6.4280 | 6.6222 | 6.3667 | 6.2525 | 6.4084 | 6.3371 | 6.6367 | 6.3712 | 5.8610 | 5.3237 | 5.5596 | 5.7737 |
| ASXL2_S600 | ASXL2 | Q76L83 | | 600 | QQPFQVSPQPFLN | 6.8821 | 6.8771 | 6.0786 | 7.1736 | 6.8945 | 5.4110 | 6.9668 | 7.2645 | 6.8173 | 6.5968 | 8.0368 | 7.0477 | 4.5627 | 5.0622 | 4.0620 | 4.2663 |
| ASXL2_S648 | ASXL2 | Q76L83 | | 1154 | DRFCLSSPTEALK | 7.2417 | 7.3949 | 6.6578 | 7.4570 | 5.8836 | 6.4236 | 6.8611 | 6.0251 | 6.6416 | 7.0908 | 6.8461 | 6.8623 | 4.4584 | 4.8064 | 4.6091 | 4.7404 |
| ASXL3_S929 | ASXL3 | Q9C0F0 | | 185 | VPRVVLTPLKVSD | 7.8779 | 7.5783 | 6.6568 | 6.9847 | 6.2630 | 6.2614 | 6.9538 | 6.3722 | 7.5855 | 6.6149 | 6.3097 | 6.4471 | 4.5612 | 4.8286 | 4.1006 | 4.6042 |
| ASXL3_T185 | ASXL3 | Q9C0F0 | | 929 | KTHKQGSTQSRLE | 6.1817 | 5.8411 | 7.0342 | 7.0915 | 5.7504 | 6.5551 | 6.2974 | 6.5267 | 6.7014 | 6.6085 | 6.2868 | 6.1755 | 5.1994 | 6.3514 | 5.6680 | 5.7309 |
| ATAD2_S1131 | ATAD2 | Q6PL18 | | 327 | FYSGPASPARPRY | 6.4975 | 5.9580 | 6.2504 | 6.7957 | 6.1618 | 5.7773 | 6.2027 | 5.8013 | 6.6881 | 6.4820 | 6.2767 | 6.4985 | 6.0168 | 6.0570 | 6.0851 | 6.4512 |
| ATAD2_S1170 | ATAD2 | Q6PL18 | | 750 | LESDLAYSDDDVP | 5.9790 | 6.0212 | 6.0480 | 6.6689 | 6.0134 | 5.6242 | 5.7754 | 6.3256 | 6.5814 | 6.5196 | 6.5782 | 6.4641 | 6.2189 | 6.2395 | 6.1553 | 6.7873 |
| ATAD2_S1190 | ATAD2 | Q6PL18 | | 1170 | RKIRKKSNWYLGT | 6.0574 | 4.1952 | 5.7094 | 6.1804 | 8.9859 | 4.8145 | 4.8886 | 4.4986 | 9.4060 | 7.5156 | 7.1150 | 7.5678 | 7.2255 | 7.4007 | 4.1362 | 4.3033 |
| ATAD2_S1200 | ATAD2 | Q6PL18 | | 342 | SSAGPRSPYCKRM | 7.6529 | 7.0184 | 6.3464 | 7.1558 | 6.7666 | 6.5262 | 6.3945 | 5.0902 | 6.1378 | 6.1790 | 6.2389 | 6.4665 | 5.9754 | 5.4528 | 5.3287 | 5.2698 |
| ATAD2_S1235 | ATAD2 | Q6PL18 | | 1283 | SQIIHISDENEGK | 6.3363 | 5.8817 | 6.2298 | 7.1544 | 6.6718 | 6.1600 | 6.5314 | 5.9863 | 5.9346 | 5.4691 | 5.3403 | 5.2101 | 7.3603 | 6.2625 | 7.0051 | 6.4663 |
| ATAD2_S1243 | ATAD2 | Q6PL18 | | 1200 | IDHKIESDTEETQ | 5.8250 | 6.5013 | 5.9332 | 6.7202 | 5.5281 | 5.4137 | 5.5045 | 5.9738 | 7.4869 | 6.7754 | 6.8877 | 6.7165 | 6.2002 | 6.0091 | 5.8971 | 6.6273 |
| ATAD2_S1283 | ATAD2 | Q6PL18 | | 61 | KAGDGSSVKEVET | 6.1698 | 6.0507 | 5.1946 | 6.0938 | 6.2478 | 5.4556 | 6.3301 | 5.6385 | 5.9106 | 5.3891 | 5.4006 | 5.5568 | 8.1010 | 7.1340 | 7.7388 | 7.5884 |
| ATAD2_S170 | ATAD2 | Q6PL18 | | 1152 | KLKTPSTPVACST | 5.9501 | 6.4496 | 6.4688 | 6.6627 | 6.1541 | 6.2256 | 6.7770 | 6.2781 | 6.6536 | 6.6961 | 6.9569 | 6.5172 | 5.6586 | 5.6186 | 5.2616 | 5.6715 |
| ATAD2_S327 | ATAD2 | Q6PL18 | | 1190 | SQAKDDSQNAIDH | 6.3763 | 7.0449 | 7.1529 | 7.8177 | 6.4762 | 5.8024 | 6.8041 | 6.4364 | 7.9036 | 7.6086 | 7.5545 | 7.8606 | 3.5922 | 4.0857 | 3.5259 | 3.9580 |
| ATAD2_S336;S342 | ATAD2 | Q6PL18 | | 1243 | LELRNNSNTCNIE | 6.0590 | 6.4673 | 5.9278 | 7.2964 | 6.3195 | 5.0125 | 6.8493 | 6.5369 | 7.3088 | 6.5765 | 6.2455 | 6.3564 | 5.6761 | 5.7914 | 5.6770 | 5.8996 |
| ATAD2_S337 | ATAD2 | Q6PL18 | | 170 | YSGVNQSMLFDKL | 9.6042 | 8.5002 | 7.6270 | 10.1232 | 7.6370 | 5.0862 | 6.0888 | 5.6943 | 7.1205 | 5.1574 | 6.6404 | 5.7647 | 3.6727 | 4.1879 | 3.5032 | 3.5924 |
| ATAD2_S342 | ATAD2 | Q6PL18 | | 337 | PRYRLSSAGPRSP | 6.6840 | 6.9046 | 6.0910 | 7.0714 | 6.2754 | 5.5651 | 6.5081 | 6.4228 | 7.0829 | 6.3027 | 6.6266 | 6.3476 | 5.2379 | 5.2895 | 5.3651 | 6.2252 |
| ATAD2_S61 | ATAD2 | Q6PL18 | | 1131 | VMPKQNSTLVGDK | 6.8125 | 7.0868 | 6.4644 | 7.5694 | 6.2471 | 5.7859 | 7.7635 | 6.0815 | 6.9452 | 6.7431 | 8.7750 | 7.5444 | 3.6070 | 4.2526 | 3.9089 | 4.4129 |
| ATAD2_S696 | ATAD2 | Q6PL18 | | 696 | SQRAVTSPGQALS | 7.3983 | 6.8108 | 7.4835 | 7.4871 | 5.6616 | 5.6487 | 6.0370 | 5.7038 | 6.6897 | 6.5091 | 7.4556 | 6.4855 | 5.3334 | 5.2209 | 4.8211 | 5.2539 |
| ATAD2_T1152 | ATAD2 | Q6PL18 | | 1235 | QQNASESKLELRN | 6.9063 | 7.2008 | 6.8476 | 7.2946 | 6.4937 | 6.3012 | 7.4143 | 7.0761 | 7.2559 | 7.2716 | 7.3047 | 7.7163 | 3.1982 | 3.7967 | 3.2004 | 4.7218 |
| ATAD2_T1152;T1158 | ATAD2 | Q6PL18 | 1152;1158 | | NA | 7.0391 | 7.1473 | 7.3668 | 8.1298 | 7.0966 | 6.7083 | 6.6792 | 7.0553 | 6.9235 | 7.0833 | 6.0990 | 7.0417 | 3.9292 | 4.2784 | 3.3289 | 4.0936 |
| ATAD2_Y750 | ATAD2 | Q6PL18 | 336;342 | | NA | 6.8456 | 6.9618 | 6.7060 | 7.3473 | 6.5588 | 6.3094 | 6.1271 | 5.3475 | 6.0352 | 6.2031 | 8.2367 | 7.4337 | 4.9515 | 4.9263 | 4.8778 | 5.1323 |
| ATAD2B_S1347 | ATAD2B | Q9ULI0 | | 939 | LPLALPSPPRQLS | 5.7704 | 6.5219 | 6.5746 | 6.6554 | 6.6180 | 5.1512 | 5.7063 | 6.7864 | 6.1732 | 6.4818 | 6.6047 | 5.9614 | 6.5145 | 6.0429 | 6.1471 | 6.2901 |
| ATAD2B_S939 | ATAD2B | Q9ULI0 | | 1347 | EKLEPGSDVEVKD | 6.0553 | 6.3561 | 6.4685 | 7.0923 | 6.2694 | 6.0553 | 6.6243 | 5.8718 | 5.9946 | 5.2728 | 5.4111 | 6.6761 | 6.5473 | 6.3001 | 6.7145 | 6.2905 |
| ATAD3A_S369 | ATAD3A | Q9NVI7 | | 369 | LEGVVLSPSLEAR | 6.3603 | 6.3117 | 6.0240 | 6.8685 | 6.2428 | 5.9314 | 6.4698 | 6.1366 | 6.3150 | 6.2282 | 6.2906 | 6.1527 | 5.7213 | 6.4613 | 6.2713 | 6.2145 |
| ATAD3B_S321 | ATAD3B | Q5T9A4 | | 321 | LEGVVLSPSLEAR | 5.6543 | 5.8597 | 6.8223 | 6.9136 | 6.5246 | 6.5040 | 6.3899 | 6.3084 | 5.6011 | 4.7145 | 5.8184 | 6.3725 | 7.0940 | 7.2417 | 6.4410 | 5.7401 |
| ATAD5_S1222 | ATAD5 | Q96QE3 | | 1222 | RKVPPPSPKSSGP | 6.8168 | 6.5223 | 6.2101 | 7.0019 | 6.2529 | 5.9185 | 6.5541 | 5.9721 | 6.6173 | 6.2738 | 6.2990 | 6.3601 | 5.5949 | 5.8545 | 5.6816 | 6.0703 |
| ATAD5_S219 | ATAD5 | Q96QE3 | | 653 | VPFDSESPIRMKF | 6.2084 | 6.5144 | 6.2461 | 6.8692 | 6.4909 | 6.4722 | 6.1274 | 6.7828 | 6.4126 | 6.4773 | 6.3560 | 6.7161 | 5.5295 | 5.7254 | 5.6138 | 5.4578 |
| ATAD5_S306 | ATAD5 | Q96QE3 | | 756 | VIIIDSSPTALKH | 6.7867 | 7.0638 | 6.6760 | 7.1199 | 6.3432 | 6.1173 | 7.0360 | 6.8330 | 7.0345 | 6.4917 | 6.6667 | 7.3460 | 4.4468 | 4.8584 | 4.4610 | 4.7192 |
| ATAD5_S44 | ATAD5 | Q96QE3 | | 44 | TITKYLSPLGKTR | 6.9242 | 6.6955 | 7.0179 | 7.7069 | 6.5366 | 6.4518 | 7.1781 | 6.7409 | 7.3870 | 7.1470 | 7.2154 | 7.2074 | 3.5435 | 4.4303 | 3.6200 | 4.1975 |
| ATAD5_S653 | ATAD5 | Q96QE3 | | 70 | DYFRKTSPTNEKT | 7.2400 | 6.8366 | 6.2230 | 6.5370 | 6.6829 | 5.6015 | 6.6808 | 6.0662 | 6.6315 | 6.3388 | 6.2195 | 6.5538 | 5.3831 | 5.6668 | 5.5146 | 5.8237 |
| ATAD5_S70 | ATAD5 | Q96QE3 | | 219 | VVDLSESLPLAEE | 6.4629 | 6.4133 | 6.9113 | 7.7127 | 6.3111 | 6.8173 | 6.9382 | 6.6888 | 7.4030 | 6.5812 | 6.9036 | 7.1397 | 3.9929 | 4.9407 | 4.1469 | 4.6364 |
| ATAD5_S756 | ATAD5 | Q96QE3 | | 306 | VKSGYISESENSE | 5.1283 | 8.1723 | 7.0708 | 6.5322 | 6.1125 | 5.4200 | 6.2900 | 6.0196 | 6.3266 | 6.2199 | 6.0653 | 6.2770 | 5.9731 | 6.2054 | 5.8619 | 6.3250 |
| ATAD5_T603 | ATAD5 | Q96QE3 | | 603 | SGRISSTPTTETI | 6.3978 | 6.1987 | 6.1658 | 6.7380 | 5.7672 | 6.5787 | 6.4898 | 6.0755 | 6.6137 | 7.0327 | 6.5801 | 6.5092 | 5.5123 | 5.8081 | 5.8138 | 5.7185 |
| ATF1_T184 | ATF1 | P18846 | | 184 | TYQIRTTPSATSL | 6.9131 | 6.7250 | 6.6101 | 7.0656 | 6.5730 | 7.0423 | 6.0136 | 7.0824 | 7.4577 | 6.9710 | 8.1127 | 7.6196 | 3.5863 | 4.4521 | 3.4021 | 4.3734 |
| ATF2_S112 | ATF2 | P15336 | | 112 | KMPLDLSPLATPI | 6.5650 | 6.6007 | 6.4643 | 6.6894 | 6.2866 | 6.0122 | 6.0094 | 6.4384 | 6.4981 | 6.3267 | 6.2608 | 6.4222 | 5.8138 | 5.9748 | 5.8267 | 5.8110 |
| ATF4_S248 | ATF4 | P18848 | | 248 | PNRSLPSPGVLCG | 5.0812 | 4.8376 | 3.7962 | 4.5669 | 6.4059 | 5.8807 | 4.3566 | 4.8150 | 5.5568 | 4.8619 | 4.0083 | 4.6806 | 11.4384 | 9.3629 | 10.7742 | 9.5768 |
| ATF6_T166 | ATF6 | P18850 | | 166 | KTENGLTPKKKIQ | 6.4463 | 7.2319 | 6.8297 | 7.4464 | 6.3133 | 6.9724 | 7.3529 | 6.1761 | 6.9432 | 6.8248 | 6.5720 | 7.1824 | 4.5183 | 4.6180 | 3.6731 | 4.8993 |
| ATF7_S101 | ATF7 | P17544 | | 413 | TQGYLESPKESSE | 6.6225 | 6.4700 | 6.5317 | 6.4794 | 6.4299 | 5.7584 | 6.4376 | 5.4991 | 6.1928 | 6.1811 | 6.0782 | 5.8542 | 7.0963 | 6.1581 | 6.1354 | 6.0752 |
| ATF7_S171 | ATF7 | P17544 | | 97 | AGPLDMSLPSTPD | 5.8791 | 6.1162 | 6.3448 | 6.1391 | 6.3444 | 5.9705 | 6.2615 | 6.7134 | 6.6975 | 5.9331 | 6.5321 | 6.1416 | 6.3698 | 6.7822 | 5.3965 | 6.3783 |
| ATF7_S246 | ATF7 | P17544 | | 101 | DMSLPSTPDIKIK | 6.7219 | 6.6444 | 6.7077 | 6.7118 | 6.5847 | 6.0727 | 6.6432 | 6.4655 | 6.5457 | 6.6527 | 6.6901 | 6.5770 | 5.6827 | 5.1507 | 5.0455 | 5.1038 |
| ATF7_S413 | ATF7 | P17544 | | 246 | NSSGSISPSGHPI | 5.8281 | 5.5482 | 6.2626 | 7.1703 | 6.9721 | 6.3704 | 5.8068 | 6.6826 | 6.0876 | 6.0892 | 5.9432 | 6.2853 | 6.3782 | 6.1929 | 6.1511 | 6.2313 |
| ATF7_S73 | ATF7 | P17544 | | 73 | FNELASSFEHEFK | 6.4419 | 7.1761 | 6.9268 | 7.2265 | 6.9742 | 6.2042 | 6.2784 | 7.1634 | 6.2663 | 7.2941 | 7.6826 | 6.9253 | 4.1219 | 4.5089 | 4.3797 | 4.4297 |
| ATF7_S97 | ATF7 | P17544 | | 171 | LHPTLPSPTSVIT | 5.4942 | 5.5105 | 7.0889 | 6.5167 | 5.3398 | 5.5316 | 6.4582 | 6.4793 | 6.6890 | 7.0303 | 6.9946 | 6.6792 | 5.5479 | 5.5641 | 6.4614 | 6.6142 |
| ATF7_S97;T101 | ATF7 | P17544 | | 53 | IADQTPTPTRFLK | 7.6519 | 6.0346 | 5.9485 | 6.9376 | 5.9624 | 5.6129 | 6.8122 | 6.9756 | 5.9738 | 5.8307 | 5.9510 | 5.6572 | 6.6210 | 6.8629 | 5.4812 | 5.6864 |
| ATF7_T53 | ATF7 | P17544 | 97;101 | | NA | 6.3501 | 6.5596 | 6.1169 | 7.0749 | 6.3251 | 6.1411 | 5.7924 | 6.2158 | 6.4420 | 6.3466 | 6.3359 | 6.2035 | 6.0294 | 6.0322 | 5.9837 | 6.0509 |
| ATF7IP_S121 | ATF7IP | Q6VMQ6-4 | | 485 | MESSFGSPSKQES | 6.4333 | 6.4204 | 6.0862 | 6.5964 | 6.3332 | 6.0283 | 6.0713 | 5.9668 | 6.4727 | 6.2077 | 6.1750 | 6.3521 | 6.3808 | 6.1624 | 6.1395 | 6.1739 |
| ATF7IP_S437 | ATF7IP | Q6VMQ6-4 | | 121 | LNCEPLSPHNITP | 6.2037 | 6.5145 | 6.5748 | 6.6630 | 6.1779 | 5.8629 | 5.8963 | 6.1503 | 6.2156 | 6.3742 | 6.3612 | 6.3554 | 6.2051 | 6.1425 | 6.1893 | 6.1133 |
| ATF7IP_S453 | ATF7IP | Q6VMQ6-4 | | 681 | PPNPPVSPGKTVN | 6.3386 | 6.5546 | 6.3204 | 6.7814 | 6.3510 | 6.2327 | 6.2282 | 6.1325 | 6.4578 | 6.4550 | 6.3655 | 6.6116 | 5.9136 | 5.8990 | 5.7068 | 5.6511 |
| ATF7IP_S481;S485 | ATF7IP | Q6VMQ6-4 | | 504 | EAFLVLSDEEDIS | 6.0799 | 6.2132 | 6.2662 | 6.5613 | 6.3005 | 6.1370 | 5.9840 | 6.2893 | 6.4147 | 6.4558 | 6.2915 | 6.4655 | 5.9596 | 6.3080 | 5.9864 | 6.2872 |
| ATF7IP_S485 | ATF7IP | Q6VMQ6-4 | | 65 | GDLDPTSPLENMD | 5.9438 | 6.1803 | 5.9615 | 6.1988 | 6.4338 | 6.0629 | 6.0032 | 6.2821 | 6.1580 | 6.3091 | 6.1601 | 5.9299 | 6.6073 | 6.7128 | 6.7551 | 6.3011 |
| ATF7IP_S504 | ATF7IP | Q6VMQ6-4 | | 64 | NGDLDPTSPLENM | 5.8362 | 6.0480 | 5.9969 | 6.1149 | 5.9591 | 6.0806 | 6.1258 | 5.9799 | 6.0648 | 6.2298 | 6.1610 | 6.4161 | 6.6119 | 6.8214 | 7.0167 | 6.5370 |
| ATF7IP_S526 | ATF7IP | Q6VMQ6-4 | | 567 | RRKRSKSEDMDNV | 6.0149 | 5.4973 | 6.0087 | 6.2174 | 6.4075 | 6.2892 | 6.4560 | 6.5073 | 5.8970 | 5.8283 | 6.4608 | 6.1326 | 7.0068 | 6.8015 | 6.1788 | 6.2959 |
| ATF7IP_S567 | ATF7IP | Q6VMQ6-4 | | 453 | LEKLAPSEDELTC | 6.7171 | 6.5777 | 7.0210 | 7.1810 | 6.5531 | 6.3098 | 6.2599 | 6.1485 | 6.2343 | 6.1632 | 6.3731 | 6.2900 | 5.5680 | 5.8816 | 5.3328 | 5.3889 |
| ATF7IP_S65 | ATF7IP | Q6VMQ6-4 | | 860 | NVPSVPSPSIQRN | 6.3206 | 5.8591 | 5.8639 | 6.6816 | 6.5568 | 6.8883 | 6.6990 | 6.5773 | 6.5184 | 6.0945 | 6.5383 | 6.4614 | 6.0177 | 6.0347 | 4.9428 | 5.9458 |
| ATF7IP_S681 | ATF7IP | Q6VMQ6-4 | | 526 | SQNETCSPAEVES | 6.0780 | 5.6037 | 5.9436 | 5.9768 | 6.4050 | 5.7386 | 6.0250 | 5.9283 | 7.2104 | 6.0139 | 5.8210 | 5.8904 | 7.2857 | 6.4791 | 6.9031 | 6.6976 |

| | | | | | | | | | | | | | | | | | | | | | |
|---|---|---|---|---|---|---|---|---|---|---|---|---|---|---|---|---|---|---|---|---|---|
| ATF7IP_S860 | ATF7IP | Q6VMQ6-4 | | 437 | ILENTDSMETDEI | 5.7935 | 6.3554 | 5.9580 | 6.2187 | 6.0095 | 5.8096 | 5.9448 | 6.1068 | 6.3733 | 6.3390 | 5.7152 | 6.1931 | 7.1590 | 7.0131 | 6.9276 | 6.0834 |
| ATF7IP_S907 | ATF7IP | Q6VMQ6-4 | | 907 | GPSGLYSPSTNRG | 5.8759 | 6.3446 | 6.2403 | 6.8043 | 5.9171 | 5.8519 | 7.1545 | 6.9033 | 6.8471 | 7.1418 | 6.9663 | 6.6236 | 5.3202 | 6.1758 | 3.9867 | 5.8466 |
| ATF7IP_T64 | ATF7IP | Q6VMQ6-4 | 481;485 | | NA | 6.6126 | 6.6469 | 6.5702 | 6.8588 | 6.3280 | 6.3744 | 6.4432 | 6.3427 | 6.8344 | 6.6857 | 6.8125 | 6.7436 | 4.9935 | 5.2033 | 5.0211 | 5.5289 |
| ATG13_S394 | ATG13 | O75143-5 | | 394 | SSEGRASPHDVLE | 5.9371 | 6.2277 | 6.0563 | 5.7349 | 6.2361 | 6.5840 | 6.2938 | 7.1687 | 6.6799 | 6.0927 | 6.4030 | 6.0878 | 5.8628 | 6.1952 | 6.1367 | 6.3032 |
| ATG16L1_S269 | ATG16L1 | Q676U5 | | 269 | AATKRLSQPAGGL | 6.5608 | 5.0629 | 6.0216 | 5.8889 | 6.8458 | 5.8301 | 5.6338 | 7.3682 | 6.7944 | 6.4973 | 8.2288 | 6.7692 | 5.6245 | 6.0431 | 5.3348 | 5.4955 |
| ATG16L1_S287 | ATG16L1 | Q676U5 | | 70 | PNRHEISPGHDGT | 6.3301 | 5.7725 | 5.7328 | 5.7881 | 6.2520 | 6.2136 | 5.5994 | 7.1017 | 6.5429 | 7.4851 | 6.3013 | 7.2051 | 5.1595 | 5.7485 | 6.1523 | 6.6150 |
| ATG16L1_S70 | ATG16L1 | Q676U5 | | 287 | NIFGRRSVSSFPV | 6.0527 | 4.6598 | 5.8626 | 7.3891 | 5.8626 | 6.1002 | 5.4676 | 6.5160 | 8.1138 | 7.4307 | 7.5317 | 6.6853 | 4.7103 | 6.4388 | 4.8410 | 6.3378 |
| ATG2A_S1263 | ATG2A | Q2TAZ0 | | 1263 | GQKLSESPASLPS | 7.2105 | 6.9110 | 7.2212 | 7.4407 | 6.3468 | 6.0411 | 6.7822 | 5.9709 | 6.5920 | 7.2443 | 6.8400 | 7.1679 | 4.5552 | 5.1284 | 3.9678 | 4.5801 |
| ATG2A_S1265 | ATG2A | Q2TAZ0-3 | | 403 | MASRRLSAQAHPA | 6.1319 | 4.4711 | 5.9211 | 5.0061 | 6.8123 | 5.8569 | 6.0570 | 6.5942 | 7.9411 | 7.2749 | 7.8187 | 7.1816 | 6.6102 | 6.6613 | 4.6881 | 4.9736 |
| ATG2A_S1329 | ATG2A | Q2TAZ0-3 | | 1329 | SGAPPPSPPVGGP | 5.9700 | 6.1689 | 6.6957 | 6.3568 | 6.0971 | 6.1804 | 6.4108 | 6.9261 | 6.4516 | 6.0785 | 7.2482 | 7.3080 | 5.9354 | 5.2894 | 5.2983 | 5.5849 |
| ATG2A_S403 | ATG2A | Q2TAZ0-3 | | 1265 | KVQLSESPASLPS | 6.6936 | 7.7882 | 9.3879 | 6.4699 | 7.1850 | 7.9509 | 6.7173 | 6.6258 | 5.4226 | 7.6967 | 7.0121 | 7.9475 | 3.9043 | 3.0977 | 2.4809 | 3.6196 |
| ATG2B_S255 | ATG2B | Q96BY7 | | 255 | ETEPKLSPSWNPK | 5.9730 | 6.1003 | 7.0149 | 6.5687 | 6.1421 | 6.6761 | 6.5137 | 6.7434 | 6.2008 | 6.5785 | 6.9675 | 6.5393 | 5.5387 | 5.7457 | 5.4094 | 5.2879 |
| ATG2B_S497 | ATG2B | Q96BY7 | | 497 | LPSRSVSVDESRP | 5.3590 | 4.9619 | 5.8562 | 5.2728 | 6.3138 | 6.0363 | 5.6630 | 6.2499 | 5.6285 | 5.1638 | 5.5351 | 5.6880 | 8.5249 | 7.8349 | 8.4206 | 7.4914 |
| ATG4B_S383 | ATG4B | Q9Y4P1 | | 383 | LERFFDSEDEDFE | 5.9432 | 6.1709 | 6.1994 | 6.4637 | 6.3152 | 6.0493 | 6.1040 | 6.3984 | 6.1227 | 6.2346 | 6.3469 | 6.7135 | 5.8901 | 6.2656 | 6.3001 | 6.4824 |
| ATG4C_S451 | ATG4C | Q96DT6 | | 451 | KQLKRFSTEEFVL | 5.9093 | 6.0631 | 7.1043 | 6.5958 | 5.9396 | 4.8355 | 5.7629 | 6.5958 | 6.6479 | 5.9590 | 6.9336 | 6.2023 | 7.0147 | 6.7689 | 6.0462 | 5.6212 |
| ATG9A_S14 | ATG9A | Q7Z3C6 | | 656 | SALRSFSPLQPGQ | 6.6527 | 6.9824 | 6.8116 | 6.8882 | 6.2945 | 6.3686 | 6.6816 | 6.3606 | 7.1268 | 6.8688 | 6.9758 | 7.2067 | 4.4908 | 5.1932 | 4.2679 | 4.8297 |
| ATG9A_S18 | ATG9A | Q7Z3C6 | | 828 | PVPEEGSEDELPP | 5.9188 | 6.6166 | 6.5174 | 6.2075 | 6.4488 | 6.6107 | 6.4665 | 6.2848 | 6.4287 | 6.0668 | 6.6180 | 6.4221 | 5.9805 | 6.6325 | 5.4449 | 5.3354 |
| ATG9A_S656 | ATG9A | Q7Z3C6 | | 14 | YQRLEASYSDSPP | 6.3367 | 6.7362 | 6.8515 | 6.8587 | 6.3268 | 6.6559 | 6.4663 | 6.4096 | 6.9243 | 6.8325 | 6.9163 | 6.8839 | 4.4278 | 5.4898 | 4.6594 | 5.2244 |
| ATG9A_S828 | ATG9A | Q7Z3C6 | | 18 | EASYSDSPPGEED | 6.0859 | 5.9104 | 6.3858 | 5.7084 | 6.2332 | 6.3378 | 5.6670 | 7.0624 | 6.7459 | 6.2920 | 6.0319 | 5.8192 | 6.4153 | 6.5701 | 6.5199 | 6.2148 |
| ATIC_S387 | ATIC | P31939 | | 387 | LFGLHLSQKRNNG | 9.4034 | 6.0330 | 7.7182 | 6.6004 | 7.3756 | 8.4316 | 3.8254 | 5.3646 | 5.2410 | 7.8137 | 6.1004 | 4.2860 | 5.3814 | 5.3477 | 5.9544 | 5.1230 |
| ATL2_S34 | ATL2 | Q8NHH9 | | 34 | AAVNHVSSTTSLG | 6.6215 | 7.2340 | 5.6123 | 6.7904 | 6.4501 | 7.1818 | 5.7700 | 7.2757 | 6.6716 | 5.8298 | 6.0467 | 5.6591 | 5.2677 | 6.1391 | 5.7588 | 5.6912 |
| ATL2_S565 | ATL2 | Q8NHH9 | | 565 | RQSVTNSIKAGLT | 5.6622 | 5.5778 | 5.8481 | 6.1212 | 5.8368 | 5.3018 | 6.3943 | 6.4225 | 6.4619 | 6.1156 | 6.3436 | 5.8481 | 6.8899 | 5.9832 | 7.4220 | 7.7711 |
| ATM_S2996 | ATM | Q13315 | | 2996 | ECKRNLSDIDQSF | 4.8563 | 6.1282 | 5.9548 | 6.4675 | 5.6766 | 5.8686 | 6.4184 | 6.5318 | 6.1457 | 5.6112 | 5.5905 | 5.8512 | 6.2537 | 7.4612 | 7.4045 | 7.7797 |
| ATM_T1885 | ATM | Q13315 | | 1885 | QTSRSTTPANLDS | 6.3660 | 6.1736 | 6.3668 | 6.6458 | 6.1878 | 5.6395 | 5.9021 | 5.7185 | 5.8580 | 6.2243 | 5.9617 | 5.7245 | 6.5421 | 7.3977 | 6.3892 | 6.9024 |
| ATN1_S101 | ATN1 | P54259 | | 645 | YGKRAPSPGAYKT | 6.3327 | 6.6162 | 6.3974 | 5.9821 | 6.3014 | 6.2573 | 5.7586 | 6.4164 | 6.5894 | 6.5474 | 6.6970 | 6.4798 | 6.0757 | 5.9172 | 5.7960 | 5.8355 |
| ATN1_S132 | ATN1 | P54259 | | 632 | AGYKTASPPGPPP | 5.8470 | 6.6390 | 6.5581 | 6.5256 | 6.3185 | 6.3559 | 6.1155 | 6.4880 | 6.6296 | 6.8011 | 6.6943 | 6.6395 | 5.7771 | 5.9126 | 5.2469 | 5.4513 |
| ATN1_S168 | ATN1 | P54259 | | 168 | PPLFPPSPQPPDS | 5.6593 | 5.8996 | 6.6235 | 6.7887 | 6.5211 | 6.4062 | 6.2556 | 6.8278 | 6.4452 | 6.4916 | 6.6145 | 6.3658 | 5.8451 | 5.7090 | 5.5581 | 5.9889 |
| ATN1_S34 | ATN1 | P54259 | | 661 | PGYKPGSPPSFRT | 6.0294 | 6.3713 | 6.3895 | 6.3483 | 6.1925 | 6.2793 | 5.8061 | 6.3432 | 6.2662 | 6.3508 | 6.4168 | 6.7221 | 6.5885 | 6.0555 | 5.8349 | 6.0056 |
| ATN1_S632 | ATN1 | P54259 | | 896 | ARPHVMSPGNRNH | 6.9591 | 6.5414 | 6.5934 | 7.2156 | 6.5404 | 7.4487 | 6.0223 | 6.8052 | 7.0715 | 7.2310 | 7.5010 | 7.0596 | 3.9095 | 4.8184 | 4.0058 | 4.2769 |
| ATN1_S645 | ATN1 | P54259 | | 34 | RSRGRASPGGVST | 5.9751 | 5.9972 | 5.9718 | 5.9327 | 6.0689 | 6.2531 | 5.7296 | 6.2393 | 6.9603 | 6.8081 | 6.4575 | 6.3530 | 6.4807 | 6.1491 | 6.1813 | 6.4421 |
| ATN1_S677 | ATN1 | P54259 | | 101 | ELPRPQSPSDLDS | 6.2381 | 6.2984 | 6.2525 | 6.3754 | 6.7047 | 5.9805 | 5.7001 | 6.4900 | 7.1470 | 6.7993 | 6.8098 | 7.1731 | 5.6988 | 5.3790 | 5.5679 | 5.3855 |
| ATN1_S77 | ATN1 | P54259 | | 77 | GRSEEISESESEE | 4.8071 | 5.9798 | 6.4140 | 6.1171 | 6.2827 | 6.5565 | 5.6307 | 6.7459 | 6.4274 | 6.8101 | 7.1615 | 7.1428 | 5.6635 | 6.4334 | 5.8590 | 5.9686 |
| ATN1_S896 | ATN1 | P54259 | | 64 | RVEEASTPKVNKQ | 7.1071 | 6.4979 | 6.9413 | 6.7381 | 5.6920 | 6.8133 | 6.5600 | 6.7822 | 6.6884 | 6.6645 | 7.2705 | 7.2558 | 4.6685 | 4.9810 | 4.4200 | 4.9194 |
| ATN1_T64 | ATN1 | P54259 | | 669 | PSFRTGTPPGYRG | 5.9704 | 6.2787 | 6.4115 | 6.0816 | 6.2959 | 6.4270 | 6.5284 | 6.1005 | 6.3473 | 6.1278 | 6.1065 | 7.0575 | 6.0849 | 6.1551 | 6.0577 | 5.9693 |
| ATN1_T653;S661 | ATN1 | P54259 | | 677 | PGYRGTSPPAGPG | 6.7562 | 6.1749 | 6.3464 | 5.7488 | 5.8780 | 5.5932 | 5.7345 | 6.0238 | 6.3720 | 5.9694 | 6.1429 | 6.0268 | 7.1471 | 6.1464 | 6.5978 | 7.3420 |
| ATN1_T661 | ATN1 | P54259 | | 132 | QDNRSTSPSIYSP | 8.0240 | 6.4696 | 6.4895 | 6.6088 | 5.9724 | 6.6452 | 7.7224 | 7.5368 | 7.7854 | 6.9568 | 5.6112 | 6.2309 | 5.2963 | 3.5861 | 4.0369 | 5.0278 |
| ATN1_T669 | ATN1 | P54259 | 653;661 | | NA | 6.2026 | 6.4103 | 6.6248 | 6.1330 | 6.4921 | 6.2797 | 6.6004 | 6.2506 | 7.2675 | 7.1991 | 7.3191 | 7.0210 | 5.1115 | 5.5715 | 4.6837 | 4.8330 |
| ATP10D_S613 | ATP10D | Q9P241 | | 613 | QKIRHPSLGGLPI | 5.9163 | 5.8208 | 5.3496 | 5.7335 | 5.7087 | 5.8459 | 5.3528 | 5.7937 | 6.8678 | 6.7198 | 6.2817 | 6.3201 | 7.0492 | 7.0209 | 7.1963 | 7.0230 |
| ATP11A_S740 | ATP11A | P98196 | | 854 | RQAARNSDYAIPK | 6.5766 | 6.4324 | 5.8739 | 6.9872 | 5.6712 | 4.9685 | 5.8041 | 5.9136 | 6.4784 | 6.4100 | 5.7889 | 6.4531 | 6.4093 | 6.0415 | 6.6647 | 7.5265 |
| ATP11A_S854 | ATP11A | P98196 | | 740 | VLRHSGSLTRDNL | 6.3031 | 7.0112 | 6.0641 | 6.3290 | 6.0283 | 6.0873 | 5.7006 | 5.8717 | 6.1932 | 5.9796 | 6.0618 | 5.9616 | 6.4000 | 6.8449 | 6.7993 | 6.3644 |
| ATP13A1_S899 | ATP13A1 | Q9HD20 | | 899 | RRRPRDSPTLSNS | 6.8994 | 7.0653 | 6.9517 | 7.6541 | 6.1537 | 6.2004 | 6.8437 | 6.5566 | 7.1165 | 6.8126 | 6.8858 | 6.9110 | 4.1745 | 4.8683 | 4.3408 | 4.5658 |
| ATP13A3_S817 | ATP13A3 | Q9H7F0 | | 817 | VKLVHDSLEDLQM | 6.2487 | 6.5554 | 6.2204 | 6.3339 | 6.0844 | 5.9953 | 6.1903 | 6.2414 | 6.4137 | 6.0803 | 6.1656 | 6.1553 | 6.4285 | 6.2448 | 6.5699 | 6.0720 |
| ATP13A3_S98 | ATP13A3 | Q9H7F0 | | 98 | ETYPVSSPKSMSN | 6.7584 | 6.7729 | 7.0720 | 7.2055 | 6.6952 | 6.2001 | 6.8330 | 6.3503 | 6.9444 | 6.7247 | 7.3410 | 6.9729 | 4.0854 | 4.5921 | 4.3815 | 5.0705 |
| ATP1A1_S16 | ATP1A1 | P05023-4 | | 16 | YEPAAVSEQGDKK | 6.2408 | 6.0530 | 6.5168 | 6.7389 | 6.2020 | 6.4988 | 6.2438 | 6.4695 | 6.8309 | 6.8368 | 7.1986 | 7.1329 | 5.0169 | 5.7859 | 5.0223 | 5.2121 |
| ATP1A1_Y260 | ATP1A1 | P05023-4 | | 260 | ARGIVVYTGDRTV | 5.6006 | 4.1735 | 2.9681 | 4.5558 | 3.9945 | 6.3636 | 6.3551 | 6.0183 | 7.6529 | 7.3953 | 9.3144 | 11.0387 | 4.3394 | 6.6134 | 3.3104 | 10.3061 |
| ATP2A2_S378 | ATP2A2 | P16615 | | 663 | FDELNPSAQRDAC | 7.0449 | 4.8991 | 5.1759 | 6.1786 | 6.3951 | 5.6953 | 5.8376 | 6.5275 | 5.6391 | 6.1866 | 5.4827 | 5.5609 | 7.4760 | 7.1993 | 6.8704 | 7.8310 |
| ATP2A2_S663 | ATP2A2 | P16615 | | 378 | VEGDTCSLNEFTI | 5.9041 | 5.9328 | 6.2116 | 6.5190 | 6.0313 | 6.0700 | 6.6263 | 6.5909 | 6.6241 | 6.9471 | 5.9804 | 5.2681 | 6.3542 | 6.5810 | 6.5810 | 5.7780 |
| ATP2B1_S1178 | ATP2B1 | P20020-1 | | 1193 | EFRIEDSEPHIPL | 5.6555 | 5.8660 | 6.2903 | 6.3468 | 5.7617 | 6.0480 | 6.1288 | 6.1004 | 6.2562 | 5.9780 | 5.7820 | 5.3967 | 6.7526 | 7.3822 | 6.9054 | 7.3495 |
| ATP2B1_S1193 | ATP2B1 | P20020-1 | | 1178 | KPESRSSIHNFMT | 5.4511 | 5.4533 | 5.6491 | 5.8891 | 5.6603 | 5.6014 | 5.1428 | 6.1263 | 5.9036 | 6.1937 | 6.1750 | 6.1785 | 7.3231 | 7.8002 | 7.9841 | 7.4683 |
| ATP2B1_S1216;S1220 | ATP2B1 | P20020-1 | | 1249 | TSSSPGSPLHSLE | 6.7505 | 6.7935 | 6.0551 | 6.9119 | 6.4824 | 7.0980 | 6.3972 | 7.5884 | 6.5770 | 6.4743 | 6.9774 | 6.5676 | 4.2968 | 5.7247 | 4.4864 | 4.8187 |
| ATP2B1_S1249 | ATP2B1 | P20020-1 | 1216;1220 | | NA | 6.2212 | 6.4387 | 5.8403 | 6.6502 | 6.4839 | 6.4638 | 6.4328 | 5.5901 | 6.0692 | 6.4223 | 5.5813 | 6.2980 | 6.3882 | 6.7353 | 5.9957 | 6.3892 |
| ATP2B1_S282;T284 | ATP2B1 | P20020-1 | 276;282 | | NA | 5.6031 | 5.9506 | 7.2379 | 6.4753 | 6.0096 | 5.5966 | 6.4515 | 6.0505 | 5.7429 | 6.4976 | 6.6959 | 6.8119 | 6.2475 | 6.1308 | 6.2188 | 6.2797 |
| ATP2B1_T276;S282 | ATP2B1 | P20020-1 | 282;284 | | NA | 5.4923 | 5.9080 | 7.2199 | 6.1113 | 6.3933 | 6.2134 | 6.2452 | 5.9511 | 6.0512 | 6.4180 | 6.3813 | 6.1202 | 6.4341 | 6.3702 | 5.9960 | 6.6945 |
| ATP2B4_S1162 | ATP2B4 | P23634 | | 1181 | EEELPRTPLLDEE | 6.4721 | 6.7591 | 5.4704 | 6.9630 | 6.7455 | 6.7691 | 5.9795 | 6.8147 | 5.8247 | 5.3894 | 5.6151 | 5.8338 | 6.0529 | 6.4467 | 6.7039 | 6.1600 |
| ATP2B4_S1240 | ATP2B4 | P23634 | | 1162 | KPYNQKSIHSFMT | 5.8116 | 7.6842 | 7.8189 | 5.9141 | 4.8736 | 7.8639 | 5.7577 | 6.4855 | 7.1774 | 8.4695 | 9.2943 | 8.8504 | 3.3012 | 4.0883 | 2.2858 | 4.3237 |
| ATP2B4_T1181 | ATP2B4 | P23634 | | 1240 | LQSLETSVxxxxx | 4.9083 | 6.1647 | 5.7188 | 6.3645 | 6.0453 | 5.5054 | 5.9914 | 6.6318 | 5.5508 | 6.2017 | 6.2260 | 6.1013 | 7.0386 | 6.8653 | 7.1834 | 7.5025 |
| ATP5F1C_S146 | ATP5F1C | P36542 | | 146 | ILYRTHSDQFLVA | 5.1947 | 5.6803 | 6.5643 | 5.7728 | 6.0423 | 5.2741 | 6.1946 | 6.6945 | 5.8613 | 6.0735 | 5.9316 | 5.4837 | 7.9470 | 6.6333 | 7.4887 | 7.1632 |
| ATP6AP1_S465 | ATP6AP1 | Q15904 | | 465 | HKGPTISLTQIVx | 3.9252 | 3.7758 | 4.8386 | 4.3365 | 3.8933 | 4.3841 | 4.9800 | 4.5784 | 4.6738 | 4.6515 | 4.0557 | 4.2415 | 11.3132 | 11.6893 | 12.8175 | 11.8456 |
| ATP6V0A2_S154 | ATP6V0A2 | Q9Y487 | | 695 | TLIRKDSEEEVSL | 5.4847 | 6.2837 | 6.1827 | 6.5239 | 6.2497 | 5.6019 | 6.1190 | 5.8571 | 6.1363 | 6.5875 | 5.8616 | 6.1804 | 6.9265 | 7.4067 | 6.1469 | 6.4513 |
| ATP6V0A2_S695 | ATP6V0A2 | Q9Y487 | | 154 | TYEEFPSLESDSL | 6.5319 | 7.4348 | 7.4197 | 7.1893 | 5.8897 | 6.1050 | 6.8530 | 5.8141 | 7.3971 | 5.8765 | 6.7114 | 6.5130 | 4.7941 | 4.5995 | 6.0540 | 4.8168 |
| ATP6V1A_T136 | ATP6V1A | P38606 | | 136 | DIKWDFTPCKNLR | 6.5108 | 6.4754 | 6.9080 | 6.5644 | 6.0656 | 6.0745 | 6.3011 | 6.4477 | 6.4134 | 6.7093 | 6.6590 | 6.2936 | 5.5396 | 5.9292 | 5.4165 | 5.6918 |
| ATP7A_S1544 | ATP7A | Q04656-3 | | 1544 | VNYSRASINSLLS | 5.5460 | 5.1683 | 5.7475 | 5.5710 | 5.1860 | 5.1387 | 5.5607 | 6.2754 | 6.6501 | 6.6723 | 6.1714 | 6.8887 | 7.2140 | 7.6374 | 7.0025 | 7.5700 |
| ATP7A_S1544;S1547 | ATP7A | Q04656-3 | | 351 | EGSQQRSPSYTND | 5.5014 | 6.2532 | 5.7239 | 6.2890 | 6.2914 | 6.2919 | 6.8481 | 5.9942 | 6.2241 | 6.6424 | 5.8364 | 6.1163 | 6.1062 | 6.2133 | 7.0377 | 6.6304 |
| ATP7A_S351 | ATP7A | Q04656-3 | | 557 | FYTKGMTPVQDKE | 6.5103 | 6.2449 | 6.9984 | 7.8254 | 6.2760 | 7.3837 | 6.9478 | 6.4384 | 6.8270 | 8.5046 | 7.7935 | 7.0213 | 3.2261 | 4.6580 | 2.9045 | 4.4400 |
| ATP7A_S434;S438 | ATP7A | Q04656-3 | 1544;1547 | | NA | 5.0241 | 4.8053 | 6.1255 | 5.5609 | 6.0007 | 4.3045 | 5.4770 | 5.5042 | 6.6240 | 5.9160 | 6.0760 | 6.5844 | 6.9931 | 8.5109 | 8.4105 | 8.0830 |
| ATP7A_T557 | ATP7A | Q04656-3 | 434;438 | | NA | 5.9674 | 6.2857 | 7.2915 | 8.0786 | 7.0487 | 6.4065 | 7.0597 | 4.3723 | 6.8403 | 6.6766 | 6.5324 | 6.1037 | 5.8224 | 6.0996 | 4.9818 | 4.4328 |
| ATP8B2_S1183 | ATP8B2 | P98198-3 | | 1183 | GKNMRLSSLALSS | 6.9254 | 5.1921 | 8.4230 | 6.6137 | 7.4765 | 5.1807 | 5.5950 | 5.3936 | 5.9523 | 9.3770 | 6.5757 | 6.9406 | 4.9831 | 7.4081 | 4.3559 | 3.6071 |
| ATR_S428 | ATR | Q13535 | | 428 | SNSDGISPKRRRL | 5.9117 | 6.4092 | 6.7190 | 6.5564 | 6.5452 | 5.9866 | 6.2541 | 6.2298 | 6.7719 | 6.5826 | 6.9716 | 6.8215 | 5.4176 | 5.7678 | 5.4463 | 5.6086 |
| ATR_S435 | ATR | Q13535 | | 1989 | CFPENETPPEGKN | 4.9729 | 6.0011 | 6.3323 | 6.4430 | 6.0234 | 6.2022 | 6.0431 | 5.9691 | 6.0563 | 6.2077 | 6.0560 | 5.8097 | 6.8343 | 6.7397 | 7.6697 | 6.6396 |
| ATR_T1989 | ATR | Q13535 | | 435 | PKRRRLSSSLNPS | 5.3457 | 5.1703 | 5.2652 | 5.5560 | 7.3469 | 6.2028 | 6.1181 | 6.2856 | 5.7803 | 5.1894 | 5.6577 | 5.9174 | 7.6004 | 7.1800 | 7.6240 | 7.7602 |
| ATRIP_S224 | ATRIP | Q8WXE1 | | 224 | PSVSHVSPRKNPS | 6.2532 | 7.0928 | 6.1053 | 6.5557 | 6.7809 | 6.7406 | 5.8665 | 7.3071 | 6.3320 | 5.8237 | 6.2452 | 6.1537 | 5.3470 | 5.7610 | 6.0274 | 5.6078 |
| ATRIP_S239 | ATRIP | Q8WXE1 | | 239 | IKPEACSPQFGKT | 6.0329 | 5.8075 | 7.3214 | 6.7998 | 6.1490 | 6.2875 | 6.2340 | 6.2991 | 6.8422 | 7.0080 | 7.1766 | 7.2576 | 4.9376 | 5.7361 | 4.5688 | 5.5418 |
| ATRX_S1061 | ATRX | P46100 | | 92 | VTKYVESDDEKPL | 5.9060 | 6.1965 | 6.3021 | 6.5093 | 5.7610 | 5.4077 | 6.1668 | 5.6463 | 6.4014 | 6.2783 | 6.0062 | 6.0191 | 6.7402 | 6.9154 | 6.8831 | 6.8605 |
| ATRX_S1244;S1245 | ATRX | P46100 | | 598 | PVSLSNSPIKGAD | 6.8514 | 6.7203 | 6.2980 | 6.8169 | 6.8940 | 6.4148 | 6.7891 | 6.4132 | 6.6258 | 6.4686 | 6.1618 | 6.1749 | 5.2111 | 5.4406 | 5.3153 | 5.4042 |
| ATRX_S1245 | ATRX | P46100 | | 634 | GLGQENSDNEHLV | 5.7157 | 6.1025 | 5.8954 | 6.7097 | 6.3995 | 5.6102 | 6.1189 | 6.0867 | 5.7494 | 6.2076 | 5.9321 | 6.1556 | 6.5630 | 7.2712 | 6.4576 | 7.0249 |
| ATRX_S1527 | ATRX | P46100 | | 1527 | IEIEDASPTKCPI | 5.8783 | 6.2658 | 6.1250 | 6.8484 | 6.3122 | 6.0433 | 6.1682 | 6.1408 | 6.4664 | 6.2394 | 6.2677 | 6.4024 | 5.8596 | 6.5082 | 6.2404 | 6.2339 |
| ATRX_S1990 | ATRX | P46100 | | 677 | NPVTSNSDEECNE | 5.7213 | 5.5621 | 5.9410 | 6.1869 | 6.0350 | 5.6037 | 5.7566 | 5.8417 | 6.0782 | 6.1474 | 6.6388 | 6.7414 | 6.0640 | 7.3734 | 7.3467 | 6.9619 |
| ATRX_S1990;S1996 | ATRX | P46100 | | 729 | ETVDQNSDSDEML | 5.8863 | 6.3556 | 6.0735 | 6.6212 | 5.7138 | 6.3025 | 6.2192 | 5.6672 | 6.5569 | 6.4317 | 6.6196 | 6.5880 | 5.7701 | 6.8287 | 6.1574 | 6.2084 |
| ATRX_S1996 | ATRX | P46100 | | 84 | RSKRKPSIVTKYV | 4.4458 | 4.2500 | 4.4449 | 5.2736 | 8.0830 | 4.9933 | 4.9328 | 5.6071 | 6.7058 | 7.0762 | 6.1710 | 6.1150 | 8.5413 | 8.4325 | 7.7657 | 7.1618 |
| ATRX_S594 | ATRX | P46100 | | 594 | VKLTPVSLSNSPI | 5.2280 | 4.7230 | 5.3865 | 5.9384 | 5.7985 | 5.1308 | 5.1989 | 5.2257 | 5.1750 | 5.0497 | 5.1849 | 5.2760 | 9.3531 | 9.6260 | 9.0416 | 8.6640 |
| ATRX_S594;S598 | ATRX | P46100 | | 962 | KVQDGLSDIAEKF | 6.3514 | 6.3115 | 7.1019 | 6.8160 | 6.2999 | 5.9666 | 6.1087 | 5.6001 | 6.6381 | 5.9267 | 5.7871 | 5.9417 | 6.0547 | 6.5658 | 7.5538 | 6.7761 |
| ATRX_S598 | ATRX | P46100 | | 1996 | SSSNPSSPAPDWY | 6.2071 | 6.6456 | 6.2415 | 7.0577 | 6.9052 | 6.5325 | 7.2244 | 6.9271 | 6.6214 | 6.8927 | 6.6465 | 6.6201 | 4.7563 | 5.3334 | 4.7546 | 4.6339 |
| ATRX_S634 | ATRX | P46100 | | 1245 | TGFCQSSGDEALS | 6.0765 | 6.1616 | 6.7165 | 7.0383 | 5.7193 | 6.0464 | 5.3302 | 7.3424 | 5.8744 | 6.6235 | 6.6217 | 5.9312 | 5.9657 | 6.8788 | 6.4657 | 5.2078 |
| ATRX_S677 | ATRX | P46100 | | 784 | KRKRKSSTSGSDF | 5.8623 | 5.6632 | 5.1585 | 6.5594 | 5.2640 | 5.2859 | 5.3235 | 5.7290 | 7.0489 | 6.9890 | 7.2438 | 6.9696 | 5.9054 | 7.3843 | 6.7776 | 6.8358 |
| ATRX_S729 | ATRX | P46100 | | 1061 | KKKDELSDYAEKS | 6.4620 | 6.3192 | 5.9110 | 7.3234 | 5.7713 | 6.1129 | 6.3427 | 5.9093 | 5.2404 | 5.8704 | 5.6501 | 6.0157 | 6.8627 | 7.1071 | 6.6337 | 6.4680 |
| ATRX_S784 | ATRX | P46100 | | 1990 | EESKATSSSNPSS | 6.5018 | 6.0862 | 5.8235 | 6.0794 | 6.8885 | 6.6055 | 6.9846 | 7.3574 | 6.5722 | 6.2343 | 6.4833 | 6.1706 | 5.3903 | 5.8163 | 5.4956 | 5.5104 |
| ATRX_S84 | ATRX | P46100 | | 887 | RKQERETFSSAEG | 6.3026 | 6.2255 | 7.0804 | 8.1926 | 6.2571 | 6.4712 | 6.6599 | 6.8141 | 5.6016 | 5.0147 | 5.2865 | 5.1795 | 5.9007 | 6.2268 | 6.1859 | 6.6009 |
| ATRX_S92 | ATRX | P46100 | 594;598 | | NA | 5.9578 | 6.1528 | 5.2380 | 7.0541 | 5.4143 | 5.3242 | 4.9082 | 4.8556 | 5.4226 | 5.5849 | 4.8590 | 5.4226 | 8.9524 | 8.3192 | 8.3990 | 8.1354 |
| ATRX_S962 | ATRX | P46100 | 1244;1245 | | NA | 6.5266 | 7.1197 | 6.6609 | 7.1068 | 6.3755 | 6.3174 | 6.6248 | 5.9287 | 5.7427 | 6.4113 | 6.0807 | 6.2410 | 5.3462 | 5.7896 | 5.9969 | 5.7312 |
| ATRX_T1346;S1352 | ATRX | P46100 | 1990;1996 | | NA | 5.7854 | 6.1379 | 5.9762 | 6.1090 | 5.8998 | 5.8990 | 6.4393 | 6.0026 | 6.0778 | 5.6623 | 5.7197 | 5.5991 | 7.0552 | 7.2792 | 7.6680 | 6.6893 |
| ATRX_T887 | ATRX | P46100 | 1346;1352 | | NA | 6.5009 | 6.9053 | 5.7638 | 6.5572 | 6.1737 | 5.8769 | 5.7878 | 5.1871 | 6.2872 | 5.7598 | 5.9301 | 5.8808 | 6.6944 | 6.7252 | 6.8924 | 7.0774 |
| ATXN1_S406 | ATXN1 | P54253 | | 775 | TRKRRWSAPESRK | 6.7827 | 6.6440 | 5.5668 | 5.7166 | 6.4552 | 5.6744 | 5.8588 | 6.2829 | 5.9961 | 6.1323 | 6.1014 | 6.4012 | 7.6181 | 6.3661 | 6.2018 | 6.2016 |
| ATXN1_S612 | ATXN1 | P54253 | | 406 | ATHREASPSTLND | 6.5372 | 6.4720 | 5.6739 | 6.4251 | 6.0152 | 6.4064 | 6.0424 | 6.3241 | 6.3302 | 5.7602 | 6.1712 | 6.7426 | 6.6457 | 6.2873 | 6.0121 | 6.1545 |
| ATXN1_S775 | ATXN1 | P54253 | | 612 | NDLKIDSSTVERI | 5.3608 | 6.1666 | 7.7617 | 6.8548 | 6.2523 | 6.0861 | 6.4926 | 5.6054 | 6.3492 | 6.2688 | 6.1996 | 7.2334 | 6.1295 | 6.1398 | 5.2329 | 5.8664 |
| ATXN10_S12 | ATXN10 | Q9UBB4 | | 12 | PPPARLSGVMVPA | 6.7959 | 6.3371 | 7.0365 | 8.2149 | 6.2404 | 6.5890 | 7.3469 | 6.4136 | 7.8079 | 6.3506 | 7.4166 | 6.8993 | 3.7285 | 5.0936 | 3.3485 | 4.3807 |
| ATXN1L_S284 | ATXN1L | P0C7T5 | | 330 | APAHRGTPDTDLE | 6.5876 | 6.9885 | 6.2202 | 6.4843 | 7.2277 | 6.6833 | 5.8691 | 6.3896 | 5.9628 | 5.9485 | 6.7025 | 6.5034 | 5.2548 | 5.8193 | 5.7265 | 5.6318 |
| ATXN1L_S284;S289 | ATXN1L | P0C7T5 | | 62 | QAGARVSLGGDGA | 5.0821 | 5.4390 | 5.7228 | 6.0612 | 7.2527 | 6.3190 | 5.8331 | 5.8183 | 5.7914 | 5.5531 | 6.1446 | 6.4135 | 8.4534 | 7.0014 | 6.4877 | 6.6268 |
| ATXN1L_S62 | ATXN1L | P0C7T5 | | 284 | NLVRRESEALDSP | 6.1177 | 5.7438 | 5.8395 | 5.6564 | 6.6310 | 6.1265 | 5.1743 | 6.0068 | 5.3027 | 6.0745 | 5.4782 | 6.0182 | 8.5081 | 7.6559 | 6.5577 | 7.1087 |
| ATXN1L_T330 | ATXN1L | P0C7T5 | 284;289 | | NA | 6.9084 | 6.4778 | 6.3262 | 6.2771 | 7.7108 | 6.4060 | 6.3173 | 6.8052 | 6.6018 | 6.6166 | 5.9693 | 6.0935 | 6.0520 | 5.4634 | 5.0853 | 4.8892 |
| ATXN2_S114 | ATXN2 | Q99700 | | 624 | EGPPRMSPKAQRH | 7.2360 | 7.2088 | 7.4090 | 5.9955 | 7.1034 | 7.6089 | 6.4472 | 6.0425 | 6.2860 | 6.6538 | 6.9421 | 6.6286 | 4.9378 | 5.1527 | 4.2257 | 4.1221 |
| ATXN2_S507 | ATXN2 | Q99700 | | 741 | ASNRAVTPSSEAK | 6.4272 | 6.3770 | 6.7475 | 6.1017 | 6.4760 | 6.7657 | 6.2603 | 6.6857 | 6.5693 | 6.4989 | 6.7138 | 6.6963 | 5.4111 | 5.6875 | 5.2706 | 5.3115 |
| ATXN2_S554;S558 | ATXN2 | Q99700 | | 772 | IKPNETSPSFSKA | 6.5985 | 6.0750 | 7.2847 | 6.1681 | 6.1942 | 6.2204 | 6.5321 | 5.8994 | 6.9313 | 7.0254 | 6.4702 | 6.5239 | 5.5131 | 5.4752 | 5.0734 | 6.0151 |
| ATXN2_S624 | ATXN2 | Q99700 | | 758 | QDQRQNSPAGNKE | 6.4519 | 6.6159 | 7.4755 | 6.5579 | 6.3191 | 6.9044 | 6.4893 | 6.5778 | 6.9214 | 7.1839 | 7.2605 | 7.3928 | 4.2751 | 4.8492 | 4.2118 | 4.5134 |
| ATXN2_S642 | ATXN2 | Q99700 | | 861 | GSSKPNSPSISPS | 5.3192 | 6.3558 | 7.3199 | 6.5302 | 5.4419 | 6.0649 | 5.9568 | 6.3554 | 6.6533 | 6.6009 | 6.7690 | 6.5434 | 6.0753 | 5.5843 | 5.7559 | 6.6737 |
| ATXN2_S667 | ATXN2 | Q99700 | | 642 | VSAGRGSISSGLE | 6.2481 | 5.5593 | 5.9300 | 5.5840 | 6.7171 | 5.6950 | 5.8019 | 6.5545 | 7.5830 | 7.3898 | 6.9750 | 7.0703 | 5.6927 | 7.0020 | 4.8076 | 5.3899 |
| ATXN2_S758 | ATXN2 | Q99700 | | 667 | PPVARTSPSGGTW | 5.9158 | 6.4214 | 6.7206 | 6.4914 | 6.4207 | 6.4086 | 6.4196 | 6.4338 | 6.2182 | 6.7369 | 6.8643 | 6.6754 | 5.8290 | 5.2762 | 5.2755 | 5.8927 |
| ATXN2_S772 | ATXN2 | Q99700 | | 784 | AENKGISPVVSEH | 7.1361 | 7.6836 | 6.4617 | 8.4780 | 6.8653 | 7.1804 | 5.9628 | 7.4699 | 6.4967 | 7.2298 | 7.3643 | 7.2383 | 3.4152 | 4.4864 | 3.2637 | 3.2679 |
| ATXN2_S784 | ATXN2 | Q99700 | | 580 | LPPRAATPTRPPS | 5.8584 | 6.5462 | 6.6018 | 5.8629 | 5.8750 | 6.0310 | 6.0176 | 6.3063 | 7.6062 | 6.1283 | 5.6683 | 6.9556 | 6.1117 | 6.4929 | 6.2198 | 5.7180 |
| ATXN2_S857;S861;S865 | ATXN2 | F8WB06 | | 507 | PPVARTSPSGGTW | 6.5592 | 6.5150 | 7.9720 | 6.6785 | 6.2190 | 5.7065 | 5.9208 | 6.2433 | 6.8276 | 6.7139 | 7.2529 | 6.9447 | 4.4472 | 5.6788 | 4.6637 | 5.6568 |
| ATXN2_S861 | ATXN2 | S4R3R6 | | 114 | GRASASSPAFENx | 5.8294 | 7.2963 | 6.7312 | 7.0698 | 7.1439 | 5.4315 | 4.8409 | 6.2972 | 6.1321 | 7.1376 | 6.2528 | 5.4950 | 5.3405 | 6.7375 | 6.1999 | 6.0644 |
| ATXN2_S861;S865 | ATXN2 | Q99700 | 857;861;865 | | NA | 5.6186 | 6.0696 | 6.8970 | 6.3575 | 5.6427 | 6.5660 | 6.4031 | 6.3282 | 6.6237 | 6.2864 | 7.0615 | 7.0057 | 5.8143 | 6.0486 | 5.7876 | 5.4894 |
| ATXN2_T580 | ATXN2 | Q99700 | 861;865 | | NA | 6.0654 | 5.8129 | 6.8961 | 6.1273 | 5.7527 | 5.5096 | 5.7722 | 6.1115 | 6.4294 | 6.7040 | 6.5135 | 6.6048 | 6.3954 | 6.2463 | 6.6337 | 6.4253 |
| ATXN2_T741 | ATXN2 | Q99700 | 554;558 | | NA | 6.9118 | 6.8821 | 7.0049 | 6.3928 | 6.5039 | 7.8512 | 6.9809 | 6.7796 | 7.4763 | 7.8755 | 7.6483 | 7.2435 | 3.1020 | 4.3565 | 3.1253 | 3.8655 |
| ATXN2L_S111 | ATXN2L | Q8WWM7-3 | | 594 | TSEPMGSPVSSKT | 6.3144 | 6.6236 | 6.7357 | 6.4728 | 6.1498 | 6.5508 | 6.1187 | 6.6235 | 6.5030 | 6.9140 | 6.6480 | 6.8551 | 5.4917 | 5.5185 | 5.2098 | 5.2708 |
| ATXN2L_S238 | ATXN2L | Q8WWM7-3 | | 449 | RMYPPRSPKSAAP | 6.4777 | 6.2075 | 6.5136 | 6.1152 | 6.6008 | 6.7146 | 6.2163 | 6.6204 | 6.5743 | 6.2577 | 6.5115 | 6.4824 | 5.9932 | 5.7864 | 5.4528 | 5.4757 |
| ATXN2L_S306 | ATXN2L | Q8WWM7-3 | | 238 | EGGDSNSDDYDLE | 6.1613 | 6.0537 | 6.7582 | 6.1334 | 6.1858 | 6.2645 | 5.6301 | 6.3016 | 6.9284 | 6.8547 | 7.1038 | 7.0153 | 5.8547 | 5.7989 | 5.6094 | 5.3463 |
| ATXN2L_S32 | ATXN2L | Q8WWM7-3 | | 339 | QGSGRESPSLASR | 6.3708 | 6.5429 | 6.7328 | 6.1270 | 6.2650 | 6.2532 | 5.8608 | 6.9735 | 6.9310 | 6.6630 | 6.4972 | 6.4771 | 5.6203 | 5.6922 | 5.4819 | 5.5112 |
| ATXN2L_S339 | ATXN2L | Q8WWM7-3 | | 684 | STSTPTSPGPRTH | 5.9410 | 5.5408 | 5.1161 | 4.8692 | 5.7472 | 7.0047 | 6.5926 | 6.6677 | 5.8724 | 4.9415 | 5.3816 | 5.7049 | 7.8456 | 7.4663 | 7.5809 | 7.7275 |
| ATXN2L_S341 | ATXN2L | Q8WWM7-3 | | 559 | KLQPSSSPENSLD | 6.4525 | 6.4432 | 7.0407 | 6.7987 | 6.2007 | 6.5355 | 6.3296 | 6.4174 | 6.4129 | 6.4170 | 6.7338 | 6.7833 | 5.4972 | 5.4595 | 5.1164 | 5.3615 |
| ATXN2L_S391 | ATXN2L | Q8WWM7-3 | | 424 | GAKTLSSPSNRPS | 7.1262 | 6.6904 | 7.1294 | 7.3319 | 6.8429 | 6.9781 | 6.8604 | 7.0496 | 7.5292 | 7.0382 | 7.5482 | 7.0582 | 3.5768 | 4.0218 | 3.3932 | 3.8256 |
| ATXN2L_S424 | ATXN2L | Q8WWM7-3 | | 604 | SKTESVSDKEDKP | 6.7575 | 6.5184 | 7.2382 | 6.0934 | 6.1659 | 6.1118 | 6.1166 | 6.4537 | 5.9505 | 6.0055 | 5.8580 | 6.3878 | 6.2421 | 5.9019 | 5.8208 | 6.3780 |
| ATXN2L_S449 | ATXN2L | Q8WWM7-3 | | 306 | AREIESSPQYRLR | 7.4103 | 7.7431 | 7.1215 | 7.5528 | 6.9092 | 7.2543 | 7.1461 | 7.4292 | 7.3337 | 7.1002 | 7.5939 | 7.4346 | 2.8208 | 3.5128 | 2.5299 | 3.1077 |
| ATXN2L_S559 | ATXN2L | Q8WWM7-3 | | 32 | RPPGGTSPPNGGL | 6.8813 | 5.6449 | 6.9942 | 5.9868 | 6.3760 | 5.1881 | 6.4825 | 7.5894 | 6.8794 | 6.4392 | 6.5317 | 6.3632 | 5.9824 | 5.8305 | 5.0745 | 5.7559 |
| ATXN2L_S594 | ATXN2L | Q8WWM7-3 | | 391 | HHLDNSSPGPGSE | 6.5324 | 7.1587 | 6.7328 | 6.9500 | 6.2656 | 7.3069 | 6.4150 | 7.0561 | 7.5062 | 6.9855 | 6.9434 | 6.8728 | 4.3777 | 4.7481 | 3.6413 | 4.5075 |
| ATXN2L_S604 | ATXN2L | Q8WWM7-3 | | 111 | GKGPPQSPVFEGV | 6.4892 | 5.7101 | 6.4141 | 6.0690 | 6.3002 | 5.8950 | 5.8141 | 6.4667 | 6.0222 | 5.7812 | 5.6708 | 5.7257 | 6.4482 | 7.2603 | 6.9175 | 7.0158 |
| ATXN2L_S634 | ATXN2L | Q8WWM7-3 | | 634 | CPSQTGSPPVGLI | 6.1120 | 5.8546 | 6.7264 | 6.0911 | 5.5720 | 6.3161 | 5.5448 | 5.9278 | 6.2397 | 6.2544 | 7.5595 | 6.5234 | 6.1110 | 6.0116 | 6.7745 | 6.3810 |
| ATXN2L_S680;S684 | ATXN2L | Q8WWM7-3 | | 341 | SGRESPSLASREG | 6.1269 | 6.3384 | 6.5143 | 5.9097 | 7.1270 | 6.6840 | 6.2542 | 6.9413 | 6.5189 | 6.7218 | 6.9275 | 7.4382 | 5.8043 | 5.1928 | 4.9584 | 4.5423 |
| ATXN2L_S684 | ATXN2L | Q8WWM7-3 | 680;684 | | NA | 6.6248 | 7.2457 | 7.3205 | 6.5388 | 5.8050 | 6.1888 | 6.0236 | 6.5152 | 7.1674 | 6.7203 | 7.0210 | 6.8490 | 4.8307 | 4.9723 | 4.8960 | 5.2810 |
| ATXN2L_T632;S634 | ATXN2L | Q8WWM7-3 | 632;634 | | NA | 5.8483 | 6.3933 | 7.3608 | 6.2757 | 6.2699 | 6.5801 | 6.3320 | 6.4126 | 6.8103 | 6.5681 | 7.3755 | 6.9225 | 5.0082 | 4.9469 | 5.4909 | 5.4048 |
| ATXN7_S302 | ATXN7 | O15265-2 | | 302 | PKPTLPSPGQILN | 6.4634 | 5.7032 | 6.3475 | 5.8904 | 5.9493 | 5.8025 | 5.7704 | 5.7611 | 6.8513 | 6.0021 | 4.8302 | 5.3784 | 7.4677 | 7.4636 | 6.4396 | 7.8794 |
| ATXN7L1_S253 | ATXN7L1 | Q9ULK2 | | 38 | LDRKVPSPEAFLG | 6.4667 | 6.4002 | 5.9057 | 6.1412 | 5.8380 | 5.6830 | 5.9019 | 5.9570 | 5.9990 | 5.9590 | 5.8406 | 5.9335 | 6.6541 | 6.6421 | 7.2466 | 7.4315 |
| ATXN7L1_S38 | ATXN7L1 | Q9ULK2 | | 253 | SKSVPPSPEKILN | 6.0938 | 6.2028 | 6.6235 | 6.5114 | 6.3090 | 6.1884 | 6.2551 | 6.1760 | 6.2012 | 6.3416 | 5.8960 | 5.9263 | 6.4754 | 6.0570 | 6.4137 | 6.3288 |
| ATXN7L2_S575 | ATXN7L2 | Q5T6C5 | | 575 | SRKRKLSPGPTTL | 6.2945 | 6.1931 | 6.1540 | 6.6348 | 6.5349 | 6.3016 | 6.5325 | 6.2450 | 6.3624 | 7.0383 | 6.3397 | 6.6937 | 5.5596 | 6.1469 | 5.4211 | 5.5479 |
| ATXN7L3_S170 | ATXN7L3 | Q14CW9-2 | | 170 | PFQNPNSPRRSKS | 6.3415 | 5.7277 | 6.4469 | 6.0549 | 5.3937 | 6.7613 | 5.5329 | 6.4581 | 6.8666 | 6.5800 | 6.3092 | 6.3438 | 6.1189 | 6.3363 | 6.7192 | 6.0090 |
| AUNIP_S267 | AUNIP | Q9H7T9-2 | | 267 | SVKQSRSPVSVFS | 5.3572 | 5.6263 | 5.6635 | 6.3777 | 4.7947 | 6.0034 | 6.4802 | 6.4364 | 6.8219 | 6.6459 | 6.4262 | 6.2874 | 7.9692 | 6.6487 | 5.7315 | 6.7297 |
| AUP1_S288 | AUP1 | Q9Y679 | | 367 | PSSGPVTPQPTAL | 6.9770 | 7.2930 | 6.7602 | 7.0595 | 7.0223 | 7.2505 | 7.2886 | 6.9484 | 6.9422 | 7.0230 | 6.9288 | 6.8435 | 3.8487 | 4.4707 | 3.3317 | 4.0120 |
| AUP1_S356 | AUP1 | Q9Y679 | | 356 | QSLPTASASKFPS | 6.8418 | 7.3321 | 6.3633 | 7.3548 | 5.5072 | 6.2725 | 6.7141 | 6.7121 | 6.0046 | 5.7052 | 6.6843 | 6.6168 | 5.1769 | 5.7859 | 5.2156 | 5.7129 |
| AUP1_S363 | AUP1 | Q9Y679 | | 363 | ASKFPSSGPVTPQ | 6.6890 | 6.6633 | 6.3666 | 7.1999 | 6.6667 | 7.2263 | 7.3015 | 6.9664 | 6.8757 | 7.3699 | 6.9255 | 6.6291 | 3.8977 | 4.7716 | 3.8341 | 4.6169 |
| AUP1_S363;T367 | AUP1 | Q9Y679 | | 288 | QSSFPPSPGPSPD | 6.1528 | 6.3327 | 6.0338 | 7.0912 | 5.7818 | 6.6713 | 6.9563 | 6.3261 | 5.7322 | 6.5476 | 6.4498 | 6.5132 | 5.4340 | 6.3929 | 6.0860 | 5.4981 |
| AUP1_S367 | AUP1 | Q9Y679 | 363;367 | | NA | 6.4773 | 7.3930 | 7.2072 | 7.6319 | 5.7395 | 6.9544 | 7.6099 | 6.4385 | 6.8767 | 7.0458 | 7.5510 | 7.5333 | 3.8186 | 4.2516 | 3.4771 | 3.9941 |
| AURKA_T288 | AURKA | O14965 | | 288 | PSSRRTTLCGTLD | 7.2545 | 7.4221 | 5.8588 | 8.3363 | 5.3963 | 5.3242 | 6.5079 | 5.3221 | 6.4082 | 5.7866 | 7.0233 | 6.3360 | 5.9988 | 6.3700 | 5.1715 | 5.4833 |
| AURKB_T233 | AURKB | Q96GD4-5 | | 233 | PSLRRKTMCGTLD | 8.2562 | 7.1357 | 7.3839 | 6.9611 | 7.4928 | 6.3391 | 7.0484 | 5.8310 | 6.5358 | 5.9424 | 6.9656 | 6.3808 | 4.4090 | 4.2003 | 4.2906 | 4.8273 |
| AXIN1_S217 | AXIN1 | O15169 | | 77 | EPEGSASPTPPYL | 6.1952 | 6.4488 | 6.8512 | 6.2840 | 5.9777 | 6.3349 | 6.1148 | 6.6694 | 7.7253 | 6.7981 | 6.5808 | 6.7032 | 4.9657 | 6.0866 | 4.9018 | 5.3625 |
| AXIN1_S77 | AXIN1 | O15169 | | 217 | TRTGSESPKVCSD | 6.6469 | 5.7966 | 5.5399 | 6.3596 | 6.3435 | 6.8708 | 6.7252 | 6.0723 | 7.0992 | 6.4641 | 6.4779 | 7.3476 | 5.7789 | 5.5810 | 4.9836 | 5.9128 |
| AXL_S612 | AXL | P30530 | | 702 | KIYNGDYYRQGRI | 7.7625 | 8.2123 | 8.4221 | 8.9758 | 7.5158 | 6.6406 | 6.3232 | 7.1098 | 7.2606 | 6.5432 | 6.7712 | 7.1768 | 2.6325 | 2.9788 | 2.6224 | 3.0523 |
| AXL_S875 | AXL | P30530 | | 612 | QGSERESFPAPVV | 6.0598 | 7.1190 | 6.9525 | 7.5898 | 6.8352 | 6.8100 | 6.6983 | 6.3862 | 5.9040 | 6.5180 | 7.0166 | 6.4642 | 4.9172 | 5.2426 | 4.6018 | 4.8847 |
| AXL_S884 | AXL | P30530 | | 866 | VHPAGRYVLCPST | 6.9894 | 6.9848 | 7.9000 | 8.3222 | 8.0632 | 6.9088 | 6.1122 | 6.4101 | 6.8677 | 6.5789 | 6.4464 | 6.2113 | 4.2474 | 4.3795 | 3.6433 | 3.9346 |
| AXL_T509 | AXL | P30530 | | 509 | SYSRRTTEATLNS | 6.2834 | 6.1375 | 6.3985 | 6.5579 | 6.7707 | 5.7088 | 5.9805 | 5.3667 | 6.7971 | 6.3112 | 6.6765 | 6.3200 | 6.6009 | 6.3750 | 5.8749 | 5.8404 |
| AXL_T872 | AXL | P30530 | | 703 | IYNGDYYRQGRIA | 7.2058 | 7.1355 | 7.8274 | 8.3115 | 7.0142 | 6.3787 | 5.9078 | 6.4441 | 7.5580 | 6.9511 | 7.1433 | 7.1853 | 3.7532 | 3.8697 | 3.4645 | 3.8498 |

| | | | | | | | | | | | | | | | | | | | | | |
|---|---|---|---|---|---|---|---|---|---|---|---|---|---|---|---|---|---|---|---|---|---|
| AXL_Y702 | AXL | P30530 | | 884 | QPADRGSPAAPGQ | 6.3536 | 6.6092 | 6.6032 | 7.2078 | 6.5221 | 6.8639 | 6.5839 | 5.8751 | 6.3823 | 6.5264 | 6.2340 | 7.0251 | 4.9030 | 5.0968 | 5.4909 | 5.7228 |
| AXL_Y702;Y703 | AXL | P30530 | | 875 | CPSTTPSPAQPAD | 6.6051 | 6.9138 | 6.0320 | 7.4143 | 6.9224 | 6.4371 | 6.3553 | 7.0521 | 6.8315 | 6.7742 | 6.8860 | 6.6178 | 4.3360 | 5.2242 | 4.6684 | 4.9298 |
| AXL_Y703 | AXL | P30530 | | 872 | YVLCPSTTPSPAQ | 5.8780 | 5.6204 | 6.2067 | 8.7178 | 6.8670 | 7.2579 | 7.3023 | 6.5087 | 6.9440 | 6.7012 | 6.0497 | 7.5955 | 4.4004 | 4.7705 | 3.6216 | 5.5582 |
| AXL_Y866 | AXL | P30530 | 702;703 | | NA | 8.8765 | 7.8822 | 7.9986 | 10.3160 | 7.8176 | 5.5552 | 6.0476 | 5.8491 | 8.6490 | 6.8316 | 6.8716 | 7.4536 | 2.4082 | 2.4769 | 2.1763 | 2.7899 |
| AZI2_S353 | AZI2 | Q9H6S1 | | 353 | NSWVFPSPPKSSE | 5.8483 | 5.4511 | 7.7226 | 5.6497 | 5.7058 | 6.6603 | 6.9168 | 6.1670 | 5.7850 | 6.5383 | 7.3005 | 6.7876 | 5.7138 | 6.2498 | 4.7940 | 6.7093 |
| B2RBV5_S18 | B2RBV5 | B2RBV5 | | 18 | PREEPGSPLSPAP | 6.5399 | 6.3008 | 7.0811 | 5.3918 | 6.6760 | 6.6242 | 5.8640 | 7.2518 | 6.4561 | 7.5538 | 7.6911 | 7.5129 | 4.7158 | 4.8619 | 4.3577 | 5.1212 |
| BABAM1_S49 | BABAM1 | Q9NWV8 | | 49 | ASVGSRSEGEGEA | 6.1187 | 6.0118 | 7.1959 | 6.4591 | 7.6391 | 6.6360 | 6.3718 | 6.7975 | 5.8685 | 5.9146 | 5.8992 | 6.1655 | 6.3102 | 6.1201 | 5.3211 | 5.1708 |
| BACH1_S196 | BACH1 | O14867 | | 196 | QGNAKASPPLQDS | 6.5980 | 7.1246 | 6.7150 | 7.2824 | 6.2227 | 6.9159 | 6.6619 | 7.2564 | 6.8714 | 6.9821 | 7.0625 | 7.1438 | 3.9299 | 5.0968 | 3.8385 | 4.2982 |
| BAD_S118 | BAD | Q92934 | | 118 | RELRRMSDEFVDS | 2.4990 | 2.2308 | 2.5328 | 2.1091 | 4.8598 | 4.3687 | 4.1459 | 5.1908 | 3.2082 | 3.3246 | 3.3927 | 3.5245 | 16.3136 | 14.5563 | 14.4585 | 13.2848 |
| BAD_S134 | BAD | Q92934 | | 99 | FRGRSRSAPPNLW | 5.5050 | 5.4141 | 5.3664 | 5.4756 | 6.5357 | 5.7694 | 5.8766 | 6.3498 | 5.7740 | 5.4648 | 5.5254 | 5.5040 | 8.5798 | 8.0620 | 7.4326 | 7.3649 |
| BAD_S146 | BAD | Q92934 | | 134 | GLPRPKSAGTATQ | 2.2700 | 2.1122 | 1.9512 | 1.8279 | 5.7693 | 5.3537 | 4.9086 | 5.9087 | 2.5899 | 2.4424 | 2.4062 | 2.5831 | 17.3131 | 14.7710 | 14.6687 | 13.1239 |
| BAD_S99 | BAD | Q92934 | | 146 | QMRQSSSWTRVFQ | 2.8883 | 2.7853 | 3.1165 | 2.8636 | 5.5813 | 4.7332 | 4.8077 | 5.5834 | 3.3013 | 3.2293 | 2.9655 | 3.4346 | 14.4292 | 13.6476 | 13.5295 | 13.1036 |
| BAG2_S73 | BAG2 | O95816 | | 73 | QDMRQISDGEREE | 6.1326 | 6.3449 | 6.0127 | 6.2847 | 6.1256 | 5.9793 | 6.0940 | 6.6711 | 6.6047 | 6.2614 | 5.7673 | 6.3401 | 6.5818 | 6.3140 | 6.0127 | 6.4731 |
| BAG3_S173 | BAG3 | O95817 | | 377 | VPCPPPSPGPSAV | 6.0405 | 5.5504 | 6.5441 | 5.6394 | 6.8977 | 6.3138 | 6.0032 | 6.7927 | 6.3730 | 6.0578 | 6.4395 | 6.5938 | 6.7133 | 6.3683 | 5.7990 | 5.8737 |
| BAG3_S195 | BAG3 | O95817 | | 406 | TAPAEATPPKPGE | 6.4855 | 6.1555 | 7.4381 | 6.3609 | 6.6268 | 6.6922 | 6.3169 | 6.8714 | 6.5052 | 7.3107 | 7.1293 | 7.0368 | 4.8165 | 5.2942 | 4.3300 | 4.6302 |
| BAG3_S275 | BAG3 | O95817 | | 285 | SPARSSTPLHSPS | 6.1175 | 6.6397 | 7.2061 | 5.7492 | 6.5110 | 6.6341 | 6.1453 | 7.2877 | 6.9007 | 6.7521 | 6.9942 | 7.1477 | 5.0358 | 5.3743 | 4.8055 | 4.6990 |
| BAG3_S289 | BAG3 | O95817 | | 289 | SSTPLHSPSPIRV | 6.3997 | 5.9341 | 6.3971 | 6.2866 | 7.2759 | 6.6151 | 5.6443 | 7.1019 | 5.9199 | 5.5937 | 6.0555 | 5.8327 | 6.2977 | 6.9763 | 6.3235 | 5.3460 |
| BAG3_S377 | BAG3 | O95817 | | 275 | SVQGASSREGSPA | 6.7767 | 5.9978 | 6.9424 | 5.1769 | 6.6221 | 6.4989 | 5.6979 | 7.3876 | 6.7452 | 6.9445 | 7.2626 | 6.9412 | 5.6987 | 5.4214 | 4.7876 | 5.0986 |
| BAG3_S377;S386 | BAG3 | O95817 | | 173 | GPERSQSPAASDC | 5.5545 | 5.8962 | 7.4958 | 6.0521 | 6.3745 | 6.5511 | 5.9451 | 6.1260 | 6.9176 | 7.0088 | 7.4809 | 6.2714 | 5.5584 | 6.1389 | 5.3190 | 5.3098 |
| BAG3_S386 | BAG3 | O95817 | | 386 | PSAVPSSPKSVAT | 6.3844 | 5.9145 | 7.2449 | 6.6911 | 6.0262 | 6.2892 | 5.8692 | 6.1036 | 6.5401 | 6.5668 | 6.2539 | 6.7089 | 6.0699 | 5.9297 | 5.3910 | 6.0165 |
| BAG3_T285 | BAG3 | O95817 | | 195 | PSSGRSSLGSHQL | 7.2764 | 6.9721 | 6.5544 | 7.1279 | 6.8902 | 7.3856 | 6.4995 | 7.5407 | 6.6061 | 7.2802 | 7.7260 | 7.1709 | 3.6254 | 4.5371 | 3.2996 | 3.5078 |
| BAG3_T285;S289 | BAG3 | O95817 | 377;386 | | NA | 6.0706 | 5.7926 | 6.7160 | 5.8857 | 6.4465 | 6.3232 | 6.4522 | 6.7509 | 6.4084 | 6.5319 | 6.7483 | 6.6383 | 6.1280 | 5.9815 | 5.5331 | 5.5930 |
| BAG3_T285;S291 | BAG3 | O95817 | 285;289 | | NA | 6.5813 | 5.7895 | 6.9052 | 5.1784 | 7.1313 | 7.2483 | 6.5757 | 7.9433 | 6.4817 | 6.4432 | 6.7322 | 6.7642 | 5.2680 | 5.3469 | 4.7900 | 4.8207 |
| BAG3_T406 | BAG3 | O95817 | 285;291 | | NA | 6.3585 | 6.4845 | 8.0588 | 6.4000 | 6.4280 | 7.0461 | 6.4173 | 7.0287 | 8.0425 | 6.7144 | 8.0740 | 7.2753 | 3.5527 | 5.2423 | 3.0607 | 3.8162 |
| BAG4_S179 | BAG4 | O95429 | | 289 | SPQSPPSPPVQQP | 5.4708 | 6.6049 | 5.6794 | 6.1088 | 7.0583 | 5.8816 | 4.9463 | 5.5767 | 5.8368 | 6.2094 | 6.1733 | 7.3446 | 6.9764 | 7.3359 | 6.3853 | 6.4115 |
| BAG4_S289 | BAG4 | O95429 | | 179 | YRSSGNSPTPVSR | 6.3776 | 5.5735 | 6.2142 | 5.2665 | 6.9206 | 5.5692 | 5.3492 | 6.8134 | 5.7370 | 7.2529 | 6.7468 | 7.1616 | 6.2904 | 6.6500 | 5.9663 | 6.1107 |
| BAG6_S1003 | BAG6 | P46379-3 | | 994 | QGAERASPEPQRE | 6.0266 | 6.0446 | 6.4469 | 5.8770 | 6.4453 | 6.1175 | 6.2342 | 6.4172 | 6.3517 | 6.2203 | 6.5247 | 6.7512 | 6.1519 | 6.3837 | 5.9828 | 6.0245 |
| BAG6_S1111 | BAG6 | P46379-3 | | 1147 | QEDPNYSPQRFPN | 6.9820 | 6.8252 | 6.9576 | 6.4646 | 6.2003 | 6.2112 | 6.4579 | 7.0728 | 7.1977 | 7.0084 | 7.3386 | 7.3190 | 4.1380 | 4.8026 | 4.3499 | 4.6741 |
| BAG6_S1114 | BAG6 | P46379-3 | | 1111 | GARPLTSPESLSR | 6.6950 | 6.1850 | 7.1396 | 6.0962 | 6.8079 | 6.8246 | 6.3448 | 5.8495 | 6.3923 | 6.7493 | 7.1824 | 7.3683 | 5.3239 | 5.4174 | 4.6736 | 4.9503 |
| BAG6_S113 | BAG6 | P46379-3 | | 1003 | PQRENASPAPGTT | 5.9200 | 6.1833 | 6.8098 | 6.1096 | 6.3173 | 6.1450 | 5.8400 | 6.9422 | 6.3216 | 6.2370 | 6.5546 | 6.0385 | 6.4784 | 6.1976 | 5.7208 | 6.1844 |
| BAG6_S1147 | BAG6 | P46379-3 | | 344 | RCNLACTPPRHLH | 6.0666 | 5.8301 | 7.1475 | 6.3604 | 6.3463 | 6.5427 | 6.4524 | 6.2759 | 6.3439 | 6.7711 | 7.1323 | 6.9247 | 5.5334 | 5.5733 | 5.2296 | 5.4699 |
| BAG6_S832 | BAG6 | P46379-3 | | 538 | VVIARPTPPQARP | 6.3672 | 7.4842 | 7.0619 | 7.0043 | 6.4444 | 6.6871 | 6.6406 | 6.5876 | 6.4769 | 7.6200 | 6.3449 | 7.5251 | 4.1732 | 4.8251 | 4.0086 | 4.7489 |
| BAG6_S994 | BAG6 | P46379-3 | | 113 | ATHGGGSPPGTRG | 6.1820 | 8.1299 | 7.7429 | 7.8833 | 6.6555 | 6.8445 | 6.0254 | 7.3162 | 6.2072 | 5.0911 | 6.3783 | 6.0776 | 3.9857 | 5.1811 | 5.8310 | 4.4682 |
| BAG6_S994;S1003 | BAG6 | P46379-3 | | 1114 | PLTSPESLSRDLE | 6.0946 | 5.4480 | 6.8596 | 6.3614 | 6.7290 | 6.4889 | 5.9002 | 6.0869 | 6.8490 | 6.6321 | 7.5497 | 6.9472 | 5.3891 | 5.8439 | 5.2169 | 5.6038 |
| BAG6_T344 | BAG6 | P46379-4 | | 832 | QGAERASPEPQEW | 5.7944 | 6.1181 | 6.1204 | 6.0314 | 6.4937 | 6.9537 | 7.2534 | 6.8360 | 6.1816 | 6.6416 | 6.4201 | 6.4790 | 5.5938 | 5.7533 | 5.5605 | 5.7688 |
| BAG6_T538 | BAG6 | P46379-3 | 994;1003 | | NA | 6.1410 | 6.0758 | 7.2293 | 5.9038 | 5.8334 | 6.3614 | 5.9027 | 6.0234 | 6.1516 | 6.4379 | 7.0929 | 6.3633 | 6.2693 | 6.7851 | 5.4433 | 5.9858 |
| BAHCC1_S1141 | BAHCC1 | Q9P281 | | 1141 | GEPPPCSPRSLEE | 6.1462 | 6.5047 | 5.8650 | 7.0489 | 6.5679 | 6.3080 | 5.9713 | 6.1438 | 6.5319 | 6.2165 | 6.1818 | 5.9925 | 6.1149 | 6.2666 | 6.4754 | 5.6646 |
| BAHCC1_S2274 | BAHCC1 | Q9P281 | | 2274 | LGRPLPSPSYVHP | 5.6213 | 5.9692 | 5.5619 | 5.9947 | 7.8529 | 6.2450 | 6.7498 | 6.9747 | 7.6789 | 4.9934 | 7.2335 | 5.0316 | 5.2819 | 6.9535 | 6.3425 | 5.5153 |
| BAHD1_S101 | BAHD1 | Q8TBE0 | | 184 | GGSRDLSPEPAPD | 7.2513 | 7.7689 | 6.6972 | 7.8921 | 6.5244 | 5.9339 | 6.7567 | 6.4116 | 7.9068 | 7.4401 | 7.1475 | 6.8247 | 3.7266 | 3.9530 | 3.8247 | 3.9406 |
| BAHD1_S121 | BAHD1 | Q8TBE0 | | 8 | THTRRKSLPMLSS | 5.6700 | 6.0568 | 6.3782 | 6.9384 | 6.0463 | 6.2851 | 6.1139 | 6.0061 | 8.0948 | 7.6952 | 7.1181 | 8.2618 | 4.6848 | 4.9934 | 4.0696 | 5.5875 |
| BAHD1_S166 | BAHD1 | Q8TBE0 | | 101 | DLPKPPSPAPSSE | 6.3904 | 6.1341 | 6.4411 | 6.7990 | 6.1064 | 6.3137 | 6.3637 | 6.3180 | 6.7241 | 6.7855 | 6.2956 | 6.6328 | 5.2761 | 6.0531 | 5.5146 | 5.8519 |
| BAHD1_S184 | BAHD1 | Q8TBE0 | | 206 | APKRLASLNAAAF | 6.4303 | 5.9299 | 7.7455 | 6.4703 | 6.4303 | 7.1737 | 6.7477 | 6.6162 | 8.0315 | 7.0765 | 7.4825 | 8.3374 | 2.7134 | 4.1287 | 4.7834 | 3.9028 |
| BAHD1_S206 | BAHD1 | Q8TBE0 | | 121 | RKRRLASLNAEAL | 5.7286 | 5.1287 | 5.5776 | 6.3899 | 6.2741 | 6.6479 | 5.0696 | 6.5808 | 7.6312 | 6.9954 | 7.1128 | 6.5377 | 6.3524 | 5.8604 | 5.5840 | 6.5289 |
| BAHD1_S8 | BAHD1 | Q8TBE0 | | 166 | ATGGWSSSKKRPR | 8.9916 | 7.2595 | 6.6396 | 8.3576 | 6.0519 | 8.1599 | 7.0295 | 7.2370 | 8.3043 | 8.0631 | 8.2973 | 7.7223 | 1.0926 | 2.8247 | 1.6999 | 2.2693 |
| BAIAP2_S261 | BAIAP2 | Q9UQB8-6 | | 367 | TLPRSSSMAAGLE | 7.4870 | 7.0003 | 6.8682 | 6.1542 | 5.5789 | 6.2014 | 6.2187 | 5.7185 | 6.2948 | 6.5678 | 6.3928 | 6.3853 | 6.1149 | 6.4293 | 5.4534 | 6.0446 |
| BAIAP2_S325 | BAIAP2 | Q9UQB8-6 | | 261 | PSALSASKSNLVI | 5.8907 | 6.0491 | 6.9081 | 6.2565 | 5.9248 | 6.2539 | 5.8912 | 6.4157 | 6.2098 | 5.9417 | 6.4103 | 5.8784 | 6.5415 | 6.6043 | 6.4897 | 6.3342 |
| BAIAP2_S367 | BAIAP2 | Q9UQB8-6 | | 454 | LQQGKSSSTGNLL | 5.5457 | 7.3118 | 7.9886 | 6.3269 | 4.4216 | 6.6771 | 6.9779 | 6.7564 | 5.1211 | 7.0126 | 9.2150 | 9.2261 | 3.5316 | 4.2546 | 4.2283 | 5.4047 |
| BAIAP2_S454 | BAIAP2 | Q9UQB8-6 | | 340 | SDSYSNTLPVRKS | 10.1408 | 7.3323 | 7.5653 | 8.2133 | 6.8917 | 3.7921 | 5.4797 | 3.9942 | 6.3172 | 4.5122 | 4.5215 | 4.2899 | 7.7290 | 6.7365 | 6.7128 | 5.7714 |
| BAIAP2_T340 | BAIAP2 | Q9UQB8-6 | | 325 | AQPKSLSPPQSQS | 6.1317 | 6.7451 | 6.8950 | 6.3649 | 6.3192 | 6.7033 | 6.5821 | 6.9837 | 6.7700 | 6.4427 | 6.6513 | 6.7674 | 5.0002 | 5.9275 | 5.0886 | 4.6273 |
| BAIAP2_T521 | BAIAP2 | Q9UQB8-6 | | 521 | SGTLVSTVxxxxx | 6.2727 | 5.8621 | 6.0367 | 6.4754 | 6.2684 | 5.8173 | 6.8595 | 5.3035 | 6.5642 | 6.5275 | 7.2005 | 6.7404 | 6.1784 | 6.1630 | 5.6184 | 6.1122 |
| BAIAP2L1_S252;S261 | BAIAP2L1 | Q9UHR4 | | 414 | VTVPTPSPTPVRS | 5.9334 | 6.2697 | 6.6642 | 6.3688 | 6.2174 | 6.3000 | 6.1264 | 6.5325 | 6.5730 | 6.4641 | 6.5039 | 6.5026 | 5.8338 | 6.0535 | 5.7565 | 5.9004 |
| BAIAP2L1_S278 | BAIAP2L1 | Q9UHR4 | | 422 | TPVRSISTVNLSE | 5.7132 | 5.6805 | 5.9363 | 5.7731 | 5.7876 | 6.1144 | 6.1754 | 5.9235 | 5.5619 | 5.8620 | 5.8899 | 6.1714 | 7.7897 | 7.1542 | 7.1548 | 7.3121 |
| BAIAP2L1_S295 | BAIAP2L1 | Q9UHR4 | | 261 | SGTPQASPMIERS | 5.7529 | 6.3037 | 5.9525 | 6.5362 | 6.1631 | 5.7565 | 5.8647 | 6.0954 | 5.7897 | 5.8026 | 5.6738 | 5.8601 | 7.0133 | 7.3093 | 7.2868 | 6.8394 |
| BAIAP2L1_S331 | BAIAP2L1 | Q9UHR4 | | 331 | SLQRSVSVATGLN | 7.0919 | 5.9907 | 5.9807 | 5.9611 | 6.5841 | 5.4564 | 5.9396 | 5.2122 | 5.6008 | 4.6455 | 5.4459 | 5.4027 | 7.8976 | 8.2306 | 7.2220 | 7.3381 |
| BAIAP2L1_S354 | BAIAP2L1 | Q9UHR4 | | 295 | SGRAYTSPLIDMF | 6.5116 | 6.7252 | 6.2886 | 7.0366 | 6.1649 | 6.5341 | 6.5286 | 6.7193 | 6.6791 | 7.1527 | 6.9769 | 6.3483 | 4.8763 | 5.3682 | 4.7925 | 5.2970 |
| BAIAP2L1_S395 | BAIAP2L1 | Q9UHR4 | | 278 | KDYDTLSKCSPKM | 8.2882 | 7.9197 | 7.2728 | 7.2715 | 5.2585 | 6.2002 | 6.6946 | 5.3269 | 6.8264 | 6.4459 | 7.8032 | 7.6725 | 4.1608 | 4.9084 | 3.6572 | 4.2932 |
| BAIAP2L1_S414 | BAIAP2L1 | Q9UHR4 | | 354 | FPHTAGSNKTLLS | 4.3323 | 4.5431 | 5.2828 | 4.6790 | 5.6684 | 5.8190 | 5.5247 | 5.3874 | 5.0276 | 5.0391 | 5.1904 | 4.5653 | 9.9178 | 9.6066 | 9.6401 | 9.7764 |
| BAIAP2L1_S422 | BAIAP2L1 | Q9UHR4 | | 257 | STPVSGTPQASPM | 6.2713 | 6.0899 | 6.3956 | 5.9896 | 5.6007 | 5.8603 | 5.7385 | 6.1211 | 6.0590 | 6.4517 | 5.5655 | 6.8237 | 6.6816 | 6.5875 | 6.9087 | 6.8554 |
| BAIAP2L1_T118 | BAIAP2L1 | Q9UHR4 | | 395 | RGWFPSSYTKLLE | 5.3855 | 5.4079 | 6.0484 | 5.1443 | 6.8017 | 6.3756 | 6.0836 | 6.7671 | 5.4832 | 5.7206 | 5.1448 | 5.3799 | 8.2955 | 8.0101 | 7.8136 | 6.1383 |
| BAIAP2L1_T248;S252;S2 | BAIAP2L1 | Q9UHR4 | | 118 | VKYMNATLKRYQT | 9.9739 | 7.2863 | 9.7808 | 8.0242 | 5.8979 | 3.7656 | 5.6306 | 4.0745 | 5.6663 | 6.3018 | 6.1548 | 4.3210 | 6.7235 | 6.2706 | 5.1406 | 4.9877 |
| BAIAP2L1_T248;S252;T2 | BAIAP2L1 | Q9UHR4 | 257;261 | | NA | 5.8641 | 6.4433 | 6.5866 | 6.1222 | 6.1913 | 6.5859 | 6.3095 | 6.3056 | 6.4407 | 6.4396 | 6.4404 | 6.6156 | 5.7623 | 6.1449 | 5.6554 | 6.0927 |
| BAIAP2L1_T248;T257;S2 | BAIAP2L1 | Q9UHR4 | 252;261 | | NA | 5.8583 | 5.9742 | 6.2947 | 5.6471 | 5.8832 | 6.8007 | 6.3213 | 6.0640 | 6.4923 | 6.8383 | 6.4871 | 6.3725 | 6.2443 | 6.3295 | 6.0994 | 6.2930 |
| BAIAP2L1_T257 | BAIAP2L1 | Q9UHR4 | 248;252;257;2 | | NA | 6.3731 | 6.8253 | 7.3596 | 5.9644 | 6.2239 | 6.3446 | 6.2046 | 6.7008 | 7.7449 | 6.7934 | 7.0476 | 7.4860 | 4.3302 | 5.4641 | 4.1722 | 4.9651 |
| BAIAP2L1_T257;S261 | BAIAP2L1 | Q9UHR4 | 248;257;261 | | NA | 6.0689 | 6.9334 | 6.5324 | 6.3388 | 5.8145 | 6.0229 | 6.0876 | 6.6698 | 6.5902 | 6.4228 | 7.3932 | 6.5872 | 4.8608 | 5.9546 | 5.9360 | 5.7869 |
| BAIAP2L1_T261 | BAIAP2L1 | Q9UHR4 | 248;252;261 | | NA | 6.2021 | 5.4708 | 7.2830 | 6.7654 | 5.8679 | 6.8149 | 6.4042 | 6.1889 | 6.6926 | 7.3163 | 6.8644 | 7.0464 | 5.4336 | 5.3250 | 5.1217 | 5.2030 |
| BAP1_S292 | BAP1 | Q92560 | | 292 | KSASNKSPLVLEA | 5.9781 | 6.2947 | 6.7063 | 6.8575 | 6.7104 | 6.2319 | 6.4274 | 6.7199 | 6.3837 | 6.6477 | 6.2257 | 5.9685 | 5.5483 | 6.1673 | 5.6850 | 5.4475 |
| BAP1_S327 | BAP1 | Q92560 | | 597 | GSQGSSSPVEKEV | 5.8924 | 6.6106 | 5.7253 | 6.1920 | 6.3113 | 6.0064 | 5.8097 | 5.9558 | 5.9284 | 5.9962 | 5.8344 | 6.1859 | 6.7896 | 6.9538 | 6.9195 | 6.8888 |
| BAP1_S521 | BAP1 | Q92560 | | 521 | NPTRPSSPVTSHI | 6.5797 | 8.1831 | 5.8763 | 7.8820 | 7.4736 | 7.2895 | 6.0973 | 8.1155 | 5.5987 | 5.5540 | 6.4625 | 5.5835 | 4.2306 | 5.4479 | 5.3186 | 4.3072 |
| BAP1_S597 | BAP1 | Q92560 | | 327 | AQAPSHSPPNKPK | 6.3155 | 6.6651 | 6.4372 | 6.6848 | 6.3084 | 6.9711 | 6.4787 | 6.8107 | 6.8993 | 6.1706 | 7.2848 | 5.6500 | 5.1653 | 5.2343 | 5.6627 | 5.2617 |
| BAP18_S110 | BAP18 | Q8IXM2-3 | | 96 | IPLPAESPKKGPK | 6.3333 | 5.1338 | 5.8016 | 5.2213 | 5.5186 | 4.6853 | 5.2933 | 6.1004 | 4.9231 | 6.2971 | 5.6348 | 6.5093 | 6.2350 | 9.0120 | 8.1591 | 9.1421 |
| BAP18_S36 | BAP18 | Q8IXM2-2 | | 36 | HPVADSSPAGAQI | 6.1949 | 5.3748 | 5.6140 | 5.7172 | 5.6635 | 4.0331 | 6.3338 | 7.5734 | 4.6100 | 6.7223 | 5.1403 | 6.1354 | 4.7232 | 9.1787 | 7.3383 | 9.6471 |
| BAP18_S36 | BAP18 | Q8IXM2-3 | | 161 | IEGLGETPPAKKL | 6.6039 | 4.7869 | 6.6537 | 5.3836 | 5.7917 | 5.8631 | 6.0569 | 6.5210 | 5.3313 | 6.8220 | 7.1535 | 7.3053 | 4.6453 | 7.4621 | 6.6945 | 6.9253 |
| BAP18_S96 | BAP18 | Q8IXM2-3 | | 110 | VASGVLSPPPAAP | 7.9080 | 6.5139 | 6.6073 | 6.0998 | 6.8908 | 5.0308 | 6.4441 | 8.2343 | 5.4787 | 7.7190 | 6.6883 | 7.9575 | 3.4555 | 5.1636 | 4.1138 | 5.6947 |
| BAP18_T161 | BAP18 | Q8IXM2-3 | | 36 | HPVADSSPAGAKW | 7.0828 | 5.8849 | 6.9154 | 5.5739 | 5.8110 | 5.4080 | 6.9085 | 6.9336 | 5.3257 | 6.5041 | 5.9992 | 7.4970 | 4.9492 | 6.8499 | 6.1150 | 6.2419 |
| BARD1_S186 | BARD1 | Q99728 | | 410 | YRRVMSSPSAMKL | 7.2278 | 7.3177 | 6.4463 | 7.1124 | 6.4497 | 6.7805 | 6.5254 | 6.6484 | 6.9360 | 7.0191 | 6.6846 | 6.6195 | 4.4544 | 4.8119 | 4.4107 | 4.5555 |
| BARD1_S330 | BARD1 | Q99728 | | 364 | PLPECSSPPSCKR | 6.5691 | 6.6220 | 6.6298 | 7.2543 | 6.5970 | 6.0800 | 6.9992 | 6.4515 | 7.2114 | 6.4350 | 6.6253 | 6.6697 | 4.8974 | 4.8504 | 4.8421 | 5.2660 |
| BARD1_S364 | BARD1 | Q99728 | | 330 | HHNRLSSPISKRC | 7.4764 | 7.6745 | 6.7242 | 7.2478 | 6.5695 | 6.5405 | 6.9541 | 6.5428 | 7.0406 | 6.5502 | 6.4402 | 6.4674 | 4.1535 | 4.6570 | 4.2625 | 4.6987 |
| BARD1_S389;T394 | BARD1 | Q99728 | | 299 | SRNEVVTPEKVCK | 6.3536 | 6.5095 | 6.2796 | 6.8456 | 5.7040 | 5.5172 | 6.3281 | 5.8803 | 6.8479 | 6.7951 | 6.5095 | 7.1888 | 5.6898 | 5.6367 | 5.4905 | 6.4238 |
| BARD1_S391;T394 | BARD1 | Q99728 | | 186 | YEFVSPSPPADVS | 5.5958 | 6.1911 | 6.2712 | 6.9556 | 6.1799 | 5.9141 | 6.2584 | 5.6972 | 6.1025 | 6.2821 | 6.2537 | 6.7850 | 6.7523 | 6.0229 | 6.2424 | 6.4958 |
| BARD1_S394 | BARD1 | Q99728 | | 394 | ISLSPGTPPSTLS | 6.7857 | 6.3892 | 7.0194 | 6.7039 | 6.5225 | 5.8526 | 6.0534 | 5.9876 | 6.7925 | 6.4322 | 7.0068 | 6.7781 | 4.7187 | 6.4263 | 4.6605 | 5.8703 |
| BARD1_S410 | BARD1 | Q99728 | 391;394 | | NA | 6.7019 | 6.7765 | 6.1574 | 6.3277 | 6.6483 | 6.2083 | 6.6866 | 6.3386 | 7.2721 | 6.1279 | 6.2709 | 6.2396 | 5.6061 | 5.8346 | 5.5249 | 5.2785 |
| BARD1_T299 | BARD1 | Q99728 | 389;394 | | NA | 6.6054 | 5.2678 | 6.5481 | 6.7935 | 7.0894 | 6.3881 | 5.6307 | 6.9259 | 6.3737 | 6.4547 | 7.2217 | 6.8951 | 5.6564 | 5.9597 | 5.2254 | 4.9645 |
| BASP1_T196 | BASP1 | P80723 | | 196 | TEAPSSTPKAQGP | 6.5968 | 5.8811 | 6.7774 | 5.9974 | 6.6734 | 5.4807 | 5.9627 | 7.2325 | 6.0186 | 6.9748 | 6.4676 | 7.2515 | 5.8296 | 5.3516 | 5.4690 | 6.0354 |
| BAZ1A_S1279 | BAZ1A | Q9NRL2 | | 731 | LDQDMVTEDEDDP | 6.5956 | 6.4044 | 6.5991 | 6.6174 | 6.4500 | 5.9583 | 6.4773 | 6.3972 | 6.1405 | 5.8883 | 6.1823 | 5.9427 | 5.9862 | 6.0806 | 6.1899 | 6.0902 |
| BAZ1A_S1339;S1353 | BAZ1A | Q9NRL2 | | 1413 | RCRKRQSPEPSPV | 6.4945 | 7.1240 | 6.3005 | 6.5742 | 6.0274 | 6.1042 | 6.6922 | 6.5399 | 6.7326 | 5.9639 | 5.7998 | 6.4112 | 5.9988 | 5.9894 | 5.6897 | 5.5576 |
| BAZ1A_S1353 | BAZ1A | Q9NRL2 | | 1279 | KTRGKLSSSFSSR | 6.1167 | 6.4115 | 6.2225 | 6.5834 | 5.7520 | 6.4584 | 6.5512 | 6.4779 | 6.5274 | 7.5951 | 7.1232 | 7.3257 | 5.0718 | 5.7086 | 4.5802 | 5.4945 |
| BAZ1A_S1413 | BAZ1A | Q9NRL2 | | 1353 | VFVELLSPRRKRR | 7.2238 | 7.6885 | 6.6110 | 7.0382 | 6.8940 | 6.5942 | 6.2246 | 7.0834 | 7.3158 | 7.0006 | 6.9446 | 6.9623 | 3.7338 | 4.8434 | 3.7384 | 4.1034 |
| BAZ1A_S270 | BAZ1A | Q9NRL2 | | 270 | PPTFIFSPANRRR | 6.7719 | 6.4333 | 7.7333 | 6.7527 | 7.0466 | 7.5768 | 5.8679 | 5.5868 | 5.9382 | 7.6695 | 6.5195 | 5.6898 | 5.6187 | 5.4239 | 4.5838 | 3.8874 |
| BAZ1A_T1547 | BAZ1A | Q9NRL2 | | 1547 | NVDQVSTPPAAKK | 7.0377 | 6.6713 | 6.0085 | 7.4334 | 6.8196 | 6.7858 | 6.5299 | 6.8003 | 6.5624 | 6.6012 | 6.6869 | 6.7842 | 4.8007 | 4.9384 | 4.7228 | 4.8168 |
| BAZ1A_T731 | BAZ1A | Q9NRL2 | 1339;1353 | | NA | 7.1323 | 5.4671 | 7.9939 | 6.6991 | 5.5066 | 6.4705 | 7.7537 | 7.3438 | 5.0827 | 8.0930 | 7.1401 | 7.5474 | 4.7653 | 4.9397 | 2.3327 | 5.7322 |
| BAZ1B_S1342 | BAZ1B | Q9UIG0 | | 361 | NSKNSKSPEEHLE | 6.2427 | 6.6567 | 6.1755 | 7.1252 | 6.1418 | 5.7804 | 6.2512 | 6.4244 | 6.1132 | 6.5972 | 6.1747 | 6.4474 | 5.8520 | 6.0823 | 5.8881 | 6.0472 |
| BAZ1B_S1468 | BAZ1B | Q9UIG0 | | 349 | KKSLSGSPLKVKN | 7.3350 | 6.9011 | 6.0729 | 6.8040 | 6.9891 | 5.9001 | 6.7227 | 6.4305 | 6.9241 | 6.3578 | 5.8998 | 6.1079 | 5.3512 | 5.5693 | 5.3988 | 5.2359 |
| BAZ1B_S158 | BAZ1B | Q9UIG0 | | 833 | FEPQVDTEAEDMI | 9.5391 | 8.7722 | 7.5386 | 9.4816 | 7.8321 | 6.7189 | 6.9895 | 5.4396 | 5.3400 | 4.2194 | 4.4820 | 4.4947 | 4.3264 | 4.7456 | 5.2532 | 4.8271 |
| BAZ1B_S158;S161 | BAZ1B | Q9UIG0 | | 1468 | AEDEGDSEPEAVG | 6.2086 | 6.6366 | 6.4719 | 7.2210 | 6.5200 | 6.0538 | 6.5925 | 6.3562 | 6.4568 | 6.2094 | 6.0950 | 6.1866 | 5.5603 | 5.9790 | 5.8044 | 5.6477 |
| BAZ1B_S161 | BAZ1B | Q9UIG0 | | 330 | DNSSLSSPLNPKL | 7.3337 | 7.4814 | 6.8597 | 7.7018 | 6.0989 | 6.7788 | 7.0913 | 6.4200 | 7.1195 | 7.4715 | 7.3896 | 7.3460 | 3.3982 | 4.2876 | 3.2838 | 3.9381 |
| BAZ1B_S304 | BAZ1B | Q9UIG0 | | 1342 | RSSRRQSLELQKC | 5.7233 | 6.1247 | 5.3168 | 6.3244 | 5.9754 | 5.6883 | 6.2010 | 6.0919 | 6.5492 | 6.5663 | 6.3537 | 6.6881 | 6.3134 | 6.5500 | 6.7660 | 6.7674 |
| BAZ1B_S312 | BAZ1B | Q9UIG0 | | 947 | CKDHTVSGDEDYC | 7.3568 | 7.6318 | 7.1712 | 8.1135 | 6.1742 | 6.5592 | 7.0073 | 7.1704 | 6.1722 | 5.8641 | 6.3766 | 6.2623 | 4.2801 | 4.7883 | 4.4400 | 4.6319 |
| BAZ1B_S330 | BAZ1B | Q9UIG0 | | 161 | ACDSPSSDKENSS | 6.1726 | 6.7045 | 5.9638 | 7.0197 | 6.1668 | 6.1488 | 6.5037 | 5.8085 | 6.7752 | 6.2458 | 6.6861 | 6.5389 | 5.7669 | 6.6233 | 5.6201 | 5.2553 |
| BAZ1B_S347 | BAZ1B | Q9UIG0 | | 158 | SDGACDSPSSDKE | 6.2910 | 6.1395 | 5.5425 | 7.0169 | 6.3515 | 5.8956 | 6.3950 | 5.7677 | 6.6134 | 6.7852 | 6.6879 | 6.4405 | 6.0626 | 5.9823 | 6.0752 | 5.9532 |
| BAZ1B_S349 | BAZ1B | Q9UIG0 | | 347 | HLKKSLSGSPLKV | 4.6320 | 4.9512 | 4.8276 | 5.2939 | 5.1157 | 5.0710 | 5.1406 | 5.4431 | 5.0441 | 5.6618 | 5.1642 | 5.6318 | 9.6669 | 8.7120 | 9.7357 | 9.9084 |
| BAZ1B_S361 | BAZ1B | Q9UIG0 | | 312 | KRKNTGSPDRKPS | 6.8152 | 6.8168 | 6.4967 | 7.6739 | 5.4393 | 5.1933 | 6.1554 | 5.3971 | 6.8807 | 6.8611 | 6.3149 | 7.2515 | 5.3590 | 5.2497 | 5.7688 | 6.3266 |
| BAZ1B_S705;T710 | BAZ1B | Q9UIG0 | | 304 | YMTLNPSTKRKNT | 6.9309 | 6.1590 | 7.8687 | 7.7368 | 6.0690 | 5.5171 | 6.6569 | 5.7151 | 6.7529 | 8.4646 | 7.5488 | 6.2570 | 4.6413 | 4.8912 | 4.2793 | 4.5113 |
| BAZ1B_S947 | BAZ1B | Q9UIG0 | 705;710 | | NA | 6.1283 | 6.0076 | 5.5589 | 6.3183 | 6.2373 | 6.1257 | 6.6868 | 6.4020 | 6.0888 | 5.8881 | 6.1830 | 6.1195 | 6.1679 | 7.0014 | 6.3073 | 6.7791 |
| BAZ1B_T833 | BAZ1B | Q9UIG0 | 158;161 | | NA | 6.6395 | 6.5297 | 6.3501 | 6.9708 | 6.6041 | 6.1716 | 6.6052 | 5.3604 | 6.3996 | 6.5146 | 6.7148 | 6.5023 | 5.6918 | 5.5723 | 5.4675 | 5.9057 |
| BAZ2A_S1370 | BAZ2A | Q9UIF9 | | 1397 | PGEMPQSPTGLGQ | 6.8393 | 7.3065 | 6.7149 | 7.4113 | 6.4069 | 6.3812 | 6.9007 | 6.3866 | 6.8440 | 6.9975 | 6.7564 | 6.7907 | 4.2321 | 4.9143 | 4.4517 | 4.6658 |
| BAZ2A_S1397 | BAZ2A | Q9UIF9 | | 1783 | YSEEGLSPSKRRR | 6.4995 | 6.6489 | 6.3916 | 7.0629 | 6.6078 | 6.1695 | 6.5073 | 6.2368 | 6.6320 | 6.6779 | 6.4433 | 6.5498 | 5.1322 | 5.6005 | 5.3010 | 5.5390 |
| BAZ2A_S1747 | BAZ2A | Q9UIF9 | | 1770 | LLRGRESPAAGPR | 6.2966 | 6.7908 | 5.6319 | 6.8237 | 6.0701 | 6.0355 | 6.3372 | 6.1436 | 6.6170 | 6.3504 | 5.9049 | 5.9906 | 5.5397 | 6.7619 | 6.3822 | 6.3239 |
| BAZ2A_S1770 | BAZ2A | Q9UIF9 | | 496 | VTSPKASPVTSPA | 5.8988 | 6.6195 | 6.3058 | 6.8646 | 5.6341 | 5.8076 | 6.0400 | 5.8661 | 5.9131 | 5.8857 | 6.0148 | 6.4726 | 6.0885 | 6.8212 | 6.7189 | 7.0485 |
| BAZ2A_S1783 | BAZ2A | Q9UIF9 | | 509 | AAFPTASPANKDV | 6.3194 | 6.7823 | 6.3539 | 6.9067 | 5.8456 | 6.0581 | 5.9967 | 6.6663 | 6.2773 | 6.7174 | 5.8478 | 6.4699 | 5.6441 | 5.7962 | 5.9834 | 6.3349 |
| BAZ2A_S38 | BAZ2A | Q9UIF9 | | 1370 | SAANPCSPVQFSS | 5.1802 | 5.7380 | 6.6062 | 6.9267 | 6.4989 | 6.2768 | 6.3904 | 6.5532 | 6.8750 | 7.1425 | 6.7803 | 8.2517 | 5.1903 | 6.1279 | 3.9397 | 5.5222 |
| BAZ2A_S496 | BAZ2A | Q9UIF9 | | 1747 | RGQKRKSGYSLNF | 6.1807 | 6.7458 | 7.0493 | 7.3931 | 5.2118 | 7.9527 | 6.7263 | 6.1779 | 6.0039 | 6.2322 | 6.4201 | 5.7784 | 4.8011 | 5.5654 | 5.3371 | 6.4243 |
| BAZ2A_S496;S509 | BAZ2A | Q9UIF9 | | 38 | GLYTNGSPMNFPQ | 6.2261 | 7.5217 | 6.8636 | 7.4520 | 5.9061 | 6.4003 | 6.5449 | 6.4790 | 7.7694 | 7.7810 | 6.9836 | 7.3643 | 4.1357 | 5.0803 | 3.9060 | 3.5860 |
| BAZ2A_S509 | BAZ2A | Q9UIF9 | | 1621 | EKALLSTPNGAPE | 6.8229 | 6.4942 | 6.3425 | 6.6628 | 5.6114 | 6.8002 | 6.6855 | 6.5069 | 7.3345 | 6.6940 | 6.2419 | 8.3335 | 4.0597 | 5.6369 | 5.0134 | 4.7597 |
| BAZ2A_T1621 | BAZ2A | Q9UIF9 | 496;509 | | NA | 6.3190 | 6.0357 | 6.5098 | 6.4808 | 6.1975 | 5.9085 | 6.2003 | 6.4479 | 6.6227 | 6.2258 | 5.7752 | 5.9900 | 6.0274 | 6.4063 | 6.4646 | 6.3885 |
| BAZ2B_S1541 | BAZ2B | Q9UIF8 | | 405 | MKLIVPSPDVLKA | 7.4393 | 8.3323 | 6.3399 | 7.5106 | 6.3564 | 7.9629 | 7.6936 | 7.1084 | 7.1360 | 8.2095 | 7.0158 | 7.1972 | 2.3783 | 3.5163 | 2.2487 | 3.5549 |
| BAZ2B_S1596 | BAZ2B | Q9UIF8 | | 1541 | GPGKFYSPLPNDQ | 6.5166 | 7.0043 | 6.3406 | 6.7748 | 6.7503 | 7.3034 | 6.9301 | 6.1684 | 6.8975 | 7.6189 | 6.8675 | 7.7083 | 3.4482 | 5.3216 | 3.5860 | 4.7636 |
| BAZ2B_S405 | BAZ2B | Q9UIF8 | | 1596 | PPSKSPSPTPAPL | 5.5974 | 5.6741 | 4.6317 | 5.6834 | 7.5610 | 6.4367 | 6.0865 | 5.3425 | 4.7892 | 6.0077 | 6.2305 | 5.8824 | 6.8595 | 8.0158 | 8.3505 | 6.8512 |
| BBX_S40 | BBX | Q8WY36 | | 649 | SGKGNSSDHEGCW | 6.3473 | 7.0409 | 6.3912 | 7.0559 | 6.2962 | 7.2092 | 6.3216 | 7.0343 | 5.5175 | 5.5936 | 5.4994 | 5.4729 | 5.1145 | 6.6627 | 6.2130 | 6.2297 |
| BBX_S479 | BBX | Q8WY36 | | 844 | TADGRVSPAGGTL | 5.4853 | 5.7984 | 5.4227 | 6.2157 | 5.7234 | 5.7019 | 5.6102 | 5.7545 | 5.7428 | 5.9714 | 5.9292 | 5.9285 | 7.0069 | 8.1222 | 7.7480 | 7.8389 |
| BBX_S649 | BBX | Q8WY36 | | 704 | NSLPQYSPVTFDR | 6.9335 | 6.8922 | 6.9440 | 7.1015 | 6.7888 | 6.7713 | 6.5792 | 6.6324 | 6.5021 | 6.3513 | 6.5021 | 6.5754 | 4.2759 | 5.8792 | 4.4000 | 4.8712 |
| BBX_S704 | BBX | Q8WY36 | | 506 | IEKLGDTPRKKVR | 6.9544 | 6.7004 | 6.1945 | 6.2081 | 5.7241 | 5.9413 | 6.5000 | 5.8038 | 6.5428 | 6.3206 | 5.8338 | 6.1010 | 5.7009 | 6.7027 | 6.0058 | 6.7659 |
| BBX_S844 | BBX | Q8WY36 | | 40 | KKLLDFSEEEEEE | 6.0000 | 6.1299 | 6.2093 | 6.5645 | 5.8551 | 6.1366 | 6.4698 | 5.5833 | 6.3014 | 5.7125 | 5.9618 | 6.0883 | 5.8405 | 7.6525 | 6.3040 | 7.1905 |
| BBX_T506 | BBX | Q8WY36 | | 479 | CKKRQSSESDIES | 5.4720 | 6.3450 | 6.3640 | 6.0957 | 6.1539 | 6.9017 | 6.7933 | 4.8527 | 6.4445 | 6.1337 | 6.7273 | 7.2125 | 6.5362 | 6.3573 | 6.1036 | 5.5066 |
| BCAN_S74 | BCAN | Q96GW7 | | 74 | RRAVLGSPRVKWT | 6.7788 | 6.9895 | 6.4328 | 6.4504 | 5.6741 | 5.8890 | 6.2022 | 5.9688 | 6.4847 | 6.6715 | 5.8142 | 6.1849 | 5.6973 | 6.1295 | 6.0353 | 6.5970 |
| BCAR1_S180;S185 | BCAR1 | P56945-6 | | 685 | QSRPLPSPPKFTS | 6.5409 | 5.4196 | 6.8802 | 6.0604 | 6.4357 | 6.6743 | 6.3911 | 6.0014 | 6.5302 | 6.6190 | 6.8897 | 6.9097 | 5.8309 | 5.9126 | 5.1715 | 5.7327 |
| BCAR1_S185 | BCAR1 | P56945-6 | | 185 | PSPQFQSPPAKQT | 6.4284 | 6.7922 | 7.2972 | 6.9153 | 6.5578 | 6.9932 | 6.4969 | 7.0230 | 7.4475 | 7.0887 | 7.5560 | 7.6309 | 3.7994 | 4.2866 | 3.5726 | 4.1142 |
| BCAR1_S338 | BCAR1 | P56945-6 | | 315 | GQEVYDTPPMAVK | 7.3883 | 7.7052 | 7.5292 | 7.1860 | 5.6080 | 7.1875 | 7.8352 | 8.0711 | 7.2875 | 7.4891 | 7.3365 | 8.4055 | 2.5426 | 3.5781 | 2.1483 | 2.7018 |
| BCAR1_S474 | BCAR1 | P56945-6 | | 474 | AEGKRLSASSTGS | 6.4469 | 6.1114 | 7.1211 | 6.3968 | 6.1217 | 7.1419 | 6.3862 | 7.0665 | 6.3466 | 6.2883 | 6.6167 | 7.2549 | 5.2610 | 5.3071 | 4.8995 | 5.2335 |
| BCAR1_S685 | BCAR1 | P56945-6 | | 408 | PPAEDVYDVPPPA | 6.0918 | 6.4009 | 6.3996 | 6.1037 | 6.4103 | 6.6332 | 5.8903 | 6.2536 | 5.7512 | 6.2643 | 6.2649 | 6.6477 | 6.5602 | 6.2643 | 6.2221 | 5.8418 |
| BCAR1_S694 | BCAR1 | P56945-6 | | 295 | PGPQDIYDVPPVR | 6.2722 | 5.5542 | 6.2117 | 5.8862 | 4.8836 | 5.9712 | 5.9124 | 6.7269 | 6.8364 | 6.0121 | 6.9885 | 5.9124 | 6.8773 | 7.4972 | 5.6638 | 6.7939 |
| BCAR1_T315 | BCAR1 | P56945-6 | | 338 | VYDVPPSVEKGLP | 9.0078 | 8.9438 | 9.4270 | 7.1620 | 7.7210 | 9.0194 | 7.8200 | 3.9478 | 4.6233 | 7.4182 | 7.4648 | 9.1243 | 2.4688 | 2.4456 | 1.6013 | 1.8051 |
| BCAR1_T844;S846 | BCAR1 | P56945-6 | | 694 | KFTSQDSPDGQYE | 6.6564 | 6.6382 | 6.8940 | 6.5460 | 6.4163 | 5.7598 | 6.2278 | 6.8879 | 5.9507 | 6.1059 | 6.9613 | 6.8164 | 5.8343 | 5.4281 | 5.3560 | 5.5209 |
| BCAR1_Y295 | BCAR1 | P56945-6 | 844;846 | | NA | 6.7695 | 8.0979 | 4.5615 | 7.2939 | 6.8353 | 7.5095 | 5.3914 | 9.3086 | 5.7741 | 5.4520 | 6.0921 | 5.1940 | 3.6335 | 6.1501 | 6.6677 | 5.2690 |
| BCAR1_Y408 | BCAR1 | P56945-6 | 180;185 | | NA | 7.2754 | 6.8373 | 7.1946 | 7.7221 | 4.8028 | 7.5585 | 6.5767 | 6.7277 | 7.9779 | 8.4868 | 7.5804 | 8.5564 | 2.5198 | 3.8724 | 2.4470 | 3.8644 |
| BCAR3_S290 | BCAR3 | O75815 | | 375 | GSEPALSPAVVRR | 5.7018 | 6.1630 | 6.6701 | 5.8328 | 6.0553 | 7.4874 | 7.3283 | 7.1279 | 5.9812 | 6.3039 | 6.8262 | 6.6533 | 5.0174 | 5.8524 | 5.0756 | 5.9233 |
| BCAR3_S375 | BCAR3 | O75815 | | 290 | DVAKRLSLTMGGV | 6.3892 | 6.0763 | 6.3377 | 5.6005 | 6.2528 | 6.0283 | 6.5581 | 6.7499 | 7.4583 | 6.0795 | 6.3208 | 6.6261 | 5.7211 | 7.1102 | 5.2942 | 5.3972 |
| BCAR3_S83 | BCAR3 | O75815 | | 83 | KSPRQNSPVTQDG | 5.6852 | 5.4211 | 6.4005 | 6.1034 | 6.5398 | 6.4494 | 7.6353 | 6.5435 | 5.7671 | 6.2110 | 6.9066 | 6.3075 | 6.1413 | 5.7292 | 6.2012 | 5.9579 |
| BCAS3_S932 | BCAS3 | Q9H6U6-8 | | 932 | NFDGYRSPLPTNE | 6.3383 | 5.5256 | 6.4206 | 7.0548 | 6.0613 | 6.6907 | 6.5401 | 6.6084 | 6.4502 | 6.8789 | 6.9913 | 6.4723 | 5.1884 | 5.9839 | 5.3433 | 5.4519 |
| BCCIP_Y64 | BCCIP | Q9P287-2 | | 64 | NIEFEAYSLSDND | 5.2417 | 4.9651 | 8.6676 | 5.3079 | 5.5790 | 7.7995 | 6.7067 | 5.1332 | 5.9501 | 6.4164 | 7.0977 | 6.0549 | 7.8104 | 7.5224 | 6.6013 | 6.9459 |
| BCKDHA_S340 | BCKDHA | P12694-2 | | 350 | DSSAYRSVDEVNY | 4.5500 | 4.5802 | 4.0624 | 4.8483 | 7.2084 | 5.6184 | 5.7521 | 6.4026 | 4.6711 | 4.4796 | 4.0821 | 3.8645 | 10.7114 | 9.4518 | 10.3642 | 9.3529 |
| BCKDHA_S340;S350 | BCKDHA | P12694-2 | | 340 | CRIGHHSTSDDSS | 5.4197 | 6.2551 | 4.9116 | 5.9983 | 7.0143 | 6.2655 | 5.9461 | 7.1371 | 5.7046 | 4.6476 | 4.5505 | 5.1420 | 7.3338 | 8.3739 | 8.1203 | 7.1797 |
| BCKDHA_S350 | BCKDHA | P12694-2 | 340;350 | | NA | 5.2091 | 5.2329 | 4.5132 | 5.2790 | 8.0055 | 7.3699 | 6.0242 | 6.7387 | 4.6659 | 4.5811 | 4.5935 | 4.1481 | 9.1783 | 8.2967 | 8.7838 | 7.3802 |
| BCKDK_S31 | BCKDK | O14874 | | 31 | LALRARSTSATDT | 5.5434 | 5.9052 | 5.8522 | 6.4453 | 6.3675 | 6.2035 | 6.2679 | 6.5228 | 6.3637 | 5.8277 | 6.5423 | 6.2278 | 6.4238 | 6.4717 | 6.7602 | 6.2750 |
| BCL2L11_S77 | BCL2L11 | O43521-12 | | 77 | PQPPKRSPLFIFM | 5.2358 | 4.9876 | 4.7014 | 5.1021 | 6.2943 | 5.3750 | 5.7002 | 5.9183 | 5.0192 | 4.7582 | 4.5756 | 5.2802 | 8.9361 | 8.7205 | 9.8203 | 9.5751 |
| BCL2L12_S242 | BCL2L12 | Q9HB09 | | 242 | SKLVRLSSDSFAR | 5.9442 | 5.7209 | 5.4026 | 5.7315 | 5.6851 | 4.9153 | 5.3189 | 6.3712 | 5.4604 | 5.1073 | 5.0149 | 5.8503 | 5.4663 | 9.3516 | 7.8600 | 10.7993 |
| BCL2L13_S303 | BCL2L13 | Q9BXK5 | | 371 | ITVEKSSPATSLF | 5.9682 | 6.0792 | 6.0760 | 6.0489 | 5.9881 | 5.9224 | 6.1156 | 6.2555 | 6.1485 | 6.5434 | 6.1586 | 6.4172 | 6.1115 | 6.9988 | 6.4612 | 6.7070 |
| BCL2L13_S353 | BCL2L13 | Q9BXK5 | | 444 | EGKSRLSPAGEMK | 6.2750 | 6.4330 | 6.0740 | 6.3541 | 6.0872 | 6.1108 | 6.1259 | 5.7508 | 6.1139 | 6.0193 | 6.1690 | 6.3228 | 6.1489 | 6.8429 | 6.5755 | 6.5969 |
| BCL2L13_S371 | BCL2L13 | Q9BXK5 | | 303 | KSENNSSNSDIVH | 5.7513 | 6.3093 | 5.4446 | 6.8145 | 5.2701 | 6.1295 | 5.9383 | 6.2237 | 5.6236 | 6.0091 | 6.0522 | 6.4932 | 5.8737 | 7.0996 | 7.7901 | 7.1774 |
| BCL2L13_S420 | BCL2L13 | Q9BXK5 | | 353 | PHITATSLLGTRE | 6.7645 | 6.2018 | 6.5543 | 7.1439 | 5.8849 | 6.0185 | 6.4588 | 6.6452 | 6.9113 | 6.6563 | 5.7031 | 6.4984 | 5.7290 | 6.0672 | 5.0800 | 5.6828 |

| | | | | | | | | | | | | | | | | | | | | |
|---|---|---|---|---|---|---|---|---|---|---|---|---|---|---|---|---|---|---|---|---|
| BCL2L13_S426 | BCL2L13 | Q9BXK5 | 437 | KKPVPPSEGKSRL | 6.6704 | 7.4017 | 6.1554 | 7.5293 | 6.2911 | 6.0930 | 7.6161 | 6.7817 | 6.0539 | 7.0484 | 7.5024 | 7.3715 | 3.5420 | 4.9560 | 3.9850 | 5.0021 |
| BCL2L13_S437 | BCL2L13 | Q9BXK5 | 420 | INAREESLVEELS | 6.9306 | 6.7426 | 6.1266 | 6.8572 | 5.8217 | 6.0174 | 6.7993 | 6.3134 | 6.6832 | 6.7577 | 6.8723 | 6.9688 | 4.8625 | 6.0942 | 4.6842 | 5.4685 |
| BCL2L13_S444 | BCL2L13 | Q9BXK5 | 426 | SLVEELSPASEKK | 6.6217 | 6.7220 | 6.0139 | 6.5405 | 6.0114 | 5.7702 | 5.8013 | 5.8494 | 6.4056 | 6.8093 | 6.2035 | 6.3534 | 5.4181 | 6.5515 | 6.5856 | 6.3424 |
| BCL6_S404 | BCL6 | P41182 | 404 | AELGRLSPRAYTA | 6.4547 | 7.5524 | 7.2030 | 6.8925 | 6.0921 | 6.6622 | 6.6336 | 5.7545 | 6.4712 | 6.8232 | 6.2274 | 6.2758 | 4.9299 | 5.9719 | 5.1121 | 4.9435 |
| BCL7C_S122 | BCL7C | Q8WUZ0-2 | 122 | GGTPQPSRPVSPA | 7.3556 | 6.1769 | 5.9515 | 7.9810 | 7.8225 | 6.8047 | 7.3532 | 8.2405 | 6.5809 | 8.2902 | 7.6982 | 7.2164 | 3.3798 | 3.8273 | 3.2897 | 3.6517 |
| BCL9_S104 | BCL9 | O00512 | 687 | PVEGPLSPSRGDF | 6.1640 | 5.8866 | 6.2443 | 5.3952 | 6.5502 | 6.2254 | 6.3657 | 6.5330 | 6.0742 | 6.5216 | 6.3778 | 6.4366 | 6.9312 | 6.4377 | 6.2751 | 6.3114 |
| BCL9_S278 | BCL9 | O00512 | 278 | RPLDRESPGVENK | 5.7756 | 6.4904 | 6.0731 | 6.3634 | 6.6786 | 6.0867 | 5.6124 | 5.7117 | 5.9186 | 6.3243 | 6.4328 | 6.6196 | 8.0967 | 7.2687 | 6.3598 | 6.8877 |
| BCL9_S291 | BCL9 | O00512 | 315 | PNNRAVTPVSQGS | 5.9230 | 5.4333 | 5.5579 | 5.3212 | 6.1361 | 6.2128 | 5.9722 | 6.2452 | 6.2225 | 6.0921 | 6.0035 | 5.7960 | 7.9870 | 7.0550 | 6.9937 | 7.0485 |
| BCL9_S35 | BCL9 | O00512 | 257 | NTRLQPTPPIPAP | 5.3400 | 5.3170 | 5.5413 | 5.1400 | 6.2101 | 6.0725 | 5.8280 | 6.4415 | 5.5135 | 6.1531 | 6.3993 | 6.0037 | 8.2516 | 7.5850 | 7.0665 | 7.1369 |
| BCL9_S687 | BCL9 | O00512 | 35 | RPPTVMSPGNPQ | 6.3461 | 5.5692 | 6.7144 | 5.4411 | 6.6156 | 6.0997 | 5.7622 | 6.5427 | 6.0412 | 6.6251 | 6.8543 | 6.5581 | 7.0480 | 6.0068 | 5.8841 | 5.8915 |
| BCL9_S885 | BCL9 | O00512 | 291 | LIPSVGSPASSTP | 5.5012 | 5.0451 | 5.9632 | 5.5203 | 7.1673 | 5.8077 | 5.9031 | 6.6437 | 6.3504 | 5.2886 | 6.1362 | 5.9045 | 7.5075 | 6.9839 | 7.0323 | 7.2450 |
| BCL9_S917 | BCL9 | O00512 | 917 | LGSAAASPVHLKS | 5.5050 | 5.5298 | 5.5574 | 4.6642 | 6.4295 | 6.2661 | 5.9827 | 6.2047 | 5.7923 | 5.7013 | 5.6977 | 5.5875 | 8.6526 | 8.0940 | 7.4602 | 6.8751 |
| BCL9_T257 | BCL9 | O00512 | 885 | PSGNLKSPQTPSQ | 6.6046 | 5.9139 | 5.9080 | 5.7809 | 6.3715 | 6.0436 | 6.1962 | 6.7725 | 6.3063 | 6.6030 | 6.5839 | 6.6448 | 6.5368 | 5.9499 | 5.9695 | 5.8148 |
| BCL9_T315 | BCL9 | O00512 | 104 | KRERSISADSFDQ | 5.1327 | 4.7168 | 5.0302 | 4.8319 | 5.9544 | 5.5079 | 5.6595 | 6.2487 | 4.9267 | 5.0234 | 5.5398 | 5.5958 | 9.8491 | 8.2454 | 9.3671 | 8.3704 |
| BCL9L_S1017 | BCL9L | Q86UU0 | 1017 | PKTAMPSPGVSQN | 5.8759 | 5.9890 | 6.0616 | 5.7561 | 5.8662 | 6.2174 | 6.0115 | 6.3914 | 6.6392 | 6.3777 | 6.5657 | 6.5928 | 6.8570 | 6.5211 | 6.0705 | 6.2069 |
| BCL9L_S116 | BCL9L | Q86UU0 | 1480 | LRQRSVSLDSQMG | 5.4794 | 5.3868 | 6.1567 | 5.2193 | 6.2479 | 5.8079 | 6.6957 | 7.0499 | 5.9116 | 5.8610 | 6.5094 | 6.7539 | 6.8779 | 7.1258 | 7.0817 | 6.8351 |
| BCL9L_S1480 | BCL9L | Q86UU0 | 21 | RREAPGSPPLSPR | 6.3231 | 6.4762 | 6.1235 | 5.5697 | 6.1967 | 5.8976 | 5.4541 | 5.9600 | 6.2508 | 6.3970 | 6.2262 | 5.9144 | 7.4426 | 6.7575 | 6.7254 | 6.2851 |
| BCL9L_S21 | BCL9L | Q86UU0 | 750 | GGRGLLSPPMGQS | 6.7243 | 6.4310 | 6.5239 | 5.9536 | 6.4281 | 6.4788 | 6.0535 | 6.8126 | 6.7427 | 6.9722 | 7.1244 | 7.1388 | 5.4865 | 5.2531 | 4.8316 | 5.0449 |
| BCL9L_S21;S25 | BCL9L | Q86UU0 | 975 | PPGPLKSPQVLGS | 5.5374 | 5.2872 | 5.6338 | 5.0703 | 6.4865 | 6.2260 | 6.1534 | 6.8349 | 6.1164 | 6.0946 | 6.2527 | 6.0100 | 7.5356 | 7.2133 | 6.9006 | 6.6472 |
| BCL9L_S750 | BCL9L | Q86UU0 | 813 | MGPQGLSPEEMAR | 6.2886 | 6.4557 | 6.2574 | 6.4132 | 6.5643 | 6.5843 | 6.2793 | 6.5803 | 6.4639 | 7.1375 | 7.0965 | 6.9066 | 5.1739 | 5.5536 | 5.2158 | 5.0290 |
| BCL9L_S813 | BCL9L | Q86UU0 | 915 | GLGRRPSDLTISI | 5.9938 | 4.8046 | 5.7678 | 5.3844 | 7.0860 | 5.3325 | 4.3252 | 6.0975 | 7.3490 | 8.3604 | 8.0401 | 8.3840 | 6.9543 | 6.6878 | 7.4446 | 4.6880 |
| BCL9L_S88 | BCL9L | Q86UU0 | 88 | AKANQISPSNSSL | 6.6724 | 6.2748 | 6.8576 | 6.0192 | 6.4099 | 6.5279 | 6.4169 | 6.9637 | 6.3097 | 6.6210 | 7.0523 | 6.8194 | 5.3678 | 5.4646 | 5.0477 | 5.1751 |
| BCL9L_S915 | BCL9L | Q86UU0 | 997 | SPSRLKSPSMAVP | 5.3503 | 5.1198 | 5.9453 | 4.7935 | 6.5099 | 6.1886 | 5.6717 | 6.5668 | 5.8752 | 6.2899 | 6.4279 | 6.1205 | 8.0808 | 7.2494 | 6.8501 | 6.9603 |
| BCL9L_S934 | BCL9L | Q86UU0 | 129 | EQREAGTPSLDSE | 6.6700 | 7.0058 | 6.1889 | 6.1521 | 6.1306 | 6.5339 | 6.3572 | 5.9185 | 6.7694 | 6.9071 | 6.8556 | 7.2348 | 5.7263 | 5.8421 | 4.7292 | 4.9785 |
| BCL9L_S934;S942;S947 | BCL9L | Q86UU0 | 934 | GMGHLKSPTLSQV | 5.8466 | 5.8086 | 6.4733 | 5.7009 | 6.0813 | 6.4403 | 6.2568 | 6.5523 | 6.5067 | 6.0374 | 6.1173 | 6.3256 | 6.7487 | 6.1583 | 6.6591 | 6.2866 |
| BCL9L_S934;S947 | BCL9L | Q86UU0 | 116 | KVKRDRSVSVDSG | 4.6945 | 4.8913 | 5.1694 | 4.7796 | 6.2895 | 5.7050 | 5.1970 | 5.9848 | 5.1555 | 5.4888 | 5.4469 | 5.7673 | 9.1470 | 8.5093 | 9.2280 | 8.5461 |
| BCL9L_S954;T957 | BCL9L | Q86UU0 | 934;947 | NA | 5.5538 | 5.7540 | 6.0050 | 5.8999 | 6.7350 | 6.6228 | 5.7965 | 6.8544 | 6.0790 | 5.9516 | 6.2844 | 6.1668 | 6.9149 | 6.7829 | 6.1751 | 6.4240 |
| BCL9L_S975 | BCL9L | Q86UU0 | 21;25 | NA | 5.9265 | 6.1169 | 6.4264 | 6.3681 | 6.1770 | 6.6220 | 5.9451 | 6.9794 | 6.5265 | 7.1429 | 6.6552 | 7.0277 | 5.6533 | 5.5340 | 5.2875 | 5.6113 |
| BCL9L_S997 | BCL9L | Q86UU0 | 954;957 | NA | 5.9944 | 5.9518 | 5.1390 | 6.3813 | 6.4112 | 6.1519 | 5.9871 | 7.4394 | 5.6090 | 6.5133 | 5.6150 | 5.9587 | 6.9778 | 6.4491 | 6.9212 | 6.4999 |
| BCL9L_T129 | BCL9L | Q86UU0 | 934;942;947 | NA | 6.1572 | 5.9935 | 6.2532 | 5.8504 | 6.2688 | 6.0087 | 5.9725 | 6.8827 | 6.1876 | 6.0152 | 6.6699 | 7.0968 | 6.4420 | 6.1643 | 5.7485 | 6.2887 |
| BCLAF1_S148 | BCLAF1 | Q9NYF8 | 658 | HRRIDISPSTLRK | 6.4703 | 6.6394 | 5.9283 | 6.6410 | 6.2845 | 5.8860 | 6.1317 | 6.1877 | 6.2925 | 6.0305 | 5.9096 | 5.9212 | 6.4268 | 6.3712 | 6.4648 | 6.4142 |
| BCLAF1_S161 | BCLAF1 | Q9NYF8 | 512 | KDLFDYSPPLHKN | 6.1333 | 6.3093 | 5.8940 | 6.5835 | 6.0634 | 5.8374 | 6.0639 | 5.9382 | 6.0794 | 6.1107 | 5.8768 | 6.0788 | 6.6058 | 6.8023 | 6.8507 | 6.7725 |
| BCLAF1_S177 | BCLAF1 | Q9NYF8 | 385 | KALFDSEKSGK | 6.2308 | 6.1776 | 6.2059 | 6.5942 | 6.1560 | 5.9568 | 6.1679 | 5.9471 | 6.3491 | 6.1619 | 6.0975 | 6.3899 | 6.6510 | 6.4700 | 6.1073 | 6.3369 |
| BCLAF1_S183 | BCLAF1 | Q9NYF8 | 496 | VKKETQSPEQVKS | 6.4040 | 6.4169 | 5.6484 | 6.4233 | 6.5926 | 5.9105 | 6.3026 | 6.1828 | 6.2172 | 6.0167 | 5.7009 | 5.7136 | 6.7861 | 6.5016 | 6.6623 | 6.5207 |
| BCLAF1_S196 | BCLAF1 | Q9NYF8 | 531 | STFREESPLRIKM | 6.6203 | 6.4903 | 5.8953 | 6.3361 | 6.9547 | 5.9404 | 5.8750 | 6.1382 | 6.9212 | 5.9024 | 5.6939 | 6.0541 | 6.2492 | 6.1612 | 6.8683 | 5.8996 |
| BCLAF1_S198 | BCLAF1 | Q9NYF8 | 177 | GEPQEESPLKSKS | 6.2774 | 6.1505 | 5.8059 | 6.3082 | 6.4992 | 6.0409 | 6.2091 | 6.0888 | 6.3170 | 5.9813 | 6.0290 | 5.9673 | 6.9571 | 6.4762 | 6.5005 | 6.3914 |
| BCLAF1_S222 | BCLAF1 | Q9NYF8 | 760 | SSASPSSPSSREE | 6.4296 | 6.2983 | 6.0693 | 6.8357 | 6.4176 | 5.7669 | 6.2469 | 5.7475 | 6.5410 | 6.0648 | 6.2255 | 6.2890 | 6.3709 | 6.1865 | 6.1370 | 6.3736 |
| BCLAF1_S259;S268 | BCLAF1 | Q9NYF8 | 222 | LSAYDNSPRSPHS | 5.6952 | 6.0323 | 6.1396 | 6.7153 | 6.0797 | 6.5717 | 6.4169 | 6.3393 | 6.2993 | 6.4207 | 6.0952 | 6.2094 | 6.2360 | 6.5532 | 6.5704 | 6.5257 |
| BCLAF1_S268 | BCLAF1 | Q9NYF8 | 290 | YSPSQNSPIHHIP | 6.2972 | 6.6537 | 6.1894 | 6.9882 | 6.6269 | 6.3729 | 6.3005 | 6.6701 | 6.4227 | 5.8579 | 6.2184 | 5.8618 | 5.7378 | 5.9610 | 5.8457 | 5.9959 |
| BCLAF1_S276 | BCLAF1 | Q9NYF8 | 648 | HSTRQKSPEIHRR | 6.1161 | 6.5304 | 5.9095 | 6.1046 | 6.1293 | 6.5693 | 6.7452 | 6.3888 | 5.6839 | 5.7285 | 6.2480 | 5.4931 | 6.3408 | 7.7589 | 6.1350 | 6.1184 |
| BCLAF1_S285 | BCLAF1 | Q9NYF8 | 198 | EHDPSESIDEFNK | 6.2739 | 6.2809 | 6.2550 | 6.9505 | 6.2765 | 6.4399 | 6.9136 | 6.8227 | 6.1789 | 5.7855 | 6.1080 | 6.0324 | 5.6815 | 6.0728 | 5.8059 | 6.1221 |
| BCLAF1_S285;S290 | BCLAF1 | Q9NYF8 | 578 | ASTLVHSVKKEQE | 7.8698 | 8.2586 | 6.8626 | 8.2852 | 6.3425 | 6.8494 | 7.4118 | 6.9084 | 7.3501 | 6.7144 | 6.6924 | 6.7074 | 3.0686 | 4.2284 | 2.8406 | 3.6100 |
| BCLAF1_S290 | BCLAF1 | Q9NYF8 | 397 | KQKFNDSEGDDTE | 5.8902 | 6.0462 | 6.0207 | 6.3761 | 5.9302 | 5.7048 | 6.3125 | 5.8022 | 6.2010 | 6.0662 | 5.9165 | 6.2473 | 7.0407 | 6.5952 | 6.6587 | 7.1918 |
| BCLAF1_S385 | BCLAF1 | Q9NYF8 | 285 | NGSSRYSPSQNSP | 6.6316 | 7.3024 | 6.0874 | 7.8821 | 6.5577 | 6.6508 | 6.0308 | 6.8327 | 6.3474 | 6.1885 | 6.5677 | 6.4148 | 4.9223 | 5.4358 | 5.1589 | 5.1893 |
| BCLAF1_S385;S389 | BCLAF1 | Q9NYF8 | 268 | SHSIQHSPERSGS | 6.2290 | 6.9763 | 6.9045 | 7.6223 | 6.4146 | 6.9752 | 5.8783 | 5.8641 | 5.4723 | 6.6862 | 6.9051 | 6.4531 | 6.5220 | 6.1446 | 5.1495 | 4.7029 |
| BCLAF1_S397 | BCLAF1 | Q9NYF8 | 450 | LKGNRESDGFREE | 6.8942 | 7.1496 | 6.7262 | 7.5343 | 7.0271 | 7.2575 | 7.6000 | 6.9816 | 6.3785 | 6.3779 | 6.2347 | 5.9303 | 4.2856 | 4.6184 | 4.2459 | 4.7583 |
| BCLAF1_S414 | BCLAF1 | Q9NYF8 | 414 | YRQFRKSVLADQG | 7.3826 | 7.1431 | 5.9304 | 6.9274 | 5.7102 | 6.4526 | 6.9289 | 5.6266 | 6.1459 | 5.8827 | 6.1740 | 6.2213 | 5.9275 | 5.9218 | 5.1309 | 6.4940 |
| BCLAF1_S450 | BCLAF1 | Q9NYF8 | 196 | TFEHDPSESIDEF | 6.0485 | 5.9603 | 5.7478 | 5.9022 | 6.6232 | 5.9553 | 6.5940 | 5.7784 | 5.5255 | 6.1200 | 5.7069 | 5.3662 | 6.7501 | 7.8362 | 7.1768 | 6.9086 |
| BCLAF1_S472 | BCLAF1 | Q9NYF8 | 564 | DSNRPASLTKDRL | 6.4750 | 7.3157 | 6.5361 | 7.5602 | 6.0038 | 6.6931 | 6.9991 | 6.6164 | 6.8754 | 7.2280 | 7.2894 | 7.6919 | 4.0064 | 4.5040 | 3.7641 | 4.4415 |
| BCLAF1_S496 | BCLAF1 | Q9NYF8 | 472 | YVVERPSTTKDKH | 7.2540 | 7.1977 | 6.3710 | 7.3683 | 6.9254 | 6.5932 | 7.0054 | 6.4109 | 6.7541 | 5.8045 | 5.8926 | 6.0655 | 5.0649 | 5.3707 | 4.9245 | 4.9973 |
| BCLAF1_S512 | BCLAF1 | Q9NYF8 | 661 | IDISPSTLRKHTR | 6.0346 | 6.5002 | 6.2837 | 6.6490 | 6.1872 | 6.1268 | 6.5805 | 5.9977 | 6.1412 | 6.1660 | 6.0534 | 6.1751 | 6.2061 | 6.3965 | 6.2378 | 6.2641 |
| BCLAF1_S525 | BCLAF1 | Q9NYF8 | 573 | KDRLLASTLVHSV | 6.0376 | 6.4367 | 6.0758 | 5.9137 | 7.0763 | 5.6203 | 6.5624 | 5.5401 | 5.4544 | 6.5971 | 6.8722 | 5.8116 | 6.3838 | 5.6421 | 8.2536 | 5.7223 |
| BCLAF1_S525;S531 | BCLAF1 | Q9NYF8 | 161 | VSKRRGSQEKQTK | 6.0423 | 6.8435 | 6.1225 | 6.9166 | 6.3028 | 5.8030 | 6.4411 | 5.3259 | 5.6695 | 6.2063 | 6.0953 | 6.2040 | 6.3547 | 7.0169 | 6.3768 | 6.2787 |
| BCLAF1_S531 | BCLAF1 | Q9NYF8 | 690 | KKLRCDSADLRHD | 5.7575 | 5.3636 | 6.2724 | 6.3411 | 5.6707 | 6.1039 | 6.6528 | 5.2354 | 6.0821 | 5.7897 | 7.7260 | 6.0294 | 7.6922 | 7.7601 | 7.1338 | 7.3893 |
| BCLAF1_S564 | BCLAF1 | Q9NYF8 | 276 | ERSGSGSVGNGSS | 5.5038 | 4.8300 | 5.4615 | 6.2533 | 6.8101 | 5.9143 | 5.5981 | 5.3583 | 7.1718 | 6.5044 | 6.1988 | 5.8663 | 7.5311 | 6.9100 | 7.4920 | 6.5963 |
| BCLAF1_S573 | BCLAF1 | Q9NYF8 | 183 | SPLKSKSQEEPKD | 4.8644 | 5.3607 | 6.5431 | 7.5225 | 6.1910 | 6.5930 | 6.3857 | 5.6793 | 6.0429 | 6.2579 | 6.5622 | 7.5395 | 5.7539 | 6.9149 | 6.0506 | 7.5385 |
| BCLAF1_S578 | BCLAF1 | Q9NYF8 | 525 | LDAREKSTFREES | 7.4859 | 7.6680 | 6.0869 | 6.8360 | 6.1077 | 5.6018 | 6.9558 | 4.6972 | 5.5568 | 5.9982 | 6.4131 | 6.3174 | 5.9763 | 6.5560 | 5.6732 | 6.0697 |
| BCLAF1_S648 | BCLAF1 | Q9NYF8 | 148 | SSSRSSSPYSKSP | 6.1001 | 6.6163 | 7.1314 | 7.1425 | 5.2190 | 5.6869 | 6.4117 | 6.0097 | 7.7089 | 6.8245 | 7.9306 | 7.6595 | 4.5670 | 5.2543 | 5.0345 | 4.7032 |
| BCLAF1_S658 | BCLAF1 | Q9NYF8 | 285;290 | NA | 6.2002 | 6.4746 | 5.7204 | 6.5830 | 6.3731 | 6.3489 | 6.0585 | 6.7291 | 6.4759 | 6.0038 | 5.9901 | 6.0083 | 6.0189 | 6.3902 | 6.3922 | 6.2328 |
| BCLAF1_S658;T661 | BCLAF1 | Q9NYF8 | 385;389 | NA | 6.0040 | 5.3840 | 6.2162 | 6.5817 | 6.2032 | 6.5554 | 6.1693 | 6.0033 | 5.7186 | 5.9744 | 5.5680 | 6.0850 | 7.3094 | 6.8957 | 7.0476 | 7.1842 |
| BCLAF1_S661 | BCLAF1 | Q9NYF8 | 525;531 | NA | 6.7895 | 6.7017 | 6.2422 | 6.7451 | 6.5571 | 6.0567 | 6.0853 | 6.1463 | 6.4492 | 6.1405 | 5.9433 | 5.7453 | 6.1073 | 6.4255 | 6.5968 | 6.1683 |
| BCLAF1_S690 | BCLAF1 | Q9NYF8 | 259;268 | NA | 5.5069 | 6.0964 | 5.5957 | 6.1498 | 4.4630 | 5.0758 | 6.1051 | 4.5373 | 6.4190 | 5.0085 | 5.1863 | 5.7629 | 8.5313 | 7.8847 | 8.4424 | 8.3349 |
| BCLAF1_S760 | BCLAF1 | Q9NYF8 | 383;389 | NA | 5.1613 | 5.5809 | 5.8063 | 6.0241 | 6.1370 | 5.3194 | 6.3673 | 5.1870 | 5.7498 | 5.7732 | 5.9676 | 5.4877 | 8.4807 | 7.4314 | 7.2510 | 8.2754 |
| BCLAF1_Y383;S389 | BCLAF1 | Q9NYF8 | 658;661 | NA | 8.4479 | 6.8584 | 7.1237 | 7.9993 | 7.4636 | 5.0656 | 6.4844 | 5.0031 | 5.8906 | 4.4798 | 6.2230 | 6.0706 | 5.6825 | 5.9548 | 5.6990 | 5.5837 |
| BCLAF3_S139 | BCLAF3 | A2AJT9 | 139 | KVQGGHSPDDHRV | 5.7815 | 6.5957 | 6.1070 | 6.2086 | 5.8556 | 6.1844 | 6.4183 | 6.1553 | 5.9685 | 6.0367 | 5.9416 | 6.2156 | 6.2755 | 6.8175 | 6.5872 | 6.8509 |
| BCLAF3_S15;S17 | BCLAF3 | A2AJT9 | 402 | LPDVKPSPINLRK | 6.4492 | 6.2735 | 6.4425 | 7.1310 | 6.7042 | 6.3405 | 6.5846 | 6.2231 | 6.6243 | 6.6767 | 6.3583 | 6.3569 | 5.2412 | 5.3534 | 5.4512 | 5.7894 |
| BCLAF3_S17 | BCLAF3 | A2AJT9 | 17 | WKHRSLSPVPRNA | 6.4913 | 7.6351 | 5.6815 | 6.0666 | 5.3087 | 5.5731 | 5.4940 | 5.0947 | 6.4645 | 6.1693 | 6.3610 | 5.9423 | 6.9763 | 6.9371 | 6.7356 | 7.0688 |
| BCLAF3_S402 | BCLAF3 | A2AJT9 | 15;17 | NA | 6.3933 | 6.5132 | 5.5554 | 6.2935 | 6.0777 | 5.7937 | 6.2721 | 5.9724 | 6.3240 | 5.8957 | 6.0805 | 6.0881 | 6.5817 | 6.9427 | 6.5622 | 6.6538 |
| BCLAF3_S78;S80 | BCLAF3 | A2AJT9 | 78;80 | NA | 6.2801 | 6.8115 | 5.9161 | 6.8227 | 7.1275 | 6.8252 | 5.9395 | 5.8093 | 5.7920 | 6.4595 | 6.5380 | 6.0186 | 5.6955 | 6.6815 | 6.1443 | 6.0388 |
| BCOR_S1047 | BCOR | Q6W2J9 | 1047 | SEMCKFSPADWER | 7.5236 | 7.9675 | 6.8176 | 7.0609 | 6.5306 | 6.5061 | 7.1802 | 6.3081 | 7.5086 | 6.5844 | 6.2766 | 6.9588 | 3.9937 | 4.3885 | 4.1871 | 4.2074 |
| BCOR_S1127 | BCOR | Q6W2J9 | 488 | IPKETLSPPGNGC | 7.1421 | 6.7905 | 7.3438 | 6.9108 | 6.4798 | 6.6537 | 6.4376 | 6.8981 | 7.3730 | 6.8897 | 6.9770 | 6.6182 | 4.2118 | 4.9830 | 4.0890 | 4.2021 |
| BCOR_S367 | BCOR | Q6W2J9 | 423 | DRKDGSSPPLLEK | 6.1535 | 6.4665 | 6.8001 | 7.4377 | 6.2598 | 6.4596 | 6.0266 | 6.2341 | 6.5411 | 6.4352 | 5.9104 | 6.4734 | 5.6274 | 6.0797 | 5.3281 | 5.7667 |
| BCOR_S423 | BCOR | Q6W2J9 | 1127 | ASDMPHSPTLRVD | 6.2441 | 6.7789 | 6.7240 | 6.3181 | 7.1267 | 6.7455 | 6.2218 | 6.3746 | 6.9166 | 5.7634 | 6.0610 | 6.3117 | 6.0165 | 5.6178 | 5.2939 | 5.4857 |
| BCOR_S488 | BCOR | Q6W2J9 | 367 | YARISTSPSVALS | 6.6094 | 6.2248 | 6.7019 | 6.2850 | 5.7694 | 6.6830 | 7.2077 | 6.5689 | 6.9004 | 6.3138 | 7.1942 | 6.5402 | 4.4874 | 6.2204 | 5.5870 | 4.7066 |
| BCORL1_S1450;T1461 | BCORL1 | Q5H9F3-3 | 599 | GTSVTFSPLKSPP | 6.4897 | 6.9325 | 6.8440 | 6.8786 | 6.2046 | 6.1760 | 6.6441 | 6.4383 | 6.8686 | 6.8540 | 6.4004 | 6.6618 | 4.8624 | 5.3748 | 4.9222 | 5.4478 |
| BCORL1_S1657 | BCORL1 | Q5H9F3-3 | 603 | TFSPLKSPPQLER | 7.1760 | 6.4678 | 6.1735 | 6.3255 | 6.4419 | 5.5720 | 5.9973 | 7.8357 | 7.1178 | 7.6158 | 6.7313 | 7.6853 | 4.2962 | 4.6584 | 4.4741 | 5.4313 |
| BCORL1_S530 | BCORL1 | Q5H9F3-3 | 1461 | KPTEPCTPSKSRS | 5.7287 | 7.4364 | 5.9587 | 7.2428 | 6.3047 | 6.4535 | 6.1325 | 5.9253 | 6.4922 | 5.3052 | 6.3093 | 5.3500 | 6.1006 | 5.8926 | 6.9521 | 6.4155 |
| BCORL1_S599 | BCORL1 | Q5H9F3-3 | 530 | NRLPCTSPSGSTT | 6.4677 | 6.1208 | 5.9061 | 6.8102 | 7.1579 | 6.6103 | 6.7125 | 6.3670 | 6.5228 | 6.8639 | 6.6419 | 6.8264 | 5.3659 | 5.8547 | 4.6330 | 5.1387 |
| BCORL1_S603 | BCORL1 | Q5H9F3-3 | 1657 | HNPPGSSDQEGDD | 6.3361 | 7.1365 | 6.4367 | 7.0858 | 5.9162 | 5.7356 | 5.8594 | 6.2288 | 6.7000 | 6.7452 | 6.2089 | 6.3019 | 6.1166 | 5.8306 | 5.6029 | 5.7588 |
| BCORL1_T1461 | BCORL1 | Q5H9F3-3 | 1450;1461 | NA | 8.1783 | 7.6558 | 6.2403 | 8.5832 | 6.8374 | 6.0025 | 6.9440 | 6.0888 | 6.8835 | 5.9897 | 6.4319 | 6.5918 | 4.1763 | 4.1706 | 4.2664 | 4.9596 |
| BCR_S122 | BCR | P11274 | 698 | ITPRRQSMTVKKG | 6.8072 | 5.8388 | 6.5471 | 6.2814 | 6.8680 | 5.9188 | 5.6556 | 6.3632 | 7.4102 | 7.1513 | 7.1196 | 7.0478 | 5.5212 | 6.4854 | 7.4446 | 4.2396 |
| BCR_S698 | BCR | P11274 | 432 | ADGSFGTPPGYGC | 6.3766 | 7.2672 | 7.3736 | 7.0485 | 6.6657 | 6.0475 | 5.8118 | 6.3613 | 6.8033 | 6.4329 | 6.5255 | 7.0147 | 4.4926 | 5.1129 | 5.3189 | 5.3472 |
| BCR_T432 | BCR | P11274 | 122 | RPDGEGSPGKARP | 5.7254 | 5.6526 | 6.0013 | 5.9317 | 5.7785 | 6.7747 | 6.6085 | 5.6778 | 6.7100 | 6.7362 | 6.1405 | 6.8062 | 6.5802 | 6.7068 | 5.9020 | 6.2675 |
| BEGAIN_S246 | BEGAIN | Q9BUH8 | 246 | RRDRRPSVDAPVT | 7.4099 | 5.8927 | 6.1969 | 5.3395 | 7.1004 | 6.4524 | 6.2733 | 5.8821 | 6.0560 | 6.9582 | 6.0468 | 6.0336 | 6.3497 | 5.5239 | 6.7158 | 5.7689 |
| BEND3_S164 | BEND3 | Q5T5X7 | 348 | QREMEDSQPSGQV | 5.9598 | 6.5476 | 6.6904 | 7.4081 | 5.7685 | 6.5334 | 6.1758 | 7.4187 | 6.8403 | 5.7567 | 6.1640 | 4.7617 | 5.5337 | 6.3009 | 6.0141 | 6.1262 |
| BEND3_S348 | BEND3 | Q5T5X7 | 717 | PDFPVPSPYLLSD | 5.1772 | 6.0407 | 6.1239 | 6.4966 | 5.5173 | 6.0709 | 5.8460 | 6.5780 | 6.4465 | 6.5490 | 7.1997 | 6.6618 | 6.4954 | 6.8053 | 6.1746 | 5.8170 |
| BEND3_S717 | BEND3 | Q5T5X7 | 124 | PCNDATTPSYKKP | 5.8585 | 6.4718 | 6.3208 | 6.6978 | 6.9218 | 7.1601 | 7.6244 | 6.7687 | 7.5279 | 5.2062 | 5.5802 | 6.4256 | 5.4137 | 5.5082 | 4.4532 | 6.0609 |
| BEND3_T124 | BEND3 | Q5T5X7 | 164 | KANASNSPSSLRL | 6.1695 | 6.4163 | 5.8132 | 6.5755 | 6.1441 | 6.3434 | 6.0556 | 5.9330 | 6.2117 | 6.1363 | 5.7397 | 6.1054 | 6.4879 | 6.8591 | 6.2815 | 6.7278 |
| BEND3_Y499 | BEND3 | Q5T5X7 | 499 | PPRDDCYDSSSLP | 6.1936 | 5.9684 | 6.6924 | 7.4002 | 5.9994 | 5.8566 | 5.9311 | 6.1225 | 7.3235 | 6.6319 | 7.3559 | 6.7579 | 5.8115 | 5.7067 | 5.2803 | 4.9680 |
| BET1_S48 | BET1 | O15155 | 50 | TAIKSLSIEIGHE | 8.0238 | 6.7250 | 6.8102 | 7.4822 | 6.8461 | 5.3416 | 4.5403 | 5.4657 | 8.1740 | 8.5949 | 6.5517 | 8.0119 | 3.5320 | 5.1594 | 4.4409 | 4.3003 |
| BET1_S50 | BET1 | O15155 | 48 | KVTAIKSLSIEIG | 7.0809 | 6.4308 | 6.4056 | 6.7101 | 7.0290 | 5.8353 | 5.5185 | 5.8335 | 7.6821 | 7.5961 | 6.6476 | 6.6203 | 4.7669 | 5.8867 | 5.1309 | 4.8258 |
| BET1L_S9 | BET1L | Q9NYM9 | 9 | DWARAQSPGAVEE | 6.1617 | 5.9050 | 6.1384 | 5.7661 | 6.2914 | 5.9023 | 5.9407 | 6.7851 | 5.8645 | 6.4931 | 5.9361 | 6.0514 | 6.2668 | 7.0519 | 6.5912 | 6.8543 |
| BICC1_S836 | BICC1 | Q9H694 | 836 | FAASIGSPKRKQN | 6.5086 | 5.7320 | 6.8084 | 6.7521 | 6.0228 | 6.8604 | 6.1987 | 6.4512 | 6.8030 | 7.1841 | 6.9663 | 5.2797 | 5.9103 | 4.7244 | 4.9165 |  |
| BICD1_S610 | BICD1 | Q96G01 | 610 | ITAPPSSPVLDTS | 5.7329 | 5.7969 | 5.9748 | 6.0970 | 5.9524 | 6.0443 | 5.7258 | 6.4540 | 6.3503 | 6.8644 | 6.1150 | 6.4665 | 6.8840 | 6.7517 | 6.4367 | 6.3530 |
| BICD2_S329 | BICD2 | Q8TD16 | 582 | EARGRRSPILLPK | 6.5047 | 6.3771 | 6.3985 | 6.5771 | 6.7819 | 6.4741 | 6.1228 | 6.6862 | 6.8626 | 6.2663 | 6.4622 | 6.3209 | 6.0582 | 5.8472 | 5.6335 | 5.5267 |
| BICD2_S582 | BICD2 | Q8TD16 | 329 | EGLAPPSPSLVSD | 5.9740 | 6.2052 | 6.3669 | 6.6524 | 6.4839 | 6.9728 | 6.6249 | 7.3081 | 7.0229 | 6.5371 | 7.0756 | 7.1434 | 4.7693 | 5.1249 | 4.4971 | 5.2416 |
| BICD2_S608 | BICD2 | Q8TD16 | 823 | TKPATPSLxxxxx | 6.0361 | 6.0361 | 6.1031 | 6.1777 | 6.7857 | 6.7277 | 5.6593 | 6.5373 | 6.4516 | 6.5030 | 6.6285 | 6.6918 | 5.8890 | 6.3891 | 5.8811 | 5.5030 |
| BICD2_S823 | BICD2 | Q8TD16 | 821 | PKTKPATPSLxxx | 6.7309 | 7.3788 | 7.4891 | 7.0132 | 5.8866 | 6.9116 | 7.1009 | 7.2803 | 6.8571 | 7.3810 | 7.5929 | 7.4511 | 3.5997 | 4.2399 | 2.7755 | 4.3113 |
| BICD2_T821 | BICD2 | Q8TD16 | 608 | TGDSSPSPGSSLP | 6.8860 | 6.3862 | 5.3172 | 5.9775 | 7.0962 | 6.2422 | 6.5848 | 7.2401 | 6.7619 | 7.4387 | 6.8562 | 5.5290 | 6.3647 | 5.7160 | 4.2084 | 5.3949 |
| BICRA_S1413 | BICRA | Q9NZM4 | 1413 | GRARGGSPAPLPA | 6.7758 | 6.6197 | 5.9895 | 6.3241 | 6.1662 | 6.3394 | 5.9222 | 6.4916 | 6.2593 | 6.3179 | 6.0061 | 6.4369 | 6.4080 | 5.8946 | 5.9808 | 6.0678 |
| BICRAL_S623 | BICRAL | Q6AI39 | 623 | KCLNQTSPISAPK | 6.7760 | 6.5711 | 6.5987 | 6.8484 | 6.1916 | 6.9176 | 7.0006 | 6.1563 | 6.8751 | 6.2390 | 6.8627 | 7.1526 | 5.3186 | 5.0755 | 4.4871 | 4.9290 |
| BICRAL_S675 | BICRAL | Q6AI39 | 675 | EKVVGSSPGHPAV | 6.1367 | 6.2916 | 5.5399 | 6.9515 | 6.4635 | 5.9436 | 5.8098 | 7.0053 | 5.2743 | 5.6142 | 6.1935 | 5.6542 | 7.9277 | 6.4541 | 6.6147 | 6.1254 |
| BIN1_S296 | BIN1 | O00499 | 296 | PAKGNKSPSPPDG | 5.2022 | 5.4579 | 6.9447 | 6.2908 | 5.7008 | 5.9246 | 5.7711 | 6.6570 | 6.4397 | 7.0543 | 5.9054 | 6.2716 | 5.5364 | 7.2114 | 6.1575 | 7.4745 |
| BIRC6_S449 | BIRC6 | Q9NR09 | 453 | VDSRRPTLAWLED | 7.8335 | 4.7077 | 6.2748 | 6.5388 | 10.1098 | 4.9335 | 5.4054 | 4.8869 | 7.7861 | 5.2636 | 6.4735 | 5.7326 | 7.1362 | 7.7229 | 4.9526 | 4.2421 |
| BIRC6_S462 | BIRC6 | Q9NR09 | 449 | PSSGVDSRRPTLA | 5.5965 | 5.3521 | 7.2186 | 5.2122 | 6.8633 | 5.7122 | 6.4164 | 6.7347 | 5.3746 | 5.7846 | 7.1768 | 7.7621 | 6.1881 | 7.1592 | 4.8264 | 6.6222 |
| BIRC6_T453 | BIRC6 | Q9NR09 | 462 | WLEDSSSCSDIPK | 5.6427 | 5.2034 | 6.5470 | 5.2023 | 5.9014 | 5.7058 | 6.0959 | 6.5427 | 5.5540 | 5.5487 | 5.1029 | 4.8892 | 8.2840 | 8.6282 | 7.9462 | 7.2055 |
| BLM_S144 | BLM | P54132 | 144 | KLEFSSSPDSLST | 6.3339 | 5.9904 | 6.1621 | 6.8434 | 6.0366 | 5.3778 | 5.7827 | 5.9157 | 6.4865 | 6.2415 | 6.4284 | 6.2070 | 6.1981 | 6.4740 | 6.9092 | 6.6127 |
| BLM_S168 | BLM | P54132 | 114 | LPDFLQTPKEVVC | 6.6329 | 6.7433 | 6.4619 | 6.8961 | 4.8853 | 6.9051 | 7.3527 | 6.0444 | 6.7685 | 6.3518 | 6.7492 | 7.2135 | 4.7372 | 5.8516 | 4.9168 | 5.4898 |
| BLM_S304 | BLM | P54132 | 766 | IKLLYVTPEKICA | 5.2817 | 5.8020 | 6.1231 | 5.4799 | 5.8271 | 6.6416 | 5.5474 | 5.7620 | 6.4340 | 6.5903 | 6.1748 | 6.2447 | 6.2644 | 7.4447 | 7.2511 | 7.1313 |
| BLM_S422 | BLM | P54132 | 171 | TSKSFVTPPQSHF | 6.7683 | 6.5981 | 6.5744 | 7.6838 | 7.0375 | 6.8891 | 6.4190 | 7.2038 | 7.5334 | 7.4552 | 7.0266 | 6.3853 | 3.6548 | 4.9751 | 3.8409 | 3.9547 |
| BLM_S462 | BLM | P54132 | 125 | VCTTQNTPTVKKS | 7.3137 | 6.8302 | 7.1193 | 8.0644 | 5.6731 | 7.3056 | 7.3356 | 6.6438 | 6.6636 | 6.9651 | 7.2756 | 8.1345 | 3.4113 | 4.7034 | 2.5112 | 4.0496 |
| BLM_S480 | BLM | P54132 | 422 | FNKSDASLLGSLW | 6.1233 | 6.7993 | 6.3750 | 7.3845 | 7.1963 | 6.3039 | 7.0300 | 6.1043 | 6.6356 | 5.8572 | 5.4944 | 5.8454 | 5.5415 | 5.7806 | 5.5276 | 6.0012 |
| BLM_T114 | BLM | P54132 | 168 | TSETSKSFVTPPQ | 7.1906 | 6.5529 | 6.8705 | 8.0323 | 5.8264 | 7.5009 | 7.5230 | 6.1407 | 7.3609 | 6.7496 | 7.2079 | 7.4728 | 3.6615 | 4.7221 | 3.4196 | 4.7402 |
| BLM_T125 | BLM | P54132 | 480 | LGKTGFSATRKNL | 7.2321 | 7.3169 | 6.8143 | 7.4020 | 5.4882 | 7.2372 | 7.7312 | 6.7401 | 7.1333 | 8.0147 | 7.2407 | 8.0487 | 2.7304 | 4.1526 | 3.0260 | 3.6914 |
| BLM_T171 | BLM | P54132 | 304 | TDFVPPSPEEIIS | 6.3275 | 6.2432 | 6.1167 | 6.4195 | 6.9128 | 6.0137 | 5.6985 | 5.2474 | 6.8690 | 6.4445 | 5.4113 | 6.0231 | 6.6771 | 6.3415 | 6.8831 | 6.3712 |
| BLM_T766 | BLM | P54132 | 462 | SHLPSNSVSPGDC | 4.6213 | 6.9678 | 5.4372 | 5.5905 | 5.3034 | 5.4111 | 4.6669 | 6.0148 | 4.1415 | 5.0684 | 4.4483 | 4.9444 | 10.6361 | 10.4076 | 7.9371 | 8.4038 |
| BLOC1S1_S22 | BLOC1S1 | P78537 | 22 | RGPGVPSPQPDVT | 5.8389 | 6.3914 | 8.1488 | 6.7414 | 6.2914 | 5.7818 | 5.7913 | 7.4487 | 6.5033 | 6.3533 | 7.9392 | 5.9342 | 5.3865 | 4.0863 | 5.2841 | 6.0794 |
| BLZF1_S15 | BLZF1 | Q9H2G9 | 15 | KVTVTSSPIRGAG | 7.0634 | 6.9439 | 6.5526 | 7.4412 | 6.9301 | 6.4199 | 6.8883 | 7.2613 | 6.5416 | 6.2657 | 6.4326 | 6.0620 | 4.8257 | 4.8816 | 4.6368 | 4.8533 |
| BLZF1_S362 | BLZF1 | Q9H2G9 | 362 | NPFFESSPTTLLA | 6.5714 | 6.7828 | 6.6662 | 7.3825 | 6.3710 | 6.3761 | 6.6392 | 6.6438 | 6.8334 | 6.7403 | 7.0482 | 7.2343 | 4.2663 | 4.5305 | 4.5799 | 5.3341 |
| BMI1_S316 | BMI1 | P35226 | 316 | NRPRKSSVNGSSA | 4.7265 | 5.1188 | 5.4876 | 6.0948 | 7.3895 | 6.7204 | 5.7942 | 6.0210 | 6.1412 | 6.6627 | 7.0055 | 5.7718 | 7.5397 | 7.2392 | 6.5718 | 5.7152 |
| BMP2K_S1076 | BMP2K | Q9NSY1 | 826 | YRDRSGSGPTQDL | 5.5356 | 5.6688 | 5.9688 | 5.9488 | 5.7942 | 5.8789 | 5.8462 | 6.2541 | 6.0597 | 6.1634 | 6.4132 | 6.0965 | 7.3144 | 7.3004 | 7.2819 | 6.4752 |
| BMP2K_S689 | BMP2K | Q9NSY1 | 689 | FISHSGSPEKKAE | 6.5256 | 6.2938 | 6.4060 | 6.9164 | 6.5892 | 6.6807 | 6.4864 | 6.5568 | 6.4802 | 6.2671 | 6.4823 | 6.3002 | 5.3343 | 6.4483 | 5.6374 | 5.4953 |
| BMP2K_S826 | BMP2K | Q9NSY1 | 1076 | GAKPFHSPDLSWH | 6.6606 | 7.8693 | 7.2476 | 7.0742 | 6.2543 | 7.3417 | 7.5300 | 7.6736 | 5.8902 | 6.5574 | 6.8017 | 7.1411 | 3.7898 | 5.6648 | 3.4529 | 3.9508 |
| BMP2K_S949 | BMP2K | Q9NSY1 | 1014 | LKPTYRTPERARR | 5.6043 | 6.1043 | 6.5909 | 6.4611 | 6.0106 | 6.4393 | 6.4195 | 6.0968 | 6.2407 | 6.5521 | 6.6903 | 6.7335 | 5.4205 | 6.7197 | 5.8589 | 6.0575 |
| BMP2K_T1014 | BMP2K | Q9NSY1 | 949 | STSKSESNEDLFG | 5.1585 | 6.1489 | 6.6282 | 5.3837 | 5.2315 | 6.8971 | 6.1939 | 5.7985 | 6.6891 | 6.0369 | 6.7462 | 5.5722 | 5.4542 | 6.9299 | 7.4957 | 6.6355 |
| BMPR2_S940 | BMPR2 | Q13873 | 940 | RAQRPNSLDLSAT | 6.4895 | 6.8690 | 7.0106 | 7.3505 | 6.5899 | 6.2516 | 6.6999 | 6.4269 | 6.8239 | 6.5961 | 6.8850 | 6.8724 | 4.7486 | 5.4214 | 4.1336 | 4.8309 |
| BMS1_S552 | BMS1 | Q14692 | 639 | NFIDETSDIENLL | 6.1501 | 6.2506 | 6.0458 | 6.7578 | 6.0216 | 6.2097 | 6.6206 | 6.0918 | 6.1900 | 6.5587 | 6.5327 | 6.0880 | 6.3466 | 6.6564 | 6.4251 | 6.6544 |
| BMS1_S625 | BMS1 | Q14692 | 552 | GSKAGLSPANCQS | 6.8646 | 6.8950 | 6.1772 | 7.0331 | 6.3337 | 6.3813 | 7.0996 | 6.4691 | 6.9029 | 6.5025 | 6.2701 | 6.2777 | 5.2037 | 5.2324 | 4.8640 | 5.4929 |
| BMS1_S639 | BMS1 | Q14692 | 708 | EEEDDDTLEELGG | 6.2351 | 6.4020 | 6.0345 | 7.0567 | 6.1515 | 6.1348 | 6.5199 | 6.1031 | 6.0950 | 6.8788 | 5.5673 | 6.1928 | 5.8697 | 6.6267 | 5.6359 | 6.4962 |
| BMS1_T708 | BMS1 | Q14692 | 625 | KPSQVSSGQKLGP | 7.2949 | 6.8320 | 6.7253 | 7.8075 | 6.8056 | 6.7644 | 7.6602 | 6.5308 | 7.7479 | 7.1371 | 7.6710 | 7.3258 | 3.1844 | 3.8302 | 2.5928 | 4.0902 |
| BNC1_S338 | BNC1 | Q01954 | 338 | FERTQLSPEAKVK | 6.5466 | 6.5452 | 6.5757 | 6.9840 | 6.1382 | 6.0279 | 6.8801 | 6.1445 | 6.8101 | 6.3521 | 6.4536 | 6.6231 | 5.6225 | 5.5775 | 5.2801 | 5.4389 |
| BNC2_S403 | BNC2 | Q6ZN30 | 403 | TEPACVSPIQNSA | 5.8907 | 6.4100 | 6.3407 | 7.0191 | 6.0464 | 6.1358 | 6.0629 | 6.0392 | 6.4726 | 6.1582 | 6.1291 | 5.9528 | 6.4576 | 6.2492 | 6.6063 |  |
| BNC2_T391 | BNC2 | Q6ZN30 | 391 | NALTSITNVEPKT | 6.3726 | 6.6853 | 6.5402 | 7.2039 | 5.7256 | 6.4813 | 6.3011 | 6.4552 | 7.2150 | 6.9713 | 6.5166 | 6.9859 | 5.8771 | 5.8801 | 4.5226 | 4.2663 |
| BNIP2_S106 | BNIP2 | J3KN59 | 235 | EVIRKGSITEYTA | 6.3452 | 6.0815 | 6.3774 | 6.5798 | 6.7689 | 6.0308 | 6.0424 | 6.3191 | 6.8182 | 6.6921 | 6.4456 | 6.6480 | 6.1276 | 6.3731 | 5.6269 | 5.6234 |
| BNIP2_S235 | BNIP2 | J3KN59 | 106 | SSGDIGSLDYQEF | 6.1753 | 6.1642 | 7.2311 | 6.2396 | 6.4730 | 5.7903 | 6.5418 | 6.1871 | 5.9418 | 6.0386 | 5.8820 | 6.6223 | 6.4427 | 6.3053 | 5.6433 | 6.3216 |
| BNIP3L_S166 | BNIP3L | O60238 | 166 | HPKRSVSLSMRKS | 8.3253 | 7.3026 | 7.2721 | 7.0951 | 5.5490 | 4.5715 | 4.4161 | 5.6489 | 7.1242 | 8.4079 | 6.9320 | 7.4259 | 4.8053 | 5.4462 | 4.9599 | 4.7179 |
| BOD1L1_S1017 | BOD1L1 | Q8NFC6 | 1531 | DKDVTLSPVKAGP | 6.5117 | 6.5106 | 5.9543 | 6.4121 | 6.4509 | 6.0135 | 6.2814 | 6.1704 | 6.2110 | 6.0038 | 5.8094 | 5.7250 | 6.0050 | 6.7593 | 6.6456 | 6.5361 |
| BOD1L1_S105 | BOD1L1 | Q8NFC6 | 2203 | APPEAESPLASTS | 5.9426 | 5.9189 | 6.0192 | 6.2669 | 5.9022 | 6.0084 | 6.2127 | 5.8519 | 6.3483 | 6.2193 | 5.6446 | 5.9668 | 6.3333 | 7.2764 | 6.9361 | 7.1524 |
| BOD1L1_S1077 | BOD1L1 | Q8NFC6 | 3019 | RSKTQLSPSIKRK | 6.1107 | 6.2578 | 6.1940 | 6.5549 | 6.4336 | 6.1252 | 6.4268 | 6.1903 | 6.3134 | 6.3188 | 6.0864 | 6.1182 | 5.7547 | 6.5253 | 6.1720 | 6.4180 |
| BOD1L1_S1318 | BOD1L1 | Q8NFC6 | 3029 | KRKREVSPPGART | 6.2900 | 6.6768 | 6.1305 | 6.6081 | 5.8727 | 5.9068 | 5.9854 | 5.8829 | 6.1758 | 6.0296 | 5.9968 | 6.2365 | 6.1431 | 6.8520 | 6.4637 | 6.7492 |
| BOD1L1_S143 | BOD1L1 | Q8NFC6 | 1077 | CENRRGSLSQEMA | 5.0262 | 4.2835 | 4.6259 | 4.8482 | 6.2684 | 5.3973 | 5.1425 | 5.3072 | 8.2639 | 7.7864 | 7.3953 | 7.8124 | 7.1249 | 9.0542 | 5.1019 | 6.5619 |
| BOD1L1_S1531 | BOD1L1 | H0Y9Y2 | 164 | SEKVSNSPPLGRS | 6.5869 | 7.0566 | 6.5194 | 7.2315 | 6.0725 | 6.2354 | 6.3923 | 6.3652 | 6.5257 | 6.7664 | 6.6379 | 6.6720 | 4.6358 | 5.7116 | 5.0669 | 5.5238 |
| BOD1L1_S164 | BOD1L1 | H0Y9Y2 | 143 | GSPDTVSPNRQK | 6.7216 | 7.3675 | 6.8381 | 7.5217 | 6.7347 | 6.4067 | 6.6727 | 6.2061 | 6.9188 | 6.4499 | 6.5260 | 6.1599 | 4.2076 | 5.1272 | 4.7450 | 5.1591 |
| BOD1L1_S2025 | BOD1L1 | Q8NFC6 | 2964 | SDDAESSEPERKR | 6.2025 | 6.3206 | 6.2299 | 6.7416 | 6.1054 | 6.2986 | 6.3284 | 6.0517 | 5.9535 | 5.9837 | 5.6133 | 6.1599 | 6.3623 | 6.8063 | 6.2783 | 6.5641 |
| BOD1L1_S2203 | BOD1L1 | Q8NFC6 | 659 | EHKRRTSTPVIME | 5.0548 | 5.3660 | 5.1542 | 5.6116 | 6.4729 | 5.6720 | 5.6832 | 6.0260 | 5.7306 | 5.7566 | 5.3700 | 6.2941 | 7.5488 | 7.8600 | 8.3062 | 8.0929 |
| BOD1L1_S2475 | BOD1L1 | Q8NFC6 | 1017 | RLERKLSDGHKSR | 5.7249 | 6.1462 | 5.8814 | 5.5493 | 6.3973 | 5.8921 | 5.8933 | 6.3204 | 5.4851 | 6.1286 | 6.0697 | 5.9187 | 6.9206 | 7.4053 | 7.2996 | 6.9677 |
| BOD1L1_S2501 | BOD1L1 | Q8NFC6 | 2501 | LEGNANSPAHLRG | 6.5204 | 7.2689 | 6.4386 | 7.4735 | 6.4908 | 6.6476 | 6.4447 | 6.2186 | 7.0419 | 6.8853 | 7.1876 | 6.6600 | 4.1105 | 5.3435 | 4.3314 | 4.9366 |
| BOD1L1_S2964 | BOD1L1 | Q8NFC6 | 1318 | GSTASTSPADHSA | 5.9724 | 6.3187 | 6.1452 | 7.4080 | 6.2830 | 6.7519 | 6.5524 | 7.3220 | 6.2548 | 6.4261 | 6.3157 | 7.0328 | 6.4243 | 6.3842 | 6.4289 | 5.5796 |
| BOD1L1_S2973 | BOD1L1 | Q8NFC6 | 2986 | DKKEQESDEEEEE | 6.2270 | 6.4598 | 6.0300 | 6.3928 | 6.2751 | 5.4449 | 6.5102 | 5.9989 | 5.3906 | 6.3512 | 6.1410 | 5.7484 | 6.6422 | 6.9348 | 6.6747 | 6.7784 |

| | | | | | | | | | | | | | | | | | | | | |
|---|---|---|---|---|---|---|---|---|---|---|---|---|---|---|---|---|---|---|---|---|
| BOD1L1_S2986 | BOD1L1 | Q8NFC6 | 1354 | EGFTVDTPAKASI | 8.1192 | 8.1246 | 6.7728 | 8.0315 | 6.8597 | 7.0300 | 7.0903 | 6.8504 | 7.2727 | 5.6657 | 6.9105 | 6.5194 | 3.2485 | 4.6179 | 2.9773 | 3.9094 |
| BOD1L1_S3019 | BOD1L1 | Q8NFC6 | 2475 | LQKEDKSPETGTA | 5.9920 | 6.7213 | 5.9878 | 6.3344 | 5.3560 | 6.9468 | 6.8366 | 6.6196 | 6.6818 | 6.6877 | 6.2032 | 6.2301 | 5.9626 | 6.0862 | 4.8791 | 6.4749 |
| BOD1L1_S3029 | BOD1L1 | Q8NFC6 | 105 | LATHTWSPHLNKN | 6.8320 | 6.8371 | 6.2932 | 7.3050 | 5.2974 | 6.7956 | 7.1845 | 6.2104 | 6.6943 | 6.8683 | 6.7994 | 7.7360 | 4.2435 | 4.6828 | 5.1654 | 5.0551 |
| BOD1L1_S538 | BOD1L1 | Q8NFC6 | 635 | KPARRLSESLHVV | 4.1933 | 4.1285 | 3.9339 | 4.3408 | 6.5166 | 5.5223 | 4.8944 | 6.0313 | 5.4634 | 5.4014 | 5.1396 | 5.0499 | 8.8913 | 10.4182 | 10.1079 | 9.9670 |
| BOD1L1_S635 | BOD1L1 | Q8NFC6 | 2973 | ERKRQKSVSDPVE | 6.3125 | 6.5852 | 7.2719 | 7.8467 | 6.8526 | 6.8042 | 7.1761 | 6.6140 | 6.5787 | 6.4487 | 7.6006 | 7.3179 | 4.0605 | 4.9020 | 3.7675 | 3.8609 |
| BOD1L1_S659 | BOD1L1 | Q8NFC6 | 938 | ATKQATTPKPDKE | 6.2526 | 6.8293 | 7.2818 | 7.4516 | 6.0938 | 6.5878 | 7.2901 | 5.9013 | 7.3614 | 7.0182 | 7.0430 | 7.3143 | 4.2141 | 4.8337 | 3.8759 | 4.6510 |
| BOD1L1_T1354 | BOD1L1 | Q8NFC6 | 2025 | CEVAHTSPSEKED | 6.1675 | 5.8843 | 5.9819 | 7.5369 | 6.2053 | 5.8355 | 6.7082 | 5.4644 | 6.6239 | 5.8318 | 4.8847 | 5.0555 | 6.4055 | 6.5519 | 7.7884 | 7.0743 |
| BOD1L1_T938 | BOD1L1 | Q8NFC6 | 538 | KGQGRSSVDLEES | 4.1179 | 4.8825 | 4.6300 | 5.6898 | 5.1511 | 5.9113 | 5.2030 | 5.8969 | 4.7234 | 5.3920 | 4.7925 | 5.4980 | 8.1081 | 9.5297 | 10.4552 | 10.0187 |
| BOK_S8 | BOK | Q9UMX3 | 8 | EVLRRSSVFAAEI | 5.4102 | 5.0501 | 5.8880 | 5.7065 | 6.3247 | 5.6252 | 5.6418 | 6.0544 | 7.2300 | 7.1604 | 7.1398 | 7.1196 | 6.5839 | 7.1420 | 5.9444 | 5.9789 |
| BOP1_S126;S127 | BOP1 | Q14137 | 323 | PPEYLLSEEERLA | 6.2399 | 6.8087 | 6.5793 | 7.4072 | 6.6470 | 6.0580 | 6.9858 | 7.3237 | 5.3750 | 5.8143 | 6.0152 | 6.5391 | 5.4966 | 5.5246 | 5.9380 | 5.2476 |
| BOP1_S323 | BOP1 | Q14137 | 106 | EQVQASTPCPRTE | 7.0794 | 7.2726 | 6.5834 | 8.3687 | 6.8791 | 5.6267 | 7.5618 | 7.3783 | 7.9056 | 5.8687 | 6.3342 | 7.0089 | 4.1515 | 4.2485 | 3.5369 | 4.1956 |
| BOP1_T106 | BOP1 | Q14137 | 126;127 | NA | 6.1936 | 6.1263 | 5.9211 | 6.9588 | 5.4559 | 5.4306 | 5.5604 | 5.5060 | 6.4207 | 6.0703 | 5.5559 | 5.9810 | 5.0864 | 8.6126 | 6.4477 | 8.6728 |
| BORA_S325 | BORA | Q6PGQ7 | 325 | PYIDGCSPIKNWS | 6.8385 | 6.3793 | 7.1404 | 6.5733 | 6.8406 | 6.4697 | 5.8465 | 6.6073 | 6.9556 | 7.0822 | 7.1697 | 7.1995 | 5.0960 | 4.7552 | 4.3828 | 4.6634 |
| BORA_S558 | BORA | Q6PGQ7 | 558 | SPFQCSSPxxxxx | 6.3376 | 6.8203 | 7.2862 | 6.7008 | 5.9861 | 5.9187 | 6.2086 | 6.4864 | 6.3299 | 6.2258 | 6.7615 | 7.4224 | 5.2794 | 5.4753 | 5.1188 | 5.6422 |
| BORCS6_T196 | BORCS6 | Q96GS4 | 196 | GGGRRATISSPLE | 6.3665 | 4.8040 | 5.8368 | 5.5922 | 7.5350 | 5.1385 | 5.2336 | 5.6406 | 6.9648 | 6.4395 | 6.7926 | 6.7949 | 7.4691 | 8.0365 | 5.8309 | 5.5246 |
| BPIFB2_S371 | BPIFB2 | Q8N4F0 | 371 | RLQLSVSKVKLQG | 6.5579 | 6.6415 | 6.5838 | 5.9622 | 6.4953 | 6.3564 | 6.0503 | 6.5131 | 6.6704 | 6.6916 | 6.7280 | 6.6743 | 5.8307 | 5.5193 | 5.3739 | 5.3514 |
| BPTF_S1231 | BPTF | Q12830 | 1673 | VKKVTSSPITSEE | 5.6948 | 6.2244 | 5.8758 | 6.4950 | 5.8935 | 5.7991 | 6.0268 | 5.8415 | 6.3330 | 6.0553 | 6.0410 | 6.2153 | 6.6805 | 7.0606 | 6.9721 | 6.7912 |
| BPTF_S1300 | BPTF | Q12830 | 216 | RVHRPRSPILEEK | 6.5288 | 6.5609 | 5.9837 | 6.5882 | 6.2846 | 5.8283 | 6.1903 | 5.8456 | 6.3226 | 6.2031 | 6.0681 | 6.1053 | 6.2806 | 6.4835 | 6.3824 | 6.3440 |
| BPTF_S1673 | BPTF | Q12830 | 2370 | QRQSKLSPQMQVH | 6.1822 | 6.6193 | 6.4996 | 6.7726 | 6.2806 | 6.0126 | 6.4668 | 6.1837 | 6.3276 | 6.3598 | 6.0886 | 6.2874 | 5.7249 | 6.1899 | 6.0696 | 5.9347 |
| BPTF_S1684 | BPTF | Q12830 | 2465 | SNVQGQSPVRVQS | 5.7062 | 5.2556 | 6.7389 | 6.6169 | 5.9814 | 5.7858 | 5.9110 | 5.8966 | 6.3142 | 6.5496 | 5.8033 | 6.7085 | 6.0997 | 6.9997 | 7.1050 | 6.5276 |
| BPTF_S216 | BPTF | Q12830 | 763 | NGELSESPGAGKG | 6.8798 | 7.5053 | 6.3758 | 7.4752 | 6.1811 | 6.1438 | 6.2003 | 6.3544 | 6.5508 | 6.3262 | 6.7832 | 6.9979 | 4.7029 | 5.2369 | 5.1394 | 5.1469 |
| BPTF_S2370 | BPTF | Q12830 | 739 | GERESHTPVSIQE | 5.4839 | 5.7727 | 5.6901 | 6.1257 | 5.7210 | 5.5438 | 5.7640 | 6.1440 | 5.7876 | 6.0131 | 5.9486 | 5.5041 | 7.2850 | 7.8146 | 7.9973 | 7.4046 |
| BPTF_S2465 | BPTF | Q12830 | 572 | DIDNVKSPEETEK | 6.0277 | 6.1363 | 5.8266 | 6.7509 | 6.3841 | 6.4204 | 6.3860 | 6.1151 | 6.3277 | 6.3123 | 7.1642 | 6.8421 | 6.2260 | 6.2842 | 5.4977 | 5.2990 |
| BPTF_S2465;S2471 | BPTF | Q12830 | 1231 | SKNLSESPVITKA | 6.5355 | 7.1210 | 6.8865 | 7.8354 | 6.1481 | 6.7912 | 6.8499 | 6.6317 | 7.2515 | 7.1283 | 7.3976 | 7.2598 | 3.6826 | 4.5068 | 3.5070 | 4.4671 |
| BPTF_S2471 | BPTF | Q12830 | 2471 | SPVRVQSPSQTRI | 5.5469 | 6.2453 | 6.2270 | 6.9488 | 5.9282 | 6.1788 | 6.2814 | 6.2840 | 6.4513 | 6.6849 | 6.4430 | 6.8374 | 5.2832 | 6.6210 | 5.9286 | 6.1102 |
| BPTF_S572 | BPTF | Q12830 | 1684 | EEESNLSNDFIDE | 6.4529 | 5.7837 | 5.9049 | 6.7399 | 6.7476 | 5.4598 | 6.1382 | 6.5296 | 6.3301 | 6.5435 | 6.7844 | 6.9118 | 4.4744 | 6.3792 | 5.9065 | 6.9134 |
| BPTF_S763 | BPTF | Q12830 | 1300 | DDVSIRSPETKCP | 6.6966 | 7.1518 | 6.7164 | 7.7502 | 6.2489 | 6.3440 | 6.9541 | 6.7401 | 7.2976 | 7.1465 | 7.0923 | 7.4314 | 3.7108 | 4.4396 | 3.9394 | 4.3402 |
| BPTF_T739 | BPTF | Q12830 | 2465;2471 | NA | 6.8823 | 6.0701 | 7.1900 | 6.6774 | 6.2473 | 6.9098 | 6.5992 | 4.8517 | 6.2399 | 6.1614 | 7.7101 | 7.2166 | 5.4036 | 5.7975 | 4.8918 | 5.1514 |
| BPTF_T739;S742 | BPTF | Q12830 | 739;742 | NA | 7.9904 | 5.2542 | 4.7103 | 8.0752 | 5.8416 | 5.1394 | 6.2595 | 7.6460 | 6.1921 | 6.9302 | 6.3214 | 6.6630 | 4.8794 | 7.2494 | 6.2825 | 4.5652 |
| BRAF_S335 | BRAF | P15056 | 365 | QRDRSSSAPNVHI | 5.8891 | 5.7953 | 6.6889 | 5.6785 | 6.3446 | 6.7240 | 5.9537 | 6.5953 | 6.0165 | 6.2239 | 6.2562 | 6.1589 | 6.5272 | 6.7342 | 6.1350 | 6.2787 |
| BRAF_S365 | BRAF | P15056 | 401 | TTGLSATPPASLP | 6.4969 | 6.9719 | 7.5329 | 6.8303 | 6.2878 | 6.7579 | 6.2164 | 6.7204 | 6.8158 | 6.7878 | 6.8876 | 6.9075 | 4.3378 | 5.2531 | 4.2338 | 4.9622 |
| BRAF_S729 | BRAF | P15056 | 335 | QILTSPSPSKSIP | 5.9332 | 6.4099 | 7.2590 | 6.7510 | 6.2088 | 7.3205 | 6.2938 | 6.5795 | 6.4815 | 6.4604 | 6.8792 | 6.9602 | 4.9725 | 5.0360 | 5.0363 | 5.4180 |
| BRAF_S750 | BRAF | P15056 | 750 | SLYACASPKTPIQ | 6.0829 | 6.3935 | 7.1731 | 6.6043 | 5.9385 | 6.2854 | 6.6309 | 6.4404 | 6.5724 | 6.7332 | 6.8150 | 6.7601 | 5.3044 | 5.3623 | 5.4993 | 5.4042 |
| BRAF_T401 | BRAF | P15056 | 729 | KIHRSASEPSLNR | 6.4373 | 6.5663 | 6.9674 | 6.3025 | 5.8637 | 6.6582 | 5.8801 | 6.7648 | 6.3406 | 6.4869 | 6.5857 | 6.7028 | 5.3196 | 6.0552 | 5.6969 | 5.3719 |
| BRAP_S117 | BRAP | Q7Z569 | 117 | INAAPDSPSKQLP | 6.4152 | 6.5830 | 6.7172 | 6.5143 | 6.1472 | 6.2315 | 6.4984 | 6.5024 | 6.6455 | 6.9153 | 6.5519 | 6.8208 | 4.8340 | 5.9462 | 5.2393 | 5.4376 |
| BRAT1_S742 | BRAT1 | Q6PJG6 | 742 | LREARGSPNTASA | 6.2286 | 6.7563 | 6.4718 | 6.4486 | 6.2991 | 6.7486 | 6.3627 | 7.1153 | 6.7253 | 7.3564 | 7.5575 | 7.1592 | 3.9239 | 5.2026 | 4.7004 | 4.9437 |
| BRAT1_S798 | BRAT1 | Q6PJG6 | 798 | SDHVEKSPQSLLQ | 5.4678 | 5.8850 | 6.2502 | 6.4639 | 5.5198 | 7.1372 | 5.7935 | 6.2055 | 6.7779 | 7.3560 | 5.8008 | 5.9216 | 5.8542 | 6.3139 | 6.7559 | 6.4968 |
| BRCA1_S114 | BRCA1 | P38398 | 1524 | QNRNYPSQEELIK | 6.3046 | 6.3468 | 6.1896 | 6.3761 | 6.1407 | 5.8657 | 5.9946 | 5.8782 | 6.3183 | 6.0290 | 5.9057 | 6.0951 | 6.7477 | 6.5731 | 6.6084 | 6.6266 |
| BRCA1_S1173 | BRCA1 | P38398 | 967 | NETGLITPNKHGL | 8.9688 | 8.2643 | 7.1749 | 7.9794 | 4.3810 | 6.6012 | 7.9382 | 5.8423 | 7.6140 | 7.1923 | 8.3420 | 7.8472 | 2.7363 | 3.5228 | 2.2792 | 3.3161 |
| BRCA1_S1174 | BRCA1 | P38398 | 1174 | NDIKESSAVFSKS | 6.4382 | 6.7508 | 6.4967 | 7.0592 | 6.6392 | 6.5457 | 6.9004 | 6.6023 | 6.2487 | 6.1819 | 6.7058 | 6.3651 | 5.1545 | 5.4285 | 5.0855 | 5.3975 |
| BRCA1_S1187;S1189;S11 | BRCA1 | P38398 | 434 | KIDLLASDPHEAL | 5.6651 | 6.1130 | 6.3265 | 6.6075 | 5.7807 | 6.2828 | 6.3116 | 6.0411 | 6.2348 | 6.0322 | 6.0619 | 6.2387 | 6.5515 | 6.4831 | 6.7735 | 6.4959 |
| BRCA1_S1189;S1191 | BRCA1 | P38398 | 803 | EPNKCVSQCAAFE | 6.3575 | 6.1581 | 6.3867 | 6.0241 | 5.2809 | 6.1236 | 6.0873 | 5.4072 | 6.1968 | 5.3264 | 5.4722 | 5.2800 | 7.3575 | 7.3207 | 7.3303 | 7.8908 |
| BRCA1_S1457 | BRCA1 | P38398 | 114 | AKKENNSPEHLKD | 5.3395 | 6.4015 | 6.3231 | 6.7833 | 5.5842 | 6.1891 | 5.8494 | 6.0431 | 6.4651 | 6.6879 | 6.0794 | 6.3150 | 6.4113 | 6.2268 | 6.8511 | 6.4503 |
| BRCA1_S1524 | BRCA1 | P38398 | 1542 | EQQLEESGPHDLT | 5.2910 | 6.1629 | 5.4637 | 6.6952 | 6.7603 | 7.2585 | 6.1261 | 7.2189 | 5.7468 | 5.3391 | 5.4864 | 5.3080 | 6.4036 | 7.4142 | 5.9620 | 7.3633 |
| BRCA1_S1542 | BRCA1 | P38398 | 632 | VVSRNLSPPNCTE | 6.0986 | 6.4077 | 6.6718 | 6.8185 | 5.1429 | 5.9634 | 6.6665 | 6.8227 | 6.5135 | 7.0050 | 7.0469 | 6.5366 | 5.7287 | 6.1489 | 5.9100 | 4.5184 |
| BRCA1_S434 | BRCA1 | P38398 | 1457 | EKAVLTSQKSSEY | 6.8895 | 6.6692 | 5.7961 | 5.9169 | 6.6520 | 5.8559 | 6.2269 | 5.6795 | 6.3601 | 5.4473 | 5.0073 | 5.1014 | 7.1375 | 6.9037 | 7.5214 | 6.8354 |
| BRCA1_S632 | BRCA1 | P38398 | 1173 | ENDIKESSAVFSK | 6.5297 | 6.9188 | 6.5203 | 7.5823 | 6.7425 | 6.3053 | 7.0134 | 6.6536 | 6.4500 | 6.4655 | 6.6006 | 6.4562 | 4.5830 | 5.0674 | 4.6578 | 5.4539 |
| BRCA1_S803 | BRCA1 | P38398 | 1189;1191 | NA | 6.2626 | 6.3969 | 6.3955 | 7.2812 | 6.2586 | 6.5885 | 6.8690 | 7.0448 | 6.5621 | 6.0229 | 6.3058 | 5.9025 | 5.3780 | 5.9156 | 5.4024 | 5.4135 |
| BRCA1_T967 | BRCA1 | P38398 | 1187;1189;11 | NA | 5.9755 | 6.6762 | 5.7211 | 6.8826 | 5.7612 | 6.6511 | 5.2664 | 6.0869 | 6.2697 | 6.2794 | 6.0121 | 6.0563 | 6.6214 | 6.5880 | 6.5686 | 6.5835 |
| BRCA2_S1680 | BRCA2 | P51587 | 93 | TLPLYQSPVKELD | 6.4126 | 6.6674 | 7.0087 | 7.5948 | 6.0555 | 6.5068 | 6.5671 | 6.7966 | 6.9188 | 6.6784 | 6.4799 | 6.7631 | 4.4520 | 5.0042 | 4.7179 | 5.3762 |
| BRCA2_S1770 | BRCA2 | P51587 | 2399 | MRHLITTGRPTKV | 7.2243 | 7.9821 | 6.4742 | 8.0116 | 6.4265 | 6.7396 | 8.1307 | 7.2770 | 6.9376 | 6.8241 | 7.0515 | 6.9414 | 3.1010 | 4.1599 | 2.9537 | 3.7648 |
| BRCA2_S1818 | BRCA2 | P51587 | 3319 | KKKELNSPQMTPF | 6.5776 | 7.0323 | 7.2229 | 7.5805 | 6.5189 | 6.3825 | 6.5504 | 6.9701 | 7.2953 | 7.9354 | 7.4653 | 7.3866 | 3.3503 | 3.9025 | 3.6556 | 4.1738 |
| BRCA2_S1869 | BRCA2 | P51587 | 1770 | SKNKLDSGIEPVL | 6.8629 | 7.5309 | 7.0064 | 6.3328 | 5.7917 | 7.4757 | 5.8635 | 7.9726 | 8.0168 | 6.8684 | 7.4702 | 7.5420 | 2.1754 | 5.8359 | 4.3397 | 2.9152 |
| BRCA2_S2072 | BRCA2 | P51587 | 70 | TPQRKPSYNQLAS | 4.4985 | 3.9821 | 4.6358 | 4.5071 | 7.9227 | 5.0774 | 4.8508 | 5.4345 | 8.5103 | 9.0681 | 8.0639 | 8.3829 | 6.4716 | 7.3437 | 5.5079 | 5.7428 |
| BRCA2_S3241 | BRCA2 | P51587 | 3284 | PLPPPVSPICTFV | 6.6100 | 6.5358 | 7.8349 | 6.9850 | 5.8640 | 6.5965 | 7.0633 | 6.0502 | 7.4626 | 8.0952 | 8.1398 | 7.1995 | 4.0739 | 4.5501 | 3.1728 | 3.7664 |
| BRCA2_S3284 | BRCA2 | P51587 | 884 | DSEELFSDNENNF | 5.9777 | 6.5314 | 5.4984 | 6.5462 | 5.9562 | 6.4553 | 7.3693 | 5.9298 | 6.9346 | 7.6122 | 7.2999 | 7.3313 | 5.2340 | 5.5133 | 5.5431 | 4.2672 |
| BRCA2_S3319 | BRCA2 | P51587 | 77 | YNQLASTPIIFKE | 6.0448 | 6.4123 | 7.3406 | 6.6235 | 5.8260 | 6.5286 | 7.5346 | 5.8653 | 7.1004 | 6.6260 | 6.9397 | 7.6321 | 4.7986 | 5.0063 | 4.3114 | 5.4097 |
| BRCA2_S3319;T3323 | BRCA2 | P51587 | 1869 | KDIFTDSFSKVIK | 9.3072 | 10.6071 | 7.0472 | 8.3966 | 4.4639 | 5.9221 | 6.6282 | 5.7374 | 6.9878 | 8.9343 | 7.8720 | 7.9941 | 2.3458 | 2.9990 | 2.1478 | 2.6097 |
| BRCA2_S492 | BRCA2 | P51587 | 1818 | ELVTSSSPCKNKN | 6.3145 | 6.8617 | 6.2842 | 7.1215 | 5.7912 | 5.6489 | 5.9877 | 7.0434 | 7.0893 | 6.0896 | 7.1554 | 6.5826 | 4.9961 | 5.6911 | 5.8472 | 5.4956 |
| BRCA2_S70 | BRCA2 | P51587 | 2072 | VSILESSLHKVKG | 8.0690 | 7.9515 | 7.3069 | 8.4273 | 7.0970 | 7.2986 | 7.1612 | 7.1580 | 7.8899 | 6.5936 | 7.5740 | 7.0206 | 2.2702 | 3.2423 | 2.2189 | 2.7209 |
| BRCA2_S844 | BRCA2 | P51587 | 844 | KYMRVASPSRKVQ | 7.9902 | 7.2803 | 6.5385 | 6.7932 | 6.6159 | 7.5058 | 6.9928 | 5.2577 | 6.2046 | 6.3383 | 7.0937 | 7.1658 | 4.7365 | 4.7006 | 4.2563 | 4.5298 |
| BRCA2_S884 | BRCA2 | P51587 | 3241 | AKRKSVSTPVSAQ | 8.3406 | 8.0785 | 8.2098 | 8.7055 | 5.0793 | 6.3789 | 7.2679 | 6.5914 | 7.8579 | 8.0963 | 8.0959 | 8.2026 | 1.7997 | 2.7570 | 1.9463 | 2.5925 |
| BRCA2_S93 | BRCA2 | P51587 | 492 | QAISGTSPVASSF | 6.4973 | 7.2797 | 6.6395 | 6.7441 | 6.0165 | 7.5042 | 6.8665 | 5.8392 | 7.0145 | 6.8467 | 7.2274 | 7.4691 | 4.3923 | 4.9375 | 4.0665 | 4.6589 |
| BRCA2_T2399 | BRCA2 | P51587 | 1680 | SCSRKTSVSQTSL | 6.1495 | 5.1326 | 5.6405 | 5.9119 | 6.4468 | 5.6884 | 4.9252 | 5.4466 | 9.1329 | 8.8259 | 7.6307 | 8.1575 | 6.0207 | 6.0423 | 4.3806 | 4.4679 |
| BRCA2_T77 | BRCA2 | P51587 | 3319;3323 | NA | 9.7167 | 9.3203 | 6.7440 | 9.1514 | 5.7836 | 7.0507 | 7.0284 | 6.1337 | 6.9077 | 5.7459 | 6.7075 | 7.0351 | 3.4453 | 3.3286 | 2.5721 | 3.4191 |
| BRD1_S128 | BRD1 | O95696-2 | 128 | VRIVEYSPPSAPR | 6.1180 | 6.6072 | 6.3704 | 7.2357 | 5.9728 | 5.9593 | 6.5378 | 6.2814 | 6.3591 | 6.2079 | 6.1579 | 6.6516 | 5.5735 | 6.1622 | 6.0141 | 5.7912 |
| BRD1_S75 | BRD1 | O95696-2 | 983 | PRRRCASESSISS | 6.0124 | 6.7159 | 6.5217 | 6.8669 | 5.6234 | 5.7463 | 6.3325 | 6.3841 | 6.3966 | 6.2661 | 6.7787 | 6.7621 | 5.9581 | 5.6304 | 5.6679 | 6.3368 |
| BRD1_S983 | BRD1 | O95696-2 | 1016 | ALVRRHTLEDRSE | 5.0063 | 5.3170 | 4.5864 | 5.9659 | 6.0260 | 5.6115 | 6.4016 | 5.5353 | 4.9655 | 4.4400 | 4.5763 | 5.1392 | 9.2081 | 8.1105 | 9.5685 | 9.5421 |
| BRD1_T1016 | BRD1 | O95696-2 | 75 | EMSECNSNKENSE | 6.2027 | 4.7982 | 6.6011 | 6.5305 | 5.1345 | 7.0815 | 5.8268 | 6.2027 | 6.9826 | 5.3436 | 6.4344 | 6.1716 | 6.3722 | 6.3100 | 7.0730 | 6.9346 |
| BRD1_T1182;S1186 | BRD1 | O95696-2 | 1182;1186 | NA | 6.3353 | 6.1950 | 6.7898 | 6.3525 | 5.2086 | 5.7770 | 6.2506 | 6.3450 | 6.1251 | 6.2370 | 5.4222 | 6.3812 | 6.5292 | 6.4419 | 6.9010 | 6.7085 |
| BRD2_S298 | BRD2 | P25440-2 | 668 | LPTGYDSEEEEES | 6.0715 | 6.3105 | 6.3737 | 6.5826 | 6.0728 | 6.1271 | 6.2242 | 5.9994 | 6.4470 | 6.4703 | 6.4784 | 6.5160 | 6.2191 | 6.3442 | 5.8142 | 5.9489 |
| BRD2_S298;S301 | BRD2 | P25440-2 | 301 | APGSPASPPGSLE | 6.6754 | 6.6268 | 6.8113 | 7.2599 | 6.2914 | 5.8257 | 6.4230 | 6.1796 | 7.1215 | 6.9330 | 6.5566 | 7.1183 | 4.6599 | 5.2184 | 5.0359 | 5.2631 |
| BRD2_S301 | BRD2 | P25440-2 | 298 | AILAPGSPASPPG | 5.6837 | 6.3963 | 5.8991 | 6.5025 | 5.7444 | 5.9175 | 5.7749 | 5.9207 | 6.0283 | 5.8770 | 5.6412 | 5.8132 | 6.9342 | 7.2672 | 7.2571 | 7.3427 |
| BRD2_S320 | BRD2 | P25440-2 | 611 | FGPSGGSGTKLQA | 6.7443 | 6.7393 | 7.1867 | 7.6453 | 6.1327 | 6.8369 | 7.0027 | 6.9915 | 6.9697 | 7.4390 | 7.2308 | 6.8679 | 3.7258 | 4.7592 | 3.4605 | 4.2679 |
| BRD2_S397 | BRD2 | P25440-2 | 320 | PPMRRESGRPIKP | 4.7621 | 4.8520 | 4.6223 | 4.7556 | 5.2007 | 5.6320 | 5.4609 | 5.9622 | 5.0063 | 4.7556 | 5.4497 | 5.3862 | 9.0494 | 8.9530 | 9.4848 | 10.6672 |
| BRD2_S611 | BRD2 | P25440-2 | 6 | xMLQNVTPHNKLP | 6.9479 | 6.8334 | 7.4765 | 6.8465 | 7.1997 | 7.1782 | 6.8573 | 6.1879 | 7.0267 | 7.2343 | 8.0158 | 6.7928 | 3.6130 | 4.6045 | 3.3075 | 3.8779 |
| BRD2_S668 | BRD2 | P25440-2 | 397 | KHPMDLSTVKRKM | 6.0957 | 7.6672 | 6.6807 | 8.2455 | 6.5787 | 6.2385 | 7.2420 | 8.1400 | 6.4868 | 5.0276 | 5.8405 | 5.5140 | 4.8099 | 5.7181 | 4.5139 | 5.2010 |
| BRD2_T286 | BRD2 | P25440-2 | 286 | KRKADTTTPTPTA | 5.6604 | 5.3675 | 5.9034 | 6.2753 | 5.8503 | 5.8729 | 5.2457 | 5.1110 | 5.7479 | 7.0587 | 5.3267 | 5.9844 | 7.4500 | 8.0332 | 7.4734 | 7.6392 |
| BRD2_T286;S301 | BRD2 | P25440-2 | 298;301 | NA | 5.8741 | 5.9843 | 6.0431 | 6.6498 | 6.1277 | 5.9301 | 6.2188 | 5.8648 | 6.2882 | 6.4031 | 6.1494 | 6.1176 | 6.6140 | 6.7113 | 6.5359 | 6.4878 |
| BRD2_T6 | BRD2 | P25440-2 | 286;301 | NA | 6.5098 | 6.6515 | 6.1183 | 6.4050 | 5.7291 | 5.4513 | 5.9663 | 6.0151 | 5.5962 | 6.6023 | 6.2596 | 6.1625 | 6.0970 | 6.5591 | 7.0070 | 6.8697 |
| BRD3_S263 | BRD3 | Q15059 | 263 | TASRSESPPPLSD | 6.1862 | 6.1752 | 6.0968 | 6.0451 | 6.4077 | 5.9768 | 6.3725 | 6.1384 | 6.5202 | 5.9898 | 6.0671 | 5.7771 | 6.5671 | 6.6050 | 6.5418 | 6.5332 |
| BRD4_S1045 | BRD4 | O60885 | 1045 | VIQHHHSPRHHKS | 6.0467 | 5.4772 | 6.1176 | 6.5036 | 6.0596 | 6.1250 | 6.0487 | 5.7684 | 6.1178 | 6.3817 | 6.3240 | 6.5503 | 7.0335 | 6.5508 | 6.1834 | 6.7119 |
| BRD4_S1070;S1078 | BRD4 | O60885 | 470 | PVVAVSSPAVPPP | 5.9953 | 6.2140 | 6.2148 | 6.7280 | 6.3413 | 5.8852 | 6.0575 | 6.1897 | 6.0769 | 6.2442 | 6.2768 | 6.4238 | 6.5971 | 6.3691 | 6.1839 | 6.2023 |
| BRD4_S1078 | BRD4 | O60885 | 601 | PPPTYESEEEDKC | 6.3930 | 6.2890 | 6.5111 | 6.6156 | 6.1860 | 6.0702 | 6.6630 | 5.8278 | 6.2628 | 6.2835 | 6.0776 | 6.4091 | 6.3716 | 6.0226 | 5.9523 | 6.0649 |
| BRD4_S1083 | BRD4 | O60885 | 1126 | IRSEPFSPSLRPE | 5.9818 | 6.8047 | 6.6567 | 6.5989 | 6.0794 | 6.4372 | 6.2504 | 6.4672 | 6.3507 | 6.9256 | 7.1753 | 7.3388 | 5.0734 | 5.2592 | 5.1113 | 5.4894 |
| BRD4_S1126 | BRD4 | O60885 | 1078 | QMSQFQSLTHQSP | 6.4541 | 6.9224 | 6.1178 | 6.8119 | 6.6589 | 6.7590 | 5.9246 | 7.3092 | 6.2218 | 6.1328 | 6.0059 | 5.9334 | 5.7231 | 6.1223 | 5.8265 | 5.0763 |
| BRD4_S470 | BRD4 | O60885 | 1083 | QSLTHQSPPQQNV | 5.7932 | 7.0194 | 6.7903 | 6.8256 | 6.8808 | 6.3552 | 6.0069 | 6.6158 | 6.4728 | 6.1344 | 5.9042 | 6.3646 | 5.2958 | 6.3840 | 5.6955 | 5.4614 |
| BRD4_S601 | BRD4 | O60885 | 296 | RKADTTTPTTIDP | 5.5704 | 6.4435 | 5.7354 | 7.1802 | 5.5349 | 6.9063 | 6.3627 | 6.2373 | 5.6249 | 5.6736 | 5.3963 | 6.2556 | 6.9459 | 6.4956 | 6.7689 | 6.8685 |
| BRD4_T296 | BRD4 | O60885 | 1070;1078 | NA | 6.8805 | 6.9485 | 5.9728 | 7.4245 | 5.7965 | 7.3484 | 5.9537 | 6.8268 | 6.1910 | 6.1027 | 6.6251 | 6.3336 | 5.2970 | 5.7613 | 6.0041 | 4.5336 |
| BRD7_S279 | BRD7 | Q9NPI1-2 | 619 | RKAMGISIPSPVM | 5.7316 | 5.8382 | 5.8437 | 6.1910 | 5.9927 | 5.8312 | 6.0166 | 5.6979 | 5.9534 | 5.9475 | 5.9991 | 6.0001 | 6.7647 | 7.5434 | 7.3294 | 7.3195 |
| BRD7_S619 | BRD7 | Q9NPI1-2 | 279 | QREREDSGDAEAH | 5.6169 | 6.5402 | 5.8203 | 6.2901 | 5.8313 | 5.7959 | 5.9775 | 6.0324 | 5.8235 | 5.6073 | 5.4368 | 5.7286 | 6.7040 | 7.4759 | 8.0587 | 7.2605 |
| BRD8_S284 | BRD8 | Q9H0E9-2 | 284 | GSLLPTSPRLVNE | 6.7453 | 6.5265 | 6.0809 | 6.5599 | 6.4888 | 6.4208 | 6.3489 | 6.4711 | 6.7098 | 6.1589 | 6.3559 | 6.3857 | 6.0066 | 5.5463 | 5.4307 | 5.7638 |
| BRD8_S460 | BRD8 | Q9H0E9-2 | 460 | VGSKAPSIDGKEE | 5.7830 | 5.7704 | 5.4073 | 5.9415 | 5.8674 | 5.8583 | 5.8907 | 6.0544 | 5.6314 | 5.3756 | 5.4574 | 5.9117 | 8.1259 | 7.2539 | 7.6393 | 8.0317 |
| BRD8_S517 | BRD8 | Q9H0E9-2 | 264 | MLGQKATPPPSPL | 6.5671 | 7.0972 | 6.6987 | 6.8648 | 6.2524 | 5.8256 | 6.0683 | 6.4760 | 6.3451 | 6.5210 | 6.2592 | 6.7088 | 5.5881 | 5.7990 | 5.4543 | 5.4745 |
| BRD8_S681 | BRD8 | Q9H0E9-2 | 683 | KFEMSDSLKEESG | 7.0170 | 6.8767 | 6.4635 | 6.6502 | 6.3836 | 5.8162 | 6.7571 | 6.5250 | 6.6832 | 6.6716 | 6.7376 | 6.7579 | 5.2441 | 4.6739 | 4.7253 | 6.0172 |
| BRD8_S683 | BRD8 | Q9H0E9-2 | 681 | QHKFEMSDSLKEE | 6.8818 | 6.8407 | 6.5903 | 7.2209 | 6.5563 | 6.5852 | 6.6471 | 6.9332 | 6.7775 | 6.0782 | 6.5112 | 6.6390 | 5.1410 | 4.9635 | 4.6725 | 4.9617 |
| BRD8_S694 | BRD8 | Q9H0E9-2 | 694 | SGTIFGSQIKDAP | 6.6428 | 6.6213 | 6.2386 | 6.8225 | 6.3548 | 6.2418 | 6.2492 | 5.8439 | 6.9190 | 6.5982 | 6.5329 | 6.9073 | 5.8097 | 5.4650 | 5.2577 | 5.4955 |
| BRD8_T264 | BRD8 | Q9H0E9-2 | 517 | AVEAALSFCEEND | 6.7224 | 6.5836 | 6.5426 | 6.9518 | 5.5896 | 7.0700 | 5.9041 | 7.2213 | 6.7910 | 6.2250 | 7.6990 | 6.6073 | 4.1495 | 5.6022 | 4.8613 | 5.4793 |
| BRD8_T264;S268 | BRD8 | Q9H0E9-2 | 642 | IGKGDETPLTNVK | 5.3458 | 6.0515 | 5.6686 | 6.9651 | 6.5611 | 6.0204 | 5.9914 | 6.2667 | 5.6230 | 5.7584 | 5.5154 | 5.4045 | 6.3314 | 7.7540 | 7.1684 | 7.5744 |
| BRD8_T642 | BRD8 | Q9H0E9-2 | 264;268 | NA | 6.2842 | 6.0165 | 6.0619 | 6.7073 | 6.5009 | 6.2451 | 6.2905 | 6.4429 | 6.5665 | 6.0764 | 6.3521 | 6.2543 | 6.2786 | 5.9472 | 5.8866 | 6.0889 |
| BRD9_S566 | BRD9 | Q9H8M2 | 588 | PYEFLQSPEPAAS | 5.5057 | 6.4811 | 6.1348 | 7.1264 | 5.9591 | 6.2602 | 5.9091 | 6.2157 | 6.4769 | 6.0414 | 6.1548 | 6.2029 | 5.9912 | 6.7752 | 6.0994 | 6.6660 |
| BRD9_S588 | BRD9 | Q9H8M2 | 566 | DQHHLGSPSRLSV | 6.4506 | 6.3586 | 6.4667 | 7.1075 | 6.0330 | 6.4673 | 6.1762 | 6.1337 | 6.3601 | 6.7119 | 6.4663 | 6.8060 | 4.9240 | 5.9224 | 5.4238 | 6.1919 |
| BRD9_T103 | BRD9 | Q9H8M2 | 103 | EREHCDTEGEADD | 5.8748 | 6.9675 | 6.6344 | 7.0582 | 6.2789 | 6.1674 | 6.3203 | 6.3104 | 5.9765 | 6.2526 | 5.6393 | 5.5706 | 5.4920 | 6.7265 | 6.1253 | 6.6051 |
| BRF1_S436 | BRF1 | Q92994 | 436 | CISSQSSDPKDAS | 4.8043 | 6.0748 | 6.1472 | 5.8597 | 5.4799 | 6.4229 | 5.2491 | 5.5869 | 6.3229 | 5.7321 | 6.4496 | 5.7004 | 6.7611 | 7.7811 | 7.9415 | 7.6864 |
| BRF1_S587 | BRF1 | Q92994 | 587 | GADPVTSVGKRLR | 7.0933 | 6.8935 | 6.3763 | 7.2202 | 5.4935 | 6.2392 | 8.0523 | 6.8554 | 8.6872 | 6.6861 | 7.2101 | 7.4242 | 3.0100 | 4.3855 | 3.9106 | 4.4625 |
| BRF2_S353 | BRF2 | Q9HAW0 | 385 | TGDENISDSEIEQ | 5.6912 | 6.1566 | 6.7246 | 6.2022 | 5.7023 | 6.6280 | 6.4959 | 6.5153 | 6.5353 | 6.5431 | 6.1193 | 6.6149 | 6.0222 | 6.2796 | 6.1939 | 5.5755 |
| BRF2_S385 | BRF2 | Q9HAW0 | 353 | QGKRPASPALLLP | 4.9421 | 5.3791 | 5.9587 | 5.3608 | 5.6877 | 6.3067 | 5.6734 | 5.9490 | 6.0294 | 6.0478 | 6.3170 | 6.2186 | 7.2385 | 7.8623 | 7.5275 | 7.5016 |
| BRIP1_S1237 | BRIP1 | Q9BX63 | 505 | QKEEKISPIYGKE | 5.5464 | 6.2022 | 6.7488 | 6.7328 | 7.2807 | 7.0418 | 6.4133 | 6.4981 | 6.6440 | 6.4913 | 7.9541 | 6.9000 | 5.1374 | 5.6687 | 4.7617 | 3.9787 |
| BRIP1_S197 | BRIP1 | Q9BX63 | 1237 | IKNFKPSPSKNKG | 5.7023 | 6.3052 | 6.6183 | 6.3772 | 5.8632 | 6.1731 | 5.9595 | 5.6426 | 6.6345 | 6.9110 | 7.0191 | 7.4431 | 5.5836 | 5.8977 | 5.7104 | 6.1594 |
| BRIP1_S505 | BRIP1 | Q9BX63 | 113 | HFNYPSTPPSERN | 6.0645 | 6.5797 | 6.4950 | 6.6424 | 6.3630 | 6.2259 | 5.5945 | 6.7605 | 6.8253 | 6.6711 | 6.6899 | 6.4178 | 5.4595 | 5.7777 | 5.4347 | 5.9983 |
| BRIP1_S930 | BRIP1 | Q9BX63 | 930 | EAASHLSPENFVE | 5.6668 | 6.0519 | 6.3337 | 6.4894 | 5.6864 | 5.8849 | 5.5712 | 6.5011 | 6.5637 | 6.2344 | 6.2588 | 6.6782 | 6.3697 | 6.6402 | 6.5263 | 6.5434 |
| BRIP1_S990 | BRIP1 | Q9BX63 | 990 | VISRSTSPTFNKQ | 6.1879 | 7.0614 | 6.5379 | 6.1200 | 6.0520 | 6.1507 | 5.8788 | 6.2369 | 6.4766 | 6.7843 | 6.6739 | 7.0372 | 5.6732 | 5.5249 | 6.5554 | 5.9488 |
| BRIP1_T113 | BRIP1 | Q9BX63 | 197 | KTVKLNSPLEKIN | 7.2418 | 7.2457 | 7.2524 | 6.7610 | 6.4165 | 6.7481 | 6.6098 | 5.9954 | 7.0172 | 7.0963 | 6.6467 | 6.8195 | 4.2812 | 4.6168 | 4.3459 | 4.9056 |
| BRIX1_S261 | BRIX1 | Q8TDN6 | 261 | ENPHYQSPNMHRR | 6.1955 | 6.2686 | 6.1764 | 7.0104 | 6.3789 | 6.0348 | 6.0422 | 6.1703 | 6.1852 | 6.2830 | 6.2606 | 6.0578 | 6.3171 | 6.3781 | 6.0506 | 6.1905 |
| BRIX1_T309 | BRIX1 | Q8TDN6 | 309 | TADVFVTPAEEKP | 7.3054 | 6.9072 | 6.5241 | 7.4775 | 5.7678 | 6.9387 | 7.6011 | 7.1989 | 6.8001 | 7.1516 | 7.0156 | 7.5249 | 3.4641 | 4.7434 | 2.8221 | 4.7572 |
| BRMS1_S173 | BRMS1 | Q9HCU9 | 237 | KARAAVSPQKRKS | 6.6483 | 6.5685 | 5.8339 | 6.5254 | 6.5220 | 5.9908 | 6.3320 | 6.2597 | 6.5057 | 6.1918 | 6.0220 | 6.0667 | 6.2490 | 5.9844 | 6.1772 | 6.1227 |
| BRMS1_S237 | BRMS1 | Q9HCU9 | 173 | LEEDRQSLDLSSE | 6.5933 | 6.1221 | 5.8947 | 6.2294 | 6.3890 | 6.1359 | 5.6278 | 5.8149 | 6.0003 | 5.5781 | 5.9522 | 5.8235 | 6.7735 | 6.1625 | 7.0567 | 7.8462 |
| BRMS1L_S197 | BRMS1L | Q5PSV4 | 197 | KRKDPFSPDKKKP | 6.6158 | 6.6437 | 6.5318 | 6.9714 | 6.2363 | 6.1782 | 6.4339 | 5.9659 | 6.6773 | 6.6290 | 6.7728 | 6.7077 | 5.3252 | 5.4645 | 5.2914 | 5.5550 |
| BRPF1_S1082 | BRPF1 | P55201-2 | 120 | DNLDVVSEDEEAP | 6.2146 | 6.4472 | 6.2292 | 6.5850 | 6.0555 | 6.1285 | 6.2176 | 5.7984 | 6.5086 | 6.3567 | 6.0615 | 6.2873 | 6.4079 | 6.1363 | 6.1958 | 6.3700 |
| BRPF1_S120 | BRPF1 | P55201-2 | 238 | RKTEGVSPIPQEI | 6.0879 | 6.3513 | 6.1498 | 6.8524 | 5.9393 | 5.5891 | 6.2067 | 6.2589 | 5.9161 | 6.3931 | 5.6730 | 6.0353 | 6.4555 | 6.6634 | 6.8493 | 6.5790 |
| BRPF1_S238 | BRPF1 | P55201-2 | 852 | SSRGSLTPHPAAC | 6.4583 | 6.5690 | 5.4507 | 7.1337 | 5.8617 | 6.4271 | 6.0500 | 6.4218 | 6.0337 | 5.9852 | 6.2533 | 6.3435 | 5.9395 | 6.5085 | 6.2806 | 6.2833 |
| BRPF1_S452 | BRPF1 | P55201-2 | 452 | IHTPPGSARRLPA | 6.5648 | 7.6479 | 6.8587 | 7.4169 | 6.5605 | 6.5842 | 7.0760 | 6.5544 | 6.6167 | 7.0594 | 7.0172 | 7.0105 | 3.9281 | 4.5607 | 3.9342 | 4.6096 |
| BRPF1_S75 | BRPF1 | P55201-2 | 932 | TPSHGGSPVGPPQ | 6.2426 | 6.3399 | 6.7071 | 7.4980 | 7.3178 | 6.3128 | 6.4115 | 6.0866 | 7.5937 | 7.1172 | 5.9456 | 6.0806 | 4.7039 | 5.6402 | 4.6631 | 5.3394 |
| BRPF1_S850 | BRPF1 | P55201-2 | 1082 | RGAGWLSEDEDSP | 6.5031 | 7.1799 | 6.4869 | 6.5968 | 6.9311 | 6.2608 | 5.9216 | 6.2430 | 6.0718 | 5.8715 | 6.6000 | 5.9248 | 5.6179 | 5.8182 | 5.8796 | 6.0928 |
| BRPF1_S866 | BRPF1 | P55201-2 | 850 | GPSSRGSLTPHPA | 5.7571 | 5.5874 | 5.5191 | 6.6038 | 7.4933 | 6.0453 | 6.2339 | 5.9901 | 7.1199 | 7.5599 | 7.6875 | 7.3504 | 5.4203 | 6.1742 | 4.4972 | 4.9606 |
| BRPF1_S897 | BRPF1 | P55201-2 | 75 | RPANKQSPSPSEV | 4.6317 | 4.5820 | 6.6087 | 5.9827 | 5.3376 | 5.8142 | 5.8227 | 5.3158 | 5.8411 | 6.4486 | 6.9864 | 6.2026 | 7.9332 | 6.9826 | 8.1326 | 7.3775 |
| BRPF1_S932 | BRPF1 | P55201-2 | 866 | KDGQTDSAAEESS | 5.6183 | 5.9538 | 5.3474 | 6.1822 | 5.6372 | 6.9036 | 6.1144 | 6.1916 | 6.5523 | 6.4956 | 7.0091 | 6.9225 | 6.1853 | 6.1192 | 6.2089 | 6.5586 |
| BRPF1_T852 | BRPF1 | P55201-2 | 897 | EVGRRTSVLFSKK | 5.5846 | 4.0819 | 4.1921 | 5.1704 | 8.3081 | 5.3305 | 5.3254 | 4.9598 | 9.2842 | 7.8230 | 7.3618 | 7.0796 | 7.8661 | 8.0236 | 4.5725 | 5.0365 |
| BRPF3_S17;S19 | BRPF3 | Q9ULD4 | 740 | ENRAHLSPEVQLK | 6.2847 | 6.9691 | 6.7952 | 7.8845 | 5.7986 | 6.5623 | 6.4646 | 6.3080 | 6.6485 | 6.7495 | 6.6554 | 6.4908 | 4.5371 | 5.2714 | 5.2215 | 5.3586 |
| BRPF3_S403 | BRPF3 | Q9ULD4 | 962 | VAGSPASPASIEE | 6.4968 | 6.6539 | 6.4749 | 7.8192 | 6.2710 | 5.4698 | 5.5623 | 6.5619 | 6.7032 | 6.7559 | 5.9905 | 5.8940 | 5.6399 | 5.8348 | 5.4031 | 6.4689 |
| BRPF3_S740 | BRPF3 | Q9ULD4 | 959 | DHGVAGSPASPAS | 5.8062 | 5.4436 | 6.0134 | 6.3448 | 6.4743 | 6.3604 | 8.9035 | 7.6553 | 6.0185 | 5.0344 | 7.1736 | 6.8421 | 6.5002 | 4.5320 | 5.4747 | 5.4229 |
| BRPF3_S852 | BRPF3 | Q9ULD4 | 403 | KGDSPRSISETGD | 5.1184 | 6.3683 | 5.9316 | 8.0978 | 5.9655 | 5.3852 | 5.9301 | 6.3698 | 6.5181 | 6.5545 | 5.9593 | 5.7418 | 5.9280 | 6.3826 | 6.8105 | 6.9383 |
| BRPF3_S959 | BRPF3 | Q9ULD4 | 852 | SLSEQESPPEPPT | 6.2338 | 5.9504 | 6.6126 | 6.4570 | 5.3174 | 5.6441 | 6.1817 | 5.9864 | 6.3299 | 7.2295 | 6.3441 | 6.3460 | 6.2951 | 6.8085 | 6.1643 | 6.0992 |
| BRPF3_S962 | BRPF3 | Q9ULD4 | 17;19 | NA | 6.0142 | 6.6357 | 6.5694 | 7.1352 | 6.0484 | 5.9646 | 6.2718 | 6.0062 | 6.4870 | 6.3117 | 6.3533 | 6.1910 | 5.7819 | 6.2995 | 5.9234 | 6.0068 |
| BRWD1_S1475 | BRWD1 | Q9NSI6 | 696 | RGFRRLSLDIQSP | 5.8969 | 6.2180 | 6.2168 | 7.1775 | 5.8467 | 5.4804 | 5.6769 | 6.0246 | 6.3699 | 6.9048 | 5.4321 | 6.3296 | 6.1759 | 6.6116 | 7.1390 | 6.4994 |
| BRWD1_S696 | BRWD1 | Q9NSI6 | 1475 | IPELVGSPTQSTS | 6.3139 | 6.3516 | 6.9988 | 7.0917 | 6.3719 | 6.1710 | 6.4191 | 6.4485 | 7.0882 | 6.7589 | 7.1365 | 6.9730 | 4.6240 | 5.2583 | 4.8958 | 5.0988 |
| BSDC1_S109 | BSDC1 | Q9NW68-3 | 330 | QKLLEASLEEQGL | 7.0776 | 6.3172 | 6.8115 | 7.5996 | 5.5669 | 5.8646 | 6.4341 | 6.4991 | 6.8539 | 6.6810 | 6.9572 | 6.1511 | 5.2353 | 5.2415 | 5.1833 | 5.5262 |
| BSDC1_S231 | BSDC1 | Q9NW68-3 | 421 | DFDLDMTEEEVQM | 6.7186 | 6.0559 | 6.3989 | 6.6110 | 6.3539 | 7.7773 | 6.8586 | 6.9860 | 7.4210 | 6.3983 | 6.2692 | 7.5652 | 4.1428 | 4.9906 | 4.6451 | 4.8079 |
| BSDC1_S330 | BSDC1 | Q9NW68-3 | 109 | SDTFAPSPDKTID | 6.2388 | 6.0531 | 7.0790 | 6.2552 | 7.0288 | 6.8035 | 7.0210 | 6.6429 | 6.1218 | 7.5780 | 7.7356 | 7.0578 | 4.3619 | 5.0175 | 4.7158 | 4.2894 |
| BSDC1_S391 | BSDC1 | Q9NW68-3 | 391 | FELNSDSGKSTPS | 5.9962 | 5.8985 | 6.6603 | 7.3436 | 6.0794 | 6.4346 | 6.1594 | 5.7017 | 6.3098 | 6.8683 | 7.8653 | 6.8555 | 4.7592 | 5.7657 | 5.3897 | 5.9129 |
| BSDC1_T421 | BSDC1 | Q9NW68-3 | 231 | KQRAEQSISEEPG | 4.7762 | 6.2511 | 5.2989 | 5.8792 | 7.2378 | 5.9802 | 7.2071 | 5.8779 | 6.5259 | 6.7611 | 6.0556 | 7.5305 | 6.5489 | 6.9157 | 5.5750 | 5.5788 |
| BSG_S362 | BSG | P35613 | 362 | DDDDAGSAPLKSS | 5.9863 | 6.2421 | 5.9770 | 6.2466 | 5.9167 | 6.0820 | 6.1447 | 5.8606 | 6.0639 | 6.1081 | 5.9470 | 6.2401 | 7.0077 | 6.4885 | 6.9345 | 6.7539 |
| BSN_S93 | BSN | Q9UPA5 | 93 | GNQRAASPTPKQA | 6.4839 | 6.5401 | 5.7599 | 5.5907 | 6.6465 | 6.7456 | 6.2034 | 6.4459 | 6.0839 | 5.9830 | 5.8960 | 6.4327 | 6.1737 | 6.4630 | 6.5616 | 5.9902 |
| BTBD10_S120 | BTBD10 | Q9BSF8-2 | 120 | SSSRPSSPRPQKA | 5.9532 | 5.4646 | 6.9730 | 6.3951 | 6.3909 | 6.4989 | 5.8745 | 5.9755 | 6.5874 | 6.6199 | 7.0232 | 6.9010 | 6.0041 | 5.9712 | 5.5658 | 5.8017 |
| BTBD7_S998 | BTBD7 | Q9P203 | 998 | YLPGQTSPKKQEE | 6.6793 | 6.4000 | 7.4681 | 6.8434 | 5.8843 | 6.5018 | 6.3431 | 6.6534 | 6.7701 | 6.9482 | 7.2826 | 7.3144 | 4.4627 | 5.0351 | 4.4333 | 4.9801 |
| BTBD9_S599 | BTBD9 | Q96Q07 | 599 | GSSLPSSPGSNSR | 6.2649 | 5.9586 | 6.8352 | 6.8629 | 5.6476 | 6.4264 | 6.1662 | 7.0586 | 6.3360 | 6.7891 | 6.5676 | 6.1459 | 5.9429 | 6.3775 | 4.8964 | 5.7242 |
| BTF3_S158 | BTF3 | P20290 | 158 | NQLGADSLTSLRR | 6.0038 | 5.8938 | 6.0440 | 6.5273 | 6.4877 | 6.2391 | 6.0574 | 6.5739 | 5.9829 | 6.1162 | 6.0184 | 6.6968 | 6.0848 | 6.3969 | 6.4540 | 6.4231 |
| BTF3_S30 | BTF3 | P20290 | 30 | GGEATLSQPPPRG | 7.6941 | 7.3254 | 7.7692 | 6.8967 | 5.2260 | 5.9716 | 6.6421 | 6.1081 | 7.4315 | 6.9151 | 7.3182 | 6.7746 | 4.2777 | 4.9698 | 4.3352 | 4.3448 |
| BTF3L4_S153 | BTF3L4 | Q96K17 | 153 | ENFDEASKNEANx | 6.1355 | 6.6305 | 7.1892 | 6.4632 | 7.2811 | 6.2581 | 7.0689 | 6.4608 | 6.1780 | 6.0860 | 6.7555 | 6.6635 | 5.0748 | 5.4991 | 4.5987 | 5.6570 |

| | | | | | | | | | | | | | | | | | | | | | |
|---|---|---|---|---|---|---|---|---|---|---|---|---|---|---|---|---|---|---|---|---|---|
| BTN2A1_S526 | BTN2A1 | Q7KYR7 | | 526 | RVGTHQSLxxxxx | 4.6414 | 4.6992 | 4.4597 | 4.6341 | 5.2751 | 5.2313 | 5.4511 | 5.1882 | 5.0139 | 5.0526 | 4.5736 | 4.5438 | 10.0480 | 9.6578 | 11.1092 | 10.4211 |
| BUB1_S159 | BUB1 | O43683 | | 593 | NKTLAPSPKSPGD | 7.2586 | 7.5247 | 7.3972 | 7.2745 | 6.2494 | 6.7016 | 7.1684 | 6.6448 | 7.6109 | 7.4728 | 7.5620 | 7.5797 | 3.0200 | 3.6994 | 3.0942 | 3.7418 |
| BUB1_S307 | BUB1 | O43683 | | 655 | SRDGKFSPIQEKS | 7.3734 | 7.3811 | 7.3956 | 7.0581 | 5.9953 | 6.2189 | 6.9116 | 6.8970 | 7.7841 | 7.2612 | 6.9385 | 7.3566 | 3.2335 | 4.0455 | 3.6554 | 4.4943 |
| BUB1_S402 | BUB1 | O43683 | | 596 | LAPSPKSPGDFTS | 6.2580 | 6.7832 | 6.9831 | 7.6020 | 6.3547 | 6.6127 | 6.8935 | 6.9451 | 6.9906 | 6.6221 | 7.1333 | 7.3341 | 4.1322 | 4.6990 | 4.3254 | 4.3311 |
| BUB1_S510 | BUB1 | O43683 | | 307 | HQVVETSHEDLPA | 6.7050 | 7.1740 | 6.9328 | 7.9314 | 6.3908 | 6.8271 | 7.6567 | 6.7701 | 7.2692 | 6.6786 | 7.4341 | 7.3586 | 3.1939 | 4.3128 | 3.0544 | 4.3106 |
| BUB1_S593 | BUB1 | O43683 | | 402 | SMFAVASKDAGCV | 5.9429 | 7.4703 | 6.9125 | 6.7734 | 6.2795 | 6.5912 | 6.3156 | 6.6014 | 7.1217 | 6.9625 | 7.2396 | 7.1589 | 4.2913 | 5.5526 | 4.1227 | 4.6641 |
| BUB1_S596 | BUB1 | O43683 | | 510 | FQKNVRSSGAWGV | 6.7849 | 6.7830 | 7.5603 | 6.8483 | 5.3927 | 6.3187 | 6.4862 | 6.7262 | 8.1359 | 7.3294 | 8.0594 | 7.6906 | 4.0589 | 4.0719 | 3.6384 | 4.1150 |
| BUB1_S655 | BUB1 | O43683 | | 159 | PAQARTSEPLHNV | 4.0127 | 5.9761 | 9.3801 | 6.3936 | 6.7665 | 6.9705 | 8.2104 | 6.3554 | 6.6390 | 6.0558 | 8.1211 | 5.7625 | 4.6948 | 4.8574 | 4.2072 | 5.5968 |
| BUB1B_S1057 | BUB1B | O60566-3 | | 368 | ARQPVMTPCKIEP | 6.9434 | 7.2693 | 7.2063 | 7.5042 | 5.4512 | 7.1466 | 7.5286 | 6.7410 | 7.4334 | 7.2690 | 8.1425 | 8.0480 | 2.8546 | 3.8915 | 2.7653 | 3.8054 |
| BUB1B_S398 | BUB1B | O60566-3 | | 684 | LSIKKLSPIIEDS | 6.8932 | 7.3331 | 7.7385 | 7.2088 | 6.0627 | 6.7077 | 6.6563 | 6.7557 | 7.2282 | 7.0453 | 7.3227 | 7.3838 | 3.7002 | 4.2698 | 3.5828 | 4.1112 |
| BUB1B_S449 | BUB1B | O60566-3 | | 1057 | KVGKLTSPGALLF | 7.3599 | 7.9302 | 6.9688 | 7.4116 | 6.0961 | 6.6712 | 6.5305 | 6.2115 | 7.8405 | 7.0824 | 7.0745 | 7.0627 | 3.3657 | 4.2987 | 3.7125 | 4.3833 |
| BUB1B_S523 | BUB1B | O60566-3 | | 557 | SEKKNKSPPADPP | 6.7749 | 6.7776 | 7.1731 | 6.8717 | 6.3236 | 7.2494 | 6.7117 | 6.7433 | 7.3991 | 6.7660 | 7.6602 | 7.6044 | 3.9196 | 4.3458 | 3.4459 | 4.2338 |
| BUB1B_S557 | BUB1B | O60566-3 | | 449 | EAELLTSAEKRAE | 6.8089 | 7.3929 | 7.0331 | 7.9451 | 6.1739 | 6.7057 | 7.4756 | 6.6793 | 6.8138 | 7.0126 | 7.5454 | 7.5239 | 3.3572 | 4.2585 | 3.1656 | 4.1084 |
| BUB1B_S578 | BUB1B | O60566-3 | | 398 | PLQRVQSHQQASE | 5.8369 | 5.7577 | 8.2256 | 6.1029 | 4.8878 | 6.8717 | 6.1799 | 6.6588 | 7.7809 | 7.8731 | 9.6180 | 7.8979 | 3.6546 | 5.0717 | 3.2535 | 4.3290 |
| BUB1B_S684 | BUB1B | O60566-3 | | 523 | CCARETSLAENIW | 8.1054 | 8.6549 | 7.8067 | 10.8052 | 6.4329 | 7.4901 | 7.7828 | 6.1164 | 7.3408 | 5.4175 | 8.5832 | 7.2691 | 1.9711 | 2.7595 | 0.0000 | 3.4643 |
| BUB1B_T368 | BUB1B | O60566-3 | | 578 | LAVLKTSESITSN | 6.4106 | 5.9329 | 6.3940 | 7.2566 | 6.1337 | 6.5491 | 5.5757 | 6.4286 | 6.3718 | 6.8204 | 5.6048 | 7.6498 | 5.4165 | 6.5297 | 5.4456 | 5.4802 |
| BUB3_S211 | BUB3 | O43684 | | 211 | VEYLDPSPEVQKK | 5.6948 | 6.2672 | 6.1064 | 6.0768 | 6.1931 | 6.1906 | 5.7348 | 6.3182 | 6.3568 | 6.4975 | 6.0701 | 6.4165 | 6.4712 | 6.6526 | 6.3389 | 6.6145 |
| BUD13_S214 | BUD13 | Q9BRD0 | | 325 | EYDPDISPPRKKQ | 6.0585 | 6.2546 | 5.9750 | 6.5413 | 6.0223 | 5.6846 | 6.0945 | 5.9682 | 6.3725 | 6.2033 | 6.0679 | 6.4062 | 6.3381 | 6.5767 | 6.5150 | 6.9212 |
| BUD13_S248 | BUD13 | Q9BRD0 | | 271 | RRARHDSPDLAPN | 6.1614 | 5.9623 | 6.4151 | 6.3903 | 6.0314 | 5.8677 | 6.1164 | 6.1700 | 5.7804 | 6.1666 | 6.4226 | 6.1860 | 6.6754 | 6.5689 | 6.4861 | 6.5995 |
| BUD13_S259 | BUD13 | Q9BRD0 | | 248 | RRVHNNSPDTSRR | 6.5059 | 6.6926 | 5.7923 | 6.6371 | 6.1338 | 5.7806 | 6.1725 | 6.1466 | 6.3925 | 6.0802 | 5.9094 | 6.0265 | 6.4540 | 6.2636 | 6.4612 | 6.5512 |
| BUD13_S271 | BUD13 | Q9BRD0 | | 214 | HNSSGASPRRVRH | 6.7112 | 6.6408 | 5.6300 | 6.9165 | 5.8708 | 5.6795 | 6.4849 | 6.2733 | 5.2765 | 6.8634 | 5.9094 | 5.5768 | 7.0771 | 6.7247 | 6.1309 | 6.2342 |
| BUD13_S325 | BUD13 | Q9BRD0 | | 358 | TDSDLSSPRHKQS | 6.1174 | 6.3762 | 5.9616 | 6.6272 | 5.3249 | 5.4979 | 6.0255 | 5.5751 | 6.8514 | 6.6533 | 7.0496 | 6.8059 | 6.1159 | 6.2452 | 6.4072 | 6.3657 |
| BUD13_S358 | BUD13 | Q9BRD0 | | 259 | RRTLGSSDTQQLR | 7.3633 | 7.9451 | 6.8171 | 7.6287 | 5.9503 | 6.8146 | 6.9292 | 6.5815 | 7.3743 | 6.9908 | 7.7162 | 7.3539 | 3.2225 | 4.2327 | 3.2803 | 3.7995 |
| BUD13_S57 | BUD13 | Q9BRD0 | | 159 | RRARHDTPDPSPL | 5.8748 | 6.5647 | 6.0502 | 6.7644 | 5.9210 | 5.9414 | 6.8568 | 6.4206 | 6.2068 | 7.0197 | 5.7276 | 5.8826 | 6.7088 | 6.6924 | 5.4848 | 5.8834 |
| BUD13_T147;S151 | BUD13 | Q9BRD0 | | 57 | IVDDDVSWTAIST | 5.9351 | 6.0448 | 6.1331 | 7.0711 | 6.1814 | 6.8508 | 5.4832 | 5.3782 | 6.6993 | 6.3497 | 5.2016 | 5.4628 | 6.7448 | 7.0116 | 6.4957 | 6.9568 |
| BUD13_T159 | BUD13 | Q9BRD0 | 147;151 | | NA | 5.9507 | 5.2435 | 6.6063 | 6.7439 | 6.9350 | 6.1616 | 5.9610 | 7.5256 | 7.9025 | 4.7111 | 6.4849 | 6.4766 | 5.7738 | 4.1125 | 6.8722 | 6.5387 |
| BUD23_S257 | BUD23 | O43709-3 | | 257 | RFPLRMSRRGMVR | 6.0250 | 6.4595 | 7.0609 | 5.5794 | 4.5162 | 6.5508 | 6.7483 | 5.7944 | 7.0189 | 7.8775 | 8.9045 | 9.5304 | 4.5521 | 5.5391 | 3.4039 | 4.4390 |
| BYSL_S167 | BYSL | Q13895 | | 98 | RMPQDGSDDEDEE | 5.7973 | 6.0663 | 6.3651 | 6.3261 | 6.0754 | 5.8299 | 5.9461 | 6.1005 | 6.2263 | 6.1393 | 6.3108 | 6.4037 | 6.4430 | 6.8831 | 6.3817 | 6.7054 |
| BYSL_S423 | BYSL | Q13895 | | 167 | EVETVMSEVSGFP | 5.8844 | 6.4996 | 6.7208 | 5.7939 | 6.1497 | 6.1884 | 6.1813 | 6.5462 | 6.1371 | 6.2196 | 6.1042 | 6.5241 | 6.0138 | 6.2363 | 6.3754 | 6.4252 |
| BYSL_S98 | BYSL | Q13895 | | 423 | IRRELQSAVPRDV | 6.4467 | 6.5170 | 6.6378 | 6.3213 | 6.4159 | 6.1399 | 6.6656 | 6.1404 | 6.4536 | 6.5719 | 6.6549 | 7.3496 | 5.3363 | 5.6401 | 5.0943 | 5.6148 |
| BZW1_S423 | BZW1 | Q7L1Q6-3 | | 423 | HVAKGKSVFLEQM | 6.1233 | 5.3634 | 7.1662 | 6.0918 | 6.9000 | 6.5262 | 6.1258 | 5.9831 | 5.7724 | 5.0625 | 5.6346 | 5.1435 | 7.6506 | 6.5058 | 7.0235 | 6.9271 |
| BZW2_S412;S414 | BZW2 | Q9Y6E2 | 412;414 | | NA | 5.7364 | 5.6593 | 6.0265 | 5.9721 | 6.4726 | 7.3944 | 6.7793 | 6.4649 | 6.6019 | 6.5423 | 7.3264 | 6.0246 | 6.3771 | 6.3745 | 5.7770 | 4.4704 |
| C11orf68_S277 | C11orf68 | Q9H3H3-3 | | 277 | RFQLGGSARGSRV | 5.5766 | 5.3610 | 5.8097 | 4.6340 | 6.8590 | 6.4622 | 5.8331 | 7.6079 | 5.5575 | 4.9439 | 5.2183 | 5.4429 | 8.2781 | 7.6138 | 7.9021 | 6.9001 |
| C11orf87_S2 | C11orf87 | Q6NUJ2 | | 2 | xxxxxMSARAPKE | 7.1623 | 7.2661 | 6.8532 | 7.7817 | 5.4579 | 6.7440 | 7.0377 | 6.6648 | 7.3456 | 7.1415 | 7.5250 | 8.2497 | 3.5101 | 4.1397 | 3.0358 | 4.0850 |
| C12orf45_S15 | C12orf45 | Q8N5I9 | | 178 | KIEVLDSPASKKK | 6.7475 | 6.5293 | 7.0375 | 6.2781 | 6.6013 | 6.6643 | 5.9344 | 6.7196 | 7.2038 | 6.5043 | 6.9136 | 6.8802 | 5.1024 | 5.0118 | 4.8176 | 5.0542 |
| C12orf45_S178 | C12orf45 | Q8N5I9 | | 15 | ASPSCSSPTRDSS | 6.7057 | 6.7026 | 7.2383 | 6.3679 | 6.3828 | 6.6017 | 5.6459 | 6.8761 | 6.7726 | 6.4900 | 7.3302 | 7.3673 | 4.9548 | 4.8135 | 4.8006 | 4.9501 |
| C12orf45_S66 | C12orf45 | Q8N5I9 | | 66 | TVRIERSPLLDQV | 7.2284 | 7.1598 | 6.5191 | 6.9888 | 7.2020 | 7.7974 | 7.4628 | 7.3287 | 7.5018 | 6.9962 | 7.1197 | 6.7925 | 3.1339 | 4.1177 | 3.1783 | 3.4728 |
| C14orf93_S224 | C14orf93 | Q9H972 | | 224 | AYAKQLSPATQLA | 6.3272 | 6.5287 | 6.4885 | 7.0381 | 5.0954 | 6.5920 | 6.7195 | 5.8520 | 7.1436 | 6.5795 | 6.4625 | 6.3549 | 5.3822 | 5.8709 | 5.7827 | 5.7823 |
| C14orf93_S285 | C14orf93 | Q9H972 | | 285 | SLGLTVSPCRTRG | 6.1625 | 6.3593 | 7.1356 | 6.4819 | 6.4441 | 6.2990 | 5.7824 | 6.3267 | 5.9724 | 6.1087 | 6.7586 | 6.3953 | 5.4968 | 6.1757 | 6.3280 | 5.7731 |
| C14orf93_S378 | C14orf93 | Q9H972 | | 378 | RREYRNSLNPFKG | 7.2057 | 8.8689 | 6.3656 | 8.3742 | 6.7968 | 6.1491 | 7.7788 | 6.8006 | 8.2958 | 6.4851 | 8.0625 | 8.6916 | 2.3446 | 3.1716 | 1.8014 | 2.8076 |
| C16orf72_S167 | C16orf72 | Q14CZ0 | | 167 | PRLTVVSPNRATS | 7.2159 | 7.5926 | 7.3960 | 6.5475 | 6.7367 | 6.9441 | 6.6391 | 6.5644 | 7.1096 | 6.7701 | 6.8622 | 6.8625 | 4.0998 | 4.6912 | 3.7775 | 4.1907 |
| C16orf74_T44 | C16orf74 | Q96GX8 | | 44 | IIITPPTPTGMML | 5.9197 | 5.8640 | 6.4767 | 5.9851 | 5.9095 | 6.5648 | 6.5718 | 6.4646 | 5.6108 | 6.4753 | 6.5138 | 6.2712 | 6.0681 | 6.7758 | 6.1803 | 6.3482 |
| C17orf100_S54 | C17orf100 | A8MU93 | | 54 | SEGPSLSPSGKRL | 6.2363 | 6.5507 | 6.8142 | 5.9015 | 6.1901 | 6.4815 | 6.0265 | 6.7172 | 6.6972 | 6.6068 | 6.4898 | 6.5810 | 5.3840 | 6.1402 | 5.5489 | 5.6342 |
| C18orf21_S158 | C18orf21 | Q32NC0 | | 130 | LKSNPATPTSKLS | 7.3905 | 6.8774 | 6.4195 | 6.6051 | 6.8115 | 7.2242 | 7.8288 | 6.6231 | 6.4775 | 6.0688 | 6.8511 | 6.2592 | 4.3418 | 4.9254 | 4.3910 | 4.9050 |
| C18orf21_T130 | C18orf21 | Q32NC0 | | 158 | SGSKGKSPASVFR | 6.7369 | 7.1721 | 5.9843 | 7.2880 | 6.3946 | 6.6784 | 7.3927 | 5.9334 | 8.1489 | 7.2916 | 6.4259 | 6.3692 | 3.9006 | 4.6681 | 4.4174 | 5.1979 |
| C18orf25_S248 | C18orf25 | Q96B23-2 | | 66 | MELRRDSSESQLA | 4.1614 | 4.7509 | 5.2571 | 4.8601 | 5.7771 | 6.0096 | 5.4387 | 6.3258 | 5.4333 | 5.8947 | 6.0961 | 6.1109 | 8.8214 | 8.2172 | 8.4360 | 8.4096 |
| C18orf25_S66 | C18orf25 | Q96B23-2 | | 248 | DPGFLASDKTSAG | 4.7915 | 6.0746 | 6.5575 | 5.7830 | 6.5725 | 7.1436 | 6.7190 | 7.0298 | 5.9523 | 5.2866 | 5.1301 | 6.3200 | 6.2552 | 6.3641 | 7.6720 | 6.3484 |
| C18orf25_S66;S69 | C18orf25 | Q96B23-2 | 66;69 | | NA | 5.7183 | 6.1424 | 6.2995 | 6.4173 | 6.9829 | 5.6529 | 6.2422 | 6.3608 | 5.4693 | 5.3784 | 5.2970 | 5.7816 | 6.8256 | 6.9580 | 7.4207 | 7.0533 |
| C19orf47_S126 | C19orf47 | Q8N9M1 | | 306 | RGPAKASPQPALT | 7.4166 | 7.6856 | 6.6111 | 7.5738 | 6.6288 | 6.7783 | 7.2109 | 6.7902 | 7.1434 | 6.9339 | 6.7523 | 7.0949 | 3.5708 | 4.2756 | 3.4371 | 4.0967 |
| C19orf47_S151 | C19orf47 | Q8N9M1 | | 151 | NSLNHDSPPSTPP | 5.8388 | 6.6081 | 5.8397 | 6.6474 | 6.1805 | 6.0898 | 5.6302 | 6.3997 | 5.8991 | 5.9270 | 5.9039 | 5.7173 | 6.7875 | 6.9110 | 7.1524 | 6.4675 |
| C19orf47_S151;S154 | C19orf47 | Q8N9M1 | | 126 | SVPCSPSPLAGEI | 6.8043 | 6.9106 | 6.7819 | 7.6817 | 6.8815 | 6.4255 | 7.0183 | 6.3909 | 6.8771 | 6.7062 | 6.5781 | 6.6977 | 4.5044 | 4.6281 | 4.3598 | 4.7540 |
| C19orf47_S280 | C19orf47 | Q8N9M1 | | 280 | EDLAWDSDNDSSS | 5.7623 | 6.0917 | 5.9136 | 6.0664 | 6.0434 | 6.2268 | 6.1416 | 6.2274 | 6.2764 | 6.3705 | 6.1047 | 6.1265 | 6.3770 | 6.9177 | 6.5930 | 6.7611 |
| C19orf47_S306 | C19orf47 | Q8N9M1 | 151;154 | | NA | 5.3594 | 5.6671 | 6.7089 | 6.8276 | 7.8426 | 5.6672 | 6.0232 | 6.5823 | 5.8386 | 6.8083 | 6.2147 | 6.0284 | 6.1970 | 6.3461 | 6.5281 | 5.3605 |
| C1orf131_S280 | C1orf131 | Q8NDD1-5 | | 280 | NGTLILSPVDIKK | 7.5572 | 7.7598 | 7.2367 | 8.0507 | 6.3499 | 6.4886 | 7.4531 | 6.0902 | 7.6313 | 7.7930 | 7.5121 | 7.8309 | 2.5909 | 3.6485 | 2.5898 | 3.4173 |
| C1orf174_S189 | C1orf174 | Q8IYL3 | | 189 | VFLDDDSNQPMPV | 5.0950 | 5.5101 | 6.3290 | 6.2206 | 5.3355 | 5.4103 | 5.6932 | 5.5541 | 6.1220 | 5.9940 | 6.0969 | 6.0626 | 7.5884 | 7.9085 | 7.9735 | 7.1061 |
| C1orf198_S289 | C1orf198 | Q9H425 | | 289 | LEGKLPSPDVRQD | 5.8081 | 6.9297 | 7.1523 | 6.5915 | 6.4785 | 6.3562 | 6.0639 | 7.8583 | 6.0690 | 5.3935 | 6.2550 | 5.8982 | 6.3358 | 5.6255 | 5.2780 | 5.9067 |
| C1orf226_S241 | C1orf226 | A1L170-2 | | 266 | ECRRASSPSLIER | 6.4004 | 7.2023 | 6.3669 | 6.5020 | 5.9356 | 6.1731 | 6.2704 | 6.8831 | 6.4947 | 6.1143 | 6.2479 | 6.1163 | 5.7207 | 6.0129 | 5.6028 | 5.9567 |
| C1orf226_S266 | C1orf226 | A1L170-2 | | 292 | ESWEDGSPPPQAR | 5.6932 | 6.1392 | 6.3397 | 7.1866 | 5.7530 | 7.0552 | 6.3211 | 6.6038 | 5.9959 | 6.1897 | 6.7591 | 6.0516 | 5.3362 | 6.7193 | 6.4923 | 5.3640 |
| C1orf226_S292 | C1orf226 | A1L170-2 | | 241 | NGEVSLSVPDLIH | 5.7733 | 5.9201 | 6.1171 | 5.3556 | 5.7842 | 6.3332 | 5.7751 | 5.6265 | 6.9734 | 5.2945 | 5.6876 | 5.3538 | 7.4530 | 7.5889 | 7.1448 | 7.8187 |
| C1orf52_S158 | C1orf52 | Q8N6N3 | | 158 | GEETLESDDEKDE | 5.9601 | 5.7768 | 6.7294 | 6.0856 | 6.2841 | 5.6484 | 5.9567 | 6.2855 | 6.1369 | 6.4590 | 6.4016 | 6.4462 | 6.3413 | 6.5124 | 6.2950 | 6.6809 |
| C1QBP_S201 | C1QBP | Q07021 | | 201 | QEDEAESDIFSIR | 6.4396 | 6.3932 | 6.4238 | 6.7620 | 6.3259 | 5.7250 | 6.1790 | 6.4870 | 6.0900 | 6.0163 | 5.9911 | 5.8700 | 6.6847 | 6.3121 | 6.0153 | 6.2850 |
| C1QBP_S210 | C1QBP | Q07021 | | 87 | AFVDFLSDEIKEE | 6.3737 | 6.7873 | 6.3106 | 6.9259 | 5.9291 | 5.8439 | 6.0042 | 5.5770 | 6.2674 | 6.3654 | 6.5344 | 6.3198 | 6.5871 | 6.1217 | 5.6438 | 6.4086 |
| C1QBP_S213 | C1QBP | Q07021 | | 213 | REVSFQSTGESEW | 6.5760 | 7.2298 | 5.7494 | 6.3279 | 6.1465 | 5.8447 | 5.8199 | 6.0195 | 6.9785 | 7.2255 | 6.1180 | 6.1982 | 6.4318 | 6.2084 | 5.2710 | 5.8549 |
| C1QBP_S87 | C1QBP | Q07021 | | 210 | FSIREVSFQSTGE | 6.2301 | 6.2314 | 6.3822 | 6.2522 | 5.8710 | 6.3841 | 6.5482 | 6.6618 | 6.9155 | 6.1733 | 6.8631 | 7.1370 | 6.1746 | 5.6022 | 5.1251 | 5.4482 |
| C2CD2_S435 | C2CD2 | Q9Y426 | | 441 | SPLSSDSPVKTPI | 6.0308 | 6.3410 | 6.2545 | 6.3173 | 6.5702 | 6.1355 | 6.4482 | 6.2011 | 6.7341 | 6.2963 | 6.2373 | 6.3167 | 5.8541 | 6.1921 | 5.9313 | 6.1394 |
| C2CD2_S439;T445 | C2CD2 | Q9Y426 | | 475 | SKTLSSSDTELLV | 6.0683 | 6.3086 | 6.2730 | 6.8515 | 6.3584 | 6.0478 | 6.2613 | 6.7711 | 6.4873 | 6.7891 | 6.3842 | 6.0856 | 6.1699 | 6.0690 | 5.5928 | 5.4823 |
| C2CD2_S441 | C2CD2 | Q9Y426 | | 519 | IIISGISKTSLSQ | 6.0256 | 6.0552 | 5.5181 | 5.9514 | 5.4017 | 5.5200 | 6.6141 | 5.7316 | 5.7420 | 5.4944 | 5.6141 | 5.6655 | 7.3409 | 7.8549 | 7.7514 | 7.7190 |
| C2CD2_S475 | C2CD2 | Q9Y426 | | 435 | VDVGRASPLSSDS | 6.5945 | 6.6565 | 5.4272 | 6.5211 | 6.3137 | 6.3237 | 7.0250 | 6.6679 | 6.8761 | 6.3544 | 5.7129 | 6.3986 | 5.6068 | 5.7101 | 5.7386 | 6.0729 |
| C2CD2_S511 | C2CD2 | Q9Y426 | | 511 | KSPRKKSTIIISG | 5.9903 | 4.4535 | 5.0032 | 5.0918 | 7.0011 | 5.3258 | 4.7284 | 5.3497 | 7.1851 | 7.2533 | 7.1272 | 7.3192 | 6.7808 | 8.5072 | 7.2056 | 5.6779 |
| C2CD2_S519 | C2CD2 | Q9Y426 | 439;445 | | NA | 6.7228 | 6.1593 | 5.8793 | 5.9154 | 7.2181 | 7.5050 | 7.1064 | 6.3476 | 6.9446 | 6.7702 | 7.3769 | 6.8356 | 4.8972 | 5.9487 | 3.8366 | 4.5363 |
| C2CD2L_S464;S468 | C2CD2L | O14523-2 | | 663 | EAGLSQSHDDLSN | 6.0571 | 6.0970 | 6.5526 | 6.0100 | 6.2188 | 5.8769 | 7.5683 | 6.1881 | 6.8454 | 5.9864 | 5.4233 | 6.7000 | 5.6434 | 6.0141 | 6.8690 | 5.9496 |
| C2CD2L_S468 | C2CD2L | O14523-2 | | 468 | LDSPSRSPSKVEV | 6.3465 | 5.9779 | 5.5273 | 7.1964 | 5.7383 | 6.0795 | 5.7418 | 5.9315 | 5.4496 | 5.3532 | 5.8950 | 6.6794 | 6.5835 | 7.6366 | 6.6573 | 7.2064 |
| C2CD2L_S663 | C2CD2L | O14523-2 | | 428 | TPRPSITPTKKIE | 6.6821 | 6.9467 | 5.5822 | 6.6966 | 6.2748 | 5.8324 | 6.4630 | 5.5636 | 5.6814 | 6.7545 | 7.0460 | 7.3685 | 6.1487 | 4.7635 | 6.1694 | 6.0267 |
| C2CD2L_T422;T428 | C2CD2L | O14523-2 | 464;468 | | NA | 6.1215 | 6.9178 | 6.6598 | 6.9468 | 6.3253 | 6.5479 | 6.6411 | 5.7145 | 7.0043 | 7.0425 | 7.2916 | 6.5903 | 4.8262 | 5.4234 | 4.5773 | 5.3697 |
| C2CD2L_T428 | C2CD2L | O14523-2 | 422;428 | | NA | 8.4202 | 7.6023 | 6.2481 | 6.6251 | 7.0511 | 6.7020 | 6.9450 | 6.3960 | 6.9943 | 6.5362 | 6.3506 | 8.5066 | 4.3650 | 4.2404 | 3.2845 | 3.7327 |
| C2CD3_S1829 | C2CD3 | Q4AC94 | | 2132 | KSSFLSSPERAVN | 6.2861 | 6.4870 | 6.6168 | 6.1055 | 6.2927 | 6.2620 | 6.3202 | 6.6005 | 6.6915 | 6.3494 | 6.5768 | 6.7310 | 5.5158 | 5.7927 | 5.7906 | 5.5815 |
| C2CD3_S2132 | C2CD3 | Q4AC94 | | 1829 | KELDFSSPGRSDT | 7.2983 | 7.2331 | 7.1321 | 7.3534 | 6.5453 | 6.9080 | 7.1844 | 6.3157 | 7.1450 | 6.7106 | 6.2991 | 7.0880 | 3.6175 | 4.8316 | 4.4449 | 3.8930 |
| C2CD3_S728 | C2CD3 | Q4AC94 | | 728 | PLCAPTSPNKALP | 6.2884 | 8.5282 | 7.1754 | 6.9093 | 7.7235 | 7.7647 | 5.3982 | 6.2282 | 6.4120 | 5.2652 | 6.3328 | 5.4362 | 5.1036 | 5.4647 | 5.8100 | 4.1595 |
| C2CD5_S250 | C2CD5 | Q86YS7-3 | | 295 | NQTYSFSPSKSYS | 6.3615 | 6.2532 | 6.7208 | 6.2406 | 6.6239 | 6.5866 | 6.5153 | 6.4777 | 6.7537 | 6.7333 | 6.7831 | 6.6748 | 5.2253 | 5.4257 | 5.2387 | 5.3856 |
| C2CD5_S260 | C2CD5 | Q86YS7-3 | | 260 | FLPACNSPSKEMK | 5.9740 | 6.0361 | 6.7862 | 6.2977 | 6.3746 | 6.3553 | 6.2593 | 6.6836 | 6.5696 | 6.4188 | 6.7196 | 6.5715 | 5.8316 | 5.8398 | 5.5880 | 5.6941 |
| C2CD5_S295 | C2CD5 | Q86YS7-3 | | 643 | SEEIIGSPIPEPR | 6.4872 | 6.5267 | 6.7915 | 7.1514 | 6.7418 | 6.7689 | 6.4017 | 6.7485 | 6.6432 | 6.8677 | 7.0116 | 6.7427 | 4.2703 | 5.5692 | 4.5022 | 4.7754 |
| C2CD5_S304 | C2CD5 | Q86YS7-3 | | 323 | TGMGSGSAGKEGG | 6.3514 | 6.4043 | 6.5188 | 6.1892 | 6.3816 | 6.7604 | 6.7966 | 6.6176 | 6.5054 | 6.4059 | 6.4253 | 6.4519 | 5.5267 | 5.8694 | 5.2098 | 5.5857 |
| C2CD5_S323 | C2CD5 | Q86YS7-3 | | 817 | KSLQRASTDNEEL | 5.9424 | 6.2245 | 7.1218 | 6.4260 | 6.3362 | 6.2074 | 5.9385 | 6.2977 | 6.4583 | 6.0230 | 6.4504 | 6.0191 | 5.8843 | 6.3387 | 6.2504 | 6.0814 |
| C2CD5_S643 | C2CD5 | Q86YS7-3 | | 864 | AAQRATSVDYSSF | 5.6264 | 5.3016 | 6.0710 | 5.8632 | 6.4343 | 5.5946 | 5.4921 | 6.2734 | 7.6223 | 7.5902 | 7.4684 | 7.3311 | 6.0078 | 6.6550 | 5.3158 | 5.3527 |
| C2CD5_S659 | C2CD5 | Q86YS7-3 | | 903 | MTVEKASPVGDGN | 7.1781 | 7.3547 | 7.3883 | 7.0809 | 6.1508 | 6.4886 | 6.3826 | 6.7647 | 7.1411 | 7.2026 | 7.3309 | 6.9975 | 3.8321 | 4.6618 | 3.6557 | 4.3896 |
| C2CD5_S659;S662 | C2CD5 | Q86YS7-3 | | 304 | KSYSRQSSSSDTD | 6.5889 | 6.3631 | 6.9914 | 3.7558 | 5.6249 | 6.9561 | 6.1904 | 6.6105 | 6.5634 | 8.3716 | 4.2820 | 6.1668 | 6.3455 | 6.3219 | 6.1963 | 6.6714 |
| C2CD5_S662 | C2CD5 | Q86YS7-3 | | 876 | FADRCSSWIELIK | 3.6895 | 4.1764 | 4.0070 | 3.9340 | 4.5776 | 4.2178 | 4.4213 | 3.7022 | 4.1681 | 4.2363 | 3.8780 | 3.8137 | 13.4621 | 13.5050 | 12.7737 | 11.4372 |
| C2CD5_S817 | C2CD5 | Q86YS7-3 | | 662 | RSQSESSDEVTEL | 5.8688 | 6.5409 | 5.7627 | 6.0545 | 6.6615 | 6.7702 | 5.9263 | 6.4253 | 7.1713 | 6.0008 | 6.8251 | 6.4765 | 6.5396 | 5.8625 | 5.7949 | 5.3192 |
| C2CD5_S864 | C2CD5 | Q86YS7-3 | | 250 | TLDKLSSPAAFLP | 5.5990 | 6.5622 | 6.8023 | 7.0628 | 5.5908 | 5.5271 | 6.7847 | 5.8567 | 6.0467 | 6.6192 | 6.1783 | 6.5527 | 6.1661 | 5.9408 | 6.3044 | 6.4062 |
| C2CD5_S876 | C2CD5 | Q86YS7-3 | | 659 | RLLRSQSESSDEV | 6.4054 | 7.1505 | 6.7677 | 5.7450 | 6.0647 | 6.9727 | 6.0996 | 6.1908 | 6.9467 | 6.9844 | 6.4001 | 6.4539 | 5.2886 | 6.3973 | 5.1028 | 5.0299 |
| C2CD5_S903 | C2CD5 | Q86YS7-3 | | 278 | EDPNPNTHSSGPS | 6.2279 | 6.4130 | 7.0523 | 6.1074 | 6.0743 | 6.7262 | 5.9229 | 6.6100 | 6.2415 | 6.5812 | 6.9563 | 6.7289 | 5.7574 | 5.9815 | 5.5940 | 5.0252 |
| C2CD5_T278 | C2CD5 | Q86YS7-3 | 278;285 | | NA | 6.5350 | 7.1449 | 7.2527 | 6.9399 | 6.1761 | 7.2974 | 5.6262 | 6.8116 | 6.4601 | 6.9634 | 5.8898 | 6.8995 | 4.0032 | 5.4866 | 5.4655 | 5.0482 |
| C2CD5_T278;T285 | C2CD5 | Q86YS7-3 | 659;662 | | NA | 5.6171 | 5.9519 | 6.4132 | 6.1321 | 5.8934 | 6.0608 | 5.9368 | 5.7641 | 6.2408 | 6.4960 | 6.4173 | 6.6619 | 6.2917 | 6.7095 | 6.4759 | 6.9375 |
| C2orf49_S143 | C2orf49 | Q9BVC5 | | 182 | DLTHRKSPSGPVK | 6.5008 | 6.6295 | 6.0908 | 6.2392 | 6.2672 | 6.2862 | 6.3615 | 6.3584 | 6.5880 | 6.4025 | 6.2979 | 6.2960 | 6.0699 | 5.8938 | 5.7824 | 5.9359 |
| C2orf49_S182 | C2orf49 | Q9BVC5 | | 198 | LSPVGTTPVKLKR | 6.0698 | 5.7871 | 5.6925 | 5.7009 | 5.9303 | 6.3120 | 6.2028 | 5.8343 | 6.6982 | 5.9363 | 5.5938 | 5.5444 | 8.0160 | 7.3716 | 6.5062 | 6.8040 |
| C2orf49_S189 | C2orf49 | Q9BVC5 | | 230 | RKIQHVTWPxxxx | 5.5275 | 5.9759 | 6.0446 | 5.7683 | 6.0341 | 6.2495 | 6.1281 | 6.0344 | 5.8035 | 6.0261 | 5.7021 | 5.9988 | 6.8171 | 7.2399 | 7.3049 | 7.3454 |
| C2orf49_S189;S193 | C2orf49 | Q9BVC5 | | 189 | PSGPVKSPPLSPV | 5.5546 | 5.8290 | 5.8717 | 6.7355 | 6.1775 | 5.5809 | 5.5526 | 5.7670 | 5.9741 | 6.2431 | 6.2010 | 6.4117 | 7.1561 | 6.7813 | 7.0534 | 7.1105 |
| C2orf49_S189;T198 | C2orf49 | Q9BVC5 | | 193 | VKSPPLSPVGTTP | 5.5300 | 6.3272 | 6.1553 | 6.5457 | 6.5161 | 6.5426 | 6.8820 | 7.3733 | 6.2350 | 6.1324 | 6.6114 | 5.7081 | 6.2300 | 5.8270 | 5.8014 | 5.5824 |
| C2orf49_S193 | C2orf49 | Q9BVC5 | | 143 | NAFRKLSNSSSSV | 4.2379 | 4.7165 | 5.2069 | 5.7585 | 7.7484 | 5.7182 | 4.4731 | 6.2144 | 7.7817 | 8.6596 | 8.0047 | 7.7420 | 6.8250 | 7.1128 | 5.1952 | 4.6050 |
| C2orf49_T198 | C2orf49 | Q9BVC5 | 189;193 | | NA | 6.2801 | 6.7679 | 6.7368 | 7.1286 | 6.4035 | 6.3981 | 5.9989 | 6.4741 | 6.6635 | 6.6026 | 7.1935 | 7.1651 | 4.9502 | 4.9191 | 5.1697 | 5.1483 |
| C2orf49_T230 | C2orf49 | Q9BVC5 | 189;198 | | NA | 6.4977 | 6.2303 | 6.1307 | 6.0463 | 6.4259 | 6.0933 | 6.2258 | 6.7710 | 7.0414 | 6.6723 | 6.8067 | 6.5855 | 5.7566 | 5.6104 | 5.0096 | 6.0966 |
| C2orf74_S153 | C2orf74 | A8MZ97 | | 153 | SVTRTPSVVESQK | 7.4845 | 6.2513 | 6.8097 | 5.6835 | 6.8545 | 5.7898 | 5.0860 | 6.7899 | 6.6749 | 6.9383 | 7.1125 | 6.9178 | 6.3035 | 6.5982 | 4.2494 | 4.4562 |
| C5orf24_S137 | C5orf24 | Q7Z6I8 | | 144 | PGRPPGSIKALSR | 6.2805 | 6.5210 | 7.9377 | 7.3261 | 5.3030 | 6.2561 | 5.6205 | 5.6950 | 7.4956 | 8.2650 | 8.7200 | 9.2972 | 3.3569 | 5.0921 | 3.1903 | 3.6427 |
| C5orf24_S137 | C5orf24 | Q7Z6I8 | | 137 | AAGYKVSPGRPPG | 6.5751 | 7.1728 | 6.7553 | 7.1584 | 6.0202 | 7.0780 | 6.9438 | 6.9217 | 7.3435 | 7.6450 | 7.8380 | 7.7616 | 3.0601 | 4.6309 | 3.1372 | 3.9583 |
| C5orf24_S144 | C5orf24 | Q7Z6I8-2 | | 137 | AAGYKVSPGRPPG | 5.3831 | 6.2744 | 6.6076 | 6.4593 | 6.0691 | 6.4790 | 6.9530 | 6.5658 | 7.7973 | 6.7406 | 7.7092 | 7.9900 | 4.5161 | 4.9950 | 4.4795 | 4.9810 |
| C5orf34_S616 | C5orf34 | Q96MH7 | | 616 | VNENRVSIALKKT | 6.1044 | 7.8862 | 7.0836 | 9.1373 | 7.0980 | 8.7446 | 7.9584 | 6.5614 | 7.4652 | 7.1885 | 9.3342 | 7.0908 | 1.7475 | 3.2434 | 1.6396 | 1.7169 |
| C6orf47_S112 | C6orf47 | O95873 | | 112 | LEAGGASPKLRWD | 6.3500 | 6.4701 | 5.9056 | 6.7012 | 6.8435 | 6.4898 | 6.7350 | 6.9096 | 6.7235 | 7.1854 | 6.5719 | 7.1478 | 4.9548 | 5.1403 | 4.5993 | 5.2722 |
| C7orf50_S175 | C7orf50 | Q9BRJ6 | | 175 | ELDEEGSDPPLPG | 5.9044 | 6.3317 | 5.8522 | 6.0961 | 6.1847 | 6.2229 | 6.0724 | 6.1576 | 5.7979 | 5.6887 | 5.8262 | 5.7947 | 6.9451 | 7.0124 | 7.0919 | 7.0210 |
| C7orf50_S59 | C7orf50 | Q9BRJ6 | | 59 | DAEPELSPEEQRV | 5.8405 | 6.6411 | 6.3931 | 7.1085 | 6.7973 | 6.3017 | 6.5899 | 6.1570 | 6.1314 | 6.6786 | 6.4655 | 6.6680 | 5.3167 | 5.3891 | 5.7161 | 5.8057 |
| C8orf37_T18 | C8orf37 | Q96NL8 | | 18 | VESKFCTPDLLRR | 6.5778 | 7.1259 | 7.3910 | 6.6151 | 5.9427 | 7.3424 | 7.3683 | 6.9683 | 6.9331 | 7.5474 | 7.4950 | 7.5627 | 3.3682 | 4.3004 | 3.3326 | 4.1293 |
| C8orf37_T187 | C8orf37 | Q96NL8 | | 187 | CQCSWRTIEEVTD | 5.8535 | 6.2370 | 6.0909 | 6.0225 | 6.6748 | 5.9532 | 6.1939 | 6.6959 | 6.0219 | 6.1727 | 6.4310 | 5.6196 | 6.6875 | 7.0241 | 6.1765 | 6.1449 |
| C8orf58_S233 | C8orf58 | Q8NAV2 | | 233 | TRAPLPSPLHTPG | 6.6591 | 6.2390 | 6.3795 | 6.6737 | 6.9301 | 6.8984 | 7.6326 | 7.3034 | 6.4259 | 6.2374 | 6.8605 | 6.8186 | 4.7266 | 4.7807 | 4.4572 | 4.9773 |
| C8orf88_S70 | C8orf88 | P0DMB2 | | 70 | EQQQQQSPVKKER | 6.9393 | 6.4651 | 8.9956 | 6.5835 | 7.6606 | 7.2113 | 6.4239 | 6.8739 | 7.0646 | 7.7445 | 8.2350 | 8.3361 | 2.6235 | 3.2592 | 2.4036 | 3.1802 |
| C9orf40_S69 | C9orf40 | Q8IXQ3 | | 69 | MAEPSASPSKRRD | 6.2694 | 5.7691 | 6.9807 | 5.6017 | 5.7211 | 5.7017 | 5.6258 | 7.0963 | 5.8798 | 7.1047 | 6.6842 | 7.4879 | 5.3204 | 6.7314 | 5.9834 | 6.0422 |
| C9orf78_S261 | C9orf78 | Q9NZ63 | | 261 | KPEPERSPPNRKR | 5.7120 | 5.8507 | 6.3399 | 5.5756 | 6.2163 | 6.2783 | 5.6021 | 6.2757 | 6.4201 | 6.5836 | 6.7964 | 6.5707 | 6.5409 | 6.7257 | 6.2946 | 6.2172 |
| C9orf85_S21 | C9orf85 | Q96MD7 | | 21 | KHQNTFSFKNDKF | 8.7432 | 7.3876 | 8.1869 | 8.3047 | 4.8183 | 5.9419 | 6.3235 | 5.5909 | 7.3882 | 8.1720 | 8.6352 | 10.4068 | 2.2172 | 3.2882 | 1.8376 | 2.7577 |
| CAAP1_S120 | CAAP1 | Q9H8G2 | | 120 | LFLAEHSDLEEGG | 5.7651 | 6.1292 | 5.8825 | 6.5730 | 5.8455 | 5.7998 | 6.0770 | 6.0075 | 6.2400 | 6.1058 | 6.4444 | 6.3594 | 6.7941 | 6.4988 | 6.6422 | 6.8355 |
| CAAP1_S203 | CAAP1 | Q9H8G2 | | 312 | LGLAESSPNEPKA | 6.0425 | 6.2984 | 5.6840 | 6.7054 | 6.3227 | 6.1243 | 5.8610 | 6.0802 | 6.4073 | 6.1878 | 5.9048 | 5.6376 | 6.8216 | 6.5506 | 6.6504 | 6.7214 |
| CAAP1_S312 | CAAP1 | Q9H8G2 | | 203 | GDNGMDSDMEEEA | 6.0040 | 6.6467 | 6.0998 | 6.1424 | 6.2877 | 5.7997 | 6.1124 | 5.5580 | 6.4568 | 5.9740 | 6.3494 | 6.5567 | 6.8281 | 6.4708 | 6.7719 | 5.9416 |
| CAAP1_Y262 | CAAP1 | Q9H8G2 | | 262 | SINADAYDSDIEG | 6.2852 | 6.6780 | 5.4078 | 7.5571 | 5.8594 | 5.9079 | 6.2159 | 6.8233 | 6.4219 | 5.1015 | 6.0152 | 5.9771 | 6.9202 | 6.1277 | 5.6379 | 7.0639 |
| CABIN1_S10 | CABIN1 | Q9Y6J0 | | 2159 | PTPTLLSPKGSIS | 6.3656 | 6.8468 | 5.9137 | 5.8601 | 6.6761 | 6.4290 | 6.1185 | 6.5135 | 7.8969 | 6.7184 | 7.5588 | 7.1638 | 4.3256 | 5.3058 | 4.9937 | 5.3139 |
| CABIN1_S2159 | CABIN1 | Q9Y6J0 | | 10 | IAALNASSTIEDD | 5.0926 | 6.7680 | 3.8651 | 7.9458 | 8.2444 | 6.5358 | 9.5936 | 5.6013 | 6.2317 | 6.6298 | 8.1172 | 8.8305 | 2.9970 | 5.6677 | 3.0467 | 4.8327 |
| CABLES1_S109 | CABLES1 | Q8TDN4 | | 313 | CRTLSGSPRPKNF | 6.6543 | 7.0019 | 6.6156 | 6.0999 | 6.6317 | 6.5730 | 6.3468 | 6.2229 | 6.7757 | 6.9833 | 6.5170 | 6.4434 | 5.4990 | 5.4727 | 5.1538 | 5.0090 |
| CABLES1_S168 | CABLES1 | Q8TDN4 | | 109 | GARTRFSLLAAAE | 8.3517 | 6.6868 | 6.9053 | 7.0861 | 6.7128 | 5.7013 | 5.7034 | 5.9444 | 7.5634 | 6.5957 | 6.5896 | 5.9958 | 4.8946 | 5.9752 | 4.5932 | 4.7008 |
| CABLES1_S313 | CABLES1 | Q8TDN4 | | 168 | VARGFASPLGAGR | 7.0188 | 7.0750 | 6.6592 | 7.1628 | 5.5414 | 7.1307 | 7.4041 | 7.2274 | 6.9395 | 7.5782 | 7.7094 | 7.4192 | 3.2551 | 4.0582 | 3.2495 | 4.5714 |
| CABLES1_T309;S313 | CABLES1 | Q8TDN4 | 309;313 | | NA | 7.6187 | 7.6071 | 7.5612 | 7.3449 | 7.2476 | 6.7661 | 7.1739 | 7.1718 | 7.9670 | 6.7232 | 7.6034 | 7.0595 | 2.8479 | 3.3682 | 2.8721 | 3.0674 |
| CACNB3_S152 | CACNB3 | P54284 | | 152 | DIGNRRSPPPSLA | 6.0226 | 6.3644 | 6.6236 | 5.9943 | 5.9932 | 6.3561 | 6.1372 | 6.3059 | 6.5756 | 6.6952 | 6.5255 | 6.3802 | 5.9387 | 6.3434 | 5.8364 | 5.9077 |
| CACTIN_S73 | CACTIN | Q8WUQ7-2 | | 73 | SGMRSRSPPRPKW | 5.9279 | 6.4235 | 5.8096 | 6.0376 | 5.9460 | 5.7372 | 6.0040 | 5.9561 | 5.6714 | 6.0886 | 5.9203 | 6.0776 | 6.8519 | 7.4097 | 7.1286 | 7.0101 |
| CAD_S1038 | CAD | P27708 | | 1859 | PRIHRASDPGLPA | 6.7665 | 6.4602 | 8.6723 | 7.0024 | 6.4363 | 6.9416 | 6.4187 | 7.0613 | 7.3970 | 7.1867 | 8.0837 | 8.0428 | 3.1564 | 3.6674 | 3.0869 | 3.6198 |
| CAD_S1406 | CAD | P27708 | | 1833 | TSEMTTTPERPRR | 6.2511 | 6.5154 | 6.7972 | 6.5614 | 6.1264 | 6.5402 | 6.6619 | 6.7585 | 6.6224 | 6.7557 | 7.0941 | 6.9007 | 4.9757 | 5.5235 | 4.8686 | 5.0472 |
| CAD_S1859 | CAD | P27708 | | 1884 | EPELMGTPDGTCY | 6.1910 | 7.5350 | 6.8068 | 6.7028 | 6.0415 | 6.8012 | 7.1353 | 7.3024 | 7.0429 | 6.8152 | 6.9662 | 6.8562 | 4.2065 | 4.5582 | 4.1981 | 4.8408 |
| CAD_S1900 | CAD | P27708 | | 1406 | AGGRRLSSFVTKG | 5.5547 | 4.2655 | 5.4949 | 4.7387 | 7.6028 | 5.6430 | 4.5817 | 6.3539 | 7.5203 | 9.2326 | 7.0642 | 7.7589 | 6.9188 | 7.7491 | 4.7484 | 4.7725 |
| CAD_S1938 | CAD | P27708 | | 1900 | PVPRQASPQNLGT | 6.2520 | 6.6549 | 7.0669 | 7.3427 | 6.7973 | 7.7357 | 6.3011 | 7.4016 | 6.4829 | 6.5019 | 6.3292 | 6.7510 | 4.5830 | 5.5333 | 4.4083 | 3.8580 |
| CAD_T1833 | CAD | P27708 | | 1938 | FTKDQMSHLFNVA | 6.8038 | 6.3180 | 8.2767 | 4.9149 | 6.3154 | 6.4963 | 4.9329 | 6.9046 | 7.5041 | 7.6901 | 6.7263 | 5.3076 | 5.3877 | 5.6074 | 5.1939 | 5.6203 |
| CAD_T1884 | CAD | P27708 | | 1038 | CRVLGTSPEAIDS | 6.1285 | 6.0039 | 6.2492 | 5.7500 | 5.9296 | 5.7757 | 6.5080 | 5.6609 | 6.0354 | 5.8837 | 7.1043 | 6.9423 | 6.4859 | 6.9716 | 6.3324 | 6.2388 |
| CADPS2_S58 | CADPS2 | Q86UW7 | | 58 | GAARSVSPSPSVL | 6.6275 | 6.5488 | 6.8397 | 6.3446 | 5.5054 | 6.2995 | 6.5674 | 6.4197 | 5.9723 | 6.3437 | 6.3764 | 7.0193 | 5.4665 | 5.9873 | 5.5081 | 6.1739 |
| CALCOCO2_S379 | CALCOCO2 | Q13137-4 | | 469 | DHVFCHSLxxxxx | 7.3761 | 7.6850 | 7.4881 | 7.5271 | 6.0884 | 6.7730 | 6.8634 | 6.4884 | 6.9531 | 7.4603 | 7.5500 | 6.7806 | 3.4373 | 4.1141 | 3.6661 | 3.7489 |
| CALCOCO2_S412 | CALCOCO2 | Q13137-4 | | 379 | MGLDFNSLPYQVP | 6.8588 | 6.5851 | 7.5210 | 8.3725 | 6.3520 | 5.5985 | 7.3825 | 7.4940 | 7.0717 | 6.4094 | 5.2505 | 6.9906 | 4.8383 | 4.9194 | 5.3553 | 3.0003 |
| CALCOCO2_S469 | CALCOCO2 | Q13137-4 | | 412 | GIQESSSPSPLSI | 5.6995 | 7.1107 | 7.6493 | 5.8714 | 7.2880 | 6.7524 | 6.9130 | 6.6592 | 6.8903 | 6.9319 | 6.9751 | 6.4653 | 4.0368 | 5.0283 | 4.2549 | 5.4737 |
| CALCOCO2_Y412;S414 | CALCOCO2 | Q13137-4 | | 414 | QESSSPSPLSIKK | 5.9313 | 6.1246 | 7.8881 | 6.4536 | 6.7716 | 7.4050 | 6.1664 | 8.1582 | 7.4069 | 5.5937 | 7.0711 | 7.5358 | 4.4231 | 4.3820 | 3.9376 | 4.7509 |
| CALCOCO2_Y414 | CALCOCO2 | Q13137-4 | 412;414 | | NA | 7.0620 | 5.8920 | 6.6587 | 7.0980 | 6.8461 | 8.0060 | 7.0235 | 7.0059 | 5.7016 | 6.4444 | 8.1256 | 6.1114 | 4.2615 | 4.9206 | 4.2534 | 4.5894 |
| CALD1_S202 | CALD1 | Q05682-4 | | 534 | SVDKVTSPTKVxx | 6.2784 | 6.5700 | 6.7689 | 6.4874 | 6.8459 | 7.1911 | 6.7439 | 6.7736 | 6.9875 | 6.5213 | 7.5068 | 7.2356 | 4.5959 | 4.6797 | 4.2575 | 4.5567 |
| CALD1_S469;T475 | CALD1 | Q05682-4 | | 202 | EKPKRGSIGENQI | 5.8240 | 5.9678 | 6.1106 | 6.1128 | 6.0477 | 6.1967 | 5.9701 | 6.0133 | 6.4056 | 6.4355 | 6.7242 | 6.8517 | 6.1718 | 6.4006 | 6.1948 | 6.5728 |
| CALD1_S504 | CALD1 | Q05682-4 | | 504 | TPDGNKSPAPKPS | 5.7309 | 6.4740 | 7.8639 | 7.2706 | 7.8104 | 9.8333 | 6.8333 | 8.1131 | 7.9434 | 4.9557 | 7.3318 | 6.0872 | 2.8838 | 4.5046 | 3.3165 | 3.0474 |
| CALD1_S534 | CALD1 | Q05682-4 | | 533 | QSVDKVTSPTKVx | 5.9241 | 6.6660 | 7.2471 | 7.6467 | 5.3542 | 6.2282 | 6.6183 | 6.4654 | 5.6423 | 7.2742 | 8.6848 | 7.8808 | 4.3543 | 5.0405 | 4.3336 | 4.6393 |
| CALD1_T533 | CALD1 | Q05682-4 | 469;475 | | NA | 6.6253 | 7.3654 | 7.0915 | 7.0990 | 5.2252 | 7.5218 | 7.2195 | 6.7322 | 7.8237 | 6.9509 | 7.2871 | 7.7141 | 3.3138 | 3.7521 | 3.8737 | 4.4048 |
| CALM1_S102 | CALM1 | P0DP23 | | 102 | DGNGYISAAELRH | 5.8596 | 5.9975 | 6.3004 | 5.4015 | 5.3586 | 5.2178 | 5.6825 | 5.9738 | 5.1363 | 5.5963 | 5.4536 | 5.1489 | 7.3453 | 9.3507 | 8.2583 | 7.9190 |
| CAMK1D_T180 | CAMK1D | Q8IU85 | | 180 | KGDVMSTACGTPG | 8.1295 | 7.5909 | 8.6893 | 8.9123 | 8.0114 | 4.9850 | 5.5089 | 4.8428 | 5.7012 | 5.0469 | 5.9450 | 4.0312 | 5.6195 | 5.9507 | 6.5195 | 4.5159 |
| CAMK2D_S330 | CAMK2D | Q13557 | | 287 | MMHRQETVDCLKK | 6.8169 | 7.0328 | 8.9946 | 7.5323 | 6.9126 | 6.6559 | 6.6750 | 5.8226 | 5.2866 | 5.5038 | 6.5177 | 5.1703 | 4.1603 | 6.6177 | 5.4461 | 4.8548 |

| | | | | | | | | | | | | | | | | | | | | | |
|---|---|---|---|---|---|---|---|---|---|---|---|---|---|---|---|---|---|---|---|---|---|
| CAMK2D_T287 | CAMK2D | Q13557 | | 330 | PDGVKESTESSNT | 8.1031 | 6.5816 | 7.8112 | 7.0926 | 6.9694 | 6.0421 | 6.1800 | 6.1628 | 6.4200 | 6.5666 | 5.9786 | 5.8623 | 4.8871 | 6.1974 | 4.7536 | 4.3917 |
| CAMK2G_S320 | CAMK2G | Q13555-6 | | 287 | MMHRQETVECLRK | 7.6233 | 7.7447 | 8.6591 | 6.9464 | 6.2123 | 6.5941 | 6.6208 | 5.8047 | 5.5835 | 6.0200 | 6.2715 | 5.4139 | 4.1337 | 6.1191 | 5.4626 | 4.7902 |
| CAMK2G_S325 | CAMK2G | Q13555-6 | | 449 | GMQPQPSLCSSAM | 7.0567 | 7.4380 | 8.0177 | 7.5289 | 6.7759 | 5.8349 | 6.5494 | 7.0540 | 6.3287 | 6.2628 | 6.3048 | 5.3522 | 4.9566 | 5.4974 | 5.0676 | 3.9744 |
| CAMK2G_S449 | CAMK2G | Q13555-9 | | 325 | SLLNKKSDGGVKP | 7.8195 | 7.3500 | 7.7830 | 7.6512 | 7.3104 | 6.1741 | 6.1183 | 6.5649 | 7.5956 | 7.4964 | 6.3124 | 6.3286 | 4.1095 | 5.1736 | 3.3878 | 2.8247 |
| CAMK2G_T287 | CAMK2G | Q13555-6 | | 320 | FSVGRQSSAPASP | 7.4432 | 7.4708 | 8.0233 | 6.8182 | 6.3115 | 6.3396 | 6.1718 | 6.4174 | 7.3422 | 6.2163 | 6.1578 | 6.2033 | 5.3457 | 5.1668 | 4.4180 | 4.1540 |
| CAMK2G_T344 | CAMK2G | Q13555-9 | | 344 | SLEPQTTVVHNAT | 7.3259 | 8.1014 | 8.2494 | 8.6507 | 6.7988 | 4.9240 | 6.4658 | 6.0119 | 7.1574 | 6.5143 | 7.0011 | 5.8006 | 3.2119 | 4.4438 | 4.0852 | 5.2578 |
| CAMKK1_S52 | CAMKK1 | Q8N5S9-2 | | 52 | PRARAASVIPGST | 7.8264 | 5.5071 | 6.4795 | 5.3363 | 7.4538 | 4.9935 | 5.1691 | 6.2997 | 7.6487 | 7.6191 | 7.1617 | 7.0869 | 6.0504 | 6.8698 | 4.2232 | 4.2747 |
| CAMKK2_S100 | CAMKK2 | Q96RR4 | | 100 | LSGRKLSLQERSQ | 7.1317 | 6.0454 | 6.7091 | 5.5615 | 7.2513 | 5.9418 | 5.3834 | 6.8989 | 7.4022 | 7.2573 | 7.1057 | 6.8025 | 5.4006 | 6.1369 | 4.6893 | 4.2821 |
| CAMKK2_S129;S133;S13 | CAMKK2 | Q96RR4 | | 495 | TMIRKRSFGNPFE | 5.5374 | 6.3929 | 6.1548 | 5.9558 | 6.1914 | 6.0987 | 5.8129 | 7.4543 | 5.8988 | 7.9365 | 6.9729 | 6.6055 | 5.6318 | 6.5528 | 5.3188 | 5.4846 |
| CAMKK2_S495 | CAMKK2 | Q96RR4 | | 511 | REERSLSAPGNLL | 4.7653 | 5.1720 | 4.5702 | 4.4464 | 5.7091 | 4.4593 | 4.3946 | 6.3460 | 4.5823 | 5.2488 | 5.2035 | 5.0759 | 9.6313 | 10.6297 | 10.6306 | 9.1349 |
| CAMKK2_S511 | CAMKK2 | Q96RR4 | 129;133;137 | | NA | 5.6890 | 5.8579 | 6.6688 | 6.1804 | 5.1614 | 6.0128 | 6.8191 | 6.0175 | 6.3286 | 5.7912 | 7.0503 | 6.6945 | 6.6716 | 7.1794 | 5.6052 | 6.2721 |
| CAMLG_S55 | CAMLG | P49069 | | 55 | GFHRPGSGAEEES | 6.9517 | 7.4633 | 7.9496 | 8.1284 | 6.4197 | 7.2135 | 7.4049 | 7.1172 | 5.1599 | 6.1097 | 6.7724 | 5.7624 | 4.3191 | 4.7127 | 4.4238 | 4.0916 |
| CAMSAP1_S1063 | CAMSAP1 | Q5T5Y3-3 | | 1243 | PLRSRASLIEVDL | 5.5877 | 5.4688 | 5.7967 | 6.1449 | 5.7300 | 5.6987 | 6.0896 | 5.9427 | 6.2571 | 6.5694 | 5.8724 | 6.3357 | 6.5859 | 7.3186 | 7.6334 | 6.9685 |
| CAMSAP1_S1091 | CAMSAP1 | Q5T5Y3-3 | | 1091 | HFVEPLSPTGVAG | 6.0166 | 5.8689 | 6.6864 | 6.5904 | 6.3378 | 6.4176 | 6.4456 | 6.7328 | 6.8250 | 6.6566 | 6.4634 | 6.5874 | 5.7278 | 5.8346 | 5.3234 | 5.4859 |
| CAMSAP1_S1196 | CAMSAP1 | Q5T5Y3-3 | | 804 | DASGRSSPCLSTA | 6.1900 | 6.3713 | 6.5543 | 6.2198 | 6.3041 | 6.3044 | 5.9450 | 6.2821 | 6.5002 | 6.4213 | 6.6400 | 6.4026 | 5.8350 | 6.2476 | 5.7356 | 6.0466 |
| CAMSAP1_S1215 | CAMSAP1 | Q5T5Y3-3 | | 574 | PEFPRASPRALGL | 6.2233 | 6.1870 | 6.7876 | 5.9631 | 6.2111 | 6.6482 | 6.4331 | 6.2167 | 6.5126 | 6.2501 | 6.1172 | 6.1229 | 5.9256 | 6.5879 | 6.0321 | 5.7816 |
| CAMSAP1_S1243 | CAMSAP1 | Q5T5Y3-3 | | 586 | LTANARSPQGQLD | 5.7724 | 5.7951 | 6.2981 | 6.2776 | 5.8264 | 6.5607 | 6.2689 | 6.9197 | 6.1297 | 6.4461 | 6.7570 | 6.5065 | 5.7410 | 6.4652 | 5.9848 | 6.2509 |
| CAMSAP1_S217 | CAMSAP1 | Q5T5Y3-3 | | 217 | KQQLLESPAHQKS | 7.1007 | 7.3994 | 7.0667 | 7.4358 | 6.3335 | 7.4616 | 7.0896 | 6.7685 | 7.2147 | 6.9925 | 7.2876 | 7.2271 | 3.2971 | 4.1628 | 3.1941 | 3.9684 |
| CAMSAP1_S386 | CAMSAP1 | Q5T5Y3-3 | | 733 | YVSCSKSPNSHDS | 6.7727 | 7.7196 | 6.1587 | 6.7528 | 6.5465 | 7.7356 | 5.5159 | 8.3794 | 6.7568 | 5.6076 | 6.1448 | 6.0491 | 4.2620 | 5.7610 | 5.0394 | 4.7982 |
| CAMSAP1_S427 | CAMSAP1 | Q5T5Y3-3 | | 873 | RQKREQSPSQHGK | 6.4619 | 7.3378 | 6.1347 | 6.4115 | 6.3940 | 6.6928 | 5.9591 | 6.9598 | 6.0859 | 5.6084 | 6.8070 | 6.4420 | 4.9186 | 6.2388 | 5.9124 | 5.6352 |
| CAMSAP1_S545 | CAMSAP1 | Q5T5Y3-3 | | 386 | KRSFLGSPAAGTL | 5.9438 | 5.8396 | 6.3976 | 6.0641 | 6.7989 | 5.8840 | 6.1703 | 6.6435 | 6.6144 | 6.6087 | 6.2397 | 6.4287 | 5.8699 | 6.3931 | 5.6753 | 6.4283 |
| CAMSAP1_S574 | CAMSAP1 | Q5T5Y3-3 | | 1196 | TLTLSSSKDANIL | 6.5223 | 6.8065 | 6.4699 | 7.1318 | 6.4115 | 6.8995 | 7.0632 | 6.6499 | 7.3920 | 6.7924 | 7.2543 | 6.8548 | 4.0136 | 4.9458 | 4.0746 | 4.7178 |
| CAMSAP1_S586 | CAMSAP1 | Q5T5Y3-3 | | 640 | IVSRRPSEGPQPL | 6.3787 | 5.4679 | 5.3538 | 4.4371 | 6.5256 | 5.5620 | 5.2617 | 5.2810 | 8.3917 | 7.6771 | 7.5491 | 7.7855 | 6.9669 | 7.8377 | 4.2186 | 5.3053 |
| CAMSAP1_S640 | CAMSAP1 | Q5T5Y3-3 | | 1215 | KEVLDASVKEVGS | 8.1874 | 6.8376 | 8.4289 | 7.1431 | 6.7630 | 6.4292 | 8.5781 | 5.9035 | 7.4699 | 6.9442 | 4.5182 | 7.3243 | 3.1009 | 5.3600 | 3.6195 | 3.3922 |
| CAMSAP1_S733 | CAMSAP1 | Q5T5Y3-3 | | 838 | KLQRLNSCETKSS | 6.2786 | 5.7965 | 7.0097 | 5.3139 | 5.8603 | 6.2259 | 5.9967 | 6.0238 | 6.1099 | 6.1402 | 5.9825 | 5.6653 | 7.0207 | 6.6751 | 6.9071 | 6.9937 |
| CAMSAP1_S804 | CAMSAP1 | Q5T5Y3-3 | | 427 | KGTAAFSPSHPLL | 6.7148 | 6.3961 | 6.6097 | 6.9714 | 6.5662 | 6.8333 | 6.7118 | 6.7473 | 7.9073 | 6.6877 | 6.7403 | 6.8983 | 4.2911 | 4.8859 | 4.4071 | 4.6317 |
| CAMSAP1_S838 | CAMSAP1 | Q5T5Y3-3 | | 1063 | EQLLMKSPTVPVP | 6.3247 | 6.8682 | 6.5106 | 6.9927 | 6.4192 | 7.4227 | 7.0068 | 6.9604 | 7.5180 | 6.8091 | 7.1742 | 7.4830 | 3.7147 | 4.6300 | 3.5032 | 4.6623 |
| CAMSAP1_S873 | CAMSAP1 | Q5T5Y3-3 | | 545 | SLLSNVSIEDEEE | 6.2906 | 6.5354 | 6.8184 | 7.0872 | 6.1662 | 7.0411 | 5.9896 | 6.0880 | 6.5435 | 7.0632 | 5.4097 | 6.6438 | 5.3415 | 6.3528 | 6.4351 | 4.1938 |
| CAMSAP2_S1019 | CAMSAP2 | Q08AD1 | | 416 | GIRRSSSMSYVDG | 5.9964 | 5.9738 | 6.2681 | 5.7781 | 5.7298 | 6.2392 | 5.8636 | 6.0990 | 6.1426 | 6.3901 | 6.2716 | 6.6166 | 6.5785 | 7.0172 | 6.4523 | 6.5833 |
| CAMSAP2_S1148 | CAMSAP2 | Q08AD1 | | 599 | IKLNQSSPDNVTD | 6.0289 | 6.8701 | 6.9384 | 7.1627 | 5.9481 | 7.0681 | 6.5022 | 6.4230 | 6.6290 | 7.0751 | 7.0528 | 7.4396 | 4.4880 | 5.2567 | 4.6730 | 4.8043 |
| CAMSAP2_S1319 | CAMSAP2 | Q08AD1 | | 936 | ISPPQPSPQKQIR | 6.0429 | 6.3148 | 6.8849 | 6.5382 | 5.8790 | 6.5446 | 7.1616 | 6.8574 | 6.4795 | 7.1093 | 7.1312 | 7.0914 | 4.2740 | 5.4040 | 4.4571 | 5.8302 |
| CAMSAP2_S416 | CAMSAP2 | Q08AD1 | | 673 | NTREALSPCPSTV | 5.9787 | 6.1650 | 6.3398 | 6.1564 | 6.1394 | 6.9786 | 7.7047 | 7.0921 | 6.7010 | 6.0779 | 6.7749 | 6.4874 | 4.8665 | 5.7517 | 5.2964 | 5.4896 |
| CAMSAP2_S599 | CAMSAP2 | Q08AD1 | | 1019 | PRLRRFSPSQVPI | 5.5685 | 6.2132 | 7.4548 | 6.9141 | 5.2507 | 6.8027 | 5.9443 | 7.4815 | 6.7410 | 6.9592 | 7.1950 | 7.3802 | 4.5324 | 5.5363 | 4.4734 | 5.5528 |
| CAMSAP2_S611 | CAMSAP2 | Q08AD1 | | 877 | KQWNLASPSEETL | 6.1729 | 6.4080 | 7.2725 | 7.0985 | 6.1520 | 6.6089 | 6.8325 | 7.2351 | 6.7169 | 6.7724 | 6.3843 | 6.2677 | 4.5983 | 5.1573 | 5.0130 | 5.3097 |
| CAMSAP2_S673 | CAMSAP2 | Q08AD1 | | 1319 | SVEGFLSPSRCGS | 6.7475 | 6.8957 | 7.0240 | 6.0220 | 6.1148 | 6.6254 | 6.6670 | 6.6415 | 7.2735 | 6.5957 | 6.4193 | 6.4428 | 4.9194 | 5.2246 | 4.9645 | 5.4222 |
| CAMSAP2_S877 | CAMSAP2 | Q08AD1 | | 1148 | EKYDGESDKEQFD | 6.7860 | 6.6632 | 6.8276 | 5.9974 | 5.7458 | 5.2254 | 6.0933 | 6.1064 | 6.1867 | 6.2869 | 5.8178 | 6.0805 | 6.9466 | 6.1722 | 6.2019 | 6.8625 |
| CAMSAP2_S936 | CAMSAP2 | Q08AD1 | | 611 | DTKGALSPITDNT | 5.9571 | 5.1283 | 5.4855 | 6.4539 | 5.4603 | 5.8449 | 4.7231 | 5.7739 | 7.1476 | 8.1824 | 6.8408 | 5.9983 | 6.9713 | 6.7469 | 6.2341 | 7.0514 |
| CAMSAP3_S1101 | CAMSAP3 | Q9P1Y5-2 | | 361 | PDGHAASPRGTEA | 6.0163 | 6.1358 | 6.2481 | 5.9444 | 6.5081 | 6.5609 | 6.1592 | 5.9746 | 5.7785 | 5.6840 | 6.6435 | 6.1338 | 6.2247 | 6.3791 | 6.8916 | 6.7173 |
| CAMSAP3_S361 | CAMSAP3 | Q9P1Y5-2 | | 395 | SSGGPQSPLRGST | 6.7490 | 6.1324 | 7.1895 | 6.9074 | 6.8546 | 6.7370 | 7.4553 | 6.3199 | 6.6565 | 6.2986 | 6.8734 | 6.6423 | 4.3488 | 5.3627 | 4.5884 | 4.8843 |
| CAMSAP3_S395 | CAMSAP3 | Q9P1Y5-2 | | 574 | PAPSEGSPKAVAS | 5.7854 | 6.9337 | 7.1138 | 6.7132 | 6.0632 | 6.6607 | 5.8895 | 5.9767 | 6.7342 | 6.4694 | 6.7585 | 8.1216 | 4.7331 | 5.4954 | 4.7622 | 5.7894 |
| CAMSAP3_S574 | CAMSAP3 | Q9P1Y5-2 | | 783 | TGPKAASPSPARR | 6.4009 | 6.3900 | 6.0908 | 5.6240 | 5.5210 | 6.7976 | 6.3584 | 6.8526 | 6.3211 | 6.3491 | 6.2365 | 6.3358 | 5.8834 | 6.7290 | 5.8422 | 6.2675 |
| CAMSAP3_S783 | CAMSAP3 | Q9P1Y5-2 | | 1101 | PTSRAPSPSGLMS | 5.8183 | 5.4484 | 6.2330 | 6.2082 | 6.2720 | 6.0118 | 6.2667 | 6.3456 | 6.2004 | 5.8427 | 6.3505 | 6.5196 | 6.3315 | 6.8320 | 6.6682 | 6.6512 |
| CAND1_S335;S340 | CAND1 | Q86VP6 | | 557 | RPLDQPSSFDATP | 5.5519 | 5.9584 | 7.0038 | 5.2697 | 6.0010 | 6.7500 | 5.8057 | 5.8412 | 6.3506 | 6.1820 | 4.2616 | 6.8014 | 7.5522 | 7.9445 | 6.9878 | 5.7383 |
| CAND1_S557 | CAND1 | Q86VP6 | 335;340 | | NA | 5.8832 | 6.1454 | 6.0486 | 5.5937 | 6.1228 | 6.0437 | 6.0766 | 5.7988 | 6.0311 | 7.2319 | 5.7700 | 5.9845 | 6.4221 | 7.2342 | 7.3109 | 6.3025 |
| CAND2_S230 | CAND2 | O75155 | | 230 | GPRVPTSPTAIRT | 5.7514 | 5.9717 | 6.4960 | 5.5352 | 5.9692 | 5.9245 | 5.6393 | 6.2901 | 6.8283 | 5.9136 | 6.2331 | 5.9052 | 6.6239 | 8.1879 | 6.5381 | 6.1924 |
| CANT1_S21 | CANT1 | Q8WVQ1 | | 21 | MHSLRISVGGLPV | 2.7528 | 3.1973 | 3.2902 | 2.8849 | 3.0859 | 3.0732 | 3.5707 | 3.6689 | 3.9468 | 3.8524 | 2.7725 | 3.2843 | 15.3428 | 14.5970 | 15.5183 | 15.1619 |
| CANX_S116 | CANX | P27824-2 | | 618 | DEILNRSPRNRKP | 6.4119 | 6.5342 | 5.9624 | 6.3937 | 6.2233 | 6.0190 | 6.2174 | 5.9667 | 6.2220 | 6.0802 | 6.0480 | 6.1831 | 6.5108 | 6.4951 | 6.4456 | 6.2868 |
| CANX_S159 | CANX | P27824-2 | | 589 | LEEKQKSDAEEDG | 6.1430 | 5.9536 | 6.5497 | 5.7925 | 6.1247 | 6.1135 | 5.8727 | 5.1367 | 5.8681 | 6.0359 | 6.1835 | 6.1158 | 7.6443 | 7.4874 | 7.0689 | 6.8096 |
| CANX_S589 | CANX | P27824-2 | | 599 | EDGGTVSQEEEDR | 5.6546 | 5.1998 | 5.6262 | 6.2527 | 5.7572 | 6.1006 | 6.2854 | 5.3200 | 6.0422 | 6.1295 | 6.2966 | 6.0598 | 7.4527 | 7.3371 | 7.2475 | 7.2380 |
| CANX_S589;S599 | CANX | P27824-2 | | 159 | KGLVLMSRAKHHA | 5.6895 | 5.2347 | 4.1408 | 4.0403 | 4.9642 | 6.2472 | 6.4698 | 8.8059 | 7.4104 | 5.7852 | 7.2620 | 6.0198 | 6.5440 | 7.3195 | 5.7445 | 8.3224 |
| CANX_S599 | CANX | P27824-2 | | 116 | FDRGTLSGWILSK | 6.5239 | 5.7551 | 5.9967 | 5.9363 | 4.6458 | 7.1664 | 6.9852 | 4.5195 | 4.1186 | 6.3811 | 6.5678 | 9.9012 | 6.5294 | 5.2059 | 7.5069 | 6.2603 |
| CANX_S618 | CANX | P27824-2 | 589;599 | | NA | 6.9112 | 6.6607 | 6.5210 | 6.7894 | 6.6937 | 6.4598 | 6.8395 | 6.0381 | 6.1568 | 6.2057 | 6.1076 | 6.0490 | 5.7194 | 5.7826 | 5.4565 | 5.6093 |
| CAP1_S308 | CAP1 | Q01518 | | 308 | APKPQTSPSPKRA | 5.6863 | 4.9339 | 7.0347 | 5.8005 | 6.8849 | 6.8976 | 6.1613 | 6.3676 | 6.5903 | 6.7974 | 6.8218 | 6.8428 | 5.9130 | 6.3558 | 5.1722 | 5.7397 |
| CAP1_S34 | CAP1 | Q01518 | | 34 | HRGYADSPSKAGA | 6.8044 | 7.4709 | 6.9080 | 7.0972 | 6.3961 | 7.4953 | 7.4790 | 7.2452 | 7.2538 | 7.3667 | 7.6872 | 7.5233 | 2.8274 | 3.9494 | 2.7422 | 3.7538 |
| CAP1_T307 | CAP1 | Q01518 | | 310 | KPQTSPSPKRATK | 6.6089 | 5.2258 | 7.3839 | 5.6900 | 6.6103 | 6.7455 | 6.1652 | 6.0086 | 6.7549 | 6.5184 | 7.1601 | 7.0633 | 5.6613 | 5.6549 | 5.0951 | 5.6539 |
| CAP1_T307;S310 | CAP1 | Q01518 | | 307 | SAPKPQTSPSPKR | 6.0758 | 5.1879 | 7.3080 | 5.8061 | 6.4934 | 6.5753 | 5.9932 | 6.1214 | 6.4786 | 6.6247 | 7.3410 | 7.0005 | 5.6098 | 6.2703 | 5.3077 | 5.8065 |
| CAP1_T310 | CAP1 | Q01518 | 307;310 | | NA | 6.2542 | 6.0597 | 7.4067 | 6.9316 | 6.4742 | 6.7984 | 7.0410 | 6.1015 | 7.0852 | 7.1646 | 7.4265 | 7.5315 | 4.2655 | 4.6740 | 3.9503 | 4.8350 |
| CAP2_S301 | CAP2 | P40123 | | 301 | QGGQTQSPTKSHT | 5.8784 | 6.1031 | 7.0448 | 6.3746 | 6.4985 | 7.0025 | 6.4540 | 6.9158 | 6.6506 | 6.7147 | 6.9046 | 7.0632 | 4.7750 | 5.5423 | 4.6788 | 5.3989 |
| CAP2_S309 | CAP2 | P40123 | | 312 | HTPSPTSPKSYPS | 6.0696 | 6.1111 | 6.6993 | 6.3711 | 6.3759 | 6.8479 | 6.2793 | 6.4220 | 6.8753 | 6.5127 | 6.9877 | 7.0834 | 5.2054 | 5.8685 | 5.1523 | 5.1385 |
| CAP2_S312 | CAP2 | P40123 | | 309 | TKSHTPSPTSPKS | 5.7913 | 6.1486 | 6.8990 | 6.1586 | 6.2487 | 6.5867 | 6.0918 | 6.7168 | 6.2426 | 6.9726 | 6.7864 | 6.9855 | 5.4137 | 6.0164 | 5.2594 | 5.6818 |
| CAPN15_S296 | CAPN15 | O75808 | | 296 | LSGKRLSVLEEEA | 5.9018 | 6.1723 | 6.1779 | 6.1794 | 6.3223 | 5.7990 | 6.8552 | 6.3567 | 6.5057 | 6.3810 | 6.7513 | 6.5877 | 6.1247 | 6.5791 | 5.7478 | 5.5579 |
| CAPN15_S335 | CAPN15 | O75808 | | 335 | VRYTPASPSSPDF | 6.3047 | 6.5517 | 6.1022 | 5.9945 | 6.5344 | 6.7456 | 5.9916 | 6.1567 | 6.4540 | 5.2161 | 7.5555 | 7.8916 | 5.3827 | 5.0394 | 6.3707 | 5.7087 |
| CAPN9_T246 | CAPN9 | O14815 | | 246 | AESEARTPFGLIK | 6.2382 | 6.3453 | 6.7719 | 6.2854 | 6.1396 | 6.4281 | 6.2437 | 6.8535 | 6.4966 | 6.2562 | 7.0307 | 6.8144 | 5.7188 | 5.7497 | 5.2057 | 5.4221 |
| CAPRIN1_S335 | CAPRIN1 | Q14444 | | 335 | QQPQAASPSVPEP | 6.5883 | 6.6939 | 6.6600 | 6.7200 | 6.2785 | 6.0007 | 6.3202 | 6.3183 | 6.0033 | 6.5615 | 6.3783 | 7.1459 | 5.8441 | 6.1187 | 5.3354 | 5.0328 |
| CAPRIN1_S504 | CAPRIN1 | Q14444 | | 504 | SKPLHSSGINVNA | 5.8538 | 5.4468 | 5.8350 | 6.3886 | 6.1822 | 6.2936 | 5.2251 | 6.8390 | 5.9558 | 6.4742 | 6.0004 | 5.7224 | 7.5052 | 7.3469 | 5.7846 | 7.1463 |
| CAPZA1_T63 | CAPZA1 | P52907 | | 63 | YNMDQFTPVKIEG | 5.6812 | 6.5341 | 6.9986 | 6.5980 | 6.2631 | 6.8469 | 6.1655 | 6.2946 | 6.4122 | 6.4165 | 6.4485 | 6.4330 | 5.4410 | 5.9505 | 5.5007 | 6.0157 |
| CAPZA2_T63 | CAPZA2 | P47755 | | 63 | YNLDQFTPVKIEG | 5.6796 | 6.7778 | 6.6906 | 6.5645 | 6.3030 | 5.4840 | 5.9954 | 6.5999 | 6.0696 | 6.2829 | 6.4738 | 7.1349 | 5.7597 | 6.4184 | 6.2334 | 5.5323 |
| CARD10_S574 | CARD10 | Q9BWT7 | | 574 | SSSSSDSVWPLGK | 5.9103 | 5.4522 | 7.7688 | 5.9053 | 5.2635 | 6.6820 | 5.6534 | 6.8957 | 6.9088 | 7.4888 | 6.9926 | 7.4743 | 5.4221 | 5.8777 | 4.8495 | 5.4552 |
| CARHSP1_S30;S32 | CARHSP1 | Q9Y2V2 | | 41 | RGNVVPSPLPTRR | 6.4608 | 6.1614 | 6.7246 | 5.5724 | 6.5256 | 6.4372 | 5.7543 | 7.1740 | 6.5034 | 6.8055 | 6.8477 | 6.8038 | 5.7074 | 5.9630 | 5.2567 | 5.3020 |
| CARHSP1_S41 | CARHSP1 | Q9Y2V2 | | 52 | RRTRTFSATVRAS | 7.0058 | 6.1172 | 7.2058 | 5.4312 | 6.6895 | 6.4500 | 5.8384 | 7.1035 | 6.4894 | 6.4942 | 6.6728 | 6.7216 | 6.0419 | 5.5879 | 5.1972 | 4.9537 |
| CARHSP1_S52 | CARHSP1 | Q9Y2V2 | | 58 | SATVRASQGPVYK | 7.0683 | 7.5593 | 6.4549 | 6.5161 | 6.4591 | 6.7719 | 6.7972 | 6.6273 | 6.7188 | 5.8862 | 6.6932 | 5.7323 | 4.8360 | 5.7548 | 4.6999 | 5.4247 |
| CARHSP1_S58 | CARHSP1 | Q9Y2V2 | 30;32 | | NA | 6.1499 | 6.1195 | 6.5420 | 5.4438 | 6.2452 | 6.4941 | 5.7970 | 7.0634 | 6.1849 | 6.6451 | 6.7500 | 6.7492 | 5.8883 | 6.2191 | 5.7606 | 5.9478 |
| CARMIL1_S1067 | CARMIL1 | Q5VZK9 | | 916 | LDTCMMTPKSKRK | 6.2196 | 6.0561 | 6.4838 | 6.1775 | 6.2210 | 6.1503 | 6.1890 | 6.2448 | 6.8560 | 6.3112 | 6.3033 | 6.1770 | 6.3713 | 6.3686 | 5.9763 | 5.8941 |
| CARMIL1_S1094 | CARMIL1 | Q5VZK9 | | 1228 | NRFGLGTPEKNTK | 6.9465 | 7.0709 | 7.0801 | 6.4158 | 6.4397 | 7.0871 | 6.9132 | 6.9974 | 7.8111 | 7.3291 | 7.4544 | 7.3356 | 3.4665 | 4.4104 | 3.2632 | 3.9789 |
| CARMIL1_S122 | CARMIL1 | Q5VZK9 | | 1331 | QEVSRRSWGQQAQ | 6.1036 | 6.0697 | 6.5448 | 6.3750 | 6.6797 | 6.4820 | 6.4216 | 6.4941 | 6.9159 | 6.3968 | 6.3657 | 6.3654 | 5.8990 | 5.8840 | 5.5026 | 5.5001 |
| CARMIL1_S1254 | CARMIL1 | Q5VZK9 | | 1254 | SSSTPTSPKPLLQ | 6.0715 | 6.1318 | 6.2266 | 5.6912 | 6.1199 | 6.4045 | 6.2096 | 6.3454 | 6.7183 | 6.5868 | 6.3878 | 6.6679 | 5.9617 | 6.1687 | 5.8497 | 6.4586 |
| CARMIL1_S1261 | CARMIL1 | Q5VZK9 | | 1261 | PKPLLQSPKPSLA | 6.9897 | 5.6601 | 6.9254 | 6.0884 | 6.8662 | 6.8596 | 6.5781 | 6.2482 | 7.5268 | 6.7108 | 6.3940 | 6.5129 | 5.3444 | 5.7347 | 4.5662 | 4.9945 |
| CARMIL1_S1291 | CARMIL1 | Q5VZK9 | | 122 | KIFPGLSPVRIMK | 6.1195 | 7.1382 | 6.8409 | 5.7034 | 6.1086 | 6.8416 | 6.4468 | 6.1037 | 6.8316 | 6.8357 | 6.3763 | 6.6961 | 5.2234 | 5.7643 | 5.8570 | 5.1129 |
| CARMIL1_S1331 | CARMIL1 | Q5VZK9 | | 1094 | MTEEPSSPKGAVR | 6.0718 | 6.0985 | 6.8335 | 6.6182 | 5.9537 | 6.3607 | 6.4986 | 6.6052 | 6.4914 | 7.1394 | 6.8503 | 6.9000 | 5.3957 | 5.6814 | 5.1735 | 5.3281 |
| CARMIL1_S933 | CARMIL1 | Q5VZK9 | | 968 | RQEKRSSGFISEL | 6.2511 | 5.9914 | 7.0731 | 5.8837 | 6.3840 | 6.1818 | 6.3132 | 6.4776 | 6.9082 | 5.9463 | 6.1171 | 7.0212 | 6.1730 | 6.0412 | 5.7652 | 5.4718 |
| CARMIL1_S968 | CARMIL1 | Q5VZK9 | | 933 | RMLRPVSRAFEME | 8.0490 | 7.4352 | 6.8055 | 5.5429 | 5.5167 | 6.8308 | 6.1127 | 6.0020 | 6.8715 | 6.7504 | 6.4483 | 6.6998 | 5.7786 | 6.0362 | 3.9177 | 5.2028 |
| CARMIL1_T1228 | CARMIL1 | Q5VZK9 | | 1067 | KRDSRKSSGFLNL | 7.5210 | 7.0698 | 6.3906 | 6.8326 | 6.1646 | 6.6668 | 7.0177 | 5.8281 | 8.0909 | 7.5084 | 7.5712 | 7.5549 | 3.5364 | 5.1021 | 2.8509 | 4.2940 |
| CARMIL1_T916 | CARMIL1 | Q5VZK9 | | 1291 | IPDSPSSPKVALL | 5.9487 | 5.7084 | 6.0729 | 5.3932 | 6.5272 | 6.1359 | 6.3593 | 5.4868 | 6.6285 | 6.6760 | 5.9794 | 6.1810 | 7.0458 | 6.6725 | 6.4628 | 6.7217 |
| CARS1_S102 | CARS1 | P49589-3 | | 79 | IEALLGSPCGKGQ | 7.1527 | 7.6646 | 6.9506 | 7.2712 | 6.7903 | 7.0265 | 7.0800 | 6.4550 | 6.9895 | 7.4159 | 7.4647 | 7.0817 | 3.1708 | 4.0656 | 3.2914 | 4.1296 |
| CARS1_S79 | CARS1 | P49589-3 | | 102 | RVQPQWSPPAGTQ | 5.9469 | 6.4802 | 6.8296 | 6.9949 | 6.1722 | 5.6472 | 5.7764 | 7.2223 | 6.6766 | 6.1567 | 7.0115 | 6.2880 | 5.0943 | 6.4316 | 5.6978 | 5.5738 |
| CASC3_S148 | CASC3 | O15234 | | 443 | VPETTPTPPTKTG | 6.3947 | 6.5140 | 6.6987 | 6.7210 | 6.2511 | 6.7087 | 6.5203 | 6.5254 | 6.6554 | 6.7072 | 6.8302 | 6.5324 | 5.3154 | 5.4587 | 5.1550 | 5.0116 |
| CASC3_S363 | CASC3 | O15234 | | 363 | TSVRDPSPEADAP | 6.2318 | 6.3023 | 6.5094 | 5.9994 | 6.2683 | 6.1599 | 5.9132 | 6.1948 | 6.3105 | 6.5903 | 6.2325 | 5.9797 | 6.5004 | 6.5074 | 6.2410 | 6.0591 |
| CASC3_T443 | CASC3 | O15234 | | 148 | VTGERQSGDGQES | 6.6390 | 5.3301 | 7.3902 | 5.9451 | 5.5666 | 6.4970 | 6.1243 | 6.0311 | 6.0010 | 6.7465 | 7.5937 | 7.6195 | 5.2398 | 6.5974 | 5.4175 | 5.2613 |
| CASK_S313 | CASK | O14936 | | 313 | AVLAAVSSHKFNS | 5.5108 | 5.9026 | 5.7265 | 6.3617 | 5.2530 | 5.7757 | 5.7370 | 5.7395 | 5.4525 | 5.2131 | 5.3773 | 5.6553 | 7.7609 | 7.8056 | 8.6776 | 8.0510 |
| CASKIN1_S787 | CASKIN1 | Q8WXD9 | | 787 | TKTRPGSPQALGG | 6.3071 | 6.5132 | 6.8360 | 5.8092 | 5.0953 | 6.0427 | 6.1617 | 6.3270 | 7.0585 | 5.8455 | 6.2300 | 6.1430 | 5.9943 | 7.0486 | 6.9858 | 5.6021 |
| CASKIN2_S396 | CASKIN2 | Q8WXE0 | | 695 | PLPPARSPSQESI | 6.0654 | 6.4361 | 6.5285 | 6.6680 | 6.2618 | 6.4121 | 6.6651 | 7.4559 | 6.9445 | 7.3272 | 6.9732 | 7.2081 | 4.6417 | 5.0641 | 4.4481 | 4.9003 |
| CASKIN2_S403 | CASKIN2 | Q8WXE0 | | 858 | GTPRSQSFALRAR | 10.0007 | 6.8163 | 7.0815 | 6.5213 | 6.1082 | 5.7045 | 6.2197 | 5.4844 | 7.2626 | 5.5744 | 6.3337 | 6.4238 | 5.5404 | 6.2514 | 4.0985 | 4.5788 |
| CASKIN2_S695 | CASKIN2 | Q8WXE0 | | 396 | VGLSPDSPAGDRN | 6.6858 | 7.2273 | 6.6632 | 6.8380 | 6.1021 | 7.2580 | 7.0718 | 6.2568 | 6.7142 | 7.5592 | 6.9247 | 6.7061 | 4.0140 | 4.9728 | 4.3290 | 4.6769 |
| CASKIN2_S800 | CASKIN2 | Q8WXE0 | | 403 | PAGDRNSVGSEGS | 5.7605 | 5.6010 | 6.8632 | 5.9329 | 5.9594 | 6.0580 | 6.1625 | 6.1281 | 6.2532 | 6.6788 | 5.9129 | 6.2698 | 6.1988 | 7.0615 | 6.6791 | 6.4803 |
| CASKIN2_S858 | CASKIN2 | Q8WXE0 | | 800 | PKRRSHSLSRPGP | 6.3010 | 6.7175 | 6.6435 | 6.0698 | 5.9685 | 6.2657 | 6.0885 | 6.4569 | 7.0500 | 7.2479 | 6.4636 | 7.2412 | 5.4314 | 5.3261 | 4.9137 | 5.8146 |
| CASKIN2_S946 | CASKIN2 | Q8WXE0 | | 946 | PPSRGSSGEGLPF | 5.6134 | 5.2517 | 6.0314 | 6.1292 | 5.7905 | 6.3054 | 4.9129 | 7.4136 | 6.5744 | 6.6502 | 7.2306 | 6.1386 | 6.6408 | 6.5931 | 6.4442 | 6.2799 |
| CASP4_S16 | CASP4 | P49662 | | 83 | FNIDQISPNKKAH | 6.0207 | 6.5341 | 7.0762 | 6.4761 | 6.5794 | 6.0098 | 7.7461 | 6.6282 | 7.3079 | 6.4848 | 7.0062 | 7.0154 | 4.4398 | 5.6392 | 4.7731 | 5.1631 |
| CASP4_S83 | CASP4 | P49662 | | 16 | PLKVLESLGKDFL | 7.0134 | 6.0851 | 6.6271 | 7.1884 | 6.2880 | 5.4942 | 5.9611 | 6.1506 | 6.5474 | 5.6729 | 5.6150 | 6.1125 | 5.5311 | 6.3881 | 6.6047 | 6.7206 |
| CASP8AP2_S1368 | CASP8AP2 | Q9UKL3 | | 875 | KVLRNESPPQVPV | 6.9158 | 7.0322 | 7.1766 | 7.6738 | 6.5053 | 5.9171 | 6.2166 | 6.5111 | 6.9796 | 7.0873 | 6.5528 | 6.6812 | 4.5299 | 5.0079 | 4.4251 | 4.7878 |
| CASP8AP2_S1537 | CASP8AP2 | Q9UKL3 | | 1368 | LVSGVASPCPKMI | 6.8870 | 7.0009 | 7.3277 | 7.5858 | 6.0194 | 5.8312 | 6.5491 | 6.0973 | 6.9789 | 6.5971 | 6.8243 | 7.0124 | 4.6153 | 5.0126 | 4.4585 | 5.2026 |
| CASP8AP2_S567 | CASP8AP2 | Q9UKL3 | | 567 | KLNLTLSPAKKQP | 6.6376 | 6.8875 | 7.3730 | 7.8996 | 6.1374 | 6.1050 | 6.4881 | 6.5935 | 6.2357 | 6.7152 | 6.6589 | 7.5564 | 4.1158 | 5.0924 | 4.9013 | 4.6028 |
| CASP8AP2_S875 | CASP8AP2 | Q9UKL3 | | 1537 | AVIMEKSNDHFIV | 5.6101 | 5.7129 | 7.4870 | 7.7862 | 5.0164 | 5.8345 | 6.8555 | 6.1040 | 5.4780 | 5.9572 | 6.7480 | 5.9474 | 5.8820 | 6.5532 | 7.2205 | 5.8072 |
| CASP8AP2_S940 | CASP8AP2 | Q9UKL3 | | 1261 | EVHCPSTPKSEKN | 6.3815 | 6.8362 | 6.4147 | 7.1481 | 6.0214 | 5.7671 | 7.3790 | 6.1361 | 6.2895 | 7.8303 | 7.0360 | 6.5703 | 4.5275 | 5.2740 | 4.8057 | 5.5825 |
| CASP8AP2_T1261 | CASP8AP2 | Q9UKL3 | | 940 | EEGEIRSDSETSK | 6.3246 | 6.8594 | 6.2249 | 8.1164 | 7.0537 | 5.6987 | 6.3409 | 6.1784 | 7.1732 | 6.4338 | 7.7071 | 7.2093 | 4.4778 | 5.3608 | 4.0049 | 4.8363 |
| CAST_S11 | CAST | P20810-6 | | 326 | SDFTCGSPTAAGK | 6.4499 | 6.1906 | 7.3780 | 6.6660 | 6.1496 | 6.6465 | 6.6186 | 6.4702 | 7.2141 | 7.1889 | 7.6689 | 7.1352 | 4.5491 | 4.8548 | 4.2620 | 4.5576 |
| CAST_S216 | CAST | P20810-6 | | 11 | GQKPAASPRPRRA | 5.9099 | 5.6239 | 6.6595 | 5.7176 | 5.8435 | 7.0912 | 6.2819 | 5.8923 | 6.1233 | 6.4009 | 6.7886 | 7.0263 | 6.5531 | 6.4577 | 5.4652 | 6.1651 |
| CAST_S326 | CAST | P20810-6 | | 494 | EAVCRTSMCSIQS | 6.2732 | 6.5658 | 6.5646 | 6.9240 | 6.1459 | 6.5975 | 6.9039 | 6.4204 | 6.9496 | 6.8277 | 7.0430 | 6.8314 | 4.9754 | 5.3984 | 4.7110 | 4.8683 |
| CAST_S34 | CAST | P20810-6 | | 34 | SEKTSESPSKPGE | 6.4823 | 6.5830 | 7.3528 | 6.3399 | 6.1473 | 6.7509 | 6.7377 | 6.6432 | 7.1776 | 7.3486 | 7.1413 | 7.1007 | 4.3416 | 4.7880 | 4.1618 | 4.9034 |
| CAST_S494 | CAST | P20810-6 | | 218 | KEKKSLTPAVPVE | 6.7135 | 7.1801 | 6.9464 | 7.8383 | 5.2150 | 6.5358 | 6.6277 | 6.8588 | 6.7976 | 7.9398 | 6.2514 | 7.1188 | 3.5695 | 5.0881 | 4.4929 | 4.8264 |
| CAST_S610 | CAST | P20810-6 | | 658 | DALDKLSDSLGQR | 6.0370 | 5.6917 | 6.3369 | 5.7913 | 5.9137 | 6.5239 | 6.6548 | 6.5242 | 6.3332 | 6.0087 | 5.9321 | 6.1792 | 7.0843 | 7.0249 | 5.8777 | 6.0865 |
| CAST_S658 | CAST | P20810-6 | | 216 | KKKEKKSLTPAVP | 5.9775 | 7.2021 | 6.5875 | 6.6882 | 5.9096 | 6.6544 | 6.4371 | 6.9546 | 7.2969 | 6.9929 | 7.2972 | 7.2355 | 4.6454 | 4.7048 | 4.5131 | 4.9033 |
| CAST_S67 | CAST | P20810-6 | | 67 | YAGGTASATKVSA | 6.6099 | 7.6884 | 6.6816 | 7.0863 | 5.9953 | 7.6402 | 7.7942 | 6.2893 | 7.5844 | 8.0546 | 7.0079 | 7.9899 | 3.7368 | 4.0007 | 2.3395 | 3.5009 |
| CAST_T218 | CAST | P20810-6 | | 610 | GPQNASSLKFEDA | 6.1363 | 6.3959 | 6.5589 | 6.4034 | 5.7892 | 7.4546 | 7.0335 | 7.0908 | 6.7045 | 6.0503 | 6.9716 | 6.4261 | 4.5947 | 5.7107 | 4.5818 | 6.0978 |
| CAV1_S37 | CAV1 | Q03135 | | 37 | AMADELSEKQVYD | 5.6843 | 5.7460 | 5.5806 | 5.8235 | 6.1281 | 5.8076 | 6.2530 | 5.8423 | 5.7594 | 5.5484 | 5.5325 | 5.5309 | 7.9819 | 7.3892 | 7.7800 | 7.6123 |
| CAV1_S6 | CAV1 | Q03135 | | 9 | GGKYVDSEGHLYT | 5.7456 | 5.7664 | 5.9950 | 6.5736 | 5.8046 | 5.7147 | 6.0155 | 6.0293 | 6.0844 | 5.8541 | 5.9241 | 5.7059 | 6.9646 | 7.3774 | 7.2221 | 7.2227 |
| CAV1_S9 | CAV1 | Q03135 | | 15 | SEGHLYTVPIREQ | 7.7675 | 7.8831 | 6.7369 | 7.8412 | 7.2943 | 6.6969 | 6.4791 | 7.0542 | 7.4948 | 6.3106 | 6.5305 | 6.6101 | 3.9009 | 4.0598 | 3.2669 | 4.0731 |
| CAV1_T15 | CAV1 | Q03135-2 | | 6 | xMADELSEKQVYD | 5.1268 | 5.4239 | 6.2025 | 5.5199 | 6.3932 | 6.5437 | 5.0366 | 6.4785 | 5.6149 | 4.9162 | 6.2353 | 4.9397 | 10.3983 | 7.6062 | 6.7571 | 6.8072 |
| CAV2_S18;Y20 | CAV2 | P51636 | | 23 | DSYSHHSGLEYAD | 6.0197 | 6.6092 | 5.1472 | 6.2166 | 6.9597 | 5.5729 | 5.6592 | 7.7770 | 6.3887 | 6.0155 | 5.5250 | 6.0745 | 5.7966 | 6.8818 | 6.6966 | 6.6597 |
| CAV2_Y18;S20;S23 | CAV2 | P51636 | 20;23 | | NA | 6.3665 | 6.6242 | 5.9273 | 7.0938 | 6.4556 | 6.2335 | 6.0802 | 6.4527 | 6.2493 | 6.0942 | 6.3228 | 6.1142 | 5.6861 | 6.2309 | 6.0823 | 5.9864 |
| CAV2_Y20;S23 | CAV2 | P51636 | 18;20;23 | | NA | 5.9173 | 6.0435 | 6.1930 | 6.6986 | 6.1493 | 6.1668 | 5.8757 | 6.3696 | 6.3959 | 5.7091 | 6.7693 | 6.3762 | 6.1243 | 6.6039 | 6.3531 | 6.2545 |
| CAV2_Y20;S23;Y27 | CAV2 | P51636 | 18;20 | | NA | 6.3413 | 6.2815 | 6.2026 | 6.9754 | 6.2373 | 6.0537 | 6.4294 | 6.7714 | 6.0018 | 5.7689 | 6.2689 | 6.0020 | 6.1005 | 6.2334 | 6.2141 | 6.1179 |
| CAV2_Y20;Y27 | CAV2 | P51636 | 20;23;27 | | NA | 6.0699 | 6.4856 | 6.1250 | 7.7132 | 6.2322 | 6.4935 | 5.4334 | 5.7130 | 6.1432 | 5.9251 | 6.6567 | 7.1348 | 5.8365 | 6.8279 | 5.1104 | 6.0994 |
| CAV2_Y23 | CAV2 | P51636 | 20;27 | | NA | 6.2953 | 6.6493 | 6.5220 | 7.1059 | 6.2448 | 6.4032 | 6.1541 | 6.3719 | 6.0479 | 5.9630 | 6.5507 | 6.5756 | 5.4796 | 5.9912 | 5.5334 | 6.1121 |
| CAVIN1_S127 | CAVIN1 | Q6NZI2 | | 365 | GDLRRGSSPDVHA | 6.0663 | 6.2316 | 6.2156 | 6.7740 | 6.2209 | 5.9899 | 6.0457 | 6.0651 | 6.6198 | 6.5029 | 6.4417 | 6.5379 | 6.0170 | 6.3038 | 5.8469 | 6.1210 |
| CAVIN1_S167 | CAVIN1 | Q6NZI2 | | 300 | RDKLRKSFTPDHV | 6.9543 | 6.5309 | 7.2374 | 7.5079 | 6.3171 | 6.0816 | 6.1162 | 5.8257 | 7.3239 | 7.4894 | 8.1122 | 8.4737 | 3.4993 | 5.2445 | 3.2073 | 4.0786 |
| CAVIN1_S202 | CAVIN1 | Q6NZI2 | | 366 | DLRRGSSPDVHAL | 6.9331 | 7.4248 | 7.3764 | 7.4450 | 6.3434 | 6.6605 | 6.6895 | 6.2547 | 6.3918 | 6.4808 | 7.0257 | 6.5025 | 4.6033 | 4.7373 | 4.5851 | 4.5461 |
| CAVIN1_S300 | CAVIN1 | Q6NZI2 | | 302 | KLRKSFTPDHVVY | 7.7630 | 8.4761 | 6.9609 | 7.9777 | 6.2888 | 7.0040 | 7.2121 | 6.8821 | 7.1554 | 7.6304 | 7.9192 | 7.4097 | 2.4563 | 3.3399 | 2.4239 | 3.1004 |
| CAVIN1_S365 | CAVIN1 | Q6NZI2 | | 269 | LEKTRHTLEKRMN | 7.2034 | 7.3693 | 6.1674 | 6.5680 | 5.7635 | 6.5074 | 6.4527 | 5.9267 | 7.1514 | 6.8234 | 6.9322 | 7.3572 | 5.1316 | 5.4432 | 4.1219 | 5.0808 |
| CAVIN1_S365;S366 | CAVIN1 | Q6NZI2 | | 156 | NFKVMIYQDEVKL | 6.9358 | 7.8291 | 6.6450 | 8.0830 | 6.5035 | 6.7574 | 6.3127 | 6.6801 | 7.3927 | 7.4884 | 7.4004 | 7.6803 | 3.1179 | 4.0730 | 3.1838 | 3.9169 |
| CAVIN1_S365;T376 | CAVIN1 | Q6NZI2 | | 127 | VKTVRGSLERQAG | 6.7881 | 7.3118 | 7.4540 | 6.4989 | 6.0018 | 6.4681 | 6.7881 | 5.7798 | 6.4548 | 6.6468 | 6.6635 | 7.4041 | 5.2686 | 4.5811 | 5.2553 | 4.6352 |
| CAVIN1_S366 | CAVIN1 | Q6NZI2 | | 167 | KLPAKLSISKSLK | 5.4541 | 5.9835 | 5.6379 | 5.7518 | 7.5878 | 7.3521 | 5.8097 | 5.3382 | 5.4961 | 5.3003 | 7.2701 | 5.7698 | 6.8406 | 8.3790 | 5.6719 | 6.3571 |
| CAVIN1_T269 | CAVIN1 | Q6NZI2 | | 202 | AAALELSSDEAVE | 5.2865 | 6.4215 | 6.5304 | 6.7146 | 5.4767 | 6.2808 | 6.3501 | 6.6235 | 7.2771 | 5.7797 | 5.5777 | 5.8748 | 5.9085 | 6.7998 | 6.1976 | 6.9008 |
| CAVIN1_T302 | CAVIN1 | Q6NZI2 | 365;366 | | NA | 5.9665 | 5.9965 | 6.0976 | 6.5964 | 6.0609 | 5.5843 | 6.0565 | 5.7926 | 7.2009 | 6.7091 | 6.6939 | 7.1001 | 6.2253 | 5.8949 | 5.9113 | 6.1131 |
| CAVIN1_Y156 | CAVIN1 | Q6NZI2 | 365;376 | | NA | 5.9951 | 6.3294 | 6.4333 | 6.4297 | 5.8867 | 5.3082 | 6.0424 | 5.9695 | 7.5770 | 6.6097 | 6.3979 | 7.4118 | 6.1228 | 5.7659 | 5.3440 | 6.3767 |
| CBARP_S133 | CBARP | Q8N350 | | 299 | RLRRHASLDGASP | 6.2309 | 6.7624 | 6.1947 | 6.4478 | 6.1880 | 6.3378 | 5.9266 | 6.3242 | 6.2967 | 6.2247 | 6.2396 | 6.2670 | 6.3158 | 6.2799 | 5.8851 | 6.0789 |
| CBARP_S299 | CBARP | Q8N350 | | 154 | DKGRRYTLTEGDF | 6.8536 | 5.5172 | 5.8729 | 5.9060 | 5.9336 | 6.0356 | 5.8569 | 5.0979 | 7.5212 | 6.8056 | 7.2446 | 7.4968 | 6.3703 | 7.3051 | 4.5259 | 5.6568 |
| CBARP_S299;S304 | CBARP | Q8N350 | | 133 | STGRRVSFNEAAL | 6.8078 | 5.9320 | 5.8365 | 6.1308 | 6.5828 | 5.6858 | 6.7469 | 5.8265 | 5.9412 | 6.8828 | 7.2598 | 7.1346 | 5.8846 | 6.4159 | 4.9111 | 6.0211 |
| CBARP_T154 | CBARP | Q8N350 | 299;304 | | NA | 7.3820 | 8.6426 | 7.3689 | 7.1247 | 5.6215 | 6.5081 | 7.6495 | 6.1493 | 5.9180 | 6.9347 | 7.9375 | 7.0270 | 3.7664 | 4.2227 | 3.5253 | 4.2216 |
| CBL_S483 | CBL | P22681 | | 483 | KVERPPSPFSMAP | 5.5611 | 5.9506 | 6.7435 | 6.2588 | 6.3039 | 6.4190 | 6.2750 | 6.3562 | 6.4650 | 6.5164 | 6.6570 | 6.5670 | 5.6331 | 6.5047 | 5.7598 | 6.0290 |
| CBL_S619 | CBL | P22681 | | 619 | ELTNRHSLPFSLP | 6.3033 | 4.4934 | 5.2347 | 6.2409 | 6.5209 | 5.3717 | 5.6411 | 7.0035 | 7.7341 | 6.8406 | 7.1481 | 6.2180 | 5.9182 | 6.1328 | 6.7082 | 6.4905 |
| CBL_S900 | CBL | P22681 | | 900 | EFVSISSPAHVAT | 6.6576 | 7.5620 | 6.7329 | 7.6025 | 6.3398 | 7.3191 | 7.2451 | 6.9732 | 7.3244 | 6.7218 | 7.1149 | 7.5031 | 3.4728 | 4.4801 | 3.2124 | 3.7384 |
| CBLL1_S201 | CBLL1 | Q75N03 | | 201 | KPVTRASLENVHP | 5.5644 | 5.5967 | 6.3925 | 7.4998 | 5.7771 | 6.4161 | 6.5944 | 6.7620 | 6.3238 | 6.2722 | 7.1261 | 6.3700 | 4.9061 | 6.2110 | 5.9930 | 6.1949 |
| CBX1_S175 | CBX1 | P83916 | | 89 | GKRKADSDSEDKG | 5.9184 | 6.1786 | 6.0246 | 6.1223 | 7.1502 | 6.9270 | 6.4425 | 6.0335 | 6.6352 | 5.9239 | 6.9704 | 6.2372 | 7.0791 | 6.1483 | 5.3710 | 4.8379 |
| CBX1_S89 | CBX1 | P83916 | | 175 | TWHSYPSEDDDKK | 6.5904 | 6.6082 | 6.5790 | 7.0284 | 6.6442 | 6.2083 | 6.1975 | 5.8486 | 6.1865 | 5.8538 | 5.9435 | 6.0945 | 6.2796 | 5.8012 | 5.8321 | 6.3042 |
| CBX1_S89;S91 | CBX1 | P83916 | 89;91 | | NA | 4.9678 | 5.5001 | 6.6857 | 6.5404 | 7.1533 | 7.4236 | 6.9707 | 5.5775 | 6.3129 | 5.9838 | 8.2319 | 6.6959 | 7.3641 | 5.9998 | 4.2700 | 4.3225 |
| CBX3_S176 | CBX3 | Q13185 | | 176 | ERLTWHSCPEDEA | 6.9289 | 7.3939 | 6.4748 | 7.1715 | 6.5077 | 6.4680 | 6.2022 | 6.0383 | 6.2884 | 6.0302 | 6.2992 | 6.2707 | 5.4448 | 5.5709 | 5.2082 | 5.7022 |
| CBX3_S93 | CBX3 | Q13185 | | 93 | DGTKRKSLSDSES | 5.2586 | 5.5606 | 6.3323 | 6.4660 | 6.0886 | 6.6148 | 6.4673 | 6.1003 | 6.7710 | 7.2492 | 7.4904 | 7.5028 | 5.1597 | 5.8452 | 5.3647 | 5.7285 |
| CBX3_S95 | CBX3 | Q13185 | | 95 | TKRKSLSDSESDD | 6.4070 | 6.8368 | 6.8601 | 7.0030 | 6.4191 | 6.3821 | 5.9512 | 6.1826 | 6.9322 | 6.2413 | 6.4162 | 6.5562 | 5.2664 | 5.8095 | 5.3429 | 5.3935 |
| CBX3_S95;S102 | CBX3 | Q13185 | 95;102 | | NA | 7.1164 | 6.5751 | 7.0972 | 7.0321 | 6.7607 | 6.2080 | 6.3038 | 6.7071 | 6.2917 | 6.3435 | 6.7190 | 6.4097 | 5.3460 | 5.3534 | 4.9067 | 4.8297 |

| | | | | | | | | | | | | | | | | | | | | | |
|---|---|---|---|---|---|---|---|---|---|---|---|---|---|---|---|---|---|---|---|---|---|
| CBX4_S182 | CBX4 | O00257 | | 182 | GHKEAPSPTCPDL | 5.7414 | 7.0434 | 6.3153 | 6.8585 | 5.8147 | 6.0100 | 6.0356 | 6.6030 | 6.7157 | 6.7841 | 6.5404 | 6.9701 | 5.2011 | 5.8569 | 5.4655 | 6.0443 |
| CBX5_S11 | CBX5 | P45973 | | 92 | ESNKRKSNFSNSA | 7.0555 | 7.4302 | 6.7929 | 7.1922 | 5.9242 | 7.2552 | 7.5629 | 6.6176 | 7.6656 | 7.7730 | 7.8018 | 7.9019 | 2.8762 | 4.0010 | 2.5474 | 3.6024 |
| CBX5_S92 | CBX5 | P45973 | | 11 | TKRTADSSSSEDE | 5.8564 | 5.7978 | 5.9812 | 6.3430 | 6.1145 | 6.2960 | 6.2065 | 5.9777 | 6.6908 | 6.8180 | 6.7341 | 6.7295 | 6.0542 | 6.5878 | 5.7878 | 6.0247 |
| CBX6_S107 | CBX6 | O95503 | | 301 | LLPETVSPSAPSW | 6.2210 | 6.4335 | 6.5486 | 6.7973 | 6.1166 | 6.3627 | 6.7174 | 6.2253 | 6.8467 | 6.5323 | 6.7513 | 6.4086 | 5.8967 | 5.6116 | 5.3097 | 5.2208 |
| CBX6_S246 | CBX6 | O95503 | | 107 | PSASASSPKLHSS | 7.0689 | 6.3784 | 6.4920 | 6.8484 | 6.5100 | 6.2787 | 6.8505 | 5.4889 | 7.3464 | 7.6686 | 6.9013 | 6.9359 | 4.8953 | 5.0265 | 4.3513 | 4.9587 |
| CBX6_S276 | CBX6 | O95503 | | 276 | ARSSGSSGCPSPT | 5.9490 | 5.9857 | 6.2439 | 5.9619 | 6.2736 | 6.2637 | 6.6182 | 6.7095 | 7.4829 | 7.1096 | 6.3322 | 5.9857 | 5.3086 | 5.9371 | 6.0046 | 5.8338 |
| CBX6_S301 | CBX6 | O95503 | | 246 | APLVAPSPGKAEA | 6.2932 | 7.5231 | 6.5978 | 6.8726 | 6.0547 | 6.6489 | 7.1987 | 5.7814 | 7.1188 | 6.9980 | 6.7979 | 6.7525 | 4.9608 | 5.5793 | 4.4603 | 4.3621 |
| CBX7_S113 | CBX7 | O95931 | | 113 | CPLGSGSPEGVVK | 7.0658 | 6.5743 | 6.2225 | 7.7491 | 6.3067 | 5.9516 | 6.5541 | 7.4495 | 6.9227 | 6.6517 | 7.2240 | 7.4950 | 4.1220 | 4.8424 | 4.6034 | 4.2651 |
| CBX8_S110 | CBX8 | Q9HC52 | | 110 | IPYPGRSPQDLAS | 7.3545 | 7.9100 | 6.5905 | 7.3823 | 6.7790 | 6.6222 | 6.9817 | 6.2858 | 7.0682 | 7.5285 | 6.9883 | 6.9562 | 3.5798 | 4.5974 | 3.6143 | 3.7613 |
| CBX8_S191 | CBX8 | Q9HC52 | | 191 | VDDKPSSPGDSSK | 6.8717 | 6.6857 | 6.3725 | 6.4454 | 7.1976 | 5.9117 | 6.9864 | 6.0559 | 6.8712 | 6.7775 | 6.0874 | 5.7491 | 5.6946 | 5.4860 | 5.5018 | 5.3054 |
| CBX8_T234 | CBX8 | Q9HC52 | | 234 | GRKLDDTPSGAGK | 7.6595 | 7.4572 | 7.0131 | 7.2395 | 6.1828 | 6.9758 | 7.4668 | 6.5092 | 7.0766 | 7.1599 | 6.6249 | 6.6823 | 3.6219 | 4.4455 | 3.5756 | 4.3094 |
| CBY1_S20 | CBY1 | Q9Y3M2 | | 20 | PPRKSASLSNLHS | 6.4182 | 5.7544 | 5.9804 | 6.5168 | 6.1655 | 6.5769 | 6.3267 | 6.5762 | 6.6609 | 6.1117 | 7.1026 | 6.6700 | 5.8394 | 6.2853 | 5.4316 | 5.5834 |
| CC2D1A_S239 | CC2D1A | Q6P1N0 | | 120 | EQKASETPPPVAQ | 6.1042 | 6.3467 | 6.7748 | 6.6651 | 6.4620 | 6.8485 | 6.8266 | 7.2172 | 6.4876 | 6.8119 | 6.6876 | 6.6045 | 4.5560 | 5.6615 | 4.4422 | 5.5037 |
| CC2D1A_S253 | CC2D1A | Q6P1N0 | | 455 | VPKKQNSPVAPTA | 7.0165 | 6.7769 | 7.3444 | 7.1400 | 5.7080 | 6.3730 | 6.4743 | 6.6373 | 7.0416 | 7.0383 | 6.8139 | 7.1071 | 4.2799 | 4.8898 | 4.4621 | 4.8969 |
| CC2D1A_S324 | CC2D1A | Q6P1N0 | | 253 | MPPGPCSPGPLAQ | 6.2456 | 5.5540 | 7.3186 | 6.8314 | 7.1105 | 6.8489 | 6.3328 | 6.8916 | 7.4412 | 6.5187 | 6.4044 | 6.9180 | 5.0109 | 5.2762 | 4.2351 | 5.0621 |
| CC2D1A_S455 | CC2D1A | Q6P1N0 | | 239 | ATAPASSPGLAKP | 6.9817 | 7.2165 | 6.9418 | 6.9246 | 6.4031 | 7.4436 | 7.4169 | 6.8336 | 6.9947 | 7.3326 | 7.1400 | 6.8060 | 3.2979 | 4.4629 | 3.4795 | 4.3246 |
| CC2D1A_T120 | CC2D1A | Q6P1N0 | | 324 | LPPDPPSPPSQPP | 5.8661 | 6.3421 | 6.8182 | 6.8169 | 6.2556 | 6.2414 | 6.5382 | 6.2645 | 6.6868 | 6.3648 | 7.1280 | 6.5319 | 5.6344 | 6.1037 | 5.0303 | 5.3774 |
| CC2D1B_S241 | CC2D1B | Q5T0F9 | | 241 | QEPANRSPETDPP | 6.1817 | 5.8828 | 6.7068 | 6.3258 | 6.4953 | 6.6000 | 5.4791 | 5.8941 | 6.3772 | 6.0803 | 6.5493 | 6.4779 | 6.1517 | 6.1884 | 6.4172 | 6.1924 |
| CCAR1_S1091 | CCAR1 | Q8IX12 | | 333 | RRSRERSPQRKRS | 6.8247 | 6.5949 | 6.2635 | 6.9305 | 5.8816 | 5.8793 | 5.7998 | 5.2229 | 5.7703 | 5.8995 | 6.1145 | 6.6331 | 6.5350 | 7.0804 | 6.4817 | 6.0882 |
| CCAR1_S333 | CCAR1 | Q8IX12 | | 1091 | EVKKDLSQLQENL | 7.2924 | 5.3402 | 5.3802 | 6.5763 | 5.7669 | 7.3590 | 6.5782 | 5.9897 | 6.0468 | 5.6507 | 5.3593 | 6.4106 | 6.2296 | 7.7990 | 5.9687 | 6.2525 |
| CCAR1_S456 | CCAR1 | Q8IX12 | | 456 | MLMASPSMEDLYH | 5.2828 | 6.8502 | 6.3458 | 5.8921 | 6.4892 | 5.9599 | 5.8850 | 5.8036 | 6.0562 | 6.4792 | 6.1953 | 7.0557 | 6.2624 | 6.9758 | 5.9927 | 6.4742 |
| CCAR1_T627 | CCAR1 | Q8IX12 | | 627 | EAKEISTPTHWSK | 6.6385 | 6.8263 | 7.4027 | 8.5084 | 7.5179 | 5.4773 | 4.8369 | 6.6043 | 5.3236 | 5.6737 | 6.0195 | 5.1827 | 5.8658 | 7.8808 | 5.5627 | 4.6790 |
| CCAR2_S124 | CCAR2 | Q8N163 | | 678 | EVRSVASNQSEME | 4.2237 | 5.2560 | 5.6631 | 5.7233 | 6.5045 | 5.8946 | 5.2486 | 6.3184 | 5.5593 | 6.7005 | 6.2164 | 6.1992 | 7.5762 | 7.1483 | 9.1294 | 6.6386 |
| CCAR2_S569 | CCAR2 | Q8N163 | | 569 | LPEKVVSPPEPEK | 6.2847 | 6.2828 | 5.8581 | 6.2185 | 6.5180 | 5.9274 | 5.9208 | 5.9216 | 5.9864 | 5.8450 | 5.7555 | 5.7806 | 7.0951 | 6.9480 | 7.0289 | 6.6286 |
| CCAR2_S627 | CCAR2 | Q8N163 | | 35 | PPPGLLTPPVATE | 6.3234 | 6.8540 | 6.6303 | 5.9831 | 6.5260 | 6.9550 | 6.3710 | 6.2635 | 6.9143 | 6.8493 | 7.2531 | 6.9339 | 5.1757 | 5.5814 | 4.3520 | 5.0341 |
| CCAR2_S640 | CCAR2 | Q8N163 | | 627 | LPKPLSSGGEEEE | 6.0838 | 5.4529 | 6.1770 | 6.1239 | 6.0591 | 5.9764 | 6.5961 | 5.7159 | 6.0503 | 5.9089 | 5.7598 | 5.8513 | 7.4548 | 6.8568 | 6.6298 | 7.3032 |
| CCAR2_S675;S678 | CCAR2 | Q8N163 | | 124 | NQPLLKSPAPPLL | 6.3444 | 6.6841 | 6.1557 | 6.4404 | 5.4978 | 7.4977 | 7.4207 | 6.9396 | 7.5389 | 7.1497 | 7.7711 | 8.0077 | 3.8942 | 4.6799 | 3.3235 | 4.6546 |
| CCAR2_S675;S678;S681 | CCAR2 | Q8N163 | | 484 | SRRNAETPEATTQ | 5.9243 | 5.7455 | 6.4598 | 6.0529 | 6.5846 | 5.7349 | 6.3846 | 5.4968 | 6.8719 | 6.5105 | 5.8566 | 6.3375 | 6.8010 | 6.7169 | 5.8264 | 6.6957 |
| CCAR2_S678 | CCAR2 | Q8N163 | | 640 | KPRGEASEDLCEM | 6.6570 | 5.7115 | 5.7945 | 5.6436 | 6.7518 | 6.3702 | 6.5211 | 6.2516 | 5.3719 | 6.3044 | 6.5826 | 6.5179 | 6.6699 | 5.9756 | 6.4004 | 6.4759 |
| CCAR2_S678;S681 | CCAR2 | Q8N163 | 678;681 | | NA | 5.6789 | 6.0993 | 6.0656 | 6.4785 | 6.3803 | 5.8043 | 5.8366 | 6.1172 | 6.4392 | 5.7503 | 5.6828 | 6.0683 | 7.0610 | 7.3666 | 6.8448 | 6.3263 |
| CCAR2_T35 | CCAR2 | Q8N163 | 675;678;681 | | NA | 6.3080 | 6.0665 | 6.0336 | 6.8083 | 6.2955 | 5.8838 | 6.0639 | 6.0303 | 6.4579 | 5.9624 | 6.0965 | 6.1217 | 6.6281 | 6.3621 | 6.2024 | 6.6792 |
| CCAR2_T484 | CCAR2 | Q8N163 | 675;678 | | NA | 4.5215 | 5.7003 | 5.6827 | 5.8100 | 5.7958 | 5.8189 | 5.6840 | 5.6710 | 5.8285 | 6.8009 | 6.9770 | 6.8013 | 6.5076 | 6.6995 | 8.1534 | 6.6476 |
| CCDC102A_S12 | CCDC102A | Q96A19 | | 12 | SPRLAESPQLSKG | 6.9385 | 7.1325 | 6.6362 | 7.1310 | 6.5445 | 6.7506 | 6.5029 | 6.9828 | 6.9117 | 7.1120 | 7.4234 | 7.0208 | 4.0209 | 4.6359 | 3.9204 | 4.3360 |
| CCDC102A_S26 | CCDC102A | Q96A19 | | 28 | TILGSPSPERMGP | 6.2076 | 6.4899 | 6.4755 | 6.8323 | 5.9838 | 6.3455 | 6.1090 | 6.4539 | 6.7353 | 7.0275 | 6.6948 | 6.9390 | 5.4705 | 5.6851 | 5.1026 | 5.4478 |
| CCDC102A_S26;S28 | CCDC102A | Q96A19 | | 26 | LLTILGSPSPERM | 6.7297 | 6.4382 | 6.2951 | 6.3962 | 6.5220 | 6.0517 | 6.3209 | 6.5433 | 6.7266 | 6.4079 | 7.0774 | 6.9708 | 5.6686 | 5.4128 | 4.6900 | 5.7488 |
| CCDC102A_S28 | CCDC102A | Q96A19 | | 537 | EAEDGTSDLDEDE | 6.2124 | 6.2861 | 5.4878 | 5.6065 | 6.2554 | 6.5419 | 6.9718 | 5.2626 | 7.8213 | 5.7887 | 6.3250 | 5.9074 | 7.3628 | 6.5686 | 5.0416 | 6.5604 |
| CCDC102A_S537 | CCDC102A | Q96A19 | 26;28 | | NA | 6.5156 | 6.9921 | 6.5685 | 7.4551 | 6.1361 | 7.0800 | 6.4270 | 6.2266 | 6.5749 | 6.6446 | 7.1808 | 7.1522 | 4.7308 | 4.5886 | 4.6897 | 5.0373 |
| CCDC106_S38 | CCDC106 | Q9BWC9 | | 38 | QIFYSLSPSRRNF | 6.3972 | 6.5541 | 6.1330 | 6.7731 | 6.0482 | 5.4151 | 6.5470 | 6.5216 | 6.2404 | 6.8565 | 6.6064 | 6.2107 | 5.6907 | 6.3718 | 5.1989 | 6.4354 |
| CCDC117_S53 | CCDC117 | Q8IWD4 | | 53 | PRAVPSSPAGSAA | 6.6401 | 6.7700 | 7.0309 | 6.3153 | 5.9345 | 7.1594 | 6.7134 | 7.0626 | 6.5650 | 6.9208 | 7.0141 | 7.2919 | 4.5588 | 5.0620 | 4.1164 | 4.8450 |
| CCDC120_S393 | CCDC120 | Q96HB5-4 | | 411 | AGGGAGSPPAPLA | 5.3605 | 5.8495 | 5.9524 | 6.2161 | 6.2610 | 6.4719 | 6.7588 | 6.4103 | 6.8414 | 6.5588 | 6.9660 | 6.4777 | 6.1053 | 5.9539 | 5.2149 | 6.6016 |
| CCDC120_S411 | CCDC120 | Q96HB5-4 | | 393 | VARTRRSNSSEAL | 6.2832 | 6.2942 | 6.2162 | 6.3769 | 6.2623 | 6.6905 | 5.6247 | 6.4026 | 7.0041 | 6.0597 | 6.5000 | 6.2937 | 6.5073 | 6.3361 | 5.6331 | 5.5153 |
| CCDC124_S141 | CCDC124 | Q96CT7 | | 141 | RVLEEGSVEARTI | 5.6608 | 6.4529 | 6.4115 | 6.1613 | 6.6505 | 6.1190 | 6.1384 | 6.6723 | 6.3030 | 6.1857 | 6.1163 | 5.9087 | 6.1392 | 6.7829 | 6.2182 | 6.0794 |
| CCDC130_S286 | CCDC130 | P13994 | | 40 | ERARKLSQGILII | 5.0899 | 4.5671 | 5.2566 | 4.9568 | 5.5975 | 5.3948 | 4.7412 | 5.6276 | 8.8311 | 9.8114 | 9.7128 | 9.5984 | 5.8683 | 6.8868 | 3.6649 | 4.3950 |
| CCDC130_S306 | CCDC130 | P13994 | | 306 | SRDVPESPQHAAD | 6.0139 | 6.7608 | 6.4412 | 6.9924 | 5.7918 | 6.0884 | 6.5164 | 5.6361 | 7.3332 | 6.5740 | 6.5634 | 6.6358 | 5.8546 | 5.9005 | 5.3165 | 5.5810 |
| CCDC130_S332 | CCDC130 | P13994 | | 332 | AQDRPMSPGDCPP | 5.6559 | 6.6112 | 6.1915 | 6.8505 | 5.9580 | 6.5097 | 5.7853 | 5.6572 | 6.0039 | 6.6720 | 6.0479 | 6.2755 | 6.5996 | 6.7328 | 6.0712 | 6.3777 |
| CCDC130_S40 | CCDC130 | P13994 | | 286 | RTALATSPITVGD | 6.7955 | 6.5897 | 6.0558 | 6.4841 | 5.9067 | 6.8360 | 6.6479 | 6.3379 | 6.5680 | 6.0649 | 6.9558 | 6.4001 | 5.3146 | 5.4976 | 5.6463 | 5.8991 |
| CCDC137_S19 | CCDC137 | Q6PK04 | | 19 | VQAGPGSPRRARG | 6.7659 | 7.0946 | 6.0809 | 7.2759 | 5.9252 | 5.7110 | 6.2144 | 6.1919 | 6.6390 | 6.4650 | 6.1008 | 6.3213 | 5.3799 | 5.8759 | 6.0839 | 5.8744 |
| CCDC137_S233 | CCDC137 | Q6PK04 | | 233 | MLRMLLSPGGVSQ | 7.1975 | 7.3644 | 7.2434 | 6.8440 | 5.8891 | 6.7107 | 7.0136 | 6.5438 | 6.6322 | 6.7768 | 7.3337 | 6.8090 | 4.0225 | 4.4753 | 3.9894 | 5.1546 |
| CCDC138_S154 | CCDC138 | Q96M89 | | 154 | CSDAGDSPLKPVS | 6.0908 | 5.8617 | 7.5340 | 6.8593 | 7.0719 | 6.4398 | 6.9849 | 6.9076 | 6.8728 | 7.6867 | 7.2981 | 7.6539 | 3.2808 | 4.4475 | 4.2407 | 4.7694 |
| CCDC14_S798 | CCDC14 | Q49A88 | | 798 | PLSKRLSPQPQIR | 6.6318 | 6.6224 | 7.1730 | 7.0811 | 6.7966 | 6.3253 | 6.3759 | 6.4328 | 6.9887 | 6.7764 | 7.0386 | 6.9561 | 4.4558 | 4.9811 | 4.5261 | 4.8383 |
| CCDC18_S1355 | CCDC18 | Q5T9S5 | | 1355 | KLRRSISASDLTF | 6.4247 | 6.4835 | 7.6959 | 6.5087 | 5.6362 | 6.6961 | 5.9768 | 6.3698 | 6.6069 | 5.9122 | 7.7268 | 7.5137 | 4.9981 | 5.1592 | 5.2717 | 5.0198 |
| CCDC186_S740 | CCDC186 | Q7Z3E2 | | 740 | SSAEDRSPENTGS | 5.8423 | 6.4514 | 6.3866 | 7.0267 | 6.1206 | 6.0383 | 6.2678 | 6.7454 | 6.7109 | 6.7911 | 7.1117 | 7.2104 | 5.5388 | 5.7614 | 5.1564 | 4.8399 |
| CCDC32_S89 | CCDC32 | Q9BV29-2 | | 89 | SEVYLASLEKKLR | 9.9716 | 5.7560 | 7.2442 | 8.6861 | 4.9511 | 4.3315 | 5.8834 | 4.7252 | 6.4567 | 7.1458 | 6.3003 | 8.5182 | 4.5978 | 7.2558 | 3.4223 | 4.7542 |
| CCDC43_T139 | CCDC43 | Q96MW1 | | 139 | AQYADVTDEEDEA | 6.2853 | 6.4287 | 7.1604 | 6.6253 | 6.2362 | 5.8525 | 5.9250 | 6.4665 | 5.8881 | 6.6573 | 6.8818 | 6.2651 | 5.8320 | 6.2381 | 5.6147 | 5.6431 |
| CCDC50_S276 | CCDC50 | Q8IVM0-2 | | 276 | RHQDRLSPKSSQK | 5.9949 | 5.5124 | 7.0388 | 6.1801 | 5.4957 | 6.3111 | 6.6585 | 6.6120 | 6.4180 | 6.8499 | 6.5348 | 6.0309 | 6.1633 | 6.4311 | 6.2583 | 5.5102 |
| CCDC59_S129 | CCDC59 | Q9P031 | | 129 | LLEEQCSIDEPLF | 5.0802 | 5.8622 | 5.9851 | 6.3090 | 6.0319 | 5.4285 | 5.3864 | 5.5205 | 6.1886 | 6.5553 | 5.8806 | 6.3328 | 6.4372 | 7.2673 | 8.2558 | 7.4785 |
| CCDC6_S240;S244 | CCDC6 | Q16204 | | 52 | SGGIVISPFRLEE | 6.0145 | 6.3359 | 6.6598 | 6.2462 | 6.1721 | 6.2451 | 5.9028 | 6.8361 | 6.5405 | 6.7189 | 6.8009 | 6.9411 | 5.8148 | 5.7697 | 5.3503 | 5.6514 |
| CCDC6_S244 | CCDC6 | Q16204 | | 244 | PVSAPPSPRDISM | 5.3168 | 5.7749 | 5.9936 | 5.3324 | 5.8545 | 6.4175 | 5.8972 | 6.9838 | 6.1910 | 6.7512 | 6.4842 | 6.8440 | 6.4891 | 6.4481 | 6.7059 | 6.5158 |
| CCDC6_S323 | CCDC6 | Q16204 | | 367 | SSSRPISPGLSYA | 6.0039 | 6.2380 | 6.2562 | 6.4844 | 6.3217 | 6.2038 | 5.9141 | 6.4269 | 6.3907 | 6.0165 | 6.4260 | 6.1764 | 6.2913 | 6.5213 | 5.9629 | 6.3658 |
| CCDC6_S352 | CCDC6 | Q16204 | | 349 | QGLRPRTVSSPIP | 6.2465 | 6.4629 | 6.8286 | 5.9620 | 6.0405 | 6.5162 | 5.6996 | 6.8018 | 6.8115 | 6.6446 | 6.5618 | 6.2929 | 5.8145 | 5.8244 | 5.9278 | 5.5644 |
| CCDC6_S367 | CCDC6 | Q16204 | | 352 | RPRTVSSPIPYTP | 6.6479 | 6.1299 | 6.7681 | 6.8445 | 6.0640 | 6.2873 | 5.8645 | 6.2314 | 6.6834 | 6.5143 | 6.5509 | 6.7499 | 5.8130 | 5.6901 | 5.5583 | 5.6025 |
| CCDC6_S367;T380 | CCDC6 | Q16204 | | 419 | SPRRSNSPDKFKR | 5.8904 | 6.6615 | 6.8191 | 5.9062 | 5.7837 | 6.4943 | 5.9509 | 5.9892 | 5.5636 | 6.1311 | 6.5737 | 6.9223 | 6.0892 | 6.3651 | 6.4694 | 6.3903 |
| CCDC6_S419 | CCDC6 | Q16204 | | 323 | ALCRQLSESESSL | 5.4556 | 5.9234 | 6.0201 | 5.5630 | 5.6952 | 4.9552 | 5.3328 | 6.9596 | 6.1182 | 6.3558 | 5.9467 | 5.9487 | 7.6409 | 7.1245 | 7.5969 | 7.3633 |
| CCDC6_S52 | CCDC6 | Q16204 | 240;244 | | NA | 6.2082 | 6.5445 | 6.7734 | 6.2683 | 6.5620 | 6.7698 | 5.9851 | 6.8243 | 6.5286 | 6.0772 | 6.7120 | 6.5499 | 5.6758 | 5.9079 | 5.1180 | 5.4949 |
| CCDC6_T349 | CCDC6 | Q16204 | 367;380 | | NA | 6.9844 | 6.4830 | 7.5559 | 7.5744 | 7.5640 | 6.6049 | 6.5713 | 7.9779 | 7.1507 | 6.6213 | 8.1496 | 8.0659 | 2.6614 | 3.4553 | 3.1554 | 3.4249 |
| CCDC61_S334;S336 | CCDC61 | Q9Y6R9 | | 447 | RRGKPPSPTPWSG | 5.8673 | 5.8887 | 6.1599 | 5.8264 | 6.3722 | 6.0972 | 6.2247 | 6.4898 | 6.2071 | 6.2246 | 6.5626 | 6.6201 | 6.1996 | 6.5255 | 6.2524 | 6.4818 |
| CCDC61_S447 | CCDC61 | Q9Y6R9 | 334;336 | | NA | 5.6134 | 5.7580 | 6.7773 | 6.3007 | 6.5974 | 6.1290 | 6.6613 | 6.0968 | 7.0513 | 5.5274 | 6.5598 | 6.2633 | 6.5887 | 6.3917 | 5.4881 | 6.1957 |
| CCDC66_S23 | CCDC66 | A2RUB6 | | 23 | KTKLILSPYEHKS | 7.0925 | 6.4984 | 6.8536 | 6.2534 | 6.4543 | 7.3791 | 6.5020 | 6.4714 | 7.2346 | 6.8291 | 5.9643 | 7.4820 | 3.5168 | 4.8349 | 4.1771 | 6.4567 |
| CCDC66_S796 | CCDC66 | A2RUB6 | | 796 | SKERSPSSPVPVV | 6.0768 | 6.5259 | 7.0046 | 6.7651 | 5.7465 | 6.1789 | 5.5743 | 6.4137 | 6.5826 | 6.4117 | 6.9549 | 7.2435 | 5.4244 | 6.4072 | 5.2079 | 5.4820 |
| CCDC71_S208 | CCDC71 | Q8IV32 | | 453 | ILRVNLSPVIRLQ | 7.2625 | 7.8168 | 6.8469 | 7.3527 | 6.1127 | 6.8988 | 7.2390 | 6.9900 | 7.2102 | 7.4971 | 7.3646 | 7.4769 | 2.8018 | 4.3434 | 2.9850 | 3.8015 |
| CCDC71_S264 | CCDC71 | Q8IV32 | | 97 | PNPTATSPPASAP | 7.0713 | 6.9308 | 6.3239 | 7.5921 | 6.5325 | 6.7602 | 6.7384 | 6.7785 | 7.2439 | 7.2190 | 7.2699 | 7.7961 | 3.5820 | 4.1101 | 3.5896 | 4.4618 |
| CCDC71_S453 | CCDC71 | Q8IV32 | | 208 | QLSLADSPLKLRK | 6.1489 | 6.7336 | 5.8053 | 6.4299 | 6.5196 | 6.1736 | 6.4529 | 6.4679 | 7.0646 | 6.5377 | 6.7931 | 6.3978 | 5.5565 | 6.0007 | 5.3510 | 5.5668 |
| CCDC71_S97 | CCDC71 | Q8IV32 | | 264 | TNRATGSPSVRRM | 6.4596 | 6.9438 | 5.7552 | 6.6203 | 6.4195 | 5.9711 | 6.1006 | 6.3935 | 6.4029 | 6.1355 | 6.1391 | 6.3850 | 5.2896 | 6.7332 | 5.9703 | 6.2807 |
| CCDC8_S261 | CCDC8 | Q9H0W5 | | 273 | PKINWASFRRRRK | 9.5530 | 7.8800 | 7.4552 | 7.0858 | 6.3887 | 5.9596 | 6.4354 | 5.8868 | 7.2239 | 6.0836 | 6.3844 | 6.2400 | 4.4269 | 5.5385 | 3.4313 | 4.0270 |
| CCDC8_S273 | CCDC8 | Q9H0W5 | | 261 | VCVPQASPRRWRP | 5.5534 | 6.2122 | 6.2848 | 6.8068 | 5.9802 | 6.3665 | 6.4697 | 6.3811 | 7.2392 | 6.7681 | 6.7363 | 7.2037 | 5.1727 | 5.9359 | 5.2194 | 5.6699 |
| CCDC82_S219 | CCDC82 | Q8N4S0 | | 219 | VVEDEGSSVEMEQ | 6.6864 | 7.6102 | 7.6943 | 6.5302 | 6.9794 | 5.4920 | 6.6422 | 7.1882 | 5.3110 | 5.6167 | 5.3128 | 4.5627 | 6.4121 | 6.1948 | 6.0537 | 5.7135 |
| CCDC85C_S179 | CCDC85C | A6NKD9 | | 258 | APPHHRSIPNGLH | 6.3166 | 5.8879 | 6.1534 | 5.7093 | 6.0067 | 6.3855 | 5.9279 | 6.5784 | 6.4565 | 5.8556 | 6.5293 | 6.1994 | 7.1201 | 6.5738 | 5.5657 | 6.7339 |
| CCDC85C_S246 | CCDC85C | A6NKD9 | | 246 | AGATRRSLDDLSA | 6.5316 | 5.9688 | 6.3909 | 6.2561 | 6.1316 | 6.2561 | 5.9238 | 6.1642 | 6.4974 | 5.9178 | 6.9233 | 6.4088 | 6.4435 | 6.0926 | 5.9360 | 6.1576 |
| CCDC85C_S258 | CCDC85C | A6NKD9 | | 179 | GAGSRSSIDSQAS | 6.4036 | 5.5669 | 5.8063 | 6.1941 | 6.9303 | 6.1342 | 6.2814 | 6.1760 | 6.6498 | 6.0214 | 6.9565 | 6.9091 | 6.6909 | 5.6156 | 5.7016 | 5.9622 |
| CCDC85C_S372 | CCDC85C | A6NKD9 | | 372 | DRQLQDSCEEDLS | 6.6464 | 7.2064 | 7.3785 | 6.4353 | 7.3644 | 8.7253 | 5.8409 | 6.6652 | 5.2120 | 7.1267 | 7.3175 | 6.0864 | 4.3767 | 5.2511 | 4.3924 | 3.9747 |
| CCDC86_S102 | CCDC86 | Q9H6F5 | | 47 | ETREPGSPPSVQR | 6.3438 | 6.9824 | 6.6104 | 7.2138 | 6.2494 | 6.0772 | 6.4487 | 6.5895 | 6.8483 | 6.5824 | 6.4914 | 6.7111 | 4.8726 | 5.2608 | 5.2019 | 5.5163 |
| CCDC86_S102;S110 | CCDC86 | Q9H6F5 | | 58 | QRAGLGSPERPPK | 7.3237 | 7.4151 | 7.0138 | 7.9136 | 6.3652 | 6.1849 | 6.7673 | 6.6119 | 7.1901 | 7.0060 | 6.9678 | 7.1010 | 3.6565 | 4.1819 | 3.9577 | 4.3434 |
| CCDC86_S102;S110;S11 | CCDC86 | Q9H6F5 | | 18 | GGLRPESPESLTS | 6.1467 | 6.2420 | 6.3941 | 7.0329 | 6.3942 | 6.1888 | 6.4227 | 6.0800 | 6.5302 | 6.1752 | 6.0363 | 5.9531 | 5.7799 | 6.1136 | 6.2141 | 6.2964 |
| CCDC86_S102;S113 | CCDC86 | Q9H6F5 | | 164 | PGSPEPYPGQQAP | 6.2394 | 6.6252 | 6.5302 | 7.0802 | 6.5153 | 6.0365 | 6.3493 | 6.1802 | 6.3195 | 6.3030 | 6.9412 | 6.3763 | 5.1598 | 6.0275 | 5.5522 | 5.7642 |
| CCDC86_S113 | CCDC86 | Q9H6F5 | | 91 | PEPGAASPQRQQD | 6.0755 | 6.1660 | 6.1426 | 6.2848 | 5.8787 | 5.9936 | 6.2052 | 5.8886 | 6.4411 | 5.8344 | 6.1863 | 6.3161 | 6.5590 | 6.7490 | 6.5700 | 6.7092 |
| CCDC86_S160 | CCDC86 | Q9H6F5 | | 102 | QDLHLESPQRQPE | 6.6147 | 7.1422 | 6.6165 | 7.6524 | 6.5861 | 6.1286 | 6.1777 | 6.6537 | 7.1267 | 7.0678 | 6.9633 | 6.9242 | 4.1313 | 4.5004 | 4.5345 | 5.1801 |
| CCDC86_S160;T182 | CCDC86 | Q9H6F5 | | 69 | PKTSPGSPRLQQG | 6.4402 | 6.5030 | 6.0773 | 6.5293 | 5.9843 | 5.6018 | 6.2064 | 5.7567 | 6.3151 | 6.0489 | 6.0098 | 6.2375 | 6.4282 | 6.4417 | 6.6159 | 6.8039 |
| CCDC86_S18 | CCDC86 | Q9H6F5 | | 113 | PEYSPESPRCQPK | 5.8422 | 6.1166 | 5.4777 | 6.3612 | 6.0528 | 5.6104 | 5.9667 | 6.0322 | 5.8945 | 5.5740 | 5.4589 | 5.5783 | 7.4113 | 7.2065 | 7.7472 | 7.6693 |
| CCDC86_S18;S21 | CCDC86 | Q9H6F5 | | 217 | AKKRKGSSSQAPA | 4.8339 | 5.0360 | 5.3724 | 6.3806 | 8.3268 | 5.3638 | 5.2431 | 5.8835 | 6.9802 | 7.7538 | 6.7237 | 7.3165 | 6.6598 | 6.8427 | 5.2599 | 6.0232 |
| CCDC86_S18;S26 | CCDC86 | Q9H6F5 | | 255 | RSKKRFSQMLQDK | 7.5952 | 6.3957 | 6.5758 | 7.3355 | 6.2406 | 6.3888 | 6.8325 | 6.2324 | 8.5603 | 8.8547 | 8.3106 | 7.7979 | 3.6302 | 3.7628 | 2.0707 | 3.4161 |
| CCDC86_S182 | CCDC86 | Q9H6F5 | | 182 | QPLLELTPRAPGS | 6.1368 | 6.0243 | 6.8062 | 7.1219 | 6.4753 | 6.6118 | 5.9380 | 5.9544 | 7.2846 | 6.0046 | 5.7687 | 5.8397 | 5.8604 | 6.8183 | 6.1968 | 5.1582 |
| CCDC86_S217 | CCDC86 | Q9H6F5 | | 146 | QNKEELTPGAPQH | 6.3321 | 5.4869 | 6.5232 | 6.4938 | 6.4295 | 5.5714 | 6.9771 | 6.4754 | 5.7643 | 5.8929 | 6.3836 | 5.8654 | 6.0087 | 6.8797 | 6.6868 | 6.2292 |
| CCDC86_S255 | CCDC86 | Q9H6F5 | | 160 | LPPVPGSPEPYPG | 6.1599 | 8.2709 | 4.6886 | 5.6896 | 6.2502 | 6.5889 | 5.7686 | 6.2277 | 6.5249 | 6.9125 | 6.6792 | 6.8561 | 4.0978 | 6.6453 | 5.6933 | 6.9464 |
| CCDC86_S47 | CCDC86 | Q9H6F5 | 80;91 | | NA | 7.1662 | 6.7341 | 7.1432 | 8.4037 | 6.2065 | 6.5615 | 6.9170 | 6.6452 | 6.9876 | 6.8200 | 7.0752 | 7.2926 | 3.4546 | 4.4947 | 3.7863 | 4.3116 |
| CCDC86_S58 | CCDC86 | Q9H6F5 | 102;110;113 | | NA | 6.7377 | 6.4232 | 6.8858 | 7.8752 | 6.9458 | 6.2647 | 6.5585 | 6.1162 | 6.9204 | 6.4772 | 6.9160 | 7.2906 | 3.9562 | 5.1674 | 4.4350 | 5.0300 |
| CCDC86_S58;S69 | CCDC86 | Q9H6F5 | 102;113 | | NA | 7.3217 | 7.3961 | 6.8963 | 8.3060 | 6.8123 | 6.5082 | 6.7273 | 6.7510 | 6.7484 | 6.8112 | 7.1947 | 7.0859 | 3.6645 | 4.0943 | 3.7538 | 3.9283 |
| CCDC86_S69 | CCDC86 | Q9H6F5 | 102;110 | | NA | 6.4545 | 6.7190 | 6.5513 | 7.8616 | 5.8331 | 6.4915 | 6.4200 | 5.9390 | 6.7339 | 7.2018 | 6.8716 | 7.0371 | 4.3207 | 5.1608 | 5.1891 | 5.2152 |
| CCDC86_S80;S91 | CCDC86 | Q9H6F5 | 160;182 | | NA | 6.5799 | 7.6886 | 7.0450 | 8.3230 | 6.0536 | 6.4485 | 7.1386 | 6.4713 | 7.6399 | 6.6774 | 7.0837 | 6.4351 | 3.6496 | 4.5443 | 3.6480 | 4.5734 |
| CCDC86_S91 | CCDC86 | Q9H6F5 | 58;69 | | NA | 5.6793 | 6.2362 | 6.3000 | 6.9819 | 6.2037 | 6.0298 | 6.1978 | 6.6265 | 5.3097 | 5.3643 | 5.7706 | 5.6508 | 7.1845 | 7.2171 | 6.7035 | 6.5444 |
| CCDC86_T146 | CCDC86 | Q9H6F5 | 146;160 | | NA | 7.0876 | 6.5487 | 6.3409 | 8.0397 | 7.0562 | 6.8726 | 6.2507 | 6.9786 | 6.4333 | 7.5022 | 7.7638 | 6.3813 | 4.1392 | 4.1020 | 3.7558 | 4.7475 |
| CCDC86_T146;S160 | CCDC86 | Q9H6F5 | 146;164 | | NA | 6.4961 | 7.7220 | 6.7425 | 6.4929 | 6.2659 | 6.8138 | 6.7974 | 6.3463 | 7.4681 | 6.7817 | 7.5394 | 7.3136 | 4.3193 | 4.8849 | 4.0905 | 3.9256 |
| CCDC86_T146;S160;T18 | CCDC86 | Q9H6F5 | 146;160;182 | | NA | 5.8250 | 6.7335 | 7.3289 | 5.7719 | 5.7888 | 6.0629 | 6.2006 | 5.9508 | 6.4449 | 6.5959 | 7.3017 | 7.2348 | 5.5283 | 6.3018 | 4.8661 | 6.0642 |
| CCDC86_T146;Y164 | CCDC86 | Q9H6F5 | 146;164;182 | | NA | 5.5274 | 6.9962 | 5.5961 | 7.2216 | 6.8272 | 6.7549 | 5.9896 | 5.2162 | 6.5115 | 6.9804 | 6.2113 | 6.5704 | 5.4788 | 6.2383 | 6.3681 | 5.5121 |
| CCDC86_T146;Y164;T18 | CCDC86 | Q9H6F5 | 18;21 | | NA | 7.3329 | 7.8426 | 6.7295 | 7.6934 | 6.1538 | 5.9199 | 6.6435 | 6.6703 | 6.6134 | 6.5524 | 6.4738 | 6.7247 | 4.8438 | 4.5684 | 4.5095 | 4.7281 |
| CCDC86_Y164 | CCDC86 | Q9H6F5 | 18;26 | | NA | 7.0282 | 7.0638 | 6.8782 | 8.3107 | 5.8755 | 6.3065 | 7.4367 | 6.8225 | 6.2854 | 6.1609 | 6.5832 | 7.3041 | 3.9451 | 4.9199 | 4.3994 | 4.6798 |
| CCDC88A_S1255 | CCDC88A | Q3V6T2-2 | | 1417 | NRERQKSLTLTPT | 8.7493 | 6.7423 | 7.1252 | 6.1287 | 6.4638 | 5.9284 | 5.7646 | 6.3947 | 8.4923 | 6.9008 | 6.8085 | 6.9479 | 4.9107 | 5.3373 | 3.6073 | 3.6982 |
| CCDC88A_S1387 | CCDC88A | Q3V6T2-2 | | 1792 | GTTRRTSIHDFLT | 5.8440 | 5.4825 | 6.2194 | 5.5097 | 6.6905 | 6.2659 | 5.5767 | 6.5436 | 7.3363 | 7.3062 | 7.2019 | 7.1346 | 6.0182 | 6.7563 | 4.9845 | 5.1297 |
| CCDC88A_S1417 | CCDC88A | Q3V6T2-2 | | 1791 | EGTTRRTSIHDFL | 5.8886 | 5.3321 | 6.2536 | 5.4677 | 6.6260 | 6.1233 | 5.5782 | 6.3414 | 7.2500 | 7.3112 | 7.3293 | 7.5798 | 5.9847 | 6.6799 | 5.0834 | 5.1708 |
| CCDC88A_S1470 | CCDC88A | Q3V6T2-2 | | 1421 | QKSLTLTPTRSDS | 6.8454 | 7.1513 | 6.7709 | 6.4373 | 6.4075 | 7.2485 | 7.2250 | 6.9967 | 7.0512 | 6.6556 | 7.0011 | 6.6879 | 4.0754 | 4.8185 | 4.0815 | 4.5463 |
| CCDC88A_S1533 | CCDC88A | Q3V6T2-2 | | 1625 | KRQTRSSPVLQHK | 6.1171 | 6.4581 | 6.8801 | 6.1060 | 6.6995 | 7.0291 | 6.4254 | 6.8423 | 6.8755 | 6.8784 | 6.5558 | 6.7832 | 5.0327 | 5.5580 | 4.8993 | 4.8594 |
| CCDC88A_S1625 | CCDC88A | Q3V6T2-2 | | 1470 | MNDLVQSMVLAGQ | 6.2322 | 6.2180 | 6.3695 | 6.3800 | 6.2160 | 6.0946 | 6.2570 | 6.3569 | 5.9055 | 6.2860 | 6.7354 | 6.8163 | 5.9715 | 6.2566 | 5.7963 | 6.1081 |
| CCDC88A_S1647 | CCDC88A | Q3V6T2-2 | | 1387 | YKFYDPSPPRRRG | 6.6937 | 7.1379 | 6.6431 | 6.7992 | 6.3240 | 6.8764 | 6.9500 | 7.1731 | 6.8390 | 7.2553 | 7.2067 | 7.3426 | 3.6539 | 4.5076 | 4.0988 | 4.4986 |
| CCDC88A_S1689 | CCDC88A | Q3V6T2-2 | | 1647 | HKIKTGSPGSEVV | 6.2787 | 6.8394 | 6.9289 | 6.3610 | 6.2321 | 6.5906 | 6.7885 | 6.1826 | 6.9827 | 7.0670 | 6.7945 | 6.6162 | 5.4249 | 5.2150 | 4.5558 | 5.1420 |
| CCDC88A_S1792 | CCDC88A | Q3V6T2-2 | | 1689 | MKSLSVSSDFLGK | 4.3070 | 5.1831 | 6.3182 | 5.2506 | 4.8309 | 6.5318 | 6.4676 | 6.3495 | 5.3540 | 6.1038 | 6.5000 | 7.0506 | 6.5343 | 7.1057 | 8.0197 | 8.0932 |
| CCDC88A_S1809 | CCDC88A | Q3V6T2-2 | | 1809 | LPISVDSPPAAAD | 6.6915 | 6.8519 | 7.2028 | 7.1556 | 6.0803 | 6.9569 | 7.5373 | 6.8936 | 6.9836 | 6.8457 | 7.2278 | 6.9487 | 3.8596 | 4.5628 | 3.5258 | 4.6760 |
| CCDC88A_T1421 | CCDC88A | Q3V6T2-2 | | 1255 | RLNHTYSQLLKET | 7.1930 | 4.3158 | 7.9558 | 7.2442 | 5.9694 | 6.3073 | 7.0547 | 5.2117 | 7.3568 | 6.4404 | 5.1144 | 4.9148 | 6.1998 | 7.1418 | 7.2442 | 4.3363 |
| CCDC88A_T1791 | CCDC88A | Q3V6T2-2 | | 1533 | NNKDTTSFEDISP | 5.2835 | 6.0634 | 6.9668 | 6.6586 | 5.5368 | 7.3898 | 6.2830 | 6.4639 | 5.3022 | 6.3491 | 5.8762 | 6.3878 | 6.8021 | 6.8183 | 6.2106 | 5.6079 |
| CCDC88C_S1580 | CCDC88C | Q9P219 | | 1806 | SLSRAFSLASADL | 9.1763 | 7.9554 | 8.0623 | 6.4720 | 5.7545 | 6.0832 | 6.7541 | 5.6724 | 6.9609 | 5.6426 | 6.6059 | 6.2717 | 4.8842 | 5.6496 | 3.8286 | 4.2264 |
| CCDC88C_S1580;S1584 | CCDC88C | Q9P219 | | 227 | QAQHPPSPIKSSS | 5.7560 | 6.1481 | 6.3571 | 6.3538 | 6.0906 | 6.5839 | 6.2214 | 6.5397 | 6.3832 | 6.1992 | 6.6964 | 6.9079 | 6.0983 | 6.0091 | 5.6785 | 5.9768 |
| CCDC88C_S1601 | CCDC88C | Q9P219 | | 239 | SADSTPSPTSSLS | 6.9461 | 6.5175 | 7.3915 | 7.0133 | 5.8850 | 6.1245 | 6.4830 | 7.0503 | 6.3083 | 6.5785 | 6.9581 | 7.4227 | 4.6117 | 4.6314 | 4.5839 | 5.4943 |
| CCDC88C_S1806 | CCDC88C | Q9P219 | | 1601 | LHGRSESFSSEDL | 6.3192 | 6.3232 | 6.9191 | 6.4485 | 6.0875 | 6.1330 | 6.1574 | 6.5974 | 7.1673 | 6.4157 | 6.8829 | 6.4753 | 5.3823 | 6.2442 | 5.2730 | 5.1740 |
| CCDC88C_S2011 | CCDC88C | Q9P219 | | 1580 | ESSRNTSSNSSPL | 5.4066 | 5.3670 | 7.2930 | 6.7156 | 5.6346 | 6.3962 | 6.3089 | 5.5063 | 4.8975 | 6.4931 | 6.9586 | 6.3211 | 6.3880 | 7.4431 | 6.8166 | 6.0536 |
| CCDC88C_S227 | CCDC88C | Q9P219 | | 2011 | SKSSPASPEPGGD | 6.0113 | 5.4276 | 5.6333 | 6.4164 | 5.6320 | 6.6060 | 7.2648 | 7.4901 | 6.9729 | 6.0482 | 5.7773 | 6.4127 | 6.0630 | 6.5087 | 6.0458 | 5.6899 |
| CCDC88C_S239 | CCDC88C | Q9P219 | 1580;1584 | | NA | 7.2722 | 6.8879 | 7.5319 | 7.4132 | 5.0389 | 5.9437 | 7.3212 | 6.3521 | 6.6058 | 7.2432 | 7.1203 | 8.5064 | 4.2372 | 3.7158 | 3.5573 | 5.2529 |
| CCDC9_S349 | CCDC9 | Q9Y3X0 | | 80 | LGPSRRSPGTPRP | 6.0716 | 6.3976 | 6.3357 | 7.0487 | 6.2357 | 6.7541 | 6.4512 | 6.3501 | 6.4524 | 6.3628 | 6.2111 | 6.2816 | 5.7831 | 5.9146 | 5.3772 | 5.9725 |
| CCDC9_S386 | CCDC9 | Q9Y3X0 | | 95 | ASKGGRTPPQQGG | 6.7207 | 7.1376 | 6.5889 | 7.1275 | 6.4366 | 6.9365 | 6.7048 | 6.6331 | 9.1841 | 6.4891 | 7.0448 | 6.8956 | 3.8929 | 4.5191 | 3.5239 | 4.1648 |
| CCDC9_S521 | CCDC9 | Q9Y3X0 | | 521 | HLAGALSPGEAWP | 6.4264 | 5.9728 | 6.4880 | 6.4231 | 6.4686 | 6.4895 | 6.3491 | 7.0941 | 6.1901 | 6.8160 | 6.2051 | 6.3008 | 5.5122 | 5.8196 | 5.3652 | 6.0794 |
| CCDC9_S80 | CCDC9 | Q9Y3X0 | | 349 | SRRRRKSSRPQAK | 6.1319 | 5.6854 | 5.9548 | 5.2702 | 6.0446 | 6.4420 | 5.5048 | 5.8880 | 8.3785 | 8.6454 | 8.2755 | 7.8016 | 5.4112 | 6.5685 | 3.7586 | 4.2391 |
| CCDC9_S80;T83 | CCDC9 | Q9Y3X0 | | 83 | SRRSPGTPRPPGA | 5.9649 | 5.7242 | 6.7435 | 7.0972 | 5.7935 | 6.6212 | 6.5117 | 6.9209 | 7.4371 | 7.0736 | 7.0427 | 8.2850 | 4.4746 | 5.5356 | 4.0452 | 4.7290 |
| CCDC9_S83 | CCDC9 | Q9Y3X0 | | 386 | ETPQPTSPETSPK | 5.0543 | 5.0632 | 6.0534 | 5.4220 | 6.1262 | 5.3480 | 6.1300 | 6.1536 | 6.8214 | 6.0253 | 6.6056 | 6.2513 | 7.3360 | 6.7065 | 7.2575 | 7.6456 |
| CCDC9_T95 | CCDC9 | Q9Y3X0 | 80;83 | | NA | 6.6553 | 6.7328 | 6.7872 | 7.2655 | 5.4809 | 7.2802 | 7.1894 | 6.4166 | 8.5448 | 7.0197 | 7.1817 | 7.3995 | 5.2229 | 3.9396 | 2.9088 | 3.9751 |
| CCDC90B_S219 | CCDC90B | Q9GZT6 | | 219 | IDAEIASLKTLME | 6.9048 | 5.4539 | 6.2557 | 5.2050 | 5.7579 | 5.9406 | 5.4759 | 5.5594 | 5.4586 | 5.8666 | 5.2428 | 5.8571 | 5.2065 | 8.0910 | 9.2489 | 8.4754 |
| CCDC93_S305 | CCDC93 | Q567U6 | | 305 | ELSAEESPEKLGT | 6.0617 | 6.7093 | 6.3765 | 6.3415 | 6.3575 | 6.4845 | 6.4347 | 6.2083 | 6.8711 | 6.5486 | 6.8833 | 6.7891 | 5.6374 | 5.3878 | 5.4204 | 5.4883 |
| CCDC97_S212 | CCDC97 | Q96F63 | | 212 | QPPKPGSPGRPAC | 6.2122 | 5.8325 | 7.0853 | 6.2893 | 6.2349 | 5.9995 | 6.1562 | 5.9741 | 6.4439 | 6.6458 | 6.6554 | 6.8395 | 6.3373 | 5.5251 | 5.4664 | 6.3028 |
| CCDC97_S257 | CCDC97 | Q96F63 | | 257 | EEEEEDSDEEDQR | 5.2888 | 6.2627 | 6.5284 | 5.8406 | 5.9060 | 5.7769 | 6.0442 | 6.8822 | 6.3878 | 6.5301 | 6.9146 | 6.1279 | 6.8338 | 5.8002 | 6.2746 | 6.6012 |
| CCDC97_T202;S212 | CCDC97 | Q96F63 | 202;212 | | NA | 5.7950 | 6.3163 | 5.6985 | 5.8799 | 7.0162 | 7.3476 | 7.3291 | 5.7005 | 6.2151 | 6.1251 | 6.7656 | 6.0649 | 6.5407 | 5.6888 | 5.8153 | 5.7015 |
| CCDC9B_S193 | CCDC9B | Q6ZUT6 | | 392 | ASPALASPEGPKG | 6.3061 | 6.3538 | 6.9871 | 6.8800 | 6.2746 | 6.4396 | 6.5346 | 6.8924 | 6.5259 | 7.0455 | 7.1373 | 7.4546 | 4.5509 | 4.9343 | 4.5909 | 5.0925 |
| CCDC9B_S201 | CCDC9B | Q6ZUT6 | | 193 | NQGIEGSPGGRVT | 6.4576 | 7.1807 | 6.8340 | 6.7915 | 6.1805 | 6.3255 | 6.8681 | 6.3994 | 6.1301 | 6.6401 | 6.4245 | 6.6063 | 4.8462 | 5.5681 | 5.0138 | 5.7337 |
| CCDC9B_S392 | CCDC9B | Q6ZUT6 | | 201 | GGRVTRSPPTQVA | 5.2550 | 6.4297 | 6.1920 | 6.4275 | 6.3644 | 6.2279 | 6.0347 | 6.3571 | 6.1739 | 6.5481 | 7.1420 | 5.9422 | 5.9725 | 6.3627 | 6.4794 | 6.0909 |
| CCM2_S15 | CCM2 | Q9BSQ5 | | 280 | VDVGGASPHSKTI | 6.6006 | 6.7510 | 6.1214 | 6.9308 | 6.2553 | 6.8520 | 6.5045 | 6.5927 | 6.9155 | 7.1319 | 6.9990 | 7.2137 | 4.6929 | 5.0870 | 4.5317 | 4.8199 |
| CCM2_S181 | CCM2 | Q9BSQ5 | | 15 | KKPGIVSPFKRVF | 6.4498 | 6.4237 | 6.6451 | 6.2286 | 6.6465 | 7.2505 | 6.6056 | 6.6089 | 7.1439 | 7.1820 | 7.2078 | 7.5334 | 4.2024 | 4.9908 | 4.0464 | 4.8346 |
| CCM2_S280 | CCM2 | Q9BSQ5 | | 384 | RGIITDSFGRHRR | 6.7127 | 6.3263 | 5.9106 | 5.8651 | 6.6495 | 5.9981 | 5.9499 | 6.1972 | 6.4653 | 5.9884 | 5.4695 | 5.6655 | 7.0621 | 7.0948 | 6.5755 | 6.0697 |
| CCM2_S287 | CCM2 | Q9BSQ5 | | 287 | PHSKTISESELSA | 5.9410 | 6.5580 | 6.5558 | 6.1311 | 6.0605 | 5.9482 | 6.3535 | 6.1289 | 7.1950 | 6.1419 | 6.6839 | 6.3499 | 6.2419 | 5.6509 | 5.8611 | 6.1980 |
| CCM2_S384 | CCM2 | Q9BSQ5 | | 181 | RGLSAGSLSESAV | 5.7295 | 6.6940 | 6.3890 | 6.1767 | 5.6744 | 5.1739 | 5.8596 | 5.8203 | 5.9990 | 6.3161 | 5.9138 | 6.5939 | 6.8998 | 6.4620 | 7.4648 | 6.8334 |
| CCNB2_S92 | CCNB2 | O95067 | | 92 | LAPKGPSPTPEDV | 6.4115 | 6.3601 | 6.3579 | 6.2523 | 6.2617 | 5.7887 | 5.8264 | 6.1863 | 6.3446 | 6.5660 | 6.2875 | 6.4318 | 6.0699 | 6.2510 | 6.2200 | 6.3844 |
| CCNB3_S954 | CCNB3 | Q8WWL7 | | 954 | PLALQESPTYKED | 5.8356 | 6.5832 | 6.7202 | 6.6233 | 5.9094 | 6.2072 | 6.2638 | 5.9871 | 6.1284 | 6.0852 | 6.3215 | 6.7245 | 5.8976 | 6.1880 | 6.1696 | 6.3556 |
| CCNE2_S21 | CCNE2 | O96020 | | 21 | QPSQTESPQEAQI | 6.4444 | 7.0116 | 7.1250 | 7.1660 | 6.2400 | 5.9200 | 6.2133 | 6.4238 | 6.8298 | 6.8465 | 6.8567 | 7.1049 | 4.9042 | 5.2507 | 4.6607 | 5.0025 |
| CCNE2_S383 | CCNE2 | O96020 | | 383 | RKGGQLSPVCNGG | 6.2406 | 6.1989 | 6.4386 | 6.7166 | 6.2773 | 5.8621 | 6.0090 | 5.8845 | 6.4110 | 6.0539 | 6.4329 | 6.1396 | 6.3380 | 6.5463 | 6.1997 | 6.2512 |
| CCNE2_S383;T392 | CCNE2 | O96020 | | 74 | PCIIIETPHKEIG | 7.7035 | 7.2863 | 7.6187 | 7.6277 | 5.7085 | 6.3688 | 6.8173 | 4.9990 | 7.4871 | 6.3633 | 6.0693 | 7.6081 | 4.9929 | 4.4335 | 3.6655 | 5.2506 |

| | | | | | | | | | | | | | | | | | | | | | |
|---|---|---|---|---|---|---|---|---|---|---|---|---|---|---|---|---|---|---|---|---|---|
| CCNE2_S392 | CCNE2 | O96020 | | 67 | VLSGGISPCIIIE | 5.4814 | 6.0619 | 6.8366 | 6.5729 | 6.1465 | 6.2201 | 6.2923 | 5.9677 | 6.6065 | 6.5976 | 6.8441 | 7.0711 | 5.6927 | 6.0806 | 5.8753 | 5.6527 |
| CCNE2_S67 | CCNE2 | O96020 | | 392 | CNGGIMTPPKSTE | 6.6400 | 6.0856 | 7.0925 | 6.2716 | 5.9401 | 6.3860 | 5.5097 | 6.0117 | 6.9013 | 6.5388 | 6.0900 | 6.0029 | 6.6540 | 5.4890 | 6.5219 | 5.8648 |
| CCNE2_T74 | CCNE2 | O96020 | 383;392 | | NA | 6.5854 | 7.3231 | 7.5097 | 6.7256 | 6.6519 | 6.7394 | 5.9765 | 6.7334 | 6.3673 | 6.8877 | 7.3654 | 6.6958 | 4.5602 | 4.6148 | 4.4269 | 4.8369 |
| CCNF_S754 | CCNF | P41002 | | 774 | INLCIHSEEEDMN | 6.3456 | 7.3486 | 7.3893 | 7.1162 | 6.4331 | 6.3410 | 7.2090 | 7.1494 | 6.5129 | 7.0144 | 7.2519 | 7.3934 | 3.7177 | 4.7099 | 3.6023 | 4.4654 |
| CCNF_S774 | CCNF | P41002 | | 754 | LQCRPPSPPESSV | 6.7809 | 7.0091 | 7.4681 | 7.4334 | 6.4405 | 7.3101 | 7.1814 | 6.7887 | 7.0077 | 7.6616 | 8.0588 | 7.7489 | 3.0069 | 3.8038 | 2.9279 | 3.3723 |
| CCNH_T315 | CCNH | P51946 | | 315 | HEEEEWTDDDLVE | 5.1116 | 5.9915 | 6.5079 | 6.5920 | 6.0498 | 6.5009 | 6.2570 | 7.3073 | 6.2845 | 6.4904 | 5.9026 | 6.4514 | 5.2666 | 6.8027 | 6.3590 | 6.1247 |
| CCNK_S324;S329 | CCNK | O75909 | | 340 | KRAVVVSPKEENK | 6.4819 | 6.5378 | 6.1193 | 6.4476 | 6.0364 | 5.8728 | 6.1708 | 6.0624 | 6.0770 | 6.4575 | 5.9632 | 6.5656 | 5.5145 | 6.5703 | 6.4485 | 6.6743 |
| CCNK_S329 | CCNK | O75909 | | 329 | PSPQPSSPRQVKR | 6.3080 | 6.2602 | 5.7455 | 6.0312 | 6.2163 | 6.0772 | 5.7393 | 6.3899 | 6.6675 | 6.1089 | 6.3121 | 6.4741 | 6.6114 | 7.1490 | 6.0081 | 5.9012 |
| CCNK_S340 | CCNK | O75909 | | 9 | ENKENSSPSVTSA | 5.9420 | 6.7600 | 6.0644 | 6.7853 | 5.7333 | 6.0578 | 5.6949 | 6.1033 | 6.1166 | 6.5465 | 6.2234 | 6.2425 | 6.3479 | 6.9073 | 6.2984 | 6.1764 |
| CCNK_S9 | CCNK | O75909 | 324;329 | | NA | 6.4506 | 6.3248 | 6.0995 | 6.3533 | 6.3268 | 6.2000 | 6.0946 | 6.0801 | 6.1756 | 6.0266 | 6.1079 | 5.8537 | 6.8540 | 6.4215 | 6.4395 | 6.1913 |
| CCNL1_S335 | CCNL1 | Q9UK58 | | 352 | VKAEEKSPISINV | 6.0192 | 6.0638 | 5.4960 | 5.9940 | 6.2554 | 5.7378 | 6.1858 | 6.0377 | 5.8488 | 5.5644 | 5.3980 | 5.4229 | 7.3836 | 7.4026 | 7.8424 | 7.3476 |
| CCNL1_S335;S338 | CCNL1 | Q9UK58 | | 65 | IPEERLSPTPSMQ | 6.1653 | 6.3924 | 5.8786 | 6.5563 | 6.0890 | 6.1922 | 5.3033 | 5.9655 | 6.1698 | 6.3373 | 6.1249 | 5.3775 | 6.4984 | 7.0472 | 6.6656 | 7.2368 |
| CCNL1_S335;S338;S342 | CCNL1 | Q9UK58 | | 335 | STLGGFSPASKPS | 6.0554 | 5.9234 | 6.1895 | 6.8620 | 5.9084 | 5.8408 | 6.5526 | 6.2166 | 6.5253 | 6.0462 | 6.0414 | 6.0919 | 5.6794 | 6.2893 | 6.6018 | 7.1760 |
| CCNL1_S338;S342 | CCNL1 | Q9UK58 | | 374 | RQQASKSPYNGVR | 5.9312 | 6.3011 | 5.4480 | 6.5084 | 6.0209 | 5.7559 | 6.0671 | 5.9298 | 6.0290 | 5.7325 | 5.5322 | 5.8672 | 6.9604 | 7.8394 | 7.0617 | 7.0151 |
| CCNL1_S342 | CCNL1 | Q9UK58 | | 342 | PASKPSSPREVKA | 5.3718 | 6.1285 | 6.9603 | 7.7301 | 7.1549 | 6.9138 | 6.4137 | 5.9638 | 5.8003 | 6.1798 | 6.2848 | 5.6034 | 6.1858 | 6.4292 | 6.1189 | 4.7608 |
| CCNL1_S352 | CCNL1 | Q9UK58 | 335;338 | | NA | 5.9553 | 5.7759 | 6.0953 | 6.5320 | 5.5604 | 5.1131 | 6.3372 | 5.7431 | 6.6949 | 6.0577 | 5.8500 | 6.3856 | 6.4139 | 6.5186 | 7.1720 | 7.7952 |
| CCNL1_S374 | CCNL1 | Q9UK58 | 335;338;342 | | NA | 5.6878 | 5.9605 | 5.6719 | 6.4195 | 5.3772 | 5.6297 | 5.7547 | 5.3754 | 5.9566 | 5.6988 | 5.5946 | 6.1665 | 7.4493 | 7.1337 | 7.9097 | 8.2143 |
| CCNL1_S65 | CCNL1 | Q9UK58 | 325;330;342 | | NA | 5.5263 | 5.8582 | 6.0566 | 6.2998 | 6.0435 | 6.2659 | 6.6508 | 6.6418 | 6.5481 | 6.7193 | 6.9666 | 6.9700 | 5.5343 | 5.9968 | 6.0517 | 5.8705 |
| CCNL1_T325;S335;S342 | CCNL1 | Q9UK58 | 338;342 | | NA | 5.6350 | 5.4077 | 5.9720 | 6.4527 | 6.0324 | 6.5937 | 5.3418 | 4.5715 | 5.9966 | 5.7293 | 4.9310 | 4.7637 | 8.2666 | 8.1361 | 8.0062 | 8.1638 |
| CCNL1_T325;T330;S342 | CCNL1 | Q9UK58 | 325;335;342 | | NA | 5.5746 | 5.1996 | 4.7310 | 8.1964 | 5.7018 | 5.5441 | 5.5186 | 6.7648 | 6.0028 | 8.0315 | 7.1701 | 8.0392 | 5.9756 | 5.4134 | 4.6399 | 7.4966 |
| CCNL1_T330;S335;S342 | CCNL1 | Q9UK58 | 330;335;342 | | NA | 6.1408 | 5.9184 | 6.1073 | 5.8577 | 6.0306 | 6.0920 | 5.0771 | 5.8056 | 5.8541 | 5.6927 | 6.1284 | 5.9990 | 7.1508 | 6.3535 | 7.7062 | 8.0859 |
| CCNL2_S330 | CCNL2 | Q96S94 | | 369 | KKAKADSPVNGLP | 6.0608 | 6.2430 | 5.6975 | 5.9921 | 6.2407 | 5.9881 | 6.3136 | 5.9839 | 6.1597 | 5.8587 | 5.7908 | 5.6308 | 7.1456 | 7.0687 | 7.0625 | 6.7637 |
| CCNL2_S348 | CCNL2 | Q96S94 | | 330 | DGTSGFSPAPKLV | 6.4525 | 6.3888 | 6.5627 | 6.7187 | 6.0256 | 6.1745 | 6.5890 | 6.4725 | 6.1242 | 6.6581 | 6.4671 | 6.5486 | 5.3363 | 6.0524 | 5.6927 | 5.7365 |
| CCNL2_S369 | CCNL2 | Q96S94 | | 348 | GKGSKPSPLSVKN | 5.9560 | 6.2769 | 6.1500 | 6.4183 | 5.9496 | 6.0271 | 6.0687 | 6.0432 | 6.2049 | 6.2496 | 5.8384 | 6.3212 | 6.4158 | 6.6841 | 6.6920 | 6.7041 |
| CCNO_S81 | CCNO | P22674 | | 81 | SAARGGSPLPGPA | 6.4297 | 5.6753 | 5.8443 | 6.4868 | 6.5593 | 5.9871 | 6.1620 | 7.1899 | 6.2524 | 6.7485 | 6.2857 | 7.0971 | 5.3410 | 6.2084 | 5.9882 | 5.7443 |
| CCNT1_S340 | CCNT1 | O60563 | | 340 | WLSSQPSFKLEPT | 7.6702 | 6.8579 | 7.0930 | 7.2093 | 5.2613 | 6.7510 | 5.1678 | 5.6955 | 6.5413 | 7.0369 | 7.6983 | 7.6943 | 4.9059 | 5.5352 | 3.6554 | 5.2266 |
| CCNY_S100 | CCNY | Q8ND76 | | 326 | ARKRSASADNLTL | 6.2232 | 6.2778 | 5.9169 | 6.1245 | 5.9021 | 5.8317 | 6.0638 | 6.3077 | 6.4447 | 6.3054 | 5.9687 | 6.1346 | 6.4543 | 6.6985 | 6.6286 | 6.7176 |
| CCNY_S25 | CCNY | Q8ND76 | | 25 | AHSRLESYRPDTD | 6.9998 | 6.9971 | 7.4263 | 6.5058 | 6.3981 | 6.0641 | 6.4008 | 6.7081 | 6.6421 | 6.2937 | 6.8906 | 5.9170 | 4.6617 | 6.2210 | 4.7126 | 5.1613 |
| CCNY_S326 | CCNY | Q8ND76 | | 83 | VREKRKSLFINHH | 6.7060 | 6.1759 | 6.8324 | 6.3767 | 5.8430 | 6.2537 | 6.2864 | 6.0842 | 6.4154 | 6.5598 | 6.3086 | 6.6437 | 5.9695 | 6.4555 | 5.2467 | 5.8426 |
| CCNY_S336 | CCNY | Q8ND76 | | 336 | LTLPRWSPAIISx | 7.3937 | 5.3548 | 6.3851 | 5.8141 | 5.8122 | 5.7211 | 7.0387 | 6.3005 | 5.5042 | 6.2123 | 7.1993 | 6.2301 | 5.9906 | 6.4470 | 6.1334 | 6.4630 |
| CCNY_S83 | CCNY | Q8ND76 | | 100 | IARKYSSCSTIFL | 6.1721 | 6.9251 | 6.5930 | 7.2299 | 5.8546 | 6.3658 | 5.5148 | 5.3299 | 6.9819 | 5.5185 | 6.6498 | 6.0490 | 5.6335 | 6.2864 | 6.5941 | 6.3014 |
| CCNYL1_S105;S112 | CCNYL1 | Q8N7R7 | | 344 | AMRRSFSADNFIG | 6.3441 | 6.5143 | 5.9816 | 6.3629 | 6.0807 | 5.9918 | 6.0749 | 6.2436 | 6.2466 | 6.0629 | 6.1695 | 6.3423 | 6.1595 | 6.5491 | 6.4283 | 6.4480 |
| CCNYL1_S122 | CCNYL1 | Q8N7R7 | | 88 | SDHPRASTIFLSK | 8.1277 | 6.7835 | 6.3147 | 7.0317 | 6.0421 | 5.2530 | 6.2995 | 4.9168 | 5.8968 | 4.1452 | 4.6250 | 4.6869 | 7.9318 | 7.3329 | 7.4170 | 7.1955 |
| CCNYL1_S344 | CCNYL1 | Q8N7R7 | | 122 | LTKKYSSCSTIFL | 6.0251 | 6.4108 | 6.7525 | 6.9539 | 6.5234 | 6.5522 | 6.3091 | 5.7080 | 6.1013 | 5.6482 | 6.2898 | 6.2280 | 5.7703 | 6.6582 | 6.1458 | 5.9234 |
| CCNYL1_S88 | CCNYL1 | Q8N7R7 | | 93 | ASTIFLSKSQTDV | 5.4029 | 6.2527 | 6.6969 | 6.3251 | 6.4217 | 5.2822 | 6.8949 | 5.4174 | 6.2865 | 7.2184 | 8.6041 | 7.1025 | 7.4260 | 7.5902 | 4.9249 | 2.1534 |
| CCNYL1_S93 | CCNYL1 | Q8N7R7 | 105;112 | | NA | 6.2223 | 6.5960 | 6.5573 | 6.8787 | 6.0504 | 5.8986 | 6.4931 | 6.8011 | 6.0904 | 6.0324 | 6.2428 | 6.9443 | 5.6546 | 5.9757 | 5.6008 | 5.9615 |
| CCP110_S170 | CCP110 | O43303 | | 372 | SPEPSMSPKMHRR | 5.8981 | 5.7767 | 6.6277 | 5.7890 | 6.3111 | 6.2905 | 5.9274 | 6.2969 | 6.4565 | 6.1870 | 6.2585 | 6.1724 | 6.7428 | 6.7815 | 6.0735 | 6.4103 |
| CCP110_S176 | CCP110 | O43303 | | 170 | DSEGFNSPKQCDS | 6.4822 | 6.6554 | 6.8263 | 6.0749 | 6.1422 | 6.2717 | 6.5245 | 5.9918 | 6.9306 | 6.3122 | 6.6045 | 6.8947 | 5.5305 | 5.9206 | 5.1162 | 5.7216 |
| CCP110_S372 | CCP110 | O43303 | | 176 | SPKQCDSSNISHV | 8.7877 | 5.9441 | 5.6222 | 7.7937 | 6.4073 | 6.5400 | 8.1213 | 6.8054 | 8.1580 | 6.1446 | 8.2456 | 6.3649 | 3.0893 | 4.5464 | 2.8323 | 4.5972 |
| CCP110_S384;S400 | CCP110 | O43303 | 384;400 | | NA | 6.2765 | 6.3140 | 6.3357 | 7.0969 | 6.4567 | 6.8340 | 6.7229 | 6.1467 | 6.4665 | 5.7987 | 6.8088 | 6.7760 | 5.3944 | 5.5085 | 5.3370 | 5.7267 |
| CCS_S267 | CCS | O14618 | | 267 | GKGRKESAQPPAH | 6.1246 | 4.6085 | 5.5324 | 4.6992 | 7.2423 | 5.3686 | 4.7264 | 6.3568 | 7.1831 | 8.0946 | 6.1885 | 6.8758 | 7.0113 | 8.5493 | 5.8565 | 5.5820 |
| CCSER2_S908 | CCSER2 | Q9H7U1-3 | | 908 | RVGRNQSPPVGYM | 6.1024 | 6.0189 | 7.0739 | 7.1102 | 7.2131 | 6.7631 | 7.3379 | 6.2006 | 6.6746 | 6.8525 | 7.2320 | 6.8605 | 5.1275 | 5.1836 | 3.8448 | 4.4043 |
| CCSER2_T430 | CCSER2 | Q9H7U1-3 | | 430 | SNRTRITPEEMSL | 7.0710 | 6.8296 | 7.0359 | 6.4188 | 6.2841 | 6.4998 | 6.4204 | 6.7492 | 6.2716 | 6.1424 | 7.1406 | 6.6011 | 5.3753 | 5.4666 | 4.9833 | 4.7102 |
| CCT2_S168 | CCT2 | P78371 | | 260 | SRVRVDSTAKVAE | 6.2102 | 5.5919 | 7.1230 | 6.4081 | 6.3769 | 6.4026 | 6.0099 | 5.7009 | 6.8169 | 5.2525 | 5.7533 | 6.1646 | 7.1060 | 7.2096 | 6.1500 | 5.7235 |
| CCT2_S260 | CCT2 | P78371 | | 168 | IAGTTLSSKLLTH | 6.7958 | 6.3990 | 8.0119 | 5.7419 | 5.5397 | 6.5458 | 7.4309 | 5.4495 | 6.3204 | 6.1349 | 6.5278 | 6.7056 | 5.4624 | 5.6711 | 5.7200 | 5.5435 |
| CCT3_S170 | CCT3 | P49368 | | 244 | IVLLDSSLEYKKG | 6.0608 | 5.9885 | 6.6293 | 5.8639 | 6.0526 | 6.5811 | 6.0725 | 6.4805 | 6.6233 | 6.4069 | 6.5344 | 6.6808 | 5.9096 | 6.1107 | 5.8623 | 6.1427 |
| CCT3_S244 | CCT3 | P49368 | | 170 | AISRWSSLACNIA | 5.7014 | 6.0828 | 7.1058 | 6.1329 | 5.7402 | 6.3387 | 6.2732 | 6.3768 | 5.9235 | 6.1477 | 6.7159 | 5.9644 | 6.3937 | 6.3282 | 6.5707 | 6.2041 |
| CCT4_S381 | CCT4 | P50991 | | 381 | KITGCASPGKTVT | 6.2956 | 6.3730 | 6.7991 | 6.5086 | 6.5652 | 6.6722 | 6.5365 | 6.2884 | 6.5797 | 6.4510 | 6.5209 | 6.6172 | 5.4668 | 5.3804 | 5.4039 | 5.5415 |
| CCT6A_S205 | CCT6A | P40227 | | 246 | EKTEVNSGFFYKS | 6.7550 | 6.6140 | 6.5166 | 6.7768 | 6.6716 | 6.7355 | 6.5791 | 6.5651 | 6.4206 | 5.8181 | 6.2803 | 5.5478 | 5.7345 | 6.1554 | 5.3928 | 5.4369 |
| CCT6A_S246 | CCT6A | P40227 | | 205 | KSETDTSLIRGLV | 5.9016 | 5.8187 | 7.1173 | 5.7354 | 6.4927 | 6.5250 | 6.1245 | 6.8792 | 6.5683 | 5.7697 | 6.4351 | 5.6265 | 5.4746 | 6.6279 | 6.0393 | 5.8643 |
| CCT6A_S528 | CCT6A | P40227 | | 528 | MRAGMSSLKGxxx | 6.5535 | 6.9186 | 7.3660 | 5.5645 | 6.4218 | 6.5956 | 5.4666 | 6.9268 | 6.8774 | 6.7100 | 6.0421 | 6.6898 | 5.4895 | 5.7859 | 5.6569 | 4.9350 |
| CCT7_S221 | CCT7 | Q99832 | | 221 | AFKKTFSYAGFEM | 6.3333 | 5.4333 | 6.9860 | 5.5632 | 5.6663 | 6.9860 | 5.9615 | 5.8957 | 5.7961 | 4.3929 | 6.3368 | 5.3729 | 7.6671 | 7.3897 | 8.0300 | 6.1892 |
| CCT8_S23 | CCT8 | P50990 | | 23 | EGAKHFSGLEEAV | 7.5380 | 7.1851 | 5.8207 | 5.3065 | 6.4021 | 5.6661 | 6.3449 | 7.4876 | 5.0309 | 5.9853 | 6.7113 | 5.1653 | 6.2038 | 7.5884 | 6.2004 | 5.3636 |
| CCZ1B_S266 | CCZ1B | P86790 | | 266 | ELAGRDSPIRAEM | 6.5008 | 6.5158 | 6.7221 | 6.4094 | 5.7189 | 6.2295 | 6.2460 | 6.9559 | 6.7567 | 6.3969 | 6.2783 | 6.6823 | 5.2111 | 5.6402 | 5.6343 | 6.1019 |
| CD2AP_S189 | CD2AP | Q9Y5K6 | | 542 | KKDTCYSPKPSVY | 7.0413 | 6.7786 | 7.1433 | 6.8458 | 6.4996 | 7.2105 | 7.2562 | 6.7891 | 7.5451 | 6.8279 | 7.1845 | 7.3871 | 3.8308 | 4.2364 | 3.3779 | 4.0459 |
| CD2AP_S458 | CD2AP | Q9Y5K6 | | 458 | LPLRPKSVDFDSL | 6.5514 | 7.5461 | 8.7861 | 6.8191 | 4.2771 | 6.0528 | 6.9539 | 6.9976 | 5.8528 | 6.5744 | 9.0255 | 8.5861 | 3.5751 | 3.7632 | 4.0419 | 4.5967 |
| CD2AP_S510 | CD2AP | Q9Y5K6 | | 554 | YLSTPSSASKANT | 6.5867 | 5.9876 | 7.1107 | 7.3868 | 6.3617 | 7.3012 | 7.1231 | 6.2753 | 7.8214 | 7.8288 | 7.9582 | 7.7441 | 3.2462 | 4.3898 | 3.1790 | 3.6993 |
| CD2AP_S510;S514 | CD2AP | Q9Y5K6 | | 565 | NTTAFLTPLEIKA | 7.4087 | 7.3501 | 6.6144 | 7.0019 | 6.0665 | 7.3297 | 6.9671 | 7.4002 | 7.3287 | 7.0500 | 7.1702 | 7.0406 | 3.2779 | 4.5604 | 3.1835 | 4.2500 |
| CD2AP_S542 | CD2AP | Q9Y5K6 | | 510 | RFNGGHSPTHSPE | 6.0399 | 5.8537 | 6.4494 | 5.5823 | 5.8013 | 5.9542 | 5.6967 | 6.8775 | 6.0880 | 6.2979 | 6.2488 | 6.4501 | 6.5887 | 7.0219 | 6.6091 | 6.4403 |
| CD2AP_S86 | CD2AP | Q9Y5K6 | | 86 | SLVQRISTYGLPA | 8.3589 | 6.3584 | 6.9322 | 6.4992 | 6.4804 | 5.7773 | 6.5373 | 6.0021 | 7.6729 | 6.0319 | 7.3277 | 6.5053 | 5.7668 | 5.8353 | 3.9557 | 3.9585 |
| CD2AP_T551;S554 | CD2AP | Q9Y5K6 | | 189 | VLAGPTSPIPSLG | 5.4815 | 6.2359 | 6.7028 | 6.1605 | 5.8587 | 7.1832 | 7.0570 | 6.6067 | 5.8841 | 6.2502 | 6.4575 | 6.4733 | 5.8262 | 5.7857 | 5.9159 | 6.1208 |
| CD2AP_T554 | CD2AP | Q9Y5K6 | 551;554 | | NA | 6.8180 | 6.7393 | 7.2398 | 7.0896 | 6.5007 | 6.9084 | 6.9959 | 6.4181 | 7.5212 | 7.4854 | 8.0850 | 7.6306 | 3.4388 | 4.4229 | 2.8946 | 3.8118 |
| CD2AP_T565 | CD2AP | Q9Y5K6 | 510;514 | | NA | 6.1753 | 6.7720 | 7.0457 | 6.6384 | 6.5705 | 7.2853 | 6.1949 | 6.6795 | 7.0763 | 7.0375 | 7.8320 | 7.3160 | 3.9970 | 4.9421 | 4.2098 | 4.2277 |
| CD2BP2_S118 | CD2BP2 | O95400 | | 195 | GPGQPSSPQRLDR | 6.3347 | 6.0759 | 6.1625 | 6.2327 | 6.2877 | 6.0581 | 5.7121 | 6.5713 | 6.4154 | 6.1327 | 6.5114 | 6.6890 | 6.4711 | 6.1562 | 5.9829 | 6.2063 |
| CD2BP2_S195 | CD2BP2 | O95400 | | 243 | LGPHNPTPPPSLD | 5.2512 | 5.2977 | 5.9096 | 6.0454 | 5.9302 | 5.5936 | 5.7106 | 6.1015 | 5.2589 | 5.9671 | 5.8929 | 6.6325 | 8.2013 | 7.7692 | 7.5987 | 6.8396 |
| CD2BP2_S49 | CD2BP2 | O95400 | | 49 | GKHSLDSDEEEDD | 6.0564 | 5.9610 | 6.5423 | 6.3866 | 6.0747 | 6.0827 | 5.6922 | 6.5610 | 6.3402 | 6.7339 | 6.2659 | 6.8897 | 6.4294 | 6.1592 | 5.8688 | 5.9559 |
| CD2BP2_T243 | CD2BP2 | O95400 | | 118 | DAQIRDSWLDNID | 7.2404 | 7.6323 | 6.7808 | 6.9715 | 6.9868 | 6.0738 | 6.9303 | 6.2205 | 6.7703 | 6.2359 | 6.7039 | 6.6187 | 4.7187 | 5.4333 | 4.1445 | 4.5384 |
| CD44_S686 | CD44 | P16070 | | 706 | NGEASKSQEMVHL | 6.3181 | 6.6849 | 6.0175 | 6.1713 | 6.0834 | 6.2371 | 6.3868 | 6.0433 | 6.2464 | 6.1794 | 6.2132 | 6.2535 | 6.0312 | 6.3567 | 6.2510 | 6.5261 |
| CD44_S697 | CD44 | P16070 | | 718 | LVNKESSETPDQF | 6.7793 | 7.1092 | 6.0951 | 7.4712 | 5.6824 | 6.2006 | 6.2407 | 6.0403 | 6.9222 | 6.4232 | 6.2464 | 6.7531 | 5.2675 | 5.7897 | 5.2238 | 5.7554 |
| CD44_S706 | CD44 | P16070 | | 686 | KKLVINSGNGAVE | 6.9280 | 6.8187 | 6.2943 | 6.7210 | 6.2806 | 6.1609 | 6.6908 | 6.8830 | 6.4460 | 6.7180 | 6.6036 | 6.5990 | 5.2509 | 5.3338 | 4.9443 | 5.3270 |
| CD44_S717 | CD44 | P16070 | | 697 | VEDRKPSGLNGEA | 4.9650 | 4.2891 | 4.4147 | 4.2830 | 5.8560 | 4.7324 | 4.6285 | 5.3729 | 6.6611 | 5.6678 | 6.2437 | 6.2425 | 8.5415 | 9.5902 | 9.2772 | 8.3343 |
| CD44_S718 | CD44 | P16070 | | 717 | HLVNKESSETPDQ | 4.8434 | 5.6770 | 5.6985 | 5.9615 | 5.5171 | 6.2379 | 5.1075 | 5.8885 | 6.2118 | 5.5514 | 4.9925 | 4.9369 | 8.8598 | 8.0154 | 8.3183 | 8.1825 |
| CDAN1_S158 | CDAN1 | Q8IWY9 | | 158 | RLRGSGSPSRPSL | 5.9215 | 6.1114 | 6.2077 | 6.1910 | 5.8876 | 6.4037 | 6.1465 | 6.5583 | 6.4507 | 6.0039 | 6.6559 | 6.7338 | 5.8853 | 6.4590 | 5.9413 | 6.4425 |
| CDAN1_S265;S277 | CDAN1 | Q8IWY9 | | 71 | LPQGPPTPAKTPG | 6.1110 | 6.2435 | 6.8583 | 6.0945 | 6.0768 | 6.5268 | 5.9303 | 6.7358 | 6.8693 | 6.5959 | 6.6201 | 6.6138 | 5.4692 | 6.1024 | 5.4902 | 5.6621 |
| CDAN1_S277 | CDAN1 | Q8IWY9 | | 277 | PTPELGSPLPSRT | 4.8272 | 5.5857 | 6.0829 | 6.4656 | 5.8580 | 5.0786 | 5.8036 | 6.1234 | 5.8315 | 5.7435 | 5.5996 | 6.1905 | 8.1529 | 8.1487 | 7.8233 | 6.6849 |
| CDAN1_T71 | CDAN1 | Q8IWY9 | 265;277 | | NA | 5.6519 | 6.5570 | 7.0474 | 6.5105 | 6.1860 | 6.3882 | 5.7982 | 5.4723 | 6.5417 | 6.7385 | 6.8446 | 6.8279 | 5.7096 | 5.7051 | 5.8591 | 6.1621 |
| CDC14A_S453 | CDC14A | Q9UNH5 | | 453 | TSKMALSPSATAK | 7.3536 | 7.4468 | 7.6483 | 7.0603 | 6.5228 | 7.1453 | 6.6847 | 7.3877 | 6.7879 | 7.2449 | 7.0353 | 7.0175 | 3.4519 | 4.1736 | 3.4797 | 3.5597 |
| CDC14A_S583 | CDC14A | Q9UNH5 | | 583 | ARFLSRSIPSLQS | 6.6822 | 6.2783 | 6.4287 | 6.7811 | 6.8768 | 6.7622 | 6.1595 | 6.6980 | 7.2502 | 6.5707 | 6.8253 | 5.9964 | 5.8418 | 5.3675 | 4.3651 | 5.1161 |
| CDC16_S560 | CDC16 | Q13042 | | 560 | TLKNIISPPWDFR | 6.5081 | 6.8711 | 6.4400 | 6.8004 | 5.9465 | 6.0369 | 6.4810 | 6.4202 | 6.2645 | 6.4559 | 6.5571 | 6.6234 | 5.4816 | 5.9784 | 5.4676 | 5.6674 |
| CDC16_T581 | CDC16 | Q13042 | | 581 | AEETGLTPLETSR | 6.4150 | 6.1573 | 6.6551 | 6.8989 | 6.4705 | 5.6501 | 6.9493 | 6.5661 | 7.3325 | 6.4004 | 7.0026 | 6.8157 | 5.4807 | 5.2742 | 4.7873 | 5.1443 |
| CDC20_S41 | CDC20 | Q12834 | | 70 | SSKVQTTPSKPGG | 6.7797 | 6.3757 | 6.9098 | 5.9234 | 6.1077 | 6.6853 | 6.6611 | 6.7698 | 7.3590 | 6.6259 | 6.8194 | 6.7711 | 4.8013 | 5.2555 | 4.8451 | 5.3101 |
| CDC20_T106 | CDC20 | Q12834 | | 41 | AAGPAPSPMRAAN | 6.9248 | 6.6995 | 6.8793 | 6.1626 | 5.2100 | 6.8901 | 6.7919 | 6.9339 | 7.3246 | 7.0891 | 7.5082 | 7.6398 | 3.9909 | 4.9071 | 4.0438 | 5.0044 |
| CDC20_T59 | CDC20 | Q12834 | | 59 | GRTPGRTPGKSSS | 7.4766 | 6.8296 | 7.4176 | 5.8253 | 5.0625 | 6.6809 | 6.6154 | 6.3259 | 7.6480 | 6.9440 | 7.6052 | 8.0440 | 4.2681 | 4.6077 | 3.6781 | 4.9712 |
| CDC20_T70 | CDC20 | Q12834 | | 106 | QPENSQTPTKKEH | 6.5390 | 7.3507 | 7.3945 | 6.5480 | 5.5447 | 7.1876 | 6.7667 | 6.3200 | 8.0797 | 7.6715 | 8.0350 | 7.9399 | 3.3473 | 4.1170 | 3.1581 | 4.0002 |
| CDC23_S578 | CDC23 | Q9UJX2 | | 596 | LNLSSVTPxxxxx | 6.5524 | 6.3604 | 7.0103 | 6.5389 | 6.2138 | 6.3980 | 6.4968 | 6.6359 | 6.8068 | 6.6328 | 6.6315 | 6.8740 | 5.2517 | 5.2799 | 4.8132 | 5.5034 |
| CDC23_S588 | CDC23 | Q9UJX2 | | 588 | TPTRRVSPLNLSS | 6.3041 | 6.5819 | 5.9696 | 6.1101 | 5.7389 | 6.3599 | 6.2064 | 6.3910 | 5.5984 | 6.1015 | 6.0384 | 6.1611 | 6.1772 | 6.7909 | 6.9059 | 6.5647 |
| CDC23_S588;T596 | CDC23 | Q9UJX2 | | 578 | FLPASLSANNTPT | 6.0745 | 6.4966 | 5.8749 | 6.5600 | 6.4386 | 6.4117 | 7.2539 | 5.5627 | 5.9592 | 6.8696 | 7.2903 | 6.8014 | 5.1311 | 6.2006 | 4.9780 | 6.0968 |
| CDC23_S596 | CDC23 | Q9UJX2 | | 582 | SLSANNTPTRRVS | 5.7378 | 6.6035 | 6.9847 | 7.2013 | 6.8803 | 5.4587 | 6.6053 | 6.4625 | 6.2264 | 6.2199 | 6.4182 | 6.1946 | 5.4374 | 5.6658 | 5.9750 | 5.9284 |
| CDC23_T565;T582 | CDC23 | Q9UJX2 | 588;596 | | NA | 6.0855 | 6.7324 | 6.6570 | 6.4734 | 6.1401 | 6.5363 | 6.8896 | 6.7021 | 6.4748 | 6.2668 | 6.5269 | 6.4860 | 5.4065 | 5.8147 | 5.0395 | 5.7683 |
| CDC23_T582 | CDC23 | Q9UJX2 | 565;582 | | NA | 6.7085 | 6.5498 | 6.4271 | 7.0555 | 6.3765 | 6.2215 | 6.1393 | 6.3775 | 6.4667 | 6.3808 | 6.6538 | 6.5903 | 5.7616 | 5.9277 | 5.6672 | 4.6960 |
| CDC25A_S116 | CDC25A | P30304 | | 18 | RLLFACSPPPASQ | 6.5615 | 6.4539 | 7.7274 | 7.5401 | 5.8007 | 7.3157 | 6.6622 | 6.5881 | 7.1005 | 7.4315 | 9.1215 | 7.7964 | 2.8609 | 3.5319 | 3.3591 | 4.1487 |
| CDC25A_S18 | CDC25A | P30304 | | 116 | QKLLGCSPALKRS | 7.3187 | 8.6000 | 7.3878 | 6.0137 | 5.1351 | 6.3059 | 7.3957 | 6.9732 | 8.8409 | 7.9663 | 8.9751 | 7.5379 | 2.2152 | 3.1569 | 2.7502 | 3.4274 |
| CDC25A_S293 | CDC25A | P30304 | | 293 | STKRRKSMSGASP | 5.4236 | 4.9212 | 6.6102 | 5.0663 | 6.5861 | 6.5889 | 4.7311 | 6.7118 | 7.7672 | 6.5222 | 7.6935 | 7.5920 | 6.3029 | 6.3229 | 5.4626 | 5.6976 |
| CDC25B_S137 | CDC25B | P30305-4 | | 146 | VRLLGHSPVLRNI | 7.5861 | 7.7285 | 7.1246 | 7.6752 | 6.4412 | 7.3045 | 7.2333 | 7.1571 | 7.9723 | 7.8515 | 8.1139 | 7.8325 | 2.0929 | 3.0689 | 1.9820 | 2.8355 |
| CDC25B_S146 | CDC25B | P30305-4 | | 270 | MEVEELSPLALGR | 6.7043 | 6.8733 | 7.1434 | 6.3238 | 6.2792 | 6.3193 | 6.3817 | 6.5609 | 6.9974 | 7.2966 | 7.2726 | 7.0633 | 4.4401 | 5.0390 | 4.6367 | 4.6684 |
| CDC25B_S259 | CDC25B | P30305-4 | | 259 | PDLMCLSPDRKME | 6.6254 | 7.4009 | 8.2789 | 6.6818 | 6.4274 | 7.7312 | 7.0251 | 7.0235 | 6.8290 | 7.5619 | 9.1486 | 8.4431 | 2.3148 | 3.2851 | 2.1144 | 3.1090 |
| CDC25B_S270 | CDC25B | P30305-4 | | 42 | SHGLLGSPVRAAA | 6.6533 | 6.8279 | 7.4336 | 6.9239 | 6.3744 | 7.6404 | 6.8887 | 7.2502 | 7.7354 | 7.7073 | 7.2017 | 7.3611 | 3.0422 | 4.2556 | 3.1172 | 3.5871 |
| CDC25B_S344 | CDC25B | P30305-4 | | 137 | AIRRFQSMPVRLL | 7.8848 | 7.8311 | 8.1081 | 7.4714 | 6.4408 | 5.9053 | 5.4385 | 5.5909 | 7.7409 | 6.6437 | 7.5086 | 7.0057 | 4.1190 | 4.7398 | 3.7024 | 3.8690 |
| CDC25B_S42 | CDC25B | P30305-4 | | 344 | RLFRSPSMPCSVI | 7.0198 | 6.7316 | 7.7863 | 7.1179 | 7.0723 | 7.2056 | 5.5972 | 6.2281 | 7.4264 | 7.7109 | 7.7554 | 8.1121 | 3.4414 | 3.9082 | 2.9433 | 3.9435 |
| CDC25C_S148 | CDC25C | P30307 | | 216 | GLYRSPSMPENLN | 7.1105 | 6.8772 | 7.3588 | 6.3618 | 6.3054 | 6.8137 | 6.4490 | 6.8820 | 6.5726 | 6.6595 | 6.7860 | 7.1769 | 4.6755 | 4.9786 | 4.0977 | 4.8949 |
| CDC25C_S168 | CDC25C | P30307 | | 263 | CLKKTVSLCDITI | 6.9923 | 6.6219 | 7.0543 | 6.5868 | 6.6023 | 6.9153 | 6.4637 | 6.4783 | 6.5862 | 6.1797 | 6.8246 | 6.5810 | 4.8825 | 5.3084 | 4.5998 | 5.3230 |
| CDC25C_S216 | CDC25C | P30307 | | 48 | CPDVPRTPVGKFL | 7.5800 | 7.3144 | 7.5567 | 6.5496 | 5.6819 | 7.5766 | 7.7150 | 6.4230 | 7.6175 | 7.0017 | 7.5096 | 7.3662 | 3.1456 | 4.1108 | 2.9869 | 3.8645 |
| CDC25C_S263 | CDC25C | P30307 | | 168 | EMKYLGSPITTVP | 6.6123 | 7.2764 | 6.9954 | 6.3985 | 6.6692 | 7.3717 | 7.5891 | 7.3276 | 6.9709 | 6.8251 | 6.7533 | 6.9188 | 3.9153 | 4.6210 | 3.7202 | 4.0354 |
| CDC25C_T37;T48 | CDC25C | P30307 | | 148 | DAMCSSSANKEND | 7.5332 | 8.3917 | 6.2551 | 7.3457 | 6.0820 | 6.9516 | 9.2055 | 7.2875 | 8.1484 | 6.5184 | 8.2457 | 7.4870 | 1.8925 | 3.3166 | 2.0616 | 3.2775 |
| CDC25C_T48 | CDC25C | P30307 | 37;48 | | NA | 8.1381 | 9.1344 | 7.3143 | 7.5296 | 5.3910 | 7.5436 | 7.4644 | 6.4260 | 7.2675 | 6.5909 | 7.4312 | 8.1059 | 3.0866 | 3.5698 | 2.0629 | 2.9438 |
| CDC26_S82 | CDC26 | Q8NHZ8 | | 82 | RSSQFGSLEFxxx | 6.2346 | 5.4087 | 6.8623 | 6.8065 | 7.5312 | 4.8219 | 5.4734 | 6.1100 | 5.6281 | 6.0936 | 5.6047 | 6.6504 | 5.7804 | 7.0146 | 7.3146 | 6.6649 |
| CDC27_S219;S220 | CDC27 | P30260-2 | | 441 | ITKLDSSIISEGK | 7.0059 | 6.7356 | 7.3136 | 7.3900 | 6.5762 | 6.8339 | 7.0763 | 6.7549 | 6.6877 | 7.0987 | 7.1071 | 7.4104 | 3.6051 | 4.6550 | 3.5778 | 4.1718 |
| CDC27_S220 | CDC27 | P30260-2 | | 220 | RLNLESSNSKYSL | 7.8640 | 7.8665 | 7.1147 | 6.9603 | 6.7655 | 7.4802 | 7.2091 | 6.9560 | 8.0566 | 7.1883 | 6.9984 | 6.6953 | 2.7346 | 3.8386 | 2.8454 | 3.4265 |
| CDC27_S370 | CDC27 | P30260-2 | | 452 | GKISTITPQIQAF | 6.4091 | 7.1100 | 7.1570 | 7.1859 | 6.0520 | 7.1055 | 6.9731 | 6.7836 | 7.1265 | 7.2815 | 7.3849 | 7.2692 | 3.7211 | 4.5393 | 3.6505 | 4.2509 |
| CDC27_S375 | CDC27 | P30260-2 | | 432 | QPNINDSLEITKL | 5.9531 | 5.8907 | 6.9088 | 6.8316 | 6.5697 | 6.8339 | 6.5666 | 7.3788 | 7.2596 | 6.6476 | 7.6252 | 6.5736 | 4.3964 | 5.0972 | 4.4487 | 5.0185 |
| CDC27_S432 | CDC27 | P30260-2 | | 375 | LSPTITSPPNALP | 5.7714 | 5.8153 | 7.0091 | 5.9191 | 6.9851 | 6.7057 | 6.9752 | 6.6158 | 5.6057 | 6.0928 | 7.1169 | 8.5602 | 5.2424 | 5.3801 | 5.4500 | 4.7553 |
| CDC27_S441 | CDC27 | P30260-2 | | 370 | TTPQVLSPTITSP | 5.8702 | 5.9103 | 7.0959 | 7.1396 | 6.3197 | 6.9201 | 5.8620 | 6.0790 | 5.4455 | 6.8081 | 6.7220 | 6.8930 | 5.1813 | 5.8089 | 5.5033 | 6.4412 |
| CDC27_T452 | CDC27 | P30260-2 | 219;220 | | NA | 6.6240 | 6.3987 | 7.8618 | 7.7374 | 6.6031 | 6.1544 | 8.4479 | 6.5250 | 7.9828 | 5.9576 | 6.9253 | 5.4054 | 4.8845 | 4.8414 | 3.0181 | 4.6327 |
| CDC37_S300 | CDC37 | Q16543 | | 300 | PVEVYESLPEELQ | 7.2538 | 5.5908 | 8.7523 | 6.0803 | 6.9026 | 6.2331 | 6.6075 | 6.9510 | 6.2254 | 6.2796 | 6.7672 | 7.1715 | 4.5256 | 4.3186 | 5.1883 | 5.1525 |
| CDC42_T138 | CDC42 | P60953-1 | | 138 | NKQKPITPETAEK | 6.3079 | 6.5942 | 7.1537 | 6.3547 | 6.0524 | 5.6184 | 5.5818 | 6.1782 | 6.2891 | 6.7448 | 7.4661 | 7.0233 | 5.0909 | 6.2598 | 5.2917 | 5.9931 |
| CDC42BPA_S1700;S1703 | CDC42BPA | Q5VT25-6 | | 750 | EKTRRESQSEREE | 6.7120 | 5.8370 | 6.8535 | 5.6587 | 6.3152 | 7.0040 | 6.4211 | 6.5727 | 6.6161 | 6.2673 | 7.0887 | 5.8025 | 5.4882 | 5.8816 | 6.3096 | 5.1716 |
| CDC42BPA_S1703 | CDC42BPA | Q5VT25-6 | | 1768 | SPRKTKSLSLEST | 6.6774 | 6.8058 | 7.9142 | 6.0513 | 6.3560 | 6.6441 | 6.4775 | 6.5461 | 6.4197 | 7.3164 | 6.7245 | 5.7377 | 5.1154 | 5.7445 | 4.9429 | 4.5264 |
| CDC42BPA_S1768 | CDC42BPA | Q5VT25-6 | | 1703 | REFSGGSYSAKRQ | 7.3722 | 7.1193 | 7.1832 | 7.0141 | 5.3887 | 6.1876 | 6.0515 | 5.6431 | 5.4167 | 5.2382 | 5.8810 | 5.7179 | 6.5989 | 6.6858 | 6.4517 | 6.0501 |
| CDC42BPA_S750 | CDC42BPA | Q5VT25-6 | 1700;1703 | | NA | 6.8293 | 7.2172 | 7.0776 | 6.3562 | 6.1245 | 6.3040 | 6.1425 | 5.8633 | 5.8433 | 6.7404 | 6.3225 | 5.5503 | 5.9762 | 6.5969 | 5.4234 | 5.6325 |
| CDC42BPB_S363 | CDC42BPB | Q9Y5S2 | | 363 | YIPDVSSPSDTSN | 5.6052 | 6.0995 | 7.0533 | 6.5787 | 5.1504 | 6.8067 | 6.8916 | 6.4054 | 6.7020 | 6.6032 | 6.8323 | 7.2755 | 6.0228 | 5.4947 | 4.7514 | 5.7273 |
| CDC42EP1_S121 | CDC42EP1 | Q00587 | | 192 | GLRRSDSLLSFRL | 5.4627 | 5.8673 | 6.0778 | 5.3509 | 6.2067 | 6.4368 | 6.4156 | 7.4462 | 6.2412 | 6.4694 | 6.7271 | 6.7989 | 6.3103 | 6.1603 | 5.9307 | 6.0981 |
| CDC42EP1_S19 | CDC42EP1 | Q00587 | | 121 | IIKNAISLPQLNQ | 6.5601 | 5.8556 | 6.5475 | 6.0945 | 6.5498 | 5.8063 | 6.4178 | 6.0166 | 5.5659 | 5.9083 | 5.7726 | 6.3592 | 7.3743 | 6.5947 | 6.2130 | 6.3637 |
| CDC42EP1_S192 | CDC42EP1 | Q00587 | | 19 | MSLGKLSPVGWVS | 7.4140 | 6.4237 | 6.3464 | 6.4839 | 6.7540 | 6.0850 | 6.6714 | 7.8971 | 7.1897 | 7.7573 | 7.1753 | 7.6656 | 3.9860 | 4.0406 | 4.0770 | 4.0330 |
| CDC42EP2_S101 | CDC42EP2 | O14613 | | 101 | ELPDGPSPLLKNA | 6.9087 | 6.0576 | 6.8011 | 5.7558 | 5.9117 | 6.9149 | 7.9762 | 7.7066 | 5.6037 | 6.4881 | 5.9154 | 6.7553 | 3.5849 | 5.6272 | 4.7613 | 7.2316 |
| CDC42EP2_S141 | CDC42EP2 | O14613 | | 141 | LETPQPSPQEGGS | 5.7914 | 6.9403 | 3.0025 | 5.1251 | 5.3344 | 3.7883 | 6.4278 | 7.0257 | 5.7017 | 9.4217 | 5.4754 | 6.8036 | 5.5650 | 6.7396 | 9.2338 | 7.6236 |
| CDC42EP2_T137;S141 | CDC42EP2 | O14613 | 137;141 | | NA | 6.8463 | 6.1810 | 5.7528 | 6.4060 | 6.4759 | 5.6271 | 6.0947 | 5.3225 | 6.4146 | 6.3473 | 6.1841 | 6.5130 | 5.4968 | 6.0832 | 5.9133 | 8.3415 |
| CDC42EP3_S108 | CDC42EP3 | Q9UKI2 | | 89 | EFFRANSTSDSVF | 5.0524 | 4.9681 | 6.4003 | 5.5193 | 6.0285 | 6.0519 | 5.3822 | 6.7474 | 5.4017 | 5.8504 | 6.1089 | 6.2508 | 7.9081 | 7.3595 | 7.7269 | 7.2438 |
| CDC42EP3_S144 | CDC42EP3 | Q9UKI2 | | 144 | AKLPRLSCEPVME | 6.6679 | 6.6311 | 6.8019 | 6.2303 | 6.7738 | 6.8405 | 6.7668 | 6.4438 | 5.5240 | 6.6000 | 6.1764 | 6.3812 | 6.6369 | 4.8318 | 5.8318 | 4.8617 |
| CDC42EP3_S27 | CDC42EP3 | Q9UKI2 | | 27 | KLRDILSPDMISP | 4.9346 | 7.0682 | 8.2506 | 6.2799 | 5.4180 | 7.5779 | 5.9961 | 9.3857 | 5.9803 | 5.4706 | 4.3723 | 6.1012 | 7.0734 | 5.7701 | 4.1148 | 6.2063 |
| CDC42EP3_S89 | CDC42EP3 | Q9UKI2 | | 108 | VLKNAISLPTIGG | 5.6460 | 5.7539 | 6.6471 | 5.1511 | 6.2954 | 6.0138 | 6.0250 | 6.6837 | 4.6034 | 5.6280 | 5.6224 | 5.0611 | 8.0581 | 8.2504 | 7.1786 | 7.3820 |
| CDC42EP4_S11 | CDC42EP4 | Q9H3Q1 | | 118 | FVKNAMSLPQLNE | 6.7816 | 6.6477 | 7.3998 | 6.2851 | 5.5583 | 6.9439 | 6.0673 | 6.4225 | 6.0870 | 6.2773 | 7.0432 | 6.9990 | 5.4896 | 5.5705 | 4.8721 | 5.5549 |
| CDC42EP4_S118 | CDC42EP4 | Q9H3Q1 | | 174 | GAAGPHSPDPLLD | 6.0360 | 6.0624 | 6.6742 | 6.9774 | 6.4169 | 6.4919 | 6.1722 | 6.6289 | 6.9304 | 6.9139 | 7.1446 | 7.1407 | 5.1651 | 5.0035 | 4.9429 | 5.2989 |
| CDC42EP4_S138;S142 | CDC42EP4 | Q9H3Q1 | | 295 | AAPSPGSARSMGS | 6.2599 | 6.4400 | 6.3540 | 6.7856 | 6.3714 | 6.5345 | 6.1268 | 6.5468 | 6.2405 | 5.6332 | 6.1427 | 5.9147 | 6.7549 | 6.4050 | 5.6764 | 5.8137 |
| CDC42EP4_S142 | CDC42EP4 | Q9H3Q1 | | 292 | WAAAAPSPGSARS | 6.0575 | 6.0463 | 6.5391 | 6.8950 | 6.3335 | 6.5346 | 6.4134 | 6.6678 | 6.4559 | 6.4775 | 6.6391 | 6.0063 | 6.0716 | 5.7098 | 5.9925 | 5.1602 |
| CDC42EP4_S174 | CDC42EP4 | Q9H3Q1 | | 142 | PKSLSSSPVKKAN | 6.4776 | 6.1644 | 5.8695 | 5.9134 | 7.3219 | 6.3949 | 6.1570 | 6.6516 | 6.1960 | 6.2484 | 5.8187 | 5.7344 | 6.9655 | 6.2926 | 6.2727 | 5.5215 |
| CDC42EP4_S292 | CDC42EP4 | Q9H3Q1 | | 64 | GEPDGESLDEQPS | 6.3613 | 6.3510 | 6.5367 | 6.2260 | 5.2511 | 5.5407 | 5.9791 | 7.2442 | 5.9494 | 6.2287 | 6.0961 | 6.3624 | 6.7267 | 6.5989 | 6.8474 | 5.7004 |
| CDC42EP4_S292;S295 | CDC42EP4 | Q9H3Q1 | | 11 | KQLVSSSVHSKRR | 11.1431 | 7.1050 | 7.5777 | 7.1648 | 5.9896 | 5.5215 | 5.9648 | 6.2983 | 6.9808 | 6.4340 | 6.9918 | 6.2932 | 4.7643 | 5.2800 | 3.2343 | 3.2567 |
| CDC42EP4_S295 | CDC42EP4 | Q9H3Q1 | 138;142 | | NA | 6.8918 | 7.1268 | 7.2215 | 6.6843 | 6.4586 | 6.7789 | 6.2703 | 6.6742 | 6.8758 | 6.7401 | 7.1418 | 7.3175 | 4.5399 | 4.5610 | 4.2690 | 4.4484 |
| CDC42EP4_S64 | CDC42EP4 | Q9H3Q1 | 292;295 | | NA | 6.1726 | 6.9929 | 6.9702 | 6.1012 | 6.4245 | 6.1266 | 6.0882 | 6.5924 | 6.1615 | 6.6066 | 6.5708 | 6.5419 | 5.5943 | 6.2776 | 5.3612 | 5.4174 |
| CDC42SE1_S74 | CDC42SE1 | Q9NRR8 | | 74 | NRDRPWSNSRGLx | 7.6985 | 7.1443 | 7.4339 | 6.1193 | 5.7946 | 5.7662 | 6.8789 | 7.0057 | 6.4122 | 6.1585 | 7.3647 | 7.0767 | 4.5258 | 5.2253 | 4.3539 | 5.0417 |
| CDC45_S144;S148 | CDC45 | O75419-3 | 144;148 | | NA | 5.8184 | 5.8701 | 6.7596 | 6.4957 | 6.0991 | 6.3925 | 5.7035 | 5.8154 | 5.9618 | 7.0023 | 6.9080 | 6.3623 | 6.1808 | 6.1954 | 6.4165 | 6.0185 |
| CDC45_S144;S148;S151 | CDC45 | O75419-3 | 144;148;151 | | NA | 5.7956 | 5.8676 | 6.9057 | 6.2915 | 6.2632 | 6.0349 | 5.4031 | 6.1134 | 5.9288 | 5.9976 | 6.8511 | 6.8380 | 6.0979 | 6.9624 | 6.5510 | 6.0982 |
| CDC5L_S303 | CDC5L | Q99459 | | 227 | ALGFYDTSEENYQ | 8.1880 | 9.2237 | 8.8107 | 9.9920 | 9.1185 | 10.7420 | 9.9352 | 10.0278 | 3.5458 | 3.1988 | 3.4185 | 3.7566 | 2.3382 | 2.7884 | 2.0805 | 2.8352 |
| CDC5L_S438;T442 | CDC5L | Q99459 | | 303 | LPAPQISDAELQE | 6.7981 | 7.5316 | 7.8962 | 8.2656 | 7.6637 | 7.9428 | 7.9427 | 8.0369 | 5.3378 | 5.1717 | 5.6400 | 5.6673 | 3.6724 | 4.1766 | 3.9044 | 4.3522 |
| CDC5L_S442 | CDC5L | Q99459 | | 377 | ALTNVDTPLKGGL | 6.0445 | 6.8347 | 6.9726 | 6.9557 | 6.3794 | 6.6977 | 6.6956 | 6.7618 | 6.4135 | 6.5007 | 6.3883 | 6.8856 | 4.9255 | 5.3333 | 5.0033 | 5.2079 |
| CDC5L_T227 | CDC5L | Q99459 | | 438 | KPVINSTPGRTPL | 7.2402 | 6.2834 | 6.5767 | 7.4228 | 5.7959 | 6.8740 | 7.4530 | 6.4056 | 7.0040 | 6.3967 | 6.7843 | 7.1473 | 4.2988 | 5.0234 | 4.1832 | 5.1106 |
| CDC5L_T377 | CDC5L | Q99459 | | 424 | NGAEGLTPRSGTT | 6.7817 | 7.0736 | 6.3252 | 7.1768 | 5.8025 | 6.2955 | 6.6441 | 6.4095 | 6.4420 | 6.1815 | 6.6776 | 6.7835 | 5.1154 | 5.2766 | 5.0559 | 5.9584 |
| CDC5L_T385 | CDC5L | Q99459 | | 385 | LKGGLNTPLHESD | 5.8096 | 8.0790 | 5.9123 | 8.6900 | 6.5742 | 7.6875 | 5.5894 | 8.7336 | 6.7500 | 6.3910 | 7.0270 | 6.0814 | 3.6155 | 4.9090 | 4.2338 | 3.9169 |
| CDC5L_T404 | CDC5L | Q99459 | | 442 | NSTPGRTPLRDKL | 5.9213 | 5.9054 | 6.6713 | 6.6746 | 5.7522 | 6.9146 | 7.1375 | 6.2229 | 6.5650 | 6.7763 | 6.3605 | 7.2414 | 5.1913 | 5.8937 | 4.9822 | 5.7896 |

| | | | | | | | | | | | | | | | | | | | | | |
|---|---|---|---|---|---|---|---|---|---|---|---|---|---|---|---|---|---|---|---|---|---|
| CDC5L_T404;T407;T424 | CDC5L | Q99459 | | 404 | QRQVVQTPNTVLS | 5.9407 | 6.5364 | 6.6372 | 5.9565 | 5.7151 | 6.5477 | 7.0314 | 6.8569 | 5.6812 | 6.0505 | 6.8787 | 7.2089 | 5.3141 | 6.4988 | 5.4849 | 5.6609 |
| CDC5L_T424 | CDC5L | Q99459 | 438;442 | | NA | 6.2802 | 6.4786 | 7.5034 | 7.5339 | 5.8681 | 6.5011 | 7.1969 | 7.1255 | 7.2872 | 7.1088 | 6.5619 | 7.2920 | 3.8457 | 4.4195 | 3.9819 | 5.0152 |
| CDC5L_T438 | CDC5L | Q99459 | 404;407;424 | | NA | 6.0062 | 5.7421 | 6.5120 | 5.9866 | 5.8481 | 6.3305 | 6.0283 | 6.6497 | 6.2261 | 6.7121 | 6.3302 | 7.0396 | 6.0191 | 6.1309 | 5.5098 | 6.9287 |
| CDC6_S45 | CDC6 | Q99741 | | 54 | VKALPLSPRKRLG | 7.7167 | 7.2158 | 7.5777 | 7.5473 | 6.2150 | 6.0194 | 6.7673 | 6.3983 | 7.6090 | 6.8629 | 7.1655 | 7.2652 | 3.6860 | 3.9618 | 3.7178 | 4.2744 |
| CDC6_S54 | CDC6 | Q99741 | | 74 | PHLPPCSPPKQGK | 6.6980 | 7.0960 | 7.1196 | 7.0598 | 6.7236 | 5.9990 | 5.9178 | 6.5746 | 7.7149 | 6.8872 | 7.5596 | 7.2707 | 4.1675 | 4.5658 | 4.0114 | 4.6347 |
| CDC6_S74 | CDC6 | Q99741 | | 45 | VQTVTCSPRVKAL | 6.9833 | 6.7087 | 7.0893 | 7.2288 | 6.4596 | 6.4513 | 6.6506 | 6.6046 | 7.2082 | 6.8990 | 6.9380 | 6.7919 | 4.4153 | 4.8167 | 3.9475 | 4.8072 |
| CDC7_S27 | CDC7 | O00311 | | 27 | RFQAEGSLKKNEQ | 7.5663 | 7.1270 | 7.6018 | 7.4979 | 5.9938 | 6.5309 | 7.2786 | 6.6658 | 7.7960 | 7.7575 | 7.5063 | 7.6243 | 2.7262 | 3.7501 | 2.9109 | 3.6666 |
| CDCA2_S1000 | CDCA2 | Q69YH5 | | 756 | CQFFKISPDLNIK | 7.0625 | 7.9959 | 7.0937 | 8.3877 | 6.0295 | 6.5455 | 7.0775 | 6.4399 | 7.0971 | 7.6304 | 7.2400 | 7.4777 | 2.8747 | 4.1125 | 3.1274 | 3.8082 |
| CDCA2_S120 | CDCA2 | Q69YH5 | | 400 | TFGEDLSPEVFDE | 6.5104 | 6.5378 | 6.5655 | 7.3277 | 6.1867 | 6.1912 | 6.6703 | 6.4812 | 6.7163 | 6.6901 | 6.7928 | 6.9548 | 4.6565 | 5.5861 | 4.9287 | 5.2041 |
| CDCA2_S291 | CDCA2 | Q69YH5 | | 591 | SKKPLLSPIPELP | 6.7748 | 7.4552 | 7.2137 | 7.4621 | 6.2486 | 6.6891 | 7.1431 | 6.3522 | 6.9948 | 7.4775 | 7.3662 | 7.2264 | 3.7719 | 4.3592 | 3.3563 | 4.1088 |
| CDCA2_S309 | CDCA2 | Q69YH5 | | 470 | SFAVLSSPNKSSI | 6.9834 | 6.7550 | 6.8567 | 7.6261 | 6.1159 | 6.1950 | 7.1031 | 6.2628 | 7.0392 | 6.8138 | 6.9486 | 7.1423 | 4.1722 | 4.7504 | 4.3614 | 4.8740 |
| CDCA2_S400 | CDCA2 | Q69YH5 | | 936 | SGFESMSPIKETV | 6.9639 | 6.9666 | 6.8991 | 7.9033 | 6.2702 | 6.5845 | 6.5357 | 5.9452 | 6.9339 | 7.0741 | 7.1450 | 7.0939 | 4.1921 | 4.5623 | 4.1909 | 4.7394 |
| CDCA2_S470 | CDCA2 | Q69YH5 | | 977 | PGKRRKSFCISTL | 7.3511 | 4.7168 | 5.4578 | 4.7857 | 7.3688 | 4.8929 | 5.8493 | 5.2826 | 9.6283 | 7.8507 | 8.3914 | 7.0919 | 6.5830 | 6.7936 | 3.6653 | 4.2908 |
| CDCA2_S591 | CDCA2 | Q69YH5 | | 120 | IKNARKSPLAQDS | 5.3059 | 7.0017 | 6.5622 | 6.7338 | 7.0083 | 6.4103 | 5.3767 | 6.9422 | 7.4694 | 6.7923 | 6.9573 | 6.6885 | 5.1569 | 5.6294 | 4.5156 | 5.4493 |
| CDCA2_S614 | CDCA2 | Q69YH5 | | 614 | SIRRLGSGYFSSN | 8.4397 | 6.8602 | 7.4087 | 7.3585 | 4.9594 | 6.6553 | 7.1682 | 5.5780 | 7.5753 | 7.7167 | 7.7564 | 6.5800 | 3.6323 | 4.4928 | 3.0970 | 4.7215 |
| CDCA2_S710 | CDCA2 | Q69YH5 | | 291 | GSSDAVSPDTFTA | 5.7284 | 6.0023 | 5.9147 | 6.0886 | 6.8765 | 6.4513 | 6.5351 | 7.1292 | 6.1173 | 6.7390 | 5.7084 | 7.8471 | 4.7403 | 6.1286 | 6.0235 | 5.9698 |
| CDCA2_S726 | CDCA2 | Q69YH5 | | 309 | AVPDVRSPATPAC | 6.1451 | 7.3845 | 6.4886 | 7.6025 | 6.2698 | 6.1742 | 6.4705 | 6.9008 | 7.5020 | 7.7605 | 7.4793 | 7.7318 | 3.5640 | 4.5128 | 3.6452 | 4.3684 |
| CDCA2_S756 | CDCA2 | Q69YH5 | | 726 | ERVASDSPKPALT | 6.6718 | 7.5366 | 5.9879 | 6.5218 | 6.1007 | 6.6945 | 6.1711 | 5.5752 | 7.1694 | 7.9246 | 7.5542 | 6.3769 | 4.0864 | 5.7810 | 4.5406 | 5.3072 |
| CDCA2_S936 | CDCA2 | Q69YH5 | | 710 | PKAGTDSPVSCAS | 6.1666 | 7.1400 | 6.5562 | 6.5192 | 6.2173 | 6.0483 | 6.9737 | 6.0437 | 6.5558 | 5.8386 | 6.0054 | 6.5677 | 5.1005 | 6.4188 | 5.4403 | 6.4078 |
| CDCA2_S977 | CDCA2 | Q69YH5 | | 1000 | GYRRRSSLNGKGE | 5.9968 | 4.8461 | 4.9031 | 5.2371 | 6.6801 | 5.8647 | 5.7200 | 5.7382 | 10.2452 | 9.1004 | 8.3099 | 8.1887 | 5.7714 | 6.2769 | 2.9675 | 4.1538 |
| CDCA3_S199 | CDCA3 | Q99618 | | 87 | SSGDPPSPLVKQL | 6.3850 | 5.1956 | 5.9788 | 5.7171 | 6.9624 | 5.9379 | 6.1018 | 6.7802 | 5.9170 | 6.7126 | 6.6776 | 6.7961 | 6.0606 | 6.6425 | 5.5889 | 6.5458 |
| CDCA3_S199;S209 | CDCA3 | Q99618 | | 37 | SAGILRTPIQVES | 6.7905 | 6.7024 | 6.9236 | 6.6003 | 6.5551 | 5.6245 | 6.0827 | 7.2953 | 7.1507 | 7.5803 | 6.7672 | 7.6142 | 4.0938 | 5.3461 | 4.1207 | 4.7526 |
| CDCA3_S209 | CDCA3 | Q99618 | | 44 | PIQVESSPQPGLP | 5.8089 | 5.5951 | 6.6276 | 6.3464 | 7.5973 | 6.2126 | 6.5034 | 7.8374 | 6.6577 | 7.3728 | 7.5324 | 8.2432 | 3.4708 | 5.6126 | 3.9268 | 4.6550 |
| CDCA3_S29;T37 | CDCA3 | Q99618 | | 199 | SKVLGRSPLTILQ | 6.5623 | 6.1862 | 6.8432 | 6.8887 | 6.5366 | 5.7149 | 6.0161 | 6.9850 | 6.8994 | 7.0625 | 6.8517 | 6.9973 | 4.7208 | 5.8599 | 4.7519 | 5.1237 |
| CDCA3_S29;T37;S44 | CDCA3 | Q99618 | | 209 | ILQDDNSPGTLTL | 6.1729 | 6.1041 | 6.5218 | 6.1670 | 6.9150 | 5.4494 | 6.2067 | 7.5048 | 6.9004 | 7.0156 | 6.8270 | 7.5452 | 4.9125 | 5.9214 | 4.2870 | 5.5493 |
| CDCA3_S68 | CDCA3 | Q99618 | | 94 | PLVKQLSEVFETE | 6.0088 | 4.5216 | 5.3800 | 4.9639 | 5.2645 | 4.9532 | 5.3800 | 6.1367 | 5.5187 | 5.1997 | 5.8270 | 4.4847 | 9.1604 | 8.2311 | 8.6503 | 10.3193 |
| CDCA3_S87 | CDCA3 | Q99618 | | 68 | QDSDPRSPTLGIA | 6.1110 | 5.8106 | 5.9675 | 5.6831 | 6.7783 | 6.3149 | 6.7692 | 7.2989 | 5.4052 | 7.0978 | 7.6467 | 7.7613 | 4.7285 | 5.9700 | 4.8679 | 5.7890 |
| CDCA3_S94 | CDCA3 | Q99618 | 199;209 | | NA | 7.6064 | 6.8578 | 7.1054 | 6.9079 | 6.4992 | 5.8515 | 6.5894 | 7.0742 | 6.0820 | 7.7291 | 7.3754 | 7.8100 | 3.8234 | 5.1414 | 3.2526 | 4.2941 |
| CDCA3_T37 | CDCA3 | Q99618 | 29;37;44 | | NA | 7.6070 | 6.7248 | 6.9310 | 6.9071 | 6.3171 | 6.2315 | 6.7015 | 7.3855 | 6.6024 | 7.0972 | 7.4088 | 8.2050 | 3.2288 | 5.0628 | 2.9717 | 4.6179 |
| CDCA3_T37;S44 | CDCA3 | Q99618 | 29;37 | | NA | 7.7292 | 6.7296 | 6.3991 | 6.6863 | 7.0586 | 6.3071 | 6.1179 | 6.7452 | 5.8894 | 7.5574 | 7.2583 | 7.5231 | 3.8871 | 5.3378 | 3.8280 | 4.9458 |
| CDCA3_T44 | CDCA3 | Q99618 | 37;44 | | NA | 6.7869 | 6.3237 | 6.6513 | 6.8972 | 6.0327 | 5.2808 | 7.1669 | 7.7909 | 6.6871 | 8.5187 | 7.1384 | 7.3132 | 4.1694 | 4.4702 | 3.4357 | 5.3371 |
| CDCA5_S209 | CDCA5 | Q96FF9 | | 209 | MTLPGISPPPEKQ | 6.9392 | 6.8176 | 7.2844 | 7.5224 | 6.7836 | 7.0118 | 6.8542 | 6.3314 | 7.8519 | 7.5234 | 7.7286 | 7.9452 | 3.1759 | 3.7477 | 2.9432 | 3.5394 |
| CDCA5_S33 | CDCA5 | Q96FF9 | | 75 | EVPAVQSPRRSPR | 7.3832 | 7.5846 | 6.8453 | 7.3906 | 6.0591 | 6.9683 | 6.9665 | 6.6903 | 7.8697 | 7.1806 | 7.0413 | 6.9388 | 3.3432 | 4.4157 | 3.3429 | 3.9799 |
| CDCA5_S75 | CDCA5 | Q96FF9 | | 83 | RRSPRISFFLEKE | 7.3426 | 6.9876 | 6.5840 | 7.2546 | 5.2618 | 6.1353 | 6.9100 | 5.9368 | 7.0802 | 6.1990 | 7.1911 | 7.5461 | 4.5168 | 5.4331 | 4.2734 | 5.3474 |
| CDCA5_S83 | CDCA5 | Q96FF9 | | 111 | THSVPATPTSTPV | 7.2479 | 7.0858 | 6.1870 | 7.9320 | 6.4334 | 5.4663 | 7.1892 | 5.8733 | 6.7250 | 5.4713 | 5.9576 | 5.5787 | 5.5406 | 5.7187 | 5.7865 | 5.8066 |
| CDCA5_T111 | CDCA5 | Q96FF9 | | 33 | RRSQRKSGSELPS | 6.4898 | 7.3319 | 7.1824 | 7.2619 | 5.8519 | 7.4009 | 6.1392 | 6.6129 | 7.3346 | 7.6188 | 7.3314 | 7.5440 | 3.2728 | 4.4818 | 3.6333 | 4.5123 |
| CDCA5_T159 | CDCA5 | Q96FF9 | | 159 | GSASTSTPGRRSC | 7.3462 | 7.2455 | 7.0904 | 7.7359 | 5.6988 | 6.3556 | 7.2314 | 7.4425 | 8.2088 | 7.0913 | 7.2568 | 8.1361 | 2.7307 | 3.6459 | 2.7500 | 4.0339 |
| CDCA7_S190 | CDCA7 | Q9BWT1 | | 190 | RSRILGSLDALPM | 6.4775 | 6.7429 | 7.0293 | 7.7645 | 6.3311 | 6.5585 | 6.8976 | 6.4599 | 7.0531 | 7.2594 | 7.1509 | 7.9106 | 3.7931 | 4.4729 | 3.7346 | 4.3639 |
| CDCA7_S232 | CDCA7 | Q9BWT1 | | 232 | SRRSRSSVTLPHI | 7.1086 | 6.6596 | 5.5497 | 6.7887 | 6.1567 | 6.3305 | 6.5293 | 6.5003 | 6.1317 | 6.9282 | 7.0625 | 6.0184 | 5.2363 | 5.7907 | 4.6820 | 6.5267 |
| CDCA7L_S139 | CDCA7L | Q96GN5 | | 139 | SRSRRSSIGLRVA | 6.1214 | 6.0903 | 6.3241 | 6.4289 | 6.7290 | 6.1467 | 6.0621 | 6.5790 | 7.1946 | 7.0000 | 7.4225 | 7.2293 | 5.4398 | 5.7897 | 4.5477 | 4.8949 |
| CDCA7L_S37;S40 | CDCA7L | Q96GN5 | | 37 | DDVPMETLSSEES | 6.9417 | 6.7615 | 6.3717 | 7.5218 | 5.7221 | 5.3351 | 6.3140 | 6.1475 | 8.1320 | 8.2043 | 7.9664 | 8.1906 | 4.2591 | 4.5839 | 3.4411 | 4.1072 |
| CDCA7L_T37 | CDCA7L | Q96GN5 | 77;81 | | NA | 6.1194 | 5.7981 | 6.8318 | 7.1952 | 6.9513 | 6.0698 | 6.2826 | 6.5499 | 6.1659 | 7.7645 | 6.2137 | 6.4473 | 6.1335 | 6.1922 | 5.6989 | 5.5859 |
| CDCA7L_T77;T81 | CDCA7L | Q96GN5 | 37;40 | | NA | 7.4362 | 7.7365 | 6.7210 | 7.7776 | 7.9995 | 6.5488 | 7.7567 | 6.1932 | 6.4300 | 5.5688 | 6.5059 | 7.2942 | 3.2121 | 5.1795 | 4.0752 | 3.5646 |
| CDCA8_S219 | CDCA8 | Q53HL2 | | 204 | KTPGLRTPAAGER | 7.5836 | 8.8679 | 7.0390 | 8.4637 | 6.4814 | 6.9155 | 7.3597 | 7.0282 | 6.9047 | 6.7245 | 6.9728 | 6.9111 | 2.4980 | 4.1011 | 2.6546 | 3.4941 |
| CDCA8_T106 | CDCA8 | Q53HL2 | | 106 | TAEAIQTPLKSAK | 10.5130 | 7.9608 | 7.8572 | 7.4703 | 5.5536 | 5.8288 | 6.8642 | 5.3345 | 6.9777 | 5.8414 | 6.5461 | 6.0828 | 4.1805 | 4.8292 | 3.7114 | 4.4486 |
| CDCA8_T204 | CDCA8 | Q53HL2 | | 219 | NISGNGSPLADSK | 6.9555 | 7.4390 | 7.0948 | 7.7882 | 6.0230 | 6.4138 | 7.4604 | 6.6934 | 7.1148 | 6.9368 | 7.3390 | 6.8544 | 3.3364 | 4.3311 | 3.5601 | 4.6592 |
| CDCP1_S797 | CDCP1 | Q9H5V8 | | 797 | EEPPPRSPPESES | 6.1502 | 6.6624 | 6.3124 | 6.9699 | 6.7590 | 5.8935 | 6.7717 | 6.7689 | 6.5023 | 5.8499 | 6.6061 | 6.3025 | 5.2355 | 5.4845 | 5.7649 | 5.9664 |
| CDK1_T161 | CDK1 | P06493 | | 161 | IPIRVYTHEVVTL | 6.9384 | 6.7264 | 6.7847 | 6.2820 | 6.7490 | 6.2228 | 6.1063 | 6.5980 | 6.5311 | 6.2705 | 6.4495 | 6.2194 | 5.5951 | 5.7182 | 5.5149 | 5.2938 |
| CDK10_T196 | CDK10 | Q15131 | | 196 | VPVKPMTPKVVTL | 5.9904 | 5.8256 | 6.1199 | 6.3914 | 6.2827 | 6.0627 | 6.2352 | 5.9843 | 6.5018 | 6.3157 | 6.4235 | 6.4709 | 6.2367 | 6.5758 | 6.3179 | 6.2657 |
| CDK11A_S65 | CDK11A | Q9UQ88 | | 583 | SPLKAYTPVVVTQ | 6.6692 | 6.8179 | 6.5785 | 6.5288 | 6.4269 | 6.2297 | 6.4287 | 6.1953 | 6.5475 | 6.5225 | 6.5235 | 6.4674 | 5.5123 | 5.6456 | 5.3825 | 5.5236 |
| CDK11A_T583 | CDK11A | Q9UQ88 | | 65 | EITIRNSPYRRED | 5.6584 | 6.5326 | 6.4488 | 6.3577 | 6.4949 | 6.5833 | 6.0416 | 5.7456 | 5.9559 | 6.3373 | 5.8769 | 5.8145 | 6.9927 | 6.7743 | 6.2682 | 6.1175 |
| CDK11B_S115 | CDK11B | P21127 | | 589 | LAREYGSPLKAYT | 6.2648 | 6.4667 | 5.8033 | 6.1421 | 6.3173 | 5.8995 | 6.2303 | 6.0701 | 6.3080 | 5.9940 | 5.9181 | 5.8670 | 6.5688 | 6.6697 | 6.8366 | 6.6435 |
| CDK11B_S283 | CDK11B | P21127 | | 595 | SPLKAYTPVVVTL | 6.1429 | 6.4205 | 6.2569 | 6.2735 | 6.1631 | 6.0746 | 6.2199 | 6.0412 | 6.3903 | 6.4456 | 6.0760 | 6.2777 | 6.2017 | 6.5603 | 6.1890 | 6.2668 |
| CDK11B_S47 | CDK11B | P21127 | | 65 | EITIRNSPYRRED | 6.0819 | 6.6024 | 6.1625 | 6.4260 | 6.5188 | 6.2662 | 6.4236 | 5.6837 | 5.9026 | 6.0025 | 5.9691 | 5.6961 | 6.5452 | 6.9939 | 6.3568 | 6.3685 |
| CDK11B_S589 | CDK11B | P21127 | | 752 | RVKRGTSPRPPEG | 5.8998 | 6.2146 | 5.8392 | 6.3595 | 6.1142 | 5.6370 | 5.9114 | 6.2120 | 6.3804 | 6.1590 | 5.9708 | 6.2531 | 6.5531 | 6.9108 | 6.7140 | 6.8711 |
| CDK11B_S65 | CDK11B | P21127 | | 47 | RDSKRDSLEEGEL | 6.1733 | 6.2413 | 6.2295 | 6.2223 | 6.0862 | 6.1050 | 6.1571 | 6.3438 | 6.3519 | 6.2137 | 5.9284 | 6.1647 | 6.1150 | 6.2618 | 6.5642 | 6.8417 |
| CDK11B_S792 | CDK11B | P21127 | | 115 | RRHRSHSAEGGKH | 6.0793 | 6.6134 | 5.9639 | 6.4725 | 5.7043 | 5.5531 | 5.6725 | 5.5821 | 6.8169 | 6.9522 | 6.3555 | 7.1539 | 5.6102 | 6.6019 | 6.4045 | 6.4636 |
| CDK11B_T595 | CDK11B | P21127 | | 751 | QRVKRGTSPRPPE | 5.5779 | 6.0868 | 5.6484 | 5.8678 | 5.9580 | 6.1588 | 5.4119 | 6.3164 | 7.0819 | 6.0262 | 5.8450 | 5.8996 | 6.5991 | 7.1380 | 7.0277 | 7.3566 |
| CDK11B_T751 | CDK11B | P21127 | | 792 | AAGPGFSLKFxxx | 7.2877 | 8.4642 | 8.6748 | 9.4667 | 6.8162 | 6.8940 | 7.0314 | 6.5644 | 4.7242 | 4.4541 | 6.5873 | 6.5507 | 4.3534 | 4.8341 | 3.4333 | 3.8636 |
| CDK11B_T751;S752 | CDK11B | P21127 | | 283 | SDLQDISDSERKT | 5.6169 | 5.5529 | 5.9514 | 6.6275 | 6.1001 | 5.7673 | 5.8725 | 6.8099 | 6.1088 | 6.9348 | 6.8335 | 6.3907 | 5.6441 | 6.6257 | 6.2399 | 6.9238 |
| CDK11B_T752 | CDK11B | P21127 | 751;752 | | NA | 5.7557 | 5.9886 | 5.9803 | 6.3490 | 5.5192 | 5.9012 | 6.2765 | 5.8754 | 6.0729 | 5.8551 | 6.3567 | 6.2444 | 6.9286 | 7.0920 | 6.7972 | 7.0072 |
| CDK12_S1083 | CDK12 | Q9NYV4 | | 1244 | PQGPRRTPTMPQE | 6.1483 | 6.3664 | 6.0254 | 6.7993 | 6.2319 | 5.8323 | 6.0043 | 5.6159 | 6.2671 | 6.0917 | 6.2756 | 6.0850 | 6.5257 | 6.5134 | 6.4068 | 6.8108 |
| CDK12_S228;S231 | CDK12 | Q9NYV4 | | 692 | PEPKAITPPQQPY | 5.9276 | 6.2501 | 6.0128 | 6.6555 | 6.0360 | 5.9163 | 6.1133 | 5.9422 | 6.3162 | 6.1293 | 6.1912 | 6.0746 | 5.9893 | 7.0110 | 6.5240 | 6.9105 |
| CDK12_S249 | CDK12 | Q9NYV4 | | 57 | KDMGLVTPEAASL | 5.6876 | 6.1339 | 5.9817 | 6.3346 | 6.0220 | 6.3675 | 6.3560 | 6.0785 | 6.1555 | 6.1462 | 6.2253 | 6.0131 | 6.1072 | 6.9131 | 6.8494 | 6.6284 |
| CDK12_S265 | CDK12 | Q9NYV4 | | 681 | LPGGDLSPPDSPE | 5.4337 | 5.6549 | 5.7832 | 5.9200 | 5.8338 | 5.6658 | 5.8577 | 5.8588 | 5.7363 | 5.6222 | 5.5848 | 5.5417 | 7.4452 | 8.1263 | 7.9332 | 8.0025 |
| CDK12_S274;S276 | CDK12 | Q9NYV4 | | 1083 | EPVKNSSPAPPQP | 6.0430 | 6.2244 | 6.2341 | 6.4757 | 5.8769 | 5.7641 | 6.0951 | 5.8199 | 6.1903 | 6.2867 | 6.0469 | 5.9095 | 6.3253 | 7.2556 | 6.6629 | 6.7896 |
| CDK12_S301;S303 | CDK12 | Q9NYV4 | | 423 | GKESKGSPVFLPR | 6.3236 | 6.5803 | 5.9088 | 6.4966 | 6.4767 | 5.9128 | 6.3642 | 6.1298 | 6.3661 | 6.1633 | 6.1752 | 6.0093 | 5.6285 | 6.7712 | 6.2933 | 6.4002 |
| CDK12_S303 | CDK12 | Q9NYV4 | | 893 | EESRPYTNKVITL | 6.2246 | 6.8205 | 6.3719 | 7.1343 | 6.5613 | 5.4422 | 5.9240 | 5.8589 | 5.5539 | 5.5402 | 5.0207 | 4.3396 | 7.0105 | 7.6576 | 8.3713 | 6.1684 |
| CDK12_S323;S325 | CDK12 | Q9NYV4 | | 303 | RRQRSVSPYSRRR | 6.7754 | 6.8879 | 6.1789 | 6.7656 | 6.2688 | 5.8349 | 6.0960 | 6.0752 | 6.5631 | 6.2916 | 6.1356 | 6.5826 | 5.6071 | 6.3309 | 5.7636 | 5.8427 |
| CDK12_S332;S333;S334 | CDK12 | Q9NYV4-2 | | 1244 | PQGPRRTPTMPQE | 5.9966 | 6.2772 | 5.9653 | 6.5081 | 5.9593 | 6.0868 | 6.0543 | 5.7389 | 5.9046 | 6.1939 | 6.1701 | 6.1678 | 6.1987 | 7.2264 | 6.8059 | 6.7463 |
| CDK12_S332;S334 | CDK12 | Q9NYV4 | | 685 | DLSPPDSPEPKAI | 6.0562 | 5.9117 | 6.4986 | 6.4408 | 6.0871 | 6.3292 | 6.1499 | 6.5484 | 5.8420 | 6.3571 | 5.8290 | 5.3388 | 6.3850 | 6.7726 | 6.4508 | 7.0028 |
| CDK12_S333 | CDK12 | Q9NYV4 | | 644 | GDDDMDSPKETLP | 6.9390 | 6.6769 | 6.5801 | 7.1213 | 5.6458 | 4.7114 | 5.9135 | 6.9447 | 6.1129 | 6.9105 | 4.6260 | 5.1786 | 6.1357 | 8.2550 | 4.7570 | 7.4916 |
| CDK12_S333;S334 | CDK12 | Q9NYV4 | | 265 | YDSYKKSPGSTSR | 5.8776 | 6.7936 | 6.3616 | 6.9282 | 5.8483 | 5.2505 | 5.7761 | 6.0981 | 5.9862 | 5.8906 | 6.4969 | 6.6641 | 5.9628 | 6.7018 | 6.4618 | 6.9016 |
| CDK12_S334 | CDK12 | Q9NYV4 | | 334 | GRRRSSSPFLSKR | 6.9468 | 6.4384 | 5.8555 | 6.3129 | 5.8258 | 5.2974 | 5.6923 | 5.1385 | 6.1265 | 5.6940 | 5.3891 | 5.1991 | 6.9922 | 7.9386 | 7.8132 | 7.3396 |
| CDK12_S341;S343;S345 | CDK12 | Q9NYV4 | | 249 | GQDYDLSPSRSHT | 3.8710 | 6.3510 | 8.6534 | 6.3747 | 5.6466 | 5.1672 | 5.5579 | 4.9364 | 7.3631 | 5.9426 | 8.6830 | 6.9133 | 6.0550 | 5.8183 | 5.0843 | 7.5821 |
| CDK12_S343;S345 | CDK12 | Q9NYV4 | | 333 | YGRRRSSSPFLSK | 6.2219 | 6.8231 | 5.9245 | 7.1242 | 4.8709 | 5.0270 | 5.8674 | 5.1821 | 6.1878 | 5.9663 | 6.1098 | 6.5450 | 6.2200 | 7.5579 | 7.1768 | 7.1954 |
| CDK12_S382;S385 | CDK12 | Q9NYV4 | 383;385 | | NA | 5.9536 | 6.0724 | 5.5486 | 6.3691 | 5.7723 | 5.3628 | 5.8145 | 5.9927 | 6.4128 | 6.2267 | 6.0483 | 6.1742 | 6.6286 | 7.1220 | 6.9962 | 7.5052 |
| CDK12_S383;S385 | CDK12 | Q9NYV4 | 301;303 | | NA | 6.2857 | 6.4688 | 6.0771 | 6.1645 | 6.0778 | 5.5963 | 6.0630 | 5.9337 | 6.0772 | 6.0970 | 5.7743 | 5.4815 | 6.3820 | 7.5855 | 6.8910 | 7.0447 |
| CDK12_S423 | CDK12 | Q9NYV4 | 274;276 | | NA | 5.7610 | 6.0469 | 6.0686 | 6.3139 | 6.0438 | 5.7020 | 5.9558 | 5.6718 | 6.2221 | 6.0141 | 6.0022 | 5.9487 | 6.4134 | 7.2758 | 7.1556 | 7.4042 |
| CDK12_S644 | CDK12 | Q9NYV4 | 681;685 | | NA | 5.9878 | 6.1470 | 6.1647 | 6.4835 | 5.7318 | 5.7986 | 6.1038 | 6.0077 | 6.4971 | 6.2759 | 6.1274 | 6.2132 | 5.9744 | 7.1704 | 6.7681 | 6.5487 |
| CDK12_S681 | CDK12 | Q9NYV4 | 382;385 | | NA | 5.5431 | 5.8866 | 5.5668 | 6.5813 | 6.0253 | 5.4097 | 6.3465 | 5.7116 | 6.0850 | 6.0160 | 6.3425 | 6.2137 | 6.5936 | 7.5424 | 7.0300 | 7.1060 |
| CDK12_S681;S685 | CDK12 | Q9NYV4 | 343;345 | | NA | 6.8735 | 7.4185 | 6.7493 | 7.0446 | 5.4147 | 5.4203 | 5.3955 | 5.4550 | 6.2241 | 6.0814 | 6.5343 | 6.5685 | 5.2090 | 6.2517 | 6.3647 | 6.9950 |
| CDK12_S685 | CDK12 | Q9NYV4 | 228;231 | | NA | 5.7924 | 5.7839 | 5.7410 | 6.5117 | 4.8558 | 4.8419 | 5.7128 | 4.7678 | 5.5638 | 6.0325 | 6.0490 | 6.3249 | 7.0026 | 7.7346 | 7.6833 | 9.6018 |
| CDK12_T1244 | CDK12 | Q9NYV4 | 332;334 | | NA | 6.1579 | 6.8261 | 6.3064 | 6.7963 | 5.6363 | 5.4946 | 5.4967 | 5.8965 | 6.3686 | 6.6104 | 6.0883 | 6.7271 | 5.9255 | 6.8085 | 6.2212 | 6.6397 |
| CDK12_T1244 | CDK12 | Q9NYV4 | 341;343;345 | | NA | 6.0822 | 6.4901 | 5.2467 | 6.2806 | 5.7869 | 5.2734 | 5.7794 | 6.1571 | 6.2686 | 6.0589 | 5.9904 | 5.9442 | 6.3557 | 7.4757 | 7.3853 | 7.4249 |
| CDK12_T57 | CDK12 | Q9NYV4 | 323;325 | | NA | 5.7320 | 6.3656 | 6.3299 | 7.1221 | 5.5980 | 5.2352 | 5.5109 | 5.7877 | 6.6604 | 6.2627 | 5.7595 | 6.0679 | 5.8558 | 7.4714 | 6.8780 | 7.3628 |
| CDK12_T692 | CDK12 | Q9NYV4 | 332;333;334 | | NA | 5.7020 | 6.1012 | 5.5621 | 6.5341 | 6.2370 | 5.7692 | 6.0072 | 6.0050 | 6.2838 | 6.3022 | 6.1790 | 5.9771 | 5.9953 | 7.0361 | 6.9583 | 7.3503 |
| CDK12_T893 | CDK12 | Q9NYV4 | 333;334 | | NA | 6.0983 | 6.3082 | 5.5561 | 6.9645 | 6.3322 | 6.2465 | 7.1023 | 6.1487 | 6.0095 | 5.7363 | 6.3096 | 6.1245 | 5.1481 | 6.4500 | 6.3713 | 7.0938 |
| CDK13_S1054 | CDK13 | Q14004 | | 383 | ERGGDVSPSPYSS | 6.1468 | 6.4610 | 5.8607 | 6.7769 | 5.8755 | 5.5622 | 5.9267 | 6.0860 | 6.3564 | 5.9790 | 5.9474 | 6.0744 | 6.4092 | 7.2294 | 6.8232 | 6.4851 |
| CDK13_S1054;T1058 | CDK13 | Q14004 | | 1246 | MRILELTPEPDRP | 6.0950 | 6.5026 | 5.9710 | 6.5753 | 5.9516 | 5.7117 | 5.9335 | 5.9775 | 6.1079 | 6.1025 | 6.2114 | 6.1361 | 6.2460 | 7.0450 | 6.8992 | 6.5337 |
| CDK13_S1499 | CDK13 | Q14004 | | 494 | KASNTSTPTKGNT | 5.4224 | 5.8369 | 6.3648 | 6.6864 | 6.6502 | 6.2984 | 6.4703 | 6.2589 | 6.4272 | 6.6314 | 6.4449 | 6.3925 | 5.4288 | 6.5864 | 6.1304 | 5.9701 |
| CDK13_S315;S317 | CDK13 | Q14004 | | 1054 | SLGLDDSRTNTPQ | 5.9597 | 6.2351 | 5.7789 | 6.6148 | 6.0198 | 5.5183 | 5.4122 | 5.0849 | 5.7844 | 5.7008 | 5.7977 | 5.7240 | 7.4144 | 8.0971 | 7.6192 | 7.2387 |
| CDK13_S358;S360 | CDK13 | Q14004 | | 525 | KIEHAPSPSSGGT | 6.1188 | 6.5105 | 6.2923 | 7.2771 | 6.0720 | 6.2723 | 5.9910 | 6.4366 | 6.7707 | 6.1385 | 6.7390 | 6.5193 | 5.3395 | 6.0534 | 5.3824 | 6.0867 |
| CDK13_S383 | CDK13 | Q14004 | | 871 | EESRPYTNKVITL | 6.7307 | 7.1143 | 6.7893 | 7.0090 | 6.8252 | 5.3254 | 5.9791 | 6.1223 | 5.5433 | 5.1919 | 4.9922 | 3.8525 | 7.0708 | 7.3995 | 8.3059 | 5.7488 |
| CDK13_S400 | CDK13 | Q14004 | | 400 | RSRSPYSPVLRRS | 6.0489 | 7.0663 | 6.2607 | 6.6310 | 7.4065 | 5.4619 | 6.1590 | 5.8204 | 6.2441 | 4.7814 | 6.1173 | 6.4108 | 5.8805 | 7.0763 | 6.8945 | 5.7404 |
| CDK13_S436;S439 | CDK13 | Q14004 | | 1499 | GYRGHISTSTGRG | 6.6747 | 5.1628 | 4.8492 | 5.2507 | 7.8555 | 5.7927 | 5.2917 | 5.3649 | 7.7559 | 8.8341 | 7.3867 | 7.1640 | 5.9773 | 6.4051 | 5.4030 | 4.8317 |
| CDK13_S437;T439 | CDK13 | Q14004 | | 1147 | KQTDPSTPQQESS | 5.7162 | 5.8797 | 6.7569 | 7.4017 | 5.9188 | 5.4802 | 6.3685 | 6.0674 | 7.4872 | 6.6677 | 5.8742 | 5.4059 | 6.5655 | 6.8461 | 5.6530 | 5.9113 |
| CDK13_S525 | CDK13 | Q14004 | 437;439 | | NA | 5.8497 | 6.1328 | 5.8879 | 6.5666 | 5.4434 | 5.7092 | 5.5293 | 5.7345 | 6.3022 | 6.0284 | 6.1450 | 6.1495 | 6.6042 | 7.6735 | 7.1990 | 7.0448 |
| CDK13_T1147 | CDK13 | Q14004 | 1054;1058 | | NA | 7.4575 | 7.3985 | 6.6958 | 6.8715 | 5.7229 | 5.9340 | 5.8227 | 5.6780 | 7.2414 | 6.7447 | 6.1400 | 5.9632 | 4.8141 | 6.5224 | 5.4541 | 5.5392 |
| CDK13_T1246 | CDK13 | Q14004 | 315;317 | | NA | 5.9034 | 5.8408 | 5.6921 | 6.2424 | 5.4348 | 5.3517 | 5.9096 | 6.1445 | 5.8989 | 5.7947 | 5.9439 | 5.7833 | 6.9264 | 7.5056 | 7.8132 | 7.8147 |
| CDK13_T494 | CDK13 | Q14004 | 436;439 | | NA | 5.7551 | 6.8415 | 6.1577 | 6.0120 | 5.3525 | 6.0740 | 5.9074 | 5.5940 | 6.1667 | 6.2924 | 6.3418 | 6.3915 | 6.5959 | 6.8662 | 6.7015 | 6.9499 |
| CDK13_T871 | CDK13 | Q14004 | 358;360 | | NA | 6.1931 | 6.5079 | 5.7714 | 6.3939 | 6.3340 | 5.4414 | 6.4857 | 5.7025 | 6.1376 | 6.0240 | 5.7237 | 5.4674 | 6.2712 | 7.4189 | 6.8642 | 7.2629 |
| CDK14_S112 | CDK14 | O94921 | | 112 | SSTGKESPKVRRH | 6.1999 | 5.9473 | 7.3907 | 5.9575 | 5.6813 | 6.6071 | 6.2453 | 6.2922 | 6.4315 | 7.0421 | 6.4472 | 6.8675 | 5.6441 | 6.1138 | 5.0771 | 6.0555 |
| CDK14_T76 | CDK14 | O94921 | | 76 | KVRVQRTQSTFDP | 7.0361 | 6.2735 | 7.0066 | 5.7454 | 6.6985 | 5.7632 | 5.6210 | 5.8719 | 6.7714 | 6.4185 | 5.8556 | 6.5933 | 6.6793 | 7.0102 | 5.3516 | 5.3040 |
| CDK16_S139 | CDK16 | Q00536-2 | | 193 | DINKRLSLPADIR | 6.6893 | 5.7495 | 7.0828 | 6.0413 | 5.9005 | 5.9247 | 5.8244 | 6.3999 | 7.2012 | 6.4592 | 6.5249 | 7.0751 | 5.9377 | 6.2893 | 5.2889 | 5.6112 |
| CDK16_S193 | CDK16 | Q00536-2 | | 227 | RRLRRVSLSEIGF | 6.1305 | 5.6529 | 6.7370 | 5.9855 | 6.4104 | 6.6441 | 5.8223 | 6.6924 | 7.1935 | 7.4485 | 7.2301 | 7.5557 | 5.4575 | 5.5769 | 4.6410 | 4.8216 |
| CDK16_S212 | CDK16 | Q00536-2 | | 212 | EKLTLNSPIFDKP | 7.2582 | 6.7760 | 7.1948 | 6.5855 | 5.9970 | 6.8960 | 6.8015 | 7.5397 | 7.1234 | 6.5242 | 6.5086 | 7.2535 | 4.5391 | 4.3435 | 4.1077 | 4.5512 |
| CDK16_S227 | CDK16 | Q00536-2 | | 86 | KIKRQLSMTLRGG | 7.0278 | 6.6564 | 6.7542 | 5.4207 | 5.3741 | 6.5697 | 6.1163 | 6.5421 | 6.9723 | 6.9068 | 7.2012 | 7.2930 | 5.3268 | 5.8863 | 4.4187 | 5.5335 |
| CDK16_S229 | CDK16 | Q00536-2 | | 139 | ARGPLSSAPEIVH | 3.1366 | 3.0826 | 3.5795 | 3.4743 | 3.3577 | 3.7521 | 3.8736 | 4.2712 | 3.4723 | 3.3554 | 3.2182 | 3.3682 | 14.8805 | 13.2549 | 15.1858 | 14.7371 |
| CDK16_S86 | CDK16 | Q00536-2 | | 229 | LRRVSLSEIGFGK | 5.3222 | 4.3032 | 5.0235 | 5.0893 | 5.2550 | 4.6662 | 6.5170 | 6.2269 | 5.0449 | 4.7019 | 4.8020 | 3.9845 | 10.7487 | 8.9680 | 10.2528 | 9.0938 |
| CDK17_S122 | CDK17 | Q00537-2 | | 180 | RRSRRASLSEIGF | 6.0139 | 5.2855 | 5.9821 | 5.3599 | 5.4859 | 6.1552 | 5.7966 | 6.1836 | 7.7840 | 7.6407 | 7.7061 | 7.7468 | 6.1443 | 6.3074 | 4.6644 | 5.7434 |
| CDK17_S137 | CDK17 | Q00537-2 | | 137 | RIHRRISMEDLNK | 5.9203 | 5.4368 | 6.0420 | 5.0992 | 5.8898 | 5.8920 | 5.5258 | 6.1505 | 6.7749 | 7.0054 | 7.0695 | 6.7950 | 6.6872 | 7.0511 | 6.0582 | 6.6023 |
| CDK17_S165 | CDK17 | Q00537-2 | | 9 | KFKRRLSLTLRGS | 7.4767 | 5.9850 | 6.1347 | 5.1975 | 5.4065 | 6.0554 | 6.1701 | 5.8271 | 7.2831 | 6.8033 | 7.1056 | 7.2811 | 6.2153 | 6.6860 | 4.2655 | 6.1071 |
| CDK17_S180 | CDK17 | Q00537-2 | | 122 | SSDEVQSPTGVCL | 5.4313 | 5.7831 | 7.2915 | 6.0013 | 5.5787 | 6.3088 | 5.9698 | 6.8631 | 6.6999 | 5.7428 | 7.4645 | 7.0213 | 5.7664 | 5.6170 | 6.1398 | 6.3206 |
| CDK17_S523 | CDK17 | Q00537-2 | | 165 | EKLQINSPPFDQP | 5.6274 | 5.9844 | 6.5846 | 7.1370 | 5.4439 | 6.8774 | 7.3007 | 6.9641 | 7.1844 | 6.9820 | 7.5643 | 7.3219 | 4.6920 | 5.9784 | 4.4171 | 4.4065 |
| CDK17_S9 | CDK17 | Q00537-2 | | 523 | NHFTCRSxxxxxx | 6.2642 | 6.5881 | 5.6037 | 6.4239 | 6.6848 | 6.5087 | 7.2313 | 6.7158 | 7.0242 | 5.6064 | 7.2003 | 6.2076 | 5.5681 | 5.5060 | 5.2935 | 5.5735 |
| CDK3_T14 | CDK3 | Q00526 | | 14 | EKIGEGTYGVVYK | 6.3262 | 5.7954 | 6.4799 | 5.4641 | 6.6418 | 5.8971 | 5.9938 | 6.2314 | 6.4779 | 6.0775 | 6.1190 | 5.9003 | 7.2232 | 6.1289 | 7.0534 | 6.1899 |
| CDK3_T14;Y15 | CDK3 | Q00526 | | 160 | VPLRTYTHEVVTL | 6.4244 | 5.9397 | 6.5521 | 5.5132 | 6.4981 | 5.7910 | 5.5586 | 6.7375 | 6.5007 | 6.2515 | 6.2127 | 6.4204 | 6.2363 | 6.7387 | 6.3457 | 6.2792 |
| CDK3_T15 | CDK3 | Q00526 | | 15 | KIGEGTYGVVYKA | 5.7613 | 5.5595 | 6.7465 | 4.7183 | 6.2002 | 6.7579 | 6.0432 | 6.6353 | 7.2402 | 7.0494 | 6.8821 | 7.2914 | 5.9295 | 5.6810 | 5.3234 | 6.1806 |
| CDK3_T160 | CDK3 | Q00526 | 14;15 | | NA | 7.4397 | 6.5322 | 7.5426 | 7.0101 | 7.0716 | 5.2253 | 5.5525 | 5.7753 | 6.0982 | 5.0673 | 5.8251 | 5.0671 | 7.0653 | 6.4347 | 7.0577 | 5.2352 |
| CDK5RAP2_S1020 | CDK5RAP2 | Q96SN8 | | 1238 | NKFRDLSPPRYDS | 6.5441 | 6.6491 | 6.6834 | 6.7440 | 6.3304 | 6.5364 | 6.4325 | 6.4588 | 6.4811 | 6.8036 | 6.7193 | 7.0110 | 5.1578 | 5.3127 | 5.0164 | 5.1196 |
| CDK5RAP2_S1074 | CDK5RAP2 | Q96SN8 | | 1074 | NPEDVLSPTSVAT | 6.0569 | 6.7561 | 7.1562 | 7.0256 | 6.3609 | 6.6654 | 7.0091 | 6.6307 | 6.7059 | 6.8241 | 6.9359 | 7.0711 | 4.6723 | 4.8399 | 4.4110 | 4.8790 |
| CDK5RAP2_S1238 | CDK5RAP2 | Q96SN8 | | 841 | HFVQTNSFSKPHD | 6.2775 | 6.9106 | 6.9566 | 7.7029 | 6.0771 | 6.5061 | 7.2226 | 6.4285 | 6.9250 | 6.7730 | 6.8317 | 6.8260 | 4.5792 | 5.2617 | 3.8513 | 4.8704 |
| CDK5RAP2_S1391 | CDK5RAP2 | Q96SN8 | | 1887 | AHPGTCSPSRPGS | 6.6288 | 7.1076 | 6.9361 | 7.0109 | 6.5807 | 6.7592 | 6.0262 | 6.5910 | 7.0438 | 6.5884 | 6.7536 | 6.5163 | 4.8082 | 4.8874 | 4.7385 | 5.0234 |
| CDK5RAP2_S1490 | CDK5RAP2 | Q96SN8 | | 1001 | VMEGRPTPDKTLL | 6.7789 | 7.6869 | 7.0709 | 7.6368 | 6.2395 | 6.7855 | 7.3360 | 6.9504 | 7.6688 | 7.5066 | 7.1743 | 7.2817 | 3.1278 | 4.1341 | 3.0772 | 3.5444 |
| CDK5RAP2_S1666 | CDK5RAP2 | Q96SN8 | | 1020 | GAAYQDSPGEQKG | 6.8063 | 7.0060 | 7.0544 | 7.7512 | 6.6945 | 7.0226 | 7.1991 | 7.2679 | 7.0863 | 7.3587 | 6.8739 | 6.9390 | 3.1734 | 4.6614 | 3.3774 | 3.7280 |
| CDK5RAP2_S1887 | CDK5RAP2 | Q96SN8 | | 1490 | LSRKTVSLEHLQR | 7.1135 | 6.1813 | 6.5861 | 6.7695 | 6.3440 | 6.1385 | 6.3301 | 6.4608 | 5.8982 | 6.2860 | 6.1743 | 6.2138 | 5.9482 | 5.7611 | 5.9675 | 5.8272 |
| CDK5RAP2_S547 | CDK5RAP2 | Q96SN8 | | 547 | SKEKKQSSDYEEL | 6.4368 | 6.8510 | 7.1347 | 7.1406 | 5.3397 | 7.1168 | 6.5201 | 6.1836 | 6.3965 | 7.2642 | 7.3954 | 7.1794 | 4.2040 | 5.0373 | 4.7279 | 5.0720 |
| CDK5RAP2_S841 | CDK5RAP2 | Q96SN8 | | 1666 | PMESDNSFDLFDS | 6.1913 | 6.7712 | 7.1846 | 6.2417 | 6.1919 | 5.6839 | 6.7331 | 7.1096 | 6.6821 | 8.1164 | 7.1330 | 7.3701 | 4.1740 | 5.4910 | 4.5014 | 4.4246 |
| CDK5RAP2_T1001 | CDK5RAP2 | Q96SN8 | | 1391 | VMVNSFSQDLLME | 6.1925 | 7.1125 | 6.4727 | 6.9350 | 5.2305 | 5.6461 | 4.6514 | 7.1779 | 7.4161 | 5.9777 | 4.6887 | 5.7488 | 5.1277 | 6.2345 | 6.4961 | 8.8918 |
| CDK5RAP3_S530 | CDK5RAP3 | Q96JB5-4 | | 530 | VNLMGTSLxxxxx | 6.3784 | 5.9441 | 6.6410 | 5.6368 | 6.0124 | 6.1495 | 6.4343 | 5.4019 | 7.2848 | 6.6133 | 5.6174 | 6.7093 | 6.6064 | 5.8440 | 6.3480 | 6.3784 |
| CDK7_S164 | CDK7 | P50613 | | 170 | SPNRAYTHQVVTR | 6.1408 | 6.9478 | 6.0436 | 6.9786 | 6.3285 | 6.7259 | 5.8804 | 7.2887 | 6.1641 | 6.0107 | 6.0703 | 5.9082 | 5.3131 | 6.3455 | 6.0197 | 5.8343 |
| CDK7_T170 | CDK7 | P50613 | | 164 | LAKSFGSPNRAYT | 6.1141 | 6.4934 | 6.1829 | 6.0210 | 5.8207 | 6.1459 | 6.0532 | 6.3260 | 6.3131 | 6.3394 | 6.2729 | 6.3733 | 5.8667 | 6.6967 | 6.4296 | 6.5510 |
| CDK9_S464 | CDK9 | P50750-2 | | 303 | SQPNRYTNRVVTL | 6.8658 | 7.1275 | 7.1173 | 6.7826 | 6.3346 | 5.8597 | 6.6674 | 5.0407 | 5.7632 | 6.0395 | 6.2895 | 5.7534 | 6.6371 | 6.3906 | 5.8528 | 5.4782 |
| CDK9_T303 | CDK9 | P50750-2 | | 464 | PPRRKGSQITQQS | 4.6918 | 6.0471 | 5.7413 | 5.8347 | 6.4693 | 6.1757 | 5.2512 | 5.9330 | 6.4887 | 5.7389 | 5.9476 | 6.1890 | 6.7399 | 8.0915 | 7.0068 | 7.6535 |
| CDKAL1_T499 | CDKAL1 | Q5VV42 | | 499 | SDAKVYTPSISKP | 7.1211 | 6.8662 | 6.4374 | 7.1886 | 6.4644 | 6.8343 | 7.4199 | 6.6230 | 6.8321 | 6.8205 | 6.8629 | 7.0337 | 4.0782 | 4.8580 | 3.8011 | 4.7587 |
| CDKL5_S306 | CDKL5 | O76039-1 | | 407 | NIPHLLSPKEAKS | 5.9666 | 6.1297 | 6.9372 | 6.8710 | 6.7000 | 5.5311 | 6.1185 | 6.9344 | 6.5688 | 7.0090 | 7.0652 | 6.9851 | 4.7595 | 6.2673 | 4.6753 | 5.4813 |
| CDKL5_S407 | CDKL5 | O76039-1 | | 306 | QRLLDRSPSRSAK | 7.0053 | 6.7144 | 5.5829 | 5.9542 | 7.6954 | 7.2427 | 4.5600 | 6.5510 | 8.0468 | 8.4616 | 7.3640 | 7.6938 | 5.3961 | 4.2976 | 4.0624 | 3.3717 |
| CDKN1B_S10 | CDKN1B | P46527 | | 10 | VRVSNGSPSLERM | 6.2009 | 6.0999 | 6.1776 | 5.1813 | 6.4998 | 6.2710 | 5.8099 | 6.4844 | 5.9263 | 6.3842 | 6.3276 | 6.4844 | 6.7273 | 7.3659 | 6.0445 | 6.0150 |
| CDKN2AIP_S268 | CDKN2AIP | Q9NXV6 | | 346 | SGTSLLTPKSSSS | 5.9657 | 6.5222 | 7.4084 | 7.3242 | 5.6177 | 6.4411 | 6.3300 | 6.5347 | 7.8487 | 6.1956 | 6.7075 | 7.1088 | 4.7220 | 5.4393 | 4.6957 | 5.1385 |
| CDKN2AIP_S382 | CDKN2AIP | Q9NXV6 | | 268 | SRQQSGSPKKSAL | 5.9054 | 6.0362 | 6.2920 | 6.7421 | 5.6952 | 6.2247 | 6.4191 | 5.1964 | 6.2930 | 6.9475 | 6.5747 | 7.2134 | 6.4233 | 6.3285 | 5.5003 | 6.2083 |
| CDKN2AIP_T346 | CDKN2AIP | Q9NXV6 | | 382 | SSQTSGSLVSKST | 6.9503 | 7.8004 | 7.0387 | 8.8181 | 5.3275 | 6.9165 | 8.0696 | 6.1957 | 6.9930 | 7.7745 | 7.2415 | 7.4464 | 2.8516 | 3.8375 | 2.7680 | 3.9708 |
| CDR2_S415 | CDR2 | Q01850 | | 418 | PVSSPTTPPEYKA | 6.2384 | 6.0516 | 6.3823 | 6.3480 | 6.2192 | 6.0681 | 6.2302 | 6.5636 | 6.2343 | 6.5097 | 7.1588 | 6.0866 | 5.9331 | 6.2230 | 6.0117 | 5.7415 |
| CDR2_T418 | CDR2 | Q01850 | | 415 | NPEPVSSPTTPPE | 6.9674 | 6.0325 | 6.4569 | 5.7138 | 6.8587 | 6.9820 | 6.4592 | 6.9215 | 6.7221 | 5.8040 | 6.0156 | 6.6174 | 5.8162 | 5.4574 | 5.5946 | 5.5807 |
| CDR2L_S179 | CDR2L | Q86X02 | | 179 | AFRLHSSSLELGP | 6.6469 | 6.5404 | 6.5064 | 6.2696 | 6.4220 | 7.7922 | 5.9204 | 5.6925 | 7.2817 | 5.9781 | 6.7860 | 6.1779 | 4.9260 | 5.6866 | 5.9337 | 5.4395 |
| CDT1_S318 | CDT1 | Q9H211 | | 93 | DIPACPSPGQKIK | 6.6036 | 6.4542 | 7.0158 | 6.6201 | 6.5220 | 6.8547 | 6.3317 | 6.4187 | 7.7458 | 7.4993 | 7.9773 | 7.6591 | 3.8231 | 4.4240 | 3.8764 | 4.1744 |
| CDT1_S45 | CDT1 | Q9H211 | | 318 | AFLASLSPAMVVP | 5.9552 | 6.3242 | 6.6997 | 6.7177 | 6.1292 | 6.8793 | 6.4353 | 6.8508 | 6.3163 | 6.8262 | 7.1402 | 6.4616 | 4.9569 | 6.0141 | 5.2125 | 5.0808 |
| CDT1_S47 | CDT1 | Q9H211 | | 45 | APASATSGSRKRA | 5.2561 | 7.2356 | 7.1482 | 7.0683 | 6.6860 | 6.2355 | 6.0654 | 6.9085 | 8.2068 | 8.3356 | 7.9802 | 8.2475 | 3.3833 | 3.6857 | 3.6540 | 3.9033 |
| CDT1_S93 | CDT1 | Q9H211 | | 47 | ASATSGSRKRARP | 5.7729 | 6.2888 | 7.1099 | 7.5919 | 5.7632 | 6.2306 | 5.6671 | 6.5077 | 7.2479 | 10.2349 | 9.6362 | 10.9124 | 2.7367 | 3.2266 | 2.2386 | 2.8347 |

| | | | | | | | | | | | | | | | | | | | | | |
|---|---|---|---|---|---|---|---|---|---|---|---|---|---|---|---|---|---|---|---|---|---|
| CDT1_T29;S31 | CDT1 | Q9H211 | 29;31 | | NA | 6.4043 | 6.6777 | 7.3268 | 6.9051 | 6.5845 | 6.9343 | 6.8609 | 6.7409 | 7.3101 | 7.5324 | 8.1188 | 7.5622 | 3.5956 | 4.2164 | 3.3967 | 3.8331 |
| CDV3_S107 | CDV3 | Q9UKY7 | | 107 | QAMQISSEKEEDD | 6.8239 | 6.8069 | 7.9232 | 6.3896 | 6.5764 | 6.6852 | 6.3083 | 6.7946 | 5.6011 | 5.6200 | 5.3616 | 5.8384 | 5.7617 | 5.8753 | 5.6732 | 5.9605 |
| CDV3_S191 | CDV3 | Q9UKY7 | | 182 | LTTTRKTPQGPPE | 6.7511 | 6.7304 | 7.0696 | 7.5150 | 6.1566 | 6.9587 | 7.2040 | 6.9694 | 7.5658 | 7.1141 | 7.4900 | 7.6308 | 3.3905 | 4.2745 | 3.2657 | 3.9138 |
| CDV3_S8 | CDV3 | Q9UKY7 | | 8 | AETEERSLDNFFA | 5.9627 | 6.2458 | 7.2438 | 6.2114 | 5.4714 | 6.6148 | 6.6023 | 7.2552 | 6.5735 | 6.6668 | 7.1279 | 7.3359 | 4.8080 | 5.6228 | 5.2145 | 5.0432 |
| CDV3_T159 | CDV3 | Q9UKY7 | | 190 | QGPPEIYSDTQFP | 5.9905 | 6.0476 | 7.2519 | 6.7792 | 6.5555 | 6.8291 | 5.8473 | 6.7302 | 7.3171 | 7.3788 | 7.1672 | 7.3222 | 4.5947 | 5.2140 | 4.4572 | 4.5175 |
| CDV3_T182 | CDV3 | Q9UKY7 | | 191 | GPPEIYSDTQFPS | 6.9030 | 7.3434 | 7.9314 | 7.0971 | 6.6010 | 7.5247 | 7.7495 | 8.2137 | 5.2982 | 5.1543 | 5.9487 | 5.6020 | 4.7102 | 5.1353 | 4.3053 | 4.4823 |
| CDV3_Y190 | CDV3 | Q9UKY7 | | 159 | PVIVTETPEPAMT | 6.1697 | 5.5630 | 5.9957 | 7.9370 | 6.0411 | 7.2268 | 5.5769 | 5.9969 | 6.6377 | 6.8749 | 6.7134 | 7.1688 | 5.1177 | 5.8418 | 5.6853 | 5.4532 |
| CDYL_S129 | CDYL | Q9Y232 | | 216 | DGFQSESPEKLDP | 6.1458 | 6.3447 | 6.3896 | 6.8835 | 5.9443 | 5.5897 | 6.2742 | 5.7451 | 6.3041 | 6.1623 | 6.3768 | 6.5483 | 6.2068 | 6.1356 | 6.3078 | 6.6414 |
| CDYL_S201 | CDYL | Q9Y232 | | 88 | RWKGYDSEDDTWE | 6.1007 | 6.6440 | 6.1852 | 7.1712 | 5.9484 | 6.2943 | 6.2412 | 6.3389 | 6.0518 | 6.1662 | 6.1599 | 5.7079 | 6.0726 | 6.3172 | 6.1668 | 6.4336 |
| CDYL_S216 | CDYL | Q9Y232 | | 129 | TRTNRTSPNNARK | 6.7026 | 7.0292 | 6.9874 | 6.9111 | 6.3215 | 6.4355 | 6.0619 | 5.4307 | 6.1934 | 6.3938 | 6.6328 | 6.4624 | 6.0410 | 5.3673 | 5.2577 | 5.7718 |
| CDYL_S88 | CDYL | Q9Y232 | | 201 | KILVPKSPVKSRT | 5.6663 | 6.2153 | 8.0957 | 6.3003 | 4.7014 | 9.1490 | 4.6302 | 7.5948 | 10.0196 | 5.1804 | 5.1218 | 6.4761 | 4.6612 | 5.6686 | 5.1207 | 5.3986 |
| CEBPZ_S34 | CEBPZ | Q03701 | | 629 | LDDHPESDDEENF | 5.9967 | 6.2498 | 5.3823 | 6.9048 | 6.1956 | 5.9961 | 6.2964 | 5.5743 | 6.1464 | 5.7647 | 6.1557 | 5.7489 | 6.4821 | 6.9856 | 6.7728 | 7.3477 |
| CEBPZ_S629 | CEBPZ | Q03701 | | 835 | RDADEESIEDVDD | 5.8351 | 5.9173 | 6.1076 | 6.7437 | 6.1369 | 5.8793 | 5.9557 | 5.5681 | 6.3699 | 6.2428 | 6.0523 | 6.3160 | 6.5294 | 6.8435 | 6.7124 | 6.7899 |
| CEBPZ_S835 | CEBPZ | Q03701 | | 34 | EDEDNTSEAENGF | 6.0336 | 6.2812 | 6.3393 | 6.7350 | 6.0072 | 6.0884 | 5.9483 | 5.4829 | 6.2351 | 6.4163 | 6.2645 | 6.1181 | 6.4515 | 6.2015 | 6.1892 | 7.2079 |
| CELF1_S18;S22 | CELF1 | Q92879-4 | | 22 | HCSLNSSPVSKKM | 6.4914 | 7.8049 | 5.8130 | 7.6265 | 7.3343 | 7.5491 | 6.1177 | 7.9715 | 6.3537 | 5.8687 | 6.9689 | 6.2928 | 4.3413 | 4.5575 | 4.6959 | 4.2127 |
| CELF1_S22 | CELF1 | Q92879-4 | 18;22 | | NA | 6.1897 | 7.0776 | 7.5932 | 7.3848 | 6.3572 | 6.8186 | 6.5285 | 6.4669 | 7.3934 | 6.6382 | 7.1205 | 7.1604 | 4.0865 | 4.6140 | 3.8432 | 4.7275 |
| CELSR1_S2871 | CELSR1 | Q9NYQ6 | | 2871 | DQSLAESDSEDPS | 6.3047 | 6.2595 | 6.9205 | 6.2556 | 6.1737 | 5.6795 | 5.7574 | 6.5697 | 6.7014 | 6.3982 | 7.2822 | 6.3974 | 5.5828 | 6.5518 | 5.7458 | 5.4199 |
| CEND1_S10 | CEND1 | Q8N111 | | 10 | RGKSASSPKPDTK | 6.4206 | 5.7616 | 6.0497 | 5.9439 | 6.4833 | 5.7275 | 5.9762 | 6.8498 | 5.6129 | 7.2760 | 6.4426 | 6.8499 | 5.8975 | 6.7457 | 6.1004 | 5.8624 |
| CENPA_S17;S19 | CENPA | P49450 | | 19 | PRRRSPSPTPTPG | 6.0074 | 6.2164 | 5.1326 | 5.5355 | 5.6348 | 5.1605 | 5.9408 | 5.8474 | 6.0145 | 5.3830 | 5.1559 | 5.2421 | 8.3434 | 8.2353 | 7.8443 | 8.3061 |
| CENPA_S17;S27 | CENPA | P49450 | 17;19 | | NA | 6.0886 | 6.2639 | 5.8010 | 6.6013 | 6.1410 | 5.8261 | 6.3308 | 5.9795 | 6.4978 | 5.7991 | 6.0903 | 5.8161 | 6.6383 | 7.0087 | 6.4216 | 6.6958 |
| CENPA_S17;T21 | CENPA | P49450 | 17;21 | | NA | 6.1815 | 6.3832 | 5.8327 | 6.2221 | 5.8118 | 5.7197 | 6.1857 | 6.3480 | 6.4276 | 5.7351 | 6.0279 | 5.7523 | 6.8822 | 6.8714 | 6.5452 | 7.0736 |
| CENPA_S19 | CENPA | P49450 | 17;27 | | NA | 6.1027 | 6.0478 | 5.4107 | 6.4264 | 5.7624 | 5.4570 | 6.2017 | 6.3480 | 6.9858 | 5.8161 | 5.5945 | 5.5745 | 7.1735 | 6.9562 | 6.8244 | 7.3184 |
| CENPB_S156 | CENPB | P07199 | | 156 | TPAAPASPAAVPS | 6.2318 | 6.2573 | 6.3215 | 6.5901 | 6.4222 | 5.8980 | 6.2438 | 6.3952 | 6.3647 | 6.5540 | 6.1352 | 6.1058 | 5.9587 | 6.2490 | 5.8690 | 6.4037 |
| CENPC_S126 | CENPC | Q03188 | | 333 | LKQRTISPAESTA | 5.9203 | 6.4248 | 5.8681 | 6.8077 | 6.3707 | 6.1831 | 6.8648 | 6.4012 | 6.4059 | 6.2948 | 6.3632 | 6.4181 | 5.4671 | 6.2172 | 5.9705 | 6.0226 |
| CENPC_S126;T130 | CENPC | Q03188 | | 734 | LVLPSNTPNVRRT | 8.1219 | 8.1156 | 7.0011 | 7.7307 | 6.6728 | 6.6360 | 7.3669 | 6.4705 | 7.1086 | 6.3143 | 6.6404 | 6.6929 | 3.4318 | 4.2928 | 3.2819 | 4.1217 |
| CENPC_S127 | CENPC | Q03188 | | 90 | KSVPVSSKKKEAS | 8.1751 | 7.9630 | 6.7325 | 7.9616 | 6.4775 | 6.4356 | 7.3688 | 6.4974 | 6.6585 | 6.2443 | 6.3794 | 6.7476 | 3.6361 | 4.6296 | 3.6150 | 4.4782 |
| CENPC_S170 | CENPC | Q03188 | | 773 | VISGVLSPDTISS | 6.6119 | 5.7125 | 6.4707 | 7.0471 | 6.7442 | 5.9359 | 6.4760 | 5.9940 | 6.5701 | 6.5261 | 6.5437 | 6.7837 | 5.4888 | 5.9346 | 5.5689 | 5.5919 |
| CENPC_S177 | CENPC | Q03188 | | 73 | KDTCIQSPSKECQ | 6.6734 | 6.4713 | 6.1599 | 6.6250 | 6.1544 | 5.9702 | 6.2617 | 5.8150 | 6.2293 | 6.1060 | 6.0947 | 6.1393 | 6.2652 | 6.5106 | 6.2263 | 6.2977 |
| CENPC_S189 | CENPC | Q03188 | | 528 | RISRRPSDWWVVK | 7.2308 | 7.0010 | 5.4566 | 7.2489 | 7.3801 | 5.8369 | 6.5505 | 6.2317 | 8.3105 | 7.7971 | 7.1598 | 7.2892 | 4.1249 | 5.2712 | 2.9968 | 4.1139 |
| CENPC_S225 | CENPC | Q03188 | | 316 | QSFASRSWITIPR | 7.0391 | 7.1076 | 6.2660 | 7.3531 | 6.7521 | 5.7223 | 6.5417 | 6.4189 | 6.4193 | 5.9275 | 5.8357 | 6.0252 | 5.3100 | 6.0937 | 5.4284 | 5.7594 |
| CENPC_S225;S232 | CENPC | Q03188 | | 538 | VVKSEESPVYSNS | 6.6238 | 7.1617 | 6.3530 | 6.8207 | 5.8587 | 5.7652 | 6.2294 | 6.3660 | 6.4532 | 6.3725 | 6.1555 | 6.3515 | 5.3640 | 6.1553 | 6.2073 | 5.7621 |
| CENPC_S277 | CENPC | Q03188 | | 177 | QNVIPSSAQKRET | 6.0205 | 6.1379 | 6.0869 | 6.7254 | 6.0092 | 5.9397 | 6.4435 | 6.0080 | 6.6298 | 6.7282 | 6.3435 | 6.5660 | 5.9953 | 5.9757 | 6.1044 | 6.2859 |
| CENPC_S316 | CENPC | Q03188 | | 613 | DEISRCSLSEPLE | 5.9271 | 5.6319 | 6.4320 | 6.5777 | 6.1080 | 5.7483 | 6.3688 | 6.0605 | 6.3369 | 6.4723 | 5.9457 | 6.3629 | 6.4207 | 6.8020 | 6.2199 | 6.5856 |
| CENPC_S333 | CENPC | Q03188 | | 126 | ILATDVSSKNTPD | 7.9425 | 7.3826 | 6.5787 | 7.3377 | 7.1825 | 6.3405 | 7.2638 | 6.9736 | 6.5161 | 6.1817 | 5.5108 | 5.7257 | 4.3891 | 5.1177 | 4.3379 | 5.2191 |
| CENPC_S384 | CENPC | Q03188 | | 62 | TKSVPNSTRKIKD | 8.9144 | 8.1879 | 6.3675 | 8.0798 | 6.8422 | 6.3700 | 7.6113 | 6.1470 | 6.5867 | 5.3471 | 6.1294 | 6.6966 | 3.9961 | 4.7683 | 3.6238 | 4.3321 |
| CENPC_S439 | CENPC | Q03188 | | 127 | LATDVSSKNTPDS | 7.1653 | 7.7128 | 6.7141 | 7.5177 | 6.4198 | 6.4483 | 6.9697 | 6.6793 | 6.3467 | 6.8022 | 5.6733 | 6.8487 | 4.2461 | 4.6765 | 4.2345 | 5.5450 |
| CENPC_S447 | CENPC | Q03188 | | 189 | TYTFENSVNMLPS | 6.4433 | 6.9283 | 6.2182 | 6.7826 | 6.7191 | 5.4075 | 6.2097 | 6.5960 | 6.1195 | 5.8774 | 6.6134 | 6.2205 | 5.2618 | 6.3122 | 6.0184 | 6.2722 |
| CENPC_S528 | CENPC | Q03188 | | 447 | DENIHTSHITQDE | 6.9618 | 6.7704 | 6.9310 | 7.4168 | 7.6007 | 6.4649 | 6.5528 | 6.6929 | 6.6643 | 6.0365 | 6.7883 | 6.4115 | 4.2182 | 5.3303 | 4.3067 | 4.8531 |
| CENPC_S538 | CENPC | Q03188 | | 384 | KTVLDTSYALIGE | 6.4336 | 6.3474 | 6.0634 | 7.9542 | 6.4554 | 5.4828 | 7.1331 | 5.9582 | 5.9210 | 5.9346 | 6.1533 | 6.7548 | 5.5172 | 6.4409 | 5.9990 | 5.4510 |
| CENPC_S55 | CENPC | Q03188 | | 439 | QLDVGQSKDENIH | 6.3311 | 5.6984 | 7.3885 | 7.1545 | 7.2113 | 6.2675 | 6.7153 | 5.6926 | 6.5892 | 7.0341 | 6.2762 | 6.7153 | 5.2274 | 5.7629 | 4.5494 | 5.3863 |
| CENPC_S613 | CENPC | Q03188 | | 225 | EIDNKVSDEEDKT | 6.1666 | 6.0206 | 5.6835 | 6.4744 | 6.5317 | 5.3173 | 6.7327 | 4.8287 | 6.2423 | 5.8899 | 5.9885 | 5.7469 | 7.0234 | 7.0165 | 6.9776 | 7.3594 |
| CENPC_S613;S615 | CENPC | Q03188 | | 96 | SKKKEASLQFVVE | 5.7925 | 6.6067 | 6.2441 | 7.1094 | 6.2441 | 6.3092 | 6.7005 | 6.7351 | 6.4631 | 6.1431 | 6.6080 | 7.1794 | 4.7264 | 6.3518 | 5.3757 | 5.4110 |
| CENPC_S62 | CENPC | Q03188 | | 277 | KRKSESSPIVRHA | 9.4916 | 8.6583 | 7.1627 | 7.7742 | 6.6107 | 5.8461 | 6.7423 | 5.8883 | 7.1592 | 7.2626 | 6.8780 | 6.6863 | 3.1067 | 3.8683 | 3.1044 | 3.7604 |
| CENPC_S73 | CENPC | Q03188 | | 55 | NDFSTNSTKSVPN | 6.0452 | 6.2390 | 5.8399 | 6.4042 | 6.4213 | 6.5549 | 7.8996 | 7.0792 | 6.7667 | 6.5152 | 6.9951 | 6.3253 | 4.6527 | 6.0452 | 4.9409 | 5.2755 |
| CENPC_S763;S773 | CENPC | Q03188 | | 170 | DAKTSVSQNVIPS | 5.3253 | 7.4303 | 6.9283 | 7.2180 | 5.8976 | 6.9522 | 6.2716 | 7.6680 | 7.0014 | 7.5541 | 5.6220 | 7.1196 | 3.9205 | 5.6051 | 4.5097 | 4.9765 |
| CENPC_S773 | CENPC | Q03188 | 763;773 | | NA | 6.6803 | 6.8925 | 5.7967 | 6.8861 | 6.1959 | 6.8449 | 6.9628 | 6.3357 | 5.8540 | 7.0418 | 6.5557 | 6.7143 | 4.6863 | 5.5976 | 4.9316 | 6.0238 |
| CENPC_S90 | CENPC | Q03188 | 613;615 | | NA | 6.8844 | 6.9799 | 6.4188 | 6.1316 | 6.5765 | 6.1619 | 6.4526 | 5.7644 | 6.8972 | 5.9657 | 6.6578 | 6.1529 | 5.6602 | 6.5936 | 5.5453 | 5.1571 |
| CENPC_S96 | CENPC | Q03188 | 225;232 | | NA | 5.9106 | 5.4864 | 5.2329 | 6.5212 | 5.6780 | 5.6093 | 6.5204 | 5.1616 | 6.3919 | 5.3924 | 5.9563 | 6.2203 | 7.5171 | 7.5993 | 7.3508 | 7.4516 |
| CENPC_T734 | CENPC | Q03188 | 126;130 | | NA | 7.1779 | 6.9327 | 6.3119 | 7.5088 | 7.4888 | 6.2893 | 7.3701 | 6.5920 | 6.5384 | 6.0918 | 6.8592 | 6.8742 | 4.1506 | 4.8296 | 3.8011 | 5.1836 |
| CENPE_S2639 | CENPE | Q02224 | | 2654 | RSKSLPSPHPVRY | 7.4587 | 7.8070 | 6.9548 | 7.9543 | 6.5018 | 6.5741 | 6.9864 | 7.0767 | 7.7727 | 7.0737 | 7.5887 | 7.5023 | 2.9014 | 3.8498 | 2.7060 | 3.2916 |
| CENPE_S2654 | CENPE | Q02224 | | 2622 | SKKKQITPSQCKE | 7.8721 | 8.1609 | 7.9122 | 7.8148 | 6.5167 | 7.0826 | 7.2458 | 7.0362 | 8.0812 | 7.3276 | 7.6659 | 7.4888 | 2.0157 | 3.0802 | 2.1379 | 2.5615 |
| CENPE_S611 | CENPE | Q02224 | | 2639 | DPVPKESPKSCFF | 8.1846 | 7.8995 | 6.5685 | 7.6964 | 6.8738 | 7.2901 | 7.7077 | 6.7146 | 7.5841 | 6.6469 | 8.8160 | 7.5105 | 2.4964 | 3.3237 | 2.0753 | 2.6122 |
| CENPE_T2622 | CENPE | Q02224 | | 611 | NIKMDLSYSLESI | 7.0560 | 6.5076 | 7.2526 | 6.6582 | 5.6437 | 6.2652 | 7.1515 | 6.7762 | 5.6637 | 5.9058 | 6.9175 | 7.4678 | 5.0765 | 5.4958 | 4.6713 | 4.5908 |
| CENPF_S1248 | CENPF | P49454 | | 3023 | AQKLALSPLSLGK | 7.1445 | 7.1264 | 6.5354 | 7.0870 | 6.9412 | 7.1447 | 7.4180 | 7.1417 | 6.8956 | 6.9811 | 7.0846 | 6.8268 | 3.7594 | 4.4800 | 3.4712 | 3.9624 |
| CENPF_S1651 | CENPF | P49454 | | 1651 | LQGLDLSSRSLLG | 6.6583 | 6.7969 | 6.3876 | 7.0951 | 6.8044 | 7.0415 | 7.2438 | 7.4510 | 7.1329 | 6.8484 | 7.0482 | 6.9035 | 3.8204 | 4.5519 | 3.7736 | 4.4426 |
| CENPF_S1651;S1654 | CENPF | P49454 | | 154 | IFTTPLTPSQYYS | 7.1077 | 7.0041 | 6.6061 | 7.4555 | 6.2876 | 7.0148 | 7.4930 | 7.2908 | 7.0721 | 6.9514 | 7.3753 | 6.9246 | 3.5972 | 4.1691 | 3.4657 | 4.1850 |
| CENPF_S1654 | CENPF | P49454 | | 2998 | IPTGKTSPYILRR | 7.1701 | 7.0523 | 6.5432 | 6.6344 | 6.9163 | 7.3357 | 7.0682 | 6.8412 | 7.0236 | 6.7032 | 7.2301 | 6.8399 | 4.4059 | 4.5689 | 3.6775 | 3.9896 |
| CENPF_S1674 | CENPF | P49454 | | 1248 | RGDLETSNLQDMQ | 6.5846 | 6.8029 | 6.9293 | 7.1671 | 6.3688 | 6.9359 | 7.1010 | 6.5875 | 7.1484 | 6.3077 | 6.4902 | 6.4861 | 4.8295 | 5.4225 | 3.9071 | 4.9313 |
| CENPF_S1722;S1726 | CENPF | P49454 | | 821 | RLEADQSPKNSAI | 6.8189 | 6.6846 | 6.2443 | 5.8231 | 6.6024 | 6.3429 | 6.7913 | 5.8423 | 6.4025 | 6.2813 | 6.2006 | 5.6895 | 4.4683 | 5.8404 | 6.0178 | 5.9498 |
| CENPF_S1726 | CENPF | P49454 | | 3054 | VKVAQRSPVDSGT | 5.9262 | 6.3436 | 5.8159 | 6.2168 | 6.1071 | 6.2682 | 6.1901 | 6.6825 | 6.1816 | 6.1015 | 6.0257 | 6.0818 | 6.3893 | 6.3962 | 6.6209 | 6.6525 |
| CENPF_S1750 | CENPF | P49454 | | 2911 | PVVPGPSPIPSVT | 6.3559 | 6.4397 | 6.7704 | 7.3218 | 6.3732 | 6.6999 | 6.5359 | 6.4715 | 6.9689 | 6.7048 | 6.2921 | 6.7893 | 5.0273 | 5.3920 | 5.0602 | 4.7969 |
| CENPF_S2417 | CENPF | P49454 | | 1726 | ECSGEQSPDTNYE | 6.0222 | 6.0914 | 5.8149 | 6.2147 | 6.1897 | 5.9597 | 5.9992 | 6.2026 | 5.7337 | 5.7060 | 5.8903 | 5.9998 | 7.2549 | 6.7736 | 7.1844 | 6.9629 |
| CENPF_S2900 | CENPF | P49454 | | 2900 | KQDSRGSPLLGPV | 6.0812 | 5.6834 | 6.2691 | 6.5920 | 6.0628 | 5.8238 | 6.5077 | 6.2655 | 6.2763 | 6.8880 | 6.1942 | 6.3193 | 6.4467 | 6.3247 | 5.6847 | 6.5804 |
| CENPF_S2900;S2911 | CENPF | P49454 | | 144 | SLNPCNTPQKIFT | 7.2995 | 7.2134 | 6.7872 | 6.8415 | 6.5138 | 7.2249 | 6.6675 | 6.7987 | 7.6718 | 6.7858 | 6.7172 | 6.5966 | 4.4071 | 4.8066 | 3.4185 | 4.2497 |
| CENPF_S2911 | CENPF | P49454 | | 2949 | RGPTPATPESFSK | 7.4758 | 7.1854 | 6.5112 | 6.9932 | 6.2257 | 7.0976 | 7.3371 | 7.2376 | 8.1360 | 7.3602 | 7.2739 | 7.1715 | 3.3164 | 3.8241 | 2.8530 | 4.0013 |
| CENPF_S2923 | CENPF | P49454 | | 1541 | NLTRKETPSAPAK | 6.6491 | 6.5816 | 6.3036 | 6.7377 | 7.1577 | 7.5205 | 7.4692 | 7.4128 | 7.4291 | 7.2058 | 7.0429 | 6.6244 | 3.7001 | 4.3946 | 3.6843 | 4.0868 |
| CENPF_S2998 | CENPF | P49454 | | 2417 | LEIINSSFENILQ | 6.5288 | 7.1103 | 6.7726 | 6.9363 | 6.3882 | 6.6946 | 6.6133 | 6.3747 | 7.1623 | 7.0235 | 6.7940 | 6.7712 | 4.4732 | 5.3339 | 4.5746 | 4.4486 |
| CENPF_S3023 | CENPF | P49454 | | 3079 | NNLPERSPTDSPR | 5.9000 | 5.8431 | 6.3961 | 6.3530 | 6.1189 | 6.4396 | 6.7634 | 6.2874 | 6.3702 | 6.5754 | 7.0178 | 6.7983 | 5.2670 | 5.8610 | 5.7843 | 6.2245 |
| CENPF_S3054 | CENPF | P49454 | | 834 | LQNRVDSLEFSLE | 4.0002 | 4.0278 | 4.5927 | 5.3867 | 5.7798 | 5.1548 | 5.6119 | 5.2454 | 5.5241 | 4.4315 | 4.1452 | 5.1748 | 10.3754 | 9.4240 | 10.4870 | 10.6388 |
| CENPF_S3079 | CENPF | P49454 | | 1654 | LDLSSRSLLGIDT | 6.7262 | 6.4249 | 6.4241 | 6.5373 | 5.7310 | 7.2086 | 6.8214 | 6.5326 | 6.5201 | 6.4608 | 6.1814 | 7.0134 | 5.2962 | 5.6818 | 4.3626 | 6.0775 |
| CENPF_S821 | CENPF | P49454 | | 1674 | NESCDISKEHTSE | 7.0862 | 7.1080 | 7.4892 | 6.9735 | 5.9740 | 7.2658 | 7.4912 | 6.1990 | 6.5945 | 7.3028 | 7.6972 | 7.2685 | 3.7846 | 4.1863 | 3.2102 | 4.3691 |
| CENPF_S834 | CENPF | P49454 | | 2923 | TEKRLSSGQNKAS | 6.9036 | 8.2856 | 7.0420 | 7.2874 | 6.1022 | 6.2327 | 5.9610 | 6.5457 | 6.7367 | 6.7712 | 7.1731 | 7.7083 | 3.9328 | 4.7975 | 4.0392 | 4.4808 |
| CENPF_T144 | CENPF | P49454 | | 1750 | ECISELSFSGPNA | 6.0500 | 6.2236 | 5.3031 | 7.5694 | 6.4743 | 6.8246 | 7.1410 | 6.2565 | 7.1688 | 4.8686 | 5.7788 | 6.0068 | 6.2575 | 6.1877 | 6.0695 | 5.8199 |
| CENPF_T154 | CENPF | P49454 | 2900;2911 | | NA | 6.5719 | 7.5982 | 6.7742 | 7.5586 | 6.3182 | 6.8539 | 7.4497 | 7.3776 | 7.3545 | 7.6183 | 7.7578 | 7.1436 | 3.1119 | 4.2766 | 2.7430 | 3.4919 |
| CENPF_T1541 | CENPF | P49454 | 1722;1726 | | NA | 5.1823 | 5.4808 | 5.3557 | 5.7137 | 5.4577 | 5.3040 | 5.4937 | 5.0988 | 5.5775 | 5.7572 | 5.4690 | 5.4412 | 8.9348 | 8.0348 | 9.0833 | 8.6155 |
| CENPF_T2949 | CENPF | P49454 | 1651;1654 | | NA | 6.3997 | 6.3752 | 7.1316 | 7.3476 | 8.1145 | 7.5210 | 7.0190 | 5.9255 | 6.2835 | 6.3546 | 7.1934 | 6.8929 | 4.3180 | 5.0236 | 4.2062 | 3.8936 |
| CENPF_T2949;S2952 | CENPF | P49454 | 2949;2952 | | NA | 7.6211 | 7.9847 | 7.2128 | 7.6993 | 6.4155 | 6.5086 | 7.3025 | 6.7017 | 7.4716 | 7.0835 | 7.1930 | 7.2216 | 2.7417 | 4.2598 | 3.0496 | 3.5330 |
| CENPH_T68 | CENPH | Q9H3R5 | | 68 | DASEEKTPEQIMQ | 6.1893 | 6.2448 | 6.9715 | 6.4220 | 6.0504 | 5.3800 | 5.8990 | 6.1120 | 6.1146 | 6.6329 | 5.9674 | 6.4544 | 5.9648 | 6.3449 | 6.3529 | 6.8990 |
| CENPI_S284 | CENPI | Q92674 | | 284 | QRNRGPSPEPLKL | 6.1876 | 6.3286 | 6.2677 | 6.2309 | 6.3692 | 6.0810 | 6.5318 | 5.9944 | 5.7691 | 6.0121 | 5.7085 | 5.7577 | 6.7994 | 6.7310 | 6.8381 | 6.3928 |
| CENPI_S520 | CENPI | Q92674 | | 520 | MKPVTNSPLETTL | 5.8707 | 6.4888 | 6.8377 | 6.1570 | 5.6127 | 5.7970 | 6.0403 | 5.5353 | 6.6804 | 5.3842 | 6.9065 | 7.9718 | 5.9825 | 6.0452 | 5.9223 | 6.7677 |
| CENPJ_S469 | CENPJ | Q9HC77 | | 759 | DVDLDLSDKDYSS | 5.8597 | 4.9522 | 6.4821 | 5.9440 | 6.7922 | 6.4571 | 6.9359 | 6.3955 | 5.7138 | 6.0968 | 5.6636 | 6.5346 | 7.1433 | 6.0649 | 5.5929 | 7.3713 |
| CENPJ_S759 | CENPJ | Q9HC77 | | 469 | SNRKSLSPSGLKI | 6.3200 | 6.4840 | 6.5853 | 6.3910 | 5.9797 | 6.8254 | 6.3393 | 7.0677 | 6.6577 | 6.3607 | 6.9974 | 7.4946 | 4.6546 | 5.1653 | 5.1281 | 5.5490 |
| CENPL_S53 | CENPL | Q8N0S6-2 | | 43 | QSSFILTPPRRKI | 6.4151 | 6.6197 | 6.0664 | 6.6869 | 6.0092 | 5.7418 | 5.8612 | 5.8346 | 6.2983 | 6.1083 | 6.0069 | 6.0031 | 6.4408 | 6.6917 | 6.6400 | 6.5762 |
| CENPL_T43 | CENPL | Q8N0S6-2 | | 53 | RKIPQCSQLQEDV | 9.2614 | 6.0152 | 8.0933 | 7.5884 | 7.3942 | 6.6561 | 6.3454 | 5.3549 | 6.4786 | 6.4536 | 7.1278 | 6.2566 | 4.0980 | 3.9870 | 4.3838 | 4.5059 |
| CENPN_S282 | CENPN | Q96H22-3 | | 282 | KSGLNGSILAERE | 7.1131 | 7.1533 | 6.0168 | 7.4835 | 7.1058 | 6.3297 | 6.8762 | 6.1904 | 6.2958 | 5.8480 | 5.9703 | 6.0346 | 5.2013 | 5.6790 | 5.6239 | 5.0782 |
| CENPO_S297 | CENPO | Q9BU64 | | 297 | GEKLDMSLVSxxx | 7.3672 | 5.0195 | 6.4295 | 7.2929 | 6.8469 | 5.6281 | 6.2275 | 5.8563 | 6.2975 | 5.9168 | 5.8322 | 5.9006 | 6.0587 | 6.7058 | 6.6478 | 5.9726 |
| CENPP_S38 | CENPP | Q6IPU0 | | 38 | KSRVQKSFQAIHQ | 7.2500 | 7.5284 | 7.0027 | 7.2306 | 6.6360 | 7.6957 | 8.1440 | 6.6032 | 7.3156 | 5.8832 | 6.7966 | 7.5258 | 3.1414 | 3.7890 | 3.5342 | 3.9236 |
| CENPT_S126 | CENPT | Q96BT3 | | 47 | ALLETASPRKLSG | 8.2703 | 7.7428 | 6.4428 | 7.2808 | 5.9670 | 6.0276 | 6.7409 | 6.2274 | 7.1942 | 6.5871 | 6.4681 | 6.5927 | 5.0913 | 4.7128 | 4.0229 | 4.6311 |
| CENPT_S280 | CENPT | Q96BT3 | | 280 | GRKTQSSGPGLQK | 8.5888 | 7.6110 | 6.7022 | 8.0942 | 6.5480 | 5.7788 | 6.3417 | 6.6968 | 6.9269 | 7.5865 | 7.4492 | 7.2298 | 2.9642 | 4.2150 | 3.7188 | 3.5478 |
| CENPT_S47 | CENPT | Q96BT3 | | 126 | PSRQESSCGSLEL | 5.2606 | 6.1866 | 6.9179 | 6.6032 | 5.6812 | 6.3394 | 6.0168 | 7.1857 | 5.7360 | 6.8200 | 6.6816 | 6.3407 | 5.7517 | 6.0873 | 6.4843 | 5.9071 |
| CENPU_S141 | CENPU | Q71F23 | | 232 | AIGSDTSDIVHIW | 6.4276 | 6.9103 | 6.3746 | 7.7078 | 6.6572 | 5.9435 | 6.7525 | 6.3980 | 6.4237 | 6.1120 | 6.0062 | 6.1389 | 5.1943 | 5.6623 | 5.5210 | 5.7702 |
| CENPU_S194 | CENPU | Q71F23 | | 50 | FDFPDNSDVSSIG | 5.4244 | 6.5089 | 6.1867 | 7.1031 | 6.1233 | 5.9899 | 6.3915 | 6.5388 | 6.5581 | 6.8995 | 6.6546 | 6.7798 | 5.2909 | 6.0619 | 5.1428 | 6.3458 |
| CENPU_S229;S232 | CENPU | Q71F23 | | 194 | PLSAQPSVEKENL | 6.0126 | 6.5919 | 6.5156 | 7.3135 | 6.7373 | 7.1556 | 6.9684 | 6.2479 | 6.2657 | 7.0280 | 6.0973 | 7.5759 | 4.3741 | 5.4061 | 4.1597 | 5.5504 |
| CENPU_S232 | CENPU | Q71F23 | | 141 | ISDDSESIEESDT | 6.7799 | 5.9196 | 5.8946 | 6.5884 | 5.8224 | 6.3448 | 5.5707 | 5.2499 | 6.2876 | 7.0155 | 6.6145 | 6.1121 | 5.9818 | 6.8230 | 5.8916 | 7.1038 |
| CENPU_S50 | CENPU | Q71F23 | | 98 | GLSLSSTPPGKEA | 6.3048 | 6.6201 | 5.9704 | 7.0157 | 5.6254 | 5.6798 | 4.5980 | 6.8754 | 5.9224 | 6.0157 | 6.2689 | 5.7990 | 6.6486 | 7.1523 | 7.1555 | 6.3480 |
| CENPU_S50;S54 | CENPU | Q71F23 | 50;54 | | NA | 5.0306 | 5.6845 | 5.7705 | 6.3885 | 6.3170 | 6.1984 | 6.1949 | 6.8451 | 6.1163 | 7.1850 | 6.4069 | 6.8201 | 5.6255 | 6.1723 | 6.1031 | 7.1413 |
| CENPU_T98 | CENPU | Q71F23 | 229;232 | | NA | 6.8166 | 7.2246 | 6.3637 | 7.0708 | 7.6389 | 6.9173 | 6.9679 | 6.7704 | 5.5912 | 5.5073 | 6.0913 | 5.6067 | 4.7103 | 5.8572 | 4.9876 | 5.8782 |
| CENPV_T101 | CENPV | Q7Z7K6 | | 101 | PPPTPATPTSSAS | 6.3452 | 6.2108 | 6.2195 | 6.7433 | 6.4630 | 6.0064 | 6.3056 | 6.6611 | 5.8514 | 6.1941 | 6.1299 | 6.1698 | 5.9107 | 5.8940 | 6.5892 | 6.3061 |
| CEP112_S242 | CEP112 | Q8N8E3 | | 242 | SLRKSSSFHDDHF | 6.2348 | 6.7983 | 6.5247 | 5.8608 | 6.2164 | 6.7594 | 5.9737 | 7.1585 | 5.8746 | 6.4460 | 6.6406 | 6.2699 | 5.5880 | 5.8018 | 6.5925 | 5.2601 |
| CEP126_S412 | CEP126 | Q9P2H0 | | 412 | RPLVTESPTFKFS | 6.6999 | 6.5013 | 6.0115 | 7.0377 | 6.9242 | 6.6097 | 7.0439 | 6.4738 | 7.6045 | 7.0211 | 7.0738 | 7.3778 | 3.7445 | 5.2557 | 4.1455 | 4.4752 |
| CEP128_S1057 | CEP128 | Q6ZU80 | | 977 | SVPEKLSLLEDFK | 6.8271 | 5.8422 | 5.8338 | 6.0851 | 6.5845 | 5.4234 | 6.2633 | 5.7557 | 5.7543 | 5.5569 | 5.2700 | 5.3757 | 7.1169 | 6.8917 | 7.2622 | 8.1574 |
| CEP128_S1057;S1061 | CEP128 | Q6ZU80 | | 18 | RCRDRLSPWAARS | 6.6585 | 7.3313 | 6.8662 | 6.3678 | 6.3654 | 6.2164 | 6.3678 | 6.8039 | 6.8753 | 6.7469 | 6.1238 | 5.9439 | 5.5557 | 5.8906 | 4.3948 | 5.4915 |
| CEP128_S18 | CEP128 | Q6ZU80 | | 1057 | LSSPRFSYVNSFT | 7.7111 | 6.1907 | 8.1425 | 7.7969 | 5.7541 | 5.1719 | 6.5052 | 5.1459 | 5.6969 | 5.8502 | 6.4688 | 6.4610 | 6.3440 | 6.9236 | 4.4234 | 5.4136 |
| CEP128_S977 | CEP128 | Q6ZU80 | 1057;1061 | | NA | 7.8085 | 7.9876 | 7.4352 | 7.2266 | 5.8862 | 6.2789 | 4.8371 | 5.0150 | 6.8896 | 4.6875 | 5.1720 | 7.4905 | 5.5612 | 5.8127 | 6.2104 | 5.7009 |
| CEP131_S114 | CEP131 | Q9UPN4 | | 89 | SQPRSGSPRPTEP | 5.9306 | 5.9753 | 6.8060 | 6.1880 | 6.2072 | 5.9585 | 5.9296 | 6.3603 | 6.2193 | 6.5102 | 6.4343 | 6.7345 | 5.9785 | 6.6508 | 6.2244 | 5.8925 |
| CEP131_S14 | CEP131 | Q9UPN4 | | 381 | QPAQELSPTPGGT | 6.3657 | 6.6407 | 6.8716 | 6.7921 | 6.3556 | 6.4406 | 6.4442 | 7.0124 | 7.0926 | 6.7211 | 7.1988 | 7.0425 | 4.6216 | 5.1685 | 4.4459 | 4.7861 |
| CEP131_S308 | CEP131 | Q9UPN4 | | 78 | NLRRSNSTTQVSQ | 5.7658 | 6.0501 | 6.5387 | 5.5489 | 5.6109 | 6.4022 | 5.9325 | 6.2527 | 6.4822 | 6.3440 | 6.3432 | 6.4478 | 6.5888 | 6.9410 | 6.2900 | 6.4613 |
| CEP131_S381 | CEP131 | Q9UPN4 | | 14 | GSVPERSPAGVDL | 7.2687 | 6.9440 | 6.3667 | 7.8559 | 7.1156 | 6.5048 | 7.1234 | 7.2430 | 6.9198 | 7.8958 | 7.9826 | 7.8858 | 2.7655 | 3.6026 | 3.1079 | 3.4176 |
| CEP131_S499 | CEP131 | Q9UPN4 | | 114 | GKKRPASLSTAPS | 5.6827 | 6.5706 | 6.6289 | 6.3401 | 6.2484 | 6.5854 | 6.1714 | 7.0474 | 6.0366 | 6.6654 | 6.6940 | 6.6412 | 5.4954 | 5.9389 | 5.2271 | 6.0265 |
| CEP131_S78 | CEP131 | Q9UPN4 | | 308 | EEQRQRSGEGTLL | 7.7002 | 7.6478 | 8.0468 | 6.8785 | 6.3000 | 9.1876 | 5.7402 | 7.5580 | 7.3398 | 6.1392 | 8.4059 | 5.9110 | 3.0285 | 3.8900 | 2.6232 | 3.6032 |
| CEP131_S89 | CEP131 | Q9UPN4 | | 499 | TEDDASSLTADNL | 7.4265 | 6.4211 | 6.6351 | 7.0395 | 5.6438 | 6.4825 | 6.0844 | 5.9066 | 7.4045 | 6.2575 | 6.5486 | 7.5555 | 5.0412 | 5.8106 | 5.1278 | 4.6148 |
| CEP131_S89;S105 | CEP131 | Q9UPN4 | 89;105 | | NA | 6.4947 | 6.3526 | 6.7883 | 6.1594 | 6.1936 | 6.6597 | 6.5984 | 6.3026 | 6.6470 | 6.6609 | 6.9438 | 6.8425 | 5.2951 | 5.6674 | 5.0952 | 5.2986 |
| CEP135_S356 | CEP135 | Q66GS9 | | 356 | EIKRKLSEMQDLE | 5.5812 | 5.2447 | 5.0328 | 5.6319 | 6.2168 | 5.4044 | 5.6613 | 5.2508 | 7.4625 | 7.0776 | 6.8112 | 7.0634 | 7.3587 | 7.2203 | 6.9629 | 6.0196 |
| CEP135_T488 | CEP135 | Q66GS9 | | 488 | KSSIFRTPEKGDY | 7.6395 | 6.7523 | 6.7897 | 6.5982 | 6.0306 | 6.9001 | 6.5455 | 6.3661 | 6.8823 | 6.7561 | 6.8385 | 6.6312 | 5.2930 | 4.5383 | 4.7533 | 4.6854 |
| CEP152_S1247 | CEP152 | O94986 | | 1351 | MAKLLETPISSKS | 7.2481 | 6.5975 | 6.3394 | 7.4583 | 7.8387 | 5.7749 | 6.5709 | 5.8588 | 6.3354 | 5.1203 | 5.7333 | 5.5385 | 6.1356 | 5.7661 | 5.8874 | 5.7966 |
| CEP152_S1461 | CEP152 | O94986 | | 1461 | SLPRNVSPEFVPC | 6.1804 | 6.4935 | 6.4904 | 6.7652 | 6.1844 | 6.3625 | 5.8152 | 6.3448 | 6.5043 | 6.0815 | 5.9467 | 6.5473 | 6.2860 | 5.9239 | 5.9801 | 6.0940 |
| CEP152_S1589 | CEP152 | O94986 | | 1618 | SPSGYLSDTEESN | 6.3708 | 6.1043 | 7.0359 | 6.4652 | 6.7544 | 6.6430 | 6.6768 | 6.1325 | 5.7469 | 5.9052 | 5.6177 | 6.3329 | 6.7095 | 6.2118 | 5.4089 | 5.8843 |
| CEP152_S1618 | CEP152 | O94986 | | 1589 | FDSREASFVHGRP | 6.4091 | 6.5866 | 6.3780 | 7.0597 | 6.8672 | 5.5818 | 6.3146 | 5.6420 | 6.3327 | 5.9891 | 5.7109 | 5.4613 | 6.2021 | 6.2654 | 6.3977 | 6.8019 |
| CEP152_S85 | CEP152 | O94986 | | 85 | MLPKSQSVNGYNE | 5.9649 | 5.4519 | 6.4490 | 5.5208 | 5.1917 | 6.4972 | 6.2399 | 6.5350 | 6.4712 | 5.9323 | 6.9042 | 6.5291 | 6.2904 | 7.0043 | 6.4787 | 6.5395 |
| CEP152_T1351 | CEP152 | O94986 | | 1247 | TPPRSLSAGAIEN | 6.3799 | 4.7914 | 4.8301 | 5.3303 | 5.9750 | 5.6553 | 5.6192 | 6.7280 | 7.6358 | 5.6372 | 5.9312 | 4.7037 | 5.8229 | 8.2934 | 8.3578 | 8.3088 |
| CEP164_S1213 | CEP164 | Q9UPV0 | | 1213 | EGTLGGSPTKKAV | 7.3433 | 6.8692 | 7.8940 | 6.3755 | 6.0259 | 6.9831 | 6.4582 | 6.9706 | 6.4813 | 6.4805 | 7.2330 | 7.1067 | 3.2202 | 5.5731 | 4.6124 | 4.3731 |
| CEP164_S285 | CEP164 | Q9UPV0 | | 285 | PTPCKPSSPGADS | 6.8415 | 7.0819 | 7.9823 | 7.8788 | 5.9724 | 5.6247 | 6.4544 | 6.1183 | 7.1839 | 7.3741 | 6.9476 | 7.7797 | 3.5465 | 4.2480 | 4.5443 | 4.4214 |
| CEP170_S1059 | CEP170 | Q5SW79 | | 838 | SLVRQGSFTIEKP | 5.8770 | 5.9851 | 6.7401 | 5.8767 | 5.8926 | 6.1226 | 6.1354 | 6.1026 | 6.7063 | 6.6666 | 6.5686 | 6.8310 | 6.0901 | 6.4303 | 5.8932 | 6.0820 |
| CEP170_S1079 | CEP170 | Q5SW79 | | 1515 | PSAMLPSPPKQKS | 5.7481 | 5.9311 | 6.3130 | 6.2843 | 6.1030 | 6.6324 | 6.2316 | 6.5245 | 6.8330 | 6.6224 | 6.8095 | 6.6927 | 5.4129 | 6.2416 | 5.5543 | 6.0655 |
| CEP170_S1160 | CEP170 | Q5SW79 | | 644 | GERRRRTLPQLPN | 6.8076 | 6.2188 | 6.3167 | 6.3456 | 6.6937 | 6.2328 | 6.5679 | 6.2670 | 7.4571 | 7.0398 | 7.0699 | 6.4032 | 5.7220 | 6.1029 | 4.4232 | 4.3317 |
| CEP170_S1198 | CEP170 | Q5SW79 | | 1529 | PVNNHHSPGQTPT | 5.9988 | 6.6790 | 6.0959 | 6.2198 | 6.0216 | 6.7870 | 6.0214 | 7.4850 | 5.9838 | 6.0920 | 6.5065 | 6.5587 | 5.4696 | 6.0479 | 6.2285 | 5.8044 |
| CEP170_S122 | CEP170 | Q5SW79 | | 1260 | HTRLRTSPALKTT | 7.1300 | 7.2220 | 7.1420 | 6.7986 | 6.5852 | 7.1853 | 7.1535 | 6.9664 | 6.7365 | 6.5219 | 6.7834 | 6.7495 | 4.0918 | 4.7094 | 3.9248 | 4.2997 |
| CEP170_S1251 | CEP170 | Q5SW79 | | 359 | DSKSIKSDVPVYL | 6.5458 | 6.4486 | 6.7663 | 5.8885 | 6.3925 | 6.1288 | 6.2937 | 6.5033 | 6.2447 | 6.1916 | 5.7778 | 5.6396 | 6.1666 | 6.4310 | 6.2133 | 6.3680 |
| CEP170_S1260 | CEP170 | Q5SW79 | | 958 | STQKRKSFTSLYK | 6.4408 | 6.1719 | 6.1641 | 6.1928 | 6.4443 | 6.2255 | 6.1750 | 6.2814 | 6.9812 | 6.9485 | 6.9481 | 7.0985 | 5.7013 | 6.1410 | 5.1252 | 4.9603 |
| CEP170_S1280 | CEP170 | Q5SW79 | | 971 | DRCSTGSPSKDVT | 6.0814 | 6.0047 | 6.6886 | 5.9455 | 6.1817 | 6.3494 | 6.3809 | 6.0855 | 6.9671 | 6.5325 | 6.6428 | 6.7428 | 5.9013 | 5.9078 | 5.7351 | 5.8527 |
| CEP170_S1362 | CEP170 | Q5SW79 | | 1160 | RRTRLGSLSARSD | 6.2420 | 5.5732 | 6.2087 | 5.2909 | 5.4603 | 6.4883 | 6.6903 | 6.4677 | 7.6385 | 6.9620 | 6.8877 | 7.0880 | 6.3265 | 6.7450 | 4.4651 | 5.4659 |
| CEP170_S1515 | CEP170 | Q5SW79 | | 760 | REEAQWTPTKLSS | 6.8416 | 6.8799 | 6.8832 | 7.0260 | 5.9561 | 6.8909 | 7.2908 | 7.1804 | 7.7287 | 7.2043 | 7.4554 | 7.7426 | 3.5157 | 4.1242 | 3.2172 | 4.0629 |
| CEP170_S1522;S1529 | CEP170 | Q5SW79 | | 1059 | GRTPLTSADEHVH | 6.0124 | 6.5189 | 7.1442 | 7.1280 | 7.3178 | 8.3121 | 7.2343 | 8.2253 | 3.1804 | 3.6477 | 3.6265 | 3.6442 | 6.7292 | 6.8220 | 7.2936 | 7.1634 |
| CEP170_S1529 | CEP170 | Q5SW79 | | 1198 | VPSDKFSPRIRAN | 7.6839 | 7.6557 | 7.2040 | 6.9319 | 6.7406 | 7.6662 | 7.5797 | 6.6520 | 7.5721 | 6.8355 | 7.3627 | 6.9281 | 2.8504 | 3.9117 | 2.9672 | 3.4582 |
| CEP170_S181 | CEP170 | Q5SW79 | | 667 | KVVTQRSEIGEKQ | 6.8064 | 6.9542 | 8.2864 | 6.9290 | 6.8557 | 6.8194 | 6.0133 | 5.2217 | 5.5924 | 6.5078 | 6.9334 | 6.4791 | 5.1202 | 6.4455 | 4.7878 | 4.2478 |
| CEP170_S356;S359 | CEP170 | Q5SW79 | | 881 | IRERSESLDPDSS | 6.2160 | 5.8479 | 7.6598 | 6.3607 | 6.4543 | 5.2391 | 5.6933 | 6.7596 | 6.2942 | 6.9689 | 6.9239 | 6.0258 | 5.2080 | 6.6614 | 5.8542 | 5.8331 |
| CEP170_S359 | CEP170 | Q5SW79 | | 1079 | VTKSKTSPVVSGS | 6.1941 | 6.4314 | 6.7416 | 7.0671 | 5.2459 | 6.3369 | 6.4797 | 5.9551 | 6.8663 | 6.9539 | 7.1055 | 7.9327 | 4.8930 | 5.4822 | 4.6494 | 5.6652 |
| CEP170_S446 | CEP170 | Q5SW79 | | 1362 | DRVFDESLNFRKI | 8.0776 | 8.2870 | 7.1140 | 8.0461 | 6.9125 | 7.0983 | 7.8351 | 6.1528 | 6.4654 | 6.0647 | 6.9888 | 6.4197 | 3.7463 | 4.7516 | 2.7151 | 3.3252 |
| CEP170_S667 | CEP170 | Q5SW79 | | 446 | KLLKQKSEEPSVS | 5.5853 | 6.2963 | 7.4540 | 6.4520 | 6.6197 | 6.0053 | 6.3047 | 5.4923 | 6.3052 | 7.0920 | 6.5638 | 6.3193 | 5.8899 | 6.3104 | 6.1422 | 5.1674 |
| CEP170_S838 | CEP170 | Q5SW79 | | 122 | TIQLQLSQKSSES | 6.4219 | 6.7061 | 7.8813 | 6.5161 | 5.5070 | 6.4389 | 6.7578 | 5.7733 | 6.8011 | 7.4311 | 8.1701 | 7.5323 | 4.0391 | 4.6128 | 4.8298 | 4.5811 |
| CEP170_S879 | CEP170 | Q5SW79 | | 181 | PLYGQPSWWGDDE | 8.2502 | 7.3460 | 6.8763 | 7.4481 | 4.4861 | 6.2757 | 6.6180 | 5.6585 | 8.3209 | 6.5833 | 7.4682 | 6.8616 | 4.7136 | 5.3709 | 2.8933 | 4.8291 |
| CEP170_S881 | CEP170 | Q5SW79 | | 682 | ELQEKETPTQVYQ | 6.8411 | 7.8414 | 7.3913 | 7.6405 | 5.8741 | 6.6586 | 7.4563 | 6.1925 | 7.3948 | 7.6611 | 7.4601 | 7.4184 | 2.6027 | 3.9020 | 2.9972 | 4.6677 |
| CEP170_S958 | CEP170 | Q5SW79 | | 879 | SRIRERSESLDPD | 7.1347 | 7.8850 | 9.6972 | 8.3238 | 6.5862 | 7.2488 | 6.3712 | 6.6433 | 5.9280 | 7.6174 | 8.2975 | 8.2448 | 2.0755 | 2.3212 | 2.0974 | 3.5279 |
| CEP170_S971 | CEP170 | Q5SW79 | | 1251 | FGSNRNSPKHTRL | 5.7143 | 6.0637 | 5.6302 | 5.5192 | 6.6165 | 6.0302 | 5.5523 | 5.5419 | 8.7929 | 8.2897 | 8.1508 | 8.2154 | 4.3704 | 5.8021 | 4.5680 | 5.1425 |
| CEP170_T644 | CEP170 | Q5SW79 | | 1280 | AMPTSSSFKHRIK | 5.8852 | 5.5613 | 6.9132 | 6.5531 | 6.6026 | 6.3310 | 6.3277 | 5.8073 | 6.9789 | 6.6711 | 6.7768 | 6.5486 | 5.7271 | 6.8770 | 4.7503 | 5.6887 |
| CEP170_T682 | CEP170 | Q5SW79 | | 762 | EAQWTPTKLSSKN | 7.0675 | 7.2980 | 6.5442 | 6.9180 | 5.9919 | 6.7352 | 7.3045 | 6.8628 | 8.0027 | 7.5112 | 7.3305 | 7.7648 | 3.5917 | 4.2135 | 2.9410 | 3.9224 |
| CEP170_T760 | CEP170 | Q5SW79 | 1522;1529 | | NA | 6.3100 | 6.4927 | 6.4634 | 6.9862 | 6.3030 | 7.0874 | 6.4674 | 6.9834 | 6.4282 | 6.2671 | 6.8272 | 6.6676 | 4.8343 | 5.3776 | 4.9807 | 5.5237 |
| CEP170_T762 | CEP170 | Q5SW79 | 356;359 | | NA | 6.6229 | 6.3380 | 6.6921 | 6.6984 | 7.5454 | 5.8858 | 6.4291 | 5.9440 | 6.8950 | 5.9544 | 6.6553 | 5.3159 | 5.1220 | 6.8460 | 5.4387 | 5.6170 |
| CEP170B_S1114 | CEP170B | Q9Y4F5 | | 492 | PARPFGSVGRRSR | 6.4525 | 6.3217 | 6.3750 | 5.9542 | 6.3960 | 6.4877 | 6.2746 | 6.5862 | 6.7593 | 6.4118 | 6.3778 | 6.3208 | 5.9661 | 6.6949 | 5.0180 | 5.7034 |

| | | | | | | | | | | | | | | | | | | | | | |
|---|---|---|---|---|---|---|---|---|---|---|---|---|---|---|---|---|---|---|---|---|---|
| CEP170B_S1179 | CEP170B | Q9Y4F5 | | 421 | RKKRSQSFTHSPS | 5.7580 | 5.4925 | 6.0265 | 5.9233 | 6.2463 | 6.2645 | 5.8416 | 6.4088 | 6.0112 | 6.1794 | 6.0278 | 5.8651 | 7.0665 | 7.5390 | 6.9155 | 6.4341 |
| CEP170B_S421 | CEP170B | Q9Y4F5 | | 853 | QLRPGRSPEPDGP | 6.0111 | 5.9217 | 6.9341 | 6.4314 | 5.7930 | 7.6885 | 6.3498 | 6.0220 | 6.2267 | 6.5271 | 6.8925 | 7.0298 | 5.8142 | 5.3060 | 5.9593 | 5.0927 |
| CEP170B_S492 | CEP170B | Q9Y4F5 | | 1304 | APGPRDTDDDEEE | 5.8046 | 5.4085 | 6.6171 | 6.7475 | 7.8647 | 6.8575 | 6.7035 | 5.8452 | 5.6878 | 6.2075 | 6.0077 | 6.2650 | 5.4102 | 6.7390 | 5.9654 | 5.8689 |
| CEP170B_S518 | CEP170B | Q9Y4F5 | | 518 | SPAARPSPEKVPP | 7.0917 | 6.4558 | 5.8930 | 6.1161 | 6.9466 | 6.7049 | 6.5041 | 6.9578 | 7.3222 | 7.6941 | 6.4322 | 6.6504 | 4.7675 | 5.0439 | 4.8555 | 4.5642 |
| CEP170B_S605 | CEP170B | Q9Y4F5 | | 1179 | PRKRAGSFTGTSD | 5.6434 | 5.6186 | 5.9932 | 5.9710 | 6.3956 | 6.2401 | 6.1841 | 6.4860 | 7.3301 | 7.2463 | 6.7955 | 6.7844 | 6.0861 | 6.1568 | 5.1446 | 5.9241 |
| CEP170B_S853 | CEP170B | Q9Y4F5 | | 1114 | ALTRSNSLSTPRP | 6.9182 | 5.4876 | 7.2012 | 5.8914 | 5.8731 | 7.8082 | 6.3137 | 6.0024 | 6.8161 | 7.1102 | 6.6357 | 6.7889 | 5.6739 | 6.3133 | 4.6643 | 4.5019 |
| CEP170B_T1304 | CEP170B | Q9Y4F5 | | 605 | ELSRASSATFRPV | 3.1545 | 3.4281 | 3.3580 | 3.4638 | 4.7319 | 4.3290 | 3.9910 | 4.3460 | 3.1272 | 3.3296 | 3.8383 | 3.4354 | 14.8131 | 13.2810 | 14.3612 | 13.0116 |
| CEP170B_T975 | CEP170B | Q9Y4F5 | | 975 | SGPSPTTPQPLRA | 5.8767 | 6.3238 | 6.5931 | 6.8338 | 6.3606 | 6.8554 | 6.3667 | 6.6289 | 6.3228 | 6.6702 | 6.9163 | 6.6863 | 5.3884 | 5.5525 | 5.1661 | 5.4584 |
| CEP192_S1755 | CEP192 | Q8TEP8 | | 2110 | VKGPQGSPLLSRA | 6.7204 | 6.9051 | 6.4197 | 6.5106 | 6.3023 | 6.7474 | 6.5021 | 6.8859 | 6.7056 | 6.5690 | 6.8217 | 7.1018 | 4.7247 | 5.2213 | 4.6155 | 5.2470 |
| CEP192_S2098 | CEP192 | Q8TEP8 | | 916 | VRNPRITSLCLLK | 6.5794 | 6.4840 | 6.2641 | 7.6255 | 6.8771 | 6.1656 | 7.1319 | 6.5650 | 6.6185 | 6.5361 | 6.4151 | 6.4396 | 4.6971 | 5.3713 | 4.6534 | 5.5761 |
| CEP192_S2110 | CEP192 | Q8TEP8 | | 387 | RGGFDLTDPVKQG | 6.4026 | 7.5483 | 6.3404 | 6.9003 | 5.5005 | 5.7701 | 7.9994 | 7.7557 | 6.0138 | 6.9159 | 7.4239 | 8.8911 | 3.2661 | 5.1117 | 3.3853 | 4.7747 |
| CEP192_S444 | CEP192 | Q8TEP8 | | 874 | TNRENNSAVVDVK | 6.2261 | 6.6423 | 7.1363 | 7.4781 | 7.2863 | 6.8573 | 7.1805 | 7.5571 | 6.6888 | 7.9024 | 7.3409 | 6.4563 | 3.6059 | 3.7152 | 2.8852 | 5.0416 |
| CEP192_S51 | CEP192 | Q8TEP8 | | 580 | ASYLRLSLGEFFA | 6.8480 | 5.7255 | 5.8640 | 6.9477 | 5.6271 | 5.5895 | 7.0008 | 5.8899 | 5.7993 | 7.7505 | 7.1251 | 6.8001 | 5.3085 | 6.3690 | 5.2101 | 6.1450 |
| CEP192_S565 | CEP192 | Q8TEP8 | | 2098 | KHGGNVSLDVLPV | 6.5979 | 6.5238 | 6.6042 | 7.3768 | 6.5175 | 7.1372 | 6.4225 | 6.0705 | 7.1316 | 7.2783 | 6.8047 | 7.3991 | 4.1173 | 4.6084 | 4.6349 | 4.7753 |
| CEP192_S580 | CEP192 | Q8TEP8 | | 1755 | SGSKPLSPGPCLD | 5.8095 | 6.2271 | 8.1234 | 5.9243 | 6.4587 | 6.4389 | 7.9492 | 6.9832 | 6.7239 | 5.1088 | 6.0826 | 6.9615 | 5.4433 | 4.7564 | 5.2770 | 5.7323 |
| CEP192_S804 | CEP192 | Q8TEP8 | | 804 | ENFSRASMSDTWD | 5.9239 | 6.5216 | 6.8339 | 6.5613 | 7.1867 | 5.5694 | 5.4400 | 7.0271 | 6.1257 | 6.6234 | 6.6795 | 7.3980 | 6.1542 | 6.3811 | 4.7664 | 4.8078 |
| CEP192_S874 | CEP192 | Q8TEP8 | | 51 | LARDRSSTDNRYP | 5.4293 | 6.0728 | 7.5839 | 6.5412 | 7.5106 | 6.9892 | 5.7877 | 7.8649 | 5.9954 | 6.8304 | 7.5635 | 7.6328 | 4.4070 | 5.4782 | 2.3827 | 5.9303 |
| CEP192_T387 | CEP192 | Q8TEP8 | | 565 | LLRKSRSTSDLDK | 5.3610 | 5.3429 | 5.4465 | 5.6836 | 5.1948 | 6.1868 | 5.8751 | 6.1479 | 6.2329 | 5.6910 | 5.7708 | 6.2009 | 7.9131 | 7.2847 | 7.6804 | 7.9877 |
| CEP192_T916 | CEP192 | Q8TEP8 | | 444 | FTDAIWSPTCERR | 5.8056 | 5.9234 | 6.2837 | 5.7199 | 6.1837 | 6.8488 | 6.4467 | 6.3858 | 6.8365 | 6.4323 | 6.7461 | 6.8940 | 6.2563 | 6.5447 | 5.4281 | 5.2644 |
| CEP250_S2229 | CEP250 | Q9BV73 | | 264 | KEAHERSQELIQL | 5.4310 | 6.5279 | 6.2259 | 6.4254 | 6.3884 | 6.4004 | 7.4743 | 5.8723 | 6.0459 | 5.9397 | 7.1160 | 6.0911 | 5.9222 | 6.0939 | 5.7283 | 6.3173 |
| CEP250_S264 | CEP250 | Q9BV73 | | 2229 | EKERLHSPGATST | 6.3771 | 6.2074 | 6.5969 | 6.2218 | 6.3179 | 6.6748 | 6.3993 | 6.6402 | 6.8887 | 6.2566 | 6.8043 | 6.1790 | 5.3924 | 5.9486 | 5.5544 | 5.5405 |
| CEP250_S5 | CEP250 | Q9BV73 | | 5 | xxMETRSPGLNNM | 6.7457 | 6.5146 | 6.3397 | 6.4666 | 5.7616 | 7.0383 | 6.8769 | 6.4348 | 7.1119 | 7.0617 | 7.8169 | 7.4562 | 4.0666 | 5.2643 | 4.7517 | 4.2924 |
| CEP295_S1890 | CEP295 | Q9C0D2-3 | | 654 | GQRLKLSPNKYQP | 6.2136 | 9.4236 | 6.6536 | 7.2411 | 7.6975 | 7.2787 | 6.3155 | 7.5093 | 6.7364 | 6.1912 | 7.5333 | 5.8557 | 3.2303 | 4.6322 | 4.5276 | 2.9604 |
| CEP295_S654 | CEP295 | Q9C0D2-3 | | 938 | LQDKRLSLSQPIL | 6.7011 | 5.3738 | 5.8207 | 5.4528 | 6.7768 | 5.8557 | 6.2575 | 6.8625 | 9.6245 | 7.0623 | 6.5307 | 7.5070 | 5.7113 | 6.2892 | 3.3218 | 4.8523 |
| CEP295_S914 | CEP295 | Q9C0D2-3 | | 1890 | EQSFFGSPLAHDP | 6.3870 | 6.4301 | 7.1486 | 6.6484 | 6.5177 | 7.0672 | 6.2941 | 6.8799 | 7.6435 | 7.5916 | 7.2153 | 7.7380 | 4.0934 | 4.5227 | 3.9239 | 3.8987 |
| CEP295_S938 | CEP295 | Q9C0D2-3 | | 914 | GRIQESSPTKNNI | 6.7755 | 7.4667 | 6.4241 | 6.7320 | 6.7538 | 6.4580 | 6.7228 | 6.3727 | 7.4922 | 6.4408 | 6.7663 | 6.8493 | 4.6990 | 4.8165 | 4.3948 | 4.8357 |
| CEP350_S1061 | CEP350 | Q5VT06 | | 878 | PWTKAVTPPVKDD | 5.6393 | 5.9060 | 6.0987 | 5.7566 | 5.9381 | 5.8991 | 6.0500 | 6.0087 | 5.8575 | 5.8061 | 5.9072 | 5.8277 | 7.2500 | 7.2058 | 7.6107 | 7.2386 |
| CEP350_S1195 | CEP350 | Q5VT06 | | 1830 | LETRSPSPISISS | 5.8076 | 6.1806 | 6.7453 | 6.3260 | 5.8267 | 5.9064 | 5.9767 | 6.3390 | 6.1706 | 6.8248 | 6.6843 | 7.0814 | 5.6327 | 6.5174 | 6.1679 | 5.8127 |
| CEP350_S1830 | CEP350 | Q5VT06 | | 802 | PQPYVTSPAAYTD | 6.6825 | 6.0202 | 5.9196 | 7.3143 | 6.6386 | 7.3474 | 6.9940 | 6.2565 | 7.2239 | 7.1443 | 6.4272 | 7.1570 | 4.8681 | 5.1617 | 3.9027 | 4.9420 |
| CEP350_S2115 | CEP350 | Q5VT06 | | 939 | SSFRVRSPGPKPE | 6.3929 | 6.5345 | 6.2598 | 6.2277 | 6.1107 | 6.6137 | 6.9789 | 6.6258 | 5.8827 | 6.5780 | 6.5712 | 6.1881 | 5.4946 | 6.2715 | 5.6640 | 5.6060 |
| CEP350_S2362 | CEP350 | Q5VT06 | | 1061 | EELAKGSPHSVIN | 7.0666 | 6.8068 | 6.6997 | 6.0217 | 6.5179 | 7.7586 | 7.7901 | 6.3339 | 7.8996 | 7.0445 | 6.1253 | 7.1820 | 4.2463 | 4.3616 | 3.5508 | 4.5946 |
| CEP350_S2460 | CEP350 | Q5VT06 | | 1195 | TGNVQNSLLDEEK | 6.5307 | 6.3551 | 6.4554 | 6.4160 | 6.0181 | 5.9823 | 5.6884 | 7.4949 | 5.1436 | 6.2905 | 6.2905 | 6.7027 | 6.5916 | 6.1974 | 5.1902 | 6.6526 |
| CEP350_S453 | CEP350 | Q5VT06 | | 2460 | RLLELKSPTELMK | 6.3899 | 6.9159 | 6.3526 | 6.2086 | 5.8792 | 6.0842 | 6.2524 | 6.1431 | 6.0759 | 6.7319 | 6.3162 | 6.1362 | 5.8551 | 6.4445 | 6.2285 | 5.9859 |
| CEP350_S802 | CEP350 | Q5VT06 | | 2115 | SQDLETSPTAKPQ | 6.1953 | 6.5489 | 6.4720 | 7.1581 | 6.0416 | 6.9469 | 6.4720 | 6.5950 | 6.8798 | 7.1111 | 6.8636 | 7.0852 | 5.0902 | 4.9996 | 4.4188 | 5.1218 |
| CEP350_S939 | CEP350 | Q5VT06 | | 2362 | TKEKDLSWSEHLF | 6.2868 | 6.7365 | 7.4334 | 8.6542 | 6.0223 | 6.6426 | 7.1270 | 6.5314 | 7.7942 | 7.8955 | 5.8172 | 7.2629 | 4.3642 | 4.2480 | 3.3707 | 3.8131 |
| CEP350_T1817 | CEP350 | Q5VT06 | | 453 | EQRTASSDRGGRE | 5.9730 | 6.9180 | 5.9337 | 6.6758 | 6.5555 | 5.1533 | 6.2672 | 6.5291 | 6.1700 | 7.6626 | 5.8740 | 6.6306 | 5.9252 | 5.4987 | 6.0378 | 6.1956 |
| CEP350_T878 | CEP350 | Q5VT06 | | 1817 | TKDNKATSPGPTD | 5.9648 | 6.3294 | 5.8441 | 6.5406 | 6.2314 | 6.0795 | 5.8193 | 7.1126 | 6.6499 | 6.8460 | 6.0800 | 6.8637 | 5.6717 | 6.1253 | 6.0396 | 5.8021 |
| CEP41_S119 | CEP41 | Q9BYV8 | | 119 | GNPGEQSPSPEQF | 6.3726 | 6.9920 | 7.2477 | 8.6216 | 6.9164 | 6.0971 | 5.3966 | 7.2225 | 5.4704 | 4.9338 | 6.1871 | 6.4896 | 5.6685 | 7.0334 | 5.1175 | 4.2334 |
| CEP41_S363 | CEP41 | Q9BYV8 | | 363 | SNPRSLSSGHLQG | 5.9468 | 5.2555 | 5.5528 | 5.2096 | 5.2705 | 6.1194 | 5.2140 | 5.6798 | 6.1806 | 6.2267 | 5.9390 | 5.4646 | 7.8447 | 8.5901 | 8.0780 | 7.4278 |
| CEP43_S156;S160 | CEP43 | O95684 | | 254 | DDMEGDSFFDDPI | 6.4695 | 6.4951 | 6.8195 | 7.3452 | 6.3274 | 6.8084 | 7.1481 | 6.5107 | 7.1448 | 6.2256 | 6.8960 | 6.7206 | 4.9947 | 4.8024 | 4.1807 | 5.1112 |
| CEP43_S254 | CEP43 | O95684 | | 170 | KTSAQTTPSKIPR | 6.8567 | 6.0605 | 6.7826 | 6.0382 | 6.5518 | 6.3851 | 6.1251 | 6.7496 | 6.7704 | 6.3720 | 6.9113 | 6.8921 | 5.8918 | 5.0354 | 5.2956 | 5.2817 |
| CEP43_T170 | CEP43 | O95684 | | 55 | EKVENKTPLVNES | 6.4680 | 7.4951 | 6.6285 | 5.4724 | 5.5487 | 6.3294 | 5.7821 | 6.3206 | 6.7153 | 5.6110 | 6.4688 | 6.1829 | 6.7171 | 5.5522 | 5.9171 | 6.7908 |
| CEP43_T55 | CEP43 | O95684 | 156;160 | | NA | 5.7483 | 5.8284 | 6.7638 | 6.2018 | 6.1972 | 6.2250 | 6.0620 | 6.4310 | 6.7095 | 6.7520 | 6.7297 | 7.3245 | 6.0992 | 5.8261 | 5.5244 | 5.5769 |
| CEP55_S40 | CEP55 | Q53EZ4 | | 428 | VAASPKSPTAALN | 5.9401 | 6.7933 | 6.5830 | 6.5991 | 6.4264 | 6.9870 | 6.1418 | 7.0648 | 6.9045 | 7.2017 | 7.5594 | 8.0789 | 4.5087 | 4.3610 | 4.3201 | 4.5303 |
| CEP55_S425;S428 | CEP55 | Q53EZ4 | | 436 | TAALNESLVECPK | 7.1552 | 7.3093 | 7.9656 | 7.7559 | 7.1467 | 6.7177 | 5.8039 | 6.7207 | 7.0791 | 6.2614 | 6.9938 | 6.8899 | 3.9946 | 3.9372 | 4.0327 | 4.2363 |
| CEP55_S428 | CEP55 | Q53EZ4 | | 40 | IAHLKTSVDEITS | 7.2252 | 7.0669 | 7.5561 | 6.2528 | 5.5280 | 6.4497 | 6.9248 | 6.9552 | 6.2589 | 5.3027 | 5.9686 | 6.9715 | 5.5138 | 5.2032 | 4.3445 | 6.4781 |
| CEP55_S436 | CEP55 | Q53EZ4 | 425;428 | | NA | 7.1715 | 6.5565 | 6.5211 | 7.0146 | 6.7173 | 6.9800 | 6.8473 | 6.9902 | 7.9816 | 7.4533 | 8.4995 | 7.9717 | 3.2518 | 4.0118 | 2.8568 | 3.1751 |
| CEP57_S55 | CEP57 | Q86XR8 | | 55 | DLRRSPSKPTLAY | 5.5171 | 5.8279 | 6.3606 | 6.8317 | 6.0287 | 6.0502 | 5.5811 | 6.5947 | 6.3713 | 6.5575 | 7.0897 | 6.6411 | 5.9632 | 6.6689 | 5.7365 | 6.1797 |
| CEP63_S495 | CEP63 | Q96MT8-2 | | 495 | ECSLPVSPLGSIA | 5.9794 | 6.1028 | 7.1519 | 6.4203 | 6.7684 | 5.9480 | 6.6537 | 7.5393 | 6.3040 | 6.4164 | 6.3244 | 6.0769 | 5.5236 | 6.3166 | 4.9767 | 5.4977 |
| CEP68_S249 | CEP68 | Q76N32 | | 53 | AEGGLISPVWGAE | 5.5709 | 5.7093 | 5.6188 | 6.0392 | 5.3924 | 6.2584 | 6.0687 | 6.5612 | 5.7547 | 5.6981 | 6.5501 | 5.7412 | 7.8902 | 6.8399 | 7.5066 | 6.8002 |
| CEP68_S53 | CEP68 | Q76N32 | | 249 | GPRPQWSPQPVFS | 5.9449 | 5.8272 | 6.2686 | 6.6805 | 6.1589 | 6.1418 | 6.2694 | 6.1294 | 6.4638 | 6.0494 | 5.9728 | 6.1773 | 6.9115 | 6.2970 | 6.3348 | 6.3728 |
| CEP72_S237 | CEP72 | Q9P209 | | 237 | ESRHLLSPQLVQY | 6.3749 | 6.1698 | 6.5584 | 5.9231 | 6.5037 | 7.3438 | 6.5541 | 7.3264 | 6.8292 | 6.2544 | 6.7197 | 7.0446 | 5.0981 | 4.9713 | 4.9751 | 5.3533 |
| CEP76_S83 | CEP76 | Q8TAP6 | | 83 | EQELPSSPKQPIC | 6.2925 | 6.6119 | 7.3528 | 5.9267 | 6.1960 | 6.4948 | 6.6815 | 6.7601 | 6.5353 | 6.8474 | 6.7163 | 7.2372 | 4.7873 | 5.4979 | 4.8801 | 5.1821 |
| CEP78_S325 | CEP78 | Q5JTW2-2 | | 325 | EYQWITSPSVKEP | 6.6042 | 6.7897 | 6.6842 | 7.6653 | 6.0976 | 6.8553 | 6.2368 | 6.4116 | 6.7257 | 7.1949 | 6.7283 | 7.2005 | 4.3721 | 5.1139 | 4.1687 | 5.1513 |
| CEP85_S17 | CEP85 | Q6P2H3 | | 17 | GISHVTSPSSDVI | 7.1557 | 7.1298 | 6.8433 | 7.1010 | 6.9278 | 6.9029 | 7.4965 | 7.3721 | 7.2892 | 7.2507 | 7.7804 | 7.3857 | 3.0120 | 3.7848 | 2.9689 | 3.5992 |
| CEP85_S51 | CEP85 | Q6P2H3 | | 51 | RFSRCSSVADSGD | 6.5740 | 5.9783 | 7.3452 | 5.6758 | 6.2511 | 6.0930 | 5.8558 | 6.3762 | 6.7299 | 6.9351 | 6.7926 | 7.1029 | 5.8243 | 5.8365 | 5.1830 | 5.4465 |
| CEP85_S77 | CEP85 | Q6P2H3 | | 77 | FCSSSGSPPFQPI | 6.9196 | 6.0813 | 5.9170 | 6.7158 | 7.1452 | 6.3797 | 6.7198 | 6.6797 | 7.1498 | 6.9179 | 7.3256 | 7.4595 | 4.8429 | 4.1460 | 4.3430 | 5.2574 |
| CEP89_S114 | CEP89 | Q96ST8 | | 114 | EMGRRSSLPSFET | 6.1060 | 6.2644 | 6.7610 | 6.4008 | 6.4633 | 6.6750 | 6.3249 | 6.5701 | 6.7966 | 6.8578 | 6.8959 | 6.9503 | 5.2089 | 5.7592 | 4.8857 | 5.0802 |
| CEP89_S188 | CEP89 | Q96ST8 | | 77 | PRQRSRSESDVSS | 5.4969 | 6.0278 | 7.1750 | 6.6719 | 6.2983 | 6.3608 | 7.0197 | 6.4215 | 6.9475 | 6.8671 | 7.5214 | 7.0902 | 4.6895 | 5.0636 | 4.8692 | 5.4797 |
| CEP89_S221 | CEP89 | Q96ST8 | | 188 | QDGFPGSPPAPQR | 5.3508 | 6.1605 | 7.5872 | 6.6858 | 6.0463 | 7.4575 | 6.3731 | 6.6391 | 6.5138 | 6.6539 | 6.5046 | 7.3205 | 4.5355 | 5.7298 | 4.9815 | 5.4601 |
| CEP89_S77 | CEP89 | Q96ST8 | | 221 | CEKPPPSPDITGR | 6.1068 | 6.1521 | 6.4755 | 6.6113 | 6.3138 | 6.8019 | 6.8238 | 6.6011 | 6.7888 | 6.6464 | 6.9228 | 7.1349 | 5.1186 | 5.5724 | 4.7714 | 5.1583 |
| CEP95_S212 | CEP95 | Q96GE4 | | 212 | SKKALASPSSKSH | 6.7510 | 6.3413 | 6.7167 | 6.5678 | 6.8038 | 7.1814 | 7.2497 | 6.5323 | 6.9704 | 6.3707 | 6.6012 | 6.7160 | 4.3136 | 5.0353 | 4.6948 | 5.1541 |
| CEP95_S449;S451 | CEP95 | Q96GE4 | 449;451 | | NA | 6.0884 | 6.8965 | 6.7009 | 7.1843 | 5.4340 | 5.9338 | 6.4006 | 6.1607 | 6.7395 | 6.4694 | 6.8215 | 7.6429 | 5.3032 | 5.3992 | 5.0805 | 5.7447 |
| CEP97_S308 | CEP97 | Q8IW35-2 | | 796 | NNEQDNSLLEQYL | 6.4615 | 5.9050 | 6.2221 | 6.3156 | 6.2800 | 5.9477 | 6.1710 | 6.4659 | 6.6843 | 6.5251 | 6.9592 | 6.8560 | 5.7996 | 6.4770 | 4.9424 | 5.9877 |
| CEP97_S331 | CEP97 | Q8IW35-2 | | 331 | LIPEHSSPVQDCQ | 5.9579 | 6.6974 | 7.4455 | 6.9113 | 6.9332 | 6.1218 | 6.8225 | 5.9367 | 6.8575 | 6.5072 | 7.3054 | 7.1534 | 4.7612 | 4.6461 | 5.1097 | 4.8332 |
| CEP97_S416 | CEP97 | Q8IW35-2 | | 308 | RQLMNQSQNEELS | 5.8883 | 6.7528 | 7.0727 | 7.9435 | 5.7732 | 6.2936 | 6.2989 | 6.5575 | 6.6504 | 6.4245 | 6.6156 | 7.2869 | 4.6099 | 5.4744 | 4.4537 | 5.9041 |
| CEP97_S789;S796 | CEP97 | Q8IW35-2 | | 416 | SGLSPLSPTVELR | 4.9593 | 6.3800 | 5.8332 | 5.1966 | 5.5294 | 7.3088 | 5.9836 | 6.5652 | 5.6972 | 6.8748 | 6.7475 | 4.7915 | 6.7649 | 7.8962 | 7.2336 | 6.2382 |
| CEP97_S796 | CEP97 | Q8IW35-2 | 789;796 | | NA | 6.6209 | 4.9059 | 5.8288 | 6.5762 | 6.8502 | 5.3794 | 6.0629 | 6.4581 | 8.2284 | 8.6685 | 5.8313 | 6.0078 | 6.5738 | 4.7642 | 5.1467 | 6.0967 |
| CEPT1_T40 | CEPT1 | Q9Y6K0 | | 40 | KLFQLPTPPLSRH | 6.2319 | 6.6359 | 7.2760 | 7.3944 | 5.8553 | 5.8332 | 6.5219 | 5.8261 | 6.0913 | 6.6262 | 5.4910 | 5.9278 | 5.4203 | 6.5201 | 6.2186 | 6.1302 |
| CETN2_S20 | CETN2 | P41208 | | 20 | SQRKRMSPKPELT | 6.6633 | 6.3691 | 6.3059 | 5.9037 | 5.9248 | 6.2741 | 5.5990 | 6.2152 | 6.1000 | 6.5116 | 6.9199 | 7.0398 | 6.3969 | 6.1137 | 5.7173 | 5.9456 |
| CFAP36_S110 | CFAP36 | Q96G28-2 | | 110 | FQEACTSPLAKTH | 6.5543 | 6.5183 | 7.0940 | 6.0376 | 6.7072 | 6.7741 | 7.1999 | 6.2524 | 7.2329 | 6.2025 | 6.4723 | 6.6892 | 4.4725 | 5.3839 | 5.4409 | 4.9682 |
| CFAP97_S488 | CFAP97 | Q9P2B7 | | 488 | STLGQYSPLRASR | 6.4808 | 6.8494 | 6.9516 | 7.1030 | 6.1291 | 6.7307 | 7.1507 | 7.2108 | 7.5961 | 7.4978 | 7.3703 | 7.2806 | 3.5995 | 4.5330 | 3.2780 | 4.2386 |
| CFDP1_S116 | CFDP1 | Q9UEE9 | | 250 | SKLDWESFKEEEG | 5.1748 | 5.9732 | 8.2038 | 5.9692 | 5.4728 | 5.7752 | 5.4954 | 5.7399 | 6.3697 | 8.2149 | 7.7379 | 7.5046 | 4.2942 | 6.9072 | 5.4630 | 5.7042 |
| CFDP1_S250 | CFDP1 | Q9UEE9 | | 116 | EDELWASFLNDVG | 7.3305 | 7.4967 | 8.6867 | 6.6262 | 5.3222 | 7.8535 | 6.7719 | 6.2358 | 6.7233 | 5.7183 | 8.1562 | 6.9905 | 3.6279 | 5.4418 | 3.6951 | 3.3234 |
| CFL1_S156 | CFL1 | P23528 | | 25 | KVRKSSTPEEVKK | 6.2137 | 7.0783 | 6.9249 | 7.2465 | 6.0696 | 7.1893 | 6.6208 | 6.5676 | 6.5624 | 7.0644 | 7.1933 | 6.8001 | 4.2366 | 5.1477 | 4.3916 | 4.6932 |
| CFL1_T25 | CFL1 | P23528 | | 156 | AEKLGGSAVISLE | 6.1971 | 6.0466 | 6.1956 | 6.6210 | 6.6330 | 6.1319 | 5.9222 | 6.2301 | 5.7526 | 5.7050 | 6.0222 | 5.6683 | 7.1762 | 7.2166 | 6.0439 | 6.4376 |
| CGAS_S129;S143 | CGAS | Q8N884 | | 143 | GPWDVPSPGLPVS | 6.3001 | 6.7580 | 6.8787 | 7.5504 | 6.0127 | 6.5051 | 6.9526 | 6.8609 | 6.7938 | 6.7990 | 6.9384 | 7.3708 | 4.3548 | 5.1365 | 4.2363 | 4.5521 |
| CGAS_S143 | CGAS | Q8N884 | | 305 | KRKRGGSPAVTLL | 6.0221 | 6.8592 | 5.8258 | 7.1657 | 6.2122 | 5.9803 | 6.4784 | 6.5333 | 6.9185 | 6.8363 | 6.3729 | 6.2347 | 5.6504 | 5.5813 | 5.7219 | 5.6070 |
| CGAS_S305 | CGAS | Q8N884 | 129;143 | | NA | 7.1723 | 5.9949 | 6.2426 | 9.4675 | 5.1257 | 6.3139 | 9.2221 | 7.2642 | 5.6492 | 7.2215 | 7.9973 | 7.3156 | 4.0707 | 3.1795 | 3.1098 | 4.6534 |
| CGN_S114 | CGN | Q9P2M7 | | 149 | PSNRSNSMLELAP | 6.6960 | 7.0620 | 6.4450 | 6.8173 | 6.1361 | 6.3107 | 6.3444 | 6.1299 | 6.1034 | 6.1067 | 6.4656 | 6.4539 | 5.9593 | 5.6875 | 5.5290 | 5.7533 |
| CGN_S149 | CGN | Q9P2M7 | | 114 | PSQSSTSDEEPGA | 5.5733 | 6.7734 | 6.9706 | 6.7131 | 6.5269 | 6.1655 | 6.1584 | 6.1988 | 6.4867 | 6.6306 | 5.9235 | 6.5799 | 5.8629 | 6.1655 | 5.4969 | 5.7741 |
| CGN_S159 | CGN | Q9P2M7 | | 332 | SVRRKVSLVLEKM | 5.4106 | 5.0885 | 5.4504 | 5.9466 | 5.8822 | 5.6483 | 5.5166 | 5.7555 | 6.9763 | 7.1742 | 6.9910 | 7.7107 | 6.5213 | 6.7806 | 6.5556 | 6.5916 |
| CGN_S252 | CGN | Q9P2M7 | | 252 | AQSQNLSPLSGFS | 6.1115 | 6.3301 | 6.4492 | 6.6482 | 6.3264 | 6.8428 | 6.2428 | 6.8029 | 6.1035 | 6.1636 | 6.3046 | 6.2671 | 5.8124 | 6.2270 | 5.8419 | 5.5261 |
| CGN_S332 | CGN | Q9P2M7 | | 159 | LAPKVASPGSTID | 6.0264 | 6.3666 | 6.5930 | 6.8589 | 6.2989 | 5.8695 | 6.2031 | 5.4787 | 6.9125 | 6.9829 | 7.1398 | 7.1182 | 5.8066 | 5.5144 | 4.9891 | 5.8413 |
| CGN_S90 | CGN | Q9P2M7 | | 90 | ASGALSSDLELPE | 5.6453 | 5.3313 | 6.3431 | 6.5197 | 5.8154 | 6.6047 | 6.1403 | 6.9231 | 5.3967 | 5.9659 | 4.9541 | 3.2773 | 6.9536 | 9.0033 | 8.0831 | 7.0430 |
| CGNL1_S388 | CGNL1 | Q0VF96 | | 388 | DRKRSRSVDSAFP | 5.9613 | 6.6750 | 7.2926 | 6.5034 | 6.2366 | 6.7966 | 5.7775 | 6.6048 | 6.5011 | 7.2732 | 6.5162 | 7.2812 | 5.2118 | 5.4291 | 4.5881 | 5.3513 |
| CHAF1A_S123 | CHAF1A | Q13111 | | 65 | TSVQSKSPDLEAS | 5.7481 | 5.8005 | 5.8314 | 6.4241 | 6.0441 | 5.6958 | 6.1565 | 6.4089 | 5.9228 | 5.6637 | 5.9646 | 5.8806 | 7.1893 | 6.6441 | 7.3322 | 7.2932 |
| CHAF1A_S141 | CHAF1A | Q13111 | | 951 | KATLTASPLGASx | 6.8753 | 6.9851 | 6.2177 | 6.7639 | 6.3540 | 6.1028 | 6.3522 | 6.4462 | 6.3749 | 6.6095 | 6.1414 | 6.4659 | 5.4908 | 5.7273 | 5.4421 | 5.6509 |
| CHAF1A_S206 | CHAF1A | Q13111 | | 775 | HTGSPRSPSTTYL | 6.3444 | 6.8832 | 6.2426 | 6.9673 | 6.1889 | 6.5448 | 6.3589 | 6.3123 | 6.3711 | 6.1467 | 6.5077 | 6.4923 | 5.3128 | 5.6704 | 5.4999 | 6.1569 |
| CHAF1A_S255 | CHAF1A | Q13111 | | 259 | IKSLPATPQGKNM | 7.7296 | 7.6387 | 6.7777 | 7.9783 | 6.7290 | 6.4922 | 7.1259 | 6.9761 | 7.1339 | 6.7801 | 7.1809 | 7.0941 | 3.1778 | 3.9402 | 3.3647 | 3.8807 |
| CHAF1A_S255;T259 | CHAF1A | Q13111 | | 123 | IDLTEDSNEQPDS | 5.5168 | 6.1952 | 6.0094 | 6.6527 | 6.3150 | 6.2159 | 6.5242 | 6.7900 | 6.4626 | 5.8940 | 6.7697 | 5.7965 | 6.1144 | 7.0271 | 5.8960 | 5.8204 |
| CHAF1A_S259 | CHAF1A | Q13111 | | 206 | QECSPRSCPELTS | 6.4226 | 7.1900 | 6.4674 | 6.7810 | 5.7888 | 5.8178 | 6.1358 | 5.7227 | 6.6779 | 6.4408 | 6.5699 | 6.1887 | 5.8065 | 5.7807 | 5.9782 | 6.2312 |
| CHAF1A_S65 | CHAF1A | Q13111 | | 772 | LSNHTGSPRSPST | 6.2825 | 7.6149 | 6.6552 | 7.9829 | 6.4017 | 6.2350 | 5.5922 | 7.1968 | 6.4990 | 6.3804 | 6.8312 | 6.7231 | 4.5422 | 4.9289 | 4.6913 | 5.4426 |
| CHAF1A_S772 | CHAF1A | Q13111 | | 141 | KLNSEASPSREAI | 6.7347 | 7.2215 | 5.9898 | 6.9881 | 6.3443 | 5.7349 | 6.5701 | 6.0422 | 6.4270 | 6.3508 | 6.5396 | 6.4880 | 5.2363 | 5.7479 | 5.8102 | 5.7747 |
| CHAF1A_S775 | CHAF1A | Q13111 | | 956 | ASPLGASxxxxxx | 6.0188 | 6.9764 | 6.2994 | 6.6834 | 6.0357 | 6.2131 | 6.0928 | 7.6858 | 5.4898 | 7.3141 | 7.0952 | 6.4999 | 5.3912 | 5.5407 | 4.3965 | 6.2671 |
| CHAF1A_S803 | CHAF1A | Q13111 | | 255 | RPPQIKSLPATPQ | 6.5453 | 6.8999 | 5.7667 | 6.6636 | 6.1804 | 6.3523 | 6.5309 | 6.4198 | 6.7941 | 6.5604 | 6.3711 | 6.3995 | 5.5033 | 6.0400 | 5.1595 | 5.8133 |
| CHAF1A_S873 | CHAF1A | Q13111 | | 722 | PAQEEQTPKASKR | 6.7651 | 7.6458 | 7.8472 | 7.6590 | 6.6717 | 7.0311 | 7.8867 | 5.8515 | 7.2417 | 6.4821 | 6.8744 | 7.5076 | 3.5030 | 3.7321 | 3.1765 | 4.1244 |
| CHAF1A_S951 | CHAF1A | Q13111 | | 803 | RLKRLISENSVYE | 5.9737 | 4.7789 | 5.1268 | 5.8994 | 5.6048 | 5.8697 | 5.9422 | 6.0619 | 5.2072 | 5.1416 | 4.7160 | 5.9850 | 8.7416 | 8.4121 | 8.5440 | 7.9952 |
| CHAF1A_S956 | CHAF1A | Q13111 | | 873 | ISLKRKSAGSMCI | 6.4316 | 6.9347 | 7.0326 | 7.1025 | 6.2978 | 6.7162 | 6.5341 | 6.0237 | 6.5406 | 6.2679 | 6.4937 | 7.8189 | 4.1996 | 5.6287 | 4.4778 | 5.4995 |
| CHAF1A_T722 | CHAF1A | Q13111 | | 865 | TGPSQGTPISLKR | 7.9580 | 7.7886 | 6.6671 | 7.2685 | 5.8564 | 6.7295 | 7.7151 | 6.1562 | 6.6334 | 7.3780 | 7.3230 | 7.4282 | 3.2757 | 4.5807 | 2.9839 | 4.2577 |
| CHAF1A_T865 | CHAF1A | Q13111 | 255;259 | | NA | 7.2649 | 7.4546 | 6.5663 | 7.6466 | 5.5791 | 6.1342 | 6.1963 | 5.7827 | 7.0452 | 6.9935 | 6.3426 | 7.6270 | 4.6028 | 5.2805 | 4.4279 | 5.0558 |
| CHAF1B_S410 | CHAF1B | Q13112 | | 410 | QTHRGSSPGPRPV | 6.4829 | 6.5159 | 5.7757 | 6.0465 | 6.2651 | 6.0199 | 6.0446 | 6.1327 | 6.0634 | 6.0249 | 6.1753 | 6.2229 | 6.5972 | 6.5885 | 6.7094 | 6.3352 |
| CHAF1B_S429 | CHAF1B | Q13112 | | 538 | PGDAQGSPPELKR | 5.9402 | 5.7690 | 6.5271 | 6.5074 | 5.9373 | 6.1849 | 6.2754 | 5.7531 | 6.5336 | 6.3330 | 6.5706 | 6.4831 | 6.1370 | 6.4587 | 6.2184 | 6.3712 |
| CHAF1B_S429;T433 | CHAF1B | Q13112 | | 458 | ITPAVKSPLPGPS | 7.0694 | 7.6548 | 6.9704 | 7.0660 | 6.3616 | 6.5929 | 6.9826 | 6.5341 | 7.4744 | 6.9420 | 7.4285 | 7.3629 | 3.5748 | 4.3904 | 3.4226 | 4.1726 |
| CHAF1B_S433 | CHAF1B | Q13112 | | 503 | PRRINLTPLKTDT | 7.0774 | 7.1642 | 6.7332 | 6.9976 | 6.0912 | 6.9081 | 7.0398 | 7.2624 | 7.8443 | 7.2758 | 6.9985 | 7.1966 | 3.5493 | 4.2133 | 3.5129 | 4.1354 |
| CHAF1B_S458 | CHAF1B | Q13112 | | 429 | RTQDPSSPGTTPP | 6.0383 | 6.0853 | 6.2015 | 6.4206 | 6.9543 | 5.7377 | 6.7638 | 6.1143 | 6.0526 | 6.2400 | 6.6568 | 6.1330 | 6.3147 | 6.4905 | 6.0536 | 5.7429 |
| CHAF1B_S473 | CHAF1B | Q13112 | | 485 | HPSRRVTLNTLQA | 7.9233 | 6.6674 | 6.5337 | 6.6853 | 5.6331 | 6.2819 | 6.2565 | 5.3126 | 9.1150 | 6.5627 | 8.3823 | 8.2700 | 4.6321 | 5.4370 | 2.5175 | 3.7894 |
| CHAF1B_S538 | CHAF1B | Q13112 | | 473 | KTLQPSSQNTKAH | 6.8988 | 6.9977 | 7.0920 | 7.0323 | 5.8432 | 6.5628 | 6.6593 | 6.5324 | 7.4400 | 7.4194 | 7.9827 | 8.5177 | 3.1555 | 4.7955 | 2.9898 | 4.0809 |
| CHAF1B_T485 | CHAF1B | Q13112 | | 433 | PSSPGTTPPQARQ | 5.0059 | 6.4746 | 7.0707 | 6.1345 | 5.4693 | 7.1080 | 6.1252 | 6.1962 | 6.1083 | 6.6147 | 6.2672 | 5.9103 | 6.5063 | 5.9103 | 6.6129 | 6.4858 |
| CHAF1B_T503 | CHAF1B | Q13112 | 429;433 | | NA | 6.6616 | 6.5810 | 5.9552 | 6.8736 | 6.0460 | 6.1242 | 6.7733 | 6.1113 | 6.7407 | 6.7445 | 6.6424 | 6.5338 | 5.4981 | 5.8840 | 5.2312 | 5.5991 |
| CHAMP1_S108 | CHAMP1 | Q96JM3 | | 214 | QKPAPVSPESVKA | 6.4091 | 5.7386 | 6.2654 | 6.5037 | 6.2443 | 5.9627 | 6.2928 | 5.6123 | 7.2268 | 6.1821 | 5.8413 | 6.1564 | 6.6033 | 6.0672 | 6.1523 | 6.7419 |
| CHAMP1_S142 | CHAMP1 | Q96JM3 | | 476 | KSSRGGSPDLWKS | 6.4292 | 6.3224 | 5.7809 | 6.2257 | 6.3467 | 5.9440 | 6.2222 | 6.2140 | 6.3272 | 6.0918 | 6.0242 | 5.9969 | 6.6981 | 6.4107 | 6.6441 | 6.3220 |
| CHAMP1_S145 | CHAMP1 | Q96JM3 | | 145 | SVLSPESPKPTPL | 6.2648 | 6.1831 | 6.0833 | 6.5428 | 5.7709 | 5.4483 | 5.9954 | 5.7027 | 6.2086 | 6.2439 | 5.9414 | 6.1890 | 6.8538 | 6.6917 | 6.8277 | 7.0525 |
| CHAMP1_S184 | CHAMP1 | Q96JM3 | | 355 | KPIPSVSPGPWKP | 6.9303 | 5.8306 | 6.2713 | 6.9405 | 7.0710 | 6.5433 | 6.9140 | 6.0243 | 6.9258 | 6.6533 | 6.3854 | 6.6329 | 5.2885 | 5.3625 | 4.7578 | 5.4685 |
| CHAMP1_S204 | CHAMP1 | Q96JM3 | | 445 | LRKPSGSPDLWKL | 6.4071 | 5.9602 | 6.0583 | 6.1936 | 5.7376 | 5.2832 | 6.0069 | 5.3704 | 6.2617 | 6.2002 | 5.5832 | 6.1722 | 7.0131 | 6.6589 | 7.2259 | 7.8673 |
| CHAMP1_S204;S214 | CHAMP1 | Q96JM3 | | 416 | KAVPPVSPELRKP | 7.0186 | 7.1377 | 6.5009 | 7.0882 | 6.4470 | 6.3382 | 6.4064 | 6.2093 | 6.4895 | 6.4315 | 6.5020 | 6.4688 | 5.3930 | 5.3246 | 5.0803 | 5.1639 |
| CHAMP1_S214 | CHAMP1 | Q96JM3 | | 297 | KPFPAVSPEPRRP | 7.5279 | 6.6235 | 6.9676 | 7.2889 | 6.5810 | 6.2088 | 6.7611 | 6.0767 | 7.2586 | 6.8580 | 6.6938 | 7.1199 | 4.3100 | 4.5760 | 4.1553 | 4.9928 |
| CHAMP1_S223 | CHAMP1 | Q96JM3 | | 204 | KLAPVPSPEPQKP | 6.3868 | 6.4537 | 5.9103 | 6.3852 | 6.2526 | 5.7286 | 6.1537 | 6.0205 | 6.3239 | 6.0719 | 5.5969 | 5.7595 | 6.8459 | 6.4722 | 6.8659 | 6.7724 |
| CHAMP1_S247 | CHAMP1 | Q96JM3 | | 344 | KPAPSVSPGPWKP | 7.0099 | 6.1733 | 6.0342 | 6.5123 | 6.8373 | 6.4951 | 6.9074 | 5.9571 | 6.7023 | 6.4424 | 6.4387 | 6.6831 | 5.7439 | 5.4105 | 5.2118 | 5.4407 |
| CHAMP1_S253;S260;S26 | CHAMP1 | Q96JM3 | | 427 | KPGPPLSPEIRSP | 5.8676 | 5.7955 | 5.6750 | 6.3270 | 6.0410 | 5.6769 | 5.9854 | 5.9167 | 6.2427 | 5.7529 | 5.6089 | 5.8008 | 7.7786 | 7.0407 | 7.2947 | 7.1955 |
| CHAMP1_S260;S264 | CHAMP1 | Q96JM3 | | 603 | QEELLASPKKLLE | 6.5267 | 6.2965 | 6.7380 | 6.9568 | 6.1060 | 5.6480 | 6.3378 | 5.6479 | 6.8081 | 6.5196 | 6.5485 | 6.8239 | 5.8308 | 5.6650 | 5.4641 | 6.0822 |
| CHAMP1_S282;S286;S29 | CHAMP1 | Q96JM3 | | 542 | KTAPPASPEARKR | 6.4734 | 6.5592 | 5.9815 | 6.5053 | 6.2857 | 5.9361 | 6.2451 | 5.9370 | 6.2693 | 6.2983 | 6.2448 | 6.4157 | 6.3602 | 6.0754 | 6.1654 | 6.2476 |
| CHAMP1_S286 | CHAMP1 | Q96JM3 | | 308 | RPAPAVSPGSWKP | 6.4444 | 6.4447 | 5.9317 | 6.3937 | 5.8540 | 5.5174 | 6.1655 | 6.2089 | 6.3723 | 6.1194 | 5.6551 | 6.2791 | 6.3555 | 6.2850 | 6.9114 | 7.0621 |
| CHAMP1_S286;S297 | CHAMP1 | Q96JM3 | | 405 | KTAPTLSPEHWKA | 7.3762 | 7.0226 | 6.6430 | 7.1216 | 6.8779 | 6.1167 | 6.5191 | 6.5530 | 6.8818 | 6.4414 | 6.4258 | 6.5399 | 4.8391 | 4.9006 | 4.9171 | 4.8244 |
| CHAMP1_S297 | CHAMP1 | Q96JM3 | | 108 | ETDPVKSPPLPEH | 7.3170 | 7.1409 | 7.0493 | 7.5762 | 6.8544 | 6.5522 | 6.3688 | 6.7299 | 7.1979 | 6.9150 | 7.1827 | 7.1965 | 3.9209 | 4.4096 | 3.7631 | 3.8256 |
| CHAMP1_S308 | CHAMP1 | Q96JM3 | | 184 | KPASVSSPEPPKS | 6.5071 | 5.7206 | 6.5464 | 6.9397 | 6.5740 | 6.3361 | 6.5846 | 5.7465 | 6.9197 | 6.8224 | 6.2332 | 6.4650 | 5.8319 | 5.6144 | 5.3484 | 5.8100 |
| CHAMP1_S308;S319 | CHAMP1 | Q96JM3 | | 459 | PDQRKTSPASLDF | 6.2915 | 6.6489 | 6.4695 | 6.9552 | 6.2723 | 6.0040 | 6.4585 | 5.9722 | 6.6781 | 6.7188 | 6.6346 | 6.7319 | 5.4727 | 5.5879 | 5.3282 | 5.7757 |
| CHAMP1_S319 | CHAMP1 | Q96JM3 | | 319 | KPGPPGSPRPWKS | 6.8754 | 6.1032 | 6.3661 | 6.4970 | 6.2092 | 5.8888 | 6.1676 | 5.4425 | 6.3559 | 6.3306 | 6.0846 | 6.6294 | 6.6103 | 6.0217 | 5.8028 | 6.6150 |
| CHAMP1_S331 | CHAMP1 | Q96JM3 | | 386 | WKSPPASPESWKS | 6.8323 | 6.2711 | 6.1806 | 7.4688 | 8.8458 | 5.8989 | 6.9326 | 6.4346 | 5.9303 | 5.3363 | 5.3875 | 5.0904 | 6.0169 | 5.7900 | 6.1748 | 5.4090 |
| CHAMP1_S342;S344 | CHAMP1 | Q96JM3 | | 507 | PSGPSESPKAASD | 6.8256 | 5.8693 | 6.5834 | 6.5451 | 5.7656 | 5.1772 | 6.4608 | 4.9598 | 6.9304 | 6.6953 | 6.3055 | 7.0995 | 6.3230 | 5.6505 | 5.7168 | 7.0923 |
| CHAMP1_S344 | CHAMP1 | Q96JM3 | | 142 | KLGSVLSPESPKP | 5.7522 | 5.9989 | 5.9142 | 6.1795 | 6.5258 | 6.5687 | 5.9268 | 5.6774 | 6.5497 | 6.4060 | 6.3149 | 6.2937 | 7.1162 | 6.4187 | 6.3481 | 6.0091 |
| CHAMP1_S353 | CHAMP1 | Q96JM3 | | 627 | KDNQESSDAELSS | 6.1656 | 6.4018 | 5.9530 | 6.6116 | 6.2760 | 5.7206 | 5.7507 | 6.2846 | 6.4652 | 5.7870 | 6.2023 | 6.2476 | 6.2998 | 6.2336 | 6.7040 | 6.8966 |
| CHAMP1_S353;S355 | CHAMP1 | Q96JM3 | | 173 | LQTPLPSPEPSKP | 6.8456 | 6.0169 | 6.2267 | 6.1600 | 6.2408 | 6.4867 | 6.7663 | 5.8655 | 6.6510 | 7.0291 | 6.5812 | 7.0816 | 5.3325 | 5.5070 | 5.1898 | 6.0192 |
| CHAMP1_S355 | CHAMP1 | Q96JM3 | | 223 | SVKATLSNPKPQK | 6.9794 | 7.6025 | 7.0966 | 7.5681 | 6.3231 | 6.8945 | 7.1162 | 6.7554 | 7.2345 | 7.3078 | 7.5399 | 7.6126 | 3.0170 | 4.0424 | 3.0599 | 3.8502 |
| CHAMP1_S376 | CHAMP1 | Q96JM3 | | 331 | SNPSASSGPWKPA | 6.3067 | 7.4407 | 6.8922 | 7.4340 | 5.8778 | 6.8441 | 6.9603 | 6.4728 | 7.3103 | 7.2367 | 7.6335 | 7.5391 | 3.9643 | 4.8739 | 3.1973 | 4.0162 |
| CHAMP1_S382;S386 | CHAMP1 | Q96JM3 | | 376 | WKSSSVSPSSWKS | 5.5562 | 5.5998 | 6.0101 | 7.4241 | 6.3435 | 6.4892 | 6.4704 | 6.5148 | 7.0744 | 7.1185 | 7.0879 | 7.1692 | 5.2540 | 5.5469 | 4.9361 | 5.4048 |
| CHAMP1_S386 | CHAMP1 | Q96JM3 | | 353 | PWKPIPSVSPGPW | 6.1710 | 5.5824 | 5.0087 | 7.3804 | 7.7548 | 8.2369 | 5.9110 | 6.4955 | 6.6450 | 5.1784 | 8.5278 | 7.1514 | 6.3918 | 5.1959 | 4.6531 | 3.7158 |
| CHAMP1_S405 | CHAMP1 | Q96JM3 | | 452 | PDLWKLSPDQRKT | 6.2438 | 5.6963 | 6.3422 | 6.7532 | 6.8320 | 6.4105 | 6.7493 | 5.8112 | 6.4608 | 6.7154 | 6.5965 | 6.8197 | 6.2543 | 5.8176 | 5.3123 | 5.1851 |
| CHAMP1_S416 | CHAMP1 | Q96JM3 | | 632 | SSDAELSSSEYIK | 5.7958 | 6.6843 | 6.5288 | 5.8208 | 6.1051 | 6.3840 | 6.5235 | 5.5478 | 6.3696 | 6.6706 | 5.6335 | 5.9155 | 6.6206 | 6.0755 | 6.8745 | 6.4500 |
| CHAMP1_S427 | CHAMP1 | Q96JM3 | | 286 | KPSPSESPEPWKP | 6.5277 | 5.9073 | 6.9774 | 6.9546 | 5.4915 | 5.4372 | 6.0186 | 5.4864 | 6.6908 | 6.6457 | 6.1751 | 6.6526 | 5.2939 | 5.9066 | 6.5086 | 7.3259 |
| CHAMP1_S427;S432 | CHAMP1 | Q96JM3 | | 615 | EDTLFPSSKKLKK | 7.1154 | 8.1217 | 6.9592 | 6.4932 | 6.6955 | 7.1641 | 7.5251 | 5.5638 | 7.1487 | 6.9720 | 8.2881 | 5.8378 | 3.1724 | 6.9720 | 2.4580 | 3.5129 |
| CHAMP1_S427;S432;S43 | CHAMP1 | Q96JM3 | | 247 | SASSPESPVLAAS | 5.3564 | 5.0350 | 5.2249 | 7.1892 | 5.7739 | 6.1330 | 5.7071 | 4.7845 | 5.5422 | 6.3104 | 6.6528 | 5.8052 | 6.6402 | 7.7988 | 6.9888 | 9.0575 |
| CHAMP1_S445 | CHAMP1 | Q96JM3 | 382;386 | | NA | 6.5922 | 6.8719 | 6.5460 | 7.0751 | 6.3362 | 6.1354 | 6.2546 | 6.2025 | 6.9719 | 6.6382 | 6.7029 | 6.5192 | 5.4303 | 5.3471 | 5.0128 | 5.3637 |
| CHAMP1_S445;S452 | CHAMP1 | Q96JM3 | 204;214 | | NA | 6.1351 | 6.4373 | 6.1074 | 6.9008 | 6.6761 | 6.4789 | 6.2942 | 6.3537 | 6.6073 | 6.3815 | 7.0822 | 6.4920 | 5.9802 | 5.8011 | 5.3505 | 4.9216 |
| CHAMP1_S452 | CHAMP1 | Q96JM3 | 169;173 | | NA | 6.3856 | 5.8855 | 6.3613 | 6.7469 | 5.8670 | 5.6379 | 5.9941 | 5.2908 | 6.6019 | 5.8374 | 5.8088 | 6.5022 | 6.6846 | 6.5075 | 6.6222 | 7.2664 |
| CHAMP1_S459 | CHAMP1 | Q96JM3 | 427;432;436 | | NA | 6.8050 | 6.5662 | 6.6255 | 7.2853 | 6.7856 | 6.2814 | 6.4555 | 6.2753 | 6.9603 | 6.4988 | 6.9918 | 6.7025 | 5.0809 | 5.0164 | 4.6812 | 4.9885 |
| CHAMP1_S476 | CHAMP1 | Q96JM3 | 286;297 | | NA | 7.0955 | 7.2302 | 6.9514 | 7.5414 | 7.8980 | 7.5054 | 7.0680 | 7.0179 | 7.5001 | 6.9926 | 8.4187 | 6.8732 | 3.1171 | 3.5249 | 2.5157 | 2.7499 |
| CHAMP1_S507 | CHAMP1 | Q96JM3 | 282;286;297 | | NA | 7.3719 | 6.9169 | 6.6060 | 7.2924 | 6.9973 | 7.1705 | 6.4448 | 6.9758 | 7.0087 | 7.3353 | 7.4320 | 6.6534 | 3.7968 | 4.6264 | 3.7827 | 3.5890 |
| CHAMP1_S542 | CHAMP1 | Q96JM3 | 308;319 | | NA | 5.3610 | 5.5865 | 6.5886 | 7.2063 | 6.9683 | 6.3508 | 6.5291 | 6.6030 | 7.2174 | 6.3292 | 7.0450 | 6.5889 | 5.4997 | 5.7918 | 5.0859 | 5.2485 |

| | | | | | | | | | | | | | | | | | | | | |
|---|---|---|---|---|---|---|---|---|---|---|---|---|---|---|---|---|---|---|---|---|
| CHAMP1_S603 | CHAMP1 | Q96JM3 | 253;260;264 | NA | 6.1727 | 5.8965 | 6.6131 | 6.5647 | 5.5954 | 6.4150 | 6.4454 | 5.6167 | 5.8097 | 6.2917 | 6.1710 | 6.2358 | 6.3263 | 6.8681 | 6.7906 | 6.1873 |
| CHAMP1_S615 | CHAMP1 | Q96JM3 | 427;432 | NA | 5.8577 | 6.0324 | 6.4544 | 7.2854 | 6.7396 | 6.1066 | 6.8545 | 6.1272 | 6.5230 | 6.4471 | 6.5766 | 6.3017 | 6.2135 | 6.1980 | 4.9262 | 5.3559 |
| CHAMP1_S627 | CHAMP1 | Q96JM3 | 342;344 | NA | 7.6513 | 6.3538 | 6.7221 | 7.9394 | 7.3618 | 7.4841 | 7.6200 | 5.7172 | 7.1813 | 7.8171 | 7.8879 | 5.9110 | 2.9595 | 3.9125 | 3.9263 | 3.5549 |
| CHAMP1_S632 | CHAMP1 | Q96JM3 | 445;452 | NA | 7.2398 | 5.9810 | 7.1890 | 6.0614 | 6.4799 | 5.5379 | 6.7916 | 5.8043 | 6.3711 | 6.3386 | 7.6579 | 6.4096 | 5.6294 | 5.6407 | 4.8423 | 6.0256 |
| CHAMP1_T169;S173 | CHAMP1 | Q96JM3 | 353;355 | NA | 7.5133 | 7.2391 | 7.0829 | 7.7559 | 6.1549 | 5.1824 | 6.6379 | 5.3714 | 7.1863 | 7.9947 | 8.4804 | 7.7676 | 3.5100 | 4.5383 | 3.1216 | 4.4631 |
| CHAMP1_T173 | CHAMP1 | Q96JM3 | 260;264 | NA | 5.7034 | 7.3775 | 6.4438 | 6.1969 | 4.7392 | 7.7633 | 7.6898 | 7.5690 | 7.0388 | 6.4328 | 5.8479 | 7.0660 | 4.6329 | 5.4361 | 4.6570 | 5.4056 |
| CHCHD3_S29 | CHCHD3 | Q9NX63 | 50 | SKSQRYSGAYGAS | 6.0886 | 6.0407 | 6.1935 | 6.7594 | 6.6868 | 5.8882 | 6.0421 | 6.3318 | 6.2606 | 5.8316 | 6.4282 | 6.7256 | 6.6588 | 5.9483 | 5.8723 | 6.2434 |
| CHCHD3_S50 | CHCHD3 | Q9NX63 | 29 | VKGIRLSENVIDR | 6.6856 | 6.3872 | 5.9896 | 5.8242 | 5.7537 | 6.3410 | 5.9266 | 6.2773 | 6.4496 | 6.0133 | 6.5938 | 5.9003 | 7.2018 | 6.5220 | 6.2255 | 5.9085 |
| CHCHD6_S13 | CHCHD6 | Q9BRQ6 | 13 | SEGRRVSFGVDEE | 4.6145 | 4.4299 | 5.2131 | 6.0245 | 7.9911 | 5.3405 | 5.2169 | 6.2088 | 5.8874 | 6.3601 | 7.0396 | 6.2590 | 7.7165 | 7.8982 | 6.8490 | 6.9509 |
| CHCHD6_S74 | CHCHD6 | Q9BRQ6 | 74 | TLPRSGSSGGQQP | 3.8583 | 3.6279 | 3.7767 | 3.5640 | 4.6390 | 4.0093 | 4.5369 | 4.2899 | 3.9668 | 4.0967 | 3.7511 | 3.5105 | 14.4408 | 11.9819 | 13.5209 | 12.4292 |
| CHD1_S1328 | CHD1 | O14646 | 1689 | SPYGSRSPFEHSV | 5.7703 | 6.1496 | 5.7445 | 6.5519 | 6.2665 | 6.1302 | 5.9969 | 6.3918 | 6.6532 | 6.2410 | 5.9129 | 6.0449 | 6.5820 | 6.5895 | 6.7599 | 6.2149 |
| CHD1_S1622 | CHD1 | O14646 | 1622 | RSNLEGSLKDRSH | 5.8891 | 6.3595 | 7.3491 | 6.0809 | 5.3800 | 6.8621 | 6.1427 | 6.0569 | 7.7633 | 9.6780 | 8.1156 | 7.5373 | 4.0306 | 5.3458 | 3.3038 | 4.1053 |
| CHD1_S1683;S1689 | CHD1 | O14646 | 241 | QATVNVSYKEDEE | 6.8228 | 7.1996 | 6.8450 | 8.1502 | 6.9211 | 5.5812 | 6.6120 | 6.1472 | 6.8757 | 6.9314 | 5.7631 | 6.6887 | 4.4419 | 4.5527 | 4.7988 | 5.6687 |
| CHD1_S1689 | CHD1 | O14646 | 90 | AEFWKSSPSILAV | 7.4425 | 7.0998 | 6.2686 | 7.0040 | 6.7972 | 6.4614 | 6.6530 | 6.0966 | 7.0171 | 7.0692 | 7.3303 | 6.8809 | 4.6595 | 4.9127 | 3.8374 | 4.4698 |
| CHD1_S241 | CHD1 | O14646 | 1328 | ALSGAGSSKRRKA | 7.3169 | 7.3020 | 7.4847 | 8.0241 | 5.0369 | 7.0859 | 8.2004 | 6.8002 | 7.1080 | 7.4385 | 7.0810 | 7.7754 | 2.5842 | 3.9062 | 3.2139 | 3.6418 |
| CHD1_S90 | CHD1 | O14646 | 1683;1689 | NA | 6.2667 | 6.2228 | 6.2090 | 7.6008 | 6.5082 | 6.9956 | 6.4059 | 6.5708 | 6.1198 | 6.0113 | 6.6347 | 6.7720 | 5.1274 | 6.0934 | 5.1767 | 5.2848 |
| CHD2_S1365 | CHD2 | O14647 | 1789 | AVSDPRSPPSQKS | 6.2001 | 6.5626 | 6.3769 | 7.3689 | 6.3598 | 6.1719 | 6.6053 | 5.9067 | 6.2166 | 6.7081 | 6.8921 | 6.3677 | 5.1481 | 6.4117 | 5.3764 | 5.3271 |
| CHD2_S1365;S1369;S137 | CHD2 | O14647 | 1365 | HGIELSSPRHSDN | 5.3050 | 6.5008 | 6.0874 | 6.7527 | 6.3923 | 5.7007 | 5.8231 | 5.6309 | 6.3288 | 6.8053 | 6.0989 | 6.0066 | 6.3828 | 7.0429 | 6.5917 | 6.5501 |
| CHD2_S1789 | CHD2 | O14647 | 1365;1369;13 | NA | 5.6507 | 6.2327 | 6.2396 | 7.0540 | 5.0543 | 6.0614 | 5.6289 | 5.5689 | 6.0392 | 6.6252 | 6.0699 | 6.3387 | 6.1273 | 6.3486 | 7.2353 | 7.7255 |
| CHD3_S132 | CHD3 | Q12873-3 | 1616 | CTSKPATPAPSEK | 6.0610 | 6.1043 | 6.6781 | 6.6338 | 5.7627 | 5.5166 | 6.4376 | 5.7308 | 6.2908 | 6.6461 | 6.1184 | 6.4848 | 5.9452 | 6.3562 | 6.2279 | 7.0057 |
| CHD3_S132;S138 | CHD3 | Q12873-3 | 1660 | TEADAPSPAPSLG | 5.2694 | 5.9594 | 5.7270 | 5.9419 | 5.8072 | 6.0830 | 6.0066 | 6.3572 | 6.2304 | 6.6897 | 5.9122 | 5.9915 | 7.0429 | 7.4191 | 6.9103 | 6.6522 |
| CHD3_S1660 | CHD3 | Q12873-3 | 656 | PPLDYGSGEDDGK | 6.5705 | 6.3347 | 6.1677 | 7.0640 | 6.0804 | 5.7078 | 6.2768 | 5.4778 | 6.2029 | 6.5403 | 5.8446 | 6.2995 | 5.8798 | 7.1085 | 5.6331 | 6.8114 |
| CHD3_S371;S375 | CHD3 | Q12873-3 | 772 | GDGPPSSPTNDPT | 5.5488 | 5.7907 | 5.9893 | 6.2052 | 5.8893 | 6.3987 | 5.8262 | 5.9241 | 6.4522 | 6.2100 | 5.6659 | 5.4177 | 6.3502 | 7.6258 | 7.5807 | 7.1253 |
| CHD3_S656 | CHD3 | Q12873-3 | 1628 | KGEGIRTPLEKEE | 5.6343 | 6.3553 | 5.2741 | 5.5722 | 6.2421 | 5.5175 | 6.0603 | 5.9261 | 6.0230 | 5.9593 | 5.2741 | 5.3820 | 7.2137 | 8.2670 | 8.2287 | 7.0705 |
| CHD3_S772 | CHD3 | Q12873-3 | 132 | KRKKRDSEEEFGS | 5.3940 | 5.8505 | 6.1489 | 6.2116 | 5.3313 | 5.5124 | 6.1359 | 5.3308 | 6.3872 | 6.3917 | 6.9213 | 6.6124 | 5.9016 | 7.9722 | 6.4704 | 7.4279 |
| CHD3_T1600;S1608 | CHD3 | H7C0J3 | 371;375 | NA | 6.0624 | 6.1209 | 5.8207 | 6.3549 | 6.5864 | 5.9786 | 6.1600 | 6.3964 | 6.3104 | 6.7298 | 6.3345 | 6.0545 | 6.0979 | 6.6034 | 6.1529 | 6.2361 |
| CHD3_T1616 | CHD3 | Q12873-3 | 132;138 | NA | 6.1233 | 6.5616 | 6.3623 | 7.0358 | 6.3798 | 6.1732 | 6.9871 | 6.6861 | 5.9290 | 5.2077 | 5.7690 | 5.9379 | 6.1122 | 6.2209 | 6.2858 | 6.2282 |
| CHD3_T1628 | CHD3 | Q12873-3 | 1600;1608 | NA | 5.1264 | 6.3962 | 7.4290 | 6.8736 | 5.6795 | 6.2101 | 5.9005 | 6.0605 | 6.0227 | 5.5757 | 6.4943 | 6.6257 | 5.8072 | 5.9223 | 7.0508 | 6.8254 |
| CHD4_S103;S105;S108 | CHD4 | Q14839-2 | 1563 | KMSQPGSPSPKTP | 6.8509 | 6.9290 | 6.3582 | 6.7843 | 6.7119 | 6.1647 | 6.5211 | 6.6055 | 6.4981 | 6.7254 | 6.3170 | 6.1390 | 5.0422 | 5.7081 | 5.4108 | 5.2339 |
| CHD4_S1559 | CHD4 | Q14839-2 | 428 | EAKEDNSEGEEIL | 6.1049 | 6.1379 | 6.3166 | 6.4748 | 6.3385 | 5.8207 | 6.0968 | 5.8824 | 6.2948 | 6.3097 | 6.1502 | 6.3787 | 6.0688 | 6.7633 | 6.4677 | 6.3942 |
| CHD4_S1559;S1563 | CHD4 | Q14839-2 | 1568 | GSPSPKTPTPSTP | 5.9474 | 6.3271 | 5.9521 | 6.7276 | 6.1419 | 6.1664 | 6.4293 | 6.0822 | 6.2164 | 6.3338 | 6.6513 | 6.6318 | 5.5190 | 6.4463 | 6.2153 | 6.2120 |
| CHD4_S1559;S1565 | CHD4 | Q14839-2 | 515 | KWGQPPSPTPVPR | 5.7338 | 6.1525 | 6.2818 | 6.8233 | 5.7335 | 6.1063 | 6.1047 | 5.9843 | 6.2363 | 6.2131 | 6.6389 | 6.5727 | 5.6111 | 6.7731 | 6.6247 | 6.4098 |
| CHD4_S1563 | CHD4 | Q14839-2 | 1681 | KSAIDLTPIVVED | 6.0839 | 6.4431 | 5.7973 | 6.4356 | 6.1516 | 5.9550 | 6.4518 | 5.8376 | 5.8632 | 5.9615 | 5.9707 | 6.1058 | 6.2673 | 7.0875 | 6.6958 | 6.8923 |
| CHD4_S1565 | CHD4 | Q14839-2 | 1581 | GDTQPNTPAPVPP | 6.2562 | 6.5587 | 6.4163 | 6.6415 | 6.5688 | 6.3267 | 6.6974 | 6.3120 | 6.5901 | 6.1671 | 6.0892 | 6.5788 | 5.4841 | 6.1324 | 5.5642 | 5.6164 |
| CHD4_S1572 | CHD4 | Q14839-2 | 367 | AVDGYETDHQDYC | 6.6664 | 6.4219 | 6.1888 | 6.8974 | 6.2199 | 6.5454 | 6.1514 | 5.9608 | 6.1405 | 6.2807 | 6.8085 | 5.7937 | 5.8037 | 6.2828 | 5.5438 | 6.2941 |
| CHD4_S1598 | CHD4 | Q14839-2 | 1630 | QAPAPASEDEKVV | 5.9253 | 6.2200 | 6.3627 | 6.6114 | 6.1982 | 6.0670 | 6.2997 | 5.6902 | 6.2640 | 6.1782 | 5.5426 | 6.3070 | 6.5101 | 6.6682 | 6.6372 | 6.5183 |
| CHD4_S1604 | CHD4 | Q14839-2 | 1707 | MLQNGETPKDLND | 6.6757 | 7.2886 | 6.3487 | 6.7817 | 6.1995 | 6.4432 | 7.0902 | 6.3948 | 6.9027 | 7.2703 | 6.7079 | 6.5537 | 4.6004 | 5.3774 | 4.1856 | 5.1796 |
| CHD4_S1630 | CHD4 | Q14839-2 | 531 | ADPNTPSPKPLEG | 6.1979 | 6.1170 | 6.1306 | 6.2770 | 6.8681 | 6.2969 | 6.4484 | 6.2582 | 6.3825 | 6.3729 | 6.0059 | 6.1206 | 5.8955 | 6.6429 | 6.0875 | 5.8981 |
| CHD4_S428 | CHD4 | Q14839-2 | 1598 | IKIEENSLKEEES | 5.9363 | 5.6155 | 5.8462 | 6.5473 | 6.2581 | 5.4749 | 6.9495 | 5.8680 | 5.7544 | 5.8640 | 4.7784 | 5.4795 | 5.8949 | 6.8118 | 8.3438 | 8.5773 |
| CHD4_S515 | CHD4 | Q14839-2 | 1559 | EENKKMSQPGSPS | 5.4987 | 5.2976 | 4.6696 | 4.7485 | 5.4938 | 5.1344 | 5.4538 | 5.2076 | 5.2804 | 4.8188 | 4.4467 | 3.9774 | 10.2967 | 9.8175 | 10.7893 | 9.0693 |
| CHD4_S515;S531 | CHD4 | Q14839-2 | 1604 | SLKEEESIEGEKE | 6.8643 | 5.9573 | 5.4126 | 6.0234 | 5.9953 | 6.5399 | 7.0075 | 6.8986 | 5.7272 | 6.9745 | 6.5008 | 5.9977 | 6.1654 | 5.5619 | 5.6733 | 6.7003 |
| CHD4_S515;T529;S531 | CHD4 | Q14839-2 | 703 | LERPPETPTVDPT | 4.7431 | 4.8963 | 4.7247 | 4.9878 | 5.1954 | 5.3287 | 5.3491 | 5.4619 | 4.6729 | 4.4085 | 4.7768 | 4.7693 | 9.3458 | 10.3771 | 11.0884 | 9.8743 |
| CHD4_S531 | CHD4 | Q14839-2 | 1570 | PSPKTPTPSTPGD | 6.2096 | 6.6598 | 5.9158 | 7.6160 | 6.3126 | 6.3077 | 6.8413 | 6.4414 | 6.0765 | 6.1010 | 6.7800 | 6.9364 | 5.3644 | 6.0397 | 4.6002 | 5.7974 |
| CHD4_T1568 | CHD4 | Q14839-2 | 1565 | SQPGSPSPKTPTP | 6.9171 | 6.6962 | 7.0624 | 7.0154 | 6.3357 | 5.9417 | 5.8132 | 6.4399 | 6.5213 | 6.9006 | 6.9075 | 6.5598 | 4.4622 | 5.4823 | 5.5506 | 5.3943 |
| CHD4_T1568;T1581 | CHD4 | Q14839-2 | 1572 | PKTPTPSTPGDTQ | 6.3349 | 6.1655 | 6.5677 | 6.7400 | 6.8210 | 6.0665 | 6.2881 | 6.2268 | 6.5601 | 5.9811 | 6.3450 | 5.6866 | 5.6866 | 6.0468 | 5.7395 | 5.8617 |
| CHD4_T1570 | CHD4 | Q14839-2 | 1573 | KTPTPSTPGDTQP | 5.9075 | 6.2129 | 5.7757 | 7.6506 | 5.8246 | 7.2803 | 7.0610 | 6.5499 | 6.4631 | 6.2059 | 6.2104 | 6.3711 | 4.8023 | 6.0168 | 5.0974 | 6.5704 |
| CHD4_T1573 | CHD4 | Q14839-2 | 515;531 | NA | 5.7475 | 6.0690 | 6.2280 | 6.5690 | 5.9037 | 5.5006 | 6.2235 | 5.5128 | 6.1580 | 6.1996 | 5.7839 | 5.9316 | 6.7501 | 7.0534 | 6.8945 | 7.4747 |
| CHD4_T1573;T1581 | CHD4 | Q14839-2 | 1559;1563 | NA | 5.6651 | 6.2669 | 6.0139 | 6.3029 | 6.0668 | 5.2687 | 5.3101 | 5.1887 | 5.3269 | 6.1175 | 5.4228 | 5.8310 | 7.3278 | 7.9771 | 7.9011 | 8.0128 |
| CHD4_T1581 | CHD4 | Q14839-2 | 103;105;108 | NA | 6.0035 | 6.2188 | 5.5019 | 6.8157 | 5.7523 | 5.3138 | 6.0240 | 5.7705 | 6.1615 | 6.2673 | 6.6917 | 6.5117 | 5.7284 | 7.5330 | 6.8418 | 6.8641 |
| CHD4_T1681 | CHD4 | Q14839-2 | 1568;1581 | NA | 6.0957 | 6.7504 | 6.2080 | 6.7112 | 6.0790 | 6.4126 | 6.5017 | 5.9922 | 6.2999 | 6.6097 | 6.5218 | 6.3303 | 5.7457 | 5.5206 | 6.1397 | 6.0814 |
| CHD4_T1707 | CHD4 | Q14839-2 | 1559;1565 | NA | 5.5873 | 5.8660 | 5.4194 | 5.5533 | 6.0616 | 5.3569 | 5.3784 | 5.4550 | 5.5538 | 5.9177 | 5.3359 | 4.9980 | 7.8246 | 8.4610 | 8.9811 | 8.2500 |
| CHD4_T367 | CHD4 | Q14839-2 | 529;531 | NA | 5.4784 | 6.5160 | 6.2182 | 6.2630 | 5.6558 | 5.6677 | 5.8293 | 5.9544 | 6.9558 | 6.0596 | 5.6245 | 5.9810 | 6.5677 | 6.8121 | 6.8894 | 7.5272 |
| CHD4_T529;S531 | CHD4 | Q14839-2 | 1573;1581 | NA | 6.3836 | 5.8634 | 4.8143 | 6.7986 | 6.4144 | 5.8871 | 5.6299 | 5.8376 | 6.7250 | 7.5001 | 4.9526 | 5.6870 | 5.5003 | 6.9872 | 7.8014 | 7.2176 |
| CHD4_T703 | CHD4 | Q14839-2 | 515;529;531 | NA | 5.2803 | 5.6184 | 6.2469 | 6.5886 | 6.1209 | 6.3275 | 7.2393 | 5.3370 | 6.0690 | 6.5651 | 6.2020 | 6.7009 | 5.5128 | 6.5739 | 6.4289 | 7.1885 |
| CHD6_S1673 | CHD6 | Q8TD26 | 21 | LKVLNHSPMSDAS | 5.4825 | 5.7526 | 6.2705 | 6.6177 | 5.6578 | 5.7201 | 5.5695 | 5.6606 | 6.5173 | 6.7391 | 6.3495 | 6.5945 | 6.4643 | 7.5498 | 6.4583 | 6.5959 |
| CHD6_S21 | CHD6 | Q8TD26 | 2357 | KSIPSKSLLDWLR | 8.1762 | 9.0842 | 7.0368 | 7.9688 | 6.1049 | 7.5192 | 6.6702 | 6.0088 | 7.0019 | 7.8117 | 7.6000 | 6.9757 | 2.3747 | 3.8218 | 2.1565 | 3.6887 |
| CHD6_S2357 | CHD6 | Q8TD26 | 1673 | SRDDHLSLPDVTC | 6.1278 | 6.9454 | 6.5594 | 7.3505 | 5.0820 | 6.2044 | 6.3782 | 6.4048 | 5.8568 | 8.3492 | 7.0205 | 7.0765 | 3.9772 | 5.3162 | 5.7905 | 5.5607 |
| CHD7_S1577;S1581 | CHD7 | Q9P2D1 | 2559 | PTRNIPSPGQLDP | 6.0829 | 6.4449 | 6.1968 | 6.5851 | 6.2114 | 5.9977 | 6.1936 | 6.0145 | 6.2433 | 6.1860 | 6.3296 | 6.3470 | 5.9600 | 6.5052 | 6.1278 | 6.5742 |
| CHD7_S1874 | CHD7 | Q9P2D1 | 1874 | IDEFANSPSEDKE | 5.8642 | 6.0163 | 6.0888 | 6.8628 | 5.7207 | 6.0276 | 6.1393 | 5.6038 | 5.9678 | 6.3469 | 6.2363 | 6.2704 | 6.3211 | 7.0341 | 6.6248 | 6.8749 |
| CHD7_S2229 | CHD7 | Q9P2D1 | 557 | VPVHQHSPSEPFL | 6.1246 | 6.3416 | 6.4346 | 7.5358 | 6.1445 | 6.3292 | 6.2794 | 6.5300 | 6.3170 | 6.2030 | 6.1972 | 6.2093 | 5.3007 | 5.9504 | 6.0159 | 6.0868 |
| CHD7_S2356 | CHD7 | Q9P2D1 | 2356 | TPTTVDSPLQKRS | 6.2287 | 6.9086 | 6.3666 | 7.0187 | 6.4481 | 6.5804 | 6.3639 | 6.4220 | 6.9057 | 6.7286 | 6.6627 | 6.7525 | 4.5171 | 5.6239 | 4.8505 | 5.6220 |
| CHD7_S2559 | CHD7 | Q9P2D1 | 2551 | FEEDIETPPTRNI | 5.8381 | 6.0662 | 6.1676 | 6.6018 | 5.3136 | 6.0524 | 5.7412 | 5.8864 | 5.8542 | 6.1860 | 5.7629 | 5.9656 | 6.7909 | 7.4024 | 7.2787 | 7.0920 |
| CHD7_S557 | CHD7 | Q9P2D1 | 2229 | VGADTGSKSISEK | 5.9089 | 6.2725 | 5.7125 | 5.8846 | 5.5426 | 4.9816 | 5.5630 | 5.4755 | 5.6964 | 5.8294 | 5.1917 | 5.5236 | 7.1983 | 8.5852 | 8.5017 | 8.1325 |
| CHD7_S637 | CHD7 | Q9P2D1 | 637 | RNSLDGSQEEKKK | 6.4250 | 6.5413 | 6.3739 | 6.8021 | 5.7624 | 5.7186 | 6.4721 | 5.5407 | 6.5518 | 6.3431 | 5.6850 | 6.7149 | 5.5750 | 6.8254 | 6.2274 | 6.4413 |
| CHD7_T2551 | CHD7 | Q9P2D1 | 1577;1581 | NA | 5.5342 | 6.4842 | 6.7403 | 7.2097 | 6.7642 | 5.7329 | 6.2807 | 5.5508 | 6.6792 | 6.4529 | 5.6023 | 6.7305 | 5.9195 | 5.4181 | 6.4992 | 6.4012 |
| CHD8_S1978 | CHD8 | Q9HCK8 | 1995 | YTSRTASPLPLRP | 5.8209 | 6.2046 | 5.8294 | 5.8107 | 5.5273 | 5.9716 | 5.8931 | 5.7211 | 5.7561 | 6.1916 | 5.8488 | 5.6868 | 7.3799 | 7.6345 | 7.4911 | 7.2324 |
| CHD8_S1995 | CHD8 | Q9HCK8 | 2046 | DYEMRVSPSDTTP | 6.8721 | 7.6571 | 6.4973 | 6.7163 | 6.3077 | 6.3446 | 6.9942 | 6.3387 | 6.6750 | 6.8243 | 6.7550 | 6.6619 | 4.4834 | 5.2359 | 4.5885 | 5.0480 |
| CHD8_S2008 | CHD8 | Q9HCK8 | 1978 | APAPSLSRCSTPL | 5.9843 | 6.6108 | 6.1953 | 5.9928 | 6.0593 | 6.1788 | 5.8454 | 6.3537 | 6.2972 | 6.2392 | 6.4293 | 5.9595 | 6.2585 | 6.6576 | 6.5948 | 6.3435 |
| CHD8_S2046 | CHD8 | Q9HCK8 | 2519 | APGYPSSPVTTAS | 6.3776 | 6.0258 | 5.6797 | 6.2858 | 6.6580 | 6.5368 | 5.8433 | 6.2961 | 6.2212 | 6.1610 | 6.0341 | 6.1675 | 6.1998 | 6.9470 | 6.1521 | 6.4141 |
| CHD8_S2048 | CHD8 | Q9HCK8 | 2008 | DAPVEKSPEETAT | 5.8729 | 6.2524 | 5.9278 | 6.5217 | 6.4871 | 6.2414 | 5.8366 | 6.3718 | 6.5094 | 6.3773 | 5.7480 | 6.5656 | 6.2980 | 6.1582 | 6.2524 | 6.5795 |
| CHD8_S2519 | CHD8 | Q9HCK8 | 2048 | EMRVSPSDTTPLV | 5.4371 | 5.3634 | 6.4446 | 6.6699 | 5.9892 | 6.1299 | 6.4474 | 6.5579 | 6.0046 | 6.0742 | 6.0495 | 6.0881 | 6.1646 | 6.8518 | 6.8780 | 6.8497 |
| CHD8_T1990 | CHD8 | Q9HCK8 | 1990 | LLHQQYTSRTASP | 5.9560 | 6.6879 | 6.1439 | 6.4679 | 5.7703 | 5.4905 | 5.1828 | 5.4236 | 6.2215 | 5.2997 | 5.5374 | 5.9982 | 8.0524 | 7.7744 | 7.3005 | 6.6930 |
| CHD9_S2079 | CHD9 | Q3L8U1 | 2079 | IKSEPVSPKNGVL | 6.1916 | 6.6579 | 6.6753 | 6.5797 | 6.0797 | 6.2268 | 6.6242 | 6.1295 | 6.4230 | 6.5632 | 6.6229 | 6.4893 | 5.4384 | 5.6503 | 5.7021 | 5.9460 |
| CHD9_S2303 | CHD9 | Q3L8U1 | 2113 | ARTEPLTPNPASK | 5.6180 | 6.0101 | 6.3983 | 6.4242 | 6.5081 | 6.6376 | 6.7908 | 6.1872 | 7.2279 | 6.4907 | 6.1197 | 6.6263 | 5.9872 | 6.0897 | 5.3051 | 5.5790 |
| CHD9_S550 | CHD9 | Q3L8U1 | 550 | NIPRVMSPENFPT | 5.8025 | 6.2763 | 6.2652 | 6.3844 | 5.8944 | 6.3492 | 6.3551 | 6.1239 | 6.1259 | 6.7417 | 6.9911 | 7.0730 | 5.4297 | 6.2237 | 6.1947 | 5.7693 |
| CHD9_T2113 | CHD9 | Q3L8U1 | 2303 | TTKLLDSPGAATE | 5.8590 | 6.0532 | 6.2816 | 5.9848 | 5.8889 | 6.3508 | 6.0894 | 6.2753 | 5.6801 | 6.3850 | 6.3323 | 5.9934 | 6.2985 | 6.7766 | 6.9299 | 6.8214 |
| CHEK1_S280 | CHEK1 | O14757 | 301 | QSNLDFSPVNSAS | 6.8366 | 7.2066 | 7.6518 | 7.5083 | 6.4300 | 7.7627 | 6.9904 | 7.0571 | 7.6523 | 7.2856 | 7.5210 | 8.0584 | 2.7785 | 3.4468 | 2.4462 | 3.3677 |
| CHEK1_S301 | CHEK1 | O14757 | 280 | KRPRVTSGGVSES | 6.0632 | 5.9658 | 6.5111 | 6.7720 | 6.4661 | 5.7438 | 5.9433 | 6.0272 | 6.8183 | 6.2458 | 6.7275 | 7.1308 | 5.4053 | 6.5747 | 5.6620 | 5.9433 |
| CHEK2_S163 | CHEK2 | O96017-9 | 303 | RKFAIGSAREADP | 5.7442 | 6.1715 | 6.0058 | 4.9739 | 5.5329 | 6.1798 | 5.6661 | 5.6477 | 5.7461 | 5.9699 | 5.9171 | 5.8947 | 7.1773 | 8.0680 | 8.1566 | 7.1485 |
| CHEK2_S303 | CHEK2 | O96017-9 | 422 | KILGETSLMRTLC | 5.8851 | 6.0692 | 6.3968 | 5.2729 | 5.8660 | 6.4906 | 5.8277 | 6.0070 | 5.9151 | 5.8025 | 6.0617 | 5.5882 | 6.8275 | 7.9495 | 7.6002 | 6.4400 |
| CHEK2_S422 | CHEK2 | O96017-9 | 163 | WFGRDKSCEYCFD | 5.8670 | 6.2988 | 6.2310 | 4.5483 | 5.5634 | 6.1857 | 5.8609 | 6.2269 | 6.3427 | 5.7375 | 6.0809 | 5.8362 | 6.2646 | 7.7382 | 8.0055 | 7.2125 |
| CHEK2_S559 | CHEK2 | O96017-9 | 559 | QVLAQPSTSRKRP | 7.0239 | 6.9000 | 5.8043 | 7.7404 | 5.2841 | 6.0139 | 6.4084 | 7.2449 | 6.4865 | 5.3146 | 5.5109 | 5.9968 | 5.8043 | 6.2045 | 6.6523 | 5.6100 |
| CHERP_S186 | CHERP | Q8IWX8 | 695 | AVEAFYSPPSHDR | 5.9430 | 7.0503 | 6.7790 | 6.2834 | 5.7940 | 6.3590 | 6.4096 | 6.4174 | 6.1614 | 6.7225 | 6.9487 | 6.0489 | 5.6654 | 5.7062 | 5.5301 | 6.1810 |
| CHERP_S695 | CHERP | Q8IWX8 | 186 | MFSNAKSPPHCEL | 4.7049 | 6.8221 | 5.7165 | 6.2392 | 6.9606 | 6.7045 | 4.8565 | 6.1321 | 6.2418 | 6.4745 | 6.7934 | 4.8643 | 7.1279 | 6.7907 | 7.7239 | 5.8471 |
| CHERP_S705 | CHERP | Q8IWX8 | 705 | HDRPRNSEGWEQN | 5.7095 | 5.9979 | 6.0511 | 6.8892 | 6.0053 | 6.2975 | 6.4776 | 6.3479 | 6.6643 | 6.1127 | 6.6410 | 7.0881 | 6.5831 | 5.7673 | 5.2101 | 6.1575 |
| CHERP_S815;S817;T819 | CHERP | Q8IWX8 | 815;817;819 | NA | 6.1706 | 5.7606 | 6.5085 | 6.9335 | 6.8750 | 5.9960 | 6.1848 | 6.0129 | 6.0602 | 5.9335 | 6.0464 | 6.1910 | 6.9708 | 5.8514 | 6.2032 | 6.3016 |
| CHFR_S165 | CHFR | Q96EP1 | 165 | PRVPPSSPATQVC | 5.9079 | 6.4285 | 6.2455 | 6.1044 | 6.3555 | 6.1010 | 7.8423 | 7.0084 | 7.1644 | 5.9565 | 5.8600 | 6.3218 | 6.3272 | 5.4825 | 5.6230 | 5.2712 |
| CHFR_S205 | CHFR | Q96EP1 | 205 | GRERSSSCGSGGG | 4.6796 | 5.3847 | 5.9125 | 4.9927 | 5.6302 | 7.1268 | 5.9159 | 5.8008 | 5.8624 | 5.3513 | 6.4357 | 6.8485 | 6.5073 | 8.2988 | 8.0018 | 7.2512 |
| CHMP1A_S258 | CHMP1A | A0A6Q8PFF8 | 258 | AATLRLSPARPAW | 7.1752 | 5.4154 | 6.7550 | 5.6061 | 6.0793 | 4.9518 | 5.8007 | 8.1530 | 5.2605 | 6.5787 | 6.4116 | 7.3005 | 4.3322 | 7.3529 | 5.9148 | 6.9123 |
| CHMP2A_S162 | CHMP2A | M0R1T5 | 162 | GPATAPSPPFPCI | 6.4829 | 6.3586 | 6.2424 | 8.4556 | 6.0127 | 8.2313 | 6.4451 | 5.2047 | 7.0936 | 8.9123 | 8.3529 | 6.4721 | 2.8618 | 5.3695 | 3.9184 | 3.5860 |
| CHMP2B_S199 | CHMP2B | Q9UQN3 | 199 | TSKATISDEEIER | 6.5479 | 6.7752 | 7.0886 | 6.4516 | 6.4073 | 6.9305 | 6.6706 | 7.1494 | 6.5121 | 6.1519 | 6.1224 | 6.5496 | 4.9762 | 5.3531 | 5.0052 | 5.3086 |
| CHMP3_S229 | CHMP3 | Q9Y3E7-3 | 229 | PGAMAASEDEEEE | 6.1464 | 6.1304 | 6.6834 | 6.5553 | 6.2908 | 6.3067 | 6.4618 | 6.2769 | 6.0971 | 6.6647 | 6.6837 | 6.5228 | 5.6150 | 6.0037 | 5.7751 | 5.7864 |
| CHMP4B_S184 | CHMP4B | Q9H444 | 184 | KNLLEISGPETVP | 6.3695 | 6.3802 | 6.9427 | 6.5900 | 6.2790 | 7.4275 | 7.1961 | 7.0442 | 6.4472 | 6.5339 | 7.2113 | 6.9922 | 4.2413 | 5.2498 | 4.1164 | 4.9789 |
| CHMP7_S417 | CHMP7 | Q8WUX9 | 417 | VPNPRISDAELEA | 5.5340 | 5.8611 | 7.6520 | 6.5035 | 6.2409 | 6.4726 | 6.1551 | 6.6565 | 6.1633 | 6.5588 | 6.4355 | 6.4804 | 5.8825 | 5.9934 | 5.6677 | 5.7427 |
| CHORDC1_T199 | CHORDC1 | Q9UHD1 | 199 | SCCRRKTSDFNTF | 6.5694 | 6.3318 | 7.1953 | 6.6245 | 6.2126 | 6.1179 | 6.7413 | 6.4751 | 5.7357 | 6.6058 | 6.8981 | 7.0246 | 4.5440 | 6.0052 | 5.3814 | 5.5373 |
| CHRM3_S457 | CHRM3 | P20309 | 457 | TATLPLSFKEATL | 7.4202 | 7.6135 | 7.3499 | 7.5783 | 5.1420 | 7.0278 | 5.3821 | 6.3953 | 8.0059 | 7.8536 | 5.9561 | 5.8741 | 4.2518 | 6.7936 | 3.9180 | 3.4378 |
| CHST14_T7 | CHST14 | Q8NCH0 | 7 | MFPRPLTPLAAPN | 6.8745 | 6.2805 | 6.5275 | 7.2255 | 6.9706 | 6.0910 | 5.9175 | 6.6516 | 7.8355 | 6.2765 | 6.6135 | 6.8858 | 4.5954 | 5.2114 | 4.6869 | 5.3562 |
| CHTF18_S253 | CHTF18 | Q8WVB6-2 | 899 | SPQVDGSPPGLEG | 5.6931 | 6.3641 | 6.5148 | 6.3488 | 5.7894 | 6.1253 | 6.3520 | 6.3631 | 6.2918 | 6.2420 | 6.3730 | 6.1255 | 6.3049 | 6.5770 | 6.3720 | 6.1631 |
| CHTF18_S893 | CHTF18 | Q8WVB6-2 | 253 | LGVSLASLKKQVD | 6.0189 | 6.4755 | 6.6527 | 6.9743 | 7.0601 | 6.8938 | 6.7619 | 6.4581 | 6.2231 | 6.0997 | 6.6708 | 6.7988 | 5.4219 | 5.7211 | 4.8586 | 4.9110 |
| CHTF18_S899 | CHTF18 | Q8WVB6-2 | 893 | SARVENSPQVDGS | 6.1926 | 6.4104 | 6.1578 | 5.9314 | 5.3622 | 6.1808 | 6.3359 | 6.1578 | 6.0666 | 6.7360 | 6.6888 | 6.6395 | 6.3289 | 6.4684 | 5.9979 | 6.3450 |
| CHTOP_S5 | CHTOP | Q9Y3Y2-3 | 33 | LKNKQPTPVNIRA | 6.2036 | 7.1933 | 5.9240 | 7.2210 | 6.8028 | 6.5796 | 7.2749 | 6.8753 | 7.6375 | 6.9052 | 7.0894 | 6.7808 | 4.6500 | 4.7704 | 3.6618 | 4.4306 |
| CHTOP_T33 | CHTOP | Q9Y3Y2-3 | 5 | xxMAAQSAPKVVL | 4.8787 | 24.4635 | 5.0376 | 5.3648 | 4.7273 | 3.7497 | 3.0172 | 11.4715 | 4.5945 | 5.6759 | 4.8481 | 4.4730 | 1.0217 | 4.0497 | 9.1139 | 3.5127 |
| CIAPIN1_S183 | CIAPIN1 | Q6FI81 | 183 | SITKKSSPSVKPA | 6.1332 | 6.1767 | 7.2702 | 7.5724 | 6.6341 | 10.6883 | 7.3772 | 5.8121 | 7.9251 | 6.3236 | 9.9111 | 7.5935 | 2.9772 | 2.9960 | 1.9554 | 2.6539 |
| CIC_S1082 | CIC | I3L2J0 | 2288 | LQSLATSPRAILG | 5.8097 | 6.4399 | 6.8024 | 6.4673 | 5.7666 | 5.9830 | 5.9879 | 6.0698 | 6.3793 | 7.0581 | 6.7825 | 6.9567 | 5.7723 | 5.8290 | 5.8502 | 6.0451 |
| CIC_S1100 | CIC | I3L2J0 | 1082 | GKRRTQSLSALPK | 6.4109 | 6.1284 | 5.6633 | 6.0819 | 7.0762 | 6.2731 | 6.3359 | 6.1546 | 6.1956 | 5.7460 | 5.8241 | 5.5899 | 7.3511 | 6.4283 | 6.3882 | 6.3525 |
| CIC_S1265 | CIC | I3L2J0 | 428 | LSPPPKSPAFVGP | 5.8261 | 5.8462 | 5.6154 | 6.1130 | 5.8163 | 5.9001 | 6.2351 | 6.3659 | 6.2713 | 5.8294 | 5.6094 | 5.5954 | 7.3766 | 6.9118 | 7.2602 | 7.4277 |
| CIC_S1405 | CIC | I3L2J0 | 472 | PAARARTPLTAAQ | 5.4789 | 6.1334 | 5.2263 | 5.7806 | 5.6941 | 5.8058 | 5.7159 | 5.8999 | 5.6612 | 5.7289 | 5.3784 | 5.5756 | 8.2173 | 7.9918 | 7.7672 | 7.9444 |
| CIC_S1648 | CIC | I3L2J0 | 1813 | PKAQSVSPVQAPP | 6.0433 | 6.1334 | 6.0828 | 6.5565 | 6.1014 | 6.6461 | 6.1334 | 6.3000 | 5.9924 | 6.4079 | 6.4248 | 6.3645 | 6.4215 | 6.2268 | 6.0978 | 6.0671 |
| CIC_S18 | CIC | I3L2J0 | 1405 | FGRKVFSPVIRSS | 6.3098 | 6.3423 | 5.9173 | 6.1685 | 6.4421 | 5.9833 | 6.2114 | 6.0196 | 5.9524 | 5.8325 | 6.0333 | 5.9267 | 7.0587 | 6.5993 | 6.6422 | 6.5605 |
| CIC_S1813 | CIC | I3L2J0 | 780 | SRSRQPSPLLLLP | 5.9597 | 6.2583 | 5.8940 | 6.2516 | 6.3748 | 6.4280 | 6.2948 | 6.2647 | 6.5693 | 6.4248 | 6.3048 | 5.8871 | 6.2351 | 6.4421 | 6.1239 | 6.2869 |
| CIC_S2080 | CIC | I3L2J0 | 2153 | TPTARSSPPLPPP | 6.0546 | 6.5750 | 6.2876 | 6.1673 | 6.4981 | 6.2990 | 6.2386 | 5.9554 | 6.3425 | 6.2755 | 6.7299 | 6.2441 | 6.6164 | 5.9944 | 5.9563 | 5.7653 |
| CIC_S2153 | CIC | I3L2J0 | 2080 | PKAQRPSPKAPQK | 6.0208 | 6.1492 | 5.7199 | 6.0861 | 6.6435 | 7.0178 | 6.6498 | 6.5117 | 6.5796 | 6.5086 | 6.3293 | 6.1155 | 6.0435 | 6.0645 | 5.5467 | 6.0134 |
| CIC_S2279 | CIC | I3L2J0 | 664 | VRSRHLSASTPKA | 6.5450 | 6.5992 | 6.3819 | 6.5613 | 5.8898 | 6.5666 | 6.6052 | 6.3890 | 6.6137 | 6.6887 | 6.8409 | 6.9433 | 5.4050 | 5.6963 | 4.8436 | 5.4303 |
| CIC_S2279;S2288 | CIC | I3L2J0 | 768 | AGFRAVSPAVPFS | 6.4333 | 6.2941 | 5.5905 | 5.6794 | 6.0826 | 6.2990 | 6.4148 | 6.3268 | 6.1082 | 5.8049 | 5.3801 | 5.3977 | 7.7730 | 6.7200 | 6.8785 | 6.8172 |
| CIC_S2288 | CIC | I3L2J0 | 2279 | PEEVLPSPTLQSL | 6.2370 | 6.6915 | 6.7863 | 6.7830 | 6.1436 | 6.1094 | 6.1350 | 5.9469 | 6.1147 | 6.5901 | 6.5137 | 7.4661 | 5.6240 | 5.7338 | 5.6249 | 5.5000 |
| CIC_S2295 | CIC | I3L2J0 | 2135 | ELEGQPTPPAPPP | 5.5436 | 6.2113 | 5.8443 | 6.5097 | 6.4525 | 5.7199 | 6.2783 | 6.3284 | 5.9150 | 6.0734 | 5.9056 | 6.3005 | 6.7636 | 6.6906 | 6.6590 | 6.8043 |
| CIC_S301 | CIC | I3L2J0 | 1492 | GAGGPATPSKATR | 7.2705 | 7.0554 | 6.1489 | 6.8459 | 5.8711 | 6.4359 | 6.9940 | 6.3614 | 7.1106 | 7.2452 | 6.6661 | 6.7606 | 4.9410 | 5.0939 | 4.1121 | 5.0875 |
| CIC_S428 | CIC | I3L2J0 | 1648 | KSAAATSPAPHLV | 5.3398 | 5.4103 | 5.3032 | 6.2025 | 6.0367 | 6.7801 | 6.3714 | 6.7679 | 6.5474 | 5.9698 | 5.8717 | 5.7906 | 6.9399 | 7.0718 | 6.6123 | 6.9848 |
| CIC_S428;S442 | CIC | I3L2J0 | 809 | PAVQRDSPVIVRN | 5.3847 | 5.8354 | 5.4499 | 6.0508 | 5.5012 | 6.0141 | 5.7344 | 6.5352 | 5.9162 | 5.8571 | 5.5856 | 5.6899 | 7.5808 | 7.2497 | 7.5769 | 8.0380 |
| CIC_S565 | CIC | I3L2J0 | 674 | PKAGVLTPPDLGP | 6.1694 | 6.4176 | 6.7156 | 7.1463 | 6.0955 | 6.2877 | 6.7346 | 6.4790 | 6.4530 | 6.8307 | 6.7998 | 6.7301 | 5.3356 | 5.4044 | 4.8238 | 5.5770 |
| CIC_S645 | CIC | I3L2J0 | 423 | PPPQLLSPPPKSP | 5.4885 | 6.0478 | 6.4047 | 6.2042 | 5.9404 | 6.1824 | 6.0745 | 6.4564 | 6.1690 | 5.7956 | 5.7354 | 6.4043 | 6.6658 | 7.2950 | 6.7603 | 6.3757 |
| CIC_S650 | CIC | I3L2J0 | 2198 | NRGEPPTPPSPAP | 5.6903 | 5.7987 | 6.3518 | 6.5995 | 6.2151 | 6.4316 | 6.2371 | 5.8957 | 6.2560 | 6.5822 | 6.1653 | 6.0607 | 7.0129 | 6.4151 | 6.1700 | 6.1181 |
| CIC_S664 | CIC | I3L2J0 | 1100 | SEKDGRSPNKREK | 6.4269 | 8.0795 | 7.1626 | 6.5289 | 6.4636 | 7.0866 | 6.4179 | 6.2742 | 7.1152 | 6.2677 | 4.7530 | 4.8706 | 6.5379 | 5.7157 | 5.4985 | 4.8012 |
| CIC_S768 | CIC | I3L2J0 | 650 | FSPINASPVIQRT | 7.1286 | 6.0368 | 7.2049 | 7.9661 | 5.4127 | 5.9205 | 7.0346 | 6.3643 | 7.1479 | 7.1733 | 6.6547 | 7.8660 | 4.6992 | 4.0929 | 4.1337 | 5.1638 |
| CIC_S776;S780 | CIC | I3L2J0 | 645 | FRPANFSPINASP | 5.6245 | 4.4890 | 5.7316 | 5.5707 | 5.4107 | 5.5227 | 5.9172 | 6.2724 | 5.6986 | 4.9978 | 5.1117 | 5.3768 | 9.2145 | 8.3184 | 8.1193 | 8.6242 |
| CIC_S780 | CIC | I3L2J0 | 846 | GRGCRETPVPPGV | 5.4882 | 5.3377 | 5.1578 | 5.6520 | 4.9859 | 5.6483 | 6.0285 | 5.1084 | 5.2342 | 4.8905 | 4.9718 | 5.3806 | 9.1598 | 7.9680 | 8.4889 | 10.4996 |
| CIC_S809 | CIC | I3L2J0 | 2201 | EPPTPPSPAPAPA | 6.4087 | 5.6082 | 6.4253 | 6.0595 | 6.6517 | 6.7000 | 6.6765 | 5.9684 | 6.9374 | 6.2514 | 5.1747 | 5.7296 | 6.4060 | 5.8207 | 6.0526 | 7.1293 |
| CIC_T143 | CIC | I3L2J0 | 2295 | PRAILGSYRKKRK | 6.9591 | 6.9993 | 6.2077 | 6.9052 | 6.3527 | 6.0758 | 6.5490 | 6.9436 | 7.0192 | 6.3415 | 6.4004 | 5.7989 | 5.9011 | 6.0721 | 4.5483 | 4.9262 |
| CIC_T1492 | CIC | I3L2J0 | 143 | PEERVRTPEEASG | 7.0101 | 6.4268 | 6.9929 | 6.5843 | 4.9016 | 5.8498 | 7.4124 | 6.9337 | 6.8604 | 5.6470 | 5.4864 | 7.7353 | 5.6782 | 5.3866 | 5.7843 | 5.3102 |
| CIC_T2135 | CIC | I3L2J0 | 301 | AYKVRLSPGPSSQ | 5.2475 | 6.4733 | 7.0511 | 6.3156 | 6.1224 | 6.0743 | 6.5771 | 6.5175 | 6.8617 | 7.6078 | 5.6926 | 6.2224 | 5.4061 | 6.6319 | 5.9397 | 5.2590 |
| CIC_T2198 | CIC | I3L2J0 | 1265 | ARPRAFSHSGVHS | 5.8981 | 5.4611 | 5.8187 | 6.4242 | 6.7355 | 6.3529 | 6.1344 | 6.6160 | 6.4046 | 5.3898 | 5.8606 | 6.2182 | 7.4150 | 7.7235 | 5.8580 | 5.6894 |
| CIC_T2198;S2201 | CIC | I3L2J0 | 755 | VDSVSHTPTPSTP | 5.8980 | 5.1856 | 5.8127 | 5.8253 | 6.4191 | 6.8440 | 5.9181 | 5.7012 | 6.4077 | 4.7699 | 5.3083 | 5.8287 | 7.7333 | 7.3937 | 7.3697 | 7.5845 |
| CIC_T2201 | CIC | I3L2J0 | 18 | SGPGSGSKSPPAT | 6.0921 | 7.2951 | 6.3831 | 6.4026 | 6.4610 | 6.4733 | 6.8022 | 6.4261 | 6.4897 | 7.2418 | 7.2582 | 7.4181 | 4.4176 | 5.1923 | 4.4607 | 5.1862 |
| CIC_T2287 | CIC | I3L2J0 | 565 | DWGDETSRDSEAS | 4.7815 | 6.9656 | 7.0463 | 5.5784 | 4.8031 | 7.4044 | 7.2254 | 5.3836 | 7.1034 | 7.6819 | 7.0030 | 6.1687 | 6.2218 | 6.4265 | 4.9055 | 5.3010 |
| CIC_T410;S423 | CIC | I3L2J0 | 2287 | TLQSLATSPRAIL | 6.0823 | 6.1922 | 6.5930 | 6.5908 | 5.6199 | 5.6804 | 6.4468 | 5.6765 | 6.5518 | 6.8129 | 7.7492 | 6.3115 | 5.3395 | 6.3115 | 5.8613 | 6.1801 |
| CIC_T423 | CIC | I3L2J0 | 2279;2288 | NA | 6.1602 | 6.1055 | 6.1638 | 6.4084 | 6.2812 | 6.3637 | 6.0675 | 6.2410 | 6.4106 | 6.0913 | 6.0184 | 6.1050 | 6.6793 | 6.3562 | 6.1736 | 6.3745 |
| CIC_T472 | CIC | I3L2J0 | 428;442 | NA | 5.3992 | 5.6132 | 5.7873 | 5.3119 | 6.0727 | 7.6515 | 7.5112 | 6.2809 | 6.2481 | 5.8887 | 5.7274 | 5.4157 | 7.3047 | 7.6553 | 5.6786 | 6.4536 |
| CIC_T674 | CIC | I3L2J0 | 776;780 | NA | 5.6353 | 5.6518 | 6.0150 | 6.1087 | 5.8335 | 5.6112 | 5.7324 | 6.2073 | 6.3660 | 6.1386 | 6.0996 | 6.2419 | 6.9712 | 6.9979 | 6.7213 | 7.6683 |
| CIC_T755 | CIC | I3L2J0 | 410;423 | NA | 5.6812 | 6.5467 | 6.3684 | 5.7974 | 6.0910 | 5.2333 | 6.0880 | 6.5498 | 5.9938 | 6.1745 | 5.5893 | 6.0439 | 6.2595 | 6.8996 | 7.3429 | 7.3408 |
| CIC_T846 | CIC | I3L2J0 | 2198;2201 | NA | 5.7239 | 6.5935 | 6.5151 | 5.2850 | 6.7446 | 6.6236 | 6.6223 | 5.6048 | 6.3786 | 6.1980 | 5.4624 | 5.8501 | 5.4511 | 7.7381 | 6.7288 | 6.4800 |
| CILK1_T157;Y159 | CILK1 | Q9UPZ9 | 157;159 | NA | 6.1589 | 6.1707 | 6.8150 | 6.5640 | 6.1221 | 6.1094 | 6.0826 | 6.3712 | 6.3737 | 6.4607 | 6.1227 | 6.4406 | 5.8572 | 6.4608 | 6.0696 | 5.8207 |
| CINP_S85 | CINP | Q9BW66-3 | 85 | IELDSSSPASKEN | 6.5925 | 5.6380 | 6.6278 | 5.6939 | 5.9438 | 6.1136 | 5.4682 | 7.2710 | 6.2329 | 7.5313 | 7.0159 | 7.8332 | 4.7278 | 5.9396 | 5.3355 | 6.0352 |
| CINP_T25 | CINP | Q9BW66-3 | 25 | KTLGTVTPRKPVL | 6.6444 | 6.3708 | 6.8009 | 6.1411 | 5.8632 | 6.3313 | 7.0791 | 8.2171 | 7.2849 | 8.3443 | 7.3303 | 9.4028 | 2.5021 | 4.5531 | 2.9320 | 4.2025 |
| CIP2A_T585 | CIP2A | Q8TCG1 | 585 | TSIKCLTPHLKDG | 6.2506 | 6.4341 | 6.9596 | 7.0583 | 5.0800 | 6.0806 | 5.8592 | 6.5702 | 5.9815 | 5.9346 | 7.1471 | 7.1163 | 5.5262 | 6.0518 | 5.4291 | 6.5209 |
| CIR1_S202 | CIR1 | Q86X95 | 202 | SQEYVASEGEEDP | 6.2030 | 5.8925 | 6.4821 | 6.3075 | 5.7883 | 5.5981 | 6.1977 | 6.3827 | 6.1657 | 5.9775 | 6.0555 | 6.3323 | 6.1330 | 6.9002 | 6.6633 | 6.9207 |
| CIR1_S305 | CIR1 | Q86X95 | 353 | DSKWSHSDSDKKS | 6.9502 | 7.4648 | 7.2755 | 8.2866 | 6.2031 | 6.4045 | 6.7328 | 6.8308 | 5.0707 | 5.6060 | 5.5749 | 5.9308 | 4.2180 | 5.3114 | 5.6761 | 6.4641 |
| CIR1_S353 | CIR1 | Q86X95 | 305 | CSGHNNSDSEEKD | 3.6803 | 6.5047 | 7.8629 | 7.0666 | 5.1911 | 6.1363 | 6.7689 | 4.7148 | 6.5121 | 5.1539 | 5.7939 | 6.3893 | 7.6434 | 7.1001 | 6.7577 | 6.7242 |
| CISD2_T106 | CISD2 | Q8N5K1 | 106 | RCWRSKTFPACDG | 6.1596 | 6.0148 | 5.5298 | 5.7634 | 5.3970 | 4.7920 | 5.3526 | 4.9625 | 5.1179 | 5.2722 | 5.4745 | 5.2773 | 9.4712 | 8.9344 | 8.1221 | 8.3589 |
| CISD3_S83 | CISD3 | P0C7P0 | 83 | FQRTGLSPLKFKA | 6.7732 | 6.6225 | 6.3478 | 6.7789 | 6.0893 | 5.5632 | 6.2902 | 5.9749 | 6.4382 | 6.6431 | 6.4822 | 6.2844 | 6.0249 | 5.8519 | 6.0355 | 5.7997 |

| | | | | | | | | | | | | | | | | | | | | | |
|---|---|---|---|---|---|---|---|---|---|---|---|---|---|---|---|---|---|---|---|---|---|
| CIT_S1364 | CIT | O14578-4 | | 1474 | CRDKMNSPGLQTK | 7.1195 | 7.4631 | 6.8695 | 7.4057 | 6.4532 | 6.9403 | 7.1904 | 7.0320 | 7.2655 | 7.4319 | 7.6496 | 7.4301 | 3.0510 | 3.9091 | 3.1069 | 3.6821 |
| CIT_S1474 | CIT | O14578-4 | | 2013 | ELRRDKSPGRPLE | 6.3916 | 7.3456 | 6.9163 | 6.3296 | 5.7207 | 6.6063 | 6.3266 | 6.5641 | 6.1342 | 6.4756 | 6.4992 | 6.5680 | 5.2296 | 5.6075 | 5.5844 | 5.7007 |
| CIT_S1624 | CIT | O14578-4 | | 440 | VVSGLDSPAKTSS | 6.5211 | 6.6991 | 6.5140 | 6.9575 | 6.2251 | 6.2393 | 6.4937 | 5.8843 | 6.9779 | 6.7487 | 6.7806 | 6.7569 | 5.3773 | 5.3504 | 5.1925 | 5.2818 |
| CIT_S1981 | CIT | O14578-4 | | 1624 | AKLLGNSLLKLEG | 7.5222 | 5.5889 | 5.6917 | 7.5235 | 6.2811 | 7.3816 | 6.5055 | 6.2432 | 6.3244 | 5.9472 | 6.1824 | 6.4568 | 4.8088 | 5.4226 | 6.5312 | 5.5889 |
| CIT_S2013 | CIT | O14578-4 | | 1364 | MSAIVRSPEHQPS | 7.0707 | 7.2035 | 6.4261 | 6.6135 | 6.2447 | 6.3998 | 6.5842 | 6.4705 | 6.0785 | 7.0403 | 6.3675 | 6.6064 | 5.5486 | 5.6683 | 5.0268 | 4.6505 |
| CIT_S2068 | CIT | O14578-4 | | 2068 | KVWDQSSVxxxxx | 7.0283 | 7.2085 | 6.5321 | 7.7383 | 6.0764 | 5.6927 | 7.0359 | 6.6782 | 6.4926 | 6.3210 | 6.4195 | 6.2214 | 4.6493 | 5.6120 | 5.1180 | 5.1759 |
| CIT_S400 | CIT | O14578-4 | | 1981 | ITKRVASSPAPPE | 6.7129 | 6.5542 | 6.4653 | 6.5436 | 6.0242 | 5.9816 | 6.8217 | 6.4414 | 6.8308 | 6.2636 | 6.7029 | 6.5251 | 5.2228 | 5.7697 | 5.1947 | 5.9456 |
| CIT_S440 | CIT | O14578-4 | | 480 | RLHRRVSEVEAVL | 5.4491 | 5.6677 | 5.9925 | 5.7704 | 5.3646 | 5.8767 | 5.2592 | 6.3371 | 7.6419 | 7.3672 | 6.9946 | 7.5422 | 6.1765 | 6.0439 | 5.1929 | 6.3234 |
| CIT_S480 | CIT | O14578-4 | | 400 | NSWVSSSPCQLSP | 6.3117 | 6.6788 | 6.4126 | 6.5709 | 6.1630 | 6.5735 | 6.4979 | 6.2186 | 5.4305 | 5.9525 | 6.9528 | 7.1894 | 5.9020 | 5.9307 | 5.2061 | 6.0090 |
| CIZ1_S350 | CIZ1 | Q9ULV3 | | 838 | KAAKNPSPTTRPV | 7.0620 | 7.1963 | 6.9188 | 7.6944 | 6.4398 | 6.0592 | 6.3085 | 6.4254 | 6.6838 | 6.5360 | 6.7255 | 6.9821 | 4.8263 | 4.6344 | 4.6050 | 4.9026 |
| CIZ1_S783 | CIZ1 | Q9ULV3 | | 567 | FSTVPLTPVPRPS | 6.8796 | 6.0692 | 5.6532 | 7.2128 | 6.1978 | 5.8448 | 6.4868 | 6.0880 | 6.6211 | 5.2819 | 5.9661 | 6.1582 | 6.5508 | 6.3378 | 5.5642 | 7.0878 |
| CIZ1_S838 | CIZ1 | Q9ULV3 | | 244 | IAKEKRTPAPEPE | 5.9002 | 6.2572 | 6.2677 | 7.3580 | 6.0363 | 5.9024 | 6.0526 | 6.1339 | 6.3072 | 6.9215 | 6.5086 | 6.6641 | 5.5820 | 5.6459 | 6.0522 | 6.4103 |
| CIZ1_T244 | CIZ1 | Q9ULV3 | | 350 | QAQTQTSPEHLVL | 6.6047 | 7.0194 | 6.4490 | 7.1133 | 6.3705 | 6.4107 | 6.1267 | 5.8691 | 6.7843 | 6.6885 | 6.2804 | 6.1147 | 5.8509 | 4.9007 | 5.8461 | 5.5712 |
| CIZ1_T295 | CIZ1 | Q9ULV3 | | 783 | KGSETYSPNTAYG | 6.4676 | 6.9215 | 6.6929 | 7.0412 | 6.7187 | 6.8346 | 6.1589 | 6.7163 | 6.5206 | 6.5464 | 6.5698 | 6.6609 | 4.4824 | 5.3037 | 4.7506 | 5.3638 |
| CIZ1_T567 | CIZ1 | Q9ULV3 | | 295 | VPKQTQTPDLLPE | 5.8185 | 5.9710 | 6.3799 | 6.3824 | 7.0460 | 5.9048 | 6.1808 | 6.2489 | 7.2753 | 6.1342 | 6.9407 | 6.6496 | 5.6326 | 5.8393 | 6.3333 | 5.2627 |
| CKAP2_S178 | CKAP2 | Q8WWK9 | | 614 | QFDGTNSAFKELK | 8.1756 | 8.2235 | 7.3916 | 7.5506 | 4.9906 | 7.0388 | 8.0540 | 6.2868 | 7.2457 | 7.0303 | 7.1825 | 7.6043 | 2.7822 | 4.2132 | 2.4304 | 3.7999 |
| CKAP2_S328 | CKAP2 | Q8WWK9 | | 597 | KYNVSTTPYLQSV | 6.9061 | 7.5297 | 7.2034 | 6.9404 | 5.4457 | 7.2679 | 7.5175 | 7.3877 | 7.6215 | 7.7791 | 7.5590 | 7.9121 | 2.7026 | 4.0972 | 2.4849 | 3.6451 |
| CKAP2_S595;T597 | CKAP2 | Q8WWK9 | | 178 | KKPVLGSYRGQIV | 10.2124 | 6.5143 | 8.2937 | 7.6854 | 4.4480 | 5.0263 | 6.6791 | 5.9647 | 9.3701 | 6.9605 | 7.9085 | 8.1730 | 3.6494 | 4.4836 | 2.1778 | 2.4533 |
| CKAP2_S614 | CKAP2 | Q8WWK9 | | 328 | VIARPASLSNDKL | 6.8195 | 5.0453 | 7.1441 | 7.7033 | 5.7229 | 6.7930 | 7.7865 | 6.7995 | 6.8961 | 7.2251 | 6.9473 | 6.9208 | 4.3942 | 5.2268 | 4.0173 | 4.5583 |
| CKAP2_S651 | CKAP2 | Q8WWK9 | | 211 | TKKLSATIPKATK | 7.3394 | 8.2424 | 6.8905 | 6.9422 | 5.4382 | 7.0018 | 6.6093 | 6.6025 | 7.7447 | 6.8001 | 6.9595 | 7.0635 | 3.8928 | 4.9370 | 4.0434 | 4.4928 |
| CKAP2_T211 | CKAP2 | Q8WWK9 | | 651 | HYPCVSSLEQLTE | 6.3350 | 5.6480 | 7.2351 | 6.9769 | 6.5883 | 6.0061 | 6.6748 | 5.8989 | 6.2387 | 7.3886 | 7.0097 | 6.2022 | 5.1206 | 5.8063 | 4.5493 | 6.3216 |
| CKAP2_T597 | CKAP2 | Q8WWK9 | | 683 | YYEADTTxxxxxx | 5.7556 | 6.5221 | 6.2674 | 6.8330 | 5.8302 | 7.0419 | 7.2266 | 7.7407 | 6.9788 | 7.2369 | 7.1417 | 6.8181 | 4.4452 | 5.7854 | 3.5284 | 4.8480 |
| CKAP2_T597;S602 | CKAP2 | Q8WWK9 | | 682 | VYYEADTTxxxxx | 7.2272 | 7.5400 | 7.5139 | 6.9986 | 4.6690 | 7.3532 | 7.0405 | 6.2934 | 6.4237 | 9.1861 | 5.4421 | 6.1414 | 3.2662 | 4.5083 | 4.0740 | 3.6223 |
| CKAP2_T682 | CKAP2 | Q8WWK9 | 597;602 | | NA | 8.4920 | 7.8965 | 7.5472 | 8.1287 | 5.1924 | 5.7137 | 7.1293 | 5.6827 | 6.4116 | 6.2190 | 6.5300 | 6.2919 | 4.6386 | 5.3965 | 3.8283 | 4.9017 |
| CKAP2_T683 | CKAP2 | Q8WWK9 | 595;597 | | NA | 6.7017 | 7.1293 | 6.8858 | 7.4573 | 5.3345 | 6.8024 | 8.7524 | 7.5766 | 7.2392 | 6.9383 | 7.2340 | 8.1026 | 2.7662 | 4.7829 | 2.1674 | 4.1293 |
| CKAP2L_S175 | CKAP2L | Q8IYA6 | | 685 | QDMKFITPVRRSS | 7.7662 | 8.3671 | 7.1016 | 6.8433 | 4.5885 | 7.5608 | 8.1071 | 6.5723 | 7.5731 | 7.9070 | 8.0487 | 8.1777 | 2.2575 | 3.8079 | 1.9904 | 3.3310 |
| CKAP2L_S420 | CKAP2L | Q8IYA6 | | 499 | IKEMNISFWKSIE | 12.3980 | 8.0304 | 7.3134 | 7.9402 | 5.6812 | 5.2580 | 8.1803 | 5.5255 | 6.5953 | 4.7467 | 6.2897 | 4.7037 | 4.2236 | 4.8595 | 3.7628 | 4.4916 |
| CKAP2L_S499 | CKAP2L | Q8IYA6 | | 742 | VTLGFQTPESxxx | 6.3288 | 6.9126 | 6.6581 | 6.8151 | 6.4450 | 6.4416 | 6.7769 | 6.5302 | 6.7604 | 6.6898 | 6.9899 | 6.6847 | 4.9128 | 5.5764 | 4.5364 | 4.9414 |
| CKAP2L_T685 | CKAP2L | Q8IYA6 | | 420 | KAQTLDSKLKKAV | 8.9374 | 7.9111 | 6.9538 | 7.1813 | 5.2853 | 6.2075 | 8.1382 | 5.7934 | 7.1982 | 6.6231 | 6.5869 | 6.7913 | 3.8382 | 4.8481 | 3.2809 | 4.4253 |
| CKAP2L_T742 | CKAP2L | Q8IYA6 | | 175 | IHVENESLDNFLK | 6.1634 | 5.7165 | 6.2638 | 5.8941 | 6.0935 | 7.0193 | 7.6016 | 6.9975 | 7.0688 | 5.5942 | 7.1343 | 6.6248 | 4.9654 | 6.2362 | 5.9145 | 4.7121 |
| CKAP5_S1134 | CKAP5 | Q14008-3 | | 1428 | RSAKRPSAAPIKQ | 5.8393 | 5.2294 | 5.6079 | 5.6880 | 6.6219 | 5.9505 | 6.0952 | 6.9144 | 7.1422 | 6.8331 | 6.3585 | 7.0782 | 6.2322 | 7.3274 | 5.5617 | 5.5199 |
| CKAP5_S1428 | CKAP5 | Q14008-3 | | 1971 | PLTSLLSKPAVPT | 8.6898 | 6.3396 | 8.7528 | 7.1746 | 4.5941 | 5.6558 | 7.7666 | 7.2358 | 7.7019 | 7.3402 | 7.3887 | 6.3018 | 3.2571 | 4.9612 | 3.7951 | 3.0448 |
| CKAP5_S1911 | CKAP5 | Q14008-3 | | 1134 | KAPGLSSKAKSAQ | 6.0098 | 4.1269 | 4.2618 | 4.7337 | 4.1606 | 6.3854 | 5.8750 | 7.0741 | 4.3870 | 7.2715 | 5.9761 | 5.3068 | 6.6936 | 9.6937 | 8.9714 | 9.0725 |
| CKAP5_S1968 | CKAP5 | Q14008-3 | | 1968 | DRPPLTSLLSKPA | 9.0436 | 6.7827 | 6.3314 | 6.0568 | 4.3201 | 5.7844 | 8.4654 | 4.7860 | 7.5253 | 6.8222 | 8.3926 | 5.5098 | 4.9316 | 6.0568 | 3.6046 | 5.5868 |
| CKAP5_S1971 | CKAP5 | Q14008-3 | | 1911 | STSTGISPQMEVT | 6.3566 | 6.3747 | 6.7191 | 6.2243 | 6.7382 | 6.3937 | 5.9874 | 6.7924 | 6.4061 | 5.8665 | 6.8257 | 6.5631 | 5.5106 | 5.9027 | 5.3888 | 5.9503 |
| CKB_S199 | CKB | P12277 | | 199 | LFDKPVSPLLLAS | 6.1319 | 5.8840 | 6.5783 | 6.1576 | 6.4769 | 6.5871 | 5.8154 | 6.2129 | 6.7488 | 6.5763 | 6.4434 | 6.5743 | 5.7682 | 6.4337 | 5.6681 | 5.9431 |
| CKB_T35 | CKB | P12277 | | 35 | HMAKVLTPELYAE | 6.2948 | 5.7496 | 6.8855 | 5.9946 | 6.3051 | 6.2916 | 6.8279 | 6.9844 | 6.3922 | 6.1789 | 6.0985 | 6.7589 | 6.2642 | 6.0092 | 5.2044 | 7.5603 |
| CLASP1_S1091 | CLASP1 | Q7Z460 | | 711 | SRGPPVTPSSEKR | 6.2267 | 6.3764 | 6.6554 | 6.0740 | 5.8248 | 6.9837 | 6.9326 | 6.9300 | 6.5227 | 6.8271 | 6.7951 | 6.7277 | 4.8484 | 5.6874 | 5.0655 | 5.5224 |
| CLASP1_S1101 | CLASP1 | Q7Z460 | | 1196 | EKFSFRSQEDLNE | 5.5628 | 5.8977 | 6.5268 | 5.6122 | 5.5690 | 6.6462 | 6.4431 | 6.0238 | 5.7209 | 6.6575 | 7.0485 | 6.7823 | 5.9995 | 6.8632 | 6.1750 | 6.4716 |
| CLASP1_S1196 | CLASP1 | Q7Z460 | | 636 | PPGSYASLGRIRT | 5.7688 | 6.2433 | 6.5284 | 6.6078 | 6.1746 | 6.4354 | 6.0356 | 6.3617 | 6.3614 | 6.1812 | 6.6744 | 6.2809 | 5.7523 | 6.6671 | 5.9626 | 5.9645 |
| CLASP1_S280 | CLASP1 | Q7Z460-4 | | 691 | SQSQPGSRSSSPG | 5.7173 | 5.9215 | 6.3697 | 5.9570 | 5.9286 | 5.8104 | 6.0597 | 6.5031 | 6.5190 | 6.8870 | 5.8167 | 6.2468 | 6.8147 | 6.5155 | 6.6689 | 6.2641 |
| CLASP1_S285 | CLASP1 | Q7Z460 | | 572 | PLSAKRSPTGSTT | 6.4156 | 6.5458 | 6.3122 | 6.5371 | 6.1657 | 6.8357 | 6.5639 | 6.7055 | 6.0799 | 6.3316 | 6.5640 | 6.5578 | 5.7953 | 6.2538 | 5.7500 | 5.4862 |
| CLASP1_S556 | CLASP1 | Q7Z460 | | 796 | AMRVLSTSTDLEA | 5.6915 | 6.2373 | 7.0483 | 6.1469 | 6.0583 | 6.2829 | 6.7854 | 6.3971 | 6.2645 | 6.2261 | 7.0252 | 6.7770 | 5.7984 | 5.9197 | 5.2940 | 6.0475 |
| CLASP1_S572 | CLASP1 | Q7Z460-4 | | 1101 | CSHGGLSPSMLDY | 6.1109 | 6.1506 | 6.9312 | 6.2296 | 6.4193 | 6.9704 | 6.5221 | 7.5555 | 6.2103 | 6.2500 | 6.4778 | 6.3824 | 5.1502 | 6.1910 | 5.4826 | 4.9662 |
| CLASP1_S594 | CLASP1 | Q7Z460 | | 1091 | SNTSVGSPSNTIG | 6.6708 | 6.5742 | 7.0016 | 6.3816 | 5.7628 | 6.3874 | 6.1414 | 5.9145 | 6.8452 | 6.9966 | 6.8907 | 6.8288 | 5.1168 | 5.6900 | 5.3536 | 5.4441 |
| CLASP1_S600 | CLASP1 | Q7Z460 | | 594 | SVSTTGSLQRSRS | 6.3407 | 6.4122 | 6.4709 | 5.6012 | 5.4206 | 6.3449 | 6.0153 | 6.1936 | 6.4862 | 6.3653 | 6.6976 | 6.3861 | 6.1686 | 7.0775 | 6.0662 | 5.9532 |
| CLASP1_S636 | CLASP1 | Q7Z460-4 | | 767 | AMRVLSTSTDLEA | 6.6653 | 6.6237 | 6.6496 | 5.8392 | 5.8125 | 7.0584 | 6.1327 | 6.4382 | 5.6639 | 6.8517 | 7.5042 | 6.5667 | 5.2172 | 6.3487 | 5.9980 | 5.5301 |
| CLASP1_S687;S691 | CLASP1 | Q7Z460 | | 285 | GSSTLGSKSSAAK | 6.3240 | 8.2036 | 7.6540 | 7.8445 | 6.0968 | 5.4007 | 4.3528 | 7.1630 | 7.3279 | 8.3062 | 8.4014 | 6.7417 | 3.7519 | 5.6791 | 3.3085 | 3.4441 |
| CLASP1_S691 | CLASP1 | Q7Z460 | | 280 | TTRRLGSSTLGSK | 7.3183 | 5.7494 | 7.1201 | 6.1408 | 5.7616 | 5.9857 | 6.0614 | 6.3206 | 8.2506 | 7.5719 | 7.3339 | 7.7457 | 5.6204 | 5.6156 | 3.5983 | 3.8057 |
| CLASP1_T711 | CLASP1 | Q7Z460 | | 600 | SLQRSRSDIDVNA | 5.7822 | 5.9729 | 6.3292 | 6.1661 | 5.6144 | 6.7090 | 5.9656 | 6.0537 | 6.5573 | 7.0985 | 6.6113 | 7.1848 | 5.7422 | 6.2390 | 5.7620 | 6.2117 |
| CLASP1_T767 | CLASP1 | Q7Z460 | | 556 | PQSDRSSSSSQES | 6.6206 | 7.2770 | 7.2824 | 6.6202 | 6.1672 | 6.6294 | 6.3582 | 6.3408 | 7.5703 | 7.8522 | 7.6814 | 7.3182 | 3.6002 | 4.6909 | 3.9563 | 4.0348 |
| CLASP1_T796 | CLASP1 | Q7Z460-4 | 687;691 | | NA | 6.9597 | 7.0867 | 6.8983 | 6.1383 | 5.8506 | 6.5665 | 6.5433 | 6.2798 | 6.8843 | 6.6148 | 6.5122 | 7.0036 | 4.8161 | 5.7411 | 4.9856 | 5.1190 |
| CLASP2_S1029 | CLASP2 | O75122 | | 596 | RVLNTGSDVEEAV | 6.7901 | 6.8430 | 7.8768 | 6.8500 | 6.7821 | 7.1617 | 7.1307 | 7.5160 | 5.7546 | 5.8463 | 6.0008 | 6.1558 | 4.7276 | 5.1925 | 4.3971 | 4.9751 |
| CLASP2_S326 | CLASP2 | O75122 | | 955 | QNFSFRSQEDMNE | 6.4116 | 7.1235 | 7.0287 | 6.8228 | 6.2380 | 6.6427 | 6.5787 | 6.7958 | 6.8762 | 6.7935 | 7.3224 | 6.5799 | 4.5744 | 4.8389 | 4.5260 | 4.8269 |
| CLASP2_S359 | CLASP2 | O75122 | | 8 | AMGDDKSFDDEES | 6.2705 | 6.8241 | 6.6493 | 7.2208 | 6.1725 | 6.8091 | 6.1291 | 6.8786 | 6.4399 | 6.4169 | 7.2796 | 7.1732 | 4.4701 | 5.5831 | 4.4349 | 5.2483 |
| CLASP2_S360 | CLASP2 | O75122 | | 360 | GSSKASSLPGSLQ | 6.8271 | 6.4551 | 7.2007 | 6.5538 | 6.4073 | 6.5767 | 6.4142 | 7.2185 | 6.8538 | 6.9779 | 6.5963 | 6.6275 | 4.4438 | 5.3770 | 4.7222 | 4.7480 |
| CLASP2_S370 | CLASP2 | O75122 | | 1029 | NYSDSISPFNKSA | 6.2874 | 6.0770 | 6.6521 | 6.8087 | 5.9072 | 6.8003 | 6.3530 | 6.4331 | 7.0727 | 6.5728 | 6.6258 | 6.5082 | 5.4232 | 5.8375 | 5.0625 | 5.5786 |
| CLASP2_S455 | CLASP2 | O75122 | | 892 | TQSSMGSPLTRPT | 6.8107 | 6.5325 | 7.2308 | 6.8852 | 6.8211 | 6.4860 | 6.5045 | 6.2311 | 6.8492 | 7.1642 | 7.1057 | 7.0720 | 4.4106 | 4.9493 | 4.3328 | 4.6142 |
| CLASP2_S455;S459 | CLASP2 | O75122 | | 901 | TRPTPRSPANWSS | 6.2467 | 4.4125 | 7.4694 | 3.7533 | 5.9649 | 6.2254 | 5.1143 | 4.8698 | 6.3158 | 8.4902 | 5.5821 | 8.3785 | 5.2897 | 8.6687 | 8.9155 | 6.1031 |
| CLASP2_S529 | CLASP2 | O75122 | | 529 | SVSQGCSREASRE | 5.8661 | 6.0082 | 6.2494 | 5.5173 | 6.9921 | 5.9397 | 5.4834 | 6.4550 | 5.9694 | 6.8777 | 6.1318 | 5.6909 | 6.7949 | 6.5531 | 7.5350 | 5.9362 |
| CLASP2_S596 | CLASP2 | O75122 | | 370 | SLQRSRSDIDVNA | 5.7671 | 5.7324 | 6.7467 | 6.3886 | 5.0151 | 6.6428 | 6.0150 | 6.6639 | 6.9091 | 6.1978 | 6.6862 | 7.1720 | 6.2426 | 6.5002 | 5.0956 | 6.2249 |
| CLASP2_S8 | CLASP2 | O75122 | | 359 | AGSSKASSLPGSL | 6.4889 | 5.3760 | 7.7448 | 7.0653 | 7.2952 | 6.6687 | 4.8423 | 7.0800 | 7.9482 | 6.4683 | 6.6525 | 7.2023 | 4.0567 | 5.1740 | 4.6743 | 5.2625 |
| CLASP2_S892 | CLASP2 | O75122 | | 455 | TKMVSQSQPGSRS | 5.9057 | 6.5315 | 6.2004 | 7.7346 | 5.9650 | 6.5694 | 6.2329 | 6.9746 | 6.1790 | 6.2904 | 5.9164 | 6.0916 | 5.5201 | 6.0501 | 5.7140 | 6.1241 |
| CLASP2_S901 | CLASP2 | O75122 | | 326 | SDRSSSSSQESLN | 5.6271 | 6.0965 | 6.7682 | 5.8490 | 5.5063 | 6.5157 | 6.0774 | 6.6563 | 6.5434 | 7.0429 | 7.6393 | 7.4114 | 5.0290 | 6.0338 | 5.6918 | 5.5121 |
| CLASP2_S955 | CLASP2 | O75122 | 455;459 | | NA | 6.9157 | 7.7822 | 7.2333 | 6.6427 | 5.7623 | 6.5444 | 6.2131 | 6.0572 | 6.5868 | 5.2307 | 6.9559 | 6.6684 | 5.0715 | 5.9201 | 5.4085 | 5.5072 |
| CLASRP_S101 | CLASRP | Q8N2M8 | | 547 | LTRPAASPAVGEK | 6.0391 | 5.8517 | 6.0967 | 6.2657 | 6.6454 | 5.8378 | 6.1272 | 6.2229 | 6.8949 | 6.3415 | 6.1139 | 6.0751 | 6.5224 | 6.3256 | 6.0473 | 6.5927 |
| CLASRP_S285 | CLASRP | Q8N2M8 | | 294 | SYARRDSPTYDPY | 6.5551 | 6.3716 | 6.2643 | 6.6093 | 6.3483 | 6.5924 | 6.0017 | 6.0912 | 6.3544 | 6.3720 | 6.0728 | 6.1781 | 6.3080 | 6.3504 | 6.3991 | 6.7312 |
| CLASRP_S294 | CLASRP | Q8N2M8 | | 285 | LRGRKISPPSYAR | 6.2002 | 6.4877 | 6.0258 | 6.3200 | 6.0472 | 6.0435 | 6.0906 | 6.0474 | 6.1630 | 5.9612 | 6.1282 | 6.1145 | 6.5475 | 6.5185 | 6.5836 | 6.7212 |
| CLASRP_S454;S456 | CLASRP | Q8N2M8 | | 101 | PLLTTISPEQESD | 6.5272 | 6.6796 | 6.5372 | 5.8697 | 6.2656 | 6.6991 | 6.6966 | 6.2857 | 6.4767 | 5.7579 | 6.4641 | 6.1732 | 6.3428 | 6.0538 | 6.6551 | 6.3158 |
| CLASRP_S456 | CLASRP | Q8N2M8 | | 456 | GHRYSRSPARRGG | 7.2493 | 7.2654 | 6.3559 | 6.4235 | 5.9268 | 6.1156 | 5.3252 | 4.9426 | 5.9006 | 5.9339 | 7.1493 | 6.8788 | 6.0429 | 6.0157 | 5.4614 | 7.0131 |
| CLASRP_S547 | CLASRP | Q8N2M8 | 454;456 | | NA | 5.9204 | 6.7681 | 7.6297 | 6.2274 | 5.6808 | 6.7681 | 5.4435 | 6.2843 | 6.1603 | 6.6547 | 5.7867 | 5.6652 | 6.5160 | 6.4491 | 6.0750 | 5.9707 |
| CLCC1_S509 | CLCC1 | Q96S66 | | 482 | GILGEGTPKESST | 6.3112 | 7.2637 | 6.8976 | 7.3629 | 6.0921 | 6.1913 | 7.2692 | 6.3158 | 6.6905 | 6.9984 | 7.0918 | 7.2239 | 4.4256 | 5.2222 | 4.0786 | 4.7652 |
| CLCC1_S532 | CLCC1 | Q96S66 | | 532 | DQSSTYSPARGVA | 7.0842 | 7.7616 | 6.3998 | 6.5120 | 7.0443 | 6.9421 | 6.8676 | 6.1914 | 7.7724 | 6.8594 | 7.1760 | 6.7697 | 3.6913 | 4.5689 | 3.8724 | 4.4870 |
| CLCC1_S548 | CLCC1 | Q96S66 | | 509 | SGNTEGSPAAEKA | 6.0216 | 6.1276 | 6.4727 | 7.3645 | 5.6687 | 6.2558 | 6.4023 | 6.3685 | 6.0987 | 6.9919 | 7.1504 | 7.1504 | 5.3714 | 5.9937 | 4.9044 | 5.6577 |
| CLCC1_T482 | CLCC1 | Q96S66 | | 548 | GQDPVSSPCGxxx | 7.5907 | 8.0892 | 6.8105 | 7.7602 | 7.3526 | 6.8085 | 6.9860 | 6.4567 | 7.3553 | 7.4273 | 6.8529 | 7.2484 | 3.1291 | 3.6968 | 2.9926 | 3.4432 |
| CLDN10_S202 | CLDN10 | P78369 | | 202 | GATSVMSSRTKYH | 7.3433 | 7.2485 | 7.6510 | 7.0664 | 6.2039 | 6.1726 | 5.9584 | 6.1528 | 6.3122 | 7.5488 | 6.7835 | 7.3544 | 4.7581 | 4.9344 | 4.4631 | 4.0487 |
| CLDN12_S231 | CLDN12 | P56749 | | 231 | SARSRLSAIEIDI | 5.9241 | 5.6950 | 5.9782 | 6.1937 | 6.0477 | 5.9949 | 5.9320 | 6.3964 | 6.6824 | 6.6955 | 6.7206 | 6.5215 | 6.1881 | 6.8679 | 6.0554 | 6.1066 |
| CLDN23_S204 | CLDN23 | Q96B33 | | 204 | AGPRRSSVSTIQV | 6.0264 | 6.7254 | 6.1353 | 6.7979 | 6.3017 | 6.7396 | 6.3043 | 6.3214 | 6.9443 | 7.0277 | 6.6097 | 6.4620 | 4.7271 | 5.8533 | 5.4453 | 5.5786 |
| CLEC16A_S863 | CLEC16A | Q2KHT3 | | 980 | VETASLSPSLVPA | 6.1174 | 5.7249 | 6.8465 | 7.6035 | 6.4434 | 6.1773 | 6.6842 | 6.1853 | 6.6283 | 5.8061 | 7.4146 | 6.7873 | 5.4269 | 5.2320 | 5.3943 | 5.5280 |
| CLEC16A_S980 | CLEC16A | Q2KHT3 | | 863 | DQGRRGSSDPTVQ | 6.2272 | 6.3397 | 5.5784 | 6.2608 | 7.1439 | 5.4995 | 5.4206 | 6.2341 | 6.6159 | 7.3377 | 6.4651 | 6.3490 | 5.7386 | 7.1509 | 6.7459 | 4.8926 |
| CLINT1_S299 | CLINT1 | Q14677-3 | | 299 | YTGDKASPDQNAS | 6.5887 | 6.8894 | 7.0598 | 6.9508 | 6.0688 | 6.3136 | 5.6034 | 6.6661 | 6.2211 | 6.1706 | 6.5960 | 6.6490 | 5.6298 | 5.7424 | 5.4179 | 5.4326 |
| CLINT1_S496 | CLINT1 | Q14677-3 | | 496 | TLPSTWSDPSVNI | 2.4350 | 2.1537 | 2.2262 | 2.6918 | 2.3643 | 2.6356 | 2.5681 | 2.7200 | 1.9556 | 2.3962 | 2.2350 | 2.2118 | 15.9917 | 16.5792 | 20.2272 | 18.6085 |
| CLIP1_S1364 | CLIP1 | P30622 | | 147 | PASRATSPLCTST | 6.1335 | 5.9430 | 6.8647 | 6.0609 | 6.2163 | 6.3397 | 5.8106 | 6.8485 | 6.2950 | 6.0798 | 6.4408 | 6.5278 | 6.3558 | 6.2931 | 5.9067 | 5.8838 |
| CLIP1_S147 | CLIP1 | P30622 | | 287 | IGFPSTTPAKAKA | 7.2315 | 7.2321 | 6.9162 | 6.4185 | 6.0334 | 7.5469 | 7.5300 | 6.9210 | 7.6927 | 6.7039 | 7.1770 | 6.6372 | 4.0621 | 4.5530 | 3.2212 | 4.1234 |
| CLIP1_S147;S152 | CLIP1 | P30622 | | 306 | MATTSASLKRSPS | 6.1436 | 6.2379 | 7.1134 | 6.0056 | 5.4459 | 6.7264 | 6.3902 | 6.4605 | 7.2638 | 6.4941 | 6.8492 | 6.7390 | 5.7137 | 5.3977 | 5.2055 | 5.8134 |
| CLIP1_S200 | CLIP1 | P30622 | | 1364 | DLNNYDSDDQEKQ | 5.9565 | 6.4637 | 7.1121 | 6.5837 | 6.3589 | 6.7217 | 5.9036 | 6.5245 | 6.6025 | 6.3903 | 6.5303 | 6.5264 | 5.9473 | 5.9526 | 5.2189 | 5.2069 |
| CLIP1_S306 | CLIP1 | P30622 | | 200 | ESISNLSEAGSIK | 6.5834 | 6.6846 | 7.4784 | 6.5590 | 6.7426 | 6.8094 | 5.8666 | 6.4357 | 6.6113 | 6.7961 | 6.8822 | 6.6354 | 4.9592 | 5.0262 | 4.6261 | 5.3039 |
| CLIP1_S312 | CLIP1 | P30622 | | 182 | AKEPSATPPISNL | 6.6942 | 7.3183 | 6.8061 | 7.4091 | 6.8430 | 7.6220 | 6.8497 | 6.9471 | 6.7603 | 7.1329 | 6.9594 | 7.8400 | 3.1374 | 4.1948 | 3.2754 | 4.2102 |
| CLIP1_S348 | CLIP1 | P30622 | | 348 | RYARKISGTTALQ | 5.1754 | 4.6463 | 4.7791 | 4.4045 | 5.5270 | 5.8650 | 4.8654 | 5.4530 | 7.5660 | 7.2897 | 6.9242 | 7.6606 | 7.5093 | 7.5677 | 7.1927 | 7.5742 |
| CLIP1_T182 | CLIP1 | P30622 | | 25 | GSTALKTPTAVVA | 5.7576 | 6.0180 | 6.1770 | 6.3024 | 6.1451 | 7.0969 | 6.9935 | 6.6703 | 6.9808 | 5.7913 | 6.7085 | 6.4475 | 5.7146 | 5.9960 | 5.3108 | 5.8897 |
| CLIP1_T25 | CLIP1 | P30622 | | 312 | SLKRSPSASSLSS | 7.9064 | 6.4892 | 5.8693 | 6.9765 | 6.6367 | 5.5129 | 5.7649 | 6.6500 | 8.4318 | 6.0616 | 7.3080 | 6.7163 | 5.3803 | 5.4466 | 4.0758 | 4.7737 |
| CLIP1_T287 | CLIP1 | P30622 | 147;152 | | NA | 6.1137 | 6.9682 | 6.0497 | 6.9000 | 6.0285 | 6.3765 | 6.0669 | 7.0667 | 7.7956 | 6.7062 | 6.2854 | 6.9941 | 4.6428 | 5.1663 | 5.0359 | 5.8038 |
| CLIP2_S195 | CLIP2 | Q9UDT6 | | 294 | IGFPSTSPAKAKK | 6.6330 | 6.2199 | 6.8336 | 5.8250 | 7.0514 | 6.9022 | 6.2188 | 7.0890 | 6.6060 | 6.3764 | 6.6969 | 6.2661 | 5.6017 | 5.3372 | 5.0409 | 5.3018 |
| CLIP2_S207 | CLIP2 | Q9UDT6 | | 923 | LEANRHSPGPERD | 6.2075 | 6.1368 | 6.6958 | 6.5106 | 6.4545 | 6.7283 | 6.2192 | 6.4451 | 6.4771 | 6.8285 | 6.7228 | 6.6756 | 5.6719 | 5.8123 | 5.1761 | 5.2380 |
| CLIP2_S294 | CLIP2 | Q9UDT6 | | 42 | ASSKEGSPLHKQS | 6.7219 | 7.2507 | 7.2863 | 6.9164 | 6.7774 | 6.9588 | 6.5419 | 6.5544 | 7.6730 | 7.5536 | 7.2776 | 7.2223 | 3.7028 | 4.1153 | 3.3985 | 4.0491 |
| CLIP2_S42 | CLIP2 | Q9UDT6 | | 195 | ESVLNSSVKTGNE | 4.7165 | 5.3521 | 7.2502 | 5.6618 | 5.7245 | 6.3651 | 5.7784 | 5.7749 | 4.8970 | 6.3526 | 5.9852 | 6.2285 | 6.5632 | 7.2201 | 7.3342 | 6.9957 |
| CLIP2_S54 | CLIP2 | Q9UDT6 | | 207 | ESGSNLSDSGSVK | 5.6992 | 6.5951 | 6.8282 | 6.3898 | 6.1788 | 6.6774 | 6.0605 | 5.8314 | 6.8780 | 7.2080 | 7.2857 | 7.6585 | 5.6285 | 5.4905 | 4.7060 | 4.8846 |
| CLIP2_S923 | CLIP2 | Q9UDT6 | | 54 | SSGPSSSPAAAAA | 6.7444 | 6.7903 | 7.0231 | 8.6566 | 7.4022 | 6.5421 | 5.7416 | 7.2373 | 7.3036 | 8.6209 | 7.2305 | 9.0305 | 3.3331 | 3.4283 | 2.3388 | 2.5767 |
| CLIP3_S143 | CLIP3 | Q96DZ5 | | 143 | AAAVRLSQQLLAL | 4.9984 | 6.0744 | 6.2091 | 5.6711 | 5.7401 | 5.5282 | 4.9074 | 6.2386 | 6.0402 | 6.2063 | 6.3288 | 6.2529 | 6.5529 | 7.7275 | 8.4279 | 7.0961 |
| CLIP3_S401 | CLIP3 | Q96DZ5 | | 401 | GKKKTPSSPSLGS | 6.4611 | 7.0519 | 6.7249 | 6.9968 | 6.4498 | 6.1552 | 6.7767 | 6.3090 | 7.7349 | 6.9612 | 6.2669 | 6.8771 | 5.4139 | 4.8863 | 3.5834 | 5.3508 |
| CLK1_S182 | CLK1 | P49759-3 | | 182 | RRKRTRSVEDDEE | 5.5196 | 5.4350 | 7.0226 | 6.1619 | 6.0682 | 5.0795 | 6.5517 | 5.5367 | 6.5665 | 6.3905 | 6.4888 | 5.6956 | 6.8329 | 7.5324 | 6.4968 | 6.6214 |
| CLK1_T180;S182 | CLK1 | P49759-3 | 180;182 | | NA | 5.4561 | 6.0011 | 5.8042 | 5.9154 | 7.1374 | 5.5922 | 5.1863 | 5.9993 | 6.8243 | 5.1598 | 5.8231 | 5.8664 | 7.3241 | 8.4697 | 6.8239 | 6.6166 |
| CLK2_S142 | CLK2 | P49760 | | 142 | SSRRAKSVEDDAE | 5.9696 | 6.1755 | 6.2281 | 6.4692 | 6.0455 | 6.0337 | 5.8952 | 6.1185 | 6.3307 | 6.4070 | 6.3609 | 6.3034 | 6.3896 | 6.3380 | 6.2971 | 6.6380 |
| CLK2_S98 | CLK2 | P49760 | | 98 | DTDYRHSYEYQRE | 4.6746 | 6.4343 | 5.6785 | 6.5449 | 6.3062 | 5.7157 | 5.1882 | 6.3900 | 5.8031 | 6.3376 | 6.0907 | 5.9987 | 6.9106 | 7.8062 | 7.0923 | 7.0282 |
| CLK3_S157 | CLK3 | P49761 | | 157 | HCKRYRSPEPDPY | 5.6656 | 6.1839 | 5.7535 | 6.1884 | 6.1761 | 5.7854 | 6.2878 | 6.3555 | 6.4015 | 6.2334 | 6.1726 | 6.2186 | 6.3446 | 6.7208 | 6.7010 | 6.8115 |
| CLK3_S224 | CLK3 | P49761 | | 224 | YRCEERSPSFGED | 5.1709 | 5.5311 | 5.5152 | 5.3951 | 6.2278 | 5.6215 | 6.5084 | 5.3799 | 5.8588 | 6.2328 | 5.9449 | 5.1506 | 7.0778 | 8.1600 | 8.2518 | 7.9733 |
| CLK3_S224;S226 | CLK3 | P49761 | 224;226 | | NA | 5.7703 | 6.4486 | 7.0032 | 6.8243 | 6.3941 | 6.4804 | 6.6142 | 5.7629 | 6.5499 | 6.2592 | 6.7305 | 7.5018 | 4.7635 | 5.5767 | 5.5218 | 5.7986 |
| CLK4_S108;S111;S112 | CLK4 | Q9HAZ1 | | 138 | RRKRSRSIEDDEE | 6.1675 | 6.3140 | 5.8671 | 6.9099 | 6.2017 | 6.0193 | 6.0397 | 5.6807 | 6.3523 | 6.3287 | 5.9021 | 6.2473 | 6.5566 | 7.1769 | 6.3042 | 5.9322 |
| CLK4_S138 | CLK4 | Q9HAZ1 | 108;111;112 | | NA | 6.3992 | 6.4075 | 6.2285 | 6.4898 | 5.2814 | 5.3242 | 6.0904 | 6.0219 | 6.0732 | 6.6343 | 6.3467 | 6.1286 | 7.1260 | 6.4971 | 6.1940 | 6.7572 |
| CLMP_T359 | CLMP | Q9H6B4 | | 359 | LTKAETTPSMIPS | 7.4975 | 8.1408 | 7.1574 | 6.3477 | 6.0261 | 6.6250 | 6.5030 | 6.1333 | 6.9134 | 6.7914 | 5.3606 | 6.9688 | 4.4955 | 5.2941 | 3.7377 | 6.0076 |
| CLN3_S12 | CLN3 | Q13286 | | 12 | GSRRRFSDSEGEE | 5.7050 | 5.9950 | 5.8861 | 6.3678 | 6.6954 | 5.4212 | 5.9461 | 6.2209 | 6.7272 | 6.4526 | 6.6827 | 6.6214 | 6.2641 | 6.9589 | 5.9344 | 6.1212 |
| CLN6_S63 | CLN6 | Q9NWW5-2 | | 63 | KGKKHGSVSADEA | 5.6132 | 5.3569 | 5.7203 | 6.2098 | 7.2644 | 4.2666 | 6.6363 | 5.3506 | 6.1615 | 6.6636 | 7.1699 | 6.4808 | 6.4388 | 8.6131 | 6.5543 | 5.4998 |
| CLNS1A_S102 | CLNS1A | P54105 | | 102 | DEEEDSDDDVEP | 5.6661 | 5.6539 | 6.4872 | 5.8477 | 6.9168 | 7.0782 | 6.0148 | 6.7435 | 6.6149 | 6.5253 | 7.6413 | 7.1796 | 5.9248 | 6.2050 | 4.8747 | 4.6261 |
| CLOCK_T461 | CLOCK | O15516 | | 461 | IPTDTSTPPRQHL | 5.6721 | 6.0159 | 6.1192 | 6.3867 | 6.2065 | 5.7000 | 5.8725 | 6.1421 | 6.2066 | 6.3365 | 5.8905 | 6.3137 | 6.6892 | 6.7095 | 6.7681 | 6.9710 |
| CLPTM1_T622 | CLPTM1 | O96005 | | 622 | TAAGALTPTPAPT | 6.7809 | 6.2228 | 6.9951 | 6.9709 | 6.7555 | 6.4873 | 6.7847 | 5.9739 | 6.7884 | 6.5581 | 7.3409 | 6.9591 | 5.1209 | 5.2594 | 4.2177 | 4.7845 |
| CLSPN_S1156 | CLSPN | Q9HAW4 | | 225 | LEDENNSPLEDEE | 5.8472 | 6.2292 | 6.5641 | 6.3878 | 6.9663 | 6.5340 | 6.1297 | 6.1484 | 5.7969 | 6.1944 | 6.0357 | 6.1966 | 6.2256 | 6.5197 | 6.1648 | 6.0595 |
| CLSPN_S1289 | CLSPN | Q9HAW4 | | 1289 | FVFHTLSPVKAEA | 6.3569 | 6.1940 | 6.4946 | 6.0935 | 6.2967 | 6.0624 | 5.9910 | 6.3097 | 6.6373 | 6.2687 | 6.8195 | 6.5995 | 6.2422 | 6.5444 | 5.5953 | 5.4944 |
| CLSPN_S1314 | CLSPN | Q9HAW4 | | 1314 | GPSFMTSPSPKHL | 6.7516 | 6.9515 | 6.6955 | 6.5415 | 6.4767 | 6.4051 | 6.4092 | 6.9560 | 7.5630 | 6.6416 | 7.0976 | 6.5941 | 4.6230 | 4.9438 | 4.3074 | 5.0424 |
| CLSPN_S1316 | CLSPN | Q9HAW4 | | 823 | SLVSSASKSSGKL | 6.5173 | 7.6591 | 8.1894 | 7.6699 | 5.1854 | 6.8871 | 7.3615 | 6.8221 | 7.3021 | 7.3485 | 8.0862 | 7.3227 | 3.0065 | 4.3115 | 3.0254 | 3.3052 |
| CLSPN_S205 | CLSPN | Q9HAW4 | | 810 | GGFRSPSPGLFRA | 5.4101 | 5.4908 | 5.5411 | 5.1278 | 5.4337 | 5.7827 | 6.0979 | 5.6686 | 5.6986 | 5.5292 | 5.6921 | 5.4430 | 8.2884 | 8.1758 | 8.3483 | 8.2718 |
| CLSPN_S225 | CLSPN | Q9HAW4 | | 1156 | DLFHRDSDDDQTE | 6.1569 | 6.8782 | 6.5650 | 5.3443 | 5.9537 | 5.7906 | 5.5722 | 6.3123 | 6.4943 | 6.6745 | 6.8488 | 6.9742 | 5.9119 | 6.1168 | 6.0851 | 6.3212 |
| CLSPN_S703 | CLSPN | Q9HAW4 | | 205 | EQPFNDSGCLLVD | 5.6427 | 6.0037 | 8.4462 | 6.1909 | 7.4300 | 5.5001 | 5.4511 | 5.4644 | 7.5147 | 7.1893 | 6.2043 | 7.9426 | 5.7853 | 4.0382 | 5.1792 | 6.0171 |
| CLSPN_S720 | CLSPN | Q9HAW4 | | 1316 | SFMTSPSPKHLKT | 5.3669 | 5.1967 | 5.4472 | 6.2445 | 5.3360 | 5.5398 | 5.6333 | 5.2417 | 5.4710 | 4.9762 | 6.3927 | 6.3203 | 8.8966 | 8.0252 | 7.9388 | 7.9732 |
| CLSPN_S810 | CLSPN | Q9HAW4 | | 839 | SLPIEDSQDLYNA | 5.4348 | 5.9675 | 7.2833 | 6.8442 | 4.6505 | 6.4963 | 6.6935 | 6.6714 | 6.2046 | 6.6124 | 6.8779 | 6.5935 | 5.7072 | 6.3962 | 5.6549 | 5.9117 |
| CLSPN_S821 | CLSPN | Q9HAW4 | | 703 | KENNDGSSEIGKA | 5.6487 | 6.2019 | 6.7032 | 5.8841 | 6.0362 | 5.7541 | 5.9859 | 5.5397 | 6.4511 | 7.1185 | 6.4246 | 6.3604 | 6.3976 | 6.6525 | 6.1856 | 6.6560 |
| CLSPN_S823 | CLSPN | Q9HAW4 | | 821 | RASLVSSASKSSG | 8.5826 | 8.9404 | 7.2667 | 7.2973 | 6.5555 | 5.7685 | 7.2746 | 7.5317 | 8.3776 | 7.5713 | 7.6053 | 7.1727 | 2.2365 | 3.1424 | 2.0270 | 2.6498 |
| CLSPN_S839 | CLSPN | Q9HAW4 | | 720 | SVPKSLSSDSTLL | 5.8915 | 6.3438 | 6.3334 | 6.0271 | 5.5497 | 6.0546 | 5.9228 | 5.8882 | 6.4542 | 6.7155 | 6.2210 | 6.3850 | 6.3376 | 6.9923 | 6.7406 | 6.1428 |
| CLSPN_S839;S846 | CLSPN | Q9HAW4 | | 950 | CSGKFTSQDASTP | 4.9606 | 6.5119 | 7.2381 | 6.4176 | 5.4174 | 6.4533 | 5.9864 | 5.9995 | 6.1651 | 6.9688 | 7.0762 | 6.6199 | 5.8727 | 6.8326 | 5.7905 | 5.6894 |
| CLSPN_S950 | CLSPN | Q9HAW4 | 839;846 | | NA | 4.9548 | 5.3147 | 9.3598 | 7.5282 | 3.0538 | 7.2731 | 4.3429 | 6.8680 | 6.3127 | 6.3647 | 5.3538 | 9.2008 | 5.4840 | 7.3578 | 4.5364 | 6.6945 |
| CLTA_S206 | CLTA | P09496-2 | | 206 | DVSRMRSVLISLK | 7.8645 | 5.3913 | 6.7683 | 5.9598 | 5.4466 | 3.9272 | 5.2329 | 4.6934 | 3.7440 | 4.7472 | 4.7821 | 5.3724 | 8.0230 | 11.5300 | 9.7911 | 6.7261 |
| CLTC_S67 | CLTC | Q00610 | | 394 | PKGILRTPDTIRR | 7.3414 | 7.0710 | 6.6496 | 5.9967 | 6.5115 | 7.9477 | 6.3157 | 4.9882 | 5.5261 | 5.8528 | 6.1950 | 6.5280 | 5.5313 | 6.9320 | 5.4505 | 5.5626 |
| CLTC_T394 | CLTC | Q00610 | | 67 | PIRRPISADSAIM | 6.0300 | 6.7475 | 6.9792 | 6.8591 | 6.0083 | 6.3980 | 5.9189 | 6.2576 | 6.7575 | 6.6437 | 6.7222 | 6.9286 | 5.8702 | 5.5554 | 4.9169 | 5.4069 |
| CLUH_S664 | CLUH | O75153 | | 664 | PGQEAGSEEEGSS | 6.4364 | 5.9108 | 6.4824 | 5.9752 | 6.3430 | 6.8050 | 6.2357 | 6.5826 | 6.3031 | 6.4295 | 6.6588 | 6.6357 | 5.5237 | 6.2566 | 5.5075 | 5.9141 |
| CMC2_S89 | CMC2 | H3BTB6 | | 89 | VIVTHGSLKLKxx | 5.6337 | 6.0735 | 5.6420 | 6.9079 | 7.5161 | 4.0840 | 4.4790 | 7.0396 | 4.6589 | 6.6360 | 5.4683 | 6.2674 | 2.8765 | 10.3385 | 8.3182 | 8.0604 |
| CMSS1_S228 | CMSS1 | Q9BQ75 | | 212 | VHLGVGTPGRIKE | 5.8707 | 5.9340 | 6.8776 | 5.0901 | 6.3828 | 5.4776 | 6.3636 | 5.7863 | 6.9371 | 6.3349 | 6.1565 | 6.6705 | 6.6437 | 7.4607 | 5.8266 | 6.1872 |
| CMSS1_T212 | CMSS1 | Q9BQ75 | | 228 | QGGLNLSPLKFLV | 6.0401 | 7.2890 | 5.7873 | 6.4911 | 5.7307 | 6.3647 | 5.8530 | 5.8348 | 6.9502 | 6.6276 | 6.2990 | 5.6872 | 6.6903 | 7.2455 | 5.9410 | 5.7684 |
| CMTM7_S14 | CMTM7 | Q96FZ5 | | 14 | VRTTCSSGSALGP | 4.0265 | 4.4783 | 5.5000 | 5.2809 | 4.9418 | 4.9976 | 5.0207 | 4.8227 | 4.1613 | 4.9849 | 4.0381 | 4.7669 | 10.2816 | 10.5628 | 11.6350 | 10.5007 |
| CMTM8_T14;S27 | CMTM8 | Q8IZV2 | | 27 | ENFSTSSSSFAYD | 6.1412 | 6.6530 | 6.2457 | 6.1731 | 6.7681 | 4.8219 | 6.7220 | 6.3077 | 6.9168 | 6.2333 | 6.6282 | 6.6247 | 6.6246 | 6.0899 | 5.4488 | 5.2646 |
| CMTM8_T27 | CMTM8 | Q8IZV2 | 14;27 | | NA | 5.7082 | 6.1152 | 6.9964 | 6.5146 | 5.6017 | 6.2266 | 6.1727 | 5.8368 | 6.2235 | 7.0362 | 7.2356 | 7.2113 | 6.1751 | 6.3157 | 5.7180 | 5.8123 |
| CMTR1_S31 | CMTR1 | Q8N1G2 | | 31 | LSLSSTSDDEPPS | 6.0421 | 7.1586 | 5.4656 | 6.3906 | 6.6190 | 6.7551 | 5.6937 | 6.6294 | 5.9152 | 6.2052 | 6.2002 | 6.3498 | 5.8283 | 6.5017 | 6.2702 | 5.9752 |
| CNBP_S49 | CNBP | P62633-6 | | 49 | FQFVSSSLPDICY | 6.1448 | 5.8589 | 6.8608 | 6.3598 | 6.3871 | 6.2998 | 5.9036 | 5.9953 | 6.4242 | 6.2048 | 6.8368 | 6.1721 | 6.8695 | 6.2780 | 6.2409 | 5.1636 |
| CNKSR2_S325 | CNKSR2 | Q8WXI2 | | 390 | VHKGSESPNSFLD | 6.0193 | 5.9978 | 6.7717 | 6.3566 | 6.4973 | 6.3811 | 6.2326 | 6.4293 | 6.3215 | 6.2520 | 6.3239 | 6.5133 | 5.9046 | 6.0004 | 5.9715 | 6.0271 |
| CNKSR2_S390 | CNKSR2 | Q8WXI2 | | 908 | EEKLGDSLQDLYR | 6.6375 | 6.3783 | 6.4857 | 6.4142 | 6.2090 | 5.8192 | 6.4200 | 6.0287 | 6.4696 | 6.3773 | 6.1915 | 6.3535 | 6.2706 | 6.1576 | 6.0255 | 5.7618 |
| CNKSR2_S908 | CNKSR2 | Q8WXI2 | | 325 | QPLIPRSPTSSVA | 5.6729 | 5.9047 | 6.4007 | 6.3251 | 6.3138 | 6.7132 | 6.2345 | 6.8958 | 6.2970 | 5.7485 | 6.7382 | 6.8262 | 6.1925 | 6.3993 | 5.6127 | 5.7252 |
| CNKSR2_T809 | CNKSR2 | Q8WXI2 | | 809 | EHRRQSTLPTQKC | 7.2443 | 6.8907 | 7.7067 | 6.5980 | 5.3855 | 6.3542 | 6.4053 | 6.1681 | 6.7385 | 6.3083 | 6.5739 | 7.3337 | 4.7725 | 5.5171 | 4.5845 | 5.4186 |
| CNKSR3_S383 | CNKSR3 | Q6P9H4 | | 383 | GPKGSESPNSFLD | 5.6209 | 5.5472 | 7.2540 | 7.2798 | 6.8864 | 6.4169 | 5.6634 | 5.3144 | 6.2491 | 5.8011 | 6.8242 | 6.4757 | 6.5130 | 6.2390 | 6.2347 | 5.6802 |
| CNKSR3_S479 | CNKSR3 | Q6P9H4 | | 479 | IIEESSSPPYRFS | 5.7985 | 6.3015 | 6.8989 | 6.8787 | 6.2017 | 6.5507 | 6.6295 | 5.7701 | 7.5851 | 6.9016 | 6.6286 | 6.5480 | 5.3295 | 4.8100 | 5.3734 | 5.7940 |
| CNN3_S259 | CNN3 | Q15417 | | 259 | ASQKGMSVYGLGR | 11.0695 | 7.4401 | 9.1469 | 6.3828 | 5.2207 | 6.1296 | 5.9821 | 5.3814 | 7.8374 | 6.3666 | 7.9550 | 6.5065 | 4.1648 | 4.6432 | 2.4015 | 3.3720 |
| CNNM3_S467 | CNNM3 | Q8NE01 | | 700 | GVPVEGSPGRNPG | 5.9809 | 5.7541 | 6.3069 | 6.2727 | 6.3351 | 6.0438 | 6.2695 | 6.2256 | 7.2429 | 6.6161 | 6.4453 | 6.7094 | 6.0280 | 5.9704 | 5.5463 | 6.2531 |

| | | | | | | | | | | | | | | | | | | | | | |
|---|---|---|---|---|---|---|---|---|---|---|---|---|---|---|---|---|---|---|---|---|---|
| CNNM3_S673 | CNNM3 | Q8NE01 | | 673 | LQVIPGSQTRLLG | 6.2995 | 6.3005 | 5.6481 | 6.0711 | 5.7576 | 4.9705 | 6.9949 | 5.1126 | 6.3689 | 5.6944 | 6.0922 | 5.2367 | 7.5063 | 7.9672 | 7.3758 | 6.6035 |
| CNNM3_S700 | CNNM3 | Q8NE01 | | 467 | VKRKPASLMAPLK | 7.1362 | 5.5507 | 7.3274 | 7.6275 | 8.0392 | 5.6490 | 6.5838 | 7.8453 | 7.6435 | 5.3488 | 5.1417 | 6.1562 | 5.3196 | 5.5932 | 4.4591 | 4.5787 |
| CNNM4_S664 | CNNM4 | Q6P4Q7 | | 664 | PLSRSASLSYPDR | 7.0495 | 6.9176 | 6.1334 | 6.4651 | 6.5958 | 5.7633 | 5.9849 | 5.9556 | 6.7681 | 5.7286 | 5.6755 | 5.6969 | 6.5958 | 6.5951 | 6.2196 | 5.8553 |
| CNNM4_S764 | CNNM4 | Q6P4Q7 | | 764 | LLNERNSLLHKAS | 6.6679 | 6.4226 | 6.0951 | 7.1322 | 6.8888 | 6.4173 | 6.3782 | 5.7961 | 6.0424 | 5.5416 | 5.9456 | 5.3035 | 6.4356 | 6.5996 | 6.4028 | 5.9308 |
| CNOT1_S1703 | CNOT1 | A5YKK6 | | 1703 | DGRAYGSPWCNKQ | 5.7062 | 6.2451 | 6.9165 | 6.5043 | 6.1787 | 6.6189 | 5.7473 | 6.0027 | 6.1758 | 6.6509 | 6.3971 | 6.0630 | 6.2302 | 6.2515 | 5.9507 | 6.3611 |
| CNOT10_S581 | CNOT10 | Q9H9A5-6 | | 581 | IPAPPSSPLRKQE | 6.3706 | 7.3718 | 6.7929 | 6.7163 | 6.3961 | 5.8719 | 6.3757 | 6.5303 | 6.5297 | 7.1561 | 6.6704 | 6.6254 | 4.9058 | 5.2063 | 5.3175 | 5.1633 |
| CNOT2_S165 | CNOT2 | Q9NZN8 | | 165 | SSSGLGSPNRSSP | 7.0724 | 6.6690 | 7.2241 | 6.1053 | 6.6182 | 6.7029 | 5.9826 | 7.0577 | 6.8524 | 7.2216 | 7.3806 | 7.3213 | 4.6048 | 4.5728 | 4.2808 | 4.3337 |
| CNOT2_S165;S170 | CNOT2 | Q9NZN8 | | 170 | GSPNRSSPSIICM | 5.3776 | 5.1079 | 5.2022 | 4.4347 | 5.9529 | 5.9754 | 5.3938 | 6.1205 | 5.5178 | 5.4549 | 5.0675 | 4.9908 | 9.9943 | 8.4824 | 8.7095 | 8.2177 |
| CNOT2_S170 | CNOT2 | Q9NZN8 | | 246 | EGSGNPTPLINPL | 6.8599 | 6.6196 | 5.7018 | 4.6591 | 5.5088 | 6.9110 | 7.3084 | 6.7653 | 7.3520 | 7.0397 | 6.7899 | 7.9291 | 4.3620 | 6.6083 | 4.8143 | 4.7707 |
| CNOT2_S305 | CNOT2 | Q9NZN8 | | 305 | KSNLNTSGKTTSS | 6.5252 | 7.1626 | 6.6339 | 7.0651 | 6.6428 | 6.9896 | 7.2747 | 7.1213 | 7.2799 | 7.6170 | 7.7914 | 7.2111 | 3.6737 | 4.2072 | 3.3837 | 3.4209 |
| CNOT2_T246 | CNOT2 | Q9NZN8 | 165;170 | | NA | 5.7543 | 5.1576 | 4.8491 | 5.4831 | 5.7880 | 5.8848 | 4.8298 | 5.9826 | 5.5938 | 5.1700 | 5.4202 | 5.5433 | 9.2970 | 8.5999 | 8.0582 | 8.5884 |
| CNOT3_S299 | CNOT3 | O75175 | | 299 | DSEVSQSPAKNGS | 6.1076 | 6.3211 | 6.8512 | 6.6952 | 6.2844 | 6.5145 | 6.2439 | 7.1108 | 6.9548 | 6.9920 | 7.0049 | 7.3734 | 4.6058 | 5.3120 | 4.7446 | 4.8838 |
| CNOT3_S507 | CNOT3 | O75175 | | 507 | PVNPPSSPTPSFS | 5.0020 | 6.2909 | 5.9805 | 5.7704 | 5.9275 | 6.0638 | 5.8755 | 6.3051 | 6.1319 | 5.0550 | 5.2651 | 6.2123 | 7.4368 | 7.6516 | 7.3886 | 7.6431 |
| CNOT4_S271 | CNOT4 | O95628-9 | | 298 | SNSDTPSPPPGLS | 5.7125 | 5.4474 | 6.8018 | 6.1174 | 6.0935 | 6.5814 | 5.6996 | 5.6547 | 6.4516 | 6.7327 | 6.2873 | 6.1575 | 6.8687 | 7.1586 | 5.8611 | 6.3742 |
| CNOT4_S298 | CNOT4 | O95628-9 | | 271 | KVTPLQSPIDKPS | 6.5458 | 6.1543 | 6.1637 | 5.9188 | 5.3105 | 5.7453 | 6.2469 | 6.5974 | 4.7174 | 5.9539 | 6.0453 | 6.0406 | 7.0017 | 6.3618 | 7.6510 | 7.5455 |
| CNP_S170 | CNP | P09543 | | 9 | RGFSRKSHTFLPK | 7.5883 | 6.9087 | 7.2643 | 5.9158 | 5.0974 | 6.5553 | 6.3749 | 6.0458 | 7.8284 | 7.7565 | 8.3103 | 8.3220 | 3.8542 | 5.1284 | 2.9650 | 4.0847 |
| CNP_S22 | CNP | P09543 | | 170 | KNQWQLSADDLKK | 3.2009 | 3.3220 | 3.6984 | 3.6641 | 4.2213 | 3.7661 | 4.1253 | 4.2686 | 3.9051 | 4.1047 | 3.5419 | 3.7338 | 13.6350 | 12.9503 | 14.5679 | 13.2947 |
| CNP_S409 | CNP | P09543 | | 409 | PVPTQGSRKGGAL | 5.3321 | 6.4282 | 7.6044 | 6.4726 | 5.6301 | 5.7089 | 5.5265 | 7.0518 | 6.4736 | 9.4129 | 8.0711 | 8.2958 | 4.2835 | 5.0743 | 3.9734 | 4.6607 |
| CNP_S9 | CNP | P09543 | | 22 | IFFRKMSSSGAKD | 6.5527 | 6.3612 | 6.8888 | 6.4265 | 5.9946 | 6.2289 | 5.6218 | 6.3900 | 7.3397 | 8.4612 | 8.2976 | 8.4264 | 3.8380 | 5.3480 | 3.7016 | 4.1230 |
| CNTLN_S1085 | CNTLN | Q9NXG0-2 | | 22 | RQLGPRSPRVGRG | 6.5510 | 6.5523 | 5.9228 | 6.5646 | 6.2000 | 5.9423 | 6.3992 | 5.9290 | 6.2520 | 6.3398 | 6.1369 | 6.0631 | 6.3198 | 6.4884 | 6.2079 | 6.1309 |
| CNTLN_S22 | CNTLN | Q9NXG0-2 | | 1085 | IQVTSLSPSRSMD | 6.9228 | 6.6262 | 6.3189 | 6.8406 | 6.2444 | 6.3354 | 6.7289 | 7.0047 | 6.9586 | 6.6933 | 7.1857 | 7.4944 | 4.3605 | 4.8091 | 4.5057 | 4.9708 |
| CNTROB_S154 | CNTROB | Q8N137-2 | | 62 | LYLPSTSPPHEGL | 6.2322 | 6.4271 | 6.5185 | 6.5453 | 6.2184 | 6.6645 | 6.3211 | 6.8460 | 6.2485 | 6.7993 | 7.1829 | 6.5520 | 5.4144 | 5.6620 | 5.1575 | 5.2106 |
| CNTROB_S62 | CNTROB | Q8N137-2 | | 154 | RPLQDLSPSSSAQ | 5.1615 | 5.6843 | 5.5156 | 5.9662 | 5.5343 | 5.8190 | 6.7665 | 6.3293 | 5.6929 | 5.6312 | 6.0563 | 5.9521 | 7.9054 | 7.4619 | 7.3448 | 7.1788 |
| COASY_S19 | COASY | Q13057-2 | | 212 | ASPVAGSPKQPVR | 6.3260 | 5.2554 | 6.6633 | 5.7113 | 6.0926 | 6.4101 | 5.7231 | 6.1163 | 7.1075 | 6.5960 | 6.2721 | 7.0030 | 6.0277 | 6.3961 | 5.5414 | 6.7582 |
| COASY_S207 | COASY | Q13057-2 | | 207 | STIRPASPVAGSP | 5.3601 | 5.2401 | 6.5992 | 5.4344 | 5.6304 | 6.6369 | 6.1390 | 6.8154 | 6.8133 | 5.9595 | 6.3104 | 6.5868 | 6.4802 | 6.8702 | 6.1817 | 6.9424 |
| COASY_S212 | COASY | Q13057-2 | | 19 | AVYQAPSPPPAPV | 5.3701 | 4.8858 | 7.7822 | 7.0177 | 6.5666 | 5.3226 | 6.6331 | 6.8088 | 5.7642 | 6.3007 | 8.7508 | 6.2580 | 5.2087 | 6.7233 | 4.8763 | 5.7310 |
| COBLL1_S326 | COBLL1 | Q53SF7-3 | | 260 | TASAPATPLVNKH | 7.1458 | 6.6517 | 6.1210 | 6.6082 | 5.5260 | 6.1323 | 7.0173 | 5.6587 | 6.4226 | 5.7257 | 6.0007 | 6.1124 | 6.2198 | 6.6464 | 5.2498 | 6.7616 |
| COBLL1_S356 | COBLL1 | Q53SF7-3 | | 949 | ERDMLPSPEQTLS | 5.9954 | 6.4663 | 6.3573 | 5.9574 | 6.4369 | 5.9358 | 6.0805 | 6.3119 | 7.0316 | 6.2245 | 6.3717 | 6.6261 | 6.0439 | 5.9763 | 6.0917 | 6.0926 |
| COBLL1_S366 | COBLL1 | Q53SF7-3 | | 356 | MSAGNSSLRRTKR | 6.4976 | 6.4082 | 6.9297 | 6.4003 | 5.9568 | 6.2677 | 6.0074 | 6.1660 | 6.8073 | 6.6241 | 6.1978 | 6.7151 | 5.7298 | 5.9467 | 5.1161 | 6.2295 |
| COBLL1_S477 | COBLL1 | Q53SF7-3 | | 477 | VEAENISPKSQDI | 5.4887 | 5.7463 | 6.1171 | 6.2116 | 6.1024 | 5.8140 | 6.2233 | 5.6202 | 6.5199 | 6.4622 | 6.0891 | 6.2332 | 6.4692 | 6.7779 | 7.0008 | 7.1240 |
| COBLL1_S904 | COBLL1 | Q53SF7-3 | | 955 | SPEQTLSPLSKMP | 6.4431 | 6.4514 | 5.8722 | 6.2388 | 6.7440 | 6.8429 | 6.6693 | 6.3417 | 6.7358 | 7.1658 | 6.8889 | 6.9886 | 4.8343 | 5.8342 | 4.4884 | 5.4607 |
| COBLL1_S949 | COBLL1 | Q53SF7-3 | | 985 | QAPAEASPPPIAP | 5.6304 | 6.2638 | 6.5934 | 6.5617 | 6.7174 | 6.0321 | 6.4992 | 6.2032 | 6.5838 | 7.0010 | 6.9865 | 7.2499 | 5.2728 | 5.2478 | 5.5727 | 5.5842 |
| COBLL1_S955 | COBLL1 | Q53SF7-3 | | 366 | TKRKAPSPPSKIP | 5.9663 | 6.3944 | 5.8233 | 6.1607 | 5.6567 | 5.8309 | 5.6246 | 6.0236 | 6.2528 | 6.7038 | 6.6318 | 6.7080 | 5.9860 | 6.9418 | 6.6089 | 6.6865 |
| COBLL1_S985 | COBLL1 | Q53SF7-3 | | 326 | CIVKSMSVDETDK | 4.3762 | 5.3747 | 7.3897 | 5.5947 | 5.8357 | 5.7313 | 5.6170 | 5.7740 | 5.4893 | 6.6707 | 7.0860 | 6.1844 | 6.0964 | 7.8544 | 7.2630 | 7.6626 |
| COBLL1_T260 | COBLL1 | Q53SF7-3 | | 284 | KPYISNTLPSDAP | 6.7500 | 5.8611 | 7.5388 | 6.8246 | 6.0525 | 5.7200 | 5.7609 | 5.0854 | 6.2921 | 5.8970 | 5.9084 | 5.7629 | 6.6109 | 7.6526 | 6.3010 | 5.9818 |
| COBLL1_T284 | COBLL1 | Q53SF7-3 | | 904 | AQAKPSSFFLQMQ | 6.4293 | 7.2057 | 7.8487 | 6.5282 | 7.1118 | 5.6677 | 6.7211 | 5.6874 | 6.2117 | 6.4441 | 7.8091 | 7.1464 | 5.2473 | 5.0890 | 4.5796 | 4.2730 |
| COG4_S6 | COG4 | Q9H9E3 | | 6 | xMADLDSPPKLSG | 6.0065 | 6.4584 | 6.2306 | 6.7632 | 6.0393 | 6.1967 | 6.0337 | 6.5589 | 6.1416 | 6.6962 | 7.0934 | 6.7316 | 5.4092 | 6.2045 | 5.7426 | 5.6935 |
| COG5_S197 | COG5 | Q9UP83-2 | | 197 | GQLQGGSREITKA | 6.4484 | 6.5551 | 6.2926 | 6.1424 | 6.0981 | 6.1959 | 6.2437 | 6.2194 | 6.5171 | 5.9524 | 6.2883 | 6.1746 | 6.2053 | 6.1196 | 6.4662 | 6.0809 |
| COIL_S235 | COIL | P38432 | | 303 | KLGFSLTPSKGKT | 6.6681 | 6.5699 | 6.2734 | 6.8925 | 6.5758 | 6.3771 | 6.7440 | 6.5296 | 6.7794 | 6.5025 | 6.2150 | 6.2563 | 5.4169 | 5.5375 | 5.2226 | 5.4394 |
| COIL_S240 | COIL | P38432 | | 566 | PRLIIESPSNTSS | 6.2938 | 6.5844 | 6.3320 | 6.8884 | 6.0631 | 6.3015 | 6.0601 | 6.0427 | 6.4250 | 6.3118 | 6.0100 | 6.4735 | 6.2806 | 6.4098 | 5.6441 | 5.8791 |
| COIL_S280 | COIL | P38432 | | 122 | LEEGEETEPDCKY | 7.1335 | 6.6382 | 6.4385 | 7.8749 | 6.7933 | 5.8341 | 6.0289 | 5.4672 | 5.9870 | 5.2570 | 5.5291 | 5.2613 | 6.7892 | 6.5162 | 6.1932 | 6.2584 |
| COIL_S301;S305 | COIL | P38432 | | 280 | KLPTELSKEEPST | 6.5477 | 6.7903 | 6.5717 | 6.9718 | 6.2535 | 6.3344 | 7.0409 | 6.3891 | 6.0725 | 6.0673 | 6.0878 | 6.5247 | 5.3241 | 5.7827 | 5.5071 | 5.7342 |
| COIL_S301;T303 | COIL | P38432 | | 486 | KLLELTSSYSPDV | 5.7978 | 5.9496 | 6.0253 | 6.7806 | 5.9546 | 6.2368 | 5.9283 | 6.1860 | 6.1422 | 5.7596 | 5.7718 | 6.1919 | 6.8436 | 6.6209 | 6.9866 | 6.8245 |
| COIL_S403 | COIL | P38432 | | 403 | REENLFSWKGAKG | 7.0692 | 6.0893 | 5.8941 | 6.9513 | 6.6621 | 6.5277 | 6.0746 | 5.7467 | 6.8997 | 7.5278 | 8.3843 | 6.8334 | 3.6525 | 5.7430 | 5.1407 | 4.8037 |
| COIL_S486 | COIL | P38432 | | 456 | IQNPVETPKKDYS | 6.6608 | 6.5922 | 6.1053 | 7.1519 | 6.5748 | 6.2478 | 7.2000 | 5.8709 | 6.7740 | 6.4092 | 7.0929 | 6.1291 | 5.2198 | 5.4257 | 4.6353 | 5.9104 |
| COIL_S566 | COIL | P38432 | | 235 | KAKRKGSVSVCSK | 4.9199 | 4.8280 | 5.8082 | 6.4518 | 8.2736 | 5.1441 | 4.9208 | 4.9304 | 8.0134 | 8.9333 | 7.8112 | 7.6999 | 6.2169 | 6.4809 | 4.3479 | 5.2198 |
| COIL_T122 | COIL | P38432 | | 240 | GSVSVCSKESPSS | 5.7752 | 7.2105 | 6.9014 | 8.0759 | 6.2634 | 5.3173 | 6.3326 | 5.2559 | 6.2278 | 7.4070 | 5.6201 | 6.7210 | 5.6674 | 5.8264 | 5.6188 | 5.7794 |
| COIL_T303 | COIL | P38432 | 301;305 | | NA | 7.1940 | 6.9274 | 6.1889 | 6.9514 | 6.7214 | 6.4480 | 6.6138 | 5.9627 | 6.3923 | 6.1734 | 6.3054 | 6.2546 | 5.5326 | 5.6781 | 5.1850 | 5.4710 |
| COIL_T303;S305 | COIL | P38432 | 301;303 | | NA | 6.7586 | 6.4624 | 6.4296 | 7.2707 | 6.5232 | 6.4543 | 6.7741 | 5.8912 | 6.3800 | 6.2290 | 6.1723 | 6.0476 | 5.6640 | 5.7838 | 5.4105 | 5.7488 |
| COIL_T456 | COIL | P38432 | 303;305 | | NA | 6.8816 | 6.6014 | 6.4349 | 7.2125 | 6.7375 | 6.3755 | 7.3862 | 5.9238 | 6.6820 | 5.4907 | 5.3456 | 5.4925 | 5.9134 | 5.4101 | 6.5785 | 6.4338 |
| COMT_S131 | COMT | P21964 | | 131 | RMARLLSPGARLI | 6.4798 | 6.6055 | 6.1339 | 6.2960 | 6.0879 | 6.0463 | 6.1529 | 6.2232 | 6.1626 | 6.3128 | 6.3775 | 6.3280 | 5.8790 | 6.2667 | 6.3149 | 6.3330 |
| COPA_S173 | COPA | P53621-2 | | 173 | LRKKNLSPGAVES | 6.0925 | 6.8662 | 6.5313 | 6.9888 | 6.0914 | 6.5677 | 6.7156 | 6.5099 | 6.2693 | 6.8951 | 6.5276 | 6.6470 | 4.7840 | 5.6419 | 5.3366 | 5.5351 |
| COPA_S904 | COPA | P53621-2 | | 600 | CRPRVLTIDPTEF | 6.7267 | 6.4617 | 6.4292 | 6.1035 | 6.8500 | 6.4739 | 6.1219 | 6.2417 | 6.4054 | 5.8084 | 6.3361 | 6.1963 | 5.5943 | 6.3055 | 5.9638 | 5.9814 |
| COPA_T600 | COPA | P53621-2 | | 904 | PPELDISPGAAGG | 6.0113 | 5.9744 | 5.8230 | 5.8045 | 6.4692 | 6.9065 | 7.0261 | 6.4869 | 6.3666 | 6.9545 | 5.9589 | 5.7402 | 6.1140 | 6.4899 | 6.0645 | 5.8097 |
| COPB2_S859 | COPB2 | P35606 | | 859 | LDGKPASPTPVIV | 6.1170 | 5.6920 | 6.9783 | 6.2097 | 6.2723 | 6.3499 | 6.0999 | 5.9932 | 6.2807 | 6.3675 | 6.3859 | 6.5663 | 6.0741 | 6.1709 | 5.8058 | 6.6366 |
| COPE_S99 | COPE | O14579 | | 99 | HESRRDSIVAELD | 5.5282 | 5.1461 | 5.9256 | 4.6196 | 5.1639 | 5.9868 | 5.7094 | 6.7779 | 5.8545 | 6.7674 | 5.5705 | 6.5274 | 6.1423 | 8.3851 | 8.4371 | 7.4581 |
| COPG1_S593 | COPG1 | Q9Y678 | | 594 | EQRTESTPITAVK | 6.1901 | 6.6867 | 6.3928 | 7.0918 | 6.4369 | 6.7813 | 7.2307 | 6.6503 | 7.2232 | 7.2573 | 7.0623 | 7.1494 | 4.1359 | 5.2459 | 3.6514 | 4.8140 |
| COPG1_T594 | COPG1 | Q9Y678 | | 593 | AEQRTESTPITAV | 5.6575 | 7.8545 | 7.3289 | 7.1939 | 5.9367 | 6.5427 | 6.5179 | 6.8116 | 6.2696 | 8.3708 | 7.7082 | 7.8174 | 3.8232 | 4.3858 | 3.1822 | 4.5991 |
| COPRS_S66 | COPRS | Q9NQ92 | | 66 | QSIPNDSPARGEG | 7.0777 | 6.4274 | 6.1143 | 5.6066 | 5.2190 | 4.8343 | 6.5555 | 7.8359 | 5.5295 | 7.1311 | 6.6005 | 6.5031 | 5.1383 | 7.4301 | 6.1562 | 6.7406 |
| COQ4_S106 | COQ4 | Q9Y3A0 | | 106 | QERPRISTSTLDL | 5.5694 | 5.4802 | 4.6933 | 3.9386 | 5.2271 | 4.9770 | 5.7873 | 5.6542 | 5.0458 | 5.2388 | 4.4491 | 5.4465 | 8.6831 | 9.1029 | 10.1384 | 10.5683 |
| CORO7_S21 | CORO7 | P57737-3 | | 21 | RPPRRESWISDIR | 6.0687 | 5.6190 | 6.0220 | 5.5322 | 6.7561 | 6.3376 | 6.1364 | 6.5326 | 6.9195 | 6.3042 | 6.7632 | 6.7606 | 6.2259 | 6.2833 | 5.2647 | 6.4739 |
| CORO7_S806 | CORO7 | P57737-3 | | 806 | CLRLRQSSLEPVA | 6.0103 | 6.5524 | 7.6608 | 6.5856 | 6.7688 | 6.4579 | 6.2988 | 7.5779 | 6.2838 | 5.5054 | 6.3593 | 6.3808 | 4.9176 | 5.7218 | 5.7483 | 5.1705 |
| COX4I1_S158 | COX4I1 | P13073 | | 158 | PIQGLASKWDYEK | 6.5812 | 6.8669 | 7.3339 | 7.6935 | 6.3097 | 5.5015 | 7.1515 | 6.0023 | 5.6080 | 5.9501 | 5.9252 | 6.5714 | 5.7134 | 6.1641 | 4.7173 | 5.9100 |
| COX6C_S73 | COX6C | P09669 | | 73 | KAGIFQSVKxxxx | 7.1078 | 4.7879 | 5.8639 | 8.5713 | 10.8295 | 8.3664 | 8.3368 | 6.7869 | 5.3753 | 5.0347 | 4.5904 | 5.1951 | 4.6003 | 4.9039 | 5.4863 | 4.1635 |
| CPD_S1355 | CPD | O75976 | | 1355 | RMMSTGSKKSLLS | 3.0001 | 3.3505 | 3.5435 | 3.1913 | 3.4531 | 4.5943 | 3.7939 | 4.2315 | 3.5850 | 4.9486 | 5.6334 | 6.3384 | 11.3068 | 13.0359 | 12.2506 | 13.7433 |
| CPD_T1368 | CPD | O75976 | | 1368 | HEFQDETDTEEET | 7.0992 | 7.4210 | 6.5309 | 7.5508 | 6.0077 | 6.0077 | 6.1117 | 5.9020 | 6.3692 | 6.4453 | 6.5054 | 6.0693 | 5.1886 | 5.7836 | 5.4623 | 5.5453 |
| CPD_T1368;T1370 | CPD | O75976 | 1368;1370 | | NA | 6.2427 | 5.9290 | 5.8397 | 6.1072 | 6.8633 | 6.6714 | 5.9664 | 7.0059 | 5.9111 | 5.7448 | 5.9224 | 6.0729 | 6.4015 | 6.9051 | 6.4830 | 5.9337 |
| CPEB4_S97 | CPEB4 | Q17RY0 | | 97 | LEKQQLSPSPGQE | 6.2021 | 5.8320 | 6.5971 | 6.2661 | 6.0226 | 6.3415 | 6.4640 | 6.2764 | 6.1354 | 6.1200 | 6.3907 | 6.5332 | 6.5474 | 6.4203 | 5.7427 | 6.1088 |
| CPEB4_S99 | CPEB4 | Q17RY0 | | 99 | KQQLSPSPGQEAG | 5.8761 | 5.9517 | 6.8105 | 6.3867 | 6.0309 | 6.7044 | 5.9131 | 5.8903 | 6.6471 | 6.3364 | 6.2329 | 6.1629 | 5.9502 | 6.0989 | 6.1106 | 6.8973 |
| CPLANE1_T3071 | CPLANE1 | Q9H799 | | 3071 | AAVRKATQSPVTF | 7.8332 | 9.2348 | 7.5412 | 7.4490 | 6.2012 | 7.2936 | 6.1301 | 7.1054 | 6.9628 | 6.7157 | 6.8556 | 7.7727 | 2.6040 | 3.7215 | 3.0470 | 3.5322 |
| CPNE1_S55 | CPNE1 | Q99829 | | 55 | RVRNCSSPEFSKT | 5.9179 | 6.1502 | 6.9602 | 6.2071 | 6.1871 | 6.8470 | 5.9367 | 6.0863 | 6.1871 | 7.0413 | 6.8792 | 6.6764 | 5.7965 | 6.0300 | 5.4856 | 5.6116 |
| CPNE7_S628 | CPNE7 | Q9UBL6 | | 628 | VPAGEASPGCTPx | 6.4398 | 6.3409 | 6.4339 | 5.9161 | 5.6879 | 6.1452 | 5.7700 | 6.3660 | 7.6220 | 6.9729 | 6.5116 | 6.7265 | 5.3820 | 6.3862 | 5.6033 | 5.6957 |
| CPNE8_S260 | CPNE8 | Q86YQ8 | | 554 | PAPPPYTPPTHVL | 6.3126 | 5.7826 | 6.3938 | 6.8021 | 6.4756 | 6.6962 | 6.5515 | 6.2135 | 6.5458 | 6.2299 | 6.5829 | 6.5596 | 5.5208 | 5.8011 | 5.4497 | 6.0823 |
| CPNE8_S7 | CPNE8 | Q86YQ8 | | 260 | ELSRGQSQFNVYE | 6.2644 | 6.4323 | 6.0264 | 7.5620 | 7.2271 | 6.6348 | 7.4321 | 6.8430 | 5.6003 | 6.1571 | 7.0343 | 6.5977 | 3.9985 | 5.1347 | 4.8248 | 6.2305 |
| CPNE8_T554 | CPNE8 | Q86YQ8 | | 7 | MDSRYNSTAGIGD | 6.5784 | 6.6231 | 6.1916 | 6.8045 | 7.3509 | 6.5337 | 6.8540 | 5.4349 | 7.4766 | 7.5237 | 7.1031 | 5.9523 | 5.8157 | 5.2668 | 4.0278 | 4.4630 |
| CPSF1_S734;S756 | CPSF1 | Q10570 | | 756 | DSGSLFSPSKEEA | 6.2971 | 6.3790 | 6.5466 | 7.1348 | 6.0256 | 6.1380 | 6.0655 | 6.5539 | 6.2264 | 6.8389 | 6.6625 | 6.1430 | 5.2924 | 5.7080 | 5.7787 | 6.2096 |
| CPSF1_S756 | CPSF1 | Q10570 | | 766 | EEARRSSQPPADR | 5.0073 | 4.5695 | 4.9952 | 5.7252 | 5.6780 | 5.9028 | 5.6550 | 6.1164 | 5.2049 | 5.0930 | 5.2282 | 5.2983 | 8.5784 | 8.7443 | 9.0026 | 9.2009 |
| CPSF1_S766 | CPSF1 | Q10570 | 734;756 | | NA | 4.1908 | 7.4538 | 7.0417 | 5.1554 | 6.1331 | 6.7312 | 6.3311 | 5.9861 | 6.0110 | 5.9778 | 7.9932 | 5.1459 | 6.4667 | 7.2125 | 6.2755 | 5.8941 |
| CPSF3_T681 | CPSF3 | Q9UKF6 | | 681 | RLYEALTPVHxxx | 6.7034 | 7.1836 | 6.0676 | 6.7192 | 6.8855 | 6.6738 | 6.8927 | 6.8168 | 6.6658 | 6.3381 | 6.2002 | 6.3012 | 4.9180 | 5.3518 | 4.9530 | 5.3293 |
| CPSF4_S212 | CPSF4 | O95639 | | 212 | LTSQNSSPNQQRT | 6.1668 | 6.2366 | 6.1996 | 6.7061 | 6.4511 | 6.6218 | 5.9568 | 6.5045 | 5.9568 | 5.9700 | 6.2064 | 6.0591 | 6.2009 | 6.5813 | 6.0793 | 6.1028 |
| CPSF6_S258 | CPSF6 | Q16630-2 | | 444 | EMDTARTPLSEAE | 5.9840 | 6.2244 | 5.9553 | 6.0453 | 6.3409 | 6.2177 | 6.0586 | 6.1043 | 6.1495 | 6.2018 | 6.3513 | 6.2314 | 6.6014 | 6.6064 | 6.4707 | 6.4570 |
| CPSF6_T444 | CPSF6 | Q16630-2 | | 258 | PWHMGHSIEEIPI | 6.0018 | 6.4161 | 8.0911 | 6.4047 | 6.5859 | 6.0310 | 5.6569 | 6.7467 | 7.1431 | 9.6553 | 7.8463 | 6.3247 | 3.5279 | 5.2802 | 4.1697 | 4.1185 |
| CPSF7_S103 | CPSF7 | Q8N684-3 | | 246 | SADGRATPSENLV | 6.3246 | 6.1530 | 5.2043 | 5.9019 | 6.2546 | 5.8181 | 6.2865 | 6.1355 | 6.5787 | 5.8232 | 5.4674 | 5.4681 | 7.1758 | 6.9571 | 7.0801 | 7.3711 |
| CPSF7_S91 | CPSF7 | Q8N684-3 | | 103 | PVRQEPSPKPNNK | 6.3785 | 6.4292 | 5.8422 | 6.0665 | 6.6876 | 6.4135 | 6.7597 | 6.4436 | 6.3181 | 5.9880 | 6.4234 | 6.2098 | 6.1800 | 6.0924 | 5.9395 | 5.8280 |
| CPSF7_T246 | CPSF7 | Q8N684-3 | | 91 | SDDRSSSTEPPPP | 6.5264 | 6.4921 | 6.0085 | 6.8289 | 6.3463 | 5.6635 | 5.6975 | 6.5229 | 6.3530 | 6.6472 | 6.5290 | 6.2764 | 6.0375 | 6.4194 | 5.8653 | 5.7861 |
| CPT1A_S429 | CPT1A | P50416 | | 429 | TEEGYRSEDPDTS | 5.2916 | 6.7148 | 6.4079 | 6.3220 | 6.4284 | 6.0956 | 6.2640 | 6.2238 | 6.8921 | 6.0124 | 6.3486 | 6.3936 | 6.3015 | 6.2361 | 6.2102 | 5.8576 |
| CRACD_S1017 | CRACD | Q6ZU35 | | 1017 | SDRRPPSPPGPEE | 6.1924 | 6.2842 | 6.0227 | 6.5300 | 6.0800 | 6.2521 | 6.1904 | 6.1890 | 6.2224 | 6.2003 | 5.6561 | 6.1163 | 6.5331 | 6.8970 | 6.3076 | 6.3265 |
| CRACD_S1072 | CRACD | Q6ZU35 | | 556 | FPKVNLSPVTPAK | 7.1231 | 7.1341 | 6.5272 | 7.3323 | 7.3612 | 6.6691 | 6.8351 | 6.6031 | 7.1354 | 7.1515 | 6.9292 | 6.5233 | 3.9168 | 4.5109 | 3.9640 | 4.2839 |
| CRACD_S1201 | CRACD | Q6ZU35 | | 1072 | MLQSRHSLDGSKL | 7.2276 | 6.3852 | 6.9716 | 6.9020 | 6.3694 | 5.2551 | 5.4068 | 6.3030 | 7.4147 | 7.2251 | 6.7332 | 6.1166 | 5.2863 | 5.9122 | 5.3972 | 5.0939 |
| CRACD_S556 | CRACD | Q6ZU35 | | 1201 | EVTKRFSTPDAAP | 6.4236 | 4.7715 | 6.4073 | 5.1519 | 7.5213 | 4.0052 | 4.3095 | 6.1573 | 8.4072 | 9.8092 | 6.3040 | 6.2497 | 6.8529 | 5.7225 | 7.1953 | 4.7117 |
| CRACD_S969;S975 | CRACD | Q6ZU35 | 969;975 | | NA | 6.1665 | 7.1664 | 5.3191 | 6.4318 | 7.1736 | 4.9244 | 6.0727 | 5.8027 | 6.8927 | 6.1949 | 6.1528 | 6.5632 | 6.2635 | 5.9896 | 5.8325 | 7.0534 |
| CRAMP1_S533 | CRAMP1 | Q96RY5 | | 533 | PAEGRDSPTREPG | 5.8847 | 6.3301 | 6.3595 | 7.0545 | 6.7614 | 5.7017 | 6.1982 | 5.8783 | 6.7934 | 5.8642 | 6.2418 | 6.7703 | 5.0669 | 6.9956 | 4.9683 | 7.1312 |
| CREB1_S142 | CREB1 | P16220-1 | | 142 | KILNDLSSDAPGV | 6.4600 | 5.9729 | 7.0869 | 6.5198 | 6.2433 | 5.7623 | 6.5451 | 6.6220 | 6.3263 | 6.9856 | 6.0559 | 6.5937 | 5.1222 | 5.9597 | 5.6144 | 6.1299 |
| CREB1_S271 | CREB1 | P16220-1 | | 271 | GVVMASSPALPTQ | 6.0503 | 6.6508 | 7.7937 | 7.0856 | 4.9590 | 6.7046 | 6.4088 | 6.3662 | 6.7203 | 6.3259 | 6.0637 | 6.9937 | 4.5153 | 6.8928 | 5.0441 | 5.4251 |
| CREB3L1_S244 | CREB3L1 | Q96BA8 | | 244 | STAISTSPLLTAP | 7.0624 | 7.0330 | 6.5761 | 7.9026 | 7.1432 | 7.5681 | 7.0267 | 6.8297 | 6.9118 | 7.4642 | 7.0830 | 7.0877 | 3.3318 | 4.1372 | 3.0905 | 3.7519 |
| CREB3L2_S249 | CREB3L2 | Q70SY1 | | 249 | PSALSSSPLLTAP | 6.2513 | 6.0238 | 6.4732 | 6.6203 | 6.8813 | 7.2347 | 7.2008 | 7.0872 | 6.7854 | 6.3057 | 6.5731 | 5.8647 | 5.1727 | 5.4933 | 4.9720 | 5.0604 |
| CREBBP_S121 | CREBBP | Q92793 | | 974 | IDNRVPTPSSVAS | 5.5191 | 5.1364 | 5.5942 | 5.5173 | 5.3856 | 5.6989 | 5.7555 | 6.0007 | 5.7590 | 5.9565 | 5.2571 | 5.8553 | 8.0805 | 7.5052 | 7.9170 | 9.0618 |
| CREBBP_S2063 | CREBBP | Q92793 | | 2063 | QPPRSISPSALQD | 5.4049 | 4.6386 | 4.1691 | 5.5853 | 6.6304 | 6.7609 | 5.9025 | 6.8760 | 5.7465 | 5.0467 | 5.9857 | 5.1721 | 7.3852 | 8.8423 | 8.4342 | 7.4197 |
| CREBBP_T974 | CREBBP | Q92793 | | 121 | LSAMGKSPLSQGD | 5.8120 | 5.9109 | 5.9014 | 6.0773 | 5.6588 | 6.7418 | 5.9068 | 5.7964 | 6.2734 | 6.0666 | 5.7037 | 6.1053 | 6.6005 | 7.8051 | 7.0586 | 6.5812 |
| CRIM1_S1016 | CRIM1 | Q9NZV1 | | 1016 | EPDARFSGFYSMQ | 8.2014 | 7.6021 | 8.2175 | 8.1471 | 7.0510 | 6.1591 | 6.6577 | 6.1312 | 7.2631 | 5.9605 | 6.4959 | 6.3673 | 4.2289 | 4.1829 | 3.5014 | 3.8328 |
| CRIM1_S1020 | CRIM1 | Q9NZV1 | | 1020 | RFSGFYSMQKQNH | 7.2233 | 6.8737 | 7.8116 | 8.8134 | 5.3474 | 5.8608 | 4.9034 | 5.6582 | 5.7387 | 7.9004 | 9.5516 | 7.0652 | 2.5003 | 5.6138 | 4.6343 | 4.5038 |
| CRIP2_S189 | CRIP2 | P52943-2 | | 189 | GPSRASSVTTFTG | 7.4033 | 6.4043 | 6.7901 | 5.6976 | 6.3499 | 6.7407 | 6.5833 | 6.2954 | 7.1248 | 6.7949 | 6.8783 | 6.4762 | 5.2843 | 5.8963 | 4.5434 | 4.7373 |
| CRIPT_T15 | CRIPT | Q9P021 | | 15 | KLGTVITPDTWKD | 6.2278 | 7.4983 | 7.5683 | 7.2182 | 6.0585 | 7.1380 | 8.0546 | 6.8847 | 8.3920 | 7.7517 | 7.8026 | 7.0171 | 2.5181 | 4.4010 | 2.3959 | 3.0732 |
| CRK_S194 | CRK | P46108-2 | | 194 | EKYRPASASVSAL | 6.5957 | 5.8514 | 6.1586 | 5.4373 | 6.8231 | 6.3931 | 7.0133 | 6.2034 | 6.5254 | 6.0445 | 5.5389 | 6.0522 | 6.2861 | 6.9436 | 5.8727 | 6.2607 |
| CRKL_S107 | CRKL | P46109 | | 107 | PAPRYPSPPMGSV | 6.2574 | 6.2176 | 6.4565 | 6.0186 | 6.1038 | 6.6345 | 6.2570 | 6.2594 | 6.3484 | 6.2476 | 6.5318 | 6.5312 | 6.0965 | 6.2039 | 5.6590 | 6.1767 |
| CRKL_S42 | CRKL | P46109 | | 207 | PEPAHAYAQPQTT | 5.6414 | 6.0383 | 7.3967 | 6.4066 | 6.6079 | 6.4392 | 5.6099 | 6.9688 | 6.9705 | 6.9997 | 7.0543 | 7.0717 | 4.8273 | 6.0721 | 4.9184 | 4.9774 |
| CRKL_Y198 | CRKL | P46109 | | 198 | NRNSNSYGIPEPA | 5.8409 | 5.7661 | 7.3169 | 7.0088 | 7.0621 | 5.3742 | 5.9411 | 6.1959 | 6.3228 | 6.2467 | 7.9180 | 6.7045 | 5.9892 | 5.2893 | 4.9900 | 6.0336 |
| CRKL_Y207 | CRKL | P46109 | | 42 | FLVRDSSTCPGDY | 5.4150 | 5.7807 | 6.7522 | 5.8394 | 7.2391 | 5.9985 | 6.3694 | 6.0156 | 6.8399 | 6.5582 | 6.8280 | 6.7083 | 6.2304 | 6.2349 | 5.1325 | 6.0579 |
| CRKL_Y207;T212 | CRKL | P46109 | 207;212 | | NA | 6.4586 | 7.2013 | 6.9402 | 6.9982 | 7.1230 | 6.3272 | 7.5144 | 7.6096 | 8.4895 | 8.5504 | 7.2137 | 8.5070 | 3.1842 | 2.6641 | 2.4144 | 2.8042 |
| CROCC_S1451 | CROCC | Q5TZA2 | | 2009 | RSSAPFSPPSGPP | 6.5615 | 6.3368 | 6.0752 | 5.7795 | 6.8112 | 6.1012 | 6.5798 | 6.7615 | 6.3847 | 6.2731 | 5.8222 | 5.6075 | 6.7347 | 6.3935 | 6.1353 | 5.6420 |
| CROCC_S1460 | CROCC | Q5TZA2 | | 1460 | PRPVPGSPARDAP | 6.2103 | 5.3989 | 6.3150 | 5.8783 | 6.3192 | 5.8452 | 5.9265 | 5.6688 | 6.4312 | 6.2212 | 6.1605 | 5.6745 | 6.9633 | 6.2700 | 6.2753 | 6.6417 |
| CROCC_S2009 | CROCC | Q5TZA2 | | 512 | SPRRGPSPACSDS | 6.2282 | 5.4583 | 6.2024 | 6.1938 | 7.8685 | 6.1015 | 5.8275 | 6.0532 | 6.2598 | 5.6640 | 6.5931 | 5.5908 | 6.9403 | 6.7624 | 6.1704 | 6.0857 |
| CROCC_S512 | CROCC | Q5TZA2 | | 1451 | GLGRAPSPAPRPV | 5.9832 | 6.4929 | 6.7679 | 6.7911 | 6.8633 | 6.0983 | 5.8938 | 6.5247 | 6.2488 | 6.3803 | 6.7043 | 6.6656 | 6.6012 | 6.2866 | 4.6466 | 5.0514 |
| CRTC1_S139 | CRTC1 | Q6UUV9 | | 64 | GQYYGGSLPNVNQ | 6.0797 | 6.0125 | 6.7082 | 5.8974 | 6.1230 | 6.5568 | 6.4364 | 6.3087 | 6.6502 | 6.1460 | 6.4233 | 6.4552 | 6.1095 | 6.5166 | 5.8550 | 5.7215 |
| CRTC1_S64 | CRTC1 | Q6UUV9 | | 139 | YGTMYLSPPADTS | 6.3517 | 5.9734 | 7.1848 | 5.0916 | 6.8181 | 6.1892 | 7.4702 | 6.4515 | 7.0873 | 7.3101 | 6.7856 | 6.4190 | 5.1426 | 5.8899 | 5.5510 | 4.2840 |
| CRTC2_S136 | CRTC2 | Q53ET0 | | 456 | QLPKQFSPTMSPT | 6.0742 | 5.6104 | 6.5393 | 5.5455 | 6.4980 | 6.5521 | 5.8240 | 6.8293 | 6.3837 | 6.2512 | 6.2375 | 6.4286 | 6.8275 | 6.0803 | 6.0713 | 6.2470 |
| CRTC2_S433 | CRTC2 | Q53ET0 | | 433 | HRRVPLSPLSLLA | 5.8285 | 5.4496 | 6.4748 | 5.9637 | 6.3706 | 7.1342 | 6.3964 | 6.5637 | 6.1694 | 5.8454 | 6.7090 | 6.1751 | 6.3634 | 6.4221 | 5.6961 | 6.4380 |
| CRTC2_S456 | CRTC2 | Q53ET0 | | 613 | HCSRHGSGPNIIL | 2.1377 | 2.1080 | 2.5988 | 2.1784 | 3.9750 | 4.0667 | 2.9806 | 3.3005 | 2.3931 | 2.6116 | 2.6752 | 2.4084 | 18.9033 | 15.4365 | 17.5177 | 14.7085 |
| CRTC2_S456;S460 | CRTC2 | Q53ET0 | | 70 | SSHYGGSLPNVNQ | 5.6184 | 6.0015 | 6.8771 | 5.8086 | 6.0935 | 6.4678 | 6.2197 | 7.4757 | 6.8990 | 6.3670 | 6.8298 | 5.8901 | 5.9594 | 6.6518 | 5.2897 | 5.5509 |
| CRTC2_S490;T501 | CRTC2 | Q53ET0 | | 136 | YSPAYLSPPPESS | 6.1155 | 6.6479 | 6.2974 | 6.2840 | 7.1540 | 6.3709 | 5.8414 | 7.0525 | 6.2018 | 5.4898 | 6.2875 | 6.9248 | 6.2560 | 6.0438 | 5.4053 | 5.6274 |
| CRTC2_S613 | CRTC2 | Q53ET0 | | 624 | ILTGDSSPGFSKE | 7.0445 | 7.6958 | 7.3552 | 7.0205 | 5.7395 | 7.4031 | 6.2984 | 7.3804 | 8.0269 | 6.4616 | 7.0433 | 7.7930 | 4.1742 | 4.0470 | 2.7216 | 3.7951 |
| CRTC2_S624 | CRTC2 | Q53ET0 | 456;460 | | NA | 6.4261 | 6.7133 | 6.5433 | 6.3340 | 6.0599 | 6.5783 | 6.8205 | 6.7398 | 6.8190 | 6.4467 | 7.0153 | 6.9896 | 5.0859 | 5.5725 | 4.7502 | 5.1057 |
| CRTC2_S70 | CRTC2 | Q53ET0 | 490;501 | | NA | 7.5171 | 6.1576 | 6.2839 | 6.5924 | 6.1564 | 7.0520 | 6.2825 | 6.3336 | 6.6073 | 7.1597 | 7.1134 | 7.5537 | 4.6016 | 5.2447 | 4.2474 | 5.0968 |
| CRTC3_S370 | CRTC3 | Q6UUV7 | | 410 | GFSRQLSSTSPLA | 6.8680 | 5.6932 | 6.2833 | 6.0146 | 5.9395 | 6.3832 | 6.3238 | 6.9860 | 6.1078 | 5.7507 | 6.2306 | 6.5750 | 6.7723 | 5.8935 | 6.4861 | 5.6925 |
| CRTC3_S391;S396 | CRTC3 | Q6UUV7 | | 370 | LRLFSLSNPSLST | 6.2354 | 6.5632 | 7.1524 | 5.9748 | 6.0760 | 6.4919 | 6.1883 | 6.5067 | 6.7921 | 6.2666 | 6.4812 | 6.4518 | 5.8429 | 5.7001 | 5.4691 | 5.8075 |
| CRTC3_S410 | CRTC3 | Q6UUV7 | | 62 | TQYHGGSLPNVSQ | 7.2789 | 6.9311 | 7.5805 | 6.6002 | 5.9769 | 7.1763 | 6.1050 | 6.5503 | 6.1511 | 6.7868 | 6.4212 | 6.3816 | 5.2761 | 5.3770 | 4.3566 | 5.0504 |
| CRTC3_S410;T412 | CRTC3 | Q6UUV7 | 391;396 | | NA | 6.1683 | 6.0349 | 6.8373 | 5.6723 | 6.1584 | 7.0732 | 6.6423 | 7.3666 | 6.4696 | 6.4151 | 6.7503 | 5.9001 | 5.6650 | 5.9963 | 5.1696 | 5.6808 |
| CRTC3_S62 | CRTC3 | Q6UUV7 | 410;412 | | NA | 5.6503 | 4.6994 | 6.5756 | 5.6773 | 6.0244 | 7.1213 | 5.8027 | 7.1763 | 5.7574 | 5.5741 | 6.7362 | 6.2942 | 5.6734 | 7.4862 | 7.5882 | 6.1628 |
| CRY2_S558 | CRY2 | Q49AN0 | | 558 | LPSGPASPKRKLE | 6.3461 | 5.8558 | 6.7472 | 6.1585 | 6.0157 | 6.0576 | 6.2976 | 5.9335 | 7.7078 | 6.7472 | 6.9476 | 6.3720 | 5.4891 | 5.9776 | 5.2559 | 6.0907 |
| CRYBG1_S103 | CRYBG1 | Q9Y4K1 | | 329 | KKVMPNSPQNGVL | 6.0983 | 6.5297 | 6.4441 | 6.6792 | 5.9907 | 6.4685 | 6.2577 | 6.4417 | 6.5759 | 6.8452 | 6.6785 | 6.7870 | 5.2734 | 5.9995 | 5.3909 | 5.5397 |
| CRYBG1_S141 | CRYBG1 | Q9Y4K1 | | 533 | PLSTERSPEAVGS | 5.8172 | 6.1635 | 6.4944 | 6.2331 | 5.9784 | 6.4346 | 6.0192 | 6.3164 | 6.3057 | 6.2905 | 6.2330 | 6.3278 | 6.0968 | 6.6939 | 6.3203 | 6.2752 |
| CRYBG1_S329 | CRYBG1 | Q9Y4K1 | | 103 | QPPPASSPTKRKG | 6.4285 | 6.1640 | 5.7220 | 5.9945 | 5.8029 | 5.9577 | 5.9949 | 6.3790 | 6.0956 | 6.1752 | 7.0648 | 7.3597 | 5.6981 | 7.1648 | 5.9356 | 6.0626 |
| CRYBG1_S461 | CRYBG1 | Q9Y4K1 | | 484 | KDTCVQSPISSFP | 5.3521 | 5.3786 | 5.7224 | 5.6590 | 6.2296 | 6.1667 | 6.2667 | 5.8478 | 5.6139 | 5.7372 | 5.9225 | 5.7696 | 7.6737 | 7.2857 | 7.6453 | 7.7293 |
| CRYBG1_S484 | CRYBG1 | Q9Y4K1 | | 141 | KRVPDPSPVTKGT | 5.3941 | 5.4905 | 5.2435 | 5.1515 | 5.6671 | 5.6883 | 5.5950 | 5.4205 | 5.3552 | 5.5376 | 5.2820 | 5.0838 | 8.5549 | 9.0336 | 9.1594 | 8.3431 |
| CRYBG1_S533 | CRYBG1 | Q9Y4K1 | | 725 | KARMPNSPAPHFA | 6.4238 | 6.5573 | 6.5303 | 6.4641 | 5.9313 | 7.4752 | 6.9062 | 6.7183 | 6.7457 | 6.9125 | 6.4762 | 7.0399 | 4.7661 | 5.3669 | 4.7607 | 4.9256 |
| CRYBG1_S725 | CRYBG1 | Q9Y4K1 | | 461 | QSPAESSPGPSLS | 6.7961 | 6.4081 | 6.8813 | 7.3962 | 6.0657 | 6.4184 | 7.0288 | 6.3704 | 6.5085 | 7.2267 | 7.3061 | 6.7883 | 4.1831 | 6.0412 | 4.2007 | 4.3805 |
| CRYBG1_S75;S81 | CRYBG1 | Q9Y4K1 | | 525 | ELGGETTPPLSTE | 6.0578 | 6.7476 | 6.1865 | 6.6618 | 6.4616 | 5.2108 | 6.3366 | 6.1954 | 6.2597 | 6.3544 | 6.3419 | 5.1840 | 6.0453 | 7.2568 | 6.6993 | 6.0006 |
| CRYBG1_S81 | CRYBG1 | Q9Y4K1 | | 81 | SPESKPSPGTKGQ | 5.7944 | 5.6663 | 6.4522 | 6.1948 | 6.1010 | 5.7388 | 6.4841 | 5.3264 | 6.3212 | 6.6379 | 6.5123 | 6.8351 | 6.9886 | 6.0274 | 6.0122 | 6.9071 |
| CRYBG1_T524;S533 | CRYBG1 | Q9Y4K1 | 525;533 | | NA | 5.8500 | 6.0278 | 6.4835 | 6.2356 | 6.1622 | 5.7873 | 6.0707 | 6.1489 | 6.2674 | 5.9018 | 6.8342 | 6.5154 | 6.0192 | 6.8264 | 6.5093 | 6.3604 |
| CRYBG1_T525 | CRYBG1 | Q9Y4K1 | 524;533 | | NA | 6.3177 | 5.9481 | 5.6418 | 6.8743 | 6.6517 | 5.8212 | 6.0809 | 5.8648 | 6.6659 | 6.7495 | 6.4212 | 6.5296 | 6.0213 | 6.3715 | 6.0014 | 6.0393 |
| CRYBG1_T525;S533 | CRYBG1 | Q9Y4K1 | 75;81 | | NA | 6.3269 | 7.1685 | 6.5585 | 7.3022 | 6.5427 | 6.6840 | 7.3583 | 6.7802 | 7.3477 | 7.4070 | 7.4218 | 7.5399 | 3.4065 | 4.7221 | 3.4017 | 4.0322 |
| CRYBG3_S177 | CRYBG3 | Q68DQ2 | | 395 | HRKVPCSPDFQRV | 6.5043 | 6.8916 | 6.7429 | 6.4529 | 6.1261 | 6.7480 | 6.3265 | 6.9424 | 7.0706 | 6.7222 | 7.0697 | 7.1224 | 4.5670 | 5.0615 | 4.6875 | 4.9646 |
| CRYBG3_S1934 | CRYBG3 | Q68DQ2 | | 931 | EVDALGSPPALLK | 6.0914 | 5.9643 | 6.5049 | 6.3990 | 6.4470 | 6.4562 | 6.2084 | 6.6856 | 6.4496 | 6.1323 | 7.0654 | 6.9874 | 5.4234 | 5.8552 | 5.4869 | 5.8429 |
| CRYBG3_S395 | CRYBG3 | Q68DQ2 | | 1934 | YFRGYESPTLSKD | 6.8604 | 6.5451 | 6.9582 | 5.9159 | 6.0698 | 6.3027 | 6.2600 | 6.7040 | 7.1180 | 6.5266 | 6.7569 | 6.5230 | 5.3983 | 5.5611 | 5.3297 | 5.1704 |
| CRYBG3_S716 | CRYBG3 | Q68DQ2 | | 9 | SGRRRGSAPWHSF | 6.3468 | 5.4721 | 5.8190 | 5.0425 | 6.5190 | 5.8922 | 5.8302 | 6.2237 | 8.5207 | 6.6378 | 8.3864 | 7.6193 | 5.9404 | 6.8031 | 4.5294 | 4.4174 |
| CRYBG3_S9 | CRYBG3 | Q68DQ2 | | 716 | EAAGRKSPPPSFC | 5.5871 | 5.7500 | 5.8196 | 6.2007 | 6.5492 | 6.3934 | 6.5795 | 6.8946 | 6.2477 | 5.8594 | 7.2669 | 6.4213 | 5.8469 | 6.7115 | 6.2239 | 5.6483 |
| CRYBG3_S931 | CRYBG3 | Q68DQ2 | | 177 | REIFSGSLRTQTH | 7.6367 | 6.3869 | 6.8822 | 6.4588 | 6.3417 | 5.8966 | 5.7172 | 6.6048 | 7.4583 | 6.7298 | 6.9824 | 6.4733 | 5.9657 | 5.6614 | 4.3635 | 4.4406 |
| CSNK1A1_S156 | CSNK1A1 | P48729-2 | | 156 | CNKCLESPVGKRK | 6.6916 | 6.9258 | 6.6083 | 6.7752 | 6.4494 | 6.5628 | 6.6986 | 6.4198 | 6.8815 | 6.9010 | 6.9611 | 6.8465 | 4.4599 | 5.1055 | 4.5840 | 5.1291 |
| CSNK1D_S331 | CSNK1D | P48730-2 | | 331 | GLPSTASGRLRGT | 7.2694 | 7.0305 | 7.1232 | 7.1375 | 5.8419 | 6.6166 | 7.0551 | 6.7982 | 7.0508 | 6.7144 | 6.7869 | 6.7943 | 4.1317 | 4.9871 | 4.0626 | 4.5998 |

| | | | | | | | | | | | | | | | | | | | | | |
|---|---|---|---|---|---|---|---|---|---|---|---|---|---|---|---|---|---|---|---|---|---|
| CSNK1D_S356 | CSNK1D | P48730-2 | | 382 | GAPVNISSSDLTG | 6.0432 | 6.1051 | 6.3480 | 6.4639 | 5.6540 | 6.1536 | 6.5245 | 6.2067 | 6.3918 | 6.1028 | 6.4330 | 6.3820 | 6.0394 | 6.6593 | 6.1360 | 6.3571 |
| CSNK1D_S382 | CSNK1D | P48730-2 | | 356 | SHTANTSPRPVSG | 5.9669 | 5.9988 | 6.4493 | 7.1617 | 6.2991 | 6.4502 | 6.8578 | 6.6652 | 6.4662 | 6.4256 | 6.5629 | 6.5068 | 5.1267 | 6.3010 | 5.5388 | 5.2230 |
| CSNK1D_S396 | CSNK1D | P48730-2 | | 347 | APPTPLTPTSHTA | 5.8802 | 5.9228 | 6.5465 | 6.8837 | 6.4547 | 6.7361 | 7.1894 | 7.1276 | 6.8933 | 6.1469 | 6.7324 | 6.7904 | 5.0493 | 5.1521 | 4.9009 | 5.5937 |
| CSNK1D_T347 | CSNK1D | P48730-2 | | 396 | QDTSRMSTSQNSI | 7.3648 | 6.3061 | 7.7903 | 6.7055 | 5.4100 | 6.5001 | 5.8760 | 5.8836 | 7.1412 | 7.3197 | 7.0245 | 7.5247 | 4.5116 | 5.3353 | 4.3691 | 4.9375 |
| CSNK1E_S363 | CSNK1E | P49674 | | 363 | QPAGNTSPRAISR | 7.3454 | 7.1196 | 6.9033 | 6.8161 | 6.2732 | 6.4353 | 6.6515 | 6.6200 | 6.9426 | 6.6047 | 6.7017 | 6.7066 | 4.5005 | 4.9648 | 4.5814 | 4.8332 |
| CSNK1G3_S346 | CSNK1G3 | Q9Y6M4-4 | | 346 | QDPALSSNREAHQ | 6.0866 | 6.2939 | 6.3694 | 6.4026 | 5.9051 | 6.3372 | 5.9677 | 6.5961 | 6.4137 | 5.4204 | 6.2091 | 6.6541 | 6.4809 | 6.4127 | 6.2017 | 6.2487 |
| CSNK1G3_S367 | CSNK1G3 | Q9Y6M4-4 | | 367 | KNQVVSSTNGELN | 6.1721 | 7.1311 | 7.4856 | 6.2918 | 5.9948 | 5.4964 | 5.7812 | 6.5766 | 7.1462 | 5.7282 | 6.0736 | 6.1903 | 6.8447 | 5.5373 | 5.8039 | 5.7464 |
| CSNK2A3_S287 | CSNK2A3 | Q8NEV1 | | 287 | WERFVHSENQHLV | 5.5914 | 6.4955 | 5.8489 | 5.2134 | 7.8294 | 5.8230 | 5.7359 | 6.6937 | 6.8030 | 5.6692 | 5.8211 | 5.5691 | 6.4066 | 7.2995 | 6.7511 | 6.4492 |
| CSNK2B_S209 | CSNK2B | P67870 | | 209 | AASNFKSPVKTIR | 4.7371 | 5.4158 | 4.7878 | 6.2474 | 6.7891 | 8.0529 | 7.0746 | 8.6542 | 6.9803 | 5.0004 | 6.0179 | 6.8682 | 5.6408 | 5.9156 | 6.5071 | 5.3108 |
| CSPP1_S459 | CSPP1 | Q1MSJ5 | | 966 | RLLHMDSDDEIPI | 6.4646 | 6.5887 | 7.0816 | 6.4395 | 6.6218 | 6.9467 | 6.5675 | 6.6509 | 6.0334 | 6.0468 | 6.1671 | 6.2463 | 5.8314 | 5.7176 | 5.3329 | 5.2632 |
| CSPP1_S882 | CSPP1 | Q1MSJ5 | | 901 | KLQRPPSVDSIIR | 5.2503 | 5.4968 | 5.3510 | 5.3024 | 5.3809 | 5.8786 | 5.8011 | 6.1576 | 5.5330 | 5.6818 | 5.7279 | 5.7271 | 8.5341 | 7.6114 | 8.3851 | 8.1809 |
| CSPP1_S901 | CSPP1 | Q1MSJ5 | | 920 | SMSRAQSPPVPAR | 6.3796 | 6.4260 | 6.5203 | 6.2137 | 6.0031 | 6.3921 | 6.2370 | 6.7234 | 6.3936 | 5.9878 | 6.9492 | 6.8081 | 5.8303 | 6.1769 | 5.3543 | 5.6044 |
| CSPP1_S901;S904 | CSPP1 | Q1MSJ5 | | 882 | KHMRQPSPIVPAL | 6.5093 | 6.0692 | 6.8086 | 6.1690 | 6.4125 | 6.4005 | 6.8007 | 7.0342 | 6.6769 | 6.1600 | 5.2948 | 6.3885 | 5.3377 | 6.3017 | 5.9953 | 5.6411 |
| CSPP1_S920 | CSPP1 | Q1MSJ5 | | 459 | AQDPEKSPDRLKQ | 5.9736 | 6.1810 | 6.3337 | 5.8118 | 7.3386 | 7.4312 | 6.8595 | 6.3332 | 6.4226 | 6.1674 | 7.3110 | 6.3976 | 5.9286 | 5.7237 | 5.0904 | 4.6959 |
| CSPP1_S966 | CSPP1 | Q1MSJ5 | 901;904 | | NA | 9.3257 | 7.1716 | 6.3149 | 7.8518 | 5.1672 | 5.9210 | 6.8156 | 5.2154 | 5.1571 | 3.9918 | 4.6992 | 3.9330 | 8.4367 | 7.8587 | 6.2484 | 5.8918 |
| CSPP1_T1186;S1190 | CSPP1 | Q1MSJ5 | 1186;1190 | | NA | 7.0582 | 6.4785 | 6.9023 | 7.9115 | 6.9049 | 6.7630 | 7.6898 | 7.6261 | 4.1060 | 5.6753 | 6.0067 | 5.9717 | 7.2141 | 5.4016 | 3.6795 | 4.6107 |
| CSRNP1_T68 | CSRNP1 | Q96S65 | | 68 | CGPRSFTPLSILK | 6.4273 | 6.7126 | 6.5730 | 6.0639 | 6.4776 | 6.6250 | 5.7743 | 6.5669 | 7.0934 | 7.4533 | 6.6875 | 7.0162 | 5.5566 | 5.1984 | 4.4621 | 5.3120 |
| CSRP1_S192 | CSRP1 | P21291 | | 192 | AGALVHSExxxxx | 6.6278 | 6.9097 | 7.1608 | 6.5656 | 6.5453 | 6.9203 | 6.7316 | 7.2460 | 6.3421 | 6.0326 | 6.4489 | 6.3060 | 4.9218 | 5.3535 | 4.8402 | 5.0479 |
| CSTF2_S113 | CSTF2 | P33240 | | 524 | SQPGGFSPGQNQV | 6.5099 | 4.9400 | 4.7817 | 4.7023 | 5.6837 | 4.1733 | 5.4303 | 6.9964 | 4.3870 | 6.1442 | 4.9761 | 6.6785 | 7.8233 | 9.0869 | 8.8806 | 8.8058 |
| CSTF2_S120 | CSTF2 | P33240 | | 113 | GAPVIESPYGETI | 5.9634 | 6.1089 | 6.2568 | 5.3712 | 6.4533 | 4.5592 | 5.2128 | 6.9642 | 6.2089 | 6.9887 | 5.7237 | 7.5207 | 6.0949 | 6.8864 | 6.7812 | 6.9057 |
| CSTF2_S524 | CSTF2 | P33240 | | 120 | PYGETISPEDAPE | 5.3451 | 5.2001 | 5.3118 | 5.0184 | 6.2396 | 5.1711 | 5.0445 | 6.5153 | 5.5226 | 5.7807 | 5.2107 | 6.0380 | 7.6800 | 9.3205 | 7.8242 | 8.7774 |
| CSTF3_S691 | CSTF3 | Q12996 | | 691 | KRPNEDSDEDEEK | 5.8754 | 5.9713 | 5.9166 | 6.4772 | 5.9023 | 5.7632 | 6.1289 | 5.7420 | 6.0676 | 6.2088 | 6.2343 | 6.3047 | 6.5499 | 6.9024 | 6.7226 | 7.2329 |
| CTBP2_S905 | CTBP2 | P56545-2 | | 905 | KEFFVTSAPWSVI | 5.9694 | 5.8126 | 5.3577 | 5.5666 | 6.3699 | 5.8831 | 6.0519 | 5.8677 | 5.2516 | 5.3671 | 5.4410 | 5.4807 | 7.6079 | 7.6499 | 8.4810 | 7.8419 |
| CTCF_S150 | CTCF | P49711 | | 289 | RHMKSHTDERPHK | 7.2142 | 8.1971 | 7.4112 | 8.1434 | 5.7929 | 7.1283 | 7.1764 | 6.5926 | 7.9023 | 7.7844 | 8.2044 | 8.0290 | 2.1541 | 3.2046 | 2.0205 | 3.0445 |
| CTCF_S224 | CTCF | P49711 | | 224 | GKDVDVSVYDFEE | 7.1939 | 7.3056 | 6.1953 | 7.4776 | 6.5226 | 6.2570 | 7.7599 | 6.1114 | 6.1258 | 6.3502 | 5.9213 | 6.1365 | 4.7253 | 5.3015 | 5.2431 | 5.3729 |
| CTCF_T289 | CTCF | P49711 | | 150 | KEGLAESEPMICH | 6.2577 | 6.5813 | 5.7681 | 6.4943 | 6.1445 | 5.5714 | 6.4479 | 6.5398 | 5.9528 | 6.6381 | 6.4360 | 5.5907 | 6.2295 | 6.7631 | 6.7853 | 5.7997 |
| CTDP1_S415 | CTDP1 | Q9Y5B0 | | 839 | RRKRQPSMSETMP | 5.0921 | 5.3816 | 5.8040 | 5.5191 | 5.5482 | 5.7510 | 5.6736 | 6.3086 | 6.0412 | 6.4966 | 6.5631 | 6.5434 | 7.3121 | 7.4991 | 7.3317 | 7.1345 |
| CTDP1_S839 | CTDP1 | Q9Y5B0 | | 904 | TLGAPASSERSAA | 7.0317 | 7.3907 | 6.8822 | 7.0561 | 6.0839 | 6.8400 | 7.3972 | 6.8470 | 7.0812 | 7.2327 | 7.3687 | 7.3934 | 3.4256 | 4.3812 | 3.4046 | 4.1838 |
| CTDP1_S869;S872 | CTDP1 | Q9Y5B0 | | 415 | PAQAPTSSQELAG | 6.3566 | 6.0763 | 5.7658 | 6.0891 | 5.5268 | 6.8869 | 7.0060 | 5.6522 | 6.6185 | 6.6698 | 7.0380 | 6.2110 | 5.1292 | 6.2732 | 6.0342 | 6.6662 |
| CTDP1_S904 | CTDP1 | Q9Y5B0 | 869;872 | | NA | 5.4035 | 5.9569 | 6.1685 | 5.8329 | 6.1228 | 5.5129 | 6.5203 | 5.9965 | 5.8760 | 5.7690 | 6.3845 | 7.3825 | 6.8496 | 6.4511 | 7.1153 | 6.6577 |
| CTDSP1_T73 | CTDSP1 | Q9GZU7-3 | | 73 | NGAIPKTPVQYLL | 5.8900 | 5.7105 | 6.5231 | 6.7132 | 6.3946 | 6.3516 | 6.3023 | 6.2643 | 6.5340 | 6.5590 | 6.2590 | 6.2897 | 5.9281 | 6.0404 | 6.3565 | 5.8837 |
| CTDSP2_S55 | CTDSP2 | O14595 | | 55 | HVGQSSSSTELAA | 6.8046 | 7.2351 | 6.2141 | 6.7821 | 6.5410 | 6.3328 | 6.3088 | 7.7139 | 6.9437 | 5.9061 | 6.5070 | 6.4919 | 4.5148 | 5.3310 | 5.4476 | 4.9255 |
| CTDSPL_S33 | CTDSPL | O15194 | | 33 | ASQCNVSLKKQRS | 6.9269 | 6.7949 | 6.4023 | 6.7021 | 6.4590 | 5.5700 | 6.5451 | 6.1992 | 6.4225 | 7.0809 | 6.7470 | 7.2838 | 4.9037 | 4.8562 | 5.2533 | 5.8530 |
| CTDSPL2_S104 | CTDSPL2 | Q05D32 | | 104 | SRVRRKSQVNGEA | 5.2902 | 5.2391 | 5.5618 | 6.3474 | 7.1540 | 5.7505 | 5.9655 | 6.2107 | 6.9285 | 6.9186 | 6.6218 | 6.7197 | 6.4843 | 6.7825 | 5.7883 | 6.2372 |
| CTDSPL2_S134 | CTDSPL2 | Q05D32 | | 86 | NNLITSTPRAGEK | 6.5006 | 6.7310 | 6.7231 | 7.6658 | 6.1739 | 6.6318 | 6.8006 | 6.5718 | 7.1119 | 6.9138 | 6.4382 | 6.5280 | 4.7116 | 5.1507 | 4.4337 | 4.9134 |
| CTDSPL2_S51 | CTDSPL2 | Q05D32 | | 134 | DNPSSGSPPRTTL | 7.5136 | 7.9387 | 6.8676 | 8.0035 | 6.5996 | 6.9501 | 6.9746 | 7.0061 | 7.3557 | 7.1959 | 7.2645 | 6.9222 | 2.9711 | 3.8385 | 3.0567 | 3.5416 |
| CTDSPL2_T86 | CTDSPL2 | Q05D32 | | 51 | ETGLLSSIKKFIK | 6.9386 | 6.7368 | 6.7285 | 7.2555 | 6.4285 | 6.3348 | 7.0451 | 6.7493 | 6.9230 | 6.8771 | 6.6982 | 6.5224 | 4.7005 | 5.2132 | 4.2665 | 4.5819 |
| CTNNA1_S641 | CTNNA1 | P35221-2 | | 641 | PEELDDSDFETED | 5.7369 | 6.3320 | 6.4128 | 6.3732 | 5.7819 | 6.1882 | 6.2119 | 6.2288 | 6.7525 | 6.0028 | 6.0718 | 6.5525 | 6.0786 | 6.4081 | 6.3749 | 6.4933 |
| CTNNA1_S655 | CTNNA1 | P35221-2 | | 655 | DVRSRTSVQTEDD | 5.7041 | 6.2620 | 6.6173 | 5.8350 | 5.9534 | 6.5246 | 6.5037 | 6.3435 | 7.2247 | 6.4160 | 6.1527 | 6.1673 | 5.9966 | 6.8687 | 5.7396 | 5.6908 |
| CTNNAL1_S538 | CTNNAL1 | Q9UBT7 | | 538 | EKYGYLSLPKPMK | 6.6100 | 6.6311 | 7.0017 | 6.3081 | 5.5076 | 6.6511 | 6.8531 | 6.4070 | 6.4847 | 6.8785 | 7.1806 | 7.0951 | 4.7066 | 5.5645 | 4.6432 | 5.4769 |
| CTNNB1_S191 | CTNNB1 | P35222 | | 675 | DYKKRLSVELTSS | 6.1160 | 6.7455 | 6.0249 | 5.9692 | 6.2120 | 6.1539 | 6.0902 | 6.0496 | 6.4034 | 6.0227 | 5.9802 | 6.2808 | 6.3897 | 6.7050 | 6.5573 | 6.2996 |
| CTNNB1_S552 | CTNNB1 | P35222 | | 191 | RHAIMRSPQMVSA | 6.6124 | 6.3689 | 6.0229 | 6.4669 | 6.3134 | 6.0096 | 6.2192 | 6.2727 | 6.3840 | 6.0608 | 5.6719 | 5.9606 | 6.3579 | 6.3595 | 6.4576 | 6.4616 |
| CTNNB1_S675 | CTNNB1 | P35222 | | 552 | DTQRRTSMGGTQQ | 6.0170 | 5.7837 | 5.7734 | 6.0434 | 6.8085 | 5.9080 | 5.9462 | 5.9997 | 6.8607 | 7.0431 | 6.8159 | 6.8144 | 5.9811 | 6.5832 | 5.8529 | 5.7689 |
| CTNNBL1_S545 | CTNNBL1 | Q8WYA6 | | 545 | NIGDGRSPEFREN | 6.0957 | 6.0588 | 5.9080 | 6.4450 | 5.7715 | 6.0323 | 6.2473 | 5.8780 | 6.1270 | 6.2143 | 5.8520 | 6.2453 | 7.2300 | 6.4482 | 6.6751 | 6.7717 |
| CTNND1_S230 | CTNND1 | O60716-3 | | 288 | LEDDQRSMGYDDL | 6.0517 | 6.3440 | 6.2886 | 6.5208 | 5.5051 | 6.4425 | 6.3156 | 6.2719 | 5.7973 | 6.4542 | 7.0675 | 7.1264 | 5.6550 | 6.0380 | 5.8815 | 6.2399 |
| CTNND1_S244 | CTNND1 | O60716-3 | | 268 | QVRVGGSSVDLHR | 6.9060 | 7.0324 | 6.2642 | 6.4797 | 5.6404 | 6.1962 | 6.4706 | 5.8709 | 5.9620 | 5.7030 | 6.0853 | 6.0951 | 6.2207 | 6.5424 | 6.0428 | 6.4882 |
| CTNND1_S252 | CTNND1 | O60716-3 | | 47 | LERVRVSPQDANP | 6.2125 | 6.6243 | 7.0795 | 6.7747 | 7.0100 | 6.8566 | 6.6884 | 6.3139 | 6.5931 | 6.3652 | 6.4998 | 6.3761 | 5.0830 | 5.3774 | 4.8568 | 5.2887 |
| CTNND1_S268 | CTNND1 | O60716-3 | | 352 | SLASLDSLRKGGP | 8.2570 | 6.1543 | 5.7211 | 6.8268 | 8.3011 | 5.1755 | 5.4573 | 5.1377 | 7.2831 | 5.3859 | 4.2870 | 4.6883 | 8.1584 | 7.8654 | 5.8450 | 5.4560 |
| CTNND1_S288 | CTNND1 | O60716-3 | | 933 | EPLKGTTPLMQKI | 7.0736 | 7.3277 | 6.5298 | 7.2810 | 6.5403 | 7.0687 | 7.3091 | 6.8148 | 6.9677 | 7.0370 | 6.9446 | 6.9942 | 3.9102 | 4.4458 | 3.5959 | 4.1594 |
| CTNND1_S320 | CTNND1 | O60716-3 | | 349 | ERGSLASLDSLRK | 4.9292 | 5.8158 | 5.6534 | 5.3061 | 5.4738 | 6.7331 | 5.8480 | 6.6139 | 5.7366 | 7.2781 | 6.2139 | 6.5236 | 6.8580 | 6.9270 | 7.1008 | 6.9886 |
| CTNND1_S346 | CTNND1 | O60716-3 | | 346 | AQHERGSLASLDS | 7.2033 | 6.1002 | 5.2208 | 5.7622 | 6.1423 | 5.5821 | 5.8757 | 5.6267 | 6.5939 | 5.1812 | 4.9391 | 5.4391 | 7.8918 | 7.9008 | 7.4314 | 7.1095 |
| CTNND1_S346;S352 | CTNND1 | O60716-3 | | 320 | PRRRLRSYEDMIG | 5.6744 | 5.9887 | 6.7413 | 6.2850 | 4.9284 | 7.1007 | 6.4221 | 6.3154 | 5.7383 | 6.2624 | 7.4800 | 8.1634 | 5.0067 | 5.6347 | 5.6815 | 6.5770 |
| CTNND1_S349 | CTNND1 | O60716-3 | | 310 | TARRTGTPSDPRR | 6.7266 | 7.1349 | 6.7536 | 6.5650 | 5.5376 | 6.6705 | 7.3408 | 6.0668 | 6.1041 | 6.7103 | 7.1440 | 7.3862 | 5.0321 | 5.1849 | 4.3734 | 5.2694 |
| CTNND1_S349;S352 | CTNND1 | O60716-3 | | 230 | GSDNYGSLSRVTR | 7.3406 | 6.7341 | 6.9881 | 6.9224 | 6.2026 | 6.3134 | 6.4461 | 6.1194 | 6.3077 | 6.1819 | 6.2756 | 6.3542 | 5.7879 | 6.0173 | 4.9739 | 5.0346 |
| CTNND1_S352 | CTNND1 | O60716-3 | | 920 | NRTLDRSGDLGDM | 7.0211 | 7.2778 | 6.5132 | 6.9553 | 6.7900 | 6.6822 | 6.5414 | 6.4768 | 6.8296 | 7.4278 | 6.9488 | 6.6567 | 4.2966 | 5.0796 | 4.0548 | 4.4485 |
| CTNND1_S47 | CTNND1 | O60716-3 | | 252 | EGYRAPSRQDVYG | 6.9068 | 7.6177 | 7.6260 | 6.4679 | 4.2376 | 7.3305 | 6.9004 | 6.1153 | 5.3360 | 6.5809 | 7.4754 | 7.1648 | 4.8299 | 5.1319 | 4.4287 | 5.8500 |
| CTNND1_S920 | CTNND1 | O60716-3 | | 906 | LDNNYSTPNERGD | 7.0316 | 7.8753 | 6.3456 | 7.3200 | 6.0699 | 6.7904 | 7.1247 | 6.9282 | 7.3350 | 7.4246 | 7.4007 | 7.8458 | 3.1996 | 4.1189 | 3.2482 | 3.9416 |
| CTNND1_T201 | CTNND1 | O60716-3 | | 244 | EERYRPSMEGYRA | 7.5529 | 7.7720 | 6.8280 | 7.2974 | 5.5504 | 6.8748 | 6.7655 | 7.1110 | 7.3788 | 7.1226 | 7.6625 | 7.2820 | 3.5913 | 4.3241 | 2.9841 | 3.9028 |
| CTNND1_T310 | CTNND1 | O60716-3 | | 201 | GQAGTATLPRNFH | 8.7651 | 7.1949 | 7.6838 | 7.1979 | 6.6561 | 5.4288 | 6.6092 | 5.6427 | 5.6136 | 5.7507 | 6.3260 | 5.9124 | 5.9117 | 5.5987 | 4.3743 | 5.3341 |
| CTNND1_T906 | CTNND1 | O60716-3 | | 96 | DHSHLLYSTIPRM | 5.7612 | 5.5793 | 6.4164 | 6.4865 | 6.1687 | 6.2280 | 6.4150 | 5.3667 | 6.6698 | 6.9912 | 6.9140 | 6.9538 | 6.5111 | 6.0471 | 5.5544 | 5.9368 |
| CTNND1_T933 | CTNND1 | O60716-3 | 349;352 | | NA | 6.4398 | 6.4686 | 6.3364 | 7.3902 | 5.7185 | 5.9525 | 6.2988 | 6.0529 | 5.7359 | 6.0602 | 5.7698 | 5.5582 | 6.4100 | 6.9672 | 6.3121 | 6.5290 |
| CTNND1_Y96 | CTNND1 | O60716-3 | 346;352 | | NA | 6.6338 | 7.2077 | 5.9758 | 7.3253 | 6.0171 | 6.2044 | 6.5479 | 7.1314 | 5.4531 | 5.0520 | 5.7824 | 6.0479 | 5.7639 | 6.1876 | 6.7923 | 5.8775 |
| CTPS1_S562 | CTPS1 | P17812 | | 562 | QKGCRLSPRDTYS | 6.9580 | 6.8225 | 7.6626 | 6.3992 | 6.5174 | 7.3139 | 5.9562 | 5.4818 | 5.8742 | 6.3704 | 6.7543 | 6.6236 | 5.7797 | 5.7090 | 4.8354 | 4.9418 |
| CTPS1_S575 | CTPS1 | P17812 | | 575 | DRSGSSSPDSEIT | 5.3411 | 5.3657 | 6.1566 | 5.4707 | 5.7955 | 5.8754 | 5.8206 | 5.9103 | 5.7826 | 5.7186 | 6.0578 | 5.9606 | 7.9014 | 7.4788 | 7.7260 | 7.6383 |
| CTR9_S970 | CTR9 | Q6PD62 | | 925 | EFVNDDTDDDLPI | 6.1676 | 6.2442 | 5.9604 | 6.6745 | 6.0445 | 6.1655 | 6.2114 | 6.1084 | 6.2016 | 6.2408 | 6.1794 | 6.4730 | 6.4090 | 6.4903 | 6.1724 | 6.2570 |
| CTR9_T925 | CTR9 | Q6PD62 | | 970 | PKGEEGSDDDETE | 5.9455 | 6.4407 | 5.6024 | 7.1585 | 6.8790 | 7.2310 | 6.5068 | 5.6348 | 6.1199 | 6.3534 | 7.1800 | 6.3312 | 5.9811 | 6.3814 | 5.3357 | 4.9185 |
| CTTN_S381 | CTTN | Q14247-2 | | 381 | EERLPSSPVYEDA | 5.8020 | 5.8439 | 6.2625 | 6.3994 | 6.3047 | 6.5660 | 6.1470 | 6.4405 | 6.1013 | 6.4011 | 6.3959 | 6.3432 | 6.3176 | 6.2135 | 6.3292 | 6.1323 |
| CTTN_T362 | CTTN | Q14247-2 | | 364 | ARAKTQTPPVSPA | 6.6897 | 6.9106 | 6.5685 | 6.9624 | 6.5468 | 6.2446 | 5.8962 | 7.1499 | 6.6847 | 6.3522 | 6.1486 | 6.4783 | 5.7268 | 5.2774 | 5.1819 | 5.1816 |
| CTTN_T364 | CTTN | Q14247-2 | | 384 | LPSSPVYEDAASF | 5.8431 | 6.2827 | 6.6868 | 6.0470 | 6.5478 | 6.6702 | 6.1977 | 6.2219 | 6.7274 | 6.7206 | 5.9551 | 5.8944 | 6.4237 | 6.5691 | 5.7698 | 5.4428 |
| CTTN_T364;S368 | CTTN | Q14247-2 | | 362 | EQARAKTQTPPVS | 6.6655 | 7.2213 | 6.1903 | 7.2886 | 6.1897 | 6.6168 | 6.7773 | 7.5169 | 6.7280 | 6.3290 | 6.2954 | 6.2768 | 4.6684 | 5.3473 | 5.5276 | 4.3614 |
| CTTN_Y384 | CTTN | Q14247-2 | | 409 | TEPEPVYSMEAAD | 6.7364 | 5.9413 | 7.7987 | 6.7445 | 6.8336 | 6.7283 | 6.9404 | 7.4408 | 7.3226 | 6.8400 | 8.3136 | 7.1914 | 4.0661 | 4.0758 | 3.0298 | 3.9965 |
| CTTN_Y409 | CTTN | Q14247-2 | 364;368 | | NA | 6.2971 | 5.9362 | 6.5673 | 6.2873 | 5.9585 | 6.3935 | 6.5000 | 6.3846 | 6.5353 | 6.1127 | 6.4212 | 6.5149 | 5.7864 | 6.2723 | 5.8790 | 6.1537 |
| CTTNBP2NL_S441 | CTTNBP2NL | Q9P2B4 | | 523 | GPIKPVSPNSSPF | 6.2626 | 6.4343 | 6.9684 | 6.2994 | 6.5493 | 6.8348 | 6.7050 | 6.2770 | 6.8468 | 6.6986 | 6.8898 | 6.8746 | 5.2141 | 5.4204 | 4.7796 | 4.9455 |
| CTTNBP2NL_S481;S488;S | CTTNBP2NL | Q9P2B4 | | 557 | TPGKVSSPLSPLS | 6.3993 | 6.4579 | 7.1959 | 6.3371 | 6.9326 | 6.8810 | 5.7768 | 6.5588 | 7.0446 | 6.4157 | 6.5356 | 6.2410 | 5.5792 | 5.6053 | 5.1368 | 4.9022 |
| CTTNBP2NL_S523 | CTTNBP2NL | Q9P2B4 | | 563 | SPLSPLSPGIKSP | 5.4618 | 5.5247 | 6.2358 | 5.4003 | 7.4902 | 6.7400 | 6.3559 | 6.3465 | 6.7928 | 5.0711 | 6.5389 | 6.1519 | 6.5425 | 6.3776 | 6.2271 | 6.7429 |
| CTTNBP2NL_S557 | CTTNBP2NL | Q9P2B4 | | 441 | KRLLGSSASSPGY | 5.7520 | 6.4665 | 6.6640 | 6.4390 | 6.2251 | 6.4104 | 7.2906 | 6.4592 | 6.5463 | 5.8368 | 5.9221 | 6.0123 | 6.0727 | 6.4348 | 6.1397 | 5.3284 |
| CTTNBP2NL_S557;S560;S | CTTNBP2NL | Q9P2B4 | 557;563 | | NA | 7.2751 | 6.1623 | 7.3202 | 6.4162 | 6.5585 | 6.6546 | 6.7036 | 7.1945 | 6.5394 | 6.7733 | 7.0102 | 7.7586 | 4.1789 | 4.2750 | 4.6454 | 4.5342 |
| CTTNBP2NL_S557;S563 | CTTNBP2NL | Q9P2B4 | 557;560;563 | | NA | 8.3347 | 8.1571 | 6.0551 | 8.2591 | 5.6743 | 8.1388 | 7.2873 | 6.8831 | 6.0578 | 8.2679 | 6.5608 | 8.1335 | 2.2019 | 3.4165 | 3.0466 | 3.5256 |
| CTTNBP2NL_S563 | CTTNBP2NL | Q9P2B4 | 481;488;490 | | NA | 6.2612 | 7.0416 | 6.3458 | 6.0938 | 6.0706 | 6.5073 | 6.0384 | 6.1241 | 7.6305 | 6.7727 | 7.1587 | 6.5596 | 5.9333 | 5.7318 | 4.9614 | 4.7692 |
| CTU2_S419 | CTU2 | Q2VPK5 | | 435 | PPGPCCSPGVGWA | 6.5433 | 6.1093 | 6.8895 | 6.6219 | 6.1886 | 6.8255 | 6.7010 | 6.9193 | 6.7245 | 6.7731 | 7.1291 | 6.7886 | 5.0914 | 5.3864 | 4.8198 | 4.4886 |
| CTU2_T428;S435 | CTU2 | Q2VPK5 | | 419 | RLSQMQSPIPLTE | 6.3132 | 6.2016 | 7.9312 | 6.5301 | 7.5386 | 5.5744 | 5.8978 | 7.7628 | 6.4216 | 6.3989 | 6.5260 | 6.4785 | 4.0555 | 5.6395 | 5.4638 | 5.2665 |
| CTU2_T435 | CTU2 | Q2VPK5 | 428;435 | | NA | 6.3610 | 6.7366 | 5.8729 | 7.3087 | 6.0982 | 5.8847 | 7.4876 | 6.6494 | 6.1363 | 6.3355 | 6.5049 | 5.9911 | 5.4116 | 6.4488 | 4.8512 | 5.9216 |
| CUEDC1_S286 | CUEDC1 | Q9NWM3 | | 286 | NDFGFSSPVPGTG | 6.6793 | 7.2814 | 6.2212 | 6.7978 | 7.2983 | 7.7731 | 5.9514 | 6.2100 | 7.0460 | 6.5752 | 7.1517 | 6.6857 | 4.6445 | 4.1024 | 4.2249 | 5.3572 |
| CUEDC2_S110 | CUEDC2 | Q9H467 | | 110 | QGQVPISPEPLQR | 6.2018 | 6.1217 | 6.8588 | 6.1092 | 6.2012 | 6.6487 | 6.5020 | 7.1290 | 6.7583 | 6.9779 | 7.3134 | 7.1035 | 4.8541 | 5.2897 | 4.6870 | 5.2435 |
| CUL3_S737 | CUL3 | Q13618 | | 737 | KARFLPSPVVIKK | 6.1490 | 6.4901 | 6.4244 | 6.1739 | 6.1177 | 6.6303 | 6.3766 | 6.3644 | 6.3318 | 6.8049 | 6.4302 | 6.5661 | 5.4980 | 6.2199 | 5.7669 | 5.6557 |
| CUL4A_S10 | CUL4A | Q13619 | | 10 | EAPRKGSFSALVG | 5.6653 | 4.5240 | 5.8380 | 5.2714 | 7.5818 | 5.5309 | 5.4078 | 6.5101 | 7.8841 | 7.9352 | 7.4555 | 7.2118 | 6.4687 | 7.1628 | 4.9353 | 4.6172 |
| CUL4B_S122 | CUL4B | Q13620-1 | | 31 | DGNTSTTPPTSAK | 6.0003 | 5.7088 | 5.9175 | 6.2078 | 5.8737 | 6.0252 | 5.5506 | 5.8365 | 5.9655 | 6.1627 | 6.1805 | 5.7173 | 6.9713 | 7.7117 | 7.2142 | 6.9564 |
| CUL4B_S13 | CUL4B | Q13620-1 | | 122 | MAEESSSSSSSSS | 6.1712 | 6.1788 | 6.1605 | 6.9570 | 5.9013 | 6.3667 | 6.9111 | 5.8560 | 6.7512 | 6.9814 | 6.4717 | 6.4212 | 4.7453 | 6.0235 | 6.0276 | 6.0755 |
| CUL4B_S29 | CUL4B | K4DI93 | | 13 | VTSGFSSPSPSAA | 6.6667 | 6.5088 | 6.4165 | 6.6040 | 6.6420 | 6.1607 | 6.6505 | 6.8457 | 7.1129 | 6.6928 | 6.3291 | 5.9563 | 5.4707 | 6.2791 | 4.5883 | 5.0760 |
| CUL4B_S29;S35 | CUL4B | Q13620-1 | | 106 | KLRFEDTLEFVGF | 7.3242 | 7.4208 | 7.1366 | 8.1439 | 6.2022 | 6.4117 | 6.0346 | 6.6084 | 6.3843 | 5.7286 | 6.7049 | 6.9964 | 4.6976 | 5.3734 | 4.2113 | 4.6211 |
| CUL4B_S49 | CUL4B | Q13620-1 | | 8 | FPTGFSSPSPSAA | 5.3240 | 5.4987 | 5.9081 | 5.6040 | 5.7268 | 7.0375 | 6.1150 | 6.0587 | 6.3475 | 6.0894 | 6.6194 | 6.2634 | 6.7663 | 7.2298 | 6.1457 | 7.2656 |
| CUL4B_S8 | CUL4B | Q13620-1 | | 29 | ATDGNTSTTPPTS | 4.9927 | 5.9141 | 5.9818 | 6.0654 | 5.5156 | 6.1040 | 5.8755 | 5.7996 | 5.3562 | 6.4348 | 6.4827 | 5.9102 | 7.1962 | 7.3087 | 7.5534 | 7.5091 |
| CUL4B_T106 | CUL4B | Q13620-1 | | 49 | SSSSSSSNSSNER | 5.8142 | 6.1227 | 5.5679 | 6.3926 | 6.5386 | 6.6489 | 6.3382 | 6.3833 | 7.5158 | 7.2795 | 6.8693 | 6.6703 | 4.6846 | 5.9395 | 5.6567 | 5.5779 |
| CUL4B_T31 | CUL4B | Q13620-1 | 29;35 | | NA | 6.0282 | 6.4567 | 6.5050 | 6.4619 | 6.5287 | 7.2712 | 7.0238 | 6.9290 | 7.0192 | 6.7062 | 7.4705 | 6.8478 | 4.6330 | 5.2537 | 4.2010 | 4.6642 |
| CUL7_S1208 | CUL7 | Q14999-2 | | 423 | PQLADVSPGLPAA | 6.1479 | 6.4876 | 6.3559 | 7.0664 | 6.0937 | 6.0470 | 6.1042 | 6.6628 | 6.5676 | 6.2362 | 6.8937 | 6.8867 | 5.1622 | 6.0672 | 5.7635 | 5.4574 |
| CUL7_S410 | CUL7 | Q14999-2 | | 1208 | PKGRNRSHDWSSL | 6.7693 | 6.9996 | 7.1626 | 6.9938 | 6.5088 | 6.3193 | 5.9340 | 6.1629 | 6.7361 | 5.9380 | 6.3436 | 6.8303 | 5.2285 | 6.0388 | 5.1657 | 4.8687 |
| CUL7_S423 | CUL7 | Q14999-2 | | 410 | PRSSARSPGSIFQ | 5.7938 | 5.7371 | 6.5130 | 6.0683 | 6.6595 | 5.8802 | 6.3173 | 6.5883 | 5.7777 | 6.8365 | 6.6366 | 6.4367 | 6.7383 | 6.0920 | 5.4059 | 6.5189 |
| CUL9_S976 | CUL9 | Q8IWT3 | | 976 | VLCREGSPGGAVR | 6.1000 | 6.6999 | 6.9162 | 6.4797 | 6.0444 | 7.2412 | 7.0974 | 6.6369 | 6.9910 | 7.1307 | 7.0770 | 7.0420 | 4.3652 | 5.2095 | 4.0125 | 4.9564 |
| CUSTOS_S179 | CUSTOS | Q96C57 | | 211 | SAVAATTPTSMAT | 6.6779 | 6.8314 | 6.9617 | 6.9595 | 6.0785 | 6.7280 | 6.9599 | 6.7116 | 6.9825 | 7.2539 | 7.4538 | 7.0063 | 4.1757 | 4.8519 | 3.8370 | 4.5306 |
| CUSTOS_S248 | CUSTOS | Q96C57 | | 179 | QESAIHSPGTVEK | 6.0932 | 6.5132 | 6.2334 | 7.0844 | 6.4925 | 6.4012 | 6.3828 | 6.4423 | 6.6925 | 6.5515 | 6.6529 | 6.6145 | 5.4396 | 5.8968 | 5.1723 | 5.3370 |
| CUSTOS_T211 | CUSTOS | Q96C57 | | 248 | KKASETSPFPPAK | 6.7032 | 6.5273 | 6.9575 | 7.9452 | 6.8416 | 6.4274 | 6.8155 | 6.8801 | 7.4192 | 6.9447 | 7.2459 | 7.2239 | 4.2612 | 4.2005 | 3.6346 | 3.9721 |
| CUX1_S1080 | CUX1 | P39880-3 | | 1080 | SSESVKSLTELVQ | 6.1152 | 6.0679 | 5.9294 | 6.4973 | 6.1257 | 5.9724 | 6.1235 | 6.1345 | 6.0533 | 5.7757 | 5.9653 | 6.2289 | 7.2755 | 6.6581 | 6.6005 | 6.4770 |
| CUX1_S1348 | CUX1 | Q13948 | | 584 | RLDPFSSFSKRER | 4.6756 | 4.8084 | 4.7748 | 4.9676 | 4.9850 | 4.9371 | 5.5108 | 5.4063 | 4.8960 | 4.9445 | 5.2706 | 5.0885 | 9.7572 | 10.1352 | 10.1211 | 9.7212 |
| CUX1_S1465 | CUX1 | P39880-3 | | 1465 | SAPRRPSSLQSLF | 4.7740 | 4.3702 | 4.6657 | 5.2670 | 8.5335 | 5.0951 | 4.7749 | 5.2393 | 8.0389 | 7.6764 | 7.3007 | 7.5608 | 7.9451 | 7.6372 | 5.2035 | 5.9177 |
| CUX1_S436 | CUX1 | P39880-3 | | 925 | NSPLPSSPIVPMS | 6.5710 | 6.3118 | 6.4739 | 6.4751 | 6.2879 | 6.5883 | 6.5258 | 6.2237 | 6.4093 | 6.6920 | 6.8196 | 6.2196 | 5.8068 | 5.7109 | 5.4961 | 5.3883 |
| CUX1_S584 | CUX1 | P39880-3 | | 760 | PKLLSTSPMPTVS | 6.5774 | 6.4869 | 6.4254 | 6.5449 | 6.6922 | 6.8435 | 6.3956 | 6.6856 | 7.0660 | 6.6127 | 6.9234 | 7.3838 | 4.5252 | 5.2477 | 4.9006 | 4.6890 |
| CUX1_S760 | CUX1 | Q13948 | | 594 | RERQRKYLSLSPW | 6.6779 | 6.4070 | 6.4665 | 7.6564 | 7.5019 | 7.2124 | 8.3029 | 8.0933 | 7.5366 | 6.8857 | 7.6032 | 7.5099 | 3.0015 | 3.5680 | 2.3888 | 3.1880 |
| CUX1_S925 | CUX1 | P39880-3 | | 436 | ESRRPGSLPAPPP | 5.9629 | 6.1656 | 6.5703 | 6.5424 | 6.4926 | 6.3243 | 7.0859 | 6.3480 | 6.5778 | 6.1257 | 6.8619 | 6.8179 | 5.8889 | 5.8602 | 5.1095 | 5.2661 |
| CUX1_Y594 | CUX1 | P39880-3 | | 1348 | PSSEGDSCDGVEA | 5.5068 | 5.7235 | 6.2058 | 5.6451 | 5.9488 | 5.4480 | 6.4703 | 6.5854 | 5.9977 | 5.2656 | 6.7066 | 6.0638 | 7.2183 | 7.3530 | 6.7396 | 7.1216 |
| CWC22_S61 | CWC22 | Q9HCG8 | | 829 | RGERRNSFSENEK | 5.9446 | 6.2304 | 5.6396 | 6.3517 | 6.0135 | 5.7747 | 6.0867 | 5.9113 | 6.2853 | 6.2615 | 6.2586 | 6.4674 | 6.7690 | 6.6943 | 6.7170 | 6.5943 |
| CWC22_S829 | CWC22 | Q9HCG8 | | 61 | HSRRGRSYDSSME | 6.0177 | 6.2404 | 6.0870 | 6.7441 | 5.8380 | 5.9343 | 6.4116 | 6.4404 | 6.5987 | 6.4938 | 6.2128 | 6.1821 | 6.7140 | 6.4459 | 5.5018 | 6.1373 |
| CWC25_S217 | CWC25 | Q9NXE8 | | 218 | KKMANSSPVLSKV | 5.9825 | 6.0730 | 6.0725 | 6.3659 | 6.5300 | 6.0797 | 6.2905 | 6.1634 | 6.6580 | 6.1703 | 5.9258 | 5.8273 | 6.2065 | 6.5029 | 6.7982 | 6.3537 |
| CWC25_S218 | CWC25 | Q9NXE8 | | 217 | QKKMANSSPVLSK | 6.0984 | 6.7110 | 6.2152 | 6.5501 | 6.1976 | 5.8611 | 6.1390 | 6.4287 | 6.3019 | 6.4946 | 6.1315 | 6.2297 | 5.8701 | 6.2619 | 6.5340 | 5.9752 |
| CWF19L2_S479 | CWF19L2 | Q2TBE0 | | 479 | KSTFAGSPERESI | 5.8109 | 6.5072 | 5.7728 | 5.9754 | 5.8557 | 5.7466 | 6.0031 | 6.2757 | 6.6034 | 6.3426 | 5.9893 | 6.1506 | 5.9600 | 6.9609 | 7.0697 | 6.9760 |
| CWF19L2_T562 | CWF19L2 | Q2TBE0 | | 562 | RVWPVNTPGKSLE | 7.4933 | 7.2094 | 7.2729 | 7.4162 | 5.2334 | 6.2237 | 6.9274 | 6.2183 | 6.7080 | 6.4423 | 6.6535 | 6.5411 | 4.5651 | 5.0094 | 4.4590 | 5.6270 |
| CXADR_S323 | CXADR | P78310 | | 332 | FERTPQSPTLPPA | 6.2030 | 6.8653 | 6.2872 | 6.6995 | 5.7634 | 6.7162 | 6.7677 | 6.5683 | 6.3515 | 6.7708 | 6.9884 | 6.8935 | 4.8989 | 5.6810 | 4.9087 | 5.6367 |
| CXADR_S332 | CXADR | P78310 | | 323 | QYNQVPSEDFERT | 5.2930 | 5.6598 | 6.6753 | 6.9283 | 6.1045 | 6.5133 | 5.8734 | 5.9986 | 6.4459 | 6.0415 | 7.2610 | 6.8915 | 6.0038 | 6.5554 | 5.6974 | 6.0572 |
| CXorf38_T314 | CXorf38 | Q8TB03 | | 314 | QEPGRQTPDRKAx | 6.3352 | 6.7879 | 7.2673 | 6.9312 | 6.2202 | 7.3909 | 6.5200 | 6.7037 | 6.7004 | 6.9760 | 7.5573 | 7.7331 | 4.2463 | 4.4435 | 3.9423 | 4.2447 |
| CXXC1_S138 | CXXC1 | Q9P0U4-2 | | 19 | AGEDSKSENGENA | 5.8535 | 6.4912 | 6.3411 | 6.1813 | 5.8982 | 6.3682 | 6.1060 | 5.9483 | 5.6388 | 6.1747 | 5.9185 | 6.0901 | 6.2063 | 6.6981 | 6.8414 | 7.2441 |
| CXXC1_S19 | CXXC1 | Q9P0U4-2 | | 138 | LARGSASPHKSSP | 6.3987 | 7.0189 | 5.8132 | 6.8311 | 6.0564 | 6.0959 | 5.8289 | 6.4661 | 6.1383 | 6.0145 | 6.2530 | 6.4126 | 5.9008 | 6.3546 | 6.4222 | 5.9946 |
| CXXC1_S215 | CXXC1 | A0A669KB29 | | 215 | QLRARLSPVTPSE | 5.7364 | 6.1974 | 6.5477 | 6.4518 | 6.0801 | 6.1489 | 5.9007 | 6.4503 | 6.7943 | 6.0805 | 6.0827 | 6.3273 | 6.2080 | 6.3531 | 6.4685 | 6.1723 |
| CXXC1_S221 | CXXC1 | Q9P0U4-2 | | 221 | SYKYFPSSLSPVT | 6.0116 | 6.4474 | 6.1681 | 7.3168 | 6.2695 | 6.3075 | 6.6588 | 6.9748 | 6.0528 | 6.6709 | 6.5462 | 6.1542 | 5.8981 | 5.1999 | 5.8333 | 5.4900 |
| CYB5B_S23 | CYB5B | O43169 | | 23 | GQEVETSVTYYRL | 7.0088 | 6.0262 | 6.0859 | 6.2030 | 5.8579 | 5.5658 | 7.1637 | 7.0046 | 5.8756 | 7.1912 | 7.0091 | 7.3305 | 4.1528 | 6.4037 | 5.4949 | 5.6262 |
| CYB5R1_S169 | CYB5R1 | Q9UHQ9 | | 169 | IQPNKKSPPEPRV | 5.1729 | 5.3629 | 6.2288 | 6.5750 | 6.2083 | 6.1800 | 6.0804 | 6.3219 | 6.6663 | 6.5562 | 6.6812 | 6.3605 | 6.2083 | 6.5500 | 6.4757 | 6.3716 |
| CYBRD1_S232 | CYBRD1 | Q53TN4 | | 248 | EQGARGSMPAYSG | 5.8522 | 6.4614 | 5.6395 | 6.5963 | 6.5069 | 5.6227 | 5.5750 | 5.7547 | 6.4937 | 6.8962 | 6.8390 | 6.6044 | 5.2370 | 7.0905 | 7.1235 | 5.7070 |
| CYBRD1_S248 | CYBRD1 | Q53TN4 | | 232 | RPKEPNSTILHPN | 5.3828 | 6.2162 | 5.4220 | 6.0480 | 6.4224 | 6.4523 | 5.7355 | 5.8046 | 5.9944 | 6.1378 | 5.0734 | 6.2471 | 6.9639 | 7.6219 | 6.8948 | 7.5827 |
| CYCS_T29 | CYCS | P99999 | | 29 | KGGKHKTGPNLHG | 6.3672 | 6.4699 | 5.9550 | 5.9446 | 6.2526 | 7.0814 | 6.5011 | 6.1871 | 7.1543 | 6.0859 | 5.6395 | 5.8092 | 6.1708 | 6.0428 | 6.7362 | 5.6023 |
| CYLD_S399 | CYLD | Q9NQC7 | | 399 | TDFDRSSPPLQPP | 6.0104 | 6.2158 | 6.3325 | 5.9457 | 6.3829 | 6.4492 | 6.1688 | 7.0058 | 6.2334 | 6.1023 | 6.2732 | 6.5463 | 6.0654 | 6.9489 | 5.6737 | 5.6457 |
| CYLD_S418 | CYLD | Q9NQC7 | | 418 | TENRFHSLPFSLT | 6.5626 | 6.4914 | 7.6693 | 6.8438 | 5.5765 | 6.7486 | 6.7073 | 7.1407 | 6.5910 | 6.3785 | 7.0430 | 7.3778 | 4.3215 | 5.6973 | 4.1434 | 4.7073 |
| CYLD_S418;S422 | CYLD | Q9NQC7 | 418;422 | | NA | 6.8417 | 6.1757 | 7.9146 | 7.4747 | 4.5209 | 6.0287 | 7.7540 | 6.5699 | 7.6509 | 4.5322 | 6.1951 | 7.8628 | 4.7030 | 5.6295 | 5.2907 | 4.8557 |
| CYP27C1_S58 | CYP27C1 | Q4G0S4 | | 58 | GAGRPGSPPGGGR | 6.4323 | 6.3901 | 5.8016 | 6.5430 | 5.8701 | 6.2417 | 6.3420 | 5.6475 | 6.4409 | 6.4168 | 5.9526 | 6.0245 | 6.5676 | 6.7804 | 6.3842 | 6.1648 |
| DAB2_S401 | DAB2 | P98082 | | 401 | PNPFVGSPPKGLS | 6.1517 | 6.3673 | 6.7554 | 6.5892 | 5.9905 | 6.1310 | 6.3986 | 6.1706 | 6.4576 | 7.1236 | 6.6238 | 6.1298 | 5.6979 | 6.4274 | 5.3990 | 5.5866 |
| DAB2IP_S1108 | DAB2IP | Q5VWQ8 | | 1108 | IISRLMSVEEELK | 5.3950 | 5.0129 | 6.8745 | 3.8686 | 5.5135 | 5.4017 | 5.8353 | 5.1358 | 5.2118 | 5.4911 | 5.0888 | 6.2175 | 8.8278 | 7.0555 | 11.0605 | 8.0098 |
| DAB2IP_S702 | DAB2IP | Q5VWQ8 | | 978 | SSSKGDSPELKPR | 6.4501 | 6.6374 | 6.5019 | 6.6509 | 5.7118 | 5.7419 | 6.3319 | 5.7556 | 6.5603 | 6.5592 | 6.4816 | 6.7275 | 5.5900 | 6.2899 | 5.7845 | 6.2254 |
| DAB2IP_S978 | DAB2IP | Q5VWQ8 | | 702 | DFTRLPSPTPENK | 6.6729 | 6.2380 | 6.3580 | 6.2156 | 6.6651 | 6.9402 | 6.2428 | 6.8526 | 6.5941 | 5.9353 | 6.6546 | 6.2001 | 5.8674 | 5.8424 | 5.5267 | 5.1940 |
| DAG1_S812 | DAG1 | Q14118 | | 790 | TLEDQATFIKKGV | 8.1649 | 7.6581 | 8.4505 | 8.4468 | 6.8802 | 4.6778 | 5.7548 | 6.0939 | 5.7678 | 5.4253 | 6.1832 | 6.2059 | 4.4209 | 6.7067 | 4.5875 | 4.5759 |
| DAG1_T790 | DAG1 | Q14118 | | 812 | DSKPPPSSSMPLI | 5.5845 | 5.4976 | 5.3197 | 5.9055 | 6.1159 | 5.4710 | 5.5065 | 5.4013 | 5.2577 | 5.9563 | 5.1814 | 5.5331 | 7.8702 | 7.8336 | 8.5441 | 9.0216 |
| DAGLB_S570 | DAGLB | Q8NCG7 | | 570 | SLLTRWSPAYSFS | 6.0559 | 6.5026 | 6.4857 | 7.5814 | 5.0719 | 7.2738 | 6.1486 | 6.2982 | 6.3930 | 7.0863 | 6.6606 | 7.5498 | 5.0065 | 6.1528 | 4.5177 | 5.2151 |
| DAGLB_S579;S584 | DAGLB | Q8NCG7 | 579;584 | | NA | 5.4186 | 5.9001 | 6.1899 | 6.0627 | 5.8288 | 5.9388 | 6.2354 | 6.2473 | 5.4031 | 5.9871 | 5.8099 | 5.5468 | 7.5728 | 7.0874 | 7.2962 | 7.4751 |
| DAP_S51 | DAP | P51397 | | 51 | QEWESPSPPKPTV | 5.7981 | 4.7714 | 6.7063 | 4.9068 | 5.9644 | 4.9098 | 4.6480 | 6.1792 | 5.0666 | 6.5717 | 6.4133 | 6.9595 | 7.1608 | 8.5038 | 7.6044 | 7.8358 |
| DAPK3_S312 | DAPK3 | O43293 | | 407 | GALLGTSGLKRRF | 6.3218 | 6.8731 | 8.3984 | 8.4719 | 5.5499 | 4.0154 | 5.3524 | 6.0737 | 6.3034 | 7.5577 | 5.7751 | 7.9252 | 4.6816 | 4.9848 | 4.4565 | 7.2590 |
| DAPK3_S407 | DAPK3 | O43293 | | 312 | TIKSHSSLPPNNS | 6.1958 | 6.6605 | 6.2016 | 6.5257 | 6.4990 | 6.4026 | 6.6512 | 5.0331 | 6.9520 | 5.6824 | 5.9113 | 6.0855 | 6.3724 | 6.6163 | 6.4990 | 5.7115 |
| DARS1_S249 | DARS1 | P14868 | | 249 | NAYLAQSPQLYKQ | 6.0839 | 5.6992 | 6.6836 | 6.1180 | 6.2063 | 6.4420 | 6.3553 | 6.1083 | 6.5209 | 6.3918 | 6.6965 | 6.2347 | 6.1599 | 6.3078 | 6.1585 | 5.8334 |
| DARS2_S242 | DARS2 | Q6PI48 | | 242 | FYSLPQSPQQFKQ | 6.7035 | 6.4875 | 6.5972 | 6.3941 | 5.7394 | 5.4126 | 6.3992 | 6.6486 | 6.1694 | 6.9603 | 6.4263 | 6.7586 | 5.7196 | 6.1284 | 5.5918 | 5.8635 |
| DAXX_S495 | DAXX | Q9UER7 | | 671 | TLPSPPSPLASLA | 5.9954 | 6.0666 | 5.8812 | 6.3477 | 6.1709 | 6.0313 | 6.1263 | 6.6231 | 6.3231 | 6.0551 | 6.1909 | 6.5415 | 6.4446 | 6.4801 | 6.2586 | 6.4635 |
| DAXX_S618 | DAXX | Q9UER7 | | 495 | GKDGDKSPMSSLQ | 6.1801 | 6.4103 | 6.0991 | 6.4420 | 6.1397 | 5.8741 | 6.0289 | 6.1454 | 6.2386 | 6.0618 | 6.1113 | 5.9571 | 6.6519 | 6.4114 | 6.5167 | 6.7317 |
| DAXX_S668 | DAXX | Q9UER7 | | 688 | SSTRVDSPSHGLV | 5.8617 | 5.9936 | 5.9679 | 6.1190 | 6.1894 | 6.4972 | 5.8127 | 5.9863 | 6.4104 | 6.5240 | 5.9492 | 6.3050 | 6.7205 | 6.6723 | 6.5674 | 6.4233 |
| DAXX_S668;S671 | DAXX | Q9UER7 | | 618 | SFNGGVSPHNWGD | 5.8244 | 6.3484 | 6.6628 | 6.2682 | 6.0377 | 6.0382 | 5.6455 | 6.7009 | 6.2975 | 6.1792 | 6.1400 | 6.2689 | 6.3504 | 6.5869 | 6.3492 | 6.3019 |
| DAXX_S671 | DAXX | Q9UER7 | | 668 | KICTLPSPPSPLA | 5.4613 | 8.5387 | 6.8768 | 6.7564 | 6.7335 | 7.9140 | 5.1289 | 6.8080 | 7.5014 | 4.6934 | 4.9742 | 8.8252 | 4.6705 | 5.6848 | 4.7106 | 4.7221 |

| | | | | | | | | | | | | | | | | | | | | |
|---|---|---|---|---|---|---|---|---|---|---|---|---|---|---|---|---|---|---|---|---|
| DAXX_S688 | DAXX | Q9UER7 | 668;671 | NA | 6.1819 | 6.9854 | 6.1989 | 6.4600 | 6.4101 | 6.2590 | 6.9152 | 7.0705 | 6.8523 | 6.7887 | 6.1744 | 6.6302 | 5.2421 | 5.4795 | 4.9439 | 5.4077 |
| DAXX_S688;S702 | DAXX | Q9UER7 | 688;702 | NA | 6.2356 | 6.7510 | 6.2844 | 7.0438 | 6.2831 | 6.6493 | 6.3197 | 6.5999 | 7.0040 | 6.5505 | 6.4178 | 6.3929 | 4.8024 | 6.0269 | 5.1851 | 5.4535 |
| DBF4_S359 | DBF4 | Q9UBU7 | 508 | SQPKQKSDTVLFP | 5.8496 | 6.1556 | 6.5231 | 6.4306 | 5.3396 | 5.7240 | 6.0620 | 5.8256 | 6.2172 | 6.2775 | 5.9916 | 6.3629 | 6.1984 | 6.5953 | 7.1744 | 7.2727 |
| DBF4_S413;S420 | DBF4 | Q9UBU7 | 359 | YSVGSLSPVSASV | 6.4628 | 6.5410 | 6.6208 | 6.6274 | 6.3863 | 6.1468 | 6.1832 | 6.2174 | 6.7038 | 6.1330 | 6.4183 | 6.2497 | 5.8439 | 5.9330 | 5.9055 | 5.6272 |
| DBF4_S420 | DBF4 | Q9UBU7 | 420 | EPIPHPSNELRGL | 5.6186 | 5.7815 | 5.4059 | 6.7457 | 6.3194 | 6.0031 | 6.0378 | 5.7181 | 6.9447 | 5.7965 | 6.6959 | 6.5495 | 6.6147 | 6.7415 | 6.4260 | 6.6011 |
| DBF4_S508 | DBF4 | Q9UBU7 | 553 | AKAPFHTPPEEPN | 6.1893 | 6.1396 | 6.9834 | 6.6342 | 6.2924 | 5.6030 | 6.3240 | 6.8565 | 6.7399 | 6.1852 | 6.5275 | 6.3302 | 6.1157 | 5.7387 | 5.5941 | 5.7464 |
| DBF4_S617 | DBF4 | Q9UBU7 | 273 | VKLRIQTDGDKYG | 8.1653 | 7.2771 | 6.6623 | 6.5265 | 5.3361 | 6.0293 | 6.9767 | 5.4762 | 7.3969 | 6.3113 | 6.6969 | 6.3437 | 5.5781 | 5.5845 | 4.2249 | 5.4142 |
| DBF4_T273 | DBF4 | Q9UBU7 | 617 | CSSPVQSLLDLFQ | 6.4314 | 5.8403 | 6.6974 | 7.0816 | 6.9693 | 6.2009 | 5.6807 | 6.3782 | 6.4964 | 4.5575 | 6.1358 | 5.3851 | 7.3890 | 5.6393 | 5.8935 | 7.2235 |
| DBF4_T553 | DBF4 | Q9UBU7 | 413;420 | NA | 6.4263 | 5.8639 | 6.7582 | 7.0858 | 5.8173 | 6.1307 | 6.6440 | 6.8300 | 6.2621 | 6.0258 | 6.2352 | 6.9552 | 5.5063 | 5.5476 | 5.8426 | 6.0689 |
| DBF4B_S278 | DBF4B | Q8NFT6-3 | 278 | SGFMELSSGPDLA | 7.2307 | 9.2543 | 6.8908 | 6.5940 | 5.2935 | 3.6855 | 6.0490 | 6.2163 | 6.7937 | 6.5346 | 8.0617 | 6.5670 | 5.3151 | 5.2126 | 3.1135 | 7.1876 |
| DBN1_S136 | DBN1 | Q16643-2 | 339 | TPIPTRSPSDSST | 6.2845 | 6.4982 | 6.1818 | 6.6828 | 6.3527 | 6.1656 | 6.2689 | 6.3313 | 6.2356 | 6.2247 | 6.5065 | 6.5937 | 6.3170 | 5.9290 | 5.6428 | 5.7847 |
| DBN1_S144 | DBN1 | Q16643-2 | 144 | GLARLSSPVLHRL | 6.8537 | 6.9003 | 5.9942 | 6.0573 | 7.0810 | 6.6137 | 6.1820 | 6.7761 | 6.6434 | 6.2405 | 5.9886 | 5.9760 | 6.1969 | 5.6136 | 5.4714 | 5.4113 |
| DBN1_S314 | DBN1 | Q16643-2 | 333 | HRRMAPTPIPTRS | 7.4679 | 6.9159 | 6.2541 | 7.2883 | 6.9197 | 6.7321 | 7.0402 | 6.9258 | 6.8945 | 6.3498 | 6.4895 | 6.6238 | 4.6933 | 4.8899 | 3.9408 | 4.5743 |
| DBN1_S339 | DBN1 | Q16643-2 | 314 | GSCDVPSPFNHRP | 7.0859 | 6.9777 | 6.1083 | 7.8232 | 7.3016 | 7.5156 | 6.8559 | 7.2336 | 6.8031 | 6.4789 | 6.8218 | 6.6898 | 4.4104 | 4.3955 | 3.7129 | 3.7857 |
| DBN1_S339;S341 | DBN1 | Q16643-2 | 136 | AIGQRLSNGLARL | 6.5576 | 6.8067 | 6.3545 | 6.8880 | 6.8021 | 6.7110 | 7.0373 | 7.6166 | 6.5735 | 6.6094 | 6.8820 | 6.8346 | 4.5402 | 4.8649 | 4.2332 | 4.6884 |
| DBN1_S383 | DBN1 | Q16643-2 | 603 | EGYFSQSQEEEFA | 6.1818 | 6.1176 | 6.4516 | 6.1984 | 6.5066 | 6.5771 | 6.0783 | 5.8221 | 6.2869 | 7.4415 | 6.2148 | 6.3098 | 6.2888 | 5.8152 | 5.8653 | 5.8441 |
| DBN1_S603 | DBN1 | Q16643-2 | 383 | QETQEPSPILDSE | 5.5680 | 6.1328 | 6.6275 | 6.1417 | 6.2106 | 5.4086 | 6.0627 | 7.3109 | 5.6534 | 5.5884 | 6.8200 | 5.6865 | 6.6198 | 7.3377 | 6.3432 | 6.4885 |
| DBN1_T333 | DBN1 | Q16643-2 | 348 | DSSTASTPVAEQI | 5.6190 | 7.1817 | 6.2330 | 6.8569 | 5.7078 | 6.4100 | 6.1962 | 6.6951 | 7.2571 | 6.9884 | 6.1652 | 6.8505 | 5.6815 | 5.2918 | 5.2679 | 5.5979 |
| DBN1_T348 | DBN1 | Q16643-2 | 339;341 | NA | 5.2781 | 6.0282 | 6.7203 | 6.4679 | 7.2148 | 6.5867 | 6.7895 | 6.2557 | 5.4102 | 5.3662 | 5.8534 | 6.1402 | 7.0073 | 6.0622 | 6.0631 | 6.7561 |
| DBNL_S178 | DBNL | Q9UJU6 | 269 | QKERAMSTTSISS | 7.2947 | 5.4713 | 6.9454 | 5.4823 | 6.4707 | 5.9523 | 5.5897 | 6.5799 | 7.4446 | 7.0505 | 6.7455 | 6.4643 | 6.5897 | 6.8274 | 4.5091 | 4.5828 |
| DBNL_S232 | DBNL | Q9UJU6 | 283 | QPGKLRSPFLQKQ | 6.9890 | 7.1844 | 6.8354 | 7.1633 | 6.8154 | 7.6093 | 7.0445 | 7.4070 | 7.5040 | 7.4403 | 7.7058 | 7.6055 | 3.0636 | 3.6967 | 2.4919 | 3.4439 |
| DBNL_S269 | DBNL | Q9UJU6 | 275 | STTSISSPQPGKL | 6.4406 | 7.0683 | 6.9900 | 6.5886 | 6.5359 | 7.7176 | 7.2933 | 7.2696 | 6.8960 | 7.4798 | 7.5532 | 7.0553 | 3.6403 | 4.1081 | 3.5241 | 3.8394 |
| DBNL_S269;S275 | DBNL | Q9UJU6 | 178 | KRVGKDSFWAKAE | 7.8929 | 6.5736 | 8.1611 | 6.6513 | 6.7386 | 5.8933 | 6.2404 | 6.6494 | 7.9458 | 5.3244 | 6.5405 | 6.5080 | 5.7563 | 6.0149 | 3.7458 | 3.3636 |
| DBNL_S275 | DBNL | Q9UJU6 | 232 | EQGGEASPQRTWE | 7.1663 | 7.4746 | 7.3439 | 6.6125 | 6.6398 | 7.1463 | 6.8500 | 7.5421 | 6.9535 | 7.3410 | 8.1137 | 7.9056 | 3.0822 | 3.3371 | 2.9588 | 3.5325 |
| DBNL_S283 | DBNL | Q9UJU6 | 291 | FLQKQLTQPETHF | 9.0762 | 7.6882 | 11.2094 | 8.4540 | 6.3741 | 5.9739 | 5.7909 | 6.5839 | 6.1182 | 6.7283 | 5.6001 | 5.7386 | 3.8852 | 5.2735 | 2.7354 | 2.7701 |
| DBNL_T291 | DBNL | Q9UJU6 | 269;275 | NA | 8.0754 | 6.9812 | 7.3159 | 7.1504 | 6.7222 | 6.2385 | 6.9873 | 6.8671 | 7.8765 | 8.0754 | 7.6393 | 6.9538 | 3.1521 | 4.2402 | 2.7145 | 3.0102 |
| DCAF1_S1000 | DCAF1 | Q9Y4B6 | 1000 | LDRHLPSPPTLDS | 5.4571 | 5.8733 | 5.7428 | 6.2973 | 6.0338 | 5.8741 | 6.0908 | 6.3560 | 5.9783 | 5.8182 | 6.2794 | 6.0511 | 6.8668 | 7.3138 | 6.7996 | 7.1676 |
| DCAF1_S255 | DCAF1 | Q9Y4B6 | 706 | GKFISGTPRRKLP | 7.8012 | 7.7520 | 7.0298 | 7.3870 | 6.3656 | 6.7771 | 7.4667 | 6.9574 | 7.2665 | 6.5333 | 6.7486 | 6.9448 | 3.2000 | 4.2948 | 3.4061 | 4.0691 |
| DCAF1_T706 | DCAF1 | Q9Y4B6 | 255 | TSSRVNSTTKPED | 5.6464 | 6.1201 | 6.4024 | 6.3305 | 6.3184 | 6.3244 | 7.2279 | 6.4588 | 7.2605 | 6.7512 | 6.4572 | 6.6010 | 5.6432 | 5.5553 | 5.1013 | 5.8012 |
| DCAF10_S63 | DCAF10 | Q5QP82 | 63 | PGAPSLSPAPRSG | 5.4836 | 5.9614 | 6.5244 | 6.0889 | 5.8041 | 6.4131 | 6.0169 | 6.4644 | 5.9437 | 6.4023 | 6.4706 | 6.3147 | 6.0890 | 6.9763 | 6.5019 | 6.5447 |
| DCAF10_S92 | DCAF10 | Q5QP82 | 92 | GEPSPPSPPCRRP | 6.3975 | 5.6656 | 6.0612 | 6.0381 | 6.0870 | 6.4273 | 5.9017 | 6.4560 | 6.3136 | 6.3558 | 6.5295 | 6.0059 | 6.1121 | 6.6142 | 6.7096 | 6.3248 |
| DCAF12_S15 | DCAF12 | Q5T6F0 | 15 | KRKAPASPGAGSD | 6.0348 | 6.7535 | 5.8363 | 6.6767 | 5.8381 | 7.1266 | 6.7204 | 6.6296 | 5.6652 | 6.8589 | 6.6400 | 6.9136 | 5.7478 | 5.7868 | 5.2438 | 5.5278 |
| DCAF15_S310 | DCAF15 | Q66K64 | 310 | PARSSGSPEPSPA | 5.2344 | 5.7590 | 6.1695 | 5.9614 | 5.9824 | 6.5048 | 6.1661 | 6.5484 | 6.1054 | 6.3209 | 6.2364 | 6.1486 | 6.2990 | 7.0578 | 6.7168 | 6.7891 |
| DCAF15_S50 | DCAF15 | Q66K64 | 50 | KISGQLSPRLFRK | 5.6703 | 6.3857 | 6.6180 | 5.7452 | 5.5762 | 6.2867 | 6.3215 | 6.2313 | 6.4984 | 6.6245 | 7.2801 | 7.0840 | 5.3487 | 6.3350 | 5.8210 | 6.1734 |
| DCAF5_S648 | DCAF5 | Q96JK2 | 794 | RAEEPPSPPVPKA | 7.1285 | 7.0612 | 7.4471 | 6.9837 | 6.4773 | 5.9508 | 6.3578 | 6.1857 | 6.9281 | 7.6509 | 7.3824 | 7.2725 | 4.1426 | 4.8083 | 3.8889 | 4.3340 |
| DCAF5_S794 | DCAF5 | Q96JK2 | 648 | IQPSRASPTSDIE | 6.0816 | 6.9166 | 6.9082 | 7.1633 | 7.3152 | 5.5755 | 5.5776 | 5.7674 | 6.4232 | 7.6378 | 5.7948 | 6.8491 | 5.4616 | 6.2903 | 4.9281 | 5.3098 |
| DCAF6_S336 | DCAF6 | Q58WW2-3 | 336 | ERDGEQSPNVSLM | 5.6625 | 5.9093 | 6.8367 | 5.8336 | 6.2540 | 6.1367 | 6.7824 | 6.4830 | 5.8936 | 5.5848 | 5.7090 | 6.6066 | 6.6974 | 6.1268 | 7.1283 | 6.3552 |
| DCBLD1_T602 | DCBLD1 | Q8N8Z6 | 602 | PEPEYATPIVERH | 6.5652 | 6.7688 | 5.8978 | 7.2497 | 6.2519 | 5.6691 | 6.1187 | 6.4853 | 6.9223 | 7.2873 | 5.8038 | 5.9777 | 5.9417 | 5.6785 | 5.2274 | 6.1547 |
| DCBLD2_S734 | DCBLD2 | Q96PD2-2 | 764 | APDELVYQVPQST | 6.1074 | 6.2048 | 6.0009 | 5.6883 | 5.3646 | 6.5027 | 6.4991 | 5.9358 | 6.5855 | 6.8056 | 6.9206 | 7.1332 | 5.9553 | 6.4744 | 5.6541 | 6.1677 |
| DCBLD2_T652 | DCBLD2 | Q96PD2-2 | 734 | TYNTLLSRTDSCS | 5.5258 | 6.2024 | 6.4573 | 6.3869 | 5.7554 | 6.6856 | 6.5632 | 7.1464 | 6.9377 | 7.4543 | 6.8065 | 6.3782 | 5.2563 | 6.7985 | 4.8655 | 4.7801 |
| DCBLD2_Y764 | DCBLD2 | Q96PD2-2 | 652 | TLHQRSTFKPEEG | 9.6245 | 7.6207 | 8.0289 | 7.5537 | 6.0041 | 5.8346 | 6.4024 | 5.1244 | 7.7860 | 6.3838 | 7.1858 | 7.1212 | 3.6967 | 5.1338 | 2.6792 | 3.8201 |
| DCK_S74 | DCK | P27707 | 74 | FEELTMSQKNGGN | 4.8601 | 5.1449 | 6.0237 | 5.0134 | 5.8173 | 5.7699 | 5.5774 | 5.8803 | 5.7330 | 5.5480 | 5.4762 | 5.3023 | 8.2087 | 8.9222 | 8.9225 | 7.8000 |
| DCLRE1B_S335 | DCLRE1B | Q9H816 | 379 | AKKEKLSPWPADL | 7.4996 | 7.3104 | 7.6129 | 7.5822 | 5.7386 | 6.5915 | 6.9185 | 6.4512 | 8.2603 | 6.7939 | 7.6243 | 8.2168 | 3.0311 | 3.9924 | 2.6387 | 3.7374 |
| DCLRE1B_S379 | DCLRE1B | Q9H816 | 335 | SSSRKPSLLWLLE | 5.9063 | 4.0886 | 4.2476 | 5.0064 | 7.1931 | 4.9705 | 4.5047 | 4.9249 | 12.1111 | 8.8103 | 9.0812 | 8.2920 | 5.8427 | 7.4004 | 3.4265 | 4.1937 |
| DCP1A_S226 | DCP1A | Q9NPI6 | 525 | RKASSPSPLTIGT | 6.1818 | 5.9981 | 6.2762 | 5.5501 | 6.2825 | 6.0056 | 5.8901 | 6.3869 | 6.4018 | 6.0514 | 6.4389 | 6.8777 | 6.4471 | 6.4259 | 6.5128 | 6.2731 |
| DCP1A_S315 | DCP1A | Q9NPI6 | 315 | TYTIPLSPVLSPT | 6.2489 | 6.7627 | 6.0264 | 5.9390 | 5.8352 | 5.8255 | 6.0615 | 5.9248 | 5.9868 | 6.3087 | 6.8546 | 7.1488 | 6.3878 | 6.1571 | 6.7000 | 5.8322 |
| DCP1A_S315;T321 | DCP1A | Q9NPI6 | 62 | VYRRSASPYHGFT | 6.0188 | 5.1072 | 6.4486 | 6.1589 | 6.3598 | 6.6451 | 5.7064 | 7.4680 | 6.4317 | 7.9659 | 6.5006 | 7.3051 | 4.8351 | 6.1427 | 5.8281 | 5.0779 |
| DCP1A_S422 | DCP1A | Q9NPI6 | 226 | EELFGTSLPKEQP | 6.2187 | 6.0490 | 7.2953 | 5.9103 | 6.4762 | 6.7079 | 6.2121 | 6.6083 | 7.4030 | 6.6275 | 7.0754 | 7.4908 | 4.7740 | 5.5377 | 4.7179 | 4.8957 |
| DCP1A_S525 | DCP1A | Q9NPI6 | 422 | AMVASFSPAAGQL | 6.4919 | 7.1425 | 7.0864 | 6.2657 | 5.3976 | 4.9417 | 6.5199 | 7.0934 | 7.4494 | 7.3143 | 7.1179 | 8.4787 | 4.3875 | 5.7080 | 4.8628 | 3.7422 |
| DCP1A_S62 | DCP1A | Q9NPI6 | 315;321 | NA | 5.7089 | 5.9176 | 6.9721 | 6.1527 | 6.4314 | 6.1228 | 5.6341 | 6.8568 | 5.6681 | 6.2224 | 5.8262 | 6.2603 | 6.1881 | 6.5774 | 5.9895 | 6.5717 |
| DCP1B_S147 | DCP1B | Q8IZD4 | 448 | GSSGVISPQELLK | 6.5165 | 6.5525 | 6.7568 | 6.1875 | 6.2229 | 6.5944 | 5.9850 | 6.4794 | 6.8549 | 6.8755 | 7.4166 | 7.1605 | 4.9480 | 5.6090 | 4.6728 | 5.1676 |
| DCP1B_S275 | DCP1B | Q8IZD4 | 511 | PLFQVISPQRIPA | 6.0641 | 6.1613 | 6.7746 | 5.8261 | 6.1956 | 6.5126 | 6.6864 | 6.5303 | 7.1412 | 6.8199 | 6.9864 | 6.8384 | 5.1654 | 5.9329 | 5.0167 | 5.3481 |
| DCP1B_S283 | DCP1B | Q8IZD4 | 283 | EEPRRHSPPIEKQ | 5.9219 | 5.7582 | 6.5102 | 5.8069 | 6.4787 | 6.3008 | 5.6372 | 6.6732 | 6.5703 | 6.7025 | 6.7051 | 6.8343 | 6.2883 | 6.3131 | 5.7567 | 5.7425 |
| DCP1B_S448 | DCP1B | Q8IZD4 | 392 | RAPTSVTPVAPGK | 7.0555 | 6.9202 | 7.1047 | 6.0954 | 6.3181 | 7.2118 | 6.9645 | 7.0787 | 7.4222 | 6.9922 | 7.7338 | 7.5007 | 3.6686 | 4.2029 | 3.3348 | 4.3960 |
| DCP1B_S511 | DCP1B | Q8IZD4 | 275 | GVVRSLSYEEPRR | 5.9945 | 5.4829 | 6.3512 | 4.9644 | 6.4833 | 6.0456 | 5.2174 | 6.6994 | 5.8691 | 5.8870 | 6.2571 | 5.9242 | 7.6739 | 7.2528 | 7.1266 | 6.7705 |
| DCP1B_T362 | DCP1B | Q8IZD4 | 147 | GTGAGISPVILNS | 5.8845 | 6.2120 | 6.0426 | 5.2949 | 6.1819 | 6.8654 | 5.7323 | 6.3551 | 6.3598 | 7.1890 | 6.5421 | 6.9800 | 6.1702 | 6.0823 | 5.9328 | 6.1750 |
| DCP1B_T392 | DCP1B | Q8IZD4 | 362 | FEKLQSTPGAANK | 6.8076 | 6.8281 | 7.3008 | 6.6643 | 5.6996 | 7.9024 | 6.9480 | 6.9908 | 7.1860 | 7.8391 | 8.7706 | 9.3848 | 2.7000 | 3.5642 | 2.2656 | 3.1481 |
| DCP2_S246;S247;S249 | DCP2 | Q8IU60 | 284 | QLFPDGSPGDQWV | 5.9074 | 7.0881 | 7.7504 | 6.7212 | 5.8294 | 7.0276 | 6.7703 | 6.4537 | 6.7670 | 7.0076 | 7.4877 | 7.8876 | 4.0637 | 4.9531 | 3.8092 | 4.4759 |
| DCP2_S284 | DCP2 | Q8IU60 | 246;247;249 | NA | 5.5357 | 6.3445 | 6.1445 | 6.1942 | 6.6962 | 6.9233 | 6.4131 | 6.3969 | 5.2307 | 6.0708 | 6.7940 | 6.4317 | 5.9596 | 7.0952 | 5.9646 | 5.8049 |
| DCTN1_T108 | DCTN1 | Q14203 | 950 | KLEDRETVIKELK | 7.6138 | 6.9328 | 7.3221 | 7.1065 | 6.7137 | 6.3304 | 6.2966 | 7.0315 | 7.3321 | 6.4101 | 7.0606 | 6.9862 | 4.2203 | 4.6613 | 3.5713 | 4.4104 |
| DCTN1_T950 | DCTN1 | Q14203 | 108 | DTTSPETPDSSAS | 6.2963 | 6.1418 | 6.8285 | 6.4556 | 5.6760 | 6.0601 | 6.7840 | 6.5349 | 6.2583 | 6.3489 | 6.3068 | 6.4589 | 5.8992 | 6.3384 | 5.4560 | 6.1564 |
| DCTN2_T203 | DCTN2 | Q13561-2 | 203 | GGKTTGTPPDSSL | 6.4689 | 5.4841 | 6.6360 | 5.6793 | 6.8140 | 5.6536 | 5.5492 | 6.9830 | 5.8095 | 7.2429 | 6.6913 | 7.4190 | 5.8325 | 6.5447 | 5.3319 | 5.8601 |
| DCTN6_T186 | DCTN6 | O00399 | 186 | TMKGSSTPVKNxx | 7.4481 | 7.5501 | 7.1405 | 7.1137 | 6.4728 | 6.9209 | 6.5536 | 7.1418 | 7.2520 | 7.1999 | 7.6205 | 7.7944 | 3.1104 | 3.8234 | 2.9844 | 3.8735 |
| DCUN1D3_S168 | DCUN1D3 | Q8IWE4 | 17 | PSSTLGSKNGDRE | 6.5814 | 5.2379 | 7.1084 | 6.3223 | 5.5825 | 5.4406 | 7.4197 | 7.4805 | 6.2789 | 5.9966 | 6.3484 | 7.1634 | 5.8243 | 5.9662 | 5.3595 | 5.8894 |
| DCUN1D3_S17 | DCUN1D3 | Q8IWE4 | 168 | ICARFPSLLTEAK | 5.4527 | 7.5055 | 4.7945 | 6.9633 | 7.6658 | 5.8448 | 6.3734 | 9.0537 | 5.5993 | 6.4007 | 5.7528 | 4.0307 | 6.1279 | 5.6095 | 6.7690 | 6.0563 |
| DCUN1D5_S9 | DCUN1D5 | Q9BTE7 | 9 | VKKKRKSPGVAAA | 5.6570 | 5.7201 | 6.5758 | 6.1826 | 5.9008 | 6.6253 | 6.2570 | 6.5584 | 6.3738 | 6.3645 | 6.6142 | 6.5622 | 5.7326 | 6.5328 | 6.0629 | 6.2800 |
| DDA1_S33 | DDA1 | Q9BW61 | 33 | ASNRRPSVYLPTR | 4.8423 | 4.2205 | 4.9837 | 4.4722 | 7.5219 | 4.8110 | 4.3913 | 5.6894 | 6.8149 | 8.2538 | 7.5916 | 8.0444 | 7.6137 | 8.5924 | 5.5319 | 6.6247 |
| DDB1_S720 | DDB1 | Q16531 | 720 | TVPLYESPRKICY | 6.5866 | 7.2441 | 6.6236 | 6.7597 | 5.9715 | 6.6868 | 6.7218 | 6.5353 | 6.6468 | 7.1109 | 7.2602 | 7.6310 | 3.6727 | 5.0941 | 4.7562 | 4.6987 |
| DDB2_S26 | DDB2 | Q92466 | 26 | RNKRSRSPLELEP | 5.8285 | 6.3623 | 6.1391 | 6.0619 | 6.1784 | 6.2125 | 6.0237 | 6.2006 | 6.3158 | 6.5103 | 6.2785 | 6.2254 | 6.3138 | 6.7515 | 6.3526 | 6.2451 |
| DDI2_S106 | DDI2 | Q5TDH0-3 | 121 | TQQSHSSPGEITS | 5.9357 | 6.4609 | 6.7084 | 6.5333 | 6.4545 | 6.3054 | 6.2604 | 6.8109 | 6.7814 | 6.4859 | 6.5984 | 6.5971 | 5.2227 | 5.9744 | 5.4096 | 5.4608 |
| DDI2_S121 | DDI2 | Q5TDH0-3 | 106 | AVPGTSSPRQRQP | 6.2383 | 6.1762 | 6.8430 | 7.0738 | 6.6447 | 6.8422 | 6.6070 | 6.8310 | 6.8753 | 6.6034 | 7.0945 | 7.5925 | 4.1299 | 5.3271 | 4.3328 | 4.7883 |
| DDI2_S121;S128 | DDI2 | Q5TDH0-3 | 121;128 | NA | 6.7897 | 7.2300 | 7.5083 | 7.4930 | 6.5524 | 7.1105 | 7.1093 | 6.7780 | 6.3671 | 5.4714 | 6.5029 | 6.4454 | 3.8967 | 5.2260 | 4.5194 | 4.9998 |
| DDRGK1_S114 | DDRGK1 | Q96HY6 | 114 | PAETHLSGKIGAK | 6.9954 | 6.8679 | 6.8390 | 7.0077 | 7.0904 | 7.3512 | 7.8121 | 6.6385 | 6.4987 | 6.8921 | 6.2662 | 6.6035 | 4.1689 | 4.5780 | 3.7012 | 4.6891 |
| DDX1_S481 | DDX1 | Q92499 | 481 | TRPGANSPEMWSE | 5.8348 | 6.1030 | 6.9214 | 5.4095 | 5.5670 | 6.5858 | 6.1959 | 6.1208 | 6.0161 | 5.7037 | 6.1159 | 6.1285 | 6.0811 | 6.8374 | 7.3348 | 7.0443 |
| DDX1_S632 | DDX1 | Q92499 | 632 | VWYHVCSSRGKGC | 6.4136 | 6.0763 | 6.5521 | 6.1324 | 6.5500 | 6.0922 | 5.1565 | 6.4364 | 6.6872 | 6.5251 | 6.2792 | 6.1525 | 6.3014 | 6.0444 | 6.8458 | 5.7549 |
| DDX1_T170 | DDX1 | Q92499 | 170 | GFGFGGTGKKSHN | 9.0587 | 7.2550 | 5.3028 | 6.7983 | 6.3074 | 4.6635 | 7.7173 | 5.3713 | 4.8005 | 7.1294 | 5.7081 | 6.7413 | 5.3485 | 6.3417 | 4.8861 | 6.5700 |
| DDX10_S539 | DDX10 | Q13206 | 539 | IEPRAPSLTNDEV | 5.8227 | 5.8881 | 5.6711 | 6.1548 | 6.7459 | 5.8614 | 6.1326 | 6.2697 | 6.1784 | 6.1915 | 6.1081 | 6.0063 | 6.8051 | 6.7598 | 6.8100 | 6.5944 |
| DDX10_S7 | DDX10 | Q13206 | 780 | EAFLDWSDDDDDD | 5.4936 | 5.8538 | 5.7305 | 6.6206 | 5.9536 | 5.7326 | 6.1558 | 5.8534 | 6.4363 | 6.1952 | 5.7069 | 6.3877 | 7.1181 | 6.7598 | 7.0028 | 6.9992 |
| DDX10_S780 | DDX10 | Q13206 | 7 | MGKTANSPGSGAR | 6.4893 | 6.1681 | 6.1439 | 7.4561 | 6.2555 | 6.4825 | 6.7112 | 6.3961 | 6.4769 | 6.3800 | 6.3218 | 6.6434 | 5.4739 | 5.5984 | 4.9874 | 6.0155 |
| DDX10_S831 | DDX10 | Q13206 | 831 | MKKRSNSEVEDVG | 4.8593 | 4.9945 | 6.1140 | 6.2462 | 5.5935 | 5.9731 | 5.8188 | 6.3066 | 6.3124 | 5.8437 | 6.6430 | 6.4101 | 7.3398 | 8.0021 | 6.8155 | 6.7274 |
| DDX11_S44 | DDX11 | Q96FC9 | 44 | KIGIFESPTGTGK | 7.3747 | 7.5463 | 6.9850 | 7.0658 | 6.5309 | 7.6569 | 7.3887 | 6.9583 | 7.6368 | 7.6170 | 7.6363 | 7.4935 | 2.7223 | 3.4207 | 2.5233 | 3.4436 |
| DDX17_S599 | DDX17 | Q92841 | 599 | DGGRRDSASYRDR | 4.3186 | 4.4152 | 4.6337 | 5.1444 | 7.8299 | 5.4348 | 4.9704 | 5.5009 | 7.3398 | 8.0395 | 7.1862 | 7.2983 | 8.0150 | 7.7179 | 5.8838 | 6.2716 |
| DDX17_S64 | DDX17 | Q92841 | 64 | EPQALPSPAIRAP | 6.3126 | 6.9233 | 6.1878 | 7.0594 | 6.5944 | 7.0936 | 6.9212 | 6.7568 | 6.6041 | 6.2195 | 6.8152 | 6.3322 | 4.9692 | 5.5424 | 4.6706 | 4.9976 |
| DDX17_T301 | DDX17 | Q92841 | 301 | VEICIATPGRLID | 5.8495 | 6.6837 | 6.5771 | 6.7587 | 6.0820 | 6.3562 | 6.3631 | 5.9998 | 6.3646 | 6.2863 | 6.4610 | 6.5447 | 5.4466 | 5.6732 | 6.1547 | 6.3988 |
| DDX17_T523 | DDX17 | Q92841 | 523 | TAYTFFTPGNLKQ | 5.9118 | 6.5956 | 7.1806 | 7.1977 | 6.8011 | 6.7919 | 6.1687 | 6.4836 | 6.6206 | 6.0462 | 7.3334 | 6.8090 | 5.4810 | 6.0633 | 3.5178 | 4.9974 |
| DDX20_S187 | DDX20 | Q9UHI6 | 714 | PEKYQESPGIQMK | 6.6571 | 6.6264 | 6.6994 | 6.5004 | 6.4901 | 7.0505 | 6.6884 | 6.9632 | 6.9249 | 6.7874 | 7.2624 | 7.0176 | 4.3204 | 4.9523 | 4.2906 | 4.7690 |
| DDX20_S268 | DDX20 | Q9UHI6 | 500 | ASSRNNSVSGLSV | 5.6439 | 4.6125 | 5.8890 | 4.8285 | 5.8866 | 5.6009 | 5.6499 | 5.7237 | 7.8255 | 7.5935 | 7.0481 | 7.0327 | 7.5122 | 8.0383 | 5.6318 | 5.4830 |
| DDX20_S48 | DDX20 | Q9UHI6 | 48 | TAQDLSSPRTRTG | 7.3524 | 7.9197 | 7.0085 | 7.7334 | 6.3983 | 6.6832 | 7.3253 | 6.6474 | 7.3959 | 7.2564 | 7.3982 | 7.4026 | 2.8632 | 3.8415 | 2.9994 | 3.7745 |
| DDX20_S500 | DDX20 | Q9UHI6 | 703 | EQPPNGSDTPNPE | 6.1681 | 5.9368 | 6.9865 | 5.6476 | 6.3668 | 6.3458 | 5.8101 | 6.6678 | 6.1345 | 5.7092 | 6.1558 | 5.4623 | 6.5894 | 6.7314 | 6.5844 | 6.7036 |
| DDX20_S672 | DDX20 | Q9UHI6 | 187 | CHIAVGSPGRIKQ | 5.4512 | 6.9710 | 7.0531 | 6.3218 | 6.3577 | 7.1748 | 5.5517 | 6.0608 | 6.5191 | 6.3826 | 7.9006 | 5.8902 | 5.3903 | 5.3433 | 5.7177 | 5.9141 |
| DDX20_S703 | DDX20 | Q9UHI6 | 672 | QSKGNKSYLEGSS | 6.5868 | 6.7507 | 7.2282 | 6.9854 | 5.9976 | 6.3197 | 6.2772 | 6.3886 | 7.0708 | 7.0968 | 7.3217 | 6.8750 | 4.4200 | 5.2822 | 4.4965 | 4.9029 |
| DDX20_S714 | DDX20 | Q9UHI6 | 705 | PPNGSDTPNPEKY | 7.5779 | 7.0414 | 6.8973 | 7.5282 | 6.4652 | 4.8383 | 5.3375 | 6.4180 | 8.0696 | 6.2838 | 6.1994 | 5.3971 | 4.9526 | 5.4940 | 5.9038 | 5.5958 |
| DDX20_T705 | DDX20 | Q9UHI6 | 268 | TFVRLNSSDPSLI | 4.4901 | 4.8017 | 5.3320 | 4.5592 | 5.7331 | 5.6543 | 5.7757 | 5.9330 | 5.0738 | 4.9890 | 4.8138 | 5.3324 | 9.4737 | 9.1607 | 9.9600 | 8.9175 |
| DDX21_S121 | DDX21 | Q9NR30 | 89 | PSQNDISPKTKSL | 6.3600 | 6.6007 | 6.6130 | 7.3184 | 6.0878 | 5.8045 | 6.5783 | 6.0039 | 6.6686 | 6.4808 | 6.3410 | 6.9127 | 4.9713 | 5.8176 | 5.4152 | 6.0262 |
| DDX21_S13 | DDX21 | Q9NR30 | 71 | SEVDMNSPKSKKA | 7.9153 | 7.8459 | 7.6838 | 8.0627 | 6.5720 | 6.1585 | 7.3385 | 6.3724 | 7.6411 | 7.2889 | 7.2278 | 7.3255 | 2.7209 | 3.4485 | 2.7960 | 3.6022 |
| DDX21_S168 | DDX21 | Q9NR30 | 121 | TKNEEPSEEEIDA | 6.0085 | 6.3534 | 6.3851 | 7.3235 | 6.0687 | 6.2547 | 6.5024 | 5.6988 | 6.0198 | 6.0351 | 6.1140 | 6.3752 | 6.0555 | 6.5266 | 5.9977 | 6.2809 |
| DDX21_S168;S171 | DDX21 | Q9NR30 | 171 | EAASEESNSEIEQ | 5.9046 | 5.6897 | 5.7759 | 6.5571 | 5.7490 | 5.7529 | 5.9636 | 5.8801 | 6.1560 | 5.6807 | 6.1863 | 6.2915 | 6.7301 | 7.3556 | 7.0829 | 7.2441 |
| DDX21_S168;S171;S173 | DDX21 | Q9NR30 | 315 | IDILVGTPGRIKD | 6.3726 | 6.8042 | 6.2115 | 7.2073 | 6.2698 | 5.5193 | 6.2620 | 6.1465 | 6.1375 | 6.1453 | 6.1131 | 6.3724 | 5.7358 | 6.4443 | 6.0356 | 6.2228 |
| DDX21_S171 | DDX21 | Q9NR30 | 13 | SDAGLESDTAMKK | 6.6760 | 7.3082 | 6.8245 | 7.8633 | 7.1764 | 6.4230 | 7.1984 | 6.5463 | 5.3066 | 4.9175 | 5.1663 | 5.0169 | 5.4262 | 6.1422 | 5.7568 | 6.2515 |
| DDX21_S171;S173 | DDX21 | Q9NR30 | 567 | KRIGVPSATEIIK | 5.9099 | 6.4585 | 6.4044 | 6.8084 | 6.0061 | 6.1846 | 6.5083 | 6.6126 | 6.5242 | 6.2980 | 6.3715 | 6.3482 | 5.2359 | 6.1376 | 5.8576 | 6.3341 |
| DDX21_S567 | DDX21 | Q9NR30 | 168 | NPSEAASEESNSE | 6.0244 | 6.1617 | 6.0157 | 6.2458 | 6.5609 | 5.4948 | 6.7300 | 5.9674 | 7.1021 | 5.7838 | 6.0863 | 6.3076 | 5.4609 | 6.8972 | 6.3850 | 6.7764 |
| DDX21_S71 | DDX21 | Q9NR30 | 168;171;173 | NA | 6.0473 | 6.1539 | 6.5223 | 6.8517 | 6.0049 | 6.0151 | 6.5972 | 5.7328 | 5.3617 | 6.0171 | 5.8619 | 6.0399 | 6.3526 | 6.9207 | 6.7075 | 6.8133 |
| DDX21_S89 | DDX21 | Q9NR30 | 171;173 | NA | 5.5918 | 5.6385 | 6.0350 | 6.3872 | 6.0380 | 5.9194 | 5.9012 | 5.5563 | 6.1047 | 5.4597 | 5.9993 | 6.2910 | 6.7029 | 7.3801 | 7.1974 | 7.7977 |
| DDX21_T315 | DDX21 | Q9NR30 | 168;171 | NA | 5.2983 | 6.0972 | 6.3368 | 7.0087 | 6.0159 | 6.2962 | 6.7503 | 6.1502 | 6.1564 | 6.1271 | 6.4792 | 6.4389 | 6.4466 | 6.4445 | 5.5842 | 6.3697 |
| DDX23_S106;S109 | DDX23 | Q9BUQ8 | 109 | RKRSSLSPGRGKD | 7.0184 | 7.4726 | 6.5960 | 7.3121 | 5.3979 | 5.4474 | 5.7667 | 6.0070 | 6.7595 | 6.8409 | 6.4939 | 7.1036 | 4.8628 | 5.6870 | 5.5857 | 5.6483 |
| DDX23_S107;S109 | DDX23 | Q9BUQ8 | 14 | KKDRDASPSKEER | 5.1902 | 5.8974 | 6.1703 | 6.7243 | 5.5610 | 6.1428 | 5.9797 | 6.0655 | 6.3790 | 6.4119 | 6.2158 | 6.4114 | 6.2865 | 7.0650 | 6.8871 | 6.6123 |
| DDX23_S109 | DDX23 | Q9BUQ8 | 106;109 | NA | 5.8201 | 6.1380 | 5.5446 | 6.0666 | 5.9310 | 5.7956 | 6.1338 | 5.9892 | 5.6852 | 5.9556 | 5.7646 | 5.7132 | 7.1524 | 7.5547 | 7.5307 | 7.2246 |
| DDX23_S14 | DDX23 | Q9BUQ8 | 107;109 | NA | 5.8549 | 6.1975 | 5.3695 | 5.9866 | 5.9744 | 5.8265 | 6.0032 | 6.0950 | 5.8451 | 5.7828 | 5.7129 | 5.6585 | 7.1852 | 7.6307 | 7.4506 | 7.4267 |
| DDX24_S82 | DDX24 | Q9GZR7 | 444 | PEIVVATPGRLWE | 6.4308 | 6.4697 | 7.6672 | 9.4479 | 5.6765 | 6.6252 | 7.4533 | 6.2558 | 6.6563 | 6.4852 | 5.9681 | 7.6322 | 3.9969 | 5.3149 | 3.8647 | 4.0552 |
| DDX24_T444 | DDX24 | Q9GZR7 | 82 | RKAQAVSEEEEEE | 4.9276 | 6.1928 | 6.1494 | 6.4505 | 6.4042 | 6.1870 | 6.3868 | 6.6734 | 6.0654 | 6.8674 | 5.0869 | 5.3474 | 6.4505 | 6.4563 | 7.2003 | 7.1540 |
| DDX27_T346 | DDX27 | Q96GQ7 | 346 | PDILIATPGRLID | 6.7144 | 6.9826 | 6.2365 | 7.1289 | 5.6197 | 5.5305 | 6.0862 | 6.6419 | 6.1722 | 5.8529 | 6.3317 | 6.2107 | 5.8434 | 6.4086 | 6.5542 | 5.6855 |
| DDX39A_S426 | DDX39A | O00148 | 171 | PHVVVGTPGRILA | 7.1737 | 7.0867 | 7.4984 | 6.7046 | 6.2076 | 5.9417 | 6.2451 | 5.9404 | 5.7800 | 6.1679 | 6.3373 | 6.1468 | 5.7071 | 5.5977 | 5.5430 | 5.9219 |
| DDX39A_T171 | DDX39A | O00148 | 426 | STYIEQSRxxxxx | 5.4798 | 6.1859 | 6.8225 | 6.7484 | 6.3341 | 6.4097 | 6.9042 | 7.8039 | 6.4415 | 6.1436 | 6.8286 | 6.8165 | 4.7253 | 6.1844 | 5.0216 | 5.1501 |
| DDX39B_S41 | DDX39B | Q13838-2 | 187 | PHIVVGTPGRILA | 6.2359 | 6.2089 | 7.0382 | 6.6564 | 5.7164 | 6.5386 | 6.8582 | 6.3205 | 6.7127 | 6.6908 | 6.6602 | 6.2234 | 5.3901 | 5.5178 | 5.3239 | 5.9081 |
| DDX39B_S436 | DDX39B | Q13838-2 | 41 | VKGSYVSIHSSGF | 6.3576 | 6.8386 | 6.1901 | 7.1636 | 6.6333 | 6.6510 | 6.2827 | 6.3733 | 6.1847 | 6.3999 | 6.5451 | 6.3004 | 5.2605 | 5.9445 | 5.5742 | 5.3004 |
| DDX39B_T187 | DDX39B | Q13838-2 | 436 | PDEIDISSYIEQT | 6.4132 | 6.0936 | 5.9224 | 6.7948 | 6.0228 | 6.5725 | 6.0267 | 6.7318 | 7.2521 | 6.9846 | 5.8359 | 6.1486 | 5.4927 | 5.8310 | 7.5503 | 6.1270 |
| DDX3X_S131 | DDX3X | A0A2R8Y4A4 | 593 | RSKSRFSGGFGAR | 7.6816 | 8.1175 | 7.0112 | 7.6562 | 6.1080 | 6.9733 | 7.5012 | 6.6705 | 7.5105 | 7.4015 | 7.3963 | 7.2097 | 3.0552 | 3.8238 | 2.5719 | 3.3115 |
| DDX3X_S492 | DDX3X | O00571 | 594 | SKSSRFSGGFGAR | 7.8258 | 7.9542 | 6.9625 | 7.5754 | 5.9469 | 6.8839 | 7.4096 | 6.6139 | 7.7902 | 7.5014 | 7.3101 | 7.2537 | 3.0604 | 3.8485 | 2.6521 | 3.4114 |
| DDX3X_S593 | DDX3X | O00571 | 323 | CHLLVATPGRLVD | 6.6572 | 7.1340 | 7.3216 | 7.4771 | 6.1759 | 6.4793 | 6.1917 | 6.0546 | 6.8254 | 6.3017 | 6.5875 | 6.1951 | 4.8657 | 5.4589 | 5.0286 | 5.2456 |
| DDX3X_S594 | DDX3X | O00571 | 82 | SDSRGKSSFFSDR | 8.0587 | 7.4489 | 8.6214 | 6.2416 | 4.8670 | 6.4840 | 7.2154 | 5.8307 | 7.3059 | 7.4349 | 8.3667 | 7.1475 | 3.4430 | 4.7850 | 2.7167 | 4.0328 |
| DDX3X_S606 | DDX3X | O00571 | 492 | QFRSGKSPILVAT | 7.9904 | 8.2851 | 6.9772 | 7.7355 | 6.7316 | 7.3395 | 6.5888 | 7.2504 | 7.1107 | 8.0671 | 7.4162 | 6.7976 | 2.2966 | 2.9812 | 3.0242 | 3.4080 |
| DDX3X_S61 | DDX3X | O00571 | 131 | SRWCDKSDEDDWS | 7.8296 | 7.8505 | 8.0096 | 7.6304 | 7.6140 | 7.4578 | 7.2483 | 6.4471 | 6.9221 | 6.2196 | 6.2141 | 5.7522 | 3.7397 | 3.8624 | 3.5771 | 3.6254 |
| DDX3X_S82 | DDX3X | O00571 | 86 | GKSSFFSDRGSGS | 6.2397 | 6.7102 | 6.9479 | 6.8662 | 6.6675 | 6.5684 | 6.6045 | 6.9138 | 6.2222 | 6.7073 | 6.4899 | 6.2842 | 4.9898 | 5.4189 | 4.9369 | 5.4326 |
| DDX3X_S86 | DDX3X | O00571 | 90 | FFSDRGSGSRGRF | 7.6418 | 7.3243 | 8.2962 | 7.5948 | 7.0089 | 7.7935 | 6.4653 | 6.1517 | 6.0042 | 7.0770 | 7.1605 | 6.9944 | 4.3163 | 4.9293 | 2.4395 | 2.8022 |
| DDX3X_S90 | DDX3X | O00571 | 606 | RDYRQSSGASSSS | 6.6087 | 6.6875 | 7.9321 | 6.6263 | 4.2542 | 6.4729 | 6.5590 | 6.8325 | 7.1735 | 9.1570 | 8.8430 | 8.2471 | 3.6158 | 4.6552 | 2.9815 | 3.3536 |
| DDX3X_T323 | DDX3X | O00571 | 61 | KDSSGWSSSKDKD | 7.4061 | 7.7074 | 6.8287 | 7.0526 | 4.7321 | 7.9551 | 8.2562 | 6.7483 | 7.4358 | 8.2747 | 9.0471 | 8.6432 | 1.8686 | 3.1446 | 1.9594 | 2.9402 |
| DDX41_S21;S23 | DDX41 | Q9UJV9 | 23 | PAGGSRSEAEDED | 5.6285 | 5.7786 | 6.1794 | 6.1132 | 5.8352 | 5.7750 | 6.1530 | 5.4720 | 6.1271 | 6.1060 | 6.4615 | 6.5880 | 7.4375 | 6.9392 | 6.6962 | 6.7097 |
| DDX41_S23 | DDX41 | Q9UJV9 | 320 | VHMMVATPGRLMD | 6.3013 | 7.2626 | 5.2056 | 7.5803 | 5.5477 | 6.5314 | 6.0467 | 5.4317 | 8.3827 | 6.1139 | 6.4764 | 5.2850 | 6.5579 | 6.5457 | 5.0895 | 5.6414 |
| DDX41_T147 | DDX41 | Q9UJV9 | 147 | PIKTSWTPPRYVL | 6.9119 | 6.7682 | 6.9503 | 6.5614 | 5.8190 | 6.5709 | 7.0569 | 6.5368 | 6.7716 | 7.0055 | 6.9434 | 7.3733 | 4.5363 | 4.9463 | 4.1577 | 5.0905 |
| DDX41_T320 | DDX41 | Q9UJV9 | 21;23 | NA | 6.5863 | 6.5393 | 6.4912 | 6.5497 | 6.5118 | 6.4443 | 6.5088 | 6.2055 | 6.2240 | 5.9705 | 5.9916 | 6.1779 | 5.9818 | 6.0634 | 5.8459 | 5.9081 |
| DDX42_S109;S111 | DDX42 | Q86XP3 | 96 | YIPAENSPTRQQF | 5.9859 | 6.1653 | 6.3225 | 6.2987 | 6.0034 | 6.1124 | 6.1514 | 6.0763 | 6.1340 | 6.4924 | 6.3283 | 6.3320 | 6.2955 | 6.7505 | 6.1489 | 6.4024 |
| DDX42_S51 | DDX42 | Q86XP3 | 169 | YMAENPTAGVVQE | 6.1515 | 5.9131 | 6.2445 | 6.2787 | 6.0720 | 5.7914 | 6.0938 | 6.0226 | 6.1287 | 5.9923 | 5.9766 | 6.0233 | 6.1984 | 7.2108 | 6.9994 | 6.9028 |
| DDX42_S96 | DDX42 | Q86XP3 | 51 | SSGFGKSAPPQLP | 5.7344 | 6.5489 | 5.6547 | 6.7547 | 6.1852 | 5.6856 | 6.5773 | 5.9850 | 5.3715 | 6.2348 | 5.6164 | 6.2185 | 6.9142 | 6.5985 | 6.5545 | 7.3657 |
| DDX42_T169 | DDX42 | Q86XP3 | 109;111 | NA | 5.9939 | 6.3642 | 6.3635 | 6.5987 | 6.6668 | 6.2698 | 6.0635 | 5.9056 | 6.2991 | 6.4235 | 6.6306 | 6.1378 | 6.5470 | 6.4946 | 5.8448 | 5.3966 |
| DDX46_S295;S296 | DDX46 | Q7L014 | 804 | ALGLQDSDDEDAA | 6.0070 | 6.2122 | 6.0305 | 6.2216 | 6.1930 | 6.0968 | 6.1013 | 6.1972 | 6.2500 | 6.2142 | 6.2006 | 6.1472 | 6.4400 | 6.6922 | 6.4412 | 6.5550 |
| DDX46_S804 | DDX46 | Q7L014 | 928 | DGGQNESFKRYEE | 6.1481 | 6.8240 | 6.2475 | 6.7084 | 6.4575 | 6.7902 | 7.4469 | 6.3642 | 6.5302 | 6.5221 | 6.8019 | 6.8783 | 4.6138 | 5.2515 | 4.9239 | 5.4916 |
| DDX46_S928 | DDX46 | Q7L014 | 295;296 | NA | 5.6796 | 6.0515 | 6.3269 | 6.2072 | 6.0579 | 6.0696 | 5.8049 | 5.8478 | 6.3439 | 6.1206 | 6.3196 | 6.2360 | 6.5839 | 6.8626 | 6.7929 | 6.6951 |
| DDX46_Y294;S295 | DDX46 | Q7L014 | 294;295 | NA | 5.4273 | 6.5314 | 5.5714 | 6.6191 | 5.7138 | 6.3123 | 5.9296 | 6.4804 | 6.6761 | 6.1991 | 6.0251 | 5.7992 | 6.3855 | 7.1499 | 6.4234 | 6.7566 |
| DDX5_S30 | DDX5 | P17844 | 480 | LVEDRGSGRSRGR | 6.8339 | 6.9308 | 6.5457 | 6.9381 | 6.7077 | 6.8475 | 6.6811 | 6.5308 | 6.2381 | 6.5447 | 6.8614 | 6.4570 | 5.0094 | 5.3203 | 4.4835 | 5.0701 |
| DDX5_S480 | DDX5 | P17844 | 498 | DRRDRYSAGKRGG | 7.2321 | 7.6145 | 6.6801 | 7.8120 | 6.6241 | 7.6309 | 7.7681 | 7.1151 | 6.8840 | 6.9452 | 7.1008 | 7.4964 | 3.0336 | 3.7695 | 2.7191 | 3.5745 |
| DDX5_S498 | DDX5 | P17844 | 30 | SRAGPLSGKKFGN | 6.8137 | 7.3399 | 6.7674 | 7.3509 | 6.0722 | 6.9809 | 7.2762 | 7.1430 | 7.2520 | 7.8573 | 8.3138 | 9.1456 | 2.2677 | 3.6980 | 2.4403 | 3.2813 |
| DDX5_S520 | DDX5 | P17844 | 520 | YDRGYSSLLKRDF | 6.1403 | 6.0527 | 6.7041 | 7.8788 | 6.2151 | 5.9279 | 7.0801 | 6.0528 | 6.4091 | 7.2056 | 8.9774 | 8.7240 | 3.5670 | 5.4361 | 3.4076 | 4.2213 |
| DDX50_S117 | DDX50 | Q9BQ39 | 82 | EGFNRLSDEFSKS | 5.9522 | 6.7611 | 6.3710 | 6.9105 | 6.1094 | 6.8329 | 7.2220 | 6.5738 | 6.4381 | 6.3237 | 6.0018 | 6.5142 | 4.7317 | 5.9779 | 5.4121 | 5.8678 |
| DDX50_S41 | DDX50 | Q9BQ39 | 117 | VSSLDTSTHKSSD | 6.9644 | 7.6438 | 6.5759 | 7.9079 | 6.6117 | 7.1573 | 7.4411 | 7.6643 | 7.2899 | 6.7798 | 7.0855 | 7.2352 | 2.9654 | 4.0185 | 2.8727 | 3.7865 |
| DDX50_S45 | DDX50 | Q9BQ39 | 41 | SRHHYDSDEKSET | 6.7486 | 6.8140 | 6.7249 | 7.7727 | 6.0461 | 5.3977 | 6.6448 | 6.2825 | 5.9607 | 5.5615 | 4.9697 | 5.8562 | 5.7738 | 6.4834 | 6.4992 | 6.4643 |
| DDX50_S683 | DDX50 | Q9BQ39 | 45 | YDSDEKSETRENG | 6.2554 | 7.3012 | 6.8508 | 6.4504 | 6.0228 | 6.1610 | 6.4595 | 5.3611 | 5.4508 | 6.2645 | 6.0288 | 6.1207 | 6.2337 | 6.5306 | 6.4169 | 6.0919 |
| DDX50_S82 | DDX50 | Q9BQ39 | 683 | RQRSGWSSGRSGR | 6.3322 | 6.5440 | 6.1941 | 6.6381 | 6.9023 | 6.1191 | 6.6862 | 6.6834 | 6.6336 | 6.5110 | 6.8349 | 6.7005 | 4.4585 | 5.9097 | 5.1751 | 5.6772 |
| DDX51_S103 | DDX51 | Q8N8A6 | 83 | NDAEPGSPEAPQG | 6.7569 | 6.7641 | 6.5122 | 6.9043 | 6.0267 | 6.4946 | 6.2291 | 5.7361 | 5.8420 | 5.9345 | 6.7131 | 6.9715 | 5.4022 | 5.9029 | 6.0026 | 5.8071 |
| DDX51_S83 | DDX51 | Q8N8A6 | 103 | EDAGAESNEEAPG | 6.5067 | 6.6745 | 6.8693 | 8.0669 | 6.5087 | 6.9098 | 6.2232 | 5.5502 | 5.3824 | 5.7218 | 5.7160 | 6.2155 | 6.0208 | 5.9648 | 5.4403 | 6.2290 |

| | | | | | | | | | | | | | | | | | | | | | |
|---|---|---|---|---|---|---|---|---|---|---|---|---|---|---|---|---|---|---|---|---|---|
| DDX52_S22 | DDX52 | Q9Y2R4 | | 22 | FDTRRFSADAARF | 4.2858 | 4.2155 | 4.8326 | 5.4370 | 7.9585 | 5.1061 | 4.7328 | 5.5230 | 7.9784 | 8.8406 | 7.8343 | 7.7350 | 7.5315 | 7.6650 | 4.8132 | 5.5106 |
| DDX52_S258 | DDX52 | Q9Y2R4 | | 565 | KPLERESISTTPK | 5.8589 | 6.2545 | 6.0160 | 7.4339 | 7.2295 | 6.4074 | 6.6268 | 5.8238 | 7.4741 | 6.4538 | 7.7818 | 6.9358 | 4.9932 | 5.6368 | 4.3861 | 4.6876 |
| DDX52_S565 | DDX52 | Q9Y2R4 | | 258 | RELIKISEGTGFR | 7.6981 | 6.4861 | 6.5677 | 6.1093 | 6.7888 | 6.3070 | 5.7994 | 6.6336 | 7.1341 | 5.9137 | 5.9676 | 6.2605 | 6.8471 | 6.2967 | 4.5366 | 4.6536 |
| DDX54_S39;S41 | DDX54 | Q8TDD1-2 | | 75 | PTSECTSDVEPDT | 6.0600 | 6.5650 | 6.5004 | 7.1891 | 6.5684 | 6.9766 | 7.0162 | 6.7736 | 5.5503 | 5.5931 | 5.5523 | 5.5330 | 5.7003 | 6.2304 | 5.9446 | 6.2466 |
| DDX54_S527 | DDX54 | Q8TDD1-2 | | 815 | RPHAPGTPAGRVR | 7.0177 | 6.1697 | 6.9605 | 7.4800 | 5.9978 | 6.9242 | 6.6336 | 6.4914 | 8.2036 | 7.9601 | 7.8252 | 7.6637 | 3.2993 | 4.0974 | 3.1757 | 4.1002 |
| DDX54_S644 | DDX54 | Q8TDD1-2 | | 782 | KIDDRDSDEEGAS | 5.7142 | 5.7996 | 6.6157 | 6.7273 | 6.4531 | 5.3677 | 5.5878 | 6.3004 | 7.4449 | 6.0393 | 5.7947 | 6.4120 | 6.1756 | 6.2971 | 6.2987 | 6.9720 |
| DDX54_S75 | DDX54 | Q8TDD1-2 | | 527 | RSRPAPSPESIKR | 7.0735 | 6.6766 | 6.7544 | 5.6520 | 6.8128 | 8.5494 | 9.6362 | 6.7129 | 6.0735 | 6.6572 | 7.3601 | 7.5909 | 2.8714 | 4.7649 | 3.2047 | 3.6094 |
| DDX54_S782 | DDX54 | Q8TDD1-2 | | 644 | EEEAGESVEDIFS | 6.0697 | 7.0854 | 7.4996 | 5.9218 | 6.7945 | 6.7945 | 7.8004 | 6.1782 | 5.4731 | 5.3843 | 6.9720 | 3.8706 | 6.2028 | 6.2522 | 5.2463 | 6.4543 |
| DDX54_T815 | DDX54 | Q8TDD1-2 | 39;41 | | NA | 6.1729 | 6.7654 | 5.1723 | 6.3850 | 6.9000 | 6.3222 | 7.4214 | 6.0828 | 5.1924 | 6.8769 | 5.9340 | 5.6267 | 6.5292 | 6.0075 | 6.6031 | 6.0083 |
| DDX55_S544 | DDX55 | Q8NHQ9 | | 544 | RKREEGSDIEDED | 5.2674 | 6.9499 | 5.8592 | 6.6710 | 6.3859 | 5.9738 | 6.7980 | 6.3766 | 5.3727 | 5.9274 | 6.3859 | 5.5649 | 6.4665 | 6.2837 | 6.7237 | 6.9932 |
| DDX55_S594 | DDX55 | Q8NHQ9 | | 594 | TVDLGISDLEDDC | 5.8582 | 5.9929 | 6.6125 | 6.2980 | 6.0627 | 6.5285 | 6.5224 | 5.3525 | 5.8421 | 6.3179 | 5.9336 | 6.6879 | 6.2325 | 6.6248 | 6.9522 | 6.1813 |
| DDX56_S517 | DDX56 | Q9NY93 | | 517 | RKKLSSSCRKAKR | 4.4757 | 5.7937 | 6.5353 | 7.3390 | 6.1714 | 6.3629 | 6.5240 | 6.7129 | 7.0547 | 8.6447 | 8.9384 | 9.1508 | 3.7690 | 4.9480 | 3.0922 | 4.4873 |
| DDX6_T222 | DDX6 | P26196 | | 222 | VHVVIATPGRILD | 6.3481 | 7.6745 | 7.5744 | 7.3108 | 5.9944 | 6.3731 | 4.9851 | 7.2474 | 6.9054 | 5.1602 | 6.4315 | 6.8837 | 5.0818 | 5.3388 | 4.3344 | 6.3565 |
| DEAF1_S176 | DEAF1 | O75398 | | 176 | LTPGPQSPPTPLA | 6.6942 | 7.2137 | 6.1435 | 7.2949 | 5.8208 | 5.8931 | 6.1082 | 5.9284 | 6.6516 | 7.1071 | 6.5507 | 6.5983 | 5.3916 | 5.5504 | 5.4133 | 5.6400 |
| DEAF1_T171 | DEAF1 | O75398 | | 312 | ENELPTTPVKKDS | 6.8671 | 7.0849 | 6.7400 | 6.7476 | 5.9828 | 6.2301 | 5.9170 | 5.7982 | 7.1704 | 6.9253 | 6.5622 | 7.0986 | 4.9434 | 5.1552 | 5.5909 | 5.1862 |
| DEAF1_T312 | DEAF1 | O75398 | | 171 | GPAAPLTPGPQSP | 6.1211 | 6.8440 | 6.2338 | 6.7592 | 6.8323 | 6.6943 | 5.9811 | 6.3337 | 6.8990 | 6.9606 | 6.0199 | 6.2755 | 5.6826 | 5.3118 | 5.1486 | 5.9024 |
| DECR2_S289 | DECR2 | Q9NUI1 | | 289 | PDFASFSAKLxxx | 6.9740 | 6.9672 | 6.2732 | 5.6204 | 4.9608 | 4.7204 | 5.6960 | 4.4902 | 7.1767 | 5.8128 | 5.8541 | 5.6823 | 8.7158 | 6.0774 | 7.7573 | 7.2214 |
| DEDD_S182 | DEDD | O75618-2 | | 182 | QRKRRKSVTPDPK | 5.2776 | 5.2609 | 5.4392 | 5.9268 | 6.9394 | 5.7657 | 5.5726 | 5.8690 | 8.2336 | 8.4095 | 7.5388 | 7.8098 | 6.2065 | 6.0649 | 4.3779 | 5.3077 |
| DEDD_T184 | DEDD | O75618-2 | | 184 | KRRKSVTPDPKEK | 6.9801 | 7.7107 | 7.3939 | 7.2834 | 6.2464 | 6.9217 | 5.6252 | 5.9920 | 7.3647 | 7.0103 | 6.8686 | 7.7003 | 3.7376 | 4.4807 | 4.1451 | 4.5391 |
| DEDD2_S111 | DEDD2 | Q8WXF8 | | 111 | KRRRPVSPERYSY | 6.6306 | 7.2434 | 6.3239 | 7.2316 | 6.2463 | 5.9197 | 5.9009 | 5.8329 | 6.9680 | 6.4045 | 6.2325 | 7.3022 | 5.3099 | 5.6706 | 5.3359 | 5.4471 |
| DEK_S210 | DEK | P35659 | | 32 | PGPREESEEEEDE | 5.5740 | 5.9168 | 6.0631 | 6.2569 | 5.7369 | 5.8292 | 6.0382 | 5.5917 | 6.3446 | 6.3159 | 6.2401 | 6.4170 | 6.8230 | 7.0303 | 6.8510 | 6.9712 |
| DEK_S243;S244 | DEK | P35659 | | 51 | EEEKEKSLIVEGK | 6.9717 | 7.3305 | 6.4111 | 6.8976 | 7.1149 | 7.0002 | 7.4326 | 7.1282 | 6.9982 | 6.7517 | 6.6943 | 6.3945 | 3.8168 | 4.7478 | 3.8827 | 4.4272 |
| DEK_S301;S303;S307 | DEK | P35659 | | 210 | SKKERNSSGMARK | 7.4268 | 6.6224 | 6.7136 | 7.7849 | 6.1469 | 6.7661 | 8.1714 | 7.1482 | 6.9054 | 7.2233 | 7.4808 | 7.9205 | 3.1409 | 3.9956 | 2.8252 | 3.7280 |
| DEK_S32 | DEK | P35659 | 301;303;306;3 | | NA | 5.6781 | 5.7713 | 6.0728 | 6.7255 | 5.9936 | 4.7919 | 6.6291 | 5.0598 | 6.0009 | 7.1879 | 6.8526 | 5.9365 | 6.8343 | 7.1837 | 6.3478 | 6.9343 |
| DEK_S51 | DEK | P35659 | 243;244 | | NA | 6.0886 | 6.4080 | 7.4024 | 6.3615 | 5.9002 | 5.3127 | 7.1941 | 5.3707 | 7.6225 | 7.1259 | 7.6696 | 8.0453 | 4.9614 | 4.8578 | 4.8502 | 4.8292 |
| DENND1A_S473 | DENND1A | Q8TEH3 | | 473 | LPKTAPSPLVEAK | 6.4374 | 6.3100 | 7.1956 | 6.0915 | 6.3681 | 6.8554 | 6.6791 | 6.6979 | 6.6881 | 6.2963 | 6.5750 | 6.3623 | 5.3904 | 5.6768 | 5.0837 | 5.2923 |
| DENND1A_S523 | DENND1A | Q8TEH3 | | 523 | RRTSVPSPEQPQP | 6.1091 | 6.6602 | 7.5449 | 6.1914 | 6.0648 | 6.3949 | 6.0803 | 6.1256 | 6.3585 | 6.9098 | 6.5503 | 7.1895 | 5.0007 | 5.9623 | 5.2045 | 5.6533 |
| DENND1A_S656 | DENND1A | Q8TEH3 | | 656 | SKPPAASPEKPSA | 6.1605 | 6.0741 | 6.2690 | 7.0144 | 6.1904 | 6.9758 | 6.4366 | 7.3938 | 6.8572 | 6.6845 | 6.7765 | 7.3665 | 4.6454 | 5.1933 | 4.9788 | 4.9832 |
| DENND2B_S539 | DENND2B | P78524 | | 539 | QDRKYNSPPTQLS | 6.3924 | 6.9697 | 6.8840 | 7.0564 | 6.1415 | 6.4826 | 6.7673 | 6.5467 | 6.4357 | 8.1805 | 6.8454 | 6.8008 | 4.0625 | 5.3193 | 4.2895 | 4.8258 |
| DENND2B_S673 | DENND2B | P78524 | | 673 | RLVHIQSMLKRAP | 8.2102 | 5.8824 | 5.8874 | 6.8778 | 8.1322 | 5.3670 | 4.4696 | 6.1664 | 5.5686 | 6.1187 | 4.5249 | 5.8044 | 7.0538 | 7.2574 | 7.5490 | 4.1302 |
| DENND2B_T364;S368 | DENND2B | P78524 | 364;368 | | NA | 7.5052 | 6.4317 | 6.0021 | 8.4079 | 6.1210 | 5.1210 | 7.4690 | 7.4126 | 7.1712 | 5.7214 | 6.4927 | 7.2273 | 4.0355 | 4.4499 | 5.2731 | 5.1585 |
| DENND2C_S504 | DENND2C | Q68D51 | | 504 | SLQKKPSGISYIP | 6.7902 | 7.2227 | 6.5569 | 6.9061 | 6.6413 | 6.2168 | 5.6908 | 6.4218 | 6.4303 | 6.2680 | 6.3807 | 5.5803 | 6.0480 | 5.4463 | 5.5947 | 5.8051 |
| DENND3_S472 | DENND3 | A2RUS2 | | 472 | HRRMVVSMPNLQD | 5.9025 | 5.8054 | 6.1106 | 6.7315 | 5.7522 | 6.3199 | 5.5487 | 7.1905 | 5.8562 | 6.4355 | 6.5361 | 6.9338 | 5.5984 | 6.5685 | 6.2598 | 6.4505 |
| DENND4A_S1058 | DENND4A | Q7Z401-2 | | 731 | KSSSPNSPLPMFR | 6.8096 | 6.9407 | 6.5564 | 6.4121 | 6.6220 | 6.6481 | 6.2220 | 6.7693 | 6.9848 | 6.8913 | 6.7333 | 6.8928 | 5.0479 | 5.3137 | 4.6270 | 4.5291 |
| DENND4A_S1160 | DENND4A | Q7Z401-2 | | 1630 | IFPMARSISTSGP | 6.8158 | 7.0066 | 7.8654 | 7.9155 | 6.5470 | 6.4059 | 6.5158 | 6.6468 | 6.7677 | 6.4281 | 7.1172 | 7.2034 | 3.8561 | 4.6326 | 4.0244 | 4.2520 |
| DENND4A_S1194 | DENND4A | Q7Z401-2 | | 1632 | PMARSISTSGPLD | 6.5127 | 6.4291 | 6.4188 | 6.0258 | 6.3558 | 6.1036 | 6.0438 | 6.2253 | 6.4067 | 6.2713 | 6.0440 | 6.1253 | 6.2404 | 6.6025 | 5.9653 | 6.2296 |
| DENND4A_S1268 | DENND4A | Q7Z401-2 | | 1299 | SSLPPNSPKPVRL | 6.9486 | 7.5292 | 6.9490 | 7.1268 | 6.0883 | 7.4560 | 7.2345 | 6.8027 | 5.4804 | 5.3349 | 5.4993 | 5.5535 | 5.3139 | 4.8687 | 5.8226 | 5.9916 |
| DENND4A_S1283 | DENND4A | Q7Z401-2 | | 1283 | DKLNKKSPPLVKA | 6.4560 | 6.4069 | 6.9932 | 6.4503 | 6.6013 | 6.3272 | 6.0048 | 6.7476 | 6.7927 | 6.4308 | 6.5009 | 6.9844 | 5.3535 | 5.6399 | 4.9400 | 5.3707 |
| DENND4A_S1299 | DENND4A | Q7Z401-2 | | 1160 | LADKVDSPVIFDL | 6.4845 | 6.1911 | 6.8885 | 7.0004 | 6.3104 | 6.3536 | 6.1358 | 5.9284 | 7.0901 | 6.5240 | 6.9112 | 6.3953 | 5.0870 | 5.7791 | 5.1097 | 5.8108 |
| DENND4A_S1324 | DENND4A | Q7Z401-2 | | 1058 | FRKRHKSDNETNL | 6.2013 | 6.2430 | 6.6394 | 6.8627 | 5.9709 | 7.4807 | 6.9072 | 6.1528 | 6.3120 | 7.1140 | 7.2775 | 6.6190 | 4.9660 | 5.3047 | 5.0137 | 4.9352 |
| DENND4A_S1554 | DENND4A | Q7Z401-2 | | 1268 | LTSRTPSIDLQRA | 5.2724 | 5.8273 | 6.0054 | 5.8154 | 5.7721 | 5.8430 | 5.6021 | 6.2165 | 5.5404 | 6.1878 | 5.7883 | 6.1401 | 7.2986 | 7.8994 | 7.3716 | 7.4198 |
| DENND4A_S1630 | DENND4A | Q7Z401-2 | | 1194 | RIQRMNSSFSVKP | 7.6207 | 6.9751 | 7.7343 | 5.4088 | 5.8704 | 5.8857 | 5.3628 | 6.3208 | 6.8395 | 6.8499 | 6.9524 | 7.0230 | 5.6770 | 6.5479 | 4.1550 | 4.7767 |
| DENND4A_S1632 | DENND4A | Q7Z401-2 | | 1324 | PRDRLWSSPAFSP | 5.1857 | 6.8116 | 6.7348 | 5.6745 | 5.9567 | 6.0972 | 5.1161 | 6.1516 | 6.4904 | 5.7678 | 6.8446 | 6.2625 | 6.5011 | 6.2980 | 6.8671 | 7.2402 |
| DENND4A_S731 | DENND4A | Q7Z401-2 | | 1554 | FLKSSPSTENMHF | 5.1935 | 5.8402 | 6.4819 | 6.6457 | 6.4951 | 6.9334 | 6.6965 | 6.3274 | 7.1714 | 6.5602 | 5.9531 | 5.8097 | 6.2277 | 6.7545 | 5.3552 | 5.5545 |
| DENND4B_S1211 | DENND4B | O75064 | | 1211 | SRPSVPSPKSAGA | 5.7115 | 5.4846 | 6.2689 | 6.0783 | 5.5767 | 5.8575 | 6.5567 | 6.1707 | 6.6862 | 6.4005 | 6.3077 | 7.1518 | 6.0341 | 6.8438 | 5.6517 | 7.2193 |
| DENND4C_S1017;S1022 | DENND4C | Q5VZ89-7 | | 741 | GNSITKSPPLMAK | 6.5874 | 6.5066 | 6.6665 | 5.8462 | 6.3840 | 6.4502 | 6.0703 | 6.6734 | 6.6438 | 6.4879 | 6.3441 | 6.3431 | 5.7916 | 6.0538 | 5.5997 | 5.5513 |
| DENND4C_S1038 | DENND4C | Q5VZ89-7 | | 1038 | SIVKVPSGIFDVN | 3.5001 | 3.4262 | 3.6213 | 3.3577 | 6.1381 | 6.2783 | 5.9093 | 6.1132 | 3.8941 | 3.6128 | 3.7991 | 3.7615 | 12.1230 | 11.5221 | 11.7965 | 11.1469 |
| DENND4C_S1049 | DENND4C | Q5VZ89-7 | | 1175 | NPEHRSSPVPEML | 5.9675 | 6.1015 | 6.4481 | 5.9642 | 6.2261 | 6.1074 | 6.2051 | 6.2867 | 6.6750 | 6.3655 | 6.5966 | 6.2989 | 6.0122 | 6.5494 | 6.2246 | 5.9712 |
| DENND4C_S1095 | DENND4C | Q5VZ89-7 | | 1274 | SEDKLFSPVIARN | 6.4977 | 7.1138 | 7.1784 | 6.6988 | 6.2926 | 7.1123 | 7.4670 | 7.0642 | 7.3460 | 6.9756 | 7.6837 | 7.2595 | 3.4160 | 4.3549 | 3.3395 | 4.2001 |
| DENND4C_S1175 | DENND4C | Q5VZ89-7 | | 1374 | HKERSTSLSALVR | 5.9739 | 5.5216 | 6.0565 | 5.2311 | 5.9096 | 5.9986 | 5.6510 | 6.1760 | 6.5929 | 6.6067 | 6.6261 | 6.6063 | 6.9109 | 7.6781 | 6.3574 | 6.1033 |
| DENND4C_S1233 | DENND4C | Q5VZ89-7 | | 1265 | ECTGGKTPDSEDK | 6.3162 | 6.2359 | 6.3584 | 6.1044 | 6.5826 | 6.5526 | 6.7019 | 6.4101 | 6.8142 | 7.2009 | 7.2273 | 6.7311 | 4.9101 | 5.3849 | 4.9972 | 5.4721 |
| DENND4C_S1274 | DENND4C | Q5VZ89-7 | | 1233 | LRNKRSSLYGIAK | 6.3920 | 5.6277 | 6.6648 | 6.1626 | 6.7654 | 5.8781 | 5.8323 | 6.1309 | 7.9590 | 6.6829 | 6.4758 | 6.8648 | 6.1830 | 6.5343 | 4.8783 | 4.9681 |
| DENND4C_S1346 | DENND4C | Q5VZ89-7 | | 1848 | NSEKKSSLLSEEQ | 4.4032 | 4.9498 | 5.5644 | 5.3336 | 5.8567 | 6.3504 | 6.1056 | 6.1337 | 5.0056 | 4.7972 | 5.5242 | 5.3811 | 9.2010 | 8.9766 | 8.9769 | 7.4399 |
| DENND4C_S1374 | DENND4C | Q5VZ89-7 | | 1672 | QTMKISSVPNSLS | 3.3788 | 3.3425 | 3.1579 | 3.2899 | 3.6969 | 3.8140 | 3.9334 | 4.0743 | 3.5846 | 3.2576 | 3.2524 | 3.3375 | 15.3327 | 14.0294 | 15.7034 | 12.8146 |
| DENND4C_S1672 | DENND4C | Q5VZ89-7 | | 1346 | PESEKSSPAVSRS | 6.8274 | 6.8806 | 7.1912 | 6.7841 | 6.2772 | 7.1577 | 6.3388 | 6.6520 | 6.8816 | 7.2918 | 6.7182 | 6.7358 | 4.3042 | 5.0379 | 4.1015 | 4.8199 |
| DENND4C_S1848 | DENND4C | Q5VZ89-7 | | 1709 | PFHGISTVSLPNS | 5.2501 | 5.0451 | 6.4724 | 6.2648 | 6.7200 | 7.3229 | 6.1331 | 8.4108 | 6.9842 | 5.7377 | 5.8676 | 5.7797 | 5.5888 | 6.4736 | 5.9434 | 6.0057 |
| DENND4C_S741 | DENND4C | Q5VZ89-7 | | 1049 | VNSRKSSTGSISN | 5.1421 | 5.3719 | 7.4291 | 5.8266 | 6.3091 | 6.4878 | 5.7122 | 6.2345 | 7.4043 | 8.4746 | 7.4922 | 8.2583 | 4.4115 | 5.1689 | 4.8582 | 5.4187 |
| DENND4C_T1265 | DENND4C | Q5VZ89-7 | | 1095 | FPERSCSFSSESR | 6.4135 | 6.3567 | 6.8225 | 5.8474 | 5.9816 | 6.3302 | 5.4807 | 5.9447 | 7.7637 | 7.1142 | 6.9560 | 6.7411 | 5.9794 | 6.0926 | 5.0497 | 5.1260 |
| DENND4C_T1709 | DENND4C | Q5VZ89-7 | 1017;1022 | | NA | 5.7614 | 5.5677 | 6.1958 | 5.8691 | 6.2755 | 6.5096 | 6.1568 | 6.6183 | 6.0336 | 6.6721 | 6.5587 | 6.5767 | 6.1245 | 6.6411 | 6.2846 | 6.1545 |
| DENND5B_S1111 | DENND5B | Q6ZUT9-2 | | 1111 | TTARRLSITSLTG | 6.1170 | 5.2146 | 6.6380 | 5.7683 | 6.5231 | 6.3689 | 5.6824 | 6.4475 | 7.3821 | 7.2767 | 7.1512 | 7.0227 | 5.5122 | 6.5713 | 5.0110 | 5.3127 |
| DENND5B_S1111;S1114 | DENND5B | Q6ZUT9-2 | 1111;1114 | | NA | 6.6917 | 5.4314 | 6.5058 | 5.7894 | 5.9656 | 6.1105 | 6.0752 | 6.1347 | 7.2972 | 6.9680 | 7.2365 | 7.5945 | 5.3467 | 7.2225 | 4.6129 | 5.0175 |
| DENND6A_S127 | DENND6A | Q8IWF6 | | 127 | SSGRRVSLHCLLD | 6.0608 | 5.9648 | 6.1485 | 6.1485 | 5.8792 | 6.0965 | 5.6690 | 6.2624 | 7.4956 | 6.5166 | 7.9441 | 7.3887 | 5.8851 | 5.9871 | 5.1342 | 5.4189 |
| DENR_S73 | DENR | O43583 | | 73 | KLTVENSPKQEAG | 6.7405 | 7.0793 | 7.0880 | 6.2971 | 6.6391 | 6.8083 | 6.9816 | 6.7251 | 6.8099 | 6.9866 | 6.4939 | 6.3178 | 4.6896 | 5.1453 | 4.3892 | 4.8088 |
| DEPDC1_S110 | DEPDC1 | Q5TB30 | | 110 | FRFPATSPLKTLP | 6.6935 | 6.9709 | 6.5873 | 6.8097 | 6.4674 | 6.7442 | 6.8134 | 6.5278 | 6.7389 | 6.8242 | 6.6411 | 6.5601 | 4.4856 | 5.2340 | 4.7606 | 5.1412 |
| DEPDC1_S341 | DEPDC1 | Q5TB30 | | 523 | SSTRRNSYINTPV | 6.1613 | 5.1046 | 6.8148 | 5.9316 | 6.0863 | 6.3815 | 6.0362 | 5.7311 | 8.0856 | 7.6575 | 7.8117 | 8.0437 | 4.9732 | 5.8902 | 4.1576 | 5.1331 |
| DEPDC1_S471 | DEPDC1 | Q5TB30 | | 341 | NLNSFKSTECLLL | 6.6679 | 6.2243 | 6.4763 | 6.9080 | 5.9061 | 6.4756 | 6.7676 | 6.5267 | 6.2416 | 6.3138 | 6.4862 | 6.4854 | 5.5067 | 6.0034 | 5.2909 | 5.7194 |
| DEPDC1_S523 | DEPDC1 | Q5TB30 | | 471 | FLLNLHSEENIQK | 7.1374 | 7.1728 | 7.3258 | 7.0990 | 6.3353 | 6.7860 | 6.0020 | 6.3168 | 8.1108 | 7.8422 | 7.6970 | 7.6307 | 3.4422 | 4.0611 | 3.1788 | 3.8620 |
| DEPDC1B_S160 | DEPDC1B | Q8WUY9 | | 160 | MWYKRHSIAIGEV | 8.4487 | 6.9618 | 7.2928 | 7.4776 | 6.3010 | 6.3551 | 6.6410 | 5.7552 | 7.4991 | 6.0793 | 6.6016 | 6.3523 | 4.5511 | 4.8468 | 4.1251 | 4.7115 |
| DEPTOR_S265 | DEPTOR | Q8TB45 | | 265 | TSFMSVSPSKEIK | 5.9369 | 5.8630 | 7.2378 | 6.4872 | 6.3346 | 6.5457 | 5.9785 | 7.5059 | 7.3426 | 7.4135 | 5.7027 | 6.6690 | 5.3374 | 5.2588 | 4.9551 | 5.4314 |
| DERL1_S201 | DERL1 | Q9BUN8 | | 201 | GGRNFLSTPQFLY | 2.7840 | 2.7238 | 3.0848 | 3.0374 | 2.9679 | 2.9939 | 3.2061 | 3.1229 | 2.7684 | 2.8428 | 3.1348 | 2.8446 | 15.8851 | 15.6552 | 17.0670 | 15.8812 |
| DFFA_S107 | DFFA | O00273 | | 315 | ISASKASPPGDLQ | 6.4676 | 6.5773 | 6.9293 | 6.4085 | 6.1164 | 6.6062 | 6.0538 | 6.8563 | 6.7097 | 6.8691 | 7.0071 | 7.4223 | 4.9304 | 5.3538 | 4.8227 | 4.8695 |
| DFFA_S315 | DFFA | O00273 | | 107 | GGTAWISQESFDV | 5.9295 | 6.4839 | 8.4901 | 6.2061 | 6.1187 | 6.3462 | 5.9633 | 6.5670 | 6.2677 | 8.0779 | 7.9815 | 7.0234 | 3.9739 | 5.7577 | 4.3589 | 4.4542 |
| DGCR2_S545 | DGCR2 | P98153 | | 545 | GRHSRSSLNTVVx | 5.6100 | 6.1351 | 5.6266 | 5.9754 | 5.9240 | 5.9267 | 6.6825 | 6.6103 | 6.0101 | 7.2618 | 6.5866 | 7.4869 | 5.8767 | 6.1546 | 5.9837 | 6.1490 |
| DGCR8_S377 | DGCR8 | Q8WYQ5 | | 377 | TPSGDVSPVKPLS | 6.4512 | 6.5355 | 6.4877 | 6.8784 | 6.3536 | 6.0399 | 6.6494 | 6.2504 | 6.9703 | 6.5991 | 6.5232 | 6.3346 | 5.3225 | 5.3464 | 5.8749 | 5.3827 |
| DGCR8_S95 | DGCR8 | Q8WYQ5 | | 95 | PNCSGHSPRTARH | 6.9315 | 7.2985 | 6.8851 | 7.0470 | 6.7232 | 6.5026 | 6.1566 | 7.0387 | 7.0573 | 6.6932 | 6.7342 | 6.7940 | 4.2184 | 4.7756 | 4.5471 | 4.5971 |
| DGKD_S44 | DGKD | Q16760 | | 44 | KLIRKVSTSGQIR | 6.2202 | 5.3186 | 5.9257 | 5.9559 | 6.7510 | 6.4182 | 5.7149 | 5.9343 | 8.0113 | 6.8914 | 7.1315 | 6.8587 | 6.3914 | 6.6511 | 5.0153 | 4.8104 |
| DGKD_S668 | DGKD | Q16760 | | 668 | PLSHSESFGVPKG | 6.5545 | 6.4912 | 7.0952 | 5.5577 | 5.7487 | 6.2453 | 6.9329 | 5.9099 | 6.7562 | 6.3157 | 6.0424 | 6.0818 | 7.5178 | 7.2014 | 5.0385 | 4.5109 |
| DGKQ_S17 | DGKQ | P52824 | | 17 | AWLGGGSPRPGSP | 5.8616 | 5.6492 | 6.5717 | 5.9550 | 6.4950 | 6.5820 | 5.9588 | 7.0874 | 6.5154 | 6.1009 | 6.6370 | 6.7777 | 5.6799 | 6.8353 | 5.1668 | 6.1264 |
| DGUOK_S42 | DGUOK | Q16854 | | 42 | RGPRRLSIEGNIA | 5.1716 | 4.9577 | 6.8350 | 5.7398 | 7.0578 | 5.3744 | 5.5125 | 7.5279 | 7.4577 | 7.2572 | 7.5770 | 6.8194 | 5.1393 | 7.0377 | 5.2407 | 5.2942 |
| DHCR7_S14 | DHCR7 | Q9UBM7 | | 14 | NIPKAKSLDGVTN | 3.6132 | 3.0390 | 2.8968 | 3.2510 | 3.2832 | 2.6457 | 3.3011 | 3.3421 | 3.0925 | 3.0704 | 2.0759 | 2.3333 | 15.2655 | 15.4034 | 17.5547 | 15.8321 |
| DHX16_S103;S106 | DHX16 | O60231 | | 160 | QTEKPESEDEWER | 5.5498 | 6.2782 | 6.0388 | 5.9824 | 6.1124 | 5.8500 | 5.9248 | 5.9187 | 6.2550 | 6.5640 | 6.1638 | 6.2930 | 6.3736 | 6.9031 | 6.9154 | 6.8772 |
| DHX16_S117 | DHX16 | O60231 | | 117 | VSRAGSSLQKKRK | 7.5336 | 7.3509 | 7.0238 | 7.2577 | 6.4622 | 7.4885 | 7.9966 | 7.3191 | 7.7477 | 7.6048 | 7.4651 | 7.6469 | 2.3594 | 3.4829 | 2.2770 | 2.9837 |
| DHX16_S160 | DHX16 | O60231 | | 712 | MESLTVTPCSKAS | 6.0455 | 6.7597 | 6.2482 | 6.6818 | 5.7978 | 6.7964 | 6.0288 | 5.9382 | 6.4341 | 6.2583 | 6.4435 | 6.3868 | 5.3911 | 6.2369 | 6.2027 | 6.3504 |
| DHX16_T712 | DHX16 | O60231 | 103;106 | | NA | 4.8237 | 5.7700 | 6.1036 | 5.5319 | 5.5693 | 6.1649 | 6.7068 | 5.7333 | 6.2489 | 6.4843 | 6.7343 | 6.4355 | 6.4195 | 7.1039 | 7.1726 | 6.9974 |
| DHX29_S200 | DHX29 | Q7Z478 | | 200 | QIQATISPPLQPK | 6.6215 | 6.8569 | 6.8111 | 6.5155 | 6.5116 | 6.9831 | 6.1865 | 6.5976 | 6.9846 | 6.6256 | 6.5241 | 6.8646 | 4.6812 | 5.6492 | 4.7601 | 4.8268 |
| DHX29_S523 | DHX29 | Q7Z478 | | 523 | SEDPEESWENLVS | 6.2311 | 5.8566 | 6.2351 | 6.0339 | 5.7420 | 6.4244 | 6.4085 | 6.2222 | 6.1315 | 6.0578 | 7.0510 | 6.6814 | 5.7330 | 6.8647 | 6.1983 | 6.1285 |
| DHX30_S408 | DHX30 | Q7L2E3-2 | | 408 | PELRLQSDDILPL | 7.8247 | 7.1953 | 5.6094 | 6.5926 | 5.7081 | 6.0290 | 6.3478 | 6.8436 | 7.2652 | 5.4284 | 6.3766 | 7.4380 | 4.7598 | 6.2450 | 5.9159 | 4.4204 |
| DHX34_S20 | DHX34 | Q14147 | | 20 | DHHRAPSEEEALE | 5.9961 | 6.3762 | 6.0641 | 6.7166 | 6.0895 | 6.2271 | 6.0737 | 6.4145 | 6.2155 | 6.4682 | 7.0657 | 7.0244 | 5.7715 | 5.9378 | 5.5340 | 6.0252 |
| DHX37_S117 | DHX37 | Q8IY37 | | 117 | RLFYTTSKLGTGN | 6.4314 | 6.9566 | 8.3495 | 7.5042 | 3.6383 | 5.9805 | 6.5549 | 6.4031 | 7.1159 | 8.6426 | 8.6411 | 7.9268 | 3.2410 | 5.9522 | 2.8675 | 3.7945 |
| DHX38_S1194 | DHX38 | Q92620 | | 1194 | QEQEKRSPLGSVR | 6.2257 | 7.1628 | 6.4693 | 6.5489 | 6.2454 | 6.8863 | 6.9849 | 6.6738 | 6.8495 | 7.0152 | 7.2707 | 7.0762 | 4.2263 | 4.9667 | 4.4656 | 4.9327 |
| DHX38_S234 | DHX38 | Q92620 | | 265 | GKYSDDTPLPTPS | 7.6612 | 6.8610 | 7.3718 | 7.2760 | 5.6972 | 7.7914 | 7.0891 | 5.4010 | 7.7638 | 7.2847 | 6.9224 | 7.7233 | 3.5341 | 4.0907 | 3.4362 | 4.0961 |
| DHX38_T265 | DHX38 | Q92620 | | 234 | SQWESPSPTPSYR | 7.4477 | 7.4636 | 6.4635 | 6.9355 | 5.9359 | 6.3157 | 6.8441 | 6.6432 | 6.8717 | 6.9708 | 6.2706 | 6.7073 | 5.2423 | 4.9014 | 4.3747 | 4.6122 |
| DHX38_T265;T269 | DHX38 | Q92620 | 265;269 | | NA | 7.5726 | 6.7602 | 6.5837 | 7.4621 | 5.6122 | 5.3659 | 6.8324 | 5.8252 | 6.6977 | 7.0095 | 5.9900 | 6.3697 | 4.7195 | 4.9870 | 5.5219 | 6.6903 |
| DHX8_S460 | DHX8 | Q14562 | | 460 | KQSMDMSPIKIVK | 6.3682 | 6.7076 | 6.2060 | 6.6788 | 6.2069 | 5.9662 | 6.2898 | 6.0149 | 6.3339 | 6.3117 | 6.2300 | 6.1308 | 5.8138 | 6.3082 | 6.3503 | 6.0828 |
| DHX9_S125 | DHX9 | Q08211 | | 87 | PAFGVASPPPLTD | 5.9305 | 6.2731 | 6.0652 | 6.5066 | 6.2659 | 6.0996 | 6.5606 | 6.2092 | 6.2531 | 6.2682 | 6.0982 | 6.2447 | 6.2682 | 6.3927 | 6.1539 | 6.4103 |
| DHX9_S87 | DHX9 | Q08211 | | 125 | LKAENNSEVGASG | 6.3299 | 6.1815 | 6.4872 | 6.5406 | 6.3114 | 6.1583 | 6.4234 | 6.3133 | 6.5841 | 6.0629 | 6.5066 | 6.6739 | 6.0699 | 6.1474 | 5.7609 | 5.4485 |
| DIAPH3_S1091;S1093 | DIAPH3 | Q9NSV4 | | 1093 | DVRQSLSPMSQRP | 6.6496 | 6.4290 | 7.0116 | 6.1439 | 6.2802 | 6.4759 | 6.4121 | 7.3923 | 7.1208 | 6.6253 | 6.9773 | 7.1351 | 4.7727 | 4.7522 | 4.5779 | 5.2440 |
| DIAPH3_S1093 | DIAPH3 | Q9NSV4 | 1091;1093 | | NA | 7.0881 | 7.7224 | 7.1369 | 8.2416 | 6.3440 | 7.5923 | 8.0933 | 5.8465 | 6.9778 | 7.3457 | 7.7602 | 7.0335 | 2.6633 | 3.6125 | 2.8909 | 3.6511 |
| DICER1_S1016 | DICER1 | Q9UPY3 | | 1255 | KSTSDGSPVMAVM | 6.0601 | 6.2008 | 6.4134 | 6.3892 | 6.2796 | 6.4274 | 6.3310 | 6.7678 | 6.3599 | 6.7230 | 6.5772 | 6.4752 | 5.6400 | 6.1083 | 5.5701 | 5.6770 |
| DICER1_S1160 | DICER1 | Q9UPY3 | | 1016 | KALPLSSAEKRKA | 6.5219 | 7.0759 | 6.6897 | 6.9908 | 6.4858 | 7.0141 | 6.9597 | 7.0742 | 6.9506 | 7.1083 | 7.2098 | 7.0095 | 3.6673 | 4.4642 | 4.1006 | 4.6776 |
| DICER1_S1255 | DICER1 | Q9UPY3 | | 1160 | RTLLSESPGKLHV | 6.4565 | 7.0136 | 6.8335 | 7.0085 | 6.0260 | 6.8227 | 6.6807 | 6.6585 | 7.1365 | 7.2355 | 7.3687 | 7.8492 | 3.7324 | 4.7149 | 3.7860 | 4.6770 |
| DICER1_S1470 | DICER1 | Q9UPY3 | | 1470 | VKKISLSPFSTTD | 6.2192 | 6.5894 | 7.0916 | 6.2972 | 6.1711 | 7.0221 | 6.5252 | 6.5190 | 7.1929 | 6.9269 | 7.1087 | 6.8921 | 4.5938 | 5.3368 | 4.6243 | 4.8898 |
| DICER1_T1001 | DICER1 | Q9UPY3 | | 1001 | SRLNLLTPRHLNQ | 6.9531 | 7.7638 | 6.6800 | 7.0252 | 4.8830 | 6.6248 | 6.4635 | 7.0450 | 6.7366 | 6.9248 | 6.6135 | 5.5296 | 4.6821 | 5.6060 | 4.8787 | 5.5905 |
| DIDO1_S1030 | DIDO1 | Q9BTC0 | | 1040 | STSESRSPPEGDT | 5.8439 | 6.1281 | 6.3097 | 6.4427 | 6.1113 | 5.9166 | 6.1677 | 6.0258 | 6.2270 | 6.2671 | 6.0043 | 6.0485 | 6.5417 | 6.7735 | 6.5519 | 6.6401 |
| DIDO1_S1040 | DIDO1 | Q9BTC0 | | 1456 | ADVRRNSVERPAE | 4.7021 | 4.8360 | 4.7436 | 5.7795 | 6.5263 | 5.7939 | 5.8958 | 5.8686 | 5.5917 | 5.7005 | 5.5751 | 5.7340 | 8.3030 | 8.2812 | 8.6003 | 8.0684 |
| DIDO1_S1237 | DIDO1 | Q9BTC0 | | 1617 | EKEPASSPWASGE | 6.0830 | 6.5471 | 6.1458 | 6.6929 | 5.9018 | 6.1994 | 6.4387 | 6.1343 | 6.6712 | 6.5225 | 6.1064 | 6.4947 | 5.8515 | 6.3628 | 5.8355 | 6.0125 |
| DIDO1_S1456 | DIDO1 | Q9BTC0 | | 1030 | YLSVPPSPNISTS | 6.1197 | 6.3921 | 6.0268 | 5.7070 | 6.3921 | 5.8206 | 5.7616 | 6.6195 | 6.4028 | 6.4610 | 5.9957 | 6.2937 | 6.1655 | 6.9782 | 6.4205 | 6.4431 |
| DIDO1_S1617 | DIDO1 | Q9BTC0 | | 1662 | RRVLLPTPPCGAL | 6.3840 | 5.6970 | 5.3464 | 6.7603 | 5.8332 | 6.4662 | 6.3782 | 5.9221 | 5.8000 | 5.8308 | 5.8490 | 5.9529 | 7.1740 | 6.7371 | 6.9082 | 6.9605 |
| DIDO1_S1714 | DIDO1 | Q9BTC0 | | 523 | EKTAAPSPSLLYK | 6.1541 | 7.0124 | 6.2145 | 7.1563 | 5.9718 | 6.2613 | 6.9611 | 6.0340 | 6.6223 | 7.1486 | 6.6754 | 6.8153 | 4.9216 | 5.7328 | 4.9233 | 5.3952 |
| DIDO1_S523 | DIDO1 | Q9BTC0 | | 1237 | VATVPQSEKKPSK | 8.8608 | 8.0453 | 8.1461 | 7.3803 | 6.3237 | 6.9536 | 8.0205 | 7.0237 | 6.2389 | 5.9803 | 6.8308 | 7.1582 | 2.9434 | 4.4852 | 1.4746 | 4.1345 |
| DIDO1_S805;S809 | DIDO1 | Q9BTC0 | | 941 | DGHPEPSPLEDLS | 5.2778 | 5.8840 | 6.5881 | 5.8300 | 6.2916 | 6.7460 | 6.5473 | 5.8698 | 5.8127 | 5.6150 | 6.1938 | 5.7974 | 6.2213 | 6.8306 | 7.9187 | 6.5759 |
| DIDO1_S941 | DIDO1 | Q9BTC0 | | 947 | SPLEDLSPCPASC | 5.7882 | 5.9626 | 5.5159 | 5.8043 | 7.0237 | 6.6347 | 5.9250 | 6.0404 | 7.1243 | 5.8298 | 5.1618 | 6.1960 | 6.3784 | 6.8734 | 6.8989 | 6.8426 |
| DIDO1_S947 | DIDO1 | Q9BTC0 | | 1714 | SAGRSSSPAGETE | 5.9432 | 7.4229 | 8.6256 | 6.8670 | 6.5553 | 7.2486 | 6.0511 | 4.8209 | 5.3354 | 5.6868 | 6.5518 | 6.5069 | 6.0589 | 6.3187 | 5.2896 | 4.7173 |
| DIDO1_T1255 | DIDO1 | Q9BTC0 | | 1255 | ADAAVSTTPPGSP | 5.7618 | 4.9707 | 6.1523 | 6.0619 | 6.5568 | 5.7419 | 5.8919 | 5.8532 | 5.8522 | 5.3434 | 5.4984 | 5.5928 | 7.6201 | 8.5075 | 6.9284 | 7.6668 |
| DIDO1_T1255;S1260 | DIDO1 | Q9BTC0 | 1255;1260 | | NA | 5.5609 | 6.8033 | 6.3260 | 6.2424 | 5.9904 | 6.6867 | 6.2588 | 5.9040 | 6.1369 | 6.3506 | 6.2356 | 6.5450 | 5.9447 | 6.5101 | 6.2746 | 6.2301 |
| DIDO1_T1662 | DIDO1 | Q9BTC0 | 805;809 | | NA | 6.3866 | 6.0983 | 5.8731 | 5.7821 | 5.6098 | 6.5321 | 6.4881 | 6.7273 | 6.3821 | 5.2135 | 6.1215 | 6.1275 | 7.2220 | 6.5722 | 6.5342 | 6.3297 |
| DIEXF_S153 | DIEXF | Q68CQ4 | | 153 | PGTSQTSPEEFTD | 6.0511 | 5.3191 | 5.8204 | 6.9428 | 5.5203 | 5.3559 | 5.7995 | 5.9247 | 6.3107 | 5.8517 | 6.3279 | 6.6501 | 6.4856 | 7.0354 | 7.2864 | 7.3183 |
| DIP2A_S94 | DIP2A | Q14689 | | 94 | RDERFRSDVHTEA | 8.6420 | 7.9269 | 8.1678 | 7.7324 | 5.7214 | 7.6521 | 7.5163 | 6.6256 | 5.7475 | 5.3622 | 5.4800 | 6.2326 | 4.5599 | 4.7688 | 3.8362 | 4.0284 |
| DIP2B_S100 | DIP2B | Q9P265 | | 283 | IQQLLNTLKRPKR | 12.5282 | 8.1384 | 7.3900 | 7.4506 | 5.8657 | 4.8755 | 5.7469 | 4.2650 | 5.1810 | 4.1319 | 4.5849 | 4.5622 | 7.1697 | 6.0625 | 6.4166 | 5.6309 |
| DIP2B_S203 | DIP2B | Q9P265 | | 100 | RDERYRSDIHTEA | 8.0629 | 8.2829 | 8.3788 | 7.1809 | 6.0199 | 7.8575 | 7.8792 | 7.0015 | 5.8734 | 5.5197 | 5.9108 | 5.7386 | 4.1143 | 4.6876 | 3.4113 | 4.0808 |
| DIP2B_S53 | DIP2B | Q9P265 | | 53 | KLLSPYSPQTQET | 6.6417 | 7.1536 | 6.6803 | 6.8039 | 6.1771 | 6.3878 | 7.0110 | 6.4675 | 7.2852 | 7.8119 | 7.0321 | 7.2183 | 4.2191 | 4.6302 | 3.7971 | 4.6832 |
| DIP2B_S9 | DIP2B | Q9P265 | | 203 | VKGTSGSLADVFA | 6.0632 | 6.2987 | 6.4810 | 6.0378 | 5.8236 | 6.2401 | 6.1652 | 6.0918 | 6.3088 | 6.1623 | 5.9515 | 5.8104 | 6.5892 | 7.0600 | 6.8187 | 6.0978 |
| DIP2B_T283 | DIP2B | Q9P265 | | 9 | ERGLEPSPAAVAA | 6.2370 | 6.6325 | 6.8498 | 6.6192 | 5.9625 | 5.8067 | 7.0207 | 6.6353 | 7.2225 | 6.7090 | 6.8505 | 7.1193 | 4.5726 | 5.6845 | 4.6463 | 5.4315 |
| DIP2C_S89 | DIP2C | Q9Y2E4 | | 264 | IQQLVNTLKRPKR | 8.7785 | 8.0319 | 6.6548 | 8.3883 | 6.9775 | 4.7957 | 5.7564 | 4.7947 | 5.8334 | 4.2759 | 4.9634 | 4.9982 | 6.6780 | 6.0727 | 6.0738 | 6.9268 |
| DIP2C_T264 | DIP2C | Q9Y2E4 | | 89 | RDERYRSDVHTEA | 7.5677 | 6.3816 | 6.7325 | 6.6333 | 5.9147 | 6.9127 | 6.4450 | 5.7994 | 5.2596 | 5.7258 | 5.8097 | 5.9023 | 6.6406 | 6.4421 | 5.2480 | 6.5851 |
| DIS3L2_S139 | DIS3L2 | Q8IYB7 | | 31 | PHDIGASPGDKKS | 6.9023 | 6.8377 | 7.1762 | 7.0425 | 6.0616 | 7.1740 | 7.2790 | 6.9110 | 7.3436 | 7.3857 | 7.3859 | 7.9332 | 3.3250 | 4.3175 | 3.1169 | 3.8078 |
| DIS3L2_S198 | DIS3L2 | Q8IYB7 | | 139 | TEAAYESDIPEEL | 6.4996 | 6.7682 | 7.0810 | 5.8807 | 7.0118 | 6.6952 | 6.7271 | 7.1761 | 6.1812 | 5.6295 | 5.3660 | 5.4503 | 6.1334 | 5.9922 | 5.7470 | 5.6605 |
| DIS3L2_S31 | DIS3L2 | Q8IYB7 | | 15 | NLRPLGTPRGVSA | 8.1637 | 8.6532 | 6.8328 | 7.3224 | 5.0591 | 6.9708 | 7.3982 | 8.0180 | 7.1223 | 7.5633 | 6.6483 | 7.9636 | 2.5237 | 4.1751 | 1.9370 | 3.6486 |
| DIS3L2_T15 | DIS3L2 | Q8IYB7 | | 198 | KLSVCVSEKGRED | 7.4292 | 5.8637 | 7.0075 | 5.7481 | 9.1508 | 7.6372 | 5.4535 | 5.1184 | 4.6332 | 7.5217 | 6.0196 | 6.9902 | 5.5055 | 5.2455 | 3.6626 | 7.0133 |
| DISP2_S107 | DISP2 | A7MBM2 | | 1373 | GGPGDGSPVVLPN | 6.5458 | 5.9863 | 6.4380 | 6.6746 | 5.8149 | 6.2062 | 6.3916 | 5.9583 | 6.3868 | 6.4822 | 6.5485 | 6.1264 | 5.7465 | 6.3366 | 6.3356 | 6.0219 |
| DISP2_S1175 | DISP2 | A7MBM2 | | 1175 | ARRRSPSFDTSTA | 6.0529 | 6.2287 | 6.1443 | 6.4998 | 5.8506 | 5.9035 | 5.8743 | 5.6002 | 6.3310 | 6.2354 | 6.1530 | 6.7266 | 6.6059 | 7.0168 | 6.3585 | 6.4183 |
| DISP2_S1235 | DISP2 | A7MBM2 | | 1235 | PALQTSSPYKQAG | 6.5425 | 6.8307 | 6.3775 | 7.1374 | 6.4181 | 6.3381 | 6.1552 | 6.6222 | 6.5351 | 6.4075 | 6.8231 | 6.7436 | 5.0465 | 5.6443 | 5.0143 | 5.3640 |
| DISP2_S1243 | DISP2 | A7MBM2 | | 1243 | YKQAGPSPKTRAR | 6.2749 | 6.7723 | 6.2198 | 6.5754 | 5.8154 | 5.7695 | 6.5030 | 5.5001 | 6.6112 | 6.3269 | 5.9545 | 6.4593 | 6.0598 | 6.7094 | 5.8613 | 6.5870 |
| DISP2_S1373 | DISP2 | A7MBM2 | | 107 | EYQGGSSLPGLGD | 5.8263 | 6.5517 | 6.1104 | 6.2066 | 5.5571 | 6.2989 | 6.3647 | 6.2434 | 6.0708 | 5.8292 | 5.1139 | 6.2124 | 6.3067 | 7.4971 | 6.8924 | 6.9185 |
| DKC1_S21 | DKC1 | O60832 | | 21 | KKKERKSLPEEDV | 6.4596 | 7.0423 | 7.0469 | 7.9852 | 6.0660 | 6.5879 | 7.0322 | 6.4999 | 6.8334 | 7.2803 | 7.0784 | 7.2944 | 3.5427 | 4.6609 | 3.9184 | 4.6715 |
| DKC1_S451 | DKC1 | O60832 | | 513 | KEVELVSExxxxx | 6.1573 | 6.4797 | 6.1054 | 6.7370 | 5.9417 | 5.7095 | 6.3506 | 6.2302 | 5.8816 | 5.8555 | 5.6501 | 5.9221 | 6.0932 | 6.9286 | 6.8570 | 7.1005 |
| DKC1_S451;S453 | DKC1 | O60832 | | 494 | EPGDGDSDTTKKK | 5.8987 | 6.0021 | 5.7694 | 6.3021 | 6.1173 | 5.8069 | 6.2507 | 6.0812 | 6.0927 | 5.9856 | 5.8107 | 5.6830 | 6.8721 | 7.3748 | 7.1110 | 6.8416 |
| DKC1_S451;S453;S455 | DKC1 | O60832 | | 485 | AKAGLESGAEPGD | 7.1451 | 6.9448 | 7.4864 | 7.7273 | 6.7715 | 6.7224 | 7.5597 | 7.7659 | 5.3812 | 4.9099 | 4.9127 | 5.2108 | 4.7412 | 5.5588 | 5.2903 | 5.8720 |
| DKC1_S451;S453;T458 | DKC1 | O60832 | | 451 | AKRKRESESESDE | 4.3684 | 5.6252 | 5.2232 | 5.9800 | 4.6331 | 5.3918 | 5.3448 | 5.2526 | 6.7486 | 7.7917 | 7.4192 | 7.8838 | 6.4388 | 6.7604 | 6.2918 | 8.8465 |
| DKC1_S451;S455 | DKC1 | O60832 | | 453 | RKRESESESDETP | 6.1023 | 5.6058 | 5.8775 | 6.4429 | 6.8359 | 5.9092 | 6.1602 | 6.1878 | 7.4565 | 5.9437 | 6.8925 | 6.4126 | 5.1893 | 7.5061 | 6.1423 | 5.3355 |
| DKC1_S451;S455;T458 | DKC1 | O60832 | 451;453 | | NA | 5.7640 | 6.1453 | 6.1881 | 7.0917 | 5.8572 | 5.7858 | 6.3380 | 6.3723 | 6.1066 | 5.9661 | 6.1033 | 6.4177 | 5.7727 | 6.6371 | 6.7897 | 6.6643 |
| DKC1_S451;T458 | DKC1 | O60832 | 451;458 | | NA | 5.7792 | 5.9322 | 5.8640 | 6.5125 | 6.1771 | 5.6940 | 6.1314 | 5.7386 | 6.3053 | 6.0483 | 6.2315 | 6.1940 | 6.2250 | 7.3330 | 6.6995 | 7.1345 |
| DKC1_S453 | DKC1 | O60832 | 451;455;458 | | NA | 5.9872 | 6.1444 | 6.0369 | 6.3612 | 5.6342 | 5.7668 | 6.3214 | 5.9498 | 5.9437 | 6.0777 | 5.9291 | 6.1975 | 6.7912 | 6.9040 | 6.9366 | 7.0183 |
| DKC1_S485 | DKC1 | O60832 | 451;453;455 | | NA | 5.5346 | 6.0080 | 5.9142 | 6.3699 | 6.0389 | 5.7474 | 6.1331 | 5.7189 | 6.0514 | 5.8912 | 5.7615 | 6.0905 | 6.6823 | 7.7180 | 7.2173 | 7.1228 |
| DKC1_S494 | DKC1 | O60832 | 451;455 | | NA | 5.8695 | 5.8985 | 6.2680 | 6.7229 | 5.7678 | 5.8532 | 6.0729 | 5.8488 | 6.1909 | 5.9693 | 6.2239 | 6.5024 | 6.1672 | 6.8838 | 6.9284 | 6.8324 |
| DKC1_S513 | DKC1 | O60832 | 451;453;458 | | NA | 5.5441 | 5.8498 | 5.1238 | 6.7880 | 6.0309 | 6.0702 | 5.4059 | 6.3852 | 5.8157 | 5.9241 | 5.4832 | 7.2496 | 6.4288 | 7.3215 | 7.2387 | 7.3406 |
| DLC1_S517 | DLC1 | Q96QB1 | | 557 | EEFDVFSPKQDLV | 7.4271 | 7.5970 | 7.1393 | 7.3972 | 6.6903 | 7.1446 | 8.4897 | 6.6683 | 6.7335 | 6.8382 | 7.0284 | 7.1049 | 3.0969 | 3.9050 | 3.3011 | 3.4385 |
| DLC1_S557 | DLC1 | Q96QB1 | | 986 | IPERRDSGVGASL | 6.0250 | 6.1484 | 6.4955 | 5.7574 | 6.5898 | 5.6000 | 6.1184 | 6.9711 | 7.4249 | 6.8409 | 7.0174 | 7.2395 | 5.9793 | 5.7813 | 4.9167 | 5.0944 |
| DLC1_S642 | DLC1 | Q96QB1 | | 517 | VMKLEISPHRKRS | 7.1453 | 7.1228 | 7.0320 | 7.0763 | 6.5122 | 7.7378 | 6.6598 | 6.4206 | 7.1323 | 7.5350 | 6.5428 | 6.5558 | 4.3635 | 4.2254 | 3.9790 | 3.9594 |

| | | | | | | | | | | | | | | | | | | | | | |
|---|---|---|---|---|---|---|---|---|---|---|---|---|---|---|---|---|---|---|---|---|---|
| DLC1_S766 | DLC1 | Q96QB1 | | 642 | SFGSLPSPKELSS | 4.9308 | 5.6478 | 7.9989 | 6.4185 | 5.9985 | 7.4195 | 6.4704 | 6.8289 | 6.1197 | 6.4046 | 6.2565 | 6.2453 | 6.3475 | 6.3180 | 5.0425 | 5.5525 |
| DLC1_S986 | DLC1 | Q96QB1 | | 766 | TRTRSLSACNKRV | 6.6734 | 5.6538 | 7.1136 | 5.8838 | 5.6852 | 6.4339 | 6.3802 | 6.7339 | 6.1513 | 5.8508 | 6.5624 | 5.8969 | 7.0940 | 6.6972 | 5.5035 | 5.6861 |
| DLG1_S158 | DLG1 | Q12959-2 | | 670 | TRDKGQSFNDKRK | 7.0929 | 7.0587 | 7.5030 | 5.7844 | 4.7373 | 6.8508 | 6.3099 | 6.5756 | 6.9747 | 7.8730 | 7.9175 | 8.7253 | 4.1456 | 5.2008 | 3.1124 | 4.1383 |
| DLG1_S301 | DLG1 | Q12959-2 | | 301 | ALKEAGSIVRLYV | 6.9321 | 7.4388 | 6.0578 | 6.2531 | 6.0896 | 5.7511 | 6.1280 | 5.8318 | 6.0842 | 5.4946 | 5.6356 | 5.5270 | 7.1693 | 6.3283 | 7.5280 | 5.7508 |
| DLG1_S670 | DLG1 | Q12959-2 | | 158 | VSHSHISPIKPTE | 5.9014 | 7.2259 | 6.5496 | 8.3479 | 6.1331 | 6.2445 | 6.3247 | 6.5475 | 6.3763 | 6.3580 | 6.9089 | 7.0900 | 4.6009 | 5.4313 | 5.1725 | 4.7873 |
| DLG4_S338 | DLG4 | P78352-2 | | 338 | TSPRRYSPVAKDL | 6.6768 | 6.8262 | 6.6861 | 6.6440 | 5.9980 | 6.4399 | 5.7573 | 6.2338 | 7.0305 | 7.3200 | 7.0644 | 6.9704 | 4.8127 | 5.4603 | 5.1172 | 4.9624 |
| DLG5_S1000 | DLG5 | Q8TDM6 | | 984 | TFKRPQTPPKIDY | 6.3504 | 6.4431 | 6.5873 | 6.8963 | 5.8067 | 5.8636 | 6.3141 | 6.4086 | 6.4032 | 6.1918 | 5.9003 | 6.4400 | 5.5724 | 6.1270 | 6.1852 | 6.5101 |
| DLG5_S1209 | DLG5 | Q8TDM6 | | 1666 | EFSRRLSMSEVKD | 6.1632 | 6.1353 | 6.1245 | 6.1257 | 6.1473 | 5.6293 | 5.6959 | 6.2711 | 7.3270 | 7.1891 | 6.7284 | 6.9726 | 5.9380 | 6.3405 | 5.6857 | 5.5265 |
| DLG5_S1232 | DLG5 | Q8TDM6 | | 1000 | GPGPAHSPQPSKR | 6.4794 | 6.5696 | 6.1950 | 6.6647 | 5.9208 | 5.9464 | 6.4671 | 6.5398 | 6.0160 | 6.3454 | 6.2040 | 6.3352 | 5.8236 | 6.0827 | 6.1333 | 6.2769 |
| DLG5_S1263 | DLG5 | Q8TDM6 | | 841 | LIQHNNSTQTDIF | 6.5352 | 7.0399 | 6.7648 | 6.3614 | 6.1163 | 5.9591 | 6.9055 | 6.4702 | 5.4355 | 6.0547 | 5.5231 | 6.0813 | 6.4928 | 6.4298 | 5.9615 | 5.8689 |
| DLG5_S1666 | DLG5 | Q8TDM6 | | 801 | VFPQSSSWSGQNI | 5.9517 | 6.2521 | 6.2408 | 6.5120 | 6.3293 | 5.9153 | 6.1184 | 6.2666 | 5.8683 | 5.9881 | 5.8926 | 5.5629 | 7.1705 | 6.8425 | 6.7426 | 6.3462 |
| DLG5_S1795 | DLG5 | Q8TDM6 | | 264 | RNLLQQSWEDMKR | 6.2097 | 6.5018 | 5.8329 | 6.0565 | 6.1053 | 5.8359 | 6.4640 | 6.0683 | 5.7530 | 6.1245 | 5.5858 | 5.6709 | 7.1061 | 7.0271 | 6.8576 | 6.8006 |
| DLG5_S264 | DLG5 | Q8TDM6 | | 1209 | RVGPCSSPPAARD | 6.8644 | 7.1261 | 6.5794 | 6.8251 | 6.3276 | 6.1811 | 6.7393 | 6.3075 | 6.2719 | 6.3831 | 6.4611 | 6.6159 | 5.4012 | 5.4556 | 5.0426 | 5.4182 |
| DLG5_S295 | DLG5 | Q8TDM6 | | 1263 | SSARLGSSSNLQF | 6.5767 | 6.4516 | 6.4696 | 6.4638 | 5.6280 | 5.9271 | 6.7009 | 5.8126 | 6.6341 | 6.2517 | 6.5415 | 7.0522 | 5.9844 | 5.8664 | 5.2354 | 6.4039 |
| DLG5_S801 | DLG5 | Q8TDM6 | | 1011 | KRAGPLTPPKPPR | 6.7864 | 6.9397 | 6.6337 | 7.1359 | 6.2234 | 6.2796 | 5.9925 | 7.1407 | 7.1342 | 6.2657 | 6.8820 | 6.9181 | 4.7482 | 4.9232 | 4.8469 | 5.1498 |
| DLG5_S841 | DLG5 | Q8TDM6 | | 1795 | VDYKRRSGHFDVT | 6.6366 | 6.7114 | 7.0333 | 6.9467 | 6.2815 | 6.5730 | 6.4145 | 5.7801 | 6.3411 | 6.0721 | 7.1879 | 6.9687 | 4.8766 | 5.5159 | 5.7884 | 4.8722 |
| DLG5_S972 | DLG5 | Q8TDM6 | | 1232 | QHQGRLSLDLSHR | 6.1914 | 8.8498 | 6.4473 | 5.8475 | 5.4223 | 5.6965 | 6.1864 | 6.2419 | 7.0722 | 7.3752 | 6.9375 | 6.9262 | 4.7728 | 5.5967 | 5.1131 | 5.3231 |
| DLG5_T1011 | DLG5 | Q8TDM6 | | 972 | KPKQRKSIFDPNT | 9.3068 | 5.2147 | 6.4255 | 5.0527 | 6.5347 | 5.6923 | 6.3746 | 5.7527 | 8.6723 | 6.9407 | 6.8794 | 6.9568 | 5.3848 | 6.8738 | 3.2095 | 4.7286 |
| DLG5_T946 | DLG5 | Q8TDM6 | | 946 | GSNSGGTWPKAML | 6.6421 | 7.0336 | 7.4780 | 7.0869 | 5.7396 | 6.8107 | 6.3539 | 6.1234 | 7.1028 | 6.3069 | 6.2383 | 7.5177 | 4.7476 | 5.2577 | 4.8009 | 4.7599 |
| DLG5_T984 | DLG5 | Q8TDM6 | | 295 | VLKHNGSSEILNK | 6.2328 | 6.2398 | 5.9381 | 5.9264 | 6.5639 | 6.2585 | 5.6565 | 5.9562 | 6.1171 | 5.6063 | 5.8992 | 5.9326 | 7.0489 | 6.8497 | 7.1551 | 6.6191 |
| DLGAP4_S665 | DLGAP4 | Q9Y2H0 | | 915 | SWQLVETPEKRKE | 6.2866 | 6.2280 | 7.0831 | 6.4613 | 6.4491 | 6.7784 | 6.3797 | 6.8865 | 6.8251 | 7.1682 | 6.7504 | 6.9597 | 4.8231 | 5.3196 | 4.8107 | 4.7904 |
| DLGAP4_S763 | DLGAP4 | Q9Y2H0 | | 665 | APKRKLSSIGIQV | 5.9854 | 5.5295 | 6.0020 | 6.5732 | 6.2290 | 6.0106 | 5.5066 | 6.8767 | 7.5180 | 7.2923 | 6.9011 | 7.0561 | 5.2970 | 5.6559 | 5.6004 | 5.9662 |
| DLGAP4_S973 | DLGAP4 | Q9Y2H0 | | 973 | ASVRQNSATESAD | 5.9080 | 5.9748 | 6.9013 | 6.5886 | 5.9821 | 6.3865 | 6.5226 | 5.9322 | 6.2050 | 6.5222 | 6.8115 | 6.8602 | 5.7969 | 5.9607 | 6.1000 | 5.5474 |
| DLGAP4_T915 | DLGAP4 | Q9Y2H0 | | 763 | QIKRNLSYGDNSD | 5.4663 | 5.5597 | 7.0627 | 6.2650 | 6.2178 | 5.6810 | 6.1094 | 6.4980 | 6.4647 | 6.4475 | 5.5544 | 6.8093 | 6.4432 | 7.1593 | 6.3123 | 5.9494 |
| DLGAP5_S202 | DLGAP5 | Q15398 | | 806 | ECHLLDSPGLNCS | 6.1903 | 6.3764 | 6.7490 | 6.4783 | 6.7750 | 6.7797 | 6.4376 | 7.1349 | 6.7453 | 6.5540 | 7.0410 | 6.5407 | 5.0603 | 5.5554 | 4.6750 | 4.9071 |
| DLGAP5_S662 | DLGAP5 | Q15398 | | 662 | IPQQTTSPENAGP | 6.2217 | 6.6864 | 6.9272 | 7.1198 | 6.1300 | 6.6241 | 6.8459 | 6.5353 | 6.9854 | 7.1838 | 7.5057 | 7.2694 | 4.0923 | 4.8710 | 4.1988 | 4.8032 |
| DLGAP5_S690 | DLGAP5 | Q15398 | | 690 | IPESRSSIEDAQC | 5.9449 | 6.6729 | 6.6153 | 7.0024 | 5.9830 | 6.7108 | 7.4612 | 6.3925 | 6.9058 | 6.7575 | 6.6099 | 6.7665 | 5.0098 | 5.1232 | 4.6310 | 5.4134 |
| DLGAP5_S725 | DLGAP5 | Q15398 | | 329 | YQVTPMTPRSANA | 7.2235 | 7.8904 | 7.3752 | 7.5809 | 6.8409 | 7.3111 | 7.9800 | 6.7363 | 7.3099 | 6.3463 | 7.2241 | 6.7635 | 3.1286 | 3.9591 | 2.7373 | 3.5929 |
| DLGAP5_S767 | DLGAP5 | Q15398 | | 344 | TPSYTWTPLKTEV | 6.9497 | 7.6855 | 7.0643 | 7.3355 | 6.4476 | 7.0729 | 7.1886 | 6.9061 | 7.1031 | 7.0563 | 6.9949 | 7.4660 | 3.4639 | 4.4181 | 2.9481 | 3.8995 |
| DLGAP5_S806 | DLGAP5 | Q15398 | | 767 | QDVLMSSPEKNTA | 5.9269 | 5.7339 | 6.1216 | 5.6837 | 6.4209 | 6.6879 | 6.0200 | 6.3713 | 6.5561 | 6.4853 | 6.5435 | 6.5188 | 6.3796 | 6.2919 | 6.0643 | 6.1944 |
| DLGAP5_S806;S812 | DLGAP5 | Q15398 | | 725 | LSSERMSLPLLAG | 6.3942 | 7.2388 | 7.2576 | 8.0765 | 6.0100 | 7.0683 | 7.1728 | 6.9799 | 6.9308 | 7.2930 | 7.2275 | 7.3672 | 3.6057 | 4.3591 | 2.9174 | 4.1011 |
| DLGAP5_T329 | DLGAP5 | Q15398 | | 202 | QPVMPTSLRMTRS | 6.7432 | 7.4408 | 7.8259 | 7.2519 | 4.9702 | 7.6019 | 7.9809 | 6.8946 | 6.2358 | 6.4489 | 7.6128 | 7.4456 | 3.5436 | 5.1712 | 2.9538 | 3.8790 |
| DLGAP5_T344 | DLGAP5 | Q15398 | 806;812 | | NA | 6.7104 | 6.7117 | 6.6313 | 6.9812 | 6.6075 | 6.7079 | 6.6121 | 7.5608 | 6.2911 | 6.9063 | 7.6784 | 7.1986 | 3.7352 | 5.3914 | 3.7649 | 4.5112 |
| DLX1_S209 | DLX1 | P56177 | | 209 | RALSAGSPPVPPG | 6.8871 | 6.0165 | 6.0306 | 6.8236 | 6.7285 | 6.5875 | 9.6715 | 5.9249 | 6.7778 | 9.2979 | 7.1514 | 6.6897 | 4.6666 | 3.1475 | 4.2436 | 3.3554 |
| DLX2_S225 | DLX2 | Q07687 | | 225 | EQHPGASASPPCA | 5.6988 | 6.8664 | 6.2955 | 7.0287 | 6.3339 | 5.6902 | 6.4249 | 6.4610 | 6.3010 | 6.0430 | 5.9929 | 6.1183 | 6.0070 | 6.3434 | 6.6429 | 5.7521 |
| DMAP1_S41 | DMAP1 | Q9NPF5 | | 445 | VVGAPLTPNSRKR | 6.3234 | 6.2802 | 6.5004 | 6.7618 | 6.1105 | 5.9317 | 6.4696 | 6.4744 | 6.2005 | 6.3293 | 6.0718 | 6.7463 | 5.8107 | 6.2375 | 5.8906 | 5.8614 |
| DMAP1_S454 | DMAP1 | Q9NPF5 | | 454 | SRKRRESASSSSS | 5.8852 | 6.2544 | 5.8074 | 6.4598 | 7.3042 | 6.4866 | 6.1939 | 5.2965 | 6.0039 | 6.3299 | 6.9102 | 7.0430 | 6.6401 | 6.8616 | 4.8599 | 5.6634 |
| DMAP1_T409 | DMAP1 | Q9NPF5 | | 409 | VLGGPATPASGPG | 5.3671 | 5.4864 | 5.6677 | 6.1170 | 6.2499 | 6.1756 | 6.6137 | 6.7206 | 5.8220 | 5.9526 | 6.2578 | 6.3231 | 6.8997 | 6.3997 | 6.5224 | 7.4245 |
| DMAP1_T445 | DMAP1 | Q9NPF5 | | 41 | KKKSKKSSETLTF | 6.3765 | 7.4550 | 6.6151 | 7.6150 | 6.1909 | 6.3819 | 6.7681 | 6.5272 | 6.6651 | 7.1896 | 7.1686 | 6.9315 | 3.8791 | 5.0499 | 4.6413 | 4.5453 |
| DMD_S291 | DMD | P11532 | | 291 | GYERTSSPKPRFK | 5.7528 | 6.2820 | 5.9211 | 6.4472 | 5.6053 | 5.6274 | 5.9496 | 6.0321 | 6.1899 | 5.9120 | 6.1006 | 6.2089 | 6.6924 | 7.0542 | 6.8788 | 7.3458 |
| DMD_Y1509 | DMD | P11532 | | 1509 | NHCVNLYKSLSEV | 6.2343 | 7.1077 | 7.3576 | 6.3654 | 5.2908 | 6.6412 | 6.3485 | 5.7623 | 6.8938 | 7.5433 | 7.4323 | 8.1619 | 4.7541 | 4.5420 | 4.5261 | 5.0386 |
| DMTN_S333 | DMTN | Q08495 | | 333 | RMDRGNSLPCVLE | 5.5151 | 5.7944 | 6.2603 | 5.6681 | 6.6481 | 5.3497 | 6.3084 | 6.6356 | 5.3818 | 6.8117 | 6.4115 | 6.6160 | 6.7548 | 7.9659 | 7.1140 | 4.7646 |
| DMWD_S495 | DMWD | Q09019 | | 495 | SLSRSNSLPHPAG | 5.9086 | 6.4405 | 8.2053 | 6.2849 | 6.4647 | 6.8592 | 6.6784 | 6.3671 | 7.0242 | 6.8254 | 7.1429 | 7.4572 | 4.5198 | 5.2816 | 4.2494 | 4.2910 |
| DMXL1_S1258 | DMXL1 | Q9Y485 | | 1908 | KDCSPSSPLKLDA | 6.1501 | 6.1815 | 6.6846 | 6.0970 | 6.2413 | 6.4743 | 6.3425 | 6.5355 | 6.3447 | 6.3043 | 6.3344 | 6.4039 | 5.9927 | 6.1963 | 5.8062 | 5.9108 |
| DMXL1_S1285 | DMXL1 | Q9Y485 | | 1901 | SSFLDTSKDCSPS | 7.0522 | 7.5576 | 7.6099 | 7.1221 | 6.4586 | 6.8580 | 6.5655 | 6.9078 | 7.2942 | 7.1567 | 7.9792 | 7.8100 | 3.2273 | 4.0455 | 2.9691 | 3.3862 |
| DMXL1_S1896 | DMXL1 | Q9Y485 | | 1896 | PFYRASSFLDTSK | 4.0072 | 3.6260 | 4.0693 | 3.7842 | 4.8853 | 4.9678 | 4.6451 | 5.1248 | 3.8417 | 3.8805 | 3.7209 | 3.9244 | 13.0215 | 12.2104 | 13.0828 | 11.2080 |
| DMXL1_S1901 | DMXL1 | Q9Y485 | | 1258 | TDSYSGSTPSITS | 4.6598 | 4.9419 | 5.1690 | 4.7181 | 6.8995 | 6.8227 | 6.8511 | 7.1068 | 5.0388 | 4.8370 | 5.2406 | 5.2734 | 8.2055 | 8.2215 | 8.1629 | 7.8515 |
| DMXL1_S1908 | DMXL1 | Q9Y485 | | 1285 | KKTLTRSMTSLAQ | 3.7755 | 3.9834 | 4.1121 | 4.1442 | 5.2140 | 5.1986 | 4.7218 | 4.5765 | 4.4400 | 3.9867 | 4.3611 | 3.8892 | 11.6910 | 11.5061 | 12.3790 | 12.0207 |
| DMXL1_S574 | DMXL1 | Q9Y485 | | 574 | GLTRSTSMLISSG | 2.8364 | 3.8433 | 2.6043 | 3.0036 | 6.3042 | 7.0210 | 6.8264 | 6.0209 | 3.1470 | 1.8568 | 3.1538 | 2.6896 | 12.7176 | 12.2261 | 13.7245 | 12.0247 |
| DMXL2_S1140;S1143 | DMXL2 | Q8TDJ6-3 | | 1400 | HLSRTISVSGSTA | 7.5287 | 6.8522 | 7.3561 | 6.3195 | 6.9099 | 6.1587 | 6.2965 | 6.3029 | 7.0749 | 6.2358 | 6.8018 | 6.5147 | 5.2428 | 5.8355 | 4.1672 | 4.4026 |
| DMXL2_S1151 | DMXL2 | Q8TDJ6-3 | | 2400 | VKPRRQSENISAP | 6.2710 | 5.7235 | 6.1144 | 5.8655 | 6.7066 | 5.1613 | 6.1114 | 6.1078 | 7.7219 | 7.3411 | 6.8123 | 7.2820 | 6.0851 | 6.5445 | 5.4594 | 4.6920 |
| DMXL2_S1400 | DMXL2 | Q8TDJ6-3 | | 1857 | IRRNLASPEGTLA | 6.4901 | 6.6469 | 7.0368 | 7.3468 | 5.5768 | 6.8619 | 6.7636 | 6.3488 | 6.4262 | 6.9135 | 6.8484 | 7.3599 | 4.5734 | 5.4445 | 4.2261 | 5.1361 |
| DMXL2_S1857 | DMXL2 | Q8TDJ6-3 | | 1151 | LFVYSKSDALLSK | 1.8810 | 1.6408 | 2.0593 | 2.0255 | 4.0333 | 3.6387 | 3.6728 | 3.6256 | 1.8122 | 1.8100 | 1.7830 | 1.9690 | 18.1523 | 17.2790 | 18.0914 | 16.5261 |
| DMXL2_S2400 | DMXL2 | Q8TDJ6-3 | 1140;1143 | | NA | 5.1351 | 4.1277 | 4.5100 | 4.6671 | 7.2061 | 6.1575 | 6.4888 | 6.4320 | 5.8237 | 4.2430 | 5.0815 | 3.9620 | 9.0041 | 8.8371 | 8.3192 | 10.0052 |
| DNAAF5_T334 | DNAAF5 | Q86Y56 | | 334 | LDFAPPTPPHYPP | 5.8435 | 6.3130 | 6.4682 | 6.6617 | 5.7448 | 5.7994 | 6.3624 | 6.1470 | 6.2464 | 6.0669 | 5.9133 | 6.0909 | 6.1935 | 6.6818 | 6.6392 | 6.8278 |
| DNAH8_S4275 | DNAH8 | Q96JB1 | | 4275 | ISILRSSLSDLKL | 6.3507 | 4.7580 | 6.3006 | 4.7211 | 7.5821 | 5.4988 | 4.8803 | 6.8468 | 6.7187 | 7.6577 | 7.8913 | 7.7943 | 6.1473 | 6.8872 | 5.0957 | 4.8693 |
| DNAH9_S351;S354 | DNAH9 | Q9NYC9 | 3036;3040 | | NA | 5.8809 | 6.6853 | 6.0465 | 6.8936 | 5.9537 | 5.8661 | 6.4294 | 6.2286 | 6.0736 | 6.1580 | 6.1897 | 6.0826 | 6.5173 | 6.5272 | 6.2679 | 6.1996 |
| DNAH9_Y3036;T3040 | DNAH9 | Q9NYC9 | 351;354 | | NA | 6.5559 | 6.6943 | 6.2969 | 6.9220 | 6.2012 | 6.0074 | 5.9334 | 5.8467 | 6.1827 | 6.1610 | 6.4484 | 6.0556 | 6.7440 | 5.8711 | 6.0066 | 6.0728 |
| DNAJA1_S112 | DNAJA1 | P31689 | | 335 | PENGFLSPDKLSL | 6.7252 | 6.9032 | 7.4951 | 7.5515 | 6.2717 | 6.3009 | 7.0861 | 6.3027 | 6.9103 | 7.5453 | 7.3781 | 7.3301 | 3.8505 | 4.2668 | 3.9590 | 4.1235 |
| DNAJA1_S335 | DNAJA1 | P31689 | | 83 | AGGGFGSPMDIFD | 6.0012 | 7.3570 | 7.1849 | 7.7920 | 6.1626 | 6.8537 | 7.1907 | 6.5486 | 7.0083 | 7.3354 | 6.9993 | 7.0087 | 3.6565 | 4.2995 | 4.1422 | 4.4595 |
| DNAJA1_S83 | DNAJA1 | P31689 | | 112 | NVVHQLSVTLEDL | 6.1144 | 6.0948 | 6.3473 | 6.2361 | 5.2554 | 6.5817 | 5.9990 | 6.5453 | 6.5502 | 5.9276 | 6.3578 | 5.8290 | 6.5747 | 6.1557 | 6.6299 | 6.8013 |
| DNAJA2_S123 | DNAJA2 | O60884 | | 382 | FDSTRGSGGGQRR | 6.1483 | 6.5337 | 7.0881 | 6.4223 | 6.4890 | 5.9995 | 6.2563 | 6.2858 | 5.9774 | 6.0343 | 5.7928 | 6.4473 | 5.7854 | 6.3417 | 5.8173 | 6.5809 |
| DNAJA2_S382 | DNAJA2 | O60884 | | 123 | MHPLKVSLEDLYN | 6.1990 | 6.2552 | 6.4031 | 6.0997 | 5.4968 | 5.1722 | 6.2281 | 6.0468 | 6.6384 | 6.1335 | 6.0612 | 6.3751 | 6.5900 | 6.3572 | 6.7612 | 7.1826 |
| DNAJB12_S24 | DNAJB12 | J3KPS0 | | 24 | PFARPASPSLVPR | 6.1863 | 5.9017 | 6.8319 | 6.7412 | 5.9528 | 6.7209 | 6.7277 | 5.7405 | 7.7746 | 6.6321 | 6.7740 | 6.4719 | 4.9531 | 4.9328 | 5.5379 | 6.1207 |
| DNAJB14_S60 | DNAJB14 | Q8TBM8 | | 60 | GSTAGNSPHCRKP | 7.7211 | 7.9880 | 5.6262 | 5.1039 | 6.8468 | 5.8363 | 7.3010 | 8.0107 | 7.1023 | 7.6871 | 6.5062 | 7.2386 | 3.4688 | 4.3091 | 5.0982 | 4.1558 |
| DNAJB14_S67 | DNAJB14 | Q8TBM8 | | 67 | PHCRKPSGSGDQS | 5.4610 | 5.7610 | 6.2707 | 6.6438 | 4.7480 | 5.3840 | 5.4543 | 5.3098 | 5.0860 | 5.4650 | 6.4120 | 6.1838 | 7.4337 | 7.9260 | 8.3117 | 8.1491 |
| DNAJB2_S313 | DNAJB2 | P25686 | | 313 | ATKRSPSPEEKAS | 6.1171 | 6.3090 | 6.4237 | 6.1934 | 5.8688 | 5.8818 | 5.9467 | 5.9426 | 6.4244 | 6.2636 | 6.5517 | 6.3704 | 6.3213 | 6.6622 | 6.2500 | 6.4733 |
| DNAJB4_S148 | DNAJB4 | Q9UDY4 | | 148 | YPRDRNSVGPSRL | 7.1575 | 6.7673 | 6.3451 | 7.4155 | 5.8767 | 6.9791 | 6.9749 | 6.6252 | 6.8775 | 7.0459 | 7.3871 | 7.0452 | 4.0337 | 4.9456 | 3.3598 | 5.1638 |
| DNAJB6_S15 | DNAJB6 | O75190 | | 277 | HAPHCLSEEEGEQ | 5.9195 | 7.1937 | 7.8527 | 7.3237 | 6.3777 | 7.3322 | 6.9433 | 7.0484 | 5.8487 | 6.1812 | 6.6038 | 6.4708 | 4.2335 | 4.6902 | 4.6214 | 5.3594 |
| DNAJB6_S277 | DNAJB6 | O75190 | | 15 | GVQRHASPEDIKK | 5.3302 | 5.6479 | 5.8754 | 5.8799 | 6.3195 | 6.2513 | 6.1495 | 6.5656 | 6.1664 | 6.7122 | 6.8673 | 7.1278 | 5.3332 | 6.8152 | 6.3252 | 6.6333 |
| DNAJB6_S85 | DNAJB6 | O75190 | | 85 | GGSHFDSPFEFGF | 6.3427 | 6.9848 | 6.4383 | 6.6302 | 6.5298 | 6.2530 | 5.8972 | 6.1959 | 7.0067 | 6.5862 | 6.9224 | 6.3130 | 5.8954 | 5.7956 | 4.8072 | 5.4018 |
| DNAJC1_S479;S480 | DNAJC1 | Q96KC8 | 479;480 | | NA | 6.3068 | 6.5223 | 6.1608 | 6.8418 | 6.0609 | 6.0366 | 6.1463 | 6.0661 | 5.8724 | 6.3644 | 6.0403 | 6.0453 | 6.1519 | 6.4558 | 6.4306 | 6.4977 |
| DNAJC17_T170 | DNAJC17 | Q9NVM6 | | 170 | NTEGQGTPKLKLK | 6.7616 | 7.6048 | 7.3357 | 6.5782 | 6.1507 | 7.8096 | 6.8790 | 6.4531 | 7.0050 | 7.7667 | 8.1138 | 7.1121 | 3.2004 | 4.2167 | 3.3401 | 3.6726 |
| DNAJC19_S70 | DNAJC19 | Q96DA6 | | 70 | ALILGVSPTANKG | 6.3847 | 6.4651 | 5.8624 | 6.9803 | 6.3083 | 5.6283 | 5.8400 | 6.2916 | 6.0820 | 6.9943 | 6.0377 | 6.8410 | 6.0917 | 6.2837 | 6.2437 | 5.6652 |
| DNAJC2_S47 | DNAJC2 | Q99543 | | 63 | KELSEESEDEELQ | 5.3215 | 5.9113 | 6.2207 | 6.0211 | 6.0972 | 5.9430 | 6.5668 | 6.3084 | 6.5294 | 6.5073 | 6.9513 | 7.1361 | 6.3264 | 6.5756 | 5.5740 | 6.0100 |
| DNAJC2_S60;S63 | DNAJC2 | Q99543 | | 47 | RRNRNASASFQEL | 5.9731 | 5.8552 | 5.8915 | 5.8447 | 6.6456 | 6.3248 | 6.2221 | 6.2160 | 6.6348 | 6.4872 | 6.3556 | 6.1665 | 6.1896 | 6.6106 | 6.2703 | 6.3125 |
| DNAJC2_S63 | DNAJC2 | Q99543 | 60;63 | | NA | 5.7502 | 6.1058 | 6.5440 | 6.1317 | 6.4159 | 6.4885 | 6.3004 | 6.5898 | 6.5580 | 6.6640 | 6.6863 | 6.4870 | 5.5290 | 6.2750 | 5.6566 | 5.8178 |
| DNAJC21_S283;Y311 | DNAJC21 | Q5F1R6-2 | | 370 | NPLDDNSEEEMED | 5.9638 | 6.5920 | 6.9933 | 6.2930 | 6.3760 | 6.1231 | 6.8992 | 6.2393 | 6.3787 | 6.8566 | 6.4517 | 6.2207 | 5.2642 | 6.0790 | 5.3110 | 5.9586 |
| DNAJC21_S370 | DNAJC21 | Q5F1R6-2 | 283;311 | | NA | 6.2371 | 5.7136 | 7.1668 | 5.9074 | 5.8359 | 6.2370 | 6.2664 | 5.6578 | 6.8600 | 6.5263 | 6.7230 | 6.6811 | 6.1599 | 6.3517 | 5.4885 | 6.1877 |
| DNAJC5_S10 | DNAJC5 | Q9H3Z4 | | 10 | QRQRSLSTSGESL | 6.3383 | 6.8466 | 6.0091 | 6.5730 | 6.5358 | 6.2575 | 5.8797 | 6.6894 | 6.2404 | 5.9326 | 6.0186 | 5.9111 | 5.8544 | 6.6565 | 6.3310 | 5.9262 |
| DNAJC5_S15 | DNAJC5 | Q9H3Z4 | | 15 | LSTSGESLYHVLG | 6.4643 | 7.4397 | 6.6080 | 6.4590 | 6.2302 | 5.7473 | 5.9337 | 5.8919 | 5.8492 | 5.7046 | 5.5162 | 5.5383 | 6.4903 | 6.3186 | 7.3014 | 6.4073 |
| DNAJC9_S109 | DNAJC9 | Q8WXX5 | | 109 | LLFKKISLEDIQA | 5.7885 | 6.3015 | 6.4789 | 6.8992 | 5.7172 | 6.8308 | 6.9085 | 6.0778 | 5.7966 | 6.8007 | 6.8186 | 6.2256 | 4.9856 | 6.3954 | 5.8673 | 6.1079 |
| DNAL1_S56 | DNAL1 | Q4LDG9 | | 56 | CEKLSLSTNCIEK | 6.7520 | 5.3218 | 7.2389 | 5.5495 | 6.4791 | 5.3058 | 5.3361 | 6.4278 | 5.5809 | 6.8353 | 5.9994 | 7.1986 | 6.3848 | 6.3821 | 6.8976 | 6.3104 |
| DNM1_S847 | DNM1 | A0A0D9SFE4 | | 849 | KGPASPTRPAAPR | 6.5314 | 6.0944 | 6.3974 | 5.8744 | 6.1693 | 5.7493 | 6.8253 | 6.4589 | 6.3811 | 6.8444 | 6.2348 | 6.3041 | 5.9639 | 6.0092 | 5.9352 | 6.2270 |
| DNM1_T849 | DNM1 | A0A0D9SFE4 | | 847 | SRKGPASPTRPAA | 5.2937 | 5.4371 | 6.6322 | 6.3380 | 5.8731 | 6.4528 | 5.8234 | 6.6595 | 6.4297 | 6.7215 | 7.0694 | 6.5422 | 5.4767 | 6.6802 | 5.9415 | 6.6289 |
| DNM1L_S629 | DNM1L | O00429-6 | | 629 | IPIMPASPQKGHA | 6.8214 | 5.9379 | 7.1389 | 6.2430 | 6.3675 | 6.5806 | 6.5551 | 6.9199 | 6.9683 | 6.7292 | 6.5677 | 6.6659 | 4.8315 | 5.4056 | 4.7785 | 5.4889 |
| DNMBP_S135 | DNMBP | Q6XZF7 | | 387 | AGGPPRSPGVEWE | 5.6297 | 5.7976 | 6.2294 | 5.8838 | 6.2534 | 6.2789 | 6.2517 | 6.8064 | 6.5763 | 5.9253 | 6.2746 | 6.3543 | 6.2816 | 6.3841 | 6.4567 | 6.6160 |
| DNMBP_S1379 | DNMBP | Q6XZF7 | | 659 | QRTNAVSPKLLSR | 6.5621 | 6.1131 | 6.7643 | 5.9159 | 6.4058 | 6.5800 | 6.4859 | 6.3388 | 6.6006 | 6.4854 | 6.5220 | 6.6123 | 5.8440 | 5.8370 | 5.3825 | 5.5502 |
| DNMBP_S1407 | DNMBP | Q6XZF7 | | 1407 | KQPQDASPPPKEC | 6.5374 | 6.8903 | 6.9456 | 6.7639 | 6.0544 | 6.5520 | 6.2827 | 6.6420 | 6.7264 | 6.3980 | 6.9003 | 7.0363 | 4.9302 | 5.2855 | 4.9357 | 5.1193 |
| DNMBP_S1430 | DNMBP | Q6XZF7 | | 689 | RSLDQTSPCPLVL | 6.1126 | 6.6116 | 6.8894 | 7.3753 | 6.7430 | 6.8921 | 7.3173 | 6.6149 | 7.0418 | 6.8300 | 6.7516 | 6.5032 | 4.4928 | 4.5498 | 4.2983 | 4.9764 |
| DNMBP_S387 | DNMBP | Q6XZF7 | | 996 | SKLNIHSIIKKSN | 9.0675 | 7.3169 | 7.9997 | 6.7292 | 6.4830 | 6.3320 | 5.7825 | 5.7934 | 8.9805 | 5.7512 | 6.5606 | 7.0200 | 4.8698 | 4.9881 | 3.2782 | 3.3653 |
| DNMBP_S495 | DNMBP | Q6XZF7 | | 578 | KILRHFSIMDFNS | 8.2076 | 6.3323 | 7.6436 | 7.7031 | 6.1739 | 6.2104 | 6.8254 | 5.3702 | 6.6138 | 6.3772 | 6.5492 | 6.7034 | 5.4557 | 5.9373 | 4.0996 | 3.7973 |
| DNMBP_S578 | DNMBP | Q6XZF7 | | 1379 | RFPRQNSGSTLTF | 5.1548 | 5.0309 | 5.3605 | 5.3640 | 5.4844 | 5.1908 | 5.2605 | 5.4382 | 5.6853 | 5.4424 | 5.7675 | 5.3819 | 8.6435 | 8.7464 | 9.4228 | 8.6260 |
| DNMBP_S659 | DNMBP | Q6XZF7 | | 135 | QSRQWHSQSALFQ | 7.5067 | 6.3609 | 6.1019 | 5.1797 | 4.5715 | 6.4276 | 5.8586 | 4.6774 | 5.1523 | 6.7729 | 8.2915 | 5.6545 | 6.2274 | 7.5773 | 6.0548 | 7.5852 |
| DNMBP_S689 | DNMBP | Q6XZF7 | | 1430 | PSNSESSPSRCPS | 6.2038 | 6.5155 | 6.1617 | 6.5952 | 5.6908 | 6.7304 | 6.3984 | 6.8250 | 7.1788 | 6.4637 | 6.7431 | 7.1518 | 5.3865 | 5.8079 | 5.1927 | 4.9546 |
| DNMBP_S996 | DNMBP | Q6XZF7 | | 495 | VVKPRQSSPQLHN | 6.8439 | 7.1265 | 6.4302 | 8.0066 | 5.6154 | 7.8682 | 7.4138 | 6.7785 | 7.3632 | 6.6643 | 7.7216 | 7.0779 | 3.5333 | 4.4084 | 3.5870 | 3.5613 |
| DNMT1_S127 | DNMT1 | P26358-2 | | 127 | GMADANSPPKPLS | 6.3482 | 6.8991 | 7.1075 | 6.8005 | 5.9627 | 6.3255 | 6.4753 | 6.3354 | 7.2478 | 7.3268 | 7.2248 | 7.5114 | 3.9570 | 4.9877 | 4.4673 | 5.0228 |
| DNMT1_S154 | DNMT1 | P26358-2 | | 730 | NIPEMPSPKKMHQ | 5.7524 | 6.3370 | 6.6164 | 6.5333 | 5.9930 | 6.3361 | 6.1683 | 5.7741 | 6.3308 | 6.6590 | 6.9015 | 6.7414 | 5.7314 | 6.2442 | 5.8380 | 6.0431 |
| DNMT1_S170 | DNMT1 | P26358-2 | | 410 | IFDANESGFESYE | 6.9667 | 6.4211 | 6.8475 | 7.4584 | 6.2530 | 6.4003 | 6.8471 | 6.3211 | 6.9665 | 6.2975 | 6.4297 | 6.8812 | 4.3539 | 5.6333 | 4.7883 | 5.1343 |
| DNMT1_S205 | DNMT1 | P26358 | | 154 | KPEPSPSPRITRK | 6.3337 | 6.8785 | 6.7372 | 6.7832 | 6.2906 | 6.7836 | 6.4694 | 6.1609 | 6.9329 | 6.9768 | 7.2341 | 7.2726 | 4.4873 | 5.0611 | 4.5504 | 5.0477 |
| DNMT1_S410 | DNMT1 | P26358-2 | | 748 | QNKNRISWVGEAV | 5.7207 | 6.1883 | 6.7905 | 6.7107 | 6.3985 | 6.0683 | 6.2400 | 5.7497 | 6.3784 | 6.3195 | 6.8167 | 6.7058 | 5.6502 | 6.2457 | 6.0616 | 5.9554 |
| DNMT1_S730 | DNMT1 | P26358-2 | | 205 | ESERAKSDESIKE | 5.7733 | 6.8913 | 6.6028 | 6.9643 | 5.0499 | 6.1280 | 5.8518 | 6.0908 | 6.0895 | 6.7193 | 6.4421 | 7.1059 | 5.2284 | 6.2693 | 6.4472 | 6.3460 |
| DNMT1_S748 | DNMT1 | P26358-2 | | 170 | RAEPSPSPRITRK | 6.4531 | 7.2005 | 6.6005 | 6.7874 | 6.0063 | 7.0147 | 6.9461 | 6.5561 | 7.6027 | 7.5323 | 7.6104 | 7.8471 | 3.3220 | 4.4953 | 3.5528 | 4.4726 |
| DNMT1_T182 | DNMT1 | P26358-2 | | 182 | KSTRQTTITSHFA | 7.6247 | 7.6724 | 9.8356 | 7.7807 | 5.2471 | 5.5835 | 6.7435 | 5.0827 | 7.6755 | 7.3003 | 8.6797 | 6.6448 | 3.4700 | 4.9421 | 2.4441 | 3.2733 |
| DNMT3A_S105 | DNMT3A | Q9Y6K1 | | 105 | PQPEEGSPAGGQK | 6.0876 | 6.5563 | 6.2518 | 6.5887 | 5.8571 | 5.4423 | 6.2601 | 6.1439 | 6.7560 | 6.5079 | 6.0921 | 6.4574 | 6.7588 | 6.3112 | 5.8122 | 6.1166 |
| DNMT3A_S75 | DNMT3A | Q9Y6K1 | | 75 | PAVISKSPSMAQD | 6.3373 | 6.7232 | 6.2831 | 7.3722 | 6.2466 | 5.5796 | 6.0561 | 6.5301 | 6.5342 | 6.5847 | 6.4612 | 6.9494 | 5.7053 | 5.7234 | 5.3300 | 5.5837 |
| DNMT3A_T257 | DNMT3A | Q9Y6K1 | | 257 | TDPASPTVATTPE | 6.3270 | 6.0464 | 5.5256 | 6.7496 | 6.4351 | 6.1500 | 6.4376 | 6.4212 | 6.8458 | 5.7065 | 6.0922 | 5.7648 | 6.8373 | 6.2557 | 5.8405 | 6.5647 |
| DNPH1_S169 | DNPH1 | O43598 | | 169 | PGQVAASPDPTTx | 6.1341 | 5.9041 | 6.4368 | 6.5946 | 6.4582 | 6.1598 | 6.1061 | 6.5581 | 6.2944 | 6.5368 | 6.6371 | 6.5708 | 5.8944 | 6.1038 | 5.4979 | 6.1131 |
| DNPH1_S28 | DNPH1 | O43598 | | 28 | ALYFCGSIRGGRE | 6.3226 | 5.9522 | 6.9510 | 6.2005 | 6.3218 | 6.6527 | 5.6305 | 6.2364 | 6.3151 | 6.7664 | 6.3526 | 6.5281 | 6.2101 | 6.3685 | 5.4242 | 5.7672 |
| DNTTIP1_S161 | DNTTIP1 | Q9H147 | | 161 | ECAHRGSPLPKKR | 6.6343 | 7.0766 | 6.5712 | 7.3155 | 6.1664 | 6.1326 | 6.4589 | 5.6796 | 6.1408 | 6.6636 | 6.6965 | 6.7218 | 5.7110 | 6.5675 | 5.1975 | 5.1763 |
| DNTTIP2_S117 | DNTTIP2 | Q5QJE6 | | 117 | QILIACSPVSSVR | 7.0246 | 6.9230 | 6.5833 | 7.0698 | 6.9322 | 6.5198 | 6.6744 | 6.3314 | 6.8136 | 6.3798 | 6.4540 | 6.3462 | 4.7436 | 4.9968 | 5.0022 | 5.2052 |
| DNTTIP2_S117;S121 | DNTTIP2 | Q5QJE6 | | 610 | LQKAVITPDFEKN | 7.4815 | 7.8749 | 6.9302 | 7.5837 | 6.2849 | 6.4624 | 6.9532 | 6.9693 | 7.0180 | 6.9387 | 7.2351 | 7.0043 | 3.2381 | 3.9819 | 3.6337 | 4.4103 |
| DNTTIP2_S184 | DNTTIP2 | Q5QJE6 | | 481 | LSGDTGSLSCDNA | 5.4955 | 6.5539 | 6.4762 | 6.7192 | 6.3012 | 6.2928 | 6.6095 | 5.8899 | 6.1055 | 5.8926 | 5.9579 | 6.0668 | 6.3273 | 6.5585 | 6.3426 | 6.4109 |
| DNTTIP2_S253 | DNTTIP2 | Q5QJE6 | | 434 | TSAPNTSQGKDNS | 7.0125 | 7.1582 | 6.6307 | 7.4275 | 7.3155 | 6.9713 | 7.6108 | 7.3551 | 6.9623 | 6.4919 | 6.8622 | 6.6906 | 3.3424 | 4.1717 | 3.7209 | 4.2764 |
| DNTTIP2_S273 | DNTTIP2 | Q5QJE6 | | 184 | SHTEAISDAETSS | 6.3617 | 6.8135 | 6.4538 | 6.8981 | 6.1565 | 6.5862 | 5.7149 | 6.0428 | 6.2136 | 6.1193 | 6.1628 | 6.5784 | 5.9954 | 6.3695 | 5.8833 | 5.6503 |
| DNTTIP2_S434 | DNTTIP2 | Q5QJE6 | | 512 | EEEDKASEVAIEE | 5.4723 | 6.6572 | 6.7502 | 7.3325 | 7.0092 | 5.0538 | 6.6634 | 4.9071 | 5.2860 | 5.7533 | 4.8860 | 5.3448 | 6.6532 | 6.6344 | 7.2108 | 8.3858 |
| DNTTIP2_S481 | DNTTIP2 | Q5QJE6 | | 341 | VSQRHSTPQNKNA | 7.5267 | 7.2643 | 6.8198 | 7.7923 | 6.2750 | 6.1598 | 6.8745 | 5.9112 | 6.7980 | 6.2265 | 6.4226 | 7.1712 | 4.6235 | 4.8700 | 4.1653 | 5.0991 |
| DNTTIP2_S512 | DNTTIP2 | Q5QJE6 | | 273 | DDFSHRSSENILT | 6.0570 | 6.2838 | 6.2241 | 7.0926 | 6.1430 | 6.0876 | 6.1215 | 6.4011 | 6.3962 | 6.1877 | 6.5848 | 6.0967 | 5.8524 | 6.4546 | 5.7474 | 6.2696 |
| DNTTIP2_S69 | DNTTIP2 | Q5QJE6 | | 232 | EKQIVGTPVNSED | 6.9070 | 6.8575 | 6.5182 | 6.7766 | 6.2115 | 5.6770 | 6.4575 | 6.7406 | 6.7430 | 6.5047 | 5.9436 | 6.4581 | 4.9437 | 5.9725 | 5.9165 | 5.3720 |
| DNTTIP2_T232 | DNTTIP2 | Q5QJE6 | | 69 | KARKRKSRTTGSL | 7.5249 | 7.8473 | 6.4372 | 8.1103 | 6.1299 | 6.6948 | 8.1506 | 6.6237 | 6.1325 | 7.2004 | 7.4550 | 7.8752 | 2.8096 | 4.6879 | 2.8527 | 3.4680 |
| DNTTIP2_T341 | DNTTIP2 | Q5QJE6 | | 253 | LQARSLSEINKPN | 6.5805 | 5.8182 | 5.7567 | 6.3879 | 6.5850 | 6.7724 | 6.8021 | 5.6959 | 5.4092 | 6.3864 | 5.9915 | 6.2182 | 6.0730 | 6.9858 | 6.1901 | 6.3471 |
| DNTTIP2_T610 | DNTTIP2 | Q5QJE6 | 117;121 | | NA | 7.8551 | 7.6631 | 7.2926 | 8.3210 | 6.0111 | 6.0913 | 8.1115 | 6.3518 | 7.0421 | 6.4293 | 6.9146 | 7.0067 | 3.4314 | 4.1199 | 3.1373 | 4.2212 |
| DOCK1_S1734 | DOCK1 | Q14185 | | 1844 | PPLPSKTPPPPPP | 6.5923 | 7.1788 | 6.3562 | 7.0824 | 6.4275 | 6.8801 | 6.8212 | 7.4122 | 6.9161 | 6.8569 | 6.9613 | 6.7513 | 4.0497 | 5.2974 | 3.9003 | 4.5162 |
| DOCK1_S1743 | DOCK1 | Q14185 | | 1743 | RPQRPKSQVMNVI | 8.0211 | 6.1201 | 7.3235 | 7.1149 | 5.7389 | 6.5564 | 5.9427 | 6.8536 | 7.7790 | 5.5328 | 6.1441 | 6.2496 | 5.3842 | 5.4177 | 4.7825 | 5.0390 |
| DOCK1_S1756 | DOCK1 | Q14185 | | 1756 | GSERRFSVSPSSP | 6.7080 | 4.7508 | 7.1454 | 3.6899 | 5.7668 | 4.7259 | 5.5789 | 3.9676 | 8.5255 | 6.5733 | 7.1553 | 7.7939 | 7.7689 | 10.4029 | 3.5835 | 5.8633 |
| DOCK1_S1761 | DOCK1 | Q14185 | | 1734 | ILSETISPLRPQR | 6.5265 | 6.7456 | 6.6104 | 5.9342 | 7.0511 | 7.1467 | 6.2000 | 5.7780 | 6.3982 | 6.7410 | 7.9441 | 5.8223 | 4.3906 | 5.4375 | 5.0878 | 6.1860 |
| DOCK1_T1844 | DOCK1 | Q14185 | | 1761 | FSVSPSSPSSQQT | 8.1251 | 9.0437 | 5.5004 | 6.8641 | 4.5761 | 8.4275 | 6.6815 | 7.8569 | 5.5917 | 5.8313 | 5.3863 | 6.2935 | 6.1794 | 4.3136 | 3.9827 | 5.3463 |
| DOCK10_S12 | DOCK10 | Q96BY6 | | 12 | TRRFTRSLLRPGQ | 8.0649 | 7.4236 | 7.8835 | 6.2574 | 5.4469 | 6.9684 | 5.8705 | 6.0726 | 6.9568 | 7.2622 | 7.2224 | 7.6554 | 4.0926 | 5.3709 | 3.5632 | 3.8889 |
| DOCK10_S1232 | DOCK10 | Q96BY6 | | 1318 | KIPRPLSLIGSTL | 6.3500 | 5.5878 | 6.1808 | 5.7022 | 6.2618 | 6.3476 | 5.8392 | 6.1227 | 6.9297 | 5.5409 | 5.5502 | 5.8702 | 6.7883 | 6.8494 | 7.5061 | 6.5730 |
| DOCK10_S1292 | DOCK10 | Q96BY6 | | 1292 | SRASLASLDSNPS | 5.9489 | 5.8117 | 6.1709 | 5.6836 | 5.9269 | 6.4087 | 5.9264 | 6.3697 | 5.9718 | 6.2425 | 6.2289 | 6.0847 | 7.1011 | 6.7001 | 6.4514 | 6.9727 |
| DOCK10_S1318 | DOCK10 | Q96BY6 | | 1232 | PFTVNTSNQGSRD | 6.0001 | 6.3602 | 6.4898 | 6.6034 | 6.7107 | 5.7872 | 5.9158 | 6.1580 | 6.6327 | 6.3186 | 6.6934 | 6.1676 | 6.1220 | 5.8698 | 6.0754 | 6.0953 |
| DOCK10_S302 | DOCK10 | Q96BY6 | | 302 | PLQGRRSTELTDL | 5.9572 | 5.6196 | 6.5846 | 5.5692 | 6.2109 | 6.1446 | 6.0527 | 6.0967 | 6.5208 | 7.0824 | 7.0300 | 6.6677 | 6.4496 | 6.2450 | 6.1827 | 5.5865 |
| DOCK10_T1406 | DOCK10 | Q96BY6 | | 1440 | KQHRSQTLPIIRG | 9.2973 | 7.1463 | 6.7790 | 6.2644 | 6.6466 | 5.8129 | 6.0860 | 5.3674 | 7.3745 | 6.1038 | 5.9204 | 5.8254 | 5.9902 | 6.1340 | 4.9043 | 4.3477 |
| DOCK10_T1440 | DOCK10 | Q96BY6 | | 1406 | STQNNGTLKGSNP | 8.7030 | 6.8694 | 7.1039 | 5.6540 | 6.6475 | 5.6018 | 6.4308 | 5.6018 | 7.8951 | 6.8149 | 6.6114 | 6.1519 | 5.6094 | 5.9847 | 4.1294 | 4.1909 |
| DOCK4_S1640 | DOCK4 | Q8N1I0-3 | | 1908 | VRKESKTPPPYSV | 5.8128 | 7.4155 | 6.8957 | 6.5296 | 6.2481 | 6.5840 | 5.8968 | 6.6543 | 6.4736 | 6.4160 | 6.1683 | 6.1408 | 5.2682 | 5.9110 | 5.7594 | 5.8260 |
| DOCK4_S1709 | DOCK4 | Q8N1I0-3 | | 1640 | RVIPRRSPLSYPA | 6.9121 | 7.4882 | 7.1187 | 7.2724 | 6.5386 | 5.5188 | 6.2128 | 7.0181 | 7.0949 | 6.4035 | 7.1571 | 6.3254 | 3.8170 | 5.7069 | 4.5361 | 4.8791 |
| DOCK4_T1908 | DOCK4 | Q8N1I0-3 | | 1709 | PSSARASPLLSDK | 6.2440 | 6.6588 | 6.2334 | 6.0017 | 6.1735 | 6.6606 | 6.2592 | 6.8144 | 6.7397 | 6.6993 | 6.4364 | 6.3884 | 5.6672 | 6.1080 | 5.4134 | 5.5022 |
| DOCK5_S1287 | DOCK5 | Q9H7D0 | | 1802 | KATRTLSSPSLQT | 6.2960 | 6.3796 | 6.5147 | 6.0482 | 6.3552 | 6.5828 | 6.3359 | 6.4507 | 6.5403 | 6.1038 | 6.5366 | 6.7156 | 5.9433 | 5.8925 | 5.7714 | 5.5335 |
| DOCK5_S1656 | DOCK5 | Q9H7D0 | | 1766 | KAQRPKSLQLMDN | 6.2341 | 6.0437 | 7.7531 | 5.8883 | 5.7223 | 5.9920 | 5.6075 | 6.7608 | 6.4976 | 5.9458 | 6.4933 | 6.8913 | 6.2856 | 6.1441 | 5.9108 | 5.8296 |
| DOCK5_S1742 | DOCK5 | Q9H7D0 | | 1742 | DWSLSKSQVIAEK | 7.4835 | 6.7030 | 7.8344 | 6.4810 | 5.7834 | 6.1600 | 6.3406 | 6.3931 | 6.8460 | 5.3531 | 6.1027 | 6.7782 | 5.6259 | 5.7152 | 5.1794 | 5.2204 |
| DOCK5_S1756 | DOCK5 | Q9H7D0 | | 1834 | EGSQRNSTELAPP | 6.5025 | 6.5762 | 7.6441 | 6.4702 | 6.1594 | 6.0975 | 6.1363 | 6.3938 | 6.8853 | 6.3267 | 6.9669 | 6.8327 | 5.4971 | 5.1917 | 5.0066 | 5.3129 |
| DOCK5_S1766 | DOCK5 | Q9H7D0 | | 1287 | HLLQKDSYYVYTQ | 6.1197 | 5.4111 | 6.6377 | 6.3305 | 6.0767 | 6.1570 | 5.8167 | 6.7216 | 6.8610 | 6.2134 | 6.4647 | 5.8571 | 6.8646 | 6.0948 | 6.3802 | 5.9933 |
| DOCK5_S1802 | DOCK5 | Q9H7D0 | | 1756 | PEPDLMSPTRKAQ | 6.7371 | 6.6967 | 5.9575 | 7.0603 | 6.5351 | 6.9264 | 6.3861 | 7.1863 | 6.1350 | 7.2623 | 7.6339 | 7.4881 | 4.0752 | 4.9427 | 4.5901 | 4.3873 |
| DOCK5_S1834 | DOCK5 | Q9H7D0 | | 1656 | KQSRTGSIVLPYI | 5.9857 | 5.5351 | 7.3469 | 6.0748 | 6.5463 | 5.8092 | 7.0836 | 6.3783 | 6.9745 | 6.2227 | 6.5672 | 6.6593 | 6.1050 | 6.4271 | 5.3346 | 4.9498 |
| DOCK6_S1230 | DOCK6 | Q96HP0 | | 2029 | PQLMAPTPPGLRN | 6.4184 | 6.8682 | 6.3939 | 6.4147 | 5.8678 | 6.3387 | 6.1078 | 6.6156 | 7.2768 | 6.2196 | 6.6059 | 6.2947 | 5.2532 | 5.9972 | 5.5338 | 5.7938 |
| DOCK6_S1232 | DOCK6 | Q96HP0 | | 1230 | APGSRASISQGPP | 7.3747 | 6.1648 | 7.5741 | 6.4971 | 6.6558 | 5.9586 | 6.3099 | 6.6300 | 7.1495 | 5.4513 | 6.1105 | 6.3818 | 6.5675 | 6.4646 | 4.5127 | 5.1068 |
| DOCK6_S1308 | DOCK6 | Q96HP0 | | 1308 | AFERINSLTFKKS | 5.3864 | 5.0428 | 5.3817 | 5.6085 | 6.1394 | 5.1528 | 4.9967 | 5.5882 | 6.1051 | 5.4500 | 4.7341 | 5.3493 | 9.0510 | 9.0350 | 8.6231 | 8.3559 |
| DOCK6_T1264 | DOCK6 | Q96HP0 | | 1264 | LWVLKNTEPALLQ | 1.1992 | 1.0528 | 0.9519 | 1.1501 | 3.4703 | 3.0727 | 3.3343 | 2.9485 | 1.1271 | 1.1429 | 0.8426 | 1.1655 | 20.7631 | 18.3368 | 20.3537 | 19.0884 |
| DOCK6_T2029 | DOCK6 | Q96HP0 | | 1232 | GSRASISQGPPTA | 5.3705 | 5.5624 | 5.9833 | 5.5296 | 5.8381 | 5.9290 | 5.4462 | 5.8531 | 6.5877 | 5.9931 | 6.0753 | 6.3073 | 7.5007 | 7.1732 | 7.2572 | 7.5932 |

| | | | | | | | | | | | | | | | | | | | | | |
|---|---|---|---|---|---|---|---|---|---|---|---|---|---|---|---|---|---|---|---|---|---|
| DOCK7_S1306 | DOCK7 | Q96N67 | | 1383 | VFERMNSLTFKKS | 5.7336 | 5.1526 | 5.5205 | 5.5657 | 6.1817 | 5.6569 | 5.8590 | 5.8266 | 6.0617 | 5.3927 | 5.2028 | 5.2788 | 8.4167 | 8.2504 | 8.1970 | 7.7032 |
| DOCK7_S1383 | DOCK7 | Q96N67 | | 1432 | PPERSPSGSAFGS | 6.7298 | 6.1803 | 6.3082 | 5.6228 | 6.2870 | 5.8840 | 5.8191 | 6.0436 | 6.6230 | 6.4676 | 6.0555 | 6.0758 | 6.8870 | 7.2243 | 6.1238 | 5.6682 |
| DOCK7_S1430 | DOCK7 | Q96N67 | | 2129 | VTCHRDSFSRMSL | 6.2974 | 5.7948 | 6.3784 | 5.9933 | 6.7294 | 5.7745 | 5.8019 | 5.8892 | 7.0094 | 6.3792 | 6.0180 | 5.6880 | 7.0755 | 7.6628 | 6.1391 | 5.3690 |
| DOCK7_S1432 | DOCK7 | Q96N67 | | 864 | TYPNSSSPGPGGL | 6.2775 | 6.2749 | 6.3410 | 6.4575 | 6.8461 | 6.6770 | 6.2771 | 7.5312 | 6.5585 | 6.0138 | 6.5856 | 5.9790 | 5.4872 | 5.9554 | 5.4026 | 5.3355 |
| DOCK7_S180 | DOCK7 | Q96N67 | | 440 | SERRNSSIVGRRS | 6.7671 | 6.3621 | 6.5252 | 5.9932 | 5.9069 | 5.9411 | 6.0914 | 5.9458 | 6.9097 | 6.7648 | 6.7392 | 6.9595 | 5.8729 | 6.8552 | 4.8358 | 5.5302 |
| DOCK7_S190 | DOCK7 | Q96N67 | | 180 | DDLKRRSMSIDDT | 6.2854 | 6.2604 | 6.1958 | 5.9336 | 6.2498 | 6.1019 | 5.8882 | 6.4479 | 6.5516 | 6.2889 | 6.1413 | 6.1040 | 6.4432 | 6.6942 | 6.1270 | 6.2868 |
| DOCK7_S2129 | DOCK7 | Q96N67 | | 929 | GLDRSNSWVNTGG | 5.7011 | 5.7829 | 6.2825 | 5.7755 | 6.0061 | 6.0892 | 5.9174 | 6.1420 | 6.3405 | 6.1954 | 6.3168 | 6.0991 | 6.7912 | 7.1028 | 6.5792 | 6.8784 |
| DOCK7_S30 | DOCK7 | Q96N67 | | 452 | SLERTTSGDDACN | 5.6597 | 5.1773 | 5.9647 | 5.5083 | 6.6495 | 6.2402 | 5.6044 | 5.7649 | 5.8817 | 5.4557 | 5.5930 | 5.8172 | 7.6005 | 8.0073 | 7.5890 | 7.4864 |
| DOCK7_S440 | DOCK7 | Q96N67 | | 190 | DDTPRGSWACSIF | 6.1261 | 5.3095 | 5.4897 | 5.6561 | 6.7044 | 5.5946 | 5.5658 | 5.8157 | 6.3119 | 5.5563 | 4.9675 | 4.8540 | 8.1069 | 8.3553 | 8.1452 | 7.4410 |
| DOCK7_S452 | DOCK7 | Q96N67 | | 1306 | QLTRPGSFLLTST | 6.2285 | 5.3376 | 6.5899 | 5.8662 | 6.1537 | 5.2134 | 6.1310 | 6.2448 | 7.1188 | 5.5374 | 5.8101 | 5.6060 | 8.1566 | 7.1653 | 6.4654 | 6.3754 |
| DOCK7_S864 | DOCK7 | Q96N67 | | 30 | SGQYSGSPQLLKN | 6.6056 | 6.5680 | 6.6915 | 6.8632 | 6.6684 | 6.7573 | 7.1134 | 6.6935 | 7.5086 | 6.9395 | 6.6895 | 6.5101 | 4.4330 | 5.3165 | 3.9988 | 4.6430 |
| DOCK7_S888 | DOCK7 | Q96N67 | | 1430 | ASPPERSPSGSAF | 5.9533 | 5.8991 | 6.2589 | 6.7377 | 5.2262 | 6.1257 | 6.3127 | 5.8908 | 6.6816 | 6.4997 | 6.1775 | 6.7646 | 5.7426 | 6.3534 | 6.6448 | 6.7314 |
| DOCK7_S898 | DOCK7 | Q96N67 | | 898 | NRSRSLSNSNPDI | 6.3628 | 6.3534 | 5.8745 | 5.7385 | 5.6301 | 5.5668 | 7.2092 | 6.2263 | 6.5908 | 5.6148 | 6.0338 | 5.8681 | 6.8120 | 7.1004 | 6.5894 | 6.4291 |
| DOCK7_S929 | DOCK7 | Q96N67 | | 142 | TGFNPNTLDKQKE | 6.3062 | 6.1201 | 5.7431 | 6.3991 | 6.2075 | 6.6728 | 6.0619 | 6.0977 | 6.3193 | 6.2966 | 5.9975 | 6.4520 | 6.2784 | 6.4820 | 6.7518 | 5.8142 |
| DOCK7_T142 | DOCK7 | Q96N67 | | 888 | SAVRPASLNLNRS | 6.5701 | 5.8386 | 6.6193 | 6.1988 | 5.8410 | 6.0182 | 5.7847 | 6.2146 | 6.4161 | 5.1081 | 5.6999 | 5.8435 | 7.4961 | 7.4523 | 6.5998 | 6.2989 |
| DOCK9_S1221 | DOCK9 | Q9BZ29-4 | | 927 | AEPYVASEYKTVH | 6.0248 | 6.3116 | 6.9371 | 6.3341 | 6.8129 | 5.7192 | 5.9289 | 6.2156 | 5.8130 | 5.9271 | 5.7098 | 5.7179 | 6.9029 | 6.8161 | 6.9724 | 5.8568 |
| DOCK9_S443 | DOCK9 | Q9BZ29-4 | | 1221 | GSTLDNSLHKDLL | 6.7452 | 6.3534 | 7.6250 | 7.7497 | 6.0260 | 7.2191 | 6.9730 | 6.8374 | 6.6245 | 6.9808 | 7.5825 | 6.8351 | 4.3769 | 4.4406 | 3.6283 | 4.0026 |
| DOCK9_S927 | DOCK9 | Q9BZ29-4 | | 443 | MNGSGQSPSVLKG | 6.1422 | 5.6199 | 5.8662 | 6.4346 | 5.9550 | 6.4276 | 5.9954 | 6.6734 | 6.8496 | 6.4527 | 6.2268 | 6.6506 | 6.5981 | 6.0842 | 6.3974 | 5.6264 |
| DOK1_S281 | DOK1 | Q99704 | | 291 | PQELLDSPPALYA | 5.8859 | 6.5649 | 6.4839 | 6.5716 | 6.0511 | 6.9529 | 6.2637 | 6.6756 | 6.8953 | 6.9692 | 6.5400 | 6.8137 | 5.0458 | 5.5770 | 5.1992 | 5.5102 |
| DOK1_S291 | DOK1 | Q99704 | | 281 | AEGKLPSPPGPQE | 6.3218 | 6.6991 | 6.0238 | 5.9984 | 6.4970 | 7.1089 | 6.0365 | 6.1459 | 7.4790 | 6.6222 | 6.9496 | 6.7110 | 4.4838 | 6.5422 | 4.6312 | 5.7496 |
| DOT1L_S1001;S1009 | DOT1L | Q8TEK3-1 | | 834 | QDPRPLSPGALQL | 6.4936 | 5.9705 | 6.7305 | 7.2452 | 5.9291 | 5.8261 | 6.6367 | 6.2814 | 6.6881 | 6.3431 | 6.7527 | 6.9460 | 5.2961 | 5.7505 | 5.1549 | 5.9553 |
| DOT1L_S1104 | DOT1L | Q8TEK3-1 | | 1104 | SLSAGVSPKRRAL | 6.2241 | 6.5247 | 6.6467 | 6.6826 | 5.7654 | 6.0883 | 6.0480 | 6.2229 | 6.3232 | 6.4687 | 6.1726 | 5.8658 | 6.1770 | 6.3958 | 6.1254 | 6.2687 |
| DOT1L_S1153 | DOT1L | Q8TEK3-1 | | 775 | PDYTRLSPAKIVL | 6.5652 | 6.3439 | 6.3188 | 6.7141 | 6.5859 | 6.5212 | 6.9415 | 6.0547 | 6.4210 | 6.0805 | 5.8020 | 5.9995 | 6.0931 | 5.9725 | 5.7474 | 5.8388 |
| DOT1L_S1213 | DOT1L | Q8TEK3-1 | | 1246 | KWKSTFSPISDIG | 6.3258 | 6.1365 | 6.1975 | 6.9499 | 5.8804 | 6.2354 | 6.2272 | 6.4826 | 6.2462 | 6.1690 | 6.1138 | 6.1800 | 6.1717 | 6.4541 | 6.0336 | 6.1964 |
| DOT1L_S1246 | DOT1L | Q8TEK3-1 | | 297 | MRVVELSPLKGSV | 6.4627 | 6.9696 | 6.4121 | 7.4817 | 5.8864 | 5.9829 | 6.4377 | 6.3120 | 6.8554 | 6.5675 | 6.4096 | 6.8401 | 5.0206 | 5.3369 | 5.3337 | 5.6908 |
| DOT1L_S1349 | DOT1L | Q8TEK3-1 | | 471 | PSVQRHSPNPLLV | 6.4740 | 5.4955 | 6.5659 | 6.3809 | 5.6413 | 5.0844 | 6.4489 | 5.8013 | 5.5641 | 6.0948 | 6.1664 | 6.0304 | 6.9679 | 7.2693 | 7.0402 | 6.9746 |
| DOT1L_S297 | DOT1L | Q8TEK3-1 | | 385 | EKAGAATVKKPSP | 8.6219 | 8.5989 | 7.9737 | 7.9168 | 5.8034 | 7.9137 | 7.6359 | 6.1788 | 7.6824 | 7.0389 | 8.2340 | 7.9637 | 1.7304 | 2.7590 | 1.4467 | 2.5019 |
| DOT1L_S447 | DOT1L | Q8TEK3-1 | | 1213 | ISLESKSPPKTLE | 5.1616 | 5.9585 | 6.1833 | 8.5648 | 6.2756 | 6.5370 | 6.7855 | 5.8342 | 6.6338 | 6.2527 | 7.3796 | 7.9142 | 6.0270 | 4.9029 | 4.7074 | 4.8819 |
| DOT1L_S471 | DOT1L | Q8TEK3-1 | | 1349 | EAAGLSSPLSFPS | 6.4872 | 6.6410 | 6.3060 | 6.6077 | 6.4166 | 6.2486 | 7.0160 | 6.6821 | 6.5633 | 7.3466 | 6.6618 | 6.5020 | 4.6414 | 5.4576 | 5.1401 | 5.2819 |
| DOT1L_S775 | DOT1L | Q8TEK3-1 | | 902 | RARSTPSPVLQPR | 6.6442 | 6.3983 | 6.1018 | 6.3819 | 5.7955 | 6.1548 | 6.4987 | 6.0809 | 5.9667 | 6.0430 | 6.2771 | 6.2883 | 6.5185 | 6.2434 | 6.4015 | 6.2055 |
| DOT1L_S834 | DOT1L | Q8TEK3-1 | | 984 | VGSRSSTPQHPLL | 6.7702 | 5.2376 | 5.5567 | 6.1003 | 5.7249 | 5.2159 | 6.2476 | 4.6853 | 6.0187 | 5.9581 | 5.5408 | 5.3993 | 8.3693 | 7.6423 | 7.9722 | 7.5608 |
| DOT1L_S902 | DOT1L | Q8TEK3-1 | | 1153 | ETSLKSSPVPYQD | 5.3637 | 6.0808 | 5.4724 | 6.0600 | 6.0950 | 5.8419 | 5.7719 | 5.8061 | 5.7711 | 5.4865 | 5.0410 | 5.1957 | 7.9051 | 7.7362 | 8.5084 | 7.8642 |
| DOT1L_T385 | DOT1L | Q8TEK3-1 | | 447 | VSQTAASSPQDAY | 5.2858 | 5.8834 | 5.3417 | 6.7215 | 6.1529 | 5.2696 | 6.8355 | 6.4000 | 5.7957 | 6.1753 | 5.6755 | 5.2920 | 7.1396 | 6.6169 | 7.7205 | 7.6942 |
| DOT1L_T984 | DOT1L | Q8TEK3-1 | 1001;1009 | | NA | 6.0115 | 6.3862 | 5.8954 | 6.7171 | 6.2198 | 6.1612 | 6.5586 | 6.1083 | 6.4671 | 5.9753 | 6.0803 | 6.0147 | 6.4395 | 6.5636 | 6.0072 | 6.3942 |
| DPF2_S200 | DPF2 | Q92785 | | 176 | EDYEEDTPKRRGK | 6.1736 | 6.0080 | 6.8257 | 6.5909 | 6.4585 | 6.4547 | 7.0895 | 6.2393 | 6.6137 | 6.4911 | 6.7003 | 6.7655 | 5.4451 | 5.5710 | 5.1065 | 5.4665 |
| DPF2_S280 | DPF2 | Q92785 | | 200 | RKKLDASILEDRD | 7.4843 | 6.9155 | 6.8253 | 7.5556 | 6.3792 | 6.6307 | 7.5421 | 6.4462 | 7.5974 | 6.5267 | 7.7371 | 7.2988 | 3.4278 | 4.6158 | 2.6879 | 4.3297 |
| DPF2_T125 | DPF2 | Q92785 | | 280 | DFCLGDSKINKKT | 5.6134 | 4.2570 | 6.5017 | 6.6291 | 7.5938 | 4.6232 | 6.5941 | 4.1297 | 5.5529 | 7.1385 | 5.7312 | 4.7442 | 6.4317 | 8.3198 | 8.1383 | 8.0014 |
| DPF2_T176 | DPF2 | Q92785 | | 125 | LEALLRTDPLEKR | 7.5477 | 6.7212 | 6.3557 | 7.8996 | 7.4285 | 6.8864 | 7.5307 | 7.2539 | 7.0759 | 5.6464 | 6.5094 | 5.7614 | 4.0964 | 4.9939 | 3.7579 | 4.5350 |
| DPF2_T248 | DPF2 | Q92785 | | 248 | EDSQPPTPVSQRS | 6.4916 | 6.1461 | 6.0366 | 6.8042 | 6.7252 | 6.3543 | 7.3032 | 5.8192 | 6.5787 | 5.5038 | 5.9801 | 5.9555 | 5.8971 | 5.6551 | 6.4457 | 6.3036 |
| DPF3_S403 | DPF3 | Q92784-3 | | 403 | RRSGRGSPTADKK | 5.9642 | 6.2105 | 5.6625 | 5.9915 | 5.8434 | 5.9326 | 5.8762 | 6.1809 | 6.0108 | 6.0458 | 5.9372 | 6.0629 | 6.7467 | 7.2891 | 7.2216 | 7.0240 |
| DPH1_S212 | DPH1 | Q9BZG8 | | 212 | PQCKPLSPGEILG | 5.9966 | 6.0614 | 7.1572 | 6.3891 | 5.6938 | 6.2087 | 6.1434 | 5.7712 | 6.6000 | 6.5619 | 6.3030 | 6.0654 | 6.1579 | 6.0128 | 6.5781 | 6.2995 |
| DPH7_S353 | DPH7 | Q9BTV6 | | 353 | SLQRAPSWSFPSN | 3.1140 | 2.9308 | 3.4307 | 3.1642 | 4.5229 | 4.2430 | 4.2765 | 4.4574 | 3.3167 | 3.5098 | 3.5187 | 3.4140 | 13.9051 | 13.3106 | 15.1729 | 13.7125 |
| DPP9_S321 | DPP9 | Q86TI2-2 | | 321 | EVIHVPSPALEER | 6.3472 | 5.3383 | 5.6163 | 7.6060 | 5.8693 | 5.7506 | 5.9630 | 6.5159 | 6.2067 | 6.6096 | 5.2540 | 7.5061 | 7.0688 | 6.2379 | 7.1562 | 4.9541 |
| DPY19L1_S23;S25 | DPY19L1 | A0A1B0GW05 | | 46 | PERAPLSPGRKGA | 6.7196 | 6.9117 | 6.1492 | 6.1678 | 6.9992 | 6.8482 | 7.2149 | 6.4147 | 6.2226 | 6.4845 | 6.4935 | 6.2328 | 5.1772 | 5.6694 | 5.1797 | 5.1150 |
| DPY19L1_S25 | DPY19L1 | A0A1B0GW05 | | 25 | PPRASQSPLRGAS | 6.1708 | 6.4773 | 5.6057 | 6.6491 | 5.5995 | 5.9632 | 6.3430 | 6.1918 | 6.2819 | 6.7730 | 6.7133 | 6.7423 | 5.5074 | 6.5818 | 6.1106 | 6.2892 |
| DPY19L1_S46 | DPY19L1 | A0A1B0GW05 | 23;25 | | NA | 6.0912 | 6.7967 | 6.3467 | 7.0525 | 6.1099 | 5.8441 | 6.0606 | 5.9769 | 7.8323 | 8.0943 | 8.4609 | 8.1435 | 3.8617 | 4.7865 | 4.1876 | 4.3546 |
| DPYD_S905 | DPYD | Q12882 | | 905 | EQNVAFSPLKRNC | 6.8399 | 7.3897 | 7.7968 | 6.1375 | 6.9178 | 6.8141 | 5.9593 | 7.0600 | 5.9421 | 6.9430 | 6.7320 | 6.2605 | 4.3672 | 4.9428 | 4.3532 | 5.5442 |
| DPYSL2_S27 | DPYSL2 | A0A1C7CYX9 | | 27 | KNLGSGSPKPRQK | 6.0621 | 5.7239 | 6.8840 | 5.4686 | 6.3360 | 6.4073 | 6.2998 | 6.3302 | 6.5206 | 6.4845 | 6.4731 | 6.7050 | 6.2040 | 6.3253 | 5.7906 | 5.9853 |
| DPYSL2_T614 | DPYSL2 | A0A1C7CYX9 | | 619 | VTPKTVTPASSAK | 6.4263 | 6.2302 | 7.1119 | 6.1152 | 6.8708 | 6.8757 | 6.3037 | 7.0416 | 6.8857 | 6.8061 | 6.9537 | 7.0892 | 4.7677 | 5.2686 | 4.4394 | 4.8141 |
| DPYSL2_T619 | DPYSL2 | A0A1C7CYX9 | | 614 | VCEVSVTPKTVTP | 7.1799 | 6.9521 | 7.6597 | 7.1649 | 6.8398 | 6.7741 | 6.4276 | 6.0975 | 7.2338 | 6.6774 | 7.1583 | 7.4161 | 4.0610 | 4.5262 | 3.5027 | 4.3286 |
| DPYSL3_S10 | DPYSL3 | Q14195-2 | | 636 | AGSARGSPTRPNP | 6.2951 | 6.3723 | 6.8623 | 5.6985 | 6.1465 | 6.8372 | 6.0916 | 6.8792 | 6.5471 | 6.5497 | 6.7662 | 6.8379 | 5.3846 | 5.7610 | 5.4896 | 5.4814 |
| DPYSL3_S102 | DPYSL3 | Q14195-2 | | 623 | VFDLTTTPKGGTP | 6.5970 | 6.7084 | 7.3792 | 6.6877 | 6.4990 | 6.7650 | 6.2821 | 6.6920 | 7.3311 | 7.6787 | 7.7332 | 7.7500 | 3.7501 | 4.2598 | 3.6612 | 4.2256 |
| DPYSL3_S516 | DPYSL3 | Q14195-2 | | 102 | REPAPASPAPAGV | 6.5368 | 6.9137 | 6.6824 | 6.4246 | 6.0627 | 7.1069 | 6.6674 | 6.9474 | 7.7259 | 6.9779 | 7.0583 | 7.0886 | 4.2760 | 4.8207 | 3.9534 | 4.7574 |
| DPYSL3_S586 | DPYSL3 | Q14195-2 | | 628 | TTPKGGTPAGSAR | 7.3063 | 7.1264 | 7.6968 | 6.6062 | 6.1313 | 6.7473 | 6.3521 | 6.8197 | 7.5464 | 7.3362 | 7.9742 | 7.9052 | 3.4075 | 4.1848 | 3.2632 | 3.5963 |
| DPYSL3_S636 | DPYSL3 | Q14195-2 | | 586 | GRFIPCSPFSDYV | 6.4261 | 6.2247 | 6.3428 | 7.1670 | 6.7561 | 6.7877 | 5.7354 | 6.6227 | 6.6963 | 5.9651 | 7.0433 | 6.2660 | 5.1094 | 6.0193 | 4.9039 | 5.9344 |
| DPYSL3_T623 | DPYSL3 | Q14195-2 | | 516 | PRKGRISVGSDSD | 6.4400 | 7.0580 | 7.4688 | 7.5226 | 5.9183 | 6.3928 | 6.1009 | 6.1074 | 5.5042 | 5.8710 | 6.5705 | 7.3498 | 4.6124 | 5.8319 | 4.8732 | 6.3781 |
| DPYSL3_T628 | DPYSL3 | Q14195-2 | | 10 | GRRGWDSSHEDDL | 7.4586 | 8.0334 | 9.0236 | 6.7491 | 8.5835 | 10.6491 | 7.4115 | 9.5804 | 5.6086 | 4.4029 | 4.2367 | 4.1941 | 3.4060 | 3.2354 | 2.9906 | 4.4366 |
| DROSHA_S373 | DROSHA | Q9NRR4 | | 807 | RHLLANSPKVKQT | 6.6577 | 7.2162 | 6.8251 | 6.8453 | 6.9542 | 6.9203 | 7.3339 | 6.4793 | 7.1419 | 6.8312 | 6.8831 | 6.6522 | 4.2122 | 4.6642 | 3.8707 | 4.5126 |
| DROSHA_S807 | DROSHA | Q9NRR4 | | 373 | EEKDRWSDNQSSG | 7.0093 | 7.9452 | 7.3754 | 7.5358 | 7.2374 | 7.0967 | 7.7407 | 7.8914 | 5.4002 | 5.4952 | 5.3986 | 4.9338 | 4.3375 | 5.0086 | 4.7393 | 4.8548 |
| DSG2_S700 | DSG2 | Q14126 | | 703 | KGSSSASIVKGQH | 8.2443 | 7.6329 | 8.8188 | 7.6196 | 6.4739 | 6.4870 | 6.4947 | 7.0394 | 7.0825 | 7.4847 | 7.6184 | 7.4343 | 2.7556 | 3.8061 | 2.4506 | 2.5573 |
| DSG2_S700;S703 | DSG2 | Q14126 | | 782 | FTDKAASYTEEDE | 5.5846 | 6.3642 | 5.4908 | 6.2962 | 5.9379 | 6.3732 | 6.0063 | 6.1838 | 6.4158 | 5.5803 | 6.1584 | 6.6299 | 6.4670 | 6.6141 | 6.6227 | 7.2750 |
| DSG2_S703 | DSG2 | Q14126 | | 922 | QAQKVATPLPDPM | 6.5526 | 6.7247 | 6.3069 | 7.5614 | 5.3076 | 6.6167 | 6.2096 | 6.8113 | 7.4380 | 6.6986 | 6.3508 | 6.8149 | 4.4970 | 5.5592 | 5.0227 | 5.5283 |
| DSG2_S782 | DSG2 | Q14126 | | 700 | ATMKGSSSASIVK | 2.6528 | 3.1011 | 3.0168 | 2.8830 | 4.2054 | 5.4495 | 4.7625 | 5.4009 | 3.3026 | 4.0644 | 3.8226 | 4.6671 | 12.1300 | 11.6714 | 13.3462 | 15.5236 |
| DSG2_T922 | DSG2 | Q14126 | 700;703 | | NA | 4.0219 | 3.5363 | 3.6775 | 3.8676 | 5.5066 | 4.8436 | 5.2871 | 4.8144 | 3.7217 | 3.2551 | 3.7757 | 3.7561 | 13.2903 | 11.8644 | 12.7815 | 12.0002 |
| DSN1_S109 | DSN1 | Q9H410 | | 58 | RIHLGSSPKKGGN | 6.3279 | 6.2410 | 6.0092 | 6.1478 | 6.5413 | 6.0525 | 5.9418 | 6.3433 | 6.4199 | 6.1723 | 6.2426 | 5.9523 | 6.5808 | 6.2266 | 6.4320 | 6.3686 |
| DSN1_S123 | DSN1 | Q9H410 | | 109 | ETNRRKSLHPIHQ | 6.4152 | 6.0231 | 5.7296 | 6.6119 | 7.1369 | 6.1483 | 5.9430 | 5.7656 | 8.1921 | 7.6408 | 7.3193 | 7.4555 | 5.4703 | 6.1930 | 3.6453 | 4.3100 |
| DSN1_S27 | DSN1 | Q9H410 | | 331 | MQQLDPSPARKLL | 6.5732 | 6.7661 | 6.7500 | 6.9745 | 6.7225 | 6.6155 | 6.2739 | 6.7042 | 7.1208 | 7.0082 | 7.0849 | 6.9524 | 4.4158 | 4.7987 | 4.2372 | 5.0022 |
| DSN1_S331 | DSN1 | Q9H410 | | 123 | ITELSRSISVDLA | 5.9649 | 5.8152 | 6.8900 | 6.2942 | 6.2455 | 6.3152 | 6.0125 | 6.0652 | 6.3316 | 6.6667 | 6.4195 | 6.2562 | 6.2761 | 6.7177 | 5.4819 | 6.2477 |
| DSN1_S58 | DSN1 | Q9H410 | | 81 | SKSLHLSPQEQSA | 6.2104 | 6.6178 | 5.9966 | 6.2943 | 6.2831 | 6.4074 | 6.9017 | 7.2071 | 6.2603 | 5.9033 | 6.3734 | 6.3308 | 5.9794 | 5.9106 | 5.7269 | 5.5970 |
| DSN1_S81 | DSN1 | Q9H410 | | 27 | HDHQLESSLSPVE | 6.2328 | 6.1223 | 6.4963 | 6.6923 | 6.8044 | 6.1732 | 6.0138 | 6.7090 | 6.3453 | 6.7720 | 6.7288 | 6.5402 | 5.7871 | 5.8594 | 5.5959 | 5.1274 |
| DST_S106 | DST | Q03001 | | 3968 | KLKRQKSFSEDVI | 5.6929 | 6.1037 | 6.4866 | 5.7164 | 6.0960 | 6.5664 | 5.8375 | 6.4820 | 6.1809 | 6.7918 | 6.2229 | 6.4042 | 6.2374 | 6.4809 | 6.5846 | 6.1158 |
| DST_S140 | DST | Q03001 | | 7421 | ASPNRSTSVSSQA | 4.6811 | 4.8165 | 5.0270 | 4.5972 | 5.4878 | 5.8936 | 5.8930 | 4.9416 | 6.0749 | 5.2208 | 5.8881 | 6.1205 | 9.6780 | 9.2780 | 7.8962 | 8.5058 |
| DST_S170 | DST | Q03001 | | 2671 | NVVQLASPSENNL | 6.1044 | 7.3865 | 7.0570 | 7.0632 | 5.9467 | 6.5937 | 6.5613 | 6.1086 | 6.6400 | 7.0645 | 7.6063 | 7.5798 | 4.2985 | 5.1211 | 4.1023 | 4.7663 |
| DST_S2229 | DST | Q03001-11 | | 77 | SEGFRASPRHLRR | 7.1719 | 7.0660 | 6.6278 | 6.8011 | 6.1086 | 6.7710 | 6.7829 | 5.7774 | 6.1346 | 6.5558 | 6.7156 | 6.4762 | 5.1613 | 5.4959 | 5.2219 | 5.1320 |
| DST_S2671 | DST | Q03001-11 | | 140 | ASIRRPSSGNASY | 5.9663 | 4.6026 | 5.0604 | 5.0991 | 6.3963 | 5.4406 | 4.7497 | 5.7796 | 8.7754 | 8.7787 | 7.8175 | 7.6437 | 6.6864 | 7.4505 | 4.7172 | 5.0359 |
| DST_S2919 | DST | Q03001-8 | | 5023 | KPWLTNSKMSTPC | 8.6831 | 6.2146 | 7.7248 | 7.6402 | 4.6324 | 6.3119 | 7.1114 | 7.0670 | 8.8967 | 7.0416 | 7.2130 | 8.6640 | 2.8788 | 4.0422 | 2.5002 | 3.3780 |
| DST_S3968 | DST | Q03001 | | 2919 | QIIVSKSPVQFEN | 5.7407 | 6.6611 | 6.3340 | 5.9601 | 5.3798 | 7.5426 | 5.8264 | 6.7403 | 6.9260 | 6.4404 | 5.6070 | 6.6897 | 5.1292 | 5.5395 | 6.6910 | 6.7922 |
| DST_S5023 | DST | Q03001 | | 2229 | RLSALLSHDKLMH | 4.7447 | 4.6505 | 5.2268 | 5.1552 | 4.7969 | 5.0538 | 5.2473 | 5.1798 | 5.2227 | 4.7017 | 5.4336 | 4.8266 | 9.4290 | 9.9664 | 10.1353 | 10.2295 |
| DST_S5853 | DST | Q03001 | | 7510 | ASSRRGSDASDFD | 5.6406 | 5.2745 | 5.8510 | 6.1871 | 6.5020 | 6.3171 | 6.0351 | 6.3361 | 7.0295 | 6.3727 | 5.7998 | 6.8892 | 6.6065 | 6.3697 | 6.5655 | 6.2236 |
| DST_S7458 | DST | Q03001 | | 7494 | TPSRPGSRAGSKA | 6.3025 | 6.2199 | 6.5406 | 6.6266 | 5.6366 | 5.8390 | 5.8114 | 6.8181 | 6.5438 | 6.3050 | 6.4009 | 6.6014 | 5.5670 | 6.5639 | 6.3891 | 5.8341 |
| DST_S7494 | DST | Q03001-8 | | 5045 | VPSAEGTPIQGSK | 7.1799 | 6.7294 | 6.3084 | 6.7798 | 5.4570 | 7.1417 | 7.5216 | 6.7885 | 7.2253 | 7.2592 | 7.3926 | 7.2030 | 4.3173 | 5.3309 | 3.7579 | 3.6073 |
| DST_S7510 | DST | Q03001 | | 5853 | TEKKLMSLGDIRL | 6.6373 | 6.7088 | 5.4781 | 6.2553 | 5.9631 | 6.1582 | 6.2612 | 5.6538 | 6.3102 | 6.0861 | 5.7367 | 6.2648 | 6.6922 | 6.5370 | 6.7016 | 6.5555 |
| DST_S7510;S7513 | DST | Q03001 | | 7458 | RLPGYLSGKGFHS | 7.6797 | 8.5549 | 6.9030 | 9.1565 | 3.7851 | 8.1063 | 6.4599 | 6.0989 | 9.7911 | 7.1765 | 6.4927 | 6.9248 | 5.0159 | 3.1233 | 2.7240 | 2.0074 |
| DST_S77 | DST | Q03001-11 | | 106 | VEVHHQSEQETSV | 5.8254 | 6.5765 | 6.2060 | 6.0400 | 6.0222 | 6.0269 | 6.3853 | 5.4916 | 5.1391 | 5.6093 | 5.1425 | 5.0616 | 7.5073 | 7.9237 | 7.6515 | 7.3909 |
| DST_T5045 | DST | Q03001-11 | | 170 | QKSGSASPAPGDT | 6.9353 | 8.2352 | 6.0471 | 5.8681 | 6.0643 | 6.3705 | 6.9806 | 6.2346 | 7.4247 | 6.3058 | 6.6270 | 7.2845 | 4.9282 | 5.1136 | 4.1219 | 5.4585 |
| DST_T7421 | DST | Q03001 | 7510;7513 | | NA | 6.1740 | 6.1226 | 5.9963 | 6.8903 | 6.4801 | 6.3412 | 5.7580 | 6.7378 | 6.3900 | 5.9344 | 6.7159 | 6.4534 | 5.7952 | 6.4198 | 6.0364 | 5.7548 |
| DTL_S444 | DTL | Q9NZJ0 | | 429 | VTSSQSTPAKAPR | 6.7558 | 6.7813 | 6.8696 | 6.0574 | 5.8427 | 6.7837 | 6.2258 | 6.0497 | 7.0502 | 6.6800 | 6.6636 | 6.8611 | 5.6307 | 5.7409 | 4.7640 | 5.2436 |
| DTL_S444;T464 | DTL | Q9NZJ0 | | 478 | SPAKARSPINRRG | 6.3164 | 6.3277 | 6.6418 | 5.6294 | 5.8092 | 6.2337 | 5.9229 | 5.6846 | 6.3213 | 6.1383 | 6.5506 | 6.5602 | 6.3620 | 6.6077 | 6.4864 | 6.4079 |
| DTL_S478 | DTL | Q9NZJ0 | | 444 | CNPSNSSPSSAAC | 5.4190 | 5.5520 | 6.5942 | 6.2521 | 5.9890 | 6.2720 | 5.9979 | 5.5303 | 6.1532 | 6.5665 | 7.1002 | 6.3291 | 6.7528 | 6.4984 | 6.5491 | 6.4440 |
| DTL_S485;S488 | DTL | Q9NZJ0 | | 557 | VKRRLDSSCLESV | 4.1735 | 3.9970 | 5.2806 | 4.3607 | 5.0548 | 5.2775 | 5.2570 | 5.2570 | 4.7102 | 4.5110 | 5.2715 | 5.0283 | 9.6315 | 9.8132 | 12.0392 | 10.3370 |
| DTL_S488 | DTL | Q9NZJ0 | | 617 | IEGAGTSISEPPS | 6.6260 | 5.8005 | 6.4615 | 6.3408 | 6.3647 | 7.1606 | 6.5562 | 6.2812 | 6.0215 | 6.9570 | 6.8108 | 5.7908 | 5.5892 | 5.8199 | 5.6106 | 5.8087 |
| DTL_S488;S490 | DTL | Q9NZJ0 | | 490 | GSVSSVSPKPPSS | 7.9739 | 9.0596 | 8.3682 | 7.3469 | 4.5701 | 6.3954 | 6.9625 | 6.3737 | 4.3201 | 4.4701 | 4.7630 | 4.6179 | 5.0811 | 4.5950 | 7.7687 | 7.3338 |
| DTL_S490 | DTL | Q9NZJ0 | | 488 | RRGSVSSVSPKPP | 8.1953 | 9.5660 | 8.4926 | 7.6306 | 4.3649 | 6.5546 | 6.9350 | 5.7559 | 4.0752 | 4.3059 | 5.1003 | 4.8198 | 4.6540 | 4.2895 | 7.9000 | 7.3602 |
| DTL_S557 | DTL | Q9NZJ0 | 488;490 | | NA | 7.8582 | 8.7228 | 8.6141 | 6.8705 | 4.9173 | 7.4975 | 7.8401 | 7.0470 | 5.2960 | 5.6354 | 6.6853 | 6.4682 | 3.3658 | 4.1189 | 4.0184 | 5.0447 |
| DTL_S617 | DTL | Q9NZJ0 | 485;488 | | NA | 4.9335 | 4.8925 | 5.5930 | 4.6828 | 5.4315 | 5.6601 | 4.8881 | 5.3652 | 6.8987 | 6.7280 | 6.7796 | 7.2282 | 7.8276 | 8.4028 | 6.9767 | 7.7117 |
| DTL_S617;Y628 | DTL | Q9NZJ0 | 444;464 | | NA | 6.2730 | 7.0037 | 7.8404 | 6.0567 | 7.0567 | 7.6607 | 6.3527 | 6.7919 | 6.5894 | 7.2501 | 6.7043 | 7.3220 | 3.6533 | 4.6759 | 4.0200 | 4.7493 |
| DTL_T429 | DTL | Q9NZJ0 | 617;628 | | NA | 5.3085 | 5.5321 | 6.2758 | 5.9874 | 5.5861 | 6.6064 | 5.2043 | 5.7143 | 6.7388 | 6.3425 | 5.8297 | 5.5836 | 7.4494 | 7.0914 | 7.9352 | 6.8143 |
| DTNA_S458 | DTNA | Q9Y4J8 | | 662 | FEDLVPSPTSEKA | 5.9427 | 6.1449 | 6.5463 | 6.6155 | 6.3885 | 6.0726 | 6.3903 | 6.0226 | 6.3460 | 6.4199 | 6.1791 | 6.1747 | 6.1801 | 6.1058 | 6.3877 | 6.0832 |
| DTNA_S662 | DTNA | Q9Y4J8 | | 458 | QPPQQRSAPDISF | 5.3438 | 5.0873 | 6.5358 | 5.5254 | 5.7142 | 7.5948 | 5.0731 | 5.2310 | 6.2627 | 4.6409 | 5.1182 | 6.8421 | 7.9236 | 7.7634 | 7.5318 | 7.8120 |
| DTNBP1_S316;S321 | DTNBP1 | Q96EV8 | 316;321 | | NA | 5.8064 | 5.6903 | 6.2642 | 5.6948 | 6.4698 | 5.9515 | 5.7021 | 5.3680 | 7.0270 | 6.2979 | 6.6453 | 6.7992 | 6.5805 | 7.2413 | 6.1213 | 6.3405 |
| DTX2_S236 | DTX2 | Q86UW9 | | 360 | RAPQPTSPPASRL | 6.4750 | 7.0906 | 6.3969 | 6.8762 | 6.1677 | 6.8204 | 6.5251 | 6.8183 | 6.5143 | 6.6192 | 7.0268 | 6.8034 | 4.6343 | 5.1839 | 4.8059 | 5.2420 |
| DTX2_S360 | DTX2 | Q86UW9 | | 236 | PPHRTASVFGTHQ | 8.8474 | 7.3971 | 8.3165 | 6.6202 | 6.9297 | 4.7784 | 5.9653 | 4.9803 | 6.8857 | 5.7654 | 5.8966 | 5.6496 | 5.3031 | 7.0568 | 4.8071 | 4.8009 |
| DTX3L_S202 | DTX3L | Q8TDB6 | | 539 | DDSKAASPPLKGS | 6.4509 | 7.1765 | 6.9876 | 6.8313 | 5.9656 | 6.7190 | 6.4849 | 6.7330 | 6.6600 | 7.0501 | 7.1577 | 7.3471 | 4.0162 | 4.9772 | 4.5176 | 4.9252 |
| DTX3L_S539 | DTX3L | Q8TDB6 | | 202 | EQKQQFSPSMTER | 6.5840 | 6.8661 | 6.6339 | 6.8637 | 6.1949 | 6.5144 | 6.4123 | 7.2206 | 6.6757 | 6.7477 | 6.6119 | 6.8866 | 4.7416 | 5.1650 | 5.0251 | 4.8564 |
| DUS2_S445 | DUS2 | Q9NX74 | | 445 | GRLGEESPSLHKR | 7.2006 | 7.1960 | 6.5131 | 7.4441 | 6.5785 | 7.6783 | 6.9339 | 7.6566 | 6.5900 | 5.9330 | 7.2217 | 7.3636 | 3.1488 | 4.7696 | 3.8252 | 3.9470 |
| DUS3L_S236 | DUS3L | Q96G46 | | 236 | QALRRFSQGPTPA | 5.9049 | 5.7310 | 6.3110 | 7.1293 | 6.0687 | 6.8910 | 7.4580 | 6.7493 | 7.8692 | 6.7825 | 7.4610 | 6.5955 | 3.9326 | 4.7448 | 5.0142 | 5.3570 |
| DUS3L_T273;S277 | DUS3L | Q96G46 | 273;277 | | NA | 5.3222 | 4.9766 | 6.9631 | 6.0306 | 4.8624 | 6.4659 | 5.3571 | 6.2032 | 6.3996 | 6.1298 | 5.9055 | 6.1235 | 7.1551 | 7.5130 | 7.8801 | 6.7123 |
| DUSP16_S446 | DUSP16 | Q9BY84 | | 446 | NKLCQFSPVQELS | 6.8418 | 6.0426 | 6.9825 | 6.1981 | 6.9195 | 6.3685 | 6.1065 | 6.7547 | 5.1711 | 4.9720 | 6.2111 | 6.5575 | 6.5380 | 5.8685 | 6.2222 | 6.2454 |
| DUSP16_S501 | DUSP16 | Q9BY84 | | 501 | AQRSLLSPLHRSG | 6.9349 | 6.3369 | 6.6862 | 6.1067 | 6.6614 | 6.6367 | 6.2771 | 7.4541 | 6.2166 | 5.8076 | 6.1928 | 6.3127 | 5.8242 | 5.5165 | 5.3502 | 5.6853 |
| DUSP22_S58 | DUSP22 | Q9NRW4-2 | | 58 | CIPAADSPSQNLT | 5.1932 | 6.1034 | 7.8974 | 5.3220 | 6.1151 | 7.9706 | 6.5512 | 6.9492 | 6.9565 | 7.1467 | 6.0083 | 7.2418 | 4.0036 | 6.3361 | 5.4361 | 4.7689 |
| DUSP7_S369 | DUSP7 | Q16829 | | 369 | RKKSNISPNFNFM | 5.3510 | 5.4329 | 6.5849 | 6.6314 | 6.4020 | 7.6196 | 6.4157 | 7.1337 | 6.8361 | 6.0253 | 5.8751 | 6.5849 | 7.2047 | 5.7714 | 5.6021 | 4.5292 |
| DUT_S243 | DUT | P33316 | | 243 | DDTERGSGGFGST | 6.1259 | 6.4821 | 6.8186 | 6.6174 | 6.3395 | 7.0527 | 6.5891 | 7.1650 | 6.6676 | 6.8961 | 7.2191 | 7.0764 | 4.4999 | 5.1644 | 4.4049 | 4.8813 |
| DUT_S88 | DUT | P33316 | | 88 | LPKAGGSPAPGPE | 6.4757 | 6.2570 | 5.8529 | 6.3907 | 6.5524 | 6.1206 | 5.9974 | 6.4436 | 6.2626 | 6.2818 | 6.2662 | 6.4055 | 6.5059 | 6.4762 | 5.6927 | 6.0190 |
| DUT_S99 | DUT | P33316 | | 99 | PETPAISPSKRAR | 6.5362 | 6.7155 | 6.1914 | 7.0119 | 6.9844 | 5.8100 | 6.4721 | 6.9556 | 6.6174 | 6.9275 | 6.6324 | 7.7257 | 4.3614 | 5.3939 | 4.6297 | 5.0347 |
| DVL1_S676;S679 | DVL1 | O14640 | 676;679 | | NA | 6.6299 | 6.7896 | 6.7064 | 6.8489 | 6.0035 | 6.1536 | 6.6398 | 5.8288 | 6.2709 | 6.0324 | 6.4172 | 6.4858 | 5.6746 | 6.1654 | 5.4784 | 5.8750 |
| DVL2_S720 | DVL2 | O14641 | | 720 | LTASRQSFHMAMG | 6.4682 | 6.4550 | 6.8359 | 6.8344 | 5.2666 | 6.0108 | 6.6462 | 5.9255 | 7.2771 | 7.4139 | 6.8241 | 6.0755 | 5.1460 | 5.4520 | 5.2490 | 6.1197 |
| DYNC1H1_S4368 | DYNC1H1 | Q14204 | | 4368 | KKTRTDSTSDGRP | 5.4547 | 5.2677 | 6.2272 | 5.4014 | 5.9389 | 5.9024 | 6.1245 | 6.2302 | 5.7538 | 5.4043 | 5.7984 | 5.9931 | 7.9504 | 7.5587 | 7.7273 | 7.2670 |
| DYNC1I1_S80 | DYNC1I1 | O14576-2 | | 80 | LVPTPMSPSSKSV | 5.8558 | 6.5028 | 7.0618 | 5.9862 | 6.7309 | 5.7486 | 6.0046 | 6.6736 | 6.8495 | 6.8259 | 6.3340 | 6.7136 | 5.2328 | 6.0725 | 5.3937 | 6.0136 |
| DYNC1I2_S73;S81 | DYNC1I2 | Q13409-2 | | 81 | IVPPPMSPSSKSV | 5.9160 | 6.0530 | 6.4949 | 6.2988 | 6.3471 | 6.6969 | 6.3859 | 6.6331 | 6.4208 | 6.3241 | 6.5711 | 6.6517 | 5.6342 | 6.0964 | 5.6642 | 5.8116 |
| DYNC1I2_T70;S81 | DYNC1I2 | Q13409-2 | | 89 | SSKSVSTPSEAGS | 6.0223 | 7.5959 | 6.1103 | 6.5189 | 5.9332 | 6.5576 | 6.6792 | 6.4298 | 5.8159 | 7.2669 | 6.6299 | 7.5027 | 4.4706 | 5.8724 | 4.5691 | 6.0254 |
| DYNC1I2_T81 | DYNC1I2 | Q13409-2 | 73;81 | | NA | 6.2430 | 6.4711 | 6.6674 | 6.4664 | 6.6885 | 6.4739 | 6.9706 | 6.8572 | 6.4332 | 6.4243 | 6.7321 | 6.3910 | 5.1559 | 5.6580 | 4.9790 | 5.3886 |
| DYNC1I2_T89 | DYNC1I2 | Q13409-2 | 70;81 | | NA | 6.0769 | 7.0166 | 6.0251 | 7.1467 | 5.7156 | 6.6778 | 6.6670 | 7.2357 | 6.5778 | 6.3409 | 6.3080 | 6.5338 | 5.1407 | 5.3141 | 5.5267 | 5.6966 |
| DYNC1LI1_S207 | DYNC1LI1 | Q9Y6G9 | | 207 | GEDFPASPQRRNT | 6.0079 | 6.4024 | 6.6439 | 6.3175 | 6.3699 | 6.2670 | 6.1990 | 6.6162 | 6.4243 | 6.3951 | 6.5910 | 6.8156 | 5.3250 | 6.1914 | 5.8089 | 5.6248 |
| DYNC1LI1_S398 | DYNC1LI1 | Q9Y6G9 | | 516 | SPTTPTSPTEGEA | 5.2124 | 5.3345 | 5.5705 | 5.9820 | 6.1756 | 6.0430 | 5.8075 | 6.4322 | 6.2518 | 5.8151 | 5.7487 | 5.9698 | 7.5714 | 7.2867 | 7.4475 | 7.3514 |
| DYNC1LI1_S405 | DYNC1LI1 | Q9Y6G9 | | 405 | PRVPGGSPRTPNR | 7.1097 | 7.4718 | 7.1732 | 6.7953 | 6.5057 | 7.1417 | 6.8466 | 7.5609 | 7.0020 | 7.0969 | 7.2556 | 6.8755 | 3.3822 | 4.4712 | 3.4371 | 3.8745 |
| DYNC1LI1_S405;T408 | DYNC1LI1 | Q9Y6G9 | | 421 | SNVASVSPIPAGS | 6.9750 | 7.1551 | 6.8826 | 7.1304 | 6.6319 | 7.1347 | 7.1654 | 7.0066 | 7.5202 | 7.3006 | 7.8163 | 7.3992 | 3.2930 | 3.8251 | 2.9792 | 3.7846 |
| DYNC1LI1_S414 | DYNC1LI1 | Q9Y6G9 | | 21 | PGLSSTYTGGPLG | 6.3342 | 5.4081 | 7.1791 | 5.5454 | 6.0059 | 5.6852 | 6.7735 | 5.6502 | 6.3903 | 6.9632 | 6.5576 | 6.8060 | 6.3167 | 6.3179 | 5.3332 | 6.7336 |
| DYNC1LI1_S421 | DYNC1LI1 | Q9Y6G9 | | 398 | GRPVDASPRVPGG | 7.7033 | 8.1429 | 6.9602 | 7.6950 | 7.1100 | 7.1187 | 7.3352 | 6.6183 | 8.1529 | 7.2014 | 7.7994 | 7.8608 | 2.1633 | 2.9770 | 2.0482 | 3.1135 |
| DYNC1LI1_S516 | DYNC1LI1 | Q9Y6G9 | | 414 | TPNRSVSSNVASV | 7.7948 | 6.5887 | 7.9361 | 6.8493 | 7.0365 | 6.1470 | 5.8632 | 6.2904 | 7.8980 | 6.8607 | 6.1330 | 6.3215 | 4.7029 | 5.5710 | 4.0924 | 3.9145 |
| DYNC1LI1_Y21 | DYNC1LI1 | Q9Y6G9 | 405;408 | | NA | 7.9812 | 8.1055 | 7.5042 | 7.3460 | 6.5656 | 7.9736 | 8.0553 | 6.5429 | 6.9914 | 7.0327 | 7.4819 | 8.1221 | 2.1909 | 3.0259 | 2.1165 | 2.9642 |
| DYNC1LI2_S194 | DYNC1LI2 | O43237 | | 194 | EEGCQGSPQRRGP | 5.9605 | 6.1729 | 6.8264 | 6.0679 | 6.1600 | 6.6945 | 6.3112 | 6.3462 | 6.4413 | 6.6540 | 6.7004 | 6.5367 | 5.8414 | 6.2383 | 5.3655 | 5.6828 |
| DYNC1LI2_S391 | DYNC1LI2 | O43237 | | 391 | ARGPSGSPRTQGR | 7.5056 | 6.9884 | 7.4837 | 7.3474 | 6.9805 | 6.3456 | 6.7039 | 7.5581 | 7.6858 | 7.1001 | 7.1041 | 7.3055 | 3.2406 | 3.9080 | 2.9827 | 3.7598 |
| DYNC1LI2_S407 | DYNC1LI2 | O43237 | | 407 | ASVPSSSPGTSVK | 6.8340 | 6.4859 | 7.0090 | 7.1907 | 6.7902 | 6.7505 | 7.1469 | 7.1489 | 8.1119 | 7.2633 | 7.2563 | 7.2238 | 3.4588 | 4.4469 | 2.9619 | 3.9209 |
| DYNC1LI2_T376 | DYNC1LI2 | O43237 | | 376 | LAKQPATPTRASE | 6.6256 | 7.4638 | 6.9485 | 8.3726 | 5.9767 | 7.6224 | 7.0498 | 6.5228 | 8.0570 | 5.9620 | 8.3227 | 8.1128 | 3.2034 | 3.6732 | 2.1596 | 3.9272 |
| DYNC2I1_S197 | DYNC2I1 | Q8WVS4 | | 434 | ERIGELSLKLFQK | 6.3002 | 5.9842 | 6.3591 | 5.3012 | 6.1776 | 6.1669 | 6.1901 | 6.0610 | 6.8029 | 7.4175 | 6.9523 | 6.8273 | 6.2401 | 6.4721 | 5.4851 | 5.2625 |
| DYNC2I1_S247 | DYNC2I1 | Q8WVS4 | | 197 | KLQYGDSKDNPLK | 5.4731 | 6.4132 | 6.1332 | 6.2135 | 5.1976 | 6.7453 | 6.2964 | 6.5698 | 5.6785 | 7.1974 | 6.8488 | 7.0378 | 4.9922 | 6.7551 | 5.9818 | 6.4663 |
| DYNC2I1_S434 | DYNC2I1 | Q8WVS4 | | 247 | SKEKSNSFSDKGE | 5.8952 | 6.7844 | 7.2646 | 5.6308 | 6.1105 | 6.1422 | 6.1167 | 6.3891 | 6.4845 | 7.1799 | 7.0119 | 6.8099 | 4.1465 | 6.6607 | 5.6408 | 5.7323 |
| DYRK1A_S97 | DYRK1A | Q13627 | | 321 | GQRIYQYIQSRFY | 6.3897 | 6.4004 | 6.1423 | 5.9554 | 6.1930 | 6.3306 | 6.1035 | 6.6778 | 6.1580 | 6.2274 | 6.0087 | 6.0776 | 6.2411 | 6.6629 | 6.4074 | 6.0241 |
| DYRK1A_Y321 | DYRK1A | Q13627 | | 97 | APLRKLSVDLIKT | 6.5815 | 4.5724 | 5.1130 | 3.8821 | 6.9650 | 5.7588 | 5.3654 | 6.3118 | 7.2619 | 9.9266 | 6.1683 | 5.3703 | 7.6862 | 9.3563 | 4.8384 | 4.8421 |
| DYRK2_S30 | DYRK2 | Q92630 | | 30 | VRQLQASPGLGAG | 5.9668 | 6.1205 | 6.3541 | 6.9020 | 6.7361 | 6.9416 | 6.6380 | 7.2315 | 6.7726 | 6.6836 | 5.9188 | 6.9652 | 5.0688 | 5.1738 | 5.2621 | 5.2644 |
| DYRK2_S48 | DYRK2 | Q92630 | | 48 | VGTGPPSPIALPP | 6.4672 | 6.7545 | 6.7679 | 7.1143 | 6.3060 | 6.5698 | 6.0472 | 6.5861 | 6.7407 | 6.2744 | 6.5833 | 6.6690 | 5.0531 | 5.5641 | 5.0784 | 5.4239 |
| DYRK2_T381 | DYRK2 | Q92630 | | 381 | EHQRVYTYIQSRF | 6.9447 | 7.3658 | 6.8267 | 6.9912 | 6.5780 | 6.8384 | 6.3794 | 6.9036 | 7.0916 | 6.2316 | 6.5292 | 6.7415 | 4.5639 | 4.5412 | 4.6133 | 4.8598 |
| DZIP3_S612 | DZIP3 | Q86Y13 | | 612 | NEEEELSPPLMEY | 6.6562 | 5.8695 | 6.1686 | 5.9025 | 6.3383 | 7.3384 | 6.7482 | 6.1240 | 5.4614 | 6.3169 | 5.5543 | 7.3697 | 5.8891 | 6.4758 | 6.4213 | 5.3658 |
| E2F1_S375 | E2F1 | Q01094 | | 375 | VDEDRLSPLVAAD | 6.1176 | 5.9473 | 5.9565 | 6.0261 | 6.0497 | 5.5387 | 5.7415 | 6.1854 | 6.3889 | 5.9848 | 6.3527 | 6.2221 | 7.2121 | 6.5732 | 6.8510 | 6.8525 |
| E2F3_S163 | E2F3 | O00716 | | 163 | GRAALRSPDSPKT | 7.3504 | 7.7055 | 7.9510 | 7.7682 | 6.2933 | 6.1660 | 6.4277 | 6.7382 | 7.1974 | 6.5345 | 7.4057 | 7.1829 | 3.9138 | 3.6864 | 3.7478 | 3.9311 |
| E2F5_T261 | E2F5 | Q15329 | | 261 | QSLTPVTPQKSSM | 5.9277 | 5.4848 | 6.1800 | 5.9768 | 6.1957 | 6.3172 | 5.9105 | 6.2973 | 6.8358 | 6.8543 | 6.8710 | 6.0515 | 6.5413 | 6.9104 | 5.6656 | 5.9801 |
| E2F7_S410 | E2F7 | Q96AV8 | | 825 | QPSLNLSPVMSRS | 6.8907 | 6.9605 | 7.9882 | 7.4231 | 6.7320 | 6.4109 | 7.0005 | 7.2768 | 6.5619 | 7.5943 | 6.7505 | 6.8847 | 3.6017 | 3.8589 | 3.8058 | 4.2594 |
| E2F7_S825 | E2F7 | Q96AV8 | | 410 | KLARHGSFNTVQA | 6.3848 | 6.0494 | 6.2952 | 5.6617 | 6.3910 | 5.5613 | 5.5721 | 6.4173 | 6.9282 | 7.2983 | 6.5775 | 6.5456 | 6.2103 | 6.5752 | 5.1028 | 6.4292 |
| E2F7_S856 | E2F7 | Q96AV8 | | 68 | PERNPITPVKFVD | 6.7184 | 7.2775 | 6.5770 | 6.2220 | 5.7671 | 6.4565 | 5.9349 | 5.5972 | 7.0158 | 7.0524 | 6.8355 | 7.2214 | 5.1802 | 5.5047 | 4.9785 | 5.6608 |

| | | | | | | | | | | | | | | | | | | | | | |
|---|---|---|---|---|---|---|---|---|---|---|---|---|---|---|---|---|---|---|---|---|---|
| E2F7_S95 | E2F7 | Q96AV8 | | 856 | VVKLHQSPVPVTP | 6.3066 | 6.4770 | 7.0029 | 6.3955 | 7.0282 | 7.0067 | 6.9499 | 6.9942 | 7.2143 | 6.7182 | 7.0539 | 7.2748 | 4.2386 | 4.9627 | 4.0389 | 4.3376 |
| E2F7_T68 | E2F7 | Q96AV8 | | 95 | MLISAASPDIRDR | 7.5321 | 7.9444 | 6.7443 | 6.0436 | 6.1267 | 6.7614 | 7.1070 | 7.0456 | 7.4685 | 7.2083 | 6.4722 | 6.8464 | 3.5821 | 4.4941 | 4.2595 | 4.3638 |
| E2F7_T82 | E2F7 | Q96AV8 | | 82 | QQAEPWTPTANLK | 7.2797 | 7.3731 | 6.7930 | 7.5174 | 6.8291 | 6.6346 | 6.5539 | 6.3933 | 7.7692 | 6.7541 | 6.8722 | 7.0158 | 3.6791 | 4.6907 | 3.2709 | 4.5739 |
| E2F8_S358 | E2F8 | A0AVK6 | | 71 | MLISAVSPEIRNR | 6.2234 | 6.4812 | 6.4087 | 6.2575 | 6.2271 | 6.5295 | 6.2748 | 6.3801 | 6.4312 | 6.7835 | 6.7806 | 6.7574 | 5.4161 | 5.9541 | 5.4966 | 5.5982 |
| E2F8_S413;S417 | E2F8 | A0AVK6 | | 417 | INSAPSSPIKTNK | 5.7151 | 6.0362 | 5.6043 | 6.0666 | 6.7387 | 6.7141 | 6.3125 | 6.2121 | 6.1339 | 5.9981 | 5.9144 | 6.2888 | 6.3903 | 6.4370 | 6.5627 | 6.8753 |
| E2F8_S417 | E2F8 | A0AVK6 | | 44 | DFGPLTTPTKPKE | 6.1304 | 6.8743 | 6.0916 | 6.8743 | 6.3500 | 6.4148 | 6.6405 | 6.4361 | 6.6275 | 6.6410 | 6.4910 | 6.3301 | 5.1402 | 6.2116 | 5.4173 | 5.3293 |
| E2F8_S71 | E2F8 | A0AVK6 | | 58 | SQGEPWTPTANLK | 6.6189 | 6.9371 | 6.8534 | 5.9940 | 6.5774 | 6.5760 | 6.7918 | 6.8133 | 7.1177 | 6.4794 | 7.0703 | 6.8749 | 4.7167 | 5.4623 | 4.2565 | 4.8603 |
| E2F8_T44 | E2F8 | A0AVK6 | | 358 | PNTSGSSPVIHFT | 6.4890 | 7.6552 | 5.8162 | 6.5306 | 6.9342 | 5.7580 | 5.9557 | 7.5290 | 6.0371 | 6.2083 | 7.3977 | 6.3727 | 5.4806 | 5.2862 | 5.3959 | 5.1533 |
| E2F8_T58 | E2F8 | A0AVK6 | 413;417 | | NA | 5.8353 | 7.0244 | 7.1934 | 5.4226 | 6.1499 | 4.4400 | 5.5683 | 4.2303 | 7.0746 | 6.7285 | 6.4977 | 6.4441 | 8.6474 | 4.2322 | 7.4861 | 7.0252 |
| EAF1_S165 | EAF1 | Q96JC9 | | 165 | PLKDNPSPEPQLD | 5.4759 | 6.4862 | 5.5876 | 5.9788 | 6.2250 | 6.5417 | 5.7029 | 6.2481 | 6.0148 | 6.0064 | 6.7539 | 6.3833 | 6.1476 | 7.6836 | 6.4329 | 6.3314 |
| EAF1_T69 | EAF1 | Q96JC9 | | 69 | PHIPGSTPPMTVF | 6.5880 | 6.7672 | 7.4043 | 6.7294 | 6.2317 | 6.8681 | 6.9325 | 6.5535 | 6.0578 | 7.0300 | 6.8057 | 7.6791 | 4.1412 | 4.9827 | 4.3204 | 4.9084 |
| EAF2_S154 | EAF2 | Q96CJ1 | | 154 | DKMSPASPIDDIE | 6.2643 | 6.5030 | 6.8747 | 6.1990 | 6.5293 | 6.2350 | 6.1040 | 6.3417 | 5.9105 | 5.9651 | 6.7505 | 6.6019 | 5.5475 | 6.0153 | 5.6469 | 6.5113 |
| EAPP_S109;S111 | EAPP | Q56P03 | | 89 | LGTGSSSGNGKVA | 6.9295 | 7.7730 | 6.9804 | 7.8461 | 5.7530 | 7.0943 | 7.1193 | 6.9488 | 7.6449 | 7.7290 | 8.3154 | 7.0304 | 3.0736 | 3.5511 | 2.6822 | 3.5290 |
| EAPP_S89 | EAPP | Q56P03 | 109;111 | | NA | 6.0873 | 6.0334 | 7.0273 | 6.2740 | 6.6015 | 6.2495 | 5.5258 | 5.8796 | 6.4183 | 6.5221 | 6.9724 | 6.8613 | 5.9738 | 6.8582 | 5.6111 | 5.1044 |
| EBAG9_S36 | EBAG9 | O00559-2 | | 36 | GRGRKLSGDQITL | 5.8741 | 6.1104 | 6.1688 | 6.6107 | 6.0899 | 6.0019 | 6.0115 | 6.2292 | 6.0825 | 6.3307 | 6.1246 | 6.2079 | 6.4166 | 6.6604 | 6.5410 | 6.5399 |
| EBAG9_S86 | EBAG9 | O00559-2 | | 86 | VATQQNSLEQLEP | 5.3788 | 6.3081 | 6.8467 | 8.3452 | 6.1038 | 7.5666 | 8.2533 | 6.0987 | 7.2858 | 6.9233 | 6.9105 | 6.8876 | 3.6097 | 5.0725 | 3.0787 | 5.3303 |
| EBAG9_T99 | EBAG9 | O00559-2 | | 99 | DYFKDMTPTIRKT | 7.9751 | 7.3584 | 6.5701 | 7.0544 | 6.3457 | 6.7655 | 7.4214 | 6.9777 | 7.1702 | 7.0010 | 6.8852 | 7.2087 | 3.9007 | 4.3601 | 3.0444 | 3.9615 |
| EBNA1BP2_S264 | EBNA1BP2 | Q99848 | | 270 | SYDDVSSFRAKTA | 6.0800 | 6.9992 | 7.7020 | 6.4604 | 5.0539 | 6.3472 | 6.6148 | 7.3134 | 6.8830 | 9.1329 | 7.4335 | 7.3705 | 3.8730 | 4.7839 | 3.4658 | 4.4866 |
| EBNA1BP2_S270 | EBNA1BP2 | Q99848 | | 264 | KWNTRESYDDVSS | 5.6536 | 6.1547 | 6.0793 | 6.9683 | 6.2826 | 5.9683 | 6.5085 | 6.2400 | 6.2518 | 6.6683 | 5.7014 | 5.9143 | 6.0540 | 6.8978 | 6.2592 | 6.3980 |
| ECE1_S34 | ECE1 | P42892 | | 36 | DLVDSLSEGDAYP | 6.0996 | 6.3599 | 6.0143 | 6.4366 | 5.9419 | 6.8135 | 6.1592 | 6.0262 | 6.1710 | 6.4332 | 6.5527 | 6.7465 | 5.7763 | 6.8736 | 5.8279 | 5.7678 |
| ECE1_S36 | ECE1 | P42892 | | 34 | EEDLVDSLSEGDA | 5.6013 | 6.0672 | 6.4353 | 6.5925 | 6.6871 | 5.7481 | 6.1995 | 6.5431 | 5.7590 | 6.3327 | 6.8575 | 5.5989 | 6.2061 | 6.7935 | 6.6391 | 5.9392 |
| ECE1_T25 | ECE1 | P42892 | | 25 | STYKRATLDEEDL | 5.5672 | 5.0488 | 7.0087 | 6.7850 | 8.0170 | 7.0513 | 7.1862 | 5.9897 | 7.5484 | 4.4914 | 6.8347 | 5.0950 | 5.6311 | 5.7447 | 6.5436 | 5.4571 |
| ECPAS_S1826 | ECPAS | Q5VYK3 | | 1826 | ATMEPDSRPELQE | 5.8905 | 6.2445 | 6.5364 | 6.3278 | 6.2490 | 6.4796 | 6.0705 | 6.0111 | 6.3246 | 6.0071 | 6.2104 | 6.1530 | 6.3474 | 6.5129 | 6.1512 | 6.4840 |
| ECPAS_S549 | ECPAS | Q5VYK3 | | 833 | PIPSEGSGFTKLH | 6.1981 | 5.9346 | 6.3226 | 5.7174 | 6.2365 | 5.8537 | 6.2588 | 6.1258 | 6.5955 | 6.2776 | 6.0240 | 6.4017 | 6.7568 | 6.5256 | 6.4811 | 6.2899 |
| ECPAS_S833 | ECPAS | Q5VYK3 | | 549 | GRNRKESTSEQMP | 6.4910 | 5.8978 | 7.4342 | 6.4530 | 6.8491 | 6.1085 | 6.0506 | 5.9828 | 7.4884 | 6.7596 | 7.3284 | 7.1249 | 4.5712 | 5.0098 | 5.3787 | 5.0722 |
| ECT2_S15 | ECT2 | Q9H8V3 | | 359 | LYEKANTPELKKS | 6.2383 | 6.4051 | 6.3009 | 5.7986 | 6.4723 | 6.1844 | 6.9377 | 6.2294 | 6.8473 | 6.3315 | 6.4943 | 6.5096 | 5.7626 | 5.7462 | 5.6994 | 6.0424 |
| ECT2_S367 | ECT2 | Q9H8V3 | | 373 | SMLSLNTPNSNRK | 6.6744 | 6.1819 | 6.5601 | 6.0810 | 5.4591 | 6.4701 | 6.7467 | 6.5012 | 6.5694 | 7.0392 | 6.9096 | 7.2924 | 4.8360 | 5.5534 | 5.1089 | 6.0163 |
| ECT2_S397 | ECT2 | Q9H8V3 | | 40 | LLIGSTSYVEEEM | 6.2756 | 6.3303 | 6.6152 | 6.9096 | 6.4020 | 6.2203 | 6.6733 | 6.2291 | 6.3529 | 6.4186 | 6.5201 | 6.5794 | 5.4231 | 5.9755 | 5.3836 | 5.6915 |
| ECT2_S40 | ECT2 | Q9H8V3 | | 716 | VIGTFRSPHGQTR | 6.8107 | 7.2899 | 6.8171 | 7.1481 | 6.3294 | 7.4401 | 7.3635 | 7.3864 | 7.4220 | 7.3434 | 7.6346 | 7.4583 | 3.1854 | 4.1118 | 2.8598 | 3.3994 |
| ECT2_S406 | ECT2 | Q9H8V3 | | 406 | PPRKRPSAEHSLS | 5.9402 | 6.1653 | 5.8501 | 6.5155 | 5.9453 | 5.9082 | 6.2953 | 6.7752 | 6.4806 | 5.5924 | 6.1454 | 6.4712 | 6.1420 | 6.4816 | 6.8116 | 6.4801 |
| ECT2_S716 | ECT2 | Q9H8V3 | | 397 | SRETDVSPFPPRK | 7.2140 | 7.3710 | 6.9658 | 7.1081 | 6.4092 | 7.2912 | 6.8438 | 6.7322 | 7.7282 | 6.8743 | 6.7710 | 7.4820 | 3.5911 | 4.4429 | 3.2739 | 3.9012 |
| ECT2_S861 | ECT2 | Q9H8V3 | | 367 | ELKKSVSMLSLNT | 6.6168 | 5.0904 | 5.7871 | 6.3709 | 5.9200 | 5.7846 | 6.2831 | 6.3062 | 6.5329 | 6.0211 | 6.0898 | 6.4862 | 6.5083 | 6.8434 | 5.4687 | 7.8905 |
| ECT2_S889 | ECT2 | Q9H8V3 | | 444 | KSSKSSTPVPSKQ | 7.6163 | 7.5586 | 6.9958 | 7.0724 | 6.8333 | 7.2591 | 8.0822 | 7.2663 | 7.7100 | 6.9251 | 7.5327 | 7.0394 | 2.6796 | 3.5298 | 2.4316 | 3.4680 |
| ECT2_T359 | ECT2 | Q9H8V3 | | 861 | LMTSHGSVEGRSP | 6.2360 | 7.2614 | 7.9070 | 6.4184 | 6.0920 | 5.8505 | 6.2410 | 6.5716 | 7.2920 | 8.1771 | 7.9426 | 7.2464 | 3.4383 | 4.7694 | 4.1103 | 4.4459 |
| ECT2_T373 | ECT2 | Q9H8V3 | | 889 | SLAGIPSPSLVSL | 6.5416 | 7.7268 | 7.6496 | 7.6375 | 4.6443 | 6.3847 | 7.9103 | 6.4113 | 6.6696 | 7.7534 | 6.4427 | 6.8820 | 4.0722 | 5.2116 | 3.7898 | 4.2726 |
| ECT2_T373;S376 | ECT2 | Q9H8V3 | | 15 | STTGRTSLADSSI | 5.9368 | 6.0942 | 7.1339 | 6.9710 | 6.5671 | 6.8333 | 7.5000 | 6.7229 | 7.3151 | 7.3426 | 7.5408 | 7.2646 | 4.0782 | 4.7956 | 3.5376 | 4.3662 |
| ECT2_T444 | ECT2 | Q9H8V3 | 373;376 | | NA | 6.5848 | 7.0373 | 7.3302 | 6.7689 | 6.2416 | 7.2206 | 7.2173 | 7.0135 | 6.9286 | 7.4312 | 7.6073 | 7.6878 | 3.4298 | 4.4561 | 3.2208 | 3.8243 |
| EDC3_S131 | EDC3 | Q96F86 | | 173 | RHPNQATPKKSGL | 7.1119 | 6.9207 | 7.4289 | 6.3841 | 5.5013 | 6.9245 | 6.3583 | 6.5110 | 8.3142 | 7.0962 | 7.6401 | 7.6431 | 3.9421 | 4.2336 | 3.6599 | 4.3300 |
| EDC3_S150 | EDC3 | Q96F86 | | 161 | FRRRHNSWSSSSR | 7.3167 | 7.4077 | 6.2032 | 6.5606 | 6.7364 | 7.3791 | 5.5886 | 8.0540 | 6.2487 | 6.0836 | 7.1838 | 7.3223 | 4.4409 | 4.9383 | 4.1039 | 4.4322 |
| EDC3_S161 | EDC3 | Q96F86 | | 131 | SQDVAVSPQQQQC | 6.0317 | 7.4322 | 7.9167 | 7.4315 | 7.0632 | 7.2515 | 5.7967 | 6.7567 | 6.2942 | 7.4556 | 7.4384 | 8.7214 | 3.7639 | 3.8539 | 3.3674 | 3.4251 |
| EDC3_T173 | EDC3 | Q96F86 | | 150 | RHMESLSQSKSFR | 7.1281 | 7.0419 | 7.7588 | 7.7697 | 5.5237 | 7.3695 | 7.3777 | 6.1753 | 8.2570 | 6.9131 | 8.1146 | 8.3868 | 2.5037 | 3.5019 | 2.9139 | 3.2642 |
| EDC4_S33 | EDC4 | Q6P2E9 | | 729 | SPSRTRSPDVISS | 5.7814 | 5.9576 | 6.5760 | 6.0896 | 6.1073 | 6.3382 | 6.0700 | 6.3800 | 6.2638 | 6.4293 | 6.4939 | 6.5377 | 6.2020 | 6.5335 | 6.0028 | 6.2369 |
| EDC4_S676 | EDC4 | Q6P2E9 | | 780 | SALHLLSPRPRPG | 5.4219 | 5.7116 | 5.9447 | 5.5908 | 6.0392 | 6.8712 | 6.2597 | 6.7363 | 6.5278 | 5.7856 | 6.0314 | 5.7276 | 6.7165 | 7.0734 | 6.6543 | 6.9082 |
| EDC4_S723 | EDC4 | Q6P2E9 | | 693 | APADKLTPKGPGQ | 6.7539 | 6.8293 | 6.7445 | 7.1465 | 6.6194 | 7.2324 | 7.6670 | 6.6519 | 7.7673 | 7.0786 | 7.2888 | 6.9963 | 3.5542 | 4.4002 | 3.1792 | 4.0904 |
| EDC4_S729 | EDC4 | Q6P2E9 | | 723 | LGLPQASPSRTRS | 5.5682 | 6.0375 | 6.8390 | 5.9122 | 6.7387 | 6.5965 | 6.1054 | 6.4984 | 6.3633 | 6.0339 | 6.6801 | 5.9699 | 6.1762 | 6.3229 | 6.2300 | 5.9279 |
| EDC4_S780 | EDC4 | Q6P2E9 | | 33 | GGPSAESPRPSSA | 5.8966 | 6.2883 | 6.3937 | 6.4952 | 6.2958 | 7.0593 | 6.2810 | 6.5372 | 6.4847 | 6.6726 | 7.0455 | 6.2735 | 5.2256 | 6.5584 | 5.3979 | 5.0949 |
| EDC4_S844 | EDC4 | Q6P2E9 | | 844 | CSTLAESPRNGLQ | 7.0840 | 7.2808 | 7.0203 | 6.7335 | 6.3132 | 6.9589 | 6.5857 | 6.8601 | 7.0796 | 7.2379 | 7.3848 | 7.0537 | 3.8613 | 4.4722 | 3.7488 | 4.3252 |
| EDC4_S879 | EDC4 | Q6P2E9 | | 676 | SGSLQASPRGLLP | 6.8157 | 6.4552 | 6.2720 | 6.4081 | 6.4900 | 6.3020 | 6.4108 | 6.8633 | 6.0621 | 6.4429 | 7.1563 | 6.1285 | 5.3133 | 5.7744 | 5.1371 | 5.9683 |
| EDC4_S914 | EDC4 | Q6P2E9 | | 879 | DASAEQSDHDDEV | 5.8171 | 5.8162 | 6.7734 | 6.0310 | 6.4726 | 6.2488 | 5.8497 | 6.7615 | 6.2688 | 6.4899 | 6.3040 | 6.2247 | 6.1288 | 6.3845 | 6.2840 | 6.1450 |
| EDC4_T693 | EDC4 | Q6P2E9 | | 914 | DWKTKGSPRTSPK | 5.7781 | 6.0748 | 5.9364 | 6.4459 | 5.7220 | 5.8499 | 5.7475 | 5.6673 | 5.8892 | 5.4896 | 5.8184 | 6.6447 | 7.1074 | 7.0830 | 7.4743 | 7.2716 |
| EEA1_S52 | EEA1 | Q15075 | | 52 | CMKSLGSADELFK | 3.4083 | 3.4661 | 3.2245 | 3.7663 | 3.5776 | 3.3410 | 3.7799 | 3.6843 | 3.5326 | 3.2183 | 2.8372 | 3.1820 | 14.7528 | 13.7447 | 15.3101 | 15.1743 |
| EED_S34 | EED | O75530-2 | | 34 | NSNPDLSGDENDD | 5.3429 | 6.3749 | 6.1273 | 6.6889 | 6.3270 | 6.1598 | 6.2670 | 6.3467 | 6.8405 | 6.7495 | 7.0705 | 7.2003 | 5.7514 | 5.4318 | 5.3140 | 6.0074 |
| EED_T179 | EED | O75530-2 | | 179 | RIINPITMQCIKH | 5.7943 | 5.8552 | 5.7398 | 7.0597 | 6.5942 | 5.2769 | 6.2623 | 7.6315 | 6.9560 | 5.7904 | 6.0263 | 6.5709 | 5.9434 | 6.1949 | 6.4944 | 5.8098 |
| EEF1A1P5_T239 | EEF1A1P5 | Q5VTE0 | | 239 | DCILPPTRPTDKP | 6.9807 | 6.2762 | 6.6368 | 6.6257 | 6.1379 | 6.0790 | 6.1379 | 6.4581 | 6.4159 | 6.2173 | 6.5626 | 6.0247 | 5.8442 | 6.0980 | 5.5796 | 5.9254 |
| EEF1B2_S106 | EEF1B2 | P24534 | | 106 | DIDLFGSDDEEES | 6.1288 | 5.7114 | 6.7098 | 5.6141 | 6.4065 | 5.9123 | 5.7504 | 6.0871 | 6.6538 | 6.6906 | 6.9794 | 7.0797 | 6.7295 | 6.1155 | 5.6150 | 5.8160 |
| EEF1B2_S128 | EEF1B2 | P24534 | | 141 | ALVAKSSILLDVK | 6.6392 | 6.4218 | 7.6701 | 3.6185 | 5.6715 | 5.5242 | 4.6564 | 6.4095 | 5.6364 | 8.7325 | 6.8864 | 6.8654 | 6.8391 | 5.4190 | 6.5621 | 6.4481 |
| EEF1B2_S141 | EEF1B2 | P24534 | | 128 | RLAQYESKKAKKP | 5.2637 | 6.0867 | 7.2681 | 5.0227 | 7.1271 | 6.7555 | 5.6519 | 6.0179 | 6.4992 | 7.3611 | 7.0632 | 6.6729 | 6.3846 | 6.4388 | 4.9199 | 5.4665 |
| EEF1B2_S43 | EEF1B2 | P24534 | | 95 | GSGATDSKDDDDI | 6.6251 | 6.9758 | 6.5018 | 6.2812 | 6.8906 | 6.0218 | 5.1293 | 6.1184 | 5.6809 | 6.9066 | 5.8163 | 6.6578 | 6.0796 | 5.9183 | 6.3580 | 6.0385 |
| EEF1B2_S95 | EEF1B2 | P24534 | | 43 | VFEAVSSPPPADL | 7.0621 | 6.0845 | 6.3305 | 6.4503 | 5.3157 | 7.4445 | 5.5678 | 6.7108 | 6.5496 | 6.2809 | 7.4982 | 6.9443 | 6.5806 | 5.8531 | 4.4084 | 4.9189 |
| EEF1B2_T95;S106 | EEF1B2 | P24534 | 95;106 | | NA | 6.3430 | 5.5571 | 6.9464 | 5.3642 | 6.0516 | 6.0363 | 6.3372 | 6.2785 | 6.3746 | 6.8145 | 7.1523 | 6.3642 | 7.4174 | 6.1617 | 5.0201 | 5.7809 |
| EEF1D_S162 | EEF1D | P29692-2 | | 499 | PQTQHVSPMRQVE | 6.4510 | 6.8280 | 6.7641 | 6.4103 | 6.5411 | 6.8177 | 6.0328 | 6.9234 | 6.7178 | 6.7063 | 6.9145 | 6.9507 | 5.1755 | 5.1748 | 4.6962 | 4.8958 |
| EEF1D_S499 | EEF1D | P29692-2 | | 528 | DIDLFGSDNEEED | 5.9993 | 5.6445 | 6.7445 | 5.7856 | 6.2482 | 6.3468 | 5.8235 | 6.2676 | 6.6344 | 6.4670 | 6.8002 | 6.9527 | 6.6019 | 6.1703 | 5.6305 | 5.8831 |
| EEF1D_S563 | EEF1D | P29692-2 | | 563 | ALVAKSSILLDVK | 7.7572 | 6.8380 | 7.4567 | 3.4840 | 5.0265 | 4.9144 | 6.2506 | 6.6496 | 4.6229 | 7.3401 | 7.0801 | 5.4031 | 7.1115 | 5.1251 | 7.7706 | 7.1698 |
| EEF1D_T513;S528 | EEF1D | E9PL12 | | 162 | DIDLFGSDNEEED | 6.1307 | 5.6730 | 7.1323 | 5.8950 | 6.4884 | 6.7823 | 5.4241 | 6.1631 | 6.4681 | 6.2845 | 6.6988 | 6.6626 | 6.2131 | 6.1280 | 5.7082 | 6.1477 |
| EEF1D_T528 | EEF1D | P29692-2 | 513;528 | | NA | 6.5254 | 6.3461 | 7.1551 | 6.3552 | 6.1431 | 6.3147 | 5.9043 | 6.1419 | 6.9441 | 7.0303 | 7.0202 | 7.5657 | 5.3181 | 5.4095 | 4.6950 | 5.1314 |
| EEF1G_S437 | EEF1G | P26641-2 | | 96 | FGQTNRTPEFLRK | 6.3447 | 6.9272 | 6.8490 | 6.7121 | 6.5325 | 6.9205 | 6.0315 | 6.6995 | 6.4082 | 6.4685 | 7.0178 | 6.4680 | 5.0055 | 5.5463 | 5.0967 | 4.9720 |
| EEF1G_T96 | EEF1G | P26641-2 | | 437 | ELAFPLSPDWQVD | 6.5270 | 6.9132 | 5.5731 | 6.0594 | 6.4519 | 6.2668 | 6.3017 | 5.9360 | 6.3821 | 6.5100 | 6.6691 | 6.0085 | 5.9718 | 6.7835 | 5.9691 | 5.6768 |
| EEF2_S279 | EEF2 | P13639 | | 435 | IMGPNYTPGKKED | 6.5388 | 6.6065 | 7.0634 | 6.4089 | 6.1357 | 6.3673 | 6.0694 | 6.5478 | 6.5670 | 6.8316 | 6.6828 | 6.6924 | 5.3304 | 5.3836 | 5.3832 | 5.3912 |
| EEF2_S33 | EEF2 | P13639 | | 57 | AGETRFTDTRKDE | 6.6693 | 6.4408 | 7.4755 | 6.2562 | 7.3198 | 7.0192 | 5.8438 | 5.6189 | 6.0579 | 5.1173 | 6.4673 | 6.6536 | 5.7472 | 7.0745 | 4.6908 | 5.5480 |
| EEF2_S422 | EEF2 | P13639 | | 500 | MRVMKFSVSPVVR | 4.7498 | 6.1173 | 6.3659 | 5.0171 | 5.2275 | 6.2756 | 5.6181 | 6.9144 | 6.1089 | 6.9291 | 7.3459 | 7.0448 | 6.3953 | 6.3603 | 6.6908 | 6.8392 |
| EEF2_S500 | EEF2 | P13639 | | 450 | LKPIQRTILMMGR | 6.3144 | 6.7057 | 5.8249 | 6.4586 | 6.0348 | 5.9086 | 6.1582 | 6.3018 | 6.2422 | 5.8527 | 5.9949 | 6.2554 | 6.5350 | 6.2600 | 6.3710 | 6.7818 |
| EEF2_S502 | EEF2 | P13639 | | 422 | VFSGLVSTGLKVR | 5.4122 | 7.1944 | 6.9025 | 5.7425 | 5.4122 | 6.6260 | 6.1228 | 5.2278 | 6.5760 | 7.4287 | 5.7271 | 9.0766 | 6.5453 | 6.2111 | 4.3750 | 5.4198 |
| EEF2_T435 | EEF2 | P13639 | | 502 | VMKFSVSPVVRVA | 5.0308 | 5.2541 | 6.1662 | 5.7243 | 6.8977 | 6.4627 | 6.5758 | 6.9197 | 5.6314 | 5.3948 | 5.3787 | 5.5662 | 7.4649 | 7.2318 | 6.9439 | 7.3570 |
| EEF2_T450 | EEF2 | P13639 | | 279 | FSKSATSPEGKKL | 5.9969 | 6.8372 | 6.9157 | 7.0213 | 5.7182 | 6.2847 | 6.2915 | 6.8999 | 6.7332 | 6.4497 | 6.9222 | 7.0936 | 4.8212 | 5.4846 | 4.9523 | 5.5778 |
| EEF2_T57 | EEF2 | P13639 | | 33 | HVDHGKSTLTDSL | 5.5678 | 5.8010 | 6.4281 | 5.6663 | 5.5236 | 5.8754 | 5.9839 | 6.2392 | 6.7638 | 7.0734 | 7.0673 | 6.6251 | 6.7397 | 6.0905 | 5.5518 | 7.0030 |
| EEF2K_S18 | EEF2K | O00418 | | 18 | GVDGGQSPRAGHD | 6.8184 | 7.0851 | 7.0942 | 6.3972 | 6.1968 | 6.9014 | 6.4642 | 7.0829 | 7.0146 | 7.1790 | 7.2171 | 6.9457 | 4.0168 | 4.9407 | 4.1882 | 4.4578 |
| EEF2K_S71;S74 | EEF2K | O00418 | | 74 | RYSSSGSPANSFH | 7.1413 | 8.8805 | 5.4198 | 5.9805 | 7.6817 | 8.2451 | 6.8385 | 9.2267 | 6.1842 | 6.3539 | 6.9716 | 7.1427 | 3.0928 | 3.9834 | 3.5177 | 3.3399 |
| EEF2K_S71;S74;S78 | EEF2K | O00418 | 71;74 | | NA | 7.1082 | 7.9469 | 8.2127 | 7.3821 | 6.5865 | 8.6468 | 6.2222 | 9.6796 | 6.5665 | 6.8276 | 7.7224 | 7.1206 | 1.8807 | 2.7741 | 2.6063 | 2.7170 |
| EEF2K_S74 | EEF2K | O00418 | 71;74;78 | | NA | 7.8646 | 8.4499 | 9.8403 | 9.8853 | 5.6100 | 9.1510 | 7.5361 | 7.9584 | 5.8686 | 5.7230 | 7.4946 | 6.2812 | 1.4123 | 2.5472 | 2.1138 | 2.2636 |
| EEFSEC_S537 | EEFSEC | P57772 | | 537 | HIPGGLSPESKKI | 6.4318 | 6.5390 | 6.4051 | 6.8710 | 6.3033 | 6.9485 | 6.5686 | 6.2823 | 6.9279 | 6.9971 | 6.9572 | 6.7919 | 4.1745 | 5.8241 | 4.8592 | 5.1184 |
| EEPD1_S102 | EEPD1 | Q7L9B9 | | 102 | FEICVSSKGSSAQ | 6.7675 | 6.7635 | 6.6958 | 6.8486 | 6.6319 | 6.3095 | 6.1005 | 5.8011 | 6.9183 | 6.3480 | 6.3101 | 6.6664 | 5.4755 | 4.9432 | 5.9083 | 5.5118 |
| EEPD1_S173 | EEPD1 | Q7L9B9 | | 25 | SHSRKFSAACNFS | 5.8385 | 4.3807 | 6.8159 | 6.1061 | 6.9357 | 5.9021 | 5.4304 | 5.3314 | 7.2909 | 8.8029 | 7.6087 | 7.7967 | 5.4739 | 6.4962 | 4.7433 | 5.0464 |
| EEPD1_S200;S205 | EEPD1 | Q7L9B9 | | 173 | EHGPFRSVEDLVR | 5.9622 | 5.9450 | 5.0193 | 5.2513 | 6.6508 | 5.8967 | 6.5923 | 6.2774 | 5.5909 | 5.5312 | 5.2063 | 5.0763 | 7.8603 | 7.5874 | 8.0971 | 7.4554 |
| EEPD1_S200;T205;T207 | EEPD1 | Q7L9B9 | 25;31 | | NA | 4.3586 | 4.2042 | 4.8034 | 4.8867 | 5.6651 | 4.6516 | 4.8525 | 4.9150 | 5.6008 | 5.8539 | 5.9230 | 5.9513 | 9.6058 | 9.0812 | 10.5028 | 9.1441 |
| EEPD1_S25 | EEPD1 | Q7L9B9 | 200;205 | | NA | 5.5308 | 5.4977 | 5.3385 | 6.3364 | 6.3064 | 8.3943 | 6.3078 | 6.8648 | 6.7450 | 7.9103 | 6.8330 | 6.5761 | 4.6563 | 5.7602 | 5.7995 | 5.1428 |
| EEPD1_S25;S31 | EEPD1 | Q7L9B9 | 200;205;207 | | NA | 7.6785 | 7.5950 | 6.6077 | 7.7811 | 6.9299 | 5.3481 | 6.3846 | 6.8272 | 6.4389 | 4.8495 | 5.4466 | 5.6448 | 5.9646 | 5.7859 | 5.2980 | 5.4196 |
| EFHD2_S74 | EFHD2 | Q96C19 | | 74 | GIGEPQSPSRRVF | 7.0498 | 6.6944 | 7.4067 | 6.6218 | 6.6109 | 7.1044 | 6.8444 | 7.3572 | 7.2759 | 7.0163 | 7.3136 | 7.1772 | 3.6778 | 4.4275 | 3.4254 | 3.9968 |
| EFNB1_S281 | EFNB1 | P98172 | | 281 | QRAAALSLSTLAS | 6.7220 | 6.4671 | 7.3821 | 6.6482 | 6.7077 | 6.1453 | 6.6777 | 7.0976 | 6.3563 | 6.5921 | 6.0993 | 5.9473 | 5.7544 | 5.8860 | 4.9367 | 4.5802 |
| EFR3A_S220 | EFR3A | Q14156 | | 694 | TDEDRLSRRKSIV | 6.2601 | 6.2977 | 6.3789 | 6.6548 | 6.1359 | 6.3286 | 6.3109 | 6.1214 | 6.1407 | 6.1138 | 5.8702 | 6.1463 | 6.3655 | 6.3301 | 6.2625 | 6.2826 |
| EFR3A_S694 | EFR3A | Q14156 | | 220 | RIGPPSSPSATDK | 5.8759 | 6.3630 | 6.4335 | 7.3545 | 7.1457 | 7.8190 | 6.7541 | 7.3844 | 7.1045 | 6.2738 | 7.4804 | 6.7321 | 4.1637 | 5.1175 | 3.9329 | 4.0650 |
| EFR3B_S216 | EFR3B | Q9Y2G0 | | 689 | TDEDRLSKRRSIG | 6.0868 | 6.4345 | 6.1751 | 6.6209 | 6.1446 | 6.2149 | 6.3274 | 5.8335 | 6.2240 | 6.2386 | 6.1661 | 5.9990 | 6.0335 | 6.6991 | 6.4476 | 6.3546 |
| EFR3B_S689 | EFR3B | Q9Y2G0 | | 216 | AESRSPSPLQAPE | 5.8892 | 6.3729 | 5.9784 | 6.2550 | 5.7844 | 6.4038 | 6.5338 | 6.7210 | 6.7389 | 7.6489 | 7.4912 | 7.8436 | 4.7296 | 5.0588 | 4.8812 | 5.6695 |
| EFR3B_S768 | EFR3B | Q9Y2G0 | | 768 | HCGARASLLQSKL | 7.5381 | 6.3439 | 6.5222 | 7.1586 | 6.2987 | 6.3105 | 6.3765 | 6.3665 | 7.2316 | 6.4265 | 7.1921 | 7.4073 | 4.6185 | 5.3584 | 4.1903 | 4.6602 |
| EFTUD2_S948 | EFTUD2 | Q15029 | | 79 | YGPEVETIVQEED | 5.5398 | 5.7887 | 6.1709 | 6.5911 | 6.5750 | 6.3689 | 6.0699 | 5.9172 | 6.2347 | 5.6448 | 6.1079 | 5.8953 | 5.9875 | 7.3516 | 7.9536 | 5.8032 |
| EFTUD2_T79 | EFTUD2 | Q15029 | | 948 | GLSEDVSISKFFD | 5.4258 | 6.0239 | 6.5296 | 7.6076 | 5.9360 | 6.2503 | 5.9721 | 5.6995 | 5.9146 | 6.1016 | 6.8326 | 5.1994 | 5.8910 | 7.4059 | 6.9475 | 6.2627 |
| EGFR_S1026 | EGFR | P00533 | | 1026 | QQGFFSSPSTSRT | 5.6545 | 6.3347 | 6.4039 | 6.3804 | 6.0285 | 6.5154 | 6.2695 | 6.6205 | 6.2031 | 6.3836 | 6.6609 | 6.4879 | 5.5993 | 6.3095 | 5.9616 | 6.1866 |
| EGFR_S1064 | EGFR | P00533 | | 693 | ELVEPLTPSGEAP | 6.1426 | 6.4887 | 6.2028 | 6.5453 | 6.2902 | 6.2995 | 6.5520 | 6.5081 | 6.0490 | 6.0093 | 6.2291 | 6.3128 | 5.9091 | 6.2254 | 5.9524 | 6.2839 |
| EGFR_S1070 | EGFR | P00533 | | 991 | ERMHLPSPTDSNF | 5.9846 | 6.1760 | 6.4275 | 6.5520 | 6.0015 | 6.7525 | 6.7114 | 6.6146 | 6.1755 | 6.4726 | 6.5562 | 6.6934 | 5.4973 | 5.7437 | 5.5360 | 6.1053 |
| EGFR_S1071 | EGFR | P00533 | | 1070 | SFLQRYSSDPTGA | 6.3968 | 6.7990 | 6.5440 | 7.2049 | 6.1079 | 6.0761 | 6.3668 | 6.4071 | 6.3988 | 6.6568 | 6.6354 | 6.7180 | 5.2989 | 5.9682 | 4.9005 | 5.5209 |
| EGFR_S991 | EGFR | P00533 | | 1071 | FLQRYSSDPTGAL | 5.8699 | 6.2250 | 6.2738 | 7.0649 | 5.9806 | 6.2152 | 6.2461 | 6.6498 | 6.7517 | 6.1681 | 6.6067 | 6.7474 | 5.7496 | 5.8683 | 5.8565 | 5.7264 |
| EGFR_T693 | EGFR | P00533 | | 1064 | CPIKEDSFLQRYS | 6.2670 | 6.2682 | 6.0191 | 6.5399 | 5.9917 | 5.3584 | 6.5906 | 5.7430 | 5.9760 | 5.4482 | 5.3514 | 5.4849 | 7.2476 | 6.9773 | 7.0647 | 7.6721 |
| EGFR_Y1172 | EGFR | P00533 | | 1172 | SLDNPDYQQDFFP | 6.0442 | 6.8640 | 6.6179 | 9.0295 | 11.9860 | 5.4449 | 5.0379 | 5.6707 | 6.3999 | 5.6484 | 5.9949 | 4.8948 | 5.5460 | 5.5860 | 5.3563 | 3.8787 |
| EGLN1_S125 | EGLN1 | Q9GZT9 | | 125 | DPAAAASPCRAAA | 6.0931 | 6.4367 | 7.0863 | 6.3304 | 6.1571 | 6.8701 | 6.6792 | 6.7187 | 6.7008 | 7.0860 | 6.9073 | 7.1965 | 4.6522 | 5.3416 | 4.8086 | 4.9354 |
| EHBP1_S171 | EHBP1 | Q8NDI1 | | 335 | FVTIKDSPPQSTK | 5.6976 | 5.9521 | 6.2082 | 6.0647 | 6.4636 | 6.5878 | 6.0272 | 6.5544 | 6.2075 | 6.5074 | 6.9300 | 6.5075 | 6.0353 | 6.3652 | 6.1024 | 5.7891 |
| EHBP1_S171;S174 | EHBP1 | Q8NDI1 | | 408 | PAPPVLSPKTGVL | 7.0131 | 7.2909 | 6.9892 | 7.1104 | 6.3822 | 6.7546 | 7.7815 | 6.9856 | 7.7300 | 7.2421 | 7.6845 | 7.5828 | 2.9275 | 4.0099 | 2.9093 | 3.6062 |
| EHBP1_S295 | EHBP1 | Q8NDI1 | | 171 | TDEDMQSLASLMS | 4.8084 | 5.6331 | 6.4379 | 5.6495 | 6.0263 | 7.0964 | 6.2350 | 7.4880 | 6.5321 | 7.4525 | 7.1132 | 6.0176 | 5.9399 | 6.2249 | 5.7704 | 5.5748 |
| EHBP1_S335 | EHBP1 | Q8NDI1 | | 378 | FYEPKSTPPPNNL | 6.6544 | 6.1964 | 7.0180 | 6.7126 | 5.9039 | 6.7104 | 7.2931 | 6.0898 | 7.2787 | 6.6083 | 7.1767 | 6.5290 | 4.7006 | 5.5086 | 4.7149 | 4.9047 |
| EHBP1_S408 | EHBP1 | Q8NDI1 | | 751 | CRSDPESPIKKTS | 6.1092 | 5.9360 | 6.3712 | 5.9748 | 5.9268 | 6.2145 | 6.3017 | 6.4543 | 6.8317 | 6.1384 | 6.2534 | 6.7213 | 6.1704 | 6.1928 | 6.3141 | 6.0894 |
| EHBP1_S428;S436 | EHBP1 | Q8NDI1 | | 295 | PITETASPRKTED | 6.2712 | 5.6790 | 6.0533 | 5.9971 | 6.7969 | 7.2872 | 6.2384 | 5.8744 | 7.1610 | 7.7506 | 6.8808 | 6.6206 | 5.2674 | 5.3383 | 5.1092 | 5.6746 |
| EHBP1_S751 | EHBP1 | Q8NDI1 | | 759 | IKKTSLSPTSKLG | 6.8882 | 6.5817 | 6.9685 | 6.4301 | 6.5824 | 7.3210 | 6.7791 | 6.8140 | 7.2780 | 6.6463 | 7.0656 | 6.9122 | 4.4071 | 4.6976 | 4.1576 | 4.4707 |
| EHBP1_S759 | EHBP1 | Q8NDI1 | 171;174 | | NA | 5.8498 | 7.2082 | 6.7588 | 6.2618 | 5.6371 | 6.7881 | 6.3365 | 7.0297 | 5.8935 | 6.1305 | 6.7165 | 5.6838 | 5.9948 | 5.8205 | 6.2098 | 5.6807 |
| EHBP1_T378 | EHBP1 | Q8NDI1 | 428;436 | | NA | 5.9792 | 5.2661 | 6.0927 | 5.5211 | 6.9039 | 5.9303 | 6.0620 | 6.4741 | 5.1068 | 5.7071 | 6.2119 | 5.9058 | 7.0320 | 7.7242 | 6.9641 | 7.1188 |
| EHBP1L1_S1168 | EHBP1L1 | Q8N3D4 | | 1257 | VRLRRPSVNGEPG | 5.9162 | 5.9805 | 6.3409 | 6.2117 | 6.5918 | 6.0309 | 5.7961 | 6.3593 | 7.2840 | 6.8529 | 6.7544 | 6.7959 | 5.8198 | 6.2387 | 5.3908 | 5.6360 |
| EHBP1L1_S1257 | EHBP1L1 | Q8N3D4 | | 191 | LDDFAESDEDEAH | 5.5076 | 6.3057 | 6.0732 | 6.2333 | 6.5043 | 6.8329 | 6.5730 | 6.9127 | 5.9445 | 6.8484 | 6.8593 | 7.1256 | 5.5294 | 5.7558 | 5.3374 | 5.6571 |
| EHBP1L1_S1273 | EHBP1L1 | Q8N3D4 | | 170 | TDDDMQSLASLMS | 6.2906 | 6.3139 | 6.8745 | 6.3401 | 5.9749 | 6.1332 | 6.1733 | 6.5871 | 7.0982 | 6.5717 | 5.9574 | 6.4822 | 5.6709 | 5.8886 | 5.9053 | 5.7381 |
| EHBP1L1_S170 | EHBP1L1 | Q8N3D4 | | 1273 | PPRAHGSFSHVRD | 6.6490 | 6.0768 | 6.4747 | 6.2882 | 6.7880 | 5.8444 | 5.5454 | 6.4512 | 6.9739 | 6.8606 | 6.5013 | 6.7139 | 6.0161 | 6.4438 | 5.2786 | 5.0941 |
| EHBP1L1_S170;S173 | EHBP1L1 | Q8N3D4 | | 285 | PRPPETSPEMRSS | 6.7208 | 6.7197 | 6.2739 | 6.7589 | 6.5127 | 6.8752 | 6.7110 | 6.6221 | 7.1286 | 6.8538 | 6.4480 | 6.8171 | 4.8366 | 5.0811 | 4.7436 | 4.8969 |
| EHBP1L1_S170;S191 | EHBP1L1 | Q8N3D4 | | 964 | EMKVLESPENKSG | 6.5920 | 7.0917 | 6.9162 | 7.3190 | 6.0706 | 6.5185 | 7.0293 | 7.1049 | 7.0173 | 7.2963 | 7.7304 | 7.9044 | 3.3411 | 4.1850 | 3.6521 | 4.2312 |
| EHBP1L1_S191 | EHBP1L1 | Q8N3D4 | | 1168 | VGSAQPSPPDDLD | 6.2623 | 5.7816 | 5.8780 | 5.9923 | 6.6466 | 6.0035 | 6.3307 | 6.1424 | 7.3593 | 6.4080 | 6.4102 | 6.8674 | 6.6444 | 5.9800 | 6.8965 | 4.3967 |
| EHBP1L1_S237 | EHBP1L1 | Q8N3D4 | | 237 | PQQAVASPSNAED | 5.7578 | 5.7410 | 6.7809 | 6.7298 | 6.4035 | 6.9323 | 7.7825 | 6.4576 | 7.3776 | 7.0345 | 6.9372 | 6.3200 | 4.6677 | 4.8899 | 4.5537 | 5.6339 |
| EHBP1L1_S285 | EHBP1L1 | Q8N3D4 | 170;191 | | NA | 6.1736 | 6.4463 | 6.5249 | 6.4345 | 6.6708 | 6.1815 | 5.8586 | 6.5867 | 5.8877 | 6.7483 | 7.9331 | 6.2965 | 4.8805 | 6.4478 | 5.5626 | 5.3667 |
| EHBP1L1_S964 | EHBP1L1 | Q8N3D4 | 170;173 | | NA | 7.1554 | 5.9758 | 6.8267 | 6.0864 | 5.3187 | 4.8968 | 5.0066 | 8.1821 | 7.9157 | 5.2830 | 6.0613 | 4.6795 | 6.1130 | 8.4100 | 5.4349 | 6.6540 |
| EHD1_S456 | EHD1 | Q9H4M9 | | 456 | EIFYTLSPVNGKI | 6.6745 | 7.1715 | 7.2351 | 7.5234 | 6.5913 | 7.0717 | 6.8448 | 6.8852 | 7.2728 | 7.4754 | 7.8672 | 7.5016 | 3.0605 | 3.9675 | 3.1060 | 3.7516 |
| EHD2_S438 | EHD2 | Q9NZN4 | | 438 | EDGEEGSDDEAEW | 5.8359 | 6.0631 | 6.2498 | 6.1691 | 6.4582 | 6.4569 | 5.7595 | 6.2396 | 6.5279 | 6.5427 | 6.3741 | 6.9046 | 6.1166 | 6.6188 | 5.7337 | 5.9495 |
| EHD2_S468 | EHD2 | Q9NZN4 | | 468 | PADGKLSGSKAKT | 6.3131 | 6.1721 | 6.5421 | 6.6171 | 6.3722 | 5.8038 | 6.4712 | 6.2165 | 6.3208 | 6.2204 | 6.6781 | 6.0966 | 5.5711 | 6.4210 | 5.7351 | 6.4487 |
| EHD4_S459 | EHD4 | Q9H223 | | 459 | ELFYTLSPINGKI | 6.7300 | 7.2223 | 7.2025 | 7.4626 | 6.2793 | 6.8944 | 7.0028 | 7.0918 | 7.1283 | 7.4075 | 7.6063 | 7.4881 | 3.3223 | 4.2080 | 3.1596 | 3.7943 |
| EHMT1_S1048 | EHMT1 | Q9H9B1 | | 1048 | SQNCVTSPMNIDR | 6.4609 | 6.1527 | 5.6049 | 6.9871 | 5.8692 | 6.3850 | 6.9817 | 6.3584 | 6.0447 | 6.4994 | 6.6533 | 5.9954 | 5.7819 | 6.4077 | 5.7040 | 6.1138 |
| EHMT1_S435 | EHMT1 | Q9H9B1 | | 435 | RKGKTDSPWIKPA | 7.4715 | 7.4350 | 6.7352 | 6.6060 | 5.7853 | 6.2726 | 6.8881 | 5.9058 | 6.9265 | 6.9142 | 6.3703 | 6.9698 | 4.8709 | 4.9318 | 4.6055 | 5.3117 |
| EHMT2_S119 | EHMT2 | Q96KQ7 | | 140 | MTGAGKSPPSVQS | 6.6806 | 7.0165 | 6.5396 | 7.0670 | 6.3866 | 6.3311 | 6.2395 | 6.3829 | 6.6030 | 6.6497 | 6.7143 | 6.7444 | 5.0885 | 5.2561 | 5.0266 | 5.2734 |
| EHMT2_S123 | EHMT2 | Q96KQ7 | | 173 | PPPATTSPEGQPK | 6.3031 | 6.9554 | 6.8804 | 7.1197 | 6.0949 | 6.3810 | 6.2981 | 6.7143 | 6.7119 | 6.6283 | 6.7893 | 6.9751 | 4.6478 | 5.2855 | 4.9449 | 5.2704 |
| EHMT2_S140 | EHMT2 | Q96KQ7 | | 119 | TKSFPSSPSKGGS | 7.0967 | 7.2416 | 7.0519 | 7.3307 | 6.2005 | 6.2733 | 6.5376 | 6.5107 | 7.1885 | 7.0512 | 7.2111 | 7.3579 | 3.9361 | 4.5053 | 4.0338 | 4.4730 |
| EHMT2_S173 | EHMT2 | Q96KQ7 | | 555 | PRGDGVTPPAGTA | 5.5671 | 5.6162 | 6.4739 | 6.1794 | 5.7819 | 5.4555 | 6.3328 | 6.0837 | 6.2383 | 6.2052 | 5.4518 | 5.3438 | 7.2948 | 6.2163 | 6.9304 | 8.8287 |
| EHMT2_S232 | EHMT2 | Q96KQ7 | | 66 | EPAGPSSPASVTV | 6.5394 | 5.9653 | 6.6749 | 6.7077 | 6.1679 | 6.0156 | 5.8444 | 6.0396 | 6.4986 | 6.5482 | 5.8677 | 6.5336 | 6.1037 | 5.5450 | 5.8735 | 7.0749 |
| EHMT2_S246 | EHMT2 | A0A0G2JK64 | | 123 | EPAGPSSPASVTV | 6.6363 | 6.3681 | 6.4809 | 6.2327 | 6.0334 | 5.4693 | 5.5771 | 6.3455 | 6.3856 | 5.9017 | 5.9845 | 5.9644 | 6.5837 | 6.5411 | 6.4684 | 7.0274 |
| EHMT2_S66 | EHMT2 | Q96KQ7 | | 232 | KRRKLNSGGGLSE | 6.0355 | 6.1559 | 6.7371 | 7.0598 | 6.0580 | 6.6797 | 6.4349 | 5.6779 | 6.5586 | 7.1971 | 6.9239 | 6.3511 | 6.1991 | 5.4472 | 5.0124 | 5.4717 |
| EHMT2_T555 | EHMT2 | Q96KQ7 | | 246 | LGSARRSGEVTLT | 6.1895 | 7.3694 | 6.7311 | 7.2935 | 5.4389 | 6.8565 | 7.3109 | 6.4640 | 7.0253 | 7.3394 | 7.3417 | 7.7985 | 3.6549 | 4.3724 | 3.6757 | 5.1384 |
| EI24_S326 | EI24 | O14681 | | 46 | QRRRRASSVLAQR | 5.8371 | 4.2155 | 5.2014 | 5.0171 | 7.1339 | 5.1318 | 4.8021 | 5.3914 | 8.6480 | 7.8661 | 7.5829 | 7.4879 | 6.8498 | 8.4040 | 5.3155 | 5.1156 |
| EI24_S326;S330 | EI24 | O14681 | | 326 | SAEKFPSPHPSPA | 6.6258 | 7.0155 | 5.8219 | 7.6141 | 6.3439 | 6.5080 | 6.3382 | 6.5337 | 6.5863 | 5.9337 | 6.5045 | 6.6170 | 5.0370 | 5.4558 | 5.2058 | 5.8587 |
| EI24_S330 | EI24 | O14681 | | 330 | FPSPHPSPAKLKA | 6.2983 | 6.3594 | 6.0655 | 7.2362 | 7.2324 | 6.7213 | 6.3589 | 6.6999 | 6.3046 | 5.9540 | 6.2095 | 5.8589 | 5.8460 | 5.8044 | 5.6486 | 5.4021 |
| EI24_S46 | EI24 | O14681 | 326;330 | | NA | 7.0867 | 6.8323 | 6.5004 | 6.9469 | 6.3988 | 6.1917 | 6.4928 | 6.2918 | 6.8400 | 6.4897 | 6.4456 | 6.5339 | 4.9629 | 5.5032 | 5.0417 | 5.4416 |
| EIF2A_S506 | EIF2A | Q9BY44 | | 506 | EARSDKSPDLAPT | 5.9218 | 6.4146 | 6.7911 | 6.0119 | 6.0446 | 6.1462 | 6.2085 | 6.2206 | 6.6472 | 6.6215 | 6.3811 | 6.4831 | 5.7082 | 6.3066 | 5.9045 | 6.1886 |
| EIF2A_T518 | EIF2A | Q9BY44 | | 518 | TPAPQSTPRNTVS | 7.0377 | 7.1195 | 6.6409 | 6.8258 | 5.8127 | 6.5770 | 7.4423 | 6.7328 | 6.9861 | 6.2464 | 7.4221 | 6.7474 | 4.3076 | 5.5151 | 4.0436 | 4.5429 |
| EIF2AK1_S258 | EIF2AK1 | Q9BQI3 | | 258 | PSLEVLSDQEEDR | 6.7425 | 6.6543 | 7.4235 | 7.2168 | 6.4874 | 6.8701 | 6.8565 | 7.2079 | 6.9396 | 7.5049 | 6.9441 | 7.4352 | 3.7686 | 4.2689 | 3.6410 | 4.0388 |
| EIF2AK2_S83 | EIF2AK2 | P19525 | | 83 | KEKKAVSPLLLTT | 6.2537 | 6.5005 | 6.6560 | 6.4549 | 6.3682 | 6.2057 | 6.3755 | 6.3750 | 6.6806 | 6.5931 | 6.7775 | 6.7034 | 5.4071 | 5.7647 | 5.5107 | 5.3733 |
| EIF2AK3_S1094 | EIF2AK3 | Q9NZJ5 | | 1094 | LRQRSRSLSSSGT | 5.8902 | 5.7015 | 5.7401 | 6.2795 | 5.9670 | 5.9405 | 5.8812 | 6.0352 | 6.3952 | 6.5475 | 6.4262 | 6.6052 | 6.7118 | 6.9435 | 6.5843 | 6.3512 |
| EIF2AK3_S715 | EIF2AK3 | Q9NZJ5 | | 715 | IEIIAPSPQRSRS | 6.5470 | 6.8721 | 7.0458 | 7.2090 | 6.2689 | 6.4399 | 6.3427 | 6.4981 | 6.3884 | 6.1370 | 6.6796 | 6.2369 | 4.9390 | 5.1505 | 5.3235 | 5.9216 |
| EIF2AK3_S845 | EIF2AK3 | Q9NZJ5 | | 845 | AFKPTSSKSSSEA | 4.1840 | 4.7676 | 4.4907 | 5.8676 | 5.4989 | 5.2717 | 6.1966 | 5.4591 | 4.7912 | 5.7410 | 6.3779 | 5.8055 | 8.5469 | 8.7555 | 9.3936 | 8.8523 |
| EIF2AK4_S230 | EIF2AK4 | Q9P2K8 | | 667 | RPAGPGTPPPDSG | 6.3814 | 6.0933 | 6.4793 | 6.0189 | 6.3067 | 6.7162 | 6.2908 | 6.2256 | 6.5909 | 6.5362 | 6.6364 | 6.6398 | 5.5777 | 6.3781 | 5.5938 | 5.5347 |
| EIF2AK4_S254 | EIF2AK4 | Q9P2K8 | | 254 | RRERQYSVCNSED | 6.3405 | 6.8832 | 7.8701 | 6.1617 | 6.1850 | 6.5008 | 5.9845 | 5.2430 | 7.0219 | 7.2769 | 7.9150 | 6.4271 | 5.2526 | 5.7064 | 3.9395 | 5.2919 |

| | | | | | | | | | | | | | | | | | | | | | |
|---|---|---|---|---|---|---|---|---|---|---|---|---|---|---|---|---|---|---|---|---|---|
| EIF2AK4_S261 | EIF2AK4 | Q9P2K8 | | 230 | AILHGGSPDFVGN | 6.0093 | 6.9940 | 7.3882 | 7.2020 | 6.4237 | 7.3019 | 6.7063 | 6.4453 | 6.9718 | 7.3289 | 7.7203 | 7.8518 | 3.5909 | 4.5078 | 3.5435 | 4.0142 |
| EIF2AK4_S264 | EIF2AK4 | Q9P2K8 | | 261 | VCNSEDSPGSCEI | 4.9243 | 5.0903 | 5.8339 | 6.1212 | 6.5774 | 7.2693 | 5.8701 | 5.6148 | 5.1657 | 5.8366 | 6.0347 | 6.0361 | 7.5176 | 7.3879 | 6.8745 | 7.8454 |
| EIF2AK4_T667 | EIF2AK4 | Q9P2K8 | | 264 | SEDSPGSCEILYF | 4.6577 | 5.3787 | 6.0603 | 5.3032 | 5.8852 | 6.7745 | 5.6551 | 5.6084 | 5.2465 | 6.9547 | 6.8861 | 4.6285 | 7.3359 | 9.1780 | 6.7934 | 6.7539 |
| EIF2B1_S7 | EIF2B1 | H0YGG4 | | 7 | XRGRLGSPGAPQQ | 6.3193 | 6.4763 | 6.9626 | 6.3905 | 5.7833 | 5.8672 | 6.6452 | 5.6468 | 7.6439 | 6.9140 | 6.9408 | 7.5257 | 5.0202 | 5.7968 | 5.0553 | 5.0122 |
| EIF2B4_S150;T157 | EIF2B4 | Q9UI10-2 | | 106 | SGIQLGTPREKVP | 7.0481 | 6.9910 | 6.5490 | 6.9722 | 6.9715 | 6.9883 | 7.3847 | 6.7060 | 8.0867 | 6.8562 | 7.1278 | 7.1001 | 3.4360 | 4.3528 | 3.5342 | 3.8954 |
| EIF2B4_T106 | EIF2B4 | Q9UI10-2 | 150;157 | | NA | 7.3840 | 8.5788 | 7.7731 | 8.5460 | 5.3179 | 7.2832 | 7.6208 | 7.1426 | 7.1284 | 6.5395 | 7.5824 | 6.3083 | 2.6957 | 4.4318 | 2.8367 | 2.8310 |
| EIF2B5_S544 | EIF2B5 | Q13144 | | 544 | PDSRGGSPQMDDI | 5.2921 | 6.5503 | 6.8767 | 6.2511 | 6.4986 | 6.1056 | 5.9200 | 6.5828 | 5.8890 | 7.1593 | 6.6872 | 6.5281 | 5.4052 | 6.3011 | 5.9880 | 5.9648 |
| EIF2D_S361 | EIF2D | P41214 | | 361 | FVIPEPSPTSQTI | 4.5326 | 5.9909 | 6.3542 | 7.2943 | 6.5465 | 6.1859 | 6.4797 | 7.3451 | 6.1966 | 6.9525 | 7.2329 | 8.5817 | 4.0972 | 5.5609 | 4.4738 | 6.1752 |
| EIF2S2_S105 | EIF2S2 | P20042 | | 105 | KDLKIESDVQEPT | 6.4647 | 5.5521 | 6.0233 | 6.0296 | 6.4773 | 6.7575 | 5.9163 | 6.8136 | 6.0002 | 6.0328 | 5.0452 | 5.5232 | 7.0451 | 6.9637 | 6.7963 | 6.5591 |
| EIF2S2_T111 | EIF2S2 | P20042 | | 111 | SDVQEPTEPEDDL | 5.4335 | 6.6586 | 6.7318 | 6.6428 | 6.5163 | 6.6769 | 6.0011 | 6.6128 | 7.0272 | 6.5487 | 4.4024 | 5.9187 | 6.0660 | 6.7326 | 5.6766 | 6.3540 |
| EIF2S2_T31 | EIF2S2 | P20042 | | 31 | LDEEGDTQTEETQ | 6.6570 | 6.5738 | 7.2798 | 7.3466 | 7.1104 | 6.7848 | 6.6119 | 6.8251 | 5.7201 | 5.1739 | 5.6211 | 5.1566 | 5.4318 | 6.1364 | 5.5100 | 6.0607 |
| EIF3A_S1188 | EIF3A | Q14152 | | 1336 | VPPPALSRDRERD | 6.0464 | 6.0764 | 6.8314 | 6.3477 | 6.1166 | 6.3516 | 6.3118 | 7.0396 | 6.7359 | 6.3080 | 6.8703 | 6.7110 | 5.2146 | 5.8671 | 5.5204 | 5.6512 |
| EIF3A_S1336 | EIF3A | Q14152 | | 882 | RRLGDSSLSRKDS | 6.7344 | 6.7840 | 7.9523 | 7.0270 | 6.3922 | 6.4371 | 6.6567 | 7.0040 | 7.1114 | 6.1765 | 6.9580 | 6.5367 | 4.3221 | 4.8889 | 4.4187 | 4.6001 |
| EIF3A_S584 | EIF3A | Q14152 | | 584 | RKERLESLNIQRE | 8.8721 | 8.2953 | 9.7335 | 9.1097 | 7.5229 | 5.7806 | 6.7367 | 6.4251 | 7.1680 | 5.9561 | 7.6997 | 7.2382 | 2.2056 | 2.3505 | 2.4708 | 2.4351 |
| EIF3A_S882 | EIF3A | Q14152 | | 1188 | EKAREESWGPPRE | 6.6034 | 6.5250 | 7.4635 | 6.9886 | 6.3113 | 6.4026 | 6.5336 | 6.5918 | 6.0579 | 6.5828 | 7.3543 | 6.5557 | 4.8442 | 5.6795 | 4.7161 | 4.7897 |
| EIF3B_S152 | EIF3B | P55884-2 | | 154 | ADEPSFSDPEDFV | 6.0926 | 6.0322 | 6.5905 | 6.1661 | 6.1457 | 6.3125 | 6.3009 | 6.2323 | 6.7395 | 6.7853 | 6.2580 | 6.6848 | 5.7379 | 6.3044 | 5.7892 | 5.8282 |
| EIF3B_S154 | EIF3B | P55884-2 | | 239 | IFLEYASPAHAVD | 6.2907 | 6.9121 | 7.1214 | 7.1929 | 6.5089 | 7.4385 | 6.7979 | 6.6391 | 6.3575 | 6.8991 | 6.7949 | 6.8693 | 4.2665 | 4.8020 | 4.1800 | 4.9290 |
| EIF3B_S239 | EIF3B | P55884-2 | | 83 | ASGPSESPSPPAA | 5.8446 | 6.1337 | 5.7662 | 6.6084 | 6.1968 | 5.9867 | 6.9495 | 5.8195 | 6.3285 | 5.8795 | 7.0702 | 7.5448 | 6.1349 | 5.7056 | 6.1576 | 5.8734 |
| EIF3B_S738 | EIF3B | P55884-2 | | 738 | EQKDRLSQSKASK | 6.6159 | 7.5553 | 7.5936 | 7.3274 | 5.9862 | 6.9386 | 7.6847 | 6.5748 | 6.6524 | 6.5860 | 7.4914 | 7.2838 | 3.5306 | 4.4413 | 3.6526 | 4.0855 |
| EIF3B_S83 | EIF3B | P55884-2 | | 152 | GDADEPSFSDPED | 6.6088 | 6.2706 | 8.5238 | 6.0466 | 5.3108 | 5.4113 | 6.6179 | 5.2331 | 6.8099 | 8.4186 | 5.2331 | 5.4799 | 6.2340 | 6.3528 | 5.2651 | 6.1837 |
| EIF3CL_S166 | EIF3CL | B5ME19 | | 39 | KQPLLLSEDEEDT | 6.3078 | 6.8603 | 7.1498 | 6.3083 | 6.0621 | 6.8253 | 6.2488 | 6.3614 | 6.5164 | 6.3232 | 6.5658 | 6.1768 | 5.4635 | 5.9564 | 5.5342 | 5.3400 |
| EIF3CL_S39 | EIF3CL | B5ME19 | | 166 | KQNPEQSADEDAE | 6.3405 | 6.5321 | 7.1040 | 6.4719 | 5.8422 | 5.9642 | 6.2603 | 6.4662 | 6.3505 | 6.5471 | 6.2823 | 6.6954 | 5.4042 | 6.0651 | 5.7119 | 5.9619 |
| EIF3D_S526;T548 | EIF3D | O15371 | 526;548 | | NA | 5.9775 | 5.9211 | 6.6381 | 6.1550 | 6.1348 | 6.4992 | 6.5754 | 6.5282 | 6.7616 | 6.5956 | 6.7067 | 6.4020 | 5.3967 | 6.0611 | 5.5386 | 6.1084 |
| EIF3E_S399 | EIF3E | P60228 | | 399 | MGNNAVSPYQQVI | 6.8028 | 7.2353 | 7.1850 | 7.3241 | 5.8226 | 6.7446 | 6.9163 | 6.9040 | 6.9869 | 6.7784 | 7.7892 | 7.4160 | 3.7209 | 4.5198 | 3.7198 | 4.1343 |
| EIF3F_S258 | EIF3F | O00303 | | 258 | IMKTCFSPNRVIG | 6.2849 | 5.6072 | 6.5680 | 5.4965 | 6.0748 | 6.0208 | 6.1487 | 6.5472 | 6.2761 | 6.6843 | 6.5396 | 7.2000 | 5.2156 | 6.4880 | 6.3058 | 6.5423 |
| EIF3G_S264 | EIF3G | O75821 | | 42 | LATGDTSPEPELL | 6.0194 | 6.1155 | 6.5554 | 6.2938 | 6.2210 | 6.3140 | 6.1597 | 6.6122 | 6.4126 | 6.4541 | 6.2799 | 6.3008 | 5.9638 | 6.2428 | 5.8981 | 6.1570 |
| EIF3G_S42 | EIF3G | O75821 | | 264 | LFRPFGSISRIYL | 6.5083 | 6.3475 | 6.5975 | 5.8478 | 6.1378 | 6.7234 | 6.2449 | 7.2517 | 6.2051 | 5.8494 | 6.4839 | 6.1617 | 6.3143 | 6.0407 | 5.9081 | 5.3780 |
| EIF3G_T27 | EIF3G | O75821 | | 38 | KGIPLATGDTSPE | 6.4940 | 6.2296 | 6.3062 | 6.2092 | 6.0272 | 6.4770 | 6.0811 | 6.7143 | 6.1475 | 6.6336 | 6.1879 | 6.4523 | 5.8641 | 6.4614 | 5.6587 | 6.0560 |
| EIF3G_T38 | EIF3G | O75821 | | 27 | EDDKCVTSELLKG | 6.4898 | 6.8886 | 7.3674 | 5.9026 | 5.2482 | 8.5701 | 7.1269 | 6.8235 | 6.2363 | 5.6556 | 7.1616 | 8.4726 | 3.8224 | 4.9340 | 3.6382 | 5.6621 |
| EIF3H_S183 | EIF3H | O15372 | | 183 | CKEKDFSPEALKK | 6.5532 | 7.0283 | 7.1848 | 6.7338 | 6.1101 | 6.2479 | 6.5770 | 6.4734 | 6.9014 | 6.9268 | 7.1358 | 6.8522 | 4.5045 | 5.1313 | 4.6705 | 4.9690 |
| EIF3J_T109 | EIF3J | O75822 | | 109 | EEPKVLTPEEQLA | 6.4733 | 6.8013 | 7.1076 | 6.2671 | 6.4095 | 6.4004 | 6.6424 | 6.3277 | 6.8083 | 6.6770 | 6.6133 | 6.6919 | 5.1843 | 5.6180 | 4.7593 | 5.2187 |
| EIF3K_S217 | EIF3K | Q9UBQ5 | | 217 | SSIMASSQxxxxx | 6.4538 | 6.1437 | 6.5430 | 5.9538 | 6.3218 | 6.0612 | 6.2842 | 6.8548 | 6.1397 | 6.4502 | 6.3263 | 6.7996 | 5.5280 | 6.4274 | 5.6983 | 6.0141 |
| EIF3M_S152 | EIF3M | Q7L2H7 | | 152 | QVRKWISDWNLTT | 6.4065 | 6.4441 | 6.9040 | 5.9216 | 6.5930 | 6.6795 | 5.8808 | 6.5479 | 6.0249 | 7.5482 | 6.4158 | 6.7208 | 5.1512 | 5.5682 | 5.7628 | 5.4307 |
| EIF4A3_T163 | EIF4A3 | P38919 | | 163 | QHVVAGTPGRVFD | 6.5503 | 6.7185 | 7.1204 | 6.7154 | 6.4510 | 6.3637 | 6.2217 | 6.4030 | 6.6647 | 6.5030 | 6.4347 | 6.3990 | 5.0404 | 5.8510 | 5.4319 | 5.1315 |
| EIF4B_S16 | EIF4B | P23588 | | 445 | RRESEKSLENETL | 6.8483 | 6.5533 | 8.0964 | 6.9898 | 6.3597 | 6.2801 | 6.0070 | 6.4320 | 7.7427 | 7.5722 | 8.2968 | 8.1973 | 3.9265 | 4.1904 | 3.1878 | 3.3197 |
| EIF4B_S219 | EIF4B | P23588 | | 406 | PRERHPSWRSEET | 6.2241 | 6.6362 | 7.1251 | 6.3691 | 6.3308 | 6.6993 | 6.3574 | 6.2281 | 6.1161 | 6.6728 | 6.6763 | 6.9619 | 5.3807 | 5.7692 | 5.1870 | 5.2658 |
| EIF4B_S283 | EIF4B | P23588 | | 93 | RSRLPKSPPYTAF | 5.6786 | 6.6368 | 6.7520 | 6.2173 | 6.0771 | 6.3060 | 6.1711 | 6.5090 | 6.4878 | 6.4908 | 6.7621 | 6.5282 | 5.9151 | 5.8264 | 5.6813 | 5.9603 |
| EIF4B_S359 | EIF4B | P23588 | | 459 | KEEDCHSPTSKPP | 6.2010 | 6.9672 | 7.0852 | 6.9907 | 6.5078 | 6.7101 | 6.0831 | 6.5109 | 6.8249 | 6.7001 | 7.0775 | 7.0899 | 4.8368 | 5.0625 | 4.7049 | 4.6474 |
| EIF4B_S406 | EIF4B | P23588 | | 283 | GRRAFGSGYRRDD | 6.4898 | 6.3733 | 6.8670 | 6.2186 | 6.1956 | 6.1085 | 5.6164 | 6.8905 | 6.9111 | 7.4135 | 6.8677 | 6.5311 | 5.7606 | 6.1084 | 4.8547 | 4.7931 |
| EIF4B_S406;S409 | EIF4B | P23588 | | 497 | PPARSQSSDTEQQ | 5.9213 | 5.7366 | 6.8326 | 6.0702 | 5.9811 | 6.3853 | 5.3261 | 6.7206 | 6.0171 | 5.8156 | 6.1337 | 6.6600 | 7.0157 | 6.4434 | 6.4681 | 6.4727 |
| EIF4B_S445 | EIF4B | P23588 | | 488 | NAWVKRSSNPPAR | 6.1677 | 6.8422 | 6.7289 | 7.0030 | 6.4113 | 7.3563 | 6.8667 | 6.7972 | 7.0474 | 6.4883 | 6.9768 | 6.4138 | 4.6391 | 4.8233 | 4.5152 | 4.9228 |
| EIF4B_S459 | EIF4B | P23588 | | 504 | SDTEQQSPTSGGG | 6.8168 | 7.3013 | 7.1925 | 6.5683 | 6.1143 | 6.4616 | 6.2836 | 6.7748 | 7.0706 | 7.2867 | 7.1248 | 7.1110 | 4.4347 | 4.7554 | 4.3452 | 4.3586 |
| EIF4B_S488 | EIF4B | P23588 | | 16 | KKGKTISLTDFLA | 5.8889 | 5.7577 | 6.5661 | 6.2131 | 6.3482 | 6.1213 | 5.6528 | 5.5325 | 5.8831 | 5.9318 | 5.8540 | 5.5928 | 7.2804 | 7.4997 | 7.1357 | 6.7420 |
| EIF4B_S497 | EIF4B | P23588 | | 359 | QSTRAASIFGGAK | 9.0875 | 6.1982 | 6.7862 | 5.0940 | 6.0320 | 5.6750 | 7.3506 | 5.5854 | 8.1648 | 7.1656 | 6.9499 | 6.5198 | 5.8835 | 6.0214 | 3.0661 | 4.4200 |
| EIF4B_S497;S504 | EIF4B | P23588 | | 219 | PRRGDDSFGDKYR | 6.8490 | 7.6022 | 6.3324 | 7.3908 | 6.1951 | 7.0472 | 7.3278 | 6.8850 | 7.3738 | 7.5016 | 7.7283 | 7.6494 | 3.0834 | 3.9394 | 3.0228 | 4.0719 |
| EIF4B_S504 | EIF4B | P23588 | 497;504 | | NA | 6.3464 | 7.1958 | 7.1822 | 6.7605 | 6.4995 | 6.8867 | 6.3096 | 7.1140 | 5.6698 | 6.0516 | 6.4550 | 6.1558 | 5.4350 | 5.4464 | 5.3343 | 5.1573 |
| EIF4B_S93 | EIF4B | P23588 | 406;409 | | NA | 6.3657 | 6.3111 | 8.4479 | 7.6481 | 6.3473 | 6.0248 | 6.1573 | 6.3616 | 7.1606 | 6.7679 | 7.3321 | 7.4617 | 4.3786 | 4.6652 | 4.3286 | 4.2415 |
| EIF4EBP1_S101 | EIF4EBP1 | Q13541 | | 101 | QSHLRNSPEDKRA | 5.6075 | 5.0142 | 5.2631 | 5.2446 | 5.0895 | 4.0797 | 4.5464 | 6.0052 | 3.9934 | 5.2158 | 4.1687 | 4.5645 | 5.1652 | 14.6491 | 9.1316 | 12.2614 |
| EIF4EBP1_S65 | EIF4EBP1 | Q13541 | | 37 | PGDYSTTPGGTLF | 5.9702 | 5.7034 | 6.4994 | 5.9211 | 5.7549 | 6.2490 | 6.1499 | 6.1834 | 5.8346 | 6.2706 | 5.8942 | 6.4122 | 5.5350 | 7.1217 | 5.8272 | 8.6733 |
| EIF4EBP1_T37 | EIF4EBP1 | Q13541 | | 46 | GTLFSTTPGGTRI | 5.0756 | 5.1003 | 5.7648 | 5.1862 | 5.5523 | 5.0509 | 4.6881 | 6.0268 | 4.8191 | 5.6435 | 5.5941 | 4.6945 | 5.4610 | 11.3128 | 9.3225 | 10.7073 |
| EIF4EBP1_T70;T82 | EIF4EBP1 | Q13541 | | 65 | LMECRNSPVTKTP | 7.2316 | 8.0056 | 7.7732 | 7.3901 | 6.2779 | 6.2419 | 6.2185 | 5.7367 | 4.9482 | 6.2571 | 6.7453 | 6.2638 | 3.1469 | 7.6084 | 4.3204 | 5.8343 |
| EIF4EBP1_T82 | EIF4EBP1 | Q13541 | | 82 | PTIPGVTSPSSDE | 6.1107 | 6.1394 | 5.7306 | 5.5313 | 4.7510 | 5.3895 | 6.2323 | 6.8690 | 6.3167 | 6.1022 | 5.4520 | 5.0095 | 5.5888 | 10.0003 | 6.8200 | 7.9567 |
| EIF4EBP1_Y37;T46 | EIF4EBP1 | Q13541 | 37;46 | | NA | 6.1816 | 6.0003 | 6.4083 | 6.3703 | 5.8540 | 5.6289 | 5.9034 | 6.3096 | 6.0230 | 6.2311 | 6.0748 | 6.3639 | 4.9756 | 8.0118 | 5.9048 | 7.7585 |
| EIF4EBP1_Y46 | EIF4EBP1 | Q13541 | 70;82 | | NA | 6.4934 | 6.1454 | 6.2951 | 7.1517 | 7.1098 | 6.2425 | 6.5096 | 6.4319 | 6.3666 | 6.2852 | 6.1072 | 6.2107 | 5.3272 | 6.9776 | 4.7381 | 5.6079 |
| EIF4EBP2_T36;T46 | EIF4EBP2 | Q13542 | 36;46 | | NA | 6.2680 | 6.3896 | 6.5054 | 6.3274 | 5.5330 | 5.6750 | 5.5871 | 6.6593 | 5.0996 | 5.8045 | 5.6173 | 6.0533 | 3.7252 | 9.8616 | 5.4800 | 9.4135 |
| EIF4ENIF1_S120 | EIF4ENIF1 | Q9NRA8-3 | | 587 | LRPRIPSPIGFTP | 6.4327 | 6.0279 | 6.3720 | 6.2243 | 5.9907 | 6.1726 | 6.3337 | 6.4160 | 6.5628 | 6.5617 | 6.5279 | 6.7119 | 6.1265 | 6.5845 | 5.2974 | 5.6573 |
| EIF4ENIF1_S138 | EIF4ENIF1 | Q9NRA8-3 | | 952 | PSQRSSSPVGLAK | 6.1954 | 6.4046 | 6.4230 | 5.8450 | 6.1395 | 6.3570 | 5.6709 | 6.3719 | 6.6629 | 6.7329 | 6.7752 | 6.7542 | 5.8827 | 6.1493 | 5.8771 | 5.7584 |
| EIF4ENIF1_S345 | EIF4ENIF1 | Q9NRA8-3 | | 345 | RWFSNPSRSGSRS | 7.4954 | 7.2404 | 6.8560 | 6.5256 | 5.9050 | 6.1793 | 6.0833 | 6.1408 | 7.0002 | 6.6112 | 7.0269 | 6.8536 | 5.0435 | 5.2547 | 4.4918 | 5.2922 |
| EIF4ENIF1_S374 | EIF4ENIF1 | Q9NRA8-3 | | 78 | PASGRSSPVESLK | 5.5064 | 5.9063 | 6.3342 | 5.8806 | 6.0440 | 6.0453 | 5.7011 | 6.5495 | 6.4473 | 6.7432 | 6.6350 | 7.0303 | 6.1717 | 6.2964 | 6.2602 | 6.4486 |
| EIF4ENIF1_S417 | EIF4ENIF1 | Q9NRA8-3 | | 682 | HRGNSSSPAPAAS | 5.8286 | 6.2977 | 6.3282 | 5.8458 | 6.2451 | 6.5096 | 6.0494 | 6.7417 | 6.3225 | 6.0516 | 6.2765 | 6.8681 | 5.9486 | 6.4226 | 6.1204 | 6.1438 |
| EIF4ENIF1_S499 | EIF4ENIF1 | Q9NRA8-3 | | 564 | LGQRAPSPPLSQV | 6.3239 | 6.0856 | 6.6081 | 6.2130 | 6.4699 | 6.4215 | 5.6231 | 6.4997 | 6.3540 | 6.7035 | 6.4864 | 6.4832 | 6.1700 | 6.2360 | 5.7570 | 5.5650 |
| EIF4ENIF1_S564 | EIF4ENIF1 | Q9NRA8-3 | | 374 | LEQAILSPGQNSG | 6.5565 | 6.1770 | 7.2362 | 6.4945 | 6.2169 | 5.9799 | 5.7957 | 6.8320 | 6.5430 | 6.7991 | 7.0891 | 6.8256 | 5.0531 | 5.7925 | 5.3915 | 5.2173 |
| EIF4ENIF1_S587 | EIF4ENIF1 | Q9NRA8-3 | | 138 | SSRRSGSPLEKDS | 6.5560 | 6.5431 | 6.4365 | 5.8182 | 6.2158 | 6.4466 | 5.9870 | 6.3181 | 6.8502 | 6.6309 | 6.8871 | 6.9456 | 5.7959 | 5.6845 | 5.4021 | 5.4824 |
| EIF4ENIF1_S671 | EIF4ENIF1 | Q9NRA8-3 | | 671 | QQRVTKSPAPVHR | 6.0257 | 5.7473 | 5.8811 | 5.7691 | 5.9338 | 6.4357 | 5.9935 | 6.7690 | 6.4411 | 6.3106 | 6.5572 | 6.2862 | 6.6612 | 6.5418 | 6.3229 | 6.3239 |
| EIF4ENIF1_S682 | EIF4ENIF1 | Q9NRA8-3 | | 120 | LSPQRRSFGGGCH | 6.0230 | 7.0474 | 6.6735 | 6.7413 | 6.2361 | 6.9847 | 6.6285 | 7.6661 | 6.5975 | 7.3401 | 7.8859 | 6.9257 | 4.1232 | 5.1837 | 3.4868 | 4.4566 |
| EIF4ENIF1_S753 | EIF4ENIF1 | Q9NRA8-3 | | 417 | PLLSSLSANKEKL | 6.9136 | 4.8094 | 6.6168 | 7.6458 | 5.6534 | 8.3317 | 7.1294 | 7.7961 | 6.3085 | 9.9773 | 8.7325 | 6.2122 | 3.3951 | 3.5107 | 3.0714 | 3.8961 |
| EIF4ENIF1_S78 | EIF4ENIF1 | Q9NRA8-3 | | 793 | KATGRKTPTLASP | 5.4176 | 5.4807 | 6.5578 | 6.7789 | 6.5090 | 5.7398 | 5.9550 | 6.9002 | 6.2748 | 6.4627 | 6.6897 | 6.1547 | 5.4069 | 6.9025 | 7.1784 | 5.5912 |
| EIF4ENIF1_S952 | EIF4ENIF1 | Q9NRA8-3 | | 455 | QQVKNSTPFMAEH | 7.1799 | 5.9578 | 6.6059 | 6.2911 | 7.1197 | 5.4023 | 6.8466 | 6.7586 | 6.3281 | 4.6431 | 7.2678 | 6.6892 | 6.1754 | 6.9253 | 4.6523 | 5.1569 |
| EIF4ENIF1_T455 | EIF4ENIF1 | Q9NRA8-3 | | 499 | ASGTLPSQPKVSR | 5.8272 | 5.5129 | 6.2395 | 6.6168 | 6.1289 | 5.8916 | 6.2812 | 5.5652 | 6.6271 | 6.4214 | 6.7273 | 6.9390 | 6.6892 | 6.5182 | 5.6527 | 6.3618 |
| EIF4ENIF1_T793 | EIF4ENIF1 | Q9NRA8-3 | | 753 | SADRDSSPTTNSK | 6.7264 | 6.5335 | 6.5636 | 6.3976 | 6.1107 | 6.7848 | 5.6809 | 6.5471 | 6.7940 | 6.5796 | 7.3496 | 7.0523 | 5.5774 | 5.4674 | 4.8944 | 4.9404 |
| EIF4G1_S1084 | EIF4G1 | Q04637-9 | | 1238 | AALPPVSPLKAAL | 6.2657 | 6.5137 | 6.8079 | 6.4629 | 6.5501 | 7.0559 | 6.7124 | 7.1404 | 7.1461 | 6.7737 | 6.9112 | 6.8063 | 4.4839 | 4.9813 | 4.4936 | 4.8950 |
| EIF4G1_S1099 | EIF4G1 | Q04637-9 | | 211 | SGARTASTPTPPQ | 6.5326 | 6.2933 | 6.9987 | 6.5773 | 6.4500 | 6.7553 | 6.5007 | 7.0971 | 7.5241 | 6.8343 | 7.3074 | 6.7853 | 4.7032 | 4.8813 | 4.1154 | 4.6441 |
| EIF4G1_S1216 | EIF4G1 | Q04637-9 | | 654 | LDKANKTPLRPLD | 6.4706 | 6.8511 | 6.2793 | 6.7610 | 5.8401 | 7.5031 | 7.3968 | 6.7758 | 7.0914 | 6.9537 | 6.6254 | 6.8056 | 5.1303 | 5.0274 | 3.9447 | 4.5435 |
| EIF4G1_S1238 | EIF4G1 | Q04637-9 | | 1186 | RLDRARTPATKRS | 7.1004 | 7.3945 | 7.0602 | 6.7980 | 5.3923 | 7.6012 | 7.5482 | 6.5766 | 6.8374 | 6.8397 | 7.1592 | 7.6577 | 3.4594 | 4.6409 | 3.5397 | 4.3944 |
| EIF4G1_S1603 | EIF4G1 | Q04637-9 | | 1084 | KITKPGSIDSNNQ | 6.7259 | 6.2826 | 8.6207 | 7.6295 | 6.6899 | 6.6526 | 6.5181 | 6.6490 | 6.7536 | 8.0272 | 5.3478 | 7.4420 | 4.3278 | 5.0366 | 3.4014 | 3.8952 |
| EIF4G1_S211 | EIF4G1 | Q04637-9 | | 1216 | GLRKAASLTEDRD | 6.4347 | 6.7357 | 5.9050 | 5.3815 | 6.3463 | 5.8386 | 5.7822 | 5.7828 | 5.9722 | 7.5202 | 5.8526 | 6.0190 | 6.2861 | 7.1563 | 6.8791 | 6.1077 |
| EIF4G1_T1186 | EIF4G1 | Q04637-9 | | 1099 | APGGRLSWGKGSS | 6.8057 | 7.5113 | 6.3501 | 7.1745 | 7.4046 | 7.4168 | 6.8758 | 5.4022 | 5.9691 | 6.7829 | 7.1761 | 6.4690 | 5.0670 | 5.2407 | 3.8357 | 4.5184 |
| EIF4G1_T654 | EIF4G1 | Q04637-9 | | 1603 | REAEEESDHNxxx | 6.2808 | 6.5221 | 6.2373 | 6.4976 | 6.7486 | 7.2127 | 5.9635 | 6.3830 | 5.5299 | 6.1803 | 6.5242 | 5.8803 | 5.7858 | 7.0480 | 5.6565 | 5.5494 |
| EIF4G2_S17 | EIF4G2 | P78344 | | 395 | VIQDRFSPTMGRH | 6.7588 | 6.7714 | 6.6437 | 6.3382 | 6.4291 | 6.5721 | 6.5511 | 6.8347 | 6.7424 | 6.5328 | 6.4248 | 6.4338 | 5.0179 | 5.6663 | 5.0677 | 5.2151 |
| EIF4G2_S395 | EIF4G2 | P78344 | | 508 | QPPRTQTPPLGQT | 6.3979 | 6.2957 | 6.9211 | 6.3195 | 6.6129 | 6.7067 | 6.6603 | 6.8094 | 6.5821 | 6.8264 | 6.9764 | 6.7026 | 5.1153 | 5.4267 | 4.6822 | 4.9650 |
| EIF4G2_T508 | EIF4G2 | P78344 | | 17 | SRFSASSGGGGSR | 6.6129 | 6.6410 | 7.4275 | 7.0890 | 6.3065 | 6.7271 | 7.0000 | 6.4788 | 6.7201 | 6.5053 | 7.1690 | 6.7832 | 4.4438 | 5.1132 | 4.2017 | 4.7810 |
| EIF4G3_S1080 | EIF4G3 | O43432-3 | | 501 | NLNSRRSPVPAQI | 5.7496 | 5.9923 | 6.4585 | 6.1600 | 6.3042 | 6.6311 | 6.0871 | 6.6477 | 6.2107 | 6.4306 | 6.6373 | 6.3725 | 6.0602 | 6.4857 | 5.9586 | 5.8137 |
| EIF4G3_S1415 | EIF4G3 | O43432-3 | | 311 | DRCELSSPREDTI | 5.5812 | 5.7401 | 6.1943 | 5.7619 | 5.6288 | 5.8300 | 6.4617 | 6.1997 | 6.1246 | 6.0595 | 6.2276 | 6.5125 | 6.7402 | 6.7875 | 7.1802 | 6.9700 |
| EIF4G3_S236 | EIF4G3 | O43432-3 | | 1080 | PKAQLGSWGKGSS | 7.9478 | 7.2265 | 5.9399 | 6.6222 | 7.0413 | 6.8717 | 8.5112 | 5.7216 | 6.3590 | 6.5949 | 6.6846 | 6.3396 | 4.7995 | 5.0939 | 3.9651 | 4.2810 |
| EIF4G3_S246 | EIF4G3 | O43432-3 | | 246 | VLRLVLSGEKKEQ | 6.3135 | 6.5890 | 6.3622 | 6.0197 | 5.7734 | 6.0892 | 6.2390 | 6.4262 | 6.0323 | 6.4027 | 6.1534 | 6.4804 | 5.7047 | 6.9844 | 6.2059 | 6.2239 |
| EIF4G3_S311 | EIF4G3 | O43432-3 | | 236 | PDPVLKSPSPVLR | 6.3903 | 6.5298 | 6.4460 | 6.3391 | 6.1811 | 6.6346 | 6.7022 | 6.9452 | 6.8391 | 6.3978 | 6.5280 | 6.6130 | 4.9844 | 5.7913 | 5.2738 | 5.4042 |
| EIF4G3_S501 | EIF4G3 | O43432-3 | | 1415 | DFIESDSPCSSEA | 5.7492 | 6.4203 | 6.5327 | 5.9029 | 5.7987 | 6.1837 | 6.0692 | 6.2129 | 6.2487 | 6.2784 | 6.7558 | 6.6729 | 6.5026 | 6.4407 | 5.9747 | 6.2567 |
| EIF4G3_S591 | EIF4G3 | O43432-3 | | 591 | NGAESVSEGEGID | 6.3923 | 6.4791 | 6.8612 | 5.8115 | 6.4579 | 6.1048 | 7.8818 | 7.9455 | 4.7696 | 7.0908 | 4.0171 | 6.8631 | 6.0778 | 5.7112 | 5.6803 | 5.8559 |
| EIF4G3_T1118;T1122 | EIF4G3 | O43432-3 | 1118;1122 | | NA | 6.1978 | 6.0841 | 6.8751 | 6.5966 | 5.4085 | 8.1188 | 7.1499 | 6.7424 | 6.0417 | 6.0874 | 6.4920 | 7.1022 | 5.6174 | 6.0295 | 4.5818 | 4.8748 |
| EIF4H_S13 | EIF4H | Q15056 | | 220 | KPRTVATPLNQVA | 6.9774 | 6.8160 | 6.6531 | 5.7618 | 6.3007 | 6.3471 | 6.6440 | 6.7330 | 7.4721 | 6.6272 | 6.8152 | 6.7209 | 4.9238 | 4.7396 | 4.3187 | 6.1492 |
| EIF4H_T220 | EIF4H | Q15056 | | 13 | YDDRAYSSFGGGR | 8.1135 | 7.4626 | 8.1731 | 7.1026 | 5.4045 | 6.1071 | 6.7446 | 6.0129 | 7.9276 | 6.4803 | 7.3166 | 7.1340 | 4.3577 | 4.7013 | 3.3479 | 3.6137 |
| EIF5_S176 | EIF5 | P55010 | | 176 | NGSVSSSETPPPP | 5.3583 | 6.9520 | 5.2599 | 6.8647 | 6.6476 | 4.8682 | 5.8910 | 7.7018 | 6.0522 | 7.1892 | 7.3011 | 7.3437 | 6.7953 | 6.2939 | 5.2711 | 4.2101 |
| EIF5_S389;S390 | EIF5 | P55010 | 389;390 | | NA | 6.0956 | 5.9277 | 6.6444 | 6.1474 | 6.6890 | 6.2497 | 6.2017 | 6.2699 | 6.3377 | 6.5498 | 6.3935 | 6.1595 | 5.8805 | 6.1601 | 6.0442 | 6.2492 |
| EIF5A_S74 | EIF5A | P63241-2 | | 76 | IVEMSTSKTGKHG | 4.8422 | 3.9667 | 4.9531 | 3.8220 | 5.8600 | 6.0492 | 5.4369 | 6.9069 | 5.0838 | 6.7552 | 6.1013 | 7.0659 | 8.3321 | 8.3323 | 8.1169 | 8.3755 |
| EIF5A_S76 | EIF5A | P63241-2 | | 74 | CKIVEMSTSKTGK | 7.5946 | 6.7639 | 6.9503 | 7.4090 | 6.9494 | 5.7996 | 6.3431 | 6.0948 | 6.3997 | 6.0721 | 6.7219 | 6.4447 | 5.0496 | 5.3761 | 4.9067 | 5.1245 |
| EIF5B_S107;S113 | EIF5B | O60841 | | 214 | PGPNIESGNEDDD | 6.0962 | 6.0223 | 7.0535 | 5.8718 | 6.1442 | 6.2899 | 6.2063 | 6.0845 | 6.3654 | 6.3734 | 6.2900 | 6.4678 | 5.8176 | 6.5810 | 5.9294 | 6.4066 |
| EIF5B_S113 | EIF5B | O60841 | | 113 | SFDDNDSEELEDK | 6.2265 | 6.0452 | 6.6681 | 6.2207 | 6.1856 | 6.5839 | 6.0845 | 6.1691 | 6.4984 | 6.0678 | 6.7064 | 6.5998 | 5.5162 | 6.6990 | 5.7445 | 5.9842 |
| EIF5B_S1168 | EIF5B | O60841 | | 164 | NKKWDGSEEDEDN | 6.3507 | 6.3420 | 6.8777 | 5.9764 | 6.2854 | 6.4203 | 6.0348 | 6.1120 | 6.3777 | 6.2080 | 6.7570 | 6.6239 | 5.3078 | 7.2102 | 5.4134 | 5.7027 |
| EIF5B_S135;S137 | EIF5B | O60841 | | 511 | DWEAMASDEETEK | 6.2638 | 6.0503 | 6.4992 | 5.9284 | 6.1668 | 5.6682 | 5.4315 | 6.1903 | 6.2662 | 6.5244 | 6.8886 | 7.1680 | 5.6883 | 6.5371 | 5.7820 | 6.9471 |
| EIF5B_S164 | EIF5B | O60841 | | 1168 | EPIPGESPKMFGR | 6.2105 | 6.5243 | 6.0815 | 6.5390 | 6.3347 | 5.7002 | 5.8635 | 5.8175 | 7.2204 | 7.6388 | 5.9065 | 4.8683 | 5.7687 | 6.6074 | 6.1294 | 6.7893 |
| EIF5B_S214 | EIF5B | O60841 | | 575 | VSDEKDSGKTLDK | 6.2097 | 6.1755 | 7.4945 | 7.2468 | 6.3362 | 6.4622 | 6.1276 | 6.3128 | 8.2897 | 7.0367 | 8.6919 | 7.9650 | 3.9241 | 4.4863 | 3.2899 | 3.9509 |
| EIF5B_S511 | EIF5B | O60841 | 135;137 | | NA | 6.0558 | 5.6088 | 6.6568 | 5.9702 | 6.6468 | 6.4134 | 6.3305 | 6.1497 | 6.4400 | 6.1167 | 6.8217 | 6.6864 | 5.9485 | 6.7894 | 5.6340 | 5.7312 |
| EIF5B_S575 | EIF5B | O60841 | 107;113 | | NA | 6.2635 | 5.0286 | 7.4673 | 6.4311 | 6.2009 | 5.4197 | 4.9949 | 6.9597 | 8.1393 | 5.6427 | 5.5198 | 5.9874 | 5.6112 | 7.4347 | 6.2226 | 6.6767 |
| EIF6_S175 | EIF6 | P56537 | | 239 | ATSMRDSLIDSLT | 6.4139 | 6.8270 | 6.6225 | 6.4598 | 6.4506 | 6.9503 | 6.2122 | 6.9393 | 6.8356 | 6.9755 | 7.0154 | 7.0246 | 4.7599 | 5.0178 | 4.5235 | 4.9721 |
| EIF6_S230 | EIF6 | P56537 | | 175 | DQDELSSLLQVPL | 6.2279 | 5.8471 | 6.5785 | 6.5527 | 6.0866 | 5.9404 | 6.5382 | 6.1931 | 6.3731 | 6.3334 | 6.6401 | 6.0333 | 6.2748 | 6.4121 | 5.8156 | 6.1531 |
| EIF6_S239 | EIF6 | P56537 | | 231 | NEAQPSTIATSMR | 6.3644 | 5.6969 | 5.9230 | 5.7317 | 6.4150 | 7.2191 | 6.6054 | 6.5615 | 6.8224 | 6.2352 | 6.5334 | 6.3007 | 6.7727 | 5.5653 | 5.1653 | 6.0878 |
| EIF6_T231 | EIF6 | P56537 | | 230 | LNEAQPSTIATSM | 6.0625 | 5.8640 | 6.7031 | 6.6261 | 7.2071 | 6.3690 | 6.3541 | 6.7939 | 6.2392 | 6.4643 | 6.5426 | 6.7373 | 5.5692 | 5.7439 | 5.1294 | 5.5944 |
| ELAC2_S199 | ELAC2 | Q9BQ52 | | 199 | KHQPWQSPERPLS | 5.7695 | 6.2440 | 6.7317 | 6.0963 | 5.9681 | 6.5537 | 6.1309 | 6.3508 | 6.5632 | 6.7891 | 7.1656 | 7.2484 | 5.0119 | 6.0790 | 5.5534 | 5.7445 |
| ELAC2_S736 | ELAC2 | Q9BQ52 | | 736 | AKVPLFSPNFSEK | 6.2976 | 6.4306 | 7.3143 | 6.9009 | 5.7617 | 6.2086 | 6.3158 | 6.0429 | 6.1899 | 6.2695 | 6.3972 | 6.5096 | 5.3547 | 6.0932 | 6.0405 | 5.8730 |
| ELAVL1_S229 | ELAVL1 | Q15717-2 | | 229 | LSQLYHSPARRFG | 7.2345 | 7.6766 | 7.0423 | 6.8339 | 6.3296 | 6.7498 | 6.9693 | 7.1615 | 7.2646 | 6.9131 | 7.1531 | 6.7372 | 3.8887 | 4.4775 | 3.5735 | 3.9947 |
| ELF1_S163;S168 | ELF1 | P32519 | | 334 | RSRVSSSPGVKGG | 6.8132 | 7.0499 | 6.1685 | 6.3217 | 6.3359 | 6.6318 | 6.7563 | 6.7363 | 7.1776 | 6.9089 | 6.6462 | 6.5941 | 4.8570 | 5.3455 | 4.4441 | 5.2132 |
| ELF1_S168 | ELF1 | P32519 | | 168 | DSPGASSPEQPKR | 7.4996 | 6.8028 | 5.3922 | 5.9638 | 7.0076 | 7.0900 | 5.7676 | 4.1152 | 5.9468 | 6.2852 | 6.3904 | 6.4729 | 7.6987 | 6.0975 | 5.6652 | 5.8046 |
| ELF1_S334 | ELF1 | P32519 | 163;168 | | NA | 7.2831 | 5.8361 | 4.3317 | 7.1441 | 7.6740 | 6.4719 | 5.8711 | 5.9507 | 6.8049 | 7.4224 | 7.2217 | 6.0697 | 4.9189 | 5.8519 | 6.2543 | 4.8934 |
| ELF2_S107 | ELF2 | Q15723 | | 107 | ALLHMESPTCLRD | 6.4226 | 6.8638 | 6.8475 | 6.6501 | 6.1429 | 6.1036 | 6.4238 | 6.8026 | 6.3191 | 6.5820 | 6.8629 | 6.2170 | 5.6185 | 5.6323 | 4.9132 | 5.5980 |
| ELF2_S185 | ELF2 | Q15723 | | 191 | SPISNGSPELGIK | 6.4434 | 6.3829 | 5.7112 | 5.2811 | 5.9372 | 6.7348 | 5.8448 | 6.0701 | 6.4924 | 6.7988 | 6.4995 | 6.7490 | 5.9777 | 6.5557 | 6.6545 | 5.8668 |
| ELF2_S191 | ELF2 | Q15723 | | 521 | QQASGQTPPRVIS | 6.2219 | 6.4029 | 6.5190 | 6.6366 | 5.5804 | 6.0768 | 6.4518 | 6.1043 | 6.7893 | 6.7626 | 6.6916 | 6.2776 | 5.8370 | 5.9996 | 6.0691 | 5.5796 |
| ELF2_S324 | ELF2 | Q15723 | | 324 | KSLERVSLSAESL | 6.4776 | 5.9863 | 7.3555 | 8.4212 | 6.8569 | 6.2066 | 6.4559 | 7.5975 | 7.4241 | 6.4957 | 6.5210 | 5.5744 | 4.5990 | 5.9429 | 3.1431 | 4.9422 |
| ELF2_S372 | ELF2 | Q15723 | | 372 | HDASSRSPTTTAS | 7.3897 | 7.5731 | 5.9023 | 6.6775 | 6.7590 | 6.2674 | 6.3755 | 6.7183 | 7.5540 | 6.4168 | 6.5571 | 6.2779 | 4.6151 | 5.3835 | 4.7553 | 4.7776 |
| ELF2_S424 | ELF2 | Q15723 | | 185 | KPKTQQSPISNGS | 6.0100 | 6.2259 | 6.2355 | 6.7561 | 6.3754 | 6.6491 | 6.2310 | 6.4191 | 6.2644 | 6.4196 | 6.5998 | 6.1081 | 5.2407 | 6.3069 | 6.1545 | 6.0038 |
| ELF2_T521 | ELF2 | Q15723 | | 424 | PLITSTSPTTATS | 5.9701 | 5.9580 | 5.9880 | 5.1388 | 5.3650 | 5.4849 | 5.7196 | 7.5786 | 6.2021 | 5.4399 | 6.1221 | 5.9073 | 7.6600 | 7.0677 | 7.3867 | 7.0113 |
| ELF4_S188 | ELF4 | Q99607 | | 188 | KGNRSTSPVTDPS | 6.1628 | 6.4646 | 6.2237 | 6.7524 | 6.0797 | 5.9110 | 6.1129 | 6.3395 | 6.7692 | 6.3362 | 6.3142 | 6.2795 | 5.7350 | 6.0452 | 5.9161 | 6.5582 |
| ELF4_S641 | ELF4 | Q99607 | | 641 | GSLLTRSPTPAPF | 7.5099 | 7.2768 | 6.6252 | 7.5301 | 6.1416 | 6.5905 | 6.8322 | 6.8382 | 7.1163 | 6.5784 | 6.9173 | 6.5544 | 4.0052 | 4.7017 | 4.1762 | 4.6059 |
| ELFN1_S461 | ELFN1 | P0C7U0 | | 735 | GLGRKASILEPLT | 7.4127 | 6.6163 | 6.2819 | 6.1306 | 6.0653 | 6.0906 | 6.3638 | 6.2768 | 6.9239 | 6.4091 | 6.2935 | 6.3729 | 5.7010 | 6.7387 | 4.7534 | 5.5693 |
| ELFN1_S623 | ELFN1 | P0C7U0 | | 461 | SAAAAGSLKKTII | 6.8901 | 6.8298 | 6.8997 | 5.9518 | 6.7023 | 6.2307 | 6.3525 | 6.7384 | 6.4900 | 6.6639 | 6.4778 | 6.2280 | 5.1775 | 6.1678 | 5.4465 | 4.7531 |
| ELFN1_S629 | ELFN1 | P0C7U0 | | 517 | LVESADTPKASKG | 6.5221 | 6.7263 | 6.2660 | 6.5076 | 6.4704 | 6.5487 | 6.9408 | 6.2408 | 6.3845 | 6.6122 | 6.0739 | 6.5175 | 5.0728 | 5.7430 | 5.1135 | 6.2600 |
| ELFN1_S645 | ELFN1 | P0C7U0 | | 629 | SLQRHHSVEAAGP | 6.7329 | 5.9053 | 6.5469 | 6.1660 | 5.2853 | 6.3893 | 5.5486 | 6.4145 | 6.8595 | 6.9168 | 5.9642 | 5.9406 | 6.2511 | 6.8432 | 5.8560 | 6.3798 |
| ELFN1_S648 | ELFN1 | P0C7U0 | | 623 | KDAFGHSLQRHHS | 7.0016 | 6.5906 | 7.5292 | 6.1067 | 5.5227 | 7.0274 | 6.6762 | 6.0551 | 6.0778 | 7.2495 | 7.8377 | 7.3231 | 4.2633 | 5.7473 | 4.2038 | 4.7881 |
| ELFN1_S735 | ELFN1 | P0C7U0 | | 648 | SSGSVRSPRAFRA | 7.2009 | 6.4129 | 5.0063 | 5.3292 | 6.4666 | 7.7330 | 6.8439 | 5.5859 | 4.7490 | 5.9764 | 6.7100 | 6.1451 | 5.7672 | 6.5231 | 6.9122 | 6.6382 |
| ELFN1_T517 | ELFN1 | P0C7U0 | | 645 | STSSSGSVRSPRA | 4.3387 | 3.1453 | 4.4108 | 4.2028 | 5.4489 | 5.8071 | 5.5731 | 5.8541 | 4.1433 | 4.1893 | 4.0339 | 4.0851 | 11.3439 | 11.4667 | 10.7395 | 11.2173 |
| ELFN2_T493 | ELFN2 | Q5R3F8 | | 493 | LEAGLDTPKVATK | 6.4123 | 6.7825 | 6.5898 | 6.4662 | 5.3451 | 6.1506 | 7.3210 | 5.5395 | 6.9978 | 6.0821 | 5.7641 | 6.1059 | 5.8732 | 5.8140 | 5.8175 | 6.9386 |
| ELK1_S196 | ELK1 | P19419 | | 196 | AAPPSGSRSTSPS | 6.3831 | 5.1217 | 6.3119 | 5.7055 | 6.2633 | 6.5610 | 6.1004 | 6.7342 | 7.0792 | 7.1106 | 6.9847 | 6.4823 | 5.2654 | 6.6835 | 5.2291 | 5.9841 |
| ELK3_S284 | ELK3 | P41970 | | 396 | SLDRAASPVLLSS | 5.6945 | 5.4639 | 6.0778 | 5.2830 | 6.0929 | 6.1403 | 6.6609 | 6.6644 | 6.4913 | 6.2321 | 6.4454 | 5.7216 | 7.3134 | 7.4231 | 6.1872 | 6.1081 |
| ELK3_S396 | ELK3 | P41970 | | 284 | SGSKTKSPSLPPK | 5.6360 | 6.2604 | 5.7136 | 5.6033 | 6.4379 | 6.7548 | 6.5785 | 6.6698 | 6.5220 | 6.5482 | 6.3787 | 6.4974 | 6.0608 | 6.3610 | 5.8021 | 6.1755 |
| ELK4_S180 | ELK4 | P28324 | | 194 | VIKFVTTPSKKPP | 7.2939 | 7.0878 | 6.4236 | 6.9358 | 5.9741 | 6.3807 | 6.8635 | 6.4149 | 6.9995 | 6.7592 | 6.8144 | 7.0404 | 4.6937 | 4.8998 | 4.4246 | 4.9940 |
| ELK4_T194 | ELK4 | P28324 | | 180 | KLAEKKSPQEPTP | 5.2223 | 5.5712 | 6.5089 | 5.7544 | 5.6912 | 6.0411 | 5.6489 | 6.3808 | 5.7014 | 5.8129 | 5.5637 | 6.5111 | 6.8111 | 7.0914 | 7.7751 | 7.9144 |
| ELL_S309 | ELL | P55199 | | 309 | GDPAASSPPGERG | 6.1843 | 6.1534 | 6.4508 | 6.6466 | 6.0942 | 6.3360 | 6.1653 | 6.6355 | 6.4134 | 6.2979 | 6.7679 | 6.4361 | 5.3917 | 6.2495 | 5.8860 | 5.8916 |
| ELL_S319 | ELL | P55199 | | 319 | ERGRSASPPQKRL | 5.9421 | 6.4887 | 6.1733 | 6.2835 | 5.9599 | 5.9446 | 5.8806 | 5.8611 | 5.8558 | 6.1090 | 5.9911 | 6.1836 | 6.3196 | 6.8595 | 7.1276 | 7.0200 |
| ELL_S442 | ELL | P55199 | | 180 | PSRKRATPINLAS | 5.2512 | 5.4835 | 5.6979 | 5.1682 | 5.3212 | 5.0598 | 6.1327 | 4.8703 | 5.2791 | 4.3121 | 5.0226 | 4.3573 | 9.2530 | 9.8018 | 9.4481 | 9.5410 |
| ELL_T180 | ELL | P55199 | | 442 | PSRPHGSPSRSKP | 6.1330 | 6.6223 | 6.3307 | 6.0766 | 5.8605 | 6.1342 | 6.7609 | 7.1353 | 6.2759 | 6.1836 | 6.0267 | 6.2229 | 5.6881 | 6.2654 | 5.8816 | 6.4024 |
| ELL2_S580 | ELL2 | O00472 | | 580 | AQRKRLSPGSKEY | 6.9909 | 7.2099 | 7.1535 | 7.3188 | 6.2428 | 6.8927 | 6.6267 | 6.7839 | 7.1719 | 7.2190 | 7.4875 | 7.5165 | 3.4406 | 4.2673 | 3.4956 | 4.1822 |
| ELMO2_S517 | ELMO2 | Q96JJ3 | | 517 | RQSERMSQDDFQS | 6.1899 | 6.0783 | 7.0637 | 5.6541 | 5.7501 | 6.7718 | 6.7189 | 5.5317 | 6.5230 | 7.4115 | 6.5014 | 6.5602 | 5.8324 | 5.9823 | 5.3749 | 6.0557 |
| ELOA_S151 | ELOA | Q14241 | | 420 | LPKVEETDMEDEF | 8.1988 | 7.1818 | 6.5280 | 7.8723 | 6.9435 | 6.0769 | 6.1281 | 5.8866 | 5.4924 | 5.0628 | 5.4581 | 5.0143 | 5.9042 | 6.7199 | 5.5481 | 5.9843 |
| ELOA_S322 | ELOA | Q14241 | | 151 | TGSRSYSPDHRQK | 6.0806 | 6.5843 | 5.6826 | 6.4542 | 6.0760 | 6.0635 | 5.8987 | 6.3569 | 6.0755 | 5.8632 | 5.8244 | 5.7284 | 6.3414 | 7.2962 | 6.8083 | 6.8658 |
| ELOA_S410 | ELOA | Q14241 | | 410 | TNLDRKSLGSLPK | 7.4908 | 7.9618 | 7.1056 | 7.1829 | 5.9533 | 7.1954 | 7.8054 | 6.9670 | 7.6327 | 7.6991 | 7.9385 | 7.6365 | 2.1712 | 3.8908 | 2.1308 | 3.2380 |
| ELOA_S542 | ELOA | Q14241 | | 322 | REKPPSSGVKKEK | 6.7550 | 7.2350 | 7.0226 | 7.6445 | 5.8157 | 7.0910 | 7.3962 | 6.9248 | 7.1471 | 7.4485 | 7.7462 | 7.8898 | 2.9186 | 4.3356 | 2.9444 | 3.6849 |
| ELOA_T420 | ELOA | Q14241 | | 542 | KRKAFSSPQEEEE | 6.1701 | 6.4342 | 6.6736 | 6.2752 | 5.7751 | 6.1139 | 6.2922 | 6.0949 | 6.6488 | 6.2314 | 6.2024 | 6.7442 | 6.0475 | 6.1567 | 5.7398 | 6.3999 |
| ELOB_S95 | ELOB | Q15370 | | 95 | CIEPFSSPPELPD | 5.3949 | 6.0734 | 6.4812 | 5.4426 | 5.9646 | 6.1040 | 6.1482 | 6.5509 | 5.8618 | 6.5423 | 6.1818 | 5.7510 | 6.2419 | 7.4450 | 6.4711 | 7.3452 |
| ELOVL5_S300 | ELOVL5 | Q9NYP7-2 | | 312 | GHTNSFSPLENNV | 6.5512 | 7.9495 | 6.6542 | 8.6022 | 6.5635 | 7.1212 | 7.0899 | 6.7897 | 6.8332 | 7.0101 | 7.4903 | 7.5210 | 2.9921 | 4.1592 | 3.0465 | 3.6260 |
| ELOVL5_S312 | ELOVL5 | Q9NYP7-2 | | 300 | KDHQNGSMAAVNG | 3.1190 | 3.5506 | 3.5727 | 3.8489 | 3.5289 | 3.2512 | 3.2282 | 3.9351 | 3.0827 | 3.2493 | 3.2498 | 3.6950 | 13.5730 | 15.6043 | 15.8528 | 13.6586 |

| | | | | | | | | | | | | | | | | | | | | |
|---|---|---|---|---|---|---|---|---|---|---|---|---|---|---|---|---|---|---|---|---|
| ELP2_S122 | ELP2 | Q6IA86-6 | 122 | VYQRRTSDPALCT | 4.9672 | 6.6541 | 6.9348 | 5.9487 | 6.3794 | 6.7981 | 6.3244 | 6.2710 | 7.1129 | 7.3090 | 7.7262 | 7.7351 | 4.6494 | 5.3444 | 4.5217 | 5.3236 |
| EMC4_S36 | EMC4 | Q5J8M3 | 36 | GRSDRGSGQGDSL | 5.2518 | 5.2067 | 4.7332 | 5.7868 | 5.3320 | 5.0584 | 4.7947 | 5.6704 | 5.9549 | 7.4942 | 5.4287 | 8.4138 | 6.4208 | 8.2919 | 7.8227 | 8.3391 |
| EMD_S141 | EMD | P50402 | 49 | TQRRRLSPPSSSA | 6.0627 | 6.7051 | 6.2263 | 6.8814 | 6.4895 | 6.0549 | 6.3582 | 6.2989 | 6.7735 | 6.9828 | 6.8730 | 6.8558 | 5.2864 | 5.8357 | 4.9874 | 5.3285 |
| EMD_S141;S142 | EMD | P50402 | 171 | THYRPVSASRSSL | 6.3059 | 7.1951 | 6.8868 | 7.3401 | 6.6447 | 7.4130 | 7.1132 | 7.2308 | 6.4137 | 6.4671 | 7.4629 | 6.7178 | 3.5493 | 4.9404 | 4.1034 | 4.2157 |
| EMD_S141;S142;S143 | EMD | P50402 | 141 | HDDDLLSSSEEEC | 6.1283 | 6.0836 | 6.7611 | 6.5586 | 6.1198 | 6.5767 | 6.3183 | 6.4818 | 6.1891 | 5.7861 | 6.4107 | 6.2160 | 5.9785 | 6.3551 | 5.9882 | 6.0481 |
| EMD_S143 | EMD | P50402 | 87 | DALLYQSKGYNDD | 7.3639 | 6.6163 | 6.4297 | 7.1451 | 7.5280 | 6.3995 | 6.8503 | 7.3664 | 7.1881 | 5.6904 | 7.2418 | 5.9070 | 4.4065 | 4.9917 | 4.2203 | 4.6549 |
| EMD_S171 | EMD | P50402 | 176 | VSASRSSLDLSYY | 5.8662 | 5.6614 | 6.2604 | 6.3302 | 6.5816 | 5.4832 | 6.0975 | 5.9221 | 5.8099 | 6.2646 | 6.4363 | 7.0898 | 6.5480 | 6.9427 | 6.3232 | 6.3828 |
| EMD_S176 | EMD | P50402 | 143 | DDLLSSSEEECKD | 6.6679 | 6.0934 | 6.2093 | 6.6224 | 6.5454 | 5.6192 | 5.9152 | 6.4865 | 5.6978 | 6.0294 | 5.7474 | 5.6322 | 7.4080 | 6.9528 | 6.5238 | 5.8493 |
| EMD_S49 | EMD | P50402 | 59 | SSAASSYSFSDLN | 8.3646 | 6.8611 | 6.8245 | 8.8861 | 7.4163 | 6.1811 | 6.7392 | 5.2213 | 7.1516 | 5.6223 | 7.7556 | 6.1173 | 3.9481 | 4.6826 | 4.0177 | 4.2106 |
| EMD_S60 | EMD | P50402 | 66 | SFSDLNSTRGDAD | 7.0790 | 6.1433 | 6.8756 | 6.7444 | 7.0576 | 7.1848 | 6.6996 | 6.3213 | 6.1738 | 5.5178 | 6.1321 | 6.7027 | 5.1252 | 5.5279 | 4.9878 | 5.7273 |
| EMD_S66 | EMD | P50402 | 60 | SAASSYSFSDLNS | 3.2935 | 3.3943 | 3.4124 | 3.9122 | 4.2569 | 3.4756 | 4.3381 | 3.9347 | 4.4826 | 2.5920 | 3.4681 | 4.1515 | 13.9989 | 14.0816 | 13.8995 | 13.3079 |
| EMD_S87 | EMD | P50402 | 67 | FSDLNSTRGDADM | 7.0475 | 7.0353 | 6.5547 | 8.3642 | 8.2920 | 6.5274 | 7.9775 | 6.8351 | 7.3784 | 4.9725 | 6.6105 | 6.2633 | 3.3509 | 4.8091 | 3.5456 | 4.4360 |
| EMD_T67 | EMD | P50402 | 141;142;143 | NA | 6.5884 | 7.2268 | 6.9792 | 8.5340 | 5.8442 | 6.4136 | 6.9605 | 6.7536 | 7.1089 | 6.4995 | 6.7711 | 7.0254 | 4.1032 | 4.6746 | 4.1129 | 4.4040 |
| EMD_Y59 | EMD | P50402 | 141;142 | NA | 6.3374 | 7.0885 | 7.6740 | 8.0851 | 6.7281 | 6.7388 | 7.6122 | 7.1180 | 5.4950 | 6.5650 | 6.1769 | 6.4390 | 3.9397 | 4.8018 | 3.9596 | 5.2411 |
| EMG1_S14 | EMG1 | Q92979 | 14 | FKPRERSGGEQAQ | 6.1857 | 6.5856 | 6.2922 | 6.1246 | 6.4888 | 7.0672 | 7.3770 | 6.4846 | 6.5390 | 6.3696 | 6.5639 | 6.2880 | 4.8958 | 5.8681 | 5.1953 | 5.6745 |
| EML1_S113 | EML1 | O00423-3 | 113 | PTGSLPSPSGVRK | 6.4691 | 6.5167 | 6.4585 | 6.2286 | 6.1556 | 6.8628 | 6.3104 | 7.0677 | 6.4721 | 6.7455 | 6.2678 | 6.0987 | 5.4148 | 6.2535 | 5.4277 | 5.2505 |
| EML2_S159 | EML2 | O95834-3 | 159 | VGGYATSPSSPKK | 6.4875 | 6.7393 | 6.3658 | 6.1572 | 5.9957 | 6.7906 | 6.3760 | 6.4813 | 5.8070 | 6.1046 | 5.7057 | 6.0843 | 6.0376 | 6.6877 | 6.1953 | 5.9844 |
| EML2_S159;S162 | EML2 | O95834-3 | 162 | YATSPSSPKKEAT | 5.8233 | 6.4359 | 6.8197 | 6.6358 | 6.5890 | 6.7337 | 6.1095 | 3.5558 | 6.4573 | 5.5629 | 6.8995 | 6.9659 | 5.5636 | 6.0739 | 5.2593 | 5.7149 |
| EML2_S162 | EML2 | O95834-3 | 158;162 | NA | 6.8844 | 6.9416 | 7.4088 | 5.6739 | 5.9489 | 6.9229 | 6.9445 | 6.9692 | 7.4776 | 7.0003 | 6.6098 | 6.6811 | 4.2618 | 4.7946 | 4.7542 | 4.7264 |
| EML2_T158;S162 | EML2 | O95834-3 | 159;162 | NA | 5.6009 | 6.3916 | 6.8754 | 6.2454 | 6.0361 | 7.1036 | 7.1441 | 6.4288 | 6.4519 | 7.2045 | 7.0064 | 6.3034 | 5.0454 | 5.5393 | 4.6925 | 5.9305 |
| EML3_S156 | EML3 | Q32P44 | 176 | LSRKAISSANLLV | 7.7214 | 7.0020 | 7.2697 | 6.3171 | 6.6093 | 6.0390 | 5.6896 | 6.1115 | 6.7531 | 6.5960 | 6.6994 | 6.4967 | 5.1064 | 6.0378 | 4.6736 | 4.8774 |
| EML3_S176 | EML3 | Q32P44 | 881 | AGPAPATPSRTPS | 6.5240 | 6.9197 | 7.3682 | 6.5808 | 6.5682 | 6.7788 | 6.8825 | 7.2380 | 7.1071 | 6.8682 | 6.9378 | 6.7102 | 4.1410 | 5.0069 | 3.8480 | 4.5206 |
| EML3_S198 | EML3 | Q32P44 | 198 | GKDPLSSPGGPGS | 6.7463 | 7.0093 | 7.0265 | 6.5048 | 6.4505 | 7.1091 | 6.7841 | 6.5413 | 7.0880 | 6.5761 | 7.1649 | 6.7034 | 4.5615 | 4.7547 | 4.1466 | 4.8329 |
| EML3_S889 | EML3 | Q32P44 | 889 | SRTPSLSPASSLD | 6.4946 | 6.3699 | 6.7133 | 6.0632 | 6.0803 | 6.5349 | 6.1800 | 6.3350 | 6.2297 | 6.3625 | 6.4386 | 6.4751 | 5.7801 | 6.1457 | 5.7854 | 6.0118 |
| EML3_T881 | EML3 | Q32P44 | 156 | DTPRRNSSSSSSP | 5.3513 | 6.2165 | 5.5973 | 6.0894 | 5.6189 | 6.7303 | 6.9845 | 4.6293 | 6.2219 | 7.5929 | 5.0944 | 6.7546 | 6.8817 | 7.4712 | 6.1814 | 6.5843 |
| EML4_S144;S146 | EML4 | Q9HC35 | 176 | KSIKRPSPAEKSH | 6.4188 | 7.1115 | 6.9187 | 6.7824 | 5.9646 | 6.8370 | 6.4276 | 6.6678 | 6.5387 | 7.0443 | 6.9970 | 6.8277 | 4.4257 | 5.3643 | 4.6619 | 5.0120 |
| EML4_S146 | EML4 | Q9HC35 | 146 | QIRASPSPQPSSQ | 6.0242 | 6.0442 | 6.5346 | 6.0295 | 6.4965 | 6.4716 | 5.8871 | 6.6899 | 6.2251 | 6.3931 | 6.1846 | 5.8911 | 6.4461 | 6.7337 | 5.8491 | 6.0994 |
| EML4_S146;T160 | EML4 | Q9HC35 | 160 | LQIHRQTPESKNA | 6.3980 | 6.5177 | 7.5101 | 6.8032 | 6.1144 | 6.4750 | 6.3612 | 6.0823 | 6.9391 | 6.3597 | 7.6044 | 7.7333 | 4.7525 | 4.9216 | 4.2608 | 5.1669 |
| EML4_S160 | EML4 | Q9HC35 | 981 | QDPSPSSxxxxxx | 5.5386 | 6.4627 | 6.6741 | 6.2550 | 5.8529 | 6.5344 | 6.3919 | 7.3244 | 5.8775 | 6.2012 | 6.5033 | 6.1786 | 6.3805 | 6.9761 | 5.1960 | 6.5628 |
| EML4_S176 | EML4 | Q9HC35 | 919;926 | NA | 6.4544 | 6.3209 | 6.1088 | 6.2581 | 6.0709 | 6.4629 | 5.8819 | 6.4631 | 6.3644 | 6.2710 | 6.0598 | 6.5368 | 6.2684 | 6.3868 | 5.9486 | 6.1431 |
| EML4_S919;S926 | EML4 | Q9HC35 | 144;146 | NA | 5.2853 | 6.2992 | 6.4813 | 6.8916 | 6.2288 | 6.1998 | 6.7110 | 6.2051 | 6.6462 | 7.0947 | 6.2479 | 6.4085 | 5.8346 | 5.9997 | 5.1220 | 6.3443 |
| EML4_S919;S926;S934 | EML4 | Q9HC35 | 146;160 | NA | 6.9850 | 7.2945 | 7.3290 | 6.0828 | 6.3945 | 7.1453 | 7.1564 | 6.9742 | 5.9347 | 7.0691 | 7.6245 | 7.1824 | 4.1312 | 4.9228 | 3.6565 | 4.1172 |
| EML4_S981 | EML4 | Q9HC35 | 919;926;934 | NA | 5.5242 | 6.5050 | 6.4302 | 5.5882 | 5.8387 | 6.6019 | 5.2608 | 7.2079 | 6.3687 | 6.4145 | 6.1580 | 5.2093 | 6.3375 | 7.4271 | 7.0855 | 6.0428 |
| EMSY_S1151 | EMSY | Q7Z589-7 | 253 | TVPVSGSPKMSNI | 6.4762 | 6.4873 | 6.5855 | 7.1728 | 6.5807 | 6.4362 | 6.4895 | 6.1714 | 6.9396 | 6.9253 | 6.5787 | 6.8564 | 4.8950 | 5.4259 | 4.7387 | 5.2407 |
| EMSY_S188 | EMSY | Q7Z589-7 | 188 | KSPRPASPASNVV | 5.1581 | 5.1845 | 5.4595 | 6.0382 | 5.7954 | 5.4146 | 5.8541 | 5.5123 | 5.9830 | 6.4541 | 5.6616 | 5.4229 | 8.4812 | 7.7421 | 7.7464 | 8.0918 |
| EMSY_S224 | EMSY | Q7Z589-7 | 224 | KRRRTNSSSSSPV | 6.5095 | 6.0506 | 6.3360 | 6.6170 | 5.9744 | 6.0444 | 6.2867 | 5.7487 | 6.1684 | 6.0330 | 6.2572 | 6.5104 | 6.5886 | 6.6670 | 6.0105 | 6.1975 |
| EMSY_S253 | EMSY | Q7Z589-7 | 836 | KRHSPESPSIAVV | 5.1382 | 6.2686 | 7.0004 | 6.8196 | 5.6251 | 5.7561 | 5.6407 | 6.6729 | 6.4508 | 6.0992 | 6.9748 | 6.6274 | 5.8813 | 5.9397 | 6.9107 | 6.1946 |
| EMSY_S299 | EMSY | Q7Z589-7 | 1151 | SPVTSISPIQASE | 6.5047 | 6.7957 | 6.3842 | 6.8076 | 6.3357 | 6.2082 | 6.8727 | 5.8252 | 6.6921 | 6.4044 | 6.2184 | 6.3809 | 5.4977 | 5.8747 | 5.4443 | 5.7535 |
| EMSY_S836 | EMSY | Q7Z589-7 | 299 | VIIVTTSPSSTFV | 6.4126 | 7.2001 | 6.4670 | 6.7794 | 6.0783 | 6.3857 | 5.9912 | 5.9349 | 6.8761 | 7.5689 | 6.7333 | 6.7487 | 5.0263 | 5.2581 | 4.7817 | 5.7575 |
| EMSY_T222;S224 | EMSY | Q7Z589-7 | 222;224 | NA | 5.2404 | 5.5798 | 5.9412 | 7.1566 | 5.0763 | 6.2342 | 6.1500 | 6.3185 | 6.1670 | 5.0894 | 6.1186 | 5.9014 | 7.9699 | 6.7483 | 7.1290 | 7.1795 |
| ENAH_S129 | ENAH | A0A0U1RRM6 | 740 | TMNGSKSPVISRP | 6.6860 | 6.6432 | 6.5916 | 5.8371 | 7.0023 | 7.0072 | 6.1105 | 6.6957 | 6.5966 | 6.5282 | 6.6732 | 6.5890 | 5.5630 | 5.5987 | 4.9749 | 4.9028 |
| ENAH_S365;S369 | ENAH | A0A0U1RRM6 | 329 | PTSTPPTPPLRHS | 5.8796 | 6.1763 | 6.0438 | 5.7802 | 6.6132 | 6.6848 | 6.5194 | 6.6452 | 6.1834 | 6.4173 | 6.4735 | 6.1221 | 6.3318 | 6.2804 | 5.7818 | 6.0671 |
| ENAH_S369 | ENAH | A0A0U1RRM6 | 369 | KNSRPSSPVNTPS | 5.8883 | 5.7889 | 5.5382 | 5.7588 | 7.3366 | 6.0087 | 5.8522 | 5.4216 | 6.6674 | 5.3937 | 5.7520 | 5.1272 | 8.6856 | 7.2538 | 7.3720 | 6.1548 |
| ENAH_S740 | ENAH | A0A0U1RRM6 | 129 | TLPRQNSQLPAQV | 6.8889 | 6.3061 | 7.3730 | 6.7528 | 5.3055 | 6.5819 | 6.1452 | 5.1682 | 6.3586 | 6.5744 | 5.4839 | 6.1165 | 6.4272 | 6.8364 | 5.7958 | 5.8857 |
| ENAH_T329 | ENAH | A0A0U1RRM6 | 365;369 | NA | 7.5701 | 5.7743 | 8.3150 | 5.5908 | 6.2823 | 5.0689 | 5.5352 | 6.3341 | 7.2850 | 5.9198 | 6.8171 | 7.0985 | 5.4575 | 5.9455 | 4.5309 | 6.4749 |
| ENO1_S263 | ENO1 | P06733 | 419 | IEEELGSKAKFAG | 5.9925 | 5.5931 | 6.7516 | 5.2640 | 6.3732 | 6.6161 | 6.3213 | 6.5309 | 6.5255 | 6.0885 | 6.2047 | 6.3198 | 6.8489 | 6.3167 | 6.2400 | 6.0131 |
| ENO1_S27 | ENO1 | P06733 | 79 | IAPALVSKKLNVT | 6.0362 | 6.0211 | 6.6140 | 5.5351 | 6.6750 | 6.9053 | 5.5920 | 7.0582 | 6.0622 | 5.8929 | 6.7838 | 6.8969 | 5.9280 | 5.7654 | 6.4532 | 5.7805 |
| ENO1_S272 | ENO1 | P06733 | 272 | DPSRYISPDQLAD | 6.3434 | 6.2119 | 6.9423 | 5.8475 | 6.6086 | 6.7280 | 5.8136 | 6.5853 | 6.1282 | 6.8407 | 6.6898 | 6.3364 | 5.8516 | 5.9112 | 5.2638 | 5.8978 |
| ENO1_S419 | ENO1 | P06733 | 263 | YDLDFKSPDDPSR | 6.0524 | 5.8099 | 7.1598 | 6.2267 | 6.5165 | 6.4163 | 6.3954 | 6.4038 | 6.7347 | 6.4868 | 7.0675 | 7.0830 | 5.5792 | 6.0401 | 4.7758 | 5.2520 |
| ENO1_S79 | ENO1 | P06733 | 27 | EVDLFTSKGLFRA | 5.9802 | 5.0807 | 7.1754 | 5.7095 | 5.9892 | 7.3072 | 6.8716 | 6.3575 | 6.4460 | 6.5376 | 6.0878 | 7.3444 | 5.8262 | 6.7468 | 4.9469 | 5.5928 |
| ENSA_S132 | ENSA | O43768-4 | 132 | LPQRKSSLVTSKL | 6.2480 | 4.2256 | 6.5470 | 4.5036 | 8.7368 | 5.5779 | 4.9598 | 7.1531 | 6.3149 | 11.3010 | 5.0066 | 7.0820 | 4.4746 | 7.6946 | 5.0620 | 5.1126 |
| ENSA_S43 | ENSA | O43768-4 | 90 | EQKYFDSGDYNMA | 10.4543 | 7.2156 | 8.1990 | 5.6637 | 5.4979 | 6.3137 | 6.7820 | 6.3810 | 7.7316 | 7.2074 | 7.8182 | 8.3658 | 2.8249 | 3.9140 | 2.1914 | 3.4394 |
| ENSA_S90 | ENSA | O43768-4 | 43 | LKAKYPSLGQKPG | 7.4062 | 5.4642 | 6.8091 | 5.7464 | 6.8949 | 5.3755 | 5.0850 | 6.8920 | 6.9464 | 6.8727 | 6.2087 | 6.9625 | 6.5017 | 7.0412 | 4.7688 | 5.0247 |
| ENTPD4_S597 | ENTPD4 | Q9Y227 | 597 | RTPRSSSAAALWM | 5.3757 | 5.1469 | 5.3820 | 5.3139 | 6.1447 | 6.6206 | 6.4840 | 6.1253 | 5.2420 | 5.4732 | 5.1754 | 5.7455 | 9.0104 | 7.3292 | 7.3842 | 8.0469 |
| ENTR1_S170 | ENTR1 | Q96C92-4 | 18 | PLSRARSLAIPDD | 6.7205 | 6.5393 | 7.7598 | 5.7361 | 5.8898 | 7.0526 | 5.6697 | 6.4238 | 6.1956 | 7.5924 | 8.1216 | 8.1478 | 4.2359 | 5.7002 | 3.6460 | 4.5690 |
| ENTR1_S170;S174 | ENTR1 | Q96C92-4 | 170 | SRVSPASPAGSPS | 6.1305 | 5.6375 | 5.7523 | 5.5379 | 6.3674 | 6.6817 | 5.5655 | 5.7913 | 6.4787 | 6.3010 | 6.9293 | 6.5866 | 6.8318 | 6.6015 | 6.2585 | 6.5486 |
| ENTR1_S18 | ENTR1 | Q96C92-4 | 170;174 | NA | 5.9089 | 6.1907 | 6.1823 | 5.9012 | 6.2082 | 6.2576 | 6.2766 | 6.5765 | 6.9627 | 5.6180 | 6.7496 | 5.8652 | 6.4563 | 6.5760 | 6.1962 | 6.0741 |
| EP300_T887 | EP300 | Q09472 | 887 | PPRQTPTPPTTQL | 5.3941 | 5.5309 | 5.8173 | 5.3109 | 6.7355 | 6.2680 | 6.6533 | 5.8363 | 5.6152 | 5.7626 | 5.2666 | 5.8689 | 7.7747 | 7.4693 | 7.4893 | 7.2072 |
| EP400_S165 | EP400 | Q96L91 | 3152 | PAVRLKTPTKPPC | 6.9773 | 6.5526 | 6.6101 | 7.2470 | 6.8345 | 6.3992 | 6.9252 | 6.6858 | 7.1553 | 6.5848 | 6.4217 | 6.5497 | 4.6566 | 4.9742 | 4.3908 | 5.0354 |
| EP400_S1732 | EP400 | Q96L91 | 736 | ATNKALSPVTSRT | 7.2701 | 7.2323 | 6.7018 | 7.4109 | 6.6495 | 6.0708 | 6.8076 | 6.4396 | 6.7885 | 6.0928 | 6.4074 | 6.1861 | 4.9119 | 4.8137 | 5.1179 | 5.0991 |
| EP400_S315;S321 | EP400 | Q96L91 | 1732 | TASKPASPIGGPT | 5.6605 | 5.5462 | 5.5052 | 5.8957 | 6.3620 | 5.8985 | 6.5883 | 5.7246 | 5.8006 | 5.8570 | 6.1973 | 5.7266 | 7.6059 | 7.5468 | 7.1473 | 6.9375 |
| EP400_S321 | EP400 | Q96L91 | 722 | PSPARSSPVNRPS | 6.6164 | 6.6353 | 6.4996 | 7.1429 | 6.3722 | 6.7608 | 6.6677 | 6.1036 | 6.2668 | 6.0174 | 6.0259 | 6.3068 | 5.5440 | 5.7421 | 5.7284 | 5.5700 |
| EP400_S722 | EP400 | Q96L91 | 321 | SPPPPTSPSRTAV | 5.7371 | 6.3736 | 5.6316 | 5.9948 | 5.7423 | 5.6834 | 5.4776 | 7.0446 | 6.2577 | 6.3943 | 5.8996 | 6.6157 | 6.1971 | 7.3923 | 6.4081 | 7.1501 |
| EP400_S736 | EP400 | Q96L91 | 165 | LGLCSSSPTGGFV | 6.8999 | 5.7717 | 5.6642 | 6.4803 | 5.7280 | 6.8168 | 6.3333 | 6.1076 | 6.5497 | 6.8143 | 6.9887 | 6.5797 | 5.8224 | 6.1676 | 5.8343 | 5.4415 |
| EP400_S941;T945 | EP400 | Q96L91 | 941;945 | NA | 6.5652 | 6.5333 | 5.7937 | 7.2856 | 6.7987 | 5.9220 | 5.9830 | 6.9516 | 6.3353 | 6.1289 | 6.4685 | 6.4290 | 5.1000 | 5.9414 | 6.0473 | 5.7165 |
| EP400_T3152 | EP400 | Q96L91 | 315;321 | NA | 5.8199 | 6.1416 | 7.4357 | 5.9676 | 5.7835 | 5.7928 | 5.7479 | 7.2066 | 6.3106 | 6.5837 | 6.6347 | 6.6912 | 5.4928 | 6.4908 | 6.1481 | 5.7525 |
| EP400P1_S154 | EP400P1 | Q6ZTU2 | 154 | LGLCSSSPTGDFV | 6.2917 | 5.7613 | 6.0599 | 5.8241 | 5.8046 | 5.6790 | 6.2944 | 5.4202 | 6.1506 | 6.6085 | 5.7374 | 6.5780 | 5.6507 | 7.6401 | 7.1690 | 7.3305 |
| EPB41_S555 | EPB41 | P11171 | 709 | EWDKRLSTHSPFR | 7.1792 | 6.8121 | 6.8491 | 6.6915 | 6.1812 | 5.9570 | 6.0803 | 6.4891 | 6.7014 | 6.8782 | 6.6355 | 6.7617 | 5.0772 | 6.2457 | 4.5965 | 4.8644 |
| EPB41_S664 | EPB41 | P11171 | 85 | LSRLFSSFLKRPK | 9.3414 | 7.9592 | 7.0732 | 7.1148 | 6.5080 | 5.6433 | 6.1562 | 5.3049 | 6.3464 | 5.1823 | 5.9915 | 5.7706 | 5.3941 | 6.6186 | 4.6273 | 4.9681 |
| EPB41_S684 | EPB41 | P11171 | 555 | VDSADRSPRPTSA | 5.9399 | 6.3360 | 6.3771 | 6.2273 | 6.2300 | 6.4816 | 6.4193 | 6.2495 | 6.6346 | 6.6975 | 6.3886 | 6.4102 | 5.6938 | 6.0355 | 5.6409 | 6.2383 |
| EPB41_S709 | EPB41 | P11171 | 664 | NIYIRHSNLMLED | 6.3956 | 6.9488 | 6.5067 | 6.3089 | 5.5335 | 5.2680 | 6.9035 | 5.8037 | 6.2098 | 5.0651 | 5.3547 | 5.3965 | 6.9337 | 7.0893 | 6.7126 | 7.5697 |
| EPB41_S84 | EPB41 | P11171 | 684 | IKKHHASISELKK | 5.8268 | 5.7000 | 5.4405 | 6.5619 | 6.0471 | 5.6075 | 5.7950 | 5.2047 | 5.9097 | 5.1157 | 5.0567 | 4.6235 | 8.7647 | 8.1850 | 8.4434 | 7.7178 |
| EPB41_S85 | EPB41 | P11171 | 84 | GLSRLFSSFLKRP | 10.0464 | 7.4413 | 7.2966 | 7.0954 | 6.2328 | 6.1969 | 7.1563 | 5.2930 | 6.4732 | 4.8492 | 5.5116 | 6.2121 | 5.2788 | 5.9337 | 3.8931 | 5.0896 |
| EPB41L1_S1498 | EPB41L1 | A0A1B0GTW6 | 1498 | TEIRSLSPIIGKD | 6.2994 | 6.8405 | 6.4788 | 6.7967 | 6.1210 | 6.4526 | 6.4834 | 6.8791 | 6.4856 | 6.5089 | 6.8268 | 6.7443 | 5.2906 | 5.3010 | 5.2385 | 5.2528 |
| EPB41L1_S544 | EPB41L1 | A0A1B0GTW6 | 866 | PKEGAGTPKNHGG | 7.5953 | 7.4020 | 6.7704 | 7.4083 | 5.6712 | 6.8083 | 7.3570 | 6.3593 | 7.5813 | 6.5067 | 7.2198 | 7.7828 | 3.3654 | 4.1644 | 3.6279 | 4.3798 |
| EPB41L1_T866 | EPB41L1 | A0A1B0GTW6 | 544 | LPSSPASPSPKGT | 6.2661 | 6.1505 | 6.5481 | 6.1616 | 6.2248 | 6.7833 | 6.6668 | 6.5343 | 6.2106 | 6.5659 | 6.3951 | 5.9491 | 5.8526 | 6.0020 | 5.6605 | 6.0287 |
| EPB41L2_S386 | EPB41L2 | O43491 | 58 | AAESQSSLRRQKR | 6.6761 | 6.8720 | 7.4078 | 6.8309 | 5.9794 | 6.3868 | 6.0902 | 6.6875 | 6.3940 | 6.8979 | 6.7978 | 6.9930 | 4.5974 | 5.9991 | 4.3569 | 5.0332 |
| EPB41L2_S39 | EPB41L2 | O43491 | 598 | GRREVRSPTKAPH | 6.7889 | 7.0476 | 6.6046 | 6.6950 | 6.7991 | 7.3983 | 7.2131 | 6.5163 | 6.6078 | 6.7369 | 6.9403 | 7.0312 | 4.1275 | 4.8570 | 4.1654 | 4.4711 |
| EPB41L2_S402 | EPB41L2 | O43491 | 715 | MNGKEISPGSGPG | 6.7725 | 7.1642 | 6.7648 | 6.9737 | 6.7165 | 6.5021 | 6.6600 | 6.3758 | 6.6508 | 6.5611 | 6.4950 | 6.4283 | 4.8460 | 5.3983 | 4.7800 | 4.9109 |
| EPB41L2_S499 | EPB41L2 | O43491 | 683 | LQTQGSSHETLNI | 5.5550 | 6.6677 | 5.8839 | 6.1921 | 6.1013 | 6.8332 | 6.2454 | 6.8649 | 5.8455 | 6.1670 | 6.4713 | 6.0759 | 5.7621 | 6.6230 | 6.6378 | 6.0739 |
| EPB41L2_S550 | EPB41L2 | O43491 | 550 | SKRVSRSLDGAPI | 6.2620 | 6.5173 | 6.5859 | 6.5934 | 5.9718 | 6.0328 | 6.4148 | 6.3741 | 6.8247 | 6.5937 | 7.1447 | 7.1456 | 5.0894 | 5.6417 | 5.2219 | 5.5862 |
| EPB41L2_S562 | EPB41L2 | O43491 | 87 | WLKKQKSYTLVVA | 6.9226 | 7.1296 | 7.7647 | 6.5651 | 6.1155 | 6.5662 | 5.7948 | 6.8131 | 7.0011 | 8.0299 | 7.7588 | 7.7188 | 3.6759 | 4.9737 | 3.2612 | 3.9090 |
| EPB41L2_S58 | EPB41L2 | O43491 | 499 | TFYRLVSPEQPPK | 6.1970 | 6.5902 | 6.0460 | 6.5006 | 6.7217 | 6.6885 | 6.7590 | 6.4808 | 6.2745 | 6.4362 | 6.5696 | 6.5910 | 5.1731 | 5.7570 | 5.5773 | 5.6374 |
| EPB41L2_S598 | EPB41L2 | O43491 | 798 | EAVPEASPVTQAG | 6.4400 | 6.8561 | 6.6288 | 6.9500 | 6.2091 | 6.3659 | 6.9467 | 6.4437 | 6.2487 | 6.7269 | 6.4036 | 6.9007 | 5.3954 | 5.3254 | 5.0710 | 5.0880 |
| EPB41L2_S613 | EPB41L2 | O43491 | 647 | ILKHQASISELKR | 6.9460 | 5.9850 | 6.5232 | 6.1793 | 5.4493 | 5.4591 | 5.6449 | 5.6521 | 6.4132 | 5.7824 | 5.9698 | 5.8578 | 6.8306 | 8.0912 | 6.7060 | 6.5100 |
| EPB41L2_S614 | EPB41L2 | O43491 | 623 | VEGDNIYVRHSNL | 7.2850 | 6.9535 | 7.5478 | 7.6505 | 7.4700 | 7.1909 | 6.2905 | 7.7669 | 7.5114 | 7.2173 | 6.6442 | 6.8570 | 3.4604 | 3.9853 | 2.7432 | 3.4260 |
| EPB41L2_S627 | EPB41L2 | O43491 | 614 | IEGKKNSLRVEGD | 7.7947 | 6.9523 | 8.1393 | 7.1060 | 6.4521 | 6.4661 | 5.8216 | 6.6178 | 7.2121 | 7.1770 | 7.4278 | 7.2256 | 4.1861 | 5.3724 | 2.9025 | 3.1466 |
| EPB41L2_S647 | EPB41L2 | O43491 | 659 | RNFMESTPEPRPN | 7.1008 | 8.2396 | 7.2305 | 7.1761 | 5.4389 | 7.3291 | 7.6426 | 6.7154 | 7.5745 | 7.7621 | 7.3563 | 7.9501 | 2.5247 | 3.8194 | 2.5379 | 3.6020 |
| EPB41L2_S682 | EPB41L2 | O43491 | 39 | NQQNQSSDPEEEK | 6.1823 | 5.9264 | 6.0107 | 5.9334 | 6.3314 | 6.3273 | 6.2188 | 6.0839 | 6.3676 | 6.4338 | 6.1898 | 5.9979 | 6.6984 | 6.7879 | 6.1476 | 6.3629 |
| EPB41L2_S682;S683 | EPB41L2 | O43491 | 88 | LKKQKSYTLVVAK | 6.8258 | 7.3235 | 8.1832 | 8.1085 | 6.7135 | 6.7205 | 6.5710 | 6.9975 | 6.8989 | 7.3239 | 7.1392 | 7.1953 | 3.4179 | 4.1781 | 2.9691 | 3.4342 |
| EPB41L2_S683 | EPB41L2 | O43491-4 | 613 | LIEGKSSHETLNI | 6.0382 | 7.4722 | 6.0655 | 6.8672 | 6.1602 | 6.5952 | 5.6541 | 6.8934 | 6.4213 | 6.1846 | 6.7853 | 6.8882 | 4.9499 | 6.0291 | 5.7553 | 5.2403 |
| EPB41L2_S715 | EPB41L2 | O43491 | 627 | NIYVRHSNLMLEE | 5.0331 | 4.7300 | 5.5556 | 4.7113 | 5.2674 | 5.5912 | 5.6151 | 5.3832 | 5.6147 | 4.8941 | 4.9281 | 4.8382 | 9.2154 | 8.9383 | 10.4909 | 9.1934 |
| EPB41L2_S798 | EPB41L2 | O43491 | 854 | KVNGEVSHVDIDV | 6.7142 | 6.1550 | 6.4925 | 6.8857 | 5.5475 | 5.8368 | 6.8621 | 7.0442 | 7.0892 | 5.8036 | 6.2707 | 6.5225 | 5.7264 | 6.2975 | 5.2379 | 5.5143 |
| EPB41L2_S854 | EPB41L2 | O43491 | 562 | IGVMDQSLMKDFP | 8.4778 | 5.8505 | 6.5429 | 7.5242 | 6.7728 | 6.4637 | 7.0735 | 6.4061 | 7.0825 | 6.4255 | 6.6165 | 6.0130 | 4.7740 | 4.9997 | 4.2087 | 4.7685 |
| EPB41L2_S87 | EPB41L2 | O43491 | 386 | KTHRGLSPAQADS | 6.7546 | 6.3168 | 6.2315 | 7.3127 | 6.4394 | 5.7470 | 5.3465 | 5.7715 | 6.3425 | 6.4195 | 7.0979 | 6.5608 | 5.3710 | 5.9525 | 5.8416 | 6.4942 |
| EPB41L2_T134 | EPB41L2 | O43491 | 402 | ENAKRLSMYGVDL | 5.7662 | 6.4461 | 6.2358 | 6.6100 | 6.1534 | 6.2443 | 6.0686 | 5.9078 | 6.4118 | 5.7622 | 6.3504 | 6.7133 | 6.4017 | 6.2403 | 6.2081 | 6.4799 |
| EPB41L2_T659 | EPB41L2 | O43491 | 682 | SLQTQGSSHETLN | 5.0971 | 5.4891 | 5.6550 | 6.6582 | 6.0431 | 7.2712 | 5.9215 | 7.1110 | 5.4396 | 6.1112 | 6.7247 | 6.5180 | 6.0036 | 6.4529 | 7.1310 | 6.3726 |
| EPB41L2_T673;S683 | EPB41L2 | Q6ZSX4 | 134 | TETKTITYESPQT | 6.7239 | 7.6299 | 7.4942 | 6.6734 | 7.0270 | 5.8021 | 6.0294 | 6.4619 | 7.1943 | 6.3009 | 5.8905 | 8.4159 | 4.1259 | 5.7043 | 3.3746 | 5.1518 |
| EPB41L2_Y623 | EPB41L2 | O43491 | 673;683 | NA | 5.7531 | 6.2067 | 6.0763 | 7.0316 | 6.3054 | 6.6921 | 6.0441 | 6.5957 | 6.8054 | 6.0717 | 6.6350 | 6.4392 | 5.4193 | 6.0362 | 6.0948 | 5.7936 |
| EPB41L2_Y88 | EPB41L2 | O43491 | 682;683 | NA | 6.8082 | 8.1012 | 5.8843 | 6.6713 | 5.5742 | 6.6169 | 5.8631 | 5.8251 | 6.0027 | 5.9469 | 6.5680 | 6.3123 | 5.1914 | 6.5910 | 6.4719 | 5.5714 |
| EPB41L4A_S389 | EPB41L4A | Q9HCS5 | 389 | IKIIAPSPVKSFK | 6.6099 | 6.8051 | 5.9172 | 6.3306 | 6.4896 | 6.1939 | 6.6493 | 6.6254 | 6.3062 | 6.0633 | 5.9363 | 6.0114 | 5.7691 | 6.2139 | 6.1358 | 5.9429 |
| EPB41L5_S418 | EPB41L5 | Q9HCM4 | 517 | QLEMENSPLLSPR | 6.4465 | 6.3934 | 6.1898 | 6.4865 | 6.0392 | 6.2502 | 6.5862 | 6.2939 | 6.2689 | 6.3604 | 6.3775 | 6.2548 | 5.8659 | 6.4046 | 5.9127 | 5.8695 |
| EPB41L5_S418;S436 | EPB41L5 | Q9HCM4 | 550 | LNNVIESPGLNVM | 6.0545 | 6.5615 | 6.5620 | 6.6944 | 6.6108 | 6.7988 | 6.8267 | 6.2857 | 6.5245 | 6.6106 | 6.9451 | 6.6044 | 4.8629 | 5.6892 | 4.7847 | 5.5840 |
| EPB41L5_S436 | EPB41L5 | Q9HCM4 | 436 | IENLPQSPGTDQH | 6.0336 | 6.2134 | 6.5326 | 7.2411 | 6.5744 | 6.1471 | 5.8127 | 6.4058 | 7.1741 | 6.9167 | 6.4456 | 7.0381 | 5.2182 | 5.5493 | 5.1008 | 5.5964 |
| EPB41L5_S517 | EPB41L5 | Q9HCM4 | 582 | KVTKEDSLLSHKN | 6.5190 | 5.5264 | 5.6757 | 5.9012 | 6.4067 | 5.6411 | 6.3306 | 5.8265 | 6.1739 | 5.4619 | 5.2626 | 5.0645 | 8.5554 | 8.0587 | 7.0924 | 6.5035 |
| EPB41L5_S550 | EPB41L5 | Q9HCM4 | 612 | VPLPKESLETLML | 5.9724 | 6.2958 | 4.9806 | 7.1459 | 6.4560 | 4.5155 | 6.3543 | 7.1952 | 6.6932 | 6.1603 | 6.1634 | 5.8708 | 6.9704 | 5.4272 | 6.2558 | 7.5433 |
| EPB41L5_S582 | EPB41L5 | Q9HCM4 | 418 | RSALPVSPSISSA | 4.8745 | 6.7100 | 5.6357 | 5.8290 | 6.8243 | 6.0264 | 6.2770 | 6.6623 | 7.1784 | 6.0591 | 6.7113 | 7.3840 | 5.1128 | 6.2402 | 5.8903 | 6.5847 |
| EPB41L5_S612 | EPB41L5 | Q9HCM4 | 418;436 | NA | 5.4655 | 6.9198 | 7.5727 | 5.0158 | 6.0197 | 5.5401 | 5.6274 | 6.5793 | 7.1988 | 6.2036 | 5.7824 | 6.6912 | 5.0716 | 6.6764 | 6.8540 | 6.7818 |
| EPC1_T352 | EPC1 | Q9H2F5 | 352 | PSSAAATPQQTSP | 5.4696 | 6.0850 | 5.8773 | 6.2746 | 6.7792 | 6.0699 | 6.2361 | 7.1050 | 5.6129 | 5.5166 | 5.0885 | 5.9694 | 6.3114 | 6.9159 | 7.3530 | 7.3356 |
| EPC2_S119 | EPC2 | Q52LR7 | 754 | TRTSAPSPTALKL | 6.3486 | 6.8091 | 6.3280 | 7.0921 | 5.9097 | 6.8033 | 6.4788 | 6.1039 | 6.6548 | 6.9726 | 6.8261 | 6.9956 | 4.7348 | 5.7708 | 5.1113 | 5.0605 |
| EPC2_S754 | EPC2 | Q52LR7 | 119 | PDYDMDSEDETLL | 6.5994 | 6.9779 | 6.5691 | 6.2092 | 6.3208 | 6.2689 | 6.4888 | 5.2437 | 6.1118 | 6.3938 | 6.3184 | 6.2733 | 6.1421 | 6.1274 | 6.0951 | 5.8601 |
| EPHA2_S570 | EPHA2 | P29317 | 897 | VSIRLPSTSGSEG | 5.5824 | 5.7203 | 5.5052 | 6.1727 | 6.6715 | 6.5786 | 6.8027 | 6.7947 | 6.1359 | 5.8842 | 5.7241 | 6.1559 | 6.3778 | 6.6112 | 6.3658 | 6.9170 |
| EPHA2_S892 | EPHA2 | P29317 | 901 | LPSTSGSEGVPFR | 5.3175 | 5.9724 | 5.7690 | 6.2867 | 6.0574 | 5.8201 | 6.1673 | 6.1789 | 5.6886 | 5.5490 | 5.9012 | 5.6432 | 7.0795 | 7.4445 | 7.4129 | 7.7118 |
| EPHA2_S892;S897 | EPHA2 | P29317 | 892 | DFDPRVSIRLPST | 8.0802 | 8.7024 | 9.0804 | 7.5288 | 6.6754 | 8.4620 | 6.3858 | 4.9550 | 5.1147 | 5.8478 | 6.4154 | 5.6913 | 4.4297 | 5.5157 | 3.6555 | 3.4599 |
| EPHA2_S892;S897;S901 | EPHA2 | P29317 | 594 | YVDPHTYEDPNQA | 7.5841 | 6.5977 | 7.1887 | 7.0951 | 7.8578 | 7.7028 | 6.9816 | 6.8357 | 7.1498 | 5.7742 | 5.7156 | 6.2967 | 4.9368 | 4.4536 | 3.5303 | 4.2997 |
| EPHA2_S897 | EPHA2 | P29317 | 570 | NQRARQSPEDVYF | 5.8400 | 6.3135 | 6.3506 | 7.1371 | 6.1494 | 6.0267 | 6.2764 | 5.9171 | 5.4451 | 5.9302 | 6.1832 | 6.7071 | 6.4309 | 6.2142 | 6.5823 | 6.4964 |
| EPHA2_S901 | EPHA2 | P29317 | 575 | QSPEDVYFSKSEQ | 7.0152 | 6.6077 | 6.8769 | 6.8181 | 6.8983 | 7.0312 | 6.4273 | 6.6482 | 7.6230 | 6.8956 | 6.7081 | 6.5859 | 4.7289 | 4.8383 | 4.1184 | 4.1789 |
| EPHA2_Y575 | EPHA2 | P29317 | 960 | HQKRIAYSLLGLK | 6.5884 | 7.2257 | 6.5034 | 7.0557 | 7.0136 | 7.6351 | 7.2888 | 6.6171 | 7.0270 | 6.8830 | 7.7370 | 7.5495 | 3.0877 | 4.2164 | 3.7133 | 3.8583 |
| EPHA2_Y594 | EPHA2 | P29317 | 772 | DDPEATYTTSGGK | 5.5318 | 5.5641 | 6.8900 | 6.1670 | 5.8231 | 7.1563 | 6.2548 | 7.0067 | 6.9959 | 7.4872 | 7.2226 | 7.9190 | 5.4531 | 5.7058 | 4.0779 | 4.7448 |
| EPHA2_Y772 | EPHA2 | P29317 | 892;897 | NA | 6.3245 | 5.9962 | 7.3580 | 6.7526 | 7.4772 | 7.2949 | 7.2502 | 7.2620 | 6.2467 | 6.1683 | 6.9217 | 6.7093 | 4.2119 | 5.3073 | 3.7955 | 4.9237 |
| EPHA2_Y960 | EPHA2 | P29317 | 892;897;901 | NA | 7.6293 | 7.2538 | 7.4278 | 7.9912 | 7.7189 | 7.3684 | 7.8254 | 6.7075 | 4.4188 | 5.7833 | 6.4504 | 6.2746 | 3.6867 | 5.1577 | 3.8576 | 4.4484 |
| EPHB2_S776 | EPHB2 | P29323 | 389 | PRQLGLTEPRIYI | 7.2939 | 8.1285 | 4.8136 | 7.6744 | 6.9348 | 7.1676 | 5.3724 | 9.2422 | 6.2420 | 6.0210 | 5.5018 | 5.4629 | 3.7534 | 5.6983 | 6.0607 | 4.6326 |
| EPHB2_T389 | EPHB2 | P29323 | 776 | FLEDDTSDPTYTS | 5.6160 | 6.1755 | 6.4949 | 5.5979 | 6.0354 | 6.0751 | 6.7086 | 6.4963 | 6.1774 | 5.3961 | 5.7102 | 6.0392 | 6.9176 | 7.3398 | 6.6584 | 5.5614 |
| EPN1_S521 | EPN1 | Q9Y6I3-1 | 521 | GEVPARSPGAFDM | 5.8875 | 5.8716 | 6.4004 | 6.0922 | 6.4330 | 6.2849 | 6.3600 | 7.2322 | 6.3818 | 6.6624 | 6.6338 | 6.5625 | 5.6301 | 5.9406 | 5.7647 | 5.8623 |
| EPN1_S622 | EPN1 | Q9Y6I3-1 | 622 | LNQLRLSPVPPVP | 4.9921 | 4.6534 | 5.6232 | 5.1082 | 5.6982 | 5.4441 | 5.9242 | 7.0255 | 5.9209 | 6.6465 | 5.9203 | 6.1900 | 7.1324 | 8.1613 | 7.6443 | 7.9155 |
| EPN1_T556 | EPN1 | Q9Y6I3-1 | 580 | VSRPGPTPPGAKA | 6.6790 | 6.0286 | 6.8161 | 6.3615 | 6.3138 | 6.5031 | 6.8264 | 6.8867 | 7.0135 | 7.6902 | 7.7703 | 8.2781 | 3.6581 | 4.6645 | 3.6351 | 4.8750 |
| EPN1_T580 | EPN1 | Q9Y6I3-1 | 556 | TPPTRKTPESFLG | 5.8494 | 5.6247 | 5.7615 | 5.4985 | 5.9814 | 6.4191 | 6.1357 | 5.8484 | 6.2759 | 6.6234 | 6.2881 | 7.2077 | 6.4705 | 6.5162 | 6.6234 | 6.8763 |
| EPN1_Y222 | EPN1 | Q9Y6I3-1 | 222 | TLKDFQYVDRDGK | 5.8991 | 6.0178 | 6.5797 | 6.3296 | 6.0746 | 6.3434 | 6.3349 | 6.4061 | 6.3208 | 6.0059 | 6.1633 | 6.6234 | 5.9354 | 6.3624 | 6.0731 | 6.5304 |
| EPN2_S172 | EPN2 | O95208 | 173 | TFGRGSSQPNLST | 5.4323 | 6.0943 | 6.1797 | 6.0420 | 6.3727 | 6.3449 | 5.6156 | 6.7578 | 5.4153 | 5.9381 | 6.5831 | 6.3832 | 6.7531 | 6.8714 | 7.0099 | 6.2067 |
| EPN2_S173 | EPN2 | O95208 | 172 | ITFGRGSSQPNLS | 6.1054 | 6.1700 | 5.9330 | 6.5326 | 6.5896 | 6.4204 | 5.9468 | 7.4629 | 5.8139 | 5.5025 | 6.3119 | 5.8569 | 6.3264 | 6.4961 | 6.5245 | 6.0071 |
| EPN2_S426 | EPN2 | O95208 | 426 | SAGKRASDAWGAV | 6.7974 | 5.1795 | 7.9356 | 6.9110 | 8.4163 | 5.8150 | 7.0156 | 6.2433 | 7.2730 | 5.5656 | 5.9145 | 6.0412 | 5.5153 | 5.2579 | 4.8084 | 5.3102 |
| EPOP_S15 | EPOP | A6NHQ4 | 15 | RLAVPASPRGSPC | 6.6731 | 6.6844 | 6.9227 | 7.2890 | 6.4964 | 6.8292 | 6.9097 | 6.8110 | 6.8913 | 6.2283 | 6.5205 | 6.6944 | 5.0694 | 4.3845 | 4.4540 | 5.1422 |
| EPRS1_S1336 | EPRS1 | P07814 | 886 | SQSSDSSPTRNSE | 5.9149 | 6.3059 | 6.4167 | 6.2981 | 6.2412 | 6.1806 | 6.0403 | 6.3666 | 6.6003 | 6.3898 | 6.5777 | 6.4251 | 5.9322 | 6.4071 | 5.9395 | 5.9640 |
| EPRS1_S434 | EPRS1 | P07814 | 1336 | YRRRLLSVNIRVR | 6.9004 | 6.6522 | 6.2402 | 6.6168 | 6.2526 | 6.4991 | 5.8611 | 6.1425 | 6.4902 | 5.9420 | 5.7058 | 5.7328 | 6.8362 | 6.3433 | 6.2978 | 5.4872 |
| EPRS1_S882;S886 | EPRS1 | P07814 | 434 | LNNTVLSKRKLTW | 5.2911 | 6.3303 | 7.5416 | 5.8340 | 5.0622 | 7.1365 | 4.7117 | 6.5896 | 6.9319 | 5.6557 | 5.4856 | 7.8171 | 6.1885 | 5.6010 | 6.0082 | 7.8151 |
| EPRS1_S886 | EPRS1 | P07814 | 954 | VEYKPVSATGAED | 6.3682 | 6.5575 | 7.6492 | 6.9490 | 5.9999 | 6.1177 | 6.6807 | 6.5207 | 6.3141 | 6.4424 | 7.4878 | 6.7706 | 5.0880 | 5.0771 | 5.1782 | 4.7990 |
| EPRS1_S954 | EPRS1 | P07814 | 882;886 | NA | 7.4431 | 5.9029 | 7.5532 | 4.8813 | 6.1788 | 6.4396 | 7.3280 | 7.0478 | 8.1977 | 5.6763 | 5.3370 | 6.1817 | 5.2317 | 6.2778 | 4.6048 | 5.7185 |
| EPS15_S108 | EPS15 | P42566 | 796 | SINKLDSPDPFKL | 5.6349 | 5.5327 | 5.8869 | 4.8474 | 6.3681 | 6.1644 | 5.6946 | 5.8753 | 6.1880 | 6.2333 | 5.9628 | 6.0380 | 8.4400 | 7.0923 | 7.0780 | 6.9634 |
| EPS15_S563 | EPS15 | P42566 | 563 | ESPARSSPELLPS | 5.8668 | 5.7684 | 6.2371 | 5.5386 | 6.0974 | 5.9529 | 5.7983 | 6.4682 | 6.4595 | 6.0473 | 5.8007 | 6.1383 | 7.0413 | 6.9157 | 7.0885 | 6.7808 |
| EPS15_S796 | EPS15 | P42566 | 108 | RFHDTSSPLLISG | 5.4139 | 5.6715 | 5.8912 | 5.5319 | 6.1629 | 6.0166 | 6.0564 | 6.8658 | 5.9377 | 6.1115 | 6.6026 | 6.5557 | 7.2257 | 7.0664 | 6.2765 | 6.6137 |
| EPS15_S814 | EPS15 | P42566 | 814 | PFPGNDSPKEKDP | 6.3357 | 6.6438 | 7.2999 | 5.9266 | 5.7546 | 6.5750 | 5.7856 | 5.7151 | 6.5072 | 6.2930 | 6.5278 | 7.0745 | 6.3735 | 5.6158 | 5.2833 | 6.2885 |
| EPS15_S847 | EPS15 | P42566 | 847 | SNFANFSAYPSEE | 5.4861 | 4.6488 | 5.2808 | 5.4989 | 5.6651 | 5.5325 | 4.8642 | 5.4834 | 5.7587 | 5.5628 | 5.4807 | 4.7424 | 9.3313 | 9.3218 | 8.9611 | 8.3816 |
| EPS15_T777 | EPS15 | P42566 | 777 | LPPKIGTPTRPCP | 6.0630 | 5.9205 | 6.7612 | 6.1307 | 6.4480 | 6.4318 | 6.0498 | 6.3084 | 6.8594 | 6.5496 | 6.7532 | 7.0294 | 5.8995 | 5.5500 | 5.3498 | 5.8957 |
| EPS15L1_S107 | EPS15L1 | Q9UBC2-2 | 107 | PKFHDTSSPLMVT | 5.6859 | 5.6457 | 5.9078 | 5.8073 | 6.5715 | 6.3625 | 5.9124 | 6.6507 | 6.3598 | 6.3651 | 6.5029 | 6.4840 | 6.4476 | 6.6705 | 6.5568 | 6.0693 |
| EPS15L1_S229 | EPS15L1 | Q9UBC2-2 | 593 | SLAERGSFGAMDD | 4.6529 | 4.5034 | 4.8358 | 4.5660 | 5.0866 | 4.7613 | 5.0326 | 5.7752 | 5.3516 | 5.0597 | 5.2272 | 5.6500 | 10.0750 | 9.0935 | 10.5288 | 9.8004 |
| EPS15L1_S244;S255 | EPS15L1 | Q9UBC2-2 | 575 | DGAHGASLTDLAN | 5.4540 | 5.4233 | 6.2521 | 5.4752 | 5.9331 | 5.9221 | 5.7253 | 6.4518 | 6.2326 | 5.7828 | 6.0763 | 6.0593 | 7.3094 | 7.2814 | 7.4520 | 7.1693 |
| EPS15L1_S255 | EPS15L1 | Q9UBC2-2 | 229 | VPVLPASPPPKDS | 6.2190 | 5.8865 | 6.6920 | 5.7608 | 7.0450 | 6.3844 | 5.8746 | 6.6728 | 6.2536 | 6.8043 | 6.6152 | 6.9052 | 5.8071 | 5.6293 | 5.5148 | 5.9353 |
| EPS15L1_S49 | EPS15L1 | Q9UBC2-2 | 366 | PPSERGTPGPDSS | 4.7535 | 4.3626 | 5.1480 | 5.0115 | 5.4392 | 5.6918 | 5.2304 | 5.6778 | 6.2411 | 5.3614 | 5.7641 | 5.4931 | 8.9827 | 8.2479 | 9.0400 | 9.5548 |
| EPS15L1_S575 | EPS15L1 | Q9UBC2-2 | 255 | NSTGSLSPKHSLK | 6.2172 | 7.4804 | 5.7129 | 6.2877 | 7.2271 | 6.7037 | 5.4246 | 8.9773 | 6.3137 | 6.5672 | 6.7776 | 6.3253 | 4.7671 | 5.7484 | 4.9585 | 4.5113 |

| | | | | | | | | | | | | | | | | | | | | | |
|---|---|---|---|---|---|---|---|---|---|---|---|---|---|---|---|---|---|---|---|---|---|
| EPS15L1_S593 | EPS15L1 | Q9UBC2-2 | | 49 | LKKSGLSDIILGK | 6.3826 | 6.4442 | 6.2917 | 6.0913 | 6.3196 | 6.2724 | 6.1473 | 6.3647 | 7.1949 | 6.4369 | 6.4665 | 6.3404 | 5.9662 | 5.9820 | 5.6292 | 5.6701 |
| EPS15L1_S672 | EPS15L1 | Q9UBC2-2 | | 797 | PPSGKSTPVSQLG | 5.5554 | 5.8593 | 6.4465 | 5.3531 | 6.0736 | 5.6754 | 5.3104 | 6.0084 | 6.0757 | 6.1546 | 5.5479 | 6.3689 | 8.0677 | 7.2343 | 6.9702 | 7.2986 |
| EPS15L1_T366 | EPS15L1 | Q9UBC2-2 | | 672 | SDPFRGSATDDFF | 4.7367 | 4.4282 | 5.2042 | 4.9110 | 6.2755 | 4.9650 | 4.5564 | 5.6077 | 5.4302 | 5.2611 | 5.4943 | 5.3937 | 9.9113 | 9.7438 | 9.4481 | 8.6328 |
| EPS15L1_T797 | EPS15L1 | Q9UBC2-2 | 244;255 | | NA | 5.2813 | 6.6892 | 6.0255 | 5.9605 | 6.1467 | 7.0634 | 5.2354 | 7.0571 | 5.5389 | 5.4764 | 6.5284 | 6.3495 | 6.8876 | 6.7818 | 6.7202 | 6.2580 |
| EPS8_S625 | EPS8 | Q12929 | | 625 | DTPPAPSPPPTPA | 6.0679 | 5.5999 | 6.5511 | 6.3307 | 6.6223 | 6.4056 | 5.9427 | 5.9427 | 6.7562 | 6.7473 | 6.5185 | 6.4408 | 6.0064 | 6.6524 | 5.6285 | 5.7869 |
| EPS8_S659 | EPS8 | Q12929 | | 685 | PVDRRKSQMEEVQ | 5.7228 | 5.8802 | 6.7025 | 6.3089 | 6.2746 | 6.6930 | 6.2437 | 5.7049 | 6.7562 | 6.5905 | 6.3435 | 6.8557 | 6.0574 | 6.4864 | 5.3856 | 5.9941 |
| EPS8_S685 | EPS8 | Q12929 | | 659 | NITRQNSSSSDSG | 6.0261 | 6.3885 | 7.2495 | 5.1788 | 6.2040 | 6.2996 | 6.1760 | 5.5335 | 6.8234 | 6.6356 | 6.6494 | 6.2287 | 6.1683 | 6.7663 | 6.1826 | 5.4896 |
| EPS8_T655 | EPS8 | Q12929 | | 655 | KVPANITRQNSSS | 9.3006 | 7.1372 | 7.3378 | 6.9287 | 4.9337 | 4.8359 | 6.3240 | 5.0886 | 7.9064 | 5.8447 | 7.2485 | 6.9049 | 5.3406 | 5.8345 | 4.5212 | 4.5127 |
| EPS8L2_S233 | EPS8L2 | Q9H6S3-3 | | 586 | KYWGPASPTHKLP | 6.3311 | 6.4155 | 6.5987 | 6.0619 | 6.4774 | 6.7987 | 6.1409 | 7.0671 | 6.3858 | 6.2085 | 6.5067 | 6.5209 | 5.5251 | 5.8556 | 5.6035 | 5.5027 |
| EPS8L2_S465 | EPS8L2 | Q9H6S3-3 | | 233 | QHRGGDSPEAKNR | 6.6975 | 6.9555 | 6.9788 | 6.4514 | 6.0783 | 6.6515 | 6.1106 | 6.5644 | 6.6124 | 6.7101 | 6.8100 | 6.5726 | 5.1753 | 5.5664 | 4.8137 | 5.2516 |
| EPS8L2_S586 | EPS8L2 | Q9H6S3-3 | | 465 | APIEEVSPVSRQS | 5.3683 | 7.0442 | 6.5365 | 6.9047 | 5.9334 | 6.6319 | 6.7252 | 6.6924 | 6.9057 | 6.1088 | 6.9929 | 6.8862 | 5.1006 | 5.1724 | 5.8052 | 5.1918 |
| ERAL1_S173 | ERAL1 | O75616 | | 173 | DTPGIISPGKQKR | 6.7519 | 6.2747 | 6.7464 | 7.1361 | 6.2791 | 6.1952 | 6.2637 | 6.5637 | 6.6095 | 6.7585 | 6.6010 | 6.8080 | 4.9574 | 5.4397 | 5.2178 | 5.3972 |
| ERBB2_S1083 | ERBB2 | P04626 | | 1083 | PSEGAGSDVFDGD | 6.1553 | 6.6188 | 6.4824 | 6.3620 | 6.0855 | 5.7833 | 6.0954 | 6.0098 | 6.6412 | 7.1248 | 6.9492 | 6.9503 | 5.9154 | 5.9583 | 5.2500 | 5.6184 |
| ERBB2_S1151 | ERBB2 | P04626 | | 1151 | VRPQPPSPREGPL | 6.7366 | 6.6263 | 5.7177 | 6.5173 | 5.9006 | 5.8856 | 5.9006 | 5.3780 | 6.5123 | 5.6399 | 6.7842 | 5.8869 | 7.5500 | 6.8456 | 5.8831 | 6.2353 |
| ERBB2_S998 | ERBB2 | P04626 | | 998 | EDLGPASPLDSTF | 6.0750 | 6.9695 | 6.1740 | 7.0887 | 6.0503 | 6.0503 | 6.3472 | 6.9994 | 6.6936 | 6.8510 | 7.2386 | 6.9702 | 5.3403 | 5.1439 | 4.8357 | 5.1724 |
| ERBB3_S982 | ERBB3 | P21860 | | 982 | LVIKRESGPGIAP | 4.9338 | 4.5666 | 5.2463 | 5.3219 | 5.8240 | 4.9785 | 5.6255 | 5.4813 | 6.7116 | 7.3774 | 6.9541 | 6.8432 | 7.0114 | 8.7199 | 7.8820 | 6.5226 |
| ERBIN_S1106 | ERBIN | Q96RT1-8 | | 440 | EDVMFISDNESFN | 6.1189 | 6.4297 | 5.8554 | 6.1704 | 6.4632 | 6.2983 | 6.4521 | 6.2889 | 6.3833 | 5.9823 | 6.3057 | 6.5678 | 6.1541 | 6.1605 | 5.9491 | 6.4202 |
| ERBIN_S1133 | ERBIN | Q96RT1-8 | | 1158 | IPERTMSVSDFNY | 7.7376 | 6.6426 | 7.2812 | 6.7659 | 5.2925 | 5.7297 | 6.6844 | 5.4804 | 6.5681 | 5.6704 | 6.7083 | 6.5910 | 5.4964 | 6.4758 | 4.9411 | 5.9347 |
| ERBIN_S1148 | ERBIN | Q96RT1-8 | | 715 | GDILNSSTEEKFK | 6.3792 | 6.7155 | 6.7041 | 6.9690 | 6.3167 | 6.3338 | 6.5176 | 6.4115 | 6.6001 | 6.9519 | 7.2875 | 7.6506 | 4.6213 | 5.0434 | 4.4010 | 5.0970 |
| ERBIN_S1158 | ERBIN | Q96RT1-8 | | 857 | KSTEDLSPQKSGP | 6.7134 | 6.6553 | 6.2840 | 6.9148 | 6.9684 | 6.5749 | 7.0260 | 6.8859 | 6.8855 | 6.2495 | 6.5731 | 6.6040 | 4.8547 | 4.9279 | 4.7840 | 5.0987 |
| ERBIN_S1179 | ERBIN | Q96RT1-8 | | 1116 | FHSAGRTPPMMPG | 6.4748 | 6.4485 | 6.3260 | 6.7677 | 6.1251 | 6.4732 | 6.8654 | 7.0361 | 6.7263 | 6.3325 | 6.6741 | 6.9938 | 5.1166 | 5.3333 | 4.6468 | 5.6599 |
| ERBIN_S1293 | ERBIN | Q96RT1-8 | | 569 | EPEAEISPGSLPV | 6.1870 | 6.7234 | 6.3437 | 7.1502 | 6.4525 | 6.3435 | 6.6117 | 6.4813 | 6.3589 | 6.6612 | 6.8070 | 7.0793 | 5.3067 | 5.2695 | 5.1314 | 5.0928 |
| ERBIN_S440 | ERBIN | Q96RT1-8 | | 1293 | SVARHPSREQLID | 5.0418 | 5.9323 | 6.3970 | 5.8115 | 4.8305 | 6.6493 | 6.5326 | 6.8858 | 4.9850 | 5.6267 | 5.9119 | 6.6165 | 6.1351 | 6.8490 | 7.2404 | 8.5546 |
| ERBIN_S444 | ERBIN | Q96RT1-8 | | 485 | NLKRYPTPYPDEL | 6.9049 | 5.1148 | 5.1825 | 5.6798 | 7.4100 | 5.0082 | 5.2773 | 5.1826 | 5.2317 | 4.5782 | 4.2360 | 4.5947 | 9.4254 | 8.8339 | 8.7226 | 8.6174 |
| ERBIN_S569 | ERBIN | Q96RT1-8 | | 1179 | PNARVGSEHSLLD | 5.0592 | 5.2901 | 6.7257 | 5.4025 | 4.1133 | 5.9269 | 6.7198 | 5.5851 | 5.2786 | 6.1102 | 7.4529 | 7.8444 | 6.0775 | 6.1956 | 6.5738 | 9.6443 |
| ERBIN_S598;S603 | ERBIN | Q96RT1-8 | | 1106 | PEGDYLSYREFHS | 6.1972 | 6.2887 | 6.5570 | 6.0269 | 5.7849 | 6.4731 | 6.3459 | 6.4723 | 6.6075 | 6.5466 | 6.5728 | 7.2554 | 5.7274 | 6.0286 | 5.3085 | 5.8073 |
| ERBIN_S602;S603 | ERBIN | Q96RT1-8 | | 444 | FISDNESFNPSLW | 5.8330 | 5.9007 | 5.8279 | 5.8368 | 6.9202 | 6.1891 | 6.2920 | 6.4510 | 6.3161 | 6.2976 | 6.4292 | 5.4543 | 7.0162 | 6.8177 | 5.9605 | 6.4575 |
| ERBIN_S620 | ERBIN | Q96RT1-8 | | 620 | PPLIETSINQPKV | 8.0143 | 7.4459 | 7.4273 | 8.6538 | 7.6981 | 6.8592 | 7.3383 | 6.6388 | 7.3618 | 6.3004 | 6.5189 | 5.7288 | 3.7797 | 4.0167 | 2.5470 | 3.6709 |
| ERBIN_S682 | ERBIN | Q96RT1-8 | | 915 | NIVRSKSATLLYD | 5.5282 | 5.4688 | 6.0852 | 5.7135 | 5.7147 | 5.8316 | 5.6480 | 5.9431 | 5.7068 | 5.9095 | 6.2621 | 6.3568 | 7.0738 | 7.6358 | 7.2111 | 7.9110 |
| ERBIN_S715 | ERBIN | Q96RT1-8 | | 872 | SVVKSHSITNMEI | 4.8867 | 4.9278 | 5.2313 | 4.9382 | 5.2556 | 6.0449 | 5.7307 | 5.6154 | 4.6497 | 4.8437 | 5.0513 | 4.8832 | 8.6074 | 8.9931 | 9.6818 | 10.6593 |
| ERBIN_S794 | ERBIN | Q96RT1-8 | | 794 | DIVLGTSFLSINS | 6.2311 | 6.4282 | 5.9486 | 6.4634 | 6.6308 | 6.8672 | 6.9681 | 5.6949 | 6.5891 | 6.0390 | 6.3933 | 6.3831 | 6.1077 | 6.0837 | 5.3118 | 5.8600 |
| ERBIN_S800 | ERBIN | Q96RT1-8 | | 852 | DLGISKSTEDLSP | 5.2997 | 6.2430 | 5.8563 | 6.0110 | 4.9003 | 5.8920 | 6.3417 | 6.1574 | 5.1653 | 5.8473 | 6.7033 | 6.8987 | 6.6337 | 6.6930 | 7.4591 | 7.8982 |
| ERBIN_S852 | ERBIN | Q96RT1-8 | | 1148 | PQSARPSINEIPE | 6.8254 | 5.3637 | 6.6899 | 5.7554 | 5.5015 | 6.8160 | 6.2638 | 7.2038 | 6.9884 | 7.2954 | 6.5802 | 7.4482 | 5.3292 | 7.1709 | 4.0061 | 4.7621 |
| ERBIN_S857 | ERBIN | Q96RT1-8 | | 1300 | REQLIDYLMLKVA | 6.7571 | 5.7372 | 7.2636 | 6.7694 | 7.7126 | 7.2171 | 5.8673 | 7.5935 | 7.8509 | 6.3848 | 6.4792 | 6.6941 | 4.8830 | 4.8351 | 2.9159 | 5.0391 |
| ERBIN_S872 | ERBIN | Q96RT1-8 | | 682 | QDTSLCSPVKQTH | 6.1014 | 6.8225 | 7.3934 | 6.2092 | 6.9789 | 6.5477 | 6.5477 | 6.1804 | 7.2491 | 6.7967 | 6.2532 | 7.1656 | 5.4608 | 5.0235 | 4.4922 | 4.7776 |
| ERBIN_S915 | ERBIN | Q96RT1-8 | | 800 | SFLSINSKEETEH | 6.3079 | 6.1969 | 5.5421 | 6.9072 | 5.0075 | 5.8996 | 6.7193 | 6.0013 | 5.3326 | 7.2477 | 6.2478 | 6.7547 | 6.4327 | 5.9936 | 6.7778 | 6.6314 |
| ERBIN_T1116 | ERBIN | Q96RT1-8 | | 1133 | LSARTYSIDGPNA | 6.1718 | 6.1865 | 7.6050 | 5.6067 | 5.1951 | 5.8887 | 6.8319 | 6.1960 | 6.6555 | 7.6176 | 7.0937 | 7.1613 | 4.5964 | 6.2272 | 5.3639 | 5.6028 |
| ERBIN_T485 | ERBIN | Q96RT1-8 | | 1104 | QIPEGDYLSYREF | 6.0960 | 6.0528 | 6.0212 | 6.3691 | 6.3955 | 7.1777 | 6.9354 | 6.0092 | 6.6766 | 6.4443 | 6.5623 | 7.0946 | 5.6321 | 6.1315 | 5.1493 | 5.2525 |
| ERBIN_Y1104 | ERBIN | Q96RT1-8 | 602;603 | | NA | 6.1892 | 5.9853 | 6.3442 | 6.7954 | 6.2002 | 5.4696 | 6.3976 | 6.3104 | 6.3623 | 6.0717 | 6.3394 | 6.1644 | 6.5611 | 5.9321 | 6.4652 | 6.4119 |
| ERBIN_Y1300 | ERBIN | Q96RT1-8 | 598;603 | | NA | 5.8427 | 6.1770 | 6.3237 | 6.6490 | 5.5949 | 5.6847 | 6.3804 | 6.0721 | 6.7234 | 6.4927 | 6.1763 | 6.0605 | 6.6655 | 6.2837 | 6.6696 | 6.2040 |
| ERC1_S17 | ERC1 | Q8IUD2 | | 191 | SIKTFWSPELKKE | 6.5978 | 6.9172 | 7.0218 | 6.7277 | 6.6077 | 7.0528 | 6.4016 | 6.8254 | 6.8067 | 6.8502 | 7.0105 | 6.8178 | 4.6569 | 4.8735 | 4.2846 | 4.5476 |
| ERC1_S17;S21 | ERC1 | Q8IUD2 | | 21 | SQSPGRSPRLPRS | 5.8974 | 6.1102 | 6.2093 | 5.6054 | 5.7233 | 6.5424 | 6.1823 | 6.3135 | 5.9494 | 6.1564 | 6.4240 | 6.2150 | 6.9454 | 6.9122 | 6.5063 | 6.3075 |
| ERC1_S191 | ERC1 | Q8IUD2 | | 17 | VEPSSQSPGRSPR | 5.4766 | 7.2617 | 7.6595 | 7.0093 | 6.5901 | 6.6222 | 5.3124 | 5.0160 | 6.2657 | 6.7704 | 8.0160 | 6.8879 | 5.4967 | 6.1055 | 5.2390 | 4.2710 |
| ERC1_S21 | ERC1 | Q8IUD2 | | 37 | GHRRTNSTGGSSG | 6.3088 | 5.7003 | 8.0103 | 6.3728 | 6.8103 | 5.5237 | 5.4310 | 7.0128 | 6.9019 | 6.6371 | 6.4594 | 7.3041 | 6.0826 | 5.0366 | 5.3891 | 5.0194 |
| ERC1_S37 | ERC1 | Q8IUD2 | | 55 | GSGKTLSMENIQS | 5.8621 | 7.2451 | 6.5266 | 6.6546 | 6.5479 | 7.2482 | 5.4467 | 7.2012 | 5.4435 | 6.3660 | 7.1955 | 6.6301 | 5.4166 | 5.5333 | 5.9048 | 4.7778 |
| ERC1_S55 | ERC1 | Q8IUD2 | 17;21 | | NA | 6.9878 | 7.0219 | 7.3707 | 5.8024 | 5.8985 | 7.0871 | 6.3877 | 7.0966 | 6.5730 | 6.1038 | 7.0948 | 6.7432 | 5.2848 | 5.1480 | 4.8791 | 4.5206 |
| ERCC2_T122 | ERCC2 | P18074 | | 122 | CIHPEVTPLRFGK | 6.4628 | 6.7161 | 8.6748 | 7.7775 | 5.9552 | 6.5120 | 7.0863 | 6.3849 | 5.9829 | 6.0926 | 6.6351 | 6.4074 | 4.4682 | 4.8856 | 4.6722 | 5.2865 |
| ERCC5_S156;S157 | ERCC5 | P28715 | | 384 | VDEGSISPRTLSA | 5.9959 | 6.1512 | 6.2044 | 6.1041 | 6.1094 | 6.0916 | 6.1771 | 5.9549 | 6.5678 | 6.1995 | 6.0995 | 5.7662 | 6.8076 | 6.5059 | 6.5231 | 6.7419 |
| ERCC5_S341 | ERCC5 | P28715 | | 526 | RELTPASPTCTNS | 6.0291 | 5.8643 | 6.0145 | 6.2816 | 6.2718 | 6.4400 | 6.3400 | 6.2175 | 6.4495 | 6.6595 | 5.9887 | 6.2628 | 6.2575 | 6.3203 | 6.3580 | 6.2448 |
| ERCC5_S384 | ERCC5 | P28715 | | 341 | PDATPPSPRTLLA | 6.3367 | 6.1823 | 5.5396 | 5.8091 | 6.0063 | 6.3123 | 6.2128 | 6.2341 | 6.1021 | 6.1429 | 6.0568 | 5.6184 | 6.7589 | 6.9632 | 6.8499 | 6.8747 |
| ERCC5_S526 | ERCC5 | P28715 | | 563 | DSSLLSSDDETKC | 5.1556 | 6.1756 | 6.1239 | 5.7672 | 5.5618 | 5.8753 | 5.8047 | 5.6875 | 5.9185 | 6.6979 | 6.1318 | 6.2013 | 7.1781 | 8.0753 | 6.5352 | 7.1104 |
| ERCC5_S563 | ERCC5 | P28715 | 156;157 | | NA | 5.7461 | 5.9937 | 6.2451 | 6.1691 | 5.8885 | 5.9297 | 6.3677 | 5.5392 | 6.1946 | 6.0523 | 6.3303 | 6.4083 | 6.9723 | 6.3518 | 6.5680 | 7.2433 |
| ERCC6_S158 | ERCC6 | Q03468 | | 158 | KIIEQLSPQAATS | 6.9878 | 7.2700 | 6.4351 | 7.0414 | 6.0920 | 6.7696 | 6.7611 | 6.8137 | 7.3416 | 7.5593 | 7.3863 | 7.3101 | 3.8273 | 4.5745 | 3.7176 | 4.1127 |
| ERCC6_T303 | ERCC6 | Q03468 | | 303 | KAPAPVTPPAPVQ | 6.3146 | 5.9654 | 6.9245 | 7.0386 | 7.0243 | 6.9060 | 6.9556 | 6.7748 | 6.5934 | 6.5144 | 6.8729 | 6.7016 | 4.4906 | 5.2201 | 4.8022 | 4.9009 |
| ERCC6L_S1000 | ERCC6L | Q2NKX8 | | 820 | LPKGFGSVEELCT | 5.6804 | 5.8664 | 6.4027 | 6.0583 | 5.7259 | 6.0694 | 5.8631 | 6.1626 | 6.3357 | 6.2288 | 6.3709 | 6.4762 | 6.4134 | 6.9000 | 6.5891 | 6.8573 |
| ERCC6L_S1000;S1004 | ERCC6L | Q2NKX8 | | 1098 | SLASRRSLINMVL | 6.5418 | 6.6917 | 6.6132 | 6.4839 | 5.7829 | 6.1884 | 6.3533 | 6.5917 | 7.2089 | 6.6371 | 6.7024 | 7.0737 | 5.3973 | 5.9846 | 4.7124 | 5.0366 |
| ERCC6L_S1004 | ERCC6L | Q2NKX8 | | 1069 | TPKNDISPPGRFF | 6.0408 | 6.7126 | 7.2755 | 6.6479 | 6.2435 | 6.5168 | 6.3577 | 6.3886 | 6.8416 | 6.6654 | 6.9537 | 6.7561 | 5.0404 | 5.5205 | 4.8019 | 5.2369 |
| ERCC6L_S1028 | ERCC6L | Q2NKX8 | | 14 | PEAEALSPEQAAH | 6.2862 | 6.6136 | 7.1860 | 7.1596 | 6.5396 | 7.2147 | 6.4844 | 6.5339 | 6.4266 | 6.8225 | 7.3163 | 6.9182 | 4.3934 | 5.2222 | 4.2659 | 4.6169 |
| ERCC6L_S1069 | ERCC6L | Q2NKX8 | | 1028 | KARRIVSDGEDED | 6.6812 | 6.5747 | 7.2491 | 6.7149 | 6.1696 | 6.5920 | 6.6758 | 6.7003 | 6.5873 | 6.5097 | 5.5170 | 6.1767 | 5.4257 | 5.8057 | 5.3134 | 5.3069 |
| ERCC6L_S1077;S1082 | ERCC6L | Q2NKX8 | | 946 | EEPSASSPQYACD | 5.7533 | 6.2121 | 6.5750 | 5.8617 | 5.9431 | 6.1774 | 5.7405 | 7.0658 | 6.5747 | 6.6222 | 6.1816 | 6.4379 | 6.0924 | 6.0662 | 6.2276 | 6.4686 |
| ERCC6L_S1082 | ERCC6L | Q2NKX8 | | 1188 | GEQLVGSPQDKAA | 6.4243 | 6.0907 | 7.4440 | 7.1077 | 6.1541 | 7.2037 | 6.5039 | 6.8462 | 6.9653 | 7.0700 | 7.3178 | 6.9951 | 3.7254 | 4.8440 | 4.4243 | 4.8832 |
| ERCC6L_S1098 | ERCC6L | Q2NKX8 | | 864 | QEDPLESFNYVLS | 8.4069 | 7.5866 | 7.0421 | 7.9736 | 6.8304 | 6.4922 | 7.3234 | 6.5052 | 7.5419 | 5.8343 | 6.9878 | 6.5918 | 3.5644 | 4.3034 | 3.1029 | 3.9131 |
| ERCC6L_S1118 | ERCC6L | Q2NKX8 | | 791 | EKQDLSSIKVNVT | 7.5864 | 8.0775 | 7.2976 | 8.3607 | 5.8816 | 6.7861 | 7.1963 | 6.3732 | 7.4474 | 6.8094 | 7.7478 | 7.6804 | 2.7011 | 3.9231 | 2.5504 | 3.5811 |
| ERCC6L_S1173;S1188 | ERCC6L | Q2NKX8 | | 995 | RAGFVHSKTCLSW | 7.5692 | 8.2930 | 7.2108 | 7.5174 | 5.6315 | 7.1983 | 7.6433 | 6.9399 | 7.7501 | 7.3583 | 8.1098 | 7.8419 | 2.2698 | 3.2508 | 2.1200 | 3.2959 |
| ERCC6L_S1188 | ERCC6L | Q2NKX8 | | 1000 | HSKTCLSWEFSEK | 6.6176 | 7.2349 | 6.5417 | 7.0437 | 6.1186 | 6.7114 | 6.5763 | 5.9275 | 7.9901 | 8.7668 | 8.1964 | 8.2045 | 3.1155 | 3.9445 | 3.3058 | 3.7046 |
| ERCC6L_S14 | ERCC6L | Q2NKX8 | | 971 | QNFSSQSLEHVEK | 6.5142 | 6.8853 | 7.3138 | 7.1409 | 6.2382 | 6.8151 | 6.5050 | 7.2566 | 7.3961 | 7.0179 | 7.4342 | 7.4754 | 3.6943 | 4.4552 | 3.6618 | 4.1960 |
| ERCC6L_S739 | ERCC6L | Q2NKX8 | | 739 | EPVFPSSTKKKCP | 8.8877 | 8.1626 | 6.8895 | 7.1503 | 4.8429 | 7.3614 | 8.2290 | 6.5651 | 8.2437 | 6.9748 | 7.8980 | 7.5440 | 2.3410 | 3.5129 | 2.0025 | 3.3943 |
| ERCC6L_S774 | ERCC6L | Q2NKX8 | | 774 | ISSKMASVVIDDL | 7.6912 | 7.7770 | 6.6051 | 7.5964 | 5.7196 | 6.8415 | 7.3449 | 6.5800 | 7.6026 | 6.4180 | 6.9717 | 7.0834 | 3.8211 | 4.6956 | 3.1289 | 4.1230 |
| ERCC6L_S791 | ERCC6L | Q2NKX8 | | 1004 | CLSWEFSEKDDEP | 6.8945 | 7.0661 | 8.3076 | 7.2066 | 7.0948 | 7.5675 | 7.4827 | 7.9613 | 4.8586 | 5.0890 | 5.0593 | 5.6219 | 4.2923 | 4.9571 | 5.3609 | 5.1798 |
| ERCC6L_S807 | ERCC6L | Q2NKX8 | | 1082 | SSQIPSSVNKSMN | 7.4777 | 7.3938 | 7.0836 | 6.8678 | 5.8269 | 6.9403 | 7.4983 | 6.7995 | 7.6939 | 7.3435 | 7.7067 | 7.9560 | 3.0764 | 4.1321 | 2.5495 | 3.6540 |
| ERCC6L_S820 | ERCC6L | Q2NKX8 | | 1063 | KQFDASTPKNDIS | 8.1598 | 8.0049 | 7.5294 | 7.0695 | 6.2674 | 6.7800 | 7.3019 | 7.4030 | 7.9751 | 7.4532 | 7.9596 | 7.6230 | 2.5916 | 3.5499 | 1.9451 | 2.3865 |
| ERCC6L_S864 | ERCC6L | Q2NKX8 | | 943 | KLEEEPSASSPQY | 5.8707 | 6.0014 | 5.8640 | 6.3072 | 6.5764 | 6.0369 | 6.4291 | 7.3298 | 6.4047 | 5.7732 | 5.7832 | 6.5543 | 5.8629 | 5.7588 | 6.9863 | 6.4612 |
| ERCC6L_S943 | ERCC6L | Q2NKX8 | | 807 | DGKGTGSADSIAT | 5.5659 | 5.7331 | 6.5734 | 7.4385 | 5.4756 | 6.6886 | 6.8117 | 6.5562 | 5.9768 | 7.3000 | 7.4249 | 7.9353 | 4.5948 | 5.6662 | 4.8150 | 5.4439 |
| ERCC6L_S946 | ERCC6L | Q2NKX8 | | 1118 | ERLDDSSEAKGPE | 7.0531 | 7.7998 | 7.4569 | 7.0631 | 6.2381 | 6.7305 | 6.6908 | 6.8307 | 7.5248 | 7.4929 | 6.8539 | 6.4907 | 3.7372 | 3.9649 | 3.8038 | 4.2685 |
| ERCC6L_S971 | ERCC6L | Q2NKX8 | 1077;1082 | | NA | 8.4900 | 7.7667 | 7.6216 | 8.1453 | 6.1119 | 7.3275 | 8.1075 | 6.6343 | 6.5617 | 6.5931 | 7.3917 | 6.7248 | 2.5533 | 4.0354 | 2.3857 | 3.5495 |
| ERCC6L_S995 | ERCC6L | Q2NKX8 | 1173;1188 | | NA | 6.6063 | 6.4613 | 7.2455 | 6.8499 | 6.0065 | 6.6573 | 6.6558 | 6.8070 | 6.4047 | 6.6295 | 6.9151 | 6.9027 | 4.7673 | 5.5633 | 4.6139 | 4.9138 |
| ERCC6L_T1063 | ERCC6L | Q2NKX8 | 1000;1004 | | NA | 6.3062 | 8.0584 | 7.7153 | 7.1171 | 6.0415 | 5.7306 | 6.4690 | 7.4347 | 6.4165 | 8.2278 | 6.7794 | 7.3736 | 4.2229 | 4.2336 | 3.5187 | 4.3547 |
| ERCC6L2_S1027 | ERCC6L2 | Q5T890 | | 1370 | DAETKKSPVSSTQ | 6.1615 | 6.1134 | 6.0689 | 6.4634 | 5.7014 | 6.1626 | 6.3424 | 5.8910 | 5.9198 | 6.9782 | 6.4937 | 6.9848 | 5.9092 | 6.4353 | 6.2472 | 6.1272 |
| ERCC6L2_S1320 | ERCC6L2 | Q5T890 | | 1027 | KKELHNSPKTMNK | 5.9164 | 6.8859 | 7.8364 | 6.5375 | 5.9589 | 7.1949 | 5.9932 | 5.8689 | 7.3990 | 7.2414 | 6.6889 | 7.2987 | 4.4758 | 5.4627 | 4.3147 | 4.9265 |
| ERCC6L2_S1370 | ERCC6L2 | Q5T890 | | 1320 | LIKPRLSDSETLS | 9.8611 | 6.7192 | 6.8782 | 6.3114 | 5.4680 | 7.7838 | 8.6306 | 5.2537 | 5.4853 | 5.3712 | 5.2260 | 4.0543 | 5.8620 | 5.5060 | 5.6443 | 5.9450 |
| ERF_S185 | ERF | P50548 | | 21 | AYKPESSPGSRQI | 5.0918 | 5.2661 | 5.4049 | 5.2342 | 5.7322 | 5.9467 | 5.6343 | 6.1890 | 5.6166 | 5.4605 | 5.5194 | 5.5355 | 8.1005 | 8.2370 | 8.6067 | 8.4246 |
| ERF_S20 | ERF | P50548 | | 327 | VYNYHLSPRAFLH | 6.7838 | 6.7821 | 6.5505 | 6.2764 | 6.7565 | 6.8701 | 6.1496 | 7.3743 | 6.6459 | 6.0635 | 6.6227 | 6.2453 | 5.1035 | 6.1482 | 4.6802 | 4.9473 |
| ERF_S21 | ERF | P50548 | | 526 | EAGGPLTPRRVSS | 5.6789 | 5.3990 | 6.0514 | 5.3271 | 6.5614 | 6.4182 | 5.8846 | 6.6282 | 7.1276 | 6.1227 | 5.7092 | 6.0619 | 6.6745 | 7.3421 | 6.5099 | 6.5033 |
| ERF_S327 | ERF | P50548 | | 489 | RFKRRWSEDCRLE | 6.0675 | 5.4725 | 6.2864 | 5.4538 | 6.4076 | 6.2400 | 5.4223 | 6.2980 | 6.7229 | 6.9934 | 6.6513 | 7.1437 | 6.2767 | 6.8889 | 5.7764 | 5.8985 |
| ERF_S373 | ERF | P50548 | | 20 | WAYKPESSPGSRQ | 6.1803 | 5.8357 | 6.7874 | 5.9925 | 6.2464 | 6.4654 | 5.8009 | 6.9285 | 6.2041 | 6.4024 | 6.7565 | 6.6259 | 5.8062 | 6.0376 | 5.8080 | 6.1221 |
| ERF_S489 | ERF | P50548 | | 373 | SSSSSSSPFKFKL | 6.1891 | 6.4822 | 6.7950 | 5.7568 | 6.2159 | 5.7571 | 6.4239 | 5.9861 | 7.3609 | 6.7386 | 6.9299 | 6.5791 | 5.9136 | 5.7402 | 5.5871 | 5.5446 |
| ERF_S531 | ERF | P50548 | | 531 | LTPRRVSSDLQHA | 6.1136 | 6.0799 | 5.1791 | 6.4984 | 7.2139 | 7.2950 | 5.6805 | 7.6871 | 6.5262 | 6.2452 | 6.0394 | 6.2042 | 6.0207 | 6.5312 | 5.4004 | 5.2852 |
| ERF_T148;S154;S161 | ERF | P50548 | | 3 | xxxxMKTPADTGF | 5.5087 | 5.2656 | 6.7717 | 5.8420 | 5.9494 | 6.6506 | 6.1125 | 6.0107 | 6.4089 | 6.8828 | 6.1831 | 6.1478 | 6.8049 | 6.9384 | 6.5113 | 6.0115 |
| ERF_T3 | ERF | P50548 | | 185 | RRLGRGSVSDCSD | 6.4520 | 5.3276 | 7.1800 | 6.3205 | 6.0408 | 5.2924 | 5.9593 | 6.3150 | 6.6428 | 6.7762 | 6.1371 | 6.8911 | 5.9148 | 6.8855 | 5.4721 | 6.3928 |
| ERF_T3;S21 | ERF | P50548 | 3;21 | | NA | 5.1616 | 5.2591 | 5.7968 | 6.4679 | 5.9945 | 6.5479 | 5.8691 | 6.7031 | 5.2299 | 5.3037 | 5.6291 | 5.9255 | 8.0785 | 7.4938 | 7.5142 | 7.0250 |
| ERF_T526 | ERF | P50548 | 148;154;161 | | NA | 6.4908 | 6.1587 | 6.0984 | 6.2831 | 7.5286 | 6.6867 | 6.2068 | 5.8622 | 5.9501 | 6.3487 | 6.5522 | 6.5557 | 5.9574 | 6.4517 | 5.3895 | 5.4793 |
| ERICH1_S254 | ERICH1 | Q86X53 | | 339 | TNEKAHSILNFLK | 5.4134 | 6.0593 | 6.1448 | 5.0502 | 5.8314 | 6.0154 | 6.0338 | 6.6708 | 5.6346 | 5.4145 | 5.4742 | 5.0728 | 7.3669 | 8.1971 | 8.3862 | 7.2347 |
| ERICH1_S339 | ERICH1 | Q86X53 | | 254 | EEGADASEEDPTP | 5.3319 | 5.9028 | 5.9579 | 5.0520 | 5.9767 | 5.8492 | 5.7756 | 6.0326 | 6.0092 | 5.9888 | 5.9688 | 6.2793 | 7.6599 | 7.4493 | 7.5082 | 7.2577 |
| ERLIN1_S326 | ERLIN1 | O75477 | | 326 | RTGRESSLPSKEA | 5.3109 | 5.5238 | 6.4843 | 4.9326 | 5.2717 | 5.4578 | 5.7453 | 5.3650 | 6.2743 | 6.1797 | 6.4756 | 6.5531 | 7.3447 | 7.9243 | 7.2164 | 7.9404 |
| ERMAP_S418 | ERMAP | Q96PL5 | | 418 | CSELHKSEESIVP | 6.2576 | 6.0356 | 6.8609 | 6.2822 | 6.0890 | 5.3037 | 6.3393 | 5.9826 | 6.1851 | 6.1466 | 6.8683 | 6.1967 | 5.9381 | 5.5048 | 7.1061 | 6.9036 |
| ERMP1_S53 | ERMP1 | Q7Z2K6 | | 53 | GRTRKRSPGGSGG | 6.0683 | 6.2904 | 5.4605 | 5.9222 | 6.3132 | 6.3670 | 6.3931 | 5.8412 | 5.9115 | 5.8168 | 6.0187 | 6.2176 | 7.1528 | 7.2431 | 6.5605 | 6.4231 |
| ERRFI1_S136 | ERRFI1 | Q9UJM3 | | 251 | RLRRSHSGPAGSF | 5.9658 | 5.4161 | 6.0961 | 5.1975 | 6.7099 | 6.6216 | 5.7599 | 6.8684 | 5.9569 | 5.4077 | 6.1742 | 5.9586 | 8.0688 | 6.7304 | 6.8667 | 6.2015 |
| ERRFI1_S251 | ERRFI1 | Q9UJM3 | | 337 | RTPSPKSLPSYLN | 6.4540 | 6.1613 | 7.5350 | 6.6248 | 6.6854 | 6.7675 | 6.1719 | 6.9587 | 6.5902 | 6.2935 | 7.0974 | 6.7089 | 5.1879 | 5.1632 | 4.8133 | 4.7869 |
| ERRFI1_S256 | ERRFI1 | Q9UJM3 | | 302 | PDYRRWSAEVTSS | 5.7086 | 5.3526 | 6.8389 | 5.7454 | 6.3229 | 6.2688 | 5.8801 | 6.7387 | 5.7143 | 5.5219 | 6.2590 | 6.2867 | 8.0005 | 6.7166 | 6.5488 | 6.0963 |
| ERRFI1_S273 | ERRFI1 | Q9UJM3 | | 326 | PPREPLSPSNSRT | 5.5309 | 5.1820 | 5.8337 | 4.9644 | 6.2031 | 6.4324 | 5.7108 | 6.4830 | 5.8165 | 5.9049 | 5.5614 | 5.5436 | 8.6389 | 7.3399 | 7.5920 | 7.2624 |
| ERRFI1_S302 | ERRFI1 | Q9UJM3 | | 131 | ASTPPLTPIKNSP | 5.2545 | 4.8272 | 5.8224 | 4.8985 | 5.4602 | 6.5553 | 5.6998 | 5.8213 | 5.8835 | 6.1850 | 5.9421 | 6.9201 | 8.2314 | 7.1363 | 7.4788 | 7.8835 |
| ERRFI1_S326 | ERRFI1 | Q9UJM3 | | 273 | CCIHRASPNSDED | 5.7535 | 5.4936 | 6.1383 | 5.7407 | 6.2239 | 6.1029 | 6.2986 | 6.4561 | 6.4315 | 5.7276 | 6.3503 | 6.2671 | 7.5697 | 6.5571 | 6.5376 | 6.3514 |
| ERRFI1_S337 | ERRFI1 | Q9UJM3 | | 127 | NGVCASTPPLTPI | 6.3102 | 6.0361 | 6.5457 | 5.7817 | 6.6154 | 6.6438 | 6.1533 | 6.1562 | 6.0822 | 5.8726 | 5.7313 | 6.7380 | 7.3609 | 6.1990 | 6.0155 | 5.7581 |
| ERRFI1_S369 | ERRFI1 | Q9UJM3 | | 256 | HSGPAGSFNKPAI | 6.2300 | 5.6531 | 6.5479 | 6.6852 | 6.2014 | 6.7337 | 6.7153 | 6.1830 | 6.0262 | 6.5615 | 6.4896 | 5.8166 | 5.6623 | 6.2582 | 5.2489 | 6.9870 |
| ERRFI1_S461 | ERRFI1 | Q9UJM3 | | 461 | HLSYVVSPxxxxx | 7.1089 | 6.6060 | 6.2922 | 6.1846 | 6.4310 | 6.4294 | 6.6873 | 7.1256 | 6.7976 | 6.5379 | 6.1704 | 6.3596 | 5.6773 | 5.4893 | 4.8151 | 5.2878 |
| ERRFI1_T127 | ERRFI1 | Q9UJM3 | | 136 | LTPIKNSPSLFPC | 6.9624 | 5.8855 | 5.9784 | 5.9092 | 6.3869 | 6.7459 | 7.0525 | 6.7124 | 6.2710 | 6.2703 | 6.4714 | 7.2334 | 5.3861 | 5.9379 | 5.2045 | 5.5921 |
| ERRFI1_T127;T131 | ERRFI1 | Q9UJM3 | | 369 | ALQRQNSEGSASK | 3.8902 | 4.0005 | 6.6656 | 6.0625 | 6.1760 | 6.0872 | 6.1012 | 6.6015 | 5.3484 | 5.6437 | 6.7024 | 7.1947 | 6.9535 | 7.6509 | 7.1867 | 7.7351 |
| ERRFI1_T131 | ERRFI1 | Q9UJM3 | 127;131 | | NA | 5.8129 | 4.8709 | 6.5927 | 5.3722 | 6.5985 | 6.6180 | 6.0681 | 7.1972 | 6.3216 | 5.8930 | 5.5006 | 5.7050 | 7.5839 | 6.1236 | 6.8312 | 6.9108 |
| ESCO1_S412 | ESCO1 | Q5FWF5 | | 523 | ESKLENSPVENVT | 5.6561 | 6.6738 | 6.6080 | 6.5121 | 6.1367 | 5.5272 | 6.8656 | 6.2138 | 6.7374 | 7.2986 | 6.4468 | 6.0744 | 5.5884 | 6.1132 | 5.3919 | 6.1561 |
| ESCO1_S523 | ESCO1 | Q5FWF5 | | 412 | KLDSQVSPKLGLL | 6.3361 | 6.9258 | 6.3507 | 7.5399 | 7.2524 | 6.8582 | 6.5259 | 5.8677 | 7.4951 | 7.6108 | 7.0545 | 7.7631 | 3.8176 | 4.7576 | 3.6767 | 4.1679 |
| ESCO2_S29 | ESCO2 | Q56NI9 | | 29 | TENLFPSPNKKHC | 6.8563 | 6.8691 | 6.3480 | 7.0684 | 6.2411 | 6.1752 | 6.3252 | 6.1052 | 6.6578 | 6.2520 | 6.1021 | 6.4822 | 5.8277 | 5.9372 | 5.1702 | 5.5824 |
| ESCO2_S512 | ESCO2 | Q56NI9 | | 75 | PSANQGSPFKSAL | 7.5103 | 7.4021 | 6.6001 | 7.0290 | 6.2359 | 5.9292 | 6.3295 | 6.1592 | 6.9113 | 6.4480 | 6.2807 | 6.2393 | 5.4370 | 5.5626 | 4.9491 | 4.9767 |
| ESCO2_S75 | ESCO2 | Q56NI9 | | 512 | EPIGPESPSSTEC | 6.4817 | 6.2140 | 6.4712 | 7.0294 | 6.2262 | 6.6641 | 6.5754 | 5.8708 | 6.9224 | 6.3442 | 6.2217 | 6.2062 | 5.7683 | 5.5837 | 5.7899 | 5.6308 |
| ESF1_S153 | ESF1 | Q9H501 | | 153 | KIDSNISPKKDSK | 5.9428 | 6.0389 | 6.0271 | 6.6527 | 6.0201 | 5.7214 | 6.3273 | 5.8564 | 6.2846 | 6.1044 | 6.0638 | 6.1820 | 6.5775 | 6.5603 | 6.6686 | 6.9721 |
| ESF1_S198 | ESF1 | Q9H501 | | 663 | SEEELPSDVDLND | 5.5075 | 5.2822 | 5.5411 | 6.0962 | 5.9548 | 5.4928 | 5.5542 | 5.4786 | 6.3672 | 6.5941 | 6.3725 | 6.4130 | 7.1603 | 7.6006 | 7.0898 | 7.4950 |
| ESF1_S657 | ESF1 | Q9H501 | | 735 | VEHQNLSKKKKKQ | 6.3969 | 7.1635 | 7.0734 | 7.5286 | 5.2642 | 6.8881 | 7.3814 | 6.9524 | 7.3100 | 7.5219 | 7.8231 | 8.5460 | 3.0558 | 4.0877 | 3.1636 | 3.8433 |
| ESF1_S657;S663 | ESF1 | Q9H501 | | 823 | KESQRKSIDPALS | 6.7548 | 6.4609 | 7.1595 | 6.1414 | 5.9019 | 6.4826 | 6.5593 | 7.0117 | 6.6688 | 6.6926 | 7.2192 | 6.6705 | 5.1079 | 5.1224 | 4.8024 | 5.2442 |
| ESF1_S663 | ESF1 | Q9H501 | | 657 | ALAEEASEEELPS | 5.1387 | 5.1938 | 6.1641 | 6.0957 | 5.3777 | 6.3531 | 6.0365 | 6.0150 | 7.6186 | 6.6973 | 6.9506 | 7.2080 | 5.4502 | 6.6227 | 6.3541 | 6.7239 |
| ESF1_S735 | ESF1 | Q9H501 | | 198 | TSEIVKSPRIECS | 5.7598 | 6.3996 | 6.0998 | 6.4293 | 6.8942 | 7.9512 | 6.7297 | 7.0722 | 6.9827 | 6.1360 | 7.4939 | 6.4302 | 4.8796 | 5.6109 | 4.7723 | 4.3586 |
| ESF1_S823 | ESF1 | Q9H501 | 657;663 | | NA | 6.0242 | 6.2917 | 6.4373 | 6.7813 | 6.2709 | 5.7101 | 6.2207 | 5.9358 | 5.7162 | 6.0049 | 5.9011 | 6.0208 | 6.5129 | 6.6417 | 6.8674 | 6.6630 |
| ESPL1_S1126 | ESPL1 | Q14674 | | 1508 | FEILRGSDGEDSA | 6.4775 | 6.5742 | 7.1832 | 6.4884 | 6.7350 | 6.7629 | 6.3022 | 6.3429 | 6.6218 | 6.5734 | 7.1278 | 6.9303 | 4.8574 | 5.0752 | 4.8419 | 5.1060 |
| ESPL1_S1508 | ESPL1 | Q14674 | | 1346 | GRGLPCTPKPPDR | 6.8943 | 7.8003 | 7.1462 | 7.2295 | 4.8894 | 8.8627 | 8.5702 | 7.3046 | 7.0670 | 8.2776 | 7.9079 | 7.6581 | 2.0496 | 3.6787 | 1.2594 | 3.4045 |
| ESPL1_S1545 | ESPL1 | Q14674 | | 1545 | SKKKLPSPCPDKE | 7.0366 | 7.2334 | 7.1630 | 8.1512 | 6.0735 | 7.5350 | 7.8297 | 6.8201 | 8.0178 | 7.3289 | 7.9780 | 7.7261 | 2.2318 | 3.6128 | 2.4341 | 2.8281 |
| ESPL1_T1346 | ESPL1 | Q14674 | | 1126 | APSTNSSPVLKTK | 6.9583 | 7.2435 | 7.1830 | 7.6582 | 5.8343 | 6.7808 | 6.4446 | 6.3897 | 6.7274 | 7.4312 | 6.4054 | 6.8303 | 4.4912 | 4.9334 | 3.8715 | 4.8172 |
| ESPNL_T252 | ESPNL | Q6ZVH7 | | 252 | AARGGHTPILDRL | 6.2853 | 6.7497 | 6.3541 | 5.9069 | 5.8074 | 5.7591 | 5.8820 | 6.1188 | 6.2449 | 6.1377 | 5.8407 | 6.2129 | 6.1553 | 6.4869 | 6.7372 | 7.3210 |
| ESRRA_S19 | ESRRA | P11474 | | 19 | IKAEPASPDSPKG | 7.5889 | 7.6537 | 7.5537 | 7.4853 | 6.5240 | 5.6320 | 6.7191 | 6.8094 | 7.0711 | 7.1573 | 7.4590 | 6.8164 | 3.5167 | 4.2869 | 3.6155 | 4.1111 |
| ESRRA_S27 | ESRRA | P11474 | | 27 | DSPKGSSETETEP | 6.1684 | 5.7384 | 5.9548 | 6.8550 | 5.9368 | 6.0588 | 4.8701 | 5.4236 | 6.5485 | 6.3418 | 6.5873 | 5.4901 | 7.0534 | 7.4612 | 6.2711 | 7.2406 |
| ESS2_S292 | ESS2 | Q96DF8 | | 110 | TPATFETPEVHAG | 5.8221 | 6.9841 | 7.2462 | 7.5984 | 6.4203 | 6.1251 | 6.4441 | 6.3591 | 6.3384 | 6.8607 | 6.7589 | 7.3759 | 4.7708 | 5.0817 | 4.7139 | 5.1003 |
| ESS2_S391;S395 | ESS2 | Q96DF8 | | 386 | ENLASLTPKGLSP | 6.1826 | 6.1187 | 7.2450 | 7.6626 | 5.9869 | 6.4709 | 6.5536 | 6.4260 | 7.2736 | 6.4695 | 6.4195 | 6.9939 | 5.4585 | 5.0592 | 4.4704 | 5.2093 |
| ESS2_S395 | ESS2 | Q96DF8 | | 421 | RASYTPSPARSTH | 6.4996 | 7.4264 | 7.7497 | 8.2421 | 5.2356 | 6.9539 | 7.0045 | 7.0913 | 6.8669 | 7.0772 | 7.8538 | 8.0308 | 3.0206 | 4.0646 | 3.1145 | 3.7686 |
| ESS2_S421 | ESS2 | Q96DF8 | | 437 | PASGLQTPTSTPA | 5.5513 | 6.3492 | 6.7560 | 7.0095 | 5.6048 | 6.1585 | 6.3565 | 7.1680 | 7.0906 | 7.3532 | 7.5435 | 7.4057 | 4.9320 | 4.8227 | 4.6229 | 5.2757 |
| ESS2_T110 | ESS2 | Q96DF8 | | 292 | ELIPQESPRVGGF | 5.9994 | 6.6752 | 7.4011 | 7.8955 | 5.8762 | 6.6280 | 6.5555 | 6.8313 | 6.3078 | 6.5423 | 7.5931 | 6.8000 | 4.7282 | 4.7129 | 4.6700 | 4.7835 |
| ESS2_T386 | ESS2 | Q96DF8 | | 395 | GLSPAMSPALQRL | 7.2242 | 7.0138 | 6.8426 | 7.2385 | 6.2968 | 5.9201 | 6.7266 | 5.9095 | 6.7340 | 6.9110 | 7.3132 | 7.0615 | 4.0336 | 5.2858 | 4.3017 | 5.1872 |
| ESS2_T437 | ESS2 | Q96DF8 | 391;395 | | NA | 6.6860 | 7.7433 | 7.7532 | 7.2477 | 5.6680 | 6.1075 | 6.5657 | 6.2739 | 5.2656 | 6.3807 | 6.9365 | 6.1506 | 5.1140 | 5.7775 | 4.7687 | 5.5612 |
| ESYT1_S1044 | ESYT1 | Q9BSJ8-2 | | 1044 | QKKRTLSPEFNER | 6.4641 | 6.4697 | 6.0048 | 6.0619 | 6.2165 | 6.0876 | 6.1401 | 5.7372 | 6.0558 | 6.1120 | 6.0046 | 6.0818 | 6.6845 | 6.7362 | 6.7130 | 6.4302 |
| ESYT1_S1077 | ESYT1 | Q9BSJ8-2 | | 973 | RLTHVDSPLEAPA | 5.9167 | 6.8920 | 6.2633 | 6.8768 | 5.7381 | 6.6660 | 6.5509 | 6.6428 | 6.3234 | 7.1498 | 6.7153 | 6.4212 | 5.3846 | 6.0157 | 5.2354 | 5.2078 |
| ESYT1_S123 | ESYT1 | Q9BSJ8-2 | | 4 | xxxMERSPGEGPS | 5.8675 | 6.7640 | 6.4527 | 6.8514 | 6.6869 | 6.8426 | 6.8433 | 6.4931 | 7.3030 | 6.2970 | 7.0196 | 6.8206 | 4.8439 | 4.8454 | 5.2140 | 4.8549 |
| ESYT1_S4 | ESYT1 | Q9BSJ8-2 | | 123 | AKTLYMSHRELPA | 5.6256 | 5.8592 | 5.1532 | 5.9489 | 5.7070 | 6.2941 | 5.2783 | 5.1104 | 5.9906 | 6.5580 | 5.1865 | 6.7415 | 7.6593 | 7.2912 | 7.8053 | 7.7907 |
| ESYT1_S830 | ESYT1 | Q9BSJ8-2 | | 830 | KGTKHLSPYATLT | 7.2681 | 9.2110 | 4.9735 | 4.4269 | 7.3006 | 6.8460 | 4.4431 | 5.6716 | 5.8989 | 5.3036 | 10.2500 | 6.7919 | 3.8370 | 5.1683 | 6.6620 | 5.9476 |
| ESYT1_S973 | ESYT1 | Q9BSJ8-2 | | 1077 | SNSSFMSRERELL | 6.2524 | 6.0461 | 6.0390 | 5.5419 | 6.1976 | 5.9544 | 6.4858 | 5.9518 | 5.7069 | 5.6911 | 5.8616 | 6.1716 | 7.1282 | 7.2806 | 7.1888 | 6.5024 |
| ESYT2_S712 | ESYT2 | A0FGR8-6 | | 779 | PTPSIASDISLPI | 5.6474 | 5.5665 | 5.6854 | 5.7688 | 5.8468 | 6.4194 | 5.9110 | 6.8724 | 6.7607 | 6.8688 | 6.6052 | 6.6774 | 6.5451 | 5.9914 | 6.3166 | 6.5169 |
| ESYT2_S776;S779 | ESYT2 | A0FGR8-6 | | 712 | SIKSHMSGSPGPG | 5.0014 | 5.4318 | 5.2337 | 6.1496 | 6.0679 | 6.1131 | 5.4432 | 5.6092 | 5.6462 | 6.0043 | 6.3607 | 5.8704 | 8.0792 | 7.3859 | 8.3006 | 7.3026 |
| ESYT2_S779 | ESYT2 | A0FGR8-6 | 776;779 | | NA | 5.2410 | 6.0124 | 6.3142 | 6.1098 | 5.8855 | 5.5885 | 6.5866 | 6.3756 | 6.4864 | 6.3057 | 6.2526 | 6.6650 | 6.7567 | 6.3186 | 6.6105 | 6.4909 |
| ETHE1_T216 | ETHE1 | O95571 | | 216 | TLNPRLTLSCEEF | 5.6248 | 6.4195 | 6.0850 | 6.9438 | 6.4924 | 5.6994 | 6.0946 | 6.4731 | 5.4974 | 5.8557 | 6.4942 | 6.7682 | 7.0079 | 6.1543 | 6.0437 | 6.3458 |
| ETS1_S26 | ETS1 | P14921 | | 38 | ADVPLLTPSSKEM | 5.5452 | 5.3300 | 5.6656 | 5.3171 | 6.3891 | 6.6460 | 5.6193 | 6.3913 | 6.1963 | 5.4555 | 5.4345 | 5.4996 | 8.4379 | 7.8223 | 7.3836 | 6.8667 |
| ETS1_S282 | ETS1 | P14921 | | 282 | SLQRVPSYDSFDS | 4.6397 | 4.5924 | 5.8401 | 5.0019 | 5.5080 | 5.6493 | 4.8839 | 6.1050 | 4.8637 | 4.8991 | 5.7686 | 5.6073 | 8.7404 | 8.9972 | 9.6726 | 9.2305 |
| ETS1_S282;S285 | ETS1 | P14921 | | 26 | DLELFPSPDMECA | 5.2101 | 5.2323 | 5.2762 | 5.6270 | 6.4104 | 6.2495 | 6.1390 | 6.5327 | 5.7025 | 5.3056 | 5.5243 | 5.7963 | 7.9582 | 7.8702 | 8.0471 | 7.1186 |
| ETS1_S285 | ETS1 | P14921 | | 285 | RVPSYDSFDSEDY | 6.6477 | 5.9688 | 7.3035 | 6.8643 | 6.1352 | 6.9132 | 5.8721 | 7.5249 | 6.8900 | 6.4251 | 8.5855 | 7.4833 | 4.0371 | 5.3118 | 4.2230 | 3.8144 |
| ETS1_T38 | ETS1 | P14921 | 282;285 | | NA | 4.7823 | 4.5747 | 6.6674 | 4.8376 | 5.6237 | 4.8166 | 5.7730 | 6.8190 | 9.1804 | 5.6143 | 4.9449 | 5.4710 | 7.4724 | 8.1733 | 8.0738 | 7.1757 |

| | | | | | | | | | | | | | | | | | | | | | |
|---|---|---|---|---|---|---|---|---|---|---|---|---|---|---|---|---|---|---|---|---|---|
| ETV1_S216 | ETV1 | P50549 | | 216 | MYQRQMSEPNIPF | 4.9142 | 6.3658 | 5.5521 | 5.5848 | 6.3675 | 6.3298 | 6.1000 | 6.4231 | 4.7883 | 5.3288 | 4.8929 | 5.3166 | 8.0137 | 7.8739 | 8.0513 | 8.0971 |
| ETV3_S139 | ETV3 | P41162 | | 159 | PPLDTHSPTNDVQ | 5.8292 | 5.9420 | 6.4292 | 6.1816 | 6.1710 | 5.9358 | 6.2277 | 6.7859 | 5.6497 | 6.0255 | 6.5329 | 6.6980 | 6.3870 | 6.5876 | 6.2457 | 6.3713 |
| ETV3_S159 | ETV3 | P41162 | | 139 | SGVVPQSAPPVPT | 5.7933 | 6.2537 | 6.5263 | 5.8223 | 6.5161 | 6.2500 | 6.2560 | 5.7846 | 6.5799 | 6.2110 | 6.1929 | 5.9097 | 6.2260 | 6.5331 | 6.5400 | 6.6053 |
| ETV3_S245;S250 | ETV3 | P41162 | 245;250 | | NA | 5.6408 | 6.3018 | 6.4841 | 5.3380 | 5.8563 | 6.4041 | 6.3406 | 5.5563 | 6.8191 | 6.5837 | 5.9483 | 6.2547 | 6.6545 | 7.6222 | 5.3400 | 6.8552 |
| ETV5_S290 | ETV5 | P41161-2 | | 290 | SYHRQMSEPIVPA | 4.2426 | 4.6157 | 5.1170 | 4.5446 | 5.9180 | 6.2291 | 5.3705 | 5.9877 | 4.8696 | 4.7996 | 5.0640 | 5.4102 | 9.9182 | 9.0347 | 9.4029 | 9.4757 |
| ETV6_S203 | ETV6 | P41212 | | 323 | HIMVSVSPPEEHA | 6.2995 | 6.3563 | 6.4162 | 7.2031 | 6.4890 | 5.9887 | 6.3967 | 6.2933 | 6.3978 | 6.3864 | 6.1581 | 6.5169 | 6.2273 | 5.5734 | 5.6782 | 5.6190 |
| ETV6_S213 | ETV6 | P41212 | | 271 | VIQLMPSPIMHPL | 6.1741 | 6.3266 | 6.2924 | 6.4133 | 6.2689 | 6.6903 | 6.4563 | 5.9766 | 6.7022 | 6.8254 | 6.6818 | 6.5948 | 5.8684 | 5.8166 | 5.4343 | 5.4780 |
| ETV6_S271 | ETV6 | P41212 | | 18 | QERISYTPPESPV | 5.7194 | 6.3121 | 5.7209 | 6.3849 | 6.3823 | 6.3882 | 6.0607 | 6.7964 | 6.7065 | 6.1922 | 6.5294 | 6.2727 | 5.8364 | 5.9448 | 6.2137 | 6.5394 |
| ETV6_S323 | ETV6 | P41212 | | 203 | EQRPLRSPLDNMI | 6.3224 | 5.4228 | 5.7840 | 6.2506 | 5.8640 | 5.9544 | 6.2717 | 6.1110 | 6.1617 | 6.2359 | 5.2836 | 6.3673 | 7.3542 | 6.5672 | 6.6073 | 7.4419 |
| ETV6_T18 | ETV6 | P41212 | | 213 | NMIRRLSPAERAQ | 6.4709 | 6.4838 | 5.8978 | 6.5303 | 6.3169 | 6.0128 | 5.7474 | 6.4237 | 6.1542 | 6.1676 | 6.1600 | 6.6233 | 6.5612 | 6.0379 | 6.2578 | 6.1543 |
| ETV6_T18;S22 | ETV6 | P41212 | 18;22 | | NA | 6.1836 | 6.3704 | 6.1552 | 6.7779 | 6.4478 | 6.1866 | 5.8478 | 6.3975 | 6.3572 | 6.2022 | 6.1538 | 6.3673 | 6.4893 | 6.3306 | 6.2178 | 5.5148 |
| EVA1B_S152 | EVA1B | Q9NVM1 | | 152 | TGTLGPSPTATGT | 6.6797 | 6.2561 | 6.6569 | 5.9383 | 6.5782 | 6.3237 | 6.5229 | 6.3895 | 7.0264 | 6.3938 | 6.8506 | 6.8985 | 4.7340 | 6.3631 | 4.5539 | 5.8344 |
| EVA1B_T94 | EVA1B | Q9NVM1 | | 94 | RLGPDDTLPGPEL | 5.3412 | 6.5177 | 6.5902 | 4.9712 | 5.6327 | 6.1538 | 6.5886 | 6.1230 | 5.3088 | 7.3581 | 6.2309 | 5.6419 | 5.3566 | 7.7682 | 6.7166 | 7.7004 |
| EXD2_S407 | EXD2 | Q9NVH0 | | 407 | PAGRPESPGDYYL | 6.2255 | 6.8116 | 6.3144 | 7.3139 | 6.3311 | 6.1062 | 6.8235 | 6.4116 | 6.8891 | 6.5194 | 6.8331 | 6.5947 | 4.9335 | 5.4834 | 5.3234 | 5.0856 |
| EXO1_S376 | EXO1 | Q9UQ84 | | 815 | PCKKPLSPVRDNI | 5.8936 | 6.0247 | 6.3105 | 6.5528 | 5.4974 | 5.5522 | 5.4462 | 6.2860 | 7.1170 | 6.5695 | 6.2512 | 6.5268 | 6.1456 | 6.2657 | 6.5420 | 7.0188 |
| EXO1_S598 | EXO1 | Q9UQ84 | | 598 | KFTRTISPPTLGT | 6.2987 | 6.3138 | 6.0884 | 6.1419 | 5.8156 | 5.8273 | 6.0077 | 6.4051 | 9.2090 | 6.6715 | 6.2547 | 6.5832 | 5.7907 | 5.4950 | 5.5450 | 5.5523 |
| EXO1_S772 | EXO1 | Q9UQ84 | | 772 | LSKKPASIQKRKH | 7.1301 | 7.6638 | 7.1042 | 6.9968 | 5.1583 | 6.1527 | 6.5581 | 5.9502 | 8.1135 | 8.0658 | 8.6368 | 8.8970 | 3.0348 | 4.8633 | 2.3474 | 3.3273 |
| EXO1_S815 | EXO1 | Q9UQ84 | | 376 | IWHRNYSPRPESG | 5.9571 | 6.9467 | 7.6671 | 6.0267 | 5.3310 | 6.3172 | 6.2420 | 5.7168 | 6.5486 | 6.1845 | 7.0537 | 6.9126 | 6.3200 | 6.0323 | 5.2312 | 6.4126 |
| EXOC1_S448 | EXOC1 | Q9NV70-2 | | 448 | KFGLHGSSGKLTG | 5.5850 | 6.0736 | 6.3254 | 6.0034 | 5.7540 | 6.3798 | 6.0207 | 6.1502 | 6.0366 | 6.7906 | 6.7678 | 6.9239 | 5.7519 | 6.6863 | 6.3885 | 6.3623 |
| EXOC1_S470 | EXOC1 | Q9NV70 | | 470 | SGKLTGSTSSLNK | 5.6293 | 6.2322 | 6.2745 | 5.1030 | 6.3273 | 6.7212 | 6.0466 | 6.9811 | 6.5629 | 6.5802 | 6.2446 | 6.0752 | 6.1687 | 5.8231 | 6.6091 | 6.6209 |
| EXOC1_S470;S473 | EXOC1 | Q9NV70 | | 445 | ESKKFATLPRKES | 7.7366 | 6.6660 | 8.1120 | 6.4440 | 6.3558 | 6.8810 | 7.1972 | 5.4214 | 6.3427 | 6.3777 | 7.0089 | 6.7720 | 4.8311 | 6.0402 | 3.8085 | 4.0051 |
| EXOC1_T445 | EXOC1 | Q9NV70 | 470;473 | | NA | 6.6720 | 6.1822 | 6.7236 | 6.1553 | 6.4352 | 6.2314 | 6.3488 | 6.2074 | 6.6877 | 6.4833 | 6.2766 | 6.1325 | 5.8402 | 6.2671 | 5.6838 | 5.6729 |
| EXOC2_S432 | EXOC2 | Q96KP1 | | 432 | SLKRGSSFQSGRD | 6.7415 | 6.2771 | 6.0601 | 5.4557 | 6.2001 | 6.1426 | 5.9045 | 6.2745 | 6.6774 | 6.1450 | 6.2417 | 6.3199 | 6.4726 | 6.7477 | 6.1729 | 6.1668 |
| EXOC2_S461 | EXOC2 | Q96KP1 | | 461 | LTKLVLSQLPNFW | 6.0482 | 6.0626 | 6.4902 | 6.2584 | 6.1973 | 6.7521 | 6.0437 | 6.2174 | 6.0974 | 6.4874 | 6.2797 | 6.6751 | 6.0958 | 6.4878 | 5.9643 | 5.8423 |
| EXOC4_T237 | EXOC4 | Q96A65 | | 237 | DVTNLPTPRKFLD | 5.8838 | 6.0330 | 6.4428 | 6.6835 | 6.8459 | 6.4297 | 6.9238 | 6.7609 | 6.4972 | 6.8384 | 6.5100 | 6.8574 | 5.2812 | 5.5106 | 5.1023 | 5.3994 |
| EXOC7_S250 | EXOC7 | Q9UPT5 | | 250 | SSGVPYSPAIPNK | 6.6405 | 6.6896 | 6.9527 | 6.3975 | 6.6585 | 7.0070 | 6.6425 | 7.0214 | 6.9297 | 7.1080 | 7.0429 | 6.6615 | 4.6189 | 5.1360 | 4.2378 | 4.2556 |
| EXOC7_S283 | EXOC7 | Q9UPT5 | | 283 | NLLKQYSQHGLDG | 8.2818 | 6.8855 | 7.3985 | 7.2067 | 6.2161 | 6.5579 | 5.4913 | 7.2355 | 7.6383 | 6.6307 | 7.1801 | 6.9459 | 4.0924 | 5.1482 | 3.5335 | 3.5577 |
| EXOSC10_S16 | EXOSC10 | Q01780 | | 217 | PLPQALSKERRER | 6.3148 | 6.1254 | 5.4200 | 7.3239 | 6.2397 | 6.2747 | 8.1325 | 6.5503 | 6.7126 | 6.3960 | 7.4251 | 8.0994 | 5.0172 | 4.8038 | 4.2506 | 4.9140 |
| EXOSC10_S217 | EXOSC10 | Q01780 | | 846 | FDPNKQTPSGKKC | 8.3166 | 7.6983 | 6.8192 | 7.6681 | 5.5557 | 7.0191 | 7.9225 | 6.2743 | 7.3300 | 6.9943 | 7.2560 | 6.8498 | 3.7523 | 4.1245 | 2.7346 | 3.6847 |
| EXOSC10_T846 | EXOSC10 | Q01780 | | 16 | RVLSATSATKSDG | 5.6636 | 6.6685 | 6.4573 | 5.8160 | 7.4545 | 6.7262 | 6.6619 | 7.0071 | 6.8645 | 6.9428 | 6.2275 | 5.9379 | 5.2520 | 4.8732 | 5.8748 | 5.5721 |
| EXOSC2_S124 | EXOSC2 | Q13868 | | 124 | GELRRRSAEDELA | 5.6938 | 6.1265 | 5.8122 | 6.2428 | 5.9363 | 5.9825 | 6.1301 | 6.0593 | 6.0200 | 6.0040 | 5.9109 | 6.3672 | 7.0000 | 6.7658 | 7.2092 | 6.7392 |
| EXOSC9_S306 | EXOSC9 | Q06265-2 | | 306 | KAPIDTSDVEEKA | 6.3766 | 6.7029 | 6.4149 | 6.7863 | 6.1649 | 6.1203 | 6.5355 | 6.0733 | 6.0592 | 5.9058 | 5.6893 | 5.9767 | 6.0461 | 6.3889 | 6.2972 | 6.4621 |
| EXPH5_S643 | EXPH5 | Q8NEV8 | | 643 | DRRNPQSPNLQNP | 6.5343 | 7.0496 | 4.8533 | 6.9217 | 6.7144 | 6.0308 | 7.7438 | 7.1195 | 6.5604 | 7.7071 | 7.0318 | 6.7985 | 3.9661 | 5.2940 | 5.2360 | 4.4387 |
| EYA3_S438 | EYA3 | Q99504 | | 438 | NVGGLLSPQRKEA | 6.4321 | 6.7996 | 7.0458 | 7.3494 | 6.1831 | 6.7107 | 6.4282 | 6.4258 | 6.9640 | 7.0366 | 7.0784 | 7.1140 | 4.2541 | 5.0061 | 4.3363 | 4.8357 |
| EYA3_T269 | EYA3 | Q99504 | | 269 | PSLSQTTPSKDTD | 5.7728 | 6.4231 | 6.5113 | 6.0306 | 6.4033 | 6.5130 | 6.7017 | 6.0503 | 6.6604 | 6.8556 | 6.5906 | 6.9383 | 5.4256 | 5.6919 | 5.4301 | 6.0014 |
| EZH2_S368 | EZH2 | Q15910-2 | | 492 | PAEDVDTPPRKKK | 6.3241 | 7.3155 | 6.1156 | 7.1649 | 6.1216 | 6.4063 | 6.8851 | 6.0745 | 7.0240 | 6.0804 | 6.0107 | 6.3865 | 5.4131 | 5.5783 | 5.5194 | 5.5801 |
| EZH2_S368;T372 | EZH2 | Q15910-2 | | 372 | NSSRPSTPTINVL | 5.4083 | 5.4623 | 5.8197 | 6.3625 | 5.6268 | 5.0955 | 6.4649 | 5.9202 | 6.1454 | 6.3910 | 6.0107 | 6.5098 | 7.0883 | 6.8798 | 7.3423 | 7.4725 |
| EZH2_T372 | EZH2 | Q15910-2 | | 368 | RLPNNSSRPSTPT | 5.5974 | 5.4579 | 5.2483 | 6.1150 | 6.0194 | 5.4264 | 5.8790 | 5.3465 | 4.8465 | 5.9053 | 6.0290 | 5.6264 | 8.4352 | 8.1474 | 7.4852 | 8.4352 |
| EZH2_T492 | EZH2 | Q15910-2 | 368;372 | | NA | 5.4203 | 6.2172 | 5.3017 | 5.9648 | 4.7787 | 5.3710 | 5.2457 | 5.3874 | 6.0555 | 6.2388 | 5.5745 | 5.6953 | 7.9362 | 8.2870 | 8.0986 | 8.4273 |
| F11R_S284 | F11R | Q9Y624 | | 296 | EFKQTSSFLVxxx | 5.8290 | 6.2292 | 6.3200 | 6.1375 | 6.0761 | 5.6639 | 5.3178 | 6.2857 | 5.7147 | 6.1304 | 6.5546 | 6.6715 | 6.4623 | 6.8075 | 6.9364 | 6.8634 |
| F11R_S296 | F11R | Q9Y624 | | 284 | VIYSQPSARSEGE | 7.4076 | 7.6621 | 7.6923 | 5.9292 | 5.9720 | 6.7604 | 6.8864 | 6.9235 | 6.2422 | 6.6679 | 6.7949 | 6.3341 | 4.2120 | 5.7898 | 4.0045 | 4.7209 |
| F2R_S418 | F2R | P25116 | | 418 | SSNLNNSIYKKLL | 6.7319 | 6.8322 | 6.9960 | 7.1442 | 6.5686 | 6.2218 | 6.7981 | 6.3694 | 6.5054 | 5.8193 | 5.8889 | 6.0984 | 5.0055 | 6.6081 | 5.0801 | 5.3321 |
| F2RL1_S373 | F2RL1 | P55085 | | 373 | VKQMQVSLTSKKH | 6.3574 | 6.4918 | 6.6757 | 5.9127 | 6.3379 | 6.2340 | 6.1203 | 6.3668 | 6.3606 | 6.5529 | 5.9757 | 5.9537 | 5.8014 | 7.1687 | 6.2434 | 5.4470 |
| F2RL1_S373;S376 | F2RL1 | P55085 | | 375 | QMQVSLTSKKHSR | 5.8042 | 6.6864 | 7.2247 | 5.5774 | 5.4904 | 6.3680 | 6.4341 | 5.9756 | 5.3532 | 6.2252 | 7.3618 | 7.1809 | 5.2251 | 6.7775 | 5.9687 | 6.3467 |
| F2RL1_S376 | F2RL1 | P55085 | | 376 | MQVSLTSKKHSRK | 6.0570 | 6.7750 | 6.4363 | 5.5543 | 5.4942 | 6.8111 | 6.5120 | 6.4307 | 5.1761 | 6.3567 | 6.6279 | 6.2001 | 5.9719 | 7.2195 | 6.2487 | 6.1285 |
| F2RL1_S384 | F2RL1 | P55085 | | 384 | KHSRKSSSYSSSS | 5.8255 | 6.8618 | 6.1822 | 5.2451 | 4.6485 | 6.4918 | 6.0731 | 6.3084 | 5.6135 | 6.7791 | 6.9489 | 6.7454 | 5.8481 | 7.4520 | 5.9982 | 6.9785 |
| F2RL1_S384;S390 | F2RL1 | P55085 | 373;376 | | NA | 7.7426 | 6.1319 | 7.5790 | 6.5509 | 6.9971 | 5.7930 | 6.2124 | 6.7550 | 7.3332 | 6.9209 | 5.7766 | 6.0266 | 5.3330 | 6.0861 | 4.7851 | 3.9765 |
| F2RL1_T375 | F2RL1 | P55085 | 384;390 | | NA | 5.8786 | 5.6614 | 6.1130 | 6.6815 | 6.3101 | 6.5252 | 5.3506 | 5.8731 | 6.7751 | 7.1932 | 7.5891 | 7.5060 | 5.3966 | 7.1832 | 5.8028 | 4.1604 |
| FAAP100_S667 | FAAP100 | Q0VG06 | | 667 | PHTRAPSPLGPTR | 6.1529 | 6.0704 | 6.1441 | 6.0943 | 6.6087 | 6.3719 | 6.1391 | 6.2565 | 6.1036 | 6.3002 | 6.0008 | 6.0512 | 6.5893 | 6.7248 | 6.1392 | 6.2530 |
| FAAP20_S216 | FAAP20 | Q6NZ36-2 | | 216 | AGGHLESPARSLP | 7.1763 | 6.6407 | 6.8131 | 7.3001 | 5.8849 | 5.6904 | 6.6058 | 7.1192 | 6.9903 | 7.2862 | 7.5330 | 8.1934 | 3.0711 | 4.8077 | 4.0410 | 4.8467 |
| FADS1_S35 | FADS1 | A0A0A0MR51 | | 35 | QQIHSWSPRTPST | 7.1392 | 8.2194 | 7.2094 | 7.9700 | 5.3835 | 7.1354 | 7.8447 | 6.5903 | 8.6004 | 8.1680 | 7.1492 | 7.2282 | 3.0376 | 3.3384 | 1.8196 | 3.1667 |
| FAF1_S270 | FAF1 | Q9UNN5 | | 320 | SSALRKSPMMPEN | 5.8843 | 5.5398 | 6.0503 | 6.2349 | 6.1201 | 6.2292 | 6.4996 | 6.3368 | 6.4099 | 6.2605 | 6.4648 | 6.5478 | 6.2563 | 6.4435 | 6.5265 | 6.1957 |
| FAF1_S320 | FAF1 | Q9UNN5 | | 582 | LRIRTPSGEFLER | 5.9416 | 6.0487 | 6.1145 | 5.7130 | 5.7461 | 5.9646 | 5.8729 | 6.1943 | 5.8291 | 5.8318 | 5.9168 | 6.1918 | 7.0368 | 7.0931 | 7.1267 | 7.3782 |
| FAF1_S582 | FAF1 | Q9UNN5 | | 270 | TVGRRSSPAQTRE | 6.0758 | 6.4528 | 6.4990 | 6.4055 | 5.7545 | 6.1182 | 6.0617 | 6.2660 | 6.2460 | 6.3481 | 6.2246 | 6.3101 | 6.2658 | 6.7919 | 5.8516 | 6.3283 |
| FAM110A_S78 | FAM110A | Q9BQ89 | | 78 | SKQPLFSPETRRT | 6.6990 | 7.2427 | 7.1151 | 6.5360 | 6.1041 | 7.2353 | 7.2933 | 7.0196 | 6.8373 | 7.0700 | 7.8252 | 7.8798 | 3.7647 | 4.2242 | 3.0951 | 4.0586 |
| FAM111A_S12 | FAM111A | Q96PZ2 | | 607 | QDVEMMSDEDLxx | 5.9500 | 6.2173 | 6.6365 | 5.9177 | 5.6926 | 6.3090 | 6.4205 | 6.1564 | 6.0802 | 6.7827 | 6.5383 | 6.7877 | 6.2856 | 6.3880 | 6.2212 | 5.6163 |
| FAM111A_S26 | FAM111A | Q96PZ2 | | 26 | KIEHYFSPVSKEQ | 7.2340 | 7.6032 | 7.4565 | 7.3573 | 7.3943 | 7.5893 | 6.8041 | 6.9181 | 8.2245 | 7.5202 | 7.7587 | 7.2183 | 2.2496 | 3.1365 | 2.4556 | 3.0797 |
| FAM111A_S607 | FAM111A | Q96PZ2 | | 12 | QRSRKHSVNEKCN | 6.6126 | 5.6344 | 6.9521 | 6.7141 | 6.2152 | 5.7917 | 5.5590 | 6.1001 | 8.2680 | 8.5937 | 8.9216 | 8.7165 | 3.9415 | 4.9606 | 3.3401 | 3.6788 |
| FAM114A1_S261 | FAM114A1 | Q8IWE2 | | 261 | LMERTVSLSQMLR | 5.1183 | 4.9996 | 5.5538 | 5.2085 | 5.8443 | 4.9534 | 4.3167 | 4.7869 | 5.2946 | 5.4781 | 5.9082 | 5.7452 | 10.0041 | 10.0505 | 8.6522 | 8.0857 |
| FAM114A2_S146 | FAM114A2 | Q9NRY5 | | 87 | RWGYWGSWGKSIL | 5.5883 | 5.9924 | 5.9363 | 5.9742 | 5.2881 | 7.3202 | 6.2662 | 6.0039 | 5.5355 | 5.5256 | 5.6658 | 5.5635 | 7.0117 | 8.3115 | 7.5973 | 6.4196 |
| FAM114A2_S87 | FAM114A2 | Q9NRY5 | | 146 | KENENSSPVAGAF | 4.3627 | 5.0896 | 6.0740 | 5.4431 | 6.3083 | 6.2644 | 6.2541 | 6.5905 | 6.2169 | 6.1937 | 6.1478 | 5.9275 | 7.5754 | 6.5275 | 7.6208 | 7.4036 |
| FAM117A_S201 | FAM117A | Q9C073 | | 67 | GGGRAASVPCSVA | 5.8387 | 5.7783 | 6.5805 | 6.0267 | 6.6285 | 6.7297 | 6.3799 | 6.1409 | 6.7491 | 7.4743 | 6.9275 | 6.4193 | 5.8383 | 5.8252 | 5.1776 | 5.4856 |
| FAM117A_S207 | FAM117A | Q9C073 | | 213 | SPCLHRSLEGLNQ | 6.4378 | 5.9200 | 7.0537 | 6.3711 | 6.4035 | 6.2505 | 5.9398 | 6.6764 | 6.2324 | 7.5895 | 6.8304 | 6.7610 | 5.4103 | 5.9389 | 4.8943 | 5.2905 |
| FAM117A_S213 | FAM117A | Q9C073 | | 428 | PEASKASPLPFEP | 6.7853 | 6.7611 | 6.1930 | 4.3190 | 6.6611 | 6.4642 | 5.5840 | 8.5578 | 5.5719 | 6.9262 | 6.6475 | 7.3004 | 5.6385 | 5.6628 | 5.8840 | 5.0432 |
| FAM117A_S306 | FAM117A | Q9C073 | | 207 | SPVLRLSPCLHRS | 6.3281 | 5.9636 | 6.5087 | 5.5509 | 6.2346 | 6.5870 | 5.2951 | 6.6218 | 7.1477 | 6.8691 | 6.5198 | 7.0932 | 5.8285 | 6.2519 | 5.5788 | 5.6212 |
| FAM117A_S355 | FAM117A | Q9C073 | | 201 | PSFPSGSPVLRLS | 6.8608 | 6.2828 | 7.5930 | 6.2273 | 6.5051 | 5.5937 | 6.4163 | 7.1144 | 6.9854 | 7.5544 | 6.8510 | 6.5803 | 4.8176 | 4.8732 | 4.7881 | 4.9565 |
| FAM117A_S428 | FAM117A | Q9C073 | | 355 | VFEEATSPGPDLA | 6.4125 | 6.5229 | 7.1020 | 7.4744 | 7.1119 | 7.5867 | 5.9850 | 7.3641 | 6.5701 | 6.0619 | 7.1139 | 7.1237 | 3.9854 | 4.5607 | 3.9342 | 5.0906 |
| FAM117A_S67 | FAM117A | Q9C073 | | 306 | PNDKASSPGHPAF | 7.1857 | 6.3017 | 6.3624 | 7.4550 | 7.1477 | 5.9868 | 6.4307 | 7.0112 | 6.2296 | 7.2046 | 7.3488 | 6.9618 | 4.9586 | 4.8069 | 4.7614 | 3.8470 |
| FAM117B_S10 | FAM117B | Q6P1L5 | | 210 | TRQPSSSPSSIIR | 6.8206 | 6.6439 | 6.5469 | 5.9050 | 6.5219 | 6.6770 | 6.2705 | 6.7382 | 6.5962 | 6.4591 | 6.5386 | 6.4754 | 5.5936 | 5.5824 | 5.1422 | 5.4883 |
| FAM117B_S106 | FAM117B | Q6P1L5 | | 220 | IIRRTSSLDTLAA | 6.0584 | 6.0730 | 6.8402 | 6.7344 | 6.0511 | 6.0662 | 6.1258 | 6.0598 | 6.3028 | 6.2526 | 6.2660 | 6.4880 | 6.3419 | 6.3537 | 5.7432 | 6.2431 |
| FAM117B_S210 | FAM117B | Q6P1L5 | | 457 | LPKYATSPKPNNS | 5.8826 | 6.4338 | 7.0543 | 6.3517 | 6.5416 | 7.0589 | 5.9806 | 7.0554 | 6.6154 | 6.2034 | 6.8480 | 6.7840 | 5.8318 | 5.6102 | 4.9276 | 4.8206 |
| FAM117B_S220 | FAM117B | Q6P1L5 | | 345 | GSRFRNSVEGLNQ | 5.6209 | 5.7623 | 7.0771 | 5.9398 | 6.0669 | 6.5534 | 6.5084 | 6.5336 | 5.9233 | 7.5965 | 5.3229 | 6.9971 | 6.0494 | 7.1713 | 5.4193 | 5.4577 |
| FAM117B_S273 | FAM117B | Q6P1L5 | | 10 | RVRRNGSPTPAGS | 6.2613 | 6.5129 | 6.8934 | 6.1240 | 6.6059 | 6.0213 | 6.2722 | 6.3935 | 6.5266 | 6.2842 | 6.7265 | 6.3721 | 5.7035 | 5.8175 | 5.6926 | 5.7926 |
| FAM117B_S307 | FAM117B | Q6P1L5 | | 273 | SHKRSASWGSTDQ | 6.5404 | 6.4087 | 6.7137 | 5.8191 | 6.3578 | 6.3076 | 5.9218 | 6.4392 | 6.8919 | 6.3913 | 6.3455 | 6.4953 | 6.1183 | 5.9354 | 5.8017 | 5.5125 |
| FAM117B_S345 | FAM117B | Q6P1L5 | | 484 | KVFEECSPKQLHE | 7.5094 | 7.7650 | 7.4055 | 7.7474 | 5.6474 | 5.8797 | 6.9709 | 5.8831 | 6.7852 | 7.1067 | 7.6582 | 8.0377 | 3.6087 | 3.9291 | 3.4457 | 4.6204 |
| FAM117B_S457 | FAM117B | Q6P1L5 | | 307 | RDKERQSPFHGNH | 6.3125 | 6.2594 | 6.7151 | 5.8742 | 7.0377 | 7.7541 | 5.8798 | 5.8077 | 6.8332 | 5.3456 | 6.5320 | 6.9870 | 5.7688 | 5.8449 | 5.0215 | 6.0264 |
| FAM117B_S484 | FAM117B | Q6P1L5 | | 106 | NAAARTSPTVATQ | 5.6489 | 6.2377 | 6.6007 | 5.5761 | 5.9634 | 6.4383 | 5.6144 | 6.1285 | 6.3954 | 7.1689 | 6.7808 | 6.5932 | 6.4271 | 6.5643 | 5.7488 | 6.1136 |
| FAM120A_S1072 | FAM120A | Q9NZB2-6 | | 383 | VAPQVPSPGGAPG | 5.8062 | 5.9174 | 6.7190 | 6.7355 | 6.7858 | 6.5865 | 6.3579 | 5.9426 | 6.3262 | 6.7418 | 6.6327 | 6.4397 | 5.3795 | 6.2838 | 5.5849 | 5.7606 |
| FAM120A_S383 | FAM120A | Q9NZB2-6 | | 1019 | PRGVISTPVIRTF | 11.0480 | 8.8992 | 7.5796 | 8.4422 | 6.3140 | 6.6942 | 7.6380 | 5.9598 | 8.2012 | 6.3103 | 6.4368 | 6.2565 | 2.3092 | 3.2597 | 1.9445 | 2.7068 |
| FAM120A_T1019 | FAM120A | Q9NZB2-6 | | 1072 | SKSSAMSSDGSLA | 6.2345 | 6.5556 | 7.2238 | 5.7229 | 6.3265 | 5.4840 | 5.6099 | 7.2383 | 5.9149 | 6.8623 | 6.1457 | 7.1576 | 6.3716 | 5.6309 | 5.9795 | 5.5421 |
| FAM126A_S415 | FAM126A | Q9BYI3 | | 485 | NRFSACSLQEEKL | 6.9633 | 6.1514 | 6.3020 | 6.8783 | 6.6942 | 5.7864 | 5.8871 | 5.9930 | 6.4402 | 5.9258 | 6.0743 | 5.4640 | 6.6158 | 6.5953 | 6.0619 | 6.1670 |
| FAM126A_S422 | FAM126A | Q9BYI3 | | 453 | ATVFSKSFEQVSG | 5.9865 | 6.6140 | 6.8703 | 5.9642 | 5.5953 | 6.4996 | 6.4387 | 6.4835 | 6.6514 | 6.5334 | 6.7753 | 7.2824 | 5.1063 | 6.0857 | 5.6100 | 5.5035 |
| FAM126A_S453 | FAM126A | Q9BYI3 | | 415 | QTQRAQSENLELL | 6.0780 | 6.1307 | 6.6973 | 5.8477 | 6.1523 | 6.8101 | 5.9185 | 5.6916 | 5.9721 | 6.6528 | 6.6273 | 6.3122 | 6.0002 | 6.8611 | 6.1640 | 6.0840 |
| FAM126A_S485 | FAM126A | Q9BYI3 | | 422 | ENLELLSLKRLTL | 7.3967 | 6.1924 | 7.3070 | 7.4848 | 5.9853 | 5.7043 | 6.3674 | 5.5119 | 6.8160 | 7.0959 | 6.4656 | 5.6970 | 5.7822 | 5.9876 | 4.9944 | 5.2116 |
| FAM126B_S430 | FAM126B | Q8IXS8 | | 430 | PMKKHLSLPAGQV | 6.1126 | 6.4064 | 6.8948 | 6.9462 | 6.1096 | 5.8965 | 6.1847 | 6.3061 | 6.0153 | 5.8583 | 5.7770 | 6.2830 | 6.4359 | 6.6300 | 5.9983 | 6.1453 |
| FAM126B_S444 | FAM126B | Q8IXS8 | | 508 | EGKELLSPGAPLT | 7.5044 | 7.1895 | 6.6622 | 7.4950 | 5.9004 | 7.0157 | 6.7705 | 7.4870 | 7.3128 | 6.7851 | 6.4574 | 7.7535 | 3.3662 | 4.6736 | 3.4085 | 4.2183 |
| FAM126B_S508 | FAM126B | Q8IXS8 | | 521 | KQSRSPSFNMQLI | 6.4070 | 6.4161 | 7.1121 | 6.1689 | 5.3051 | 6.1182 | 6.0870 | 5.6447 | 6.8961 | 6.7504 | 7.2474 | 6.3459 | 5.1165 | 6.7726 | 5.7305 | 5.8814 |
| FAM126B_S521 | FAM126B | Q8IXS8 | | 444 | PKINSLSLIRTAS | 5.1838 | 5.1206 | 5.2556 | 5.0497 | 4.7546 | 5.0782 | 5.1381 | 5.0301 | 4.7261 | 4.4114 | 4.3995 | 4.4996 | 10.0425 | 10.1735 | 10.9410 | 10.1959 |
| FAM126B_T308 | FAM126B | Q8IXS8 | | 308 | KVEVTPTVPRISR | 7.1331 | 6.4872 | 7.6781 | 5.6115 | 6.7294 | 7.4033 | 6.1037 | 6.3148 | 6.4049 | 5.4189 | 6.0742 | 5.9950 | 5.1845 | 6.1922 | 5.4158 | 5.8537 |
| FAM133B_S82 | FAM133B | Q5BKY9 | | 86 | LLSGSESSSKKRQ | 6.0901 | 6.6241 | 6.8359 | 6.4704 | 6.4819 | 6.4124 | 6.8167 | 6.2316 | 8.2061 | 6.7020 | 7.0989 | 6.6037 | 4.3844 | 5.0339 | 4.7457 | 5.2622 |
| FAM133B_S86 | FAM133B | Q5BKY9 | | 82 | HREKLLSGSESSS | 5.9804 | 7.1870 | 8.7829 | 7.6084 | 7.5677 | 8.7358 | 7.6067 | 8.6093 | 4.6149 | 5.8204 | 5.5706 | 5.4628 | 2.8165 | 4.9251 | 4.0011 | 4.7104 |
| FAM135A_S953 | FAM135A | Q9P2D6 | | 953 | MCKGFQSPDKSNN | 6.4437 | 6.5908 | 7.2356 | 6.5137 | 6.2180 | 6.5202 | 6.8922 | 6.7484 | 7.0352 | 7.3250 | 7.9043 | 7.9691 | 3.8584 | 4.4191 | 3.8308 | 4.4956 |
| FAM13B_S760 | FAM13B | Q9NYF5 | | 760 | ITPVLGSPSTKRR | 6.9469 | 6.7410 | 7.1406 | 6.8210 | 5.7495 | 5.8329 | 6.1308 | 6.4205 | 6.7016 | 6.9743 | 7.0598 | 6.8029 | 5.1602 | 5.5697 | 5.1066 | 4.8417 |
| FAM156A_S11 | FAM156A | Q8NDB6 | | 114 | GREARISPLGDRS | 7.3199 | 7.6369 | 7.1700 | 6.8866 | 6.1097 | 6.7882 | 6.7361 | 6.4433 | 7.4059 | 7.0713 | 7.7453 | 7.7132 | 3.7346 | 3.9260 | 3.5060 | 3.8067 |
| FAM156A_S114 | FAM156A | Q8NDB6 | | 11 | QKRNPASPSKSSP | 6.4398 | 6.5611 | 4.6661 | 6.3215 | 6.2933 | 6.3571 | 5.9209 | 6.1668 | 6.2276 | 6.4431 | 6.6515 | 6.5861 | 6.0980 | 5.4961 | 6.0119 | 5.9593 |
| FAM160A2_S524 | FAM160A2 | Q8N612-2 | | 873 | VKSRRPSLGELLL | 5.6546 | 5.1847 | 6.2151 | 5.3547 | 6.7076 | 6.3319 | 5.5675 | 6.4467 | 7.1048 | 7.2942 | 7.6941 | 6.8218 | 6.1490 | 6.8969 | 4.9797 | 5.5966 |
| FAM160A2_S596 | FAM160A2 | Q8N612-2 | | 524 | LFLRQQSLGGSES | 5.6688 | 5.8442 | 7.5149 | 5.2328 | 4.4101 | 5.8788 | 6.5845 | 6.0755 | 6.5978 | 8.0306 | 6.2164 | 7.9814 | 5.4029 | 6.8224 | 5.0454 | 6.6935 |
| FAM160A2_S873 | FAM160A2 | Q8N612-2 | | 596 | YDGERPSPEPSPF | 6.9290 | 7.1607 | 6.5295 | 6.2172 | 6.2069 | 6.0567 | 6.3682 | 6.6988 | 6.2876 | 5.9833 | 6.5141 | 6.5649 | 6.1116 | 6.1043 | 5.5684 | 5.6087 |
| FAM161A_T428 | FAM161A | Q3B820-3 | | 428 | HKVRCPTPDFEDL | 6.4664 | 6.0130 | 6.1699 | 6.3490 | 5.7927 | 6.1454 | 6.3783 | 6.5400 | 7.1820 | 6.3921 | 6.6694 | 6.0975 | 5.7190 | 6.2969 | 5.8334 | 5.9550 |
| FAM167A_S45 | FAM167A | Q96KS9 | | 45 | LETRRPSYLEWQA | 6.6582 | 5.1510 | 5.2380 | 5.9021 | 6.0573 | 5.4079 | 5.4618 | 5.5015 | 7.3145 | 6.9101 | 7.0029 | 7.3762 | 6.1917 | 8.2636 | 5.1533 | 6.4098 |
| FAM169A_S447 | FAM169A | Q9Y6X4 | | 448 | TEEEDSTSEVLDE | 6.0861 | 5.8844 | 6.0773 | 6.2997 | 6.1540 | 6.1517 | 5.9498 | 5.9101 | 6.0205 | 6.2820 | 5.8682 | 6.0722 | 6.7061 | 6.9220 | 6.7406 | 6.8751 |
| FAM169A_T448 | FAM169A | Q9Y6X4 | | 447 | ITEEEDSTSEVLD | 6.3997 | 6.0384 | 6.0168 | 6.5898 | 5.8265 | 5.6625 | 6.1930 | 6.2193 | 6.0964 | 5.9474 | 6.5971 | 6.0393 | 6.8817 | 6.7098 | 6.3874 | 6.3947 |
| FAM171A1_S358 | FAM171A1 | Q5VUB5 | | 358 | KRDQSTSMSHINL | 5.9427 | 6.1317 | 6.4376 | 6.6313 | 6.3695 | 6.2835 | 6.3213 | 6.5018 | 6.7947 | 6.3012 | 6.4906 | 6.3302 | 5.6641 | 6.3794 | 5.5400 | 5.8805 |
| FAM171A1_S422 | FAM171A1 | Q5VUB5 | | 511 | GPASTGSKLTIQE | 6.1815 | 6.1746 | 6.4798 | 6.6579 | 5.5243 | 5.6702 | 6.1636 | 5.8782 | 6.8512 | 6.3761 | 6.5035 | 6.3488 | 6.0364 | 6.7144 | 5.9523 | 6.4872 |
| FAM171A1_S511 | FAM171A1 | Q5VUB5 | | 525 | LYPAPSSPEKEQL | 5.9851 | 6.5644 | 6.7018 | 6.5047 | 6.1415 | 6.4662 | 6.4725 | 6.7802 | 6.6876 | 6.5130 | 6.5966 | 6.7165 | 5.3861 | 5.5441 | 5.3736 | 5.5661 |
| FAM171A1_S525 | FAM171A1 | Q5VUB5 | | 422 | KLSYSTSQEFSSR | 6.5414 | 6.6303 | 6.1248 | 6.4641 | 5.5772 | 6.0601 | 6.1791 | 6.0328 | 6.8632 | 5.9212 | 6.3225 | 6.7786 | 6.3782 | 6.0899 | 5.9469 | 6.0896 |
| FAM171A1_S564 | FAM171A1 | Q5VUB5 | | 448 | ISFDNLTPSGTLG | 5.9285 | 6.7700 | 6.8383 | 6.7163 | 5.9053 | 6.1473 | 6.2809 | 7.2369 | 6.0230 | 6.2209 | 6.3333 | 7.0943 | 5.3518 | 6.1178 | 5.3913 | 5.6440 |
| FAM171A1_S598 | FAM171A1 | Q5VUB5 | | 598 | GDHSYVSQPLVVP | 5.7749 | 5.7240 | 6.8638 | 6.0667 | 6.8046 | 6.6325 | 5.8437 | 5.9277 | 5.6799 | 5.8079 | 6.8060 | 6.5596 | 5.8864 | 7.0689 | 5.7074 | 6.8459 |
| FAM171A1_T448 | FAM171A1 | Q5VUB5 | | 564 | GQLICCSSVDQVN | 5.7174 | 5.3009 | 6.9189 | 6.4375 | 6.0947 | 6.3887 | 6.0523 | 7.0631 | 6.7035 | 6.3494 | 5.4563 | 6.5889 | 5.6053 | 6.3222 | 6.2301 | 6.7707 |
| FAM171A2_S789 | FAM171A2 | A8MVW0 | | 789 | SELRRDSLTSPED | 6.2317 | 6.3090 | 7.4988 | 6.4348 | 6.5926 | 5.8620 | 5.6843 | 5.7538 | 6.9624 | 5.8598 | 6.1776 | 6.5154 | 6.2317 | 6.2748 | 5.7008 | 5.9105 |
| FAM171B_S436 | FAM171B | Q6P995 | | 794 | DFEANTSPTKRRG | 5.0334 | 6.4542 | 6.3306 | 6.4702 | 5.6353 | 6.0326 | 6.6572 | 7.0496 | 6.8129 | 6.1524 | 6.3643 | 6.0878 | 6.2936 | 6.4134 | 5.8241 | 6.3885 |
| FAM171B_S794 | FAM171B | Q6P995 | | 436 | AKNSSYSPQKKEP | 6.4959 | 6.6252 | 6.1663 | 6.7272 | 5.6258 | 6.4283 | 6.2054 | 5.7397 | 6.5426 | 6.5250 | 6.9532 | 7.0736 | 5.6167 | 5.6613 | 5.8846 | 5.7292 |
| FAM172BP_S92 | FAM172BP | A6NC97 | | 92 | DKEEPRSFFFMSE | 6.3444 | 6.7853 | 5.8190 | 6.2742 | 5.7751 | 5.9535 | 6.3929 | 6.2956 | 6.0704 | 6.0985 | 6.3925 | 6.1529 | 5.8039 | 6.8475 | 6.5088 | 6.4855 |
| FAM174C_S117;S120 | FAM174C | Q9BVV8 | | 120 | EMASLDSDEETVF | 5.9983 | 5.9149 | 6.0991 | 6.1351 | 6.3606 | 5.7758 | 5.4949 | 6.3006 | 5.7924 | 6.2338 | 5.9523 | 6.3404 | 6.9561 | 7.2659 | 6.5215 | 6.8586 |
| FAM174C_T113;S120 | FAM174C | Q9BVV8 | 113;120 | | NA | 6.3235 | 6.9628 | 6.1653 | 5.6805 | 6.7817 | 5.3207 | 6.0286 | 5.5237 | 6.2079 | 5.7278 | 6.3034 | 8.0929 | 5.8886 | 6.1695 | 6.0687 | 6.7546 |
| FAM174C_T120 | FAM174C | Q9BVV8 | 117;120 | | NA | 6.8874 | 6.1372 | 6.1221 | 6.3503 | 6.2104 | 5.2890 | 7.5791 | 6.3712 | 5.8405 | 6.4989 | 5.9947 | 7.4698 | 6.3152 | 7.1121 | 5.7899 | 5.8521 |
| FAM177A1_S93;T94 | FAM177A1 | Q8N128-2 | 92;93 | | NA | 6.4065 | 6.4729 | 6.2288 | 6.5921 | 6.2459 | 6.1705 | 6.3820 | 6.5033 | 6.2240 | 6.3077 | 6.3472 | 6.6244 | 6.0069 | 5.9645 | 5.5901 | 5.9332 |
| FAM177A1_Y92;S93 | FAM177A1 | Q8N128-2 | 93;94 | | NA | 6.1237 | 5.6762 | 6.0185 | 6.7898 | 6.2545 | 6.3483 | 5.9305 | 6.8820 | 6.4867 | 6.7510 | 5.5262 | 7.3151 | 6.0237 | 5.9512 | 5.9798 | 5.9428 |
| FAM177A1_Y92;T94 | FAM177A1 | Q8N128-2 | 92;94 | | NA | 6.7215 | 6.5968 | 6.3812 | 6.5020 | 7.0483 | 5.7474 | 6.1798 | 6.4142 | 6.1253 | 5.7407 | 6.5572 | 6.5837 | 6.0100 | 5.8104 | 5.7393 | 5.8421 |
| FAM189B_S359 | FAM189B | P81408-4 | | 359 | LQGSVRSVDYVLF | 6.6979 | 7.4185 | 5.7361 | 7.3773 | 5.3971 | 5.4887 | 7.6247 | 5.9712 | 5.8964 | 6.2750 | 6.6414 | 7.6445 | 5.1452 | 5.2658 | 5.4490 | 5.9712 |
| FAM193A_S1129 | FAM193A | P78312 | | 393 | EESKADSPPPSYP | 5.9960 | 6.5259 | 6.4751 | 6.3577 | 6.1805 | 6.5020 | 6.1578 | 6.4058 | 6.2857 | 6.0913 | 6.2447 | 6.5797 | 5.8184 | 6.4442 | 6.1148 | 5.8204 |
| FAM193A_S1144 | FAM193A | P78312 | | 1129 | TPMEPTSPGEHQQ | 6.2033 | 6.4274 | 6.6804 | 6.1326 | 6.5775 | 6.5433 | 6.1610 | 6.6246 | 6.2505 | 6.3423 | 6.7617 | 6.3722 | 5.4937 | 6.0908 | 5.5314 | 5.8072 |
| FAM193A_S1249 | FAM193A | P78312 | | 287 | ILTDSGSAPTFCS | 5.4087 | 6.0205 | 5.4418 | 6.0106 | 5.4362 | 5.9394 | 5.7727 | 6.1209 | 6.1281 | 6.2948 | 6.3533 | 6.1532 | 6.8104 | 7.6221 | 7.3031 | 7.1841 |
| FAM193A_S287 | FAM193A | P78312 | | 1144 | KLVLAESPQPKGK | 6.2603 | 6.9450 | 6.3324 | 7.1209 | 5.5317 | 6.2440 | 6.1170 | 6.0470 | 5.9888 | 6.6852 | 7.5596 | 7.8525 | 4.4607 | 5.6871 | 5.5308 | 5.6368 |
| FAM193A_S393 | FAM193A | P78312 | | 1249 | QTRQRLSINWSNF | 6.8002 | 4.3161 | 6.2277 | 5.3480 | 6.8462 | 5.5867 | 6.0070 | 6.3533 | 7.4397 | 6.8979 | 8.5149 | 9.4727 | 5.3186 | 6.5279 | 4.6051 | 3.7380 |
| FAM193B_S785 | FAM193B | Q96PV7 | | 785 | GPGWAGSPKTEKE | 6.7606 | 6.3081 | 7.5364 | 6.9116 | 7.2169 | 6.3641 | 6.0662 | 6.0604 | 7.3548 | 6.2771 | 6.5618 | 6.6198 | 4.7963 | 5.0130 | 4.9376 | 5.2153 |
| FAM207A_S203 | FAM207A | Q9NSI2 | | 203 | FQELLASPAYRAS | 7.1942 | 7.5609 | 6.9263 | 7.2121 | 5.8304 | 7.1553 | 7.3466 | 6.4305 | 7.3519 | 6.8997 | 7.0957 | 7.7771 | 3.5817 | 4.1826 | 3.3154 | 4.1397 |
| FAM207A_S209 | FAM207A | Q9NSI2 | | 34 | PPAPEATPPPASA | 6.3516 | 6.4532 | 6.4014 | 7.1589 | 6.6635 | 6.8027 | 7.2703 | 6.8332 | 6.6347 | 6.7038 | 6.6596 | 7.1534 | 4.6344 | 5.1118 | 4.1099 | 5.0579 |
| FAM207A_T34 | FAM207A | Q9NSI2 | | 209 | SPAYRASPLVAIG | 6.4522 | 5.8411 | 5.6359 | 6.2041 | 5.6083 | 6.8044 | 6.1060 | 7.0066 | 7.7922 | 7.1367 | 7.8443 | 5.7063 | 5.7523 | 5.5042 | 5.1412 | 5.4643 |
| FAM216A_S33 | FAM216A | Q8WUB2 | | 33 | WTERSSSAEPPAV | 6.4205 | 5.9400 | 7.6034 | 6.8911 | 6.9879 | 7.0268 | 6.2445 | 5.9343 | 6.4290 | 6.1703 | 7.0550 | 6.4664 | 5.0319 | 5.4383 | 4.8744 | 5.4863 |
| FAM219A_S131 | FAM219A | Q8IW50 | | 47 | AMNYKPSPLQVKL | 7.0786 | 7.1727 | 6.7005 | 6.2260 | 5.9878 | 7.0107 | 6.7818 | 6.7388 | 6.8660 | 7.7112 | 6.7127 | 6.8376 | 3.6134 | 5.6068 | 3.7406 | 5.2147 |
| FAM219A_S28 | FAM219A | Q8IW50 | | 28 | PAAASISDGDCDA | 6.4772 | 6.2014 | 6.5818 | 6.5197 | 5.3164 | 6.1484 | 6.5757 | 7.4123 | 5.7043 | 6.2302 | 6.4969 | 5.2466 | 5.3649 | 6.4030 | 6.4166 | 6.9046 |
| FAM219A_S47 | FAM219A | Q8IW50 | | 72 | KNGSMGSPVNQQP | 7.1229 | 6.8786 | 6.4256 | 6.7082 | 5.9632 | 5.6412 | 6.4719 | 6.2528 | 6.2892 | 6.8919 | 7.2016 | 7.2755 | 4.7304 | 5.9076 | 4.7608 | 5.4787 |
| FAM219A_S72 | FAM219A | Q8IW50 | | 131 | SSSGYSSAEQINQ | 6.0068 | 5.4001 | 5.2280 | 5.2181 | 5.6256 | 5.8861 | 5.4482 | 6.1551 | 5.5880 | 5.6526 | 5.4258 | 6.1340 | 6.9577 | 8.8318 | 7.8545 | 8.5878 |
| FAM219B_S66 | FAM219B | Q5XKK7 | | 66 | MVTRAPSIQAKLQ | 6.1872 | 6.7745 | 6.3265 | 5.6654 | 4.5428 | 6.8509 | 6.6420 | 5.6699 | 7.2032 | 8.5475 | 8.9487 | 9.2448 | 3.5388 | 5.5083 | 3.2542 | 5.0951 |
| FAM219B_S91 | FAM219B | Q5XKK7 | | 91 | KGMLGASPNRPDS | 7.2732 | 7.1993 | 6.7028 | 6.4857 | 6.8574 | 6.7853 | 6.8290 | 7.0730 | 6.8361 | 7.5284 | 6.6665 | 6.3142 | 4.2732 | 4.7391 | 4.1145 | 4.3222 |
| FAM32A_S44 | FAM32A | Q9Y421 | | 44 | LEAMGTSKKNEEE | 6.4741 | 6.0563 | 6.8378 | 6.2936 | 5.8391 | 6.4476 | 6.5361 | 6.9981 | 7.7721 | 7.7175 | 6.9849 | 7.2883 | 4.5198 | 5.3427 | 4.2199 | 4.6721 |
| FAM50A_S121 | FAM50A | Q14320 | | 121 | RKISSLSFTLEEE | 6.6677 | 6.6328 | 5.9493 | 6.4584 | 6.4405 | 6.2741 | 5.8756 | 6.8759 | 6.6766 | 6.1555 | 6.8779 | 7.1379 | 5.4900 | 5.3425 | 5.3774 | 5.7679 |
| FAM50B_S118 | FAM50B | Q9Y247 | | 118 | ERKRKISCLSFAL | 5.2841 | 5.0824 | 5.6842 | 4.8713 | 6.5432 | 5.9177 | 5.1676 | 6.1167 | 8.5822 | 9.3388 | 8.4795 | 8.5239 | 5.4174 | 6.4959 | 3.8630 | 4.6320 |
| FAM53A_S346 | FAM53A | Q6NSI3 | | 346 | QRGLRTSPVHPNL | 6.6885 | 6.7334 | 7.5648 | 5.6927 | 7.0638 | 6.0375 | 6.1479 | 6.0797 | 6.8896 | 6.2458 | 6.8555 | 7.1697 | 5.9216 | 4.9151 | 4.5452 | 5.4494 |
| FAM53B_S167 | FAM53B | Q14153 | | 167 | TMQRSSSFSLPSR | 6.1563 | 6.3392 | 6.9277 | 5.6565 | 6.0668 | 6.5352 | 5.6647 | 6.4521 | 6.2301 | 6.0855 | 6.6548 | 7.0395 | 6.2370 | 6.3519 | 5.9643 | 5.6382 |
| FAM53C_S232;S234 | FAM53C | Q9NYF3 | | 234 | QRRFSLSPSLGPQ | 6.9065 | 6.6619 | 7.6752 | 6.7899 | 6.3277 | 8.0743 | 6.3229 | 6.5254 | 6.0332 | 6.0338 | 7.1072 | 6.6426 | 4.6135 | 4.9471 | 4.4260 | 4.9129 |
| FAM53C_S234 | FAM53C | Q9NYF3 | | 299 | PRRLRPSLDFDKM | 6.1507 | 5.8033 | 6.7567 | 5.7392 | 7.0933 | 6.3598 | 5.7359 | 6.4519 | 7.1042 | 7.1443 | 7.2757 | 6.0917 | 5.5993 | 5.9661 | 5.0064 | 5.7217 |
| FAM53C_S299 | FAM53C | Q9NYF3 | 232;234 | | NA | 6.5223 | 5.8994 | 6.8280 | 6.1974 | 6.2714 | 6.5916 | 6.0702 | 6.6471 | 6.3617 | 5.9664 | 7.2192 | 6.9126 | 5.8637 | 5.6661 | 5.4080 | 5.5750 |
| FAM76A_S340 | FAM76A | Q8TAV0-3 | | 340 | KSGAITSPxxxxx | 6.6949 | 7.5989 | 6.8487 | 6.8260 | 6.0202 | 6.6545 | 6.8041 | 6.4527 | 6.4897 | 6.5707 | 6.8063 | 7.3708 | 4.2838 | 4.9847 | 4.9266 | 4.6674 |
| FAM76B_S193 | FAM76B | Q5HYJ3 | | 193 | HKISNLSPEEEQG | 5.7237 | 5.8155 | 5.6921 | 6.0606 | 6.4696 | 5.8088 | 6.3845 | 6.2459 | 6.1074 | 6.0789 | 5.7064 | 5.6678 | 6.9652 | 7.0467 | 7.2342 | 6.9925 |
| FAM76B_S227 | FAM76B | Q5HYJ3 | | 338 | SGSILTSPxxxxx | 7.3486 | 7.4651 | 6.8458 | 6.5751 | 6.0926 | 6.4995 | 6.7038 | 6.6752 | 6.4375 | 6.4695 | 7.2931 | 7.0029 | 4.1327 | 5.0016 | 4.8017 | 4.6555 |
| FAM76B_S338 | FAM76B | Q5HYJ3 | | 215 | ATIQNETPKKKPK | 6.5059 | 6.2641 | 5.9870 | 6.5799 | 6.1881 | 5.5542 | 6.3397 | 5.4744 | 6.3536 | 5.9409 | 5.8470 | 5.7195 | 7.8229 | 6.7127 | 6.3919 | 6.3183 |
| FAM76B_T215 | FAM76B | Q5HYJ3 | | 227 | KLESKPSNGDSSS | 4.1269 | 2.4311 | 3.3586 | 3.4354 | 4.5549 | 6.0970 | 6.3604 | 4.7690 | 4.2805 | 3.9787 | 4.1982 | 6.3440 | 9.4337 | 9.2086 | 13.3794 | 14.0435 |

| | | | | | | | | | | | | | | | | | | | | | |
|---|---|---|---|---|---|---|---|---|---|---|---|---|---|---|---|---|---|---|---|---|---|
| FAM83G_S116 | FAM83G | A6ND36 | | 356 | ALVKAKSVDEIAK | 6.8747 | 7.1415 | 7.5820 | 6.7172 | 6.2269 | 5.8488 | 5.9513 | 6.3043 | 6.0423 | 6.5028 | 6.1046 | 6.1667 | 5.0000 | 6.1353 | 6.1585 | 5.2432 |
| FAM83G_S124;S127 | FAM83G | A6ND36 | | 760 | LLPDPGSPRLAQN | 6.6098 | 7.6139 | 6.4953 | 5.8553 | 6.1505 | 6.5400 | 6.9556 | 8.7126 | 6.7191 | 6.5355 | 6.0512 | 6.9670 | 4.5327 | 5.1118 | 4.5667 | 4.5830 |
| FAM83G_S127 | FAM83G | A6ND36 | | 127 | PQKSDRSIPQLDL | 5.7037 | 5.5602 | 5.8650 | 6.1203 | 6.1086 | 5.6688 | 6.2408 | 6.3995 | 6.1857 | 5.9753 | 5.9233 | 6.5375 | 6.5257 | 6.3984 | 6.8914 | 6.9957 |
| FAM83G_S356 | FAM83G | A6ND36 | | 650 | PPRRQLSAPHITR | 4.8487 | 5.1891 | 5.6615 | 5.7388 | 5.8610 | 4.9410 | 4.6811 | 4.7021 | 4.6538 | 4.7772 | 5.0984 | 5.0611 | 9.2225 | 9.1516 | 10.3626 | 10.0495 |
| FAM83G_S650 | FAM83G | A6ND36 | | 116 | EAEPLPSLEYWPQ | 5.7326 | 5.0048 | 5.3382 | 5.5313 | 6.1099 | 6.0063 | 5.5909 | 6.2583 | 6.4537 | 5.7691 | 6.2777 | 6.4723 | 6.7534 | 7.5654 | 7.1262 | 8.0098 |
| FAM83G_S760 | FAM83G | A6ND36 | 124;127 | | NA | 6.2322 | 5.6092 | 5.1663 | 5.5733 | 5.8900 | 5.1559 | 6.2926 | 5.6882 | 6.2897 | 5.9574 | 6.2753 | 5.5210 | 7.7943 | 7.2412 | 7.7174 | 7.5960 |
| FAM83H_S1068 | FAM83H | Q6ZRV2 | | 892 | FSTRRGSPTTGFI | 6.0892 | 5.9855 | 6.0420 | 6.2278 | 5.9568 | 6.2229 | 6.0024 | 6.2426 | 6.4734 | 6.1985 | 6.0037 | 6.6348 | 6.7359 | 6.8254 | 6.2216 | 6.1375 |
| FAM83H_S516 | FAM83H | Q6ZRV2 | | 516 | YVPSSASREVRHG | 5.7164 | 6.2956 | 5.7126 | 5.6829 | 5.9724 | 6.2005 | 6.2533 | 6.5361 | 6.1610 | 5.6971 | 6.0232 | 6.0532 | 6.6896 | 6.8457 | 7.1058 | 7.0546 |
| FAM83H_S870 | FAM83H | Q6ZRV2 | | 870 | HNESKGSPTSAYP | 6.7478 | 6.7902 | 5.8659 | 5.4628 | 6.0645 | 5.7030 | 6.9429 | 6.8849 | 6.5269 | 5.5396 | 5.9355 | 5.6598 | 6.5064 | 6.6270 | 6.5051 | 6.2379 |
| FAM83H_S881 | FAM83H | Q6ZRV2 | | 881 | YPERKGSPTPGFS | 6.2125 | 6.7476 | 6.1453 | 5.8115 | 6.3712 | 5.8992 | 5.9272 | 6.4133 | 6.4619 | 6.1498 | 6.1701 | 6.0253 | 6.4173 | 6.3796 | 6.3840 | 6.4840 |
| FAM83H_S892 | FAM83H | Q6ZRV2 | | 936 | VPERRGSLTLTIS | 5.9929 | 5.8837 | 6.2879 | 6.2591 | 5.6105 | 5.2765 | 5.2105 | 5.8212 | 6.8427 | 6.2851 | 6.3420 | 6.9274 | 7.0183 | 7.1079 | 6.6826 | 6.4519 |
| FAM83H_S936 | FAM83H | Q6ZRV2 | | 1068 | PGPTHNSPELGRP | 6.1372 | 6.3429 | 6.6728 | 6.7406 | 6.3200 | 5.8474 | 6.6947 | 6.6040 | 5.9204 | 6.5611 | 6.6675 | 6.4284 | 5.2860 | 6.1801 | 6.1099 | 5.4870 |
| FAM91A1_S671 | FAM91A1 | Q658Y4 | | 671 | LADRKLSDASDER | 5.7320 | 5.9229 | 6.1988 | 6.1915 | 5.9947 | 6.0250 | 5.9707 | 6.1468 | 7.5385 | 5.9765 | 5.9973 | 5.9603 | 6.9119 | 7.0981 | 7.0575 | 7.0778 |
| FAM91A1_S828 | FAM91A1 | Q658Y4 | | 828 | WSGRSPSSLLIAN | 5.5936 | 5.7935 | 6.3302 | 6.3440 | 5.5987 | 6.0288 | 6.3429 | 6.2746 | 6.2606 | 5.8154 | 5.8788 | 6.0349 | 6.9818 | 6.9901 | 7.1028 | 6.6294 |
| FAM9C_S52 | FAM9C | Q8IZT9 | | 52 | EHGERGSFAETDE | 5.4587 | 6.0447 | 6.5516 | 5.7743 | 6.3739 | 6.7725 | 6.3809 | 6.8456 | 6.4314 | 6.7260 | 6.8104 | 6.2829 | 6.0181 | 6.3852 | 5.6826 | 5.4611 |
| FAN1_S317 | FAN1 | Q9Y2M0 | | 317 | EAKIQLSDSEAKS | 6.2852 | 6.9834 | 7.3164 | 6.5843 | 6.4959 | 6.1720 | 5.9042 | 6.6694 | 7.1347 | 5.2671 | 6.1505 | 6.2753 | 4.9919 | 5.9984 | 5.6158 | 6.1554 |
| FANCB_S456 | FANCB | Q8NB91 | | 456 | CGEEENSVHILDE | 5.7609 | 4.8544 | 6.2846 | 6.6299 | 7.0174 | 7.3767 | 6.2635 | 5.1761 | 6.2377 | 5.7844 | 5.7680 | 6.1155 | 6.5265 | 5.5402 | 7.7666 | 6.8976 |
| FANCD2_S319 | FANCD2 | Q9BXW9-1 | | 319 | PSRLQASQVKLKS | 6.9814 | 5.9860 | 5.2047 | 6.8782 | 5.6853 | 5.2303 | 6.4064 | 4.7539 | 5.5159 | 4.2353 | 4.4893 | 4.3107 | 9.6157 | 8.7812 | 7.4256 | 8.5001 |
| FANCE_S249 | FANCE | Q9HB96 | | 249 | ADGGSASPIKDQP | 5.4044 | 5.7380 | 6.5388 | 6.1303 | 6.0117 | 6.6946 | 6.0352 | 6.3707 | 6.2899 | 6.8549 | 6.8327 | 7.4820 | 5.8951 | 5.8481 | 5.6690 | 6.2046 |
| FANCI_S730 | FANCI | Q9NVI1 | | 730 | DKSADFSQSTSIG | 5.3730 | 5.5830 | 5.2135 | 6.0017 | 6.1051 | 5.6069 | 5.5513 | 5.3284 | 5.3408 | 5.2593 | 5.2838 | 5.3801 | 8.2702 | 8.0283 | 9.0772 | 8.5974 |
| FANCM_S1437 | FANCM | Q8IYD8 | | 1437 | KGNVLNSPEDQKN | 6.1675 | 6.2176 | 7.0889 | 6.6644 | 6.0892 | 6.3900 | 6.2476 | 5.8107 | 7.0198 | 7.0816 | 7.1760 | 7.2269 | 5.2095 | 5.7280 | 4.6503 | 5.2318 |
| FANCM_S661 | FANCM | Q8IYD8 | | 661 | NLQRKSSIFSYRD | 5.5500 | 4.8301 | 5.5732 | 5.4185 | 7.9034 | 5.3276 | 4.9440 | 5.5061 | 8.9339 | 7.9918 | 7.9523 | 8.6750 | 5.9993 | 6.3834 | 3.8747 | 5.1366 |
| FARP1_S1073 | FARP1 | Q9Y4F1-2 | | 24 | ENSGISTLERGQK | 6.6677 | 6.9345 | 6.2311 | 6.6638 | 6.2023 | 6.2047 | 6.1580 | 6.1801 | 6.5683 | 6.2768 | 6.2235 | 6.0067 | 5.8282 | 6.3293 | 5.8460 | 5.6790 |
| FARP1_S427 | FARP1 | Q9Y4F1-2 | | 427 | EPGSHPSPAPRRS | 6.5106 | 6.6565 | 5.9958 | 6.5707 | 6.3967 | 6.0326 | 6.1230 | 6.2208 | 6.2669 | 6.3237 | 6.0226 | 6.1321 | 5.9790 | 6.4998 | 6.1764 | 6.0928 |
| FARP1_S433 | FARP1 | Q9Y4F1-2 | | 903 | PEFLASSPPDNKS | 5.6358 | 6.1066 | 5.9610 | 6.2755 | 6.3719 | 6.2981 | 6.8018 | 6.2670 | 5.9102 | 5.9068 | 5.9542 | 5.9358 | 6.4545 | 6.6123 | 6.9865 | 6.5221 |
| FARP1_S481 | FARP1 | A0A1B0GV68 | | 583 | RLPRHSSLIDEMF | 5.3950 | 5.4321 | 4.5842 | 4.7663 | 5.4396 | 5.5010 | 5.4229 | 5.3796 | 5.2871 | 5.2836 | 5.3940 | 5.0538 | 9.0342 | 9.2302 | 9.3132 | 9.4831 |
| FARP1_S510 | FARP1 | Q9Y4F1-2 | | 510 | PDTKQASPLISPL | 6.9831 | 5.6142 | 6.2011 | 7.0306 | 6.6799 | 6.4080 | 6.3249 | 6.3705 | 7.2043 | 6.5012 | 6.8668 | 6.7412 | 5.1192 | 5.9205 | 4.9579 | 5.0767 |
| FARP1_S583 | FARP1 | A0A1B0GV68 | | 713 | RSLANGTPPMGAA | 6.7728 | 7.0312 | 5.5818 | 6.3307 | 6.3232 | 5.8605 | 6.1967 | 6.4782 | 6.2305 | 6.2841 | 6.4187 | 6.4091 | 6.4737 | 6.3125 | 5.5590 | 5.7373 |
| FARP1_S903 | FARP1 | A0A1B0GV68 | | 481 | SRAARGSSTSIPF | 6.9831 | 5.8856 | 7.7495 | 5.2197 | 6.2800 | 5.6773 | 5.5024 | 5.8708 | 6.6967 | 4.0031 | 4.7063 | 4.9146 | 9.5279 | 7.5784 | 6.3098 | 7.0948 |
| FARP1_T24 | FARP1 | Q9Y4F1-2 | | 596 | PLHKFHTNFLKEI | 5.8430 | 6.6326 | 7.6915 | 6.6521 | 5.8869 | 7.1806 | 5.4360 | 6.9456 | 6.6375 | 6.4139 | 7.7697 | 6.7550 | 5.9217 | 4.9934 | 4.2675 | 4.9731 |
| FARP1_T34 | FARP1 | Q9Y4F1-2 | | 433 | SPAPRRSPAGNKQ | 7.0966 | 6.8573 | 5.7407 | 6.5551 | 6.4038 | 6.1345 | 6.4376 | 6.0298 | 6.1915 | 6.2958 | 6.2346 | 6.3675 | 5.6685 | 5.8340 | 6.0390 | 6.1139 |
| FARP1_T596 | FARP1 | Q9Y4F1-2 | | 34 | GQKPPPTPSGKLV | 6.7613 | 7.0834 | 6.3299 | 6.8874 | 6.5118 | 6.8102 | 6.5138 | 6.2462 | 6.4849 | 6.5645 | 7.1149 | 6.4435 | 4.5911 | 5.7521 | 4.7720 | 5.1331 |
| FARP1_T713 | FARP1 | Q9Y4F1-2 | | 1073 | VLSHKESLVYxxx | 6.9687 | 6.0947 | 6.5358 | 6.9497 | 6.3701 | 6.3464 | 5.9245 | 6.6790 | 7.5016 | 7.3846 | 6.8963 | 6.3641 | 5.7890 | 6.1129 | 3.8880 | 4.1944 |
| FARP2_S399 | FARP2 | O94887 | | 399 | GLRTPASPSSANA | 6.0972 | 6.6071 | 6.9081 | 6.6767 | 6.7934 | 6.7395 | 6.8859 | 7.1490 | 6.3675 | 6.0436 | 7.0686 | 7.0108 | 4.9209 | 4.7757 | 5.0176 | 4.9382 |
| FARSA_S184 | FARSA | Q9Y285 | | 184 | GSAFSTSISKQET | 4.4014 | 5.9169 | 8.7361 | 4.8489 | 4.9941 | 8.0608 | 5.6468 | 9.0428 | 4.8771 | 10.7823 | 8.5989 | 8.2885 | 3.2423 | 4.1520 | 4.0229 | 4.3882 |
| FAS_S212 | FAS | P25445 | | 212 | NQGSHESPTLNPE | 6.1723 | 6.5313 | 6.6151 | 7.2682 | 6.0878 | 6.2319 | 6.4592 | 6.3347 | 6.6338 | 6.3491 | 6.7022 | 6.9045 | 5.5612 | 5.6236 | 5.0195 | 5.5055 |
| FAS_S333 | FAS | P25445 | | 333 | FRNEIQSLVxxxx | 6.6492 | 5.8346 | 6.3865 | 6.1744 | 6.9938 | 6.1623 | 6.8347 | 6.1431 | 5.8129 | 6.2009 | 6.6901 | 5.9286 | 5.7647 | 6.4636 | 5.8394 | 6.1214 |
| FASN_S207 | FASN | P49327 | | 207 | LRLGMLSPEGTCK | 6.3685 | 6.3280 | 7.0846 | 6.1954 | 6.3083 | 6.8035 | 6.3083 | 6.5557 | 6.7909 | 6.3624 | 6.5828 | 6.8073 | 5.3467 | 5.5257 | 5.3157 | 5.3162 |
| FASN_S2236 | FASN | P49327 | | 974 | LFDHPESPTPNPT | 5.9769 | 6.4551 | 6.8987 | 6.0553 | 6.3333 | 6.7321 | 6.1351 | 6.5381 | 6.4282 | 6.4775 | 6.7166 | 6.7872 | 5.5916 | 5.7621 | 5.7793 | 5.3329 |
| FASN_S974 | FASN | P49327 | | 2204 | SELACPTPKEDGL | 5.6571 | 5.8890 | 6.2496 | 5.6665 | 6.1442 | 6.0864 | 6.1761 | 6.0301 | 6.2944 | 5.9819 | 5.5198 | 5.7552 | 7.1581 | 7.0071 | 7.3800 | 7.0043 |
| FASN_T2204 | FASN | P49327 | | 2236 | TLMRLNSVQSSER | 7.2531 | 8.3578 | 6.3883 | 6.5849 | 6.3613 | 6.6739 | 5.7201 | 5.5395 | 8.1423 | 5.0411 | 6.1431 | 6.8732 | 4.8821 | 4.8202 | 5.6446 | 5.5746 |
| FASTKD5_S95 | FASTKD5 | Q7L8L6 | | 95 | STLQLGSPRATGV | 8.0087 | 5.9703 | 6.5911 | 7.6207 | 5.8410 | 6.8395 | 6.9274 | 7.3051 | 5.7271 | 5.9289 | 5.7168 | 6.0117 | 3.2283 | 5.3546 | 7.1757 | 5.7530 |
| FAT1_S4344 | FAT1 | Q14517 | | 4344 | DLDPCLSKKPLEE | 9.2885 | 7.9926 | 6.9158 | 7.8688 | 6.5469 | 5.9342 | 6.6942 | 4.7101 | 7.4634 | 5.2328 | 7.1140 | 6.8611 | 4.1704 | 5.0816 | 3.3647 | 4.7609 |
| FBF1_S156 | FBF1 | Q8TES7-6 | | 417 | PTEGAGSPAKASQ | 6.1138 | 6.2673 | 6.3630 | 6.2562 | 6.7662 | 6.7420 | 6.0836 | 7.5190 | 7.1455 | 6.8937 | 6.9469 | 6.3612 | 4.9281 | 5.7302 | 4.8156 | 5.0676 |
| FBF1_S417 | FBF1 | Q8TES7-6 | | 156 | HQNRRFSSEDLED | 5.6620 | 4.5098 | 6.4233 | 5.0242 | 7.7351 | 6.2202 | 5.6959 | 7.1704 | 7.0339 | 7.5112 | 7.4609 | 6.8340 | 5.9122 | 5.9406 | 5.2230 | 5.6435 |
| FBRS_S230;S237 | FBRS | J3KNZ9 | | 309 | VPRPPVSPPAPLP | 6.0930 | 6.1887 | 6.4101 | 6.1309 | 6.2105 | 6.3495 | 6.2459 | 6.5385 | 5.7942 | 6.2663 | 6.8326 | 6.3323 | 6.0813 | 6.3924 | 6.0559 | 6.0778 |
| FBRS_S309 | FBRS | J3KNZ9 | | 967 | VPAPRPSSPPRGP | 6.4062 | 6.1023 | 7.0388 | 6.7846 | 5.6359 | 6.0361 | 5.9338 | 5.7796 | 5.8435 | 7.2329 | 6.5341 | 6.4588 | 6.0782 | 6.3054 | 6.0308 | 5.7991 |
| FBRS_S340 | FBRS | J3KNZ9 | | 948 | PHLLSKTPPGALL | 6.1811 | 5.3769 | 6.9439 | 6.8724 | 6.4592 | 5.8582 | 6.2723 | 6.5108 | 6.1415 | 6.8363 | 6.8397 | 6.3886 | 5.0971 | 6.1819 | 6.1251 | 5.9150 |
| FBRS_S59 | FBRS | J3KNZ9 | | 801 | RPQLRVSPATPKA | 5.9774 | 7.0071 | 5.8146 | 6.0867 | 6.6065 | 6.8464 | 6.5284 | 6.3867 | 6.6877 | 6.4801 | 6.5847 | 6.1538 | 5.5725 | 6.0302 | 5.5268 | 5.7104 |
| FBRS_S801 | FBRS | J3KNZ9 | | 340 | PFGLRTSPYGSSL | 5.5987 | 6.0301 | 6.5931 | 6.0540 | 6.0570 | 6.3105 | 6.0794 | 5.7718 | 6.0978 | 6.6537 | 6.0372 | 5.7426 | 6.8956 | 6.8590 | 6.4376 | 6.7819 |
| FBRS_S967 | FBRS | J3KNZ9 | | 59 | GSSSSSSPPPPAR | 5.5012 | 5.4933 | 5.7478 | 5.0572 | 6.2530 | 5.4578 | 5.1697 | 5.0197 | 8.1472 | 7.2119 | 6.7088 | 7.2257 | 6.2569 | 6.9258 | 6.7561 | 7.0679 |
| FBRS_T948 | FBRS | J3KNZ9 | 230;237 | | NA | 5.6509 | 6.4204 | 6.4866 | 6.5163 | 6.3596 | 6.5455 | 6.4559 | 6.2022 | 5.6542 | 6.5189 | 6.3383 | 6.0181 | 6.3668 | 6.6641 | 5.9038 | 5.8985 |
| FBRSL1_S790 | FBRSL1 | Q9HCM7 | | 790 | RLLSRASPATPAG | 6.0104 | 6.0968 | 6.8467 | 6.3167 | 5.8550 | 6.5094 | 6.4405 | 6.8227 | 6.5633 | 6.1769 | 7.0389 | 6.8446 | 5.5764 | 5.8422 | 5.0849 | 5.9746 |
| FBRSL1_S830 | FBRSL1 | Q9HCM7 | | 830 | RVKESRSPAKEEA | 6.4659 | 5.7206 | 6.9634 | 5.4167 | 6.1594 | 6.8953 | 6.0639 | 5.7302 | 6.7414 | 7.0692 | 7.7305 | 7.9391 | 5.5740 | 5.8422 | 4.5784 | 5.1097 |
| FBXL20_T417 | FBXL20 | Q96IG2 | | 417 | AYFAPVTPPPSVG | 5.8075 | 6.0616 | 6.6167 | 6.3391 | 5.7777 | 6.2839 | 6.8525 | 6.4335 | 6.1589 | 6.1725 | 6.0459 | 6.4233 | 6.2002 | 6.7922 | 6.1904 | 5.8440 |
| FBXL20_T417;S421 | FBXL20 | Q96IG2 | 417;421 | | NA | 5.7261 | 6.3779 | 5.9878 | 6.0464 | 6.0890 | 6.2449 | 6.7724 | 7.1281 | 6.2637 | 6.4508 | 5.3850 | 6.4260 | 7.3163 | 6.0527 | 5.4637 | 6.2693 |
| FBXL4_S267 | FBXL4 | Q9UKA2 | | 267 | SLNKKFSSAVLGE | 1.1254 | 1.2786 | 1.7366 | 1.4334 | 2.1456 | 2.1825 | 2.0209 | 1.9524 | 1.3529 | 1.5129 | 1.5560 | 1.8492 | 20.9587 | 16.8543 | 23.0985 | 18.9420 |
| FBXO10_S111 | FBXO10 | Q9UK96 | | 111 | RERRTLSVGPGRE | 6.3397 | 6.5252 | 5.8451 | 6.5018 | 5.9970 | 5.8364 | 6.0610 | 5.4087 | 5.8586 | 6.1305 | 5.6799 | 5.8593 | 6.4046 | 7.4893 | 6.9050 | 7.1580 |
| FBXO10_S326 | FBXO10 | Q9UK96 | | 326 | TSPASSSPKPGSK | 6.1325 | 6.4618 | 6.8087 | 6.5675 | 7.1013 | 7.6108 | 6.6230 | 6.6027 | 6.5187 | 6.4943 | 6.0797 | 5.8643 | 4.7667 | 6.0675 | 5.4618 | 4.8386 |
| FBXO28_S344 | FBXO28 | Q9NVF7 | | 344 | SGQNEESPRKRKK | 6.5982 | 6.7580 | 6.5659 | 7.5454 | 6.4185 | 6.0213 | 6.2592 | 6.2725 | 6.8290 | 6.5425 | 6.5726 | 6.5406 | 5.1898 | 5.3733 | 5.0853 | 5.4282 |
| FBXO4_S12 | FBXO4 | Q9UKT5 | | 12 | PRSGTNSPPPPFS | 6.1639 | 6.3269 | 6.2804 | 5.8894 | 6.2547 | 6.4546 | 5.8970 | 6.7533 | 6.3603 | 6.2451 | 6.2908 | 6.8282 | 6.1898 | 6.2168 | 5.8667 | 5.9821 |
| FBXO4_S48 | FBXO4 | Q9UKT5 | | 48 | RVARTTSREEVDE | 5.0225 | 5.1986 | 5.9581 | 5.1195 | 6.0086 | 4.9125 | 5.9719 | 6.0290 | 6.7739 | 5.5628 | 4.5368 | 5.9508 | 8.1365 | 8.4820 | 8.2807 | 8.0558 |
| FBXO42_S365 | FBXO42 | Q6P3S6 | | 488 | LAPRRGSLPDQKD | 5.8128 | 5.2661 | 6.8831 | 5.7147 | 6.0112 | 5.7257 | 5.7569 | 6.6568 | 7.1531 | 6.8568 | 7.1051 | 7.4863 | 5.7147 | 6.6253 | 5.2979 | 5.9336 |
| FBXO42_S488 | FBXO42 | Q6P3S6 | | 587 | LGSSPGSPGSQSL | 5.9227 | 5.6988 | 6.1442 | 4.9011 | 6.5064 | 6.2254 | 6.5201 | 6.9493 | 6.5529 | 6.3092 | 7.1429 | 6.2558 | 5.7999 | 7.0827 | 5.7794 | 6.2093 |
| FBXO42_S499 | FBXO42 | Q6P3S6 | | 499 | KDLRLGSIDLNWD | 7.4770 | 7.2689 | 7.4863 | 5.4734 | 6.4115 | 6.9831 | 5.1255 | 7.2130 | 6.6072 | 7.2409 | 7.1695 | 7.4242 | 6.7501 | 4.9609 | 3.6655 | 2.7429 |
| FBXO42_S587 | FBXO42 | Q6P3S6 | | 365 | SGRAPLSPSLNSR | 5.6573 | 6.5996 | 6.9117 | 5.9376 | 5.7808 | 6.3780 | 6.2634 | 6.8562 | 7.2564 | 6.6012 | 6.0989 | 7.1132 | 5.2153 | 6.2441 | 5.3304 | 5.7559 |
| FBXO46_S240 | FBXO46 | Q6PJ61 | | 240 | FEAQRDSPPTKGL | 6.1457 | 6.5113 | 6.8470 | 6.7524 | 6.1712 | 6.4760 | 6.4716 | 5.7648 | 6.2264 | 7.3317 | 6.9066 | 6.5024 | 4.8341 | 6.1882 | 5.5407 | 5.3299 |
| FBXO46_S313 | FBXO46 | Q6PJ61 | | 313 | DLYQLISPSRDAL | 6.5233 | 6.7987 | 6.5498 | 6.7676 | 5.4735 | 5.6936 | 7.1018 | 6.6213 | 7.4890 | 6.4288 | 7.8024 | 6.6789 | 4.6381 | 6.1523 | 4.8836 | 4.3973 |
| FBXO5_S98 | FBXO5 | Q9UKT4 | | 98 | RLSCIGSPIVSPR | 5.9862 | 5.7621 | 6.4245 | 6.3431 | 5.7577 | 6.3601 | 5.9045 | 6.6983 | 6.7279 | 6.9488 | 6.4556 | 6.1988 | 5.4770 | 6.6457 | 6.1561 | 6.1535 |
| FBXO6_S258 | FBXO6 | Q9NRD1 | | 258 | NSSIVVSPKMTRN | 6.1673 | 5.9854 | 7.1289 | 6.4178 | 6.5766 | 6.1988 | 6.5369 | 6.4813 | 6.7280 | 6.3537 | 6.7230 | 6.4317 | 5.6161 | 6.1215 | 5.5562 | 4.9767 |
| FBXW11_S541 | FBXW11 | Q9UKB1 | | 65 | VSRKRPSEGNYQK | 6.0910 | 7.4155 | 7.1174 | 8.0815 | 5.9209 | 6.3241 | 6.3101 | 6.7675 | 6.9792 | 6.8360 | 7.9161 | 7.6405 | 3.2286 | 4.4271 | 4.5524 | 4.3920 |
| FBXW11_S65 | FBXW11 | Q9UKB1 | | 541 | RTYTYISRxxxxx | 5.7203 | 6.5882 | 6.8628 | 6.1907 | 6.5882 | 6.6183 | 6.3020 | 6.4670 | 6.9780 | 6.0783 | 6.9401 | 7.0059 | 4.9149 | 5.9380 | 5.1562 | 5.6512 |
| FBXW5_S275 | FBXW5 | Q969U6 | | 275 | AGDPATSPCRIFD | 5.9918 | 6.5421 | 6.9792 | 6.8865 | 5.9733 | 6.0886 | 6.2188 | 6.3254 | 6.3858 | 6.8803 | 6.9505 | 6.4422 | 5.4922 | 5.7315 | 5.6141 | 5.4978 |
| FBXW8_S249 | FBXW8 | Q8N3Y1 | | 85 | VASRSRSPLAREG | 6.6094 | 6.4675 | 6.6094 | 5.9827 | 6.1452 | 6.0759 | 5.9625 | 6.7529 | 6.0727 | 6.5748 | 7.0619 | 7.1736 | 5.3221 | 6.3379 | 5.3896 | 5.4617 |
| FBXW8_S85 | FBXW8 | Q8N3Y1 | | 249 | VAPFLESEDEEDE | 7.0030 | 6.4990 | 7.1084 | 7.0960 | 6.7913 | 5.7710 | 6.2371 | 6.2832 | 6.7197 | 6.0798 | 5.9753 | 6.3343 | 5.3354 | 6.3821 | 5.6352 | 4.7491 |
| FCHO2_S312 | FCHO2 | Q0JRZ9 | | 488 | EIPRPFSPPVTSN | 5.4913 | 5.9379 | 6.1712 | 5.7268 | 6.0439 | 6.4580 | 6.0798 | 6.3084 | 6.3913 | 6.2843 | 6.3104 | 6.1253 | 6.6910 | 6.8763 | 6.5180 | 6.5862 |
| FCHO2_S387 | FCHO2 | Q0JRZ9 | | 387 | IGNITLSPAISRH | 6.3471 | 6.0950 | 6.1824 | 5.8129 | 6.1385 | 6.8675 | 6.4179 | 7.0446 | 6.7974 | 6.1901 | 6.2126 | 6.5630 | 5.5278 | 6.1433 | 5.5566 | 6.1033 |
| FCHO2_S394 | FCHO2 | Q0JRZ9 | | 394 | PAISRHSPVQMNR | 6.0195 | 6.3092 | 6.2095 | 5.6488 | 6.2587 | 5.8479 | 5.9281 | 6.2585 | 6.2207 | 6.4571 | 6.2747 | 6.2027 | 6.8341 | 6.5905 | 6.5529 | 6.3871 |
| FCHO2_S403 | FCHO2 | Q0JRZ9 | | 403 | QMNRNLSNEELTK | 5.2381 | 5.3355 | 5.9870 | 5.2437 | 6.0915 | 6.3185 | 5.9401 | 5.9859 | 5.8895 | 5.6554 | 5.4652 | 5.6307 | 8.0783 | 7.9248 | 7.6076 | 7.6083 |
| FCHO2_S468 | FCHO2 | Q0JRZ9 | | 312 | ECPDADSLNIPDV | 6.7103 | 6.3252 | 7.7460 | 6.0087 | 6.2134 | 6.7485 | 5.9712 | 5.4361 | 6.8639 | 6.8377 | 6.9946 | 7.0340 | 5.3287 | 5.6946 | 5.1033 | 4.9839 |
| FCHO2_S478;S488 | FCHO2 | Q0JRZ9 | | 468 | PPPRPASRPKLTS | 6.0742 | 4.6389 | 6.2045 | 5.4307 | 6.0201 | 6.2887 | 6.3298 | 5.3605 | 5.9579 | 5.8687 | 6.4381 | 6.0351 | 7.1387 | 7.0335 | 7.4073 | 7.7732 |
| FCHO2_S488 | FCHO2 | Q0JRZ9 | | 496 | PVTSNTSPPPAAP | 5.4203 | 5.5670 | 6.7714 | 6.7478 | 6.0883 | 6.9159 | 6.1804 | 6.4684 | 7.3366 | 6.4641 | 6.3870 | 6.2982 | 5.5424 | 6.8570 | 4.7994 | 6.1558 |
| FCHO2_S488;S496 | FCHO2 | Q0JRZ9 | 488;496 | | NA | 5.6175 | 6.1642 | 6.2230 | 6.3844 | 6.2610 | 6.3922 | 6.5286 | 6.3639 | 6.3656 | 6.6636 | 6.7125 | 6.7513 | 5.8184 | 5.9319 | 5.6628 | 6.1591 |
| FCHO2_S496 | FCHO2 | Q0JRZ9 | 478;488 | | NA | 7.8231 | 7.1742 | 8.1140 | 6.2351 | 6.5701 | 5.3094 | 5.0737 | 6.1639 | 8.7333 | 7.2448 | 6.9008 | 6.6566 | 4.4822 | 5.0998 | 4.0489 | 4.3701 |
| FCHSD2_S644 | FCHSD2 | O94868 | | 644 | MREIQISPSPKPH | 6.3452 | 6.8493 | 6.6444 | 6.3707 | 6.4581 | 6.5721 | 6.4266 | 6.5365 | 6.8829 | 6.6036 | 6.7549 | 6.9961 | 5.0499 | 5.1601 | 4.9651 | 5.3845 |
| FCHSD2_S675 | FCHSD2 | O94868 | | 681 | SLYFPRSPSANEK | 5.7840 | 5.4736 | 6.4728 | 6.1889 | 6.4478 | 6.6951 | 6.8088 | 6.6220 | 6.7196 | 6.3742 | 6.4596 | 6.0651 | 5.8001 | 6.2486 | 5.8482 | 5.9917 |
| FCHSD2_S681 | FCHSD2 | O94868 | | 675 | SPDKRSSLYFPRS | 8.1031 | 6.7293 | 7.3596 | 7.6047 | 5.5546 | 6.7144 | 6.3004 | 6.0534 | 7.7062 | 6.5800 | 7.0484 | 6.6268 | 4.3651 | 5.7470 | 3.6352 | 3.8718 |
| FEN1_S352 | FEN1 | P39748 | | 352 | VTGSLSSAKRKEP | 6.8622 | 7.4573 | 7.1281 | 7.1084 | 5.2739 | 6.6874 | 7.0636 | 6.7843 | 6.9913 | 7.3370 | 7.8258 | 7.5942 | 3.7918 | 4.7549 | 3.5131 | 3.8267 |
| FERMT2_S159 | FERMT2 | Q96AC1-3 | | 159 | KKLDDQSEDEALE | 6.5421 | 6.8399 | 6.8312 | 6.3727 | 6.2915 | 6.7202 | 6.4188 | 6.8640 | 6.0379 | 6.1971 | 6.1670 | 6.2176 | 5.4343 | 6.1704 | 5.5810 | 5.3145 |
| FERMT2_S159;S181 | FERMT2 | Q96AC1-3 | | 181 | SGSIYSSPGLYSK | 6.4784 | 6.5064 | 6.9624 | 6.9921 | 6.3551 | 6.8072 | 6.6472 | 6.9221 | 7.2391 | 7.1880 | 7.5212 | 7.5244 | 3.8844 | 4.8211 | 3.7118 | 4.4390 |
| FERMT2_S181 | FERMT2 | Q96AC1-3 | | 339 | ENHLNNSDKEVDE | 5.8282 | 7.4737 | 6.9616 | 6.9885 | 6.1236 | 7.3830 | 6.3347 | 6.7922 | 5.8551 | 4.5929 | 6.1264 | 5.5254 | 5.2448 | 6.2690 | 6.2977 | 6.2032 |
| FERMT2_S339 | FERMT2 | Q96AC1-3 | 159;181 | | NA | 7.4188 | 7.5898 | 7.2219 | 7.8342 | 6.3562 | 7.9411 | 7.5195 | 7.6254 | 7.0812 | 7.0412 | 7.2056 | 7.5955 | 2.6193 | 3.5819 | 2.6331 | 2.7353 |
| FEZ2_S195 | FEZ2 | Q9UHY8-2 | | 29 | QENCNASPEPGAE | 5.7815 | 6.6611 | 6.6717 | 6.2511 | 6.2626 | 6.5089 | 6.5241 | 6.3438 | 6.9180 | 6.7616 | 7.0785 | 6.6450 | 5.3904 | 5.9213 | 5.1930 | 5.0874 |
| FEZ2_S29 | FEZ2 | Q9UHY8-2 | | 195 | DRLSMLSQEIQTL | 6.8590 | 6.6548 | 7.2769 | 6.7828 | 6.0512 | 6.3591 | 6.6935 | 6.8572 | 6.6572 | 6.7633 | 6.9718 | 6.9090 | 4.8874 | 5.1347 | 4.6515 | 4.4906 |
| FGD1_S116 | FGD1 | P98174 | | 48 | LLARRGSGSALGG | 5.9378 | 6.4775 | 6.7040 | 5.3719 | 6.3824 | 6.0119 | 5.4267 | 5.8643 | 6.0603 | 7.1212 | 6.0270 | 6.4646 | 6.2315 | 7.3006 | 6.0978 | 6.5204 |
| FGD1_S48 | FGD1 | P98174 | | 82 | VLPCGPSPQHHRA | 6.2529 | 5.7502 | 6.1387 | 6.3964 | 6.0382 | 6.5562 | 6.4222 | 7.0318 | 6.4473 | 6.0028 | 6.3169 | 6.5632 | 5.7041 | 6.1829 | 5.6694 | 6.5269 |
| FGD1_S745 | FGD1 | P98174 | | 116 | ILVKSLSLDPGQS | 6.8873 | 6.8304 | 7.7730 | 6.3044 | 6.1020 | 6.7510 | 6.0547 | 6.8034 | 6.0407 | 6.3916 | 7.0624 | 6.8836 | 4.3109 | 6.3687 | 5.2340 | 4.2020 |
| FGD1_S816 | FGD1 | P98174 | | 450 | MEEWDRYPRIGDI | 7.3359 | 8.4146 | 5.0227 | 8.1984 | 6.9866 | 7.7916 | 5.8759 | 9.8619 | 6.1289 | 6.1263 | 6.7384 | 6.4644 | 2.6426 | 4.5069 | 4.2374 | 3.6676 |
| FGD1_S82 | FGD1 | P98174 | | 816 | ILEKQASVAAENS | 6.9984 | 7.4391 | 8.9542 | 5.9981 | 7.0148 | 7.9963 | 6.7903 | 6.9137 | 6.3683 | 6.1850 | 6.3613 | 5.4374 | 3.2453 | 5.0860 | 4.7592 | 4.4524 |
| FGD1_T711 | FGD1 | P98174 | | 711 | NREDEDTPPNSPN | 5.9850 | 6.5001 | 7.5455 | 5.5294 | 5.9880 | 6.6199 | 6.3329 | 7.1704 | 6.0483 | 6.0618 | 6.1469 | 6.8842 | 5.8841 | 6.4165 | 5.3931 | 5.4940 |
| FGD1_Y450 | FGD1 | P98174 | | 745 | CQEPFNSITKRRH | 6.6341 | 7.4202 | 6.8228 | 5.9291 | 6.5563 | 7.4054 | 6.8389 | 6.4963 | 6.3577 | 7.1666 | 7.2159 | 6.6987 | 4.6473 | 5.3420 | 3.9042 | 4.5645 |
| FGD6_S1197 | FGD6 | Q6ZV73 | | 554 | NRGVSSSFDMPKR | 6.9108 | 7.2791 | 9.2042 | 7.0394 | 5.5766 | 6.4341 | 6.0577 | 6.5897 | 6.6780 | 7.7820 | 8.0431 | 7.1501 | 2.8081 | 5.1226 | 3.6140 | 3.7103 |
| FGD6_S1293 | FGD6 | Q6ZV73 | | 1293 | HSPRIGSPGNHKS | 7.1051 | 6.9186 | 7.6805 | 6.9500 | 6.4707 | 6.6228 | 7.2260 | 6.8364 | 7.0524 | 6.9918 | 7.0921 | 6.7723 | 4.0189 | 4.4780 | 3.6879 | 4.0965 |
| FGD6_S515 | FGD6 | Q6ZV73 | | 605 | TKPRAKSLSAMDV | 5.6428 | 5.5809 | 6.9559 | 6.4152 | 6.0337 | 6.3422 | 5.9824 | 6.6300 | 6.2896 | 6.3622 | 6.6663 | 7.0483 | 5.8385 | 6.4331 | 5.8669 | 5.9119 |
| FGD6_S554 | FGD6 | Q6ZV73 | | 515 | VLKKAASEELLEK | 6.4621 | 6.8884 | 7.6377 | 6.4879 | 5.7564 | 6.5640 | 6.3023 | 6.6618 | 5.7998 | 6.4945 | 7.3366 | 6.7513 | 4.6728 | 5.6536 | 5.5026 | 5.0283 |
| FGD6_S605 | FGD6 | Q6ZV73 | | 692 | KPIKAYSTENYSL | 8.0002 | 7.7348 | 11.2279 | 8.8023 | 6.1794 | 6.0081 | 5.4104 | 6.8021 | 6.2012 | 7.5241 | 6.7254 | 6.3348 | 2.7398 | 4.9220 | 3.1650 | 2.2222 |
| FGD6_S692 | FGD6 | Q6ZV73 | | 1197 | TFCPSRSLDEADS | 7.2371 | 7.5314 | 9.1376 | 7.4702 | 5.8126 | 5.6993 | 5.7316 | 6.2468 | 6.0964 | 6.5434 | 7.3407 | 6.4653 | 4.1296 | 5.6295 | 4.9240 | 4.0045 |
| FGD6_S721 | FGD6 | Q6ZV73 | | 721 | NGLRAESLDDQML | 7.0733 | 7.2868 | 7.9869 | 7.2177 | 6.5856 | 6.2434 | 5.6220 | 6.6615 | 6.3777 | 7.1815 | 7.0110 | 5.7421 | 4.6120 | 5.4912 | 4.5367 | 4.3707 |
| FGFR3_S410 | FGFR3 | P22607-2 | | 410 | PKKGLGSPTVHKI | 6.1437 | 6.6487 | 7.2011 | 6.1842 | 5.5959 | 5.4698 | 5.4217 | 6.1040 | 6.9956 | 7.2944 | 6.9322 | 6.1666 | 5.5187 | 6.4722 | 5.8870 | 5.9642 |
| FHOD1_S358 | FHOD1 | Q9Y613 | | 523 | EPLIPASPKAEPI | 5.7240 | 5.6264 | 6.0802 | 5.6255 | 6.0722 | 6.5104 | 6.5107 | 6.3136 | 6.3765 | 6.0706 | 6.2194 | 5.9737 | 6.6930 | 6.7719 | 6.5964 | 6.8354 |
| FHOD1_S367 | FHOD1 | Q9Y613 | | 367 | GKRSRRSLEGGGC | 6.9042 | 7.3503 | 6.7992 | 7.0839 | 6.2842 | 6.7134 | 6.6040 | 6.6352 | 6.9244 | 7.0103 | 7.0392 | 7.1500 | 3.9704 | 4.8250 | 4.0650 | 4.6413 |
| FHOD1_S486 | FHOD1 | Q9Y613 | | 498 | AARTPQSPAPCVL | 6.3144 | 6.8677 | 6.6725 | 6.6815 | 6.1923 | 7.0788 | 7.1162 | 6.5121 | 7.3015 | 7.0023 | 6.2361 | 6.9537 | 4.6028 | 4.5947 | 4.6437 | 5.2297 |
| FHOD1_S498 | FHOD1 | Q9Y613 | | 358 | ERRKPSSEEGKRS | 6.8299 | 7.2282 | 6.8401 | 6.6229 | 6.4678 | 6.4997 | 6.8795 | 6.6155 | 6.8244 | 6.8461 | 6.6387 | 7.1652 | 4.5131 | 4.9892 | 4.3117 | 4.7280 |
| FHOD1_S523 | FHOD1 | Q9Y613 | | 486 | ARQLWDSPETAPA | 6.5969 | 7.7244 | 7.0939 | 7.2814 | 5.9498 | 6.7673 | 6.9230 | 6.5234 | 7.0034 | 7.4595 | 7.2785 | 7.5048 | 3.6059 | 4.3516 | 3.6930 | 4.2431 |
| FHOD3_S955 | FHOD3 | Q2V2M9-4 | | 955 | LGRGSISPDAEPN | 6.1779 | 5.1724 | 6.6726 | 6.2002 | 6.3331 | 6.8662 | 5.7305 | 6.1793 | 6.6408 | 6.0524 | 6.3149 | 6.8649 | 5.7514 | 6.4323 | 5.9849 | 6.6260 |
| FIGN_S521 | FIGN | Q5HY92 | | 521 | LTALPRSILLFGP | 7.5094 | 5.1435 | 5.3940 | 5.4471 | 6.2601 | 4.5279 | 5.7748 | 4.4606 | 8.4738 | 7.0019 | 6.6020 | 7.3124 | 7.3812 | 8.4300 | 4.6871 | 5.5940 |
| FIGN_T400 | FIGN | Q5HY92 | | 400 | QSSRALTPPSYST | 5.8701 | 5.4013 | 6.0190 | 5.4633 | 5.4485 | 6.0559 | 5.7040 | 6.0038 | 6.1965 | 6.3858 | 6.5120 | 6.4608 | 7.0266 | 7.5560 | 7.0440 | 6.8524 |
| FIP1L1_S14 | FIP1L1 | Q6UN15 | | 492 | ERERDHSPTPSVF | 5.8413 | 6.3183 | 5.9086 | 6.5055 | 5.9553 | 5.6889 | 6.2662 | 5.8596 | 6.4702 | 6.4723 | 6.2861 | 6.2427 | 6.2335 | 6.6177 | 6.5959 | 6.7380 |
| FIP1L1_S149 | FIP1L1 | Q6UN15 | | 259 | TKAEFTSPPSLFK | 7.1794 | 7.4059 | 6.7457 | 7.5309 | 6.4438 | 6.6953 | 7.0669 | 6.7090 | 7.2572 | 6.9956 | 7.2651 | 7.1843 | 3.4326 | 4.3801 | 3.5048 | 4.2036 |
| FIP1L1_S259 | FIP1L1 | Q6UN15 | | 258 | STKAEFTSPPSLF | 6.8506 | 7.6661 | 6.6455 | 7.2867 | 6.2621 | 6.9824 | 7.0271 | 6.8278 | 6.9534 | 6.8835 | 7.0086 | 7.1301 | 3.7155 | 4.5908 | 3.7292 | 4.4406 |
| FIP1L1_S304 | FIP1L1 | Q6UN15 | | 14 | RLVSELSGGTGGD | 6.3966 | 5.9876 | 6.3471 | 6.2193 | 6.4558 | 5.8058 | 6.1511 | 6.8660 | 5.7031 | 5.8062 | 5.7541 | 6.2152 | 6.6418 | 6.4843 | 6.5882 | 6.5780 |
| FIP1L1_S492 | FIP1L1 | H0Y8P7 | | 149 | ERERDHSPTPSVF | 5.8128 | 6.4751 | 6.2302 | 6.3320 | 6.0233 | 6.0351 | 6.4372 | 6.0981 | 6.1774 | 6.2059 | 6.1283 | 6.1356 | 6.3750 | 6.6064 | 6.3487 | 6.5790 |
| FIP1L1_S492;S496 | FIP1L1 | Q6UN15 | | 304 | RYGRAESPDLRRL | 6.6394 | 6.9154 | 6.0849 | 6.4813 | 5.8813 | 6.1039 | 6.1179 | 5.9640 | 7.1948 | 5.8441 | 6.4474 | 6.0733 | 6.1211 | 6.5301 | 5.8548 | 5.7461 |
| FIP1L1_S492;S500 | FIP1L1 | Q6UN15-5 | | 27 | EEEEWLYGDENEV | 6.5009 | 6.4733 | 6.3242 | 6.7973 | 6.4004 | 6.0823 | 6.2288 | 6.3745 | 6.3636 | 5.1798 | 6.4189 | 6.5503 | 6.2589 | 6.7596 | 5.3196 | 5.9676 |
| FIP1L1_T258 | FIP1L1 | Q6UN15 | 492;500 | | NA | 5.5221 | 5.9707 | 4.4818 | 6.1010 | 5.6698 | 5.6541 | 5.5839 | 6.4037 | 6.3506 | 5.3559 | 6.1697 | 7.6235 | 7.9613 | 7.1720 | 7.0391 | 6.9408 |
| FIP1L1_Y27 | FIP1L1 | Q6UN15 | 492;496 | | NA | 10.5413 | 7.7105 | 8.0625 | 8.0728 | 5.4096 | 3.9527 | 6.3666 | 3.4435 | 5.3883 | 3.8935 | 6.5403 | 5.8531 | 6.8448 | 6.9260 | 5.7222 | 5.2724 |
| FKBP15_S1065 | FKBP15 | Q5T1M5 | | 311 | RDSAAPSPIPGAD | 5.7963 | 6.1180 | 6.3651 | 5.7435 | 6.3860 | 6.3409 | 6.1233 | 6.2628 | 5.9659 | 6.7831 | 6.5685 | 6.2936 | 5.9474 | 6.5814 | 6.4184 | 6.3056 |
| FKBP15_S1097 | FKBP15 | Q5T1M5 | | 1114 | LALGPESPGEPQP | 6.3592 | 6.4499 | 6.6106 | 6.2486 | 6.1744 | 6.4664 | 6.2462 | 6.4797 | 6.9414 | 7.1293 | 6.8483 | 7.1300 | 5.1215 | 5.6514 | 5.1609 | 4.9823 |
| FKBP15_S1097;T1114 | FKBP15 | Q5T1M5 | | 1097 | ESSTRLSLTSDPE | 5.3790 | 5.6661 | 5.5924 | 5.4349 | 6.3603 | 6.1370 | 5.7093 | 6.1387 | 6.1077 | 5.7642 | 5.9901 | 5.7784 | 7.3622 | 7.7596 | 7.5429 | 7.2771 |
| FKBP15_S1114 | FKBP15 | Q5T1M5 | | 956 | ERPRRPSQEQSAS | 5.3542 | 5.6847 | 6.1345 | 5.6430 | 5.3055 | 5.7217 | 5.9587 | 5.5842 | 7.2500 | 6.4529 | 6.7582 | 6.5295 | 6.9007 | 7.2372 | 6.7427 | 6.7424 |
| FKBP15_S1162 | FKBP15 | Q5T1M5 | | 979 | NRERPESPMVPSE | 6.3841 | 6.2356 | 7.0056 | 6.7057 | 6.5095 | 6.5272 | 5.6421 | 7.3653 | 5.8458 | 5.9746 | 5.8499 | 6.5981 | 5.7218 | 5.4301 | 6.4032 | 5.8015 |
| FKBP15_S307;S311 | FKBP15 | Q5T1M5 | | 61 | ATGNQATPKTAPA | 7.0120 | 7.6097 | 6.6720 | 7.1756 | 6.0139 | 7.0656 | 7.7237 | 6.6268 | 7.0601 | 7.1434 | 7.1611 | 7.3525 | 3.2348 | 4.4987 | 3.1606 | 4.4894 |
| FKBP15_S311 | FKBP15 | Q5T1M5 | | 1065 | EESLAASPMAAKP | 6.3713 | 6.1557 | 6.4810 | 6.1911 | 6.1548 | 6.1231 | 6.1883 | 6.2640 | 7.1210 | 6.9063 | 7.5623 | 6.7247 | 5.5496 | 5.4232 | 5.4042 | 5.3793 |
| FKBP15_S311;S326 | FKBP15 | Q5T1M5 | | 344 | PALRTKSNSLSEQ | 5.8784 | 7.1244 | 7.1005 | 6.4647 | 5.5452 | 5.5220 | 5.2130 | 5.5215 | 7.2522 | 8.2718 | 7.7980 | 8.4053 | 4.7664 | 4.9635 | 5.0445 | 5.1287 |
| FKBP15_S344 | FKBP15 | Q5T1M5 | | 356 | QLAINTSPDAVKA | 6.7737 | 6.9749 | 7.0581 | 6.7832 | 6.2850 | 6.1535 | 7.3140 | 6.0972 | 6.6319 | 7.0597 | 6.8284 | 7.3615 | 5.0207 | 5.2694 | 4.2745 | 4.1145 |
| FKBP15_S346 | FKBP15 | Q5T1M5 | | 346 | LRTKSNSLSEQLA | 5.3729 | 4.8851 | 5.9766 | 6.6089 | 5.8130 | 6.0148 | 5.8757 | 5.6159 | 6.1229 | 7.1046 | 6.9738 | 6.8024 | 7.1063 | 6.3761 | 6.8568 | 6.4943 |
| FKBP15_S356 | FKBP15 | Q5T1M5 | | 1162 | HHSQRSSLSGDEE | 6.7303 | 7.1806 | 6.4977 | 5.9586 | 7.1131 | 6.9086 | 6.0089 | 6.3944 | 6.2511 | 6.5420 | 6.7816 | 6.9182 | 5.0519 | 5.7501 | 4.7151 | 5.1978 |
| FKBP15_S956 | FKBP15 | Q5T1M5 | 311;326 | | NA | 6.2252 | 6.4588 | 6.8082 | 7.3404 | 6.3282 | 6.7481 | 6.7363 | 6.7251 | 7.1473 | 7.3709 | 7.7950 | 7.2559 | 4.0618 | 4.5766 | 3.9114 | 4.5108 |
| FKBP15_S979 | FKBP15 | Q5T1M5 | 1097;1114 | | NA | 5.4114 | 5.8104 | 6.1487 | 6.5792 | 5.5508 | 6.6206 | 5.4903 | 6.2110 | 6.1757 | 5.8301 | 6.2958 | 6.5519 | 6.3068 | 6.7778 | 6.5531 | 7.6862 |
| FKBP15_T61 | FKBP15 | Q5T1M5 | 307;311 | | NA | 6.3355 | 7.0740 | 6.3160 | 6.9395 | 7.4314 | 6.4922 | 5.1886 | 6.6910 | 5.7716 | 6.1399 | 7.2913 | 6.5608 | 5.7997 | 5.7726 | 4.7708 | 5.4251 |
| FKBP3_S152 | FKBP3 | Q00688 | | 152 | DTNIQTSAKKKKN | 6.4039 | 6.9288 | 6.9885 | 7.1813 | 6.4967 | 7.2067 | 7.1051 | 6.7645 | 6.8290 | 6.9137 | 7.2810 | 6.8936 | 3.8785 | 5.0200 | 3.8224 | 4.2864 |
| FKBP3_S36 | FKBP3 | Q00688 | | 36 | QEHGSDSFLAEHK | 5.8412 | 6.9414 | 6.4800 | 6.3334 | 6.3951 | 6.7663 | 6.0431 | 6.7764 | 6.0094 | 6.4080 | 6.6240 | 6.2794 | 5.6985 | 5.9087 | 5.8042 | 5.6909 |
| FKBP4_S118 | FKBP4 | Q02790 | | 118 | AYGSAGSPPKIPP | 6.5046 | 6.4865 | 6.5626 | 6.0572 | 5.7182 | 6.6522 | 6.3163 | 5.9758 | 5.9178 | 7.0491 | 5.9148 | 6.5151 | 5.4335 | 5.8614 | 6.4323 | 6.6025 |
| FKBP5_S13 | FKBP5 | Q13451 | | 13 | AKNNEESPTATVA | 5.8556 | 6.7279 | 6.0082 | 6.2984 | 5.7744 | 6.0882 | 5.6587 | 6.9063 | 5.8651 | 6.0173 | 5.9902 | 6.5473 | 6.2954 | 6.6853 | 6.4497 | 6.8319 |
| FKBP5_S445 | FKBP5 | Q13451 | | 445 | NEKGTDSQAMEEE | 5.7708 | 6.0501 | 5.8793 | 5.9036 | 6.1105 | 6.0855 | 5.3383 | 5.8805 | 5.9137 | 5.6858 | 5.7831 | 6.0128 | 7.4219 | 7.8115 | 7.3179 | 7.0345 |
| FLAD1_S563 | FLAD1 | Q8NFF5 | | 563 | PALKCLSPGGHPT | 5.7735 | 6.6117 | 6.9294 | 6.5769 | 7.2401 | 7.0720 | 6.5793 | 7.5206 | 7.5856 | 6.0181 | 6.6604 | 6.1178 | 4.5388 | 5.2391 | 4.9550 | 4.5817 |
| FLI1_S241 | FLI1 | Q01543 | | 241 | NSGLNKSPPLGGA | 6.0027 | 6.0110 | 6.3947 | 5.8546 | 6.2777 | 6.1794 | 6.0119 | 6.5475 | 6.4982 | 6.1584 | 6.6537 | 6.8190 | 6.1928 | 6.2285 | 6.1191 | 6.0509 |
| FLII_S352 | FLII | Q13045-2 | | 352 | LRLAGASPATVAA | 5.5190 | 6.0341 | 7.1784 | 7.5838 | 6.3297 | 6.6211 | 6.3128 | 8.3988 | 7.7105 | 7.3854 | 6.4775 | 6.2833 | 4.5689 | 5.8019 | 3.6906 | 4.1044 |
| FLII_S801 | FLII | Q13045-2 | | 801 | AEAVLQSPGLSGK | 6.2117 | 6.4042 | 6.8645 | 6.1068 | 6.7939 | 6.4062 | 7.0723 | 6.2632 | 6.4839 | 6.8815 | 7.0130 | 6.1007 | 5.1123 | 6.1698 | 4.6333 | 5.4826 |

| | | | | | | | | | | | | | | | | | | | | |
|---|---|---|---|---|---|---|---|---|---|---|---|---|---|---|---|---|---|---|---|---|
| FLNA_S1055 | FLNA | P21333 | 1459 | CSGPGLSPGMVRA | 6.1766 | 6.3951 | 6.6794 | 6.2378 | 6.0211 | 6.3765 | 6.1865 | 6.1738 | 6.6138 | 6.6455 | 6.6146 | 6.6081 | 5.7332 | 6.2011 | 5.6224 | 5.7144 |
| FLNA_S1084 | FLNA | P21333 | 2152 | RRRRAPSVANVGS | 5.5026 | 5.3406 | 6.0893 | 5.6626 | 6.0901 | 5.9506 | 5.7412 | 6.0600 | 6.6412 | 6.3049 | 6.7020 | 6.6366 | 6.9694 | 7.1716 | 6.8413 | 6.2960 |
| FLNA_S1338 | FLNA | P21333 | 2327 | FVVPVASPSGDAR | 6.2437 | 6.0156 | 6.6547 | 6.7115 | 5.8204 | 6.3645 | 5.8848 | 6.3012 | 6.6981 | 6.5343 | 6.5126 | 6.7801 | 5.5699 | 6.2178 | 5.7293 | 5.9616 |
| FLNA_S1436 | FLNA | P21333 | 1533 | DEEVPRSPFKVKV | 6.6821 | 7.4306 | 6.8259 | 6.9745 | 6.5406 | 7.4527 | 7.3293 | 7.0070 | 7.5062 | 7.2371 | 7.4277 | 6.8751 | 3.3731 | 4.3990 | 3.1520 | 3.7871 |
| FLNA_S1459 | FLNA | P21333 | 1084 | QGGSAGSPARFTI | 6.5944 | 7.0387 | 7.0925 | 7.2497 | 5.9424 | 6.8838 | 6.7252 | 6.9453 | 6.9754 | 7.0718 | 7.2214 | 6.9012 | 3.8915 | 4.8835 | 4.0025 | 4.5807 |
| FLNA_S1533 | FLNA | P21333 | 2510 | PYHIGGSPFKAKV | 7.0280 | 8.0111 | 7.1459 | 7.7303 | 6.6966 | 6.2111 | 6.1574 | 7.4585 | 7.1677 | 7.0071 | 7.0345 | 7.2982 | 3.5407 | 4.0177 | 3.5482 | 3.9470 |
| FLNA_S1630 | FLNA | P21333 | 2284 | PSKAEISFEDRKD | 6.5756 | 6.7822 | 7.4239 | 7.0536 | 6.4027 | 6.6383 | 6.0748 | 5.9454 | 6.8228 | 6.4452 | 7.2269 | 7.4915 | 4.7629 | 5.2559 | 4.6746 | 4.4235 |
| FLNA_S1734 | FLNA | P21333 | 2599 | GVHGPRTPCEEIL | 5.9356 | 6.1256 | 6.4413 | 5.8538 | 6.4827 | 6.3542 | 6.0434 | 6.9526 | 6.6077 | 6.0567 | 6.2116 | 6.3665 | 5.9763 | 6.1860 | 6.3426 | 6.0634 |
| FLNA_S2053 | FLNA | P21333 | 1436 | GHQVPGSPFKVPV | 6.3997 | 6.5243 | 6.1377 | 6.5353 | 6.8501 | 7.3332 | 6.6812 | 6.2032 | 7.1655 | 6.3078 | 7.2488 | 5.6437 | 4.7825 | 5.4920 | 5.2278 | 5.4673 |
| FLNA_S2152 | FLNA | P21333 | 2053 | ASRVRVSGQGLHE | 6.7226 | 6.9354 | 6.3445 | 6.5833 | 6.0148 | 5.6341 | 7.2961 | 6.4937 | 6.8831 | 7.2128 | 5.5321 | 6.8048 | 5.1079 | 5.9066 | 5.0420 | 5.4861 |
| FLNA_S2158 | FLNA | P21333 | 1734 | GEHVPNSPFQVTA | 6.4218 | 6.4663 | 7.0572 | 6.4207 | 6.0787 | 5.9167 | 5.9846 | 6.0358 | 6.5443 | 6.4992 | 5.9484 | 5.9233 | 6.6896 | 6.0770 | 6.3204 | 5.6159 |
| FLNA_S2284 | FLNA | P21333 | 1055 | GVPVPGSPFPLEA | 6.7371 | 6.7320 | 6.2437 | 6.4348 | 6.5186 | 7.0374 | 6.6837 | 6.5714 | 6.7888 | 6.2742 | 6.6823 | 6.2008 | 4.9425 | 5.5268 | 4.8711 | 5.7547 |
| FLNA_S2327 | FLNA | P21333 | 1630 | GDEIPFSPYRVRA | 5.9398 | 6.6110 | 6.5446 | 6.7161 | 5.8141 | 6.7414 | 5.7454 | 7.0297 | 6.3826 | 6.1500 | 6.5488 | 6.3814 | 6.1412 | 5.6697 | 5.8476 | 5.7366 |
| FLNA_S2370 | FLNA | P21333 | 2615 | VGSRLYSVSYLLK | 5.9848 | 5.9028 | 6.0293 | 6.1184 | 6.9282 | 6.5004 | 5.9660 | 5.9391 | 6.9176 | 5.2981 | 5.8641 | 6.0680 | 6.8321 | 6.5649 | 6.5485 | 6.5379 |
| FLNA_S2510 | FLNA | P21333 | 2158 | SVANVGSHCDLSL | 5.0854 | 5.4954 | 6.3332 | 5.5847 | 6.1190 | 6.0202 | 5.4842 | 6.2809 | 6.0449 | 5.3592 | 5.6648 | 5.4176 | 7.4111 | 8.3663 | 8.0772 | 7.2560 |
| FLNA_S2615 | FLNA | P21333 | 2336 | GDARRLTVSSLQE | 6.9668 | 6.8633 | 6.8053 | 6.9780 | 7.4962 | 5.8707 | 6.2970 | 6.5964 | 7.3880 | 6.0089 | 6.5653 | 6.0178 | 5.3085 | 5.5976 | 4.7204 | 4.5197 |
| FLNA_T2336 | FLNA | P21333 | 2370 | IDAKVHSPSGALE | 6.1667 | 6.3752 | 7.9837 | 4.1821 | 6.1441 | 6.4522 | 5.4237 | 6.5111 | 5.6774 | 7.2179 | 8.9850 | 5.5369 | 4.4540 | 7.2768 | 6.5338 | 5.0793 |
| FLNA_T2599 | FLNA | P21333 | 1338 | DVTYDGSPVPSSP | 5.8790 | 7.0331 | 6.1768 | 7.1085 | 6.2536 | 6.3052 | 5.6871 | 7.3071 | 5.9849 | 6.3674 | 7.0318 | 6.7790 | 5.2239 | 6.1609 | 5.1220 | 5.5799 |
| FLNB_S1316 | FLNB | O75369-8 | 983 | LDVTILSPSRKVV | 6.8949 | 7.7769 | 6.7303 | 7.8417 | 6.4642 | 6.9531 | 7.5428 | 7.1844 | 7.3002 | 7.1097 | 7.1826 | 6.8155 | 3.1485 | 4.0834 | 3.1527 | 3.8192 |
| FLNB_S1536 | FLNB | O75369-8 | 1536 | DEEIPRSPFKVKV | 6.5830 | 6.6512 | 6.6741 | 7.0425 | 6.3075 | 6.8955 | 6.7665 | 6.7393 | 6.6375 | 6.2781 | 6.8820 | 6.9707 | 4.5881 | 5.3236 | 4.5770 | 5.0833 |
| FLNB_S2138 | FLNB | O75369-8 | 2509 | TGQRLVSPGSANE | 5.6680 | 6.0010 | 6.4767 | 6.3024 | 6.5168 | 6.4348 | 5.8920 | 6.3842 | 6.4627 | 5.8861 | 6.3886 | 6.3488 | 6.6710 | 6.2516 | 6.1484 | 6.1670 |
| FLNB_S2400 | FLNB | O75369-8 | 2400 | GSHVVGSPFKVRV | 6.8010 | 8.2506 | 7.2011 | 7.3519 | 5.9473 | 7.6020 | 7.5278 | 6.5512 | 7.4713 | 7.2265 | 7.6874 | 6.9849 | 2.8071 | 4.1113 | 2.9326 | 3.5462 |
| FLNB_S2496 | FLNB | O75369-8 | 2138 | RTSRAPSVATVGS | 5.2104 | 5.2497 | 5.5707 | 5.9207 | 5.7884 | 5.9872 | 5.5429 | 5.9288 | 5.5538 | 5.4572 | 6.7563 | 6.4864 | 7.4750 | 7.4574 | 7.9229 | 7.6922 |
| FLNB_S2509 | FLNB | O75369-8 | 2496 | PNHIVGSPFKAKV | 6.5921 | 6.2799 | 7.2108 | 7.1973 | 6.4274 | 6.6981 | 6.8552 | 6.6625 | 5.9282 | 6.9005 | 7.6772 | 7.2599 | 3.8102 | 4.6389 | 4.9357 | 4.9260 |
| FLNB_S983 | FLNB | O75369-8 | 1316 | DVPIPNSPFKVAV | 6.2865 | 6.1532 | 6.6879 | 7.6168 | 6.1604 | 6.3300 | 6.4763 | 6.2169 | 7.2951 | 6.8952 | 8.2617 | 7.1995 | 4.6200 | 5.0591 | 3.6229 | 5.1185 |
| FLNB_T994 | FLNB | O75369-8 | 994 | VVPCLVTPVTGRE | 6.2315 | 6.7798 | 6.7205 | 6.3786 | 5.9039 | 6.2671 | 8.0522 | 8.1780 | 7.8243 | 7.0101 | 6.2433 | 6.3834 | 4.9591 | 4.7644 | 3.9715 | 4.3323 |
| FLNC_S1050 | FLNC | Q14315 | 2233 | GRERLGSFGSITR | 6.1746 | 5.4342 | 6.3659 | 4.9109 | 6.5703 | 6.2984 | 6.0874 | 6.7288 | 6.3253 | 6.2831 | 5.8147 | 4.0825 | 7.1395 | 8.0394 | 7.0496 | 6.6953 |
| FLNC_S1527 | FLNC | Q14315 | 2718 | DESVPGSPFKVKV | 6.9367 | 7.2463 | 7.4125 | 6.9107 | 6.3925 | 7.2578 | 7.4229 | 6.5816 | 7.7105 | 7.2361 | 7.3668 | 7.1943 | 3.2991 | 4.1737 | 3.1255 | 3.7329 |
| FLNC_S1540 | FLNC | Q14315 | 1050 | GHPVPGSPFAVEG | 6.2674 | 6.2830 | 6.5659 | 6.9617 | 6.7853 | 6.7528 | 6.6648 | 6.9875 | 6.1394 | 6.1247 | 6.8202 | 6.7183 | 4.9882 | 5.3566 | 5.2239 | 5.3604 |
| FLNC_S1728 | FLNC | Q14315 | 2624 | SSSRGSSYSSIPK | 4.7918 | 5.2325 | 5.2728 | 5.3830 | 5.0017 | 5.4129 | 5.1694 | 5.6072 | 5.1995 | 5.4019 | 5.7615 | 5.6576 | 8.5547 | 9.4305 | 9.2446 | 8.8784 |
| FLNC_S2122 | FLNC | Q14315 | 1527 | DQEVPRSPFKIKV | 6.2741 | 6.4644 | 7.7844 | 6.7299 | 5.8711 | 6.1634 | 5.8797 | 6.0934 | 6.8211 | 6.3936 | 6.4175 | 7.2504 | 5.3615 | 5.7028 | 5.2248 | 5.5678 |
| FLNC_S2233 | FLNC | Q14315 | 2586 | PQHIVGSPFKAKV | 7.5041 | 8.1874 | 7.3747 | 8.6729 | 6.2295 | 7.5553 | 7.7470 | 6.3764 | 7.8584 | 6.9575 | 7.9326 | 7.2379 | 2.2874 | 3.2386 | 1.8888 | 2.9515 |
| FLNC_S2233;S2236 | FLNC | Q14315 | 2714 | VKWGDESVPGSPF | 5.9307 | 7.4900 | 6.8933 | 7.3853 | 6.2299 | 6.5548 | 6.1866 | 6.3338 | 6.1333 | 7.3108 | 6.4906 | 6.4657 | 4.8447 | 5.5814 | 4.9794 | 5.1896 |
| FLNC_S2236 | FLNC | Q14315 | 2236 | RLGSFGSITRQQE | 7.0043 | 7.2854 | 8.2497 | 6.3461 | 5.7673 | 6.5717 | 6.4391 | 6.7920 | 7.1562 | 7.7896 | 8.0134 | 7.3623 | 3.4222 | 4.4351 | 3.4312 | 3.9344 |
| FLNC_S2490 | FLNC | Q14315 | 1540 | LPAHDASKVRASG | 7.1205 | 6.5143 | 6.0939 | 6.6002 | 6.5232 | 5.8659 | 7.0917 | 6.0436 | 6.6261 | 6.1724 | 5.7697 | 6.1035 | 5.8363 | 6.2590 | 5.7786 | 5.6010 |
| FLNC_S2586 | FLNC | Q14315 | 2490 | GAHIPGSPFKIRV | 6.9003 | 7.6858 | 7.7741 | 8.0478 | 6.4141 | 6.4457 | 7.1843 | 7.0188 | 6.4035 | 6.8384 | 7.3143 | 8.1024 | 3.1233 | 4.0849 | 3.1655 | 3.4968 |
| FLNC_S2598 | FLNC | Q14315 | 1728 | GEHIPNSPFHVLA | 5.6164 | 5.8209 | 6.9019 | 7.0053 | 6.8525 | 5.9063 | 6.2546 | 6.3400 | 4.7466 | 4.6545 | 7.2952 | 7.2660 | 5.7580 | 7.0929 | 5.7085 | 6.7805 |
| FLNC_S2624 | FLNC | Q14315 | 2412 | DDARRLTVTSLQE | 7.5391 | 7.7990 | 7.2531 | 8.0357 | 6.9755 | 5.7919 | 6.1411 | 6.4547 | 6.7057 | 5.6070 | 6.6298 | 5.8935 | 4.7277 | 5.2633 | 4.4787 | 4.7041 |
| FLNC_S2714 | FLNC | Q14315 | 2122 | DKHVPGSPFTVKV | 6.6189 | 7.3198 | 7.0647 | 7.2744 | 6.0651 | 6.0759 | 5.4413 | 6.1169 | 6.9195 | 6.1018 | 7.0565 | 6.3114 | 4.8780 | 5.3087 | 5.8334 | 5.6136 |
| FLNC_S2718 | FLNC | Q14315 | 2598 | VTGPRLSGGHSLH | 6.3278 | 7.4938 | 6.6643 | 8.1758 | 5.8615 | 7.1191 | 6.0473 | 7.7877 | 5.9683 | 6.0090 | 7.0669 | 6.7572 | 4.2224 | 5.4169 | 4.5194 | 4.5625 |
| FLNC_T2412 | FLNC | Q14315 | 2233;2236 | NA | 5.8206 | 5.8981 | 7.2855 | 5.9835 | 6.6790 | 6.2980 | 6.4341 | 6.5779 | 7.0518 | 6.8714 | 7.1837 | 6.4890 | 5.2800 | 5.7157 | 5.3537 | 5.0779 |
| FLOT1_S19 | FLOT1 | O75955 | 19 | VSGFCRSPPVMVA | 6.1352 | 8.0007 | 7.4834 | 6.5012 | 5.6598 | 5.9441 | 6.3904 | 6.6697 | 6.6426 | 6.5929 | 6.2177 | 6.0009 | 5.2241 | 5.6768 | 5.8453 | 5.0152 |
| FLOT1_S385 | FLOT1 | O75955 | 238 | AQADLAYQLQVAK | 5.9454 | 5.7703 | 7.0549 | 6.0919 | 6.6876 | 6.9769 | 5.3611 | 5.6523 | 6.9617 | 6.4040 | 6.9483 | 7.9342 | 5.1537 | 6.4116 | 4.8625 | 5.7836 |
| FLOT1_Y238 | FLOT1 | O75955 | 385 | TLVSSGSGTMGAA | 5.9053 | 6.2011 | 5.5197 | 5.8813 | 5.7935 | 6.0376 | 6.3102 | 6.3162 | 6.3412 | 5.7072 | 5.4535 | 5.3534 | 7.5728 | 7.5009 | 7.2187 | 6.8874 |
| FLYWCH1_S503 | FLYWCH1 | Q4VC44 | 503 | NTAQRGSPGGPEF | 6.9822 | 7.4013 | 6.3736 | 7.1856 | 6.4942 | 6.5383 | 6.9981 | 6.7923 | 6.8332 | 6.8371 | 6.8936 | 6.6986 | 4.1715 | 4.8740 | 4.2925 | 4.6338 |
| FLYWCH1_S591 | FLYWCH1 | Q4VC44 | 591 | NLAQWDSPDPLRP | 6.5878 | 6.7524 | 7.5373 | 7.5553 | 6.5878 | 6.3922 | 7.7355 | 8.1060 | 7.3649 | 6.4925 | 8.5950 | 7.1796 | 2.8847 | 4.1225 | 3.4380 | 2.6686 |
| FMN1_S168 | FMN1 | Q68DA7-5 | 397 | GEVKPKSPETALE | 5.8831 | 6.0860 | 6.5078 | 6.3234 | 5.7398 | 6.4180 | 6.4994 | 6.5371 | 6.5251 | 6.2824 | 6.4842 | 6.6347 | 5.6719 | 6.0925 | 6.1387 | 6.1760 |
| FMN1_S397 | FMN1 | Q68DA7-5 | 88 | EAGSRVSPDLGND | 4.4863 | 5.2052 | 7.1958 | 6.3723 | 6.6112 | 6.4815 | 5.0619 | 6.1289 | 6.3245 | 7.8283 | 5.5510 | 7.0684 | 7.5143 | 6.0856 | 6.1516 | 5.9332 |
| FMN1_S88 | FMN1 | Q68DA7-5 | 168 | STNGPKSPSGVLS | 6.0877 | 6.0178 | 6.5077 | 5.8982 | 6.3002 | 6.8815 | 6.2213 | 6.3064 | 6.2021 | 7.3720 | 6.5015 | 6.6278 | 5.5143 | 5.5487 | 5.9901 | 6.0228 |
| FMNL1_S1031 | FMNL1 | O95466-2 | 184 | NKPLEQSVEDLSK | 5.5188 | 6.7467 | 6.7355 | 6.4259 | 4.8926 | 6.1503 | 6.9888 | 6.6152 | 5.3138 | 6.4764 | 7.1091 | 7.4145 | 5.2348 | 6.0508 | 5.8149 | 6.5118 |
| FMNL1_S1103 | FMNL1 | O95466-2 | 1031 | EPPAPKSPPKARR | 5.3262 | 6.2071 | 6.7224 | 6.5662 | 7.3924 | 7.7714 | 6.6841 | 5.7937 | 6.9286 | 6.3788 | 7.6455 | 6.6054 | 5.7498 | 6.0501 | 4.5833 | 3.5947 |
| FMNL1_S184 | FMNL1 | O95466-2 | 624 | ALGRRDSELGPGV | 5.7174 | 5.3591 | 5.2906 | 5.5341 | 6.3507 | 5.0345 | 5.6967 | 6.1803 | 6.2748 | 6.2720 | 6.2239 | 6.3989 | 7.4741 | 8.1077 | 7.3443 | 6.7409 |
| FMNL1_S624 | FMNL1 | O95466-2 | 1103 | QVTSDLSLxxxxx | 6.2461 | 5.9291 | 6.3108 | 6.9114 | 6.1238 | 5.7153 | 6.3848 | 6.5343 | 6.8225 | 6.5934 | 6.1788 | 6.4362 | 5.7472 | 6.3252 | 5.6681 | 6.0729 |
| FMNL2_S171 | FMNL2 | Q96PY5-3 | 183 | RGSNLPSPVGNSV | 6.6956 | 7.1781 | 6.4972 | 6.7840 | 7.0889 | 7.0501 | 7.5524 | 7.2314 | 6.6346 | 6.6216 | 6.7083 | 6.5195 | 3.8705 | 4.8619 | 4.0763 | 4.6296 |
| FMNL2_S183 | FMNL2 | Q96PY5-3 | 171 | SKPWSRSIEDLHR | 6.1969 | 6.0815 | 6.1079 | 6.4429 | 5.8062 | 6.1624 | 6.2010 | 6.2522 | 6.3400 | 6.1678 | 6.0852 | 6.0377 | 6.2425 | 6.8821 | 6.5236 | 6.4699 |
| FMNL2_S679 | FMNL2 | Q96PY5-3 | 679 | AIDLSSSKQKIPQ | 6.0476 | 5.2980 | 5.4327 | 5.2666 | 7.2372 | 5.5382 | 6.4977 | 5.1667 | 5.4854 | 5.7244 | 5.3418 | 5.6313 | 7.5761 | 7.4201 | 8.0497 | 8.2865 |
| FMNL3_S174 | FMNL3 | Q8IVF7 | 174 | LRSWSRSIEDLQP | 6.3284 | 6.2869 | 6.3378 | 6.1000 | 5.9185 | 6.2397 | 6.3010 | 6.3391 | 6.0599 | 6.4827 | 6.8445 | 6.6717 | 5.6970 | 6.5779 | 5.7866 | 6.0285 |
| FMNL3_S93 | FMNL3 | Q8IVF7 | 93 | QSFLDPSVTRKKF | 6.3347 | 6.2567 | 6.5865 | 6.5886 | 5.7928 | 6.0808 | 6.3404 | 7.1376 | 6.3759 | 5.2715 | 6.1950 | 6.5943 | 6.3432 | 6.3503 | 5.9765 | 5.7751 |
| FMNL3_T205 | FMNL3 | Q8IVF7 | 205 | SVLRYSTLPGRRA | 8.2977 | 7.3057 | 8.0932 | 6.6501 | 5.6572 | 5.7642 | 6.2459 | 5.7219 | 7.4983 | 7.0941 | 7.2310 | 7.2250 | 4.0584 | 5.4210 | 3.4454 | 4.2910 |
| FNBP1_S296 | FNBP1 | Q96RU3 | 296 | PMKRTVSDNSLSN | 6.1205 | 6.2667 | 6.9153 | 6.1588 | 6.0161 | 6.8369 | 6.0913 | 6.7909 | 6.3591 | 6.8648 | 6.7003 | 6.7004 | 5.4851 | 5.6980 | 5.5800 | 5.4158 |
| FNBP1_S359 | FNBP1 | Q96RU3 | 359 | VPNGPQSPKQQKE | 6.6866 | 6.6699 | 6.4480 | 6.2728 | 5.8345 | 6.9339 | 5.5666 | 7.6623 | 6.1936 | 6.0327 | 7.1170 | 7.1756 | 4.8382 | 5.3605 | 5.5024 | 5.7053 |
| FNBP1_S497 | FNBP1 | Q96RU3 | 497 | EQARRQSGLYDSQ | 5.9455 | 5.2944 | 5.8783 | 7.0444 | 5.8995 | 6.5702 | 6.4976 | 6.8887 | 6.2198 | 7.1240 | 7.6726 | 8.2229 | 5.4979 | 7.1116 | 4.5494 | 3.5832 |
| FNBP1L_S295 | FNBP1L | Q5T0N5 | 295 | HIYRTISDGTISA | 6.3243 | 5.9387 | 6.3513 | 5.7658 | 6.1486 | 6.2379 | 6.0446 | 6.4078 | 6.2633 | 5.9695 | 6.1123 | 6.0615 | 7.0125 | 6.8271 | 6.4608 | 6.0739 |
| FNBP4_S118 | FNBP4 | Q8N3X1-2 | 501 | EKIDENSDKEMEV | 6.3659 | 6.3393 | 6.3576 | 6.5729 | 5.8337 | 5.6703 | 6.1655 | 5.5251 | 6.2208 | 6.1107 | 6.0598 | 6.3042 | 6.9298 | 6.2580 | 6.4518 | 6.8345 |
| FNBP4_S18 | FNBP4 | Q8N3X1-2 | 18 | RPILQLSPPGPRG | 6.4824 | 6.8605 | 6.2843 | 6.4219 | 6.5940 | 6.0699 | 6.1612 | 6.0980 | 6.2922 | 6.0790 | 6.2813 | 6.2154 | 5.9639 | 6.1367 | 5.9743 | 6.0849 |
| FNBP4_S434 | FNBP4 | Q8N3X1-2 | 510 | EMEVEESPEKIKV | 6.3457 | 6.4141 | 6.0699 | 6.1783 | 6.2916 | 5.8403 | 6.2005 | 5.9985 | 6.3523 | 6.1994 | 5.9144 | 5.9703 | 6.9856 | 6.3280 | 6.5440 | 6.3670 |
| FNBP4_S466 | FNBP4 | Q8N3X1-2 | 466 | MFVRATSPESTSR | 6.5954 | 6.8328 | 5.8324 | 6.5367 | 5.9919 | 5.7773 | 6.1211 | 6.1395 | 6.3578 | 5.8621 | 6.1411 | 6.2863 | 6.6141 | 6.2836 | 6.3174 | 6.3106 |
| FNBP4_S501 | FNBP4 | Q8N3X1-2 | 118 | LGAYADSDDDDND | 6.3817 | 6.0499 | 6.0905 | 6.6211 | 6.3511 | 5.6095 | 6.1707 | 5.7311 | 6.5099 | 6.4147 | 6.0168 | 6.0850 | 6.8442 | 6.4015 | 6.2489 | 6.4735 |
| FNBP4_S510 | FNBP4 | Q8N3X1-2 | 481 | SKTGRDTPENGET | 6.0377 | 6.2631 | 5.6792 | 6.2873 | 6.4279 | 5.7782 | 6.2285 | 5.8048 | 6.4848 | 6.0122 | 5.3443 | 6.0858 | 7.6996 | 6.3181 | 6.8586 | 6.6898 |
| FNBP4_T481 | FNBP4 | Q8N3X1-2 | 434 | GSVSGSSPRSDIS | 5.5814 | 6.1090 | 6.3815 | 6.4380 | 6.0595 | 6.1254 | 6.1934 | 5.8160 | 6.5747 | 6.3294 | 6.0378 | 6.2361 | 6.5088 | 6.9680 | 6.4502 | 6.1908 |
| FNDC3A_S202;S207 | FNDC3A | Q9Y2H6 | 203 | QKDKMSSPPSSPQ | 6.0944 | 6.1302 | 6.1715 | 6.3578 | 6.0249 | 5.7603 | 5.7532 | 5.6142 | 5.9617 | 6.2179 | 6.0252 | 6.0984 | 7.0510 | 6.9393 | 7.0277 | 6.7724 |
| FNDC3A_S203 | FNDC3A | Q9Y2H6 | 213 | SPQKCPSPINEHN | 5.7210 | 5.8608 | 6.0144 | 6.5664 | 6.1262 | 6.1168 | 6.3737 | 5.9930 | 6.1360 | 6.4911 | 6.1647 | 6.1556 | 6.5853 | 6.6332 | 6.4974 | 6.5645 |
| FNDC3A_S203;S207 | FNDC3A | Q9Y2H6 | 203;207 | NA | 6.4836 | 6.2878 | 6.4234 | 6.3114 | 6.3376 | 6.2686 | 6.4367 | 6.1189 | 6.1121 | 6.3816 | 6.4697 | 5.8865 | 6.0571 | 6.0151 | 6.1689 | 6.2411 |
| FNDC3A_S213 | FNDC3A | Q9Y2H6 | 202;207 | NA | 6.6898 | 6.9476 | 6.6716 | 6.5608 | 6.2418 | 5.8655 | 6.1407 | 5.7825 | 6.5701 | 6.4307 | 6.3037 | 6.5958 | 5.4514 | 5.9402 | 5.7065 | 6.1011 |
| FNDC3B_S208 | FNDC3B | Q53EP0 | 208 | RQNRLNSPPSSIY | 6.2845 | 6.2926 | 6.0615 | 6.1065 | 6.5156 | 6.3325 | 6.9221 | 6.0477 | 6.4906 | 6.3592 | 6.2201 | 6.3294 | 6.1514 | 6.0816 | 5.8588 | 5.9460 |
| FNIP1_S170 | FNIP1 | Q8TF40 | 763 | DSMSPDSDTELRS | 6.7380 | 6.4299 | 7.3247 | 7.3993 | 6.8189 | 6.2686 | 5.2465 | 6.7906 | 6.1998 | 7.0125 | 7.4195 | 6.3166 | 4.9292 | 5.2783 | 4.7939 | 5.0338 |
| FNIP1_S232 | FNIP1 | Q8TF40 | 170 | TARTGSSICGSLN | 6.9285 | 5.7119 | 6.8468 | 6.2263 | 6.9226 | 6.4831 | 6.0118 | 7.2848 | 6.7155 | 7.6040 | 6.9775 | 7.2440 | 4.4582 | 5.3148 | 4.5116 | 4.7588 |
| FNIP1_S760 | FNIP1 | Q8TF40 | 760 | LIGDSMSPDSDTE | 6.0915 | 5.5920 | 6.4553 | 6.5751 | 5.8551 | 6.2349 | 5.8564 | 6.1966 | 7.2046 | 6.4935 | 6.5273 | 7.4397 | 5.7233 | 5.9036 | 5.4327 | 6.4184 |
| FNIP1_S763 | FNIP1 | Q8TF40 | 232 | RLIRSASFFAVHS | 7.1202 | 5.6097 | 6.5625 | 6.3686 | 5.7081 | 6.7027 | 6.8726 | 6.6221 | 7.4721 | 6.6907 | 7.2484 | 5.8766 | 4.3541 | 5.7110 | 4.7508 | 6.3299 |
| FNIP1_S96 | FNIP1 | Q8TF40 | 96 | LKPGGDSSSSLDS | 6.7160 | 6.7252 | 6.6074 | 6.3666 | 6.7396 | 7.0301 | 6.7461 | 5.7790 | 6.6271 | 6.7959 | 6.5852 | 7.3455 | 4.9598 | 6.0486 | 4.4010 | 4.5267 |
| FOSL1_S260 | FOSL1 | P15407 | 74 | LTYPQYSPPQPRP | 5.8079 | 4.7983 | 5.9353 | 5.7235 | 7.4326 | 6.6841 | 6.6524 | 6.7403 | 6.1315 | 5.6839 | 6.0494 | 6.3100 | 6.9151 | 6.2422 | 6.2276 | 6.6660 |
| FOSL1_S265 | FOSL1 | P15407 | 265 | SSDPLGSPTLLAL | 5.6707 | 5.6467 | 5.9622 | 6.0719 | 6.8212 | 6.4177 | 6.8185 | 6.7216 | 5.7328 | 5.8793 | 6.0810 | 6.4696 | 6.2991 | 6.4407 | 6.2768 | 6.6902 |
| FOSL1_S74 | FOSL1 | P15407 | 260 | SSGDPSSDPLGSP | 6.0892 | 5.3394 | 6.6078 | 6.0773 | 6.5549 | 5.9077 | 6.2221 | 7.1361 | 6.2200 | 6.6165 | 5.9034 | 6.4360 | 5.5695 | 6.6554 | 5.9315 | 6.7331 |
| FOSL2_S200 | FOSL2 | P15408 | 200 | SPEERRSPPAPGL | 5.5925 | 5.6344 | 5.8204 | 6.1269 | 6.3590 | 6.1376 | 6.0181 | 6.1319 | 5.8326 | 5.5880 | 5.7445 | 5.9416 | 7.8710 | 6.8674 | 7.3531 | 6.9812 |
| FOSL2_S230 | FOSL2 | P15408 | 230 | EPLEEDSPSSSSA | 6.3775 | 5.5259 | 6.5646 | 6.2184 | 7.4585 | 5.3099 | 6.3455 | 6.5263 | 6.4984 | 5.9197 | 5.5208 | 6.5677 | 6.4571 | 5.2769 | 6.5842 | 6.8488 |
| FOXC1_S9 | FOXC1 | Q12948 | 232 | IKTENGTCPSPPQ | 6.3254 | 6.1842 | 6.1219 | 6.1617 | 6.4698 | 5.9075 | 6.2677 | 5.7336 | 6.6612 | 6.2602 | 6.3855 | 5.9730 | 7.0756 | 6.2268 | 6.0095 | 6.2364 |
| FOXC1_T232 | FOXC1 | Q12948 | 9 | ARYSVSSPNSLGV | 6.2319 | 5.7996 | 5.6621 | 6.0534 | 5.4132 | 6.2433 | 6.1320 | 7.4681 | 6.6199 | 8.1116 | 6.6871 | 7.0178 | 5.4361 | 7.1734 | 4.9580 | 4.9924 |
| FOXJ2_S164 | FOXJ2 | Q9P0K8 | 164 | DDLSQDSPEQEAS | 4.8506 | 5.1795 | 7.2068 | 4.5294 | 6.1662 | 5.7297 | 5.6298 | 6.6627 | 6.3384 | 6.8025 | 6.4075 | 5.9019 | 7.5772 | 7.5910 | 6.7641 | 6.6627 |
| FOXJ2_S46 | FOXJ2 | Q9P0K8 | 46 | RKCSPGSPTDPNA | 6.1555 | 5.4522 | 6.6798 | 6.4068 | 5.6070 | 6.5783 | 5.8297 | 5.8575 | 6.3606 | 6.5338 | 6.4834 | 7.2682 | 6.1609 | 6.0257 | 5.4297 | 7.1709 |
| FOXK1_S101 | FOXK1 | P85037 | 213 | KEEAPASPLRPLY | 5.7058 | 5.8727 | 6.3278 | 6.0808 | 5.9878 | 6.0134 | 5.6955 | 6.2886 | 5.8198 | 6.3067 | 6.1986 | 6.1991 | 6.9644 | 6.8604 | 7.1165 | 6.5620 |
| FOXK1_S213 | FOXK1 | P85037 | 223 | PLYPQISPLKIHI | 4.8377 | 5.4450 | 5.7864 | 5.2571 | 5.7980 | 5.8265 | 5.3637 | 5.5543 | 5.7991 | 5.8372 | 5.9439 | 5.7459 | 8.5863 | 8.0500 | 8.3121 | 7.8568 |
| FOXK1_S213;S223 | FOXK1 | P85037 | 420 | SRSAPASPTHPGL | 5.2160 | 5.8449 | 5.8587 | 6.2238 | 6.7599 | 6.5568 | 5.7630 | 6.9710 | 5.9087 | 6.1951 | 5.9970 | 5.4106 | 6.7736 | 6.6897 | 7.7920 | 6.0394 |
| FOXK1_S223 | FOXK1 | P85037 | 445 | CLSREGSPIPHDP | 5.3652 | 5.6868 | 6.0655 | 5.6069 | 6.3013 | 5.8455 | 5.9528 | 5.8767 | 6.2564 | 6.2747 | 5.8872 | 6.0639 | 7.4717 | 6.8857 | 7.3180 | 7.1416 |
| FOXK1_S239;S243 | FOXK1 | P85037 | 299 | DTSGGDSPKDESK | 5.0121 | 5.4704 | 5.9860 | 5.4854 | 5.7345 | 6.1495 | 5.3133 | 5.1447 | 6.3888 | 6.9752 | 6.6049 | 6.8025 | 8.0639 | 7.3756 | 6.9332 | 6.5599 |
| FOXK1_S239;S257 | FOXK1 | P85037 | 101 | AASVRQSPGPALA | 6.3175 | 6.4435 | 6.4345 | 6.0895 | 6.5813 | 6.7355 | 6.4437 | 6.6025 | 6.5343 | 6.6422 | 7.0023 | 6.8428 | 5.3835 | 5.4657 | 5.3247 | 5.1566 |
| FOXK1_S295;S299 | FOXK1 | P85037 | 441 | QTPECLSREGSPI | 5.8377 | 6.1566 | 6.1262 | 6.0751 | 6.3193 | 6.3905 | 6.3229 | 6.6652 | 6.4335 | 6.5451 | 6.7137 | 6.3998 | 6.1822 | 6.0140 | 5.8986 | 5.9195 |
| FOXK1_S299 | FOXK1 | P85037 | 436 | RSGGLQTPECLSR | 5.2997 | 5.8302 | 5.5740 | 6.0030 | 6.1243 | 5.5178 | 5.6283 | 5.6376 | 5.7942 | 6.3790 | 5.5135 | 5.4803 | 8.2403 | 7.7233 | 8.1475 | 7.1068 |
| FOXK1_S416 | FOXK1 | P85037 | 468 | EYRYSQSAPGSPV | 5.6328 | 5.9234 | 5.3088 | 5.8610 | 6.2413 | 5.9794 | 5.6994 | 6.0739 | 5.9211 | 6.4653 | 5.6787 | 5.8342 | 8.0530 | 7.8520 | 7.2519 | 6.2237 |
| FOXK1_S416;S420 | FOXK1 | P85037 | 416 | GPLSSRSAPASPT | 5.2644 | 6.9515 | 5.8427 | 7.2084 | 6.0661 | 6.1259 | 5.8248 | 6.5620 | 5.9371 | 5.8463 | 6.0446 | 7.2168 | 6.0363 | 5.9741 | 6.2752 | 6.8237 |
| FOXK1_S416;S420;S428 | FOXK1 | P85037 | 213;223 | NA | 5.9500 | 6.0646 | 6.4567 | 5.8541 | 6.6749 | 6.7328 | 6.2267 | 6.5900 | 6.6951 | 6.4200 | 6.8158 | 6.5582 | 5.8752 | 5.8263 | 5.5809 | 5.6788 |
| FOXK1_S420 | FOXK1 | P85037 | 416;420 | NA | 6.2053 | 6.0814 | 5.8365 | 5.1885 | 5.5126 | 6.0766 | 5.4652 | 5.4625 | 6.1705 | 6.0878 | 6.1900 | 6.1223 | 7.8958 | 7.8237 | 6.7328 | 7.1484 |
| FOXK1_S420;S428 | FOXK1 | P85037 | 436;441 | NA | 5.4085 | 5.5924 | 5.8125 | 5.4311 | 4.9173 | 5.2782 | 5.1382 | 5.6199 | 6.0544 | 6.5208 | 5.7849 | 5.5909 | 8.4319 | 7.9082 | 7.9867 | 8.5242 |
| FOXK1_S441;S445 | FOXK1 | P85037 | 420;428 | NA | 5.2211 | 6.2023 | 6.1246 | 6.5990 | 6.9863 | 6.5282 | 6.2826 | 7.2740 | 6.4422 | 6.1889 | 6.2101 | 6.7198 | 5.8260 | 5.8220 | 6.0681 | 5.5048 |
| FOXK1_S445 | FOXK1 | P85037 | 416;420;428 | NA | 6.9194 | 6.7641 | 6.3377 | 6.0795 | 5.6932 | 6.2882 | 6.4615 | 6.0641 | 6.8887 | 6.7558 | 6.1653 | 6.8675 | 5.6980 | 5.9406 | 4.7263 | 6.3501 |
| FOXK1_S468 | FOXK1 | P85037 | 468;472 | NA | 5.6735 | 5.5656 | 6.5296 | 5.9415 | 5.7108 | 6.1454 | 6.2848 | 5.6386 | 6.1061 | 6.5448 | 6.7122 | 6.9318 | 6.6279 | 6.9078 | 6.4835 | 6.1962 |
| FOXK1_S468;S472 | FOXK1 | P85037 | 436;441;445 | NA | 5.6728 | 6.1595 | 5.9425 | 5.3942 | 5.4690 | 6.0450 | 5.4946 | 5.2956 | 6.1345 | 5.9408 | 6.3705 | 6.2466 | 7.7186 | 7.9971 | 6.8384 | 7.2804 |
| FOXK1_T436 | FOXK1 | P85037 | 441;445 | NA | 5.7352 | 5.9030 | 6.7489 | 6.1923 | 6.6497 | 6.9685 | 6.6325 | 6.8417 | 6.0492 | 6.2295 | 7.1100 | 6.6650 | 6.0505 | 5.5583 | 5.5300 | 5.1359 |
| FOXK1_T436;S441 | FOXK1 | P85037 | 295;299 | NA | 5.9192 | 5.4970 | 6.1337 | 5.3927 | 6.6334 | 6.9894 | 6.2057 | 5.9101 | 5.6697 | 6.1524 | 6.1504 | 6.7906 | 6.8483 | 6.7550 | 6.3072 | 6.6451 |
| FOXK1_T436;S441;S445 | FOXK1 | P85037 | 239;243 | NA | 6.6122 | 5.8245 | 5.9792 | 6.5951 | 7.1878 | 6.4496 | 6.1899 | 7.1017 | 6.3132 | 5.5362 | 7.7463 | 6.8320 | 4.7449 | 6.0577 | 5.5526 | 5.2772 |
| FOXK1_T441 | FOXK1 | P85037 | 239;257 | NA | 6.4768 | 5.1252 | 6.0718 | 6.4124 | 7.1465 | 5.9341 | 7.0771 | 7.6463 | 6.8268 | 5.4720 | 6.6402 | 6.1621 | 4.8586 | 6.0908 | 5.5413 | 6.5180 |
| FOXK2_S170 | FOXK2 | Q01167 | 252 | EASGGDSPKDDSK | 5.5619 | 5.8327 | 6.3381 | 6.0815 | 6.4481 | 6.3741 | 5.9971 | 5.5212 | 6.6896 | 6.3758 | 7.3938 | 6.9969 | 6.6413 | 6.3955 | 5.6893 | 5.6633 |
| FOXK2_S252 | FOXK2 | Q01167 | 398 | SLSREGSPAPLEP | 5.7387 | 6.3976 | 6.3918 | 5.9005 | 6.4228 | 5.7157 | 6.3588 | 6.0883 | 6.5270 | 6.3239 | 6.6162 | 5.8471 | 6.7834 | 6.1789 | 6.1676 | 6.5418 |
| FOXK2_S398 | FOXK2 | Q01167 | 634 | ELKRIKTEDGEGI | 5.8913 | 6.6987 | 6.7384 | 5.7601 | 5.8524 | 6.4246 | 5.9607 | 6.2874 | 6.6080 | 6.1317 | 7.1822 | 6.6481 | 5.6173 | 6.0607 | 6.1642 | 5.9744 |
| FOXK2_S428 | FOXK2 | Q01167 | 428 | AQSAPGSPLSSQP | 6.3406 | 5.5908 | 5.9758 | 7.4090 | 6.2944 | 6.1169 | 6.2894 | 6.5936 | 7.0496 | 6.7116 | 6.2090 | 6.8115 | 5.7666 | 6.1029 | 5.1773 | 5.5611 |
| FOXK2_T634 | FOXK2 | Q01167 | 170 | KQEASESPVKAVQ | 6.7332 | 7.0324 | 6.3134 | 4.4649 | 6.1797 | 6.1263 | 6.2963 | 6.1439 | 6.5609 | 6.3338 | 6.4489 | 6.5086 | 5.6934 | 5.8620 | 5.7222 | 5.5805 |
| FOXM1_S35 | FOXM1 | Q08050-3 | 519 | RPIKVESPPLEEW | 5.8902 | 5.9369 | 5.6479 | 5.7011 | 5.8136 | 5.7240 | 5.9251 | 5.9293 | 6.1770 | 6.1356 | 6.2614 | 6.1133 | 7.3396 | 6.9272 | 7.1815 | 7.2963 |
| FOXM1_S519 | FOXM1 | Q08050-3 | 638 | VGGPFKTPIKETL | 6.3953 | 6.6842 | 7.0635 | 6.4862 | 6.1900 | 6.8980 | 6.7165 | 6.4100 | 7.2140 | 7.3568 | 7.4658 | 7.5546 | 4.2018 | 5.0349 | 4.0968 | 4.2317 |
| FOXM1_S519;S527 | FOXM1 | Q08050-3 | 649 | TLPISSTPSKSVL | 7.2187 | 7.3753 | 7.1032 | 6.6351 | 6.1682 | 6.6124 | 6.9452 | 6.6772 | 7.7302 | 7.1300 | 7.2112 | 7.2597 | 3.6720 | 4.3292 | 3.4610 | 4.4713 |
| FOXM1_S560 | FOXM1 | Q08050-3 | 560 | SYSGLRSPTRCVS | 6.2621 | 6.6115 | 6.8059 | 6.7339 | 6.1802 | 6.6071 | 6.8267 | 6.1593 | 7.1482 | 7.0545 | 7.3148 | 7.0781 | 4.2375 | 5.2653 | 4.6350 | 5.0798 |
| FOXM1_T638 | FOXM1 | Q08050-3 | 665 | PESWRLTPPAKVG | 6.5884 | 6.0637 | 6.6228 | 6.2154 | 6.2045 | 6.3012 | 6.2953 | 5.7477 | 6.1869 | 6.6069 | 6.4914 | 6.0364 | 6.3574 | 6.3365 | 5.7586 | 6.1869 |
| FOXM1_T649 | FOXM1 | Q08050-3 | 35 | EEEPKRSPAQQES | 5.6301 | 6.0729 | 6.3100 | 6.0769 | 5.8463 | 5.8842 | 6.0269 | 6.1834 | 6.0406 | 6.6858 | 6.9511 | 6.0342 | 6.9479 | 6.5648 | 6.1342 | 6.6108 |
| FOXM1_T665 | FOXM1 | Q08050-3 | 519;527 | NA | 6.7602 | 7.0174 | 6.2050 | 7.0228 | 6.6337 | 6.2674 | 6.5724 | 6.1979 | 6.4755 | 8.6184 | 7.4155 | 7.4152 | 4.1780 | 4.5103 | 4.4414 | 4.2688 |
| FOXO3_S253 | FOXO3 | O43524 | 43 | RPELQASPAKPSG | 6.6683 | 7.1613 | 7.4562 | 6.8205 | 6.1338 | 6.4962 | 5.9998 | 6.6209 | 6.5649 | 7.0689 | 6.8910 | 7.0868 | 4.5605 | 5.0592 | 4.4220 | 4.9895 |
| FOXO3_S294 | FOXO3 | O43524 | 294 | LSKWPGSPTSRSS | 6.3666 | 6.5708 | 7.4373 | 6.1901 | 6.0904 | 6.5761 | 6.4180 | 6.9120 | 6.7482 | 6.8747 | 7.2388 | 7.1828 | 4.7793 | 5.1500 | 4.5671 | 4.8977 |
| FOXO3_S413 | FOXO3 | O43524 | 425 | KGSGLGSPTSSFN | 6.7484 | 6.6664 | 6.3299 | 7.0449 | 7.0155 | 6.6811 | 6.3089 | 7.4886 | 6.8451 | 5.4677 | 6.9335 | 6.4540 | 5.1144 | 4.4730 | 5.1039 | 5.3247 |
| FOXO3_S425 | FOXO3 | O43524 | 253 | PRRRAVSMDNSNK | 5.4506 | 4.9243 | 6.4735 | 5.4072 | 6.2207 | 6.7411 | 5.8665 | 6.5094 | 6.1036 | 5.4674 | 5.1725 | 5.4160 | 8.2645 | 7.4111 | 7.2859 | 7.2859 |
| FOXO3_S43 | FOXO3 | O43524 | 413 | LMQRSSSFPYTTK | 6.2695 | 5.7798 | 5.6029 | 4.9286 | 5.3420 | 5.5141 | 4.7240 | 5.3882 | 5.5102 | 6.0491 | 5.8264 | 5.8613 | 8.7388 | 8.5315 | 8.1681 | 7.7656 |
| FOXP2_S616 | FOXP2 | O15409-4 | 616 | SQKITGSPTLVKN | 6.6299 | 7.0754 | 6.7701 | 6.2781 | 6.6512 | 6.9271 | 6.4900 | 6.7773 | 6.8532 | 6.4138 | 6.8094 | 7.4024 | 4.3496 | 5.0604 | 4.5837 | 4.9286 |
| FOXP4_S554 | FOXP4 | Q8IVH2 | 554 | PPKMTGSPTLVKN | 6.5947 | 6.6556 | 6.5207 | 6.6111 | 6.2742 | 6.8401 | 6.4060 | 6.4938 | 6.8692 | 7.2040 | 7.1018 | 7.2443 | 4.6550 | 5.1055 | 4.5385 | 4.8853 |
| FOXP4_T253 | FOXP4 | Q8IVH2 | 253 | QEGLDLTGTAATA | 6.2974 | 6.1814 | 5.8601 | 6.7887 | 6.2717 | 5.8618 | 5.9471 | 6.0668 | 6.0149 | 5.8908 | 6.0178 | 6.2319 | 5.9995 | 7.2709 | 6.5587 | 6.7407 |
| FRMD4A_S357 | FRMD4A | Q9P2Q2 | 694 | TRSVDISPTRLHS | 6.4814 | 6.4390 | 6.2341 | 5.6698 | 6.5267 | 6.3410 | 6.3840 | 6.5995 | 6.3766 | 5.9303 | 5.8122 | 6.0774 | 6.2728 | 6.2654 | 6.3940 | 6.1959 |
| FRMD4A_S694 | FRMD4A | Q9P2Q2 | 357 | KLANMGSKGKIIS | 5.7817 | 5.5285 | 9.0003 | 6.1114 | 6.4897 | 5.9898 | 6.3787 | 5.2560 | 6.4571 | 4.8644 | 6.0097 | 4.9974 | 6.5429 | 6.9450 | 6.3554 | 7.2920 |
| FRMD4A_S710 | FRMD4A | Q9P2Q2 | 800 | GSASSGSMPNLAA | 5.9116 | 7.2680 | 6.2206 | 5.8771 | 6.0435 | 6.7784 | 6.1985 | 5.6159 | 6.8218 | 6.7713 | 6.8740 | 6.6509 | 5.8797 | 6.8058 | 5.0626 | 5.2201 |
| FRMD4A_S800 | FRMD4A | Q9P2Q2 | 710 | HFRHRSSSLESQG | 6.3033 | 6.5421 | 7.1928 | 6.6449 | 6.3186 | 5.9257 | 5.4810 | 6.4015 | 6.6134 | 6.0493 | 6.7205 | 7.0446 | 5.5604 | 5.3633 | 5.8161 | 6.0226 |
| FRMD4A_T906;S908 | FRMD4A | Q9P2Q2 | 906;908 | NA | 6.3824 | 5.9679 | 7.8212 | 5.3784 | 6.1324 | 6.8980 | 6.1119 | 7.7184 | 7.8958 | 5.5430 | 5.1887 | 5.9355 | 5.8341 | 6.0137 | 5.1106 | 6.0679 |
| FRMD5_S375 | FRMD5 | Q7Z6J6 | 375 | RRAVHISIMEGLE | 6.0936 | 5.4548 | 5.9141 | 6.5064 | 6.7468 | 5.2400 | 6.8169 | 5.7261 | 5.3153 | 5.3199 | 5.4155 | 5.4738 | 8.0283 | 7.3895 | 7.8455 | 6.7134 |
| FRMD6_S349 | FRMD6 | Q96NE9 | 375 | SGSSAGSMKHKRL | 7.0857 | 6.8898 | 7.4296 | 6.1895 | 5.6760 | 6.7833 | 6.3089 | 6.5288 | 6.5421 | 6.6594 | 6.8382 | 7.0224 | 4.7574 | 6.0099 | 4.4693 | 4.8098 |
| FRMD6_S375 | FRMD6 | Q96NE9 | 544 | TDRHSLSLDDIRL | 6.3407 | 6.6075 | 6.7529 | 6.5151 | 5.9188 | 6.3873 | 5.9755 | 6.3064 | 6.1748 | 6.2574 | 6.1430 | 6.0665 | 5.8842 | 6.2682 | 6.1540 | 6.2476 |
| FRMD6_S522;S525 | FRMD6 | Q96NE9 | 609 | RTSKYFSLDLTHD | 6.1630 | 6.1846 | 6.6663 | 6.6288 | 5.4302 | 6.5836 | 6.5471 | 6.1738 | 6.1491 | 6.1250 | 6.4439 | 6.3612 | 5.6741 | 6.2698 | 5.9735 | 6.6262 |
| FRMD6_S525 | FRMD6 | Q96NE9 | 349 | KKQYRESYISDNL | 7.7339 | 6.6032 | 5.9793 | 7.4454 | 6.9744 | 6.0532 | 6.7684 | 5.4799 | 8.3641 | 4.9669 | 6.3911 | 5.3578 | 5.6414 | 6.5181 | 4.4453 | 5.2776 |
| FRMD6_S542 | FRMD6 | Q96NE9 | 542 | TSTDRHSLSLDDI | 6.1846 | 6.5171 | 6.3988 | 6.1041 | 6.7597 | 6.3013 | 6.7049 | 6.9278 | 5.2487 | 5.4523 | 6.0486 | 5.9817 | 6.1751 | 6.5311 | 6.6414 | 6.0229 |
| FRMD6_S544 | FRMD6 | Q96NE9 | 525 | RGQSTDSLPQTIC | 6.4182 | 6.5535 | 6.5028 | 6.4506 | 6.6902 | 5.9070 | 6.4288 | 6.2984 | 6.8666 | 6.2090 | 6.3047 | 6.5351 | 5.5533 | 6.5356 | 4.9469 | 5.7992 |
| FRMD6_S609 | FRMD6 | Q96NE9 | 522;525 | NA | 5.7732 | 6.2411 | 6.9416 | 6.8946 | 6.5479 | 7.0885 | 6.9537 | 6.9186 | 5.6393 | 5.9871 | 6.9822 | 7.1255 | 5.3037 | 6.0137 | 4.9253 | 4.6640 |
| FRMD8_S21 | FRMD8 | Q9BZ67 | 439 | KPKRTTSFFSRQL | 7.2428 | 6.6957 | 7.3691 | 5.8186 | 6.8658 | 6.2818 | 5.6601 | 6.4851 | 7.0403 | 7.0447 | 5.9814 | 5.9718 | 5.9733 | 6.0650 | 4.8772 | 4.6273 |
| FRMD8_S408 | FRMD8 | Q9BZ67 | 446 | FFSRQLSLGQGSY | 5.8586 | 6.7628 | 7.6482 | 6.7183 | 6.5984 | 5.4938 | 6.7559 | 6.8981 | 7.2731 | 6.4357 | 6.3912 | 6.7833 | 5.3756 | 5.1890 | 4.9115 | 4.9064 |
| FRMD8_S439 | FRMD8 | Q9BZ67 | 408 | KLRRQGSVVSSRI | 6.4747 | 6.6511 | 7.0806 | 5.9503 | 5.4495 | 6.3962 | 6.3264 | 5.8669 | 6.3020 | 7.2627 | 6.8036 | 6.5149 | 6.0272 | 5.9653 | 5.3549 | 5.5737 |
| FRMD8_S446 | FRMD8 | Q9BZ67 | 21 | ERSHRSSVSSVGA | 6.0511 | 6.6316 | 6.1124 | 5.7109 | 5.2521 | 5.8718 | 5.6895 | 6.6251 | 6.4998 | 7.0653 | 6.9610 | 7.0748 | 5.9271 | 6.0836 | 5.8776 | 6.5662 |
| FRS2_S177 | FRS2 | Q8WU20 | 428 | FDIRRPSLEHRQL | 5.9461 | 5.6188 | 5.9104 | 6.1426 | 6.2394 | 5.9513 | 5.7564 | 6.3751 | 7.1280 | 6.9567 | 6.9740 | 7.1633 | 6.0884 | 6.7693 | 5.4758 | 5.5045 |
| FRS2_S211 | FRS2 | Q8WU20 | 211 | ERKNRTSVHVPLE | 6.0581 | 5.9355 | 6.8620 | 6.2758 | 6.5633 | 5.8095 | 5.8808 | 6.0652 | 6.8851 | 6.9279 | 6.6102 | 6.9296 | 6.0329 | 6.2620 | 5.3906 | 5.5115 |

| | | | | | | | | | | | | | | | | | | | | | |
|---|---|---|---|---|---|---|---|---|---|---|---|---|---|---|---|---|---|---|---|---|---|
| FRS2_S221 | FRS2 | Q8WU20 | | 221 | PLEARVSNAESST | 6.9900 | 5.7135 | 5.8144 | 6.4638 | 6.9886 | 5.4603 | 5.6775 | 6.0559 | 7.2078 | 6.1886 | 5.3683 | 5.9947 | 6.7135 | 7.0230 | 6.2774 | 6.0625 |
| FRS2_S292 | FRS2 | Q8WU20 | | 177 | GSARLPSVGEEST | 6.2286 | 6.3900 | 6.8871 | 6.9098 | 6.1877 | 6.1159 | 5.8710 | 6.9965 | 6.1066 | 6.3596 | 5.9906 | 6.4731 | 5.5827 | 6.3081 | 5.8479 | 5.7449 |
| FRS2_S428 | FRS2 | Q8WU20 | | 292 | WDTGYDSDERRDA | 6.5192 | 6.8806 | 7.0624 | 6.9592 | 6.4618 | 7.1176 | 6.8372 | 6.0124 | 6.4559 | 6.0160 | 6.3691 | 5.6464 | 5.1631 | 5.7238 | 5.6734 | 5.1021 |
| FRS2_T135 | FRS2 | Q8WU20 | | 463 | TTPLPQTPTRRTE | 6.8812 | 6.0787 | 6.7748 | 6.4265 | 6.6832 | 5.6509 | 5.6120 | 5.4946 | 6.5060 | 6.1210 | 5.8308 | 5.6230 | 7.2245 | 6.3305 | 6.4029 | 6.3596 |
| FRS2_T135;T137 | FRS2 | Q8WU20 | | 135 | VPRTPRTPTTPGF | 5.8723 | 6.0254 | 6.3211 | 6.6954 | 6.2077 | 6.1990 | 5.4660 | 5.8916 | 5.9256 | 7.4361 | 6.0313 | 6.5305 | 6.1030 | 6.1592 | 6.1146 | 7.0211 |
| FRS2_T463 | FRS2 | Q8WU20 | 135;137 | | NA | 6.0458 | 6.3780 | 5.6294 | 6.2168 | 5.5785 | 6.1542 | 5.9004 | 6.1907 | 6.2509 | 5.8241 | 6.1753 | 5.8805 | 7.2727 | 6.6853 | 6.6126 | 7.2050 |
| FRYL_S1914 | FRYL | O94915 | | 1959 | DRRRSNTLDIMDG | 6.5004 | 6.1734 | 6.5219 | 5.8421 | 6.1392 | 6.3881 | 6.0226 | 6.5809 | 6.2881 | 6.2197 | 6.3802 | 6.3175 | 6.1558 | 6.4093 | 6.1741 | 5.8868 |
| FRYL_S1923 | FRYL | O94915 | | 2454 | WGVRRRSLDSIDK | 6.4757 | 6.2149 | 6.3757 | 5.4788 | 6.6862 | 6.6866 | 6.1786 | 6.6443 | 6.5980 | 6.4622 | 6.2451 | 6.1514 | 5.9177 | 6.2304 | 5.8731 | 5.7814 |
| FRYL_S1941 | FRYL | O94915 | | 2272 | YGGDTGSPEISFT | 6.1916 | 6.5783 | 6.9412 | 6.7225 | 6.0040 | 7.0119 | 6.7652 | 6.8437 | 6.5791 | 6.7644 | 6.7959 | 6.9478 | 4.6727 | 5.3583 | 4.6601 | 5.1633 |
| FRYL_S1945 | FRYL | O94915 | | 1914 | YAANRKSTGQLNL | 5.8257 | 6.0543 | 6.9016 | 6.0241 | 6.1058 | 6.6584 | 5.9439 | 6.3833 | 5.7742 | 6.5984 | 6.3458 | 5.8739 | 6.1830 | 6.7975 | 5.4534 | 7.0766 |
| FRYL_S1957 | FRYL | O94915 | | 1957 | RGDRRRSNTLDIM | 5.5803 | 4.9242 | 5.7184 | 5.2308 | 6.5660 | 5.2348 | 5.7702 | 6.1051 | 7.5208 | 6.9694 | 7.0074 | 7.2711 | 6.8719 | 7.6398 | 5.8223 | 5.7676 |
| FRYL_S2272 | FRYL | O94915 | | 1945 | SNSLRLSLIGDRR | 6.9179 | 6.8039 | 6.6682 | 6.3436 | 6.0606 | 6.4060 | 6.3213 | 6.7164 | 7.2749 | 6.9386 | 7.1984 | 6.9226 | 4.8840 | 5.6127 | 4.0545 | 4.8762 |
| FRYL_S2332 | FRYL | O94915 | | 1923 | QLNLSTSPINSSS | 6.8464 | 6.6085 | 6.6768 | 6.0678 | 6.1563 | 6.9476 | 6.8955 | 7.0471 | 7.5641 | 7.0760 | 6.5274 | 6.7789 | 4.8164 | 5.2322 | 4.5903 | 4.1687 |
| FRYL_S2454 | FRYL | O94915 | | 1941 | SNARSNSLRLSLI | 5.3946 | 7.1543 | 7.6641 | 7.0326 | 6.8930 | 6.6397 | 6.3735 | 6.5277 | 7.8751 | 7.4141 | 6.3784 | 6.8930 | 4.8881 | 4.0471 | 4.3540 | 4.4709 |
| FRYL_T1959 | FRYL | O94915 | | 2332 | AVTRSTSSTSSGS | 5.8821 | 5.4195 | 6.9581 | 6.1806 | 6.4501 | 6.2009 | 6.4945 | 6.4259 | 7.1716 | 6.5495 | 6.8480 | 6.2839 | 5.0988 | 6.1091 | 5.4610 | 6.4665 |
| FSCN1_S38 | FSCN1 | Q16658 | | 39 | VNASASSLKKKQI | 6.5040 | 6.3991 | 6.8910 | 5.9350 | 6.4361 | 6.7118 | 6.3096 | 6.4379 | 6.6775 | 6.8068 | 6.3999 | 6.1275 | 5.8365 | 5.7611 | 5.4963 | 5.2698 |
| FSCN1_S39 | FSCN1 | Q16658 | | 38 | KVNASASSLKKKQ | 6.4881 | 5.9483 | 6.3455 | 5.9544 | 6.7086 | 6.3070 | 7.6727 | 6.9577 | 7.1797 | 6.7592 | 6.6452 | 6.6905 | 4.7887 | 5.1609 | 5.3361 | 5.0575 |
| FTO_S256 | FTO | Q9C0B1 | | 256 | EGPEEESEDDSHL | 6.5907 | 6.6080 | 6.9768 | 6.2988 | 6.5329 | 7.0310 | 5.9461 | 5.8584 | 5.7262 | 6.2089 | 6.8646 | 6.6951 | 5.8072 | 5.9255 | 5.5943 | 5.3356 |
| FTSJ1_S271 | FTSJ1 | Q9UET6 | | 328 | DNEMSCSPxxxxx | 6.1115 | 6.3528 | 7.2994 | 6.5937 | 6.1042 | 6.4065 | 6.4645 | 6.2308 | 6.5226 | 7.0801 | 6.8374 | 6.9891 | 5.2956 | 5.4945 | 4.8824 | 5.3348 |
| FTSJ1_S328 | FTSJ1 | Q9UET6 | | 271 | PTQPPISPPYQEA | 6.3996 | 6.8094 | 6.6815 | 6.4475 | 6.0882 | 6.8717 | 7.1933 | 6.5134 | 6.8284 | 6.4935 | 6.8930 | 6.8833 | 4.8633 | 5.3190 | 4.5216 | 5.1933 |
| FTSJ3_S335;S336 | FTSJ3 | Q8IY81 | | 584 | LKTEIMSPLYQDE | 6.6093 | 6.8466 | 6.4463 | 7.2956 | 6.1590 | 6.5489 | 7.3688 | 6.4575 | 7.4605 | 7.2125 | 7.3290 | 7.2315 | 3.9613 | 4.5781 | 3.9718 | 4.5233 |
| FTSJ3_S441 | FTSJ3 | Q8IY81 | | 688 | RDLIDNSFNRYTF | 6.6215 | 7.3552 | 6.1890 | 6.8702 | 6.7257 | 7.1223 | 7.7467 | 7.1721 | 7.2869 | 6.8978 | 7.1939 | 6.4862 | 3.5350 | 4.9936 | 3.2465 | 4.5575 |
| FTSJ3_S549 | FTSJ3 | Q8IY81 | | 573 | KQQLPQTPPSCLK | 6.5086 | 7.2280 | 6.3337 | 7.0637 | 6.3480 | 6.4847 | 6.7380 | 6.2304 | 7.1153 | 7.0556 | 7.0223 | 6.6542 | 4.5971 | 5.0239 | 4.4637 | 5.1327 |
| FTSJ3_S584 | FTSJ3 | Q8IY81 | | 441 | VTQGDMSAADTFL | 6.1380 | 7.2544 | 6.5093 | 7.2610 | 6.3144 | 6.5207 | 7.0989 | 6.9877 | 6.1715 | 6.6319 | 6.2839 | 5.7398 | 5.2943 | 5.4480 | 5.1550 | 5.1914 |
| FTSJ3_S599 | FTSJ3 | Q8IY81 | | 676 | LGAVIASSKKAKR | 7.4712 | 7.4828 | 7.2866 | 7.0665 | 5.7497 | 7.1930 | 7.5535 | 6.6685 | 7.0547 | 6.4676 | 6.8494 | 6.7284 | 3.8103 | 4.9377 | 3.4353 | 4.2450 |
| FTSJ3_S644 | FTSJ3 | Q8IY81 | | 644 | RSRGPKSDDDGFE | 6.5029 | 6.8599 | 7.3518 | 7.8401 | 7.3482 | 7.5613 | 7.6838 | 6.8616 | 6.4337 | 5.4428 | 5.0521 | 4.8319 | 4.8515 | 5.3611 | 5.0823 | 4.9349 |
| FTSJ3_S676 | FTSJ3 | Q8IY81 | | 549 | DEALEISQAQLLF | 7.3883 | 6.8808 | 5.7330 | 7.9392 | 7.3694 | 6.1659 | 6.7923 | 6.1257 | 6.5996 | 5.9782 | 5.9044 | 5.8798 | 5.4223 | 5.8970 | 5.1231 | 4.8009 |
| FTSJ3_S688 | FTSJ3 | Q8IY81 | | 599 | KGTEASSGTEAAT | 5.9777 | 6.7349 | 6.5339 | 6.8513 | 5.7279 | 6.1110 | 7.2085 | 6.1403 | 6.4987 | 5.9399 | 5.9269 | 6.2496 | 6.4545 | 6.3426 | 5.4026 | 5.8996 |
| FTSJ3_T573 | FTSJ3 | Q8IY81 | 335;336 | | NA | 5.6368 | 6.0981 | 5.9797 | 6.7471 | 5.7954 | 5.6214 | 5.9497 | 5.9522 | 5.5643 | 6.6498 | 6.1684 | 6.4869 | 5.9534 | 7.1893 | 6.6499 | 6.5578 |
| FUBP1_S630 | FUBP1 | Q96AE4 | | 630 | AYYAQTSPQGMPQ | 6.3953 | 6.0984 | 6.1178 | 5.9955 | 6.5910 | 6.5807 | 6.0954 | 6.5554 | 6.3634 | 6.4335 | 6.6793 | 6.4771 | 6.1719 | 6.1388 | 5.7012 | 5.6052 |
| FUBP1_T153 | FUBP1 | Q96AE4 | | 153 | SCMLTGTPESVQS | 5.9714 | 5.9163 | 6.6490 | 6.1868 | 6.3110 | 6.0409 | 6.0657 | 6.6707 | 6.6982 | 6.4644 | 6.8472 | 6.3972 | 6.1192 | 6.3126 | 5.7298 | 5.6196 |
| FUBP1_T629 | FUBP1 | Q96AE4 | | 629 | AAYYAQTSPQGMP | 6.2111 | 5.8324 | 5.7027 | 6.0613 | 6.7383 | 6.8153 | 6.2777 | 5.7477 | 6.2548 | 6.1091 | 6.9110 | 5.9933 | 6.3977 | 5.9586 | 5.6936 | 7.2953 |
| FUBP3_S296 | FUBP3 | Q96I24 | | 539 | AAPQASSPPDYTM | 6.5443 | 6.3911 | 6.2694 | 6.6540 | 6.4625 | 6.2031 | 6.1770 | 6.6296 | 6.2502 | 6.2533 | 6.3501 | 6.5384 | 5.9797 | 5.8276 | 5.6305 | 5.8392 |
| FUBP3_S539 | FUBP3 | Q96I24 | | 296 | KPDDGISPERAAQ | 6.4568 | 6.6712 | 6.2166 | 6.5898 | 6.5463 | 6.2981 | 6.5163 | 6.1426 | 6.0365 | 6.4131 | 6.5302 | 6.2539 | 6.2659 | 5.4532 | 5.4570 | 6.1525 |
| FUBP3_S569 | FUBP3 | Q96I24 | | 569 | GQAQAHSQEQxxx | 6.0601 | 6.2756 | 5.9177 | 6.6789 | 5.9377 | 6.4788 | 6.4549 | 6.5134 | 5.7129 | 6.6220 | 6.2156 | 5.4882 | 6.2410 | 6.1447 | 6.7428 | 6.5157 |
| FUNDC1_S13 | FUNDC1 | Q8IVP5 | | 13 | PPQDYESDDDSYE | 6.0530 | 6.9173 | 6.7835 | 6.6927 | 6.2629 | 6.1702 | 6.3537 | 5.7391 | 6.2216 | 7.0028 | 5.8887 | 6.3283 | 5.5695 | 5.9499 | 6.0547 | 6.0120 |
| FUNDC2_S151 | FUNDC2 | Q9BWH2 | | 151 | QLKIRKSNQIPTE | 7.6984 | 7.4985 | 6.7161 | 7.5889 | 5.6715 | 4.4005 | 4.8876 | 5.0669 | 9.5282 | 8.2128 | 7.8544 | 8.6629 | 3.3455 | 4.3394 | 4.0697 | 4.4587 |
| FUS_S277 | FUS | P35637 | | 277 | QGSRHDSEQDNSD | 5.4139 | 5.9514 | 6.4363 | 6.1113 | 6.3138 | 5.8671 | 6.1111 | 6.1995 | 5.9354 | 5.5640 | 6.1066 | 6.1179 | 7.0924 | 6.5075 | 6.9112 | 7.3607 |
| FUS_S462 | FUS | P35637 | | 462 | GGGPGGSHMGGNY | 5.5915 | 6.4576 | 6.0897 | 6.3523 | 5.0336 | 7.2968 | 6.2438 | 6.1770 | 6.7838 | 7.0964 | 7.7545 | 7.5407 | 5.7238 | 5.4072 | 5.2185 | 5.2326 |
| FXR1_S406;S409 | FXR1 | P51114 | | 409 | SELSNPSETESER | 7.3130 | 7.2225 | 7.4360 | 7.2452 | 7.0784 | 7.4228 | 6.8890 | 7.3729 | 5.3507 | 5.1066 | 5.6499 | 5.4127 | 4.8433 | 5.3216 | 4.9771 | 5.3583 |
| FXR1_S409 | FXR1 | P51114 | | 464 | PRGGKSSISSVLK | 6.4714 | 5.8944 | 6.2681 | 6.0449 | 6.6512 | 6.2272 | 5.8624 | 6.0246 | 7.1907 | 7.0823 | 7.0925 | 7.0270 | 5.5234 | 6.3791 | 4.9856 | 5.2753 |
| FXR1_S420 | FXR1 | P51114 | | 420 | ERKDELSDWSLAG | 6.0660 | 7.3290 | 7.9174 | 6.8190 | 6.9694 | 8.2564 | 7.7445 | 8.5707 | 4.8636 | 4.9629 | 4.9973 | 4.7737 | 4.9325 | 5.3012 | 4.9072 | 5.5893 |
| FXR1_S464 | FXR1 | P51114 | | 587 | NGPTSASGDDISK | 5.3983 | 6.1486 | 6.3058 | 5.7715 | 6.3469 | 6.2567 | 6.0175 | 6.0624 | 5.4166 | 5.5191 | 5.2185 | 5.2682 | 7.4611 | 8.2252 | 7.5375 | 7.0462 |
| FXR1_S587 | FXR1 | P51114 | 406;409 | | NA | 5.4802 | 6.8293 | 6.2078 | 6.3849 | 6.5556 | 6.4695 | 6.6259 | 6.4562 | 5.5811 | 6.1426 | 6.0234 | 5.8825 | 6.1465 | 6.5994 | 6.4652 | 6.1500 |
| FXR2_S453 | FXR2 | P51116 | | 601 | GNRTDGSISGDRQ | 6.4330 | 6.8113 | 7.2932 | 6.5420 | 6.2867 | 6.4384 | 6.4928 | 5.6540 | 6.6449 | 6.2481 | 6.5252 | 6.6282 | 5.4990 | 5.7525 | 5.3077 | 5.4431 |
| FXR2_S566 | FXR2 | P51116 | | 566 | MDGGLESDGPNMT | 7.1430 | 7.1234 | 6.9514 | 6.8049 | 7.1685 | 6.6092 | 6.5394 | 7.1072 | 5.9563 | 5.3114 | 5.7127 | 6.0977 | 5.6081 | 5.8224 | 4.9994 | 5.0450 |
| FXR2_S601 | FXR2 | P51116 | | 455 | VSTASETESEKRE | 7.6141 | 7.8892 | 7.7763 | 6.8305 | 6.8989 | 7.4566 | 7.1321 | 7.0211 | 6.0368 | 5.0882 | 5.3535 | 5.8647 | 4.5070 | 4.8382 | 4.6508 | 5.0418 |
| FXR2_S601;S603 | FXR2 | P51116 | | 453 | SDVSTASETESEK | 6.9810 | 7.4528 | 8.4159 | 8.0414 | 7.5501 | 8.7807 | 7.5003 | 6.1189 | 6.4898 | 6.7160 | 5.0050 | 5.3577 | 4.0104 | 4.2134 | 3.5580 | 3.8085 |
| FXR2_T455 | FXR2 | P51116 | 601;603 | | NA | 5.8045 | 6.0220 | 6.5209 | 5.9830 | 5.9132 | 5.4809 | 5.8322 | 6.2690 | 5.9326 | 6.3153 | 6.6674 | 6.7146 | 6.6157 | 6.7016 | 6.2719 | 6.9551 |
| FYCO1_S196 | FYCO1 | Q9BQS8-4 | | 196 | YLWKPPSRSSSMS | 6.5481 | 6.3558 | 6.0992 | 6.2139 | 5.4064 | 5.5586 | 5.7483 | 5.6353 | 6.6036 | 6.6872 | 6.4160 | 6.9531 | 6.5736 | 6.6669 | 6.0270 | 6.5071 |
| FYTTD1_S118 | FYTTD1 | Q96QD9 | | 118 | GIRKGISPMNRPP | 7.3517 | 7.3503 | 6.8034 | 7.7060 | 6.7422 | 6.3132 | 6.8358 | 6.8128 | 7.0830 | 6.6867 | 6.7237 | 6.8940 | 4.0989 | 4.3379 | 3.8813 | 4.3792 |
| FYTTD1_S16 | FYTTD1 | Q96QD9 | | 16 | GATATSSPPPKAR | 7.1549 | 6.6911 | 6.3846 | 6.6238 | 6.6614 | 5.9085 | 6.7538 | 6.4289 | 6.9530 | 6.3549 | 5.8319 | 5.5906 | 6.5509 | 5.6635 | 5.6959 | 5.6522 |
| FYTTD1_T12 | FYTTD1 | Q96QD9 | | 12 | TRLVGATATSSPP | 7.1903 | 6.6432 | 6.1249 | 7.0072 | 6.7749 | 6.0964 | 6.8184 | 6.6273 | 7.1234 | 5.8951 | 5.9269 | 5.5822 | 5.7828 | 5.5000 | 5.4651 | 5.4419 |
| FZD6_S629 | FZD6 | O60353 | | 653 | RSEGRISPKSDIT | 6.3527 | 6.6795 | 6.3132 | 6.9317 | 6.0589 | 6.0785 | 6.3012 | 5.7330 | 6.2649 | 6.6192 | 6.4954 | 6.7124 | 5.7028 | 6.1248 | 5.6828 | 5.9489 |
| FZD6_S653 | FZD6 | O60353 | | 629 | ASISRLSGEQVDG | 6.2151 | 6.2526 | 6.2084 | 6.0180 | 5.4596 | 5.2160 | 6.0014 | 6.1691 | 6.9943 | 6.4115 | 5.9534 | 5.8501 | 6.9003 | 6.6919 | 6.7595 | 6.8989 |
| FZD6_S683 | FZD6 | O60353 | | 683 | PSSLKGSTSLLVH | 5.7335 | 5.4728 | 5.7464 | 4.9199 | 4.7882 | 4.4630 | 6.1732 | 5.3754 | 4.4774 | 5.2250 | 4.8455 | 5.2708 | 9.7239 | 8.9175 | 9.1653 | 9.7024 |
| FZR1_S133 | FZR1 | Q9UM11 | | 121 | RRLQPSTPEKKGL | 6.2863 | 6.7498 | 7.0153 | 6.9642 | 5.4732 | 6.4887 | 6.6341 | 6.5582 | 7.3914 | 7.2601 | 7.3198 | 8.1024 | 4.1156 | 4.9063 | 4.0869 | 4.6477 |
| FZR1_S138 | FZR1 | Q9UM11 | | 133 | LFTYSLSTKRSSP | 7.7628 | 6.2446 | 8.6529 | 6.4291 | 5.1254 | 6.4146 | 6.4343 | 6.1078 | 8.6187 | 8.5441 | 8.9390 | 7.9607 | 3.1358 | 3.8498 | 2.8871 | 2.8933 |
| FZR1_S36 | FZR1 | Q9UM11 | | 40 | ASSPVSSPSKHGD | 6.7227 | 6.2732 | 7.3520 | 7.4084 | 8.1109 | 6.8401 | 6.3339 | 7.0844 | 7.1829 | 6.0379 | 7.0359 | 6.6305 | 3.7453 | 4.8289 | 4.2834 | 4.1296 |
| FZR1_S36;S40 | FZR1 | Q9UM11 | | 36 | TLTPASSPVSSPS | 7.5939 | 7.4552 | 6.1015 | 5.2298 | 8.4033 | 6.1904 | 6.7739 | 6.0410 | 7.5512 | 6.1780 | 5.7350 | 7.3342 | 4.0860 | 5.4629 | 5.5180 | 4.3457 |
| FZR1_S40 | FZR1 | Q9UM11 | | 138 | LSTKRSSPDDGND | 7.1696 | 6.7390 | 7.1449 | 8.3432 | 6.2565 | 6.7629 | 6.2751 | 7.1719 | 6.5262 | 7.5512 | 7.7955 | 7.1465 | 3.5367 | 4.5670 | 3.0470 | 3.9668 |
| FZR1_T121 | FZR1 | Q9UM11 | 32;35 | | NA | 6.4999 | 6.0782 | 9.3992 | 6.0672 | 5.8991 | 6.8923 | 3.4497 | 6.2472 | 7.2357 | 9.7416 | 8.2195 | 5.9538 | 4.0503 | 3.4234 | 4.8783 | 5.9645 |
| FZR1_T32;S35 | FZR1 | Q9UM11 | 36;40 | | NA | 7.0728 | 6.9245 | 6.4722 | 7.2293 | 7.8673 | 6.3108 | 6.8481 | 6.7300 | 7.1853 | 6.4278 | 6.7555 | 6.4738 | 4.2675 | 4.3367 | 4.1489 | 4.9495 |
| G3BP1_S149 | G3BP1 | Q13283 | | 149 | TEPQEESEEEVEE | 5.9157 | 6.2947 | 6.6127 | 6.2799 | 6.3574 | 6.1496 | 6.0453 | 6.3257 | 6.3736 | 6.4950 | 6.3229 | 6.5092 | 5.9993 | 6.3276 | 5.8680 | 6.1234 |
| G3BP1_S230 | G3BP1 | Q13283 | | 232 | DAQKSSSPAPADI | 6.2978 | 6.3537 | 6.6518 | 6.1746 | 6.2488 | 6.6358 | 6.1147 | 6.5775 | 6.8213 | 6.6806 | 6.7330 | 6.7129 | 5.4963 | 5.7488 | 5.3609 | 5.3916 |
| G3BP1_S232 | G3BP1 | Q13283 | | 250 | EDLRTFSWASVTS | 4.9062 | 5.0538 | 5.4042 | 5.4075 | 4.9813 | 5.2419 | 5.2993 | 4.7089 | 5.0446 | 4.7590 | 5.2076 | 5.0755 | 10.5831 | 10.4351 | 9.3458 | 8.5464 |
| G3BP1_S250 | G3BP1 | Q13283 | | 67 | IHRKVMSQNFTNC | 4.8277 | 5.8210 | 4.7426 | 6.1258 | 3.4237 | 5.5887 | 4.8256 | 7.2685 | 4.7261 | 3.8551 | 4.8360 | 4.9106 | 10.1924 | 9.8067 | 9.9540 | 9.0954 |
| G3BP1_S253 | G3BP1 | Q13283 | | 253 | RTFSWASVTSKNL | 7.7359 | 6.6774 | 7.6427 | 7.8600 | 6.5739 | 6.3197 | 6.1171 | 5.9220 | 7.9517 | 5.8289 | 5.9368 | 5.6648 | 4.8044 | 6.1644 | 4.9094 | 3.8908 |
| G3BP1_S67 | G3BP1 | Q13283 | | 230 | PEDAQKSSSPAPA | 6.6787 | 7.2568 | 6.7449 | 7.8661 | 6.8215 | 4.6661 | 5.6237 | 6.8633 | 4.4362 | 6.4489 | 6.9260 | 6.2226 | 7.5736 | 6.3479 | 5.0665 | 4.4571 |
| G3BP2_S141;S149 | G3BP2 | Q9UN86-2 | | 227 | LEEKSTTPPPAEP | 6.2043 | 6.5065 | 6.5846 | 6.4103 | 5.9963 | 6.2534 | 6.0600 | 6.2673 | 6.3184 | 6.6216 | 6.4019 | 6.6768 | 5.8252 | 5.9658 | 5.8074 | 6.1003 |
| G3BP2_S149 | G3BP2 | Q9UN86-2 | | 163 | QEERQPSPEPVQE | 5.8462 | 6.4731 | 6.4783 | 6.8203 | 7.2247 | 6.4862 | 5.4480 | 7.6186 | 6.9660 | 5.9762 | 6.4513 | 6.3963 | 4.9869 | 5.8087 | 5.2988 | 5.7202 |
| G3BP2_S163 | G3BP2 | Q9UN86-2 | | 149 | PELDEESEDEVEE | 5.5080 | 6.3689 | 6.0180 | 6.6586 | 6.1623 | 6.0810 | 6.5729 | 6.8639 | 6.2565 | 6.8535 | 6.9861 | 7.2024 | 6.2558 | 5.4255 | 4.9032 | 5.8835 |
| G3BP2_T227 | G3BP2 | Q9UN86-2 | 141;149 | | NA | 5.9319 | 6.4284 | 6.5015 | 6.4699 | 6.7985 | 6.1133 | 6.2416 | 6.0627 | 6.1644 | 6.4170 | 5.7840 | 6.4825 | 6.0171 | 6.4041 | 6.2243 | 5.9585 |
| GAB1_S274 | GAB1 | Q13480-2 | | 581 | LQAPVRSPITRSF | 5.8691 | 5.5069 | 6.2499 | 5.8050 | 6.7004 | 6.9226 | 6.2807 | 6.4514 | 7.0106 | 6.2778 | 6.1275 | 6.3094 | 6.4903 | 6.4353 | 5.6414 | 5.9217 |
| GAB1_S454 | GAB1 | Q13480-2 | | 454 | VPMNPNSPPRQHS | 5.4896 | 5.7554 | 5.7856 | 6.1458 | 6.6606 | 6.1515 | 6.4532 | 7.0246 | 6.3193 | 6.3872 | 6.7530 | 7.1396 | 6.7021 | 5.6593 | 6.1383 | 5.4349 |
| GAB1_S581 | GAB1 | Q13480-2 | | 274 | DVLPKVSPSSTEA | 5.1834 | 6.6366 | 6.8562 | 6.0581 | 6.4738 | 6.1483 | 6.7896 | 6.0718 | 6.5425 | 6.6444 | 5.6854 | 5.1912 | 6.7543 | 6.9131 | 6.3130 | 5.7384 |
| GAB2_S623 | GAB2 | Q9UQC2 | | 631 | SPHRKPSTSSVTS | 6.5924 | 5.5171 | 6.2816 | 5.3715 | 6.5443 | 6.0919 | 5.2993 | 6.6020 | 7.1666 | 7.0490 | 7.2496 | 6.5628 | 6.4105 | 7.0658 | 5.3443 | 4.8512 |
| GAB2_S631 | GAB2 | Q9UQC2 | | 287 | SLTGSETDNEDVY | 7.2203 | 5.5897 | 9.5457 | 7.6310 | 8.7768 | 9.4963 | 8.8509 | 9.6970 | 5.2006 | 3.7738 | 4.9134 | 3.2890 | 4.0703 | 4.3421 | 3.8140 | 3.7893 |
| GAB2_T287 | GAB2 | Q9UQC2 | | 623 | LDFQPSSPSPHRK | 6.5862 | 5.5919 | 6.3955 | 6.5148 | 6.9053 | 5.8558 | 7.0075 | 6.5247 | 5.7157 | 6.1360 | 6.8131 | 7.1675 | 5.3939 | 6.3919 | 5.7410 | 5.2592 |
| GAB3_S345 | GAB3 | Q8WWW8-2 | | 345 | LVPRRISLSGLDN | 6.7530 | 6.5692 | 6.7831 | 5.7143 | 6.6682 | 6.0969 | 6.2084 | 6.3419 | 6.9381 | 6.4766 | 6.7710 | 6.7968 | 6.0322 | 6.0967 | 4.7993 | 4.9542 |
| GAB3_S483 | GAB3 | Q8WWW8-2 | | 483 | SRTSFLSPERNGI | 6.6000 | 6.6842 | 6.6774 | 6.5508 | 6.3872 | 6.4870 | 7.4280 | 6.7153 | 6.4890 | 6.3629 | 6.3191 | 6.6530 | 5.2613 | 5.3041 | 4.9015 | 5.1790 |
| GAGE12F_S33 | GAGE12F | P0CL80 | | 40 | DEVEPATPEEGEP | 4.9553 | 4.8873 | 5.1811 | 4.8670 | 4.5010 | 4.3439 | 4.3478 | 5.1590 | 4.1454 | 4.9986 | 4.4823 | 4.7420 | 5.2721 | 14.1779 | 10.3846 | 13.5547 |
| GAGE12F_T40 | GAGE12F | P0CL80 | | 33 | MRPEQFSDEVEPA | 6.8000 | 6.4666 | 6.4052 | 6.1357 | 4.8045 | 4.3198 | 4.6557 | 6.5795 | 5.1223 | 6.3913 | 5.7323 | 5.8857 | 4.2680 | 10.0713 | 6.9364 | 9.4255 |
| GAGE13_S33 | GAGE13 | Q4V321 | | 40 | DEVEPATPEEGEP | 5.2539 | 4.9093 | 5.2061 | 5.0040 | 4.5421 | 4.2863 | 4.6224 | 5.3558 | 4.2187 | 5.2827 | 4.7010 | 4.6553 | 5.4053 | 13.5001 | 10.5930 | 12.4639 |
| GAGE13_T40 | GAGE13 | Q4V321 | | 33 | MRPEQFSDEVEPA | 6.2425 | 6.2883 | 6.1134 | 6.3820 | 5.1083 | 5.2661 | 4.9873 | 7.0521 | 5.2838 | 6.3625 | 6.0200 | 6.0080 | 4.2445 | 9.7792 | 7.0183 | 7.8437 |
| GAGE5_T40 | GAGE5 | Q13069 | | 40 | DEVEPATPEEGEP | 5.1116 | 4.7598 | 5.1441 | 4.8430 | 4.1913 | 4.1630 | 4.5519 | 5.4312 | 4.0648 | 4.9314 | 4.6997 | 4.3649 | 5.2171 | 14.5961 | 10.0209 | 13.9093 |
| GAK_S906 | GAK | O14976 | | 906 | ACKAPSSNTDLLS | 5.3457 | 5.2423 | 3.7036 | 4.2596 | 4.8673 | 5.1573 | 5.2145 | 5.4399 | 5.8740 | 5.6726 | 5.2718 | 5.6486 | 8.6078 | 10.5159 | 9.9285 | 9.2506 |
| GAPDH_S151 | GAPDH | P04406 | | 83 | FQERDPSKIKWGD | 6.1354 | 6.1500 | 7.2002 | 5.8285 | 6.7888 | 6.9628 | 6.0903 | 6.1518 | 6.2420 | 6.7726 | 6.7403 | 6.6363 | 5.8261 | 5.9869 | 5.4272 | 5.0609 |
| GAPDH_S210 | GAPDH | P04406 | | 184 | VHAITATQKTVDG | 6.1766 | 6.2880 | 6.4606 | 6.8391 | 6.0792 | 5.1600 | 6.3487 | 4.8125 | 6.4530 | 6.0665 | 5.8385 | 5.5912 | 7.1377 | 6.4117 | 7.6337 | 6.7032 |
| GAPDH_S25 | GAPDH | P04406 | | 151 | KIISNASCTTNCL | 5.1847 | 6.1752 | 6.7115 | 6.4997 | 5.9403 | 5.9677 | 6.1456 | 6.2165 | 6.3117 | 6.2952 | 6.7733 | 5.6144 | 6.3413 | 6.7763 | 7.1809 | 5.8659 |
| GAPDH_S83 | GAPDH | P04406 | | 210 | QNIIPASTGAAKA | 6.1951 | 5.9570 | 6.5805 | 6.2098 | 5.6662 | 6.4919 | 6.5653 | 6.4257 | 6.3732 | 6.7384 | 6.8317 | 6.8024 | 5.7608 | 5.9254 | 6.0488 | 5.4277 |
| GAPDH_T184 | GAPDH | P04406 | | 25 | TRAAFNSGKVDIV | 6.2831 | 6.1223 | 6.3042 | 5.7970 | 5.9495 | 6.8747 | 6.1658 | 6.6003 | 6.9608 | 6.8172 | 6.5470 | 6.7224 | 5.2751 | 5.9222 | 5.6986 | 5.9600 |
| GAPVD1_S1046 | GAPVD1 | Q14C86-6 | | 758 | DVREVSSRPSTPG | 5.8561 | 6.0954 | 6.5349 | 6.0094 | 6.0901 | 6.2547 | 6.0125 | 5.9269 | 6.4750 | 6.3726 | 6.2939 | 6.8113 | 6.3871 | 6.5377 | 5.8567 | 6.4859 |
| GAPVD1_S1114 | GAPVD1 | Q14C86-6 | | 1046 | TLTDDPSPRLSAQ | 6.2867 | 6.6144 | 6.2817 | 6.5146 | 6.3241 | 7.5329 | 7.4495 | 6.7431 | 6.9551 | 6.7222 | 6.4312 | 6.5261 | 4.5634 | 5.4315 | 4.5505 | 5.0729 |
| GAPVD1_S466 | GAPVD1 | Q14C86-6 | | 941 | SVRRPMSDPSWNR | 6.2749 | 6.3118 | 6.6649 | 6.0785 | 5.8614 | 6.5053 | 5.7260 | 6.4896 | 6.0847 | 6.0649 | 6.7840 | 6.4477 | 6.4455 | 6.2382 | 5.9452 | 6.0774 |
| GAPVD1_S566;S569 | GAPVD1 | Q14C86-6 | | 466 | YNTPQLSPATTPA | 6.2625 | 6.3414 | 7.0731 | 6.2382 | 6.6693 | 6.6380 | 6.1763 | 6.3907 | 6.5314 | 6.9588 | 6.9565 | 7.1718 | 5.6167 | 5.5795 | 4.6172 | 4.7786 |
| GAPVD1_S758 | GAPVD1 | Q14C86-6 | | 929 | RLVRSRSSDIVSS | 5.3468 | 5.5211 | 6.4183 | 5.0919 | 5.3831 | 5.3700 | 5.1415 | 5.6545 | 6.1631 | 6.0490 | 5.8339 | 5.8222 | 7.7238 | 8.4191 | 8.4623 | 7.5993 |
| GAPVD1_S758;T762 | GAPVD1 | Q14C86-6 | | 762 | VSSRPSTPGLSVV | 5.0909 | 5.5084 | 6.2475 | 5.2882 | 6.4584 | 6.8428 | 5.9061 | 7.4157 | 6.0281 | 7.4412 | 6.9556 | 7.1861 | 5.3766 | 5.8657 | 6.3887 | 5.9999 |
| GAPVD1_S762 | GAPVD1 | Q14C86-6 | | 935 | SSDIVSSVRRPMS | 8.6882 | 7.2113 | 8.1007 | 6.0898 | 5.0968 | 6.3999 | 7.2342 | 6.0341 | 8.2000 | 6.5893 | 7.8341 | 7.0578 | 3.6991 | 4.9952 | 2.8929 | 3.8766 |
| GAPVD1_S791 | GAPVD1 | Q14C86-6 | | 791 | LRSECSSDFGGKD | 6.2050 | 6.4026 | 7.2829 | 5.8540 | 5.7213 | 6.6684 | 5.9771 | 5.9233 | 6.4345 | 6.7127 | 6.0930 | 6.7071 | 5.4039 | 6.5448 | 5.8415 | 6.2278 |
| GAPVD1_S929 | GAPVD1 | Q14C86-6 | | 1114 | DLPDSASQAAHPQ | 6.1903 | 7.0342 | 7.1120 | 7.6574 | 6.5062 | 6.3633 | 6.2992 | 5.7878 | 6.3303 | 5.8043 | 7.2685 | 6.1173 | 5.4776 | 5.4339 | 5.3775 | 5.2404 |
| GAPVD1_S935 | GAPVD1 | Q14C86-6 | 758;762 | | NA | 6.4952 | 6.4878 | 6.9895 | 6.1603 | 6.2620 | 6.3009 | 6.4457 | 6.5264 | 6.6374 | 5.5644 | 6.6617 | 6.2803 | 5.5286 | 5.9245 | 5.4570 | 5.2782 |
| GAPVD1_S941 | GAPVD1 | Q14C86-6 | 566;569 | | NA | 6.4136 | 6.4840 | 7.5651 | 7.2008 | 6.5423 | 5.9820 | 5.7818 | 6.5990 | 6.8868 | 6.6415 | 6.6790 | 7.0305 | 5.5212 | 4.8479 | 4.8909 | 4.9335 |
| GAREM1_S739 | GAREM1 | Q9H706 | | 739 | VEEKVASETSPLP | 7.4761 | 7.8701 | 6.9174 | 7.2611 | 6.7067 | 6.7621 | 6.9037 | 7.3072 | 7.6373 | 6.4329 | 6.8449 | 6.7186 | 3.4084 | 4.5215 | 3.3265 | 3.9056 |
| GARNL3_S424 | GARNL3 | Q5VVW2 | | 426 | LNRRSFSDVLPES | 5.9186 | 5.4385 | 5.5455 | 5.3180 | 4.8965 | 5.1658 | 3.7623 | 4.0968 | 5.7195 | 4.8195 | 4.5134 | 5.3883 | 9.0401 | 10.4052 | 9.3713 | 10.6009 |
| GARNL3_S426 | GARNL3 | Q5VVW2 | | 424 | NMLNRRSFSDVLP | 5.7567 | 5.9292 | 7.4435 | 4.6645 | 6.3975 | 6.5223 | 5.6011 | 7.1569 | 7.7007 | 6.3359 | 6.6640 | 7.2571 | 4.9695 | 6.6979 | 4.9002 | 6.0031 |
| GARRE1_S22 | GARRE1 | O15063 | | 22 | RFLLGGSKQKVQQ | 6.9058 | 7.9530 | 8.0898 | 6.1183 | 5.6164 | 6.8548 | 5.9622 | 7.1190 | 6.3823 | 7.0729 | 7.6465 | 7.9384 | 3.5227 | 4.8975 | 4.0238 | 3.8964 |
| GARRE1_S723 | GARRE1 | O15063 | | 946 | QRRKHSSGEQDTS | 6.2456 | 5.7262 | 6.8786 | 6.4237 | 6.3497 | 6.2429 | 5.7576 | 6.7989 | 6.9181 | 6.8030 | 6.6070 | 6.7456 | 5.6035 | 6.2530 | 5.3508 | 5.2957 |
| GARRE1_S946 | GARRE1 | O15063 | | 723 | LAPQQQSPKQQQP | 6.1527 | 6.6062 | 6.6332 | 6.2424 | 6.7127 | 6.7980 | 6.7023 | 6.3725 | 6.8502 | 6.8143 | 7.1394 | 6.9082 | 5.2268 | 5.0952 | 4.6556 | 5.0903 |
| GARRE1_S946;T951 | GARRE1 | O15063 | | 817 | QGPRNNTWPNRDQ | 6.5089 | 6.5927 | 7.4603 | 5.8671 | 5.9521 | 6.2386 | 5.9694 | 6.1804 | 7.2123 | 7.0110 | 7.2710 | 6.9047 | 5.6087 | 5.6126 | 4.7586 | 4.8515 |
| GARRE1_T817 | GARRE1 | O15063 | 946;951 | | NA | 5.6273 | 7.3955 | 7.0173 | 7.3774 | 6.5776 | 5.7265 | 6.9457 | 7.1414 | 5.7183 | 7.1734 | 7.3412 | 8.3324 | 4.8266 | 4.9205 | 3.4076 | 4.4713 |
| GAS2L1_S306 | GAS2L1 | Q99501 | | 492 | SRVSSPSPELGTT | 5.5814 | 6.2942 | 6.5228 | 6.3446 | 5.8631 | 6.3755 | 6.2861 | 7.0338 | 5.9548 | 6.2045 | 6.4162 | 6.4826 | 5.8737 | 6.3099 | 6.6524 | 5.8043 |
| GAS2L1_S394 | GAS2L1 | Q99501 | | 306 | TSPRPASPVPGSE | 5.8930 | 5.6661 | 6.2055 | 5.6960 | 5.7631 | 6.1278 | 5.6575 | 5.7679 | 6.3608 | 6.2208 | 6.1042 | 6.2858 | 7.1547 | 7.1028 | 6.9497 | 7.0443 |
| GAS2L1_S429 | GAS2L1 | Q99501 | | 479 | QTPRARSPAAPRL | 7.5442 | 7.1239 | 6.4191 | 6.7548 | 6.1911 | 6.0095 | 5.9262 | 5.8178 | 6.5548 | 6.3964 | 7.1412 | 6.1917 | 5.2541 | 5.9440 | 5.5958 | 5.1356 |
| GAS2L1_S479 | GAS2L1 | Q99501 | | 394 | TTGTPASPRRPPA | 6.3775 | 6.7417 | 6.7893 | 6.0108 | 5.1268 | 6.6807 | 6.1943 | 6.5908 | 6.5886 | 6.8857 | 6.6652 | 6.6611 | 5.5054 | 5.8412 | 5.6705 | 5.6703 |
| GAS2L1_S490 | GAS2L1 | Q99501 | | 429 | AQLSVPSPARRAR | 6.0649 | 6.7278 | 6.7208 | 6.0729 | 5.0512 | 7.0047 | 6.4779 | 6.8180 | 6.9808 | 6.6096 | 7.0215 | 6.9132 | 4.9587 | 5.4450 | 5.5337 | 5.5990 |
| GAS2L1_S490;S492 | GAS2L1 | Q99501 | | 490 | RLSRVSSPSPELG | 6.1887 | 5.2949 | 5.7074 | 5.8680 | 5.6521 | 6.3393 | 5.6134 | 6.2977 | 6.4734 | 5.9319 | 6.3106 | 7.1247 | 6.4655 | 7.0666 | 6.6190 | 7.0465 |
| GAS2L1_S492 | GAS2L1 | Q99501 | 490;492 | | NA | 6.8358 | 6.8592 | 6.9264 | 6.3435 | 5.6313 | 6.5195 | 6.1729 | 7.2869 | 6.0523 | 6.6705 | 7.9763 | 6.7858 | 4.8934 | 5.3208 | 4.7585 | 4.9668 |
| GAS2L3_S195 | GAS2L3 | Q86XJ1 | | 570 | VSVSSVSPVKATQ | 6.2885 | 6.2540 | 6.4350 | 6.0912 | 5.9167 | 6.3754 | 6.7211 | 5.9748 | 6.5519 | 6.7742 | 6.3321 | 6.6511 | 5.4005 | 6.4582 | 5.8394 | 5.9358 |
| GAS2L3_S431 | GAS2L3 | Q86XJ1 | | 431 | APCISESPRKCIS | 6.1202 | 6.4879 | 6.4034 | 6.3297 | 6.0934 | 6.4119 | 6.2778 | 6.1885 | 6.6700 | 7.0150 | 6.8996 | 6.4531 | 4.8901 | 6.3150 | 5.3362 | 6.1084 |
| GAS2L3_S518 | GAS2L3 | Q86XJ1 | | 195 | ETLLNTSGPEDSI | 5.8190 | 6.2524 | 6.3647 | 7.2680 | 5.8660 | 5.9510 | 6.9865 | 6.3173 | 6.8371 | 7.0234 | 6.9062 | 6.9816 | 5.1923 | 6.0422 | 5.0719 | 5.1203 |
| GAS2L3_S570 | GAS2L3 | Q86XJ1 | | 518 | KPSFQSSAKMTKT | 8.6959 | 6.5053 | 7.7519 | 7.0776 | 5.1226 | 5.8665 | 6.7912 | 6.2176 | 8.4840 | 8.8438 | 7.9203 | 7.4157 | 3.2241 | 4.4085 | 2.3969 | 3.2781 |
| GATA6_S266 | GATA6 | Q92908 | | 266 | SARFPYSPSPPMA | 5.7188 | 5.5436 | 6.8280 | 6.9500 | 6.8147 | 6.4309 | 5.5325 | 7.1231 | 6.9212 | 6.4487 | 4.9735 | 5.2996 | 6.3910 | 7.2073 | 6.7792 | 5.0378 |
| GATAD2A_S100 | GATAD2A | Q86YP4-3 | | 49 | DGDMRVTPEPGAG | 6.2203 | 6.5096 | 6.5116 | 6.5984 | 5.9743 | 6.0625 | 6.4907 | 6.2118 | 6.2697 | 6.4004 | 6.1798 | 6.6424 | 5.8580 | 6.0167 | 5.8892 | 6.1646 |
| GATAD2A_S100;S107;S1 | GATAD2A | Q86YP4-3 | | 189 | VGSTVTTPPPLVR | 6.6168 | 6.6000 | 6.6478 | 7.0662 | 6.8147 | 6.8235 | 7.1373 | 6.5625 | 6.9645 | 6.5295 | 7.0878 | 7.0910 | 4.5135 | 4.7091 | 4.1730 | 4.6628 |
| GATAD2A_S340 | GATAD2A | Q86YP4-3 | | 340 | VVTSAESPASRQA | 6.1299 | 6.0918 | 6.1115 | 6.4523 | 6.9428 | 6.6362 | 6.7877 | 6.1003 | 6.4622 | 6.3084 | 6.2024 | 6.2991 | 6.0145 | 5.9787 | 5.6131 | 5.8692 |
| GATAD2A_S547 | GATAD2A | Q86YP4-3 | | 547 | GVLHTFSPSPKLQ | 7.7947 | 8.0416 | 7.1893 | 6.7177 | 5.5060 | 7.3229 | 7.1994 | 7.4129 | 8.0160 | 7.9324 | 7.2645 | 7.5316 | 2.8230 | 3.7602 | 2.3560 | 3.1318 |
| GATAD2A_S549 | GATAD2A | Q86YP4-3 | | 100 | SERRPPSPDVIVL | 5.5546 | 5.8435 | 5.8881 | 6.4394 | 5.8993 | 5.6192 | 6.1030 | 5.7314 | 6.1971 | 6.2995 | 6.3373 | 6.2786 | 6.9950 | 6.8482 | 6.8980 | 7.0677 |
| GATAD2A_T189 | GATAD2A | Q86YP4-3 | | 549 | LHTFSPSPKLQNS | 5.5242 | 6.8100 | 5.9300 | 5.9244 | 6.4199 | 5.4301 | 6.8963 | 6.5242 | 7.8279 | 7.4456 | 7.0521 | 7.1777 | 4.9559 | 6.0881 | 5.2014 | 4.7922 |
| GATAD2A_T49 | GATAD2A | Q86YP4-3 | 545;549 | | NA | 7.0466 | 6.3686 | 7.1682 | 6.4060 | 5.8488 | 7.0977 | 6.5610 | 6.1201 | 7.5355 | 5.9942 | 5.9511 | 6.9198 | 5.0094 | 5.3445 | 4.9854 | 5.6430 |
| GATAD2A_T545;S549 | GATAD2A | Q86YP4-3 | 100;107;114 | | NA | 5.6211 | 5.9555 | 7.0132 | 6.2028 | 7.2579 | 6.5558 | 7.1586 | 6.3512 | 6.2335 | 5.8121 | 6.5893 | 5.7957 | 5.6734 | 6.3573 | 5.4919 | 5.9307 |
| GATAD2B_S129 | GATAD2B | Q8WXI9 | | 120 | PERGRLTPSPDII | 5.6881 | 6.2466 | 5.8685 | 6.0577 | 6.4831 | 5.9765 | 5.5553 | 5.9125 | 6.3510 | 6.3047 | 5.7624 | 6.5125 | 6.8655 | 7.0286 | 6.5549 | 6.8322 |
| GATAD2B_S135 | GATAD2B | Q8WXI9 | | 338 | VSSPLPSPSAMTD | 5.9524 | 6.5642 | 6.4610 | 6.4178 | 5.6773 | 6.4042 | 6.2323 | 6.3905 | 6.3959 | 6.9580 | 6.4692 | 6.9670 | 5.7507 | 5.9740 | 5.2464 | 6.1389 |
| GATAD2B_S333 | GATAD2B | Q8WXI9 | | 122 | RGRLTPSPDIIVL | 6.7511 | 6.5874 | 5.5536 | 6.4321 | 6.2349 | 6.3706 | 5.7187 | 5.9845 | 7.3848 | 6.4811 | 6.6867 | 6.3860 | 6.7553 | 5.7802 | 5.1717 | 5.7215 |
| GATAD2B_S333;S338 | GATAD2B | Q8WXI9 | | 129 | PDIIVLSDNEASS | 6.4929 | 7.1066 | 7.2414 | 7.3098 | 6.3291 | 7.0631 | 5.9974 | 6.3136 | 6.4691 | 6.8713 | 6.0813 | 5.2478 | 5.6335 | 5.3453 | 5.2675 | 5.2302 |
| GATAD2B_S338 | GATAD2B | Q8WXI9 | | 135 | SDNEASSPRSSSR | 5.4451 | 6.2044 | 7.4317 | 6.7787 | 5.9795 | 6.4009 | 5.7726 | 7.0549 | 6.7606 | 6.4038 | 6.4645 | 6.1693 | 5.4926 | 5.9804 | 5.7906 | 5.8703 |
| GATAD2B_T120 | GATAD2B | Q8WXI9 | | 333 | GTVNRVSSPLPSP | 6.1202 | 6.0515 | 6.4194 | 6.3052 | 5.7472 | 6.3729 | 6.6349 | 7.3758 | 6.0887 | 6.7894 | 5.8717 | 6.5512 | 5.8510 | 6.0557 | 6.0784 | 5.6867 |
| GATAD2B_T120;S122 | GATAD2B | Q8WXI9 | 333;338 | | NA | 5.9686 | 6.4556 | 5.9650 | 6.9658 | 5.7731 | 6.3978 | 6.5280 | 7.0187 | 6.6207 | 6.9372 | 7.5567 | 7.1993 | 5.0168 | 5.8839 | 4.8337 | 4.8792 |
| GATAD2B_T120;S122;S1 | GATAD2B | Q8WXI9 | 120;122;129 | | NA | 6.5195 | 6.8574 | 5.6781 | 6.8062 | 6.5361 | 6.1365 | 5.7132 | 5.4460 | 5.8532 | 5.8855 | 6.2760 | 6.5231 | 6.5390 | 7.2745 | 6.0528 | 5.9027 |
| GATAD2B_T120;S122;S1 | GATAD2B | Q8WXI9 | 120;122 | | NA | 5.4026 | 5.8990 | 5.9641 | 6.1230 | 6.0074 | 5.6173 | 5.6297 | 5.6457 | 6.5790 | 6.0165 | 6.1248 | 6.0582 | 8.2267 | 7.1962 | 6.6827 | 6.8271 |
| GATAD2B_T122 | GATAD2B | Q8WXI9 | 120;122;135 | | NA | 5.9484 | 6.5086 | 6.3433 | 5.6668 | 5.7966 | 4.9358 | 6.4480 | 5.8630 | 7.3068 | 7.6124 | 5.5141 | 5.6401 | 6.0444 | 7.6820 | 6.8449 | 5.8449 |
| GBF1_S1298 | GBF1 | Q92538 | | 314 | DLEQPGSPGYSTA | 5.6131 | 6.0816 | 6.1629 | 6.0068 | 6.2242 | 6.3233 | 5.9214 | 6.3002 | 5.8524 | 6.0366 | 6.2295 | 5.9986 | 6.2990 | 6.9793 | 6.9386 | 7.0324 |
| GBF1_S1318 | GBF1 | Q92538 | | 1318 | LDRGYTSDSEVYT | 5.8030 | 6.3035 | 6.5372 | 6.1977 | 6.3213 | 6.7620 | 6.1016 | 6.7315 | 6.0048 | 5.8356 | 6.1888 | 6.0977 | 5.9620 | 6.4786 | 6.4690 | 6.2057 |
| GBF1_S1773 | GBF1 | Q92538 | | 1298 | PDAGAQSDSELPS | 6.2328 | 7.2593 | 6.3726 | 6.5157 | 6.7215 | 6.5729 | 6.1613 | 7.8863 | 5.9336 | 5.6767 | 6.3255 | 5.9432 | 5.0442 | 6.1483 | 5.8423 | 5.3638 |
| GBF1_S1780;S1784 | GBF1 | Q92538 | | 599 | QAKVLNSLTQQEK | 6.2889 | 7.0599 | 6.5537 | 7.1492 | 6.4911 | 7.3165 | 7.2433 | 6.3996 | 7.3648 | 7.0310 | 7.1617 | 7.3254 | 3.6946 | 4.6537 | 3.6690 | 4.5973 |
| GBF1_S314 | GBF1 | Q92538 | | 1337 | KIHRSATDADVVN | 2.7670 | 2.4896 | 2.3418 | 2.6624 | 3.7870 | 3.0295 | 3.1620 | 2.8924 | 2.3454 | 2.4175 | 2.0929 | 2.2075 | 17.4804 | 16.5046 | 18.0674 | 15.7525 |
| GBF1_S599 | GBF1 | Q92538 | | 1773 | DFEKPESPRAASS | 5.9203 | 6.2974 | 6.8121 | 6.5155 | 6.8749 | 6.5228 | 6.6665 | 6.4351 | 6.9236 | 6.5411 | 6.1847 | 7.1380 | 5.5707 | 5.3087 | 4.7855 | 5.5031 |
| GBF1_T1337 | GBF1 | Q92538 | 1780;1784 | | NA | 5.0846 | 6.0898 | 5.6628 | 4.8246 | 6.9890 | 5.8712 | 5.4683 | 6.6961 | 5.2868 | 5.1186 | 4.3506 | 5.1229 | 7.6003 | 9.1551 | 7.6706 | 9.0087 |
| GCC2_S1483 | GCC2 | Q8IWJ2 | | 1483 | NEPTTRSPVSSQQ | 6.2697 | 6.1155 | 6.1735 | 6.2581 | 6.2507 | 6.3884 | 6.3248 | 7.0728 | 6.6034 | 6.3313 | 6.3904 | 6.2247 | 6.0127 | 5.8875 | 5.9497 | 5.7468 |

| | | | | | | | | | | | | | | | | | | | | | |
|---|---|---|---|---|---|---|---|---|---|---|---|---|---|---|---|---|---|---|---|---|---|
| GCC2_S1542 | GCC2 | Q8IWJ2 | | 591 | LEGKINSLTEEKD | 5.9102 | 4.7764 | 6.4756 | 5.4950 | 8.2810 | 7.2913 | 7.0735 | 6.3270 | 5.3433 | 5.8875 | 5.3798 | 5.2834 | 7.7308 | 6.5417 | 5.9992 | 6.2042 |
| GCC2_S236 | GCC2 | Q8IWJ2 | | 1542 | LEQLLNSPETKLE | 6.2598 | 5.4156 | 6.3777 | 6.3601 | 6.0167 | 6.3317 | 5.5298 | 6.3041 | 6.2008 | 5.8787 | 6.7169 | 6.6810 | 6.6379 | 6.3058 | 6.5245 | 6.4589 |
| GCC2_S591 | GCC2 | Q8IWJ2 | | 713 | DLEVFLSQKEDVI | 5.3381 | 6.4725 | 5.5661 | 4.9368 | 9.5099 | 8.0458 | 7.7613 | 6.0523 | 4.6881 | 5.4738 | 4.3734 | 3.7440 | 9.1161 | 6.6835 | 6.9982 | 5.2401 |
| GCC2_S713 | GCC2 | Q8IWJ2 | | 236 | YQKNINSLQEELL | 5.8437 | 5.3584 | 6.5231 | 6.4850 | 6.8506 | 6.9595 | 5.9974 | 6.9491 | 5.7087 | 6.5455 | 5.9786 | 6.4874 | 5.5306 | 6.6992 | 6.0751 | 6.0082 |
| GCC2_T1500 | GCC2 | Q8IWJ2 | | 1500 | LRERRNTDLPLLD | 7.7152 | 5.9216 | 6.0818 | 7.0544 | 7.8182 | 5.2665 | 5.5153 | 5.7914 | 7.3018 | 5.6884 | 5.2550 | 7.1016 | 7.1745 | 6.6553 | 4.5828 | 5.0762 |
| GCFC2_S429;S430 | GCFC2 | P16383 | 429;430 | | NA | 5.6632 | 6.1329 | 6.2433 | 5.9897 | 5.9046 | 6.0874 | 5.5941 | 6.4469 | 6.0829 | 6.2384 | 6.2620 | 6.0023 | 6.9345 | 6.8014 | 6.8401 | 6.7762 |
| GCFC2_S96;T97 | GCFC2 | P16383 | 428;430 | | NA | 5.6147 | 6.0017 | 6.4731 | 6.1612 | 5.8670 | 6.3138 | 5.9415 | 6.4267 | 6.2810 | 6.0393 | 6.2062 | 6.4128 | 6.5028 | 6.5764 | 6.6578 | 6.5238 |
| GCFC2_T428;S429 | GCFC2 | P16383 | 96;97 | | NA | 5.7074 | 5.7681 | 6.8109 | 6.1163 | 6.0759 | 6.2527 | 6.2036 | 5.6079 | 6.4296 | 6.6703 | 6.3015 | 6.4106 | 6.0920 | 6.3101 | 6.5250 | 6.7181 |
| GCFC2_T428;S430 | GCFC2 | P16383 | 428;429 | | NA | 4.2405 | 5.2032 | 6.4228 | 6.0707 | 6.3941 | 6.9970 | 6.1346 | 6.2009 | 7.4958 | 6.2634 | 5.8996 | 6.1480 | 6.0167 | 6.9687 | 6.0630 | 7.4808 |
| GDAP1_S11 | GDAP1 | Q8TB36 | | 11 | QEEQRGSPPLRAE | 5.9303 | 7.1531 | 6.7745 | 6.1949 | 6.1113 | 6.5423 | 6.1910 | 5.3262 | 5.8026 | 5.8779 | 5.5687 | 5.8838 | 6.4603 | 6.9902 | 6.8333 | 6.3597 |
| GDI2_S61 | GDI2 | P50395 | | 61 | RFKIPGSPPESMG | 6.4236 | 6.6733 | 6.7566 | 6.4246 | 6.3641 | 6.5102 | 6.0017 | 6.6374 | 6.4957 | 6.8237 | 6.9614 | 6.7540 | 5.1152 | 5.5294 | 5.3504 | 5.1787 |
| GEM_S23 | GEM | P55040 | | 23 | PQQQRWSIPADGR | 7.4409 | 6.8159 | 8.1645 | 6.8421 | 6.3080 | 6.3208 | 6.5418 | 7.0446 | 7.0990 | 6.2729 | 7.4176 | 7.4486 | 4.1242 | 4.1650 | 3.6881 | 4.3059 |
| GEMIN2_S81 | GEMIN2 | O14893 | | 81 | KLKRKQSVNISLS | 5.9065 | 4.6293 | 5.6385 | 4.9944 | 7.3140 | 5.9483 | 4.8457 | 5.0915 | 6.7350 | 7.1407 | 7.1640 | 7.7123 | 6.8825 | 8.3539 | 5.1235 | 6.5199 |
| GEMIN4_S205 | GEMIN4 | P57678 | | 205 | PPKRLRSDPDACP | 3.4879 | 3.0431 | 4.8134 | 4.5884 | 5.5758 | 5.6593 | 5.5308 | 5.9627 | 4.3994 | 3.6165 | 4.6077 | 4.1436 | 12.0013 | 10.2837 | 11.4974 | 10.7890 |
| GEMIN4_T84 | GEMIN4 | P57678 | | 84 | LQPHPVTPSDTET | 5.4365 | 4.2803 | 5.8748 | 5.0891 | 5.9291 | 4.8285 | 5.0606 | 5.7364 | 5.8179 | 3.6818 | 4.5422 | 5.7039 | 9.5906 | 8.6705 | 10.7170 | 9.0409 |
| GEMIN5_S778 | GEMIN5 | Q8TEQ6 | | 778 | PVENGVSDQEGEE | 6.8053 | 7.0974 | 6.5359 | 6.9649 | 5.9245 | 6.3296 | 5.4950 | 6.7719 | 6.3180 | 6.3694 | 5.6695 | 7.2549 | 5.4945 | 6.1881 | 5.3248 | 5.4563 |
| GEMIN8_S117 | GEMIN8 | Q9NWZ8 | | 117 | KEDQALSKEEEME | 6.4684 | 6.1025 | 6.8443 | 5.5410 | 6.7190 | 5.9115 | 5.9559 | 6.8416 | 5.8530 | 6.0982 | 5.6578 | 5.7187 | 7.0882 | 6.4644 | 6.3576 | 6.3780 |
| GEN1_S245 | GEN1 | Q17RS7 | | 245 | NRWNETSCNSSPQ | 6.2219 | 7.2252 | 6.8038 | 7.0091 | 6.2491 | 7.0406 | 6.9015 | 6.5886 | 6.9785 | 6.9864 | 6.4889 | 6.7248 | 4.8421 | 4.7202 | 4.3950 | 4.8244 |
| GFPT1_S261 | GFPT1 | Q06210 | | 261 | NLSRVDSTTCLFP | 5.9195 | 5.3655 | 6.3039 | 5.5085 | 6.7916 | 6.1585 | 6.1785 | 5.9803 | 5.8061 | 5.8002 | 5.8305 | 5.5716 | 7.5427 | 7.5776 | 6.8867 | 6.7785 |
| GFPT2_S244 | GFPT2 | O94808 | | 244 | RMKRLDSSACLHA | 6.6892 | 5.0504 | 6.6582 | 5.9507 | 6.7141 | 6.2550 | 6.5379 | 5.2760 | 6.1466 | 5.6327 | 5.5940 | 6.0215 | 7.4750 | 6.8956 | 6.8684 | 6.2349 |
| GGA2_S177 | GGA2 | Q9UJY4 | | 177 | KILPPPSPWPKSS | 8.0090 | 7.1939 | 6.8816 | 6.8867 | 6.6385 | 6.1233 | 7.1596 | 7.4332 | 6.7174 | 6.3518 | 5.7490 | 6.2890 | 4.6354 | 4.9740 | 4.0851 | 4.8725 |
| GGA3_S159 | GGA3 | Q9NZ52 | | 159 | DRTLIPSPPPRPK | 6.7334 | 6.5028 | 6.7691 | 6.2204 | 7.0482 | 6.7338 | 6.2511 | 7.0076 | 6.4288 | 6.4348 | 6.8427 | 6.8403 | 5.1793 | 5.3055 | 4.8601 | 4.8421 |
| GGA3_S538 | GGA3 | Q9NZ52 | | 538 | LVKPTTSPLIPTT | 7.5228 | 7.7677 | 4.9880 | 6.7408 | 7.4197 | 6.8139 | 5.4777 | 8.2231 | 7.8794 | 6.4831 | 8.5797 | 7.6259 | 3.7506 | 5.0653 | 3.3511 | 2.3114 |
| GGCX_S11 | GGCX | P38435 | | 11 | AGSARTSPSSDKV | 6.0357 | 7.2331 | 7.1002 | 6.8047 | 5.5903 | 6.0916 | 7.3459 | 6.4371 | 10.8291 | 6.9449 | 6.6777 | 6.7474 | 3.8828 | 4.3695 | 3.9955 | 3.9144 |
| GGT7_S72 | GGT7 | Q9UJ14 | | 72 | RLQRLPSSSSEMG | 4.8860 | 5.3953 | 5.5754 | 5.3839 | 5.7857 | 5.0783 | 5.1996 | 5.6921 | 5.6339 | 5.3594 | 5.1747 | 5.0298 | 9.0352 | 8.7018 | 9.0470 | 9.0218 |
| GHDC_S87 | GHDC | Q8N2G8 | | 87 | HCSLRRSTDISTF | 6.5234 | 6.8377 | 6.1983 | 6.3659 | 5.7953 | 5.8681 | 5.7720 | 6.0465 | 6.0181 | 6.2682 | 6.4455 | 5.6919 | 6.2079 | 5.9880 | 7.3338 | 6.6393 |
| GIGYF1_S230 | GIGYF1 | O75420 | | 538 | PFAPGPSPPPLLG | 7.1400 | 6.1121 | 6.1989 | 6.2686 | 6.1372 | 6.0133 | 5.8621 | 6.2000 | 6.2175 | 6.5913 | 5.8620 | 6.4205 | 6.2380 | 6.3864 | 6.1255 | 6.2265 |
| GIGYF1_S24 | GIGYF1 | O75420 | | 406 | IRGIQLSPGVGSS | 6.3172 | 6.2796 | 6.5862 | 6.4025 | 6.3323 | 6.8372 | 6.4755 | 6.7964 | 6.3651 | 6.8505 | 6.8164 | 7.0263 | 5.4719 | 5.5709 | 4.7759 | 5.0960 |
| GIGYF1_S24;S28 | GIGYF1 | O75420 | | 756 | VPPAPSSPPPLWA | 6.7909 | 6.3193 | 6.6894 | 6.2586 | 6.5817 | 6.7776 | 6.0505 | 6.5058 | 6.9598 | 6.6874 | 6.6801 | 6.5158 | 5.4657 | 5.2444 | 5.0756 | 5.3975 |
| GIGYF1_S368 | GIGYF1 | O75420 | | 368 | QEEKSSSPSPLPT | 7.0057 | 6.0310 | 7.0317 | 6.3108 | 6.7252 | 6.8314 | 5.6482 | 6.8995 | 6.5735 | 6.9174 | 6.8225 | 6.5865 | 5.6393 | 5.6052 | 4.6937 | 4.6783 |
| GIGYF1_S406 | GIGYF1 | O75420 | | 230 | DRWRSASPDGGPR | 7.0479 | 6.7790 | 6.8138 | 6.3608 | 6.3838 | 7.0164 | 5.6030 | 6.0931 | 7.1725 | 6.9115 | 6.3575 | 6.4815 | 5.5800 | 5.2096 | 4.7706 | 5.4189 |
| GIGYF1_S538 | GIGYF1 | O75420 | | 24 | GGGSVASPPPSPA | 6.1656 | 6.3987 | 6.4521 | 5.5354 | 6.7851 | 6.5129 | 5.7563 | 7.2796 | 7.1094 | 7.0999 | 6.7426 | 7.0337 | 6.6824 | 5.7468 | 4.1248 | 4.5747 |
| GIGYF1_S756 | GIGYF1 | O75420 | 24;28 | | NA | 7.1358 | 7.4095 | 6.6139 | 5.9914 | 6.3342 | 6.9084 | 6.5963 | 6.7121 | 7.3008 | 7.3303 | 6.5377 | 7.2678 | 4.7253 | 4.8792 | 3.9028 | 4.3547 |
| GIGYF2_S160 | GIGYF2 | Q6Y7W6-3 | | 258 | ERWRPHSPDGPRS | 6.5876 | 6.7488 | 6.5396 | 6.1847 | 6.6346 | 6.6928 | 5.9167 | 6.5942 | 6.6306 | 6.4740 | 6.6836 | 6.6463 | 5.7349 | 5.6583 | 5.0838 | 5.1894 |
| GIGYF2_S23 | GIGYF2 | Q6Y7W6-3 | | 614 | WGRVPFSPGPAPP | 6.2433 | 6.4882 | 6.5789 | 6.6931 | 6.1423 | 7.0396 | 6.5415 | 6.5505 | 6.5702 | 6.5739 | 6.6582 | 6.5489 | 5.2366 | 5.5624 | 5.3483 | 5.2242 |
| GIGYF2_S258 | GIGYF2 | Q6Y7W6-3 | | 26 | SGGSITSPPLSPA | 6.1844 | 6.1899 | 6.8631 | 6.2935 | 6.7028 | 6.7392 | 6.2137 | 6.8458 | 6.5858 | 6.9978 | 7.3289 | 6.9900 | 4.8404 | 5.1707 | 4.9097 | 5.1440 |
| GIGYF2_S26 | GIGYF2 | Q6Y7W6-3 | | 306 | IDDDRDSLPEWCL | 6.3818 | 5.8770 | 6.6274 | 6.6034 | 5.8460 | 6.3956 | 6.2923 | 6.1436 | 6.7959 | 6.6485 | 7.1444 | 7.0888 | 5.4153 | 5.7107 | 5.0911 | 5.9381 |
| GIGYF2_S26;S30 | GIGYF2 | Q6Y7W6-3 | | 160 | EMHRSQSWEERGD | 5.4277 | 5.3706 | 6.0315 | 5.1761 | 5.9832 | 5.9664 | 5.4344 | 6.6136 | 5.9676 | 5.4793 | 5.9255 | 6.4277 | 7.9558 | 7.5620 | 7.4596 | 7.2188 |
| GIGYF2_S306 | GIGYF2 | Q6Y7W6-3 | | 409 | TPSDHQSQEASQF | 6.5490 | 6.8951 | 7.3126 | 5.9983 | 5.1096 | 5.9674 | 4.3866 | 8.5912 | 5.8456 | 6.9016 | 7.3386 | 6.8529 | 5.5206 | 5.2379 | 5.8521 | 5.6409 |
| GIGYF2_S357;S359;S362 | GIGYF2 | Q6Y7W6-3 | | 23 | ALSSGGSITSPPL | 5.5534 | 6.6047 | 6.1165 | 5.9483 | 6.4903 | 6.2785 | 5.9742 | 7.2263 | 7.3434 | 6.7109 | 6.6326 | 7.2107 | 5.3947 | 5.9687 | 5.0461 | 5.5009 |
| GIGYF2_S409 | GIGYF2 | Q6Y7W6-3 | 26;30 | | NA | 6.7116 | 6.6577 | 6.9436 | 6.6262 | 6.5421 | 7.6279 | 7.1419 | 7.8482 | 7.7707 | 7.2836 | 7.2067 | 7.0469 | 3.5336 | 4.1123 | 3.1554 | 3.7916 |
| GIGYF2_S614 | GIGYF2 | Q6Y7W6-3 | 357;359;362 | | NA | 6.1231 | 5.6651 | 5.9616 | 7.3133 | 6.5475 | 5.6965 | 6.7658 | 6.1899 | 6.2552 | 6.9371 | 6.4089 | 6.2900 | 6.0012 | 6.2247 | 6.3317 | 5.2883 |
| GIPC1_S232 | GIPC1 | O14908 | | 232 | AGGRPGSGPQLGT | 5.7205 | 5.7231 | 6.7509 | 7.0307 | 6.4190 | 6.2570 | 6.4468 | 6.5110 | 5.9756 | 5.8238 | 6.7483 | 6.3178 | 5.9780 | 6.8334 | 5.6396 | 5.8246 |
| GIT1_S371 | GIT1 | Q9Y2X7-3 | | 419 | RSNRARSMDSSDL | 6.3626 | 6.0442 | 6.2074 | 6.1169 | 6.2418 | 5.9355 | 5.6678 | 6.4461 | 6.6381 | 6.8576 | 6.4349 | 6.6229 | 6.1233 | 6.4366 | 6.0507 | 5.8136 |
| GIT1_S419 | GIT1 | Q9Y2X7-3 | | 580 | VPFTPSSPLLSCS | 6.3541 | 6.4036 | 6.7457 | 7.0182 | 6.3296 | 6.6255 | 6.8609 | 6.4069 | 7.1110 | 7.2602 | 7.1985 | 6.6733 | 5.0770 | 4.8114 | 4.2548 | 4.8694 |
| GIT1_S570 | GIT1 | Q9Y2X7-3 | | 601 | KLSRHGSGADSDY | 6.3194 | 4.2864 | 7.0696 | 5.8118 | 8.0791 | 5.4967 | 5.6361 | 4.0158 | 6.3696 | 6.4226 | 6.7628 | 6.1074 | 6.3500 | 8.0122 | 6.8911 | 6.3696 |
| GIT1_S580 | GIT1 | Q9Y2X7-3 | | 570 | RIRKGVSASAVPF | 6.2883 | 7.7242 | 6.3776 | 6.5802 | 6.0987 | 6.5410 | 5.5518 | 5.7109 | 7.1228 | 6.3101 | 5.4429 | 6.5192 | 5.2860 | 6.4234 | 5.3034 | 6.7197 |
| GIT1_S601 | GIT1 | Q9Y2X7-3 | | 371 | QGKSLSSPTDNLE | 6.7327 | 6.1653 | 6.7204 | 7.1603 | 6.1304 | 6.4581 | 6.1733 | 6.8068 | 6.1647 | 7.2965 | 5.9528 | 6.3739 | 5.2341 | 6.1009 | 4.7744 | 5.7554 |
| GIT2_S415 | GIT2 | Q14161 | | 614 | RKGRQRSMVWPGD | 6.2461 | 5.8214 | 6.1565 | 5.7053 | 5.9175 | 6.1172 | 6.7861 | 6.4706 | 7.3354 | 6.1358 | 6.7919 | 6.7881 | 6.8071 | 6.2234 | 5.2663 | 5.4311 |
| GIT2_S421 | GIT2 | Q14161 | | 421 | SLDSDLSDGPVTV | 5.7307 | 5.6359 | 6.7875 | 6.1433 | 5.9948 | 6.9425 | 6.0309 | 6.5285 | 6.2964 | 6.1391 | 5.8894 | 7.0715 | 6.0739 | 6.2247 | 6.1664 | 6.3445 |
| GIT2_S514 | GIT2 | Q14161 | | 514 | RGSRPMSMYETGS | 8.0666 | 6.3693 | 7.7356 | 6.5982 | 6.3503 | 5.9632 | 5.9174 | 6.0480 | 6.5500 | 5.4384 | 6.5308 | 6.2169 | 5.8948 | 6.0971 | 5.0221 | 5.2012 |
| GIT2_S556 | GIT2 | Q14161 | | 415 | KTNRQKSLDSDLS | 6.9702 | 6.6542 | 6.9539 | 5.9955 | 6.9399 | 4.6161 | 5.9220 | 5.8112 | 7.3060 | 4.6665 | 6.9084 | 6.3452 | 5.8870 | 6.7825 | 4.9146 | 5.5267 |
| GIT2_S614 | GIT2 | Q14161 | | 556 | RSALVTSSSSLPS | 6.8391 | 6.1044 | 5.2626 | 5.5356 | 5.4565 | 5.8009 | 6.4029 | 7.2141 | 5.8902 | 5.7499 | 5.9463 | 6.1988 | 5.7601 | 8.0483 | 6.1019 | 7.6886 |
| GLCCI1_S105;S108 | GLCCI1 | Q86VQ1 | | 223 | SHQRSASWGSADQ | 6.5809 | 6.3895 | 7.1268 | 6.0816 | 6.3687 | 6.6224 | 5.8321 | 6.6458 | 6.5290 | 6.6066 | 6.8357 | 6.5927 | 5.9689 | 5.5391 | 5.2769 | 5.0031 |
| GLCCI1_S148 | GLCCI1 | Q86VQ1 | | 406 | LPKYASSPKPNNS | 5.8543 | 6.2071 | 6.7781 | 6.1350 | 6.9758 | 7.7461 | 6.2877 | 7.1368 | 6.4809 | 6.6662 | 7.4991 | 7.1137 | 5.4731 | 5.5538 | 4.1663 | 3.9260 |
| GLCCI1_S171 | GLCCI1 | Q86VQ1 | | 303 | CNVEGISPELEKV | 6.1494 | 5.8419 | 6.9839 | 5.9776 | 6.3842 | 6.6187 | 5.8772 | 6.7052 | 6.9231 | 6.4387 | 6.5079 | 6.7955 | 5.8952 | 5.9680 | 5.3330 | 5.6006 |
| GLCCI1_S223 | GLCCI1 | Q86VQ1 | | 30 | RRSAAGSPPAVAA | 6.2368 | 6.1198 | 5.8921 | 7.2677 | 5.8084 | 6.1531 | 6.3721 | 6.4269 | 6.8973 | 6.1542 | 6.3764 | 6.3131 | 5.6301 | 6.4817 | 6.0317 | 5.8385 |
| GLCCI1_S30 | GLCCI1 | Q86VQ1 | | 148 | ERRSPGSPVCRAD | 6.9493 | 6.4805 | 7.1589 | 5.2089 | 6.7212 | 7.4140 | 6.4328 | 6.7295 | 5.7711 | 6.0283 | 6.8083 | 6.7709 | 6.7543 | 5.0326 | 4.5928 | 5.1467 |
| GLCCI1_S303 | GLCCI1 | Q86VQ1 | | 171 | STIRRTSSLDTIT | 5.8591 | 5.9707 | 7.1822 | 6.0525 | 6.3261 | 6.4886 | 5.8463 | 5.9422 | 6.2643 | 6.6943 | 6.9195 | 6.4320 | 6.1313 | 5.9123 | 6.1128 | 5.8658 |
| GLCCI1_S394;S398 | GLCCI1 | Q86VQ1 | 105;108 | | NA | 6.2942 | 6.3259 | 6.6704 | 7.0110 | 7.1405 | 6.0689 | 6.9078 | 6.7976 | 6.9128 | 6.5421 | 6.1091 | 6.3500 | 5.2919 | 5.2181 | 4.6431 | 5.7166 |
| GLCCI1_S406 | GLCCI1 | Q86VQ1 | 394;398 | | NA | 5.6591 | 6.7742 | 7.2687 | 6.3539 | 5.9308 | 6.3324 | 5.9055 | 7.2158 | 6.9883 | 7.0857 | 7.2118 | 7.3864 | 5.1617 | 4.8182 | 4.7542 | 5.1532 |
| GLE1_S112 | GLE1 | Q53GS7 | | 112 | TKGKDESQHTESM | 7.5985 | 7.5945 | 5.6014 | 8.5931 | 6.1166 | 6.3851 | 6.3374 | 6.8347 | 7.6681 | 7.4613 | 9.1898 | 9.1142 | 2.1343 | 2.7371 | 3.1170 | 3.5168 |
| GLE1_S99 | GLE1 | Q53GS7 | | 99 | SAFSPASPATPNG | 6.1301 | 5.7255 | 5.9760 | 6.4890 | 6.2291 | 5.5981 | 6.6375 | 6.4830 | 6.2360 | 5.9876 | 6.2674 | 6.3384 | 6.2837 | 6.8639 | 6.3259 | 6.4288 |
| GLG1_S1201 | GLG1 | Q92896-2 | | 1201 | SQDVTGSPAxxxx | 7.0095 | 7.2601 | 6.7565 | 8.0669 | 5.5977 | 6.3257 | 7.7425 | 6.4916 | 6.8963 | 7.2143 | 6.3656 | 7.0327 | 3.8634 | 4.8986 | 3.8210 | 4.6574 |
| GLI4_S86 | GLI4 | P10075 | | 86 | PEQAPRSPGSQAP | 6.3066 | 5.9078 | 6.6936 | 6.8416 | 6.8433 | 6.1804 | 6.0742 | 6.3651 | 7.0147 | 6.2907 | 6.0951 | 6.2748 | 5.9923 | 6.2347 | 5.2583 | 5.6269 |
| GLIS3_S85 | GLIS3 | Q8NEA6-2 | | 85 | IHLPALSPRRQML | 6.3601 | 6.4850 | 6.1567 | 6.8652 | 6.0881 | 6.0732 | 6.7305 | 7.0310 | 6.9409 | 5.6969 | 6.0466 | 6.8950 | 5.2960 | 5.7707 | 5.5033 | 6.0609 |
| GLMN_S585;T586 | GLMN | Q92990 | 585;586 | | NA | 6.0700 | 6.8703 | 6.0032 | 6.4189 | 6.2243 | 5.9424 | 6.4118 | 6.0692 | 5.8569 | 6.2492 | 6.9732 | 6.0452 | 6.0129 | 6.0241 | 6.3185 | 6.5098 |
| GLUD1_T410 | GLUD1 | P00367 | | 410 | GANGPTTPEADKI | 5.2536 | 5.8408 | 5.5624 | 6.4892 | 6.6695 | 5.9293 | 6.5704 | 5.9190 | 6.1006 | 6.6259 | 5.8225 | 5.6618 | 7.1891 | 7.4566 | 6.4529 | 6.4563 |
| GLYR1_S132 | GLYR1 | Q49A26 | | 167 | KRAQEQSPRKRGR | 6.5723 | 6.7857 | 5.8061 | 6.5758 | 6.2340 | 6.9216 | 5.9859 | 5.6933 | 5.9136 | 6.0588 | 7.8156 | 6.5567 | 5.8928 | 6.0761 | 5.8801 | 5.2318 |
| GLYR1_S148 | GLYR1 | Q49A26 | | 132 | KRKLSLSEGKVKK | 6.1408 | 6.8992 | 7.6699 | 8.3142 | 6.5833 | 6.9490 | 7.3009 | 6.0012 | 7.8968 | 7.6643 | 7.7476 | 7.7121 | 2.9945 | 3.9902 | 3.0596 | 3.0765 |
| GLYR1_S167 | GLYR1 | Q49A26 | | 148 | EGKKRVSSGSSER | 6.6242 | 7.4941 | 5.8843 | 8.0338 | 6.9724 | 5.8968 | 7.0175 | 6.7977 | 6.8636 | 5.9012 | 6.9571 | 6.4642 | 4.6151 | 5.1771 | 4.3300 | 4.9709 |
| GMEB1_S129 | GMEB1 | Q9Y692 | | 129 | FNDQLISPKHFVH | 6.3546 | 8.1992 | 5.2294 | 6.7097 | 6.2670 | 7.6920 | 7.9087 | 6.7466 | 4.8651 | 6.3869 | 7.6043 | 7.8072 | 4.5054 | 4.8190 | 4.4132 | 4.4916 |
| GMIP_S19 | GMIP | Q9P107 | | 437 | ESRSLDSPTSSPG | 5.9837 | 6.3276 | 6.4216 | 6.4543 | 6.1355 | 6.3556 | 6.1428 | 6.2944 | 6.2317 | 6.5056 | 6.4667 | 6.1138 | 6.0158 | 6.4428 | 5.9048 | 6.2032 |
| GMIP_S437 | GMIP | Q9P107 | | 19 | EGRKRYSDIFRSL | 6.9153 | 7.1756 | 7.3276 | 7.3093 | 6.1924 | 6.9739 | 7.0151 | 6.5441 | 8.0190 | 7.5068 | 7.1961 | 7.3115 | 3.5545 | 4.2990 | 3.0735 | 3.5863 |
| GMIP_S437 | GMIP | Q9P107 | | 919 | PSPAAASPEGSPL | 6.2639 | 6.6551 | 6.6752 | 6.7349 | 6.4034 | 5.3493 | 5.8756 | 6.5278 | 6.4653 | 5.9159 | 6.6213 | 6.9262 | 5.7663 | 6.1122 | 5.6549 | 6.0525 |
| GMIP_S919 | GMIP | Q9P107-2 | | 437 | ESRSLDSPTSSPD | 5.9492 | 5.7746 | 6.1338 | 5.9081 | 5.9435 | 6.0934 | 6.5133 | 6.2021 | 7.0546 | 5.9372 | 5.5628 | 6.2116 | 7.0040 | 7.0882 | 6.2856 | 6.3381 |
| GMPS_S332 | GMPS | P49915 | | 332 | TLNMTTSPEEKRK | 6.1730 | 6.0808 | 6.6859 | 5.6793 | 5.8963 | 6.4776 | 6.0898 | 6.2974 | 6.1848 | 6.4957 | 6.4842 | 6.2796 | 5.9104 | 6.8502 | 6.4041 | 6.0110 |
| GMPS_T318 | GMPS | P49915 | | 318 | ISDEDRTPRKRIS | 6.0945 | 6.7398 | 6.9974 | 6.5927 | 7.0164 | 6.9662 | 6.7813 | 6.8174 | 7.2312 | 6.6703 | 6.4436 | 7.0636 | 4.3061 | 5.3976 | 4.3855 | 4.4965 |
| GNAI2_S253 | GNAI2 | P04899-4 | | 253 | SMKLFDSICNNKW | 5.3737 | 4.2138 | 7.7460 | 7.7034 | 6.5533 | 6.4681 | 6.7564 | 5.4294 | 6.4779 | 5.6096 | 4.8789 | 5.4327 | 6.3534 | 7.8476 | 6.5893 | 6.5664 |
| GNAI2_S282 | GNAI2 | P04899-4 | | 282 | EEKITHSPLTICF | 4.1578 | 5.4048 | 7.4030 | 5.8493 | 5.6998 | 6.1600 | 7.6548 | 6.8799 | 5.6841 | 5.9122 | 7.1159 | 6.7658 | 6.9782 | 5.3418 | 7.1238 | 5.8689 |
| GNAS_S995 | GNAS | Q5JWF2 | | 995 | LRISTASGDGRHY | 6.2850 | 6.4481 | 5.5988 | 6.0833 | 5.9306 | 5.9551 | 5.9595 | 6.1408 | 6.1754 | 6.3428 | 5.8364 | 6.1106 | 6.6810 | 6.9559 | 6.8743 | 6.6224 |
| GNG12_S26 | GNG12 | Q9UBI6 | | 49 | CEEHARSDPLLIG | 5.6480 | 5.5280 | 5.8926 | 5.6290 | 6.1276 | 6.4070 | 5.8703 | 6.2812 | 5.5756 | 6.4401 | 6.2355 | 6.6046 | 5.8233 | 7.1289 | 7.1201 | 7.6882 |
| GNG12_S49 | GNG12 | Q9UBI6 | | 26 | QLRLEASIERIKV | 6.0561 | 5.3675 | 5.1163 | 5.0069 | 5.7765 | 5.2362 | 5.5527 | 6.2283 | 4.9215 | 6.2144 | 5.5456 | 5.5871 | 5.2469 | 10.0691 | 8.2668 | 9.8080 |
| GNG12_S7 | GNG12 | Q9UBI6 | | 7 | MSSKTASTNNIAQ | 4.8799 | 4.4578 | 4.3793 | 4.2526 | 5.1178 | 4.5878 | 4.9153 | 5.1916 | 4.4550 | 4.4802 | 4.5864 | 4.8701 | 8.7716 | 11.9267 | 11.1522 | 11.9757 |
| GNL1_S324 | GNL1 | P36915 | | 324 | ATWGNGSGEEEEE | 6.0313 | 5.8263 | 6.5281 | 6.1693 | 5.9802 | 6.5227 | 5.9689 | 6.3497 | 6.5996 | 6.2895 | 6.0054 | 6.4832 | 6.0757 | 6.6630 | 5.9381 | 6.5691 |
| GNL3L_S465 | GNL3L | Q9NVN8 | | 465 | EIKLLHSPMTKIA | 7.4067 | 7.3240 | 6.5926 | 7.2602 | 7.0313 | 6.6663 | 7.2280 | 6.3917 | 7.3155 | 6.7938 | 6.8217 | 6.6903 | 3.6611 | 4.5064 | 3.9768 | 4.3336 |
| GOLGA1_S194 | GOLGA1 | Q92805 | | 194 | LLKKEESLGKMEQ | 5.8519 | 5.7164 | 6.0509 | 5.7183 | 6.0310 | 5.9953 | 5.8362 | 6.2204 | 6.7499 | 5.9529 | 6.9933 | 6.6894 | 6.9893 | 6.6533 | 6.0912 | 6.4603 |
| GOLGA3_S1391;T1396 | GOLGA3 | Q08378 | | 983 | KMRRLGSDLTSAQ | 2.3987 | 2.5517 | 3.1768 | 2.6971 | 2.7794 | 2.5877 | 2.9723 | 3.2180 | 2.8707 | 2.4334 | 2.5775 | 2.8038 | 18.3496 | 15.6538 | 17.4429 | 15.4866 |
| GOLGA3_S983 | GOLGA3 | Q08378 | 1391;1396 | | NA | 7.0717 | 7.2931 | 6.8276 | 6.3523 | 6.2138 | 7.1747 | 6.9097 | 5.8205 | 6.8920 | 6.4813 | 6.7605 | 6.7495 | 4.5384 | 5.5062 | 4.4991 | 4.9097 |
| GOLGA4_S2224 | GOLGA4 | Q13439-5 | | 93 | LQLRVPSVESLFR | 6.0092 | 6.3507 | 6.1553 | 6.3032 | 6.0760 | 5.7689 | 5.8884 | 6.1386 | 5.9917 | 6.0016 | 6.2174 | 6.1184 | 6.9847 | 6.6794 | 6.7732 | 6.5433 |
| GOLGA4_S288 | GOLGA4 | Q13439-5 | | 288 | KEENPESDGEPVV | 6.1705 | 5.9703 | 6.0736 | 6.7430 | 6.5237 | 6.5914 | 6.2956 | 6.1175 | 6.3162 | 6.5028 | 6.0969 | 5.8004 | 5.9700 | 6.5953 | 6.1940 | 6.0386 |
| GOLGA4_S40 | GOLGA4 | Q13439-5 | | 40 | RMRSRTSSFTEQL | 6.4994 | 6.2776 | 5.7543 | 6.1353 | 6.5739 | 6.0026 | 5.9345 | 5.7405 | 6.4738 | 6.1996 | 6.4678 | 6.5431 | 6.9404 | 6.1602 | 6.2663 | 6.0308 |
| GOLGA4_S63 | GOLGA4 | Q13439-5 | | 2169 | YATTVGTPYKGGN | 7.5793 | 7.7812 | 7.3809 | 8.1044 | 5.8196 | 7.2049 | 7.0207 | 6.3782 | 7.7674 | 6.6996 | 7.0761 | 7.1149 | 3.5659 | 4.4664 | 2.7046 | 3.3360 |
| GOLGA4_S93 | GOLGA4 | Q13439-5 | | 63 | STHASKSPDSVNG | 5.4154 | 6.5987 | 6.4953 | 6.8962 | 6.3365 | 7.0698 | 6.5798 | 6.6605 | 6.5756 | 6.9819 | 7.2572 | 7.1832 | 5.0842 | 5.1712 | 4.6871 | 5.0073 |
| GOLGA4_T2169 | GOLGA4 | Q13439-5 | | 1830 | EEKNKKYSLIVAQ | 7.8679 | 7.1113 | 7.8208 | 7.7679 | 7.4278 | 8.2633 | 7.4819 | 7.5478 | 5.5226 | 5.9615 | 5.4708 | 5.8450 | 3.7386 | 4.4247 | 3.8210 | 3.9269 |
| GOLGA4_Y1830 | GOLGA4 | Q13439 | | 2224 | ARLMFTSPRSGIF | 7.1534 | 7.7039 | 7.0734 | 6.9675 | 6.3554 | 7.0508 | 7.4813 | 6.8768 | 7.4233 | 7.6287 | 8.4468 | 8.1301 | 2.4866 | 3.5919 | 2.5306 | 3.0993 |
| GOLGA5_S116 | GOLGA5 | Q8TBA6 | | 116 | FVRRKKSEPDDEL | 5.1721 | 5.4506 | 5.4099 | 5.6434 | 6.0616 | 5.6868 | 5.8487 | 5.8502 | 6.4241 | 6.3675 | 6.3187 | 6.3141 | 7.4242 | 7.4822 | 7.4889 | 7.0568 |
| GOLGB1_S138 | GOLGB1 | Q14789-2 | | 17 | VVLHELSGDDDTD | 6.2400 | 6.4157 | 6.4593 | 7.2506 | 6.3806 | 6.5999 | 6.6453 | 7.0473 | 6.5532 | 6.3172 | 6.1232 | 6.0447 | 5.1883 | 5.6164 | 5.2935 | 5.8247 |
| GOLGB1_S17 | GOLGB1 | Q14789-2 | | 3042 | RTQLNDSLKEIHQ | 5.9595 | 6.2544 | 6.2072 | 6.7479 | 6.5881 | 7.2942 | 7.3665 | 6.8623 | 6.8323 | 6.5065 | 7.0082 | 6.6907 | 4.8896 | 5.6076 | 4.0290 | 5.1560 |
| GOLGB1_S3015 | GOLGB1 | Q14789-2 | | 565 | QKHKELSVLLLEM | 2.6328 | 2.2431 | 4.0110 | 4.4720 | 4.1821 | 3.9587 | 1.8487 | 2.4712 | 2.5045 | 3.2079 | 3.4930 | 2.5758 | 13.5586 | 16.3340 | 17.6647 | 14.8417 |
| GOLGB1_S3042 | GOLGB1 | Q14789-2 | | 3015 | QYQRQASPETSAS | 5.7230 | 5.6635 | 5.8061 | 5.6328 | 6.4922 | 5.7558 | 6.4943 | 5.8933 | 5.9579 | 5.7958 | 5.9343 | 5.3435 | 7.7568 | 7.2840 | 7.9548 | 6.5117 |
| GOLGB1_S565 | GOLGB1 | Q14789-2 | | 138 | LSKHDKSSTEEEM | 6.7503 | 7.0363 | 6.5935 | 6.5804 | 6.7282 | 6.4903 | 7.4412 | 6.5404 | 6.6064 | 6.6329 | 6.9949 | 7.2620 | 4.4291 | 5.1287 | 3.8553 | 4.9300 |
| GOLIM4_Y506 | GOLIM4 | O00461 | | 506 | QGEEGAYERDNQH | 5.3129 | 6.4201 | 6.3806 | 6.3458 | 6.1810 | 6.0203 | 5.4435 | 6.1494 | 6.5039 | 6.0098 | 6.3053 | 6.0914 | 6.8636 | 6.7904 | 6.3321 | 6.8499 |
| GOLM2_Y389 | GOLM2 | Q6P4E1 | | 389 | DGNVGEYEADKQA | 6.0125 | 5.8824 | 6.2943 | 6.6727 | 6.6423 | 5.8939 | 5.9251 | 6.1022 | 5.9712 | 6.6081 | 6.6076 | 6.8357 | 6.0398 | 6.3329 | 6.0081 | 6.1712 |
| GOLPH3_S9 | GOLPH3 | Q9H4A6 | | 9 | SLTQRSSGLVQRR | 5.4075 | 5.4218 | 5.0219 | 5.4346 | 5.7176 | 4.6480 | 5.2827 | 5.0905 | 5.6706 | 5.2852 | 5.0166 | 5.5361 | 9.7277 | 9.3740 | 8.6236 | 8.7416 |
| GON4L_S1255 | GON4L | Q3T8J9 | | 346 | IPTWNISPIKKAN | 6.5691 | 6.6304 | 6.3952 | 7.1562 | 6.2675 | 6.2181 | 6.5173 | 6.4991 | 6.2506 | 6.3236 | 6.3343 | 6.2669 | 5.4305 | 5.7775 | 5.6404 | 5.7233 |
| GON4L_S1268 | GON4L | Q3T8J9 | | 1255 | LEPQELSPLSATV | 6.4828 | 6.5484 | 6.1957 | 6.8995 | 6.2301 | 6.4060 | 6.6461 | 6.3318 | 6.5223 | 6.3216 | 6.5220 | 6.4251 | 5.6244 | 5.9634 | 5.4395 | 5.4413 |
| GON4L_S1339 | GON4L | Q3T8J9 | | 1977 | DGPPPHSPETPQF | 6.2231 | 6.5306 | 6.7087 | 7.0446 | 6.0813 | 6.3312 | 6.5756 | 6.3855 | 6.1177 | 6.3719 | 6.4297 | 6.8133 | 5.8167 | 5.6682 | 5.2545 | 5.6474 |
| GON4L_S1426 | GON4L | Q3T8J9 | | 1339 | REEISGSPERDIC | 7.0583 | 6.4565 | 6.3034 | 6.5593 | 6.2277 | 5.3054 | 6.4576 | 6.2207 | 6.4055 | 6.0611 | 6.6022 | 6.5802 | 6.3672 | 6.1268 | 5.7027 | 5.5655 |
| GON4L_S1977 | GON4L | Q3T8J9 | | 1426 | PEVRLSSPPGKPE | 6.0068 | 6.2704 | 6.1088 | 7.0642 | 5.9617 | 6.3792 | 6.4948 | 6.2400 | 6.3667 | 6.9560 | 6.3739 | 6.8112 | 5.6604 | 5.6687 | 5.4355 | 6.2018 |
| GON4L_S2053 | GON4L | Q3T8J9 | | 1268 | FPKVEHSPGPPLA | 6.5652 | 5.5528 | 6.4603 | 6.7708 | 6.1449 | 5.1186 | 6.6960 | 6.7798 | 5.6906 | 6.2776 | 6.3395 | 6.0320 | 5.8454 | 6.4074 | 6.8667 | 6.4524 |
| GON4L_S206 | GON4L | Q3T8J9 | | 206 | QPDVCASPQEKPL | 6.4506 | 6.5603 | 6.4922 | 6.9597 | 6.4145 | 6.1906 | 6.3608 | 5.8978 | 6.5129 | 5.8672 | 6.2033 | 6.4719 | 5.9793 | 6.0354 | 5.8001 | 5.8032 |
| GON4L_S346 | GON4L | Q3T8J9 | | 2053 | PPASFLSPVSSKT | 6.9590 | 7.2351 | 6.8535 | 7.9859 | 6.5649 | 6.9787 | 7.8317 | 6.2523 | 6.3573 | 7.1926 | 7.4670 | 7.5893 | 3.7530 | 4.1190 | 3.1539 | 3.7068 |
| GON4L_T433 | GON4L | Q3T8J9 | | 433 | EAEKVTTPAIRHI | 7.8178 | 8.5335 | 6.7179 | 7.1246 | 6.5253 | 7.1482 | 6.8930 | 7.5884 | 7.8178 | 6.7854 | 6.7695 | 6.7553 | 2.8090 | 4.2185 | 3.1531 | 3.3425 |
| GOPC_S376 | GOPC | Q9HD26 | | 376 | VAPEVDSDDENVE | 6.6956 | 6.2507 | 6.7780 | 6.3700 | 6.0435 | 6.0419 | 6.2452 | 6.4777 | 6.2927 | 6.2876 | 6.8336 | 6.1618 | 6.6465 | 6.0050 | 5.2837 | 5.5865 |
| GORAB_S335 | GORAB | Q5T7V8 | | 335 | CQEQAVSPKVDDQ | 5.7733 | 5.8657 | 5.4035 | 5.7959 | 6.1053 | 6.1521 | 6.0582 | 6.0546 | 5.5830 | 5.7808 | 6.1838 | 5.6772 | 7.6649 | 7.3655 | 7.0029 | 7.5332 |
| GORASP1_T216 | GORASP1 | Q9BQQ3 | | 216 | HKKPPGTPPPSAL | 7.1594 | 5.9717 | 5.3763 | 7.1292 | 6.2464 | 6.5560 | 6.4798 | 5.4779 | 6.5576 | 6.5322 | 5.0730 | 6.2004 | 5.3588 | 6.0622 | 7.1515 | 6.6672 |
| GORASP1_T216;S241 | GORASP1 | Q9BQQ3 | 216;241 | | NA | 6.6730 | 5.7484 | 5.8614 | 7.2797 | 5.5199 | 5.9894 | 7.2629 | 6.7601 | 5.1434 | 5.6275 | 7.2335 | 6.8476 | 5.9660 | 6.1944 | 5.7328 | 6.1603 |
| GORASP1_T216;T237;S2 | GORASP1 | Q9BQQ3 | 216;237;241 | | NA | 5.5167 | 6.2195 | 6.5068 | 6.7229 | 7.1006 | 7.5254 | 6.2458 | 6.9996 | 6.7288 | 5.7695 | 6.9437 | 5.4638 | 5.8864 | 5.5618 | 5.5226 | 5.2861 |
| GORASP2_S463 | GORASP2 | Q9H8Y8-3 | | 237 | MAGTPITPLKDGF | 6.3211 | 6.2973 | 6.1912 | 6.3003 | 6.6988 | 6.8136 | 7.1073 | 7.3970 | 7.2131 | 7.2875 | 7.1012 | 7.5986 | 3.7897 | 4.9438 | 4.1215 | 4.8179 |
| GORASP2_T234 | GORASP2 | Q9H8Y8-3 | | 463 | DANASESPxxxxx | 6.0072 | 5.7032 | 6.5457 | 5.8847 | 5.9744 | 6.5764 | 6.2846 | 6.4569 | 6.9192 | 6.9198 | 6.8161 | 7.2912 | 5.5559 | 5.7976 | 5.2141 | 6.0531 |
| GORASP2_T237 | GORASP2 | Q9H8Y8-3 | | 445 | DRVGDSTPVSEKP | 5.3535 | 5.8306 | 6.0015 | 5.6649 | 5.4369 | 5.8124 | 6.0910 | 5.8257 | 5.8742 | 6.3271 | 6.1974 | 6.4467 | 6.6984 | 7.5591 | 7.1963 | 7.6843 |
| GORASP2_T427 | GORASP2 | Q9H8Y8-3 | | 234 | PGQMAGTPITPLK | 6.5481 | 5.7069 | 5.9215 | 6.2061 | 6.3144 | 6.6239 | 6.9818 | 7.3747 | 6.0654 | 6.7588 | 6.5266 | 6.4277 | 5.3843 | 5.9262 | 5.5264 | 5.7073 |
| GORASP2_T445 | GORASP2 | Q9H8Y8-3 | | 427 | SLTVDVTPPTAKA | 5.7926 | 5.6031 | 5.8362 | 5.9373 | 5.9238 | 6.5397 | 6.5296 | 7.0842 | 6.2928 | 7.0177 | 6.9637 | 7.3797 | 5.1452 | 6.2366 | 5.8082 | 5.9094 |
| GOSR1_S49 | GOSR1 | E9PCW1 | | 49 | RDGRRDSSDTTPL | 6.1369 | 5.7184 | 5.8383 | 6.1983 | 6.5805 | 5.5581 | 5.4526 | 5.9820 | 6.5473 | 6.6101 | 6.8289 | 7.2016 | 6.4210 | 6.2382 | 6.6927 | 5.9950 |
| GOSR2_S114 | GOSR2 | O14653-2 | | 114 | TFTTNDSDTTIPM | 6.5393 | 6.5779 | 6.4690 | 7.3043 | 6.1386 | 6.2763 | 7.2298 | 6.3245 | 6.3960 | 6.5909 | 6.9974 | 7.1708 | 4.6284 | 4.9213 | 5.0813 | 5.3543 |
| GOT2_S360 | GOT2 | P00505 | | 360 | MRTQLVSNLKKEG | 5.8010 | 5.5734 | 7.0865 | 6.1732 | 7.1589 | 5.9775 | 7.5779 | 5.5507 | 5.5386 | 5.4762 | 6.5881 | 5.4287 | 6.9518 | 5.6280 | 7.0786 | 6.7809 |
| GPALPP1_S105 | GPALPP1 | Q8IXQ4 | | 105 | FKKQDDSPPRPII | 5.6376 | 5.8412 | 6.0522 | 5.8897 | 5.9190 | 6.0853 | 5.8710 | 6.0567 | 6.1135 | 6.3085 | 6.4671 | 6.4138 | 6.9402 | 7.0243 | 6.7789 | 6.6011 |
| GPAM_S688;S695 | GPAM | Q9HCL2 | 688;695 | | NA | 6.6701 | 7.2409 | 6.8276 | 7.6795 | 5.8092 | 5.5023 | 5.9176 | 5.3544 | 6.8773 | 6.4252 | 5.9619 | 6.0758 | 6.0507 | 6.3390 | 5.4082 | 5.8603 |
| GPAT2_T666;S668 | GPAT2 | Q6NUI2-4 | | 668 | SGDFTDSDSDDFG | 7.2488 | 8.1219 | 7.4794 | 6.4470 | 8.6766 | 8.5118 | 9.8133 | 4.7831 | 8.0450 | 6.7326 | 11.0324 | 4.9643 | 1.3289 | 2.8775 | 0.0000 | 3.9374 |
| GPAT2_T668 | GPAT2 | Q6NUI2-4 | 666;668 | | NA | 9.6134 | 8.8585 | 8.9086 | 9.4198 | 6.4162 | 5.9023 | 7.3377 | 7.6028 | 4.8890 | 7.6931 | 6.5845 | 7.1315 | 2.7293 | 3.2765 | 1.7598 | 1.8769 |
| GPAT3_S68 | GPAT3 | Q53EU6 | | 68 | ILKNSASVGIIQR | 1.3933 | 1.3146 | 1.3033 | 1.4919 | 3.5339 | 3.2013 | 3.5971 | 3.2877 | 1.4792 | 1.4138 | 1.2114 | 1.4280 | 19.5781 | 16.9597 | 19.8556 | 18.9512 |
| GPATCH1_S200;Y210 | GPATCH1 | Q9BRR8 | | 715 | PPKQQSSPLVNKE | 6.1935 | 6.3966 | 5.9713 | 6.1737 | 6.0776 | 6.2061 | 6.3777 | 6.4419 | 6.5903 | 6.6866 | 6.4789 | 6.3984 | 5.5260 | 6.4530 | 5.8360 | 6.1923 |
| GPATCH1_S715 | GPATCH1 | Q9BRR8 | | 210 | EGEDDDYLPDNVT | 4.9314 | 5.2613 | 5.3678 | 5.4130 | 5.7019 | 5.7099 | 5.6454 | 5.0627 | 6.9746 | 6.4017 | 6.6370 | 6.0908 | 7.3700 | 7.9572 | 7.5397 | 7.9356 |
| GPATCH1_Y210 | GPATCH1 | Q9BRR8 | 200;210 | | NA | 5.5729 | 6.0151 | 5.9921 | 6.5379 | 5.8896 | 6.0700 | 5.7735 | 5.6534 | 6.3862 | 6.1180 | 5.8048 | 6.0990 | 6.9008 | 7.2527 | 6.9112 | 7.0227 |
| GPATCH11_S141 | GPATCH11 | Q8N954 | | 141 | GIGHEASLKRKAE | 6.3417 | 6.7862 | 7.8715 | 7.5119 | 6.6821 | 5.8868 | 5.6129 | 6.3408 | 7.7971 | 8.7309 | 8.0143 | 8.6725 | 3.4083 | 4.4915 | 2.5738 | 3.2777 |
| GPATCH2_S115;S117 | GPATCH2 | Q9NW75 | | 195 | RDQDMDSDRAYQY | 6.1715 | 6.4775 | 8.1162 | 6.9144 | 7.0214 | 8.0242 | 7.3836 | 6.3601 | 5.2694 | 5.4268 | 4.8175 | 6.1136 | 5.6553 | 6.3867 | 4.8812 | 4.9807 |
| GPATCH2_S195 | GPATCH2 | Q9NW75 | | 284 | QGDDEQSDWFYEK | 7.4489 | 7.6427 | 8.5202 | 7.7280 | 8.5564 | 7.1309 | 7.6789 | 6.9901 | 4.9806 | 4.2233 | 4.5127 | 4.2155 | 4.6614 | 5.6202 | 5.3424 | 4.7479 |
| GPATCH2_S284 | GPATCH2 | Q9NW75 | 115;117 | | NA | 5.7107 | 6.3769 | 6.8512 | 6.4099 | 6.8696 | 6.6575 | 6.0542 | 6.6029 | 5.8521 | 6.6110 | 6.0808 | 6.4925 | 5.9120 | 5.9952 | 5.6616 | 5.8619 |
| GPATCH2L_S137 | GPATCH2L | Q9NWQ4 | | 31 | WEEMALSPRQQRR | 6.4818 | 7.0154 | 6.9351 | 6.7934 | 6.8208 | 6.7344 | 6.7150 | 6.6220 | 6.3022 | 6.6548 | 6.2993 | 6.3942 | 4.8130 | 5.5987 | 4.7049 | 5.1150 |
| GPATCH2L_S248 | GPATCH2L | Q9NWQ4 | | 248 | QGDDEQSDWFYEG | 7.2631 | 8.0559 | 9.2872 | 8.1481 | 7.2988 | 6.7455 | 7.6801 | 6.1419 | 4.4130 | 4.9843 | 4.7510 | 5.5789 | 4.9306 | 5.6051 | 3.8542 | 5.2623 |
| GPATCH2L_S31 | GPATCH2L | Q9NWQ4 | | 447 | ASQAPKSPSSEWL | 6.1918 | 6.4048 | 6.8640 | 6.3617 | 6.4713 | 6.7464 | 6.8400 | 7.1275 | 6.0021 | 5.8394 | 6.2622 | 6.0769 | 5.2881 | 6.0415 | 5.6990 | 5.7833 |
| GPATCH2L_S447 | GPATCH2L | Q9NWQ4 | | 137 | KVKRVTSEVAASL | 7.3345 | 6.5887 | 6.8333 | 5.3437 | 5.9709 | 6.4902 | 5.8234 | 6.1955 | 7.7323 | 7.3907 | 6.6269 | 6.9424 | 5.5165 | 6.3634 | 4.0616 | 4.7859 |
| GPATCH4_S135 | GPATCH4 | Q5T3I0-3 | | 135 | KDLESCSDDDNQG | 5.7698 | 5.8132 | 5.6780 | 7.0104 | 6.2779 | 5.7606 | 5.9556 | 5.4012 | 6.6658 | 6.6515 | 6.2854 | 6.5555 | 6.3570 | 6.4868 | 6.1338 | 7.1974 |
| GPATCH8_S1009;S1014 | GPATCH8 | Q9UKJ3 | | 740 | EPPGSGSPAPPRR | 5.7787 | 5.9230 | 5.8554 | 6.1148 | 5.9577 | 5.6667 | 5.7198 | 5.7695 | 5.7022 | 5.6468 | 5.6315 | 5.5122 | 6.9650 | 8.2955 | 7.7865 | 7.6746 |
| GPATCH8_S1033;S1035 | GPATCH8 | Q9UKJ3 | | 758 | DDSQRRSLPAEEG | 6.3553 | 6.8485 | 5.9469 | 5.6036 | 6.1358 | 5.8797 | 6.4314 | 6.2243 | 6.2444 | 5.9928 | 5.9501 | 6.0490 | 6.0059 | 6.9425 | 6.6886 | 6.7012 |
| GPATCH8_S1035 | GPATCH8 | Q9UKJ3 | | 1035 | KIYRSQSPHYFRS | 6.1002 | 6.8623 | 5.3135 | 6.4862 | 6.4346 | 6.2248 | 5.6175 | 6.5513 | 5.7355 | 5.5340 | 5.6978 | 5.4575 | 6.3988 | 7.6625 | 7.1638 | 6.7595 |
| GPATCH8_S1081 | GPATCH8 | Q9UKJ3 | | 1107 | KVERKPSVSEEVQ | 4.5193 | 5.0579 | 5.4513 | 5.8075 | 6.5058 | 5.4529 | 5.8100 | 6.3550 | 5.6471 | 5.0337 | 5.0989 | 4.8843 | 8.1323 | 8.9728 | 8.6891 | 8.5820 |
| GPATCH8_S1107 | GPATCH8 | Q9UKJ3 | | 439 | ESKKGSSPKPKSC | 5.8789 | 6.8152 | 5.5576 | 6.4538 | 7.0966 | 6.6839 | 6.1968 | 6.1534 | 6.0740 | 6.1276 | 6.7850 | 5.5888 | 6.7334 | 7.0004 | 5.7944 | 5.0601 |

| | | | | | | | | | | | | | | | | | | | | | |
|---|---|---|---|---|---|---|---|---|---|---|---|---|---|---|---|---|---|---|---|---|---|
| GPATCH8_S1107;T1115 | GPATCH8 | Q9UKJ3 | | 1175 | GEEQEQSETEEGP | 5.3043 | 6.0237 | 6.1806 | 6.2273 | 6.2859 | 6.2260 | 5.7626 | 6.3742 | 6.5154 | 5.8786 | 6.2040 | 6.1561 | 5.4171 | 6.9650 | 6.8685 | 7.6106 |
| GPATCH8_S1175 | GPATCH8 | Q9UKJ3 | | 1081 | GSGDCSPEDKNS | 5.8622 | 7.2374 | 6.8396 | 7.1638 | 5.5323 | 4.8489 | 6.2272 | 6.1509 | 7.2166 | 5.8339 | 6.4733 | 6.2973 | 5.1726 | 6.5966 | 5.7855 | 7.7616 |
| GPATCH8_S439 | GPATCH8 | Q9UKJ3 | 1009;1014 | | NA | 6.1503 | 6.5041 | 5.6944 | 5.8562 | 6.1077 | 6.5506 | 5.7334 | 5.9496 | 5.9067 | 5.5316 | 5.5350 | 5.6898 | 6.8890 | 7.8887 | 7.4177 | 7.4954 |
| GPATCH8_S738;S740 | GPATCH8 | Q9UKJ3 | 1107;1115 | | NA | 7.0671 | 6.5356 | 6.5651 | 7.2788 | 6.9062 | 6.5106 | 7.0383 | 6.2839 | 7.1306 | 5.8252 | 6.2176 | 6.0246 | 5.0922 | 5.7377 | 4.3605 | 5.4259 |
| GPATCH8_S740 | GPATCH8 | Q9UKJ3 | 738;740 | | NA | 5.5181 | 6.7906 | 6.4574 | 7.8306 | 5.5941 | 6.4325 | 6.3214 | 6.2355 | 6.9044 | 6.4791 | 6.8408 | 6.6935 | 5.1407 | 6.1886 | 4.6688 | 5.5036 |
| GPATCH8_S758 | GPATCH8 | Q9UKJ3 | 1033;1035 | | NA | 6.8500 | 7.2562 | 7.1292 | 7.2515 | 6.3676 | 6.3983 | 6.2886 | 5.7293 | 6.0969 | 6.3262 | 6.5953 | 6.1052 | 5.0353 | 5.7892 | 5.4391 | 5.3419 |
| GPBP1L1_T273 | GPBP1L1 | Q9HC44 | | 273 | SRESAFTSPISVT | 6.1105 | 6.1681 | 5.6331 | 6.3506 | 6.2284 | 6.7594 | 6.5969 | 6.8387 | 6.2587 | 6.4126 | 6.3851 | 6.6623 | 5.8867 | 6.2571 | 5.4068 | 6.0449 |
| GPHN_S188;S194;S200 | GPHN | Q9NQX3-2 | | 338 | ILRASHSAVDITK | 3.4098 | 3.4747 | 3.4676 | 3.2973 | 4.8713 | 4.7630 | 4.3518 | 4.5858 | 4.3020 | 2.8382 | 3.4694 | 3.8652 | 14.2088 | 12.0893 | 14.0307 | 12.9750 |
| GPHN_S194;S200 | GPHN | Q9NQX3-2 | | 319 | STASCPTPKVQSR | 8.5122 | 8.2425 | 7.3994 | 7.4346 | 6.2558 | 7.2269 | 7.3827 | 6.2148 | 8.0523 | 6.5509 | 7.5462 | 6.8112 | 3.1246 | 3.5598 | 2.4539 | 3.2324 |
| GPHN_S303 | GPHN | Q9NQX3-2 | | 303 | STTPSESPRAQAT | 7.6695 | 7.6962 | 6.8897 | 6.6383 | 6.1946 | 7.3424 | 7.4534 | 7.1315 | 7.5757 | 7.1996 | 8.1875 | 7.4567 | 2.8576 | 3.4037 | 2.8366 | 3.4670 |
| GPHN_S338 | GPHN | Q9NQX3-2 | 188;194;200 | | NA | 6.2394 | 5.9959 | 6.6195 | 6.3337 | 6.8702 | 6.2843 | 6.2371 | 6.7506 | 6.0428 | 6.7293 | 6.8076 | 6.7976 | 5.6648 | 5.9487 | 5.3436 | 5.3348 |
| GPHN_T319 | GPHN | Q9NQX3-2 | 194;200 | | NA | 6.3537 | 6.6155 | 7.2383 | 6.3915 | 6.5092 | 6.6786 | 5.5508 | 7.1721 | 6.7742 | 6.1608 | 7.0368 | 6.3538 | 5.1570 | 5.2255 | 5.3418 | 5.4406 |
| GPI_S146 | GPI | P06744-2 | | 148 | LRNRSNTPILVDG | 6.3584 | 6.1587 | 6.7892 | 5.4974 | 6.3789 | 6.4856 | 6.0536 | 6.3651 | 6.6068 | 6.5613 | 6.5487 | 6.5024 | 6.0625 | 6.3634 | 5.7964 | 5.4715 |
| GPI_T148 | GPI | P06744-2 | | 146 | VALRNRSNTPILV | 5.1905 | 5.3375 | 7.9449 | 6.4079 | 7.0315 | 5.4845 | 6.6162 | 6.4207 | 6.0625 | 6.9916 | 5.7985 | 7.1685 | 5.8085 | 6.2566 | 4.7524 | 6.7276 |
| GPKOW_T216 | GPKOW | Q92917 | | 473 | YMGPSDTDDDxxx | 5.6552 | 6.1243 | 6.4649 | 6.0603 | 5.9383 | 6.3648 | 5.7046 | 6.2253 | 6.4758 | 6.1136 | 6.0368 | 6.4709 | 6.6705 | 6.6751 | 6.4676 | 6.5519 |
| GPKOW_T473 | GPKOW | Q92917 | | 216 | TEAQALTPTGPSR | 6.4515 | 7.3021 | 6.9778 | 6.4060 | 6.5984 | 6.2843 | 7.6298 | 6.7357 | 7.0696 | 6.6344 | 7.1750 | 7.8535 | 3.7218 | 5.0306 | 4.0529 | 4.0767 |
| GPN1_S328 | GPN1 | Q9HCN4-5 | | 328 | TAKDSLSPVLHPS | 5.9158 | 7.4344 | 6.3054 | 7.2722 | 7.0511 | 7.1338 | 6.6473 | 7.6061 | 6.2879 | 5.8165 | 5.9866 | 5.9627 | 4.8266 | 5.3514 | 5.2735 | 5.1288 |
| GPR161_S380 | GPR161 | Q8N6U8-6 | | 380 | RTSRLFSISNRIT | 6.9119 | 6.2257 | 6.2611 | 6.4032 | 6.4730 | 5.8186 | 5.4843 | 5.8188 | 7.1506 | 6.6871 | 6.7186 | 6.9666 | 5.9313 | 6.9026 | 4.8113 | 5.4352 |
| GPR161_S450 | GPR161 | Q8N6U8-6 | | 450 | PPKRRSSVTFEDE | 6.3582 | 6.3134 | 6.2273 | 6.4274 | 6.1649 | 5.9237 | 6.1669 | 6.1463 | 6.2795 | 7.0619 | 6.1176 | 7.0636 | 5.5950 | 6.2527 | 6.0122 | 5.8894 |
| GPR176_S367 | GPR176 | Q14439 | | 367 | PSIRSGSQLLEMF | 5.0046 | 5.2141 | 5.5317 | 5.5965 | 5.2542 | 5.3172 | 5.3474 | 5.2760 | 5.4192 | 5.1392 | 5.5930 | 5.2463 | 8.8679 | 8.7296 | 9.5322 | 8.9309 |
| GPR176_S397 | GPR176 | Q14439 | | 397 | EAKYIGSADFQAK | 5.9242 | 6.1393 | 6.1921 | 6.3160 | 5.5307 | 6.2084 | 6.6238 | 6.0863 | 6.6141 | 6.2448 | 6.6564 | 6.8150 | 5.8964 | 6.0934 | 5.9621 | 6.6970 |
| GPR39_S384 | GPR39 | O43194 | | 396 | RPLLFASRRQSSA | 8.4208 | 6.7613 | 7.2582 | 6.2668 | 5.5166 | 6.5375 | 6.5603 | 5.8185 | 6.7104 | 6.1996 | 6.4242 | 6.4370 | 5.0654 | 5.6565 | 4.1490 | 5.3179 |
| GPR39_S396 | GPR39 | O43194 | | 384 | AHSTTDSARFVQR | 6.3487 | 7.5371 | 6.5383 | 6.2818 | 5.2337 | 7.5728 | 6.2260 | 7.9156 | 5.9470 | 6.9343 | 6.5288 | 6.7789 | 4.1324 | 6.0941 | 4.7351 | 5.1953 |
| GPR39_S425 | GPR39 | O43194 | | 425 | PQSKSQSLSLESL | 6.3919 | 6.2731 | 6.4522 | 6.6152 | 6.4882 | 6.2216 | 5.9817 | 5.8197 | 7.0555 | 6.6731 | 6.6570 | 6.5388 | 5.2397 | 6.3237 | 5.2037 | 6.0649 |
| GPR39_S427 | GPR39 | O43194 | | 427 | SKSQSLSLESLEP | 7.0090 | 6.5638 | 6.6385 | 6.0915 | 6.9522 | 6.3052 | 6.0431 | 6.3009 | 6.3679 | 6.4576 | 6.4906 | 6.5309 | 7.5085 | 6.7767 | 6.0923 | 6.0711 |
| GPR39_S427;S430 | GPR39 | O43194 | 427;430 | | NA | 6.1875 | 6.6948 | 6.0810 | 5.4509 | 5.9951 | 6.5884 | 5.8859 | 6.7964 | 7.0561 | 6.3909 | 6.2915 | 5.4642 | 5.9114 | 6.0662 | 6.1517 | 6.9880 |
| GPRASP2_S512 | GPRASP2 | Q96D09 | | 512 | VGFRSTSPFGIPE | 6.1458 | 6.3440 | 6.6249 | 6.2791 | 6.4309 | 6.7304 | 6.1404 | 6.3786 | 6.3748 | 6.4368 | 6.3054 | 6.7577 | 5.5839 | 6.2484 | 5.4199 | 5.7992 |
| GPRASP2_S534 | GPRASP2 | Q96D09 | | 534 | PKNLELSPEGEEQ | 5.8320 | 5.5426 | 6.8868 | 6.3659 | 6.8627 | 4.9798 | 7.1956 | 7.3081 | 7.3708 | 5.9751 | 7.2277 | 6.7373 | 5.1117 | 5.2885 | 5.7967 | 5.5185 |
| GPRC5A_S301 | GPRC5A | Q8NFJ5 | | 345 | RAHAWPSPYKDYE | 6.4426 | 6.6742 | 6.6684 | 6.5779 | 6.2457 | 6.9156 | 6.9242 | 6.5795 | 6.7626 | 6.0412 | 6.4829 | 6.2750 | 4.8611 | 5.6712 | 5.3253 | 5.5525 |
| GPRC5A_S345 | GPRC5A | Q8NFJ5 | | 301 | VENRAYSQEEITQ | 4.7777 | 4.8060 | 4.4980 | 5.6561 | 4.6475 | 5.9119 | 4.8481 | 4.9018 | 4.8035 | 5.0310 | 5.2154 | 5.4367 | 9.2012 | 9.1521 | 10.5680 | 10.5452 |
| GPRC5A_S59 | GPRC5A | Q8NFJ5 | | 59 | VCKVQDSNRRKML | 6.4011 | 6.3644 | 5.8568 | 6.3668 | 6.2508 | 5.7708 | 6.3530 | 6.2972 | 6.4427 | 6.1630 | 5.8085 | 5.9061 | 6.2887 | 6.6126 | 6.6759 | 6.4415 |
| GPRIN1_S100;S104 | GPRIN1 | Q7ZZK8 | | 615 | LPLEKGSPVTTTK | 6.6700 | 6.7925 | 6.0537 | 6.3733 | 6.7205 | 6.6460 | 6.4973 | 6.5660 | 6.6226 | 6.6181 | 6.4135 | 6.1808 | 5.4593 | 5.6770 | 5.3336 | 5.3760 |
| GPRIN1_S104 | GPRIN1 | Q7ZZK8 | | 895 | VRVRPGSALAAAV | 6.4943 | 5.9585 | 6.3073 | 6.7453 | 6.7808 | 6.1883 | 6.4259 | 6.3769 | 6.3142 | 5.5104 | 5.6936 | 5.6804 | 6.6142 | 5.6870 | 6.2222 | 6.1008 |
| GPRIN1_S286 | GPRIN1 | Q7ZZK8 | | 452 | GKAGTVSPGKEDP | 6.9521 | 7.4234 | 6.8115 | 7.0947 | 6.1375 | 6.2349 | 6.5506 | 6.6593 | 6.8005 | 6.9491 | 6.8042 | 6.9963 | 4.3793 | 4.7769 | 4.4130 | 5.0169 |
| GPRIN1_S452 | GPRIN1 | Q7ZZK8 | | 877 | VATGPMTPQAAAP | 5.6155 | 5.7138 | 5.8151 | 5.7660 | 6.0382 | 6.5327 | 6.4317 | 6.6130 | 5.9935 | 5.7645 | 5.5839 | 6.1799 | 6.7658 | 6.8330 | 7.2739 | 7.0795 |
| GPRIN1_S615 | GPRIN1 | Q7ZZK8 | | 704 | PSRKTESPSLGKV | 6.6009 | 6.6631 | 6.1649 | 6.4955 | 6.5380 | 6.3221 | 6.8072 | 6.5771 | 6.7094 | 6.8178 | 6.6564 | 6.8534 | 5.3115 | 5.4216 | 4.8816 | 5.1796 |
| GPRIN1_S704 | GPRIN1 | Q7ZZK8 | | 95 | GSLACPSPTCFSP | 6.2885 | 6.2089 | 5.9734 | 5.7419 | 6.1412 | 6.3185 | 6.0695 | 6.2809 | 5.5490 | 5.7580 | 5.7194 | 5.9100 | 6.9343 | 6.8042 | 7.3547 | 6.9477 |
| GPRIN1_S799 | GPRIN1 | Q7ZZK8 | | 578 | IGKVVSTPGKTVP | 7.5402 | 7.8214 | 6.6683 | 7.6643 | 6.4176 | 7.1262 | 7.2150 | 6.9661 | 7.0130 | 7.3276 | 7.2574 | 7.2453 | 3.0646 | 4.0835 | 2.8765 | 3.7132 |
| GPRIN1_S850 | GPRIN1 | Q7ZZK8 | | 799 | NFTKAPSWEASAP | 4.4529 | 4.3457 | 4.2354 | 4.4371 | 5.1537 | 4.8313 | 5.0064 | 5.1691 | 4.9852 | 4.3812 | 4.1480 | 4.3142 | 11.5691 | 10.7076 | 11.5224 | 10.7408 |
| GPRIN1_S895 | GPRIN1 | Q7ZZK8 | | 850 | AAPRAPSPPSRRD | 6.6887 | 7.2598 | 6.5925 | 6.6895 | 6.5082 | 7.0381 | 6.1679 | 5.5281 | 5.7567 | 6.5156 | 6.9638 | 6.4409 | 5.9029 | 5.7225 | 5.2005 | 5.0244 |
| GPRIN1_S95 | GPRIN1 | Q7ZZK8 | | 286 | KVDLVLSGKRDPG | 8.9832 | 8.3560 | 5.7330 | 7.1717 | 6.0970 | 5.5356 | 7.5050 | 7.3077 | 7.5007 | 6.4611 | 5.7593 | 5.1627 | 4.0661 | 7.3603 | 3.2941 | 3.7065 |
| GPRIN1_S95;S104 | GPRIN1 | Q7ZZK8 | | 993 | FRALLQSVRRPRC | 8.0974 | 7.0389 | 7.4987 | 7.4918 | 6.6091 | 6.2156 | 6.3157 | 7.1415 | 6.9582 | 6.6748 | 7.8065 | 6.5941 | 4.4688 | 4.1316 | 3.4591 | 3.4985 |
| GPRIN1_S993 | GPRIN1 | Q7ZZK8 | | 120 | GASISGTPEATTS | 6.1977 | 7.0856 | 6.9181 | 7.6867 | 6.5415 | 6.7555 | 6.5086 | 6.9460 | 6.7030 | 7.7140 | 6.9986 | 7.8372 | 3.8888 | 4.3207 | 3.8472 | 4.0508 |
| GPRIN1_T120 | GPRIN1 | Q7ZZK8 | | 104 | CFSPQESPSKETL | 5.8020 | 6.5599 | 6.9297 | 6.6825 | 6.5930 | 6.7662 | 6.4307 | 6.3132 | 6.3959 | 5.8574 | 5.9101 | 6.1878 | 5.9317 | 6.1347 | 6.1281 | 6.2770 |
| GPRIN1_T578 | GPRIN1 | Q7ZZK8 | 95;104 | | NA | 6.0136 | 5.9204 | 6.6482 | 7.5518 | 7.4835 | 7.1884 | 6.4405 | 6.5027 | 6.1683 | 6.6356 | 6.2564 | 6.4739 | 4.6655 | 5.0665 | 6.2452 | 4.7394 |
| GPRIN1_T877 | GPRIN1 | Q7ZZK8 | 100;104 | | NA | 6.6122 | 6.8796 | 6.6665 | 6.9283 | 6.5833 | 7.0105 | 6.5122 | 7.5899 | 7.5191 | 7.0695 | 7.7813 | 7.3338 | 3.4870 | 4.2157 | 3.3251 | 4.4861 |
| GPS1_S276 | GPS1 | Q13098-7 | | 276 | QRGERDSQTQAIL | 6.4683 | 5.8335 | 5.7231 | 5.7388 | 7.1820 | 6.4513 | 5.7461 | 6.0933 | 5.5568 | 6.0471 | 5.8894 | 7.3872 | 5.7910 | 7.1808 | 6.8592 | 6.0520 |
| GPS1_T515 | GPS1 | Q13098-7 | | 515 | GSQGELTPANSQS | 5.3837 | 6.1054 | 5.8539 | 6.2169 | 6.4012 | 7.1830 | 7.1001 | 6.4084 | 6.1211 | 5.6824 | 7.6960 | 6.4741 | 4.8677 | 6.5241 | 5.9710 | 6.0111 |
| GPSM1_S413 | GPSM1 | Q86YR5 | | 413 | KRTQRLSAETWDL | 7.1305 | 6.7293 | 7.0515 | 7.2750 | 7.2320 | 6.3428 | 6.4111 | 6.7977 | 6.5429 | 6.1769 | 6.6239 | 6.8064 | 4.9284 | 4.7283 | 4.4550 | 4.7683 |
| GPSM1_S445 | GPSM1 | Q86YR5 | | 445 | RGPSRDSLPLPVR | 4.6772 | 5.8137 | 6.2950 | 6.5271 | 5.4014 | 5.9304 | 6.0156 | 6.4535 | 6.7713 | 5.6944 | 5.8733 | 6.8514 | 6.8880 | 7.6607 | 6.9417 | 7.1052 |
| GPSM2_S408 | GPSM2 | P81274 | | 408 | KLGRRHSMENMEL | 6.6742 | 6.9208 | 7.3952 | 6.2699 | 5.9362 | 6.6220 | 6.0036 | 6.3051 | 6.9939 | 7.1351 | 7.6099 | 7.4500 | 4.5136 | 5.0277 | 4.5882 | 4.5546 |
| GRAMD1A_S16 | GRAMD1A | Q96CP6 | | 550 | DARGLLSGLRRRK | 7.6647 | 8.1989 | 7.0091 | 7.8243 | 6.5501 | 6.9215 | 7.5183 | 7.0913 | 7.2003 | 7.1897 | 7.1663 | 6.8463 | 2.7628 | 3.6022 | 2.9227 | 3.5314 |
| GRAMD1A_S263;S267 | GRAMD1A | Q96CP6 | | 527 | KGGVPGTPSTQSL | 7.9580 | 7.6610 | 7.0896 | 7.9380 | 6.3408 | 7.1420 | 8.3019 | 6.9158 | 6.7901 | 6.5126 | 6.7201 | 6.3111 | 3.4091 | 3.7710 | 2.9331 | 4.2020 |
| GRAMD1A_S267 | GRAMD1A | Q96CP6 | | 267 | DRSQEPSPVGSRR | 5.8767 | 6.3758 | 5.5855 | 7.0206 | 6.3976 | 5.9700 | 6.0800 | 5.9150 | 6.2980 | 5.9912 | 6.4063 | 5.7709 | 6.6275 | 6.8424 | 6.4531 | 6.3894 |
| GRAMD1A_S550 | GRAMD1A | Q96CP6 | | 83 | SWYSMLSPTYKQR | 6.7034 | 7.9378 | 8.0758 | 8.8757 | 6.4676 | 7.4210 | 7.3242 | 7.5425 | 6.2370 | 7.9975 | 8.2168 | 7.6207 | 2.2146 | 2.9126 | 2.0117 | 2.4411 |
| GRAMD1A_S83 | GRAMD1A | Q96CP6 | | 16 | RSTPSSSPSLRKR | 5.9438 | 6.1143 | 5.0424 | 6.7396 | 7.4847 | 6.4408 | 5.8297 | 5.6848 | 5.2543 | 4.8028 | 5.5735 | 6.0103 | 7.6557 | 7.0009 | 7.6570 | 6.7653 |
| GRAMD1A_T57 | GRAMD1A | Q96CP6 | | 230 | TSEDEDYVSPLQL | 6.7032 | 6.6144 | 6.7495 | 8.0908 | 6.1783 | 6.9005 | 6.4573 | 6.2180 | 6.7462 | 6.0559 | 6.0648 | 6.1679 | 6.4641 | 5.4418 | 5.2251 | 5.7423 |
| GRAMD1A_Y230 | GRAMD1A | Q96CP6 | 263;267 | | NA | 5.4691 | 8.0071 | 6.7031 | 6.2545 | 6.2130 | 6.0510 | 7.3448 | 6.9838 | 6.6772 | 7.3475 | 7.1454 | 5.3609 | 4.9881 | 6.6311 | 4.2695 | 4.5536 |
| GRAMD1B_S21 | GRAMD1B | Q3KR37-4 | | 557 | AEMHRQSPKEKAS | 6.4699 | 6.3555 | 6.5318 | 6.4985 | 6.8632 | 6.6022 | 6.7356 | 6.1796 | 6.2237 | 6.7575 | 6.2788 | 6.0095 | 5.5488 | 6.1641 | 5.3172 | 5.4642 |
| GRAMD1B_S55 | GRAMD1B | Q3KR37-4 | | 594 | SPVTTPTDEDVGH | 6.0653 | 6.3000 | 7.2482 | 5.0627 | 5.9860 | 5.1980 | 6.7601 | 5.3518 | 6.4290 | 6.7726 | 6.4664 | 5.6394 | 7.3601 | 6.4088 | 6.6451 | 6.3063 |
| GRAMD1B_S557 | GRAMD1B | Q3KR37-4 | | 21 | RSTPACSPILRKR | 6.2132 | 6.1852 | 6.4637 | 6.7255 | 6.1147 | 6.1336 | 6.3421 | 6.6379 | 6.3939 | 6.5841 | 6.0307 | 6.6870 | 5.7382 | 5.8482 | 5.7453 | 6.1565 |
| GRAMD1B_S95 | GRAMD1B | Q3KR37-4 | | 95 | SWYNVLSPTYKQR | 6.3965 | 6.8021 | 6.3743 | 6.9239 | 6.8630 | 6.8601 | 7.1619 | 7.0435 | 6.9453 | 7.4563 | 6.5263 | 6.4465 | 4.3585 | 4.9552 | 3.9530 | 4.9337 |
| GRAMD1B_T594 | GRAMD1B | Q3KR37-4 | | 55 | SSDKSPSTPEQGV | 6.7115 | 6.7865 | 6.2307 | 6.9282 | 6.2771 | 6.3186 | 6.7994 | 6.5281 | 6.6002 | 6.5827 | 6.6466 | 6.7476 | 5.2099 | 5.3013 | 4.8647 | 5.4669 |
| GRAMD2B_S267 | GRAMD2B | Q96HH9-3 | | 267 | DFNDEFSDLDGVV | 6.4633 | 5.9781 | 5.7272 | 6.9400 | 6.2485 | 6.1509 | 6.0305 | 6.3444 | 5.9915 | 5.5224 | 5.4062 | 5.7785 | 7.1831 | 7.2376 | 6.3291 | 6.6688 |
| GRIP1_S43 | GRIP1 | Q9Y3R0 | | 43 | LAVRRQSIPEEFK | 6.5953 | 6.5536 | 6.5916 | 6.9153 | 6.1709 | 6.2687 | 6.1871 | 6.4250 | 6.6845 | 6.9438 | 6.0199 | 6.1650 | 6.6176 | 6.8167 | 6.3490 | 6.4962 |
| GRIP1_S433 | GRIP1 | Q9Y3R0 | | 847 | KQRGSLSPVTKPR | 6.0860 | 6.1355 | 6.4102 | 5.0854 | 6.3846 | 6.2670 | 5.9879 | 6.4835 | 6.1835 | 5.8486 | 5.9255 | 5.8482 | 7.1265 | 6.9710 | 6.7891 | 6.4674 |
| GRIP1_S847 | GRIP1 | Q9Y3R0 | | 996 | EIKEIMSPTPVEL | 5.5838 | 6.8730 | 7.4803 | 7.1439 | 6.1802 | 5.5838 | 6.5966 | 6.3093 | 6.4075 | 7.3894 | 7.7239 | 8.2439 | 4.0347 | 5.4402 | 4.0238 | 4.9856 |
| GRIP1_S996 | GRIP1 | Q9Y3R0 | | 433 | RSLYSTSPRGTMM | 6.1514 | 6.4723 | 6.3143 | 5.8335 | 5.5264 | 6.5951 | 6.5942 | 6.6432 | 6.7198 | 6.2245 | 5.8630 | 6.2396 | 6.0347 | 6.4728 | 6.1930 | 6.0222 |
| GRIPAP1_S655 | GRIPAP1 | Q4V328 | | 655 | VLSEMNSPSRTQT | 6.2903 | 6.5504 | 6.8021 | 6.7810 | 6.0725 | 6.8560 | 6.6202 | 6.4816 | 7.0733 | 7.1144 | 7.0012 | 7.0522 | 4.3724 | 5.1445 | 4.5888 | 5.1990 |
| GRIPAP1_S666 | GRIPAP1 | Q4V328 | | 688 | SAVPARSLSSSPQ | 6.3342 | 6.0563 | 6.7721 | 6.7679 | 6.6173 | 6.6431 | 6.0766 | 7.1592 | 7.0380 | 6.8144 | 7.0424 | 7.1650 | 5.1925 | 5.0259 | 4.6799 | 4.6151 |
| GRIPAP1_S688 | GRIPAP1 | Q4V328 | | 666 | QTGDSSSISSFSY | 5.3166 | 5.4097 | 6.4931 | 5.6121 | 6.3307 | 6.5319 | 5.7893 | 6.7775 | 6.3547 | 5.9680 | 6.1722 | 6.4112 | 6.8445 | 6.5326 | 6.7415 | 6.7145 |
| GRIPAP1_S688;S691 | GRIPAP1 | Q4V328 | | 691 | PARSLSSSPQAQP | 6.5999 | 6.2772 | 6.2688 | 6.5069 | 6.3804 | 5.7294 | 5.6550 | 6.4242 | 6.7888 | 7.3096 | 8.1076 | 7.3970 | 4.6663 | 5.4559 | 5.2588 | 5.1741 |
| GRIPAP1_S691 | GRIPAP1 | Q4V328 | 688;691 | | NA | 6.5502 | 6.5999 | 7.2104 | 5.6371 | 6.0244 | 5.4042 | 6.4400 | 6.8587 | 6.4091 | 6.0146 | 6.0365 | 6.4858 | 5.4712 | 6.9380 | 6.0061 | 5.9138 |
| GRK2_S670 | GRK2 | P25098 | | 670 | MKNKPRSPVVELS | 5.7724 | 5.8141 | 6.4274 | 6.4998 | 5.9427 | 6.5043 | 6.8349 | 6.5363 | 6.4278 | 6.3582 | 6.2160 | 6.3436 | 5.9472 | 6.3310 | 5.6430 | 6.3958 |
| GRK5_S484 | GRK5 | P34947 | | 484 | LDIEQFSTVKGVN | 7.2001 | 7.5868 | 6.8295 | 7.8614 | 6.5295 | 6.0207 | 6.5393 | 5.9932 | 6.1695 | 5.4702 | 5.9457 | 6.0535 | 5.1911 | 5.6864 | 5.7630 | 5.1601 |
| GRPR_S354 | GRPR | P30550 | | 360 | STTCMTSLKSTNP | 6.6345 | 6.9023 | 6.9144 | 6.6103 | 6.9653 | 6.3916 | 6.5475 | 6.4185 | 6.8300 | 6.7285 | 6.3804 | 6.2851 | 4.7958 | 5.5755 | 4.9696 | 5.0508 |
| GRPR_S360 | GRPR | P30550 | | 354 | SHSTGRSTTCMTS | 4.7257 | 5.3337 | 6.2403 | 6.6776 | 5.9819 | 6.2870 | 5.7548 | 6.6333 | 5.7484 | 6.1373 | 5.9400 | 6.6180 | 6.3651 | 6.6244 | 6.9618 | 7.9707 |
| GRSF1_S244 | GRSF1 | Q12849 | | 244 | SLQVKSSPVVNDG | 6.2414 | 5.8003 | 6.8347 | 7.0082 | 6.6771 | 6.8344 | 6.6129 | 6.7381 | 6.1484 | 6.3273 | 6.4304 | 6.4751 | 5.7304 | 4.9852 | 5.4757 | 5.6802 |
| GRWD1_S119;S122 | GRWD1 | Q9BQ67 | | 344 | RQFKSGSPVATFK | 6.1315 | 6.4414 | 6.5055 | 6.5912 | 6.2682 | 6.2314 | 6.1003 | 6.4439 | 6.6865 | 6.5325 | 6.9322 | 7.0928 | 5.3508 | 5.5310 | 5.4057 | 5.7553 |
| GRWD1_S344 | GRWD1 | Q9BQ67 | | 10 | RKGRRRTCETGEP | 6.0259 | 6.3614 | 6.3648 | 6.3135 | 6.4949 | 6.9616 | 6.6957 | 5.1382 | 5.7372 | 7.4819 | 7.1670 | 5.7418 | 6.0476 | 6.2598 | 6.4441 | 6.5645 |
| GRWD1_T10 | GRWD1 | Q9BQ67 | 119;122 | | NA | 6.0399 | 5.7412 | 6.5159 | 7.0436 | 6.4639 | 5.7768 | 5.4296 | 7.3428 | 6.5035 | 6.2383 | 6.7978 | 5.9017 | 5.8569 | 5.6066 | 6.7204 | 6.0212 |
| GSE1_S710 | GSE1 | Q14687 | | 710 | GPLKPGSPYRPPV | 6.3069 | 5.9162 | 6.7628 | 6.8218 | 6.7827 | 7.0692 | 6.9937 | 6.7219 | 7.2247 | 6.7749 | 7.1135 | 7.0675 | 4.1771 | 5.1825 | 4.1059 | 4.9785 |
| GSE1_S826;S828 | GSE1 | Q14687 | | 909 | ADSLTNSPRDSPA | 6.7301 | 7.0090 | 7.9798 | 7.5707 | 5.8824 | 6.5296 | 6.4421 | 5.7745 | 6.4838 | 6.5907 | 6.8339 | 6.8197 | 4.6957 | 5.3369 | 4.2265 | 5.0946 |
| GSE1_S909 | GSE1 | Q14687 | | 433 | KPSEQLTPTRAEK | 7.3860 | 7.2482 | 7.6538 | 7.8441 | 5.8065 | 6.2481 | 6.9011 | 6.7308 | 7.0978 | 6.5806 | 7.3876 | 7.8297 | 3.8652 | 3.6850 | 3.4698 | 4.2655 |
| GSE1_S95 | GSE1 | Q14687 | | 95 | PATNHSSPASTPK | 5.4799 | 5.8678 | 6.9929 | 6.3430 | 5.9357 | 6.7068 | 5.6496 | 7.2159 | 4.9852 | 5.4944 | 4.5294 | 6.1976 | 7.3226 | 9.1848 | 5.9308 | 6.1636 |
| GSE1_T433 | GSE1 | Q14687 | 826;828 | | NA | 5.9484 | 6.1095 | 5.8418 | 6.1990 | 7.4883 | 5.5811 | 6.7209 | 5.9719 | 6.7166 | 7.3553 | 7.0610 | 7.0114 | 6.0695 | 6.2954 | 4.7106 | 4.9192 |
| GSK3A_S21 | GSK3A | P49840 | | 278 | RGEPNVSYICSRY | 6.4984 | 6.4892 | 6.8519 | 6.0733 | 6.1802 | 6.4662 | 6.0509 | 6.2754 | 6.5205 | 6.3708 | 6.6690 | 6.6980 | 5.6154 | 5.8971 | 5.5848 | 5.7590 |
| GSK3A_S278 | GSK3A | P49840 | | 21 | GRARTSSFAEPGG | 5.9685 | 5.2144 | 6.1926 | 4.9554 | 6.5315 | 6.2686 | 5.7525 | 6.5136 | 7.2747 | 6.6615 | 6.8944 | 6.5603 | 7.0017 | 7.1720 | 5.7342 | 6.2041 |
| GSK3B_S25 | GSK3B | P49841-2 | | 9 | GRPRTTSFAESCK | 6.1971 | 5.5738 | 6.9330 | 5.5410 | 5.8197 | 6.1749 | 6.0342 | 5.8798 | 6.8605 | 6.2072 | 6.6369 | 6.4659 | 6.5964 | 7.2512 | 6.0294 | 5.7990 |
| GSK3B_S9 | GSK3B | P49841-2 | | 25 | QPSAFGSMKVSRD | 6.8127 | 6.0187 | 6.9531 | 6.9558 | 6.1866 | 5.5766 | 5.9434 | 7.5557 | 8.0978 | 7.6287 | 7.0822 | 7.6873 | 4.7283 | 5.4835 | 3.6003 | 5.4894 |
| GTF2A1_S316;S321 | GTF2A1 | P52655 | 316;321 | | NA | 5.6623 | 5.6994 | 6.0718 | 5.3956 | 5.5925 | 6.1653 | 5.6834 | 5.8555 | 6.6953 | 6.5712 | 6.7048 | 6.5162 | 6.8762 | 7.0785 | 6.6517 | 6.7804 |
| GTF2A1_S321;T331 | GTF2A1 | P52655 | 321;331 | | NA | 5.4062 | 5.2459 | 6.0764 | 5.3678 | 6.1161 | 6.0376 | 5.8867 | 6.1182 | 6.1339 | 6.2092 | 6.5477 | 6.4918 | 7.2753 | 7.3828 | 6.7627 | 6.9417 |
| GTF2B_S76 | GTF2B | Q00403 | | 76 | SGNPLLSDGDLST | 6.2401 | 6.7612 | 7.1062 | 7.1588 | 6.8418 | 7.1916 | 7.1215 | 7.0270 | 5.1221 | 5.4830 | 5.9199 | 5.9859 | 5.0956 | 5.7633 | 5.2852 | 5.8969 |
| GTF2E1_S268 | GTF2E1 | P29083 | | 268 | EDLHRASLEGKSA | 5.2970 | 6.2165 | 6.4775 | 6.6000 | 5.7839 | 6.3302 | 6.3514 | 6.0163 | 6.4335 | 6.1395 | 6.6865 | 6.1725 | 6.0882 | 6.4401 | 6.2121 | 6.7547 |
| GTF2F1_S377 | GTF2F1 | P35269 | | 437 | SLSGKSTPQPPSG | 6.2872 | 6.5118 | 6.0904 | 5.2145 | 4.3232 | 5.9800 | 5.5852 | 5.4347 | 7.1292 | 5.7575 | 6.8241 | 7.5450 | 6.0212 | 7.1390 | 5.8775 | 6.4596 |
| GTF2F1_S381 | GTF2F1 | P35269 | | 433 | TGPQSLSGKSTPQ | 6.0102 | 6.5732 | 6.2031 | 5.9856 | 6.5929 | 6.5760 | 5.8236 | 6.1131 | 7.0732 | 7.2567 | 6.7278 | 6.8964 | 5.4111 | 6.4681 | 5.5016 | 5.6875 |
| GTF2F1_S385 | GTF2F1 | P35269 | | 331 | EEKKAPTPQEKKR | 5.7789 | 6.5439 | 6.3686 | 5.9792 | 5.3620 | 6.0657 | 5.5895 | 6.2634 | 6.2855 | 6.5710 | 6.1844 | 6.9160 | 5.6494 | 6.8319 | 6.5831 | 7.0277 |
| GTF2F1_S385;T389 | GTF2F1 | P35269 | | 427 | KRLRLDTGPQSLS | 5.6586 | 5.8070 | 6.7332 | 6.2456 | 5.2820 | 5.9368 | 6.2108 | 5.7427 | 6.4672 | 5.4524 | 6.7132 | 6.4026 | 6.8037 | 6.5273 | 7.0784 | 6.9385 |
| GTF2F1_S433 | GTF2F1 | P35269 | | 377 | KRERKPSGGSSRG | 6.0043 | 6.4965 | 6.6928 | 6.0213 | 5.8764 | 6.4967 | 5.7315 | 6.1715 | 6.6549 | 6.7013 | 6.9559 | 6.6283 | 5.4507 | 6.2695 | 6.1867 | 5.6619 |
| GTF2F1_S442 | GTF2F1 | P35269 | | 446 | PPSGKTTPNSGDV | 5.5540 | 5.4432 | 6.0828 | 6.1192 | 5.7575 | 6.0399 | 6.4031 | 5.5899 | 5.8423 | 6.0499 | 6.5328 | 6.0653 | 7.1464 | 7.7306 | 6.3138 | 7.3295 |
| GTF2F1_T331 | GTF2F1 | P35269 | | 442 | STPQPPSGKTTPN | 5.4565 | 5.6596 | 5.1569 | 5.5700 | 5.4283 | 5.2128 | 5.3618 | 5.9545 | 5.1328 | 5.7367 | 5.8020 | 5.9470 | 7.7421 | 8.6947 | 8.7735 | 8.3709 |
| GTF2F1_T389 | GTF2F1 | P35269 | | 385 | GSSRGNSRPGTPS | 4.8845 | 5.2929 | 5.8652 | 5.3269 | 6.2735 | 5.4866 | 5.9464 | 6.2242 | 5.7811 | 5.9290 | 5.5422 | 6.2707 | 6.9048 | 8.0925 | 8.1244 | 8.0549 |
| GTF2F1_T427 | GTF2F1 | P35269 | | 389 | GNSRPGTPSAEGG | 5.1553 | 5.2506 | 5.8299 | 5.0951 | 6.5200 | 6.1428 | 5.3264 | 7.4142 | 6.2303 | 6.3003 | 6.7221 | 5.3711 | 6.7338 | 6.8718 | 6.6777 | 6.3586 |
| GTF2F1_T437 | GTF2F1 | P35269 | | 381 | KPSGGSSRGNSRP | 4.6371 | 5.1577 | 4.5486 | 4.1542 | 5.0239 | 5.1591 | 4.4560 | 4.9030 | 4.9081 | 5.7954 | 5.4639 | 5.4442 | 9.8364 | 11.3536 | 10.2472 | 8.9116 |
| GTF2F1_T446 | GTF2F1 | P35269 | 385;389 | | NA | 4.8808 | 5.5703 | 5.3685 | 6.2010 | 5.8773 | 5.7877 | 5.7541 | 5.4451 | 6.2640 | 6.1545 | 5.9804 | 6.2986 | 7.6351 | 7.8793 | 7.0373 | 7.8659 |
| GTF2I_S103 | GTF2I | P78347 | | 831 | PRKINSSPNVNTT | 6.1318 | 6.4027 | 6.2744 | 6.7777 | 6.1984 | 6.3644 | 6.3462 | 6.5173 | 6.2563 | 6.2816 | 6.3280 | 6.7314 | 5.6517 | 6.1422 | 5.7033 | 5.8924 |
| GTF2I_S210 | GTF2I | P78347 | | 210 | ADRSILSPGGSCG | 6.7603 | 7.5719 | 6.9894 | 7.4726 | 6.4327 | 7.2033 | 7.2732 | 7.1129 | 7.3051 | 7.5938 | 7.5312 | 7.3254 | 2.9895 | 3.9562 | 2.9132 | 3.5694 |
| GTF2I_S210;S214 | GTF2I | P78347 | | 412 | IPFRRPSTYGIPR | 4.9444 | 4.9453 | 5.2024 | 5.2905 | 6.5033 | 5.3649 | 5.3137 | 5.6948 | 5.4100 | 5.3937 | 5.3249 | 5.1472 | 9.0936 | 8.4577 | 9.3469 | 8.5667 |
| GTF2I_S412 | GTF2I | P78347 | | 674 | SPKRPRSPGSNSK | 6.4846 | 6.4250 | 5.7046 | 6.1111 | 6.0605 | 6.2629 | 6.0357 | 6.5385 | 5.8788 | 6.3090 | 5.7607 | 5.7608 | 6.1716 | 7.0660 | 6.7131 | 6.7171 |
| GTF2I_S515 | GTF2I | P78347 | | 703 | QTSAVRTPTQTNG | 6.5361 | 6.6780 | 6.6263 | 7.1489 | 6.5194 | 6.7811 | 7.3188 | 7.3528 | 7.5847 | 7.1451 | 7.2932 | 6.9456 | 3.7959 | 4.5979 | 3.6905 | 3.9857 |
| GTF2I_S517 | GTF2I | P78347 | | 517 | IPFRSPSWYGIPR | 5.1276 | 5.3429 | 5.2316 | 5.2394 | 5.9844 | 5.7313 | 5.7863 | 5.6720 | 5.2476 | 4.8678 | 5.3417 | 5.3253 | 8.8774 | 8.4666 | 9.2518 | 8.5063 |
| GTF2I_S674 | GTF2I | P78347 | | 784 | VPFRRPSTFGIPR | 4.4208 | 5.1527 | 5.4119 | 4.7311 | 5.7766 | 6.1762 | 5.4415 | 6.4616 | 5.5412 | 8.2902 | 6.2103 | 4.7393 | 8.1885 | 7.7333 | 8.2780 | 7.4467 |
| GTF2I_S722 | GTF2I | P78347 | | 515 | ENIPFRSPSWYGI | 6.7995 | 6.8495 | 6.1393 | 6.7178 | 6.3814 | 6.6594 | 6.2518 | 6.1408 | 6.2987 | 6.6142 | 6.3695 | 6.4255 | 5.0993 | 5.8964 | 5.3857 | 5.6854 |
| GTF2I_S784 | GTF2I | P78347 | | 722 | PRGREFSFEAWNA | 5.3891 | 4.9018 | 5.1115 | 4.6601 | 6.9850 | 5.7664 | 5.3282 | 4.9875 | 5.5793 | 5.8966 | 5.1318 | 5.0101 | 9.2991 | 8.3806 | 8.5046 | 9.0683 |
| GTF2I_S823 | GTF2I | P78347 | | 103 | TQANRMSVDAVEI | 5.6003 | 6.0541 | 5.8257 | 6.3068 | 6.1228 | 5.7485 | 6.5315 | 6.1232 | 6.0443 | 6.0047 | 6.0409 | 6.0705 | 6.4862 | 6.9688 | 6.8635 | 7.2081 |
| GTF2I_S831 | GTF2I | P78347 | | 823 | ESTSSKSPPRKIN | 5.8357 | 7.1761 | 6.6126 | 6.7123 | 6.6771 | 7.9077 | 6.5934 | 7.3882 | 7.0437 | 5.5543 | 6.0257 | 6.5588 | 6.0343 | 5.8965 | 4.9197 | 3.9639 |
| GTF2I_T703 | GTF2I | P78347 | 210;214 | | NA | 6.2763 | 7.5833 | 6.0010 | 5.3907 | 5.3668 | 7.8243 | 5.2961 | 7.1947 | 6.1813 | 6.5473 | 8.0160 | 6.6014 | 4.6738 | 3.7224 | 7.3990 | 5.9256 |
| GTF2IRD2B_S516 | GTF2IRD2B | Q6EKJ0 | | 516 | QVFANPSPTQKSP | 5.6893 | 6.1210 | 6.4575 | 7.2105 | 6.1468 | 6.6845 | 6.5137 | 6.2918 | 6.1215 | 7.1213 | 7.3139 | 6.3389 | 5.1212 | 6.3440 | 5.0710 | 5.4531 |
| GTF3A_S175 | GTF3A | Q92664 | | 175 | CGKHFASPSKLKR | 5.8985 | 7.0924 | 6.5460 | 6.5264 | 6.6530 | 6.4585 | 6.8883 | 6.4759 | 6.5617 | 7.0500 | 6.9746 | 6.6359 | 4.5259 | 5.2787 | 5.1496 | 5.2845 |
| GTF3C1_S1062;S1068 | GTF3C1 | Q12789 | | 1653 | LMRGYYSPGIVST | 6.3941 | 7.0273 | 6.1899 | 6.8049 | 6.6283 | 6.7661 | 6.7548 | 7.1933 | 6.7897 | 6.4442 | 6.5728 | 6.4439 | 4.5942 | 5.4219 | 4.7831 | 5.1916 |
| GTF3C1_S1068 | GTF3C1 | Q12789 | | 1865 | GTKRRASWASENG | 5.2648 | 5.1763 | 6.5547 | 6.6993 | 5.7816 | 6.2999 | 5.3328 | 5.4932 | 6.5318 | 7.3911 | 7.0735 | 7.0869 | 6.0399 | 6.8609 | 6.7673 | 6.5460 |
| GTF3C1_S1202 | GTF3C1 | Q12789 | | 1856 | QAPPSHSPRGTKR | 6.0110 | 5.8274 | 6.2091 | 6.7579 | 6.1795 | 5.7629 | 6.1036 | 5.3364 | 6.9019 | 6.3257 | 6.5737 | 6.6705 | 6.4005 | 6.2232 | 6.0058 | 6.6840 |
| GTF3C1_S1653 | GTF3C1 | Q12789 | | 1682 | KFQLRCTPVPARL | 5.9035 | 7.1001 | 5.8141 | 6.2059 | 6.7133 | 6.4400 | 6.2582 | 7.2206 | 7.1262 | 7.9219 | 6.1165 | 6.6580 | 5.5820 | 4.8516 | 4.6778 | 5.4102 |
| GTF3C1_S1856 | GTF3C1 | Q12789 | | 1202 | EVDREPSLDRNRR | 6.5335 | 6.6530 | 5.8798 | 6.4178 | 5.9591 | 6.3318 | 6.0010 | 6.1686 | 5.8849 | 6.2157 | 7.0301 | 6.6229 | 5.8048 | 6.1988 | 6.2958 | 6.9022 |
| GTF3C1_S1865 | GTF3C1 | Q12789 | | 1068 | SSTDQGSDEEGSL | 6.4941 | 7.5120 | 6.7397 | 6.4354 | 6.4925 | 5.8638 | 6.1959 | 4.4535 | 6.7675 | 6.6439 | 7.1181 | 6.6640 | 6.2809 | 6.2716 | 4.5137 | 5.5533 |
| GTF3C1_S1865;S1868 | GTF3C1 | Q12789 | | 514 | TQPHHSTPTKGGW | 6.6184 | 7.2692 | 5.8341 | 8.2553 | 7.1830 | 6.3503 | 5.7602 | 7.4444 | 6.4337 | 5.5499 | 6.0719 | 6.1600 | 4.5411 | 5.4988 | 6.0037 | 5.0261 |
| GTF3C1_S846 | GTF3C1 | Q12789 | | 846 | GVRPSSSGSAWEA | 5.2798 | 6.9247 | 5.7053 | 6.6214 | 4.6345 | 7.3285 | 8.4597 | 7.0562 | 5.4639 | 6.4868 | 6.1324 | 5.4283 | 5.7657 | 6.1308 | 6.4744 | 6.1076 |
| GTF3C1_T1682 | GTF3C1 | Q12789 | 1865;1868 | | NA | 5.5189 | 5.6968 | 5.5068 | 6.2168 | 6.0446 | 5.7548 | 6.5219 | 6.3533 | 5.8937 | 6.0039 | 5.8595 | 5.8366 | 7.0642 | 6.9248 | 7.4387 | 7.3647 |
| GTF3C1_T514 | GTF3C1 | Q12789 | 1062;1068 | | NA | 5.5902 | 5.5418 | 5.9767 | 6.7806 | 6.9805 | 5.6943 | 6.6198 | 6.2140 | 6.3563 | 5.7069 | 6.7222 | 6.3868 | 6.6197 | 6.7228 | 5.8747 | 6.2128 |
| GTF3C2_S167 | GTF3C2 | Q8WUA4 | | 893 | AMFQPSSPTRRPG | 6.1781 | 6.6565 | 6.5326 | 6.9992 | 6.4762 | 5.9129 | 6.6556 | 6.3938 | 6.7731 | 6.0780 | 6.1387 | 6.4059 | 5.2926 | 5.8374 | 5.6036 | 6.0658 |
| GTF3C2_S893 | GTF3C2 | Q8WUA4 | | 167 | DRPESQSPKRPPE | 6.1538 | 6.6687 | 6.1293 | 7.0342 | 6.0342 | 5.8019 | 6.4765 | 5.8965 | 6.4787 | 6.6434 | 6.4823 | 6.5548 | 5.8081 | 6.1599 | 5.7372 | 5.9404 |
| GTF3C2_S901 | GTF3C2 | Q8WUA4 | | 901 | TRRPGFSPTSHRL | 7.0591 | 7.6499 | 6.8410 | 7.6004 | 6.4727 | 6.5904 | 6.7913 | 6.8502 | 7.2614 | 7.0221 | 6.9976 | 6.7170 | 3.6324 | 4.3209 | 4.0939 | 4.0998 |
| GTF3C3_S43 | GTF3C3 | Q9Y5Q9 | | 43 | QEKGKLSAEENPD | 6.1941 | 6.4395 | 6.3194 | 6.5559 | 6.3451 | 6.1306 | 6.3498 | 6.3454 | 5.9790 | 6.0157 | 5.9097 | 5.9371 | 6.0257 | 6.4985 | 6.4257 | 6.5286 |
| GTF3C4_S19 | GTF3C4 | Q9UKN8 | | 611 | KILLVDSPGMGNA | 6.0180 | 5.8862 | 5.7856 | 6.3993 | 6.1817 | 5.7940 | 5.8370 | 6.0332 | 6.2932 | 6.1851 | 6.0410 | 6.0824 | 7.0334 | 6.9560 | 6.7351 | 6.7388 |
| GTF3C4_S611 | GTF3C4 | Q9UKN8 | | 19 | DDGPAPSGEEEGE | 5.8685 | 7.0013 | 6.6285 | 6.6947 | 6.3789 | 6.4158 | 6.9026 | 6.1311 | 5.8944 | 6.4704 | 5.8805 | 5.8585 | 5.4449 | 6.1738 | 5.9117 | 6.3444 |
| GTF3C5_S358 | GTF3C5 | Q9Y5Q8-3 | | 477 | PALFSSSAKADGG | 5.8658 | 6.7507 | 7.3252 | 5.0566 | 4.9117 | 6.9110 | 7.0380 | 6.9623 | 6.5853 | 8.1037 | 8.5346 | 7.1393 | 4.0204 | 5.3259 | 3.7960 | 5.6735 |
| GTF3C5_S477 | GTF3C5 | Q9Y5Q8-3 | | 358 | TVKKTSSQLVTMH | 6.1012 | 6.2481 | 7.6411 | 5.6225 | 3.8927 | 5.1996 | 5.4402 | 5.7466 | 7.5018 | 7.0282 | 6.6508 | 8.5680 | 6.1569 | 8.2717 | 4.5259 | 5.4047 |
| GTF3C6_S160 | GTF3C6 | Q969F1 | | 160 | ASAPDKSLELEEE | 6.0260 | 6.0096 | 6.2262 | 6.4677 | 6.5119 | 6.2183 | 5.9464 | 6.0686 | 6.1169 | 6.4971 | 6.4513 | 6.8535 | 5.8485 | 6.5545 | 5.8046 | 6.3989 |
| GTF3C6_S199 | GTF3C6 | Q969F1 | | 199 | PLIDIPSETEGSV | 5.8548 | 6.2766 | 6.7769 | 6.1134 | 6.1422 | 6.3096 | 6.2598 | 6.5562 | 5.5993 | 5.9190 | 5.9790 | 6.0918 | 6.5808 | 7.2173 | 6.5292 | 5.7942 |
| GTPBP1_S12;S25 | GTPBP1 | O00178 | | 25 | FAPEPSSPGAARA | 6.0058 | 6.5879 | 6.7142 | 6.5367 | 6.2458 | 6.2824 | 6.2077 | 6.6064 | 6.4755 | 6.6047 | 6.4175 | 6.2050 | 5.6583 | 5.8513 | 5.5200 | 6.0808 |
| GTPBP1_S25 | GTPBP1 | O00178 | | 580 | LQTTNNSPMNSKP | 5.8304 | 6.5951 | 7.0838 | 6.8517 | 6.1890 | 6.5670 | 6.5475 | 6.4763 | 7.0727 | 6.9678 | 6.9007 | 7.0765 | 4.4270 | 5.4693 | 4.8928 | 5.4223 |
| GTPBP1_S44;S47 | GTPBP1 | O00178 | 44;47 | | NA | 5.9063 | 5.7611 | 6.4705 | 6.3598 | 6.2803 | 6.2922 | 6.2406 | 6.1899 | 6.5643 | 6.5280 | 6.2051 | 6.3688 | 5.5973 | 6.4540 | 6.2776 | 6.5039 |
| GTPBP1_S580 | GTPBP1 | O00178 | 12;25 | | NA | 6.1787 | 6.5832 | 6.5740 | 6.8430 | 7.2409 | 6.1247 | 5.9123 | 6.1770 | 6.3328 | 6.9589 | 6.6173 | 6.7002 | 5.1632 | 5.7545 | 5.5956 | 5.2434 |
| GTPBP4_T30 | GTPBP4 | Q9BZE4 | | 30 | SKTQRKTPTVIHK | 7.4927 | 7.8117 | 6.7309 | 8.2988 | 6.4990 | 7.3587 | 7.3762 | 7.0800 | 7.3751 | 7.0598 | 7.1522 | 6.9175 | 2.8480 | 3.7053 | 2.7088 | 3.5853 |
| GTSE1_S152 | GTSE1 | Q9NYZ3 | | 575 | SRLVDVSPDRGSP | 6.1271 | 6.5777 | 6.6969 | 5.7455 | 6.0370 | 6.1314 | 5.9874 | 6.6711 | 6.8620 | 6.5121 | 6.1489 | 6.4647 | 5.6643 | 6.0267 | 5.9690 | 5.9084 |
| GTSE1_S168 | GTSE1 | Q9NYZ3 | | 446 | KTKVMPTPTNQFK | 8.4716 | 8.3637 | 6.8951 | 7.5764 | 6.5777 | 6.8387 | 7.3518 | 6.7257 | 7.4472 | 6.7505 | 7.1499 | 7.1526 | 3.0622 | 4.1534 | 2.9064 | 3.4772 |
| GTSE1_S326 | GTSE1 | Q9NYZ3 | | 705 | SPLIQLSPEADKE | 5.3925 | 4.8728 | 6.1711 | 5.7851 | 5.3586 | 5.9424 | 5.6689 | 5.6450 | 6.0996 | 6.0565 | 6.0388 | 6.4307 | 7.6154 | 7.8616 | 7.3697 | 7.6913 |
| GTSE1_S352 | GTSE1 | Q9NYZ3 | | 592 | PQALNFSPEESDS | 6.7621 | 6.8948 | 7.1847 | 7.1594 | 6.5518 | 6.6859 | 7.0404 | 6.7852 | 6.7884 | 7.2060 | 7.2665 | 7.2153 | 3.9062 | 4.6080 | 3.8169 | 4.1284 |
| GTSE1_S435 | GTSE1 | Q9NYZ3 | | 152 | TYYLSDSPLLGPP | 5.9935 | 6.4522 | 7.5590 | 7.0288 | 6.0136 | 6.7395 | 6.5387 | 7.2716 | 7.9405 | 6.9309 | 7.0815 | 6.8744 | 3.8149 | 4.7078 | 4.2328 | 4.8201 |

| | | | | | | | | | | | | | | | | | | | | |
|---|---|---|---|---|---|---|---|---|---|---|---|---|---|---|---|---|---|---|---|---|
| GTSE1_S461 | GTSE1 | Q9NYZ3 | 477 | RAQRPQSCTSVGR | 7.5933 | 6.7313 | 7.2159 | 6.2313 | 6.0487 | 6.2135 | 6.0526 | 5.8310 | 7.8790 | 6.8408 | 7.3327 | 7.9138 | 4.7725 | 5.5682 | 3.6526 | 4.1228 |
| GTSE1_S477 | GTSE1 | Q9NYZ3 | 715 | DKENVDSPLLKFx | 6.3223 | 6.2037 | 7.4728 | 8.0179 | 6.3204 | 7.0864 | 7.1427 | 6.7928 | 7.0211 | 7.4911 | 7.8929 | 7.1076 | 3.5518 | 4.6060 | 3.2459 | 3.7243 |
| GTSE1_S575 | GTSE1 | Q9NYZ3 | 435 | SIRRRDSCLNSKT | 6.9410 | 5.8632 | 5.9599 | 5.8311 | 5.3261 | 5.9335 | 6.7208 | 5.4471 | 8.6770 | 7.1912 | 7.9314 | 7.1244 | 6.4253 | 6.7418 | 3.1018 | 4.7844 |
| GTSE1_S575;S580 | GTSE1 | Q9NYZ3 | 352 | RPLSNISKSGRMG | 9.0091 | 6.7661 | 7.1622 | 8.2672 | 4.4077 | 7.4907 | 8.6224 | 6.2672 | 7.9345 | 6.3973 | 8.5206 | 7.5416 | 2.6076 | 3.4743 | 1.7666 | 3.7650 |
| GTSE1_S580 | GTSE1 | Q9NYZ3 | 326 | ASSACTSPAVGKA | 7.1601 | 7.4635 | 6.8376 | 6.7945 | 5.5196 | 7.4265 | 6.8219 | 7.0622 | 7.5066 | 7.2716 | 7.4570 | 7.8074 | 3.3389 | 4.8558 | 2.9250 | 3.7516 |
| GTSE1_S592 | GTSE1 | Q9NYZ3 | 580 | VSPDRGSPPSRVP | 6.0269 | 5.8221 | 5.8115 | 6.1147 | 5.7668 | 5.3441 | 5.4437 | 5.8407 | 7.0008 | 6.5145 | 6.3607 | 5.6786 | 7.0471 | 6.9945 | 7.0601 | 7.1732 |
| GTSE1_S705 | GTSE1 | Q9NYZ3 | 489 | RVTVHSTPVRRSS | 7.1513 | 7.4528 | 6.6178 | 7.2767 | 4.8545 | 7.1227 | 7.2638 | 6.6624 | 8.0099 | 7.2341 | 8.3458 | 8.7257 | 2.8248 | 3.8449 | 2.6525 | 3.9603 |
| GTSE1_S715 | GTSE1 | Q9NYZ3 | 461 | KFSIGDSPDSSTP | 6.1991 | 6.9207 | 6.1143 | 7.5268 | 6.7251 | 7.0423 | 6.7079 | 6.3161 | 6.9651 | 7.1637 | 7.4432 | 6.8615 | 3.9413 | 5.0238 | 4.2267 | 4.8223 |
| GTSE1_T446 | GTSE1 | Q9NYZ3 | 168 | PRLLASSPALPSS | 6.2045 | 6.7102 | 7.2716 | 6.2800 | 5.4002 | 6.5706 | 6.1277 | 6.5984 | 7.2480 | 7.2595 | 7.2389 | 7.3549 | 4.4787 | 5.7392 | 4.3784 | 5.1392 |
| GTSE1_T489 | GTSE1 | Q9NYZ3 | 575;580 | NA | 6.5831 | 6.5586 | 6.4029 | 5.9785 | 6.0273 | 6.5010 | 6.3470 | 6.3562 | 6.6514 | 6.6531 | 6.4932 | 6.8406 | 5.4468 | 5.9786 | 5.5283 | 5.6536 |
| GULP1_T16 | GULP1 | Q9UBP9 | 16 | DKTWMHTPEALSK | 6.9064 | 7.5346 | 6.7577 | 6.9737 | 6.7408 | 7.2564 | 7.4230 | 7.5428 | 6.8799 | 7.3421 | 7.4710 | 7.0025 | 3.3331 | 4.2906 | 2.8451 | 3.7005 |
| GULP1_T208 | GULP1 | Q9UBP9 | 208 | PPAGSMTPKSPST | 6.9102 | 6.5814 | 6.9453 | 7.1414 | 5.7224 | 6.0543 | 6.6203 | 5.9018 | 6.8335 | 6.1548 | 6.3010 | 6.9958 | 5.1037 | 5.5519 | 5.1435 | 6.0389 |
| GYS1_S698 | GYS1 | P13807 | 698 | EWPRRASCTSSTS | 5.9157 | 5.6347 | 6.1767 | 5.0216 | 7.1690 | 6.6061 | 5.8047 | 6.8152 | 7.0261 | 7.4091 | 7.0152 | 7.0420 | 5.9937 | 6.8306 | 4.5952 | 4.9442 |
| GZF1_S613 | GZF1 | Q9H116 | 613 | LVIVDGSPKNDDG | 6.1448 | 6.7153 | 6.7052 | 6.9064 | 6.0436 | 5.9970 | 6.2339 | 6.0876 | 6.3030 | 6.2560 | 6.3585 | 6.4702 | 6.0094 | 6.0164 | 5.8493 | 5.9034 |
| H1-10_S31 | H1-10 | Q92522 | 31 | GGSAALSPSKKRK | 6.2799 | 6.4822 | 5.4657 | 6.2376 | 6.4161 | 6.1409 | 6.3261 | 6.1893 | 6.2749 | 6.1051 | 5.6915 | 5.6495 | 6.1375 | 7.2107 | 6.7177 | 6.6753 |
| H1-2_T146 | H1-2 | P16403 | 146 | KAAGGATPKKSAK | 7.7268 | 7.6115 | 6.1528 | 6.1904 | 6.7124 | 6.6101 | 6.9012 | 7.3149 | 5.9657 | 6.9187 | 6.4191 | 6.5303 | 4.0079 | 5.5586 | 4.1652 | 5.2145 |
| H1-2_T165 | H1-2 | P16403 | 165 | KKPAAATVTKKVA | 10.5183 | 9.5983 | 8.0394 | 11.2504 | 5.4944 | 4.4841 | 8.0200 | 4.9475 | 6.0540 | 5.3640 | 5.7349 | 5.8700 | 3.5996 | 4.2816 | 2.9481 | 3.7954 |
| H1-3_S37 | H1-3 | P16402 | 147 | KVAGAATPKKSIK | 7.5041 | 7.1521 | 6.1640 | 6.6308 | 7.2276 | 7.0546 | 7.5893 | 7.0566 | 6.9690 | 6.3815 | 6.0531 | 5.9958 | 4.0687 | 5.2235 | 4.0314 | 4.8978 |
| H1-3_S56 | H1-3 | P16402 | 37 | AGKRKASGPPVSE | 5.2457 | 5.8264 | 5.2323 | 5.8330 | 6.3626 | 5.9394 | 5.8941 | 6.3883 | 6.3177 | 7.4776 | 6.5011 | 6.3212 | 6.1343 | 7.2029 | 6.4692 | 6.8541 |
| H1-3_S59 | H1-3 | P16402 | 59 | KERSGVSLAALKK | 3.6478 | 3.9338 | 4.9103 | 4.9393 | 4.4956 | 5.6494 | 5.7282 | 6.5933 | 5.2904 | 6.7789 | 5.8709 | 6.2628 | 7.4845 | 8.7676 | 9.7511 | 9.8962 |
| H1-3_T147 | H1-3 | P16402 | 56 | AASKERSGVSLAA | 4.5672 | 5.5168 | 4.3360 | 4.8800 | 6.0104 | 6.3258 | 7.1800 | 5.7873 | 4.4106 | 4.5902 | 4.7058 | 4.4548 | 8.5199 | 9.6592 | 9.4663 | 9.5897 |
| H1-4_T146 | H1-4 | P10412 | 146 | KATGAATPKKSAK | 7.6144 | 7.4334 | 6.4776 | 6.8507 | 7.0314 | 7.5974 | 7.1053 | 7.0635 | 7.1323 | 6.6468 | 6.4654 | 6.2610 | 3.5871 | 4.8193 | 3.5292 | 4.3851 |
| H1-5_S105 | H1-5 | P16401 | 138 | KKPAGATPKKAKK | 6.8756 | 6.4887 | 6.4859 | 7.1490 | 6.7733 | 6.7628 | 7.2865 | 6.3296 | 6.9570 | 6.9820 | 6.7370 | 6.9976 | 3.9399 | 5.2360 | 3.9319 | 5.0671 |
| H1-5_S107 | H1-5 | P16401 | 107 | GTGASGSFKLNKK | 5.4287 | 7.0502 | 7.7212 | 7.7914 | 6.5805 | 6.2001 | 6.0818 | 8.5309 | 6.5370 | 9.5612 | 8.0266 | 8.2409 | 2.3227 | 3.9752 | 2.7672 | 3.1845 |
| H1-5_T138 | H1-5 | P16401 | 105 | TKGTGASGSFKLN | 5.4616 | 6.1062 | 5.2822 | 6.1873 | 5.7495 | 6.0385 | 6.0653 | 5.9642 | 5.5720 | 6.4252 | 5.6533 | 6.0335 | 6.3988 | 7.9183 | 7.4783 | 7.6658 |
| H1-5_T39 | H1-5 | P16401 | 39 | AAKRKATGPPVSE | 5.6663 | 5.6616 | 5.9420 | 6.1919 | 6.0620 | 6.5431 | 7.4714 | 6.8979 | 6.7288 | 6.8021 | 6.9066 | 6.3327 | 5.0009 | 5.7834 | 5.1177 | 5.0916 |
| H2AX_S122 | H2AX | P16104 | 140 | KKATQASQEYxxx | 5.9686 | 5.9276 | 6.0922 | 7.3596 | 6.3603 | 5.4442 | 6.2174 | 5.8305 | 6.0822 | 5.8674 | 5.7119 | 6.2053 | 6.4237 | 6.4943 | 6.8551 | 7.1599 |
| H2AX_S140 | H2AX | P16104 | 122 | LLPKKTSATVGPK | 7.7262 | 7.3844 | 7.0302 | 7.6865 | 7.6635 | 5.6048 | 7.3666 | 6.5576 | 6.8365 | 5.6717 | 6.5471 | 5.2100 | 4.7195 | 5.2991 | 4.7317 | 4.8645 |
| H2AX_T121;T124 | H2AX | P16104 | 121;124 | NA | 7.7391 | 7.3482 | 7.0506 | 8.6569 | 5.7942 | 5.7025 | 6.9537 | 5.8107 | 6.3289 | 5.9926 | 6.7745 | 6.2749 | 4.6788 | 5.4165 | 4.5534 | 4.9246 |
| H2BC18_S113 | H2BC18 | Q5QNW6-2 | 39 | SRKESYSVYVYKV | 5.7079 | 6.1010 | 6.1605 | 6.7947 | 6.4408 | 4.9434 | 6.5040 | 5.5516 | 5.7165 | 5.1941 | 5.0677 | 4.8780 | 7.5556 | 7.0311 | 9.2981 | 7.0550 |
| H2BC18_S37 | H2BC18 | Q5QNW6-2 | 113 | LAKHAVSEGTKAV | 5.4268 | 5.7863 | 5.4369 | 6.0267 | 5.6741 | 5.8677 | 6.1341 | 6.0500 | 5.7129 | 6.1152 | 5.9137 | 6.2712 | 7.2077 | 7.6476 | 7.2470 | 7.4820 |
| H2BC18_S39 | H2BC18 | Q5QNW6-2 | 37 | KRSRKESYSVYVY | 6.1324 | 6.1749 | 6.0715 | 7.1430 | 6.0920 | 4.9395 | 5.6806 | 5.3810 | 5.3611 | 4.8109 | 4.9466 | 4.5369 | 8.0403 | 7.2069 | 9.5217 | 7.0607 |
| H2BC18_S56 | H2BC18 | Q5QNW6-2 | 57 | PDTGISSKAMGIM | 4.0157 | 4.8262 | 4.6393 | 5.4542 | 4.5450 | 5.5268 | 6.2175 | 5.4858 | 5.1209 | 7.2255 | 6.7586 | 7.2780 | 7.0374 | 8.0413 | 8.1446 | 9.6831 |
| H2BC18_S57 | H2BC18 | Q5QNW6-2 | 56 | HPDTGISSKAMGI | 5.3196 | 5.3394 | 4.7782 | 6.4277 | 5.1480 | 6.8547 | 6.8924 | 5.7459 | 7.1012 | 7.0740 | 8.1160 | 8.1759 | 5.8842 | 6.0218 | 4.9369 | 7.1843 |
| H2BC3_S37 | H2BC3 | P33778 | 37 | KRSRKESYSIYVY | 5.4633 | 5.2809 | 5.5658 | 6.1410 | 6.4363 | 5.5613 | 6.2950 | 5.6997 | 5.4564 | 5.7687 | 5.5020 | 4.8997 | 7.7366 | 7.6625 | 9.2349 | 7.2960 |
| H2BC3_S39 | H2BC3 | P33778 | 39 | SRKESYSIYVYKV | 6.0235 | 6.2814 | 6.4242 | 7.4791 | 6.2393 | 4.5534 | 6.2438 | 5.0273 | 5.3405 | 4.7265 | 4.6775 | 4.3076 | 8.1453 | 7.4383 | 10.1839 | 6.9084 |
| H3-2_S29 | H3-2 | Q5TEC6 | 29 | TKAARKSAPATGG | 7.9945 | 8.3870 | 6.8234 | 8.2293 | 6.7456 | 7.7871 | 6.8864 | 7.4512 | 6.7053 | 7.2223 | 6.9132 | 6.7768 | 2.3906 | 4.0688 | 2.2317 | 3.3869 |
| H3-2_S58 | H3-2 | Q5TEC6 | 58 | IRRYQKSTELLIR | 6.2693 | 6.2167 | 5.4833 | 6.1890 | 6.1780 | 6.0328 | 6.2089 | 6.5360 | 5.7766 | 5.8475 | 5.5891 | 5.8761 | 6.6412 | 7.1324 | 7.0964 | 6.9267 |
| H3-3A_S29 | H3-3A | P84243 | 29 | TKAARKSAPSTGG | 7.8142 | 8.3226 | 6.8086 | 8.0643 | 6.5577 | 6.8135 | 8.0743 | 7.3442 | 6.9266 | 7.1859 | 7.1585 | 7.0128 | 2.4550 | 3.7853 | 2.4230 | 3.2537 |
| H3BNC9_S103 | H3BNC9 | H3BNC9 | 562 | KLLDFGSLSNLQV | 7.0269 | 6.6692 | 6.4747 | 6.8068 | 7.2628 | 6.1113 | 6.5112 | 6.0810 | 6.6618 | 5.4026 | 5.7727 | 5.3518 | 5.8833 | 6.5358 | 5.8635 | 5.5846 |
| H3BNC9_S562 | H3BNC9 | H3BNC9 | 103 | TRPILDSRSSSPS | 7.6445 | 7.3237 | 7.1910 | 6.8765 | 6.4376 | 6.2665 | 6.9575 | 6.3219 | 6.8477 | 6.0777 | 6.9844 | 6.7706 | 4.6772 | 5.0644 | 4.0498 | 4.5089 |
| H4C1_S48 | H4C1 | P62805 | 48 | GGVKRISGLIYEE | 6.8309 | 6.5293 | 5.9651 | 6.7845 | 6.5934 | 5.9745 | 6.6433 | 6.2957 | 6.2697 | 6.0073 | 5.7333 | 5.9759 | 6.0369 | 6.2517 | 6.0087 | 6.0997 |
| H4C1_T81 | H4C1 | P62805 | 81 | EHAKRKTVTAMDV | 6.4754 | 6.2199 | 6.0073 | 6.9458 | 6.0988 | 6.3055 | 6.3670 | 6.6452 | 5.8682 | 6.3530 | 6.2115 | 6.3095 | 5.7674 | 5.9581 | 6.1875 | 6.2799 |
| HABP4_S108 | HABP4 | Q5JVS0 | 108 | VAQRPDSPGGLQ | 6.2874 | 6.1811 | 7.1644 | 6.1881 | 6.4580 | 6.5515 | 6.4390 | 6.5990 | 6.8767 | 6.8660 | 6.5249 | 5.4710 | 5.4819 | 5.2305 | 5.2920 |
| HABP4_S7 | HABP4 | Q5JVS0 | 7 | MKGALGSPVAAAG | 6.0695 | 6.3893 | 6.5124 | 6.6597 | 5.8874 | 6.0018 | 5.6636 | 6.3733 | 6.1066 | 6.1724 | 6.6355 | 6.9874 | 6.1665 | 5.9322 | 5.9034 | 6.5389 |
| HABP4_S97;S108 | HABP4 | Q5JVS0 | 97;108 | NA | 6.6198 | 7.3503 | 7.3662 | 7.4791 | 5.7096 | 6.5342 | 7.6217 | 6.8516 | 7.7131 | 6.8825 | 7.5853 | 7.7519 | 3.4937 | 4.1333 | 3.3921 | 3.5155 |
| HACD3_S114 | HACD3 | Q9P035 | 114 | DRWLDESDAEMEL | 6.3157 | 6.5039 | 6.1576 | 6.5152 | 6.0987 | 6.0257 | 6.3455 | 6.2628 | 6.0846 | 6.1668 | 6.0696 | 6.0528 | 6.2173 | 6.4535 | 6.4468 | 6.2836 |
| HACD3_S135 | HACD3 | Q9P035 | 135 | NKLRLESEGSPET | 5.7819 | 6.5434 | 7.2922 | 6.8188 | 5.6373 | 6.1893 | 5.6690 | 5.7210 | 5.8759 | 7.2236 | 7.4508 | 6.3454 | 5.1715 | 6.8429 | 5.6715 | 5.7654 |
| HACE1_S385 | HACE1 | Q8IYU2 | 385 | MKNKRDSTEITSI | 6.2436 | 6.6466 | 6.9713 | 5.8521 | 6.0697 | 6.2214 | 6.2864 | 6.8854 | 6.7051 | 6.3374 | 6.4695 | 6.8773 | 5.5575 | 5.7091 | 5.5010 | 5.6667 |
| HADHA_S756 | HADHA | P40939 | 756 | LADHANSPNKKFY | 6.2231 | 6.1726 | 6.0829 | 6.9185 | 6.1677 | 6.1995 | 6.0748 | 6.0618 | 6.3771 | 6.2992 | 6.1141 | 6.0083 | 6.3772 | 6.4644 | 6.3191 | 6.1398 |
| HARS2_S67 | HARS2 | P49590 | 67 | KGTRDLSPQHMVV | 6.6020 | 6.5367 | 6.0509 | 6.4564 | 6.0367 | 6.0345 | 6.0818 | 6.2885 | 6.2549 | 6.1541 | 6.0256 | 6.1759 | 6.5565 | 6.4117 | 6.2351 | 6.0989 |
| HASPIN_S147 | HASPIN | Q8TF76 | 147 | RDSGRLSPDLSVC | 6.1674 | 6.2667 | 6.1047 | 6.8196 | 6.2021 | 6.0128 | 6.2755 | 5.9691 | 6.7437 | 6.4813 | 6.3142 | 6.6012 | 5.6145 | 6.1140 | 6.1680 | 6.1452 |
| HASPIN_T112 | HASPIN | Q8TF76 | 97 | RPSLTVTPKRWKL | 6.7849 | 6.9151 | 6.4198 | 7.2080 | 6.7576 | 6.3926 | 7.1373 | 6.8763 | 7.0875 | 6.7849 | 6.4282 | 6.8676 | 4.2539 | 4.8279 | 4.2536 | 5.0050 |
| HASPIN_T97 | HASPIN | Q8TF76 | 112 | RPSLTVTPRRLGL | 8.8598 | 7.1785 | 6.6083 | 7.2511 | 5.5902 | 6.7730 | 6.3900 | 7.3600 | 8.8373 | 8.6415 | 7.1790 | 7.2986 | 2.8805 | 3.4936 | 2.4796 | 3.1791 |
| HAT1_S361 | HAT1 | O14929 | 361 | IKRRLISPYKKKQ | 5.7641 | 6.6721 | 6.6189 | 6.0499 | 7.2691 | 7.8411 | 6.7539 | 7.0309 | 6.5501 | 6.8825 | 6.9878 | 6.0571 | 5.2791 | 5.6584 | 4.4806 | 4.1042 |
| HAUS6_S387 | HAUS6 | Q7Z4H7 | 584 | RNQIPRTPENLIT | 6.5813 | 6.7732 | 6.6882 | 6.6098 | 6.0391 | 6.4027 | 6.7483 | 6.5074 | 6.6335 | 6.4289 | 7.0554 | 6.5072 | 5.0608 | 5.3848 | 5.1936 | 5.3857 |
| HAUS6_S406 | HAUS6 | Q7Z4H7 | 805 | FKLEPNSPMHGGT | 6.4942 | 6.7968 | 6.6938 | 7.3241 | 6.4037 | 6.5751 | 7.0368 | 7.1980 | 6.7967 | 6.8079 | 6.8420 | 7.1933 | 4.1021 | 4.7688 | 4.2648 | 4.7018 |
| HAUS6_S459 | HAUS6 | Q7Z4H7 | 459 | LHDLANSPASFLS | 6.2726 | 6.6557 | 6.8849 | 7.2171 | 5.9584 | 6.7375 | 6.5727 | 6.7181 | 7.0545 | 6.6351 | 7.1699 | 7.4805 | 4.4077 | 5.1596 | 4.1768 | 4.8988 |
| HAUS6_S507 | HAUS6 | Q7Z4H7 | 552 | RAVLSDSPQLSEG | 6.7741 | 7.3004 | 6.7731 | 7.1994 | 6.2567 | 6.9543 | 6.6436 | 7.0606 | 7.0779 | 6.7795 | 7.2430 | 7.3428 | 3.8950 | 4.6199 | 3.6558 | 4.4239 |
| HAUS6_S552 | HAUS6 | Q7Z4H7 | 406 | DLLPPMSPLSFDP | 5.8795 | 5.6410 | 6.3080 | 6.4355 | 6.2894 | 6.2052 | 6.0499 | 6.4737 | 6.4858 | 6.2979 | 6.3925 | 6.4311 | 6.4553 | 6.0406 | 5.9176 | 6.6972 |
| HAUS6_S660 | HAUS6 | Q7Z4H7 | 507 | EFEVENSPLSDVA | 6.5196 | 7.7033 | 7.0625 | 7.7752 | 5.9027 | 7.6848 | 7.5917 | 7.1152 | 7.2305 | 7.3292 | 7.9113 | 7.7471 | 2.7677 | 3.8499 | 2.5726 | 3.2368 |
| HAUS6_S805 | HAUS6 | Q7Z4H7 | 906 | IGERKRSLSPLIK | 7.2322 | 4.7088 | 4.9390 | 4.1283 | 5.8753 | 5.3990 | 4.8630 | 5.1078 | 10.2492 | 7.6320 | 7.8985 | 8.1076 | 7.3105 | 8.3190 | 3.7402 | 4.4896 |
| HAUS6_S906 | HAUS6 | Q7Z4H7 | 387 | KEFLGLSPFSLIK | 6.6570 | 7.2582 | 7.1320 | 7.4242 | 6.2657 | 6.6218 | 6.7189 | 6.8519 | 6.5238 | 7.0035 | 7.3875 | 7.3035 | 3.7753 | 4.4837 | 4.0236 | 4.5694 |
| HAUS6_S908 | HAUS6 | Q7Z4H7 | 908 | ERKRSLSPLIKFS | 6.2756 | 6.2415 | 6.3097 | 5.7555 | 6.1094 | 6.8169 | 6.2775 | 6.4614 | 6.9578 | 6.5125 | 6.6215 | 6.7007 | 5.6858 | 5.9898 | 5.6063 | 5.6780 |
| HAUS6_T584 | HAUS6 | Q7Z4H7 | 660 | TEEKVISDCECVP | 6.6291 | 6.9792 | 7.8423 | 7.2080 | 6.5563 | 7.0087 | 6.5037 | 7.4356 | 6.0577 | 6.5279 | 5.6648 | 6.2512 | 4.9513 | 5.6787 | 4.5509 | 5.1547 |
| HAUS8_S131 | HAUS8 | Q9BT25 | 21 | GPTNSSSAKKKDK | 7.5981 | 6.7733 | 7.4452 | 8.4178 | 6.0313 | 7.1985 | 7.3644 | 6.3247 | 7.7746 | 8.3115 | 8.0897 | 8.2539 | 2.6242 | 3.1342 | 1.9398 | 2.7186 |
| HAUS8_S21 | HAUS8 | Q9BT25 | 131 | KKPESTSFSAPRK | 6.5611 | 6.2999 | 6.4952 | 6.9824 | 5.8748 | 6.5308 | 6.8251 | 6.8585 | 6.6121 | 6.1630 | 5.7627 | 5.7568 | 5.7518 | 6.9025 | 5.1070 | 5.5165 |
| HBS1L_S127 | HBS1L | Q9Y450 | 33 | EDDYCISPSTAAQ | 5.6899 | 6.1894 | 6.2271 | 6.1942 | 5.8792 | 5.5194 | 6.2262 | 6.3301 | 6.3825 | 6.5460 | 6.6029 | 6.6185 | 6.2665 | 6.8607 | 6.1418 | 6.3255 |
| HBS1L_S246 | HBS1L | Q9Y450 | 231 | SEEQSSTPAPVKK | 7.5802 | 7.8172 | 6.9794 | 7.5455 | 6.2843 | 7.8181 | 7.5954 | 7.2462 | 7.3667 | 6.9189 | 7.2424 | 7.0177 | 2.6820 | 3.9564 | 2.5699 | 3.3797 |
| HBS1L_S33 | HBS1L | Q9Y450-2 | 246 | PSVRLSSTDSLES | 4.9009 | 4.7493 | 5.3485 | 5.3423 | 5.5752 | 5.6554 | 5.4036 | 5.4790 | 5.3225 | 5.3347 | 5.3658 | 5.3074 | 9.0926 | 9.1632 | 9.6414 | 8.3180 |
| HBS1L_T231 | HBS1L | Q9Y450 | 127 | EQDRVQSLKDKNE | 6.9702 | 7.3661 | 9.5446 | 7.6516 | 5.8535 | 7.3387 | 6.7379 | 6.2477 | 6.2690 | 7.0638 | 8.4661 | 7.2454 | 3.3030 | 3.8397 | 2.8266 | 3.2762 |
| HCFC1_S1507 | HCFC1 | P51610-4 | 2064 | NKRPMSSPEMKSA | 7.3654 | 7.2624 | 6.5451 | 7.0249 | 7.0351 | 6.6167 | 6.7465 | 6.7539 | 7.3185 | 6.7875 | 7.1319 | 7.0615 | 4.0518 | 4.3095 | 3.7554 | 4.2339 |
| HCFC1_S1551 | HCFC1 | P51610 | 1507 | PEELQVSPGPRQQ | 6.4563 | 6.1174 | 6.3662 | 6.9208 | 6.5058 | 6.0198 | 6.2037 | 6.8496 | 6.7162 | 6.4511 | 6.1980 | 6.5983 | 5.5170 | 5.8447 | 5.4259 | 5.8091 |
| HCFC1_S1882 | HCFC1 | P51610-4 | 598 | TVKVASSPVMVSN | 6.3349 | 7.0120 | 6.9952 | 6.2223 | 6.7643 | 6.9685 | 6.3610 | 6.7047 | 6.8624 | 6.8754 | 6.6508 | 6.7226 | 4.5934 | 5.7153 | 4.4956 | 4.7216 |
| HCFC1_S2063 | HCFC1 | P51610-4 | 666 | TITLVKSPISVPG | 6.3297 | 6.2953 | 6.5521 | 6.8406 | 6.7086 | 6.2638 | 6.8267 | 6.5408 | 6.9848 | 6.3958 | 6.3900 | 6.2832 | 5.4633 | 5.4271 | 5.3211 | 5.3770 |
| HCFC1_S2063;S2064 | HCFC1 | P51610-4 | 2063 | ANKRPMSSPEMKS | 6.9792 | 7.1514 | 6.6022 | 6.9938 | 6.7992 | 6.8900 | 6.4340 | 6.5671 | 7.0009 | 6.8520 | 7.3250 | 7.7767 | 4.3579 | 4.5349 | 3.6408 | 4.0952 |
| HCFC1_S2064 | HCFC1 | P51610-4 | 794 | SSPGIKSPITIIT | 6.4577 | 6.8476 | 6.3927 | 6.6427 | 6.4134 | 6.4875 | 7.5537 | 6.5562 | 6.5176 | 6.4681 | 5.9877 | 6.6230 | 4.9058 | 5.4146 | 5.1589 | 5.5727 |
| HCFC1_S598 | HCFC1 | P51610-4 | 1551 | PEELQVSPGPRQQ | 5.8331 | 6.3337 | 6.6052 | 6.7021 | 5.9702 | 6.4398 | 6.3700 | 6.0753 | 6.4420 | 6.8895 | 6.2136 | 6.9458 | 5.7981 | 6.0921 | 5.4030 | 5.8865 |
| HCFC1_S666 | HCFC1 | P51610-4 | 405 | KYDIPATAATATS | 6.4471 | 6.0891 | 6.6423 | 7.3035 | 5.8311 | 6.4973 | 6.2432 | 6.1738 | 6.9689 | 7.0697 | 6.6625 | 6.8482 | 5.3421 | 5.4668 | 5.0346 | 5.3797 |
| HCFC1_S794 | HCFC1 | P51610-4 | 1882 | PDDAVPSDDDLGT | 6.1227 | 6.3085 | 6.7705 | 6.3283 | 6.5808 | 5.9378 | 6.7626 | 7.1198 | 6.0545 | 5.7298 | 6.3490 | 6.4831 | 5.6306 | 6.0505 | 5.8370 | 5.9346 |
| HCFC1_T405 | HCFC1 | P51610-4 | 2063;2064 | NA | 7.4105 | 7.7147 | 8.0262 | 7.6513 | 7.0730 | 6.8601 | 7.2555 | 7.1847 | 6.5353 | 6.3915 | 6.8344 | 6.6527 | 3.5712 | 3.8314 | 3.2721 | 3.7354 |
| HDAC1_S393 | HDAC1 | Q13547 | 393 | DAIPEESGDEDED | 6.0706 | 6.3170 | 6.4169 | 6.4044 | 6.1624 | 6.0810 | 6.2009 | 6.1674 | 6.3545 | 6.3704 | 6.2653 | 6.4459 | 6.4197 | 6.3628 | 6.0648 | 5.8960 |
| HDAC1_S406 | HDAC1 | Q13547 | 409 | KRISICSSDKRIA | 6.1584 | 6.1778 | 6.2265 | 6.8675 | 6.1915 | 6.1566 | 6.9348 | 6.5065 | 6.2240 | 6.1808 | 6.5378 | 6.7022 | 5.7253 | 5.6903 | 5.8094 | 5.9106 |
| HDAC1_S409 | HDAC1 | Q13547 | 406 | DPDKRISICSSDK | 6.2086 | 6.2244 | 6.3145 | 7.4955 | 6.2576 | 7.4876 | 7.3453 | 5.9463 | 7.0619 | 6.8085 | 6.7303 | 6.7727 | 4.1095 | 5.2149 | 4.5030 | 5.5193 |
| HDAC1_S410 | HDAC1 | Q13547 | 410 | RISICSSDKRIAC | 5.9609 | 6.5130 | 6.1496 | 6.5165 | 5.5622 | 5.3956 | 5.9848 | 6.5387 | 5.9937 | 6.0885 | 6.0265 | 7.5488 | 6.2799 | 6.5378 | 7.0384 | 5.8652 |
| HDAC1_S421;S423 | HDAC1 | Q13547 | 421;423 | NA | 5.6817 | 5.7993 | 6.0700 | 6.0561 | 6.3738 | 5.9476 | 6.5665 | 6.0562 | 5.8698 | 6.1735 | 6.1111 | 6.0481 | 7.1694 | 7.3916 | 6.6811 | 6.9141 |
| HDAC2_S394 | HDAC2 | Q92769 | 394 | DAVHEDSGDEDGE | 6.1118 | 6.6604 | 6.6342 | 6.6723 | 6.1691 | 6.2737 | 6.2367 | 5.9876 | 6.2263 | 6.2689 | 6.3469 | 6.1545 | 6.2537 | 6.0047 | 6.1833 | 5.8159 |
| HDAC2_S422;S424 | HDAC2 | Q92769 | 422;424 | NA | 6.2705 | 5.9819 | 6.5000 | 6.8787 | 6.6568 | 6.1897 | 6.1362 | 5.8372 | 6.6348 | 6.3562 | 6.2258 | 6.7041 | 6.4099 | 5.5865 | 5.9467 | 5.6851 |
| HDAC3_S425 | HDAC3 | O15379-2 | 425 | HDNDKESDVEIxx | 6.0997 | 6.1469 | 6.5613 | 6.5081 | 6.0237 | 5.9339 | 6.1383 | 6.4803 | 6.5781 | 5.8195 | 5.9346 | 5.9837 | 6.3187 | 6.5423 | 6.1483 | 6.7825 |
| HDAC3_S75 | HDAC3 | O15379-2 | 75 | DFLQRVSPTNMQG | 6.3450 | 5.8674 | 6.2413 | 6.4665 | 6.3305 | 7.0861 | 6.4097 | 7.2783 | 6.1026 | 6.3457 | 6.5788 | 6.3807 | 5.8278 | 5.6178 | 5.5636 | 5.5583 |
| HDAC4_S467 | HDAC4 | P56524 | 467 | PLGRTQSAPLPQN | 5.5431 | 5.9363 | 6.7235 | 5.9168 | 6.2901 | 6.8857 | 5.8615 | 6.6269 | 6.7486 | 6.9971 | 6.6480 | 6.6803 | 5.9403 | 6.3623 | 5.2228 | 6.6165 |
| HDAC4_S565 | HDAC4 | P56524 | 632 | PLSRAQSSPASAT | 6.2039 | 5.7171 | 6.7477 | 6.3086 | 6.4188 | 6.4240 | 5.9645 | 5.9934 | 6.6097 | 6.6130 | 6.6633 | 6.7987 | 5.3642 | 6.6082 | 5.5003 | 6.0648 |
| HDAC4_S632 | HDAC4 | P56524 | 565 | KQEPIESDEEEAE | 6.5351 | 6.5878 | 6.9453 | 5.4995 | 6.4510 | 7.2002 | 6.6476 | 6.1149 | 6.8863 | 6.8584 | 7.5203 | 6.5265 | 6.5037 | 5.9881 | 4.8472 |
| HDAC5_S260 | HDAC5 | Q9UQL6-3 | 260 | PLRKTASEPNLKV | 6.1582 | 6.0149 | 6.7713 | 5.7709 | 6.1420 | 6.5319 | 6.1486 | 6.1988 | 5.8617 | 6.0724 | 6.0846 | 6.1707 | 6.5399 | 6.7616 | 6.5045 | 6.2679 |
| HDAC5_S280 | HDAC5 | Q9UQL6-3 | 499 | PLSRTQSSPLPQS | 5.9501 | 4.7202 | 7.2937 | 6.6284 | 5.7203 | 6.9654 | 6.3440 | 6.2652 | 5.7947 | 7.3243 | 6.1886 | 6.1339 | 6.3834 | 6.2958 | 5.0813 | 6.9107 |
| HDAC5_S499 | HDAC5 | Q9UQL6-3 | 280 | VAERRSSPLLRRK | 6.7155 | 6.8448 | 7.5005 | 6.3829 | 5.9231 | 6.9332 | 6.3694 | 6.8431 | 6.3145 | 6.9202 | 6.2463 | 6.5520 | 5.1515 | 5.6157 | 4.9888 | 4.6983 |
| HDAC5_S662 | HDAC5 | Q9UQL6-3 | 662 | ALGRTQSSPAAPG | 5.5962 | 5.9333 | 6.3487 | 5.8669 | 7.2236 | 6.3622 | 5.8371 | 6.4373 | 5.6114 | 6.1177 | 6.3248 | 5.7856 | 7.4554 | 6.4843 | 6.4531 | 6.1624 |
| HDAC6_S22 | HDAC6 | Q9UBN7 | 22 | SRQNPQSPPQDSS | 6.7299 | 6.3837 | 7.4733 | 6.4930 | 6.0748 | 6.9380 | 6.3758 | 6.4413 | 6.9334 | 7.0499 | 6.6029 | 6.9053 | 4.5448 | 5.2496 | 4.5088 | 5.2954 |
| HDAC7_S165 | HDAC7 | Q8WUI4-8 | 237 | PLLRKESAPPSLR | 6.0523 | 5.9422 | 6.7099 | 6.1402 | 6.4859 | 6.2922 | 5.7010 | 6.7525 | 6.6881 | 6.5951 | 6.9047 | 7.0276 | 5.8616 | 5.9720 | 5.4978 | 5.3769 |
| HDAC7_S211 | HDAC7 | Q8WUI4-8 | 414 | PLSRTRSEPLPPS | 5.9189 | 6.1989 | 6.2902 | 6.4422 | 6.1052 | 6.0651 | 6.3482 | 6.6638 | 7.0040 | 6.7009 | 6.8985 | 6.8019 | 5.9166 | 5.6212 | 5.3774 | 5.6470 |
| HDAC7_S237 | HDAC7 | Q8WUI4-8 | 211 | PLRKTVSEPNLKL | 6.2881 | 6.2882 | 6.7440 | 5.8839 | 6.1987 | 6.4125 | 6.0922 | 6.5979 | 6.3854 | 6.3199 | 6.4670 | 6.5441 | 5.9051 | 6.2293 | 5.8387 | 5.8050 |
| HDAC7_S414 | HDAC7 | Q8WUI4-8 | 542 | PLSRAQSSPAAPA | 5.7915 | 6.4672 | 6.4014 | 5.8944 | 6.5072 | 6.5589 | 5.7235 | 6.4023 | 6.3225 | 6.5218 | 6.7134 | 6.3771 | 6.1518 | 6.3498 | 5.9282 | 5.8888 |
| HDAC7_S542 | HDAC7 | Q8WUI4-8 | 165 | RTVHPNSPGIPYR | 6.9114 | 7.6440 | 7.1485 | 6.8489 | 5.7363 | 7.3964 | 7.1370 | 6.7524 | 7.1994 | 7.2897 | 7.3419 | 7.1710 | 3.4353 | 4.4918 | 3.3312 | 4.1648 |
| HDAC7_S651 | HDAC7 | Q8WUI4-8 | 651 | YGTNPLSRLKLDN | 6.2660 | 5.4622 | 7.2636 | 6.2366 | 5.9590 | 5.3014 | 6.1924 | 5.7319 | 6.3221 | 6.0712 | 5.8775 | 6.4632 | 7.3134 | 6.7673 | 6.4632 | 6.3091 |
| HDAC9_S454 | HDAC9 | Q9UKV0-7 | 454 | PLNRTQSAPLPQS | 5.3617 | 5.7345 | 6.1496 | 6.0479 | 6.2138 | 6.1240 | 5.4822 | 6.0393 | 6.0557 | 6.1957 | 6.5873 | 6.8716 | 6.8337 | 7.0874 | 6.7108 | 6.5046 |
| HDGF_S119 | HDGF | P51858-3 | 181 | GDLLEDSPKRPKE | 5.5974 | 5.5690 | 6.4809 | 5.2649 | 5.8443 | 5.8110 | 5.6713 | 5.7264 | 6.3755 | 6.4229 | 6.5270 | 6.3927 | 7.3812 | 7.2718 | 6.8168 | 6.8469 |
| HDGF_S149 | HDGF | P51858-3 | 119 | ASGYQSSQKKSCV | 5.4572 | 5.8984 | 6.6585 | 5.7861 | 5.4201 | 6.3178 | 5.3199 | 5.8638 | 6.2959 | 6.7355 | 6.9371 | 7.0894 | 6.7087 | 7.1812 | 6.1232 | 6.2072 |
| HDGF_S181 | HDGF | P51858-3 | 149 | GNAEGSSDEEGKL | 5.8521 | 5.3471 | 7.2281 | 5.2325 | 4.6169 | 6.5549 | 4.3030 | 4.4016 | 5.9848 | 6.5220 | 7.3568 | 8.5216 | 6.7432 | 6.5320 | 8.4592 | 7.2442 |
| HDGF_T216 | HDGF | P51858-3 | 216 | EVEKNSTPSEPGS | 6.3271 | 6.8043 | 6.6630 | 6.1612 | 5.7628 | 7.0340 | 6.3029 | 5.7915 | 6.4620 | 7.5545 | 6.9009 | 6.9529 | 4.8996 | 5.8891 | 4.8526 | 5.6416 |
| HDGFL2_S137 | HDGFL2 | Q7Z4V5-4 | 454 | ERTRKRSEGFSMD | 6.6835 | 7.1022 | 7.7265 | 7.4775 | 6.4052 | 6.6453 | 6.8314 | 6.8463 | 7.3995 | 7.4401 | 8.3174 | 7.7539 | 3.1431 | 3.7600 | 3.0092 | 3.4590 |
| HDGFL2_S149 | HDGFL2 | Q7Z4V5-4 | 240 | SDSKADSDGAPKE | 5.2140 | 6.1040 | 6.2300 | 6.0302 | 6.0774 | 5.7221 | 5.6811 | 5.5722 | 6.2232 | 6.8296 | 6.3056 | 6.2013 | 6.6671 | 7.1219 | 6.9222 | 7.0982 |
| HDGFL2_S230;S232;S234 | HDGFL2 | Q7Z4V5-4 | 490 | FALKVDSPDVKRC | 6.0829 | 7.0644 | 7.0143 | 6.4429 | 5.9080 | 6.2081 | 6.3801 | 5.9650 | 6.7144 | 7.1359 | 7.0126 | 7.2251 | 4.9154 | 5.3683 | 4.8112 | 5.7513 |
| HDGFL2_S232 | HDGFL2 | Q7Z4V5-4 | 656 | DLDRPGSDRQERE | 5.4944 | 5.9650 | 6.0637 | 5.8055 | 5.5344 | 6.0304 | 5.8864 | 5.9611 | 6.2984 | 5.8420 | 5.9499 | 6.0249 | 6.8070 | 7.2064 | 7.5511 | 7.5796 |
| HDGFL2_S240 | HDGFL2 | Q7Z4V5-4 | 630 | HEEGRDSEEGPRC | 5.5987 | 5.9334 | 6.2813 | 5.3399 | 6.2250 | 5.2499 | 6.7229 | 6.1556 | 6.8495 | 5.7993 | 5.4627 | 5.8865 | 6.7454 | 7.2368 | 7.2114 | 7.3015 |
| HDGFL2_S366;S369;S370 | HDGFL2 | Q7Z4V5-4 | 232 | KKAPSASDSDSKA | 5.6608 | 6.8195 | 7.3096 | 6.6040 | 5.9662 | 6.6865 | 5.7175 | 6.0307 | 8.3387 | 8.2611 | 9.5824 | 8.7307 | 3.7897 | 4.7255 | 2.9120 | 2.8651 |
| HDGFL2_S454 | HDGFL2 | Q7Z4V5-4 | 137 | VTATAASDRMESD | 5.2210 | 6.0179 | 5.4509 | 6.7868 | 6.4033 | 6.5916 | 6.4332 | 6.4777 | 6.0517 | 6.2767 | 6.2091 | 6.0836 | 6.5124 | 6.9471 | 6.1386 | 6.3985 |
| HDGFL2_S490 | HDGFL2 | Q7Z4V5-4 | 149 | DSDSDKSSDNSGL | 6.2846 | 7.5824 | 7.6966 | 6.6316 | 5.2843 | 5.3231 | 5.6309 | 5.2066 | 6.3476 | 7.0175 | 6.9925 | 6.9303 | 5.5375 | 6.0164 | 5.3292 | 6.1889 |
| HDGFL2_S613 | HDGFL2 | Q7Z4V5-4 | 639 | GPRCGSSEDLHDV | 5.3681 | 5.9988 | 6.2415 | 6.0133 | 6.3107 | 5.1717 | 5.7415 | 5.5361 | 6.6586 | 6.2803 | 6.3245 | 6.6426 | 6.9884 | 6.3218 | 6.9006 | 7.5016 |
| HDGFL2_S630 | HDGFL2 | Q7Z4V5-4 | 613 | VNGEATSQKGESA | 4.8641 | 5.5864 | 5.9388 | 5.8229 | 6.1236 | 6.8563 | 6.1209 | 5.1405 | 6.1591 | 6.0220 | 6.8778 | 6.4791 | 7.4941 | 7.7768 | 6.3106 | 6.4270 |
| HDGFL2_S639 | HDGFL2 | Q7Z4V5-4 | 366;369;370 | NA | 6.0202 | 5.6520 | 6.6060 | 6.1466 | 5.6410 | 5.6033 | 6.6494 | 5.2667 | 6.4452 | 7.0227 | 7.2590 | 7.0833 | 6.7728 | 6.3919 | 5.2230 | 6.2172 |
| HDGFL2_S656 | HDGFL2 | Q7Z4V5-4 | 230;232;234 | NA | 4.9195 | 5.7369 | 6.0894 | 5.2135 | 5.5805 | 5.0789 | 5.4907 | 4.9091 | 7.0831 | 6.6825 | 7.0490 | 6.7472 | 7.6744 | 8.1597 | 6.6691 | 6.9166 |
| HDGFL3_S121;S122 | HDGFL3 | Q9Y3E1 | 121;122 | NA | 5.4212 | 5.8906 | 5.9079 | 6.3561 | 5.7465 | 5.1816 | 5.5818 | 5.6575 | 6.3117 | 6.3162 | 6.4349 | 6.8630 | 6.7255 | 7.4794 | 7.1224 | 7.0038 |
| HDLBP_S1237 | HDLBP | Q00341 | 28 | QQIKVATLNSEEE | 7.1039 | 7.4174 | 7.1584 | 6.8167 | 6.6357 | 7.2812 | 6.7678 | 6.9080 | 6.5240 | 6.1881 | 5.9399 | 5.9892 | 4.7002 | 5.0184 | 4.5418 | 4.7093 |
| HDLBP_S31 | HDLBP | Q00341 | 31 | KVATLNSEEESDP | 6.3935 | 7.2139 | 6.6841 | 6.8037 | 6.5121 | 6.0640 | 6.3037 | 6.0887 | 6.5706 | 6.1821 | 6.8698 | 6.1695 | 5.3258 | 5.9696 | 5.4202 | 5.4287 |
| HDLBP_T28 | HDLBP | Q00341 | 437 | LINRMDYVEINID | 5.4757 | 5.3082 | 5.6621 | 5.5843 | 6.2708 | 6.4407 | 6.0679 | 4.7161 | 5.6556 | 5.0629 | 5.9806 | 5.6621 | 8.2738 | 6.8370 | 8.0260 | 6.9762 |
| HDLBP_Y437 | HDLBP | Q00341 | 1237 | APWTASSSEKAPD | 6.3557 | 7.1434 | 6.8166 | 7.3969 | 6.5410 | 6.3735 | 6.5393 | 6.6693 | 7.2136 | 7.3065 | 7.1717 | 6.9296 | 3.6750 | 3.5755 | 3.9958 | 4.4966 |
| HEATR1_S516 | HEATR1 | Q9H583 | 516 | KEGVDESFIKEAV | 6.5944 | 6.8660 | 7.8461 | 7.3474 | 6.0142 | 6.0833 | 6.3352 | 5.8093 | 5.5476 | 6.7796 | 5.4167 | 5.4982 | 5.3131 | 6.3944 | 6.3253 | 5.8291 |
| HEATR3_T340 | HEATR3 | Q7Z4Q2 | 340 | RRVRRKTFVSDLL | 6.4520 | 6.2121 | 7.2039 | 6.8352 | 6.4640 | 6.1190 | 6.2995 | 5.9046 | 6.5814 | 5.7692 | 6.7526 | 6.6310 | 5.4458 | 5.8530 | 5.5341 | 5.9427 |
| HEATR5A_S1704 | HEATR5A | Q86XA9 | 1704 | NPKLTGSPGVKAT | 6.2923 | 6.4711 | 6.1294 | 6.1579 | 5.9122 | 6.7364 | 6.1254 | 6.5056 | 6.3267 | 6.5287 | 6.3855 | 6.8754 | 5.6887 | 6.4188 | 5.6991 | 5.7469 |
| HEATR5B_S1737 | HEATR5B | Q9P2D3 | 1737 | STKVSDSPSHIAT | 6.1014 | 6.3599 | 6.5377 | 6.2499 | 6.2668 | 6.7917 | 6.1224 | 6.5051 | 6.1902 | 6.2492 | 7.0394 | 6.5800 | 5.5203 | 6.3127 | 5.3942 | 5.7790 |
| HEATR6_S336 | HEATR6 | Q6AI08 | 643 | EETSVSSPKGSSE | 5.7860 | 5.9947 | 6.5005 | 6.1436 | 6.1010 | 6.6529 | 6.3447 | 6.7940 | 6.6549 | 6.0035 | 6.4354 | 7.0974 | 5.9548 | 6.1590 | 5.1036 | 6.2738 |
| HEATR6_S643 | HEATR6 | Q6AI08 | 336 | EEEEKESSGEIEA | 6.1172 | 5.7120 | 7.0983 | 6.3509 | 6.6411 | 6.6753 | 6.7309 | 6.5925 | 5.8994 | 5.5260 | 6.6634 | 6.2584 | 6.8996 | 5.9891 | 5.3913 | 6.4547 |
| HEBP2_S181 | HEBP2 | Q9Y5Z4 | 181 | YTAGYNSPVKLLN | 7.2681 | 6.4684 | 6.6060 | 5.7060 | 6.3230 | 6.7959 | 5.9211 | 7.0394 | 6.3921 | 7.0411 | 6.4982 | 6.9166 | 4.9747 | 5.8899 | 4.8342 | 5.3254 |
| HECTD1_S1488 | HECTD1 | Q9ULT8 | 1772 | VLKRQFSALVPAF | 6.5838 | 5.9464 | 6.9931 | 6.5196 | 6.2580 | 6.0113 | 6.0381 | 6.1925 | 6.4010 | 6.1450 | 6.1784 | 6.4781 | 6.1431 | 6.2482 | 6.0283 | 5.8350 |
| HECTD1_S1772 | HECTD1 | Q9ULT8 | 632 | TLAGPSSDDENEE | 6.3819 | 6.3695 | 7.2262 | 6.4030 | 6.0746 | 6.0810 | 5.9893 | 6.3353 | 6.2245 | 6.7164 | 6.4808 | 6.7097 | 5.8151 | 6.0107 | 5.4356 | 5.7463 |
| HECTD1_S632 | HECTD1 | Q9ULT8 | 1760 | KGGRRRTWDDDYV | 6.6122 | 6.6063 | 7.4759 | 6.5224 | 6.0335 | 6.5536 | 6.5934 | 6.3937 | 6.0821 | 5.9648 | 6.5678 | 6.8592 | 5.8548 | 5.2908 | 5.1877 | 5.4017 |
| HECTD1_T1760 | HECTD1 | Q9ULT8 | 1488 | GIVSVSSPDVSSV | 6.0256 | 6.0207 | 6.5689 | 6.1512 | 6.4275 | 6.1516 | 6.2300 | 6.0126 | 6.4905 | 6.6393 | 6.3213 | 6.1753 | 5.6821 | 6.8399 | 6.3599 | 5.9037 |
| HEG1_S1332 | HEG1 | Q9ULI3 | 1332 | MTDVYYSPTSVRN | 6.6111 | 6.6502 | 6.8747 | 6.8154 | 6.8095 | 6.4613 | 6.3965 | 7.0102 | 6.5210 | 6.8703 | 7.1014 | 6.7575 | 5.3063 | 5.0221 | 4.0810 | 4.7116 |
| HELB_S1058 | HELB | Q8NG08 | 1058 | IFMVGESPQVSSR | 6.0701 | 6.3378 | 7.3988 | 6.4669 | 6.6011 | 6.8065 | 6.6170 | 6.5503 | 6.9234 | 6.1964 | 6.5914 | 6.7295 | 5.1353 | 5.3409 | 4.9642 | 5.2705 |
| HELB_S967 | HELB | Q8NG08 | 967 | PPADFPSPRKSSG | 6.2221 | 5.9592 | 6.8144 | 6.3918 | 5.9905 | 6.6310 | 5.9309 | 6.6174 | 6.7782 | 6.9673 | 6.2929 | 6.6599 | 5.7239 | 5.8764 | 5.3407 | 5.8034 |
| HELLS_S115 | HELLS | Q9NRZ9 | 115 | VKKGKNSIDASEE | 5.4642 | 6.4330 | 7.3706 | 7.0640 | 6.6459 | 6.3403 | 6.5155 | 6.5737 | 6.7339 | 6.6230 | 7.3664 | 6.9697 | 4.4763 | 5.3656 | 5.2282 | 4.8297 |
| HELLS_S503 | HELLS | Q9NRZ9 | 503 | KETIELSPTGRPK | 7.3490 | 7.3585 | 7.8833 | 7.6373 | 6.3418 | 7.1298 | 7.2762 | 7.3258 | 7.9670 | 7.8819 | 8.0874 | 7.9892 | 1.9691 | 2.9052 | 2.2020 | 2.6964 |
| HELZ_S1126 | HELZ | P42694 | 1763 | LYQRRISSSSVQP | 5.3689 | 5.0832 | 6.0925 | 5.7411 | 6.4478 | 6.0550 | 5.8301 | 6.0737 | 7.0228 | 6.9486 | 6.5926 | 6.9389 | 6.4636 | 6.9200 | 6.3606 | 6.0605 |
| HELZ_S1763 | HELZ | P42694 | 1126 | TNKQQQSPPKGKS | 6.3602 | 6.6098 | 6.7929 | 6.4000 | 5.6070 | 6.1996 | 6.2874 | 6.2777 | 6.2740 | 6.3645 | 6.9722 | 6.3289 | 5.6289 | 6.1312 | 5.9068 | 5.8590 |
| HELZ_T1163 | HELZ | P42694 | 1163 | NPIRAYTPPPPLG | 6.3877 | 6.1330 | 6.6613 | 6.3102 | 6.5041 | 6.6691 | 6.4262 | 6.4720 | 6.4912 | 6.3344 | 6.6034 | 6.4185 | 5.4834 | 6.0592 | 5.3442 | 5.7021 |
| HERC1_S1342 | HERC1 | Q15751 | 2720 | EVGRRQSLTSPDS | 4.5694 | 4.5133 | 10.6051 | 9.4881 | 11.0609 | 7.1442 | 3.8445 | 4.5669 | 5.0370 | 8.8961 | 5.5338 | 5.3526 | 5.0578 | 5.5541 | 4.3646 | 4.4116 |
| HERC1_S1428 | HERC1 | Q15751 | 1502 | SRLVHTSPNYRLI | 6.0998 | 7.0269 | 7.1926 | 7.1078 | 5.9413 | 7.2732 | 6.5097 | 6.3651 | 7.1055 | 6.8098 | 7.0031 | 6.9983 | 4.3019 | 4.8219 | 4.0387 | 5.4042 |
| HERC1_S1502 | HERC1 | Q15751 | 1428 | QQERRVSTDLPEG | 5.8831 | 5.5012 | 6.1262 | 6.0474 | 6.0164 | 5.8700 | 6.0699 | 6.7643 | 6.6761 | 6.6742 | 7.2251 | 6.2116 | 6.7155 | 7.0477 | 5.9394 | 5.2318 |
| HERC1_S2720 | HERC1 | Q15751 | 1342 | VDPQEHSFTRTID | 5.5960 | 6.4426 | 8.6268 | 7.5643 | 5.7442 | 6.1971 | 7.7294 | 7.9030 | 5.5748 | 7.0775 | 5.9854 | 6.4511 | 4.4616 | 5.3717 | 4.1949 | 5.0796 |

| | | | | | | | | | | | | | | | | | | | | | |
|---|---|---|---|---|---|---|---|---|---|---|---|---|---|---|---|---|---|---|---|---|---|
| HERC2_S1577 | HERC2 | O95714 | | 2454 | VKRRKQSPVPALP | 6.3402 | 6.0445 | 6.4781 | 6.3505 | 5.8126 | 6.5304 | 6.3964 | 6.0918 | 6.3385 | 6.1486 | 6.0915 | 6.2812 | 6.3049 | 6.4518 | 6.3514 | 5.9877 |
| HERC2_S1577;S1588 | HERC2 | O95714 | | 1588 | ACILPHSPINVDK | 6.3051 | 6.2417 | 6.8200 | 6.3703 | 6.0428 | 6.2718 | 6.2763 | 6.7738 | 6.9220 | 6.8813 | 6.7163 | 6.9358 | 5.4369 | 5.5338 | 5.0836 | 5.3885 |
| HERC2_S1588 | HERC2 | O95714 | | 2928 | AVPFLASDNEEEE | 5.8400 | 6.1979 | 7.1772 | 6.5388 | 6.1378 | 6.8746 | 6.3892 | 5.5805 | 6.5159 | 6.3070 | 5.7766 | 6.6737 | 6.1031 | 6.2532 | 5.7007 | 5.9340 |
| HERC2_S2454 | HERC2 | O95714 | | 1577 | KIGNEESDLEEAC | 6.3187 | 6.5463 | 6.4652 | 6.2956 | 6.0321 | 6.2542 | 6.1534 | 6.4076 | 5.8689 | 5.8214 | 5.5916 | 5.9351 | 6.9058 | 6.5172 | 6.4316 | 6.4554 |
| HERC2_S2928 | HERC2 | O95714 | | 3462 | DIEDRLSPNPWQE | 5.4379 | 5.9470 | 6.1124 | 6.1196 | 5.8967 | 5.6356 | 6.7258 | 7.2685 | 7.4985 | 6.1591 | 7.4003 | 5.8236 | 5.6308 | 6.3628 | 5.6344 | 6.3472 |
| HERC2_S3462 | HERC2 | O95714 | 1577;1588 | | NA | 5.9863 | 5.6684 | 6.8407 | 6.6712 | 6.8902 | 7.0791 | 6.4739 | 6.5246 | 6.2474 | 6.2362 | 6.8155 | 6.4911 | 4.9356 | 5.6261 | 5.6479 | 5.8659 |
| HERC4_S830 | HERC4 | Q5GLZ8 | | 830 | LLKKKPSLDDLKE | 4.2752 | 4.0457 | 4.7675 | 4.0120 | 4.8359 | 5.0538 | 4.7194 | 5.1417 | 4.5581 | 4.3860 | 4.1935 | 4.8184 | 11.1247 | 10.6997 | 11.3536 | 12.0148 |
| HERPUD2_T90 | HERPUD2 | Q9BSE4 | | 90 | LVCTSRTPPSSPK | 6.6749 | 6.3226 | 6.1128 | 6.2475 | 5.9697 | 5.6314 | 5.8418 | 5.8465 | 6.1381 | 6.4338 | 5.8023 | 6.2002 | 6.5829 | 6.7688 | 6.7724 | 6.6544 |
| HERPUD2_T90;S94 | HERPUD2 | Q9BSE4 | 90;94 | | NA | 6.8852 | 6.2315 | 6.2946 | 6.2871 | 6.3661 | 6.0645 | 6.8135 | 6.4895 | 6.2686 | 6.6466 | 6.5112 | 6.6165 | 5.5162 | 5.6816 | 5.4506 | 5.8766 |
| HEXD_S477 | HEXD | Q8WVB3 | | 477 | PPLPPTSPGRDVA | 5.4737 | 6.0835 | 7.2874 | 5.6936 | 5.7529 | 5.8992 | 5.8265 | 5.8336 | 6.4972 | 7.2819 | 5.8154 | 6.9117 | 6.1951 | 6.8334 | 6.2236 | 6.3912 |
| HEXIM2_S76 | HEXIM2 | Q96MH2 | | 76 | RSPRTQSPGGCSA | 6.0165 | 6.3970 | 5.9453 | 5.9552 | 6.2985 | 6.2205 | 5.8682 | 5.9928 | 6.7372 | 6.6980 | 6.6428 | 6.5032 | 6.0133 | 6.5104 | 5.8087 | 6.3922 |
| HGS_Y237 | HGS | O14964 | | 237 | TELPPEYLTSPLS | 6.8457 | 7.2102 | 6.8193 | 7.4998 | 6.2833 | 6.9862 | 7.4366 | 6.9140 | 7.1609 | 7.3082 | 7.6754 | 7.5887 | 3.4843 | 3.9295 | 3.0406 | 3.8172 |
| HGS_Y308 | HGS | O14964 | | 308 | PPASSLYSSPVNS | 6.0915 | 6.6397 | 6.2944 | 6.5875 | 6.6280 | 6.3046 | 6.4677 | 6.6358 | 6.8567 | 6.3806 | 6.4314 | 6.9324 | 5.4133 | 5.5651 | 5.0785 | 5.6927 |
| HIBCH_S258 | HIBCH | Q6NVY1 | | 258 | KIDRDKSFILEEH | 6.8692 | 7.0289 | 6.5918 | 6.8596 | 6.1327 | 6.5803 | 6.2733 | 6.0428 | 5.5168 | 6.7582 | 6.0562 | 5.6000 | 6.2838 | 6.2446 | 5.8209 | 5.3408 |
| HIC1_S704 | HIC1 | Q14526 | | 704 | TAELGLSPDKAAE | 6.5337 | 6.5535 | 7.3138 | 5.7273 | 6.4711 | 6.6492 | 5.4833 | 5.2920 | 6.8619 | 7.3253 | 6.7828 | 6.3820 | 5.9236 | 6.2237 | 5.2310 | 5.2458 |
| HID1_S670 | HID1 | Q8IV36 | | 670 | REQRRPSTSSASG | 4.8825 | 4.3756 | 6.3643 | 5.4613 | 6.2555 | 5.8553 | 5.9025 | 5.8697 | 6.6126 | 7.1360 | 7.0108 | 6.7932 | 8.1827 | 7.5321 | 5.9456 | 5.8204 |
| HIPK1_T351 | HIPK1 | Q86Z02 | | 351 | SKAVCSTYLQSRY | 6.1711 | 6.4156 | 6.3998 | 6.1205 | 6.3388 | 6.3391 | 5.9925 | 6.2888 | 7.0072 | 6.3324 | 6.6303 | 6.3255 | 6.0181 | 6.0448 | 5.8500 | 5.7257 |
| HIRA_S549 | HIRA | P54198 | | 555 | SPSVLTTPSKIEP | 6.4342 | 6.4376 | 6.3433 | 7.0250 | 6.6109 | 6.2723 | 6.5178 | 6.0031 | 6.7975 | 6.4445 | 6.2042 | 6.3422 | 5.7973 | 5.6897 | 5.3460 | 5.7345 |
| HIRA_S549;T555 | HIRA | P54198 | | 586 | PALTSMTPTAVER | 6.2197 | 6.3393 | 6.4772 | 6.7775 | 6.6859 | 6.5972 | 6.6009 | 6.5200 | 7.0376 | 6.4593 | 6.4744 | 6.4979 | 5.7431 | 5.4800 | 5.0539 | 5.0361 |
| HIRA_S555 | HIRA | P54198 | | 549 | STPAALSPSVLTT | 5.7579 | 5.4968 | 5.5609 | 6.1402 | 5.6137 | 5.8698 | 6.1018 | 5.8904 | 5.5863 | 5.9382 | 6.0219 | 5.5083 | 7.6551 | 7.4153 | 7.8898 | 7.5533 |
| HIRA_S661 | HIRA | P54198 | | 661 | VSLSVQSPAALTA | 5.6458 | 6.1261 | 5.7183 | 6.9974 | 6.0410 | 6.6768 | 6.3269 | 5.6805 | 6.5097 | 7.0899 | 6.4362 | 6.1850 | 5.6269 | 5.9464 | 6.0410 | 6.9522 |
| HIRA_T586 | HIRA | P54198 | 549;555 | | NA | 6.4903 | 6.2741 | 5.4044 | 6.6181 | 6.6446 | 5.9810 | 6.8910 | 6.0915 | 6.6598 | 6.0147 | 5.7894 | 6.3866 | 6.3893 | 6.5829 | 6.0602 | 5.7220 |
| HIRIP3_S125 | HIRIP3 | Q9BW71 | | 530 | YRRTLDSDEERPR | 5.1973 | 5.7869 | 5.9649 | 5.3998 | 5.8022 | 6.2716 | 5.6107 | 5.9336 | 5.9103 | 6.5820 | 6.7589 | 6.3078 | 6.9270 | 7.6342 | 7.1975 | 6.7154 |
| HIRIP3_S223;S227 | HIRIP3 | Q9BW71 | | 125 | GVAAEVSPAKEEN | 5.0442 | 5.6715 | 6.2604 | 5.2451 | 5.4251 | 5.8974 | 5.8539 | 5.5416 | 6.4165 | 6.5224 | 6.6820 | 6.5465 | 6.9210 | 7.5510 | 7.1012 | 7.3203 |
| HIRIP3_S227 | HIRIP3 | Q9BW71 | | 84 | KGKRPPTPCSDPE | 4.3871 | 5.2415 | 5.1141 | 4.1897 | 5.1724 | 5.8377 | 5.2119 | 4.7682 | 5.5843 | 6.8534 | 7.2965 | 6.3997 | 8.4423 | 9.4330 | 8.3797 | 7.6885 |
| HIRIP3_S289;S291 | HIRIP3 | Q9BW71 | | 87 | RPPTPCSDPERKR | 5.7727 | 6.3130 | 6.4062 | 5.7890 | 6.0861 | 6.7917 | 6.4908 | 6.8821 | 5.8640 | 6.4091 | 6.2389 | 5.9967 | 6.4158 | 6.9099 | 5.7429 | 5.8909 |
| HIRIP3_S530 | HIRIP3 | Q9BW71 | | 227 | KESEQESEEEILA | 5.5352 | 5.8104 | 5.5536 | 5.2314 | 5.3720 | 6.0202 | 5.1584 | 6.5209 | 7.1325 | 6.9700 | 6.8455 | 6.5724 | 6.1392 | 7.5692 | 6.9123 | 6.6569 |
| HIRIP3_T471 | HIRIP3 | Q9BW71 | | 471 | ALGMKGTPSLGKC | 7.8602 | 8.0552 | 7.2090 | 7.1818 | 6.6815 | 7.1262 | 6.9451 | 6.4617 | 8.4967 | 6.8742 | 7.4008 | 6.5660 | 3.0646 | 3.9906 | 2.5909 | 3.4956 |
| HIRIP3_T84 | HIRIP3 | Q9BW71 | 223;227 | | NA | 5.2410 | 5.9001 | 6.5659 | 5.3504 | 6.0030 | 5.7653 | 5.9173 | 6.0756 | 5.6157 | 6.0078 | 6.2170 | 5.5443 | 7.2431 | 7.6650 | 7.5632 | 7.3254 |
| HIRIP3_T84;S87 | HIRIP3 | Q9BW71 | 84;87 | | NA | 5.3518 | 5.9512 | 6.0504 | 5.4589 | 5.7662 | 6.6633 | 5.9153 | 6.1153 | 5.8355 | 6.1905 | 5.9350 | 6.0742 | 6.8854 | 7.4810 | 7.4116 | 6.9145 |
| HIRIP3_T87 | HIRIP3 | Q9BW71 | 289;291 | | NA | 5.1528 | 5.3631 | 6.3071 | 6.0473 | 5.3689 | 6.2936 | 5.8963 | 6.3578 | 6.1389 | 6.4138 | 5.6453 | 6.4200 | 7.0632 | 6.6816 | 7.3552 | 7.4951 |
| HIVEP1_S1158 | HIVEP1 | P15822 | | 1753 | MLSPANSLDIAME | 5.8391 | 7.0552 | 6.4453 | 6.9376 | 6.0672 | 6.2867 | 6.1181 | 6.2278 | 5.6647 | 6.3620 | 6.7862 | 6.5921 | 4.7308 | 6.4494 | 6.4015 | 6.0365 |
| HIVEP1_S130 | HIVEP1 | P15822 | | 779 | SLSRRGSIDSPKS | 6.5187 | 6.8496 | 6.6276 | 6.6663 | 6.1083 | 6.0470 | 6.1997 | 5.8920 | 6.7925 | 6.4897 | 6.3789 | 6.4391 | 5.6993 | 5.9195 | 5.6443 | 5.7276 |
| HIVEP1_S1387 | HIVEP1 | P15822 | | 65 | LKKIPKSPLRNPL | 7.0306 | 7.1421 | 6.1803 | 6.8843 | 6.6772 | 6.4665 | 6.8376 | 6.0881 | 6.5950 | 5.8864 | 5.9146 | 5.6236 | 5.8629 | 5.7135 | 5.6443 | 5.4529 |
| HIVEP1_S1735 | HIVEP1 | P15822 | | 1158 | EPFERASPVSFQE | 5.9565 | 6.0428 | 5.6691 | 6.5257 | 6.0419 | 5.8400 | 6.8903 | 6.1716 | 6.5679 | 5.9040 | 5.6862 | 5.5765 | 6.6345 | 6.8876 | 6.7197 | 6.8855 |
| HIVEP1_S1749;S1753 | HIVEP1 | P15822 | | 2327 | AVAITQSPSSVRL | 6.6525 | 6.9617 | 6.3891 | 6.8537 | 6.5274 | 6.5674 | 7.0914 | 6.1436 | 6.8090 | 6.8750 | 5.7964 | 6.3157 | 5.2615 | 5.4309 | 4.9578 | 5.3668 |
| HIVEP1_S1753 | HIVEP1 | P15822 | | 1735 | ISVGRLSPQQESS | 6.5481 | 6.2192 | 6.0534 | 6.9106 | 6.8462 | 6.8275 | 6.6999 | 5.9683 | 6.2980 | 6.0994 | 5.7654 | 6.3640 | 6.5887 | 6.1642 | 5.3218 | 5.3253 |
| HIVEP1_S1884 | HIVEP1 | P15822 | | 534 | LLKSSFTPSSPEN | 6.5041 | 6.2765 | 6.2325 | 7.2119 | 6.5827 | 5.9895 | 6.2516 | 6.5093 | 6.0680 | 6.7214 | 6.0643 | 6.0467 | 6.3052 | 5.9263 | 5.6136 | 5.6965 |
| HIVEP1_S232 | HIVEP1 | P15822 | | 681 | YFSRSESADQTVS | 6.0662 | 7.1202 | 6.9926 | 8.3695 | 6.3469 | 6.2433 | 5.7058 | 6.5752 | 5.6577 | 6.7929 | 7.1097 | 6.3409 | 4.1652 | 6.5587 | 4.9881 | 4.9670 |
| HIVEP1_S2327 | HIVEP1 | P15822 | | 791 | SYIFKDSFQFDLK | 6.0373 | 7.7914 | 9.2001 | 9.8820 | 6.1875 | 3.9528 | 5.4816 | 5.4426 | 6.7732 | 7.4580 | 6.1995 | 6.6020 | 4.1871 | 5.0611 | 4.5625 | 5.1813 |
| HIVEP1_S479 | HIVEP1 | P15822 | | 1884 | SENTHISPLKCTD | 6.5574 | 6.7206 | 6.0048 | 6.8851 | 6.5597 | 6.2595 | 6.3950 | 6.2110 | 6.2979 | 6.1960 | 6.0254 | 6.7187 | 5.7584 | 5.6628 | 6.1395 | 5.6083 |
| HIVEP1_S571 | HIVEP1 | P15822 | | 1387 | SDLRSKSFDCGSI | 5.6407 | 6.4239 | 6.2470 | 7.2271 | 6.3113 | 6.2040 | 5.6855 | 5.3674 | 6.0868 | 6.6570 | 6.9797 | 6.8137 | 6.1081 | 6.7919 | 5.3461 | 6.1098 |
| HIVEP1_S65 | HIVEP1 | P15822 | | 479 | LFLSHESPKALSI | 6.0993 | 5.7430 | 6.2296 | 6.2644 | 6.2727 | 6.7669 | 6.2727 | 6.2100 | 5.9765 | 5.8923 | 5.6088 | 6.2866 | 6.7201 | 6.2809 | 6.6511 | 6.7251 |
| HIVEP1_S681 | HIVEP1 | P15822 | | 130 | KSEESVSPKKPLF | 6.3910 | 6.3021 | 6.3639 | 6.3512 | 5.9829 | 5.5197 | 6.5036 | 5.5300 | 6.9782 | 6.6064 | 5.7717 | 5.8809 | 6.5061 | 6.3220 | 6.7603 | 6.2299 |
| HIVEP1_S779 | HIVEP1 | P15822 | | 232 | PNKTARSPPKLKN | 6.7213 | 6.9028 | 6.4429 | 6.6269 | 6.2263 | 6.4685 | 6.3310 | 5.3046 | 6.7224 | 6.4419 | 6.2839 | 6.9817 | 5.6905 | 5.8915 | 5.6170 | 5.3467 |
| HIVEP1_S791 | HIVEP1 | P15822 | | 571 | VHHVTVSPLRTDS | 6.2549 | 6.8490 | 6.2195 | 6.7285 | 6.3982 | 6.3670 | 6.5975 | 6.2889 | 6.1912 | 6.1929 | 6.1358 | 5.9260 | 5.8973 | 5.7841 | 5.8752 | 6.2941 |
| HIVEP1_S866 | HIVEP1 | P15822 | | 866 | PLRGSQSFDDKIG | 5.4189 | 6.7415 | 6.6697 | 6.9822 | 5.8119 | 5.7929 | 6.6546 | 6.2225 | 5.7465 | 6.6259 | 6.7609 | 7.1586 | 4.7429 | 6.9495 | 5.7034 | 6.0181 |
| HIVEP1_T534 | HIVEP1 | P15822 | 1749;1753 | | NA | 7.3501 | 6.6752 | 7.7671 | 7.3902 | 5.3655 | 6.1620 | 6.5655 | 6.6295 | 6.6766 | 7.6881 | 7.1805 | 8.1605 | 3.2577 | 5.3032 | 3.4989 | 4.3294 |
| HIVEP2_S1050 | HIVEP2 | P31629 | | 1085 | FLVRQASFSGSPE | 5.7941 | 6.0540 | 6.4885 | 6.2302 | 6.0993 | 6.7355 | 6.0575 | 6.1304 | 6.1464 | 6.2132 | 5.7159 | 6.3799 | 6.1403 | 6.7440 | 6.3252 | 6.7457 |
| HIVEP2_S1085 | HIVEP2 | P31629 | | 629 | LKEKKLSPGDRVG | 6.6757 | 6.6137 | 6.4110 | 6.5965 | 6.3921 | 7.0649 | 7.0321 | 7.0361 | 6.7542 | 6.2272 | 6.0590 | 6.3346 | 4.9706 | 5.3404 | 4.8834 | 5.6084 |
| HIVEP2_S1443 | HIVEP2 | P31629 | | 1447 | MLSPASSLELFME | 4.8168 | 5.6083 | 5.9976 | 4.8192 | 7.0655 | 6.6157 | 5.9312 | 6.6472 | 5.3782 | 6.5678 | 6.9553 | 7.2086 | 4.5611 | 7.6290 | 7.4244 | 6.7741 |
| HIVEP2_S1443;S1447 | HIVEP2 | P31629 | | 296 | EASDKMSPGPPIP | 5.8987 | 5.9880 | 7.0158 | 5.7676 | 5.3103 | 6.1891 | 6.3366 | 6.5302 | 6.9682 | 6.8177 | 5.9687 | 6.2219 | 6.6688 | 6.7224 | 5.4994 | 6.0968 |
| HIVEP2_S1447 | HIVEP2 | P31629 | | 769 | PQLQPGSPSLVSE | 5.9964 | 5.2460 | 5.9746 | 6.0316 | 6.6164 | 6.4029 | 6.3674 | 6.4471 | 6.2581 | 5.7454 | 6.1398 | 5.7531 | 6.3808 | 7.2494 | 6.7799 | 6.6113 |
| HIVEP2_S296 | HIVEP2 | P31629 | | 819 | SFERSESAELVAC | 5.7170 | 6.0882 | 6.7145 | 5.9751 | 6.7224 | 6.2747 | 6.3623 | 6.5161 | 6.2439 | 6.5538 | 6.9341 | 5.9091 | 5.2460 | 6.4147 | 6.0962 | 6.2322 |
| HIVEP2_S565 | HIVEP2 | P31629 | | 1050 | PEVRSKSFDYGNL | 5.7673 | 5.4370 | 6.4733 | 6.2324 | 7.4011 | 6.1573 | 6.3290 | 6.1465 | 6.7357 | 5.7291 | 5.9975 | 6.0452 | 5.8687 | 7.0398 | 5.7446 | 6.8955 |
| HIVEP2_S622 | HIVEP2 | P31629 | | 1443 | GSKRMLSPASSLE | 5.7809 | 5.7860 | 5.3204 | 6.2686 | 5.7587 | 6.6422 | 7.5291 | 6.5639 | 6.3818 | 5.5349 | 5.3936 | 5.8362 | 6.8550 | 6.5920 | 6.7086 | 7.0482 |
| HIVEP2_S629 | HIVEP2 | P31629 | | 565 | SLRGSHSFDERMT | 5.9443 | 5.9897 | 6.7725 | 6.5007 | 6.3349 | 6.0818 | 5.6670 | 6.4399 | 5.9293 | 6.6985 | 6.1820 | 7.1660 | 4.6657 | 7.0052 | 5.7444 | 6.8782 |
| HIVEP2_S769 | HIVEP2 | P31629 | | 622 | KGISSSSLKEKKL | 3.9657 | 4.8299 | 6.2838 | 5.4916 | 5.8128 | 4.6357 | 4.1144 | 19.9093 | 5.1487 | 4.6888 | 5.2726 | 5.0530 | 5.6032 | 6.5412 | 6.8700 | 5.7791 |
| HIVEP2_S819 | HIVEP2 | P31629 | 1443;1447 | | NA | 6.9533 | 7.0550 | 8.5065 | 7.7946 | 6.4875 | 5.8683 | 5.5226 | 6.5467 | 6.6808 | 7.3779 | 6.9879 | 6.0847 | 4.2472 | 5.4120 | 4.0201 | 4.4550 |
| HJURP_S123 | HJURP | Q8NCD3 | | 412 | TLKWLISPVKIVS | 6.6862 | 6.9072 | 6.1758 | 6.4091 | 6.5318 | 6.3581 | 6.7082 | 6.2247 | 6.5538 | 6.4255 | 6.1839 | 6.1866 | 5.3307 | 5.7682 | 5.6024 | 5.9480 |
| HJURP_S123;S140 | HJURP | Q8NCD3 | | 473 | YRGGPASPGGLQG | 6.5649 | 6.8062 | 6.3209 | 6.5563 | 6.2074 | 6.1735 | 6.6783 | 6.2778 | 6.4743 | 6.5994 | 6.6958 | 6.6529 | 5.2442 | 5.8243 | 5.2085 | 5.7151 |
| HJURP_S140 | HJURP | Q8NCD3 | | 595 | QKYCLKSPGQMTV | 6.7407 | 7.0076 | 6.7550 | 7.1644 | 6.3638 | 6.9791 | 6.9884 | 5.5632 | 6.9116 | 7.0769 | 7.1565 | 7.3172 | 3.9757 | 4.7438 | 3.7407 | 4.5153 |
| HJURP_S185 | HJURP | Q8NCD3 | | 140 | APAVPQSPLKNEL | 6.1418 | 6.1924 | 6.3806 | 6.1627 | 5.9665 | 6.2403 | 5.8895 | 5.4460 | 6.9647 | 6.4659 | 6.9982 | 6.6803 | 5.9145 | 6.2224 | 5.9653 | 6.3689 |
| HJURP_S201 | HJURP | Q8NCD3 | | 123 | GEVDATSDQEESV | 5.8464 | 6.6387 | 6.6005 | 6.8460 | 6.4762 | 6.4642 | 6.2676 | 6.2551 | 6.1932 | 6.2222 | 6.1248 | 5.9199 | 5.9865 | 6.1436 | 6.0071 | 6.0080 |
| HJURP_S412 | HJURP | Q8NCD3 | | 642 | FQKLPSSPLGCRK | 6.8123 | 6.9105 | 6.3758 | 6.4139 | 6.5392 | 6.9623 | 6.9121 | 6.7514 | 6.6805 | 6.1756 | 6.1555 | 6.2845 | 4.9952 | 5.4145 | 4.9644 | 5.6523 |
| HJURP_S448 | HJURP | Q8NCD3 | | 201 | SRISRKSPGDPAK | 7.0774 | 7.6524 | 6.7051 | 7.3362 | 6.0234 | 6.9778 | 7.0305 | 6.3747 | 7.1940 | 7.1934 | 7.0821 | 7.2804 | 3.4905 | 4.5903 | 3.6753 | 4.3165 |
| HJURP_S473 | HJURP | Q8NCD3 | | 448 | HREYCLSPRNQPR | 6.7152 | 7.1762 | 6.6670 | 6.4933 | 6.5247 | 6.6307 | 6.3967 | 6.1826 | 6.7542 | 6.3660 | 6.7408 | 6.3668 | 4.8447 | 5.2328 | 5.1392 | 5.7690 |
| HJURP_S486 | HJURP | Q8NCD3 | | 486 | LETRRLSLPSSKA | 5.1339 | 4.7559 | 5.5140 | 6.8436 | 6.6844 | 6.2936 | 5.9214 | 5.5345 | 6.7508 | 8.2768 | 6.9052 | 8.8210 | 5.6078 | 6.9628 | 4.8663 | 5.1281 |
| HJURP_S496 | HJURP | Q8NCD3 | | 185 | PLPSLASPAVPAP | 7.0223 | 6.5774 | 8.1929 | 6.6173 | 5.9683 | 6.8950 | 6.4853 | 6.4163 | 6.3396 | 6.6495 | 6.0879 | 8.0579 | 4.1624 | 5.5755 | 3.5104 | 5.4420 |
| HJURP_S595 | HJURP | Q8NCD3 | | 496 | SKAKAKSLSEAFE | 6.3480 | 6.8037 | 6.3300 | 7.2569 | 6.2898 | 6.0130 | 6.2278 | 6.4222 | 6.3291 | 6.9026 | 6.5860 | 6.7663 | 5.4192 | 5.0362 | 5.2844 | 5.9849 |
| HJURP_S642 | HJURP | Q8NCD3 | 123;140 | | NA | 5.6785 | 6.3946 | 6.1780 | 6.2869 | 6.2901 | 6.0332 | 6.0182 | 6.2221 | 6.2448 | 6.6699 | 6.5723 | 6.8704 | 6.0599 | 6.2537 | 6.1284 | 6.0989 |
| HLA-A_S343 | HLA-A | A0A0G2JPD3 | | 352 | QAASSDSAQGSDM | 5.2657 | 5.6022 | 5.1698 | 5.3345 | 5.9135 | 6.1295 | 5.8921 | 6.0123 | 5.7153 | 5.9292 | 5.3685 | 5.2046 | 8.4593 | 7.9017 | 8.1424 | 7.9592 |
| HLA-A_S352 | HLA-A | A0A0G2JPD3 | | 359 | AQGSDMSLTACKG | 6.4873 | 7.2076 | 6.4160 | 7.1597 | 6.8486 | 6.2087 | 6.2937 | 6.4021 | 6.3553 | 5.3595 | 6.0452 | 5.7985 | 6.1206 | 5.9368 | 5.9217 | 5.4387 |
| HLA-A_S356 | HLA-A | A0A140T9X5 | | 365 | AQGSDVSLTACKV | 5.2005 | 5.9447 | 5.6231 | 5.3272 | 5.8931 | 5.9408 | 5.7299 | 6.2962 | 5.3869 | 6.0882 | 5.7716 | 5.6603 | 7.4947 | 7.8692 | 7.9512 | 7.8225 |
| HLA-A_S358 | HLA-A | A0A0G2JPD3 | | 343 | SDRKGGSYSQAAS | 6.2118 | 5.5721 | 7.1742 | 6.5431 | 5.9087 | 5.9854 | 6.0638 | 6.9123 | 6.0576 | 7.6094 | 7.1837 | 7.6436 | 5.8328 | 5.9256 | 4.6134 | 4.7627 |
| HLA-A_S358;S365 | HLA-A | A0A0G2JPD3 | | 356 | SDSAQGSDMSLTA | 4.0557 | 4.2003 | 3.7693 | 3.1036 | 4.6704 | 6.5815 | 6.2721 | 6.0395 | 5.8858 | 6.2567 | 5.3961 | 6.3091 | 9.0376 | 8.4941 | 9.0865 | 10.8419 |
| HLA-A_S359 | HLA-A | A0A140T9X5 | | 358 | QAASSDSAQGSDV | 4.9862 | 5.1575 | 5.1289 | 5.9961 | 6.9666 | 5.4136 | 5.5363 | 6.1481 | 5.6713 | 5.5795 | 6.1288 | 7.5785 | 8.2473 | 7.9642 | 7.2181 | 8.0992 |
| HLA-A_S365 | HLA-A | A0A140T9X5 | 358;365 | | NA | 6.2506 | 5.9325 | 5.9893 | 6.4785 | 5.5969 | 5.8790 | 5.8723 | 5.2481 | 6.5578 | 7.1243 | 6.2952 | 6.8524 | 7.0730 | 6.6625 | 6.1401 | 6.0474 |
| HLCS_S124 | HLCS | P50747 | | 124 | ESVKFASAENIPD | 6.1163 | 6.3638 | 6.2042 | 5.3131 | 6.7458 | 6.9141 | 5.0406 | 6.4430 | 6.6340 | 5.8884 | 5.9817 | 6.7252 | 7.0432 | 7.1192 | 5.9101 | 5.5573 |
| HLCS_S147 | HLCS | P50747 | | 147 | SVADETSPEREGR | 5.7639 | 7.0315 | 6.1304 | 5.3823 | 5.9623 | 7.2431 | 6.6499 | 6.9597 | 5.8659 | 6.0775 | 7.2582 | 6.7859 | 5.9944 | 5.4578 | 6.0454 | 5.3917 |
| HLTF_S398 | HLTF | Q14527 | | 635 | GLRRLQSLIKNIT | 6.0124 | 6.2076 | 5.7714 | 5.3815 | 6.4354 | 6.0900 | 5.7294 | 4.9394 | 6.4583 | 6.0162 | 6.1940 | 6.4155 | 7.0044 | 7.0565 | 6.9704 | 7.3175 |
| HLTF_S51 | HLTF | Q14527 | | 51 | PDDFLTSDEEVDS | 6.5012 | 6.3841 | 6.0956 | 6.4150 | 5.8690 | 5.8085 | 5.6051 | 5.7686 | 7.3137 | 6.5553 | 6.9918 | 6.6532 | 6.2811 | 6.3879 | 5.7119 | 5.6579 |
| HLTF_S635 | HLTF | Q14527 | | 398 | VQYIESSDSEEIE | 4.1041 | 4.4936 | 4.1606 | 4.4188 | 4.6443 | 4.6170 | 4.5690 | 5.1384 | 6.5544 | 5.6900 | 5.1528 | 5.1047 | 10.1757 | 11.3117 | 9.8962 | 9.9686 |
| HM13_S368 | HM13 | Q8TCT9-2 | | 368 | FPTVSGSPASLAD | 5.8099 | 6.5013 | 5.8932 | 6.6701 | 7.0849 | 7.3066 | 8.4128 | 6.1495 | 5.4255 | 7.0178 | 6.6762 | 7.7621 | 3.9817 | 5.9786 | 4.1769 | 5.1530 |
| HMBOX1_S170 | HMBOX1 | Q6NT76-5 | | 230 | WYQLEKTNPGATL | 6.5770 | 6.8628 | 5.7810 | 6.3748 | 6.3694 | 5.4051 | 6.0163 | 6.7504 | 6.5337 | 5.9832 | 5.7150 | 5.9030 | 6.1716 | 6.2785 | 6.4580 | 6.8202 |
| HMBOX1_T230 | HMBOX1 | Q6NT76-5 | | 170 | ELMRRDSSVIKEE | 4.6663 | 5.6765 | 5.2286 | 5.5768 | 6.1858 | 5.9234 | 6.0818 | 6.1947 | 6.0749 | 6.3485 | 6.4788 | 6.5738 | 7.2375 | 6.8662 | 7.6763 | 7.2102 |
| HMG20A_S105 | HMG20A | Q9NP66 | | 105 | DSNAPKSPLTGYV | 7.2178 | 7.4083 | 6.4265 | 7.2596 | 6.5663 | 6.1963 | 6.5579 | 6.5911 | 6.5643 | 6.6896 | 6.4963 | 6.2871 | 4.9018 | 5.0166 | 4.7710 | 5.0495 |
| HMGA1_S102;S103 | HMGA1 | P17096 | | 53 | EPSEVPTPKRPRG | 6.5721 | 6.7727 | 6.4664 | 6.7260 | 6.6034 | 6.6829 | 7.0175 | 6.5322 | 7.2959 | 6.9813 | 6.7487 | 6.7447 | 4.2014 | 5.4862 | 4.1466 | 5.0221 |
| HMGA1_S14 | HMGA1 | P17096-2 | | 42 | EPSEVPTPKRPRG | 6.5821 | 6.7809 | 6.4616 | 6.7294 | 6.6073 | 6.7144 | 7.0304 | 6.5359 | 7.3030 | 6.9815 | 6.7831 | 6.7740 | 4.1863 | 5.4591 | 4.1135 | 4.9575 |
| HMGA1_S36 | HMGA1 | P17096 | | 36 | RKQPPVSPGTALV | 7.1317 | 7.6554 | 6.6983 | 6.9966 | 6.6033 | 7.3558 | 7.6410 | 6.8487 | 6.9111 | 7.1006 | 7.4635 | 7.0385 | 3.1989 | 4.5574 | 3.0586 | 3.7405 |
| HMGA1_S36;S44;T53 | HMGA1 | P17096 | | 44 | GTALVGSQKEPSE | 6.0437 | 5.9739 | 6.1886 | 6.4231 | 6.4882 | 5.6807 | 5.4872 | 6.0117 | 6.0483 | 6.5816 | 6.0324 | 6.0505 | 6.2117 | 7.2947 | 6.5819 | 6.9017 |
| HMGA1_S38 | HMGA1 | P17096 | | 78 | TRKTTTTPGRKPR | 7.9692 | 8.1369 | 6.8473 | 6.8162 | 5.2477 | 7.3914 | 8.1479 | 6.9412 | 7.4794 | 7.5272 | 7.5868 | 7.4339 | 2.3578 | 4.1092 | 2.2393 | 3.7686 |
| HMGA1_S38;T42 | HMGA1 | P17096 | | 49 | GSQKEPSEVPTPK | 5.8961 | 6.3254 | 6.2250 | 6.2244 | 6.2039 | 6.1382 | 6.1361 | 6.1402 | 6.4799 | 6.1700 | 6.1959 | 6.1977 | 6.6916 | 6.6034 | 6.1078 | 6.2644 |
| HMGA1_S42 | HMGA1 | P17096 | | 14 | SSQPLASKQEKDG | 4.3534 | 4.9458 | 4.7567 | 4.6095 | 5.4869 | 5.6983 | 5.6265 | 5.6460 | 5.0784 | 5.5333 | 5.2814 | 4.9948 | 8.8841 | 9.7970 | 9.7865 | 9.5215 |
| HMGA1_S44 | HMGA1 | P17096 | | 8 | SESSSKSSQPLAS | 5.3672 | 5.3967 | 4.8621 | 4.9508 | 5.9729 | 5.4612 | 5.7510 | 5.6748 | 5.1253 | 4.9799 | 4.9062 | 4.7255 | 8.6777 | 9.5880 | 9.4726 | 9.0880 |
| HMGA1_S44;T53 | HMGA1 | P17096 | | 9 | ESSSKSSQPLASK | 5.5414 | 5.6105 | 5.1020 | 5.9096 | 5.9555 | 5.4346 | 6.0080 | 5.3907 | 5.6491 | 5.6074 | 5.3693 | 5.2065 | 7.7511 | 8.7344 | 8.2039 | 8.5259 |
| HMGA1_S49 | HMGA1 | P17096-2 | | 38 | QPPKEPSEVPTPK | 6.0978 | 6.4732 | 6.1213 | 6.1377 | 5.7997 | 6.1735 | 6.0693 | 6.1002 | 6.2679 | 6.1552 | 6.2528 | 6.2331 | 6.4784 | 6.5941 | 6.3636 | 6.6821 |
| HMGA1_S53 | HMGA1 | P17096 | | 39 | PPVSPGTALVGSQ | 5.4877 | 6.1811 | 6.4769 | 7.2491 | 6.2768 | 7.2296 | 6.4129 | 6.6737 | 6.1932 | 6.1232 | 6.7458 | 7.7579 | 5.0309 | 5.2296 | 5.0807 | 5.8509 |
| HMGA1_S8 | HMGA1 | P17096 | 102;103 | | NA | 5.3307 | 5.9803 | 8.1044 | 6.3498 | 4.2426 | 5.3720 | 5.7735 | 5.6183 | 5.8877 | 5.3741 | 6.2873 | 5.7284 | 6.8172 | 7.5676 | 6.9807 | 8.5853 |
| HMGA1_S9 | HMGA1 | P17096-2 | 38;42 | | NA | 6.7196 | 6.5789 | 6.6560 | 6.5243 | 6.6373 | 6.4570 | 7.2113 | 5.8694 | 6.7916 | 6.4025 | 6.5572 | 6.2730 | 5.2929 | 5.9111 | 4.6057 | 5.5122 |
| HMGA1_T39 | HMGA1 | P17096 | 44;53 | | NA | 6.9285 | 6.4139 | 6.5923 | 7.0024 | 7.4700 | 6.6532 | 6.6582 | 6.0345 | 7.2162 | 6.8004 | 7.2671 | 6.5425 | 4.4503 | 5.6710 | 4.1447 | 4.1546 |
| HMGA1_T78 | HMGA1 | P17096 | 36;44;53 | | NA | 6.4530 | 7.5008 | 6.0428 | 8.1219 | 6.4318 | 7.5700 | 7.9414 | 6.2197 | 7.2216 | 7.1778 | 8.1755 | 6.4861 | 3.6533 | 4.7570 | 2.8324 | 3.4147 |
| HMGA2_S101 | HMGA2 | P52926 | | 44 | EPTGEPSPKRPRG | 6.0519 | 6.2424 | 6.1895 | 6.1574 | 6.3528 | 6.2529 | 6.4535 | 5.9943 | 6.7844 | 6.4654 | 6.3721 | 6.3786 | 5.6617 | 6.6741 | 5.8658 | 6.1033 |
| HMGA2_S101;S102;S105 | HMGA2 | P52926 | | 12 | EGAGQPSTSAQGQ | 4.9983 | 5.3641 | 5.3330 | 5.8266 | 5.5470 | 6.6324 | 5.6290 | 6.0086 | 5.1976 | 5.7791 | 5.1286 | 5.3132 | 7.8542 | 9.0949 | 8.6264 | 7.6670 |
| HMGA2_S101;S105 | HMGA2 | H0YFY4 | | 7 | EPTGEPSPKRPRG | 5.8862 | 6.2682 | 6.0625 | 6.6151 | 5.8450 | 5.8763 | 6.3041 | 6.2457 | 6.2930 | 5.9997 | 6.3457 | 5.7305 | 6.1609 | 6.9629 | 6.6602 | 6.7439 |
| HMGA2_S102;S105 | HMGA2 | P52926 | | 105 | ETSSQESAEEDxx | 5.0037 | 5.1999 | 5.0343 | 4.9918 | 4.4748 | 4.5977 | 5.3318 | 5.0277 | 6.9296 | 7.0207 | 6.0461 | 6.5661 | 7.9653 | 8.2172 | 9.3700 | 8.2232 |
| HMGA2_S105 | HMGA2 | P52926 | | 100 | QEETEETSSQESA | 5.0020 | 4.0702 | 5.0384 | 4.0762 | 5.7790 | 5.7911 | 5.4512 | 3.5421 | 6.5196 | 7.7579 | 6.0582 | 5.9520 | 8.2314 | 9.5092 | 8.1039 | 9.1177 |
| HMGA2_S12 | HMGA2 | P52926 | | 101 | EETEETSSQESAE | 4.9874 | 4.7705 | 3.9575 | 4.9767 | 5.0941 | 4.8471 | 5.0078 | 4.6689 | 6.0592 | 7.2999 | 6.1307 | 6.1909 | 7.8619 | 9.2578 | 10.0958 | 8.7938 |
| HMGA2_S44 | HMGA2 | P52926 | 101;105 | | NA | 5.2034 | 5.7088 | 5.0417 | 5.0903 | 5.3970 | 5.3024 | 5.3041 | 5.2979 | 6.5687 | 6.6135 | 6.0404 | 5.9106 | 7.4445 | 8.6114 | 8.4829 | 7.9824 |
| HMGA2_S7 | HMGA2 | P52926 | 100;105 | | NA | 4.9639 | 5.6618 | 5.2698 | 5.6276 | 5.0698 | 5.6977 | 5.2217 | 5.3928 | 6.5045 | 6.2811 | 6.3038 | 6.0666 | 7.6913 | 8.4759 | 7.8036 | 7.9682 |
| HMGA2_T100 | HMGA2 | P52926 | 102;105 | | NA | 4.8878 | 5.4591 | 5.1968 | 5.2301 | 5.2172 | 5.3282 | 5.4929 | 4.5316 | 6.1712 | 6.5512 | 5.8085 | 6.1494 | 7.9038 | 8.6266 | 8.6148 | 8.8308 |
| HMGA2_T100;S101;S102 | HMGA2 | P52926 | 101;102;105 | | NA | 8.0690 | 6.0747 | 5.9120 | 6.3539 | 6.6011 | 5.3492 | 6.5771 | 6.4532 | 5.9172 | 5.8360 | 5.8579 | 6.2328 | 5.6579 | 6.7500 | 5.7346 | 6.6236 |
| HMGA2_T100;S101;S105 | HMGA2 | P52926 | 100;101;102 | | NA | 7.2385 | 6.5263 | 5.5555 | 5.7157 | 7.1997 | 5.5534 | 5.5483 | 6.4853 | 6.2910 | 5.8692 | 6.6845 | 5.6905 | 5.7663 | 7.0912 | 5.8366 | 6.9480 |
| HMGA2_T100;S105 | HMGA2 | P52926 | 100;101;105 | | NA | 6.6500 | 7.5718 | 5.0363 | 7.3016 | 4.0491 | 6.7859 | 5.4923 | 4.5622 | 6.5628 | 5.2229 | 4.4495 | 6.1946 | 6.3000 | 8.1396 | 7.5644 | 8.1169 |
| HMGCR_S524 | HMGCR | P04035-3 | | 524 | LLSKKLSEPSSLQ | 2.6635 | 2.8965 | 2.8650 | 2.9750 | 5.2448 | 5.0797 | 4.9199 | 5.2763 | 2.9584 | 2.9383 | 3.3074 | 3.3728 | 14.2870 | 12.5254 | 14.9187 | 13.7714 |
| HMGCS1_S495 | HMGCS1 | Q01581 | | 495 | ATEHIPSPAKKVP | 6.3936 | 6.1659 | 7.5599 | 6.6479 | 5.7121 | 6.6877 | 6.0567 | 7.0061 | 6.9109 | 6.7054 | 7.2900 | 7.5189 | 4.5072 | 5.3158 | 4.7845 | 4.7373 |
| HMGN1_S25 | HMGN1 | P05114 | | 89 | TEESPASDEAGEK | 6.1368 | 5.9236 | 6.1789 | 6.3819 | 5.5757 | 5.6440 | 6.2907 | 5.7864 | 6.0617 | 6.2674 | 5.9534 | 6.3552 | 6.5453 | 7.0596 | 6.8010 | 7.0382 |
| HMGN1_S86 | HMGN1 | P05114 | | 25 | RRSARLSAKPPAK | 8.2233 | 8.0296 | 7.5686 | 7.3919 | 5.8799 | 7.8410 | 6.2922 | 4.5463 | 6.4630 | 7.3203 | 8.6069 | 7.9181 | 3.1140 | 4.5920 | 2.6878 | 3.5250 |
| HMGN1_S86;S89 | HMGN1 | P05114 | | 86 | ETKTEESPASDEA | 5.9088 | 6.8951 | 5.2661 | 6.9075 | 5.0864 | 5.2255 | 5.2189 | 4.9051 | 6.5671 | 6.1812 | 6.6052 | 6.6011 | 6.2168 | 7.3348 | 7.2503 | 7.8300 |
| HMGN1_S89 | HMGN1 | P05114 | 86;89 | | NA | 5.6544 | 5.5746 | 5.3055 | 6.4737 | 6.0233 | 5.1278 | 5.9560 | 5.6444 | 5.5159 | 6.2166 | 6.0436 | 6.3418 | 7.1228 | 7.8202 | 7.6262 | 7.5531 |
| HMGN3_S106 | HMGN3 | A0A087WZE9 | | 6 | xMPKRKSPENTEG | 4.7620 | 5.4068 | 5.9618 | 6.2562 | 5.4013 | 5.7923 | 5.9754 | 5.7472 | 5.8115 | 6.4185 | 5.8223 | 6.0600 | 6.4566 | 7.9332 | 8.0635 | 8.1313 |
| HMGN3_S6 | HMGN3 | A0A087WZE9 | | 103 | VSTSRGTPPSTLS | 6.6013 | 6.2048 | 6.1072 | 6.7060 | 5.8245 | 6.4488 | 6.1905 | 6.9998 | 6.4050 | 6.4711 | 6.3207 | 6.1244 | 4.8241 | 6.6470 | 6.3959 | 5.7290 |
| HMGN3_T103 | HMGN3 | A0A087WZE9 | | 106 | SRGTPPSTLSVKG | 5.6800 | 6.0757 | 5.8345 | 6.2110 | 7.0552 | 7.4249 | 6.3495 | 6.3574 | 6.5068 | 6.2784 | 7.5101 | 6.6040 | 6.1794 | 6.8526 | 4.8593 | 4.2213 |
| HMGN5_T11 | HMGN5 | P82970 | | 31 | AMLVPVTPEVKPK | 6.7876 | 7.4666 | 6.8536 | 7.0264 | 6.5329 | 7.8040 | 7.0603 | 7.0030 | 7.6419 | 7.2513 | 7.7690 | 7.3915 | 3.0679 | 4.0668 | 2.7172 | 3.5598 |
| HMGN5_T31 | HMGN5 | Q5JSK9 | | 11 | KMLVPVTPEVKPK | 7.2404 | 6.2274 | 8.0370 | 6.4296 | 6.4527 | 6.1866 | 6.6461 | 7.4001 | 6.5627 | 6.4953 | 5.7413 | 7.5527 | 4.9890 | 5.3421 | 3.6158 | 5.0813 |
| HMGXB4_S102 | HMGXB4 | Q9UGU5 | | 497 | SSVGVLSPQKKSP | 6.7760 | 6.7578 | 6.2676 | 6.8550 | 6.2225 | 5.7395 | 6.4108 | 6.1054 | 6.5083 | 6.1783 | 6.0180 | 6.0359 | 5.6163 | 6.1181 | 5.9620 | 6.4285 |
| HMGXB4_S118 | HMGXB4 | Q9UGU5 | | 118 | LLKAITSPLAAGS | 7.1328 | 7.4490 | 6.7237 | 7.2753 | 6.7564 | 6.8788 | 7.3506 | 7.0191 | 7.2406 | 7.2026 | 7.0516 | 6.8416 | 3.0374 | 4.4712 | 3.4502 | 4.1190 |
| HMGXB4_S480 | HMGXB4 | Q9UGU5 | | 512 | TMLLPASPAKAPE | 6.1656 | 6.0841 | 5.9331 | 6.2360 | 6.2938 | 5.8793 | 6.0691 | 5.9951 | 6.1426 | 6.0333 | 5.8568 | 5.8321 | 6.6349 | 7.1331 | 6.8801 | 6.8311 |
| HMGXB4_S497 | HMGXB4 | Q9UGU5 | | 102 | KKKKKSSPQSTDT | 5.9707 | 5.8892 | 5.8978 | 6.5979 | 6.2345 | 5.8100 | 6.1996 | 6.3128 | 6.3635 | 6.1811 | 6.0106 | 6.1241 | 6.2217 | 6.9975 | 6.4882 | 6.7007 |
| HMGXB4_S502;S512 | HMGXB4 | Q9UGU5 | | 480 | TVKRKASSSEGSM | 4.4148 | 4.1038 | 4.2108 | 4.8509 | 7.1491 | 5.4790 | 5.6734 | 5.4569 | 5.1939 | 4.8404 | 4.9220 | 4.4147 | 9.5444 | 9.8121 | 10.3646 | 9.5693 |
| HMGXB4_S512 | HMGXB4 | Q9UGU5 | 502;512 | | NA | 6.0783 | 6.3323 | 6.1419 | 6.7871 | 6.1002 | 5.9707 | 6.3724 | 6.0535 | 6.2349 | 6.2505 | 6.1116 | 6.3803 | 5.7954 | 6.8328 | 6.2687 | 6.2893 |
| HMMR_S20 | HMMR | O75330-3 | | 20 | PSGCAPSPGAYDV | 7.4209 | 7.7866 | 7.4076 | 7.0782 | 5.0431 | 8.0140 | 8.1804 | 6.8301 | 7.7041 | 7.5697 | 7.5346 | 8.0415 | 2.4371 | 3.4340 | 2.2032 | 3.3148 |
| HMOX2_S179 | HMOX2 | P30519 | | 179 | RALKLPSTGEGTQ | 5.5004 | 5.4302 | 5.9500 | 5.7809 | 6.1949 | 6.1225 | 6.4906 | 6.5847 | 6.1217 | 6.4759 | 6.0332 | 6.4845 | 6.7421 | 6.5960 | 6.3449 | 7.1474 |
| HNRNPA0_S68 | HNRNPA0 | Q13151 | | 68 | DAAMAASPHAVDG | 7.0806 | 6.3699 | 6.5521 | 7.0394 | 6.2400 | 6.4729 | 6.5251 | 5.8187 | 6.2294 | 7.0067 | 6.9053 | 5.8245 | 4.8962 | 6.1207 | 5.9195 | 4.9992 |
| HNRNPA1_S22 | HNRNPA1 | P09651-2 | | 6 | xMSKSESPKEPEQ | 5.8044 | 5.7712 | 5.9511 | 6.2764 | 6.3387 | 5.9315 | 6.4088 | 5.7980 | 6.3223 | 6.1410 | 6.2187 | 5.7815 | 7.2342 | 6.6337 | 6.7728 | 6.6155 |
| HNRNPA1_S286 | HNRNPA1 | P09651-2 | | 316 | SSSSYGSGRRFxx | 5.5605 | 5.9445 | 6.9787 | 6.9734 | 5.8505 | 6.4027 | 6.9078 | 6.0172 | 6.4281 | 7.5459 | 7.8817 | 7.4918 | 4.9257 | 5.2782 | 4.3938 | 5.4193 |
| HNRNPA1_S316 | HNRNPA1 | P09651-2 | | 286 | NFGGRSSGPYGGG | 6.1651 | 6.9285 | 6.4549 | 7.1740 | 6.1902 | 6.7069 | 7.1506 | 6.8670 | 7.0803 | 7.7089 | 7.7256 | 7.2435 | 3.9853 | 4.5786 | 3.5265 | 4.5143 |
| HNRNPA1_S4;S6 | HNRNPA1 | P09651-2 | | 95 | AVSREDSQRPGAH | 6.0397 | 7.2827 | 7.7313 | 6.9802 | 5.7415 | 7.2369 | 6.4173 | 6.7747 | 7.1519 | 8.4703 | 8.3709 | 7.3990 | 3.4866 | 4.3749 | 2.8333 | 3.7089 |
| HNRNPA1_S6 | HNRNPA1 | P09651-2 | | 260 | FGGGGSYNDFGNY | 6.2627 | 6.8388 | 6.8442 | 8.2882 | 6.5315 | 6.2222 | 6.8881 | 5.5835 | 7.0649 | 6.7220 | 7.0874 | 7.2813 | 4.3812 | 5.3105 | 4.1090 | 4.5847 |
| HNRNPA1_S95 | HNRNPA1 | P09651-2 | | 22 | LFIGGLSFETTDE | 6.0297 | 6.4956 | 6.7625 | 6.1833 | 5.6466 | 7.0103 | 5.5618 | 6.5577 | 5.9718 | 7.2926 | 6.4471 | 7.1656 | 5.5907 | 6.1047 | 5.4777 | 5.7024 |
| HNRNPA1_Y260 | HNRNPA1 | P09651-2 | 4;6 | | NA | 6.3002 | 6.4666 | 6.6072 | 7.0482 | 6.0470 | 5.2130 | 6.0505 | 5.2864 | 6.4381 | 6.8086 | 6.3940 | 6.2067 | 6.5731 | 5.8889 | 5.9177 | 6.7538 |
| HNRNPA2B1_S212 | HNRNPA2B1 | P22626 | | 259 | GGNFGGSPGYGGG | 6.8397 | 7.6439 | 6.9079 | 7.8890 | 6.5951 | 7.0096 | 7.0884 | 7.0909 | 6.8618 | 7.2497 | 7.4458 | 7.1092 | 3.3600 | 4.0131 | 3.1385 | 3.7574 |
| HNRNPA2B1_S259 | HNRNPA2B1 | P22626 | | 344 | PGGSGGSGGYGGR | 6.1531 | 6.2050 | 5.9143 | 6.7327 | 6.5701 | 6.2891 | 6.5024 | 6.1858 | 6.3593 | 6.1617 | 6.2154 | 5.8946 | 6.4350 | 6.2488 | 5.9932 | 6.1394 |
| HNRNPA2B1_S341;S344 | HNRNPA2B1 | P22626 | | 212 | NFGFGDSRGGGGN | 4.6769 | 5.3503 | 5.9037 | 5.5601 | 6.3130 | 6.0052 | 6.1894 | 6.7107 | 6.2153 | 6.7054 | 6.6118 | 6.5552 | 7.1785 | 6.5985 | 6.7403 | 6.6857 |
| HNRNPA2B1_S344 | HNRNPA2B1 | P22626 | | 85 | MAARPHSIDGRVV | 6.0478 | 5.6584 | 5.4190 | 6.3459 | 6.1085 | 5.8120 | 6.2506 | 6.1845 | 6.6990 | 6.0693 | 5.8744 | 6.7181 | 6.8161 | 6.2056 | 6.4267 | 7.3640 |
| HNRNPA2B1_S85 | HNRNPA2B1 | P22626 | | 4 | xxxMEKTLETVPL | 8.0219 | 7.8341 | 6.5015 | 7.6200 | 7.3344 | 6.5706 | 7.3438 | 5.7198 | 6.4464 | 6.0601 | 6.6069 | 6.3123 | 4.3062 | 5.2916 | 3.8441 | 4.1862 |
| HNRNPA2B1_T4 | HNRNPA2B1 | P22626 | 341;344 | | NA | 7.3384 | 6.7043 | 6.3117 | 7.0943 | 7.1152 | 6.2799 | 6.6541 | 6.7633 | 5.7435 | 6.7478 | 6.8865 | 6.0771 | 5.5673 | 5.0771 | 4.5088 | 5.1306 |
| HNRNPA3_S14 | HNRNPA3 | P51991 | | 358 | GGRSSGSPYGGGY | 5.7485 | 6.2454 | 5.9459 | 6.9976 | 6.3550 | 6.7451 | 6.4842 | 6.5528 | 6.5247 | 6.6315 | 6.6547 | 6.6096 | 5.3188 | 6.0051 | 5.4491 | 5.7320 |
| HNRNPA3_S17 | HNRNPA3 | P51991 | | 14 | GRPQPDSGRRRRR | 6.4361 | 6.7184 | 6.5036 | 7.8067 | 5.8472 | 6.9432 | 6.8042 | 6.8300 | 7.7288 | 7.1435 | 7.6312 | 6.8616 | 3.3879 | 4.7786 | 3.6759 | 4.9029 |

| | | | | | | | | | | | | | | | | | | | | | |
|---|---|---|---|---|---|---|---|---|---|---|---|---|---|---|---|---|---|---|---|---|---|
| HNRNPA3_S358 | HNRNPA3 | A0A7I2V4G0 | | 17 | SLCRRESEPGIGG | 5.0592 | 4.8959 | 5.0807 | 5.9612 | 7.1089 | 5.8734 | 5.2790 | 5.4468 | 6.0360 | 6.4583 | 6.3476 | 6.5614 | 7.5300 | 7.4160 | 7.0306 | 7.9150 |
| HNRNPAB_S221 | HNRNPAB | Q99729-4 | | 256 | NRGNRGSGGGGGG | 5.6548 | 6.7342 | 6.1736 | 7.0285 | 6.7946 | 7.3974 | 7.0471 | 6.8296 | 7.1355 | 6.9307 | 7.1202 | 6.9491 | 4.3801 | 5.2588 | 3.8507 | 4.7151 |
| HNRNPAB_S243 | HNRNPAB | Q99729-4 | | 243 | QQQQYGSGGRGNR | 6.7377 | 7.7517 | 6.7302 | 7.0158 | 6.2055 | 6.4081 | 6.8092 | 6.4831 | 7.5959 | 6.4093 | 7.2825 | 7.3910 | 3.7988 | 4.8388 | 4.3586 | 4.1838 |
| HNRNPAB_S256 | HNRNPAB | Q99729-4 | | 82 | GLSWDTSKKDLKD | 2.3565 | 2.1935 | 2.3139 | 2.3957 | 2.9488 | 2.9486 | 3.0227 | 2.8928 | 2.5783 | 2.6381 | 2.3965 | 2.7028 | 17.0311 | 17.0236 | 18.5073 | 16.0499 |
| HNRNPAB_S82 | HNRNPAB | Q99729-4 | | 221 | KKFHTVSGSKCEI | 5.0484 | 5.2154 | 4.7699 | 5.4680 | 5.7324 | 6.0351 | 6.4010 | 6.2069 | 5.2808 | 5.9165 | 6.0382 | 6.8465 | 8.1433 | 7.1329 | 7.2717 | 8.4930 |
| HNRNPC_S107 | HNRNPC | P07910 | | 260 | EGGADDSAEEGDL | 5.7999 | 5.9442 | 5.6866 | 6.2968 | 6.5021 | 6.8345 | 6.4256 | 6.1690 | 6.6144 | 6.1780 | 6.9455 | 6.3732 | 6.8250 | 6.5105 | 5.7252 | 5.1695 |
| HNRNPC_S113 | HNRNPC | P07910 | | 113 | SVTEHPSPSPLLS | 6.4592 | 7.0782 | 6.5211 | 7.6509 | 6.5728 | 6.6618 | 6.7529 | 7.0669 | 6.6586 | 6.3846 | 6.5173 | 6.7416 | 4.5821 | 5.0419 | 4.3953 | 4.9148 |
| HNRNPC_S115 | HNRNPC | P07910 | | 299 | GEDDRDSANGEDD | 6.5181 | 6.7167 | 6.3252 | 6.3910 | 6.2850 | 5.5537 | 6.1004 | 6.9140 | 6.6394 | 5.9069 | 6.1419 | 6.2083 | 6.2479 | 5.9197 | 6.0448 | 6.0870 |
| HNRNPC_S138 | HNRNPC | P07910 | | 241 | EEQSSSSVKKDET | 6.5069 | 6.5941 | 5.7818 | 6.8311 | 6.2909 | 6.0424 | 6.6858 | 6.5722 | 6.1396 | 6.1140 | 6.1700 | 6.1199 | 5.8031 | 6.1099 | 6.0922 | 6.1460 |
| HNRNPC_S166 | HNRNPC | B4DY08 | | 247 | EGGADDSAEEGDL | 6.1575 | 6.0172 | 6.3883 | 6.8598 | 5.9648 | 5.4472 | 6.2145 | 5.7876 | 6.2603 | 6.4167 | 6.2207 | 6.3244 | 6.1226 | 6.9835 | 6.2381 | 6.5969 |
| HNRNPC_S182 | HNRNPC | P07910 | | 209 | IKQKVDSLLENLE | 6.1559 | 5.8414 | 6.2147 | 6.5629 | 6.2763 | 5.6953 | 6.1194 | 6.0866 | 6.3491 | 6.5685 | 6.1643 | 6.1240 | 6.2181 | 6.5134 | 6.2683 | 6.8418 |
| HNRNPC_S209 | HNRNPC | P07910 | | 233 | EMKNDKSEEEQSS | 5.9324 | 6.3677 | 6.1446 | 6.5377 | 5.8791 | 5.3115 | 6.1398 | 5.5547 | 5.9545 | 6.1027 | 5.7352 | 5.7710 | 7.2438 | 6.7517 | 7.1351 | 7.4387 |
| HNRNPC_S233 | HNRNPC | B4DY08 | | 107 | AAEMYGSSFDLDY | 7.3096 | 7.1039 | 7.3903 | 7.6401 | 6.6127 | 6.1292 | 6.6390 | 6.5672 | 6.5321 | 6.7969 | 6.6997 | 6.6059 | 4.3560 | 4.8111 | 4.4735 | 4.3327 |
| HNRNPC_S233;S238;S24 | HNRNPC | P07910 | | 253 | TNVKMESEGGADD | 5.7896 | 5.9709 | 5.9426 | 6.5160 | 6.6301 | 6.9700 | 6.3920 | 6.2279 | 6.4294 | 6.3645 | 7.1081 | 6.2816 | 6.9121 | 6.5057 | 5.2404 | 4.7189 |
| HNRNPC_S233;S241 | HNRNPC | P07910 | | 182 | KSGQRGSSKSGKL | 6.8745 | 7.4707 | 7.1449 | 6.8022 | 6.7031 | 8.1348 | 6.8987 | 4.7700 | 4.7929 | 6.0446 | 7.0297 | 6.8120 | 5.6804 | 5.8020 | 4.6835 | 4.3562 |
| HNRNPC_S238;S241 | HNRNPC | P07910 | | 247 | SVKKDETNVKMES | 8.0171 | 8.0697 | 7.9046 | 10.1174 | 6.9796 | 5.8256 | 6.7942 | 5.1663 | 6.6534 | 5.7066 | 5.6754 | 5.7200 | 4.1540 | 5.0607 | 4.0683 | 4.0871 |
| HNRNPC_S241 | HNRNPC | P07910 | | 138 | YYDRMYSYPARVP | 5.7767 | 6.4492 | 6.0196 | 6.4521 | 5.7430 | 5.5630 | 5.4212 | 5.3598 | 5.5735 | 5.4020 | 5.4456 | 5.5798 | 8.0677 | 7.8001 | 7.7617 | 7.5849 |
| HNRNPC_S247 | HNRNPC | P07910 | | 166 | RVSGNTSRRGKSG | 5.7888 | 6.9661 | 6.2869 | 6.7629 | 6.5770 | 6.2562 | 6.9415 | 6.8369 | 7.6125 | 6.3790 | 6.8749 | 6.3583 | 4.4092 | 5.1952 | 5.0984 | 6.6561 |
| HNRNPC_S253 | HNRNPC | P07910 | | 115 | TEHPSPSPLLSSS | 6.8690 | 7.4693 | 5.7550 | 7.5984 | 7.0624 | 6.4297 | 5.9229 | 7.7430 | 5.8590 | 6.3248 | 6.8037 | 6.8010 | 4.2825 | 5.2664 | 4.9376 | 4.8755 |
| HNRNPC_S253;S260 | HNRNPC | P07910 | 253;260 | | NA | 5.9019 | 5.9746 | 5.9329 | 6.5499 | 6.6020 | 6.7524 | 6.7805 | 6.2051 | 6.2715 | 5.7587 | 6.2492 | 5.9097 | 6.6844 | 6.4945 | 6.3134 | 5.6195 |
| HNRNPC_S260 | HNRNPC | P07910 | 238;241 | | NA | 6.4900 | 6.2786 | 6.0484 | 6.7218 | 6.0808 | 5.8131 | 6.4658 | 5.7809 | 6.7803 | 6.5411 | 6.3912 | 6.3539 | 6.0760 | 6.0086 | 5.8666 | 6.3030 |
| HNRNPC_S299 | HNRNPC | P07910 | 233;238;241 | | NA | 5.6426 | 5.4679 | 6.5608 | 6.4942 | 6.2025 | 5.9999 | 7.3155 | 5.0441 | 5.8199 | 5.8480 | 5.5970 | 5.9425 | 7.9256 | 6.7189 | 6.5296 | 6.8909 |
| HNRNPC_T247 | HNRNPC | P07910 | 233;241 | | NA | 6.1187 | 5.7117 | 5.9933 | 6.3632 | 6.5101 | 6.2677 | 6.9849 | 5.2027 | 6.3268 | 5.8333 | 6.3113 | 6.1036 | 7.1617 | 6.5969 | 5.9054 | 6.6088 |
| HNRNPD_S250 | HNRNPD | Q14103 | | 193 | GGLSPDTPEEKIR | 6.1053 | 6.4611 | 6.3805 | 6.4911 | 6.0934 | 6.2854 | 6.0988 | 6.0104 | 6.2761 | 6.6509 | 6.5233 | 6.5928 | 5.9867 | 6.1590 | 5.8096 | 6.0756 |
| HNRNPD_S83 | HNRNPD | Q14103 | | 250 | YHNVGLSKCEIKV | 4.4802 | 4.8767 | 4.3910 | 5.1822 | 5.2889 | 6.6159 | 5.4396 | 6.4218 | 6.8754 | 7.5084 | 6.5881 | 7.7942 | 7.9779 | 6.2526 | 5.5920 | 8.7151 |
| HNRNPD_T193 | HNRNPD | Q14103 | | 83 | EGHSNSSPRHSEA | 5.5079 | 5.9912 | 6.4956 | 6.4034 | 6.5497 | 6.2168 | 5.3806 | 5.8759 | 5.8408 | 6.4445 | 5.3846 | 6.9348 | 6.7332 | 7.0361 | 6.9017 | 6.3031 |
| HNRNPDL_S241 | HNRNPDL | O14979 | | 241 | VFVGGLSPDTSEE | 6.2957 | 6.0279 | 6.2113 | 6.6620 | 6.2924 | 6.0342 | 5.8296 | 6.0380 | 6.6952 | 6.2169 | 6.2115 | 6.2079 | 6.7933 | 6.4120 | 5.9298 | 6.1423 |
| HNRNPH1_S104 | HNRNPH1 | P31943 | | 104 | KHTGPNSPDTAND | 5.8541 | 6.2444 | 5.7263 | 6.2759 | 6.3499 | 6.0500 | 6.1127 | 6.2371 | 6.2816 | 6.0778 | 6.1594 | 6.3449 | 6.7966 | 6.4561 | 6.6282 | 6.4051 |
| HNRNPH1_S123 | HNRNPH1 | P31943 | | 272 | YCFSGMSDHRYGD | 6.2219 | 6.3394 | 6.0618 | 6.8337 | 6.2394 | 6.2552 | 6.8235 | 6.5983 | 6.4790 | 6.0037 | 6.0612 | 6.5552 | 5.9445 | 6.1075 | 5.6541 | 5.8216 |
| HNRNPH1_S272 | HNRNPH1 | P31943 | | 63 | AFVELESEDEVKL | 6.9780 | 7.0476 | 6.3956 | 6.9512 | 7.8178 | 6.9323 | 6.7583 | 5.8816 | 5.6259 | 5.2141 | 6.0191 | 5.6292 | 6.3403 | 5.8766 | 5.3504 | 5.1818 |
| HNRNPH1_S310 | HNRNPH1 | P31943 | | 310 | DIYNFFSPLNPVR | 6.1573 | 6.2749 | 6.0024 | 6.5903 | 6.2931 | 6.4695 | 6.0159 | 5.5577 | 6.0061 | 6.8064 | 6.5129 | 6.6715 | 6.3304 | 6.1307 | 6.2082 | 5.9725 |
| HNRNPH1_S63 | HNRNPH1 | P31943 | | 123 | GLPFGCSKEEIVQ | 4.4755 | 4.6371 | 4.6627 | 5.7989 | 5.8742 | 5.2800 | 6.0917 | 5.1232 | 5.8803 | 5.7794 | 5.0126 | 4.3139 | 8.9741 | 9.5817 | 9.6133 | 8.9012 |
| HNRNPH2_S310 | HNRNPH2 | P55795 | | 310 | DIYNFFSPLNPMR | 6.3272 | 6.6016 | 6.5698 | 6.3403 | 6.5255 | 6.3325 | 6.1682 | 6.0250 | 6.3644 | 6.0050 | 6.4821 | 6.0427 | 6.4437 | 5.8543 | 5.8920 | 6.0257 |
| HNRNPH3_S216 | HNRNPH3 | P31942 | | 216 | DIANFFSPLNPIR | 6.0419 | 6.6905 | 5.8890 | 6.8144 | 6.4032 | 5.2927 | 6.0708 | 6.1368 | 6.4081 | 6.5364 | 6.6658 | 6.4285 | 5.9506 | 6.4661 | 6.4420 | 5.7632 |
| HNRNPH3_T270 | HNRNPH3 | P31942 | | 270 | ELFLNSTPGGGSG | 6.0206 | 6.1550 | 6.2335 | 6.0399 | 5.6500 | 6.6294 | 6.4717 | 6.6454 | 6.6813 | 5.8456 | 6.2901 | 6.8204 | 6.2661 | 6.2588 | 5.8875 | 6.1044 |
| HNRNPK_S216 | HNRNPK | P61978-2 | | 284 | RDYDDMSPRRGPP | 6.8600 | 7.0219 | 6.5924 | 6.8102 | 6.4525 | 6.4796 | 6.2535 | 6.7081 | 7.0114 | 6.9302 | 7.1993 | 7.1585 | 4.6837 | 4.7885 | 4.3867 | 4.6634 |
| HNRNPK_S284 | HNRNPK | P61978-2 | | 379 | YAGGRGSYGDLGG | 5.8057 | 6.1984 | 5.9852 | 6.4723 | 6.4528 | 6.1561 | 6.1756 | 6.5929 | 6.5130 | 6.7603 | 6.7264 | 6.4920 | 5.8706 | 6.2418 | 5.4509 | 6.1060 |
| HNRNPK_S379 | HNRNPK | P61978-2 | | 216 | LDLISESPIKGRA | 6.0079 | 6.4178 | 6.3578 | 6.2829 | 6.1062 | 6.3356 | 5.8644 | 6.1889 | 6.2069 | 6.5308 | 6.5468 | 6.4507 | 6.0212 | 6.4642 | 6.1740 | 6.0437 |
| HNRNPK_S401 | HNRNPK | P61978-2 | | 401 | PKDLAGSIIGKGG | 4.9238 | 4.6775 | 5.9729 | 6.8096 | 6.3095 | 6.6399 | 6.3670 | 6.0483 | 6.3729 | 6.3295 | 6.5817 | 6.7838 | 5.5412 | 6.9977 | 7.2690 | 6.3756 |
| HNRNPL_S101 | HNRNPL | P14866 | | 101 | PHKTPASPVVHIR | 6.4975 | 6.6748 | 6.0302 | 6.9761 | 6.1724 | 6.1858 | 6.1466 | 6.4847 | 6.2909 | 6.2616 | 6.2903 | 6.2933 | 6.1362 | 6.0095 | 5.6445 | 5.9057 |
| HNRNPL_S454 | HNRNPL | P14866 | | 487 | RNNRFSTPEQAAK | 6.2604 | 6.7013 | 5.7924 | 6.6275 | 5.9953 | 6.2340 | 6.5844 | 6.6524 | 6.4208 | 6.1556 | 6.5311 | 7.0796 | 5.6281 | 5.5809 | 5.4875 | 6.2686 |
| HNRNPL_S52 | HNRNPL | P14866 | | 454 | KLNVCVSKQPAIM | 2.7392 | 3.8505 | 5.0671 | 4.1937 | 3.1889 | 5.7980 | 5.2306 | 7.0803 | 6.0458 | 8.8953 | 7.3950 | 8.7763 | 7.1484 | 7.0406 | 7.6688 | 9.8814 |
| HNRNPL_S589 | HNRNPL | P14866 | | 52 | RYYGGGSEGGRAP | 6.4294 | 6.5264 | 5.6354 | 6.8078 | 6.8639 | 5.7210 | 6.8779 | 6.7184 | 6.1658 | 5.5808 | 6.5563 | 6.0174 | 5.8846 | 6.1898 | 5.7073 | 6.3179 |
| HNRNPL_T487 | HNRNPL | P14866 | | 589 | STAQHASxxxxxx | 5.6674 | 5.7702 | 6.9105 | 7.5150 | 5.7235 | 6.6675 | 7.9605 | 5.6954 | 7.4963 | 5.5521 | 6.6737 | 6.0880 | 5.3932 | 5.9883 | 5.4867 | 5.4119 |
| HNRNPM_S377 | HNRNPM | P52272 | | 618 | GGGGGASFDRAIE | 4.9339 | 5.9420 | 7.3487 | 5.7486 | 5.5879 | 6.3070 | 5.6822 | 6.6039 | 6.4179 | 10.3406 | 9.5897 | 8.2897 | 3.2279 | 5.2908 | 4.0247 | 4.6644 |
| HNRNPM_S468 | HNRNPM | P52272 | | 575 | ERMGANSLERMGL | 5.8333 | 6.4476 | 6.0293 | 6.5637 | 6.1378 | 5.9071 | 6.0807 | 5.9348 | 6.1360 | 6.6085 | 6.3569 | 6.2560 | 6.3284 | 6.4447 | 6.2979 | 6.6373 |
| HNRNPM_S481 | HNRNPM | P52272 | | 528 | IERMGLSMERMVP | 5.4512 | 6.5572 | 7.7157 | 5.7187 | 5.7177 | 6.3954 | 5.8374 | 6.9827 | 7.0859 | 10.2895 | 8.9719 | 7.9135 | 3.0798 | 4.5723 | 3.5755 | 4.1356 |
| HNRNPM_S528 | HNRNPM | P52272 | | 468 | LDHMASSIERMGQ | 5.2259 | 6.2230 | 6.0894 | 6.4155 | 6.3668 | 6.3576 | 6.3169 | 6.4169 | 6.5126 | 6.7772 | 6.5030 | 6.7579 | 6.2170 | 6.1456 | 5.8108 | 5.8639 |
| HNRNPM_S575 | HNRNPM | P52272 | | 377 | RINEILSNALKRG | 3.4685 | 5.2414 | 5.9801 | 5.0637 | 5.5877 | 5.2630 | 6.0442 | 6.3620 | 5.8191 | 6.9599 | 6.1271 | 5.3905 | 8.2659 | 7.2087 | 10.1447 | 7.0735 |
| HNRNPM_S618 | HNRNPM | P52272 | | 701 | GVVKFESPEVAER | 6.2596 | 6.8427 | 6.0094 | 6.8232 | 5.9234 | 6.2452 | 6.6967 | 6.2291 | 6.1012 | 6.2519 | 6.7111 | 6.4868 | 5.0611 | 6.3297 | 6.0094 | 6.0194 |
| HNRNPM_S633 | HNRNPM | P52272 | | 633 | RGNFGGSFAGSFG | 6.1384 | 6.1253 | 6.8957 | 6.5106 | 5.8126 | 6.1503 | 6.5195 | 5.7706 | 7.4186 | 6.9453 | 6.8497 | 7.2412 | 5.5387 | 5.8811 | 4.7416 | 5.4609 |
| HNRNPM_S637 | HNRNPM | P52272 | | 481 | TMERIGSGVERMG | 4.6413 | 5.0354 | 5.2082 | 4.6914 | 4.7871 | 4.7927 | 5.1172 | 4.6478 | 5.1832 | 6.5161 | 6.0782 | 6.1432 | 9.0144 | 8.9454 | 9.3200 | 9.8783 |
| HNRNPM_S701 | HNRNPM | P52272 | | 637 | GGSFAGSFGGAGG | 5.8268 | 6.0630 | 7.1619 | 6.1553 | 6.4482 | 7.0536 | 4.8583 | 6.1639 | 8.1033 | 8.4356 | 7.7022 | 7.7034 | 4.2086 | 5.2988 | 4.1224 | 4.6946 |
| HNRNPU_S252 | HNRNPU | Q00839-2 | | 252 | KYSRAKSPQPPVE | 6.0558 | 6.4194 | 6.1561 | 6.7413 | 6.1741 | 6.1236 | 6.2097 | 6.0285 | 6.1967 | 6.1650 | 6.1281 | 5.9625 | 6.2504 | 6.4004 | 6.6922 | 6.2960 |
| HNRNPU_S311 | HNRNPU | Q00839-2 | | 59 | MEPGNGSLDLGGD | 5.7628 | 6.2749 | 6.2430 | 6.7394 | 6.2032 | 6.3254 | 6.5537 | 6.3693 | 6.2866 | 6.2409 | 6.2495 | 6.2826 | 5.8483 | 6.3464 | 6.1344 | 6.1397 |
| HNRNPU_S59 | HNRNPU | Q00839-2 | | 66 | LDLGGDSAGRSGA | 5.8872 | 6.0076 | 6.0724 | 6.5186 | 6.5106 | 5.8693 | 6.4115 | 6.5643 | 5.7953 | 6.2541 | 6.3704 | 6.2183 | 6.0594 | 6.8261 | 6.3334 | 6.3014 |
| HNRNPU_S66 | HNRNPU | Q00839-2 | | 216 | AGGDGKTEQKGGD | 5.7682 | 6.3018 | 6.7014 | 6.4860 | 5.5784 | 5.6353 | 6.5206 | 5.3953 | 5.7012 | 6.2549 | 6.4112 | 5.3294 | 6.6501 | 7.4974 | 7.1334 | 6.6356 |
| HNRNPU_T191 | HNRNPU | Q00839-2 | | 311 | RASYGVSKGKVCF | 6.7701 | 6.8997 | 5.4121 | 6.7649 | 6.3325 | 6.0793 | 7.1503 | 6.0506 | 6.0471 | 5.6174 | 5.0336 | 5.8488 | 6.3516 | 6.0906 | 6.9328 | 6.6187 |
| HNRNPU_T216 | HNRNPU | Q00839-2 | | 191 | GKSSGPTSLFAVT | 7.0623 | 7.0138 | 6.2406 | 6.8807 | 6.3674 | 5.5530 | 6.8962 | 5.4023 | 6.7700 | 6.3114 | 6.4410 | 6.5364 | 5.8254 | 6.0276 | 5.0887 | 5.5833 |
| HNRNPUL1_S194 | HNRNPUL1 | Q9BUJ2 | | 194 | EDRRGRSPQPPAE | 5.9547 | 6.2976 | 6.1077 | 6.4309 | 6.1306 | 6.1871 | 6.0678 | 5.9619 | 6.1817 | 6.2323 | 6.1861 | 6.1113 | 6.5272 | 6.6108 | 6.5561 | 6.4564 |
| HNRNPUL1_S512 | HNRNPUL1 | Q9BUJ2 | | 718 | PPPPSYSPARNPP | 5.7279 | 6.0649 | 6.3783 | 6.4805 | 6.0838 | 6.1647 | 6.4111 | 6.7030 | 6.6435 | 6.5071 | 6.3775 | 6.3496 | 5.8526 | 6.2967 | 5.7085 | 6.2503 |
| HNRNPUL1_S718 | HNRNPUL1 | Q9BUJ2 | | 512 | QTNVYGSAQRRKM | 5.7745 | 6.4006 | 6.2584 | 6.2349 | 6.6221 | 6.0905 | 6.2870 | 5.9277 | 6.5251 | 6.2552 | 6.0850 | 6.0863 | 6.8717 | 6.1447 | 6.1674 | 6.2688 |
| HNRNPUL2_S161 | HNRNPUL2 | Q1KMD3 | | 228 | YHSRSKSPLPPEE | 6.1276 | 6.4001 | 5.7689 | 6.6809 | 6.2402 | 5.9174 | 5.9952 | 6.0097 | 6.0181 | 5.9372 | 6.0060 | 5.7379 | 6.6087 | 6.9819 | 7.0152 | 6.5548 |
| HNRNPUL2_S193 | HNRNPUL2 | Q1KMD3 | | 193 | KSKPAGSDGERRG | 6.2366 | 6.6063 | 5.8107 | 6.5686 | 6.2607 | 5.8893 | 6.2638 | 6.1880 | 5.8593 | 5.7025 | 5.7345 | 6.0862 | 6.8580 | 6.8353 | 6.5874 | 6.5127 |
| HNRNPUL2_S228 | HNRNPUL2 | Q1KMD3 | | 161 | DEPEERSGDETPG | 5.7824 | 6.1676 | 6.0693 | 6.5096 | 5.9661 | 5.5825 | 6.1157 | 6.0656 | 6.5748 | 6.0860 | 5.6426 | 6.0250 | 6.4461 | 6.7851 | 6.8257 | 7.3559 |
| HOMER3_S256 | HOMER3 | Q9NSC5 | | 256 | GLGQGQSLEQLEA | 5.9732 | 6.3755 | 6.4553 | 5.7834 | 5.9498 | 6.4112 | 5.5199 | 6.3107 | 6.6767 | 6.6684 | 6.6086 | 6.6364 | 6.2965 | 6.4553 | 6.0477 | 5.8313 |
| HOMEZ_T453 | HOMEZ | Q8IX15-3 | | 453 | LNERAETPPLPIP | 5.7749 | 6.0872 | 6.5513 | 6.6183 | 6.4371 | 6.1145 | 6.3995 | 6.3837 | 6.1102 | 6.1281 | 6.1740 | 6.1249 | 5.9552 | 6.6029 | 6.3238 | 6.2146 |
| HORMAD1_S242 | HORMAD1 | Q86X24 | | 242 | IDSTILSPKQIKT | 6.5816 | 6.8648 | 6.9375 | 7.2312 | 6.2422 | 6.9422 | 6.7330 | 7.0317 | 6.9610 | 7.4845 | 7.7277 | 7.3213 | 3.5061 | 4.1513 | 3.9750 | 4.3090 |
| HOXA4_S109 | HOXA4 | Q00056 | | 109 | GYRGGASPGRPPQ | 6.4969 | 6.5964 | 6.3014 | 7.1894 | 6.3583 | 5.9666 | 6.0979 | 5.8289 | 6.5495 | 7.2095 | 6.6843 | 6.1287 | 5.2327 | 6.0689 | 5.6041 | 5.6866 |
| HOXB4_S116;S130 | HOXB4 | P17483 | | 46 | GGQRRESSFQPEA | 5.6243 | 4.3818 | 5.2093 | 4.7372 | 7.0265 | 5.7628 | 5.4743 | 5.1782 | 9.1137 | 7.9452 | 7.4687 | 7.9430 | 7.3263 | 7.8531 | 3.9773 | 4.9782 |
| HOXB4_S130 | HOXB4 | P17483 | | 130 | QNPLHPSPSHSAC | 6.8036 | 6.3592 | 4.5917 | 5.1517 | 8.2315 | 6.8176 | 6.2507 | 5.4002 | 6.2787 | 6.6846 | 5.2917 | 5.4177 | 7.1851 | 7.8256 | 5.3407 | 6.3696 |
| HOXB4_S46 | HOXB4 | P17483 | 116;130 | | NA | 6.0973 | 6.7342 | 6.5339 | 6.9616 | 6.4701 | 6.2738 | 6.6323 | 6.4038 | 7.0120 | 6.7948 | 6.7460 | 6.6050 | 5.1032 | 5.3412 | 5.0888 | 5.2020 |
| HOXC10_S189 | HOXC10 | Q9NYD6 | | 189 | RAEHLESPQLGGK | 5.7778 | 6.6832 | 6.3061 | 6.5308 | 5.9562 | 7.1257 | 6.1637 | 6.2970 | 6.7482 | 7.1491 | 6.9059 | 6.9903 | 4.9010 | 5.5278 | 5.6020 | 5.3353 |
| HOXC10_T201 | HOXC10 | Q9NYD6 | | 201 | KVSFPETPKSDSQ | 8.1068 | 7.4900 | 7.1289 | 6.7535 | 6.4605 | 6.7019 | 6.6420 | 6.0293 | 7.7427 | 6.6833 | 6.3883 | 6.5069 | 3.7125 | 4.7482 | 4.0952 | 4.8100 |
| HP1BP3_S156 | HP1BP3 | Q5SSJ5 | | 156 | QTPMASSPRPKMD | 6.4952 | 6.9124 | 5.9852 | 6.9202 | 6.0469 | 6.0201 | 6.4859 | 6.1552 | 5.9961 | 6.2796 | 5.9537 | 6.0829 | 5.4776 | 6.9787 | 6.0925 | 6.1178 |
| HP1BP3_S441;S442;S446 | HP1BP3 | Q5SSJ5 | | 91 | TPPATSSEAEQPK | 5.7229 | 5.9881 | 5.4761 | 5.9094 | 6.0012 | 5.9348 | 6.2509 | 5.5816 | 5.7823 | 5.8335 | 5.5764 | 5.9080 | 7.2853 | 7.6498 | 7.4532 | 7.6465 |
| HP1BP3_S74 | HP1BP3 | Q5SSJ5 | | 74 | DISSEESVSTVEE | 5.4166 | 5.6477 | 5.7401 | 5.7787 | 5.8726 | 5.9278 | 6.2394 | 5.7500 | 5.9855 | 5.9119 | 5.5476 | 6.1402 | 6.7406 | 8.3068 | 7.3465 | 7.6478 |
| HP1BP3_S91 | HP1BP3 | Q5SSJ5 | | 51 | VNSTRETPPKSKL | 6.3717 | 6.6954 | 5.6700 | 6.5912 | 5.9525 | 6.1686 | 6.5337 | 6.0017 | 5.7388 | 6.4911 | 5.6247 | 5.9820 | 6.5920 | 7.1736 | 6.5751 | 6.7380 |
| HP1BP3_T51 | HP1BP3 | Q5SSJ5 | 441;442;446 | | NA | 6.6729 | 6.3298 | 5.5595 | 6.0398 | 6.1986 | 5.6257 | 6.5069 | 6.1945 | 5.6894 | 5.1100 | 5.7133 | 5.8082 | 6.9532 | 7.3861 | 6.9560 | 7.2561 |
| HPRT1_S5 | HPRT1 | P00492 | | 5 | xxMATRSPGVVIS | 6.3398 | 6.3616 | 6.4426 | 6.5288 | 6.4784 | 6.2546 | 6.4538 | 6.5804 | 6.4369 | 6.2660 | 6.7846 | 6.6714 | 5.3748 | 6.1291 | 5.4101 | 5.4871 |
| HPS4_S368 | HPS4 | Q9NQG7-4 | | 368 | VLGLSSSLGKELV | 7.2376 | 6.8597 | 7.6120 | 6.5698 | 5.9661 | 6.9604 | 6.0447 | 7.1372 | 6.9696 | 6.1421 | 7.5273 | 6.5936 | 4.2881 | 5.0660 | 4.5748 | 4.4509 |
| HPS5_S534 | HPS5 | Q9UPZ3 | | 534 | LPFRSPSPLVSLQ | 6.1033 | 5.7835 | 6.1270 | 5.5007 | 6.5512 | 7.5592 | 6.6712 | 6.7099 | 6.3379 | 5.2453 | 5.6689 | 5.7491 | 6.8592 | 6.6959 | 6.1541 | 6.2835 |
| HRH1_S230 | HRH1 | P35367 | | 233 | INRSLPSFSEIKL | 7.6045 | 7.3509 | 7.3844 | 6.4441 | 6.5122 | 6.6686 | 6.1427 | 6.2630 | 6.7204 | 7.1230 | 6.7993 | 6.6653 | 4.3800 | 5.4610 | 4.3122 | 4.1683 |
| HRH1_S230;S233 | HRH1 | P35367 | | 230 | RELINRSLPSFSE | 5.4898 | 6.0713 | 5.0912 | 5.0635 | 5.0198 | 5.9949 | 5.7125 | 5.8608 | 5.0045 | 4.8047 | 5.6901 | 6.3009 | 7.6484 | 7.7676 | 8.6293 | 9.8508 |
| HRH1_S233 | HRH1 | P35367 | | 380 | GKLRSGSNTGLDY | 5.3650 | 4.8184 | 5.8141 | 5.2853 | 5.8749 | 5.1234 | 4.8462 | 5.1323 | 6.7526 | 6.0479 | 6.1062 | 6.2638 | 8.1193 | 8.9193 | 8.2585 | 7.2728 |
| HRH1_S380 | HRH1 | P35367 | 230;233 | | NA | 3.2119 | 3.7029 | 3.8610 | 3.3606 | 4.0302 | 3.5561 | 3.8283 | 3.8091 | 3.6050 | 3.6602 | 3.3371 | 3.1614 | 12.9423 | 13.0752 | 16.4339 | 14.4248 |
| HS1BP3_S249 | HS1BP3 | Q53T59 | | 249 | GPGRKLSPQDPSE | 5.6685 | 5.6471 | 6.7535 | 5.5378 | 6.3093 | 6.5322 | 5.8017 | 7.0773 | 6.1164 | 6.2590 | 6.5193 | 6.1519 | 6.4074 | 6.8255 | 6.2281 | 6.1650 |
| HSD17B4_T290 | HSD17B4 | P51659-2 | | 290 | QKNHPMTPEAVKA | 6.5445 | 6.6036 | 6.3026 | 6.4414 | 6.0191 | 6.1123 | 6.2132 | 5.8232 | 6.4024 | 6.1757 | 6.2434 | 6.2893 | 6.2488 | 6.3989 | 6.2070 | 5.9747 |
| HSF1_S292 | HSF1 | Q00613 | | 314 | PRVEEASPGRPSS | 5.5086 | 5.7213 | 6.1718 | 5.5004 | 5.9476 | 6.1681 | 5.6353 | 6.7226 | 6.7502 | 6.8172 | 6.8570 | 6.8499 | 5.9407 | 6.7913 | 6.4451 | 6.1728 |
| HSF1_S303 | HSF1 | Q00613 | | 363 | TEGRPPSPPPTST | 4.9104 | 5.5946 | 5.4593 | 5.1580 | 5.4130 | 5.8885 | 5.2259 | 6.3171 | 5.2962 | 5.7927 | 5.9039 | 6.0597 | 8.0137 | 8.6569 | 8.4429 | 7.8672 |
| HSF1_S314 | HSF1 | Q00613 | | 292 | ERPLSSSPLVRVK | 6.1800 | 5.4295 | 6.3536 | 5.9445 | 6.1740 | 6.0270 | 6.0203 | 6.5143 | 7.2283 | 6.7241 | 6.7481 | 7.1834 | 5.1466 | 6.3976 | 5.8212 | 6.1075 |
| HSF1_S314;S326 | HSF1 | Q00613 | | 303 | VKEEPPSPPQSPR | 5.9126 | 5.4224 | 5.6221 | 4.9837 | 6.0363 | 6.6531 | 5.1387 | 5.7114 | 5.6810 | 6.1658 | 6.0515 | 5.7904 | 7.7847 | 8.2577 | 7.7911 | 6.9976 |
| HSF1_S363 | HSF1 | Q00613 | 363;369 | | NA | 5.7314 | 6.3538 | 6.6002 | 5.8653 | 6.1229 | 7.0406 | 5.9234 | 6.3169 | 7.1962 | 6.5528 | 7.3603 | 7.1389 | 5.5359 | 5.7700 | 4.9999 | 5.4916 |
| HSF1_S363;T369 | HSF1 | Q00613 | 314;326 | | NA | 6.8393 | 6.4626 | 6.6091 | 5.8291 | 5.8617 | 6.7446 | 6.2538 | 7.2240 | 7.5902 | 6.9342 | 6.4515 | 6.9656 | 4.6997 | 5.8855 | 4.6595 | 4.9898 |
| HSP90AA1_S353 | HSP90AA1 | P07900-2 | | 385 | EIEDVGSDEEEEK | 5.6091 | 5.6612 | 6.6411 | 5.6133 | 6.2383 | 6.5874 | 6.2582 | 6.0554 | 6.4593 | 6.4205 | 6.8949 | 6.7106 | 6.6187 | 6.5377 | 5.8196 | 5.8747 |
| HSP90AA1_S385 | HSP90AA1 | P07900-2 | | 353 | ERDKEVSDDEAEE | 5.3490 | 5.6419 | 6.6516 | 5.7421 | 6.3870 | 6.5076 | 6.2045 | 6.4099 | 6.4614 | 6.5192 | 6.3366 | 6.2880 | 6.3564 | 6.6923 | 6.2819 | 6.1704 |
| HSP90AA1_S437 | HSP90AA1 | P07900-2 | | 848 | EGDDDTSRMEEVD | 6.5653 | 6.4578 | 6.3264 | 7.2223 | 6.1703 | 6.3950 | 6.8561 | 6.5675 | 6.5317 | 6.1880 | 6.2641 | 6.4855 | 5.4296 | 5.8791 | 5.1994 | 5.4618 |
| HSP90AA1_S452 | HSP90AA1 | P07900-2 | | 598 | SGDEMVSLKDYCT | 5.6870 | 5.8248 | 6.7130 | 5.8871 | 6.1404 | 6.3662 | 6.2983 | 6.1987 | 6.6493 | 6.1017 | 6.5745 | 6.2760 | 6.3660 | 6.4933 | 6.1322 | 6.2917 |
| HSP90AA1_S528 | HSP90AA1 | P07900-2 | | 847 | LEGDDDTSRMEEV | 6.1079 | 6.1588 | 6.1456 | 6.5900 | 6.2166 | 6.4470 | 6.7043 | 6.5455 | 6.2784 | 5.9481 | 6.6634 | 6.3122 | 5.6147 | 6.3577 | 5.6120 | 6.2976 |
| HSP90AA1_S564 | HSP90AA1 | P07900-2 | | 717 | SNRLVTSPCCIVT | 6.5542 | 5.7695 | 6.7257 | 7.1882 | 6.5462 | 7.0498 | 6.0317 | 6.4446 | 6.0638 | 6.0199 | 6.0686 | 6.3317 | 5.7813 | 6.0166 | 5.5111 | 5.8971 |
| HSP90AA1_S575 | HSP90AA1 | P07900-2 | | 452 | LAVKHFSVEGQLE | 6.3320 | 5.9332 | 6.2225 | 5.5835 | 6.8494 | 6.3847 | 6.3261 | 6.8728 | 6.0817 | 5.4050 | 6.1177 | 6.1362 | 7.3435 | 6.7566 | 5.9003 | 5.7548 |
| HSP90AA1_S598 | HSP90AA1 | P07900-2 | | 763 | EINPDHSIIETLR | 5.6774 | 5.9089 | 6.2851 | 6.4393 | 6.4811 | 6.4608 | 6.7452 | 6.6694 | 6.1716 | 6.1128 | 6.2282 | 5.9459 | 5.9125 | 6.1411 | 6.4169 | 6.4038 |
| HSP90AA1_S717 | HSP90AA1 | P07900-2 | | 437 | YGEFYKSLTNDWE | 6.2258 | 6.8065 | 6.8453 | 6.2564 | 6.2728 | 6.3847 | 5.6919 | 6.6202 | 6.2416 | 6.4808 | 6.3210 | 6.4938 | 6.0395 | 5.8496 | 6.1326 | 5.3374 |
| HSP90AA1_S763 | HSP90AA1 | P07900-2 | | 439 | EFYKSLTNDWEDH | 6.0525 | 6.4385 | 6.4465 | 6.4171 | 6.5340 | 6.0525 | 6.0867 | 6.6870 | 6.2043 | 5.7949 | 6.7084 | 6.4767 | 5.9515 | 6.1678 | 6.3674 | 5.6143 |
| HSP90AA1_S848 | HSP90AA1 | P07900-2 | | 528 | REMLQQSKILKVI | 5.9565 | 5.5136 | 6.8737 | 5.5188 | 5.9253 | 5.4198 | 8.8280 | 6.6496 | 7.0665 | 6.4099 | 6.4360 | 7.1708 | 6.0972 | 6.8894 | 5.0862 | 4.1586 |
| HSP90AA1_T439 | HSP90AA1 | P07900-2 | | 575 | LGIHEDSQNRKKL | 5.8681 | 5.6549 | 7.6227 | 6.5641 | 6.0161 | 6.2613 | 5.8930 | 6.5076 | 6.6053 | 6.4003 | 6.8247 | 6.4463 | 5.6827 | 6.2575 | 5.2640 | 6.1315 |
| HSP90AA1_T847 | HSP90AA1 | P07900-2 | | 564 | KFYEQFSKNIKLG | 6.3128 | 5.7612 | 5.5384 | 6.5920 | 6.0202 | 6.4323 | 8.0901 | 5.3424 | 6.2210 | 6.5107 | 6.0298 | 7.0996 | 5.3405 | 6.2172 | 5.4925 | 6.9992 |
| HSP90AB1_S226 | HSP90AB1 | P08238 | | 255 | KIEDVGSDEEDDS | 5.8582 | 5.6882 | 6.9830 | 5.9229 | 6.5500 | 6.7876 | 6.3406 | 6.2030 | 6.4125 | 6.3911 | 6.7214 | 6.5449 | 6.1029 | 6.1454 | 5.5646 | 5.7838 |
| HSP90AB1_S255 | HSP90AB1 | P08238 | | 226 | EREKEISDDEAEE | 5.8871 | 6.0799 | 6.8466 | 5.9092 | 6.2017 | 6.3906 | 6.1885 | 6.4003 | 6.4148 | 6.3130 | 6.4961 | 6.5656 | 6.0943 | 6.3158 | 5.8837 | 6.0128 |
| HSP90AB1_S255;S261 | HSP90AB1 | P08238 | | 718 | EGDEDASRMEEVD | 6.2259 | 6.2421 | 6.1850 | 7.1981 | 6.0859 | 5.9959 | 6.4410 | 6.2215 | 6.7088 | 6.4124 | 6.3041 | 6.3261 | 6.0191 | 6.2861 | 5.4621 | 5.8859 |
| HSP90AB1_S434 | HSP90AB1 | P08238 | | 452 | TNRRRLSELLRYH | 6.5286 | 6.6128 | 7.3342 | 6.4978 | 6.4173 | 6.4430 | 5.7314 | 6.6501 | 8.2059 | 7.7514 | 8.3113 | 7.7863 | 3.6992 | 4.0954 | 3.8273 | 4.1081 |
| HSP90AB1_S452 | HSP90AB1 | P08238 | | 434 | KFYEAFSKNLKLG | 5.9221 | 4.3880 | 5.9451 | 5.8211 | 5.9345 | 7.1585 | 5.5996 | 4.8007 | 6.9389 | 7.8618 | 6.9531 | 9.6298 | 5.1249 | 5.8849 | 3.7095 | 8.3277 |
| HSP90AB1_S532 | HSP90AB1 | P08238 | | 532 | KEFDGKSLVSVTK | 6.3542 | 6.8509 | 7.4668 | 6.3078 | 6.3682 | 6.7372 | 6.0797 | 6.4482 | 6.4187 | 5.8757 | 6.5119 | 6.1239 | 5.7244 | 5.8022 | 5.7185 | 5.2116 |
| HSP90AB1_S587 | HSP90AB1 | P08238 | | 587 | SNRLVSSPCCIVT | 7.0025 | 6.6236 | 5.9626 | 6.5035 | 6.4793 | 5.8723 | 6.4132 | 6.9034 | 5.6795 | 6.1741 | 5.9582 | 6.1257 | 6.3151 | 6.2843 | 5.7864 | 5.9164 |
| HSP90AB1_S718 | HSP90AB1 | P08238 | 255;261 | | NA | 5.9485 | 5.3213 | 7.0766 | 5.5227 | 6.4266 | 6.8076 | 7.1727 | 5.2789 | 5.5259 | 6.1716 | 6.9607 | 7.3630 | 6.9907 | 5.8690 | 5.6605 | 5.9035 |
| HSP90B1_S306 | HSP90B1 | P14625 | | 786 | EEMDVGTDEEEET | 5.8686 | 6.3869 | 6.0925 | 6.4804 | 5.9582 | 6.3043 | 6.2543 | 5.7595 | 5.6816 | 6.5507 | 6.2996 | 5.8150 | 7.0074 | 6.7394 | 6.2326 | 6.5689 |
| HSP90B1_T786 | HSP90B1 | P14625 | | 306 | KEEKEESDDEAAV | 5.4500 | 6.4070 | 6.3786 | 6.1715 | 6.6187 | 5.3341 | 6.3845 | 6.2933 | 5.2831 | 5.8994 | 5.3142 | 5.5155 | 6.9092 | 6.8916 | 7.0220 | 8.1270 |
| HSPA4_S415 | HSPA4 | P34932 | | 647 | SEDDRNSFTLKLE | 5.9959 | 5.8995 | 6.6118 | 5.8867 | 6.0390 | 6.1755 | 6.0116 | 6.2075 | 6.3032 | 6.2127 | 6.4065 | 6.4606 | 6.0343 | 6.8471 | 6.5030 | 6.4052 |
| HSPA4_S647 | HSPA4 | P34932 | | 76 | FHGRAFSDPFVEA | 5.6285 | 5.3249 | 6.3914 | 5.9027 | 6.2481 | 6.1268 | 5.6211 | 5.6217 | 6.1358 | 6.2763 | 6.9129 | 6.6138 | 6.5743 | 7.4992 | 6.4501 | 6.6725 |
| HSPA4_S76 | HSPA4 | P34932 | | 415 | PAEEGSSDCEVFS | 5.3666 | 5.8741 | 7.4009 | 6.1750 | 5.4478 | 5.9559 | 6.1495 | 6.0189 | 5.8486 | 6.2656 | 6.3858 | 6.0215 | 7.3243 | 6.5607 | 6.4436 | 6.7611 |
| HSPA8_S254 | HSPA8 | P11142 | | 637 | PPSGGASSGPTIE | 6.1899 | 6.3169 | 6.4760 | 6.3439 | 6.1965 | 6.3727 | 6.3743 | 6.5857 | 6.3036 | 6.3168 | 6.2696 | 6.4544 | 5.8408 | 6.1887 | 5.7124 | 6.0578 |
| HSPA8_S329 | HSPA8 | P11142 | | 541 | KVSSKNSLESYAF | 6.1126 | 6.0774 | 6.2806 | 6.1225 | 6.2967 | 5.9714 | 6.2165 | 6.0229 | 6.6075 | 6.4815 | 6.2059 | 6.2196 | 6.7457 | 6.5438 | 6.0201 | 6.0753 |
| HSPA8_S541 | HSPA8 | P11142 | | 329 | DAKLDKSQIHDIV | 5.6308 | 5.7754 | 7.2096 | 5.7350 | 6.3370 | 5.9382 | 6.2439 | 6.7081 | 6.1344 | 6.3168 | 6.1354 | 6.6337 | 6.2615 | 6.2668 | 6.3912 | 6.2822 |
| HSPA8_S637 | HSPA8 | P11142 | | 254 | KHKKDISENKRAV | 5.2973 | 6.3401 | 6.5856 | 5.5543 | 6.1940 | 7.1131 | 7.1749 | 7.3352 | 6.1370 | 5.6832 | 6.0536 | 6.0302 | 6.4818 | 7.1992 | 5.4117 | 5.4090 |
| HSPA9_S148 | HSPA9 | P38646 | | 162 | AHGKLYSPSQIGA | 6.2276 | 6.0066 | 6.7347 | 5.8564 | 6.1025 | 6.0291 | 5.6871 | 6.8346 | 5.5417 | 6.7796 | 6.7993 | 5.8122 | 5.6953 | 6.9692 | 6.5559 | 6.3683 |
| HSPA9_S162 | HSPA9 | P38646 | | 336 | PYLTMDSSGPKHL | 6.3487 | 6.1267 | 5.8195 | 6.1675 | 6.4213 | 5.9719 | 5.9269 | 6.8023 | 6.4033 | 6.8984 | 5.6275 | 6.6325 | 6.2257 | 6.5005 | 6.1273 | 5.9995 |
| HSPA9_S336 | HSPA9 | P38646 | | 148 | FKIVRASNGDAWV | 5.6244 | 6.1226 | 6.5458 | 6.7055 | 7.2197 | 7.4577 | 5.9901 | 5.7202 | 6.6896 | 5.9294 | 5.8943 | 6.8221 | 6.5554 | 5.2747 | 5.1150 | 6.3334 |
| HSPB1_S15 | HSPB1 | P04792 | | 82 | ALSRQLSSGVSEI | 5.0258 | 4.3006 | 5.4349 | 4.0906 | 5.5017 | 5.0187 | 4.7503 | 6.5991 | 4.9265 | 5.7237 | 5.3303 | 5.8933 | 7.8529 | 9.8946 | 9.9869 | 9.6701 |
| HSPB1_S82 | HSPB1 | P04792 | | 15 | SLLRGPSWDPFRD | 2.8814 | 3.2046 | 3.8317 | 3.2153 | 5.3109 | 4.3012 | 4.9490 | 7.6209 | 4.5285 | 5.9356 | 5.0301 | 5.9260 | 6.8882 | 10.0097 | 12.7232 | 13.6438 |
| HSPD1_S70 | HSPD1 | P10809 | | 70 | IEQSWGSPKVTKD | 5.9852 | 6.0858 | 6.0878 | 6.7643 | 6.4325 | 6.2512 | 6.2190 | 5.9962 | 6.3180 | 6.4585 | 6.2115 | 6.6285 | 6.3818 | 5.9478 | 6.0491 | 6.1829 |
| HSPD1_T164 | HSPD1 | P10809 | | 164 | QSKPVTTPEEIAQ | 6.2438 | 5.5476 | 5.9997 | 6.2074 | 6.4981 | 6.3216 | 6.1557 | 6.0730 | 6.3480 | 6.6072 | 6.3911 | 6.3368 | 6.5974 | 6.2196 | 6.1589 | 6.2941 |
| HSPH1_S559 | HSPH1 | Q92598-4 | | 811 | PKPKIESPKLERT | 5.7118 | 5.6809 | 6.9112 | 6.0385 | 7.5835 | 8.2877 | 6.7470 | 5.6849 | 6.2728 | 6.5665 | 8.0553 | 6.6949 | 6.3464 | 5.5774 | 4.1260 | 3.7151 |
| HSPH1_S811 | HSPH1 | Q92598-4 | | 817 | SPKLERTPNGPNI | 6.6557 | 7.3719 | 6.7528 | 6.8655 | 6.7508 | 7.2324 | 7.3651 | 6.9334 | 6.8734 | 6.9999 | 7.1703 | 6.9270 | 3.7601 | 4.3767 | 3.7612 | 4.2038 |
| HSPH1_T817 | HSPH1 | Q92598-4 | | 559 | TSQSPPSPELTSE | 5.7553 | 5.4908 | 6.1320 | 5.6131 | 5.9425 | 6.1841 | 6.1182 | 6.2750 | 6.0030 | 6.4477 | 6.1760 | 6.5427 | 7.0516 | 6.9377 | 6.5154 | 6.8150 |
| HTATSF1_S452;S453 | HTATSF1 | O43719 | | 579 | DLDEEGSEKELHE | 5.6323 | 6.0486 | 6.1916 | 6.3437 | 6.0388 | 5.7785 | 6.2329 | 5.6548 | 5.9142 | 6.1010 | 6.1976 | 5.9930 | 7.0619 | 6.9629 | 6.8545 | 6.9937 |
| HTATSF1_S481 | HTATSF1 | O43719 | | 702 | LFEDDDSNEKLFD | 5.8373 | 6.0484 | 5.9074 | 6.2950 | 5.6862 | 5.7691 | 6.0106 | 5.8084 | 5.9097 | 6.1729 | 6.1937 | 6.2747 | 6.7759 | 7.1330 | 7.0513 | 7.1262 |
| HTATSF1_S498 | HTATSF1 | O43719 | | 714 | DEEEDSSEKLFDD | 6.0105 | 6.0570 | 6.2878 | 6.2520 | 6.0095 | 6.0445 | 5.9892 | 5.7216 | 6.5073 | 6.3091 | 6.4089 | 6.3462 | 6.5793 | 6.5593 | 6.4022 | 6.5158 |
| HTATSF1_S579 | HTATSF1 | O43719 | | 642 | KVFDDESDEKEDE | 5.9296 | 6.7718 | 6.8622 | 6.5783 | 6.0617 | 5.5965 | 6.1494 | 5.1693 | 6.0148 | 6.0666 | 5.6860 | 5.8079 | 6.8506 | 6.5396 | 6.7642 | 7.1515 |
| HTATSF1_S616;S624 | HTATSF1 | O43719 | | 481 | GFEGSCSQKESEE | 6.5827 | 6.8954 | 6.5246 | 6.9328 | 6.4157 | 6.3922 | 5.9148 | 5.8025 | 6.7162 | 6.4246 | 6.1831 | 6.0484 | 5.9118 | 5.8238 | 5.7353 | 5.6961 |
| HTATSF1_S642 | HTATSF1 | O43719 | | 498 | RGSEEDSPKKESK | 5.4118 | 5.9689 | 6.4413 | 6.6683 | 5.8066 | 5.8389 | 6.0796 | 5.4035 | 6.4246 | 6.3960 | 6.4667 | 7.0178 | 6.6248 | 6.4524 | 6.1918 | 6.8068 |
| HTATSF1_S702 | HTATSF1 | O43719 | | 721 | EKLFDDSDERGTL | 5.7704 | 6.0907 | 5.5482 | 6.1952 | 5.8968 | 5.6921 | 6.0065 | 6.6236 | 5.7708 | 5.7924 | 5.5234 | 5.8690 | 6.9794 | 6.8317 | 7.4233 | 7.9866 |
| HTATSF1_S713;S714 | HTATSF1 | O43719 | 616;624 | | NA | 5.6837 | 6.1423 | 6.4372 | 6.4584 | 5.9565 | 5.9807 | 6.1588 | 5.2486 | 6.0175 | 6.3239 | 6.2415 | 6.3638 | 6.8623 | 6.7009 | 6.4179 | 7.0059 |
| HTATSF1_S714 | HTATSF1 | O43719 | 452;453 | | NA | 6.1476 | 6.5192 | 7.3529 | 6.8716 | 6.0595 | 5.7810 | 6.7798 | 5.4346 | 5.4737 | 6.6482 | 6.1387 | 6.4584 | 6.0439 | 6.3752 | 5.4841 | 6.4316 |
| HTATSF1_S721 | HTATSF1 | O43719 | 713;714 | | NA | 5.3303 | 6.4277 | 6.6738 | 6.0139 | 6.1061 | 5.9157 | 6.2807 | 5.6111 | 6.1734 | 5.8943 | 5.8441 | 6.5449 | 6.5344 | 6.8849 | 6.8598 | 6.9048 |
| HTT_S1179 | HTT | P42858 | | 1199 | QASVPLSPKKGSE | 6.2161 | 6.4538 | 6.8111 | 6.5192 | 6.1050 | 6.4031 | 6.0953 | 6.5951 | 7.0939 | 6.7140 | 7.0431 | 7.1749 | 4.9099 | 5.3530 | 5.0562 | 5.4562 |
| HTT_S1199 | HTT | P42858 | | 1874 | LLSPQMSGEEEDS | 4.8139 | 5.0268 | 5.3022 | 4.5570 | 5.7239 | 5.5831 | 5.3648 | 5.8053 | 5.1347 | 4.7922 | 5.0561 | 4.9364 | 9.7052 | 9.2848 | 10.0243 | 8.8893 |
| HTT_S1864 | HTT | P42858 | | 1864 | PKRHSLSSTKLLS | 6.3746 | 6.5023 | 6.6010 | 5.6713 | 5.8951 | 6.1167 | 6.1688 | 6.6971 | 6.2789 | 6.0183 | 6.2441 | 6.1416 | 5.9572 | 6.4560 | 6.4418 | 6.4352 |
| HTT_S1874 | HTT | P42858 | | 1179 | TNPPSLSPIRRKG | 6.7772 | 6.7547 | 6.6517 | 6.3078 | 6.5242 | 6.2623 | 6.3425 | 6.8565 | 6.6932 | 6.7599 | 6.9318 | 6.9566 | 4.6145 | 5.6960 | 4.7812 | 5.0899 |
| HTT_S419 | HTT | P42858 | | 419 | GRSRSGSIVELIA | 5.2490 | 6.6732 | 6.0139 | 6.1133 | 6.5849 | 6.0391 | 6.2016 | 6.4540 | 6.5028 | 6.7710 | 6.0250 | 5.7474 | 6.3451 | 6.1574 | 6.6117 | 6.5107 |

| | | | | | | | | | | | | | | | | | | | | |
|---|---|---|---|---|---|---|---|---|---|---|---|---|---|---|---|---|---|---|---|---|
| HTT_S419;S432 | HTT | P42858 | 419;432 | NA | 5.7889 | 5.9405 | 6.5446 | 6.4036 | 6.3378 | 6.3971 | 5.9973 | 5.9925 | 6.7040 | 6.9430 | 6.1455 | 6.7076 | 5.9187 | 6.4633 | 5.8298 | 5.8858 |
| HTT_S640;S643 | HTT | P42858 | 640;643 | NA | 6.4121 | 6.0691 | 6.6582 | 5.5501 | 5.7136 | 5.6678 | 6.4794 | 6.2405 | 5.6945 | 5.9110 | 6.8472 | 6.2500 | 6.9265 | 6.4923 | 6.8881 | 6.1998 |
| HUWE1_S1084 | HUWE1 | Q7Z6Z7 | 3816 | MDVDQPSPSAQDT | 5.7613 | 5.8946 | 6.2073 | 5.8414 | 6.0196 | 6.3988 | 6.2353 | 5.8132 | 6.3394 | 6.4451 | 5.9103 | 6.0822 | 6.5553 | 7.1346 | 6.4365 | 6.9250 |
| HUWE1_S134 | HUWE1 | Q7Z6Z7 | 3919 | DEPPPLSPAPLTP | 5.9564 | 5.8468 | 6.7231 | 6.1531 | 6.1473 | 6.3668 | 6.2235 | 6.7834 | 6.4665 | 6.1105 | 6.3430 | 6.5265 | 6.2324 | 6.3259 | 5.6728 | 6.1218 |
| HUWE1_S1368 | HUWE1 | Q7Z6Z7 | 3373 | RGNKACSPCSSQS | 6.1930 | 6.4478 | 6.6811 | 6.7762 | 6.3964 | 6.7491 | 6.6804 | 6.4939 | 6.5738 | 6.6999 | 6.9461 | 6.8612 | 4.7437 | 5.5201 | 4.9119 | 5.3254 |
| HUWE1_S1395 | HUWE1 | Q7Z6Z7 | 649 | MRRRRSSDPLGDT | 6.1848 | 6.4414 | 6.5489 | 6.0249 | 6.1804 | 5.7220 | 6.1692 | 6.2562 | 6.1183 | 7.0952 | 6.6422 | 6.3712 | 6.0374 | 6.1212 | 6.3412 | 5.7454 |
| HUWE1_S1907 | HUWE1 | Q7Z6Z7 | 1907 | RGSGTASDDEFEN | 6.2472 | 6.5765 | 6.4833 | 6.0818 | 6.1786 | 6.3822 | 6.2163 | 6.3927 | 5.7975 | 5.7585 | 5.7756 | 5.7178 | 6.4862 | 6.7103 | 6.6499 | 6.5455 |
| HUWE1_S2362 | HUWE1 | Q7Z6Z7 | 1395 | MDQRAESPEEVAC | 6.6796 | 7.9101 | 7.1267 | 6.5936 | 7.2787 | 8.6103 | 5.7696 | 5.0434 | 4.7187 | 5.5826 | 7.1750 | 6.5290 | 6.0652 | 6.1063 | 4.4228 | 4.3884 |
| HUWE1_S2362;T2365 | HUWE1 | Q7Z6Z7 | 3127 | LSAILRSPAFTSR | 7.3383 | 7.9911 | 7.0407 | 7.4708 | 6.5708 | 6.9941 | 7.3740 | 7.2992 | 7.2780 | 7.0036 | 7.4370 | 7.4685 | 2.8239 | 3.7525 | 2.6866 | 3.4708 |
| HUWE1_S2365 | HUWE1 | Q7Z6Z7 | 1368 | GVVRDLSMSEEDQ | 6.2185 | 6.2071 | 6.9388 | 6.1890 | 5.7400 | 6.5892 | 6.5006 | 6.1123 | 6.2722 | 6.3978 | 6.7735 | 6.0616 | 5.9542 | 6.2592 | 5.7083 | 6.0778 |
| HUWE1_S3072 | HUWE1 | Q7Z6Z7 | 3757 | GSSGLGSASSIQA | 4.5999 | 4.7063 | 4.8943 | 5.2616 | 5.3782 | 5.2934 | 5.1445 | 5.2446 | 5.2353 | 5.3958 | 5.3326 | 5.6266 | 9.2758 | 9.4917 | 9.6500 | 9.4694 |
| HUWE1_S3117 | HUWE1 | Q7Z6Z7 | 2889 | PRAGSSTPGDAPP | 6.0916 | 6.6662 | 6.3737 | 6.3509 | 5.9313 | 6.7207 | 5.8777 | 5.8827 | 5.8731 | 6.6139 | 5.6837 | 6.3042 | 6.6094 | 6.3605 | 6.3596 | 6.3010 |
| HUWE1_S3127 | HUWE1 | Q7Z6Z7 | 3072 | PSDLRRSVLEDME | 5.3669 | 7.2068 | 5.9041 | 4.9900 | 5.9824 | 6.8939 | 6.9337 | 6.4042 | 6.3837 | 6.8170 | 5.8785 | 5.9926 | 6.8029 | 6.4324 | 5.8426 | 6.1683 |
| HUWE1_S3373 | HUWE1 | Q7Z6Z7 | 1084 | PVRQRRSHHAAST | 7.1712 | 6.7027 | 5.6116 | 8.1762 | 7.0631 | 5.6997 | 6.9109 | 8.6687 | 7.5075 | 7.1952 | 6.6887 | 7.7277 | 3.2793 | 3.9319 | 2.7487 | 4.9169 |
| HUWE1_S3662 | HUWE1 | Q7Z6Z7 | 2362 | LERDGGSGNSTII | 4.9173 | 5.8062 | 6.1533 | 5.4339 | 5.4498 | 6.8375 | 6.1847 | 7.1680 | 6.1042 | 8.4267 | 7.9351 | 8.1570 | 4.6333 | 5.2607 | 5.4741 | 6.0584 |
| HUWE1_S3757 | HUWE1 | Q7Z6Z7 | 3924 | LSPAPLTPATPSS | 5.5652 | 5.4127 | 5.2397 | 5.4394 | 6.8458 | 7.0976 | 6.4379 | 6.1523 | 5.7141 | 5.9066 | 6.1208 | 7.1557 | 7.2416 | 6.4028 | 6.3423 | 6.9257 |
| HUWE1_S3816 | HUWE1 | Q7Z6Z7 | 3927 | APLTPATPSSLDP | 5.9233 | 6.3272 | 6.3420 | 6.0246 | 6.4752 | 5.9573 | 6.1127 | 6.3635 | 6.6239 | 5.1850 | 5.8242 | 5.5482 | 7.3977 | 6.1615 | 6.7222 | 7.0115 |
| HUWE1_S3919 | HUWE1 | Q7Z6Z7 | 3117 | RLFGHSSTSALSA | 5.4628 | 6.5175 | 4.8306 | 5.0335 | 6.5411 | 4.7326 | 5.6352 | 6.6054 | 4.3878 | 3.8267 | 4.5365 | 4.0599 | 7.6736 | 9.9182 | 11.3109 | 8.9277 |
| HUWE1_S3919;T3924 | HUWE1 | Q7Z6Z7 | 2365 | DGGSGNSTIIVSR | 5.6093 | 6.5929 | 6.2818 | 6.7239 | 6.0388 | 6.5460 | 6.7119 | 6.0119 | 7.0762 | 6.6661 | 6.7079 | 6.3510 | 5.5733 | 5.7546 | 5.2388 | 6.1154 |
| HUWE1_S3919;T3924;T3 | HUWE1 | Q7Z6Z7 | 648 | AMRRRRSSDPLGD | 5.4652 | 5.9898 | 7.2585 | 4.1407 | 6.0619 | 5.9078 | 5.6324 | 6.7143 | 6.0357 | 6.8782 | 6.9700 | 7.6290 | 6.9241 | 7.3733 | 6.3209 | 4.6981 |
| HUWE1_S3919;T3927 | HUWE1 | Q7Z6Z7 | 3662 | AQCETLSPDGLPE | 4.7393 | 5.3158 | 6.7492 | 6.6838 | 5.3102 | 7.1694 | 7.4088 | 5.4909 | 6.1838 | 6.4455 | 7.2714 | 5.7869 | 6.0519 | 6.8900 | 6.3390 | 6.1639 |
| HUWE1_S3924 | HUWE1 | A0A087X1S3 | 134 | NISSGPSPGPSPG | 6.0247 | 6.5983 | 7.6479 | 6.4885 | 5.2437 | 5.9556 | 7.1556 | 6.6634 | 6.4356 | 6.3258 | 3.9460 | 5.7969 | 5.7969 | 5.9230 | 6.7895 | 7.2085 |
| HUWE1_S3927 | HUWE1 | Q7Z6Z7 | 3919;3924 | NA | 5.7207 | 6.1073 | 6.6006 | 5.8209 | 6.1185 | 6.0883 | 6.6056 | 6.5871 | 6.2473 | 6.2746 | 5.9522 | 5.8541 | 6.2663 | 6.7251 | 6.5630 | 6.4684 |
| HUWE1_S648 | HUWE1 | Q7Z6Z7 | 3919;3927 | NA | 5.8353 | 6.1908 | 6.2952 | 6.3371 | 6.6778 | 5.6637 | 6.2987 | 6.0643 | 6.4007 | 5.8547 | 5.9315 | 5.7464 | 7.1738 | 6.3834 | 6.3532 | 6.7935 |
| HUWE1_S649 | HUWE1 | Q7Z6Z7 | 2362;2365 | NA | 6.8138 | 6.9964 | 6.8785 | 6.4078 | 6.2637 | 6.3861 | 6.3531 | 6.2049 | 6.6217 | 6.4020 | 5.9580 | 6.3966 | 5.4091 | 6.1338 | 4.9540 | 5.8204 |
| HUWE1_T2889 | HUWE1 | Q7Z6Z7 | 3919;3924;39 | NA | 6.9018 | 5.3909 | 5.7169 | 6.9518 | 5.8026 | 5.8201 | 6.0726 | 5.6300 | 6.1500 | 6.8666 | 6.0456 | 5.8513 | 7.0592 | 6.3658 | 7.2909 | 6.0839 |
| HYLS1_S297 | HYLS1 | Q96M11 | 297 | KLPFPLSPSxxxx | 6.5545 | 6.6887 | 6.1800 | 7.3433 | 6.6170 | 7.5428 | 7.2957 | 8.0756 | 7.5864 | 7.0364 | 7.7333 | 7.5193 | 2.9492 | 4.3216 | 3.0830 | 3.4733 |
| IARS1_S1047 | IARS1 | P41252 | 1047 | LKPYPVSPSDKVL | 6.2070 | 5.7024 | 6.7857 | 6.1915 | 6.1722 | 6.1354 | 6.1070 | 5.5321 | 6.7037 | 6.9087 | 6.2093 | 6.3331 | 6.5107 | 6.0907 | 6.2563 | 6.1542 |
| IBTK_S1004 | IBTK | Q9P2D0 | 1045 | SPRDLQSPDFTTG | 5.9466 | 6.4342 | 6.3680 | 5.5208 | 6.4121 | 6.7080 | 6.2182 | 6.8784 | 6.5361 | 6.8536 | 6.9823 | 6.5156 | 5.5617 | 6.0355 | 5.6098 | 5.4191 |
| IBTK_S1045 | IBTK | Q9P2D0 | 992 | PRKRSDSSGGYNL | 6.3676 | 5.6549 | 5.6629 | 5.8419 | 6.1270 | 6.4133 | 7.0004 | 6.2983 | 5.5626 | 6.5255 | 6.1183 | 6.0595 | 6.3740 | 7.2548 | 5.7844 | 5.9547 |
| IBTK_S992 | IBTK | Q9P2D0 | 1004 | LSDIIQSPSSTGL | 6.4702 | 6.9879 | 7.1866 | 6.9489 | 6.8455 | 6.7119 | 6.1180 | 6.5855 | 6.5394 | 7.4999 | 5.9796 | 6.4829 | 4.7789 | 5.0310 | 4.3694 | 5.4643 |
| IBTK_S992;S1004 | IBTK | Q9P2D0 | 992;1004 | NA | 6.4260 | 6.9305 | 8.0501 | 6.5812 | 6.1405 | 7.0536 | 6.3043 | 6.7517 | 6.7506 | 7.4693 | 7.2687 | 6.9699 | 4.2511 | 4.7704 | 4.0019 | 4.2803 |
| ICAM1_T521 | ICAM1 | P05362 | 521 | QQAQKGTPMKPNT | 7.0741 | 7.1988 | 6.2749 | 7.4565 | 6.5370 | 6.7259 | 6.9461 | 6.3839 | 6.2405 | 6.4276 | 6.6637 | 6.5273 | 4.7922 | 5.2169 | 4.6058 | 4.9288 |
| ICE1_S1470 | ICE1 | Q9Y2F5 | 533 | KSELCSSPLGKRP | 6.7388 | 6.7973 | 6.1550 | 6.8498 | 6.2566 | 5.9149 | 6.2464 | 6.1863 | 6.5808 | 5.8801 | 6.3455 | 6.3107 | 6.0328 | 5.7511 | 5.8716 | 6.0822 |
| ICE1_S1617 | ICE1 | Q9Y2F5 | 1642 | LAPLIATPPRTSQ | 6.6266 | 6.4259 | 6.3066 | 7.0255 | 6.1916 | 5.6982 | 6.0561 | 6.7888 | 6.3908 | 6.9699 | 6.1989 | 6.3902 | 5.5244 | 5.5152 | 5.7904 | 6.1010 |
| ICE1_S1712 | ICE1 | Q9Y2F5 | 960 | NSRLSFSPENILI | 5.7591 | 6.6602 | 6.1563 | 6.7025 | 6.7536 | 6.3326 | 6.3900 | 6.4144 | 6.3013 | 6.0067 | 6.3511 | 6.4341 | 5.7823 | 6.0560 | 5.5486 | 6.3511 |
| ICE1_S1854 | ICE1 | Q9Y2F5 | 1712 | SERVVPSPLQFCA | 6.2461 | 6.1086 | 6.2075 | 6.5494 | 6.5087 | 6.0751 | 6.3015 | 6.3280 | 6.3004 | 6.2023 | 5.7069 | 5.6267 | 6.8259 | 6.4348 | 6.2827 | 6.2954 |
| ICE1_S1891;S1903 | ICE1 | Q9Y2F5 | 516 | IERLSASPAQEKE | 7.5395 | 6.9184 | 6.3844 | 7.1806 | 6.6714 | 6.1268 | 6.1921 | 6.1609 | 6.7605 | 6.4811 | 6.1356 | 6.3620 | 5.4542 | 5.1409 | 5.1783 | 5.3134 |
| ICE1_S1903 | ICE1 | Q9Y2F5 | 1903 | VSQLPLSPKETVE | 6.0550 | 6.0668 | 6.4359 | 6.8034 | 5.6782 | 5.5499 | 6.8149 | 6.2320 | 6.6848 | 6.2493 | 6.1303 | 6.7486 | 6.2640 | 6.6206 | 5.8995 | 5.7667 |
| ICE1_S516 | ICE1 | Q9Y2F5 | 1854 | LRLDTGSPEPETR | 6.8859 | 6.8319 | 6.2784 | 7.1885 | 6.2523 | 5.9407 | 6.1416 | 6.2655 | 6.2861 | 5.8678 | 6.0989 | 6.3398 | 5.7271 | 5.8428 | 5.9189 | 6.1341 |
| ICE1_S533 | ICE1 | Q9Y2F5 | 707 | NSLCALSPELGAS | 5.9435 | 6.4223 | 5.7863 | 7.6265 | 6.5304 | 6.5627 | 5.7888 | 6.3341 | 6.7676 | 6.3838 | 6.3291 | 6.0745 | 6.4167 | 6.1360 | 5.9037 | 4.9939 |
| ICE1_S707 | ICE1 | Q9Y2F5 | 832 | LQNRGPTPKPDLL | 6.0860 | 5.7585 | 4.6610 | 4.9655 | 4.4991 | 5.5307 | 5.7806 | 5.1995 | 5.0610 | 4.3272 | 4.6179 | 4.9223 | 9.7017 | 9.4160 | 8.8155 | 10.6574 |
| ICE1_S925 | ICE1 | Q9Y2F5 | 939 | RKLDFNSPGGSSP | 7.2298 | 7.0907 | 6.9615 | 8.3925 | 6.7379 | 5.4807 | 8.8000 | 4.6609 | 7.8957 | 8.0248 | 6.5143 | 7.0509 | 4.1689 | 3.8708 | 3.9752 | 3.1453 |
| ICE1_S939 | ICE1 | Q9Y2F5 | 1470 | VTSAEKSPEASHT | 6.5281 | 6.4448 | 6.3366 | 6.2319 | 6.3556 | 5.9437 | 5.9771 | 5.5528 | 5.9193 | 6.4892 | 7.1569 | 6.6827 | 6.0908 | 6.0913 | 5.6984 | 6.5012 |
| ICE1_S960 | ICE1 | Q9Y2F5 | 925 | AAVKSISPEVSAS | 6.7238 | 6.8557 | 5.7164 | 6.8800 | 5.9577 | 5.6258 | 6.2840 | 6.0772 | 6.1806 | 6.2525 | 5.9399 | 5.9137 | 6.3120 | 6.2058 | 6.5349 | 6.5401 |
| ICE1_T1642 | ICE1 | Q9Y2F5 | 1617 | STPTDCSPDTLSK | 7.1331 | 8.7492 | 6.6586 | 8.6158 | 6.4418 | 6.3129 | 6.9512 | 6.1962 | 7.1452 | 5.9930 | 6.5676 | 6.3508 | 3.9524 | 4.0600 | 4.6361 | 4.2359 |
| ICE1_T832 | ICE1 | Q9Y2F5 | 1891;1903 | NA | 6.4377 | 6.8939 | 6.1320 | 6.7282 | 6.0384 | 5.8431 | 5.9154 | 5.5202 | 6.7504 | 6.5009 | 6.5340 | 6.3203 | 5.7434 | 6.7244 | 6.4344 | 5.4834 |
| ICE2_S17 | ICE2 | Q659A1 | 416 | TTKVSKSPSPAST | 6.0918 | 6.3637 | 6.4805 | 7.1007 | 5.9800 | 6.3455 | 6.4513 | 6.2924 | 6.4945 | 6.3949 | 6.2394 | 6.6695 | 5.5982 | 5.8623 | 5.5972 | 6.0381 |
| ICE2_S326 | ICE2 | Q659A1 | 605 | LSSRPASPNSSSG | 5.5824 | 5.5873 | 6.1272 | 6.1752 | 6.4108 | 6.2885 | 6.1552 | 6.2619 | 6.7581 | 6.9120 | 6.6632 | 6.5436 | 5.7346 | 6.0759 | 6.1220 | 6.6021 |
| ICE2_S416 | ICE2 | Q659A1 | 326 | KVIYINSPLPQKK | 6.6351 | 6.9849 | 6.3534 | 7.4815 | 7.0365 | 6.6412 | 7.0496 | 6.3951 | 7.1285 | 6.8314 | 7.1355 | 6.7446 | 4.2161 | 4.4089 | 4.1794 | 4.7783 |
| ICE2_S601;S605 | ICE2 | Q659A1 | 17 | GLNWDISPKNGLK | 6.7903 | 6.6026 | 6.6789 | 6.5852 | 5.7531 | 6.3072 | 6.8902 | 5.8445 | 6.2440 | 6.9766 | 6.7903 | 7.1165 | 5.2523 | 5.7740 | 4.9754 | 5.4188 |
| ICE2_S605 | ICE2 | Q659A1 | 601;605 | NA | 6.4552 | 5.9307 | 5.6870 | 6.6174 | 5.1167 | 5.3493 | 5.7365 | 6.1943 | 6.1182 | 6.5005 | 6.0611 | 6.0884 | 7.3886 | 7.4118 | 6.0206 | 7.3237 |
| IER3_S31 | IER3 | P46695 | 31 | PGPRRGSGPEIFT | 5.6291 | 5.3787 | 5.3036 | 6.4945 | 6.1105 | 4.5634 | 7.4629 | 7.0566 | 6.2497 | 5.9185 | 6.3025 | 6.3164 | 8.2448 | 6.5780 | 4.3603 | 8.0305 |
| IER5_S284 | IER5 | Q5VY09 | 284 | SGLLRKSPGGGRE | 6.8714 | 7.0816 | 6.7773 | 7.1046 | 6.3943 | 6.9159 | 6.8398 | 7.4391 | 6.4201 | 7.0100 | 7.3245 | 7.2644 | 3.7881 | 4.6641 | 3.6585 | 4.4464 |
| IFFO1_S158 | IFFO1 | Q0D2I5-5 | 158 | SARVLGSPARSPA | 7.4967 | 7.3851 | 6.7928 | 6.9053 | 6.5323 | 7.3019 | 6.5759 | 6.2224 | 7.6643 | 7.2744 | 7.2970 | 7.4198 | 3.9455 | 3.7068 | 3.6143 | 3.8655 |
| IFI16_S153 | IFI16 | Q16666 | 153 | EQTQPPSPAGAGM | 5.6153 | 5.8952 | 6.1611 | 5.8366 | 5.2548 | 6.1548 | 5.8217 | 5.9132 | 5.8593 | 6.3777 | 6.0501 | 5.8514 | 8.0316 | 7.4928 | 6.5404 | 7.1440 |
| IFIH1_S301 | IFIH1 | Q9BYX4 | 301 | NVAARASPEPELQ | 6.3137 | 6.4628 | 6.9241 | 6.1694 | 6.4268 | 6.7437 | 6.1512 | 6.8391 | 6.2629 | 6.7811 | 6.8953 | 6.6006 | 5.4167 | 5.7897 | 5.1350 | 5.0880 |
| IFIH1_S88 | IFIH1 | Q9BYX4 | 88 | ALRRTGSPLAARY | 6.8423 | 8.0754 | 7.1327 | 6.6860 | 6.0799 | 6.2633 | 6.9129 | 6.3148 | 6.6792 | 7.0054 | 6.7498 | 7.3289 | 4.1749 | 5.0602 | 4.0939 | 4.6004 |
| IFIT3_S203 | IFIT3 | O14879 | 203 | KQAIELSPDNQYV | 7.4129 | 7.3932 | 6.8166 | 7.1336 | 5.8629 | 6.8845 | 6.5204 | 7.0136 | 6.6312 | 6.9645 | 7.3075 | 6.7193 | 3.8553 | 4.7931 | 4.4502 | 4.2411 |
| IFNAR1_T494 | IFNAR1 | P17181 | 494 | KNLLLSTSEEQIE | 6.5257 | 6.6277 | 6.1712 | 6.6237 | 6.2909 | 6.0178 | 6.5014 | 5.7691 | 6.3638 | 6.4902 | 6.4516 | 6.3129 | 5.8324 | 6.0560 | 6.1580 | 5.8077 |
| IFNGR1_S286 | IFNGR1 | P15260 | 286 | SIILPKSLISVVR | 7.8994 | 8.0557 | 7.7493 | 7.7402 | 5.4554 | 5.9910 | 6.8163 | 6.6615 | 7.4595 | 6.9619 | 7.9192 | 8.0436 | 3.0691 | 3.6592 | 2.6625 | 3.8564 |
| IFNGR1_S351 | IFNGR1 | P15260 | 351 | HTEELSSITEVVT | 5.7939 | 5.9445 | 7.4487 | 6.2535 | 5.1445 | 5.1347 | 6.2614 | 6.7895 | 5.7958 | 6.8501 | 6.9538 | 6.8990 | 6.1910 | 6.5254 | 6.1694 | 5.8447 |
| IFT140_S360 | IFT140 | Q96RY7 | 360 | VPDFLGSPGAEGK | 6.5309 | 6.3649 | 6.7882 | 7.4039 | 5.7170 | 6.7904 | 6.1758 | 6.7087 | 7.4200 | 6.5831 | 6.4342 | 6.6338 | 4.6327 | 5.7988 | 4.9754 | 5.0422 |
| IFT22_S137 | IFT22 | Q9H7X7 | 137 | KGSLSLSPPLNKL | 6.3482 | 6.1738 | 5.9592 | 5.4531 | 5.1925 | 5.3074 | 5.5589 | 7.6931 | 5.8070 | 7.0378 | 6.1966 | 7.2928 | 4.0211 | 8.3608 | 6.1304 | 7.4672 |
| IFT43_S78 | IFT43 | Q96FT9 | 78 | KFRRKASEEIEDF | 6.8931 | 5.4944 | 6.5098 | 7.0034 | 7.7124 | 5.5213 | 5.2027 | 5.7540 | 8.0316 | 6.8130 | 8.0246 | 7.0553 | 6.0393 | 5.3206 | 4.6335 | 3.9911 |
| IFTAP_S193 | IFTAP | Q86VG3 | 193 | MLTSKFSPAEIEN | 6.8651 | 6.7368 | 8.9015 | 6.9007 | 6.9002 | 6.8793 | 6.2697 | 6.4973 | 7.2922 | 7.6348 | 7.0162 | 7.0547 | 3.6974 | 3.9706 | 3.2741 | 4.1094 |
| IGF2BP1_S181 | IGF2BP1 | Q9NZI8 | 181 | GQPRQGSPVAAGA | 6.1615 | 6.2438 | 6.3154 | 5.5224 | 6.4850 | 6.1006 | 6.3397 | 6.0613 | 6.5846 | 6.3631 | 6.2091 | 6.1884 | 6.4746 | 6.5196 | 6.3417 | 6.0892 |
| IGF2BP1_S314 | IGF2BP1 | Q9NZI8 | 446 | KIAPPETPDSKVR | 6.8645 | 6.8006 | 6.5997 | 6.9832 | 7.1484 | 6.6054 | 6.4007 | 6.2437 | 6.9561 | 6.0336 | 6.0629 | 5.8908 | 5.5486 | 5.6000 | 5.1998 | 5.0620 |
| IGF2BP1_T446 | IGF2BP1 | Q9NZI8 | 314 | TKITISSLQDLTL | 5.9806 | 7.3491 | 7.8986 | 6.5595 | 6.8506 | 6.1936 | 5.9341 | 5.9659 | 7.6901 | 6.1211 | 5.4538 | 5.2748 | 6.1618 | 5.5931 | 5.1468 | 5.8265 |
| IGF2BP2_S11 | IGF2BP2 | Q9Y6M1 | 164 | EEVSSPSPPQRAQ | 6.1506 | 6.1757 | 6.6015 | 6.3468 | 6.0874 | 6.2307 | 6.1368 | 6.6914 | 6.5716 | 6.3317 | 6.5855 | 7.0044 | 5.8114 | 6.0147 | 5.6574 | 5.6023 |
| IGF2BP2_S162 | IGF2BP2 | Q9Y6M1 | 162 | PDEEVSSPSPPQR | 6.1847 | 6.4992 | 7.0215 | 7.1883 | 6.3067 | 6.6302 | 5.9668 | 6.6127 | 6.5535 | 6.5576 | 6.5501 | 6.4982 | 5.1330 | 5.6106 | 5.0660 | 5.6210 |
| IGF2BP2_S164 | IGF2BP2 | Q9Y6M1 | 11 | LYIGNLSPAVTAD | 3.8972 | 5.7195 | 6.8911 | 6.1037 | 4.6552 | 5.5170 | 5.4132 | 6.3719 | 5.5205 | 5.1969 | 6.3477 | 5.8268 | 7.6439 | 8.6251 | 7.6110 | 8.6597 |
| IGF2BP3_S184 | IGF2BP3 | O00425 | 184 | GSSRQGSPGSVSK | 5.7565 | 5.8670 | 6.3965 | 5.9918 | 6.1770 | 6.1735 | 6.2650 | 6.2772 | 6.2929 | 6.5366 | 6.5618 | 6.5098 | 6.2980 | 6.6295 | 6.1311 | 6.1358 |
| IGF2BP3_S184;S187 | IGF2BP3 | O00425 | 184;187 | NA | 6.7699 | 6.2814 | 6.5952 | 6.2800 | 7.0356 | 5.4885 | 6.1446 | 5.9580 | 6.3720 | 5.3307 | 6.9021 | 6.3060 | 6.4139 | 6.3393 | 5.7745 | 6.0082 |
| IGF2R_S2347 | IGF2R | P11717 | 2409 | HGDDQDSEDEVLT | 6.1176 | 6.8327 | 5.9272 | 6.7241 | 6.6124 | 6.8170 | 6.1921 | 7.1103 | 6.1804 | 6.0065 | 6.4564 | 6.1325 | 5.5448 | 6.2235 | 5.6194 | 5.5032 |
| IGF2R_S2409 | IGF2R | P11717 | 2484 | VSFHDDSDEDLLH | 6.5118 | 7.4179 | 6.3871 | 6.8205 | 6.9328 | 6.6429 | 6.1668 | 6.9365 | 6.2107 | 6.4759 | 6.3357 | 6.3399 | 4.6824 | 5.8939 | 5.1803 | 5.0648 |
| IGF2R_S2484 | IGF2R | P11717 | 2347 | TCCRRSSNVSYKY | 5.3249 | 4.8374 | 5.0589 | 5.1477 | 6.2999 | 5.3874 | 5.1359 | 5.8631 | 7.9580 | 7.7786 | 7.4194 | 7.8151 | 6.3561 | 7.7224 | 5.9651 | 5.9303 |
| IGF2R_T2362 | IGF2R | P11717 | 2362 | VNKEEETDENETE | 5.9050 | 6.0551 | 6.0459 | 7.3486 | 6.1949 | 6.4732 | 6.8439 | 6.2168 | 6.3889 | 5.2213 | 6.5713 | 5.3991 | 5.9304 | 6.9259 | 6.3346 | 6.1452 |
| IGSF1_S60 | IGSF1 | Q8N6C5-4 | 60 | ITLWCRSPSRISS | 6.3686 | 7.8440 | 7.0264 | 7.5546 | 6.3162 | 6.5163 | 6.0835 | 5.8542 | 6.6099 | 5.8706 | 5.9859 | 5.5414 | 4.9353 | 5.9767 | 5.8370 | 5.6795 |
| IK_S382 | IK | Q13123 | 460 | DDMAVDSDEEVDY | 6.0797 | 6.2394 | 5.9519 | 6.5839 | 6.1885 | 5.8992 | 6.1645 | 6.0024 | 6.2393 | 6.0133 | 6.1030 | 5.9615 | 6.5249 | 6.7027 | 6.6326 | 6.7132 |
| IK_S460 | IK | Q13123 | 449 | NSYAECYPATMDD | 6.1124 | 5.9301 | 6.2090 | 6.5036 | 6.1994 | 5.9927 | 6.3307 | 5.9369 | 6.3839 | 5.9748 | 6.5959 | 6.0539 | 6.0342 | 6.9565 | 6.5725 | 6.2135 |
| IK_Y449 | IK | Q13123 | 382 | EEKKRHSYFEKPK | 7.2122 | 6.9984 | 5.9304 | 7.0549 | 7.5180 | 6.4374 | 6.9148 | 6.1816 | 6.9121 | 5.6945 | 5.9463 | 5.6264 | 5.5061 | 5.5466 | 4.9180 | 5.6023 |
| IKZF2_S56 | IKZF2 | Q9UKS7 | 56 | VKLEMQSDEECDR | 7.3120 | 7.2173 | 6.8752 | 6.8765 | 5.0607 | 5.1988 | 6.9712 | 5.7725 | 6.0120 | 6.1594 | 4.9449 | 6.2620 | 6.3160 | 6.3515 | 6.4475 | 6.2225 |
| IL10RB_S261 | IL10RB | Q08334 | 261 | AFSPRNSLPQHLK | 5.1956 | 5.9703 | 6.7234 | 6.2829 | 5.6925 | 5.7774 | 5.9762 | 5.1996 | 6.0611 | 6.8561 | 6.9589 | 7.7344 | 5.7837 | 7.1295 | 6.1491 | 6.5094 |
| IL10RB_S296 | IL10RB | Q08334 | 296 | DVFDKLSVIAEDS | 6.8458 | 6.8580 | 6.9179 | 7.4683 | 5.8109 | 5.2392 | 6.8232 | 5.9857 | 6.1194 | 6.7276 | 6.4728 | 6.4510 | 5.0644 | 5.4198 | 5.7166 | 6.0796 |
| IL17RA_S629 | IL17RA | Q96F46 | 708 | ACPLLGSPGAGRN | 6.3127 | 6.4874 | 6.9421 | 6.4445 | 6.1455 | 6.2239 | 6.5737 | 6.7494 | 6.8273 | 6.2543 | 6.0901 | 6.6027 | 5.8995 | 5.5332 | 5.1763 | 5.7373 |
| IL17RA_S708 | IL17RA | Q96F46 | 629 | LVREPGSQACLAI | 6.7819 | 7.0787 | 6.6075 | 6.4555 | 6.0662 | 5.4673 | 6.1948 | 5.3648 | 6.6291 | 5.7766 | 6.2775 | 5.3855 | 6.2370 | 7.1129 | 6.6588 | 5.9061 |
| IL17RA_S801 | IL17RA | Q96F46 | 801 | GYISRSSPQPPEG | 5.2628 | 5.9522 | 5.7713 | 6.3259 | 5.8805 | 6.3895 | 5.9766 | 6.6828 | 6.4242 | 5.9351 | 5.9581 | 6.3841 | 7.0569 | 7.1936 | 6.6684 | 6.1382 |
| IL1RAP_S555 | IL1RAP | Q9NPH3 | 555 | KKSPRRSSSDEQG | 5.8258 | 6.1084 | 5.9112 | 5.7673 | 6.0582 | 5.2101 | 5.4814 | 5.8949 | 6.3740 | 6.7471 | 7.3398 | 7.1224 | 6.2746 | 7.3910 | 6.4708 | 6.0229 |
| IL31RA_T630 | IL31RA | Q8NI17-2 | 740 | EGAPNPYLKNSVT | 5.5140 | 5.5336 | 6.9668 | 6.2438 | 6.1808 | 5.4868 | 5.9534 | 6.7744 | 6.5581 | 6.5947 | 6.1578 | 6.0761 | 6.0446 | 6.8476 | 6.9515 | 6.1161 |
| IL31RA_Y740 | IL31RA | Q8NI17-2 | 630 | ILKPCSTPSDKLV | 4.6252 | 5.8552 | 6.6510 | 5.6033 | 7.7882 | 5.4706 | 6.9228 | 6.3924 | 6.3658 | 6.0608 | 6.3725 | 6.2995 | 7.1649 | 8.0137 | 4.9103 | 5.5038 |
| IL4R_S336 | IL4R | P24394 | 336 | EMPFQGSGKSAWC | 7.2369 | 6.7934 | 8.3970 | 6.9985 | 6.3688 | 6.4319 | 6.5203 | 7.6567 | 6.6908 | 7.5131 | 8.3007 | 7.0727 | 3.1710 | 4.2285 | 3.0400 | 3.5798 |
| IL6ST_S667 | IL6ST | P40189 | 667 | SHIAQWSPHTPPR | 7.0206 | 6.8857 | 6.7036 | 6.8264 | 6.3809 | 6.1235 | 6.2691 | 6.4409 | 7.0031 | 6.9699 | 6.7031 | 6.4878 | 4.9995 | 5.3089 | 4.7687 | 5.1083 |
| IL6ST_S667;T670 | IL6ST | P40189 | 670 | AQWSPHTPPRHNF | 7.2349 | 7.5078 | 7.1350 | 7.7956 | 6.7234 | 6.3073 | 6.5489 | 6.6734 | 7.3520 | 6.5250 | 7.0641 | 6.7424 | 3.8512 | 4.9085 | 3.8128 | 3.8176 |
| IL6ST_S670 | IL6ST | P40189 | 839 | RSKQVSSVNEEDF | 6.6072 | 6.9665 | 7.0892 | 7.6313 | 6.7711 | 5.8787 | 6.4102 | 6.9402 | 7.5375 | 6.7505 | 6.9265 | 6.8949 | 4.4791 | 4.0150 | 4.2089 | 4.8931 |
| IL6ST_S839 | IL6ST | P40189 | 783 | FSRSESTQPLLDS | 5.8459 | 6.0827 | 6.6541 | 6.4063 | 5.5589 | 5.4785 | 5.4675 | 6.5508 | 5.6348 | 5.0095 | 6.4219 | 6.4411 | 7.9421 | 7.7474 | 6.4411 | 6.3176 |
| IL6ST_T783 | IL6ST | P40189 | 667;670 | NA | 6.7042 | 6.7533 | 6.6900 | 7.7323 | 6.6846 | 6.5983 | 6.7983 | 6.6810 | 7.4227 | 6.8910 | 7.1309 | 7.1156 | 3.7756 | 4.5956 | 3.6942 | 4.7326 |
| ILF2_S385 | ILF2 | Q12905 | 385 | QGEEEESMETQEx | 6.3492 | 6.4225 | 6.0404 | 6.2408 | 5.9290 | 6.1929 | 6.4541 | 5.9385 | 5.9166 | 6.5617 | 6.3400 | 6.3451 | 6.3622 | 6.0978 | 6.0304 | 6.7788 |
| ILF2_T51 | ILF2 | Q12905 | 51 | KPAPDETSFSEAL | 5.4214 | 6.6261 | 5.2551 | 7.0881 | 6.0348 | 4.9373 | 7.4743 | 6.1125 | 5.1405 | 5.9221 | 6.9125 | 6.7167 | 7.0253 | 6.1956 | 6.2584 | 6.8793 |
| ILF3_S341 | ILF3 | Q12906-7 | 368 | STTYAITPMKRPM | 6.8231 | 6.8769 | 6.9104 | 7.2501 | 6.8447 | 6.5624 | 7.1243 | 6.2607 | 7.1220 | 6.7920 | 6.8326 | 6.9351 | 4.2374 | 4.7957 | 3.9738 | 4.6587 |
| ILF3_S382 | ILF3 | Q12906-7 | 596 | EKLFPDTPLALDA | 6.1524 | 6.7087 | 6.3165 | 7.0131 | 6.1714 | 6.5649 | 6.9551 | 6.8416 | 6.5203 | 6.5042 | 6.3498 | 6.3133 | 4.8904 | 5.5442 | 5.3843 | 5.7697 |
| ILF3_S476 | ILF3 | Q12906-7 | 62 | SSEQAESDNMDVP | 5.8388 | 6.2318 | 6.0177 | 6.7398 | 6.0365 | 5.8056 | 5.9615 | 5.7342 | 5.9765 | 5.8703 | 5.9606 | 5.9515 | 6.8662 | 7.1858 | 7.0429 | 6.7804 |
| ILF3_S476;S477 | ILF3 | Q12906-7 | 482 | SSKGEDSAEETEA | 5.8854 | 5.9077 | 5.7908 | 6.4248 | 6.3037 | 5.8723 | 6.1284 | 5.9344 | 6.1821 | 6.1008 | 5.8786 | 5.8340 | 6.9974 | 6.8227 | 7.0714 | 6.8654 |
| ILF3_S476;S477;S482 | ILF3 | Q12906-7 | 476 | GAEGRDSSKGEDS | 6.0714 | 6.5101 | 6.0461 | 6.4580 | 6.1120 | 5.6635 | 6.2261 | 5.6690 | 5.9209 | 6.0639 | 5.6573 | 5.5956 | 6.8258 | 6.5065 | 7.3097 | 7.3642 |
| ILF3_S476;S482 | ILF3 | Q12906-7 | 864 | SQSNYNSPGSGQN | 6.3411 | 6.8614 | 6.6965 | 7.5466 | 6.2014 | 6.8784 | 7.2183 | 6.4778 | 7.2683 | 6.4224 | 7.3863 | 6.6806 | 4.3526 | 4.9387 | 4.0464 | 4.6832 |
| ILF3_S477;S482 | ILF3 | Q12906-7 | 816 | KFNYSGSGGRSGG | 6.7118 | 8.2241 | 6.8419 | 7.6647 | 6.2030 | 6.8849 | 7.6560 | 7.1560 | 6.5682 | 7.1700 | 7.2138 | 6.9942 | 3.2325 | 4.2909 | 3.2183 | 3.9696 |
| ILF3_S482 | ILF3 | Q12906-7 | 766 | QQSYNQSPYSNYG | 7.3364 | 7.1540 | 6.2560 | 8.1337 | 6.7142 | 8.2710 | 6.9349 | 7.4536 | 6.8872 | 6.9242 | 7.3241 | 6.8511 | 3.1794 | 3.9158 | 3.3898 | 3.2747 |
| ILF3_S503 | ILF3 | Q12906-7 | 503 | PVVEAVSTPSAAF | 6.1895 | 6.2169 | 6.1218 | 6.9145 | 6.2041 | 6.4932 | 6.3718 | 5.7964 | 6.4546 | 5.9950 | 5.6173 | 6.0307 | 6.3101 | 6.6891 | 6.1552 | 6.4398 |
| ILF3_S506 | ILF3 | Q12906-7 | 382 | EDGEEKSPSKKKK | 5.5146 | 5.9562 | 6.3426 | 6.0589 | 6.4763 | 9.9469 | 6.0435 | 6.7158 | 8.3234 | 5.1002 | 7.0682 | 6.6869 | 6.5565 | 5.4286 | 4.1805 | 3.6007 |
| ILF3_S62 | ILF3 | Q12906-7 | 793 | SYSYSNSYNSPGG | 7.2070 | 7.7968 | 5.8476 | 7.7708 | 6.6080 | 7.7651 | 6.5395 | 7.2546 | 6.0714 | 6.6471 | 7.6461 | 6.2805 | 3.5754 | 4.8039 | 4.3602 | 3.8259 |
| ILF3_S766 | ILF3 | Q12906-7 | 341 | GMDPLPSKMPKKP | 4.2190 | 4.4913 | 5.3157 | 5.6328 | 6.5131 | 5.1758 | 4.8568 | 6.8488 | 5.0005 | 4.2619 | 4.8028 | 5.0415 | 11.3084 | 9.1336 | 9.7585 | 7.6397 |
| ILF3_S793 | ILF3 | Q12906 | 506 | EAVSTPSAAFPSD | 6.0347 | 5.2910 | 7.0064 | 7.1254 | 5.4466 | 6.2393 | 6.2326 | 5.3010 | 5.8702 | 7.7024 | 6.3060 | 6.2182 | 6.4294 | 6.5195 | 5.6690 | 6.6084 |
| ILF3_S816 | ILF3 | Q12906-7 | 476;482 | NA | 6.0169 | 5.9379 | 6.1744 | 6.4318 | 6.0959 | 5.8330 | 6.3961 | 5.6611 | 6.0073 | 5.9665 | 5.8206 | 6.1050 | 7.3651 | 6.7778 | 6.7569 | 6.6538 |
| ILF3_S864 | ILF3 | Q12906-7 | 476;477;482 | NA | 5.8653 | 6.1640 | 6.1261 | 6.0820 | 5.8249 | 5.8085 | 6.2816 | 5.1914 | 6.2018 | 6.1305 | 5.8727 | 6.2145 | 7.7032 | 6.5865 | 6.7038 | 7.2431 |
| ILF3_T368 | ILF3 | Q12906-7 | 477;482 | NA | 5.2438 | 4.3078 | 5.7547 | 5.6367 | 6.3703 | 5.6756 | 6.9366 | 5.0153 | 5.9266 | 6.6057 | 6.3304 | 6.1613 | 6.7998 | 8.4262 | 7.4796 | 7.3295 |
| ILF3_T596 | ILF3 | Q12906-7 | 476;477 | NA | 5.5976 | 6.8297 | 6.2209 | 6.4363 | 5.8158 | 5.7181 | 7.1284 | 5.0363 | 6.3445 | 6.4954 | 5.6222 | 5.9949 | 6.9273 | 6.6205 | 6.2842 | 6.9280 |
| ILKAP_S82 | ILKAP | Q9H0C8 | 82 | GAKRKTSEEEKNG | 4.4251 | 4.2069 | 4.7978 | 4.0850 | 5.0808 | 5.0476 | 5.0849 | 5.2890 | 4.4757 | 4.3032 | 4.7190 | 4.6852 | 10.4017 | 11.0376 | 11.7585 | 10.6020 |
| ILRUN_S215 | ILRUN | Q9H6K1 | 215 | NFNPFASPQKNRQ | 6.8265 | 6.3267 | 7.2981 | 6.4522 | 6.2868 | 6.5667 | 6.3230 | 7.0121 | 7.1528 | 6.7340 | 7.1310 | 7.5224 | 4.5895 | 4.8545 | 4.1657 | 4.7579 |
| ILRUN_S272 | ILRUN | Q9H6K1 | 272 | QNSVNLSPSSHAN | 5.8482 | 6.6411 | 6.4162 | 6.9624 | 6.6968 | 7.4447 | 6.6063 | 7.5236 | 5.9503 | 6.1065 | 7.7035 | 6.8569 | 4.2115 | 5.6363 | 5.0291 | 4.3668 |
| IMMT_S103 | IMMT | Q16891 | 34 | RPCRRYSTSGSSG | 6.5731 | 6.0827 | 6.3708 | 6.6300 | 6.4165 | 6.0945 | 6.2063 | 5.5162 | 7.0530 | 6.4500 | 6.0516 | 6.9473 | 6.0087 | 6.1726 | 5.5878 | 5.8393 |
| IMMT_S113 | IMMT | Q16891 | 103 | VPLPKKSIQSGPL | 4.9588 | 5.3840 | 4.8938 | 5.8471 | 7.0732 | 5.2157 | 5.8249 | 5.8698 | 5.2212 | 5.9994 | 5.2243 | 5.2126 | 8.6563 | 8.1409 | 8.6849 | 7.7929 |
| IMMT_S34 | IMMT | Q16891 | 390 | WKGMSVSDLADKL | 5.9093 | 6.1996 | 5.6325 | 6.5933 | 5.9386 | 6.6248 | 7.4812 | 7.2245 | 5.2334 | 6.1659 | 5.5515 | 6.5982 | 6.2839 | 5.9120 | 5.9984 | 6.6531 |
| IMMT_S390 | IMMT | Q16891 | 113 | GPLKISSVSEVMK | 5.1843 | 5.3299 | 6.3401 | 7.0254 | 6.4236 | 7.0217 | 7.0912 | 6.5250 | 5.2012 | 5.7561 | 5.3834 | 6.1147 | 6.2208 | 6.1129 | 7.2573 | 7.0123 |
| IMPDH2_S122 | IMPDH2 | P12268 | 416 | KYRGMGSLDAMDK | 6.0745 | 6.1247 | 6.7812 | 5.6876 | 6.4443 | 6.4831 | 5.8014 | 6.6865 | 6.5173 | 6.7787 | 6.7040 | 6.5455 | 5.9347 | 6.1470 | 5.7613 | 5.5282 |
| IMPDH2_S160 | IMPDH2 | P12268 | 160 | LVGIISSRDIDFL | 6.4966 | 6.0105 | 6.8819 | 5.8963 | 6.4871 | 6.5121 | 5.7839 | 6.6094 | 6.5404 | 6.3922 | 6.2723 | 6.6524 | 6.0213 | 6.2626 | 5.6780 | 5.5029 |
| IMPDH2_S416 | IMPDH2 | P12268 | 122 | TDPVVLSPKDRVR | 6.1797 | 6.0181 | 6.7191 | 6.6412 | 6.3876 | 6.9470 | 6.4292 | 6.6092 | 6.7686 | 6.7003 | 6.8904 | 7.1656 | 5.0645 | 5.5585 | 4.7454 | 5.1756 |
| IMUP_S29 | IMUP | Q9GZP8 | 29 | GGDPKLSPHKVQG | 6.7910 | 6.2375 | 7.9433 | 5.8676 | 7.7965 | 8.4824 | 6.3203 | 5.3502 | 5.7352 | 6.3201 | 7.5985 | 6.9148 | 5.7151 | 5.1771 | 3.8941 | 3.8561 |
| INA_S30 | INA | Q16352 | 440 | ATTSKVSSTGLSL | 2.1422 | 1.8347 | 1.7230 | 2.0344 | 2.1191 | 1.9665 | 2.1614 | 1.7591 | 2.1017 | 1.7299 | 1.3360 | 1.5657 | 20.3824 | 19.2168 | 18.1278 | 19.7993 |
| INA_S437 | INA | Q16352 | 30 | RLSARLSGAGGAG | 6.8270 | 6.4699 | 5.8183 | 7.0342 | 6.5897 | 6.2461 | 6.6699 | 6.1428 | 7.2139 | 7.0914 | 7.1025 | 7.5217 | 4.6542 | 5.3985 | 4.2267 | 4.9932 |
| INA_S440 | INA | Q16352 | 437 | ILSATTSKVSSTG | 6.7568 | 5.5986 | 5.7821 | 5.8025 | 5.9520 | 5.2715 | 5.5986 | 5.3959 | 6.0748 | 5.3169 | 6.0094 | 5.7936 | 7.7478 | 7.2952 | 8.1879 | 7.4164 |
| INA_S445 | INA | Q16352 | 445 | VSSTGLSLKKEEE | 6.9265 | 6.0746 | 5.1954 | 6.3775 | 6.1450 | 6.7995 | 6.9265 | 5.7913 | 4.7558 | 5.1426 | 6.4498 | 4.5213 | 7.8331 | 6.4342 | 7.6886 | 6.9383 |
| INCENP_S119 | INCENP | Q9NQS7 | 481 | RSKTPSSPCPASK | 6.8974 | 6.5763 | 6.1735 | 7.0964 | 6.6343 | 6.3961 | 6.6830 | 6.2401 | 6.7583 | 6.5487 | 6.3477 | 6.2642 | 5.0669 | 5.5672 | 5.1318 | 5.6180 |
| INCENP_S143 | INCENP | Q9NQS7 | 263 | LRIAQVSPGPRDS | 7.5143 | 7.4148 | 6.7383 | 7.7340 | 7.0070 | 6.5113 | 7.1042 | 6.8425 | 7.2128 | 6.8596 | 6.6106 | 6.6269 | 3.5356 | 4.1847 | 3.7634 | 4.3401 |
| INCENP_S143;S148 | INCENP | Q9NQS7 | 275 | SPAFPDSPWRERV | 7.1874 | 7.3953 | 6.6910 | 7.6064 | 6.4402 | 6.2074 | 6.8314 | 6.6482 | 7.1891 | 7.0411 | 6.9112 | 6.6817 | 3.9247 | 4.5993 | 4.0205 | 4.6250 |
| INCENP_S148 | INCENP | Q9NQS7 | 306 | SQSVRHSPIAPSS | 6.3787 | 6.6410 | 6.2435 | 7.2127 | 6.1800 | 6.0956 | 6.6955 | 6.5249 | 6.8021 | 6.5722 | 6.6170 | 6.7422 | 5.1270 | 5.3698 | 5.1670 | 5.6308 |
| INCENP_S197 | INCENP | Q9NQS7 | 292 | LPDNFSTPTGSRT | 6.7006 | 6.9692 | 6.7079 | 6.9038 | 6.2386 | 6.6049 | 6.2495 | 6.5706 | 6.6984 | 6.6871 | 6.6588 | 6.5514 | 5.0432 | 5.2889 | 4.9727 | 5.1544 |
| INCENP_S197;S214;T219 | INCENP | Q9NQS7 | 119 | VVGENGSVLRRVT | 8.5172 | 8.0995 | 7.0652 | 8.7260 | 6.8159 | 6.1132 | 7.2331 | 6.1923 | 6.9677 | 5.9083 | 6.0734 | 5.8392 | 3.7288 | 4.7678 | 3.7595 | 4.1928 |
| INCENP_S197;T219 | INCENP | Q9NQS7 | 239 | VSSLMATPQDPKG | 7.6097 | 7.4524 | 6.6403 | 7.3271 | 6.3372 | 6.7598 | 6.9514 | 6.5391 | 7.1113 | 6.5577 | 6.7254 | 6.5539 | 4.2348 | 4.6498 | 3.8811 | 4.6689 |
| INCENP_S214 | INCENP | Q9NQS7 | 143 | ALAAPSSPTPESP | 7.0083 | 6.3421 | 6.3567 | 6.8945 | 6.5806 | 6.5417 | 6.1238 | 5.9573 | 6.7378 | 6.5476 | 6.1654 | 6.4320 | 5.7120 | 5.7122 | 5.4053 | 5.4825 |
| INCENP_S214;T219 | INCENP | Q9NQS7 | 214 | SQGIPTSDEESTP | 5.7244 | 5.5951 | 5.7818 | 6.0213 | 5.7425 | 5.3956 | 6.0823 | 5.4075 | 6.3287 | 5.6857 | 6.1111 | 6.1456 | 7.5641 | 6.9634 | 7.4090 | 8.0418 |
| INCENP_S219 | INCENP | Q9NQS7 | 148 | SSPTPESPTMLTK | 6.2420 | 6.1263 | 5.9655 | 6.1648 | 6.0608 | 6.1376 | 5.8391 | 6.1077 | 6.0636 | 6.0885 | 6.1662 | 6.1176 | 6.2510 | 6.7559 | 6.9006 | 7.0129 |
| INCENP_S263 | INCENP | Q9NQS7 | 899 | SSAVWNSPPLQGA | 6.0896 | 6.2390 | 6.5268 | 6.4419 | 7.5547 | 5.9425 | 6.2155 | 5.5455 | 6.3987 | 6.9325 | 7.0268 | 6.9871 | 6.2426 | 5.6817 | 5.7492 | 6.2260 |
| INCENP_S263;S269 | INCENP | Q9NQS7 | 914 | PSSLAYSLKKHxx | 6.9487 | 7.5170 | 7.0731 | 7.2001 | 5.2284 | 6.2335 | 7.6909 | 6.3402 | 7.2019 | 7.1536 | 8.3513 | 7.3714 | 2.9667 | 4.5308 | 3.5691 | 4.6233 |
| INCENP_S263;S269;S275 | INCENP | Q9NQS7 | 478 | QPPRSKTPSSPCP | 7.5570 | 6.8615 | 6.0270 | 6.9788 | 6.8266 | 6.3715 | 6.9778 | 6.2628 | 6.7215 | 6.2053 | 6.3491 | 5.7755 | 5.2541 | 5.3352 | 5.0773 | 5.4190 |
| INCENP_S275 | INCENP | Q9NQS7 | 197 | PLPRTLSPTPASA | 5.9626 | 5.8972 | 5.3561 | 6.5968 | 6.0047 | 5.6851 | 5.3297 | 6.0019 | 6.6748 | 6.4387 | 6.3795 | 6.6738 | 6.6324 | 6.7785 | 6.7650 | 6.8230 |
| INCENP_S306 | INCENP | Q9NQS7 | 72 | RKKRRISYVQDEN | 5.4244 | 6.0308 | 5.9042 | 6.8154 | 5.9111 | 6.1451 | 6.2959 | 6.1984 | 6.9514 | 7.1922 | 7.3623 | 7.3306 | 5.6510 | 6.6913 | 4.6206 | 5.4753 |
| INCENP_S306;S312 | INCENP | Q9NQS7 | 219 | TSDEESTPKKSKA | 5.0297 | 4.9286 | 4.7819 | 5.5750 | 5.9180 | 5.1864 | 5.3827 | 4.9306 | 6.0720 | 5.6048 | 5.4541 | 5.2571 | 9.2095 | 8.3087 | 9.1176 | 9.2432 |
| INCENP_S420 | INCENP | Q9NQS7 | 909 | QGARVPSSLAYSL | 6.1414 | 7.8816 | 7.9889 | 8.0793 | 5.3194 | 5.8165 | 6.8081 | 7.4550 | 5.6442 | 7.0793 | 7.1273 | 6.7657 | 3.6865 | 5.2064 | 4.2261 | 4.7741 |
| INCENP_S480 | INCENP | Q9NQS7 | 480 | PRSKTPSSPCPAS | 6.6737 | 7.2519 | 7.0402 | 7.0353 | 5.9558 | 6.0085 | 6.4346 | 5.8160 | 6.4326 | 6.8950 | 7.0257 | 6.8413 | 4.9126 | 5.5622 | 4.6625 | 5.4520 |

| | | | | | | | | | | | | | | | | | | | | |
|---|---|---|---|---|---|---|---|---|---|---|---|---|---|---|---|---|---|---|---|---|
| INCENP_S481 | INCENP | Q9NQS7 | 894 | YHKRTSSAVWNSP | 4.8142 | 5.7590 | 6.4141 | 6.1192 | 6.0096 | 5.2422 | 5.7968 | 5.9821 | 7.0248 | 7.6265 | 8.0871 | 8.3194 | 5.6402 | 6.7273 | 4.8846 | 5.5528 |
| INCENP_S72 | INCENP | Q9NQS7 | 420 | NPKPAASSPETPS | 6.3964 | 6.3564 | 7.0383 | 6.5034 | 6.2066 | 6.0843 | 6.6300 | 5.8954 | 6.2486 | 6.7888 | 6.6295 | 7.1254 | 5.0867 | 5.3159 | 5.6081 | 6.0863 |
| INCENP_S798;T807 | INCENP | Q9NQS7 | 143;148 | NA | 6.3478 | 6.5795 | 6.5894 | 7.2717 | 6.5757 | 6.2113 | 6.3364 | 6.2012 | 7.0879 | 6.3792 | 6.5583 | 6.7599 | 5.1193 | 5.3011 | 5.2278 | 5.4535 |
| INCENP_S894 | INCENP | Q9NQS7 | 214;219 | NA | 5.7662 | 6.0885 | 6.1422 | 6.8624 | 5.6646 | 6.3344 | 6.6069 | 5.7492 | 6.5381 | 6.1315 | 6.4317 | 6.7708 | 6.0181 | 6.2387 | 5.9311 | 6.7256 |
| INCENP_S894;S899 | INCENP | Q9NQS7 | 197;219 | NA | 5.4105 | 5.2365 | 5.0356 | 5.8584 | 5.5767 | 5.2460 | 5.9959 | 5.5922 | 6.1298 | 5.9774 | 5.8710 | 5.8748 | 8.3276 | 7.5343 | 8.1763 | 8.1571 |
| INCENP_S899 | INCENP | Q9NQS7 | 306;312 | NA | 6.6850 | 7.2737 | 6.9443 | 7.8666 | 5.7116 | 6.4358 | 6.6851 | 6.7438 | 7.2906 | 7.5747 | 7.2813 | 7.7370 | 3.6281 | 4.3489 | 3.7657 | 4.0279 |
| INCENP_S909 | INCENP | Q9NQS7 | 478;481 | NA | 6.7579 | 7.4853 | 6.9226 | 7.0512 | 6.2526 | 6.7845 | 7.1597 | 6.8183 | 6.1326 | 6.5866 | 6.4740 | 6.1269 | 4.8499 | 5.0453 | 4.5871 | 4.9655 |
| INCENP_S914 | INCENP | Q9NQS7 | 197;214;219 | NA | 5.5787 | 6.3404 | 7.1058 | 7.1999 | 5.8494 | 6.9964 | 7.5138 | 6.5594 | 6.8984 | 7.6661 | 6.6317 | 6.5997 | 4.3545 | 5.0825 | 4.6324 | 4.9910 |
| INCENP_T213;T219 | INCENP | Q9NQS7 | 798;807 | NA | 7.1157 | 7.7462 | 5.8830 | 7.1122 | 6.2885 | 6.7723 | 7.0694 | 6.8461 | 6.6531 | 6.7757 | 6.5317 | 6.8424 | 4.7302 | 5.1660 | 3.4935 | 4.9740 |
| INCENP_T239 | INCENP | Q9NQS7 | 263;269 | NA | 6.6944 | 7.1363 | 7.1862 | 7.5136 | 6.1673 | 6.4370 | 5.5680 | 7.4344 | 6.4334 | 6.7599 | 8.0777 | 6.7155 | 4.4339 | 4.8083 | 3.9392 | 4.6948 |
| INCENP_T292 | INCENP | Q9NQS7 | 213;219 | NA | 6.3888 | 5.7894 | 5.6627 | 5.8975 | 6.5018 | 5.8415 | 6.5672 | 6.1100 | 6.5613 | 6.0922 | 6.1758 | 5.9800 | 6.8659 | 6.6192 | 5.9735 | 6.9733 |
| INCENP_T478 | INCENP | Q9NQS7 | 263;269;275 | NA | 7.2589 | 8.0284 | 7.1195 | 7.7440 | 6.0766 | 6.8673 | 7.2823 | 7.0033 | 7.4907 | 7.2275 | 7.6084 | 7.5192 | 2.6153 | 4.3034 | 2.4800 | 3.3753 |
| INCENP_T478;S481 | INCENP | Q9NQS7 | 894;899 | NA | 5.5108 | 5.6621 | 6.0671 | 6.3905 | 5.8325 | 5.7845 | 5.9633 | 6.9083 | 6.6784 | 6.7721 | 6.7239 | 6.8363 | 5.8974 | 6.1422 | 6.1863 | 6.6442 |
| INF2_S1077 | INF2 | Q27J81-2 | 1229 | KKRPSRSQEGLRP | 6.2904 | 6.7768 | 7.1789 | 6.2693 | 6.3230 | 6.8237 | 5.8781 | 5.7924 | 6.1221 | 6.6206 | 6.7112 | 6.5772 | 5.6751 | 5.9396 | 5.2958 | 5.7258 |
| INF2_S1194 | INF2 | Q27J81-2 | 372 | QSQRGSSPQNTTT | 6.1736 | 6.4034 | 6.3609 | 5.9021 | 5.9917 | 6.3117 | 6.2145 | 6.2532 | 6.3610 | 6.4638 | 6.3938 | 6.5293 | 5.9658 | 6.5586 | 6.0502 | 6.0665 |
| INF2_S1229 | INF2 | Q27J81-2 | 589 | MWASLSSPDAEAV | 5.4468 | 6.0425 | 7.2370 | 5.5006 | 7.4620 | 4.4233 | 4.7845 | 6.9075 | 5.3644 | 7.8644 | 8.7231 | 4.5089 | 5.3739 | 7.7693 | 6.2706 | 6.3213 |
| INF2_S372 | INF2 | Q27J81-2 | 1194 | SLDKSFSEDAVTD | 5.6469 | 6.4166 | 6.1858 | 6.2337 | 6.5522 | 6.7142 | 6.3624 | 6.0648 | 6.2643 | 6.3770 | 5.9835 | 6.1225 | 6.5105 | 6.3610 | 5.6622 | 6.5424 |
| INF2_S589 | INF2 | Q27J81-2 | 1077 | PLERRSSWYVDAS | 6.5449 | 5.4201 | 6.4524 | 5.6659 | 6.2341 | 6.1358 | 5.5864 | 5.4100 | 6.8153 | 8.0485 | 5.8958 | 7.1970 | 5.0038 | 6.7011 | 6.6418 | 6.2471 |
| ING5_S118 | ING5 | Q8WYH8 | 152 | RTSEEDTPKKKKH | 7.1286 | 7.0341 | 7.5218 | 8.0160 | 5.9386 | 5.6608 | 6.3974 | 5.3608 | 8.1371 | 7.7111 | 7.7459 | 7.6985 | 3.8679 | 4.1713 | 3.4595 | 4.1507 |
| ING5_S148 | ING5 | Q8WYH8 | 118 | KDKMEGSDFESSG | 6.5896 | 6.4989 | 6.7230 | 7.4563 | 5.8950 | 6.4026 | 5.8000 | 6.4746 | 5.9617 | 6.2289 | 5.8588 | 5.6503 | 6.4128 | 6.2554 | 5.8468 | 5.9455 |
| ING5_T152 | ING5 | Q8WYH8 | 148 | GRGRRTSEEDTPK | 5.1548 | 5.6674 | 5.5230 | 6.2351 | 5.0461 | 5.3197 | 5.7361 | 5.7056 | 5.6447 | 6.1024 | 5.7916 | 6.1798 | 7.8569 | 7.9906 | 8.2521 | 7.7941 |
| INO80B_S132 | INO80B | Q9C086 | 351 | GPEGPGSPLLATx | 6.0773 | 6.1978 | 6.1163 | 6.8360 | 5.9126 | 6.1593 | 6.2870 | 6.2263 | 6.1766 | 6.1810 | 6.3885 | 5.9519 | 6.0773 | 6.5454 | 6.4290 | 6.4378 |
| INO80B_S351 | INO80B | Q9C086 | 132 | DSNLSPSPLRDLS | 6.0623 | 6.6498 | 6.9278 | 5.8019 | 6.3479 | 5.5808 | 6.3755 | 5.7692 | 6.8566 | 6.9454 | 6.8306 | 6.7850 | 5.7956 | 6.2989 | 5.4125 | 5.5603 |
| INO80B_S9 | INO80B | Q9C086 | 9 | KLWRRGSTSGAME | 5.7471 | 6.5153 | 5.8939 | 5.3128 | 6.9034 | 5.4536 | 6.6179 | 6.0186 | 6.7204 | 6.4932 | 6.9356 | 6.4248 | 5.4073 | 6.1855 | 7.2955 | 6.0749 |
| INO80B_S97;S99 | INO80B | Q9C086 | 97;99 | NA | 6.5953 | 6.2427 | 6.8066 | 6.5798 | 6.8145 | 6.2618 | 6.3859 | 6.7499 | 6.5218 | 6.4084 | 6.7873 | 7.5088 | 5.5136 | 4.8613 | 4.8078 | 5.1546 |
| INO80D_S21 | INO80D | Q53TQ3 | 232 | QGSVCKSPQPQNT | 7.6530 | 6.3221 | 5.0308 | 5.4991 | 6.4185 | 5.9330 | 5.7040 | 6.1603 | 5.5594 | 5.2030 | 5.4044 | 5.7522 | 7.8630 | 7.3706 | 6.8937 | 7.2329 |
| INO80D_S232 | INO80D | Q53TQ3 | 21 | KPLCSYSPKLCKQ | 6.8367 | 5.8880 | 6.8275 | 7.0164 | 6.4198 | 5.9079 | 7.2369 | 5.2452 | 9.7107 | 7.7396 | 6.8652 | 6.7770 | 4.2428 | 4.4413 | 4.1033 | 4.7418 |
| INO80E_S154 | INO80E | Q8NBZ0 | 90 | TPKLSDTPAPKRK | 7.9979 | 7.2232 | 6.4129 | 7.1804 | 7.1999 | 6.0656 | 6.7356 | 5.8682 | 6.7863 | 5.7445 | 5.9315 | 5.4446 | 5.8852 | 5.4660 | 5.0815 | 4.9767 |
| INO80E_S227 | INO80E | Q8NBZ0 | 154 | LQLPEPSPLRPKR | 8.2473 | 7.2006 | 6.6941 | 7.7030 | 5.6126 | 6.9987 | 7.0843 | 6.1690 | 7.5286 | 7.4701 | 6.8384 | 7.2183 | 3.8376 | 4.5573 | 3.1074 | 3.7326 |
| INO80E_S98;S106 | INO80E | Q8NBZ0 | 227 | MFSDAGSGDDALD | 5.9239 | 6.0984 | 6.0175 | 6.4844 | 6.4667 | 6.1261 | 6.6426 | 6.2893 | 6.2637 | 6.8512 | 6.3900 | 6.1247 | 6.3709 | 6.6540 | 5.1739 | 6.1226 |
| INO80E_T90 | INO80E | Q8NBZ0 | 98;106 | NA | 6.7467 | 6.9056 | 5.7231 | 5.8754 | 6.6732 | 5.9304 | 5.6556 | 6.3634 | 6.9332 | 5.9732 | 6.2943 | 5.9137 | 6.0400 | 7.2458 | 5.7501 | 5.9763 |
| INPP5F_S1069 | INPP5F | Q9Y2H2 | 878 | DRQLANSLESVGP | 6.0481 | 6.8294 | 5.7476 | 5.7654 | 6.0352 | 6.4649 | 6.7557 | 6.4552 | 6.2193 | 7.0454 | 6.2490 | 5.5199 | 5.7191 | 6.4999 | 6.2775 | 6.3685 |
| INPP5F_S1103 | INPP5F | Q9Y2H2 | 942 | PLKKSPSAGDVHI | 6.0504 | 5.9611 | 6.4301 | 5.9489 | 6.3613 | 6.4393 | 6.1528 | 6.6310 | 6.1215 | 6.1962 | 6.0444 | 6.1793 | 6.3487 | 6.7056 | 6.1445 | 6.2851 |
| INPP5F_S878 | INPP5F | Q9Y2H2 | 940 | ESPLKKSPSAGDV | 6.7078 | 6.7998 | 7.0178 | 7.2168 | 6.2935 | 6.4426 | 6.1777 | 6.4037 | 6.3181 | 6.7127 | 7.1077 | 6.7252 | 4.7064 | 5.2985 | 4.8671 | 5.2045 |
| INPP5F_S935 | INPP5F | Q9Y2H2 | 1069 | SSSRAVSPFAKIR | 6.2252 | 5.8098 | 5.4138 | 5.1040 | 5.9443 | 7.3459 | 7.5735 | 6.8895 | 6.2568 | 5.6873 | 5.7501 | 5.7597 | 5.9614 | 6.7015 | 6.3929 | 7.1845 |
| INPP5F_S940 | INPP5F | Q9Y2H2 | 1103 | VSKVQKSPPEPEI | 5.8504 | 6.7861 | 6.6707 | 6.8422 | 6.3102 | 7.3217 | 6.2697 | 6.1105 | 5.7789 | 7.1855 | 7.0878 | 7.0552 | 4.9132 | 5.5583 | 5.0825 | 5.1770 |
| INPP5F_S942 | INPP5F | Q9Y2H2 | 935 | LGKGQESPLKKSP | 6.0379 | 6.8324 | 7.2055 | 6.6884 | 6.1572 | 7.5707 | 6.6929 | 6.6424 | 6.5999 | 6.5826 | 8.2930 | 7.2855 | 4.4822 | 4.9416 | 3.9789 | 4.0090 |
| INPPL1_S1003 | INPPL1 | O15357 | 886 | GTRERLYEWISID | 7.3607 | 7.4326 | 7.1486 | 6.4106 | 7.0349 | 7.5322 | 5.8987 | 6.6129 | 7.9576 | 6.6812 | 6.8668 | 7.0337 | 4.1598 | 4.2744 | 3.5367 | 4.0586 |
| INPPL1_Y886 | INPPL1 | O15357 | 1003 | LPPEPPSPARAPV | 6.5967 | 5.6924 | 6.1322 | 7.0505 | 6.6659 | 7.0289 | 5.3417 | 7.4534 | 6.3630 | 5.3232 | 6.4845 | 8.3071 | 5.4463 | 6.1230 | 4.8495 | 5.1417 |
| INSYN2B_S17 | INSYN2B | A6NMK8 | 17 | VLLKRNSLESVEF | 6.7423 | 6.5441 | 7.0911 | 6.7363 | 7.3635 | 6.4209 | 6.0305 | 6.0008 | 6.9835 | 7.2403 | 6.0208 | 6.2287 | 4.6329 | 6.0988 | 4.8904 | 4.9751 |
| INTS1_S104 | INTS1 | Q8N201 | 104 | AEKRAISPSIKEP | 5.8913 | 5.9704 | 4.9801 | 5.5021 | 5.9383 | 5.4725 | 5.6412 | 5.8258 | 5.6587 | 5.7293 | 5.2227 | 5.2055 | 8.2175 | 7.9504 | 8.5881 | 8.2062 |
| INTS1_S13 | INTS1 | Q8N201 | 1327 | APKPKSSPEQPIG | 6.1318 | 6.3531 | 5.6519 | 6.3581 | 6.1570 | 6.0493 | 6.1274 | 6.3024 | 6.3150 | 6.0460 | 6.2139 | 5.9955 | 6.4803 | 6.5075 | 6.7345 | 6.5763 |
| INTS1_S1318 | INTS1 | Q8N201 | 307 | IAEEKLSPEQEGQ | 6.0297 | 6.2266 | 6.3155 | 6.4948 | 5.7746 | 5.7396 | 6.2352 | 5.6954 | 6.1591 | 6.0969 | 5.5283 | 6.1161 | 6.7968 | 6.5138 | 6.8510 | 7.4264 |
| INTS1_S1327 | INTS1 | Q8N201 | 83 | RPKLSSTPPLSAL | 6.1204 | 6.3669 | 5.9069 | 6.2269 | 6.2384 | 6.4064 | 6.3001 | 6.5021 | 6.4182 | 6.1877 | 6.4805 | 6.1191 | 5.9358 | 6.3576 | 6.3128 | 6.1201 |
| INTS1_S284;S307 | INTS1 | Q8N201 | 13 | TTVRRPSAAAKPS | 4.9715 | 3.8479 | 3.8441 | 4.1187 | 5.0098 | 5.1363 | 5.1941 | 4.6834 | 9.3681 | 7.4881 | 7.5520 | 8.5719 | 8.7595 | 9.2848 | 5.1379 | 7.0319 |
| INTS1_S307 | INTS1 | Q8N201 | 1318 | LPPRRDSTEAPKP | 4.9067 | 4.9807 | 5.5697 | 6.1405 | 6.0538 | 6.0960 | 6.3275 | 5.9804 | 5.7412 | 5.4287 | 6.2607 | 6.6547 | 7.9799 | 7.6542 | 7.1692 | 7.0561 |
| INTS1_S82 | INTS1 | Q8N201 | 82 | KRPKLSSTPPLSA | 5.8436 | 6.4380 | 5.5886 | 6.2971 | 6.0888 | 6.6557 | 6.2402 | 6.2614 | 6.4673 | 6.6504 | 6.8579 | 6.5421 | 5.8633 | 6.2703 | 5.7269 | 6.2084 |
| INTS1_T83 | INTS1 | Q8N201 | 284;307 | NA | 6.9272 | 6.5088 | 5.2484 | 6.1496 | 5.7882 | 7.1289 | 4.7325 | 6.8990 | 4.7907 | 6.1252 | 6.4493 | 5.3384 | 6.5091 | 7.8612 | 7.3216 | 6.2220 |
| INTS10_S231 | INTS10 | Q9NVR2 | 231 | DTSDLMSPSKRSS | 7.2842 | 6.9990 | 6.8653 | 7.5218 | 6.2917 | 5.9426 | 6.0054 | 6.9028 | 6.5446 | 6.5569 | 6.6655 | 7.2486 | 4.6849 | 4.7350 | 4.3213 | 5.4304 |
| INTS12_S378 | INTS12 | Q96CB8 | 378 | GLSRSVSCDNVSK | 5.0774 | 6.0217 | 5.9355 | 6.2538 | 5.6628 | 6.1616 | 6.0025 | 6.3384 | 6.2629 | 6.9819 | 6.6521 | 6.7140 | 5.9189 | 6.7057 | 6.4315 | 6.8795 |
| INTS12_S389 | INTS12 | Q96CB8 | 389 | SKVGLPSPSSLVP | 5.6231 | 5.3409 | 6.3672 | 6.3876 | 6.2905 | 5.6443 | 6.1976 | 6.2694 | 6.2081 | 6.4911 | 7.2408 | 6.1046 | 6.3179 | 6.3940 | 4.6608 | 6.6622 |
| INTS12_T125 | INTS12 | Q96CB8 | 125 | RLEKPETQSSPIT | 6.0280 | 6.0117 | 6.9312 | 6.4585 | 5.8673 | 5.9752 | 6.8168 | 5.3410 | 6.8253 | 6.7855 | 6.4820 | 6.7263 | 5.9278 | 6.2459 | 5.2047 | 6.3729 |
| INTS13_S623 | INTS13 | Q9NVM9 | 626 | IKDSPDSPEPPNK | 5.5588 | 5.9890 | 5.9577 | 6.4265 | 6.7931 | 6.3730 | 6.1627 | 5.9797 | 6.2923 | 6.5948 | 6.5643 | 6.3524 | 6.5017 | 6.4410 | 6.1895 | 5.8235 |
| INTS13_S626 | INTS13 | Q9NVM9 | 623 | AEIIKDSPDSPEP | 5.9386 | 5.9651 | 6.1841 | 5.8403 | 6.4658 | 6.0243 | 6.2934 | 6.3131 | 6.6495 | 6.3433 | 6.2297 | 5.9635 | 6.5271 | 6.3346 | 6.3495 | 6.5783 |
| INTS3_S502 | INTS3 | Q68E01 | 502 | EFCSSPSPPVEVK | 5.8240 | 5.7988 | 5.8018 | 6.1190 | 6.3200 | 5.8468 | 5.8069 | 5.5535 | 5.8657 | 5.9416 | 5.6919 | 5.4950 | 7.7051 | 7.0923 | 7.6115 | 7.5261 |
| INTS3_S537 | INTS3 | Q68E01 | 537 | NAEAAFSDDEEDL | 5.7789 | 6.1507 | 8.0206 | 6.1720 | 6.5235 | 6.3376 | 6.5897 | 5.8933 | 6.3043 | 5.7691 | 6.0063 | 6.1222 | 6.0954 | 5.8798 | 6.1214 | 6.2353 |
| INTS5_S279 | INTS5 | Q6P9B9 | 279 | TDPFPGSPAIPAE | 6.4567 | 6.2868 | 5.7155 | 6.6700 | 6.9853 | 7.1451 | 6.2327 | 5.5128 | 6.4699 | 5.6481 | 6.0559 | 6.3025 | 5.6576 | 6.2843 | 6.9576 | 5.6192 |
| INTS6_S150 | INTS6 | Q9UL03 | 150 | LHLPLNSPLPGSE | 6.0426 | 6.5530 | 5.8896 | 7.1002 | 6.0720 | 6.1767 | 6.2067 | 5.9873 | 6.4312 | 6.1324 | 5.9650 | 6.6150 | 6.1623 | 6.5524 | 6.1093 | 6.0043 |
| INTS9_S167 | INTS9 | Q9NV88 | 564 | RPAQPTSGKKRKR | 6.1290 | 6.4391 | 6.3525 | 7.1134 | 5.9677 | 5.9687 | 6.9313 | 6.4346 | 6.5963 | 6.7800 | 6.6302 | 7.4459 | 5.0664 | 5.5452 | 5.0142 | 5.5855 |
| INTS9_S564 | INTS9 | Q9NV88 | 167 | IQRLLPSPLKDAV | 6.4829 | 6.2690 | 6.2566 | 6.7648 | 5.6628 | 5.9867 | 6.4296 | 6.3059 | 6.3055 | 6.6149 | 6.3703 | 6.5370 | 5.7046 | 5.9750 | 5.9767 | 6.3577 |
| IPO5_S670 | IPO5 | O00410-3 | 670 | QDMENMSDDDGWE | 5.4205 | 6.4088 | 5.8832 | 6.2512 | 6.6661 | 6.3839 | 6.2433 | 6.7888 | 6.3669 | 5.2090 | 6.3101 | 6.0528 | 6.0727 | 6.7320 | 6.9494 | 6.2612 |
| IPO7_S886;Y915 | IPO7 | O95373 | 886;915 | NA | 5.9949 | 6.1539 | 6.6570 | 6.5765 | 5.9968 | 6.8268 | 5.8642 | 5.9945 | 6.4129 | 6.5511 | 6.7622 | 6.2263 | 5.8262 | 5.9811 | 6.2354 | 5.9399 |
| IQCB1_S472 | IQCB1 | Q15051 | 472 | VRRHLGSPMSDVV | 7.4349 | 7.3103 | 6.4466 | 7.4244 | 6.1334 | 6.7702 | 6.2987 | 7.0077 | 6.7399 | 7.6665 | 7.2533 | 7.7853 | 3.5842 | 3.8298 | 3.8670 | 4.4479 |
| IQCB1_S572 | IQCB1 | Q15051 | 572 | KKLGEESGDEIDV | 6.1309 | 6.0858 | 7.2231 | 7.7814 | 6.3236 | 6.3613 | 5.9138 | 7.1020 | 5.7565 | 6.6493 | 6.3753 | 7.6499 | 4.8799 | 5.8052 | 4.5978 | 5.3643 |
| IQCC_S518 | IQCC | Q4KMZ1-3 | 518 | IPWRSKSPEILSS | 5.8680 | 6.1820 | 6.6632 | 6.3795 | 6.1832 | 6.5503 | 6.0869 | 6.5960 | 6.4090 | 6.2799 | 6.9001 | 6.7392 | 5.5867 | 6.0678 | 5.5828 | 5.9253 |
| IQCE_S130 | IQCE | Q6IPM2 | 590 | PVPRVPSPIAQAT | 5.2414 | 6.2454 | 6.1164 | 6.3639 | 5.6340 | 6.9590 | 6.1910 | 6.0706 | 7.3048 | 7.5636 | 6.5167 | 6.3170 | 5.5977 | 5.8986 | 5.9617 | 6.0181 |
| IQCE_S590 | IQCE | Q6IPM2 | 130 | HLRRSASNGHVPG | 5.1320 | 5.9444 | 9.1426 | 5.9893 | 10.2715 | 6.5392 | 5.4668 | 6.3074 | 5.9735 | 5.6188 | 6.0158 | 5.4601 | 5.0124 | 5.9045 | 6.1147 | 5.1071 |
| IQCJ-SCHIP1_S139 | IQCJ-SCHIP1 | B3KU38 | 139 | ECKSSSSPGGSLD | 6.1096 | 6.7464 | 7.2089 | 6.2992 | 6.2498 | 6.6732 | 6.0296 | 6.4219 | 7.0056 | 6.7418 | 6.5401 | 7.3048 | 5.0593 | 5.6679 | 4.6875 | 5.2546 |
| IQCJ-SCHIP1_S406 | IQCJ-SCHIP1 | B3KU38 | 406 | SLDTPLSPMSKQS | 6.1275 | 6.5747 | 6.7287 | 6.1984 | 6.2477 | 6.9890 | 6.1322 | 6.6512 | 6.2562 | 6.1974 | 6.7104 | 6.9768 | 6.3158 | 5.4985 | 5.0231 | 5.3726 |
| IQGAP1_S1227 | IQGAP1 | P46940 | 1227 | QRRNLGSIAKMLQ | 4.5856 | 5.4191 | 5.3535 | 5.4436 | 5.8789 | 5.9478 | 5.4707 | 5.5330 | 6.0472 | 5.0176 | 5.1454 | 5.9226 | 8.1978 | 8.8306 | 8.4012 | 8.8054 |
| IQGAP1_S86 | IQGAP1 | P46940 | 86 | KLGNFFSPKVVSL | 7.0460 | 8.0895 | 8.9160 | 9.1706 | 7.7931 | 7.1921 | 6.7538 | 5.0507 | 6.9249 | 8.1229 | 6.2946 | 6.6327 | 1.8658 | 4.0030 | 2.8384 | 3.3059 |
| IQGAP1_T435 | IQGAP1 | P46940 | 435 | YQKELATLQRQSP | 5.7808 | 5.8317 | 6.8375 | 6.3131 | 6.2092 | 6.8734 | 6.0864 | 5.8677 | 5.9306 | 5.8147 | 7.0262 | 6.7326 | 5.5790 | 6.9543 | 6.3880 | 5.7748 |
| IQGAP1_T898 | IQGAP1 | P46940 | 898 | MREEVITLIRSNQ | 7.2563 | 8.4015 | 7.9150 | 8.0991 | 6.7196 | 6.6094 | 6.8349 | 6.1651 | 7.3243 | 6.2567 | 6.5953 | 6.4719 | 3.6049 | 4.0033 | 3.6374 | 4.1054 |
| IQGAP3_S1343 | IQGAP3 | Q86VI3 | 1424 | PLRRHRSLTAHSL | 7.3095 | 6.4244 | 6.2499 | 6.6331 | 7.3764 | 6.3216 | 6.1988 | 6.2395 | 7.3397 | 6.1237 | 6.1131 | 5.7214 | 5.9924 | 6.0624 | 4.8849 | 5.0091 |
| IQGAP3_S1424 | IQGAP3 | Q86VI3 | 1415 | QACTAQTPEPLRR | 6.8854 | 6.6020 | 7.0725 | 6.4361 | 5.6291 | 6.9751 | 6.6795 | 6.1382 | 6.6910 | 6.3273 | 6.9268 | 6.7574 | 4.7676 | 5.5204 | 5.0643 | 5.5273 |
| IQGAP3_T1415 | IQGAP3 | Q86VI3 | 1343 | LSKLEVSLTLTNK | 4.9601 | 6.4856 | 7.0026 | 5.2470 | 6.4771 | 5.3436 | 6.5453 | 7.0225 | 6.9686 | 5.3351 | 5.7640 | 5.7640 | 8.5168 | 7.9401 | 5.0794 | 5.5481 |
| IQSEC1_S166 | IQSEC1 | Q6DN90-3 | 166 | NMRMQFSFEGPEK | 5.8062 | 6.8322 | 8.1829 | 6.3460 | 6.3172 | 6.4784 | 6.0040 | 6.6776 | 6.2787 | 7.7356 | 7.6461 | 6.7644 | 3.9710 | 5.3715 | 4.8081 | 4.7803 |
| IQSEC1_S248 | IQSEC1 | Q6DN90-3 | 403 | LGGQQGSPKHGPH | 6.3933 | 6.2954 | 6.5278 | 5.9515 | 6.3069 | 6.4230 | 6.7227 | 6.3041 | 6.2492 | 6.2000 | 5.9164 | 6.1545 | 6.3241 | 6.1928 | 5.9372 | 6.1009 |
| IQSEC1_S403 | IQSEC1 | Q6DN90-3 | 911 | RSALSSSLRDLSE | 6.3687 | 6.3024 | 6.1791 | 5.8967 | 5.8858 | 6.1742 | 6.0941 | 6.5291 | 6.6369 | 6.3738 | 6.3516 | 6.3676 | 6.1394 | 6.4379 | 6.0385 | 6.2242 |
| IQSEC1_S498 | IQSEC1 | Q6DN90-3 | 75 | KLQHSTSILRKQA | 8.6039 | 7.1962 | 6.6508 | 6.8514 | 6.1396 | 6.1935 | 6.5000 | 6.5500 | 6.8585 | 5.4488 | 6.2170 | 6.0118 | 5.2343 | 6.1680 | 4.5450 | 4.8313 |
| IQSEC1_S75 | IQSEC1 | Q6DN90-3 | 498 | HKEARNSWDSPAF | 6.5436 | 6.3118 | 6.4830 | 6.2250 | 6.1243 | 6.2905 | 6.1024 | 6.4226 | 6.3153 | 6.3292 | 6.5960 | 6.3806 | 5.7934 | 5.7043 | 5.9938 | 6.3843 |
| IQSEC1_S911 | IQSEC1 | Q6DN90-3 | 248 | DALNCRSLHTEEA | 7.0726 | 6.7410 | 7.5572 | 7.9735 | 7.1526 | 6.2235 | 6.6210 | 5.5438 | 4.6641 | 7.0115 | 7.6230 | 6.4258 | 4.6030 | 4.5959 | 6.0024 | 4.1890 |
| IQSEC2_S1143 | IQSEC2 | Q5JU85 | 393 | NMRMQFSFEEYEK | 6.0919 | 6.7635 | 7.7643 | 6.0418 | 6.1394 | 6.5002 | 6.2020 | 6.7631 | 6.4280 | 7.1049 | 7.0734 | 6.8356 | 4.7152 | 5.6217 | 5.1589 | 4.7961 |
| IQSEC2_S393 | IQSEC2 | Q5JU85 | 1143 | RGALSSSLRDLSD | 6.1378 | 6.1527 | 6.3555 | 5.6520 | 5.5184 | 6.2554 | 5.8810 | 6.4361 | 6.7929 | 6.4794 | 6.3185 | 6.2071 | 6.3856 | 6.6688 | 6.1303 | 6.6286 |
| IQSEC2_S720 | IQSEC2 | Q5JU85 | 720 | GSSSRDSLREPPA | 8.4486 | 6.7983 | 8.3828 | 6.9418 | 6.7043 | 5.9268 | 5.5859 | 6.3242 | 8.2139 | 7.0384 | 7.0189 | 6.5021 | 5.1138 | 4.5870 | 3.1955 | 3.2176 |
| IRAG2_S378 | IRAG2 | Q12912 | 378 | RRVTIASLPRNIG | 6.3050 | 6.1928 | 6.1914 | 6.2854 | 5.7138 | 5.4198 | 5.8771 | 5.6315 | 5.8903 | 6.1607 | 5.5330 | 6.0099 | 6.5161 | 6.7921 | 7.2970 | 8.1840 |
| IRAK1_S131 | IRAK1 | P51617 | 131 | AEAEAWSPRKLPS | 6.1007 | 5.8589 | 6.9775 | 6.6223 | 6.3502 | 6.7961 | 7.0318 | 6.5465 | 8.0054 | 6.7092 | 7.2329 | 6.7974 | 4.2003 | 5.6975 | 4.0223 | 5.0509 |
| IRAK2_S144 | IRAK2 | O43187 | 144 | FPGPGSSPARAHQ | 6.4925 | 6.4556 | 6.9432 | 6.1035 | 6.3752 | 6.8617 | 6.2363 | 6.7305 | 6.7586 | 6.4784 | 6.5554 | 6.4341 | 5.3262 | 5.6574 | 5.2805 | 5.3107 |
| IRAK3_S467 | IRAK3 | Q9Y616 | 467 | KSFRCPSPLFLEN | 5.3514 | 6.1046 | 6.2878 | 6.2589 | 5.6124 | 6.1079 | 6.5427 | 6.2789 | 6.5372 | 6.4327 | 6.3717 | 6.1316 | 6.1246 | 6.8397 | 6.3591 | 6.6588 |
| IRAK4_S152 | IRAK4 | Q9NWZ3 | 152 | MPPDSSSPENKSL | 5.7753 | 6.3512 | 6.9324 | 6.4712 | 6.0001 | 6.6194 | 6.7852 | 6.7273 | 6.6244 | 7.1453 | 6.8354 | 7.0356 | 5.1747 | 5.4525 | 4.9171 | 5.1529 |
| IRF2BP1_S186 | IRF2BP1 | Q8IU81 | 66 | VLPEGRSPGPPAL | 6.0242 | 6.3763 | 6.2678 | 6.2160 | 5.7803 | 6.6476 | 6.1780 | 6.6820 | 6.6172 | 6.9275 | 6.9683 | 6.8717 | 5.6971 | 5.9148 | 5.4626 | 5.3685 |
| IRF2BP1_S384 | IRF2BP1 | Q8IU81 | 421 | ETPGVPSPIAALK | 6.2190 | 6.3599 | 6.4139 | 6.3659 | 6.7784 | 6.2198 | 6.1188 | 6.0404 | 6.5858 | 6.4564 | 6.5868 | 5.9968 | 6.3013 | 6.0502 | 5.7298 | 5.7769 |
| IRF2BP1_S421 | IRF2BP1 | Q8IU81 | 384 | PRRRKASPEPEGE | 5.8839 | 5.9318 | 6.2528 | 5.9760 | 6.3444 | 6.3389 | 6.0002 | 5.9437 | 6.5636 | 6.5797 | 6.4340 | 6.2019 | 6.6765 | 6.2657 | 6.2185 | 6.3883 |
| IRF2BP1_S436 | IRF2BP1 | Q8IU81 | 436 | AEALGHSPKDPGG | 6.5461 | 6.8168 | 6.5002 | 6.4052 | 6.4227 | 6.6540 | 6.5798 | 6.7032 | 6.7516 | 6.9919 | 6.9840 | 6.7561 | 4.9037 | 5.5264 | 4.4534 | 5.0050 |
| IRF2BP1_S478 | IRF2BP1 | Q8IU81 | 478 | NGEAEVSPTAGAE | 5.3017 | 5.9856 | 6.3518 | 6.3653 | 6.7465 | 5.9915 | 5.9463 | 6.4859 | 6.4380 | 6.3825 | 6.2425 | 5.6291 | 6.6814 | 6.4331 | 7.0670 | 5.9517 |
| IRF2BP1_S66 | IRF2BP1 | Q8IU81 | 186 | TLAPGLSPARPLF | 6.3349 | 6.4142 | 6.8569 | 6.3777 | 5.7939 | 6.9376 | 6.7231 | 6.6278 | 6.9728 | 7.0781 | 7.0286 | 7.5798 | 4.5161 | 5.6973 | 4.2993 | 4.7619 |
| IRF2BP2_S175 | IRF2BP2 | Q7Z5L9 | 175 | PELNRQSPNPRRG | 6.2267 | 6.1487 | 6.1054 | 5.9016 | 6.5111 | 6.5967 | 6.1535 | 6.2064 | 6.4054 | 6.3226 | 6.0064 | 6.1273 | 6.9350 | 6.3172 | 6.0423 | 5.9937 |
| IRF2BP2_S240 | IRF2BP2 | Q7Z5L9 | 406 | PPPPTASPHSNRT | 5.8532 | 6.3561 | 6.2481 | 6.3460 | 6.1976 | 5.8420 | 6.2582 | 7.2301 | 6.4434 | 5.8711 | 6.7088 | 6.4707 | 5.9592 | 6.1160 | 5.8393 | 6.2602 |
| IRF2BP2_S360 | IRF2BP2 | Q7Z5L9 | 460 | NSNSPPSPSSMNQ | 6.4047 | 6.5471 | 6.5147 | 6.2484 | 5.6430 | 6.6974 | 6.5867 | 6.1919 | 6.2371 | 5.5358 | 5.9139 | 6.1706 | 6.4767 | 6.0416 | 6.2272 | 6.5631 |
| IRF2BP2_S406 | IRF2BP2 | Q7Z5L9 | 360 | ARKRKPSPEPEGE | 5.8554 | 6.1447 | 6.0761 | 6.1029 | 6.5839 | 6.2677 | 5.5750 | 6.9845 | 6.5488 | 5.9702 | 6.1190 | 5.8992 | 6.9181 | 6.2386 | 6.4751 | 6.2406 |
| IRF2BP2_S455;S460 | IRF2BP2 | Q7Z5L9 | 147 | SLAQPPTPQPPPV | 5.2373 | 5.4252 | 6.1768 | 6.2379 | 5.5469 | 6.5914 | 6.3621 | 5.7373 | 5.8437 | 5.7934 | 5.7405 | 6.4195 | 7.8213 | 7.2997 | 7.1927 | 6.5742 |
| IRF2BP2_S460 | IRF2BP2 | Q7Z5L9 | 240 | AHKRPASVSSSAA | 5.2586 | 6.1528 | 5.7560 | 5.3583 | 5.7262 | 6.8085 | 6.0702 | 6.7336 | 5.1275 | 5.8616 | 6.9193 | 5.4128 | 8.0204 | 7.9735 | 7.0231 | 5.7977 |
| IRF2BP2_T147 | IRF2BP2 | Q7Z5L9 | 455;460 | NA | 4.7379 | 4.9448 | 4.1585 | 4.7792 | 6.6021 | 5.7834 | 5.6567 | 6.0026 | 6.0791 | 6.8899 | 7.3271 | 7.1341 | 8.1985 | 7.2521 | 6.7281 | 7.7260 |
| IRF2BPL_S215 | IRF2BPL | Q9H1B7 | 215 | PELNRQSPNSSSA | 5.8593 | 5.9115 | 6.0426 | 5.9609 | 6.2906 | 6.3430 | 5.9619 | 5.8997 | 6.2364 | 6.2308 | 6.3396 | 6.1640 | 7.0767 | 6.6737 | 6.5034 | 6.5061 |
| IRF2BPL_S334 | IRF2BPL | Q9H1B7 | 69 | CFQDGRSPGPPPP | 6.4848 | 6.4292 | 6.1444 | 6.1194 | 6.4965 | 6.4906 | 6.3326 | 6.7846 | 6.7786 | 6.3982 | 6.6236 | 6.4563 | 5.9353 | 5.6267 | 5.3018 | 5.5971 |
| IRF2BPL_S519 | IRF2BPL | Q9H1B7 | 639 | TLGTAHSPKDGSS | 6.0503 | 6.2757 | 6.0804 | 6.2065 | 6.5381 | 6.3497 | 5.9684 | 5.9445 | 6.1548 | 6.2652 | 6.4208 | 5.9181 | 6.9222 | 6.5514 | 6.5420 | 5.8120 |
| IRF2BPL_S547 | IRF2BPL | Q9H1B7 | 547 | LRKRKASPEPPDS | 5.7202 | 6.1324 | 6.2172 | 5.9271 | 5.9774 | 5.8121 | 5.9207 | 6.2756 | 6.3332 | 6.4544 | 5.6596 | 6.2329 | 7.2795 | 6.4339 | 6.7352 | 6.8887 |
| IRF2BPL_S657 | IRF2BPL | Q9H1B7 | 657 | ASARRNSSSPVSP | 4.6508 | 4.4515 | 4.8248 | 5.0217 | 7.4999 | 5.2398 | 4.7581 | 5.5316 | 7.5214 | 8.5245 | 7.4521 | 7.1543 | 8.2636 | 7.9052 | 5.4622 | 5.7384 |
| IRF2BPL_S657;S659 | IRF2BPL | Q9H1B7 | 659 | ARRNSSSPVSPAS | 6.9740 | 6.7436 | 6.3311 | 6.1697 | 6.3449 | 5.9643 | 5.6139 | 5.6922 | 6.9517 | 6.9503 | 6.8212 | 6.5832 | 6.2055 | 6.0508 | 5.2288 | 5.3751 |
| IRF2BPL_S657;S662 | IRF2BPL | Q9H1B7 | 519 | SLSRAPSAPPGTG | 5.5349 | 5.8692 | 6.5970 | 6.3563 | 5.5306 | 6.7254 | 6.1381 | 5.9601 | 6.3520 | 6.3277 | 6.8015 | 6.2733 | 6.2182 | 6.7620 | 5.8525 | 6.7011 |
| IRF2BPL_S659 | IRF2BPL | Q9H1B7 | 662 | NSSSPVSPASVPG | 7.6589 | 7.9892 | 7.5387 | 7.0782 | 5.8030 | 6.5882 | 6.8707 | 6.5126 | 5.8256 | 5.6829 | 6.3722 | 6.1009 | 4.9264 | 5.0147 | 4.8840 | 5.1539 |
| IRF2BPL_S659;S662 | IRF2BPL | Q9H1B7 | 334 | GGKRPGSVSSTDQ | 5.1936 | 5.3539 | 4.9918 | 4.9965 | 6.1532 | 6.4839 | 5.9736 | 5.9628 | 4.8339 | 5.2363 | 5.5151 | 5.0423 | 9.4874 | 8.1184 | 8.3533 | 8.3039 |
| IRF2BPL_S662 | IRF2BPL | Q9H1B7 | 657;659 | NA | 5.8387 | 5.5325 | 5.4498 | 5.7707 | 6.3890 | 5.6442 | 6.2562 | 6.3827 | 6.0784 | 5.7602 | 6.2978 | 5.3767 | 7.9724 | 7.1297 | 7.2480 | 6.8728 |
| IRF2BPL_S69 | IRF2BPL | Q9H1B7 | 659;662 | NA | 5.6299 | 6.7353 | 7.3270 | 7.2590 | 6.5365 | 6.7819 | 6.3269 | 7.3082 | 6.9112 | 6.3849 | 6.6797 | 6.9339 | 5.6428 | 5.1816 | 3.8935 | 4.4677 |
| IRF2BPL_T611;S639 | IRF2BPL | Q9H1B7 | 657;662 | NA | 6.7927 | 6.2783 | 6.6825 | 6.6152 | 6.5361 | 6.5443 | 6.2405 | 5.1826 | 7.8310 | 7.4136 | 6.9867 | 6.6638 | 5.2907 | 4.6680 | 4.9980 | 5.2760 |
| IRF2BPL_T639 | IRF2BPL | Q9H1B7 | 611;639 | NA | 6.1985 | 6.0100 | 6.5697 | 6.7361 | 6.3956 | 5.9046 | 6.0051 | 5.7337 | 6.1127 | 6.3277 | 6.3630 | 6.8595 | 6.7881 | 6.0406 | 5.6689 | 6.2860 |
| IRF3_S175 | IRF3 | Q14653-4 | 175 | QPLRSPSLDNPTP | 5.1554 | 4.9834 | 5.6603 | 5.6057 | 5.6538 | 5.7519 | 5.4689 | 5.9597 | 6.7511 | 7.0928 | 5.9861 | 6.3519 | 7.5578 | 7.9300 | 7.1774 | 6.9138 |
| IRS1_S1078 | IRS1 | P35568 | 1078 | FTRVNLSPNRNQS | 6.2472 | 6.5496 | 7.0314 | 6.1873 | 5.8193 | 6.6637 | 6.2642 | 6.6913 | 6.8087 | 6.6965 | 6.8369 | 7.0271 | 5.2030 | 5.5717 | 5.1966 | 5.2055 |
| IRS1_S1101 | IRS1 | P35568 | 766 | PVLSYYSLPRSFK | 6.4458 | 5.6833 | 6.6559 | 6.2585 | 5.9867 | 6.0853 | 6.1310 | 5.4542 | 6.1893 | 6.1111 | 6.0687 | 6.2672 | 6.8397 | 7.1301 | 5.9599 | 6.7334 |
| IRS1_S312 | IRS1 | P35568 | 629 | PSGRKGSGDYMPM | 5.2829 | 4.7118 | 5.8430 | 5.2154 | 5.9232 | 6.2747 | 5.3153 | 6.0275 | 6.3577 | 8.1197 | 7.0031 | 6.8147 | 6.3221 | 7.7480 | 6.7957 | 6.2455 |
| IRS1_S629 | IRS1 | P35568 | 636 | GDYMPMSPKSVSA | 5.7763 | 6.0325 | 6.9428 | 6.6200 | 6.0113 | 6.5861 | 6.7184 | 7.4645 | 7.2578 | 6.8508 | 7.0787 | 6.3942 | 4.7930 | 5.2623 | 4.7137 | 5.4974 |
| IRS1_S636 | IRS1 | P35568 | 1101 | CRRRHSSETFSST | 4.4112 | 7.2034 | 9.0098 | 5.1070 | 5.1873 | 7.5335 | 7.1632 | 6.0660 | 6.3113 | 6.6726 | 7.8992 | 7.8323 | 4.5272 | 5.7047 | 5.8296 | 3.5415 |
| IRS1_S766 | IRS1 | P35568 | 312 | ESITATSPASMVG | 6.3458 | 6.2016 | 7.7301 | 7.2860 | 6.5010 | 5.8964 | 6.6069 | 6.7748 | 6.1042 | 6.9886 | 7.3292 | 7.4416 | 4.6018 | 5.1786 | 4.4075 | 4.6059 |
| ISCA2_S29 | ISCA2 | Q86U28 | 29 | GRLLTASLGPQAR | 7.2095 | 5.6795 | 6.9152 | 5.2024 | 5.1311 | 6.2462 | 6.5829 | 7.0860 | 7.7522 | 8.3758 | 8.7460 | 9.6255 | 3.3905 | 5.2705 | 2.9271 | 3.8597 |
| ISCU_S14 | ISCU | Q9H1K1 | 14 | RLRRAASALLLRS | 6.2275 | 5.9544 | 6.4941 | 5.3490 | 6.9797 | 6.7074 | 5.8472 | 7.1649 | 6.3181 | 6.4927 | 6.5920 | 6.5137 | 6.0360 | 6.0892 | 5.6277 | 5.6063 |
| ISCU_S29 | ISCU | Q9H1K1 | 29 | LPARELSAPARLY | 6.4677 | 6.4712 | 6.6508 | 5.4029 | 6.6306 | 6.6226 | 6.0109 | 7.3369 | 6.1602 | 6.6177 | 6.5094 | 6.4539 | 5.9543 | 5.9284 | 5.4992 | 5.2833 |
| ISL1_S148 | ISL1 | P61371 | 148 | GAGDPLSPLHPAR | 6.2678 | 6.9303 | 6.1897 | 6.2808 | 5.7720 | 6.0531 | 6.1572 | 6.1702 | 5.9346 | 6.6661 | 6.6596 | 6.5021 | 6.4630 | 6.5398 | 5.3750 | 6.0388 |
| IST1_S227 | IST1 | P53990-5 | 277 | GLPMGTYQAFPNI | 6.2907 | 6.4714 | 7.2129 | 6.1756 | 6.1139 | 6.6817 | 6.2284 | 7.2812 | 6.6489 | 6.7804 | 7.3296 | 7.0609 | 4.9829 | 5.4678 | 4.4598 | 4.8138 |
| IST1_T291 | IST1 | P53990-5 | 291 | PPQIPATPPSYES | 6.0903 | 6.3267 | 7.4596 | 6.5972 | 6.3070 | 6.5641 | 6.4799 | 7.1614 | 7.0007 | 6.9553 | 7.2410 | 7.0207 | 4.5564 | 5.1424 | 4.3924 | 4.7049 |
| IST1_Y277 | IST1 | P53990-5 | 227 | GPGRGGSGGFTAP | 6.8768 | 6.3892 | 5.5904 | 5.8134 | 6.1410 | 7.1780 | 6.6790 | 6.5455 | 6.9423 | 7.4098 | 6.2632 | 6.5039 | 4.3216 | 5.9230 | 5.8978 | 5.5249 |
| ISY1_S222 | ISY1 | Q9ULR0-1 | 222 | AVTEEESDEEGSQ | 6.2054 | 6.4749 | 6.3000 | 6.2091 | 6.0112 | 5.8332 | 5.9525 | 5.2625 | 6.0156 | 6.5528 | 6.1726 | 6.2091 | 7.5834 | 6.3952 | 6.0786 | 6.7439 |
| ITCH_S240 | ITCH | Q96J02 | 240 | RVSGNNSPSLSNG | 6.2296 | 6.1566 | 7.0171 | 6.4986 | 6.3543 | 6.4899 | 6.5075 | 6.7466 | 6.8100 | 7.0023 | 6.5418 | 6.6977 | 4.7825 | 5.7029 | 4.8817 | 5.5811 |
| ITCH_S240;S244 | ITCH | Q96J02 | 240;244 | NA | 6.2397 | 5.9681 | 8.1379 | 6.5049 | 6.1883 | 7.0486 | 7.2428 | 6.9703 | 6.2557 | 6.7990 | 7.3814 | 6.0325 | 4.7090 | 5.4759 | 5.2531 | 3.7927 |
| ITGA4_S1021 | ITGA4 | P13612 | 1021 | EENRRDSWSYINS | 6.0305 | 6.2880 | 5.9802 | 6.3419 | 6.1663 | 6.0102 | 6.0617 | 6.1997 | 6.4958 | 6.4708 | 6.3780 | 6.6433 | 6.3518 | 6.2428 | 6.0342 | 6.3047 |
| ITGA6_S1103 | ITGA6 | P23229-6 | 1103 | EIHAQPSDKERLT | 6.7280 | 7.0606 | 7.6129 | 7.2173 | 6.2049 | 6.0637 | 5.7859 | 6.6532 | 7.2198 | 7.0576 | 7.5964 | 7.9677 | 3.9428 | 4.8102 | 3.8661 | 4.2129 |
| ITGA7_S27 | ITGA7 | G3V2I4 | 27 | RGFSIDSGKGLVR | 5.9093 | 6.4790 | 6.4419 | 5.6173 | 6.7878 | 7.6849 | 5.1871 | 7.1512 | 6.6963 | 5.8211 | 7.3579 | 6.6238 | 6.9790 | 5.5929 | 4.7323 | 4.9383 |
| ITGB1_Y783 | ITGB1 | P05556 | 783 | TGENPIYKSAVTT | 6.3640 | 6.3023 | 6.6695 | 6.2761 | 5.4926 | 7.1481 | 6.3616 | 7.3122 | 8.1568 | 8.4324 | 7.4260 | 7.7669 | 3.9535 | 4.9753 | 3.2696 | 4.0932 |
| ITGB3BP_S110 | ITGB3BP | Q13352-5 | 72 | KSVITYSPTTGTC | 6.1125 | 6.1681 | 5.9087 | 6.4948 | 6.1808 | 6.0477 | 5.8366 | 6.2818 | 6.0132 | 6.0028 | 5.7858 | 5.9305 | 6.9740 | 6.4727 | 6.9482 | 6.8419 |
| ITGB3BP_S72 | ITGB3BP | Q13352-5 | 110 | KKLNHPSLTESKE | 6.4670 | 6.2204 | 6.8242 | 7.1110 | 7.6877 | 6.7258 | 7.1305 | 7.7157 | 6.9622 | 7.0107 | 6.5952 | 6.9169 | 4.1875 | 4.3312 | 3.6637 | 4.4504 |
| ITGB4_S1069 | ITGB4 | P16144 | 1474 | AYGTHLSPHVPHR | 5.7654 | 6.2609 | 5.7128 | 6.0280 | 5.9413 | 6.5032 | 6.2958 | 6.3962 | 5.8136 | 6.0910 | 6.1144 | 6.0110 | 6.6457 | 7.1694 | 6.8121 | 6.4393 |
| ITGB4_S1457 | ITGB4 | P16144 | 1069 | ELQEVDSLLRGRQ | 6.5290 | 6.8239 | 6.1381 | 6.5861 | 6.7711 | 6.0186 | 6.3212 | 5.7572 | 6.1804 | 5.7636 | 5.7979 | 5.5669 | 6.6040 | 6.8771 | 6.2114 | 6.0535 |
| ITGB4_S1474 | ITGB4 | P16144 | 1457 | FPGSTNSLHRMTT | 6.4048 | 6.9866 | 6.2391 | 6.4004 | 6.1618 | 6.0984 | 6.3848 | 6.2733 | 6.2823 | 6.1803 | 5.9487 | 5.7248 | 6.5022 | 6.7214 | 5.8404 | 5.8508 |
| ITGB4_S1483 | ITGB4 | P16144 | 1530 | TAGVPDTPTRLVF | 6.2541 | 6.6100 | 6.0848 | 6.4515 | 6.2135 | 6.0812 | 6.5149 | 6.1307 | 6.2733 | 6.2156 | 6.0908 | 6.0148 | 6.0426 | 6.3977 | 6.3235 | 6.3010 |
| ITGB4_S1494 | ITGB4 | P16144 | 1494 | LTRDYNSLTRSEH | 6.4568 | 6.6777 | 5.9903 | 6.1777 | 5.7280 | 5.9890 | 6.1374 | 6.3625 | 6.1604 | 6.3467 | 6.1365 | 6.1205 | 6.3428 | 6.7820 | 6.2498 | 6.3419 |
| ITGB4_T1530 | ITGB4 | P16144 | 1483 | VPHRVLSTSSTLT | 6.8855 | 6.8850 | 6.5505 | 6.0770 | 4.7443 | 6.8304 | 6.3909 | 6.0834 | 7.2834 | 7.2385 | 7.3839 | 7.5683 | 4.8941 | 5.6920 | 4.4002 | 5.0926 |
| ITGB5_S770 | ITGB5 | P18084 | 779 | LYRKPISTHTVDF | 7.1704 | 5.8011 | 8.2417 | 6.7167 | 5.8730 | 6.1162 | 5.4604 | 6.3642 | 7.3472 | 7.3273 | 6.6524 | 7.1135 | 5.0141 | 5.5139 | 4.5019 | 4.7858 |
| ITGB5_S779 | ITGB5 | P18084 | 770 | ARYEMASNPLYRK | 6.5989 | 8.1874 | 7.6341 | 6.6546 | 5.5531 | 6.7255 | 6.8133 | 6.8884 | 6.3580 | 7.2503 | 6.6071 | 7.3677 | 3.9840 | 4.8109 | 4.2402 | 4.3265 |
| ITM2C_T37 | ITM2C | Q9NQX7 | 37 | ATEILLTPAREEQ | 6.6658 | 6.6658 | 7.4008 | 6.6279 | 6.2849 | 6.4868 | 6.8566 | 6.1582 | 6.3287 | 5.8714 | 6.4482 | 7.0219 | 5.4598 | 5.5166 | 4.8659 | 5.3409 |
| ITPKA_S180 | ITPKA | P23677 | 20 | GGARPCSPGLERA | 6.1695 | 5.9093 | 6.6881 | 6.3682 | 6.7239 | 6.3675 | 6.4671 | 5.5564 | 6.2628 | 6.3864 | 6.6370 | 6.4300 | 6.5038 | 6.3414 | 5.8400 | 5.3487 |

| | | | | | | | | | | | | | | | | | | | | | |
|---|---|---|---|---|---|---|---|---|---|---|---|---|---|---|---|---|---|---|---|---|---|
| ITPKA_S20 | ITPKA | P23677 | | 95 | PDVPPTSPGPPER | 5.1986 | 6.2844 | 5.7533 | 6.2708 | 6.1792 | 6.6992 | 6.2300 | 6.0678 | 6.4726 | 6.2089 | 6.1012 | 6.1173 | 7.3876 | 7.0025 | 6.2300 | 5.7966 |
| ITPKA_S95 | ITPKA | P23677 | | 180 | VNLPVISPFKKRY | 7.1645 | 6.6001 | 7.0273 | 7.5251 | 6.0347 | 6.9419 | 8.4905 | 7.3485 | 7.9815 | 7.6124 | 6.3559 | 6.0328 | 3.2784 | 4.4786 | 3.7301 | 3.3977 |
| ITPKB_S43 | ITPKB | P27987 | | 43 | PRRAVLSPGSVFS | 5.7605 | 6.2380 | 7.2552 | 7.0839 | 5.3246 | 7.3071 | 5.9059 | 5.8747 | 6.7051 | 6.4871 | 6.7673 | 8.1582 | 3.5238 | 6.8296 | 3.4304 | 7.3486 |
| ITPKC_S322 | ITPKC | Q96DU7 | | 322 | YSHLKCSPLCPVP | 6.8221 | 7.2271 | 7.2350 | 6.9088 | 6.1856 | 6.8229 | 7.4969 | 6.9989 | 7.5749 | 6.7587 | 7.9658 | 7.6596 | 3.1837 | 3.8920 | 2.8630 | 4.4050 |
| ITPR1_S1764 | ITPR1 | Q14643 | | 1764 | PSGRRESLTSFGN | 5.4136 | 5.7500 | 5.8067 | 6.1873 | 5.7183 | 5.3125 | 5.3648 | 5.7720 | 7.0378 | 7.1913 | 6.7931 | 6.7893 | 6.0091 | 6.9260 | 6.7204 | 7.2076 |
| ITPR2_S1855 | ITPR2 | Q14571 | | 1855 | RMRVRDSTLHLKE | 1.7365 | 1.9762 | 2.3205 | 2.1341 | 3.1848 | 2.1484 | 2.2209 | 2.2943 | 2.4210 | 2.0985 | 1.7426 | 1.5952 | 17.3952 | 17.7179 | 20.1378 | 18.8762 |
| ITPR3_S1832 | ITPR3 | Q14573 | | 934 | VLSRKQSVFSAPS | 5.5941 | 4.8712 | 4.9425 | 5.7446 | 6.6140 | 5.2911 | 5.3979 | 5.6963 | 6.8244 | 7.1340 | 6.8746 | 7.0949 | 6.9817 | 7.7938 | 6.8832 | 6.2616 |
| ITPR3_S2670 | ITPR3 | Q14573 | | 1832 | TKGRVASFSIPGS | 5.3981 | 4.8173 | 5.1452 | 5.3842 | 5.7685 | 5.1347 | 5.4497 | 5.3356 | 6.8346 | 6.6380 | 6.5628 | 6.5388 | 7.6660 | 8.1401 | 7.9810 | 7.2053 |
| ITPR3_S934 | ITPR3 | Q14573 | | 2670 | DVQNCISRxxxxx | 6.0303 | 6.1621 | 6.3136 | 5.9880 | 6.6054 | 6.3850 | 5.6369 | 6.2016 | 6.0327 | 5.5406 | 6.1266 | 5.6559 | 6.7316 | 7.0306 | 7.2721 | 6.2870 |
| ITPRID2_S1055 | ITPRID2 | P28290 | | 466 | MTQQKDSFEMEEV | 6.3116 | 6.5429 | 6.8793 | 6.4304 | 5.9707 | 6.1706 | 5.9794 | 6.3339 | 6.2401 | 6.7403 | 6.6142 | 6.7238 | 5.2587 | 6.2490 | 5.7989 | 5.7564 |
| ITPRID2_S1134 | ITPRID2 | P28290 | | 767 | GKETRCSPPSFTY | 6.4204 | 6.7985 | 6.8087 | 6.9784 | 6.1369 | 6.5283 | 6.5535 | 5.8103 | 6.4673 | 6.6695 | 6.8961 | 6.8364 | 5.0708 | 5.3614 | 4.9993 | 5.6641 |
| ITPRID2_S1161 | ITPRID2 | P28290 | | 739 | PLRRSQSLPTTLL | 6.1793 | 6.0533 | 6.0522 | 6.0751 | 6.2545 | 6.0778 | 6.2614 | 6.3173 | 6.2432 | 6.3026 | 6.2666 | 6.1235 | 6.4584 | 6.3865 | 6.6548 | 6.2934 |
| ITPRID2_S153 | ITPRID2 | P28290 | | 153 | FHQKGRSMNSTGS | 5.5238 | 6.1365 | 4.3641 | 5.4349 | 4.7418 | 6.6362 | 6.3476 | 5.7519 | 5.4565 | 5.4937 | 6.8659 | 6.9890 | 7.3753 | 6.9755 | 7.3027 | 8.6046 |
| ITPRID2_S209 | ITPRID2 | P28290 | | 1134 | GVPSTASVGKSKT | 6.8025 | 6.9634 | 6.4224 | 7.1301 | 6.6803 | 5.8896 | 6.5677 | 6.2029 | 6.6660 | 6.1074 | 6.1685 | 6.1758 | 5.7804 | 5.8204 | 5.2465 | 5.3763 |
| ITPRID2_S270 | ITPRID2 | P28290 | | 92 | RTPLGASLDEQSS | 5.8246 | 6.3861 | 6.6420 | 6.7244 | 6.1138 | 6.8530 | 6.1337 | 6.1014 | 5.6847 | 6.7087 | 6.6382 | 6.8062 | 5.2076 | 6.4210 | 5.8729 | 5.8816 |
| ITPRID2_S4 | ITPRID2 | P28290 | | 668 | ILKSLASIEAKCS | 7.0168 | 7.0920 | 8.2211 | 7.5793 | 7.2283 | 5.9316 | 5.9346 | 6.7207 | 6.2710 | 7.2854 | 6.3847 | 5.6249 | 3.8927 | 5.6191 | 5.2530 | 3.9448 |
| ITPRID2_S466 | ITPRID2 | P28290 | | 803 | KSTIFISPSSVKK | 7.6197 | 7.5739 | 7.2197 | 7.6011 | 6.7820 | 6.5288 | 6.9207 | 7.7109 | 7.8530 | 5.9767 | 6.8874 | 6.9000 | 3.2013 | 4.2784 | 3.0251 | 3.9211 |
| ITPRID2_S593 | ITPRID2 | P28290 | | 1161 | LTPTAPSRTGSVQ | 6.6140 | 6.7674 | 6.4034 | 7.1774 | 6.6082 | 6.3878 | 5.9760 | 6.1008 | 7.0165 | 6.0648 | 6.0996 | 6.0215 | 5.5741 | 6.0996 | 5.4667 | 5.6223 |
| ITPRID2_S668 | ITPRID2 | P28290 | | 1156 | RASVALTPTAPSR | 7.1343 | 7.4957 | 6.2493 | 6.8852 | 6.2246 | 6.8808 | 7.0766 | 7.0547 | 7.1946 | 7.6735 | 7.9551 | 7.4798 | 3.2647 | 4.1256 | 3.2417 | 4.0638 |
| ITPRID2_S737 | ITPRID2 | P28290 | | 1055 | GMCGSRSADNLSC | 5.4891 | 5.8093 | 6.0509 | 5.8064 | 5.5772 | 5.8255 | 5.3103 | 5.6461 | 5.2355 | 5.5572 | 5.3724 | 5.2382 | 8.7429 | 8.1283 | 8.3722 | 7.8385 |
| ITPRID2_S737;S739 | ITPRID2 | P28290 | | 270 | PLQRIGSMSSVTS | 6.6335 | 7.6408 | 8.2440 | 7.7960 | 6.3835 | 5.0464 | 5.4078 | 5.5255 | 5.9419 | 6.1615 | 6.1346 | 6.3085 | 6.2060 | 5.7416 | 5.3797 | 5.4488 |
| ITPRID2_S739 | ITPRID2 | P28290 | | 209 | RFFNSSSFAKGID | 6.2385 | 9.2030 | 9.7667 | 7.6741 | 7.8754 | 6.0880 | 5.8729 | 5.2764 | 5.5138 | 6.6750 | 5.9429 | 6.8509 | 3.7412 | 5.4608 | 4.3705 | 3.4498 |
| ITPRID2_S739;S746 | ITPRID2 | P28290 | | 593 | DKRKSGSQDFPQC | 6.3145 | 6.0289 | 7.5389 | 7.1482 | 7.4340 | 6.0965 | 5.7114 | 5.9486 | 7.5251 | 7.5302 | 6.6387 | 6.3698 | 4.5747 | 5.4207 | 4.7889 | 4.9312 |
| ITPRID2_S767 | ITPRID2 | P28290 | | 737 | GYPLRRSQSLPTT | 7.2900 | 6.6245 | 6.7112 | 7.3587 | 5.9820 | 6.5482 | 6.5814 | 5.8769 | 7.0955 | 6.9191 | 5.8167 | 6.7291 | 4.7013 | 5.7085 | 4.7516 | 5.3053 |
| ITPRID2_S803 | ITPRID2 | P28290 | | 1140 | SVGKSKTPLVARK | 6.5406 | 7.0257 | 6.8247 | 7.2142 | 6.0121 | 7.4109 | 6.8642 | 6.6012 | 6.3532 | 6.2017 | 6.8600 | 6.8845 | 4.3930 | 5.2651 | 4.5049 | 5.0441 |
| ITPRID2_S92 | ITPRID2 | P28290 | | 1168 | RTGSVQTPPDLES | 7.1825 | 6.0371 | 6.8384 | 6.5198 | 5.4331 | 6.7757 | 5.6544 | 5.5331 | 6.7364 | 6.8308 | 6.7060 | 6.9053 | 5.2903 | 5.9213 | 5.2290 | 6.4067 |
| ITPRID2_T1140 | ITPRID2 | H7BZ26 | | 4 | xxxGSRSADNLSC | 5.8203 | 6.4609 | 5.7460 | 5.5193 | 5.5641 | 6.2064 | 6.2172 | 6.0353 | 5.1753 | 5.9574 | 6.1294 | 6.0434 | 7.6810 | 6.8264 | 8.1603 | 6.4573 |
| ITPRID2_T1156 | ITPRID2 | P28290 | 739;746 | | NA | 6.5760 | 6.4588 | 6.3218 | 7.1038 | 6.0555 | 6.5712 | 6.4938 | 7.0234 | 6.5725 | 6.5077 | 7.0788 | 6.3829 | 5.1706 | 5.7729 | 5.0538 | 4.8565 |
| ITPRID2_T1168 | ITPRID2 | P28290 | 737;739 | | NA | 7.8683 | 7.6848 | 10.2078 | 7.7421 | 7.4985 | 5.0552 | 5.9574 | 6.5251 | 6.3398 | 6.4417 | 5.7915 | 5.6703 | 3.9543 | 5.5504 | 4.3428 | 3.3699 |
| ITPRIPL2_S78 | ITPRIPL2 | Q3MIP1 | | 78 | QRFLPGSPRLEGH | 6.6989 | 7.2327 | 6.4432 | 7.4428 | 6.0320 | 6.3356 | 6.6501 | 6.7389 | 6.7955 | 6.9073 | 6.8945 | 6.8409 | 4.1970 | 5.0577 | 4.8261 | 4.9069 |
| ITSN1_S1137 | ITSN1 | Q15811 | | 904 | ATGSSPSPVLGQG | 5.9061 | 6.0647 | 6.0222 | 5.7343 | 6.1256 | 5.9480 | 5.5719 | 6.1677 | 6.3219 | 6.6891 | 6.1468 | 6.3013 | 6.8557 | 6.7910 | 6.9154 | 6.4382 |
| ITSN1_S203 | ITSN1 | Q15811 | | 315 | RRVRSGSGISVIS | 5.2595 | 5.8353 | 5.9509 | 5.7589 | 6.0393 | 5.9267 | 5.9040 | 5.9947 | 6.1065 | 5.7495 | 5.5572 | 5.5440 | 8.1467 | 7.4765 | 7.7817 | 6.9687 |
| ITSN1_S315 | ITSN1 | Q15811 | | 897 | SAFTPATATGSSP | 6.0952 | 5.9997 | 6.1641 | 6.1147 | 5.8503 | 6.2763 | 5.8683 | 6.4967 | 6.1921 | 6.1899 | 6.0822 | 6.7980 | 7.2477 | 6.5521 | 5.8758 | 6.1969 |
| ITSN1_S904 | ITSN1 | Q15811 | | 203 | KLQKAQSFDVASV | 5.8170 | 6.7304 | 7.1276 | 5.6800 | 5.7598 | 6.6303 | 6.2346 | 6.5905 | 6.4774 | 7.1124 | 6.1236 | 7.6546 | 5.3132 | 6.2867 | 4.7732 | 5.6887 |
| ITSN1_S986 | ITSN1 | Q15811 | | 1137 | NYVKLLSPGTSKI | 6.1906 | 6.1371 | 5.9362 | 5.9316 | 6.3613 | 6.5378 | 6.2561 | 6.2105 | 6.6821 | 6.8317 | 6.2431 | 6.2633 | 6.4252 | 6.2042 | 6.1110 | 5.6782 |
| ITSN1_S989 | ITSN1 | Q15811 | | 989 | SSESPASLKRVAS | 5.6294 | 5.3668 | 5.8986 | 5.9490 | 6.6014 | 5.8203 | 6.7804 | 7.9130 | 6.0140 | 6.4860 | 5.8893 | 7.1213 | 6.0391 | 6.1638 | 6.0604 | 6.2672 |
| ITSN1_T1144 | ITSN1 | Q15811 | | 1144 | PGTSKITPTEPPK | 5.4634 | 5.3463 | 5.7569 | 4.9837 | 6.2115 | 5.7532 | 5.8773 | 5.4291 | 5.4426 | 5.1794 | 5.5931 | 5.7918 | 8.9848 | 7.6103 | 8.1841 | 8.3923 |
| ITSN1_T897 | ITSN1 | Q15811 | | 986 | DSGSSESPASLKR | 6.0489 | 6.6194 | 5.9655 | 6.4686 | 6.0414 | 5.3696 | 6.1041 | 6.4049 | 6.3965 | 7.0785 | 6.9745 | 6.7815 | 5.6731 | 6.1950 | 6.1407 | 5.7378 |
| ITSN1_T897;S904 | ITSN1 | Q15811 | 897;904 | | NA | 5.6940 | 5.3370 | 5.4583 | 5.4145 | 6.0874 | 6.0798 | 5.6511 | 6.0577 | 6.0277 | 5.6946 | 5.9367 | 5.8907 | 8.3226 | 7.9779 | 7.5395 | 6.8307 |
| ITSN2_S110 | ITSN2 | Q9NZM3 | | 889 | VSPGSVSPIHGQG | 5.9970 | 6.3313 | 5.6912 | 5.9863 | 5.9964 | 6.3604 | 6.5115 | 5.8732 | 6.5578 | 6.2578 | 6.1861 | 6.2759 | 6.2437 | 6.8317 | 6.6945 | 6.2054 |
| ITSN2_S210 | ITSN2 | Q9NZM3 | | 884 | AFTRTVSPGSVSP | 5.9981 | 6.3910 | 6.1228 | 6.3625 | 6.4528 | 6.5769 | 5.7395 | 7.0542 | 6.6507 | 5.9645 | 6.2505 | 6.0359 | 5.8688 | 6.0796 | 6.4634 | 5.9889 |
| ITSN2_S884 | ITSN2 | Q9NZM3 | | 210 | SIQKAQSLIDLGS | 6.0333 | 5.7171 | 5.8995 | 5.5217 | 5.8843 | 6.7006 | 6.6087 | 6.4668 | 7.7036 | 5.6049 | 6.4647 | 6.2736 | 6.6724 | 6.3576 | 5.3343 | 6.7570 |
| ITSN2_S884;S889 | ITSN2 | Q9NZM3 | | 1123 | PSSERATPAFHPV | 5.2355 | 5.1203 | 5.8312 | 5.0342 | 5.8910 | 5.4540 | 6.1341 | 5.3547 | 5.4535 | 5.6557 | 5.7360 | 6.3467 | 7.8383 | 8.2930 | 7.9230 | 8.6987 |
| ITSN2_S889 | ITSN2 | Q9NZM3 | | 110 | KQPPMFSPLISAR | 6.3598 | 6.1870 | 6.9851 | 6.1964 | 5.4375 | 6.4713 | 6.2090 | 6.8641 | 5.8775 | 6.2372 | 8.2734 | 7.3464 | 5.1956 | 6.3472 | 4.5122 | 5.5004 |
| ITSN2_T1123 | ITSN2 | Q9NZM3 | 884;889 | | NA | 5.5336 | 5.3139 | 5.5184 | 5.4657 | 5.9284 | 6.1720 | 6.1979 | 6.0647 | 6.2510 | 5.9256 | 5.9766 | 6.1489 | 7.3804 | 7.3288 | 7.6202 | 7.1740 |
| IVNS1ABP_S269 | IVNS1ABP | Q9Y6Y0 | | 269 | GHKQISSSSTGCL | 6.2582 | 6.4163 | 6.0542 | 7.5653 | 6.1864 | 7.3105 | 5.8808 | 6.4800 | 8.5572 | 6.3856 | 7.4629 | 7.1928 | 4.8334 | 4.6834 | 3.5787 | 5.1543 |
| IWS1_S157 | IWS1 | Q96ST2 | | 377 | KKQKMDSDEDEKE | 5.9200 | 6.2406 | 6.3716 | 6.1604 | 5.7592 | 5.5521 | 6.1849 | 5.4889 | 6.1025 | 6.3356 | 6.0683 | 6.3597 | 7.0613 | 6.6730 | 6.7473 | 6.9745 |
| IWS1_S157;S159 | IWS1 | Q96ST2 | | 527 | KHMDFLSDFEMML | 8.0083 | 9.1968 | 7.4069 | 8.9015 | 8.4114 | 7.0627 | 6.8148 | 7.1575 | 4.2639 | 3.8349 | 4.2452 | 5.3928 | 5.1348 | 5.1283 | 4.2107 | 4.8294 |
| IWS1_S235;S237 | IWS1 | Q96ST2 | | 725 | NSTGGQTPRRDLE | 7.4340 | 7.6667 | 6.5931 | 6.5948 | 5.0707 | 6.1486 | 6.3229 | 5.6770 | 6.4079 | 6.6937 | 6.9601 | 6.4708 | 5.6334 | 5.3867 | 5.0827 | 5.8570 |
| IWS1_S248;S250 | IWS1 | Q96ST2 | | 720 | QRRRMNSTGGQTP | 5.0129 | 4.9988 | 5.5819 | 5.9700 | 6.5883 | 5.3042 | 5.6501 | 6.2991 | 6.0762 | 5.9374 | 5.5578 | 5.9532 | 8.1036 | 7.7172 | 7.7978 | 7.4516 |
| IWS1_S300 | IWS1 | Q96ST2 | | 157 | VGKHPASDSEIEE | 5.2540 | 6.5589 | 6.5759 | 6.2642 | 5.3012 | 6.1761 | 5.6828 | 5.7082 | 6.3623 | 7.5025 | 7.4451 | 7.7956 | 5.0347 | 5.7332 | 6.4450 | 6.1604 |
| IWS1_S300;S302 | IWS1 | Q96ST2 | | 300 | LPKPRVSDSESEG | 5.0639 | 5.6860 | 6.1327 | 5.4024 | 4.9809 | 5.9125 | 5.8221 | 5.7722 | 6.9736 | 8.2237 | 7.5629 | 7.9293 | 6.3228 | 6.5137 | 5.5276 | 6.1738 |
| IWS1_S313;S315 | IWS1 | Q96ST2 | | 400 | AAVLSDSEDEEKA | 4.8817 | 5.5201 | 5.3871 | 5.0934 | 5.2279 | 4.9065 | 4.7477 | 5.5487 | 7.0457 | 6.4598 | 6.0330 | 6.0789 | 9.1849 | 8.1427 | 7.8504 | 7.8912 |
| IWS1_S329;S333 | IWS1 | Q96ST2 | 363;365 | | NA | 4.7625 | 5.0636 | 5.4847 | 5.5751 | 5.7014 | 5.6156 | 5.5021 | 5.3185 | 6.0731 | 6.1060 | 6.1766 | 6.1296 | 8.2314 | 8.1728 | 7.9657 | 8.1213 |
| IWS1_S363;S365 | IWS1 | Q96ST2 | 235;237 | | NA | 5.6065 | 6.3593 | 6.4630 | 6.7437 | 5.9485 | 5.1885 | 6.1159 | 6.0167 | 5.9627 | 6.4126 | 5.8678 | 6.1158 | 6.4374 | 6.3055 | 6.8184 | 7.6375 |
| IWS1_S377 | IWS1 | Q96ST2 | 329;333 | | NA | 5.5455 | 6.1227 | 6.2506 | 6.4168 | 5.1246 | 5.8246 | 5.8949 | 5.6451 | 5.6653 | 6.4912 | 6.0392 | 6.5776 | 6.5540 | 6.6761 | 7.0376 | 8.1343 |
| IWS1_S400 | IWS1 | Q96ST2 | 300;302 | | NA | 5.8690 | 5.8437 | 6.3111 | 6.6550 | 6.2678 | 5.5138 | 5.8527 | 5.4806 | 5.9214 | 6.3891 | 6.5007 | 6.1480 | 6.6154 | 6.3951 | 6.8814 | 7.3552 |
| IWS1_S438;S440 | IWS1 | Q96ST2 | 438;440 | | NA | 5.5453 | 6.1160 | 6.7617 | 6.2952 | 6.0881 | 6.0812 | 6.5639 | 6.0797 | 6.4006 | 6.2858 | 5.8793 | 6.2976 | 5.8223 | 6.9443 | 6.2600 | 6.5791 |
| IWS1_S527 | IWS1 | Q96ST2 | 248;250 | | NA | 6.1244 | 5.9015 | 6.0885 | 6.7641 | 6.2594 | 6.5729 | 6.2813 | 6.8037 | 5.9321 | 5.6329 | 5.5828 | 5.7312 | 6.2555 | 6.5477 | 6.6297 | 6.8922 |
| IWS1_S720 | IWS1 | Q96ST2 | 313;315 | | NA | 6.0699 | 6.3839 | 6.0543 | 4.9418 | 6.6056 | 5.5705 | 6.5568 | 6.6608 | 5.7179 | 6.2693 | 5.0624 | 7.0239 | 7.7217 | 7.2325 | 6.0711 | 6.0576 |
| IWS1_T725 | IWS1 | Q96ST2 | 157;159 | | NA | 5.7241 | 6.6402 | 6.8959 | 6.8536 | 6.2587 | 4.5585 | 6.3929 | 5.6803 | 5.7961 | 5.8326 | 5.0888 | 5.2270 | 6.0551 | 6.3510 | 7.2941 | 8.4511 |
| JADE1_S743 | JADE1 | Q6IE81 | | 743 | VPTTPASPVKNWG | 6.4870 | 6.5149 | 6.4313 | 7.1119 | 6.5018 | 6.2511 | 6.4927 | 6.3372 | 6.4597 | 6.2465 | 6.2109 | 6.1316 | 5.5849 | 5.6761 | 5.7042 | 5.8581 |
| JADE1_T702 | JADE1 | Q6IE81 | | 702 | DNTRAATSPGVGQ | 6.8319 | 6.3217 | 5.4833 | 6.3471 | 6.6160 | 5.8411 | 6.2540 | 6.3069 | 7.3295 | 6.7536 | 6.5207 | 6.3260 | 5.9639 | 6.1968 | 5.7310 | 5.1764 |
| JADE2_S117 | JADE2 | Q9NQC1 | | 117 | GPPAQASPSSTML | 5.8758 | 6.3457 | 6.2119 | 6.5486 | 5.9868 | 6.2669 | 6.2426 | 6.3782 | 6.5227 | 6.6645 | 6.4224 | 6.0915 | 6.0043 | 6.4038 | 6.1177 | 5.9166 |
| JAK1_T688 | JAK1 | P23458 | | 688 | KSDVLTTPWKFKV | 6.9964 | 6.4399 | 8.3656 | 8.2275 | 7.0629 | 6.2686 | 6.2852 | 6.6188 | 7.1894 | 6.4898 | 6.3052 | 7.0413 | 4.1133 | 4.3862 | 3.9220 | 4.2880 |
| JCAD_S1321 | JCAD | Q9P266 | | 757 | KGHRSLSPSSNSA | 6.6368 | 6.7079 | 6.2488 | 6.4108 | 6.1568 | 6.1437 | 6.2052 | 6.2786 | 6.4930 | 6.4045 | 6.8072 | 6.2951 | 5.9434 | 5.9288 | 5.6810 | 5.6583 |
| JCAD_S757 | JCAD | Q9P266 | | 931 | PRAWPPSPGRFRV | 5.4188 | 6.8053 | 6.6554 | 6.0652 | 6.6685 | 6.0071 | 6.9028 | 6.6910 | 6.7885 | 6.7435 | 5.9116 | 6.5149 | 5.5855 | 6.0390 | 5.1883 | 6.0146 |
| JCAD_S931 | JCAD | Q9P266 | | 1321 | GHSLSVSKDSISR | 4.5119 | 6.5553 | 5.2588 | 5.5351 | 6.6282 | 7.5695 | 6.9440 | 6.2547 | 5.4653 | 5.8357 | 6.7405 | 7.2506 | 7.0351 | 6.7345 | 5.0584 | 6.6221 |
| JMJD1C_S1225 | JMJD1C | Q15652 | | 1225 | RKEGSYSSLSPPT | 6.1051 | 6.6549 | 6.7008 | 6.9156 | 6.4327 | 6.4280 | 6.7290 | 6.4226 | 6.4677 | 6.4497 | 6.7446 | 6.7785 | 5.0454 | 5.5524 | 5.2711 | 5.3019 |
| JMJD1C_S2007 | JMJD1C | Q15652 | | 617 | DKLHKRSPPPETI | 6.1833 | 6.4420 | 6.0363 | 6.3740 | 6.1985 | 6.2791 | 6.3115 | 6.1049 | 5.9918 | 6.2677 | 6.3989 | 6.3747 | 6.2859 | 6.2696 | 6.3391 | 6.1427 |
| JMJD1C_S2053 | JMJD1C | Q15652 | | 1978 | ESQQQNTPPKSEK | 6.0399 | 6.6796 | 6.7315 | 6.7486 | 6.6957 | 5.9796 | 6.0925 | 6.7716 | 6.3710 | 6.4922 | 6.9020 | 6.6608 | 5.1132 | 5.3824 | 5.4735 | 5.8658 |
| JMJD1C_S317;S320 | JMJD1C | Q15652 | | 652 | KPKITHSPDSVKS | 6.3024 | 6.3228 | 5.7791 | 6.2642 | 7.0331 | 7.1731 | 6.8041 | 6.9822 | 5.8532 | 5.8057 | 6.1422 | 6.4493 | 5.7346 | 6.0961 | 5.5753 | 5.6825 |
| JMJD1C_S617 | JMJD1C | Q15652 | | 943 | KITAHSSPPLTKT | 5.9075 | 6.6304 | 6.6410 | 6.6407 | 6.9484 | 6.4937 | 6.8784 | 6.7260 | 6.4050 | 6.6760 | 6.3175 | 6.0939 | 4.9230 | 5.6929 | 5.3335 | 5.6922 |
| JMJD1C_S639;S641 | JMJD1C | Q15652 | | 2007 | TPPESQSPLHWLA | 6.5300 | 6.3709 | 5.8005 | 7.1857 | 6.2634 | 6.1689 | 6.1517 | 6.6843 | 6.0108 | 6.5123 | 5.9482 | 6.1299 | 5.8408 | 6.7218 | 5.8530 | 5.8276 |
| JMJD1C_S641 | JMJD1C | Q15652 | | 2053 | SPNGRTSPLVSQN | 5.4457 | 6.3456 | 6.2109 | 5.9755 | 6.0024 | 5.9428 | 6.2772 | 6.2029 | 5.9403 | 6.3338 | 6.3443 | 6.6955 | 6.7005 | 6.8885 | 6.4085 | 6.2856 |
| JMJD1C_S652 | JMJD1C | Q15652 | | 641 | KIKSSPSPEVVKP | 6.2191 | 6.8577 | 5.7805 | 6.8778 | 6.2890 | 6.0319 | 6.5630 | 6.2051 | 6.8347 | 6.2939 | 6.2373 | 6.4600 | 5.3403 | 6.0604 | 6.0704 | 5.8789 |
| JMJD1C_S943 | JMJD1C | Q15652 | | 505 | FVSRPPTPKCVID | 5.5946 | 5.3621 | 7.3436 | 6.4921 | 5.9559 | 5.7441 | 6.4506 | 6.0594 | 6.2021 | 5.6245 | 7.3413 | 5.4565 | 5.2770 | 6.9362 | 7.2193 | 6.9408 |
| JMJD1C_T1978 | JMJD1C | Q15652 | 317;320 | | NA | 5.8763 | 6.3474 | 6.7916 | 5.7043 | 5.1751 | 5.7354 | 6.9487 | 6.1000 | 5.8416 | 5.7422 | 5.8824 | 5.8037 | 7.4208 | 6.4907 | 6.6731 | 7.4667 |
| JMJD1C_T505 | JMJD1C | Q15652 | 639;641 | | NA | 6.0500 | 6.7494 | 6.8137 | 7.2251 | 5.9889 | 6.5305 | 6.6500 | 6.1249 | 6.3934 | 7.0749 | 7.2440 | 7.1507 | 4.6549 | 5.5261 | 4.8556 | 4.9680 |
| JMY_S130 | JMY | Q8N9B5 | | 130 | GDPRLRSPGSKGA | 6.6443 | 6.7107 | 6.8779 | 6.6660 | 6.5000 | 7.2192 | 6.7548 | 6.7630 | 7.2184 | 7.0484 | 7.0778 | 7.1956 | 3.9426 | 4.8259 | 3.9379 | 4.6175 |
| JMY_S713 | JMY | Q8N9B5 | | 713 | RRKGAASPVLQED | 5.8016 | 5.9390 | 6.1307 | 6.2150 | 6.2255 | 6.3784 | 5.9781 | 6.4980 | 6.4264 | 6.3021 | 6.4341 | 6.1187 | 6.2888 | 6.5680 | 6.3671 | 6.3286 |
| JMY_S888 | JMY | Q8N9B5 | | 888 | LQRRRVSSPMDEV | 5.2865 | 5.0709 | 4.8755 | 4.8695 | 6.9356 | 5.8773 | 4.6793 | 6.1516 | 7.8956 | 7.7977 | 7.3827 | 7.4754 | 7.0592 | 7.5759 | 5.2988 | 5.7686 |
| JPH1_S157;S162 | JPH1 | Q9HDC5 | | 216 | GLFRRGSLLGSMK | 5.5524 | 5.1761 | 5.2240 | 5.7010 | 6.1764 | 5.7843 | 6.0890 | 5.6983 | 7.2542 | 6.5456 | 6.4219 | 6.9645 | 7.0541 | 7.3063 | 5.8005 | 7.2513 |
| JPH1_S162 | JPH1 | Q9HDC5 | | 448 | VPEKPPTPKESPH | 5.8612 | 7.0443 | 6.0778 | 6.6983 | 5.7452 | 5.3018 | 6.3665 | 6.0993 | 6.5097 | 6.3314 | 5.8570 | 6.1470 | 6.1900 | 6.2249 | 6.8466 | 6.6990 |
| JPH1_S162;S165 | JPH1 | Q9HDC5 | | 452 | PPTPKESPHFYRK | 6.1505 | 6.3739 | 6.1148 | 6.6916 | 5.9574 | 6.6555 | 6.0404 | 6.5263 | 6.3414 | 5.9215 | 5.8542 | 5.9882 | 6.3639 | 6.3509 | 6.6037 | 7.0657 |
| JPH1_S216 | JPH1 | Q9HDC5 | | 162 | RSPLRTSLASLRS | 5.7398 | 5.7959 | 5.8005 | 5.5952 | 5.5601 | 5.5570 | 5.7448 | 5.8559 | 6.4557 | 6.4950 | 6.2294 | 6.5527 | 7.0697 | 6.8862 | 7.2949 | 7.3672 |
| JPH1_S216;S220 | JPH1 | Q9HDC5 | | 220 | RGSLLGSMKLRKS | 9.2087 | 9.5925 | 7.0682 | 9.6414 | 5.7339 | 5.1672 | 7.5730 | 6.4525 | 4.4717 | 4.9225 | 5.2007 | 5.1620 | 3.8973 | 4.4485 | 6.4165 | 5.0435 |
| JPH1_S220 | JPH1 | Q9HDC5 | | 461 | FYRKGTTPPRSPE | 6.1640 | 5.7637 | 5.7201 | 6.6860 | 6.0484 | 5.2103 | 5.6475 | 5.6651 | 6.2890 | 5.9946 | 5.6361 | 5.4881 | 7.9481 | 7.7330 | 7.1088 | 6.8972 |
| JPH1_S452 | JPH1 | Q9HDC5 | | 594 | KWSPSKSVTKPVA | 7.0402 | 7.2205 | 6.2814 | 7.0117 | 7.4925 | 6.3298 | 7.0132 | 6.9124 | 6.6578 | 5.9626 | 6.2534 | 5.8180 | 7.5915 | 5.8200 | 4.3380 | 4.0569 |
| JPH1_S594 | JPH1 | Q9HDC5 | 216;220 | | NA | 7.8717 | 7.7139 | 8.4329 | 7.6128 | 6.6995 | 6.4078 | 6.7620 | 5.8487 | 6.7248 | 6.3689 | 6.6965 | 6.4731 | 4.2936 | 4.7149 | 3.5694 | 3.8094 |
| JPH1_T448 | JPH1 | Q9HDC5 | 461;465 | | NA | 6.5210 | 6.4640 | 6.4561 | 6.7219 | 5.5561 | 5.5223 | 6.1227 | 5.9775 | 6.0299 | 5.2996 | 6.2276 | 5.7360 | 6.6789 | 6.8436 | 7.0102 | 6.8327 |
| JPH1_T461 | JPH1 | Q9HDC5 | 157;162 | | NA | 6.6787 | 6.4691 | 5.9823 | 7.2769 | 6.3357 | 7.1380 | 6.5369 | 7.2638 | 6.8930 | 7.2241 | 6.7511 | 7.5174 | 3.4455 | 5.0116 | 3.6159 | 5.8601 |
| JPH1_T461;S465 | JPH1 | Q9HDC5 | 162;165 | | NA | 8.6777 | 7.4304 | 7.6128 | 7.2711 | 6.5373 | 6.4853 | 6.2255 | 5.4003 | 5.2190 | 6.0235 | 7.3238 | 6.5322 | 5.1902 | 5.7396 | 3.8479 | 4.4835 |
| JPT1_S131 | JPT1 | Q9UK76 | | 87 | GLQRRNSSEASSG | 5.9604 | 4.9516 | 6.5376 | 5.2613 | 6.3325 | 5.0797 | 5.2144 | 7.1959 | 5.3132 | 7.0181 | 6.4273 | 7.7675 | 5.5781 | 7.7452 | 6.4999 | 7.1172 |
| JPT1_S28 | JPT1 | Q9UK76 | | 131 | PAAPVPSPVAPAP | 6.5757 | 5.7705 | 6.7609 | 6.3027 | 6.4502 | 5.9908 | 5.9442 | 7.0757 | 6.1853 | 7.2627 | 7.0712 | 7.6854 | 4.3815 | 5.9361 | 4.7365 | 5.8707 |
| JPT1_S87 | JPT1 | Q9UK76 | | 54 | ASNIFGTPEENQA | 7.1515 | 6.5309 | 7.1813 | 6.6115 | 6.3130 | 5.6392 | 6.2618 | 7.4113 | 6.0480 | 8.2099 | 7.7092 | 8.8920 | 3.0482 | 5.0578 | 3.3179 | 4.6166 |
| JPT1_T54 | JPT1 | Q9UK76 | | 28 | RPPGGGSNFSLGF | 5.6871 | 4.8779 | 8.4976 | 5.9538 | 6.9248 | 4.6432 | 4.2032 | 7.1095 | 5.1264 | 6.2615 | 6.3550 | 7.9096 | 4.6409 | 8.3267 | 5.5777 | 7.9050 |
| JPT2_S125 | JPT2 | Q9H910-3 | | 125 | TSDIFGSPVTATS | 7.4563 | 6.5009 | 7.5990 | 6.5111 | 6.5352 | 6.8385 | 6.4622 | 7.5798 | 7.2645 | 7.6846 | 8.2416 | 8.1088 | 2.9921 | 3.7679 | 2.7756 | 3.6820 |
| JPT2_S14 | JPT2 | Q9H910-3 | | 104 | GIFDESTPVQTRQ | 7.6227 | 6.6515 | 7.1131 | 6.3905 | 5.6792 | 6.7119 | 6.7498 | 7.8409 | 7.2704 | 7.6327 | 8.0981 | 8.0578 | 3.6675 | 3.9324 | 2.7695 | 3.8121 |
| JPT2_S58 | JPT2 | Q9H910-3 | | 58 | SSNLFGSPEEATP | 6.5835 | 6.5751 | 7.5438 | 6.7004 | 6.3777 | 6.5666 | 6.4819 | 8.4004 | 7.8693 | 8.3217 | 9.0336 | 8.9620 | 2.4877 | 3.0602 | 2.1094 | 2.9268 |
| JPT2_T104 | JPT2 | Q9H910-3 | | 89 | QNIPKRTNPPGGK | 5.9769 | 5.7261 | 7.4701 | 5.6678 | 5.8307 | 6.8031 | 5.9474 | 6.8734 | 7.4376 | 7.4443 | 8.5930 | 9.1169 | 3.9790 | 4.8239 | 3.9849 | 4.3249 |
| JPT2_T89 | JPT2 | Q9H910-2 | | 14 | SSNLFGSPEEATP | 7.4870 | 6.3462 | 7.6269 | 5.4939 | 6.9012 | 7.5046 | 6.8001 | 6.3480 | 8.7369 | 7.5926 | 8.3569 | 8.3077 | 3.2193 | 3.4902 | 2.7645 | 3.0240 |
| JRK_S557 | JRK | O75564-2 | | 557 | CGATAQSPLPCSS | 5.3899 | 7.2018 | 5.9372 | 7.2764 | 7.8444 | 6.2109 | 7.6164 | 5.2987 | 4.3949 | 6.8079 | 5.4729 | 7.3676 | 4.9587 | 6.1860 | 6.3187 | 5.7175 |
| JUN_S243 | JUN | P05412 | | 243 | GETPPLSPIDMES | 5.8646 | 5.9002 | 6.4931 | 6.3643 | 6.2819 | 5.7636 | 5.9653 | 6.3827 | 6.3798 | 5.8510 | 6.2301 | 6.0036 | 6.7486 | 7.1390 | 6.4869 | 6.1453 |
| JUN_S63 | JUN | P05412 | | 63 | NSDLLTSPDVGLL | 6.8899 | 6.6923 | 8.1236 | 7.4800 | 6.4241 | 6.9700 | 7.3116 | 6.7018 | 5.9179 | 5.9905 | 6.5711 | 6.4403 | 4.9072 | 4.3888 | 4.7796 | 4.4111 |
| JUNB_S251;T255;S259 | JUNB | P17275 | | 28 | RAPGGLSLHDYKL | 7.9928 | 7.3521 | 6.5997 | 5.6459 | 7.0134 | 6.0376 | 6.2720 | 5.8015 | 7.1525 | 6.2189 | 7.3667 | 7.3356 | 4.8495 | 4.3076 | 4.1117 | 5.9424 |
| JUNB_S28 | JUNB | P17275 | 251;255;259 | | NA | 5.6277 | 4.7552 | 6.5586 | 6.8161 | 6.5713 | 5.4975 | 6.1806 | 5.4022 | 4.3715 | 6.4604 | 5.1673 | 5.4693 | 7.8607 | 7.8579 | 7.7670 | 7.6368 |
| JUND_S100 | JUND | P17535 | | 251 | TVPDVPSFGESPP | 6.3904 | 5.9147 | 6.2009 | 6.3537 | 6.4872 | 5.7919 | 6.0372 | 6.3079 | 6.5380 | 5.9449 | 5.8119 | 6.0180 | 6.7529 | 6.1964 | 6.8294 | 6.4245 |
| JUND_S251 | JUND | P17535 | | 116 | SNGLVTTTPTSSQ | 6.4602 | 6.9734 | 6.1456 | 6.6676 | 6.4794 | 6.9226 | 7.2834 | 6.5496 | 7.2080 | 6.9641 | 7.4182 | 7.0877 | 3.7400 | 5.1010 | 4.5071 | 4.4921 |
| JUND_S251;S259 | JUND | P17535 | | 100 | GLLKLASPELERL | 6.9397 | 6.8488 | 6.9089 | 7.0535 | 6.7435 | 6.5265 | 7.0895 | 7.1006 | 5.8916 | 5.6888 | 6.1029 | 5.7764 | 5.4503 | 4.9911 | 5.5262 | 5.3619 |
| JUND_S255 | JUND | P17535 | | 90 | PDGLLASPDLGLL | 7.0423 | 6.9446 | 6.4318 | 7.1530 | 7.0778 | 6.8771 | 7.2117 | 7.0407 | 7.4412 | 6.9567 | 7.3043 | 7.7770 | 3.5872 | 4.0371 | 3.2351 | 3.8824 |
| JUND_S255;S259 | JUND | P17535 | | 259 | GESPPLSPIDMDT | 6.4436 | 6.1489 | 6.3924 | 7.1014 | 6.5589 | 6.7417 | 6.8816 | 6.9151 | 6.6604 | 5.8585 | 6.3528 | 5.9290 | 5.3013 | 6.3192 | 5.2622 | 5.1331 |
| JUND_S259 | JUND | P17535 | | 255 | VPSFGESPPLSPI | 6.2699 | 6.1544 | 6.0966 | 6.0599 | 5.9103 | 5.8064 | 5.8061 | 6.1219 | 6.1268 | 6.4251 | 6.2544 | 6.2333 | 6.9844 | 6.7448 | 6.2375 | 6.7682 |
| JUND_S90 | JUND | P17535 | | 117 | NGLVTTTPTSSQF | 6.8351 | 7.4676 | 6.9648 | 7.1375 | 6.9049 | 6.8172 | 7.2113 | 7.3105 | 6.7810 | 7.2089 | 6.7031 | 6.9943 | 3.7184 | 4.1768 | 3.8518 | 3.9171 |
| JUND_T116 | JUND | P17535 | 255;259 | | NA | 6.4883 | 6.1978 | 6.4830 | 5.9804 | 6.7251 | 6.7081 | 6.6132 | 6.1905 | 5.8682 | 6.2770 | 6.4590 | 6.5091 | 5.9516 | 6.1317 | 5.8209 | 5.5960 |
| JUND_T117 | JUND | P17535 | 251;259 | | NA | 6.3023 | 6.2439 | 5.9212 | 6.8711 | 6.6311 | 5.9061 | 6.2159 | 6.2281 | 6.7359 | 5.5138 | 6.4048 | 6.4051 | 6.3721 | 5.8803 | 6.6781 | 5.6903 |
| KANK2_S323 | KANK2 | Q63ZY3-2 | | 375 | MPGRPESPPVFRS | 6.0653 | 6.0882 | 6.0323 | 5.7191 | 6.5782 | 6.0422 | 5.8644 | 6.3354 | 6.5405 | 6.1399 | 6.1459 | 6.0960 | 6.6181 | 6.9398 | 6.5328 | 6.2620 |
| KANK2_S375 | KANK2 | Q63ZY3-2 | | 548 | PRERVPSVAEAPQ | 5.4160 | 4.7374 | 5.7761 | 5.2553 | 6.7486 | 5.4095 | 5.4020 | 6.2019 | 6.7222 | 6.3846 | 6.1551 | 6.2230 | 8.0327 | 7.7009 | 7.2724 | 6.5623 |
| KANK2_S406 | KANK2 | Q63ZY3-2 | | 323 | AWPPPDSPVRVDT | 5.7986 | 5.6458 | 6.4787 | 5.6987 | 6.6400 | 6.2713 | 5.8287 | 6.2437 | 6.4168 | 6.2699 | 6.2354 | 6.1000 | 6.9186 | 6.8516 | 6.4459 | 6.1560 |
| KANK2_S548 | KANK2 | Q63ZY3-2 | | 83 | PGSWWTSTESLCS | 5.8009 | 5.6408 | 5.8885 | 5.2118 | 5.4367 | 5.7435 | 6.0996 | 6.2989 | 6.5630 | 6.2174 | 5.7094 | 6.8949 | 6.0085 | 8.0169 | 6.7365 | 7.7325 |
| KANK2_S83 | KANK2 | Q63ZY3-2 | | 406 | HMVKKISITERSC | 6.0389 | 6.0134 | 6.5865 | 6.4377 | 6.4602 | 5.7814 | 5.6111 | 6.1754 | 7.0034 | 6.1677 | 6.9006 | 6.7400 | 6.3108 | 6.6773 | 6.2125 | 4.8832 |
| KANK2_S83;S86 | KANK2 | Q63ZY3-2 | 83;86 | | NA | 6.0711 | 9.8729 | 7.1355 | 6.1557 | 3.7050 | 7.6115 | 5.1368 | 4.1803 | 5.5183 | 7.4603 | 3.1390 | 6.2499 | 8.4241 | 6.0378 | 5.5194 | 7.7825 |
| KANSL1_S1045 | KANSL1 | Q7Z3B3 | | 1082 | TEAAPTSPPIVPL | 7.0439 | 7.2084 | 6.7524 | 7.3234 | 6.3365 | 6.3482 | 6.6267 | 6.5437 | 6.9618 | 6.8946 | 6.8794 | 7.1418 | 4.1619 | 4.7247 | 4.2955 | 4.7570 |
| KANSL1_S1082 | KANSL1 | Q7Z3B3 | | 1045 | TFPLAHSPQAECE | 5.8265 | 6.2574 | 6.6589 | 7.1869 | 6.0817 | 6.2769 | 6.3355 | 6.1268 | 6.2210 | 6.3323 | 6.9282 | 6.3922 | 6.0032 | 6.0210 | 5.5708 | 5.7807 |
| KANSL1_S249 | KANSL1 | Q7Z3B3 | | 991 | HGQSPRSPISPEL | 5.9255 | 6.1348 | 6.1131 | 7.5198 | 6.1513 | 6.1490 | 6.3027 | 6.2891 | 6.5785 | 6.3986 | 6.6919 | 6.4169 | 5.4960 | 6.2659 | 5.4235 | 6.1434 |
| KANSL1_S268 | KANSL1 | Q7Z3B3 | | 249 | QGSSRLSPGTDSS | 6.3062 | 6.6271 | 5.7784 | 6.8499 | 6.2115 | 5.9383 | 6.2185 | 5.7606 | 6.5297 | 6.1122 | 6.3865 | 6.0392 | 6.0924 | 6.2476 | 6.3811 | 6.5208 |
| KANSL1_S564 | KANSL1 | Q7Z3B3 | | 718 | PGSLPDSARKDRH | 7.4464 | 7.8849 | 7.0442 | 7.7339 | 5.3204 | 7.3964 | 7.4645 | 6.6066 | 7.4652 | 8.2852 | 8.3237 | 8.1361 | 2.2330 | 3.8389 | 2.1238 | 2.6970 |
| KANSL1_S718 | KANSL1 | Q7Z3B3 | | 268 | KLEGKKSPLSSIL | 5.9813 | 7.0150 | 5.6049 | 6.4579 | 6.2370 | 6.3467 | 6.5771 | 6.1557 | 6.0681 | 6.2914 | 6.5763 | 5.7256 | 6.2180 | 6.4761 | 6.5810 | 5.6878 |
| KANSL1_S991 | KANSL1 | Q7Z3B3 | | 564 | DHIPGDSSDAEEQ | 5.0027 | 8.9305 | 6.2833 | 6.4291 | 6.6894 | 7.6759 | 7.4026 | 7.0668 | 5.7628 | 5.7524 | 6.3068 | 5.6196 | 4.8830 | 4.0969 | 5.7498 | 6.3484 |
| KANSL1_T828 | KANSL1 | Q7Z3B3 | | 828 | LVRQLSTSSDSPA | 5.6905 | 5.5299 | 6.6672 | 6.7135 | 5.8047 | 5.7462 | 5.7149 | 5.5463 | 6.3190 | 6.0986 | 6.8794 | 6.8998 | 6.5597 | 6.9624 | 5.9136 | 6.9543 |
| KANSL2_S149 | KANSL2 | Q9H9L4 | | 149 | LDEDSWSDGEQEP | 6.7851 | 6.9496 | 5.1880 | 7.4239 | 6.0591 | 4.4282 | 5.4058 | 5.7349 | 6.4560 | 7.4045 | 7.2836 | 6.1946 | 5.7204 | 6.2866 | 7.1626 | 5.5171 |
| KANSL3_S511;S515 | KANSL3 | Q9P2N6 | | 773 | GLPANPSPGPAPQ | 6.4745 | 6.3111 | 5.9571 | 6.8464 | 6.3134 | 6.6766 | 7.4734 | 6.5195 | 7.4145 | 7.1610 | 6.0108 | 6.3174 | 5.0038 | 5.6401 | 4.4836 | 5.3965 |
| KANSL3_S536;S540 | KANSL3 | Q9P2N6 | | 559 | KSSQIGSSQLLKR | 7.5023 | 7.2050 | 7.1314 | 7.0513 | 5.7341 | 7.0467 | 7.9069 | 6.7471 | 7.4013 | 7.3175 | 7.6606 | 7.1550 | 3.1414 | 4.4141 | 2.8312 | 3.7539 |
| KANSL3_S559 | KANSL3 | Q9P2N6 | 536;540 | | NA | 6.3698 | 6.1102 | 7.2011 | 6.9139 | 6.2523 | 6.1513 | 6.1264 | 5.7147 | 7.5511 | 6.6940 | 6.4564 | 7.6325 | 5.0865 | 4.9217 | 4.6603 | 6.1580 |
| KANSL3_S773 | KANSL3 | Q9P2N6 | 511;515 | | NA | 6.7281 | 6.7064 | 6.3978 | 6.7149 | 5.6270 | 5.3952 | 5.9733 | 5.3658 | 7.0353 | 7.1355 | 6.8854 | 6.8611 | 6.1039 | 6.1348 | 4.9410 | 5.9944 |
| KAT5_S119;S123 | KAT5 | Q92993-3 | | 123 | PGSRPGSPEREVP | 6.3149 | 6.3719 | 6.5362 | 7.1197 | 6.7132 | 6.2690 | 7.2698 | 6.1327 | 6.3913 | 6.5118 | 6.7017 | 6.4893 | 5.4935 | 5.0332 | 5.1674 | 5.4845 |
| KAT5_S123 | KAT5 | Q92993-3 | 119;123 | | NA | 6.3789 | 6.7982 | 5.5978 | 6.2241 | 5.4337 | 5.3986 | 5.7131 | 5.3395 | 5.7866 | 6.4369 | 5.5213 | 6.5840 | 7.3542 | 7.4886 | 6.5514 | 7.3931 |
| KAT6A_S1089 | KAT6A | Q92794 | | 941 | LPKRRLSEGVEPW | 5.1713 | 5.1193 | 4.7349 | 5.3195 | 6.5523 | 5.3531 | 6.1238 | 6.2052 | 5.7190 | 5.0133 | 5.2906 | 5.4646 | 8.4751 | 8.1651 | 8.9468 | 8.3462 |
| KAT6A_S420 | KAT6A | Q92794 | | 1089 | EYFRRLSSQDVLR | 4.5867 | 4.4311 | 4.5854 | 5.7761 | 7.9188 | 5.0847 | 4.5387 | 5.8406 | 8.1261 | 8.5266 | 7.0128 | 7.2703 | 7.0532 | 7.8218 | 5.7399 | 5.6873 |
| KAT6A_S448 | KAT6A | Q92794 | | 473 | EERLFGSQEIMTE | 7.0993 | 7.6154 | 6.3918 | 7.5945 | 5.5577 | 6.1777 | 5.8880 | 5.8657 | 6.7468 | 6.6840 | 7.1041 | 6.5559 | 4.7852 | 5.5210 | 5.2426 | 5.1703 |
| KAT6A_S473 | KAT6A | Q92794 | | 420 | TKFFTPSPDGRKA | 7.3813 | 7.4748 | 6.0849 | 7.1311 | 6.5641 | 6.4694 | 6.1474 | 6.7315 | 7.6856 | 6.6093 | 6.5313 | 6.8702 | 4.1234 | 5.0754 | 4.4465 | 4.6738 |
| KAT6A_S941 | KAT6A | Q92794 | | 448 | RGNRKSSTSDWPT | 5.1325 | 5.9393 | 5.2227 | 6.5965 | 6.6416 | 6.0468 | 6.3720 | 5.3964 | 5.6151 | 6.4075 | 6.4248 | 5.7801 | 6.5869 | 6.7663 | 7.6652 | 7.4062 |
| KAT6B_S1301 | KAT6B | Q8WYB5 | | 998 | ISNAAVSEEEREA | 7.1406 | 7.0683 | 6.2518 | 6.6170 | 6.4544 | 6.0242 | 6.5745 | 5.7510 | 5.5613 | 6.0723 | 5.8514 | 5.9108 | 6.1911 | 5.9308 | 6.5620 | 6.0384 |
| KAT6B_S358 | KAT6B | Q8WYB5 | | 1275 | FKLNLYTPPETPM | 4.9816 | 6.0725 | 5.2262 | 5.1362 | 6.0367 | 5.7113 | 6.7418 | 6.3959 | 5.7696 | 5.5076 | 5.4339 | 5.2313 | 7.9698 | 7.9053 | 7.8613 | 8.0189 |
| KAT6B_S371 | KAT6B | Q8WYB5 | | 647 | PQMGTPSPGKGSL | 6.7527 | 5.8634 | 6.3690 | 6.8508 | 6.7037 | 6.3900 | 7.0002 | 6.0742 | 6.0230 | 6.0255 | 5.7003 | 6.1822 | 5.9205 | 6.0255 | 5.9859 | 6.1331 |
| KAT6B_S445 | KAT6B | Q8WYB5 | | 445 | TKFFTPSPDGRRS | 7.4248 | 7.5245 | 5.9531 | 6.9536 | 6.4883 | 6.2745 | 5.9641 | 6.8307 | 7.8854 | 6.5334 | 6.5577 | 7.0872 | 4.0171 | 5.1264 | 4.5910 | 4.7882 |
| KAT6B_S647 | KAT6B | Q8WYB5 | | 358 | RLLSVTSDEGSMN | 5.6576 | 6.1449 | 6.6647 | 6.4565 | 5.9560 | 6.5119 | 5.8609 | 6.0955 | 7.1303 | 7.0088 | 6.0522 | 6.3458 | 4.9801 | 6.4204 | 6.9426 | 5.7719 |
| KAT6B_S998 | KAT6B | Q8WYB5 | | 1301 | TSEGKTSPSPIRI | 6.2035 | 6.5098 | 5.7491 | 6.7316 | 5.8190 | 5.5641 | 6.3610 | 5.7180 | 6.0626 | 6.6568 | 5.9832 | 6.1626 | 6.4718 | 6.4923 | 6.7179 | 6.7967 |
| KAT6B_T1275 | KAT6B | Q8WYB5-3 | | 371 | AFTGRGSPDTEIK | 6.0313 | 6.6000 | 6.1145 | 6.4778 | 5.7903 | 5.4811 | 5.8065 | 6.1753 | 6.2638 | 6.1643 | 6.3175 | 6.1749 | 6.4373 | 6.3519 | 7.0833 | 6.7304 |

| | | | | | | | | | | | | | | | | | | | | | |
|---|---|---|---|---|---|---|---|---|---|---|---|---|---|---|---|---|---|---|---|---|---|
| KAT7_S102 | KAT7 | O95251 | | 88 | QQPTPVTPKKYPL | 6.2584 | 6.2494 | 6.0567 | 6.4613 | 6.0970 | 6.1170 | 6.2585 | 5.9535 | 6.7266 | 6.3088 | 6.1638 | 6.2528 | 6.4993 | 6.3135 | 5.9813 | 6.3022 |
| KAT7_S162 | KAT7 | O95251 | | 57 | QSSQDSSPVRNLQ | 7.4435 | 7.6900 | 6.7128 | 7.2580 | 6.1553 | 5.9751 | 5.9833 | 5.7907 | 7.4328 | 7.6276 | 7.1738 | 6.7942 | 4.2732 | 5.0600 | 4.3358 | 4.2937 |
| KAT7_S164 | KAT7 | O95251 | | 178 | RRRFHESYNFNMK | 6.3238 | 6.4336 | 7.1570 | 7.9773 | 6.6310 | 6.5761 | 7.3964 | 6.7245 | 7.0795 | 6.9806 | 8.3340 | 7.3088 | 3.2225 | 5.0609 | 2.9781 | 3.8160 |
| KAT7_S178 | KAT7 | O95251 | | 506 | VEEKVGSPERPLS | 6.8002 | 7.1435 | 7.2311 | 6.5834 | 6.9102 | 5.9375 | 7.1023 | 5.8801 | 7.1682 | 7.8375 | 7.1971 | 6.7742 | 4.5721 | 4.6189 | 3.8017 | 4.4421 |
| KAT7_S506 | KAT7 | O95251 | | 162 | DMSLKDSGSDLSH | 6.2688 | 6.4850 | 6.8527 | 7.5279 | 6.9254 | 7.3096 | 7.0208 | 6.9960 | 6.0343 | 6.1639 | 6.8932 | 6.8493 | 4.6740 | 5.4265 | 4.4227 | 4.1501 |
| KAT7_S57 | KAT7 | O95251 | | 164 | SLKDSGSDLSHRP | 6.6074 | 6.7297 | 6.3814 | 6.8179 | 5.0178 | 6.4812 | 6.3228 | 5.8586 | 6.9351 | 6.4419 | 6.3029 | 6.2443 | 5.9056 | 6.1490 | 5.5058 | 6.2984 |
| KAT7_T85;T88 | KAT7 | O95251 | | 102 | QTRSSGSETEQVV | 6.3852 | 6.6982 | 6.0573 | 6.8207 | 5.6324 | 5.7634 | 6.1801 | 5.4472 | 6.0732 | 6.7515 | 6.0587 | 6.1625 | 6.6929 | 6.4561 | 6.3651 | 6.4554 |
| KAT7_T88 | KAT7 | O95251 | 85;88 | | NA | 6.6265 | 7.2048 | 6.6265 | 6.4110 | 6.9139 | 6.7545 | 7.5603 | 6.5262 | 6.6185 | 6.3472 | 6.5749 | 6.5323 | 4.8466 | 4.9487 | 4.5259 | 4.9822 |
| KAT8_S348 | KAT8 | Q9H7Z6-2 | | 45 | GRVSPPTPARGEP | 6.5975 | 6.5025 | 6.7738 | 7.3947 | 6.6555 | 6.3606 | 6.7763 | 6.8852 | 6.8742 | 6.5609 | 6.5466 | 6.8806 | 4.1713 | 5.5411 | 4.3580 | 5.1212 |
| KAT8_S42 | KAT8 | Q9H7Z6-2 | | 42 | PSPGRVSPPTPAR | 5.9863 | 5.2034 | 6.4763 | 6.2069 | 5.8457 | 5.2222 | 6.1962 | 6.5349 | 6.4590 | 6.1957 | 6.1732 | 6.0454 | 6.3673 | 6.7162 | 7.0942 | 7.2771 |
| KAT8_T45 | KAT8 | Q9H7Z6-2 | | 348 | LESTVGSPEKPLS | 6.9712 | 7.1864 | 6.8330 | 7.6400 | 7.1118 | 7.7818 | 7.3725 | 6.5098 | 7.3030 | 7.5342 | 8.1301 | 7.0627 | 3.0857 | 3.6732 | 2.7422 | 3.0625 |
| KATNA1_T81 | KATNA1 | O75449 | | 81 | SFKLDSTPLKAAQ | 6.2554 | 6.6843 | 6.4888 | 6.8581 | 6.1480 | 6.7408 | 6.7485 | 6.5406 | 6.9712 | 6.8122 | 6.8567 | 6.8346 | 4.6809 | 5.4775 | 4.7038 | 5.1983 |
| KATNIP_S233 | KATNIP | O60303 | | 777 | PKPLWLSPEKPLA | 5.9615 | 5.7920 | 7.0304 | 6.2838 | 6.4505 | 6.7484 | 6.8305 | 6.3898 | 6.7954 | 6.6284 | 7.3604 | 6.5514 | 5.2761 | 5.5712 | 5.1508 | 5.1796 |
| KATNIP_S758 | KATNIP | O60303 | | 758 | PSWLQPSPTGKDR | 6.2043 | 5.9013 | 6.5435 | 6.5427 | 6.0521 | 6.6342 | 6.3927 | 7.6574 | 7.2053 | 6.4223 | 7.3906 | 7.1646 | 5.2218 | 5.4218 | 4.5796 | 4.6658 |
| KATNIP_S777 | KATNIP | O60303 | | 1164 | AMRRPSTADGEGD | 5.9343 | 5.2339 | 6.5294 | 5.4546 | 6.8173 | 6.0451 | 6.3904 | 6.3109 | 7.8684 | 5.9809 | 7.5125 | 7.4796 | 7.9807 | 5.4531 | 4.4380 | 4.5709 |
| KATNIP_T1164 | KATNIP | O60303 | | 233 | RHDRPPSSGDWTQ | 5.9694 | 6.1376 | 7.0726 | 6.3966 | 5.9270 | 6.4097 | 6.6871 | 6.4497 | 6.6983 | 6.4414 | 6.1952 | 6.3082 | 5.8185 | 5.9663 | 5.5415 | 5.9809 |
| KCNK12_S5 | KCNK12 | Q9HB15 | | 5 | xxMSSRSPRPPPR | 6.2008 | 6.4564 | 5.8213 | 6.6097 | 5.7506 | 5.6399 | 6.1765 | 5.8676 | 5.9715 | 6.1188 | 6.1244 | 6.3898 | 6.5269 | 6.7163 | 6.6016 | 7.0279 |
| KCNMA1_S153 | KCNMA1 | Q12791-7 | | 981 | PGMDRSSPDNSPV | 6.2007 | 6.3299 | 5.9388 | 7.1341 | 5.8427 | 5.9702 | 5.4338 | 6.6181 | 6.3764 | 6.0240 | 6.1182 | 6.4037 | 6.0345 | 6.5444 | 6.7172 | 6.3134 |
| KCNMA1_S768 | KCNMA1 | Q12791-7 | | 768 | EQPSTLSPKKKQR | 5.9461 | 6.3400 | 6.4547 | 6.7001 | 5.6874 | 5.8660 | 6.6538 | 5.8072 | 6.9706 | 6.5972 | 6.5752 | 6.5392 | 5.8925 | 5.6962 | 5.9968 | 6.2768 |
| KCNMA1_S781;S785 | KCNMA1 | A0A1W2PQ93 | | 153 | EAQKSSSALEVHK | 5.4179 | 4.9812 | 5.2815 | 6.1323 | 5.9006 | 5.5451 | 5.3785 | 6.4023 | 6.2932 | 4.7321 | 5.0776 | 5.4249 | 8.0535 | 8.5976 | 8.4350 | 8.3467 |
| KCNMA1_S981 | KCNMA1 | Q12791-7 | 781;785 | | NA | 6.1685 | 5.8888 | 6.5083 | 5.9622 | 5.8500 | 5.8288 | 6.4973 | 5.1829 | 6.7649 | 6.2786 | 6.1497 | 6.1126 | 6.6275 | 6.3595 | 6.9963 | 6.8244 |
| KCNQ5_S476 | KCNQ5 | Q9NR82-6 | | 850 | DLGKSLSVQNLIR | 5.3803 | 5.5028 | 5.0711 | 5.7017 | 4.8657 | 5.5909 | 6.0638 | 5.3992 | 5.1204 | 4.9692 | 5.5349 | 5.9483 | 7.9394 | 8.1616 | 8.9169 | 9.8341 |
| KCNQ5_S485 | KCNQ5 | Q9NR82-6 | | 485 | KVQKSWSFNDRTR | 6.4570 | 6.5968 | 6.1039 | 5.9373 | 5.6395 | 5.9655 | 6.1514 | 6.1020 | 5.9993 | 5.8939 | 6.0410 | 6.0040 | 6.7217 | 6.8109 | 6.7383 | 6.8375 |
| KCNQ5_S696 | KCNQ5 | Q9NR82-6 | | 78 | GGGLRESRRGKQG | 6.9092 | 7.0564 | 7.2652 | 6.2364 | 5.6949 | 6.7648 | 6.1786 | 6.5277 | 6.5498 | 6.8535 | 6.5290 | 6.5828 | 4.7929 | 5.8277 | 4.7741 | 5.4571 |
| KCNQ5_S78 | KCNQ5 | Q9NR82-6 | | 826 | DRSMRKSFDMGGE | 5.8827 | 5.4937 | 6.9766 | 6.2053 | 6.3335 | 6.1428 | 6.2747 | 5.9701 | 6.3825 | 6.9705 | 6.2445 | 6.4760 | 5.9927 | 6.7300 | 5.6595 | 6.2649 |
| KCNQ5_S826 | KCNQ5 | Q9NR82-6 | | 696 | QNSGCLSRSTSAN | 4.7746 | 5.2725 | 4.8782 | 5.0111 | 5.9147 | 5.6190 | 4.9018 | 5.7104 | 5.3754 | 5.4097 | 4.9068 | 5.3582 | 8.4642 | 8.6342 | 9.6657 | 10.1036 |
| KCNQ5_S850 | KCNQ5 | Q9NR82-6 | | 921 | AAFASDSLRTGRS | 6.8240 | 6.7792 | 7.0086 | 6.8735 | 6.5684 | 5.8403 | 6.5287 | 6.4932 | 6.6841 | 5.9688 | 6.2392 | 6.1542 | 5.4602 | 5.8227 | 5.3604 | 5.3946 |
| KCNQ5_S921 | KCNQ5 | Q9NR82-6 | | 476 | DITAEGSPTKVQK | 5.8252 | 5.3622 | 6.1734 | 6.4872 | 5.6092 | 6.2118 | 6.9092 | 5.7997 | 6.2407 | 6.1670 | 5.7458 | 5.9148 | 6.7066 | 6.6799 | 7.4674 | 6.6997 |
| KCTD1_S12 | KCTD1 | Q719H9 | | 12 | ITRSPASPLNNQG | 6.3233 | 5.9687 | 6.3355 | 6.7490 | 6.0639 | 4.8540 | 7.2374 | 7.0327 | 7.6245 | 7.0753 | 5.9443 | 5.7356 | 5.6545 | 7.2739 | 5.2613 | 4.8661 |
| KCTD12_S176 | KCTD12 | Q96CX2 | | 176 | ASAGAPSPTLELA | 6.2268 | 6.1342 | 6.4151 | 5.9083 | 6.0086 | 6.2784 | 6.0741 | 6.5521 | 6.3305 | 5.9515 | 6.2408 | 6.1181 | 6.6041 | 6.8211 | 6.0208 | 6.3152 |
| KCTD15_S31;S35 | KCTD15 | Q96SI1 | | 35 | RLSLTRSPVSPLA | 5.2029 | 5.8030 | 6.3057 | 5.9350 | 5.7905 | 6.1033 | 6.3429 | 6.8036 | 6.7410 | 6.7802 | 6.8046 | 6.8808 | 5.4405 | 6.4827 | 5.9885 | 6.5947 |
| KCTD15_S35 | KCTD15 | Q96SI1 | 31;35 | | NA | 7.1058 | 6.4204 | 6.6667 | 6.4998 | 6.8014 | 6.3748 | 6.5249 | 7.2287 | 6.8503 | 7.3403 | 7.1339 | 7.1418 | 4.2873 | 4.9153 | 4.0280 | 4.6805 |
| KCTD16_T341 | KCTD16 | Q68DU8 | | 341 | RQTNIQTLDRPIK | 7.8435 | 7.3665 | 8.1031 | 6.9660 | 6.2612 | 5.4688 | 7.0013 | 5.7025 | 6.8432 | 5.4091 | 6.7098 | 5.3683 | 5.4966 | 5.7831 | 5.3946 | 4.2824 |
| KCTD2_S33 | KCTD2 | Q14681 | | 33 | PVRGPPSPRPAGP | 7.4315 | 6.2148 | 6.0811 | 5.6784 | 6.7084 | 7.0638 | 7.6067 | 6.5332 | 5.5638 | 5.4466 | 5.9182 | 5.5634 | 6.5773 | 6.3599 | 5.9533 | 5.2995 |
| KCTD20_S374 | KCTD20 | Q7Z5Y7 | | 374 | QCVRSKSLTNLVA | 6.1033 | 6.8244 | 6.5482 | 6.4110 | 5.4712 | 6.2318 | 5.9424 | 6.8358 | 6.1749 | 6.9075 | 6.0290 | 6.9171 | 5.3611 | 6.8131 | 5.1373 | 6.2921 |
| KCTD3_S653 | KCTD3 | Q9Y597 | | 653 | MRPYRESPLLARA | 6.3138 | 5.2055 | 6.4800 | 6.6035 | 6.1418 | 6.8463 | 6.8032 | 6.6080 | 6.9375 | 6.8169 | 6.0742 | 6.6259 | 5.8197 | 5.9681 | 5.0717 | 5.6838 |
| KCTD8_S410 | KCTD8 | Q6ZWB6 | | 410 | PDKRRNSELFQTL | 5.6974 | 6.2532 | 5.8341 | 7.3379 | 6.4285 | 5.1505 | 5.5101 | 5.7271 | 8.7645 | 8.3217 | 6.6919 | 5.9916 | 4.8117 | 5.6558 | 4.9752 | 6.6187 |
| KCTD9_S11 | KCTD9 | Q7L273 | | 11 | TLFLNGSPKNGKV | 4.1877 | 6.8995 | 7.9252 | 5.6537 | 5.5886 | 7.5199 | 7.9502 | 5.7037 | 5.1283 | 9.9765 | 7.4899 | 4.5179 | 7.4498 | 3.5173 | 4.2678 | 6.2240 |
| KDELR3_S209 | KDELR3 | O43731 | | 209 | LKGKKLSLPMPIx | 5.5739 | 4.7458 | 4.3979 | 5.3890 | 6.4201 | 5.2894 | 6.0668 | 5.4072 | 7.2953 | 8.7631 | 6.7571 | 6.1374 | 6.7716 | 5.9453 | 7.9549 | 7.0851 |
| KDM1A_S131 | KDM1A | O60341 | | 166 | PEEENESEPEEPS | 6.0039 | 6.4509 | 6.2593 | 6.2960 | 6.0700 | 5.8074 | 6.0412 | 6.3252 | 6.3412 | 6.2011 | 6.2401 | 6.2193 | 6.3748 | 6.7631 | 6.1977 | 6.4087 |
| KDM1A_S131;S137 | KDM1A | O60341 | | 69 | KEPPRASPPGGLA | 5.8899 | 5.8358 | 6.0350 | 6.5105 | 6.3117 | 6.1901 | 6.1085 | 6.2592 | 6.2850 | 6.1845 | 6.0918 | 5.9420 | 6.2818 | 6.9759 | 6.5893 | 6.5092 |
| KDM1A_S166 | KDM1A | O60341 | | 131 | ESLANLSEDEYYS | 5.6519 | 5.9744 | 4.6445 | 6.5872 | 5.4754 | 8.3384 | 4.8380 | 5.3313 | 6.3463 | 5.8094 | 6.0522 | 5.4451 | 7.9096 | 7.7047 | 6.7257 | 7.1659 |
| KDM1A_S69 | KDM1A | O60341 | | 849 | GVPAQQSPSMxxx | 6.0732 | 6.4977 | 7.0065 | 6.9330 | 6.4460 | 6.0913 | 6.2628 | 5.9707 | 6.9167 | 6.2120 | 6.1022 | 8.0477 | 5.2360 | 5.7875 | 4.7762 | 5.6406 |
| KDM1A_S849 | KDM1A | O60341 | 131;137 | | NA | 5.6260 | 6.2898 | 6.6145 | 6.6625 | 6.1766 | 5.8234 | 6.2987 | 6.3066 | 6.2116 | 6.3372 | 6.2892 | 6.2115 | 6.0185 | 6.7772 | 6.3125 | 6.0441 |
| KDM1B_S17 | KDM1B | Q8NB78 | | 17 | KASFDHSPDSLPL | 6.8095 | 7.6951 | 6.6349 | 7.3405 | 6.6525 | 7.4178 | 6.5548 | 7.2136 | 6.9229 | 6.6855 | 7.1937 | 6.3440 | 3.7504 | 4.6001 | 4.0674 | 4.1171 |
| KDM2A_S558 | KDM2A | Q9Y2K7 | | 558 | VRPAAASPIVSGA | 5.9291 | 6.4312 | 6.0636 | 6.4768 | 6.0734 | 5.9844 | 6.2535 | 5.7602 | 6.0790 | 6.1072 | 6.2038 | 6.3185 | 6.1448 | 6.8264 | 6.4457 | 6.9023 |
| KDM2A_S692 | KDM2A | Q9Y2K7 | | 718 | LTPPPHSPTSMLQ | 6.1313 | 5.9839 | 6.1711 | 7.1645 | 5.9438 | 6.0428 | 6.1153 | 6.2563 | 6.1768 | 6.1816 | 5.8938 | 6.0834 | 6.2783 | 6.7104 | 6.3875 | 6.4792 |
| KDM2A_S718 | KDM2A | Q9Y2K7 | | 692 | KRKMEESDEEAVQ | 6.5854 | 7.1764 | 6.6339 | 7.8408 | 6.0815 | 6.3048 | 6.7182 | 6.4388 | 6.5798 | 5.7225 | 6.1494 | 6.0344 | 5.3776 | 5.5167 | 5.4416 | 5.3980 |
| KDM2A_S718;S731 | KDM2A | Q9Y2K7 | | 550 | KPAPRLTPVRPAA | 7.2496 | 7.1154 | 6.8291 | 8.1353 | 5.8181 | 7.1303 | 8.4484 | 7.0140 | 6.4086 | 7.3778 | 6.5816 | 6.5875 | 4.4464 | 3.6710 | 3.2207 | 3.9662 |
| KDM2A_T550 | KDM2A | Q9Y2K7 | 713;718;731 | | NA | 6.2084 | 6.5540 | 6.6755 | 6.9797 | 6.4002 | 6.5733 | 6.4378 | 6.6745 | 6.6857 | 6.5291 | 6.5621 | 6.7858 | 5.1473 | 5.3105 | 5.3198 | 5.1562 |
| KDM2A_T550;S558 | KDM2A | Q9Y2K7 | 718;731 | | NA | 6.4458 | 6.5605 | 6.6876 | 6.9330 | 6.3025 | 6.5562 | 6.5930 | 6.1534 | 6.8475 | 6.5904 | 6.3512 | 6.7855 | 5.1264 | 5.9206 | 4.8660 | 5.2804 |
| KDM2A_T713;S718 | KDM2A | Q9Y2K7 | 713;731 | | NA | 6.4652 | 6.3964 | 6.9615 | 7.4078 | 6.1205 | 6.3101 | 6.8036 | 6.5502 | 6.6529 | 6.4945 | 6.4641 | 6.3843 | 5.1208 | 5.3858 | 5.2408 | 5.2415 |
| KDM2A_T713;S718;S731 | KDM2A | Q9Y2K7 | 713;718 | | NA | 5.8546 | 6.1981 | 6.3797 | 6.2977 | 6.2340 | 6.1681 | 5.9023 | 6.2948 | 5.8895 | 5.8872 | 5.7657 | 5.6782 | 6.5752 | 7.0443 | 6.7800 | 7.0504 |
| KDM2A_T713;S731 | KDM2A | Q9Y2K7 | 550;558 | | NA | 6.7695 | 7.1432 | 7.4139 | 7.3178 | 6.5026 | 7.5408 | 7.6368 | 6.7929 | 7.3433 | 7.8372 | 7.3865 | 7.0698 | 3.0974 | 3.8874 | 2.8372 | 3.4236 |
| KDM2A_T713;T720;S731 | KDM2A | Q9Y2K7 | 713;720;731 | | NA | 6.1690 | 6.7613 | 6.8156 | 7.3807 | 7.0174 | 6.5760 | 5.8494 | 6.1288 | 6.3222 | 6.6533 | 6.4389 | 7.0437 | 5.3227 | 5.6284 | 4.8849 | 5.0078 |
| KDM2B_S1018 | KDM2B | Q8NHM5 | | 483 | SEESVKSTTLAVD | 6.2918 | 6.4997 | 6.2529 | 6.7155 | 5.7657 | 5.6301 | 5.9868 | 5.6693 | 6.4941 | 6.3129 | 5.6700 | 5.8973 | 6.3396 | 6.5401 | 6.7690 | 7.1655 |
| KDM2B_S1031 | KDM2B | Q8NHM5 | | 1031 | RPPPSVSPPKCIQ | 6.5869 | 6.3685 | 6.0075 | 6.8484 | 6.0684 | 5.6300 | 6.4536 | 6.0949 | 6.2776 | 5.8859 | 6.2065 | 6.0075 | 6.2585 | 6.5641 | 6.6435 | 6.0982 |
| KDM2B_S474 | KDM2B | Q8NHM5 | | 1018 | LGPSLRSPPRVIS | 8.0897 | 7.9585 | 7.2228 | 8.2161 | 6.9635 | 7.2179 | 6.9385 | 6.3100 | 7.0000 | 6.9646 | 6.5230 | 6.7980 | 3.2974 | 3.8545 | 3.3127 | 3.3328 |
| KDM2B_S483 | KDM2B | Q8NHM5 | | 493 | AVDYPKTPTGSPA | 6.3062 | 6.5496 | 5.7458 | 6.4601 | 5.5631 | 6.2510 | 6.3853 | 5.9690 | 5.9868 | 6.4436 | 6.1443 | 5.8427 | 5.6379 | 7.0861 | 6.6661 | 6.9623 |
| KDM2B_T444 | KDM2B | Q8NHM5 | | 474 | FLKRTLSNESEES | 6.6810 | 6.7654 | 6.2046 | 7.0654 | 5.7976 | 5.6644 | 6.0625 | 5.4681 | 7.3691 | 6.4339 | 6.2478 | 6.8450 | 5.1774 | 5.8489 | 6.0358 | 6.3330 |
| KDM2B_T493 | KDM2B | Q8NHM5 | | 444 | PPTDGSTSPTSTP | 5.5775 | 6.1938 | 5.9571 | 6.6265 | 6.0534 | 6.2667 | 6.0289 | 6.6805 | 6.7683 | 6.4567 | 5.6087 | 6.0043 | 6.0829 | 6.7417 | 6.9061 | 6.0469 |
| KDM2B_T493;S497 | KDM2B | Q8NHM5 | 493;497 | | NA | 6.5547 | 6.6924 | 5.7010 | 6.0660 | 5.7620 | 5.7445 | 5.9720 | 6.1142 | 5.9316 | 6.0383 | 5.8332 | 5.6346 | 6.6985 | 6.9499 | 7.3649 | 6.9420 |
| KDM3A_S264 | KDM3A | Q9Y4C1 | | 308 | LGCTAATPPSKDP | 7.1907 | 7.6057 | 7.0053 | 7.6779 | 6.5887 | 7.1304 | 6.8403 | 6.8259 | 7.4983 | 7.0830 | 7.3074 | 7.2409 | 3.5508 | 4.1599 | 2.9530 | 3.3416 |
| KDM3A_S325 | KDM3A | Q9Y4C1 | | 325 | TPQAANSPPNLGA | 6.5047 | 6.9619 | 6.4542 | 7.2595 | 6.2522 | 6.7833 | 6.9925 | 6.4190 | 6.5335 | 7.0110 | 6.5890 | 7.1679 | 4.4350 | 4.9315 | 4.4080 | 5.2967 |
| KDM3A_S892 | KDM3A | Q9Y4C1 | | 892 | RNLLNSSTGKTEN | 6.6640 | 6.9862 | 7.2759 | 7.1157 | 7.2011 | 7.5205 | 8.7625 | 7.6605 | 6.9556 | 7.4102 | 7.1733 | 7.6269 | 2.8054 | 4.2508 | 1.8118 | 2.7795 |
| KDM3A_T308 | KDM3A | Q9Y4C1 | | 264 | GAVKRKSSENNGT | 4.5913 | 5.0335 | 4.9211 | 4.9121 | 5.2213 | 6.0944 | 5.2764 | 5.1477 | 4.9026 | 4.6320 | 5.1902 | 5.5568 | 8.9958 | 9.1993 | 10.5567 | 9.7689 |
| KDM3B_S1267 | KDM3B | Q7LBC6 | | 727 | GNGRSSSPTSSLT | 5.8049 | 6.9257 | 6.4550 | 6.1391 | 5.8852 | 6.1720 | 6.0320 | 6.1114 | 6.7424 | 7.0755 | 7.0263 | 6.8759 | 5.4872 | 6.2507 | 5.4020 | 5.6147 |
| KDM3B_S727 | KDM3B | Q7LBC6 | | 798 | EAVKRFSLDERSL | 4.4614 | 4.7094 | 4.9891 | 4.8365 | 6.1244 | 5.2602 | 5.3657 | 5.4397 | 5.0761 | 5.1366 | 5.1099 | 4.8825 | 9.5458 | 9.0621 | 10.4061 | 9.5946 |
| KDM3B_S727;S743 | KDM3B | Q7LBC6 | | 1267 | NSTAKVSPLTPKL | 7.1929 | 7.5137 | 6.7615 | 7.1548 | 6.4662 | 7.1464 | 7.3384 | 6.8870 | 7.3026 | 7.3565 | 7.4480 | 6.6962 | 3.3049 | 4.1845 | 3.2901 | 3.9564 |
| KDM3B_S727;S744 | KDM3B | Q7LBC6 | | 743 | EMPTLSSSPTEER | 6.1011 | 6.5797 | 7.3279 | 6.9460 | 5.9944 | 6.3677 | 5.9149 | 6.0676 | 7.5337 | 6.8712 | 6.8564 | 6.7208 | 4.3296 | 5.6101 | 5.1916 | 5.5875 |
| KDM3B_S743 | KDM3B | Q7LBC6 | | 744 | MPTLSSSPTEERP | 6.6181 | 7.0088 | 6.8135 | 6.6762 | 5.8288 | 6.1808 | 6.2284 | 6.3468 | 6.6930 | 7.2765 | 7.4896 | 7.0685 | 4.7855 | 4.9840 | 5.1029 | 4.8986 |
| KDM3B_S744 | KDM3B | Q7LBC6 | 727;744 | | NA | 5.0434 | 5.5475 | 5.6543 | 5.6911 | 5.8017 | 6.2482 | 5.6888 | 5.7410 | 5.6715 | 5.8395 | 5.9567 | 5.9460 | 7.4994 | 8.1444 | 7.9981 | 7.5283 |
| KDM3B_S773;S778 | KDM3B | Q7LBC6 | 727;743 | | NA | 5.1553 | 5.2538 | 5.6369 | 5.4111 | 5.4314 | 6.2892 | 5.8330 | 5.3705 | 6.2763 | 6.1925 | 5.5123 | 5.2589 | 7.6793 | 8.3187 | 8.6038 | 7.7771 |
| KDM3B_S798 | KDM3B | Q7LBC6 | 773;778 | | NA | 7.0377 | 7.9816 | 6.7892 | 7.3776 | 6.3498 | 7.6138 | 7.9328 | 8.2884 | 7.4122 | 7.0006 | 6.7914 | 6.7167 | 2.7568 | 4.0353 | 2.5437 | 3.3725 |
| KDM4A_S410 | KDM4A | O75164 | | 616 | VLQECVSDDETSE | 5.9585 | 5.9670 | 6.4303 | 6.6182 | 5.5942 | 6.0720 | 5.7307 | 6.5231 | 7.1567 | 6.7003 | 6.4203 | 6.6803 | 5.6325 | 6.1848 | 6.2554 | 6.0759 |
| KDM4A_S616 | KDM4A | O75164 | | 410 | EIPQEVSQSELFP | 6.1044 | 7.1313 | 7.1049 | 6.8110 | 5.8202 | 5.9723 | 6.5249 | 6.3762 | 6.9566 | 6.3329 | 6.8098 | 7.7665 | 5.1995 | 6.0122 | 4.9289 | 5.1483 |
| KDM4B_S622 | KDM4B | O94953 | | 622 | GRPPTRSPLSVVK | 6.3608 | 6.7462 | 6.2395 | 6.8459 | 6.2887 | 6.4327 | 6.3720 | 6.6204 | 6.6715 | 6.8640 | 6.5695 | 6.6106 | 4.9550 | 5.8973 | 5.1744 | 5.3517 |
| KDM5A_S1111 | KDM5A | P29375 | | 1666 | KKQGPVSPGPAPP | 6.2888 | 6.3111 | 6.0378 | 6.6157 | 6.2470 | 5.9644 | 6.4133 | 6.1595 | 6.3847 | 6.2199 | 6.0409 | 6.0741 | 6.0367 | 6.5059 | 6.4701 | 6.2298 |
| KDM5A_S1331 | KDM5A | P29375 | | 1331 | VSSVSSSPRQTMD | 7.0435 | 6.9688 | 6.2300 | 7.0805 | 6.1727 | 5.8421 | 6.9419 | 6.5591 | 6.4928 | 6.3462 | 6.3437 | 6.4303 | 5.1029 | 5.3185 | 5.4430 | 5.6840 |
| KDM5A_S1365 | KDM5A | P29375 | | 1488 | HIMEDDSMEEKPL | 5.9251 | 5.7784 | 5.8961 | 7.2540 | 5.9297 | 6.1984 | 6.5913 | 5.4719 | 6.0077 | 6.3547 | 6.2481 | 6.3692 | 5.9855 | 6.5451 | 6.8466 | 6.5982 |
| KDM5A_S1365;S1368 | KDM5A | P29375 | | 1111 | LDLEPLSDLEEGL | 5.9232 | 6.2921 | 6.3179 | 7.3051 | 6.0055 | 6.1257 | 6.1636 | 6.3762 | 6.5463 | 5.5202 | 5.7541 | 6.6803 | 6.8051 | 6.2865 | 6.0545 | 5.8437 |
| KDM5A_S1367 | KDM5A | P29375 | | 1365 | DTASVKSSSSLEP | 5.3445 | 6.1213 | 5.6244 | 7.0381 | 6.5158 | 5.4246 | 5.5159 | 6.5473 | 6.8201 | 5.5260 | 5.3273 | 4.9338 | 6.4874 | 7.1395 | 7.8829 | 7.7510 |
| KDM5A_S140 | KDM5A | P29375 | | 204 | STDTQTSPEPGTR | 6.4366 | 5.7187 | 6.5166 | 6.0382 | 5.9689 | 6.4379 | 6.5320 | 6.4793 | 6.7335 | 6.8649 | 6.4846 | 6.8622 | 5.5445 | 5.7100 | 5.4765 | 6.1957 |
| KDM5A_S1488 | KDM5A | P29375 | | 1367 | ASVKSSSSLEPNL | 8.1330 | 8.3563 | 5.5376 | 6.4506 | 6.1914 | 5.5057 | 6.8253 | 6.2034 | 8.2805 | 7.3875 | 6.4346 | 5.8605 | 5.2266 | 5.1429 | 3.2971 | 5.1668 |
| KDM5A_S1666 | KDM5A | P29375 | | 140 | TKEKKWSKVGSRL | 1.3899 | 1.7027 | 1.3494 | 1.6261 | 1.5221 | 1.4435 | 1.4959 | 1.5455 | 1.4099 | 1.9839 | 1.4346 | 1.6421 | 19.0168 | 18.8317 | 23.7344 | 19.8714 |
| KDM5A_S204 | KDM5A | P29375 | 1365;1368 | | NA | 5.3600 | 6.6159 | 6.5524 | 5.3875 | 5.2230 | 5.3006 | 6.8586 | 5.8734 | 5.6699 | 6.5016 | 5.9479 | 6.4967 | 7.0571 | 7.1512 | 7.0149 | 6.9892 |
| KDM5C_S1359 | KDM5C | P41229 | | 287 | VKVESTSPKTFLE | 5.7660 | 6.5831 | 6.4247 | 6.0763 | 5.8937 | 6.2917 | 6.1923 | 6.0111 | 6.1419 | 6.5250 | 6.5007 | 6.5047 | 6.0030 | 6.5710 | 6.2514 | 6.2634 |
| KDM5C_S287 | KDM5C | P41229 | | 301 | KEELSHSPEPCTK | 6.5013 | 6.9797 | 5.9695 | 5.9183 | 6.5304 | 6.3334 | 6.1398 | 6.1544 | 6.2379 | 6.0939 | 6.3231 | 6.2464 | 5.9211 | 6.5577 | 5.9460 | 6.1471 |
| KDM5C_S301 | KDM5C | P41229 | | 897 | LASLPSSPGLLQS | 5.2010 | 6.4400 | 5.7696 | 6.6726 | 6.4251 | 6.8612 | 6.6577 | 6.2250 | 6.2479 | 6.8182 | 6.6024 | 6.7963 | 6.0583 | 5.9425 | 5.6117 | 5.6705 |
| KDM5C_S317 | KDM5C | P41229 | | 317 | RLRRNHSNAQFIE | 6.0087 | 7.5710 | 5.9541 | 4.7119 | 6.1180 | 5.6963 | 5.0728 | 7.5965 | 6.0200 | 6.0115 | 5.9072 | 5.9763 | 6.9029 | 6.5817 | 6.9462 | 6.9248 |
| KDM5C_S897 | KDM5C | P41229 | | 1359 | NGDSVTSPEKVAP | 6.1007 | 6.8896 | 7.0132 | 6.7158 | 5.9491 | 6.4590 | 6.5642 | 6.6012 | 7.3859 | 6.7827 | 6.8114 | 5.9580 | 4.9619 | 5.6738 | 5.0808 | 5.0527 |
| KDM6A_S769 | KDM6A | O15550 | | 769 | SHGDSKSPGLLSS | 5.9403 | 6.0036 | 5.9394 | 6.2624 | 6.1294 | 5.9288 | 6.3591 | 6.0669 | 5.8663 | 6.0092 | 5.9010 | 5.8623 | 7.1311 | 6.9672 | 6.9442 | 6.6886 |
| KDM6A_S829 | KDM6A | O15550 | | 829 | ISTATPSPKSTEQ | 6.3371 | 6.3255 | 6.0330 | 6.3017 | 6.4244 | 6.3039 | 6.4767 | 5.8211 | 6.4854 | 6.2137 | 6.4041 | 6.8408 | 5.9288 | 6.0657 | 5.8896 | 6.1484 |
| KHDC4_S571;S572 | KHDC4 | Q7Z7F0 | 571;572 | | NA | 6.2658 | 6.3759 | 7.1629 | 6.7150 | 5.6961 | 6.3889 | 6.2872 | 6.5233 | 5.8077 | 5.7984 | 6.6351 | 6.0292 | 5.7726 | 6.4737 | 5.8822 | 6.1861 |
| KHDRBS1_S20 | KHDRBS1 | Q07666 | | 58 | RGGARASPATQPP | 6.5990 | 6.6214 | 6.5536 | 6.7741 | 6.1471 | 6.0302 | 6.6433 | 6.4859 | 6.5093 | 6.5952 | 7.0296 | 6.5381 | 5.1150 | 5.7016 | 5.3472 | 5.3093 |
| KHDRBS1_S58 | KHDRBS1 | Q07666 | | 20 | SSGRSGSMDPSGA | 5.9805 | 5.9762 | 5.2834 | 6.2442 | 6.2280 | 5.6110 | 6.4282 | 6.6232 | 9.2195 | 6.6157 | 6.8766 | 6.0571 | 5.8585 | 6.4574 | 5.1377 | 5.4030 |
| KHNYN_S10 | KHNYN | O15037 | | 10 | WGARPASPDRFAV | 6.1027 | 5.5644 | 6.8874 | 6.0547 | 6.3429 | 6.9322 | 6.0543 | 4.8085 | 5.9335 | 7.0843 | 7.3696 | 7.3463 | 6.3990 | 6.3025 | 5.0662 | 5.7515 |
| KHNYN_S353;S359 | KHNYN | O15037 | 353;359 | | NA | 6.7574 | 6.7162 | 6.4464 | 6.5350 | 6.4208 | 6.7486 | 5.7178 | 6.4295 | 6.5233 | 6.6557 | 6.9900 | 6.8522 | 5.4386 | 5.6849 | 4.9589 | 5.1245 |
| KHSRP_S125 | KHSRP | Q92945 | | 181 | GCKVQISPDSGGL | 6.4257 | 6.7004 | 5.9489 | 5.8431 | 6.6160 | 6.2772 | 5.6748 | 6.8559 | 6.0747 | 6.2941 | 6.3024 | 6.2326 | 6.1551 | 6.2469 | 6.1663 | 6.1859 |
| KHSRP_S181 | KHSRP | Q92945 | | 480 | LFIIRGSPQQIDH | 6.2461 | 6.0501 | 5.9492 | 5.3017 | 6.7406 | 5.9362 | 5.6914 | 6.7260 | 6.0094 | 6.6005 | 6.3422 | 6.5030 | 7.2017 | 6.5716 | 6.0384 | 6.0919 |
| KHSRP_S193 | KHSRP | Q92945 | | 100 | TTVNNSTPDFGFG | 6.2173 | 6.1163 | 6.5519 | 6.4579 | 6.3367 | 6.0747 | 6.2916 | 6.9030 | 6.7573 | 6.7388 | 6.5753 | 7.3687 | 5.4107 | 5.6747 | 4.9825 | 5.5426 |
| KHSRP_S480 | KHSRP | Q92945 | | 54 | GGPGGGSAGGPSQ | 5.2260 | 6.3896 | 5.1164 | 4.4660 | 6.0762 | 5.7197 | 4.8343 | 7.4356 | 5.8568 | 7.6510 | 7.0869 | 6.8401 | 7.9762 | 7.6275 | 6.7813 | 4.9166 |
| KHSRP_S54 | KHSRP | Q92945 | | 125 | ESKKLASQGDSIS | 6.3198 | 5.9603 | 5.5164 | 6.0410 | 6.8191 | 5.9286 | 6.1144 | 6.1036 | 6.4135 | 7.1146 | 6.3875 | 6.2132 | 6.2636 | 6.3962 | 6.3933 | 6.0150 |
| KHSRP_T100 | KHSRP | Q92945 | | 193 | LPERSVSLTGAPE | 7.0265 | 6.1498 | 6.9231 | 6.1365 | 7.3423 | 5.9449 | 5.9771 | 6.3747 | 7.0312 | 6.7206 | 7.2793 | 5.6356 | 7.6340 | 7.1063 | 3.6267 | 3.0912 |
| KIAA0100_S2094 | KIAA0100 | Q14667 | | 2094 | RRSFRKSPEHPVD | 6.4597 | 6.5448 | 6.5984 | 6.6916 | 5.9078 | 6.2202 | 6.2615 | 6.1596 | 6.7959 | 6.8380 | 6.0989 | 6.4652 | 5.5529 | 5.8742 | 5.6352 | 5.8961 |
| KIAA0232_S1294 | KIAA0232 | Q92628 | | 1294 | WEKALYSPLFPAS | 5.5128 | 5.4501 | 6.0841 | 5.2672 | 6.3261 | 6.3563 | 6.1503 | 6.2973 | 5.5359 | 6.0761 | 5.8276 | 6.0387 | 6.8181 | 8.2753 | 7.5060 | 6.4781 |
| KIAA0753_S287 | KIAA0753 | Q2KHM9 | | 287 | EELDKLSPHKIKH | 5.8811 | 6.0103 | 6.3717 | 6.4696 | 6.1739 | 6.4952 | 6.8439 | 6.3782 | 6.7103 | 6.6985 | 6.4949 | 7.0035 | 5.2017 | 6.0181 | 5.5519 | 5.6971 |
| KIAA0753_S700 | KIAA0753 | Q2KHM9 | | 700 | KAQRVNSTTEANI | 5.4485 | 5.4899 | 5.0126 | 5.8698 | 6.0651 | 5.3417 | 6.0277 | 5.5280 | 5.9432 | 5.1705 | 4.5349 | 5.1488 | 9.1046 | 8.5936 | 8.2431 | 8.4781 |
| KIAA0930_S309 | KIAA0930 | Q6ICG6-2 | | 298 | SFSTPPTPERNNR | 5.9629 | 6.1657 | 6.6081 | 6.3548 | 6.2088 | 6.4142 | 6.0912 | 6.5703 | 6.6423 | 6.5286 | 6.8374 | 7.1386 | 5.5535 | 5.6338 | 5.5452 | 5.7447 |
| KIAA0930_S311 | KIAA0930 | Q6ICG6-2 | | 367 | GRSLVGSWLKLNR | 5.9401 | 5.7796 | 6.2202 | 5.5421 | 6.1572 | 6.2094 | 6.2301 | 6.3426 | 6.0489 | 5.9286 | 5.9948 | 5.8497 | 7.2231 | 6.9530 | 6.8087 | 6.7721 |
| KIAA0930_S329 | KIAA0930 | Q6ICG6-2 | | 309 | NRPAFFSPSLKRK | 6.6387 | 6.1196 | 7.0212 | 6.0985 | 5.8597 | 6.2511 | 7.0643 | 6.4257 | 6.6156 | 6.3702 | 6.9656 | 7.0247 | 5.2337 | 5.0364 | 5.2771 | 5.9979 |
| KIAA0930_S367 | KIAA0930 | Q6ICG6-2 | | 329 | EMKKSHSANDSEE | 5.7904 | 5.2336 | 6.8369 | 5.4496 | 6.6026 | 6.2000 | 6.1397 | 6.0219 | 5.6754 | 5.6558 | 6.0962 | 5.2714 | 7.1735 | 8.6997 | 7.3629 | 5.7904 |
| KIAA0930_T298 | KIAA0930 | Q6ICG6-2 | | 311 | PAFFSPSLKRKVP | 7.4394 | 6.5824 | 9.0567 | 7.1545 | 5.5580 | 5.3266 | 5.5948 | 5.7335 | 8.0771 | 8.6291 | 9.0285 | 10.7689 | 2.2083 | 3.6409 | 2.2832 | 2.9180 |
| KIAA1107_S254 | KIAA1107 | Q9UPP5 | | 694 | CFSGQLSEKNSPK | 5.5906 | 6.2333 | 7.4392 | 6.1549 | 5.7383 | 7.8165 | 6.9938 | 5.0194 | 7.5672 | 6.7272 | 5.8122 | 5.5329 | 5.4706 | 6.5091 | 5.7152 | 5.6795 |
| KIAA1107_S694 | KIAA1107 | Q9UPP5 | | 254 | KLSFCDSPGQMMK | 7.1514 | 6.8686 | 6.8809 | 8.1998 | 5.5281 | 5.8048 | 6.4074 | 6.6441 | 7.2652 | 7.1576 | 6.6687 | 8.0215 | 4.1476 | 4.8332 | 4.0338 | 4.3874 |
| KIAA1109_S1305 | KIAA1109 | Q2LD37 | | 1323 | RLGSTKSLTAAFY | 6.9216 | 6.8120 | 6.3542 | 7.0603 | 6.8168 | 5.8536 | 5.9879 | 5.6269 | 6.2692 | 5.2797 | 5.7013 | 5.5365 | 6.5060 | 6.9053 | 6.4329 | 5.9358 |
| KIAA1109_S1323 | KIAA1109 | Q2LD37 | | 3653 | IRVDAASPGPRVT | 6.6253 | 6.3845 | 6.4137 | 7.1287 | 6.2990 | 5.8576 | 6.6258 | 6.3058 | 6.7061 | 5.7903 | 6.2625 | 6.0843 | 5.7987 | 6.3152 | 5.4370 | 5.9655 |
| KIAA1109_S1529 | KIAA1109 | Q2LD37 | | 2287 | GMPVKDSPRKESL | 6.6917 | 6.6758 | 6.1166 | 6.9861 | 6.5434 | 6.8635 | 6.8816 | 5.6323 | 6.7437 | 7.0463 | 7.0063 | 6.7015 | 5.3013 | 5.4222 | 4.7623 | 4.6255 |
| KIAA1109_S1682 | KIAA1109 | Q2LD37 | | 3996 | LETIPITPVERNY | 5.8784 | 5.9749 | 7.0308 | 6.4359 | 5.4679 | 5.8838 | 6.3657 | 6.0638 | 6.3415 | 6.1549 | 6.4721 | 5.8498 | 6.9034 | 6.6005 | 6.0046 | 6.5720 |
| KIAA1109_S2227 | KIAA1109 | Q2LD37 | | 3196 | RLQTKASPGSSSY | 7.3217 | 5.3130 | 7.5224 | 6.3607 | 5.9223 | 5.7439 | 6.2245 | 7.0889 | 7.5199 | 5.6622 | 6.8808 | 4.8498 | 6.4325 | 6.3359 | 5.5210 | 5.3006 |
| KIAA1109_S2287 | KIAA1109 | Q2LD37 | | 1305 | GQSPLRSPLKRQA | 6.0959 | 6.4637 | 6.3582 | 6.4701 | 6.0740 | 7.0992 | 6.8512 | 6.0852 | 6.1799 | 6.4305 | 6.3573 | 5.7924 | 5.4199 | 6.0968 | 6.3005 | 5.9254 |
| KIAA1109_S2601;S2603 | KIAA1109 | Q2LD37 | | 1682 | FSENLSSKQDIRG | 6.2879 | 6.0018 | 5.8385 | 5.6831 | 5.9164 | 5.3738 | 6.0526 | 5.8165 | 5.8623 | 6.0374 | 5.4206 | 5.7397 | 7.5109 | 7.5190 | 7.4488 | 7.4908 |
| KIAA1109_S2603 | KIAA1109 | Q2LD37 | | 1529 | KRTSKSSLHRPLD | 5.9573 | 7.3197 | 5.8137 | 5.5254 | 6.7311 | 6.4724 | 5.8232 | 6.6599 | 6.4581 | 6.4724 | 6.9281 | 5.2073 | 6.1009 | 6.3762 | 6.3252 | 5.8292 |
| KIAA1109_S3196 | KIAA1109 | Q2LD37 | | 2603 | YTAGSASPTPTFK | 6.0670 | 5.9630 | 6.1858 | 6.3625 | 6.3916 | 5.8701 | 6.3222 | 6.0920 | 6.9405 | 6.6869 | 6.1975 | 6.4557 | 5.9886 | 6.3070 | 6.2010 | 5.9685 |
| KIAA1109_S3653 | KIAA1109 | Q2LD37 | | 2227 | NRPPPGSSGPVTG | 6.6425 | 6.8974 | 6.8322 | 7.3242 | 6.2990 | 6.6536 | 6.4858 | 6.8407 | 6.9233 | 6.8881 | 7.3645 | 7.4633 | 4.1498 | 4.4959 | 3.9527 | 4.7871 |
| KIAA1109_T3996 | KIAA1109 | Q2LD37 | 2601;2603 | | NA | 6.3177 | 6.3057 | 5.5472 | 6.4150 | 6.2972 | 6.0958 | 6.1589 | 5.9849 | 7.5338 | 6.0649 | 7.1135 | 6.2698 | 5.3829 | 6.1355 | 6.3872 | 5.9900 |
| KIAA1143_S115 | KIAA1143 | Q96AT1 | | 115 | YSGLTASSKKKKP | 8.2305 | 8.0247 | 7.3309 | 7.2427 | 5.9850 | 7.4188 | 7.2193 | 7.0512 | 7.9956 | 7.4650 | 8.0313 | 8.2369 | 2.0840 | 3.0527 | 1.8341 | 2.7974 |
| KIAA1143_S50 | KIAA1143 | Q96AT1 | | 50 | DEDGDHSDKEDEQ | 6.0855 | 5.6908 | 6.9915 | 5.8854 | 5.7186 | 6.3733 | 5.7875 | 6.2555 | 6.1256 | 6.5256 | 6.5356 | 7.2233 | 6.1004 | 6.4040 | 6.0949 | 6.2026 |
| KIAA1191_S183 | KIAA1191 | Q96A73 | | 183 | STTPHSSPKQRPR | 6.1401 | 5.8812 | 4.5959 | 5.9797 | 6.4735 | 6.0389 | 5.2789 | 6.6959 | 5.2278 | 5.3481 | 5.3569 | 5.5443 | 8.3798 | 8.3067 | 7.9371 | 6.8152 |
| KIAA1217_S1806 | KIAA1217 | Q5T5P2 | | 361 | PVSRPISPSPSAI | 5.9698 | 5.6733 | 5.2475 | 5.9502 | 6.2058 | 5.8545 | 5.8523 | 6.2444 | 6.1225 | 6.3060 | 5.8679 | 6.2635 | 7.0836 | 7.4388 | 7.1233 | 6.7966 |
| KIAA1217_S361 | KIAA1217 | Q5T5P2 | | 1806 | SKIPALSPSSGKS | 6.2509 | 6.8190 | 6.9448 | 6.8656 | 6.4901 | 7.4958 | 7.1263 | 7.0591 | 7.4361 | 7.4588 | 7.1319 | 7.0646 | 4.0385 | 4.6512 | 2.9556 | 4.2117 |
| KIAA1522_S1038 | KIAA1522 | Q9P206-2 | | 604 | LAGPPASPGKAQP | 6.4772 | 6.4421 | 6.3380 | 6.3939 | 6.2043 | 6.0805 | 6.3449 | 6.1021 | 6.5218 | 6.4058 | 6.3399 | 6.4672 | 6.0043 | 6.3926 | 5.7533 | 5.7320 |
| KIAA1522_S401 | KIAA1522 | Q9P206-2 | | 401 | RFSSVSSPQPRSR | 6.1537 | 6.6526 | 6.2647 | 6.1593 | 6.0358 | 6.4656 | 6.5016 | 6.7709 | 6.8594 | 6.6252 | 6.6398 | 6.7593 | 5.2743 | 5.7893 | 5.5463 | 5.5022 |
| KIAA1522_S604 | KIAA1522 | Q9P206-2 | | 671 | KVPAPFSPPPSKP | 5.5772 | 5.8945 | 6.3944 | 6.9932 | 6.2959 | 6.0765 | 6.8769 | 6.8134 | 6.7964 | 7.5077 | 6.6168 | 6.8402 | 5.0853 | 5.3914 | 5.1240 | 5.7161 |
| KIAA1522_S671 | KIAA1522 | Q9P206-2 | | 1038 | GVPPPASPSYPRA | 5.8599 | 6.0371 | 6.3433 | 6.3092 | 5.6556 | 6.3957 | 6.6285 | 5.9358 | 6.4808 | 6.3663 | 6.1857 | 6.4482 | 6.2705 | 6.7211 | 5.9080 | 6.4542 |
| KIAA1522_S965 | KIAA1522 | Q9P206-2 | | 988 | QLERPVSPETQAD | 6.2651 | 6.9787 | 7.7743 | 5.5771 | 5.7500 | 6.3048 | 6.8403 | 6.4483 | 5.7513 | 7.1504 | 6.5315 | 6.4470 | 5.4835 | 6.3163 | 4.7558 | 5.6257 |
| KIAA1522_S988 | KIAA1522 | Q9P206-2 | | 965 | EPRPPQSPASTAS | 5.6216 | 6.2712 | 5.9226 | 5.9004 | 6.8843 | 6.0631 | 5.9166 | 6.3924 | 6.6681 | 6.3540 | 6.1045 | 6.1418 | 6.3308 | 6.3651 | 5.8398 | 7.2237 |
| KIAA1549L_S1310 | KIAA1549L | Q6ZVL6 | | 1310 | PSSGRRSPQNVMA | 6.1195 | 6.6111 | 6.0794 | 6.6578 | 6.5573 | 6.3102 | 5.9163 | 6.1897 | 6.3319 | 6.0670 | 6.2236 | 6.2278 | 6.1396 | 6.3807 | 6.1029 | 6.0853 |
| KIAA1549L_S1653 | KIAA1549L | Q6ZVL6 | | 1653 | GPVAVASLRRSTS | 6.9925 | 6.4932 | 6.1858 | 7.1033 | 5.8644 | 5.8900 | 6.7593 | 5.5761 | 6.1101 | 5.7228 | 5.9458 | 5.8629 | 6.6388 | 6.5310 | 5.9423 | 6.3815 |
| KIAA1549L_S1659 | KIAA1549L | Q6ZVL6 | | 1659 | SLRRSTSDIGSKT | 6.7290 | 6.2513 | 5.6688 | 6.2859 | 5.9112 | 5.6750 | 5.7269 | 5.7520 | 6.4752 | 6.1083 | 6.1387 | 6.4115 | 6.7454 | 7.2245 | 6.2324 | 6.6640 |
| KIAA1671_S1019 | KIAA1671 | Q9BY89 | | 284 | RKPRPLSMDLTAR | 6.9409 | 6.0778 | 6.9823 | 7.0903 | 6.1677 | 6.1252 | 6.2845 | 5.9099 | 6.9302 | 6.4782 | 6.6296 | 6.8203 | 5.4940 | 5.5423 | 4.8958 | 5.6311 |
| KIAA1671_S1111 | KIAA1671 | Q9BY89 | | 1366 | VRVSPKSPPTDQK | 6.3282 | 6.5054 | 5.7422 | 6.2442 | 6.3174 | 6.0776 | 6.3997 | 6.2683 | 6.1849 | 6.1205 | 5.9746 | 5.9876 | 6.3964 | 6.4631 | 6.6264 | 6.3634 |
| KIAA1671_S1224 | KIAA1671 | Q9BY89 | | 1760 | QPKSPKSPFQPGV | 6.5070 | 7.3255 | 6.5612 | 6.8104 | 6.7072 | 6.7870 | 7.1065 | 6.5960 | 6.9302 | 7.0650 | 7.3323 | 7.0257 | 3.8075 | 4.9230 | 4.0575 | 4.4579 |
| KIAA1671_S128 | KIAA1671 | Q9BY89 | | 1224 | EAQERRSPTVEPS | 5.7558 | 5.9755 | 5.8755 | 6.0826 | 6.0972 | 6.4247 | 6.4259 | 6.4874 | 5.8270 | 5.5688 | 6.0245 | 5.6101 | 7.2135 | 7.1102 | 6.8365 | 6.6848 |
| KIAA1671_S1312 | KIAA1671 | Q9BY89 | | 465 | PLATPASPSAAPE | 5.3911 | 5.7292 | 5.8017 | 5.8729 | 6.4156 | 6.4546 | 6.0218 | 6.1107 | 6.0864 | 5.5637 | 5.7043 | 6.2508 | 7.1323 | 7.2459 | 7.3213 | 6.8977 |
| KIAA1671_S1366 | KIAA1671 | Q9BY89 | | 1506 | FCKDRRSGPFVDQ | 6.7391 | 8.2502 | 6.8850 | 7.4949 | 6.0726 | 7.0905 | 7.6821 | 6.6741 | 6.9241 | 7.0126 | 7.2398 | 7.1341 | 3.4939 | 4.1359 | 3.1485 | 4.0226 |
| KIAA1671_S141 | KIAA1671 | Q9BY89 | | 1312 | ERYPGGSPIPADP | 6.5159 | 6.4479 | 6.3433 | 6.6825 | 6.4488 | 6.8514 | 7.0558 | 6.5393 | 6.6084 | 6.6704 | 6.6019 | 6.3771 | 4.8672 | 5.6943 | 5.1136 | 5.1821 |
| KIAA1671_S1441 | KIAA1671 | Q9BY89 | | 1488 | RPLQQVSPVASVP | 6.2408 | 6.5847 | 5.6173 | 6.2673 | 6.1308 | 6.4067 | 6.8331 | 6.7848 | 6.2478 | 6.0664 | 6.3356 | 5.6813 | 6.2427 | 6.6136 | 5.8754 | 6.0715 |
| KIAA1671_S1488 | KIAA1671 | Q9BY89 | | 1786 | ERSDEPSPQWLKE | 7.0973 | 8.1164 | 6.4560 | 7.1657 | 6.5681 | 7.1979 | 8.2478 | 7.0644 | 6.8725 | 6.7835 | 7.2852 | 6.8265 | 3.1271 | 3.9726 | 3.2122 | 4.0068 |

| | | | | | | | | | | | | | | | | | | | | |
|---|---|---|---|---|---|---|---|---|---|---|---|---|---|---|---|---|---|---|---|---|
| KIAA1671_S1506 | KIAA1671 | Q9BY89 | 635 | GRCLSTTPPGDMA | 6.5648 | 6.9794 | 6.4456 | 7.2982 | 6.4933 | 6.7620 | 6.6047 | 6.1635 | 6.4521 | 6.5525 | 7.0302 | 6.3668 | 4.9259 | 5.2950 | 4.8084 | 5.2576 |
| KIAA1671_S1675 | KIAA1671 | Q9BY89 | 128 | EGPRTSSPLFNKA | 6.5280 | 6.3358 | 5.6652 | 5.4394 | 7.0795 | 6.2943 | 6.8985 | 6.2106 | 5.9547 | 5.6722 | 5.4854 | 4.9778 | 7.3912 | 6.7600 | 6.9449 | 6.3624 |
| KIAA1671_S1701 | KIAA1671 | Q9BY89 | 951 | DRWRRRTLPPNVK | 9.3990 | 7.5415 | 7.6627 | 6.3326 | 6.3106 | 6.9818 | 6.8025 | 6.8889 | 7.7293 | 6.4989 | 6.2307 | 6.0494 | 4.2539 | 4.5595 | 2.9645 | 3.7942 |
| KIAA1671_S1744 | KIAA1671 | Q9BY89 | 508 | FQARPLSADLTKL | 7.5116 | 6.9897 | 8.2069 | 7.6577 | 5.7004 | 5.9258 | 6.6693 | 5.9342 | 7.5708 | 6.9287 | 6.8870 | 7.2793 | 4.1853 | 4.2492 | 3.8303 | 4.4739 |
| KIAA1671_S1760 | KIAA1671 | Q9BY89 | 1675 | SESESRSPLEDET | 5.6724 | 5.7659 | 5.6581 | 5.6496 | 5.7071 | 5.9027 | 6.1080 | 5.9910 | 5.7227 | 5.9594 | 5.9818 | 5.7086 | 7.2584 | 8.0754 | 7.5113 | 7.3278 |
| KIAA1671_S1786 | KIAA1671 | Q9BY89 | 1019 | PAVKQGSPVEPKA | 5.6900 | 5.6952 | 5.7902 | 5.9867 | 5.9086 | 5.9336 | 6.3688 | 5.9340 | 5.8637 | 5.3916 | 5.4506 | 5.6374 | 8.1347 | 7.2262 | 7.6873 | 7.3014 |
| KIAA1671_S1800 | KIAA1671 | Q9BY89 | 366 | SKPEMGSPRALVG | 6.4508 | 7.2139 | 6.0154 | 6.6332 | 6.8362 | 7.4333 | 6.8953 | 7.0368 | 6.4958 | 6.5788 | 6.8780 | 6.3084 | 5.0995 | 5.1530 | 4.4338 | 4.5379 |
| KIAA1671_S244 | KIAA1671 | Q9BY89 | 558 | GACIPRSPWKPGT | 5.9370 | 6.2414 | 5.5309 | 6.1518 | 6.3126 | 7.0475 | 7.2442 | 6.8553 | 6.3791 | 5.8824 | 6.2134 | 6.3807 | 6.1270 | 6.1316 | 5.5783 | 5.9868 |
| KIAA1671_S284 | KIAA1671 | Q9BY89 | 1800 | KSKKRQSLYENQV | 6.5865 | 5.7239 | 5.8587 | 5.8548 | 6.7765 | 5.6317 | 6.1170 | 6.2464 | 7.4362 | 6.9755 | 6.6023 | 6.8425 | 6.1175 | 6.8546 | 5.0568 | 5.3190 |
| KIAA1671_S366 | KIAA1671 | Q9BY89 | 764 | TLRSLRSWLSLKD | 2.1043 | 2.5588 | 2.0231 | 2.0338 | 1.9963 | 2.2847 | 2.7757 | 1.7276 | 2.2838 | 1.5642 | 1.9981 | 1.7472 | 17.5391 | 18.7122 | 19.5336 | 19.1176 |
| KIAA1671_S404 | KIAA1671 | Q9BY89 | 404 | KAAESPSPRLGRG | 5.7492 | 6.0591 | 5.2588 | 5.8820 | 5.9324 | 5.7804 | 6.2852 | 6.1611 | 6.3023 | 6.1425 | 6.4684 | 5.6773 | 7.9760 | 6.8522 | 6.9584 | 6.5147 |
| KIAA1671_S465 | KIAA1671 | Q9BY89 | 244 | LKRRPVSAIFTES | 8.2906 | 7.0969 | 7.7749 | 6.7167 | 6.7359 | 5.8580 | 6.1838 | 6.9618 | 7.1285 | 6.7735 | 6.9146 | 6.6643 | 3.8137 | 4.8467 | 3.8792 | 4.3609 |
| KIAA1671_S508 | KIAA1671 | Q9BY89 | 774 | LKDRQLSQEVTPA | 6.0188 | 6.8713 | 6.7799 | 6.1233 | 5.6676 | 5.9303 | 6.1330 | 6.0135 | 6.4625 | 7.2642 | 6.7271 | 6.6052 | 5.5079 | 6.4248 | 5.8490 | 5.6216 |
| KIAA1671_S529 | KIAA1671 | Q9BY89 | 1701 | SPRKEESDEEETA | 5.5717 | 5.5202 | 5.8536 | 6.1639 | 6.1287 | 5.9778 | 5.8778 | 5.7364 | 5.9532 | 6.0418 | 5.9909 | 6.0844 | 7.3566 | 7.1664 | 7.5495 | 7.0272 |
| KIAA1671_S558 | KIAA1671 | Q9BY89 | 529 | EVKYEKSAELSGE | 7.7041 | 7.0933 | 6.3465 | 5.3437 | 5.9256 | 6.0246 | 6.5280 | 7.1016 | 7.5101 | 7.2584 | 5.1622 | 5.4634 | 3.8170 | 8.2281 | 5.7605 | 4.7330 |
| KIAA1671_S764 | KIAA1671 | Q9BY89 | 600 | PCPSDVTPEDDRS | 5.0498 | 5.2859 | 5.0722 | 5.8346 | 5.9518 | 6.0516 | 5.8412 | 5.6799 | 5.7614 | 5.6741 | 4.9043 | 5.8246 | 7.7492 | 8.5424 | 8.4501 | 8.3270 |
| KIAA1671_S774 | KIAA1671 | Q9BY89 | 1744 | KRPEVDSPGETPS | 7.0422 | 7.0461 | 5.8154 | 7.4523 | 6.1297 | 6.2963 | 6.4994 | 5.8337 | 6.7904 | 6.8552 | 7.1136 | 6.5541 | 4.9444 | 5.7484 | 4.1309 | 5.7479 |
| KIAA1671_S88 | KIAA1671 | Q9BY89 | 1441 | QPKGRPSLTGENL | 5.8482 | 4.8412 | 5.3430 | 5.1626 | 7.0716 | 5.4045 | 5.5923 | 6.6500 | 7.6654 | 7.3162 | 7.4327 | 6.4737 | 6.7415 | 7.3826 | 6.0195 | 5.0550 |
| KIAA1671_S981 | KIAA1671 | Q9BY89 | 981 | RRTDYVSPTASAL | 6.5817 | 6.8470 | 7.2629 | 6.8553 | 6.1061 | 7.0369 | 6.7805 | 6.5561 | 6.4221 | 6.8297 | 7.3110 | 6.6497 | 4.1447 | 5.2690 | 4.5627 | 4.7846 |
| KIAA1671_T1059;S1063 | KIAA1671 | Q9BY89 | 1111 | PTDRPSSLGAWSL | 6.9892 | 6.8402 | 6.8846 | 6.7421 | 5.9421 | 7.0633 | 7.4280 | 6.5874 | 6.1572 | 7.0640 | 7.0306 | 6.4137 | 4.2659 | 5.6784 | 4.0552 | 4.8581 |
| KIAA1671_T600 | KIAA1671 | Q9BY89 | 88 | PSLRPSSTGPSPS | 5.1033 | 6.2129 | 5.6313 | 6.3914 | 6.2874 | 5.4144 | 6.1318 | 5.8876 | 5.2172 | 5.5732 | 5.8372 | 5.5009 | 7.5021 | 7.9830 | 7.8494 | 7.4769 |
| KIAA1671_T635 | KIAA1671 | Q9BY89 | 141 | VFLRPSSSTMILF | 4.4127 | 5.6036 | 5.2502 | 5.6617 | 4.2795 | 4.7176 | 5.7512 | 4.3570 | 5.3930 | 4.5192 | 5.6084 | 4.1706 | 9.3869 | 9.4401 | 10.2897 | 11.1587 |
| KIAA1671_T951 | KIAA1671 | Q9BY89 | 1059;1063 | NA | 5.6554 | 6.2471 | 5.0495 | 6.9005 | 6.3329 | 6.4376 | 6.1443 | 6.2481 | 5.9987 | 5.0699 | 5.7462 | 5.2101 | 7.2027 | 7.7970 | 7.4501 | 6.5098 |
| KIAA1958_S291 | KIAA1958 | Q8N8K9-3 | 291 | ARVSLASPNRGPP | 7.0639 | 6.9440 | 6.6146 | 7.0148 | 5.7477 | 6.2416 | 6.7889 | 6.7275 | 7.4784 | 7.3153 | 6.8874 | 6.5680 | 4.0465 | 4.6505 | 4.7263 | 5.1847 |
| KIAA1958_S97 | KIAA1958 | Q8N8K9-3 | 97 | PSETQTSPVERYP | 5.4634 | 6.2769 | 5.8750 | 7.5116 | 5.8052 | 5.1481 | 6.3251 | 6.8786 | 5.8220 | 7.2059 | 6.6067 | 5.7619 | 6.2071 | 6.3419 | 4.5272 | 8.2433 |
| KIDINS220_S1329 | KIDINS220 | Q9ULH0 | 918 | TITRQMSFDLTKL | 6.3404 | 6.7977 | 6.3220 | 6.2689 | 5.9930 | 5.8238 | 6.1337 | 5.9627 | 6.4442 | 6.3601 | 6.2320 | 6.1799 | 6.1641 | 6.2020 | 6.4629 | 6.3125 |
| KIDINS220_S1555 | KIDINS220 | Q9ULH0 | 1555 | VERVPKSPEHSAE | 6.5127 | 7.1232 | 6.2033 | 6.9008 | 6.2839 | 6.2650 | 6.4606 | 6.5388 | 6.6560 | 6.2954 | 6.3308 | 6.4285 | 5.3617 | 5.5369 | 5.4243 | 5.6781 |
| KIDINS220_S1644 | KIDINS220 | Q9ULH0 | 1681 | NLNRTPSTVTLNN | 6.7261 | 6.3521 | 6.4424 | 6.7942 | 6.2765 | 5.8160 | 6.2554 | 6.1945 | 7.0295 | 6.2813 | 6.0842 | 6.5706 | 5.6859 | 5.9300 | 5.6888 | 5.8726 |
| KIDINS220_S1681 | KIDINS220 | Q9ULH0 | 885 | RRVSQNSLGEMTK | 5.7283 | 5.8078 | 6.0149 | 6.3820 | 5.9437 | 6.0863 | 6.2837 | 5.9290 | 5.9331 | 6.7742 | 6.5007 | 6.1444 | 6.1683 | 7.0707 | 6.4789 | 6.7540 |
| KIDINS220_S1718 | KIDINS220 | Q9ULH0 | 1329 | PYTLNFSFEELNT | 6.5610 | 7.1772 | 6.6624 | 7.0412 | 6.6846 | 6.5342 | 6.6282 | 6.5833 | 5.6663 | 6.0204 | 5.7268 | 5.9351 | 5.8188 | 5.9527 | 5.4792 | 5.5288 |
| KIDINS220_S882;S885 | KIDINS220 | Q9ULH0 | 1718 | ILRPSSSPNPTTI | 6.5415 | 6.5869 | 6.3039 | 7.3878 | 6.1616 | 6.7577 | 6.7801 | 6.8628 | 6.3244 | 6.4179 | 6.4648 | 6.7626 | 5.0256 | 5.3901 | 5.1825 | 5.0498 |
| KIDINS220_S885 | KIDINS220 | Q9ULH0 | 1644 | PIIARMSICSEDK | 8.4817 | 6.4932 | 7.7392 | 6.2955 | 5.4740 | 5.8092 | 7.3147 | 6.6169 | 6.6187 | 5.9329 | 5.8590 | 6.0429 | 5.7679 | 5.3417 | 5.2557 | 4.9567 |
| KIDINS220_S918 | KIDINS220 | Q9ULH0 | 882;885 | NA | 5.4009 | 5.5756 | 6.2710 | 6.6456 | 6.1953 | 5.7465 | 5.7912 | 6.2431 | 7.0659 | 6.3111 | 6.5377 | 6.0867 | 6.4706 | 6.9233 | 6.5449 | 6.1906 |
| KIF11_T926 | KIF11 | P52732 | 926 | DIPTGTTPQRKSY | 7.0261 | 7.1845 | 6.9104 | 6.7872 | 6.4060 | 6.8501 | 6.9705 | 6.9652 | 7.5213 | 7.1055 | 7.1744 | 6.9826 | 3.5946 | 4.6029 | 3.6634 | 4.2552 |
| KIF13A_S1287 | KIF13A | Q9H1H9 | 1460 | SPFKAFSPQPPKF | 6.9603 | 7.4079 | 7.3254 | 7.0620 | 6.2478 | 7.1974 | 6.7755 | 7.0083 | 7.3634 | 7.2442 | 7.2426 | 7.7067 | 3.1848 | 4.1391 | 3.3164 | 3.8183 |
| KIF13A_S1384 | KIF13A | Q9H1H9 | 1384 | HIRRSLSTPNVHN | 6.1774 | 6.0496 | 6.7887 | 6.1913 | 6.3910 | 6.4694 | 5.5681 | 6.8210 | 6.2409 | 6.4237 | 6.6559 | 6.7963 | 5.9088 | 6.5719 | 5.5590 | 5.3870 |
| KIF13A_S1460 | KIF13A | Q9H1H9 | 1287 | KQSFTQSLKRRIS | 6.8159 | 7.4638 | 8.5776 | 8.1270 | 4.8755 | 5.1238 | 6.2190 | 6.0285 | 7.2614 | 5.6646 | 7.9246 | 6.7682 | 5.7870 | 5.3925 | 3.6477 | 4.3228 |
| KIF13A_S1698 | KIF13A | Q9H1H9 | 1698 | SLCRTGSCSELDA | 5.9640 | 6.2006 | 7.1198 | 5.2670 | 5.5651 | 6.1024 | 5.9203 | 5.6222 | 6.4491 | 6.9194 | 5.8767 | 6.4541 | 6.9322 | 6.6698 | 6.4868 | 6.4506 |
| KIF13B_S1391 | KIF13B | Q9NQT8 | 1644 | EAPAPGSPFRVRR | 6.5933 | 6.8901 | 7.0230 | 6.3451 | 5.6817 | 7.1569 | 6.8493 | 6.6872 | 6.6344 | 6.9149 | 6.9373 | 6.6699 | 4.4930 | 5.3779 | 4.5969 | 5.1488 |
| KIF13B_S1497 | KIF13B | Q9NQT8 | 1793 | EARRSATLSGSAT | 7.6837 | 6.3642 | 7.3118 | 6.4181 | 6.6779 | 6.3317 | 6.5197 | 5.9742 | 6.2583 | 5.8827 | 6.3004 | 6.3442 | 5.9138 | 5.8308 | 4.7957 | 5.3929 |
| KIF13B_S1644 | KIF13B | Q9NQT8 | 1497 | IMVQSASPDIRVT | 6.5429 | 8.7296 | 6.6574 | 6.6644 | 5.6335 | 7.0600 | 7.6605 | 7.4662 | 6.3381 | 6.6817 | 7.1121 | 9.3197 | 3.1239 | 4.7310 | 3.0719 | 3.2072 |
| KIF13B_T1793 | KIF13B | Q9NQT8 | 1391 | VNRLSGSRQDLIP | 7.0254 | 6.3845 | 6.0381 | 5.8581 | 6.4581 | 5.9018 | 6.1309 | 6.9190 | 6.0954 | 6.5072 | 6.3899 | 8.1545 | 4.7509 | 7.3636 | 5.0400 | 4.9827 |
| KIF14_S101 | KIF14 | Q15058 | 1200 | MKNRRISGCLHDI | 6.4995 | 6.6512 | 5.7766 | 6.8452 | 4.7314 | 6.6215 | 6.0160 | 6.6403 | 8.8135 | 7.6150 | 8.5460 | 8.1767 | 4.4592 | 5.2102 | 2.8946 | 4.5030 |
| KIF14_S1200 | KIF14 | Q15058 | 1466 | AMGLIRSLENIFA | 6.5938 | 6.8139 | 5.9403 | 6.0628 | 5.4479 | 6.9455 | 5.9720 | 6.4848 | 6.0083 | 5.9062 | 5.4910 | 5.2097 | 7.3516 | 5.5659 | 6.7503 | 7.4560 |
| KIF14_S1466 | KIF14 | Q15058 | 298 | SLPQLKSPAPSIL | 6.4419 | 7.2107 | 7.0310 | 6.9492 | 4.5590 | 7.1568 | 7.8774 | 6.8199 | 6.4749 | 7.3568 | 8.1708 | 8.7089 | 3.1474 | 4.5736 | 3.0619 | 4.4597 |
| KIF14_S298 | KIF14 | Q15058 | 101 | TRNKESSLLVSEL | 5.5753 | 6.4052 | 6.5424 | 6.6355 | 5.4860 | 6.4833 | 7.2787 | 6.6329 | 6.8810 | 6.5432 | 6.9935 | 6.6243 | 4.7561 | 5.8276 | 5.1991 | 6.1360 |
| KIF15_S568 | KIF15 | Q9NS87 | 568 | KNQQGFSPKAQKE | 6.4136 | 6.8199 | 7.2352 | 6.4388 | 6.3154 | 6.6185 | 6.8624 | 6.5336 | 6.7488 | 7.0196 | 7.0761 | 7.3426 | 4.4086 | 5.1040 | 4.2485 | 4.8146 |
| KIF15_T1144 | KIF15 | Q9NS87 | 661 | TLKIITTPTKAYQ | 7.0858 | 7.5281 | 7.0302 | 7.2720 | 6.1559 | 7.4910 | 7.3626 | 7.2666 | 7.7782 | 7.5080 | 8.0644 | 8.0529 | 2.5025 | 3.5700 | 2.2723 | 3.0596 |
| KIF15_T1171 | KIF15 | Q9NS87 | 1144 | DPQSPKTPPHFQT | 6.4548 | 6.5203 | 6.8255 | 7.2595 | 6.1094 | 6.8094 | 6.7430 | 6.4755 | 6.1612 | 7.2052 | 6.9216 | 6.9723 | 4.2516 | 5.1357 | 4.6771 | 5.4779 |
| KIF15_T661 | KIF15 | Q9NS87 | 1171 | DGRASKTSLEHLV | 6.0251 | 7.4353 | 7.6173 | 5.4488 | 8.7798 | 5.8482 | 7.6729 | 7.7891 | 4.0538 | 6.8490 | 6.8136 | 6.1211 | 5.5702 | 4.6907 | 4.5036 | 4.7816 |
| KIF16B_S398 | KIF16B | Q96L93-2 | 398 | QIALLDSPTALSM | 6.5225 | 6.4349 | 6.7108 | 6.4643 | 6.3595 | 6.2953 | 6.6641 | 6.6052 | 6.3995 | 6.8054 | 7.2050 | 6.3609 | 4.9014 | 5.7715 | 4.9348 | 5.5650 |
| KIF16B_S582 | KIF16B | Q96L93-2 | 582 | MTDLSKSRENLSA | 5.7901 | 6.4901 | 6.7024 | 6.7089 | 6.3482 | 6.8806 | 6.3053 | 5.8834 | 6.2269 | 6.1760 | 6.9772 | 6.3841 | 5.4751 | 5.7616 | 5.5320 | 6.3580 |
| KIF18A_S674 | KIF18A | Q8NI77 | 674 | RRKLMPSPLKGQH | 6.3729 | 6.6060 | 5.8897 | 6.5307 | 6.3240 | 5.9945 | 6.5893 | 5.9987 | 6.3904 | 6.0883 | 6.1157 | 6.1519 | 5.9848 | 6.2947 | 6.1397 | 6.5287 |
| KIF18A_S684 | KIF18A | Q8NI77 | 684 | GQHTLKSPPSQSV | 6.7338 | 7.4702 | 6.7243 | 7.2328 | 6.9175 | 6.9758 | 6.6883 | 6.5896 | 6.8990 | 6.9305 | 6.8591 | 7.0396 | 3.6147 | 4.7282 | 4.1111 | 4.4856 |
| KIF18A_S695 | KIF18A | Q8NI77 | 695 | SVQLNDSLSKELQ | 5.3645 | 5.4847 | 5.2735 | 6.6751 | 6.3671 | 6.2812 | 6.4565 | 6.6050 | 6.5407 | 6.0113 | 5.4905 | 5.9076 | 6.4736 | 6.9275 | 7.0078 | 7.1336 |
| KIF18B_S404 | KIF18B | Q86Y91 | 579 | ESIPVPSPLCPEP | 6.1684 | 6.7658 | 6.5396 | 7.2881 | 6.1087 | 6.4785 | 7.0997 | 6.9379 | 6.1260 | 6.8069 | 6.0458 | 6.1031 | 5.5053 | 5.9628 | 4.7367 | 5.3267 |
| KIF18B_S417 | KIF18B | Q86Y91 | 682 | KGERASSPCHSPR | 6.7442 | 7.0386 | 6.0479 | 7.2038 | 6.2239 | 5.8152 | 5.9245 | 6.0650 | 6.3797 | 5.8420 | 5.9233 | 6.0562 | 5.9132 | 6.1198 | 6.4850 | 6.2178 |
| KIF18B_S417;S425;T429 | KIF18B | Q86Y91 | 404 | PQDLPGSPKSGPP | 8.1553 | 8.1011 | 7.1892 | 7.5412 | 6.6167 | 6.0570 | 6.8116 | 6.1226 | 7.6993 | 7.0175 | 6.9663 | 6.9214 | 3.5585 | 4.1618 | 3.2090 | 3.8716 |
| KIF18B_S579 | KIF18B | Q86Y91 | 662 | PCLRRGSLPDTQP | 4.3186 | 3.6875 | 4.6574 | 5.2356 | 7.5132 | 5.1072 | 4.3861 | 5.6298 | 8.1530 | 8.7628 | 7.2563 | 7.9796 | 8.1230 | 7.8556 | 5.6492 | 5.6850 |
| KIF18B_S601 | KIF18B | Q86Y91-6 | 417 | PEHLPSSPLPPHP | 6.8473 | 7.9999 | 7.2645 | 7.9579 | 6.3322 | 5.8674 | 6.5534 | 6.5338 | 6.7326 | 7.0816 | 7.1665 | 6.7587 | 4.0420 | 3.9505 | 4.6980 | 4.2137 |
| KIF18B_S633 | KIF18B | Q86Y91 | 639 | SALEADSPMAPKR | 8.2805 | 8.5429 | 7.3696 | 8.4610 | 6.0834 | 7.2549 | 7.0322 | 7.2286 | 7.3191 | 6.8974 | 7.1377 | 6.8084 | 2.8743 | 3.2682 | 2.6036 | 2.8382 |
| KIF18B_S639 | KIF18B | Q86Y91 | 601 | TMARRLSGPLHTL | 4.4958 | 4.4638 | 5.8004 | 6.1112 | 8.1257 | 6.2826 | 4.9933 | 4.9243 | 7.5156 | 8.7396 | 8.0016 | 6.8518 | 6.9119 | 6.7175 | 5.2965 | 4.7682 |
| KIF18B_S662 | KIF18B | Q86Y91 | 633 | KKRRRPSALEADS | 5.0555 | 4.6982 | 5.6955 | 5.8889 | 6.4668 | 6.3272 | 5.7209 | 5.1696 | 6.8051 | 6.6833 | 6.4455 | 7.9786 | 7.6048 | 7.1843 | 5.4831 | 6.7927 |
| KIF18B_S662;T674 | KIF18B | Q86Y91 | 780 | SSLPGTSACKKKR | 8.1208 | 7.1133 | 7.6893 | 7.6457 | 4.8233 | 7.4631 | 8.3424 | 5.4927 | 6.8551 | 6.7795 | 7.6194 | 8.4156 | 3.5609 | 3.9886 | 2.2736 | 3.8167 |
| KIF18B_S682 | KIF18B | Q86Y91 | 822 | LPELPLSPLCPSN | 6.9551 | 7.0542 | 5.8553 | 6.3811 | 5.8831 | 6.2022 | 5.8082 | 6.5635 | 7.5002 | 6.0705 | 6.1357 | 6.9503 | 5.3780 | 5.8481 | 5.6970 | 5.7176 |
| KIF18B_S780 | KIF18B | Q86Y91-6 | 417;425;429 | NA | 6.2000 | 6.7213 | 7.1650 | 7.4719 | 6.2665 | 5.8972 | 6.2798 | 7.2153 | 7.3603 | 6.6557 | 7.5248 | 7.2329 | 4.4825 | 4.6152 | 4.0962 | 4.8154 |
| KIF18B_S822 | KIF18B | Q86Y91 | 662;674 | NA | 6.4864 | 6.7788 | 6.2805 | 7.2793 | 5.6380 | 5.6052 | 6.8586 | 6.1862 | 7.7706 | 8.2126 | 7.2402 | 8.0193 | 4.9863 | 4.6853 | 3.6907 | 4.2820 |
| KIF1B_S1053 | KIF1B | O60333-4 | 1659 | ANSRASSPCPEFE | 6.4593 | 6.3568 | 6.7013 | 6.1295 | 6.5093 | 7.0366 | 6.3586 | 7.1409 | 6.2566 | 6.1078 | 6.6109 | 6.6359 | 5.6881 | 6.1309 | 5.1733 | 4.7040 |
| KIF1B_S1057 | KIF1B | O60333-4 | 1487 | WRPRGDSLILEHQ | 5.8895 | 5.5258 | 6.4063 | 5.8181 | 6.0881 | 6.2168 | 6.0617 | 6.6104 | 6.2243 | 6.3907 | 6.1605 | 6.4579 | 6.8690 | 6.4460 | 6.5191 | 6.3159 |
| KIF1B_S1141 | KIF1B | O60333-3 | 1141 | RMRRQFSAPNLKA | 6.4652 | 6.1323 | 6.6543 | 5.4995 | 5.9814 | 6.2267 | 6.0018 | 6.5502 | 6.3316 | 6.2145 | 6.1329 | 6.4245 | 6.2423 | 6.4715 | 6.0966 | 6.5745 |
| KIF1B_S1454 | KIF1B | O60333-4 | 1057 | MTRSGLSLEELRI | 6.0070 | 6.0875 | 6.6591 | 6.0293 | 6.5894 | 6.4624 | 5.9985 | 6.3225 | 6.3634 | 6.4152 | 6.0960 | 6.4060 | 6.2822 | 6.2895 | 5.7275 | 6.2645 |
| KIF1B_S1487 | KIF1B | O60333-4 | 1613 | CKLSDISPIGRDP | 6.2771 | 6.8156 | 6.5044 | 6.4840 | 6.3087 | 6.2859 | 6.5345 | 7.2839 | 6.7165 | 6.4351 | 6.3853 | 6.4301 | 4.9502 | 5.5793 | 5.2448 | 5.7647 |
| KIF1B_S1613 | KIF1B | O60333-3 | 1053 | LRWRSNSLNNGQP | 5.8146 | 5.3721 | 6.5761 | 5.8006 | 6.1596 | 6.1770 | 6.0833 | 6.3944 | 6.6325 | 6.4197 | 6.5870 | 6.6732 | 6.5752 | 6.6727 | 6.1704 | 5.8917 |
| KIF1B_S1659 | KIF1B | O60333-4 | 1454 | KMSDTGSPGMQRR | 7.3228 | 5.6000 | 7.3252 | 6.0838 | 5.6864 | 6.5737 | 5.5396 | 6.8057 | 7.2931 | 7.0254 | 5.8888 | 6.3713 | 5.2064 | 5.8197 | 5.0990 | 6.3590 |
| KIF1C_S1033 | KIF1C | O43896 | 1083 | RYPPYTTPPRMRR | 6.7335 | 6.8786 | 7.3528 | 6.4085 | 6.0921 | 6.8437 | 6.4646 | 6.8229 | 6.8518 | 6.6684 | 6.7686 | 6.9358 | 4.6363 | 4.9093 | 4.6048 | 5.0284 |
| KIF1C_S295 | KIF1C | O43896 | 1033 | HHPRRNSLDGGGR | 6.0840 | 6.1615 | 6.7423 | 5.5457 | 6.1229 | 6.4106 | 5.9497 | 6.9987 | 6.7254 | 6.7348 | 6.8927 | 7.1674 | 5.5383 | 5.8571 | 5.6137 | 5.4551 |
| KIF1C_S915 | KIF1C | O43896 | 915 | PSARPPSPPLSSW | 6.0269 | 6.2969 | 6.7436 | 6.4195 | 6.5266 | 6.2986 | 6.8063 | 6.5287 | 6.8978 | 7.0553 | 6.2071 | 6.6235 | 5.3322 | 5.7991 | 5.3907 | 5.0473 |
| KIF1C_T1083 | KIF1C | O43896 | 295 | QSKKRKSDFIPYR | 7.3586 | 7.6743 | 7.2352 | 7.6333 | 6.0405 | 7.0237 | 7.1264 | 6.3766 | 7.4411 | 7.0682 | 7.8694 | 7.6672 | 2.9972 | 3.8212 | 3.0703 | 3.5968 |
| KIF20A_S240 | KIF20A | O95235 | 825 | KLQGQVSAKKRLG | 7.4502 | 8.0620 | 6.8624 | 8.3768 | 5.9351 | 6.7968 | 7.4880 | 6.7720 | 7.0224 | 6.8460 | 7.2309 | 7.5429 | 2.9172 | 4.0108 | 2.7112 | 3.9752 |
| KIF20A_S532 | KIF20A | O95235 | 532 | EHSLQVSPSLEKG | 6.9653 | 7.5383 | 6.6883 | 7.7020 | 6.1174 | 6.7517 | 7.2459 | 7.0793 | 6.9380 | 7.3933 | 7.4336 | 6.7594 | 3.3382 | 4.5263 | 3.3548 | 4.1683 |
| KIF20A_S825 | KIF20A | O95235 | 240 | EEELSTSLKRSVY | 6.9039 | 6.7249 | 6.5241 | 7.3862 | 6.8561 | 6.1695 | 6.6183 | 6.2932 | 6.4189 | 6.8308 | 7.6315 | 5.5116 | 4.8913 | 5.9222 | 4.2717 | 5.0457 |
| KIF20B_S1621 | KIF20B | Q96Q89-2 | 1773 | GDFLQHSPSILQS | 7.0889 | 6.9273 | 6.1861 | 7.4352 | 6.4561 | 6.2283 | 6.6903 | 6.5703 | 6.9521 | 6.6279 | 6.5238 | 6.7578 | 4.7733 | 5.2910 | 4.5864 | 4.9052 |
| KIF20B_S1691 | KIF20B | Q96Q89-2 | 1661 | ELEIQFTPLQPNK | 8.3995 | 7.7641 | 6.8682 | 8.5217 | 6.7782 | 6.7587 | 7.9492 | 6.6358 | 7.3759 | 5.9795 | 6.2843 | 6.5565 | 3.3593 | 4.2270 | 2.8665 | 3.6755 |
| KIF20B_S1773 | KIF20B | Q96Q89-2 | 1789 | KIIETMSSSKLSN | 9.1712 | 8.3595 | 7.4495 | 8.7909 | 6.5027 | 6.7072 | 8.1839 | 6.4972 | 7.6167 | 5.9794 | 6.8505 | 7.0320 | 2.4563 | 3.2045 | 2.2134 | 2.9851 |
| KIF20B_S1789 | KIF20B | Q96Q89-2 | 1621 | TEPLSTSFEISRN | 6.4079 | 6.7186 | 6.8158 | 7.8086 | 6.6616 | 6.3991 | 7.2410 | 6.9029 | 6.6531 | 6.5527 | 7.0146 | 6.8323 | 4.1571 | 4.9575 | 4.1843 | 4.6929 |
| KIF20B_S303 | KIF20B | Q96Q89-2 | 1676 | VKHPGCTTPVTVK | 6.8023 | 6.5532 | 6.5573 | 7.2323 | 6.4937 | 6.3391 | 6.8375 | 6.3178 | 6.7591 | 6.2420 | 6.5551 | 6.1056 | 5.1413 | 5.9127 | 4.9937 | 5.1571 |
| KIF20B_T1661 | KIF20B | Q96Q89-2 | 1677 | KHPGCTTPVTVKI | 7.6762 | 6.9541 | 6.3711 | 7.5410 | 6.3840 | 6.4895 | 7.1045 | 6.2797 | 7.1709 | 6.3944 | 6.5274 | 6.5961 | 4.5971 | 4.8559 | 4.4290 | 4.6294 |
| KIF20B_T1676 | KIF20B | Q96Q89-2 | 1691 | KARKRKSNEMEED | 6.4341 | 6.0980 | 6.2652 | 7.2024 | 5.7788 | 5.6072 | 6.9085 | 6.1358 | 6.4654 | 7.5501 | 5.7750 | 7.0351 | 5.4004 | 5.6872 | 5.7662 | 5.8907 |
| KIF20B_T1677 | KIF20B | Q96Q89-2 | 303 | RKMLRLSQDVKGY | 7.0078 | 7.5279 | 7.3297 | 8.8829 | 5.9858 | 6.1045 | 6.5583 | 6.5870 | 7.2093 | 7.2248 | 7.2093 | 7.1232 | 3.5398 | 4.1355 | 3.6347 | 3.9395 |
| KIF21A_S1212 | KIF21A | Q7Z4S6-4 | 1212 | AREKELSPPPGLP | 6.4770 | 7.0233 | 7.0362 | 6.8132 | 5.9770 | 6.5367 | 6.6700 | 6.8815 | 6.4100 | 6.7568 | 6.9852 | 7.0275 | 4.5506 | 5.2000 | 4.7369 | 4.9181 |
| KIF21A_S1229 | KIF21A | Q7Z4S6-4 | 1239 | KKIPEPSPVTRRK | 6.0978 | 6.3058 | 6.7026 | 6.3857 | 5.8883 | 6.6405 | 6.4331 | 6.9030 | 6.4742 | 6.6338 | 6.5745 | 6.4006 | 5.3010 | 6.1810 | 5.5331 | 5.5451 |
| KIF21A_S1239 | KIF21A | Q7Z4S6-4 | 524 | SGSSTFSPTILSS | 6.1556 | 6.4011 | 6.4766 | 6.6758 | 6.4988 | 6.6385 | 6.3080 | 6.3689 | 6.5956 | 6.6852 | 6.6964 | 6.8605 | 5.2237 | 5.7155 | 5.1989 | 5.5007 |
| KIF21A_S524 | KIF21A | Q7Z4S6-4 | 1229 | SISRQSSLSEKKI | 5.9872 | 6.2347 | 6.9661 | 5.7740 | 5.8271 | 6.2026 | 5.7172 | 6.2993 | 6.2563 | 6.3606 | 6.0037 | 5.8885 | 6.7403 | 7.3057 | 6.4466 | 5.9901 |
| KIF21A_S853 | KIF21A | Q7Z4S6-4 | 853 | KVTRKLSSSDAPA | 5.6876 | 5.3564 | 6.7478 | 6.0630 | 6.4565 | 6.2408 | 6.0539 | 6.6668 | 7.2387 | 6.7584 | 6.0779 | 6.3037 | 5.7499 | 6.7318 | 6.1413 | 5.7254 |
| KIF22_S412 | KIF22 | Q14807 | 543 | IQEQAASPNAEIH | 6.7161 | 7.0900 | 6.4477 | 7.4507 | 6.5023 | 6.9243 | 7.0394 | 6.7311 | 7.1048 | 7.3209 | 6.8826 | 7.2524 | 3.7308 | 4.7590 | 3.6959 | 4.3522 |
| KIF22_S543 | KIF22 | Q14807 | 463 | GAPLLSTPKRERM | 8.5783 | 7.5381 | 7.2198 | 7.8198 | 5.7719 | 6.4700 | 6.9108 | 5.9316 | 8.1404 | 7.3894 | 7.1350 | 7.4783 | 3.3035 | 3.9691 | 2.6594 | 3.6845 |
| KIF22_S581 | KIF22 | Q14807 | 581 | CWELQISPELLAH | 6.1294 | 6.4620 | 5.9714 | 6.1144 | 6.1253 | 5.9960 | 6.5585 | 6.8000 | 6.8958 | 6.2417 | 6.9273 | 6.4114 | 5.5376 | 6.5030 | 5.2995 | 6.0268 |
| KIF22_T463 | KIF22 | Q14807 | 412 | EEEEIGSPEPMAA | 5.9220 | 7.3108 | 6.8563 | 7.6629 | 6.9114 | 6.4480 | 6.7025 | 6.2747 | 6.9942 | 7.1439 | 7.1800 | 6.4557 | 4.3190 | 5.1433 | 4.4533 | 4.2220 |
| KIF23_S160 | KIF23 | Q02241-2 | 684 | VTQRSVSPSPVPL | 5.9781 | 6.5066 | 6.4132 | 6.3684 | 6.2469 | 6.2087 | 6.1312 | 6.3725 | 6.4318 | 6.2708 | 6.2907 | 6.5688 | 6.0469 | 6.3315 | 5.8088 | 6.0251 |
| KIF23_S684 | KIF23 | Q02241 | 160 | KSNDRNSMDIQCE | 6.1265 | 6.3812 | 6.4269 | 7.1371 | 5.9655 | 6.4496 | 6.5340 | 6.2673 | 6.2767 | 6.5544 | 6.4362 | 6.2420 | 5.2570 | 5.9342 | 6.0751 | 5.9363 |
| KIF23_S684 | KIF23 | Q02241 | 450 | KAICGLTPGRRYR | 7.5331 | 7.7784 | 7.3826 | 7.7350 | 5.8295 | 6.9145 | 7.5323 | 6.5538 | 8.3522 | 6.6153 | 7.3478 | 6.9956 | 2.9434 | 3.9731 | 2.7484 | 3.7651 |
| KIF23_S684;S686 | KIF23 | Q02241 | 867 | THQELASDGEIET | 6.6004 | 7.3095 | 7.3578 | 8.1333 | 6.8176 | 6.8856 | 6.7976 | 6.8632 | 6.0720 | 6.3961 | 6.2455 | 6.3246 | 4.1712 | 4.8604 | 4.3790 | 4.7860 |
| KIF23_S686 | KIF23 | Q02241 | 911 | GSRKRRSSTVAPA | 5.5016 | 5.0584 | 5.3950 | 5.6327 | 6.3739 | 5.9946 | 5.7973 | 5.7726 | 7.1775 | 7.0703 | 6.5743 | 6.8107 | 6.8915 | 6.9896 | 6.5289 | 6.4313 |
| KIF23_S716 | KIF23 | Q02241 | 738 | KIPTYNTPLKVTS | 7.1018 | 6.7409 | 6.2009 | 6.3126 | 6.5991 | 6.7565 | 7.3578 | 6.7069 | 6.1797 | 5.6360 | 5.8382 | 5.7759 | 5.4522 | 5.8729 | 5.3156 | 6.1532 |
| KIF23_S867 | KIF23 | Q02241 | 716 | LHRRSNSCSSISV | 5.4584 | 5.5363 | 5.9665 | 6.1342 | 6.1053 | 6.2151 | 6.2457 | 6.0224 | 5.7071 | 5.5712 | 5.8508 | 5.9198 | 7.0501 | 6.8910 | 7.8337 | 7.4924 |
| KIF23_S911 | KIF23 | Q02241 | 684 | VTQRSVSPSPVPL | 5.6841 | 6.0167 | 6.4809 | 6.4212 | 6.1658 | 6.4324 | 6.2240 | 7.2192 | 6.3377 | 5.7963 | 6.1356 | 6.4287 | 6.0282 | 6.4824 | 5.8906 | 6.2561 |
| KIF23_S911;T913 | KIF23 | Q02241-2 | 686 | QRSVSPSPVPLLF | 6.8948 | 5.4199 | 6.0979 | 5.9428 | 4.9728 | 6.9056 | 5.4776 | 8.0379 | 5.1819 | 5.4055 | 5.2324 | 6.0871 | 6.0294 | 7.7927 | 8.4598 | 6.0618 |
| KIF23_T450 | KIF23 | Q02241 | 911;913 | NA | 5.9477 | 6.2165 | 6.2643 | 6.5289 | 5.5441 | 6.3582 | 6.7047 | 5.7151 | 6.9601 | 6.7663 | 7.3081 | 6.9677 | 4.8133 | 6.0133 | 5.2981 | 6.5935 |
| KIF23_T738 | KIF23 | Q02241-2 | 684;686 | NA | 4.4732 | 7.2190 | 5.8539 | 3.4809 | 6.9064 | 6.7951 | 5.5162 | 6.9773 | 5.4254 | 8.3294 | 8.3628 | 6.8562 | 3.9857 | 7.6318 | 6.2550 | 5.9316 |
| KIF24_S102 | KIF24 | Q5T7B8 | 102 | RQLNFDSPADNKD | 6.6722 | 6.8630 | 7.0067 | 6.5540 | 6.1780 | 6.7371 | 6.5529 | 7.0925 | 7.5949 | 6.8826 | 7.4446 | 7.6300 | 3.9349 | 4.7500 | 3.8060 | 4.3007 |
| KIF24_S1268 | KIF24 | Q5T7B8 | 574 | PKRIQSSPGALSE | 6.6434 | 6.6279 | 6.6784 | 6.6694 | 6.0736 | 6.7449 | 5.7794 | 6.7104 | 7.4912 | 6.7095 | 6.9962 | 7.1474 | 4.2721 | 5.1609 | 4.9598 | 5.3356 |
| KIF24_S574 | KIF24 | Q5T7B8 | 854 | LENSEDSFFLHQT | 7.1389 | 7.3770 | 8.7653 | 8.9831 | 8.0559 | 3.9266 | 6.9109 | 5.6797 | 6.8146 | 4.3877 | 5.5632 | 4.7677 | 6.5917 | 8.0610 | 3.9824 | 2.9944 |
| KIF24_S854 | KIF24 | Q5T7B8 | 1268 | CLARPSSPLVPSC | 5.3366 | 4.4461 | 6.1132 | 4.4066 | 6.1713 | 6.0533 | 6.4102 | 5.2851 | 6.3982 | 6.4695 | 5.9952 | 6.0539 | 7.4360 | 7.4264 | 8.0743 | 7.9240 |
| KIF2A_S624 | KIF2A | O00139-4 | 624 | QWGVGSSPQRDDL | 5.6435 | 6.3097 | 6.1724 | 6.7333 | 5.9979 | 6.1789 | 6.1814 | 6.2355 | 6.7218 | 6.8966 | 6.5975 | 6.8317 | 5.2348 | 6.0125 | 6.0255 | 6.2271 |
| KIF2A_T78 | KIF2A | O00139-4 | 78 | IEPSPETPPPPAS | 6.3448 | 5.8979 | 6.6225 | 5.7880 | 6.1295 | 6.0494 | 5.9172 | 5.7167 | 6.4978 | 6.3924 | 6.7472 | 6.4265 | 6.4250 | 6.3552 | 5.9603 | 6.7294 |
| KIF2C_S187 | KIF2C | Q99661 | 621 | LIPGNLSKEEEEL | 5.8374 | 7.4411 | 6.4699 | 6.6015 | 6.5279 | 6.7034 | 6.2338 | 6.8457 | 6.3292 | 6.7965 | 6.1134 | 6.6336 | 5.2156 | 5.6003 | 5.2518 | 5.3989 |
| KIF2C_S621 | KIF2C | Q99661 | 187 | SANPVNSVRRKSC | 7.2211 | 7.0546 | 7.9189 | 6.8537 | 4.6169 | 6.2596 | 6.5645 | 6.4271 | 7.3237 | 7.2755 | 8.5861 | 8.3804 | 3.4393 | 4.7801 | 3.2965 | 4.0021 |
| KIF3A_S687 | KIF3A | Q9Y496 | 633 | NNMRKQTPVPDKK | 6.7788 | 6.6767 | 6.5784 | 5.9370 | 6.6258 | 6.4135 | 6.3629 | 7.4646 | 6.1752 | 6.1548 | 6.7085 | 6.8096 | 5.4575 | 5.7235 | 4.9165 | 5.2167 |
| KIF3A_T633 | KIF3A | Q9Y496 | 687 | TGRRKRSAKPETV | 6.3068 | 5.7545 | 6.7259 | 6.5683 | 6.9201 | 5.8442 | 5.5827 | 6.2338 | 6.9470 | 7.8339 | 7.1175 | 6.7823 | 5.3693 | 6.6042 | 4.6080 | 4.8016 |
| KIF4A_S1001 | KIF4A | O95239 | 1181 | QVLSKKTPPAPSP | 6.2402 | 6.9339 | 6.6761 | 6.8776 | 6.5815 | 6.8146 | 6.8966 | 6.7100 | 6.8004 | 7.1632 | 7.1206 | 6.8470 | 4.0839 | 5.0795 | 4.3513 | 4.8237 |
| KIF4A_S1013 | KIF4A | O95239 | 1225 | SFFSGCSPIEEEA | 6.4523 | 7.0058 | 7.1540 | 7.7878 | 5.8604 | 6.3996 | 6.6315 | 6.5217 | 7.0146 | 7.4401 | 7.1114 | 6.8829 | 3.9905 | 4.9250 | 4.1200 | 4.7024 |
| KIF4A_S1013;S1017 | KIF4A | O95239 | 1161 | FNPVCATPNSKIL | 7.1904 | 7.2681 | 7.1353 | 7.0911 | 5.7681 | 6.9963 | 7.2217 | 6.0993 | 7.3023 | 6.5650 | 6.5129 | 6.8323 | 4.1747 | 5.3002 | 3.8708 | 4.6713 |
| KIF4A_S1017 | KIF4A | O95239 | 1038 | EKFLEQSMDIEDL | 6.4311 | 6.6132 | 7.1240 | 6.9868 | 5.6905 | 6.9343 | 7.1122 | 6.6608 | 6.6756 | 6.8070 | 7.1125 | 7.0563 | 4.3229 | 5.3933 | 4.1021 | 4.9773 |
| KIF4A_S1038 | KIF4A | O95239 | 799 | PKLRRRTFSLTEV | 7.9346 | 6.9152 | 6.0781 | 8.2849 | 6.7685 | 6.2927 | 7.0384 | 5.9996 | 7.6288 | 7.0133 | 7.1120 | 8.1065 | 3.3654 | 4.5131 | 2.7449 | 4.2041 |
| KIF4A_S1048;S1051 | KIF4A | O95239 | 801 | LRRRTFSLTEVRG | 4.7499 | 4.9618 | 5.7828 | 4.8421 | 5.1542 | 5.6309 | 5.5487 | 5.8447 | 5.1974 | 5.5398 | 5.5564 | 5.8085 | 8.5010 | 8.6599 | 9.1280 | 9.0937 |
| KIF4A_S1225 | KIF4A | O95239 | 953 | QLEESVSEKEQQL | 6.2463 | 6.5483 | 6.5002 | 6.0036 | 5.8794 | 5.8404 | 6.4622 | 5.3316 | 6.5296 | 6.0329 | 5.7740 | 6.3269 | 6.3172 | 7.1620 | 6.4505 | 6.5949 |
| KIF4A_S394 | KIF4A | O95239 | 961 | KEQQLLSTLKCQD | 8.1123 | 7.9089 | 7.0659 | 8.7953 | 6.7641 | 6.3608 | 7.4496 | 5.6228 | 6.2593 | 5.1262 | 6.1211 | 5.7028 | 4.7470 | 5.0998 | 4.1788 | 4.6853 |
| KIF4A_S508 | KIF4A | O95239 | 394 | LMEKNQSLVEENE | 5.9656 | 6.4700 | 6.9318 | 6.7273 | 5.9518 | 6.2056 | 6.5689 | 6.5091 | 6.3152 | 6.5842 | 6.3770 | 6.6008 | 5.4772 | 6.1254 | 5.3606 | 5.8296 |
| KIF4A_S801 | KIF4A | O95239 | 1001 | TLLQVASRQKHLP | 6.9954 | 8.1702 | 6.2219 | 7.6916 | 5.2163 | 6.1639 | 8.3394 | 8.2620 | 6.4346 | 8.3926 | 7.7399 | 7.7931 | 3.6113 | 3.1182 | 2.6493 | 3.2004 |
| KIF4A_S815 | KIF4A | O95239 | 1013 | PKDTLLSPDSSFE | 6.3434 | 6.8817 | 6.7185 | 7.3422 | 6.2592 | 6.4426 | 6.9537 | 6.4873 | 7.5382 | 7.1987 | 7.1780 | 7.4145 | 3.9089 | 4.6827 | 4.2023 | 4.4480 |
| KIF4A_S951 | KIF4A | O95239 | 815 | VSESEDSITKQIE | 5.7434 | 6.6816 | 6.1834 | 7.0361 | 6.3537 | 6.0315 | 6.4321 | 6.5585 | 6.4191 | 6.3137 | 6.3618 | 6.4782 | 5.5357 | 6.0896 | 5.9954 | 5.7862 |
| KIF4A_S953 | KIF4A | O95239 | 951 | EKQLEESVSEKEQ | 7.5972 | 8.2446 | 6.2711 | 8.2314 | 6.1052 | 6.4183 | 7.5110 | 6.3454 | 6.4993 | 6.5521 | 6.6296 | 6.8055 | 3.9810 | 5.0566 | 3.4136 | 4.3381 |
| KIF4A_S961 | KIF4A | O95239 | 1017 | LLSPDSSFEYVPP | 5.4296 | 6.4192 | 6.1424 | 6.6316 | 5.4829 | 7.0503 | 6.8930 | 6.6402 | 6.8293 | 7.1801 | 5.7941 | 6.3479 | 4.8243 | 6.2902 | 6.7072 | 5.3376 |
| KIF4A_T1161 | KIF4A | O95239 | 508 | PETSRSSDAFTTQ | 6.6605 | 7.4340 | 5.8516 | 7.2019 | 6.9960 | 6.2029 | 7.2245 | 6.4772 | 7.3051 | 6.4045 | 6.8951 | 6.4766 | 4.8275 | 4.8055 | 4.0680 | 5.1691 |
| KIF4A_T1181 | KIF4A | O95239 | 1181;1186 | NA | 6.1751 | 6.9639 | 6.7596 | 7.1887 | 5.5744 | 6.7752 | 6.6274 | 6.1885 | 6.7752 | 7.1045 | 6.5174 | 6.9386 | 4.7063 | 5.8504 | 4.4523 | 5.4026 |
| KIF4A_T1181;S1186 | KIF4A | O95239 | 1048;1051 | NA | 6.0110 | 7.2319 | 7.5552 | 7.6719 | 5.6731 | 6.8864 | 6.7619 | 7.3771 | 6.6336 | 6.8258 | 6.5212 | 7.2382 | 3.7283 | 4.8785 | 4.0905 | 4.9155 |
| KIF4A_T799 | KIF4A | O95239 | 1013;1017 | NA | 6.1095 | 10.8669 | 6.2222 | 8.3957 | 4.8385 | 10.3115 | 8.4712 | 5.5777 | 6.0349 | 8.5243 | 8.2532 | 7.2341 | 1.8359 | 2.4201 | 3.3665 | 1.5377 |
| KIF5B_S933 | KIF5B | P33176 | 933 | GQHPAASPTHPSA | 7.1888 | 6.5595 | 6.7399 | 7.4636 | 7.3882 | 8.0012 | 5.9439 | 8.0350 | 6.3932 | 6.0507 | 6.8686 | 6.8293 | 3.6953 | 4.9990 | 3.7641 | 4.0797 |
| KIFC1_S31 | KIFC1 | Q9BW19 | 6 | xMDPQRSPLLEVK | 6.1286 | 6.9385 | 6.0538 | 6.9091 | 5.9761 | 6.2216 | 6.3466 | 6.7120 | 6.6554 | 6.6590 | 6.3128 | 6.3040 | 5.0609 | 6.2576 | 5.5271 | 5.9369 |
| KIFC1_S6 | KIFC1 | Q9BW19 | 359 | SDERRGTLSGAPA | 7.8779 | 6.4510 | 6.3863 | 6.1653 | 6.6747 | 5.5166 | 6.2549 | 5.3354 | 7.4149 | 6.3935 | 6.6963 | 6.0371 | 5.4512 | 6.6211 | 5.1955 | 5.5283 |
| KIFC1_S71 | KIFC1 | Q9BW19 | 71 | SHPRVPSLTTVPQ | 5.9760 | 6.7606 | 6.4204 | 6.8530 | 5.8743 | 6.4901 | 6.7146 | 6.5990 | 6.7358 | 7.0032 | 7.0225 | 7.2308 | 4.6074 | 5.4410 | 4.9460 | 5.3254 |
| KIFC1_T359 | KIFC1 | Q9BW19 | 31 | PSQLPLSGSRLKR | 7.2152 | 7.8422 | 6.5021 | 7.5494 | 5.4666 | 7.1168 | 7.8190 | 7.0365 | 7.2315 | 7.4668 | 7.5956 | 7.8117 | 2.9804 | 3.8039 | 2.6071 | 3.9552 |
| KIN_S208 | KIN | O60870 | 238 | TIGSSASVKRKES | 6.9904 | 6.0875 | 7.6841 | 6.0269 | 4.7801 | 5.8168 | 6.5803 | 5.8579 | 8.6326 | 8.3272 | 8.8979 | 8.3831 | 3.6742 | 5.5040 | 2.5623 | 4.1947 |
| KIN_S238 | KIN | O60870 | 208 | KVTFNLSKGACSS | 5.2421 | 7.0861 | 5.2471 | 6.9862 | 6.8662 | 6.6663 | 5.3770 | 7.3360 | 6.3215 | 7.9606 | 7.3060 | 6.3515 | 4.1177 | 6.1916 | 5.0622 | 5.8818 |
| KIRREL1_S589 | KIRREL1 | Q96J84-2 | 638 | SSRAVLYADYRAP | 6.5126 | 6.5346 | 6.9723 | 6.3932 | 6.0530 | 6.7663 | 6.6118 | 6.7756 | 6.7697 | 6.4666 | 6.3883 | 6.7947 | 5.1200 | 5.5938 | 4.9863 | 5.2612 |
| KIRREL1_S590 | KIRREL1 | Q96J84-2 | 590 | MKAIYSSFKDDVD | 6.9254 | 6.4834 | 7.5804 | 6.7714 | 5.7991 | 6.1869 | 5.9821 | 6.7407 | 7.4755 | 7.7882 | 6.7031 | 7.1028 | 4.3202 | 5.4561 | 3.8397 | 4.8449 |
| KIRREL1_S699 | KIRREL1 | Q96J84-2 | 737 | GLERTPYEAYDPI | 6.7123 | 6.4772 | 6.6053 | 5.9177 | 6.4075 | 6.0570 | 6.4979 | 6.3940 | 6.8691 | 6.6244 | 6.4971 | 6.1457 | 5.7979 | 5.9563 | 5.0248 | 6.0156 |
| KIRREL1_S753 | KIRREL1 | Q96J84-2 | 753 | ATATRFSYTSQHS | 6.1807 | 6.1427 | 5.5940 | 6.2951 | 6.1827 | 6.4520 | 5.8699 | 6.8287 | 7.3564 | 6.1701 | 5.7674 | 6.7554 | 5.8089 | 6.7209 | 6.0037 | 5.8714 |
| KIRREL1_Y638 | KIRREL1 | Q96J84-2 | 699 | DTTSQLSYENYEK | 5.6676 | 6.1505 | 6.2095 | 6.7938 | 6.2676 | 6.1462 | 6.1973 | 5.9138 | 5.1292 | 7.2984 | 6.2866 | 7.1458 | 6.1470 | 6.3291 | 5.9702 | 6.3473 |

| | | | | | | | | | | | | | | | | | | | | |
|---|---|---|---|---|---|---|---|---|---|---|---|---|---|---|---|---|---|---|---|---|
| KIRREL1_Y737 | KIRREL1 | Q96J84-2 | 589 | VMKAIYSSFKDDV | 6.5162 | 6.8638 | 6.2746 | 5.6261 | 5.4318 | 6.4228 | 6.7438 | 6.0003 | 6.7609 | 6.4197 | 6.7152 | 6.7262 | 6.2527 | 5.5646 | 5.5312 | 6.1502 |
| KIZ_S276 | KIZ | Q2M2Z5 | 276 | KSAELNSPLRERL | 6.5340 | 6.5466 | 8.0901 | 6.4433 | 6.1640 | 6.6968 | 7.0050 | 7.5346 | 7.4611 | 6.1687 | 7.6129 | 6.4433 | 3.8256 | 4.5524 | 4.1378 | 4.7840 |
| KLC1_S162 | KLC1 | Q07866-4 | 600 | GMKRASSLNVLNV | 6.1035 | 6.0272 | 6.9680 | 5.6522 | 5.8445 | 6.7575 | 6.2525 | 6.7185 | 6.4656 | 6.8254 | 7.0378 | 7.1461 | 5.5262 | 5.7923 | 5.1681 | 5.7148 |
| KLC1_S460 | KLC1 | Q07866-4 | 631 | SRALSASHTDLAH | 6.1533 | 6.4090 | 6.6458 | 6.2567 | 6.0280 | 6.5656 | 6.2290 | 6.7347 | 6.2863 | 6.2555 | 6.7957 | 7.0378 | 5.4587 | 5.8581 | 5.5081 | 5.7776 |
| KLC1_S521 | KLC1 | Q07866-4 | 460 | KACKVDSPTVTTT | 6.5061 | 7.1400 | 7.3327 | 7.1100 | 6.3456 | 6.9947 | 6.9882 | 6.5264 | 6.9539 | 7.3279 | 7.2538 | 7.1462 | 3.8293 | 4.5449 | 3.6936 | 4.3066 |
| KLC1_S521;S524 | KLC1 | Q07866-4 | 599 | PGMKRASSLNVLN | 6.0842 | 5.9104 | 7.0871 | 5.9365 | 5.6921 | 6.7998 | 6.0600 | 6.5813 | 6.4482 | 7.0396 | 7.1571 | 7.4836 | 5.2870 | 5.4684 | 5.1242 | 5.8403 |
| KLC1_S524 | KLC1 | Q07866-4 | 524 | KRRSRESLNVDVV | 6.7080 | 6.5718 | 7.4428 | 6.7595 | 6.2772 | 6.6018 | 6.9588 | 6.5927 | 6.4165 | 6.7219 | 7.0461 | 6.8158 | 4.6984 | 5.3846 | 4.0743 | 4.9297 |
| KLC1_S561 | KLC1 | Q07866-4 | 162 | KYDDDISPSEDKD | 5.8815 | 6.1272 | 6.8128 | 5.8670 | 6.1993 | 6.4143 | 6.0294 | 6.7032 | 6.3223 | 6.4259 | 6.5431 | 6.9527 | 6.1343 | 6.4566 | 5.3404 | 5.7898 |
| KLC1_S599 | KLC1 | Q07866-4 | 521 | NMEKRRSRESLNV | 7.1727 | 7.0706 | 7.5036 | 6.5253 | 6.0531 | 5.9583 | 5.9880 | 6.1447 | 6.8715 | 7.0330 | 6.7487 | 7.2378 | 5.1579 | 4.8349 | 5.0943 | 4.6057 |
| KLC1_S600 | KLC1 | Q07866-4 | 561 | SLKRSGSFSKLRA | 6.2853 | 6.5276 | 7.5667 | 5.8452 | 7.6039 | 8.1104 | 7.4592 | 4.8501 | 4.6083 | 5.7590 | 6.5283 | 6.3984 | 6.3109 | 5.8987 | 5.0019 | 5.2462 |
| KLC1_S631 | KLC1 | Q07866-4 | 521;524 | NA | 7.4553 | 6.6893 | 7.3629 | 6.3294 | 6.4374 | 6.5355 | 6.7094 | 6.5481 | 7.3351 | 6.6123 | 7.5419 | 6.4842 | 4.3775 | 4.6053 | 4.1046 | 4.8716 |
| KLC2_S581 | KLC2 | Q9H0B6 | 610 | SRTLSSSSMDLSR | 6.6620 | 6.8522 | 7.1068 | 6.4847 | 6.3566 | 6.7060 | 6.4809 | 6.4606 | 6.6015 | 6.2670 | 6.6720 | 6.4989 | 5.4719 | 5.2406 | 4.7085 | 5.4297 |
| KLC2_S582 | KLC2 | Q9H0B6 | 582 | RMKRASSLNFLNK | 6.0627 | 5.8048 | 6.8897 | 5.8767 | 6.0909 | 6.9871 | 5.6060 | 6.6127 | 7.0710 | 6.6200 | 6.7781 | 7.3831 | 5.8293 | 5.5600 | 4.8999 | 5.9279 |
| KLC2_S610 | KLC2 | Q9H0B6 | 581 | PRMKRASSLNFLN | 6.3355 | 6.1339 | 6.9605 | 5.5225 | 5.8664 | 6.8951 | 6.1529 | 6.9190 | 6.7000 | 6.8963 | 6.8245 | 7.0374 | 5.3176 | 5.5399 | 5.2662 | 5.6323 |
| KLC3_S480 | KLC3 | Q6P597-3 | 480 | GMKRAMSLNTLNV | 6.4636 | 5.7002 | 6.7601 | 5.4713 | 6.0656 | 6.2119 | 5.8606 | 6.9281 | 6.1070 | 6.2089 | 7.1073 | 6.9199 | 6.3669 | 6.2518 | 5.5494 | 6.0275 |
| KLC4_S181 | KLC4 | Q9NSK0-3 | 608 | NMKRAASLNYLNQ | 6.2050 | 5.3987 | 7.0447 | 5.9236 | 6.1023 | 6.9611 | 6.0710 | 6.7165 | 6.5119 | 6.5574 | 6.7225 | 6.4175 | 6.1274 | 6.1840 | 5.4249 | 5.6314 |
| KLC4_S478 | KLC4 | Q9NSK0-3 | 629 | SRGLSASTMDLSS | 6.4225 | 6.3765 | 6.8864 | 6.5271 | 6.4694 | 6.4943 | 6.1674 | 6.4745 | 6.5624 | 6.0055 | 6.2981 | 6.3920 | 5.8022 | 6.1562 | 5.3425 | 5.6231 |
| KLC4_S537 | KLC4 | Q9NSK0-3 | 478 | KACKVSSPTVNTT | 7.2919 | 7.6170 | 7.0685 | 7.3326 | 6.5375 | 7.0786 | 6.8742 | 7.0985 | 7.1610 | 7.5334 | 7.5593 | 7.0697 | 3.3232 | 3.8341 | 2.9815 | 3.6392 |
| KLC4_S584 | KLC4 | Q9NSK0-3 | 584 | DVLRRSSELLVRK | 6.2600 | 5.4082 | 6.1049 | 5.7969 | 7.2564 | 6.0764 | 5.3896 | 6.4750 | 6.9500 | 6.5177 | 7.3458 | 7.0484 | 6.8849 | 7.3057 | 4.6259 | 4.5544 |
| KLC4_S608 | KLC4 | Q9NSK0-3 | 537 | SDGRRTSQEGPGD | 5.6443 | 6.4496 | 6.2524 | 6.6344 | 5.4992 | 6.2872 | 6.4109 | 5.7502 | 6.9060 | 7.3939 | 7.3160 | 7.3730 | 5.6789 | 5.8120 | 4.8738 | 5.7181 |
| KLC4_S629 | KLC4 | Q9NSK0-3 | 181 | DEDGHTSEEKEGD | 5.9635 | 5.9529 | 7.0986 | 5.6378 | 6.3702 | 6.5354 | 6.0987 | 6.9452 | 6.5511 | 6.2856 | 6.7973 | 5.8693 | 6.4781 | 6.0906 | 5.0786 | 6.2472 |
| KLF10_S140 | KLF10 | Q13118 | 140 | FKEEEKSPVSAPK | 6.7868 | 6.9172 | 6.5793 | 6.9468 | 5.8928 | 6.6295 | 6.5678 | 6.2198 | 7.1193 | 6.9679 | 7.1202 | 6.7705 | 4.5341 | 5.0449 | 4.7649 | 5.1382 |
| KLF10_S206 | KLF10 | Q13118 | 206 | IPCAAVSPNRSKC | 6.1716 | 5.8993 | 6.3394 | 6.6533 | 5.1685 | 6.7193 | 6.7373 | 6.0819 | 6.2998 | 6.5185 | 6.4298 | 6.6184 | 6.2370 | 5.9182 | 5.7731 | 6.4347 |
| KLF10_S249 | KLF10 | Q13118 | 249 | SQPAPVSPQQKSV | 6.2119 | 6.0043 | 6.5248 | 6.8123 | 5.8169 | 5.7321 | 6.0122 | 6.1460 | 6.7686 | 6.5319 | 6.8712 | 6.6433 | 5.7189 | 5.7990 | 5.9939 | 6.4127 |
| KLF11_S124 | KLF11 | O14901 | 124 | STRTPVSPQVTDS | 6.2228 | 6.4117 | 7.1321 | 6.8457 | 5.7677 | 6.2414 | 6.0789 | 5.9437 | 6.1653 | 6.3933 | 6.1900 | 6.1850 | 6.0472 | 6.3286 | 5.9109 | 6.1356 |
| KLF11_T107;S111 | KLF11 | O14901 | 107;111 | NA | 5.4760 | 8.5817 | 7.8748 | 6.6559 | 6.5664 | 4.8714 | 4.3897 | 4.4306 | 6.7959 | 6.3465 | 8.1740 | 4.9497 | 5.3753 | 5.4986 | 8.7238 | 5.2896 |
| KLF13_S162 | KLF13 | Q9Y2Y9 | 162 | SRADLESPQRKHK | 5.8584 | 6.1628 | 6.6191 | 5.9347 | 6.5764 | 6.6882 | 6.5367 | 6.6296 | 6.7984 | 7.2320 | 6.9449 | 6.6066 | 5.6218 | 5.5442 | 5.0431 | 5.2031 |
| KLF16_S238 | KLF16 | Q9BXK1 | 238 | PCSLAGSPAPSPA | 6.5472 | 6.0138 | 6.6845 | 6.3424 | 6.0699 | 5.9546 | 6.4614 | 6.2706 | 6.3931 | 6.6110 | 6.2905 | 6.0429 | 6.3336 | 6.3252 | 5.7830 | 5.8762 |
| KLF16_S99 | KLF16 | Q9BXK1 | 99 | AAPGGASPASSSS | 7.7722 | 6.7341 | 5.3646 | 6.4245 | 4.4095 | 6.0450 | 5.5216 | 6.5379 | 8.1472 | 5.3428 | 6.7690 | 7.2619 | 6.8955 | 7.1921 | 6.0843 | 3.4979 |
| KLF16_T152 | KLF16 | Q9BXK1 | 152 | SHLRTHTGERPFA | 7.2114 | 7.4169 | 7.8161 | 7.9182 | 6.6336 | 7.5486 | 6.7324 | 6.9320 | 7.6885 | 7.1726 | 8.4718 | 8.8013 | 2.1399 | 2.6717 | 1.8661 | 2.9789 |
| KLF2_T173 | KLF2 | Q9Y5W3 | 173 | PPPPPDTPPLSPD | 6.0288 | 5.4223 | 7.4333 | 5.9233 | 6.0927 | 5.9594 | 6.8185 | 6.0940 | 5.7859 | 5.8775 | 5.4501 | 5.9317 | 6.9351 | 5.9025 | 7.1294 | 7.2154 |
| KLF2_T244;S248 | KLF2 | Q9Y5W3 | 244;248 | NA | 5.3062 | 5.5240 | 5.5765 | 5.6114 | 6.3419 | 6.4005 | 6.3713 | 6.2872 | 5.3707 | 5.8188 | 5.7079 | 5.8325 | 7.3603 | 7.2724 | 7.8418 | 7.3766 |
| KLF3_S111 | KLF3 | P57682 | 250 | RPLPVESPDTQRK | 5.9834 | 6.2505 | 6.6017 | 5.8650 | 5.9590 | 6.3401 | 5.9733 | 6.4811 | 6.4969 | 6.7271 | 6.8482 | 6.5672 | 5.9165 | 6.2247 | 5.7318 | 6.0335 |
| KLF3_S250 | KLF3 | P57682 | 111 | KKYSPPSPGVQPF | 5.9533 | 4.9450 | 4.9849 | 6.6969 | 6.5079 | 5.6886 | 7.0162 | 5.9239 | 4.3526 | 8.5749 | 7.4363 | 7.5330 | 6.1213 | 6.7305 | 6.5688 | 4.9660 |
| KLF3_S92;S101 | KLF3 | P57682 | 92;101 | NA | 5.9899 | 6.4233 | 6.3469 | 6.0165 | 6.2817 | 5.7337 | 6.2297 | 6.4348 | 6.5696 | 6.3921 | 6.8614 | 6.9095 | 6.0278 | 5.9342 | 6.0136 | 5.8351 |
| KLF4_S254 | KLF4 | O43474 | 254 | ISVSKGSPDGSHP | 5.4934 | 5.8700 | 6.3568 | 5.6676 | 5.7266 | 5.9233 | 5.6150 | 6.7431 | 6.1390 | 6.9658 | 6.6617 | 6.6792 | 6.9771 | 7.1148 | 6.2226 | 5.8440 |
| KLF5_S153 | KLF5 | Q13887 | 153 | RTGLYKSQRPCVT | 5.0705 | 5.4928 | 5.9735 | 4.2750 | 5.0516 | 5.3207 | 4.6070 | 5.3346 | 5.9786 | 4.8165 | 4.8399 | 5.7685 | 9.9533 | 8.7589 | 9.4581 | 9.3004 |
| KLF6_S171 | KLF6 | Q99612 | 171 | VPGELPSPGKVRS | 5.5116 | 5.8881 | 6.0176 | 6.1453 | 6.1077 | 5.9998 | 5.6468 | 6.1044 | 6.5448 | 6.0237 | 6.0836 | 6.4384 | 7.3385 | 7.1307 | 6.5502 | 6.4688 |
| KLF7_S65 | KLF7 | O75840 | 65 | DCFLHASPPPCIE | 5.0127 | 5.5104 | 5.5451 | 5.3183 | 5.9964 | 5.7445 | 5.9933 | 5.9031 | 5.4646 | 5.4921 | 5.7897 | 5.3841 | 8.7388 | 7.2453 | 8.6997 | 8.1619 |
| KLHL22_T605 | KLHL22 | Q53GT1 | 605 | LEPPRGTPDRSQA | 6.6627 | 6.6665 | 6.7187 | 6.3100 | 5.9625 | 6.3316 | 5.9891 | 6.7251 | 6.3213 | 6.1852 | 6.6853 | 6.2650 | 5.6511 | 6.2201 | 5.6197 | 5.6860 |
| KLHL36_T329 | KLHL36 | Q8N4N3 | 329 | EQWVKETPLPARR | 6.0898 | 7.2526 | 7.1536 | 6.4934 | 5.3176 | 6.8355 | 7.0029 | 7.5544 | 6.0515 | 7.0870 | 7.0175 | 7.5044 | 3.7806 | 4.8720 | 4.6128 | 5.3744 |
| KMT2A_S1115;S1119 | KMT2A | Q03164-3 | 337 | RKDKEGTPPLTKE | 6.7104 | 6.3483 | 6.4687 | 6.6319 | 5.6167 | 5.1922 | 6.3551 | 5.3702 | 6.3749 | 6.4207 | 5.9323 | 6.4594 | 6.1531 | 6.5752 | 6.1554 | 7.2355 |
| KMT2A_S142 | KMT2A | Q03164-3 | 518 | PLPISQSPENESN | 6.7213 | 6.4589 | 6.8275 | 6.9181 | 6.1999 | 6.0216 | 6.2108 | 6.2763 | 6.4794 | 5.9271 | 6.2791 | 6.3707 | 5.6404 | 5.7964 | 5.5301 | 6.3425 |
| KMT2A_S153 | KMT2A | Q03164-3 | 1840 | GPGEPDSPTPLHP | 6.2375 | 6.3592 | 6.1410 | 6.9358 | 6.1832 | 6.1869 | 6.3179 | 6.0303 | 6.4954 | 5.9351 | 6.1280 | 6.1497 | 5.8442 | 6.5670 | 6.3251 | 6.1637 |
| KMT2A_S161;S165 | KMT2A | Q03164-3 | 2869 | MQALGESPESSSS | 5.9494 | 6.0173 | 6.0494 | 6.3362 | 6.4024 | 5.8499 | 6.0919 | 5.8232 | 6.4281 | 5.9387 | 6.2837 | 5.9936 | 6.5287 | 6.8816 | 6.6559 | 6.7699 |
| KMT2A_S1688 | KMT2A | Q03164-3 | 2528 | TVKVTLTPLKMEN | 7.2795 | 7.2029 | 6.1941 | 7.0589 | 6.9117 | 6.5295 | 7.1146 | 6.7879 | 6.9567 | 6.3618 | 6.1322 | 6.2323 | 4.3955 | 5.2766 | 4.6281 | 4.9378 |
| KMT2A_S1840 | KMT2A | Q03164-3 | 3530 | SGQRSASPSVPGP | 6.0036 | 6.1821 | 5.7883 | 6.4240 | 5.9771 | 5.9472 | 6.2054 | 6.0405 | 6.0907 | 6.1553 | 6.0622 | 6.0333 | 6.7516 | 6.6911 | 6.8224 | 6.8253 |
| KMT2A_S1840;T1848 | KMT2A | Q03164-3 | 1688 | SIPSRSSPEGPDP | 5.7083 | 5.8453 | 5.4016 | 6.3327 | 6.0777 | 5.6562 | 5.9296 | 6.0765 | 5.9270 | 5.8166 | 5.8932 | 5.9167 | 6.8658 | 7.5520 | 7.6588 | 7.3421 |
| KMT2A_S187 | KMT2A | Q03164-3 | 351 | KTVVRQSPRRIKP | 7.1147 | 7.5396 | 6.6221 | 7.1564 | 6.4529 | 7.0269 | 6.3512 | 5.0044 | 5.4096 | 6.1824 | 6.6540 | 6.4196 | 5.4635 | 6.2671 | 5.1372 | 5.1984 |
| KMT2A_S197 | KMT2A | Q03164-3 | 3039 | GLQVPVSPTVPIQ | 5.8571 | 6.0778 | 6.3362 | 6.4164 | 6.3270 | 6.0729 | 6.5482 | 6.0185 | 6.1996 | 6.2373 | 6.4515 | 6.1771 | 6.2327 | 6.3598 | 6.2425 | 6.4455 |
| KMT2A_S2154 | KMT2A | Q03164-3 | 458 | SAPSCGSSEKSSA | 7.1154 | 7.0808 | 6.9759 | 8.0430 | 6.2451 | 6.7431 | 6.8612 | 6.5917 | 7.8293 | 7.3820 | 7.7702 | 7.4188 | 2.9638 | 4.0838 | 3.2575 | 3.6384 |
| KMT2A_S2159;S2170 | KMT2A | Q03164-3 | 2958 | LAVISDSGEKRVT | 6.6716 | 6.3672 | 7.1387 | 8.3735 | 6.4255 | 6.6676 | 7.5063 | 6.7292 | 7.1543 | 7.2740 | 6.6587 | 6.8928 | 3.3653 | 4.4469 | 3.6672 | 4.6611 |
| KMT2A_S2170 | KMT2A | Q03164-3 | 2941 | SVLTTRSPTVPSQ | 6.8771 | 7.3138 | 6.2194 | 7.3495 | 6.3344 | 6.3688 | 6.6095 | 6.5109 | 6.8437 | 6.4393 | 6.5361 | 6.6587 | 4.8443 | 5.2228 | 4.8776 | 4.9940 |
| KMT2A_S2199 | KMT2A | Q03164-3 | 2170 | PLPSAGSPTPTTH | 5.4241 | 5.5320 | 5.6313 | 6.5337 | 6.0749 | 5.7034 | 6.3813 | 5.8101 | 6.5563 | 6.8282 | 6.2302 | 6.4326 | 5.9771 | 7.1013 | 6.6431 | 7.1404 |
| KMT2A_S261 | KMT2A | Q03164-3 | 2154 | IRTPSYSPTQRSP | 6.7189 | 6.6801 | 6.0920 | 7.2607 | 6.3215 | 6.1458 | 6.6130 | 6.3647 | 6.5231 | 6.6110 | 6.3790 | 6.8345 | 4.7825 | 5.7106 | 5.4134 | 5.5492 |
| KMT2A_S279 | KMT2A | Q03164-3 | 2199 | IGSRRHSTSSLSP | 4.4761 | 4.6748 | 3.9591 | 5.3249 | 6.9387 | 5.4651 | 4.4819 | 5.7394 | 6.7219 | 6.2086 | 6.5519 | 7.2749 | 8.0785 | 9.4098 | 7.1794 | 7.5149 |
| KMT2A_S2869 | KMT2A | Q03164-3 | 680 | ASARLFSPLHSGT | 8.4908 | 7.8463 | 6.8882 | 7.1199 | 5.8872 | 6.7732 | 7.3332 | 6.9772 | 7.9035 | 7.4715 | 6.9262 | 7.8355 | 2.5048 | 3.9889 | 2.3559 | 3.6977 |
| KMT2A_S2941 | KMT2A | Q03164-3 | 153 | QFLGFGSDEEVRV | 6.6702 | 6.9296 | 6.8939 | 7.0197 | 6.1181 | 6.3215 | 6.3136 | 6.1815 | 6.9531 | 6.0856 | 6.3261 | 6.6912 | 4.6952 | 6.1779 | 5.3621 | 5.2606 |
| KMT2A_S2958 | KMT2A | Q03164-3 | 3514 | VQASPTSPGGSPS | 6.2125 | 5.9557 | 5.9785 | 6.2784 | 6.8704 | 6.1906 | 6.1434 | 5.9433 | 6.3530 | 5.5456 | 5.2875 | 6.0423 | 6.5709 | 6.6330 | 6.9400 | 7.0551 |
| KMT2A_S3039 | KMT2A | Q03164-3 | 187 | SGSDRNSAILSDP | 6.8161 | 5.7643 | 4.8083 | 6.8528 | 6.0456 | 5.2895 | 5.8969 | 6.3589 | 7.7889 | 5.7738 | 5.9481 | 7.1086 | 5.7914 | 7.4906 | 6.7793 | 6.4868 |
| KMT2A_S308 | KMT2A | Q03164-3 | 3070 | NAAVQTTPPHLKP | 6.1009 | 6.6273 | 6.7098 | 6.6997 | 6.3422 | 5.6021 | 6.3063 | 6.4821 | 5.9927 | 5.4441 | 6.2179 | 6.1546 | 6.0703 | 5.8837 | 6.8003 | 6.5660 |
| KMT2A_S351 | KMT2A | Q03164-3 | 261 | KIKRTPSATFQQA | 6.0227 | 5.8119 | 5.6724 | 5.8864 | 5.0799 | 6.1754 | 5.9616 | 6.4066 | 6.7834 | 7.2873 | 7.0348 | 6.8370 | 5.9924 | 6.5202 | 5.3261 | 7.2021 |
| KMT2A_S3514 | KMT2A | Q03164-3 | 279 | LRAGKLSPLKSKF | 6.8935 | 6.5164 | 6.0233 | 6.5589 | 6.6765 | 5.9197 | 6.4306 | 6.4957 | 6.2784 | 6.2463 | 5.9869 | 6.0491 | 5.5051 | 6.3487 | 5.9330 | 6.1379 |
| KMT2A_S3514;S3521 | KMT2A | Q03164-3 | 627 | SLKHSRSEPQYFS | 8.3870 | 6.7138 | 7.7094 | 5.9190 | 5.7475 | 5.6555 | 6.1742 | 5.6346 | 7.3747 | 7.5128 | 10.0310 | 6.3833 | 4.3587 | 4.6432 | 3.5974 | 4.1580 |
| KMT2A_S3521 | KMT2A | Q03164-3 | 532 | RRSRRYSVSERSF | 6.2447 | 5.6514 | 5.1475 | 6.0622 | 6.2719 | 6.0129 | 5.5872 | 5.8475 | 7.4830 | 6.8434 | 6.2570 | 7.5820 | 6.4505 | 7.2168 | 5.0217 | 6.3204 |
| KMT2A_S3530 | KMT2A | Q03164-3 | 197 | SDPSVFSPLNKSE | 6.7017 | 6.7766 | 7.1433 | 6.7744 | 7.0785 | 6.6129 | 7.0218 | 6.4267 | 7.0298 | 6.8268 | 7.2015 | 6.9890 | 4.2180 | 4.8255 | 3.9707 | 4.4028 |
| KMT2A_S3662 | KMT2A | Q03164-3 | 3662 | EQKRKESITEKKP | 5.7709 | 6.1106 | 5.4952 | 6.1683 | 6.7204 | 5.3871 | 5.7274 | 6.1199 | 7.0557 | 7.0365 | 7.8612 | 6.8552 | 5.7448 | 6.1261 | 5.6684 | 6.1522 |
| KMT2A_S458 | KMT2A | Q03164-3 | 142 | SGGGGGSGEDEQF | 5.8880 | 6.8147 | 6.6636 | 6.1449 | 7.2580 | 4.3367 | 3.4502 | 4.0193 | 5.7167 | 6.7845 | 6.2809 | 5.6513 | 7.7465 | 7.8221 | 8.8244 | 6.5982 |
| KMT2A_S518 | KMT2A | Q03164-3 | 3521 | PGGSPSSPSSGQR | 5.8457 | 6.2752 | 6.7788 | 5.8797 | 5.2340 | 5.5800 | 5.9214 | 5.1660 | 7.0646 | 6.8082 | 6.1454 | 6.0898 | 6.5177 | 7.0507 | 6.6985 | 6.9441 |
| KMT2A_S532 | KMT2A | Q03164-3 | 926 | DVATSSSAKKATG | 6.8449 | 7.5862 | 6.2288 | 7.0421 | 6.4217 | 6.3475 | 7.7301 | 7.3232 | 6.9287 | 7.8144 | 6.5470 | 6.6547 | 3.6751 | 5.0625 | 3.6490 | 4.1442 |
| KMT2A_S627 | KMT2A | Q03164-3 | 750 | IRSEPRSPSHSMR | 6.6078 | 7.1304 | 5.9346 | 7.1760 | 5.6592 | 5.6603 | 6.1462 | 6.0927 | 6.3122 | 6.2813 | 5.8463 | 6.1161 | 5.6736 | 6.8613 | 6.1073 | 6.3949 |
| KMT2A_S680 | KMT2A | Q03164-3 | 314 | STERIKTPSGLLI | 5.8840 | 6.0883 | 6.1981 | 6.4594 | 6.2506 | 6.2838 | 6.5968 | 6.3117 | 6.5026 | 6.2737 | 6.2901 | 6.1870 | 5.9707 | 6.2112 | 5.8043 | 6.6876 |
| KMT2A_S680;S684 | KMT2A | Q03164-3 | 829 | PAEPFSSSSPTPL | 5.3269 | 5.8088 | 6.9308 | 6.1596 | 6.5989 | 7.2041 | 6.4140 | 5.8878 | 7.7667 | 6.1217 | 6.4646 | 6.7648 | 4.3471 | 5.7109 | 6.0869 | 6.4061 |
| KMT2A_S750 | KMT2A | Q03164-3 | 308 | RRGRPPSTERIKT | 6.1322 | 6.1006 | 6.2301 | 7.0658 | 5.7056 | 5.8986 | 6.9062 | 6.4940 | 6.3382 | 5.8666 | 6.6146 | 6.4943 | 5.5333 | 6.0863 | 5.9487 | 6.5848 |
| KMT2A_S829 | KMT2A | Q03164-3 | 1540 | KSCGSTTPGKGWD | 5.6721 | 6.8909 | 5.9475 | 6.8017 | 6.5671 | 6.6272 | 6.6136 | 7.0237 | 6.9811 | 6.7552 | 7.6210 | 6.3402 | 4.2032 | 5.3909 | 5.0196 | 5.5451 |
| KMT2A_S926 | KMT2A | Q03164-3 | 1288 | ESKQATTPASRKS | 7.8610 | 8.0891 | 6.9775 | 7.5700 | 6.8828 | 6.5265 | 7.3517 | 6.8779 | 7.3558 | 6.3037 | 6.7718 | 6.9485 | 3.1065 | 4.1248 | 3.3053 | 3.9472 |
| KMT2A_T1288 | KMT2A | Q03164-3 | 3514;3521 | NA | 6.2587 | 6.4885 | 6.1910 | 7.0030 | 6.1275 | 6.3367 | 6.0867 | 6.3455 | 6.2429 | 6.2163 | 6.3406 | 6.0320 | 5.7290 | 6.1554 | 5.9701 | 6.4762 |
| KMT2A_T1540 | KMT2A | Q03164-3 | 1840;1848 | NA | 5.5578 | 6.1793 | 5.4800 | 5.8121 | 6.1077 | 5.9233 | 6.0980 | 6.2509 | 5.5451 | 5.7392 | 5.7163 | 5.5337 | 6.8490 | 7.7851 | 7.6177 | 7.8049 |
| KMT2A_T2150;T2154 | KMT2A | Q03164-3 | 161;165 | NA | 7.6729 | 7.4542 | 6.7451 | 7.7739 | 6.3961 | 6.5058 | 6.8112 | 6.2426 | 6.9668 | 7.0107 | 6.9779 | 7.4271 | 3.5449 | 4.5542 | 3.6824 | 4.2342 |
| KMT2A_T2528 | KMT2A | Q03164-3 | 2159;2170 | NA | 5.6161 | 6.7847 | 6.5566 | 6.7684 | 6.4492 | 6.1221 | 7.1074 | 6.8802 | 6.5416 | 6.8838 | 6.5212 | 7.1121 | 4.6366 | 5.5920 | 4.8771 | 5.5507 |
| KMT2A_T3031;S3039 | KMT2A | Q03164-3 | 1115;1119 | NA | 5.8436 | 5.9219 | 5.4825 | 6.7584 | 5.4696 | 6.7626 | 6.1118 | 5.9295 | 5.8115 | 6.0471 | 6.2433 | 6.2754 | 6.8191 | 7.3748 | 6.6446 | 6.5043 |
| KMT2A_T3070 | KMT2A | Q03164-3 | 2150;2154 | NA | 6.6671 | 7.0666 | 6.3009 | 6.9599 | 5.9129 | 6.5151 | 6.7639 | 7.0577 | 7.2645 | 6.7819 | 7.2699 | 7.0181 | 3.9347 | 5.3429 | 4.5143 | 4.6295 |
| KMT2A_T314 | KMT2A | Q03164-3 | 680;684 | NA | 7.9728 | 7.5151 | 6.3325 | 8.2683 | 5.4237 | 6.7936 | 7.1991 | 6.3777 | 7.3186 | 7.4404 | 7.2668 | 6.9197 | 3.3106 | 4.5194 | 2.9263 | 4.4153 |
| KMT2A_T337 | KMT2A | Q03164-3 | 3031;3039 | NA | 6.0210 | 6.0060 | 6.6686 | 6.6551 | 6.0462 | 5.5920 | 6.6020 | 6.2010 | 6.2529 | 6.5253 | 6.4755 | 6.9680 | 6.0157 | 6.1632 | 5.3830 | 6.4245 |
| KMT2B_S1032;S1035 | KMT2B | Q9UMN6 | 2083 | QPRGQGTPPSGPG | 6.3143 | 6.8133 | 6.0520 | 6.9607 | 6.2741 | 6.0976 | 6.3987 | 6.1972 | 6.6019 | 6.4190 | 6.1237 | 6.4803 | 5.5979 | 5.9249 | 5.8151 | 5.9293 |
| KMT2B_S1092;S1095 | KMT2B | Q9UMN6 | 821 | AASMPLSPGGQME | 6.3553 | 6.7629 | 6.2459 | 6.9427 | 6.2629 | 6.2209 | 6.6859 | 6.4532 | 6.9435 | 6.4992 | 6.6598 | 6.5777 | 5.1439 | 5.5563 | 5.2258 | 5.4640 |
| KMT2B_S1164 | KMT2B | Q9UMN6 | 2288 | STFSGRSPPAPPP | 7.0580 | 7.3969 | 6.7454 | 7.5941 | 6.5215 | 6.2596 | 6.7462 | 6.7765 | 7.2781 | 6.9644 | 7.0488 | 7.1516 | 3.7236 | 4.3079 | 3.9893 | 4.4380 |
| KMT2B_S1836 | KMT2B | Q9UMN6 | 713 | FAPVVTTPVKAEV | 7.4243 | 6.7509 | 5.8062 | 6.9388 | 5.5126 | 6.3233 | 7.3528 | 5.7890 | 7.5002 | 5.5599 | 6.1404 | 6.3100 | 5.5072 | 6.2161 | 4.5855 | 6.2828 |
| KMT2B_S1873 | KMT2B | Q9UMN6 | 1930 | PPSRWASPPLKTS | 7.1405 | 7.6914 | 6.5466 | 7.3317 | 6.6198 | 6.5996 | 7.3884 | 6.7486 | 7.8131 | 7.0189 | 7.3169 | 7.1222 | 3.1617 | 4.2716 | 3.3373 | 3.8918 |
| KMT2B_S1917 | KMT2B | Q9UMN6 | 632 | PPPPAPSPPPAPA | 6.5547 | 6.0386 | 6.1704 | 6.1757 | 6.4682 | 5.4329 | 6.0633 | 6.6884 | 6.5981 | 6.0136 | 6.2361 | 6.6846 | 6.1841 | 6.7364 | 6.0528 | 5.9021 |
| KMT2B_S1930 | KMT2B | Q9UMN6 | 351 | GSGQGGSPCWKKQ | 6.5798 | 7.0053 | 5.9191 | 6.6408 | 6.5118 | 6.2228 | 6.6318 | 6.3595 | 7.1868 | 6.5188 | 6.4253 | 6.2142 | 5.3361 | 5.5820 | 5.1702 | 5.6958 |
| KMT2B_S2270 | KMT2B | Q9UMN6 | 861 | RPSGPESPVQGPR | 6.0178 | 5.9360 | 5.9944 | 6.8417 | 6.3834 | 5.5281 | 6.5388 | 6.2840 | 7.0971 | 6.5113 | 6.4659 | 6.3179 | 6.0944 | 6.3184 | 5.5718 | 6.0989 |
| KMT2B_S2288 | KMT2B | Q9UMN6 | 588 | SPPRAPTPPSTPV | 5.5866 | 6.1483 | 5.9430 | 6.2990 | 5.6575 | 5.6772 | 6.3557 | 6.2333 | 6.7107 | 6.2370 | 6.2279 | 6.3416 | 6.4537 | 6.8585 | 6.5936 | 6.6763 |
| KMT2B_S2348 | KMT2B | Q9UMN6 | 493 | AGPEGTSPPTPTP | 6.2583 | 6.6877 | 6.5354 | 7.3784 | 6.1289 | 6.1492 | 6.6708 | 6.1657 | 6.6414 | 7.3220 | 6.9254 | 6.6944 | 4.7866 | 5.7488 | 4.6574 | 5.2495 |
| KMT2B_S2455 | KMT2B | Q9UMN6 | 2348 | EPAGEESPGPLQE | 6.4422 | 5.9141 | 6.2351 | 8.1404 | 6.3944 | 6.3300 | 6.2013 | 6.5058 | 5.6971 | 6.3878 | 7.5678 | 6.5470 | 5.0930 | 6.1336 | 4.9948 | 5.4157 |
| KMT2B_S351 | KMT2B | Q9UMN6 | 1917 | RRSRRPSPLAPRP | 6.2839 | 6.4916 | 5.6551 | 7.0110 | 6.3062 | 6.1683 | 6.7353 | 6.7527 | 6.6981 | 6.3850 | 6.4589 | 6.5260 | 5.3567 | 5.9872 | 5.1257 | 6.0583 |
| KMT2B_S493 | KMT2B | Q9UMN6 | 653 | LRAPQFTPSEAHL | 7.4975 | 8.3448 | 6.6989 | 7.8211 | 6.3248 | 7.0360 | 7.7728 | 6.7988 | 6.7474 | 7.5143 | 7.1054 | 6.7352 | 3.0006 | 4.1965 | 2.7883 | 3.6177 |
| KMT2B_S570 | KMT2B | Q9UMN6 | 172 | LADVAPTPPKTPA | 5.9909 | 6.5576 | 6.3121 | 6.4141 | 6.4311 | 6.1250 | 5.9509 | 6.3681 | 6.9459 | 7.2954 | 6.2321 | 6.8354 | 5.0634 | 5.9466 | 5.5154 | 6.0161 |
| KMT2B_S570;S582;T588 | KMT2B | Q9UMN6 | 1164 | WTGKQKSPDGVHR | 6.5243 | 7.5841 | 6.3621 | 7.0930 | 5.6968 | 5.7550 | 6.1223 | 6.0441 | 6.2754 | 6.6517 | 6.2191 | 6.3538 | 4.9174 | 5.8759 | 6.1087 | 6.4162 |
| KMT2B_S582;T588 | KMT2B | Q9UMN6 | 1836 | GAPERHSPIQNLD | 6.8005 | 7.5695 | 6.4951 | 6.7144 | 6.6314 | 5.5585 | 4.8490 | 5.6102 | 6.3792 | 6.0895 | 6.0832 | 7.7481 | 5.8999 | 5.4943 | 5.5898 | 6.4873 |
| KMT2B_S632 | KMT2B | Q9UMN6 | 844 | DRGPVRSEDESVE | 6.6955 | 7.2747 | 6.8915 | 7.3799 | 6.8206 | 6.4607 | 7.2226 | 6.9975 | 6.3478 | 5.9908 | 5.5683 | 5.9536 | 4.8671 | 5.3809 | 4.9840 | 5.1645 |
| KMT2B_S821 | KMT2B | Q9UMN6 | 2270 | LSSGPASPPRQAI | 5.6697 | 6.2346 | 6.4742 | 6.9751 | 6.2722 | 6.1699 | 6.1329 | 6.2241 | 6.6209 | 6.0584 | 5.8286 | 6.4287 | 6.0473 | 6.3289 | 6.0757 | 6.4588 |
| KMT2B_S844 | KMT2B | Q9UMN6 | 2455 | ARLRHLSFSGMSG | 8.1559 | 6.3395 | 5.4412 | 6.3672 | 6.1021 | 6.4741 | 5.3067 | 6.9925 | 6.4860 | 3.8227 | 8.3102 | 5.5046 | 6.7076 | 7.2418 | 6.0625 | 4.6854 |
| KMT2B_S861 | KMT2B | Q9UMN6 | 475 | RGRPPLTPSQRAE | 7.3639 | 7.4847 | 6.1259 | 7.4799 | 6.1516 | 6.1001 | 7.2723 | 6.6068 | 7.3282 | 7.3989 | 6.5341 | 6.7137 | 3.9335 | 4.9051 | 4.0756 | 4.5256 |
| KMT2B_T172 | KMT2B | Q9UMN6 | 1873 | VPNYSPSRRPLGG | 8.0509 | 8.4287 | 7.5343 | 8.3098 | 6.4207 | 6.5504 | 7.2427 | 7.3605 | 8.1367 | 7.4282 | 7.4593 | 7.5109 | 2.0386 | 2.8736 | 2.0192 | 2.6355 |
| KMT2B_T2083 | KMT2B | Q9UMN6 | 570 | RPPITTSPPVPQE | 6.0349 | 6.2333 | 7.2405 | 6.8410 | 5.4202 | 6.2417 | 6.0968 | 6.3641 | 6.6019 | 6.8410 | 5.8422 | 6.1151 | 6.1742 | 6.0392 | 5.8169 | 6.0968 |
| KMT2B_T475 | KMT2B | Q9UMN6 | 582;588 | NA | 6.0058 | 6.3268 | 5.6837 | 5.8623 | 6.1987 | 5.4234 | 6.2544 | 6.4123 | 5.9483 | 5.7262 | 6.4526 | 6.0010 | 7.1025 | 7.0206 | 6.5545 | 7.0270 |
| KMT2B_T569;S582;T588 | KMT2B | Q9UMN6 | 570;582;588 | NA | 6.1193 | 6.0981 | 5.8846 | 6.8891 | 6.1494 | 6.1504 | 6.5292 | 6.4431 | 6.9908 | 5.9072 | 6.2059 | 7.0181 | 5.8958 | 6.1908 | 5.5944 | 5.9338 |
| KMT2B_T588 | KMT2B | Q9UMN6 | 1032;1035 | NA | 6.0164 | 6.0227 | 6.4465 | 6.1805 | 6.0036 | 6.3621 | 6.8303 | 6.5039 | 6.3973 | 7.1105 | 6.1003 | 5.9721 | 5.1414 | 6.0331 | 6.5863 | 6.2931 |
| KMT2B_T653 | KMT2B | Q9UMN6 | 569;582;588 | NA | 5.5786 | 5.6589 | 5.6554 | 7.2599 | 5.8828 | 5.0410 | 5.9097 | 6.7683 | 6.0611 | 7.0660 | 6.1989 | 7.2882 | 5.8094 | 6.3380 | 7.3379 | 6.1459 |
| KMT2B_T713 | KMT2B | Q9UMN6 | 1092;1095 | NA | 6.3155 | 6.4087 | 6.5958 | 7.3735 | 5.6160 | 6.6098 | 6.7228 | 6.6223 | 6.8720 | 5.9721 | 6.3384 | 5.6889 | 5.1891 | 6.2322 | 5.5592 | 5.8839 |
| KMT2C_S1493 | KMT2C | Q8NEZ4-3 | 4091 | PPEPVPSPIIPIL | 5.8720 | 6.2022 | 6.1100 | 6.7614 | 5.9598 | 6.1915 | 6.5639 | 6.2797 | 6.1979 | 6.1997 | 5.9134 | 5.9832 | 6.2639 | 6.4054 | 6.5101 | 6.5859 |
| KMT2C_S2068 | KMT2C | Q8NEZ4-3 | 1493 | QLDGILSPELDKM | 5.6878 | 6.1933 | 6.1787 | 6.4377 | 6.0293 | 6.0126 | 4.9745 | 5.5635 | 7.2763 | 5.7076 | 5.7797 | 6.2528 | 6.9264 | 6.5912 | 7.0163 | 7.3724 |
| KMT2C_S3758 | KMT2C | Q8NEZ4-3 | 2068 | QASRRLSVDPYER | 5.0354 | 4.7190 | 5.0354 | 5.6875 | 6.9178 | 6.3391 | 5.7874 | 6.4390 | 6.7785 | 8.5468 | 7.2461 | 7.7229 | 6.4690 | 6.9710 | 4.8745 | 5.4306 |
| KMT2C_S4091 | KMT2C | Q8NEZ4-3 | 4324 | IPSLPQSPMRETP | 6.5542 | 6.1478 | 5.7451 | 5.9354 | 6.4822 | 7.3726 | 6.3677 | 5.8338 | 6.1423 | 6.9108 | 6.4748 | 6.1700 | 5.3849 | 6.2993 | 6.3732 | 5.8061 |
| KMT2C_S4324 | KMT2C | Q8NEZ4-3 | 3758 | PVSSAQSPPHSAG | 5.9997 | 5.9080 | 6.1949 | 6.5432 | 6.0266 | 5.8681 | 6.5411 | 6.0503 | 6.7848 | 6.1366 | 6.4429 | 6.0277 | 6.0827 | 6.5012 | 5.8163 | 7.0760 |
| KMT2C_S4357;S4361 | KMT2C | Q8NEZ4-3 | 4357;4361 | NA | 6.0921 | 6.2880 | 6.1324 | 6.5055 | 6.0504 | 5.9229 | 6.4325 | 6.0240 | 6.0777 | 6.1935 | 6.2193 | 6.0947 | 6.2615 | 6.9155 | 6.4322 | 6.3578 |
| KMT2D_S1151 | KMT2D | O14686-3 | 4362 | PPPGRVSPAAAQL | 5.9289 | 5.6550 | 6.2434 | 6.5738 | 6.4073 | 6.0580 | 6.3037 | 5.7194 | 6.1582 | 6.2172 | 6.0056 | 6.2139 | 6.7243 | 6.7965 | 6.3872 | 6.6079 |
| KMT2D_S1151;T1164 | KMT2D | O14686-3 | 4741 | QEDRALSPVIPLI | 6.4144 | 6.6557 | 5.9145 | 6.7496 | 6.1110 | 5.8190 | 6.2271 | 6.1666 | 6.0619 | 6.2243 | 6.1273 | 5.9498 | 6.2342 | 6.4257 | 6.5759 | 6.3432 |
| KMT2D_S1606 | KMT2D | O14686-3 | 2643 | QRSGITSPVEKRE | 6.3261 | 6.4716 | 6.0318 | 6.5049 | 6.3080 | 6.1152 | 6.3729 | 6.0995 | 6.1404 | 6.0667 | 6.1725 | 6.1793 | 6.1457 | 6.3237 | 6.4044 | 6.3373 |
| KMT2D_S1671 | KMT2D | O14686-3 | 4852 | KGLEGKSPDTGPD | 6.1362 | 6.1349 | 5.9986 | 6.3852 | 7.5015 | 6.5699 | 6.0351 | 5.7844 | 5.9956 | 6.0507 | 5.6269 | 5.8183 | 7.1991 | 6.7286 | 7.3913 | 7.3537 |
| KMT2D_S2254 | KMT2D | O14686-3 | 1671 | KLEGPVSPDVEPG | 5.9298 | 6.4032 | 6.3107 | 6.7981 | 6.3439 | 5.9072 | 6.3134 | 6.0500 | 6.2386 | 6.2101 | 5.8175 | 5.9446 | 6.1950 | 6.6775 | 6.2656 | 6.5949 |
| KMT2D_S2263 | KMT2D | O14686-3 | 1151 | ASELKGSPVLLDP | 5.9184 | 6.2071 | 6.3204 | 6.8346 | 6.2985 | 5.9270 | 5.7394 | 5.9950 | 6.2021 | 6.4846 | 6.6847 | 6.2140 | 6.1154 | 6.3478 | 6.1983 | 6.5127 |
| KMT2D_S2277 | KMT2D | O14686-3 | 2263 | NLAVPESPGVGGG | 5.6188 | 5.8561 | 5.6726 | 6.2877 | 6.0309 | 6.0948 | 6.0632 | 6.1443 | 6.4098 | 6.0478 | 5.8454 | 6.1376 | 6.7545 | 6.6285 | 7.1417 | 7.2665 |
| KMT2D_S2417 | KMT2D | O14686-3 | 4825 | EVLFPESPARAGT | 5.8949 | 6.1035 | 6.0749 | 6.7689 | 6.2768 | 6.2210 | 6.2603 | 6.1221 | 6.4636 | 6.1528 | 5.9434 | 6.3344 | 6.2817 | 6.4335 | 6.2242 | 6.4439 |
| KMT2D_S2643 | KMT2D | O14686-3 | 3133 | EGGRHPSPCQFTI | 5.3517 | 5.9722 | 5.4010 | 6.2348 | 6.0236 | 5.8910 | 6.0871 | 5.6437 | 6.0181 | 6.0770 | 5.9244 | 6.0007 | 7.4450 | 7.5160 | 7.1699 | 7.2438 |
| KMT2D_S3133 | KMT2D | O14686-3 | 654 | PEVSRLSPLPVVS | 5.5396 | 5.8004 | 5.5583 | 6.2541 | 5.6479 | 5.0137 | 5.8268 | 6.7529 | 6.6595 | 5.9381 | 5.4432 | 5.8940 | 7.3675 | 7.2101 | 7.6226 | 7.4713 |
| KMT2D_S3202 | KMT2D | O14686-3 | 2243 | PRHQPSTPDPFLK | 5.1366 | 5.2793 | 5.2092 | 5.6253 | 5.4861 | 5.8725 | 6.2735 | 6.1586 | 5.4148 | 5.5382 | 5.4318 | 5.6912 | 8.1270 | 7.9321 | 8.5565 | 8.2673 |
| KMT2D_S3232 | KMT2D | O14686-3 | 1806 | PSFGLGTPKAKGD | 6.4030 | 6.1476 | 6.6275 | 5.9251 | 6.5568 | 6.0468 | 6.1848 | 5.9798 | 7.1603 | 6.6332 | 6.8325 | 6.2633 | 5.6561 | 6.1698 | 5.8273 | 5.5861 |
| KMT2D_S3623 | KMT2D | O14686-3 | 2277 | ASEPLLSPPPFGE | 5.9720 | 6.2069 | 5.8957 | 6.5499 | 6.1286 | 6.2591 | 6.2508 | 6.1646 | 6.1548 | 5.6925 | 6.0345 | 5.9093 | 6.5306 | 6.7966 | 6.6617 | 6.7924 |
| KMT2D_S373 | KMT2D | O14686-3 | 2417 | FSRVPASPQSQSS | 6.0817 | 6.5362 | 7.0303 | 6.4027 | 6.1267 | 6.4000 | 6.0409 | 6.0764 | 6.9043 | 6.3423 | 6.4223 | 6.4318 | 6.2703 | 5.4842 | 5.8529 | 5.5970 |
| KMT2D_S3989 | KMT2D | O14686-3 | 4685 | LLAALPTPPHNQT | 6.5752 | 6.6193 | 6.2494 | 6.8534 | 6.2571 | 5.9340 | 6.1619 | 6.3299 | 6.2978 | 6.1790 | 6.3806 | 6.5946 | 5.6751 | 6.2858 | 5.7943 | 5.8125 |
| KMT2D_S4362 | KMT2D | O14686-3 | 1606 | LRNLTMSPLHKRR | 7.2064 | 7.1192 | 6.6410 | 7.0677 | 6.4933 | 6.4543 | 6.6726 | 6.3356 | 6.6180 | 6.5544 | 6.7980 | 6.2051 | 5.0435 | 5.3391 | 4.7870 | 4.6650 |
| KMT2D_S4741 | KMT2D | O14686-3 | 2232 | PGEFHTTPPGTPR | 6.2077 | 5.6349 | 6.0647 | 6.7541 | 6.0391 | 5.8202 | 6.2296 | 5.7524 | 6.5642 | 6.1948 | 6.0345 | 6.1749 | 6.9743 | 6.8055 | 6.4135 | 6.3355 |
| KMT2D_S4825 | KMT2D | O14686-3 | 1846 | LEGKADTPGPEDG | 5.8829 | 5.8788 | 5.7096 | 6.2271 | 6.5541 | 5.4928 | 6.2737 | 5.5276 | 6.2570 | 6.2323 | 5.8608 | 5.5984 | 7.5770 | 6.7870 | 6.8747 | 7.2661 |
| KMT2D_S4852 | KMT2D | O14686-3 | 2254 | LKPRCPSLDNLAV | 6.0358 | 5.6292 | 6.5673 | 6.3231 | 5.9482 | 6.5079 | 6.7992 | 6.4925 | 5.9707 | 5.5553 | 5.5328 | 5.8114 | 6.5568 | 6.8318 | 6.4820 | 6.9559 |
| KMT2D_S654 | KMT2D | O14686-3 | 3232 | PGGPAASGDELDK | 6.2185 | 7.0647 | 7.2681 | 7.4009 | 5.9301 | 6.5651 | 6.3508 | 7.1679 | 6.8084 | 5.8668 | 6.0847 | 5.9057 | 5.2022 | 5.5845 | 5.3407 | 5.2411 |
| KMT2D_T1806 | KMT2D | O14686-3 | 373 | PVCSRFSPPEPGD | 5.5754 | 6.3732 | 5.9766 | 6.2205 | 5.6850 | 6.0776 | 6.0263 | 6.1783 | 6.5556 | 6.3873 | 6.1855 | 6.4795 | 6.4886 | 6.4373 | 6.4978 | 6.8554 |
| KMT2D_T1846 | KMT2D | O14686-3 | 3989 | QSRTLLSPQQQQQ | 6.0322 | 7.0102 | 6.0615 | 7.2515 | 5.3117 | 6.4012 | 7.2773 | 6.3392 | 6.6314 | 6.9746 | 6.6175 | 6.6126 | 5.0857 | 6.5121 | 4.8492 | 5.0320 |
| KMT2D_T1989 | KMT2D | O14686-3 | 3623 | SAVLALSPSQSPR | 5.7356 | 6.8845 | 5.0540 | 6.2778 | 6.3039 | 6.4138 | 7.2593 | 6.1122 | 6.7799 | 6.1261 | 5.4026 | 6.2970 | 6.1470 | 6.6822 | 5.9535 | 6.5707 |
| KMT2D_T2232 | KMT2D | O14686-3 | 3202 | AHLLTPSPLSGPG | 5.8109 | 5.8925 | 6.0397 | 6.0807 | 5.2649 | 6.4981 | 6.9919 | 6.0602 | 6.8736 | 6.2186 | 5.5994 | 6.5191 | 6.2163 | 6.7068 | 6.8349 | 6.3924 |
| KMT2D_T2243 | KMT2D | O14686-3 | 1989 | TPSTPTTPTTEGE | 6.1716 | 6.4207 | 6.6865 | 7.1069 | 6.1792 | 7.1797 | 6.9140 | 6.6840 | 5.8215 | 6.1741 | 6.4731 | 6.3364 | 5.2439 | 5.4394 | 5.1711 | 5.9978 |
| KMT2D_T4685 | KMT2D | O14686-3 | 1151;1164 | NA | 5.4025 | 5.7840 | 5.5753 | 6.3878 | 6.4205 | 6.0491 | 6.0777 | 5.8430 | 5.6854 | 5.8163 | 6.2646 | 5.9758 | 7.1000 | 7.1435 | 7.2404 | 7.2342 |
| KMT2E_S1002 | KMT2E | Q8IZD2 | 1359 | KMSKPGSPGSVIP | 6.0661 | 6.0253 | 6.1674 | 6.6317 | 6.3576 | 5.6487 | 6.4670 | 5.3603 | 5.9887 | 6.6142 | 6.0431 | 6.3520 | 6.9611 | 6.5583 | 6.1560 | 6.6026 |
| KMT2E_S1359 | KMT2E | Q8IZD2 | 854 | VNGENKSPLLLND | 5.8221 | 6.0982 | 5.6971 | 6.2358 | 6.1001 | 5.6953 | 5.9511 | 5.6840 | 6.1871 | 6.3859 | 6.1247 | 6.4458 | 6.5544 | 6.8739 | 6.9455 | 7.1991 |
| KMT2E_S1444 | KMT2E | Q8IZD2 | 1444 | KLHCPPSPHLENP | 5.8625 | 5.7932 | 5.7107 | 6.2980 | 6.0940 | 6.5394 | 6.2290 | 5.9194 | 6.1337 | 6.1374 | 5.8626 | 5.8872 | 7.0195 | 7.1157 | 6.7766 | 6.6212 |
| KMT2E_S522 | KMT2E | Q8IZD2 | 522 | TTNKMKSPETKQR | 7.4628 | 7.4368 | 6.5704 | 7.6364 | 6.4738 | 6.2742 | 6.8017 | 6.6336 | 6.9425 | 6.5685 | 6.9057 | 7.0252 | 4.1642 | 4.6799 | 3.9848 | 4.4398 |
| KMT2E_S795;S799 | KMT2E | Q8IZD2 | 641 | ENASKPTPAKVNR | 10.6380 | 8.6001 | 5.8647 | 8.2216 | 6.9606 | 6.9514 | 7.2668 | 5.2321 | 6.9329 | 5.1901 | 5.8241 | 5.9218 | 4.1929 | 4.7010 | 3.3441 | 4.1578 |
| KMT2E_S854 | KMT2E | Q8IZD2 | 1002 | KTIGYTSPRSRTE | 6.3741 | 7.0346 | 6.2524 | 6.8498 | 6.1049 | 5.8659 | 6.5200 | 6.1504 | 6.1208 | 6.1087 | 6.2101 | 6.2568 | 5.7817 | 6.0478 | 6.1971 | 6.1250 |

| | | | | | | | | | | | | | | | | | | | | |
|---|---|---|---|---|---|---|---|---|---|---|---|---|---|---|---|---|---|---|---|---|
| KMT2E_T641 | KMT2E | Q8IZD2 | 795;799 | NA | 7.0246 | 9.3808 | 6.0273 | 7.8010 | 7.2476 | 6.7981 | 8.3182 | 5.7593 | 5.6080 | 7.6481 | 8.1951 | 7.1931 | 2.8850 | 3.8284 | 2.9336 | 3.3516 |
| KMT5A_S100 | KMT5A | Q9NQR1 | 181 | AAEPPKTPPSSCD | 6.0870 | 6.6812 | 6.9151 | 7.8551 | 5.8599 | 6.9724 | 7.0865 | 6.5461 | 5.8058 | 6.9210 | 7.0518 | 7.0379 | 4.1416 | 5.2721 | 4.0705 | 4.6961 |
| KMT5A_T181 | KMT5A | Q9NQR1 | 100 | KIYSYMSPNKCSG | 6.6024 | 6.6766 | 6.7708 | 7.0312 | 5.8899 | 7.1144 | 7.7295 | 6.9850 | 6.8461 | 6.8463 | 7.4228 | 7.3851 | 3.5473 | 4.9289 | 3.5091 | 4.7146 |
| KMT5B_S635 | KMT5B | Q4FZB7 | 635 | STPVHDSPGKDDA | 7.0922 | 7.4018 | 6.5644 | 6.2328 | 4.8573 | 5.7675 | 7.4357 | 6.1143 | 6.2328 | 5.0417 | 7.0296 | 5.7269 | 4.6898 | 6.6693 | 7.4255 |
| KNL1_S1675 | KNL1 | Q8NG31 | 1017 | TPTVICTPTEESV | 5.6693 | 6.1733 | 5.8657 | 6.4808 | 6.2353 | 6.2846 | 6.0625 | 6.2232 | 6.0623 | 5.5204 | 5.4620 | 5.7686 | 6.7920 | 7.2370 | 6.9745 | 7.1886 |
| KNL1_S1690 | KNL1 | Q8NG31 | 60 | KNSRRVSFADTIK | 5.7171 | 4.7230 | 4.8294 | 4.8934 | 7.0682 | 5.8603 | 4.9618 | 6.0275 | 8.7183 | 8.2175 | 7.3935 | 7.6031 | 6.8101 | 7.2333 | 4.6627 | 5.2807 |
| KNL1_S1732 | KNL1 | Q8NG31 | 1732 | AGKLNLSPSQYIN | 5.0042 | 5.8378 | 5.8996 | 6.1184 | 6.1093 | 5.8940 | 6.0209 | 6.0531 | 5.7292 | 5.9635 | 6.0450 | 5.8704 | 7.1744 | 7.3225 | 7.6539 | 7.3038 |
| KNL1_S1732;Y1745 | KNL1 | Q8NG31 | 1675 | DKVRRCSLGIFLP | 6.8775 | 4.5786 | 5.0951 | 5.3222 | 7.1148 | 4.9754 | 5.5352 | 4.0752 | 8.7364 | 7.1472 | 7.5401 | 7.2160 | 8.1720 | 8.2291 | 4.1085 | 5.2767 |
| KNL1_S1831 | KNL1 | Q8NG31 | 32 | ILKPPRSPLQDLR | 6.3212 | 6.7577 | 5.9157 | 6.5089 | 6.1176 | 5.9576 | 6.3673 | 6.7873 | 6.8168 | 6.4319 | 6.3129 | 6.1919 | 5.5469 | 5.8793 | 5.8240 | 6.2633 |
| KNL1_S32 | KNL1 | Q8NG31 | 1831 | EEDIDKSANSVLI | 6.3371 | 7.4459 | 6.5289 | 6.8448 | 6.5377 | 7.5338 | 7.1662 | 7.2550 | 7.2940 | 7.8092 | 7.6428 | 7.6529 | 3.0024 | 4.0130 | 2.9849 | 3.9514 |
| KNL1_S60 | KNL1 | Q8NG31 | 767 | PLRKSLSNPTPDY | 6.5147 | 5.9697 | 6.5420 | 7.3796 | 6.9548 | 6.1605 | 6.1096 | 6.4441 | 8.0238 | 6.9319 | 7.6111 | 7.1663 | 4.7715 | 5.2364 | 4.0535 | 4.5807 |
| KNL1_S659 | KNL1 | Q8NG31 | 539 | QIAAAPTPEKEMM | 6.3970 | 6.5982 | 6.4133 | 7.3943 | 6.7962 | 6.1773 | 6.6454 | 7.1367 | 6.6724 | 6.9595 | 6.6531 | 6.6786 | 4.8326 | 4.6631 | 4.3034 | 5.6790 |
| KNL1_S767 | KNL1 | Q8NG31 | 1690 | PNKRNCSVTGIDD | 5.9059 | 5.1204 | 6.8178 | 6.7040 | 6.1462 | 5.3252 | 6.4272 | 5.7231 | 6.6998 | 8.0985 | 6.6873 | 6.2914 | 5.6939 | 6.9317 | 6.1452 | 5.2823 |
| KNL1_T1017 | KNL1 | Q8NG31 | 659 | DIIAKNSLTDTWN | 6.9350 | 6.1406 | 6.4000 | 6.8999 | 6.5053 | 6.2904 | 6.5969 | 7.4324 | 7.4281 | 6.9008 | 7.4204 | 7.2886 | 4.2333 | 4.7889 | 3.9764 | 4.7632 |
| KNL1_T539 | KNL1 | Q8NG31 | 1732;1745 | NA | 5.9026 | 6.1636 | 6.6375 | 6.6320 | 5.9337 | 7.1162 | 6.8115 | 6.5122 | 5.7807 | 5.8663 | 7.7707 | 6.6675 | 5.1445 | 6.4764 | 5.2710 | 5.3135 |
| KNOP1_S185 | KNOP1 | Q1ED39 | 238 | SRSMESSPRKGSK | 6.5281 | 6.7328 | 6.0004 | 7.0192 | 6.2150 | 6.0150 | 6.0388 | 5.7929 | 5.6280 | 5.9618 | 6.2482 | 6.3685 | 5.9686 | 6.7165 | 6.4520 | 6.3142 |
| KNOP1_S238 | KNOP1 | Q1ED39 | 387 | GGFKNLSPSFSRP | 7.1669 | 7.4006 | 7.2882 | 7.5833 | 6.7302 | 6.6934 | 7.5277 | 6.6130 | 7.6269 | 7.3318 | 7.5311 | 7.3326 | 2.8235 | 3.8933 | 2.7471 | 3.7103 |
| KNOP1_S387 | KNOP1 | Q1ED39 | 310 | WKEETDTDLEVVL | 6.5971 | 6.6909 | 6.7524 | 7.3376 | 6.6114 | 5.7182 | 6.6970 | 6.4636 | 5.8064 | 5.1186 | 5.6536 | 5.5735 | 5.9460 | 6.4727 | 6.1261 | 6.4347 |
| KNOP1_T310 | KNOP1 | Q1ED39 | 185 | DVGDTCSVGKKDE | 6.5048 | 6.6652 | 6.2745 | 7.0120 | 5.4631 | 6.5505 | 6.5031 | 6.5327 | 6.3432 | 6.4189 | 7.4994 | 6.8391 | 5.0888 | 6.1032 | 5.4525 | 5.6491 |
| KNTC1_T1035 | KNTC1 | P50748 | 1035 | KHKPGSTPEPIAA | 6.6396 | 7.3982 | 7.3406 | 7.9605 | 5.8198 | 7.3667 | 6.9632 | 7.1361 | 7.1203 | 7.2875 | 7.2595 | 7.0220 | 3.7910 | 4.1399 | 2.9270 | 3.8282 |
| KPNA1_S244 | KPNA1 | P52294 | 244 | NLCRGKSPPPEFA | 5.5709 | 5.7092 | 6.0995 | 5.9281 | 6.8297 | 6.4581 | 5.7846 | 5.7591 | 5.9800 | 5.8364 | 6.4053 | 6.4077 | 7.0573 | 6.6823 | 7.2522 | 6.2392 |
| KPNA2_S24 | KPNA2 | P52292 | 62 | FPDDATSPLQENR | 6.2921 | 7.1948 | 7.0124 | 7.2056 | 6.4738 | 6.0764 | 6.7298 | 6.4538 | 6.9042 | 6.8528 | 6.6083 | 6.7729 | 4.5809 | 5.1000 | 4.6003 | 5.1419 |
| KPNA2_S490 | KPNA2 | P52292 | 490 | VYKASLSLIEKYF | 6.2212 | 5.8191 | 5.6420 | 6.4688 | 5.8788 | 6.0780 | 6.6732 | 6.3175 | 5.7912 | 5.6235 | 5.6544 | 5.8943 | 6.0720 | 7.1731 | 7.2728 | 7.4201 |
| KPNA2_S62 | KPNA2 | P52292 | 24 | KNKGKDSTEMRRR | 6.5784 | 6.8521 | 6.1455 | 6.9297 | 6.8449 | 6.5163 | 7.5560 | 7.1562 | 7.1581 | 6.5971 | 7.4331 | 7.6473 | 3.9883 | 4.3603 | 3.4603 | 4.5765 |
| KPNA2_Y515 | KPNA2 | P52292 | 515 | ETTSEGYTFQVQD | 6.1096 | 6.3281 | 6.2884 | 6.2370 | 6.5069 | 6.0757 | 5.9338 | 6.5023 | 7.2115 | 6.4619 | 5.8829 | 6.2113 | 5.6825 | 6.0161 | 5.8777 | 6.6740 |
| KPNA3_S60 | KPNA3 | O00505 | 60 | EESLEDSDVDADF | 5.9762 | 6.2870 | 6.7568 | 5.7839 | 6.3940 | 6.1268 | 5.9200 | 6.1727 | 6.4551 | 5.9707 | 6.3493 | 6.4799 | 6.3275 | 6.3493 | 6.3777 | 6.2731 |
| KPNA4_S60 | KPNA4 | O00629 | 60 | EDICEDSDIDGDY | 5.9834 | 6.0213 | 7.3813 | 6.8995 | 6.0536 | 7.1391 | 6.8603 | 5.8576 | 6.2483 | 5.3524 | 5.8386 | 5.4517 | 6.4032 | 5.9954 | 5.9221 | 6.5922 |
| KPNA4_S72 | KPNA4 | O00629 | 72 | YRVQNTSLEAIVQ | 6.0968 | 6.5236 | 6.4636 | 6.5875 | 6.1432 | 6.0800 | 6.0928 | 6.7345 | 7.1117 | 5.5158 | 7.1117 | 7.4754 | 5.6045 | 6.0489 | 4.9652 | 5.4447 |
| KPNB1_S12 | KPNB1 | Q14974 | 12 | ILEKTVSPDRLEL | 5.8690 | 6.3817 | 6.6973 | 6.8307 | 6.5512 | 6.4279 | 6.4571 | 6.9196 | 6.6621 | 6.5768 | 6.8402 | 6.8982 | 4.6903 | 5.7753 | 5.0553 | 5.3671 |
| KRBA1_S253 | KRBA1 | A5PL33 | 253 | ADLGPGSPPWAVK | 6.1663 | 6.4593 | 5.9993 | 7.6050 | 7.4838 | 6.0395 | 6.1178 | 5.8823 | 5.9571 | 6.9760 | 7.1873 | 5.9024 | 5.0213 | 5.8940 | 5.8954 | 5.4133 |
| KRI1_S171 | KRI1 | Q8N9T8 | 639 | EEEAPVSPHKKPA | 6.1125 | 6.2210 | 6.3172 | 7.0183 | 6.1465 | 6.0450 | 6.4975 | 6.1671 | 6.6818 | 6.2784 | 6.2296 | 6.4033 | 5.9271 | 6.0377 | 5.7450 | 6.1720 |
| KRI1_S309 | KRI1 | Q8N9T8 | 171 | RAFVEDSEDEDGA | 6.1865 | 6.1731 | 6.1548 | 6.6963 | 6.1087 | 5.9730 | 6.1062 | 6.0599 | 6.5321 | 5.9617 | 6.0408 | 6.0156 | 6.4834 | 6.6780 | 6.3475 | 6.4824 |
| KRI1_S480 | KRI1 | Q8N9T8 | 628 | SLMGPESPPAQEE | 5.8610 | 6.0835 | 5.8330 | 6.6638 | 6.0134 | 5.8363 | 6.2169 | 6.0986 | 6.0211 | 6.2304 | 5.8860 | 5.8253 | 6.5789 | 7.2609 | 6.9944 | 6.5965 |
| KRI1_S628 | KRI1 | Q8N9T8 | 480 | GKKKRKSPFAAAV | 7.5522 | 8.1048 | 7.5507 | 8.4012 | 5.4578 | 6.0400 | 6.2172 | 5.8839 | 7.5557 | 7.8202 | 7.4529 | 7.3875 | 3.2504 | 4.2912 | 3.0807 | 3.9537 |
| KRI1_S628;S639 | KRI1 | Q8N9T8 | 309 | EEPDSASVKTYPR | 6.7088 | 6.1845 | 6.2929 | 7.2722 | 5.9502 | 5.5127 | 7.1960 | 5.5488 | 7.1784 | 5.2832 | 5.4493 | 6.0198 | 6.5233 | 6.6132 | 5.8356 | 6.4312 |
| KRI1_S639 | KRI1 | Q8N9T8 | 628;639 | NA | 5.9328 | 6.2763 | 6.0354 | 6.6523 | 5.9848 | 5.8958 | 6.2599 | 6.0978 | 6.3582 | 6.2626 | 6.1304 | 6.1968 | 6.4417 | 6.3671 | 6.4767 | 6.6314 |
| KRTCAP2_T150 | KRTCAP2 | Q8N6L1-2 | 150 | AAAPVLTPAKVTG | 7.1286 | 7.4224 | 6.6060 | 6.8288 | 6.5047 | 6.9837 | 6.4948 | 7.0235 | 7.0484 | 6.8316 | 6.7247 | 6.8413 | 4.0285 | 4.9850 | 4.1683 | 4.3798 |
| KSR1_S334 | KSR1 | Q8IVT5 | 334 | FDLSHGSPQMVRR | 6.6204 | 6.6011 | 7.3349 | 6.3231 | 6.1954 | 6.6752 | 6.7177 | 6.8921 | 7.2034 | 6.7892 | 6.9243 | 6.0876 | 5.2277 | 5.4383 | 4.5117 | 4.4580 |
| KTI12_S200 | KTI12 | Q96EK9 | 200 | KSAKHGSGAFYSP | 5.1265 | 6.0059 | 6.2308 | 4.1971 | 4.3084 | 5.8718 | 4.6356 | 4.8492 | 5.0969 | 4.0812 | 3.7585 | 5.1833 | 9.8825 | 11.0619 | 10.0529 | 9.6576 |
| KTN1_S1084 | KTN1 | Q86UP2 | 75 | NGNLHESDSESVP | 6.2448 | 6.3304 | 6.4589 | 6.6089 | 6.1647 | 6.2253 | 6.2662 | 6.2112 | 6.2087 | 6.1476 | 6.3215 | 6.1420 | 5.5896 | 6.8116 | 5.9808 | 6.2876 |
| KTN1_S109 | KTN1 | Q86UP2 | 153 | PVTKQPTPPSEAA | 6.3817 | 6.8094 | 5.7748 | 6.8100 | 5.9250 | 5.8961 | 6.0446 | 6.3546 | 6.1244 | 6.1913 | 6.2096 | 6.1829 | 5.9615 | 6.6536 | 6.5153 | 6.1652 |
| KTN1_S110 | KTN1 | Q86UP2 | 109 | NVVETSSSVRERK | 5.8552 | 6.5861 | 6.6813 | 7.1433 | 6.5175 | 6.8631 | 6.2870 | 5.7032 | 6.5157 | 6.0479 | 6.6281 | 6.0249 | 5.8647 | 6.2175 | 5.3964 | 5.6680 |
| KTN1_S1313 | KTN1 | Q86UP2 | 206 | TKQESGSGKKKAS | 6.8310 | 7.9958 | 6.8672 | 8.2259 | 6.3896 | 7.0260 | 7.8669 | 6.8884 | 6.4443 | 7.3305 | 8.0740 | 7.5079 | 2.8514 | 3.9425 | 2.7198 | 3.0388 |
| KTN1_S206 | KTN1 | Q86UP2 | 243 | MDNADSSPVVDKR | 6.3525 | 6.5969 | 6.4316 | 6.8239 | 6.3503 | 6.6018 | 6.1357 | 6.6977 | 6.0628 | 6.1388 | 6.2636 | 6.8626 | 5.0493 | 5.9037 | 6.1250 | 5.6038 |
| KTN1_S243 | KTN1 | Q86UP2 | 1313 | IENSDVSPETESS | 5.9809 | 6.0973 | 6.3158 | 6.1970 | 5.8859 | 5.2318 | 6.3091 | 5.6145 | 5.7700 | 6.4938 | 6.0749 | 6.7008 | 6.8010 | 7.0482 | 6.7915 | 6.6875 |
| KTN1_S75 | KTN1 | Q86UP2 | 1084 | KLFPKVSVPSNLS | 3.4998 | 3.5743 | 3.0588 | 3.9545 | 4.3720 | 4.2045 | 4.2926 | 3.7808 | 3.0687 | 4.2119 | 3.8187 | 3.5699 | 13.3275 | 14.2704 | 14.9627 | 12.0329 |
| KTN1_S75;S79 | KTN1 | Q86UP2 | 110 | VVETSSSVRERKK | 6.0654 | 6.3292 | 5.4009 | 6.4144 | 6.7452 | 6.1980 | 6.3683 | 7.6749 | 5.5293 | 5.9998 | 6.3167 | 6.7089 | 5.1315 | 6.0640 | 6.6629 | 6.3906 |
| KTN1_T153 | KTN1 | Q86UP2 | 75;79 | NA | 6.9156 | 7.1704 | 5.7439 | 7.0536 | 6.2309 | 5.9537 | 7.0532 | 6.1887 | 6.3505 | 6.0238 | 6.1369 | 5.8554 | 5.8413 | 5.5288 | 5.9911 | 5.9622 |
| L1RE1_S18 | L1RE1 | Q9UN81 | 27 | PKERSSSPATEQS | 6.0651 | 6.3879 | 6.2709 | 6.3572 | 6.2408 | 5.9078 | 5.9951 | 6.3040 | 6.5501 | 7.0273 | 6.5780 | 6.3704 | 5.6986 | 6.5100 | 5.8615 | 5.8754 |
| L1RE1_S27 | L1RE1 | Q9UN81 | 18 | SKTQSASPPPKER | 6.3264 | 6.6378 | 6.1770 | 6.2240 | 6.0437 | 6.2949 | 6.0514 | 6.1414 | 6.4788 | 6.6488 | 6.3758 | 6.5264 | 5.7260 | 6.3720 | 6.1085 | 5.8669 |
| L1RE1_S33 | L1RE1 | Q9UN81 | 33 | SPATEQSWMENDF | 5.7484 | 6.2461 | 6.7270 | 6.6510 | 6.5865 | 6.2092 | 6.0813 | 6.3273 | 6.5974 | 6.5439 | 6.4557 | 6.1637 | 5.5030 | 6.3112 | 5.4938 | 6.3544 |
| L1RE1_T203 | L1RE1 | Q9UN81 | 203 | IQEIQRTPQRYSS | 7.1572 | 7.7346 | 6.7155 | 6.9227 | 5.9968 | 6.6180 | 6.2761 | 5.6116 | 6.3152 | 6.9514 | 6.5535 | 6.8177 | 5.4501 | 5.3027 | 4.5916 | 4.9853 |
| L3MBTL2_S661 | L3MBTL2 | Q969R5 | 67 | AGELPTSPLHLLS | 6.0553 | 6.2344 | 6.1842 | 6.8866 | 6.2085 | 5.7627 | 6.2727 | 6.5669 | 6.5564 | 6.4756 | 6.3467 | 5.8737 | 5.8374 | 5.9635 | 6.4646 | 6.3107 |
| L3MBTL2_S67 | L3MBTL2 | Q969R5 | 79 | SPGTPRSLDGSGS | 6.1541 | 6.0858 | 6.8328 | 6.4685 | 7.0667 | 5.5947 | 6.5555 | 6.2183 | 7.0078 | 6.9703 | 7.0292 | 5.1705 | 5.7058 | 5.6108 | 5.1920 | 5.1371 |
| L3MBTL2_S67;S73 | L3MBTL2 | Q969R5 | 689 | SPDKASSPELPVS | 5.8167 | 6.0332 | 5.2649 | 6.4034 | 5.7550 | 5.6738 | 6.0954 | 5.9697 | 6.6954 | 6.5741 | 6.2968 | 6.1482 | 6.5067 | 6.6765 | 7.1211 | 6.9688 |
| L3MBTL2_S683;S688;S68 | L3MBTL2 | Q969R5 | 661 | QGARKISSEPVPG | 7.0470 | 5.1675 | 4.3083 | 5.7004 | 4.4157 | 5.5599 | 6.4398 | 6.2126 | 6.0391 | 6.2497 | 5.2377 | 7.0015 | 8.2077 | 7.8855 | 5.4319 | 9.0958 |
| L3MBTL2_S688;S689 | L3MBTL2 | Q969R5 | 688;689 | NA | 5.5162 | 6.0474 | 6.0477 | 6.2200 | 5.7442 | 6.1408 | 6.3139 | 5.5210 | 5.5138 | 5.7650 | 5.6826 | 5.8690 | 7.3863 | 7.4342 | 7.3677 | 7.4304 |
| L3MBTL2_S689 | L3MBTL2 | Q969R5 | 683;688;689 | NA | 6.5557 | 6.5743 | 6.7399 | 7.1364 | 5.4100 | 5.8032 | 7.4751 | 5.8714 | 5.8159 | 7.4835 | 5.5681 | 6.8679 | 6.0549 | 6.2334 | 4.5629 | 5.8475 |
| L3MBTL2_S79 | L3MBTL2 | Q969R5 | 67;73 | NA | 6.0839 | 6.8110 | 6.8025 | 7.4171 | 6.6325 | 6.4704 | 6.5675 | 6.9352 | 6.8930 | 6.3520 | 5.7464 | 6.9705 | 4.7302 | 5.1762 | 4.9577 | 5.4539 |
| L3MBTL3_S608 | L3MBTL3 | Q96JM7 | 678 | VFLSFKSPIPCLP | 7.6335 | 7.9336 | 6.9226 | 7.6882 | 6.2238 | 7.0324 | 7.0758 | 6.9445 | 7.1738 | 7.3276 | 7.4106 | 7.1556 | 2.9102 | 3.7678 | 2.9988 | 3.8013 |
| L3MBTL3_S670 | L3MBTL3 | Q96JM7 | 670 | QQAQRRSAVFLSF | 7.1002 | 7.5471 | 6.4195 | 8.1204 | 8.0024 | 6.8885 | 7.5328 | 7.0177 | 6.8754 | 7.4247 | 7.4132 | 6.5886 | 2.8049 | 3.8998 | 2.7716 | 3.5934 |
| L3MBTL3_S678 | L3MBTL3 | Q96JM7 | 608 | GEMPPASPSFPRN | 6.4266 | 6.0097 | 6.1830 | 6.4334 | 6.3700 | 6.1927 | 6.2386 | 6.5773 | 6.4889 | 6.1462 | 6.5718 | 6.1291 | 6.2081 | 6.3930 | 6.2702 | 5.7614 |
| LAMTOR1_S27 | LAMTOR1 | Q6IAA8 | 27 | LLLDPSSPPTKAL | 7.0672 | 7.1424 | 6.7311 | 6.9542 | 6.6133 | 7.1271 | 7.6169 | 7.2576 | 7.3174 | 7.2673 | 7.1927 | 7.1336 | 2.8538 | 4.3738 | 3.0814 | 4.2703 |
| LAMTOR1_S56 | LAMTOR1 | Q6IAA8 | 98 | LAVLSSSLTHWKK | 7.0227 | 7.0675 | 6.6169 | 7.2426 | 6.1910 | 6.5615 | 6.5534 | 6.9192 | 6.6481 | 6.4840 | 6.7653 | 6.9619 | 4.1721 | 4.8256 | 4.6580 | 5.3100 |
| LAMTOR1_S98 | LAMTOR1 | Q6IAA8 | 56 | EQALLSSILAKTA | 7.0487 | 6.3121 | 6.4291 | 6.7479 | 6.8982 | 5.0679 | 6.3716 | 5.4226 | 5.4712 | 4.9133 | 5.7476 | 5.6946 | 6.6964 | 7.2184 | 7.7705 | 7.1897 |
| LAP3_S238 | LAP3 | P28838 | 238 | VHIRPKSWIEEQA | 6.0572 | 5.8994 | 6.7647 | 6.9579 | 6.6047 | 6.5030 | 5.6914 | 7.1237 | 7.0116 | 6.2126 | 7.2791 | 6.7144 | 5.7988 | 4.9930 | 5.3016 | 5.0867 |
| LARP1_S1040 | LARP1 | Q6PKG0 | 548 | PKGLSASLPDLDS | 5.8505 | 5.8396 | 6.0325 | 5.5424 | 6.0650 | 6.4918 | 5.9939 | 6.1928 | 6.1624 | 6.2853 | 5.9997 | 6.0859 | 6.7783 | 7.1529 | 6.7649 | 6.7623 |
| LARP1_S1056 | LARP1 | Q6PKG0 | 824 | RKTRHSSNPPLES | 5.2175 | 5.2742 | 5.7527 | 5.2915 | 5.7940 | 6.2220 | 5.8349 | 6.1243 | 5.5613 | 5.4099 | 5.4151 | 5.7332 | 7.9562 | 8.2085 | 8.3925 | 7.8121 |
| LARP1_S143 | LARP1 | Q6PKG0 | 1071 | MISQPPTPPTGQP | 6.0954 | 6.1875 | 6.3242 | 7.2928 | 6.5043 | 6.9991 | 6.3744 | 6.3580 | 6.0334 | 6.4930 | 6.7745 | 6.4143 | 5.2606 | 6.0753 | 5.4937 | 5.3194 |
| LARP1_S225;S228 | LARP1 | Q6PKG0 | 526 | GSPRAVTPVPTKT | 6.5571 | 6.5514 | 6.4789 | 5.8567 | 6.8867 | 6.7577 | 6.5335 | 6.8016 | 6.6979 | 6.1799 | 6.1297 | 5.9397 | 5.5674 | 5.9352 | 5.6025 | 5.5240 |
| LARP1_S324 | LARP1 | Q6PKG0 | 774 | PTTVPESPNYRNT | 6.9042 | 7.1576 | 6.9452 | 6.3466 | 6.8316 | 6.6674 | 6.8621 | 7.3618 | 7.0055 | 7.0244 | 6.9031 | 6.7441 | 3.9553 | 4.8669 | 3.9095 | 4.5146 |
| LARP1_S324;S327 | LARP1 | Q6PKG0 | 1040 | GNHKRHSVVAGGG | 6.5279 | 4.1093 | 4.4795 | 4.3077 | 6.8105 | 5.0704 | 5.1336 | 5.2195 | 6.9188 | 6.4996 | 5.6118 | 5.8636 | 9.6239 | 10.5650 | 7.7533 | 6.4059 |
| LARP1_S548 | LARP1 | Q6PKG0 | 649 | ILIVTQTPHYMRR | 8.1256 | 7.3164 | 6.6358 | 8.0779 | 7.7638 | 6.3398 | 7.2741 | 6.0750 | 7.1193 | 5.8936 | 6.2622 | 6.3166 | 3.8751 | 4.8974 | 3.6108 | 4.4166 |
| LARP1_S627 | LARP1 | Q6PKG0 | 143 | TTVNGQSPPEHSA | 6.3759 | 6.6745 | 6.8587 | 6.6196 | 6.3348 | 6.6523 | 6.4720 | 5.9714 | 6.4045 | 6.4308 | 6.7799 | 6.7694 | 5.4687 | 5.6989 | 5.1896 | 5.2988 |
| LARP1_S627;S631 | LARP1 | Q6PKG0 | 90 | AEGPAISDGEEGG | 6.0955 | 6.0869 | 6.7503 | 6.1864 | 6.7327 | 6.5659 | 5.9822 | 6.8474 | 6.4306 | 6.0185 | 6.3462 | 6.0928 | 5.7899 | 6.0926 | 6.0277 | 5.9544 |
| LARP1_S631 | LARP1 | Q6PKG0 | 376 | GVEGPRTPKYMNN | 7.1379 | 7.3242 | 6.8388 | 6.8751 | 6.4476 | 7.3386 | 6.9150 | 6.4032 | 6.9402 | 6.9345 | 6.5246 | 6.9218 | 4.2914 | 4.8894 | 3.8964 | 4.3213 |
| LARP1_S75 | LARP1 | Q6PKG0 | 766 | PSTIARSLPTTVP | 7.1060 | 6.5519 | 7.5289 | 6.9550 | 6.7182 | 6.5325 | 6.4072 | 6.2137 | 7.0470 | 6.2829 | 7.1142 | 6.7948 | 4.6080 | 5.1677 | 4.4381 | 4.5340 |
| LARP1_S766 | LARP1 | Q6PKG0 | 324 | DQDETSSVKSDGA | 5.7659 | 5.9023 | 6.6682 | 6.1884 | 5.8264 | 5.5620 | 5.3562 | 5.6155 | 6.3095 | 6.2731 | 5.4455 | 6.0193 | 6.9079 | 7.0701 | 7.4750 | 7.6148 |
| LARP1_S766;S774 | LARP1 | Q6PKG0 | 627 | NTFTAWSDEESDY | 6.0364 | 6.3654 | 7.2632 | 6.6166 | 5.6813 | 5.9445 | 5.8799 | 6.2716 | 7.3981 | 8.0022 | 8.1521 | 8.1603 | 4.3488 | 4.8118 | 4.4523 | 4.6155 |
| LARP1_S774 | LARP1 | Q6PKG0 | 847 | HRPRTASISSSPS | 5.6062 | 5.3964 | 6.2616 | 5.6326 | 6.1718 | 5.9292 | 6.0909 | 6.2086 | 6.4610 | 6.4902 | 6.2774 | 6.6318 | 6.9011 | 7.6163 | 6.0687 | 6.2563 |
| LARP1_S824 | LARP1 | Q6PKG0 | 850 | RTASISSSPSEGT | 6.0910 | 6.1828 | 7.1524 | 5.9650 | 5.9374 | 6.7299 | 6.4284 | 6.5462 | 6.4386 | 6.6285 | 7.1893 | 6.8221 | 5.1302 | 5.5841 | 5.6586 | 5.5156 |
| LARP1_S847 | LARP1 | Q6PKG0 | 75 | GQQERESPRPLQL | 4.6605 | 5.9354 | 6.4435 | 6.4736 | 5.4666 | 7.2818 | 6.2407 | 7.2356 | 7.0539 | 6.5649 | 6.6975 | 5.8361 | 5.1443 | 6.7416 | 5.1544 | 5.2698 |
| LARP1_S847;S850 | LARP1 | Q6PKG0 | 724 | EQFDTLTPEPPVD | 5.1332 | 7.1890 | 7.8638 | 8.2252 | 5.6826 | 6.2029 | 6.8025 | 7.7489 | 6.3219 | 6.7398 | 8.0581 | 6.2238 | 3.8504 | 4.3309 | 4.2432 | 5.3839 |
| LARP1_S850 | LARP1 | Q6PKG0 | 178 | DAINWPTPGEIAH | 6.7721 | 6.7692 | 6.5502 | 7.7345 | 6.6646 | 8.1457 | 8.6409 | 7.2837 | 6.8445 | 7.0104 | 6.4147 | 6.5922 | 3.9969 | 2.7028 | 2.1421 | 4.8075 |
| LARP1_S850;T865 | LARP1 | Q6PKG0 | 631 | AWSDEESDYEIDD | 6.5525 | 6.3333 | 6.6711 | 6.7620 | 5.9749 | 6.1855 | 5.9876 | 5.9200 | 5.9608 | 7.0389 | 8.3985 | 7.5900 | 5.1897 | 5.8097 | 5.1295 | 5.3957 |
| LARP1_S865 | LARP1 | Q6PKG0 | 788 | TPRTPRTPQLKDS | 8.1257 | 8.0227 | 8.1007 | 6.5206 | 7.2721 | 8.6916 | 7.1438 | 5.3966 | 6.3700 | 7.0986 | 7.6484 | 7.4967 | 2.9492 | 3.5656 | 2.6476 | 2.9501 |
| LARP1_S89 | LARP1 | Q6PKG0 | 1056 | GRKRCPSQSSSRP | 4.8398 | 4.7120 | 5.0456 | 5.1040 | 4.9763 | 5.4640 | 5.9176 | 5.5895 | 6.2930 | 5.1018 | 5.2680 | 4.5898 | 10.0252 | 9.2446 | 9.2193 | 8.6094 |
| LARP1_S90 | LARP1 | Q6PKG0 | 303 | VPVAPPTPAWQPE | 7.6557 | 7.6451 | 6.6662 | 7.4833 | 6.6963 | 6.5947 | 7.5758 | 6.4735 | 6.9444 | 6.2429 | 7.5349 | 6.0832 | 4.4689 | 5.5324 | 3.7931 | 4.3096 |
| LARP1_T1071 | LARP1 | Q6PKG0 | 865 | VGSYGCTPQSLPK | 7.1060 | 6.8077 | 7.6337 | 6.8302 | 6.8019 | 6.7368 | 7.2825 | 6.1163 | 6.3818 | 7.5016 | 7.0867 | 7.2542 | 3.0044 | 5.0191 | 4.1976 | 4.2395 |
| LARP1_T178 | LARP1 | H0YAN4 | 89 | ETSSVKSDGAAHF | 5.7349 | 6.2257 | 6.1488 | 5.6886 | 5.8320 | 6.7328 | 5.6807 | 6.2621 | 6.0972 | 6.2406 | 5.9601 | 5.7983 | 6.6978 | 7.0565 | 6.7569 | 7.0872 |
| LARP1_T303 | LARP1 | Q6PKG0 | 627;631 | NA | 5.9552 | 6.0002 | 6.8879 | 5.9221 | 6.3378 | 6.3015 | 6.1810 | 6.0684 | 6.2202 | 6.0684 | 6.1689 | 6.1664 | 6.4495 | 6.6851 | 6.2851 | 6.3023 |
| LARP1_T376 | LARP1 | Q6PKG0 | 847;850 | NA | 4.9944 | 4.8158 | 5.3351 | 4.9157 | 6.0762 | 6.2992 | 5.6021 | 5.9432 | 5.9270 | 5.8745 | 5.3505 | 5.7284 | 8.5192 | 8.0866 | 8.0014 | 8.5307 |
| LARP1_T526 | LARP1 | Q6PKG0 | 766;774 | NA | 7.0176 | 7.1334 | 7.2293 | 6.8567 | 6.1421 | 7.0836 | 7.2663 | 6.7844 | 6.8143 | 6.6151 | 7.0155 | 6.6315 | 4.0847 | 4.7325 | 4.1155 | 4.4775 |
| LARP1_T649 | LARP1 | Q6PKG0 | 324;327 | NA | 5.7799 | 5.3714 | 6.3314 | 5.5666 | 6.1108 | 5.9552 | 5.6307 | 5.6302 | 6.1102 | 6.1101 | 6.4358 | 6.3053 | 7.4833 | 7.7696 | 6.8795 | 6.5300 |
| LARP1_T724 | LARP1 | Q6PKG0 | 225;228 | NA | 6.5195 | 6.2042 | 6.9657 | 6.2118 | 5.9110 | 6.0450 | 6.7049 | 5.6028 | 5.8586 | 6.7485 | 6.2723 | 6.5004 | 5.8380 | 6.5424 | 5.8720 | 6.2030 |
| LARP1_T788 | LARP1 | Q6PKG0 | 850;865 | NA | 6.4743 | 6.2370 | 6.9642 | 6.5322 | 5.9507 | 6.5516 | 7.0342 | 7.1184 | 6.5984 | 7.5489 | 7.7674 | 7.4986 | 4.1126 | 4.6293 | 4.2543 | 4.7277 |
| LARP1B_S1101 | LARP1B | A0A3B3ISF0 | 564 | SRGLSTSLPDLDS | 5.8965 | 5.7258 | 5.9396 | 5.6376 | 6.0614 | 6.2220 | 5.7404 | 6.1788 | 5.8387 | 5.6519 | 5.9230 | 5.7026 | 7.0268 | 7.8534 | 7.3265 | 7.2751 |
| LARP1B_S112 | LARP1B | A0A3B3ISF0 | 655 | ILIVTQTPPYVKK | 6.3156 | 6.6730 | 6.6655 | 6.0952 | 6.3396 | 6.8780 | 6.2356 | 6.7725 | 6.8082 | 6.7270 | 6.8283 | 7.1078 | 4.8692 | 5.6040 | 4.9648 | 5.1157 |
| LARP1B_S159 | LARP1B | A0A3B3ISF0 | 112 | PPPPPPSPLPPLL | 5.5818 | 5.8342 | 6.3871 | 5.8902 | 6.4649 | 6.1146 | 6.3993 | 6.4975 | 6.3595 | 6.5863 | 5.6643 | 5.3251 | 7.0489 | 7.2522 | 6.3365 | 6.9475 |
| LARP1B_S541 | LARP1B | A0A3B3ISF0 | 544 | RIGSPLSPKKNSE | 5.0872 | 5.2530 | 5.7535 | 5.1323 | 6.0124 | 6.0086 | 5.8164 | 6.1381 | 5.8159 | 5.7402 | 5.5861 | 5.6480 | 7.8434 | 8.0969 | 8.1091 | 7.9587 |
| LARP1B_S541;S544 | LARP1B | A0A3B3ISF0 | 541 | NSPRIGSPLSPKK | 6.0759 | 5.8824 | 6.9618 | 6.0451 | 6.0385 | 6.0108 | 6.1586 | 5.6065 | 6.6066 | 7.1015 | 5.9679 | 5.8813 | 6.5000 | 6.9295 | 6.0533 | 6.1803 |
| LARP1B_S564 | LARP1B | A0A3B3ISF0 | 536 | TESAPNSPRIGSP | 5.9067 | 5.8362 | 5.8218 | 6.0809 | 6.4372 | 6.4067 | 6.3388 | 6.6545 | 6.3854 | 6.8065 | 6.4411 | 6.7229 | 6.2077 | 6.8148 | 6.8567 | 6.2821 |
| LARP1B_S655 | LARP1B | A0A3B3ISF0 | 159 | KPPQHLSPDTTGP | 6.2260 | 6.4632 | 6.8440 | 6.2723 | 6.5794 | 6.5369 | 6.2670 | 6.5497 | 6.8519 | 6.8471 | 6.3670 | 6.4350 | 5.4742 | 6.0918 | 5.3092 | 5.8851 |
| LARP1B_S794 | LARP1B | A0A3B3ISF0 | 1101 | LQVPINSPRRNIS | 6.4838 | 5.7000 | 7.2552 | 6.8361 | 6.8936 | 7.6574 | 6.4638 | 6.7632 | 6.9589 | 6.7940 | 6.3682 | 6.5228 | 4.1345 | 6.1114 | 3.8161 | 5.2409 |
| LARP1B_T530;S536 | LARP1B | A0A3B3ISF0 | 794 | PTAVPESPRIHPT | 5.9089 | 6.2474 | 6.6637 | 6.3262 | 5.2708 | 6.0081 | 6.0704 | 6.4487 | 6.5975 | 6.4458 | 6.8222 | 6.3592 | 6.8300 | 6.4672 | 5.9712 | 5.5626 |
| LARP1B_T536 | LARP1B | A0A3B3ISF0 | 541;544 | NA | 6.1786 | 5.9066 | 7.6498 | 5.7528 | 6.6151 | 7.0276 | 6.5353 | 4.9392 | 6.2291 | 7.2850 | 6.0871 | 5.6746 | 5.9947 | 5.3878 | 6.2753 | 6.4614 |
| LARP1B_T655 | LARP1B | A0A3B3ISF0 | 530;536 | NA | 7.4370 | 6.1164 | 6.3366 | 7.5528 | 5.8578 | 6.3178 | 5.9644 | 5.9000 | 7.0582 | 6.8014 | 6.5836 | 7.1768 | 4.4906 | 5.0874 | 5.6889 | 5.6303 |
| LARP4_S589 | LARP4 | Q71RC2-4 | 728 | QYVPPRSPKxxxx | 6.0396 | 6.1924 | 6.6071 | 6.4143 | 6.2432 | 6.2829 | 6.2007 | 6.1709 | 6.5506 | 6.3019 | 6.4753 | 6.6273 | 6.1175 | 6.0336 | 5.8428 | 5.8998 |
| LARP4_S653 | LARP4 | Q71RC2-4 | 653 | LRSNVVSPTKNED | 6.3492 | 6.2714 | 6.6217 | 6.2934 | 6.6299 | 6.5576 | 6.5691 | 6.6507 | 6.9374 | 6.5519 | 6.8828 | 6.6916 | 5.1576 | 5.2643 | 5.2026 | 5.3690 |
| LARP4_S728 | LARP4 | Q71RC2-4 | 601 | KPSRASTASPCNN | 6.1836 | 6.6383 | 6.8553 | 6.3292 | 6.2634 | 6.3806 | 6.5806 | 6.4035 | 6.7699 | 6.8575 | 6.6087 | 6.7095 | 5.2873 | 5.5150 | 5.1499 | 5.4677 |
| LARP4_T601 | LARP4 | Q71RC2-4 | 589 | QTTIPVSPPSTTK | 5.8263 | 6.1304 | 6.5228 | 6.6326 | 6.3042 | 6.2591 | 6.3793 | 6.6707 | 6.5030 | 6.7711 | 6.3795 | 6.7742 | 5.6722 | 5.6387 | 5.7488 | 5.7869 |
| LARP4B_S398 | LARP4B | Q92615 | 664 | SKEPPSSPLQPQK | 6.6172 | 6.9243 | 6.7992 | 6.3672 | 6.5202 | 7.0502 | 6.8014 | 6.8466 | 7.2666 | 6.6653 | 7.6018 | 7.0198 | 5.2426 | 4.2916 | 3.8458 | 4.1401 |
| LARP4B_S434 | LARP4B | Q92615 | 601 | TYERSPSPAHLPD | 6.0979 | 5.9881 | 6.1406 | 6.4349 | 6.4509 | 6.2638 | 6.0656 | 6.2983 | 6.3225 | 6.3608 | 6.6570 | 6.4672 | 6.0317 | 6.2792 | 6.0185 | 6.1485 |
| LARP4B_S451 | LARP4B | Q92615 | 516 | TSSQTQSPTPPKP | 6.1071 | 6.4202 | 6.3820 | 6.7430 | 6.2246 | 6.0556 | 6.1259 | 6.5425 | 6.4905 | 6.7076 | 6.5444 | 6.9734 | 5.6822 | 5.7395 | 5.2382 | 6.0233 |
| LARP4B_S488 | LARP4B | Q92615 | 451 | LINGVRSPQTRQA | 7.4644 | 7.3178 | 7.6488 | 7.0252 | 5.6660 | 8.3961 | 7.0817 | 5.8921 | 6.4277 | 7.0192 | 7.6518 | 7.2850 | 3.2696 | 4.3432 | 2.9815 | 3.6299 |
| LARP4B_S498 | LARP4B | Q92615 | 398 | AFKPAASPLTSLR | 7.6820 | 6.5719 | 6.9543 | 7.2597 | 5.8498 | 6.8915 | 7.1372 | 6.4297 | 6.7563 | 7.1553 | 7.1569 | 7.7896 | 3.4477 | 4.2288 | 3.2917 | 4.4976 |
| LARP4B_S516 | LARP4B | Q92615 | 524 | TPPKPPSPSFELG | 5.6304 | 5.8989 | 6.3771 | 6.0073 | 5.8460 | 5.8890 | 5.7444 | 6.4154 | 6.1763 | 5.8979 | 6.0339 | 6.2368 | 6.9875 | 7.1432 | 7.1729 | 6.5430 |
| LARP4B_S516;T518;S524 | LARP4B | Q92615 | 649 | TELRKPSYAEICQ | 5.2563 | 4.3015 | 4.9980 | 4.9386 | 6.3423 | 5.2885 | 4.5368 | 5.3929 | 6.6451 | 4.6804 | 6.2505 | 6.2270 | 9.0450 | 9.4012 | 7.9344 | 6.9615 |
| LARP4B_S518 | LARP4B | Q92615 | 498 | GRGRKNSFGYRKK | 6.3487 | 6.7116 | 6.0000 | 5.4588 | 6.8927 | 6.4090 | 4.9177 | 6.0221 | 6.9775 | 6.9357 | 7.0076 | 6.6989 | 6.0835 | 7.4518 | 6.1232 | 3.9612 |
| LARP4B_S524 | LARP4B | Q92615 | 434 | GPGLLESPSIFNF | 7.9295 | 6.3165 | 7.5064 | 6.8448 | 5.8934 | 7.7448 | 7.0225 | 6.6157 | 7.5297 | 6.6765 | 7.0069 | 7.1978 | 4.1518 | 4.5749 | 3.3920 | 3.8970 |
| LARP4B_S562 | LARP4B | Q92615 | 488 | PGSLESSPGLGRG | 6.8637 | 7.0621 | 6.8768 | 6.1271 | 6.4084 | 6.4068 | 7.1402 | 6.9101 | 7.4418 | 6.1092 | 6.7190 | 6.4475 | 4.8157 | 5.3588 | 4.5262 | 4.7864 |
| LARP4B_S599;S601 | LARP4B | Q92615 | 562 | SLIGPSKERTLS | 6.7700 | 7.4141 | 8.1168 | 6.3601 | 3.7206 | 7.5222 | 5.1169 | 6.3376 | 4.3782 | 7.5943 | 7.4771 | 7.0087 | 5.0268 | 6.2249 | 5.8916 | 5.0403 |
| LARP4B_S601 | LARP4B | Q92615 | 518 | SQTQSPTPPKPPS | 5.8667 | 6.0877 | 6.4686 | 5.5208 | 5.7802 | 5.6134 | 5.5626 | 6.5098 | 6.0870 | 5.8420 | 5.9889 | 6.7836 | 7.5029 | 7.6422 | 7.0321 | 5.9916 |
| LARP4B_S649 | LARP4B | Q92615 | 599;601 | NA | 6.3668 | 6.5066 | 7.5248 | 6.4310 | 7.0062 | 6.4212 | 5.9870 | 6.6500 | 6.6209 | 6.2683 | 6.1384 | 6.6411 | 5.8954 | 4.8643 | 5.1945 | 5.4838 |
| LARP4B_S664 | LARP4B | Q92615 | 731;736 | NA | 6.1935 | 6.0464 | 7.1342 | 6.4080 | 6.6619 | 6.2622 | 6.6518 | 6.2427 | 6.8065 | 6.3500 | 6.9157 | 6.5346 | 5.3242 | 5.3264 | 5.7745 | 5.3673 |
| LARP4B_S731;S736 | LARP4B | Q92615 | 516;518;524 | NA | 6.3062 | 6.5887 | 6.2764 | 6.0286 | 6.9949 | 6.3829 | 6.4345 | 6.7875 | 5.6313 | 5.3975 | 7.2350 | 7.1419 | 5.6243 | 5.6482 | 5.4263 | 6.0956 |
| LARP6_S409 | LARP6 | Q9BRS8 | 409 | RLNCSTSPEIFRK | 6.3833 | 6.3068 | 6.9526 | 5.9239 | 6.4201 | 7.0576 | 6.5167 | 6.4327 | 6.4733 | 6.8073 | 6.6939 | 6.3836 | 5.4016 | 5.6663 | 5.3407 | 5.2394 |
| LARP6_S56 | LARP6 | Q9BRS8 | 202 | KLWALATPQKNGR | 7.7271 | 8.4011 | 7.7706 | 7.4205 | 3.8516 | 6.7748 | 8.4478 | 5.6582 | 7.7641 | 7.7140 | 7.6727 | 8.5946 | 2.6471 | 3.4385 | 2.3003 | 2.9069 |
| LARP6_T202 | LARP6 | Q9BRS8 | 56 | LPSGWGSAEEEP | 6.1628 | 6.5950 | 6.3123 | 6.0718 | 5.9125 | 6.3118 | 5.8694 | 6.4512 | 6.6845 | 6.6076 | 5.5209 | 6.7334 | 6.1536 | 6.2949 | 6.1642 | 6.1540 |
| LARP6_T70 | LARP6 | Q9BRS8 | 70 | RGHSGTTASGGEN | 6.1813 | 6.5340 | 7.8021 | 6.4397 | 6.2650 | 7.6194 | 7.1902 | 6.8605 | 5.6974 | 6.2697 | 6.2597 | 6.5804 | 4.8271 | 5.4461 | 5.1491 | 4.8780 |
| LARP7_S265;S268 | LARP7 | Q4G0J3-3 | 306 | GKRKRSSSEDAES | 5.7265 | 5.9512 | 5.7617 | 5.0057 | 6.0693 | 5.7686 | 6.2197 | 6.2309 | 5.8115 | 5.5982 | 5.6122 | 5.9865 | 6.8973 | 7.8046 | 7.2143 | 7.2919 |
| LARP7_S268 | LARP7 | Q4G0J3-3 | 345 | RDIEISTEEEKDT | 6.0980 | 6.1522 | 6.0937 | 6.1898 | 6.0175 | 6.1001 | 6.2902 | 5.8674 | 6.1016 | 5.8533 | 5.7762 | 5.8773 | 6.5702 | 7.0516 | 7.0815 | 6.8795 |
| LARP7_S306 | LARP7 | Q4G0J3-3 | 268 | RPTSEGSDIESTE | 5.5362 | 6.0034 | 6.0429 | 6.6075 | 5.7316 | 5.7041 | 5.3252 | 5.7678 | 5.9837 | 6.4944 | 6.1057 | 6.5643 | 6.1311 | 7.7278 | 7.2302 | 7.0442 |
| LARP7_S344;T345 | LARP7 | Q4G0J3-3 | 265;268 | NA | 5.4382 | 6.3374 | 5.9614 | 6.8911 | 6.0113 | 6.2553 | 5.9720 | 5.6180 | 5.5287 | 6.1555 | 5.6791 | 5.9777 | 6.4634 | 7.8721 | 7.2049 | 6.6340 |
| LARP7_S345 | LARP7 | Q4G0J3-3 | 344;345 | NA | 6.6523 | 6.6219 | 5.9929 | 6.5920 | 5.7639 | 5.9773 | 6.2295 | 5.8982 | 6.1624 | 6.3854 | 6.2933 | 5.8209 | 5.7086 | 6.6009 | 6.5930 | 6.7074 |
| LAS1L_S235;S238 | LAS1L | Q9Y4W2 | 523 | QEGSEASPIGKSP | 6.1169 | 6.6860 | 6.3878 | 6.8066 | 6.7190 | 6.7205 | 6.6041 | 6.2992 | 6.1575 | 6.2423 | 6.8174 | 6.1572 | 5.8988 | 5.9995 | 5.2781 | 5.1091 |
| LAS1L_S523 | LAS1L | Q9Y4W2 | 617 | FSTGQESPTAENA | 6.0315 | 6.3194 | 6.0056 | 6.7533 | 6.7256 | 6.1447 | 6.4577 | 6.5471 | 6.2111 | 6.2341 | 6.0589 | 6.1404 | 5.8715 | 6.2812 | 6.0264 | 6.1913 |
| LAS1L_S560 | LAS1L | Q9Y4W2 | 560 | QQEEQGSVNDVKE | 5.7488 | 6.0709 | 5.6416 | 6.1426 | 6.0771 | 5.6036 | 6.2330 | 5.4220 | 5.7298 | 5.9518 | 5.4493 | 5.8738 | 7.3981 | 7.2939 | 7.8883 | 7.4753 |
| LAS1L_S617 | LAS1L | Q9Y4W2 | 235;238 | NA | 6.0904 | 6.0343 | 6.0831 | 6.1212 | 6.1534 | 5.5305 | 6.4791 | 5.4703 | 6.4764 | 6.0873 | 6.3752 | 6.1798 | 7.1696 | 6.5523 | 6.0170 | 7.1861 |
| LASP1_S146 | LASP1 | Q14847-2 | 146 | EPERRDSQDGSSY | 6.1641 | 5.9016 | 6.4533 | 6.1810 | 6.6694 | 6.0697 | 5.4013 | 6.4158 | 7.1124 | 7.9325 | 7.5300 | 7.4597 | 5.6866 | 5.7531 | 4.7601 | 4.5093 |
| LASP1_T104 | LASP1 | Q14847-2 | 104 | FSVVADTPELQRI | 7.2853 | 7.4618 | 7.2588 | 7.0597 | 5.7323 | 7.1952 | 7.3108 | 7.5286 | 7.5576 | 7.5236 | 8.0038 | 7.8238 | 2.6837 | 3.6509 | 2.4956 | 3.4284 |
| LASP1_T68 | LASP1 | Q14847-2 | 68 | FTMVADTPENLRL | 6.9119 | 6.9018 | 6.4804 | 6.7761 | 6.0772 | 6.1452 | 5.9323 | 6.1881 | 6.6278 | 6.4364 | 6.6846 | 6.9711 | 5.6792 | 5.2652 | 5.3158 | 5.6069 |
| LAT2_S165 | LAT2 | Q9GZY6 | 59 | TGSRTYSLVGQAW | 6.1117 | 7.9024 | 6.7017 | 7.2037 | 6.4214 | 6.2752 | 5.4469 | 6.6703 | 6.2340 | 5.9255 | 6.0423 | 6.7723 | 5.3295 | 6.3213 | 5.2699 | 5.3719 |
| LAT2_S59 | LAT2 | Q9GZY6 | 165 | LCRGDLSLSLALK | 8.1581 | 6.2274 | 8.0340 | 8.1678 | 7.4942 | 6.0815 | 5.9234 | 6.0304 | 6.0280 | 5.0553 | 6.3149 | 5.8821 | 5.8918 | 6.2249 | 5.1185 | 3.3678 |
| LAT2_S86 | LAT2 | Q9GZY6 | 86 | LLQFYPSLEDPAS | 5.4187 | 5.8125 | 7.7413 | 5.9161 | 6.2351 | 6.2879 | 6.4963 | 6.7764 | 6.5182 | 6.6418 | 6.8761 | 6.8603 | 5.1456 | 5.4855 | 6.2451 | 5.5184 |
| LATS1_S278 | LATS1 | O95835 | 613 | KKQITTSPITVRK | 6.4412 | 6.7738 | 7.0426 | 6.4582 | 6.0905 | 6.5375 | 6.3419 | 6.7636 | 6.6244 | 6.5164 | 6.7997 | 6.8438 | 5.1154 | 5.5291 | 4.9122 | 5.2095 |
| LATS1_S464 | LATS1 | O95835 | 464 | IPVRSNFNNPLG | 6.8432 | 6.3100 | 6.8150 | 6.0965 | 5.8636 | 6.6315 | 6.2442 | 6.5558 | 6.9160 | 6.5080 | 6.5783 | 6.6726 | 5.6910 | 5.9311 | 5.2062 | 5.1370 |
| LATS1_S613 | LATS1 | O95835 | 278 | SQTKRYSGNMEYV | 6.9957 | 6.6326 | 6.9802 | 6.4846 | 6.5722 | 6.7610 | 6.5442 | 7.0077 | 6.7189 | 5.9954 | 6.5709 | 6.5481 | 5.2341 | 5.3248 | 4.8383 | 4.7913 |
| LATS1_S635 | LATS1 | O95835 | 246 | PQVRSVTPPPPPR | 6.7193 | 6.6805 | 6.8924 | 6.4914 | 5.7483 | 6.4214 | 6.7692 | 6.6220 | 6.9136 | 6.8476 | 6.7946 | 6.6967 | 4.7027 | 5.4086 | 5.0475 | 5.2444 |

| | | | | | | | | | | | | | | | | | | | | | |
|---|---|---|---|---|---|---|---|---|---|---|---|---|---|---|---|---|---|---|---|---|---|
| LATS1_T246 | LATS1 | O95835 | | 262 | PPPRGTTPPPPSW | 6.3196 | 6.1762 | 6.9850 | 6.2745 | 6.2585 | 6.9196 | 6.3792 | 7.0819 | 6.8941 | 6.3763 | 6.8156 | 7.0739 | 4.8645 | 5.3665 | 5.0202 | 5.1943 |
| LATS1_T255 | LATS1 | O95835 | | 255 | PPPRGQTPPPPRGT | 5.9976 | 5.7208 | 8.1171 | 7.0173 | 6.2800 | 6.2015 | 7.1614 | 6.9048 | 5.8827 | 6.0114 | 6.9655 | 7.3451 | 5.5695 | 5.2255 | 4.0011 | 5.2684 |
| LATS1_T262 | LATS1 | O95835 | | 635 | SRIQSYSPQAFKF | 6.4151 | 7.6599 | 6.1082 | 6.3463 | 6.5228 | 7.4847 | 7.2803 | 8.3249 | 7.2913 | 6.5452 | 6.8619 | 6.2690 | 3.7651 | 5.2395 | 3.4493 | 4.4466 |
| LATS2_S576 | LATS2 | Q9NRM7 | | 576 | KKQIQTSPVPVRK | 6.7279 | 6.3904 | 6.6089 | 6.4047 | 6.0924 | 6.3890 | 6.3605 | 6.7428 | 6.8303 | 6.4032 | 6.1685 | 6.4117 | 5.6077 | 5.5960 | 5.6409 | 5.6252 |
| LATS2_T1026 | LATS2 | Q9NRM7 | | 279 | PSFQSKTPPETGG | 6.7273 | 7.0054 | 7.3080 | 7.1619 | 5.9302 | 6.4018 | 6.2052 | 6.2867 | 6.9269 | 6.1721 | 7.3977 | 6.4773 | 5.2021 | 5.9759 | 4.4067 | 4.4149 |
| LATS2_T279 | LATS2 | Q9NRM7 | | 1026 | KAWDTLTSPNNKH | 6.2923 | 7.0766 | 7.4965 | 7.7374 | 5.8926 | 7.1303 | 7.4325 | 7.3354 | 7.7461 | 7.1294 | 7.6689 | 8.2683 | 2.7632 | 3.7542 | 2.9542 | 3.6920 |
| LAYN_S286 | LAYN | Q6UX15 | | 299 | NVIRKQSEADLAE | 5.2585 | 5.4055 | 5.8523 | 5.8150 | 5.7132 | 6.0268 | 4.9755 | 6.0021 | 7.1660 | 6.9193 | 8.5366 | 7.5058 | 5.9317 | 6.6924 | 6.2161 | 5.9833 |
| LAYN_S299 | LAYN | Q6UX15 | | 286 | SPHQGNSPDLEVY | 6.4146 | 6.1332 | 5.5292 | 8.1256 | 6.9360 | 6.5262 | 5.9620 | 6.7387 | 6.1164 | 5.7632 | 6.8458 | 5.8060 | 5.6485 | 7.0644 | 4.7188 | 5.6714 |
| LBR_S86 | LBR | Q14739 | | 86 | SRSRSRSPGRPPK | 5.4244 | 5.9010 | 6.5336 | 5.7832 | 5.2527 | 6.0581 | 6.1366 | 6.8095 | 6.2190 | 6.4573 | 5.9372 | 6.1843 | 6.8764 | 6.8280 | 6.7873 | 7.7113 |
| LBR_S99 | LBR | Q14739 | | 118 | EVEVKLTPLILKP | 7.9781 | 8.0882 | 6.7294 | 7.8708 | 6.0431 | 6.4162 | 7.6767 | 7.0390 | 7.1342 | 7.0441 | 6.5180 | 6.9394 | 3.0361 | 4.2778 | 3.0471 | 4.1618 |
| LBR_T118 | LBR | Q14739 | | 99 | SARRSASASHQAD | 5.8214 | 6.3706 | 6.2392 | 6.5480 | 5.7795 | 6.0949 | 6.1359 | 6.3577 | 6.9736 | 7.1479 | 6.9362 | 6.9611 | 5.1349 | 6.3884 | 5.5477 | 5.5631 |
| LBR_T200 | LBR | Q14739 | | 200 | VRTFEVTPIRAKD | 5.7587 | 7.2164 | 6.5907 | 5.3427 | 6.2468 | 5.7653 | 6.8925 | 5.9462 | 6.0538 | 6.3755 | 5.5845 | 5.8075 | 6.9579 | 6.0948 | 7.5625 | 5.8042 |
| LCA5_S41 | LCA5 | Q86VQ0 | | 41 | RSSLVSSSPASVR | 5.6092 | 6.8782 | 6.9674 | 6.4758 | 5.8818 | 6.9826 | 6.9902 | 6.2935 | 6.5161 | 6.9433 | 6.9082 | 7.3830 | 4.4378 | 5.2816 | 4.3789 | 5.1727 |
| LCN10_S54 | LCN10 | E9PQQ7 | | 54 | RGKGEQSGPAPRA | 6.0470 | 5.9913 | 6.0422 | 6.2801 | 5.7720 | 5.7183 | 5.6000 | 5.5555 | 5.9679 | 6.2486 | 6.3178 | 6.5064 | 7.0627 | 7.1559 | 6.8065 | 6.9279 |
| LCOR_S1406 | LCOR | Q96JN0-3 | | 37 | QSLPKASPVTTSP | 6.2153 | 6.3778 | 6.0992 | 6.8437 | 6.4729 | 5.7971 | 6.0847 | 6.0971 | 6.3368 | 6.2570 | 6.1815 | 6.5844 | 6.1559 | 6.1941 | 5.9634 | 6.3388 |
| LCOR_S249 | LCOR | Q96JN0-3 | | 583 | EGGGDVSPRKEPQ | 6.1855 | 6.8525 | 6.4208 | 6.8218 | 5.8883 | 5.8883 | 6.3037 | 6.5487 | 6.1817 | 5.8462 | 6.3521 | 6.4328 | 5.3811 | 6.7307 | 5.7016 | 6.4643 |
| LCOR_S328 | LCOR | Q96JN0-3 | | 375 | TLQCPKTPLRQDL | 7.4366 | 7.2127 | 6.8774 | 7.1571 | 6.5503 | 6.5166 | 7.1206 | 6.2947 | 6.6571 | 6.3074 | 7.4044 | 7.1771 | 4.2480 | 4.9885 | 3.7824 | 5.1689 |
| LCOR_S37 | LCOR | Q96JN0-3 | | 1406 | KRTEGSSPPDSKN | 6.2344 | 7.2497 | 6.2630 | 7.0184 | 5.7408 | 5.6701 | 6.2473 | 6.2676 | 5.8389 | 5.9217 | 6.1910 | 6.8103 | 5.3086 | 5.3608 | 5.7227 | 7.1548 |
| LCOR_S37;T40 | LCOR | Q96JN0 | | 328 | GQPYPTSDQEGDP | 6.1573 | 6.4874 | 6.5694 | 6.5679 | 5.9901 | 6.2649 | 6.4226 | 6.0178 | 5.4792 | 6.7695 | 5.2724 | 5.6271 | 6.6917 | 6.8949 | 6.6149 | 6.1729 |
| LCOR_S583 | LCOR | Q96JN0 | | 249 | KDVSHSSPVDLKI | 7.1134 | 7.2288 | 7.2303 | 7.6867 | 6.5838 | 6.7208 | 6.5511 | 6.8518 | 6.6170 | 7.1255 | 6.9967 | 6.8853 | 4.0456 | 3.9768 | 4.3874 | 3.9990 |
| LCOR_T375 | LCOR | Q96JN0-3 | 37;40 | | NA | 5.4207 | 6.1665 | 5.9075 | 7.0459 | 5.7315 | 5.5182 | 7.4984 | 6.5225 | 6.6194 | 6.0023 | 6.1056 | 6.4242 | 5.7752 | 6.7163 | 6.1779 | 6.3677 |
| LCORL_S282 | LCORL | Q8N3X6-3 | | 282 | TASCPLSPIKMCL | 5.9327 | 5.9327 | 7.4438 | 6.0387 | 5.8479 | 5.6648 | 7.5306 | 6.6593 | 5.3178 | 6.0753 | 7.3147 | 5.8922 | 5.3545 | 5.7303 | 7.2993 | 5.9655 |
| LDB1_S265 | LDB1 | Q86U70 | | 302 | RRKRKMSGGSTMS | 5.8297 | 6.0433 | 6.2706 | 6.1223 | 5.5639 | 6.3230 | 5.9172 | 6.0528 | 6.0401 | 7.1029 | 6.4947 | 6.6620 | 6.3377 | 6.7396 | 6.2274 | 6.2728 |
| LDB1_S302 | LDB1 | Q86U70 | | 265 | HKTYSLSPRDCLK | 6.1796 | 6.6287 | 6.4220 | 6.1266 | 5.7940 | 6.3917 | 6.4499 | 6.3100 | 6.5857 | 6.8628 | 6.2113 | 6.5697 | 5.5292 | 6.5024 | 5.8538 | 6.4825 |
| LDHB_S275 | LDHB | P07195 | | 302 | LNARGLTSVINQK | 4.9040 | 5.5605 | 5.7376 | 5.7513 | 6.1579 | 7.0505 | 5.4424 | 5.6400 | 6.5690 | 7.5980 | 7.0187 | 8.4656 | 5.4196 | 5.0426 | 6.2987 | 7.3436 |
| LDHB_T302 | LDHB | P07195 | | 275 | SRIHPVSTMVKGM | 4.2764 | 4.0392 | 4.4077 | 3.7071 | 4.1614 | 6.6316 | 6.1427 | 6.5441 | 7.3833 | 7.7810 | 8.1787 | 10.2713 | 5.4987 | 4.6577 | 6.0150 | 10.3041 |
| LDLRAP1_S14 | LDLRAP1 | Q5SW96 | | 14 | GRALIRSPSLAKQ | 7.1502 | 6.8122 | 7.3549 | 6.8948 | 5.7625 | 7.9219 | 6.9654 | 5.7503 | 6.9847 | 6.9502 | 7.1708 | 7.0366 | 3.9376 | 5.0010 | 3.9712 | 4.3357 |
| LEMD2_S166 | LEMD2 | Q8NC56 | | 499 | TKPSSFSDSERxx | 6.7648 | 6.8349 | 6.7258 | 7.3821 | 6.1259 | 6.0428 | 6.4924 | 5.9622 | 6.4357 | 6.5363 | 6.3631 | 6.6865 | 5.2144 | 5.4637 | 5.3655 | 5.6040 |
| LEMD2_S175 | LEMD2 | Q8NC56 | | 166 | RRWWAASPAPARL | 6.9923 | 7.4005 | 6.4474 | 7.1490 | 6.5539 | 6.8477 | 7.2263 | 6.9842 | 6.7334 | 6.8488 | 7.2905 | 7.2439 | 3.8661 | 4.6342 | 3.8474 | 4.8342 |
| LEMD2_S499 | LEMD2 | Q8NC56 | | 82 | ARPAAASPRAEPW | 6.5187 | 7.6907 | 7.3520 | 7.4361 | 6.4765 | 7.3959 | 8.1788 | 6.8594 | 6.0721 | 6.3936 | 7.4270 | 7.6075 | 3.1205 | 4.2152 | 3.2392 | 4.0168 |
| LEMD2_S82 | LEMD2 | Q8NC56 | | 175 | PARLPSSLLGPDP | 6.4978 | 7.1354 | 6.8335 | 7.2787 | 6.0948 | 6.0057 | 6.8357 | 6.7293 | 6.6952 | 7.2999 | 7.4541 | 7.4406 | 4.0620 | 4.8810 | 4.1236 | 4.6327 |
| LEMD3_S144 | LEMD3 | Q9Y2U8 | | 209 | AGPELQTPPGKDG | 7.0020 | 7.8413 | 6.9747 | 7.7406 | 6.7837 | 6.7618 | 7.1083 | 6.9805 | 7.2826 | 7.1846 | 7.3730 | 7.3684 | 3.0803 | 4.0220 | 3.0156 | 3.4805 |
| LEMD3_S27 | LEMD3 | Q9Y2U8 | | 402 | GAFSVDSPRIYSN | 6.6591 | 7.2861 | 7.0546 | 7.5465 | 6.3484 | 6.8156 | 7.1861 | 6.8537 | 7.0586 | 7.0758 | 7.4172 | 7.3444 | 3.3791 | 4.4464 | 3.3895 | 4.1392 |
| LEMD3_S352 | LEMD3 | Q9Y2U8 | | 144 | GFSSDESDVEASP | 5.5648 | 6.3347 | 6.2982 | 6.8726 | 6.7104 | 6.4349 | 6.3373 | 6.2432 | 6.2283 | 5.7957 | 5.3430 | 5.9065 | 6.0389 | 6.3308 | 6.1549 | 6.4058 |
| LEMD3_S402 | LEMD3 | Q9Y2U8 | | 883 | QALTSNTPLKPSN | 6.4065 | 7.0072 | 6.4073 | 7.5124 | 6.2146 | 6.2838 | 6.8115 | 6.2244 | 6.6797 | 6.9758 | 6.4509 | 6.4317 | 4.4391 | 5.5475 | 5.0817 | 5.5260 |
| LEMD3_T209 | LEMD3 | Q9Y2U8 | | 27 | LRRYGLSPGPVTE | 6.9869 | 7.6191 | 6.8969 | 7.8665 | 6.4578 | 7.1528 | 7.3996 | 7.1386 | 7.0488 | 7.2821 | 7.2329 | 7.4697 | 3.0229 | 4.1280 | 2.7507 | 3.5467 |
| LEMD3_T883 | LEMD3 | Q9Y2U8 | | 352 | CDQVDSSPVPRYR | 6.7294 | 7.3090 | 6.6325 | 6.6735 | 6.2066 | 6.6935 | 6.8113 | 6.2534 | 6.4872 | 6.9574 | 6.1498 | 6.7236 | 4.5205 | 5.4742 | 4.7610 | 5.6170 |
| LENG1_S59 | LENG1 | Q96BZ8 | | 59 | KARHQNSLPELEA | 6.6151 | 6.5247 | 6.7655 | 6.2008 | 6.4959 | 6.2321 | 6.4006 | 5.5572 | 6.5576 | 6.0841 | 7.0861 | 6.1145 | 6.2855 | 5.9016 | 5.8679 | 5.3107 |
| LENG8_S351 | LENG8 | Q96PV6 | | 351 | REPVAESPKKKRW | 6.6718 | 6.8274 | 6.4791 | 6.5183 | 6.4173 | 5.8755 | 6.6911 | 5.7274 | 6.4384 | 6.0443 | 6.1809 | 6.2314 | 5.9789 | 5.9361 | 5.8717 | 6.1103 |
| LEO1_S151;S154 | LEO1 | Q8WVC0 | | 179 | EDKLQNSDDDEKM | 5.8463 | 5.8838 | 6.1875 | 6.2909 | 6.5245 | 6.1261 | 6.4101 | 5.9066 | 6.3797 | 6.2236 | 5.9721 | 5.7680 | 6.7225 | 6.5253 | 6.5375 | 6.6955 |
| LEO1_S162 | LEO1 | Q8WVC0 | | 630 | KAKKLTSDEEGEP | 6.4326 | 6.7825 | 6.5224 | 7.1362 | 6.1778 | 6.1605 | 6.1714 | 6.4845 | 6.2931 | 5.8525 | 6.2422 | 6.2730 | 5.7374 | 6.0549 | 5.7506 | 5.9284 |
| LEO1_S179 | LEO1 | Q8WVC0 | | 205 | DERQQLSEEEKAN | 5.2400 | 5.5764 | 6.0516 | 6.3482 | 6.1825 | 5.6597 | 6.2098 | 6.1667 | 6.1972 | 6.2699 | 5.8402 | 5.8554 | 6.8927 | 6.9039 | 7.2730 | 7.3326 |
| LEO1_S205 | LEO1 | Q8WVC0 | | 162 | DEKIQNSDDEERA | 5.6662 | 6.0464 | 5.7484 | 6.7194 | 5.9603 | 5.6511 | 6.4725 | 6.3591 | 6.5983 | 6.5265 | 5.9747 | 6.2079 | 6.2160 | 6.5967 | 6.5453 | 6.7113 |
| LEO1_S212;S220 | LEO1 | Q8WVC0 | | 658 | HKKYVISDEEEED | 5.4922 | 6.0544 | 6.0262 | 6.5168 | 5.9339 | 6.3872 | 6.5212 | 6.2954 | 6.1312 | 6.5997 | 6.3971 | 6.1876 | 6.4480 | 6.2354 | 6.4488 | 6.3247 |
| LEO1_S630 | LEO1 | Q8WVC0 | | 629 | LKAKKLTSDEEGE | 6.3194 | 6.2254 | 6.7559 | 7.2440 | 5.9136 | 5.0087 | 6.3053 | 6.2757 | 6.5861 | 6.1623 | 6.6247 | 7.0000 | 5.7080 | 5.6615 | 5.6253 | 6.5842 |
| LEO1_S658 | LEO1 | Q8WVC0 | 151;154 | | NA | 5.5719 | 5.8040 | 6.3959 | 6.2471 | 5.9781 | 5.9530 | 6.4693 | 5.0549 | 6.2682 | 6.3765 | 6.6632 | 6.9003 | 7.0599 | 6.3641 | 6.1100 | 6.7837 |
| LEO1_T629 | LEO1 | Q8WVC0 | 212;220 | | NA | 4.9802 | 6.3467 | 6.7012 | 7.2231 | 6.0949 | 4.7397 | 6.8715 | 5.2266 | 6.9759 | 5.5709 | 5.1147 | 6.8660 | 6.8418 | 6.5723 | 6.2565 | 7.6181 |
| LETM1_S190 | LETM1 | O95202 | | 190 | RILNGHSLTRRER | 6.2328 | 6.2464 | 5.8934 | 6.6101 | 5.6941 | 5.9017 | 6.0162 | 5.9132 | 5.9297 | 6.4482 | 6.4288 | 6.5264 | 6.3057 | 6.3115 | 6.8885 | 6.6533 |
| LIG1_S141 | LIG1 | P18858 | | 183 | DGDQPTTPPKPLK | 5.3272 | 5.8522 | 6.2415 | 5.6950 | 6.6681 | 6.1780 | 5.9026 | 5.7308 | 6.1227 | 6.2898 | 6.5475 | 6.4065 | 6.7572 | 7.4690 | 6.8379 | 6.9739 |
| LIG1_S51 | LIG1 | P18858 | | 91 | LDCSQVSPPRPAT | 4.8697 | 4.6426 | 5.7541 | 5.3560 | 5.4776 | 5.7919 | 5.4906 | 5.1276 | 5.9080 | 6.1484 | 5.8765 | 5.7966 | 8.6667 | 8.3749 | 8.2400 | 8.4788 |
| LIG1_S66 | LIG1 | P18858 | | 51 | VVSESDSPVKRPG | 5.6050 | 6.0124 | 6.2939 | 5.5857 | 5.9547 | 6.2474 | 5.7135 | 5.7977 | 6.3517 | 6.5298 | 6.5377 | 6.2375 | 6.6862 | 7.0098 | 6.7284 | 6.7085 |
| LIG1_S66;S76 | LIG1 | P18858 | | 233 | TLSSFFTPRKPAV | 8.3877 | 7.4028 | 7.2246 | 6.5805 | 4.8539 | 7.2529 | 7.2764 | 6.6460 | 8.0996 | 6.8090 | 7.4967 | 8.0274 | 2.8132 | 4.4633 | 2.6965 | 3.9695 |
| LIG1_S76 | LIG1 | P18858 | | 819 | KALVLPSPRPVVR | 5.8603 | 6.4572 | 6.6943 | 6.4307 | 5.6513 | 6.1986 | 6.1352 | 6.2287 | 6.5334 | 5.9903 | 5.7460 | 5.8616 | 5.9607 | 7.1875 | 6.4868 | 6.5773 |
| LIG1_S819 | LIG1 | P18858 | | 66 | AARVLGSEGEEED | 5.6781 | 6.3018 | 6.7685 | 6.1001 | 5.2458 | 5.6875 | 5.0046 | 6.2228 | 5.6802 | 6.6978 | 5.8403 | 6.2456 | 7.1665 | 6.6926 | 6.8516 | 7.8162 |
| LIG1_S91 | LIG1 | P18858 | | 76 | EEDEALSPAKGQK | 5.5361 | 5.3855 | 5.8763 | 5.4070 | 5.6723 | 5.9663 | 5.7653 | 6.0882 | 6.5394 | 6.2235 | 6.7867 | 6.2061 | 7.2500 | 7.4340 | 6.9736 | 6.8895 |
| LIG1_S91;S98 | LIG1 | P18858 | | 141 | EVLEEQSEDEDRE | 5.8026 | 6.0897 | 6.6060 | 6.5123 | 5.6142 | 5.4723 | 6.3382 | 6.7888 | 5.6239 | 6.0667 | 6.7493 | 6.9004 | 6.0552 | 6.8425 | 5.7819 | 6.7562 |
| LIG1_T183 | LIG1 | P18858 | 91;98 | | NA | 4.8834 | 4.7788 | 5.3710 | 5.8981 | 4.9350 | 5.6706 | 5.2672 | 5.4007 | 5.8473 | 6.2826 | 6.2820 | 6.3119 | 7.9993 | 8.4000 | 8.7392 | 7.9329 |
| LIG1_T233 | LIG1 | P18858 | 66;76 | | NA | 6.2775 | 6.8569 | 6.6476 | 6.1617 | 5.8025 | 5.9583 | 5.8545 | 6.8111 | 6.6383 | 6.6690 | 6.9722 | 7.1179 | 5.1350 | 5.6318 | 5.2667 | 6.1989 |
| LIG3_S210 | LIG3 | P49916 | | 210 | TTGQVTSPVKGAS | 5.7597 | 6.3567 | 6.4583 | 6.5984 | 6.2364 | 6.2561 | 6.4369 | 6.2981 | 6.5447 | 6.4422 | 6.4819 | 6.5791 | 5.7643 | 6.0411 | 5.7550 | 5.9912 |
| LIG3_S242 | LIG3 | P49916 | | 242 | SGEAPSSPTPKRS | 6.0209 | 5.5186 | 6.6125 | 6.7447 | 6.2018 | 6.0571 | 6.3155 | 5.5372 | 6.4018 | 6.5994 | 6.4625 | 6.8196 | 6.2842 | 6.0831 | 5.7506 | 6.5905 |
| LIG3_T244 | LIG3 | P49916 | | 244 | EAPSSPTPKRSLS | 5.5395 | 5.7515 | 6.1216 | 6.2125 | 6.4189 | 6.2654 | 6.1716 | 5.9241 | 6.6516 | 6.7252 | 6.4892 | 6.5347 | 6.3610 | 6.6455 | 6.0277 | 6.1600 |
| LIMA1_S132 | LIMA1 | Q9UHB6-4 | | 225 | ASGRKISENSYSL | 6.8614 | 7.0108 | 8.0426 | 8.0118 | 6.6256 | 6.5009 | 7.0064 | 6.5765 | 7.3427 | 6.6337 | 8.1080 | 7.2627 | 3.3337 | 3.8307 | 3.3158 | 3.5368 |
| LIMA1_S15 | LIMA1 | Q9UHB6-4 | | 132 | PRSRLRSPPEALV | 6.2903 | 6.8030 | 6.2290 | 6.8350 | 6.0600 | 6.1874 | 6.6616 | 6.0475 | 6.3263 | 6.4430 | 6.0319 | 6.1303 | 5.8767 | 6.0307 | 5.7598 | 6.2876 |
| LIMA1_S215 | LIMA1 | Q9UHB6-4 | | 610 | KSPKTVSPPIRKG | 6.1519 | 6.5488 | 6.2517 | 7.0072 | 6.1263 | 5.9784 | 6.3402 | 6.8867 | 6.6216 | 6.5217 | 6.3384 | 6.5225 | 5.3102 | 5.7949 | 5.5844 | 6.0151 |
| LIMA1_S230 | LIMA1 | Q9UHB6-4 | | 375 | SSLSESSPPKAMK | 5.8669 | 6.4473 | 6.5432 | 7.1364 | 6.3323 | 6.4173 | 6.6024 | 6.1677 | 6.6569 | 6.5711 | 6.8165 | 7.1192 | 5.1518 | 5.2932 | 5.1419 | 5.7360 |
| LIMA1_S363 | LIMA1 | Q9UHB6-4 | | 687 | VENGADSDEDDNS | 5.9829 | 6.2448 | 6.0517 | 6.6064 | 6.2043 | 6.0548 | 5.7862 | 6.0624 | 5.9142 | 5.7385 | 6.1272 | 6.6501 | 6.6279 | 6.6800 | 6.6602 | 6.6085 |
| LIMA1_S375 | LIMA1 | Q9UHB6-4 | | 491 | ARETPHSPGVEDA | 5.9761 | 6.1554 | 5.7093 | 6.1307 | 6.5137 | 6.0615 | 6.1057 | 6.3355 | 5.9651 | 5.5807 | 5.6687 | 5.8158 | 7.2642 | 6.7853 | 7.1207 | 6.8116 |
| LIMA1_S491 | LIMA1 | Q9UHB6-4 | | 699 | SFLKQQSPQEPKS | 5.4032 | 5.8817 | 6.0724 | 6.3916 | 5.7310 | 5.8913 | 6.3472 | 6.0466 | 5.9179 | 5.8403 | 6.1740 | 6.0980 | 6.9552 | 6.9153 | 7.0467 | 7.2875 |
| LIMA1_S610 | LIMA1 | Q9UHB6-4 | | 15 | RQWTSLSLRVTAK | 7.4007 | 7.7445 | 8.1074 | 6.1269 | 6.4897 | 6.8060 | 6.5966 | 6.7820 | 6.6239 | 7.7781 | 7.1037 | 6.3298 | 4.2279 | 4.7549 | 3.4227 | 3.7053 |
| LIMA1_S687 | LIMA1 | Q9UHB6-4 | | 230 | ISENSYSLDDLEI | 6.3517 | 6.5627 | 7.2139 | 6.8451 | 5.9700 | 6.1725 | 6.2434 | 5.9231 | 7.1002 | 7.0142 | 6.8306 | 6.8077 | 5.1992 | 5.5493 | 5.0616 | 5.1548 |
| LIMA1_S699 | LIMA1 | Q9UHB6-4 | | 742 | EVVKELSVEEQIK | 5.5693 | 6.0002 | 6.0839 | 6.6259 | 6.0748 | 6.6356 | 6.9445 | 6.4493 | 6.6863 | 5.6827 | 6.4800 | 6.5137 | 6.0458 | 6.2521 | 5.9922 | 5.9637 |
| LIMA1_S705 | LIMA1 | Q9UHB6-4 | | 705 | SPQEPKSLNWSSF | 5.8994 | 6.6143 | 7.3545 | 6.7699 | 5.8676 | 6.6331 | 6.3644 | 7.1868 | 6.2512 | 6.0158 | 6.6510 | 6.5712 | 5.4027 | 5.7561 | 5.0102 | 5.6519 |
| LIMA1_S710 | LIMA1 | Q9UHB6-4 | | 363 | VHPKPLSPDSRAS | 6.7877 | 6.2364 | 6.7847 | 6.8671 | 6.5904 | 6.2199 | 6.2292 | 5.9270 | 6.8807 | 6.3487 | 6.1710 | 6.4381 | 5.9995 | 5.6281 | 5.2733 | 5.6180 |
| LIMA1_S742 | LIMA1 | Q9UHB6-4 | | 710 | KSLNWSSFVDNTF | 7.3977 | 7.1726 | 8.0257 | 7.8037 | 6.8031 | 6.1827 | 5.9516 | 6.8518 | 7.0814 | 5.4650 | 7.0647 | 6.0808 | 4.2394 | 5.5958 | 4.1299 | 4.1542 |
| LIMA1_S93 | LIMA1 | Q9UHB6-4 | | 93 | TDSLRNSSTEIRH | 6.5192 | 6.6598 | 6.2490 | 7.0206 | 6.2645 | 7.0140 | 6.7788 | 6.0666 | 5.8698 | 6.1211 | 6.2918 | 5.8198 | 6.3574 | 5.6074 | 5.7297 | 5.6307 |
| LIMD1_S272 | LIMD1 | Q9UGP4 | | 384 | GTGPKLSPTSLVH | 6.2049 | 6.2650 | 6.7518 | 6.3553 | 6.4328 | 6.9454 | 5.8705 | 7.2496 | 6.3332 | 5.8905 | 6.4619 | 6.2323 | 5.8886 | 5.8538 | 5.7447 | 5.5199 |
| LIMD1_S316 | LIMD1 | Q9UGP4 | | 272 | ASSQRVSPGLPSP | 5.6436 | 6.0453 | 6.2974 | 6.1044 | 5.9274 | 6.7536 | 6.2608 | 6.5994 | 6.4088 | 6.1315 | 6.2088 | 6.7296 | 6.4405 | 6.3011 | 5.9813 | 6.1666 |
| LIMD1_S353 | LIMD1 | Q9UGP4 | | 316 | GLPRSNSGLGGEV | 5.5354 | 5.2911 | 5.5326 | 5.0811 | 5.5935 | 5.0297 | 4.5116 | 5.1540 | 5.5072 | 5.0324 | 4.9841 | 5.0479 | 10.4971 | 9.1284 | 9.7824 | 8.2916 |
| LIMD1_S384 | LIMD1 | Q9UGP4 | | 424 | LLDSPSSPRVRLP | 5.4780 | 5.8809 | 6.2095 | 5.0995 | 7.0842 | 6.8822 | 6.2166 | 6.7776 | 6.4680 | 6.3135 | 5.5274 | 6.8691 | 6.1935 | 6.4900 | 6.3961 | 6.1139 |
| LIMD1_S421 | LIMD1 | Q9UGP4 | | 421 | GSVLLDSPSSPRV | 6.7210 | 6.9075 | 7.4497 | 6.3764 | 6.4123 | 5.7923 | 5.5712 | 6.8247 | 7.6337 | 6.8161 | 6.5320 | 6.9569 | 5.6144 | 4.2979 | 4.9833 | 5.1105 |
| LIMD1_S424 | LIMD1 | Q9UGP4 | | 353 | SSAPSSSPAGLDG | 6.1854 | 6.8617 | 7.8521 | 6.5418 | 6.0022 | 6.9483 | 6.6069 | 6.1735 | 7.1148 | 6.8982 | 7.4420 | 7.3900 | 4.5403 | 4.4328 | 3.8771 | 5.1330 |
| LIMK1_S310 | LIMK1 | P53667 | | 310 | GAGSLGSPASQRK | 6.2300 | 6.2937 | 7.0440 | 6.4813 | 6.1232 | 6.8038 | 6.3014 | 6.8014 | 6.4335 | 7.0396 | 6.9586 | 6.6744 | 5.0018 | 5.5852 | 4.9844 | 5.2438 |
| LIN28B_S211 | LIN28B | A0A1B0GVD3 | | 211 | GGHGCTSPPFPQE | 6.2675 | 6.9046 | 6.5996 | 6.7381 | 6.1544 | 6.6410 | 6.5813 | 6.2740 | 6.6507 | 6.3928 | 6.7878 | 6.4750 | 5.1805 | 5.6828 | 5.3647 | 5.3053 |
| LIN28B_S229 | LIN28B | A0A1B0GVD3 | | 3 | xxxxMRSFNQVSS | 8.1489 | 7.4130 | 7.9985 | 6.8993 | 5.5818 | 5.8826 | 5.7599 | 6.3083 | 7.7259 | 8.3939 | 6.7457 | 6.9271 | 3.3558 | 5.0957 | 3.7603 | 3.8232 |
| LIN28B_S3 | LIN28B | A0A1B0GVD3 | | 229 | SERSGRSPQEASS | 6.1525 | 6.9848 | 6.0556 | 6.6940 | 5.8095 | 5.7512 | 6.4818 | 6.9472 | 6.4577 | 6.5596 | 6.4383 | 6.1993 | 5.1902 | 6.0687 | 5.8510 | 6.3584 |
| LIN28B_S9 | LIN28B | A0A1B0GVD3 | | 9 | SFNQVSSAPGGAS | 6.4571 | 7.1731 | 7.8886 | 7.5441 | 7.2382 | 10.5392 | 8.6729 | 8.2229 | 5.2171 | 4.8986 | 4.4082 | 5.6703 | 4.2470 | 4.6186 | 3.3919 | 3.8122 |
| LIN37_S138 | LIN37 | Q96GY3 | | 182 | CRSRIPSPLQPEM | 5.8064 | 5.8451 | 5.8966 | 6.4367 | 6.1199 | 6.0830 | 6.5147 | 6.0984 | 6.2227 | 5.6846 | 6.6620 | 5.9673 | 6.4362 | 6.8070 | 6.8250 | 6.5943 |
| LIN37_S182 | LIN37 | Q96GY3 | | 138 | RECSPSSPLPPLP | 5.8593 | 6.0219 | 6.2611 | 6.5294 | 5.9845 | 6.2774 | 6.0882 | 6.1441 | 6.3006 | 6.3601 | 6.3102 | 6.2356 | 6.3165 | 6.4905 | 6.4542 | 6.3664 |
| LIN37_S182;S202 | LIN37 | Q96GY3 | | 167 | YKLPPPTPPGPPG | 5.8014 | 6.2809 | 6.4524 | 6.5895 | 6.1519 | 6.9631 | 6.6607 | 6.0188 | 6.8097 | 7.5570 | 6.7598 | 6.3490 | 5.4056 | 5.7420 | 5.5577 | 5.9007 |
| LIN37_T167 | LIN37 | Q96GY3 | 182;202 | | NA | 6.1057 | 5.7398 | 6.4125 | 6.5779 | 6.1026 | 6.5709 | 6.2317 | 7.0527 | 6.0997 | 6.9008 | 5.0979 | 7.1754 | 6.1172 | 6.0478 | 6.0694 | 5.6982 |
| LIN52_S53 | LIN52 | Q52LA3 | | 53 | KSPITSSPPKWMA | 4.9546 | 4.9057 | 4.8148 | 5.7319 | 5.5255 | 5.3055 | 4.6776 | 4.5816 | 4.3145 | 4.8216 | 4.7639 | 4.9291 | 5.0574 | 13.0797 | 8.0733 | 14.4632 |
| LIN54_S264 | LIN54 | Q6MZP7 | | 310 | PNKIAISPLKSPN | 6.3429 | 6.5135 | 6.0128 | 6.3661 | 6.1227 | 5.8034 | 5.9280 | 6.0304 | 6.2677 | 6.2492 | 5.8987 | 6.1477 | 6.7007 | 6.4425 | 6.7147 | 6.4590 |
| LIN54_S310 | LIN54 | Q6MZP7 | | 635 | CKNFEESPERKTL | 6.5250 | 6.9662 | 6.4950 | 6.6709 | 6.9060 | 6.4286 | 6.2880 | 6.1990 | 6.6362 | 6.7986 | 6.6675 | 6.5611 | 5.2599 | 5.4137 | 4.9900 | 5.1943 |
| LIN54_S635 | LIN54 | Q6MZP7 | | 280 | SQSTPGTPSKTIT | 7.4117 | 7.5449 | 6.7844 | 6.7375 | 6.1011 | 6.3959 | 6.7058 | 6.6300 | 7.2200 | 6.6349 | 6.8244 | 7.0030 | 4.4025 | 4.7074 | 4.2530 | 4.6434 |
| LIN54_T235 | LIN54 | Q6MZP7 | | 264 | GQTTQVSPPVIAG | 5.5478 | 5.8264 | 5.3944 | 6.6829 | 6.4806 | 5.8073 | 5.7899 | 6.3106 | 6.0477 | 5.7493 | 5.7708 | 6.7318 | 6.7368 | 6.5311 | 6.7948 | 7.7981 |
| LIN54_T280 | LIN54 | Q6MZP7 | | 235 | IAKKPRTPTSGPV | 6.2912 | 6.4658 | 6.0354 | 6.3177 | 6.5752 | 6.6601 | 6.8858 | 6.9163 | 6.2434 | 5.9728 | 6.2084 | 6.3393 | 5.8226 | 5.9157 | 5.7720 | 6.4783 |
| LIN9_S194 | LIN9 | Q5TKA1-2 | | 313 | PSRFFMTPPRLHY | 6.8823 | 7.0812 | 6.9269 | 7.1508 | 6.3787 | 6.5307 | 6.8516 | 6.7804 | 6.9814 | 6.8043 | 6.7485 | 7.1050 | 4.3240 | 4.7929 | 4.1061 | 4.5552 |
| LIN9_S223 | LIN9 | Q5TKA1-2 | | 112 | FTATMSTPDKKAS | 6.7179 | 6.9546 | 6.9398 | 7.0666 | 6.7864 | 6.3272 | 6.8605 | 6.5576 | 6.9462 | 6.6727 | 6.5464 | 6.6412 | 4.4877 | 4.6968 | 4.6318 | 5.1665 |
| LIN9_S325 | LIN9 | Q5TKA1-2 | | 223 | RKVADVSQFKDLP | 7.5480 | 6.8827 | 7.1591 | 6.9575 | 6.6783 | 6.3947 | 7.9059 | 6.6025 | 7.1751 | 7.2154 | 7.0163 | 6.5015 | 3.3639 | 4.5105 | 3.5518 | 4.5367 |
| LIN9_S325;S337 | LIN9 | Q5TKA1-2 | | 337 | DPLLGQSPWRSKI | 6.9071 | 6.8852 | 6.6030 | 6.6379 | 6.1747 | 6.4518 | 6.5209 | 6.5418 | 6.3821 | 5.9000 | 6.4060 | 5.7249 | 5.2736 | 6.5160 | 5.3766 | 5.6985 |
| LIN9_S337 | LIN9 | Q5TKA1-2 | | 325 | YTPPLQSPIIDND | 5.9051 | 6.0800 | 6.6906 | 6.2167 | 6.2279 | 6.9876 | 6.4068 | 5.8840 | 6.5341 | 5.9183 | 6.0886 | 6.0602 | 6.5678 | 6.3507 | 6.1665 | 5.9150 |
| LIN9_T112 | LIN9 | Q5TKA1-2 | | 194 | KPRRCSSAFFEEE | 5.4040 | 5.8177 | 6.2874 | 5.9777 | 6.4402 | 5.6817 | 6.1896 | 6.1381 | 6.4962 | 6.5038 | 6.3456 | 6.0617 | 6.6571 | 7.1587 | 6.3220 | 6.5184 |
| LIN9_T313 | LIN9 | Q5TKA1-2 | 325;337 | | NA | 6.4822 | 6.5729 | 7.0257 | 6.3888 | 6.5371 | 6.8394 | 6.3413 | 6.4381 | 6.6542 | 6.5052 | 6.4995 | 6.2565 | 5.3142 | 5.6525 | 5.1932 | 5.2992 |
| LIPE_S855 | LIPE | Q05469 | | 897 | TPEMSLSAETLSP | 5.8104 | 6.0950 | 6.5348 | 5.9061 | 6.2406 | 5.8583 | 6.8018 | 6.5207 | 6.2201 | 6.4524 | 6.2453 | 6.4160 | 5.9163 | 6.5264 | 5.9642 | 6.4918 |
| LIPE_S897 | LIPE | Q05469 | | 950 | GFHPRRSSQGATQ | 5.8792 | 5.5594 | 6.4978 | 6.0807 | 6.1914 | 6.0376 | 5.4679 | 6.8109 | 7.2054 | 6.8977 | 7.2650 | 7.6849 | 5.7140 | 6.6835 | 5.0571 | 6.9674 |
| LIPE_S950 | LIPE | Q05469 | | 855 | PMRRSVSEAALAQ | 6.4026 | 6.4096 | 6.2124 | 6.2302 | 6.6155 | 6.1795 | 6.3266 | 6.2693 | 6.7148 | 5.9293 | 6.0753 | 6.1503 | 6.2565 | 6.0766 | 5.8410 | 6.3105 |
| LLGL1_S1044 | LLGL1 | Q15334 | | 663 | KKSLRQSFRRIRK | 6.8025 | 7.1190 | 7.4900 | 5.9972 | 5.6083 | 6.9582 | 6.6033 | 6.1137 | 6.3826 | 7.0942 | 6.9169 | 7.1405 | 4.6339 | 6.0056 | 4.3334 | 4.8006 |
| LLGL1_S488 | LLGL1 | Q15334 | | 488 | DCEHADSLAQAAE | 6.2588 | 5.8943 | 6.2419 | 6.0369 | 6.7378 | 5.4686 | 6.2554 | 5.6010 | 5.5705 | 5.9690 | 5.0440 | 5.2246 | 6.9880 | 8.1128 | 7.5150 | 7.0814 |
| LLGL1_S659;S663 | LLGL1 | Q15334 | | 1044 | LGSSEESEKNLRN | 5.1154 | 5.3976 | 5.3273 | 6.1019 | 5.2536 | 4.9409 | 5.3768 | 5.5168 | 5.1090 | 5.9666 | 5.4753 | 5.7258 | 8.5243 | 8.7661 | 8.6561 | 8.7466 |
| LLGL1_S663 | LLGL1 | Q15334 | | 961 | RDATQASYRIRES | 9.2079 | 7.3049 | 7.7313 | 8.5839 | 7.0162 | 5.5389 | 5.7366 | 5.9420 | 6.7084 | 5.7494 | 6.5524 | 6.4411 | 4.1738 | 5.1748 | 4.0146 | 4.1236 |
| LLGL1_S682;S683 | LLGL1 | Q15334 | 682;683 | | NA | 7.1644 | 7.4384 | 6.1752 | 7.1788 | 6.2535 | 6.7508 | 7.3616 | 6.6630 | 6.9407 | 6.8758 | 6.2117 | 6.6319 | 3.9459 | 5.1875 | 3.8617 | 5.3590 |
| LLGL1_S961 | LLGL1 | Q15334 | 659;663 | | NA | 7.6679 | 7.2316 | 7.1381 | 6.5393 | 5.9225 | 6.9163 | 6.6068 | 6.6758 | 7.2682 | 6.7854 | 6.8464 | 7.0504 | 4.3301 | 5.3262 | 3.2163 | 4.4787 |
| LLGL2_S480 | LLGL2 | Q6P1M3 | | 480 | DPNENFSAQGEDE | 6.3737 | 7.0693 | 7.1312 | 6.6689 | 6.2932 | 6.1198 | 8.0105 | 4.0393 | 6.8464 | 6.8464 | 7.4697 | 6.2003 | 5.0073 | 5.5687 | 5.5151 | 4.8401 |
| LMNA_S107 | LMNA | P02545 | | 390 | EERLRLSPSPTSQ | 6.3478 | 6.3309 | 5.7347 | 6.2397 | 6.3510 | 5.9743 | 6.1338 | 6.1569 | 6.4067 | 6.0389 | 6.5516 | 6.5322 | 6.8551 | 6.8201 | 6.8579 | 6.4683 |
| LMNA_S143 | LMNA | P02545 | | 22 | ASSTPLSPTRITR | 6.8623 | 7.4611 | 7.2537 | 7.1889 | 6.2559 | 6.6383 | 6.8331 | 6.6631 | 7.5078 | 7.5122 | 8.1123 | 8.0024 | 3.0183 | 3.8622 | 3.0972 | 3.7309 |
| LMNA_S19;S22 | LMNA | P02545 | | 392 | RLRLSPSPTSQRS | 6.2805 | 6.3430 | 5.7086 | 6.1893 | 6.3820 | 6.0264 | 6.2799 | 6.3059 | 6.1734 | 5.9242 | 5.7234 | 5.6934 | 6.7828 | 6.7372 | 6.8274 | 6.6226 |
| LMNA_S212 | LMNA | P02545 | | 458 | VRLRNKSNEDQSM | 5.9030 | 6.1048 | 5.7560 | 5.9638 | 6.0597 | 6.0030 | 6.3112 | 6.0943 | 6.2524 | 6.0144 | 5.9205 | 5.8195 | 7.3682 | 6.9254 | 6.7977 | 6.7060 |
| LMNA_S22 | LMNA | P02545 | | 628 | VTRSYRSVGGSGG | 6.7097 | 6.9185 | 6.5200 | 6.8416 | 6.4776 | 6.8821 | 7.1861 | 6.3331 | 6.9478 | 6.8065 | 6.6993 | 6.5635 | 4.6366 | 5.4327 | 4.3772 | 4.6676 |
| LMNA_S301 | LMNA | P02545 | | 107 | RLQLELSKVREEF | 6.6353 | 6.4339 | 5.7452 | 6.2397 | 6.8066 | 5.7233 | 6.3940 | 6.0863 | 6.1941 | 5.6749 | 5.0803 | 4.8202 | 7.6579 | 6.6322 | 7.4680 | 6.4081 |
| LMNA_S390 | LMNA | P02545 | | 616 | PISSGSSASSVTV | 5.4920 | 5.8002 | 5.5221 | 5.7775 | 6.0671 | 6.0943 | 6.0502 | 6.5544 | 6.0425 | 5.5483 | 5.6097 | 6.1704 | 7.8464 | 7.2355 | 7.6019 | 7.4877 |
| LMNA_S390;S392 | LMNA | P02545 | | 652 | YLLGNSSPRTQSP | 6.6249 | 6.9702 | 6.0985 | 6.3532 | 5.9797 | 6.2768 | 6.3858 | 6.1741 | 6.5925 | 7.0493 | 6.9358 | 7.0696 | 5.0330 | 5.5277 | 5.4161 | 5.5125 |
| LMNA_S392 | LMNA | P02545 | | 423 | STESRSSFSQHAR | 6.0790 | 6.2741 | 5.1458 | 6.1444 | 6.4314 | 6.5375 | 6.0466 | 6.8046 | 5.6610 | 5.3254 | 5.5780 | 5.3737 | 7.1889 | 7.1499 | 7.6890 | 6.5708 |
| LMNA_S404 | LMNA | P02545 | | 423 | KKRKLESTESRSS | 3.8557 | 3.7818 | 3.7417 | 4.1635 | 5.3972 | 5.0458 | 5.2450 | 5.1810 | 3.7939 | 3.8073 | 3.7333 | 4.1363 | 12.6204 | 10.7409 | 12.9325 | 11.8235 |
| LMNA_S404;S407 | LMNA | P02545 | | 94 | ARKTLDSVAKERA | 6.2615 | 6.8942 | 6.4417 | 7.1306 | 5.8019 | 5.7394 | 6.1952 | 5.9981 | 6.0076 | 6.2577 | 6.0278 | 5.7649 | 6.4565 | 6.3622 | 6.6421 | 6.0187 |
| LMNA_S404;S407;S414 | LMNA | P02545 | | 19 | GAQASSTPLSPTR | 6.0535 | 6.2447 | 5.4591 | 6.5295 | 6.5988 | 5.9799 | 6.2299 | 5.8950 | 6.1265 | 5.6885 | 5.9234 | 6.1078 | 6.8831 | 6.7391 | 6.8433 | 6.6978 |
| LMNA_S407 | LMNA | P02545 | | 533 | RTALINSTGEEVA | 5.8273 | 6.3562 | 6.1349 | 6.2174 | 5.9996 | 6.2302 | 6.0066 | 6.0561 | 6.5514 | 6.0386 | 5.9069 | 6.5220 | 6.4754 | 6.4267 | 6.6100 | 6.4007 |
| LMNA_S423 | LMNA | P02545 | | 301 | SRIRIDSLSAQLS | 5.7797 | 6.0269 | 5.8131 | 6.5501 | 6.5711 | 6.7836 | 6.4969 | 5.0110 | 4.9672 | 5.8655 | 6.3933 | 5.5309 | 7.5668 | 7.8887 | 6.3182 | 6.4370 |
| LMNA_S429 | LMNA | P02545 | | 404 | SRGRASSHSSQTQ | 5.2870 | 5.6570 | 3.8264 | 5.6498 | 7.0161 | 6.6096 | 5.7882 | 7.1352 | 4.7593 | 4.3787 | 5.0254 | 4.7057 | 8.3920 | 8.6942 | 9.5796 | 7.4961 |
| LMNA_S458 | LMNA | P02545 | | 143 | LEALLNSKEAALS | 7.1213 | 5.3797 | 7.5388 | 8.0596 | 7.8608 | 3.6740 | 7.0934 | 6.2704 | 6.6481 | 4.8748 | 6.1352 | 5.0736 | 5.7416 | 5.8330 | 6.1551 | 6.5408 |
| LMNA_S463 | LMNA | P02545 | | 212 | FQKNIYSEELRET | 6.2957 | 6.4113 | 5.9329 | 5.9718 | 6.4600 | 5.8286 | 6.4521 | 5.8644 | 5.5989 | 5.9520 | 6.3381 | 6.1029 | 6.8240 | 6.8945 | 6.4719 | 6.6009 |
| LMNA_S533 | LMNA | P02545 | | 407 | RASSHSSQTQGGG | 4.9862 | 5.3539 | 4.0601 | 5.5967 | 6.4539 | 6.2996 | 5.9231 | 5.2477 | 4.7463 | 4.7463 | 4.4966 | 4.5192 | 10.3735 | 7.6985 | 10.2761 | 9.2223 |
| LMNA_S568 | LMNA | P02545 | | 463 | KSNEDQSMGNWQI | 6.0220 | 6.3895 | 5.2252 | 6.5294 | 5.7302 | 5.9744 | 5.7742 | 6.0223 | 6.3663 | 5.9696 | 5.8565 | 6.2216 | 7.1219 | 7.1363 | 7.2515 | 6.4091 |
| LMNA_S616 | LMNA | P02545-2 | | 568 | LHHHHVSGSRRxx | 5.2840 | 5.5636 | 4.4756 | 6.4371 | 6.6018 | 7.9195 | 6.9919 | 6.7535 | 6.3525 | 6.6625 | 5.3014 | 6.3959 | 6.6733 | 7.5468 | 5.1236 | 5.9169 |
| LMNA_S628 | LMNA | P02545 | 390;392 | | NA | 6.3850 | 6.8388 | 6.5851 | 7.0308 | 6.1292 | 6.4036 | 6.4811 | 6.6033 | 6.9335 | 7.0120 | 6.8208 | 6.7769 | 4.7286 | 5.3065 | 4.8481 | 5.1168 |
| LMNA_S652 | LMNA | P02545 | 404;407 | | NA | 4.9629 | 5.1438 | 4.8363 | 5.5045 | 6.6261 | 6.0884 | 6.1903 | 6.5450 | 5.1617 | 4.7708 | 4.9824 | 4.7811 | 8.7904 | 8.9572 | 8.9457 | 7.7134 |
| LMNA_S94 | LMNA | P02545 | 19;22 | | NA | 6.9620 | 7.2395 | 6.9026 | 7.1969 | 6.3075 | 6.8618 | 7.0549 | 6.6773 | 7.5992 | 7.3051 | 7.4619 | 7.1799 | 3.2074 | 4.3116 | 3.4492 | 4.2831 |
| LMNA_T19 | LMNA | P02545 | 404;407;414 | | NA | 5.4574 | 5.3822 | 5.8910 | 5.8700 | 7.3916 | 7.7980 | 7.9388 | 7.7632 | 5.7195 | 5.3473 | 5.1735 | 5.0629 | 6.4266 | 6.2788 | 5.8565 | 6.6426 |
| LMNB1_S158 | LMNB1 | P20700 | | 575 | LFHQQGTPRASNR | 5.9552 | 6.5110 | 6.4287 | 6.2202 | 5.9865 | 6.0427 | 6.3601 | 6.5053 | 6.1949 | 6.3037 | 6.3049 | 6.1242 | 6.1193 | 6.4013 | 6.2584 | 6.2837 |
| LMNB1_S200 | LMNB1 | P20700 | | 23 | GPTTPLSPTRLSR | 7.3848 | 7.7310 | 6.6383 | 7.5816 | 6.5905 | 6.7667 | 7.4710 | 7.0282 | 7.3618 | 7.3000 | 7.5670 | 7.2995 | 2.8290 | 3.7865 | 2.9844 | 3.6796 |
| LMNB1_S210 | LMNB1 | P20700 | | 20 | RAGGPTTPLSPTR | 6.3982 | 6.4059 | 5.7935 | 6.5729 | 6.4814 | 6.2299 | 6.6877 | 6.5354 | 6.2076 | 5.9518 | 6.1010 | 6.2000 | 5.9931 | 6.0506 | 6.0998 | 6.2913 |
| LMNB1_S23 | LMNB1 | P20700 | | 158 | LENRCQSLTEDLE | 5.8672 | 6.0732 | 6.3812 | 6.5442 | 6.2313 | 5.9632 | 6.2455 | 5.4690 | 5.6488 | 6.3726 | 6.0656 | 6.0778 | 6.6565 | 6.7659 | 6.9685 | 6.6694 |
| LMNB1_S375 | LMNB1 | P20700 | | 158 | ALGDKKSLEGDLE | 6.6210 | 6.7330 | 5.5639 | 7.1994 | 5.8690 | 5.5269 | 6.1176 | 5.9389 | 5.2023 | 5.8402 | 5.7139 | 6.1433 | 6.8881 | 7.0340 | 6.0467 | 7.5620 |
| LMNB1_T19;S23 | LMNB1 | P20700 | | 375 | ALDMEISAYRKLL | 6.8832 | 7.4664 | 6.3777 | 6.9853 | 6.4675 | 5.9105 | 5.8645 | 6.0744 | 5.2781 | 5.1797 | 5.9227 | 5.1752 | 6.8959 | 6.3908 | 7.0751 | 6.0531 |
| LMNB1_T20 | LMNB1 | P20700 | | 210 | DLEFRKSMYEEEI | 6.3249 | 7.3580 | 6.7507 | 7.2208 | 6.3821 | 6.4962 | 6.3597 | 6.3526 | 6.4980 | 5.9226 | 6.0093 | 6.0488 | 5.3878 | 5.6383 | 5.1653 | 6.0847 |
| LMNB1_T20;S23 | LMNB1 | P20700 | 19;23 | | NA | 7.3376 | 7.6814 | 6.7252 | 7.9386 | 6.5140 | 7.3233 | 7.4520 | 7.2923 | 7.3808 | 7.4952 | 7.5107 | 7.3400 | 2.3462 | 3.8080 | 2.4389 | 3.4158 |
| LMNB1_T575 | LMNB1 | P20700 | 20;23 | | NA | 7.4071 | 8.3786 | 6.5566 | 7.4261 | 6.5748 | 7.0319 | 7.7279 | 7.1714 | 6.9248 | 7.0421 | 8.0356 | 7.1413 | 2.7067 | 3.8663 | 2.4567 | 3.5522 |
| LMNB2_S214 | LMNB2 | Q03252 | | 407 | RLKLSPSPSSRVT | 6.9473 | 7.1917 | 6.3941 | 7.2433 | 6.3266 | 6.1429 | 6.3115 | 6.3577 | 6.7869 | 6.6231 | 6.1968 | 6.5255 | 5.0919 | 5.0937 | 5.4404 | 5.3268 |
| LMNB2_S316 | LMNB2 | Q03252 | | 37 | GPATPLSPTRLSR | 7.3661 | 7.7488 | 6.6983 | 7.6012 | 6.7055 | 6.7500 | 7.2300 | 6.9953 | 7.3181 | 7.3128 | 7.5078 | 7.4242 | 2.8828 | 3.7248 | 3.0511 | 3.6831 |
| LMNB2_S37 | LMNB2 | Q03252 | | 34 | RAGGPATPLSPTR | 5.9815 | 6.1403 | 5.6494 | 6.4544 | 6.3685 | 5.8063 | 6.2215 | 6.2993 | 5.9836 | 5.5211 | 6.5659 | 5.9149 | 7.0876 | 6.9959 | 7.0047 | 6.9110 |

| | | | | | | | | | | | | | | | | | | | | | |
|---|---|---|---|---|---|---|---|---|---|---|---|---|---|---|---|---|---|---|---|---|---|
| LMNB2_S405 | LMNB2 | Q03252 | | 405 | EERLKLSPSPSSR | 6.8825 | 6.9659 | 6.3497 | 7.2024 | 7.0863 | 6.5943 | 6.5969 | 6.9208 | 6.5066 | 6.4748 | 6.9291 | 6.4119 | 4.8762 | 5.0782 | 4.6476 | 4.4769 |
| LMNB2_S405;S407 | LMNB2 | Q03252 | | 420 | VSRATSSSSGSLS | 7.2716 | 7.5477 | 6.3100 | 7.5111 | 6.4354 | 5.9556 | 6.6697 | 6.2956 | 6.6871 | 6.7389 | 6.1756 | 6.5021 | 4.8140 | 5.1567 | 4.7484 | 5.1805 |
| LMNB2_S407 | LMNB2 | Q03252 | | 518 | EIAYKFTPKYILR | 6.3937 | 6.5327 | 6.4684 | 6.7391 | 6.3122 | 6.4815 | 6.3778 | 5.7795 | 6.2176 | 6.3041 | 6.5479 | 6.3764 | 6.2833 | 6.1005 | 5.5882 | 5.4971 |
| LMNB2_S420 | LMNB2 | Q03252 | | 424 | TSSSSGSLSATGR | 6.9911 | 6.9985 | 5.8692 | 7.0675 | 6.7711 | 6.1009 | 6.4529 | 6.4543 | 6.4031 | 6.2186 | 6.0445 | 6.4355 | 5.5728 | 5.5096 | 5.5664 | 5.5440 |
| LMNB2_S420;S426 | LMNB2 | Q03252 | | 214 | LENRCQSLQEELD | 6.1310 | 7.1473 | 6.5121 | 7.2665 | 7.1544 | 5.9812 | 6.4558 | 4.6952 | 5.4609 | 7.0445 | 6.5178 | 6.0233 | 6.2195 | 6.3123 | 5.7271 | 5.3510 |
| LMNB2_S424 | LMNB2 | Q03252 | | 426 | SSSGSLSATGRLG | 7.0174 | 6.6660 | 6.4935 | 6.2351 | 5.9220 | 6.1461 | 6.4185 | 6.6286 | 7.5163 | 6.7621 | 6.7238 | 6.1539 | 5.5129 | 5.5355 | 5.2591 | 5.0093 |
| LMNB2_S426 | LMNB2 | Q03252 | | 497 | NSDKDQSLGNWRI | 6.6736 | 6.7128 | 6.1218 | 7.0679 | 6.4550 | 5.6012 | 6.4933 | 6.4081 | 6.5339 | 5.9020 | 5.5501 | 6.1660 | 6.4318 | 6.0646 | 6.2529 | 5.5650 |
| LMNB2_S497 | LMNB2 | Q03252 | | 541 | GAGVAHSPPSTLV | 5.9896 | 7.1431 | 6.9448 | 8.3701 | 6.5291 | 6.4112 | 6.6835 | 6.9772 | 6.1854 | 6.8623 | 7.5967 | 7.1961 | 3.9462 | 4.5043 | 4.0387 | 4.6217 |
| LMNB2_S541 | LMNB2 | Q03252 | | 316 | ARMRLESLSYQLS | 6.6766 | 6.0021 | 6.9901 | 6.3271 | 7.0474 | 5.5557 | 6.8950 | 5.4878 | 6.0840 | 5.6243 | 5.5275 | 4.7261 | 6.4327 | 6.7409 | 8.1226 | 5.7600 |
| LMNB2_T34 | LMNB2 | Q03252 | 405;407 | | NA | 7.0388 | 7.9868 | 7.1340 | 8.2939 | 6.1595 | 6.6807 | 6.9212 | 6.6123 | 7.1827 | 7.5956 | 7.5927 | 7.6860 | 2.9209 | 3.7279 | 2.8896 | 3.5773 |
| LMNB2_T34;S37 | LMNB2 | Q03252 | 34;37 | | NA | 7.7840 | 8.1185 | 6.9010 | 7.8673 | 6.4410 | 7.0529 | 7.4775 | 7.2281 | 7.3301 | 7.5547 | 7.5939 | 7.7473 | 2.2792 | 3.3294 | 2.1967 | 3.0985 |
| LMNB2_T420;S424 | LMNB2 | Q03252 | 420;426 | | NA | 7.6459 | 8.0163 | 6.9953 | 7.9838 | 6.5263 | 6.3029 | 7.3630 | 6.3415 | 7.5609 | 6.3682 | 7.7651 | 6.6344 | 3.7622 | 3.7686 | 3.0910 | 3.8747 |
| LMNB2_T518 | LMNB2 | Q03252 | 420;424 | | NA | 5.4887 | 5.9061 | 6.8252 | 7.2144 | 7.0179 | 6.9512 | 6.4924 | 5.7026 | 6.8696 | 6.4611 | 7.1072 | 5.9969 | 5.4170 | 5.1973 | 5.7945 | 5.5578 |
| LMO7_S1089 | LMO7 | Q8WWI1-3 | | 531 | TEIQLPSQSPVEE | 5.6723 | 5.4105 | 5.8384 | 6.6645 | 6.0289 | 5.8183 | 6.0162 | 6.4622 | 6.1785 | 6.5343 | 5.9766 | 6.1049 | 7.0667 | 6.4817 | 6.8394 | 6.9065 |
| LMO7_S1159 | LMO7 | Q8WWI1-3 | | 714 | RISINQTPGKSLD | 7.5623 | 7.6231 | 7.2166 | 8.0598 | 6.5930 | 6.3748 | 7.2790 | 7.0137 | 7.1477 | 6.9727 | 6.8969 | 6.7970 | 3.3783 | 3.8436 | 3.4031 | 3.8385 |
| LMO7_S116 | LMO7 | Q8WWI1-3 | | 417 | WKDRRKSYTSDLQ | 6.3164 | 5.2014 | 5.7522 | 5.0296 | 5.6451 | 5.8154 | 5.4599 | 5.9357 | 8.3461 | 8.2853 | 7.9259 | 8.2841 | 6.1993 | 6.8219 | 4.0819 | 4.8999 |
| LMO7_S1176 | LMO7 | Q8WWI1-3 | | 246 | HHKREDSFESLDS | 5.5627 | 6.0044 | 6.2146 | 5.8138 | 5.6508 | 6.3785 | 5.6245 | 5.7165 | 5.6812 | 6.3664 | 6.0547 | 5.9418 | 7.1271 | 7.2862 | 7.1547 | 7.4222 |
| LMO7_S1252 | LMO7 | Q8WWI1-3 | | 654 | DVTRLPSPTSPFS | 6.2475 | 6.4730 | 5.6732 | 6.5955 | 6.5371 | 5.7570 | 6.2483 | 6.1330 | 5.8834 | 6.0695 | 6.0603 | 6.2266 | 6.6449 | 6.6432 | 6.5019 | 6.3056 |
| LMO7_S246 | LMO7 | Q8WWI1-3 | | 692 | SGEGEISPQREVS | 5.5883 | 6.1839 | 6.5165 | 6.5246 | 5.9146 | 6.0262 | 6.1042 | 6.0171 | 6.6463 | 6.4515 | 6.1191 | 7.0844 | 6.4232 | 6.6467 | 5.9661 | 5.7872 |
| LMO7_S276 | LMO7 | Q8WWI1-3 | | 471 | SRRRMYSFDDVLE | 4.4235 | 7.3800 | 9.3523 | 5.9976 | 5.2105 | 7.5545 | 6.2760 | 7.1076 | 5.8892 | 8.5882 | 8.7957 | 6.5743 | 3.1634 | 4.6767 | 5.0102 | 4.0002 |
| LMO7_S300 | LMO7 | Q8WWI1-3 | | 863 | KNLKRRSQFFEQG | 6.4006 | 6.4446 | 6.2384 | 5.7052 | 6.0114 | 6.7953 | 6.4196 | 6.7712 | 7.1627 | 6.4275 | 6.4562 | 6.7323 | 5.6464 | 5.6969 | 5.4745 | 5.6172 |
| LMO7_S322 | LMO7 | Q8WWI1-3 | | 1176 | IMRRGESLDNLDS | 4.8832 | 6.0415 | 7.0491 | 6.3868 | 4.8351 | 6.3304 | 6.6241 | 6.7721 | 5.3210 | 6.8436 | 7.1938 | 7.2316 | 5.2117 | 5.4216 | 6.5633 | 7.2909 |
| LMO7_S342 | LMO7 | Q8WWI1-3 | | 342 | HEDNRRSWASPVY | 6.2775 | 6.4356 | 6.5469 | 6.5010 | 5.4297 | 6.5076 | 6.8699 | 6.0780 | 7.2760 | 6.7218 | 5.8336 | 6.7630 | 6.1463 | 5.9541 | 5.3702 | 5.2886 |
| LMO7_S417 | LMO7 | Q8WWI1-3 | | 795 | RYGKAGSPETKWI | 6.4062 | 6.5374 | 6.0905 | 6.7336 | 6.0443 | 5.9682 | 6.3825 | 6.0080 | 6.1065 | 6.3660 | 6.2349 | 6.4182 | 6.0059 | 6.2082 | 6.0989 | 6.3907 |
| LMO7_S471 | LMO7 | Q8WWI1-3 | | 561 | STRVSASLPRSYR | 7.7936 | 7.8700 | 6.9815 | 7.8393 | 5.8275 | 6.2816 | 6.7700 | 6.2045 | 6.7466 | 6.8772 | 6.7151 | 6.7462 | 3.9520 | 4.8838 | 3.8570 | 4.6540 |
| LMO7_S531 | LMO7 | Q8WWI1-3 | | 1252 | GVATTQSPTPRSH | 6.5084 | 6.5362 | 6.5703 | 6.9187 | 6.6936 | 6.9058 | 7.1628 | 6.2670 | 6.1169 | 6.7741 | 6.0998 | 6.1082 | 5.1831 | 5.7860 | 4.7282 | 5.6411 |
| LMO7_S539 | LMO7 | Q8WWI1-3 | | 598 | SLPRSYTMDDAWK | 7.5896 | 8.1692 | 9.0876 | 7.1765 | 5.7763 | 6.3232 | 6.1312 | 6.0393 | 6.0314 | 5.9997 | 6.2657 | 5.6352 | 4.7971 | 5.2868 | 4.7703 | 4.9210 |
| LMO7_S561 | LMO7 | Q8WWI1-3 | | 1159 | IRQRSASVNKEPV | 5.5191 | 5.8886 | 6.2768 | 6.5837 | 6.9638 | 6.6273 | 6.4780 | 6.3435 | 6.1654 | 6.7004 | 6.8756 | 6.4556 | 6.3733 | 6.4947 | 5.3408 | 4.9133 |
| LMO7_S592 | LMO7 | E9PMS6 | | 721 | ATKIIGSPETKWI | 7.1889 | 7.5622 | 6.1901 | 7.8927 | 6.4706 | 6.4028 | 7.3368 | 6.8925 | 6.9147 | 7.0914 | 6.9029 | 7.0132 | 3.5003 | 4.4507 | 3.8495 | 4.3407 |
| LMO7_S654 | LMO7 | Q8WWI1-3 | | 300 | MSYRRISAVEPKT | 6.4085 | 5.0573 | 6.0267 | 5.5048 | 5.5010 | 6.4796 | 6.4024 | 5.5994 | 8.1957 | 7.9369 | 7.9042 | 7.1170 | 5.6841 | 7.2476 | 3.9948 | 4.9401 |
| LMO7_S654;S657 | LMO7 | Q8WWI1-3 | | 322 | NKSRQPSYVPAPL | 8.7254 | 6.3933 | 8.6267 | 7.0574 | 4.4544 | 5.5977 | 6.2951 | 4.8882 | 7.9684 | 7.7885 | 7.6066 | 7.9865 | 4.3869 | 5.2544 | 3.3991 | 3.5713 |
| LMO7_S692 | LMO7 | Q8WWI1-3 | | 579 | RLTSVVTPRPFGS | 6.8034 | 8.2612 | 7.3646 | 7.9047 | 5.0055 | 7.0486 | 6.6093 | 9.2437 | 8.2913 | 6.7522 | 7.3345 | 8.3816 | 2.5908 | 3.4559 | 1.8987 | 3.0541 |
| LMO7_S721 | LMO7 | Q8WWI1-3 | | 276 | GREGFESDTDSEF | 5.8138 | 6.3364 | 6.7035 | 7.2946 | 6.1636 | 6.3144 | 5.8325 | 6.1624 | 6.1416 | 6.2372 | 5.5945 | 5.9555 | 6.0906 | 6.4284 | 6.1217 | 6.8093 |
| LMO7_S795 | LMO7 | Q8WWI1-3 | | 116 | KKINRLSTPIAGL | 6.7605 | 7.3205 | 7.1409 | 7.9267 | 6.1497 | 6.1151 | 6.9676 | 5.8314 | 7.2262 | 6.3076 | 6.9855 | 7.1391 | 4.3137 | 5.3102 | 3.9476 | 4.5577 |
| LMO7_S863 | LMO7 | Q8WWI1-3 | | 539 | SPVEEQSPASLSS | 6.6318 | 6.3394 | 6.5160 | 7.5537 | 6.2505 | 5.5928 | 6.1434 | 5.2031 | 7.2066 | 6.4113 | 6.7998 | 6.7280 | 4.9948 | 6.7791 | 5.4516 | 5.3980 |
| LMO7_T579 | LMO7 | Q8WWI1-3 | | 626 | VSTPAPSPDASQL | 4.7506 | 5.7336 | 5.1433 | 6.1502 | 6.3184 | 5.6493 | 6.1570 | 6.5844 | 5.5303 | 5.8040 | 5.8563 | 6.0831 | 7.2464 | 7.6574 | 8.3555 | 6.9801 |
| LMO7_T598 | LMO7 | Q8WWI1-3 | | 592 | QTRGISSLPRSYT | 7.3996 | 7.3607 | 6.7407 | 6.7377 | 6.0838 | 6.2940 | 6.7397 | 5.6333 | 6.3375 | 6.7690 | 6.8732 | 6.4701 | 4.8149 | 5.4129 | 4.5282 | 4.8748 |
| LMO7_T622;S626 | LMO7 | Q8WWI1-3 | | 1089 | DRNKSRSTTELDD | 5.5164 | 6.4649 | 5.7232 | 6.2011 | 5.5740 | 6.1601 | 6.1135 | 6.4525 | 6.3803 | 6.3786 | 6.7582 | 6.5091 | 5.7122 | 6.6657 | 6.4076 | 6.9824 |
| LMO7_T626 | LMO7 | Q8WWI1-3 | 654;657 | | NA | 5.8980 | 5.6514 | 6.0919 | 6.4083 | 5.9175 | 5.8146 | 6.4196 | 6.4456 | 6.3243 | 6.0270 | 6.1320 | 5.8488 | 6.5913 | 6.7280 | 6.8304 | 6.8714 |
| LMO7_T714 | LMO7 | Q8WWI1-3 | 622;626 | | NA | 6.6957 | 6.2386 | 5.9249 | 6.6776 | 6.2044 | 6.1969 | 6.9577 | 5.7475 | 6.0037 | 6.0682 | 5.6787 | 6.0504 | 6.2205 | 6.7360 | 6.4118 | 6.1875 |
| LMTK2_S1107 | LMTK2 | Q8IWU2 | | 630 | VTSGPESPFNNIF | 6.1628 | 6.3766 | 6.3229 | 6.4247 | 6.1042 | 6.3769 | 6.2246 | 6.5795 | 6.4278 | 6.9067 | 6.7634 | 6.4413 | 5.3236 | 5.9622 | 5.6275 | 5.9753 |
| LMTK2_S1307;S1308;S13 | LMTK2 | Q8IWU2 | | 617 | TDPKDSSLPGDLH | 5.6811 | 5.9004 | 6.1634 | 5.8955 | 6.2738 | 6.2318 | 5.9474 | 6.2283 | 5.8330 | 5.8826 | 5.8123 | 6.1300 | 6.9908 | 7.3300 | 7.0919 | 6.6077 |
| LMTK2_S1397 | LMTK2 | Q8IWU2 | | 1397 | PAPASGSPYLSRC | 5.5736 | 6.0275 | 5.9027 | 5.7226 | 5.3218 | 5.7721 | 6.2159 | 6.0229 | 7.4326 | 7.2891 | 6.0490 | 6.7162 | 6.8330 | 6.2927 | 6.1418 | 6.6864 |
| LMTK2_S617 | LMTK2 | Q8IWU2 | | 1107 | RDSAYFSDNDSEP | 6.0204 | 6.5255 | 6.3289 | 6.7323 | 5.9809 | 6.5822 | 6.7132 | 6.2470 | 5.9630 | 5.7972 | 6.0764 | 6.1126 | 5.9399 | 6.5284 | 5.8070 | 6.6451 |
| LMTK2_S617;S630 | LMTK2 | Q8IWU2 | 617;630 | | NA | 5.7001 | 5.7715 | 5.5722 | 8.5606 | 5.9667 | 6.5423 | 6.7408 | 6.0205 | 6.2138 | 6.9540 | 7.9998 | 6.2799 | 4.9944 | 5.7327 | 4.7161 | 6.2349 |
| LMTK2_S630 | LMTK2 | Q8IWU2 | 1307;1308;13 | | NA | 5.9858 | 6.7603 | 5.4890 | 5.3312 | 6.0386 | 5.7843 | 5.9955 | 6.8914 | 6.4300 | 6.3066 | 6.0637 | 6.4332 | 5.8064 | 7.1580 | 6.7815 | 6.7443 |
| LNPK_S177 | LNPK | Q9C0E8-4 | | 194 | PPQVPVSPGPPKD | 5.4451 | 6.1775 | 5.9055 | 6.3119 | 6.1988 | 6.5533 | 7.2643 | 6.5808 | 6.2391 | 6.4398 | 6.5391 | 6.6954 | 5.6213 | 6.0649 | 6.0466 | 5.9167 |
| LNPK_S177;S182;S194 | LNPK | Q9C0E8-4 | | 415 | QVIEKASDSEEPE | 6.1678 | 6.4506 | 6.1044 | 6.6998 | 5.5718 | 6.0985 | 6.1975 | 5.8490 | 6.1861 | 6.2647 | 5.7523 | 6.2430 | 6.4780 | 6.4097 | 6.5026 | 7.0243 |
| LNPK_S182;S194 | LNPK | Q9C0E8-4 | | 177 | AAQRNLSPTPASP | 6.7653 | 5.9959 | 6.3279 | 6.3400 | 6.3596 | 6.7592 | 5.9662 | 5.8817 | 6.4063 | 7.2169 | 7.4454 | 5.8283 | 5.2834 | 5.8087 | 5.4240 | 6.1913 |
| LNPK_S194 | LNPK | Q9C0E8-4 | 182;194 | | NA | 7.5305 | 7.5510 | 7.2250 | 8.0586 | 7.2830 | 6.7963 | 7.3150 | 7.1091 | 7.0058 | 7.4990 | 7.4147 | 7.4900 | 2.5290 | 3.0879 | 2.2456 | 3.8595 |
| LNPK_S415 | LNPK | Q9C0E8-4 | 177;182;194 | | NA | 7.0190 | 6.9636 | 5.8626 | 7.3901 | 5.9488 | 7.0734 | 7.1112 | 6.6243 | 7.1498 | 8.1191 | 8.1463 | 8.1657 | 3.3872 | 5.0271 | 2.9407 | 3.0711 |
| LONP2_S177 | LONP2 | Q86WA8 | | 177 | LRRLLDSLPREAL | 6.2495 | 6.1269 | 6.8518 | 5.5031 | 6.5739 | 6.3823 | 5.9679 | 6.1738 | 6.2760 | 6.0898 | 5.8407 | 5.6136 | 6.5193 | 6.9679 | 6.7895 | 6.0741 |
| LPGAT1_S233 | LPGAT1 | Q92604 | | 233 | AQQKNGSPAGGDA | 5.4441 | 5.8148 | 6.6866 | 6.5456 | 5.4235 | 5.6688 | 6.5368 | 5.1321 | 6.7303 | 6.8283 | 6.6872 | 6.4039 | 6.3920 | 6.5624 | 6.1874 | 6.9562 |
| LPIN1_S155 | LPIN1 | Q14693-7 | | 211 | RKRRRKSQLDSLK | 5.3485 | 5.0363 | 5.0903 | 4.8650 | 5.7657 | 5.2348 | 4.9877 | 5.6319 | 6.1671 | 6.1589 | 5.9380 | 5.9381 | 8.4301 | 8.7587 | 8.6715 | 7.9774 |
| LPIN1_S211 | LPIN1 | Q14693-7 | | 337 | RPSTPKSDSELVS | 5.8169 | 6.7653 | 6.7304 | 6.2326 | 5.4024 | 5.9754 | 5.7893 | 5.8325 | 5.6672 | 6.6533 | 6.3396 | 6.6469 | 6.1215 | 6.7876 | 6.5260 | 6.7130 |
| LPIN1_S337 | LPIN1 | Q14693-7 | | 379 | HKMKESSPLSSRK | 6.1882 | 6.5179 | 6.8172 | 5.9595 | 5.7246 | 6.1402 | 6.3541 | 6.1691 | 6.0339 | 6.6276 | 6.8904 | 6.9957 | 5.8258 | 6.1673 | 5.7969 | 5.7917 |
| LPIN1_S379 | LPIN1 | Q14693-7 | | 523 | ARSANQSPQSVGS | 5.8345 | 5.4454 | 5.8563 | 5.4846 | 5.3045 | 5.3234 | 7.1763 | 6.1961 | 5.8156 | 5.9507 | 6.1961 | 6.7261 | 6.3195 | 7.7135 | 7.3926 | 7.2648 |
| LPIN1_S523 | LPIN1 | Q14693-7 | | 155 | PMHLATSPILSEG | 5.0677 | 5.8692 | 6.4208 | 6.2915 | 6.4983 | 5.7097 | 5.3445 | 6.9777 | 6.7267 | 6.9045 | 6.4919 | 6.9271 | 6.5360 | 5.3876 | 6.0696 | 6.7773 |
| LPIN2_S106 | LPIN2 | Q92539 | | 106 | PAYLATSPIPTED | 6.3984 | 6.4341 | 7.1548 | 6.2846 | 6.2906 | 6.9222 | 6.5072 | 6.7609 | 6.5538 | 6.7509 | 6.6908 | 6.6754 | 5.1399 | 5.5708 | 4.7979 | 5.0676 |
| LPIN2_S203 | LPIN2 | Q92539 | | 203 | RGSSNASLKEEEC | 6.0834 | 6.0548 | 5.9769 | 5.9237 | 5.0122 | 6.2985 | 5.7701 | 7.0216 | 6.9519 | 5.0265 | 5.5919 | 6.4378 | 6.9458 | 6.3989 | 7.5767 | 6.9294 |
| LPP_S116 | LPP | Q93052 | | 333 | KREPGYTPPGAGN | 6.8084 | 7.2705 | 6.6831 | 7.5295 | 6.8777 | 6.9500 | 6.9054 | 7.5375 | 7.1273 | 7.5883 | 7.9830 | 7.6254 | 3.2062 | 3.6816 | 2.7518 | 3.4743 |
| LPP_T156 | LPP | Q93052 | | 116 | LEERRSSLDAEID | 5.6733 | 5.7655 | 6.3031 | 5.4372 | 6.7301 | 6.1058 | 5.9504 | 7.1182 | 6.6126 | 5.9205 | 6.9068 | 6.4764 | 7.0272 | 6.8714 | 5.6062 | 5.4952 |
| LPP_T333 | LPP | Q93052 | | 156 | ASPPVSTPVTGHK | 7.7722 | 6.4648 | 6.7459 | 6.5964 | 5.7351 | 6.9213 | 6.6350 | 6.3823 | 7.2720 | 6.1116 | 6.5499 | 6.9445 | 4.7809 | 6.0110 | 4.2575 | 4.8196 |
| LRBA_S1081 | LRBA | P50851 | | 1118 | ELKVEGSPTEEAN | 5.8196 | 5.8710 | 6.8545 | 6.2997 | 6.4147 | 6.3065 | 6.2967 | 6.2626 | 6.4995 | 6.5340 | 6.4668 | 6.1502 | 5.9897 | 6.4631 | 6.0176 | 5.7540 |
| LRBA_S1100 | LRBA | P50851 | | 1767 | SDMGGESPGSRSS | 5.5569 | 6.9018 | 6.4082 | 7.2223 | 5.9103 | 6.6428 | 6.2136 | 5.4296 | 6.7115 | 6.6958 | 7.0148 | 7.2308 | 4.5583 | 6.0991 | 5.7300 | 5.6742 |
| LRBA_S1118 | LRBA | P50851 | | 1261 | RLELKASPNVEAP | 5.5975 | 6.9279 | 6.2691 | 6.8601 | 6.9065 | 6.2477 | 7.4068 | 6.4768 | 6.0029 | 5.4991 | 7.2548 | 6.4190 | 5.6040 | 5.5383 | 5.3799 | 5.6097 |
| LRBA_S1261 | LRBA | P50851 | | 1100 | PEFLDKSIVEEEE | 6.1686 | 6.3516 | 7.4025 | 7.5903 | 7.0026 | 6.6977 | 6.8097 | 6.3484 | 6.4640 | 6.1307 | 7.2426 | 6.6538 | 4.6597 | 4.7777 | 4.4603 | 5.2397 |
| LRBA_S1767 | LRBA | P50851 | | 2201 | RRISLASPRQLFK | 6.3338 | 6.9136 | 6.6498 | 6.9754 | 6.1187 | 7.0490 | 6.5732 | 6.8142 | 6.5128 | 6.6925 | 6.4097 | 6.2613 | 4.8645 | 5.6185 | 5.3559 | 4.8571 |
| LRBA_S2201 | LRBA | P50851 | | 1081 | VSEVTASISSPSE | 5.7604 | 6.3390 | 5.6400 | 6.3029 | 5.6881 | 6.3089 | 5.5979 | 6.3856 | 6.4910 | 6.3435 | 6.0794 | 6.4150 | 6.8446 | 7.1847 | 6.2600 | 6.3593 |
| LRBA_T1079 | LRBA | P50851 | | 1079 | MTVSEVTASISSP | 5.2899 | 6.4802 | 6.1682 | 6.8145 | 6.1552 | 6.4284 | 5.1969 | 6.8051 | 6.5991 | 6.7156 | 5.1333 | 6.3448 | 6.1340 | 7.2231 | 6.3719 | 6.1399 |
| LRCH1_S335 | LRCH1 | Q9Y2L9-3 | | 335 | NGDKRLSATEPSD | 6.0262 | 5.7214 | 6.4319 | 6.1997 | 6.5580 | 6.0214 | 6.1593 | 6.6315 | 5.7214 | 5.6658 | 6.4289 | 5.9419 | 6.8133 | 6.7441 | 6.9968 | 5.9384 |
| LRCH1_S532;S536 | LRCH1 | Q9Y2L9-3 | | 536 | ENSPAVSPTTNST | 5.8843 | 6.0363 | 6.1181 | 6.1367 | 6.0903 | 6.0014 | 5.5156 | 5.9450 | 6.9218 | 5.9632 | 5.9737 | 5.6600 | 6.7516 | 6.9753 | 6.7616 | 7.2651 |
| LRCH1_S536 | LRCH1 | Q9Y2L9-3 | 532;536 | | NA | 5.8943 | 5.9095 | 6.6859 | 6.2323 | 6.0428 | 5.7520 | 6.1561 | 6.5295 | 6.4356 | 6.2854 | 6.1259 | 6.3491 | 6.4431 | 6.4851 | 6.2545 | 6.4189 |
| LRCH2_S521 | LRCH2 | Q5VUJ6 | | 521 | EVNSPLSPLTWQP | 5.4636 | 5.8256 | 5.9348 | 6.4692 | 6.2549 | 6.0478 | 6.3631 | 7.2601 | 6.4777 | 6.2858 | 6.5800 | 6.4887 | 6.0686 | 6.2310 | 6.0067 | 6.2423 |
| LRCH3_S324 | LRCH3 | Q96II8-2 | | 419 | RISHEGSPVKPVA | 7.0142 | 6.6410 | 6.6111 | 6.2455 | 6.6965 | 7.1008 | 6.4813 | 7.2349 | 7.3816 | 6.8317 | 6.5962 | 6.7399 | 4.6219 | 4.8212 | 4.2533 | 4.7290 |
| LRCH3_S419 | LRCH3 | Q96II8-2 | | 516 | KQKASQSPQKQHP | 6.7222 | 6.6506 | 6.9027 | 6.4533 | 5.9619 | 6.6963 | 6.1709 | 6.8133 | 6.9559 | 6.8159 | 7.5057 | 7.8027 | 4.4049 | 4.9737 | 4.3847 | 4.7853 |
| LRCH3_S516 | LRCH3 | Q96II8-2 | | 324 | SGDKRWSGNEPTD | 5.7087 | 5.9279 | 6.1191 | 5.6702 | 6.0840 | 6.6760 | 5.7998 | 5.8125 | 6.0681 | 6.4459 | 5.9487 | 5.9746 | 7.2655 | 7.1059 | 6.6558 | 6.7373 |
| LRCH4_S281 | LRCH4 | O75427 | | 281 | SRPPSFSPCPAED | 6.3440 | 6.5998 | 6.2472 | 6.5388 | 5.9697 | 6.5336 | 7.1705 | 6.3801 | 6.6288 | 6.2830 | 6.6663 | 6.9508 | 5.2930 | 5.2349 | 5.3668 | 5.7926 |
| LRCH4_S313 | LRCH4 | O75427 | | 506 | SIQRPNSFLFRSS | 4.1416 | 3.5153 | 3.6837 | 3.8380 | 4.6315 | 4.0175 | 4.1475 | 3.8639 | 4.1962 | 4.0665 | 4.5079 | 4.4270 | 12.7067 | 11.7617 | 12.7769 | 13.7179 |
| LRCH4_S506 | LRCH4 | O75427 | | 589 | PFIHVPSPAVPKL | 7.2791 | 7.8006 | 7.0609 | 7.1579 | 6.8337 | 6.9663 | 6.5394 | 6.8806 | 6.8765 | 6.9582 | 6.3729 | 6.2961 | 3.9460 | 4.7617 | 3.5514 | 4.7188 |
| LRCH4_S521 | LRCH4 | O75427 | | 313 | SGSKRWSGNESTD | 6.2165 | 6.3864 | 6.1975 | 5.7538 | 6.8591 | 5.5470 | 6.4679 | 6.5361 | 6.3048 | 6.4154 | 6.4869 | 6.4378 | 5.7236 | 5.9963 | 6.7641 | 5.9069 |
| LRCH4_S589 | LRCH4 | O75427 | | 521 | SGSGPSSPDSVLR | 6.6628 | 6.7818 | 6.0550 | 7.0736 | 7.0051 | 6.5954 | 6.4100 | 7.2969 | 6.1419 | 6.6322 | 6.7423 | 6.9551 | 4.5496 | 4.9652 | 5.0563 | 5.0769 |
| LRFN1_S674 | LRFN1 | Q9P244 | | 674 | GPRRSRSGALEPP | 6.2555 | 6.4354 | 6.4825 | 6.6245 | 5.8408 | 5.8821 | 5.8854 | 6.5933 | 6.5837 | 6.2655 | 6.1100 | 6.3083 | 7.1093 | 6.4508 | 5.3772 | 5.7957 |
| LRIF1_T732 | LRIF1 | Q5T3J3 | | 732 | EDIFPVTPPELEE | 6.1555 | 6.7538 | 6.4028 | 7.3243 | 5.6352 | 6.2077 | 6.1230 | 6.7335 | 6.9286 | 5.9563 | 5.9062 | 6.7206 | 6.2280 | 6.2456 | 5.4692 | 5.2097 |
| LRP1_S4520 | LRP1 | Q07954 | | 4520 | GHGSRHSLASTDE | 6.4094 | 5.6101 | 5.6654 | 5.7414 | 7.0441 | 5.8480 | 5.8530 | 5.7987 | 7.2777 | 7.6296 | 6.7921 | 6.3122 | 6.3605 | 6.9368 | 5.3707 | 5.3503 |
| LRP6_S1490 | LRP6 | O75581 | | 1490 | ILNPPPSPATERS | 6.3011 | 7.5121 | 5.8179 | 5.5643 | 5.7239 | 6.3861 | 6.5293 | 5.9536 | 5.9670 | 6.8827 | 5.2616 | 5.7806 | 6.0371 | 7.5926 | 5.9103 | 6.7798 |
| LRRC1_S37 | LRRC1 | Q9BTT6 | | 456 | RAVNRVSAIRFVE | 8.1699 | 7.5813 | 6.9167 | 6.8567 | 6.5760 | 6.3870 | 6.4605 | 6.5028 | 6.9618 | 5.8190 | 5.8862 | 6.3938 | 5.1705 | 5.1206 | 4.3380 | 4.8592 |
| LRRC1_S456 | LRRC1 | Q9BTT6 | | 37 | IYRYARSLEELLL | 6.1520 | 6.5879 | 6.0686 | 6.0184 | 6.2134 | 6.1008 | 6.2341 | 5.8800 | 6.2928 | 6.0540 | 6.5153 | 5.8128 | 6.3610 | 6.7578 | 6.6221 | 6.3291 |
| LRRC1_T480 | LRRC1 | Q9BTT6 | | 480 | TLLRRATPHPGEL | 7.3370 | 6.6360 | 6.9024 | 6.0490 | 5.7604 | 6.1459 | 5.7047 | 6.1698 | 6.9732 | 7.8316 | 8.2243 | 9.0912 | 4.5565 | 5.6117 | 3.2032 | 3.8031 |
| LRRC41_S357 | LRRC41 | Q15345 | | 357 | APGTKRSPSAPAA | 5.6349 | 6.1466 | 6.2026 | 5.9126 | 5.1442 | 6.5321 | 7.2479 | 6.6346 | 6.1168 | 6.3043 | 6.6863 | 6.5960 | 5.5385 | 7.0806 | 6.0730 | 6.1492 |
| LRRC47_S195 | LRRC47 | Q8N1G4 | | 520 | KEEGSLSDTEADA | 5.9477 | 6.1470 | 6.6254 | 6.2061 | 6.2343 | 6.1764 | 6.1043 | 6.3100 | 6.3534 | 6.4543 | 6.4928 | 6.5926 | 6.0448 | 6.3425 | 5.9600 | 6.0084 |
| LRRC47_S315 | LRRC47 | Q8N1G4 | | 431 | KQKKRQSVSGLHR | 7.3108 | 8.3785 | 7.1721 | 8.1368 | 6.3747 | 7.1884 | 7.2577 | 7.0545 | 7.2072 | 7.0857 | 7.6939 | 7.5306 | 2.4043 | 3.5841 | 2.4382 | 3.1822 |
| LRRC47_S431 | LRRC47 | Q8N1G4 | | 561 | VVDLEGSLKVVYP | 6.4955 | 7.2042 | 6.4450 | 6.8523 | 5.9792 | 6.3558 | 6.1933 | 6.3409 | 5.8444 | 6.3350 | 6.1110 | 6.1596 | 6.2815 | 6.5104 | 5.6759 | 5.2161 |
| LRRC47_S520 | LRRC47 | Q8N1G4 | | 195 | NCLRELSPDIAHL | 7.0503 | 6.4340 | 7.3401 | 6.2142 | 6.7788 | 6.0227 | 5.8344 | 7.2252 | 6.2675 | 5.9211 | 6.5273 | 7.8131 | 4.8901 | 5.5580 | 5.1666 | 4.9567 |
| LRRC47_S561 | LRRC47 | Q8N1G4 | | 315 | LRVLHVSENPVPL | 6.1126 | 6.8675 | 7.1849 | 5.9539 | 5.3577 | 6.6952 | 6.1339 | 7.1771 | 8.0578 | 6.5868 | 6.3158 | 4.7790 | 5.4254 | 5.8493 | 6.0235 | 5.4796 |
| LRRC59_S23 | LRRC59 | Q96AG4 | | 23 | GNELDLSLSDLNE | 6.6251 | 6.4472 | 6.3363 | 7.3425 | 6.7458 | 6.7693 | 7.5148 | 6.6042 | 6.9072 | 7.0175 | 6.8002 | 6.9951 | 4.0514 | 4.9161 | 4.0560 | 4.8713 |
| LRRC8A_S193 | LRRC8A | Q8IWT6 | | 217 | MLQRTKSRIEQGI | 6.8603 | 6.9110 | 6.9453 | 6.6992 | 6.8002 | 6.4380 | 6.3235 | 6.7223 | 6.5332 | 6.4486 | 6.3043 | 6.7760 | 4.7267 | 5.9079 | 4.9267 | 4.6768 |
| LRRC8A_S199 | LRRC8A | Q8IWT6 | | 199 | SMDKKSSTVSEDV | 5.7192 | 4.1520 | 5.2563 | 5.1999 | 7.0466 | 5.1680 | 5.2610 | 5.5495 | 6.7204 | 6.9899 | 6.5112 | 6.3408 | 8.2413 | 8.3724 | 7.1735 | 6.2980 |
| LRRC8A_S199;S202 | LRRC8A | Q8IWT6 | | 193 | FSKMNGSMDKKSS | 6.3162 | 6.3310 | 6.3887 | 6.3656 | 5.8037 | 5.7579 | 5.7116 | 5.5001 | 6.6232 | 6.9210 | 6.6061 | 6.7943 | 6.3531 | 6.6891 | 6.3318 | 5.5066 |
| LRRC8A_S217 | LRRC8A | Q8IWT6 | 199;202 | | NA | 5.1767 | 5.4003 | 5.2285 | 5.3608 | 5.4583 | 5.3488 | 5.3589 | 4.7181 | 6.5261 | 6.9455 | 6.7851 | 7.2075 | 7.7635 | 7.7553 | 7.7776 | 7.1891 |
| LRRFIP1_S120 | LRRFIP1 | Q32MZ4 | | 120 | TSSRRGSGDTSIS | 6.2246 | 6.2170 | 6.7007 | 5.9797 | 6.5743 | 6.3912 | 5.8053 | 6.3954 | 6.5766 | 6.7011 | 6.8463 | 6.9824 | 5.8964 | 6.0635 | 5.2310 | 5.4144 |
| LRRFIP1_S16 | LRRFIP1 | Q32MZ4 | | 521 | KTEVPGSPAGTEG | 6.0913 | 5.8051 | 7.0963 | 6.4049 | 6.3750 | 6.6311 | 6.1778 | 6.3075 | 6.7213 | 6.5989 | 7.3260 | 7.0715 | 5.8588 | 5.3622 | 4.9533 | 5.2190 |
| LRRFIP1_S467 | LRRFIP1 | Q32MZ4 | | 16 | REIDCLSPEAQKL | 6.7790 | 7.0524 | 7.1780 | 7.4564 | 7.0264 | 7.3507 | 6.9618 | 7.0321 | 7.5173 | 7.3401 | 8.0013 | 7.1093 | 3.2613 | 3.7082 | 2.6482 | 3.5774 |
| LRRFIP1_S521 | LRRFIP1 | Q32MZ4 | | 733 | TLPEHESPSQDIS | 6.1925 | 6.7549 | 7.0396 | 7.1059 | 5.8718 | 6.6852 | 6.0023 | 5.7274 | 6.7765 | 7.1930 | 7.1795 | 6.6921 | 5.3313 | 5.4253 | 4.7546 | 5.2681 |
| LRRFIP1_S565 | LRRFIP1 | Q32MZ4 | | 714 | DGATQSSPAEPKS | 6.3954 | 6.6914 | 6.8210 | 6.6089 | 6.3768 | 6.5897 | 6.3815 | 6.5719 | 6.8211 | 7.0075 | 7.1194 | 7.2741 | 4.7895 | 5.1290 | 4.4819 | 4.9409 |
| LRRFIP1_S714 | LRRFIP1 | Q32MZ4 | | 565 | GEALDSSQKKTKN | 6.1902 | 6.3025 | 7.2880 | 7.9635 | 6.3119 | 7.2441 | 6.9879 | 6.7227 | 7.2824 | 7.7950 | 8.2040 | 7.9363 | 3.5369 | 3.8298 | 2.8180 | 3.5867 |
| LRRFIP1_S733 | LRRFIP1 | Q32MZ4 | | 467 | PQELETSTGHSLE | 6.2042 | 6.9149 | 6.9755 | 6.6702 | 6.8931 | 6.5987 | 6.3846 | 6.7397 | 6.4945 | 6.4043 | 7.8319 | 7.2010 | 4.1246 | 4.9923 | 4.1614 | 5.4091 |
| LRRFIP2_S309 | LRRFIP2 | Q9Y608 | | 328 | NSSRRGSGDTSSL | 6.0248 | 6.3127 | 6.3122 | 6.0749 | 6.2990 | 6.2334 | 5.7110 | 6.0238 | 6.1300 | 6.5153 | 6.2603 | 6.4177 | 6.7798 | 6.4675 | 6.4205 | 6.0171 |
| LRRFIP2_S320;S324 | LRRFIP2 | Q9Y608 | | 309 | NYTRPSSRNSASA | 6.5544 | 6.3954 | 6.9414 | 6.2653 | 6.7764 | 6.7626 | 6.6340 | 6.7364 | 7.1855 | 6.7256 | 6.9577 | 7.1573 | 4.6964 | 4.8795 | 4.4697 | 4.8623 |
| LRRFIP2_S324 | LRRFIP2 | Q9Y608 | | 324 | PLSGNSSRRGSGD | 6.0870 | 6.3870 | 7.3568 | 6.0110 | 5.8341 | 6.3312 | 5.9109 | 6.7280 | 6.6328 | 6.1559 | 6.2511 | 6.4477 | 6.1896 | 6.2552 | 5.1061 | 6.3156 |
| LRRFIP2_S328 | LRRFIP2 | Q9Y608 | 320;324 | | NA | 6.7769 | 6.2251 | 6.0946 | 6.8135 | 6.1925 | 6.0676 | 6.7460 | 5.1184 | 6.3319 | 7.0074 | 7.3240 | 6.4376 | 4.6713 | 5.7535 | 5.9614 | 6.4782 |
| LRRK2_S973 | LRRK2 | Q5S007 | | 973 | HMRHSDSISSLAS | 5.5761 | 6.3580 | 5.7866 | 6.1595 | 5.6763 | 7.2844 | 6.1776 | 7.7997 | 5.8888 | 5.9650 | 5.4457 | 6.2558 | 6.7671 | 6.5164 | 5.5460 | 6.7971 |
| LRWD1_S212 | LRWD1 | Q9UFC0 | | 259 | SAQVEGSPVAGSD | 5.9558 | 6.4623 | 6.3649 | 7.1760 | 6.4530 | 6.2466 | 6.4847 | 6.2396 | 6.6474 | 6.4234 | 6.7593 | 6.6698 | 5.2805 | 5.8753 | 5.2496 | 5.7119 |
| LRWD1_S243 | LRWD1 | Q9UFC0 | | 251 | SKRACASPSAQVE | 5.2581 | 5.9414 | 5.8572 | 5.8983 | 6.0359 | 6.0258 | 6.2488 | 6.1412 | 5.7564 | 5.8776 | 6.0376 | 6.2896 | 6.9006 | 7.2204 | 7.4167 | 7.0942 |
| LRWD1_S251 | LRWD1 | Q9UFC0 | | 243 | DVPLSLSPSKRAC | 6.0595 | 6.1683 | 6.2957 | 6.2656 | 6.2098 | 6.0398 | 6.1356 | 6.1608 | 6.9047 | 6.4230 | 6.4262 | 6.3766 | 5.9535 | 6.2990 | 6.0882 | 6.1936 |
| LRWD1_S251;S259 | LRWD1 | Q9UFC0 | | 212 | QVQKANSPEKPPE | 5.7218 | 6.6741 | 6.7789 | 6.2328 | 6.1020 | 6.3932 | 6.2186 | 6.1182 | 6.4853 | 6.3438 | 6.4645 | 5.9868 | 6.1971 | 6.5084 | 5.6044 | 6.1701 |
| LRWD1_S259 | LRWD1 | Q9UFC0 | 251;259 | | NA | 5.3572 | 6.2795 | 6.1544 | 6.1342 | 6.5002 | 6.4415 | 6.0435 | 6.7670 | 7.5261 | 6.6470 | 5.9692 | 6.3538 | 5.2970 | 6.0620 | 6.0464 | 6.4211 |
| LSG1_S252 | LSG1 | Q9H089 | | 252 | IPLNGDSEEEANR | 7.5611 | 6.9166 | 6.0771 | 7.3184 | 6.7354 | 5.5428 | 6.3431 | 6.0008 | 6.2106 | 6.0983 | 5.8407 | 6.6495 | 5.8004 | 5.6965 | 5.5216 | 5.6870 |
| LSM11_S15;S21 | LSM11 | P83369 | | 21 | SPARPPSPRLDVS | 6.3851 | 6.3839 | 6.7489 | 7.0809 | 6.3024 | 5.7996 | 6.0212 | 6.0840 | 6.1938 | 6.5380 | 6.1289 | 6.5595 | 5.9159 | 5.9345 | 5.9695 | 5.9539 |
| LSM11_S21 | LSM11 | P83369 | 15;21 | | NA | 6.4311 | 6.6875 | 6.2720 | 6.9903 | 6.1158 | 5.5345 | 5.7818 | 5.8383 | 6.4322 | 6.1886 | 5.9455 | 6.3082 | 5.9303 | 6.1723 | 6.9142 | 6.4572 |
| LSM12_T78 | LSM12 | Q3MHD2-2 | | 78 | INDRTETPPPLAS | 5.8072 | 5.2686 | 6.4775 | 5.5189 | 5.9243 | 5.9969 | 5.7078 | 6.1882 | 6.7353 | 6.9795 | 6.7664 | 7.1509 | 6.3422 | 6.4513 | 6.2457 | 6.4394 |
| LSM14A_S178 | LSM14A | Q8ND56-2 | | 192 | LDPLRKSPTMEQA | 6.1090 | 6.3498 | 6.4073 | 6.4521 | 6.3518 | 6.4164 | 5.9345 | 6.5917 | 6.4604 | 6.3769 | 6.6915 | 6.3641 | 5.8683 | 6.0722 | 5.7540 | 5.8000 |
| LSM14A_S178;S182 | LSM14A | Q8ND56-2 | | 183 | LSQGRSSPQLDPL | 6.4359 | 6.3264 | 6.3201 | 5.9069 | 6.3552 | 6.2213 | 5.9607 | 6.7234 | 6.9193 | 6.5880 | 6.5978 | 6.5366 | 5.7837 | 5.9995 | 5.6952 | 5.6301 |
| LSM14A_S178;S183 | LSM14A | Q8ND56-2 | | 216 | AAVGRRSPVSTRP | 6.2682 | 6.1569 | 6.3522 | 5.7847 | 6.5767 | 6.4318 | 5.9777 | 6.6659 | 6.4806 | 6.5445 | 6.6230 | 6.2548 | 6.0935 | 6.2468 | 5.8424 | 5.7002 |
| LSM14A_S182 | LSM14A | Q8ND56-2 | | 178 | SLKTQLSQGRSSP | 4.4910 | 4.4958 | 5.1366 | 4.3428 | 5.4630 | 5.2756 | 4.8869 | 5.9838 | 4.9766 | 5.1313 | 4.8566 | 4.9574 | 10.2005 | 9.8497 | 10.3115 | 9.6408 |
| LSM14A_S183 | LSM14A | Q8ND56-2 | | 182 | QLSQGRSSPQLDP | 6.2869 | 6.3618 | 6.4771 | 5.8714 | 6.4668 | 6.2135 | 6.0091 | 6.6224 | 6.7862 | 6.6452 | 6.7998 | 6.6976 | 5.6712 | 6.0471 | 5.5426 | 5.5011 |
| LSM14A_S192 | LSM14A | Q8ND56-2 | | 347 | GVDTQNSEGNADE | 5.6829 | 5.8758 | 6.4102 | 6.0940 | 6.5163 | 6.3555 | 6.0777 | 6.0348 | 6.3555 | 5.5799 | 5.9282 | 6.7986 | 6.4553 | 6.8357 | 6.3424 | 6.6572 |
| LSM14A_S216 | LSM14A | Q8ND56-2 | 178;183 | | NA | 5.1136 | 5.1646 | 5.9941 | 5.6499 | 5.7070 | 5.7997 | 6.1799 | 6.0149 | 5.6966 | 5.6927 | 5.7098 | 5.4330 | 8.8772 | 8.2029 | 7.6340 | 7.1300 |
| LSM14A_S347 | LSM14A | Q8ND56-2 | 178;182 | | NA | 4.5746 | 4.5385 | 5.9207 | 4.9390 | 6.0873 | 6.4310 | 5.1092 | 5.0607 | 4.7779 | 5.2389 | 5.3549 | 5.5143 | 10.3317 | 9.8071 | 8.9033 | 7.4109 |
| LSM14B_S154 | LSM14B | Q9BX40 | | 154 | SIPVGKSPMVEQA | 5.9058 | 6.9480 | 6.7442 | 6.6496 | 6.3392 | 6.8020 | 7.2446 | 6.4219 | 6.4907 | 5.2962 | 6.7331 | 6.8562 | 5.4330 | 5.7488 | 5.2888 | 5.0978 |
| LSR_S332 | LSR | Q86X29 | | 336 | HLSPAKTPPPPAM | 5.9416 | 6.5514 | 5.8821 | 6.3231 | 6.2737 | 6.6011 | 6.2990 | 6.9001 | 6.2880 | 6.2098 | 6.4871 | 6.4050 | 5.9849 | 6.2084 | 5.7480 | 5.8967 |
| LSR_S432 | LSR | Q86X29 | | 464 | EEWGGHSPRSPRG | 5.9935 | 6.1683 | 5.3432 | 6.0433 | 5.6908 | 6.1088 | 6.2840 | 6.3742 | 5.8822 | 6.1358 | 5.8657 | 5.8971 | 6.8810 | 7.0199 | 7.0501 | 7.2622 |
| LSR_S436 | LSR | Q86X29 | | 332 | STYAHLSPAKTPP | 5.7292 | 6.2485 | 6.1516 | 6.2895 | 6.2043 | 6.2986 | 5.9289 | 7.0495 | 6.2674 | 6.3440 | 6.5579 | 6.3741 | 5.8826 | 6.4532 | 5.9085 | 6.3122 |
| LSR_S464 | LSR | Q86X29 | | 493 | RRPRARSVDALDD | 4.3524 | 7.2872 | 6.5123 | 7.1669 | 4.9950 | 5.9109 | 7.6068 | 6.3473 | 5.8645 | 6.3937 | 9.1412 | 7.7305 | 4.4589 | 4.7957 | 4.8369 | 6.5999 |
| LSR_S464;S467 | LSR | Q86X29 | | 512 | AESGSRSPTSNGG | 6.1042 | 6.5346 | 6.7795 | 6.4843 | 5.7942 | 6.2266 | 5.8554 | 5.6214 | 6.1111 | 6.9295 | 5.8870 | 6.2661 | 5.6984 | 7.0905 | 6.4576 | 6.1595 |
| LSR_S493 | LSR | Q86X29 | | 436 | AMSEVTSLHEDDW | 6.9508 | 6.4612 | 5.9387 | 6.8618 | 5.7312 | 6.0098 | 5.7147 | 6.6423 | 6.7773 | 6.0157 | 6.8632 | 6.8501 | 5.7061 | 6.1342 | 5.3300 | 6.0129 |
| LSR_S512 | LSR | Q86X29 | | 453 | SRGPALTPIRDEE | 6.7141 | 8.2107 | 5.5560 | 8.6082 | 5.3052 | 5.0239 | 7.0995 | 7.2218 | 6.8589 | 7.6826 | 6.1860 | 6.0555 | 4.3836 | 5.0422 | 4.7241 | 5.3277 |
| LSR_S646 | LSR | Q86X29 | | 432 | RVERAMSEVTSLH | 8.9292 | 7.7190 | 6.4614 | 6.3104 | 6.5379 | 6.3038 | 5.8692 | 5.7268 | 7.2357 | 6.5834 | 6.6184 | 6.7153 | 4.4745 | 6.4384 | 4.4692 | 3.6073 |
| LSR_T336 | LSR | Q86X29 | | 501 | DALDDLTPPSTAE | 6.0087 | 6.0120 | 6.0518 | 6.4508 | 6.2642 | 6.1099 | 6.6307 | 7.0612 | 6.5909 | 5.4912 | 6.7170 | 6.3695 | 6.3380 | 6.5984 | 5.9548 | 5.3510 |
| LSR_T453 | LSR | Q86X29 | | 646 | LALSRESLVVxxx | 8.1591 | 7.2827 | 6.5809 | 7.0391 | 4.9590 | 6.0622 | 6.9177 | 5.2832 | 6.5592 | 5.2288 | 5.4358 | 6.2312 | 6.2110 | 6.5718 | 5.4267 | 6.0516 |
| LSR_T501 | LSR | Q86X29 | 464;467 | | NA | 5.7241 | 6.4864 | 5.9806 | 6.2984 | 5.9244 | 7.3071 | 7.4424 | 6.4868 | 6.0357 | 6.5299 | 6.4148 | 6.1807 | 5.2618 | 5.9867 | 5.9470 | 5.9931 |
| LTBP2_S250 | LTBP2 | Q14767 | | 250 | PNLRRSSAAGEGT | 5.4948 | 4.8527 | 4.7187 | 5.7384 | 7.1252 | 5.4762 | 5.5275 | 5.0427 | 7.9841 | 7.6030 | 7.1124 | 8.3780 | 7.1462 | 8.2417 | 4.6091 | 4.9494 |
| LTV1_S17 | LTV1 | Q96GA3 | | 17 | EKKKAVSFHLVHR | 10.3914 | 7.6666 | 10.2823 | 8.9343 | 4.8476 | 5.6685 | 5.9226 | 7.2180 | 8.1902 | 6.7013 | 7.3427 | 8.6802 | 2.0665 | 2.9334 | 1.4351 | 1.7193 |
| LUC7L2_S18 | LUC7L2 | Q9Y383 | | 18 | DQLMGTSRDGDTT | 4.8654 | 5.7488 | 7.2041 | 6.1699 | 5.2033 | 6.2325 | 5.8627 | 6.8751 | 5.9521 | 7.4170 | 7.5468 | 6.6019 | 5.0753 | 7.0540 | 6.6411 | 5.5500 |
| LUC7L2_S347 | LUC7L2 | Q9Y383 | | 347 | RDQDLASCDRDRS | 5.8172 | 5.7149 | 6.4398 | 6.5898 | 6.1718 | 6.2431 | 5.3022 | 6.4883 | 6.9164 | 6.8342 | 6.2273 | 6.4416 | 6.1749 | 6.7585 | 6.0752 | 5.8050 |
| LUC7L2_S370 | LUC7L2 | Q9Y383 | | 370 | RKDKKRSYESANG | 6.8321 | 6.8819 | 5.6610 | 7.2373 | 6.3343 | 6.8349 | 7.3534 | 6.2002 | 6.8722 | 6.6883 | 6.9317 | 6.8045 | 4.3709 | 5.5254 | 4.4718 | 5.0000 |
| LUC7L3_S425;S431 | LUC7L3 | O95232 | | 238 | EKLRKRTEEPDRD | 5.9327 | 6.3401 | 6.7178 | 6.8801 | 5.9956 | 6.3831 | 5.8929 | 6.3144 | 6.4959 | 6.7694 | 6.4655 | 6.7781 | 5.5990 | 6.2586 | 5.5758 | 5.6011 |
| LUC7L3_T238 | LUC7L3 | O95232 | 425;431 | | NA | 5.1214 | 5.9400 | 6.1401 | 6.4158 | 5.2200 | 6.7318 | 6.5593 | 5.7949 | 6.0683 | 5.6969 | 5.7789 | 5.3955 | 7.1036 | 7.4033 | 7.4431 | 7.1871 |
| LUZP1_S1023 | LUZP1 | Q86V48 | | 394 | HTARELSPQHKRE | 6.0354 | 6.3153 | 5.8936 | 6.3059 | 6.0477 | 5.8572 | 6.1696 | 6.1076 | 6.3216 | 6.1284 | 5.8955 | 6.0066 | 6.6638 | 6.8211 | 6.7747 | 6.6561 |

| | | | | | | | | | | | | | | | | | | | | | |
|---|---|---|---|---|---|---|---|---|---|---|---|---|---|---|---|---|---|---|---|---|---|
| LUZP1_S394 | LUZP1 | Q86V48 | | 659 | GREKPDSDDDLDI | 6.5765 | 6.7230 | 6.7344 | 7.1276 | 5.8799 | 6.3023 | 6.3363 | 6.0154 | 6.2059 | 6.5246 | 6.3766 | 6.7419 | 5.6067 | 5.7609 | 5.4526 | 5.6354 |
| LUZP1_S423 | LUZP1 | Q86V48 | | 423 | SNRQVSSPSFTNR | 7.3010 | 7.5667 | 6.5728 | 6.9639 | 5.9023 | 6.9625 | 7.5041 | 6.7436 | 7.8073 | 7.1336 | 7.1774 | 7.2627 | 3.9845 | 4.1698 | 3.1463 | 3.8016 |
| LUZP1_S570 | LUZP1 | Q86V48 | | 745 | KSQRPFSPREALR | 7.1599 | 7.2765 | 6.5439 | 6.7453 | 5.3966 | 6.7667 | 6.8074 | 7.0119 | 7.0209 | 6.5335 | 6.4648 | 6.6316 | 4.2413 | 5.0241 | 4.6775 | 4.6981 |
| LUZP1_S574 | LUZP1 | Q86V48 | | 995 | SSRRTQSSLTVSE | 7.2986 | 6.7356 | 7.1500 | 7.2307 | 6.1262 | 6.4610 | 6.6086 | 5.9525 | 7.2883 | 6.7358 | 6.6193 | 7.0677 | 4.7507 | 5.0481 | 4.3455 | 4.5815 |
| LUZP1_S639 | LUZP1 | Q86V48 | | 1023 | AWNHSASMEEEGE | 7.0578 | 6.3876 | 8.4094 | 7.4703 | 6.9476 | 5.8815 | 5.4547 | 6.2302 | 6.7068 | 7.4081 | 6.0572 | 6.0140 | 4.4899 | 6.1802 | 4.8016 | 4.5029 |
| LUZP1_S659 | LUZP1 | Q86V48 | | 679 | LVNTTITPEPEPK | 6.7054 | 7.0913 | 6.9404 | 7.5117 | 6.2020 | 6.5522 | 7.1336 | 6.9225 | 6.9111 | 6.9933 | 6.8363 | 7.2932 | 3.7122 | 4.5819 | 3.8186 | 4.7943 |
| LUZP1_S703 | LUZP1 | Q86V48 | | 639 | VVMEDSSPHEALR | 6.4431 | 6.7400 | 6.5970 | 7.0004 | 6.4715 | 6.7211 | 6.3898 | 6.5938 | 7.1087 | 6.9828 | 6.2734 | 6.4914 | 4.8479 | 5.6060 | 4.6331 | 5.1001 |
| LUZP1_S745 | LUZP1 | Q86V48 | | 574 | ASSRRSSSEGLSK | 5.5347 | 5.6250 | 5.6230 | 6.0055 | 6.1184 | 5.3953 | 5.3940 | 5.7980 | 6.6781 | 7.1705 | 6.6654 | 6.8512 | 6.3296 | 7.3899 | 7.0520 | 6.3693 |
| LUZP1_S805 | LUZP1 | Q86V48 | | 703 | RGAPRTSLFENDK | 7.0017 | 6.8871 | 6.8943 | 7.1757 | 5.9244 | 6.4142 | 7.1283 | 6.1116 | 7.2718 | 7.4344 | 7.5653 | 7.6874 | 4.0730 | 4.8057 | 3.4261 | 4.1989 |
| LUZP1_S957 | LUZP1 | Q86V48 | | 570 | IGALASSRRSSSE | 6.8964 | 6.4540 | 6.0660 | 6.2399 | 4.7582 | 5.7733 | 6.5222 | 5.6990 | 7.4218 | 7.1348 | 7.5025 | 7.4908 | 5.2219 | 6.0376 | 5.1156 | 5.6659 |
| LUZP1_S995 | LUZP1 | Q86V48 | | 958 | YTQRSSTDFSELE | 7.1112 | 6.8950 | 7.2600 | 7.6716 | 6.5954 | 6.5635 | 6.8436 | 6.5286 | 6.8954 | 5.9999 | 6.6874 | 6.1022 | 4.6083 | 5.0289 | 4.6000 | 4.6090 |
| LUZP1_T679 | LUZP1 | Q86V48 | | 805 | YPSDSSSPRAAPG | 6.9929 | 6.7174 | 6.8282 | 8.1277 | 6.4180 | 6.6904 | 7.1031 | 6.1560 | 7.7279 | 6.7439 | 6.7211 | 7.0278 | 4.0103 | 4.7177 | 3.7722 | 4.2452 |
| LUZP1_T958 | LUZP1 | Q86V48 | | 957 | AYTQRSSTDFSEL | 6.2761 | 7.3777 | 6.6660 | 7.0971 | 6.2046 | 6.1270 | 6.5705 | 6.5209 | 6.6575 | 6.9877 | 7.0618 | 7.2261 | 4.4316 | 5.0460 | 4.5930 | 5.1565 |
| LYAR_S276 | LYAR | Q9NX58 | | 276 | KRKRRHSEVETDS | 4.9442 | 5.4555 | 5.8426 | 6.1239 | 7.3874 | 5.4242 | 6.0238 | 6.6521 | 7.2376 | 7.9138 | 7.4988 | 7.8645 | 4.7993 | 6.1560 | 4.7637 | 5.9125 |
| LYN_S13 | LYN | P07948 | | 508 | TATEGQYQQQPxx | 5.8937 | 6.8393 | 6.2825 | 6.4806 | 6.5056 | 6.6456 | 6.1447 | 6.3650 | 6.2144 | 6.2614 | 6.0891 | 6.1185 | 5.8476 | 6.1661 | 5.9707 | 6.1752 |
| LYN_Y508 | LYN | P07948 | | 13 | KGKDSLSDDGVDL | 6.6573 | 7.0629 | 6.6478 | 6.6094 | 5.9824 | 6.0736 | 6.2612 | 6.2378 | 6.5930 | 6.3470 | 6.6267 | 6.4180 | 5.4702 | 6.8507 | 5.5265 | 5.6355 |
| LYSMD2_S24 | LYSMD2 | Q8IV50 | | 24 | PRPSAPSPPPRSR | 5.8993 | 5.1789 | 6.3984 | 5.7934 | 6.1055 | 5.9477 | 5.6486 | 6.0062 | 6.4589 | 7.0217 | 7.1900 | 7.0499 | 6.6731 | 6.6092 | 5.6623 | 6.3567 |
| LYSMD3_S134 | LYSMD3 | Q7Z3D4 | | 134 | QTSRHSSVQYSSE | 6.0653 | 4.8703 | 4.5185 | 4.7026 | 7.4280 | 4.8785 | 5.5083 | 4.8046 | 7.6977 | 7.9581 | 9.2398 | 7.2838 | 6.8029 | 7.5347 | 5.5270 | 5.1799 |
| LYSMD3_S53;S55 | LYSMD3 | Q7Z3D4 | 53;55 | | NA | 5.9743 | 6.7391 | 5.8685 | 6.5540 | 5.8296 | 6.2652 | 6.2954 | 5.3738 | 7.0042 | 7.4984 | 7.6697 | 7.4962 | 5.4415 | 5.7707 | 4.7073 | 5.5121 |
| LYST_S2105 | LYST | Q99698 | | 2105 | HMLRSRSLPAFPT | 6.0917 | 6.4488 | 5.7986 | 6.0997 | 6.4086 | 6.1890 | 6.5076 | 6.6295 | 6.3510 | 6.3554 | 6.5196 | 6.2050 | 5.8621 | 6.3950 | 6.0973 | 6.0411 |
| LYST_S2264 | LYST | Q99698 | | 2264 | AVGRWPSLVDRNT | 4.9036 | 5.1545 | 5.8004 | 5.7496 | 5.9580 | 5.9437 | 5.7927 | 6.0153 | 5.6940 | 5.0101 | 5.1361 | 5.2210 | 8.5816 | 8.5388 | 8.6283 | 7.8722 |
| LZTS2_S346 | LZTS2 | Q9BRK4 | | 570 | AAGVGGSLRAQVE | 6.4468 | 6.4205 | 5.4588 | 5.9863 | 6.5959 | 5.6971 | 5.9680 | 6.1678 | 5.7334 | 5.4200 | 5.2880 | 5.1267 | 8.0123 | 7.3811 | 7.6567 | 6.6406 |
| LZTS2_S570 | LZTS2 | Q9BRK4 | | 346 | LQQLRDSLDENEA | 5.9784 | 5.9534 | 6.0083 | 6.4242 | 7.0142 | 6.6502 | 6.3111 | 5.3865 | 5.7761 | 6.3355 | 6.2699 | 6.5515 | 6.2381 | 6.5384 | 5.7024 | 6.8618 |
| LZTS3_S316 | LZTS3 | O60299 | | 316 | AACSPPSPSALIQ | 5.4649 | 5.9393 | 5.9600 | 7.3004 | 6.4637 | 5.4186 | 6.1796 | 7.3369 | 5.8039 | 5.3198 | 6.8784 | 6.3479 | 6.9198 | 6.5820 | 5.9552 | 6.1296 |
| MACF1_S1376 | MACF1 | Q9UPN3 | | 4496 | SMDAMSSPTKTET | 6.1482 | 6.4626 | 6.6594 | 6.5358 | 6.1284 | 6.5143 | 6.2856 | 6.6003 | 6.5059 | 6.7234 | 6.7137 | 6.7541 | 5.1504 | 5.7285 | 5.5331 | 5.5565 |
| MACF1_S200 | MACF1 | Q9UPN3 | | 4521 | AELEQNSPKIQKV | 5.9800 | 6.3464 | 6.9456 | 6.6307 | 6.0357 | 6.5282 | 6.3783 | 6.3051 | 6.4822 | 6.9631 | 6.8315 | 6.9541 | 5.1368 | 5.8461 | 5.1851 | 5.4512 |
| MACF1_S213 | MACF1 | Q9UPN3 | | 6967 | RSGGRKSLSQPTP | 6.7845 | 6.0894 | 6.2986 | 6.7074 | 6.6703 | 6.1023 | 6.1471 | 6.4644 | 7.4740 | 7.2987 | 7.4012 | 6.9879 | 4.9853 | 5.8191 | 4.2931 | 4.4767 |
| MACF1_S291 | MACF1 | A0A0A6YYJ5 | | 802 | EMERRDSDSFLNI | 5.0690 | 5.3402 | 4.9237 | 5.4274 | 5.6703 | 5.1047 | 5.2251 | 5.3710 | 6.7171 | 6.2111 | 6.0818 | 6.0028 | 8.3479 | 7.8815 | 8.4518 | 8.1747 |
| MACF1_S295 | MACF1 | Q9UPN3 | | 3927 | LLKRQGSFSEDVI | 5.8232 | 5.9256 | 6.4667 | 5.8794 | 5.8678 | 6.0756 | 5.8418 | 5.8723 | 6.0680 | 6.1135 | 6.3792 | 6.5724 | 6.7565 | 6.9619 | 6.7839 | 6.6121 |
| MACF1_S342 | MACF1 | A0A0A6YYJ5 | | 200 | LEHVSKSIGIPEV | 6.8048 | 7.4744 | 6.8234 | 6.9523 | 6.1764 | 6.2264 | 6.3428 | 6.4003 | 6.4656 | 6.0367 | 6.6531 | 6.3357 | 5.1554 | 5.6101 | 5.2132 | 5.3294 |
| MACF1_S37 | MACF1 | A0A0A6YYJ5 | | 390 | RTFGHLSLGQICC | 5.6439 | 6.3929 | 5.8199 | 6.5925 | 5.3937 | 5.9117 | 6.1984 | 5.0583 | 5.1583 | 4.6161 | 5.7624 | 5.0332 | 8.0887 | 7.8190 | 8.3948 | 8.1163 |
| MACF1_S390 | MACF1 | Q9UPN3 | | 5808 | LIAQLPSPAIDHE | 6.1028 | 5.9532 | 6.3460 | 6.2678 | 5.9550 | 6.0639 | 5.9692 | 6.1206 | 6.2413 | 6.6296 | 6.3224 | 6.5131 | 6.3508 | 6.2642 | 6.3632 | 6.5369 |
| MACF1_S3927 | MACF1 | A0A0A6YYJ5 | | 213 | QDFKNLSGDCQDF | 6.4799 | 6.9468 | 6.5597 | 7.1856 | 6.6600 | 6.4806 | 7.1264 | 6.2289 | 6.6284 | 6.7213 | 6.7579 | 6.7145 | 4.8838 | 4.9722 | 4.6347 | 5.0193 |
| MACF1_S4496 | MACF1 | A0A0A6YYJ5 | | 617 | SHEEKLSGFIASE | 3.5903 | 3.6035 | 2.8326 | 3.3984 | 3.7038 | 2.6883 | 3.5419 | 3.2787 | 3.1837 | 3.3632 | 3.1063 | 2.5105 | 14.0389 | 14.3197 | 16.5064 | 16.3339 |
| MACF1_S4521 | MACF1 | A0A0A6YYJ5 | | 258 | SRFSSATWPRATK | 6.8779 | 6.8977 | 6.1705 | 6.8830 | 6.2029 | 6.0131 | 6.3405 | 5.8127 | 6.3140 | 5.8763 | 5.8435 | 5.9017 | 6.3000 | 6.5992 | 5.8510 | 6.1161 |
| MACF1_S5718 | MACF1 | A0A0A6YYJ5 | | 68 | MEKMEGSTISSNG | 5.7451 | 6.0133 | 5.9973 | 6.3253 | 5.9204 | 5.9266 | 6.7230 | 6.1653 | 6.3154 | 6.6783 | 6.5260 | 6.4635 | 6.2204 | 6.6078 | 6.0819 | 6.2903 |
| MACF1_S5808 | MACF1 | Q9UPN3 | | 7330 | ASSRRGSDASDFD | 5.7173 | 5.9668 | 6.8846 | 6.7616 | 6.1088 | 5.9903 | 5.6813 | 5.8496 | 6.8165 | 6.0885 | 6.7249 | 6.1795 | 6.4846 | 6.1815 | 6.2810 | 6.2830 |
| MACF1_S611 | MACF1 | Q9UPN3 | | 7267 | VIPSSGSKLKRPT | 7.3852 | 6.5165 | 6.5629 | 7.0583 | 6.1325 | 5.6880 | 5.8672 | 5.8548 | 7.4785 | 7.1052 | 6.3839 | 6.6863 | 4.7120 | 6.5559 | 4.9580 | 5.0549 |
| MACF1_S617 | MACF1 | Q9UPN3 | | 7265 | TKVIPSSGSKLKR | 7.4417 | 7.3271 | 6.7983 | 7.3239 | 6.4193 | 6.9045 | 7.2558 | 6.5455 | 7.8098 | 7.0551 | 6.9903 | 6.6517 | 3.7032 | 4.3428 | 3.3375 | 4.0935 |
| MACF1_S68 | MACF1 | Q9UPN3 | | 7314 | SASRPGSRAGSRA | 7.0095 | 6.9208 | 6.6668 | 6.3746 | 5.4256 | 6.3382 | 5.9744 | 6.2329 | 6.6030 | 6.8264 | 6.5098 | 6.4154 | 5.5475 | 5.9719 | 5.3025 | 5.8806 |
| MACF1_S6967 | MACF1 | Q9UPN3 | | 7279 | TPTFHSSRTSLAG | 5.6486 | 6.3779 | 5.5984 | 5.8286 | 5.1546 | 6.2366 | 6.4661 | 5.5321 | 5.9078 | 5.6687 | 6.4636 | 6.1169 | 6.9450 | 7.3951 | 6.5462 | 8.1138 |
| MACF1_S7265 | MACF1 | A0A0A6YYJ5 | | 342 | WEEEKLSDREKTF | 5.8629 | 6.1970 | 6.1555 | 6.5713 | 6.3350 | 5.8658 | 6.5970 | 6.0652 | 6.2819 | 6.0610 | 6.7284 | 5.9615 | 6.2840 | 6.5920 | 6.2264 | 6.2152 |
| MACF1_S7267 | MACF1 | A0A0A6YYJ5 | | 611 | NSDHRVSHEEKLS | 4.0515 | 4.8864 | 3.8393 | 5.0083 | 5.3048 | 5.0798 | 4.8718 | 5.6082 | 5.2371 | 4.3419 | 4.5630 | 4.4407 | 10.8587 | 10.6757 | 11.1333 | 10.0996 |
| MACF1_S7279 | MACF1 | A0A0A6YYJ5 | | 37 | CYVPQRSIYDTMR | 6.6365 | 7.0650 | 6.7548 | 6.9536 | 5.7098 | 6.2812 | 6.4564 | 6.4173 | 6.3670 | 6.4576 | 6.6791 | 6.6340 | 5.1557 | 5.9138 | 5.2592 | 5.2589 |
| MACF1_S7314 | MACF1 | A0A0A6YYJ5 | | 804 | ERRDSDSFLNIFP | 7.9131 | 7.2658 | 7.5836 | 8.2662 | 6.8405 | 6.0990 | 7.9064 | 8.1073 | 7.2658 | 6.4353 | 6.5899 | 5.8233 | 3.8460 | 3.5224 | 2.9112 | 3.6241 |
| MACF1_S7330 | MACF1 | A0A0A6YYJ5 | | 295 | PLSPCLSEELLDP | 7.2959 | 5.9358 | 5.8262 | 6.7479 | 7.5624 | 6.0529 | 6.4166 | 6.9272 | 6.1750 | 6.8674 | 4.5235 | 5.4327 | 6.7479 | 6.4714 | 5.4601 | 5.5572 |
| MACF1_S7455 | MACF1 | A0A0A6YYJ5 | | 291 | VEMPPLSPCLSEE | 6.0005 | 5.6036 | 7.9715 | 6.6194 | 8.3448 | 5.9399 | 5.4388 | 6.5554 | 7.5275 | 5.9567 | 4.7896 | 5.2504 | 5.5800 | 7.8235 | 4.6853 | 5.9130 |
| MACF1_S802 | MACF1 | Q9UPN3 | | 5718 | QVQKAFSIDIIRH | 5.5593 | 6.2865 | 6.7555 | 6.0261 | 5.9835 | 6.4301 | 5.4965 | 8.2862 | 5.8690 | 7.1618 | 7.0922 | 7.1842 | 4.5471 | 5.5705 | 5.3910 | 6.3605 |
| MACF1_S804 | MACF1 | Q9UPN3 | | 1376 | KRRRMLSSSDAIT | 5.9935 | 6.2882 | 6.9064 | 5.8409 | 6.1078 | 6.0732 | 6.5147 | 6.5771 | 5.8774 | 5.9993 | 5.6307 | 6.5810 | 6.1020 | 5.5453 | 6.9285 | 7.0340 |
| MACF1_T258 | MACF1 | H3BQK9 | | 7455 | PIRDSHSPDLQLP | 6.0163 | 5.5258 | 6.4369 | 7.5299 | 6.7328 | 6.2295 | 6.4743 | 7.4542 | 6.4497 | 5.5455 | 6.1450 | 6.2590 | 5.5582 | 5.3312 | 6.5775 | 5.7342 |
| MACIR_S167 | MACIR | Q96GV9 | | 25 | PGAEANSPGKAEA | 6.2232 | 6.4858 | 6.8513 | 6.2455 | 6.0540 | 6.3407 | 6.1822 | 6.5279 | 6.8728 | 6.7433 | 6.7351 | 6.5975 | 5.2316 | 5.6895 | 5.4889 | 5.7307 |
| MACIR_S25 | MACIR | Q96GV9 | | 167 | KRAHSKSLDYLNL | 6.5989 | 7.0723 | 7.8199 | 7.1001 | 5.6149 | 6.9890 | 6.4634 | 7.0033 | 6.5509 | 6.6420 | 7.5094 | 7.3297 | 3.8062 | 4.3695 | 4.2167 | 4.9137 |
| MACO1_S305 | MACO1 | Q8N5G2 | | 332 | SGVVNSSPRSHSA | 6.5283 | 6.8940 | 6.3560 | 6.7484 | 6.2245 | 6.0196 | 6.4909 | 6.0310 | 6.4175 | 6.0969 | 6.2826 | 6.3537 | 5.8096 | 5.9845 | 5.7216 | 6.0411 |
| MACO1_S332 | MACO1 | Q8N5G2 | | 305 | NNDLVGSTENLLK | 5.5750 | 6.0131 | 5.8572 | 6.2585 | 5.6454 | 5.6933 | 5.6357 | 5.7446 | 5.6278 | 6.1118 | 5.8742 | 6.0198 | 6.9694 | 6.7692 | 8.2136 | 7.9914 |
| MACO1_S648 | MACO1 | Q8N5G2 | | 648 | SKFVETSPSGLDP | 7.0143 | 7.1941 | 6.7282 | 7.4876 | 6.5133 | 6.4117 | 7.2975 | 6.5985 | 7.1215 | 7.2408 | 7.0048 | 6.9261 | 3.4715 | 4.6753 | 3.8017 | 4.5128 |
| MACROH2A1_S138 | MACROH2A1 | O75367 | | 129 | KLEAIITPPPAKK | 7.2013 | 7.4352 | 6.5645 | 8.0225 | 6.2984 | 6.3185 | 7.2946 | 6.9161 | 6.8461 | 7.2045 | 6.7724 | 7.0639 | 3.2677 | 4.6200 | 3.7660 | 4.4084 |
| MACROH2A1_S173 | MACROH2A1 | O75367 | | 138 | PAKKAKSPSQKKP | 7.2957 | 6.7570 | 6.0401 | 7.1177 | 7.5596 | 7.4228 | 7.7089 | 6.8918 | 7.0543 | 5.9531 | 7.2402 | 6.5720 | 4.3193 | 4.9255 | 3.6256 | 3.5164 |
| MACROH2A1_T129 | MACROH2A1 | O75367 | | 173 | KAASADSTTEGTP | 5.6803 | 6.0134 | 5.3167 | 6.3496 | 5.7749 | 5.5771 | 6.3126 | 5.8370 | 5.9649 | 5.6643 | 5.2938 | 6.3222 | 7.3784 | 8.0426 | 7.2957 | 7.1765 |
| MACROH2A2_S129 | MACROH2A2 | Q9P0M6 | | 129 | KSETILSPPPEKR | 7.2352 | 7.0363 | 6.8918 | 8.2925 | 6.1699 | 6.2119 | 7.7090 | 7.1573 | 7.2529 | 7.4685 | 6.9994 | 6.3449 | 3.1292 | 4.6628 | 3.1650 | 4.2734 |
| MAD1L1_S428 | MAD1L1 | Q9Y6D9 | | 428 | LTPAEYSPQLTRR | 6.3361 | 6.8916 | 6.4381 | 6.8659 | 6.8348 | 6.2561 | 6.5528 | 6.8648 | 6.8392 | 6.1336 | 6.5350 | 5.8965 | 5.3595 | 5.8260 | 5.0960 | 5.2740 |
| MAD2L1BP_S134 | MAD2L1BP | Q15013-3 | | 134 | HFYRKPSPQAEEM | 6.1769 | 6.1186 | 6.2739 | 6.4121 | 6.1510 | 6.2161 | 5.9214 | 6.2727 | 6.2690 | 6.6245 | 6.8549 | 6.8213 | 6.1427 | 5.9170 | 5.7776 | 6.0504 |
| MADD_S1239 | MADD | Q8WXG6 | | 1270 | KEKLAGSPIRTSE | 6.9323 | 7.2430 | 6.7862 | 6.6439 | 6.4330 | 7.6851 | 7.2361 | 7.6751 | 7.5631 | 7.0751 | 7.1827 | 6.5328 | 3.2136 | 4.0891 | 3.5352 | 4.1737 |
| MADD_S1270 | MADD | Q8WXG6 | | 1239 | SSRGTLSDSEIET | 6.4533 | 6.2549 | 7.0111 | 5.7579 | 6.7173 | 6.9138 | 6.3890 | 6.5453 | 6.5298 | 6.3575 | 6.3019 | 6.4556 | 5.4060 | 6.0844 | 5.3988 | 5.4235 |
| MAF1_S60;S68 | MAF1 | Q9H063 | | 68 | PQTSGLSPSRLSK | 5.6681 | 6.0598 | 5.8886 | 5.9919 | 6.6114 | 6.5378 | 6.2239 | 6.6876 | 6.0917 | 6.1030 | 6.1430 | 6.3024 | 6.4918 | 6.8007 | 6.3214 | 6.0769 |
| MAF1_S68 | MAF1 | Q9H063 | | 75 | PSRLSKSQGGEEE | 6.8735 | 6.4536 | 7.6905 | 6.8624 | 5.9184 | 6.7465 | 5.6937 | 6.2441 | 6.5679 | 6.8595 | 7.1066 | 7.4489 | 4.5754 | 4.8878 | 5.1339 | 4.9374 |
| MAF1_S75 | MAF1 | Q9H063 | 60;68 | | NA | 6.1686 | 6.5267 | 7.5314 | 6.8775 | 6.2854 | 7.0172 | 4.6605 | 7.2139 | 6.9427 | 7.0055 | 7.4330 | 7.4972 | 4.0165 | 4.9197 | 4.2859 | 3.8184 |
| MAGEA10_S116 | MAGEA10 | P43363 | | 116 | SSQKEESPSTLQV | 5.7261 | 6.0496 | 6.2666 | 5.7021 | 6.6390 | 5.5646 | 5.6296 | 5.8675 | 6.2195 | 6.7157 | 5.6191 | 6.4817 | 6.3526 | 6.7304 | 7.3015 | 7.1346 |
| MAGEA10_S325 | MAGEA10 | P43363 | | 325 | NGSDPRSFPLWYE | 5.7380 | 6.0816 | 6.4212 | 5.6402 | 6.0968 | 6.2751 | 5.4942 | 6.1562 | 5.8437 | 5.7235 | 5.9963 | 6.0862 | 7.4492 | 7.3844 | 7.0818 | 6.5315 |
| MAGEA4_S304 | MAGEA4 | P43358 | | 99 | EEGPSTSPDAESL | 6.2730 | 7.3601 | 6.7690 | 6.5899 | 6.0766 | 6.9697 | 6.5212 | 6.5389 | 6.8989 | 6.7588 | 7.2222 | 6.9996 | 4.6758 | 5.2206 | 4.1287 | 4.9970 |
| MAGEA4_S88;S90 | MAGEA4 | P43358 | | 90 | PNEGSSSQEEEGP | 6.5521 | 6.1897 | 6.7934 | 6.8276 | 6.1833 | 6.1476 | 6.1600 | 6.9957 | 6.0830 | 6.6562 | 6.4705 | 6.7329 | 5.6976 | 6.1296 | 5.3932 | 4.9876 |
| MAGEA4_S90 | MAGEA4 | P43358 | | 304 | VRIAYPSLREAAL | 6.9090 | 6.6551 | 7.0583 | 6.1406 | 6.9313 | 6.4786 | 6.2419 | 6.8781 | 6.3242 | 5.9731 | 6.4926 | 6.3384 | 5.5237 | 5.7001 | 5.1995 | 5.1556 |
| MAGEA4_S90;S99 | MAGEA4 | P43358 | 90;99 | | NA | 6.4203 | 6.8471 | 6.4815 | 7.5637 | 5.6382 | 6.9808 | 7.5626 | 6.5828 | 7.2944 | 6.4856 | 6.9444 | 7.5272 | 4.0405 | 4.7711 | 3.8694 | 4.9906 |
| MAGEA4_S99 | MAGEA4 | P43358 | 88;90 | | NA | 7.3570 | 7.9108 | 7.7197 | 6.9574 | 7.6213 | 7.3850 | 6.8620 | 7.5996 | 5.8327 | 5.7302 | 5.8419 | 7.6470 | 4.1783 | 3.3102 | 3.8058 | 4.2409 |
| MAGED2_S19 | MAGED2 | Q9UNF1 | | 247 | ALLSLRSPKARRG | 6.7286 | 7.3079 | 7.0498 | 6.4937 | 6.5088 | 7.3447 | 7.1137 | 6.8395 | 6.2885 | 7.2882 | 7.3133 | 7.2460 | 3.8157 | 4.7934 | 3.6659 | 4.2025 |
| MAGED2_S247 | MAGED2 | Q9UNF1 | | 72 | ATEVSKTPEAREA | 6.5010 | 7.2258 | 6.9170 | 6.9747 | 6.3416 | 7.3409 | 7.2284 | 6.9076 | 7.1578 | 7.5329 | 7.8934 | 7.2465 | 3.5567 | 4.4595 | 3.0484 | 3.6676 |
| MAGED2_T72 | MAGED2 | Q9UNF1 | | 19 | RFQAEASEKDSSS | 5.2088 | 7.8934 | 8.4504 | 4.7170 | 7.0805 | 6.8396 | 4.6768 | 3.6632 | 7.9085 | 6.2776 | 6.5134 | 5.8260 | 5.6002 | 7.2411 | 4.8776 | 7.2260 |
| MAGI1_S1071 | MAGI1 | Q96QZ7 | | 1071 | CSITNKSHSDIVN | 6.5183 | 6.0737 | 7.4733 | 7.2555 | 5.9208 | 6.9034 | 5.8933 | 5.0372 | 6.7185 | 5.7536 | 5.2364 | 5.5929 | 5.9043 | 6.2839 | 7.6802 | 5.7547 |
| MAGI3_S1475 | MAGI3 | Q5TCQ9-1 | | 684 | LRGGPPSPTKTAK | 8.7382 | 5.7068 | 5.7838 | 5.3253 | 5.4717 | 5.7152 | 10.2285 | 8.4933 | 7.0326 | 5.5084 | 5.5844 | 5.5721 | 4.9433 | 5.7963 | 5.1449 | 4.9551 |
| MAGI3_S684 | MAGI3 | Q5TCQ9-1 | | 701 | KKENAGSLEAINE | 6.1117 | 6.7323 | 6.3783 | 6.8092 | 5.8948 | 6.3136 | 6.4130 | 6.7052 | 6.3572 | 5.6643 | 6.1659 | 5.6462 | 6.3030 | 5.9776 | 5.9279 | 6.5998 |
| MAGI3_S701 | MAGI3 | Q5TCQ9-1 | | 1475 | FKPESSSPVKKTL | 6.5444 | 6.5906 | 6.6821 | 5.8478 | 6.7671 | 6.6078 | 6.2512 | 6.7296 | 6.8957 | 6.8338 | 5.7353 | 6.2268 | 5.7185 | 6.0285 | 5.6468 | 4.8941 |
| MAGOHB_S108 | MAGOHB | Q96A72 | | 108 | TTSKIGSLIDVNQ | 6.0837 | 5.4330 | 6.6954 | 5.0565 | 5.5387 | 5.9582 | 6.1039 | 5.6447 | 6.2768 | 6.6677 | 6.2097 | 6.9340 | 7.0211 | 7.0122 | 6.5756 | 7.7888 |
| MAK16_S229;S232 | MAK16 | Q9BXY0 | 229;232 | | NA | 5.6432 | 6.0988 | 6.2115 | 6.8668 | 6.0458 | 5.4097 | 6.2433 | 5.7033 | 6.3551 | 6.0526 | 5.7907 | 6.0465 | 6.7005 | 6.6379 | 6.8161 | 7.3782 |
| MAML1_S120 | MAML1 | Q92585 | | 451 | PMEKPASPSSYKQ | 4.5080 | 4.1305 | 5.1817 | 4.5051 | 4.8847 | 5.7967 | 5.3584 | 5.1599 | 5.1511 | 5.5775 | 5.8722 | 6.0451 | 9.5638 | 9.3344 | 9.4059 | 9.5250 |
| MAML1_S45 | MAML1 | Q92585 | | 45 | ARYEAVSPERLEL | 5.9859 | 6.5534 | 6.4208 | 6.1096 | 6.5102 | 6.3787 | 6.3021 | 6.3656 | 6.5461 | 6.4183 | 6.8805 | 6.5970 | 5.6161 | 6.3868 | 5.4387 | 5.4900 |
| MAML1_S451 | MAML1 | Q92585 | | 120 | NLDSATSPQNGDQ | 5.6697 | 5.7858 | 6.1894 | 5.7827 | 6.1863 | 5.3678 | 6.0877 | 6.0419 | 6.0243 | 5.5077 | 7.0970 | 6.1900 | 7.0593 | 7.5703 | 6.8497 | 6.5905 |
| MAML2_S1089 | MAML2 | Q8IZL2 | | 1089 | TPSNFPSPNQSSR | 6.8061 | 7.0470 | 6.0917 | 6.4223 | 6.0049 | 5.9292 | 6.5427 | 7.4994 | 6.7417 | 5.8284 | 6.2710 | 6.6954 | 5.5272 | 5.7121 | 5.1574 | 5.7233 |
| MAML2_T352 | MAML2 | Q8IZL2 | | 352 | QQSQRSTPRPSLP | 6.8503 | 7.2914 | 6.8651 | 6.3894 | 6.3830 | 7.2061 | 6.3733 | 6.1759 | 7.6376 | 6.8510 | 7.9704 | 7.6601 | 4.0294 | 5.0623 | 3.1799 | 4.0749 |
| MAP1A_S1069 | MAP1A | P78559-2 | | 2022 | ELSSPISPKSLQS | 6.2637 | 6.1982 | 6.5647 | 7.5551 | 6.1500 | 6.3576 | 5.9812 | 6.1767 | 6.8005 | 6.9106 | 6.6676 | 6.5452 | 5.8552 | 6.2875 | 5.8611 | 5.6250 |
| MAP1A_S1146 | MAP1A | P78559-2 | | 1172 | QEPTPKSPCGLTE | 5.6730 | 5.4424 | 5.8954 | 5.3922 | 6.1439 | 6.7705 | 6.4915 | 6.7318 | 6.3410 | 5.8702 | 5.8433 | 5.7145 | 6.8362 | 7.0379 | 6.8403 | 6.9759 |
| MAP1A_S1172 | MAP1A | P78559-2 | | 2617 | GKAKPASPARRLD | 6.1603 | 6.3288 | 6.2208 | 5.5679 | 6.2433 | 6.4687 | 5.5909 | 6.7062 | 6.5556 | 7.0641 | 6.4777 | 6.4754 | 6.1275 | 6.5737 | 5.8801 | 5.5591 |
| MAP1A_S1218 | MAP1A | P78559-2 | | 2235 | SLPQLPSPSSPGA | 5.7304 | 5.9525 | 6.9607 | 6.3131 | 6.4457 | 6.0493 | 6.1368 | 6.5172 | 6.5404 | 6.0573 | 6.3728 | 5.5589 | 5.9966 | 6.5178 | 6.2076 | 6.6431 |
| MAP1A_S1326 | MAP1A | P78559-2 | | 2182 | APPQLPSPAEPRS | 6.3626 | 5.5570 | 6.6096 | 5.9519 | 6.2091 | 6.5649 | 6.3037 | 6.0612 | 6.5660 | 6.0431 | 6.8731 | 6.4595 | 6.0442 | 6.5592 | 5.5514 | 6.2833 |
| MAP1A_S1762 | MAP1A | P78559-2 | | 1326 | DSSFSKSPESLPG | 5.9247 | 6.1826 | 6.4033 | 6.3014 | 6.1856 | 6.5803 | 6.3092 | 5.8677 | 6.4361 | 6.6905 | 6.0234 | 6.4389 | 5.7510 | 6.8834 | 6.1861 | 5.8360 |
| MAP1A_S1776 | MAP1A | P78559-2 | | 1146 | YPDRSLSPEDAES | 5.4260 | 6.0672 | 6.1040 | 6.1565 | 5.9732 | 5.9133 | 6.1899 | 6.0551 | 6.1511 | 6.1167 | 6.4953 | 5.9561 | 6.6745 | 6.5999 | 6.7655 | 7.3558 |
| MAP1A_S2022 | MAP1A | P78559-2 | | 1776 | ERWLAESPVGLPP | 5.4643 | 6.4768 | 5.8954 | 5.3201 | 5.4936 | 6.9468 | 6.2602 | 6.7917 | 7.1689 | 6.7603 | 7.1292 | 7.3199 | 5.9692 | 5.8100 | 5.2730 | 5.9207 |
| MAP1A_S2182 | MAP1A | P78559-2 | | 1069 | KVPPPRSPQAQEA | 5.4809 | 5.5868 | 6.7859 | 6.0621 | 5.5301 | 6.9195 | 6.2840 | 6.5917 | 6.3874 | 6.3168 | 6.4694 | 7.1099 | 5.8944 | 6.5526 | 6.3017 | 5.7268 |
| MAP1A_S2235 | MAP1A | P78559-2 | | 644 | TEAREESEPEVKE | 6.3027 | 6.3705 | 6.6646 | 6.3936 | 6.5045 | 6.5695 | 6.3244 | 6.8117 | 5.9334 | 6.7642 | 6.3682 | 6.7351 | 5.4244 | 6.4708 | 4.8195 | 5.5427 |
| MAP1A_S2449 | MAP1A | P78559-2 | | 1218 | VSSKQLSPESLGT | 5.4607 | 6.6718 | 5.6650 | 6.5316 | 6.5870 | 6.0754 | 6.0300 | 6.1501 | 6.6718 | 6.7147 | 6.5799 | 6.4522 | 6.3279 | 7.2276 | 5.1528 | 5.7015 |
| MAP1A_S2617 | MAP1A | P78559-2 | | 2629 | DLRGKRSPTPGKG | 5.8951 | 6.7987 | 6.4842 | 6.3287 | 6.8035 | 6.6454 | 5.6863 | 7.0177 | 6.5366 | 7.0305 | 6.8331 | 7.5934 | 4.7569 | 5.4607 | 5.2517 | 4.8774 |
| MAP1A_S2629 | MAP1A | P78559-2 | | 2449 | PHRGELSPSFLNP | 5.7605 | 4.8835 | 7.8716 | 6.7046 | 5.0603 | 6.8249 | 5.5483 | 6.8744 | 6.1777 | 5.8595 | 5.8948 | 5.5483 | 6.2060 | 9.9438 | 5.1593 | 5.6827 |
| MAP1A_S644 | MAP1A | P78559-2 | | 1762 | KDFQESSPQKGLE | 5.9322 | 6.8325 | 7.0386 | 6.2288 | 6.0974 | 7.0012 | 6.1611 | 6.3137 | 6.5262 | 7.0384 | 7.1334 | 7.6307 | 4.7407 | 5.5765 | 4.5904 | 5.1581 |
| MAP1A_S896 | MAP1A | P78559-2 | | 896 | PSAGTISPTSSLE | 5.8643 | 6.0709 | 6.2932 | 5.5383 | 5.6049 | 6.3179 | 5.6673 | 6.3310 | 7.0502 | 6.3962 | 6.2129 | 6.7119 | 6.7764 | 6.7119 | 6.1127 | 6.3399 |
| MAP1B_S1016 | MAP1B | P46821 | | 1625 | ISPPDFSPKTAKS | 5.9760 | 5.7048 | 6.9859 | 5.8549 | 6.4959 | 6.5837 | 6.1615 | 5.8380 | 6.4668 | 6.5381 | 7.0077 | 6.8236 | 5.7255 | 6.5223 | 5.4121 | 5.9033 |
| MAP1B_S1154 | MAP1B | P46821 | | 1501 | RKLGDVSPTQIDV | 5.6609 | 5.8427 | 6.4061 | 5.6998 | 6.0245 | 6.3413 | 6.0378 | 6.2227 | 6.1060 | 6.4272 | 6.1446 | 6.2067 | 6.2656 | 7.3227 | 6.5636 | 6.7280 |
| MAP1B_S1208 | MAP1B | P46821 | | 1653 | IEFGQESPEQSLA | 6.0157 | 6.1530 | 6.2732 | 6.2649 | 5.9275 | 6.1004 | 6.0254 | 6.5660 | 6.4038 | 6.5583 | 6.4334 | 6.5908 | 5.9833 | 6.7061 | 6.0674 | 5.9308 |
| MAP1B_S1256 | MAP1B | P46821 | | 1400 | VLSPLRSPPLIGS | 5.7268 | 5.8785 | 6.3267 | 5.8196 | 5.9759 | 6.2350 | 5.8247 | 6.1105 | 5.8820 | 6.1673 | 6.3177 | 6.3694 | 6.3820 | 7.5367 | 6.6263 | 6.8208 |
| MAP1B_S1265 | MAP1B | P46821 | | 2271 | SKPLAASPKPAGL | 6.6655 | 6.2447 | 7.0809 | 6.6716 | 6.3447 | 6.9247 | 6.9418 | 6.2779 | 7.0771 | 7.2221 | 7.0739 | 7.2288 | 4.1083 | 4.9237 | 4.1035 | 5.1109 |
| MAP1B_S1276;T1282 | MAP1B | P46821 | | 1427 | SGRGAESPFEEKS | 6.1349 | 6.2969 | 6.8075 | 5.9291 | 6.1706 | 6.5441 | 6.4103 | 6.4008 | 6.3912 | 6.3493 | 6.4774 | 6.2160 | 5.3289 | 6.6953 | 5.8819 | 5.9658 |
| MAP1B_S1282 | MAP1B | P46821 | | 1438 | KSGKQGSPDQVSP | 5.6615 | 6.1046 | 6.7672 | 6.0821 | 6.0019 | 6.2779 | 6.2921 | 6.3394 | 6.0295 | 6.6866 | 6.6562 | 6.6611 | 5.7778 | 6.5557 | 5.5423 | 6.5642 |
| MAP1B_S1298 | MAP1B | P46821 | | 1282 | SVNFSLTPNEIKV | 6.8089 | 7.3646 | 6.8962 | 7.3923 | 6.2625 | 6.8891 | 7.0835 | 7.0236 | 7.0577 | 7.3824 | 7.3977 | 7.5585 | 3.1268 | 4.3363 | 3.2673 | 4.1525 |
| MAP1B_S1298;S1312 | MAP1B | P46821 | | 614 | PSKEEPSPVKAEV | 6.0464 | 6.7779 | 6.7034 | 6.3803 | 5.9894 | 7.0732 | 6.3180 | 6.6090 | 6.7678 | 6.6697 | 6.6230 | 6.9665 | 4.4989 | 5.9504 | 4.9957 | 5.6301 |
| MAP1B_S1312 | MAP1B | P46821 | | 2098 | HPKTELSPSFINP | 4.8616 | 5.6654 | 6.4599 | 6.0643 | 6.0095 | 6.1540 | 6.1839 | 6.1264 | 6.3052 | 6.3518 | 6.3745 | 6.6513 | 6.2943 | 7.0189 | 6.6294 | 6.8497 |
| MAP1B_S1322 | MAP1B | P46821 | | 2072 | YTAEKKSPSEARQ | 6.0616 | 6.5955 | 6.6494 | 6.3386 | 5.9540 | 6.5924 | 6.0844 | 6.4622 | 6.3968 | 6.7722 | 6.6690 | 6.7882 | 4.9803 | 6.3435 | 5.4817 | 5.8302 |
| MAP1B_S1378 | MAP1B | P46821 | | 2305 | AAKPTTTPEVKAA | 6.8170 | 6.7621 | 6.9978 | 7.3946 | 6.3139 | 6.9594 | 7.4846 | 6.4685 | 7.0444 | 7.5643 | 7.6358 | 7.7335 | 3.0415 | 4.2476 | 3.1673 | 4.3676 |
| MAP1B_S1389 | MAP1B | P46821 | | 1881 | PEETTRSPDEEDY | 5.7931 | 6.0971 | 6.5445 | 6.0330 | 6.0554 | 6.4650 | 5.6811 | 6.0501 | 6.0906 | 6.5723 | 6.7341 | 6.6068 | 5.7498 | 7.3812 | 6.2245 | 5.9214 |
| MAP1B_S1396;S1400 | MAP1B | P46821 | | 1154 | NNEETESPSQEFV | 6.0401 | 5.9538 | 6.4433 | 6.1792 | 6.0536 | 5.9753 | 5.9238 | 6.1264 | 6.5858 | 6.5144 | 6.4094 | 6.6823 | 6.0939 | 6.8343 | 5.9944 | 6.1901 |
| MAP1B_S1396;S1408 | MAP1B | P46821 | | 1887 | SPDEEDYDYESYE | 5.6025 | 6.2314 | 6.3601 | 6.2692 | 5.8304 | 6.3818 | 6.1152 | 6.5962 | 5.9193 | 6.7222 | 6.7727 | 6.7356 | 5.6460 | 7.2544 | 5.7059 | 5.8572 |
| MAP1B_S1396;Y1410 | MAP1B | P46821 | | 832 | ERSLMSSPEDLTK | 5.6495 | 5.8686 | 6.3144 | 5.7015 | 6.1024 | 6.2073 | 5.5819 | 6.6056 | 6.5033 | 6.6691 | 6.9039 | 7.1531 | 6.0046 | 6.4891 | 5.8277 | 6.4182 |
| MAP1B_S1400 | MAP1B | P46821 | | 1378 | NERASVSPMDEPV | 5.8403 | 6.0862 | 6.3461 | 6.0205 | 6.3031 | 6.5725 | 5.9095 | 5.7594 | 5.8665 | 6.3469 | 6.5877 | 6.0107 | 6.0092 | 7.6393 | 5.9900 | 6.7120 |
| MAP1B_S1427 | MAP1B | P46821 | | 1620 | PRPMSISPPDFSP | 4.7260 | 5.1634 | 5.7311 | 4.5132 | 5.7645 | 6.0066 | 4.9410 | 5.0925 | 4.9177 | 5.7021 | 5.4072 | 5.2129 | 9.1274 | 10.3717 | 9.2230 | 8.0996 |
| MAP1B_S1438 | MAP1B | P46821 | | 1016 | KGEAEQSEEEADE | 5.7231 | 5.5021 | 6.2777 | 5.3961 | 6.0304 | 7.0489 | 6.4509 | 6.0838 | 7.1692 | 6.4008 | 7.8944 | 7.3142 | 6.0711 | 6.7597 | 5.0694 | 4.8081 |
| MAP1B_S1438;S1443 | MAP1B | P46821 | | 1265 | LSPSPPSPLEKTP | 5.3777 | 5.3719 | 6.0796 | 5.2566 | 6.0946 | 6.5000 | 5.5875 | 6.2604 | 6.0176 | 5.9068 | 6.1125 | 6.1398 | 6.6270 | 8.1331 | 7.4241 | 7.1109 |
| MAP1B_S1443 | MAP1B | P46821 | | 1666 | MDFSRQSPDHPTV | 5.5073 | 6.7326 | 6.5225 | 7.0467 | 6.3121 | 7.2212 | 5.3514 | 7.5807 | 6.4382 | 6.1435 | 6.3079 | 6.5139 | 5.3189 | 6.0023 | 5.6947 | 5.3060 |
| MAP1B_S1501 | MAP1B | P46821 | | 1443 | GSPDQVSPVSEMT | 5.5759 | 5.9044 | 5.8954 | 6.2059 | 5.8015 | 6.1264 | 5.7559 | 6.3769 | 6.2853 | 6.1826 | 6.5051 | 6.3375 | 6.4886 | 7.1251 | 6.5472 | 6.8864 |
| MAP1B_S1618;S1620 | MAP1B | P46821 | | 1633 | KTAKSRTPVQDHR | 6.3017 | 7.5148 | 7.0810 | 7.1258 | 6.2843 | 7.1393 | 6.9080 | 6.9556 | 6.8854 | 7.0187 | 7.2139 | 7.2660 | 3.3634 | 4.5540 | 3.8601 | 4.5279 |
| MAP1B_S1618;S1620;S1 | MAP1B | P46821 | | 1256 | KVSPSKSPSLSPS | 5.7670 | 6.1081 | 6.6467 | 6.0946 | 6.4164 | 6.3521 | 5.3583 | 6.1159 | 7.2596 | 6.7761 | 6.9794 | 7.0844 | 5.1966 | 6.3464 | 5.4568 | 6.0415 |
| MAP1B_S1618;S1625 | MAP1B | P46821 | | 1067 | QYGFLTTPTKQLG | 6.2166 | 6.1736 | 6.4221 | 6.5836 | 6.1322 | 6.8909 | 6.4765 | 6.6673 | 6.6850 | 6.3494 | 7.2296 | 6.7570 | 4.6254 | 6.1727 | 5.0654 | 5.5527 |
| MAP1B_S1620 | MAP1B | P46821 | | 1298 | AEVAPVSPEVTQE | 5.7054 | 6.2937 | 6.5631 | 6.0962 | 6.3652 | 6.4746 | 5.7162 | 5.9976 | 6.8876 | 6.7737 | 6.0495 | 6.0930 | 6.0688 | 6.7074 | 6.3026 | 5.9052 |
| MAP1B_S1620;S1625 | MAP1B | P46821 | | 1797 | ESSPLYSPTFSDS | 5.8314 | 6.0779 | 6.2114 | 5.8621 | 5.8002 | 6.3760 | 6.0105 | 6.5722 | 6.6206 | 6.5973 | 6.4836 | 6.5907 | 5.4934 | 6.7736 | 6.1952 | 6.5039 |
| MAP1B_S1625 | MAP1B | P46821 | | 744 | DAKKSSTPLSEAK | 7.1695 | 7.5198 | 6.7468 | 7.5728 | 6.1357 | 7.1083 | 7.0706 | 6.3861 | 7.8331 | 7.4664 | 7.6901 | 7.6325 | 2.9956 | 4.2260 | 2.9033 | 3.5435 |
| MAP1B_S1653 | MAP1B | P46821 | | 1793 | LTPRESSPLYSPT | 5.8796 | 5.8028 | 6.3423 | 5.8617 | 6.1974 | 5.9918 | 5.6237 | 6.2630 | 6.4981 | 6.2876 | 6.1916 | 6.6857 | 6.2660 | 7.1247 | 6.3982 | 6.5859 |
| MAP1B_S1666 | MAP1B | P46821 | | 937 | AGFEESSETGDYE | 5.2298 | 6.3264 | 6.9110 | 5.3021 | 6.0567 | 6.4242 | 5.7216 | 6.0065 | 6.4161 | 6.3066 | 6.2088 | 6.4906 | 5.8212 | 7.4678 | 6.7571 | 6.5534 |
| MAP1B_S1779;S1782;T1 | MAP1B | P46821 | | 1963 | YDISEKTTSPPEV | 5.9968 | 6.2575 | 7.0135 | 6.3983 | 5.8530 | 6.5066 | 6.1514 | 6.6448 | 6.4872 | 6.6067 | 6.9161 | 6.6430 | 4.9947 | 6.2800 | 5.7754 | 5.4750 |
| MAP1B_S1785 | MAP1B | P46821 | | 995 | KRESVASGDDRAE | 5.7261 | 6.4314 | 6.8356 | 5.4203 | 5.9970 | 6.2249 | 5.1066 | 5.8048 | 6.1470 | 7.0544 | 6.9226 | 6.9238 | 5.7451 | 6.8859 | 6.0469 | 6.7276 |
| MAP1B_S1793 | MAP1B | P46821 | | 2209 | APVQDRSPSPRHP | 6.1082 | 6.5720 | 6.4456 | 6.9105 | 6.5983 | 5.8498 | 6.2269 | 6.1642 | 6.1687 | 6.3639 | 6.5871 | 6.9128 | 5.2142 | 6.2319 | 5.4335 | 6.2123 |
| MAP1B_S1793;S1797 | MAP1B | P46821 | | 948 | YEEKAETEEAEEP | 5.8803 | 5.8396 | 6.0662 | 5.4717 | 6.0802 | 6.1919 | 6.2329 | 6.2392 | 6.4710 | 6.1108 | 6.2333 | 5.7445 | 6.4311 | 7.3353 | 6.9859 | 6.6862 |
| MAP1B_S1797 | MAP1B | P46821 | | 1788 | SDISPLTPRESSP | 5.9646 | 6.1718 | 6.5353 | 5.6555 | 5.6938 | 6.3914 | 6.2712 | 6.7783 | 6.3325 | 5.4139 | 6.1493 | 5.9494 | 6.0232 | 7.1378 | 6.8404 | 6.6915 |
| MAP1B_S1819 | MAP1B | P46821 | | 2087 | DLCLVSSCEYKHP | 6.0075 | 7.4635 | 7.2923 | 7.1925 | 5.5254 | 7.2324 | 7.0442 | 6.2426 | 7.1902 | 6.8863 | 7.3412 | 7.2543 | 3.9758 | 4.8795 | 3.9158 | 4.5564 |
| MAP1B_S1881 | MAP1B | P46821 | | 1322 | KTLEVVSPSQSVT | 5.8872 | 6.7066 | 5.9541 | 5.5989 | 6.0613 | 6.9535 | 5.3973 | 7.5542 | 5.7675 | 5.6889 | 6.1520 | 6.5344 | 5.4856 | 7.3559 | 6.7566 | 6.1459 |
| MAP1B_S1899 | MAP1B | P46821 | | 2034 | STKTTRTPDTSTY | 7.3673 | 8.0064 | 7.2796 | 7.0130 | 6.8080 | 7.4033 | 7.3380 | 5.6705 | 6.3845 | 7.3795 | 7.5870 | 7.4061 | 3.4331 | 4.6301 | 2.9660 | 3.3275 |
| MAP1B_S1915 | MAP1B | P46821 | | 1208 | ASASTISPPSSME | 5.7431 | 5.8261 | 6.6411 | 6.3770 | 6.1063 | 6.9393 | 6.2691 | 6.2527 | 6.0158 | 6.1099 | 6.3302 | 6.5346 | 5.2641 | 6.9166 | 5.9933 | 6.6807 |
| MAP1B_S2072 | MAP1B | P46821 | | 1819 | TCHSSSSPPIDAA | 6.1841 | 7.0085 | 5.2841 | 6.3480 | 6.0903 | 8.2690 | 5.4467 | 8.4169 | 5.8536 | 6.6589 | 6.5150 | 6.3484 | 4.3910 | 6.8451 | 5.7339 | 4.6063 |
| MAP1B_S2087 | MAP1B | P46821 | | 1853 | LNRDLSTPGLEKD | 5.3152 | 5.4998 | 6.2426 | 5.5432 | 5.4669 | 5.4735 | 5.3955 | 5.6412 | 6.5909 | 5.7783 | 5.8051 | 5.7942 | 7.3201 | 8.6264 | 7.8364 | 7.6707 |
| MAP1B_S2098 | MAP1B | P46821 | | 1899 | EKTTRTSDVGGYY | 6.8015 | 6.8183 | 6.9011 | 7.2920 | 6.4906 | 6.4996 | 7.1096 | 6.5372 | 6.9404 | 6.9342 | 7.4350 | 6.8898 | 3.8920 | 5.1647 | 3.7788 | 4.5152 |
| MAP1B_S2209 | MAP1B | P46821 | | 1312 | VEEHCASPEDKTL | 6.0335 | 6.1592 | 6.7628 | 6.1571 | 6.0590 | 5.4990 | 5.5272 | 5.7493 | 6.2328 | 6.2824 | 6.4594 | 6.5694 | 6.6047 | 7.0600 | 6.5848 | 6.2594 |
| MAP1B_S2209;S2211 | MAP1B | P46821 | | 2051 | AEKITRTPQASTY | 5.1532 | 6.9781 | 6.5836 | 6.8852 | 6.4825 | 6.1510 | 6.2587 | 7.0560 | 7.2283 | 6.0465 | 7.0195 | 7.8980 | 4.7288 | 5.1648 | 4.7454 | 5.6206 |
| MAP1B_S2271 | MAP1B | P46821 | | 1389 | PVPDSESPIEKVL | 6.2068 | 5.8031 | 6.5470 | 5.0932 | 5.6611 | 5.8963 | 5.6197 | 5.7750 | 7.3219 | 5.7277 | 5.8297 | 7.0439 | 6.2039 | 7.8928 | 6.8871 | 6.4908 |
| MAP1B_S339 | MAP1B | P46821 | | 1785 | SPKSDISPLTPRE | 6.5942 | 7.0271 | 6.4136 | 6.1107 | 5.7954 | 6.0265 | 6.0127 | 6.3974 | 6.7292 | 6.1400 | 6.6192 | 6.8419 | 5.0723 | 6.7166 | 5.4602 | 6.0431 |
| MAP1B_S614 | MAP1B | P46821 | | 1915 | IERTTKSPSDSGY | 6.2621 | 6.5561 | 6.9068 | 7.5074 | 5.9285 | 6.5036 | 6.4729 | 6.5417 | 6.2898 | 6.8446 | 6.9256 | 6.8838 | 4.2101 | 5.7828 | 4.6558 | 5.7284 |

| | | | | | | | | | | | | | | | | | | | | |
|---|---|---|---|---|---|---|---|---|---|---|---|---|---|---|---|---|---|---|---|---|
| MAP1B_S831;S832 | MAP1B | P46821 | 339 | EEQSQGSTTNSDW | 4.8930 | 5.5705 | 6.1999 | 6.2169 | 5.0886 | 5.7661 | 6.7640 | 6.8632 | 5.9561 | 4.7172 | 6.8094 | 5.9305 | 6.6563 | 8.1304 | 7.4274 | 7.0106 |
| MAP1B_S832 | MAP1B | P46821 | 1618;1620 | NA | 5.8864 | 5.8209 | 6.6740 | 5.6783 | 6.0102 | 6.0634 | 5.6136 | 6.0818 | 6.2542 | 6.3983 | 6.5066 | 6.3348 | 6.3288 | 7.1662 | 6.6298 | 6.5527 |
| MAP1B_S936;S937 | MAP1B | P46821 | 1396;1400 | NA | 5.5664 | 5.8592 | 6.4580 | 6.1685 | 5.9905 | 6.3672 | 6.3911 | 6.3471 | 6.2192 | 6.3922 | 6.2788 | 6.3628 | 5.6062 | 7.1516 | 6.3478 | 6.4934 |
| MAP1B_S937 | MAP1B | P46821 | 1618;1620;16 | NA | 5.9994 | 6.5394 | 7.5737 | 6.4999 | 5.7650 | 6.3583 | 6.4870 | 6.1795 | 7.2340 | 6.9259 | 6.7429 | 6.8936 | 4.8791 | 5.6315 | 4.8119 | 5.4791 |
| MAP1B_S992;S995 | MAP1B | P46821 | 1620;1625 | NA | 6.0012 | 6.3453 | 6.5491 | 5.9060 | 6.3509 | 6.3032 | 5.8747 | 5.4465 | 6.3310 | 7.1124 | 6.2976 | 6.2422 | 6.1030 | 7.3683 | 5.7977 | 5.9709 |
| MAP1B_S995 | MAP1B | P46821 | 1276;1282 | NA | 7.2267 | 8.2795 | 7.0885 | 8.2311 | 6.1666 | 7.0716 | 7.8931 | 6.5949 | 7.4888 | 7.0043 | 7.5585 | 6.8939 | 2.5175 | 4.1071 | 2.2809 | 3.5970 |
| MAP1B_T1067 | MAP1B | P46821 | 2209;2211 | NA | 6.3793 | 6.1901 | 6.0340 | 6.3407 | 6.3955 | 6.6729 | 6.0417 | 6.0522 | 6.2197 | 6.6693 | 6.7035 | 7.0731 | 5.3175 | 6.0393 | 5.7678 | 6.1033 |
| MAP1B_T1633 | MAP1B | P46821 | 1793;1797 | NA | 5.9554 | 6.3474 | 6.1888 | 5.9050 | 5.9570 | 6.6986 | 6.3213 | 6.0268 | 6.6105 | 6.4475 | 6.3006 | 6.4470 | 5.5786 | 6.3009 | 6.3340 | 6.5807 |
| MAP1B_T1788 | MAP1B | P46821 | 1396;1410 | NA | 5.1322 | 5.7416 | 6.2682 | 5.9175 | 5.8331 | 6.1914 | 6.1649 | 6.2935 | 6.3508 | 6.5732 | 6.7244 | 6.6370 | 5.6440 | 7.1264 | 6.4357 | 6.9661 |
| MAP1B_T1853 | MAP1B | P46821 | 1779;1782;17 | NA | 5.5269 | 6.5045 | 7.0182 | 6.1565 | 5.9679 | 6.4842 | 6.5476 | 5.9513 | 6.5034 | 6.8331 | 7.1063 | 6.9815 | 5.3251 | 6.3110 | 5.2838 | 5.4987 |
| MAP1B_T1963 | MAP1B | P46821 | 2051;2067 | NA | 11.3941 | 7.3292 | 7.7173 | 8.3420 | 7.1984 | 8.3530 | 8.0369 | 6.2057 | 6.1630 | 6.9733 | 5.4868 | 7.4876 | 2.7865 | 3.3221 | 1.2883 | 1.9159 |
| MAP1B_T2034 | MAP1B | P46821 | 936;937 | NA | 5.5103 | 6.9363 | 7.8869 | 6.8709 | 8.5233 | 5.9716 | 6.1447 | 6.5929 | 5.6286 | 6.4060 | 5.4223 | 4.1384 | 5.8758 | 7.6264 | 3.9797 | 6.4859 |
| MAP1B_T2051 | MAP1B | P46821 | 1396;1408 | NA | 5.9941 | 5.4894 | 6.0646 | 6.4003 | 5.9833 | 6.0996 | 5.9073 | 6.3068 | 6.4216 | 5.7098 | 6.8570 | 6.3440 | 5.7592 | 7.2040 | 6.4329 | 7.0261 |
| MAP1B_T2051;T2067 | MAP1B | P46821 | 1618;1625 | NA | 5.6480 | 5.5380 | 6.9180 | 6.1940 | 6.3613 | 6.2531 | 5.9941 | 5.9692 | 6.4017 | 7.1732 | 6.6114 | 6.4542 | 5.8450 | 6.5215 | 5.7585 | 6.3590 |
| MAP1B_T2305 | MAP1B | P46821 | 992;995 | NA | 4.8056 | 5.0858 | 6.0323 | 5.2404 | 5.8801 | 6.0742 | 5.5777 | 6.1642 | 6.5595 | 6.4731 | 7.3677 | 5.8901 | 6.2533 | 7.5991 | 7.4101 | 7.5869 |
| MAP1B_T744 | MAP1B | P46821 | 1438;1443 | NA | 5.8889 | 6.0010 | 6.4999 | 5.8847 | 5.6493 | 5.9588 | 6.0388 | 6.0538 | 6.4445 | 6.2200 | 6.2269 | 6.9047 | 6.0480 | 7.4907 | 6.2204 | 6.4696 |
| MAP1B_T948 | MAP1B | P46821 | 1298;1312 | NA | 6.2758 | 6.0776 | 6.1399 | 6.3857 | 6.3516 | 6.7303 | 6.5988 | 5.3714 | 6.0598 | 6.6828 | 6.1450 | 6.8156 | 5.3443 | 7.1247 | 5.6869 | 6.2099 |
| MAP1B_Y1887 | MAP1B | P46821 | 831;832 | NA | 6.5418 | 5.9107 | 6.3567 | 5.6888 | 6.1164 | 6.3026 | 5.7318 | 5.9571 | 6.6671 | 6.0764 | 6.2089 | 6.2932 | 6.3129 | 6.7490 | 6.5651 | 6.5215 |
| MAP1S_S546 | MAP1S | Q66K74 | 731 | RARRSASPHDVDL | 5.8984 | 5.9256 | 6.3366 | 5.8054 | 6.5471 | 6.5380 | 5.8088 | 6.6666 | 5.9815 | 6.1097 | 6.3506 | 6.1315 | 6.3359 | 6.9196 | 6.4073 | 6.2375 |
| MAP1S_S592 | MAP1S | Q66K74 | 657 | SERLSLSPLRGGE | 6.4609 | 6.2221 | 6.5351 | 5.4569 | 6.3165 | 6.2581 | 5.7246 | 6.5485 | 6.5107 | 6.0695 | 5.8344 | 5.8680 | 6.5032 | 6.8398 | 6.6060 | 6.2457 |
| MAP1S_S592;S600 | MAP1S | Q66K74 | 759 | MAPAPASPGSSND | 6.1817 | 6.2665 | 6.7914 | 6.2827 | 6.3576 | 6.2997 | 6.0638 | 6.5277 | 6.6138 | 6.6500 | 6.3859 | 6.6123 | 5.8313 | 5.7740 | 5.7236 | 5.6380 |
| MAP1S_S655;S657 | MAP1S | Q66K74 | 941 | RPGVSATPPKSPV | 6.3555 | 6.5875 | 7.0548 | 6.8603 | 6.2458 | 6.7041 | 6.7819 | 6.1069 | 6.2734 | 6.6337 | 6.9232 | 6.8085 | 5.1265 | 5.4847 | 4.7845 | 5.2687 |
| MAP1S_S657 | MAP1S | Q66K74 | 809 | APGAADSDEDTEG | 5.6087 | 5.3944 | 6.9904 | 6.1199 | 5.8250 | 6.6005 | 6.1247 | 6.3512 | 6.5389 | 6.3352 | 6.7916 | 6.5869 | 6.3643 | 6.5340 | 5.8236 | 6.0108 |
| MAP1S_S729;S731 | MAP1S | Q66K74 | 592 | LRCGEASPPSAAC | 6.0783 | 6.1424 | 6.7952 | 6.7899 | 6.0667 | 5.1583 | 6.3036 | 6.1896 | 6.2894 | 7.3643 | 6.0444 | 6.3170 | 5.4130 | 6.7890 | 5.9019 | 6.3570 |
| MAP1S_S731 | MAP1S | Q66K74 | 546 | EVRRAASSVPNLK | 7.2177 | 7.0268 | 6.3529 | 5.1433 | 5.5485 | 6.8868 | 5.7827 | 5.3340 | 6.2389 | 6.4985 | 6.2974 | 6.1785 | 6.2028 | 7.3805 | 5.7949 | 6.1156 |
| MAP1S_S759 | MAP1S | Q66K74 | 608 | ASQLVATPSLELG | 5.9953 | 6.3224 | 6.5487 | 5.7984 | 5.8143 | 6.5352 | 5.7817 | 6.7488 | 6.3661 | 5.9318 | 6.3550 | 6.4526 | 5.6745 | 6.6511 | 6.6646 | 6.3597 |
| MAP1S_S809 | MAP1S | Q66K74 | 592;600 | NA | 6.4650 | 6.5603 | 6.9935 | 6.9512 | 6.3538 | 6.5475 | 6.7851 | 6.1201 | 6.7234 | 6.8067 | 6.6618 | 7.3189 | 4.9740 | 5.4799 | 4.3781 | 4.8806 |
| MAP1S_T608 | MAP1S | Q66K74 | 655;657 | NA | 6.2860 | 6.8282 | 6.7352 | 7.0072 | 5.9004 | 6.0950 | 6.5288 | 7.2950 | 7.7722 | 7.4115 | 6.3617 | 6.9354 | 4.1443 | 4.7904 | 4.7325 | 5.1764 |
| MAP1S_T941 | MAP1S | Q66K74 | 729;731 | NA | 6.1080 | 6.2800 | 6.5208 | 6.4540 | 6.2570 | 6.8004 | 6.7688 | 6.4040 | 6.0763 | 6.5768 | 6.2955 | 7.1538 | 5.3420 | 5.7803 | 5.1300 | 6.0524 |
| MAP2_S1155 | MAP2 | P11137 | 1605 | TAITPGTPPSYSS | 5.9853 | 5.8726 | 6.1683 | 5.9089 | 6.1115 | 6.0411 | 6.0694 | 6.0123 | 6.3595 | 6.3010 | 6.2290 | 6.3333 | 6.6717 | 7.1949 | 6.2284 | 6.5128 |
| MAP2_S1480 | MAP2 | P11137 | 285 | GLVAPISPGPLTP | 5.8485 | 5.8658 | 6.7622 | 5.8376 | 6.2706 | 6.5140 | 5.9249 | 6.5334 | 6.3226 | 6.7245 | 6.7652 | 6.6372 | 6.0270 | 6.4874 | 5.9699 | 5.5092 |
| MAP2_S1555 | MAP2 | P11137 | 1155 | EGKKETSPESSLI | 5.6302 | 6.6549 | 6.8731 | 6.2263 | 6.2150 | 6.4166 | 5.8686 | 6.4421 | 6.1592 | 6.4408 | 6.7054 | 6.7380 | 5.6896 | 6.3149 | 5.9318 | 5.6936 |
| MAP2_S1782 | MAP2 | P11137 | 1649 | KVAIIRTPPKSPA | 6.7536 | 6.6722 | 6.7675 | 5.5190 | 6.7444 | 6.3903 | 5.4864 | 6.1899 | 6.1318 | 6.8108 | 6.8578 | 6.5080 | 5.8760 | 6.2162 | 5.5635 | 5.5126 |
| MAP2_S1790 | MAP2 | P11137 | 1782 | AEIITQSPGRSSV | 5.9797 | 6.6328 | 6.8203 | 5.8394 | 6.5576 | 6.6808 | 6.0678 | 6.8819 | 6.6158 | 6.6400 | 6.6488 | 6.3571 | 5.4134 | 6.0495 | 5.2865 | 5.5286 |
| MAP2_S285 | MAP2 | P11137 | 311 | EGKQFDSPMPSPF | 6.7062 | 6.2441 | 7.0650 | 5.8730 | 6.0681 | 6.2371 | 6.3206 | 7.0555 | 6.3185 | 6.3512 | 6.0916 | 5.6361 | 6.2330 | 6.4570 | 6.1693 | 5.1736 |
| MAP2_S285;T290 | MAP2 | P11137 | 1790 | GRSSVASPRRLSN | 6.2513 | 6.6811 | 6.5010 | 5.9296 | 6.9395 | 6.6165 | 5.8837 | 6.8847 | 6.3553 | 7.0853 | 6.6497 | 6.2879 | 5.4547 | 5.8486 | 5.5066 | 5.1245 |
| MAP2_S311 | MAP2 | P11137 | 821 | TRSRLASVSADAE | 5.9632 | 5.6411 | 5.7157 | 6.3919 | 6.5733 | 5.9465 | 6.4198 | 6.3492 | 6.7288 | 6.4768 | 5.8059 | 5.6008 | 6.4937 | 7.5672 | 6.1742 | 6.1519 |
| MAP2_S821 | MAP2 | P11137 | 1616 | SSRTPGTPGTPSY | 6.0353 | 6.2959 | 7.2163 | 5.9008 | 5.1909 | 6.6054 | 6.4880 | 6.7128 | 6.1979 | 6.8853 | 7.0149 | 6.3420 | 5.7715 | 6.6142 | 5.4045 | 5.3244 |
| MAP2_S881 | MAP2 | P11137 | 881 | YTVPLPSPVQDSE | 5.9362 | 6.1459 | 6.8585 | 6.2107 | 6.0801 | 6.6336 | 5.2654 | 6.3657 | 6.4297 | 7.2706 | 6.7153 | 6.4110 | 5.6384 | 6.5466 | 6.0326 | 5.4596 |
| MAP2_T1592;T1605 | MAP2 | P11137 | 1555 | PPRRGVSGDRDEN | 5.2860 | 5.6191 | 5.3842 | 4.5214 | 5.7809 | 6.3127 | 5.7422 | 6.4752 | 5.6840 | 6.1115 | 6.2648 | 6.0399 | 8.4113 | 8.5514 | 7.1997 | 6.6157 |
| MAP2_T1605 | MAP2 | P11137 | 1631 | TPHTPGTPKSAIL | 6.0980 | 6.3687 | 7.3958 | 5.8953 | 5.6176 | 7.3959 | 6.3937 | 7.1607 | 6.8897 | 7.2717 | 7.0621 | 6.7513 | 4.8271 | 5.8453 | 4.4432 | 4.5837 |
| MAP2_T1616 | MAP2 | P11137 | 1480 | TEVQAHSPSRKFI | 6.0041 | 5.9900 | 6.0867 | 5.5553 | 5.9561 | 6.7323 | 6.0874 | 6.4857 | 6.4639 | 6.5072 | 6.4771 | 6.0121 | 6.6697 | 6.9681 | 5.9908 | 6.0135 |
| MAP2_T1631 | MAP2 | P11137 | 285;290 | NA | 4.9476 | 4.6860 | 4.8979 | 4.9825 | 6.0354 | 6.7056 | 6.7520 | 7.0308 | 5.5267 | 5.2010 | 4.8204 | 4.8415 | 8.4564 | 8.9382 | 8.1831 | 7.9950 |
| MAP2_T1649 | MAP2 | P11137 | 1649;1653 | NA | 5.7631 | 5.9138 | 7.5163 | 5.9580 | 5.7499 | 7.0770 | 6.1532 | 6.5477 | 6.4589 | 6.9033 | 6.3303 | 6.7626 | 5.8521 | 6.0888 | 5.0181 | 5.9069 |
| MAP2_T1649;S1653 | MAP2 | P11137 | 1592;1605 | NA | 5.6482 | 6.3703 | 6.0524 | 5.3630 | 6.3123 | 6.1003 | 6.6234 | 5.2099 | 5.9124 | 6.2877 | 6.3164 | 6.2108 | 6.3612 | 7.8324 | 6.5039 | 6.8953 |
| MAP2K1_T286 | MAP2K1 | Q02750 | 386 | GLNQPSTPTHAAG | 5.9186 | 5.8120 | 6.8268 | 6.1523 | 6.2895 | 6.5588 | 6.5560 | 6.6999 | 6.1256 | 5.8932 | 6.0711 | 6.1373 | 6.2504 | 6.3515 | 5.9697 | 6.3873 |
| MAP2K1_T386 | MAP2K1 | Q02750 | 286 | EGDAAETPPRPRT | 5.1318 | 6.4870 | 6.1709 | 6.4794 | 6.7354 | 6.5837 | 6.2107 | 6.0082 | 5.9260 | 6.4777 | 6.8464 | 7.7702 | 5.7014 | 5.9548 | 5.9455 | 5.5709 |
| MAP2K2_S226 | MAP2K2 | P36507 | 295 | GEPHSISPRPRPP | 6.2372 | 6.4019 | 6.7225 | 6.5934 | 6.3692 | 6.7580 | 6.7639 | 7.4186 | 7.0598 | 6.6195 | 6.9625 | 7.1607 | 4.6068 | 4.9900 | 4.4520 | 4.8840 |
| MAP2K2_S23 | MAP2K2 | P36507 | 394 | RLNQPGTPTRTAV | 6.4557 | 6.4169 | 6.5966 | 5.8392 | 6.5741 | 6.8049 | 6.1720 | 7.0484 | 6.6012 | 6.1973 | 6.1803 | 6.3160 | 5.9666 | 5.7584 | 5.5942 | 5.4782 |
| MAP2K2_S295 | MAP2K2 | P36507 | 306 | PPGRPVSGHGMDS | 6.3233 | 4.6837 | 7.8438 | 6.5538 | 6.8667 | 7.2196 | 5.9412 | 6.4776 | 7.1974 | 5.9051 | 6.5687 | 6.8204 | 6.0641 | 5.2532 | 4.6868 | 5.5946 |
| MAP2K2_S306 | MAP2K2 | P36507 | 226 | IDSMANSFVGTRS | 6.3304 | 6.2219 | 6.1656 | 5.7596 | 6.6479 | 6.4389 | 6.9011 | 7.1101 | 7.6326 | 4.9518 | 5.6752 | 6.3746 | 5.8441 | 5.7717 | 6.0129 | 6.1616 |
| MAP2K2_T394 | MAP2K2 | P36507 | 23 | TIAEGPSPTSEGA | 4.9094 | 4.6270 | 6.3196 | 5.6256 | 6.2296 | 6.9363 | 7.0537 | 5.9282 | 6.0043 | 5.6055 | 5.7039 | 6.1894 | 7.6736 | 7.3520 | 6.7966 | 7.0452 |
| MAP2K3_S223 | MAP2K3 | P46734-3 | 223 | SGYLVDSVAKTMD | 5.3464 | 5.0149 | 5.3801 | 5.2510 | 6.2142 | 5.6128 | 5.3678 | 5.2591 | 4.8478 | 4.8150 | 4.8670 | 4.3537 | 10.0859 | 9.5405 | 9.6134 | 8.4303 |
| MAP2K4_S101 | MAP2K4 | P45985-2 | 101 | SGKLKISPEQHWD | 5.6830 | 5.4825 | 6.3240 | 5.8247 | 6.3798 | 6.4747 | 6.0925 | 6.2444 | 6.5301 | 5.7068 | 6.9510 | 6.0010 | 6.7365 | 6.9647 | 6.3253 | 6.2789 |
| MAP2K4_S268 | MAP2K4 | P45985-2 | 268 | SGQLVDSIAKTRD | 6.1654 | 6.4411 | 6.7283 | 6.7084 | 6.8536 | 6.0846 | 6.0289 | 6.5342 | 6.8214 | 5.7675 | 5.6692 | 6.3448 | 6.1502 | 6.1224 | 5.7739 | 5.8061 |
| MAP2K7_S35 | MAP2K7 | O14733-2 | 35 | DLNLDISPQRPRP | 6.2792 | 5.9857 | 7.3722 | 6.3133 | 5.7670 | 6.3317 | 6.4086 | 7.3559 | 7.9281 | 6.5445 | 7.2504 | 7.3158 | 4.3125 | 4.9545 | 4.7192 | 5.1613 |
| MAP2K7_T83 | MAP2K7 | O14733-2 | 83 | LPSTLFTPRSMES | 5.0934 | 7.8857 | 7.3150 | 5.5738 | 6.2891 | 7.9002 | 5.8158 | 8.0049 | 7.3113 | 8.9766 | 7.5425 | 6.2927 | 3.8652 | 5.3462 | 3.6195 | 3.1680 |
| MAP3K1_S1157 | MAP3K1 | Q13233 | 1455 | WNAEKHSNHLALI | 7.4300 | 7.0540 | 7.4349 | 7.1982 | 6.6416 | 6.3396 | 6.5000 | 6.3961 | 7.3147 | 6.7871 | 7.4395 | 7.1602 | 4.2465 | 3.9275 | 4.0928 | 4.0371 |
| MAP3K1_S1455 | MAP3K1 | Q13233 | 275 | VRRKRVSPVPFQS | 6.2243 | 6.6681 | 6.4107 | 6.9757 | 6.5068 | 6.2793 | 5.8668 | 6.5571 | 7.3385 | 6.9299 | 6.6115 | 7.1205 | 4.9764 | 5.5771 | 4.8921 | 5.0653 |
| MAP3K1_S275 | MAP3K1 | Q13233 | 292 | PPRRAPSPDGFSP | 6.2835 | 5.9648 | 7.1124 | 6.5195 | 6.1839 | 6.7531 | 6.5670 | 6.4498 | 6.9232 | 6.8788 | 6.5118 | 6.1939 | 5.2018 | 5.5925 | 5.7005 | 5.1635 |
| MAP3K1_S292 | MAP3K1 | Q13233 | 1157 | SEVAVLSPEKAEN | 5.8744 | 6.2864 | 7.5593 | 6.4990 | 5.6614 | 6.3337 | 6.1321 | 6.0922 | 7.1328 | 6.9842 | 6.6954 | 7.7035 | 4.8904 | 5.4746 | 4.6440 | 6.0366 |
| MAP3K1_S923 | MAP3K1 | Q13233 | 923 | ATKLSASSEDISE | 5.9693 | 6.4004 | 6.5067 | 5.7331 | 6.0822 | 5.8268 | 6.0534 | 6.1178 | 6.8020 | 6.5935 | 6.8760 | 6.8893 | 5.4828 | 6.3555 | 6.0636 | 6.2477 |
| MAP3K11_S507 | MAP3K11 | Q16584 | 507 | RITVQASPGLDRR | 6.4650 | 6.6060 | 6.5967 | 6.4779 | 5.9389 | 6.7214 | 6.6773 | 6.7798 | 6.3096 | 6.4333 | 6.5094 | 6.4231 | 5.1905 | 5.7949 | 5.3259 | 5.7502 |
| MAP3K11_S570 | MAP3K11 | Q16584 | 758 | TPGTPRSPPLGLI | 6.4600 | 6.4738 | 6.3890 | 5.8351 | 6.6216 | 5.7323 | 5.6826 | 6.9831 | 8.3512 | 6.0444 | 5.5437 | 5.7842 | 6.4450 | 6.1357 | 5.9699 | 5.5484 |
| MAP3K11_S705 | MAP3K11 | Q16584 | 748 | PPGTSRSAPGTPG | 6.3170 | 7.1979 | 6.6863 | 6.4719 | 6.0931 | 6.5313 | 5.5191 | 6.5187 | 6.6581 | 6.6686 | 6.1183 | 6.4186 | 5.1680 | 6.7109 | 5.4583 | 5.4638 |
| MAP3K11_S748 | MAP3K11 | Q16584 | 793 | RPSPLPSPQPAPR | 7.1587 | 6.5481 | 7.4949 | 6.4419 | 5.8756 | 4.7031 | 5.2562 | 7.1542 | 4.0793 | 7.4462 | 4.9951 | 7.0879 | 5.2783 | 7.5082 | 7.0525 | 5.9198 |
| MAP3K11_S758 | MAP3K11 | Q16584 | 570 | WAWGPSSPKPGEA | 6.3869 | 7.3210 | 6.7945 | 5.9537 | 5.4738 | 6.9570 | 5.8965 | 6.7253 | 5.5385 | 7.3014 | 6.9660 | 6.5193 | 6.5238 | 4.9985 | 5.4874 | 5.1564 |
| MAP3K11_S793 | MAP3K11 | Q16584 | 705 | SLKTPDSPPTPAP | 6.1564 | 6.0296 | 6.3052 | 5.4999 | 6.7272 | 5.5150 | 6.2900 | 7.0847 | 6.7720 | 5.9272 | 5.8240 | 6.7917 | 6.0509 | 5.9333 | 6.3758 | 6.7173 |
| MAP3K12_S728 | MAP3K12 | Q12852-2 | 728 | GGSRAGSQHLTPA | 7.7515 | 7.3104 | 4.7606 | 5.2448 | 7.2673 | 8.5853 | 5.1748 | 10.2743 | 6.4228 | 9.2361 | 4.8359 | 8.8865 | 2.8456 | 5.2448 | 3.9430 | 2.2162 |
| MAP3K14_S733 | MAP3K14 | Q99558 | 733 | SPPLTLSKEESGM | 6.8558 | 8.7981 | 9.9139 | 9.3902 | 5.8584 | 6.1768 | 6.1400 | 6.7847 | 6.2479 | 9.8271 | 11.3825 | 8.2770 | 0.3316 | 1.5133 | 1.0738 | 1.4291 |
| MAP3K15_S994 | MAP3K15 | Q6ZN16 | 994 | DRGLASSPEDRDQ | 6.2188 | 6.6608 | 7.3365 | 6.3229 | 6.8062 | 7.0586 | 5.7350 | 6.4481 | 6.8054 | 7.3349 | 6.7037 | 7.6100 | 4.4694 | 5.0346 | 4.8733 | 4.5820 |
| MAP3K2_S153 | MAP3K2 | Q9Y2U5 | 153 | ERKKRLSIIGPTS | 6.3320 | 5.5459 | 7.1465 | 5.8210 | 6.5158 | 5.9849 | 5.2681 | 6.4295 | 7.7761 | 7.3751 | 7.0501 | 7.3159 | 5.9019 | 6.2669 | 4.6868 | 4.5835 |
| MAP3K2_S164 | MAP3K2 | Q9Y2U5 | 164 | TSRDRSSPPPGYI | 6.1067 | 6.2431 | 7.0637 | 6.2553 | 6.0956 | 6.5332 | 6.4606 | 6.6364 | 6.4097 | 6.8192 | 6.5174 | 6.2929 | 5.4923 | 6.1214 | 5.5490 | 5.4035 |
| MAP3K2_S239 | MAP3K2 | Q9Y2U5 | 239 | RMPRAQSYPDNHQ | 5.8643 | 5.7372 | 6.0802 | 5.5771 | 6.2665 | 6.6630 | 5.7669 | 6.8078 | 5.6361 | 5.8751 | 5.6999 | 5.7726 | 7.3631 | 6.9579 | 7.2095 | 6.7230 |
| MAP3K2_S331 | MAP3K2 | Q9Y2U5 | 331 | RIRRRGSDIDNPT | 6.5748 | 5.7515 | 5.6756 | 6.1688 | 6.7342 | 5.7542 | 6.0488 | 6.4316 | 6.6922 | 7.3410 | 7.0287 | 6.0358 | 6.4472 | 6.1683 | 6.0294 | 5.1178 |
| MAP3K20_S599 | MAP3K20 | Q9NYL2 | 727 | ALNPHQSPDFKRS | 5.7115 | 6.0732 | 6.6821 | 6.3181 | 6.4185 | 6.6532 | 6.6374 | 6.7636 | 6.3109 | 6.4455 | 7.1364 | 6.9528 | 5.2169 | 6.2761 | 5.1801 | 5.2237 |
| MAP3K20_S633 | MAP3K20 | Q9NYL2 | 599 | SNPILGSPFFSHF | 6.2611 | 6.1709 | 6.1516 | 6.2645 | 6.3210 | 6.9950 | 6.8717 | 6.6232 | 6.8270 | 5.2290 | 5.3612 | 6.1743 | 6.4490 | 6.9206 | 5.5044 | 5.8755 |
| MAP3K20_S727 | MAP3K20 | Q9NYL2 | 633 | ITPVNQSRSSSPT | 6.8139 | 7.2417 | 6.5532 | 6.3189 | 6.0269 | 5.8564 | 5.6726 | 5.8956 | 6.6562 | 6.0933 | 6.6486 | 5.9903 | 6.3038 | 6.2182 | 5.2399 | 6.4706 |
| MAP3K20_T226 | MAP3K20 | Q9NYL2 | 226 | EKNERLTIPSSCP | 5.8899 | 6.7724 | 6.2490 | 5.7006 | 6.2964 | 5.5814 | 6.1900 | 7.9790 | 6.4406 | 5.9333 | 6.5425 | 6.6828 | 6.0374 | 6.3877 | 5.9795 | 5.3375 |
| MAP3K21_T592 | MAP3K21 | Q5TCX8 | 592 | FKKKGCTWGPNSI | 6.0884 | 5.1980 | 7.6600 | 7.1658 | 6.9940 | 6.2451 | 6.1662 | 5.9449 | 5.9277 | 6.0631 | 6.3734 | 5.9176 | 6.1683 | 5.1192 | 6.5938 | 6.3744 |
| MAP3K3_S197 | MAP3K3 | Q99759-2 | 197 | PRSRHLSVSSQNP | 6.6248 | 5.4204 | 6.2897 | 5.4099 | 5.0947 | 6.3570 | 6.6253 | 7.5425 | 7.1755 | 6.7862 | 6.9819 | 6.5287 | 5.8650 | 6.4715 | 5.3885 | 5.4383 |
| MAP3K3_S347 | MAP3K3 | Q99759-2 | 386 | RNVPTKSPSAPIN | 7.4116 | 7.2155 | 7.3628 | 6.8229 | 6.3971 | 6.7551 | 6.3269 | 6.9670 | 7.2747 | 6.9647 | 7.0614 | 7.2365 | 3.7431 | 4.3960 | 3.8241 | 4.2406 |
| MAP3K3_S386 | MAP3K3 | Q99759-2 | 347 | PSSRSLSTNGENM | 6.1274 | 6.1348 | 6.4098 | 6.3097 | 5.7683 | 5.8362 | 6.4938 | 5.8667 | 7.1229 | 6.3924 | 6.7433 | 6.8466 | 6.5374 | 5.9494 | 5.5344 | 5.9270 |
| MAP3K4_S1135 | MAP3K4 | J3KNB8 | 447 | EYEGDDTEGELKE | 6.3450 | 6.3044 | 6.9354 | 6.2721 | 6.3937 | 6.6584 | 6.5091 | 6.1546 | 6.1763 | 5.2770 | 5.8859 | 5.9427 | 6.5219 | 6.3885 | 5.8729 | 6.3621 |
| MAP3K4_S1155 | MAP3K4 | J3KNB8 | 1155 | KVPRCHSDPPNPH | 6.2006 | 5.9107 | 6.4814 | 5.8661 | 5.9129 | 6.2383 | 6.3191 | 5.7076 | 6.7086 | 5.6185 | 5.9688 | 6.2224 | 6.9630 | 7.0475 | 6.4263 | 6.4081 |
| MAP3K4_S431 | MAP3K4 | J3KNB8 | 499 | KLERLESEDDSLG | 5.5130 | 5.1633 | 7.3487 | 5.6693 | 5.7115 | 6.3701 | 4.9673 | 5.9261 | 6.4929 | 4.8619 | 5.4659 | 6.3986 | 8.2739 | 7.3896 | 7.6637 | 6.7843 |
| MAP3K4_S499 | MAP3K4 | J3KNB8 | 1135 | VIGKPHSPVTAIH | 5.6200 | 5.3759 | 7.2545 | 6.6388 | 5.6562 | 6.1487 | 5.5871 | 5.7640 | 6.9394 | 6.3759 | 6.2681 | 6.7103 | 6.2024 | 6.0733 | 6.3382 | 7.0472 |
| MAP3K4_T447 | MAP3K4 | J3KNB8 | 431 | PVFEIPSPRPSKG | 6.0816 | 6.4552 | 7.2286 | 6.3049 | 5.9173 | 7.2204 | 6.1766 | 6.6382 | 5.8221 | 6.8391 | 6.7629 | 6.4640 | 5.3045 | 6.2043 | 5.2990 | 5.2815 |
| MAP3K7_S367 | MAP3K7 | O43318 | 367 | SESGRLSLGASRG | 7.1838 | 7.7309 | 7.0382 | 7.4201 | 6.2638 | 6.6768 | 7.2174 | 7.2495 | 8.0939 | 7.6934 | 7.5006 | 7.5373 | 2.7557 | 3.6638 | 2.5959 | 3.3791 |
| MAP3K7_S389 | MAP3K7 | O43318 | 389 | SEGKRMSADMSEI | 5.8011 | 6.0429 | 6.7319 | 6.4253 | 6.2160 | 6.3076 | 6.2577 | 6.7981 | 6.9146 | 6.3019 | 6.7266 | 6.9686 | 5.6247 | 5.6100 | 5.3559 | 5.9170 |
| MAP3K7_S417 | MAP3K7 | O43318 | 439 | GQPRRRSIQDLTV | 6.6043 | 6.4743 | 6.5917 | 5.6729 | 6.7848 | 5.5896 | 6.2509 | 6.8050 | 6.8428 | 6.4857 | 6.4769 | 6.2875 | 5.3789 | 5.5051 | 5.2691 | 6.9804 |
| MAP3K7_S439 | MAP3K7 | O43318 | 417 | GHRKTASFGNILD | 6.3432 | 5.9578 | 6.6581 | 6.6327 | 5.8331 | 6.4760 | 6.2940 | 6.2243 | 6.1332 | 6.9738 | 6.3752 | 5.9750 | 5.9873 | 6.2792 | 6.0037 | 5.8536 |
| MAP3K7_S439;S455 | MAP3K7 | O43318 | 439;455 | NA | 6.6157 | 6.4431 | 7.4739 | 6.2131 | 6.6384 | 5.7874 | 6.9607 | 5.3738 | 6.2313 | 6.4155 | 6.2640 | 5.5928 | 7.1773 | 5.7007 | 5.8995 | 5.2127 |
| MAP3K9_S519 | MAP3K9 | P80192-4 | 519 | KDGNRISLPSDFQ | 6.2268 | 5.0863 | 7.1914 | 6.4905 | 6.7993 | 6.6616 | 6.2711 | 6.8227 | 6.5947 | 6.9261 | 6.5652 | 6.3411 | 5.7732 | 5.3570 | 5.3866 | 5.5064 |
| MAP3K9_S533 | MAP3K9 | P80192-4 | 533 | KFTVQASPTMDKR | 6.2349 | 6.6016 | 6.4404 | 6.3597 | 8.0564 | 6.6516 | 5.8612 | 6.4314 | 5.3498 | 6.2669 | 6.2247 | 6.2298 | 5.7466 | 6.0717 | 5.6096 | 5.8637 |
| MAP4_S1000 | MAP4 | P27816 | 280 | LAKDMESPTKLDV | 6.2966 | 6.4909 | 6.6501 | 5.9428 | 6.1118 | 6.4309 | 6.1234 | 6.5966 | 6.4010 | 6.6020 | 6.6459 | 6.6845 | 5.4980 | 6.2090 | 5.5862 | 5.7302 |
| MAP4_S1073 | MAP4 | P27816 | 687 | TGNDITTPPNKEL | 5.7578 | 6.1842 | 6.6773 | 6.6579 | 6.2201 | 6.7845 | 6.2230 | 6.6309 | 6.5385 | 7.1397 | 7.0823 | 7.1447 | 4.8341 | 5.8763 | 4.8244 | 5.4242 |
| MAP4_S116 | MAP4 | P27816 | 507 | GLLKDMSPLSETE | 5.7551 | 5.9203 | 6.5219 | 5.8920 | 6.1436 | 6.1150 | 6.0793 | 6.2504 | 6.5286 | 6.5728 | 6.6084 | 6.3093 | 6.0864 | 6.9109 | 6.1321 | 6.1738 |
| MAP4_S156 | MAP4 | P27816 | 696 | NKELPPSPEKKTK | 5.7543 | 5.7674 | 6.5347 | 5.7513 | 6.1660 | 6.5581 | 6.1827 | 6.5342 | 6.4995 | 6.3448 | 6.7120 | 6.8114 | 5.9492 | 6.5250 | 5.7505 | 6.1590 |
| MAP4_S1679 | MAP4 | P27816 | 521 | ALGKDVTPPPETE | 6.1568 | 5.9756 | 6.4653 | 5.6866 | 5.9875 | 6.3225 | 5.8455 | 6.3350 | 6.3176 | 6.6836 | 6.5712 | 6.5566 | 5.9929 | 6.9156 | 6.0059 | 6.1820 |
| MAP4_S280 | MAP4 | P27816 | 868 | KSTSTSSMKKTTT | 8.0313 | 8.3327 | 8.5046 | 7.1028 | 4.9552 | 6.1420 | 5.7327 | 7.3306 | 7.5299 | 7.9540 | 7.3358 | 7.3885 | 3.2122 | 4.6100 | 2.8398 | 2.9979 |
| MAP4_S358 | MAP4 | P27816 | 585 | LSETEATPVPIKD | 7.3848 | 7.7278 | 6.6490 | 7.7706 | 6.4182 | 7.0985 | 7.4118 | 6.6338 | 7.6151 | 6.9464 | 7.2489 | 7.0199 | 3.1602 | 4.2231 | 2.9758 | 3.8160 |
| MAP4_S507 | MAP4 | P27816 | 358 | KETERASPIKMDL | 6.6452 | 7.0730 | 6.9791 | 6.6144 | 6.2067 | 6.8228 | 6.4270 | 6.5145 | 6.5924 | 6.9212 | 7.2013 | 7.4243 | 4.2491 | 5.3239 | 4.2134 | 4.7918 |
| MAP4_S507;S510 | MAP4 | P27816 | 99 | GSDTTGSPTEFLE | 5.9560 | 5.9339 | 6.4110 | 5.9237 | 6.1177 | 6.4496 | 5.5184 | 6.5148 | 6.0352 | 6.6028 | 6.2821 | 6.3720 | 6.2933 | 7.0383 | 6.2383 | 6.3131 |
| MAP4_S580 | MAP4 | P27816 | 941 | VRSKVGSTENIKH | 5.7878 | 5.4781 | 6.2884 | 4.6191 | 5.7064 | 5.8674 | 5.9486 | 5.5258 | 5.9505 | 6.1562 | 5.6200 | 5.4138 | 8.1891 | 8.2777 | 7.5579 | 7.6131 |
| MAP4_S580;T585 | MAP4 | P27816 | 825 | TGPSSRSPSTLLP | 6.7298 | 7.5226 | 6.9489 | 7.2638 | 6.3613 | 7.0792 | 7.0748 | 7.1160 | 6.8596 | 7.1433 | 7.3162 | 7.2927 | 3.2122 | 4.4675 | 3.4739 | 4.1382 |
| MAP4_S585 | MAP4 | P27816 | 925 | RLSRLATNTSAPD | 2.6649 | 2.6731 | 2.4618 | 2.4064 | 2.7770 | 3.3166 | 2.8753 | 3.1091 | 2.7192 | 2.7062 | 2.4682 | 2.4594 | 15.3736 | 17.5785 | 17.4729 | 16.9378 |
| MAP4_S605;S624 | MAP4 | P27816 | 636 | GTGKKCSLPAEED | 6.2336 | 6.7915 | 6.8077 | 7.4872 | 5.6010 | 6.4886 | 6.5152 | 6.4716 | 6.8386 | 7.3074 | 7.0484 | 7.5407 | 4.2542 | 5.1977 | 4.2371 | 5.1794 |
| MAP4_S624 | MAP4 | P27816 | 571 | TLANNVTPAKDVP | 6.5778 | 7.0209 | 6.9731 | 7.2379 | 6.3127 | 6.8734 | 7.1425 | 6.7466 | 7.4343 | 7.2911 | 7.6779 | 7.4123 | 3.4376 | 4.4755 | 3.0833 | 4.3029 |
| MAP4_S636 | MAP4 | P27816 | 629 | ISPETVTGTGKKC | 6.1772 | 7.3428 | 6.8233 | 7.4111 | 6.1509 | 7.0107 | 6.8094 | 6.9413 | 6.8477 | 6.7502 | 7.2344 | 6.9103 | 3.9295 | 5.1246 | 3.9265 | 4.6101 |
| MAP4_S696 | MAP4 | P27816 | 1073 | AQAKVGSLDNVGH | 5.8179 | 5.4395 | 6.5109 | 5.5779 | 5.8447 | 6.0019 | 6.0281 | 5.5857 | 6.2795 | 6.2519 | 6.1711 | 6.2319 | 7.0558 | 7.3978 | 7.0304 | 6.7750 |
| MAP4_S723 | MAP4 | P27816 | 723 | AKTQPTSLPKQPA | 6.7362 | 7.3060 | 6.8414 | 7.1864 | 6.0454 | 7.5972 | 7.9628 | 7.2034 | 7.1773 | 6.8813 | 7.3534 | 6.8638 | 3.2270 | 4.3414 | 2.9986 | 4.2784 |
| MAP4_S723 | MAP4 | P27816 | 624 | APTFMISPETVTG | 5.6871 | 6.9636 | 6.6231 | 7.5107 | 6.5782 | 6.3514 | 6.7721 | 6.8140 | 6.5561 | 6.6289 | 7.0127 | 7.3694 | 4.3411 | 5.3551 | 4.3679 | 5.0686 |
| MAP4_S825 | MAP4 | P27816 | 580 | KDVPPLSETEATP | 6.2402 | 7.5119 | 7.2673 | 8.3069 | 6.4741 | 7.3296 | 7.1535 | 6.8464 | 6.4898 | 6.9434 | 6.3870 | 6.9954 | 3.5815 | 4.3729 | 3.7982 | 4.3019 |
| MAP4_S859 | MAP4 | P27816 | 116 | YQEYPNSQNWPED | 5.2421 | 6.4272 | 6.3676 | 8.3789 | 9.5010 | 6.0530 | 6.0367 | 6.0108 | 6.1572 | 6.1477 | 5.9955 | 5.5492 | 5.2884 | 5.5897 | 5.5866 | 5.6683 |
| MAP4_S868 | MAP4 | P27816 | 631 | PETVTGTGKKCSL | 6.1934 | 7.0017 | 7.2425 | 7.3888 | 5.7637 | 7.0557 | 6.3263 | 7.6981 | 6.5773 | 7.3193 | 7.5479 | 7.3748 | 3.6404 | 4.8888 | 3.6620 | 4.3193 |
| MAP4_S941 | MAP4 | E7EVA0 | 723 | PELLGGSPPWKTL | 6.5819 | 6.0805 | 7.3233 | 5.8461 | 5.8873 | 5.8625 | 6.2047 | 6.0669 | 7.0528 | 6.8078 | 7.0376 | 7.7502 | 5.0770 | 5.7281 | 5.7592 | 4.9342 |
| MAP4_S99 | MAP4 | P27816 | 28 | KRDFIATLEAEAF | 5.3722 | 6.0517 | 5.2051 | 9.3790 | 13.5030 | 5.7698 | 5.7801 | 6.1357 | 6.8382 | 5.6066 | 6.4969 | 5.9124 | 4.4297 | 4.9477 | 4.8709 | 3.7011 |
| MAP4_T1462 | MAP4 | P27816 | 859 | SVPADLSRPKSTS | 5.3391 | 5.8683 | 6.3951 | 5.3188 | 6.1824 | 6.4705 | 5.5422 | 6.0209 | 6.1398 | 6.1144 | 5.3956 | 5.2998 | 6.8540 | 8.3364 | 7.1246 | 7.5982 |
| MAP4_T28 | MAP4 | P27816 | 1000 | IVSKKVSYSHIQS | 7.8948 | 6.7027 | 6.8349 | 5.8770 | 5.9550 | 6.0749 | 5.6839 | 6.1862 | 7.3840 | 6.8076 | 6.1164 | 6.7727 | 5.5587 | 6.6893 | 4.4331 | 5.0286 |
| MAP4_T521 | MAP4 | P27816 | 864 | LSRPKSTSTSSMK | 7.8706 | 6.3573 | 8.2975 | 6.9735 | 6.0111 | 7.6341 | 5.8351 | 7.5642 | 7.5537 | 7.0049 | 8.7260 | 7.6043 | 3.2967 | 4.4760 | 2.3769 | 2.4180 |
| MAP4_T571 | MAP4 | P27816 | 156 | ADLVFPSSATADT | 7.0930 | 6.8306 | 7.9300 | 7.9433 | 5.9172 | 5.8956 | 6.4038 | 6.3573 | 6.2311 | 4.4558 | 7.3023 | 6.2112 | 4.5621 | 6.2992 | 5.1616 | 5.4057 |
| MAP4_T629 | MAP4 | E7EVA0 | 1679 | VDLTLLSPKSEND | 6.6905 | 7.2483 | 7.3995 | 6.3898 | 5.9438 | 6.4217 | 6.5301 | 6.1722 | 6.5796 | 6.0513 | 7.4448 | 6.7880 | 4.3528 | 5.4473 | 4.7636 | 5.7766 |
| MAP4_T631 | MAP4 | E7EVA0 | 1462 | ACEKLPTPTPQVV | 6.3221 | 6.3084 | 6.5092 | 6.8260 | 6.4081 | 6.1541 | 6.6713 | 6.8344 | 6.4537 | 5.9207 | 6.3492 | 6.7288 | 5.1045 | 6.4088 | 5.3484 | 5.6523 |
| MAP4_T687 | MAP4 | P27816 | 94;99 | NA | 5.6560 | 5.6512 | 6.3039 | 5.8382 | 6.2273 | 6.3352 | 5.5441 | 6.1744 | 5.7877 | 6.4153 | 6.4651 | 6.2357 | 6.4672 | 7.6096 | 6.5015 | 6.7873 |
| MAP4_T76;S99 | MAP4 | P27816 | 76;99 | NA | 7.4086 | 7.5754 | 7.0394 | 7.9865 | 6.9565 | 7.8507 | 7.5351 | 6.5739 | 6.5964 | 6.6930 | 7.3237 | 6.8016 | 3.1840 | 4.3285 | 2.9494 | 3.1975 |
| MAP4_T864 | MAP4 | P27816 | 507;510 | NA | 6.3331 | 6.1039 | 6.6207 | 6.0916 | 6.1099 | 6.3798 | 6.0503 | 6.8231 | 5.6985 | 7.0124 | 6.3244 | 6.6024 | 6.1836 | 6.2271 | 5.3645 | 6.0747 |
| MAP4_T925 | MAP4 | P27816 | 580;585 | NA | 7.1299 | 7.8880 | 6.8201 | 7.2128 | 6.5030 | 6.8838 | 6.9745 | 6.5059 | 7.0793 | 6.2751 | 6.2539 | 7.2910 | 3.9102 | 4.9844 | 3.6090 | 4.6791 |
| MAP4_T94;T99 | MAP4 | P27816 | 605;624 | NA | 6.1624 | 7.3637 | 7.2327 | 6.5067 | 5.0117 | 7.8637 | 7.3804 | 6.3352 | 6.5465 | 6.1709 | 6.9593 | 6.0028 | 4.6678 | 6.4083 | 4.0341 | 5.3538 |
| MAP4K2_S328 | MAP4K2 | Q12851 | 328 | PAERTPSEIQFHQ | 4.8996 | 4.7783 | 5.5409 | 5.5065 | 6.1015 | 5.2178 | 5.3186 | 5.8533 | 5.5667 | 5.0578 | 5.7290 | 5.4918 | 8.6628 | 9.2860 | 8.8896 | 8.0997 |
| MAP4K3_S280 | MAP4K3 | Q8IVH8 | 467 | RCPMSGSPAKPSQ | 6.5315 | 7.0253 | 7.1653 | 6.1583 | 6.2103 | 6.2617 | 6.2570 | 6.6397 | 7.2096 | 7.0649 | 6.9630 | 6.8007 | 4.6944 | 5.0326 | 4.8252 | 5.1605 |
| MAP4K3_S329 | MAP4K3 | Q8IVH8 | 280 | TQHLTRSLAIELL | 6.6788 | 6.5971 | 6.3019 | 6.7891 | 6.2669 | 6.2386 | 6.0511 | 6.4549 | 6.3068 | 6.2996 | 5.7473 | 6.2404 | 6.0206 | 6.0722 | 6.0708 | 5.8640 |
| MAP4K3_S467 | MAP4K3 | Q8IVH8 | 329 | REEKTRSEITFGQ | 7.6445 | 6.2913 | 6.3274 | 6.1789 | 5.8265 | 5.5148 | 5.8614 | 5.7559 | 6.7477 | 4.8785 | 5.3698 | 5.4587 | 7.9376 | 7.0565 | 6.4407 | 6.7098 |
| MAP4K4_S264 | MAP4K4 | O95819-3 | 708 | VRTTSRSPVLSRR | 6.6214 | 6.9482 | 6.6382 | 6.2134 | 6.2049 | 6.5344 | 6.1549 | 5.7354 | 6.2002 | 6.5437 | 6.6112 | 6.9051 | 5.7101 | 6.1183 | 5.3840 | 5.4765 |
| MAP4K4_S324;S326 | MAP4K4 | O95819-3 | 933 | TLQKHKSSSSFTP | 5.5940 | 5.4224 | 5.9131 | 5.3535 | 5.2000 | 5.9206 | 5.3516 | 5.6079 | 5.3152 | 5.9855 | 5.7972 | 6.2524 | 7.5780 | 8.2038 | 7.8719 | 8.6329 |
| MAP4K4_S580 | MAP4K4 | O95819-3 | 886 | ASSLNLSNGETES | 6.0773 | 6.4021 | 6.0580 | 6.7699 | 6.7007 | 6.8768 | 6.5731 | 6.7178 | 7.0044 | 7.0694 | 7.0742 | 6.9351 | 4.7952 | 5.2042 | 4.4365 | 5.3053 |
| MAP4K4_S643 | MAP4K4 | O95819-3 | 923 | IVRQTQSASSTLQ | 6.1483 | 6.9197 | 6.7759 | 6.4512 | 5.5823 | 6.7453 | 6.4281 | 6.6739 | 6.7683 | 6.7224 | 7.0858 | 6.9656 | 4.9892 | 5.4211 | 4.9351 | 5.3879 |
| MAP4K4_S648 | MAP4K4 | O95819-3 | 981 | DPTRKGSVVNVNP | 6.4899 | 5.0287 | 6.0838 | 5.6361 | 6.9489 | 5.4652 | 5.5991 | 5.7275 | 8.2149 | 7.3141 | 6.8900 | 6.6120 | 6.8697 | 7.0158 | 4.9061 | 5.1983 |
| MAP4K4_S706;S708 | MAP4K4 | O95819-3 | 927 | TQSASSTLQKHKS | 10.1209 | 7.7081 | 7.0757 | 6.4603 | 6.1158 | 5.7722 | 6.0864 | 4.8855 | 5.6994 | 4.5231 | 5.4605 | 5.4242 | 7.2026 | 6.2712 | 5.4993 | 5.6948 |
| MAP4K4_S708 | MAP4K4 | O95819-3 | 881 | EDTRAASSLNLSN | 6.1877 | 6.0255 | 6.3883 | 5.8002 | 5.5891 | 5.2189 | 6.1142 | 5.8212 | 5.9266 | 5.6637 | 6.7203 | 6.2314 | 6.5329 | 7.2995 | 7.0309 | 7.4496 |
| MAP4K4_S716 | MAP4K4 | O95819-3 | 264 | WSKKFFSFIEGCL | 7.4802 | 6.0703 | 7.2687 | 7.6079 | 6.9795 | 6.0107 | 5.1866 | 5.9937 | 5.7895 | 5.3458 | 5.4613 | 5.4370 | 7.3149 | 6.0703 | 6.2769 | 5.7068 |
| MAP4K4_S733 | MAP4K4 | O95819-3 | 915 | TPSKEGTLIVRQT | 10.2947 | 8.1949 | 6.7717 | 7.8578 | 5.9137 | 4.5221 | 5.9672 | 4.6742 | 6.1450 | 5.0373 | 5.2231 | 5.4292 | 6.6539 | 6.1845 | 5.6664 | 5.4643 |
| MAP4K4_S881 | MAP4K4 | O95819-3 | 716 | VLSRRDSPLQGSG | 6.8194 | 8.4206 | 7.0722 | 5.9800 | 6.4078 | 6.4791 | 7.3283 | 8.0219 | 3.9737 | 6.9718 | 6.7514 | 7.9733 | 4.1844 | 5.2345 | 4.5150 | 3.8667 |
| MAP4K4_S886 | MAP4K4 | O95819-3 | 911 | EPAMTPSKEGTLI | 5.4899 | 7.5660 | 7.1521 | 6.6150 | 6.1982 | 7.4450 | 6.7523 | 5.8020 | 6.1819 | 7.7593 | 7.0702 | 8.2389 | 4.1906 | 4.4216 | 4.2261 | 4.8908 |
| MAP4K4_S911 | MAP4K4 | O95819-3 | 949 | PRLLQISPSSGTT | 6.2705 | 6.9527 | 6.5676 | 6.6204 | 6.2223 | 6.5519 | 6.7759 | 6.7842 | 6.4667 | 5.5512 | 6.7833 | 6.4519 | 5.5522 | 6.4380 | 4.8264 | 5.1847 |
| MAP4K4_S923 | MAP4K4 | O95819-3 | 996 | TRPQSDTPEIRKY | 5.6828 | 7.3147 | 6.5111 | 7.2478 | 6.7548 | 6.9364 | 7.6612 | 6.4506 | 6.7524 | 5.9153 | 6.5493 | 7.6795 | 3.5275 | 5.6318 | 4.4833 | 4.9014 |
| MAP4K4_S933 | MAP4K4 | O95819-3 | 936 | KHKSSSSFTPFID | 5.0868 | 4.8522 | 5.8677 | 6.4282 | 5.2818 | 5.0020 | 5.5667 | 6.3235 | 5.0629 | 5.3911 | 5.6824 | 5.8054 | 8.2723 | 8.0622 | 8.9209 | 8.3940 |
| MAP4K4_S933;S936 | MAP4K4 | O95819-3 | 909 | ESEPAMTPSKEGT | 6.3778 | 7.5217 | 7.3660 | 6.9422 | 6.0730 | 6.3276 | 7.2152 | 7.7411 | 7.5552 | 7.0896 | 7.0595 | 7.3794 | 2.1840 | 4.5673 | 3.6512 | 4.9492 |
| MAP4K4_S936 | MAP4K4 | O95819-3 | 580 | RKTNHSSPEAQSK | 6.1519 | 7.0025 | 6.2168 | 6.7208 | 6.0042 | 6.0637 | 6.5871 | 6.4280 | 6.3037 | 6.6449 | 6.8882 | 7.3872 | 5.0373 | 5.9872 | 4.8779 | 5.6984 |

| | | | | | | | | | | | | | | | | | | | | | |
|---|---|---|---|---|---|---|---|---|---|---|---|---|---|---|---|---|---|---|---|---|---|
| MAP4K4_S949 | MAP4K4 | O95819-3 | | 643 | PVSRSHSFSDPSP | 6.0440 | 6.2506 | 6.7719 | 6.9461 | 5.9600 | 6.1568 | 6.4704 | 6.7209 | 6.9316 | 6.5435 | 7.1631 | 7.0201 | 4.8621 | 5.0448 | 5.1094 | 6.0047 |
| MAP4K4_S981 | MAP4K4 | O95819-3 | | 733 | QAGQRNSTSSIEP | 5.8530 | 6.1429 | 6.1488 | 5.7309 | 6.0974 | 6.5761 | 6.4465 | 6.4527 | 6.5535 | 6.6402 | 6.6474 | 6.7863 | 5.9967 | 6.2594 | 5.5747 | 6.0933 |
| MAP4K4_T909 | MAP4K4 | O95819-5 | | 648 | QAGQRNSTSIEPR | 6.0446 | 6.5775 | 6.3664 | 5.8337 | 5.9109 | 6.4774 | 6.3766 | 5.7114 | 6.5803 | 6.4194 | 6.8714 | 6.3482 | 6.2945 | 6.0353 | 5.6721 | 6.4802 |
| MAP4K4_T915 | MAP4K4 | O95819-3 | 706;708 | | NA | 5.4318 | 5.3341 | 5.5752 | 5.6543 | 5.8455 | 5.7883 | 6.0599 | 6.0425 | 5.8258 | 5.8218 | 5.2683 | 5.7660 | 8.1034 | 7.8205 | 8.1448 | 7.5178 |
| MAP4K4_T927 | MAP4K4 | O95819-3 | 324;326 | | NA | 5.8612 | 6.1893 | 6.2895 | 5.6587 | 5.8400 | 5.7737 | 5.5471 | 6.7163 | 6.4679 | 6.3395 | 6.2990 | 7.2296 | 6.4780 | 6.6025 | 6.3047 | 6.4029 |
| MAP4K4_T996 | MAP4K4 | O95819-3 | 933;936 | | NA | 4.7016 | 5.1161 | 4.7731 | 5.0632 | 6.2075 | 5.7505 | 6.1021 | 6.2512 | 4.7147 | 4.7686 | 5.9935 | 6.0429 | 9.9503 | 8.6336 | 8.8879 | 7.0433 |
| MAP4K5_S335 | MAP4K5 | Q9Y4K4 | | 335 | RAERTASEINFDK | 6.1650 | 5.2787 | 6.3035 | 5.7144 | 6.0097 | 5.7981 | 5.7980 | 5.9315 | 6.2208 | 4.9288 | 5.3219 | 5.2921 | 8.1139 | 8.1673 | 7.7066 | 7.2498 |
| MAP4K5_S433 | MAP4K5 | Q9Y4K4 | | 433 | QILRRQSSPSCGP | 6.4005 | 5.5513 | 6.7424 | 5.7476 | 6.2612 | 6.5129 | 5.8076 | 6.5362 | 7.0329 | 6.7751 | 7.0757 | 6.8606 | 5.7869 | 6.4356 | 5.5016 | 4.9720 |
| MAP7_S205 | MAP7 | Q14244-7 | | 387 | AQVRPPSPGNIRP | 5.5033 | 5.5981 | 5.7378 | 5.6692 | 5.5682 | 6.6051 | 6.2346 | 6.5481 | 5.8600 | 5.9109 | 6.0990 | 6.2417 | 6.8092 | 7.4137 | 7.1420 | 7.0592 |
| MAP7_S231 | MAP7 | Q14244-7 | | 337 | GTRRAVSPSNPKA | 5.4129 | 5.6218 | 5.5362 | 5.6144 | 5.1224 | 6.4807 | 6.2787 | 6.3428 | 6.1128 | 5.9108 | 5.8099 | 6.6015 | 6.9485 | 7.4034 | 7.2158 | 7.5874 |
| MAP7_S241 | MAP7 | Q14244-7 | | 264 | FLARSKSTAALSG | 6.8245 | 6.2337 | 6.1333 | 5.9602 | 5.4536 | 6.2995 | 6.6270 | 5.4316 | 6.5688 | 6.5463 | 7.2199 | 5.9686 | 5.8545 | 6.5588 | 5.2493 | 7.0704 |
| MAP7_S264 | MAP7 | Q14244-7 | | 205 | PDRRSVSTMNLSK | 7.2645 | 7.3953 | 6.8708 | 6.0464 | 5.3302 | 6.8850 | 7.0891 | 6.3570 | 6.1836 | 6.2931 | 6.5277 | 5.9992 | 5.2862 | 5.9322 | 4.7525 | 5.7871 |
| MAP7_S315 | MAP7 | Q14244-7 | | 241 | ARRLQLSPWESSV | 6.9569 | 7.5818 | 6.8705 | 6.7198 | 6.1794 | 6.1362 | 6.9312 | 8.0110 | 8.0137 | 7.3658 | 7.5405 | 6.8457 | 3.0649 | 4.5518 | 3.3314 | 3.8994 |
| MAP7_S337 | MAP7 | Q14244-7 | | 231 | SATLLNSPDRARR | 6.0461 | 6.0296 | 6.4936 | 6.7208 | 6.7832 | 6.1808 | 6.9765 | 6.7644 | 6.6350 | 7.0389 | 6.1717 | 6.8088 | 4.9037 | 5.9401 | 5.3354 | 5.1714 |
| MAP7_S387 | MAP7 | Q14244-7 | | 253 | VVNRLLTPTHSFL | 7.5398 | 7.7323 | 7.0964 | 7.2851 | 6.1902 | 7.2539 | 7.4795 | 6.8384 | 7.2598 | 7.7427 | 8.3774 | 7.1379 | 2.7097 | 3.5855 | 2.3363 | 3.4351 |
| MAP7_T253 | MAP7 | Q14244-7 | | 315 | IIHGTASYKKERE | 8.7699 | 7.3373 | 6.9751 | 7.0980 | 5.6873 | 7.3325 | 7.3142 | 6.9117 | 7.9373 | 7.6377 | 7.4709 | 7.3731 | 2.7790 | 4.2467 | 2.1385 | 2.9906 |
| MAP7D1_S113 | MAP7D1 | Q3KQU3 | | 503 | PRGTTASPKGRVR | 6.5652 | 6.6219 | 7.1811 | 6.4860 | 6.3449 | 6.8425 | 6.7190 | 6.7419 | 7.1854 | 7.0782 | 7.2427 | 7.2960 | 4.1591 | 4.7210 | 4.1146 | 4.7004 |
| MAP7D1_S113;S116 | MAP7D1 | Q3KQU3 | | 97 | AKSRGPTPPAMGP | 6.0381 | 6.0197 | 6.1054 | 5.9343 | 5.9224 | 6.0517 | 5.7566 | 6.6281 | 6.4650 | 6.4560 | 6.5504 | 6.5617 | 6.1335 | 6.4389 | 6.6858 | 6.2524 |
| MAP7D1_S125 | MAP7D1 | Q3KQU3 | | 399 | KPNAGGSPAPVRR | 6.2964 | 6.6931 | 6.5825 | 6.1285 | 5.7019 | 6.9119 | 6.3979 | 7.0668 | 6.8818 | 6.9899 | 6.8348 | 6.7170 | 4.9113 | 5.6883 | 4.9545 | 5.2434 |
| MAP7D1_S252 | MAP7D1 | Q3KQU3 | | 585 | APAPPVTPSKPMA | 6.3011 | 6.2437 | 6.6654 | 6.2629 | 6.0496 | 6.6480 | 6.4091 | 7.0459 | 7.0321 | 6.5365 | 7.0011 | 7.0555 | 4.8506 | 5.7425 | 4.7360 | 5.4199 |
| MAP7D1_S273 | MAP7D1 | Q3KQU3 | | 446 | RQSLPASPRARLS | 6.5019 | 6.7895 | 6.7575 | 6.3917 | 6.2983 | 6.7963 | 6.6879 | 7.0617 | 7.0849 | 7.1062 | 6.9988 | 7.2200 | 4.3577 | 4.8362 | 4.3061 | 4.8052 |
| MAP7D1_S280 | MAP7D1 | Q3KQU3 | | 70 | LPLEPESPSGQVG | 6.1350 | 6.7349 | 6.5855 | 6.7358 | 5.8122 | 6.2946 | 6.4515 | 6.6978 | 6.6508 | 6.3894 | 6.5768 | 6.6751 | 5.2864 | 5.9023 | 5.3432 | 5.7285 |
| MAP7D1_S313 | MAP7D1 | Q3KQU3 | | 280 | SATLWNSPSRNRS | 7.8559 | 8.1481 | 6.9014 | 6.5165 | 5.2618 | 7.5263 | 7.4159 | 6.8968 | 7.8373 | 7.6552 | 7.4047 | 7.8023 | 2.7881 | 3.6581 | 2.6847 | 3.6469 |
| MAP7D1_S366 | MAP7D1 | Q3KQU3 | | 460 | STASELSPKSKAR | 6.5020 | 6.2869 | 6.8883 | 6.5249 | 6.3909 | 6.6706 | 6.5250 | 6.2757 | 7.3388 | 6.9624 | 6.9790 | 7.1667 | 4.6878 | 5.0687 | 4.5426 | 5.1896 |
| MAP7D1_S399 | MAP7D1 | Q3KQU3 | | 834 | KKAVVQSPQVTEV | 6.4970 | 6.7356 | 6.6543 | 6.5129 | 6.1267 | 6.5472 | 6.7199 | 6.5947 | 6.5532 | 6.6316 | 6.8778 | 6.7053 | 5.2198 | 5.9602 | 4.7225 | 4.9412 |
| MAP7D1_S410 | MAP7D1 | Q3KQU3 | | 366 | VCPRSASASPLTP | 5.9165 | 6.2986 | 5.9830 | 5.6964 | 5.6134 | 6.4358 | 5.8755 | 5.8935 | 6.3816 | 6.1224 | 6.5578 | 6.2580 | 6.8424 | 7.3882 | 6.5771 | 6.1598 |
| MAP7D1_S442;S446 | MAP7D1 | Q3KQU3 | | 252 | TSGSRCSVSAVNL | 7.3749 | 6.5291 | 6.5773 | 6.1550 | 5.9664 | 5.8293 | 6.8811 | 5.7130 | 7.3984 | 6.5299 | 7.0963 | 6.2439 | 5.6449 | 6.3755 | 4.7699 | 4.9151 |
| MAP7D1_S446 | MAP7D1 | Q3KQU3 | | 302 | IVDRLMTPTLSFL | 7.9549 | 7.9491 | 7.9457 | 7.1224 | 5.6989 | 6.8432 | 7.1011 | 6.7947 | 7.6824 | 7.3962 | 7.2047 | 6.7534 | 3.3443 | 4.3022 | 2.5812 | 3.3255 |
| MAP7D1_S460 | MAP7D1 | Q3KQU3 | | 113 | RPPRRSSQPSPTA | 4.7533 | 4.1046 | 5.3823 | 4.6238 | 6.4185 | 5.3877 | 5.2722 | 5.9704 | 8.6405 | 6.8215 | 6.5729 | 6.4128 | 7.7543 | 7.9727 | 7.2885 | 6.6242 |
| MAP7D1_S479;S496 | MAP7D1 | Q3KQU3 | | 410 | RRRPEASPVQKKE | 4.7243 | 5.2047 | 6.5417 | 5.7255 | 5.6375 | 6.5107 | 6.9021 | 5.5840 | 7.5908 | 6.4607 | 6.7316 | 6.6367 | 6.5252 | 6.5007 | 6.1688 | 6.5547 |
| MAP7D1_S503 | MAP7D1 | Q3KQU3 | | 125 | AVPASDSPPTKQE | 7.1587 | 7.0387 | 6.8345 | 6.4701 | 6.5097 | 6.5233 | 6.5867 | 5.9564 | 5.8766 | 6.2947 | 6.1567 | 6.3836 | 5.2628 | 6.2331 | 5.2435 | 5.4710 |
| MAP7D1_S544 | MAP7D1 | Q3KQU3 | | 313 | FLARSRSAVTLPR | 6.4867 | 6.1629 | 6.3297 | 6.0526 | 5.7754 | 6.1191 | 6.2757 | 6.0890 | 6.7190 | 6.4132 | 6.5693 | 6.1155 | 6.2722 | 6.8102 | 5.7134 | 6.0960 |
| MAP7D1_S70 | MAP7D1 | Q3KQU3 | | 544 | ESAAPASPAPSPA | 7.0157 | 7.0261 | 6.7229 | 7.0563 | 5.9695 | 5.8341 | 5.4121 | 6.4030 | 6.8062 | 7.0709 | 7.0928 | 5.9112 | 5.5027 | 5.4819 | 5.1297 | 5.5652 |
| MAP7D1_S742 | MAP7D1 | Q3KQU3 | | 742 | ANANGSSPEPVKA | 6.2250 | 7.1515 | 7.6297 | 7.3338 | 5.8838 | 6.5109 | 6.6475 | 6.5665 | 6.8506 | 6.5970 | 7.4514 | 7.4539 | 4.8726 | 4.3427 | 4.0227 | 4.4604 |
| MAP7D1_S834 | MAP7D1 | Q3KQU3 | | 47 | SALVPDTPPDTPP | 5.4576 | 5.3705 | 6.6956 | 5.3882 | 5.8303 | 6.7656 | 6.1031 | 6.1646 | 6.3168 | 6.0404 | 6.5081 | 6.6110 | 7.1042 | 7.1255 | 6.1323 | 6.3863 |
| MAP7D1_T302 | MAP7D1 | Q3KQU3 | | 273 | NKRLSKSSATLWN | 5.9510 | 6.2431 | 6.6762 | 5.9831 | 5.6408 | 6.8848 | 6.0643 | 5.9702 | 5.8365 | 6.4604 | 6.7110 | 6.6798 | 5.7740 | 6.1950 | 6.4312 | 6.4986 |
| MAP7D1_T47 | MAP7D1 | Q3KQU3 | 442;446 | | NA | 7.2528 | 7.3558 | 7.2829 | 7.3018 | 5.8440 | 6.8743 | 7.2258 | 6.9660 | 7.6587 | 7.1783 | 7.5139 | 7.7246 | 2.9602 | 4.0085 | 2.9256 | 3.9270 |
| MAP7D1_T585 | MAP7D1 | Q3KQU3 | 479;496 | | NA | 5.7303 | 6.0719 | 5.9094 | 5.9629 | 6.0894 | 6.4815 | 6.1717 | 6.2401 | 6.6206 | 6.5520 | 6.9392 | 6.6554 | 6.0244 | 6.4259 | 6.1986 | 5.9266 |
| MAP7D1_T97 | MAP7D1 | Q3KQU3 | 113;116 | | NA | 4.7922 | 5.1329 | 5.6899 | 4.5604 | 6.0515 | 5.3676 | 4.7268 | 5.7494 | 6.7149 | 6.1662 | 6.7727 | 6.3222 | 7.7115 | 8.7868 | 8.3963 | 7.0586 |
| MAP7D2_S331 | MAP7D2 | Q96T17-2 | | 331 | IPKRPSSPVISKT | 6.5021 | 6.5970 | 6.4760 | 6.1798 | 5.7740 | 6.4981 | 6.7637 | 6.8920 | 6.8610 | 7.1568 | 6.2830 | 6.6985 | 5.4888 | 5.5173 | 5.3849 | 4.9271 |
| MAP7D3_S165 | MAP7D3 | Q8IWC1 | | 524 | QKNCPPSPLPLIS | 6.4705 | 6.4235 | 6.8060 | 6.3613 | 5.1344 | 7.4650 | 7.1423 | 6.5634 | 7.4163 | 7.0309 | 7.2134 | 7.5059 | 4.5764 | 5.1129 | 3.8147 | 4.9632 |
| MAP7D3_S441 | MAP7D3 | Q8IWC1 | | 165 | YQQKRWSWGGSAM | 6.0710 | 7.0125 | 7.3945 | 6.7754 | 6.1910 | 6.4607 | 6.5542 | 6.7958 | 7.1357 | 6.9629 | 7.1473 | 6.9972 | 4.1606 | 5.1241 | 4.4056 | 4.8116 |
| MAP7D3_S461 | MAP7D3 | Q8IWC1 | | 461 | EASMEASPKAKAR | 7.0481 | 6.9466 | 7.3199 | 6.8304 | 6.5796 | 6.6635 | 7.0655 | 6.3629 | 7.2639 | 6.8316 | 7.1883 | 7.0997 | 4.0278 | 4.4534 | 3.9072 | 4.4116 |
| MAP7D3_S489 | MAP7D3 | Q8IWC1 | | 441 | KESMEASPEAMVK | 6.8977 | 7.1823 | 7.1791 | 7.8108 | 6.4144 | 6.2721 | 7.1546 | 6.6208 | 7.3815 | 7.1810 | 6.9322 | 7.3051 | 3.4923 | 4.4229 | 3.3425 | 4.4108 |
| MAP7D3_S500;S509 | MAP7D3 | Q8IWC1 | | 489 | IAKKRLSSYTECY | 6.3174 | 6.3585 | 6.5133 | 6.2917 | 6.4258 | 6.9990 | 6.5161 | 7.2667 | 6.8849 | 6.4916 | 6.6305 | 6.5406 | 4.8168 | 5.5942 | 4.7419 | 5.6111 |
| MAP7D3_S524 | MAP7D3 | Q8IWC1 | | 770 | ILNGTGSPTKFKM | 6.1348 | 6.2865 | 6.5169 | 5.6611 | 6.5227 | 6.1436 | 6.2415 | 6.0822 | 6.5240 | 6.2348 | 6.2718 | 6.8443 | 6.0482 | 6.0289 | 6.2462 | 6.2125 |
| MAP7D3_S770 | MAP7D3 | Q8IWC1 | 500;509 | | NA | 6.8412 | 6.1249 | 7.1317 | 7.5008 | 6.5055 | 6.5923 | 6.4936 | 7.4187 | 6.9198 | 7.6958 | 7.0538 | 6.6989 | 4.3077 | 4.6302 | 3.4563 | 4.6290 |
| MAP9_S79 | MAP9 | Q49MG5 | | 79 | MNDFHISDDEEKN | 6.2685 | 4.3405 | 7.4165 | 5.5505 | 6.5163 | 6.2284 | 5.5487 | 7.1449 | 4.9018 | 5.8730 | 6.0316 | 7.0739 | 6.2502 | 7.3764 | 6.1609 | 7.3181 |
| MAPK1_S360 | MAPK1 | P28482 | | 187 | TGFLTEYVATRWY | 7.6899 | 5.9326 | 6.7999 | 5.5321 | 7.4063 | 6.3349 | 5.7477 | 6.8135 | 7.7708 | 6.4618 | 5.3808 | 5.9606 | 6.3830 | 6.1299 | 4.6665 | 4.9896 |
| MAPK1_Y187 | MAPK1 | P28482 | | 360 | FQPGYRSxxxxxx | 6.1525 | 6.7347 | 6.7658 | 6.0247 | 6.3512 | 6.5416 | 6.3768 | 6.4443 | 6.3336 | 6.6607 | 7.1524 | 7.1305 | 5.0971 | 5.5793 | 5.0543 | 5.6006 |
| MAPK14_T180 | MAPK14 | Q16539-2 | | 180 | HTDDEMTGYVATR | 5.3569 | 6.6145 | 7.5975 | 5.4201 | 5.5423 | 6.8233 | 5.6820 | 6.7517 | 6.5609 | 6.5585 | 7.0202 | 7.6222 | 6.0016 | 5.2985 | 5.9709 | 5.1788 |
| MAPK3_Y204 | MAPK3 | P27361 | | 204 | TGFLTEYVATRWY | 7.4304 | 5.7886 | 6.6560 | 5.5095 | 7.3582 | 6.3226 | 6.0229 | 6.7219 | 8.1619 | 6.2658 | 6.0222 | 6.0284 | 6.1482 | 5.9253 | 4.7497 | 4.8883 |
| MAPK6_S189 | MAPK6 | Q16659 | | 189 | SHKGHLSEGLVTK | 4.4670 | 4.9119 | 5.1847 | 4.4691 | 6.7384 | 6.6991 | 6.7362 | 7.4385 | 5.2965 | 5.0456 | 5.1157 | 5.2668 | 8.5135 | 7.8836 | 8.2355 | 7.9978 |
| MAPK6_S451 | MAPK6 | Q16659 | | 684 | GIPQFHSPVGSPL | 5.1621 | 5.9067 | 6.2568 | 5.8133 | 7.0021 | 6.8774 | 6.6481 | 7.0661 | 5.9715 | 5.6214 | 6.0196 | 5.8495 | 6.2487 | 6.4846 | 6.5904 | 6.4816 |
| MAPK6_S684 | MAPK6 | Q16659 | | 451 | NYKTRSSSYLDNL | 5.5242 | 4.5458 | 4.4938 | 4.2414 | 7.8739 | 6.4453 | 4.8763 | 6.5650 | 7.0022 | 3.7886 | 6.1383 | 6.2007 | 8.2459 | 9.3154 | 7.3092 | 7.4341 |
| MAPK6_S684;S688 | MAPK6 | Q16659 | 684;688 | | NA | 5.1143 | 5.2837 | 5.5337 | 4.7279 | 7.4490 | 7.8231 | 6.8497 | 7.7160 | 6.3874 | 5.4244 | 5.9273 | 6.2926 | 6.7159 | 6.1610 | 5.8981 | 6.6960 |
| MAPK7_T219 | MAPK7 | Q13164 | | 733 | PPVFSGTPKGSGA | 6.5007 | 7.2976 | 7.3844 | 6.6135 | 6.8170 | 6.9360 | 7.0059 | 6.6849 | 6.9299 | 7.5318 | 7.1349 | 7.0320 | 3.7383 | 4.9053 | 3.2131 | 4.2749 |
| MAPK7_T733 | MAPK7 | Q13164 | | 219 | EHQYFMTEYVATR | 6.6832 | 6.1207 | 6.9227 | 7.8703 | 7.6691 | 6.9875 | 5.3545 | 6.7705 | 8.8800 | 6.7467 | 7.4401 | 7.7789 | 3.2304 | 3.8061 | 3.5004 | 4.2388 |
| MAPK8_S377 | MAPK8 | P45983-2 | | 377 | VIRGQPSPLAQVQ | 6.2400 | 7.0213 | 6.6251 | 6.4721 | 6.5853 | 6.6282 | 5.9399 | 7.0036 | 6.5780 | 6.4004 | 6.7531 | 6.5609 | 5.0502 | 5.5829 | 5.4696 | 5.0893 |
| MAPKAP1_S186 | MAPKAP1 | Q9BPZ7 | | 510 | KLNRRTSFSFQKE | 7.4185 | 6.3134 | 6.6430 | 5.8594 | 6.5774 | 6.3810 | 6.3159 | 6.2540 | 7.2127 | 6.5960 | 6.5325 | 6.9227 | 5.4507 | 6.2140 | 4.3616 | 4.9472 |
| MAPKAP1_S510 | MAPKAP1 | Q9BPZ7 | | 509 | RKLNRRTSFSFQK | 7.3536 | 6.4761 | 6.5403 | 5.8500 | 6.3548 | 6.4273 | 6.3893 | 6.0749 | 7.1143 | 6.7220 | 6.5678 | 6.9083 | 5.5375 | 6.2751 | 4.3126 | 5.0959 |
| MAPKAP1_T509 | MAPKAP1 | Q9BPZ7 | | 186 | YLPLHSSQDRLLP | 5.4335 | 6.0666 | 6.7507 | 6.3569 | 6.0573 | 6.5033 | 6.2257 | 6.8099 | 5.8052 | 6.8041 | 6.1688 | 6.0364 | 5.7634 | 7.1073 | 6.4615 | 5.6496 |
| MAPKAPK5_T182 | MAPKAPK5 | Q8IW41 | | 182 | DQGDLMTPQFTPY | 6.3038 | 6.0834 | 6.5071 | 6.5351 | 6.3138 | 5.8146 | 6.3172 | 6.1047 | 6.8147 | 5.5792 | 6.0747 | 6.8189 | 6.9850 | 6.4708 | 5.7207 | 5.5562 |
| MAPKBP1_S1263 | MAPKBP1 | O60336 | | 1314 | PTLAAFSPVTKGR | 6.7017 | 5.7099 | 6.8194 | 6.6080 | 6.6714 | 7.0629 | 7.3064 | 5.9587 | 6.8506 | 6.6116 | 7.9735 | 6.5465 | 4.9215 | 5.0080 | 4.2633 | 4.9866 |
| MAPKBP1_S1314 | MAPKBP1 | O60336 | | 1367 | AHPGPSSPCAQQL | 6.1068 | 4.9573 | 6.6660 | 7.0259 | 6.0435 | 6.0178 | 5.5605 | 6.4768 | 6.6495 | 6.9690 | 6.5273 | 6.4245 | 5.6422 | 7.0553 | 6.3143 | 5.5633 |
| MAPKBP1_S1367 | MAPKBP1 | O60336 | | 1263 | KISRSISVGENLG | 6.4524 | 5.9358 | 6.9037 | 6.6201 | 6.0714 | 6.3923 | 5.9079 | 5.7547 | 5.9234 | 6.2899 | 6.7122 | 5.7277 | 6.2754 | 6.9099 | 6.1077 | 6.0155 |
| MAPKBP1_S18;S20 | MAPKBP1 | O60336 | 18;20 | | NA | 7.1958 | 6.3483 | 7.6202 | 6.1161 | 5.5898 | 6.4085 | 6.3381 | 5.7043 | 7.0336 | 7.1176 | 6.8586 | 7.0208 | 6.0566 | 5.9061 | 4.2778 | 4.4079 |
| MAPRE2_S200 | MAPRE2 | Q15555 | | 217 | AAKPGSTPSRPSS | 7.0999 | 6.4654 | 7.4890 | 6.6534 | 6.0976 | 7.3979 | 7.5759 | 6.1136 | 7.8749 | 7.7199 | 7.3045 | 7.6032 | 3.4136 | 4.3179 | 2.8107 | 4.0626 |
| MAPRE2_S229 | MAPRE2 | Q15555 | | 200 | KSHHANSPTAGAA | 6.5467 | 7.6775 | 6.8209 | 7.5467 | 5.9825 | 7.1394 | 6.3151 | 7.4058 | 6.6677 | 7.3880 | 6.7275 | 6.9666 | 3.5179 | 5.0451 | 3.7295 | 4.5230 |
| MAPRE2_T217 | MAPRE2 | Q15555 | | 229 | SAKRASSSGSASK | 5.7134 | 7.0444 | 6.2479 | 6.3796 | 5.8174 | 7.2081 | 6.6446 | 6.3734 | 5.7509 | 6.4355 | 6.4294 | 6.9645 | 5.8832 | 5.7522 | 5.8746 | 5.4809 |
| MAPRE3_S162 | MAPRE3 | Q9UPY8 | | 162 | AVPQRTSPTGPKN | 5.8174 | 5.7773 | 5.9053 | 6.6843 | 5.2814 | 5.8180 | 5.6990 | 5.2130 | 6.1666 | 6.5347 | 6.6693 | 7.5968 | 6.0862 | 7.5609 | 5.8658 | 7.3238 |
| MAPT_S537 | MAPT | P10636-9 | | 537 | GYSSPGSPGTPGS | 6.5765 | 5.9537 | 6.7554 | 5.6658 | 6.2135 | 6.3876 | 5.7185 | 6.4669 | 6.5819 | 5.8947 | 6.1887 | 6.3788 | 6.2810 | 6.6815 | 6.0866 | 6.1690 |
| MAPT_T566 | MAPT | P10636-9 | | 566 | KVAVVRTPPKSPS | 6.7115 | 6.2309 | 7.1338 | 6.3323 | 6.1708 | 6.5763 | 5.8677 | 5.6350 | 6.0633 | 6.6860 | 6.7970 | 7.3121 | 5.4843 | 5.9571 | 5.1842 | 5.8578 |
| MARCHF7_S13 | MARCHF7 | Q9H992 | | 13 | RIPRRISVQPSSS | 7.8282 | 6.0575 | 6.1237 | 4.7336 | 7.7652 | 5.7894 | 5.4901 | 5.1614 | 8.3429 | 7.6574 | 7.8761 | 8.4985 | 5.5811 | 5.9194 | 3.4106 | 3.7649 |
| MARCKS_S101 | MARCKS | P29966 | | 170 | FKLSGFSFKKNKK | 7.7084 | 6.3148 | 6.5029 | 5.2409 | 5.3985 | 5.5775 | 5.7526 | 6.8830 | 6.1551 | 7.0344 | 6.3332 | 7.4742 | 5.8627 | 6.9282 | 4.7468 | 6.0868 |
| MARCKS_S118 | MARCKS | P29966 | | 27 | EAAVASSPSKANG | 6.9798 | 6.6249 | 6.3356 | 6.3746 | 6.6629 | 6.1274 | 6.3560 | 7.1239 | 7.0115 | 7.3159 | 7.2137 | 7.3342 | 4.6266 | 4.8154 | 4.0915 | 5.0061 |
| MARCKS_S118;S135 | MARCKS | P29966 | | 150 | PSPSNETPKKKKK | 6.4703 | 6.2231 | 6.2656 | 5.9659 | 5.6763 | 5.3421 | 5.7870 | 6.4381 | 5.8702 | 6.4621 | 6.1788 | 6.7470 | 7.2084 | 6.6314 | 6.1944 | 6.5395 |
| MARCKS_S135 | MARCKS | P29966 | | 167 | KKSFKLSGFSFKK | 8.0657 | 7.6817 | 6.5676 | 5.9387 | 5.4400 | 6.7008 | 7.0514 | 7.1995 | 6.2541 | 7.4960 | 7.3283 | 7.7405 | 3.8116 | 4.8006 | 3.2085 | 4.7150 |
| MARCKS_S145 | MARCKS | P29966 | | 145 | EDGATPSPSNETP | 6.6939 | 6.4999 | 6.2985 | 5.9166 | 6.8380 | 5.5771 | 5.9610 | 7.0235 | 6.6254 | 6.7842 | 6.8100 | 6.7445 | 5.5092 | 5.5230 | 5.3581 | 5.8373 |
| MARCKS_S163;S167 | MARCKS | P29966 | | 118 | EAAEPGSPTAAEG | 5.5447 | 5.5658 | 5.6759 | 5.7094 | 6.2037 | 5.7221 | 6.1765 | 6.5155 | 5.6118 | 5.6558 | 6.0265 | 5.8840 | 7.7145 | 7.6250 | 7.5883 | 6.7804 |
| MARCKS_S167 | MARCKS | P29966 | | 101 | APEAGASPVEKEA | 6.4376 | 5.8632 | 6.0752 | 6.3101 | 6.6583 | 5.7980 | 6.3289 | 6.8859 | 5.1087 | 6.4365 | 6.0252 | 6.5337 | 6.7649 | 6.0636 | 6.3423 | 6.3677 |
| MARCKS_S170 | MARCKS | P29966 | | 46 | KVNGDASPAAAES | 7.7631 | 7.7532 | 6.3741 | 6.1839 | 5.7928 | 6.3423 | 6.3196 | 6.3001 | 5.7844 | 6.9518 | 7.5729 | 7.1110 | 4.8435 | 5.3838 | 4.4316 | 5.0917 |
| MARCKS_S27 | MARCKS | P29966 | | 143 | KAEDGATPSPSNE | 6.2777 | 6.3423 | 6.6587 | 6.0400 | 6.8376 | 6.3957 | 6.0761 | 6.8024 | 6.1030 | 6.7191 | 7.6578 | 7.0433 | 6.1210 | 5.6313 | 4.9575 | 4.3366 |
| MARCKS_S46 | MARCKS | P29966 | | 135 | AASSTSSPKAEDG | 6.0718 | 6.4868 | 6.3248 | 5.8992 | 5.9529 | 6.5416 | 5.4480 | 6.1942 | 7.4078 | 8.1750 | 7.6632 | 8.7218 | 4.5119 | 5.1298 | 4.3789 | 5.0925 |
| MARCKS_S77;S81 | MARCKS | P29966 | | 81 | AGSGAASPSAAEK | 7.0678 | 7.2485 | 6.7533 | 6.4808 | 6.5616 | 6.5839 | 6.8143 | 7.5570 | 6.5877 | 7.2992 | 7.4200 | 6.7882 | 3.5980 | 4.9264 | 3.8424 | 4.4710 |
| MARCKS_S81 | MARCKS | P29966 | 77;81 | | NA | 6.7416 | 7.4429 | 6.7786 | 6.2556 | 5.4949 | 6.0509 | 6.5165 | 6.9944 | 7.0127 | 6.8178 | 8.0353 | 7.0130 | 4.4675 | 5.0431 | 4.2741 | 5.0610 |
| MARCKS_T143 | MARCKS | P29966 | 143;145 | | NA | 7.5738 | 8.0015 | 7.0065 | 7.0076 | 6.2486 | 5.4509 | 6.5183 | 7.3831 | 5.8933 | 7.5783 | 7.1514 | 7.5445 | 4.0799 | 4.8888 | 3.4866 | 4.1870 |
| MARCKS_T143;S145 | MARCKS | P29966 | 118;135 | | NA | 6.7990 | 7.1023 | 6.3781 | 6.9112 | 5.4395 | 6.4929 | 6.7443 | 6.5458 | 6.6801 | 7.4663 | 7.5786 | 7.7840 | 4.5277 | 4.6894 | 3.9357 | 4.9249 |
| MARCKS_T150 | MARCKS | P29966 | 163;167 | | NA | 8.0609 | 7.2447 | 7.1148 | 7.0466 | 4.6939 | 6.4827 | 6.8614 | 7.0996 | 7.2858 | 8.7617 | 8.6770 | 9.5299 | 2.5201 | 3.8682 | 1.6985 | 3.0545 |
| MARCKSL1_S101 | MARCKSL1 | P49006 | | 104 | FKLSGLSFKRNRK | 8.0367 | 6.0114 | 6.5416 | 5.1151 | 5.6376 | 5.3369 | 5.9196 | 6.5688 | 6.0658 | 7.0885 | 6.3202 | 7.4617 | 5.5026 | 7.0335 | 4.7331 | 6.6266 |
| MARCKSL1_S101;S104 | MARCKSL1 | P49006 | | 85 | EVPPKETPKKKKK | 6.1075 | 5.3250 | 5.9550 | 5.5590 | 6.8979 | 5.4635 | 5.8087 | 6.6029 | 5.9043 | 6.6684 | 6.2914 | 7.2817 | 6.8006 | 6.7494 | 5.9660 | 6.6186 |
| MARCKSL1_S104 | MARCKSL1 | P49006 | | 36 | ENGHVKSNGDLSP | 5.6681 | 5.6977 | 5.6593 | 5.6353 | 6.5922 | 5.6203 | 6.0298 | 6.3953 | 6.0465 | 5.9791 | 6.4896 | 6.9226 | 6.7504 | 6.7517 | 6.8448 | 6.9173 |
| MARCKSL1_S22 | MARCKSL1 | P49006 | | 101 | KKPFKLSGLSFKR | 5.6525 | 4.9779 | 7.5750 | 5.2526 | 5.6396 | 5.6367 | 5.2576 | 8.2649 | 5.9138 | 9.8299 | 7.4217 | 9.2984 | 4.5002 | 5.6863 | 4.0413 | 5.0516 |
| MARCKSL1_S36 | MARCKSL1 | P49006 | | 22 | EEAAGASPAKANG | 6.2688 | 6.2995 | 6.4072 | 6.6621 | 6.2743 | 5.7715 | 6.2488 | 6.7677 | 6.4876 | 6.7441 | 6.6201 | 7.3119 | 5.5924 | 5.6136 | 5.1779 | 5.7527 |
| MARCKSL1_S41 | MARCKSL1 | P49006 | | 41 | KSNGDLSPKGEGE | 7.3726 | 6.7340 | 6.8158 | 6.8459 | 7.0031 | 5.4533 | 6.3602 | 7.2236 | 6.6988 | 8.0134 | 7.3319 | 8.4247 | 3.8245 | 4.4716 | 3.2461 | 4.1805 |
| MARCKSL1_T85 | MARCKSL1 | P49006 | 101;104 | | NA | 8.6247 | 4.6075 | 10.8049 | 6.7055 | 7.2230 | 6.9709 | 7.0065 | 6.1249 | 4.8928 | 6.5630 | 10.0293 | 8.0713 | 3.0645 | 5.1089 | 2.3959 | 1.8064 |
| MARF1_S1093 | MARF1 | Q9Y4F3-5 | | 1571 | LSSLSLSPANHEN | 6.7094 | 7.0749 | 6.7077 | 6.2444 | 6.3433 | 6.9793 | 6.5311 | 6.1952 | 7.4052 | 7.2420 | 6.9704 | 7.0464 | 4.6063 | 4.4744 | 4.6728 | 4.7969 |
| MARF1_S1571 | MARF1 | Q9Y4F3-5 | | 715 | ARSVTSSPVEKKD | 6.0055 | 6.3147 | 6.2153 | 5.9469 | 6.3043 | 6.4428 | 5.9301 | 5.9147 | 6.2518 | 6.6348 | 6.1646 | 6.0991 | 6.7062 | 7.0387 | 6.3829 | 5.6476 |
| MARF1_S1718 | MARF1 | Q9Y4F3-5 | | 1718 | SKDPVESPAKKQP | 6.3071 | 6.5703 | 6.9639 | 6.7639 | 6.1085 | 6.4763 | 6.0066 | 6.5016 | 7.1685 | 6.5726 | 7.3321 | 7.6026 | 4.6185 | 5.4135 | 4.4728 | 5.1213 |
| MARF1_S715 | MARF1 | Q9Y4F3-5 | | 768 | NLLNRASPLAFNI | 5.7931 | 5.9450 | 6.9514 | 6.7222 | 5.7576 | 6.8297 | 6.7577 | 5.6963 | 7.0970 | 6.0267 | 6.0152 | 6.2585 | 6.2772 | 6.0081 | 5.6377 | 6.2266 |
| MARF1_S768 | MARF1 | Q9Y4F3-5 | | 1093 | WLLRSKSPVGNPQ | 5.4932 | 5.9622 | 6.6446 | 6.1664 | 6.3935 | 6.7198 | 6.3554 | 6.3418 | 6.9534 | 7.0934 | 6.1604 | 6.1838 | 6.1533 | 6.1626 | 5.5983 | 5.6179 |
| MARK2_S376 | MARK2 | Q7KZI7 | | 486 | TNRSRNSPLLERA | 5.8517 | 6.2000 | 6.1954 | 6.1573 | 5.7328 | 6.0331 | 6.1441 | 6.1354 | 6.3753 | 6.4230 | 6.4102 | 6.4989 | 6.2812 | 6.5091 | 6.2624 | 6.7899 |
| MARK2_S400 | MARK2 | Q7KZI7 | | 208 | FGNKLDTFCGSPP | 6.7704 | 6.9851 | 7.5730 | 7.3041 | 6.6220 | 4.7058 | 6.2036 | 5.0960 | 5.5463 | 5.1940 | 5.2301 | 4.2824 | 7.6035 | 6.6979 | 8.2885 | 5.8974 |
| MARK2_S409 | MARK2 | Q7KZI7 | | 400 | KVQRSVSANPKQR | 7.4600 | 6.8910 | 7.2394 | 5.9474 | 6.0282 | 6.6022 | 5.9648 | 6.5355 | 6.9849 | 6.6324 | 6.7403 | 7.1110 | 5.3008 | 5.8724 | 4.1158 | 4.5737 |
| MARK2_S456 | MARK2 | Q7KZI7 | | 456 | TAKVPASPLPGLE | 6.3078 | 6.6082 | 6.4200 | 6.3160 | 5.6361 | 6.5186 | 6.8283 | 6.3803 | 6.1312 | 6.5295 | 6.5551 | 6.4630 | 5.2862 | 5.9426 | 5.7775 | 6.2996 |
| MARK2_S486 | MARK2 | Q7KZI7 | | 571 | VPVASPSAHNISS | 7.2680 | 6.8474 | 6.8074 | 7.2203 | 6.3400 | 6.2601 | 6.1781 | 6.2956 | 6.6064 | 6.0009 | 6.3114 | 6.0502 | 4.9404 | 6.0509 | 5.1172 | 5.7057 |
| MARK2_S498 | MARK2 | Q7KZI7 | | 631 | QGRRGASGSIFSK | 7.7093 | 6.9215 | 8.0507 | 7.1664 | 5.5745 | 6.5279 | 6.1413 | 6.6909 | 7.6407 | 6.7270 | 7.3853 | 7.8489 | 3.6439 | 4.6310 | 3.1047 | 4.2360 |
| MARK2_S514 | MARK2 | Q7KZI7-14 | | 376 | PKQRRFSDQAGPA | 5.5739 | 5.8990 | 6.0892 | 6.3733 | 6.4075 | 6.0672 | 6.0471 | 6.5258 | 7.0519 | 7.8844 | 7.7750 | 7.5703 | 4.8459 | 6.1745 | 4.8887 | 4.8262 |
| MARK2_S571 | MARK2 | Q7KZI7 | | 409 | PKQRRFSDQAAGP | 5.1566 | 5.2851 | 5.8767 | 5.9718 | 6.5249 | 5.6299 | 6.2911 | 6.6664 | 7.9538 | 7.7013 | 7.6232 | 7.5943 | 4.9959 | 6.4321 | 5.4057 | 4.8911 |
| MARK2_S631 | MARK2 | Q7KZI7 | | 759 | CKLPRLSLNGVRF | 5.7864 | 6.1756 | 6.2654 | 5.5427 | 6.2833 | 6.9947 | 5.6469 | 6.2211 | 6.5007 | 5.7217 | 6.0462 | 5.9067 | 6.7132 | 7.2120 | 6.4737 | 6.5097 |
| MARK2_S722 | MARK2 | Q7KZI7 | | 722 | KVLDANSCQSELH | 6.8704 | 7.9335 | 6.4706 | 7.2406 | 6.8882 | 6.2100 | 7.7944 | 5.6326 | 5.5644 | 4.2348 | 6.2041 | 4.0541 | 7.6048 | 6.6009 | 5.1647 | 5.5319 |
| MARK2_S759 | MARK2 | Q7KZI7 | | 498 | ASLGQASIQNGKD | 6.1696 | 5.6397 | 7.8397 | 6.7752 | 5.7903 | 6.6409 | 6.7445 | 5.7994 | 7.2029 | 6.5119 | 7.2140 | 7.8704 | 5.0580 | 5.3002 | 4.4509 | 4.9923 |
| MARK2_T208 | MARK2 | Q7KZI7 | | 514 | MPGSRASTASASA | 7.5617 | 6.7670 | 6.5780 | 5.7319 | 5.6515 | 6.1294 | 6.6924 | 5.5786 | 6.9531 | 6.6860 | 6.9034 | 6.7975 | 5.2668 | 6.2258 | 4.6228 | 5.8542 |
| MARK2_T467 | MARK2 | Q7KZI7 | | 560 | EVPRPSTAPQRVP | 6.0979 | 6.6619 | 7.6258 | 7.5898 | 5.9496 | 6.1653 | 5.8642 | 5.9564 | 7.6033 | 6.8798 | 6.5832 | 6.0732 | 5.4441 | 7.0933 | 3.2602 | 5.1520 |
| MARK2_T560 | MARK2 | Q7KZI7 | | 467 | LERKKTTPTPSTN | 5.2209 | 6.7707 | 7.8338 | 6.7273 | 5.8619 | 6.7845 | 6.4877 | 6.8107 | 6.0037 | 7.7360 | 8.5911 | 8.3522 | 3.8329 | 5.1314 | 3.4026 | 4.4525 |
| MARK3_S374 | MARK3 | P27448 | | 469 | KGIAPASPMLGNA | 5.8212 | 6.2556 | 6.3493 | 6.3891 | 5.7349 | 6.7478 | 6.8762 | 6.5564 | 6.3202 | 6.4822 | 6.5419 | 6.6693 | 5.8219 | 5.5912 | 5.6412 | 6.2018 |
| MARK3_S419 | MARK3 | P27448 | | 601 | QTRSRGSTNLFSK | 6.8359 | 6.3181 | 6.8785 | 6.0468 | 5.3420 | 6.7819 | 6.4374 | 6.2495 | 8.4096 | 7.4936 | 7.3981 | 7.9432 | 4.5794 | 5.3978 | 3.2194 | 4.6689 |
| MARK3_S42 | MARK3 | P27448 | | 42 | GARCRNSIASCAD | 7.9059 | 6.5355 | 6.7659 | 6.0833 | 5.9895 | 5.9719 | 6.4097 | 5.7188 | 7.1874 | 6.0972 | 6.9987 | 6.8757 | 5.4471 | 6.4620 | 4.3292 | 5.2221 |
| MARK3_S455 | MARK3 | P27448 | | 419 | QKQRRYSDHAGPA | 5.7473 | 6.1971 | 6.6126 | 6.1993 | 6.9919 | 5.7820 | 6.4213 | 6.6468 | 7.0502 | 7.2700 | 7.3739 | 7.4756 | 5.2286 | 5.7689 | 4.6759 | 4.5588 |
| MARK3_S456 | MARK3 | P27448 | | 507 | GMTRRNTYVCSER | 9.3087 | 7.1936 | 7.2210 | 6.4480 | 5.6872 | 5.6730 | 4.6648 | 5.4275 | 7.5254 | 6.0599 | 6.7368 | 6.6281 | 5.2292 | 6.0731 | 4.0975 | 4.2260 |
| MARK3_S469 | MARK3 | P27448 | | 540 | QRTPVASTHSISS | 6.9745 | 5.8940 | 6.3182 | 6.4136 | 6.6020 | 6.5081 | 6.0840 | 6.3604 | 7.3038 | 5.9926 | 6.0542 | 6.4938 | 5.9786 | 6.5725 | 5.2684 | 5.1811 |
| MARK3_S540 | MARK3 | P27448 | | 456 | ISSRKSSGSAVGG | 6.4453 | 6.4421 | 6.4491 | 6.2148 | 6.1490 | 6.4000 | 5.9838 | 6.1736 | 6.9948 | 6.8913 | 7.0806 | 7.1499 | 5.2519 | 5.9477 | 5.1926 | 5.2335 |
| MARK3_S583 | MARK3 | P27448 | | 583 | YNGPPASPSLSHE | 7.0306 | 6.0593 | 5.8090 | 7.2101 | 5.5209 | 7.7186 | 6.3049 | 8.1106 | 5.8200 | 6.3364 | 6.2262 | 6.5096 | 4.6378 | 5.4233 | 5.3178 | 5.9649 |
| MARK3_S583;S587 | MARK3 | P27448 | | 455 | GISSRKSSGSAVG | 7.2647 | 7.8953 | 6.9972 | 7.1323 | 5.6101 | 6.6136 | 7.3828 | 6.8251 | 7.2429 | 7.6226 | 7.6921 | 7.8912 | 3.1045 | 3.9010 | 3.2102 | 3.6144 |
| MARK3_S601 | MARK3 | P27448 | | 374 | SSELDASDSSSSS | 6.3678 | 6.2237 | 6.1730 | 6.2137 | 6.6161 | 6.1497 | 5.9018 | 5.7489 | 6.2551 | 6.4028 | 6.3767 | 6.0318 | 6.1223 | 7.0137 | 6.1384 | 6.2644 |
| MARK3_T507 | MARK3 | P27448 | 583;587 | | NA | 6.3626 | 6.3726 | 7.8342 | 7.2130 | 5.9606 | 6.4877 | 7.3559 | 7.9279 | 6.9116 | 7.0477 | 7.2183 | 7.2931 | 3.4337 | 4.6727 | 3.1149 | 4.7935 |
| MARK3_T576;S587 | MARK3 | P27448 | 576;587 | | NA | 6.2715 | 6.3390 | 6.7147 | 6.4682 | 7.4757 | 6.7255 | 6.0463 | 6.7408 | 5.9621 | 7.3832 | 6.6613 | 6.9252 | 4.7117 | 5.2502 | 4.9043 | 5.4205 |
| MARK4_S423 | MARK4 | Q96L34 | | 423 | HRQRRHSDFCGPS | 6.1468 | 5.5517 | 6.4521 | 5.7694 | 6.3401 | 6.4845 | 5.9190 | 6.1070 | 8.4672 | 7.8331 | 7.8322 | 7.8392 | 4.9203 | 5.8037 | 3.6808 | 4.8529 |
| MARS1_S229 | MARS1 | P56192 | | 227 | EEEELATLSEEEI | 5.8274 | 6.2389 | 6.3810 | 6.0903 | 6.0793 | 6.5899 | 6.0558 | 6.9361 | 6.2245 | 6.3207 | 6.2633 | 6.2723 | 6.2228 | 6.4761 | 6.1240 | 5.8976 |
| MARS1_S825 | MARS1 | P56192 | | 229 | EELATLSEEEIAM | 6.5050 | 5.3721 | 5.8478 | 6.7313 | 6.0965 | 6.8911 | 5.8076 | 6.3652 | 5.9683 | 6.6695 | 6.1204 | 6.5622 | 6.4733 | 5.8339 | 6.8209 | 5.9351 |
| MARS1_T227 | MARS1 | P56192 | | 825 | GGQAKTSPKPAVV | 5.9498 | 6.2106 | 6.7671 | 5.3072 | 7.4078 | 8.2775 | 6.6071 | 6.6979 | 6.5324 | 6.5063 | 7.7665 | 6.1270 | 5.5931 | 5.7294 | 4.6253 | 3.8950 |
| MAST2_S1381 | MAST2 | Q6P0Q8 | | 900 | RSWVIGSPEILRK | 6.8960 | 7.0537 | 7.1587 | 6.2839 | 6.2658 | 6.6186 | 6.3729 | 6.7854 | 7.2012 | 6.8220 | 7.1182 | 6.7414 | 4.4012 | 5.0590 | 4.2989 | 4.9232 |
| MAST2_S144 | MAST2 | Q6P0Q8 | | 1381 | PTKLHLSPPLGRQ | 6.2336 | 6.0726 | 8.8569 | 6.5518 | 6.7542 | 7.0326 | 7.4588 | 6.5366 | 6.4899 | 7.0662 | 6.9928 | 7.1325 | 3.6720 | 4.4319 | 4.0381 | 4.6796 |
| MAST2_S299 | MAST2 | Q6P0Q8 | | 657 | LTDFGLSKIGLMS | 6.8414 | 5.3849 | 5.4746 | 5.9273 | 7.2507 | 6.5883 | 6.3829 | 5.7740 | 5.8101 | 5.7754 | 5.4818 | 5.9707 | 7.3172 | 6.4104 | 7.0294 | 6.5810 |
| MAST2_S657 | MAST2 | Q6P0Q8 | | 299 | MRPRSRSLSPGRS | 6.6764 | 6.5947 | 6.1956 | 6.6337 | 5.4453 | 6.1742 | 6.3834 | 6.3959 | 7.1286 | 7.0453 | 5.7096 | 6.2869 | 5.7921 | 6.1588 | 5.4092 | 5.9703 |
| MAST2_S900 | MAST2 | Q6P0Q8 | | 144 | VRMRNQSLGQSAP | 6.8107 | 5.9407 | 7.4662 | 6.7392 | 6.3922 | 5.9393 | 6.2492 | 5.8596 | 6.9467 | 7.0374 | 6.6526 | 6.9763 | 5.5641 | 5.6631 | 4.9931 | 4.7697 |
| MAST3_S710 | MAST3 | O60307 | | 710 | FSKVYSSSEFLAV | 6.4366 | 6.4945 | 6.6007 | 7.7510 | 6.1696 | 6.7391 | 6.4672 | 7.0206 | 7.4115 | 5.6065 | 5.3813 | 6.8871 | 4.7909 | 5.7979 | 4.7683 | 5.6773 |
| MAST3_S774 | MAST3 | O60307 | | 774 | QPERGPSPSLLNT | 5.9699 | 6.7077 | 6.3624 | 6.1337 | 5.8209 | 5.9862 | 6.7542 | 7.2802 | 7.1924 | 6.7320 | 6.4657 | 6.0238 | 5.1031 | 7.0116 | 5.1628 | 5.2934 |
| MAST3_S889 | MAST3 | O60307 | | 889 | GGSGGGSGGRVPK | 7.3017 | 7.0741 | 7.7894 | 7.2529 | 5.9439 | 7.0285 | 7.3370 | 6.6620 | 7.7835 | 7.1216 | 7.3191 | 7.6431 | 3.8826 | 3.7609 | 2.8127 | 3.2870 |
| MASTL_S453 | MASTL | Q96GX5 | | 453 | FELVDSSPCKKII | 5.8036 | 6.3009 | 7.0937 | 5.5611 | 5.2559 | 6.5343 | 6.1284 | 6.6484 | 7.1603 | 6.9758 | 6.9797 | 7.2761 | 5.1228 | 5.7728 | 5.1991 | 6.1873 |
| MASTL_S512 | MASTL | Q96GX5 | | 512 | KENIVNSFTDKQQ | 6.3259 | 7.4916 | 7.3166 | 7.6791 | 6.1875 | 6.6722 | 7.5143 | 6.9593 | 7.5502 | 7.2705 | 7.7918 | 7.5407 | 2.9817 | 4.5092 | 2.4560 | 3.7535 |
| MAT2B_S282 | MAT2B | Q9NZL9 | | 282 | LRPITDSPVLGAQ | 5.0571 | 4.9731 | 6.7216 | 5.6039 | 5.4090 | 5.9385 | 5.5530 | 5.6213 | 6.6750 | 5.9707 | 6.0546 | 7.0409 | 7.2093 | 6.9535 | 7.3015 | 7.9171 |
| MATR3_S11 | MATR3 | A8MXP9 | | 598 | SRKRSYSPDGKES | 6.5065 | 6.4879 | 5.9757 | 6.5662 | 6.2700 | 5.8824 | 6.3396 | 5.7602 | 6.1865 | 6.0443 | 5.9772 | 6.0595 | 6.7349 | 6.3815 | 6.4365 | 6.3910 |
| MATR3_S188 | MATR3 | A8MXP9 | | 188 | RHFRRDSFDDRGP | 4.9238 | 5.3565 | 4.9380 | 5.9475 | 6.4816 | 6.0941 | 5.8446 | 6.1201 | 6.0585 | 6.4598 | 5.9578 | 5.6574 | 7.6909 | 7.4938 | 7.4223 | 7.5533 |
| MATR3_S206 | MATR3 | A8MXP9 | | 206 | LDYDHGSRSQESG | 6.3277 | 6.3716 | 6.2979 | 7.8621 | 6.2088 | 5.8728 | 6.2115 | 6.6765 | 6.3624 | 6.1622 | 6.1866 | 6.1403 | 5.5994 | 6.3342 | 5.6359 | 5.7501 |
| MATR3_S234 | MATR3 | A8MXP9 | | 814 | ENADGQSDENKDD | 6.0417 | 6.0963 | 5.7568 | 6.2021 | 6.8612 | 6.2273 | 6.6898 | 5.8873 | 6.2725 | 6.2914 | 6.9053 | 6.1107 | 7.0811 | 6.6767 | 5.5551 | 5.3447 |
| MATR3_S41 | MATR3 | A8MXP9 | | 604 | SPDGKESPSDKKS | 6.0328 | 6.2710 | 5.8453 | 6.5964 | 6.3158 | 6.2335 | 6.2436 | 6.0707 | 6.1787 | 6.4959 | 6.2453 | 6.4153 | 6.4949 | 6.3023 | 6.1980 | 6.0605 |
| MATR3_S596;S598 | MATR3 | A8MXP9 | | 722 | KRRFPGSMEGFVT | 5.6676 | 5.7946 | 5.5468 | 6.0179 | 6.2741 | 5.9871 | 5.9945 | 6.1665 | 5.8499 | 5.6127 | 5.6450 | 6.1980 | 7.4207 | 7.3622 | 7.3764 | 7.0860 |
| MATR3_S596;S598;S604 | MATR3 | A8MXP9 | | 234 | ERCRDDSFFGETS | 8.1175 | 7.9666 | 6.6148 | 8.3249 | 6.9095 | 6.4835 | 7.2329 | 6.5348 | 7.2626 | 6.2967 | 6.9672 | 6.6283 | 3.4715 | 4.2071 | 3.1227 | 3.8593 |

| | | | | | | | | | | | | | | | | | | | | | |
|---|---|---|---|---|---|---|---|---|---|---|---|---|---|---|---|---|---|---|---|---|---|
| MATR3_S596;S604 | MATR3 | A8MXP9 | | 11 | FQQSSLSRDSQGH | 4.1537 | 5.6889 | 6.0791 | 5.9735 | 4.9672 | 6.1150 | 6.3528 | 6.4170 | 6.1324 | 7.2336 | 7.0458 | 6.2896 | 6.2903 | 7.3896 | 6.9206 | 6.9507 |
| MATR3_S598 | MATR3 | A8MXP9 | | 9 | KSFQQSSLSRDSQ | 4.5116 | 6.1412 | 6.7260 | 6.7018 | 5.5412 | 6.3404 | 6.3451 | 6.9573 | 6.6409 | 9.1647 | 8.0612 | 8.5657 | 4.3161 | 5.1928 | 3.9219 | 4.8721 |
| MATR3_S598;S604 | MATR3 | A8MXP9 | | 41 | SLSMPASLGRMNQ | 6.0697 | 6.9548 | 5.6026 | 5.8457 | 6.6324 | 6.6379 | 6.6078 | 8.3780 | 6.0998 | 4.9279 | 4.7995 | 7.0859 | 6.3238 | 7.2935 | 5.2065 | 5.5343 |
| MATR3_S598;S606 | MATR3 | A8MXP9 | | 150 | QLKRRRTEEGPTL | 4.8229 | 5.6792 | 6.6825 | 5.8491 | 6.0429 | 6.0005 | 6.2541 | 6.4002 | 6.2478 | 6.1943 | 5.9825 | 5.7469 | 6.5511 | 7.0033 | 7.3874 | 7.1552 |
| MATR3_S604 | MATR3 | A8MXP9 | | 705 | AVKKDGSASAAAK | 6.1525 | 6.5079 | 6.1085 | 6.7975 | 5.6164 | 6.1835 | 6.5670 | 5.7065 | 5.9332 | 6.2564 | 6.3388 | 6.8047 | 5.9489 | 6.5098 | 6.0770 | 6.4914 |
| MATR3_S705 | MATR3 | A8MXP9 | 598;604 | | NA | 6.2823 | 6.8976 | 6.7950 | 7.1183 | 6.7015 | 6.2629 | 6.7928 | 5.6033 | 6.1824 | 6.5083 | 6.5610 | 6.6536 | 5.7426 | 5.5863 | 5.1753 | 5.1368 |
| MATR3_S722 | MATR3 | A8MXP9 | 596;598 | | NA | 6.1677 | 5.7468 | 5.7652 | 5.8918 | 6.1744 | 6.1262 | 6.4068 | 5.5075 | 5.7841 | 5.6358 | 6.1003 | 5.8408 | 7.9024 | 7.2068 | 7.0985 | 6.6449 |
| MATR3_S814 | MATR3 | A8MXP9 | 598;606 | | NA | 6.9837 | 6.4498 | 6.0941 | 6.3394 | 6.6251 | 5.5908 | 6.7912 | 5.9923 | 6.7845 | 6.4330 | 6.1786 | 6.1366 | 6.2555 | 5.9518 | 5.7429 | 5.6508 |
| MATR3_S9 | MATR3 | A8MXP9 | 596;604 | | NA | 7.0251 | 6.6258 | 6.4395 | 6.6809 | 6.3520 | 5.0713 | 6.4174 | 6.1176 | 6.7954 | 6.9052 | 7.0170 | 6.6632 | 5.2812 | 5.3609 | 5.1883 | 6.0592 |
| MATR3_T150 | MATR3 | A8MXP9 | 596;598;604 | | NA | 5.6731 | 6.0980 | 5.4864 | 7.2172 | 6.8993 | 6.8943 | 6.4841 | 5.6179 | 6.3806 | 6.2182 | 6.6157 | 6.8917 | 6.1293 | 6.2444 | 5.6570 | 5.4927 |
| MAVS_S152 | MAVS | Q7Z434 | | 222 | PSRGPVSPSVSFQ | 5.7241 | 5.8930 | 5.9281 | 6.2140 | 6.1736 | 5.9091 | 6.3211 | 6.3823 | 6.2590 | 6.1769 | 6.0476 | 6.3190 | 6.9548 | 6.5238 | 6.5978 | 6.5756 |
| MAVS_S165 | MAVS | Q7Z434 | | 152 | ETQAPESPGENSE | 5.6383 | 6.3171 | 6.1088 | 5.7852 | 6.3629 | 6.1541 | 6.2338 | 6.1652 | 5.5064 | 6.0708 | 6.2336 | 6.1234 | 7.2718 | 6.9477 | 6.7982 | 6.2828 |
| MAVS_S222 | MAVS | Q7Z434 | | 419 | GLGSELSKPGVLA | 5.7149 | 5.5128 | 5.7810 | 6.0083 | 5.2362 | 5.5141 | 5.9991 | 6.0168 | 5.3782 | 5.5554 | 6.0843 | 6.4472 | 7.7083 | 8.2200 | 7.5650 | 7.2584 |
| MAVS_S246;T250 | MAVS | Q7Z434 | | 165 | QALQTLSPRAIPR | 6.0622 | 5.8326 | 6.5629 | 6.5764 | 5.8102 | 5.7161 | 6.2415 | 6.8015 | 7.2227 | 5.0071 | 6.3804 | 5.3946 | 6.5170 | 7.5666 | 6.5708 | 5.7374 |
| MAVS_S419 | MAVS | Q7Z434 | 246;250 | | NA | 7.3254 | 7.7033 | 6.5797 | 7.8081 | 7.5604 | 5.8979 | 7.2008 | 7.1883 | 6.7317 | 7.0898 | 6.9191 | 7.7746 | 3.7956 | 3.5555 | 3.2642 | 3.6056 |
| MAZ_S361 | MAZ | P56270-2 | | 361 | HVRQVHSTERPFK | 7.1386 | 7.9558 | 6.9368 | 8.3517 | 6.3752 | 7.5275 | 7.0252 | 7.7443 | 7.6255 | 7.6688 | 8.1350 | 7.7338 | 1.9658 | 3.2746 | 1.9326 | 2.6088 |
| MBD1_S115 | MBD1 | Q9UIS9-12 | | 311 | PRALAPSPPAEFI | 5.9323 | 6.3273 | 6.3989 | 6.6168 | 6.3094 | 6.1015 | 6.3170 | 6.3329 | 6.8722 | 6.4724 | 5.9862 | 6.2919 | 5.8833 | 6.3392 | 6.1195 | 5.6992 |
| MBD1_S17 | MBD1 | K7EN65 | | 17 | PRALAPSPPAEFI | 6.0706 | 6.3004 | 6.5482 | 6.9638 | 6.4326 | 6.3059 | 6.3878 | 6.2394 | 6.1888 | 6.3862 | 6.3641 | 6.1610 | 5.8488 | 5.9634 | 5.8858 | 5.9529 |
| MBD1_S262 | MBD1 | Q9UIS9-12 | | 297 | PPPPSQSPEPTEP | 6.6805 | 6.4359 | 6.3481 | 7.1475 | 6.4607 | 5.8816 | 6.0843 | 6.2258 | 6.5378 | 6.4091 | 6.3377 | 6.6197 | 5.7481 | 5.5312 | 5.6050 | 5.9469 |
| MBD1_S297 | MBD1 | Q9UIS9-12 | | 513 | DEWTPGTAVLTSP | 7.1351 | 7.3385 | 6.6640 | 7.5350 | 6.3244 | 6.6888 | 6.8700 | 7.0001 | 6.7259 | 7.0441 | 7.1081 | 6.9842 | 3.9650 | 4.4737 | 3.9322 | 4.2109 |
| MBD1_S311 | MBD1 | Q9UIS9-12 | | 399 | SEDGAGSPPPYRR | 5.5720 | 5.8362 | 5.5337 | 6.2893 | 5.8191 | 5.5720 | 6.1338 | 6.1662 | 5.9523 | 5.7813 | 5.7938 | 6.3143 | 6.9463 | 7.4562 | 7.1390 | 7.6944 |
| MBD1_S391 | MBD1 | Q9UIS9-12 | | 570 | EAGGAGTPVITEI | 5.8634 | 6.6524 | 6.2086 | 6.6211 | 6.0178 | 6.0930 | 6.6188 | 6.4367 | 6.8397 | 6.9869 | 6.4240 | 6.3579 | 5.5816 | 6.0439 | 5.6033 | 5.6508 |
| MBD1_S391;S399 | MBD1 | Q9UIS9-10 | | 262 | PRALAPSPPAEFI | 6.0213 | 6.2698 | 5.9462 | 7.1095 | 5.9268 | 5.7658 | 6.1106 | 6.4348 | 6.3904 | 6.3205 | 6.5156 | 7.2016 | 5.8000 | 6.5019 | 5.8648 | 5.8204 |
| MBD1_S399 | MBD1 | Q9UIS9-12 | | 391 | LLPSVWSESEDGA | 6.4578 | 6.2162 | 6.6210 | 6.9412 | 6.5630 | 6.2436 | 7.0578 | 6.7102 | 6.1590 | 4.9781 | 4.9248 | 5.7397 | 6.0805 | 6.0271 | 6.3564 | 6.9236 |
| MBD1_T513 | MBD1 | K7EN65 | | 115 | LREGASSPVQVPG | 5.9390 | 6.7029 | 5.8588 | 7.0098 | 5.5662 | 6.0854 | 6.6446 | 6.2052 | 6.3269 | 6.2419 | 6.2934 | 6.6554 | 5.8532 | 6.1349 | 6.2930 | 6.1896 |
| MBD1_T570 | MBD1 | Q9UIS9-12 | 391;399 | | NA | 5.8599 | 6.7259 | 6.4797 | 6.9055 | 6.2349 | 5.8685 | 6.1315 | 6.8862 | 6.4725 | 6.0402 | 6.1789 | 6.8234 | 6.1109 | 5.8453 | 5.3254 | 6.1114 |
| MBD2_S181 | MBD2 | Q9UBB5 | | 181 | SDVYYFSPSGKKF | 7.6388 | 7.8633 | 6.8692 | 7.6944 | 6.6037 | 7.0629 | 7.2667 | 6.8436 | 7.7665 | 7.5801 | 7.1935 | 7.2442 | 2.7786 | 3.6416 | 2.5721 | 3.3809 |
| MBD3_S144 | MBD3 | O95983 | | 56 | ARYLGGSMDLSTF | 7.1036 | 7.0370 | 7.2632 | 7.0301 | 6.4392 | 6.2847 | 5.9843 | 6.2759 | 6.9733 | 7.6337 | 7.0057 | 6.8372 | 4.2940 | 5.1549 | 4.3539 | 4.3293 |
| MBD3_S37 | MBD3 | O95983 | | 37 | RDVFYYSPSGKKF | 7.2189 | 7.2493 | 6.6137 | 7.1099 | 6.8500 | 6.7996 | 6.5405 | 6.9153 | 7.1962 | 7.0505 | 6.8535 | 7.1006 | 3.9575 | 4.4323 | 3.7999 | 4.3124 |
| MBD3_S56 | MBD3 | O95983 | | 85 | QRVRYDSSNQVKG | 6.9879 | 6.7954 | 7.1706 | 7.1151 | 5.8920 | 5.9089 | 6.7667 | 5.7593 | 6.0765 | 6.3139 | 6.7272 | 6.8800 | 5.4832 | 5.3451 | 5.2247 | 5.5533 |
| MBD3_S85 | MBD3 | O95983 | | 144 | FWEKKLSGLNAFD | 6.5650 | 6.4041 | 5.6943 | 6.5461 | 6.5556 | 5.8070 | 7.0451 | 6.0144 | 6.6906 | 6.5349 | 7.2078 | 6.2295 | 5.6220 | 5.6177 | 5.2580 | 6.2079 |
| MBD3_S86 | MBD3 | O95983 | | 86 | RVRYDSSNQVKGK | 7.0614 | 6.7236 | 7.3532 | 7.7869 | 6.5044 | 5.6697 | 6.4219 | 6.1125 | 6.1301 | 6.3401 | 6.9807 | 6.9767 | 5.4534 | 4.5623 | 4.8726 | 5.0506 |
| MBD5_S300 | MBD5 | Q9P267 | | 300 | RTNIPLSPTLTTK | 5.7815 | 6.2197 | 6.4768 | 5.8648 | 6.5679 | 6.7489 | 5.7883 | 6.5105 | 6.4245 | 6.2833 | 6.4522 | 6.4566 | 6.2781 | 6.2215 | 5.7314 | 6.1943 |
| MBOAT2_T474 | MBOAT2 | Q6ZWT7 | | 474 | TQRRKNTHENIQL | 3.3732 | 3.2778 | 4.0811 | 4.0636 | 4.3782 | 3.7986 | 4.2840 | 4.0175 | 4.0670 | 4.0509 | 4.0465 | 3.6705 | 13.5728 | 12.5615 | 13.8838 | 12.8729 |
| MBOAT7_S285 | MBOAT7 | Q96N66 | | 285 | QCPPPSSPEKAAS | 5.9229 | 5.9882 | 5.9885 | 6.4868 | 6.1499 | 5.8226 | 5.9340 | 6.3630 | 6.4146 | 6.2519 | 6.0974 | 5.9413 | 6.9829 | 6.8265 | 5.9904 | 6.8392 |
| MBP_S26 | MBP | P02686 | | 26 | ETNRGESEKKRNL | 6.9920 | 5.5130 | 7.9495 | 6.6236 | 6.9795 | 5.0520 | 5.0996 | 7.5010 | 5.9436 | 8.0118 | 8.8163 | 9.7586 | 3.7543 | 5.3736 | 3.2261 | 3.4056 |
| MBP_S40 | MBP | P02686 | | 40 | ELSRTTSEDNEVF | 6.1155 | 6.7627 | 6.7404 | 5.9986 | 5.9824 | 6.0701 | 5.8977 | 5.8423 | 6.1040 | 6.9166 | 6.4964 | 6.2071 | 6.3510 | 6.3402 | 6.7242 | 5.4506 |
| MBTD1_S612 | MBTD1 | Q05BQ5 | | 612 | NFLQGASDQESNG | 6.3492 | 6.1576 | 5.9850 | 6.4546 | 6.7149 | 6.3186 | 6.3993 | 6.5415 | 5.5876 | 6.1685 | 5.8768 | 5.6868 | 6.4670 | 6.3450 | 6.7344 | 6.2129 |
| MCC_S484 | MCC | P23508-2 | | 484 | QATGPSSPGRLTS | 6.5186 | 6.5670 | 6.4666 | 6.4967 | 6.4232 | 6.7834 | 6.4726 | 6.9491 | 6.7692 | 6.5875 | 6.8643 | 6.7127 | 5.2607 | 5.3193 | 4.6044 | 5.2046 |
| MCL1_T163 | MCL1 | Q07820 | | 163 | DGSLPSTPPPAEE | 5.8465 | 6.7575 | 6.3951 | 5.9481 | 6.8528 | 5.6509 | 6.5212 | 6.9782 | 6.0340 | 5.8202 | 6.1437 | 5.9688 | 6.0835 | 6.3375 | 5.5851 | 7.0767 |
| MCM10_S143 | MCM10 | Q7L590 | | 548 | ASGIMGSPKPAIK | 8.1963 | 7.5379 | 6.7199 | 7.4531 | 6.0018 | 5.9999 | 6.9503 | 6.1157 | 8.6831 | 7.3242 | 7.0368 | 7.4722 | 3.5697 | 4.1619 | 3.0480 | 3.7291 |
| MCM10_S155 | MCM10 | Q7L590 | | 196 | ARTPKASPPDPKS | 6.9740 | 7.1390 | 6.5314 | 7.1205 | 6.3963 | 6.3653 | 6.5187 | 6.2216 | 7.0258 | 6.4387 | 6.8515 | 6.8555 | 4.6556 | 5.0636 | 4.8178 | 5.0247 |
| MCM10_S196 | MCM10 | Q7L590 | | 155 | KSPVEKSPRPPLK | 7.2902 | 7.0019 | 6.7006 | 7.2328 | 6.7523 | 6.5573 | 6.9415 | 6.6043 | 6.9814 | 6.5796 | 6.5237 | 6.6629 | 4.5222 | 4.7553 | 4.1329 | 4.7612 |
| MCM10_S548 | MCM10 | Q7L590 | | 644 | VLFYDESPPPRPK | 6.3320 | 6.8197 | 6.8840 | 7.6212 | 6.8290 | 6.5176 | 6.9092 | 6.9959 | 7.4146 | 6.4667 | 6.8971 | 7.4127 | 4.0176 | 4.4214 | 4.1351 | 4.3262 |
| MCM10_S593 | MCM10 | Q7L590 | | 680 | TKTNPNSIKKKQK | 7.0261 | 6.8899 | 7.1805 | 7.1658 | 5.6616 | 6.2495 | 6.7666 | 6.0608 | 7.4747 | 7.4735 | 7.6514 | 8.1210 | 3.7840 | 4.6269 | 3.6963 | 4.1715 |
| MCM10_S644 | MCM10 | Q7L590 | | 593 | SSSEVESPAVPSS | 6.4653 | 6.8215 | 6.4344 | 6.8498 | 6.1501 | 6.5382 | 6.5062 | 7.4939 | 6.6835 | 7.5590 | 6.3536 | 7.3040 | 4.0452 | 5.2316 | 4.3987 | 5.1650 |
| MCM10_S680 | MCM10 | Q7L590 | | 143 | TIKQTASPARLQK | 7.9395 | 6.1547 | 4.8761 | 6.4443 | 6.0796 | 6.3864 | 8.1369 | 6.1568 | 8.1926 | 6.6953 | 7.0321 | 6.5065 | 5.0799 | 5.7878 | 3.6212 | 4.9104 |
| MCM10_T85 | MCM10 | Q7L590 | | 85 | GDMEDLTDEEEVP | 5.7107 | 6.7176 | 6.5524 | 5.9018 | 6.4440 | 6.0825 | 5.7947 | 5.7275 | 6.6957 | 6.7086 | 6.6944 | 6.9151 | 6.0012 | 7.2585 | 6.2374 | 4.5580 |
| MCM2_S108 | MCM2 | P49736 | | 41 | TDALTSSPGRDLP | 5.8110 | 6.2432 | 6.0706 | 5.9794 | 5.9990 | 5.6620 | 5.7649 | 6.3082 | 6.4601 | 6.5226 | 6.1571 | 6.0670 | 6.7304 | 6.7557 | 6.8699 | 6.5990 |
| MCM2_S139 | MCM2 | P49736 | | 108 | VEELTASQREAAE | 5.7130 | 6.1525 | 6.2313 | 6.3901 | 6.1345 | 6.0387 | 6.2296 | 6.4466 | 6.2116 | 6.2310 | 6.3155 | 6.3059 | 6.1953 | 6.6541 | 6.3000 | 6.4503 |
| MCM2_S26;S27 | MCM2 | P49736 | | 139 | RGLLYDSDEEDEE | 5.4053 | 5.8879 | 6.1432 | 5.8884 | 5.9424 | 6.2514 | 5.9415 | 6.1176 | 6.2071 | 6.3015 | 6.1906 | 6.2460 | 6.6983 | 7.0088 | 6.9780 | 6.7922 |
| MCM2_S27 | MCM2 | P49736 | | 27 | NDPLTSSPGRSSR | 5.6574 | 6.1200 | 5.7925 | 5.6906 | 5.9660 | 5.9992 | 5.8418 | 5.9683 | 6.1470 | 6.2187 | 6.2061 | 6.0506 | 7.1155 | 7.1798 | 7.2014 | 6.8452 |
| MCM2_S381 | MCM2 | P49736 | | 381 | RIRIQESPGKVAA | 5.8057 | 6.4580 | 6.4906 | 6.5298 | 5.6361 | 6.3173 | 6.0897 | 6.2167 | 6.2218 | 6.6670 | 6.8224 | 7.0125 | 5.6462 | 5.9454 | 5.7858 | 6.3552 |
| MCM2_S40 | MCM2 | P49736 | | 40 | RTDALTSSPGRDL | 6.0707 | 6.5303 | 6.3101 | 6.4900 | 6.0173 | 6.0847 | 5.8509 | 6.0705 | 6.2566 | 6.3393 | 6.4593 | 6.5992 | 6.4301 | 6.4448 | 5.9692 | 6.0770 |
| MCM2_S40;S41 | MCM2 | P49736 | | 53 | PPFEDESEGLLGT | 5.4284 | 6.4752 | 5.8863 | 6.2077 | 6.5548 | 6.4114 | 5.6672 | 6.4574 | 6.4044 | 6.3145 | 6.0297 | 6.5721 | 6.1800 | 6.2334 | 6.8618 | 6.3159 |
| MCM2_S40;S53 | MCM2 | P49736 | 40;41 | | NA | 5.7155 | 5.8981 | 6.3871 | 5.9061 | 5.7444 | 5.6971 | 5.6287 | 6.0449 | 6.1621 | 6.3544 | 5.9342 | 5.9942 | 7.0490 | 7.4962 | 6.8927 | 7.0952 |
| MCM2_S41 | MCM2 | P49736 | 26;27 | | NA | 5.7869 | 6.4761 | 6.1106 | 5.8511 | 6.0168 | 6.2166 | 5.9745 | 5.8760 | 6.3807 | 5.7992 | 6.1985 | 6.2343 | 6.7239 | 6.7457 | 6.7831 | 6.8262 |
| MCM2_S41;S53 | MCM2 | P49736 | 39;41 | | NA | 5.7727 | 6.2252 | 6.7393 | 6.1057 | 5.7390 | 5.5937 | 6.1350 | 6.2933 | 6.4603 | 6.7891 | 6.3761 | 6.6650 | 6.0523 | 6.3029 | 6.3774 | 6.3730 |
| MCM2_S53 | MCM2 | P49736 | 39;53 | | NA | 5.6248 | 6.2343 | 6.3681 | 5.6673 | 5.6331 | 6.2509 | 5.9329 | 6.0439 | 6.8447 | 6.1828 | 6.1016 | 6.2672 | 6.7855 | 6.8033 | 6.2457 | 7.0139 |
| MCM2_T39;S41 | MCM2 | P49736 | 41;53 | | NA | 6.1784 | 6.3342 | 6.8987 | 5.1628 | 5.1496 | 5.8256 | 5.4928 | 6.4556 | 5.6411 | 6.5545 | 5.9333 | 6.4307 | 7.9177 | 7.6026 | 6.4750 | 5.9473 |
| MCM2_T39;S53 | MCM2 | P49736 | 40;53 | | NA | 5.4856 | 7.1390 | 5.9739 | 5.6082 | 6.3547 | 6.0569 | 5.6912 | 6.2177 | 7.0338 | 5.8527 | 6.4577 | 6.0367 | 6.8494 | 6.9072 | 5.6861 | 6.6493 |
| MCM3_S205 | MCM3 | P25205-2 | | 205 | TIERRYSDLTTLV | 4.1021 | 4.5820 | 5.4578 | 4.8562 | 6.5687 | 5.8542 | 5.0343 | 5.9411 | 7.0045 | 9.0811 | 7.7090 | 7.6246 | 7.4730 | 7.6723 | 5.2874 | 5.7517 |
| MCM3_S713;S717;T719 | MCM3 | P25205-2 | | 756 | YDPYDFSDTEEEM | 5.7436 | 6.1553 | 6.4039 | 6.6116 | 5.7307 | 5.9831 | 5.8189 | 5.9024 | 6.6176 | 6.5000 | 6.3293 | 6.7577 | 6.3314 | 6.2884 | 6.5206 | 6.3054 |
| MCM3_S717;T719 | MCM3 | P25205-2 | | 773 | TPKTADSQETKES | 5.6001 | 6.0518 | 5.8584 | 5.2694 | 5.5855 | 5.8345 | 5.9391 | 5.4627 | 5.9642 | 6.2059 | 6.2039 | 6.1150 | 7.0561 | 7.4506 | 7.5868 | 7.8159 |
| MCM3_S756 | MCM3 | P25205-2 | | 758 | PYDFSDTEEEMPQ | 5.7082 | 6.3454 | 6.7971 | 6.8308 | 5.6704 | 5.5739 | 5.5543 | 5.5631 | 6.7102 | 6.5763 | 7.1357 | 6.7701 | 5.9802 | 6.5422 | 5.7153 | 6.5268 |
| MCM3_S756;T758 | MCM3 | P25205-2 | | 574 | AVDILATDDPNFS | 5.3900 | 5.9617 | 6.4965 | 6.6507 | 6.1815 | 5.7087 | 5.3900 | 6.3644 | 6.8151 | 5.6283 | 6.1206 | 6.1289 | 6.4679 | 6.6415 | 6.6803 | 7.3739 |
| MCM3_S756;T758;T767 | MCM3 | P25205-2 | 758;767 | | NA | 5.1292 | 5.7766 | 5.7363 | 5.9310 | 5.7249 | 5.8780 | 5.8550 | 5.8531 | 5.8504 | 6.2747 | 6.5730 | 6.3680 | 7.0854 | 7.4828 | 7.2327 | 7.2490 |
| MCM3_S756;T767 | MCM3 | P25205-2 | 713;717;719 | | NA | 5.4747 | 5.2897 | 6.1607 | 5.7413 | 5.8893 | 5.8512 | 5.9662 | 5.7458 | 5.8591 | 5.6541 | 5.9560 | 6.3418 | 7.6786 | 7.2823 | 7.3045 | 7.8048 |
| MCM3_S758 | MCM3 | P25205-2 | 756;767 | | NA | 5.3928 | 5.4605 | 5.7877 | 5.6026 | 5.6935 | 6.0363 | 5.5686 | 5.7655 | 6.0694 | 6.3301 | 6.4254 | 5.9601 | 7.1939 | 7.8012 | 7.7039 | 7.2085 |
| MCM3_S773 | MCM3 | P25205-2 | 756;758;767 | | NA | 5.5216 | 5.7343 | 6.2187 | 5.5697 | 5.5626 | 6.2194 | 6.5857 | 5.9652 | 5.5905 | 5.6611 | 5.6704 | 5.6052 | 7.4174 | 7.6099 | 7.4778 | 7.5905 |
| MCM3_T574 | MCM3 | P25205-2 | 717;719 | | NA | 5.5729 | 6.4323 | 5.8970 | 5.9519 | 5.4809 | 5.3013 | 6.2440 | 5.8263 | 6.2156 | 6.0462 | 5.6449 | 6.4809 | 7.2524 | 6.8079 | 7.6610 | 7.1845 |
| MCM3_T758;T767 | MCM3 | P25205-2 | 756;758 | | NA | 6.0806 | 6.4431 | 7.5393 | 6.8624 | 6.2737 | 6.5789 | 6.4011 | 6.2601 | 6.5830 | 6.8569 | 6.2762 | 6.0557 | 5.5706 | 5.6522 | 5.3264 | 5.2398 |
| MCM3AP_S114 | MCM3AP | O60318 | | 527 | PGDGEVSPSTEDA | 5.3952 | 5.5263 | 4.8245 | 5.6254 | 5.8035 | 5.3586 | 5.8398 | 5.6255 | 5.0787 | 4.9865 | 5.2336 | 5.2141 | 8.9615 | 8.1373 | 9.6297 | 8.7598 |
| MCM3AP_S300 | MCM3AP | O60318 | | 300 | KEDQDRSPRRHGH | 6.7944 | 6.5981 | 5.8119 | 6.9273 | 7.4226 | 7.0884 | 6.6571 | 6.6984 | 6.5705 | 5.8984 | 6.1304 | 4.8646 | 5.6840 | 5.9367 | 5.7636 | 5.1536 |
| MCM3AP_S430 | MCM3AP | O60318 | | 425 | RSESTDSLGGLSP | 3.9810 | 4.9536 | 5.3775 | 5.2166 | 5.1408 | 6.0889 | 5.4142 | 5.4405 | 5.5176 | 5.4796 | 5.4264 | 6.3483 | 8.2880 | 8.7406 | 9.3352 | 9.2514 |
| MCM3AP_S527 | MCM3AP | O60318 | | 430 | DSLGGLSPSEVTA | 6.3855 | 6.3051 | 7.4280 | 7.0935 | 5.3191 | 4.7305 | 6.3855 | 5.1301 | 6.3833 | 7.5257 | 7.6343 | 7.9297 | 5.2039 | 5.2191 | 5.1822 | 6.1444 |
| MCM3AP_S527;S538 | MCM3AP | O60318 | | 114 | TGFSFKSPTSVGA | 6.4300 | 6.4672 | 6.1391 | 6.9394 | 6.4308 | 7.0093 | 7.1721 | 5.9939 | 6.7171 | 6.8230 | 6.3475 | 6.1291 | 4.4959 | 5.9935 | 5.3539 | 5.5582 |
| MCM3AP_T423 | MCM3AP | O60318 | | 423 | SNRSESTDSLGGL | 5.5209 | 3.9717 | 5.2705 | 4.2100 | 5.5370 | 5.2557 | 5.5182 | 4.4172 | 5.7833 | 4.6299 | 4.7268 | 4.9166 | 9.7079 | 9.1925 | 11.4253 | 9.9166 |
| MCM3AP_T423;S425 | MCM3AP | O60318 | 527;538 | | NA | 5.8076 | 5.9190 | 6.1713 | 6.5006 | 7.2905 | 6.0946 | 6.3952 | 6.5672 | 6.3809 | 6.3017 | 6.5115 | 6.5553 | 6.0327 | 6.1161 | 5.5820 | 5.7738 |
| MCM3AP_T425 | MCM3AP | O60318 | 423;425 | | NA | 3.6649 | 3.9562 | 4.2067 | 4.0309 | 5.9686 | 3.6993 | 4.3804 | 4.1183 | 2.3767 | 3.1094 | 2.8127 | 1.6786 | 15.5546 | 12.0513 | 14.3890 | 14.0023 |
| MCM4_S120 | MCM4 | P33991 | | 120 | QRPDLGSAQKGLQ | 7.2231 | 7.9080 | 7.5007 | 7.4797 | 5.7776 | 7.1191 | 6.9698 | 6.8119 | 7.6914 | 7.6077 | 8.0581 | 8.5297 | 2.2909 | 3.5073 | 2.2830 | 3.2422 |
| MCM4_S131 | MCM4 | P33991 | | 326 | GRIAEPSVCGRCH | 5.9484 | 6.3351 | 6.0304 | 5.9839 | 5.9491 | 6.2148 | 5.8509 | 5.9864 | 6.0212 | 6.1396 | 6.1975 | 6.2484 | 6.6320 | 6.6212 | 7.1161 | 6.7251 |
| MCM4_S326 | MCM4 | P33991 | | 131 | LQVDLQSDGAAAE | 6.1417 | 6.6022 | 6.8314 | 6.6394 | 6.0097 | 6.3520 | 6.2739 | 6.1121 | 6.3605 | 6.7696 | 6.5442 | 6.5875 | 5.7947 | 5.7434 | 5.6553 | 5.5826 |
| MCM4_T110 | MCM4 | P33991 | | 110 | RSGVRGTPVRQRP | 7.0354 | 7.7583 | 7.6684 | 6.2830 | 5.5814 | 7.9511 | 6.6198 | 5.5528 | 6.5374 | 6.8349 | 7.7616 | 8.0596 | 4.0047 | 4.6343 | 3.6163 | 4.1008 |
| MCM5_S604 | MCM5 | P33992 | | 604 | RDSDRRSSIPITV | 6.5826 | 7.6137 | 6.8992 | 5.5364 | 5.9875 | 5.8813 | 4.9280 | 6.6452 | 6.3476 | 5.6634 | 5.2559 | 5.5421 | 6.3173 | 6.4007 | 7.1531 | 7.2460 |
| MCM6_S271 | MCM6 | Q14566 | | 699 | VNGINGYNEDINQ | 5.3527 | 5.8094 | 5.9961 | 6.2068 | 6.3640 | 6.2520 | 5.9568 | 6.2189 | 5.9852 | 5.7776 | 6.4968 | 5.8089 | 6.7070 | 6.9137 | 7.2412 | 6.9130 |
| MCM6_S762 | MCM6 | Q14566 | | 271 | ETNSRVSGVDGYE | 6.0428 | 6.5749 | 5.9385 | 6.4467 | 6.1309 | 6.0061 | 6.8581 | 6.8463 | 6.1964 | 6.5102 | 6.6250 | 6.5817 | 5.2898 | 5.8900 | 5.6896 | 6.3729 |
| MCM6_S802 | MCM6 | Q14566 | | 762 | IESEIDSEEELIN | 5.9170 | 6.0186 | 6.3151 | 6.4390 | 6.5254 | 6.3199 | 5.8331 | 6.0974 | 6.2407 | 6.2608 | 6.6330 | 6.6436 | 6.3117 | 6.4617 | 6.2492 | 5.7337 |
| MCM6_Y699 | MCM6 | Q14566 | | 802 | KGSTEGSESYEED | 6.5390 | 6.3811 | 6.0111 | 6.4503 | 6.0053 | 6.4215 | 6.4768 | 6.4491 | 7.0623 | 6.3650 | 6.5148 | 6.3915 | 5.2757 | 6.3189 | 5.9454 | 5.3921 |
| MCM7_S121 | MCM7 | P33993 | | 365 | VGGVDQSPRGMKI | 5.9871 | 7.0050 | 6.2251 | 6.6900 | 5.9160 | 7.0541 | 6.3555 | 6.8195 | 6.1819 | 6.2954 | 6.8737 | 7.2218 | 5.6391 | 5.1775 | 4.8032 | 5.7551 |
| MCM7_S365 | MCM7 | P33993 | | 500 | RYNPRRSLEQNIQ | 6.6576 | 6.2845 | 6.1167 | 6.7475 | 6.1753 | 6.5793 | 6.9403 | 6.8280 | 6.1700 | 6.3538 | 6.7614 | 6.8186 | 5.2216 | 5.9436 | 5.2569 | 5.1446 |
| MCM7_S500 | MCM7 | P33993 | | 121 | DPGMVRSPQNQYP | 6.2347 | 6.8061 | 6.4245 | 7.3823 | 5.8434 | 6.2167 | 6.4356 | 6.8772 | 6.6675 | 6.8469 | 6.8721 | 6.9631 | 4.8162 | 5.1895 | 4.9013 | 5.5229 |
| MCM7_S678 | MCM7 | P33993 | | 678 | GRSVRFSEAEQRC | 6.4836 | 6.2182 | 6.7266 | 6.7830 | 6.2953 | 6.2322 | 6.0943 | 6.0978 | 5.3850 | 5.2918 | 6.0497 | 6.1639 | 7.1197 | 6.1289 | 6.7226 | 6.2073 |
| MCM9_S1088 | MCM9 | Q9NXL9 | | 1088 | GERGPSSPPTTTA | 6.4511 | 7.0240 | 6.5104 | 6.5820 | 6.3421 | 6.3838 | 6.3157 | 5.9511 | 6.7067 | 6.4943 | 6.6964 | 6.8308 | 5.1341 | 5.5797 | 5.4310 | 5.5667 |
| MCM9_S802 | MCM9 | Q9NXL9 | | 915 | NVKPPGSPVAKLA | 5.8891 | 6.0611 | 6.4008 | 6.3764 | 6.4329 | 6.4733 | 6.1985 | 5.9836 | 7.0272 | 6.2896 | 6.1378 | 6.9918 | 5.9600 | 5.9583 | 5.4626 | 6.3570 |
| MCM9_S883 | MCM9 | Q9NXL9 | | 952 | TKIAVHSPKISQR | 7.4811 | 7.2025 | 6.8757 | 6.8289 | 6.4525 | 7.0997 | 6.9124 | 7.1183 | 8.0307 | 7.4523 | 7.5509 | 7.7607 | 2.9084 | 4.1637 | 2.5561 | 3.6062 |
| MCM9_S915 | MCM9 | Q9NXL9 | | 883 | QSTPVHSPDRMLD | 6.4554 | 6.3262 | 6.3869 | 6.6754 | 7.4318 | 5.6855 | 7.1269 | 7.3952 | 6.8201 | 5.7386 | 5.8833 | 6.3927 | 6.0097 | 5.6334 | 5.3276 | 4.7111 |
| MCM9_S952 | MCM9 | Q9NXL9 | | 802 | DIGLLPSPGETGV | 6.2804 | 6.5848 | 6.2593 | 7.1823 | 6.3819 | 6.0314 | 5.9352 | 7.4564 | 6.8246 | 6.5967 | 6.9676 | 6.8404 | 5.0425 | 5.3746 | 5.0405 | 5.2013 |
| MCMBP_S154 | MCMBP | Q9BTE3 | | 154 | ANQARVSPSTSYT | 5.4337 | 5.7080 | 6.2246 | 5.6921 | 5.8238 | 6.0604 | 5.4668 | 6.2428 | 6.3119 | 6.1100 | 6.3594 | 6.7302 | 6.7631 | 7.2047 | 7.0505 | 6.8178 |
| MCMBP_S298 | MCMBP | Q9BTE3 | | 298 | EEQRVHSPPASLV | 5.8099 | 5.8644 | 5.9859 | 5.3373 | 5.7771 | 6.0898 | 5.6066 | 6.1034 | 6.0191 | 5.8953 | 6.3941 | 6.2230 | 7.1284 | 7.3172 | 7.4971 | 6.9514 |
| MCMBP_T160 | MCMBP | Q9BTE3 | | 160 | SPSTSYTPSRHKR | 6.8324 | 6.7769 | 7.4612 | 6.4413 | 6.7715 | 6.6173 | 6.8018 | 6.7125 | 7.3864 | 6.7355 | 7.2227 | 7.4659 | 4.0300 | 4.4581 | 3.7039 | 4.5826 |
| MCOLN1_S547 | MCOLN1 | Q9GZU1 | | 547 | IAQCQDSPTSGKF | 6.1081 | 6.0401 | 6.3273 | 6.8818 | 6.3884 | 6.2091 | 5.9084 | 6.2740 | 6.5044 | 6.2912 | 6.6126 | 6.4990 | 5.3692 | 5.9080 | 5.9537 | 6.7248 |
| MCOLN1_T34 | MCOLN1 | Q9GZU1 | | 34 | PSPAPPTPPEEED | 5.3877 | 6.9795 | 7.4801 | 7.7358 | 6.4609 | 5.0600 | 7.4153 | 5.6542 | 6.4933 | 9.0611 | 5.5317 | 6.6590 | 3.5834 | 5.6182 | 6.3457 | 4.5342 |
| MCPH1_S254 | MCPH1 | Q8NEM0 | | 312 | IAGKVVTPDQKQA | 6.8616 | 7.5303 | 6.7369 | 7.1680 | 6.8941 | 6.7329 | 6.7353 | 7.0816 | 7.3499 | 6.7100 | 6.9670 | 7.2930 | 3.7707 | 4.1905 | 3.5866 | 4.3916 |
| MCPH1_S296 | MCPH1 | Q8NEM0 | | 254 | ERKLEGSINDIKS | 7.0625 | 6.7169 | 7.0429 | 6.2930 | 5.3605 | 8.3733 | 7.0136 | 5.2203 | 7.4733 | 7.9657 | 7.8385 | 8.6993 | 3.7758 | 4.5747 | 2.8824 | 3.7073 |
| MCPH1_S438 | MCPH1 | Q8NEM0 | | 296 | KFLSNLSKEEINL | 7.3581 | 6.0925 | 6.4222 | 7.9644 | 7.4935 | 6.4810 | 8.5531 | 5.6510 | 7.4935 | 6.9814 | 6.1220 | 5.6569 | 3.1493 | 6.1691 | 3.8321 | 4.5797 |
| MCPH1_T312 | MCPH1 | Q8NEM0 | | 438 | ESQLPSSPAQLSC | 6.0706 | 7.0261 | 6.4397 | 5.7089 | 6.7575 | 6.4930 | 6.2797 | 7.1871 | 6.2170 | 5.5291 | 7.2582 | 6.9739 | 6.6759 | 5.1800 | 5.1267 | 5.0765 |
| MCRIP1_S21 | MCRIP1 | C9JLW8 | | 21 | RTSSPRSPPSSSE | 6.2619 | 5.8359 | 7.1611 | 5.8238 | 6.2659 | 5.5340 | 5.8986 | 6.2909 | 6.5251 | 6.8542 | 6.1728 | 6.6206 | 5.5466 | 6.4672 | 6.2206 | 6.5208 |
| MCRS1_S295 | MCRS1 | Q96EZ8-2 | | 295 | DQVLNFSDAEDLI | 6.2302 | 6.1572 | 6.3847 | 6.2285 | 6.1504 | 6.1433 | 6.3779 | 6.1823 | 6.6065 | 6.4309 | 6.2936 | 6.2409 | 5.9904 | 6.2065 | 6.1721 | 6.2048 |
| MCRS1_T116 | MCRS1 | Q96EZ8-2 | | 116 | VSKAPSTPVPPSP | 5.8211 | 6.0997 | 6.1512 | 6.6848 | 6.0687 | 6.0881 | 6.3433 | 6.1260 | 6.3770 | 6.4646 | 6.2766 | 6.2791 | 6.1583 | 6.5340 | 6.0483 | 6.4792 |
| MDC1_S1033 | MDC1 | Q14676 | | 168 | LLLAEDSEEEVDF | 6.1930 | 6.3030 | 6.4009 | 6.9773 | 6.3511 | 5.9164 | 6.0908 | 6.2177 | 6.1478 | 6.2585 | 6.2574 | 6.3325 | 5.8205 | 6.4357 | 5.9699 | 6.3274 |
| MDC1_S1068 | MDC1 | Q14676 | | 793 | GDQHPESPVHTEP | 6.2077 | 6.1898 | 6.1843 | 6.7039 | 6.2222 | 5.9500 | 6.2614 | 6.0843 | 6.1713 | 6.3069 | 5.9850 | 5.9184 | 5.8940 | 6.7977 | 6.4169 | 6.7062 |
| MDC1_S108 | MDC1 | Q14676 | | 780 | AYGPCLSPPRAIP | 6.1362 | 6.4045 | 6.0907 | 6.7333 | 6.1805 | 5.9851 | 6.3829 | 5.9900 | 6.2429 | 6.2293 | 6.0443 | 6.0963 | 5.8965 | 6.4664 | 6.3392 | 6.7821 |
| MDC1_S1086 | MDC1 | Q14676 | | 1033 | SRGDQESPDACLP | 5.9462 | 6.0122 | 6.1577 | 6.7272 | 5.8392 | 5.7555 | 5.9517 | 5.9530 | 5.9976 | 5.8401 | 5.8451 | 5.9048 | 6.2218 | 7.1395 | 7.2509 | 7.4575 |
| MDC1_S1400;T1403 | MDC1 | Q14676 | | 1466 | STDQPVTPEPTSQ | 5.9751 | 6.4517 | 5.9149 | 7.1387 | 6.2103 | 6.2681 | 5.9099 | 6.2686 | 6.3389 | 6.7628 | 5.9284 | 6.6000 | 5.7633 | 6.3141 | 5.9180 | 6.2370 |
| MDC1_S168 | MDC1 | Q14676 | | 1321 | NMSSVKTPETVVP | 6.3339 | 6.9709 | 6.6981 | 7.8742 | 6.0834 | 6.0465 | 6.7652 | 6.6733 | 6.4813 | 6.8335 | 6.7447 | 7.0615 | 4.3203 | 5.3090 | 4.6554 | 5.1487 |
| MDC1_S1711 | MDC1 | Q14676 | | 1485 | DRSSVKTPETVVP | 6.1625 | 6.5293 | 6.3572 | 7.0430 | 6.2203 | 5.9682 | 6.3492 | 6.5970 | 6.4568 | 6.7712 | 6.9948 | 6.3632 | 5.1014 | 5.8233 | 5.4499 | 5.8128 |
| MDC1_S1732 | MDC1 | Q14676 | | 505 | VEADKSSPGIHLE | 6.3601 | 7.0425 | 5.8100 | 7.2225 | 5.8436 | 6.0008 | 6.0405 | 6.4877 | 6.2489 | 6.1704 | 6.1075 | 6.3450 | 5.1222 | 6.5197 | 6.2428 | 6.4357 |
| MDC1_S1775 | MDC1 | Q14676 | | 376 | QESQAGSDTDVEE | 5.5374 | 5.7169 | 5.6505 | 6.2711 | 5.9005 | 5.7896 | 6.1235 | 5.9244 | 6.4755 | 6.6167 | 6.4349 | 6.6892 | 6.0454 | 7.3118 | 6.6450 | 6.8678 |
| MDC1_S1814 | MDC1 | Q14676 | | 964 | QKGQASSPTPEPG | 5.9461 | 5.9880 | 6.4871 | 6.6705 | 6.0984 | 6.0232 | 6.4237 | 5.8941 | 6.5424 | 6.2231 | 6.4987 | 6.0690 | 6.2110 | 6.3269 | 6.0327 | 6.5651 |
| MDC1_S1820 | MDC1 | Q14676 | | 329 | PFGFIDSDTDAEE | 4.7664 | 5.1898 | 5.7311 | 5.6237 | 5.1658 | 6.0011 | 5.6861 | 6.3149 | 6.3564 | 7.4593 | 6.8020 | 7.5024 | 6.2128 | 7.0101 | 6.7612 | 7.4169 |
| MDC1_S190 | MDC1 | Q14676 | | 1239 | NRSSVKTPEPVVP | 5.8279 | 6.3764 | 6.4754 | 6.7850 | 6.0752 | 5.9792 | 6.3148 | 6.4855 | 6.5507 | 6.4671 | 6.0323 | 6.5112 | 5.6376 | 6.3839 | 5.8213 | 6.2764 |
| MDC1_S190;S218 | MDC1 | Q14676 | | 1567 | NRSSVKTPESIVP | 6.5539 | 6.6905 | 6.6969 | 7.3559 | 6.4929 | 6.6425 | 6.9478 | 6.8419 | 7.1426 | 6.8815 | 6.9854 | 6.8030 | 4.1067 | 4.8038 | 4.2554 | 4.7993 |
| MDC1_S299;T301 | MDC1 | Q14676 | | 1814 | SQSRKRSLATMDS | 4.3444 | 3.5088 | 3.9652 | 4.0341 | 6.4568 | 5.1328 | 4.5959 | 5.1242 | 9.3847 | 9.0416 | 7.9376 | 8.5890 | 8.1195 | 9.8980 | 4.3261 | 5.5414 |
| MDC1_S329 | MDC1 | Q14676 | | 1384 | STEQPVTPEPTSR | 6.0548 | 6.1650 | 6.2145 | 6.7191 | 6.1799 | 6.1478 | 6.2348 | 6.0527 | 6.4802 | 6.6096 | 6.6274 | 6.9323 | 5.6780 | 6.4634 | 5.3415 | 6.0992 |
| MDC1_S329;T331 | MDC1 | Q14676 | | 1820 | SLATMDSPPHQKQ | 7.6692 | 8.0660 | 7.1660 | 8.3493 | 6.8788 | 6.9034 | 7.2133 | 7.0452 | 6.4263 | 6.5202 | 6.6609 | 6.5377 | 3.1862 | 4.0661 | 3.4051 | 3.9063 |
| MDC1_S331 | MDC1 | Q14676 | | 1548 | STDQPVTPEPTSR | 6.3356 | 6.4726 | 6.3732 | 6.9979 | 6.5576 | 6.2669 | 6.1314 | 6.4203 | 6.5028 | 6.9014 | 6.4298 | 6.6778 | 5.0914 | 5.9698 | 5.2428 | 5.6288 |
| MDC1_S376 | MDC1 | Q14676 | | 1697 | AMPVPTTPEFQSP | 5.8193 | 6.3570 | 6.5617 | 7.2471 | 5.8585 | 5.7132 | 6.4339 | 6.2667 | 6.3779 | 6.4752 | 6.6964 | 6.4211 | 5.6068 | 6.0321 | 6.0763 | 6.0570 |
| MDC1_S376;T378 | MDC1 | Q14676 | | 1068 | APPPLLSPLLPSI | 7.3221 | 7.5881 | 7.1415 | 7.8026 | 5.9969 | 6.8487 | 7.4483 | 6.8738 | 7.7598 | 7.9125 | 7.1520 | 7.7560 | 2.6333 | 3.7354 | 2.5935 | 3.4354 |
| MDC1_S378 | MDC1 | Q14676 | | 1198 | SRSSVKTPETVVP | 5.6636 | 6.1475 | 5.8996 | 6.2905 | 6.5281 | 6.3883 | 6.7198 | 6.4178 | 6.4453 | 6.2552 | 6.4538 | 6.1170 | 5.8628 | 7.0334 | 6.0811 | 5.6963 |
| MDC1_S402;T404 | MDC1 | Q14676 | | 1858 | KEEDVVTPKPGKR | 7.1672 | 7.3598 | 6.9175 | 7.2928 | 5.8314 | 6.3286 | 7.1052 | 6.6052 | 7.1393 | 7.1597 | 7.0108 | 7.1185 | 3.6885 | 4.8060 | 3.8209 | 4.6487 |
| MDC1_S422 | MDC1 | Q14676 | | 331 | GFIDSDTDAEEER | 4.7188 | 4.9284 | 5.7259 | 5.5747 | 5.5929 | 6.1027 | 5.8302 | 6.4632 | 6.5449 | 7.6102 | 6.8440 | 7.4414 | 6.3928 | 6.3804 | 6.8912 | 6.9582 |
| MDC1_S453 | MDC1 | Q14676 | | 1086 | KTRQDGSQEAPEA | 5.6814 | 5.7723 | 5.9722 | 6.3787 | 5.9940 | 5.4832 | 5.7119 | 5.9913 | 6.2017 | 6.2420 | 5.8347 | 6.1763 | 6.7097 | 7.0702 | 7.4603 | 7.3199 |
| MDC1_S453;T455 | MDC1 | Q14676 | | 1157 | NRSSVKTPEPVVP | 6.0934 | 6.6410 | 6.5518 | 6.1227 | 6.0955 | 6.2760 | 6.2926 | 5.9813 | 6.6008 | 6.1879 | 6.0261 | 6.3247 | 6.0232 | 6.2002 | 6.4636 | 6.1191 |
| MDC1_S495 | MDC1 | Q14676 | | 1302 | STDRPVTPKPTSR | 5.0077 | 4.7611 | 4.9674 | 5.4437 | 5.2283 | 5.0295 | 5.4587 | 5.4349 | 5.6091 | 5.4360 | 4.9641 | 5.3210 | 8.2824 | 9.3587 | 9.3668 | 10.3306 |
| MDC1_S505 | MDC1 | Q14676 | | 1343 | STDQPVTPKPTSR | 6.6382 | 6.7747 | 6.5725 | 6.9945 | 6.3282 | 6.4489 | 6.7750 | 5.6262 | 7.3235 | 6.8748 | 6.6859 | 6.6504 | 4.6995 | 5.3803 | 4.6570 | 5.5702 |
| MDC1_S513 | MDC1 | Q14676 | | 1526 | NRSSVKTPETVVP | 5.9666 | 6.6351 | 6.3780 | 6.7365 | 6.2541 | 6.4635 | 6.1782 | 6.3353 | 6.4682 | 6.5863 | 6.7524 | 6.8703 | 5.4782 | 6.0475 | 5.1311 | 5.7185 |
| MDC1_S641 | MDC1 | Q14676 | | 955 | GEPEGGSQDQKGQ | 6.4966 | 6.5373 | 5.9358 | 6.6920 | 6.1146 | 5.5697 | 5.9502 | 6.0768 | 6.2422 | 5.7598 | 6.1071 | 6.0735 | 6.2150 | 6.7943 | 6.5023 | 6.9327 |
| MDC1_S780 | MDC1 | Q14676 | | 513 | GIHLERSQASTTV | 6.4823 | 6.2575 | 6.0524 | 6.6623 | 6.0533 | 5.4014 | 6.3182 | 5.5701 | 6.3046 | 5.7658 | 5.4700 | 5.3295 | 6.8305 | 7.5790 | 6.7023 | 7.2207 |
| MDC1_S793 | MDC1 | Q14676 | | 1630 | STDQPVTPKLTSR | 6.5978 | 6.9519 | 5.8877 | 7.5849 | 7.2950 | 6.8223 | 6.8633 | 7.3248 | 6.3370 | 6.4792 | 6.3183 | 6.2028 | 4.0312 | 5.5482 | 4.8615 | 4.8939 |
| MDC1_S955 | MDC1 | Q14676 | | 1775 | AIPEPASPQLLET | 6.5957 | 7.3312 | 5.9318 | 7.4494 | 6.4112 | 6.6278 | 5.7781 | 7.5372 | 6.2785 | 5.6825 | 6.5092 | 6.3515 | 4.6959 | 5.8960 | 5.5606 | 5.3633 |
| MDC1_S964 | MDC1 | Q14676 | | 1403 | NRSSGKTPETLVP | 6.3373 | 7.1168 | 6.9043 | 8.5045 | 5.9357 | 6.0670 | 6.7986 | 6.5138 | 6.8965 | 7.2172 | 6.7688 | 7.0886 | 3.7630 | 4.8066 | 4.2471 | 5.0342 |
| MDC1_S964;S998 | MDC1 | Q14676 | | 1589 | SRNQLVTPEPTSR | 6.9199 | 7.4460 | 7.0227 | 6.7269 | 5.9719 | 6.9399 | 7.4122 | 6.6163 | 7.3060 | 7.1182 | 7.2356 | 7.3137 | 3.6642 | 4.3149 | 3.5354 | 4.4561 |
| MDC1_S995 | MDC1 | Q14676 | | 1444 | NRSSVKTPETVVP | 5.9861 | 6.7163 | 6.8934 | 6.9339 | 6.3967 | 6.1619 | 5.6172 | 6.2430 | 5.8471 | 6.2049 | 7.1319 | 6.5541 | 5.1230 | 6.8946 | 5.0431 | 6.2528 |
| MDC1_S998 | MDC1 | Q14676 | | 995 | QSGGRGSPVSPRR | 4.9279 | 5.5563 | 5.2065 | 5.5462 | 5.3104 | 4.9328 | 4.8485 | 5.3475 | 5.2278 | 5.6804 | 4.8429 | 5.4428 | 8.5384 | 9.5181 | 9.5159 | 9.5577 |
| MDC1_T1157 | MDC1 | Q14676 | | 1425 | STDQPVTPEPTSQ | 6.3381 | 7.1417 | 6.6030 | 7.3743 | 5.6652 | 6.3731 | 7.1968 | 6.7135 | 6.7853 | 6.6790 | 8.0215 | 7.7200 | 3.6700 | 4.7841 | 4.2075 | 4.7268 |
| MDC1_T1198 | MDC1 | Q14676 | | 1671 | PTDQSVTPEAIAQ | 5.9420 | 6.0430 | 6.7057 | 7.5440 | 6.0383 | 6.5878 | 7.2966 | 6.1881 | 7.0871 | 7.3769 | 7.5278 | 7.1478 | 4.1081 | 5.0630 | 4.2742 | 5.0695 |
| MDC1_T1239 | MDC1 | Q14676 | | 1732 | AAGNPGSLAAPID | 6.7873 | 5.9880 | 5.1506 | 7.0745 | 5.4578 | 6.4245 | 6.2197 | 6.6380 | 6.4983 | 6.4931 | 6.6892 | 5.5411 | 5.2000 | 6.7083 | 6.2293 | 6.9001 |
| MDC1_T1280;T1302 | MDC1 | Q14676 | | 422 | LAHLKESQPAIWN | 6.2103 | 6.6367 | 5.7997 | 8.4488 | 9.1002 | 4.5047 | 6.2774 | 4.3744 | 4.9666 | 5.5509 | 4.3705 | 5.3693 | 6.6406 | 6.3958 | 6.9051 | 8.4488 |

| | | | | | | | | | | | | | | | | | | | | |
|---|---|---|---|---|---|---|---|---|---|---|---|---|---|---|---|---|---|---|---|---|
| MDC1_T1302 | MDC1 | Q14676 | 341 | EERIPATPVVIPM | 9.2970 | 6.8105 | 7.1376 | 7.6342 | 5.6563 | 5.0569 | 6.4372 | 6.8619 | 7.2095 | 4.6853 | 6.0433 | 4.9765 | 5.1854 | 7.0725 | 5.3515 | 4.5843 |
| MDC1_T1321 | MDC1 | Q14676 | 1608 | NRSSVKTPEPVVP | 5.8302 | 7.4491 | 5.4112 | 6.8722 | 6.4581 | 7.0350 | 5.8402 | 6.3599 | 6.1808 | 6.3462 | 5.5492 | 6.1858 | 5.6076 | 6.9331 | 5.5492 | 6.3922 |
| MDC1_T1343 | MDC1 | Q14676 | 108 | RPPKVLSPGVSHR | 6.0004 | 6.3130 | 6.7892 | 7.0060 | 5.7830 | 5.7737 | 6.3107 | 6.2306 | 6.0337 | 6.2996 | 6.1926 | 5.8245 | 6.2744 | 6.3346 | 5.9151 | 6.9189 |
| MDC1_T1384 | MDC1 | Q14676 | 998 | GRGSPVSPRRHQK | 7.6973 | 7.8545 | 7.4455 | 8.0944 | 6.4448 | 5.9169 | 6.4993 | 6.3683 | 6.6747 | 6.3796 | 6.5358 | 7.1699 | 3.6925 | 4.5623 | 4.0350 | 4.6292 |
| MDC1_T1403 | MDC1 | Q14676 | 378 | SQAGSDTDVEEGK | 7.4262 | 7.7747 | 7.1066 | 7.8739 | 6.5299 | 6.6315 | 6.4995 | 6.1697 | 5.9384 | 5.3227 | 6.5018 | 5.2594 | 4.4874 | 5.7446 | 4.6437 | 6.0900 |
| MDC1_T1425 | MDC1 | Q14676 | 453 | TTTERDSDTDVEE | 3.3095 | 5.2646 | 4.5068 | 4.8373 | 6.0628 | 6.7320 | 4.7930 | 4.7003 | 4.6640 | 9.3320 | 6.9779 | 7.8123 | 6.6997 | 9.8400 | 6.9335 | 7.5342 |
| MDC1_T1444 | MDC1 | Q14676 | 495 | QPPFGDSDDSVEA | 5.9907 | 5.0182 | 5.1470 | 7.0469 | 6.1491 | 6.0989 | 5.4304 | 6.1234 | 6.3063 | 5.8838 | 6.0409 | 6.4441 | 6.3333 | 7.6317 | 6.7442 | 7.6111 |
| MDC1_T1466 | MDC1 | Q14676 | 1800 | AGRSRFTPELQPK | 7.5655 | 8.0495 | 6.6299 | 7.1103 | 6.5843 | 7.1812 | 7.5004 | 7.0422 | 7.0201 | 7.5975 | 7.5547 | 7.0792 | 2.8348 | 3.9900 | 2.7876 | 3.4729 |
| MDC1_T1485 | MDC1 | Q14676 | 1711 | TTDQPISPEPITQ | 6.4517 | 6.2769 | 6.6480 | 7.3397 | 6.1630 | 6.3236 | 5.9379 | 6.4823 | 6.0846 | 6.9496 | 6.6086 | 6.3231 | 6.0067 | 5.5832 | 5.2549 | 5.5662 |
| MDC1_T1526 | MDC1 | Q14676 | 190 | SRTTSSSVIVPES | 5.3497 | 6.5892 | 6.4243 | 6.6900 | 5.4666 | 5.5903 | 5.7805 | 7.0887 | 6.3005 | 6.3830 | 8.1793 | 6.9627 | 5.1435 | 6.4724 | 4.9213 | 6.6580 |
| MDC1_T1548 | MDC1 | Q14676 | 641 | EQDLPISRENLTD | 7.0806 | 7.0588 | 5.9530 | 7.6909 | 5.4457 | 6.1135 | 5.6597 | 6.8963 | 6.2820 | 7.2847 | 5.2653 | 5.9966 | 6.0858 | 6.5019 | 4.2765 | 6.4088 |
| MDC1_T1567 | MDC1 | Q14676 | 299;301 | NA | 6.0051 | 6.4210 | 5.8560 | 6.8656 | 5.7992 | 5.8049 | 6.0881 | 5.9232 | 6.1579 | 6.3406 | 6.4136 | 6.5343 | 5.9229 | 6.8901 | 6.2341 | 6.7435 |
| MDC1_T1589 | MDC1 | Q14676 | 1697;1705 | NA | 6.3661 | 6.1886 | 6.2744 | 7.4677 | 6.1111 | 7.4013 | 7.0305 | 7.2883 | 6.4078 | 7.2944 | 7.6913 | 7.5114 | 3.9748 | 4.6309 | 4.4178 | 3.9436 |
| MDC1_T1608 | MDC1 | Q14676 | 1400;1403 | NA | 7.1093 | 7.9871 | 7.6397 | 7.4006 | 6.2586 | 7.4827 | 6.4974 | 6.6618 | 6.3085 | 7.9416 | 7.9872 | 7.8618 | 2.4844 | 3.9379 | 2.2135 | 4.2278 |
| MDC1_T1630 | MDC1 | Q14676 | 329;331 | NA | 5.7022 | 5.6228 | 6.1207 | 6.0800 | 6.3230 | 6.3500 | 7.0476 | 6.1457 | 6.4833 | 6.2574 | 6.2074 | 6.2486 | 5.3295 | 7.3605 | 5.9275 | 6.7938 |
| MDC1_T1671 | MDC1 | Q14676 | 1280;1302 | NA | 6.1000 | 6.9344 | 6.0574 | 6.3348 | 6.2768 | 5.7084 | 5.8175 | 6.0838 | 6.7210 | 6.2697 | 5.7998 | 6.8428 | 4.4605 | 6.4921 | 5.7336 | 6.3674 |
| MDC1_T1697 | MDC1 | Q14676 | 964;998 | NA | 6.0205 | 6.4709 | 4.8298 | 6.1247 | 5.6732 | 5.6279 | 5.8138 | 4.6842 | 4.6793 | 5.3105 | 5.3922 | 4.6919 | 8.2378 | 9.6251 | 7.9059 | 8.9125 |
| MDC1_T1697;S1711 | MDC1 | Q14676 | 402;404 | NA | 6.9734 | 6.2832 | 6.3940 | 6.4874 | 5.3867 | 6.3203 | 5.9911 | 5.2792 | 6.3273 | 5.9333 | 6.2220 | 6.3609 | 5.8303 | 6.9724 | 6.3179 | 6.9208 |
| MDC1_T1697;T1705 | MDC1 | Q14676 | 190;218 | NA | 4.7309 | 4.8093 | 5.9235 | 6.4191 | 5.4983 | 5.0048 | 7.6961 | 7.0479 | 6.9376 | 5.6752 | 5.9323 | 6.4032 | 6.8585 | 7.1822 | 6.0657 | 7.8154 |
| MDC1_T1800 | MDC1 | Q14676 | 453;455 | NA | 5.6775 | 5.9464 | 6.3440 | 6.2862 | 6.0701 | 5.1260 | 6.0468 | 5.9624 | 6.1989 | 5.9327 | 6.2764 | 6.7473 | 6.2185 | 7.5502 | 7.1523 | 6.4644 |
| MDC1_T1858 | MDC1 | Q14676 | 376;378 | NA | 8.2269 | 7.1352 | 6.0085 | 7.5386 | 7.0054 | 6.1556 | 6.2866 | 5.9811 | 5.5836 | 5.5083 | 5.6933 | 5.8111 | 5.7332 | 6.3150 | 5.1526 | 5.8651 |
| MDC1_T341 | MDC1 | Q14676 | 1697;1711 | NA | 6.4815 | 7.4938 | 7.6595 | 7.9740 | 6.1537 | 7.6623 | 7.2446 | 6.5814 | 7.9975 | 7.0303 | 7.3447 | 8.4440 | 1.7968 | 3.6946 | 3.0394 | 3.4019 |
| MDFIC_S237 | MDFIC | Q9P1T7-1 | 237 | KLSAPVSQKMHRK | 6.3276 | 6.7730 | 6.7376 | 6.6933 | 5.5713 | 6.5747 | 7.5218 | 6.5801 | 5.9900 | 6.2166 | 6.6763 | 6.6939 | 4.8718 | 6.4580 | 4.9075 | 5.4065 |
| MDFIC_S252 | MDFIC | Q9P1T7-1 | 252 | SSLSVNSDISKKS | 5.9295 | 6.4664 | 6.4535 | 6.6822 | 5.7975 | 6.0457 | 6.0507 | 5.9535 | 6.3827 | 6.5058 | 6.5447 | 6.6419 | 6.2030 | 6.4896 | 5.7327 | 6.1206 |
| MDH1_S259 | MDH1 | P40925-3 | 259 | IKARKLSSAMSAA | 6.0042 | 6.1521 | 6.3767 | 6.8375 | 4.9973 | 6.0667 | 5.6884 | 5.2647 | 6.1890 | 8.0463 | 7.3921 | 8.1857 | 5.3870 | 4.2151 | 6.3910 | 6.8062 |
| MDM1_S263 | MDM1 | Q8TC05 | 314 | YRAKFLSPAQYLY | 7.7228 | 6.8651 | 5.8219 | 6.7724 | 5.7784 | 6.2268 | 6.9639 | 6.2014 | 7.4724 | 6.5393 | 7.1708 | 6.5341 | 4.8130 | 5.4820 | 4.4828 | 5.1530 |
| MDM1_S314 | MDM1 | Q8TC05 | 263 | RNLETVSPERKSN | 6.4453 | 6.5671 | 6.6156 | 6.6347 | 6.3667 | 6.2305 | 6.8016 | 6.6494 | 6.7905 | 5.8731 | 6.2501 | 6.2411 | 5.9435 | 5.5744 | 5.3817 | 5.6348 |
| MDM1_S356 | MDM1 | Q8TC05 | 584 | AVSLLTSPAAGIK | 6.5504 | 6.6609 | 6.4493 | 7.0160 | 6.3121 | 6.7740 | 6.6141 | 6.7367 | 7.0399 | 6.7996 | 6.6920 | 6.3884 | 4.7918 | 5.1647 | 4.9560 | 5.0540 |
| MDM1_S543 | MDM1 | Q8TC05 | 356 | VQGTHFSRDHLNQ | 6.1121 | 6.3890 | 5.8586 | 6.2042 | 5.6377 | 6.0506 | 6.8150 | 6.2021 | 6.1924 | 7.0006 | 6.5531 | 7.1619 | 5.6084 | 6.2680 | 5.6880 | 6.2583 |
| MDM1_S580;S584 | MDM1 | Q8TC05 | 543 | GGAVLVSPSKMKP | 5.6955 | 6.0194 | 6.5134 | 7.0135 | 5.9760 | 6.6792 | 6.1299 | 6.3829 | 7.0895 | 6.2706 | 7.0631 | 6.4436 | 5.7172 | 5.6226 | 5.1700 | 6.2137 |
| MDM1_S584 | MDM1 | Q8TC05 | 580;584 | NA | 6.3020 | 6.2810 | 6.4824 | 7.9371 | 5.8849 | 5.8313 | 7.0918 | 6.6016 | 5.9442 | 5.9007 | 6.7788 | 6.6126 | 6.0558 | 5.2208 | 4.8574 | 6.2176 |
| MDN1_S2492 | MDN1 | Q9NU22 | 3992 | RSSLVESDKEEQP | 5.7421 | 6.0435 | 6.2837 | 6.4249 | 5.9936 | 5.5009 | 6.1246 | 5.9333 | 6.5411 | 6.3576 | 5.9480 | 5.9801 | 6.8975 | 6.3029 | 6.3842 | 7.5422 |
| MDN1_S3992 | MDN1 | Q9NU22 | 4538 | ENTDQASPQEDYA | 5.9698 | 6.2732 | 6.6236 | 6.0841 | 6.0516 | 6.1589 | 6.1488 | 5.8806 | 6.0694 | 6.0456 | 5.8934 | 6.1666 | 6.4372 | 6.7755 | 6.9357 | 6.4861 |
| MDN1_S4538 | MDN1 | Q9NU22 | 2492 | LEKIMQSPSPENL | 6.0566 | 6.1924 | 7.6673 | 6.5766 | 6.5941 | 6.3531 | 5.5902 | 5.7278 | 6.3928 | 6.1822 | 6.2053 | 6.4833 | 5.5339 | 7.0615 | 5.7343 | 5.6484 |
| MDN1_S4752 | MDN1 | Q9NU22 | 5015 | EEEREDSDTEEQV | 6.3168 | 6.2459 | 6.3733 | 7.0169 | 6.4488 | 6.1316 | 5.9024 | 5.4034 | 6.1436 | 6.6071 | 6.1179 | 6.3019 | 6.2516 | 6.5934 | 5.8658 | 6.2796 |
| MDN1_S4752;S4754 | MDN1 | Q9NU22 | 4212 | LNAALATPAKEMG | 6.4251 | 6.2047 | 6.7959 | 8.1870 | 5.4316 | 5.9423 | 6.6210 | 6.2490 | 6.4554 | 5.8583 | 6.9393 | 6.0811 | 4.6165 | 6.5743 | 6.2863 | 5.3324 |
| MDN1_S5015 | MDN1 | Q9NU22 | 4898 | KNGGEDTDNEEGE | 6.0188 | 6.4589 | 6.0979 | 6.6829 | 6.3366 | 6.0569 | 5.9581 | 6.8405 | 6.0377 | 6.4172 | 6.5666 | 5.8200 | 6.1493 | 6.0403 | 6.1031 | 6.4153 |
| MDN1_T4212 | MDN1 | Q9NU22 | 4752 | EEDDEKSDSEGGD | 5.8677 | 6.4772 | 6.3770 | 6.3681 | 6.4764 | 5.9501 | 6.8006 | 6.1183 | 7.3398 | 6.3269 | 6.3681 | 6.8119 | 5.0988 | 6.0253 | 5.8272 | 5.7666 |
| MDN1_T4898 | MDN1 | Q9NU22 | 4752;4754 | NA | 6.3441 | 6.2398 | 6.0906 | 6.3077 | 5.9027 | 5.8031 | 6.6349 | 5.5173 | 6.4484 | 6.1391 | 6.6701 | 6.4709 | 6.5165 | 6.0627 | 6.0951 | 6.7572 |
| ME2_T400 | ME2 | P23368 | 400 | GAGRLFTPDVIRA | 6.1724 | 7.1116 | 5.8433 | 7.0663 | 5.2363 | 5.4839 | 6.6647 | 6.6254 | 6.4714 | 6.2449 | 6.0607 | 6.9998 | 7.1810 | 6.0365 | 4.3455 | 6.4563 |
| ME3_S469 | ME3 | Q16798 | 469 | GIFASGSPFKSVT | 6.0070 | 6.6766 | 7.0490 | 7.1317 | 5.7437 | 6.5562 | 6.8665 | 7.2446 | 7.0320 | 6.0597 | 6.3343 | 7.8257 | 3.4963 | 6.5054 | 4.6247 | 4.8466 |
| MEAF6_S118;S122 | MEAF6 | Q9HAF1-3 | 118;122 | NA | 6.0783 | 6.4404 | 5.9272 | 7.1592 | 6.1117 | 5.8213 | 6.0483 | 6.2977 | 5.9998 | 5.8546 | 6.1821 | 6.5325 | 6.6606 | 6.6193 | 6.2697 | 5.9974 |
| MEAK7_S425 | MEAK7 | Q6P9B6 | 425 | LAKGNKSILDADP | 6.3446 | 7.0680 | 6.8821 | 6.0079 | 6.1707 | 7.3548 | 6.9514 | 7.0856 | 6.2660 | 6.6425 | 6.1143 | 6.4566 | 4.9544 | 5.2624 | 5.1236 | 5.3151 |
| MECOM_S1048 | MECOM | Q03112-3 | 526 | QTNKSQSPLMTHP | 6.4317 | 5.9822 | 5.6883 | 5.6349 | 6.5673 | 6.7243 | 6.2567 | 6.5540 | 5.9521 | 6.4097 | 6.7918 | 6.0229 | 6.2627 | 6.3095 | 6.2801 | 6.1318 |
| MECOM_S526 | MECOM | Q03112-3 | 726 | LKMEPQSPGEVKK | 6.6756 | 6.0918 | 5.9276 | 6.2525 | 6.6061 | 6.1073 | 6.3630 | 6.5279 | 6.1840 | 5.8658 | 6.3440 | 6.3927 | 5.6161 | 6.4915 | 6.1842 | 6.3699 |
| MECOM_S726 | MECOM | Q03112-3 | 1048 | SNHGSQSPRNVEE | 6.6751 | 8.4765 | 5.2617 | 7.1505 | 7.5668 | 7.3843 | 5.5970 | 7.7569 | 6.4631 | 5.8502 | 6.1996 | 5.5842 | 4.7323 | 4.6603 | 5.6202 | 5.0214 |
| MECOM_S759 | MECOM | Q03112-3 | 759 | PLTPVPSKPPVTP | 5.7457 | 5.4518 | 6.5430 | 7.2474 | 6.2395 | 5.9766 | 7.2944 | 5.8932 | 6.6382 | 6.7151 | 7.3414 | 7.1758 | 5.5608 | 6.3592 | 5.0329 | 4.7849 |
| MECP2_S161 | MECP2 | P51608-2 | 241 | KMPFQTSPGGKAE | 7.2448 | 7.6098 | 6.6373 | 7.6901 | 6.4361 | 6.3595 | 6.9397 | 6.5156 | 7.0229 | 6.7421 | 6.7601 | 6.8786 | 3.7019 | 4.6910 | 4.1189 | 4.6516 |
| MECP2_S241 | MECP2 | P51608-2 | 372 | RSSSASSPPKKEH | 6.5519 | 6.5881 | 6.1612 | 6.6469 | 5.6549 | 5.3228 | 6.0053 | 5.8413 | 6.1138 | 6.2798 | 5.5458 | 6.1092 | 6.4210 | 6.6574 | 7.0220 | 7.0786 |
| MECP2_S25 | MECP2 | P51608-2 | 92 | VPEASASPKQRRS | 5.9314 | 6.1999 | 6.1493 | 6.2620 | 6.2291 | 5.8969 | 6.6752 | 5.7192 | 7.8560 | 5.8884 | 6.2478 | 5.8498 | 6.6236 | 6.0772 | 6.3300 | 6.0640 |
| MECP2_S368;S372 | MECP2 | P51608-2 | 25 | ERLEEKSEDQDLQ | 5.2656 | 5.7737 | 7.1064 | 6.2962 | 6.6715 | 6.7821 | 6.7331 | 5.2935 | 5.2367 | 6.6898 | 5.4686 | 6.1201 | 6.2664 | 6.9197 | 6.6407 | 6.7360 |
| MECP2_S372 | MECP2 | P51608-2 | 435 | KMPRGGSLESDGC | 5.9696 | 6.1490 | 5.5719 | 6.8214 | 6.0151 | 6.2725 | 6.3077 | 6.2187 | 6.0639 | 6.4270 | 6.0909 | 5.7464 | 5.7677 | 7.2902 | 6.4761 | 6.8118 |
| MECP2_S435 | MECP2 | P51608-2 | 161 | EKVGDTSLDPNDF | 6.4394 | 7.5227 | 6.9856 | 6.6531 | 5.8475 | 6.0540 | 7.1044 | 6.2896 | 5.5443 | 7.4405 | 5.7416 | 7.0770 | 4.3149 | 6.1033 | 4.6309 | 6.2513 |
| MECP2_S92 | MECP2 | P51608-2 | 368;372 | NA | 5.9332 | 6.6996 | 6.0539 | 7.6896 | 5.8435 | 5.8565 | 6.6562 | 6.5278 | 6.8979 | 5.9602 | 6.9637 | 6.8157 | 5.2110 | 5.8504 | 5.3515 | 5.6894 |
| MED1_S1021 | MED1 | Q15648 | 1215 | MKPVPGTPPSSKA | 5.7302 | 5.4710 | 5.9900 | 6.4566 | 6.0544 | 5.4965 | 5.8068 | 5.4194 | 5.8986 | 6.0759 | 5.8939 | 6.0407 | 7.0095 | 7.3751 | 7.4360 | 7.8454 |
| MED1_S1156 | MED1 | Q15648 | 805 | DCPAIGTPLRDSS | 5.6455 | 6.2297 | 6.1382 | 6.2132 | 5.8397 | 5.9523 | 6.1281 | 5.8609 | 6.2640 | 5.9646 | 6.0454 | 6.1905 | 6.6702 | 7.0073 | 6.7570 | 7.0934 |
| MED1_S1207;T1215 | MED1 | Q15648 | 1433 | SSKNYGSPLISGS | 5.5452 | 5.8785 | 5.9534 | 6.1303 | 5.5711 | 5.7556 | 6.2747 | 5.9602 | 6.1063 | 5.9123 | 6.0075 | 6.3817 | 6.5997 | 7.5963 | 6.8383 | 7.4888 |
| MED1_S1401 | MED1 | Q15648 | 1051 | SAGRSQTPPGVAT | 5.5043 | 6.0279 | 5.3795 | 6.1449 | 5.9455 | 5.8734 | 6.3580 | 5.9928 | 5.8036 | 5.7127 | 5.4721 | 5.6353 | 6.9786 | 8.1541 | 7.6477 | 7.3695 |
| MED1_S1433 | MED1 | Q15648 | 1156 | GGKPGSSPITKHG | 5.7857 | 5.4304 | 5.8255 | 5.6588 | 5.3071 | 5.1881 | 6.0791 | 5.0194 | 5.8310 | 5.6533 | 5.0447 | 5.6907 | 7.7202 | 8.0108 | 8.3679 | 9.3875 |
| MED1_S1433;T1440 | MED1 | Q15648 | 1440 | PLISGSTPKHERG | 4.7896 | 5.2426 | 5.3868 | 4.9193 | 4.9902 | 6.0451 | 6.3441 | 4.9462 | 5.5118 | 5.4378 | 5.4795 | 5.7574 | 8.3117 | 8.8681 | 8.8718 | 9.0980 |
| MED1_S1440 | MED1 | Q15648 | 1561 | DRPSRLSPDFMIG | 5.6757 | 6.4314 | 6.8773 | 6.2511 | 6.2231 | 6.0395 | 5.3334 | 5.7178 | 5.1413 | 5.8441 | 6.8093 | 6.2069 | 7.6471 | 6.6895 | 6.2252 | 6.8871 |
| MED1_S1447 | MED1 | Q15648 | 1447 | PKHERGSPSHSKS | 5.5082 | 6.1904 | 5.4466 | 6.1996 | 5.2852 | 5.1088 | 5.7981 | 5.7713 | 5.7215 | 5.6317 | 5.7691 | 6.0401 | 6.9941 | 7.9044 | 8.1006 | 8.5303 |
| MED1_S1465 | MED1 | Q15648 | 1482 | YQNSPSSDDGIRP | 6.0051 | 6.2464 | 5.6461 | 6.6162 | 5.8894 | 5.8210 | 6.6629 | 5.4774 | 5.5019 | 6.4683 | 5.8911 | 5.7410 | 6.1279 | 7.1590 | 7.1561 | 7.5900 |
| MED1_S1479;S1482 | MED1 | Q15648 | 1021 | ADTEGKSPSHSSS | 5.4457 | 7.1173 | 5.5937 | 6.7389 | 5.7338 | 6.2585 | 4.8873 | 8.4087 | 4.6657 | 5.5095 | 6.1025 | 5.5698 | 5.5671 | 8.0267 | 8.1136 | 6.2611 |
| MED1_S1482 | MED1 | Q15648 | 1401 | SKHDGGSPSIKAK | 6.4915 | 6.9148 | 5.5404 | 6.4725 | 6.7043 | 6.5880 | 6.5567 | 6.3289 | 6.6652 | 5.9566 | 5.8197 | 6.5280 | 5.6766 | 6.1417 | 5.5445 | 6.0705 |
| MED1_S1561 | MED1 | Q15648 | 1465 | QNLDSESESGSSI | 6.7024 | 6.6822 | 7.0920 | 7.9852 | 6.9270 | 6.5293 | 6.3410 | 6.4105 | 6.2118 | 6.3333 | 6.2827 | 6.3791 | 5.0321 | 5.9294 | 4.6304 | 4.5318 |
| MED1_S545 | MED1 | Q15648-3 | 545 | NLPPASSPGSKNP | 5.6312 | 6.9296 | 7.1855 | 6.5045 | 6.0330 | 6.7075 | 7.4244 | 6.4559 | 5.8363 | 5.4134 | 5.7687 | 5.5234 | 5.0243 | 7.1326 | 6.7794 | 5.6502 |
| MED1_S771;S774 | MED1 | Q15648 | 1479;1482 | NA | 6.1724 | 6.1390 | 6.1211 | 6.7857 | 5.9742 | 6.1040 | 6.0402 | 6.2237 | 6.5404 | 6.4553 | 6.1577 | 6.4746 | 5.5002 | 6.7485 | 6.2927 | 6.2704 |
| MED1_T1051 | MED1 | Q15648 | 1207;1215 | NA | 5.3983 | 6.3631 | 5.0583 | 7.5323 | 6.2910 | 5.9809 | 6.6434 | 6.3779 | 6.3212 | 5.0765 | 6.8175 | 7.0177 | 5.7869 | 6.8460 | 5.0425 | 7.4466 |
| MED1_T1215 | MED1 | Q15648 | 1433;1440 | NA | 6.0889 | 5.6577 | 5.2642 | 5.8282 | 5.9939 | 6.4605 | 6.7893 | 5.9703 | 5.9970 | 6.6135 | 6.3578 | 5.6818 | 6.7601 | 6.9036 | 6.9223 | 6.7109 |
| MED1_T805 | MED1 | Q15648 | 771;774 | NA | 5.4842 | 5.8970 | 6.2859 | 6.0325 | 6.5182 | 5.6126 | 5.5480 | 6.7777 | 6.3265 | 5.5477 | 5.7642 | 5.3554 | 6.8727 | 7.3403 | 7.2147 | 7.4224 |
| MED12_S2166 | MED12 | Q93074-2 | 688 | PDFSLFSPTMPCE | 5.9484 | 5.6038 | 6.7628 | 6.4788 | 5.7085 | 5.4306 | 5.8481 | 5.4825 | 6.3130 | 6.5388 | 5.5861 | 6.6423 | 6.6791 | 6.6974 | 6.6014 | 7.6786 |
| MED12_S635 | MED12 | Q93074-2 | 635 | PGPRPPSPFDDPA | 5.6873 | 5.8939 | 5.9908 | 6.9353 | 6.2434 | 6.2360 | 6.2971 | 5.8613 | 6.2598 | 6.2107 | 6.5812 | 6.8890 | 5.9310 | 6.3357 | 5.8850 | 6.7626 |
| MED12_S688 | MED12 | Q93074-2 | 698 | PCEGKGSPSPEKP | 6.0534 | 6.3544 | 5.7589 | 7.8977 | 6.4047 | 5.6143 | 5.8531 | 6.5878 | 6.0877 | 6.5600 | 6.8460 | 7.4564 | 5.0681 | 5.5340 | 5.7375 | 6.1862 |
| MED12_S698 | MED12 | Q93074-2 | 2166 | QLQQQLSNTQPQP | 3.3832 | 3.2319 | 5.3127 | 6.0098 | 4.3615 | 7.5717 | 5.5937 | 5.1992 | 9.5336 | 4.5344 | 5.8963 | 6.6584 | 9.3877 | 6.9070 | 6.0639 | 10.3551 |
| MED12_S698;S700 | MED12 | Q93074-2 | 698;700 | NA | 5.9447 | 5.9096 | 6.3553 | 6.3690 | 5.7953 | 5.4291 | 6.0624 | 5.7032 | 6.4567 | 6.5197 | 5.9733 | 6.1069 | 6.3245 | 6.9625 | 7.0091 | 7.0787 |
| MED13_S395 | MED13 | Q9UHV7 | 395 | QNKRKYSASSGGL | 5.6391 | 5.3865 | 5.5709 | 5.7077 | 5.6406 | 6.0107 | 6.2908 | 5.9429 | 5.4939 | 5.5967 | 5.9476 | 5.3728 | 7.8356 | 7.4110 | 7.8924 | 7.3608 |
| MED13_S530;S537 | MED13 | Q9UHV7 | 890 | VDEGFCSPKPSEI | 5.9041 | 6.0514 | 6.3278 | 6.0400 | 5.8059 | 5.5877 | 5.7678 | 6.2961 | 6.3232 | 6.1941 | 6.1484 | 5.9257 | 6.6873 | 6.5012 | 7.0753 | 7.3640 |
| MED13_S749 | MED13 | Q9UHV7 | 789 | LDNLFNSDEDELT | 6.2504 | 5.8112 | 6.3707 | 7.4826 | 6.5304 | 6.2121 | 6.0803 | 5.7481 | 5.8654 | 6.0777 | 6.9823 | 6.4343 | 6.1251 | 6.3551 | 5.9061 | 5.7682 |
| MED13_S789 | MED13 | Q9UHV7 | 749 | DAMSLFSPSIKQD | 6.5909 | 6.3039 | 6.1930 | 6.9633 | 6.6630 | 6.2586 | 6.5634 | 6.5569 | 6.7601 | 6.0619 | 6.0894 | 6.5737 | 5.7890 | 5.5532 | 5.4241 | 5.6555 |
| MED13_S848 | MED13 | Q9UHV7 | 474 | PQKRPLTPFHHRV | 6.0285 | 6.8276 | 5.9493 | 6.3640 | 6.5585 | 6.1776 | 7.0068 | 6.2153 | 6.0273 | 5.7848 | 5.4330 | 5.9447 | 6.4667 | 6.6197 | 6.3825 | 6.2137 |
| MED13_S890 | MED13 | Q9UHV7 | 848 | QHIMGFSPMNMNN | 5.8299 | 7.0625 | 5.5270 | 6.7150 | 7.1813 | 6.8397 | 6.9348 | 5.3874 | 5.1083 | 6.4299 | 5.5745 | 6.1685 | 6.0467 | 6.5605 | 6.6585 | 5.9755 |
| MED13_S925;Y927 | MED13 | Q9UHV7 | 925;927 | NA | 6.3974 | 6.1397 | 5.8683 | 6.2752 | 5.8881 | 5.3063 | 5.3835 | 5.2198 | 6.2012 | 5.4908 | 5.0064 | 5.4283 | 8.0718 | 6.9679 | 8.2687 | 8.0867 |
| MED13_T474 | MED13 | Q9UHV7 | 530;537 | NA | 6.6788 | 6.4664 | 5.6101 | 7.1599 | 6.7400 | 5.8853 | 5.9605 | 7.0606 | 5.7192 | 6.4566 | 5.7337 | 6.1661 | 6.2195 | 6.4514 | 6.0460 | 5.6458 |
| MED13L_S878 | MED13L | Q71F56 | 923 | VEDGLGSPKPEEI | 5.8935 | 6.0493 | 6.1068 | 6.4525 | 5.9205 | 5.4394 | 6.4578 | 5.8551 | 6.3869 | 6.1744 | 6.1910 | 6.4597 | 6.5016 | 6.7356 | 6.2883 | 7.0875 |
| MED13L_S923 | MED13L | Q71F56 | 878 | EQHPAFSPVMNYK | 5.8373 | 5.9374 | 5.7173 | 6.1113 | 6.2929 | 6.3175 | 5.6453 | 6.2245 | 5.7512 | 6.0234 | 5.7421 | 5.6505 | 7.3764 | 7.0429 | 7.2732 | 7.0568 |
| MED14_S617 | MED14 | O60244 | 986 | EDDNPPSPIGGDM | 6.2516 | 6.3307 | 6.6157 | 6.7952 | 6.0082 | 6.0321 | 6.3307 | 6.3488 | 6.3781 | 6.4239 | 6.2415 | 6.3439 | 5.8797 | 6.2333 | 5.7559 | 6.0307 |
| MED14_S986 | MED14 | O60244 | 617 | NAKRKLSDDPCPV | 5.1105 | 5.4185 | 5.9574 | 6.4946 | 5.6050 | 5.6874 | 5.7105 | 5.6050 | 5.3344 | 5.6104 | 6.4071 | 6.6305 | 7.2222 | 6.9320 | 7.5949 | 7.6795 |
| MED15_S505;T511 | MED15 | Q96RN5 | 603 | NDMAVPTPPPPPV | 6.5310 | 5.8473 | 5.9944 | 5.4839 | 6.4000 | 5.8017 | 5.9284 | 6.2139 | 6.0666 | 6.8008 | 6.0348 | 6.4308 | 7.1324 | 6.3775 | 6.4187 | 6.5377 |
| MED15_S511 | MED15 | Q96RN5 | 511 | SPGPLNTPVNPSS | 5.4088 | 5.8790 | 5.9157 | 6.1598 | 6.0992 | 6.6479 | 6.1410 | 7.0541 | 6.3432 | 7.1420 | 6.2425 | 6.6530 | 6.3722 | 5.8833 | 5.5863 | 6.4721 |
| MED15_T497;S505;S511 | MED15 | Q96RN5 | 505 | QNFSVPSPGPLNT | 5.1155 | 4.5878 | 6.2057 | 4.8003 | 6.6039 | 5.7108 | 5.8736 | 6.3431 | 6.0565 | 6.3239 | 6.4426 | 5.9162 | 8.2373 | 7.3972 | 7.5027 | 6.8830 |
| MED15_T505 | MED15 | Q96RN5 | 497;505;511 | NA | 6.6800 | 5.7603 | 6.3444 | 6.5056 | 6.8881 | 5.6683 | 5.8101 | 6.4480 | 5.6510 | 6.4796 | 6.4888 | 7.5061 | 5.3506 | 5.5695 | 6.3619 | 6.4878 |
| MED15_T603 | MED15 | Q96RN5 | 505;511 | NA | 6.5836 | 6.0847 | 5.4723 | 5.6308 | 8.2280 | 6.9130 | 6.2330 | 6.6166 | 6.5963 | 6.8215 | 8.0824 | 6.8921 | 5.0899 | 4.3925 | 4.7287 | 5.6346 |
| MED16_T571 | MED16 | Q9Y2X0 | 571 | RPHFLNTPDKSPG | 6.6587 | 6.6066 | 6.6773 | 6.9282 | 6.5548 | 6.8041 | 6.2973 | 5.5043 | 5.5037 | 6.2365 | 6.6909 | 6.3603 | 5.6889 | 5.8329 | 5.5754 | 6.0799 |
| MED17_S10 | MED17 | Q9NVC6 | 10 | VRAVRISIESACE | 4.7546 | 4.6023 | 4.6494 | 5.1073 | 5.4238 | 5.2185 | 5.5824 | 6.0340 | 4.8337 | 4.6072 | 4.8765 | 4.7887 | 9.7819 | 8.9991 | 10.4437 | 10.2969 |
| MED19_S194 | MED19 | A0JLT2 | 226 | KKKNRHSPDHPGM | 6.0866 | 7.7725 | 5.5997 | 7.5774 | 6.9079 | 6.8495 | 4.7275 | 6.4901 | 5.5046 | 6.5535 | 6.5660 | 5.0844 | 5.3228 | 7.6999 | 6.2708 | 4.9868 |
| MED19_S226 | MED19 | A0JLT2 | 194 | VPPETPSDSDHKK | 5.9145 | 6.3581 | 6.7843 | 6.6720 | 6.8960 | 6.1472 | 6.6742 | 6.7814 | 6.1000 | 4.7974 | 5.2236 | 6.2051 | 5.2186 | 6.9793 | 6.6307 | 6.6176 |
| MED24_S873 | MED24 | O75448 | 873 | DANILSSPTDRSM | 6.0876 | 6.2288 | 6.6055 | 6.9180 | 6.4796 | 6.2512 | 6.8808 | 6.3455 | 6.7732 | 6.5510 | 6.5530 | 7.2870 | 5.1532 | 5.3138 | 5.4162 | 5.1557 |
| MED26_S337 | MED26 | O95402 | 373 | LSRAGFSPDSSKA | 6.4737 | 6.9024 | 6.2622 | 6.6877 | 6.0196 | 6.2550 | 6.3339 | 6.2556 | 6.6726 | 6.5505 | 6.6251 | 6.6037 | 5.6169 | 5.3599 | 5.6502 | 5.7308 |
| MED26_S373 | MED26 | O95402 | 323 | PLAQPSTPPVRRL | 6.2839 | 5.9281 | 5.8213 | 6.3742 | 6.3138 | 5.3123 | 6.3907 | 5.8835 | 5.9769 | 5.8646 | 7.0241 | 6.0617 | 6.9472 | 6.1447 | 7.1193 | 6.5536 |
| MED26_S535 | MED26 | O95402 | 337 | LLPSAESPVCWLE | 5.5772 | 6.9881 | 6.3477 | 7.6591 | 6.6998 | 5.9548 | 6.2406 | 5.5568 | 7.2560 | 7.1897 | 6.0135 | 6.4294 | 5.1129 | 6.2865 | 5.2405 | 5.4471 |
| MED26_T323 | MED26 | O95402 | 535 | HVLVPQSPPTDLP | 6.6718 | 6.3216 | 6.2523 | 6.2283 | 6.1641 | 5.9802 | 6.0003 | 5.4977 | 6.3694 | 6.9464 | 6.9817 | 5.8164 | 6.4916 | 6.0595 | 6.1477 | 6.0709 |
| MED8_S82 | MED8 | Q96G25-2 | 82 | IIPLVLSPDRDED | 5.2890 | 5.4358 | 6.2008 | 5.8291 | 7.1596 | 5.4886 | 6.3828 | 7.3553 | 6.6763 | 6.1480 | 5.3712 | 7.1146 | 5.9112 | 6.5002 | 5.4416 | 7.6958 |
| MEF2A_S408 | MEF2A | Q02078 | 408 | IKSEPISPPRDRM | 4.8086 | 6.1514 | 4.9567 | 5.9376 | 5.0312 | 5.3394 | 5.5132 | 6.1333 | 15.5130 | 5.6170 | 4.9813 | 5.1499 | 6.2911 | 6.4385 | 6.1317 | 6.0062 |
| MEF2A_S479 | MEF2A | Q02078 | 494 | NTEDRESPSVKRM | 5.2142 | 5.5074 | 5.4744 | 4.9232 | 6.1439 | 6.2021 | 5.6919 | 5.8901 | 5.6798 | 5.7366 | 5.6037 | 5.7288 | 8.0812 | 8.5209 | 7.8273 | 7.7746 |
| MEF2A_S479;S494 | MEF2A | Q02078 | 479 | PRGDFHSPIVLGR | 4.4767 | 6.1145 | 6.2395 | 4.9260 | 5.7481 | 6.0302 | 6.0838 | 6.1489 | 6.4591 | 6.2893 | 5.9753 | 5.4519 | 7.6897 | 7.5058 | 7.2850 | 7.5760 |
| MEF2A_S494 | MEF2A | Q02078 | 479;494 | NA | 6.4057 | 6.0260 | 6.1747 | 5.8585 | 6.4580 | 6.5563 | 6.7807 | 7.0592 | 7.7021 | 6.1305 | 6.5323 | 6.5011 | 5.5282 | 5.6780 | 5.3500 | 5.2585 |
| MEF2C_S240 | MEF2C | Q06413-5 | 240 | YGNPRNSPGLLVS | 5.1165 | 4.5414 | 4.7748 | 4.1775 | 5.8225 | 6.0385 | 5.7514 | 5.4428 | 5.0132 | 5.5332 | 4.9215 | 5.1052 | 9.7813 | 10.4534 | 9.3848 | 8.1419 |
| MEF2D_S121 | MEF2D | Q14814 | 251 | KVIPAKSPPPPTH | 6.0891 | 5.7470 | 5.3028 | 5.7168 | 6.4179 | 7.1145 | 5.7475 | 7.3664 | 5.8668 | 5.4298 | 5.4824 | 5.6603 | 6.1093 | 7.2028 | 7.8632 | 6.8834 |
| MEF2D_S180 | MEF2D | Q14814 | 121 | DKYRRASEELDGL | 4.1236 | 4.3063 | 4.8154 | 5.3568 | 8.6450 | 5.5788 | 4.8806 | 5.3098 | 6.6557 | 6.9193 | 5.6062 | 5.9433 | 7.9973 | 8.1805 | 8.1026 | 7.5788 |
| MEF2D_S231 | MEF2D | Q14814 | 180 | TDPRLLSPQQPAL | 6.3573 | 6.5522 | 6.1431 | 6.8811 | 5.8329 | 6.2571 | 7.0052 | 6.4626 | 6.6637 | 6.0303 | 6.5384 | 5.9394 | 5.8022 | 6.0127 | 5.4695 | 6.0522 |
| MEF2D_S251 | MEF2D | Q14814 | 231 | YVSARASPGLLPV | 4.9299 | 5.4578 | 5.5572 | 4.9018 | 5.4212 | 5.7461 | 6.5545 | 6.2627 | 5.7989 | 6.0809 | 5.3056 | 5.4712 | 8.1593 | 8.7858 | 7.6469 | 7.9203 |
| MEF2D_S444 | MEF2D | Q14814 | 98 | GFNGCDSPEPDGE | 5.4444 | 5.3436 | 6.2870 | 6.2691 | 6.0125 | 5.8174 | 6.6820 | 6.1729 | 6.8812 | 5.6060 | 6.2743 | 6.4214 | 7.2831 | 6.3745 | 6.4932 | 6.6374 |
| MEF2D_S472 | MEF2D | Q14814 | 444 | IKSEPVSPSRERS | 5.3899 | 5.2815 | 5.5074 | 5.3876 | 5.6112 | 6.1573 | 6.0901 | 6.5969 | 6.2620 | 5.6287 | 5.4644 | 5.4499 | 7.2863 | 8.1071 | 7.9944 | 7.7852 |
| MEF2D_S98 | MEF2D | Q14814 | 472 | PGDGLSSPAGGSY | 5.6883 | 4.8600 | 6.4101 | 6.5531 | 6.0979 | 6.0076 | 6.2682 | 5.4896 | 5.7927 | 6.7406 | 6.2915 | 6.7528 | 7.0376 | 6.7680 | 6.2489 | 6.9930 |
| MEIOC_S130 | MEIOC | A2RUB1 | 130 | ERNDYGSETDLYG | 6.4838 | 6.3550 | 6.4295 | 6.3686 | 5.3173 | 6.3766 | 6.1505 | 6.1843 | 5.9633 | 6.6549 | 6.9415 | 6.5352 | 6.1197 | 6.5756 | 6.1436 | 5.4006 |
| MELK_S335 | MELK | Q14680 | 498 | LMTGVISPERRCR | 6.9962 | 7.3371 | 7.1478 | 7.3751 | 6.2536 | 6.9046 | 6.8841 | 6.8371 | 7.1311 | 7.3142 | 7.6118 | 7.6606 | 3.2712 | 4.0451 | 3.2547 | 3.9758 |
| MELK_S356 | MELK | Q14680 | 505 | PERRCRSVELDLN | 6.4335 | 6.3868 | 7.5051 | 6.5046 | 6.0936 | 6.6031 | 6.4431 | 6.6676 | 6.8250 | 6.8830 | 7.0418 | 7.1582 | 4.6559 | 5.1031 | 4.6151 | 5.0807 |
| MELK_S498 | MELK | Q14680 | 529 | GAKVFGSLERGLD | 7.3498 | 7.1379 | 7.4373 | 6.2549 | 5.7682 | 6.5924 | 6.5657 | 6.7441 | 7.1883 | 7.1328 | 7.2047 | 7.1644 | 4.0966 | 4.6928 | 4.0447 | 4.6254 |
| MELK_S505 | MELK | Q14680 | 356 | IKSNNWSLEDVTA | 6.5227 | 6.5740 | 7.0350 | 5.9740 | 5.8731 | 6.4511 | 6.2543 | 6.3398 | 6.3847 | 6.7404 | 6.8776 | 6.6520 | 5.5060 | 5.7160 | 5.3369 | 5.7625 |
| MELK_S529 | MELK | Q14680 | 335 | PVRLRLSSFSCGQ | 6.2638 | 6.5233 | 7.1609 | 6.6749 | 6.6967 | 6.3772 | 6.3300 | 6.4902 | 7.2335 | 7.9718 | 6.8160 | 7.1781 | 4.5363 | 5.0717 | 3.6950 | 4.9805 |
| MELK_T409 | MELK | Q14680 | 460 | KREILTTPNRYTT | 7.6925 | 7.7764 | 6.6984 | 6.9128 | 5.6352 | 7.6727 | 7.5920 | 6.7643 | 8.8240 | 8.1159 | 7.9912 | 7.9152 | 2.0538 | 3.1930 | 1.8761 | 3.2866 |
| MELK_T460 | MELK | Q14680 | 409 | VESKSLTPALCRT | 6.9847 | 7.3934 | 7.4854 | 6.9877 | 4.9295 | 7.1984 | 7.4779 | 6.9297 | 7.8342 | 7.6712 | 7.7148 | 7.4744 | 3.0250 | 4.2522 | 2.6550 | 3.9866 |
| MELK_T542 | MELK | Q14680 | 542 | KVITVLTRSKRKG | 10.5212 | 8.7839 | 8.2309 | 7.2296 | 4.3975 | 5.2793 | 7.7601 | 7.9469 | 7.0876 | 8.3019 | 8.9931 | 5.6828 | 2.0998 | 3.0674 | 2.4173 | 2.2006 |
| MEPCE_S101 | MEPCE | Q7L2J0 | 213 | RTLNAETPKSSPL | 6.0736 | 6.2126 | 6.1994 | 5.8800 | 5.4846 | 5.4583 | 5.9463 | 5.5205 | 6.5969 | 6.7427 | 6.3905 | 6.6391 | 6.6196 | 6.6154 | 6.5065 | 7.1141 |
| MEPCE_S152 | MEPCE | Q7L2J0 | 60 | CAPSAGSPAAAVG | 5.9592 | 6.2499 | 5.8738 | 6.0929 | 5.8090 | 5.7093 | 6.1303 | 5.9744 | 6.3095 | 6.3108 | 6.1344 | 6.0350 | 6.7266 | 6.8026 | 6.9658 | 6.9165 |
| MEPCE_S216;S217 | MEPCE | Q7L2J0 | 217 | AETPKSSPLPAKG | 6.1191 | 6.4167 | 5.8589 | 5.9828 | 6.0977 | 5.8023 | 6.1548 | 6.0727 | 6.3064 | 6.5390 | 6.0964 | 6.0768 | 6.3399 | 6.7010 | 6.7947 | 6.6410 |
| MEPCE_S217 | MEPCE | Q7L2J0 | 254 | GHVVLASPLKTGR | 6.0356 | 6.5717 | 6.3790 | 6.9363 | 6.3154 | 6.1996 | 6.5614 | 6.4485 | 6.6595 | 6.6570 | 6.6054 | 6.7690 | 5.4226 | 5.6639 | 5.1176 | 5.6576 |
| MEPCE_S254 | MEPCE | Q7L2J0 | 152 | GGKRRNSCNVGGG | 4.7448 | 3.8705 | 4.5511 | 4.7154 | 5.7747 | 5.3111 | 5.4004 | 4.8563 | 8.3741 | 7.5610 | 7.0744 | 7.6672 | 8.3129 | 8.6103 | 5.5946 | 7.5812 |
| MEPCE_S330 | MEPCE | Q7L2J0 | 330 | CRDEVVSPLPSAL | 5.7470 | 5.9237 | 5.8914 | 5.9640 | 5.7337 | 6.3721 | 6.1102 | 6.3750 | 6.3828 | 6.8449 | 6.4384 | 6.6266 | 6.2488 | 6.2780 | 6.2574 | 6.8061 |
| MEPCE_S60 | MEPCE | Q7L2J0 | 69 | AAVGRESPGAAAT | 5.3858 | 6.4625 | 6.3318 | 6.8140 | 5.8968 | 5.9095 | 6.5244 | 5.2695 | 6.2647 | 6.0402 | 6.1988 | 6.3354 | 6.4900 | 6.6386 | 6.8867 | 6.5512 |
| MEPCE_S60;S69 | MEPCE | Q7L2J0 | 101 | HGEARLSDPPGRA | 6.3768 | 7.2479 | 6.5065 | 7.0043 | 6.4151 | 6.5563 | 7.6016 | 7.0242 | 6.5981 | 7.1562 | 6.5705 | 6.7958 | 3.7256 | 5.1768 | 4.0346 | 5.2096 |
| MEPCE_S69 | MEPCE | Q7L2J0 | 60;69 | NA | 5.7980 | 6.1837 | 6.0614 | 5.8533 | 5.9443 | 6.9459 | 5.8782 | 5.9990 | 5.8684 | 6.3367 | 6.8250 | 6.2336 | 6.1623 | 6.8155 | 6.3768 | 6.7179 |
| MEPCE_T213 | MEPCE | Q7L2J0 | 216;217 | NA | 5.0480 | 6.9739 | 7.2728 | 5.8228 | 6.1052 | 5.1997 | 5.9899 | 6.3796 | 6.4705 | 7.5867 | 7.0164 | 5.6069 | 4.6298 | 6.3328 | 7.0080 | 6.5569 |
| MERTK_Y753 | MERTK | Q12866 | 929 | KPHEGRYILNGGS | 6.1269 | 6.6521 | 7.3165 | 6.6861 | 5.3217 | 7.0042 | 6.8169 | 6.8526 | 6.7432 | 7.9991 | 6.7813 | 8.0479 | 4.2953 | 4.5778 | 4.0666 | 4.7120 |
| MERTK_Y929 | MERTK | Q12866 | 753 | KIYSGDYYRQGRI | 7.4896 | 7.6865 | 7.9901 | 7.6089 | 6.6040 | 6.3231 | 6.0003 | 6.5948 | 7.3563 | 6.8061 | 6.6346 | 6.9519 | 4.1110 | 3.9586 | 3.5334 | 4.3507 |
| MET_S1008 | MET | P08581-2 | 1008 | VSARSVSPTTEMV | 5.8547 | 6.4529 | 6.3263 | 6.6964 | 6.1129 | 6.3290 | 6.0583 | 6.9473 | 6.9794 | 6.8397 | 6.6697 | 6.9202 | 5.2301 | 5.7242 | 5.5654 | 5.2936 |
| METAP2_S91 | METAP2 | F8VSC4 | 91 | DEGYKISLPSPRL | 7.3928 | 7.3693 | 6.7297 | 6.7144 | 5.8304 | 5.3957 | 6.2469 | 6.5185 | 6.9592 | 6.7555 | 6.0484 | 6.3321 | 5.0056 | 5.2453 | 5.6151 | 5.8410 |
| METTL1_S27 | METTL1 | Q9UBP6 | 27 | YRQRAHSNPMADH | 3.8727 | 4.6996 | 4.4887 | 4.4594 | 5.3187 | 5.6373 | 5.4460 | 6.0025 | 4.8216 | 4.6303 | 4.5512 | 4.1988 | 11.5607 | 10.3068 | 11.1774 | 8.8283 |
| METTL16_S329 | METTL16 | Q86W50 | 329 | ELSLKASPLRSET | 6.1048 | 6.6975 | 6.7238 | 6.0689 | 6.0029 | 6.6978 | 5.6575 | 5.8319 | 5.8964 | 5.8026 | 6.4795 | 6.4270 | 6.2933 | 6.9018 | 6.4836 | 5.9305 |
| METTL16_S498 | METTL16 | Q86W50 | 498 | ASEQFGSPVAERG | 6.3051 | 6.4719 | 7.2922 | 6.8353 | 5.4034 | 6.4475 | 5.9539 | 6.0963 | 6.5261 | 5.9851 | 6.9505 | 7.0007 | 5.2569 | 6.5220 | 5.3261 | 5.6271 |
| METTL2B_T154 | METTL2B | Q6P1Q9 | 154 | LEHKTQTPPVEEN | 6.7340 | 6.4999 | 7.1557 | 7.4647 | 6.2905 | 6.1645 | 6.5492 | 5.7864 | 7.7508 | 6.1853 | 7.2097 | 7.5670 | 4.6748 | 4.8198 | 4.4881 | 4.6596 |
| METTL3_S43 | METTL3 | Q86U44 | 43 | NPEAALSPTFRSD | 7.2185 | 7.2540 | 6.7281 | 7.0829 | 6.2179 | 6.8132 | 7.3211 | 6.7410 | 7.4332 | 6.8527 | 7.6196 | 7.3405 | 3.6356 | 4.0450 | 3.4297 | 4.2669 |
| METTL3_S50 | METTL3 | Q86U44 | 50 | PTFRSDSPVPTAP | 6.0934 | 5.8326 | 5.5863 | 5.0723 | 5.1517 | 6.5875 | 5.8036 | 6.1313 | 6.1345 | 6.5865 | 6.4932 | 6.1294 | 6.7929 | 7.5020 | 6.7473 | 7.3554 |
| MEX3C_S545 | MEX3C | Q5U5Q3 | 545 | PSTPRLSPTFPES | 5.9373 | 6.1570 | 7.1708 | 6.8296 | 6.5180 | 6.2878 | 6.4415 | 7.4707 | 6.2940 | 7.6096 | 7.0061 | 6.3532 | 5.0994 | 5.5832 | 4.8112 | 4.4305 |
| MEX3D_S514 | MEX3D | Q86XN8-3 | 514 | AGTPRHSPTLPEP | 6.0387 | 6.2260 | 6.8740 | 6.2097 | 6.5863 | 6.1630 | 5.8459 | 6.6862 | 6.4305 | 6.4183 | 6.6385 | 6.9232 | 5.6124 | 5.7266 | 5.7525 | 5.8682 |
| MFAP1_S116 | MFAP1 | P55081 | 267 | ALDALNTDDENDE | 5.9542 | 6.3263 | 6.1632 | 6.4798 | 6.0933 | 6.0615 | 5.9723 | 6.1583 | 6.4634 | 6.2063 | 6.1914 | 6.3655 | 6.4595 | 6.3788 | 6.3681 | 6.3583 |
| MFAP1_S116;S118 | MFAP1 | P55081 | 116 | PEVVGESDSEVEG | 5.6028 | 5.7757 | 5.6155 | 5.3425 | 5.4463 | 6.1215 | 6.2859 | 6.5880 | 6.6311 | 7.5448 | 6.7682 | 6.3781 | 6.5067 | 6.6900 | 6.4691 | 6.2337 |
| MFAP1_S52;S53 | MFAP1 | P55081 | 53 | YAPMESSDEEDEE | 4.3085 | 4.0490 | 4.9119 | 5.6082 | 5.3107 | 5.1841 | 4.2621 | 4.0384 | 7.5535 | 7.7139 | 8.3532 | 6.5577 | 7.7962 | 8.4545 | 8.3490 | 7.5493 |

| | | | | | | | | | | | | | | | | | | | | |
|---|---|---|---|---|---|---|---|---|---|---|---|---|---|---|---|---|---|---|---|---|
| MFAP1_S53 | MFAP1 | P55081 | 52;53 | NA | 6.0805 | 6.2432 | 5.9707 | 6.4767 | 6.2941 | 6.1460 | 5.9914 | 6.0392 | 6.4961 | 6.0154 | 6.1198 | 6.0626 | 6.6663 | 6.3967 | 6.5657 | 6.4353 |
| MFAP1_T267 | MFAP1 | P55081 | 116;118 | NA | 6.1662 | 6.5239 | 6.3816 | 6.8086 | 6.1503 | 5.8306 | 6.0369 | 6.0619 | 6.6099 | 6.3325 | 6.1721 | 6.3134 | 6.1686 | 6.4040 | 6.0260 | 6.0136 |
| MFF_S131 | MFF | Q9GZY8-5 | 131 | KRERSMSENAVRQ | 7.4900 | 6.3867 | 6.9186 | 6.6397 | 6.1728 | 5.3721 | 5.8592 | 6.9698 | 6.3650 | 7.0334 | 7.0217 | 7.6019 | 6.4641 | 5.3734 | 5.1378 | 5.0137 |
| MFF_S146 | MFF | Q9GZY8-5 | 112 | LERPPTTPQNEEI | 6.4879 | 5.6880 | 6.4351 | 6.4666 | 6.1404 | 5.7800 | 6.4967 | 6.9093 | 6.2283 | 6.4573 | 6.1989 | 7.0123 | 5.1743 | 6.5217 | 5.7899 | 6.2133 |
| MFF_S146 | MFF | Q9GZY8-5 | 89 | KPLALKTPPRVLT | 7.3840 | 5.3998 | 6.3242 | 6.7538 | 7.8223 | 7.3581 | 7.2652 | 6.9304 | 6.6226 | 6.9065 | 7.9978 | 7.7491 | 3.7967 | 4.9026 | 3.3865 | 3.4004 |
| MFF_T112 | MFF | Q9GZY8-4 | 146 | QLVRNDSLYGISN | 5.7058 | 5.5512 | 5.3150 | 5.9522 | 5.8630 | 6.0402 | 6.2640 | 6.0882 | 5.9756 | 6.2220 | 5.9798 | 6.3825 | 6.9212 | 7.5771 | 7.2228 | 6.9394 |
| MFF_T89 | MFF | Q9GZY8-5 | 146 | QLVRNDSLPVLRG | 5.2446 | 4.6150 | 5.0273 | 4.2887 | 6.7096 | 5.3297 | 6.3297 | 6.7821 | 4.8527 | 5.4816 | 5.0895 | 6.1878 | 8.6547 | 8.1893 | 8.3188 | 8.8988 |
| MFHAS1_S1038 | MFHAS1 | Q9Y4C4 | 1038 | PTPTVISPCSKKN | 6.0492 | 5.7762 | 5.9463 | 5.1757 | 6.4439 | 6.5702 | 6.1090 | 6.7533 | 6.2381 | 5.6420 | 7.1663 | 5.9236 | 6.2637 | 7.1685 | 5.9791 | 6.7949 |
| MFN2_S10 | MFN2 | O95140 | 10 | LFSRCNSIVTVKK | 8.7326 | 6.4275 | 8.5424 | 6.6966 | 4.9876 | 5.4984 | 6.3807 | 4.9043 | 7.6038 | 6.3343 | 7.6390 | 6.6765 | 4.9724 | 6.0100 | 3.8399 | 4.7542 |
| MGA_S1195 | MGA | Q8IWI9 | 924 | ILPYPVSPKQKYS | 6.4319 | 6.5217 | 6.0102 | 6.9529 | 6.3545 | 6.0966 | 6.4738 | 6.3042 | 6.4551 | 6.1939 | 6.1997 | 6.2376 | 5.6753 | 6.0329 | 6.0269 | 6.0328 |
| MGA_S1328 | MGA | Q8IWI9 | 2963 | EPESVSSPPTLHM | 6.7078 | 7.1301 | 6.9837 | 7.5904 | 6.4183 | 6.3662 | 6.5313 | 6.5404 | 6.5866 | 7.0403 | 7.3046 | 6.9117 | 4.4396 | 4.7217 | 4.1928 | 4.5345 |
| MGA_S1422 | MGA | Q8IWI9 | 56 | LASSVSSPVKSKG | 6.3713 | 6.2201 | 6.7125 | 7.1427 | 6.0786 | 6.2501 | 6.6890 | 6.3196 | 6.7750 | 6.4950 | 6.5023 | 6.5326 | 5.2795 | 5.5590 | 5.2732 | 5.7996 |
| MGA_S1430 | MGA | Q8IWI9 | 2910 | LEDDDFSENEKQL | 6.3758 | 6.7425 | 6.3718 | 6.9999 | 6.5956 | 6.2275 | 6.8160 | 6.4823 | 5.8894 | 5.9575 | 6.0135 | 6.0217 | 5.7736 | 6.0407 | 5.8583 | 5.8339 |
| MGA_S1457 | MGA | Q8IWI9 | 1457 | PFYAGLSPAGKLV | 7.4374 | 7.1732 | 6.9540 | 7.5429 | 6.2979 | 6.7929 | 7.0980 | 6.6680 | 6.9244 | 6.5370 | 6.9510 | 6.9985 | 3.6807 | 5.0672 | 3.8487 | 4.0283 |
| MGA_S2541 | MGA | Q8IWI9 | 449 | SKWLPSSPSGVAK | 6.8730 | 6.9185 | 5.9984 | 6.8664 | 6.6316 | 6.1464 | 6.5283 | 6.3680 | 6.4486 | 6.2267 | 5.9677 | 5.7644 | 5.6052 | 5.8380 | 6.0347 | 5.7843 |
| MGA_S2620 | MGA | Q8IWI9 | 1195 | PQPEVLSPTVKGK | 6.9912 | 6.4652 | 6.9830 | 7.0980 | 6.4794 | 6.8876 | 6.7186 | 6.2948 | 7.2296 | 6.9239 | 7.1193 | 7.3223 | 4.1082 | 4.7009 | 4.1583 | 4.5197 |
| MGA_S2910 | MGA | Q8IWI9 | 851 | SSNAPTSPVVYQL | 5.5858 | 6.2750 | 6.5366 | 6.9092 | 6.0393 | 6.5243 | 6.3717 | 6.7569 | 6.1624 | 6.5013 | 6.4805 | 6.5890 | 5.5954 | 6.3325 | 5.7229 | 5.6172 |
| MGA_S2921 | MGA | Q8IWI9 | 369 | DDSRVASPLDQNG | 5.8419 | 5.9451 | 6.0631 | 6.7487 | 5.4649 | 6.2177 | 6.1152 | 5.9203 | 6.5306 | 6.4824 | 5.9715 | 6.6408 | 6.0919 | 6.0534 | 6.8734 | 7.0391 |
| MGA_S2963 | MGA | Q8IWI9 | 376 | PLDQNGSFNVVIK | 6.5463 | 7.3500 | 7.2291 | 6.8385 | 6.5810 | 7.4317 | 6.2239 | 6.8071 | 7.0713 | 7.5370 | 7.2045 | 7.0064 | 4.1496 | 4.6881 | 3.3985 | 3.9370 |
| MGA_S369 | MGA | Q8IWI9 | 2921 | QLAEPASEPDVLK | 5.3674 | 6.0260 | 6.1943 | 6.7322 | 5.4774 | 5.6848 | 6.1036 | 5.7160 | 6.2465 | 5.6679 | 5.6474 | 5.8909 | 6.8541 | 7.0397 | 7.1666 | 8.1853 |
| MGA_S369;S376 | MGA | Q8IWI9 | 607 | NANQNASPNVPGK | 7.0100 | 6.8464 | 7.1026 | 7.3286 | 6.9110 | 6.4481 | 6.9346 | 6.4438 | 7.2511 | 6.2457 | 7.0466 | 6.8722 | 4.0820 | 4.3102 | 4.3533 | 4.8140 |
| MGA_S376 | MGA | Q8IWI9 | 1422 | TPDGPLSPGKMED | 6.8499 | 6.1031 | 6.7608 | 6.8284 | 7.1130 | 6.2130 | 7.2071 | 6.7129 | 7.1041 | 6.0734 | 6.7231 | 6.5452 | 4.9552 | 5.0262 | 5.0272 | 4.7574 |
| MGA_S449 | MGA | Q8IWI9 | 2541 | SDEFDISPRISKQ | 6.8514 | 7.2562 | 6.2232 | 7.0386 | 6.5591 | 6.3403 | 6.3631 | 6.5833 | 6.9363 | 6.8601 | 6.5215 | 6.6648 | 5.0401 | 4.9494 | 4.4665 | 5.3460 |
| MGA_S56 | MGA | Q8IWI9 | 1430 | GKMEDISPVQTDA | 5.6491 | 6.9916 | 6.3306 | 7.4054 | 5.5929 | 6.3868 | 6.1883 | 4.7129 | 7.8960 | 7.2912 | 5.4075 | 6.8249 | 6.3288 | 6.4317 | 4.1362 | 6.4261 |
| MGA_S569;S572 | MGA | Q8IWI9 | 1328 | SYMHQRSPGGPTK | 7.0862 | 7.3604 | 6.7172 | 7.7986 | 6.6890 | 6.2439 | 6.8721 | 7.0150 | 6.7628 | 6.2709 | 6.3550 | 6.6777 | 4.0781 | 4.9046 | 4.6076 | 4.5609 |
| MGA_S607 | MGA | Q8IWI9 | 2620 | GPVVAVSPDLLES | 6.7915 | 6.8330 | 6.0317 | 6.7654 | 6.0847 | 5.8421 | 6.2778 | 5.2968 | 6.3415 | 6.8821 | 6.1144 | 6.5105 | 5.7282 | 5.7859 | 5.9814 | 6.7329 |
| MGA_S851 | MGA | Q8IWI9 | 369;376 | NA | 6.5780 | 6.4175 | 6.4911 | 6.6585 | 6.5023 | 6.2979 | 6.9412 | 6.5780 | 7.0614 | 6.5880 | 6.6269 | 6.8451 | 4.8880 | 4.9555 | 4.7265 | 5.8440 |
| MGA_S924 | MGA | Q8IWI9 | 1416;1422 | NA | 6.0799 | 6.2427 | 4.4655 | 6.2175 | 6.3657 | 6.0733 | 6.4783 | 5.9165 | 7.3435 | 5.9646 | 6.9037 | 7.3687 | 5.8397 | 5.2652 | 6.0118 | 7.4632 |
| MGA_T1416;S1422 | MGA | Q8IWI9 | 569;572 | NA | 5.5940 | 6.6069 | 7.3278 | 6.3379 | 5.4996 | 6.0661 | 7.2848 | 5.7925 | 8.3735 | 7.0045 | 7.2671 | 6.7689 | 5.4128 | 5.2456 | 4.4736 | 4.9445 |
| MGMT_S201 | MGMT | P16455 | 201 | AGATSGSPPAGRN | 5.7518 | 5.1885 | 7.0371 | 5.3089 | 6.3271 | 5.2437 | 6.2720 | 7.8382 | 5.6756 | 6.6627 | 6.1368 | 7.4700 | 4.5284 | 7.3909 | 6.5659 | 6.6023 |
| MGRN1_S524 | MGRN1 | O60291-2 | 524 | NVLQDSSPEHCGR | 5.9523 | 12.1336 | 6.5414 | 6.8492 | 5.6652 | 5.7037 | 6.3500 | 5.7402 | 6.0599 | 5.9267 | 5.1749 | 5.6317 | 5.8320 | 5.7126 | 4.8819 | 5.8448 |
| MIA2_S1130 | MIA2 | Q96PC5 | 1255 | RSFNMPSLDKMDG | 7.5074 | 6.9461 | 8.7168 | 7.5156 | 6.7616 | 5.5367 | 6.1366 | 6.2079 | 6.6529 | 7.5346 | 6.3626 | 6.0886 | 3.5655 | 5.8104 | 4.6085 | 4.0481 |
| MIA2_S1144 | MIA2 | Q96PC5 | 1144 | ETRAFLSPPTLLE | 6.0777 | 6.6582 | 5.9564 | 6.1360 | 6.2944 | 6.1531 | 5.7700 | 6.3000 | 6.3333 | 6.4686 | 6.4405 | 6.5510 | 6.4241 | 6.3437 | 6.0180 | 6.0749 |
| MIA2_S1144;S1156 | MIA2 | Q96PC5 | 745 | CEKLNRSNSELED | 5.9994 | 6.0577 | 5.9522 | 6.3818 | 5.6960 | 5.9630 | 6.1799 | 5.6446 | 5.9415 | 6.0898 | 5.8844 | 6.1695 | 7.1141 | 7.0262 | 6.9956 | 6.9045 |
| MIA2_S1156 | MIA2 | Q96PC5 | 1243 | SNSGRLSGPAELR | 5.9084 | 6.0917 | 5.3830 | 5.7188 | 5.9348 | 5.8138 | 6.1246 | 6.0645 | 5.9565 | 5.6833 | 5.1391 | 5.3208 | 8.1057 | 7.3118 | 7.7282 | 7.7151 |
| MIA2_S1204 | MIA2 | Q96PC5 | 1130 | HSPYGPSPLGWPS | 5.4121 | 5.4684 | 5.5216 | 5.5594 | 5.9481 | 6.7797 | 6.6156 | 5.7717 | 6.1573 | 5.3543 | 5.4493 | 5.8267 | 6.6247 | 7.4649 | 7.4327 | 8.6135 |
| MIA2_S1243 | MIA2 | Q96PC5 | 1156 | EGPLRLSPLLPGG | 4.6194 | 3.9266 | 2.7859 | 3.8394 | 4.3805 | 7.0416 | 7.2854 | 6.0802 | 3.8602 | 2.6699 | 3.2596 | 3.0961 | 10.4905 | 10.7997 | 12.7382 | 13.1267 |
| MIA2_S1255 | MIA2 | Q96PC5 | 1204 | SDTGSLSPPWDQD | 6.2995 | 6.0537 | 6.7891 | 6.4347 | 6.1906 | 6.5436 | 6.2252 | 5.9051 | 5.9086 | 6.5165 | 6.7373 | 5.8040 | 6.5005 | 6.3257 | 5.8588 | 5.9071 |
| MIA2_S745 | MIA2 | Q96PC5 | 1144;1156 | NA | 4.2244 | 4.0939 | 3.4929 | 3.9840 | 5.0592 | 6.5207 | 6.6095 | 5.8781 | 5.0255 | 4.0855 | 4.1021 | 4.1526 | 10.2921 | 10.2759 | 10.8801 | 11.3234 |
| MIA3_S1678 | MIA3 | Q5JRA6 | 1741 | TMMNSSSRGSSPT | 6.2792 | 6.3013 | 6.6356 | 6.7839 | 6.5273 | 6.6164 | 6.1947 | 6.2772 | 6.1120 | 6.3171 | 6.6919 | 6.5244 | 6.3832 | 5.7843 | 5.3006 | 5.2709 |
| MIA3_S1706 | MIA3 | Q5JRA6 | 1906 | SQALKQSPxxxxx | 6.3312 | 6.6587 | 6.6607 | 6.2021 | 6.3907 | 6.4086 | 6.5634 | 6.5270 | 6.4424 | 6.9333 | 6.2544 | 6.4209 | 5.5846 | 5.8936 | 5.3632 | 5.3653 |
| MIA3_S1741 | MIA3 | Q5JRA6 | 1678 | VSGGECSPPLTVE | 6.1094 | 6.4633 | 6.1295 | 6.1734 | 6.3573 | 5.8496 | 6.1931 | 6.7200 | 6.4529 | 6.6441 | 6.2220 | 5.9672 | 5.9733 | 6.3719 | 6.0539 | 6.3191 |
| MIA3_S1906 | MIA3 | Q5JRA6 | 1706 | PRSEFGSVDGPLP | 5.7895 | 6.4720 | 6.3753 | 6.7235 | 6.8099 | 6.5914 | 5.5141 | 6.4756 | 6.1609 | 7.2358 | 7.0047 | 6.7024 | 5.2234 | 6.4262 | 5.7169 | 5.6783 |
| MIB1_S416 | MIB1 | Q86YT6 | 805 | GQVGSRSPSMISN | 5.7746 | 6.2717 | 6.5635 | 6.2417 | 5.9484 | 6.4572 | 6.0183 | 6.5771 | 6.3368 | 5.9323 | 6.3107 | 6.7757 | 5.8621 | 6.8532 | 6.0492 | 6.0274 |
| MIB1_S805 | MIB1 | Q86YT6 | 416 | ASGERLSQLLKKL | 6.6591 | 6.3690 | 7.3522 | 5.5963 | 6.1449 | 7.1377 | 5.7200 | 6.3250 | 5.9978 | 6.6069 | 6.5959 | 6.3924 | 6.0459 | 5.9359 | 4.9761 | 6.1449 |
| MICA_S338 | MICA | A0A140T986 | 338 | ALGSTGSTEGAxx | 6.1487 | 6.2530 | 6.1714 | 6.7830 | 6.5282 | 6.0082 | 5.8527 | 6.5090 | 6.5310 | 6.2514 | 6.1860 | 6.3842 | 6.0175 | 5.9247 | 5.7720 | 6.6790 |
| MICAL1_S1076 | MICAL1 | Q8TDZ2-4 | 1076 | QEERRLSELALGT | 5.5245 | 5.3635 | 6.2643 | 5.7738 | 6.2610 | 6.3419 | 6.6476 | 6.5920 | 7.0971 | 7.7870 | 7.5189 | 6.2217 | 5.4714 | 6.8617 | 4.7407 | 5.5327 |
| MICAL1_S786 | MICAL1 | Q8TDZ2-4 | 494 | LNLRAVTPNQVRD | 6.6660 | 6.8692 | 6.9619 | 7.5498 | 6.1808 | 5.5632 | 5.9005 | 6.6715 | 6.2137 | 5.9584 | 6.2175 | 6.4169 | 5.7866 | 6.2715 | 5.6195 | 5.9529 |
| MICAL1_S877 | MICAL1 | Q8TDZ2-4 | 877 | WGLPVQSPQALVA | 5.9639 | 6.8458 | 6.4502 | 6.8888 | 7.3747 | 6.6946 | 6.8756 | 7.0433 | 7.1630 | 6.8234 | 7.1887 | 6.6209 | 4.1485 | 5.3936 | 4.4298 | 4.0951 |
| MICAL1_T494 | MICAL1 | Q8TDZ2-4 | 786 | SDRGPESPELPTP | 5.5258 | 5.8965 | 7.1206 | 7.1967 | 6.6738 | 5.6704 | 6.1542 | 6.2836 | 6.4488 | 6.0759 | 5.9422 | 5.7704 | 6.1433 | 6.1042 | 5.8835 | 7.1098 |
| MICAL2_S515 | MICAL2 | O94851-4 | 515 | NLSRKESDIRPSK | 4.9575 | 5.5703 | 5.6899 | 5.5888 | 6.0077 | 5.7077 | 5.5468 | 5.0383 | 5.9139 | 5.7079 | 5.5615 | 5.9902 | 8.4398 | 8.1130 | 7.7334 | 8.4334 |
| MICAL2_T866 | MICAL2 | O94851-4 | 866 | EAPRRDTPTESSC | 5.8875 | 6.1996 | 6.2132 | 6.8116 | 5.9312 | 5.8520 | 5.8724 | 6.6594 | 6.1364 | 6.1296 | 6.7275 | 6.3921 | 6.2117 | 6.4903 | 6.4553 | 6.0304 |
| MICAL3_S1134 | MICAL3 | Q7RTP6 | 1310 | TKDRLGSPLAVDE | 6.1127 | 6.3972 | 6.5325 | 6.4204 | 6.0635 | 6.0618 | 6.5663 | 6.5724 | 7.3619 | 6.8071 | 6.8536 | 6.9545 | 5.0797 | 5.5096 | 5.0910 | 5.6158 |
| MICAL3_S1156 | MICAL3 | Q7RTP6 | 1649 | KERRPDSPTRPTL | 6.1048 | 6.3335 | 6.3168 | 5.9186 | 6.0606 | 6.2715 | 5.9268 | 6.2455 | 6.3748 | 6.6658 | 6.6610 | 6.7603 | 5.8582 | 6.4470 | 5.9777 | 6.0769 |
| MICAL3_S1221 | MICAL3 | Q7RTP6 | 1134 | ELELRVSEDEEKL | 5.5315 | 5.9496 | 6.6732 | 6.2047 | 6.0078 | 6.3011 | 6.1897 | 6.2176 | 6.4249 | 6.4132 | 6.5052 | 6.4363 | 6.3613 | 6.5350 | 6.1773 | 6.0715 |
| MICAL3_S1310 | MICAL3 | Q7RTP6 | 1321 | DEALRRSDLVEEF | 6.7124 | 6.8295 | 6.6221 | 7.7031 | 6.2152 | 6.5227 | 7.6452 | 7.4628 | 7.7328 | 7.3558 | 7.3683 | 7.6474 | 3.1328 | 4.1043 | 3.0520 | 3.8936 |
| MICAL3_S1321 | MICAL3 | Q7RTP6 | 1704 | KRSSLFSPRRNKK | 6.6957 | 6.7971 | 6.8404 | 6.4740 | 6.2975 | 6.5710 | 6.4999 | 6.6220 | 7.2199 | 6.7635 | 6.9794 | 7.0341 | 4.5967 | 5.1397 | 4.5829 | 4.8861 |
| MICAL3_S1371 | MICAL3 | Q7RTP6 | 1371 | SYSVEKSPQDEGL | 7.4994 | 7.3991 | 6.3952 | 7.1348 | 6.7348 | 6.6118 | 7.1554 | 7.0048 | 7.4326 | 7.1409 | 7.1382 | 7.3006 | 3.3116 | 4.1933 | 3.4020 | 4.1453 |
| MICAL3_S1512 | MICAL3 | Q7RTP6 | 649 | LIASTRSPISFLS | 6.9546 | 7.8815 | 6.8661 | 7.5702 | 6.5541 | 7.1622 | 7.3801 | 6.7576 | 7.1460 | 7.5604 | 7.5316 | 7.6572 | 2.9896 | 3.6687 | 2.8181 | 3.5018 |
| MICAL3_S1636 | MICAL3 | Q7RTP6 | 984 | EEELEASKSFGPG | 6.8107 | 7.2958 | 7.3391 | 6.9645 | 6.2840 | 6.3386 | 7.7444 | 6.5958 | 7.1058 | 6.5650 | 7.2786 | 7.4776 | 3.7691 | 4.5956 | 3.6864 | 4.1489 |
| MICAL3_S1649 | MICAL3 | Q7RTP6 | 2002 | GFSLNWSxxxxxx | 6.9122 | 7.8978 | 6.3108 | 6.4885 | 7.1307 | 5.8326 | 6.7284 | 7.5214 | 6.6800 | 6.2343 | 6.4704 | 7.2430 | 4.1806 | 5.2559 | 4.5504 | 5.5630 |
| MICAL3_S1704 | MICAL3 | Q7RTP6 | 1156 | GPSQATSPIRSPQ | 5.9751 | 5.9946 | 6.5273 | 5.7672 | 5.9617 | 6.1408 | 5.9302 | 6.5696 | 6.7092 | 6.6480 | 6.4555 | 6.5579 | 5.9354 | 6.3916 | 6.3683 | 6.0676 |
| MICAL3_S1726 | MICAL3 | Q7RTP6 | 1176 | PVHSPSTEGPQLP | 5.8003 | 6.0209 | 6.2625 | 5.6357 | 5.6636 | 5.8316 | 6.0483 | 6.2446 | 5.7903 | 6.4151 | 5.9702 | 6.3977 | 7.1194 | 6.7993 | 7.3216 | 6.6790 |
| MICAL3_S1745 | MICAL3 | Q7RTP6 | 1221 | DLKAVHSPIRSQP | 6.0253 | 6.3983 | 6.5811 | 6.1475 | 6.0331 | 6.5537 | 6.3280 | 6.1949 | 6.3832 | 6.5683 | 6.1512 | 7.2622 | 5.6962 | 5.9373 | 5.6469 | 6.0928 |
| MICAL3_S1998 | MICAL3 | Q7RTP6 | 1512 | REEVRKSFVESVE | 6.3714 | 5.8720 | 6.2368 | 6.1082 | 6.1183 | 5.8834 | 6.6130 | 6.6413 | 6.8325 | 6.5760 | 5.3571 | 7.3016 | 6.3263 | 6.6689 | 5.2319 | 5.8612 |
| MICAL3_S2002 | MICAL3 | Q7RTP6 | 1341 | RRSLGLTPVDRSK | 7.3051 | 7.7274 | 6.8242 | 6.1964 | 6.1431 | 6.8383 | 7.4900 | 6.7972 | 7.3870 | 6.6805 | 6.8129 | 7.2074 | 3.9774 | 4.7409 | 3.8041 | 4.0682 |
| MICAL3_S649 | MICAL3 | Q7RTP6 | 1745 | LWKSVFSGYKKDK | 7.8876 | 8.2361 | 6.8164 | 7.1710 | 6.2092 | 7.0420 | 7.1857 | 6.8096 | 7.6192 | 7.0996 | 7.4902 | 7.6569 | 2.8157 | 3.8535 | 2.5139 | 3.5934 |
| MICAL3_S984 | MICAL3 | Q7RTP6 | 1545 | LQEKFFTPPSCWP | 6.5163 | 7.3176 | 7.3918 | 7.5549 | 6.3901 | 7.1727 | 7.7210 | 6.7411 | 7.7657 | 7.9662 | 8.2283 | 7.6038 | 2.7090 | 3.3122 | 2.2523 | 3.3571 |
| MICAL3_T1176 | MICAL3 | Q7RTP6 | 1998 | MLSKGFSLNWSxx | 7.9056 | 7.1244 | 6.9198 | 7.3148 | 7.0995 | 5.3540 | 6.0426 | 5.4519 | 6.5373 | 6.4856 | 6.9483 | 7.5675 | 4.6138 | 5.1814 | 3.9181 | 5.5355 |
| MICAL3_T1341 | MICAL3 | Q7RTP6 | 1636 | PRPRKASSAPSQG | 6.3165 | 5.1799 | 5.4264 | 4.8792 | 6.5567 | 6.3478 | 5.8258 | 6.1859 | 7.9317 | 7.0932 | 7.2370 | 7.3337 | 6.7782 | 6.5303 | 4.8522 | 5.5257 |
| MICAL3_T1454;S1458 | MICAL3 | Q7RTP6 | 1726 | PPEKPSSNLLEEA | 6.4025 | 6.6894 | 7.8742 | 7.3725 | 5.8409 | 6.3638 | 7.1964 | 5.6121 | 6.0487 | 7.7756 | 6.0363 | 7.9692 | 4.4731 | 4.7865 | 4.3041 | 5.2547 |
| MICAL3_T1545 | MICAL3 | Q7RTP6 | 1454;1458 | NA | 6.5470 | 5.8733 | 6.7982 | 5.9959 | 6.5424 | 5.7057 | 5.6019 | 6.5630 | 6.6121 | 6.5361 | 6.9817 | 6.1494 | 5.5842 | 6.1019 | 6.2175 | 6.1898 |
| MICALL1_S273 | MICALL1 | Q8N3F8 | 578 | EQMPQASPGLAPR | 6.4638 | 6.4898 | 6.4068 | 6.3047 | 6.3196 | 6.5145 | 6.5410 | 6.7689 | 7.7666 | 6.5634 | 6.6412 | 6.5426 | 5.2853 | 5.4116 | 4.9816 | 4.9986 |
| MICALL1_S295 | MICALL1 | Q8N3F8 | 295 | KLQELASPPAGRP | 6.4280 | 7.1754 | 7.1339 | 6.4773 | 6.3299 | 6.9488 | 6.9850 | 7.0328 | 6.8940 | 6.9580 | 7.3907 | 6.8465 | 4.1977 | 4.4864 | 4.1397 | 4.5759 |
| MICALL1_S391 | MICALL1 | Q8N3F8 | 391 | PLPPSSSPGPPSQ | 6.1568 | 6.4582 | 6.8765 | 6.6566 | 6.4084 | 6.6801 | 6.5793 | 6.7321 | 6.8331 | 7.0057 | 7.1447 | 7.0045 | 5.0751 | 5.3667 | 4.3907 | 4.6313 |
| MICALL1_S414 | MICALL1 | Q8N3F8 | 640 | PFNRKPSPAASPA | 5.9726 | 6.1836 | 6.1887 | 5.9478 | 6.0027 | 6.0174 | 5.7700 | 6.3143 | 6.1215 | 6.1807 | 6.3975 | 6.4972 | 6.6348 | 7.0070 | 6.5931 | 6.4712 |
| MICALL1_S578 | MICALL1 | Q8N3F8 | 414 | EEVAQPSPTASLE | 6.1976 | 5.8959 | 6.3120 | 6.9095 | 6.4324 | 6.2975 | 6.2110 | 6.3180 | 6.2416 | 6.4562 | 6.8360 | 7.0736 | 5.5264 | 5.6257 | 5.7923 | 5.8743 |
| MICALL1_S589 | MICALL1 | Q8N3F8 | 621 | PSPGSSSPQLQVK | 5.3478 | 5.8741 | 5.8489 | 5.5282 | 6.1352 | 6.5228 | 5.7326 | 6.2401 | 5.8288 | 5.8008 | 6.0515 | 5.8959 | 7.5216 | 6.7213 | 7.6562 | 7.2944 |
| MICALL1_S612;S621 | MICALL1 | Q8N3F8 | 273 | ADGPKASPEARPQ | 6.2704 | 7.3582 | 8.2477 | 7.4766 | 6.0879 | 7.3777 | 6.1978 | 5.3440 | 5.9177 | 7.5298 | 7.8198 | 7.2112 | 3.8305 | 4.7940 | 4.1268 | 4.4098 |
| MICALL1_S621 | MICALL1 | Q8N3F8 | 644 | KPSPAASPATKKA | 6.6268 | 5.8585 | 6.9861 | 6.9180 | 6.2796 | 5.8252 | 5.4316 | 5.4320 | 6.6956 | 7.0823 | 6.3592 | 6.7985 | 6.2575 | 6.2654 | 5.9383 | 5.2452 |
| MICALL1_S640 | MICALL1 | Q8N3F8 | 589 | PRTRGSSGPQPAK | 5.7657 | 6.3508 | 6.4501 | 5.9920 | 5.9308 | 6.3585 | 5.7800 | 6.2589 | 6.1945 | 5.9753 | 6.3523 | 6.2021 | 6.7492 | 6.9266 | 6.3198 | 6.3933 |
| MICALL1_S640;S644 | MICALL1 | Q8N3F8 | 197 | CRRCSSTLLPGAY | 5.7121 | 6.1701 | 6.8929 | 7.5047 | 6.0703 | 6.2498 | 6.5704 | 5.8596 | 6.8306 | 5.7011 | 7.1640 | 6.4568 | 5.8871 | 5.1378 | 5.5280 | 6.2645 |
| MICALL1_S644 | MICALL1 | Q8N3F8 | 612;621 | NA | 5.9673 | 5.5327 | 6.7974 | 6.1868 | 6.0491 | 6.7603 | 6.3560 | 6.1907 | 6.4547 | 6.7206 | 6.5199 | 6.8500 | 6.0134 | 6.3836 | 5.4304 | 5.7871 |
| MICALL1_T197 | MICALL1 | Q8N3F8 | 467;471 | NA | 6.6543 | 6.5765 | 6.8952 | 7.1530 | 6.2312 | 7.3550 | 5.8528 | 7.2150 | 7.4572 | 6.4924 | 6.6506 | 6.1997 | 4.3282 | 5.7800 | 4.3033 | 4.8556 |
| MICALL1_T467;S471 | MICALL1 | Q8N3F8 | 640;644 | NA | 6.4160 | 6.5993 | 6.9964 | 6.4045 | 6.2989 | 6.9641 | 6.5184 | 6.8376 | 7.3353 | 7.0242 | 6.8234 | 7.1173 | 4.5985 | 5.3013 | 4.1863 | 4.5784 |
| MICALL2_S143 | MICALL2 | Q8IY33 | 272 | DSRTSCSPQKAQE | 6.4966 | 6.3212 | 6.4860 | 6.0666 | 6.3667 | 6.1459 | 6.1924 | 6.1620 | 6.8222 | 6.5334 | 6.8376 | 6.4144 | 5.7904 | 5.8126 | 5.8138 | 5.7380 |
| MICALL2_S153 | MICALL2 | Q8IY33 | 143 | QAAKLPSPAPARK | 6.2073 | 6.1474 | 6.0004 | 5.7007 | 6.1804 | 6.3407 | 5.9453 | 6.2253 | 6.7444 | 6.2864 | 5.9455 | 5.8820 | 6.8505 | 6.8069 | 6.2778 | 6.4589 |
| MICALL2_S249 | MICALL2 | Q8IY33 | 712 | KPGRPLSPANVPA | 5.2926 | 4.9582 | 5.5565 | 4.9974 | 6.1663 | 6.2349 | 6.2410 | 5.5824 | 6.2793 | 5.3355 | 5.4663 | 5.3511 | 8.2346 | 8.1596 | 7.5696 | 8.5745 |
| MICALL2_S272 | MICALL2 | Q8IY33 | 249 | PAAASASPKLTGL | 6.1875 | 6.2599 | 6.1588 | 5.9839 | 6.3605 | 6.7487 | 6.3356 | 6.3612 | 6.8397 | 6.7620 | 6.2598 | 6.2682 | 6.0636 | 6.4513 | 5.6051 | 5.3543 |
| MICALL2_S494 | MICALL2 | Q8IY33 | 153 | ARKPPLSPAQTNP | 5.7563 | 5.9233 | 6.4786 | 5.9585 | 6.2446 | 6.4146 | 6.8678 | 6.2888 | 6.8728 | 6.3676 | 6.7276 | 6.3770 | 5.3967 | 6.2804 | 5.9433 | 6.1021 |
| MICALL2_S712 | MICALL2 | Q8IY33 | 494 | TEAPQASPLAKPL | 7.0368 | 7.1758 | 6.3164 | 6.4253 | 6.9326 | 7.0105 | 6.8919 | 6.9640 | 7.3678 | 7.0131 | 6.9131 | 6.4241 | 4.3312 | 4.9920 | 4.0643 | 4.1412 |
| MICB_T380 | MICB | A0A140T9M6 | 380 | TGSTGSTEGAxxx | 5.5379 | 6.9707 | 5.8327 | 6.6689 | 6.6802 | 5.8980 | 6.3687 | 5.6615 | 6.8492 | 6.4924 | 6.2699 | 6.3812 | 5.7587 | 6.1695 | 6.6176 | 5.8429 |
| MIDEAS_S461 | MIDEAS | Q6PJG2 | 461 | RRRRRASQEANLL | 5.3661 | 6.0057 | 5.7775 | 6.1724 | 6.4428 | 5.6239 | 6.0780 | 6.1560 | 6.3723 | 6.1413 | 6.2713 | 6.2100 | 6.9990 | 6.7388 | 6.9727 | 6.6722 |
| MIDEAS_S636 | MIDEAS | Q6PJG2 | 704 | TNSAEVTPPVLSV | 6.7169 | 6.1215 | 6.4250 | 6.1712 | 6.2481 | 6.4766 | 6.6370 | 6.8237 | 6.5469 | 6.7455 | 6.7680 | 6.5207 | 5.3529 | 5.8691 | 5.3497 | 5.2271 |
| MIDEAS_T655 | MIDEAS | Q6PJG2 | 687 | PAPPPITPKSAHR | 5.9376 | 6.4648 | 6.4765 | 6.5021 | 5.7539 | 5.8722 | 5.7756 | 6.3535 | 5.6032 | 6.5541 | 6.0091 | 6.4367 | 6.8396 | 6.6212 | 6.5077 | 6.2920 |
| MIDEAS_T687 | MIDEAS | Q6PJG2 | 661 | TPPPILSPVREGS | 6.1679 | 6.9184 | 5.2515 | 6.4014 | 6.4711 | 5.6306 | 6.5118 | 6.5118 | 6.5674 | 6.3698 | 5.6742 | 7.1315 | 5.9077 | 6.7383 | 5.7417 | 6.0048 |
| MIDEAS_T704 | MIDEAS | Q6PJG2 | 655 | FELPPYTPPPILS | 6.4154 | 6.4683 | 6.2363 | 6.4636 | 6.4710 | 5.4880 | 5.3760 | 6.5219 | 6.9378 | 5.5165 | 5.9988 | 7.5760 | 5.7777 | 6.7316 | 7.3924 | 6.4487 |
| MIDEAS_T704;T715 | MIDEAS | Q6PJG2 | 636 | YQSHLRSPVRLAD | 5.4722 | 7.0848 | 6.0463 | 7.1088 | 6.3584 | 6.7967 | 6.3667 | 6.2508 | 4.9525 | 6.2780 | 6.9334 | 5.9878 | 6.5525 | 5.9743 | 5.2771 | 6.5598 |
| MIDEAS_Y655;S661 | MIDEAS | Q6PJG2 | 655;661 | NA | 5.5756 | 5.7901 | 5.9603 | 5.8830 | 6.1324 | 6.0999 | 5.8448 | 6.4142 | 5.8943 | 5.7716 | 5.9431 | 6.0322 | 6.7384 | 7.2135 | 7.4404 | 7.2662 |
| MIDEAS_Y661 | MIDEAS | Q6PJG2 | 704;715 | NA | 6.0424 | 6.6734 | 6.7284 | 6.6316 | 5.7788 | 6.0674 | 6.0148 | 6.2576 | 6.2927 | 5.8209 | 6.4272 | 6.3545 | 6.6359 | 6.7660 | 5.4621 | 6.0462 |
| MIEF1_S59 | MIEF1 | Q9NQG6 | 94 | KTGLSRSLQTLPT | 5.8083 | 6.3023 | 6.3819 | 6.4490 | 5.7597 | 5.8424 | 6.1039 | 6.0160 | 6.3888 | 6.4425 | 6.2192 | 6.1465 | 6.5342 | 6.3389 | 6.5977 | 6.6686 |
| MIEF1_S94 | MIEF1 | Q9NQG6 | 59 | AISAPTSPTRLSH | 5.5347 | 5.7778 | 5.3785 | 6.1478 | 6.2200 | 5.8868 | 6.1568 | 5.8984 | 5.7811 | 5.3808 | 5.4080 | 5.4282 | 7.6368 | 7.3824 | 8.1003 | 7.8816 |
| MIER1_S213;S219 | MIER1 | Q8N108-12 | 219 | SEVEEESEEDEDY | 5.3999 | 5.1815 | 5.5753 | 6.0584 | 5.7706 | 6.0088 | 5.3080 | 5.9141 | 6.9138 | 7.0023 | 6.6776 | 6.7873 | 7.0828 | 7.4458 | 7.2170 | 6.6568 |
| MIER1_S219 | MIER1 | Q8N108-12 | 430 | ASSRAPSPPPTAS | 5.7951 | 6.3186 | 6.6717 | 7.1550 | 7.2911 | 5.8719 | 5.9078 | 6.3261 | 6.6617 | 6.0564 | 6.4588 | 6.1023 | 5.6157 | 5.8795 | 6.4972 | 5.3911 |
| MIER1_S430 | MIER1 | Q8N108-12 | 541 | NSNGKESPGSSEF | 6.1393 | 6.9239 | 7.1944 | 7.0311 | 5.3232 | 5.2118 | 5.4152 | 5.6371 | 7.2124 | 7.1430 | 6.4703 | 6.6147 | 5.4167 | 6.1563 | 5.8743 | 6.2365 |
| MIER1_S536;S541 | MIER1 | Q8N108-12 | 213;219 | NA | 6.4498 | 6.0615 | 6.1395 | 6.5310 | 5.8858 | 5.8576 | 5.9388 | 6.0764 | 5.9671 | 6.0617 | 6.2686 | 6.5048 | 6.7915 | 6.2326 | 6.5166 | 6.7165 |
| MIER1_S541 | MIER1 | Q8N108-12 | 536;541 | NA | 5.4219 | 5.3650 | 5.9429 | 5.8117 | 6.2058 | 5.7726 | 6.0571 | 5.6465 | 6.1540 | 6.1936 | 6.1672 | 6.4132 | 7.7524 | 7.2112 | 6.9267 | 6.9580 |
| MIGA1_S140 | MIGA1 | Q8NAN2 | 140 | NLTLSLSSTKDKG | 5.6242 | 6.6930 | 6.4702 | 7.0357 | 6.4699 | 6.6074 | 6.1935 | 6.0646 | 7.1881 | 7.0299 | 6.9200 | 6.8931 | 4.6206 | 5.7105 | 5.0072 | 5.4722 |
| MIGA1_S147 | MIGA1 | Q8NAN2 | 147 | STKDKGSQVCNYA | 2.0427 | 2.1244 | 2.0560 | 2.1567 | 2.3267 | 2.2199 | 2.4606 | 2.3437 | 1.7781 | 2.1705 | 1.6130 | 2.3615 | 18.5393 | 16.2927 | 21.7774 | 17.7370 |
| MIGA1_S289 | MIGA1 | Q8NAN2 | 289 | EATLGASDPNSLA | 5.9188 | 6.0710 | 6.2742 | 5.9320 | 5.9591 | 5.5130 | 6.4377 | 6.4556 | 6.2696 | 6.2246 | 6.9010 | 5.9797 | 5.9538 | 6.3530 | 6.8560 | 6.9010 |
| MIGA2_S111 | MIGA2 | Q7L4E1 | 276 | LPLTEGSLRLRAD | 6.7148 | 6.7483 | 6.0662 | 6.7120 | 6.3396 | 5.8834 | 6.3356 | 6.0712 | 5.9507 | 5.9221 | 5.7921 | 5.8830 | 6.5846 | 6.4937 | 6.2613 | 6.2415 |
| MIGA2_S276 | MIGA2 | Q7L4E1 | 111 | SSRRVQSPSKSSN | 6.9613 | 7.3495 | 6.6085 | 6.9454 | 6.1123 | 6.4893 | 6.2044 | 6.4977 | 6.3180 | 6.0555 | 5.6431 | 6.2491 | 5.1469 | 5.9801 | 5.8703 | 5.5686 |
| MIIP_S303 | MIIP | Q5JXC2 | 303 | HIHRRKSFDASDT | 6.4722 | 6.8027 | 6.9531 | 6.2433 | 6.7376 | 6.7583 | 6.0506 | 6.6706 | 6.7230 | 7.3955 | 7.4681 | 7.0798 | 4.5651 | 4.9154 | 4.2911 | 4.4925 |
| MINDY1_S489 | MINDY1 | Q8NBJ2-3 | 489 | MRTRVLSLQGRGA | 8.0354 | 6.3247 | 6.7709 | 5.9705 | 5.6958 | 5.9188 | 5.8159 | 6.4133 | 7.8002 | 7.0088 | 6.5634 | 7.1471 | 5.2024 | 6.6169 | 4.3811 | 4.3348 |
| MINDY2_S587 | MINDY2 | Q8NBR6 | 587 | GQPAQASPSSGRQ | 5.9117 | 6.4148 | 6.4193 | 6.6956 | 6.6681 | 6.6821 | 6.4895 | 6.5741 | 6.8418 | 6.8256 | 6.5417 | 6.1083 | 5.9243 | 5.7627 | 5.4801 | 5.5602 |
| MINDY2_S62 | MINDY2 | Q8NBR6 | 62 | AAAARRSLPDSAS | 4.9636 | 5.1048 | 5.1973 | 6.2976 | 5.9081 | 6.7407 | 7.7144 | 6.3414 | 5.6355 | 6.4242 | 6.5946 | 7.9311 | 6.9938 | 6.6920 | 4.8419 | 6.6190 |
| MINDY4_S524 | MINDY4 | Q4G0A6 | 524 | SRTQQFSPTGKYK | 6.3364 | 6.5172 | 7.4314 | 6.7758 | 5.8650 | 6.8911 | 6.0778 | 6.3210 | 7.9874 | 7.1064 | 7.0845 | 6.9879 | 4.6575 | 4.9439 | 4.4597 | 4.5572 |
| MINK1_S535 | MINK1 | Q8N4C8 | 187 | TVGRRNTFIGTPY | 6.7337 | 6.7747 | 6.3681 | 6.4238 | 6.4196 | 6.5539 | 6.7298 | 6.5365 | 6.4409 | 6.0540 | 6.1417 | 5.9126 | 6.2071 | 6.5804 | 5.6367 | 6.0866 |
| MINK1_S641 | MINK1 | Q8N4C8 | 641 | AVIRQNSDPTSEG | 5.8485 | 5.4913 | 5.4368 | 5.5288 | 6.4977 | 6.0291 | 6.1965 | 6.0911 | 6.3531 | 6.5071 | 6.1407 | 6.3588 | 6.9656 | 7.3946 | 6.4436 | 7.1677 |
| MINK1_S733 | MINK1 | Q8N4C8 | 733 | GPPNASSNPDLRR | 5.3523 | 6.2819 | 6.5489 | 6.4626 | 5.5907 | 5.8554 | 5.8994 | 5.8527 | 6.8184 | 6.6770 | 6.7099 | 6.8798 | 5.8545 | 6.4934 | 6.1648 | 6.5584 |
| MINK1_S778 | MINK1 | Q8N4C8 | 778 | SSKPDSSPVLSPG | 6.1431 | 5.8431 | 5.8565 | 5.5203 | 6.5202 | 6.3152 | 6.0047 | 6.5073 | 6.7098 | 6.3324 | 6.0125 | 5.6409 | 6.9396 | 7.1474 | 6.4041 | 6.1029 |
| MINK1_S782 | MINK1 | Q8N4C8 | 535 | MNKQQNSPLAKSK | 6.0345 | 6.2831 | 6.5736 | 6.7879 | 5.9169 | 6.4131 | 6.2200 | 5.7544 | 7.0826 | 7.7200 | 6.8882 | 7.8772 | 4.8845 | 5.4770 | 4.9158 | 5.1710 |
| MINK1_S927 | MINK1 | Q8N4C8 | 993 | YDVRKGSVVNVNP | 6.8517 | 5.3237 | 5.9903 | 5.7639 | 6.0013 | 5.1565 | 5.4674 | 5.7861 | 7.9613 | 6.8737 | 6.1897 | 6.3865 | 6.7410 | 7.4806 | 5.6707 | 6.3556 |
| MINK1_S993 | MINK1 | Q8N4C8 | 782 | DSSPVLSPGNKAK | 6.3965 | 5.6020 | 7.9818 | 5.8488 | 5.8911 | 6.5586 | 6.7831 | 6.9758 | 7.2731 | 8.4401 | 7.7773 | 8.4036 | 3.7218 | 4.6572 | 3.3751 | 4.3141 |
| MINK1_T187 | MINK1 | Q8N4C8 | 927 | ENSKGQSPPSKDG | 6.6872 | 7.2180 | 6.7878 | 6.7335 | 6.2065 | 6.8165 | 6.4612 | 6.7328 | 7.0919 | 7.1171 | 7.0468 | 7.2995 | 3.9232 | 4.8556 | 4.0415 | 4.9811 |
| MIOS_S766 | MIOS | Q9NXC5 | 766 | QYGVSGSPTKSKV | 5.9421 | 6.3977 | 6.5433 | 6.3903 | 6.3192 | 6.4585 | 6.0595 | 6.5183 | 6.5126 | 6.7796 | 6.7900 | 6.8611 | 5.3692 | 6.0774 | 5.6555 | 5.3256 |
| MIOS_T342 | MIOS | Q9NXC5 | 342 | NRMIVVTPNRTMS | 6.1553 | 5.6172 | 6.0607 | 6.0540 | 5.4146 | 5.7036 | 7.5007 | 6.2135 | 7.7465 | 7.1868 | 6.6254 | 5.8896 | 5.3598 | 7.0256 | 5.4063 | 6.0407 |
| MIS18A_S233 | MIS18A | Q9NYP9 | 233 | SFATCKSxxxxxx | 5.5540 | 7.1929 | 6.6795 | 5.1898 | 6.0155 | 6.4833 | 6.7486 | 7.5712 | 6.0830 | 8.3828 | 8.4095 | 7.6528 | 3.8366 | 5.3672 | 4.4284 | 4.4049 |
| MIS18BP1_S1008 | MIS18BP1 | Q6P0N0 | 653 | KAYILVTPLKSRK | 7.2464 | 6.9135 | 5.6785 | 7.1905 | 6.8878 | 6.0970 | 6.7354 | 5.9642 | 6.7854 | 5.9885 | 5.7931 | 5.9532 | 5.4657 | 5.8597 | 5.7520 | 5.6889 |
| MIS18BP1_S110 | MIS18BP1 | Q6P0N0 | 218 | APLHNLTYELPTL | 6.4014 | 6.5898 | 7.5112 | 6.8493 | 5.9907 | 5.8257 | 6.5976 | 6.6148 | 7.2661 | 5.8390 | 6.6493 | 6.0669 | 5.5978 | 5.4344 | 5.2355 | 5.5306 |
| MIS18BP1_S172 | MIS18BP1 | Q6P0N0 | 702 | MGTLENTFEGHKS | 6.6735 | 6.9994 | 6.6607 | 7.8358 | 6.4133 | 6.1616 | 6.7039 | 6.3576 | 6.7702 | 6.6025 | 6.3029 | 6.6840 | 4.5138 | 5.1293 | 4.7637 | 5.4278 |
| MIS18BP1_S365 | MIS18BP1 | Q6P0N0 | 161 | KKKLQHTYLCEEK | 10.2311 | 8.8968 | 7.1082 | 10.3428 | 7.6556 | 5.0894 | 6.7495 | 4.8834 | 6.0548 | 4.5044 | 5.6265 | 5.4419 | 2.6114 | 4.0427 | 4.8241 | 3.7470 | 4.7986 |
| MIS18BP1_S541 | MIS18BP1 | Q6P0N0 | 824 | EPETEESENEFYI | 6.2511 | 6.3891 | 6.0800 | 6.9830 | 6.5178 | 5.8988 | 6.2522 | 6.0162 | 6.7017 | 6.0601 | 6.6720 | 6.6323 | 5.5127 | 6.0091 | 5.7832 | 6.2406 |
| MIS18BP1_T1023 | MIS18BP1 | Q6P0N0 | 821 | VILEPETEESENE | 5.7131 | 6.6408 | 6.5784 | 7.2491 | 5.8583 | 5.9504 | 6.3586 | 6.5926 | 6.3855 | 5.9915 | 5.7882 | 5.7920 | 6.4233 | 6.3614 | 6.0183 | 6.2984 |
| MIS18BP1_T161 | MIS18BP1 | Q6P0N0 | 1008 | LPSFQDSEDDDDI | 5.9690 | 6.6429 | 6.8066 | 7.3214 | 6.2747 | 5.9788 | 6.6542 | 6.2360 | 6.2308 | 6.3862 | 6.7926 | 6.4497 | 5.1810 | 5.9798 | 5.3658 | 5.7304 |
| MIS18BP1_T218 | MIS18BP1 | Q6P0N0 | 1023 | NMDKNPTTPSSVI | 6.6221 | 7.4348 | 6.6509 | 7.2794 | 6.0396 | 7.0519 | 7.3691 | 6.7127 | 6.8375 | 7.6697 | 7.3598 | 7.3191 | 3.2624 | 4.3811 | 3.4967 | 4.5131 |
| MIS18BP1_T653 | MIS18BP1 | Q6P0N0 | 110 | NKANYESPGKIFL | 6.3187 | 7.3036 | 6.1628 | 6.9661 | 6.8318 | 6.0138 | 6.2843 | 6.0667 | 6.1023 | 6.4383 | 6.2941 | 6.6078 | 5.0455 | 5.6498 | 5.5559 | 6.3584 |
| MIS18BP1_T702 | MIS18BP1 | Q6P0N0 | 365 | RNISKLSPPRIFQ | 6.5031 | 7.4739 | 6.4669 | 7.1616 | 6.9036 | 7.3961 | 6.8547 | 6.1003 | 6.5779 | 6.3082 | 7.0675 | 6.3851 | 4.8560 | 5.4171 | 4.3105 | 4.2175 |
| MIS18BP1_T821 | MIS18BP1 | Q6P0N0 | 172 | EKENNKSFQSDDS | 6.5263 | 7.0920 | 6.0313 | 6.4413 | 5.8564 | 6.3598 | 6.3630 | 6.0197 | 6.2418 | 6.7000 | 6.7467 | 6.4942 | 5.3036 | 5.9384 | 5.4671 | 6.4184 |
| MIS18BP1_T821;S824 | MIS18BP1 | Q6P0N0 | 541 | SNKHSESPGATEL | 6.2571 | 7.1167 | 6.5740 | 7.9314 | 5.9098 | 6.5138 | 6.8369 | 6.2566 | 6.7668 | 6.8245 | 6.4432 | 6.3679 | 4.8085 | 5.0605 | 4.7359 | 5.5965 |
| MIS18BP1_T824 | MIS18BP1 | Q6P0N0 | 821;824 | NA | 6.7512 | 7.2776 | 7.1429 | 7.3325 | 6.2570 | 7.2159 | 6.6917 | 5.9522 | 6.5369 | 5.5579 | 6.0123 | 5.7821 | 5.7922 | 5.0661 | 5.3419 | 5.2894 |
| MKI67_S1071 | MKI67 | P46013 | 1193 | IKAFMGTPVQKLD | 7.7579 | 7.9375 | 6.7275 | 8.0567 | 6.6954 | 6.9609 | 7.9334 | 7.0397 | 7.2924 | 6.9931 | 6.6859 | 6.7105 | 2.6679 | 4.1766 | 2.7696 | 3.5951 |
| MKI67_S1131 | MKI67 | P46013 | 584 | ISPPAPSPRKTPV | 6.4158 | 6.5112 | 6.2754 | 6.6464 | 5.8463 | 6.0888 | 6.2941 | 6.3423 | 6.5617 | 6.6468 | 6.5122 | 6.7325 | 5.5238 | 6.0781 | 5.5254 | 5.9992 |
| MKI67_S1207 | MKI67 | P46013 | 507 | LKRRRVSFGGHLR | 5.1552 | 4.9511 | 5.1276 | 5.6594 | 6.2308 | 5.8800 | 5.5905 | 5.9832 | 8.0042 | 8.0676 | 7.7807 | 7.8339 | 6.4133 | 7.4894 | 4.3971 | 5.4360 |
| MKI67_S1253 | MKI67 | P46013 | 2223 | TKIACRSPQPDPV | 6.6794 | 7.1184 | 6.7760 | 7.7185 | 6.2672 | 6.3091 | 6.6691 | 6.3790 | 6.9397 | 7.0263 | 6.8780 | 7.1350 | 3.9994 | 4.8767 | 4.3567 | 4.8714 |
| MKI67_S1291 | MKI67 | P46013 | 827 | SDKCSASPPLRRQ | 7.1905 | 7.2432 | 6.5405 | 7.7091 | 6.4417 | 6.2902 | 6.8302 | 6.2533 | 6.7962 | 6.9413 | 6.8242 | 7.0082 | 3.9390 | 5.0407 | 4.1897 | 4.7621 |
| MKI67_S1329 | MKI67 | P46013 | 1937 | LGNLPGSKRRPQT | 7.6962 | 8.1395 | 7.1453 | 8.3534 | 6.6254 | 6.6960 | 7.7346 | 7.1778 | 7.3727 | 7.0446 | 6.7571 | 6.7594 | 2.5004 | 4.1191 | 2.5808 | 3.2977 |
| MKI67_S1376 | MKI67 | P46013 | 1815 | PGNLPGSNRRLQT | 7.5101 | 7.7818 | 7.0546 | 7.7073 | 6.4291 | 7.1134 | 7.6054 | 6.7881 | 7.2069 | 6.9525 | 6.9453 | 7.1337 | 2.9776 | 4.2850 | 2.9410 | 3.5682 |
| MKI67_S1447 | MKI67 | P46013 | 1557 | IYAFMGTPVQKLD | 7.5404 | 7.9594 | 6.9347 | 7.9709 | 6.1141 | 7.1722 | 7.5142 | 6.9565 | 7.2356 | 7.5057 | 7.0440 | 7.1327 | 2.6778 | 4.1023 | 2.7030 | 3.4366 |

| | | | | | | | | | | | | | | | | | | | | |
|---|---|---|---|---|---|---|---|---|---|---|---|---|---|---|---|---|---|---|---|---|
| MKI67_S1451 | MKI67 | P46013 | 2466 | TEVSCKSPQPESF | 7.7706 | 8.0754 | 6.6660 | 7.8180 | 6.6329 | 6.6896 | 7.4150 | 6.9285 | 7.1943 | 7.1783 | 7.0578 | 6.8991 | 2.7525 | 4.1508 | 3.1201 | 3.6508 |
| MKI67_S1679 | MKI67 | P46013 | 1017 | QPEPINTPTHTKQ | 7.5787 | 7.5828 | 6.8224 | 7.7159 | 5.6707 | 7.4803 | 7.6673 | 6.9994 | 6.9842 | 7.4385 | 7.1544 | 7.2505 | 2.8730 | 4.3164 | 2.7328 | 3.7328 |
| MKI67_S1740;T1747 | MKI67 | P46013 | 1679 | MKAFMESPKQILD | 7.8698 | 7.9034 | 7.1122 | 7.8390 | 6.5861 | 6.2854 | 7.7339 | 6.4610 | 7.3805 | 7.3894 | 7.0490 | 7.0821 | 2.7400 | 3.9125 | 2.9392 | 3.7166 |
| MKI67_S1815 | MKI67 | P46013 | 1315 | IIIFVGTPVQKLD | 7.7741 | 8.1473 | 6.6933 | 8.0533 | 6.5211 | 6.8230 | 7.4170 | 7.1078 | 7.2777 | 6.8252 | 6.7874 | 6.8540 | 2.9952 | 4.2182 | 2.7704 | 3.7350 |
| MKI67_S1861 | MKI67 | P46013 | 1167 | LALRKLTPSAGKA | 7.6157 | 8.0271 | 6.7767 | 7.9428 | 6.6000 | 6.9485 | 7.7120 | 7.3427 | 7.2202 | 7.4061 | 7.1502 | 7.1162 | 2.4487 | 3.9767 | 2.4991 | 3.2172 |
| MKI67_S1861;S1864 | MKI67 | P46013 | 3041 | PRARGKSSEPVVI | 5.3209 | 5.7469 | 5.3821 | 6.0121 | 5.3725 | 5.7585 | 6.1224 | 6.0771 | 4.8707 | 5.4273 | 5.3879 | 5.8768 | 7.2502 | 7.6751 | 8.8539 | 8.8656 |
| MKI67_S1937 | MKI67 | P46013 | 1291 | CRNLMPSAGKAMH | 7.6903 | 8.0988 | 6.8608 | 8.2704 | 6.4194 | 7.0933 | 7.6335 | 6.7391 | 6.9281 | 7.1039 | 7.1350 | 6.9712 | 2.7253 | 4.1048 | 2.6251 | 3.6010 |
| MKI67_S1983 | MKI67 | P46013 | 2638 | THKEPASGDEGIK | 7.2987 | 8.2081 | 7.7750 | 8.3256 | 7.2588 | 7.5337 | 8.2640 | 8.0581 | 4.9628 | 4.9714 | 5.1721 | 5.2595 | 3.5926 | 4.5223 | 4.2008 | 4.5966 |
| MKI67_S2022 | MKI67 | P46013 | 2588 | TKIPCKSPPPELT | 6.7058 | 7.3941 | 6.8206 | 7.9377 | 6.1339 | 6.7735 | 6.8436 | 6.6080 | 6.5306 | 7.1017 | 7.1630 | 7.0767 | 3.6591 | 4.8498 | 3.8017 | 4.6003 |
| MKI67_S2105 | MKI67 | P46013 | 1923 | INTFVGTPVEKLD | 7.4271 | 7.8126 | 6.9095 | 8.4702 | 6.4519 | 6.8871 | 7.4951 | 7.0307 | 7.1426 | 7.3041 | 7.1392 | 7.0068 | 2.6168 | 4.0696 | 2.6680 | 3.5686 |
| MKI67_S2105;T2113 | MKI67 | P46013 | 605 | CKTAPASSSKSQT | 7.9798 | 8.1498 | 7.2309 | 7.9569 | 5.6380 | 6.9572 | 7.8186 | 6.5322 | 7.0061 | 7.1284 | 7.5549 | 7.6461 | 2.4410 | 4.1149 | 2.3413 | 3.5037 |
| MKI67_S2135 | MKI67 | P46013 | 2299 | PGNLPGSKRWPQT | 7.5033 | 7.8599 | 7.5177 | 8.3830 | 6.3202 | 6.9778 | 7.7752 | 6.9168 | 6.8356 | 6.9644 | 7.0412 | 7.0794 | 2.8491 | 4.0701 | 2.6661 | 3.2401 |
| MKI67_S2223 | MKI67 | P46013 | 374 | TTGRRESVNLGKS | 5.6668 | 4.6974 | 4.4009 | 4.9402 | 6.2528 | 5.7740 | 5.2806 | 5.4408 | 9.1572 | 8.8000 | 7.8914 | 9.1176 | 6.4212 | 7.8199 | 3.2958 | 5.0434 |
| MKI67_S2223;T2231 | MKI67 | P46013 | 1983 | TEVSCKSPQPDPV | 7.1704 | 7.9010 | 6.8246 | 7.8202 | 6.3139 | 6.0842 | 6.9072 | 6.7105 | 7.0683 | 7.5432 | 6.9404 | 7.1095 | 3.2851 | 4.3976 | 3.6718 | 4.2520 |
| MKI67_S2261 | MKI67 | P46013 | 2105 | TKIACKSPPPESM | 6.1478 | 6.8404 | 6.2909 | 7.3264 | 6.1645 | 6.2109 | 6.4203 | 6.4707 | 6.8251 | 6.5065 | 6.5173 | 6.4790 | 4.9969 | 5.8312 | 5.3769 | 5.5953 |
| MKI67_S2299 | MKI67 | P46013 | 2344 | TKIACKSPQPDPV | 6.9212 | 7.2950 | 6.6423 | 7.5107 | 6.4683 | 6.6037 | 7.0156 | 6.7427 | 7.1012 | 7.2575 | 6.7734 | 7.0638 | 3.6366 | 4.4387 | 3.9948 | 4.5344 |
| MKI67_S2344 | MKI67 | P46013 | 308 | AVAEPASPEQELD | 6.6634 | 7.6638 | 5.9259 | 8.0892 | 6.7207 | 6.8333 | 5.7176 | 7.7910 | 5.7520 | 6.1254 | 6.6463 | 5.5563 | 3.7501 | 5.9307 | 5.6275 | 5.2069 |
| MKI67_S235 | MKI67 | P46013 | 1801 | INTFLGTPVQKLD | 7.6903 | 7.9462 | 7.0689 | 7.5840 | 6.6421 | 7.1238 | 7.5373 | 6.9716 | 7.1266 | 7.2004 | 6.7359 | 6.9823 | 2.8432 | 4.2078 | 2.7959 | 3.5437 |
| MKI67_S2466 | MKI67 | P46013 | 2406 | INTFVETPVQKLD | 7.3398 | 7.9034 | 6.9493 | 7.8896 | 6.5825 | 6.8148 | 7.5785 | 6.9191 | 7.1740 | 7.0492 | 6.6447 | 7.0152 | 3.0175 | 4.4035 | 2.9224 | 3.7965 |
| MKI67_S2528 | MKI67 | P46013 | 1841 | FRELFQTPCTDNP | 7.2729 | 7.6483 | 6.5870 | 7.7751 | 6.2970 | 6.8219 | 6.9465 | 6.5687 | 6.8134 | 6.9077 | 7.0619 | 6.4748 | 3.8875 | 4.9733 | 3.6863 | 4.2778 |
| MKI67_S2588 | MKI67 | P46013 | 2528 | IKAFKESPKQILD | 6.5828 | 7.0268 | 6.3849 | 7.4573 | 6.4638 | 6.4660 | 7.0476 | 6.5867 | 6.5845 | 7.0365 | 6.6498 | 6.6894 | 4.1865 | 5.3130 | 4.3766 | 5.1478 |
| MKI67_S2638 | MKI67 | P46013 | 543 | KSLVMHTPPVLKK | 6.8314 | 7.9530 | 6.2129 | 7.7335 | 7.8852 | 6.7804 | 6.7848 | 7.9028 | 6.9064 | 6.8878 | 6.5316 | 5.8791 | 3.4911 | 4.4610 | 3.7685 | 3.9905 |
| MKI67_S264 | MKI67 | P46013 | 2708 | TKIPCESPPLEVV | 6.7301 | 7.0063 | 6.3493 | 6.9406 | 6.2116 | 6.4493 | 6.5757 | 6.4750 | 6.7003 | 6.4359 | 6.3505 | 6.2737 | 4.9517 | 5.7801 | 5.1547 | 5.6152 |
| MKI67_S2689 | MKI67 | P46013 | 357 | YSQQQNSPQKHKN | 6.2611 | 6.1670 | 6.0963 | 6.5593 | 6.2300 | 5.7721 | 6.1119 | 6.2868 | 6.4569 | 6.1697 | 5.9213 | 6.2176 | 6.0621 | 6.8153 | 6.5571 | 6.3154 |
| MKI67_S2708 | MKI67 | P46013 | 1207 | AGTLPGSKRQLQT | 7.1890 | 7.8371 | 6.7454 | 7.9922 | 6.1883 | 6.7546 | 7.6201 | 6.5810 | 6.5309 | 6.6503 | 6.9854 | 7.0922 | 3.4859 | 4.4623 | 3.5832 | 4.3020 |
| MKI67_S2828 | MKI67 | P46013 | 1355 | FKELFQTPGHTEE | 7.3873 | 7.9779 | 6.8233 | 8.2512 | 6.3461 | 6.7400 | 7.3774 | 6.7824 | 6.9540 | 7.1249 | 7.0463 | 7.2440 | 2.8909 | 4.4470 | 2.8937 | 3.7135 |
| MKI67_S3041 | MKI67 | P46013 | 1476 | LKELFQTPVCTDK | 8.2522 | 8.4128 | 7.1068 | 8.2469 | 6.3779 | 6.2172 | 7.3147 | 6.6630 | 7.0089 | 7.0208 | 6.7501 | 6.9209 | 2.6636 | 4.0050 | 2.9953 | 4.0439 |
| MKI67_S308 | MKI67 | P46013 | 1329 | TENLTGSKRRPQT | 7.8336 | 7.8852 | 7.2011 | 8.5398 | 7.0425 | 6.8416 | 7.7852 | 6.7255 | 7.0942 | 6.6205 | 6.5302 | 6.3709 | 2.9057 | 4.3412 | 2.7207 | 3.5622 |
| MKI67_S357 | MKI67 | P46013 | 1963 | FKELFQTPGHTEE | 7.5256 | 7.4732 | 7.2357 | 8.1460 | 6.2669 | 6.9376 | 7.5693 | 6.6730 | 6.7855 | 7.2301 | 7.6520 | 6.9431 | 3.1145 | 4.2667 | 2.7426 | 3.4380 |
| MKI67_S374 | MKI67 | P46013 | 2259 | LALRKRTPSVGKA | 8.2009 | 8.8292 | 7.3406 | 8.4899 | 5.8899 | 6.2625 | 6.9811 | 6.3391 | 7.9385 | 8.1687 | 7.7646 | 7.8586 | 1.7640 | 3.2265 | 2.0109 | 2.9349 |
| MKI67_S507 | MKI67 | P46013 | 1376 | KMPCESSPPESAD | 6.8394 | 6.5176 | 6.4730 | 7.3487 | 5.9141 | 6.5514 | 6.5057 | 6.1208 | 6.7237 | 6.5990 | 6.6360 | 6.6658 | 4.8619 | 5.6840 | 5.3040 | 5.2550 |
| MKI67_S507;T525 | MKI67 | P46013 | 1327 | DLTENLTGSKRRP | 8.4730 | 8.3138 | 7.4468 | 8.6200 | 7.1075 | 6.7146 | 7.9226 | 6.6941 | 7.0028 | 6.4153 | 6.7315 | 6.3068 | 2.6438 | 3.9380 | 2.5346 | 3.1347 |
| MKI67_S538 | MKI67 | P46013 | 1719 | FIELFQTPSHTKE | 7.9317 | 7.7975 | 6.4211 | 8.4519 | 6.8859 | 6.6111 | 7.0767 | 6.1944 | 7.4713 | 6.6881 | 6.5874 | 6.4044 | 3.4697 | 4.5066 | 3.5938 | 3.9085 |
| MKI67_S538;T543 | MKI67 | P46013 | 1213 | SKRQLQTPKEKAQ | 6.7968 | 7.3808 | 7.1684 | 7.7727 | 6.3490 | 7.4334 | 7.7672 | 6.6461 | 5.9121 | 6.7882 | 7.1402 | 7.7389 | 3.5160 | 4.6095 | 2.9311 | 4.0497 |
| MKI67_S543 | MKI67 | P46013 | 538 | APTKRKSLVMHTP | 4.8235 | 4.2974 | 4.6849 | 5.6207 | 8.7436 | 4.7215 | 4.6295 | 5.0863 | 8.4150 | 9.9494 | 8.6962 | 10.0296 | 5.4732 | 7.4272 | 3.0265 | 4.3757 |
| MKI67_S579 | MKI67 | P46013 | 2135 | DIVEELSALKQLT | 8.4909 | 8.2159 | 7.3766 | 8.8594 | 7.9218 | 6.6537 | 8.5819 | 6.4775 | 7.0570 | 6.2834 | 6.2795 | 5.6528 | 2.3368 | 4.5145 | 2.2373 | 3.0610 |
| MKI67_S579;S584 | MKI67 | P46013 | 1861 | KKILCKSPQSDPA | 7.1676 | 7.9747 | 7.2724 | 8.0571 | 6.8976 | 6.1827 | 6.7419 | 7.6654 | 7.1072 | 7.3953 | 7.0027 | 7.6779 | 2.7400 | 3.8577 | 3.2906 | 3.8694 |
| MKI67_S584 | MKI67 | P46013 | 1335 | SKRRPQTPKEEAQ | 6.8649 | 7.4351 | 6.5477 | 7.7485 | 6.6824 | 7.2674 | 7.2177 | 6.8241 | 6.6046 | 6.8899 | 7.4407 | 6.3249 | 3.9831 | 4.8800 | 3.5965 | 3.6926 |
| MKI67_S605 | MKI67 | P46013 | 1131 | TKIACKSPPPESV | 6.4824 | 6.7447 | 6.5396 | 7.3108 | 5.9702 | 6.1393 | 6.3288 | 6.4722 | 6.3427 | 6.7620 | 6.5045 | 6.5303 | 4.6577 | 5.8379 | 5.6912 | 5.6857 |
| MKI67_S621 | MKI67 | P46013 | 621 | KRGGRKSGNLPSK | 7.8623 | 8.4944 | 6.7690 | 7.8608 | 5.4514 | 6.8549 | 7.7592 | 6.6036 | 7.1532 | 7.6044 | 7.7924 | 8.2924 | 2.1182 | 3.6788 | 2.1273 | 3.5777 |
| MKI67_S648 | MKI67 | P46013 | 1253 | TKIPCDSPQSDPV | 6.0329 | 6.8287 | 6.0542 | 8.6347 | 5.7513 | 6.5463 | 6.4131 | 6.5279 | 6.5843 | 6.1177 | 6.5075 | 7.1617 | 4.3464 | 5.7192 | 5.1209 | 5.6531 |
| MKI67_S827 | MKI67 | P46013 | 2085 | FKELFQTPDHTEE | 7.1815 | 8.1410 | 6.8398 | 7.8064 | 6.4869 | 6.6768 | 7.3289 | 6.5950 | 7.3224 | 7.0315 | 7.1103 | 7.1611 | 3.2937 | 4.4525 | 2.8578 | 3.7145 |
| MKI67_S859 | MKI67 | P46013 | 648 | ICSKRRSGASEAN | 7.1503 | 7.7832 | 6.7016 | 8.0658 | 5.9536 | 6.0998 | 6.9763 | 6.5372 | 6.7303 | 6.9683 | 7.4350 | 7.7324 | 3.1693 | 4.3694 | 3.7221 | 4.6052 |
| MKI67_T1017 | MKI67 | P46013 | 1747 | QPDLVDTPTSSKP | 7.5784 | 7.7981 | 7.4861 | 8.4147 | 6.0775 | 6.7234 | 7.2105 | 6.9874 | 8.0716 | 6.9900 | 7.2786 | 6.9512 | 2.7563 | 3.8440 | 2.6197 | 3.2125 |
| MKI67_T1167 | MKI67 | P46013 | 2022 | GKLTQTSGKTTQT | 7.2796 | 8.2084 | 6.6324 | 8.8000 | 6.3255 | 7.0622 | 7.8197 | 7.1200 | 7.1261 | 7.3989 | 7.4124 | 7.3222 | 2.0437 | 3.7357 | 2.2026 | 3.5106 |
| MKI67_T1176 | MKI67 | P46013 | 1261 | QSDPVDTPTSTKQ | 7.7659 | 8.3222 | 6.5014 | 7.6971 | 6.1886 | 6.5771 | 6.2930 | 6.5974 | 7.2675 | 6.5776 | 6.3529 | 5.9733 | 4.2554 | 4.9864 | 4.1282 | 4.5157 |
| MKI67_T1193 | MKI67 | P46013 | 328 | DVESVQTPSKAVG | 6.9186 | 7.3807 | 6.7255 | 7.1831 | 5.4378 | 6.8232 | 7.4888 | 6.5153 | 6.7977 | 7.0525 | 7.0501 | 7.0093 | 4.0107 | 4.9209 | 3.8775 | 4.8082 |
| MKI67_T1213 | MKI67 | P46013 | 859 | SLETKTSDTETEP | 6.7969 | 6.5241 | 6.3429 | 7.2056 | 6.0963 | 5.7571 | 6.8547 | 6.4102 | 5.4928 | 5.8087 | 5.5691 | 5.8044 | 6.0518 | 6.3030 | 6.1766 | 6.8058 |
| MKI67_T1233 | MKI67 | P46013 | 2426 | SKRQPQTPKEKAE | 7.2849 | 8.8071 | 6.2642 | 8.8222 | 6.0100 | 7.0508 | 7.5988 | 7.0284 | 7.2359 | 7.5881 | 7.1810 | 7.5449 | 2.0416 | 4.0336 | 2.4249 | 3.0837 |
| MKI67_T1261 | MKI67 | P46013 | 2268 | VGKAMDTPKPAGG | 7.2323 | 7.6988 | 7.0013 | 9.7177 | 5.5341 | 7.0570 | 7.2493 | 6.2082 | 7.0591 | 7.8898 | 7.7425 | 8.5463 | 2.1921 | 3.6159 | 2.0712 | 3.1845 |
| MKI67_T1315 | MKI67 | P46013 | 1176 | AGKAMLTPKPAGG | 7.1084 | 7.1562 | 6.6319 | 7.7623 | 6.5177 | 7.2156 | 6.6066 | 6.2770 | 6.4387 | 6.1432 | 7.1928 | 6.9303 | 4.9301 | 4.8328 | 4.0900 | 4.1664 |
| MKI67_T1327 | MKI67 | P46013 | 235 | NSKKNESPFWKLY | 7.4339 | 8.7158 | 6.7691 | 7.0370 | 5.4935 | 6.9796 | 7.0193 | 6.5813 | 7.4679 | 7.4133 | 7.0108 | 6.7146 | 3.0246 | 5.1832 | 3.0175 | 4.1386 |
| MKI67_T1335 | MKI67 | P46013 | 761 | IAEMFKTPVKEQP | 6.3321 | 7.5111 | 6.4511 | 7.4257 | 7.6484 | 7.4024 | 7.5476 | 7.1273 | 7.0653 | 6.6460 | 7.3666 | 6.3648 | 3.8381 | 4.2955 | 3.3173 | 3.6606 |
| MKI67_T1355 | MKI67 | P46013 | 2203 | LKELFQTPICTDK | 8.8216 | 8.0658 | 6.6821 | 8.7619 | 6.5053 | 6.0721 | 7.0654 | 6.8073 | 7.2058 | 6.5460 | 6.3123 | 6.6512 | 3.2537 | 4.4623 | 2.7403 | 4.0470 |
| MKI67_T1476 | MKI67 | P46013 | 264 | LQYCRKSGLQTDY | 7.6008 | 7.5432 | 6.4592 | 7.5619 | 5.7529 | 7.0130 | 7.7153 | 6.7861 | 6.7165 | 6.7735 | 7.2166 | 7.2520 | 3.5121 | 4.6808 | 2.9670 | 4.4492 |
| MKI67_T1503 | MKI67 | P46013 | 1451 | AASVTGSKRHPKT | 6.9375 | 7.6671 | 6.1948 | 6.8939 | 5.9309 | 5.8608 | 7.2656 | 6.5347 | 7.2656 | 5.7576 | 6.4223 | 6.3939 | 5.1398 | 5.3826 | 5.1411 | 5.2119 |
| MKI67_T1531 | MKI67 | P46013 | 579 | AQSLVISPPAPSP | 6.4468 | 6.6238 | 6.7271 | 7.2946 | 5.7986 | 7.5158 | 6.2993 | 6.2069 | 6.7022 | 7.6828 | 7.1548 | 7.5818 | 3.9609 | 5.2341 | 4.1449 | 4.6255 |
| MKI67_T1557 | MKI67 | P46013 | 2949 | TSAPKQTPDSGKP | 7.1917 | 8.3591 | 6.6697 | 8.2025 | 6.2806 | 6.9117 | 7.5075 | 7.1163 | 6.8579 | 7.4959 | 7.2282 | 7.4889 | 2.4595 | 4.2035 | 2.6439 | 3.3831 |
| MKI67_T1691 | MKI67 | P46013 | 1071 | IRTFKESPKQILD | 6.2894 | 6.6371 | 6.9836 | 7.0956 | 6.9403 | 6.7733 | 7.4882 | 6.3830 | 7.0133 | 6.8529 | 6.3979 | 6.4810 | 4.1267 | 5.2518 | 4.2079 | 5.0781 |
| MKI67_T1719 | MKI67 | P46013 | 2325 | FKELFQTPGTDKP | 7.8813 | 8.4448 | 7.2850 | 8.4350 | 5.7570 | 7.0763 | 7.8104 | 6.4538 | 7.2615 | 7.5104 | 7.3484 | 7.3636 | 2.2574 | 3.7039 | 2.2751 | 3.1361 |
| MKI67_T1747 | MKI67 | P46013 | 588 | APSPRKTPVASDQ | 6.9480 | 7.0899 | 6.7198 | 7.5318 | 6.2363 | 6.7213 | 7.6064 | 6.8056 | 6.4539 | 7.3196 | 7.8559 | 7.2803 | 3.3292 | 4.3664 | 3.3384 | 4.3974 |
| MKI67_T1775 | MKI67 | P46013 | 1233 | FKELFQTPGHTEE | 7.0176 | 7.1619 | 7.5497 | 8.9699 | 6.1811 | 7.0944 | 8.7978 | 6.4815 | 6.5861 | 7.3961 | 7.3445 | 7.4133 | 2.2673 | 3.8847 | 2.5532 | 3.3010 |
| MKI67_T1801 | MKI67 | P46013 | 1447 | KLDPAASVTGSKR | 7.2156 | 7.6572 | 7.6109 | 8.2476 | 6.1279 | 6.4376 | 7.4669 | 6.0565 | 6.3324 | 6.2550 | 6.3733 | 6.3487 | 3.7509 | 5.2153 | 3.4145 | 5.4895 |
| MKI67_T1841 | MKI67 | P46013 | 2261 | LRKRTPSVGKAMD | 4.9486 | 5.4617 | 5.5078 | 5.9441 | 5.6116 | 5.9591 | 6.2836 | 5.2892 | 4.6623 | 4.7400 | 5.7345 | 5.8684 | 7.1306 | 7.5408 | 9.4846 | 9.8331 |
| MKI67_T1923 | MKI67 | P46013 | 1503 | QPDPVDTPTSSKP | 7.8122 | 8.6328 | 7.0342 | 8.0506 | 6.4378 | 7.2552 | 7.4952 | 6.5623 | 7.2426 | 6.6814 | 8.1619 | 7.4469 | 2.8517 | 3.9563 | 2.2721 | 2.1067 |
| MKI67_T1963 | MKI67 | P46013 | 1775 | LAFRKQTPSAGKA | 7.7456 | 8.5441 | 6.9033 | 8.1437 | 6.1280 | 6.9420 | 7.7667 | 7.1243 | 7.0257 | 7.4189 | 7.4145 | 7.5958 | 2.0952 | 3.7930 | 2.2784 | 3.0807 |
| MKI67_T2085 | MKI67 | P46013 | 2828 | KIPCKSSPELEDT | 5.6025 | 6.8422 | 6.7714 | 7.2356 | 6.3460 | 6.3787 | 5.9069 | 5.9693 | 6.5606 | 6.5956 | 7.2120 | 6.8543 | 4.9777 | 6.1306 | 4.8773 | 5.7395 |
| MKI67_T2203 | MKI67 | P46013 | 2689 | TELSETSGHTQES | 6.2643 | 7.1293 | 6.2591 | 6.8561 | 6.3718 | 7.2175 | 6.6114 | 6.3149 | 6.3673 | 5.9956 | 5.5808 | 6.4970 | 5.4231 | 6.3018 | 5.2370 | 5.5728 |
| MKI67_T2259 | MKI67 | P46013 | 1691 | DSAASLTGSKRQL | 7.9645 | 7.7365 | 6.3852 | 7.6198 | 6.7961 | 6.8848 | 8.0723 | 6.7616 | 6.5710 | 6.2038 | 6.7603 | 6.5369 | 3.6766 | 4.6662 | 3.4806 | 3.8838 |
| MKI67_T2268 | MKI67 | P46013 | 1531 | FALRKRTPSAGKA | 8.3007 | 8.8292 | 7.1624 | 8.4075 | 6.0788 | 6.7860 | 7.3050 | 6.7123 | 7.7457 | 7.7763 | 7.7444 | 7.6794 | 1.6549 | 3.1738 | 1.9101 | 2.7334 |
| MKI67_T2325 | MKI67 | P46013 | 2223;2231 | NA | 7.1093 | 7.8408 | 7.0001 | 8.4572 | 6.3277 | 7.1463 | 7.3948 | 6.6344 | 6.8850 | 7.5194 | 7.3577 | 7.3275 | 2.6399 | 4.1634 | 2.7336 | 3.4629 |
| MKI67_T2406 | MKI67 | P46013 | 538;543 | NA | 7.6983 | 7.8251 | 6.7930 | 7.9220 | 6.0171 | 7.2142 | 7.3566 | 6.3379 | 7.2220 | 7.1381 | 7.5753 | 7.5458 | 2.8104 | 4.2564 | 2.6417 | 3.6461 |
| MKI67_T2426 | MKI67 | P46013 | 507;525 | NA | 7.2698 | 7.1611 | 6.8373 | 7.2845 | 5.5100 | 6.7726 | 7.6042 | 6.2572 | 7.1332 | 6.5581 | 7.9687 | 7.7692 | 3.3349 | 4.6288 | 3.3801 | 4.5301 |
| MKI67_T2949 | MKI67 | P46013 | 347;357 | NA | 7.1356 | 7.8506 | 7.1329 | 7.6415 | 5.7245 | 6.9216 | 7.2088 | 6.6187 | 7.4450 | 7.3403 | 7.2552 | 7.6399 | 2.9987 | 4.1334 | 2.5236 | 4.4297 |
| MKI67_T328 | MKI67 | P46013 | 2105;2113 | NA | 7.2419 | 7.3998 | 6.5643 | 7.3483 | 5.9278 | 6.0313 | 7.2106 | 6.7928 | 6.6663 | 5.9665 | 6.9757 | 6.3978 | 4.3444 | 5.5624 | 4.0725 | 5.4976 |
| MKI67_T347;S357 | MKI67 | P46013 | 1861;1864 | NA | 8.1476 | 7.2152 | 8.1651 | 9.8151 | 5.1790 | 5.6422 | 9.6588 | 7.0225 | 6.1181 | 7.5856 | 5.1287 | 8.1318 | 1.5779 | 3.7469 | 3.3207 | 3.5449 |
| MKI67_T588 | MKI67 | P46013 | 579;584 | NA | 7.3731 | 6.9999 | 7.3023 | 8.2721 | 6.4469 | 6.1001 | 7.0304 | 7.4326 | 6.5408 | 6.6550 | 6.8873 | 7.3664 | 3.2971 | 4.7868 | 3.2563 | 4.2530 |
| MKI67_T761 | MKI67 | P46013 | 1740;1747 | NA | 5.8747 | 7.0097 | 6.3894 | 8.7378 | 6.0696 | 6.3941 | 7.5090 | 7.1887 | 7.1157 | 6.8634 | 7.1426 | 7.4450 | 3.3587 | 5.2228 | 3.2818 | 4.3970 |
| MKNK2_S437 | MKNK2 | Q9HBH9 | 437 | SRCLQLSPPSQSK | 7.3156 | 7.1750 | 6.9931 | 5.7678 | 5.5737 | 6.1846 | 5.2913 | 6.0611 | 8.2163 | 9.0279 | 7.4405 | 7.1781 | 3.9123 | 5.5254 | 4.1956 | 4.1417 |
| MKRN2_S139 | MKRN2 | Q9H000 | 139 | CSDPQPSPEMKPH | 6.3553 | 6.5716 | 6.4651 | 6.7952 | 6.7210 | 6.5694 | 6.2055 | 6.7263 | 6.3414 | 6.3430 | 6.6290 | 6.7418 | 5.6194 | 5.4474 | 5.3894 | 5.0791 |
| MLF1_S34 | MLF1 | P58340 | 34 | QMIRSFSEPFGRD | 5.0844 | 4.8444 | 5.2785 | 5.0022 | 6.1196 | 5.6926 | 5.9927 | 6.5025 | 4.8218 | 4.4277 | 4.6359 | 4.7383 | 9.2544 | 9.0570 | 9.2705 | 9.2777 |
| MLF1_S8 | MLF1 | P58340 | 8 | FRMLNSSFEDDPF | 6.3971 | 6.5694 | 6.9682 | 6.5098 | 6.6821 | 6.6535 | 7.1238 | 7.1692 | 6.6487 | 6.8883 | 6.6416 | 6.6139 | 4.6538 | 5.2922 | 4.2148 | 4.9737 |
| MLF2_S238 | MLF2 | Q15773 | 238 | IQGPEDSPSRQSR | 6.6324 | 6.9492 | 6.6799 | 6.4699 | 6.7409 | 6.7568 | 6.5410 | 6.9159 | 6.6352 | 6.7320 | 6.7058 | 6.6981 | 4.8531 | 5.1939 | 4.5740 | 4.9219 |
| MLLT1_S267 | MLLT1 | Q03111 | 267 | TKLESTSPKGGPP | 6.7738 | 6.7772 | 6.4893 | 6.6813 | 6.3266 | 6.0025 | 6.5535 | 5.9637 | 6.7248 | 6.6352 | 6.2780 | 6.3216 | 5.6686 | 5.6346 | 5.5577 | 5.6116 |
| MLLT1_S292 | MLLT1 | Q03111 | 475 | KVSGRRSPESCSK | 6.1905 | 6.6243 | 6.2570 | 6.8705 | 6.1401 | 6.1404 | 6.2820 | 6.1537 | 6.3754 | 6.6428 | 6.7502 | 6.5937 | 5.8360 | 5.9789 | 5.4087 | 5.7558 |
| MLLT1_S315 | MLLT1 | Q03111 | 315 | RSAPGTSPRTSSS | 6.3662 | 6.8468 | 5.8661 | 6.8459 | 6.2046 | 5.9486 | 5.9791 | 6.0524 | 6.3366 | 6.6214 | 6.3289 | 6.3693 | 5.9942 | 6.1742 | 6.1113 | 5.9544 |
| MLLT1_S475 | MLLT1 | Q03111 | 292 | RPATADSPKPSAK | 6.2289 | 6.8710 | 6.4847 | 6.8441 | 6.0829 | 6.2663 | 6.0205 | 5.9313 | 6.3856 | 6.7032 | 6.5692 | 6.4332 | 5.9080 | 6.0458 | 5.6617 | 5.5635 |
| MLLT1_Y158 | MLLT1 | Q03111 | 158 | SRPSPDYPMLPTI | 5.9645 | 5.9494 | 7.1940 | 6.7085 | 5.9064 | 6.8273 | 6.2882 | 6.6118 | 6.5751 | 6.1067 | 5.7686 | 6.4209 | 6.3710 | 6.2869 | 5.2054 | 5.8153 |
| MLLT10_S686;S689 | MLLT10 | P55197 | 689 | GSLSPRSPVSSLQ | 6.2973 | 6.5414 | 5.9343 | 6.5740 | 5.9947 | 5.7666 | 6.3562 | 6.5578 | 6.5131 | 5.9806 | 5.8525 | 5.9132 | 5.9786 | 6.4064 | 6.4106 | 6.9228 |
| MLLT10_S689 | MLLT10 | P55197 | 686;689 | NA | 6.2178 | 6.9091 | 6.0338 | 6.7911 | 6.7704 | 6.5958 | 6.2140 | 5.8885 | 5.9969 | 6.4459 | 6.2603 | 6.6210 | 5.5485 | 6.1153 | 5.8099 | 5.7815 |
| MLLT3_S364 | MLLT3 | P42568 | 364 | IVDPNDSDVEENI | 6.1488 | 6.8959 | 7.0481 | 6.8836 | 5.4982 | 5.4359 | 6.1693 | 6.2125 | 6.1693 | 7.5876 | 6.6320 | 6.9569 | 5.4434 | 5.5832 | 4.9039 | 6.4312 |
| MLLT6_S222 | MLLT6 | P55198 | 423 | GRARAPSPGDYKS | 6.2897 | 6.3536 | 5.6050 | 6.2604 | 6.0302 | 5.7936 | 6.1050 | 5.9785 | 5.9893 | 5.7228 | 5.5362 | 5.7413 | 6.9605 | 7.2185 | 7.3331 | 7.0823 |
| MLLT6_S258 | MLLT6 | P55198 | 258 | HKKRPESPPSILT | 6.5224 | 6.6636 | 5.9767 | 6.4867 | 5.7817 | 5.7972 | 6.0135 | 6.2185 | 5.9564 | 6.1335 | 5.9315 | 6.2713 | 6.2357 | 6.6243 | 6.6143 | 6.7728 |
| MLLT6_S382 | MLLT6 | P55198 | 382 | APSAPPSPSAPEP | 5.5829 | 5.3449 | 6.2880 | 6.6188 | 6.2900 | 6.0533 | 6.1801 | 6.0032 | 6.2219 | 6.4943 | 6.9446 | 6.6863 | 6.2402 | 6.1097 | 6.3691 | 6.5727 |
| MLLT6_S423 | MLLT6 | P55198 | 222 | RRSRSASPSTQQE | 5.8861 | 5.8048 | 6.4305 | 6.5435 | 5.3555 | 5.4575 | 6.0280 | 5.6738 | 6.2279 | 7.2160 | 6.6758 | 6.4966 | 5.4465 | 8.0112 | 6.0887 | 6.6578 |
| MMACHC_S245 | MMACHC | Q9Y4U1 | 245 | PSEKPSSPSPDLP | 6.5595 | 6.1795 | 7.4082 | 6.4868 | 6.6214 | 6.9118 | 6.6059 | 7.1334 | 6.8803 | 6.7338 | 7.3836 | 8.0346 | 4.1184 | 4.6444 | 3.9324 | 4.3659 |
| MMACHC_S275 | MMACHC | Q9Y4U1 | 275 | WLSPRVSPPASPG | 6.5941 | 5.8033 | 6.3779 | 5.9078 | 5.8712 | 6.2795 | 5.6662 | 6.8594 | 6.4285 | 6.1190 | 6.4520 | 6.8973 | 5.8547 | 6.6515 | 5.7417 | 6.4958 |
| MMACHC_S275;S279 | MMACHC | Q9Y4U1 | 275;279 | NA | 5.7018 | 6.2371 | 7.3733 | 6.7595 | 6.4169 | 6.8272 | 6.3325 | 6.6399 | 6.4285 | 6.7731 | 6.2744 | 7.0030 | 5.1558 | 5.7186 | 4.7373 | 5.6211 |
| MME_S6 | MME | P08473 | 6 | xMGKSESQMDITD | 5.4185 | 5.9667 | 5.8387 | 5.8796 | 5.4756 | 6.0057 | 5.8658 | 6.2394 | 5.7078 | 5.7650 | 5.8699 | 6.1474 | 7.0967 | 7.3140 | 7.6922 | 7.7168 |
| MMS19_S1048 | MMS19 | Q96T76-8 | 1048 | EWFLLGSPGSxxx | 6.6526 | 7.9235 | 7.1075 | 6.3910 | 6.0503 | 6.9598 | 6.8374 | 7.1360 | 7.3294 | 7.0466 | 6.9516 | 6.8521 | 3.8622 | 5.0733 | 3.6464 | 4.1801 |
| MMTAG2_S177 | MMTAG2 | Q9BU76 | 220 | ATSSPTSPERPRH | 5.9214 | 5.9043 | 5.6220 | 6.5230 | 5.8993 | 5.6405 | 5.7871 | 5.9545 | 6.1762 | 6.0238 | 5.9381 | 5.6651 | 6.6862 | 7.6636 | 7.2687 | 7.3262 |
| MMTAG2_S217;S220 | MMTAG2 | Q9BU76 | 177 | AEDQTESSCESHR | 6.9308 | 8.4125 | 7.8210 | 9.2724 | 7.1267 | 6.9597 | 7.3478 | 6.1249 | 4.4850 | 5.4062 | 5.5327 | 5.4550 | 4.7157 | 5.0794 | 4.5124 | 4.8177 |
| MMTAG2_S220 | MMTAG2 | Q9BU76 | 231 | RHHHHDSDSNSPC | 6.9745 | 5.1774 | 5.5807 | 6.0060 | 7.3806 | 5.2542 | 5.3201 | 6.1075 | 6.9279 | 4.6835 | 6.9718 | 6.2913 | 6.7386 | 7.4190 | 7.1940 | 5.9731 |
| MMTAG2_S231 | MMTAG2 | Q9BU76 | 217;220 | NA | 5.8847 | 6.0834 | 6.0040 | 5.7367 | 5.3007 | 6.0284 | 6.1291 | 5.9971 | 6.0639 | 6.3453 | 5.5848 | 6.3024 | 6.7472 | 6.8865 | 7.0183 | 7.8876 |
| MNT_S162 | MNT | Q99583 | 162 | TIPPNGSPKPLQP | 6.6247 | 6.2346 | 6.3616 | 6.4255 | 6.5750 | 6.2893 | 6.4250 | 6.1525 | 6.5380 | 6.2528 | 6.4362 | 6.4163 | 6.2891 | 5.7822 | 5.4752 | 5.7220 |
| MOB1B_T40 | MOB1B | Q7L9L4-2 | 40 | LKHAEATLGSGNL | 7.6013 | 7.4055 | 7.1807 | 7.4320 | 7.2442 | 4.5719 | 6.6912 | 4.4687 | 6.1065 | 4.5269 | 5.6038 | 5.2836 | 5.0773 | 7.4796 | 6.6250 | 6.7018 |
| MOCS2_S20 | MOCS2 | O96007 | 20 | ETKLPLSPPLVED | 6.0183 | 6.4096 | 5.7191 | 7.0405 | 6.2115 | 5.4412 | 6.3918 | 6.6509 | 6.4760 | 6.4199 | 6.8878 | 7.1322 | 5.6269 | 6.1906 | 5.7983 | 5.5855 |
| MON1A_S153 | MON1A | Q86VX9 | 153 | MFVHARSYEDLTE | 5.6809 | 6.7778 | 6.1443 | 5.9816 | 6.1680 | 6.5388 | 5.9828 | 6.3027 | 6.4065 | 6.8175 | 6.6723 | 6.8391 | 6.1087 | 6.2650 | 5.4656 | 5.8485 |
| MON1B_S59 | MON1B | Q7L1V2 | 59 | DKDQPPSPSPPPQ | 5.4624 | 6.1929 | 5.9628 | 6.1431 | 5.8513 | 5.8798 | 5.7832 | 6.8421 | 7.3241 | 5.9409 | 6.2717 | 6.2314 | 6.0737 | 6.2670 | 6.9032 | 6.8705 |
| MON2_S1214 | MON2 | Q7Z3U7 | 1214 | PFVRTDSIGEKLG | 6.0632 | 5.7323 | 7.1856 | 6.0286 | 5.7677 | 5.7942 | 5.6281 | 5.6920 | 6.4613 | 6.4877 | 6.6815 | 6.7984 | 6.4714 | 6.8960 | 6.0057 | 6.3062 |
| MON2_S205 | MON2 | Q7Z3U7 | 205 | GNSNRRSVSTLKP | 5.8818 | 6.8602 | 6.7689 | 7.8533 | 7.5182 | 8.4725 | 7.8242 | 6.8412 | 6.6015 | 7.4721 | 8.5642 | 7.2374 | 3.0273 | 3.9395 | 2.4567 | 2.6810 |
| MORC2_S615 | MORC2 | Q9Y6X9 | 615 | RPQRPRSPPLPAV | 5.9174 | 6.1270 | 5.8754 | 6.3550 | 6.2762 | 5.9749 | 6.0274 | 5.8860 | 6.1405 | 6.1207 | 5.9332 | 6.1455 | 6.6450 | 6.8421 | 6.9395 | 6.7942 |
| MORC2_S696 | MORC2 | Q9Y6X9 | 856 | RLMKPPSPEHQSL | 6.2340 | 6.7965 | 5.5417 | 6.7810 | 6.4040 | 6.7950 | 5.1279 | 6.7728 | 5.8489 | 5.8049 | 6.2648 | 5.6686 | 5.7915 | 6.5133 | 6.8916 | 5.8636 |
| MORC2_S725 | MORC2 | Q9Y6X9 | 733 | IKLSPATPSRKRS | 6.7658 | 8.1096 | 6.6779 | 6.3622 | 5.9840 | 6.7264 | 7.0431 | 6.9892 | 6.5009 | 6.2700 | 6.5237 | 6.4039 | 4.3511 | 5.2436 | 4.8171 | 5.2314 |
| MORC2_S725;S730 | MORC2 | Q9Y6X9 | 696 | PLVQQLSPSLLPN | 6.3702 | 6.4136 | 6.7979 | 7.1179 | 6.5069 | 7.0405 | 6.7060 | 6.3665 | 7.5347 | 7.8027 | 7.2979 | 7.5439 | 3.8502 | 4.4101 | 3.7773 | 4.4638 |
| MORC2_S730 | MORC2 | Q9Y6X9 | 743 | KRSVAVSDEEEVE | 6.7236 | 6.1327 | 6.6523 | 6.9918 | 6.1807 | 5.5162 | 6.3353 | 5.7636 | 5.6403 | 6.9429 | 5.7500 | 6.6363 | 6.4441 | 6.3064 | 5.9109 | 6.0727 |
| MORC2_S743 | MORC2 | Q9Y6X9 | 730 | ESPIKLSPATPSR | 6.1208 | 7.0065 | 6.5425 | 7.1948 | 6.1509 | 5.9027 | 6.2533 | 6.5162 | 6.4576 | 6.5807 | 6.6127 | 6.5033 | 4.6247 | 5.7830 | 5.5233 | 6.2270 |
| MORC2_S777;S779 | MORC2 | Q9Y6X9 | 717 | SPKVIKTPVVKKT | 6.4672 | 7.0903 | 7.3547 | 7.3716 | 5.7322 | 8.0565 | 7.9608 | 5.9929 | 7.2132 | 7.3186 | 8.2340 | 7.8790 | 2.5129 | 4.3578 | 2.8790 | 3.5794 |
| MORC2_S856 | MORC2 | Q9Y6X9 | 725 | VVKKTESPIKLSP | 5.1959 | 5.7702 | 5.6931 | 6.8129 | 5.5900 | 5.9857 | 5.9083 | 5.8433 | 5.6227 | 6.0126 | 6.0587 | 6.6845 | 7.0041 | 7.2490 | 7.1518 | 7.4173 |
| MORC2_T650 | MORC2 | Q9Y6X9 | 650 | APVISSTPKLPAL | 7.4474 | 7.6943 | 6.1352 | 6.7018 | 5.1600 | 6.2883 | 8.2918 | 7.1265 | 7.0857 | 7.0475 | 6.9833 | 6.4031 | 3.7502 | 4.6539 | 4.2514 | 4.9798 |
| MORC2_T717 | MORC2 | Q9Y6X9 | 725;730 | NA | 5.7206 | 6.2111 | 6.1135 | 6.6100 | 7.0802 | 6.7084 | 6.9582 | 6.6592 | 6.7814 | 6.8643 | 7.2700 | 6.9841 | 4.9326 | 5.3957 | 4.7050 | 5.0057 |
| MORC2_T733 | MORC2 | Q9Y6X9 | 777;779 | NA | 5.5721 | 5.8385 | 6.6872 | 6.2497 | 5.6690 | 6.0140 | 6.2175 | 5.4273 | 5.6703 | 6.3767 | 6.6007 | 6.2614 | 7.2597 | 7.4262 | 6.5675 | 6.1621 |
| MORC3_S560 | MORC3 | Q14149 | 560 | AKNRRLSSQFENS | 4.4112 | 4.4214 | 4.8973 | 5.0166 | 6.6871 | 5.9611 | 5.3172 | 5.8623 | 6.5132 | 6.7267 | 6.5887 | 6.5698 | 8.2883 | 8.5132 | 7.1314 | 7.0947 |
| MORC3_S765 | MORC3 | Q14149 | 765 | INGKSESPDHMVS | 6.6622 | 6.4283 | 6.9582 | 5.1729 | 5.4196 | 6.4867 | 5.5238 | 4.8038 | 7.2907 | 6.6950 | 6.0117 | 5.2533 | 5.9276 | 6.7426 | 7.2980 | 7.3255 |
| MORF4L2_S71 | MORF4L2 | Q15014 | 71 | ENPPSGSVRKTRK | 6.3405 | 6.6036 | 7.1264 | 6.1616 | 4.1820 | 6.1448 | 6.5628 | 6.0153 | 8.0350 | 8.1219 | 9.2959 | 9.9130 | 2.8344 | 5.5356 | 2.9278 | 4.1995 |
| MOSPD2_S261 | MOSPD2 | Q8NHP6 | 261 | ENGPITSEDETSS | 5.8868 | 7.3238 | 7.5342 | 6.6674 | 6.6379 | 4.6097 | 5.5165 | 6.6190 | 6.6716 | 6.6926 | 7.2691 | 6.1877 | 5.8321 | 6.1140 | 5.2241 | 5.2135 |
| MOV10_S815 | MOV10 | Q9HCE1 | 815 | LLLAPSSKKGKAR | 5.6314 | 5.7664 | 6.2834 | 5.2988 | 7.0008 | 6.1629 | 7.0196 | 5.9581 | 6.1513 | 6.1237 | 4.7717 | 4.8719 | 6.0496 | 7.0327 | 8.2220 | 7.6557 |
| MOV10_S969 | MOV10 | Q9HCE1 | 254 | AQLKPMTPFKRTR | 6.1472 | 6.7820 | 6.5711 | 7.1398 | 6.1543 | 6.4378 | 6.4632 | 6.1468 | 6.4802 | 6.5582 | 6.8447 | 6.7617 | 4.9898 | 5.7213 | 5.2149 | 5.5870 |
| MOV10_T254 | MOV10 | Q9HCE1 | 969 | QGLSKLSPSTSGP | 6.8597 | 6.7324 | 5.9858 | 7.8436 | 6.7549 | 6.4844 | 5.5283 | 7.9789 | 6.3458 | 6.6031 | 6.4486 | 5.4753 | 4.7605 | 5.2824 | 5.0622 | 5.8539 |
| MPDZ_S1078 | MPDZ | O75970 | 483 | TKDADLSPVNASI | 6.1792 | 6.5321 | 6.6193 | 6.8259 | 6.2611 | 6.4465 | 6.5348 | 6.2544 | 6.4107 | 6.6835 | 7.2073 | 7.1289 | 5.2077 | 5.2993 | 4.9469 | 5.4623 |
| MPDZ_S1194 | MPDZ | O75970 | 230 | SLPQLVSPIVSRS | 6.1726 | 6.2930 | 6.4269 | 6.1575 | 6.2793 | 6.4013 | 6.4108 | 6.5907 | 6.3657 | 6.6415 | 6.8888 | 7.1583 | 5.5664 | 5.5239 | 5.7691 | 5.3542 |
| MPDZ_S1283 | MPDZ | O75970 | 790 | AKPLPLSPEEGYV | 5.7311 | 5.2044 | 6.4341 | 6.6138 | 6.2885 | 6.3286 | 6.1826 | 5.7414 | 6.2052 | 6.2581 | 6.9830 | 6.9552 | 6.4429 | 6.1698 | 5.8520 | 6.6094 |
| MPDZ_S1595 | MPDZ | O75970 | 802 | VSAKEDSFLYPPH | 6.7184 | 6.9112 | 7.2926 | 7.4728 | 7.0319 | 6.9989 | 6.9782 | 5.9870 | 7.5116 | 6.0966 | 6.9998 | 6.6993 | 3.8933 | 4.7997 | 3.7290 | 4.8796 |
| MPDZ_S1818 | MPDZ | O75970 | 1818 | HSERRPSQSSQVS | 5.7179 | 5.8276 | 5.8173 | 6.2048 | 6.6743 | 6.1525 | 5.5176 | 6.5859 | 6.9296 | 7.2911 | 7.6896 | 7.0107 | 5.7531 | 6.4752 | 5.5146 | 4.8381 |
| MPDZ_S1818;S1821 | MPDZ | O75970 | 354 | GITLSSSPTSTPE | 6.6545 | 5.2184 | 6.5568 | 6.1563 | 6.0781 | 6.0026 | 6.8608 | 6.6343 | 6.7887 | 7.7623 | 7.0496 | 7.5216 | 5.3074 | 5.3405 | 5.4268 | 4.6413 |
| MPDZ_S230 | MPDZ | O75970 | 1595 | MVPQSGSPEPESI | 5.8772 | 5.9278 | 7.0681 | 6.1893 | 5.7433 | 6.1571 | 6.6404 | 6.0181 | 6.4894 | 6.5854 | 7.0460 | 6.4060 | 5.5986 | 6.0547 | 5.7566 | 6.4420 |
| MPDZ_S230;S234 | MPDZ | O75970 | 1078 | AMLRRHSLIGPDI | 6.2372 | 5.5602 | 5.8927 | 6.3152 | 6.9517 | 5.9709 | 5.3317 | 6.0987 | 7.2832 | 7.8207 | 7.2799 | 7.1059 | 6.1449 | 6.1810 | 5.1214 | 4.7045 |
| MPDZ_S354 | MPDZ | O75970 | 1194 | KHVLEDSPAGKNG | 6.1118 | 6.4800 | 6.9390 | 6.5965 | 6.0693 | 7.5145 | 6.5809 | 6.8474 | 7.3189 | 6.6370 | 7.2748 | 7.1163 | 3.9972 | 5.2343 | 4.1640 | 5.1182 |
| MPDZ_S483 | MPDZ | O75970 | 797 | PEEGYVSAKEDSF | 6.0436 | 5.6752 | 6.1674 | 6.6061 | 6.3537 | 7.4168 | 7.1123 | 6.7903 | 5.4369 | 6.0521 | 6.2089 | 5.7111 | 5.6070 | 6.2047 | 6.4310 | 6.1829 |
| MPDZ_S790 | MPDZ | O75970 | 1283 | NADKAPSQSESEP | 6.2637 | 5.4709 | 6.1059 | 7.3085 | 5.5135 | 6.3072 | 6.1803 | 6.0883 | 6.1377 | 5.7039 | 6.5991 | 6.2478 | 6.0379 | 7.1812 | 6.4795 | 6.3747 |
| MPDZ_S797 | MPDZ | O75970 | 1609 | NTSRSSTPAIFAS | 5.1996 | 5.3252 | 4.9996 | 5.7894 | 5.7189 | 6.2716 | 6.4331 | 5.9529 | 5.0072 | 4.7790 | 5.5285 | 4.7386 | 8.6848 | 9.0861 | 8.5354 | 7.9500 |
| MPDZ_S802 | MPDZ | O75970 | 1818;1821 | NA | 5.7186 | 6.3186 | 6.0830 | 5.6643 | 6.0191 | 6.2409 | 6.0060 | 6.6259 | 6.5365 | 6.8586 | 6.1976 | 7.3186 | 6.0559 | 5.7383 | 6.3519 | 6.2664 |
| MPDZ_T1609 | MPDZ | O75970 | 230;234 | NA | 6.0676 | 7.1660 | 7.3780 | 6.2761 | 6.4328 | 6.0446 | 6.0413 | 6.5746 | 6.5745 | 6.6229 | 6.1072 | 6.0691 | 6.0758 | 5.3400 | 5.9433 | 5.2863 |
| MPG_S78 | MPG | P29372 | 78 | RSIYFSSPKGHLT | 7.3602 | 7.8784 | 7.0941 | 7.7261 | 5.9223 | 6.9707 | 8.0366 | 6.7850 | 7.5559 | 8.0316 | 8.2075 | 7.3875 | 2.1961 | 3.5376 | 2.1038 | 3.2067 |
| MPG_T67 | MPG | P29372 | 67 | CLGPPTTPGPYRS | 8.5790 | 7.9399 | 6.6144 | 6.3021 | 5.2109 | 6.8242 | 8.8002 | 7.1527 | 8.5091 | 6.0387 | 7.2519 | 6.2631 | 3.1649 | 4.0334 | 3.0267 | 4.2887 |
| MPHOSPH10_S163;S167 | MPHOSPH10 | O00566 | 16 | TLERCLTEVGKAT | 8.2007 | 8.5533 | 7.3186 | 8.1020 | 6.1304 | 6.7616 | 7.7928 | 7.0330 | 7.3446 | 6.6203 | 7.7659 | 6.9706 | 2.6013 | 3.2467 | 2.3385 | 3.2198 |
| MPHOSPH10_T16 | MPHOSPH10 | O00566 | 163;167;171 | NA | 5.5172 | 5.9084 | 5.6818 | 6.5670 | 6.4890 | 5.0584 | 5.3206 | 7.0921 | 7.2357 | 6.2744 | 6.2135 | 6.6283 | 6.2923 | 7.3357 | 7.9254 | 4.4600 |
| MPHOSPH6_T147 | MPHOSPH6 | Q99547 | 147 | DENGDITPIKAKK | 6.6054 | 7.0252 | 6.5532 | 6.8419 | 6.4254 | 6.2432 | 6.5426 | 6.5793 | 6.8043 | 6.4603 | 6.3323 | 6.7526 | 5.1733 | 5.2609 | 4.8440 | 5.5561 |
| MPHOSPH8_S136;S138 | MPHOSPH8 | Q99549-2 | 85 | RWKGYTSDDDTWE | 5.9447 | 6.3287 | 6.2154 | 6.8348 | 6.2346 | 5.9503 | 6.0658 | 5.9013 | 5.8479 | 6.0988 | 5.9097 | 5.7704 | 6.0903 | 6.8645 | 7.1423 | 6.8004 |

| | | | | | | | | | | | | | | | | | | | | | |
|---|---|---|---|---|---|---|---|---|---|---|---|---|---|---|---|---|---|---|---|---|---|
| MPHOSPH8_S149 | MPHOSPH8 | Q99549-2 | | 496 | LKERRNTRDETDT | 5.6000 | 5.6297 | 5.9296 | 5.8117 | 5.9510 | 5.9495 | 5.9620 | 6.0582 | 6.5557 | 6.0448 | 5.8983 | 6.0742 | 7.0147 | 7.3489 | 7.1287 | 7.0429 |
| MPHOSPH8_S188;S189 | MPHOSPH8 | Q99549-2 | | 189 | KPDLESSLESLVF | 6.2579 | 5.9138 | 6.1048 | 6.3283 | 6.4230 | 6.3929 | 6.4104 | 6.9700 | 6.7394 | 7.0860 | 6.4865 | 6.7090 | 4.8209 | 6.2936 | 5.3797 | 5.6837 |
| MPHOSPH8_S189 | MPHOSPH8 | Q99549-2 | | 403 | GLWSTDSAEEDKE | 6.3566 | 6.5683 | 6.1075 | 6.6337 | 6.2814 | 6.0017 | 6.5451 | 6.0177 | 5.9868 | 6.0070 | 5.8638 | 6.1817 | 6.0172 | 6.7617 | 6.0705 | 6.5991 |
| MPHOSPH8_S319 | MPHOSPH8 | Q99549-2 | | 392 | PKGRRLSGEERGL | 5.7159 | 5.6967 | 5.2309 | 6.1755 | 5.6614 | 5.5523 | 5.7721 | 5.9386 | 5.7935 | 5.4722 | 5.5567 | 5.5960 | 7.1379 | 8.2250 | 8.2485 | 8.2267 |
| MPHOSPH8_S392 | MPHOSPH8 | Q99549-2 | | 51 | AEAFGDSEEDGED | 6.5543 | 6.2791 | 5.7364 | 6.3127 | 5.9372 | 5.7384 | 6.3598 | 6.0159 | 6.0198 | 6.3310 | 5.7340 | 6.2172 | 6.2220 | 7.3669 | 6.6455 | 6.5298 |
| MPHOSPH8_S403 | MPHOSPH8 | Q99549-2 | | 385 | MPVSAQTPKGRRL | 6.9852 | 7.0786 | 6.9913 | 7.3864 | 6.4228 | 6.8438 | 7.6217 | 6.5157 | 6.6371 | 5.9011 | 6.5764 | 6.6264 | 3.7136 | 5.4102 | 4.2383 | 5.0513 |
| MPHOSPH8_S51 | MPHOSPH8 | Q99549-2 | | 319 | PLDSAMSAEEDTD | 6.3912 | 5.8983 | 6.6869 | 7.2646 | 6.6074 | 5.7985 | 6.0232 | 6.3147 | 6.4814 | 6.0734 | 5.6731 | 6.3625 | 6.4336 | 6.0686 | 5.9688 | 5.9539 |
| MPHOSPH8_S85 | MPHOSPH8 | Q99549-2 | | 149 | ETKEDTSPKKKKK | 6.0728 | 6.3405 | 6.4879 | 7.4293 | 4.9935 | 5.2507 | 5.6181 | 5.0058 | 5.5865 | 6.3518 | 6.6139 | 7.7367 | 5.8721 | 6.8953 | 6.7388 | 7.0065 |
| MPHOSPH8_T385 | MPHOSPH8 | Q99549-2 | 136;138 | | NA | 6.4652 | 5.2259 | 6.9205 | 7.1433 | 5.1049 | 4.9721 | 5.6424 | 6.8562 | 6.7998 | 6.9555 | 5.4696 | 6.5910 | 6.1312 | 7.4403 | 6.5508 | 6.6315 |
| MPHOSPH8_T496 | MPHOSPH8 | Q99549-2 | 188;189 | | NA | 4.7018 | 6.0817 | 5.2040 | 5.9178 | 6.1228 | 7.5726 | 7.1521 | 7.9941 | 4.9020 | 6.7056 | 6.0507 | 5.8676 | 5.6048 | 6.7595 | 6.9166 | 6.4461 |
| MPHOSPH9_S1007 | MPHOSPH9 | Q99550 | | 994 | SPKENLSPGFSHL | 6.7919 | 7.0232 | 6.3649 | 6.3752 | 6.7724 | 7.0536 | 6.7569 | 6.9429 | 7.0319 | 6.4375 | 6.9311 | 6.8555 | 4.7687 | 5.1039 | 4.0016 | 4.7889 |
| MPHOSPH9_S267 | MPHOSPH9 | Q99550 | | 267 | KEDVSISPGEFEH | 6.1502 | 5.9120 | 7.0200 | 6.8310 | 6.6111 | 6.6968 | 5.8853 | 6.4651 | 6.4079 | 6.9717 | 6.4195 | 6.6766 | 5.5334 | 5.2908 | 5.4060 | 5.7227 |
| MPHOSPH9_S516 | MPHOSPH9 | Q99550 | | 516 | PSHTKASPVDSWK | 6.2678 | 6.8079 | 6.5911 | 6.7300 | 6.0689 | 7.1056 | 6.7246 | 6.5775 | 7.0118 | 6.7745 | 6.9879 | 7.3149 | 4.3907 | 5.1269 | 4.5264 | 4.9936 |
| MPHOSPH9_S994 | MPHOSPH9 | Q99550 | | 1096 | CKPVSVTPQGNDF | 5.3963 | 6.6290 | 7.6635 | 8.5729 | 5.4136 | 7.4350 | 8.5169 | 7.9048 | 7.5083 | 8.9220 | 7.1247 | 8.1290 | 2.7197 | 4.7067 | 0.0000 | 3.3576 |
| MPHOSPH9_T1096 | MPHOSPH9 | Q99550 | | 1007 | LSKNESSPIRFDI | 5.8670 | 6.0299 | 6.5689 | 6.3841 | 6.4290 | 7.0959 | 7.3769 | 6.4782 | 6.3896 | 6.9560 | 7.6054 | 7.2457 | 4.8753 | 6.0190 | 4.3264 | 4.3527 |
| MPLKIP_S115 | MPLKIP | Q8TAP9 | | 133 | VREKRMSNELENY | 5.5965 | 4.9959 | 6.1766 | 5.0800 | 5.9112 | 5.5144 | 5.4625 | 6.3247 | 5.6202 | 6.0449 | 6.2358 | 6.4933 | 7.4143 | 7.6210 | 7.9036 | 7.6051 |
| MPLKIP_S133 | MPLKIP | Q8TAP9 | | 115 | QTHPQGSPRTSTP | 5.9254 | 5.2714 | 6.7145 | 4.6439 | 6.8618 | 7.9360 | 5.5385 | 6.6840 | 7.4902 | 6.4930 | 6.7781 | 7.1717 | 5.8636 | 6.0290 | 5.6858 | 4.9130 |
| MPLKIP_S40 | MPLKIP | Q8TAP9 | | 40 | GGPRPPSPRDGYG | 5.6228 | 5.5470 | 5.9901 | 5.6613 | 6.5423 | 6.5308 | 5.7536 | 7.3451 | 6.3552 | 5.9560 | 6.2399 | 6.4501 | 6.2258 | 6.7668 | 6.5395 | 6.4735 |
| MPLKIP_T120 | MPLKIP | Q8TAP9 | | 120 | GSPRTSTPFGSGR | 8.3120 | 6.8577 | 7.6196 | 6.3983 | 5.7932 | 6.0358 | 6.5443 | 7.7444 | 7.1061 | 8.5696 | 7.8838 | 9.7048 | 2.0727 | 3.4188 | 2.1763 | 3.7626 |
| MPND_S214 | MPND | Q8N594 | | 214 | RRPLGKSPSEPAH | 6.2838 | 6.1154 | 6.2910 | 6.3465 | 5.1675 | 6.9987 | 6.6888 | 6.1172 | 6.5132 | 7.2281 | 5.9918 | 6.3985 | 4.5099 | 7.0830 | 6.2247 | 6.0419 |
| MPP1_S110 | MPP1 | Q00013 | | 110 | MIHRQGSLHVGDE | 6.0073 | 6.7674 | 8.5158 | 7.1552 | 6.7320 | 6.2031 | 5.7844 | 6.9104 | 5.8681 | 8.1596 | 7.8819 | 6.6734 | 4.1957 | 5.1021 | 4.0875 | 3.9561 |
| MPP2_T162 | MPP2 | Q14168-4 | | 162 | ASKTYETPPPSPG | 6.1033 | 6.9036 | 6.3599 | 5.8843 | 6.1137 | 6.7145 | 6.1449 | 6.6021 | 6.3870 | 6.2331 | 6.2665 | 6.0670 | 6.1812 | 6.1754 | 5.9500 | 5.9137 |
| MPP2_T162;S166 | MPP2 | Q14168-4 | 162;166 | | NA | 6.3834 | 5.9938 | 5.5181 | 6.9105 | 6.4820 | 4.9024 | 5.2934 | 6.6997 | 4.2935 | 6.1647 | 6.6759 | 6.3086 | 7.5969 | 6.5318 | 7.6173 | 6.6281 |
| MPP5_S84 | MPP5 | Q8N3R9 | | 84 | ELDLNSSMRLKKL | 7.7460 | 6.9015 | 7.5292 | 7.2808 | 6.7424 | 5.8800 | 6.1315 | 6.2081 | 6.6807 | 5.7479 | 5.9168 | 5.9201 | 5.6660 | 6.0152 | 4.8282 | 4.8057 |
| MPP6_S106 | MPP6 | Q9NZW5 | | 106 | ASKCYDSPPSSPE | 5.9031 | 6.0117 | 6.1191 | 6.1631 | 6.2822 | 6.5813 | 6.3425 | 6.5422 | 6.5471 | 5.9400 | 6.7519 | 6.2212 | 5.7211 | 6.3849 | 5.9775 | 6.5112 |
| MPRIP_S217;S224 | MPRIP | Q6WCQ1-2 | | 993 | DIMKSKSNPDFLK | 5.8699 | 6.0720 | 5.8870 | 5.4989 | 5.9316 | 5.5080 | 5.7006 | 5.9431 | 6.2268 | 6.2140 | 5.7591 | 6.2311 | 7.3246 | 7.2485 | 7.2880 | 7.2966 |
| MPRIP_S269 | MPRIP | Q6WCQ1-2 | | 542 | REGRSKTFDWAEF | 8.1210 | 7.2286 | 7.8366 | 6.3712 | 6.9889 | 7.6275 | 6.6561 | 6.5382 | 6.3648 | 6.0500 | 6.1496 | 5.9415 | 4.9456 | 5.2172 | 3.8925 | 4.0707 |
| MPRIP_S365 | MPRIP | Q6WCQ1-2 | | 619 | RMEVDRSPGLPMS | 5.8485 | 6.3497 | 5.9774 | 6.3341 | 6.1497 | 5.9955 | 6.1899 | 6.6453 | 6.0596 | 6.1060 | 6.0432 | 6.0707 | 6.6398 | 6.2893 | 6.8930 | 6.4082 |
| MPRIP_S377 | MPRIP | Q6WCQ1-2 | | 377 | KSLDRRSTEPSVT | 6.4380 | 6.7170 | 6.9811 | 7.0031 | 6.4306 | 6.3377 | 6.7464 | 5.9817 | 7.5115 | 7.1956 | 7.0108 | 7.3264 | 4.7338 | 5.2535 | 4.3312 | 4.0017 |
| MPRIP_S619 | MPRIP | Q6WCQ1-2 | | 365 | ASASPLSPHRRAK | 5.5211 | 5.8413 | 6.1031 | 6.4493 | 6.3655 | 6.1083 | 6.0142 | 6.7061 | 7.0283 | 6.0245 | 6.3476 | 6.5258 | 6.1479 | 6.3282 | 6.0257 | 6.4631 |
| MPRIP_S891 | MPRIP | Q6WCQ1-2 | | 977 | EALGEKSPDSATV | 5.4765 | 6.4883 | 5.9037 | 5.9241 | 5.8187 | 6.1179 | 5.7653 | 5.6430 | 6.1578 | 6.3369 | 6.2012 | 5.8078 | 7.1132 | 7.2247 | 6.9207 | 7.1002 |
| MPRIP_S977 | MPRIP | Q6WCQ1-2 | | 269 | VESGYFSLEKTKQ | 6.4756 | 7.4525 | 6.1998 | 5.6520 | 6.3318 | 6.2477 | 6.5226 | 6.1939 | 6.2972 | 7.1954 | 6.6551 | 6.1202 | 5.3233 | 6.2977 | 5.6446 | 5.3905 |
| MPRIP_S993 | MPRIP | Q6WCQ1-2 | | 891 | GGEATGSPLAQGK | 6.1301 | 6.3233 | 6.9244 | 6.8491 | 6.5805 | 5.9748 | 6.4608 | 6.1272 | 6.4660 | 6.0018 | 7.1355 | 6.3348 | 5.5014 | 6.1266 | 5.4818 | 5.5819 |
| MPRIP_T542 | MPRIP | Q6WCQ1-2 | 217;224 | | NA | 5.4272 | 6.5505 | 7.0004 | 6.2638 | 5.1548 | 5.6148 | 5.9031 | 6.3095 | 5.4060 | 6.7246 | 5.9086 | 5.7244 | 7.7054 | 6.6440 | 6.5216 | 7.1412 |
| MPST_S15 | MPST | P25325-2 | | 15 | SETRARSPSVAAM | 6.0300 | 5.9740 | 6.8143 | 5.5464 | 6.2595 | 6.2079 | 5.9408 | 6.5665 | 6.4806 | 6.8473 | 6.9363 | 6.9075 | 5.9614 | 5.8488 | 7.5513 | 5.9276 |
| MPV17L2_S14 | MPV17L2 | Q567V2 | | 14 | RLRRLLSAGQLLF | 5.3526 | 5.9358 | 6.5506 | 4.5163 | 5.0136 | 5.2034 | 6.5642 | 8.3816 | 4.9955 | 6.4467 | 8.0967 | 5.3526 | 6.2523 | 6.3201 | 8.1691 | 6.8490 |
| MPZL1_S210 | MPZL1 | O95297 | | 260 | KINKSESVVYADI | 6.8208 | 6.4535 | 7.0434 | 5.9168 | 5.6885 | 5.5365 | 5.3770 | 5.3349 | 5.8445 | 6.0823 | 6.0212 | 5.8592 | 7.2576 | 7.3355 | 6.7531 | 6.6753 |
| MPZL1_S221 | MPZL1 | O95297 | | 263 | KSESVVYADIRKN | 6.9366 | 7.2045 | 7.4098 | 7.1516 | 6.3400 | 6.5684 | 6.0811 | 6.0529 | 6.0825 | 6.6391 | 7.1796 | 6.8985 | 4.9742 | 5.4298 | 4.3854 | 4.6660 |
| MPZL1_S260 | MPZL1 | O95297 | | 210 | STSESLSPVKQAP | 6.2464 | 6.3010 | 6.7464 | 6.1276 | 5.8534 | 6.1452 | 6.1887 | 5.9720 | 6.2694 | 6.5026 | 6.5616 | 6.8589 | 5.7560 | 6.1709 | 5.8752 | 6.4248 |
| MPZL1_Y263 | MPZL1 | O95297 | | 221 | APRKSPSDTEGLV | 2.9487 | 3.4834 | 3.5417 | 3.7656 | 3.6760 | 3.4548 | 3.8197 | 3.7545 | 3.5151 | 3.8500 | 3.3472 | 3.7989 | 13.5216 | 13.3710 | 16.4210 | 13.7308 |
| MRE11_S168 | MRE11 | P49959-3 | | 652 | VIEVDESDVEEDI | 6.1896 | 6.3889 | 6.3331 | 6.7829 | 6.3888 | 6.1677 | 6.3955 | 6.1131 | 5.9314 | 6.1879 | 6.1799 | 6.0476 | 6.1972 | 6.3520 | 6.0369 | 6.3075 |
| MRE11_S278 | MRE11 | P49959-3 | | 622 | SASRNMSIIDAFK | 6.8336 | 3.9531 | 5.7407 | 5.6222 | 5.9270 | 4.9067 | 6.0095 | 4.6118 | 10.4114 | 7.7329 | 7.7642 | 6.5064 | 8.4020 | 7.7381 | 3.2772 | 4.5632 |
| MRE11_S539 | MRE11 | P49959-3 | | 684 | MSQSQVSKGVDFE | 5.6194 | 6.5569 | 7.0422 | 6.0344 | 4.8928 | 6.4800 | 6.1906 | 6.2260 | 6.5172 | 7.3363 | 7.0888 | 6.5698 | 5.7860 | 6.5004 | 5.5849 | 5.5745 |
| MRE11_S622 | MRE11 | P49959-3 | | 674 | RWSSTSSSKIMSQ | 7.1771 | 7.7253 | 7.1365 | 7.2331 | 5.4528 | 6.9181 | 7.3683 | 6.2302 | 7.1970 | 6.7222 | 7.1445 | 7.4060 | 3.9070 | 4.5785 | 3.6912 | 4.1122 |
| MRE11_S652 | MRE11 | P49959-3 | | 168 | VEKIDISPVLLQK | 8.6945 | 5.9287 | 5.1796 | 5.2948 | 6.6329 | 8.1311 | 6.5753 | 7.2284 | 5.0067 | 5.2756 | 5.9735 | 7.3980 | 5.0868 | 6.1208 | 5.7750 | 5.6982 |
| MRE11_S674 | MRE11 | P49959-3 | | 278 | SVVTSLSPGEAVK | 6.2558 | 6.2999 | 6.8642 | 6.8191 | 6.5264 | 5.8700 | 5.5216 | 6.1176 | 5.9813 | 5.9842 | 6.3959 | 6.5149 | 6.7356 | 6.2558 | 6.2721 | 5.5859 |
| MRE11_S679 | MRE11 | P49959-3 | | 681 | SKIMSQSQVSKGV | 7.8081 | 7.6655 | 6.5299 | 7.9617 | 6.2579 | 5.7597 | 6.0223 | 5.2206 | 6.4647 | 5.1804 | 5.3168 | 5.3254 | 6.4402 | 6.4672 | 5.2976 | 6.2820 |
| MRE11_S681 | MRE11 | P49959-3 | | 679 | SSSKIMSQSQVSK | 7.0373 | 6.4412 | 7.0459 | 7.9307 | 7.2266 | 5.6819 | 6.4412 | 5.3828 | 6.3106 | 5.7524 | 5.2075 | 5.2933 | 6.5700 | 6.3782 | 5.0658 | 6.2347 |
| MRE11_S684 | MRE11 | P49959-3 | | 539 | RSQSEESASAFSA | 6.0851 | 5.5912 | 7.3763 | 6.2250 | 5.8646 | 6.8328 | 6.6123 | 7.2591 | 7.2583 | 6.5041 | 6.2014 | 6.5301 | 5.5245 | 6.2169 | 4.4155 | 5.5025 |
| MRE11_S691;S692 | MRE11 | P49959-3 | 691;692 | | NA | 5.8539 | 5.8282 | 5.7163 | 6.1840 | 6.6047 | 5.8898 | 6.4877 | 5.8749 | 5.9227 | 6.4228 | 6.5355 | 6.3399 | 6.4330 | 6.6769 | 6.6134 | 6.6163 |
| MREG_S213 | MREG | Q8N565 | | 213 | HYLPFPSPxxxxx | 6.8077 | 7.2763 | 6.7941 | 6.8622 | 6.2694 | 6.2929 | 6.2252 | 6.4131 | 7.0513 | 6.3862 | 6.8424 | 6.9984 | 4.6169 | 5.3761 | 4.7019 | 5.0859 |
| MRGBP_S23 | MRGBP | Q9NV56 | | 195 | ANSNPSSPSAAKR | 7.3528 | 7.1359 | 6.3807 | 6.5441 | 6.8243 | 6.1500 | 6.0351 | 6.7322 | 7.3583 | 6.7300 | 6.4918 | 6.4894 | 5.4778 | 5.1272 | 4.2351 | 4.9354 |
| MRGBP_T188 | MRGBP | Q9NV56 | | 23 | PGEAATSPAEETV | 5.7065 | 5.7035 | 6.5893 | 6.2846 | 5.9755 | 6.2417 | 5.8570 | 6.1319 | 5.7472 | 6.2002 | 6.1421 | 5.9566 | 6.8918 | 7.0889 | 6.9354 | 6.5478 |
| MRGBP_T188;S195 | MRGBP | Q9NV56 | | 188 | VTDKVLTANSNPS | 5.3463 | 6.1612 | 4.8786 | 5.3707 | 6.4820 | 5.2825 | 4.9718 | 6.1871 | 5.8110 | 5.4295 | 5.0712 | 5.5095 | 9.2550 | 7.4489 | 8.9738 | 7.8209 |
| MRGBP_T195 | MRGBP | Q9NV56 | 188;195 | | NA | 6.7812 | 6.8323 | 6.1526 | 7.2659 | 5.4984 | 6.2196 | 5.9989 | 6.0310 | 6.4802 | 7.2230 | 6.4099 | 7.1021 | 5.8768 | 5.9562 | 4.6360 | 5.5359 |
| MRPL23_S76 | MRPL23 | Q16540 | | 76 | TRVQHGSNKRRDH | 6.1831 | 6.0214 | 5.7678 | 6.6094 | 5.9324 | 6.0385 | 5.8485 | 6.2617 | 5.9845 | 5.9766 | 6.4634 | 7.3294 | 6.1970 | 6.2687 | 6.3975 | 6.7201 |
| MRPL34_S71 | MRPL34 | Q9BQ48 | | 71 | GWVRRLSTPAGVQ | 6.0589 | 3.7865 | 6.5206 | 4.2307 | 6.4833 | 6.1448 | 5.7453 | 4.9439 | 10.4919 | 9.7696 | 9.6866 | 6.6868 | 6.3166 | 6.6614 | 2.2188 | 4.2544 |
| MRPL9_S264 | MRPL9 | Q9BYD2 | | 264 | KAMAPTSPQIxxx | 6.2879 | 5.9463 | 6.4402 | 6.3651 | 6.4148 | 5.9109 | 6.7937 | 7.0545 | 6.0809 | 6.2668 | 5.6846 | 6.2432 | 6.6116 | 6.1744 | 5.6392 | 6.0859 |
| MRPS18B_S38 | MRPS18B | Q9Y676 | | 38 | LCTKAPSEEDSLS | 6.1218 | 6.4095 | 6.1767 | 6.9691 | 6.5029 | 6.1620 | 6.1523 | 6.0511 | 6.8832 | 6.5326 | 6.1972 | 6.9369 | 5.7000 | 5.7887 | 5.3904 | 6.0259 |
| MRPS36_S61 | MRPS36 | P82909 | | 61 | HSKGSKSPDLLMY | 6.1271 | 6.0392 | 6.1628 | 6.1544 | 6.6396 | 5.9113 | 5.9287 | 6.3748 | 6.2089 | 6.6584 | 6.0139 | 6.7659 | 6.4189 | 6.3619 | 6.1538 | 6.0807 |
| MRTFA_S156 | MRTFA | Q969V6 | | 156 | LSATSASPTQVVS | 5.4591 | 5.8391 | 6.1475 | 5.7579 | 6.0835 | 5.8880 | 6.1864 | 5.8051 | 6.1503 | 5.7380 | 5.8352 | 6.0700 | 7.1601 | 7.3302 | 7.2431 | 7.3066 |
| MRTFB_S222 | MRTFB | Q9ULH7-5 | | 222 | KVSESPSPVTTNT | 5.4140 | 5.5433 | 5.8494 | 5.6518 | 6.1469 | 5.9183 | 5.7778 | 6.1452 | 5.7774 | 5.8047 | 5.9614 | 5.9173 | 7.7754 | 7.5510 | 7.4404 | 7.3257 |
| MRTFB_S236 | MRTFB | Q9ULH7-5 | | 236 | AQFASVSPTVPEF | 5.7789 | 6.2396 | 6.7736 | 6.4832 | 6.2917 | 6.6974 | 6.3122 | 6.5438 | 5.9097 | 6.6717 | 7.1291 | 6.9693 | 5.5714 | 5.5359 | 5.3767 | 5.7157 |
| MRTFB_S554 | MRTFB | Q9ULH7-5 | | 554 | NNEDSLSPTSSTL | 6.0347 | 6.1067 | 6.6540 | 6.7445 | 6.3309 | 6.3101 | 6.2786 | 6.2745 | 6.2904 | 6.4173 | 6.7282 | 6.2817 | 5.9408 | 6.1582 | 5.4211 | 6.0283 |
| MRTFB_S77 | MRTFB | Q9ULH7-5 | | 77 | IMPPLKSPAAFHE | 5.6123 | 4.5827 | 5.4414 | 5.9822 | 7.1697 | 6.3738 | 6.0560 | 7.2975 | 6.6539 | 5.5168 | 4.8316 | 5.3130 | 8.0198 | 7.7633 | 6.4839 | 6.9022 |
| MSANTD2_S27 | MSANTD2 | Q6P1R3 | | 436 | CIERPLSPHMEQS | 5.7658 | 6.3090 | 6.0678 | 6.2304 | 6.1880 | 6.0752 | 5.7933 | 6.0673 | 5.9932 | 6.1259 | 6.2809 | 5.9215 | 6.6211 | 6.7995 | 6.9759 | 6.7852 |
| MSANTD2_S436 | MSANTD2 | Q6P1R3 | | 27 | EVLSPASPGGLSD | 5.5109 | 6.2454 | 6.5626 | 6.5563 | 6.1303 | 5.6539 | 6.1267 | 6.4873 | 6.3330 | 6.7453 | 6.1846 | 6.3428 | 5.8165 | 6.4623 | 6.5436 | 6.2986 |
| MSANTD2_S48 | MSANTD2 | Q6P1R3 | | 48 | STPRGASPLGPGS | 6.5616 | 6.3785 | 6.4758 | 5.4804 | 5.5691 | 6.2155 | 6.0596 | 6.0753 | 5.5419 | 6.2198 | 6.0496 | 7.0865 | 6.5616 | 7.5956 | 6.5959 | 5.5333 |
| MSANTD3_S274 | MSANTD3 | Q96H12 | | 98 | KVKRSVSPLLSTH | 6.4228 | 6.8033 | 6.0001 | 6.8355 | 7.0738 | 6.8059 | 5.9632 | 6.9479 | 6.0548 | 5.6728 | 5.7612 | 5.5783 | 6.1878 | 6.0199 | 6.1687 | 5.7041 |
| MSANTD3_S98 | MSANTD3 | Q96H12 | | 274 | NRPFPNSPxxxxx | 6.5903 | 6.6821 | 6.4843 | 6.6042 | 6.4719 | 6.0645 | 6.2155 | 5.9826 | 6.1121 | 6.7068 | 5.9273 | 6.0432 | 6.1294 | 6.1868 | 6.0354 | 5.7636 |
| MSANTD4_S137 | MSANTD4 | Q8NCY6 | | 137 | FRDAGGSLTEVKV | 6.1797 | 5.8538 | 6.6533 | 6.8182 | 6.3337 | 5.4066 | 6.8983 | 7.1044 | 7.8105 | 6.0062 | 6.8337 | 6.4977 | 5.6213 | 5.2044 | 5.3639 | 5.4144 |
| MSH2_Y863 | MSH2 | P43246 | | 863 | IGESQGYDIMEPA | 5.9880 | 5.5325 | 6.5414 | 5.7166 | 6.4072 | 6.3249 | 5.7480 | 6.0164 | 6.3024 | 6.2113 | 6.3964 | 6.2221 | 6.3974 | 7.1193 | 6.6815 | 6.3945 |
| MSH3_S33 | MSH3 | P20585 | | 33 | FFQSTGSLKSTSS | 5.6111 | 6.4197 | 7.8329 | 6.8706 | 5.7313 | 6.0455 | 5.6948 | 7.1945 | 6.7550 | 9.7302 | 8.4105 | 9.5493 | 3.2216 | 4.5459 | 2.8087 | 3.5783 |
| MSH6_S137 | MSH6 | P52701 | | 309 | MVTGNGSLKRKSS | 4.5523 | 5.9722 | 7.1764 | 5.7597 | 6.0663 | 6.4415 | 5.7624 | 7.0284 | 6.7546 | 9.6874 | 8.8759 | 8.3181 | 4.1078 | 5.2584 | 3.7211 | 4.5176 |
| MSH6_S14 | MSH6 | P52701 | | 830 | KIHNVGSPLKSQN | 5.9795 | 6.7952 | 6.5702 | 6.3276 | 5.9500 | 6.3760 | 5.9724 | 6.0517 | 6.5494 | 6.7997 | 6.6981 | 6.6262 | 5.3494 | 6.2152 | 5.8843 | 5.8551 |
| MSH6_S200 | MSH6 | P52701 | | 200 | AVCDEPSEPEEEE | 5.0454 | 5.6850 | 6.0401 | 5.6783 | 5.7592 | 6.1753 | 5.4777 | 5.2886 | 6.4908 | 6.8472 | 6.3921 | 6.5699 | 6.8743 | 7.3413 | 7.2624 | 7.0725 |
| MSH6_S261 | MSH6 | P52701 | | 137 | VQFFDDSPTRGWV | 5.9360 | 6.3402 | 6.5373 | 6.2093 | 6.3487 | 6.4175 | 6.1418 | 6.2056 | 6.4485 | 6.5049 | 6.4229 | 6.4933 | 6.0010 | 6.1644 | 5.9082 | 5.9205 |
| MSH6_S309 | MSH6 | P52701 | | 79 | NGGLRRSVAPAAP | 7.0060 | 6.3657 | 6.4063 | 6.5678 | 5.6935 | 6.5608 | 6.8612 | 5.8944 | 7.2419 | 6.6482 | 7.2669 | 6.9678 | 4.9607 | 5.4898 | 4.6924 | 5.3767 |
| MSH6_S41 | MSH6 | P52701 | | 41 | AAAPGASPSPGGD | 6.3104 | 6.3872 | 6.9436 | 6.5161 | 6.0035 | 6.7137 | 6.5454 | 6.4341 | 6.6975 | 6.3027 | 6.7124 | 6.7685 | 5.4820 | 6.0026 | 5.0114 | 5.1692 |
| MSH6_S43 | MSH6 | P52701 | | 43 | APGASPSPGGDAA | 6.2886 | 6.1313 | 6.2379 | 5.8806 | 6.1488 | 5.7179 | 6.9488 | 6.1790 | 7.0518 | 6.1112 | 5.6432 | 6.5653 | 6.3216 | 6.7988 | 5.6568 | 6.3186 |
| MSH6_S5 | MSH6 | P52701 | | 14 | YSFFPKSPALSDA | 6.3746 | 7.9802 | 6.2983 | 5.2673 | 5.5432 | 6.9567 | 6.8550 | 6.9107 | 6.7344 | 6.2246 | 7.3512 | 6.9387 | 4.5186 | 5.7285 | 4.8612 | 5.4569 |
| MSH6_S79 | MSH6 | P52701 | | 261 | ESDIGGSDVEFKP | 4.5336 | 5.5962 | 5.6472 | 5.0527 | 4.0098 | 4.3119 | 5.0707 | 4.3938 | 7.4466 | 9.0014 | 8.2883 | 7.8267 | 6.0799 | 7.7118 | 7.5390 | 7.4904 |
| MSH6_S830 | MSH6 | P52701 | | 5 | xxMSRQSTLYSFF | 6.1729 | 5.5577 | 5.7688 | 6.2181 | 6.0030 | 6.0422 | 6.2060 | 6.2040 | 8.2144 | 6.7669 | 7.7419 | 8.2084 | 5.7859 | 7.1268 | 3.1000 | 4.8832 |
| MSI2_S274 | MSI2 | A0A6Q8PF05 | | 274 | NFGAPASPAGSNP | 5.2516 | 6.2341 | 6.3814 | 5.8805 | 6.9452 | 6.3362 | 6.2742 | 7.4080 | 6.6577 | 7.0971 | 6.8868 | 6.4484 | 5.4218 | 5.8825 | 5.1404 | 5.7540 |
| MSL1_S126 | MSL1 | Q68DK7 | | 442 | WHQPPPSPLPLRE | 5.7819 | 6.5111 | 6.0173 | 6.8349 | 6.0273 | 6.2475 | 6.1705 | 6.0297 | 6.6908 | 6.2207 | 6.4090 | 6.1806 | 6.1908 | 6.4233 | 5.9416 | 6.3230 |
| MSL1_S442 | MSL1 | Q68DK7 | | 126 | AAGAGCSPRPKYQ | 6.7305 | 6.7192 | 6.3756 | 7.1084 | 6.6570 | 6.3593 | 6.6732 | 6.5541 | 6.8517 | 6.6416 | 6.6887 | 6.4228 | 4.9102 | 5.0982 | 5.0214 | 5.1881 |
| MSL2_S211 | MSL2 | Q9HCI7 | | 211 | FGINIPSPEHSNT | 5.9795 | 6.2908 | 6.1037 | 6.4420 | 6.3277 | 6.3943 | 6.4580 | 6.4220 | 6.4289 | 6.2693 | 6.2333 | 5.7331 | 5.9952 | 6.3480 | 6.4390 | 6.1352 |
| MSL3_S311 | MSL3 | Q8N5Y2 | | 405 | SSPIPLTPSKEGS | 6.9143 | 6.8868 | 7.1470 | 7.8659 | 6.3498 | 6.3867 | 6.6010 | 5.9018 | 6.7372 | 6.8470 | 6.9173 | 6.6949 | 4.5077 | 5.1750 | 4.2234 | 4.8441 |
| MSL3_S367 | MSL3 | Q8N5Y2 | | 367 | LSESSASPQPKRR | 6.6047 | 6.5838 | 6.5852 | 7.2059 | 6.3207 | 6.3949 | 6.2276 | 6.1953 | 6.8306 | 6.6848 | 6.6318 | 6.5590 | 5.5299 | 5.2678 | 5.0285 | 5.3495 |
| MSL3_S398;T405 | MSL3 | Q8N5Y2 | | 311 | QEELSPSPPLLNP | 5.7100 | 6.2868 | 5.7391 | 5.9608 | 5.8045 | 7.3968 | 5.8687 | 5.4773 | 6.6007 | 6.2214 | 5.8760 | 6.9472 | 6.3341 | 7.3350 | 6.3001 | 6.1414 |
| MSL3_T405 | MSL3 | Q8N5Y2 | 398;405 | | NA | 6.5953 | 6.5081 | 6.1178 | 6.5628 | 5.9229 | 5.9435 | 5.9418 | 6.0513 | 6.5808 | 6.2352 | 5.7295 | 6.2418 | 6.7129 | 6.2650 | 6.0949 | 6.4963 |
| MSL3P1_S292 | MSL3P1 | P0C860 | | 292 | LSKSSTSPQPKRW | 7.1485 | 7.3390 | 6.0605 | 7.1332 | 6.3780 | 6.0112 | 6.4239 | 6.2185 | 6.8147 | 7.0895 | 6.6350 | 6.8852 | 4.7419 | 5.1011 | 4.5567 | 5.4633 |
| MSN_S56 | MSN | P26038 | | 576 | RIDEFESMxxxxx | 5.9247 | 6.7827 | 6.8566 | 5.9768 | 6.8514 | 6.9131 | 6.5853 | 6.0607 | 5.9848 | 6.8074 | 6.7902 | 6.3923 | 5.7407 | 5.8264 | 5.3844 | 5.1224 |
| MSN_S576 | MSN | P26038 | | 74 | QDVRKESPLLFKF | 6.1349 | 6.7266 | 7.7227 | 6.1221 | 6.1366 | 5.9603 | 5.1026 | 6.6488 | 6.2123 | 6.0401 | 6.8675 | 6.7752 | 6.0497 | 5.9894 | 5.8842 | 5.6269 |
| MSN_S74 | MSN | P26038 | | 56 | QDTKGFSTWLKLN | 6.1561 | 6.9780 | 7.8670 | 6.2309 | 6.7277 | 5.3567 | 8.2032 | 7.7699 | 6.8734 | 6.0329 | 5.4165 | 5.8984 | 3.5973 | 6.3093 | 6.5932 | 3.9895 |
| MSTO1_S495 | MSTO1 | Q9BUK6 | | 495 | GMVLDGSPKGAAV | 6.8940 | 6.3028 | 6.4121 | 5.3896 | 6.0355 | 6.6044 | 5.6043 | 9.2810 | 6.0578 | 6.9709 | 6.6713 | 7.5540 | 4.9159 | 4.8248 | 5.4038 | 5.0778 |
| MTA1_S386 | MTA1 | Q13330 | | 576 | PVINNGSPTILGK | 7.1518 | 7.2276 | 6.8337 | 7.4866 | 6.8376 | 6.4489 | 6.9546 | 6.4644 | 7.3606 | 6.8965 | 6.8306 | 6.6312 | 4.1093 | 4.5275 | 3.7820 | 4.4570 |
| MTA1_S449 | MTA1 | Q13330 | | 564 | SVLSSLTPAKVAP | 7.3804 | 7.3546 | 6.8662 | 7.9349 | 6.4344 | 6.2189 | 7.0820 | 6.6996 | 7.6116 | 6.7443 | 6.9562 | 7.0452 | 3.8968 | 4.2251 | 3.3781 | 4.1719 |
| MTA1_S460 | MTA1 | Q13330 | | 522 | LPEASQSPLVLKQ | 6.5816 | 6.6403 | 6.0986 | 6.7483 | 6.2732 | 6.2027 | 6.5271 | 5.8784 | 6.5565 | 6.9102 | 6.3747 | 6.0117 | 5.6195 | 6.0334 | 5.5355 | 6.0082 |
| MTA1_S522 | MTA1 | Q13330 | | 449 | PNRSNMSPHGLPA | 7.1451 | 7.6413 | 6.7317 | 7.8884 | 6.1587 | 6.0410 | 6.6266 | 6.5928 | 7.1320 | 7.4075 | 7.5779 | 7.1826 | 3.6033 | 4.1505 | 3.7350 | 4.3858 |
| MTA1_S558 | MTA1 | Q13330 | | 639 | RLIRGGSLPPVKR | 5.6544 | 5.9870 | 5.6377 | 6.2316 | 6.7013 | 5.7263 | 5.9720 | 6.3403 | 6.5047 | 6.5977 | 6.3422 | 6.5628 | 6.5393 | 6.3367 | 6.3379 | 6.5279 |
| MTA1_S576 | MTA1 | Q13330 | | 386 | TGAPGQSPGAGRA | 7.0851 | 7.0497 | 6.4427 | 6.9413 | 6.0108 | 6.4799 | 7.4946 | 6.8010 | 6.9189 | 7.3408 | 6.8818 | 6.4757 | 4.0378 | 4.8446 | 4.2476 | 4.9477 |
| MTA1_S639 | MTA1 | Q13330 | | 460 | PARSSGSPKFAMK | 6.3031 | 6.9191 | 5.8345 | 7.0928 | 6.2355 | 5.5571 | 6.0042 | 5.9996 | 6.6096 | 6.5381 | 5.6447 | 6.9061 | 5.9420 | 6.9084 | 6.1648 | 5.3405 |
| MTA1_T564 | MTA1 | Q13330 | | 558 | PVKSVSSVLSSLT | 6.6053 | 6.0045 | 5.2006 | 4.4795 | 5.6357 | 4.5583 | 5.2006 | 7.0943 | 6.3318 | 3.6880 | 4.4795 | 5.2834 | 8.3499 | 6.4147 | 9.0088 | 11.6650 |
| MTA2_S54 | MTA2 | O94776 | | 54 | RRDISSSLNSLAD | 6.2136 | 6.3584 | 6.4162 | 6.0291 | 6.3633 | 5.8759 | 6.6056 | 5.8535 | 6.3199 | 6.0024 | 6.1848 | 6.1986 | 6.5924 | 6.6111 | 5.7569 | 6.6182 |
| MTA3_S430 | MTA3 | Q9BTC8 | | 519 | HAELSGSPLKSKS | 6.4479 | 6.7617 | 6.2373 | 6.7686 | 6.3569 | 6.1485 | 6.5527 | 6.2112 | 6.5320 | 6.3521 | 6.2789 | 6.3377 | 5.5855 | 5.8809 | 5.9102 | 5.6380 |
| MTA3_S519 | MTA3 | Q9BTC8 | | 430 | EEKLSPSPTTEDP | 6.3250 | 6.5324 | 6.4921 | 7.3396 | 6.1470 | 5.7553 | 6.4870 | 6.6046 | 6.5119 | 6.6460 | 6.3118 | 6.6671 | 5.2439 | 5.7042 | 5.5321 | 5.6999 |
| MTBP_S539 | MTBP | Q96DY7 | | 639 | VLTPELSPGKLQV | 6.9977 | 7.1636 | 6.6447 | 6.8662 | 6.6331 | 6.3768 | 6.6954 | 6.5776 | 6.8409 | 6.9594 | 6.6015 | 6.4616 | 4.6246 | 4.8726 | 4.5280 | 5.1564 |
| MTBP_S597 | MTBP | Q96DY7 | | 539 | NILNDFSPVEPNS | 6.0189 | 6.3237 | 6.5030 | 6.9636 | 5.7567 | 6.0808 | 6.3216 | 6.3418 | 6.0278 | 6.4147 | 6.8512 | 6.2173 | 6.0676 | 6.2935 | 5.8364 | 5.9815 |
| MTBP_S639 | MTBP | Q96DY7 | | 597 | KEGPRDSITLLDA | 4.6376 | 5.2357 | 5.3941 | 4.6633 | 5.9031 | 6.6290 | 5.4704 | 6.1052 | 7.9778 | 9.9871 | 9.3375 | 9.2177 | 5.2832 | 6.5894 | 3.5968 | 3.9721 |
| MTBP_S703 | MTBP | Q96DY7 | | 703 | PVPTVLSPLPSPV | 6.0026 | 6.8207 | 6.3147 | 7.2175 | 6.2325 | 6.1323 | 6.1978 | 6.0026 | 6.8245 | 7.4166 | 8.0716 | 5.8601 | 5.0112 | 5.9846 | 4.6143 | 5.2963 |
| MTBP_T386;S388 | MTBP | Q96DY7 | 386;388 | | NA | 6.0714 | 6.8424 | 6.5207 | 7.1690 | 6.7488 | 6.8955 | 8.1174 | 6.8596 | 7.0735 | 6.4258 | 6.4516 | 7.0581 | 3.8958 | 5.2072 | 3.7213 | 4.9418 |
| MTCH1_S374 | MTCH1 | Q9NZJ7 | | 381 | LLFRRVSSGSCFA | 6.1502 | 6.4982 | 5.6733 | 6.1581 | 6.2961 | 6.1184 | 6.4195 | 4.9928 | 7.3757 | 7.2335 | 6.7244 | 6.2430 | 6.1464 | 5.8135 | 6.2280 | 5.9289 |
| MTCH1_S381 | MTCH1 | Q9NZJ7 | | 374 | QLFRGSSLLFRRV | 5.7514 | 4.7558 | 4.3967 | 6.3426 | 5.9573 | 5.7371 | 5.8089 | 5.3685 | 5.6916 | 3.9827 | 4.0880 | 3.6428 | 10.2535 | 9.0089 | 9.6790 | 9.5354 |
| MTCH2_S283 | MTCH2 | Q9Y6C9 | | 283 | NMSRGNSLFFRKV | 5.3151 | 4.9375 | 4.8134 | 5.0640 | 5.9004 | 5.6378 | 5.5250 | 5.6064 | 5.2735 | 5.0680 | 4.7248 | 4.6642 | 9.7166 | 9.2751 | 9.4181 | 9.0601 |
| MTCL1_S1385 | MTCL1 | Q9Y4B5 | | 1399 | QRLMDISPFLPEK | 7.3753 | 7.3550 | 7.7926 | 6.7491 | 5.9720 | 7.3478 | 7.2624 | 7.0996 | 7.4176 | 7.5110 | 8.0328 | 7.3546 | 2.5567 | 3.3359 | 2.8518 | 3.9857 |
| MTCL1_S1399 | MTCL1 | Q9Y4B5 | | 1514 | GLRVLHSPPAVRR | 6.5921 | 6.5184 | 7.1086 | 6.1282 | 6.1391 | 6.2812 | 6.4988 | 6.7097 | 6.9754 | 6.2873 | 6.8226 | 7.0074 | 5.2027 | 5.3625 | 4.9224 | 5.4435 |
| MTCL1_S1514 | MTCL1 | Q9Y4B5 | | 1438 | NKSWDYTPNRGHN | 6.8163 | 7.0705 | 6.8207 | 6.8511 | 6.2349 | 6.4819 | 7.1185 | 6.7240 | 6.5143 | 6.6471 | 6.7576 | 6.7126 | 4.5440 | 5.1050 | 4.4833 | 5.1182 |
| MTCL1_S1561 | MTCL1 | Q9Y4B5 | | 1561 | SRWPCTSPRHSRD | 6.5906 | 6.8130 | 6.6267 | 6.0499 | 6.0807 | 7.0988 | 6.8125 | 7.1765 | 6.6183 | 6.4223 | 6.9485 | 7.3438 | 4.6413 | 4.7930 | 4.7465 | 5.2376 |
| MTCL1_S1661 | MTCL1 | Q9Y4B5 | | 263 | RGAPPGSPEPPAL | 5.5535 | 6.7944 | 7.7919 | 6.8132 | 5.4621 | 5.3801 | 5.2820 | 6.9826 | 6.1988 | 5.8184 | 7.5271 | 6.4704 | 6.4314 | 5.1328 | 5.4352 | 6.9261 |
| MTCL1_S1730 | MTCL1 | Q9Y4B5 | | 43 | RLHRAPSPARPFL | 7.3696 | 6.1658 | 7.2810 | 5.3955 | 5.1822 | 6.2075 | 6.4496 | 5.9150 | 7.4980 | 6.4078 | 6.4057 | 6.5887 | 6.4613 | 6.1325 | 4.7633 | 5.7764 |
| MTCL1_S1812;S1814 | MTCL1 | Q9Y4B5 | | 756 | DAGKKESDGEESR | 6.4477 | 6.6672 | 7.6669 | 6.5631 | 6.5858 | 6.8094 | 6.6736 | 5.6792 | 5.4509 | 6.3925 | 6.6580 | 6.7957 | 5.3524 | 5.7566 | 5.1271 | 5.3739 |
| MTCL1_S263 | MTCL1 | Q9Y4B5 | | 1385 | NLQRAVSVSSMSE | 6.1892 | 5.7889 | 6.8995 | 5.9947 | 5.6332 | 6.0637 | 6.1960 | 6.5416 | 6.6427 | 6.4022 | 6.6964 | 6.8304 | 5.3719 | 6.3480 | 5.8607 | 6.5407 |
| MTCL1_S43 | MTCL1 | Q9Y4B5 | | 622 | APSIPTSPFGDSL | 5.4001 | 5.3622 | 6.4091 | 6.3703 | 6.1996 | 6.5997 | 6.5057 | 7.3155 | 6.6506 | 6.5109 | 6.8541 | 7.0576 | 5.5873 | 6.1263 | 5.3424 | 5.7089 |
| MTCL1_S622 | MTCL1 | Q9Y4B5 | | 87 | AGKAPPSPGSLAA | 6.8749 | 7.0904 | 6.7692 | 6.6150 | 5.5733 | 7.0678 | 6.7976 | 6.5808 | 7.1656 | 6.5742 | 6.8657 | 7.1914 | 4.4379 | 4.9596 | 4.4182 | 5.0185 |
| MTCL1_S756 | MTCL1 | Q9Y4B5 | | 1661 | GSGVTSSPHKCLT | 6.3722 | 6.9466 | 6.8043 | 6.9516 | 6.8290 | 7.5627 | 6.3743 | 8.4443 | 7.2727 | 6.5500 | 7.8871 | 7.3277 | 3.2752 | 4.2696 | 3.3101 | 3.8227 |
| MTCL1_S87 | MTCL1 | Q9Y4B5 | | 1730 | QPNNRTSPGMAQK | 7.6792 | 7.5299 | 7.4798 | 7.5365 | 6.3178 | 6.7502 | 7.1847 | 6.8217 | 7.0902 | 7.6472 | 7.8497 | 8.0747 | 2.8891 | 3.0891 | 2.7356 | 3.3246 |
| MTCL1_T1417;S1421 | MTCL1 | Q9Y4B5 | 1812;1814 | | NA | 6.3043 | 6.4787 | 6.4620 | 6.7241 | 6.1245 | 5.9156 | 6.1363 | 6.7009 | 6.7781 | 6.1712 | 6.2468 | 6.8858 | 5.6269 | 5.6205 | 5.9918 | 5.8326 |
| MTCL1_T1438 | MTCL1 | Q9Y4B5 | 1417;1421 | | NA | 6.0945 | 6.5023 | 6.3733 | 6.3866 | 6.0231 | 5.6598 | 5.9244 | 6.2619 | 6.2643 | 6.5257 | 6.4994 | 7.0538 | 5.5286 | 6.3103 | 6.1273 | 6.4648 |
| MTDH_S180 | MTDH | Q86UE4 | | 568 | SETSWESPKQIKK | 6.0505 | 6.2931 | 6.5587 | 6.6262 | 6.7603 | 6.3609 | 7.0718 | 6.1237 | 6.5604 | 6.7493 | 6.5931 | 6.5347 | 5.0986 | 5.8625 | 5.0805 | 5.6756 |
| MTDH_S298 | MTDH | Q86UE4 | | 426 | PDDQKVSDDDKEK | 5.8512 | 5.9245 | 6.2248 | 6.4135 | 5.4876 | 5.4677 | 6.5056 | 5.1037 | 6.0611 | 6.3121 | 6.1553 | 6.2924 | 6.9069 | 7.2263 | 6.5982 | 7.4690 |
| MTDH_S308 | MTDH | Q86UE4 | | 308 | EKWNSVSPASAGK | 6.5346 | 7.0099 | 6.4682 | 7.0831 | 6.7496 | 6.8296 | 7.1757 | 7.0221 | 6.8790 | 6.9677 | 6.7700 | 6.6312 | 3.9730 | 5.0787 | 4.1688 | 4.6589 |
| MTDH_S308;S311 | MTDH | Q86UE4 | | 298 | KLSSQISAGEEKW | 3.6313 | 3.6587 | 3.9933 | 4.0545 | 5.0638 | 4.7577 | 4.9700 | 4.8517 | 3.7658 | 3.6162 | 3.3816 | 3.8606 | 12.1126 | 11.9208 | 13.4657 | 12.8956 |
| MTDH_S362 | MTDH | Q86UE4 | | 143 | EGEAVRTPQSVTA | 6.6839 | 7.2822 | 6.5569 | 7.7230 | 6.6786 | 6.9573 | 7.3841 | 6.9998 | 6.5708 | 6.7438 | 7.4552 | 7.0871 | 3.2825 | 4.8810 | 3.6368 | 4.0773 |
| MTDH_S426 | MTDH | Q86UE4 | | 478 | DLPVNTSKTRPKQ | 6.3730 | 6.8276 | 6.2185 | 7.3519 | 6.2390 | 7.0874 | 7.3439 | 6.6829 | 5.9502 | 6.0488 | 6.3994 | 6.0367 | 4.9022 | 6.2294 | 4.6131 | 5.6961 |
| MTDH_S478 | MTDH | Q86UE4 | | 494 | FSLKTISTSDPAE | 6.4727 | 6.8600 | 6.7100 | 7.6684 | 6.6149 | 5.8802 | 6.9250 | 5.9069 | 6.3877 | 6.2119 | 6.3728 | 6.2521 | 4.6573 | 5.8864 | 5.4919 | 5.7019 |
| MTDH_S494 | MTDH | Q86UE4 | | 180 | KAVQNSSRHDGKE | 6.0534 | 8.5400 | 5.8184 | 7.4644 | 5.9435 | 6.7101 | 8.0482 | 6.8542 | 7.4199 | 6.6684 | 6.3406 | 6.8295 | 3.4162 | 5.0688 | 3.8625 | 4.9618 |
| MTDH_S568 | MTDH | Q86UE4 | | 362 | VSQSTTSDYQWDV | 5.9485 | 6.7808 | 7.0499 | 7.7841 | 7.4708 | 5.5566 | 4.7823 | 5.9195 | 6.6510 | 5.7594 | 7.1631 | 6.1473 | 6.2356 | 5.5662 | 5.7084 | 5.4765 |
| MTDH_T143 | MTDH | Q86UE4 | 308;311 | | NA | 6.6826 | 7.5298 | 6.2121 | 7.4408 | 6.4231 | 6.8239 | 7.5113 | 6.9194 | 6.8008 | 7.0686 | 6.9817 | 6.8226 | 3.5853 | 5.0222 | 3.6564 | 4.5194 |
| MTF2_T576 | MTF2 | Q9Y483 | | 576 | VLARRVTLDGKVQ | 6.8805 | 7.2989 | 6.6871 | 7.4995 | 6.0202 | 7.0528 | 7.3586 | 6.7256 | 7.2191 | 7.2077 | 7.3757 | 8.1912 | 3.2488 | 4.2244 | 2.9717 | 4.0381 |
| MTFR1_S119 | MTFR1 | Q15390 | | 119 | APSRQISLPDLSQ | 4.8169 | 5.1499 | 4.9778 | 5.3246 | 6.7403 | 6.7235 | 6.3359 | 6.6368 | 5.7671 | 5.0881 | 5.2139 | 5.3622 | 8.2317 | 7.9573 | 8.0932 | 7.5808 |
| MTFR1L_S103 | MTFR1L | Q9H019 | | 238 | SLSKASSFADMMG | 6.4368 | 6.2071 | 6.0989 | 6.5951 | 7.2766 | 6.3113 | 6.6521 | 6.8807 | 6.3930 | 5.9540 | 6.3883 | 6.4862 | 5.7033 | 5.9247 | 5.0518 | 5.6402 |
| MTFR1L_S234 | MTFR1L | Q9H019 | | 103 | VMQRNASVPNLRG | 6.7708 | 6.3581 | 6.2489 | 6.4678 | 7.2841 | 6.4757 | 6.9144 | 6.5484 | 6.7334 | 6.5086 | 6.5728 | 6.7634 | 5.3600 | 5.4960 | 4.3135 | 5.1840 |
| MTFR1L_S238 | MTFR1L | Q9H019 | | 41 | KPCARASFETLPN | 3.7035 | 4.7025 | 4.5363 | 5.2859 | 5.5245 | 5.1766 | 5.7446 | 6.0602 | 4.8595 | 5.3136 | 5.9601 | 5.2875 | 9.9330 | 8.5846 | 9.4805 | 9.8468 |
| MTFR1L_S41 | MTFR1L | Q9H019 | | 234 | DDCVSLSKASSFA | 6.4647 | 5.9851 | 6.4409 | 6.3898 | 7.0043 | 5.4181 | 6.0822 | 6.9598 | 6.0326 | 5.4796 | 6.0592 | 5.9584 | 7.0850 | 6.7648 | 6.0259 | 5.8495 |
| MTFR2_S119 | MTFR2 | Q6P444 | | 328 | AFQEDDSFEKENR | 7.2850 | 7.8315 | 7.0265 | 8.7342 | 6.6144 | 6.5645 | 7.2378 | 6.7851 | 7.4436 | 8.0892 | 8.0733 | 8.1619 | 2.1862 | 3.1689 | 1.8911 | 2.9067 |
| MTFR2_S291 | MTFR2 | Q6P444 | | 291 | LRAIERSPGGRPI | 7.8232 | 9.4124 | 6.6014 | 8.6922 | 6.9278 | 5.2499 | 5.9028 | 5.7876 | 7.1271 | 5.6628 | 6.1357 | 7.2015 | 4.4337 | 3.5671 | 4.2801 | 5.1947 |
| MTFR2_S305 | MTFR2 | Q6P444 | | 119 | LVRDPLSPAVRQK | 8.8768 | 5.4458 | 6.1785 | 6.1310 | 5.6714 | 5.4661 | 10.6558 | 9.7960 | 7.7913 | 5.4661 | 5.9020 | 5.2660 | 3.3784 | 4.3298 | 4.7996 | 4.8454 |
| MTFR2_S328 | MTFR2 | Q6P444 | | 305 | KRKRQNSHWDPVS | 6.1347 | 5.5915 | 6.4989 | 6.2941 | 6.9305 | 5.7441 | 5.4023 | 5.3165 | 8.6610 | 6.6594 | 7.3215 | 7.8838 | 5.4803 | 6.4975 | 4.4149 | 5.1692 |

| | | | | | | | | | | | | | | | | | | | | | |
|---|---|---|---|---|---|---|---|---|---|---|---|---|---|---|---|---|---|---|---|---|---|
| MTHFD1L_S357 | MTHFD1L | Q6UB35 | | 62 | QARSSCSPGGRTP | 6.7350 | 7.2529 | 6.9235 | 6.8145 | 5.6082 | 5.7664 | 6.3876 | 5.9455 | 6.3637 | 6.6339 | 6.2467 | 6.2265 | 5.8371 | 5.6147 | 5.3662 | 6.2778 |
| MTHFD1L_S62 | MTHFD1L | Q6UB35 | | 357 | LKLQPLSPVPSDI | 7.2589 | 7.1892 | 6.5879 | 7.4681 | 6.6533 | 6.1392 | 6.3527 | 8.7709 | 7.3417 | 5.2808 | 5.8255 | 5.9126 | 5.5815 | 4.7405 | 4.3310 | 4.5662 |
| MTM1_S588;S591 | MTM1 | Q13496 | | 591 | PPTSPSSPSQMMP | 5.9459 | 6.1847 | 6.6435 | 6.3814 | 7.0665 | 6.0607 | 6.6077 | 6.1039 | 6.3097 | 5.5467 | 6.2279 | 6.4837 | 6.1955 | 6.6429 | 5.7969 | 5.8026 |
| MTM1_S591 | MTM1 | Q13496 | 588;591 | | NA | 5.5610 | 7.4294 | 7.0201 | 7.1106 | 6.1658 | 5.2638 | 8.8097 | 7.7620 | 6.5950 | 6.0927 | 6.3944 | 5.6374 | 6.3562 | 5.5019 | 3.7757 | 4.5242 |
| MTMR1_S49 | MTMR1 | Q13613 | | 49 | SVETLDSPTGSHV | 5.5833 | 5.8769 | 6.1369 | 6.2114 | 6.5400 | 7.0527 | 6.8844 | 6.8088 | 5.9654 | 6.2216 | 6.6345 | 7.0725 | 5.6513 | 5.7000 | 5.5786 | 6.0817 |
| MTMR10_S569 | MTMR10 | Q9NXD2 | | 607 | LLPRRNSLILKPK | 7.0332 | 6.4478 | 6.0261 | 6.1302 | 6.2495 | 6.2347 | 6.1502 | 6.0334 | 6.7223 | 6.1737 | 6.2313 | 6.5320 | 6.1337 | 6.2097 | 5.3956 | 6.2964 |
| MTMR10_S607 | MTMR10 | Q9NXD2 | | 755 | RRSILGTPLSKFL | 7.9746 | 8.1714 | 6.8132 | 8.2164 | 6.7804 | 7.0497 | 7.8379 | 6.5212 | 7.2899 | 6.9827 | 6.7942 | 6.9631 | 2.5973 | 3.5259 | 2.8198 | 3.6622 |
| MTMR10_S625 | MTMR10 | Q9NXD2 | | 751 | GNLCRRSILGTPL | 7.4340 | 6.9088 | 6.5145 | 7.0945 | 6.7947 | 5.9753 | 6.4509 | 5.9434 | 5.9289 | 5.8847 | 6.2082 | 6.1981 | 5.7629 | 5.8260 | 5.3924 | 5.6827 |
| MTMR10_S751 | MTMR10 | Q9NXD2 | | 769 | GAKIWLSTETLAN | 5.7382 | 6.1139 | 6.4602 | 5.7041 | 6.0061 | 6.1753 | 6.4182 | 6.0921 | 5.9573 | 6.2716 | 6.6655 | 6.5684 | 5.4152 | 7.1292 | 7.0915 | 6.1931 |
| MTMR10_S769 | MTMR10 | Q9NXD2 | | 569 | PCIQNGSVKSFKR | 5.4309 | 5.4683 | 6.4489 | 6.1205 | 7.0079 | 6.2068 | 7.2777 | 5.8433 | 6.6659 | 5.9075 | 5.6716 | 5.7649 | 6.3551 | 7.1386 | 5.8723 | 6.8199 |
| MTMR10_T755 | MTMR10 | Q9NXD2 | | 625 | QTDSQNSDTEQYF | 6.3420 | 5.7249 | 6.7281 | 7.0334 | 6.3301 | 7.0851 | 6.7778 | 6.1037 | 6.4699 | 5.6043 | 6.0917 | 5.8990 | 5.8215 | 5.9394 | 5.8341 | 6.2149 |
| MTMR12_S564 | MTMR12 | Q9C0I1 | | 564 | KHQRQLSLPLTQS | 6.9594 | 6.4241 | 7.0956 | 5.8402 | 4.9420 | 6.3047 | 6.2475 | 6.3920 | 6.7262 | 6.8172 | 6.9258 | 7.0607 | 5.3703 | 6.1176 | 5.2686 | 5.5080 |
| MTMR14_S624 | MTMR14 | Q8NCE2 | | 624 | LRAVAPSPSGAIG | 6.8422 | 5.9136 | 6.5298 | 6.7383 | 6.4218 | 6.2816 | 6.0242 | 6.8556 | 6.7545 | 6.8184 | 6.9175 | 6.5124 | 5.2778 | 5.5609 | 6.2283 | 5.0232 |
| MTMR14_S647 | MTMR14 | Q8NCE2 | | 647 | GLRSISSNALxxx | 4.8021 | 5.5250 | 5.3278 | 3.8675 | 4.5101 | 5.6856 | 3.8091 | 3.4829 | 5.6540 | 3.9916 | 4.4954 | 5.2913 | 8.9933 | 11.7461 | 11.2544 | 11.5635 |
| MTMR2_S631 | MTMR2 | Q13614 | | 631 | SSERASSPAQCVT | 5.7193 | 5.4388 | 8.0737 | 6.7669 | 7.5911 | 6.4364 | 7.2305 | 6.0631 | 5.9088 | 6.2461 | 6.5168 | 5.7542 | 5.5389 | 5.7604 | 5.5282 | 5.4270 |
| MTMR3_S613 | MTMR3 | Q13615 | | 647 | WQEHRRSLELSSL | 6.0726 | 6.0500 | 6.1197 | 6.0861 | 6.2460 | 6.2776 | 6.0540 | 6.5112 | 6.4373 | 5.9416 | 6.4615 | 6.0911 | 6.2003 | 6.9081 | 6.5513 | 5.9915 |
| MTMR3_S633 | MTMR3 | Q13615 | | 613 | RLPKTRSYDNLTT | 6.0958 | 6.2307 | 7.2628 | 6.7093 | 6.6299 | 6.4128 | 6.4895 | 6.3094 | 6.1767 | 6.0846 | 6.5555 | 6.5258 | 5.7865 | 6.0722 | 5.1081 | 5.5504 |
| MTMR3_S647 | MTMR3 | Q13615 | | 633 | LASRRCSDPSLNE | 6.2381 | 6.3140 | 6.1581 | 6.6265 | 5.8941 | 5.7722 | 6.6977 | 6.2707 | 6.0989 | 6.5541 | 6.6730 | 6.3750 | 6.1833 | 6.3519 | 6.0362 | 5.7563 |
| MTMR4_S1002 | MTMR4 | Q9NYA4 | | 1002 | VPLNCPSPVPPLY | 5.7992 | 6.3226 | 6.5381 | 6.3732 | 6.5695 | 6.3959 | 7.2151 | 5.8053 | 5.9344 | 6.0932 | 5.7695 | 6.2956 | 6.1098 | 6.5093 | 5.5278 | 6.7414 |
| MTMR4_S699 | MTMR4 | Q9NYA4 | | 699 | KSHKSCSPSYKLL | 5.8380 | 6.6303 | 6.3585 | 6.9693 | 5.8035 | 7.3095 | 7.0377 | 7.1916 | 6.5844 | 6.8184 | 6.9087 | 7.7805 | 4.9611 | 5.2248 | 3.9366 | 4.6473 |
| MTOR_S1261 | MTOR | P42345 | | 1261 | MKKLHVSTINLQK | 2.0161 | 1.9987 | 2.0234 | 2.2398 | 2.8370 | 2.4099 | 2.5880 | 2.7478 | 2.1774 | 2.0224 | 1.7731 | 2.4053 | 17.5056 | 17.9292 | 19.4584 | 17.8680 |
| MTOR_S567 | MTOR | P42345 | | 567 | LAHQLASPGLTTL | 6.0562 | 6.7581 | 6.6279 | 5.5938 | 5.2429 | 6.4722 | 5.8215 | 7.0486 | 6.4792 | 6.3886 | 6.9870 | 6.1446 | 6.3479 | 5.6891 | 6.3700 | 5.9726 |
| MTREX_S40 | MTREX | P42285 | | 40 | WKGPPGSADKAGK | 7.2313 | 7.7276 | 6.6250 | 7.7549 | 6.2744 | 6.9363 | 7.0461 | 6.5137 | 7.7159 | 7.0513 | 7.5627 | 7.6210 | 3.2503 | 3.8208 | 3.0239 | 3.8449 |
| MTSS2_S428 | MTSS2 | Q765P7 | | 579 | TVRRALSSAGPIP | 6.9442 | 6.8575 | 7.0015 | 5.7332 | 6.5637 | 6.4154 | 6.5412 | 6.3802 | 6.7688 | 6.5908 | 6.6481 | 6.6941 | 5.4107 | 6.2861 | 4.5048 | 4.6597 |
| MTSS2_S579 | MTSS2 | Q765P7 | | 428 | APRPRMSPATIAA | 6.9483 | 7.2316 | 7.1702 | 7.0529 | 6.4080 | 7.5293 | 7.1514 | 6.6234 | 7.2892 | 6.8686 | 7.6384 | 7.1660 | 3.2705 | 4.4743 | 3.2225 | 3.9553 |
| MTSS2_S612 | MTSS2 | Q765P7 | | 612 | GPTRAGSEECVFY | 5.8495 | 6.0083 | 6.7785 | 6.3676 | 6.5882 | 5.8597 | 5.9367 | 6.7977 | 6.5673 | 6.5919 | 6.4478 | 6.3746 | 5.9052 | 6.7838 | 5.4271 | 5.7161 |
| MTX3_S249 | MTX3 | Q5HYI7 | | 311 | NSFQRLSPxxxxx | 6.1422 | 6.4964 | 6.9469 | 6.7589 | 6.0787 | 6.1528 | 6.1615 | 6.0394 | 6.1702 | 6.1907 | 6.0777 | 6.5954 | 6.0737 | 6.2365 | 5.8075 | 6.0715 |
| MTX3_S284 | MTX3 | Q5HYI7 | | 249 | LSLGGISPAGQET | 6.9034 | 6.1365 | 6.3318 | 6.4719 | 6.4986 | 5.8363 | 6.5656 | 7.0913 | 6.1455 | 6.9724 | 7.1422 | 7.5822 | 5.2423 | 4.9185 | 4.9154 | 5.2460 |
| MTX3_S311 | MTX3 | Q5HYI7 | | 284 | DDNLRQSPQLPPR | 6.2003 | 6.9600 | 6.3076 | 5.5628 | 6.3132 | 6.3436 | 6.7045 | 6.7424 | 5.7613 | 7.7179 | 6.8856 | 7.6844 | 5.0487 | 5.1430 | 5.0115 | 5.6131 |
| MVB12A_S170 | MVB12A | Q96EY5 | | 170 | RDMQGLSLDAASQ | 7.7052 | 6.9796 | 9.5714 | 7.2653 | 6.4902 | 5.9387 | 5.5369 | 6.4897 | 7.0603 | 7.2501 | 6.8677 | 6.3073 | 3.9990 | 5.4339 | 3.6821 | 3.4226 |
| MVB12A_S232 | MVB12A | Q96EY5 | | 232 | FEGKSCSPLAFSA | 5.9277 | 6.1537 | 6.6897 | 7.0107 | 6.9817 | 6.4378 | 6.2887 | 6.2047 | 6.1718 | 6.6897 | 7.0610 | 6.5500 | 5.6546 | 5.6036 | 5.1131 | 5.4615 |
| MVD_S96 | MVD | P53602 | | 96 | ARKRRNSRDGDPL | 5.8986 | 4.9647 | 7.0123 | 5.4332 | 7.2866 | 7.4543 | 5.1569 | 5.6005 | 6.5739 | 6.8949 | 7.4652 | 7.6548 | 5.3496 | 6.7386 | 5.5309 | 4.9850 |
| MVP_S445 | MVP | Q14764 | | 873 | SPGEGISPQSAQA | 5.6107 | 5.3715 | 6.4565 | 5.9903 | 6.0558 | 6.5705 | 5.7372 | 6.9660 | 6.4735 | 6.2258 | 6.4360 | 6.3431 | 6.6698 | 6.5587 | 6.2341 | 6.3006 |
| MVP_S873 | MVP | Q14764 | | 445 | EKDTAKSLQPLAP | 6.4325 | 6.7767 | 6.3853 | 5.7419 | 6.7379 | 6.3406 | 6.2403 | 6.0528 | 5.9898 | 5.4852 | 5.7096 | 5.6053 | 6.6537 | 7.1890 | 6.8921 | 5.7675 |
| MXRA7_S128 | MXRA7 | P84157 | | 128 | GEKGPSSEGPEEE | 6.9125 | 6.1172 | 5.5708 | 6.6075 | 5.8644 | 5.5765 | 5.7767 | 6.3679 | 5.5768 | 5.8395 | 5.6468 | 6.5341 | 5.4811 | 7.7282 | 6.8845 | 7.5153 |
| MXRA7_S144 | MXRA7 | P84157 | | 144 | GFSFKYSPGKLRG | 6.1892 | 5.9414 | 4.8196 | 5.6076 | 5.4206 | 4.8632 | 5.6448 | 5.4238 | 4.2240 | 5.1669 | 5.2102 | 6.1274 | 5.7173 | 12.1952 | 8.7042 | 8.7446 |
| MXRA8_S429 | MXRA8 | Q9BRK3-2 | | 429 | RAELAHSPLPAKY | 7.6625 | 6.9996 | 6.3813 | 6.9512 | 6.0784 | 7.1345 | 6.9270 | 6.8978 | 7.1389 | 7.5513 | 6.9234 | 7.0104 | 3.7568 | 5.1562 | 3.7124 | 4.6181 |
| MYBBP1A_S1159;S1163 | MYBBP1A | Q9BQG0-2 | | 775 | ALGGEDSENEEEL | 5.8995 | 5.8798 | 5.7137 | 6.4709 | 6.0681 | 6.1185 | 6.3672 | 6.3768 | 6.2985 | 6.0831 | 5.9940 | 6.5479 | 6.0473 | 6.9472 | 6.5771 | 6.6104 |
| MYBBP1A_S1163 | MYBBP1A | Q9BQG0-2 | | 1163 | IPSATQSPISKKR | 6.5740 | 6.9214 | 6.2081 | 7.0035 | 6.5742 | 6.4338 | 6.9838 | 6.4896 | 7.2029 | 6.3628 | 6.5077 | 6.5954 | 4.6746 | 5.3230 | 4.8225 | 5.3227 |
| MYBBP1A_S1241 | MYBBP1A | Q9BQG0-2 | | 1244 | PTRSPSTPAKSPK | 7.4735 | 7.9508 | 7.1276 | 7.1805 | 6.2383 | 7.3807 | 7.5686 | 7.1737 | 7.1856 | 7.2680 | 7.6164 | 7.8584 | 2.4183 | 3.3931 | 2.4424 | 3.7240 |
| MYBBP1A_S1241;T1244 | MYBBP1A | Q9BQG0-2 | | 1241 | PGAPTRSPSTPAK | 7.2312 | 6.9403 | 6.5033 | 7.6427 | 6.8284 | 7.0110 | 7.4755 | 6.4515 | 7.0564 | 6.9946 | 6.9475 | 6.6914 | 3.7238 | 4.6036 | 3.7035 | 4.1953 |
| MYBBP1A_S1244 | MYBBP1A | Q9BQG0-2 | | 1290 | KGVLGKSPLSALA | 7.7768 | 8.0763 | 6.5676 | 7.7454 | 6.4248 | 7.0755 | 7.5216 | 7.2282 | 7.9575 | 7.4460 | 7.4716 | 7.3483 | 2.3591 | 3.3704 | 2.3470 | 3.2841 |
| MYBBP1A_S1267 | MYBBP1A | Q9BQG0-2 | | 1310 | LVIRSPSLLQSGA | 5.6310 | 6.2088 | 6.5657 | 5.5495 | 4.8836 | 6.9232 | 6.7941 | 6.6310 | 7.7261 | 8.4430 | 7.4576 | 7.4732 | 4.6954 | 5.5784 | 3.8940 | 5.5454 |
| MYBBP1A_S1290 | MYBBP1A | Q9BQG0-2 | | 1227 | LKLGFSPTPTKSP | 7.7362 | 7.8494 | 6.7873 | 7.7858 | 6.9418 | 7.0897 | 7.5515 | 7.1655 | 7.9277 | 7.1614 | 7.0774 | 7.3974 | 2.4228 | 3.4609 | 2.4212 | 3.2241 |
| MYBBP1A_S1303 | MYBBP1A | Q9BQG0-2 | | 1267 | VNGAPGSPTEPAG | 5.7814 | 5.2729 | 5.9624 | 6.8692 | 6.2143 | 5.7298 | 6.5852 | 5.2396 | 6.7255 | 6.5149 | 6.1074 | 6.0265 | 7.0607 | 6.6918 | 6.4741 | 6.7444 |
| MYBBP1A_S1303;S1308 | MYBBP1A | Q9BQG0-2 | | 1303 | RKKARLSLVIRSP | 5.5892 | 6.4605 | 6.4634 | 6.6627 | 5.3283 | 6.3907 | 6.7438 | 6.2601 | 7.7071 | 8.5064 | 7.7413 | 8.4048 | 4.5503 | 5.3944 | 3.3091 | 4.4879 |
| MYBBP1A_S1303;S1308; | MYBBP1A | Q9BQG0-2 | | 1308 | LSLVIRSPSLLQS | 5.6683 | 7.0660 | 6.4572 | 7.2998 | 6.9880 | 7.2386 | 6.5337 | 6.6431 | 6.7678 | 6.8144 | 6.4673 | 6.8445 | 4.2503 | 5.0945 | 4.1124 | 4.8539 |
| MYBBP1A_S1308 | MYBBP1A | Q9BQG0-2 | 1308;1314 | | NA | 8.4895 | 7.7477 | 6.6240 | 7.6991 | 5.5803 | 6.6184 | 7.8742 | 6.5506 | 8.0499 | 7.0136 | 7.7020 | 7.2655 | 2.6581 | 4.0231 | 2.5924 | 3.5115 |
| MYBBP1A_S1308;S1314 | MYBBP1A | Q9BQG0-2 | 1159;1163 | | NA | 6.3832 | 6.5869 | 6.2559 | 7.2158 | 5.7614 | 6.0829 | 6.3774 | 6.2261 | 7.0244 | 6.7444 | 6.8041 | 6.6211 | 4.9842 | 5.8702 | 5.1744 | 5.8878 |
| MYBBP1A_S1310 | MYBBP1A | Q9BQG0-2 | 1241;1244 | | NA | 6.9282 | 7.5439 | 6.5328 | 8.1940 | 6.3470 | 6.1184 | 7.3291 | 7.5941 | 8.2956 | 7.4687 | 7.3654 | 7.4078 | 2.8819 | 3.7675 | 2.5731 | 3.6525 |
| MYBBP1A_S775 | MYBBP1A | Q9BQG0-2 | 1303;1308 | | NA | 6.6834 | 6.7209 | 6.8361 | 7.2026 | 6.0651 | 8.2573 | 7.8892 | 5.2524 | 6.9778 | 7.8440 | 8.9728 | 9.1116 | 2.7485 | 3.4194 | 2.1758 | 3.8431 |
| MYBBP1A_T1227 | MYBBP1A | Q9BQG0-2 | 1303;1308;13 | | NA | 10.0423 | 9.0365 | 6.6806 | 6.9723 | 5.8479 | 7.3744 | 8.9578 | 6.1558 | 6.8382 | 5.7568 | 6.5835 | 8.0160 | 2.4524 | 3.4300 | 1.8681 | 3.9876 |
| MYBL1_S434 | MYBL1 | P10243 | | 458 | KMRVGHSPGSELR | 6.5955 | 7.1638 | 6.6245 | 7.3453 | 6.8826 | 6.8875 | 6.6744 | 7.3001 | 6.8429 | 6.3803 | 6.9919 | 6.7067 | 4.0458 | 4.6402 | 4.3190 | 4.5997 |
| MYBL1_S458 | MYBL1 | P10243 | | 541 | ENVGFRTPTIRRS | 7.5068 | 7.8580 | 7.3776 | 7.9732 | 6.2881 | 7.0853 | 7.3682 | 6.6860 | 7.2257 | 7.0704 | 7.1597 | 7.5142 | 2.9175 | 3.8292 | 2.6646 | 3.4755 |
| MYBL1_S487 | MYBL1 | P10243 | | 522 | QKALITTPLHKET | 7.4615 | 7.5557 | 7.0262 | 8.3026 | 6.2261 | 7.2715 | 7.6107 | 6.6549 | 7.3891 | 6.9829 | 7.2221 | 7.5330 | 2.9346 | 3.7662 | 2.5178 | 3.5451 |
| MYBL1_S626 | MYBL1 | P10243 | | 626 | GKKVRKSLVLDNW | 6.2226 | 6.8324 | 6.6073 | 8.2875 | 6.0942 | 6.5395 | 6.3661 | 6.4863 | 7.1935 | 6.5120 | 6.9238 | 7.7230 | 4.5541 | 4.9077 | 4.4150 | 4.3350 |
| MYBL1_T442 | MYBL1 | P10243 | | 487 | LKTLPFSPSQFFN | 6.1140 | 6.2642 | 7.6787 | 7.7260 | 7.2311 | 6.4208 | 6.2696 | 7.7141 | 6.1540 | 7.0658 | 7.4256 | 8.0631 | 3.9467 | 4.3832 | 4.0159 | 4.4372 |
| MYBL1_T522 | MYBL1 | P10243 | | 442 | NIAKFSTPPAILR | 7.0389 | 7.0902 | 7.1074 | 7.2508 | 6.6892 | 7.1537 | 7.4269 | 7.0766 | 7.3356 | 6.8620 | 7.0248 | 6.9923 | 3.6761 | 3.9280 | 3.3149 | 4.0326 |
| MYBL1_T541 | MYBL1 | P10243 | | 434 | EAVPLTSPNIAKF | 6.4384 | 7.3869 | 6.4665 | 6.8602 | 6.8837 | 6.5181 | 7.0975 | 7.1084 | 6.7410 | 6.1221 | 6.2057 | 6.5164 | 4.8904 | 5.1254 | 4.3746 | 5.2647 |
| MYBL2_S241 | MYBL2 | P10244 | | 266 | DLDAVRTPEPLEE | 6.0369 | 6.2488 | 5.9248 | 6.2407 | 6.5637 | 5.7700 | 5.7416 | 5.9054 | 6.4053 | 6.1772 | 6.1860 | 6.3480 | 6.8233 | 6.5569 | 6.5791 | 6.4923 |
| MYBL2_S282 | MYBL2 | P10244 | | 405 | GGSGIGTPPSVLK | 6.1842 | 6.8999 | 6.4149 | 6.1515 | 6.5345 | 6.2673 | 6.4794 | 6.0042 | 6.9264 | 6.7244 | 6.3268 | 6.7038 | 5.7029 | 5.9102 | 5.3403 | 5.4295 |
| MYBL2_S393 | MYBL2 | P10244 | | 487 | QKVVVTTPLHRDK | 8.2759 | 8.0743 | 7.1008 | 7.5863 | 7.0771 | 7.1331 | 6.9368 | 6.3821 | 7.7071 | 6.8464 | 6.8979 | 7.2229 | 3.0149 | 3.9712 | 2.5807 | 3.1924 |
| MYBL2_S393;T405 | MYBL2 | P10244 | | 505 | KHAAFVTPDQKYS | 7.6703 | 8.1431 | 6.6942 | 7.1524 | 7.4099 | 7.3974 | 7.3688 | 7.5714 | 7.2823 | 7.1536 | 7.0811 | 7.1052 | 3.1382 | 3.8697 | 2.7899 | 3.1725 |
| MYBL2_S405 | MYBL2 | P10244 | | 282 | REDQEGSPPETSL | 6.0256 | 6.3932 | 6.0923 | 6.1364 | 6.3699 | 6.1326 | 5.9009 | 5.9745 | 6.3781 | 6.2027 | 6.3244 | 6.2349 | 6.2196 | 6.6365 | 6.6681 | 6.3103 |
| MYBL2_S452 | MYBL2 | P10244 | | 538 | LKPLPQTPHLEED | 7.6428 | 6.3658 | 7.5329 | 7.3781 | 6.1932 | 7.2648 | 7.1235 | 6.3071 | 7.3642 | 7.8939 | 7.3699 | 7.9609 | 3.2339 | 4.1402 | 2.7590 | 3.4698 |
| MYBL2_T266 | MYBL2 | P10244 | | 241 | IKEEENSEEELAA | 6.2714 | 6.3956 | 6.1631 | 6.2077 | 6.0711 | 5.7403 | 6.3386 | 6.1661 | 6.1839 | 5.8374 | 5.9876 | 6.3676 | 6.3021 | 6.9038 | 6.8302 | 6.2334 |
| MYBL2_T440 | MYBL2 | P10244 | | 440 | DSCNSLTPKSTPV | 6.6640 | 6.9997 | 6.3211 | 6.4417 | 5.6620 | 6.2590 | 6.3377 | 6.0923 | 7.1860 | 6.5853 | 6.2099 | 6.5189 | 5.8468 | 5.9833 | 5.1790 | 5.7133 |
| MYBL2_T487 | MYBL2 | P10244 | | 452 | VKTLPFSPSQFLN | 6.4372 | 6.1023 | 6.4088 | 8.2688 | 6.7778 | 6.7513 | 6.3294 | 7.0938 | 6.9576 | 5.4438 | 5.8828 | 6.9065 | 5.7976 | 4.8648 | 3.5005 | 6.4769 |
| MYBL2_T505 | MYBL2 | P10244 | | 393 | GELIPISPSTEVG | 5.2803 | 5.9535 | 6.5086 | 6.6446 | 6.1054 | 6.2352 | 6.4204 | 6.4738 | 6.1830 | 6.5321 | 6.0826 | 6.0474 | 6.0978 | 6.4518 | 6.3473 | 6.6362 |
| MYBL2_T538 | MYBL2 | P10244 | 393;405 | | NA | 6.4559 | 6.6818 | 6.3848 | 6.8947 | 7.3771 | 6.4693 | 6.1376 | 6.8124 | 6.8543 | 7.1291 | 7.1311 | 6.7428 | 4.8289 | 4.8945 | 4.5345 | 4.6712 |
| MYC_S308 | MYC | P01106-2 | | 77 | LPTPPLSPSRRSG | 5.8685 | 5.4561 | 6.2412 | 6.2947 | 6.9428 | 6.0973 | 6.6581 | 6.9397 | 6.1267 | 5.3000 | 6.2035 | 5.8548 | 7.0902 | 6.1594 | 6.7087 | 6.0583 |
| MYC_S77 | MYC | P01106-2 | | 308 | HSKPPHSPLVLKR | 4.9201 | 4.2743 | 4.9535 | 5.1574 | 5.3873 | 4.9675 | 5.8402 | 5.0713 | 4.7647 | 4.2095 | 4.5727 | 4.9653 | 10.5382 | 9.0453 | 10.3505 | 10.9823 |
| MYC_T73 | MYC | P01106-2 | | 73 | KFELLPTPPLSPS | 4.2609 | 4.2943 | 4.2058 | 4.4571 | 4.7341 | 4.4567 | 5.4381 | 4.8290 | 4.5926 | 4.1520 | 4.1540 | 4.2601 | 11.9722 | 10.7492 | 12.4438 | 11.0001 |
| MYC_T73;S77 | MYC | P01106-2 | 73;77 | | NA | 4.9183 | 4.9721 | 5.1796 | 5.4928 | 5.5248 | 5.2561 | 5.4748 | 5.3610 | 5.0542 | 4.5503 | 4.7782 | 5.1739 | 10.0198 | 8.8551 | 9.9491 | 9.4399 |
| MYCBP2_S2334 | MYCBP2 | O75592 | | 2789 | SHSRSLPNHNTL | 6.6359 | 6.9617 | 6.3340 | 7.1664 | 6.6561 | 7.0927 | 5.7574 | 7.1821 | 6.1311 | 6.4969 | 6.5639 | 5.9776 | 4.9918 | 5.8892 | 5.2796 | 4.8836 |
| MYCBP2_S2789 | MYCBP2 | O75592 | | 2839 | SPSGASSPRSSSP | 7.2839 | 6.2988 | 6.3726 | 6.4024 | 6.2812 | 6.4233 | 5.7236 | 6.3631 | 6.5733 | 5.7837 | 6.1446 | 6.2478 | 5.9301 | 6.3801 | 6.1750 | 5.6165 |
| MYCBP2_S2839 | MYCBP2 | O75592 | | 2920 | SENRAPSPHVVQE | 6.5671 | 6.1100 | 7.0547 | 6.6864 | 6.1628 | 6.3870 | 6.0452 | 6.5919 | 6.4286 | 7.3189 | 5.7073 | 5.7666 | 5.4367 | 6.4607 | 6.1885 | 5.0877 |
| MYCBP2_S2871 | MYCBP2 | O75592 | | 2334 | PQRIPGSPAVTAA | 5.8506 | 6.7829 | 7.1254 | 7.7644 | 6.1364 | 7.0098 | 7.0377 | 5.8913 | 6.4586 | 7.5364 | 7.4401 | 6.2980 | 4.4324 | 5.5798 | 3.8747 | 4.7813 |
| MYCBP2_S2896;S2898 | MYCBP2 | O75592 | | 2871 | PRERSKSDSYTLD | 5.7720 | 5.8224 | 5.9257 | 6.2953 | 6.2099 | 6.7196 | 6.2142 | 6.5291 | 5.5861 | 6.1385 | 6.6182 | 5.8799 | 7.0004 | 7.0462 | 6.5906 | 5.6520 |
| MYCBP2_S2920 | MYCBP2 | O75592 | | 3478 | VFQRSYSVVASEY | 7.8362 | 6.1155 | 7.0978 | 6.9138 | 6.2004 | 5.1390 | 5.6697 | 5.1884 | 5.9686 | 5.5804 | 6.3156 | 5.8854 | 6.9026 | 7.3241 | 6.2588 | 5.6036 |
| MYCBP2_S3460 | MYCBP2 | O75592 | | 3460 | PPSLETSPITDTD | 5.9767 | 5.4238 | 5.9006 | 7.3127 | 8.1194 | 5.8607 | 6.6493 | 6.5042 | 7.3055 | 5.9441 | 6.9230 | 6.0873 | 5.5272 | 6.2106 | 5.2462 | 5.0087 |
| MYCBP2_S3478 | MYCBP2 | O75592 | | 2683 | LDQNSQTPPPSPF | 5.9969 | 6.1650 | 6.5540 | 6.6135 | 6.8651 | 6.4145 | 6.2693 | 6.0171 | 5.5625 | 6.7037 | 7.0748 | 5.7620 | 5.6623 | 5.5042 | 6.1331 | 5.7021 |
| MYCBP2_T2683 | MYCBP2 | O75592 | 2896;2898 | | NA | 6.1762 | 6.7380 | 7.3796 | 6.1704 | 5.8394 | 5.6887 | 6.3183 | 5.5480 | 6.1949 | 6.9425 | 6.2373 | 6.0367 | 6.2533 | 6.8625 | 6.0411 | 5.5731 |
| MYEF2_S17 | MYEF2 | Q9P2K5 | | 17 | GATGGDSPHLQPA | 6.1472 | 6.2301 | 6.0003 | 6.4504 | 5.9914 | 6.0881 | 6.1471 | 6.1284 | 6.1868 | 6.0991 | 6.1242 | 6.0616 | 6.6650 | 6.6629 | 6.6196 | 6.3981 |
| MYH10_S1176 | MYH10 | P35580-4 | | 1987 | GASLELSDDDTES | 5.5296 | 6.0357 | 6.1526 | 5.8850 | 6.5170 | 6.1679 | 5.9187 | 6.0347 | 6.4879 | 6.3467 | 6.4351 | 6.1626 | 6.4582 | 7.0679 | 6.5109 | 6.2895 |
| MYH10_S1970 | MYH10 | P35580-4 | | 1970 | PISFSSSRSGRRQ | 5.5831 | 5.9610 | 6.6168 | 5.8264 | 5.2853 | 6.2131 | 6.2073 | 6.2596 | 6.3759 | 6.3273 | 6.8647 | 6.3118 | 6.2529 | 7.3713 | 6.1072 | 6.4363 |
| MYH10_S1983 | MYH10 | P35580-4 | | 1176 | KQKRDLSEELEAL | 6.2501 | 6.7471 | 6.4455 | 4.7278 | 6.5345 | 7.7505 | 5.7359 | 5.1060 | 8.0943 | 7.8052 | 4.8638 | 5.2123 | 6.6377 | 6.3361 | 4.8294 | 6.9237 |
| MYH10_S1983;S1987 | MYH10 | P35580-4 | | 1983 | LHLEGASLELSDD | 6.2866 | 6.8953 | 5.9304 | 6.8867 | 6.2232 | 7.0662 | 5.7797 | 7.9821 | 6.4695 | 6.2943 | 5.5665 | 6.6529 | 4.5431 | 6.3509 | 5.8507 | 5.2219 |
| MYH10_S1987 | MYH10 | P35580-4 | 1983;1987 | | NA | 7.1981 | 6.8482 | 8.2613 | 8.5226 | 5.9576 | 5.9692 | 6.4585 | 6.1504 | 6.5156 | 7.5930 | 6.1466 | 5.1497 | 4.3169 | 6.1382 | 4.7439 | 4.0303 |
| MYH9_S1057 | MYH9 | P35579 | | 1943 | KGAGDGSDEEVDG | 5.9302 | 6.0653 | 6.7399 | 6.1478 | 6.3809 | 6.3995 | 6.0898 | 6.5236 | 6.4016 | 6.5025 | 6.4684 | 6.5392 | 5.9037 | 6.2097 | 5.8181 | 5.8798 |
| MYH9_S1943 | MYH9 | P35579 | | 725 | QRYEILTPNSIPK | 7.1755 | 6.3823 | 7.3210 | 6.9494 | 5.6604 | 6.0627 | 5.7638 | 7.3510 | 6.8117 | 6.5035 | 5.9999 | 5.7788 | 5.3037 | 5.9008 | 5.8416 | 5.1939 |
| MYH9_T725 | MYH9 | P35579 | | 1057 | RKLEGDSTDLSDQ | 5.8021 | 5.7999 | 6.0946 | 6.6819 | 5.6202 | 6.2830 | 6.2744 | 6.1456 | 6.6481 | 6.3284 | 5.8941 | 6.7938 | 5.8194 | 6.7657 | 6.1417 | 7.1071 |
| MYL12B_S20 | MYL12B | O14950 | | 20 | RPQRATSNVFAMF | 8.1774 | 8.0549 | 8.8313 | 7.6026 | 6.4094 | 7.3029 | 8.2194 | 7.1986 | 6.9379 | 7.9153 | 6.0728 | 4.8862 | 4.0367 | 3.7758 | 2.7187 | 3.8602 |
| MYL12B_T135 | MYL12B | O14950 | | 19 | KRPQRATSNVFAM | 10.1757 | 7.7294 | 8.4803 | 7.6610 | 7.3180 | 6.5720 | 7.2503 | 6.5859 | 6.7164 | 5.4368 | 5.3006 | 4.7544 | 4.5213 | 4.1001 | 3.4498 | 3.9481 |
| MYL12B_T19 | MYL12B | O14950 | | 135 | TMGDRFTDEEVDE | 7.4025 | 7.0416 | 8.7442 | 8.0814 | 7.4749 | 8.0788 | 7.4920 | 8.6666 | 5.4440 | 5.3912 | 6.0386 | 5.8197 | 3.2071 | 3.8979 | 3.1440 | 4.0753 |
| MYNN_S584 | MYNN | Q9NPC7 | | 584 | PVTDTQSPTSDTL | 5.4638 | 7.9221 | 7.4203 | 5.9891 | 6.3326 | 6.0598 | 5.9336 | 5.0041 | 6.7721 | 6.2097 | 5.9066 | 6.2804 | 5.4604 | 6.6727 | 6.2333 | 6.3393 |
| MYO18A_S1067 | MYO18A | Q92614 | | 2007 | GSLKSSSPTSYWK | 6.1918 | 6.2661 | 5.8544 | 5.7424 | 6.5363 | 6.1925 | 6.0134 | 6.0439 | 6.6408 | 6.1629 | 5.9516 | 5.9733 | 6.7000 | 6.7081 | 6.5899 | 6.4326 |
| MYO18A_S140 | MYO18A | Q92614 | | 140 | MIVKRFSFSQRSR | 6.7668 | 6.4108 | 6.5136 | 5.7642 | 6.6107 | 6.1807 | 6.1243 | 5.8567 | 6.8421 | 6.1391 | 5.9196 | 5.7006 | 6.6019 | 6.9329 | 5.9296 | 5.7066 |
| MYO18A_S198 | MYO18A | Q92614 | | 1998 | GPSKAASDDGSLK | 5.4589 | 6.4582 | 5.8113 | 5.6033 | 5.5921 | 5.8928 | 6.0413 | 6.2455 | 6.4131 | 6.6114 | 6.3865 | 7.0070 | 5.8173 | 7.3103 | 6.6709 | 6.9801 |
| MYO18A_S1998 | MYO18A | Q92614 | | 1067 | ASSRRVSSSSELD | 5.8428 | 6.5382 | 6.5194 | 6.3907 | 6.6753 | 6.9781 | 5.4724 | 6.2727 | 6.9781 | 6.2422 | 6.7842 | 6.9830 | 6.5606 | 6.0739 | 5.6811 | 5.8173 |
| MYO18A_S2007 | MYO18A | A0A1B0GV45 | | 198 | SLSRTLSPSLSRA | 6.2660 | 6.1544 | 6.4281 | 6.7511 | 5.9473 | 5.8176 | 6.3347 | 6.3140 | 7.2113 | 6.1603 | 6.7325 | 6.2264 | 5.5511 | 6.2210 | 6.0008 | 5.8833 |
| MYO19_S963 | MYO19 | Q96H55 | | 963 | RLIHVTSSAFTGL | 3.7152 | 3.6765 | 7.3482 | 6.4496 | 3.8021 | 2.9035 | 5.9713 | 5.3046 | 5.3433 | 5.8988 | 3.5799 | 4.5703 | 13.2615 | 10.0391 | 9.3290 | 8.8072 |
| MYO1C_S389 | MYO1C | O00159-3 | | 6 | xMRYRASALGSDG | 8.4211 | 6.4658 | 6.6829 | 6.2577 | 5.9950 | 6.1803 | 6.4869 | 5.7796 | 7.5430 | 6.2115 | 6.6792 | 7.3835 | 5.3172 | 6.1590 | 3.9491 | 4.4882 |
| MYO1C_S391 | MYO1C | O00159-3 | | 389 | ASKDVESPSWRST | 5.8153 | 6.2633 | 6.0070 | 6.4910 | 6.3127 | 6.0618 | 6.0770 | 6.2297 | 6.3290 | 6.5192 | 6.5124 | 6.4760 | 6.0668 | 6.4323 | 6.0781 | 6.3283 |
| MYO1C_S6 | MYO1C | O00159-3 | | 717 | LEVRRQSLATKIQ | 5.7437 | 6.3641 | 5.6995 | 6.4286 | 5.5932 | 5.9407 | 5.5752 | 5.6220 | 6.3287 | 7.2064 | 7.1624 | 6.8937 | 6.0052 | 6.5213 | 6.5520 | 6.3635 |
| MYO1C_S717 | MYO1C | O00159-3 | | 391 | KDVESPSWRSTTV | 5.5063 | 5.7982 | 5.0687 | 5.7203 | 7.0085 | 6.6773 | 6.4203 | 6.4223 | 6.2922 | 5.4609 | 5.7243 | 5.2912 | 7.0081 | 6.8737 | 7.5930 | 7.1348 |
| MYO1C_S845 | MYO1C | O00159-3 | | 845 | KYCRSIRSPEWKQ | 5.8836 | 6.7559 | 6.5175 | 6.7821 | 6.2110 | 6.6511 | 6.8476 | 7.0912 | 5.9019 | 5.7945 | 5.1527 | 5.2470 | 6.6223 | 7.1017 | 5.8888 | 5.5509 |
| MYO5A_S1677 | MYO5A | Q9Y4I1-3 | | 600 | DDEKAISPTSATS | 6.0148 | 6.4455 | 5.9965 | 5.8521 | 6.0769 | 6.0723 | 6.3059 | 5.9935 | 6.5231 | 6.4700 | 6.3900 | 6.3824 | 6.1254 | 6.5596 | 6.3472 | 6.4448 |
| MYO5A_S600 | MYO5A | Q9Y4I1-3 | | 1677 | LRKRTSSIADEGT | 6.0768 | 5.0666 | 6.3614 | 6.3668 | 6.2533 | 6.4967 | 6.2625 | 5.8755 | 6.8857 | 7.0359 | 6.3841 | 6.3812 | 6.0254 | 6.3126 | 5.8656 | 6.3499 |
| MYO6_S1155 | MYO6 | Q9UM54 | | 1155 | APFLNNSPQQNPA | 4.5877 | 7.8499 | 7.0283 | 5.9474 | 6.0501 | 8.5738 | 6.3191 | 9.0189 | 6.6321 | 5.6246 | 7.7570 | 9.2341 | 2.9395 | 3.9079 | 4.0106 | 4.5192 |
| MYO9A_S1358 | MYO9A | B2RTY4-4 | | 1358 | GDLQFSPSKIGSS | 6.5065 | 6.2227 | 7.0227 | 7.2423 | 6.0127 | 5.9874 | 6.5831 | 6.7242 | 7.1640 | 7.0033 | 6.8463 | 7.0411 | 4.7204 | 5.4080 | 4.7108 | 5.1046 |
| MYO9A_S2344 | MYO9A | B2RTY4-4 | | 2365 | NYPGPSSPVVVRL | 6.2671 | 6.0071 | 6.4853 | 5.7523 | 6.6438 | 6.0317 | 6.7369 | 7.2394 | 6.8835 | 5.9374 | 6.0606 | 6.6269 | 5.6650 | 6.1573 | 5.4723 | 6.0336 |
| MYO9A_S2365 | MYO9A | B2RTY4-4 | | 2344 | KAKTRLSLIRRSM | 6.4412 | 6.2330 | 6.5793 | 6.0418 | 6.6828 | 6.5281 | 6.6949 | 5.8742 | 6.7456 | 6.5265 | 6.3274 | 5.7864 | 5.7333 | 6.7612 | 6.7674 | 5.8411 |
| MYO9B_S1043 | MYO9B | Q13459 | | 1242 | KTLPSGSPRPGQL | 6.7008 | 7.0183 | 7.0158 | 6.7606 | 6.2649 | 6.7379 | 6.6402 | 7.1083 | 7.0397 | 6.9902 | 6.8185 | 6.7969 | 4.0674 | 5.1228 | 4.2578 | 4.6600 |
| MYO9B_S1114;S1115 | MYO9B | Q13459 | | 1354 | TEERRTSFSTSDV | 5.9674 | 5.7587 | 6.5807 | 5.8005 | 6.5651 | 5.9680 | 5.5869 | 6.4804 | 7.2507 | 7.5063 | 7.2985 | 7.0628 | 5.6309 | 6.4586 | 5.0136 | 5.0708 |
| MYO9B_S1115 | MYO9B | Q13459 | | 1122 | PEKEAPSPEKTLP | 6.4297 | 6.7210 | 7.0147 | 6.4966 | 6.1383 | 6.4792 | 6.2350 | 6.6525 | 6.5426 | 6.6443 | 6.6670 | 6.8124 | 4.9277 | 5.6667 | 5.1327 | 5.4395 |
| MYO9B_S1122 | MYO9B | Q13459 | | 1281 | KSKPCGSPRVQEK | 6.6907 | 6.6211 | 6.9255 | 6.2296 | 6.5002 | 7.0169 | 6.5491 | 6.8773 | 6.3349 | 6.3339 | 6.5163 | 6.5024 | 5.0513 | 5.6203 | 5.2148 | 5.0156 |
| MYO9B_S1242 | MYO9B | Q13459 | | 1267 | SPPAPGSAPETPE | 6.2320 | 5.9984 | 6.9774 | 6.0071 | 6.4032 | 6.2635 | 6.1841 | 6.7183 | 6.6906 | 6.7072 | 6.4919 | 6.4177 | 5.7135 | 6.2749 | 5.6494 | 5.2707 |
| MYO9B_S1253 | MYO9B | Q13459 | | 2141 | GAKRRYSDPPTYC | 6.9033 | 5.5097 | 6.1722 | 6.2567 | 6.5122 | 6.5219 | 6.3376 | 6.5782 | 6.1113 | 6.3381 | 6.0601 | 5.6382 | 6.1282 | 6.2331 | 6.2777 | 6.4215 |
| MYO9B_S1261 | MYO9B | Q13459 | | 1323 | KLVAAASPSAMLS | 6.4757 | 6.3035 | 6.7719 | 6.4734 | 5.9225 | 6.4966 | 6.5002 | 6.6558 | 6.8476 | 6.5759 | 6.6780 | 6.9079 | 5.1511 | 5.4783 | 5.3269 | 5.4349 |
| MYO9B_S1267 | MYO9B | Q13459 | | 1405 | LPDAGLSPGSQVD | 6.4547 | 7.1796 | 8.0674 | 8.0207 | 5.9604 | 6.3764 | 6.1237 | 6.9881 | 7.2021 | 7.7836 | 7.6519 | 7.3231 | 3.2071 | 4.2712 | 3.0911 | 4.2991 |
| MYO9B_S1281 | MYO9B | Q13459 | | 1926 | LKLGFSSPYEGVL | 7.0993 | 6.6602 | 6.3659 | 7.0406 | 5.7979 | 6.9546 | 7.9670 | 6.8329 | 7.2923 | 6.5786 | 7.3969 | 7.0882 | 3.7720 | 5.0433 | 3.5862 | 4.5241 |
| MYO9B_S1290 | MYO9B | Q13459 | | 1261 | ALDSRVSPPAPGS | 5.5368 | 6.0726 | 6.4514 | 5.9899 | 6.4268 | 6.1203 | 5.4943 | 6.3776 | 6.6087 | 7.0157 | 6.3195 | 7.1565 | 6.1050 | 6.9989 | 5.9647 | 5.7016 |
| MYO9B_S1323 | MYO9B | Q13459 | | 1115 | SPLEHSSPEKEAP | 5.5218 | 6.2955 | 7.0369 | 6.5633 | 5.4900 | 6.8790 | 6.1659 | 5.5621 | 6.6132 | 6.3265 | 7.0349 | 6.9682 | 5.5568 | 5.3502 | 5.6216 | 6.0143 |
| MYO9B_S1354 | MYO9B | Q13459 | | 1290 | VQEKPDSPGGSTQ | 6.0134 | 6.3202 | 7.2013 | 6.6303 | 5.8690 | 6.6440 | 5.6241 | 5.9969 | 7.1569 | 6.3622 | 7.4505 | 7.3103 | 5.1522 | 6.5415 | 4.6250 | 5.1021 |
| MYO9B_S1405 | MYO9B | Q13459 | | 1043 | GHLQRKSFSQMIS | 5.8003 | 4.7015 | 5.7655 | 5.4625 | 6.4098 | 5.1595 | 5.2083 | 5.3812 | 6.1608 | 5.7777 | 5.4816 | 5.3754 | 10.0874 | 9.8814 | 7.1533 | 6.1938 |
| MYO9B_S1926 | MYO9B | Q13459 | | 1346 | ATGAALTPTEERR | 6.8014 | 6.6831 | 6.9775 | 6.5804 | 5.9796 | 6.4893 | 6.5376 | 6.6523 | 6.5597 | 6.2844 | 6.4242 | 6.1331 | 5.2590 | 5.7296 | 5.5577 | 5.3512 |
| MYO9B_S2141 | MYO9B | Q13459 | | 1253 | QLERPTSLALDSR | 7.0310 | 5.9185 | 7.6694 | 6.6276 | 6.2272 | 5.9516 | 7.0446 | 6.2386 | 7.6721 | 6.2617 | 5.9521 | 5.7417 | 6.3102 | 5.7521 | 4.5784 | 5.0231 |
| MYO9B_T1346 | MYO9B | Q13459 | 1114;1115 | | NA | 6.9040 | 6.8771 | 6.9753 | 6.7317 | 6.5422 | 6.7049 | 6.5431 | 6.7384 | 5.7988 | 6.0246 | 6.4953 | 6.6944 | 5.1437 | 5.5717 | 5.0729 | 5.1821 |
| MYOF_S174 | MYOF | Q9NZM1 | | 174 | GPVGTVSEAQLAR | 7.1176 | 7.1138 | 6.8323 | 7.1824 | 6.6780 | 5.7902 | 6.1534 | 5.8451 | 5.8476 | 5.3242 | 5.5177 | 5.0344 | 6.3228 | 6.7365 | 6.8489 | 5.6550 |
| MYOF_S193 | MYOF | Q9NZM1 | | 1781 | GPPFNITPRKAKK | 6.1496 | 6.6022 | 6.4143 | 7.1449 | 6.1663 | 6.1068 | 6.7887 | 6.4824 | 6.7436 | 6.5106 | 6.5591 | 6.9201 | 5.2231 | 5.7435 | 5.1526 | 5.2920 |
| MYOF_S729 | MYOF | Q9NZM1 | | 731 | LRSRSLSQIHEAA | 6.4400 | 6.7955 | 6.0932 | 6.6399 | 6.0728 | 6.1249 | 6.0498 | 6.2771 | 6.3747 | 6.6136 | 6.1135 | 6.3801 | 5.6996 | 5.8755 | 6.1731 | 6.2768 |
| MYOF_S731 | MYOF | Q9NZM1 | | 729 | RKLRSRSLSQIHE | 6.1883 | 6.4785 | 6.6295 | 6.5555 | 5.8844 | 6.0547 | 6.3686 | 6.2613 | 6.3583 | 6.3167 | 6.4764 | 6.6411 | 5.7593 | 6.0599 | 5.7997 | 6.1677 |
| MYOF_S963 | MYOF | Q9NZM1 | | 963 | NGDKAASPSELTC | 6.4280 | 6.6076 | 6.9268 | 6.8670 | 5.8772 | 5.8025 | 6.7311 | 6.2360 | 6.2551 | 7.3856 | 6.6076 | 6.3039 | 5.4395 | 5.7599 | 5.6753 | 5.0968 |
| MYOF_T1781 | MYOF | Q9NZM1 | | 193 | NSRRMLSNKPQDF | 6.8648 | 5.9030 | 6.7166 | 5.7560 | 6.2510 | 6.8746 | 7.5399 | 5.9386 | 6.4066 | 5.5857 | 5.7756 | 6.6823 | 4.9155 | 7.1687 | 5.6874 | 5.9337 |
| MYPN_S418 | MYPN | Q86TC9 | | 644 | KTPEPSSPVKEPP | 6.2135 | 5.8451 | 6.2775 | 5.5783 | 6.5246 | 6.6115 | 6.2954 | 6.1372 | 6.5746 | 5.8987 | 6.2204 | 6.2337 | 6.5533 | 6.4553 | 6.4187 | 6.1622 |
| MYPN_S644 | MYPN | Q86TC9 | | 418 | TIQQCQSPTNYLQ | 7.1522 | 7.4390 | 6.9236 | 7.5203 | 6.6144 | 5.8387 | 6.9316 | 7.1417 | 6.8292 | 6.6184 | 7.9861 | 7.7621 | 4.0198 | 4.2841 | 3.2435 | 3.6954 |
| MYSM1_S110 | MYSM1 | Q5VVJ2 | | 110 | AKIMVHSPTKPAS | 6.3968 | 6.7383 | 6.4139 | 6.4425 | 6.3692 | 6.2092 | 6.0790 | 5.9949 | 6.7124 | 6.4931 | 6.0898 | 6.2965 | 6.0616 | 5.7535 | 6.0671 | 5.8821 |
| MYSM1_S218 | MYSM1 | Q5VVJ2 | | 218 | VKIEKLSDDEEVD | 6.1425 | 5.8065 | 5.8634 | 6.0479 | 6.3076 | 5.8611 | 6.2413 | 5.7716 | 6.3155 | 6.1510 | 6.3535 | 6.1011 | 6.7198 | 6.9575 | 6.7993 | 6.5604 |
| MYSM1_S340 | MYSM1 | Q5VVJ2 | | 340 | DARQLPSPEPCEI | 6.2367 | 6.2015 | 6.3100 | 6.6323 | 6.1472 | 6.4958 | 6.1127 | 6.3537 | 6.4435 | 6.4260 | 6.2557 | 6.7762 | 5.6231 | 5.9707 | 5.9770 | 6.0377 |
| MYSM1_S41 | MYSM1 | Q5VVJ2 | | 41 | DHYLDSSWRTENG | 6.9167 | 7.3529 | 7.3820 | 6.8368 | 6.8877 | 5.6046 | 5.8372 | 7.5128 | 6.1825 | 6.0735 | 6.3461 | 7.6037 | 4.2416 | 6.0008 | 4.9904 | 4.2307 |
| MZF1_S111 | MZF1 | P28698 | | 111 | QGQRPGSPEEAAA | 5.8695 | 6.9631 | 6.9161 | 4.4897 | 6.3523 | 5.8439 | 4.3530 | 7.3732 | 7.3690 | 5.3142 | 5.9123 | 5.5705 | 8.8556 | 6.7709 | 6.2497 | 5.7969 |
| N4BP1_S258 | N4BP1 | O75113 | | 242 | ARNKAGTPVSELT | 5.4973 | 5.5104 | 6.5964 | 5.5232 | 6.2263 | 5.9946 | 6.0989 | 6.1361 | 6.4955 | 6.5948 | 6.4147 | 6.5658 | 6.8657 | 6.7523 | 6.2714 | 6.4566 |
| N4BP1_S288 | N4BP1 | O75113 | | 258 | DTVLSSSPDVLFD | 6.3222 | 6.3424 | 6.3497 | 6.5915 | 6.4430 | 6.1044 | 6.3108 | 6.4533 | 6.5371 | 6.3268 | 6.9454 | 6.9115 | 5.3323 | 5.8941 | 5.4896 | 5.6461 |
| N4BP1_S335 | N4BP1 | O75113 | | 335 | ADSENLSPDIKET | 5.8240 | 6.1351 | 5.9113 | 5.6668 | 5.9016 | 6.2591 | 6.0707 | 5.7506 | 6.9470 | 5.9667 | 5.8033 | 5.9771 | 7.0745 | 7.1708 | 7.0128 | 6.5286 |
| N4BP1_T242 | N4BP1 | O75113 | | 288 | CQKRRFSDSEERH | 5.5656 | 6.2231 | 5.7181 | 5.7062 | 6.4106 | 6.4954 | 5.8208 | 6.9200 | 6.0781 | 6.5073 | 6.5422 | 6.8315 | 6.1272 | 6.9163 | 6.3518 | 5.7858 |
| N4BP2_S607 | N4BP2 | Q86UW6 | | 1210 | AEMRAVTPENHES | 6.4226 | 6.6108 | 6.1470 | 6.3567 | 6.9530 | 7.1007 | 5.4697 | 6.2630 | 6.1303 | 6.7338 | 5.9477 | 6.3741 | 6.2717 | 5.8087 | 5.8286 | 5.5816 |
| N4BP2_S626 | N4BP2 | Q86UW6 | | 724 | SSMERVSPSTCCS | 5.8324 | 5.4791 | 5.8049 | 5.9521 | 6.3494 | 5.9479 | 6.2372 | 6.0327 | 7.1691 | 5.9762 | 6.6288 | 5.9080 | 6.6038 | 7.1633 | 6.6371 | 6.2780 |
| N4BP2_S724 | N4BP2 | Q86UW6 | | 626 | ENILSLSLKHLEF | 7.2956 | 5.6994 | 6.7295 | 5.8482 | 8.7810 | 6.4464 | 6.4639 | 6.5569 | 8.1507 | 6.2392 | 7.2840 | 5.5826 | 6.0962 | 4.9231 | 4.0564 | 4.5466 |

| | | | | | | | | | | | | | | | | | | | | | |
|---|---|---|---|---|---|---|---|---|---|---|---|---|---|---|---|---|---|---|---|---|---|
| N4BP2_S751 | N4BP2 | Q86UW6 | 751 | | LQNEKSSPGEIVE | 6.9222 | 6.9855 | 6.9606 | 6.6211 | 6.2698 | 6.6887 | 6.4028 | 6.5079 | 6.7736 | 6.7177 | 6.9158 | 6.5960 | 4.6176 | 5.2001 | 4.8120 | 5.0089 |
| N4BP2_T1210 | N4BP2 | Q86UW6 | 607 | | CEDRSTSPRDDED | 6.2505 | 5.9535 | 6.5780 | 6.0506 | 6.4531 | 6.5415 | 6.1942 | 6.5148 | 6.4935 | 6.5105 | 6.1501 | 6.4038 | 6.2815 | 6.0849 | 5.9862 | 5.5534 |
| NAA10_S186 | NAA10 | P41227 | 186 | | VESKGNSPPSSGE | 6.5337 | 6.4032 | 6.8562 | 6.0420 | 6.1883 | 6.2755 | 5.7500 | 6.9144 | 6.6715 | 6.5294 | 6.6450 | 6.9272 | 5.5813 | 5.7992 | 5.3986 | 5.4843 |
| NAA10_S205 | NAA10 | P41227 | 205 | | GLAAEDSGGDSKD | 6.4024 | 6.0357 | 6.7652 | 5.4826 | 6.7977 | 6.6084 | 6.0946 | 6.8445 | 6.1194 | 6.0454 | 5.9920 | 5.9544 | 6.3793 | 6.5183 | 6.0916 | 5.8685 |
| NAA15_S855 | NAA15 | Q9BXJ9 | 855 | | ITVNGDSSAEAEE | 5.8928 | 6.1856 | 6.4784 | 6.2105 | 6.1809 | 6.6097 | 6.3127 | 6.2262 | 6.2969 | 6.5266 | 6.2765 | 6.5530 | 6.0658 | 6.4484 | 5.8890 | 5.8471 |
| NAA30_S39 | NAA30 | Q147X3 | 89 | | QLNGLISPELRHL | 6.1464 | 6.5186 | 6.5378 | 7.0634 | 6.6093 | 6.6615 | 6.7885 | 6.8044 | 7.2763 | 6.7856 | 6.8995 | 6.7159 | 4.7288 | 5.3324 | 4.6867 | 4.4451 |
| NAA30_S39;S55 | NAA30 | Q147X3 | 55 | | EGGGSRSPAGGES | 6.2444 | 6.4499 | 6.6849 | 6.2901 | 6.7508 | 5.9351 | 6.0132 | 7.6426 | 6.7089 | 7.0025 | 6.5723 | 6.0162 | 5.2676 | 5.7744 | 5.4362 | 5.2111 |
| NAA30_S55 | NAA30 | Q147X3 | 39 | | AALACCSEDEEDD | 5.2339 | 5.7335 | 6.2007 | 5.5982 | 6.6191 | 6.6563 | 6.1891 | 6.0646 | 6.8705 | 6.3817 | 6.2579 | 6.8976 | 6.0932 | 6.7166 | 6.3515 | 6.1357 |
| NAA30_S89 | NAA30 | Q147X3 | 117 | | VAATTATPDGGPR | 6.1312 | 6.2431 | 6.2081 | 6.6894 | 5.5271 | 6.1147 | 6.6902 | 5.8702 | 6.3926 | 6.1669 | 6.0370 | 6.4507 | 6.2109 | 6.7224 | 6.1183 | 6.4272 |
| NAA30_T117 | NAA30 | Q147X3 | | 39;55 | NA | 6.2546 | 6.3448 | 6.4951 | 6.0959 | 6.2761 | 7.0225 | 6.3649 | 6.3647 | 6.8644 | 7.5173 | 7.0541 | 6.8457 | 4.5103 | 5.3320 | 5.5481 | 5.1093 |
| NAA50_T12 | NAA50 | Q9GZZ1 | 12 | | IELGDVTPHNIKQ | 6.8474 | 7.1638 | 7.7572 | 7.2340 | 6.8118 | 6.5504 | 7.1368 | 7.2244 | 6.9391 | 7.5311 | 7.6784 | 7.5043 | 3.2623 | 3.8894 | 2.7777 | 3.6920 |
| NAA80_T29 | NAA80 | Q93015-2 | 29 | | MELILSTSPAELT | 6.1610 | 6.7344 | 6.7733 | 6.6521 | 6.6026 | 6.6223 | 5.7472 | 7.0443 | 6.1994 | 6.5852 | 7.2136 | 6.8640 | 5.0936 | 5.5774 | 5.0305 | 5.0991 |
| NAB1_S395 | NAB1 | Q13506 | 395 | | HAERRLSAGLYRQ | 5.1147 | 5.4336 | 5.2840 | 5.5978 | 6.5107 | 5.1157 | 5.5537 | 6.3599 | 6.2498 | 7.2207 | 7.0392 | 7.0346 | 7.0266 | 7.1783 | 6.6981 | 6.5464 |
| NAB2_S159;S162 | NAB2 | Q15742 | 171 | | ELGEKLSPLPGGP | 5.2098 | 6.2478 | 6.3717 | 5.4302 | 5.3697 | 5.2820 | 5.8321 | 5.8793 | 6.1030 | 5.7013 | 5.8331 | 5.7299 | 7.6630 | 7.6977 | 7.7166 | 7.9329 |
| NAB2_S171 | NAB2 | Q15742 | 434 | | GSCPRLTPPPADL | 5.8086 | 6.3031 | 6.0035 | 5.4171 | 5.4208 | 6.0999 | 6.7093 | 5.8449 | 6.0977 | 6.4639 | 7.0096 | 6.5833 | 6.0784 | 7.0889 | 6.6545 | 6.4165 |
| NAB2_T434 | NAB2 | Q15742 | | 159;162 | NA | 4.9998 | 5.4783 | 6.0017 | 5.1427 | 5.2512 | 5.7429 | 5.6023 | 5.7623 | 5.5586 | 6.0599 | 6.1423 | 6.0357 | 8.0512 | 8.5553 | 8.3369 | 7.2790 |
| NACA_S1304 | NACA | E9PAV3 | 2024 | | ENTQTPTVQEESE | 6.1981 | 6.2390 | 6.5627 | 6.4380 | 6.2043 | 6.2846 | 6.0867 | 6.3817 | 6.5785 | 6.6649 | 6.7104 | 6.6896 | 5.7443 | 5.9525 | 5.5241 | 5.7409 |
| NACA_S2029 | NACA | E9PAV3 | 2022 | | IQENTQTPTVQEE | 5.3414 | 5.9044 | 7.1340 | 6.5627 | 6.7689 | 6.3750 | 6.1867 | 5.9052 | 6.2609 | 6.9077 | 6.3651 | 6.7748 | 6.5290 | 5.7458 | 5.7932 | 6.3452 |
| NACA_T2020;T2022 | NACA | E9PAV3 | 2029 | | PTVQEESEEEEVD | 6.5317 | 5.9775 | 6.6222 | 6.3568 | 6.5657 | 6.1995 | 6.4592 | 6.7631 | 6.3638 | 6.2454 | 7.0023 | 7.1039 | 6.5564 | 5.5442 | 5.3435 | 5.2648 |
| NACA_T2022 | NACA | E9PAV3 | 1304 | | KGGPATSPPKGAP | 6.4431 | 6.7511 | 6.4233 | 6.1783 | 6.2643 | 6.2781 | 5.6464 | 6.3385 | 6.1916 | 6.7900 | 6.5305 | 6.3007 | 5.7748 | 6.3451 | 5.9313 | 5.8128 |
| NACA_T2024 | NACA | E9PAV3 | | 2020;2022 | NA | 6.3708 | 5.6788 | 7.6421 | 6.5721 | 5.7742 | 6.6321 | 5.8479 | 6.6545 | 6.6451 | 6.5013 | 8.0742 | 6.1627 | 5.6310 | 5.3168 | 5.5046 | 4.9917 |
| NACC1_S141 | NACC1 | Q96RE7 | 188 | | AKRLWDSGQKEAG | 7.3503 | 7.1511 | 7.0284 | 7.7223 | 5.1680 | 6.2453 | 7.3108 | 7.0452 | 7.6518 | 7.6877 | 8.1211 | 8.4121 | 2.9039 | 3.6501 | 2.8366 | 3.7152 |
| NACC1_S188 | NACC1 | Q96RE7 | 141 | | AEEAPSSEPQSPV | 6.0776 | 3.9774 | 3.8210 | 6.1724 | 7.6041 | 7.1964 | 4.7644 | 7.0020 | 7.3860 | 5.2242 | 6.8266 | 6.8835 | 5.5656 | 7.7889 | 6.6370 | 7.0731 |
| NAF1_S315 | NAF1 | Q96HR8 | 315 | | PEALDFSDDEKEK | 5.8107 | 6.0500 | 6.9202 | 6.4480 | 5.8627 | 6.1723 | 6.3219 | 6.8991 | 5.3476 | 6.3102 | 5.7365 | 7.1817 | 5.9746 | 6.5976 | 6.5846 | 6.7824 |
| NAF1_S50 | NAF1 | Q96HR8 | 50 | | LQSFEGSPDAGQT | 5.6658 | 6.0167 | 6.6198 | 6.1658 | 5.8637 | 5.2400 | 5.5019 | 5.7732 | 6.2622 | 6.6124 | 5.8165 | 6.3985 | 6.5811 | 7.0741 | 6.9996 | 7.4086 |
| NAF1_S98 | NAF1 | Q96HR8 | 40 | | SAPVPGTQPPLQS | 5.8977 | 5.8591 | 7.1547 | 5.9704 | 5.8414 | 5.6861 | 6.1884 | 6.1575 | 6.5744 | 6.2833 | 5.4990 | 6.5040 | 5.8914 | 6.6747 | 6.1702 | 7.6479 |
| NAF1_T40 | NAF1 | Q96HR8 | 98 | | CGDCVTSPGAAEP | 6.0104 | 4.2967 | 5.9340 | 6.7351 | 6.1384 | 6.3717 | 6.1322 | 6.1941 | 6.3304 | 7.8789 | 7.6229 | 6.6835 | 5.0152 | 6.1342 | 5.5334 | 6.9890 |
| NAGK_S122 | NAGK | Q9UJ70-2 | 122 | | SLGLSLSGGDQED | 6.0894 | 6.0845 | 6.4186 | 6.1063 | 6.2351 | 6.1717 | 6.5328 | 6.4037 | 6.5702 | 6.2564 | 6.6192 | 6.6497 | 5.7414 | 6.1738 | 5.7886 | 6.1584 |
| NAMPT_S398 | NAMPT | P43490 | 398 | | RDLLNCSFKCSYV | 6.9904 | 6.6623 | 7.1408 | 5.3989 | 6.1930 | 5.6723 | 7.0121 | 6.6282 | 6.0198 | 7.6877 | 6.4971 | 5.1232 | 5.4946 | 5.5379 | 6.1690 | 5.7726 |
| NANS_S248 | NANS | Q9NR45 | 275 | | VERALGSPTKQLL | 6.0296 | 5.9614 | 6.2260 | 5.5585 | 6.2974 | 6.5450 | 5.8062 | 6.8151 | 6.2752 | 6.6086 | 6.4701 | 6.5539 | 5.6026 | 6.6955 | 6.3888 | 6.1661 |
| NANS_S275 | NANS | Q9NR45 | 248 | | DKTWKGSDHSASL | 4.5040 | 6.8672 | 6.6625 | 4.6820 | 7.0720 | 8.2870 | 4.6108 | 5.1271 | 7.6950 | 7.7485 | 7.7752 | 5.4787 | 4.8645 | 5.3140 | 8.0689 | 5.2428 |
| NAP1L1_S10 | NAP1L1 | P55209 | 62 | | LDGLVETPTGYIE | 6.7517 | 6.4197 | 6.8369 | 6.7062 | 6.8547 | 6.7453 | 6.2706 | 6.9798 | 6.5819 | 6.3721 | 6.4754 | 6.6813 | 5.5815 | 4.9788 | 4.8168 | 4.9472 |
| NAP1L1_S143 | NAP1L1 | P55209 | 69 | | PTGYIESLPRVVK | 6.4160 | 5.9909 | 6.9676 | 6.6695 | 6.7984 | 6.1079 | 6.1513 | 6.6332 | 6.4395 | 6.4565 | 6.1788 | 6.4204 | 5.5643 | 6.0497 | 5.5062 | 5.6500 |
| NAP1L1_S69 | NAP1L1 | P55209 | 10 | | IDNKEQSELDQDL | 7.1276 | 6.6789 | 6.9369 | 6.4178 | 6.4742 | 6.5284 | 6.5466 | 6.7443 | 7.0634 | 6.1795 | 7.1365 | 6.7949 | 5.2893 | 4.6309 | 4.4124 | 5.0383 |
| NAP1L1_T62 | NAP1L1 | P55209 | 143 | | DEEDEISEELKEK | 5.9314 | 5.2494 | 6.7382 | 5.6576 | 6.2195 | 6.5077 | 6.2231 | 5.6306 | 6.5221 | 6.6134 | 5.8383 | 7.1170 | 7.4472 | 6.1391 | 5.7591 | 6.4062 |
| NAP1L4_S125 | NAP1L4 | Q99733-2 | 125 | | AESEWHSENEEEE | 6.1533 | 6.4763 | 6.8830 | 6.1580 | 6.5621 | 6.6745 | 5.8977 | 6.5216 | 6.3623 | 6.3605 | 6.6975 | 6.5404 | 5.9661 | 5.8168 | 5.4396 | 5.4902 |
| NAP1L4_T51 | NAP1L4 | Q99733-2 | 51 | | LDNVPHTPSSYIE | 7.0683 | 7.8377 | 7.0340 | 7.4753 | 6.4521 | 7.0154 | 7.0108 | 7.8967 | 7.0082 | 6.9082 | 7.1507 | 6.1965 | 3.7825 | 4.0383 | 3.3008 | 3.8246 |
| NAPG_T287 | NAPG | Q99747 | 287 | | KKKSPATPQAKPD | 6.3613 | 6.1915 | 6.5546 | 6.5259 | 6.6321 | 6.3853 | 6.2280 | 6.4886 | 6.7351 | 7.1372 | 7.2985 | 7.2092 | 5.4566 | 5.1595 | 5.0112 | 5.5253 |
| NARF_S29 | NARF | Q9UHQ1-2 | 29 | | VSADAPSPAQENG | 5.9294 | 6.8782 | 5.3982 | 7.4594 | 5.5707 | 6.3561 | 5.9793 | 5.8567 | 6.4787 | 6.3516 | 7.1733 | 7.3913 | 5.0894 | 7.1189 | 6.2063 | 4.7626 |
| NASP_S141 | NASP | P49321-3 | 423 | | RAKLVPSQEETKL | 5.7925 | 6.3207 | 6.1534 | 5.7852 | 5.6653 | 6.0180 | 5.6823 | 6.0016 | 6.2163 | 6.1765 | 6.5204 | 6.6129 | 6.7646 | 6.6612 | 6.7383 | 6.8908 |
| NASP_S191 | NASP | P49321-2 | 141 | | IDETEGSEEDDKE | 4.8806 | 4.8725 | 5.7938 | 4.5882 | 5.3035 | 5.5904 | 5.1342 | 5.1926 | 6.2576 | 6.3435 | 6.2112 | 6.2826 | 8.7890 | 8.5628 | 7.9957 | 8.2018 |
| NASP_S346 | NASP | P49321-3 | 125 | | EEEGEKTEDESLV | 5.7883 | 5.9722 | 6.3435 | 5.5254 | 5.8066 | 6.3179 | 6.0762 | 6.1498 | 6.3072 | 6.3416 | 6.5375 | 6.5416 | 7.0317 | 6.5498 | 6.1093 | 6.6012 |
| NASP_S423 | NASP | P49321-3 | 499 | | EEMPNDSVLENKS | 6.4465 | 6.6646 | 7.0609 | 7.1772 | 6.5064 | 6.5228 | 6.0139 | 6.4068 | 6.6627 | 5.4864 | 7.2103 | 6.1962 | 5.2244 | 5.9256 | 5.1239 | 5.3714 |
| NASP_S499 | NASP | P49321-3 | 322 | | PEEKVVTSENEAG | 5.6146 | 6.0162 | 6.3543 | 6.0706 | 6.3654 | 6.2623 | 6.3522 | 6.3818 | 5.5227 | 5.7310 | 5.6999 | 5.9040 | 6.6880 | 6.9737 | 6.9108 | 7.1526 |
| NASP_S505 | NASP | P49321-3 | 346 | | VPPAEESPEVTTE | 5.7298 | 5.8889 | 5.9621 | 5.8857 | 5.9515 | 6.0854 | 5.5923 | 6.3781 | 5.4560 | 6.1087 | 6.3643 | 6.6294 | 6.9394 | 6.8598 | 7.3049 | 6.8637 |
| NASP_S664 | NASP | P49321-3 | 505 | | SVLENKSLQENEE | 5.0897 | 6.1569 | 5.8811 | 5.4540 | 5.7630 | 5.9197 | 5.2447 | 6.2043 | 5.7875 | 6.1623 | 6.3654 | 6.2354 | 7.6387 | 7.5181 | 7.2128 | 7.3663 |
| NASP_S728 | NASP | P49321-3 | 728 | | RKPEEESPRKDDA | 5.5336 | 6.1339 | 6.0356 | 6.5715 | 6.4486 | 6.0356 | 6.5738 | 6.7152 | 6.4024 | 6.8778 | 6.5809 | 6.5144 | 6.9497 | 6.7101 | 7.1270 | 6.8900 |
| NASP_T125 | NASP | P49321-3 | 191 | | REDMDISKSAEEP | 6.5512 | 6.7300 | 6.4445 | 6.7413 | 7.1539 | 7.0510 | 6.9315 | 7.4759 | 7.0559 | 6.5109 | 6.9420 | 6.9627 | 4.1574 | 4.8298 | 3.7854 | 4.6766 |
| NASP_T125;S129 | NASP | P49321-3 | 685 | | SSTSGFTPGGGGS | 6.3089 | 6.7485 | 6.6768 | 6.6578 | 6.5752 | 6.9556 | 6.6278 | 6.8458 | 6.8236 | 6.5550 | 6.6500 | 6.9248 | 5.0763 | 5.0771 | 5.0242 | 5.1725 |
| NASP_T129 | NASP | P49321-3 | 664 | | AKESQRSGNVAEL | 6.0487 | 7.3324 | 6.1015 | 5.0224 | 4.7716 | 5.8837 | 6.4216 | 7.3522 | 5.0323 | 6.1807 | 4.8904 | 5.7385 | 7.6459 | 7.4083 | 6.6889 | 7.4809 |
| NASP_T322 | NASP | P49321-3 | 129 | | EKTEDESLVENND | 6.2137 | 6.3273 | 7.2834 | 6.3938 | 6.4043 | 7.2135 | 5.8432 | 5.4097 | 5.4184 | 6.9181 | 5.6824 | 6.3623 | 5.9603 | 6.4899 | 5.9079 | 6.1718 |
| NASP_T479;S482 | NASP | P49321-3 | | 125;129 | NA | 5.3843 | 6.3066 | 7.1471 | 6.5470 | 6.2394 | 5.9077 | 6.4102 | 6.8483 | 6.7297 | 6.3182 | 6.6951 | 6.4810 | 6.2113 | 5.9834 | 5.0888 | 5.7018 |
| NASP_T685 | NASP | P49321-3 | | 479;482 | NA | 5.2297 | 5.1737 | 6.3991 | 5.9408 | 5.6391 | 5.5278 | 5.8617 | 5.7820 | 6.1323 | 6.0405 | 6.2510 | 6.5383 | 7.6967 | 7.4165 | 6.6638 | 7.7071 |
| NAT10_S987 | NAT10 | Q9H0A0 | 987 | | PNASIISLKSDKK | 6.0885 | 6.6449 | 6.8456 | 6.9626 | 6.0657 | 6.7620 | 7.2280 | 7.0422 | 6.5607 | 6.3902 | 7.2032 | 6.7705 | 4.5842 | 5.5890 | 4.3567 | 5.4043 |
| NAV1_S1253 | NAV1 | Q8NEY1 | 391 | | LKSGYMSDSDLMG | 6.3754 | 6.8852 | 7.3055 | 6.4480 | 6.1394 | 7.0374 | 6.4770 | 6.9749 | 5.7407 | 6.3164 | 5.9499 | 6.1570 | 5.6167 | 5.8023 | 5.1628 | 5.6114 |
| NAV1_S142 | NAV1 | Q8NEY1 | 452 | | TASRRNSTIVLRT | 6.0601 | 5.6344 | 6.5010 | 5.4705 | 6.0819 | 6.1053 | 5.5296 | 6.0822 | 7.3844 | 7.4912 | 7.2025 | 7.3241 | 6.0270 | 7.0932 | 4.7270 | 5.2856 |
| NAV1_S1862 | NAV1 | Q8NEY1 | 142 | | SKTLSKSEHSLFQ | 7.0196 | 7.0069 | 7.4288 | 6.1040 | 5.3730 | 7.0736 | 6.7780 | 6.3668 | 6.4294 | 7.2143 | 7.3819 | 7.6251 | 4.4547 | 4.9521 | 4.0355 | 4.7562 |
| NAV1_S312 | NAV1 | Q8NEY1 | 760 | | SLKSIGSPESTPK | 6.1564 | 6.4663 | 6.7993 | 6.4479 | 6.0161 | 6.7637 | 6.3281 | 6.6098 | 6.3378 | 6.9201 | 7.0473 | 7.2292 | 4.9705 | 5.3880 | 5.0009 | 5.5187 |
| NAV1_S322 | NAV1 | Q8NEY1 | 90 | | NLRKQKSLTNLSF | 6.5294 | 6.1132 | 6.9814 | 5.7320 | 5.3659 | 7.2069 | 6.1667 | 6.1971 | 6.8820 | 6.8729 | 6.9976 | 7.9330 | 5.1068 | 5.8145 | 4.3575 | 5.7430 |
| NAV1_S391 | NAV1 | Q8NEY1 | 312 | | SLSNRSSPLSWRY | 5.9584 | 5.9941 | 6.4982 | 6.0079 | 6.4593 | 6.4945 | 6.0726 | 6.7860 | 6.5120 | 6.0796 | 6.1305 | 6.4293 | 6.1091 | 6.3751 | 6.2810 | 5.8125 |
| NAV1_S452 | NAV1 | Q8NEY1 | 672 | | SNIQYRSLPRPAK | 6.6629 | 7.0698 | 6.3845 | 6.3043 | 5.8444 | 6.2746 | 5.9277 | 6.2390 | 6.6325 | 7.1078 | 7.0732 | 7.7606 | 4.8054 | 5.9543 | 4.7081 | 5.2508 |
| NAV1_S648 | NAV1 | Q8NEY1 | 884 | | LSGLHRSMESLQM | 4.8392 | 4.7596 | 6.7166 | 5.6187 | 6.5711 | 5.8328 | 6.4614 | 7.4632 | 5.7532 | 6.7441 | 6.8237 | 5.8794 | 7.6389 | 6.1265 | 6.2912 | 6.4806 |
| NAV1_S672 | NAV1 | Q8NEY1 | 1253 | | DSSAPSSPKLQHG | 6.2950 | 4.7489 | 7.0121 | 5.7951 | 5.8200 | 4.4233 | 6.0406 | 6.4943 | 7.1740 | 5.6011 | 6.8946 | 6.2381 | 7.0459 | 6.4018 | 7.0548 | 6.9605 |
| NAV1_S69 | NAV1 | Q8NEY1 | 1862 | | AANYIESPDRETI | 6.3108 | 6.1527 | 6.3745 | 6.8798 | 5.7819 | 7.2660 | 6.1681 | 7.1606 | 6.1663 | 7.0542 | 7.3078 | 6.8589 | 4.3287 | 4.9503 | 5.2584 | 5.9809 |
| NAV1_S760 | NAV1 | Q8NEY1 | 322 | | WRYGQSSPRLQAG | 5.7845 | 6.1952 | 6.1611 | 6.4296 | 5.9686 | 6.3964 | 6.9321 | 5.8818 | 6.1710 | 6.7346 | 6.3746 | 6.4829 | 5.8946 | 6.4422 | 5.6738 | 6.4771 |
| NAV1_S884 | NAV1 | Q8NEY1 | 648 | | VNGRKTSLDVSNS | 6.3839 | 5.3465 | 6.9362 | 5.5481 | 6.7126 | 6.4049 | 5.8369 | 6.1697 | 6.0542 | 7.6849 | 6.6412 | 6.4459 | 5.7445 | 6.7661 | 5.5586 | 5.7655 |
| NAV1_S90 | NAV1 | Q8NEY1 | 69 | | ELKVFKSGSVDSR | 6.2230 | 6.6809 | 6.2707 | 6.4942 | 6.3914 | 5.9198 | 6.2416 | 5.6619 | 6.3970 | 6.1430 | 5.5821 | 5.8063 | 6.1065 | 6.6856 | 6.5995 | 6.7965 |
| NAV3_S1113 | NAV3 | Q8IVL0 | 1245 | | KYPDIASPTFRRL | 6.2827 | 6.2299 | 6.7167 | 6.1914 | 6.4663 | 6.5686 | 6.5699 | 6.8129 | 6.5354 | 6.1861 | 6.3590 | 6.4138 | 5.5539 | 5.4900 | 5.7126 | 5.9110 |
| NAV3_S1170 | NAV3 | Q8IVL0 | 275 | | LNRRSQSFNSIDK | 6.4680 | 5.8442 | 6.6164 | 5.7655 | 5.9596 | 6.4946 | 6.0537 | 6.7800 | 6.6940 | 5.8470 | 6.1785 | 6.5988 | 6.1343 | 6.5352 | 5.8607 | 6.1695 |
| NAV3_S1245 | NAV3 | Q8IVL0 | 1170 | | SNVSSKSAGATTS | 6.2732 | 6.0492 | 7.1112 | 6.4488 | 6.1358 | 6.4890 | 5.8700 | 6.2070 | 6.7104 | 6.1603 | 6.4990 | 6.7819 | 5.7521 | 6.0231 | 5.5111 | 5.9779 |
| NAV3_S275 | NAV3 | Q8IVL0 | 542 | | TVKQTISPGSTAS | 6.9216 | 7.5467 | 7.4166 | 7.5339 | 6.3495 | 6.9900 | 7.2759 | 6.7805 | 6.4286 | 7.4250 | 7.3687 | 7.6914 | 3.1332 | 3.4747 | 3.1982 | 4.4654 |
| NAV3_S542 | NAV3 | Q8IVL0 | 1113 | | NAGRKTSLDGSQN | 6.0993 | 6.0207 | 5.0494 | 6.1387 | 7.5701 | 7.8257 | 5.1241 | 7.7864 | 7.2752 | 4.0269 | 5.9932 | 5.1359 | 6.5673 | 8.4235 | 6.4415 | 4.5224 |
| NBAS_S473 | NBAS | A2RRP1 | 473 | | DEALVDSDSDYEI | 6.3463 | 6.6059 | 5.5057 | 6.2815 | 6.3774 | 5.9198 | 6.0630 | 5.6458 | 6.7634 | 6.5508 | 6.0231 | 5.8071 | 7.1339 | 6.8051 | 6.9726 | 6.0985 |
| NBEAL1_S734 | NBEAL1 | Q6ZS30 | 734 | | PDPPFSSPITPHR | 6.7633 | 6.8860 | 6.7874 | 6.4601 | 5.9392 | 6.1308 | 6.4340 | 6.4173 | 6.1569 | 6.5233 | 6.3410 | 5.6156 | 4.7302 | 6.0248 | 6.2443 | 5.6456 |
| NBEAL2_S1312 | NBEAL2 | Q6ZNJ1 | 1312 | | SPESPTSPKPAPP | 7.1394 | 6.0511 | 6.1907 | 7.2424 | 5.8383 | 6.2056 | 6.5593 | 6.9971 | 7.3711 | 7.3481 | 7.5934 | 7.3372 | 3.5480 | 5.1052 | 3.9546 | 5.5185 |
| NBEAL2_T1298;S1309 | NBEAL2 | Q6ZNJ1 | | 1298;1309 | NA | 5.7116 | 5.5105 | 6.2746 | 6.5716 | 6.2527 | 6.4029 | 6.3495 | 6.6214 | 6.6451 | 6.5942 | 6.5624 | 6.9422 | 6.5593 | 5.8838 | 6.2155 | 5.8028 |
| NBN_S343 | NBN | O60934 | 432 | | IPNYQLSPTKLPS | 6.7122 | 6.5569 | 6.1075 | 6.6491 | 6.7479 | 6.0299 | 6.4926 | 6.2978 | 6.6687 | 6.0030 | 5.8601 | 5.7384 | 6.3122 | 5.9909 | 5.9920 | 5.8408 |
| NBN_S397 | NBN | O60934 | 397 | | QKFRMLSQDAPTV | 6.2587 | 6.7424 | 6.3194 | 6.9119 | 5.9637 | 6.6398 | 5.3907 | 5.6642 | 6.3001 | 6.4751 | 6.1939 | 6.2680 | 6.0382 | 6.7321 | 6.6128 | 6.4891 |
| NBN_S432 | NBN | O60934 | 486 | | SARIETSCSLLEQ | 7.3052 | 7.4561 | 7.2059 | 7.0123 | 6.3613 | 6.6047 | 6.6429 | 6.3438 | 6.8098 | 6.2347 | 6.1717 | 6.3029 | 4.8774 | 5.2757 | 4.3334 | 5.0621 |
| NBN_S486 | NBN | O60934 | 343 | | TPGPSLSQGVSVD | 5.7632 | 6.0960 | 6.3429 | 6.9245 | 5.8713 | 5.5163 | 6.0386 | 5.9624 | 6.6072 | 6.0242 | 5.4478 | 6.1175 | 6.5935 | 6.8430 | 6.7353 | 7.1162 |
| NBN_S58 | NBN | O60934 | 58 | | FSVTNLSQTDEIP | 5.5428 | 5.9437 | 5.8519 | 6.8964 | 5.7259 | 5.6619 | 6.0528 | 5.7611 | 5.8597 | 6.4174 | 6.3723 | 6.3162 | 6.9426 | 6.9358 | 7.1299 | 6.5895 |
| NBR1_S625 | NBR1 | Q14596 | 625 | | LPDSMVSVKRKAE | 9.4520 | 7.7628 | 9.4158 | 8.2500 | 6.6289 | 6.2013 | 7.3586 | 5.9780 | 5.6120 | 6.8778 | 6.7119 | 6.1758 | 3.6123 | 4.5058 | 2.8805 | 2.5763 |
| NCAPD2_S1320 | NCAPD2 | Q15021 | 585 | | EKNPRESTGNMVT | 6.4189 | 7.2710 | 7.0697 | 7.6570 | 6.3793 | 6.8337 | 7.5651 | 6.3343 | 6.7983 | 7.3265 | 7.4479 | 6.9828 | 3.5958 | 4.3341 | 3.4687 | 4.5168 |
| NCAPD2_S1333 | NCAPD2 | Q15021 | 1389 | | ETPKKTTPILRAS | 7.4309 | 7.7553 | 6.9353 | 7.3418 | 6.8691 | 7.0928 | 7.3433 | 7.4699 | 7.2879 | 7.3498 | 7.4252 | 7.2792 | 2.6685 | 3.7366 | 2.6559 | 3.3586 |
| NCAPD2_S585 | NCAPD2 | Q15021 | 1333 | | PLASTASDNDFVT | 7.1054 | 7.2044 | 6.3232 | 6.8727 | 6.4965 | 7.6499 | 6.3018 | 6.4206 | 5.4338 | 5.9321 | 6.6318 | 6.1434 | 4.3645 | 5.6005 | 5.4404 | 6.0790 |
| NCAPD2_T1389 | NCAPD2 | Q15021 | 1320 | | PSAKKPSTGSRYQ | 6.8048 | 7.3779 | 6.4690 | 7.3904 | 6.5752 | 7.1601 | 7.5049 | 7.1418 | 6.6678 | 6.8597 | 6.9159 | 6.6386 | 3.6914 | 4.8088 | 3.5737 | 4.4200 |
| NCAPD3_S1348 | NCAPD3 | P42695 | 1415 | | TKRAISTPEKSIS | 7.2986 | 7.6124 | 7.3037 | 7.9728 | 6.4466 | 6.5174 | 6.8396 | 6.5078 | 7.6228 | 7.4708 | 7.3673 | 7.4519 | 3.1453 | 3.8536 | 2.9574 | 3.6319 |
| NCAPD3_S1348;S1357 | NCAPD3 | P42695 | 1384 | | FTLNSGSPEKTCS | 6.8874 | 7.2427 | 7.0177 | 6.7355 | 6.7137 | 6.5673 | 6.5282 | 5.9098 | 6.0745 | 6.1732 | 6.7768 | 6.1018 | 5.5265 | 5.5473 | 5.1477 | 5.0498 |
| NCAPD3_S1372 | NCAPD3 | P42695 | 1372 | | SRHRSRSLGVLPF | 5.7745 | 5.1994 | 6.1250 | 5.4629 | 6.6042 | 5.9520 | 5.6762 | 5.8845 | 6.8664 | 7.4895 | 7.4001 | 7.0207 | 7.1900 | 6.9257 | 4.8059 | 5.6231 |
| NCAPD3_S1384 | NCAPD3 | P42695 | 1481 | | PARNKDTPACSRR | 8.0538 | 8.5436 | 7.5696 | 8.2339 | 5.4096 | 7.1582 | 7.7935 | 6.0941 | 8.0901 | 6.7150 | 8.1726 | 7.7350 | 2.4087 | 3.3305 | 1.7250 | 2.9668 |
| NCAPD3_S531 | NCAPD3 | P42695 | 531 | | EINIDSSGETVGS | 5.6675 | 6.4966 | 6.3900 | 6.4103 | 5.9676 | 6.7836 | 6.2875 | 6.2272 | 7.1429 | 6.3254 | 6.7184 | 7.1976 | 5.4147 | 5.2259 | 5.2007 | 5.6441 |
| NCAPD3_T1415 | NCAPD3 | P42695 | 1348 | | PKARPMSLSTIAI | 4.9847 | 6.0996 | 5.6571 | 6.5302 | 6.3886 | 8.9724 | 8.8544 | 8.1996 | 7.0906 | 7.5861 | 7.8870 | 9.0196 | 3.9346 | 2.9967 | 2.4481 | 3.3506 |
| NCAPD3_T1481 | NCAPD3 | P42695 | | 1348;1357 | NA | 7.2859 | 6.8748 | 6.3971 | 8.6289 | 6.6442 | 6.1387 | 7.8563 | 6.4223 | 7.0539 | 6.8985 | 6.8504 | 5.1957 | 3.8123 | 6.3204 | 3.8089 | 4.8118 |
| NCAPG_S1015 | NCAPG | Q9BPX3 | 931 | | NKEVYMTPLRGVK | 8.1357 | 7.8128 | 7.1333 | 7.6949 | 6.4044 | 6.8942 | 7.6031 | 6.8013 | 7.6625 | 6.5999 | 6.9272 | 6.9402 | 3.0549 | 4.0674 | 2.7495 | 3.5186 |
| NCAPG_S390 | NCAPG | Q9BPX3 | 390 | | EHRGDFSYIGNLM | 7.2554 | 7.4883 | 6.7889 | 8.2037 | 7.1127 | 6.3493 | 7.0773 | 6.3585 | 6.6332 | 5.9996 | 6.3628 | 6.1476 | 4.2041 | 4.9651 | 4.3695 | 4.6840 |
| NCAPG_S674 | NCAPG | Q9BPX3 | 1015 | | FLNEDLSxxxxxx | 6.2185 | 6.8745 | 7.1903 | 6.2402 | 6.5135 | 6.5169 | 6.5397 | 6.3040 | 6.5864 | 6.1830 | 6.2525 | 6.6436 | 5.2394 | 6.0866 | 5.5098 | 5.7910 |
| NCAPG_T931 | NCAPG | Q9BPX3 | 674 | | EGTEINSDDEQES | 6.1945 | 6.6609 | 6.9293 | 6.7208 | 6.3053 | 6.6677 | 5.8778 | 6.5180 | 6.6341 | 6.1798 | 6.3577 | 6.8810 | 5.3202 | 5.6499 | 5.3074 | 5.7955 |
| NCAPH_S201 | NCAPH | Q15003 | 49 | | PLNIPGTPVLEDF | 6.6221 | 6.5546 | 6.7311 | 7.0996 | 6.2781 | 6.5681 | 6.7516 | 6.7044 | 7.0357 | 6.7778 | 7.0561 | 7.1943 | 4.5714 | 5.1499 | 4.0408 | 4.8643 |
| NCAPH_S27 | NCAPH | Q15003 | 81 | | LQFSTDSPRLLAS | 6.1885 | 6.2284 | 6.3333 | 6.7569 | 6.5498 | 6.1894 | 6.5239 | 5.8301 | 6.7785 | 6.9457 | 6.4966 | 6.8160 | 5.4817 | 5.7380 | 5.8545 | 5.9886 |
| NCAPH_S359 | NCAPH | Q15003 | 87 | | SPRLLASPSSRSI | 7.7680 | 8.3085 | 7.1090 | 7.7404 | 6.4843 | 6.8507 | 7.4358 | 7.3168 | 7.7711 | 7.5847 | 7.7735 | 7.6107 | 2.2859 | 3.0834 | 2.0529 | 2.8243 |
| NCAPH_S432 | NCAPH | Q15003 | 70 | | RLQRRRSRVFDLQ | 5.2091 | 5.4139 | 6.1193 | 5.6587 | 6.7663 | 6.5409 | 6.2521 | 6.1829 | 7.8737 | 8.0873 | 7.5117 | 7.5019 | 6.0583 | 5.7729 | 4.0436 | 5.0073 |
| NCAPH_S70 | NCAPH | Q15003 | 359 | | NAEVDESDCGDFP | 6.4151 | 6.3476 | 6.7416 | 5.9315 | 6.4534 | 5.9842 | 6.2003 | 6.5799 | 6.1814 | 7.0020 | 6.9223 | 7.0796 | 5.7145 | 5.8341 | 5.7930 | 5.8193 |
| NCAPH_S81 | NCAPH | Q15003 | 27 | | GHPHSASSPSERV | 7.0088 | 7.4588 | 6.8387 | 6.7851 | 6.8512 | 6.9015 | 6.4966 | 7.0350 | 6.9342 | 6.9748 | 7.2940 | 6.8315 | 3.8226 | 5.0661 | 3.6028 | 4.0980 |
| NCAPH_S87 | NCAPH | Q15003 | 432 | | GEYSYFSPRTMSM | 6.4209 | 6.3395 | 7.2291 | 7.4193 | 6.6968 | 6.9624 | 7.2352 | 6.5599 | 7.9256 | 7.1888 | 7.7857 | 7.9013 | 3.5604 | 4.1788 | 2.7580 | 3.8382 |
| NCAPH_T49 | NCAPH | Q15003 | 201 | | GHVADGSATEMGT | 5.7675 | 6.5975 | 6.3387 | 6.3481 | 6.1359 | 7.1884 | 6.7888 | 6.9933 | 7.1747 | 7.2010 | 6.7603 | 7.4938 | 4.1652 | 5.3547 | 4.9277 | 4.7643 |
| NCAPH2_S208 | NCAPH2 | Q6IBW4-4 | 284 | | EPKESRSPQQSAA | 6.2340 | 6.7066 | 6.3776 | 6.5837 | 6.3556 | 6.2213 | 6.0909 | 6.2010 | 6.2260 | 6.2707 | 6.4985 | 6.5655 | 5.8741 | 6.2588 | 5.7595 | 5.7761 |
| NCAPH2_S232 | NCAPH2 | Q6IBW4-4 | 208 | | MEPAGVSPMPGTQ | 6.2158 | 6.7143 | 6.5820 | 6.5321 | 6.0870 | 6.4595 | 6.0519 | 6.2146 | 6.2764 | 6.5268 | 6.4363 | 6.8543 | 5.8186 | 5.9650 | 5.6164 | 5.6490 |
| NCAPH2_S232;S246 | NCAPH2 | Q6IBW4-4 | 232 | | EVSVCRSPVPALG | 5.7280 | 5.9389 | 6.4204 | 6.6938 | 6.0314 | 6.0872 | 5.8710 | 6.3666 | 6.1985 | 6.6319 | 6.1169 | 6.2196 | 6.2679 | 6.4333 | 6.5199 | 6.4748 |
| NCAPH2_S246 | NCAPH2 | Q6IBW4-4 | 246 | | SQEPGPSPEGPMP | 5.9485 | 6.9435 | 6.8539 | 6.4360 | 5.9710 | 6.1051 | 6.3420 | 5.6295 | 6.5043 | 7.1647 | 6.2493 | 6.7418 | 6.0230 | 5.8131 | 5.7379 | 5.5362 |
| NCAPH2_S284 | NCAPH2 | Q6IBW4-4 | 325 | | SLDPFDSLESKPF | 5.7630 | 7.1940 | 6.4153 | 6.8997 | 6.1889 | 6.4212 | 6.4955 | 6.0766 | 6.4802 | 6.0865 | 5.7004 | 7.2589 | 5.7460 | 5.7402 | 5.8402 | 5.6934 |
| NCAPH2_S308 | NCAPH2 | Q6IBW4-4 | 308 | | GAPEPASCVKETP | 6.9149 | 6.9625 | 6.8289 | 7.5550 | 6.2875 | 6.3273 | 6.6370 | 5.9283 | 6.8022 | 7.0724 | 6.6451 | 6.2592 | 4.9026 | 5.4042 | 4.1255 | 5.3473 |
| NCAPH2_S325 | NCAPH2 | Q6IBW4-4 | | 232;246 | NA | 5.0525 | 6.1694 | 6.2014 | 6.1385 | 6.2093 | 5.9840 | 5.5894 | 6.2550 | 6.4232 | 6.4242 | 5.9176 | 6.5184 | 6.4241 | 6.9785 | 7.2398 | 6.4748 |
| NCBP1_S22 | NCBP1 | Q09161 | 22 | | HKRRKTSDANETE | 5.1560 | 5.2284 | 5.5594 | 5.9001 | 6.3007 | 6.2647 | 6.4402 | 6.0734 | 5.4056 | 5.3815 | 5.7199 | 5.7746 | 7.7750 | 7.3595 | 7.6830 | 7.9781 |
| NCBP1_S7 | NCBP1 | Q09161 | 7 | | MSRRRHSDENDGG | 4.9885 | 5.1304 | 5.1225 | 5.0639 | 5.8313 | 5.7196 | 5.9174 | 6.1120 | 5.1588 | 4.6049 | 5.5763 | 5.2611 | 8.8267 | 8.3159 | 9.3878 | 8.9829 |
| NCBP1_T21 | NCBP1 | Q09161 | 21 | | PHKRRKTSDANET | 4.0388 | 3.9112 | 4.4273 | 4.4505 | 5.3424 | 5.1441 | 5.4460 | 5.5905 | 4.5004 | 4.1263 | 4.4362 | 4.6694 | 11.2056 | 10.5640 | 11.3196 | 10.8275 |
| NCBP3_S25 | NCBP3 | Q53F19 | 415 | | KMISTPSPKKSMK | 6.3667 | 6.1941 | 5.9363 | 6.4751 | 6.1500 | 5.6001 | 6.3662 | 5.8062 | 6.3313 | 5.7924 | 5.4716 | 5.6939 | 7.0041 | 6.6679 | 7.0415 | 7.1026 |
| NCBP3_S500 | NCBP3 | Q53F19 | 25 | | PALGLPSPEAESG | 5.7283 | 6.1565 | 6.5141 | 6.8044 | 6.1402 | 6.0627 | 6.6836 | 6.0539 | 5.9475 | 6.3088 | 6.3167 | 6.1885 | 5.8189 | 6.3084 | 6.4212 | 6.5462 |
| NCBP3_T413;S415 | NCBP3 | Q53F19 | 500 | | LGKRPHSPEKAFS | 6.5273 | 6.5416 | 5.8463 | 6.7488 | 5.7662 | 5.7822 | 6.1465 | 6.1277 | 6.1847 | 6.1761 | 6.0038 | 6.3065 | 6.4289 | 6.2829 | 6.5490 | 6.5816 |
| NCBP3_T415 | NCBP3 | Q53F19 | | 413;415 | NA | 7.0710 | 7.3585 | 6.8842 | 7.1376 | 6.5822 | 6.3605 | 7.0544 | 6.3293 | 7.1429 | 6.9982 | 6.9382 | 6.9132 | 4.3271 | 4.4695 | 4.0178 | 4.4155 |
| NCK1_S166 | NCK1 | P16333 | 166 | | VTEEGDSPLGDHV | 5.5574 | 6.1400 | 6.2133 | 5.9520 | 6.2395 | 6.5149 | 6.0416 | 6.5910 | 6.0944 | 6.2379 | 6.1111 | 6.2207 | 6.4090 | 6.9875 | 6.4251 | 6.2647 |
| NCK1_S85 | NCK1 | P16333 | 85 | | KVKRKPSVPDSAS | 6.0623 | 4.8033 | 5.5199 | 4.7955 | 7.3547 | 5.6552 | 5.5055 | 6.5637 | 6.2243 | 6.7102 | 6.5970 | 6.9731 | 7.8128 | 7.8982 | 6.1400 | 5.3845 |
| NCK1_S85;S96 | NCK1 | P16333 | 91 | | SVPDSASPADDSF | 6.3787 | 8.4905 | 7.2279 | 6.1892 | 6.0743 | 7.4488 | 6.7936 | 6.7429 | 6.9205 | 5.6803 | 6.1668 | 6.7250 | 4.8057 | 4.6311 | 5.7922 | 3.9326 |
| NCK1_S91 | NCK1 | P16333 | 105 | | DPGERLYDLNMPA | 7.8100 | 7.1029 | 8.4878 | 7.3852 | 6.4470 | 6.6601 | 6.0032 | 6.4182 | 7.1058 | 6.6888 | 6.8025 | 6.9400 | 4.2349 | 4.8903 | 3.5990 | 4.4244 |
| NCK1_S96 | NCK1 | P16333 | 96 | | ASPADDSFVDPGE | 7.0124 | 8.4790 | 8.3209 | 8.7819 | 5.8312 | 7.3198 | 7.1090 | 7.2539 | 6.9377 | 6.8499 | 10.0026 | 6.1957 | 1.8135 | 2.6873 | 1.9320 | 3.4733 |
| NCK1_Y105 | NCK1 | P16333 | | 85;96 | NA | 5.6606 | 3.6783 | 6.4993 | 6.6218 | 8.4758 | 5.3502 | 6.4478 | 6.7803 | 9.6582 | 6.8281 | 8.3851 | 9.4903 | 3.7039 | 4.9367 | 4.1237 | 3.3598 |
| NCK2_S90 | NCK2 | O43639 | 90 | | TSARDASPTPSTD | 5.1230 | 6.8345 | 6.1018 | 5.5649 | 6.8960 | 6.1409 | 5.3412 | 6.2640 | 3.9821 | 6.4206 | 4.8826 | 6.2136 | 9.4519 | 7.1532 | 7.3098 | 6.3199 |
| NCKAP5L_S440 | NCKAP5L | Q9HCH0 | 477 | | PGGPQLSPQLPRN | 4.8876 | 5.4925 | 5.5881 | 5.5188 | 5.3471 | 6.4800 | 6.2632 | 6.2498 | 5.5902 | 6.2046 | 6.3202 | 6.3445 | 7.0399 | 7.3938 | 7.6143 | 7.6655 |
| NCKAP5L_S440;S451 | NCKAP5L | Q9HCH0 | 735 | | VALGTNSLKQQEP | 6.0497 | 6.2007 | 6.8099 | 6.1259 | 5.4660 | 6.0109 | 6.0698 | 6.1040 | 6.6927 | 6.7731 | 6.3997 | 6.2295 | 6.4391 | 7.0255 | 6.2646 | 5.3388 |
| NCKAP5L_S451 | NCKAP5L | Q9HCH0 | 630 | | DKAGSESPHPGRR | 6.3159 | 7.6731 | 6.2014 | 5.9980 | 5.1051 | 6.4782 | 6.0269 | 7.1052 | 6.4121 | 6.7068 | 6.5490 | 7.0189 | 5.3309 | 5.8640 | 6.0220 | 5.1926 |
| NCKAP5L_S477 | NCKAP5L | Q9HCH0 | 451 | | QGPLLPSPARGLK | 6.1998 | 6.0752 | 5.9616 | 6.0341 | 6.4917 | 7.1221 | 7.1487 | 6.1762 | 6.2753 | 5.4085 | 6.6833 | 6.1230 | 6.2693 | 6.1738 | 5.7458 | 6.1115 |
| NCKAP5L_S571 | NCKAP5L | Q9HCH0 | 571 | | STFRGPSPEPPPS | 6.7730 | 6.4952 | 6.1077 | 6.5756 | 5.7035 | 6.2403 | 6.3373 | 6.7322 | 6.8105 | 6.4223 | 6.5240 | 7.0527 | 5.3408 | 6.0060 | 5.7544 | 5.1246 |
| NCKAP5L_S571;S577 | NCKAP5L | Q9HCH0 | 440 | | LQIGPPSPGEAQG | 6.2631 | 7.2694 | 6.7168 | 5.8594 | 5.7629 | 6.3870 | 6.7538 | 7.0027 | 5.9153 | 6.2440 | 6.5610 | 6.1678 | 6.2393 | 6.2869 | 5.8653 | 5.6152 |
| NCKAP5L_S577 | NCKAP5L | Q9HCH0 | 617 | | ESYPYGSPQEKSL | 7.4476 | 7.2865 | 6.7013 | 7.0131 | 6.1721 | 6.3731 | 6.5254 | 7.0279 | 5.9756 | 7.7299 | 8.6448 | 7.5068 | 2.9397 | 4.6498 | 3.6300 | 4.3764 |
| NCKAP5L_S617 | NCKAP5L | Q9HCH0 | 577 | | SPEPPPSPLQVPT | 5.6278 | 5.3923 | 5.8983 | 5.5276 | 6.0054 | 6.1202 | 6.8871 | 6.5464 | 6.1942 | 5.9781 | 5.5081 | 5.6472 | 7.2773 | 7.1936 | 7.5216 | 6.6749 |
| NCKAP5L_S630 | NCKAP5L | Q9HCH0 | | 440;451 | NA | 6.6072 | 7.0717 | 6.2705 | 6.4621 | 5.5109 | 7.4333 | 6.4142 | 7.0931 | 7.4178 | 7.9298 | 6.8744 | 7.1558 | 4.9091 | 4.9749 | 3.6223 | 4.2530 |
| NCKAP5L_S735 | NCKAP5L | Q9HCH0 | | 571;577 | NA | 5.6945 | 6.5708 | 6.9982 | 6.0293 | 6.4764 | 6.3948 | 5.8903 | 6.7309 | 6.7330 | 6.1595 | 6.2486 | 6.0821 | 5.7452 | 5.8909 | 6.7919 | 5.5635 |
| NCKIPSD_T181 | NCKIPSD | Q9NZQ3-2 | 181 | | RRAAPTTPPPPVK | 6.1610 | 5.9525 | 6.5000 | 6.2584 | 6.8974 | 6.5780 | 6.3224 | 6.9466 | 6.7557 | 6.6401 | 6.9022 | 6.5093 | 5.2630 | 5.9522 | 5.1260 | 5.2353 |
| NCL_S145;S153 | NCL | P19338 | 67 | | AKKVVVSPTKKVA | 6.3336 | 6.5196 | 5.9180 | 6.2252 | 6.7734 | 6.3962 | 6.2803 | 6.2980 | 6.4235 | 6.0855 | 6.0234 | 5.9036 | 6.3029 | 6.2226 | 6.2244 | 6.0697 |
| NCL_S184;S206 | NCL | P19338 | 84 | | AKKAAVTPGKKAA | 8.2248 | 8.4484 | 7.2550 | 7.1394 | 5.6030 | 7.1350 | 7.5531 | 7.1284 | 7.9836 | 7.7839 | 7.3012 | 7.6124 | 2.1785 | 3.2421 | 2.1049 | 3.3063 |
| NCL_S28;S34;S41 | NCL | P19338 | 92 | | GKKAAATPAKKTV | 8.5071 | 8.6193 | 7.1382 | 7.1419 | 5.9662 | 7.5661 | 7.9566 | 7.2530 | 8.0013 | 7.6041 | 7.5463 | 7.4109 | 1.8535 | 2.9461 | 1.6661 | 2.8234 |
| NCL_S356 | NCL | P19338 | 106 | | PAKAVTTPGKKGA | 7.6235 | 8.2247 | 7.3310 | 7.5044 | 5.8644 | 7.2698 | 7.4742 | 7.1649 | 8.0237 | 7.8875 | 7.7410 | 8.1395 | 1.9135 | 3.0162 | 1.8275 | 2.9944 |
| NCL_S460 | NCL | P19338 | 121 | | GKALVATPGKKGA | 7.8592 | 8.3669 | 7.2679 | 7.2236 | 5.8126 | 7.2979 | 7.8017 | 7.0806 | 7.9501 | 7.8480 | 7.5945 | 7.8071 | 2.0272 | 3.0304 | 1.9388 | 3.0936 |
| NCL_S563 | NCL | P19338 | 76 | | KKVAVATPAKKAA | 8.6188 | 8.7113 | 7.3863 | 7.2392 | 5.8260 | 7.5384 | 7.9134 | 7.1477 | 8.0470 | 7.5074 | 7.3273 | 7.4797 | 1.8376 | 2.9257 | 1.6536 | 2.8403 |
| NCL_S60 | NCL | P19338 | 563 | | LQGPRGSPNARSQ | 5.3683 | 6.8939 | 7.0574 | 6.6484 | 5.8727 | 6.9684 | 5.6292 | 7.0344 | 7.9998 | 5.9608 | 6.2172 | 6.5460 | 4.6965 | 6.3469 | 5.3735 | 5.3865 |
| NCL_S619 | NCL | P19338 | 619 | | GFVDFNSEEDAKA | 6.7084 | 6.9635 | 6.4104 | 6.6770 | 6.1159 | 6.2032 | 6.3490 | 5.5417 | 6.3718 | 6.6499 | 6.2260 | 6.5139 | 5.8184 | 5.9851 | 5.6680 | 5.7980 |
| NCL_S67 | NCL | P19338 | 356 | | GYVDFESAEDLEK | 6.1145 | 6.7999 | 6.0197 | 6.5618 | 6.5162 | 5.7649 | 5.8677 | 5.6636 | 5.7997 | 6.1471 | 5.9343 | 7.0156 | 6.2064 | 6.4988 | 6.6704 | 6.4192 |
| NCL_T106 | NCL | P19338 | 60 | | KKAAATSAKKVVV | 7.3802 | 8.6845 | 7.5956 | 7.6212 | 5.0013 | 7.2300 | 7.7756 | 7.0221 | 7.5142 | 7.4609 | 7.7433 | 8.0601 | 2.0041 | 3.4120 | 2.1185 | 3.3766 |
| NCL_T121 | NCL | P19338 | 460 | | IDGRSISLYYTGE | 5.6133 | 6.9961 | 8.0165 | 5.9478 | 5.5312 | 6.5981 | 6.5176 | 5.2184 | 5.9121 | 6.1769 | 6.3086 | 5.2587 | 6.2110 | 5.9199 | 6.2931 | 7.4808 |
| NCL_T76 | NCL | P19338 | | 184;206 | NA | 5.6547 | 6.1184 | 5.7639 | 6.2245 | 6.1084 | 5.8368 | 5.9575 | 5.9609 | 6.0900 | 6.0517 | 5.8969 | 5.9708 | 6.8539 | 7.0955 | 7.2130 | 7.2032 |

| | | | | | | | | | | | | | | | | | | | | |
|---|---|---|---|---|---|---|---|---|---|---|---|---|---|---|---|---|---|---|---|---|
| NCL_T84 | NCL | P19338 | 145;153 | NA | 5.6684 | 5.9545 | 5.8871 | 5.9689 | 5.8412 | 5.8291 | 6.1587 | 5.5614 | 6.3627 | 6.2381 | 6.2149 | 6.2826 | 7.1400 | 6.9686 | 6.7998 | 7.1240 |
| NCL_T92 | NCL | P19338 | 28;34;41 | NA | 6.5121 | 6.3246 | 6.8530 | 6.4615 | 6.1917 | 6.3948 | 6.3133 | 6.0171 | 7.2545 | 6.7341 | 6.6865 | 7.1286 | 5.1589 | 5.5012 | 5.1463 | 5.3218 |
| NCOA1_S395 | NCOA1 | Q15788 | 698 | RLLQEGSPSDITT | 5.9684 | 6.4209 | 6.0499 | 6.2104 | 6.5769 | 6.1208 | 5.8562 | 6.0119 | 6.1114 | 6.2954 | 6.4502 | 6.4203 | 6.7132 | 6.4064 | 6.3217 | 6.0662 |
| NCOA1_S698 | NCOA1 | Q15788 | 395 | SVNPSISPAHGVA | 5.7155 | 7.0188 | 6.4850 | 5.5320 | 5.6543 | 6.9360 | 6.8264 | 7.5832 | 4.9663 | 6.9245 | 6.8647 | 6.5359 | 5.3918 | 5.9945 | 6.2009 | 5.3702 |
| NCOA2_S29 | NCOA2 | Q15596 | 736 | IKQEPVSPKKKEN | 5.8375 | 6.1932 | 6.0386 | 5.4644 | 6.1848 | 5.9926 | 6.0048 | 5.8033 | 6.0601 | 5.9792 | 6.0532 | 5.8984 | 7.4266 | 7.2133 | 7.1381 | 6.7120 |
| NCOA2_S487 | NCOA2 | Q15596 | 29 | PDQLGPSPKRNTE | 5.9049 | 6.3658 | 6.2557 | 6.0987 | 6.5141 | 6.2125 | 6.2751 | 6.1422 | 6.6543 | 6.4928 | 6.5437 | 6.2572 | 6.1765 | 6.2796 | 5.9260 | 5.9008 |
| NCOA2_S493 | NCOA2 | Q15596 | 565 | MGNLQNSPVNMNP | 5.0543 | 5.2144 | 5.3998 | 6.2368 | 5.9401 | 5.7965 | 5.7333 | 5.7614 | 5.4034 | 5.6726 | 5.9945 | 5.4172 | 8.6135 | 7.8840 | 8.1200 | 7.7582 |
| NCOA2_S565 | NCOA2 | Q15596 | 487 | QPTSMLSPRHRMS | 7.3379 | 6.9424 | 7.5814 | 6.9649 | 7.0916 | 7.4646 | 6.3667 | 7.5519 | 5.0196 | 6.3526 | 7.0916 | 6.1921 | 3.6570 | 5.5348 | 4.7493 | 4.1018 |
| NCOA2_S699 | NCOA2 | Q15596 | 699 | LLQDSSSPVDLAK | 6.3023 | 6.9189 | 6.2620 | 6.5384 | 6.0793 | 6.7624 | 7.0192 | 6.3351 | 6.4514 | 7.2199 | 6.7875 | 6.6524 | 4.8608 | 5.5696 | 4.9902 | 5.2505 |
| NCOA2_S736 | NCOA2 | Q15596 | 763 | IGLPEITPKLERL | 7.2080 | 6.4359 | 6.5831 | 7.9072 | 6.7506 | 6.6560 | 7.4064 | 7.0111 | 7.1878 | 7.1816 | 7.0498 | 6.6653 | 4.2296 | 4.0947 | 3.4079 | 4.2249 |
| NCOA2_T763 | NCOA2 | Q15596 | 493 | SPRHRMSPGVAGS | 4.7725 | 6.2577 | 6.3902 | 5.7189 | 6.6155 | 5.7012 | 5.9739 | 5.9673 | 5.8812 | 6.0358 | 6.3748 | 6.5824 | 7.4967 | 6.8452 | 6.8496 | 6.5371 |
| NCOA3_S1330 | NCOA3 | Q9Y6Q9 | 1330 | AFGRVSSPPNAMM | 6.0453 | 6.4428 | 6.1605 | 5.6625 | 6.0200 | 6.5972 | 5.8270 | 6.5274 | 6.2170 | 6.9432 | 6.6992 | 6.4941 | 6.4115 | 6.3561 | 5.9007 | 5.6953 |
| NCOA3_S214 | NCOA3 | Q9Y6Q9 | 728 | VKQEQLSPKKKEN | 5.5694 | 5.9691 | 6.2503 | 5.5735 | 5.8760 | 6.0159 | 5.8351 | 5.9715 | 6.3124 | 6.1962 | 6.2702 | 6.3446 | 7.2643 | 6.8190 | 6.9122 | 6.8203 |
| NCOA3_S551 | NCOA3 | Q9Y6Q9 | 551 | GPKLDNSPNMNIT | 5.5561 | 6.9075 | 6.4808 | 6.5565 | 6.8975 | 6.5227 | 6.4871 | 7.2021 | 7.0990 | 6.6039 | 7.1501 | 6.1735 | 5.4631 | 5.5506 | 4.5731 | 4.7765 |
| NCOA3_S569 | NCOA3 | Q9Y6Q9 | 569 | SNQDSKSPLGFYC | 4.6448 | 5.3354 | 6.3255 | 6.2391 | 6.4285 | 7.0327 | 6.0453 | 5.7886 | 5.8855 | 5.8497 | 6.7516 | 6.6049 | 7.5959 | 7.0100 | 6.3447 | 6.1177 |
| NCOA3_S728 | NCOA3 | Q9Y6Q9 | 857 | PYNRAVSLDSPVS | 5.1542 | 4.9618 | 6.2538 | 5.2171 | 5.7512 | 6.1228 | 5.5063 | 6.4877 | 5.5872 | 6.5210 | 6.5255 | 5.8932 | 7.7256 | 7.5556 | 7.5700 | 7.1668 |
| NCOA3_S857 | NCOA3 | Q9Y6Q9 | 214 | LEDINASPEMRQR | 5.4062 | 6.2175 | 6.1102 | 6.0468 | 6.3958 | 6.3803 | 5.9809 | 7.0763 | 6.1911 | 6.3608 | 5.7572 | 6.1961 | 6.4013 | 6.9600 | 6.1427 | 6.3768 |
| NCOA5_S29 | NCOA5 | Q9HCD5 | 378 | ERLMRSSTDSLPG | 5.5893 | 5.4539 | 5.7050 | 6.1467 | 6.3413 | 5.9651 | 6.4720 | 6.3195 | 5.9633 | 5.8879 | 5.7430 | 5.7898 | 7.0668 | 6.9808 | 7.3835 | 7.1921 |
| NCOA5_S29;S34 | NCOA5 | Q9HCD5 | 29 | DSRRDRSPIRGSP | 6.1631 | 6.3448 | 5.8899 | 7.3325 | 5.7118 | 5.2878 | 5.5068 | 6.2153 | 6.5389 | 6.3430 | 6.2834 | 6.4079 | 6.5560 | 6.3905 | 6.4337 | 6.5946 |
| NCOA5_S378 | NCOA5 | Q9HCD5 | 274 | VNIMFGTPQEHRN | 6.1307 | 6.8617 | 7.2028 | 6.9812 | 6.9345 | 5.6419 | 6.1046 | 5.9992 | 6.0231 | 5.2454 | 5.6929 | 5.5767 | 6.3685 | 6.2252 | 5.8526 | 7.1593 |
| NCOA5_T274 | NCOA5 | Q9HCD5 | 29;34 | NA | 6.4550 | 6.4268 | 5.6826 | 6.4843 | 5.9621 | 5.7567 | 6.1788 | 6.0777 | 6.2404 | 6.0648 | 6.1898 | 6.2349 | 6.6532 | 6.4838 | 6.6131 | 6.4960 |
| NCOA6_S1343 | NCOA6 | Q14686 | 1481 | ENKNLVSPAMREA | 5.9742 | 6.1210 | 5.5107 | 6.1022 | 6.1867 | 6.2992 | 6.2494 | 6.3857 | 6.0390 | 5.7985 | 5.4935 | 5.5647 | 7.5252 | 6.8912 | 6.9426 | 6.9162 |
| NCOA6_S1481 | NCOA6 | Q14686 | 1892 | LLKMTSSPVGPGT | 5.9462 | 5.7901 | 6.5879 | 6.9070 | 4.9576 | 7.3094 | 6.3641 | 5.6453 | 6.1976 | 7.0822 | 6.9547 | 6.6373 | 6.2688 | 6.1291 | 5.9219 | 5.3010 |
| NCOA6_S1745 | NCOA6 | Q14686 | 1809 | SSGNRRSPVSSSK | 6.7567 | 7.1034 | 6.0947 | 6.6578 | 6.3468 | 6.2754 | 6.0642 | 6.2201 | 6.5274 | 6.2209 | 6.4456 | 6.3241 | 5.7877 | 5.7301 | 5.7325 | 5.7127 |
| NCOA6_S1809 | NCOA6 | Q14686 | 1745 | TPVQLPSPPCTSS | 5.8070 | 6.3923 | 6.2758 | 6.4273 | 5.8626 | 5.6255 | 6.2155 | 6.4814 | 6.3392 | 6.2367 | 6.1658 | 6.3430 | 6.5405 | 6.7245 | 6.2039 | 6.3589 |
| NCOA6_S1892 | NCOA6 | Q14686 | 1343 | SRKTTPSPGRQNS | 6.7558 | 7.0279 | 6.0946 | 6.4863 | 6.4205 | 6.6112 | 6.5335 | 6.2793 | 6.7651 | 6.4722 | 6.4524 | 6.7275 | 5.5831 | 5.4795 | 5.0989 | 5.2121 |
| NCOA6_T1880 | NCOA6 | Q14686 | 1880 | LDSKTPTPPAPTL | 6.3507 | 6.4340 | 6.3075 | 6.3799 | 6.7720 | 6.5995 | 6.0007 | 5.8005 | 6.1429 | 6.3091 | 6.1450 | 6.4445 | 6.1654 | 6.2278 | 6.0762 | 5.8444 |
| NCOA7_S162 | NCOA7 | Q8NI08-2 | 162 | FVPDANSPSSTLR | 6.7740 | 6.8031 | 6.7914 | 6.6769 | 6.0881 | 6.7480 | 6.6841 | 6.3714 | 6.2918 | 7.0488 | 6.8242 | 6.9828 | 4.8809 | 5.4093 | 4.8676 | 4.7575 |
| NCOA7_S179 | NCOA7 | Q8NI08-2 | 89 | ELKRYYSIDDNQN | 6.0207 | 6.4222 | 8.1827 | 5.2760 | 5.0318 | 7.2601 | 6.1080 | 6.2961 | 6.4109 | 8.1930 | 8.0181 | 7.2551 | 4.3180 | 6.1243 | 4.4765 | 4.6065 |
| NCOA7_S211 | NCOA7 | Q8NI08-2 | 347 | EKSTGHTPTKPSG | 6.5798 | 6.6187 | 6.8156 | 5.8722 | 5.9267 | 6.6888 | 6.3056 | 6.6240 | 6.8234 | 6.7933 | 6.5389 | 6.8732 | 4.8595 | 6.0029 | 5.3407 | 5.3369 |
| NCOA7_S295 | NCOA7 | Q8NI08-2 | 179 | SPGATVSPSSSDA | 5.1738 | 5.3223 | 5.7381 | 4.5976 | 4.7419 | 5.1713 | 4.9637 | 6.0182 | 7.2769 | 7.3140 | 7.1917 | 7.0955 | 7.3334 | 7.5215 | 8.0800 | 6.4603 |
| NCOA7_S585 | NCOA7 | Q8NI08-2 | 134 | ALKFNITPNKLVE | 6.4505 | 7.4880 | 7.0582 | 7.3659 | 5.7618 | 7.2347 | 6.8978 | 6.9318 | 6.7405 | 7.5292 | 7.9131 | 7.9050 | 2.9099 | 4.2463 | 3.2160 | 4.3513 |
| NCOA7_S89 | NCOA7 | Q8NI08-2 | 295 | IKDALPSPGEWED | 4.9498 | 4.9665 | 6.5034 | 4.4692 | 8.2510 | 7.2474 | 5.2955 | 5.5152 | 5.5152 | 4.3022 | 4.9254 | 6.2297 | 7.3798 | 8.2536 | 8.2266 | 7.9696 |
| NCOA7_T134 | NCOA7 | Q8NI08-2 | 585 | PVKLNSSTEANVI | 6.8395 | 8.4527 | 7.6595 | 7.7135 | 6.4107 | 6.6085 | 7.1369 | 6.4953 | 7.2117 | 7.1252 | 6.9046 | 6.7972 | 3.1048 | 3.7230 | 3.3430 | 4.4740 |
| NCOA7_T347 | NCOA7 | Q8NI08-2 | 211 | RVLSSTSEEDEPG | 5.9392 | 6.2570 | 6.5298 | 6.4120 | 6.1691 | 6.4699 | 5.9339 | 6.0604 | 6.4907 | 6.1527 | 5.8827 | 6.5308 | 5.8069 | 7.0731 | 6.0991 | 6.1927 |
| NCOR1_S1111 | NCOR1 | O75376 | 224 | EPEKPVSPPPVEQ | 6.0798 | 6.2513 | 6.2879 | 6.5809 | 6.1465 | 6.0141 | 6.2504 | 5.7208 | 6.2660 | 6.2529 | 6.1947 | 6.1420 | 6.6900 | 6.3538 | 6.2879 | 6.4811 |
| NCOR1_S1196 | NCOR1 | O75376 | 999 | PCGTSKSPNREWE | 6.2877 | 6.4644 | 6.4086 | 6.5529 | 6.5863 | 5.9009 | 6.0718 | 6.1105 | 6.2743 | 6.4942 | 6.3148 | 6.3951 | 5.9519 | 6.2925 | 6.0004 | 5.8937 |
| NCOR1_S1263 | NCOR1 | O75376 | 992 | ECRSSTSPCGTSK | 6.2231 | 6.5400 | 6.2032 | 6.4860 | 6.2184 | 6.3613 | 6.3632 | 6.1320 | 6.4093 | 6.4840 | 6.6302 | 6.3049 | 5.7929 | 6.1662 | 5.8759 | 5.8094 |
| NCOR1_S13 | NCOR1 | O75376 | 1533 | ESIPAKSPVPGVD | 6.0770 | 6.3500 | 5.9731 | 5.9957 | 5.8160 | 6.6969 | 6.6595 | 6.0596 | 6.3048 | 5.6601 | 5.8909 | 5.7536 | 6.6461 | 6.5553 | 6.4398 | 7.1219 |
| NCOR1_S1533 | NCOR1 | O75376 | 2120 | RPGSRVSPENLVD | 6.6298 | 6.7173 | 6.7856 | 6.5921 | 6.3539 | 6.4079 | 6.5358 | 6.1160 | 6.4103 | 6.3476 | 6.3531 | 6.1644 | 6.0679 | 5.6169 | 5.4596 | 5.4418 |
| NCOR1_S158;S172 | NCOR1 | O75376 | 2184 | QRNDARSPGSISY | 6.0520 | 5.7625 | 5.9315 | 6.4406 | 6.3348 | 6.3131 | 6.4750 | 6.5071 | 6.1199 | 5.8457 | 6.4014 | 6.2247 | 6.4455 | 6.5245 | 6.3443 | 6.2773 |
| NCOR1_S1977 | NCOR1 | O75376 | 1977 | DAIEVISPASSPA | 6.5970 | 6.4280 | 6.2432 | 6.7266 | 6.3547 | 5.9170 | 6.1872 | 6.1934 | 6.0414 | 6.0003 | 6.0164 | 6.5319 | 6.2888 | 6.1416 | 6.0639 | 6.2685 |
| NCOR1_S2102 | NCOR1 | O75376 | 2202 | TKLENTSPMVKSK | 7.5566 | 7.8155 | 6.6112 | 7.0278 | 6.6292 | 7.3419 | 7.0829 | 7.1210 | 7.3869 | 7.4075 | 7.1723 | 7.1733 | 3.1226 | 4.0979 | 2.7019 | 3.7515 |
| NCOR1_S2120 | NCOR1 | O75376 | 2136 | GSRPGKSPERSHV | 6.0071 | 5.8498 | 6.1041 | 6.6327 | 6.4011 | 6.3493 | 6.3395 | 6.2958 | 6.5582 | 6.5309 | 6.8613 | 6.5152 | 6.6155 | 6.0970 | 5.4447 | 5.3978 |
| NCOR1_S2136 | NCOR1 | O75376 | 2337 | NSRKSKSPIPGQG | 6.3220 | 6.7014 | 5.7336 | 6.4841 | 5.6544 | 5.2887 | 6.2608 | 5.9635 | 6.2796 | 6.1304 | 6.0462 | 5.9010 | 6.1100 | 6.7953 | 6.8321 | 7.4970 |
| NCOR1_S2151 | NCOR1 | O75376 | 1263 | GMSMRESPVSAPL | 6.4202 | 6.7127 | 5.9143 | 6.6418 | 6.0537 | 6.1078 | 6.4697 | 6.8817 | 6.5397 | 6.2548 | 6.1481 | 6.8158 | 5.9725 | 5.9705 | 5.0828 | 6.0139 |
| NCOR1_S2184 | NCOR1 | O75376 | 1367 | TQESRKTPEVVQS | 5.7479 | 6.3370 | 6.2860 | 6.7346 | 6.9629 | 6.4835 | 6.4946 | 5.8788 | 6.4123 | 7.1082 | 5.9851 | 6.1566 | 6.1972 | 5.9016 | 5.6954 | 5.6183 |
| NCOR1_S2202 | NCOR1 | O75376 | 1196 | MPIEDSSPEKGRE | 6.4507 | 6.5766 | 6.0114 | 6.6856 | 6.4441 | 6.0609 | 6.6231 | 5.9981 | 6.3707 | 6.2188 | 6.1958 | 6.5349 | 6.2535 | 5.7735 | 5.6584 | 6.1439 |
| NCOR1_S224 | NCOR1 | O75376 | 1522 | ERKSTLTPTQRES | 6.4543 | 6.8189 | 6.0992 | 6.4522 | 6.4614 | 6.7834 | 7.3695 | 6.7437 | 7.0278 | 6.6622 | 6.6213 | 6.1906 | 5.1827 | 5.3939 | 4.6554 | 5.0834 |
| NCOR1_S2337 | NCOR1 | O75376 | 913 | LLNPTGSILVSSP | 6.5431 | 6.1453 | 6.9752 | 6.2901 | 6.5686 | 6.0981 | 5.6567 | 5.7008 | 6.7744 | 6.1801 | 7.1486 | 6.1484 | 6.1932 | 5.9862 | 5.3956 | 6.1956 |
| NCOR1_S70 | NCOR1 | O75376 | 1111 | IKQEEFSPRSQNS | 6.0361 | 6.6649 | 6.4448 | 6.6088 | 6.5335 | 6.0685 | 6.7325 | 6.3898 | 6.8813 | 6.7276 | 6.5787 | 6.9387 | 5.6427 | 5.2614 | 5.0883 | 5.4026 |
| NCOR1_S913 | NCOR1 | O75376 | 2151 | EPYEPISPPQVPV | 6.7899 | 7.1236 | 6.6359 | 6.4842 | 6.5254 | 6.8723 | 6.5659 | 7.2690 | 6.2640 | 5.6861 | 6.2049 | 5.9552 | 4.9464 | 6.1925 | 5.6037 | 4.8810 |
| NCOR1_S992 | NCOR1 | O75376 | 70 | QLRRRPSLLSEFH | 4.8892 | 4.7566 | 5.4396 | 5.6259 | 6.8649 | 6.3467 | 5.6281 | 7.2539 | 7.7760 | 7.1893 | 6.8031 | 7.2924 | 6.2442 | 6.6769 | 5.5086 | 5.7046 |
| NCOR1_S999 | NCOR1 | O75376 | 2102 | KTSNRYSPESQAQ | 6.1057 | 7.3119 | 7.5404 | 7.4658 | 6.5699 | 6.2837 | 6.4352 | 6.2368 | 6.0576 | 5.6956 | 6.9078 | 5.8664 | 5.6920 | 6.2031 | 4.5856 | 5.0426 |
| NCOR1_T1367 | NCOR1 | O75376 | 757 | VELEPTTETAPST | 5.5310 | 5.6694 | 6.3124 | 6.9290 | 5.7018 | 5.8735 | 5.6985 | 5.6624 | 5.9754 | 6.3054 | 5.5995 | 5.9782 | 6.7906 | 6.4263 | 7.6382 | 7.9085 |
| NCOR1_T1522 | NCOR1 | H0Y459 | 13 | PCGTSKSPNREWE | 6.5968 | 6.7529 | 6.5200 | 6.4851 | 6.3518 | 6.2169 | 6.1874 | 6.1893 | 6.3752 | 6.2964 | 6.2577 | 6.5868 | 6.2838 | 5.7727 | 5.6899 | 5.4371 |
| NCOR1_T757 | NCOR1 | O75376 | 158;172 | NA | 6.1401 | 5.8379 | 6.5889 | 6.8558 | 6.1850 | 6.3747 | 6.1205 | 6.5120 | 6.6389 | 5.7549 | 6.6576 | 6.6216 | 6.3736 | 5.9893 | 5.2991 | 6.0502 |
| NCOR2_S1018 | NCOR2 | Q9Y618 | 215 | EPEKPVSPPPIES | 6.3502 | 6.2601 | 6.2409 | 6.5812 | 5.8726 | 5.7645 | 6.0959 | 5.6491 | 6.2578 | 6.7565 | 6.2274 | 6.3617 | 6.3578 | 6.5704 | 6.1944 | 6.4595 |
| NCOR2_S1025 | NCOR2 | Q9Y618 | 2223 | DGIEPVSPPEGMT | 5.9016 | 6.3741 | 6.4947 | 6.2239 | 6.2877 | 6.1898 | 6.1901 | 6.1393 | 6.3521 | 6.1107 | 6.3382 | 6.0891 | 6.1170 | 6.5264 | 6.3493 | 6.3161 |
| NCOR2_S1173 | NCOR2 | Q9Y618 | 2258 | SRMGSKSPGNTSQ | 5.9664 | 6.2603 | 6.3448 | 6.4352 | 6.3001 | 6.3785 | 6.1711 | 6.1787 | 6.3747 | 6.3974 | 6.2258 | 6.1439 | 6.4935 | 6.1625 | 6.1128 | 6.0543 |
| NCOR2_S1251 | NCOR2 | Q9Y618 | 67 | PQRRRPSLLSEFQ | 5.0506 | 4.9031 | 5.1418 | 5.4209 | 6.3263 | 5.8778 | 5.5306 | 5.8989 | 7.7216 | 7.7486 | 6.9866 | 7.2921 | 7.0143 | 7.2727 | 5.6827 | 6.1314 |
| NCOR2_S1341 | NCOR2 | Q9Y618 | 939 | DKNRLLSPRPSLL | 6.5395 | 6.9243 | 6.8474 | 7.2356 | 6.8814 | 6.9661 | 6.4555 | 6.1541 | 6.3125 | 6.8251 | 6.8513 | 6.4214 | 5.0601 | 5.2590 | 4.7190 | 4.5476 |
| NCOR2_S1479 | NCOR2 | Q9Y618 | 1025 | PRGKSRSPAPPAD | 6.1055 | 6.3240 | 6.0665 | 6.2665 | 5.5511 | 6.3578 | 6.2133 | 6.1028 | 6.1665 | 6.1096 | 6.1881 | 6.1608 | 6.7298 | 6.5511 | 6.4126 | 6.6939 |
| NCOR2_S149;S152 | NCOR2 | Q9Y618 | 1479 | VRSLIGSPGRTFP | 6.8056 | 6.6152 | 6.6171 | 7.2109 | 6.7037 | 6.6621 | 6.6162 | 6.7074 | 7.2851 | 6.8771 | 6.8202 | 6.8393 | 4.4520 | 4.8505 | 4.3625 | 4.5750 |
| NCOR2_S152 | NCOR2 | Q9Y618 | 956 | DPRANASPQKPLD | 7.2316 | 7.1113 | 6.4371 | 6.5555 | 6.4810 | 6.2367 | 6.4660 | 6.2665 | 6.8158 | 6.5283 | 6.4203 | 6.3488 | 5.4260 | 5.3360 | 5.0772 | 5.2621 |
| NCOR2_S1539 | NCOR2 | Q9Y618 | 2005 | LAPHHASPDPPAP | 6.1437 | 7.0502 | 6.4497 | 7.0406 | 6.6816 | 6.3371 | 6.3917 | 6.1364 | 7.2783 | 7.0288 | 7.0353 | 7.0853 | 4.8007 | 5.4100 | 4.2193 | 4.9114 |
| NCOR2_S1971 | NCOR2 | Q9Y618 | 1971 | GLEPASSPSKGSE | 6.8297 | 6.7640 | 6.5143 | 6.3997 | 6.3144 | 6.2710 | 6.3807 | 6.1780 | 6.7802 | 6.3790 | 6.4075 | 6.5089 | 5.7378 | 5.6024 | 5.4334 | 5.4988 |
| NCOR2_S2005 | NCOR2 | Q9Y618 | 2203 | SEGGKRSPEPNKT | 6.5685 | 6.6893 | 6.1571 | 6.3434 | 5.8570 | 5.5732 | 5.7524 | 5.7172 | 6.4083 | 6.4362 | 5.8588 | 6.2676 | 6.2874 | 6.3504 | 6.7808 | 6.9523 |
| NCOR2_S215 | NCOR2 | Q9Y618 | 1341 | IPPERHSPHHLKE | 6.2993 | 6.0683 | 5.7201 | 6.2057 | 6.3860 | 6.6531 | 6.3772 | 6.3444 | 6.4253 | 5.9485 | 6.2324 | 5.8250 | 6.7941 | 6.7008 | 6.1382 | 5.8816 |
| NCOR2_S2194 | NCOR2 | Q9Y618 | 54 | DYASHLSPGSIIQ | 6.2334 | 6.2907 | 5.8653 | 6.6958 | 5.9382 | 6.0948 | 6.5996 | 6.3831 | 6.5944 | 6.6580 | 6.3525 | 6.6886 | 5.8970 | 6.0360 | 5.5760 | 6.0966 |
| NCOR2_S2203 | NCOR2 | Q9Y618 | 152 | EPVSPPSPPHTDP | 6.8459 | 7.0051 | 6.6929 | 7.0522 | 6.4049 | 6.0167 | 6.5735 | 6.5928 | 6.8890 | 7.1457 | 6.8703 | 6.6848 | 4.4778 | 5.4029 | 4.5836 | 4.7619 |
| NCOR2_S2223 | NCOR2 | Q9Y618 | 946 | PRPSLLTPTGDPR | 5.3803 | 5.3843 | 6.3157 | 6.5510 | 6.3925 | 5.9787 | 6.0160 | 6.0067 | 6.3792 | 6.3188 | 6.2234 | 6.2358 | 6.8587 | 6.3814 | 6.3490 | 7.2285 |
| NCOR2_S2258 | NCOR2 | Q9Y618 | 1018 | PQQPGSSPRGKSR | 5.6535 | 5.8145 | 6.5524 | 6.7777 | 6.3199 | 6.4700 | 5.7713 | 6.6211 | 6.0505 | 6.2252 | 6.6187 | 6.6516 | 6.4509 | 6.2953 | 5.9111 | 5.8164 |
| NCOR2_S2453 | NCOR2 | Q9Y618 | 2453 | WEDRPSSAGSTPF | 7.3678 | 6.5312 | 6.8023 | 7.0416 | 6.2293 | 7.3145 | 5.7385 | 5.7806 | 7.7688 | 6.7088 | 6.2508 | 6.3396 | 5.4852 | 6.0479 | 4.4373 | 4.1560 |
| NCOR2_S54 | NCOR2 | Q9Y618 | 1436 | REELRHTPELPLA | 6.7414 | 6.9917 | 6.4907 | 6.6369 | 6.1413 | 6.5298 | 6.2806 | 6.4188 | 7.7152 | 7.3178 | 7.2350 | 7.2761 | 4.5298 | 5.0001 | 4.1393 | 4.5556 |
| NCOR2_S67 | NCOR2 | Q9Y618 | 1173 | VKQEQLSPRGQAG | 5.2329 | 5.3242 | 5.6591 | 5.7266 | 5.2955 | 6.7783 | 6.3563 | 6.0497 | 6.1839 | 6.0354 | 5.7256 | 5.3699 | 7.1709 | 7.6313 | 7.3289 | 8.1314 |
| NCOR2_S939 | NCOR2 | Q9Y618 | 1383 | LLKREGTPPPPPP | 7.7193 | 7.2728 | 6.5258 | 6.1949 | 6.2783 | 6.1741 | 5.0136 | 6.2653 | 6.6318 | 6.7386 | 6.4598 | 5.7093 | 5.7085 | 6.4303 | 5.9656 | 4.9120 |
| NCOR2_S956 | NCOR2 | Q9Y618 | 1455 | GSITQGTPLKYDT | 8.0936 | 7.9515 | 7.4739 | 7.0263 | 5.2324 | 6.4242 | 8.2892 | 6.8318 | 8.4961 | 7.8068 | 7.6186 | 7.9464 | 2.5464 | 3.4976 | 2.1492 | 2.6159 |
| NCOR2_T1383 | NCOR2 | Q9Y618 | 2194 | GAPARGSPHSEGG | 5.5889 | 5.8765 | 5.8035 | 6.8640 | 6.7253 | 6.4361 | 5.9782 | 6.0379 | 6.3411 | 6.2443 | 6.3166 | 6.0688 | 7.2566 | 7.0356 | 6.0568 | 5.3698 |
| NCOR2_T1436 | NCOR2 | Q9Y618 | 1539 | LGKPRQSPLTYED | 5.6174 | 5.4359 | 6.2964 | 6.3577 | 7.1490 | 6.3577 | 7.4245 | 5.4711 | 5.6866 | 6.4609 | 6.4282 | 7.0563 | 6.8121 | 6.0587 | 5.4528 | 5.9347 |
| NCOR2_T1455 | NCOR2 | Q9Y618 | 1251 | RIIGEDSPSRLDR | 7.0834 | 7.1984 | 6.1717 | 6.9526 | 6.6970 | 6.9947 | 6.8327 | 6.2300 | 6.3301 | 6.9988 | 7.3369 | 7.4965 | 4.6865 | 4.6784 | 4.2138 | 4.0984 |
| NCOR2_T946 | NCOR2 | Q9Y618 | 149;152 | NA | 6.3297 | 6.2882 | 6.1073 | 6.4850 | 6.3626 | 5.8542 | 6.1738 | 5.6882 | 6.4085 | 6.7927 | 5.8982 | 6.2379 | 6.1409 | 6.5043 | 6.2313 | 6.4973 |
| NDC1_S406 | NDC1 | Q9BTX1 | 500 | TEFSNPSPSTSIS | 6.4068 | 6.6762 | 6.3863 | 7.1528 | 6.5928 | 6.7120 | 6.6645 | 6.2879 | 6.7123 | 6.7378 | 6.6013 | 6.6475 | 4.8378 | 5.5473 | 4.8485 | 5.1881 |
| NDC1_S436 | NDC1 | Q9BTX1 | 504 | NPSPSTSISAEGK | 6.3158 | 6.2812 | 6.2240 | 6.9288 | 6.4983 | 5.9991 | 6.5124 | 6.5737 | 6.5426 | 6.7048 | 6.3565 | 6.4093 | 5.5034 | 5.9502 | 5.5746 | 5.6255 |
| NDC1_S471;S474 | NDC1 | Q9BTX1 | 414 | EETAFQTPKSSQM | 7.1344 | 6.9270 | 6.8312 | 7.4613 | 6.4889 | 6.0391 | 7.3021 | 7.0371 | 7.1566 | 6.6227 | 7.0062 | 6.9070 | 4.0268 | 4.4901 | 3.6902 | 4.8793 |
| NDC1_S474 | NDC1 | Q9BTX1 | 474 | CYGSPQSPQLIRR | 6.2032 | 6.5078 | 6.7425 | 6.1149 | 5.8885 | 6.2035 | 7.2084 | 6.6127 | 6.6079 | 6.0868 | 6.2098 | 6.7250 | 5.5508 | 6.0323 | 5.6301 | 5.6757 |
| NDC1_S500 | NDC1 | Q9BTX1 | 406 | EPKKLNSPEETAF | 6.8447 | 6.9177 | 5.2674 | 6.2592 | 6.7635 | 5.6008 | 7.1990 | 6.1655 | 5.7196 | 6.2280 | 6.5155 | 7.3115 | 5.1487 | 5.5612 | 5.7654 | 6.7322 |
| NDC1_S500;S504 | NDC1 | Q9BTX1 | 436 | KTSLFSSKLSTPD | 7.8167 | 8.5306 | 7.0720 | 8.2166 | 6.4172 | 6.4532 | 9.3252 | 6.8992 | 6.8670 | 6.4705 | 7.1594 | 6.8305 | 2.3934 | 3.8625 | 2.3497 | 3.3363 |
| NDC1_S504 | NDC1 | Q9BTX1 | 471;474 | NA | 6.7962 | 8.1590 | 7.2950 | 7.0013 | 6.7620 | 7.0529 | 6.9882 | 6.5951 | 7.0043 | 7.0280 | 7.2972 | 6.2565 | 3.6122 | 4.1676 | 3.2460 | 4.7385 |
| NDC1_T414 | NDC1 | Q9BTX1 | 500;504 | NA | 6.3007 | 6.6278 | 6.4967 | 7.4482 | 6.3454 | 6.4206 | 7.2441 | 5.8910 | 7.4415 | 6.7111 | 6.4532 | 6.8293 | 4.7201 | 5.7266 | 4.5318 | 4.8119 |
| NDC80_S5 | NDC80 | O14777 | 69 | GHGSRNSQLGIFS | 6.9311 | 6.3776 | 6.3216 | 5.2594 | 4.1553 | 6.4924 | 7.0589 | 6.0976 | 7.4145 | 6.5614 | 7.8065 | 7.5004 | 6.1396 | 5.9837 | 4.3531 | 5.5469 |
| NDC80_S50 | NDC80 | O14777 | 31 | NKQGLYTPQTKEK | 7.0795 | 6.8594 | 7.2061 | 7.3836 | 4.9879 | 6.8599 | 7.7351 | 6.5256 | 7.0186 | 6.2958 | 7.2210 | 7.0853 | 4.3784 | 4.9448 | 3.5270 | 4.8920 |
| NDC80_S69 | NDC80 | O14777 | 5 | xxMKRSSVSSGGA | 5.8523 | 5.5758 | 6.5463 | 6.1163 | 5.7029 | 5.0449 | 6.2891 | 6.1758 | 9.1634 | 8.5385 | 7.6205 | 7.8210 | 6.1398 | 5.9753 | 3.1950 | 4.2430 |
| NDC80_T31 | NDC80 | O14777 | 50 | SINKPTSERKVSL | 7.2717 | 8.0208 | 7.7401 | 7.2464 | 4.2134 | 6.9196 | 8.4371 | 7.8562 | 7.9567 | 7.4792 | 7.7825 | 8.6353 | 2.1984 | 3.3374 | 1.8173 | 3.0878 |
| NDE1_S211 | NDE1 | Q9NXR1-1 | 282 | NLVYDQSPNRTGG | 6.9521 | 6.7404 | 6.6166 | 5.9594 | 6.9195 | 6.7528 | 6.5824 | 6.8408 | 7.1512 | 7.0466 | 7.3454 | 7.1576 | 4.4796 | 4.7001 | 4.0762 | 4.6794 |
| NDE1_S282 | NDE1 | Q9NXR1-1 | 228 | PSSSLNTPGSFRR | 7.5453 | 6.8766 | 6.6139 | 6.5661 | 6.5867 | 7.0719 | 7.1207 | 7.2661 | 7.0365 | 7.2690 | 7.7218 | 7.4303 | 3.8075 | 4.2085 | 3.0780 | 3.8012 |
| NDE1_S306 | NDE1 | Q9NXR1-1 | 211 | AVQATGSVPSTPI | 6.6365 | 6.7217 | 6.0459 | 6.0975 | 7.0623 | 6.5995 | 5.6324 | 6.9211 | 7.5767 | 6.5928 | 7.1161 | 7.4333 | 4.4042 | 5.6413 | 4.4198 | 5.0990 |
| NDE1_T228 | NDE1 | Q9NXR1-1 | 306 | GGERRPSSTSVPL | 5.6954 | 5.2434 | 6.0239 | 5.7708 | 6.5830 | 5.9791 | 4.6384 | 5.9583 | 8.1739 | 8.0980 | 8.2608 | 8.9702 | 5.4008 | 5.8211 | 4.3165 | 5.0663 |
| NDEL1_S194;S198 | NDEL1 | Q9GZM8-2 | 231 | TENTFPSPKAIPN | 6.6045 | 6.5534 | 6.8842 | 6.6130 | 6.8382 | 6.8962 | 6.9516 | 6.9189 | 6.9168 | 6.8079 | 7.4145 | 7.7045 | 4.2472 | 4.5956 | 3.6428 | 4.4108 |
| NDEL1_S231 | NDEL1 | Q9GZM8-2 | 194;198 | NA | 6.5717 | 7.0060 | 7.1074 | 6.7167 | 5.6265 | 6.1615 | 6.5263 | 5.7066 | 6.2451 | 6.6791 | 7.0223 | 6.8750 | 5.5794 | 5.8481 | 5.2906 | 5.0377 |
| NDRG2_S332 | NDRG2 | Q9UN36 | 332 | SRSRTASLTSAAS | 5.2613 | 5.3118 | 5.9923 | 5.7483 | 5.9713 | 5.7011 | 5.8548 | 7.0028 | 6.0268 | 7.1369 | 6.6709 | 7.0353 | 6.3042 | 7.4656 | 6.2119 | 6.3046 |
| NDRG3_S331 | NDRG3 | Q9UGV2 | 331 | ARSRTHSTSSSLG | 5.9766 | 6.0982 | 6.3346 | 5.7227 | 6.9095 | 6.4698 | 5.5476 | 8.1633 | 6.7091 | 5.5194 | 7.0126 | 6.2587 | 6.0437 | 6.5933 | 5.1848 | 5.4562 |
| NDRG3_S352 | NDRG3 | Q9UGV2 | 352 | SVTSNQSDGTQES | 6.6074 | 6.1127 | 7.0347 | 5.1561 | 6.3493 | 5.7691 | 7.5426 | 5.8332 | 5.1347 | 5.9910 | 7.4177 | 6.7372 | 6.4973 | 5.2333 | 6.1258 | 6.4578 |
| NDUFAF1_S45 | NDUFAF1 | Q9Y375 | 45 | LQKPVASPGKASS | 5.9019 | 5.9143 | 6.2485 | 7.4919 | 6.4311 | 6.0980 | 5.7258 | 5.8050 | 5.8701 | 6.4195 | 6.4989 | 7.5366 | 6.1584 | 6.3642 | 5.4129 | 6.1230 |
| NDUFAF2_S134 | NDUFAF2 | Q8N183 | 134 | QIKGHASAPYFGK | 2.8077 | 2.7610 | 2.3543 | 2.6827 | 3.9800 | 3.3698 | 3.3141 | 3.6446 | 2.9201 | 2.6130 | 2.6122 | 2.5861 | 16.7297 | 13.4773 | 17.0118 | 17.1358 |
| NDUFB10_S161 | NDUFB10 | O96000-2 | 161 | QNHCKSSLPSLVL | 5.9810 | 8.2836 | 8.2712 | 8.2465 | 5.9315 | 4.5763 | 4.9347 | 6.3805 | 6.4094 | 5.3919 | 5.9109 | 7.2414 | 6.5618 | 5.6720 | 4.0285 | 6.1787 |
| NDUFB4_S26 | NDUFB4 | O95168 | 26 | PAEYNISPETRRA | 6.5364 | 5.1864 | 6.1471 | 6.4684 | 6.3442 | 5.8307 | 5.9897 | 6.2001 | 7.0432 | 5.8175 | 5.9318 | 5.6435 | 7.6661 | 7.0333 | 5.9814 | 6.1802 |
| NDUFV3_S100 | NDUFV3 | P56181-2 | 470 | PSSGRESPRHxxx | 6.2715 | 6.6107 | 5.8719 | 6.4501 | 6.4666 | 6.2249 | 5.7563 | 6.0322 | 6.1699 | 6.1231 | 6.1187 | 6.3263 | 6.6849 | 6.5891 | 6.1001 | 6.2036 |
| NDUFV3_S470 | NDUFV3 | P56181-2 | 98 | KGRKVASPSPSGS | 6.3891 | 6.0533 | 5.4074 | 6.5329 | 6.5888 | 5.8664 | 6.3848 | 6.1911 | 6.2033 | 6.2571 | 6.3065 | 6.4261 | 6.5416 | 6.2164 | 6.4789 | 6.1561 |
| NDUFV3_S98 | NDUFV3 | P56181-2 | 100 | RKVASPSPSGSVL | 6.5167 | 5.2594 | 5.9679 | 6.6067 | 5.9282 | 5.7221 | 6.1015 | 5.6863 | 6.0579 | 7.3597 | 5.7889 | 6.3870 | 6.5690 | 8.0818 | 6.1934 | 5.7734 |
| NEB_T34;T35 | NEB | P20929-2 | 34;35 | NA | 7.2181 | 7.9120 | 8.1678 | 8.5708 | 6.7721 | 6.5663 | 6.2160 | 6.8279 | 6.9846 | 6.5909 | 6.8541 | 7.3774 | 3.2328 | 3.6180 | 3.4022 | 3.6892 |
| NEB_T35;T36 | NEB | P20929-2 | 35;36 | NA | 7.3371 | 8.0408 | 8.3262 | 9.0706 | 7.2554 | 6.5403 | 6.0357 | 6.9220 | 7.1888 | 6.9116 | 6.7251 | 7.1815 | 2.9395 | 3.1807 | 2.9615 | 3.3833 |
| NECAP2_S181 | NECAP2 | Q9NVZ3-2 | 181 | PRVRPASTGGLSL | 6.9018 | 6.6999 | 6.9727 | 6.9226 | 6.1603 | 6.7703 | 6.8788 | 6.9370 | 7.7904 | 6.6237 | 7.0243 | 6.9385 | 4.5109 | 5.0833 | 3.7396 | 4.0459 |
| NECTIN2_S433 | NECTIN2 | Q92692-2 | 470 | ESQLDGSLISRRA | 5.8370 | 6.4448 | 5.9427 | 6.4457 | 6.2512 | 6.5007 | 6.6580 | 6.5812 | 6.2129 | 6.4042 | 6.1193 | 6.3493 | 5.8104 | 6.2363 | 6.0342 | 6.1721 |
| NECTIN2_S465 | NECTIN2 | Q92692 | 433 | LGASEHSPLKTPY | 6.2361 | 7.0926 | 6.3667 | 7.0193 | 6.7499 | 7.1667 | 6.5047 | 7.0624 | 6.4829 | 6.4572 | 7.0179 | 6.6233 | 4.4716 | 5.3172 | 4.6937 | 4.7378 |
| NECTIN2_S465;S470 | NECTIN2 | Q92692-2 | 465 | LEDDMESQLDGSL | 6.3711 | 6.4452 | 6.2010 | 6.4853 | 6.3645 | 5.9085 | 6.2279 | 6.0146 | 6.1374 | 5.7997 | 5.8252 | 5.9839 | 6.2144 | 6.7311 | 6.8005 | 6.4898 |
| NECTIN2_S470 | NECTIN2 | Q92692 | 410 | PSYKPPTPKAKLE | 6.1391 | 6.3880 | 6.7099 | 6.7263 | 6.1361 | 6.2451 | 6.1326 | 6.0759 | 6.6701 | 6.7027 | 6.5389 | 6.8042 | 5.5958 | 5.8335 | 5.3716 | 5.9303 |
| NECTIN2_S476 | NECTIN2 | Q92692 | 520 | DALSYSSPSDSYQ | 5.9955 | 6.4592 | 6.3409 | 6.6149 | 7.0780 | 7.1760 | 6.5944 | 7.1050 | 6.8142 | 7.0495 | 6.4840 | 6.8394 | 4.4515 | 5.4417 | 4.5849 | 4.9708 |
| NECTIN2_S476;S491 | NECTIN2 | Q92692 | 513 | DKINPIYDALSYS | 5.9231 | 6.4964 | 6.0519 | 5.9705 | 5.7705 | 6.3227 | 5.6614 | 6.4534 | 7.0669 | 7.1285 | 7.1202 | 6.9983 | 5.5077 | 6.3523 | 5.2391 | 5.9373 |
| NECTIN2_S517 | NECTIN2 | Q92692 | 476 | GATSLGSPIPVPP | 6.5048 | 7.2831 | 6.1707 | 6.5638 | 6.3520 | 7.1894 | 7.3160 | 7.0634 | 6.5851 | 5.9212 | 6.7245 | 7.3300 | 4.2471 | 5.2956 | 4.4059 | 5.0473 |
| NECTIN2_S517;S520 | NECTIN2 | Q92692 | 517 | PIYDALSYSSPSD | 7.2595 | 7.3352 | 8.0654 | 7.8413 | 6.3827 | 6.3923 | 6.1715 | 6.7163 | 6.5471 | 6.2360 | 7.6044 | 6.9420 | 3.8715 | 5.3237 | 3.3896 | 3.9215 |
| NECTIN2_S517;S520;S52 | NECTIN2 | Q92692 | 524 | YSSPSDSYQGKGF | 6.1819 | 6.0435 | 6.4146 | 6.5816 | 6.2619 | 6.1605 | 7.7292 | 6.4098 | 6.9216 | 7.1649 | 6.2535 | 6.5553 | 4.9973 | 5.1608 | 4.4831 | 6.6806 |
| NECTIN2_S519;S520 | NECTIN2 | Q92692 | 517;520 | NA | 7.2639 | 8.1839 | 7.1565 | 6.8921 | 5.5615 | 7.8698 | 7.9417 | 6.2037 | 5.8804 | 6.5777 | 8.0987 | 7.7666 | 2.7875 | 3.8064 | 3.2631 | 4.7465 |
| NECTIN2_S520 | NECTIN2 | Q92692 | 520;524 | NA | 5.7051 | 6.8568 | 5.9919 | 6.7835 | 6.9963 | 6.2348 | 6.1793 | 6.3783 | 6.9432 | 6.7885 | 5.6641 | 6.2417 | 6.0773 | 6.4769 | 5.4547 | 5.2276 |
| NECTIN2_S520;S524 | NECTIN2 | Q92692 | 476;491 | NA | 7.4173 | 7.9014 | 7.2132 | 7.2604 | 7.8923 | 7.6395 | 8.2179 | 6.4429 | 6.8530 | 5.2228 | 7.8023 | 6.9497 | 3.4136 | 3.8612 | 2.6897 | 3.2228 |
| NECTIN2_S524 | NECTIN2 | Q92692-2 | 465;470 | NA | 6.4721 | 5.7050 | 6.8760 | 6.1365 | 6.3341 | 6.4789 | 6.4521 | 5.4476 | 5.6453 | 6.6275 | 6.5936 | 7.0071 | 6.3010 | 6.0436 | 5.9442 | 5.9353 |
| NECTIN2_T410 | NECTIN2 | Q92692 | 513;524 | NA | 6.4055 | 5.2887 | 6.5870 | 6.5745 | 6.1093 | 6.0339 | 6.6802 | 6.1610 | 6.2700 | 7.1145 | 5.6925 | 6.9505 | 5.7971 | 6.8072 | 5.6780 | 5.8502 |
| NECTIN2_Y513 | NECTIN2 | Q92692 | 517;520;524 | NA | 5.8986 | 7.5601 | 6.3249 | 6.2841 | 8.5031 | 7.7265 | 9.6680 | 4.2461 | 7.0396 | 5.0306 | 6.2265 | 5.9952 | 4.7164 | 6.5526 | 3.5354 | 4.6922 |
| NECTIN2_Y513;S524 | NECTIN2 | Q92692 | 519;520 | NA | 6.9820 | 7.6074 | 6.6771 | 8.1074 | 6.4487 | 6.7980 | 7.0465 | 7.4757 | 7.0840 | 7.1531 | 6.4599 | 7.5895 | 3.2336 | 3.7098 | 3.4807 | 4.1466 |
| NECTIN3_S397 | NECTIN3 | Q9NQS3-3 | 397 | PLYEERSPPLPQK | 6.5935 | 6.6869 | 6.4256 | 6.1426 | 6.1143 | 6.1025 | 6.5381 | 6.6215 | 6.7835 | 6.4593 | 6.3231 | 6.3280 | 5.5922 | 5.7831 | 5.4614 | 6.0445 |
| NEDD1_S412 | NEDD1 | Q8NHV4-3 | 557 | PINGSSTPNPKIA | 6.1759 | 6.5829 | 7.0813 | 6.5601 | 6.4775 | 6.9408 | 6.7681 | 6.6324 | 7.4882 | 7.0309 | 6.9324 | 6.7636 | 4.2220 | 5.1579 | 4.3296 | 4.8564 |
| NEDD1_T557 | NEDD1 | Q8NHV4-3 | 412 | DDTGKSSLGDMFS | 5.1641 | 5.3144 | 6.2618 | 5.3921 | 6.3501 | 6.3017 | 6.0678 | 6.6490 | 5.8227 | 5.9289 | 6.0895 | 5.8934 | 6.7241 | 7.1353 | 6.9611 | 7.9441 |
| NEDD4_S747 | NEDD4 | P46934 | 747 | HSSRRGSLQAYTF | 6.1965 | 5.5398 | 6.3982 | 6.5236 | 6.0751 | 6.7191 | 5.7913 | 6.8007 | 6.7883 | 6.8526 | 6.9178 | 6.6353 | 5.6691 | 6.2449 | 5.2219 | 5.6258 |
| NEDD4L_S303 | NEDD4L | Q96PU5 | 449 | RPRSLSSPTVTLS | 6.7992 | 7.1857 | 7.0368 | 6.7935 | 6.3982 | 6.9139 | 6.9518 | 6.3642 | 7.4344 | 6.8507 | 6.9161 | 7.3599 | 4.0526 | 4.4833 | 3.9062 | 4.5533 |
| NEDD4L_S446 | NEDD4L | Q96PU5 | 448 | RRPRSLSSPTVTL | 5.7845 | 5.6312 | 6.5253 | 6.0598 | 5.9176 | 6.1133 | 5.7526 | 6.2991 | 6.5299 | 6.4346 | 6.7495 | 6.6773 | 6.7184 | 6.4488 | 5.8176 | 6.5404 |
| NEDD4L_S448 | NEDD4L | Q96PU5 | 318 | SRRLQITPDSNGE | 5.5597 | 6.0915 | 6.9339 | 8.2903 | 6.4244 | 7.7272 | 7.6356 | 7.1060 | 6.8557 | 6.8691 | 6.6568 | 7.4099 | 3.2871 | 5.9194 | 3.2692 | 3.9642 |
| NEDD4L_S448;S449 | NEDD4L | Q96PU5 | 446 | QIRRPRSLSSPTV | 6.8748 | 7.4368 | 7.4653 | 6.8517 | 6.4545 | 6.7554 | 6.7685 | 7.3535 | 8.0367 | 7.5517 | 6.6857 | 6.3581 | 3.1012 | 4.2885 | 3.6166 | 4.4012 |
| NEDD4L_S449 | NEDD4L | Q96PU5 | 303 | SPGSRTSPQELSE | 5.7303 | 6.4009 | 6.9405 | 6.3887 | 6.0391 | 5.8779 | 5.4920 | 6.0142 | 6.5946 | 6.3652 | 6.8761 | 6.6012 | 5.9094 | 6.5133 | 5.6998 | 6.5570 |
| NEDD4L_T318 | NEDD4L | Q96PU5 | 448;449 | NA | 6.0744 | 6.0875 | 5.8553 | 6.3734 | 6.2319 | 6.6576 | 6.1231 | 6.7752 | 6.5725 | 6.5090 | 6.5528 | 6.2946 | 6.1672 | 5.9082 | 5.6990 | 6.1184 |
| NEIL2_S68 | NEIL2 | Q969S2 | 68 | MGPPGSSPTPEPP | 5.3369 | 6.0042 | 6.6025 | 6.3074 | 6.0440 | 6.0171 | 6.0156 | 5.8751 | 6.5604 | 6.7440 | 6.8887 | 6.5705 | 6.0985 | 6.2645 | 6.3146 | 6.3561 |
| NEIL3_S474 | NEIL3 | Q8TAT5 | 474 | KTAQYSSPELKSC | 7.8039 | 8.1024 | 7.6355 | 8.7585 | 5.4697 | 5.9715 | 6.9102 | 6.8168 | 8.5420 | 8.0421 | 8.1336 | 8.3412 | 1.9609 | 3.0297 | 1.9750 | 2.5071 |
| NEK1_S1080 | NEK1 | Q96PY6-3 | 1080 | KMLRTCSLPDLSK | 6.2237 | 6.0683 | 6.7779 | 5.8013 | 6.2585 | 6.3258 | 6.0966 | 6.0593 | 6.3785 | 6.3394 | 6.5452 | 6.1570 | 6.2771 | 6.4447 | 6.1944 | 6.0522 |
| NEK1_S826 | NEK1 | Q96PY6-3 | 826 | KLGPNGSPRRAWG | 6.4486 | 6.0732 | 6.8925 | 6.5482 | 5.9044 | 6.6129 | 6.2700 | 6.7821 | 6.9546 | 6.3960 | 6.7035 | 7.0118 | 5.0273 | 5.5747 | 5.1595 | 5.6408 |
| NEK1_S865 | NEK1 | Q96PY6-3 | 852 | AELQLQTELLENT | 6.1710 | 6.4000 | 5.8769 | 6.9208 | 6.2644 | 6.3699 | 5.8867 | 5.8959 | 5.9742 | 6.2336 | 5.9995 | 6.1626 | 6.6903 | 6.4206 | 6.7105 | 6.0232 |
| NEK1_S896 | NEK1 | Q96PY6-3 | 896 | PSAIVDSPVETKS | 5.8234 | 5.4328 | 6.6122 | 6.0468 | 5.8621 | 6.4876 | 6.0218 | 6.5764 | 6.1352 | 5.8827 | 6.1143 | 6.1091 | 6.7107 | 6.7901 | 6.8692 | 6.5257 |
| NEK1_S909 | NEK1 | Q96PY6-3 | 909 | PEFSEASPQMSLK | 5.7623 | 6.7647 | 6.7685 | 6.2723 | 5.3113 | 6.3844 | 7.1897 | 6.7706 | 6.2739 | 6.6011 | 6.6032 | 7.1531 | 5.0919 | 6.1225 | 5.5158 | 5.4149 |
| NEK1_T852 | NEK1 | Q96PY6-3 | 865 | TIRSEISPEGEKY | 6.1725 | 6.7156 | 6.2821 | 6.0417 | 5.9008 | 6.3605 | 6.5294 | 6.2718 | 6.1159 | 6.3982 | 6.5243 | 6.5050 | 5.6943 | 6.3784 | 5.9819 | 6.1277 |
| NEK3_S355 | NEK3 | P51956 | 479 | GLDEEDTDFEEED | 5.9886 | 6.3960 | 6.8211 | 6.3051 | 6.3318 | 6.5819 | 6.1519 | 6.5029 | 6.5694 | 6.4855 | 6.5768 | 6.4474 | 5.6382 | 5.9320 | 5.3399 | 5.9315 |
| NEK3_T479 | NEK3 | P51956 | 355 | HLRKASSPNLHRR | 5.5542 | 6.3230 | 6.4164 | 6.8079 | 6.6020 | 5.8173 | 5.8581 | 6.2588 | 6.2738 | 6.5394 | 5.8164 | 6.2438 | 6.9297 | 6.3309 | 5.6918 | 6.5365 |

| | | | | | | | | | | | | | | | | | | | | | |
|---|---|---|---|---|---|---|---|---|---|---|---|---|---|---|---|---|---|---|---|---|---|
| NEK4_S461 | NEK4 | P51957 | | 461 | DQSLALSPKLECS | 7.1975 | 7.0614 | 7.6651 | 6.9492 | 6.3475 | 6.5486 | 6.9608 | 7.0726 | 7.4525 | 7.1744 | 7.4000 | 8.1637 | 3.2238 | 3.6947 | 3.0605 | 4.0278 |
| NEK4_S661 | NEK4 | P51957 | | 661 | LPARRLSSDCSVT | 6.3235 | 5.8578 | 6.7788 | 6.4556 | 6.0842 | 6.0961 | 6.1470 | 5.6759 | 7.5460 | 7.7469 | 7.0836 | 6.7534 | 3.3586 | 6.2688 | 4.4907 | 5.3332 |
| NEK9_S29 | NEK9 | Q8TD19 | | 827 | FIPMPDSPSPLSA | 6.2180 | 6.0101 | 6.8410 | 6.3437 | 6.4445 | 6.8166 | 6.5233 | 6.8430 | 6.6936 | 6.4836 | 6.8810 | 6.6512 | 5.1587 | 5.8122 | 5.0196 | 5.2599 |
| NEK9_S793 | NEK9 | Q8TD19 | | 829 | PMPDSPSPLSAAF | 6.2773 | 6.4164 | 6.8061 | 6.0268 | 6.2744 | 6.7005 | 6.2201 | 6.6793 | 6.6859 | 6.7590 | 6.6298 | 6.8403 | 5.3248 | 5.8248 | 5.1499 | 5.3846 |
| NEK9_S827 | NEK9 | Q8TD19 | | 978 | TDSCRPSLxxxxx | 6.0369 | 6.4399 | 7.1079 | 6.7558 | 5.9745 | 6.9287 | 6.7631 | 6.9199 | 6.5315 | 6.7296 | 6.8826 | 6.7530 | 5.0626 | 5.3332 | 4.6851 | 5.0956 |
| NEK9_S829 | NEK9 | Q8TD19 | | 793 | GMEGLISPTEAMG | 6.3219 | 6.8048 | 6.8369 | 6.1018 | 6.2291 | 6.3731 | 6.2604 | 7.0280 | 6.0606 | 6.5669 | 5.9586 | 6.5967 | 5.9790 | 6.0149 | 5.7995 | 6.0677 |
| NEK9_S869 | NEK9 | Q8TD19 | | 869 | PQVEASSPRLNPA | 6.4329 | 6.5966 | 6.9578 | 6.8997 | 6.5848 | 7.2054 | 6.7694 | 6.8523 | 6.8632 | 6.8082 | 6.9444 | 6.5452 | 4.2942 | 5.2277 | 4.3950 | 4.6232 |
| NEK9_S978 | NEK9 | Q8TD19 | | 29 | GGCGDSSPGPSAS | 6.0526 | 7.3040 | 6.7439 | 7.3055 | 6.2708 | 8.2074 | 8.0975 | 7.8106 | 5.7565 | 7.2765 | 6.4463 | 6.3151 | 3.7787 | 4.4074 | 4.2273 | 3.9999 |
| NEK9_T333 | NEK9 | Q8TD19 | | 886 | GKGTPLTPPACAC | 5.7372 | 6.2608 | 6.8547 | 6.4073 | 6.3493 | 7.2890 | 6.1733 | 6.5695 | 5.7485 | 6.4446 | 5.7476 | 5.4000 | 6.2496 | 6.6579 | 5.9158 | 6.1950 |
| NEK9_T886 | NEK9 | Q8TD19 | | 333 | KRPRSSTVTEAPI | 5.3740 | 5.0377 | 5.8993 | 5.2915 | 5.6548 | 5.5261 | 5.7287 | 6.1644 | 5.5726 | 5.0064 | 5.0473 | 5.0449 | 8.4137 | 8.5707 | 9.9969 | 7.6609 |
| NELFA_S225 | NELFA | Q9H3P2 | | 277 | AKRRRKTLDAEVV | 4.1907 | 4.4098 | 4.3870 | 4.2178 | 4.9192 | 5.5869 | 5.4624 | 5.6878 | 4.4599 | 4.4672 | 4.4439 | 4.8104 | 10.7215 | 10.0942 | 11.5533 | 10.5881 |
| NELFA_S233 | NELFA | Q9H3P2 | | 363 | EEPSAPSPTLPAQ | 5.7679 | 6.1406 | 5.9840 | 5.9221 | 6.3281 | 6.5307 | 6.2736 | 6.8483 | 6.1947 | 6.3818 | 6.3982 | 6.0379 | 6.2319 | 6.3702 | 6.4214 | 6.1687 |
| NELFA_S363 | NELFA | Q9H3P2 | | 157 | TLAGPLTPPVKHF | 5.8085 | 6.5307 | 6.6101 | 6.3880 | 6.2712 | 6.5820 | 6.5251 | 6.1908 | 6.7958 | 6.7102 | 6.7639 | 6.5441 | 5.5279 | 6.1412 | 5.0958 | 5.5146 |
| NELFA_T157 | NELFA | Q9H3P2 | | 233 | TAPSVFSPTGNRT | 6.2253 | 6.8596 | 6.9095 | 6.5036 | 6.3278 | 6.7011 | 6.2648 | 7.0603 | 6.8259 | 6.9332 | 6.8101 | 7.1248 | 4.4536 | 5.3989 | 4.5192 | 5.0823 |
| NELFA_T277 | NELFA | Q9H3P2 | | 225 | KQAPFRSPTAPSV | 6.0823 | 6.7655 | 7.1596 | 6.4234 | 6.1273 | 6.6049 | 6.2934 | 6.8333 | 7.7617 | 6.6632 | 6.4847 | 6.5287 | 5.0481 | 5.7536 | 4.5305 | 4.9398 |
| NELFB_S581 | NELFB | Q8WX92-2 | | 605 | LDHRKPSPAQAAE | 5.4438 | 5.7049 | 5.7748 | 5.6134 | 6.0088 | 5.9864 | 5.8289 | 6.0157 | 6.0510 | 6.2620 | 5.8904 | 6.0964 | 7.2794 | 7.4325 | 7.2790 | 7.3325 |
| NELFB_S605 | NELFB | Q8WX92-2 | | 581 | LEPTGQSGEAVKE | 5.7191 | 6.8989 | 6.1439 | 6.6045 | 6.6743 | 7.0112 | 5.8360 | 6.6664 | 6.1193 | 6.1750 | 5.6167 | 6.0682 | 5.4095 | 6.7525 | 6.4783 | 5.8262 |
| NELFE_S122 | NELFE | P18615-3 | | 188 | GAHSSASPPRSRS | 6.1414 | 6.9743 | 5.4263 | 6.2688 | 6.3288 | 6.6845 | 5.5639 | 7.1157 | 5.8117 | 5.6275 | 6.0698 | 6.0416 | 6.2480 | 6.7649 | 6.9948 | 5.9379 |
| NELFE_S138 | NELFE | P18615-3 | | 138 | RRPQRKSLYESFV | 5.8405 | 6.1013 | 7.1500 | 6.2414 | 6.2747 | 6.5891 | 5.5682 | 6.4375 | 6.9032 | 7.0470 | 7.1418 | 7.1695 | 5.0034 | 5.7824 | 4.6594 | 5.0905 |
| NELFE_S188 | NELFE | P18615-3 | | 360 | SLAVQNSPKGCHR | 6.1787 | 6.8355 | 7.1067 | 6.5464 | 6.1539 | 6.9050 | 6.7755 | 6.3603 | 7.4423 | 7.9261 | 7.5550 | 7.5543 | 3.8778 | 4.7785 | 3.5751 | 4.4290 |
| NELFE_S258 | NELFE | P18615-3 | | 58 | GVKRSLSEQPVMD | 4.7053 | 4.5851 | 5.4171 | 5.0665 | 5.3952 | 5.6781 | 4.9002 | 6.0260 | 5.7548 | 5.6127 | 5.3354 | 5.6183 | 8.6340 | 8.7120 | 9.8757 | 8.6838 |
| NELFE_S360 | NELFE | P18615-3 | | 122 | PFQRSIADDDLQ | 3.9483 | 4.5829 | 4.7849 | 4.5947 | 5.1375 | 5.5982 | 5.0823 | 5.2416 | 4.6498 | 4.7273 | 4.5933 | 4.8927 | 10.3473 | 10.6505 | 11.5242 | 9.6455 |
| NELFE_S8 | NELFE | P18615-3 | | 258 | PFRRSDSFPERRA | 5.0111 | 5.6280 | 6.6741 | 5.3671 | 5.7200 | 5.7311 | 5.7120 | 5.9195 | 5.6388 | 7.1799 | 6.7373 | 6.7044 | 6.7721 | 7.2375 | 6.9564 | 7.0107 |
| NEMF_S417 | NEMF | O60524 | | 417 | RNPYLLSEEEDDD | 5.8458 | 6.2545 | 6.9762 | 6.0928 | 5.8877 | 6.4578 | 6.0434 | 6.2015 | 6.7443 | 6.1112 | 6.2858 | 6.8364 | 5.8804 | 5.9988 | 6.0609 | 6.3226 |
| NEMF_S747;S748 | NEMF | O60524 | | 376 | ANQIDWTEIGLIV | 5.8670 | 5.1147 | 9.7727 | 5.9701 | 7.6594 | 6.5902 | 9.0183 | 5.6133 | 5.3379 | 6.2687 | 6.1934 | 6.3894 | 4.9946 | 5.0502 | 4.8114 | 5.3487 |
| NEMF_T376 | NEMF | O60524 | 747;748 | | NA | 5.5859 | 5.6640 | 5.5143 | 6.4307 | 5.9057 | 5.8667 | 6.3307 | 6.2126 | 6.3808 | 6.2792 | 5.7855 | 6.0621 | 6.7168 | 7.1650 | 6.8203 | 7.2797 |
| NEPRO_S465 | NEPRO | Q6NW34 | | 265 | FLINEQSPRASEE | 7.4086 | 7.9546 | 7.1181 | 7.4961 | 6.1281 | 6.8955 | 7.1660 | 6.7043 | 8.0139 | 7.2626 | 7.2420 | 7.3756 | 2.9323 | 3.7993 | 3.0078 | 3.4952 |
| NES_S768 | NES | P48681 | | 768 | TQQRRRSLGEQDQ | 5.9265 | 6.3211 | 6.3128 | 6.5620 | 5.5547 | 6.5713 | 5.7709 | 6.4521 | 6.4362 | 7.0533 | 6.2092 | 6.6651 | 6.6082 | 6.4163 | 5.9114 | 6.1288 |
| NET1_S106 | NET1 | Q7Z628 | | 539 | TAQRRASTVSSVT | 5.9912 | 4.9835 | 6.2397 | 5.6594 | 6.4504 | 5.9583 | 6.3615 | 5.9033 | 7.8876 | 7.2905 | 7.3998 | 7.6259 | 6.3148 | 6.7246 | 4.2185 | 4.9911 |
| NET1_S153 | NET1 | Q7Z628 | | 106 | SLANLISPVRNGA | 5.9705 | 7.4167 | 6.1882 | 5.9553 | 6.3260 | 6.7317 | 6.3724 | 6.8276 | 6.9615 | 6.4516 | 5.8974 | 7.1396 | 5.2322 | 5.5785 | 5.4468 | 5.5039 |
| NET1_S50 | NET1 | Q7Z628 | | 50 | CSLRRGSSFTFLT | 7.5631 | 4.2409 | 4.9431 | 5.0413 | 7.2611 | 5.1279 | 4.8988 | 4.5809 | 11.3868 | 5.7489 | 7.2980 | 5.4109 | 8.5525 | 8.3709 | 4.1268 | 5.4479 |
| NET1_S539 | NET1 | Q7Z628 | | 153 | PAKRRSSALWSEM | 5.5918 | 7.6030 | 5.1924 | 5.8080 | 6.8371 | 4.2667 | 4.9293 | 5.9349 | 8.6744 | 9.4826 | 7.0203 | 8.4536 | 5.7422 | 6.0946 | 4.9951 | 3.3739 |
| NEU3_S450 | NEU3 | Q9UQ49-2 | | 450 | LQGDCTSPGRNPS | 6.6350 | 6.7480 | 5.9179 | 7.1612 | 6.1554 | 6.8161 | 6.7105 | 8.1600 | 6.9167 | 6.2274 | 6.1642 | 6.2613 | 4.4656 | 5.7178 | 4.3615 | 5.5815 |
| NEURL4_S502 | NEURL4 | Q96JN8 | | 902 | NTATEKSFPLHSP | 6.2799 | 6.1921 | 6.4604 | 6.8856 | 6.2511 | 6.0011 | 6.1095 | 6.2823 | 6.4925 | 6.2985 | 6.1334 | 6.6360 | 6.1589 | 6.1485 | 5.5978 | 6.0725 |
| NEURL4_S902 | NEURL4 | Q96JN8 | | 502 | AILRALSPEGALR | 6.4350 | 6.7282 | 6.6328 | 6.3333 | 6.1647 | 6.6972 | 6.7358 | 6.8812 | 6.7601 | 6.4969 | 6.8441 | 6.7294 | 4.9687 | 4.9593 | 5.1707 | 5.4626 |
| NEXN_S160 | NEXN | Q0ZGT2 | | 160 | NNTGTESAEEGD | 5.7274 | 6.2837 | 6.6873 | 6.6358 | 6.9756 | 6.1376 | 5.9660 | 6.9707 | 6.3402 | 6.2500 | 5.8509 | 6.2527 | 5.8774 | 6.3125 | 5.9483 | 5.7839 |
| NF1_S2496 | NF1 | P21359 | | 2496 | ETQPWSSPKGSEG | 6.7830 | 6.9905 | 6.9279 | 7.0805 | 6.2313 | 6.7847 | 7.2441 | 6.7224 | 7.2523 | 6.9695 | 7.0139 | 7.2370 | 3.8268 | 4.5589 | 3.9327 | 4.4445 |
| NF1_S2515 | NF1 | P21359 | | 2597 | PHLRKVSVSESNV | 5.9433 | 6.1444 | 6.0956 | 6.0046 | 6.1818 | 6.3944 | 6.1427 | 6.1717 | 6.4335 | 6.2251 | 6.0456 | 6.0458 | 6.6187 | 6.3480 | 6.9419 | 6.2630 |
| NF1_S2523 | NF1 | P21359 | | 2565 | MESGITTPPKMRR | 6.1734 | 6.4895 | 6.3576 | 6.6695 | 6.2399 | 6.5372 | 5.9488 | 6.5128 | 6.2641 | 6.4518 | 6.5409 | 6.3545 | 5.6204 | 6.4243 | 5.7251 | 5.6903 |
| NF1_S2543 | NF1 | P21359 | | 864 | SGLATYSPPMGPV | 6.0450 | 6.6142 | 6.5525 | 6.8345 | 6.5975 | 6.1929 | 6.4642 | 6.8494 | 6.5851 | 6.6464 | 6.7472 | 6.2287 | 5.4303 | 5.7985 | 4.9486 | 5.4649 |
| NF1_S2597 | NF1 | P21359 | | 2543 | LLGTRKSFDHLIS | 6.3646 | 5.9340 | 6.3811 | 5.4832 | 6.3618 | 5.5510 | 5.0432 | 6.5193 | 6.8101 | 7.3410 | 6.5959 | 6.5625 | 5.7890 | 6.9987 | 6.3044 | 5.9603 |
| NF1_S864 | NF1 | P21359 | | 2515 | PTVGQTSPRARKS | 5.9631 | 5.7060 | 5.7460 | 6.1313 | 6.1546 | 6.3550 | 6.5694 | 5.7661 | 6.6991 | 6.1830 | 5.9498 | 6.5810 | 6.6005 | 6.6183 | 6.6378 | 6.3389 |
| NF1_T2565 | NF1 | P21359 | | 2523 | RARKSMSLDMGQP | 6.1284 | 5.3173 | 5.7251 | 4.9106 | 6.7510 | 5.4496 | 5.5666 | 5.5490 | 7.3427 | 7.7591 | 7.6260 | 6.6136 | 5.9736 | 7.7436 | 5.5120 | 6.5317 |
| NF2_S13 | NF2 | P35240 | | 518 | TDMKRLSMEIEKE | 6.1850 | 6.1690 | 6.3293 | 6.3319 | 6.2293 | 6.0583 | 6.2970 | 6.0888 | 6.6592 | 6.1811 | 6.3604 | 6.4008 | 6.1589 | 6.0117 | 6.3246 | 6.2146 |
| NF2_S518 | NF2 | P35240 | | 13 | SRMSFSSLKRKQP | 6.2762 | 6.0749 | 7.5123 | 6.4879 | 5.7806 | 6.2473 | 6.3261 | 6.2773 | 7.1393 | 7.6247 | 7.9000 | 8.0017 | 4.4878 | 5.5952 | 3.6193 | 4.6492 |
| NF2_T581 | NF2 | P35240 | | 581 | NTIKKLTLQSAKS | 12.6178 | 6.1703 | 5.2422 | 4.7443 | 8.5345 | 4.3672 | 5.1244 | 4.4356 | 9.6439 | 4.1424 | 4.1056 | 4.0566 | 9.3356 | 7.1278 | 5.5327 | 4.8190 |
| NFAT5_S452 | NFAT5 | O94916-5 | | 452 | TLQTPSSPILCTQ | 6.1353 | 5.9303 | 6.3664 | 6.3111 | 5.9018 | 6.2599 | 5.8582 | 5.9416 | 6.5337 | 6.3100 | 6.4366 | 6.0703 | 6.3485 | 6.5717 | 6.3532 | 6.6715 |
| NFAT5_S579 | NFAT5 | O94916-5 | | 579 | VKKEISSPARPCS | 6.5570 | 6.6714 | 6.6278 | 6.6176 | 6.8079 | 6.8227 | 6.9053 | 6.8407 | 7.2817 | 6.5903 | 6.8410 | 6.6611 | 4.9960 | 5.2883 | 4.0458 | 4.4454 |
| NFATC1_S233 | NFATC1 | O95644 | | 233 | ACTLLGSPRHSPS | 6.2342 | 6.2656 | 6.5169 | 5.8509 | 5.3096 | 6.6737 | 6.5998 | 6.7455 | 6.6498 | 6.5857 | 7.1595 | 7.4734 | 5.2252 | 5.7256 | 5.2424 | 5.7423 |
| NFATC1_S903 | NFATC1 | O95644 | | 903 | EVHEDGSPNLAPI | 5.9887 | 7.0161 | 7.2320 | 7.5730 | 7.3500 | 7.0651 | 7.3399 | 7.0147 | 7.3313 | 6.4046 | 7.4881 | 6.7758 | 3.4448 | 4.0304 | 3.3886 | 4.5570 |
| NFATC2IP_S127 | NFATC2IP | Q8NCF5 | | 204 | SPLSPPSPRTKSR | 5.5614 | 5.6619 | 6.0792 | 5.3563 | 5.6957 | 5.9585 | 5.8235 | 6.3297 | 6.2144 | 6.2853 | 6.1809 | 6.4935 | 6.6882 | 7.4502 | 7.1907 | 7.0306 |
| NFATC2IP_S170 | NFATC2IP | Q8NCF5 | | 234 | DLRSCLSPKPPQG | 5.9938 | 6.0242 | 7.3065 | 6.2966 | 5.6396 | 7.1414 | 6.4544 | 6.3909 | 6.8877 | 7.2912 | 7.7886 | 8.1139 | 4.1468 | 5.0051 | 4.4413 | 5.0779 |
| NFATC2IP_S204 | NFATC2IP | Q8NCF5 | | 127 | PLVPVYSGKVKSS | 6.1110 | 6.2492 | 8.1683 | 7.8790 | 4.0504 | 6.3167 | 5.4841 | 6.5546 | 8.1362 | 6.2524 | 8.5026 | 8.0397 | 4.0086 | 5.2366 | 5.4455 | 3.5650 |
| NFATC2IP_S234 | NFATC2IP | Q8NCF5 | | 369 | VSLSRDSPLKTLM | 6.0614 | 7.1776 | 6.8947 | 6.7896 | 6.5668 | 7.2415 | 5.4545 | 7.5241 | 6.7516 | 6.1959 | 7.2845 | 7.1925 | 4.1484 | 5.2526 | 4.7576 | 4.7067 |
| NFATC2IP_S300 | NFATC2IP | Q8NCF5 | | 314 | FGETELSPTATPR | 5.9547 | 6.3161 | 6.4615 | 5.9616 | 6.6526 | 6.1901 | 5.5032 | 6.2967 | 6.6332 | 6.0392 | 6.3009 | 6.7662 | 6.3327 | 6.8257 | 5.7650 | 6.0004 |
| NFATC2IP_S314 | NFATC2IP | Q8NCF5 | | 170 | YHEGSPSPGSPWK | 5.3955 | 6.5660 | 5.9425 | 6.0755 | 6.5869 | 5.8071 | 4.9530 | 7.1703 | 5.7537 | 5.8241 | 6.7045 | 6.2844 | 5.9440 | 7.3351 | 7.1943 | 6.4631 |
| NFATC2IP_S369 | NFATC2IP | Q8NCF5 | | 300 | ATHLGVSPSRILL | 7.4436 | 7.1259 | 6.5001 | 4.7411 | 5.7320 | 7.6664 | 6.8035 | 5.5519 | 5.2532 | 6.5380 | 7.1828 | 5.1157 | 5.4665 | 5.4523 | 6.7561 | 6.6708 |
| NFATC3_S1063 | NFATC3 | Q12968 | | 416 | PIFRTSSLPPLDW | 5.7029 | 5.4423 | 6.2658 | 6.0003 | 6.0899 | 6.5287 | 5.9037 | 5.8555 | 5.7287 | 5.7609 | 5.5855 | 6.0885 | 7.3871 | 7.2715 | 7.2885 | 6.8001 |
| NFATC3_S261;S269 | NFATC3 | Q12968 | | 1063 | RQAPLPSPESLDL | 6.2885 | 6.2935 | 6.9158 | 6.6982 | 6.1416 | 6.8943 | 5.4651 | 6.5528 | 6.8055 | 6.4139 | 6.9981 | 6.7027 | 5.4601 | 5.7544 | 5.1466 | 5.4691 |
| NFATC3_S344 | NFATC3 | Q12968 | | 98 | IPESKYSPLGGPK | 5.6863 | 5.6330 | 6.7412 | 6.8613 | 5.7795 | 6.5283 | 6.3374 | 7.1316 | 6.8508 | 7.0511 | 7.4736 | 7.3658 | 4.3826 | 5.2716 | 4.7809 | 6.1252 |
| NFATC3_S416 | NFATC3 | Q12968 | | 344 | LKTRKTSEDQAAI | 5.4551 | 4.7939 | 5.9124 | 5.2871 | 5.7288 | 6.3332 | 5.2959 | 6.3467 | 6.7749 | 6.3603 | 6.6387 | 7.4463 | 7.5987 | 7.1509 | 6.3162 | 5.5608 |
| NFATC3_S98 | NFATC3 | Q12968 | 261;269 | | NA | 4.6780 | 5.5543 | 5.7683 | 5.5196 | 4.8878 | 7.1014 | 6.2876 | 6.4637 | 5.7308 | 6.2793 | 7.7044 | 7.2968 | 5.9992 | 6.0395 | 6.8061 | 7.8832 |
| NFATC4_T289 | NFATC4 | Q14934-2 | | 289 | ASPRPWTPEDPWS | 5.9409 | 5.9235 | 7.2864 | 5.1584 | 4.9215 | 7.4977 | 7.5961 | 6.8311 | 5.8168 | 5.9686 | 6.0035 | 5.8948 | 5.6281 | 5.8004 | 6.5460 | 7.1859 |
| NFIA_S258 | NFIA | Q12857-2 | | 258 | SSYYSMSPGAMRR | 5.7497 | 5.8782 | 6.2425 | 6.2787 | 5.7751 | 6.2082 | 6.1811 | 6.2542 | 6.4653 | 6.6593 | 6.5821 | 6.4538 | 5.8010 | 6.1773 | 5.9458 | 6.3477 |
| NFIA_S265 | NFIA | Q12857-2 | | 265 | PGAMRRSLPSTSS | 5.9913 | 7.2211 | 6.7388 | 8.2489 | 7.5809 | 5.7991 | 7.2816 | 7.0013 | 7.1667 | 6.6118 | 6.9983 | 7.2328 | 3.9992 | 4.6357 | 3.6987 | 3.7937 |
| NFIA_S472 | NFIA | Q12857-2 | | 472 | GAASPTSPILVPG | 5.4152 | 4.9569 | 5.8357 | 6.5588 | 6.4183 | 6.4093 | 6.6695 | 6.1869 | 5.9033 | 6.1239 | 5.6863 | 6.1680 | 6.8757 | 7.1198 | 7.1360 | 6.5363 |
| NFIB_S194 | NFIB | O00712-5 | | 295 | FYPSPSSPAAGSR | 5.9410 | 6.2441 | 6.1788 | 6.3452 | 6.1910 | 6.0127 | 6.0826 | 6.1118 | 5.8939 | 6.3670 | 6.0505 | 6.1298 | 6.2552 | 6.7304 | 6.7209 | 6.7452 |
| NFIB_S264 | NFIB | O00712-5 | | 194 | DSGQSGSPSHNDP | 5.6364 | 6.4898 | 6.0377 | 6.1640 | 6.4197 | 6.4005 | 6.2155 | 5.9330 | 6.3255 | 6.2679 | 6.5423 | 5.9683 | 6.5929 | 6.3417 | 6.4815 | 6.1832 |
| NFIB_S264;S265 | NFIB | O00712-5 | | 312 | RDQDMSSPTTMKK | 6.7220 | 7.2150 | 6.7694 | 7.0879 | 6.4745 | 6.2911 | 6.8506 | 6.5152 | 7.2010 | 7.1250 | 7.2536 | 7.2522 | 4.0264 | 4.6799 | 4.0232 | 4.5132 |
| NFIB_S265 | NFIB | O00712-5 | | 265 | LQRSLSSPPSSKR | 6.9474 | 7.0030 | 6.6798 | 7.3802 | 6.5601 | 6.6066 | 6.7941 | 6.7673 | 7.0955 | 7.2230 | 7.2773 | 7.2950 | 3.7559 | 4.7115 | 3.7740 | 4.1295 |
| NFIB_S284;S295 | NFIB | O00712-5 | | 264 | NLQRSLSSPPSSK | 4.5961 | 4.8231 | 4.9862 | 4.7900 | 5.0658 | 5.6353 | 5.3081 | 5.6685 | 4.9704 | 5.4113 | 5.2496 | 5.3214 | 9.6267 | 9.4841 | 9.8266 | 9.2368 |
| NFIB_S292 | NFIB | O00712-5 | | 478 | NRYVGLSPRDPSF | 6.2779 | 6.9008 | 6.0113 | 6.1698 | 5.5309 | 6.4613 | 6.5875 | 6.6888 | 6.2201 | 6.1889 | 6.5303 | 6.7572 | 5.3142 | 6.3989 | 5.8072 | 6.1549 |
| NFIB_S295 | NFIB | O00712-5 | | 328 | PLFSSASPQDSSP | 5.7925 | 5.9741 | 6.6261 | 6.2830 | 6.0672 | 6.0252 | 7.0972 | 5.2794 | 6.0911 | 6.1105 | 6.3786 | 6.8025 | 6.6694 | 6.3773 | 5.9838 | 6.4420 |
| NFIB_S312 | NFIB | O00712-5 | | 292 | TGDFYPSPSSPAA | 6.7566 | 6.3458 | 5.4519 | 6.7066 | 6.8676 | 6.7622 | 6.3402 | 6.4457 | 7.2563 | 3.9085 | 6.3902 | 6.6567 | 6.1626 | 4.3971 | 6.2236 | 7.3284 |
| NFIB_S328 | NFIB | O00712-5 | 284;295 | | NA | 5.7518 | 5.9354 | 5.6827 | 6.2454 | 5.6497 | 6.1200 | 5.5273 | 6.9953 | 6.5834 | 5.6502 | 6.4345 | 6.0992 | 6.7527 | 6.7375 | 7.0150 | 6.8200 |
| NFIB_S478 | NFIB | O00712-5 | 264;265 | | NA | 5.5674 | 5.9433 | 4.7472 | 5.9246 | 6.9987 | 6.6494 | 5.8957 | 7.2141 | 5.7475 | 4.9500 | 5.0816 | 4.2679 | 7.8941 | 8.6233 | 7.7296 | 7.6658 |
| NFIC_S277 | NFIC | P08651 | | 323 | MEGGISSPVKKTE | 6.0244 | 6.1991 | 6.4475 | 6.7719 | 6.0657 | 6.1773 | 6.2836 | 6.0182 | 6.5945 | 6.6268 | 6.5423 | 6.7431 | 5.8268 | 6.0244 | 6.5479 | 6.0066 |
| NFIC_S305 | NFIC | P08651 | | 339 | SPFNSPSPQDSPR | 5.9827 | 6.3303 | 6.0190 | 6.2775 | 5.9782 | 5.7769 | 5.7948 | 5.8077 | 6.2446 | 6.2647 | 6.0130 | 6.1297 | 6.8425 | 6.9723 | 6.7918 | 6.7743 |
| NFIC_S323 | NFIC | P08651 | | 305 | YYTSPSSPTSSSR | 6.0560 | 6.8665 | 6.9711 | 6.7064 | 6.2566 | 5.9171 | 5.9204 | 5.8746 | 6.5168 | 6.1854 | 6.2559 | 6.3123 | 5.9636 | 6.4130 | 5.9685 | 5.8158 |
| NFIC_S333 | NFIC | P08651 | | 333 | KTEMDKSPFNSPS | 6.0770 | 6.2676 | 5.8617 | 6.1237 | 5.8738 | 5.7806 | 6.3359 | 6.0292 | 6.2492 | 6.0755 | 5.9518 | 6.1990 | 6.8716 | 6.6324 | 6.7219 | 6.9491 |
| NFIC_S333;S339 | NFIC | P08651 | | 277 | LPSTSSSGSKRHK | 6.8825 | 7.6165 | 6.9845 | 7.6402 | 6.7452 | 6.6793 | 7.2920 | 6.9701 | 7.6132 | 7.4489 | 7.6957 | 7.5410 | 2.7722 | 3.7822 | 2.7090 | 3.6275 |
| NFIC_S333;S343 | NFIC | P08651 | | 487 | TSPANRSFVGLGP | 6.9119 | 7.1284 | 6.4914 | 7.1624 | 5.2764 | 6.9128 | 7.8168 | 6.7448 | 8.0383 | 6.9107 | 7.5994 | 8.4800 | 2.7749 | 4.4259 | 2.8975 | 4.4285 |
| NFIC_S339 | NFIC | P08651 | | 343 | SPSPQDSPRLSSF | 5.6181 | 5.4479 | 6.3908 | 7.0719 | 6.6786 | 7.3282 | 6.0463 | 6.9976 | 6.8977 | 6.8423 | 6.4163 | 6.1373 | 4.9847 | 5.5078 | 5.7929 | 5.8417 |
| NFIC_S343 | NFIC | P08651 | 333;339 | | NA | 5.3670 | 6.2715 | 5.6755 | 6.3846 | 6.3603 | 5.8848 | 6.2430 | 5.9215 | 6.9711 | 5.7198 | 6.8376 | 6.5224 | 5.9594 | 6.2147 | 6.0156 | 6.2110 |
| NFIC_S487 | NFIC | P08651 | 333;343 | | NA | 5.2597 | 6.2346 | 7.4218 | 7.1995 | 6.0583 | 6.5331 | 6.3777 | 5.5589 | 6.9287 | 6.5508 | 6.8356 | 6.2146 | 5.9987 | 5.4948 | 5.8612 | 6.3718 |
| NFIL3_S301 | NFIL3 | Q16649 | | 353 | ATQKLSSPIDMTS | 6.3381 | 6.4067 | 5.8576 | 6.7966 | 6.0477 | 6.1819 | 6.4694 | 6.2040 | 5.8147 | 5.9921 | 6.1993 | 6.5318 | 6.1203 | 6.2668 | 6.3009 | 6.4721 |
| NFIL3_S353 | NFIL3 | Q16649 | | 301 | PKGPIHSPVELKH | 6.5119 | 6.7916 | 6.0504 | 6.9009 | 6.8990 | 6.5219 | 6.5828 | 6.5364 | 6.4304 | 6.1694 | 5.9164 | 6.0796 | 5.7219 | 5.5128 | 5.6952 | 5.6790 |
| NFIX_S301 | NFIX | Q14938 | | 301 | YPGTGRSPAAGSS | 6.1359 | 6.5620 | 6.2243 | 6.6182 | 6.3326 | 6.0169 | 5.7388 | 6.1930 | 6.1992 | 6.8574 | 6.3765 | 6.3177 | 5.9223 | 6.5228 | 5.9690 | 6.0132 |
| NFIX_T347 | NFIX | Q14938 | | 347 | SPRMAFTHHPLPV | 2.4974 | 2.8168 | 23.3075 | 3.5073 | 37.4353 | 8.0385 | 0.4418 | 2.2815 | 2.5890 | 1.9830 | 2.4934 | 2.0716 | 2.9879 | 3.1939 | 2.2278 | 2.1273 |
| NFKB1_S894 | NFKB1 | P19838-2 | | 908 | AHSLPLSPASTRQ | 6.5540 | 6.8496 | 6.6215 | 6.4856 | 6.7643 | 7.0087 | 6.1101 | 7.1882 | 6.7972 | 6.3684 | 6.8985 | 6.6475 | 4.9242 | 5.2242 | 4.7543 | 4.8037 |
| NFKB1_S908 | NFKB1 | P19838-2 | | 894 | VIQAASSPVKTTS | 6.1375 | 6.3273 | 6.5689 | 6.1396 | 6.7307 | 6.8821 | 6.3395 | 6.6350 | 6.8127 | 6.1496 | 6.7350 | 6.9001 | 5.3887 | 5.7678 | 5.0077 | 5.4779 |
| NFKB1_S938 | NFKB1 | P19838-2 | | 938 | TSFRKLSFTESLT | 6.3069 | 5.4335 | 5.8321 | 5.5055 | 6.6496 | 6.1951 | 5.4410 | 7.5014 | 6.4916 | 6.5475 | 6.3660 | 7.0535 | 6.6635 | 6.5582 | 5.5388 | 5.9158 |
| NFKB2_S22 | NFKB2 | Q00653 | | 222 | PIHDSKSPGASNL | 6.1928 | 6.2733 | 6.5562 | 6.0959 | 6.3203 | 6.4553 | 6.3131 | 6.5897 | 6.2469 | 6.1788 | 6.3332 | 6.1983 | 6.0270 | 6.2871 | 6.1780 | 5.7542 |
| NFKB2_S222 | NFKB2 | Q00653 | | 23 | DFKLNSSIVEPKE | 8.2275 | 7.9830 | 7.3220 | 8.1015 | 6.6811 | 6.9019 | 7.5063 | 6.5257 | 7.6801 | 6.5911 | 7.5309 | 7.1475 | 2.6597 | 3.5391 | 2.3906 | 3.2120 |
| NFKB2_S23 | NFKB2 | Q00653 | | 812 | TYRQTTSPSGSLL | 6.8966 | 7.2619 | 7.2558 | 7.4001 | 6.3852 | 7.1576 | 7.6380 | 7.2642 | 6.9158 | 7.0249 | 7.1027 | 7.7659 | 3.1616 | 4.4031 | 3.3738 | 3.9928 |
| NFKB2_S277 | NFKB2 | Q00653 | | 22 | DDFKLNSSIVEPK | 7.6707 | 7.5051 | 7.2870 | 8.1763 | 6.4587 | 6.7734 | 7.4739 | 6.6979 | 8.2581 | 6.9905 | 7.8351 | 7.3641 | 2.4812 | 3.5547 | 2.2703 | 3.2028 |
| NFKB2_S812 | NFKB2 | Q00653 | | 277 | QAFGDFSPTDVHK | 6.3168 | 6.4293 | 6.0585 | 6.9232 | 5.5873 | 6.7381 | 6.6114 | 5.8777 | 6.8285 | 6.8861 | 6.7317 | 6.5396 | 6.4314 | 5.4265 | 5.3311 | 5.2827 |
| NFKBIE_S183 | NFKBIE | O00221 | | 183 | SGPSDGSPQPCTH | 6.4450 | 5.4164 | 6.3605 | 6.1365 | 5.2255 | 6.0444 | 6.2973 | 7.1613 | 6.1878 | 8.0634 | 7.2638 | 8.6506 | 4.0308 | 6.1404 | 4.8400 | 7.3363 |
| NFKBIL1_S150 | NFKBIL1 | Q9UBC1 | | 150 | WGPPWDSAEEEEE | 6.0939 | 6.1137 | 6.6561 | 6.6885 | 6.1639 | 6.1281 | 6.4352 | 5.9580 | 6.4201 | 6.3005 | 6.4705 | 6.4559 | 6.0332 | 6.1073 | 5.8204 | 6.1547 |
| NFRKB_S251 | NFRKB | Q6P4R8-2 | | 323 | NAGRKGSLAALYD | 4.9565 | 4.7387 | 5.1577 | 5.8055 | 7.6593 | 5.2546 | 5.3664 | 5.6514 | 7.3274 | 7.3085 | 6.9660 | 7.0625 | 6.9945 | 6.7717 | 6.4500 | 6.5293 |
| NFRKB_S323 | NFRKB | Q6P4R8-2 | | 376 | PLSQAPSPLAIPA | 5.9473 | 6.2738 | 5.8587 | 6.4838 | 6.3483 | 6.3908 | 6.5221 | 6.8265 | 6.3370 | 6.4519 | 6.3063 | 6.5819 | 6.1630 | 5.7952 | 5.8483 | 5.8651 |
| NFRKB_S372;S376 | NFRKB | Q6P4R8-2 | | 251 | PSSPARSPSPAVP | 5.9237 | 6.6662 | 6.0498 | 6.7785 | 6.3535 | 5.5366 | 6.3336 | 7.3845 | 5.6304 | 6.6705 | 5.9417 | 6.4502 | 5.8190 | 5.9029 | 6.5312 | 6.0275 |
| NFRKB_S376 | NFRKB | Q6P4R8-2 | 372;376 | | NA | 5.5105 | 5.7433 | 6.1303 | 6.3084 | 5.3534 | 5.2278 | 5.2633 | 5.5369 | 6.0708 | 6.1232 | 6.4405 | 5.9406 | 7.9083 | 8.0141 | 7.7973 | 7.6313 |
| NFX1_S326 | NFX1 | Q12986 | | 326 | QDPQVVSPFSRGK | 5.8551 | 6.3946 | 7.1219 | 6.8264 | 6.3486 | 7.1554 | 7.0848 | 6.7896 | 7.2047 | 6.5577 | 7.2466 | 7.0608 | 4.2695 | 5.0124 | 4.1041 | 4.9679 |
| NFXL1_S331 | NFXL1 | Q6ZNB6 | | 50 | AVPSGTSPGGVAT | 6.8541 | 6.5998 | 6.3436 | 7.8314 | 6.6248 | 6.3897 | 7.0849 | 7.1363 | 7.2147 | 6.8142 | 7.3464 | 5.9773 | 3.7621 | 5.8427 | 3.8376 | 4.3404 |
| NFXL1_S50 | NFXL1 | Q6ZNB6 | | 331 | NPCHAGSCQPCPR | 6.0576 | 6.4270 | 6.0269 | 6.8379 | 6.6207 | 5.1488 | 6.3057 | 4.8972 | 7.2199 | 6.2224 | 5.6376 | 6.7655 | 5.9382 | 6.8977 | 6.4777 | 6.5193 |
| NFXL1_S65 | NFXL1 | Q6ZNB6 | | 65 | AAGSRHSPAGSQA | 6.7948 | 6.3816 | 6.2588 | 7.5830 | 6.4368 | 6.7228 | 6.4473 | 6.5415 | 6.9964 | 6.7146 | 6.5021 | 7.0847 | 4.7700 | 5.2269 | 4.5385 | 5.0002 |
| NFYC_S375 | NFYC | Q13952 | | 375 | PQQSSTSPPPSDA | 6.0701 | 6.4969 | 6.0208 | 5.6521 | 6.3987 | 5.4496 | 5.7507 | 6.3419 | 6.5378 | 6.5315 | 6.4685 | 6.2689 | 6.6518 | 6.7023 | 6.5791 | 6.0793 |
| NGDN_S143 | NGDN | Q8NEJ9 | | 204 | RRALSSSVIRELK | 7.7499 | 7.8542 | 7.9936 | 7.8494 | 5.6472 | 7.0245 | 7.8569 | 6.7640 | 6.8234 | 6.9847 | 7.4803 | 7.2307 | 2.6463 | 3.7925 | 2.6839 | 3.6185 |
| NGDN_S204 | NGDN | Q8NEJ9 | | 143 | MMSKLSSEDEEED | 6.0505 | 5.9806 | 5.2132 | 6.2771 | 5.7285 | 6.0028 | 6.2406 | 5.9832 | 6.5893 | 6.4274 | 6.1367 | 6.3967 | 6.3621 | 6.7330 | 6.9342 | 6.9440 |
| NHSL1_S1187 | NHSL1 | Q5SYE7-2 | | 1187 | TPLPDSSPSRKPP | 6.2936 | 5.6495 | 7.1350 | 6.2184 | 6.2990 | 6.4524 | 6.0310 | 6.3581 | 7.0807 | 6.9045 | 7.2323 | 7.2322 | 5.3734 | 5.2977 | 5.0350 | 5.4073 |
| NHSL1_S1272 | NHSL1 | Q5SYE7-2 | | 209 | KVKRRKTITGVPD | 9.1596 | 7.7042 | 7.3929 | 6.7828 | 6.5057 | 6.5968 | 6.6643 | 6.2835 | 7.0052 | 5.8364 | 6.0953 | 6.4153 | 4.7024 | 5.2030 | 3.5870 | 4.0657 |
| NHSL1_S1288 | NHSL1 | Q5SYE7-2 | | 155 | KSLPLPTPEEKMR | 7.5829 | 7.6465 | 7.4567 | 6.8939 | 6.7221 | 7.1429 | 7.8595 | 7.4614 | 8.3027 | 7.0632 | 7.0378 | 8.1698 | 2.4614 | 2.9682 | 2.0814 | 3.3586 |
| NHSL1_S1382;S1384 | NHSL1 | Q5SYE7-2 | | 194 | RRSLIYTDTLVRR | 6.5868 | 5.9742 | 6.8589 | 6.8752 | 6.7168 | 6.2840 | 6.6797 | 6.3153 | 7.1792 | 6.2967 | 6.0693 | 6.5799 | 5.3574 | 5.5442 | 5.4601 | 5.2222 |
| NHSL1_S1463 | NHSL1 | Q5SYE7-2 | | 1288 | MVQPDVSPAPKQE | 5.4232 | 6.3845 | 6.2020 | 6.6412 | 5.8472 | 6.6732 | 6.7261 | 7.1587 | 6.2945 | 5.9077 | 6.1878 | 6.9620 | 5.6092 | 6.1593 | 5.5615 | 6.2620 |
| NHSL1_S1463;S1468 | NHSL1 | Q5SYE7-2 | | 635 | FGNPRHSVINVFV | 9.2440 | 5.2234 | 6.0436 | 5.6208 | 7.3230 | 6.7289 | 5.9889 | 4.1080 | 8.0411 | 6.2003 | 7.1371 | 5.9379 | 6.2003 | 5.3656 | 4.3778 | 6.4591 |
| NHSL1_S1520 | NHSL1 | Q5SYE7-2 | | 937 | PPPPVPSPPFPCP | 6.0144 | 6.8888 | 7.2412 | 6.1362 | 4.9679 | 6.2848 | 6.2228 | 6.3738 | 6.0472 | 6.3867 | 6.8865 | 5.6913 | 6.9790 | 6.0952 | 5.9980 | 5.7862 |
| NHSL1_S324 | NHSL1 | Q5SYE7-2 | | 324 | SDAGFHSLPRSGA | 6.7977 | 6.7617 | 6.6244 | 5.5765 | 5.3181 | 6.4742 | 6.4318 | 5.7644 | 6.1127 | 6.8435 | 6.7594 | 6.7689 | 5.7661 | 6.1313 | 5.2570 | 5.7023 |
| NHSL1_S635 | NHSL1 | Q5SYE7-2 | | 1463 | SPDAPESPSSCSP | 4.9817 | 5.0941 | 5.0346 | 5.6317 | 5.8608 | 6.1583 | 4.8473 | 5.8719 | 6.2464 | 4.9861 | 4.1511 | 6.7532 | 9.8312 | 7.9628 | 8.5819 | 8.0069 |
| NHSL1_S719 | NHSL1 | Q5SYE7-2 | | 847 | KSHRVISPSSGYS | 5.6704 | 7.0363 | 6.8464 | 6.7153 | 5.9707 | 6.5588 | 6.9473 | 7.2239 | 6.8289 | 7.8850 | 6.6549 | 6.8504 | 4.5142 | 5.6306 | 4.2965 | 4.3704 |
| NHSL1_S847 | NHSL1 | Q5SYE7-2 | | 1502 | RYGRSRTPPSAAS | 8.1770 | 9.3530 | 5.5026 | 6.1526 | 4.7534 | 6.8622 | 8.8072 | 5.6168 | 8.0778 | 6.4106 | 9.3728 | 5.8202 | 4.2622 | 3.9397 | 3.1904 | 3.7015 |
| NHSL1_S926;S937 | NHSL1 | Q5SYE7-2 | | 1272 | GAAGSMSPSRVEA | 6.4384 | 6.2486 | 5.8610 | 7.3772 | 6.0649 | 7.6255 | 5.4572 | 7.7002 | 6.5858 | 5.2149 | 6.3617 | 5.8892 | 5.9034 | 6.4485 | 5.3623 | 5.4612 |
| NHSL1_S937 | NHSL1 | Q5SYE7-2 | | 719 | PGKSGSSPSQSPC | 5.7492 | 6.7386 | 6.9740 | 5.8145 | 6.0935 | 5.5813 | 7.5247 | 7.0066 | 5.9191 | 6.5533 | 6.2068 | 7.1832 | 6.6623 | 5.8145 | 6.0434 | 6.1349 |
| NHSL1_T1502 | NHSL1 | Q5SYE7-2 | | 1520 | RNRIQSSPMTVIS | 6.5153 | 6.1243 | 5.7097 | 6.0862 | 5.9675 | 6.6494 | 6.7402 | 6.6886 | 6.4503 | 6.5163 | 6.1089 | 6.1770 | 5.4167 | 6.2368 | 5.8530 | 6.7598 |
| NHSL1_T155 | NHSL1 | Q5SYE7-2 | 1463;1468 | | NA | 5.5107 | 5.2771 | 7.8877 | 5.9554 | 6.3754 | 8.2358 | 6.3715 | 5.8858 | 7.5621 | 5.9030 | 6.1660 | 7.7297 | 5.0721 | 5.7126 | 5.3081 | 5.0469 |
| NHSL1_T194 | NHSL1 | Q5SYE7-2 | 926;937 | | NA | 6.5241 | 6.8837 | 6.4835 | 6.2737 | 6.0317 | 5.7078 | 6.3602 | 7.1733 | 6.6942 | 7.3545 | 7.1931 | 7.0391 | 5.3701 | 4.9422 | 5.3132 | 4.6554 |
| NHSL1_T209 | NHSL1 | Q5SYE7-2 | 1382;1384 | | NA | 5.3635 | 5.8705 | 5.9320 | 6.4234 | 6.6612 | 6.2207 | 6.0194 | 6.2318 | 6.6091 | 6.1922 | 6.4186 | 6.1322 | 6.6461 | 6.4292 | 6.0204 | 6.8295 |
| NIBAN1_S579;S582 | NIBAN1 | Q9BZQ8 | | 602 | SPARRASAILPGV | 5.3629 | 5.2144 | 6.0042 | 5.2673 | 6.0886 | 5.6637 | 5.7236 | 6.1205 | 6.7634 | 6.6329 | 6.5124 | 6.7791 | 7.0328 | 7.6722 | 6.3947 | 6.7673 |
| NIBAN1_S579;S582;S596 | NIBAN1 | Q9BZQ8 | | 582 | PSESVSSLTDLKP | 5.8072 | 5.3216 | 5.6367 | 5.2837 | 5.8854 | 5.9917 | 5.6880 | 6.0878 | 5.8826 | 5.6220 | 5.9194 | 6.1964 | 7.8936 | 7.5936 | 7.5968 | 7.5933 |
| NIBAN1_S579;S592;S596 | NIBAN1 | Q9BZQ8 | | 596 | TGSNQASPARRAS | 5.8160 | 5.9314 | 6.5769 | 5.9728 | 6.5955 | 6.5527 | 5.7925 | 6.4247 | 6.1870 | 6.0991 | 6.7745 | 6.1479 | 6.5266 | 6.5955 | 5.9219 | 6.0850 |
| NIBAN1_S579;S596 | NIBAN1 | Q9BZQ8 | | 633 | QPEVPSSLAKGES | 7.3052 | 7.4802 | 8.1078 | 7.5680 | 6.2032 | 6.1265 | 6.2541 | 5.8128 | 7.5453 | 6.2201 | 7.3102 | 6.8653 | 4.1723 | 4.7039 | 4.1099 | 4.2151 |
| NIBAN1_S582 | NIBAN1 | Q9BZQ8 | | 641 | AKGESLSLPGPSP | 6.2664 | 6.1137 | 7.2443 | 6.2135 | 6.1919 | 5.6947 | 6.8681 | 6.3140 | 5.8948 | 6.1584 | 6.0452 | 6.5989 | 5.9610 | 6.2977 | 5.5522 | 6.4796 |
| NIBAN1_S582;T590;S592 | NIBAN1 | Q9BZQ8 | | 646 | LSLPGPSPPPDGT | 4.3669 | 7.2317 | 5.6276 | 6.5615 | 6.7694 | 5.9506 | 6.0249 | 7.3951 | 6.6060 | 6.6097 | 6.6469 | 7.0553 | 6.3052 | 6.0100 | 4.6305 | 6.2087 |
| NIBAN1_S582;T590;S596 | NIBAN1 | Q9BZQ8 | | 719 | RKLLTASVEVPVD | 5.9454 | 5.9372 | 6.7906 | 5.7460 | 5.9856 | 5.7020 | 6.4868 | 6.7358 | 6.7780 | 6.0277 | 5.4869 | 6.3453 | 6.6258 | 6.4950 | 6.5975 | 6.3145 |
| NIBAN1_S592;S596 | NIBAN1 | Q9BZQ8 | 582;596 | | NA | 5.5809 | 6.0435 | 6.6780 | 6.3676 | 6.3058 | 6.0331 | 5.9821 | 6.8957 | 7.2166 | 6.6591 | 6.9775 | 6.9705 | 5.3768 | 5.8496 | 5.4428 | 5.6205 |
| NIBAN1_S596 | NIBAN1 | Q9BZQ8 | 592;596 | | NA | 5.7928 | 6.5654 | 6.1181 | 6.9231 | 6.3353 | 7.0493 | 5.6985 | 6.8582 | 7.4994 | 6.6904 | 7.2785 | 6.7103 | 5.1733 | 5.2703 | 5.1359 | 4.9013 |
| NIBAN1_S602 | NIBAN1 | Q9BZQ8 | 579;582 | | NA | 5.4851 | 5.7489 | 5.7987 | 5.5766 | 6.1090 | 6.4477 | 4.9717 | 5.8156 | 5.9760 | 5.4005 | 5.4689 | 6.2572 | 7.7279 | 7.2275 | 8.2733 | 7.7152 |
| NIBAN1_S633 | NIBAN1 | Q9BZQ8 | 579;582;596 | | NA | 6.1084 | 6.2159 | 6.8955 | 6.5519 | 6.2173 | 6.0533 | 6.6810 | 6.7222 | 6.3125 | 6.5157 | 6.3096 | 6.8787 | 5.4626 | 5.7786 | 5.6738 | 5.6229 |
| NIBAN1_S641 | NIBAN1 | Q9BZQ8 | 579;596 | | NA | 6.1921 | 6.1553 | 6.9445 | 5.8466 | 5.7967 | 6.9915 | 6.6203 | 6.3500 | 6.7036 | 5.8532 | 5.4861 | 6.7036 | 6.6270 | 6.7855 | 5.5657 | 5.3782 |
| NIBAN1_S646 | NIBAN1 | Q9BZQ8 | 582;590;592;5 | | NA | 6.8635 | 5.3218 | 6.4303 | 6.2067 | 6.0721 | 6.4281 | 6.2844 | 6.6398 | 6.8396 | 6.4814 | 6.8468 | 7.1422 | 5.7829 | 5.7371 | 5.3974 | 5.5259 |
| NIBAN1_S719 | NIBAN1 | Q9BZQ8 | 582;592;596 | | NA | 6.9771 | 6.9731 | 6.5495 | 7.1598 | 5.7086 | 6.3351 | 6.4713 | 6.4343 | 7.9270 | 7.2605 | 7.2307 | 5.9201 | 4.2443 | 5.0583 | 4.5748 | 5.1755 |
| NIBAN1_T582;S592;S596 | NIBAN1 | Q9BZQ8 | 582;590;596 | | NA | 6.7066 | 6.8964 | 6.2960 | 6.7596 | 6.7269 | 5.9458 | 6.6609 | 7.2634 | 6.9277 | 7.3363 | 7.0287 | 7.1061 | 3.5408 | 5.4411 | 4.6956 | 4.6683 |
| NIBAN1_T582;S596 | NIBAN1 | Q9BZQ8 | 579;582;596 | | NA | 6.2673 | 6.2779 | 5.2999 | 5.9733 | 5.8534 | 6.5983 | 6.9632 | 6.2456 | 6.5170 | 6.2182 | 5.6473 | 5.7061 | 7.0670 | 6.8558 | 5.9206 | 6.5892 |
| NIBAN2_S568;S574 | NIBAN2 | Q96TA1 | | 665 | QGLRPESPPPAGP | 6.1305 | 5.7968 | 6.5790 | 5.7863 | 6.4232 | 6.2026 | 5.9867 | 6.5880 | 6.3234 | 6.3889 | 6.4320 | 6.4665 | 6.2460 | 6.6038 | 6.0165 | 6.0299 |
| NIBAN2_S568;Y593 | NIBAN2 | Q96TA1 | | 646 | ESPPPASPDGVTE | 5.8499 | 5.9910 | 6.7182 | 6.1872 | 6.1884 | 6.3874 | 6.0705 | 6.3078 | 6.3529 | 6.2533 | 6.3874 | 6.5019 | 5.9467 | 6.6094 | 6.2493 | 5.9986 |
| NIBAN2_S574 | NIBAN2 | Q96TA1 | | 692 | AAPEASSPPASPL | 6.1673 | 6.3287 | 6.8412 | 6.5588 | 6.2994 | 6.6770 | 6.0200 | 7.2413 | 6.4296 | 6.0884 | 6.7155 | 6.6713 | 5.2552 | 5.9406 | 5.5528 | 5.2130 |
| NIBAN2_S574;S593 | NIBAN2 | Q96TA1 | | 641 | FEASPESPPPASP | 6.0753 | 6.1217 | 6.8093 | 6.6243 | 5.6953 | 6.4076 | 6.2216 | 6.2014 | 6.1245 | 6.4263 | 6.0590 | 6.3659 | 6.2143 | 6.1797 | 5.9615 | 6.5124 |
| NIBAN2_S574;T606 | NIBAN2 | Q96TA1 | | 574 | SMVMHNSDPNLHL | 4.4788 | 5.4095 | 6.4341 | 5.4255 | 6.0988 | 6.4623 | 5.1822 | 6.7142 | 5.0452 | 6.1446 | 6.5588 | 5.4007 | 7.2440 | 8.1628 | 8.3501 | 6.8885 |
| NIBAN2_S624 | NIBAN2 | Q96TA1 | | 624 | RAKQVVSVVQDEE | 4.8614 | 5.2796 | 7.4498 | 7.0256 | 6.4937 | 5.5058 | 7.2298 | 6.2382 | 6.3714 | 6.8862 | 4.8492 | 7.0158 | 6.6380 | 6.6673 | 5.5278 | 5.9606 |

| | | | | | | | | | | | | | | | | | | | | | |
|---|---|---|---|---|---|---|---|---|---|---|---|---|---|---|---|---|---|---|---|---|---|
| NIBAN2_S638 | NIBAN2 | Q96TA1 | | 638 | GLPFEASPESPPP | 5.3264 | 6.8003 | 6.3440 | 6.2760 | 6.2840 | 5.4044 | 6.2529 | 6.8653 | 6.0908 | 6.6102 | 6.3600 | 6.6252 | 5.4744 | 7.5317 | 5.7366 | 6.0178 |
| NIBAN2_S638;S646 | NIBAN2 | Q96TA1 | 692;696 | | NA | 6.2061 | 6.6871 | 6.5873 | 6.5628 | 6.3085 | 6.6307 | 6.4420 | 6.6238 | 6.4844 | 6.6762 | 7.0345 | 7.0458 | 4.8896 | 5.4794 | 4.9572 | 5.3847 |
| NIBAN2_S641 | NIBAN2 | Q96TA1 | 665;681 | | NA | 6.2142 | 6.4192 | 6.9971 | 6.6658 | 6.3726 | 7.2217 | 6.8137 | 7.1016 | 7.0754 | 7.0918 | 7.2811 | 7.2368 | 3.9924 | 4.9597 | 4.0445 | 4.5122 |
| NIBAN2_S641;S646 | NIBAN2 | Q96TA1 | 641;646 | | NA | 6.0651 | 5.9053 | 6.6110 | 6.7253 | 6.1683 | 6.4023 | 6.6052 | 6.3149 | 6.8351 | 5.8173 | 6.1942 | 7.0132 | 5.5934 | 6.0859 | 5.7242 | 5.9393 |
| NIBAN2_S646 | NIBAN2 | Q96TA1 | 574;593 | | NA | 4.8802 | 6.1279 | 6.7009 | 6.0019 | 6.9282 | 6.5975 | 5.8865 | 6.8467 | 5.5239 | 6.4097 | 6.7610 | 6.3275 | 5.8967 | 6.5785 | 6.4439 | 6.0890 |
| NIBAN2_S665 | NIBAN2 | Q96TA1 | 574;606 | | NA | 5.0116 | 5.9228 | 6.3000 | 6.0488 | 6.4298 | 6.3775 | 6.1554 | 7.3785 | 5.5149 | 6.4232 | 7.0256 | 6.5765 | 5.7449 | 6.2879 | 6.2917 | 6.5108 |
| NIBAN2_S665;S681 | NIBAN2 | Q96TA1 | 568;574 | | NA | 6.4848 | 6.7674 | 6.1125 | 3.6676 | 5.7418 | 7.0668 | 5.9459 | 6.5499 | 5.5217 | 5.2954 | 6.6302 | 7.3631 | 6.7569 | 6.8789 | 6.2335 | 6.9837 |
| NIBAN2_S692 | NIBAN2 | Q96TA1 | 568;593 | | NA | 6.9985 | 6.1643 | 6.1039 | 4.6132 | 4.8452 | 7.0070 | 5.2926 | 5.7221 | 5.4920 | 6.5451 | 6.5711 | 7.7125 | 7.7517 | 6.8127 | 5.7827 | 6.5853 |
| NIBAN2_S692;S696 | NIBAN2 | Q96TA1 | 638;646 | | NA | 6.0976 | 6.4283 | 5.4903 | 7.2456 | 6.4220 | 6.9045 | 5.1639 | 6.5225 | 5.9900 | 6.8581 | 6.3143 | 5.5689 | 5.8507 | 6.8598 | 6.6633 | 5.6203 |
| NIFK_S145 | NIFK | Q9BYG3 | | 223 | VSGTLDTPEKTVD | 7.2094 | 7.0590 | 6.6128 | 7.1656 | 6.8076 | 6.2151 | 6.8568 | 6.6352 | 6.9630 | 6.3500 | 6.3268 | 6.2232 | 4.6952 | 4.9728 | 4.9062 | 5.0013 |
| NIFK_S218;T223 | NIFK | Q9BYG3 | | 247 | FLERRKSQVAELN | 4.9464 | 4.2682 | 5.2032 | 5.0484 | 6.8594 | 5.0217 | 4.8713 | 5.5555 | 8.2932 | 8.7262 | 7.3209 | 7.4623 | 7.6937 | 7.8492 | 5.2460 | 5.6345 |
| NIFK_S223 | NIFK | Q9BYG3 | | 238 | GPTPVCTPTFLER | 6.1612 | 6.9252 | 6.6295 | 7.1730 | 6.9113 | 6.7232 | 7.1907 | 7.0074 | 6.9951 | 6.9474 | 7.3673 | 6.8338 | 4.1016 | 4.6507 | 3.9854 | 4.3973 |
| NIFK_S230 | NIFK | Q9BYG3 | | 279 | EIQETQTPTHSRK | 6.6670 | 6.5487 | 6.9430 | 6.8398 | 5.6261 | 6.4593 | 6.8359 | 6.7728 | 6.9706 | 6.4342 | 6.7482 | 6.7862 | 4.6742 | 5.6179 | 5.0836 | 4.9925 |
| NIFK_S230;T234;T238 | NIFK | Q9BYG3 | | 230 | PEKTVDSQGPTPV | 6.0422 | 5.6998 | 6.4638 | 6.2679 | 6.3668 | 6.8151 | 6.1956 | 6.5650 | 6.4727 | 6.8976 | 7.0557 | 6.4312 | 5.4204 | 5.9923 | 5.7600 | 5.5538 |
| NIFK_S230;T238 | NIFK | Q9BYG3 | | 268 | VFKQPISCVKEEI | 8.4844 | 8.8041 | 7.2165 | 8.7064 | 5.8987 | 6.7705 | 8.2972 | 6.5160 | 7.4094 | 6.7654 | 7.4458 | 7.4310 | 2.1860 | 3.0949 | 1.9876 | 2.9862 |
| NIFK_S234 | NIFK | Q9BYG3 | | 234 | VDSQGPTPVCTPT | 5.7864 | 6.8381 | 5.9840 | 7.2147 | 5.7302 | 6.5794 | 6.6846 | 5.5138 | 6.6302 | 6.6000 | 6.4331 | 6.2724 | 5.9774 | 6.5867 | 5.9435 | 5.2255 |
| NIFK_S238 | NIFK | Q9BYG3 | | 145 | KQPSYPSVKRYNR | 5.0821 | 5.8055 | 7.0629 | 6.9871 | 4.7455 | 6.0689 | 6.5776 | 7.1220 | 7.0989 | 8.7290 | 8.3260 | 8.5342 | 3.9878 | 5.6561 | 3.3969 | 4.8195 |
| NIFK_S247 | NIFK | Q9BYG3 | 230;238 | | NA | 5.9093 | 6.3979 | 6.3458 | 6.5199 | 6.3605 | 6.3345 | 6.1760 | 6.1921 | 6.5310 | 6.8456 | 6.4610 | 6.8312 | 5.4290 | 5.8247 | 5.4842 | 6.3572 |
| NIFK_S268 | NIFK | Q9BYG3 | 218;223 | | NA | 6.8959 | 7.5738 | 6.5494 | 7.7077 | 6.5140 | 7.1313 | 8.0807 | 7.1451 | 6.6353 | 6.9724 | 7.2686 | 6.9706 | 3.1634 | 4.0611 | 3.1724 | 4.1583 |
| NIFK_T279 | NIFK | Q9BYG3 | 230;234;238 | | NA | 7.4751 | 7.0881 | 7.0914 | 7.2173 | 6.2866 | 6.3582 | 7.2572 | 5.5616 | 7.0091 | 6.7894 | 6.2166 | 6.6939 | 3.7675 | 5.1514 | 5.1612 | 4.8754 |
| NIM1K_S381 | NIM1K | Q8IY84 | | 381 | QGRDARSSITGVY | 6.4294 | 6.4634 | 6.1665 | 6.8185 | 5.8613 | 5.4754 | 6.1691 | 5.2747 | 6.4311 | 6.3433 | 5.9664 | 6.1243 | 6.6465 | 6.1964 | 6.5737 | 7.0601 |
| NIPBL_S2658 | NIPBL | Q6KC79 | | 350 | MYDIISSPSKDST | 6.7741 | 6.7909 | 6.1085 | 6.9195 | 6.2606 | 6.2081 | 6.5597 | 6.3615 | 6.3613 | 6.2200 | 6.2319 | 6.3605 | 5.5422 | 5.8358 | 5.6844 | 5.7812 |
| NIPBL_S274 | NIPBL | Q6KC79 | | 318 | PDILLDSPERKQK | 6.2910 | 6.5840 | 6.0808 | 6.7565 | 6.2601 | 6.0806 | 6.3282 | 6.1196 | 6.3514 | 6.2845 | 6.1183 | 6.2918 | 6.1185 | 6.2441 | 6.1254 | 5.9653 |
| NIPBL_S318 | NIPBL | Q6KC79 | | 2658 | SLLGGGSPKNNTA | 6.1718 | 6.4253 | 5.7656 | 6.3615 | 6.1101 | 5.8259 | 6.2978 | 5.6470 | 6.1742 | 5.9706 | 5.8347 | 6.0112 | 6.9072 | 6.8742 | 6.7631 | 6.8600 |
| NIPBL_S350 | NIPBL | Q6KC79 | | 367 | RLSRVRSSDMDQQ | 4.9072 | 5.1306 | 5.2184 | 5.9623 | 5.7408 | 5.5816 | 6.3593 | 6.0358 | 5.4860 | 5.3517 | 5.2652 | 5.4391 | 8.5505 | 8.1753 | 8.3348 | 8.4612 |
| NIPBL_S367 | NIPBL | Q6KC79 | | 702 | TKSRPETPKQKGE | 5.6365 | 5.7138 | 5.2448 | 5.7921 | 5.7507 | 5.7078 | 6.2349 | 5.6848 | 5.6246 | 5.7683 | 5.5529 | 5.6496 | 7.8338 | 8.0833 | 7.8373 | 7.8847 |
| NIPBL_S768 | NIPBL | Q6KC79 | | 274 | ASFPLRSPQPVCS | 5.6632 | 6.1163 | 7.0613 | 6.7571 | 5.5417 | 6.6039 | 6.7864 | 6.1282 | 6.4585 | 6.2928 | 6.8563 | 7.2517 | 5.5104 | 5.8709 | 5.2877 | 5.8134 |
| NIPBL_S892 | NIPBL | Q6KC79 | | 406 | TSKTPITPQDINR | 5.4585 | 4.9099 | 5.2047 | 6.3937 | 5.7080 | 5.7760 | 5.3813 | 5.0444 | 5.9094 | 5.7404 | 5.3743 | 5.3057 | 8.4532 | 8.7048 | 8.4758 | 8.1600 |
| NIPBL_T1734 | NIPBL | Q6KC79 | | 892 | QKSRPDSPRVKQG | 6.1284 | 6.4761 | 5.6389 | 6.3542 | 5.9600 | 5.9120 | 6.0323 | 6.1195 | 5.6802 | 5.5739 | 5.4119 | 5.6844 | 6.7539 | 7.4688 | 7.8293 | 6.9762 |
| NIPBL_T406 | NIPBL | Q6KC79 | | 1734 | RSIIKTTPSQFST | 6.0913 | 7.0559 | 6.3146 | 6.7984 | 6.4894 | 6.4113 | 5.9407 | 5.8088 | 6.2173 | 5.8357 | 5.9905 | 6.8176 | 6.2808 | 6.8377 | 6.0151 | 5.0948 |
| NIPBL_T702 | NIPBL | Q6KC79 | | 713 | GESRPETPKQKSD | 5.5650 | 6.1102 | 6.0073 | 6.8977 | 5.6387 | 5.9333 | 6.2273 | 5.6675 | 5.7500 | 5.8922 | 5.9054 | 6.0870 | 7.0301 | 7.0493 | 6.8831 | 7.3561 |
| NIPBL_T713 | NIPBL | Q6KC79 | | 768 | HDNRRDSGKPSTE | 3.9657 | 3.8628 | 4.0523 | 5.5900 | 7.1952 | 5.2251 | 4.9378 | 5.3144 | 7.5925 | 8.3772 | 7.0702 | 7.5095 | 7.8362 | 8.2571 | 6.5658 | 6.6482 |
| NIPBL_Y1146 | NIPBL | Q6KC79 | | 1146 | SSGGGRYRNRSPS | 7.7347 | 7.4093 | 6.9151 | 7.2139 | 8.1691 | 6.2917 | 5.9903 | 5.8943 | 5.8859 | 6.1271 | 6.0932 | 6.0395 | 5.0496 | 5.5259 | 4.7173 | 4.9429 |
| NISCH_S1284 | NISCH | Q9Y2I1 | | 1284 | SLERTPSPEPVDK | 6.0400 | 6.3418 | 6.3878 | 6.9189 | 6.2902 | 6.2706 | 6.0902 | 6.3474 | 6.2964 | 6.2171 | 6.5168 | 6.0840 | 5.8736 | 6.3458 | 6.2566 | 5.7228 |
| NKAP_S149 | NKAP | Q8N5F7 | | 149 | PEVWGLSPKNPEP | 5.8032 | 5.9365 | 6.1306 | 6.1464 | 5.8950 | 5.7883 | 6.1234 | 5.6824 | 6.0079 | 6.3141 | 5.9677 | 6.1079 | 6.6133 | 7.2998 | 6.8985 | 7.2850 |
| NKAPD1_S138 | NKAPD1 | Q6ZUT1-2 | | 138 | TNGKKTSPQVKSS | 6.7805 | 6.6388 | 5.8493 | 6.2885 | 6.9438 | 5.8365 | 6.3223 | 6.0753 | 6.1165 | 5.9083 | 5.3680 | 5.2687 | 6.0362 | 7.1583 | 6.9201 | 6.4890 |
| NKRF_S444 | NKRF | O15226-2 | | 444 | QCHTGSSPRGSGK | 6.1849 | 7.0751 | 6.3675 | 7.4682 | 6.6271 | 6.5073 | 5.5772 | 6.7840 | 5.8160 | 5.8417 | 6.1972 | 5.9201 | 5.8631 | 6.2579 | 6.0274 | 5.4853 |
| NKTR_S1146 | NKTR | P30414 | | 613 | VIPLSDSPPPSRW | 5.7414 | 6.6227 | 6.1188 | 6.8248 | 5.7749 | 6.0160 | 6.1185 | 6.0407 | 6.1032 | 6.4041 | 6.0661 | 6.2101 | 6.1485 | 6.7968 | 6.3767 | 6.6368 |
| NKTR_S379 | NKTR | P30414 | | 1146 | AKVEETSPLGNAR | 6.3807 | 6.5115 | 5.9967 | 6.5853 | 5.9526 | 5.8116 | 6.2448 | 6.0108 | 6.1676 | 6.0382 | 5.7651 | 5.8035 | 6.0677 | 7.1235 | 5.6692 | 6.9712 |
| NKTR_S463 | NKTR | P30414 | | 1155 | GNARLDTPDINIV | 6.0284 | 6.2779 | 6.3729 | 6.7783 | 5.5978 | 5.4686 | 6.3158 | 5.2635 | 6.5241 | 6.4294 | 6.2691 | 6.3787 | 6.1583 | 6.6568 | 6.2048 | 7.2756 |
| NKTR_S463;S467 | NKTR | P30414 | | 463 | RRILIPSDIESSK | 6.8197 | 6.9963 | 6.6640 | 7.2959 | 5.8758 | 5.9543 | 5.8597 | 5.6467 | 6.5103 | 6.4237 | 5.7511 | 5.8777 | 5.8484 | 6.4014 | 6.1743 | 5.9008 |
| NKTR_S572;S575 | NKTR | P30414 | | 379 | RAYRPPSGEKWSK | 5.7877 | 5.9696 | 5.9405 | 6.4026 | 5.9076 | 5.8485 | 5.9704 | 5.7275 | 5.7976 | 5.8636 | 5.7274 | 5.7716 | 6.6326 | 7.6399 | 7.4114 | 7.6015 |
| NKTR_S613 | NKTR | P30414 | | 866 | TLKENLSDHLRNG | 5.2589 | 5.4018 | 5.3301 | 6.1949 | 6.1648 | 5.8805 | 6.3153 | 5.6601 | 5.6353 | 5.0014 | 4.6262 | 5.1397 | 7.9322 | 8.1058 | 8.3246 | 9.0285 |
| NKTR_S775;S776 | NKTR | P30414 | | 343 | GTIRYHTPPRSRS | 6.2857 | 5.9053 | 6.8126 | 6.8950 | 6.0926 | 6.0474 | 6.2969 | 6.4463 | 6.6111 | 6.0626 | 6.1761 | 6.7405 | 5.8761 | 5.7264 | 6.1449 | 5.8807 |
| NKTR_S866 | NKTR | P30414 | 463;467 | | NA | 6.4334 | 6.6250 | 6.9217 | 7.0616 | 6.4327 | 6.4727 | 6.2202 | 5.2683 | 6.1044 | 6.2686 | 5.9197 | 6.4993 | 5.4325 | 6.1432 | 5.8323 | 6.3647 |
| NKTR_S887;S889;S891 | NKTR | P30414 | 887;889;891 | | NA | 3.7661 | 6.2695 | 6.7294 | 6.9755 | 4.8282 | 5.4783 | 6.9987 | 6.4627 | 4.8842 | 4.7603 | 5.7149 | 4.3925 | 8.1938 | 9.6023 | 7.2185 | 7.7252 |
| NKTR_T1155 | NKTR | P30414 | 572;575 | | NA | 5.6710 | 5.2530 | 6.3178 | 6.1695 | 4.9882 | 5.2951 | 5.5801 | 5.4556 | 6.2402 | 6.4523 | 6.0019 | 6.4665 | 7.2066 | 8.1922 | 6.9484 | 7.7614 |
| NKTR_T343 | NKTR | P30414 | 775;776 | | NA | 6.1579 | 6.2813 | 5.6003 | 6.6126 | 6.4082 | 6.2211 | 6.1160 | 6.0158 | 5.8965 | 5.6460 | 5.7571 | 5.8321 | 6.6605 | 7.5813 | 6.6833 | 6.5301 |
| NKX2-4_S271 | NKX2-4 | Q9H2Z4 | | 271 | PPPPPPSPRRVAV | 5.8861 | 5.5040 | 6.1438 | 6.8400 | 6.6169 | 5.9774 | 6.0895 | 6.2506 | 6.2355 | 6.5716 | 6.0066 | 5.8770 | 6.4661 | 6.5696 | 6.5175 | 6.4477 |
| NKX2-6_T204 | NKX2-6 | A6NCS4 | | 204 | LAGHPLTPRRVAV | 6.3428 | 7.0768 | 7.1384 | 7.2286 | 5.7402 | 6.4148 | 6.7789 | 7.0157 | 6.7669 | 6.6522 | 7.1561 | 6.7635 | 3.9333 | 5.5969 | 4.4721 | 4.9229 |
| NLN_S53 | NLN | Q9BYT8 | | 53 | VLRWDLSPEQIKT | 5.9379 | 6.6528 | 6.6595 | 6.3310 | 7.0489 | 6.3269 | 5.8807 | 6.3700 | 6.4534 | 6.8739 | 6.4044 | 6.2684 | 5.7260 | 6.1125 | 5.4942 | 5.4594 |
| NMD3_S258 | NMD3 | Q96D46 | | 258 | DNVVCLSPKLAQS | 4.9014 | 6.5774 | 6.9423 | 6.1043 | 6.7767 | 7.4701 | 6.0391 | 6.4666 | 5.8958 | 6.8588 | 5.8410 | 6.1773 | 5.6025 | 6.2229 | 5.3745 | 6.7494 |
| NME2_S235 | NME2 | P22392-2 | | 235 | RNIIHGSDSVKSA | 6.3096 | 5.6258 | 7.4947 | 5.8935 | 6.0906 | 6.7506 | 6.2628 | 7.2235 | 6.8978 | 6.6537 | 6.8324 | 6.9481 | 4.9973 | 5.4518 | 5.2207 | 5.3470 |
| NME2_S237 | NME2 | P22392-2 | | 237 | IIHGSDSVKSAEK | 7.0399 | 6.7896 | 6.7010 | 5.5321 | 6.0619 | 6.8893 | 5.9625 | 6.9131 | 7.1772 | 6.2722 | 6.6273 | 6.5568 | 5.2635 | 6.1843 | 5.4561 | 4.5732 |
| NMNAT1_S117 | NMNAT1 | Q9HAN9 | | 117 | CDHQQNSPTLERP | 6.1860 | 6.6464 | 6.4773 | 6.9938 | 6.4680 | 6.6368 | 6.2976 | 6.5108 | 6.7652 | 7.0155 | 7.0185 | 6.7736 | 5.0063 | 5.1967 | 5.0198 | 4.9877 |
| NMT1_S47 | NMT1 | P30419 | | 47 | YNRGGLSPANDTG | 6.4965 | 6.6109 | 6.7672 | 6.3399 | 6.5614 | 6.5529 | 6.6879 | 6.7864 | 6.8791 | 6.7421 | 6.8404 | 6.5785 | 4.7296 | 5.5283 | 4.8517 | 5.0474 |
| NMT1_T71 | NMT1 | P30419 | | 71 | KEKGSETDSAQDQ | 5.5920 | 6.6863 | 6.9841 | 5.4022 | 6.5905 | 6.5189 | 6.0043 | 6.6023 | 5.7870 | 5.7864 | 5.9068 | 5.3441 | 6.9136 | 7.2243 | 6.7814 | 5.8757 |
| NMT2_S38 | NMT2 | O60551 | | 38 | TEHAKGSPGGYLG | 5.5512 | 6.2286 | 6.7978 | 6.3223 | 6.0906 | 6.8477 | 6.3982 | 6.7114 | 6.6267 | 6.9859 | 7.4872 | 7.7241 | 4.8464 | 5.7591 | 4.4845 | 5.1383 |
| NOB1_S184 | NOB1 | Q9ULX3 | | 201 | EDRKDDSDDDGGG | 5.4316 | 5.4340 | 7.2644 | 5.7079 | 6.3519 | 6.2251 | 6.0953 | 6.0016 | 4.4608 | 6.2972 | 6.3986 | 6.8661 | 6.3680 | 6.3499 | 6.0710 | 6.6765 |
| NOB1_S201 | NOB1 | Q9ULX3 | | 184 | RGEDVPSEEEEEE | 5.9451 | 6.5616 | 7.5425 | 6.3182 | 5.8320 | 5.9153 | 6.5214 | 6.4112 | 5.9107 | 6.6402 | 6.2918 | 6.5851 | 6.1185 | 6.0599 | 5.7150 | 5.6317 |
| NOB1_S352 | NOB1 | Q9ULX3 | | 352 | FPQLRLSQKARQK | 7.3542 | 7.1450 | 10.3952 | 6.8527 | 7.0313 | 9.6429 | 6.3866 | 4.9542 | 6.7463 | 7.1658 | 7.3273 | 7.1512 | 4.3707 | 3.6514 | 1.7578 | 2.0673 |
| NOC2L_S49 | NOC2L | Q9Y3T9 | | 56 | PDKPGGSPSASRR | 6.2911 | 6.0100 | 6.1793 | 6.6826 | 5.5278 | 5.3173 | 5.9394 | 5.1658 | 6.0420 | 6.1160 | 5.6053 | 6.3104 | 6.9632 | 6.5785 | 7.1145 | 8.1569 |
| NOC2L_S49;S56 | NOC2L | Q9Y3T9 | | 691 | GILRPLSTRHGVE | 5.4874 | 6.6349 | 7.7626 | 7.1183 | 5.0397 | 6.1823 | 6.3679 | 5.8428 | 7.5134 | 7.3784 | 7.3903 | 7.7189 | 5.3097 | 5.1439 | 4.0639 | 5.0457 |
| NOC2L_S49;S60 | NOC2L | Q9Y3T9 | | 49 | AREAARSPDKPGG | 5.5712 | 5.9371 | 5.3678 | 6.9291 | 5.5784 | 5.2593 | 5.5565 | 6.1149 | 6.0425 | 6.0708 | 5.6628 | 6.1627 | 7.2560 | 7.2312 | 7.6237 | 7.6359 |
| NOC2L_S56 | NOC2L | Q9Y3T9 | 672;673 | | NA | 5.9272 | 6.3638 | 6.0917 | 6.4222 | 6.3086 | 6.1885 | 6.5096 | 6.1472 | 6.1693 | 6.1616 | 5.9161 | 6.4039 | 6.1129 | 6.4411 | 6.1033 | 6.7330 |
| NOC2L_S672;T673 | NOC2L | Q9Y3T9 | 49;56 | | NA | 6.4895 | 5.7657 | 6.4930 | 6.8823 | 6.3123 | 6.5203 | 6.3576 | 6.0742 | 6.8820 | 7.3832 | 6.5838 | 6.3664 | 5.6634 | 5.2365 | 5.3567 | 5.6331 |
| NOC2L_S691 | NOC2L | Q9Y3T9 | 49;60 | | NA | 6.5311 | 6.7714 | 6.1136 | 7.1950 | 6.7866 | 5.9269 | 6.4064 | 6.8894 | 7.1149 | 6.5620 | 6.3782 | 6.1924 | 5.0772 | 4.9441 | 5.3632 | 5.7476 |
| NOL12_S134 | NOL12 | Q9UGY1 | | 134 | GGAGDRSEEEASS | 6.2641 | 6.5550 | 6.4234 | 7.1651 | 5.9668 | 6.0951 | 6.6307 | 6.3174 | 6.1016 | 6.1017 | 5.4751 | 6.0508 | 5.5318 | 6.2131 | 6.6456 | 6.4627 |
| NOL12_S134;S139;S140 | NOL12 | Q9UGY1 | | 124 | ARLLGLTPPEGGA | 6.3455 | 7.3539 | 5.1693 | 6.6287 | 5.9964 | 5.1019 | 6.1852 | 6.1522 | 6.6287 | 6.1687 | 6.9778 | 7.6666 | 6.5103 | 6.6617 | 5.1334 | 6.2197 |
| NOL12_T124 | NOL12 | Q9UGY1 | 134;139;140 | | NA | 5.8696 | 5.9930 | 6.5670 | 7.0195 | 5.2175 | 5.9746 | 6.3250 | 5.6551 | 6.1708 | 6.0885 | 6.5893 | 6.6141 | 5.6230 | 6.7167 | 6.0820 | 7.4945 |
| NOL7_S126 | NOL7 | Q9UMY1 | | 126 | EKLTTASQTNIKK | 6.5869 | 6.7363 | 6.0868 | 7.0798 | 6.3643 | 6.3799 | 6.5871 | 6.1703 | 6.1112 | 5.9762 | 5.8298 | 6.2236 | 5.6516 | 6.1332 | 5.7451 | 6.3378 |
| NOL8_S1036 | NOL8 | Q76FK4 | | 660 | QDRKAVSPSSSEK | 6.7371 | 6.7481 | 6.2254 | 7.1316 | 6.1959 | 5.9033 | 6.4991 | 6.1664 | 6.5045 | 6.1496 | 6.1229 | 6.1621 | 5.6050 | 5.8486 | 5.9188 | 6.0815 |
| NOL8_S1099 | NOL8 | Q76FK4 | | 1099 | TDHRNSSPGEASL | 6.1896 | 6.1879 | 6.2443 | 7.0249 | 6.1308 | 5.8618 | 6.3855 | 6.1276 | 6.3294 | 6.4925 | 6.2831 | 6.8754 | 5.8934 | 5.9491 | 6.0274 | 5.9973 |
| NOL8_S268 | NOL8 | Q76FK4 | | 263 | RTCDSITPSKSSP | 7.2267 | 7.1584 | 6.6520 | 7.5817 | 5.7563 | 6.1103 | 6.3841 | 6.2859 | 7.1665 | 6.6270 | 6.6008 | 6.8285 | 4.8179 | 5.0698 | 4.3726 | 5.3616 |
| NOL8_S331 | NOL8 | Q76FK4 | | 268 | ITPSKSSPVPVSD | 6.7090 | 7.3076 | 6.4036 | 7.4002 | 6.2344 | 6.3263 | 6.6801 | 6.2173 | 6.2648 | 6.0967 | 6.4173 | 6.8254 | 4.8234 | 5.3867 | 5.3167 | 5.5905 |
| NOL8_S378;T381 | NOL8 | Q76FK4 | | 331 | SINESESDPFEVV | 6.5523 | 6.8211 | 7.3315 | 7.0856 | 6.0748 | 6.4833 | 6.6422 | 5.7767 | 5.6720 | 5.9549 | 6.2157 | 5.8176 | 6.3120 | 5.7976 | 5.4786 | 5.9842 |
| NOL8_S432 | NOL8 | Q76FK4 | | 1036 | DPAALTSDAEQPS | 6.2673 | 6.3386 | 6.3349 | 6.8846 | 6.2648 | 6.7359 | 6.6558 | 6.4505 | 6.3208 | 6.1344 | 5.9467 | 6.0166 | 5.7854 | 6.0424 | 5.8403 | 5.9812 |
| NOL8_S660 | NOL8 | Q76FK4 | | 432 | KLQKRKSNVESAL | 6.6371 | 7.0669 | 6.9318 | 6.6592 | 6.2445 | 7.0271 | 6.4128 | 6.8634 | 7.7525 | 7.9662 | 8.4937 | 8.1191 | 3.4929 | 4.4466 | 2.8626 | 3.0237 |
| NOL8_S681 | NOL8 | Q76FK4 | | 681 | PLEGKKSLSLSAK | 6.1342 | 7.4559 | 7.3044 | 8.4212 | 5.6668 | 6.0929 | 6.2779 | 5.5851 | 6.7332 | 7.3461 | 8.1075 | 8.2779 | 3.4578 | 5.4468 | 3.7404 | 3.9519 |
| NOL8_S795;S803 | NOL8 | Q76FK4 | 795;803 | | NA | 5.8919 | 6.3130 | 5.9104 | 6.9900 | 6.2085 | 6.0515 | 6.1313 | 5.7978 | 6.3131 | 5.8927 | 6.0166 | 6.0566 | 6.5655 | 6.5166 | 7.0129 | 6.3315 |
| NOL8_S795;S803;T807 | NOL8 | Q76FK4 | 888;890 | | NA | 6.3546 | 6.2343 | 6.2791 | 6.9444 | 6.2511 | 5.3588 | 6.4111 | 6.1958 | 6.9543 | 6.2760 | 5.9079 | 6.0364 | 6.1125 | 6.1709 | 6.3113 | 6.2015 |
| NOL8_T263 | NOL8 | Q76FK4 | 795;803;807 | | NA | 6.0134 | 6.0308 | 6.1133 | 6.1575 | 7.2964 | 7.1771 | 6.0610 | 6.1041 | 6.2182 | 6.6220 | 6.2191 | 5.1597 | 6.5427 | 6.7865 | 5.7535 | 5.7446 |
| NOL8_T888;S890 | NOL8 | Q76FK4 | 378;381 | | NA | 6.2914 | 6.7767 | 6.3149 | 7.2456 | 5.9470 | 5.5113 | 6.0914 | 5.1525 | 6.2504 | 5.8511 | 6.2256 | 6.2228 | 7.0160 | 6.3014 | 6.4477 | 6.3542 |
| NOL9_S487 | NOL9 | Q5SY16 | | 487 | ADEEKESPVEFTG | 5.5283 | 5.9848 | 5.7481 | 6.1188 | 5.0012 | 5.5096 | 5.9420 | 5.5145 | 12.2885 | 5.9023 | 4.6652 | 5.2902 | 5.8266 | 6.1010 | 6.8935 | 7.6855 |
| NOLC1_S548 | NOLC1 | Q14978-2 | | 653 | SRVADNSFDAKRG | 6.7591 | 6.8466 | 6.3945 | 7.2454 | 7.0391 | 6.6919 | 7.4083 | 7.0564 | 6.9055 | 6.5016 | 6.7577 | 6.7421 | 4.0451 | 4.5768 | 4.1388 | 4.8910 |
| NOLC1_S573 | NOLC1 | Q14978-2 | | 633 | GEKRASSPFRRVR | 6.3380 | 6.7176 | 6.3931 | 6.9266 | 6.4286 | 6.1079 | 7.0095 | 4.7486 | 4.8971 | 5.7220 | 5.8923 | 6.5992 | 6.6394 | 6.5683 | 6.2151 | 6.7968 |
| NOLC1_S588 | NOLC1 | Q14978-2 | | 548 | KLKGKGSPRPQAP | 6.3044 | 6.7737 | 6.1696 | 7.1852 | 6.0537 | 5.9841 | 6.2757 | 6.0988 | 6.2006 | 6.2051 | 6.2459 | 6.5063 | 5.7191 | 6.0970 | 5.9589 | 6.2219 |
| NOLC1_S632;S633 | NOLC1 | Q14978-2 | | 588 | KAAVVVSKSGSLK | 5.9096 | 6.4752 | 5.7255 | 6.8003 | 6.1327 | 6.3948 | 6.9661 | 6.2276 | 6.0209 | 5.7731 | 6.2781 | 6.4020 | 5.8030 | 6.0109 | 6.2148 | 6.8654 |
| NOLC1_S633 | NOLC1 | Q14978-2 | | 573 | GKAAKNSEEEEEE | 5.8147 | 5.8899 | 6.4263 | 6.7174 | 6.1365 | 6.2148 | 7.0175 | 5.7806 | 6.1139 | 5.7724 | 6.5081 | 6.3180 | 6.7634 | 6.6159 | 5.8220 | 6.0886 |
| NOLC1_S653 | NOLC1 | Q14978-2 | | 188 | NQKPKITPVTVKA | 7.3897 | 7.9197 | 7.2628 | 7.9252 | 6.6000 | 7.2350 | 7.7175 | 6.8068 | 7.5427 | 7.5598 | 7.4554 | 7.5316 | 2.1875 | 3.2197 | 2.2719 | 3.3748 |
| NOLC1_S696 | NOLC1 | Q14978-2 | | 617 | KKIKLQTPNTFPK | 6.2415 | 7.0883 | 6.3550 | 6.4081 | 4.8815 | 7.1658 | 7.0591 | 7.4038 | 7.7056 | 8.3674 | 8.3895 | 9.5142 | 2.8517 | 3.6376 | 2.9294 | 4.0015 |
| NOLC1_S708 | NOLC1 | Q14978-2 | | 620 | KLQTPNTFPKRKK | 7.9707 | 7.8837 | 7.4640 | 8.3673 | 6.8824 | 6.9146 | 7.2736 | 6.6919 | 7.7122 | 7.2299 | 7.7907 | 7.7994 | 2.0513 | 3.0054 | 2.1375 | 2.8254 |
| NOLC1_T188 | NOLC1 | Q14978-2 | | 708 | NSIKFDSExxxxx | 6.2952 | 6.5648 | 6.2341 | 6.7473 | 6.6614 | 7.1655 | 6.9489 | 5.5859 | 5.7649 | 5.8122 | 6.3396 | 6.0611 | 6.5061 | 6.3911 | 5.6343 | 5.2875 |
| NOLC1_T607 | NOLC1 | Q14978-2 | | 607 | AAKEAETPQAKKI | 7.3498 | 7.2282 | 7.1569 | 8.0129 | 7.0072 | 7.5390 | 7.8330 | 7.1809 | 7.6978 | 7.1294 | 7.3019 | 6.8115 | 2.7491 | 3.5331 | 2.3960 | 3.0733 |
| NOLC1_T617 | NOLC1 | Q14978-2 | | 696 | GSYRGGSISVQVN | 6.0975 | 4.8186 | 5.6025 | 6.2823 | 7.2421 | 5.5408 | 5.5692 | 5.1889 | 9.1172 | 8.0877 | 7.1314 | 6.8057 | 7.2176 | 6.4788 | 4.0693 | 4.7503 |
| NOLC1_T617;T620 | NOLC1 | Q14978-2 | 617;620 | | NA | 7.4926 | 7.9191 | 7.2868 | 7.9654 | 6.4309 | 6.8494 | 7.5769 | 6.6504 | 7.5927 | 7.2350 | 7.8745 | 7.6595 | 2.4955 | 3.3190 | 2.3707 | 3.2818 |
| NOLC1_T620 | NOLC1 | Q14978-2 | 632;633 | | NA | 7.0711 | 8.4401 | 6.7551 | 7.5252 | 6.6987 | 7.9966 | 7.1837 | 6.8097 | 6.8854 | 7.2888 | 7.6980 | 7.1975 | 2.5043 | 3.7149 | 2.8584 | 3.3728 |
| NOM1_S139 | NOM1 | Q5C9Z4 | | 139 | APSRDPSPPRKPR | 6.1663 | 6.0234 | 6.1174 | 6.6473 | 6.1065 | 5.8663 | 6.5945 | 5.6394 | 6.0641 | 5.8552 | 6.1305 | 6.4861 | 6.7694 | 6.5618 | 5.9340 | 7.0379 |
| NONO_S147 | NONO | Q15233 | | 450 | IGAIGGTPPAFNR | 6.6763 | 6.8839 | 6.4937 | 6.9342 | 6.4599 | 6.5758 | 6.9011 | 6.7376 | 7.2146 | 6.7054 | 7.0327 | 7.0003 | 4.3334 | 4.9712 | 4.2914 | 4.7886 |
| NONO_S196 | NONO | Q15233 | | 428 | DGTLGLTPPTTER | 6.4580 | 6.7965 | 6.4398 | 6.6560 | 6.2126 | 6.8180 | 6.7952 | 6.3810 | 6.5027 | 6.8737 | 6.8474 | 6.4532 | 5.2707 | 5.3515 | 5.0515 | 5.0922 |
| NONO_T15 | NONO | Q15233 | | 410 | APVPAGTPAPPGP | 6.5075 | 6.1039 | 6.2970 | 6.7476 | 6.1927 | 6.0009 | 7.0113 | 6.4249 | 6.1806 | 6.0817 | 6.2560 | 6.0734 | 6.3107 | 6.4272 | 5.5340 | 5.8506 |
| NONO_T410 | NONO | Q15233 | | 15 | LEKQNHTPRKHHQ | 7.7625 | 7.7012 | 6.6656 | 7.2647 | 6.7629 | 7.0729 | 6.9383 | 5.5030 | 6.2077 | 6.3934 | 6.3962 | 6.1679 | 5.3465 | 5.2912 | 4.0014 | 4.5247 |
| NONO_T428 | NONO | Q15233 | | 196 | KGIVEFSGKPAAR | 3.7783 | 5.5088 | 5.3611 | 5.5357 | 4.7988 | 5.9848 | 5.3951 | 6.5065 | 6.8478 | 7.7867 | 6.3634 | 5.7233 | 6.9125 | 7.5225 | 8.7451 | 7.2296 |
| NONO_T450 | NONO | Q15233 | | 147 | VRFACHSASLTVR | 6.7829 | 7.1735 | 6.8473 | 7.6896 | 6.8140 | 6.7080 | 6.3807 | 6.5489 | 6.6119 | 5.7648 | 6.4633 | 6.2993 | 5.3180 | 5.2162 | 4.8813 | 4.5002 |
| NOP14_S146;S148 | NOP14 | P78316 | | 96 | EYNSNMSPEEKMM | 6.6973 | 6.7561 | 6.6808 | 7.6141 | 6.6718 | 6.0752 | 7.1941 | 6.2759 | 7.2296 | 6.9069 | 6.9929 | 7.1249 | 4.1181 | 4.7735 | 4.1084 | 4.7805 |
| NOP14_S96 | NOP14 | P78316 | 146;148 | | NA | 5.7460 | 6.0965 | 5.9971 | 6.9308 | 6.2227 | 5.8137 | 6.3195 | 5.8628 | 6.4873 | 6.1379 | 6.2054 | 6.0147 | 6.6936 | 6.4378 | 5.9794 | 7.0549 |
| NOP16_S16 | NOP16 | Q9Y3C1 | | 16 | RQKFGYSVNRKRL | 6.8825 | 8.3870 | 8.7420 | 8.8833 | 5.9334 | 6.2809 | 8.4022 | 6.3944 | 6.8809 | 6.2766 | 5.7017 | 5.2879 | 4.1238 | 4.5740 | 2.5835 | 4.6659 |
| NOP16_T144 | NOP16 | Q9Y3C1 | | 144 | KNYYQDTPKQIRS | 6.6191 | 7.2922 | 7.0108 | 7.5831 | 6.7779 | 7.5194 | 7.9526 | 6.4709 | 7.0781 | 6.9798 | 7.0224 | 6.8926 | 3.2663 | 4.2733 | 3.0887 | 4.1727 |
| NOP2_S214;T218 | NOP2 | P46087-4 | | 819 | PQPPTVSPIRSSR | 6.8683 | 7.0563 | 6.2264 | 7.0799 | 6.4915 | 6.3012 | 6.6000 | 6.4447 | 6.5885 | 6.4180 | 6.5452 | 6.6510 | 4.6125 | 5.5928 | 5.0983 | 5.4254 |
| NOP2_S36 | NOP2 | P46087-4 | | 765 | QTPAVLSPSKTQA | 6.6209 | 6.5533 | 6.4327 | 7.3340 | 6.4463 | 6.1459 | 6.6011 | 6.4221 | 6.6091 | 6.3513 | 6.4340 | 6.6417 | 4.8777 | 5.5780 | 5.3445 | 5.6073 |
| NOP2_S58 | NOP2 | P46087-4 | | 228 | EEEKEVTPESGPP | 5.8624 | 6.0719 | 5.6731 | 6.7162 | 5.8863 | 5.6248 | 6.0875 | 5.8451 | 6.0488 | 5.9916 | 5.7660 | 5.8145 | 6.4419 | 7.5942 | 7.4112 | 7.1646 |
| NOP2_S699 | NOP2 | P46087-4 | | 58 | AKRRLGSVEAPKT | 5.1380 | 5.3294 | 5.3696 | 5.5923 | 5.6179 | 5.7663 | 6.4869 | 5.7039 | 5.0206 | 5.0906 | 5.1522 | 5.5186 | 7.9175 | 8.0398 | 8.7707 | 9.4856 |
| NOP2_S765 | NOP2 | P46087-4 | | 845 | NSQLLLSxxxxxx | 6.8458 | 6.6892 | 6.6050 | 6.6381 | 6.3134 | 6.1483 | 6.7209 | 6.9378 | 7.0537 | 7.9155 | 8.0654 | 7.7806 | 3.6745 | 5.0992 | 3.5180 | 3.9945 |
| NOP2_S819 | NOP2 | P46087-4 | | 699 | ELSTVPSVTKTQA | 6.6807 | 7.1327 | 6.4950 | 7.1842 | 6.1396 | 6.7763 | 7.1519 | 6.2702 | 6.6769 | 6.6305 | 7.1125 | 7.0126 | 4.6016 | 4.9462 | 4.2487 | 4.9406 |
| NOP2_S845 | NOP2 | P46087-4 | | 634 | QTGNSETATPTNV | 5.6155 | 5.9911 | 5.9527 | 6.9665 | 5.8069 | 6.5526 | 6.3812 | 6.0043 | 7.0377 | 6.2669 | 6.1736 | 6.5713 | 5.3329 | 6.9162 | 5.9088 | 6.5219 |
| NOP2_T228 | NOP2 | P46087-4 | | 36 | RFLPAVSDENSKR | 6.3626 | 6.3634 | 6.9766 | 7.6564 | 7.2169 | 6.9293 | 7.4231 | 7.4711 | 5.3487 | 5.3766 | 6.1557 | 6.7045 | 3.8170 | 5.6999 | 4.5185 | 5.9797 |
| NOP2_T634 | NOP2 | P46087-4 | 214;218 | | NA | 5.8421 | 5.8051 | 5.6305 | 6.6976 | 5.4403 | 5.5790 | 6.2220 | 5.5623 | 6.6525 | 5.7081 | 6.2641 | 6.5025 | 7.9510 | 7.0640 | 6.2679 | 6.8110 |
| NOP53_S93 | NOP53 | Q9NZM5 | | 93 | FFVDTGSKEKGLT | 6.4553 | 8.0408 | 6.8650 | 7.1739 | 6.2723 | 6.0185 | 8.1528 | 7.0992 | 7.0535 | 7.5369 | 7.0115 | 7.4370 | 3.6406 | 3.7628 | 3.4530 | 4.0270 |
| NOP56_S314 | NOP56 | O00567 | | 520 | KEELMSSDLEETA | 6.0665 | 6.1472 | 6.1863 | 6.5188 | 6.0557 | 5.8570 | 6.2056 | 5.9979 | 6.1321 | 5.9446 | 5.8817 | 6.0035 | 6.4160 | 6.9812 | 6.7879 | 6.8180 |
| NOP56_S462 | NOP56 | O00567 | | 314 | LIAHAGSLTNLAK | 4.3874 | 6.1059 | 5.5722 | 7.5222 | 7.8825 | 5.6091 | 5.8232 | 6.2375 | 6.4560 | 6.9076 | 6.1901 | 6.0376 | 6.0024 | 5.9728 | 6.7518 | 6.5419 |
| NOP56_S462;T468 | NOP56 | O00567 | | 468 | SENSSSTPEECEE | 5.9008 | 6.2378 | 6.0641 | 7.0982 | 6.2495 | 5.8850 | 6.1687 | 6.0437 | 6.1389 | 6.2084 | 5.8470 | 5.8371 | 6.0120 | 6.7169 | 6.8469 | 6.7450 |
| NOP56_S468 | NOP56 | O00567 | | 519 | SKEELMSSDLEET | 5.9560 | 6.6994 | 6.0029 | 6.4065 | 6.0203 | 5.9154 | 6.2137 | 6.1227 | 6.1234 | 5.9263 | 5.6486 | 6.2783 | 6.6052 | 7.0847 | 6.4565 | 6.5402 |
| NOP56_S519 | NOP56 | O00567 | | 462 | ALALASSENSSST | 6.3131 | 6.3030 | 6.5814 | 6.6236 | 5.8679 | 5.7492 | 6.2139 | 6.2394 | 6.1933 | 5.9038 | 6.2060 | 6.3569 | 6.3740 | 6.7648 | 6.0219 | 6.2877 |
| NOP56_S519;S520 | NOP56 | O00567 | | 570 | KEEPVSSGPEEAV | 6.5479 | 6.6963 | 6.2763 | 7.0452 | 6.1267 | 5.7863 | 6.5754 | 6.2826 | 5.8875 | 5.6117 | 5.5966 | 5.6493 | 6.2818 | 6.5859 | 6.3913 | 6.6592 |
| NOP56_S520 | NOP56 | O00567 | | 591 | KKFHKASQEDxxx | 5.0645 | 5.4444 | 5.5717 | 6.0384 | 6.3032 | 5.6207 | 6.0149 | 5.1103 | 5.0773 | 4.6979 | 5.5014 | 4.5807 | 8.2622 | 9.0709 | 9.0943 | 8.5472 |
| NOP56_S569;S570 | NOP56 | O00567 | 462;468 | | NA | 6.3298 | 6.2267 | 6.1674 | 6.5951 | 5.9233 | 6.1038 | 6.4753 | 5.7315 | 6.1891 | 6.1864 | 6.0223 | 5.8918 | 6.0358 | 6.7851 | 6.6630 | 6.6737 |
| NOP56_S570 | NOP56 | O00567 | 569;570 | | NA | 7.5833 | 7.0218 | 6.6188 | 6.8274 | 6.3474 | 5.8232 | 7.1690 | 7.1305 | 6.2140 | 5.2583 | 5.2298 | 5.0359 | 5.4885 | 6.3356 | 5.6336 | 6.2830 |
| NOP56_S591 | NOP56 | O00567 | 519;520 | | NA | 6.9989 | 6.3013 | 7.0171 | 8.7705 | 6.0900 | 6.5482 | 6.6305 | 6.7322 | 5.4029 | 6.2475 | 6.0339 | 6.2621 | 4.5492 | 5.4957 | 4.9304 | 5.9897 |
| NOP58_S109 | NOP58 | Q9Y2X3 | | 502 | IKEEPLSEEEPCT | 5.4531 | 5.8296 | 5.6395 | 6.2158 | 5.7392 | 5.8943 | 5.8874 | 5.7541 | 5.5637 | 6.0361 | 5.6934 | 5.5678 | 7.3062 | 8.3008 | 7.7934 | 7.3256 |
| NOP58_S304 | NOP58 | Q9Y2X3 | | 34 | LWKEFETPEKANK | 6.3641 | 6.5715 | 5.7568 | 6.6977 | 5.9378 | 5.8006 | 6.3226 | 6.0111 | 6.3095 | 6.0892 | 5.9826 | 6.0445 | 5.8736 | 6.7777 | 6.6439 | 6.8168 |
| NOP58_S345 | NOP58 | Q9Y2X3 | | 514 | TSTAIASPEKKKK | 5.3922 | 5.2221 | 5.2842 | 5.6349 | 6.2666 | 5.4309 | 5.7118 | 5.2077 | 6.6889 | 6.3271 | 6.6198 | 6.1639 | 7.9954 | 7.7101 | 7.1100 | 7.2342 |
| NOP58_S483 | NOP58 | Q9Y2X3 | | 109 | NLSCIHSPVVNEL | 6.2499 | 6.5016 | 6.5258 | 6.9555 | 5.8217 | 6.1122 | 5.8704 | 6.3335 | 6.3469 | 6.5518 | 6.5258 | 6.0440 | 5.7461 | 6.5616 | 5.8152 | 6.0383 |
| NOP58_S502 | NOP58 | Q9Y2X3 | | 304 | LIAHAGSLLNLAK | 3.8521 | 5.3314 | 5.8523 | 7.3483 | 5.7950 | 5.7721 | 6.2402 | 6.2311 | 5.9851 | 7.5614 | 6.7359 | 6.2364 | 6.9436 | 6.9214 | 5.8240 | 7.3697 |
| NOP58_S502;S514 | NOP58 | Q9Y2X3 | | 483 | AEEEETSVKKKKK | 7.2245 | 7.4688 | 6.5279 | 7.1987 | 6.3335 | 6.3170 | 7.7659 | 6.0601 | 6.5540 | 7.2412 | 7.2993 | 7.3562 | 3.5835 | 4.7447 | 3.5823 | 4.7423 |
| NOP58_S514 | NOP58 | Q9Y2X3 | | 345 | GLIYHASLVGQTS | 6.9022 | 8.6854 | 5.1279 | 8.0836 | 8.0767 | 7.5789 | 4.2606 | 10.0575 | 6.0258 | 5.2243 | 5.6070 | 5.2070 | 3.1006 | 7.1386 | 5.0808 | 3.8432 |
| NOP58_T34 | NOP58 | Q9Y2X3 | 502;514 | | NA | 5.9061 | 6.2455 | 6.3733 | 6.7061 | 6.3593 | 6.7115 | 6.3507 | 5.8906 | 6.6032 | 6.0623 | 6.4175 | 6.2431 | 6.0968 | 6.3053 | 5.8544 | 5.8744 |
| NOS1AP_S266 | NOS1AP | O75052 | | 266 | EPMLTASPRMLLP | 6.5451 | 7.1716 | 7.5602 | 6.8425 | 5.6965 | 6.5784 | 6.8797 | 6.6950 | 6.3135 | 6.4618 | 6.2730 | 6.9559 | 4.3491 | 5.2096 | 4.9312 | 5.5371 |
| NOSIP_S138 | NOSIP | Q9Y314 | | 138 | KALSGTSPDDVQP | 6.2210 | 6.6679 | 6.9909 | 7.4571 | 6.3721 | 6.5992 | 6.7204 | 6.7055 | 7.1250 | 6.7162 | 6.9084 | 7.3146 | 4.1396 | 4.8081 | 4.6333 | 4.6207 |
| NOSIP_S138 | NOSIP | A0A075B6F9 | | 138 | KALSGTSPGDSDD | 6.0470 | 6.5371 | 8.0591 | 7.0802 | 6.7273 | 7.7464 | 6.9556 | 5.9312 | 6.5867 | 7.3774 | 7.5797 | 7.3714 | 3.2789 | 5.1665 | 3.6760 | 3.8796 |
| NOSIP_T168 | NOSIP | Q9Y314 | | 168 | FWIPSLTPEAKAT | 7.1536 | 6.9396 | 6.4710 | 7.5924 | 6.0166 | 6.9784 | 6.7429 | 6.9776 | 7.5982 | 7.3511 | 7.3866 | 8.2683 | 3.0616 | 4.1796 | 3.0211 | 4.2614 |
| NOTCH2_S1804 | NOTCH2 | Q04721 | | 1804 | DIRRTPSLALTPP | 6.2510 | 6.2141 | 6.2369 | 6.1673 | 6.5012 | 5.9920 | 6.0833 | 5.8206 | 6.3042 | 6.0793 | 5.7814 | 6.1941 | 6.2558 | 7.0081 | 6.5043 | 6.6065 |
| NOTCH2_S2070;S2081 | NOTCH2 | Q04721 | | 2081 | VLTSALSPVICGP | 5.7580 | 6.1489 | 5.9644 | 6.2903 | 6.0776 | 6.8216 | 6.7414 | 6.4213 | 5.9930 | 6.2302 | 6.7354 | 6.4688 | 5.6631 | 6.1167 | 5.7367 | 6.8328 |
| NOTCH2_S2081 | NOTCH2 | Q04721 | | 2115 | KSTMPTSLPNLAK | 6.7627 | 6.8768 | 6.5213 | 6.3567 | 6.1597 | 6.3235 | 6.6749 | 6.6543 | 6.4364 | 6.9141 | 6.7749 | 6.2473 | 5.2358 | 5.6317 | 5.0380 | 5.3918 |
| NOTCH2_S2115 | NOTCH2 | Q04721 | | 359 | CIDRVASFSCMCP | 5.1875 | 5.8471 | 5.6096 | 6.1558 | 9.9818 | 5.9078 | 4.8629 | 4.7178 | 7.2825 | 8.2087 | 5.1453 | 5.1875 | 6.1717 | 8.2641 | 6.9949 | 4.4751 |
| NOTCH2_S359 | NOTCH2 | Q04721 | 2070;2081 | | NA | 6.1203 | 6.9366 | 6.6074 | 7.2281 | 6.4577 | 7.0095 | 6.8774 | 6.6847 | 6.5184 | 7.2863 | 7.2310 | 7.1524 | 4.1801 | 4.5889 | 4.3334 | 4.7877 |
| NOTCH3_S2032 | NOTCH3 | Q9UM47 | | 2032 | QPSGPRSPPGPHG | 6.5352 | 6.7329 | 6.0165 | 6.3853 | 7.6360 | 6.1381 | 6.9408 | 6.0730 | 6.6795 | 7.6878 | 6.8145 | 7.1644 | 4.6340 | 4.9738 | 4.8788 | 4.7093 |
| NOVA1_S154 | NOVA1 | P51513 | | 154 | KQTLPSSPTTTKS | 6.4651 | 6.7033 | 5.8367 | 6.8809 | 5.9907 | 5.8409 | 5.9087 | 5.9776 | 5.9384 | 6.7303 | 6.5153 | 6.6656 | 6.0749 | 6.2100 | 6.0093 | 6.2523 |
| NPAT_S1100 | NPAT | Q14207 | | 1350 | SRTTSATPLKDNT | 8.9398 | 7.6662 | 6.9620 | 7.5855 | 5.2365 | 6.0586 | 7.1551 | 5.9732 | 8.0136 | 6.7227 | 7.0782 | 7.6811 | 3.5624 | 4.0456 | 3.2065 | 4.1130 |
| NPAT_S1296 | NPAT | Q14207 | | 1296 | PSSRRFSEDSSTS | 5.0138 | 5.4434 | 5.2176 | 6.3810 | 6.5566 | 5.2436 | 5.5707 | 5.5696 | 6.5739 | 6.1070 | 6.0577 | 6.1330 | 7.1681 | 7.4122 | 7.6219 | 7.9296 |

| | | | | | | | | | | | | | | | | | | | | | |
|---|---|---|---|---|---|---|---|---|---|---|---|---|---|---|---|---|---|---|---|---|---|
| NPAT_S207 | NPAT | Q14207 | | 207 | AHASLMSPGRRKS | 6.0939 | 8.9673 | 6.6563 | 6.8143 | 6.2740 | 6.1332 | 6.2962 | 8.3687 | 4.8182 | 5.4269 | 6.4612 | 6.6805 | 4.9963 | 5.3514 | 5.4299 | 5.2317 |
| NPAT_S775 | NPAT | Q14207 | | 775 | PTIILSSPTKSPT | 5.5012 | 6.2126 | 7.3689 | 7.2908 | 5.5947 | 5.7865 | 6.2244 | 6.3463 | 6.8955 | 5.8421 | 6.4765 | 6.5108 | 5.5545 | 6.2232 | 5.5119 | 6.6600 |
| NPAT_T1350 | NPAT | Q14207 | | 1100 | SFPNLDSPNVSST | 6.5079 | 7.4936 | 6.5805 | 7.3649 | 5.4359 | 6.4988 | 7.3453 | 7.0001 | 6.7413 | 7.8826 | 6.9903 | 7.0145 | 4.0272 | 4.6332 | 3.9240 | 4.5600 |
| NPHP4_S142 | NPHP4 | O75161 | | 451 | FQFSLGSEEHLDA | 6.8781 | 5.4271 | 6.6096 | 6.4605 | 7.0309 | 5.9939 | 5.6944 | 6.6896 | 6.4843 | 6.6860 | 5.7349 | 6.7349 | 5.5715 | 6.7886 | 5.7481 | 5.4676 |
| NPHP4_S451 | NPHP4 | O75161 | | 142 | FSNQPDSPISASQ | 6.2263 | 6.3804 | 6.6811 | 6.3762 | 6.8293 | 6.4634 | 6.4853 | 6.2305 | 6.0280 | 6.7501 | 6.0313 | 7.2138 | 5.5584 | 5.9816 | 5.2519 | 5.5125 |
| NPHP4_S481 | NPHP4 | O75161 | | 481 | PPTSPSSPPAPVP | 6.7133 | 6.5077 | 6.9987 | 6.3486 | 6.3595 | 6.8519 | 6.6042 | 6.7161 | 6.3742 | 6.8355 | 6.6565 | 6.3236 | 5.4210 | 5.8548 | 4.7058 | 4.7287 |
| NPHP4_S868 | NPHP4 | O75161 | | 868 | SLLTTGSSRRKHV | 7.6216 | 6.1757 | 7.3093 | 6.1159 | 5.4642 | 6.0342 | 6.5631 | 6.6038 | 7.4068 | 6.3626 | 6.9872 | 7.5897 | 4.9782 | 5.7263 | 4.1759 | 4.8855 |
| NPM1_S112 | NPM1 | P06748 | | 70 | AMNYEGSPIKVTL | 6.1863 | 6.4012 | 6.3091 | 6.7714 | 6.3202 | 6.1670 | 6.0434 | 6.2924 | 6.4129 | 6.3061 | 6.3507 | 6.5224 | 6.1697 | 5.9234 | 5.7829 | 6.0410 |
| NPM1_S125 | NPM1 | P06748 | | 125 | VEEDAESEDEEEE | 6.1824 | 6.5702 | 6.0310 | 7.0124 | 6.5772 | 6.5287 | 5.7269 | 6.6989 | 6.2639 | 5.9679 | 6.3515 | 6.2797 | 5.8553 | 5.9768 | 6.0743 | 5.9030 |
| NPM1_S137 | NPM1 | P06748 | | 254 | KAKMQASIEKGGS | 8.0424 | 8.1821 | 7.0443 | 7.7558 | 7.1238 | 7.3878 | 7.6588 | 7.2621 | 7.7849 | 7.6078 | 7.8035 | 7.2970 | 1.9691 | 2.7112 | 1.7538 | 2.6156 |
| NPM1_S139 | NPM1 | P06748 | | 199 | KKSIRDTPAKNAQ | 8.4160 | 8.3780 | 7.0757 | 7.1566 | 6.1212 | 7.9070 | 7.4504 | 7.3895 | 7.7038 | 7.4764 | 7.8463 | 7.7862 | 1.9878 | 2.8077 | 1.7095 | 2.7880 |
| NPM1_S143 | NPM1 | P06748 | | 139 | VKLLSISGKRSAP | 8.2198 | 7.9131 | 7.0395 | 6.8902 | 5.6009 | 7.3507 | 7.8042 | 7.3646 | 7.5908 | 7.0928 | 7.4238 | 7.4037 | 2.7307 | 3.4788 | 2.6806 | 3.4158 |
| NPM1_S149 | NPM1 | P06748 | | 112 | SGPVHISGQHLVA | 6.9091 | 8.0967 | 4.8261 | 7.9960 | 7.2635 | 8.0521 | 4.8799 | 9.8633 | 5.7545 | 5.5272 | 5.7633 | 5.1511 | 3.8232 | 5.6778 | 6.0536 | 4.3628 |
| NPM1_S214;T219 | NPM1 | P06748 | | 242 | KTPKGPSSVEDIK | 6.2271 | 6.4607 | 5.8381 | 6.0502 | 7.1524 | 6.4286 | 6.9582 | 6.4232 | 6.0024 | 5.8650 | 5.8329 | 5.6209 | 6.6470 | 6.2285 | 6.1032 | 6.1615 |
| NPM1_S227 | NPM1 | P06748 | | 219 | DSKPSSTPRSKGQ | 6.8400 | 7.1317 | 6.5225 | 7.0837 | 6.6645 | 7.8845 | 7.0782 | 7.2708 | 7.5236 | 7.1861 | 8.1052 | 7.6490 | 3.2944 | 3.6444 | 2.8612 | 3.2601 |
| NPM1_S242 | NPM1 | P06748 | | 88 | SVQPTVSLGGFEI | 6.2593 | 5.9227 | 6.4695 | 6.4915 | 6.3402 | 5.9300 | 6.0225 | 6.3174 | 6.2822 | 5.8867 | 6.3894 | 6.6200 | 6.4373 | 5.8120 | 6.5968 | 6.2226 |
| NPM1_S243 | NPM1 | P06748 | | 243 | TPKGPSSVEDIKA | 6.4131 | 6.7681 | 6.5207 | 6.9968 | 6.3764 | 6.5822 | 6.6091 | 6.2548 | 6.3563 | 6.1661 | 6.6875 | 6.5803 | 5.4568 | 5.4654 | 5.1446 | 5.6217 |
| NPM1_S254 | NPM1 | P06748 | | 143 | SISGKRSAPGGGS | 6.4691 | 6.6641 | 7.0509 | 6.7283 | 5.4383 | 6.6011 | 6.4082 | 6.7843 | 6.9951 | 6.3805 | 6.9646 | 7.5141 | 4.3583 | 4.9038 | 4.9218 | 5.8176 |
| NPM1_S48 | NPM1 | P06748 | | 137 | EDVKLLSISGKRS | 8.9917 | 8.5128 | 7.1233 | 7.7044 | 6.9222 | 7.1767 | 7.7599 | 6.8595 | 7.1995 | 6.4864 | 6.6290 | 6.1668 | 3.0147 | 3.6096 | 2.6027 | 3.2410 |
| NPM1_S70 | NPM1 | P06748 | | 227 | RSKGQESFKKQEK | 5.0076 | 6.7806 | 8.2938 | 6.5533 | 5.7624 | 6.9799 | 6.6843 | 7.9821 | 6.6737 | 10.3413 | 9.0210 | 7.9482 | 2.8140 | 3.8035 | 2.2935 | 3.0607 |
| NPM1_S88 | NPM1 | P06748 | | 149 | SAPGGGSKVPQKK | 6.0179 | 6.1575 | 5.6282 | 6.3123 | 6.4666 | 5.5378 | 6.0893 | 6.5394 | 6.3096 | 6.1360 | 6.0616 | 6.2882 | 7.1438 | 6.3636 | 6.3169 | 6.6314 |
| NPM1_T199 | NPM1 | P06748 | | 78 | IKVTLATLKMSVQ | 6.2108 | 5.5248 | 6.0891 | 5.5617 | 5.2261 | 6.9595 | 5.3884 | 8.5638 | 7.7967 | 5.7092 | 5.8936 | 5.3183 | 7.6197 | 5.7092 | 4.7245 | 7.7045 |
| NPM1_T219 | NPM1 | P06748 | | 48 | LSLRTVSLGAGAK | 6.0956 | 5.4368 | 6.6386 | 7.6688 | 5.0615 | 6.2500 | 7.0259 | 7.0765 | 6.3711 | 7.6980 | 8.6656 | 9.1767 | 3.8437 | 4.6569 | 3.5070 | 4.8272 |
| NPM1_T78 | NPM1 | P06748 | 214;219 | | NA | 8.2326 | 8.2010 | 8.5275 | 9.0489 | 5.4962 | 6.4620 | 6.5924 | 6.7639 | 7.7148 | 8.6110 | 10.4604 | 7.3985 | 1.2571 | 2.4716 | 0.8885 | 1.8735 |
| NPM3_S147;S151;S158 | NPM3 | O75607 | 147;151;158 | | NA | 6.2511 | 6.0208 | 6.3732 | 5.7890 | 6.6363 | 6.8614 | 6.1336 | 6.4767 | 5.4059 | 6.2533 | 6.3453 | 6.6322 | 6.6230 | 5.7793 | 5.9536 | 6.4655 |
| NPR2_S523;S526 | NPR2 | P20594 | | 526 | RGSSYGSLMTAHG | 7.9055 | 7.2118 | 7.1529 | 7.0780 | 7.0808 | 6.1848 | 5.8081 | 6.2447 | 7.0294 | 5.9938 | 5.9333 | 5.7728 | 5.4164 | 5.5585 | 5.2884 | 4.3408 |
| NPR2_S526 | NPR2 | P20594 | 523;526 | | NA | 6.7610 | 7.3944 | 6.6510 | 7.4115 | 6.2458 | 6.6276 | 6.2420 | 6.7425 | 5.9569 | 5.6776 | 5.9920 | 5.8317 | 5.5551 | 4.4208 | 6.9896 | 5.5006 |
| NPRL3_S538 | NPRL3 | Q12980 | | 538 | NENTRRSQLLMLF | 8.5250 | 7.0267 | 8.8618 | 7.3229 | 7.3810 | 7.6655 | 5.3484 | 6.5389 | 4.9768 | 4.8316 | 6.3008 | 5.6794 | 2.8862 | 6.1092 | 4.8316 | 5.7143 |
| NR2C1_S215 | NR2C1 | P13056 | | 215 | IRKDLRSPLTATP | 5.7918 | 5.8426 | 6.3130 | 6.6668 | 5.9224 | 6.3693 | 6.1733 | 6.6378 | 6.9989 | 6.2083 | 6.3237 | 5.8583 | 5.7526 | 6.5801 | 5.9417 | 6.6193 |
| NR2C1_S64 | NR2C1 | P13056 | | 64 | ILARQDSTPGKVF | 5.2489 | 5.6029 | 6.2251 | 6.1990 | 5.7161 | 5.7941 | 5.8204 | 6.0379 | 5.9816 | 6.8964 | 6.2737 | 6.4131 | 6.2040 | 7.5595 | 7.0689 | 6.9584 |
| NR2C1_T53 | NR2C1 | P13056 | | 53 | TNHDGSTPSKVIL | 8.0260 | 8.2230 | 6.7571 | 7.8589 | 6.4668 | 7.5825 | 6.7892 | 7.1161 | 7.6794 | 7.4365 | 7.6699 | 7.4785 | 2.3525 | 3.5466 | 2.1125 | 2.9042 |
| NR2C2_S19 | NR2C2 | P49116-2 | | 65 | AVDASGSPKQQFI | 6.0006 | 6.0262 | 6.3712 | 6.7624 | 5.9843 | 6.0564 | 6.5452 | 5.9884 | 6.6303 | 6.5175 | 6.3330 | 6.4797 | 5.8999 | 6.0779 | 5.8599 | 6.4669 |
| NR2C2_S65 | NR2C2 | P49116-2 | | 19 | TDSAVASPQRIQG | 6.3011 | 6.6655 | 6.4980 | 6.7537 | 6.3820 | 6.2346 | 6.7710 | 6.6069 | 6.8135 | 7.1369 | 6.3172 | 6.8562 | 4.8718 | 5.4474 | 4.7578 | 5.5863 |
| NR2C2_S74 | NR2C2 | P49116-2 | | 87 | GKVILASPETSSA | 6.3892 | 7.0948 | 6.9574 | 7.8600 | 6.3485 | 6.9149 | 7.1933 | 6.7807 | 7.1758 | 7.6086 | 7.1957 | 7.6792 | 3.3131 | 4.1397 | 3.0941 | 4.2550 |
| NR2C2_S87 | NR2C2 | P49116-2 | | 74 | QQFILTSPDGAGT | 6.3732 | 7.1242 | 7.7530 | 7.1810 | 6.0753 | 6.5220 | 7.4157 | 6.7993 | 7.1247 | 7.4496 | 7.8880 | 7.6821 | 3.4246 | 3.9156 | 3.1688 | 4.1028 |
| NR2F2_S21 | NR2F2 | P24468 | | 21 | VPGSQGSQASQAP | 5.6291 | 5.2654 | 5.5384 | 6.3524 | 5.9740 | 5.2585 | 6.0863 | 5.9336 | 4.8752 | 5.7956 | 5.6281 | 5.9888 | 8.5924 | 7.6887 | 7.6385 | 7.7548 |
| NR3C1_S134 | NR3C1 | P04150-3 | | 203 | LEFSSGSPGKETN | 5.8072 | 5.9304 | 5.8916 | 6.0482 | 6.0964 | 5.8746 | 6.3142 | 5.9669 | 6.2578 | 6.3293 | 6.5423 | 6.0547 | 6.6838 | 7.0462 | 6.3629 | 6.7934 |
| NR3C1_S203 | NR3C1 | P04150-3 | | 267 | GDLVLSSPSNVTL | 6.1894 | 6.5707 | 6.4591 | 6.6106 | 6.1672 | 6.8576 | 6.5032 | 6.5224 | 6.6148 | 6.9926 | 6.6999 | 6.9504 | 5.3526 | 5.2690 | 4.8436 | 5.3971 |
| NR3C1_S267 | NR3C1 | P04150-3 | | 8 | DSKESLTPGREEN | 6.6991 | 6.7118 | 7.2380 | 7.2498 | 5.8810 | 6.4307 | 6.9071 | 6.5459 | 6.0890 | 6.6367 | 7.1160 | 6.8661 | 5.3011 | 5.1391 | 4.6714 | 4.5172 |
| NR3C1_T8 | NR3C1 | P04150-3 | | 134 | NLNRSTSVPENPK | 3.7750 | 4.0187 | 4.4074 | 4.0109 | 5.1018 | 4.8059 | 4.4941 | 4.7888 | 4.0696 | 4.2377 | 3.8849 | 4.0636 | 12.5718 | 10.9597 | 13.1054 | 11.7048 |
| NRBF2_S113 | NRBF2 | Q96F24 | | 113 | EDAEGQSPLSQKY | 6.1229 | 6.4378 | 6.8969 | 5.8799 | 7.5958 | 6.7757 | 6.0768 | 8.2332 | 6.6885 | 7.2924 | 8.1306 | 7.7784 | 4.1898 | 4.5195 | 3.6912 | 3.6906 |
| NRBP1_S31 | NRBP1 | Q9UHY1 | | 431 | VPPSVKTPTPEPA | 6.6312 | 6.4869 | 6.9792 | 6.0378 | 6.4535 | 6.7287 | 6.1088 | 6.1207 | 6.3575 | 6.9373 | 6.4542 | 6.7675 | 5.6031 | 5.6171 | 5.3870 | 5.3294 |
| NRBP1_S422 | NRBP1 | Q9UHY1 | | 422 | QQEEVTSPVVPPS | 5.8529 | 6.5140 | 6.5114 | 6.1634 | 6.1885 | 6.8014 | 6.6265 | 6.3789 | 6.1298 | 6.4825 | 6.3931 | 6.7306 | 5.7980 | 6.0802 | 5.5214 | 5.8274 |
| NRBP1_S509 | NRBP1 | Q9UHY1 | | 509 | DQSRLTSLLEETL | 7.7969 | 7.5796 | 4.4641 | 6.0358 | 6.5987 | 6.8439 | 6.2531 | 5.5119 | 6.8105 | 4.6090 | 6.7157 | 5.8686 | 5.7515 | 6.8606 | 5.9410 | 6.3590 |
| NRBP1_T421 | NRBP1 | Q9UHY1 | | 421 | PQQEEVTSPVVPP | 5.6798 | 6.4390 | 6.8876 | 6.4098 | 5.8333 | 7.0437 | 5.8195 | 6.1592 | 5.9039 | 6.8955 | 7.7603 | 7.0745 | 5.5577 | 5.6687 | 5.5689 | 5.2987 |
| NRBP1_T431 | NRBP1 | Q9UHY1 | | 31 | PGLTSVSPPVTST | 6.4158 | 6.1306 | 8.6367 | 5.5202 | 6.5142 | 7.3014 | 5.0704 | 6.8716 | 6.1547 | 6.9660 | 6.4921 | 6.7833 | 5.3214 | 4.6125 | 5.3616 | 5.8475 |
| NRDC_S94 | NRDC | O43847-2 | | 94 | EEGRRGSLSNAGD | 5.5227 | 5.2014 | 6.1269 | 5.2910 | 6.2928 | 5.6970 | 5.7696 | 5.7967 | 7.4584 | 7.4673 | 7.3321 | 7.5901 | 5.9453 | 6.8748 | 5.7410 | 5.8928 |
| NRDE2_S341 | NRDE2 | Q9H7Z3 | | 341 | QDEVMKSPGLYAI | 6.1091 | 6.9363 | 6.5498 | 6.3570 | 6.1678 | 6.1433 | 6.4306 | 7.0994 | 6.5027 | 6.9388 | 6.6449 | 6.5651 | 5.5993 | 5.3524 | 5.2634 | 5.3401 |
| NRF1_S136 | NRF1 | Q16656-4 | | 136 | AIVLCISPSKPNP | 5.7795 | 6.6185 | 6.4964 | 6.9754 | 6.0931 | 5.8236 | 6.0002 | 5.7400 | 6.3355 | 6.3808 | 6.0499 | 6.4605 | 6.7151 | 5.6696 | 6.2461 | 6.6159 |
| NRG1_S443 | NRG1 | Q02297-2 | | 443 | TTPARMSPVDFHT | 6.1575 | 6.1189 | 5.9336 | 6.6928 | 6.2701 | 5.6829 | 5.6908 | 6.0807 | 6.2382 | 5.8037 | 5.7942 | 5.9858 | 7.1949 | 6.7569 | 6.8065 | 6.7925 |
| NRG1_S443;S452 | NRG1 | Q02297-2 | 443;452 | | NA | 7.0216 | 6.8216 | 7.0799 | 7.3481 | 6.3044 | 5.7457 | 6.4960 | 6.1216 | 6.9624 | 6.3840 | 6.4396 | 6.9416 | 5.6613 | 4.7924 | 4.6385 | 5.2415 |
| NRIP1_S807 | NRIP1 | P48552 | | 807 | FKSEPVSPQDFSF | 5.3871 | 5.7292 | 5.9339 | 6.4854 | 6.8668 | 6.2703 | 6.1580 | 6.8999 | 6.6446 | 5.3208 | 5.3492 | 5.4847 | 6.8237 | 6.9663 | 7.3112 | 6.3693 |
| NRIP3_S40 | NRIP3 | Q9NQ35 | | 40 | QFIHKDSADLLPL | 8.1464 | 6.6607 | 8.6171 | 6.2888 | 4.9775 | 6.2180 | 6.9002 | 6.1922 | 8.4899 | 6.9116 | 8.0638 | 7.9934 | 3.4023 | 3.8599 | 3.1431 | 4.1353 |
| NSD1_S1334 | NSD1 | Q96L73 | | 822 | SSDTKGSPLASIS | 6.8994 | 7.3326 | 6.4221 | 7.5005 | 6.0367 | 5.9463 | 6.2732 | 6.3132 | 6.5492 | 6.9569 | 7.2868 | 7.1587 | 4.4742 | 5.2886 | 4.7240 | 4.8376 |
| NSD1_S2369 | NSD1 | Q96L73 | | 2369 | KSIGAASPRPQSL | 6.8290 | 7.2417 | 6.3975 | 7.0831 | 6.5431 | 6.3961 | 7.0495 | 5.8803 | 6.6409 | 7.1133 | 6.9023 | 7.0973 | 4.3983 | 5.1322 | 4.4761 | 4.8192 |
| NSD1_S2397 | NSD1 | Q96L73 | | 1334 | KQVDENSLISTKE | 6.6182 | 5.5787 | 5.8535 | 6.3745 | 6.6789 | 6.3819 | 6.2941 | 6.8987 | 5.7797 | 6.6272 | 7.3565 | 6.5747 | 5.0422 | 6.2424 | 5.9307 | 5.7682 |
| NSD1_S543 | NSD1 | Q96L73 | | 543 | FISGDISDTQASN | 5.0824 | 5.2498 | 6.5310 | 7.1446 | 5.6884 | 5.3633 | 6.1905 | 7.9526 | 6.9965 | 7.3601 | 5.8712 | 6.0597 | 6.2290 | 5.5114 | 5.9827 | 6.7868 |
| NSD1_S822 | NSD1 | Q96L73 | | 1208 | EFPEHRTPSASIL | 5.9264 | 6.0292 | 6.1893 | 6.7419 | 5.9095 | 6.3836 | 6.7936 | 6.3334 | 6.1565 | 5.9597 | 6.6167 | 7.0123 | 5.3205 | 6.1962 | 6.3692 | 6.0620 |
| NSD1_T1208 | NSD1 | Q96L73 | | 2397 | RLLITSSPKPQTS | 7.1410 | 7.3314 | 5.9703 | 6.9309 | 4.8091 | 6.1305 | 6.3423 | 5.9751 | 7.3790 | 7.8383 | 6.1258 | 7.6114 | 3.9374 | 5.2905 | 4.9764 | 6.2106 |
| NSD2_S121 | NSD2 | O96028-3 | | 437 | PRRGVGSPPGRKK | 7.4961 | 7.6858 | 7.1711 | 8.0176 | 6.6990 | 6.4109 | 6.9318 | 6.0891 | 6.9833 | 7.0515 | 7.2313 | 7.2987 | 3.4259 | 4.2278 | 3.4278 | 3.8521 |
| NSD2_S194 | NSD2 | O96028-3 | | 121 | TPIKNGSPEIKLK | 6.6616 | 6.9743 | 6.6079 | 7.0695 | 5.8415 | 5.5757 | 6.6089 | 5.7709 | 6.8281 | 6.6107 | 6.5735 | 6.9175 | 5.3660 | 5.4700 | 5.3726 | 5.7511 |
| NSD2_S408 | NSD2 | O96028-3 | | 544 | DAEAEDTPRKRLR | 7.0013 | 7.0484 | 6.2210 | 7.5658 | 6.3171 | 5.4510 | 6.7752 | 6.2834 | 6.9691 | 6.2350 | 6.1637 | 6.1626 | 5.1134 | 5.8199 | 5.1620 | 5.7111 |
| NSD2_S437 | NSD2 | O96028-3 | | 8 | EFSIKQSPLSVQS | 6.8769 | 7.6760 | 6.8118 | 7.6843 | 6.3075 | 6.5057 | 7.3170 | 6.7580 | 7.5971 | 6.8990 | 7.2535 | 7.6023 | 3.2034 | 4.2024 | 3.2896 | 4.0154 |
| NSD2_S495 | NSD2 | O96028-3 | | 495 | SQWSLLSEKQRAR | 7.4514 | 8.4727 | 6.5905 | 7.6872 | 5.1044 | 7.9525 | 7.0429 | 5.8117 | 6.4846 | 6.8978 | 5.7344 | 7.0276 | 4.1700 | 4.9947 | 4.4287 | 4.1490 |
| NSD2_S618 | NSD2 | O96028-3 | | 115 | GTPPNTTPIKNGS | 7.1177 | 6.6637 | 6.3904 | 7.1552 | 7.7233 | 6.7294 | 7.4456 | 7.3493 | 6.5530 | 6.1113 | 6.6128 | 5.8423 | 4.4299 | 4.8887 | 4.6614 | 4.3261 |
| NSD2_S618 | NSD2 | O96028-3 | | 194 | LVSKISSPSDKKI | 7.3805 | 8.1534 | 7.2560 | 8.3975 | 6.1909 | 6.5849 | 7.2053 | 6.4500 | 7.3889 | 7.5197 | 7.6742 | 8.0457 | 2.3506 | 3.5201 | 2.5497 | 3.3325 |
| NSD2_S8 | NSD2 | O96028-3 | | 618 | GFSKSSSPSASLT | 6.0297 | 6.1602 | 5.6676 | 6.9331 | 5.8302 | 6.3421 | 6.3458 | 5.7425 | 6.0843 | 6.4372 | 6.2824 | 6.6279 | 6.0347 | 6.3568 | 6.5365 | 6.5889 |
| NSD2_T110 | NSD2 | O96028-3 | | 110 | EMKGIGTPPNTTP | 6.3523 | 6.7054 | 6.5923 | 7.2822 | 6.2245 | 5.5535 | 5.6616 | 5.6017 | 6.6114 | 6.6189 | 6.0305 | 6.5182 | 5.6308 | 6.5416 | 5.9031 | 6.1721 |
| NSD2_T110;T115 | NSD2 | O96028-3 | | 35 | GSANGKTPSCEVN | 7.1171 | 8.0732 | 7.6049 | 6.5707 | 6.3195 | 8.4488 | 7.7317 | 6.5659 | 7.2756 | 6.3122 | 7.3902 | 7.7146 | 2.4024 | 3.6585 | 4.0098 | 2.8049 |
| NSD2_T115 | NSD2 | O96028-3 | | 408 | RAKLCSSAETLES | 6.3697 | 6.2804 | 5.8996 | 7.2600 | 6.2644 | 7.3606 | 6.2888 | 6.6104 | 6.4618 | 6.3631 | 7.0457 | 7.3907 | 4.4207 | 5.2387 | 5.0271 | 5.7182 |
| NSD2_T35 | NSD2 | O96028-5 | | 618 | GFSKSSSPSASLT | 6.7441 | 6.8416 | 6.3589 | 6.5845 | 6.4611 | 5.9731 | 6.5456 | 6.0301 | 6.7685 | 6.0274 | 6.0022 | 6.1936 | 5.7237 | 6.0857 | 5.8417 | 5.8182 |
| NSD2_T544 | NSD2 | O96028-3 | 110;115 | | NA | 6.4975 | 6.8695 | 6.6030 | 7.3926 | 5.8932 | 5.6182 | 5.9623 | 5.8653 | 6.6504 | 6.5283 | 6.4270 | 6.7614 | 5.2433 | 5.8218 | 5.7721 | 6.0942 |
| NSD3_S478 | NSD3 | Q9BZ95 | | 498 | LQKCNMSPVVKIE | 7.3096 | 7.3433 | 7.0144 | 7.2986 | 5.3292 | 6.2680 | 7.3184 | 6.9139 | 7.5186 | 7.1185 | 7.2400 | 7.2119 | 3.4231 | 4.5883 | 3.7734 | 4.3310 |
| NSD3_S498 | NSD3 | Q9BZ95 | | 478 | TAAARKSLPASIT | 6.4115 | 4.7723 | 7.3436 | 8.1645 | 6.1137 | 6.1742 | 6.7674 | 7.2420 | 7.6536 | 7.2323 | 6.7795 | 5.9103 | 2.2905 | 6.5446 | 3.7481 | 6.8521 |
| NSF_S207 | NSF | P46459 | | 207 | TKENRQSIINPDW | 6.9323 | 6.3945 | 6.2632 | 6.2776 | 7.2171 | 6.1403 | 6.7552 | 6.3941 | 6.1612 | 5.5371 | 5.0796 | 6.0618 | 6.2318 | 6.7622 | 5.7879 | 6.0042 |
| NSFL1C_S116 | NSFL1C | Q9UNZ2-5 | | 116 | GPPRKKSPNELVD | 6.2322 | 6.0191 | 6.5213 | 5.8854 | 6.2729 | 6.0778 | 5.8444 | 6.5202 | 6.4715 | 6.2838 | 6.6651 | 6.8436 | 6.5693 | 6.1204 | 5.8516 | 5.8216 |
| NSFL1C_S142 | NSFL1C | Q9UNZ2-5 | | 142 | VERVTKSPGETSK | 7.2562 | 7.4196 | 7.1377 | 6.7991 | 6.7498 | 7.1660 | 6.7374 | 7.3295 | 7.3168 | 7.2213 | 7.5613 | 7.6130 | 3.3075 | 3.7218 | 3.1009 | 3.5620 |
| NSFL1C_S274 | NSFL1C | Q9UNZ2-5 | | 274 | QVLSTSSPAQQAE | 6.2990 | 6.7885 | 6.7768 | 6.7622 | 6.1367 | 7.2115 | 6.5952 | 6.7619 | 7.0313 | 7.1643 | 7.2730 | 7.6433 | 4.2019 | 4.9694 | 4.0205 | 4.3643 |
| NSL1_T242 | NSL1 | Q96IY1 | | 242 | ITQIETTPTETAS | 7.1238 | 7.3569 | 7.4835 | 6.6349 | 5.4278 | 6.3629 | 7.5274 | 6.5685 | 7.1351 | 7.1564 | 8.4375 | 6.8367 | 3.3306 | 5.0404 | 3.4008 | 4.1767 |
| NSMAF_T211 | NSMAF | Q92636-2 | | 211 | QSRLARTSFDKNR | 6.4411 | 6.7908 | 6.3174 | 6.8840 | 7.0772 | 5.8523 | 7.1222 | 5.7618 | 5.7648 | 5.8440 | 5.3028 | 5.1073 | 6.0927 | 6.6802 | 6.1522 | 6.8092 |
| NSMCE2_S6 | NSMCE2 | Q96MF7 | | 6 | xMPGRSSSNSGST | 6.5350 | 7.1563 | 6.7083 | 7.1932 | 6.8071 | 7.1066 | 7.0876 | 7.3681 | 6.3323 | 6.9479 | 7.0300 | 6.6972 | 3.6848 | 4.6925 | 3.8429 | 4.8101 |
| NSMCE3_S64 | NSMCE3 | Q96MG7 | | 64 | QGSQGPSPQGARR | 6.0958 | 7.1264 | 6.1835 | 7.2036 | 6.0107 | 6.5671 | 6.3092 | 6.8689 | 6.6618 | 7.3753 | 7.4782 | 6.7230 | 4.4025 | 5.3686 | 4.5190 | 5.1063 |
| NSMCE4A_S30;S32 | NSMCE4A | Q9NXX6 | 30;32 | | NA | 5.8447 | 6.2410 | 5.5802 | 6.2149 | 5.8420 | 5.3362 | 5.2238 | 6.2609 | 6.2406 | 5.9578 | 5.8400 | 5.8014 | 7.8867 | 7.3672 | 7.3525 | 7.0100 |
| NSRP1_T275 | NSRP1 | Q9H0G5 | | 275 | REKVIETPENDFK | 6.4105 | 6.7957 | 6.4473 | 6.8854 | 6.6281 | 6.6134 | 6.7111 | 6.3101 | 6.5273 | 6.4087 | 6.5691 | 6.3756 | 5.1507 | 5.4146 | 5.1799 | 5.5724 |
| NSUN2_S473 | NSUN2 | Q08J23 | | 473 | DPSKLESPSFTGT | 6.0101 | 6.5671 | 6.5997 | 6.5897 | 6.2974 | 6.4311 | 6.5405 | 6.7515 | 6.5679 | 6.6436 | 6.6928 | 6.5722 | 4.9533 | 5.9267 | 5.4614 | 5.3951 |
| NSUN2_S743 | NSUN2 | Q08J23 | | 743 | RAGEPNSPDAEEA | 5.5883 | 6.0453 | 6.0546 | 5.3284 | 5.6374 | 6.4820 | 5.5656 | 5.8381 | 6.5749 | 5.8993 | 5.6013 | 5.4602 | 7.2192 | 7.5772 | 7.4681 | 7.6599 |
| NSUN2_S743;S751 | NSUN2 | Q08J23 | | 751 | DAEEANSPDVTAG | 4.7358 | 6.1358 | 6.1634 | 5.6495 | 5.6558 | 6.0639 | 6.3149 | 6.3269 | 6.1670 | 6.1124 | 6.7960 | 5.9093 | 6.7064 | 6.9974 | 7.3260 | 6.9397 |
| NSUN2_S751 | NSUN2 | Q08J23 | 743;751 | | NA | 5.1228 | 5.7172 | 5.8883 | 5.5446 | 5.7261 | 6.3519 | 5.9142 | 5.5454 | 6.3907 | 6.2328 | 6.1516 | 6.1790 | 6.9400 | 7.5697 | 7.5910 | 7.1348 |
| NSUN4_S206 | NSUN4 | Q96CB9 | | 206 | LAANDLSPSRIAR | 6.2241 | 6.5373 | 6.2996 | 6.6427 | 6.3777 | 5.8095 | 6.2294 | 5.7174 | 6.4904 | 5.8472 | 6.0056 | 5.7189 | 6.7522 | 6.5468 | 6.3919 | 6.4095 |
| NSUN5_S327 | NSUN5 | Q96P11-2 | | 444 | APERTPSPAPKRK | 5.9770 | 5.9840 | 5.5313 | 5.5570 | 6.1944 | 5.7479 | 5.8360 | 5.9901 | 6.0569 | 5.9755 | 5.7116 | 5.8243 | 7.3435 | 7.3078 | 7.6176 | 7.3452 |
| NSUN5_T442;S444 | NSUN5 | Q96P11-2 | | 462 | AAAGACTPPCTxx | 6.1972 | 7.0201 | 6.0538 | 6.2177 | 6.2882 | 6.0637 | 6.0313 | 6.1099 | 6.2238 | 6.5025 | 6.3092 | 6.1641 | 5.7864 | 5.9395 | 6.5729 | 6.5197 |
| NSUN5_T444 | NSUN5 | Q96P11-2 | | 327 | PGAGTPSPVRLHA | 6.0033 | 6.3825 | 6.2589 | 6.9335 | 6.6131 | 6.2227 | 6.4213 | 6.6221 | 6.8100 | 6.8347 | 6.7687 | 6.5388 | 5.0275 | 5.5641 | 5.3481 | 5.6506 |
| NSUN5_T462 | NSUN5 | Q96P11-2 | 442;444 | | NA | 6.1637 | 6.3223 | 6.4718 | 6.3204 | 6.5947 | 6.6176 | 6.5538 | 6.6368 | 6.9910 | 6.8628 | 6.8011 | 6.7613 | 5.1685 | 5.2847 | 5.1591 | 5.2904 |
| NT5C2_S529 | NT5C2 | P49902 | | 529 | QLTRSISEIKPPN | 6.8024 | 6.3102 | 7.9273 | 6.4986 | 6.3119 | 6.6109 | 6.1201 | 6.0817 | 6.7026 | 6.8808 | 6.8883 | 6.8017 | 4.6636 | 5.8372 | 4.8784 | 4.6842 |
| NTHL1_S22 | NTHL1 | P78549 | | 22 | MLTRSRSLGPGAG | 6.1085 | 7.8366 | 6.8847 | 5.8378 | 4.4564 | 6.3535 | 6.4634 | 6.4269 | 7.5217 | 8.2085 | 8.2829 | 8.6236 | 3.5853 | 5.1612 | 3.4809 | 4.7680 |
| NTHL1_S71 | NTHL1 | P78549 | | 71 | RVAYEGSDSEKGE | 7.0786 | 7.4014 | 7.2490 | 7.6770 | 6.2184 | 6.8501 | 6.3655 | 6.5697 | 5.9197 | 6.1376 | 6.2968 | 6.2872 | 4.6489 | 5.0160 | 4.8352 | 5.4488 |
| NUAK1_S388 | NUAK1 | O60285 | | 455 | EVPGKLSPKQSAT | 5.8944 | 5.9540 | 5.8156 | 5.6052 | 5.7949 | 6.0624 | 6.5931 | 6.0847 | 6.2205 | 6.4568 | 6.1318 | 6.4874 | 7.6683 | 6.7352 | 6.9166 | 6.4791 |
| NUAK1_S413 | NUAK1 | O60285 | | 388 | QDAVPESPSKLSS | 5.5707 | 5.7357 | 6.5647 | 5.5132 | 6.2808 | 5.6649 | 6.3377 | 6.2283 | 6.4698 | 6.7253 | 7.0212 | 6.0639 | 5.9842 | 7.1686 | 6.1366 | 6.5343 |
| NUAK1_S455 | NUAK1 | O60285 | | 413 | SEHRSHSTGFIEG | 5.3654 | 5.6774 | 7.1212 | 5.3743 | 6.9994 | 6.7528 | 5.0267 | 5.2080 | 6.2329 | 5.5704 | 5.6863 | 5.4427 | 6.1646 | 8.0452 | 8.7968 | 6.5359 |
| NUBP1_S312 | NUBP1 | P53384 | | 312 | FCNLHQSKEENLI | 5.9976 | 5.7447 | 6.4676 | 5.6211 | 6.4778 | 6.7443 | 5.2790 | 6.1424 | 6.4372 | 6.9065 | 6.7608 | 6.4417 | 6.3935 | 6.6844 | 5.7759 | 6.1253 |
| NUCKS1_S113 | NUCKS1 | Q9H1E3 | | 181 | KATVTPSPVKGKG | 5.4961 | 5.5106 | 6.5464 | 5.2751 | 5.6854 | 6.0059 | 5.4331 | 6.2301 | 6.2778 | 6.3363 | 6.6753 | 6.6169 | 6.2785 | 7.5684 | 7.1988 | 6.8650 |
| NUCKS1_S19 | NUCKS1 | Q9H1E3 | | 113 | LMEDVGSEEEQEE | 5.6125 | 6.0478 | 6.4993 | 6.2222 | 6.0916 | 5.8190 | 5.6140 | 5.8952 | 5.9233 | 6.4319 | 6.3402 | 6.2831 | 6.0921 | 7.0728 | 6.9373 | 7.1179 |
| NUCKS1_S19;S30 | NUCKS1 | Q9H1E3 | | 223 | EKKTSTSPPPEKS | 5.4534 | 5.4457 | 5.9952 | 5.1205 | 6.1442 | 6.3275 | 5.6012 | 6.1378 | 6.0561 | 6.2464 | 6.0115 | 5.7752 | 6.9247 | 8.1088 | 7.6123 | 7.0395 |
| NUCKS1_S214 | NUCKS1 | Q9H1E3 | | 214 | EDEEPESPPEKKT | 4.9649 | 5.1972 | 6.0862 | 5.2871 | 5.8914 | 6.3231 | 5.6989 | 5.9903 | 6.2281 | 6.2833 | 6.3708 | 6.1341 | 6.9343 | 8.1864 | 7.3120 | 7.1119 |
| NUCKS1_S223 | NUCKS1 | Q9H1E3 | | 19 | YSQFQESDDADED | 5.7627 | 5.9684 | 6.3839 | 5.7096 | 6.2277 | 6.2462 | 5.6557 | 6.5396 | 6.0600 | 6.1863 | 5.6684 | 5.6166 | 6.8065 | 7.3967 | 7.0994 | 6.6724 |
| NUCKS1_S75;S79 | NUCKS1 | Q9H1E3 | | 202 | KASKEKTPSPKEE | 6.8729 | 7.6001 | 7.8157 | 7.0757 | 5.3622 | 6.7080 | 6.4016 | 6.5740 | 6.9474 | 8.0400 | 7.4843 | 7.6308 | 3.3806 | 4.2618 | 3.4592 | 4.3858 |
| NUCKS1_T179 | NUCKS1 | Q9H1E3 | | 179 | RLKATVTPSPVKG | 5.0123 | 6.5584 | 6.5350 | 5.0389 | 5.0123 | 7.5058 | 5.8457 | 7.4116 | 6.8514 | 8.7281 | 8.6554 | 8.8351 | 3.7219 | 5.0950 | 3.9136 | 5.2794 |
| NUCKS1_T179;S181 | NUCKS1 | Q9H1E3 | 179;181 | | NA | 6.5436 | 6.9306 | 6.8042 | 6.3083 | 6.3058 | 7.0504 | 6.6620 | 6.9657 | 6.9846 | 6.9184 | 7.4638 | 7.3048 | 3.8504 | 5.0045 | 4.1962 | 4.7067 |
| NUCKS1_T181 | NUCKS1 | Q9H1E3 | 202;204;214 | | NA | 5.3079 | 5.0865 | 6.4732 | 4.9607 | 5.3953 | 5.6102 | 6.0363 | 5.2485 | 6.3137 | 6.5970 | 6.5154 | 6.4360 | 7.7987 | 7.6950 | 7.2040 | 7.3216 |
| NUCKS1_T202 | NUCKS1 | Q9H1E3 | 75;79 | | NA | 5.8384 | 5.9065 | 6.6077 | 5.2289 | 5.8189 | 5.7938 | 5.7678 | 5.6519 | 6.3193 | 6.2510 | 6.2278 | 6.0751 | 6.3022 | 7.5654 | 7.2861 | 7.3592 |
| NUCKS1_T202;S204;S21[illegible] | NUCKS1 | Q9H1E3 | 19;30 | | NA | 6.0772 | 6.3968 | 6.7784 | 6.4906 | 5.9654 | 7.0600 | 7.5351 | 6.7577 | 5.6908 | 6.1521 | 6.5271 | 6.3482 | 5.2990 | 5.7490 | 5.6966 | 5.4762 |
| NUDC_S139 | NUDC | Q9Y266 | | 139 | KNGSLDSPGKQDT | 6.0679 | 6.1050 | 7.2401 | 5.8820 | 5.5929 | 6.2071 | 6.6234 | 6.0695 | 6.6062 | 7.2896 | 7.2088 | 7.4904 | 5.8973 | 5.5185 | 4.5827 | 5.6185 |
| NUDC_S260 | NUDC | Q9Y266 | | 145 | SPGKQDTEEDEEE | 5.5500 | 5.6157 | 7.4541 | 5.7959 | 5.5876 | 6.3410 | 6.8142 | 5.8142 | 6.1664 | 6.5985 | 7.0388 | 6.6574 | 6.6314 | 5.9668 | 5.6732 | 6.2948 |
| NUDC_T145 | NUDC | Q9Y266 | | 260 | WSRLVSSDPEINT | 6.1319 | 6.0219 | 7.0340 | 5.7092 | 5.9827 | 6.6804 | 5.9984 | 6.1147 | 6.4415 | 6.0392 | 7.7569 | 6.9475 | 5.8412 | 6.7134 | 5.4656 | 5.1215 |
| NUDCD1_T387 | NUDCD1 | Q96RS6 | | 387 | ERLMHLTSEELNP | 3.0615 | 3.2302 | 3.1682 | 2.8944 | 4.3356 | 5.0424 | 4.0610 | 5.0657 | 2.9745 | 3.4093 | 3.9077 | 3.9241 | 11.9660 | 14.2569 | 15.1334 | 13.5691 |
| NUDCD3_S340 | NUDCD3 | Q8IVD9 | | 355 | PAMFNISPGAVQF | 6.6987 | 6.6177 | 5.8588 | 7.0055 | 6.4946 | 7.3800 | 7.3761 | 6.4857 | 7.1537 | 7.5016 | 7.0332 | 7.5440 | 3.8286 | 4.5998 | 3.7683 | 4.6537 |
| NUDCD3_S355 | NUDCD3 | Q8IVD9 | | 340 | GWDAEGSPFRGQR | 7.2050 | 7.6496 | 7.1604 | 6.5912 | 5.2830 | 8.1800 | 7.8113 | 7.2008 | 7.7137 | 7.1161 | 8.1483 | 8.2042 | 2.2786 | 3.4487 | 2.2023 | 3.8068 |
| NUDT11_S148 | NUDT11 | Q96G61 | | 148 | KLKLGGSPTNGNS | 5.7644 | 6.0666 | 6.5059 | 5.7848 | 6.8112 | 7.3138 | 6.1266 | 6.0516 | 5.7869 | 6.1105 | 6.0173 | 6.0194 | 5.6616 | 6.6623 | 6.4577 | 6.8595 |
| NUDT19_S272 | NUDT19 | A8MXV4 | | 272 | RLANFASLSDLHK | 6.3677 | 6.6750 | 6.2867 | 6.3964 | 6.5011 | 4.8754 | 6.9620 | 7.1105 | 6.3357 | 5.7583 | 6.3964 | 6.1331 | 5.9119 | 6.0099 | 6.6581 | 5.6216 |
| NUDT19_T31 | NUDT19 | A8MXV4 | | 31 | SRPETATPPSRPP | 6.0972 | 8.2955 | 7.7613 | 5.6938 | 5.7774 | 6.4896 | 7.0928 | 7.6923 | 4.1750 | 5.8501 | 6.0863 | 6.4169 | 6.1480 | 5.4431 | 4.4911 | 6.4896 |
| NUDT9_S121 | NUDT9 | Q9BW91 | | 121 | ISESNFSPKFNEK | 6.4825 | 6.8075 | 6.5962 | 7.1427 | 7.0978 | 6.7814 | 6.4974 | 6.5414 | 6.6297 | 6.3069 | 7.0561 | 7.3257 | 5.0491 | 4.9732 | 4.3913 | 4.3212 |
| NUDT9_S68 | NUDT9 | Q9BW91 | | 68 | HNKARTSPYPGSK | 6.3306 | 6.7051 | 6.4780 | 6.6112 | 6.0987 | 7.2694 | 6.4393 | 5.8678 | 6.8329 | 7.1358 | 7.2763 | 6.9298 | 5.2973 | 5.5123 | 4.3116 | 4.9040 |
| NUF2_S247 | NUF2 | Q9BZD4 | | 247 | KTKIVDSPEKLKN | 7.0023 | 7.2909 | 6.9151 | 6.6695 | 6.5498 | 6.5879 | 6.7542 | 6.5844 | 6.9106 | 6.8243 | 6.4575 | 6.5885 | 4.5228 | 4.8930 | 4.4775 | 4.9717 |
| NUFIP1_S403 | NUFIP1 | Q9UHK0 | | 403 | DSSAPKSPSQDVK | 6.2377 | 6.4152 | 6.3094 | 6.6132 | 6.2808 | 6.0655 | 6.3985 | 6.2929 | 6.3056 | 6.0320 | 6.2856 | 5.9608 | 6.0815 | 6.0607 | 6.3579 | 6.3028 |
| NUFIP2_S112 | NUFIP2 | Q7Z417 | | 572 | ELEKRTSPQVLGS | 6.3290 | 6.6487 | 6.8835 | 7.0551 | 6.0763 | 6.4236 | 6.1836 | 6.8443 | 6.7709 | 6.9287 | 7.0238 | 7.1316 | 4.7890 | 5.2081 | 4.6587 | 5.0452 |
| NUFIP2_S212;S214 | NUFIP2 | Q7Z417 | | 652 | GLERNDSWGSFDL | 4.3186 | 4.3646 | 5.0363 | 4.4151 | 5.5564 | 5.5663 | 5.7261 | 5.5434 | 4.5939 | 4.7015 | 5.1185 | 5.0484 | 10.8993 | 9.3609 | 10.1619 | 9.5888 |
| NUFIP2_S212;T220 | NUFIP2 | Q7Z417 | | 629 | SQNPLASPTNTLL | 6.4104 | 6.3790 | 6.4674 | 6.7907 | 6.5451 | 6.4139 | 6.1350 | 6.4998 | 6.7837 | 6.7385 | 6.9413 | 6.8835 | 5.6036 | 5.4613 | 5.0398 | 4.9071 |
| NUFIP2_S399 | NUFIP2 | Q7Z417 | | 87 | LQQHQETPKKKTG | 7.5684 | 7.6301 | 7.2673 | 7.6905 | 6.1415 | 6.9084 | 7.4901 | 6.6213 | 7.9156 | 7.3124 | 7.6672 | 7.5996 | 2.9304 | 3.7013 | 2.3886 | 3.1673 |
| NUFIP2_S572 | NUFIP2 | Q7Z417 | | 399 | QSSSRLSQVPMSA | 5.8265 | 7.2597 | 6.1131 | 4.3464 | 5.9440 | 6.4562 | 8.4672 | 6.6394 | 8.8056 | 7.8517 | 6.6629 | 6.2212 | 5.5822 | 5.3237 | 4.2759 | 4.2242 |
| NUFIP2_S629 | NUFIP2 | Q7Z417 | | 112 | ISLKNLSSDEATN | 6.7932 | 7.0849 | 7.1199 | 6.3925 | 6.6067 | 6.5437 | 6.6619 | 6.7975 | 5.7338 | 5.9384 | 5.7989 | 5.8371 | 5.7872 | 5.9058 | 5.4555 | 5.5430 |
| NUFIP2_S652 | NUFIP2 | Q7Z417 | 212;214 | | NA | 7.2195 | 6.2934 | 7.2612 | 6.8962 | 6.2014 | 6.4172 | 5.5069 | 5.9749 | 5.1994 | 4.7623 | 4.6797 | 4.8543 | 7.5427 | 7.5132 | 6.6832 | 6.9945 |
| NUFIP2_T87 | NUFIP2 | Q7Z417 | 212;220 | | NA | 6.3434 | 6.7467 | 7.1177 | 6.6639 | 5.6970 | 6.8273 | 6.3641 | 6.3987 | 7.0220 | 7.2985 | 7.0283 | 7.1707 | 5.1510 | 5.1432 | 4.3749 | 4.6526 |
| NUMA1_S1225 | NUMA1 | Q14980 | | 2055 | AFSILNTPKKLGN | 8.2724 | 7.9819 | 7.2088 | 7.8700 | 5.9339 | 6.5283 | 7.1533 | 5.6131 | 6.9866 | 6.2565 | 6.5118 | 6.4513 | 4.3497 | 4.5389 | 3.8421 | 4.5012 |
| NUMA1_S169 | NUMA1 | Q14980 | | 1969 | ETLRRASMQPIQI | 5.0933 | 4.9771 | 5.0708 | 5.5386 | 7.2580 | 5.5313 | 5.5278 | 5.8027 | 8.2792 | 8.0768 | 7.0880 | 7.1595 | 7.0964 | 7.3722 | 4.8368 | 5.2914 |
| NUMA1_S1724 | NUMA1 | Q14980 | | 2047 | QADRRQSMAFSIL | 9.7004 | 5.0211 | 5.1504 | 5.8468 | 8.7751 | 4.5787 | 5.6191 | 4.2101 | 9.1755 | 6.5848 | 6.1901 | 5.5715 | 7.8878 | 8.4184 | 3.3860 | 3.8844 |
| NUMA1_S1728 | NUMA1 | Q14980 | | 1757 | VPGEPASPISQRL | 6.3537 | 6.4955 | 6.1146 | 6.9075 | 6.2122 | 6.0365 | 6.4179 | 6.1635 | 6.6128 | 6.3585 | 6.0939 | 6.2518 | 5.7431 | 6.0556 | 6.1042 | 6.0786 |
| NUMA1_S1757 | NUMA1 | Q14980 | | 169 | TFPEELSPPSHQA | 6.6237 | 6.9759 | 6.2701 | 7.1966 | 6.3305 | 6.4049 | 6.3332 | 6.2014 | 6.7568 | 7.0020 | 6.7450 | 6.5777 | 4.8497 | 5.4277 | 5.1874 | 5.1174 |
| NUMA1_S1769 | NUMA1 | Q14980 | | 1769 | LPPKVESLESLYF | 6.1004 | 6.4667 | 6.5645 | 6.9404 | 6.1743 | 6.2318 | 6.7136 | 6.6149 | 6.5403 | 7.1518 | 7.2770 | 6.6173 | 4.6189 | 5.5262 | 5.0195 | 5.4423 |
| NUMA1_S1769;T1776 | NUMA1 | Q14980 | | 1225 | EAERKNSLISSLE | 5.4634 | 5.4221 | 5.7937 | 6.5267 | 6.1796 | 5.8691 | 5.6847 | 5.9053 | 6.5388 | 7.1666 | 7.2258 | 6.6356 | 6.6637 | 6.9118 | 6.0351 | 5.9780 |
| NUMA1_S1776 | NUMA1 | Q14980 | | 1945 | PLESRPSLSLGTI | 6.0772 | 5.4565 | 5.9244 | 6.8838 | 7.2414 | 6.0446 | 6.5332 | 5.7559 | 7.8537 | 7.4128 | 6.8515 | 7.0332 | 5.6311 | 6.5283 | 4.1654 | 4.6070 |
| NUMA1_S1789;S1800 | NUMA1 | Q14980 | | 1840 | SFYSTRSAPASQA | 6.8623 | 7.9161 | 6.9385 | 7.5840 | 6.0760 | 6.3551 | 7.2342 | 6.6196 | 6.6853 | 6.6799 | 7.4117 | 7.0353 | 3.6005 | 4.7131 | 3.9324 | 4.3559 |
| NUMA1_S1800 | NUMA1 | Q14980 | | 2015 | CFPRPMTPRDRHE | 7.9039 | 7.9994 | 6.6073 | 7.1909 | 5.4477 | 6.5228 | 6.6489 | 5.8934 | 6.5814 | 6.9612 | 6.7673 | 6.6229 | 4.7221 | 5.2439 | 4.2719 | 4.6153 |
| NUMA1_S1830;S1834 | NUMA1 | Q14980 | | 2000 | PHQGPGTPESKKA | 7.4824 | 7.6755 | 6.5871 | 7.6811 | 6.3470 | 6.4965 | 6.2014 | 6.9670 | 6.9309 | 6.3050 | 6.8977 | 6.5309 | 4.2092 | 4.9459 | 4.4486 | 4.2938 |
| NUMA1_S1834 | NUMA1 | Q14980 | | 1834 | PDSANSSFYSTRS | 8.0255 | 7.8367 | 6.8932 | 7.6552 | 6.1722 | 6.9190 | 7.8499 | 6.4080 | 7.6091 | 6.3210 | 7.2979 | 6.8240 | 3.2806 | 4.3464 | 2.6528 | 3.9085 |
| NUMA1_S1840 | NUMA1 | Q14980 | | 2062 | PKKLGNSLLRRGA | 4.9747 | 5.7192 | 5.4991 | 4.9132 | 3.7604 | 5.9792 | 5.7892 | 6.2673 | 5.8169 | 9.4748 | 9.5414 | 12.9348 | 4.2923 | 6.5475 | 2.9231 | 5.5671 |
| NUMA1_S1853 | NUMA1 | Q14980 | | 1800 | GDVFLDSGRKTRS | 7.5768 | 7.8292 | 6.8325 | 7.8689 | 6.0960 | 7.2399 | 7.4217 | 6.5910 | 7.5417 | 7.1745 | 7.7118 | 7.6259 | 3.0264 | 3.4995 | 2.5455 | 3.4188 |
| NUMA1_S1862 | NUMA1 | Q14980 | | 1776 | LESLYFTPIPARS | 6.2497 | 6.5585 | 6.4782 | 7.3095 | 6.1787 | 6.0600 | 6.6294 | 6.3077 | 6.6658 | 6.6764 | 6.7096 | 6.5727 | 4.7698 | 5.9016 | 5.3602 | 5.5723 |
| NUMA1_S1887 | NUMA1 | Q14980 | | 1862 | SLARLGSPDYGNS | 6.5694 | 6.7950 | 6.0850 | 6.4545 | 6.1379 | 6.6012 | 6.8389 | 6.3181 | 6.5713 | 6.8087 | 7.1278 | 6.3863 | 5.0688 | 5.6561 | 5.2093 | 5.3717 |
| NUMA1_S1901 | NUMA1 | Q14980 | | 1812 | SARRRTTQIINIT | 9.1873 | 9.0484 | 9.3507 | 8.9975 | 6.4042 | 7.7089 | 7.1989 | 7.1891 | 5.3026 | 5.2444 | 5.1792 | 5.2322 | 3.8392 | 3.8880 | 2.6064 | 3.6230 |

| | | | | | | | | | | | | | | | | | | | | |
|---|---|---|---|---|---|---|---|---|---|---|---|---|---|---|---|---|---|---|---|---|
| NUMA1_S1945 | NUMA1 | Q14980 | 200 | NNFLSGSPASPMG | 6.1447 | 6.7354 | 6.8698 | 6.4572 | 6.5014 | 6.3100 | 5.8559 | 6.5988 | 6.6950 | 6.5186 | 6.3106 | 6.9318 | 5.0484 | 5.6983 | 5.5572 | 5.7669 |
| NUMA1_S1969 | NUMA1 | Q14980 | 1811 | RSARRRTTQIINI | 6.7478 | 6.8858 | 6.5453 | 6.1345 | 5.9796 | 6.5087 | 6.2668 | 6.5249 | 6.4178 | 6.1193 | 6.5823 | 6.5994 | 5.3409 | 6.0543 | 5.5815 | 5.7112 |
| NUMA1_S1991 | NUMA1 | Q14980 | 1853 | SLRATSSTQSLAR | 4.6072 | 4.9017 | 4.5328 | 5.1162 | 5.3844 | 4.8694 | 4.5494 | 5.0986 | 5.2439 | 5.4166 | 5.7351 | 6.4299 | 8.2709 | 10.2089 | 8.8406 | 10.7943 |
| NUMA1_S1991;T2000 | NUMA1 | H0YFY6 | 582 | TIIPDLSPNNPLS | 8.2163 | 7.9757 | 7.8069 | 7.5412 | 7.3293 | 5.1567 | 8.8196 | 7.4478 | 7.5089 | 6.3346 | 7.6777 | 6.3633 | 2.5856 | 2.6897 | 2.8764 | 3.6701 |
| NUMA1_S200 | NUMA1 | Q14980 | 1991 | QQRKRVSLEPHQG | 4.9030 | 5.3862 | 5.9067 | 5.4821 | 7.1515 | 5.7366 | 5.9272 | 5.8220 | 6.7535 | 7.0144 | 6.7449 | 7.1667 | 6.6727 | 7.1369 | 6.3287 | 5.8669 |
| NUMA1_S2047 | NUMA1 | Q14980 | 1887 | RSSARRSQAGVSS | 6.5973 | 6.8111 | 5.8489 | 6.0070 | 5.7811 | 6.0748 | 5.6516 | 6.0823 | 7.7117 | 7.9571 | 7.0656 | 7.1394 | 4.9619 | 5.7751 | 4.9679 | 5.5673 |
| NUMA1_S2047;T2055 | NUMA1 | Q14980 | 1901 | APPGRNSFYMGTC | 6.3216 | 6.9357 | 6.9055 | 6.7887 | 4.4831 | 7.2295 | 9.2337 | 7.6401 | 7.2936 | 6.6983 | 6.5702 | 5.9599 | 3.5903 | 5.2517 | 4.3701 | 4.7280 |
| NUMA1_S2062 | NUMA1 | Q14980 | 395 | QLEEHLSQLQDNP | 6.1822 | 6.3516 | 5.8501 | 6.5531 | 6.0437 | 4.9033 | 6.2143 | 5.3452 | 5.6626 | 5.4769 | 5.3041 | 5.1414 | 8.2603 | 7.5956 | 8.0284 | 7.0873 |
| NUMA1_S395 | NUMA1 | Q14980 | 1952 | LSLGTITDEEMKT | 6.2078 | 6.0431 | 7.5039 | 7.0301 | 6.6851 | 6.8545 | 6.5316 | 6.5257 | 6.2868 | 5.9279 | 6.3110 | 5.9483 | 5.4689 | 5.8174 | 4.9698 | 5.8879 |
| NUMA1_S582 | NUMA1 | Q14980 | 1724 | LDLSIDSLDLSCE | 6.1380 | 5.7201 | 6.3393 | 6.8146 | 6.2519 | 6.3419 | 6.0875 | 6.1791 | 5.8957 | 7.6214 | 6.1055 | 6.3470 | 5.9556 | 6.8429 | 5.1934 | 6.1663 |
| NUMA1_T1811 | NUMA1 | Q14980 | 1728 | IDSLDLSCEEGTP | 6.3673 | 5.9370 | 6.2396 | 7.9525 | 7.3977 | 5.8806 | 6.8699 | 5.9953 | 6.0559 | 6.3361 | 6.2070 | 6.4009 | 5.7934 | 5.6358 | 4.6773 | 6.2536 |
| NUMA1_T1812 | NUMA1 | Q14980 | 1789;1800 | NA | 7.0578 | 7.2517 | 6.6275 | 7.3018 | 6.2022 | 6.9389 | 7.6231 | 7.0835 | 6.9013 | 7.1851 | 7.2712 | 6.9522 | 3.5721 | 4.7805 | 3.2896 | 3.9615 |
| NUMA1_T1952 | NUMA1 | Q14980 | 2047;2055 | NA | 10.8195 | 6.3700 | 6.4298 | 6.2907 | 7.4106 | 5.2477 | 6.7495 | 4.5043 | 10.4518 | 7.5320 | 6.7017 | 6.4979 | 5.4508 | 4.4702 | 2.0902 | 2.9832 |
| NUMA1_T2000 | NUMA1 | Q14980 | 1991;2000 | NA | 6.3760 | 7.2571 | 7.2762 | 7.3263 | 5.6962 | 7.1202 | 7.1106 | 6.8504 | 7.0351 | 7.0219 | 7.6552 | 7.5790 | 3.4540 | 4.7614 | 3.4589 | 4.0215 |
| NUMA1_T2015 | NUMA1 | Q14980 | 1769;1776 | NA | 7.2677 | 6.3178 | 6.0351 | 6.0781 | 6.0190 | 6.4884 | 6.6580 | 6.8344 | 7.1105 | 6.6135 | 7.1137 | 7.0903 | 4.2070 | 5.8979 | 4.9906 | 5.2782 |
| NUMA1_T2055 | NUMA1 | Q14980 | 1830;1834 | NA | 7.3705 | 7.5663 | 6.5146 | 7.0946 | 5.5600 | 7.0924 | 8.1516 | 7.0529 | 7.8112 | 7.2559 | 7.3118 | 7.2470 | 3.1930 | 4.1592 | 2.7026 | 3.9162 |
| NUMB_S284 | NUMB | P49757 | 438 | GHRRTPSEADRWL | 6.1742 | 6.4734 | 6.6424 | 6.3778 | 6.1407 | 6.8633 | 5.9193 | 6.0409 | 5.6975 | 6.1292 | 6.5932 | 6.3466 | 6.4684 | 6.4012 | 5.9165 | 5.8152 |
| NUMB_S361 | NUMB | P49757 | 361 | TVVAPQSPTFQAN | 5.8758 | 5.5519 | 5.8713 | 6.4251 | 6.2586 | 6.5631 | 6.1237 | 6.4152 | 5.9476 | 5.8182 | 6.5073 | 6.5347 | 6.3911 | 6.7889 | 6.3522 | 6.5753 |
| NUMB_S361;S377 | NUMB | P49757 | 425 | QSSGAASPGLFQA | 6.0659 | 6.1688 | 6.2842 | 5.7912 | 5.7381 | 6.1409 | 6.1153 | 6.6076 | 6.2475 | 6.1501 | 6.7936 | 5.9333 | 5.9857 | 6.6676 | 6.4050 | 6.9053 |
| NUMB_S377 | NUMB | P49757 | 634 | KQRTNPSPTNPFS | 5.1032 | 5.2666 | 5.2380 | 5.3617 | 5.2708 | 5.9004 | 6.0041 | 5.6141 | 5.6653 | 5.9680 | 6.5838 | 6.3993 | 7.2267 | 7.8731 | 7.7445 | 8.7805 |
| NUMB_S425 | NUMB | P49757 | 284 | RGFPALSQKMSPF | 9.7711 | 7.6622 | 8.4982 | 7.0996 | 5.4166 | 6.1020 | 6.7026 | 5.8405 | 7.0991 | 6.2435 | 7.1801 | 7.2750 | 3.4111 | 5.3491 | 2.8914 | 3.4578 |
| NUMB_S438 | NUMB | P49757-2 | 377 | QSSGAASPGLFQA | 5.8432 | 5.3820 | 6.5222 | 7.4960 | 6.2292 | 6.8211 | 6.3618 | 6.1659 | 5.9343 | 6.4509 | 5.8828 | 5.5364 | 5.6314 | 7.4782 | 6.2549 | 6.0095 |
| NUMB_S634 | NUMB | P49757 | 436 | QAGHRRTPSEADR | 5.9820 | 7.0080 | 7.8033 | 6.8106 | 6.6517 | 6.9291 | 5.8781 | 5.1729 | 5.5472 | 5.9918 | 6.9823 | 6.1033 | 6.7604 | 5.9843 | 5.3703 | 5.0247 |
| NUMB_T243;S244 | NUMB | P49757-2 | 361;377 | NA | 6.1158 | 5.6384 | 5.8646 | 5.7448 | 5.5940 | 6.7209 | 6.9246 | 5.4749 | 6.4411 | 5.8913 | 6.9634 | 6.7189 | 6.0557 | 6.1514 | 6.5142 | 7.1859 |
| NUMB_T436 | NUMB | P49757 | 243;244 | NA | 6.9141 | 6.7476 | 4.9960 | 6.1456 | 6.7378 | 6.7006 | 6.6981 | 7.2168 | 7.3993 | 6.7867 | 6.6529 | 5.6897 | 5.5956 | 5.7069 | 5.1690 | 4.8434 |
| NUMBL_S165 | NUMBL | Q9Y6R0 | 263 | AQPGHVSPTPATT | 5.8110 | 5.8754 | 6.1016 | 5.7966 | 6.3291 | 6.5616 | 5.9042 | 6.7936 | 6.0771 | 6.2727 | 6.4040 | 6.5437 | 6.2191 | 6.2978 | 6.5014 | 6.5110 |
| NUMBL_S165;S172 | NUMBL | M0QXQ4 | 165 | AQPGHVSPTPATT | 5.6154 | 5.7536 | 6.4275 | 5.9719 | 6.1319 | 6.6935 | 6.1062 | 6.3251 | 6.3837 | 6.3481 | 6.6263 | 6.7672 | 6.2341 | 6.5223 | 5.9613 | 6.1321 |
| NUMBL_S263 | NUMBL | Q9Y6R0 | 279 | EKGEAGTPVAAGT | 6.9539 | 7.1204 | 7.4289 | 7.1871 | 6.3146 | 6.7535 | 7.0880 | 6.7153 | 7.6115 | 7.1036 | 7.3781 | 7.8296 | 3.4521 | 3.8757 | 3.2457 | 3.9420 |
| NUMBL_T279 | NUMBL | M0QXQ4 | 165;172 | NA | 6.1648 | 6.7341 | 7.1617 | 6.8996 | 5.9211 | 6.7358 | 7.5352 | 7.1730 | 6.7084 | 6.8545 | 6.3986 | 7.0019 | 4.5540 | 5.5026 | 3.8618 | 4.7931 |
| NUP107_S11 | NUP107 | P57740 | 11 | GFGEISSPVIREA | 7.1169 | 7.3171 | 6.4300 | 7.0461 | 6.3210 | 6.8062 | 7.1027 | 6.9840 | 7.4955 | 7.1835 | 7.0964 | 7.2495 | 3.7397 | 4.3856 | 3.5300 | 4.1958 |
| NUP107_S129 | NUP107 | P57740 | 129 | TNTEPHSITEDVT | 6.5420 | 6.6179 | 6.0662 | 6.6723 | 6.1734 | 6.5282 | 6.4082 | 6.5823 | 6.4608 | 6.2912 | 6.1405 | 5.9961 | 6.0205 | 6.0193 | 5.6678 | 5.8134 |
| NUP107_S311 | NUP107 | P57740 | 86 | LGTGGKSPRLTQS | 6.1107 | 5.9217 | 5.8068 | 6.8343 | 6.5674 | 6.6480 | 6.2997 | 5.8963 | 6.0462 | 5.7016 | 6.3274 | 6.5173 | 6.8154 | 6.6981 | 5.5481 | 6.2610 |
| NUP107_S4 | NUP107 | P57740 | 82 | ISCILGTGGKSPR | 8.0751 | 8.1991 | 7.8074 | 8.3425 | 6.4009 | 6.8381 | 7.3347 | 6.5351 | 7.9430 | 6.1677 | 6.5603 | 6.4646 | 2.7482 | 4.0960 | 2.7416 | 3.7456 |
| NUP107_S86 | NUP107 | P57740 | 4 | xxxMDRSGFGEIS | 7.1425 | 7.1274 | 6.7835 | 7.5153 | 6.4748 | 6.1637 | 6.6866 | 6.2241 | 7.2893 | 5.5854 | 6.7963 | 6.8961 | 4.6166 | 5.3634 | 4.1204 | 5.2146 |
| NUP107_T64 | NUP107 | P57740 | 64 | SFRQPFTPTSRSL | 8.0613 | 7.5811 | 6.3282 | 7.7628 | 6.5690 | 6.3796 | 7.1091 | 6.5696 | 7.2451 | 5.9415 | 6.4379 | 6.9148 | 4.0303 | 4.6967 | 3.8170 | 4.5560 |
| NUP107_T82 | NUP107 | P57740 | 311 | KQRQLTSYVGSVR | 7.9330 | 7.3808 | 7.2137 | 7.3262 | 6.3921 | 7.3842 | 7.4762 | 6.6342 | 7.2376 | 6.3989 | 6.7501 | 6.1330 | 2.6864 | 4.6671 | 3.4671 | 4.9194 |
| NUP133_S27;T28 | NUP133 | Q8WUM0 | 28 | GLGPGSTPRTASR | 7.3673 | 6.8422 | 6.1188 | 7.3217 | 7.0668 | 6.4381 | 7.2316 | 6.6882 | 7.0843 | 6.0041 | 6.5192 | 6.6815 | 4.5868 | 5.1212 | 3.9417 | 4.9864 |
| NUP133_S28 | NUP133 | Q8WUM0 | 57 | PVGRRSSLSSRGT | 5.5726 | 5.5286 | 5.0484 | 5.6574 | 6.6103 | 5.4260 | 5.2282 | 5.5339 | 7.5995 | 7.4369 | 6.7051 | 7.0859 | 7.2054 | 7.2578 | 5.6365 | 6.4675 |
| NUP133_S45 | NUP133 | Q8WUM0 | 480 | TSRENVSILAEDL | 5.9373 | 6.2245 | 6.5067 | 6.4690 | 6.3276 | 5.5148 | 6.2647 | 6.7672 | 6.1596 | 5.9006 | 6.6546 | 6.4438 | 6.0570 | 6.4121 | 6.3920 | 5.9685 |
| NUP133_S480 | NUP133 | Q8WUM0 | 501 | AGPNSESMIFETT | 5.8356 | 6.2937 | 5.6644 | 6.4190 | 6.6707 | 5.8518 | 5.6924 | 5.5503 | 6.1986 | 4.7186 | 5.7769 | 5.3859 | 7.7094 | 7.8845 | 7.3274 | 7.0208 |
| NUP133_S50 | NUP133 | Q8WUM0 | 594 | PGFSNTSLIILHQ | 6.1407 | 7.1455 | 5.4389 | 6.6285 | 6.5036 | 5.8954 | 5.9469 | 6.0710 | 6.3283 | 6.4809 | 6.8656 | 6.2512 | 5.8475 | 6.0257 | 5.9857 | 6.4446 |
| NUP133_S501 | NUP133 | Q8WUM0 | 50 | SSPVLFSPVGRRS | 6.8484 | 5.7569 | 6.8531 | 8.5901 | 6.7416 | 5.3041 | 6.9878 | 6.1726 | 6.7509 | 6.3282 | 6.1703 | 6.0403 | 4.5354 | 5.9520 | 5.7198 | 5.2484 |
| NUP133_S553 | NUP133 | Q8WUM0 | 45 | LGSAVSSPVLFSP | 5.9018 | 6.2078 | 6.2373 | 6.4026 | 5.8189 | 6.6563 | 6.7724 | 5.6733 | 5.4724 | 6.6754 | 6.5328 | 6.9088 | 5.9117 | 6.1722 | 6.3755 | 6.2809 |
| NUP133_S57 | NUP133 | Q8WUM0 | 553 | SHSDLDSDSELDR | 6.6242 | 6.5502 | 6.0265 | 6.8847 | 6.8056 | 6.4369 | 6.4202 | 6.6370 | 6.2703 | 6.3186 | 5.9647 | 6.0857 | 5.9422 | 5.9068 | 4.9488 | 6.1777 |
| NUP133_S594 | NUP133 | Q8WUM0 | 27;28 | NA | 5.9981 | 7.0046 | 6.5970 | 7.0050 | 6.1722 | 6.9331 | 7.0193 | 7.1467 | 7.3466 | 7.9860 | 7.6833 | 7.7617 | 3.4418 | 4.4975 | 3.2980 | 4.1090 |
| NUP153_S1078 | NUP153 | P49790-3 | 320 | SLEKMSSPLADAK | 7.5410 | 7.8302 | 6.8157 | 7.2995 | 5.6359 | 7.2659 | 7.5163 | 6.9624 | 7.7154 | 7.5271 | 7.7706 | 7.8323 | 2.7670 | 3.6460 | 2.5039 | 3.3706 |
| NUP153_S1492 | NUP153 | P49790-3 | 730 | KQTGIETPNKSGK | 7.5466 | 7.2597 | 6.9213 | 7.2680 | 6.3898 | 6.3903 | 6.8748 | 6.2348 | 7.2197 | 6.7594 | 6.7451 | 6.8311 | 4.1588 | 4.7715 | 3.9833 | 4.6459 |
| NUP153_S1494 | NUP153 | P49790-3 | 209 | SLPPLWSPEAERS | 6.7976 | 6.7253 | 6.7242 | 6.9493 | 6.5755 | 6.5739 | 6.7404 | 6.1905 | 7.0260 | 6.5947 | 6.7527 | 6.6829 | 4.8207 | 5.1924 | 4.6301 | 5.0238 |
| NUP153_S174 | NUP153 | P49790-3 | 553 | SPSSTGSPMFKFS | 6.9647 | 6.9165 | 6.4018 | 7.3071 | 6.7219 | 6.5145 | 6.5564 | 6.1182 | 7.1390 | 6.6390 | 6.5197 | 6.7005 | 4.7451 | 5.2920 | 4.5144 | 4.9492 |
| NUP153_S209 | NUP153 | P49790-3 | 334 | IPSIVSSPLNSPL | 6.3421 | 6.3409 | 6.3789 | 6.5831 | 6.5474 | 6.3675 | 6.1393 | 6.3403 | 6.3811 | 6.6265 | 6.1867 | 6.4453 | 5.9815 | 5.9040 | 5.5817 | 5.8537 |
| NUP153_S234 | NUP153 | P49790-3 | 645 | VLDILKSPGFASP | 7.1093 | 7.3772 | 6.7078 | 7.5776 | 6.4539 | 6.7313 | 7.1218 | 6.7283 | 7.2528 | 6.5941 | 6.9979 | 6.9919 | 3.8943 | 4.3409 | 3.8360 | 4.2850 |
| NUP153_S240 | NUP153 | P49790-3 | 102 | ITDGRITPEPAVS | 5.5473 | 5.9856 | 6.2261 | 6.4659 | 5.4756 | 5.9163 | 6.3236 | 6.2569 | 6.9652 | 6.2587 | 6.0494 | 6.2876 | 6.9652 | 6.3937 | 5.9757 | 6.9073 |
| NUP153_S257 | NUP153 | P49790-3 | 388 | YFKPSLTPSGEFR | 7.3727 | 6.6446 | 7.2414 | 7.2492 | 5.4267 | 7.3047 | 7.7348 | 5.9320 | 7.2208 | 7.1205 | 7.4669 | 8.1978 | 3.6071 | 4.2183 | 2.9568 | 4.3058 |
| NUP153_S320 | NUP153 | P49790-3 | 664 | AQPTATSPVVYTR | 5.5440 | 6.4791 | 6.3575 | 6.6656 | 6.3426 | 6.1278 | 6.5820 | 6.3776 | 6.9046 | 6.6022 | 6.8517 | 6.6878 | 5.5827 | 5.8262 | 5.4682 | 5.6004 |
| NUP153_S334 | NUP153 | P49790-3 | 560 | PMFKFSSPIVKST | 7.6702 | 7.7130 | 6.8738 | 7.6180 | 6.4558 | 7.0823 | 7.3886 | 6.8685 | 7.6841 | 7.3264 | 7.2273 | 7.3463 | 2.8455 | 3.6510 | 2.7538 | 3.4954 |
| NUP153_S334;S338 | NUP153 | P49790-3 | 619 | STNCKKTPPEDCE | 6.5238 | 6.7817 | 6.8693 | 7.4343 | 6.0033 | 6.0178 | 6.7294 | 6.1754 | 6.4843 | 6.7794 | 6.8727 | 6.6331 | 4.9647 | 5.3799 | 5.0526 | 5.2983 |
| NUP153_S338 | NUP153 | P49790-3 | 638 | EILKEGSVLDILK | 7.2806 | 8.6439 | 7.4671 | 7.5265 | 6.2116 | 5.7831 | 7.3146 | 6.9000 | 6.4778 | 5.7433 | 5.9469 | 6.2245 | 4.6744 | 5.1222 | 4.0228 | 4.6607 |
| NUP153_S343 | NUP153 | P49790-3 | 343 | NSPLDRSGIDITD | 7.6775 | 7.3139 | 6.6465 | 7.9852 | 6.8040 | 6.2779 | 7.7261 | 6.4461 | 7.7468 | 6.5959 | 6.7690 | 7.0257 | 3.5069 | 4.1664 | 2.8703 | 4.4418 |
| NUP153_S547 | NUP153 | P49790-3 | 1494 | SSGTSFSGRKIKT | 7.0102 | 7.2877 | 6.5720 | 7.2692 | 6.1120 | 6.9672 | 7.1660 | 6.9065 | 7.6607 | 7.0765 | 7.4876 | 7.7629 | 3.4011 | 4.1659 | 3.3138 | 3.8407 |
| NUP153_S547;S553 | NUP153 | P49790-3 | 240 | SAFGTLSPSLGNS | 6.9005 | 6.2292 | 6.5823 | 7.5074 | 6.5339 | 6.8160 | 6.9930 | 6.2348 | 7.0677 | 7.0154 | 7.2595 | 7.3785 | 4.2440 | 4.7840 | 3.9011 | 4.5529 |
| NUP153_S553 | NUP153 | P49790-3 | 650 | KSPGFASPKIDSV | 6.4594 | 6.7465 | 6.8841 | 7.6702 | 6.1733 | 6.5400 | 6.5062 | 5.9774 | 7.3292 | 7.2817 | 7.1954 | 7.2196 | 4.1235 | 5.0238 | 4.2127 | 4.6570 |
| NUP153_S560 | NUP153 | P49790-3 | 257 | TSQLGDSPFYPGK | 7.3947 | 7.5103 | 7.2241 | 7.3118 | 5.4462 | 7.1682 | 7.7968 | 6.4836 | 7.8406 | 7.9640 | 7.7060 | 7.8807 | 2.7540 | 3.6130 | 2.4066 | 3.4994 |
| NUP153_S638 | NUP153 | P49790-3 | 885 | FKKPEGSWDCELC | 4.6467 | 4.6986 | 4.3454 | 4.9346 | 4.1761 | 4.4739 | 4.3973 | 4.7450 | 4.4073 | 4.5127 | 4.3362 | 4.4090 | 11.2090 | 10.8119 | 11.9992 | 11.8971 |
| NUP153_S645 | NUP153 | P49790-3 | 338 | VSSPLNSPLDRSG | 6.3772 | 6.4359 | 6.4151 | 6.8071 | 6.6867 | 6.0103 | 6.4357 | 6.6354 | 6.8440 | 5.8915 | 6.0266 | 6.1576 | 6.2159 | 5.8025 | 5.8066 | 5.4520 |
| NUP153_S645;S650 | NUP153 | P49790-3 | 922 | FKGFDTSSSSSNS | 6.2098 | 6.1922 | 6.3441 | 6.5647 | 6.0869 | 7.1593 | 7.3225 | 7.2320 | 7.3769 | 6.8738 | 7.0177 | 6.7628 | 4.2428 | 5.4229 | 4.3093 | 4.8822 |
| NUP153_S650 | NUP153 | P49790-3 | 1078 | TSENKSSFNLGTI | 6.0680 | 6.5553 | 6.6105 | 7.0231 | 6.2523 | 6.7779 | 6.9898 | 6.8649 | 6.8246 | 6.8007 | 7.3474 | 7.3369 | 4.1885 | 4.8108 | 4.5050 | 5.0442 |
| NUP153_S664 | NUP153 | P49790-3 | 369 | PVQRLMTPKPVSI | 7.5686 | 7.0291 | 6.6850 | 7.1422 | 5.7367 | 7.3571 | 7.3174 | 5.6843 | 7.1596 | 6.2823 | 7.7096 | 7.2945 | 4.4858 | 4.9838 | 3.0826 | 4.4815 |
| NUP153_S885 | NUP153 | P49790-3 | 281 | QSKLRNTPYQAPV | 6.4461 | 6.5528 | 13.5234 | 7.0871 | 6.8544 | 6.7023 | 6.5514 | 5.5047 | 6.8474 | 5.9653 | 7.6014 | 6.7175 | 2.9330 | 3.8147 | 3.0487 | 3.8501 |
| NUP153_S922 | NUP153 | P49790-3 | 234 | KPAFNLSAFGTLS | 7.3400 | 7.3145 | 6.9505 | 8.0834 | 6.8086 | 6.1985 | 7.4965 | 5.7350 | 6.4376 | 6.2826 | 6.3034 | 6.9351 | 4.9229 | 4.7787 | 4.0005 | 4.4123 |
| NUP153_T102 | NUP153 | P49790-3 | 1492 | FSSSGTSFSGRKI | 5.7123 | 5.7196 | 7.0221 | 6.4359 | 5.3714 | 6.5277 | 6.5413 | 5.5190 | 6.6963 | 7.9517 | 7.8711 | 8.0061 | 5.1839 | 5.9265 | 4.3602 | 5.1548 |
| NUP153_T281 | NUP153 | P49790-3 | 547 | NKVQMTSPSSTGS | 5.6753 | 6.3665 | 5.8364 | 6.5605 | 5.9037 | 6.5061 | 6.6827 | 6.1361 | 5.7042 | 5.3935 | 5.4698 | 6.6258 | 6.9046 | 6.8103 | 6.7430 | 6.6817 |
| NUP153_T369 | NUP153 | P49790-3 | 174 | EIKDSTSQHDDDN | 6.3043 | 5.7117 | 6.8845 | 7.5483 | 6.1725 | 5.4016 | 5.5210 | 6.7724 | 6.0497 | 6.9852 | 6.6251 | 6.3059 | 5.5661 | 6.0301 | 6.5809 | 5.5407 |
| NUP153_T388 | NUP153 | P49790-3 | 547;553 | NA | 7.3189 | 7.3351 | 6.7232 | 7.5382 | 6.5408 | 7.3055 | 7.5509 | 6.9817 | 7.4823 | 7.5867 | 7.3602 | 7.1870 | 3.0119 | 3.9359 | 2.7445 | 3.3973 |
| NUP153_T619 | NUP153 | P49790-3 | 334;338 | NA | 6.2352 | 7.0181 | 6.5849 | 7.0558 | 5.9720 | 6.5076 | 6.6722 | 6.2967 | 6.6463 | 6.7204 | 6.7547 | 7.2434 | 4.7487 | 5.4351 | 4.7947 | 5.3141 |
| NUP153_T730 | NUP153 | P49790-3 | 645;650 | NA | 7.5491 | 7.8985 | 7.1861 | 7.6511 | 5.8911 | 7.4563 | 7.5455 | 6.6926 | 8.2221 | 7.6745 | 7.9794 | 7.9656 | 1.8747 | 3.4779 | 2.0249 | 2.9107 |
| NUP155_S992 | NUP155 | O75694 | 992 | SKAAPQSPSVPKK | 6.3254 | 6.4028 | 6.1061 | 6.4972 | 6.3313 | 6.3326 | 6.6124 | 6.3545 | 6.4684 | 6.2046 | 6.2245 | 6.1267 | 5.8951 | 6.0035 | 6.0284 | 6.0864 |
| NUP160_S1157 | NUP160 | Q12769 | 1157 | YDRPGASPKRNHD | 6.1168 | 6.2790 | 6.0886 | 6.7125 | 6.1293 | 5.8928 | 6.0332 | 5.9316 | 6.3637 | 6.2276 | 5.9552 | 6.1836 | 6.4910 | 6.4992 | 6.3485 | 6.7476 |
| NUP160_S490 | NUP160 | Q12769 | 490 | ERNLDLSWSELKK | 6.2067 | 7.0731 | 5.6373 | 6.2800 | 6.2092 | 5.7035 | 6.5022 | 6.2910 | 6.1199 | 5.7136 | 5.6970 | 6.1776 | 6.1245 | 6.3924 | 6.3628 | 7.5091 |
| NUP188_S1523 | NUP188 | Q5SRE5 | 1708 | FRRGAPSSPATGV | 5.6901 | 5.7647 | 5.3516 | 6.1177 | 6.1470 | 6.0762 | 5.8441 | 5.8425 | 5.7787 | 5.3755 | 5.9219 | 5.5733 | 7.7542 | 7.4391 | 7.6910 | 7.6324 |
| NUP188_S1708 | NUP188 | Q5SRE5 | 1523 | RVQRPPSAASAAP | 5.9447 | 5.6327 | 5.3776 | 6.2009 | 6.2877 | 5.8266 | 6.2612 | 5.8275 | 5.7426 | 5.6362 | 5.5935 | 5.7858 | 8.2717 | 7.3171 | 7.4280 | 6.8663 |
| NUP188_S1708;S1717 | NUP188 | Q5SRE5 | 1717 | ATGVLPSPQGKST | 6.6980 | 6.1534 | 6.3230 | 7.6276 | 6.8301 | 6.5607 | 6.8623 | 7.3136 | 8.3060 | 6.6616 | 5.6695 | 7.1904 | 3.9702 | 4.7801 | 3.7537 | 4.3998 |
| NUP188_S1717 | NUP188 | Q5SRE5 | 1708;1717 | NA | 6.1699 | 6.3280 | 5.9617 | 6.5084 | 6.2062 | 6.1725 | 6.7154 | 6.1352 | 6.2950 | 5.8023 | 6.3123 | 6.4958 | 6.2320 | 6.2958 | 5.9481 | 6.4212 |
| NUP205_S1153 | NUP205 | Q92621 | 1165 | IEDENRSVSGFLH | 6.9738 | 6.5391 | 6.1549 | 7.0308 | 6.5753 | 5.5252 | 6.3622 | 6.1423 | 6.3534 | 5.2494 | 5.6479 | 5.6399 | 6.3444 | 6.5123 | 6.2552 | 6.6937 |
| NUP205_S1165 | NUP205 | Q92621 | 1167 | DENRSVSGFLHFD | 7.2423 | 7.0158 | 6.4464 | 7.4688 | 5.1620 | 5.7514 | 7.3623 | 5.3785 | 6.0580 | 5.1183 | 5.9115 | 5.7314 | 6.3045 | 5.9252 | 6.3509 | 6.7729 |
| NUP205_S1165;S1167 | NUP205 | Q92621 | 1153 | MPVKPYSDGEGGI | 4.9670 | 6.7541 | 6.2539 | 6.1763 | 6.0505 | 5.5899 | 6.3125 | 6.6441 | 6.4272 | 5.9310 | 6.2302 | 5.4023 | 6.6844 | 6.8055 | 7.1466 | 6.6243 |
| NUP205_S1167 | NUP205 | Q92621 | 1942 | LQDSFASETNLDF | 6.6473 | 6.3392 | 5.8260 | 6.4675 | 6.3461 | 5.8888 | 5.6438 | 5.7775 | 6.3473 | 5.1955 | 5.6100 | 5.7494 | 7.1303 | 7.0202 | 6.7333 | 7.2778 |
| NUP205_S1942 | NUP205 | Q92621 | 1165;1167 | NA | 6.9519 | 6.4298 | 3.8774 | 7.2237 | 7.5046 | 6.1051 | 7.0147 | 6.8617 | 7.0223 | 5.0486 | 5.2069 | 5.2538 | 6.3797 | 6.8537 | 5.5903 | 6.6759 |
| NUP210_S1874 | NUP210 | Q8TEM1 | 1844 | AVPAALTPRASPG | 6.5197 | 7.1408 | 6.2762 | 7.5845 | 6.4051 | 6.4195 | 6.7886 | 6.6720 | 6.6843 | 6.6246 | 6.6877 | 6.6734 | 4.5318 | 5.0910 | 4.9431 | 4.9579 |
| NUP210_S1874;S1881 | NUP210 | Q8TEM1 | 1874 | PPARKASPPSGLW | 6.9801 | 7.1571 | 5.9287 | 6.6851 | 6.5127 | 6.2579 | 6.0992 | 6.5081 | 6.5782 | 6.0613 | 6.9081 | 6.3222 | 5.4616 | 5.4996 | 5.4164 | 5.6237 |
| NUP210_T1844 | NUP210 | Q8TEM1 | 1874;1881 | NA | 7.2556 | 7.6043 | 6.2522 | 8.2165 | 6.7629 | 7.8377 | 7.7688 | 9.2055 | 6.5268 | 6.7167 | 8.3079 | 7.3923 | 2.5131 | 3.3230 | 1.9898 | 2.3266 |
| NUP214_S1012 | NUP214 | P35658-5 | 1251 | EAIPESSPPSGIT | 6.0803 | 5.8372 | 6.3160 | 7.0391 | 6.2740 | 6.5178 | 6.4233 | 6.1965 | 6.6421 | 6.7275 | 6.9878 | 6.6624 | 5.6753 | 5.7580 | 5.1741 | 5.6886 |
| NUP214_S1070 | NUP214 | P35658-5 | 635 | KSSVLPSPSGRSA | 6.5050 | 6.4047 | 6.4835 | 6.2471 | 6.4834 | 6.5700 | 6.4528 | 6.5603 | 6.7194 | 6.3647 | 6.3051 | 6.3088 | 5.6947 | 5.8954 | 5.5686 | 5.4364 |
| NUP214_S1146 | NUP214 | P35658-5 | 645 | RSAQGSSSPVPSM | 6.0100 | 6.2030 | 6.3443 | 6.9095 | 6.6214 | 6.6756 | 6.7147 | 6.6159 | 6.8057 | 6.7987 | 6.7281 | 6.9973 | 5.1937 | 5.3028 | 4.9143 | 5.1649 |
| NUP214_S1251 | NUP214 | P35658-5 | 929 | KVPAKLSPMKQAQ | 6.0569 | 6.6065 | 6.2849 | 6.6366 | 7.1010 | 7.1157 | 6.5130 | 6.6553 | 6.3996 | 6.5516 | 7.1186 | 6.7548 | 5.4194 | 5.5586 | 4.7098 | 4.5176 |
| NUP214_S1332 | NUP214 | P35658-5 | 667 | PAAKPGSPQAKSL | 6.0726 | 5.7641 | 6.3514 | 6.4311 | 6.1345 | 6.1193 | 6.5738 | 5.8300 | 6.2224 | 6.2502 | 6.5325 | 6.5354 | 6.5251 | 6.1011 | 5.9345 | 6.6220 |
| NUP214_S1942 | NUP214 | P35658-5 | 1291 | TSSKLETPPSKLG | 6.0140 | 6.4290 | 5.9670 | 6.2986 | 5.8710 | 6.1857 | 6.8235 | 6.1461 | 6.4102 | 6.1798 | 6.3590 | 6.6825 | 6.1286 | 6.2159 | 5.8892 | 6.4000 |
| NUP214_S430 | NUP214 | P35658-5 | 1070 | VKHGAPSPSHPIS | 6.3255 | 6.4158 | 6.1062 | 6.3965 | 7.0034 | 6.8419 | 6.5222 | 6.9800 | 7.1938 | 6.2485 | 6.3135 | 6.7638 | 4.9497 | 5.5915 | 4.9464 | 5.4013 |
| NUP214_S433;T437 | NUP214 | P35658-5 | 430 | GERQPKSPGSTPT | 6.1503 | 6.3491 | 6.3899 | 6.7975 | 6.2004 | 6.0820 | 6.2262 | 6.1336 | 6.4493 | 6.2828 | 6.5773 | 6.4285 | 6.2243 | 6.1433 | 5.7344 | 5.8311 |
| NUP214_S437 | NUP214 | P35658-5 | 1942 | QGFGFSSPNKTGG | 6.1225 | 5.7580 | 6.4173 | 6.6555 | 6.8150 | 6.6272 | 6.4953 | 6.2743 | 7.1082 | 6.8792 | 6.7392 | 6.6067 | 5.1229 | 5.5892 | 5.0818 | 5.7078 |
| NUP214_S599 | NUP214 | P35658-5 | 659 | QKSPRITPPAAKP | 6.4801 | 6.8220 | 6.3893 | 6.7674 | 5.8295 | 5.8143 | 6.2567 | 6.4193 | 6.5453 | 6.5554 | 6.3321 | 6.6804 | 5.5049 | 5.8170 | 5.6653 | 6.1211 |
| NUP214_S635 | NUP214 | P35658-5 | 1012 | PVVRTPSIQPSLL | 6.1305 | 6.4409 | 6.2761 | 6.4178 | 6.8916 | 7.1444 | 5.4756 | 6.1962 | 6.8630 | 6.4325 | 7.1097 | 6.8778 | 6.0266 | 6.4234 | 4.4920 | 4.8020 |
| NUP214_S645 | NUP214 | P35658-5 | 1146 | NQAKQGSLINSLK | 6.6075 | 6.7344 | 7.6881 | 6.5077 | 5.9896 | 6.6573 | 7.6732 | 6.6043 | 7.2829 | 6.7898 | 6.4742 | 7.7519 | 4.5514 | 4.2398 | 4.1845 | 4.2632 |
| NUP214_S655;T659 | NUP214 | P35658-5 | 416 | VKSLIKTPERLSL | 6.3642 | 6.5530 | 5.9650 | 6.5299 | 6.7518 | 6.5929 | 6.6041 | 6.4390 | 6.0194 | 6.3057 | 6.9475 | 6.2813 | 6.1550 | 5.9525 | 5.5229 | 5.0159 |
| NUP214_S655;T659;S667 | NUP214 | P35658-5 | 1135 | TPHPVLTPVAANQ | 6.6418 | 6.9269 | 6.9688 | 7.8909 | 6.0359 | 7.7553 | 7.1255 | 6.8368 | 7.6441 | 7.1060 | 7.6920 | 8.5790 | 3.0390 | 3.9337 | 2.6000 | 3.2242 |
| NUP214_S667 | NUP214 | P35658-5 | 437 | PGSTPTTPTSSQA | 5.4842 | 5.9015 | 5.6844 | 5.6901 | 5.8818 | 5.8038 | 5.5726 | 5.9232 | 5.3573 | 5.1440 | 6.5867 | 6.7409 | 7.5877 | 7.5962 | 8.3932 | 6.6525 |
| NUP214_S929 | NUP214 | P35658-5 | 1332 | KPTNKASSTSLTS | 2.1554 | 2.9713 | 3.0930 | 2.7238 | 3.1161 | 2.7651 | 3.4649 | 2.6546 | 2.6176 | 2.6903 | 2.8098 | 2.2631 | 15.3426 | 16.7556 | 17.8197 | 16.7570 |
| NUP214_T1135 | NUP214 | P35658-5 | 599 | QSAPPMSPFSSAS | 6.1331 | 6.6290 | 5.7877 | 6.0637 | 6.1496 | 7.8263 | 6.4415 | 6.2328 | 7.1065 | 6.1516 | 5.8174 | 5.6490 | 5.8669 | 6.1780 | 5.6351 | 6.3318 |
| NUP214_T1291 | NUP214 | P35658-5 | 617 | GPLSHPTPLSAPP | 8.1508 | 6.4042 | 6.1682 | 7.0323 | 6.3551 | 8.3315 | 7.2009 | 6.1253 | 6.4992 | 5.9936 | 7.6207 | 7.5992 | 4.4400 | 4.5994 | 2.5341 | 4.9456 |
| NUP214_T416 | NUP214 | P35658-5 | 433;437 | NA | 5.7533 | 6.1061 | 6.1389 | 6.0479 | 5.7142 | 6.7468 | 6.3660 | 6.3899 | 6.1144 | 5.4692 | 5.5442 | 5.6796 | 7.1736 | 6.8776 | 6.9266 | 6.9519 |
| NUP214_T617 | NUP214 | P35658-5 | 655;659 | NA | 8.2019 | 8.5052 | 7.9404 | 7.6258 | 5.1952 | 7.1340 | 6.7472 | 5.6310 | 5.8471 | 7.0601 | 7.6416 | 7.1370 | 2.9495 | 4.9405 | 3.8173 | 3.6263 |
| NUP214_T659 | NUP214 | P35658-5 | 655;659;667 | NA | 7.2042 | 7.0075 | 7.2413 | 7.5134 | 6.9496 | 7.8706 | 7.4088 | 5.3726 | 6.0571 | 5.9994 | 7.3297 | 7.2260 | 4.3214 | 4.9739 | 3.5249 | 3.9997 |
| NUP35_S100 | NUP35 | Q8NFH5 | 48 | DLPAPVTPQPRSI | 5.7787 | 5.2612 | 5.7046 | 6.0708 | 6.7450 | 6.0816 | 6.3040 | 5.7709 | 5.5662 | 5.8275 | 5.5531 | 6.5528 | 7.6538 | 7.1315 | 6.4375 | 7.5609 |
| NUP35_S259 | NUP35 | Q8NFH5 | 259 | DRCALSSPSLAFT | 5.2621 | 4.8873 | 4.9132 | 5.1200 | 6.0532 | 5.2391 | 5.5180 | 5.6779 | 5.1165 | 5.3538 | 5.3258 | 5.3179 | 10.1362 | 8.2796 | 9.0355 | 8.7638 |
| NUP35_S259;T265 | NUP35 | Q8NFH5 | 265 | SPSLAFTPPIKTL | 6.2699 | 5.7577 | 6.0788 | 6.3055 | 6.6495 | 6.0508 | 6.4497 | 5.8483 | 6.4490 | 6.2107 | 6.4589 | 6.2655 | 6.7021 | 6.0897 | 6.1474 | 6.2666 |
| NUP35_S73 | NUP35 | Q8NFH5 | 73 | PLLAGGSPPQPVV | 6.1529 | 6.4076 | 6.7000 | 7.3523 | 6.5514 | 6.4184 | 6.3443 | 6.1167 | 6.7560 | 7.0025 | 6.6863 | 6.9394 | 4.9837 | 5.4571 | 4.9335 | 5.1978 |
| NUP35_T106 | NUP35 | Q8NFH5 | 273 | PIKTLGTPTQPGS | 6.0730 | 5.2194 | 5.7641 | 6.1543 | 6.7626 | 5.9438 | 6.3532 | 6.6057 | 6.4229 | 6.1490 | 6.0475 | 5.8263 | 7.1378 | 6.6694 | 6.4519 | 6.4191 |
| NUP35_T265 | NUP35 | Q8NFH5 | 106 | SPGLGSTPLTSRR | 6.4302 | 5.9638 | 6.3196 | 6.2042 | 6.5217 | 6.0550 | 6.1137 | 6.1347 | 6.5672 | 5.8886 | 6.3862 | 6.0709 | 6.2169 | 6.5099 | 6.2338 | 6.3836 |
| NUP35_T273 | NUP35 | Q8NFH5 | 100 | IYDDISSPGLGST | 6.6666 | 6.3877 | 6.8826 | 6.4328 | 6.3281 | 6.1294 | 6.1723 | 6.4343 | 6.7370 | 6.8049 | 6.9007 | 7.2034 | 5.4083 | 5.2908 | 5.0123 | 5.2088 |
| NUP35_T308 | NUP35 | Q8NFH5 | 308 | VISDRQTPKKDES | 6.6757 | 5.9769 | 5.8653 | 6.7873 | 6.3433 | 6.4406 | 6.5100 | 6.5173 | 6.3802 | 6.4274 | 6.5488 | 6.8045 | 6.1107 | 5.8373 | 4.7697 | 6.0050 |
| NUP35_T48 | NUP35 | Q8NFH5 | 259;265 | NA | 6.3884 | 6.2430 | 6.0954 | 6.5232 | 6.2383 | 6.4564 | 6.5062 | 6.6073 | 6.6422 | 6.6805 | 6.7019 | 6.9234 | 5.4150 | 5.7755 | 5.1585 | 5.6448 |
| NUP42_S106 | NUP42 | O15504 | 106 | NPFASLSPDEQKD | 6.5329 | 7.1899 | 7.2632 | 7.2202 | 6.4398 | 6.8186 | 6.4380 | 6.4979 | 7.1036 | 7.1430 | 7.5492 | 7.3210 | 3.7213 | 4.7426 | 3.7934 | 4.2255 |
| NUP42_S305;S313 | NUP42 | O15504 | 305;313 | NA | 5.6886 | 6.1752 | 6.5540 | 6.2275 | 6.4534 | 5.8798 | 6.6339 | 6.1950 | 6.9793 | 6.6246 | 6.5502 | 6.9098 | 5.6345 | 6.2719 | 5.2268 | 5.9954 |
| NUP43_S300 | NUP43 | Q8NFH3 | 300 | DVPEKSSLFHQGG | 6.4610 | 5.8150 | 4.7814 | 7.1377 | 7.6104 | 5.7478 | 5.1443 | 5.6036 | 5.1189 | 3.6641 | 4.6492 | 4.9210 | 9.0653 | 8.2505 | 7.9891 | 8.0406 |
| NUP50_S221 | NUP50 | Q9UKX7 | 221 | VAAETQSPSLFGS | 6.7947 | 7.0128 | 7.0848 | 7.5555 | 6.5613 | 6.1292 | 6.8975 | 6.2521 | 6.9594 | 7.1055 | 7.4307 | 6.8554 | 4.1418 | 4.8887 | 4.0202 | 4.3105 |
| NUP50_S315 | NUP50 | Q9UKX7 | 315 | QSKPVSSPFPTKP | 6.9256 | 5.5952 | 5.7049 | 5.4797 | 5.7599 | 7.0589 | 6.6663 | 5.4199 | 6.6402 | 5.3602 | 5.8578 | 6.4403 | 6.9353 | 6.7559 | 5.6478 | 7.7522 |
| NUP54_S289 | NUP54 | Q7Z3B4 | 289 | MTRTELSPAQIKQ | 6.2474 | 6.0224 | 6.2856 | 6.3258 | 5.8994 | 5.8092 | 6.0601 | 6.0569 | 5.9903 | 6.2086 | 6.6476 | 6.5492 | 6.7299 | 6.5144 | 6.2139 | 6.4393 |
| NUP58_T331 | NUP58 | Q9BVL2 | 331 | ALRTQKTPPGLQH | 6.3351 | 6.0507 | 6.0212 | 6.5031 | 6.2350 | 5.4568 | 5.7286 | 6.6670 | 6.2632 | 6.1880 | 6.2506 | 6.2270 | 6.0295 | 6.3907 | 6.7696 | 6.8838 |
| NUP62_S468 | NUP62 | P37198 | 468 | GAPADTSDPLQQI | 6.4087 | 5.2127 | 6.0329 | 5.8916 | 5.9264 | 6.0534 | 6.0508 | 6.3437 | 5.9838 | 6.4240 | 6.7926 | 7.9410 | 5.1871 | 6.9974 | 6.2310 | 6.5228 |
| NUP88_S167 | NUP88 | Q99567 | 517 | DVEVAESPLRVLA | 6.4863 | 6.7606 | 6.1481 | 6.7022 | 6.4904 | 6.2915 | 6.6804 | 6.6710 | 6.9578 | 6.3274 | 6.5602 | 6.5226 | 5.1777 | 5.5881 | 5.1720 | 5.4639 |
| NUP88_S35 | NUP88 | Q99567 | 50 | SSSLPSSPPPQLL | 6.1347 | 5.9939 | 6.6948 | 6.2991 | 6.0448 | 6.1193 | 6.3053 | 6.2473 | 6.8726 | 6.6123 | 6.6317 | 6.2303 | 6.1799 | 5.9976 | 5.4902 | 6.1461 |
| NUP88_S46;S50 | NUP88 | Q99567 | 525 | LRVLAETPDSFEK | 6.2040 | 6.6048 | 6.4039 | 6.6822 | 6.5227 | 6.0400 | 6.5887 | 6.4921 | 6.7377 | 6.3358 | 6.2376 | 6.6838 | 5.5052 | 6.0153 | 5.3190 | 5.6272 |
| NUP88_S50 | NUP88 | Q99567 | 35 | EGLKNQSPTEAEK | 5.5557 | 5.8363 | 5.9433 | 5.9013 | 5.6608 | 6.1781 | 6.1793 | 5.7566 | 5.8796 | 5.7550 | 6.1100 | 6.0897 | 7.4698 | 6.9402 | 7.3315 | 7.4128 |
| NUP88_S517 | NUP88 | Q99567 | 167 | AERFFTSSTSLTL | 6.1372 | 6.9176 | 6.1379 | 6.9995 | 6.0683 | 5.9684 | 6.2683 | 6.4542 | 6.2714 | 6.2598 | 6.0293 | 6.2508 | 6.2845 | 6.0985 | 5.7490 | 6.1054 |
| NUP88_S540 | NUP88 | Q99567 | 540 | RSILQRSVANPAF | 6.5791 | 7.5939 | 6.6917 | 6.8976 | 7.4388 | 6.2870 | 6.4950 | 6.1350 | 5.9197 | 5.2774 | 4.9817 | 5.4974 | 6.6839 | 6.3000 | 6.0333 | 5.1887 |
| NUP88_T525 | NUP88 | Q99567 | 46;50 | NA | 5.9642 | 5.4859 | 5.5552 | 5.8589 | 6.8989 | 5.9165 | 7.0428 | 6.3351 | 5.9629 | 5.6087 | 6.0200 | 6.0102 | 6.9962 | 6.5387 | 6.7193 | 7.0868 |
| NUP93_S52 | NUP93 | Q8N1F7 | 767 | KRLKGTSPSSSSR | 6.8686 | 7.1708 | 6.6516 | 6.8106 | 6.3810 | 6.6717 | 6.4779 | 5.6434 | 5.9118 | 6.3281 | 6.5345 | 6.3637 | 5.7511 | 5.7166 | 5.3257 | 5.3929 |
| NUP93_S61 | NUP93 | Q8N1F7 | 75 | LDISHISQRLESL | 6.7585 | 6.8430 | 6.0904 | 6.7715 | 6.8986 | 6.4325 | 6.4565 | 6.3342 | 6.3754 | 6.0519 | 5.7182 | 6.2322 | 5.6595 | 5.8570 | 5.5320 | 5.9885 |
| NUP93_S647 | NUP93 | Q8N1F7 | 66 | ASVLLGSRGLDIS | 5.8875 | 6.9137 | 5.8859 | 6.6351 | 6.4108 | 6.0657 | 6.9309 | 6.0930 | 6.8520 | 6.1515 | 5.4892 | 5.6583 | 6.7994 | 6.1644 | 5.9342 | 6.1285 |
| NUP93_S66 | NUP93 | Q8N1F7 | 61 | TADVKASVLLGSR | 9.1469 | 7.0116 | 7.0726 | 8.8854 | 6.9497 | 4.1631 | 5.7026 | 4.6659 | 6.3484 | 4.6696 | 4.6814 | 5.2526 | 6.2546 | 6.4496 | 6.5361 | 6.2100 |
| NUP93_S75 | NUP93 | Q8N1F7 | 647 | LMNKLLSPVVPQI | 7.0486 | 7.4745 | 7.1368 | 8.4414 | 4.6850 | 6.3272 | 6.4769 | 5.9704 | 6.4001 | 4.5392 | 6.8414 | 6.2543 | 5.2644 | 6.8222 | 4.7157 | 5.6020 |
| NUP93_S767 | NUP93 | Q8N1F7 | 80 | ISQRLESLSAATT | 5.7704 | 5.8332 | 5.8782 | 6.2402 | 7.0281 | 5.1321 | 7.0207 | 6.1094 | 5.5706 | 5.0327 | 5.2839 | 5.4366 | 8.1570 | 7.4686 | 6.8230 | 7.2154 |
| NUP93_S80 | NUP93 | Q8N1F7 | 159 | EDALDFTQESEPS | 5.9729 | 6.5127 | 5.7043 | 5.7030 | 6.1816 | 6.9273 | 6.2929 | 7.4671 | 5.4483 | 6.5642 | 6.6880 | 8.0223 | 5.9478 | 6.0981 | 5.3579 | 5.1117 |
| NUP93_T159 | NUP93 | Q8N1F7 | 52 | RTLTRTSQETADV | 5.9645 | 5.9510 | 5.5736 | 5.6998 | 5.8508 | 5.4805 | 6.3246 | 5.7850 | 5.9079 | 5.8322 | 5.6526 | 5.6441 | 7.7943 | 6.9996 | 7.9063 | 7.6334 |
| NUP98_S1028 | NUP98 | P52948 | 888 | KYGLQDSDEEEEE | 6.1132 | 6.5757 | 5.7426 | 7.0417 | 6.3295 | 6.4184 | 6.0202 | 6.5170 | 6.0901 | 5.8350 | 6.4160 | 5.9039 | 6.1969 | 6.4946 | 6.2495 | 6.0558 |
| NUP98_S1043 | NUP98 | P52948 | 934 | PPPQSQSPEVEQL | 6.1997 | 5.9413 | 6.3707 | 6.4540 | 6.0060 | 5.8676 | 5.9297 | 5.8980 | 6.6058 | 6.6163 | 6.2866 | 6.3933 | 6.4826 | 6.5833 | 5.9373 | 6.4278 |
| NUP98_S1060 | NUP98 | P52948 | 612 | NNSNLFSPVNRDS | 5.9648 | 5.5965 | 5.4099 | 5.7310 | 6.3506 | 6.0964 | 5.8547 | 6.0352 | 5.9603 | 5.5871 | 5.7418 | 5.7493 | 8.4523 | 6.8197 | 7.1492 | 7.5009 |
| NUP98_S1099 | NUP98 | P52948 | 670 | AKPIPQTPESAGN | 6.2810 | 5.5263 | 6.0927 | 6.6438 | 6.8027 | 6.3069 | 6.4589 | 5.9221 | 6.6480 | 6.2073 | 6.4575 | 6.6341 | 6.4770 | 5.9780 | 5.4858 | 6.0778 |
| NUP98_S1465 | NUP98 | P52948 | 1043 | HSSKTRSLVGGLL | 6.2962 | 6.7888 | 6.2998 | 6.7810 | 5.8649 | 6.4077 | 6.5703 | 6.7576 | 7.0234 | 6.7064 | 7.0919 | 6.9899 | 5.0071 | 5.3562 | 4.6240 | 5.4349 |
| NUP98_S1816 | NUP98 | P52948 | 1028 | TSQEICSPRLPIS | 6.4264 | 6.6394 | 6.2160 | 6.8517 | 6.2201 | 6.1472 | 6.1630 | 6.0636 | 6.2857 | 6.3402 | 6.1030 | 6.0388 | 6.1216 | 6.1757 | 5.9478 | 6.2598 |

| | | | | | | | | | | | | | | | | | | | | | |
|---|---|---|---|---|---|---|---|---|---|---|---|---|---|---|---|---|---|---|---|---|---|
| NUP98_S524 | NUP98 | P52948 | | 916 | PPASQTTPLQMAL | 6.4269 | 6.5289 | 6.4012 | 6.8785 | 6.2308 | 6.6072 | 6.7070 | 6.7002 | 7.0566 | 6.7663 | 7.0772 | 7.0860 | 4.8026 | 5.2487 | 4.4660 | 5.0160 |
| NUP98_S608;S612 | NUP98 | P52948 | | 546 | AQKALTTPTHYKL | 7.2421 | 7.5649 | 6.2286 | 7.2047 | 6.6586 | 6.9520 | 6.6632 | 6.7943 | 7.0778 | 6.2926 | 6.9412 | 7.3616 | 4.3846 | 4.4974 | 3.5068 | 4.6295 |
| NUP98_S612 | NUP98 | P52948 | | 524 | LFRNPMSDPKKKE | 7.1295 | 7.1744 | 6.4999 | 6.8562 | 6.4285 | 6.6699 | 6.7347 | 6.6130 | 7.4011 | 6.6336 | 7.0912 | 7.2247 | 4.8050 | 4.3913 | 3.7572 | 4.5896 |
| NUP98_S623 | NUP98 | P52948 | | 1060 | TSGAFLSPSVSVQ | 6.3780 | 6.2642 | 6.6252 | 5.9272 | 6.1471 | 6.4344 | 6.3423 | 6.4941 | 6.5543 | 6.4216 | 5.6159 | 5.5852 | 6.4534 | 5.9233 | 6.2358 | 6.5979 |
| NUP98_S623;S635 | NUP98 | P52948 | | 653 | QDGDEDSLVSHFY | 6.2299 | 5.6192 | 6.0249 | 6.7347 | 7.1964 | 6.1813 | 6.3530 | 6.1449 | 6.6338 | 5.4781 | 5.9592 | 6.3128 | 6.6731 | 6.1379 | 5.8280 | 6.4927 |
| NUP98_S635 | NUP98 | P52948 | | 681 | GNKHSNSNSVDDT | 5.7060 | 6.7194 | 6.4602 | 7.0120 | 6.5132 | 6.3366 | 6.2506 | 6.8520 | 6.4717 | 6.8096 | 6.6182 | 7.3436 | 5.6217 | 5.1085 | 4.9518 | 5.2249 |
| NUP98_S653 | NUP98 | P52948 | | 673 | IPQTPESAGNKHS | 5.9906 | 5.3535 | 6.1156 | 7.0948 | 6.8352 | 6.2409 | 6.3150 | 5.8081 | 6.1655 | 6.1051 | 6.6065 | 6.9561 | 6.7807 | 5.7409 | 5.3401 | 6.5515 |
| NUP98_S681 | NUP98 | P52948 | | 1816 | RELAVGSLxxxxx | 5.9583 | 6.6464 | 6.4371 | 6.5592 | 6.0032 | 6.5754 | 5.8083 | 6.2134 | 5.9188 | 6.0122 | 6.0310 | 6.2332 | 6.0463 | 5.6178 | 6.6904 | 7.2491 |
| NUP98_S683 | NUP98 | P52948 | | 746 | GYYTIPSMDDLAK | 6.1327 | 6.4349 | 6.0301 | 6.2472 | 5.9295 | 6.1175 | 5.8601 | 5.9545 | 6.0828 | 6.1262 | 6.3728 | 5.5133 | 6.7848 | 7.0363 | 6.9655 | 6.4120 |
| NUP98_S746 | NUP98 | P52948 | | 623 | DSENLASPSEYPE | 6.0060 | 6.4563 | 5.0212 | 6.4694 | 6.5519 | 6.4081 | 5.6750 | 5.8703 | 5.5189 | 6.5593 | 6.3452 | 5.8017 | 6.6655 | 6.6091 | 7.3782 | 6.6639 |
| NUP98_S888 | NUP98 | P52948 | | 635 | ENGERFSFLSKPV | 7.2493 | 7.0208 | 6.4005 | 7.2710 | 7.9645 | 5.9922 | 6.4152 | 5.8271 | 6.1623 | 6.0514 | 6.5423 | 6.5787 | 5.4564 | 5.3262 | 4.4716 | 5.2704 |
| NUP98_S934 | NUP98 | P52948 | | 1465 | SDRYACSPLPSYL | 5.9108 | 5.6885 | 6.3414 | 6.6948 | 6.2766 | 6.9944 | 6.0322 | 5.4829 | 6.1427 | 5.7134 | 6.9764 | 6.0640 | 6.5099 | 5.8901 | 5.7921 | 7.4899 |
| NUP98_T546 | NUP98 | P52948 | | 683 | KHSNSNSVDDTIV | 5.6709 | 5.2208 | 5.3583 | 5.5535 | 5.9827 | 5.8336 | 6.1005 | 6.6936 | 5.8544 | 6.4768 | 6.1303 | 5.5477 | 7.5772 | 7.1603 | 7.5130 | 7.3264 |
| NUP98_T670 | NUP98 | P52948 | | 1099 | SVFTMPSPAPEVP | 6.3844 | 5.9833 | 6.0692 | 6.6960 | 6.0927 | 6.4020 | 6.8136 | 6.4161 | 5.2272 | 5.7129 | 6.0421 | 6.2162 | 6.5572 | 6.8018 | 5.7423 | 6.8430 |
| NUP98_T670;S673 | NUP98 | P52948 | 608;612 | | NA | 7.4222 | 7.5516 | 6.4257 | 7.0115 | 6.0330 | 6.9368 | 7.1578 | 6.7710 | 7.3241 | 6.6456 | 6.9433 | 7.5723 | 4.0350 | 4.1755 | 3.4724 | 4.5221 |
| NUP98_T673 | NUP98 | P52948 | 623;635 | | NA | 6.8737 | 6.1616 | 5.4812 | 7.4987 | 7.6081 | 5.7191 | 6.7305 | 6.1517 | 6.2589 | 6.0106 | 5.5715 | 5.8596 | 6.7713 | 5.7258 | 5.6298 | 5.9477 |
| NUP98_T916 | NUP98 | P52948 | 670;673 | | NA | 6.2896 | 5.9532 | 7.1465 | 6.9089 | 6.5438 | 6.5584 | 6.7233 | 6.2688 | 6.7493 | 6.7642 | 7.3155 | 7.1722 | 4.9009 | 4.5082 | 4.6410 | 5.5560 |
| NUSAP1_S135 | NUSAP1 | Q9BXS6 | | 182 | KRTAITTPNFKKL | 6.8050 | 7.5616 | 7.7605 | 6.8576 | 5.5419 | 7.5025 | 7.8820 | 7.3506 | 6.9880 | 7.7652 | 7.4564 | 7.2279 | 2.9342 | 4.4688 | 2.8235 | 4.0743 |
| NUSAP1_S240 | NUSAP1 | Q9BXS6 | | 247 | VASTPISQRRSQG | 7.3141 | 8.0140 | 6.9826 | 7.0550 | 4.7456 | 7.5020 | 7.9569 | 7.1465 | 7.3315 | 7.4408 | 7.1176 | 7.9849 | 2.7777 | 4.4231 | 2.6910 | 3.5167 |
| NUSAP1_S247 | NUSAP1 | Q9BXS6 | | 338 | NSAAVITPFKLTT | 7.2808 | 7.3918 | 7.7319 | 7.4030 | 5.0656 | 7.4069 | 8.5684 | 6.5450 | 6.7078 | 6.9351 | 7.7102 | 7.4125 | 2.7933 | 4.5888 | 2.3941 | 4.0648 |
| NUSAP1_S255 | NUSAP1 | Q9BXS6 | | 135 | ATAKVPSPPDEHQ | 6.0913 | 6.6288 | 6.0624 | 7.3223 | 6.2296 | 5.9088 | 6.4299 | 6.1544 | 6.6110 | 6.6654 | 6.1501 | 6.5156 | 5.4128 | 6.1564 | 5.6651 | 5.9960 |
| NUSAP1_T182 | NUSAP1 | Q9BXS6 | | 255 | RRSQGRSCGPASQ | 5.9710 | 5.9730 | 6.5537 | 6.5902 | 5.5553 | 6.1489 | 6.0836 | 5.4023 | 6.8619 | 6.6454 | 6.7300 | 7.0360 | 6.1303 | 6.2027 | 5.6236 | 6.4922 |
| NUSAP1_T244 | NUSAP1 | Q9BXS6 | | 240 | PPRGRLSVASTPI | 5.8663 | 5.9478 | 5.8972 | 5.6306 | 5.4745 | 7.6726 | 7.6563 | 6.6859 | 8.3370 | 8.4476 | 7.9109 | 8.1552 | 4.2838 | 4.8873 | 2.5932 | 4.5538 |
| NUSAP1_T338 | NUSAP1 | Q9BXS6 | | 244 | RLSVASTPISQRR | 7.0061 | 6.7150 | 6.9969 | 6.9567 | 5.6670 | 6.7834 | 7.1616 | 6.8737 | 6.9458 | 6.6561 | 7.0722 | 6.6199 | 4.2713 | 5.4590 | 4.1762 | 4.6389 |
| NVL_S185 | NVL | O15381 | | 191 | SFFLDLSCEKSNP | 6.1435 | 7.1690 | 7.0522 | 7.1941 | 6.2499 | 7.1279 | 7.0440 | 6.8026 | 7.5752 | 7.1377 | 7.4431 | 6.7683 | 3.6334 | 4.6387 | 3.7989 | 4.2214 |
| NVL_S191 | NVL | O15381 | | 185 | PSVKKDSFFLDLS | 6.4717 | 6.5354 | 6.4080 | 8.1965 | 6.6787 | 7.6207 | 6.1002 | 5.7394 | 7.6047 | 8.5441 | 7.9550 | 6.1162 | 3.9987 | 3.4654 | 3.5423 | 5.0230 |
| NVL_S211 | NVL | O15381 | | 211 | QDSKDSSLLESDM | 6.4348 | 6.5312 | 6.7830 | 7.6699 | 7.2648 | 6.9434 | 6.8695 | 6.5115 | 6.4313 | 7.0714 | 5.8011 | 7.2894 | 4.6526 | 4.8305 | 3.6855 | 5.2300 |
| NXF1_S575 | NXF1 | Q9UBU9 | | 554 | FAMPAPTPSSSPV | 6.1228 | 6.1470 | 6.2679 | 6.6273 | 6.3235 | 6.2113 | 6.0335 | 6.1384 | 5.9875 | 6.1465 | 6.0404 | 6.4949 | 6.2317 | 6.7532 | 6.1875 | 6.2868 |
| NXF1_T554 | NXF1 | Q9UBU9 | | 575 | EMLQAFSTQSGMN | 5.7583 | 5.6967 | 5.6867 | 6.0576 | 6.3674 | 5.9275 | 6.2862 | 6.5595 | 5.9529 | 6.3659 | 6.0143 | 5.9397 | 6.5212 | 7.1060 | 6.9536 | 6.8065 |
| NXF1_T554;S575 | NXF1 | Q9UBU9 | 554;575 | | NA | 6.2011 | 6.8739 | 6.6514 | 6.0976 | 6.3855 | 5.7782 | 5.4629 | 6.2562 | 6.1185 | 5.0398 | 5.1680 | 5.6774 | 6.6684 | 6.9379 | 7.5048 | 7.1782 |
| NXF2_S33 | NXF2 | Q9GZY0 | | 34 | RKKGGSSFRDNFD | 6.2171 | 6.4678 | 6.9080 | 6.3112 | 6.1817 | 7.0035 | 6.5379 | 7.3224 | 7.2505 | 8.3302 | 8.8256 | 8.0107 | 4.0337 | 4.1969 | 3.0497 | 3.3531 |
| NXF2_S34 | NXF2 | Q9GZY0 | | 33 | GRKKGGSSFRDNF | 6.5528 | 6.7534 | 6.4515 | 7.0360 | 5.9387 | 7.3609 | 7.0859 | 6.6659 | 6.4291 | 7.3125 | 8.1471 | 7.3690 | 4.0537 | 4.7177 | 4.1861 | 3.9396 |
| OBI1_S553 | OBI1 | Q5W0B1 | | 553 | ESDNSKSPCNNGF | 6.1222 | 6.2584 | 6.2530 | 6.1590 | 6.4194 | 6.5156 | 6.2146 | 6.2971 | 6.4687 | 6.0424 | 6.5572 | 6.0969 | 6.2442 | 5.8936 | 6.3415 | 6.1163 |
| OBI1_S592 | OBI1 | Q5W0B1 | | 719 | SSLSSASPSKATK | 6.2213 | 5.6868 | 6.6244 | 6.1197 | 6.1220 | 6.4476 | 6.0654 | 6.3144 | 6.5587 | 6.2675 | 6.4731 | 6.8917 | 6.4138 | 5.9563 | 6.0204 | 5.8171 |
| OBI1_S719 | OBI1 | Q5W0B1 | | 592 | KTELNLSKGSLTN | 6.5896 | 6.4182 | 8.0996 | 7.3296 | 6.1516 | 6.1360 | 7.2404 | 6.8035 | 7.5070 | 7.9241 | 8.0673 | 7.7269 | 3.3460 | 2.9158 | 3.2592 | 4.4852 |
| OBSL1_S10 | OBSL1 | O75147 | | 10 | SSGDQGSPPCFLR | 6.2408 | 5.9708 | 6.0699 | 6.2104 | 6.0732 | 6.7322 | 6.7015 | 6.2027 | 6.4955 | 6.4845 | 6.0784 | 6.5451 | 5.7282 | 6.3709 | 6.2515 | 5.8443 |
| OCIAD1_S108 | OCIAD1 | Q9NX40 | | 108 | FKKLENSPLGEAL | 7.9175 | 8.2299 | 7.2885 | 8.2029 | 6.1121 | 5.9390 | 6.3946 | 6.7460 | 7.3373 | 7.5927 | 7.2204 | 7.6098 | 2.9647 | 3.7795 | 3.0516 | 3.6135 |
| OCIAD1_S123 | OCIAD1 | Q9NX40 | | 123 | GQARRSSPPGHYY | 6.8131 | 7.3094 | 6.1427 | 7.7646 | 6.6104 | 6.8352 | 5.8987 | 7.5819 | 6.5719 | 6.4855 | 6.9083 | 6.7227 | 4.4098 | 4.9080 | 4.4462 | 4.5916 |
| OCIAD1_T198 | OCIAD1 | Q9NX40 | | 198 | PKRKNITYEELRN | 4.3815 | 4.5455 | 4.7924 | 5.1548 | 5.0416 | 4.4333 | 5.2118 | 4.9923 | 4.2598 | 4.8523 | 4.8293 | 4.3825 | 11.7352 | 9.9136 | 11.6321 | 9.8421 |
| OCLN_S310 | OCLN | Q16625 | | 341 | RFYPESSYKSTPV | 7.4925 | 7.4580 | 8.5038 | 8.1870 | 6.1151 | 6.0142 | 6.1359 | 6.4818 | 7.1889 | 6.9812 | 7.2668 | 7.3250 | 3.7385 | 4.9444 | 2.9667 | 3.2001 |
| OCLN_S321 | OCLN | Q16625 | | 321 | YVERVDSPMAYSS | 6.5094 | 7.0157 | 7.3314 | 7.7963 | 6.5236 | 6.4586 | 7.1997 | 6.7690 | 6.7698 | 6.9553 | 7.5215 | 7.3907 | 3.5563 | 4.2435 | 3.6020 | 4.3573 |
| OCLN_S327 | OCLN | Q16625 | | 376 | SGGNFETPSKRAP | 8.1556 | 8.1743 | 7.4522 | 7.9878 | 5.8647 | 7.1512 | 7.5106 | 6.7831 | 7.4356 | 6.9758 | 7.1234 | 6.7602 | 2.6513 | 3.7364 | 2.6921 | 3.5457 |
| OCLN_S341 | OCLN | Q16625 | | 8 | SSRPLESPPPYRP | 6.0411 | 5.6506 | 6.2812 | 7.0394 | 6.2127 | 6.6266 | 6.7304 | 6.3905 | 6.1618 | 6.5800 | 6.4858 | 6.5902 | 5.6775 | 5.5147 | 5.6048 | 6.4126 |
| OCLN_S358 | OCLN | Q16625 | | 369 | FRQPRYSSGGNFE | 6.9491 | 6.8669 | 6.8426 | 7.1926 | 6.3325 | 6.1873 | 6.4690 | 6.3511 | 7.3467 | 7.0482 | 7.1561 | 7.6142 | 4.2255 | 5.3753 | 3.8816 | 4.1613 |
| OCLN_S369 | OCLN | Q16625 | | 378 | GNFETPSKRAPAK | 6.7651 | 6.0436 | 7.1625 | 7.1663 | 5.2300 | 6.5199 | 6.9295 | 6.0821 | 7.4979 | 6.7548 | 7.0798 | 7.3063 | 4.6372 | 6.4786 | 4.3770 | 3.9693 |
| OCLN_S378 | OCLN | Q16625 | | 358 | QELPLTSPVDDFR | 7.0581 | 6.7941 | 6.8770 | 7.8138 | 5.5295 | 6.6008 | 6.4764 | 6.4177 | 5.6671 | 5.7582 | 6.9236 | 6.6676 | 4.8932 | 4.8892 | 5.3414 | 6.2923 |
| OCLN_S8 | OCLN | Q16625 | | 327 | SPMAYSSNGKVND | 5.5567 | 5.8612 | 6.5253 | 6.7330 | 5.8073 | 6.5798 | 6.1923 | 6.0605 | 6.0475 | 6.1605 | 6.0553 | 6.4032 | 6.3085 | 6.3890 | 6.8856 | 6.4344 |
| OCLN_T345 | OCLN | Q16625 | | 310 | GTQDVPSPPSDYV | 6.0874 | 6.4871 | 7.6570 | 7.1014 | 5.6755 | 6.9800 | 5.9427 | 6.6227 | 6.6298 | 5.9821 | 6.2938 | 6.2179 | 4.9984 | 5.2484 | 5.4923 | 6.5833 |
| OCLN_T376 | OCLN | Q16625 | | 345 | ESSYKSTPVPEVV | 6.0673 | 6.2584 | 7.4093 | 5.8154 | 4.4995 | 5.4419 | 9.2334 | 6.0890 | 6.2063 | 8.8078 | 5.5809 | 5.7807 | 5.5331 | 6.9403 | 4.2432 | 6.0934 |
| ODF2_S124 | ODF2 | Q5BJF6-4 | | 124 | YDKKIDSLMNAVG | 7.1189 | 6.0483 | 6.2147 | 7.3534 | 7.2574 | 5.8499 | 7.1435 | 5.7441 | 6.2229 | 5.2302 | 6.2442 | 5.8606 | 6.3729 | 5.4490 | 5.7105 | 6.1794 |
| ODF2_S787 | ODF2 | Q5BJF6-4 | | 7 | MKDRSSTPPLHVH | 6.1484 | 6.7060 | 6.1765 | 7.1525 | 6.3762 | 6.6058 | 6.5041 | 7.0392 | 6.8066 | 6.9630 | 6.5540 | 6.9027 | 4.7553 | 5.3120 | 4.8644 | 5.1332 |
| ODF2_S815 | ODF2 | Q5BJF6-4 | | 787 | SESTNRSMQNYVQ | 5.8678 | 7.1255 | 7.6049 | 5.9700 | 7.1805 | 6.7049 | 6.8778 | 6.2608 | 7.1490 | 7.4320 | 4.9874 | 6.5281 | 4.1542 | 4.1424 | 5.4237 | 6.5909 |
| ODF2_T7 | ODF2 | Q5BJF6-4 | | 815 | STFLTSSPIRSRS | 6.4954 | 6.3557 | 6.7776 | 6.1649 | 5.8499 | 6.0923 | 6.8817 | 6.0998 | 6.8109 | 6.1738 | 6.1415 | 6.4701 | 5.9427 | 6.1574 | 5.6586 | 5.9277 |
| OFD1_S620 | OFD1 | O75665 | | 774 | DRMPLPSPTESRH | 6.1760 | 6.1561 | 6.5668 | 6.4598 | 6.5487 | 6.4891 | 6.0721 | 6.6083 | 6.5470 | 6.4451 | 6.8521 | 6.8056 | 5.5737 | 5.7012 | 5.5396 | 5.4590 |
| OFD1_S669 | OFD1 | O75665 | | 669 | SPLAAKSPPSLHL | 6.6657 | 7.1714 | 6.5822 | 7.1080 | 7.0228 | 6.9403 | 5.8412 | 7.3900 | 6.4934 | 6.3953 | 6.8308 | 6.7664 | 4.6926 | 4.8522 | 4.6867 | 4.5609 |
| OFD1_S669;S672 | OFD1 | O75665 | | 735 | TSSRRLSSTPLPK | 6.4636 | 4.9289 | 5.9655 | 5.2457 | 6.5325 | 6.2757 | 5.7704 | 6.0093 | 8.1296 | 7.6650 | 7.8695 | 8.5641 | 5.5606 | 6.0978 | 4.0213 | 4.9005 |
| OFD1_S735 | OFD1 | O75665 | | 620 | AWVEGSSPDSDLE | 6.8294 | 6.7299 | 6.6592 | 6.8865 | 6.2475 | 6.9360 | 6.7787 | 6.7496 | 6.9626 | 6.8389 | 7.0862 | 7.6415 | 4.2251 | 4.5973 | 4.3319 | 4.4998 |
| OFD1_S735;T737 | OFD1 | O75665 | | 899 | REERRQSNLQEVL | 6.1441 | 5.1790 | 5.9817 | 5.7744 | 7.9418 | 5.9182 | 5.3402 | 6.3061 | 7.7174 | 7.7126 | 7.9305 | 8.1247 | 5.3586 | 5.6768 | 4.3819 | 4.5119 |
| OFD1_S762 | OFD1 | O75665 | | 762 | GRSHIASPSPCPD | 5.6296 | 5.3954 | 7.1158 | 7.1257 | 6.2789 | 7.0945 | 7.2409 | 6.4414 | 7.5370 | 5.2802 | 6.7740 | 6.9228 | 4.4029 | 5.5549 | 4.5287 | 6.6771 |
| OFD1_S774 | OFD1 | O75665 | | 789 | SIPPVSSPPEQKV | 6.1201 | 6.2733 | 7.7207 | 6.5806 | 6.8086 | 7.0045 | 6.2287 | 6.7667 | 6.4274 | 6.0809 | 6.9164 | 6.7952 | 4.7760 | 5.0312 | 5.1042 | 5.3656 |
| OFD1_S789 | OFD1 | O75665 | 669;672 | | NA | 6.8049 | 7.2368 | 6.8872 | 5.5527 | 6.2593 | 6.0031 | 6.2665 | 6.9568 | 6.8916 | 6.1672 | 6.2666 | 6.4520 | 5.4292 | 6.0462 | 5.1425 | 5.6373 |
| OFD1_S899 | OFD1 | O75665 | 735;737 | | NA | 6.9009 | 6.7155 | 7.3878 | 6.8725 | 6.2322 | 6.8680 | 6.8809 | 7.3582 | 7.8232 | 6.5845 | 7.4185 | 7.6025 | 3.5852 | 4.5694 | 3.4029 | 3.7979 |
| OGA_S364 | OGA | O60502 | | 364 | KLENEGSDEDIET | 6.1232 | 6.2467 | 6.5059 | 6.1515 | 6.5923 | 6.4977 | 6.1697 | 6.4309 | 6.2559 | 6.3720 | 6.1375 | 6.1482 | 6.3065 | 6.2644 | 6.0087 | 5.7890 |
| OGA_S505 | OGA | O60502 | | 511 | SIAESKSPEMSMQ | 5.8141 | 6.2253 | 6.6083 | 6.2854 | 6.1222 | 6.4746 | 6.1718 | 6.4044 | 6.2315 | 6.5662 | 6.7341 | 6.6960 | 5.8247 | 6.2617 | 5.6343 | 5.9453 |
| OGA_S509 | OGA | O60502 | | 509 | KESIAESKSPEMS | 6.3383 | 7.2400 | 8.3139 | 7.3692 | 5.7859 | 5.4326 | 6.2685 | 6.6358 | 6.0322 | 6.8739 | 6.7516 | 6.7144 | 4.0659 | 5.3325 | 5.2545 | 5.5909 |
| OGA_S511 | OGA | O60502 | | 505 | MDTDKESIAESKS | 5.6606 | 6.3098 | 7.9503 | 6.3482 | 6.6081 | 6.1360 | 6.2710 | 6.3087 | 5.6292 | 5.9390 | 6.5090 | 6.3475 | 6.6070 | 5.9194 | 5.7412 | 5.7150 |
| OGFOD1_S393 | OGFOD1 | Q8N543 | | 393 | ITEEGTSHSPPEP | 5.5503 | 5.9912 | 6.0180 | 5.5142 | 5.7649 | 6.3512 | 6.2011 | 6.3567 | 6.1280 | 5.9063 | 7.3069 | 6.6066 | 5.6328 | 7.3210 | 7.3973 | 5.9535 |
| OGFR_S349 | OGFR | Q9NZT2 | | 349 | EGPQPRSVEPQDA | 5.9952 | 6.4650 | 6.0908 | 5.9176 | 5.6382 | 6.1173 | 6.0298 | 6.1772 | 5.9651 | 6.1965 | 6.5568 | 6.4544 | 6.3950 | 7.0573 | 6.4911 | 6.4529 |
| OGFR_S517 | OGFR | Q9NZT2 | | 514 | TGPKEGTPGSPSE | 5.0206 | 5.6153 | 6.1307 | 6.3441 | 6.1031 | 5.9203 | 5.6547 | 6.5284 | 6.7879 | 5.8927 | 6.1645 | 6.4491 | 6.9072 | 7.0269 | 7.0203 | 6.4342 |
| OGFR_S537 | OGFR | Q9NZT2 | | 577 | GDEPAESPSETPG | 5.9712 | 6.0015 | 5.3436 | 4.8461 | 6.1706 | 6.3809 | 5.0325 | 6.1912 | 6.5456 | 6.3982 | 5.3479 | 6.9662 | 6.3917 | 7.6198 | 6.6421 | 8.1509 |
| OGFR_S577 | OGFR | Q9NZT2 | | 537 | GDEPAESPSETPG | 5.9895 | 5.2826 | 5.9004 | 6.4016 | 5.6649 | 6.0627 | 5.8112 | 7.0193 | 6.5966 | 6.6816 | 5.8855 | 6.4676 | 6.0062 | 6.8540 | 6.5906 | 6.7857 |
| OGFR_T514 | OGFR | Q9NZT2 | | 517 | KEGTPGSPSETPG | 5.3864 | 6.6727 | 6.0882 | 4.8007 | 5.9476 | 6.1179 | 6.0934 | 6.5037 | 6.2366 | 6.5127 | 6.6211 | 6.6056 | 5.6160 | 6.4805 | 7.1255 | 7.1913 |
| OGT_S1045 | OGT | O15294 | | 1045 | PVEVTESAxxxxx | 5.9230 | 6.5617 | 6.3763 | 6.5003 | 6.2563 | 7.8671 | 6.1577 | 6.4897 | 6.5483 | 4.9723 | 6.0590 | 5.3069 | 7.1604 | 5.7856 | 5.2363 | 6.7990 |
| OIP5_S48 | OIP5 | O43482 | | 48 | QVVKGSSPLGPAG | 6.2882 | 6.2661 | 6.5146 | 6.3556 | 6.1425 | 6.5560 | 6.3158 | 6.4984 | 6.7845 | 6.6767 | 6.8308 | 6.6534 | 5.4320 | 5.8276 | 5.4148 | 5.4431 |
| OIP5_T14 | OIP5 | O43482 | | 14 | HRSRCATPPRGDF | 6.9257 | 7.3455 | 7.2860 | 6.2526 | 6.4621 | 6.5937 | 6.1786 | 5.8356 | 5.8960 | 6.1475 | 7.3336 | 6.3130 | 5.3244 | 5.8274 | 4.7830 | 5.4954 |
| OIP5_T221 | OIP5 | O43482 | | 221 | KILSEVTPDQSKP | 5.3929 | 7.1612 | 9.2566 | 5.6472 | 4.9811 | 5.3384 | 4.8085 | 4.6147 | 6.3073 | 7.1007 | 4.6964 | 7.1461 | 7.1067 | 5.1567 | 6.6344 | 8.6510 |
| OPA3_S169 | OPA3 | Q9H6K4 | | 169 | LCNPGRSASHAVP | 5.2946 | 6.3259 | 5.4233 | 6.1272 | 6.0692 | 6.3851 | 6.7967 | 6.4074 | 5.3088 | 5.3731 | 4.8255 | 5.0221 | 7.8254 | 7.6447 | 7.8035 | 7.3675 |
| OPHN1_S372 | OPHN1 | O60890 | | 703 | GPTKTPSFHIKRP | 7.6203 | 6.0110 | 6.5737 | 5.7866 | 6.8086 | 6.5460 | 5.9829 | 6.1911 | 7.6483 | 6.0924 | 5.8844 | 5.7592 | 6.8632 | 6.9428 | 4.8551 | 4.4344 |
| OPHN1_S652 | OPHN1 | O60890 | | 753 | PPCRAATPQKPEP | 6.6365 | 6.5510 | 7.5541 | 6.9127 | 6.7231 | 7.0110 | 7.4533 | 6.6602 | 6.9830 | 7.0893 | 7.2315 | 7.6476 | 3.7865 | 4.3605 | 3.3936 | 4.0060 |
| OPHN1_S703 | OPHN1 | O60890 | | 372 | KEPIYHSPITKQQ | 7.4979 | 8.0287 | 7.9210 | 6.8537 | 5.7769 | 5.6107 | 7.1219 | 7.0804 | 5.8166 | 6.6327 | 6.9859 | 5.4104 | 4.0975 | 5.2064 | 4.0200 | 5.9394 |
| OPHN1_T753 | OPHN1 | O60890 | | 652 | TDPGRKSPSRPIL | 6.3063 | 6.7493 | 6.6379 | 6.3724 | 6.5559 | 6.2273 | 6.6158 | 6.7417 | 6.7621 | 6.1979 | 6.3159 | 6.2243 | 5.4373 | 5.8170 | 5.5099 | 5.5292 |
| OPTN_S198 | OPTN | Q96CV9 | | 198 | VKEIKHSPGPTRT | 6.0016 | 6.6565 | 7.0463 | 6.2220 | 7.6772 | 8.0850 | 6.6490 | 7.3549 | 6.6411 | 6.2181 | 7.3311 | 6.2506 | 4.9255 | 5.0813 | 3.9666 | 3.8933 |
| ORC1_S199 | ORC1 | Q13415 | | 287 | DKLQTLSPALKAP | 8.8939 | 8.8088 | 7.6553 | 8.8595 | 6.3234 | 5.9543 | 6.2113 | 6.3059 | 8.1185 | 7.6982 | 7.6042 | 7.7233 | 1.9221 | 2.9878 | 2.1694 | 2.7641 |
| ORC1_S258 | ORC1 | Q13415 | | 258 | SCASLDSPGRIKR | 7.6199 | 7.9620 | 7.6363 | 8.0174 | 6.3343 | 6.5304 | 6.2925 | 6.0730 | 7.7165 | 7.5909 | 7.0285 | 7.1758 | 2.9882 | 4.1300 | 3.3556 | 3.5486 |
| ORC1_S273 | ORC1 | Q13415 | | 273 | AFSEITSPSKRSQ | 7.8691 | 7.8455 | 7.7406 | 8.4147 | 6.0781 | 6.2056 | 6.6218 | 6.3371 | 7.7663 | 7.4463 | 7.4510 | 8.0711 | 2.8148 | 3.5335 | 2.5390 | 3.2655 |
| ORC1_S287 | ORC1 | Q13415 | | 311 | EDDKKASPEHRII | 7.7848 | 8.2229 | 7.4142 | 8.6432 | 5.7690 | 6.0677 | 6.2736 | 6.0734 | 7.3999 | 7.1872 | 7.1613 | 7.5036 | 3.2685 | 3.9336 | 3.5561 | 3.7410 |
| ORC1_S311 | ORC1 | Q13415 | | 337 | REERTLTPISGGQ | 8.7469 | 8.5743 | 7.5582 | 8.6652 | 5.7570 | 6.1011 | 6.9449 | 6.3827 | 8.1885 | 7.8439 | 7.1225 | 7.6515 | 2.2746 | 2.9589 | 2.1053 | 3.1247 |
| ORC1_S383 | ORC1 | Q13415 | | 230 | TPTHPLTPRARKR | 7.6408 | 7.9082 | 7.6886 | 8.2761 | 6.1500 | 6.3229 | 6.7625 | 6.4644 | 7.6248 | 7.3611 | 7.0911 | 7.4891 | 2.8128 | 3.6921 | 3.0645 | 3.6511 |
| ORC1_T230 | ORC1 | Q13415 | | 199 | KSKSAESPSWTPA | 7.3023 | 7.7075 | 7.1439 | 7.8227 | 6.2679 | 6.3087 | 6.0537 | 6.3344 | 7.5280 | 7.6244 | 7.6219 | 7.3347 | 3.5222 | 4.1242 | 3.4901 | 3.8134 |
| ORC1_T337 | ORC1 | Q13415 | | 375 | QNEATSTPHRIRR | 7.6794 | 7.1512 | 7.0871 | 7.5088 | 5.5625 | 5.7370 | 5.7829 | 5.7342 | 6.5152 | 6.6688 | 6.2732 | 6.6031 | 5.6384 | 5.6724 | 4.7786 | 5.6072 |
| ORC1_T375 | ORC1 | Q13415 | | 383 | HRIRRKSSVLTMN | 7.3930 | 5.4420 | 5.7653 | 6.5113 | 6.8320 | 4.1184 | 5.9304 | 4.0291 | 9.8155 | 8.0991 | 7.5651 | 7.0664 | 7.0233 | 7.0310 | 3.1213 | 4.2567 |
| ORC2_S280 | ORC2 | Q13416 | | 280 | NLLSKVSPSFSAE | 6.5909 | 6.5720 | 6.4905 | 6.2175 | 6.4731 | 6.5448 | 6.2743 | 6.3785 | 6.9091 | 6.3475 | 6.2057 | 5.7981 | 6.0744 | 5.7373 | 5.6874 | 5.6989 |
| ORC3_S207 | ORC3 | Q9UBD5-2 | | 23 | SKKRKISLPIEDY | 5.8181 | 6.3298 | 6.6782 | 6.7666 | 6.3479 | 6.6462 | 6.5896 | 6.9933 | 7.2313 | 6.9966 | 7.1563 | 7.3584 | 4.3375 | 5.7681 | 4.1424 | 4.8396 |
| ORC3_S23 | ORC3 | Q9UBD5-2 | | 207 | SKKRTTSSQWQSP | 7.0644 | 7.0341 | 7.0956 | 7.9339 | 6.2754 | 5.8831 | 7.0133 | 6.4781 | 6.9059 | 6.7968 | 6.6548 | 6.5604 | 4.0488 | 5.3236 | 4.3935 | 4.5381 |
| ORC6_T195 | ORC6 | Q9Y5N6 | | 195 | EPGDVATPPRKRK | 6.2845 | 5.9988 | 7.6061 | 6.7669 | 5.9743 | 6.6826 | 6.4541 | 6.8913 | 6.9539 | 6.8749 | 7.1673 | 7.4961 | 4.3721 | 5.0144 | 4.3410 | 5.1216 |
| OSBP_S190;S193 | OSBP | P22059 | | 351 | SGKGDMSDEDDEN | 6.1546 | 6.3731 | 6.6039 | 6.5489 | 6.2355 | 6.5501 | 6.1864 | 6.4118 | 6.5466 | 6.9635 | 6.7719 | 6.7457 | 5.3078 | 5.8364 | 5.2516 | 5.5121 |
| OSBP_S238 | OSBP | P22059 | | 240 | ALQRSLSELESLK | 5.8356 | 6.1858 | 6.0389 | 5.8382 | 6.0619 | 6.4547 | 6.3037 | 6.3742 | 6.4087 | 6.1563 | 6.2275 | 6.1525 | 6.6335 | 6.4543 | 6.4849 | 6.3894 |
| OSBP_S240 | OSBP | P22059 | | 238 | GTALQRSLSELES | 5.7810 | 6.2999 | 6.3532 | 6.6025 | 6.1870 | 7.0453 | 6.2548 | 6.6542 | 6.3710 | 6.2332 | 6.5303 | 6.5643 | 5.6840 | 6.0101 | 5.7112 | 5.7179 |
| OSBP_S351 | OSBP | P22059 | | 332 | TVLPANTPGNVGS | 5.9587 | 6.3332 | 5.9980 | 7.0197 | 6.1067 | 6.2877 | 6.1505 | 6.3646 | 7.9373 | 7.2816 | 7.2970 | 7.2918 | 5.3109 | 5.3280 | 4.3723 | 4.9620 |
| OSBP_S379;S382 | OSBP | P22059 | | 382 | RTGSNISGASSDI | 5.9439 | 5.9986 | 6.6539 | 5.9957 | 5.8237 | 5.2765 | 5.7903 | 5.7399 | 7.3355 | 6.5572 | 6.0327 | 6.0938 | 6.9375 | 7.2445 | 6.5366 | 6.0398 |
| OSBP_S382 | OSBP | P22059 | 190;193 | | NA | 6.2918 | 6.4871 | 6.9024 | 6.4127 | 6.1869 | 6.1038 | 6.3152 | 6.1541 | 6.9560 | 6.9965 | 6.2276 | 6.4412 | 5.6124 | 6.1164 | 5.1178 | 5.6782 |
| OSBP_T332 | OSBP | P22059 | 379;382 | | NA | 5.6250 | 6.1114 | 7.7074 | 6.4635 | 7.3667 | 6.7706 | 6.9132 | 6.5679 | 5.1186 | 6.2007 | 5.8756 | 6.2741 | 5.6967 | 5.9223 | 5.8681 | 5.5182 |
| OSBPL10_S223 | OSBPL10 | Q9BXB5 | | 223 | TITHHKSPAAARR | 6.9755 | 7.1444 | 5.8504 | 6.5423 | 6.4460 | 6.0650 | 6.7056 | 6.2834 | 6.4533 | 6.0788 | 6.0534 | 6.3114 | 5.3154 | 6.0782 | 5.5045 | 6.1924 |
| OSBPL11_S172 | OSBPL11 | Q9BXB4 | | 27 | GQATAVTPNKNSS | 6.5836 | 6.8002 | 6.7655 | 6.7248 | 6.5685 | 7.0248 | 6.8508 | 6.8772 | 7.2517 | 6.9967 | 6.9932 | 7.1836 | 4.0780 | 4.8041 | 3.8341 | 4.6633 |
| OSBPL11_S174 | OSBPL11 | Q9BXB4 | | 189 | ISQRRPSQNAISF | 5.3440 | 5.1104 | 6.2559 | 5.8375 | 6.6690 | 6.0832 | 5.3062 | 6.6473 | 6.9465 | 7.8675 | 7.2762 | 7.3745 | 6.1731 | 6.5890 | 5.4979 | 5.0217 |
| OSBPL11_S181 | OSBPL11 | Q9BXB4 | | 181 | LASSSNSPISQRR | 6.8620 | 7.6037 | 6.8653 | 6.7352 | 6.6632 | 7.0682 | 6.6966 | 7.5216 | 6.8363 | 6.9337 | 6.6420 | 6.5403 | 4.0390 | 4.6675 | 3.9710 | 4.3544 |
| OSBPL11_S189 | OSBPL11 | Q9BXB4 | | 174 | LKSRSFSLASSSN | 5.8025 | 4.2146 | 6.4426 | 7.0475 | 6.3839 | 5.5118 | 6.1578 | 5.7563 | 7.1408 | 6.6715 | 6.0928 | 6.4262 | 7.3422 | 7.2086 | 5.8791 | 5.9218 |
| OSBPL11_S286 | OSBPL11 | Q9BXB4 | | 286 | ASHQKGSLPSGTT | 5.2594 | 5.7499 | 6.1760 | 5.9409 | 6.0310 | 5.5709 | 6.6560 | 6.1755 | 6.3340 | 6.5795 | 6.4900 | 6.0295 | 6.3720 | 6.6035 | 6.8360 | 7.1960 |
| OSBPL11_T27 | OSBPL11 | Q9BXB4 | | 172 | PPLKSRSFSLASS | 6.3816 | 5.0452 | 6.4555 | 5.8582 | 6.5137 | 6.6390 | 5.6101 | 5.4664 | 5.7694 | 6.5553 | 6.8311 | 6.4272 | 7.2045 | 7.1976 | 5.9667 | 6.0787 |
| OSBPL1A_S490 | OSBPL1A | Q9BXW6 | | 499 | EAVTARSFEEEGE | 6.6504 | 7.3304 | 6.7764 | 7.4801 | 6.1187 | 6.8094 | 6.6854 | 6.5669 | 7.0078 | 6.9935 | 7.2150 | 7.2962 | 4.2195 | 4.6681 | 4.0659 | 4.1162 |
| OSBPL1A_S499 | OSBPL1A | Q9BXW6 | | 490 | ESERSLSRLEAVT | 8.0776 | 7.6750 | 7.2105 | 7.4844 | 7.6293 | 7.6508 | 5.0764 | 6.5072 | 4.8965 | 7.3958 | 7.9005 | 7.3636 | 3.0308 | 4.8589 | 3.2348 | 4.0079 |
| OSBPL3_S200 | OSBPL3 | Q9H4L5 | | 304 | PVRLHSSNPNLST | 5.4684 | 5.6112 | 5.7948 | 5.7500 | 5.4160 | 5.7666 | 5.5423 | 5.6573 | 6.2302 | 6.1286 | 6.1379 | 6.2331 | 7.3624 | 7.5405 | 7.3605 | 7.9999 |
| OSBPL3_S251 | OSBPL3 | Q9H4L5 | | 372 | EQDASSSPSAQVI | 5.4470 | 5.9164 | 6.0298 | 6.2751 | 5.4761 | 5.8419 | 6.0205 | 5.9833 | 6.5330 | 6.6621 | 6.9341 | 7.1557 | 6.1437 | 6.5205 | 6.6737 | 6.3871 |
| OSBPL3_S265 | OSBPL3 | Q9H4L5-2 | | 251 | VLHRTYSAPAINA | 4.4837 | 4.5285 | 4.8093 | 4.7864 | 5.2748 | 4.9892 | 5.0842 | 5.0454 | 6.1730 | 5.7422 | 5.9491 | 6.0883 | 9.4753 | 8.8302 | 9.6052 | 9.1352 |
| OSBPL3_S304 | OSBPL3 | Q9H4L5 | | 410 | ESLLLDSPAVAKS | 5.6158 | 6.5056 | 6.1216 | 6.2713 | 6.2520 | 6.2100 | 6.4562 | 6.1216 | 6.8572 | 6.9126 | 6.7041 | 6.2125 | 5.8716 | 6.1808 | 5.6434 | 6.0638 |
| OSBPL3_S371 | OSBPL3 | Q9H4L5 | | 265 | QGGSFESPKKEKR | 6.1472 | 6.1411 | 7.0776 | 6.5167 | 6.6496 | 7.2319 | 7.1535 | 6.5208 | 6.8068 | 6.5361 | 5.9991 | 6.9153 | 5.1185 | 5.0554 | 5.1409 | 4.9897 |
| OSBPL3_S372 | OSBPL3 | Q9H4L5 | | 200 | QTGSNVSFSCGGE | 5.4813 | 5.8128 | 5.3361 | 5.5347 | 6.2429 | 6.2360 | 5.8936 | 6.0819 | 6.2127 | 6.6846 | 6.5914 | 6.4038 | 6.9722 | 6.7421 | 6.5490 | 7.2249 |
| OSBPL3_S410 | OSBPL3 | Q9H4L5 | | 371 | MEQDASSSPSAQV | 4.9580 | 5.6259 | 5.0062 | 6.6825 | 6.0386 | 6.3307 | 5.4642 | 5.4217 | 6.7122 | 5.9067 | 7.5830 | 6.5818 | 6.9250 | 7.6525 | 6.9675 | 6.1435 |
| OSBPL5_S747 | OSBPL5 | Q9H0X9 | | 747 | QTTFLGSPGPRHE | 5.8804 | 6.2594 | 5.7765 | 6.2051 | 6.1436 | 6.4461 | 6.7146 | 6.5251 | 6.2618 | 6.4848 | 6.7087 | 6.4839 | 5.8578 | 6.1380 | 5.9128 | 6.2014 |
| OSBPL6_S290 | OSBPL6 | Q9BZF3-5 | | 369 | PVRLHSSNPNLCA | 5.4302 | 5.4702 | 5.7655 | 5.7026 | 5.9169 | 6.3813 | 6.3318 | 6.2310 | 5.9170 | 5.8040 | 6.2153 | 6.1433 | 6.9267 | 6.9143 | 7.5462 | 7.3034 |
| OSBPL6_S369 | OSBPL6 | Q9BZF3-5 | | 9 | SDEKGISPAHKTS | 6.1591 | 6.7225 | 6.6078 | 7.1277 | 6.5666 | 6.8383 | 6.6628 | 6.8706 | 7.0934 | 7.8910 | 6.7680 | 7.9121 | 3.6817 | 4.3943 | 3.5764 | 4.2175 |
| OSBPL6_S9 | OSBPL6 | Q9BZF3-5 | | 290 | ILQRTQSAPNFTD | 4.2583 | 4.0463 | 4.7444 | 4.4619 | 5.3410 | 6.1423 | 5.7240 | 5.7093 | 5.2916 | 5.1658 | 5.8492 | 5.5088 | 8.9485 | 8.9548 | 9.8113 | 10.0423 |
| OSBPL8_S314 | OSBPL8 | Q9BZF1 | | 328 | KDQDMYSDKSDKE | 6.2843 | 6.6825 | 6.3060 | 6.5473 | 5.6464 | 5.3801 | 5.7590 | 5.5088 | 5.6839 | 6.2237 | 5.4482 | 5.5201 | 6.8399 | 6.8705 | 7.4469 | 7.8525 |
| OSBPL8_S328 | OSBPL8 | Q9BZF1 | | 54 | GKEAYPTPTKDLH | 6.5895 | 7.2385 | 6.0780 | 7.1598 | 6.5640 | 6.9343 | 7.2131 | 6.7873 | 6.5672 | 6.9175 | 6.8264 | 6.8399 | 4.0418 | 5.0838 | 4.3899 | 4.7691 |
| OSBPL8_S328;S331 | OSBPL8 | Q9BZF1 | | 39 | DESQLLTPGKMSQ | 6.9352 | 6.7273 | 5.9596 | 6.8691 | 7.1157 | 6.8896 | 7.9804 | 6.9639 | 6.8987 | 6.2891 | 6.2800 | 6.6507 | 4.4691 | 5.0433 | 4.6120 | 4.3162 |
| OSBPL8_S65 | OSBPL8 | Q9BZF1 | | 68 | PSLSPASPHSQGF | 6.2169 | 6.2949 | 6.0833 | 6.4893 | 6.6914 | 6.4994 | 7.0053 | 7.1706 | 6.6905 | 7.0878 | 6.5522 | 6.9024 | 4.6827 | 5.3130 | 5.2323 | 5.0880 |
| OSBPL8_S65;S68 | OSBPL8 | Q9BZF1 | | 65 | LHQPSLSPASPHS | 6.1515 | 6.5761 | 6.1084 | 6.7041 | 6.6062 | 6.2911 | 6.8233 | 6.8396 | 6.6840 | 6.7039 | 6.7300 | 6.7221 | 4.9885 | 5.2241 | 5.3274 | 5.5198 |
| OSBPL8_S68 | OSBPL8 | Q9BZF1 | | 314 | NFQLNDSEIERQH | 6.2116 | 7.4477 | 6.0475 | 7.6735 | 6.8044 | 6.7805 | 6.9047 | 6.4964 | 5.5024 | 5.2625 | 5.4709 | 5.1671 | 5.8251 | 6.5028 | 5.9051 | 5.9977 |
| OSBPL8_T39 | OSBPL8 | Q9BZF1 | 328;331 | | NA | 5.7242 | 5.9836 | 6.9238 | 6.3072 | 5.8848 | 5.1637 | 6.7126 | 4.7405 | 5.5571 | 6.1060 | 6.0806 | 5.8349 | 7.0957 | 7.1509 | 7.0522 | 7.6820 |
| OSBPL8_T54 | OSBPL8 | Q9BZF1 | 65;68 | | NA | 6.4099 | 7.0375 | 5.6770 | 7.4595 | 6.6059 | 6.9143 | 7.2350 | 6.1629 | 7.0381 | 6.3950 | 6.1794 | 7.3796 | 4.2448 | 5.8223 | 4.8647 | 4.5741 |
| OSBPL9_S621 | OSBPL9 | Q96SU4-6 | | 621 | ITAEIFSPNDKKS | 7.3182 | 6.6290 | 7.8496 | 6.3522 | 5.9371 | 6.2138 | 6.6733 | 5.7793 | 7.0276 | 5.4776 | 6.6207 | 6.4546 | 4.4175 | 6.3771 | 5.8872 | 4.9849 |
| OSER1_S82;S84 | OSER1 | Q9NX31 | | 84 | RRRRSKSPVLHPP | 6.3253 | 6.9380 | 6.5067 | 6.6466 | 6.2803 | 6.5898 | 6.4797 | 6.9703 | 5.9385 | 6.0602 | 6.7186 | 6.5759 | 5.3837 | 5.6101 | 5.3762 | 5.6002 |
| OSER1_S84 | OSER1 | Q9NX31 | 82;84 | | NA | 5.2784 | 6.0364 | 7.0240 | 6.6127 | 6.3194 | 6.8113 | 6.9239 | 6.8481 | 6.6033 | 6.3780 | 7.1171 | 6.9698 | 5.3886 | 5.5200 | 4.7022 | 5.4668 |
| OSMR_S826 | OSMR | Q99650 | | 826 | FLGTRKSLTETEL | 7.9282 | 6.6512 | 7.2925 | 7.1086 | 7.0058 | 6.0703 | 5.8093 | 6.3706 | 8.1080 | 6.5124 | 7.2928 | 6.8703 | 4.1101 | 4.6748 | 3.9594 | 4.2357 |
| OSTM1_S325 | OSTM1 | Q86WC4 | | 325 | RLKSSTSFANIQE | 6.5243 | 6.6036 | 6.2815 | 6.5952 | 7.2699 | 6.9693 | 6.6186 | 6.2348 | 6.6055 | 6.4016 | 7.2851 | 7.0568 | 4.5189 | 5.6021 | 4.5415 | 4.8912 |
| OTUB1_S16 | OTUB1 | Q96FW1 | | 16 | KQEPLGSDSEGVN | 5.9492 | 6.0511 | 6.4857 | 6.4457 | 6.4714 | 6.4299 | 6.0526 | 6.4864 | 6.5400 | 6.0669 | 6.6313 | 6.3921 | 5.7040 | 6.0126 | 5.9175 | 6.3635 |
| OTUD4_S1006 | OTUD4 | Q01804 | | 557 | KKLECPSPAEQKP | 6.5964 | 6.3225 | 6.6781 | 6.1613 | 6.3242 | 6.2629 | 6.4828 | 6.4992 | 6.6063 | 6.6380 | 6.2085 | 6.4332 | 5.3081 | 5.9819 | 5.6529 | 5.8437 |
| OTUD4_S416 | OTUD4 | Q01804 | | 893 | KGELDLSLENLDL | 6.2100 | 6.6454 | 6.9431 | 7.0126 | 6.1963 | 6.7790 | 6.8027 | 6.7014 | 6.8442 | 7.3729 | 7.2320 | 7.2761 | 4.2315 | 4.8239 | 4.3359 | 4.5930 |
| OTUD4_S443 | OTUD4 | Q01804 | | 1006 | TAADVVSPGANSV | 6.5540 | 6.7711 | 6.9187 | 6.9214 | 6.1232 | 6.5942 | 6.6204 | 6.5289 | 6.9502 | 6.9832 | 7.0594 | 6.9855 | 4.1560 | 5.2083 | 4.5128 | 5.1128 |
| OTUD4_S546 | OTUD4 | Q01804 | | 416 | NLSRTPSQIIRKP | 8.3781 | 7.4554 | 7.6210 | 5.8006 | 4.5775 | 7.0906 | 6.9059 | 6.2578 | 8.4149 | 7.6394 | 7.9430 | 7.5144 | 3.7276 | 4.6724 | 2.5385 | 3.4628 |
| OTUD4_S557 | OTUD4 | Q01804 | | 443 | SNYFGLSPEERRE | 6.1898 | 7.2211 | 7.4687 | 7.3452 | 5.7473 | 6.7721 | 6.0800 | 5.7258 | 6.8442 | 6.2905 | 6.0722 | 7.2218 | 5.1676 | 6.0156 | 4.7653 | 5.0727 |
| OTUD4_S893 | OTUD4 | Q01804 | | 546 | KYATVSSPSKSKK | 6.3043 | 6.0515 | 6.2109 | 5.8551 | 6.6636 | 6.6686 | 6.6572 | 6.5130 | 6.5415 | 6.3046 | 6.4462 | 6.6663 | 6.1643 | 6.0349 | 5.7190 | 5.1989 |
| OTUD5_S508 | OTUD5 | Q96G74 | | 64 | GARPRASPPPQGP | 6.3156 | 6.1465 | 6.8094 | 6.4910 | 6.1017 | 6.0213 | 6.1230 | 6.6408 | 6.6198 | 6.4154 | 6.7087 | 6.6992 | 5.9510 | 5.5928 | 5.4903 | 5.8735 |
| OTUD5_S64 | OTUD5 | Q96G74 | | 508 | GADRATSPLVSLY | 6.3839 | 6.3566 | 6.0110 | 6.2160 | 6.1306 | 6.3839 | 5.5967 | 6.6096 | 6.1754 | 6.9708 | 6.6299 | 6.1619 | 6.4018 | 6.0823 | 5.8245 | 6.0651 |
| OTUD7B_S449 | OTUD7B | Q6GQQ9 | | 449 | PLAQPESPTASAG | 5.8696 | 6.5989 | 6.3364 | 5.8968 | 6.1545 | 6.5996 | 5.8910 | 6.7803 | 6.6907 | 6.3967 | 6.3187 | 6.7636 | 5.7253 | 6.2816 | 5.8353 | 5.8612 |
| OTUD7B_T65 | OTUD7B | Q6GQQ9 | | 65 | GSGGSRTPEKGFS | 6.4272 | 6.6074 | 6.9539 | 6.4071 | 6.4589 | 6.9119 | 6.3499 | 6.9721 | 6.7986 | 6.2671 | 6.9259 | 7.4427 | 4.5667 | 4.8559 | 4.7627 | 5.2918 |
| OTULIN_S351 | OTULIN | Q96BN8 | | 351 | RVCEETSLxxxxx | 5.6763 | 6.1851 | 7.6190 | 5.3371 | 7.5573 | 8.4310 | 6.3804 | 6.9355 | 5.5735 | 4.9774 | 5.7277 | 6.0926 | 5.7354 | 5.6609 | 5.6609 | 6.4498 |
| OXR1_S202 | OXR1 | Q8N573 | | 202 | PARVVSSTSEEEE | 6.2984 | 5.8489 | 6.2633 | 6.1993 | 6.8644 | 7.0327 | 6.4421 | 6.4642 | 6.8023 | 5.5415 | 6.3199 | 6.5022 | 6.4523 | 5.9104 | 5.3767 | 5.6813 |
| OXSR1_S339 | OXSR1 | O95747 | | 359 | AISQLRSPRVKES | 5.8767 | 6.1571 | 5.9496 | 5.6746 | 5.2590 | 7.8231 | 7.5814 | 6.7616 | 6.2953 | 5.9164 | 6.0925 | 6.5646 | 5.3989 | 6.7304 | 5.6336 | 6.2852 |

| | | | | | | | | | | | | | | | | | | | | | |
|---|---|---|---|---|---|---|---|---|---|---|---|---|---|---|---|---|---|---|---|---|---|
| OXSR1_S359 | OXSR1 | O95747 | | 339 | DGGWEWSDDEFDE | 5.8394 | 5.9971 | 7.3869 | 6.1185 | 6.0034 | 6.2005 | 5.8444 | 6.0542 | 6.3699 | 6.0614 | 6.2734 | 6.6648 | 6.3445 | 6.2571 | 6.2268 | 6.3577 |
| P2RY1_S354 | P2RY1 | P47900 | | 354 | ANLQSKSEDMTLN | 3.9364 | 5.9872 | 5.8142 | 5.3593 | 6.7395 | 5.7170 | 6.5626 | 6.2069 | 4.1424 | 5.3846 | 5.9464 | 6.2516 | 6.8503 | 8.2305 | 9.9800 | 6.8911 |
| PA2G4_S361 | PA2G4 | Q9UQ80 | | 361 | KALLQSSASRKTQ | 6.7188 | 7.1923 | 6.5776 | 6.6054 | 6.4065 | 6.3150 | 6.4623 | 6.6503 | 6.9704 | 6.4642 | 6.4851 | 6.9995 | 5.0359 | 5.6388 | 4.5457 | 4.9324 |
| PABIR1_S143;S147 | PABIR1 | Q96E09 | | 270 | TTDSPVSPAQAAS | 5.0567 | 5.2882 | 6.2486 | 4.8988 | 5.8453 | 6.1494 | 5.2234 | 6.4817 | 5.9627 | 6.7147 | 6.5415 | 6.8514 | 6.9456 | 7.7442 | 7.1803 | 6.8675 |
| PABIR1_S158;S173 | PABIR1 | Q96E09 | | 62 | PSARRNSTTFPSR | 5.1365 | 4.8415 | 5.7400 | 4.7408 | 5.9315 | 5.6685 | 4.7751 | 5.9372 | 6.2927 | 7.0056 | 6.5364 | 7.1527 | 7.6555 | 8.4398 | 7.0178 | 7.1283 |
| PABIR1_S189 | PABIR1 | Q96E09 | | 76 | GLLLPASPVRMHS | 5.8966 | 5.2436 | 6.2504 | 4.9057 | 5.8688 | 5.8943 | 5.4632 | 6.8720 | 6.1556 | 6.5022 | 6.6462 | 7.4160 | 6.5922 | 7.2495 | 6.5942 | 6.4493 |
| PABIR1_S19 | PABIR1 | Q96E09 | | 189 | TTRRSQSPINCIR | 5.0074 | 4.5331 | 5.7631 | 4.5988 | 5.5627 | 5.3902 | 5.0746 | 6.1231 | 5.6209 | 6.2249 | 6.1071 | 5.9178 | 7.3505 | 9.1299 | 8.9150 | 8.6809 |
| PABIR1_S270 | PABIR1 | Q96E09 | | 19 | PPGTGGSPAEGGG | 5.1194 | 6.5149 | 5.3928 | 7.2771 | 5.6267 | 6.3441 | 7.0695 | 6.8067 | 4.8987 | 5.9841 | 5.1957 | 7.4348 | 5.8816 | 7.9446 | 5.1326 | 7.3769 |
| PABIR1_S270;S276 | PABIR1 | Q96E09 | 143;147 | | NA | 5.5510 | 5.6383 | 6.5618 | 4.9536 | 5.1464 | 6.3125 | 5.7507 | 6.8686 | 6.1832 | 6.8553 | 6.5361 | 7.1957 | 6.3300 | 7.0433 | 6.4541 | 6.6193 |
| PABIR1_S62 | PABIR1 | Q96E09 | 270;276 | | NA | 5.6810 | 6.4143 | 6.4259 | 5.9679 | 6.2713 | 6.1957 | 6.0007 | 6.2637 | 6.6888 | 7.2136 | 6.8083 | 6.2149 | 5.5666 | 6.5341 | 6.0024 | 5.7506 |
| PABIR1_S76 | PABIR1 | Q96E09 | 158;173 | | NA | 5.7834 | 6.2371 | 4.9762 | 5.9227 | 7.4458 | 6.2554 | 6.4864 | 7.4524 | 5.3874 | 5.8448 | 6.6830 | 6.4816 | 6.0733 | 6.5193 | 6.1114 | 6.3398 |
| PABIR2_S134;S138 | PABIR2 | Q7Z309-3 | | 50 | LRTRRNSTTIMSR | 5.0938 | 4.9316 | 5.4109 | 4.4885 | 5.6097 | 5.8930 | 4.8320 | 6.1980 | 5.9214 | 6.2752 | 6.7065 | 6.6538 | 8.1835 | 8.1852 | 7.8841 | 7.7328 |
| PABIR2_S156;S160 | PABIR2 | Q7Z309-3 | | 64 | SLLLSSSPNRIPS | 5.2584 | 5.6063 | 5.5668 | 5.1159 | 6.1462 | 6.2459 | 5.7520 | 6.5752 | 6.2500 | 5.7295 | 6.0035 | 6.1380 | 7.4297 | 7.4996 | 7.6330 | 7.0500 |
| PABIR2_S246 | PABIR2 | Q7Z309-3 | | 25 | TLRRSSSAPLIHG | 5.1818 | 5.3835 | 5.7296 | 4.7651 | 5.8247 | 6.9105 | 5.0075 | 7.2834 | 4.3021 | 6.2245 | 7.5140 | 6.6301 | 7.3113 | 8.9287 | 7.0710 | 5.9321 |
| PABIR2_S25 | PABIR2 | Q7Z309-2 | | 58 | TIMSRHSLEEGLD | 5.9526 | 6.2124 | 6.2372 | 5.8630 | 6.5042 | 6.2442 | 6.0543 | 5.8985 | 6.1624 | 6.0885 | 5.6325 | 6.2712 | 6.8407 | 6.7044 | 6.8800 | 6.4538 |
| PABIR2_S50 | PABIR2 | Q7Z309-3 | | 246 | GSATAESPVACSN | 5.4916 | 5.7257 | 5.9150 | 6.0188 | 6.3479 | 5.8662 | 5.4954 | 5.7118 | 5.5055 | 5.9846 | 6.2434 | 5.9441 | 7.4497 | 7.7548 | 8.0491 | 6.4966 |
| PABIR2_S58 | PABIR2 | Q7Z309-3 | 134;138 | | NA | 5.1813 | 5.3128 | 6.0909 | 4.9174 | 6.0360 | 5.8654 | 5.9084 | 6.2356 | 5.5159 | 6.1994 | 6.5593 | 6.4108 | 7.3908 | 7.7830 | 7.4969 | 7.0963 |
| PABIR2_S64 | PABIR2 | Q7Z309-3 | 156;160 | | NA | 5.6329 | 5.2460 | 6.4845 | 5.8459 | 5.0226 | 6.5119 | 6.1941 | 6.5555 | 7.2090 | 5.9136 | 7.2774 | 5.7014 | 5.3592 | 8.4011 | 6.2719 | 6.3731 |
| PABPC1_S315 | PABPC1 | P11940 | | 434 | IAQLRPSPRWTAQ | 7.3047 | 7.0848 | 6.7445 | 7.9575 | 7.1684 | 5.8115 | 7.0833 | 6.7375 | 7.2763 | 7.6087 | 7.6978 | 7.3231 | 3.1744 | 3.9148 | 3.1988 | 3.9138 |
| PABPC1_S434 | PABPC1 | P11940 | | 315 | RLRKEFSPFGTIT | 6.4606 | 6.7077 | 7.1362 | 6.9599 | 5.9672 | 6.5834 | 6.1036 | 6.7305 | 6.9871 | 6.6749 | 6.7737 | 6.1175 | 5.1361 | 5.2357 | 5.0912 | 5.3347 |
| PABPC1_T635 | PABPC1 | P11940 | | 635 | SATGVPTVxxxxx | 7.9680 | 8.1335 | 5.9198 | 6.5776 | 6.4914 | 6.1603 | 7.8319 | 5.0534 | 5.5955 | 6.3190 | 6.6706 | 7.6141 | 5.0330 | 5.8019 | 4.3730 | 4.4569 |
| PABPC4_S547 | PABPC4 | Q13310-3 | | 547 | YASSVRSPHPAIQ | 5.9963 | 6.3949 | 6.4279 | 6.3609 | 6.7508 | 5.0557 | 6.7395 | 7.5305 | 7.3207 | 5.5908 | 6.2461 | 6.1190 | 5.8075 | 5.8075 | 6.4010 | 5.4507 |
| PABPN1_S150 | PABPN1 | Q86U42 | | 150 | EKQMNMSPPPGNA | 6.5906 | 6.9506 | 6.6475 | 7.2620 | 6.2454 | 6.2632 | 6.5240 | 6.2538 | 6.9919 | 7.2719 | 6.8426 | 6.9169 | 4.9368 | 5.0190 | 4.7651 | 5.0637 |
| PABPN1_S162 | PABPN1 | Q86U42 | | 162 | AGPVIMSIEEKME | 6.7130 | 7.1808 | 6.6409 | 7.3768 | 6.2519 | 6.6818 | 6.1082 | 6.3757 | 7.0135 | 7.3143 | 6.6248 | 6.8287 | 4.8370 | 5.0086 | 4.3310 | 4.7131 |
| PABPN1_S304 | PABPN1 | Q86U42 | | 304 | RATSWYSPYxxxx | 7.5112 | 7.5634 | 7.1606 | 7.4970 | 6.0827 | 7.2195 | 7.4106 | 7.2571 | 7.4061 | 7.5810 | 7.3440 | 7.4228 | 2.9749 | 3.6426 | 2.6571 | 3.2695 |
| PACC1_S9 | PACC1 | Q9H813 | | 9 | RQERSTSYQELSE | 4.7575 | 5.3322 | 5.9665 | 6.8112 | 5.9589 | 5.2877 | 5.6248 | 6.3873 | 5.5471 | 6.7410 | 5.8631 | 6.4967 | 7.0389 | 7.0940 | 7.7260 | 7.3670 |
| PACRGL_S47 | PACRGL | Q8N7B6 | | 47 | SSLSTSSPESARK | 6.9326 | 6.3819 | 7.0271 | 6.1616 | 6.0856 | 6.7509 | 5.8736 | 6.9938 | 6.9676 | 6.9671 | 6.6772 | 6.8238 | 5.2116 | 5.5568 | 4.7874 | 4.8014 |
| PACS1_S409 | PACS1 | Q6VY07-2 | | 495 | DLQGSASPSKVEG | 6.1702 | 6.0974 | 6.9431 | 6.0642 | 6.4614 | 6.3926 | 6.6080 | 6.6118 | 6.8490 | 6.4554 | 6.8757 | 6.4769 | 5.4159 | 5.7471 | 5.2789 | 5.5525 |
| PACS1_S430 | PACS1 | Q6VY07-2 | | 430 | LGKDTTSPMELAA | 6.6566 | 7.2214 | 6.7947 | 7.2463 | 6.2044 | 6.7085 | 6.8574 | 6.7266 | 7.1987 | 7.2562 | 7.3786 | 7.1052 | 3.8521 | 4.6278 | 3.8254 | 4.3400 |
| PACS1_S495 | PACS1 | Q6VY07-2 | | 504 | KVEGVHTPRQKRS | 6.6874 | 6.7637 | 7.0670 | 4.4680 | 5.5814 | 7.0080 | 6.8130 | 6.8815 | 7.3307 | 6.8914 | 6.9387 | 7.1468 | 4.1921 | 5.0275 | 4.4077 | 4.7951 |
| PACS1_S781 | PACS1 | Q6VY07-2 | | 46 | QPPQQPTPPKLAQ | 6.2184 | 6.3390 | 6.7079 | 6.1311 | 6.1076 | 6.3202 | 6.1799 | 6.9937 | 6.9316 | 6.9342 | 6.8820 | 6.9646 | 5.2253 | 5.4483 | 4.7812 | 5.8350 |
| PACS1_T46 | PACS1 | Q6VY07-2 | | 781 | LTVPSTSPPSSSG | 6.8683 | 6.3563 | 6.2070 | 7.0885 | 5.3182 | 5.8691 | 6.5582 | 6.7937 | 7.0644 | 6.1646 | 5.7132 | 6.7147 | 6.2407 | 6.6759 | 4.7352 | 5.6320 |
| PACS1_T504 | PACS1 | Q6VY07-2 | | 409 | FEGMSQSSSQTEI | 4.7695 | 5.5154 | 5.5260 | 5.2758 | 5.1185 | 5.9862 | 5.8620 | 5.6751 | 5.2149 | 5.7993 | 6.1725 | 5.9542 | 8.3017 | 8.4815 | 8.3384 | 8.0090 |
| PACSIN2_S372;S375 | PACSIN2 | Q9UNF0 | | 375 | DTGSTVSEKDDTK | 5.6900 | 5.3560 | 6.7917 | 5.6299 | 5.9288 | 5.9387 | 5.8554 | 5.8044 | 6.4506 | 6.1916 | 6.9335 | 6.6989 | 7.2124 | 6.8058 | 6.0908 | 6.6214 |
| PACSIN2_S375 | PACSIN2 | Q9UNF0 | | 399 | SYPTDWSDDESNN | 5.8836 | 5.9171 | 7.2206 | 5.8479 | 5.8343 | 5.5008 | 6.2250 | 5.7343 | 6.2371 | 6.3457 | 6.2050 | 6.8892 | 6.0814 | 6.8934 | 6.8092 | 6.3757 |
| PACSIN2_S399 | PACSIN2 | Q9UNF0 | 372;375 | | NA | 5.4804 | 4.9477 | 6.8828 | 5.8298 | 6.2361 | 5.9693 | 6.0405 | 5.4216 | 6.5320 | 6.3334 | 6.6768 | 6.9404 | 7.5934 | 6.5843 | 6.0586 | 6.4728 |
| PACSIN3_S319 | PACSIN3 | Q9UKS6 | | 319 | KEKGGRSPDEVTL | 6.3754 | 6.6380 | 6.6188 | 6.7045 | 6.4630 | 6.1043 | 6.0675 | 6.4103 | 6.4706 | 6.1958 | 6.2409 | 6.3261 | 6.0458 | 5.9612 | 5.6455 | 5.7324 |
| PACSIN3_S354 | PACSIN3 | Q9UKS6 | | 354 | GQDEEWSDEESPR | 5.5683 | 6.3349 | 5.7879 | 6.3842 | 6.4401 | 6.1512 | 5.5875 | 5.8614 | 6.4443 | 5.6218 | 5.6785 | 5.7036 | 7.2493 | 7.3938 | 7.7262 | 6.0669 |
| PACSIN3_S383 | PACSIN3 | Q9UKS6 | | 383 | QEADELSFRAGEE | 6.0199 | 5.8522 | 5.7593 | 6.2761 | 6.2470 | 6.1891 | 6.5970 | 5.6937 | 5.4112 | 5.0480 | 5.8259 | 5.4336 | 8.3264 | 7.2602 | 6.9830 | 7.0774 |
| PAG1_S229 | PAG1 | Q9NWQ8 | | 298 | KRFSSLSYKSREE | 6.4804 | 6.2318 | 6.5094 | 6.4839 | 5.9160 | 6.0324 | 5.8809 | 5.9668 | 6.4811 | 6.3375 | 6.2381 | 6.3314 | 6.4779 | 6.6495 | 6.0559 | 5.9269 |
| PAG1_S295 | PAG1 | Q9NWQ8 | | 229 | EFAEYASVDRNKK | 6.6425 | 6.4047 | 6.6280 | 7.5708 | 6.0777 | 6.1771 | 5.8568 | 6.9007 | 6.4624 | 5.9916 | 5.3800 | 5.7249 | 6.8291 | 6.0476 | 5.4775 | 5.8285 |
| PAG1_S298 | PAG1 | Q9NWQ8 | | 295 | ETNKRFSSLSYKS | 4.4054 | 5.0342 | 4.9955 | 5.1655 | 5.3413 | 6.1071 | 5.9678 | 7.0436 | 6.3703 | 6.5088 | 5.5769 | 6.0600 | 7.8232 | 7.7838 | 8.1375 | 7.6792 |
| PAGR1_S237 | PAGR1 | Q9BTK6 | | 237 | EDPSPASPPLRSS | 6.3422 | 6.3211 | 6.5983 | 6.5702 | 6.2595 | 6.2435 | 6.0147 | 6.7548 | 6.5801 | 6.5126 | 6.5302 | 6.6431 | 5.7283 | 5.9693 | 5.4090 | 5.5231 |
| PAGR1_T176 | PAGR1 | Q9BTK6 | | 176 | FDDEPVTPKDSLI | 6.7306 | 6.3840 | 7.2355 | 7.3579 | 6.0383 | 5.9407 | 6.1427 | 6.1016 | 7.3742 | 7.2116 | 7.3014 | 6.6159 | 4.8163 | 4.6913 | 4.5108 | 5.5471 |
| PAICS_S114 | PAICS | P22234-2 | | 27 | VYELLDSPGKVLL | 7.2584 | 7.7703 | 7.1981 | 7.6562 | 6.0771 | 7.0401 | 7.3401 | 6.8505 | 7.5453 | 7.5427 | 7.7018 | 7.5217 | 2.5768 | 3.8930 | 2.5704 | 3.4574 |
| PAICS_S27 | PAICS | P22234-2 | | 114 | RRIATGSFLKRNP | 6.1652 | 6.4862 | 7.2682 | 5.6827 | 6.1075 | 6.6450 | 6.6364 | 6.6354 | 6.2067 | 6.0938 | 6.3136 | 6.2145 | 5.8411 | 5.8181 | 5.7594 | 6.1263 |
| PAICS_T245 | PAICS | P22234-2 | | 245 | RDLKEVTPEGLQM | 6.0514 | 7.1624 | 7.4218 | 6.9768 | 5.7535 | 6.9386 | 6.4793 | 6.4294 | 6.5763 | 6.9852 | 7.6017 | 7.2419 | 4.4062 | 5.0619 | 4.1915 | 4.7221 |
| PAIP1_S77 | PAIP1 | Q9H074 | | 77 | QCEVPASPQRPSR | 5.7087 | 5.4783 | 6.6614 | 6.3436 | 6.2267 | 6.4220 | 6.9315 | 7.0428 | 7.3335 | 6.2954 | 6.8178 | 7.1607 | 5.4820 | 5.5194 | 5.1120 | 5.4642 |
| PAK1_S144 | PAK1 | Q13153-2 | | 144 | NSQKYMSFTDKSA | 7.0158 | 6.0588 | 7.2803 | 6.3765 | 6.7471 | 6.2251 | 6.0019 | 6.5938 | 6.6744 | 5.5827 | 6.1076 | 6.1743 | 6.3004 | 5.8542 | 5.7563 | 5.2506 |
| PAK1_S174 | PAK1 | Q13153-2 | | 212 | IEPLPVTPTRDVA | 6.8365 | 7.1395 | 6.8660 | 6.8488 | 6.0606 | 6.5180 | 6.6957 | 7.2558 | 7.3620 | 6.8901 | 6.9282 | 6.7650 | 4.3004 | 4.9105 | 4.1978 | 4.4252 |
| PAK1_S220 | PAK1 | Q13153-2 | | 230 | PTENNTTPPDALT | 6.7599 | 5.4383 | 7.2173 | 6.9906 | 6.6978 | 6.2782 | 6.6762 | 6.1662 | 6.0785 | 6.1601 | 6.8827 | 6.5110 | 5.3776 | 5.5476 | 5.7412 | 5.4767 |
| PAK1_T212 | PAK1 | Q13153-2 | | 174 | PAVPPVSEDEDDD | 6.4950 | 6.0976 | 6.7694 | 5.8752 | 6.5939 | 6.0399 | 6.7409 | 6.6820 | 6.9431 | 5.6920 | 6.3942 | 6.7257 | 5.7490 | 5.5754 | 5.5519 | 6.0748 |
| PAK1_T230 | PAK1 | Q13153-2 | | 220 | TRDVATSPISPTE | 5.5349 | 5.9272 | 5.2288 | 5.8826 | 6.9201 | 6.1384 | 5.9543 | 6.3331 | 6.1606 | 6.6353 | 5.6269 | 6.2992 | 6.6547 | 7.0286 | 6.9811 | 6.6944 |
| PAK1IP1_S320 | PAK1IP1 | Q9NWT1 | | 320 | PPAAEPSPVSKEQ | 6.6194 | 7.0431 | 6.4393 | 7.2548 | 6.5008 | 6.5875 | 6.9227 | 6.7155 | 6.7368 | 6.8799 | 6.6874 | 6.6256 | 4.1785 | 5.0864 | 4.6186 | 5.1037 |
| PAK1IP1_S358 | PAK1IP1 | Q9NWT1 | | 358 | RGLTGDSKKATKE | 6.7042 | 6.3075 | 6.1560 | 8.7454 | 4.6269 | 6.9422 | 5.8603 | 5.6764 | 7.8907 | 7.9051 | 7.9772 | 12.1389 | 2.5064 | 4.3528 | 3.1952 | 3.0149 |
| PAK2_S132 | PAK2 | Q13177 | | 141 | VKQKYLSFTPPEK | 6.6346 | 5.9329 | 7.2086 | 6.0852 | 6.8667 | 6.3068 | 6.1670 | 6.7727 | 6.6028 | 5.5258 | 6.1719 | 6.2161 | 6.3394 | 5.9547 | 5.6834 | 5.5314 |
| PAK2_S141 | PAK2 | Q13177 | | 132 | VLKFYDSNTVKQK | 7.4375 | 6.7240 | 7.4918 | 6.6119 | 7.0160 | 6.4473 | 6.1755 | 6.7418 | 6.7933 | 5.7823 | 6.4491 | 6.3042 | 5.2384 | 5.1634 | 5.0792 | 4.5444 |
| PAK2_S141;T143 | PAK2 | Q13177 | | 75 | KERPEISPPSDFE | 6.6949 | 6.3277 | 5.8865 | 6.1898 | 6.3037 | 6.0235 | 6.2037 | 6.7298 | 6.6706 | 6.3747 | 6.5461 | 6.6549 | 5.8717 | 5.7628 | 6.0475 | 5.7119 |
| PAK2_S143 | PAK2 | Q13177 | | 152 | EKDGFPSGTPALN | 6.3987 | 6.0653 | 7.2839 | 6.0383 | 6.8457 | 6.2468 | 6.2355 | 7.1312 | 6.3581 | 5.6511 | 6.3459 | 6.1400 | 5.9806 | 5.7447 | 5.7955 | 5.7387 |
| PAK2_S152 | PAK2 | Q13177 | | 55 | PRHKIISIFSGTE | 8.1188 | 5.9751 | 6.3388 | 7.0917 | 6.1825 | 7.1883 | 6.1655 | 6.2820 | 6.8772 | 7.0463 | 6.5604 | 5.8913 | 5.9680 | 5.5234 | 4.2121 | 4.5786 |
| PAK2_S42 | PAK2 | Q13177 | | 58 | KIISIFSGTEKGS | 6.4579 | 6.0470 | 6.9295 | 5.7837 | 6.9377 | 6.3033 | 6.2787 | 6.7541 | 6.1224 | 5.6963 | 5.7454 | 5.6700 | 6.7295 | 6.7098 | 5.6192 | 6.2153 |
| PAK2_S55 | PAK2 | Q13177 | | 42 | SLKPLPSVPEEKK | 6.2477 | 5.7372 | 7.5452 | 6.5076 | 5.7769 | 7.1036 | 6.6490 | 6.1838 | 8.2672 | 7.4906 | 8.4563 | 8.6095 | 3.6650 | 4.3721 | 3.1700 | 4.2182 |
| PAK2_S58 | PAK2 | Q13177 | | 143 | QKYLSFTPPEKDG | 5.7462 | 5.6361 | 6.3670 | 6.1710 | 6.4730 | 6.8366 | 6.6084 | 6.2697 | 6.2010 | 6.3703 | 6.3447 | 6.2774 | 6.7158 | 5.9061 | 5.8926 | 6.1842 |
| PAK2_S75 | PAK2 | Q13177 | | 154 | DGFPSGTPALNAK | 6.7851 | 6.1919 | 6.9401 | 6.5333 | 6.5654 | 6.6608 | 6.4326 | 6.6858 | 7.0455 | 6.4817 | 6.4426 | 6.3727 | 5.5614 | 5.5518 | 4.7227 | 5.0266 |
| PAK2_T154 | PAK2 | Q13177 | 141;143 | | NA | 6.5627 | 6.1099 | 7.5274 | 6.8009 | 6.4116 | 6.3710 | 6.2639 | 6.7310 | 6.2108 | 6.1677 | 6.6697 | 7.6302 | 5.1419 | 4.4486 | 5.3229 | 5.6301 |
| PAK4_S104 | PAK4 | O96013 | | 474 | EVPRRKSLVGTPY | 6.7217 | 6.6281 | 6.5796 | 6.3951 | 6.2285 | 6.5154 | 6.4565 | 6.4945 | 6.3088 | 6.6193 | 6.2860 | 6.6568 | 5.4553 | 5.7942 | 5.3085 | 5.5519 |
| PAK4_S181 | PAK4 | O96013 | | 207 | AGRPFNTYPRADT | 7.0212 | 6.6628 | 7.5116 | 6.9400 | 6.7927 | 6.3711 | 6.6541 | 6.1495 | 6.6864 | 6.5765 | 6.5685 | 6.3910 | 4.8308 | 5.5191 | 4.4705 | 4.8542 |
| PAK4_S258;S267 | PAK4 | O96013 | | 181 | RDKRPLSGPDVGT | 6.6990 | 6.7080 | 7.2009 | 6.7628 | 6.5514 | 6.7144 | 6.4970 | 6.8410 | 6.7822 | 6.7421 | 6.7282 | 6.8553 | 4.6576 | 5.1721 | 4.4879 | 4.6002 |
| PAK4_S267 | PAK4 | O96013 | | 104 | NSLRRDSPPPPAR | 6.1451 | 6.4728 | 6.3437 | 6.3185 | 6.1164 | 6.1442 | 6.2561 | 6.3890 | 6.4338 | 6.3726 | 6.0686 | 6.3146 | 5.8059 | 6.4955 | 6.1865 | 6.1366 |
| PAK4_S291 | PAK4 | O96013 | | 267 | VLGPHASEPQLAP | 6.2070 | 6.5699 | 6.7458 | 6.3289 | 6.1019 | 6.4771 | 6.7937 | 6.2679 | 6.5358 | 6.4231 | 6.4660 | 6.0245 | 5.6897 | 6.4677 | 5.3861 | 5.5148 |
| PAK4_S474 | PAK4 | O96013 | | 291 | GPPGPRSPQREPQ | 6.9055 | 6.8473 | 7.0414 | 6.7831 | 6.4110 | 6.8059 | 6.8479 | 6.5969 | 6.1363 | 6.7062 | 7.1690 | 6.7827 | 4.4459 | 5.2533 | 4.5089 | 4.7587 |
| PAK4_S99;S104 | PAK4 | O96013 | | 187 | SGPDVGTPQPAGL | 7.5453 | 8.0731 | 7.2141 | 7.2905 | 5.8576 | 7.4835 | 8.1151 | 5.9801 | 7.3418 | 7.7221 | 7.3261 | 7.1447 | 2.4873 | 4.6510 | 2.4558 | 3.3119 |
| PAK4_T187 | PAK4 | O96013 | 99;104 | | NA | 7.5549 | 7.3804 | 7.4777 | 6.6365 | 6.8654 | 6.9953 | 6.4769 | 5.8484 | 6.1327 | 6.3523 | 6.4750 | 6.2763 | 5.1968 | 5.9099 | 4.2346 | 4.1868 |
| PAK4_T207 | PAK4 | O96013 | 258;267 | | NA | 6.2315 | 6.0580 | 6.1827 | 7.1049 | 6.1955 | 6.9753 | 6.8997 | 5.9367 | 6.2042 | 6.2807 | 6.0864 | 6.0172 | 5.6120 | 6.5164 | 5.4811 | 6.2176 |
| PALB2_S209 | PALB2 | Q86YC2 | | 809 | DSVPPGTPPPIES | 5.9135 | 6.4258 | 6.7665 | 6.5590 | 5.9620 | 6.3636 | 6.4084 | 6.2833 | 6.4807 | 6.1822 | 6.4268 | 6.6420 | 6.1056 | 5.9918 | 5.5313 | 5.9573 |
| PALB2_S387 | PALB2 | Q86YC2 | | 660 | IFPEELSPKRMDT | 6.2006 | 6.2132 | 6.3197 | 6.7221 | 6.3038 | 5.8989 | 6.5123 | 5.6711 | 6.8867 | 6.1996 | 6.2378 | 6.7019 | 6.3140 | 5.9679 | 5.8991 | 5.9514 |
| PALB2_S660 | PALB2 | Q86YC2 | | 209 | KSELPDSPEPVTE | 7.0621 | 6.8240 | 5.7302 | 6.1003 | 6.5955 | 6.5859 | 6.8254 | 5.9819 | 5.2513 | 6.8063 | 5.5262 | 6.0839 | 5.8323 | 6.4254 | 5.4024 | 6.9669 |
| PALB2_S771;T773 | PALB2 | Q86YC2 | | 387 | LSLEATSPLSAEK | 6.4929 | 6.4756 | 6.0886 | 7.2043 | 6.2171 | 6.3940 | 6.5918 | 6.3403 | 7.0490 | 6.5449 | 6.4105 | 6.8302 | 5.3406 | 5.1779 | 5.2917 | 5.5509 |
| PALB2_S781 | PALB2 | Q86YC2 | | 781 | QFDSSGSPAKPHT | 6.8952 | 6.6152 | 6.4158 | 7.2539 | 6.2387 | 6.2271 | 6.4540 | 5.8713 | 6.5044 | 6.4343 | 6.9331 | 6.4837 | 5.3869 | 5.4805 | 5.5975 | 5.2085 |
| PALB2_T809 | PALB2 | Q86YC2 | 771;773 | | NA | 6.9778 | 5.4734 | 6.3923 | 4.8516 | 6.8339 | 4.0118 | 4.4490 | 6.4383 | 6.0825 | 6.8039 | 6.0598 | 6.8661 | 6.4035 | 7.6039 | 7.1064 | 7.6457 |
| PALLD_S1104 | PALLD | Q8WX93 | | 641 | TPAVLLSPTKEPP | 6.6850 | 6.2882 | 6.7787 | 6.1938 | 6.1628 | 6.9793 | 5.9547 | 6.5910 | 6.7988 | 6.8424 | 6.9312 | 7.0246 | 5.0895 | 5.3204 | 5.1787 | 5.1809 |
| PALLD_S641 | PALLD | Q8WX93 | | 877 | GLPKGVTPAGFPK | 8.0067 | 7.5517 | 7.7652 | 7.7884 | 4.2177 | 6.5086 | 7.6401 | 6.7394 | 7.5959 | 7.2716 | 8.7424 | 8.5928 | 2.1689 | 3.6720 | 2.3163 | 3.4221 |
| PALLD_T877 | PALLD | Q8WX93 | | 1104 | RGRSPRSPSGHPH | 6.1945 | 6.5225 | 6.1159 | 5.7743 | 6.7170 | 6.7997 | 6.1708 | 7.6089 | 6.0027 | 6.6167 | 7.3643 | 6.9719 | 5.5242 | 5.9248 | 4.7164 | 4.9753 |
| PALM_S124 | PALM | O75781 | | 162 | LRTVDGSPMMKAA | 6.9284 | 6.9837 | 6.4025 | 6.1476 | 6.3263 | 6.4933 | 6.2764 | 6.7227 | 6.5073 | 6.7780 | 6.4418 | 6.4376 | 5.3120 | 5.8218 | 5.2382 | 5.1825 |
| PALM_S152 | PALM | O75781 | | 124 | VRAPAPSPAKEER | 6.2827 | 6.1303 | 6.2844 | 5.9988 | 6.2794 | 6.3188 | 6.2455 | 6.5152 | 6.3278 | 6.4908 | 6.1086 | 6.4891 | 5.9643 | 6.3345 | 6.1241 | 6.1057 |
| PALM_S162 | PALM | O75781 | | 152 | PKDKRVSNTPLRT | 7.2266 | 6.6431 | 6.2093 | 6.4818 | 6.4767 | 6.0929 | 6.1961 | 6.7024 | 6.7765 | 5.8362 | 5.9479 | 6.5467 | 5.7686 | 5.9321 | 5.5061 | 5.6570 |
| PALM2_S383 | PALM2 | Q8IXS6-2 | | 383 | DTTEPSSPEGKEE | 5.1160 | 5.5638 | 7.4472 | 5.9203 | 5.5005 | 5.5798 | 6.3705 | 6.1566 | 6.4250 | 6.1294 | 6.2495 | 7.0250 | 7.3335 | 6.6189 | 6.0789 | 6.4851 |
| PALM3_S124 | PALM3 | A6NDB9 | | 260 | DRKGAGSLELPAW | 5.9580 | 6.1901 | 5.6756 | 6.2816 | 6.2115 | 6.4488 | 6.1193 | 6.6017 | 5.7764 | 6.1193 | 5.7200 | 6.1474 | 6.9314 | 6.4408 | 6.5571 | 6.8209 |
| PALM3_S143;T151;S155 | PALM3 | A6NDB9 | | 124 | QGHRPLSQSIVEA | 6.2065 | 6.2758 | 6.3730 | 6.9097 | 6.3106 | 5.9355 | 5.8506 | 6.0372 | 6.7915 | 6.3268 | 6.4499 | 6.5442 | 6.0450 | 5.9734 | 6.1629 | 5.8073 |
| PALM3_S260 | PALM3 | A6NDB9 | | 303 | EVVQTSSPRLQER | 6.3224 | 6.7896 | 6.3425 | 6.5285 | 5.9977 | 5.8751 | 6.3814 | 6.4470 | 6.5872 | 6.5682 | 6.4852 | 6.7332 | 5.8482 | 5.8128 | 5.5045 | 5.7764 |
| PALM3_S303 | PALM3 | A6NDB9 | | 375 | RERGDESPLGAEG | 7.5219 | 5.1014 | 6.0810 | 7.4094 | 6.6747 | 6.5751 | 6.6391 | 5.5115 | 6.0796 | 6.5722 | 5.8233 | 5.6311 | 7.0606 | 5.8176 | 6.1351 | 5.3663 |
| PALM3_S375 | PALM3 | A6NDB9 | 143;151;155 | | NA | 4.9229 | 5.7980 | 5.9745 | 6.1185 | 5.9031 | 6.0743 | 6.5796 | 6.6545 | 7.6710 | 6.6421 | 6.0984 | 5.9530 | 6.3404 | 6.3529 | 6.1091 | 6.8075 |
| PALM3_T151;S155 | PALM3 | A6NDB9 | 151;155 | | NA | 5.6074 | 5.7810 | 6.1246 | 7.5426 | 6.2022 | 6.0850 | 6.4901 | 6.2683 | 6.5392 | 6.9338 | 5.8166 | 6.3249 | 6.6321 | 5.9960 | 5.7635 | 5.8928 |
| PAM_S947 | PAM | P19021-5 | | 947 | RLSTEGSDQEKED | 6.3429 | 6.9970 | 6.5721 | 6.4708 | 5.5235 | 6.2694 | 6.7792 | 5.6013 | 6.4945 | 6.1237 | 6.6893 | 6.9896 | 5.6120 | 5.8502 | 5.9667 | 5.7178 |
| PAN2_S1189 | PAN2 | Q504Q3 | | 1189 | EPEGQTSPKNAAV | 6.1435 | 6.5332 | 6.8584 | 6.3703 | 6.1638 | 6.4932 | 6.6264 | 6.2642 | 6.4808 | 6.8137 | 6.6360 | 7.0009 | 5.0868 | 5.6898 | 5.2536 | 5.5854 |
| PAN2_S791 | PAN2 | Q504Q3 | | 791 | DWKELGSPEGVLV | 6.6686 | 6.8564 | 6.7800 | 7.1704 | 6.0701 | 6.5393 | 6.9674 | 6.7500 | 6.3064 | 6.4094 | 7.1734 | 6.9887 | 4.8540 | 4.9435 | 4.5739 | 4.9486 |
| PAN3_S191 | PAN3 | Q58A45 | | 191 | RMTNSSSSPSLLN | 6.0593 | 6.4377 | 6.4468 | 6.4396 | 5.5781 | 6.4005 | 6.4651 | 6.0567 | 5.9575 | 6.1204 | 6.4553 | 6.0180 | 6.2959 | 6.6242 | 6.0914 | 6.5533 |
| PAN3_S354 | PAN3 | Q58A45 | | 354 | KITPHTSPAPRRR | 5.2614 | 4.9893 | 7.6193 | 5.9203 | 6.8080 | 7.1536 | 8.0075 | 6.0262 | 5.8025 | 5.0915 | 5.6831 | 5.4022 | 6.6591 | 6.4578 | 6.5718 | 6.5464 |
| PANK2_S189 | PANK2 | Q9BZ23 | | 189 | RRDRLGSYSGPTS | 6.0405 | 5.4628 | 6.7032 | 5.6597 | 6.3488 | 6.0722 | 5.8110 | 6.2467 | 6.5947 | 6.3667 | 6.6137 | 6.3948 | 6.9585 | 6.9468 | 6.0243 | 5.7556 |
| PANK4_S393 | PANK4 | Q9NVE7 | | 393 | SGLMSASPELGPA | 6.0636 | 6.0147 | 7.1592 | 6.1457 | 6.0286 | 6.0243 | 6.7313 | 6.5047 | 6.4222 | 6.9079 | 7.1039 | 6.8031 | 5.6661 | 5.9174 | 4.9993 | 5.5078 |
| PANX1_S182 | PANX1 | Q96RD7 | | 182 | TENLGQSLWEVSE | 6.3686 | 6.9090 | 6.3955 | 7.1639 | 6.8881 | 6.1930 | 6.1594 | 5.6391 | 5.2974 | 5.4490 | 5.5673 | 5.5519 | 6.1403 | 6.8551 | 7.0729 | 6.3495 |
| PAPOLA_S24 | PAPOLA | P51003 | | 537 | NSMSVPSPTSATK | 5.1281 | 5.4610 | 5.6854 | 5.0503 | 5.5825 | 5.7943 | 5.5614 | 5.6378 | 6.0939 | 6.1068 | 6.1439 | 6.0546 | 7.4784 | 8.3225 | 8.0671 | 7.8321 |
| PAPOLA_S526;S537 | PAPOLA | P51003 | | 545 | TSATKTSPLNSSG | 5.9843 | 6.2641 | 5.7679 | 5.2588 | 5.9692 | 6.0660 | 6.2380 | 5.5788 | 5.8372 | 6.6531 | 6.2945 | 6.1387 | 6.3944 | 7.1538 | 7.0872 | 7.3138 |
| PAPOLA_S537 | PAPOLA | P51003 | | 24 | KHYGITSPISLAA | 6.0932 | 6.3022 | 7.0655 | 6.3541 | 6.2071 | 6.5018 | 6.1036 | 6.5378 | 6.4682 | 6.3926 | 7.1461 | 6.9757 | 5.0138 | 5.8082 | 5.4778 | 5.5524 |
| PAPOLA_S537;S540 | PAPOLA | P51003 | | 540 | SVPSPTSATKTSP | 4.8297 | 5.0251 | 5.5074 | 4.7588 | 5.1223 | 5.6718 | 5.2321 | 5.3017 | 5.9143 | 5.9814 | 5.8704 | 6.2958 | 8.1987 | 8.9686 | 8.8699 | 8.4520 |
| PAPOLA_S540 | PAPOLA | P51003 | | 644 | AATKIPTPIVGVK | 7.1001 | 8.0595 | 7.2638 | 7.2080 | 6.6802 | 7.2917 | 7.4690 | 7.1327 | 7.2811 | 6.8207 | 7.7211 | 7.5774 | 2.6765 | 4.0479 | 2.4297 | 3.2406 |
| PAPOLA_S545 | PAPOLA | P51003 | 526;537 | | NA | 4.8896 | 5.3865 | 5.6010 | 5.1661 | 5.1868 | 5.4708 | 5.3795 | 6.4066 | 5.5712 | 6.2334 | 5.9075 | 5.9515 | 8.1120 | 8.5141 | 8.1636 | 8.0600 |
| PAPOLA_T644 | PAPOLA | P51003 | 537;540 | | NA | 4.9171 | 5.0907 | 5.2853 | 5.0538 | 5.6857 | 4.7133 | 5.9346 | 5.1539 | 4.7972 | 5.1642 | 5.3150 | 6.2374 | 8.9351 | 9.1018 | 9.6928 | 8.9220 |
| PAPOLG_S525 | PAPOLG | Q9BWT3 | | 525 | LQSKRLSLDSSCL | 4.1138 | 4.3193 | 4.6686 | 4.9793 | 7.0965 | 5.1386 | 5.0935 | 5.7134 | 5.7385 | 5.8418 | 5.6137 | 5.6032 | 9.1858 | 9.0397 | 9.2923 | 8.5622 |
| PAPOLG_S599 | PAPOLG | Q9BWT3 | | 599 | STVKTVSPPTVCT | 5.3879 | 7.6283 | 6.6704 | 6.9906 | 5.4417 | 5.8300 | 6.9736 | 7.3619 | 6.3090 | 6.1899 | 6.4493 | 6.1999 | 5.7818 | 6.2225 | 5.2858 | 5.2773 |
| PAPOLG_S648 | PAPOLG | Q9BWT3 | | 648 | TPKRSHSPSIDGT | 5.3947 | 5.7385 | 6.4334 | 6.2179 | 6.1331 | 6.4048 | 6.0538 | 5.9550 | 6.1810 | 5.8936 | 6.2820 | 6.6221 | 6.5378 | 6.9155 | 6.4069 | 6.8299 |
| PARD3_S1178 | PARD3 | Q8TEW0 | | 144 | LHVRRSSDPALIG | 5.8382 | 6.1401 | 6.5524 | 6.3800 | 6.4634 | 5.8702 | 6.2047 | 6.4053 | 6.2075 | 6.0683 | 6.4913 | 6.2200 | 6.3170 | 6.4578 | 6.2065 | 6.1772 |
| PARD3_S1196 | PARD3 | Q8TEW0 | | 889 | GLKKSSSLESLQT | 5.8444 | 6.2518 | 6.4013 | 5.9845 | 6.3306 | 6.5610 | 5.9419 | 6.5040 | 6.2104 | 6.6190 | 6.3831 | 6.7616 | 5.7512 | 6.1955 | 6.3062 | 5.9537 |
| PARD3_S1223 | PARD3 | Q8TEW0 | | 174 | KNPTRWSTTAGFL | 7.6256 | 6.5958 | 6.5355 | 6.3626 | 6.5207 | 5.8854 | 6.5180 | 5.8355 | 6.5979 | 5.4462 | 5.5081 | 5.7752 | 6.4251 | 6.7982 | 5.3868 | 6.1835 |
| PARD3_S1335 | PARD3 | Q8TEW0 | | 852 | KTRKSKSMDLGIA | 6.2007 | 6.3979 | 6.7908 | 6.2730 | 5.7941 | 6.4814 | 6.3736 | 6.4192 | 6.3806 | 6.6888 | 7.1373 | 6.8034 | 5.3760 | 5.6982 | 5.3784 | 5.8067 |
| PARD3_S144 | PARD3 | Q8TEW0 | | 1178 | DRRRTYSFEQPWP | 5.7360 | 5.6334 | 6.0950 | 5.9899 | 6.0030 | 5.7306 | 5.9471 | 6.1675 | 6.3374 | 6.2265 | 5.6948 | 5.7227 | 7.4153 | 7.3133 | 7.1919 | 6.7956 |
| PARD3_S174 | PARD3 | Q8TEW0 | | 383 | YYSSRFSPDSQYI | 6.6195 | 6.6195 | 6.7757 | 6.7838 | 6.4696 | 6.3731 | 6.7721 | 6.3515 | 6.8112 | 6.8410 | 6.9745 | 6.6739 | 4.9442 | 5.3227 | 4.7981 | 4.8696 |
| PARD3_S187 | PARD3 | Q8TEW0 | | 1223 | AQRQYSSLPRQSR | 7.0695 | 7.1400 | 6.7110 | 5.9618 | 5.4361 | 7.0780 | 6.2748 | 6.3569 | 6.2886 | 6.9229 | 6.5331 | 6.9248 | 5.2454 | 5.8716 | 4.5874 | 5.5981 |
| PARD3_S221 | PARD3 | Q8TEW0 | | 837 | KRTKQFSDASQLD | 5.8631 | 6.5812 | 6.6202 | 6.3947 | 6.1345 | 6.9449 | 6.6397 | 6.2497 | 6.9500 | 7.4522 | 6.8674 | 7.3331 | 4.9169 | 4.9727 | 4.6650 | 5.4147 |
| PARD3_S221;S223 | PARD3 | Q8TEW0 | | 187 | KQNTAGSPKTCDR | 6.2575 | 6.0932 | 6.2225 | 6.5459 | 6.2247 | 6.5202 | 6.3316 | 5.8141 | 6.7534 | 6.5831 | 6.4229 | 7.0612 | 5.8570 | 6.0712 | 5.7422 | 5.4993 |
| PARD3_S383 | PARD3 | Q8TEW0 | | 715 | DRERRISHSLYSG | 6.1746 | 6.0763 | 5.4750 | 6.2566 | 6.3181 | 5.5220 | 5.5194 | 5.8534 | 6.5982 | 5.8551 | 5.5271 | 6.2198 | 7.4618 | 7.5241 | 6.5863 | 7.0321 |
| PARD3_S695 | PARD3 | Q8TEW0 | | 1196 | TQSGRHSVSVEVQ | 6.1993 | 6.3331 | 7.4031 | 6.2807 | 5.0188 | 4.4081 | 6.8448 | 7.1210 | 5.7486 | 6.1906 | 7.4525 | 6.4668 | 6.6238 | 5.2427 | 6.0074 | 6.6587 |
| PARD3_S715 | PARD3 | Q8TEW0 | | 221 | RDNARSSLSASHP | 7.2498 | 7.5076 | 7.5127 | 7.1695 | 5.7678 | 7.1010 | 6.1888 | 7.0407 | 8.1721 | 7.7626 | 7.4005 | 7.0765 | 4.1466 | 4.3391 | 2.9087 | 3.6560 |
| PARD3_S728 | PARD3 | Q8TEW0 | | 1335 | RQDVPPSPSQVAR | 6.2389 | 6.6155 | 6.2915 | 6.3953 | 6.0760 | 5.9927 | 5.7698 | 6.0984 | 6.6574 | 7.7456 | 5.9239 | 5.6669 | 6.3683 | 7.1298 | 6.4405 | 6.5894 |
| PARD3_S837 | PARD3 | Q8TEW0 | | 728 | IEGLDESPSRNAA | 5.6643 | 6.1126 | 6.5092 | 6.1780 | 5.9576 | 7.0208 | 6.9449 | 6.1411 | 6.0324 | 6.1559 | 6.8172 | 6.4080 | 5.8795 | 6.1991 | 6.0356 | 5.9438 |
| PARD3_S852 | PARD3 | Q8TEW0 | | 695 | ELKSPGSPPGPEL | 6.8006 | 6.3507 | 5.9737 | 6.0073 | 5.7582 | 5.9327 | 6.1007 | 6.5064 | 6.3827 | 6.6990 | 6.3237 | 6.9564 | 5.5426 | 6.5400 | 5.5738 | 6.5515 |
| PARD3_S889 | PARD3 | Q8TEW0 | 221;223 | | NA | 6.6555 | 7.1585 | 5.5556 | 5.6052 | 5.6953 | 6.8753 | 7.4956 | 6.2720 | 6.0715 | 6.8953 | 5.2038 | 6.6074 | 6.3235 | 7.7438 | 4.7968 | 5.0449 |
| PARD3B_S363;S368 | PARD3B | Q8TEW8 | | 133 | SALKLGTPLLVRR | 6.5553 | 6.7492 | 6.6284 | 6.3900 | 5.6772 | 5.8264 | 6.7441 | 5.9618 | 6.2902 | 6.2984 | 6.0109 | 6.6496 | 6.0843 | 6.3198 | 5.8050 | 6.0092 |
| PARD3B_S364;S368 | PARD3B | Q8TEW8 | | 746 | GLKKSSSLESLQT | 5.3955 | 5.6228 | 5.7104 | 6.3590 | 5.7432 | 6.2912 | 6.2229 | 7.0495 | 5.9754 | 6.3685 | 6.2867 | 7.5185 | 6.2104 | 6.7350 | 6.1708 | 6.3402 |
| PARD3B_S746 | PARD3B | Q8TEW8 | 364;368 | | NA | 6.3660 | 5.4562 | 6.8943 | 7.3829 | 6.3037 | 8.1646 | 7.3795 | 7.0952 | 6.3361 | 7.3582 | 7.9704 | 7.1823 | 3.9391 | 4.5340 | 3.5953 | 4.0420 |
| PARD3B_T133 | PARD3B | Q8TEW8 | 363;368 | | NA | 7.0792 | 7.4393 | 6.5006 | 6.2592 | 6.6094 | 6.8888 | 6.1984 | 7.1967 | 7.8177 | 6.9314 | 7.0465 | 6.4649 | 4.2286 | 4.9241 | 4.1753 | 4.2397 |
| PARD6B_S11 | PARD6B | Q9BYG5 | | 11 | HRHGAGSGCLGTM | 5.0854 | 5.7525 | 6.4249 | 6.3631 | 6.5364 | 5.7946 | 5.3855 | 6.7552 | 6.4670 | 6.2912 | 6.4310 | 6.6499 | 6.4817 | 6.3526 | 6.7424 | 6.4867 |
| PARD6B_S283 | PARD6B | Q9BYG5 | | 283 | PQQIEPSFEPEDE | 6.6996 | 6.4946 | 5.7451 | 6.8685 | 7.4360 | 6.0337 | 5.4581 | 7.2047 | 6.0698 | 6.1666 | 6.0993 | 5.8927 | 6.2633 | 6.0173 | 5.5548 | 5.9960 |
| PARN_S163 | PARN | O95453 | | 619 | RMKKELSPAGSIS | 5.9579 | 6.5003 | 6.2602 | 6.8644 | 6.1048 | 6.0517 | 6.1305 | 6.2724 | 6.3609 | 6.4385 | 6.3600 | 6.4463 | 5.7844 | 6.2291 | 6.0190 | 6.2197 |
| PARN_S557 | PARN | O95453 | | 163 | GALSYVSPNTSKC | 6.9363 | 6.8581 | 6.8219 | 7.3852 | 6.5392 | 6.6305 | 6.6506 | 6.6109 | 7.3823 | 6.8999 | 6.5708 | 7.0383 | 4.1801 | 4.8663 | 4.0405 | 4.5889 |
| PARN_S570;S583 | PARN | O95453 | | 557 | HYYRNNSFTAPST | 5.4569 | 5.9436 | 6.0254 | 5.8063 | 5.4071 | 5.3570 | 5.8736 | 5.6843 | 5.9609 | 6.0592 | 5.6785 | 6.0005 | 8.0320 | 7.9171 | 7.3781 | 7.4196 |
| PARN_S619 | PARN | O95453 | 619;623 | | NA | 5.9392 | 6.7817 | 6.7961 | 7.9446 | 5.5089 | 6.1071 | 6.9253 | 6.9129 | 6.4720 | 6.1229 | 5.9741 | 6.3445 | 5.8749 | 4.9290 | 6.1689 | 5.1978 |
| PARN_S619;S623 | PARN | O95453 | 570;583 | | NA | 5.8887 | 7.3351 | 5.8838 | 5.9089 | 6.2387 | 6.4378 | 5.8083 | 5.7518 | 5.5990 | 6.6262 | 5.7444 | 5.8093 | 6.3183 | 7.3559 | 5.7234 | 7.5703 |
| PARP1_S185 | PARP1 | P09874 | | 504 | KSGAALSKKSKGQ | 8.3937 | 7.2789 | 7.1568 | 7.6251 | 5.8969 | 5.9008 | 5.4465 | 5.8740 | 8.6800 | 8.8212 | 8.1404 | 7.4227 | 3.0668 | 4.3724 | 3.7209 | 2.2028 |
| PARP1_S257 | PARP1 | P09874 | | 257 | ELKKVCSTNDLKE | 2.1204 | 2.3210 | 2.4761 | 2.5563 | 4.6626 | 3.9331 | 4.5987 | 4.4614 | 2.3817 | 2.5693 | 2.1862 | 2.3021 | 15.0207 | 14.1307 | 18.3045 | 15.9751 |
| PARP1_S362 | PARP1 | P09874 | | 185 | SQLKGFSLLATED | 5.8956 | 6.4799 | 6.4010 | 6.6575 | 6.2753 | 5.8314 | 6.8695 | 4.7786 | 5.5602 | 5.8931 | 5.3704 | 5.7624 | 6.3271 | 7.5821 | 6.5465 | 7.7695 |
| PARP1_S504 | PARP1 | P09874 | | 782 | SLLRGGSDDSSKD | 4.7742 | 5.5987 | 5.1530 | 5.0929 | 6.0221 | 7.5218 | 6.8131 | 5.9318 | 6.3151 | 6.5056 | 7.8460 | 6.2159 | 7.2744 | 7.4315 | 5.9675 | 5.5363 |
| PARP1_S782 | PARP1 | P09874 | | 362 | IFPPETSASVAAT | 5.2599 | 6.6913 | 6.4546 | 6.5348 | 6.3243 | 6.6269 | 6.5565 | 6.6247 | 5.8651 | 6.8292 | 6.6853 | 6.7977 | 5.3348 | 5.8322 | 5.8045 | 5.7782 |
| PARP12_S263 | PARP12 | Q9H0J9 | | 70 | RVVLAASPLRLCR | 8.5025 | 8.0158 | 8.0691 | 6.7267 | 5.5632 | 6.2286 | 7.0994 | 6.5214 | 8.6851 | 7.0081 | 8.0348 | 9.3885 | 2.2207 | 2.2435 | 2.0002 | 3.6923 |
| PARP12_S70 | PARP12 | Q9H0J9 | | 263 | DSSGSVSPNTLSQ | 5.1229 | 6.6315 | 8.3977 | 6.5497 | 5.3191 | 8.8270 | 5.9978 | 6.8809 | 4.4810 | 4.2357 | 6.0387 | 6.9586 | 6.0387 | 5.6298 | 7.4410 | 5.4499 |
| PARP14_S33 | PARP14 | Q460N5 | | 33 | LQMYFQSPKRSGG | 5.8809 | 7.0349 | 7.4574 | 6.7337 | 5.8883 | 6.4363 | 6.0395 | 6.2863 | 6.7497 | 6.9075 | 7.0340 | 7.1835 | 4.5290 | 5.6192 | 4.8267 | 5.3931 |
| PARP2_S226 | PARP2 | Q9UGN5 | | 226 | KEESLKSPLKPES | 7.8707 | 7.9343 | 6.7273 | 8.3074 | 6.2300 | 6.6753 | 6.7783 | 6.9733 | 7.1509 | 7.7151 | 7.3712 | 7.6135 | 2.5190 | 3.7666 | 2.7092 | 3.6579 |
| PARP4_S1236 | PARP4 | Q9UKK3 | | 101 | YKPLDITPPPDQK | 6.2102 | 6.8088 | 6.4898 | 5.7349 | 6.0519 | 6.1082 | 5.9182 | 6.5376 | 6.1542 | 6.4548 | 5.9049 | 6.3535 | 6.1438 | 6.6705 | 6.0439 | 6.4148 |
| PARP4_S1253 | PARP4 | Q9UKK3 | | 1507 | LEESVGSLEGSRC | 6.3092 | 6.1118 | 6.5116 | 6.1292 | 6.4135 | 6.0996 | 6.3222 | 6.6427 | 6.0377 | 6.7063 | 6.4072 | 6.3634 | 5.7943 | 6.3678 | 5.9876 | 5.7958 |

| | | | | | | | | | | | | | | | | | | | | | |
|---|---|---|---|---|---|---|---|---|---|---|---|---|---|---|---|---|---|---|---|---|---|
| PARP4_S1288 | PARP4 | Q9UKK3 | | 1335 | PTARAHSPASLSF | 6.5120 | 7.3510 | 6.0938 | 6.5255 | 6.5320 | 7.5619 | 6.0105 | 7.6523 | 6.7541 | 6.0710 | 6.6072 | 6.0247 | 4.3603 | 5.8230 | 5.2256 | 4.8952 |
| PARP4_S1335 | PARP4 | Q9UKK3 | | 1288 | EKLLDLSWTESCK | 7.1136 | 7.6571 | 7.0054 | 7.2279 | 5.5641 | 6.6004 | 7.2269 | 6.7232 | 7.4177 | 7.4911 | 8.0624 | 8.0375 | 3.0413 | 4.1874 | 3.0486 | 3.5956 |
| PARP4_S1507 | PARP4 | Q9UKK3 | | 1236 | WPELRLSKRKHRK | 6.5759 | 6.3698 | 6.9648 | 6.4726 | 5.9435 | 6.8268 | 7.0171 | 6.3220 | 6.7854 | 7.2398 | 7.2736 | 7.4097 | 4.8662 | 5.3759 | 3.9655 | 4.5913 |
| PARP4_S1713 | PARP4 | Q9UKK3 | | 1713 | QPISTVSPLHRVL | 5.7203 | 6.3659 | 6.6364 | 6.2139 | 6.6735 | 6.6506 | 6.5074 | 7.1294 | 6.3920 | 6.9444 | 6.6770 | 6.8815 | 5.1832 | 5.6240 | 4.9504 | 5.4500 |
| PARP4_S1722 | PARP4 | Q9UKK3 | | 1253 | KRKMELSQPEVSE | 5.9629 | 5.1724 | 6.9315 | 6.1130 | 8.1530 | 5.0852 | 6.7988 | 5.9612 | 5.5651 | 6.4638 | 6.2928 | 7.0047 | 5.8338 | 5.4953 | 6.3486 | 6.8180 |
| PARP4_T101 | PARP4 | Q9UKK3 | | 1722 | HRVLHYSQGxxxx | 4.4406 | 4.5105 | 4.6658 | 5.7700 | 6.7086 | 6.7240 | 6.2368 | 5.8443 | 5.0695 | 4.0003 | 4.5327 | 4.1795 | 8.8172 | 9.3325 | 10.1295 | 9.0382 |
| PARP4_T1428 | PARP4 | Q9UKK3 | | 1428 | TRPSAGTFPELDS | 6.8980 | 5.2700 | 5.5190 | 8.1939 | 6.1542 | 9.7134 | 7.4183 | 7.0257 | 7.0640 | 7.5013 | 7.6609 | 7.6130 | 2.7388 | 3.6070 | 3.4729 | 4.1496 |
| PARP8_S323 | PARP8 | Q8N3A8 | | 456 | AEGRRLSLTSGLI | 5.6568 | 4.8528 | 6.9747 | 5.9034 | 6.3128 | 6.4887 | 6.1446 | 6.3170 | 6.8208 | 7.8262 | 7.6189 | 6.9932 | 5.5807 | 6.2326 | 5.1238 | 5.1529 |
| PARP8_S435 | PARP8 | Q8N3A8 | | 323 | LLRRTCSSTVKTD | 6.5197 | 6.2084 | 7.1768 | 6.3182 | 5.7928 | 6.2712 | 6.1355 | 5.9285 | 6.8099 | 6.5366 | 7.0924 | 7.3091 | 5.7979 | 5.7207 | 5.0827 | 5.2996 |
| PARP8_S456 | PARP8 | Q8N3A8 | | 435 | LSKSYSSAPKSSK | 7.6114 | 6.9329 | 7.3791 | 7.6874 | 6.3785 | 6.0683 | 5.7748 | 5.5566 | 7.6394 | 6.4865 | 7.3125 | 7.7969 | 4.5766 | 5.2234 | 3.4314 | 4.1443 |
| PARP9_S851 | PARP9 | Q8IXQ6 | | 851 | RGFASGSPVDxxx | 6.9844 | 6.2301 | 7.5879 | 7.1255 | 6.9538 | 6.5968 | 7.0605 | 8.3595 | 7.2531 | 5.9197 | 6.7637 | 7.0838 | 3.9816 | 4.3447 | 3.7523 | 4.0025 |
| PASK_S120 | PASK | Q96RG2-2 | | 120 | LSSGWSSPLLPAP | 6.2375 | 5.6671 | 6.1242 | 6.0083 | 5.7244 | 6.5187 | 6.6268 | 7.0240 | 5.4848 | 6.1372 | 5.8012 | 6.6789 | 7.4459 | 5.9354 | 6.3364 | 6.2492 |
| PATJ_S522 | PATJ | Q8NI35 | | 522 | LEKVPDSPENELK | 6.5786 | 6.2464 | 6.3400 | 6.1718 | 6.0521 | 6.5391 | 6.1358 | 6.1526 | 6.5039 | 6.4924 | 6.6400 | 6.7460 | 5.8016 | 6.1480 | 5.7935 | 5.6582 |
| PATL1_S177 | PATL1 | Q86TB9-4 | | 179 | LPRRSTSPIIGSP | 5.2921 | 5.3922 | 5.6021 | 4.9886 | 5.8532 | 5.7031 | 5.5636 | 6.2724 | 5.8913 | 5.8840 | 5.4212 | 5.7196 | 8.3499 | 8.1072 | 8.1901 | 7.7694 |
| PATL1_S177;S179 | PATL1 | Q86TB9-4 | | 194 | RAVPIGTPPKQMA | 7.6370 | 7.3016 | 8.3467 | 7.2549 | 6.1630 | 6.8116 | 6.7991 | 7.0363 | 7.5757 | 7.3321 | 7.8597 | 7.7903 | 2.7368 | 3.4273 | 2.7464 | 3.1816 |
| PATL1_S177;S184 | PATL1 | Q86TB9-4 | | 177 | RALPRRSTSPIIG | 5.8437 | 5.6364 | 5.6274 | 5.5706 | 6.2178 | 5.4404 | 5.6570 | 6.2737 | 6.2221 | 5.9337 | 5.7765 | 6.0358 | 7.5488 | 7.5604 | 7.5520 | 7.1036 |
| PATL1_S179 | PATL1 | Q86TB9-4 | | 261 | VPGFVGSPLAAMN | 7.2428 | 6.5037 | 6.7728 | 6.4819 | 5.4731 | 6.7723 | 7.2993 | 6.3711 | 7.1930 | 7.4426 | 7.9159 | 7.5763 | 3.8813 | 5.1609 | 3.9865 | 3.9266 |
| PATL1_S179;S184 | PATL1 | Q86TB9-4 | | 233 | PYGERMSPNQLCS | 6.3908 | 5.5247 | 5.8130 | 6.6786 | 6.3755 | 6.8334 | 6.6721 | 6.9237 | 6.6490 | 6.4759 | 6.6000 | 7.1778 | 5.3374 | 5.6482 | 5.1584 | 5.7415 |
| PATL1_S233 | PATL1 | Q86TB9-4 | | 248 | LQPGRMSPSQFAR | 6.8866 | 6.7775 | 6.7226 | 5.3995 | 6.1397 | 7.5499 | 6.6130 | 7.5127 | 6.7533 | 6.5852 | 7.0846 | 7.2350 | 4.7703 | 4.9428 | 4.1701 | 4.8572 |
| PATL1_S248 | PATL1 | Q86TB9-4 | 177;184 | | NA | 7.3873 | 6.2884 | 7.3410 | 7.0760 | 5.9733 | 7.0278 | 6.6794 | 6.9710 | 7.7151 | 6.9649 | 7.2627 | 7.1525 | 3.6625 | 4.7574 | 3.6498 | 4.0911 |
| PATL1_S261 | PATL1 | Q86TB9-4 | 177;179 | | NA | 5.8390 | 5.6764 | 6.6084 | 6.5651 | 6.5577 | 6.0463 | 5.9759 | 6.0176 | 6.5340 | 7.3198 | 6.3682 | 6.1762 | 6.0399 | 6.5110 | 5.6516 | 6.1130 |
| PATL1_T194 | PATL1 | Q86TB9-4 | 179;184 | | NA | 7.0369 | 6.2020 | 7.7664 | 6.3066 | 5.7602 | 6.3543 | 6.4253 | 6.6765 | 6.9187 | 6.9712 | 7.5416 | 7.0504 | 4.5989 | 5.4653 | 4.7803 | 4.1453 |
| PATZ1_S243 | PATZ1 | Q9HBE1 | | 378 | RHKLSHSGEKPYS | 6.6157 | 7.6999 | 6.9495 | 7.8826 | 5.9256 | 6.9247 | 7.4098 | 6.8072 | 7.5953 | 8.3879 | 8.4953 | 8.8728 | 2.0584 | 3.0826 | 2.2148 | 3.0780 |
| PATZ1_S282 | PATZ1 | Q9HBE1 | | 282 | LDSMFGSPGGLRE | 5.8889 | 6.4448 | 5.9206 | 6.9490 | 6.7269 | 6.6772 | 7.0014 | 5.9875 | 6.5621 | 7.0994 | 6.1337 | 6.0227 | 5.5357 | 6.0006 | 5.5337 | 5.5157 |
| PATZ1_S378 | PATZ1 | Q9HBE1 | | 243 | MVAGPLSPQLLTS | 5.9451 | 6.3138 | 6.8792 | 6.6346 | 6.9291 | 5.8397 | 6.6854 | 6.5857 | 6.8660 | 6.3101 | 7.1285 | 6.2264 | 5.6309 | 5.3694 | 5.1690 | 5.4870 |
| PATZ1_S585 | PATZ1 | Q9HBE1 | | 585 | KQSANGSFSCDMA | 5.6470 | 6.9999 | 6.0425 | 7.5008 | 5.9821 | 7.2244 | 7.2244 | 7.4198 | 5.8859 | 6.9281 | 7.4550 | 7.0885 | 4.2635 | 5.2293 | 4.0413 | 5.0675 |
| PAWR_S10 | PAWR | Q96IZ0 | | 10 | GGYRTSSGLGGST | 6.4013 | 6.0705 | 6.1004 | 6.1379 | 6.1875 | 5.6040 | 6.3531 | 6.4504 | 6.8743 | 6.2721 | 6.1110 | 7.0086 | 6.3582 | 6.5902 | 5.7737 | 5.7068 |
| PAXBP1_S154 | PAXBP1 | Q9Y5B6 | | 295 | EIGIEGSDDDALV | 5.4292 | 6.4027 | 6.9719 | 6.5335 | 5.9312 | 5.8446 | 6.4549 | 6.2781 | 6.2644 | 6.7271 | 5.8822 | 6.6528 | 5.9648 | 6.0170 | 6.2878 | 6.3577 |
| PAXBP1_S295 | PAXBP1 | Q9Y5B6 | | 62 | LLGPGPSPPSALT | 5.9470 | 6.3296 | 6.0948 | 6.5579 | 5.7170 | 5.9592 | 6.6205 | 6.8447 | 6.0428 | 6.1964 | 5.8422 | 6.4864 | 6.5993 | 6.4328 | 5.7674 | 6.5620 |
| PAXBP1_S557;S558 | PAXBP1 | Q9Y5B6 | | 154 | IKTELNSSAESEQ | 5.6264 | 6.2918 | 7.1261 | 6.9233 | 6.8101 | 6.4403 | 6.3670 | 6.5096 | 6.3674 | 6.5010 | 6.3923 | 6.5168 | 5.7916 | 5.8654 | 4.8890 | 5.5820 |
| PAXBP1_S62 | PAXBP1 | Q9Y5B6 | 557;558 | | NA | 5.8873 | 6.1345 | 6.0729 | 6.5240 | 6.1673 | 6.0300 | 5.9191 | 6.0224 | 6.1733 | 6.3373 | 6.3808 | 6.0003 | 6.5575 | 6.5637 | 6.7770 | 6.4524 |
| PAXBP1_T558;S564 | PAXBP1 | Q9Y5B6 | 558;564 | | NA | 5.9162 | 6.5971 | 6.0992 | 6.2554 | 6.4231 | 5.6555 | 6.2690 | 6.3243 | 6.0056 | 6.2060 | 5.8825 | 6.3035 | 6.3360 | 6.7074 | 6.6429 | 6.3764 |
| PAXIP1_S235 | PAXIP1 | Q6ZW49 | | 235 | SGDQQFSPKSNTE | 6.4942 | 6.4397 | 6.1854 | 7.1158 | 8.2903 | 6.4054 | 6.1471 | 6.0522 | 6.2359 | 7.4972 | 6.3105 | 7.6970 | 4.6254 | 4.9080 | 4.4822 | 5.1138 |
| PAXX_S148 | PAXX | Q9BUH6 | | 148 | AEETAVSPRKSPR | 6.3175 | 5.8214 | 7.0535 | 5.7160 | 6.5403 | 6.7550 | 6.0025 | 6.5356 | 6.2708 | 6.4474 | 6.6452 | 6.8855 | 5.8531 | 6.0775 | 5.6557 | 5.4228 |
| PAXX_S152 | PAXX | Q9BUH6 | | 152 | AVSPRKSPRPAGP | 6.4700 | 6.3406 | 7.1919 | 6.0162 | 6.8039 | 7.4227 | 6.0561 | 5.5812 | 6.0199 | 5.8216 | 7.0585 | 7.2926 | 6.1730 | 5.9257 | 5.0181 | 4.8080 |
| PBDC1_S181 | PBDC1 | Q9BVG4 | | 197 | GEKGADSGEEKEE | 6.8143 | 6.3224 | 6.9572 | 5.8474 | 6.2400 | 6.5279 | 5.8366 | 6.6389 | 6.6977 | 6.3463 | 6.4873 | 6.8532 | 5.8163 | 5.7599 | 5.5016 | 5.3530 |
| PBDC1_S197 | PBDC1 | Q9BVG4 | | 181 | GEKRADSGEEENT | 5.7484 | 6.3030 | 7.1362 | 5.9660 | 6.1000 | 6.1423 | 5.9829 | 6.3695 | 6.5438 | 6.3948 | 6.5218 | 7.0052 | 6.1217 | 6.2068 | 5.4564 | 6.0012 |
| PBK_S275 | PBK | Q96KB5-2 | | 24 | KSVLCSTPTINIP | 5.8687 | 6.9208 | 7.5697 | 6.8045 | 5.6428 | 7.3998 | 6.5475 | 7.4130 | 6.9254 | 8.4269 | 8.1731 | 7.9308 | 3.4330 | 3.4295 | 3.4408 | 4.0737 |
| PBK_S59 | PBK | Q96KB5-2 | | 59 | PRGLSHSPWAVKK | 7.6922 | 7.8939 | 7.1607 | 7.2098 | 6.8488 | 7.5503 | 7.7167 | 7.4324 | 7.7590 | 7.4528 | 8.0223 | 7.5112 | 2.1717 | 2.9721 | 1.7553 | 2.8509 |
| PBK_T24 | PBK | Q96KB5-2 | | 275 | DKTFDESDFDDEA | 6.2333 | 6.3022 | 7.3060 | 6.7493 | 7.0131 | 6.8351 | 6.7143 | 6.1952 | 6.7350 | 7.2721 | 6.4611 | 7.1900 | 4.4206 | 5.0155 | 4.6044 | 4.9530 |
| PBK_T24;S32 | PBK | Q96KB5-2 | 24;32 | | NA | 7.0706 | 7.2972 | 7.1081 | 7.7729 | 6.6486 | 7.1249 | 7.1564 | 6.5797 | 7.7035 | 6.6923 | 7.6825 | 7.2900 | 3.1417 | 3.8982 | 2.9349 | 3.8986 |
| PBRM1_S1453 | PBRM1 | Q86U86 | | 39 | RKRRRLSNLPTVD | 5.0107 | 5.1140 | 5.1164 | 5.5740 | 6.6365 | 5.8626 | 5.4499 | 5.9206 | 5.9080 | 5.7790 | 5.4881 | 5.8559 | 8.2317 | 8.5055 | 7.5343 | 8.0128 |
| PBRM1_S353 | PBRM1 | Q86U86 | | 948 | GLHRTYSQDCSFK | 3.6684 | 3.6726 | 3.7143 | 4.1050 | 5.0241 | 4.6600 | 4.9639 | 4.6426 | 3.7945 | 3.7274 | 3.5230 | 3.8188 | 13.0236 | 11.2227 | 13.4280 | 13.0110 |
| PBRM1_S353;S355 | PBRM1 | Q86U86 | | 1453 | QQQPSASPRAGTP | 6.0718 | 6.4579 | 6.4970 | 6.8816 | 5.9972 | 5.8834 | 6.5766 | 5.1305 | 5.8294 | 6.0270 | 5.6680 | 5.4916 | 6.1394 | 6.8319 | 7.3836 | 7.1333 |
| PBRM1_S355 | PBRM1 | Q86U86 | | 355 | LQYGSESEEDAAL | 6.3238 | 5.8858 | 6.4796 | 6.0802 | 6.5342 | 6.4503 | 5.7953 | 6.6713 | 6.5794 | 6.0815 | 7.3236 | 6.8510 | 5.8086 | 5.4411 | 6.0269 | 5.6675 |
| PBRM1_S39 | PBRM1 | Q86U86 | | 301 | SSLRMRTPSNLAA | 8.1585 | 7.2256 | 7.4393 | 8.7242 | 6.1832 | 7.7119 | 7.9311 | 6.6859 | 7.4612 | 7.0599 | 6.3764 | 7.1585 | 2.9382 | 3.6519 | 2.3752 | 2.9190 |
| PBRM1_S948 | PBRM1 | Q86U86 | | 353 | MALQYGSESEEDA | 6.2458 | 6.0362 | 6.3980 | 6.3745 | 6.2799 | 7.3101 | 6.4124 | 6.0778 | 5.9552 | 6.7878 | 5.8795 | 6.2640 | 6.1664 | 6.2701 | 5.7455 | 5.7969 |
| PBRM1_T301 | PBRM1 | Q86U86 | 353;355 | | NA | 6.0119 | 6.6959 | 6.7341 | 6.3572 | 6.4394 | 6.3468 | 6.4642 | 6.5145 | 6.9468 | 5.7957 | 6.7446 | 6.1045 | 6.1265 | 5.7763 | 5.9380 | 5.0035 |
| PBX2_S334 | PBX2 | P40425 | | 334 | GHSRTSSPTPPSS | 6.3589 | 5.4331 | 5.9478 | 5.8621 | 6.0974 | 5.6624 | 6.8970 | 6.3901 | 6.2883 | 6.1765 | 6.5620 | 6.6956 | 6.8753 | 6.2210 | 6.5598 | 5.9728 |
| PBX2_S395 | PBX2 | P40425 | | 395 | GGGQMYSPREMRA | 5.7801 | 6.0588 | 6.5763 | 7.8799 | 6.9593 | 5.8402 | 6.3757 | 6.2474 | 6.7367 | 6.4800 | 7.5971 | 6.0789 | 5.0079 | 5.9907 | 5.2265 | 5.1644 |
| PBXIP1_S43 | PBXIP1 | Q96AQ6 | | 43 | ALQAPHSPSKTDG | 6.6940 | 7.2466 | 6.3910 | 6.7351 | 6.7431 | 6.6439 | 6.9331 | 6.7684 | 6.5332 | 6.6778 | 6.4448 | 6.1322 | 4.9431 | 5.2887 | 4.8381 | 4.9869 |
| PCBP1_S173 | PCBP1 | Q15365 | | 190 | QPMPASSPVICAG | 5.9123 | 6.1506 | 6.4355 | 6.4982 | 6.4999 | 6.4566 | 6.5921 | 6.4736 | 6.4651 | 6.2812 | 6.3915 | 6.3088 | 5.8184 | 6.0658 | 5.7404 | 5.9101 |
| PCBP1_S190 | PCBP1 | Q15365 | | 173 | LETLSQSPQGRVM | 6.3606 | 6.5335 | 6.5095 | 6.4321 | 6.1053 | 6.2771 | 6.3659 | 6.7639 | 6.4286 | 6.1089 | 6.6085 | 6.5507 | 5.6007 | 6.0144 | 5.7658 | 5.5743 |
| PCBP1_S264 | PCBP1 | Q15365 | | 264 | AGIDSSSPEVKGY | 7.0799 | 7.2883 | 7.3190 | 7.4104 | 6.4744 | 7.2398 | 6.7649 | 6.8140 | 7.3524 | 7.2138 | 7.9697 | 7.4839 | 3.2938 | 3.8739 | 2.9811 | 3.4407 |
| PCBP1_T255 | PCBP1 | Q15365 | | 255 | AMMHGGTGFAGID | 7.1064 | 7.2253 | 7.6394 | 6.9147 | 6.2182 | 6.8475 | 6.8039 | 6.7832 | 7.1615 | 7.5836 | 7.8562 | 7.2291 | 3.5835 | 4.0259 | 3.3912 | 3.6304 |
| PCBP2_S173 | PCBP2 | Q15366-5 | | 189 | RPKPSSSPVIFAG | 5.8328 | 5.8176 | 6.8403 | 6.4054 | 6.2253 | 6.9639 | 6.4633 | 6.2803 | 6.3939 | 6.4051 | 6.6471 | 6.4838 | 5.7528 | 6.1917 | 5.4789 | 5.8179 |
| PCBP2_S189 | PCBP2 | Q15366-2 | | 189 | RPKPSSSPVIFAG | 6.7799 | 5.8551 | 6.6931 | 6.3340 | 6.7659 | 6.6113 | 6.2168 | 6.3529 | 6.6892 | 6.7443 | 6.6258 | 6.6828 | 5.2431 | 5.7680 | 5.0960 | 5.5418 |
| PCBP2_S189 | PCBP2 | F8VZX2 | | 189 | RPKPSSSPVIFAG | 6.2986 | 5.9511 | 6.8954 | 6.7983 | 6.5993 | 6.4510 | 6.6349 | 5.9508 | 6.5883 | 6.9771 | 7.0010 | 6.8677 | 5.0959 | 5.8420 | 4.8396 | 5.2090 |
| PCBP2_S189 | PCBP2 | Q15366-2 | | 173 | LETLSQSPPKGVT | 6.5516 | 6.2192 | 6.2210 | 6.0442 | 6.7341 | 6.6239 | 6.3507 | 6.0135 | 6.2918 | 7.5598 | 6.5338 | 6.3688 | 5.4581 | 6.1673 | 5.3364 | 5.5259 |
| PCBP2_S272 | PCBP2 | Q15366-2 | | 272 | SGIESSSPEVKGY | 7.4456 | 7.1480 | 7.6164 | 7.6887 | 6.4379 | 7.2388 | 6.1634 | 7.3908 | 6.9980 | 6.7742 | 8.5011 | 7.8006 | 3.2542 | 3.6187 | 2.8506 | 3.0729 |
| PCBP2_S366 | PCBP2 | Q15366-2 | | 366 | TGGMGSSxxxxxx | 7.1413 | 7.3620 | 7.2546 | 6.9613 | 6.3954 | 6.2336 | 6.7412 | 7.2320 | 6.2036 | 6.7135 | 6.3918 | 6.3465 | 4.5636 | 4.7124 | 4.7404 | 5.0066 |
| PCDH1_S351;T352 | PCDH1 | Q08174-2 | 351;352 | | NA | 5.7610 | 5.8190 | 6.5386 | 6.2543 | 6.1739 | 6.2704 | 6.0568 | 6.0039 | 6.1455 | 6.4539 | 6.4001 | 6.3328 | 6.5164 | 6.7795 | 6.2381 | 6.2559 |
| PCDH7_S664 | PCDH7 | H0YAH0 | | 925 | TGEYKPSPVNTLT | 5.9769 | 6.3553 | 6.1065 | 6.4616 | 6.1566 | 6.3528 | 6.6864 | 6.2952 | 6.1442 | 6.9736 | 7.0059 | 6.1892 | 6.1104 | 6.4123 | 4.8776 | 5.8953 |
| PCDH7_S690;S701 | PCDH7 | H0YAH0 | | 701 | VQLHPQSPTAGKK | 5.9922 | 5.7852 | 5.4954 | 6.9436 | 5.7663 | 6.2606 | 6.2888 | 6.3621 | 6.1051 | 5.9991 | 6.0615 | 6.8460 | 6.3220 | 6.7177 | 6.4884 | 6.5657 |
| PCDH7_S701 | PCDH7 | H0YAH0 | | 664 | NEKLSDSPSMGRY | 6.3092 | 6.9927 | 6.3427 | 6.6276 | 5.8219 | 6.0693 | 6.2731 | 6.1521 | 6.0358 | 6.5501 | 6.1959 | 6.0929 | 6.0997 | 6.3664 | 6.1785 | 5.8921 |
| PCDH7_S925 | PCDH7 | H0YAH0 | 690;701 | | NA | 6.8590 | 7.3242 | 6.0169 | 6.9020 | 6.3483 | 6.5910 | 6.8921 | 7.0311 | 7.1591 | 6.8526 | 6.3265 | 6.3284 | 4.6435 | 5.2310 | 4.7217 | 4.7727 |
| PCDHAC2_S950 | PCDHAC2 | Q9Y5I4 | | 950 | KFIIPGSPAIISI | 6.0163 | 7.3429 | 6.2912 | 6.7610 | 7.0119 | 7.0065 | 7.5964 | 6.4300 | 6.5127 | 7.0599 | 6.0430 | 5.6346 | 4.3481 | 4.7911 | 4.6684 | 6.4861 |
| PCDHB11_S792 | PCDHB11 | Q9Y5F2 | | 792 | NSTFRNSFGFNFx | 8.5190 | 8.7200 | 8.6960 | 6.9975 | 5.8004 | 6.2516 | 5.8963 | 6.3041 | 7.8016 | 7.1654 | 6.5566 | 8.4961 | 3.6312 | 3.5501 | 2.9230 | 2.6912 |
| PCDHGC3_S733 | PCDHGC3 | Q9UN70 | | 780 | KKPGAASPLASRQ | 6.5078 | 7.0604 | 6.1478 | 6.5218 | 6.9946 | 7.3811 | 7.2117 | 7.0223 | 6.3759 | 6.7677 | 6.6622 | 6.6436 | 4.1422 | 5.0108 | 4.3353 | 5.2147 |
| PCDHGC3_S780 | PCDHGC3 | Q9UN70 | | 733 | YRAPVSSLYRTPG | 7.3376 | 8.2016 | 8.1514 | 6.9486 | 6.4341 | 7.3737 | 6.8236 | 7.2568 | 6.5959 | 7.1413 | 7.0387 | 7.3836 | 2.8193 | 4.6781 | 2.8067 | 3.0090 |
| PCF11_S169 | PCF11 | O94913 | | 705 | VREEQRSPFNDRF | 6.1910 | 6.5215 | 6.3125 | 7.1162 | 6.0976 | 6.1298 | 6.4956 | 6.3270 | 6.2218 | 6.4688 | 6.4544 | 6.4405 | 5.5069 | 6.0867 | 5.9102 | 5.7196 |
| PCF11_S182 | PCF11 | E9PQ01 | | 800 | ERARRLSPISGSR | 6.4770 | 6.9859 | 6.0866 | 6.7470 | 6.1091 | 5.8673 | 6.3337 | 6.3153 | 6.2333 | 6.0742 | 5.8055 | 5.9574 | 5.9965 | 6.3776 | 6.3859 | 6.2475 |
| PCF11_S370;S372 | PCF11 | O94913 | | 851 | GFRFEGSPGLRFE | 6.8755 | 7.4065 | 6.9794 | 7.5447 | 6.0864 | 7.0927 | 7.2665 | 6.9697 | 7.1545 | 6.9146 | 6.9933 | 7.1803 | 3.4865 | 4.2662 | 3.4309 | 4.3524 |
| PCF11_S489;S494 | PCF11 | O94913 | | 494 | RSPIIHSPKRRDR | 6.6068 | 6.9782 | 5.9193 | 6.9690 | 6.4300 | 5.9243 | 5.8980 | 6.5717 | 6.2506 | 5.9728 | 6.2799 | 6.3192 | 5.5443 | 6.1897 | 6.3910 | 5.7553 |
| PCF11_S494 | PCF11 | O94913 | | 182 | PGTVVSSPSISTP | 5.5394 | 5.4338 | 5.4920 | 5.9527 | 5.9730 | 5.7764 | 6.0434 | 6.0181 | 5.8764 | 5.6959 | 5.6836 | 5.9341 | 7.1591 | 7.8520 | 7.6033 | 7.9669 |
| PCF11_S509;S511 | PCF11 | O94913 | | 777 | EGPNKLSPRIDGP | 5.8628 | 6.4300 | 6.1746 | 6.7939 | 6.0736 | 6.1096 | 6.2608 | 6.5840 | 6.1972 | 6.7279 | 6.7321 | 6.6185 | 5.6261 | 6.2577 | 5.7135 | 5.8377 |
| PCF11_S509;T515 | PCF11 | O94913 | | 1530 | KEERIDTPPACTE | 6.2443 | 6.7076 | 6.6393 | 7.3426 | 6.1821 | 5.7998 | 5.9961 | 5.7768 | 5.7825 | 5.7492 | 6.1854 | 6.4249 | 6.3883 | 6.7224 | 5.9138 | 6.1447 |
| PCF11_S511 | PCF11 | O94913 | | 511 | RRQRSMSPTSTPK | 6.5494 | 6.7443 | 5.9278 | 6.3586 | 6.3725 | 4.8868 | 5.3219 | 5.4947 | 6.1827 | 6.0221 | 5.2866 | 5.4711 | 7.3758 | 7.4181 | 7.8943 | 6.6932 |
| PCF11_S645 | PCF11 | O94913 | | 645 | QQQHRLSVDANLQ | 4.8485 | 5.2517 | 4.8956 | 5.1946 | 6.3732 | 5.9353 | 5.6077 | 5.1040 | 5.9899 | 5.2951 | 5.6127 | 6.1586 | 7.8620 | 8.1064 | 9.2143 | 8.5506 |
| PCF11_S705 | PCF11 | O94913 | | 169 | PKFLNKSPEEPST | 5.7348 | 6.3852 | 6.5170 | 6.9303 | 6.1511 | 5.9857 | 6.7739 | 6.5693 | 5.5013 | 6.4815 | 6.7227 | 6.3022 | 5.9219 | 6.5832 | 5.5380 | 5.9020 |
| PCF11_S777 | PCF11 | O94913 | | 785 | RIDGPPTPASLRF | 7.2368 | 7.4764 | 6.4291 | 7.8456 | 6.1263 | 6.5505 | 7.3754 | 6.5162 | 6.9898 | 6.1218 | 6.9686 | 7.1143 | 3.7622 | 4.8643 | 4.0084 | 4.6144 |
| PCF11_S800 | PCF11 | O94913 | | 1541 | TEESIATPSEIKT | 5.8963 | 5.2606 | 6.5752 | 6.9496 | 6.2460 | 7.0340 | 6.1493 | 5.9456 | 7.1492 | 4.1620 | 5.2832 | 6.4826 | 6.7706 | 7.0957 | 6.2892 | 6.7110 |
| PCF11_S851 | PCF11 | O94913 | 509;515 | | NA | 6.5408 | 6.4625 | 6.2004 | 6.5938 | 5.9544 | 5.6539 | 6.1450 | 6.1792 | 6.3924 | 6.0358 | 6.2217 | 5.7605 | 6.3184 | 6.3428 | 6.7811 | 6.4174 |
| PCF11_T1530 | PCF11 | O94913 | 489;494 | | NA | 5.4808 | 5.8878 | 6.5223 | 6.3888 | 5.9571 | 5.9193 | 6.0889 | 6.4407 | 6.7699 | 6.4066 | 6.3388 | 6.3800 | 6.1211 | 6.5142 | 6.2778 | 6.5060 |
| PCF11_T1541 | PCF11 | O94913 | 509;511 | | NA | 6.3701 | 6.6706 | 6.1046 | 6.1423 | 6.3790 | 6.5420 | 5.9027 | 5.7775 | 5.7976 | 6.2674 | 6.1899 | 5.5628 | 6.3560 | 6.5282 | 6.7221 | 6.6870 |
| PCF11_T785 | PCF11 | O94913 | 370;372 | | NA | 4.9517 | 4.8357 | 6.2758 | 6.7213 | 5.7799 | 6.6122 | 6.5487 | 5.5755 | 6.3181 | 6.4143 | 6.1293 | 6.4001 | 6.3371 | 7.4369 | 6.6803 | 6.9831 |
| PCGF6_T35 | PCGF6 | Q9BYE7 | | 35 | VSPPALTPAPAAG | 6.2382 | 6.4742 | 6.9319 | 5.7483 | 5.6885 | 5.7494 | 6.5462 | 6.3368 | 6.1650 | 5.6619 | 5.9190 | 6.1883 | 6.6438 | 6.4986 | 6.5684 | 6.6415 |
| PCID2_S45 | PCID2 | Q5JVF3-4 | | 45 | PRLQMASPEEKCQ | 6.3384 | 6.8475 | 5.8317 | 6.6097 | 6.7677 | 5.5516 | 5.9083 | 6.3328 | 7.1579 | 5.8269 | 6.3288 | 5.7335 | 6.2482 | 6.5842 | 5.5460 | 6.3871 |
| PCIF1_S143 | PCIF1 | Q9H4Z3 | | 144 | GQSVPSSPSIPGT | 4.9855 | 5.1654 | 5.4559 | 4.8367 | 5.5656 | 6.1594 | 5.3406 | 5.7627 | 5.6464 | 5.4115 | 5.5055 | 5.3540 | 8.2800 | 8.6267 | 9.3116 | 8.5927 |
| PCIF1_S143;T150 | PCIF1 | Q9H4Z3 | | 159 | LKMWGTSPEDKQQ | 5.8174 | 6.2774 | 5.9316 | 5.8688 | 5.4972 | 6.2218 | 6.3139 | 6.1230 | 6.2525 | 6.7716 | 6.3023 | 6.7555 | 5.7667 | 6.6529 | 6.6093 | 6.8383 |
| PCIF1_S144 | PCIF1 | Q9H4Z3 | | 150 | SPSIPGTPTLKMW | 4.9111 | 5.4303 | 6.6807 | 6.5420 | 6.2761 | 6.4057 | 5.3845 | 5.8188 | 6.6876 | 6.3403 | 6.1959 | 6.5741 | 6.9019 | 6.6658 | 6.5134 | 6.6716 |
| PCIF1_S150 | PCIF1 | Q9H4Z3 | | 143 | TGQSVPSSPSIPG | 4.8117 | 6.0595 | 5.7089 | 5.9437 | 5.8657 | 5.6245 | 6.1519 | 5.9365 | 5.9115 | 6.3827 | 6.0755 | 6.0386 | 7.4736 | 7.5870 | 7.3908 | 7.0379 |
| PCIF1_S159 | PCIF1 | Q9H4Z3 | 143;150 | | NA | 5.5711 | 5.9345 | 5.8388 | 5.3035 | 5.7959 | 6.1566 | 5.7189 | 6.2444 | 5.6918 | 5.3413 | 5.4137 | 5.5078 | 8.0171 | 8.2584 | 7.9236 | 7.2826 |
| PCK2_S290 | PCK2 | Q16822 | | 290 | LILGITSPAGKKR | 6.4701 | 6.2003 | 5.8866 | 7.0235 | 6.2900 | 6.1454 | 5.5967 | 6.4226 | 6.6668 | 6.8278 | 6.6357 | 5.8281 | 5.8994 | 6.4381 | 6.1208 | 5.5482 |
| PCLAF_S72 | PCLAF | Q15004 | | 72 | GEFFRLSPKDSEK | 8.5173 | 8.8605 | 8.5369 | 7.9573 | 6.7007 | 7.4064 | 6.9246 | 6.0257 | 7.3962 | 7.8543 | 8.4861 | 7.6135 | 1.5353 | 2.6972 | 1.5318 | 1.9561 |
| PCLO_S573 | PCLO | Q9Y6V0 | | 906 | EQSRRFSLNLGSI | 6.3766 | 4.3367 | 6.7238 | 5.5032 | 7.0029 | 5.9739 | 5.4407 | 7.0821 | 6.5349 | 7.4153 | 7.5486 | 7.1557 | 5.5060 | 6.8571 | 5.3976 | 5.1449 |
| PCLO_S906 | PCLO | Q9Y6V0 | | 573 | LQPPTVSPSAKQP | 7.4104 | 5.7980 | 6.5989 | 5.9475 | 6.1944 | 6.7572 | 6.7423 | 6.7380 | 6.9475 | 7.1083 | 6.2591 | 6.9383 | 4.9036 | 5.6155 | 4.6809 | 5.3601 |
| PCM1_S1187 | PCM1 | Q15154-5 | | 93 | RYMSQMSVPEQAE | 5.1085 | 5.2112 | 5.5548 | 5.3275 | 6.2837 | 6.0881 | 5.5021 | 6.2192 | 5.3075 | 5.0201 | 5.3467 | 5.3572 | 8.8122 | 8.6261 | 8.4009 | 7.8343 |
| PCM1_S1188 | PCM1 | Q15154-5 | | 65 | RVTNDISPESSPG | 6.2551 | 6.4137 | 6.1325 | 5.8796 | 6.1113 | 6.1348 | 5.8864 | 6.4392 | 6.1612 | 6.3944 | 6.3420 | 6.4873 | 6.2055 | 6.4745 | 6.3960 | 6.2864 |
| PCM1_S1257 | PCM1 | Q15154-5 | | 1290 | AQASLASKDKTPK | 8.2619 | 7.5967 | 8.1416 | 7.1984 | 5.2007 | 6.3880 | 4.4974 | 6.2416 | 7.7417 | 6.3473 | 7.8065 | 8.3988 | 4.7431 | 5.1600 | 2.9032 | 3.3732 |
| PCM1_S1290 | PCM1 | Q15154-5 | | 861 | GLAETASPVAVSL | 6.8294 | 7.4935 | 7.6241 | 6.7801 | 6.4410 | 7.0794 | 7.0941 | 6.5285 | 6.7278 | 7.5087 | 7.0726 | 7.0925 | 3.7291 | 3.9915 | 3.6426 | 4.3651 |
| PCM1_S1722 | PCM1 | Q15154-5 | | 1883 | SAAHKESPPTVDS | 5.8793 | 6.1290 | 6.9313 | 6.8035 | 6.0505 | 6.7539 | 6.6217 | 6.5200 | 6.3790 | 6.3461 | 6.1924 | 6.5420 | 5.5291 | 5.9709 | 5.3252 | 6.0262 |
| PCM1_S1757;S1760 | PCM1 | Q15154-5 | | 1187 | KPFESSSSIGAEK | 5.8607 | 5.4120 | 6.1384 | 6.1517 | 6.4672 | 6.4014 | 6.0724 | 5.8511 | 6.0522 | 5.6786 | 5.8022 | 5.7637 | 7.3631 | 7.2225 | 6.8716 | 6.8911 |
| PCM1_S1757;S1760;S17 | PCM1 | Q15154-5 | | 1722 | ILEDHGSPAGEID | 6.5927 | 6.4450 | 7.0828 | 6.6538 | 6.1536 | 6.0710 | 6.5458 | 6.2620 | 6.6970 | 6.9170 | 6.7989 | 6.8887 | 5.1650 | 5.7014 | 5.1983 | 4.8269 |
| PCM1_S1757;S1768 | PCM1 | Q15154-5 | | 384 | SQSRKPSASERLP | 5.3309 | 4.3606 | 5.4132 | 5.3183 | 6.7000 | 5.5087 | 5.4098 | 6.5348 | 7.2354 | 7.2400 | 7.3015 | 7.5320 | 6.8181 | 7.3843 | 6.0980 | 5.8143 |
| PCM1_S1757;S1768;S17 | PCM1 | Q15154-5 | | 804 | VNQHETSTSKSVF | 5.9657 | 6.2816 | 6.2270 | 6.9254 | 7.1235 | 7.2857 | 6.3667 | 7.2810 | 6.3468 | 6.6422 | 6.9707 | 7.0231 | 4.7199 | 5.3578 | 4.7461 | 4.7368 |
| PCM1_S1760 | PCM1 | Q15154-5 | | 977 | TRQQNISMQRQEN | 6.9586 | 7.1751 | 6.7576 | 6.9198 | 6.5195 | 6.8165 | 6.6382 | 7.0423 | 6.6474 | 6.7565 | 6.9126 | 7.4350 | 4.1939 | 4.7714 | 3.8721 | 4.5834 |
| PCM1_S1883 | PCM1 | Q15154-5 | | 1557 | DGAGAETPVIENR | 6.7393 | 6.5595 | 6.2329 | 6.3044 | 5.6921 | 6.8572 | 6.5408 | 6.5090 | 6.7923 | 5.9153 | 6.8193 | 6.2665 | 5.5586 | 5.6692 | 5.6199 | 5.9237 |
| PCM1_S1950 | PCM1 | Q15154-5 | | 2015 | ETVGAQSIxxxxx | 5.4152 | 5.7518 | 6.1795 | 6.1465 | 5.5175 | 6.0437 | 6.8608 | 6.5656 | 5.8193 | 5.7094 | 6.5854 | 6.6001 | 6.4732 | 7.5171 | 6.2865 | 6.5284 |
| PCM1_S1969 | PCM1 | Q15154-5 | | 1760 | NVRSDISDQEEDE | 6.3605 | 6.2665 | 6.9083 | 7.2447 | 5.5528 | 5.9727 | 6.4988 | 6.0693 | 7.2060 | 8.0113 | 7.7125 | 7.4625 | 4.5891 | 5.3506 | 4.4191 | 4.3753 |
| PCM1_S2015 | PCM1 | Q15154-5 | | 1969 | LKLTIYSEADLRK | 3.5194 | 3.6305 | 3.1024 | 3.3016 | 3.8048 | 2.9181 | 3.2855 | 2.6169 | 2.5781 | 2.0249 | 2.3196 | 1.9418 | 16.5352 | 17.9942 | 17.1222 | 13.3047 |
| PCM1_S372 | PCM1 | Q15154-5 | | 808 | ETSTSKSVFEPED | 6.0923 | 6.1820 | 6.7826 | 6.2539 | 6.1657 | 5.5159 | 6.3630 | 6.2523 | 7.3108 | 6.6310 | 6.7789 | 6.0073 | 6.1794 | 5.8725 | 6.0121 | 5.6003 |
| PCM1_S384 | PCM1 | Q15154-5 | | 960 | KNNCPFSADENYR | 6.0977 | 6.2451 | 6.0605 | 6.5114 | 6.0699 | 6.2176 | 5.7846 | 5.8578 | 5.9322 | 6.0067 | 6.2550 | 6.2005 | 7.1290 | 6.2843 | 6.6406 | 6.7069 |
| PCM1_S644 | PCM1 | Q15154-5 | | 1257 | RQFDEESLESFSS | 5.6522 | 6.7424 | 6.4182 | 6.9871 | 6.0260 | 7.4055 | 6.1040 | 6.5629 | 6.5147 | 7.6467 | 7.0516 | 7.7240 | 4.4479 | 4.9096 | 4.5429 | 5.2642 |
| PCM1_S65 | PCM1 | Q15154-5 | | 816 | FEPEDSSIVDNEL | 6.0495 | 6.4681 | 6.4750 | 7.1063 | 6.6031 | 5.2614 | 5.9168 | 6.6237 | 6.1856 | 6.1444 | 6.1147 | 7.6073 | 4.9858 | 7.0674 | 5.9809 | 5.4101 |
| PCM1_S804 | PCM1 | Q15154-5 | | 372 | RQAESLSLTREVS | 6.4404 | 6.6644 | 6.3334 | 6.9059 | 6.6192 | 6.4005 | 6.2573 | 6.3958 | 6.1958 | 6.4726 | 6.6398 | 6.8326 | 5.3858 | 5.6149 | 5.3780 | 5.4635 |
| PCM1_S808 | PCM1 | Q15154-5 | | 1950 | GNISQKSDEEDFV | 6.6959 | 6.5204 | 6.5686 | 5.7148 | 6.5851 | 6.2680 | 6.2115 | 6.5307 | 6.3911 | 7.7806 | 6.6425 | 7.2121 | 4.5214 | 6.0627 | 4.5860 | 5.7087 |
| PCM1_S816 | PCM1 | Q15154-5 | | 991 | RWVSELSYVEEKE | 6.6802 | 6.8730 | 6.7337 | 7.6384 | 7.5656 | 6.4602 | 6.8550 | 6.1455 | 6.5225 | 5.5144 | 6.4526 | 5.3723 | 5.3920 | 5.6526 | 5.2228 | 4.9191 |
| PCM1_S861 | PCM1 | Q15154-5 | | 1188 | PFESSSSIGAEKP | 5.0235 | 4.8390 | 5.9762 | 5.2203 | 5.8058 | 7.0113 | 6.3977 | 5.6422 | 5.6954 | 6.1931 | 6.3879 | 5.9890 | 8.0770 | 7.4459 | 6.9887 | 7.3070 |
| PCM1_S93 | PCM1 | Q15154-5 | | 644 | LSSHRSSLVDEHP | 6.6352 | 5.1673 | 7.5971 | 6.6093 | 7.5317 | 5.4153 | 6.0093 | 6.0817 | 7.5383 | 8.1068 | 8.3520 | 8.0602 | 4.2040 | 5.3292 | 3.9913 | 3.3715 |
| PCM1_S960 | PCM1 | Q15154-5 | 61;69 | | NA | 7.1075 | 7.0157 | 7.4712 | 6.4177 | 6.1763 | 6.4391 | 6.4525 | 7.2971 | 6.9263 | 6.8941 | 7.0007 | 7.4450 | 4.2517 | 4.6875 | 3.7744 | 4.6431 |
| PCM1_S977 | PCM1 | Q15154-5 | 1757;1760;17 | | NA | 5.6064 | 6.3177 | 6.5098 | 5.9523 | 6.1891 | 5.7310 | 5.7921 | 6.5931 | 5.6214 | 6.3619 | 5.7898 | 5.9567 | 7.1374 | 6.8247 | 6.6048 | 7.0118 |
| PCM1_S991 | PCM1 | Q15154-5 | 1757;1768;17 | | NA | 5.6408 | 6.1281 | 6.1325 | 5.7050 | 5.6866 | 7.2837 | 5.8418 | 5.8837 | 5.9130 | 5.8085 | 5.9932 | 6.2080 | 7.3037 | 7.1598 | 6.7920 | 6.5197 |
| PCM1_T1557 | PCM1 | Q15154-5 | 1757;1768 | | NA | 6.7205 | 6.7107 | 6.8703 | 6.2127 | 5.7994 | 6.1823 | 6.7439 | 6.5238 | 6.5444 | 7.0067 | 6.5341 | 6.2915 | 5.3553 | 5.3810 | 5.2615 | 5.8619 |
| PCM1_T61;S69 | PCM1 | Q15154-5 | 1757;1760 | | NA | 5.9665 | 6.9806 | 7.0698 | 6.1467 | 5.9607 | 6.4374 | 5.6893 | 5.9812 | 5.9903 | 6.4469 | 7.2290 | 6.4683 | 5.5741 | 6.1006 | 5.8385 | 6.1199 |
| PCMTD1_S302 | PCMTD1 | Q96MG8 | | 302 | IPQPLDSEEDEKM | 6.2043 | 5.5723 | 5.1982 | 6.3455 | 6.7310 | 6.1850 | 6.8801 | 6.0538 | 5.1316 | 6.9216 | 6.8385 | 6.6056 | 5.9234 | 6.1735 | 6.4924 | 6.7432 |
| PCNP_S119 | PCNP | Q8WW12 | | 119 | FNEDEDSEPEEMP | 5.6162 | 6.0296 | 6.3128 | 6.1018 | 6.2492 | 6.1843 | 5.7688 | 6.1697 | 6.3795 | 6.3644 | 6.6755 | 6.4954 | 6.0917 | 6.7924 | 6.1351 | 6.6335 |
| PCNP_S87 | PCNP | Q8WW12 | | 87 | KKASAISIKLGSS | 6.5667 | 5.3792 | 7.5462 | 5.4452 | 4.1462 | 6.4018 | 5.7565 | 5.8522 | 8.6452 | 8.9972 | 10.0947 | 10.2092 | 3.5657 | 5.4786 | 2.4533 | 3.4621 |
| PCNP_T139 | PCNP | Q8WW12 | | 139 | KNIGRDTPTSAGP | 8.1432 | 7.9299 | 7.1808 | 6.3021 | 4.7809 | 7.9177 | 7.8169 | 6.8827 | 7.8665 | 7.8719 | 8.0229 | 7.7466 | 2.7060 | 3.4171 | 1.8944 | 3.5204 |
| PCNT_S1630 | PCNT | O95613 | | 2317 | VEDFITTSFDSQE | 5.5502 | 5.8933 | 5.7577 | 6.2523 | 6.0761 | 5.8572 | 6.0118 | 6.1936 | 5.8576 | 6.1362 | 6.2324 | 6.0299 | 6.9083 | 7.1826 | 7.1964 | 6.8644 |
| PCNT_S1632 | PCNT | O95613 | | 2177 | PDEMPDSPIQEKS | 6.1887 | 5.9643 | 6.5811 | 6.2163 | 6.1846 | 6.3082 | 6.1819 | 6.2521 | 6.2812 | 6.1976 | 6.4901 | 6.3657 | 6.2480 | 6.3104 | 6.0461 | 6.1839 |
| PCNT_S1653 | PCNT | O95613 | | 1653 | ALLRRESEVLDLK | 4.9605 | 5.0292 | 4.9841 | 5.3060 | 5.8536 | 5.6925 | 5.5288 | 5.5162 | 6.9966 | 6.7551 | 6.8310 | 6.4405 | 7.8199 | 7.4113 | 7.5035 | 7.3713 |
| PCNT_S2044 | PCNT | O95613 | | 2352 | GFGARLSPGSGGP | 6.8093 | 6.8889 | 6.1298 | 7.3191 | 5.9646 | 6.5985 | 5.7649 | 6.8806 | 6.6435 | 7.0793 | 6.4471 | 6.3011 | 5.2051 | 5.2863 | 5.4449 | 5.2369 |
| PCNT_S2177 | PCNT | O95613 | | 2327 | SQETLSSPPPGLE | 5.3200 | 6.0028 | 5.9486 | 6.0013 | 6.0126 | 6.0443 | 5.6002 | 6.3158 | 6.5715 | 6.1786 | 6.2468 | 6.2659 | 6.8280 | 7.1393 | 6.6919 | 6.8325 |
| PCNT_S2192 | PCNT | O95613 | | 3242 | RARQPQSPPRTRE | 6.1680 | 6.4031 | 6.0051 | 5.9471 | 5.9306 | 6.0213 | 5.9829 | 6.1691 | 6.3789 | 6.0457 | 5.9906 | 6.2943 | 6.5823 | 6.9102 | 6.6716 | 6.4992 |
| PCNT_S2214 | PCNT | O95613 | | 682 | EHKVQLSLLQTEL | 8.0322 | 8.2872 | 7.1997 | 9.2148 | 7.7473 | 6.6064 | 7.2382 | 6.8573 | 7.5734 | 6.0099 | 6.9003 | 5.4102 | 2.6895 | 3.9925 | 3.2171 | 3.0242 |
| PCNT_S2327 | PCNT | O95613 | | 3274 | RLAAAASPHSGGR | 5.7611 | 6.0510 | 5.8977 | 6.4828 | 6.2633 | 6.3537 | 6.3127 | 6.0072 | 6.8423 | 6.3206 | 6.3959 | 6.1768 | 6.1202 | 6.4231 | 6.2572 | 6.3346 |
| PCNT_S2352 | PCNT | O95613 | | 2155 | ACDANTTPGGVTD | 6.2686 | 6.8485 | 6.5340 | 7.3520 | 6.3850 | 6.5085 | 6.1193 | 5.9525 | 6.7074 | 6.9059 | 6.9425 | 6.6930 | 4.8146 | 5.2060 | 5.3040 | 5.4584 |
| PCNT_S3242 | PCNT | O95613 | | 1630 | RCPEPPSGSPPEG | 6.1337 | 6.4353 | 6.7907 | 6.4784 | 5.8282 | 6.0431 | 6.8343 | 6.0598 | 6.1656 | 6.6827 | 6.0705 | 6.6099 | 5.9944 | 5.7246 | 5.7398 | 6.4090 |
| PCNT_S3274 | PCNT | O95613 | | 2044 | KNEMRLSLEDGGK | 6.9731 | 5.2812 | 6.8052 | 8.1154 | 6.3278 | 7.1740 | 7.7607 | 7.1990 | 7.3291 | 7.3352 | 7.2394 | 6.8450 | 3.9602 | 4.3621 | 3.1740 | 4.1187 |
| PCNT_S3324 | PCNT | O95613 | | 2214 | EAGPRKSPVGMLD | 7.2565 | 9.6151 | 8.0187 | 6.7188 | 5.6317 | 6.1928 | 5.5412 | 7.5564 | 7.5379 | 6.9232 | 5.4876 | 5.0873 | 4.9784 | 3.9532 | 5.2950 | 4.2062 |
| PCNT_S682 | PCNT | O95613 | | 2192 | QDMSLSSPTSVLG | 6.0289 | 6.2686 | 6.6389 | 6.7559 | 6.5321 | 5.4735 | 7.3363 | 6.6434 | 6.6093 | 6.0687 | 6.3822 | 6.0233 | 5.8290 | 5.2986 | 6.1062 | 6.0051 |
| PCNT_T2155 | PCNT | O95613 | | 3282 | HSGGRATPSPNSR | 6.9953 | 6.4418 | 6.2700 | 6.0678 | 5.7486 | 5.5535 | 6.8578 | 6.0894 | 6.1092 | 6.0038 | 5.2779 | 5.6496 | 7.3273 | 7.0425 | 6.3096 | 6.2560 |
| PCNT_T2317 | PCNT | O95613 | | 3324 | GGVLPDSTSKKSC | 6.7868 | 7.6585 | 6.8205 | 8.4866 | 6.3555 | 7.5630 | 7.7849 | 6.4609 | 7.2616 | 6.2010 | 6.9420 | 6.7601 | 3.5479 | 4.1197 | 3.1553 | 4.0958 |
| PCNT_T3282 | PCNT | O95613 | | 1632 | PEPPSGSPPEGPE | 6.2104 | 6.8405 | 6.7161 | 6.7193 | 5.6934 | 5.9543 | 5.4583 | 6.0642 | 6.1353 | 6.4608 | 6.5788 | 6.2872 | 6.4374 | 5.6345 | 7.0053 | 5.8041 |
| PCNX1_S117 | PCNX1 | Q96RV3 | | 117 | STRRKDSNGPSDP | 5.6378 | 6.3826 | 7.4387 | 5.2868 | 5.0517 | 5.3417 | 4.9510 | 4.4748 | 7.9118 | 8.2690 | 8.6597 | 8.5132 | 5.6103 | 7.5089 | 4.1726 | 4.7892 |
| PCNX1_S217 | PCNX1 | Q96RV3 | | 217 | RLVSNDSFISIQP | 6.2706 | 5.8616 | 6.4536 | 6.7155 | 6.3405 | 6.3540 | 6.6894 | 6.0374 | 6.9505 | 5.8822 | 7.3281 | 6.4275 | 5.6490 | 5.4517 | 5.6759 | 5.9127 |
| PCNX3_S136 | PCNX3 | Q9H6A9 | | 246 | LLKGSLSQELSKS | 5.4429 | 5.8383 | 6.0154 | 6.6633 | 5.6773 | 6.2074 | 6.3008 | 6.1433 | 6.1346 | 5.9456 | 6.2582 | 6.0444 | 6.2878 | 6.9987 | 6.4682 | 7.5739 |
| PCNX3_S136;S139 | PCNX3 | Q9H6A9 | | 1909 | PGPPPASPIPTEG | 5.4329 | 6.0271 | 6.4675 | 6.7360 | 5.8789 | 5.7689 | 6.5605 | 6.4082 | 6.6393 | 6.1349 | 6.3580 | 6.4392 | 5.9862 | 6.5067 | 6.1085 | 6.5473 |

| | | | | | | | | | | | | | | | | | | | | | |
|---|---|---|---|---|---|---|---|---|---|---|---|---|---|---|---|---|---|---|---|---|---|
| PCNX3_S1882 | PCNX3 | Q9H6A9 | | 505 | THARVLSMDGAGG | 6.7090 | 6.7367 | 7.2409 | 7.2241 | 5.5122 | 6.1639 | 5.7349 | 6.3817 | 6.0784 | 5.9731 | 6.0177 | 6.6122 | 5.7198 | 5.8997 | 5.6903 | 6.3053 |
| PCNX3_S1909 | PCNX3 | Q9H6A9 | | 392 | LALLRPSKRQPPL | 9.0239 | 6.8712 | 7.5734 | 6.2391 | 4.9873 | 5.8070 | 6.8758 | 5.8443 | 7.6600 | 6.9266 | 8.2715 | 8.1560 | 3.8116 | 5.8266 | 2.6258 | 3.4999 |
| PCNX3_S246 | PCNX3 | Q9H6A9 | | 1882 | GWGPRSSLSGSGD | 5.5461 | 6.2949 | 5.8301 | 5.8895 | 6.1451 | 6.4446 | 6.3568 | 5.6132 | 7.7924 | 6.4524 | 7.3431 | 7.0746 | 4.9987 | 6.1399 | 5.5538 | 6.5247 |
| PCNX3_S392 | PCNX3 | Q9H6A9 | | 136 | PPVRCSSQHSVFG | 7.1213 | 7.3949 | 6.9046 | 4.6354 | 3.5293 | 8.3756 | 7.6800 | 9.3392 | 4.3275 | 5.3937 | 5.7985 | 7.1954 | 5.6047 | 6.6081 | 4.7608 | 5.3310 |
| PCNX3_S505 | PCNX3 | Q9H6A9 | 136;139 | | NA | 8.0556 | 7.2426 | 8.3674 | 8.3703 | 7.5579 | 7.2892 | 6.2593 | 3.9595 | 5.1152 | 5.7366 | 5.6559 | 6.2092 | 3.4313 | 5.9167 | 4.6596 | 6.1736 |
| PCYT1A_S233 | PCYT1A | P49585 | | 315 | RMLQAISPKQSPS | 5.7354 | 5.4879 | 6.1322 | 5.1428 | 5.9634 | 5.8651 | 5.9121 | 6.1213 | 6.3705 | 5.8890 | 6.0985 | 5.9356 | 6.7917 | 7.7982 | 7.5300 | 7.2262 |
| PCYT1A_S315 | PCYT1A | P49585 | | 343 | PFSGKTSPPCSPA | 5.5016 | 5.7745 | 6.4147 | 5.3609 | 5.6548 | 5.8822 | 5.9411 | 5.9623 | 6.4044 | 5.9892 | 6.0816 | 5.9238 | 7.1836 | 7.6151 | 7.6879 | 6.6224 |
| PCYT1A_S323 | PCYT1A | P49585 | | 233 | AKELNVSFINEKK | 7.8545 | 7.2317 | 7.1858 | 7.7322 | 7.0735 | 6.5886 | 7.1909 | 6.5142 | 7.8105 | 6.2678 | 7.5411 | 7.1996 | 2.9285 | 4.2792 | 2.9458 | 3.6559 |
| PCYT1A_S329;S331 | PCYT1A | P49585 | | 362 | AAAYDISEDEEDx | 5.8214 | 5.9905 | 6.3307 | 5.4485 | 5.3293 | 6.0978 | 5.4564 | 6.0587 | 6.3489 | 6.2446 | 6.7187 | 6.2117 | 6.8347 | 7.1832 | 7.2106 | 6.7144 |
| PCYT1A_S329;S333 | PCYT1A | P49585 | | 331 | TRERSPSPSFRWP | 5.0125 | 4.7493 | 5.0636 | 4.1851 | 4.5582 | 4.8632 | 4.0738 | 5.0870 | 5.5908 | 5.1567 | 4.8238 | 5.2833 | 9.9431 | 10.4466 | 11.4585 | 9.7046 |
| PCYT1A_S331 | PCYT1A | P49585 | | 323 | KQSPSSSPTRERS | 5.0359 | 4.9235 | 5.4382 | 4.8754 | 5.6994 | 5.5642 | 5.5596 | 6.2156 | 5.5750 | 5.5584 | 6.3109 | 5.8334 | 8.3982 | 8.3018 | 9.1840 | 7.5265 |
| PCYT1A_S343 | PCYT1A | P49585 | 343;347 | | NA | 5.8131 | 5.9805 | 7.0693 | 5.5067 | 5.8954 | 6.3070 | 5.6490 | 5.7945 | 6.4492 | 6.4243 | 6.3323 | 6.3645 | 6.7778 | 6.8396 | 6.4143 | 6.3826 |
| PCYT1A_S343;S347 | PCYT1A | P49585 | 329;331 | | NA | 5.9796 | 6.0944 | 6.0639 | 5.3297 | 6.0352 | 6.5075 | 5.6385 | 6.7048 | 5.9509 | 6.4693 | 6.4281 | 6.1934 | 6.1029 | 6.9978 | 7.0931 | 6.4108 |
| PCYT1A_S362 | PCYT1A | P49585 | 329;333 | | NA | 5.6776 | 5.8716 | 6.2616 | 5.4048 | 5.9043 | 6.0195 | 5.5482 | 6.2487 | 5.9964 | 6.2391 | 6.2648 | 6.2873 | 6.6352 | 7.4440 | 7.3545 | 6.8425 |
| PDAP1_S170 | PDAP1 | Q13442 | | 178 | KRMQSLSLNKxxx | 7.2719 | 7.0988 | 7.9331 | 6.6133 | 5.9505 | 6.3505 | 5.8488 | 6.9883 | 7.3991 | 7.2355 | 7.1845 | 7.5805 | 4.3944 | 4.7287 | 3.5304 | 3.8919 |
| PDAP1_S178 | PDAP1 | Q13442 | | 170 | KDDATLSGKRMQS | 7.9870 | 7.3176 | 6.8634 | 7.3821 | 5.7767 | 6.5260 | 7.1143 | 6.9602 | 6.8876 | 7.3429 | 6.1348 | 7.5816 | 3.8150 | 4.3979 | 3.4484 | 4.4648 |
| PDAP1_S19 | PDAP1 | Q13442 | | 19 | RARQYTSPEEIDA | 5.5973 | 7.3065 | 6.1788 | 7.1750 | 6.4670 | 6.8290 | 6.8182 | 6.4353 | 6.4694 | 6.8797 | 6.6199 | 6.8906 | 5.1202 | 5.8429 | 4.5055 | 4.8647 |
| PDAP1_S60;S63 | PDAP1 | Q13442 | 60;63 | | NA | 5.3304 | 5.8677 | 5.9670 | 4.4606 | 5.9805 | 5.4587 | 6.2632 | 6.5276 | 6.3467 | 6.7094 | 6.6881 | 7.1976 | 5.9364 | 7.1826 | 5.8813 | 6.2020 |
| PDCD11_S1360 | PDCD11 | Q14690 | | 1360 | SHVSQHSPSKKAL | 6.5765 | 8.1774 | 5.6974 | 8.2695 | 6.2762 | 6.3605 | 6.0954 | 7.3395 | 5.8437 | 5.9825 | 6.4663 | 6.7191 | 4.3003 | 5.5812 | 5.0176 | 5.2972 |
| PDCD2L_S139 | PDCD2L | Q9BRP1 | | 139 | DTEEGPSPQFTLD | 6.3016 | 6.0099 | 7.1844 | 6.0994 | 6.2620 | 6.1085 | 5.9272 | 6.2840 | 6.0378 | 6.4984 | 6.6638 | 7.5016 | 6.0300 | 5.9518 | 5.1472 | 5.9925 |
| PDCD2L_S20 | PDCD2L | Q9BRP1 | | 20 | DAPVHGSPTGPGA | 6.3103 | 5.8600 | 6.9066 | 5.9533 | 7.4495 | 6.4990 | 6.1860 | 7.0502 | 5.9314 | 6.3633 | 7.1494 | 6.9980 | 5.2700 | 5.7505 | 5.4198 | 4.9028 |
| PDCD4_S94 | PDCD4 | Q53EL6 | | 94 | GLTVPTSPKGRLL | 6.3518 | 6.4152 | 6.2871 | 5.3149 | 6.6346 | 6.7763 | 6.4244 | 6.6547 | 6.4663 | 7.0045 | 6.4728 | 6.0941 | 5.5558 | 6.3986 | 5.7273 | 5.4216 |
| PDCD4_T126 | PDCD4 | Q53EL6 | | 126 | GKGVWGTPGQVYD | 7.0975 | 7.2934 | 6.7942 | 6.3536 | 6.1338 | 6.8898 | 7.4039 | 6.6455 | 7.2269 | 7.2559 | 7.0931 | 7.0281 | 4.0001 | 4.7918 | 3.6185 | 4.3740 |
| PDCD5_S119 | PDCD5 | O14737 | | 119 | RRKVMDSDEDDDY | 6.1969 | 5.5182 | 6.6965 | 5.9352 | 6.3644 | 5.6865 | 5.8867 | 6.8109 | 5.6843 | 6.6667 | 6.3750 | 6.9107 | 5.8337 | 7.0108 | 6.0983 | 6.3253 |
| PDCD6IP_S735 | PDCD6IP | Q8WUM4-2 | | 735 | TPAYQSSPAGGHA | 6.0821 | 7.1402 | 6.6491 | 6.8312 | 6.1209 | 6.7367 | 6.6921 | 7.1936 | 6.6319 | 6.8972 | 6.8995 | 6.6294 | 4.9575 | 5.3361 | 4.5617 | 4.6409 |
| PDCD6IP_S735;T743 | PDCD6IP | Q8WUM4-2 | | 484 | KERWQRTPSNELY | 5.9318 | 6.1727 | 7.1073 | 6.4562 | 5.8708 | 6.4935 | 5.9685 | 6.7484 | 7.3349 | 7.4063 | 6.7778 | 7.1780 | 5.0374 | 5.8112 | 4.7901 | 4.9150 |
| PDCD6IP_T484 | PDCD6IP | Q8WUM4-2 | 735;743 | | NA | 6.1677 | 7.5284 | 6.4464 | 6.3374 | 5.8110 | 6.6998 | 6.8444 | 5.7747 | 5.8276 | 7.0025 | 8.2040 | 7.3132 | 4.7058 | 5.6375 | 4.3624 | 5.3372 |
| PDCL3_S43 | PDCL3 | Q9H2J4 | | 43 | QRILQQSVVKTYE | 9.0506 | 7.4419 | 7.2144 | 8.0655 | 6.9262 | 6.3565 | 7.3821 | 6.0877 | 7.7781 | 5.7868 | 6.2655 | 5.8323 | 4.0020 | 4.5093 | 3.5351 | 3.7660 |
| PDE10A_S777 | PDE10A | Q9Y233-2 | | 777 | TATWISSPSVAQK | 6.3184 | 6.5133 | 6.3302 | 7.1288 | 6.2830 | 6.8783 | 6.7782 | 6.8893 | 6.7206 | 7.0059 | 6.6498 | 6.7218 | 4.8235 | 5.5042 | 4.3356 | 5.1191 |
| PDE12_S217 | PDE12 | Q6L8Q7 | | 220 | SSLSPSSPSSSWT | 5.3822 | 6.1330 | 6.4775 | 5.8200 | 5.9092 | 6.1017 | 6.2688 | 5.5803 | 6.2620 | 6.2683 | 6.5278 | 6.7196 | 6.5797 | 7.1546 | 6.1252 | 6.6899 |
| PDE12_S220 | PDE12 | Q6L8Q7 | | 217 | GVPSSLSPSSPSS | 5.9296 | 6.2590 | 6.9176 | 6.3827 | 6.1813 | 6.1823 | 6.5152 | 6.4560 | 6.3700 | 6.3149 | 6.5649 | 6.6919 | 5.8741 | 5.6915 | 5.5720 | 6.0970 |
| PDE3B_S279 | PDE3B | Q13370 | | 318 | KIFRRPSLPCISR | 4.5583 | 4.2738 | 4.4102 | 4.8490 | 5.9410 | 5.0616 | 4.6720 | 5.0358 | 5.9583 | 5.9143 | 5.7109 | 5.9051 | 9.7257 | 8.7271 | 9.7653 | 9.4915 |
| PDE3B_S318 | PDE3B | Q13370 | | 65 | LRPPPASPQQPRR | 5.9074 | 6.7087 | 7.2137 | 6.3697 | 5.6931 | 5.8364 | 6.2788 | 5.7496 | 6.7115 | 6.8830 | 6.7783 | 6.8492 | 5.4389 | 6.2540 | 5.7179 | 5.6098 |
| PDE3B_S442 | PDE3B | Q13370 | | 279 | PLHPRLSSAAEEK | 4.9249 | 5.2361 | 5.0688 | 5.2449 | 5.9344 | 5.5334 | 5.6068 | 5.7589 | 5.1931 | 5.1712 | 4.7804 | 5.1154 | 9.1807 | 8.3590 | 9.8653 | 9.0267 |
| PDE3B_S65 | PDE3B | Q13370 | | 442 | PQLRRSSGTSGLL | 5.1611 | 4.0029 | 5.7214 | 5.2412 | 5.7827 | 5.5390 | 5.3140 | 5.5112 | 7.8128 | 6.8755 | 7.8635 | 7.6525 | 6.5687 | 8.4410 | 6.5687 | 5.9438 |
| PDE4A_S302 | PDE4A | P27815 | | 346 | KLMHSNSLNNSNI | 5.9096 | 5.9910 | 6.6961 | 6.1683 | 6.0282 | 5.8599 | 6.1366 | 6.1165 | 6.9920 | 7.5605 | 7.3922 | 6.6898 | 5.3625 | 6.5050 | 5.4191 | 5.1728 |
| PDE4A_S346 | PDE4A | P27815 | | 302 | NEVEIPSPTMKER | 7.4177 | 6.7179 | 7.3601 | 7.3860 | 7.0701 | 7.1876 | 6.7097 | 6.4232 | 6.5324 | 6.7579 | 5.1834 | 6.5794 | 4.3146 | 4.7408 | 4.8535 | 4.7655 |
| PDE4DIP_S195 | PDE4DIP | Q5VU43-2 | | 195 | KVARSISCGPSSR | 6.3455 | 6.2645 | 6.9032 | 5.9773 | 5.4258 | 6.2631 | 5.5132 | 5.6064 | 6.8448 | 6.4663 | 6.3920 | 7.0844 | 6.7216 | 6.9615 | 5.5669 | 5.6636 |
| PDE8A_S386 | PDE8A | O60658 | | 457 | DGLRRLSGNEYVL | 6.2984 | 6.1175 | 5.8225 | 5.9408 | 6.3357 | 6.2341 | 6.0933 | 6.1755 | 7.1979 | 6.1570 | 6.1655 | 6.5038 | 6.1863 | 6.5433 | 6.3287 | 5.8999 |
| PDE8A_S457 | PDE8A | O60658 | | 386 | SMARIHSMTIEAP | 6.9364 | 6.3846 | 6.0785 | 5.8978 | 6.7901 | 5.6235 | 5.7461 | 5.4470 | 6.2921 | 5.3503 | 5.0624 | 5.7028 | 7.5044 | 7.4042 | 6.9429 | 6.8369 |
| PDHA1_S331 | PDHA1 | P08559-4 | | 331 | YRYHGHSMSDPGV | 6.2583 | 6.6537 | 6.0183 | 7.5293 | 7.2626 | 4.5936 | 4.5101 | 4.9755 | 6.8148 | 5.9843 | 6.0581 | 5.5793 | 9.0707 | 6.7081 | 6.9461 | 5.0374 |
| PDHX_S162 | PDHX | O00330 | | 162 | PSEPRPSPEPQIS | 6.1814 | 5.5951 | 6.1424 | 7.5571 | 5.9919 | 5.4189 | 5.8604 | 5.6330 | 6.5894 | 6.5181 | 6.9852 | 7.4712 | 6.0711 | 5.6764 | 5.9395 | 6.3687 |
| PDIA6_S480 | PDIA6 | Q15084-2 | | 480 | EDDIDLSDVELDD | 5.9397 | 6.3559 | 7.4529 | 6.7473 | 7.0372 | 6.2753 | 6.1371 | 6.6268 | 5.6888 | 5.9087 | 5.7791 | 5.5324 | 6.1158 | 5.7983 | 6.0931 | 6.5117 |
| PDLIM1_S130 | PDLIM1 | O00151 | | 90 | SEHKVWSPLVTEE | 6.6244 | 8.9677 | 7.0464 | 6.9447 | 6.2389 | 6.5353 | 6.6453 | 7.6170 | 6.9513 | 7.6256 | 7.7015 | 7.4246 | 2.9724 | 3.8782 | 3.3155 | 3.5113 |
| PDLIM1_S90 | PDLIM1 | O00151 | | 130 | AMPFTASPASSTT | 6.8284 | 7.0549 | 7.0336 | 6.7838 | 6.1868 | 6.2815 | 6.4985 | 6.5153 | 7.6464 | 7.0145 | 7.7188 | 7.3657 | 4.5430 | 4.8355 | 3.2595 | 4.4338 |
| PDLIM2_S516 | PDLIM2 | Q96JY6-5 | | 529 | ASRALATPPKLHT | 7.0896 | 6.5256 | 6.8077 | 6.7245 | 6.4565 | 6.9132 | 6.9621 | 7.3929 | 7.3594 | 7.2739 | 7.6554 | 7.6982 | 3.6351 | 4.2105 | 3.2455 | 4.0497 |
| PDLIM2_T529 | PDLIM2 | Q96JY6-5 | | 516 | FLPSSLSPQSSLP | 6.5495 | 6.2780 | 7.8175 | 6.6949 | 6.1387 | 5.0618 | 5.6516 | 6.6918 | 6.3430 | 6.2241 | 6.9237 | 6.1651 | 6.6471 | 5.7573 | 5.5580 | 5.4980 |
| PDLIM3_S170 | PDLIM3 | Q53GG5-2 | | 89 | GETHLWSPQVSED | 6.8104 | 6.8879 | 7.3228 | 7.3352 | 6.4058 | 6.6725 | 7.1135 | 6.5112 | 6.9562 | 6.5337 | 7.7879 | 7.5057 | 3.6347 | 4.4943 | 3.3006 | 4.7276 |
| PDLIM3_S89 | PDLIM3 | Q53GG5-2 | | 170 | RGLIPSSPQNEPT | 5.5237 | 7.7309 | 6.8293 | 6.3514 | 7.2237 | 5.9863 | 5.7466 | 7.3421 | 6.1103 | 5.9612 | 7.3937 | 6.4280 | 4.4855 | 5.2004 | 6.2566 | 5.4303 |
| PDLIM5_S111 | PDLIM5 | Q96HC4 | | 111 | KPVPITSPAVSKV | 6.8070 | 6.2738 | 6.9576 | 7.1622 | 6.2754 | 6.9944 | 7.1187 | 6.3434 | 7.2847 | 6.8563 | 6.4859 | 6.9183 | 4.3078 | 4.7475 | 4.4008 | 5.0661 |
| PDLIM5_S213 | PDLIM5 | Q96HC4 | | 360 | STGVIKSPSWQRP | 5.9217 | 5.9177 | 6.1054 | 5.8808 | 6.5955 | 6.7977 | 7.2407 | 6.8352 | 7.0092 | 6.5366 | 6.2594 | 6.7279 | 5.4151 | 6.1778 | 4.9832 | 5.5960 |
| PDLIM5_S228 | PDLIM5 | Q96HC4-2 | | 213 | IPLHVFSPKYTKL | 8.7394 | 7.1041 | 7.3609 | 7.2648 | 5.1628 | 9.4982 | 8.5472 | 5.8707 | 8.3394 | 7.8668 | 7.5845 | 6.3491 | 2.0020 | 3.4550 | 1.8569 | 2.9981 |
| PDLIM5_S360 | PDLIM5 | Q96HC4 | | 228 | EGQRRGSQGDSKQ | 5.3594 | 4.7845 | 6.1579 | 5.8783 | 6.2413 | 5.6627 | 5.2942 | 6.3702 | 7.1405 | 7.5460 | 7.7163 | 8.0419 | 6.3273 | 6.8268 | 5.5303 | 5.1224 |
| PDLIM7_S111 | PDLIM7 | Q9NR12 | | 217 | PGPTAPSPTSRPP | 6.0284 | 6.1146 | 6.6871 | 6.3963 | 5.3055 | 6.2029 | 5.9196 | 6.6000 | 6.4067 | 6.6762 | 6.8254 | 6.6000 | 5.8838 | 6.5075 | 5.7667 | 6.0792 |
| PDLIM7_S217 | PDLIM7 | Q9NR12 | | 78 | ACGERLSLGLSRA | 6.3388 | 7.1362 | 6.3911 | 6.8299 | 6.3255 | 6.9286 | 7.1285 | 7.2050 | 6.4463 | 6.5609 | 6.7087 | 6.7269 | 4.2967 | 5.2667 | 4.4398 | 5.2705 |
| PDLIM7_S78 | PDLIM7 | Q9NR12 | | 111 | TFAPSVSLNKTAR | 7.8670 | 7.2657 | 8.4104 | 6.9654 | 5.5843 | 6.0838 | 6.3391 | 6.4036 | 7.7741 | 8.1228 | 7.9432 | 7.6049 | 3.4569 | 4.4831 | 2.5077 | 3.1882 |
| PDLIM7_T251 | PDLIM7 | Q9NR12 | | 279 | GSNNGKTPVCHQC | 6.4725 | 7.4421 | 7.2149 | 7.4876 | 5.7076 | 6.7140 | 7.1518 | 6.7986 | 7.5753 | 7.1656 | 6.8036 | 7.4012 | 3.5759 | 4.5393 | 3.5728 | 4.3773 |
| PDLIM7_T279 | PDLIM7 | Q9NR12 | | 251 | RHSQPATPTPLQS | 6.5897 | 7.0476 | 7.2837 | 7.0169 | 5.7002 | 6.8288 | 7.5467 | 7.2013 | 6.9787 | 7.0596 | 7.3751 | 7.1735 | 4.1256 | 4.7716 | 2.8463 | 4.4546 |
| PDPK1_S241 | PDPK1 | O15530 | | 241 | KQARANSFVGTAQ | 6.1622 | 6.3154 | 7.1456 | 5.8830 | 6.4409 | 6.7240 | 6.2112 | 6.7281 | 6.9214 | 6.2800 | 6.2759 | 6.4633 | 5.8241 | 5.6708 | 5.6332 | 5.3207 |
| PDS5A_S1305 | PDS5A | Q29RF7 | | 1208 | VRIISVTPVKNID | 6.8097 | 6.8367 | 6.1585 | 6.6328 | 7.0774 | 6.6834 | 7.0688 | 6.9456 | 6.8849 | 6.4780 | 6.5017 | 6.3414 | 4.7168 | 5.1902 | 4.5300 | 5.1439 |
| PDS5A_T1208 | PDS5A | Q29RF7 | | 1305 | AAVGQESPGGLEA | 6.2781 | 6.5592 | 5.8535 | 6.3237 | 6.2572 | 6.4545 | 6.5425 | 6.5744 | 6.6544 | 6.6277 | 6.5851 | 6.5550 | 5.3909 | 6.1351 | 5.4413 | 5.7674 |
| PDS5B_S1140 | PDS5B | Q9NTI5 | | 1283 | NEDEQNSPPKKGK | 6.0657 | 6.3268 | 5.8004 | 6.6183 | 5.9608 | 5.7265 | 6.3659 | 5.8826 | 6.0686 | 6.3214 | 6.3475 | 6.3056 | 6.3982 | 6.6258 | 6.3765 | 6.8091 |
| PDS5B_S1166 | PDS5B | Q9NTI5 | | 1182 | SSEMDHSENEDYT | 6.7165 | 6.9647 | 6.9124 | 7.2529 | 5.8795 | 6.3005 | 6.0772 | 5.9505 | 6.8495 | 6.8522 | 7.0597 | 6.6780 | 4.9310 | 5.6156 | 4.8977 | 5.0622 |
| PDS5B_S1176 | PDS5B | Q9NTI5 | | 1301 | KPLGGGTPKEEPT | 6.8583 | 6.3756 | 6.6870 | 7.1464 | 6.0342 | 6.3790 | 7.0151 | 5.8708 | 6.9104 | 7.1872 | 6.9005 | 7.1599 | 4.4284 | 5.3909 | 4.4190 | 5.2372 |
| PDS5B_S1176;Y1187 | PDS5B | Q9NTI5 | | 1140 | VNKPLSSAGKQSQ | 7.2150 | 7.0166 | 7.1608 | 7.9093 | 6.3385 | 7.2515 | 7.7560 | 6.5151 | 7.5979 | 7.6402 | 7.7027 | 7.6398 | 2.5327 | 3.8132 | 2.4218 | 3.4890 |
| PDS5B_S1182 | PDS5B | Q9NTI5 | | 1121 | EMKSFFTPGKPKT | 7.3601 | 7.4872 | 6.7945 | 7.5452 | 6.2769 | 6.4991 | 7.3072 | 6.7675 | 7.1352 | 6.6041 | 6.4236 | 7.1099 | 3.5268 | 4.7138 | 3.7635 | 4.6853 |
| PDS5B_S1257 | PDS5B | Q9NTI5 | | 1166 | SSSNPSSPGRIKG | 6.5968 | 6.6219 | 6.2715 | 6.7985 | 6.2699 | 5.9880 | 6.2229 | 6.4214 | 6.8063 | 6.4136 | 6.2766 | 6.1998 | 5.5581 | 5.8067 | 5.4574 | 6.2906 |
| PDS5B_S1283 | PDS5B | Q9NTI5 | | 1220 | PRGRKKTPVTEQE | 6.2939 | 6.6835 | 6.5597 | 6.7820 | 6.0977 | 6.1439 | 6.3053 | 6.2276 | 6.7143 | 6.1317 | 6.5856 | 6.3537 | 5.4073 | 5.9612 | 5.9531 | 5.7996 |
| PDS5B_S1358;T1370 | PDS5B | Q9NTI5 | | 1176 | IKGRLDSSEMDHS | 4.6506 | 4.7892 | 4.1155 | 4.9787 | 5.1085 | 5.1595 | 6.0131 | 5.0663 | 6.5316 | 4.9150 | 4.8585 | 4.5805 | 10.0673 | 9.7575 | 10.0561 | 9.3522 |
| PDS5B_T1121 | PDS5B | Q9NTI5 | | 1257 | KRGHTASESDEQQ | 6.1676 | 5.8535 | 6.6658 | 7.3393 | 5.8037 | 6.6482 | 6.3351 | 6.2278 | 6.1879 | 6.0712 | 6.3952 | 6.0676 | 5.6694 | 6.5870 | 6.0525 | 5.9281 |
| PDS5B_T1220 | PDS5B | Q9NTI5 | 1176;1187 | | NA | 4.6731 | 6.1021 | 5.2965 | 5.6119 | 6.2095 | 6.2352 | 5.7673 | 5.6221 | 4.4360 | 5.0319 | 5.2298 | 5.7721 | 8.1415 | 8.2193 | 8.7439 | 8.9077 |
| PDS5B_T1301 | PDS5B | Q9NTI5 | 1381;1385 | | NA | 6.5060 | 6.9597 | 6.0080 | 6.6609 | 6.3191 | 5.6741 | 6.6650 | 6.7182 | 6.3228 | 6.3835 | 6.4273 | 6.3113 | 5.1726 | 5.8154 | 6.0230 | 6.0331 |
| PDS5B_T1381;S1383 | PDS5B | Q9NTI5 | 1381;1383 | | NA | 6.2953 | 6.9693 | 5.8335 | 6.5275 | 6.2646 | 6.3007 | 6.7787 | 6.5294 | 6.3748 | 6.1517 | 6.3599 | 6.6161 | 5.1997 | 6.2164 | 6.0232 | 5.5590 |
| PDS5B_T1381;S1385 | PDS5B | Q9NTI5 | 1358;1370 | | NA | 5.7069 | 6.3430 | 6.1189 | 7.2799 | 6.2683 | 6.4487 | 6.2898 | 6.9189 | 7.2121 | 6.1486 | 6.8640 | 5.8210 | 5.1122 | 5.4601 | 6.0308 | 5.9770 |
| PDXDC1_S718;T720;S72 | PDXDC1 | Q6P996 | | 691 | GVTLPPTPSGSRT | 7.0502 | 6.2458 | 6.1317 | 6.1687 | 6.3739 | 6.0242 | 6.3035 | 6.5196 | 6.3830 | 6.1657 | 6.2215 | 5.6745 | 6.9148 | 6.5748 | 5.6842 | 5.5640 |
| PDXDC1_T691 | PDXDC1 | Q6P996 | 718;720;721 | | NA | 6.0250 | 4.5239 | 5.5604 | 5.1618 | 7.1122 | 5.3873 | 5.7082 | 6.1831 | 6.0681 | 6.4326 | 7.5750 | 7.7809 | 6.2058 | 5.2458 | 7.4591 | 7.5709 |
| PDZD8_S521 | PDZD8 | Q8NEN9 | | 538 | KPLGAISPVLNRK | 6.3170 | 5.9698 | 6.4368 | 6.4259 | 6.2522 | 6.3649 | 6.3163 | 5.7571 | 6.4618 | 6.4449 | 6.4431 | 6.4040 | 6.2529 | 6.1464 | 5.7744 | 6.2324 |
| PDZD8_S538 | PDZD8 | Q8NEN9 | | 521 | AQSLSHSPKRVPT | 6.9923 | 7.5219 | 6.7524 | 7.9542 | 6.3773 | 6.5281 | 6.1013 | 7.0552 | 6.4880 | 6.0210 | 6.7803 | 6.4363 | 4.3591 | 5.0316 | 4.8912 | 4.7100 |
| PEA15_S137 | PEA15 | Q15121-2 | | 137 | DIIRQPSEEEIIK | 6.5802 | 4.9772 | 7.3212 | 5.2875 | 5.9390 | 4.7633 | 4.7479 | 6.3653 | 5.2978 | 7.0882 | 5.5393 | 6.2612 | 5.2309 | 9.3939 | 7.7811 | 7.4262 |
| PEAK1_S281 | PEAK1 | Q9H792 | | 1151 | PNASQPTPPPLPK | 6.5908 | 6.5264 | 6.5604 | 6.2986 | 6.1069 | 6.1543 | 7.0230 | 6.3417 | 6.0440 | 6.4146 | 5.9909 | 5.9387 | 5.4767 | 6.4568 | 6.1127 | 5.9636 |
| PEAK1_S540 | PEAK1 | Q9H792 | | 854 | IKPVTPSPSKLVT | 6.5255 | 5.7890 | 6.2233 | 6.4700 | 6.3281 | 6.1413 | 6.2457 | 5.7630 | 6.6071 | 6.2114 | 6.2920 | 6.5814 | 6.0489 | 6.0821 | 5.9938 | 6.6973 |
| PEAK1_S587 | PEAK1 | Q9H792 | | 779 | SIQPPQSPPETPQ | 5.8818 | 6.9435 | 6.1059 | 6.5915 | 6.1017 | 6.2312 | 6.6687 | 6.8363 | 6.7535 | 6.8538 | 6.8136 | 6.7788 | 5.4587 | 5.1378 | 5.8307 | 5.0125 |
| PEAK1_S648 | PEAK1 | Q9H792 | | 837 | APQTTDSPTTKVQ | 6.2967 | 7.1229 | 7.1243 | 7.3134 | 6.2077 | 6.8867 | 6.8602 | 6.5508 | 6.5763 | 6.1794 | 6.8597 | 6.6280 | 4.7712 | 5.2716 | 4.4880 | 4.8632 |
| PEAK1_S779 | PEAK1 | Q9H792 | | 648 | KSFLGTSGELSVK | 6.3968 | 5.8827 | 6.3886 | 6.2909 | 5.2168 | 5.5081 | 6.2848 | 6.2955 | 5.9227 | 5.8125 | 5.9700 | 5.8583 | 6.7650 | 6.8792 | 6.9449 | 7.5833 |
| PEAK1_S837 | PEAK1 | Q9H792 | | 635 | VINPNAYDNLAIY | 5.7519 | 5.8283 | 6.6611 | 6.5888 | 6.2054 | 4.7930 | 5.7911 | 6.5040 | 6.1207 | 5.3200 | 6.0081 | 6.8119 | 7.2500 | 6.5381 | 7.2335 | 6.5939 |
| PEAK1_S861 | PEAK1 | Q9H792 | | 587 | KTIPVKSPNLSEI | 6.0550 | 5.8897 | 6.0420 | 6.3255 | 6.4124 | 7.2366 | 7.4743 | 6.7703 | 6.6642 | 5.7590 | 6.0964 | 6.3746 | 5.4817 | 5.6075 | 5.6140 | 6.1968 |
| PEAK1_T1151 | PEAK1 | Q9H792 | | 281 | FRANTLSPVRFFV | 6.6317 | 6.8231 | 6.0124 | 6.1839 | 6.1789 | 6.0349 | 6.0205 | 6.2271 | 6.0828 | 5.8895 | 5.9858 | 6.1797 | 6.4952 | 6.6481 | 6.4713 | 6.1349 |
| PEAK1_T852;S854 | PEAK1 | Q9H792 | | 540 | VWTSSTSPRQKIP | 7.0678 | 7.3262 | 6.4471 | 7.4564 | 6.8426 | 6.6748 | 7.4141 | 6.7906 | 7.2305 | 7.4243 | 7.0887 | 6.8822 | 3.3639 | 4.4282 | 3.3133 | 4.2491 |
| PEAK1_T854 | PEAK1 | Q9H792 | | 861 | PSKLVTSPQSEPP | 6.0370 | 5.7534 | 6.5110 | 5.9753 | 6.4048 | 6.1766 | 6.2069 | 6.0464 | 6.6513 | 6.0390 | 5.7532 | 6.2238 | 6.5616 | 6.9890 | 6.5286 | 6.1421 |
| PEAK1_Y635 | PEAK1 | Q9H792 | 852;854 | | NA | 6.3413 | 6.6506 | 6.7104 | 7.2253 | 5.9987 | 6.9683 | 6.9315 | 6.6037 | 6.4604 | 6.5105 | 7.1573 | 7.0383 | 4.6089 | 4.9029 | 4.6504 | 5.2413 |
| PEAR1_S953 | PEAR1 | Q5VY43 | | 953 | KGPPSGSPPRQPP | 7.1335 | 6.8148 | 6.4370 | 6.7231 | 6.3554 | 6.4746 | 6.6629 | 5.7241 | 6.4684 | 6.1509 | 6.5751 | 6.9804 | 5.7518 | 5.1594 | 4.6775 | 5.9111 |
| PEBP1_T42 | PEBP1 | P30086 | | 42 | ELGKVLTPTQVKN | 5.8779 | 6.9913 | 6.6288 | 6.3768 | 6.6607 | 7.0249 | 6.7924 | 7.2188 | 7.1314 | 7.7541 | 8.2086 | 7.7792 | 3.5117 | 4.3094 | 3.3203 | 4.4135 |
| PEG10_S35 | PEG10 | B4DSP0 | | 397 | SSPAGNSPAPLxx | 6.5944 | 7.1164 | 6.9028 | 6.6113 | 6.1099 | 6.9378 | 6.4790 | 6.9328 | 6.7273 | 6.7770 | 6.8378 | 6.5630 | 4.4748 | 5.1967 | 4.8499 | 4.8891 |
| PEG10_S397 | PEG10 | B4DSP0 | | 35 | EATAGRSPPTPTV | 5.9189 | 6.1494 | 6.2031 | 6.7639 | 5.8502 | 6.1494 | 6.0665 | 6.2698 | 6.3365 | 6.4803 | 7.2728 | 5.9215 | 5.8328 | 6.2219 | 6.1559 | 6.4071 |
| PELP1_S1033 | PELP1 | Q8IZL8 | | 745 | ILAPSGTPPPTIP | 6.0270 | 6.1387 | 6.3407 | 6.3934 | 6.3405 | 6.0598 | 6.1410 | 6.5040 | 6.2635 | 6.1530 | 6.2854 | 6.3565 | 6.2742 | 6.4335 | 6.0223 | 6.2663 |
| PELP1_S1043 | PELP1 | Q8IZL8 | | 658 | PPPEAPSPFRAPP | 6.0497 | 6.2021 | 6.3652 | 6.3691 | 6.3297 | 6.1336 | 6.4749 | 6.3604 | 6.3716 | 6.3323 | 6.4071 | 6.3780 | 6.2591 | 6.2370 | 5.6448 | 6.0851 |
| PELP1_S357 | PELP1 | Q8IZL8 | | 1043 | VEREGESPAAGPP | 5.8118 | 5.9108 | 5.9601 | 6.5556 | 6.1453 | 5.6514 | 6.1486 | 5.8435 | 6.1385 | 6.2356 | 5.9561 | 5.9237 | 6.8824 | 6.9041 | 6.8400 | 7.0925 |
| PELP1_S477;S481 | PELP1 | Q8IZL8 | | 481 | LRSPRGSPDGSLQ | 6.1641 | 6.3652 | 5.6584 | 6.3383 | 5.5934 | 5.2332 | 6.1639 | 6.3728 | 6.5290 | 5.9915 | 5.8351 | 6.1514 | 6.4880 | 6.5450 | 7.1315 | 7.4392 |
| PELP1_S477;S485 | PELP1 | Q8IZL8 | | 1033 | APEALPSQGEVER | 5.8939 | 5.9797 | 6.3620 | 6.5045 | 6.3096 | 6.2826 | 5.6306 | 6.0617 | 6.2399 | 5.8431 | 6.3868 | 6.0202 | 6.5241 | 6.9575 | 6.3662 | 6.6376 |
| PELP1_S481 | PELP1 | Q8IZL8 | | 1090 | AEEEMETETEAEA | 6.1355 | 6.1660 | 6.2003 | 6.3827 | 6.6147 | 6.0522 | 6.3224 | 6.5853 | 5.8344 | 5.7775 | 6.1576 | 5.4509 | 6.8506 | 6.5269 | 7.0379 | 5.9051 |
| PELP1_S658 | PELP1 | Q8IZL8 | | 357 | VSSKNISLHGDGP | 5.0064 | 6.2985 | 6.0429 | 6.2305 | 6.6338 | 6.2586 | 6.9176 | 7.1333 | 6.2211 | 5.8670 | 4.9244 | 5.5880 | 7.2975 | 6.7839 | 6.0570 | 6.7394 |
| PELP1_T1014 | PELP1 | Q8IZL8 | | 1014 | EVEEPGTEEERGA | 5.2311 | 6.3558 | 5.6154 | 7.4327 | 6.7922 | 5.8209 | 6.1954 | 5.3789 | 6.8457 | 5.3944 | 5.2156 | 5.6281 | 6.8401 | 7.1371 | 6.7922 | 7.3244 |
| PELP1_T1090 | PELP1 | Q8IZL8 | | 647 | MGPTCPTPAPVPP | 6.2422 | 5.6976 | 5.7477 | 6.3581 | 5.8980 | 6.0390 | 6.2180 | 6.0362 | 6.1819 | 6.0260 | 6.2357 | 5.5844 | 6.3841 | 7.1356 | 7.2692 | 6.9464 |
| PELP1_T647 | PELP1 | Q8IZL8 | 477;481 | | NA | 6.3231 | 6.7890 | 5.9624 | 6.8596 | 6.6290 | 6.9304 | 6.6794 | 5.6792 | 5.6157 | 6.2554 | 7.0501 | 6.4728 | 6.3458 | 5.9970 | 5.0937 | 5.3175 |
| PELP1_T745 | PELP1 | Q8IZL8 | 477;485 | | NA | 6.9087 | 6.9262 | 6.0119 | 7.1343 | 5.4719 | 5.2456 | 5.7149 | 5.4658 | 5.4966 | 6.0886 | 7.3790 | 5.8701 | 6.4348 | 6.7994 | 6.0603 | 6.9919 |
| PEPD_T487 | PEPD | P12955 | | 487 | GCDKAFTPFSGPK | 6.5884 | 6.6340 | 6.6234 | 6.0189 | 5.9093 | 7.7157 | 6.6354 | 6.7701 | 7.0253 | 6.4028 | 6.7259 | 6.2805 | 4.9347 | 5.7435 | 5.0864 | 4.9058 |
| PER1_S704 | PER1 | O15534 | | 704 | VVGGTLSPLALAN | 6.1784 | 5.9560 | 6.2549 | 6.3335 | 6.1265 | 6.1412 | 6.2779 | 6.2946 | 6.2256 | 6.6231 | 6.1886 | 6.4701 | 6.1005 | 6.3314 | 6.3453 | 6.1524 |
| PER1_S815 | PER1 | O15534 | | 692 | LSGEGATPRKEPV | 6.9937 | 7.2319 | 6.3524 | 6.6715 | 6.4042 | 6.1865 | 6.7719 | 6.8233 | 7.2232 | 5.9838 | 6.3552 | 6.2638 | 5.5089 | 5.1094 | 4.7590 | 5.3612 |
| PER1_S980 | PER1 | O15534 | | 815 | SSSTAPSALGERG | 7.9065 | 6.6025 | 6.7715 | 6.5000 | 5.8876 | 5.7396 | 6.1177 | 5.9421 | 5.9254 | 5.8367 | 6.6966 | 6.1752 | 5.5077 | 6.4353 | 5.8528 | 6.1027 |
| PER1_T692 | PER1 | O15534 | | 980 | FNSRCSSPLQLNL | 6.5420 | 7.0082 | 6.6283 | 4.9795 | 6.7958 | 5.1573 | 6.5006 | 7.6384 | 5.3023 | 6.7509 | 6.4919 | 6.1691 | 5.5164 | 6.4557 | 5.9515 | 6.1121 |
| PER2_S977 | PER2 | O15055 | | 977 | SAMGRASPPLFQS | 7.7844 | 7.2746 | 6.6589 | 7.3717 | 6.0300 | 5.4176 | 5.8678 | 7.5284 | 6.5652 | 5.3746 | 7.8318 | 6.0587 | 4.7247 | 5.3194 | 4.8008 | 5.3912 |
| PER3_S919 | PER3 | P56645-2 | | 919 | EKWEAQSEGHPFI | 6.7756 | 6.7986 | 6.1354 | 6.4236 | 6.7804 | 6.2519 | 5.6118 | 6.6839 | 6.9952 | 5.7841 | 6.1603 | 6.3187 | 6.3533 | 6.0875 | 5.3042 | 5.5354 |
| PES1_S457 | PES1 | O00541 | | 457 | PGNLNESEEEEEE | 6.0306 | 5.9193 | 6.4822 | 6.3991 | 6.2996 | 4.8566 | 6.0064 | 5.6061 | 6.6576 | 7.1828 | 6.5422 | 6.9855 | 6.2119 | 6.3479 | 5.7904 | 6.6817 |
| PES1_S457;S488 | PES1 | O00541 | 457;488 | | NA | 5.9218 | 5.5733 | 6.1654 | 6.6400 | 6.1624 | 5.8647 | 6.3956 | 5.5454 | 6.2324 | 6.0301 | 5.6263 | 6.2785 | 6.5444 | 6.7225 | 6.6304 | 7.6669 |
| PEX1_S354 | PEX1 | O43933 | | 438 | ITPVEVTPKIPRS | 6.8627 | 6.5468 | 6.5522 | 6.6333 | 5.9157 | 6.1976 | 6.4973 | 5.7944 | 7.1547 | 5.8106 | 6.4147 | 6.3606 | 5.7967 | 5.8863 | 5.7728 | 5.8037 |
| PEX1_T438 | PEX1 | O43933 | | 354 | TKQNVLSPEKEKQ | 7.0122 | 7.5461 | 6.9865 | 7.7413 | 6.1367 | 6.8017 | 7.1797 | 6.5975 | 7.3110 | 7.9648 | 8.0591 | 7.9854 | 2.7083 | 3.6548 | 2.7657 | 3.5494 |
| PEX13_S354 | PEX13 | Q92968 | | 354 | VSKQQQSFTNPTL | 5.8643 | 6.0696 | 5.9605 | 6.4849 | 5.1077 | 5.8993 | 5.4362 | 5.8010 | 5.8892 | 6.1786 | 7.5053 | 6.5360 | 6.4398 | 6.8840 | 7.2475 | 6.6962 |
| PEX14_S232 | PEX14 | O75381 | | 232 | RRQFPPSPSAPKI | 6.1169 | 6.2274 | 6.4370 | 6.7889 | 6.4176 | 6.3029 | 6.4723 | 6.5206 | 6.1965 | 6.2410 | 6.3725 | 6.3200 | 5.9678 | 6.0253 | 5.6799 | 5.9135 |
| PEX14_S282 | PEX14 | O75381 | | 335 | EEDDDVSHVDEED | 5.8694 | 5.9844 | 6.8212 | 6.7464 | 6.4629 | 5.8613 | 6.1457 | 6.0461 | 6.7341 | 6.5650 | 6.3796 | 6.7006 | 5.9048 | 5.8929 | 5.8813 | 6.0041 |
| PEX14_S335 | PEX14 | O75381 | | 282 | PGKEGHSPEGSTV | 6.0159 | 6.3440 | 6.3909 | 7.2571 | 6.6920 | 5.8677 | 5.9232 | 6.5126 | 6.2894 | 5.5056 | 6.4481 | 6.5177 | 6.2203 | 6.0579 | 5.8076 | 6.1500 |
| PEX19_S147 | PEX19 | P40855 | | 147 | TDLQNSSMSEEEL | 7.1122 | 6.0734 | 5.3755 | 6.4198 | 5.9813 | 5.4160 | 6.1355 | 7.3244 | 6.0969 | 6.5770 | 6.4366 | 7.9745 | 4.3047 | 7.0625 | 5.4915 | 6.2182 |
| PEX19_S54 | PEX19 | P40855 | | 54 | SGPQKRSPGDTAK | 6.1215 | 5.6905 | 5.8017 | 6.2448 | 4.5629 | 5.1155 | 5.4155 | 6.6430 | 5.4896 | 6.8370 | 5.7948 | 7.0603 | 5.0008 | 8.7448 | 6.6293 | 8.8482 |
| PEX3_S221 | PEX3 | P56589 | | 221 | LVEQHKSSSWINK | 1.6579 | 1.7402 | 1.5827 | 1.9836 | 2.4313 | 2.3402 | 2.3030 | 2.5942 | 1.7842 | 1.8411 | 1.4448 | 1.6412 | 18.7329 | 18.4314 | 20.9374 | 18.5539 |
| PFKFB2_S466 | PFKFB2 | O60825 | | 493 | RPLKPLSPLRAQD | 5.5771 | 5.3050 | 7.0503 | 6.1879 | 6.9871 | 7.9397 | 6.6908 | 6.3278 | 6.1191 | 6.6522 | 7.7341 | 6.6519 | 5.6596 | 5.8772 | 4.7676 | 4.4727 |
| PFKFB2_S493 | PFKFB2 | O60825 | | 466 | VRMRRNSFTPLSS | 5.6926 | 4.8922 | 6.3180 | 4.7022 | 5.9172 | 6.0339 | 5.7639 | 5.9453 | 6.7408 | 6.7226 | 6.8066 | 7.1470 | 6.9753 | 7.4656 | 5.9805 | 6.8962 |
| PFKL_S424 | PFKL | P17858-2 | | 424 | TQLRGGSFENNWN | 6.2947 | 6.5901 | 6.7940 | 5.0977 | 6.5350 | 6.6011 | 5.8483 | 6.8645 | 6.2760 | 6.6849 | 5.4107 | 6.6077 | 6.6386 | 5.8141 | 5.6488 | 6.2936 |
| PFKL_S822 | PFKL | P17858-2 | | 822 | VTRRTLSMDKGFx | 4.4690 | 3.9260 | 4.7514 | 4.2245 | 4.9836 | 3.9447 | 4.3814 | 4.3317 | 3.2383 | 4.0794 | 4.0728 | 3.8149 | 11.9967 | 12.4281 | 14.7831 | 10.5743 |
| PFKM_S738 | PFKM | P08237-3 | | 738 | HMQQGGSPTPFDR | 5.6547 | 6.4610 | 6.5382 | 5.9833 | 7.3990 | 6.3246 | 5.8777 | 6.0227 | 7.0419 | 7.0459 | 6.0976 | 7.0238 | 6.2521 | 5.5262 | 4.6245 | 6.1268 |
| PFKP_S386 | PFKP | Q01813 | | 386 | VRLRGRSFAGNLN | 6.1993 | 6.1982 | 7.1434 | 6.4867 | 6.8085 | 6.5034 | 6.2305 | 6.5537 | 6.1911 | 6.1600 | 6.2798 | 6.2655 | 5.7286 | 6.0819 | 5.5258 | 5.6437 |
| PFKP_S783 | PFKP | Q01813 | | 783 | EHVQPWSVxxxxx | 4.6492 | 5.3280 | 4.9940 | 5.2959 | 6.4527 | 6.8219 | 7.1273 | 7.1609 | 4.7422 | 4.9556 | 4.8649 | 5.1951 | 7.8842 | 8.5551 | 8.1460 | 7.8271 |
| PGAM1_S118 | PGAM1 | P18669 | | 118 | VKIWRRSYDVPPP | 6.0248 | 6.0407 | 7.0993 | 6.0660 | 6.5972 | 6.7299 | 6.2201 | 6.8977 | 6.4470 | 6.4289 | 6.7811 | 7.0268 | 5.2268 | 5.7434 | 5.2841 | 5.3863 |
| PGAM1_S31 | PGAM1 | P18669 | | 92 | RLNERHYGGLTGL | 6.2423 | 5.8636 | 6.5251 | 5.5174 | 6.6811 | 6.7591 | 6.0395 | 7.2258 | 6.2978 | 6.0753 | 6.3843 | 6.8470 | 5.8387 | 6.3299 | 5.7150 | 5.6580 |
| PGAM1_Y92 | PGAM1 | P18669 | | 31 | WYDADLSPAGHEE | 6.0700 | 6.7381 | 6.2837 | 6.5601 | 6.5926 | 6.6585 | 5.9446 | 6.2515 | 7.3395 | 6.2061 | 6.4209 | 6.7448 | 5.5870 | 5.7673 | 5.9410 | 4.8942 |
| PGAM5_S188 | PGAM5 | Q96HS1 | | 188 | EPDPPVSHWKPEA | 6.7029 | 6.3390 | 5.7611 | 8.0786 | 6.0550 | 4.8759 | 5.8374 | 7.2426 | 5.9947 | 6.2037 | 4.3104 | 5.9136 | 7.5352 | 6.6488 | 6.9844 | 5.5165 |
| PGD_S257 | PGD | P52209 | | 257 | LPKIRDSAGQKGT | 5.6487 | 6.2381 | 7.3095 | 6.0059 | 5.9864 | 6.5804 | 6.4199 | 6.8427 | 6.1025 | 6.5216 | 6.5369 | 6.7067 | 5.4565 | 5.9622 | 5.7639 | 5.9180 |
| PGK1_S203 | PGK1 | P00558 | | 203 | FAKALESPERPFL | 5.8127 | 5.8138 | 6.8107 | 5.5939 | 6.6472 | 6.7961 | 6.2035 | 6.9785 | 6.3671 | 6.5241 | 6.3767 | 6.5444 | 5.9019 | 6.4219 | 5.7386 | 5.4688 |
| PGK1_T329 | PGK1 | P00558 | | 329 | KYAEAVTRAKQIV | 6.2033 | 5.8477 | 7.0386 | 6.0406 | 6.2466 | 6.2653 | 5.8126 | 6.7792 | 6.5876 | 6.8059 | 7.3264 | 7.4096 | 5.3806 | 5.8627 | 5.1215 | 5.2719 |
| PGM1_S135 | PGM1 | P36871-2 | | 135 | GIILTASHNPGGP | 9.3711 | 8.3290 | 9.7455 | 7.3570 | 6.7836 | 5.0786 | 3.8404 | 4.3113 | 8.2611 | 9.5048 | 8.0580 | 8.0858 | 2.4128 | 3.9045 | 2.6427 | 2.3138 |
| PGM2_S165 | PGM2 | Q96G03 | | 165 | GIMITASHNPKQD | 7.1295 | 6.8341 | 8.3066 | 6.5951 | 5.9735 | 6.6391 | 5.6460 | 5.8047 | 7.0150 | 7.3971 | 6.8556 | 6.6333 | 4.7034 | 5.3540 | 4.8221 | 4.2909 |
| PGM2L1_S175 | PGM2L1 | Q6PCE3 | | 175 | GVMITASHNRKED | 7.1406 | 6.6963 | 6.9080 | 6.2927 | 6.5938 | 6.3644 | 6.2514 | 6.4231 | 6.5486 | 6.2746 | 6.3637 | 6.1574 | 5.5499 | 5.8676 | 5.2141 | 5.3540 |
| PGM3_S92 | PGM3 | O95394-4 | | 92 | GVMVTASHNPEED | 7.9235 | 7.8183 | 9.2140 | 6.9907 | 5.8441 | 5.5762 | 4.5013 | 5.6711 | 7.6559 | 9.1207 | 8.6832 | 8.8418 | 2.8435 | 3.9265 | 2.5672 | 2.8220 |
| PGRMC1_S181 | PGRMC1 | O00264 | | 181 | EEPTVYSDEEEPK | 6.4211 | 5.6825 | 5.9866 | 5.6542 | 5.6862 | 5.0295 | 5.7363 | 5.8706 | 5.7293 | 6.5840 | 5.9968 | 6.8226 | 6.9747 | 7.4538 | 6.9752 | 7.3965 |
| PGRMC1_S57 | PGRMC1 | O00264 | | 57 | PAASGDSDDDEPP | 6.3610 | 6.2907 | 6.4201 | 5.9948 | 6.0519 | 5.8134 | 6.1430 | 6.4725 | 6.0462 | 5.8784 | 6.1415 | 6.7935 | 6.1628 | 6.8048 | 6.2949 | 6.3305 |
| PGRMC2_S114 | PGRMC2 | O15173-2 | | 235 | EEPSEYTDEEDTK | 6.0175 | 5.7898 | 6.0502 | 5.8474 | 6.6101 | 5.8797 | 6.0464 | 6.0675 | 6.1740 | 6.2256 | 6.0729 | 6.1635 | 7.2456 | 6.8397 | 6.5711 | 6.3990 |
| PGRMC2_S128 | PGRMC2 | O15173-2 | | 128 | MKKRDFSLEQLRQ | 1.1867 | 2.2985 | 1.3818 | 1.3671 | 1.8762 | 1.9259 | 2.0676 | 1.9459 | 1.4343 | 1.4065 | 0.7917 | 1.1565 | 21.0729 | 18.8637 | 21.9877 | 19.2369 |
| PGRMC2_S232;T235 | PGRMC2 | O15173-2 | | 114 | AGAGEESPATSLP | 7.3297 | 6.3260 | 6.4158 | 5.9648 | 6.2200 | 5.7302 | 6.4356 | 6.5677 | 6.8633 | 5.9926 | 6.5479 | 6.1337 | 5.7500 | 5.8201 | 6.3188 | 5.5838 |
| PGRMC2_T235 | PGRMC2 | O15173-2 | 232;235 | | NA | 6.8070 | 6.5141 | 6.7460 | 7.0564 | 6.1743 | 5.9689 | 6.5826 | 6.3494 | 6.4887 | 6.5810 | 5.7319 | 5.7648 | 6.0677 | 6.1762 | 5.6051 | 5.3858 |
| PHACTR1_S237 | PHACTR1 | Q9C0D0 | | 237 | CLPVKLSPPLPPK | 5.5358 | 5.9566 | 6.9604 | 6.8913 | 6.0269 | 7.1436 | 6.8130 | 6.5345 | 6.7415 | 6.6457 | 6.7427 | 6.9429 | 5.2602 | 5.1859 | 5.5222 | 5.0968 |
| PHACTR1_S67 | PHACTR1 | Q9C0D0 | | 67 | RRVRSKSDTPYLA | 5.3464 | 5.8643 | 5.6692 | 6.7906 | 7.3950 | 6.1892 | 6.1912 | 7.0792 | 7.6002 | 6.1982 | 6.9082 | 8.1895 | 4.3608 | 5.7607 | 5.4419 | 5.0155 |

| | | | | | | | | | | | | | | | | | | | | | |
|---|---|---|---|---|---|---|---|---|---|---|---|---|---|---|---|---|---|---|---|---|---|
| PHACTR2_S225 | PHACTR2 | J3KP75 | | 548 | RLSRKLSLRPTVA | 8.3969 | 6.0527 | 5.8123 | 6.7123 | 7.3677 | 5.5816 | 5.9194 | 5.6797 | 7.9278 | 5.7052 | 5.5410 | 5.2556 | 6.7786 | 7.5819 | 4.5540 | 5.1333 |
| PHACTR2_S46 | PHACTR2 | J3KP75 | | 510 | KLVRRLSQRPTTE | 6.1110 | 6.1024 | 6.2696 | 5.6155 | 5.6029 | 5.8204 | 6.1227 | 5.6575 | 6.9305 | 6.4493 | 6.2358 | 6.4093 | 6.6302 | 7.3399 | 6.1660 | 6.5369 |
| PHACTR2_S510 | PHACTR2 | J3KP75 | | 93 | PTAGSQTPPFKRK | 6.2557 | 6.1239 | 7.6967 | 6.2733 | 6.5024 | 7.2299 | 6.5914 | 5.9667 | 6.4651 | 6.9415 | 6.6532 | 6.4841 | 5.0439 | 5.3028 | 4.9657 | 5.5035 |
| PHACTR2_S548 | PHACTR2 | J3KP75 | | 225 | GSKASASPSTSST | 6.8540 | 7.2143 | 7.1322 | 6.5313 | 6.8773 | 7.4175 | 6.4367 | 6.3879 | 6.4980 | 7.1422 | 7.2849 | 7.0940 | 4.1142 | 5.0477 | 4.0794 | 3.8884 |
| PHACTR2_S72 | PHACTR2 | J3KP75 | | 46 | WLPGDPSPRGRSQ | 6.6858 | 7.0025 | 6.8007 | 6.8861 | 6.1583 | 6.4883 | 6.8625 | 6.8057 | 7.2232 | 7.0885 | 7.0600 | 7.1649 | 3.9928 | 4.7929 | 4.0069 | 4.9811 |
| PHACTR2_T93 | PHACTR2 | J3KP75 | | 72 | RVHISGSVDGLDK | 6.0285 | 6.3782 | 6.8532 | 7.1424 | 6.0531 | 6.4849 | 5.8629 | 7.2749 | 6.0994 | 6.3567 | 6.7295 | 6.6406 | 4.9924 | 5.6287 | 5.8771 | 5.5975 |
| PHACTR4_S128 | PHACTR4 | Q8IZ21-2 | | 474 | TLGRTRSLPITIE | 6.2568 | 6.2218 | 6.0481 | 6.0901 | 6.1887 | 5.7525 | 5.6262 | 6.1289 | 6.8100 | 6.3243 | 6.0055 | 6.2014 | 6.6181 | 6.8357 | 6.6339 | 6.2580 |
| PHACTR4_S186 | PHACTR4 | Q8IZ21-2 | | 453 | GSHRAHSLLFENS | 5.9936 | 6.0237 | 6.6340 | 6.2645 | 5.9449 | 6.0510 | 6.4429 | 6.0805 | 6.6500 | 6.0037 | 6.3784 | 6.0941 | 6.0547 | 6.5138 | 6.2805 | 6.5897 |
| PHACTR4_S301 | PHACTR4 | Q8IZ21-2 | | 236 | TVNLSVTPSPAPR | 6.2219 | 6.4982 | 6.0090 | 5.9377 | 5.6486 | 6.8409 | 6.9300 | 6.7165 | 6.1849 | 5.8318 | 5.9767 | 6.2727 | 5.5683 | 6.2755 | 6.3228 | 7.6647 |
| PHACTR4_S352;S354;T3€ | PHACTR4 | Q8IZ21-2 | | 600 | TLIRRLSQRPTPE | 7.3111 | 6.1856 | 5.7161 | 6.1289 | 6.7586 | 6.8851 | 5.7532 | 5.9231 | 7.9155 | 6.3204 | 5.4410 | 5.5775 | 7.1674 | 7.1068 | 5.5920 | 5.2178 |
| PHACTR4_S354;T368 | PHACTR4 | Q8IZ21-2 | | 638 | RLTRKLSQRPTVA | 8.2475 | 6.5256 | 6.3651 | 7.0530 | 7.0048 | 5.7040 | 5.9945 | 6.3171 | 7.2783 | 5.7574 | 5.3175 | 6.1851 | 5.9240 | 6.2175 | 4.7844 | 5.3242 |
| PHACTR4_S437 | PHACTR4 | Q8IZ21-2 | | 128 | GNARSSSPVQVEE | 6.7751 | 6.4810 | 6.3951 | 6.2467 | 6.1546 | 5.7813 | 6.1198 | 6.4611 | 7.2998 | 6.5019 | 6.5613 | 6.4721 | 5.8081 | 6.0116 | 5.4601 | 5.4704 |
| PHACTR4_S453 | PHACTR4 | Q8IZ21-2 | | 437 | IQQALTSPLPMTP | 5.5490 | 6.3086 | 6.7892 | 6.6177 | 6.9695 | 7.1479 | 6.3798 | 6.5381 | 6.7929 | 6.6177 | 6.1068 | 6.2309 | 5.6670 | 5.5794 | 4.7064 | 5.9991 |
| PHACTR4_S474 | PHACTR4 | Q8IZ21-2 | | 557 | KVKRKDTLAMKLN | 13.7894 | 8.7215 | 9.6198 | 9.1764 | 6.1893 | 4.1224 | 5.5627 | 3.7665 | 5.7618 | 4.0994 | 5.1016 | 5.0517 | 5.7223 | 5.2250 | 4.6161 | 3.4739 |
| PHACTR4_S524;Y534 | PHACTR4 | Q8IZ21-2 | | 301 | TLLSKPSPPLPPK | 5.7912 | 5.6502 | 6.5935 | 6.3400 | 6.1089 | 6.6122 | 6.4221 | 6.2990 | 6.9451 | 7.2245 | 5.8358 | 6.3516 | 5.4029 | 6.3454 | 6.4070 | 5.6707 |
| PHACTR4_S600 | PHACTR4 | Q8IZ21-2 | | 186 | PPKRPLSSSHEAS | 6.2821 | 6.5800 | 6.7822 | 6.7207 | 7.0476 | 6.0464 | 6.0339 | 7.1922 | 6.7089 | 5.7521 | 6.3558 | 6.2801 | 5.6218 | 5.7639 | 5.9396 | 4.8927 |
| PHACTR4_S638 | PHACTR4 | Q8IZ21-2 | 354;368 | | NA | 6.0693 | 6.2653 | 6.1085 | 6.2853 | 5.9927 | 6.2233 | 5.7665 | 6.0375 | 6.3124 | 6.4871 | 6.1856 | 6.4622 | 6.6163 | 6.7189 | 6.2435 | 6.2257 |
| PHACTR4_T236 | PHACTR4 | Q8IZ21-2 | 352;354;368 | | NA | 5.9324 | 6.5048 | 7.0434 | 6.7434 | 6.5948 | 6.2817 | 7.0705 | 6.9397 | 6.8988 | 7.2204 | 7.5532 | 7.5164 | 4.2072 | 5.0206 | 3.9038 | 4.5688 |
| PHACTR4_T557 | PHACTR4 | Q8IZ21-2 | 524;534 | | NA | 6.6461 | 6.5166 | 8.1032 | 6.2586 | 5.9267 | 7.6889 | 7.3815 | 4.9967 | 6.1768 | 6.9005 | 6.3469 | 7.9049 | 5.0404 | 4.7543 | 4.5051 | 4.8526 |
| PHAX_S368 | PHAX | Q9H814 | | 368 | LASLDESQEGHAE | 5.6978 | 6.8868 | 6.3256 | 6.4752 | 6.4676 | 6.1667 | 5.9240 | 6.3576 | 6.6326 | 7.1201 | 6.9099 | 6.8107 | 5.1506 | 5.5866 | 5.3366 | 6.1517 |
| PHB2_S243 | PHB2 | Q99623 | | 243 | MLGEALSKNPGYI | 6.6111 | 6.4853 | 5.3669 | 6.9435 | 5.8136 | 5.7790 | 5.2665 | 6.2291 | 6.0952 | 7.0035 | 6.1379 | 6.6538 | 5.8459 | 6.6561 | 5.5031 | 7.6095 |
| PHB2_S261 | PHB2 | Q99623 | | 92 | RPRKISSPTGSKD | 6.3308 | 6.7273 | 5.8479 | 6.4590 | 6.3064 | 5.8953 | 6.5077 | 6.3554 | 6.1229 | 6.0243 | 5.7632 | 5.8861 | 6.2187 | 6.4775 | 6.5858 | 6.4919 |
| PHB2_S293 | PHB2 | Q99623 | | 261 | RAAQNISKTIATS | 6.0302 | 5.9008 | 4.8796 | 6.5704 | 7.7196 | 7.7276 | 6.7213 | 6.2120 | 7.2561 | 6.0854 | 5.9183 | 5.8564 | 4.8338 | 5.9062 | 5.3579 | 7.0244 |
| PHB2_S92 | PHB2 | Q99623 | | 293 | FTRGSDSLIKGKK | 6.4795 | 6.4722 | 5.8587 | 6.7175 | 6.4082 | 5.9740 | 6.1313 | 5.8240 | 6.5769 | 5.8149 | 6.0381 | 5.9683 | 6.5872 | 6.7748 | 6.8535 | 6.4210 |
| PHC1_S645 | PHC1 | P78364 | | 669 | ASPVAESPKVMDE | 6.8115 | 6.7983 | 6.3969 | 6.5573 | 6.3481 | 5.9321 | 6.6065 | 5.7578 | 6.1283 | 6.3865 | 6.1312 | 6.7546 | 6.1664 | 5.9338 | 5.4880 | 5.8026 |
| PHC1_S669 | PHC1 | P78364 | | 645 | VKRKADSEEERDD | 6.6925 | 7.2686 | 6.4270 | 7.4236 | 6.5371 | 5.9174 | 6.7820 | 6.4967 | 6.2921 | 6.0794 | 5.9227 | 6.4860 | 5.8211 | 5.6773 | 5.1261 | 5.0505 |
| PHC1_S786 | PHC1 | P78364 | | 922 | PNTAPPTPELHGI | 6.2618 | 6.8675 | 6.2990 | 7.0183 | 6.6856 | 5.8939 | 6.1351 | 6.3343 | 6.1693 | 6.7539 | 6.5265 | 6.5941 | 5.0014 | 6.1016 | 5.8152 | 5.5426 |
| PHC1_T922 | PHC1 | P78364 | | 786 | GPLGVDSPSAELD | 6.6736 | 6.7560 | 6.1022 | 6.4460 | 6.3747 | 6.8175 | 5.8519 | 5.7806 | 6.6779 | 6.2954 | 5.9337 | 6.0598 | 6.4608 | 6.7456 | 5.3741 | 5.6502 |
| PHC2_S592 | PHC2 | Q8IXK0-5 | | 752 | SYEEPLSPISASS | 5.9969 | 6.4033 | 6.3909 | 6.8548 | 5.7783 | 5.8077 | 6.8763 | 5.9186 | 7.3499 | 7.0740 | 7.0074 | 7.2338 | 5.2616 | 5.9179 | 4.7103 | 5.4182 |
| PHC2_S752 | PHC2 | Q8IXK0-5 | | 592 | FPVGRSSLLVGNL | 6.7513 | 7.0312 | 6.8604 | 7.0658 | 7.2070 | 7.0543 | 7.1544 | 5.8052 | 7.4099 | 6.5125 | 6.4753 | 5.8193 | 4.1299 | 4.5214 | 4.6755 | 5.5266 |
| PHC3_S275 | PHC3 | Q8NDX5-7 | | 621 | CVRMDRTPPPPTL | 6.2708 | 6.0487 | 6.4311 | 6.6886 | 6.1822 | 5.9143 | 6.0746 | 5.8101 | 6.7150 | 6.2500 | 6.0209 | 6.2017 | 6.5233 | 6.2677 | 5.9965 | 6.6045 |
| PHC3_S327 | PHC3 | Q8NDX5-7 | | 327 | QQJPLHSPPSKVS | 7.7230 | 7.4211 | 7.1726 | 7.6580 | 6.7214 | 6.6305 | 7.1477 | 6.9477 | 7.7988 | 7.2076 | 7.5340 | 7.2426 | 2.9550 | 3.8835 | 2.6835 | 3.2728 |
| PHC3_S774 | PHC3 | Q8NDX5-7 | | 854 | HRGRRPSGPDGAA | 5.4844 | 5.0247 | 4.9762 | 5.4139 | 6.7464 | 5.5627 | 5.2228 | 5.5749 | 8.0799 | 7.9289 | 7.4410 | 7.7947 | 7.1577 | 7.0680 | 4.6595 | 5.8643 |
| PHC3_S854 | PHC3 | Q8NDX5-7 | | 275 | ISQRDPSPESNKK | 6.3894 | 6.7737 | 6.2037 | 6.6331 | 6.0016 | 5.6926 | 5.7555 | 5.6163 | 6.5645 | 6.4733 | 6.3292 | 6.8618 | 6.2165 | 6.0732 | 6.2146 | 6.2009 |
| PHC3_T621 | PHC3 | Q8NDX5-7 | | 774 | KHADNSSDTEMED | 5.3389 | 5.7787 | 6.2123 | 6.1874 | 5.9627 | 5.7374 | 6.0397 | 6.3594 | 7.0540 | 7.0257 | 6.3979 | 6.5139 | 6.5324 | 6.1732 | 6.2462 | 6.4403 |
| PHC3_T621;S628 | PHC3 | Q8NDX5-7 | 621;628 | | NA | 6.4672 | 6.3317 | 6.0248 | 6.4070 | 6.0580 | 6.2214 | 6.1490 | 6.0208 | 6.5806 | 6.6300 | 6.0108 | 6.5353 | 6.4673 | 6.1559 | 6.0642 | 5.8761 |
| PHETA1_S192 | PHETA1 | Q8N4B1-4 | | 226 | PPRRRASAPHGPL | 6.2189 | 5.9902 | 6.5370 | 6.0014 | 6.0746 | 5.9888 | 6.0550 | 6.2113 | 7.0016 | 6.9986 | 7.0570 | 6.6797 | 6.0453 | 6.3912 | 5.3242 | 5.4252 |
| PHETA1_S226 | PHETA1 | Q8N4B1-4 | | 192 | PLPRRPSALPPKE | 6.2282 | 4.7187 | 5.4301 | 4.7178 | 6.0321 | 5.2051 | 5.4027 | 5.7850 | 7.9277 | 7.5111 | 7.2836 | 7.7609 | 6.9829 | 8.0997 | 5.4440 | 5.4705 |
| PHF1_S446 | PHF1 | O43189 | | 515 | RRSAPPSPLCRSL | 6.7845 | 6.7384 | 6.2141 | 6.7886 | 6.4699 | 6.2752 | 6.3669 | 6.2801 | 6.7084 | 6.1962 | 5.8907 | 6.0776 | 5.7258 | 5.8600 | 5.7513 | 5.8721 |
| PHF1_S515 | PHF1 | O43189 | | 446 | ARCLPSSPIRMFA | 7.2241 | 7.9277 | 7.0507 | 7.4774 | 6.1186 | 6.6341 | 7.0506 | 6.7476 | 7.5639 | 7.5323 | 7.5282 | 7.9214 | 2.9005 | 3.8932 | 3.1101 | 3.3196 |
| PHF10_S270 | PHF10 | Q8WUB8 | | 270 | EYYKRYSPDELRY | 5.9445 | 6.9520 | 6.2500 | 6.4453 | 5.8312 | 5.9962 | 6.2016 | 6.1590 | 6.2492 | 6.4547 | 6.6819 | 5.9769 | 5.8242 | 6.6437 | 6.0822 | 6.3074 |
| PHF10_S327 | PHF10 | Q8WUB8 | | 56 | SGDSSRSCETSSQ | 5.7637 | 6.1641 | 6.3307 | 6.5888 | 6.3629 | 6.0383 | 5.7984 | 6.2599 | 6.6177 | 6.4975 | 6.3634 | 6.1135 | 6.2478 | 6.2041 | 6.6659 | 5.9833 |
| PHF10_S50;S53;S56 | PHF10 | Q8WUB8 | | 327 | SSSGNVSEGESPP | 6.4683 | 6.2297 | 7.2285 | 8.0635 | 5.5708 | 5.8510 | 6.3809 | 6.0716 | 7.0135 | 6.6750 | 6.6237 | 6.1312 | 5.6846 | 5.9925 | 4.6303 | 5.3849 |
| PHF10_S56 | PHF10 | Q8WUB8 | 50;53;56 | | NA | 5.8580 | 5.3235 | 5.8473 | 5.6838 | 5.8858 | 5.9092 | 7.5380 | 5.9430 | 6.4006 | 5.4126 | 6.5345 | 6.1777 | 6.3187 | 6.6831 | 6.0722 | 7.5119 |
| PHF12_S662 | PHF12 | Q96QT6 | | 662 | DSRPLGSPPNATR | 6.4178 | 6.7767 | 6.8461 | 7.1120 | 6.0982 | 6.4107 | 6.4423 | 6.4222 | 6.5007 | 6.7054 | 7.1689 | 6.8253 | 4.5500 | 5.4680 | 4.5678 | 5.0878 |
| PHF12_S691 | PHF12 | Q96QT6 | | 880 | SEKTPPTPPSSIV | 6.0139 | 6.2984 | 6.2586 | 6.8466 | 6.2381 | 5.8616 | 6.1395 | 5.9738 | 6.3955 | 6.4364 | 6.1943 | 6.2498 | 6.1707 | 6.2573 | 6.4984 | 6.1671 |
| PHF12_S701 | PHF12 | Q96QT6 | | 586 | GWPRPLTPPAAGG | 6.2447 | 6.5862 | 6.0615 | 6.7047 | 5.8942 | 6.0760 | 6.4373 | 5.7832 | 5.8601 | 6.3534 | 6.2936 | 6.1606 | 6.5844 | 6.5588 | 6.0652 | 6.3361 |
| PHF12_T586 | PHF12 | Q96QT6 | | 701 | SSDGKVSPGTLSI | 5.4201 | 5.8865 | 6.3365 | 6.8678 | 5.5842 | 6.1795 | 6.5798 | 6.0543 | 6.1572 | 5.8772 | 6.7922 | 5.8398 | 6.4020 | 6.8721 | 6.7173 | 6.4337 |
| PHF12_T671 | PHF12 | Q96QT6 | | 671 | NATRVLTPPQAAG | 6.1377 | 7.2997 | 6.8735 | 8.0170 | 5.4081 | 4.5496 | 5.7545 | 5.9691 | 6.5025 | 5.9967 | 5.7177 | 6.5761 | 8.1060 | 5.4081 | 5.9047 | 5.7790 |
| PHF12_T880 | PHF12 | Q96QT6 | | 691 | ANQRFSSPAPSSD | 6.4626 | 6.8269 | 6.0324 | 6.9219 | 5.9272 | 6.2500 | 6.3267 | 6.0718 | 6.7280 | 7.2855 | 6.4179 | 6.3636 | 5.2014 | 5.5551 | 5.8409 | 5.7881 |
| PHF13_S54 | PHF13 | Q86YI8 | | 54 | ELPLRSSPSPANS | 6.1785 | 6.5096 | 8.0927 | 5.2015 | 6.4336 | 6.2201 | 6.0700 | 7.0108 | 6.4409 | 6.6743 | 5.9542 | 6.3613 | 5.4006 | 6.4373 | 5.4259 | 5.5887 |
| PHF14_S208 | PHF14 | O94880 | | 208 | DMDDYDSEDDNDW | 6.0447 | 6.0758 | 6.3546 | 6.5612 | 6.0197 | 5.9544 | 6.3294 | 5.8674 | 6.2946 | 6.2587 | 6.2907 | 6.4490 | 6.3891 | 6.4867 | 6.2805 | 6.3435 |
| PHF14_S298 | PHF14 | O94880 | | 835 | TRGRKRSFVPEEE | 4.4909 | 4.4980 | 4.6278 | 5.1432 | 6.6663 | 5.1594 | 5.5802 | 5.8612 | 5.6064 | 5.7351 | 5.3283 | 5.1902 | 9.6191 | 8.5279 | 9.2927 | 8.6733 |
| PHF14_S302 | PHF14 | O94880 | | 302 | SKSNEDSLLEKS | 6.3808 | 6.4165 | 6.2870 | 6.8747 | 6.5723 | 6.2165 | 6.5991 | 6.2007 | 6.3445 | 6.0513 | 6.1374 | 6.0755 | 5.9758 | 6.2740 | 5.6648 | 5.9291 |
| PHF14_S308 | PHF14 | O94880 | | 298 | TLSQSKSNEDSLI | 6.4976 | 6.5953 | 5.9546 | 6.6097 | 5.9563 | 5.5200 | 6.1521 | 5.6939 | 6.1394 | 6.2854 | 6.2281 | 6.6139 | 6.3348 | 6.3266 | 6.4699 | 6.6223 |
| PHF14_S602 | PHF14 | O94880 | | 603 | NSDTSSSVDGRRK | 6.2254 | 6.3057 | 6.2553 | 5.5575 | 7.1312 | 5.5134 | 6.5499 | 7.1721 | 6.0584 | 5.9481 | 5.9970 | 6.2978 | 5.7985 | 6.1797 | 6.0537 | 6.9563 |
| PHF14_S603 | PHF14 | O94880 | | 602 | DNSDTSSSVDGRR | 5.7047 | 5.9372 | 7.4447 | 7.1173 | 7.1220 | 5.5019 | 6.3482 | 6.5147 | 5.9595 | 5.6293 | 5.7744 | 5.7791 | 6.6430 | 6.1418 | 6.4468 | 5.9353 |
| PHF14_S835 | PHF14 | O94880 | | 152 | ASAAATTPATSPP | 5.9554 | 6.2333 | 5.6717 | 6.8502 | 5.8182 | 6.0912 | 6.3065 | 5.7934 | 5.7900 | 6.3327 | 6.0180 | 6.7052 | 6.5815 | 7.4346 | 6.2828 | 6.1353 |
| PHF14_S84 | PHF14 | O94880 | | 84 | EEQLKNSAEEEVL | 7.0458 | 5.9781 | 6.0863 | 6.3813 | 6.3080 | 5.9194 | 6.7902 | 5.5224 | 5.3652 | 6.1362 | 5.9079 | 6.7398 | 6.6908 | 6.2196 | 5.7423 | 7.1668 |
| PHF14_T152 | PHF14 | O94880 | | 308 | SLILEKSQNWSSQ | 7.3084 | 7.6183 | 6.7809 | 8.1221 | 7.0758 | 6.4450 | 7.5104 | 6.5829 | 7.4804 | 5.7882 | 6.6314 | 7.0192 | 3.5298 | 4.3909 | 3.0687 | 4.6477 |
| PHF14_T292 | PHF14 | O94880 | | 292 | LTNDSLTLSQSKS | 5.7291 | 6.1994 | 5.8908 | 6.4967 | 4.8327 | 5.4793 | 6.6618 | 7.0405 | 6.5397 | 5.8049 | 5.4748 | 5.8931 | 8.1178 | 7.1773 | 6.3893 | 6.2729 |
| PHF19_S183 | PHF19 | Q5T6S3-2 | | 183 | LPSQSLSSKQKGH | 7.4048 | 7.0373 | 6.7722 | 6.9072 | 5.2645 | 5.5117 | 6.4892 | 6.8323 | 7.2015 | 8.1790 | 7.2194 | 7.1592 | 4.4026 | 4.6807 | 4.2887 | 4.6498 |
| PHF19_S187 | PHF19 | Q5T6S3 | | 187 | EELEWDSPHRTNQ | 6.1208 | 8.1843 | 6.4658 | 6.9697 | 6.5175 | 6.5407 | 6.6557 | 6.2862 | 5.1788 | 6.8327 | 6.1195 | 6.3133 | 4.4643 | 5.9825 | 5.3145 | 6.0536 |
| PHF2_S1056;S1059 | PHF2 | O75151 | | 474 | EKEEPPSPIEATP | 5.8479 | 5.9873 | 6.0612 | 6.4154 | 6.1150 | 5.7493 | 6.1307 | 5.9619 | 6.0256 | 5.9461 | 5.9217 | 5.8438 | 6.7278 | 7.3731 | 7.0345 | 6.8586 |
| PHF2_S474 | PHF2 | O75151 | | 655 | ALRPPTSPGVFGA | 6.5269 | 6.4587 | 6.0957 | 7.0331 | 6.2552 | 5.9322 | 6.5446 | 6.3620 | 6.4575 | 6.3033 | 6.2811 | 6.0743 | 5.5054 | 6.3906 | 5.8238 | 5.9556 |
| PHF2_S537 | PHF2 | O75151 | | 882 | VYPSLESDEDNPI | 6.0068 | 6.4280 | 6.1085 | 6.7479 | 6.1299 | 5.7136 | 6.3935 | 6.3188 | 6.1024 | 6.2519 | 5.8875 | 5.8642 | 6.0582 | 6.9943 | 6.6175 | 6.3770 |
| PHF2_S537;S539 | PHF2 | O75151 | | 539 | KSRESASPTIPNL | 5.7834 | 6.3610 | 5.9376 | 6.7198 | 6.2729 | 6.1140 | 5.9917 | 6.3128 | 6.0407 | 6.3526 | 6.1299 | 6.3004 | 6.1678 | 6.7246 | 6.5908 | 6.2000 |
| PHF2_S539 | PHF2 | O75151 | | 625 | EQKLEKSPLAGNK | 7.1404 | 7.1748 | 6.3837 | 7.6461 | 6.3545 | 6.1837 | 6.9660 | 6.6027 | 6.6565 | 6.5072 | 6.5561 | 6.8694 | 4.2327 | 5.1143 | 4.5558 | 5.0562 |
| PHF2_S625 | PHF2 | O75151 | | 705 | APKRDLSFLLDKK | 7.2105 | 6.7908 | 6.4345 | 7.0124 | 6.6547 | 6.3449 | 7.2268 | 6.7966 | 7.0083 | 6.5061 | 6.4659 | 6.4481 | 4.1972 | 5.7766 | 4.4446 | 4.6819 |
| PHF2_S640 | PHF2 | O75151 | | 681 | DEYEYVSDDGELK | 6.4198 | 6.2959 | 6.1539 | 6.8299 | 5.6266 | 5.8736 | 5.6946 | 5.2323 | 6.4742 | 6.8496 | 6.5029 | 6.0587 | 6.5459 | 6.3118 | 6.7725 | 6.3578 |
| PHF2_S655 | PHF2 | O75151 | | 640 | KFSFSFSNKKLLG | 9.7680 | 9.6707 | 6.0081 | 7.2891 | 6.2081 | 6.2943 | 8.5591 | 6.4860 | 6.6528 | 5.6023 | 7.7032 | 7.6476 | 3.0846 | 3.7266 | 2.3621 | 2.9373 |
| PHF2_S681 | PHF2 | O75151 | | 537 | GKKSRESASPTIP | 7.6285 | 8.3048 | 5.3725 | 6.9453 | 6.9643 | 6.8782 | 8.5612 | 6.5857 | 7.3239 | 7.8866 | 7.5562 | 8.5664 | 1.9084 | 4.3159 | 2.1063 | 3.0958 |
| PHF2_S705 | PHF2 | O75151 | | 905 | SDDAPYSPTARVG | 7.4483 | 8.2678 | 6.6256 | 7.6209 | 6.3224 | 6.2794 | 6.3954 | 6.2591 | 7.4967 | 6.9261 | 7.3107 | 7.3530 | 3.2382 | 4.3812 | 3.6627 | 4.1124 |
| PHF2_S882 | PHF2 | O75151 | 537;539 | | NA | 6.0241 | 8.3277 | 6.1450 | 7.9685 | 6.1708 | 6.4484 | 7.0776 | 5.8600 | 6.6501 | 6.8925 | 6.9410 | 6.4768 | 4.8513 | 5.0181 | 4.2483 | 4.8997 |
| PHF2_S905 | PHF2 | O75151 | 1056;1059 | | NA | 5.3703 | 6.2306 | 5.2964 | 6.6463 | 7.6616 | 5.4593 | 6.5194 | 6.0030 | 7.0284 | 7.2941 | 7.0281 | 6.2650 | 5.6220 | 6.3477 | 5.6917 | 5.5362 |
| PHF20_S159 | PHF20 | Q9BVI0 | | 519 | KRVSASSPTTKDK | 6.6092 | 6.7201 | 6.0938 | 6.6573 | 6.3501 | 5.8298 | 6.3871 | 6.5251 | 6.7534 | 6.4580 | 6.0633 | 6.3640 | 5.2212 | 5.9839 | 5.9283 | 6.0555 |
| PHF20_S248 | PHF20 | Q9BVI0 | | 902 | SDPKPGSPKVKEY | 6.5928 | 6.9287 | 6.1850 | 6.8948 | 6.1668 | 6.0282 | 6.8222 | 5.8833 | 6.4867 | 6.3872 | 6.5639 | 6.5591 | 5.5282 | 5.8573 | 5.4486 | 5.6672 |
| PHF20_S332 | PHF20 | Q9BVI0 | | 488 | SLEPEESPGKRHV | 6.3520 | 6.8802 | 6.0403 | 6.8952 | 6.0993 | 5.7323 | 6.6712 | 5.8292 | 6.4677 | 6.0295 | 6.5120 | 6.6297 | 5.8252 | 5.9446 | 6.1547 | 5.9368 |
| PHF20_S415 | PHF20 | Q9BVI0 | | 248 | ENDIVKSPQENLR | 7.6219 | 7.8039 | 7.3927 | 9.0348 | 6.3687 | 7.1912 | 7.1587 | 5.9308 | 6.8603 | 7.7194 | 7.8270 | 6.4372 | 3.1365 | 3.7043 | 2.9013 | 2.9113 |
| PHF20_S488 | PHF20 | Q9BVI0 | | 159 | HKSLSSSPDKREK | 6.6833 | 6.5327 | 5.9019 | 6.5089 | 6.4892 | 5.6399 | 6.8589 | 6.0135 | 6.6782 | 6.3482 | 5.9023 | 6.0180 | 5.3673 | 6.4085 | 6.4182 | 6.2311 |
| PHF20_S519 | PHF20 | Q9BVI0 | | 415 | MKTEPTSPLVELQ | 6.0030 | 6.1678 | 6.4676 | 7.5389 | 5.6396 | 5.1312 | 6.7675 | 5.8381 | 5.6763 | 6.0764 | 6.5729 | 6.5699 | 5.9493 | 6.3177 | 7.3771 | 5.9066 |
| PHF20_S902 | PHF20 | Q9BVI0 | | 332 | RRSSRLSTNGTHE | 4.4277 | 4.7360 | 5.0557 | 5.8391 | 5.6995 | 6.0090 | 5.6281 | 6.1139 | 5.0102 | 5.1694 | 4.6823 | 5.2945 | 8.1205 | 8.6310 | 9.6008 | 9.9824 |
| PHF20L1_S435 | PHF20L1 | A8MW92 | | 435 | EKVSSPSPATDGK | 5.7502 | 5.6679 | 5.5258 | 6.2046 | 6.2504 | 5.6211 | 6.1462 | 6.2754 | 6.4259 | 5.8429 | 5.3437 | 5.5097 | 7.5371 | 7.3343 | 7.1126 | 7.4520 |
| PHF21A_S448 | PHF21A | Q96BD5-3 | | 479 | QPVSLPSPTSTDG | 5.7829 | 6.0843 | 6.2609 | 5.9422 | 5.8816 | 6.4452 | 6.3339 | 5.6273 | 6.6353 | 6.6966 | 6.0374 | 6.3262 | 6.2885 | 6.6172 | 6.3822 | 6.6583 |
| PHF21A_S448;S456 | PHF21A | Q96BD5-3 | | 351 | TESRTITPPAAPK | 6.1318 | 6.1435 | 5.5869 | 6.5898 | 6.1908 | 6.2472 | 6.3349 | 6.0488 | 6.2765 | 6.4986 | 6.2144 | 6.4270 | 6.2157 | 6.2254 | 6.3111 | 6.2576 |
| PHF21A_S479 | PHF21A | Q96BD5-3 | | 448 | GALTPTSPQSSHP | 7.0955 | 8.0960 | 6.7825 | 7.8157 | 6.7784 | 6.2403 | 7.2039 | 5.9825 | 7.4761 | 6.7129 | 6.6884 | 7.3472 | 3.4557 | 4.3539 | 4.9390 | 3.0322 |
| PHF21A_T351 | PHF21A | Q96BD5-3 | 448;456 | | NA | 6.1989 | 6.3873 | 5.1756 | 6.5108 | 7.0018 | 6.8809 | 6.0184 | 7.2759 | 6.1307 | 5.9808 | 6.2513 | 6.2572 | 6.1548 | 6.7099 | 5.2885 | 5.7770 |
| PHF21A_T447;S456 | PHF21A | Q96BD5-3 | 447;456 | | NA | 6.3780 | 6.8633 | 5.3845 | 6.5887 | 7.0758 | 6.7675 | 6.1125 | 6.9364 | 5.8856 | 5.5514 | 6.9302 | 5.7722 | 5.7885 | 6.1721 | 6.4424 | 5.3509 |
| PHF23_S332 | PHF23 | Q9BUL5 | | 332 | EDIMVESGDDSWD | 5.7190 | 6.4762 | 6.4366 | 5.9500 | 6.1204 | 6.7357 | 6.0040 | 6.3830 | 6.2966 | 6.4811 | 5.8469 | 6.7060 | 5.8838 | 7.0473 | 6.5070 | 5.4063 |
| PHF23_S64 | PHF23 | Q9BUL5 | | 64 | PASGSSSPLRGES | 5.8883 | 6.1206 | 6.3695 | 6.0848 | 6.0948 | 5.8683 | 5.9634 | 5.9255 | 6.3211 | 6.4217 | 6.5811 | 6.3534 | 6.3577 | 6.5699 | 6.3444 | 6.7356 |
| PHF23_T30 | PHF23 | Q9BUL5 | | 30 | PEKRRRTIEDFNK | 5.1731 | 5.7027 | 6.0820 | 6.1652 | 7.2282 | 6.2080 | 6.6762 | 6.6628 | 6.7223 | 6.4779 | 7.1706 | 6.9615 | 5.0177 | 5.7386 | 5.9752 | 6.0379 |
| PHF3_S1084 | PHF3 | Q92576 | | 1133 | PPVDDLSPKKVKV | 6.7047 | 6.5090 | 6.2316 | 6.5453 | 6.2538 | 5.6331 | 6.3352 | 5.8804 | 6.4665 | 6.2679 | 5.9773 | 6.0617 | 6.0887 | 6.1932 | 6.3364 | 6.5151 |
| PHF3_S1133 | PHF3 | Q92576 | | 1184 | SPKSTFSPAPRPE | 6.4377 | 6.7417 | 6.2692 | 6.7710 | 6.2752 | 6.2198 | 6.2043 | 6.3181 | 6.6372 | 6.7269 | 6.3789 | 6.3827 | 5.5050 | 5.8441 | 5.6656 | 5.6226 |
| PHF3_S1184 | PHF3 | Q92576 | | 283 | KAKPVGSPFLKFS | 6.5635 | 6.2371 | 6.4323 | 7.1252 | 5.5425 | 5.6916 | 6.2139 | 5.3923 | 6.4685 | 6.7283 | 5.9552 | 6.7748 | 5.6674 | 6.2596 | 6.1486 | 6.7993 |
| PHF3_S1250 | PHF3 | Q92576 | | 1642 | VANTARSPQFINL | 7.3639 | 7.6233 | 6.4508 | 7.5968 | 6.7255 | 6.5075 | 7.1461 | 6.8064 | 7.3129 | 6.9650 | 6.9528 | 6.7054 | 3.4738 | 4.5350 | 3.7026 | 4.1323 |
| PHF3_S1614 | PHF3 | Q92576 | | 680 | QRRSSKSFSLDEP | 5.3038 | 5.0904 | 5.8876 | 6.3102 | 5.8312 | 5.6173 | 5.9436 | 5.6257 | 5.4361 | 5.2075 | 5.8591 | 5.7468 | 7.3237 | 7.7716 | 8.4309 | 8.6145 |
| PHF3_S1642 | PHF3 | Q92576 | | 1758 | HPFRRGSAVATSH | 4.5533 | 4.0873 | 4.8748 | 5.1660 | 7.6264 | 5.5185 | 5.2159 | 5.5621 | 8.1798 | 8.3444 | 7.4035 | 8.0607 | 7.0547 | 8.0763 | 4.9024 | 5.3739 |
| PHF3_S1722 | PHF3 | Q92576 | | 1614 | NQTSNSSPCRSNV | 6.2144 | 6.3504 | 6.3685 | 6.7465 | 6.2312 | 6.5988 | 6.4221 | 5.9873 | 6.5123 | 6.1514 | 6.6722 | 6.4722 | 5.4681 | 6.6150 | 5.5731 | 6.6164 |
| PHF3_S1758 | PHF3 | Q92576 | | 1925 | DRQRFYSDSHHLK | 6.1400 | 6.7445 | 5.6567 | 7.2284 | 6.4306 | 6.7646 | 6.2179 | 7.2369 | 5.8133 | 5.9334 | 6.1411 | 5.7775 | 5.1852 | 6.1425 | 6.6297 | 5.9578 |
| PHF3_S1784 | PHF3 | Q92576 | | 1722 | LKVAQNSPSVENI | 6.1572 | 6.7656 | 6.1235 | 6.7386 | 6.2294 | 6.3413 | 6.1629 | 6.3895 | 6.4542 | 6.1604 | 6.4085 | 6.2034 | 5.3690 | 6.0181 | 6.2823 | 6.1962 |
| PHF3_S1796 | PHF3 | Q92576 | | 1898 | RPERRHSDPWGRQ | 4.6079 | 5.0216 | 4.6645 | 5.9252 | 6.6987 | 5.8360 | 5.4014 | 6.0097 | 6.0443 | 6.4560 | 6.7466 | 6.1784 | 7.5211 | 7.9689 | 7.7266 | 7.1929 |
| PHF3_S1898 | PHF3 | Q92576 | | 419 | STEFNKSNLEVVD | 5.4129 | 7.2740 | 7.2934 | 6.7045 | 6.6255 | 6.1792 | 7.9101 | 6.6809 | 6.3178 | 6.4827 | 7.2629 | 6.7530 | 4.7075 | 5.0096 | 4.1976 | 5.1884 |
| PHF3_S1925 | PHF3 | Q92576 | | 1084 | EPAANKSLEKPEG | 6.4497 | 7.3223 | 6.7862 | 7.6748 | 6.6397 | 6.8234 | 7.8920 | 6.6693 | 6.2160 | 6.5564 | 7.0046 | 6.3839 | 3.6640 | 4.9078 | 3.7299 | 5.0800 |
| PHF3_S256 | PHF3 | Q92576 | | 1250 | QVGGRISPQTVWD | 5.9585 | 6.1734 | 6.3232 | 7.0003 | 6.0705 | 5.9245 | 6.0977 | 6.7446 | 5.6529 | 6.9352 | 6.7627 | 5.9986 | 5.7187 | 6.8535 | 5.4191 | 6.3663 |
| PHF3_S283 | PHF3 | Q92576 | | 256 | SHELNCSLLSETC | 6.8640 | 7.7349 | 7.0132 | 8.0039 | 7.0329 | 6.5911 | 7.0565 | 5.7100 | 7.4722 | 6.8098 | 6.5809 | 7.3377 | 3.4977 | 4.4317 | 3.5104 | 4.3533 |
| PHF3_S419 | PHF3 | Q92576 | | 1784 | KSITFTSRSTSPR | 7.2579 | 7.7263 | 5.2704 | 6.0466 | 4.6956 | 6.4854 | 6.8797 | 5.7129 | 8.0959 | 7.0787 | 7.8425 | 7.9575 | 4.9342 | 5.5683 | 4.0863 | 4.3619 |
| PHF3_S680 | PHF3 | Q92576 | | 1796 | RTSTNFSPMRPQQ | 6.2259 | 6.6015 | 5.0888 | 6.4735 | 7.8788 | 8.5928 | 7.4806 | 6.6097 | 6.7996 | 7.1834 | 6.0608 | 6.3163 | 2.8333 | 4.5874 | 3.4730 | 5.7946 |
| PHF5A_S94 | PHF5A | Q7RTV0 | | 94 | IVNLGSSKTDLFY | 6.0602 | 5.7964 | 5.2585 | 6.4852 | 6.5052 | 6.0176 | 6.1988 | 5.8723 | 6.1222 | 5.5013 | 5.6376 | 5.2171 | 5.7657 | 7.1626 | 7.0080 | 7.4813 |
| PHF6_S155 | PHF6 | Q8IWS0 | | 155 | HELEPSSPKSKKK | 5.4714 | 6.7340 | 6.5224 | 6.4466 | 5.4273 | 6.2684 | 5.5206 | 7.1379 | 6.4600 | 5.9986 | 6.9097 | 5.6921 | 6.2751 | 6.9347 | 6.2243 | 5.9769 |
| PHF6_S204 | PHF6 | Q8IWS0 | | 204 | RSPHRSSPSDTRP | 5.0802 | 6.2531 | 5.4167 | 6.3447 | 5.8556 | 5.5203 | 5.6511 | 6.2859 | 6.6844 | 6.3029 | 6.1877 | 6.3267 | 6.5487 | 7.1870 | 7.1227 | 7.2320 |
| PHF8_S1021 | PHF8 | Q9UPP1 | | 1021 | LTQRRPSVGSQSN | 4.7664 | 4.8616 | 4.7606 | 5.9594 | 6.4745 | 6.1890 | 5.3043 | 6.0373 | 7.4760 | 8.2485 | 7.5366 | 8.1188 | 6.0907 | 7.3536 | 5.0745 | 5.7482 |
| PHF8_S120 | PHF8 | Q9UPP1 | | 886 | SPKARVTPTLPKQ | 6.3535 | 6.5213 | 7.4936 | 7.0492 | 7.6514 | 6.2944 | 6.4214 | 6.4234 | 7.4445 | 6.8560 | 6.9093 | 7.2043 | 3.8959 | 4.6603 | 4.0528 | 4.7689 |
| PHF8_S857 | PHF8 | Q9UPP1 | | 880 | SDDAPWSPKARVT | 6.5457 | 6.8294 | 6.9195 | 7.1876 | 6.4014 | 6.3127 | 7.0113 | 6.1437 | 7.0860 | 6.9868 | 7.1200 | 7.2568 | 4.3341 | 4.9434 | 4.1613 | 5.1204 |
| PHF8_S880 | PHF8 | Q9UPP1 | | 120 | KPVKTGSPTFVRE | 5.9469 | 6.3996 | 6.9355 | 6.8550 | 5.8251 | 5.7895 | 6.7588 | 5.9660 | 6.4702 | 6.8254 | 6.4815 | 6.9539 | 5.4805 | 5.9548 | 5.4043 | 5.9530 |
| PHF8_S969 | PHF8 | Q9UPP1 | | 969 | QLVTSSSPLPPPE | 5.4904 | 5.5098 | 5.2531 | 5.7639 | 6.5379 | 6.0153 | 6.1410 | 5.9696 | 5.9464 | 5.2858 | 5.5051 | 6.0576 | 7.7615 | 7.5183 | 7.7741 | 8.3703 |
| PHF8_T886 | PHF8 | Q9UPP1 | | 857 | IYPSLESDDDDPA | 6.0808 | 6.0355 | 5.5171 | 6.1981 | 5.9112 | 5.6028 | 6.1049 | 5.8172 | 6.8537 | 6.1375 | 6.1093 | 6.1132 | 6.6906 | 7.5010 | 6.4055 | 6.9218 |
| PHGDH_S349 | PHGDH | O43175 | | 349 | MRAWAGSPKGTIQ | 6.7510 | 6.3311 | 7.1586 | 5.2420 | 6.1461 | 6.6563 | 6.0703 | 6.6440 | 6.3780 | 6.3523 | 7.0786 | 6.7215 | 5.7670 | 5.7047 | 5.4542 | 5.9143 |
| PHIP_S1281;S1283 | PHIP | Q8WWQ0 | | 911 | EKDGPISPKKKKP | 6.5495 | 6.2875 | 5.8980 | 6.5112 | 6.1922 | 5.4357 | 6.4393 | 5.9620 | 6.5020 | 5.9156 | 5.7570 | 5.9776 | 6.3386 | 6.5201 | 6.9289 | 7.6848 |
| PHIP_S1315 | PHIP | Q8WWQ0 | | 1315 | LRNRAQSYDIQAW | 4.3043 | 4.5401 | 4.5089 | 5.4013 | 5.9135 | 5.1982 | 5.8105 | 5.8719 | 4.4402 | 4.4809 | 4.1037 | 4.4429 | 9.9385 | 9.7094 | 11.3215 | 10.0141 |
| PHIP_S1525 | PHIP | Q8WWQ0 | | 1480 | STSAFSTPTRSIP | 7.0805 | 7.2940 | 6.4979 | 7.9840 | 6.6549 | 6.6654 | 7.1104 | 6.5623 | 7.0863 | 6.9563 | 6.5354 | 7.0992 | 3.6449 | 4.5229 | 3.8275 | 4.4781 |
| PHIP_S1548 | PHIP | Q8WWQ0 | | 1783 | FYNEDDSEEEQRQ | 6.2459 | 6.8435 | 6.0752 | 6.7279 | 6.2272 | 5.7795 | 6.0495 | 5.8037 | 6.0382 | 6.0680 | 5.8716 | 5.9847 | 6.1602 | 6.7303 | 6.8103 | 6.5842 |
| PHIP_S1734 | PHIP | Q8WWQ0 | | 1548 | KTILENSVKHSKA | 7.6269 | 7.7355 | 7.0203 | 9.2670 | 6.9040 | 6.7554 | 8.1487 | 6.6829 | 6.6293 | 6.4348 | 6.4530 | 6.2021 | 2.9499 | 4.3196 | 2.7871 | 4.0834 |
| PHIP_S1783 | PHIP | Q8WWQ0 | | 692 | NVGLRRSGQIEGV | 7.2469 | 8.3227 | 6.5479 | 8.1596 | 6.2094 | 6.7368 | 7.6957 | 6.9824 | 6.8747 | 7.3507 | 7.1227 | 7.0856 | 2.7938 | 4.2839 | 2.9232 | 3.6639 |
| PHIP_S674 | PHIP | Q8WWQ0 | | 1734 | DLLVPASVKVLRR | 9.5584 | 8.9227 | 7.5451 | 10.4003 | 7.5798 | 5.6779 | 7.5523 | 5.3029 | 6.9381 | 5.3919 | 6.1666 | 5.6647 | 3.2081 | 4.3255 | 2.6935 | 3.0722 |
| PHIP_S692 | PHIP | Q8WWQ0 | | 674 | SRLSRGSISSTSE | 4.3993 | 4.1789 | 5.0117 | 5.4630 | 9.9832 | 6.2368 | 4.8439 | 6.7142 | 6.2973 | 6.4885 | 6.5818 | 6.0878 | 7.1254 | 8.3677 | 6.9091 | 5.3113 |
| PHIP_S879;S880;S881 | PHIP | Q8WWQ0 | | 1525 | EQPSTSSAAKTFI | 7.4219 | 8.2917 | 6.1076 | 8.0875 | 6.4266 | 6.4562 | 6.8592 | 6.5220 | 7.0837 | 6.7114 | 7.0590 | 7.7497 | 3.3693 | 4.5802 | 3.0232 | 4.2509 |
| PHIP_S911 | PHIP | Q8WWQ0 | 1281;1283 | | NA | 5.8637 | 6.2093 | 6.0790 | 6.6434 | 5.9553 | 6.5532 | 6.4951 | 5.8315 | 6.2937 | 6.1520 | 6.8052 | 6.5407 | 6.1499 | 6.7025 | 5.9521 | 5.7736 |
| PHIP_T1295;S1296 | PHIP | Q8WWQ0 | 1295;1296 | | NA | 6.8029 | 7.5064 | 7.6642 | 9.0527 | 7.4861 | 7.1573 | 7.3247 | 5.6903 | 6.2953 | 5.5250 | 6.3073 | 5.2865 | 3.9589 | 5.0966 | 4.0738 | 4.7719 |
| PHIP_T1480 | PHIP | Q8WWQ0 | 879;880;881 | | NA | 5.8008 | 6.8085 | 5.6169 | 6.3142 | 6.3672 | 4.8890 | 4.1117 | 4.7209 | 8.1076 | 6.8543 | 4.8854 | 6.9057 | 6.8404 | 7.1610 | 7.9050 | 6.7116 |
| PHKA1_S735 | PHKA1 | P46020 | | 758 | HLPRDQSGEVDFK | 2.6501 | 2.7482 | 3.0630 | 2.7646 | 4.8074 | 5.0437 | 5.7437 | 5.5032 | 3.2204 | 2.8597 | 3.5398 | 3.4846 | 15.0748 | 13.1613 | 13.4361 | 12.8995 |
| PHKA1_S758 | PHKA1 | P46020 | | 735 | SFNLLDSPHPRQE | 6.0569 | 5.8320 | 7.5807 | 6.3789 | 6.6145 | 6.4737 | 5.8384 | 7.2203 | 6.1211 | 7.5153 | 7.5024 | 7.7014 | 3.6894 | 5.7778 | 4.4695 | 5.2279 |
| PHKA1_S985 | PHKA1 | P46020 | | 985 | NVSPAISIHEIGA | 7.5828 | 7.6691 | 7.2548 | 6.3648 | 6.9068 | 7.5816 | 6.1675 | 6.3217 | 6.9010 | 6.9580 | 6.7635 | 7.5143 | 4.0267 | 4.4726 | 3.7902 | 3.7246 |
| PHKA2_S729 | PHKA2 | P46019 | | 729 | LSAHRKSLNLVDS | 5.3698 | 5.0294 | 5.6542 | 5.3219 | 6.3402 | 5.7306 | 5.2521 | 5.7432 | 6.5079 | 6.3700 | 6.0779 | 6.1109 | 8.4351 | 7.9761 | 7.4421 | 6.6386 |
| PHKB_S25 | PHKB | Q93100 | | 701 | KVKRQSSTPSAPE | 6.1056 | 6.3123 | 6.4913 | 6.0273 | 6.2048 | 6.4182 | 6.1369 | 7.0005 | 6.2178 | 5.9403 | 6.6272 | 6.2748 | 6.0249 | 6.7337 | 5.8334 | 5.6510 |
| PHKB_S27 | PHKB | Q93100 | | 25 | RARTKRSGSVYEP | 6.1754 | 5.5717 | 6.7232 | 5.8523 | 6.4980 | 6.1633 | 5.4530 | 5.8700 | 7.4924 | 6.8000 | 6.7803 | 7.1180 | 6.1492 | 6.8816 | 5.0163 | 5.4555 |
| PHKB_S700 | PHKB | Q93100 | | 27 | RTKRSGSVYEPLK | 6.5455 | 5.4979 | 6.9741 | 5.7793 | 6.7598 | 5.7574 | 5.2407 | 5.9641 | 7.6266 | 6.8278 | 7.0229 | 7.1511 | 6.2889 | 6.6418 | 4.8338 | 5.0882 |
| PHKB_S700;T702 | PHKB | Q93100 | | 700 | SKVKRQSSTPSAP | 6.5057 | 7.4500 | 7.6632 | 7.8688 | 5.7036 | 7.6759 | 5.7062 | 6.3687 | 7.8764 | 8.0668 | 6.8738 | 7.5998 | 3.0841 | 4.4827 | 3.4775 | 3.5968 |
| PHKB_S701 | PHKB | Q93100 | 700;702 | | NA | 6.0067 | 6.2673 | 6.5294 | 6.0905 | 5.5029 | 6.0158 | 6.0227 | 5.8412 | 6.1425 | 5.5664 | 7.1209 | 6.4546 | 5.8319 | 6.9961 | 6.1267 | 6.4846 |
| PHLDA2_S144 | PHLDA2 | Q53GA4 | | 144 | SEPSRPSPQPKPR | 6.6132 | 6.3917 | 7.2268 | 7.1947 | 6.2323 | 6.3066 | 6.6032 | 6.4496 | 6.5806 | 7.3380 | 7.3762 | 7.5098 | 5.0476 | 4.4111 | 4.0780 | 4.6407 |
| PHLDB1_S1017 | PHLDB1 | Q86UU1 | | 404 | LQDRPPSPFREPP | 5.6812 | 5.4193 | 5.5729 | 5.2328 | 5.4579 | 6.8572 | 6.2331 | 6.7120 | 6.4379 | 5.5470 | 5.6053 | 5.7701 | 6.8399 | 7.4973 | 7.4639 | 7.6721 |
| PHLDB1_S220;S223 | PHLDB1 | Q86UU1 | | 461 | RELPPLSPSLSRR | 6.1893 | 6.3224 | 6.4582 | 5.7143 | 4.9380 | 6.8383 | 6.6608 | 6.6826 | 6.3577 | 6.6172 | 6.6551 | 6.9665 | 5.6174 | 6.2038 | 5.7021 | 6.0762 |
| PHLDB1_S324 | PHLDB1 | Q86UU1 | | 563 | CVQRKLSSGDLRV | 6.1074 | 5.8382 | 6.6949 | 5.7096 | 6.6430 | 6.6950 | 5.6070 | 6.6582 | 6.9686 | 6.8675 | 7.3248 | 6.8402 | 5.7740 | 6.4140 | 5.0977 | 4.7600 |
| PHLDB1_S381 | PHLDB1 | Q86UU1 | | 324 | GHERPPSPGLRGL | 6.0476 | 6.0484 | 6.7484 | 6.1929 | 6.3462 | 6.2220 | 6.2917 | 6.4784 | 6.5575 | 6.2557 | 6.2380 | 6.4611 | 6.1902 | 6.2358 | 5.5340 | 6.1522 |
| PHLDB1_S404 | PHLDB1 | Q86UU1 | | 443 | TLQPPESPRLGRR | 6.3375 | 6.2962 | 6.7207 | 5.9257 | 5.8000 | 6.4160 | 6.3081 | 6.3540 | 6.7237 | 6.3952 | 6.2070 | 6.6245 | 5.9463 | 6.1257 | 5.6624 | 6.1571 |
| PHLDB1_S419 | PHLDB1 | Q86UU1 | | 470 | LSRRALSPLPTRT | 5.4930 | 5.9053 | 5.1119 | 4.6113 | 4.5619 | 6.5420 | 6.9562 | 7.0764 | 6.4873 | 6.0940 | 5.9218 | 6.5102 | 6.2256 | 6.8450 | 7.6021 | 8.0560 |
| PHLDB1_S443 | PHLDB1 | Q86UU1 | | 489 | NREVAESPRPRRW | 5.9504 | 6.3622 | 6.2510 | 6.0036 | 5.5519 | 6.6758 | 6.5557 | 6.4401 | 6.8264 | 6.4702 | 6.3233 | 6.4345 | 5.6948 | 6.4845 | 5.5734 | 6.4024 |
| PHLDB1_S461 | PHLDB1 | Q86UU1 | | 539 | SFSGRLSPAYSLG | 6.0073 | 5.5578 | 5.9726 | 6.4601 | 6.6358 | 5.6419 | 6.3025 | 6.8229 | 6.0058 | 5.7369 | 6.4760 | 5.8470 | 6.1536 | 7.0114 | 6.4684 | 6.8998 |
| PHLDB1_S470 | PHLDB1 | Q86UU1 | | 1017 | TQNGTGSLPRNLA | 6.2591 | 6.4796 | 7.1285 | 6.2007 | 5.5518 | 6.3096 | 6.1135 | 6.1244 | 6.5236 | 6.6344 | 6.5689 | 6.1093 | 5.9824 | 6.4194 | 5.6230 | 5.9718 |
| PHLDB1_S489 | PHLDB1 | Q86UU1 | | 381 | PTSIPGSPKFQPP | 6.2030 | 5.8181 | 6.0384 | 5.7961 | 6.4353 | 5.8574 | 5.7614 | 6.5615 | 6.3301 | 7.3451 | 7.1366 | 6.5441 | 5.9854 | 7.1421 | 5.3435 | 5.7020 |

| | | | | | | | | | | | | | | | | | | | | | |
|---|---|---|---|---|---|---|---|---|---|---|---|---|---|---|---|---|---|---|---|---|---|
| PHLDB1_S501 | PHLDB1 | Q86UU1 | | 501 | WAAHGASPEDFSL | 7.4734 | 7.2163 | 7.3875 | 7.3493 | 6.0279 | 7.2574 | 7.2049 | 6.8243 | 7.4627 | 7.9471 | 7.8749 | 8.2925 | 2.5176 | 3.6392 | 2.3720 | 3.1529 |
| PHLDB1_S539 | PHLDB1 | Q86UU1 | | 419 | ERVLTTSPSRQLV | 6.5262 | 6.9762 | 7.1435 | 6.0442 | 5.8172 | 6.3159 | 6.8807 | 6.7454 | 6.7602 | 6.3986 | 6.7756 | 6.3929 | 5.3666 | 6.0442 | 4.7179 | 5.0948 |
| PHLDB1_S539;S551 | PHLDB1 | Q86UU1 | 539;551 | | NA | 5.8596 | 5.3675 | 5.7976 | 6.4067 | 6.3744 | 6.5659 | 6.0014 | 6.1814 | 6.4112 | 6.4439 | 6.2907 | 6.5429 | 6.2513 | 6.3520 | 6.2754 | 6.8782 |
| PHLDB1_S539;T548;S555 | PHLDB1 | Q86UU1 | 539;548;555 | | NA | 5.8325 | 5.7460 | 6.2878 | 6.5096 | 5.8522 | 6.4274 | 7.2157 | 6.6739 | 6.2468 | 6.0335 | 6.3072 | 6.4491 | 6.1271 | 6.6929 | 5.7312 | 5.8672 |
| PHLDB1_S563 | PHLDB1 | Q86UU1 | 220;223 | | NA | 6.3969 | 6.7298 | 7.2225 | 7.2667 | 6.1704 | 7.3490 | 6.7877 | 6.7912 | 7.0602 | 7.3472 | 7.6695 | 7.4577 | 3.6966 | 4.1325 | 3.4081 | 4.5141 |
| PHLDB2_S157 | PHLDB2 | Q86SQ0 | | 415 | LRERKSSISSISG | 5.5239 | 5.6627 | 6.2824 | 5.6675 | 6.3295 | 6.1309 | 5.9722 | 6.4195 | 7.1492 | 7.0257 | 6.8784 | 7.0541 | 6.0463 | 6.5400 | 5.5220 | 5.7959 |
| PHLDB2_S175 | PHLDB2 | Q86SQ0 | | 204 | AASMPSSPKQARK | 5.7458 | 5.9537 | 5.9021 | 6.4304 | 6.3414 | 5.7680 | 6.0640 | 5.7230 | 6.5009 | 6.0777 | 5.8937 | 5.9415 | 6.8805 | 6.9615 | 6.8957 | 6.9201 |
| PHLDB2_S200;S204 | PHLDB2 | Q86SQ0 | | 513 | RCHRKDSLPDADL | 5.6798 | 5.8757 | 6.3993 | 5.9878 | 6.2198 | 5.9826 | 6.0121 | 6.0464 | 6.8283 | 6.6296 | 6.5430 | 6.4423 | 6.4161 | 6.7241 | 6.1746 | 6.0384 |
| PHLDB2_S204 | PHLDB2 | Q86SQ0 | | 212 | KQARKMSIQDSLA | 5.6273 | 4.9135 | 5.8921 | 5.0560 | 6.1657 | 6.0674 | 5.6347 | 6.2325 | 8.1235 | 8.0603 | 8.0016 | 8.3522 | 6.0638 | 6.8757 | 4.1628 | 4.7708 |
| PHLDB2_S212 | PHLDB2 | Q86SQ0 | | 157 | YSSRHKSHDNVYS | 6.1502 | 6.7432 | 6.7858 | 6.1739 | 5.4380 | 6.6327 | 6.5383 | 6.3619 | 6.6115 | 6.6496 | 6.7763 | 6.7632 | 5.2847 | 5.7821 | 5.3765 | 5.9320 |
| PHLDB2_S308 | PHLDB2 | Q86SQ0 | | 550 | LSDLTRTPPPPSS | 6.5731 | 6.5178 | 6.6262 | 6.2603 | 6.7202 | 6.8526 | 6.6384 | 6.3272 | 6.8977 | 6.6206 | 6.7430 | 6.3622 | 5.1356 | 5.5222 | 4.8062 | 5.3968 |
| PHLDB2_S334 | PHLDB2 | Q86SQ0 | | 489 | LEKLQLSDEESVF | 6.3452 | 6.5426 | 6.2186 | 6.6657 | 5.9841 | 5.8936 | 6.5829 | 6.4665 | 6.3357 | 6.3562 | 6.1150 | 6.4467 | 6.1797 | 5.7853 | 6.0623 | 6.0198 |
| PHLDB2_S384 | PHLDB2 | Q86SQ0 | | 73 | PAKRSPSPLGTSV | 5.6455 | 5.6968 | 5.3241 | 5.1463 | 6.1039 | 6.8914 | 6.7387 | 6.4689 | 6.2161 | 5.1606 | 5.1815 | 5.3703 | 7.4577 | 7.4604 | 7.4713 | 7.6665 |
| PHLDB2_S387 | PHLDB2 | Q86SQ0 | | 404 | LRQASGTPQPALR | 6.6948 | 6.8750 | 6.9136 | 6.4747 | 5.3925 | 6.7372 | 6.9194 | 6.3517 | 6.7979 | 6.2618 | 6.4369 | 6.6520 | 5.1117 | 5.5840 | 5.1320 | 5.6648 |
| PHLDB2_S415 | PHLDB2 | Q86SQ0 | | 387 | RNFSCGSVEFDEA | 5.9821 | 6.3679 | 6.4097 | 5.8715 | 6.1592 | 6.0591 | 6.2170 | 6.2827 | 6.1343 | 6.4101 | 6.0150 | 6.3156 | 6.3646 | 6.8077 | 6.2283 | 6.3754 |
| PHLDB2_S468 | PHLDB2 | Q86SQ0 | | 308 | LNFSSLSSGALPY | 6.6086 | 6.9630 | 7.1740 | 6.7886 | 6.0449 | 6.7558 | 6.2602 | 6.2265 | 6.6731 | 6.0297 | 6.5197 | 6.4030 | 5.1997 | 5.9301 | 5.1300 | 5.2932 |
| PHLDB2_S489 | PHLDB2 | Q86SQ0 | | 934 | PLGQSNSCGSVLP | 4.4601 | 6.3632 | 6.6885 | 6.3643 | 6.4184 | 6.1804 | 6.3226 | 6.1896 | 7.1366 | 6.3889 | 6.5096 | 6.9277 | 5.7058 | 5.9360 | 5.2573 | 5.1509 |
| PHLDB2_S513 | PHLDB2 | Q86SQ0 | | 334 | TLSVPASPRVARK | 5.8660 | 6.4396 | 6.0514 | 6.2813 | 6.2306 | 6.2327 | 6.3015 | 6.1755 | 6.6578 | 6.1744 | 6.0246 | 6.3316 | 6.3226 | 6.2057 | 6.0345 | 6.6702 |
| PHLDB2_S563 | PHLDB2 | Q86SQ0 | | 468 | KPDSRLSTGTTVE | 6.0832 | 6.4641 | 6.7274 | 7.3062 | 6.4590 | 6.8795 | 7.0869 | 6.8519 | 6.9723 | 6.5853 | 6.3213 | 6.8837 | 4.8331 | 5.0693 | 4.5851 | 4.8918 |
| PHLDB2_S71;S73 | PHLDB2 | Q86SQ0 | | 980 | FYNRTASESNVYL | 6.2596 | 7.4614 | 6.6279 | 7.2242 | 6.6124 | 7.0685 | 6.1196 | 7.0725 | 6.4288 | 5.6863 | 6.2573 | 6.5771 | 4.6778 | 6.1046 | 4.8668 | 4.9552 |
| PHLDB2_S73 | PHLDB2 | Q86SQ0 | | 384 | ATRRNFSCGSVEF | 5.6220 | 6.4787 | 6.0691 | 6.5393 | 5.4013 | 7.0498 | 7.1061 | 6.1700 | 7.0902 | 7.2618 | 6.5047 | 6.5739 | 5.3162 | 6.2998 | 5.6984 | 4.8186 |
| PHLDB2_S934 | PHLDB2 | Q86SQ0 | | 175 | GRKASGSLLAMWN | 5.9440 | 6.8744 | 5.9366 | 6.6573 | 5.5970 | 6.3103 | 6.5909 | 6.8109 | 6.9039 | 6.0046 | 7.0442 | 6.2438 | 4.6976 | 6.1729 | 6.3354 | 5.8761 |
| PHLDB2_S980 | PHLDB2 | Q86SQ0 | | 563 | TFPKASSESSYLS | 5.2476 | 4.0627 | 5.3178 | 5.3710 | 5.2355 | 5.2863 | 5.0759 | 4.2513 | 4.9478 | 5.3008 | 5.2017 | 4.5681 | 10.4082 | 9.0975 | 10.9717 | 9.6561 |
| PHLDB2_T404 | PHLDB2 | Q86SQ0 | 200;204 | | NA | 6.7751 | 6.8770 | 7.3115 | 6.7570 | 5.4017 | 6.3006 | 6.5836 | 5.8403 | 6.9974 | 6.9509 | 7.6581 | 7.4093 | 4.3327 | 5.3867 | 4.4314 | 4.9867 |
| PHLDB2_T550 | PHLDB2 | Q86SQ0 | 71;73 | | NA | 6.3001 | 7.0476 | 6.4242 | 6.5577 | 5.7281 | 6.5952 | 6.4278 | 6.8349 | 7.4610 | 5.9355 | 6.1753 | 6.4662 | 5.6534 | 5.8757 | 4.9424 | 5.5929 |
| PHLDB3_S254 | PHLDB3 | Q6NSJ2 | | 254 | EEEDRDSPGPQVP | 6.0286 | 5.9488 | 6.2049 | 6.3416 | 6.4446 | 6.7598 | 6.2749 | 6.3410 | 6.5482 | 6.1040 | 6.2993 | 7.0343 | 6.0199 | 6.0291 | 5.5111 | 6.1100 |
| PHRF1_S1124 | PHRF1 | Q9P1Y6 | | 973 | EPEAPPSPDVLQA | 6.2694 | 6.0071 | 5.9806 | 6.4254 | 6.1071 | 6.1345 | 6.2074 | 6.1372 | 6.5782 | 6.3938 | 6.0350 | 6.0567 | 6.2677 | 6.6728 | 6.2933 | 6.4339 |
| PHRF1_S1202 | PHRF1 | Q9P1Y6 | | 1202 | GAVREASPAPLAQ | 5.9462 | 6.2952 | 5.9419 | 6.6876 | 6.0769 | 5.5027 | 6.2450 | 6.0692 | 6.4055 | 6.0301 | 5.9009 | 6.1707 | 6.2081 | 6.8455 | 6.7649 | 6.9097 |
| PHRF1_S1229 | PHRF1 | Q9P1Y6 | | 867 | RTISINSPKAQTV | 6.7941 | 6.4472 | 6.0698 | 6.6566 | 6.8216 | 6.1572 | 6.7856 | 6.3126 | 6.7046 | 6.3306 | 5.8745 | 5.8028 | 5.5354 | 6.0427 | 5.6466 | 6.0181 |
| PHRF1_S1359 | PHRF1 | Q9P1Y6 | | 814 | QRKENPSPLFSIK | 6.1377 | 6.0791 | 5.4668 | 5.8903 | 6.1794 | 5.8074 | 6.2093 | 5.9162 | 5.9276 | 5.4888 | 5.8232 | 5.2968 | 7.0302 | 7.5513 | 7.4923 | 7.7040 |
| PHRF1_S1359;S1360 | PHRF1 | Q9P1Y6 | | 936 | ARLRRPSPPEPWD | 5.9744 | 5.8856 | 6.3675 | 6.7317 | 6.2400 | 6.0295 | 5.8680 | 6.1121 | 6.1949 | 6.2227 | 6.3874 | 6.2988 | 5.7876 | 6.6703 | 6.3343 | 6.8953 |
| PHRF1_S1360 | PHRF1 | Q9P1Y6 | | 1359 | EKAEAPSSPDVAP | 6.2685 | 6.0111 | 5.3140 | 5.8350 | 6.5078 | 5.5964 | 6.1513 | 5.7899 | 6.2856 | 6.0471 | 5.3887 | 5.1479 | 7.1853 | 7.3484 | 7.8734 | 7.2497 |
| PHRF1_S814 | PHRF1 | Q9P1Y6 | | 1229 | LGEAHVSPEVATA | 6.2079 | 6.5270 | 6.3541 | 6.6819 | 6.2599 | 6.1837 | 6.5797 | 5.8103 | 6.2800 | 6.3693 | 6.2516 | 5.9486 | 5.8052 | 6.1467 | 6.1333 | 6.4607 |
| PHRF1_S867 | PHRF1 | Q9P1Y6 | | 98 | KLEAAGSPNSDDD | 6.1120 | 6.7117 | 6.1577 | 5.7961 | 6.3527 | 5.4828 | 6.3137 | 6.1295 | 6.7493 | 6.1671 | 5.8969 | 6.7654 | 6.9564 | 6.3554 | 5.4895 | 6.5638 |
| PHRF1_S915 | PHRF1 | Q9P1Y6 | | 1124 | PRGRECSPTSSLE | 6.6366 | 6.9581 | 5.7482 | 6.6856 | 6.1015 | 5.7753 | 5.7187 | 6.0426 | 6.4703 | 6.0598 | 6.0127 | 5.9508 | 6.0050 | 6.5316 | 6.7538 | 6.5493 |
| PHRF1_S936 | PHRF1 | Q9P1Y6 | | 991 | VELRPPSRSRSTS | 6.9163 | 6.5734 | 5.7504 | 6.9153 | 6.1760 | 5.5066 | 6.8334 | 5.7741 | 6.2816 | 5.8373 | 5.9373 | 6.1969 | 5.9777 | 6.5509 | 6.1907 | 6.5820 |
| PHRF1_S973 | PHRF1 | Q9P1Y6 | | 1360 | KAEAPSSPDVAPA | 6.0752 | 5.6715 | 5.3781 | 6.1918 | 5.9196 | 5.6003 | 5.7156 | 6.0543 | 6.1671 | 5.7689 | 5.3241 | 5.8768 | 7.4790 | 7.6095 | 8.0393 | 7.1288 |
| PHRF1_S98 | PHRF1 | Q9P1Y6 | | 915 | VAAEGASDTEREE | 6.2937 | 5.7663 | 5.0760 | 7.0565 | 5.8654 | 6.1132 | 6.4317 | 6.0211 | 6.4282 | 6.5663 | 5.5857 | 6.3273 | 6.6371 | 6.9450 | 5.8636 | 7.0229 |
| PHRF1_S991 | PHRF1 | Q9P1Y6 | 1359;1360 | | NA | 7.2111 | 7.7288 | 7.1919 | 7.0701 | 5.2767 | 5.2502 | 6.6278 | 6.3300 | 5.5659 | 5.9399 | 5.0542 | 6.1515 | 4.9955 | 5.9809 | 6.3291 | 7.2963 |
| PHTF1_T222 | PHTF1 | Q9UMS5 | | 222 | SNKGTETDNDPSC | 8.1766 | 8.5742 | 8.5454 | 10.1563 | 7.2675 | 7.5763 | 9.4948 | 8.2331 | 3.6355 | 4.1736 | 3.2080 | 3.8740 | 4.0134 | 4.1747 | 4.5031 | 4.3937 |
| PHTF2_S279 | PHTF2 | Q8N3S3 | | 279 | TDNGYVSLDGKKT | 5.7625 | 6.0193 | 6.6493 | 7.3465 | 5.7128 | 5.5478 | 5.7399 | 6.0344 | 6.8438 | 6.5818 | 7.2108 | 7.2859 | 5.7118 | 5.7607 | 5.7256 | 6.0674 |
| PHYHIPL_S12 | PHYHIPL | Q96FC7 | | 12 | LDHALNSPTSPCE | 7.3269 | 6.8993 | 8.7641 | 7.4731 | 7.3594 | 6.5501 | 6.1576 | 8.3148 | 7.9197 | 5.8004 | 7.0752 | 6.9372 | 2.6877 | 3.6729 | 3.4402 | 3.6215 |
| PI4K2A_S462 | PI4K2A | Q9BTU6 | | 462 | SHQRSSSESYTQS | 6.2061 | 6.1157 | 6.7809 | 6.7806 | 5.9915 | 5.6493 | 5.8813 | 6.2222 | 6.5826 | 6.1193 | 6.3978 | 6.5505 | 5.9140 | 6.2033 | 6.2504 | 6.3544 |
| PI4K2A_S47;S51 | PI4K2A | Q9BTU6 | 47;51 | | NA | 6.1101 | 6.1222 | 7.8189 | 6.6173 | 5.5765 | 6.0612 | 7.3346 | 6.5531 | 6.0016 | 5.4273 | 6.9805 | 6.4406 | 6.6796 | 5.6274 | 5.0093 | 5.6398 |
| PI4KA_S1436 | PI4KA | P42356 | | 1436 | DNQDTRSNLDITV | 6.0644 | 5.8606 | 4.6009 | 5.7010 | 5.3870 | 5.5542 | 6.1145 | 5.5414 | 5.4752 | 4.6567 | 4.4516 | 5.0381 | 8.5137 | 9.2201 | 8.6981 | 9.1225 |
| PI4KA_S1466 | PI4KA | P42356 | | 1763 | RKKACLSALSEVK | 6.0765 | 6.4254 | 6.1569 | 6.6540 | 7.2566 | 6.8455 | 7.4321 | 6.6442 | 6.5435 | 4.8853 | 5.5487 | 5.3816 | 6.1297 | 6.4806 | 5.7536 | 5.7857 |
| PI4KA_S1479 | PI4KA | P42356 | | 1466 | SGMSTISKKSGMS | 5.8048 | 5.7997 | 5.3669 | 6.0748 | 5.5916 | 6.2216 | 6.1252 | 5.7981 | 5.8352 | 5.8974 | 5.4448 | 5.6966 | 7.6941 | 7.6203 | 7.3533 | 7.6756 |
| PI4KA_S1763 | PI4KA | P42356 | | 250 | TVCQEGTLKRKTS | 6.7530 | 6.6964 | 7.3330 | 6.8846 | 6.1871 | 6.0559 | 6.0823 | 5.7103 | 5.9531 | 5.7674 | 6.2480 | 5.8741 | 6.1953 | 6.4205 | 5.9945 | 5.8444 |
| PI4KA_T250 | PI4KA | P42356 | | 1479 | KKTNRGSLHKYY | 5.9921 | 6.3832 | 6.3864 | 6.6459 | 5.9794 | 5.8938 | 6.1065 | 5.9654 | 6.7179 | 6.6726 | 6.3687 | 6.9848 | 5.7816 | 6.3226 | 5.9683 | 5.8306 |
| PI4KB_S266 | PI4KB | Q9UBF8 | | 511 | DIRRRLSEQLAHT | 5.4981 | 5.4050 | 6.4433 | 5.6473 | 6.5494 | 6.0089 | 5.5371 | 6.4134 | 6.9297 | 7.3308 | 6.9247 | 7.1732 | 6.1366 | 6.8088 | 5.6495 | 5.5441 |
| PI4KB_S294 | PI4KB | Q9UBF8 | | 294 | NLKRTASNPKVEN | 6.5222 | 6.4453 | 6.8725 | 6.1425 | 6.5765 | 6.4973 | 6.4632 | 5.5207 | 5.3532 | 5.7697 | 6.2013 | 5.8196 | 6.8276 | 6.6944 | 6.2431 | 6.0509 |
| PI4KB_S511 | PI4KB | Q9UBF8 | | 266 | APDTGLSPSKRTH | 6.6120 | 6.3197 | 6.4279 | 6.5467 | 6.1090 | 6.6966 | 6.0672 | 6.3944 | 7.0766 | 6.3938 | 6.8802 | 6.5822 | 5.4750 | 5.8653 | 5.2706 | 5.2827 |
| PI4KB_S511;T517 | PI4KB | Q9UBF8 | | 341 | IGKRLATLPTKEQ | 5.9375 | 6.3745 | 6.1833 | 5.9526 | 6.0285 | 6.0205 | 5.7945 | 6.1036 | 6.8618 | 6.3716 | 7.1925 | 5.9428 | 6.2843 | 6.0370 | 6.3574 | 6.5576 |
| PI4KB_T341 | PI4KB | Q9UBF8 | 511;517 | | NA | 6.8135 | 6.8807 | 7.2315 | 7.2421 | 6.4967 | 7.1323 | 7.0359 | 7.3879 | 8.0843 | 7.5066 | 7.9756 | 7.5249 | 2.7959 | 3.7407 | 2.8765 | 3.4549 |
| PIAS1_S485;S487 | PIAS1 | O75925-2 | | 505 | SLPHQASPVSRTP | 6.3298 | 6.4662 | 6.5835 | 6.3509 | 6.3662 | 6.6632 | 6.6746 | 6.8077 | 6.8926 | 6.7998 | 6.8603 | 6.8249 | 4.8438 | 5.7623 | 4.6950 | 5.0792 |
| PIAS1_S490 | PIAS1 | O75925-2 | | 490 | PSLSPTSPLNNKG | 6.0064 | 6.0629 | 6.6174 | 6.4969 | 6.5902 | 6.1043 | 6.4513 | 6.6442 | 6.8467 | 7.0824 | 6.9926 | 6.9331 | 5.5344 | 5.3144 | 5.4050 | 5.8180 |
| PIAS1_S505 | PIAS1 | O75925-2 | | 512 | PVSRTPSLPAVDT | 5.5296 | 6.0452 | 6.0211 | 6.6947 | 6.3184 | 5.4104 | 6.5166 | 7.0349 | 5.9380 | 6.0358 | 5.4988 | 6.1617 | 6.3398 | 6.3519 | 6.8300 | 7.2732 |
| PIAS1_S512 | PIAS1 | O75925-2 | | 636 | TVTNRSSTDTASI | 5.8192 | 5.4230 | 6.4878 | 6.9831 | 8.1263 | 6.3764 | 5.9802 | 4.8246 | 8.2418 | 5.1506 | 5.1589 | 6.3475 | 6.2278 | 8.1634 | 6.4631 | 4.2262 |
| PIAS1_S636 | PIAS1 | O75925-2 | 485;487 | | NA | 6.0515 | 5.8937 | 5.9583 | 6.3572 | 6.5675 | 5.6898 | 5.9363 | 6.5314 | 6.5412 | 6.5605 | 6.6958 | 6.8400 | 6.3153 | 7.1655 | 6.8091 | 6.8968 |
| PIAS2_S499 | PIAS2 | O75928 | | 499 | MSETQSSPTKGVL | 6.0141 | 5.9009 | 6.2946 | 6.0690 | 5.8953 | 6.0031 | 5.9940 | 5.7767 | 6.4223 | 6.5532 | 6.4783 | 6.3633 | 6.6922 | 6.5515 | 6.5518 | 6.4396 |
| PIAS3_S431 | PIAS3 | Q9Y6X2 | | 431 | LDGLQYSPVQGGD | 6.1867 | 6.5864 | 6.3133 | 6.4400 | 6.2728 | 5.8876 | 6.7878 | 6.6390 | 6.8803 | 6.9532 | 7.2997 | 6.9456 | 5.2447 | 5.6773 | 4.7195 | 5.1661 |
| PIEZO1_S1646 | PIEZO1 | Q92508 | | 1646 | GLMRTASELLLDR | 6.1384 | 6.2011 | 5.7739 | 6.1302 | 6.4777 | 6.1004 | 6.2927 | 5.9962 | 5.9243 | 5.8120 | 5.6119 | 5.6948 | 7.0928 | 7.1882 | 6.8975 | 6.6678 |
| PIEZO1_T734 | PIEZO1 | Q92508 | | 734 | QDAVSGTPLLREE | 6.7376 | 7.2917 | 6.1122 | 6.6420 | 6.6572 | 6.3715 | 6.8543 | 6.5351 | 6.1905 | 6.3377 | 6.4693 | 6.1730 | 4.9598 | 6.1083 | 4.9148 | 5.6450 |
| PIGB_S39 | PIGB | Q92521 | | 39 | KLRKRKSTLYFNT | 7.7021 | 7.6659 | 6.9529 | 7.1140 | 5.2499 | 5.1794 | 5.5240 | 5.4643 | 7.0596 | 7.4095 | 6.4012 | 6.8937 | 4.9702 | 5.8470 | 4.5210 | 6.0454 |
| PIGB_S416 | PIGB | Q92521 | | 416 | GTLDVMSHIQKVC | 3.1142 | 2.8383 | 18.2433 | 5.3000 | 11.1976 | 11.1574 | 15.3957 | 4.3595 | 3.6421 | 2.2494 | 6.8501 | 3.2482 | 2.6333 | 2.7860 | 3.4525 | 3.5322 |
| PIGW_S416 | PIGW | Q7Z7B1 | | 416 | SWKLIQSPVTNKK | 6.3378 | 6.2398 | 5.7200 | 6.6019 | 6.0899 | 5.9447 | 6.1363 | 6.2245 | 6.1688 | 6.2907 | 6.5538 | 5.9826 | 6.2557 | 6.6483 | 6.8183 | 6.8869 |
| PIK3C2A_S108 | PIK3C2A | O00443 | | 108 | KLLLDDSFETKKT | 6.0643 | 7.0437 | 7.4564 | 7.4135 | 5.9966 | 7.2552 | 7.2376 | 7.5339 | 7.3482 | 8.1400 | 8.5391 | 8.2092 | 2.7997 | 3.1998 | 2.4397 | 3.3231 |
| PIK3C2A_S259 | PIK3C2A | O00443 | | 259 | VSNLQVSPKSEDI | 6.5643 | 6.8000 | 6.6352 | 6.7272 | 6.6117 | 6.8256 | 6.9693 | 6.7503 | 6.1108 | 6.0121 | 6.3881 | 5.8219 | 5.2720 | 5.4027 | 5.6331 | 5.4757 |
| PIK3C3_S244 | PIK3C3 | Q8NEB9 | | 244 | KDGEDSPILTSF | 5.7900 | 6.2548 | 6.3291 | 6.1967 | 6.7862 | 6.0534 | 6.1613 | 6.1650 | 6.5944 | 6.6162 | 6.3087 | 6.6398 | 5.9119 | 6.2203 | 6.1019 | 5.8702 |
| PIK3R1_T86 | PIK3R1 | P27986-4 | | 86 | KKISPPTPKPRPP | 8.9451 | 8.1028 | 7.1273 | 7.8907 | 6.6170 | 6.9536 | 7.5116 | 5.8248 | 7.5478 | 5.7129 | 7.0730 | 6.3134 | 3.6057 | 4.2658 | 2.2926 | 3.5159 |
| PIK3R2_S263 | PIK3R2 | O00459 | | 263 | APPPPSSPPPGGA | 5.5353 | 6.0907 | 6.0794 | 6.4693 | 6.7285 | 6.9675 | 6.2815 | 6.4871 | 6.4730 | 6.4819 | 6.0353 | 6.6327 | 5.8660 | 6.5401 | 5.8236 | 5.5080 |
| PIK3R4_S865 | PIK3R4 | Q99570 | | 894 | KPPRSESSAGICV | 5.1390 | 5.0095 | 5.2931 | 5.6987 | 5.2125 | 5.1742 | 5.2477 | 5.2806 | 5.8570 | 4.9133 | 5.4977 | 6.1869 | 8.8328 | 8.3421 | 9.2554 | 9.0597 |
| PIK3R4_S894 | PIK3R4 | Q99570 | | 865 | WKSMFGSLDPPNM | 6.3795 | 7.7440 | 7.9045 | 9.6674 | 6.2457 | 4.4888 | 4.8158 | 6.8432 | 7.8629 | 8.1840 | 6.7124 | 7.8867 | 4.0637 | 4.2867 | 3.4662 | 3.4484 |
| PIK3R4_S925 | PIK3R4 | Q99570 | | 925 | PVIPVLSSTILPS | 8.4285 | 7.3531 | 6.3619 | 5.5058 | 7.2420 | 6.1156 | 5.5869 | 6.4430 | 7.1248 | 5.5929 | 7.5844 | 7.4913 | 4.9622 | 6.7434 | 4.2743 | 3.1900 |
| PIKFYVE_S1544 | PIKFYVE | Q9Y2I7 | | 307 | ARNRSASITNLSL | 5.8998 | 6.5080 | 6.2498 | 5.7332 | 6.4375 | 5.5611 | 5.5705 | 6.2513 | 6.7168 | 6.7116 | 6.4686 | 6.6095 | 6.6951 | 6.8674 | 6.1685 | 5.5513 |
| PIKFYVE_S1549 | PIKFYVE | Q9Y2I7 | | 1544 | ISAMDASPRNISP | 6.7894 | 6.7379 | 7.2424 | 6.5331 | 6.3596 | 7.1550 | 6.8828 | 7.0943 | 6.7794 | 6.6671 | 6.7874 | 6.0101 | 4.7386 | 4.7252 | 4.6416 | 4.8560 |
| PIKFYVE_S1754 | PIKFYVE | Q9Y2I7 | | 299 | QEDAGKSPARNRS | 6.2656 | 6.1089 | 7.0767 | 7.7613 | 6.7247 | 7.4698 | 7.0358 | 6.7827 | 7.2318 | 6.2061 | 8.0264 | 6.8365 | 4.4492 | 5.1955 | 3.6974 | 4.1316 |
| PIKFYVE_S299 | PIKFYVE | Q9Y2I7 | | 1754 | RGTAGKSPDLSSQ | 6.5505 | 6.6729 | 6.7172 | 6.5739 | 6.1917 | 6.3148 | 6.1412 | 6.3916 | 6.2271 | 6.3004 | 6.5343 | 6.7377 | 5.5069 | 5.8226 | 5.5507 | 5.7666 |
| PIKFYVE_S307 | PIKFYVE | Q9Y2I7 | | 1549 | ASPRNISPGLQNG | 6.4172 | 6.6213 | 6.6219 | 6.3550 | 5.8230 | 6.6090 | 5.9815 | 6.2956 | 6.1580 | 5.8718 | 6.7730 | 6.6849 | 5.7715 | 6.2082 | 5.9963 | 5.8118 |
| PIKFYVE_S76 | PIKFYVE | Q9Y2I7 | | 76 | LSGSWTSPQLPSR | 5.9373 | 6.7206 | 7.4998 | 8.0098 | 7.4874 | 7.5516 | 6.8732 | 5.6150 | 5.0042 | 5.3238 | 5.4708 | 6.0816 | 5.1623 | 5.8607 | 5.1996 | 6.2023 |
| PIKFYVE_T283 | PIKFYVE | Q9Y2I7 | | 283 | HPDSSNTPLSTRL | 6.2162 | 7.0399 | 6.7982 | 7.3531 | 6.3428 | 7.8661 | 6.4513 | 7.4929 | 6.2319 | 6.3839 | 7.5775 | 7.2150 | 4.1784 | 5.1049 | 3.8644 | 3.8833 |
| PIMREG_S131 | PIMREG | Q9BSJ6 | | 131 | AQKGSGSPTHSLS | 6.1395 | 6.3107 | 5.1973 | 5.8567 | 6.6020 | 6.4038 | 6.3065 | 6.6779 | 6.1775 | 6.3279 | 6.2239 | 5.6918 | 6.3729 | 6.4526 | 6.7739 | 6.4851 |
| PIN1_S43 | PIN1 | Q13526 | | 43 | PSGNSSSGGKNGQ | 6.5343 | 6.7402 | 7.7734 | 7.3002 | 5.6470 | 7.0006 | 6.8556 | 6.6260 | 7.3221 | 7.6242 | 8.2297 | 8.2246 | 3.0142 | 3.9453 | 2.8406 | 4.3221 |
| PIN4_Y147 | PIN4 | Q9Y237-2 | | 147 | VKTKFGYHIIMVE | 3.6630 | 4.4794 | 7.8905 | 7.0199 | 6.5451 | 7.0080 | 6.9724 | 6.3947 | 4.2460 | 8.2110 | 8.5117 | 5.9792 | 4.5784 | 4.7169 | 6.1573 | 4.9266 |
| PINX1_S161 | PINX1 | Q96BK5 | | 161 | TPEGDASPSTPEE | 6.0113 | 6.9131 | 6.4740 | 6.9107 | 5.8857 | 6.7889 | 6.2526 | 5.8371 | 6.2875 | 5.9249 | 7.2143 | 6.6071 | 5.0401 | 6.0859 | 5.5623 | 6.2044 |
| PINX1_S163 | PINX1 | Q96BK5 | | 163 | EGDASPSTPEENE | 5.7649 | 5.6808 | 5.5720 | 6.6055 | 5.9768 | 5.9058 | 5.9044 | 5.8038 | 5.9938 | 5.7076 | 5.8567 | 5.4763 | 6.6799 | 7.6954 | 7.3675 | 8.0089 |
| PIP4K2B_T322 | PIP4K2B | P78356 | | 322 | LLCSYGTPPDSPG | 6.3747 | 5.8683 | 6.7140 | 5.7919 | 6.4044 | 6.7333 | 6.1518 | 5.8219 | 6.5806 | 6.5946 | 6.4556 | 5.9544 | 6.1583 | 6.2752 | 6.2686 | 5.8523 |
| PIP4K2B_T322;S326 | PIP4K2B | P78356 | 322;326 | | NA | 5.8914 | 5.7001 | 6.9529 | 6.9626 | 5.7357 | 5.8570 | 7.1944 | 6.1299 | 5.9479 | 6.3192 | 5.7928 | 6.5465 | 6.4808 | 7.0175 | 6.4895 | 5.8819 |
| PIP4P1_S169 | PIP4P1 | Q86T03-2 | | 169 | VHPGPLSPEPQPM | 6.2768 | 7.3597 | 7.1442 | 5.8592 | 7.0364 | 6.2768 | 7.6587 | 6.3657 | 6.8048 | 6.4222 | 5.4444 | 5.7488 | 4.6470 | 5.9616 | 5.9777 | 5.0160 |
| PIP4P2_S33 | PIP4P2 | Q8N4L2 | | 47 | PYTAIASPDASGI | 6.9367 | 5.9930 | 7.2973 | 6.3037 | 6.6672 | 7.1020 | 5.8262 | 6.2610 | 6.6815 | 7.0949 | 7.3586 | 6.1255 | 5.0493 | 5.1063 | 5.1305 | 5.0664 |
| PIP4P2_S47 | PIP4P2 | Q8N4L2 | | 33 | PYLQESSPRAELP | 6.4288 | 6.2544 | 7.0426 | 7.8444 | 6.2301 | 7.1332 | 7.3198 | 6.0300 | 6.1800 | 5.4283 | 6.4680 | 5.8109 | 5.3729 | 5.8339 | 5.1268 | 5.4959 |
| PIP5K1A_S468 | PIP5K1A | Q99755 | | 486 | CITYQPSVSGEHK | 12.7828 | 11.0648 | 8.7381 | 9.2427 | 6.1431 | 4.4452 | 5.1139 | 5.7306 | 5.4663 | 5.8588 | 5.0378 | 7.1203 | 3.2518 | 4.3210 | 3.3679 | 2.3147 |
| PIP5K1A_S486 | PIP5K1A | Q99755 | | 468 | KFRSGSSFSRRAG | 7.7785 | 6.8005 | 7.4180 | 7.0438 | 5.1339 | 6.7091 | 6.4957 | 6.3006 | 6.5731 | 6.8983 | 7.3407 | 4.5884 | 5.4184 | 3.9483 | 4.8555 |
| PIP5K1C_S476 | PIP5K1C | O60331-3 | | 476 | QPTAAFSASQIPS | 6.8642 | 5.7170 | 6.8793 | 6.1599 | 5.1241 | 6.1017 | 6.5664 | 5.5694 | 6.7461 | 6.0553 | 7.1180 | 7.3399 | 5.9785 | 6.8355 | 5.5298 | 5.4151 |
| PIP5K1C_S555 | PIP5K1C | O60331-3 | | 555 | YRRRTQSSGQDGR | 6.0324 | 6.3922 | 7.4781 | 6.4441 | 6.1524 | 6.0216 | 6.2442 | 6.9541 | 6.1578 | 6.2831 | 6.7347 | 5.8909 | 5.6132 | 6.1750 | 5.7774 | 5.6488 |
| PITPNC1_S270 | PITPNC1 | Q9UKF7 | | 299 | DRPRKKSAPETLT | 6.3878 | 6.5539 | 6.3031 | 5.8238 | 6.3762 | 6.2591 | 5.8780 | 5.9025 | 6.8900 | 6.3286 | 6.2410 | 6.5633 | 6.2380 | 6.9878 | 5.8967 | 6.2702 |
| PITPNC1_S274 | PITPNC1 | Q9UKF7 | | 274 | VRSAPSSAPSTPL | 5.7760 | 5.6917 | 6.2192 | 5.2876 | 6.1180 | 6.7709 | 5.8275 | 5.9764 | 6.1905 | 6.5937 | 6.4306 | 6.9476 | 6.4730 | 6.9969 | 6.5146 | 6.1858 |
| PITPNC1_S299 | PITPNC1 | Q9UKF7 | | 270 | LPSSVRSAPSSAP | 6.8457 | 6.4506 | 6.4623 | 5.9949 | 6.4721 | 6.9200 | 6.0340 | 4.5984 | 6.1826 | 5.9108 | 7.5264 | 7.0022 | 6.0946 | 6.7283 | 5.2810 | 5.4961 |
| PITPNM1_S300 | PITPNM1 | O00562 | | 373 | EMTKWNSNDFIDA | 5.9988 | 6.2860 | 6.1683 | 6.0862 | 5.9614 | 6.0046 | 5.9549 | 6.0395 | 6.6118 | 6.9876 | 6.5135 | 6.6771 | 5.9854 | 6.6106 | 6.1717 | 5.9425 |
| PITPNM1_S373 | PITPNM1 | O00562 | | 593 | GSSRRGSMNNELL | 6.1309 | 5.8968 | 6.7613 | 6.0924 | 5.8571 | 5.9533 | 6.0641 | 5.9802 | 6.6649 | 6.0451 | 6.8488 | 6.9256 | 6.0302 | 6.5587 | 5.9310 | 6.2597 |
| PITPNM1_S593 | PITPNM1 | O00562 | | 896 | EEPSIYSPAFPRE | 5.7529 | 6.7953 | 6.9216 | 6.5877 | 6.0040 | 6.2547 | 6.5549 | 6.0959 | 6.5892 | 6.6715 | 7.0608 | 7.1802 | 5.4543 | 5.0133 | 5.4581 | 5.6058 |
| PITPNM1_S593;S600 | PITPNM1 | O00562 | | 664 | WEPRRASTAFCPP | 5.3619 | 6.4417 | 6.3565 | 5.4366 | 6.9654 | 5.7392 | 5.6484 | 5.4276 | 7.8987 | 7.0787 | 7.0697 | 7.3967 | 6.2811 | 7.4443 | 5.6941 | 5.5596 |
| PITPNM1_S600 | PITPNM1 | O00562 | | 600 | MNNELLSPEFGPV | 5.3907 | 6.3133 | 5.6770 | 5.7488 | 5.3159 | 5.5163 | 6.1578 | 6.3671 | 5.3436 | 5.1738 | 5.9649 | 5.4901 | 7.8744 | 7.8946 | 7.8422 | 7.9297 |
| PITPNM1_S621 | PITPNM1 | O00562 | | 621 | EGLGRGSPEPSAL | 5.7986 | 6.3957 | 6.3499 | 6.0739 | 5.5493 | 6.2922 | 7.2632 | 6.8724 | 6.8379 | 7.0815 | 6.1943 | 5.7612 | 5.6127 | 6.2950 | 5.7549 | 5.8676 |
| PITPNM1_S664 | PITPNM1 | O00562 | | 300 | PPGPDASPDASFG | 5.7666 | 6.0148 | 6.3228 | 6.7180 | 6.1224 | 6.4470 | 5.5491 | 6.6104 | 6.7351 | 5.8878 | 6.7089 | 6.5534 | 5.5007 | 6.1953 | 6.2750 | 6.5927 |
| PITPNM1_S896 | PITPNM1 | O00562 | 593;600 | | NA | 4.7848 | 5.0015 | 4.9764 | 5.0049 | 5.5503 | 5.2799 | 5.5282 | 6.2556 | 6.2169 | 5.2642 | 5.3250 | 6.1336 | 8.8311 | 7.8575 | 9.4228 | 8.5673 |
| PITPNM2_S1277 | PITPNM2 | Q9BZ72 | | 400 | EFRVASSVEQLNI | 6.0632 | 6.7524 | 6.5353 | 6.5566 | 5.9473 | 6.1701 | 6.1124 | 6.3598 | 6.7151 | 6.2600 | 6.0741 | 6.4998 | 5.6904 | 6.2089 | 6.4265 | 5.6280 |
| PITPNM2_S308 | PITPNM2 | Q9BZ72 | | 368 | DITKWSSNDLMDK | 6.4828 | 7.0731 | 7.0599 | 6.1129 | 5.8950 | 6.4607 | 6.0509 | 5.7816 | 7.0901 | 6.4760 | 6.9996 | 6.8506 | 5.0104 | 5.6937 | 5.4873 | 5.4754 |
| PITPNM2_S368 | PITPNM2 | Q9BZ72 | | 1277 | MALRKGSFGLPGQ | 4.9619 | 3.5719 | 7.0806 | 6.0907 | 5.5684 | 5.3536 | 5.0082 | 6.7268 | 8.9213 | 6.9332 | 4.9703 | 6.5456 | 5.8254 | 8.2347 | 6.1960 | 8.0115 |
| PITPNM2_S400 | PITPNM2 | Q9BZ72 | | 872 | PPARKASPGLERA | 6.5832 | 7.1423 | 6.1844 | 6.6705 | 6.4295 | 5.9275 | 5.6045 | 6.3572 | 5.9523 | 5.9243 | 5.5373 | 5.9088 | 5.9742 | 6.5516 | 6.7244 | 6.5279 |
| PITPNM2_S894 | PITPNM2 | Q9BZ72 | | 308 | KQWSSNSDLLSSS | 5.7628 | 5.8238 | 5.9433 | 6.7132 | 5.6088 | 6.2955 | 6.0761 | 5.4654 | 5.6035 | 6.3203 | 6.6831 | 6.7345 | 6.3663 | 6.9760 | 7.2468 | 6.3805 |
| PJA1_S265 | PJA1 | Q8NG27 | | 265 | KIRNLASPNCVKP | 6.5984 | 6.7575 | 7.1352 | 6.5310 | 6.8268 | 7.2841 | 6.9648 | 7.3032 | 6.7312 | 6.2762 | 7.1480 | 6.9885 | 4.3258 | 4.5215 | 3.8319 | 4.7756 |
| PKD2_S166 | PKD2 | Q13563 | | 829 | SSRRRGSISSGVS | 5.7597 | 6.3191 | 6.2940 | 6.2515 | 5.9948 | 6.5606 | 6.0711 | 5.7141 | 6.5067 | 6.3627 | 6.4083 | 6.3597 | 5.7954 | 6.4938 | 6.4881 | 6.2606 |
| PKD2_S829 | PKD2 | Q13563 | | 166 | QGPPCPSPVGGGD | 5.1272 | 6.0337 | 6.1931 | 7.0346 | 6.4923 | 5.9207 | 7.1330 | 6.5987 | 5.9118 | 6.3996 | 6.1305 | 6.5151 | 5.4231 | 6.9021 | 6.7338 | 6.3508 |
| PKM_S100 | PKM | P14618-2 | | 37 | CRLDIDSPPITAR | 6.0255 | 6.1987 | 6.8076 | 5.8957 | 6.1700 | 6.4899 | 6.0137 | 6.5124 | 6.5273 | 6.4230 | 6.2303 | 6.5352 | 5.9875 | 6.3910 | 5.9177 | 5.8745 |
| PKM_S37 | PKM | P14618-2 | | 100 | ATESFASDPILYR | 5.7699 | 6.2833 | 6.6985 | 6.1478 | 6.5074 | 6.4596 | 6.0210 | 6.3365 | 5.9728 | 6.2323 | 6.3212 | 6.7201 | 6.2596 | 6.3794 | 5.9508 | 5.9397 |
| PKMYT1_S120 | PKMYT1 | Q99640 | | 40 | HAEPGFSLKRPRG | 9.4641 | 8.9729 | 7.9412 | 8.4473 | 5.9189 | 5.9530 | 8.0996 | 6.4574 | 7.3145 | 6.5829 | 7.4636 | 6.8726 | 2.1852 | 3.3203 | 2.1628 | 2.8436 |
| PKMYT1_S143 | PKMYT1 | Q99640 | | 120 | SRLGHGSYGEVFK | 6.0233 | 6.4141 | 5.6053 | 6.7583 | 5.3229 | 5.1739 | 5.7933 | 5.6511 | 5.9108 | 5.9752 | 5.9591 | 6.0057 | 6.6517 | 7.2501 | 7.8581 | 7.6471 |
| PKMYT1_S40 | PKMYT1 | Q99640 | | 479 | SEPPRGSFPSFEP | 4.9453 | 4.8235 | 4.8406 | 5.5379 | 5.1733 | 4.9615 | 5.0158 | 4.9352 | 5.0694 | 5.3807 | 4.8837 | 5.0155 | 9.8374 | 9.3447 | 10.3804 | 9.8551 |
| PKMYT1_S479 | PKMYT1 | Q99640 | | 50 | PRGLSRSLPPPPP | 7.5247 | 7.6311 | 7.0112 | 7.5694 | 5.8675 | 6.3953 | 6.9001 | 6.4405 | 7.3000 | 7.5642 | 7.7153 | 7.9950 | 3.0977 | 4.0150 | 3.0407 | 3.9322 |
| PKMYT1_S50 | PKMYT1 | Q99640 | | 60 | PPPAKGSIPISRL | 7.7721 | 8.5789 | 6.7996 | 8.0730 | 6.1940 | 6.7203 | 7.0892 | 6.6126 | 6.7521 | 7.5068 | 7.6912 | 7.4378 | 2.9097 | 3.9584 | 2.9570 | 2.9474 |
| PKMYT1_S60 | PKMYT1 | Q99640 | | 94 | ASETLQSPGYDPS | 6.4621 | 6.9917 | 7.1894 | 7.6847 | 7.0305 | 6.1973 | 6.6740 | 7.0158 | 7.1148 | 7.0109 | 7.5116 | 6.9623 | 3.8615 | 4.6250 | 3.9640 | 3.7045 |
| PKMYT1_S94 | PKMYT1 | Q99640 | | 143 | AVKRSMSPFRGPK | 8.2536 | 8.2314 | 7.4646 | 7.3879 | 5.7684 | 5.7658 | 6.7498 | 5.3449 | 5.0094 | 6.2331 | 6.7065 | 6.5206 | 4.1433 | 5.0589 | 3.4853 | 4.1484 |
| PKMYT1_T455 | PKMYT1 | Q99640 | | 455 | TVGSTSTPRSRCT | 7.7694 | 5.7733 | 6.5734 | 7.4585 | 6.6451 | 6.2942 | 6.5165 | 7.0001 | 7.3160 | 6.6169 | 6.3183 | 6.2826 | 4.3949 | 4.6062 | 4.3357 | 4.5992 |
| PKN1_S380 | PKN1 | Q16512-2 | | 779 | GYGDRTSTFCGTP | 6.8424 | 6.7818 | 7.4637 | 7.0265 | 7.0417 | 5.7635 | 6.1447 | 5.7056 | 6.0820 | 5.6546 | 5.6504 | 5.4104 | 6.6436 | 6.1765 | 6.1540 | 5.4586 |
| PKN1_S539 | PKN1 | Q16512-2 | | 922 | GEAPTLSPPRDAR | 6.1093 | 6.4663 | 6.6356 | 6.1262 | 6.2402 | 6.2825 | 6.3811 | 6.6141 | 6.2468 | 6.2695 | 6.6439 | 6.6207 | 6.0246 | 6.1308 | 5.3656 | 5.8427 |
| PKN1_S539;S543 | PKN1 | Q16512-2 | | 568 | GTDSDSSPQKSSR | 6.2144 | 6.3319 | 6.8322 | 6.6803 | 6.3817 | 6.6324 | 6.5535 | 6.7990 | 6.9922 | 7.2786 | 7.0714 | 7.3975 | 4.3927 | 5.1433 | 4.3956 | 4.9034 |
| PKN1_S568 | PKN1 | Q16512-2 | | 380 | LYSRSGSLSGRSS | 7.0047 | 6.5242 | 7.0289 | 5.9648 | 5.9779 | 6.3485 | 5.9415 | 6.2447 | 7.2871 | 7.1172 | 7.0529 | 7.1828 | 5.1322 | 5.9217 | 4.3890 | 4.8819 |
| PKN1_S614 | PKN1 | Q16512-2 | | 614 | CSPLRKSPLTLED | 7.0323 | 4.8212 | 6.3575 | 5.9422 | 6.5927 | 6.7030 | 6.2001 | 7.3162 | 5.3144 | 5.3614 | 8.0835 | 7.5125 | 5.6599 | 5.1457 | 6.2245 | 5.7329 |
| PKN1_S779 | PKN1 | Q16512-2 | | 539 | TGTGTFSPGASPG | 6.7071 | 8.0810 | 7.5331 | 6.9094 | 5.2869 | 6.5638 | 5.7905 | 5.6978 | 6.9600 | 6.6060 | 8.0936 | 7.7375 | 3.9783 | 4.6737 | 4.1090 | 5.2721 |
| PKN1_S922 | PKN1 | Q16512-2 | 539;543 | | NA | 5.7991 | 6.8810 | 6.7342 | 6.9717 | 6.2493 | 6.2752 | 7.3233 | 7.1668 | 6.0164 | 5.9998 | 6.6648 | 6.8112 | 5.1299 | 5.6933 | 4.9315 | 5.3524 |
| PKN2_S110 | PKN2 | Q16513 | | 958 | SEAPILTPPREPR | 6.0495 | 6.2875 | 6.5670 | 6.4380 | 6.2424 | 6.0617 | 6.4958 | 6.4713 | 6.4832 | 6.2565 | 6.2962 | 6.4453 | 5.8240 | 6.4682 | 5.7670 | 5.8465 |
| PKN2_S360 | PKN2 | Q16513 | | 816 | YGDRTSTFCGTPE | 7.5245 | 6.7773 | 7.5969 | 7.3818 | 6.6346 | 5.3472 | 5.9140 | 5.5323 | 5.9069 | 5.5699 | 5.8521 | 5.1755 | 6.8675 | 6.0363 | 6.7839 | 5.0993 |
| PKN2_S535 | PKN2 | Q16513 | | 360 | VALPGWSPSETRS | 6.2994 | 7.1728 | 6.7422 | 7.0947 | 6.2964 | 7.2387 | 7.0467 | 6.9852 | 7.2087 | 7.2717 | 7.3178 | 7.2272 | 3.8598 | 4.4813 | 3.3518 | 4.0755 |
| PKN2_S583 | PKN2 | Q16513 | | 535 | NHSGTFSPQAPVP | 6.9845 | 7.3502 | 7.5491 | 6.9878 | 6.2963 | 6.9373 | 7.3870 | 6.8462 | 7.8587 | 7.0977 | 7.7978 | 7.6673 | 3.0494 | 3.7510 | 2.8116 | 3.6281 |
| PKN2_T121 | PKN2 | Q16513 | | 110 | NAHIVVSPDEDIT | 6.5502 | 6.5672 | 7.1430 | 6.9453 | 6.3315 | 6.8585 | 6.5836 | 6.5368 | 6.5889 | 6.3234 | 6.5966 | 6.3743 | 5.4927 | 5.4203 | 5.0843 | 5.5035 |
| PKN2_T533 | PKN2 | Q16513 | | 583 | APPRASSLGEIDE | 5.8583 | 6.2008 | 6.2837 | 5.8458 | 6.0809 | 6.1388 | 6.3503 | 6.3011 | 5.9792 | 6.1855 | 6.0941 | 6.3650 | 6.6536 | 6.9351 | 6.3183 | 6.4093 |
| PKN2_T628 | PKN2 | Q16513 | | 533 | TVNHSGTFSPQAP | 6.3393 | 6.8261 | 6.5558 | 6.3483 | 7.2950 | 6.9906 | 6.9718 | 7.7346 | 7.3479 | 6.4647 | 6.9800 | 6.8668 | 4.6453 | 4.1763 | 3.6707 | 4.7787 |
| PKN2_T816 | PKN2 | Q16513 | | 628 | SEYKPDTPQSGLE | 6.4653 | 7.0943 | 6.8524 | 6.3469 | 5.5881 | 6.8040 | 6.2298 | 7.1159 | 5.5805 | 6.0821 | 6.4386 | 8.3560 | 4.4639 | 6.7518 | 4.9897 | 4.8407 |
| PKN2_T958 | PKN2 | Q16513 | | 121 | ITDCPRTPDTPNN | 7.3438 | 6.1480 | 6.6992 | 6.7982 | 5.7398 | 5.8237 | 5.2986 | 6.0257 | 6.0614 | 8.0393 | 5.9157 | 7.5775 | 6.2250 | 5.1941 | 4.8890 | 6.2209 |
| PKN3_S171 | PKN3 | Q6P5Z2 | | 860 | GLPPALTPPAPHS | 6.2387 | 6.3830 | 6.4740 | 6.5596 | 6.2514 | 6.3203 | 6.1650 | 6.3593 | 6.0069 | 6.8231 | 6.5097 | 6.5312 | 5.6222 | 6.3328 | 5.7559 | 5.6671 |
| PKN3_S544 | PKN3 | Q6P5Z2 | | 508 | PLGEEMTPPPKPP | 6.6456 | 6.6602 | 6.6139 | 7.4060 | 6.3891 | 6.3038 | 6.8970 | 6.6152 | 6.6861 | 6.6420 | 6.4503 | 6.8521 | 4.7775 | 5.1197 | 4.7698 | 5.1716 |
| PKN3_T508 | PKN3 | Q6P5Z2 | | 718 | FGDRTSTFCGTPE | 6.5609 | 6.3876 | 7.3449 | 7.6079 | 6.3098 | 5.9192 | 6.1905 | 5.3475 | 6.0327 | 5.9058 | 5.6925 | 5.6580 | 6.3271 | 6.2755 | 6.5903 | 5.8497 |
| PKN3_T527 | PKN3 | Q6P5Z2 | | 527 | EPTSEETPRTKRP | 6.9839 | 6.9304 | 6.7442 | 7.5788 | 5.8166 | 6.7458 | 6.8574 | 6.9578 | 6.7116 | 6.0831 | 6.1506 | 6.3805 | 4.8972 | 5.2785 | 4.7599 | 5.1239 |
| PKN3_T718 | PKN3 | Q6P5Z2 | | 544 | RTRRGPSPPPASPT | 6.6410 | 6.7432 | 7.5683 | 7.7157 | 6.0275 | 5.4925 | 6.0180 | 6.2106 | 6.4460 | 6.4484 | 5.8112 | 5.9396 | 4.9576 | 5.8278 | 5.7898 | 6.3628 |
| PKN3_T860 | PKN3 | Q6P5Z2 | | 171 | SLEASGSPEPGPE | 6.3589 | 5.2991 | 6.2697 | 6.8730 | 8.6044 | 6.9517 | 4.9790 | 5.9759 | 9.3075 | 5.9759 | 6.6369 | 7.0672 | 4.8164 | 4.4544 | 4.1448 | 6.2854 |
| PKNOX1_S41 | PKNOX1 | P55347 | | 41 | PDAEGVSPPPVES | 6.0008 | 6.1441 | 6.2982 | 5.9199 | 6.2509 | 6.1356 | 6.0429 | 5.6420 | 6.7079 | 6.3976 | 5.7985 | 6.1951 | 6.7924 | 6.7335 | 6.6322 | 6.3083 |
| PKP2_S151 | PKP2 | Q99959 | | 251 | RPGTSRSMGNLLE | 6.6648 | 6.7634 | 6.9909 | 6.0384 | 5.2604 | 6.3707 | 6.2610 | 5.9454 | 5.7284 | 6.1204 | 7.2891 | 7.4093 | 5.2350 | 5.6872 | 5.3908 | 6.8449 |
| PKP2_S183 | PKP2 | Q99959 | | 183 | TLHHQSRRAALL | 9.5906 | 8.3237 | 8.4525 | 7.2262 | 5.6387 | 5.3790 | 6.6730 | 5.7178 | 7.5401 | 5.0222 | 6.1333 | 5.5936 | 5.1871 | 5.3881 | 4.0557 | 4.0783 |

| | | | | | | | | | | | | | | | | | | | | |
|---|---|---|---|---|---|---|---|---|---|---|---|---|---|---|---|---|---|---|---|---|
| PKP2_S251 | PKP2 | Q99959 | 81 | GNLHRTSSVPEYV | 6.7740 | 6.3086 | 6.8499 | 5.9028 | 6.3189 | 6.7172 | 5.7715 | 5.8132 | 6.3730 | 6.5632 | 6.6371 | 6.6980 | 5.8967 | 6.3695 | 5.7420 | 5.2644 |
| PKP2_S329 | PKP2 | Q99959 | 151 | LRRLEISPDSSPE | 5.7892 | 5.5498 | 6.2712 | 6.2762 | 6.2906 | 5.6395 | 6.6823 | 5.6257 | 6.7819 | 6.7361 | 6.8383 | 6.8095 | 5.8770 | 6.4397 | 5.8594 | 6.5337 |
| PKP2_S81 | PKP2 | Q99959 | 329 | QAAAGGSGNLLTE | 5.2619 | 4.8211 | 4.9994 | 5.0755 | 7.9786 | 6.7868 | 6.1838 | 6.6960 | 4.5082 | 4.9274 | 4.1905 | 4.4032 | 9.5803 | 8.0076 | 8.9376 | 7.6421 |
| PKP3_S253 | PKP3 | Q9Y446-2 | 253 | PEATEVSPSIRTR | 6.3173 | 6.3086 | 6.3737 | 7.1419 | 6.0959 | 6.5662 | 5.9295 | 6.0737 | 6.4575 | 6.1167 | 7.0863 | 6.2717 | 4.8559 | 6.1992 | 5.5349 | 6.6709 |
| PKP3_S328 | PKP3 | Q9Y446-2 | 329 | TLQRLSSGFDDID | 7.7900 | 6.4399 | 7.0388 | 5.3226 | 7.8805 | 6.0407 | 6.1024 | 6.8033 | 7.7727 | 7.0446 | 6.8780 | 5.8784 | 5.1301 | 6.0177 | 4.0903 | 3.7700 |
| PKP3_S329 | PKP3 | Q9Y446-2 | 328 | RTLQRLSSGFDDI | 5.7625 | 6.2983 | 7.0070 | 6.0785 | 5.7987 | 5.8169 | 6.6174 | 6.0980 | 7.3293 | 6.3800 | 6.7463 | 6.9045 | 5.7497 | 5.9421 | 5.4201 | 6.0505 |
| PKP4_S1100 | PKP4 | Q99569 | 314 | TTARVGSPLTLTD | 6.4054 | 5.9705 | 6.4291 | 6.2932 | 6.1400 | 5.9224 | 6.7818 | 5.9064 | 6.6351 | 6.3136 | 6.0618 | 6.0585 | 6.3761 | 6.1985 | 6.0351 | 6.4725 |
| PKP4_S214 | PKP4 | Q99569 | 1100 | YISSYSSPAREQN | 6.5320 | 5.6862 | 6.7377 | 7.1565 | 6.7742 | 6.9774 | 6.9226 | 6.5307 | 6.5642 | 6.6225 | 7.0355 | 6.7042 | 5.0389 | 5.0963 | 4.5074 | 5.1138 |
| PKP4_S221 | PKP4 | Q99569 | 281 | YSQRPASPTAIRR | 6.3567 | 6.8015 | 6.9362 | 7.0236 | 6.2436 | 6.1954 | 6.0323 | 5.9925 | 6.2702 | 6.2404 | 6.5263 | 6.6161 | 5.7896 | 6.0582 | 5.5342 | 5.3834 |
| PKP4_S221;S231 | PKP4 | Q99569 | 776 | DLLGKESPSKDSE | 6.5125 | 6.4378 | 6.4550 | 6.8268 | 6.4030 | 6.2728 | 6.3133 | 6.0474 | 6.1732 | 6.0892 | 6.1612 | 6.6070 | 5.9981 | 6.1490 | 5.8746 | 6.5791 |
| PKP4_S231 | PKP4 | Q99569 | 1017 | FITPVSTLERDRF | 6.7349 | 6.6439 | 6.5614 | 6.3890 | 6.5503 | 6.2986 | 6.5000 | 6.1905 | 6.4760 | 6.3481 | 6.3553 | 5.9713 | 6.1057 | 6.2365 | 5.2704 | 5.3679 |
| PKP4_S236 | PKP4 | Q99569 | 447 | ALYRTGSVGIGNL | 7.6350 | 7.1055 | 8.0370 | 6.2590 | 5.3865 | 6.0115 | 6.4974 | 6.0081 | 7.1057 | 7.3358 | 7.3072 | 7.5151 | 4.5730 | 5.3943 | 3.4489 | 4.3800 |
| PKP4_S273;T281 | PKP4 | Q99569 | 382 | VPPRPDSLTGLRS | 5.7480 | 6.1514 | 5.7705 | 5.8295 | 5.8245 | 5.6059 | 5.7721 | 6.0252 | 6.1139 | 5.8600 | 5.2850 | 5.3937 | 8.0785 | 7.5351 | 7.9344 | 7.0723 |
| PKP4_S281 | PKP4 | Q99569 | 231 | VISTGVSPSRGSL | 6.9419 | 6.6073 | 6.1856 | 7.0161 | 6.3806 | 6.7171 | 6.8097 | 6.3734 | 6.8393 | 6.7130 | 6.9474 | 6.9802 | 4.9478 | 5.4197 | 4.2307 | 4.8902 |
| PKP4_S290 | PKP4 | Q99569 | 406 | DLRSAVSPDLHIT | 6.7271 | 6.9339 | 6.0913 | 7.6930 | 6.7152 | 6.9146 | 7.4126 | 8.1045 | 6.1467 | 6.4296 | 6.9150 | 6.4426 | 3.9308 | 4.7842 | 4.4169 | 4.3419 |
| PKP4_S314 | PKP4 | Q99569 | 422 | EGRTYYSPVYRSP | 7.3646 | 7.1676 | 6.5039 | 7.0386 | 6.7496 | 6.5488 | 7.1621 | 7.1710 | 7.3012 | 6.5743 | 6.6976 | 6.6438 | 3.8903 | 4.4685 | 4.1596 | 4.5586 |
| PKP4_S327 | PKP4 | Q99569 | 221 | VPSRAQSPSYVIS | 6.3245 | 6.1222 | 5.7911 | 6.0950 | 6.3807 | 6.0057 | 6.3754 | 5.7797 | 6.3959 | 5.8494 | 5.7311 | 5.1186 | 6.7247 | 7.0549 | 6.8784 | 7.3726 |
| PKP4_S327;S337 | PKP4 | Q99569 | 327 | AQTRVASPSQGQV | 6.6094 | 6.9076 | 6.3626 | 6.4203 | 6.3654 | 6.1959 | 5.6726 | 6.0131 | 5.9599 | 5.0870 | 5.4609 | 6.8572 | 7.3076 | 6.6013 | 6.0509 | 7.1283 |
| PKP4_S337 | PKP4 | Q99569 | 337 | GQVGSSSPKRSGM | 6.8061 | 6.6188 | 6.1357 | 6.2596 | 5.6921 | 5.6287 | 6.4534 | 5.8240 | 6.7695 | 6.0865 | 6.0007 | 5.9704 | 6.5917 | 6.5533 | 6.0485 | 6.5611 |
| PKP4_S382 | PKP4 | Q99569 | 290 | AIRRIGSVTSRQT | 8.0065 | 7.4651 | 7.4363 | 6.2955 | 5.7719 | 6.1496 | 6.9714 | 7.5457 | 6.9064 | 6.5859 | 6.4721 | 6.8752 | 4.7794 | 5.6671 | 3.7681 | 5.1038 |
| PKP4_S406 | PKP4 | Q99569 | 214 | AMRRVSSVPSRAQ | 6.7114 | 7.2657 | 7.5223 | 6.0037 | 5.7155 | 6.6021 | 5.9492 | 6.7338 | 6.4025 | 7.1686 | 7.2101 | 7.6720 | 4.5617 | 5.6565 | 3.9663 | 4.8586 |
| PKP4_S422 | PKP4 | Q99569 | 512 | GTTRSPSIDSIQK | 6.5253 | 6.4459 | 5.7186 | 6.3787 | 6.3762 | 5.7237 | 5.8352 | 6.0433 | 6.7012 | 5.9000 | 5.3587 | 5.6874 | 7.2420 | 6.7648 | 6.6537 | 6.6453 |
| PKP4_S427 | PKP4 | Q99569 | 427 | YSPVYRSPNHGTV | 4.8273 | 5.5218 | 6.7456 | 6.4531 | 7.4533 | 6.2952 | 5.8518 | 7.1671 | 6.9990 | 6.5376 | 5.7884 | 6.5493 | 5.7071 | 5.7931 | 6.8840 | 5.4264 |
| PKP4_S447 | PKP4 | Q99569 | 75 | CRLGAESPSIAST | 5.8882 | 5.7242 | 5.8394 | 6.8014 | 6.1980 | 6.3688 | 6.4601 | 6.2692 | 5.8202 | 5.8198 | 5.9431 | 5.7410 | 7.0483 | 5.9495 | 6.9050 | 7.2239 |
| PKP4_S447 | PKP4 | Q99569 | 236 | VSPSRGSLRTSLG | 6.7930 | 6.4484 | 8.6782 | 6.3293 | 6.0259 | 7.0659 | 6.0917 | 6.7798 | 6.8251 | 6.8760 | 6.9682 | 7.8723 | 4.5784 | 4.7102 | 3.6442 | 4.3135 |
| PKP4_S512 | PKP4 | A0A0D9SF60 | 447 | ALYRTGSIGNLQ | 7.5060 | 6.4988 | 7.2974 | 5.8805 | 5.6746 | 6.1671 | 6.3049 | 5.6696 | 7.2169 | 6.9513 | 6.6443 | 7.2759 | 5.6790 | 5.8345 | 4.3146 | 5.0845 |
| PKP4_S75 | PKP4 | Q99569 | 273;281 | NA | 6.3560 | 6.5053 | 5.7503 | 6.2642 | 5.9927 | 6.1682 | 6.2446 | 6.5268 | 6.2836 | 5.9419 | 6.1686 | 6.2361 | 6.2247 | 6.6214 | 6.5460 | 6.1695 |
| PKP4_S776 | PKP4 | Q99569 | 221;231 | NA | 5.6626 | 5.6176 | 6.5579 | 6.4896 | 6.2854 | 5.9268 | 5.7944 | 6.3421 | 6.0321 | 6.2327 | 6.2018 | 6.2587 | 6.5916 | 7.2397 | 5.6896 | 6.1774 |
| PKP4_T1017 | PKP4 | Q99569 | 327;337 | NA | 6.2573 | 6.6963 | 6.3864 | 7.1746 | 6.1253 | 6.3394 | 6.6044 | 5.8464 | 6.5728 | 6.6574 | 7.1412 | 6.6225 | 5.2836 | 6.5698 | 5.3395 | 5.2830 |
| PLA2G4C_S347 | PLA2G4C | Q9UP65-3 | 347 | DPERKGSLSNLMD | 2.9656 | 2.9126 | 3.1936 | 3.1212 | 3.9920 | 3.5164 | 3.6096 | 4.0421 | 4.1886 | 4.0356 | 3.5848 | 3.7056 | 14.4359 | 12.7054 | 16.2473 | 13.7435 |
| PLAA_S50 | PLAA | Q9Y263 | 50 | RLWAPDSPNRSFT | 6.2736 | 6.4876 | 6.6792 | 6.7343 | 6.1232 | 6.0378 | 5.8845 | 6.2704 | 6.5562 | 6.6943 | 6.8454 | 7.0099 | 5.2292 | 5.7993 | 5.6615 | 5.7137 |
| PLAGL1_S341 | PLAGL1 | Q9UM63 | 341 | PLQEPQSPQKLNP | 6.2577 | 6.3289 | 6.6463 | 7.1510 | 6.7623 | 6.5223 | 5.8382 | 6.2907 | 7.0075 | 6.5064 | 6.8430 | 7.1821 | 4.7337 | 5.3326 | 5.1393 | 5.4580 |
| PLAGL2_S273 | PLAGL2 | Q9UPG8 | 273 | SVKEELSPVLCMA | 7.0874 | 6.7420 | 6.3351 | 6.1538 | 6.6052 | 6.6565 | 6.5095 | 6.7369 | 7.5593 | 6.7813 | 7.1746 | 6.9882 | 3.8027 | 4.4303 | 5.6391 | 4.7979 |
| PLCB3_S474 | PLCB3 | Q01970 | 474 | KKRHRPSAGGPDS | 6.3591 | 6.9972 | 7.1275 | 7.0106 | 6.5688 | 6.5440 | 6.2083 | 6.5241 | 6.3868 | 6.0490 | 6.1651 | 6.9292 | 4.7509 | 6.1580 | 6.2118 | 4.9096 |
| PLCB3_S916 | PLCB3 | Q01970 | 926 | PPGPTTSPASTSL | 6.1973 | 7.0247 | 6.4714 | 6.7494 | 6.3720 | 5.8814 | 6.4164 | 6.2043 | 6.5595 | 6.9549 | 7.3679 | 6.6520 | 4.9347 | 5.8432 | 4.8719 | 5.4989 |
| PLCB3_S926 | PLCB3 | Q01970 | 916 | PSPLDASPRRPPG | 5.7903 | 6.4667 | 7.4045 | 7.5287 | 6.5988 | 7.4045 | 5.9439 | 6.7085 | 8.1430 | 6.9503 | 5.9733 | 7.0745 | 4.0846 | 5.0387 | 4.5813 | 5.2086 |
| PLCB3_T925 | PLCB3 | Q01970 | 925 | RPPGPTTSPASTS | 6.3252 | 6.8571 | 7.1212 | 6.7268 | 6.2143 | 6.2689 | 6.9463 | 6.9078 | 7.3802 | 7.4548 | 6.8956 | 7.4754 | 3.9126 | 5.1476 | 3.8954 | 4.4708 |
| PLCB3_T925;S934 | PLCB3 | Q01970 | 925;934 | NA | 6.9348 | 7.6104 | 6.9995 | 7.3132 | 5.7824 | 6.5407 | 7.5326 | 6.6898 | 7.5939 | 7.9294 | 7.8486 | 7.7274 | 3.1162 | 3.8530 | 2.9124 | 3.6156 |
| PLCB4_S1187 | PLCB4 | Q15147-4 | 886 | TAKANVTPQSSSE | 5.2226 | 5.3667 | 5.8570 | 5.8858 | 6.3809 | 6.2709 | 6.4536 | 6.0638 | 6.1394 | 6.0126 | 6.4251 | 6.3515 | 7.0084 | 6.5728 | 6.8124 | 7.1766 |
| PLCB4_S450 | PLCB4 | Q15147-4 | 450 | PGRALPSPNDLKR | 6.4629 | 7.5146 | 8.3380 | 8.4644 | 6.5173 | 7.1143 | 6.1258 | 5.3375 | 7.1055 | 7.5602 | 7.5743 | 8.9034 | 2.5581 | 3.5712 | 3.4255 | 3.4272 |
| PLCB4_T886 | PLCB4 | Q15147-4 | 1187 | PQTSNSSMKLQNA | 7.3495 | 7.8583 | 8.2047 | 8.2334 | 6.2163 | 6.5955 | 5.8256 | 6.5367 | 7.5165 | 7.2932 | 6.9414 | 6.9728 | 3.8769 | 4.4342 | 3.5328 | 3.5122 |
| PLCD1_S481 | PLCD1 | P51178-2 | 481 | PEATVVSDEDEAA | 5.5636 | 6.3286 | 6.1007 | 5.5462 | 6.4105 | 6.9269 | 5.7924 | 6.0020 | 6.7639 | 5.9766 | 6.4076 | 5.7127 | 6.6285 | 6.5918 | 6.0210 | 6.2270 |
| PLCD3_S105 | PLCD3 | Q8N3E9 | 105 | RIPRAPSQHIFFV | 5.6946 | 5.2293 | 6.0465 | 6.2049 | 5.6573 | 5.2791 | 6.1458 | 6.1534 | 6.0526 | 6.1230 | 5.5335 | 5.6455 | 7.5388 | 7.4628 | 7.8926 | 7.3410 |
| PLCD3_S496 | PLCD3 | Q8N3E9 | 496 | EDGRALSDREEEE | 7.1673 | 6.3492 | 5.7375 | 6.8475 | 5.8144 | 5.8279 | 6.7061 | 6.2005 | 6.7824 | 6.1893 | 6.0383 | 6.1741 | 6.1270 | 6.2386 | 5.6764 | 6.1236 |
| PLCG1_S1222 | PLCG1 | P19174-2 | 1264 | FEDFRISQEHLAD | 5.5573 | 6.3114 | 6.3704 | 6.3586 | 5.7062 | 6.8615 | 5.9688 | 6.5390 | 6.1148 | 6.5120 | 6.9663 | 6.1851 | 5.3267 | 6.5159 | 6.0189 | 6.6870 |
| PLCG1_S1264 | PLCG1 | P19174-2 | 1222 | QENGDLSPFSGTS | 6.7451 | 7.2258 | 6.5838 | 6.8693 | 6.1919 | 6.1219 | 5.9655 | 6.6065 | 6.9445 | 7.1403 | 7.0024 | 7.2596 | 4.1477 | 5.2340 | 4.8912 | 5.0704 |
| PLCH1_S1027;S1030 | PLCH1 | Q4KWH8 | 1307 | ESNLPGSPNTSRG | 6.1276 | 5.8655 | 5.7510 | 6.6453 | 6.5963 | 5.8115 | 6.1254 | 6.6345 | 6.1427 | 6.3357 | 6.0721 | 5.7208 | 6.5251 | 7.0550 | 6.1982 | 6.3933 |
| PLCH1_S1307 | PLCH1 | Q4KWH8 | 1027;1030 | NA | 5.9802 | 5.6120 | 5.8180 | 5.9475 | 5.9860 | 6.0449 | 6.6188 | 6.3933 | 6.0356 | 5.1695 | 5.2037 | 5.6777 | 7.0654 | 8.1902 | 7.0579 | 7.1994 |
| PLD1_S499 | PLD1 | Q13393 | 499 | RLTDVGSVKRVTS | 6.5219 | 6.6645 | 6.4224 | 6.7138 | 5.2864 | 5.6740 | 5.8399 | 6.1496 | 6.5794 | 6.3492 | 6.6291 | 5.8514 | 5.7735 | 7.0099 | 6.2522 | 6.2828 |
| PLD1_S505 | PLD1 | Q13393 | 629 | DTGSIRSLQTGVG | 6.2693 | 6.8792 | 6.7033 | 6.3524 | 6.1075 | 5.5015 | 6.1090 | 6.3199 | 6.3947 | 6.3612 | 6.2096 | 6.3311 | 5.5166 | 6.2519 | 6.5459 | 6.1469 |
| PLD1_S629 | PLD1 | Q13393 | 505 | SVKRVTSGPSLGS | 3.4891 | 3.9174 | 4.9202 | 6.8450 | 6.2704 | 4.2804 | 5.1448 | 4.5128 | 5.0247 | 4.6591 | 5.2205 | 5.4817 | 9.3677 | 10.9347 | 8.7070 | 11.2246 |
| PLEC_S1047 | PLEC | Q15149 | 1435 | QLEPVASPAKKPK | 6.6624 | 6.6443 | 6.1612 | 6.4057 | 6.9383 | 6.6535 | 6.6933 | 6.7356 | 6.9890 | 6.7322 | 6.6590 | 6.1941 | 4.9623 | 5.4612 | 4.9830 | 5.1259 |
| PLEC_S125 | PLEC | Q15149 | 4642 | AGSRRGSFDATGS | 5.6831 | 6.0542 | 6.1715 | 5.9432 | 6.3512 | 6.2956 | 5.9917 | 6.0310 | 6.4413 | 6.3047 | 6.4337 | 6.4542 | 6.7502 | 6.6971 | 6.2975 | 6.0997 |
| PLEC_S1435 | PLEC | Q15149-4 | 21 | GRKRTSSEDNLYL | 5.8540 | 5.8131 | 5.5988 | 5.5743 | 5.8927 | 6.3203 | 6.2092 | 5.7668 | 5.7844 | 5.8290 | 6.0283 | 6.1233 | 7.2570 | 7.1933 | 7.2330 | 7.1623 |
| PLEC_S149 | PLEC | Q15149 | 1721 | LQSKRASFAEKTA | 6.2354 | 6.1165 | 5.5995 | 5.7427 | 6.2679 | 5.9171 | 6.0477 | 6.1675 | 6.1002 | 5.7837 | 5.8469 | 5.6434 | 7.1490 | 7.2695 | 7.3328 | 6.7803 |
| PLEC_S1721 | PLEC | Q15149 | 4406 | SRTQLASWSDPTE | 5.7396 | 5.9041 | 6.4208 | 6.6663 | 6.3634 | 6.3735 | 6.1734 | 6.2501 | 6.0218 | 5.7862 | 6.3149 | 6.0945 | 6.4438 | 6.3025 | 6.6539 | 6.4911 |
| PLEC_S1732 | PLEC | Q15149 | 1047 | QLIRSWSLATFRT | 4.6761 | 4.9703 | 4.7124 | 4.4124 | 5.6986 | 4.8478 | 5.1379 | 5.1395 | 4.2414 | 3.7557 | 3.6014 | 3.0329 | 12.5624 | 10.2098 | 14.4915 | 8.5099 |
| PLEC_S20 | PLEC | Q15149 | 125 | VQGPLGSPPKRGP | 6.0948 | 6.2121 | 6.4344 | 6.5041 | 6.2398 | 6.3762 | 6.6745 | 6.1218 | 6.1684 | 6.3361 | 6.3776 | 6.5695 | 5.7418 | 6.4249 | 5.7580 | 5.9659 |
| PLEC_S20 | PLEC | Q15149 | 4386 | FRSRSSSVGSSSS | 5.6782 | 5.4365 | 5.8718 | 5.6128 | 6.1459 | 5.4901 | 5.8760 | 5.6082 | 6.2419 | 6.6070 | 5.7746 | 5.7127 | 7.7201 | 7.7452 | 7.6359 | 6.8430 |
| PLEC_S20;S21 | PLEC | Q15149 | 4626 | SGSTAGSRTGSRT | 5.6824 | 5.6289 | 6.5325 | 6.0226 | 6.5984 | 6.7175 | 5.8229 | 6.0645 | 5.9186 | 6.3470 | 6.6026 | 6.1807 | 6.5423 | 7.0883 | 6.7988 | 6.3520 |
| PLEC_S21 | PLEC | Q15149 | 4382 | NAGGFRSRSSSVG | 6.0742 | 6.6086 | 7.0168 | 6.3506 | 6.7524 | 7.4258 | 6.1575 | 5.5945 | 5.9792 | 5.9340 | 6.5505 | 6.2586 | 6.2297 | 6.3287 | 5.5836 | 5.1553 |
| PLEC_S2833 | PLEC | Q15149 | 1732 | TAQLERSLQEEHV | 6.1743 | 6.8601 | 7.4705 | 6.3855 | 6.7179 | 7.4376 | 6.5953 | 6.1129 | 7.0523 | 7.8813 | 7.8848 | 6.6349 | 3.7260 | 5.0723 | 3.5910 | 4.4033 |
| PLEC_S2834 | PLEC | Q15149 | 4030 | GLRKQITMEELVR | 5.3196 | 7.3820 | 8.9354 | 5.6817 | 5.1123 | 6.6209 | 5.1949 | 5.9348 | 6.6878 | 9.5209 | 7.3046 | 6.8595 | 4.4787 | 5.7900 | 4.8537 | 5.3232 |
| PLEC_S3441 | PLEC | Q15149 | 149 | KELEEVSPETPVV | 6.0891 | 5.6921 | 6.9246 | 6.2961 | 6.3445 | 6.1281 | 5.6330 | 6.1526 | 6.3181 | 6.2992 | 6.2080 | 6.5395 | 6.2778 | 6.9227 | 6.2583 | 5.9161 |
| PLEC_S4161 | PLEC | Q15149 | 4613 | STKGYYSPYSVSG | 5.1939 | 6.3450 | 5.3859 | 5.0301 | 6.6400 | 6.0330 | 5.0115 | 5.8291 | 6.3154 | 5.8002 | 5.8121 | 5.9855 | 7.2249 | 7.7363 | 8.4576 | 7.1997 |
| PLEC_S4382 | PLEC | Q15149 | 720 | SDEGQLSPATRGA | 6.2593 | 6.5007 | 6.1371 | 5.9185 | 6.1368 | 5.7992 | 5.9477 | 6.0222 | 6.1855 | 6.0301 | 6.1556 | 6.1559 | 6.8426 | 6.5040 | 6.7589 | 6.6457 |
| PLEC_S4386 | PLEC | Q15149 | 4618 | YSPYSVSGSGSTA | 4.9057 | 5.4977 | 5.8128 | 5.4833 | 6.1053 | 6.3028 | 5.6992 | 6.6812 | 6.2022 | 6.7205 | 6.8268 | 6.9214 | 6.2868 | 7.1406 | 6.8468 | 5.5670 |
| PLEC_S4386;S4389 | PLEC | Q15149 | 4622 | SVSGSGSTAGSRT | 5.6621 | 6.1469 | 6.2206 | 6.6161 | 6.3921 | 6.0161 | 6.1256 | 6.1022 | 5.8049 | 5.9332 | 6.3957 | 6.1387 | 6.6597 | 7.2515 | 6.4708 | 6.0639 |
| PLEC_S4386;S4396 | PLEC | Q15149 | 2834 | YLQGRSSIAGLLL | 6.2437 | 3.5739 | 5.7603 | 3.5496 | 4.7187 | 6.2336 | 4.2636 | 7.5907 | 5.4812 | 12.9667 | 6.7048 | 10.0421 | 8.0195 | 6.2032 | 4.5245 | 4.1240 |
| PLEC_S4396 | PLEC | Q15149-2 | 20 | DVSNGSSGSPSPG | 6.6189 | 6.7118 | 7.1926 | 7.0721 | 6.5493 | 6.4076 | 6.4351 | 6.7057 | 6.8804 | 6.9632 | 6.6706 | 6.5336 | 4.6806 | 5.6090 | 4.5637 | 4.4058 |
| PLEC_S4406 | PLEC | Q15149 | 4161 | YSGKLISLFQAMK | 4.0272 | 4.1474 | 6.4258 | 4.9217 | 5.1798 | 6.5326 | 4.2364 | 6.7595 | 8.2013 | 7.4938 | 6.7417 | 7.6807 | 5.9229 | 5.7894 | 8.9489 | 6.9909 |
| PLEC_S4613 | PLEC | Q15149 | 4396 | SSSYPISPAVSRT | 7.5882 | 8.2428 | 6.5666 | 6.5862 | 6.6485 | 7.0649 | 7.0963 | 6.7706 | 6.1129 | 5.4350 | 5.9997 | 6.0523 | 4.9249 | 5.0939 | 4.5892 | 5.3282 |
| PLEC_S4613;S4622 | PLEC | Q15149 | 3441 | LRQERLSFSGLRA | 6.0734 | 5.8921 | 6.1134 | 5.8755 | 7.0179 | 5.8030 | 6.4563 | 5.9006 | 6.5721 | 6.7411 | 6.4531 | 6.0579 | 6.8403 | 6.7731 | 5.5321 | 5.8979 |
| PLEC_S4618 | PLEC | Q15149 | 2833 | HYLQGRSSIAGLL | 5.7097 | 3.7250 | 4.8265 | 3.5387 | 5.2510 | 6.5748 | 4.4600 | 7.9226 | 4.8365 | 12.2304 | 8.8138 | 9.9934 | 6.9954 | 6.0822 | 5.1469 | 3.8932 |
| PLEC_S4618;S4626 | PLEC | Q15149-4 | 20 | LGRKRTSSEDNLY | 6.1783 | 5.3140 | 5.5004 | 5.2857 | 5.7484 | 6.0600 | 6.2079 | 5.2611 | 5.9777 | 5.7075 | 6.3587 | 6.3511 | 7.4474 | 7.3409 | 7.9007 | 7.3602 |
| PLEC_S4622 | PLEC | Q15149 | 4386;4389 | NA | 6.5198 | 6.1100 | 6.2737 | 5.9732 | 5.9526 | 6.3839 | 5.6076 | 6.1522 | 6.4363 | 6.3918 | 5.9236 | 6.2541 | 6.6496 | 6.7396 | 6.4475 | 6.1848 |
| PLEC_S4622;S4626 | PLEC | Q15149-4 | 19;21 | NA | 1.4962 | 1.4403 | 1.7341 | 1.6986 | 1.6255 | 1.7362 | 1.7859 | 1.8383 | 1.7036 | 1.6614 | 1.4498 | 1.6848 | 18.4154 | 18.7129 | 22.9967 | 20.0203 |
| PLEC_S4626 | PLEC | Q15149 | 4613;4622 | NA | 6.0837 | 6.6312 | 6.0494 | 6.3613 | 6.4633 | 6.5186 | 6.1119 | 5.9623 | 6.8461 | 6.5291 | 6.1726 | 6.2000 | 6.1233 | 6.7053 | 6.3588 | 5.7831 |
| PLEC_S4642 | PLEC | Q15149 | 4386;4396 | NA | 6.5016 | 5.4682 | 5.4864 | 6.0319 | 6.3344 | 5.5519 | 6.5181 | 6.4182 | 6.9063 | 7.0269 | 6.4874 | 7.1126 | 6.2304 | 6.6085 | 5.4551 | 5.8622 |
| PLEC_S720 | PLEC | Q15149 | 4618;4626 | NA | 5.7280 | 5.7633 | 6.7566 | 6.3094 | 6.0443 | 5.7586 | 6.1971 | 7.0566 | 6.4614 | 6.4256 | 6.3467 | 6.5355 | 6.1262 | 6.0925 | 5.8350 | 6.5633 |
| PLEC_T19;S21 | PLEC | Q15149-4 | 20;21 | NA | 2.7862 | 2.8359 | 3.9717 | 3.4714 | 3.5451 | 4.3887 | 4.4982 | 5.4014 | 4.6848 | 4.5371 | 4.7137 | 4.3235 | 12.2805 | 11.7844 | 14.4997 | 12.2776 |
| PLEC_T4030 | PLEC | Q15149 | 4622;4626 | NA | 6.7526 | 6.3597 | 7.0761 | 7.6593 | 5.9272 | 6.4864 | 5.9953 | 5.9588 | 6.6159 | 5.8801 | 6.1889 | 6.0943 | 5.7538 | 6.1640 | 6.6345 | 5.4533 |
| PLEKHA1_S308 | PLEKHA1 | Q9HB21 | 332 | TASRSNSLVSTFT | 7.0710 | 6.1574 | 6.4952 | 5.8173 | 6.1352 | 6.5287 | 6.2013 | 6.2911 | 7.0701 | 6.6196 | 6.5978 | 6.6608 | 6.1060 | 6.4045 | 4.8793 | 4.9648 |
| PLEKHA1_S332 | PLEKHA1 | Q9HB21-2 | 308 | RQARRLSNPCIQR | 5.8519 | 5.4828 | 6.8240 | 5.7262 | 5.8841 | 6.1646 | 5.8488 | 6.1996 | 7.0832 | 7.0936 | 7.0654 | 6.9288 | 5.7081 | 6.7451 | 5.6062 | 5.7877 |
| PLEKHA2_S184 | PLEKHA2 | Q9HB19 | 184 | ILRRSQSYIPTSG | 6.4969 | 6.4487 | 6.2261 | 5.9877 | 6.1763 | 5.9467 | 5.8933 | 5.6424 | 6.9859 | 5.7766 | 6.4155 | 6.7695 | 7.3515 | 6.9165 | 5.2673 | 5.6990 |
| PLEKHA2_S349 | PLEKHA2 | Q9HB19 | 349 | ALCKAPSVASSWQ | 8.2541 | 6.4267 | 6.6154 | 5.9848 | 6.1515 | 5.4756 | 6.8519 | 6.4965 | 7.3753 | 6.2729 | 6.6361 | 6.9152 | 5.4187 | 5.9705 | 4.4495 | 4.7054 |
| PLEKHA2_S401 | PLEKHA2 | Q9HB19 | 401 | SRIRHRSEPQHPK | 6.4271 | 6.6696 | 7.1425 | 5.9097 | 6.1055 | 6.6680 | 6.2131 | 6.1201 | 6.9605 | 6.5809 | 6.9687 | 7.1659 | 4.8769 | 5.9624 | 5.0108 | 5.2182 |
| PLEKHA4_S559 | PLEKHA4 | Q9H4M7 | 559 | PHLGLGSPRVSRA | 6.5380 | 7.0087 | 6.9435 | 5.9977 | 6.5973 | 5.4111 | 6.0107 | 5.9673 | 5.9658 | 7.0405 | 5.9456 | 5.8934 | 5.1214 | 6.3830 | 6.9102 | 6.2657 |
| PLEKHA4_S566;S567 | PLEKHA4 | Q9H4M7 | 575 | EGRHLPSPQLGTK | 6.2259 | 5.8921 | 6.2164 | 5.7874 | 6.5531 | 6.3566 | 8.0166 | 8.1250 | 7.0242 | 5.7561 | 5.5344 | 6.0025 | 4.9739 | 5.4124 | 5.8904 | 6.2329 |
| PLEKHA4_S575 | PLEKHA4 | Q9H4M7 | 566;567 | NA | 6.9089 | 6.8570 | 5.8183 | 5.7523 | 5.5512 | 4.7826 | 6.0566 | 6.2678 | 6.3032 | 7.1327 | 6.7316 | 7.2490 | 5.2990 | 7.1743 | 5.8591 | 6.2565 |
| PLEKHA5_S1021 | PLEKHA5 | Q9HAU0-6 | 1258 | VKSLSPSPESSAS | 6.0564 | 6.1877 | 6.5494 | 6.0410 | 6.0804 | 6.1539 | 6.2375 | 6.2562 | 6.0967 | 6.5040 | 6.3156 | 6.3706 | 6.4368 | 6.8813 | 6.1628 | 6.3158 |
| PLEKHA5_S1099 | PLEKHA5 | Q9HAU0-6 | 975 | DYRLYKSEPELTT | 6.0638 | 6.8065 | 6.7650 | 6.3945 | 6.0813 | 7.0078 | 6.3596 | 6.6094 | 6.6128 | 6.7642 | 7.5680 | 6.8281 | 4.9778 | 5.4774 | 4.5856 | 5.0983 |
| PLEKHA5_S1099;S1103 | PLEKHA5 | Q9HAU0-6 | 466 | YPEGYRTLPRNSK | 8.7567 | 7.7393 | 7.6889 | 7.4936 | 5.7572 | 5.9018 | 6.5034 | 5.1177 | 6.2531 | 6.5994 | 6.2560 | 5.0682 | 5.9604 | 4.2934 | 5.0195 | |
| PLEKHA5_S1256 | PLEKHA5 | Q9HAU0-6 | 1021 | VPPRAKSPTPESS | 6.2132 | 7.0127 | 7.2624 | 6.2044 | 6.1579 | 6.7972 | 6.1788 | 6.4885 | 6.9531 | 5.9728 | 6.1074 | 6.0544 | 5.6769 | 5.9603 | 5.7401 | 5.2201 |
| PLEKHA5_S1258 | PLEKHA5 | Q9HAU0-6 | 388 | NKIVNVSLADLRG | 8.2728 | 8.0869 | 6.5209 | 7.1634 | 5.3504 | 5.7673 | 7.2825 | 5.1176 | 5.7390 | 5.1584 | 6.5302 | 5.4387 | 5.4952 | 6.4169 | 5.5871 | 6.0726 |
| PLEKHA5_S140 | PLEKHA5 | Q9HAU0-6 | 444 | GVISYQTLPRNMP | 9.5950 | 7.9097 | 8.6494 | 7.6590 | 6.1862 | 5.1049 | 6.1243 | 4.9745 | 5.7467 | 5.4774 | 5.8546 | 5.3559 | 5.8047 | 5.8778 | 4.8766 | 4.8031 |
| PLEKHA5_S388 | PLEKHA5 | Q9HAU0-6 | 1099 | IGASDQSPLQSPS | 6.7264 | 5.9483 | 6.4549 | 5.8997 | 6.1477 | 6.3117 | 6.1557 | 6.3320 | 6.2781 | 5.9355 | 6.4961 | 6.1236 | 6.1706 | 6.2064 | 6.5698 | 6.2433 |
| PLEKHA5_S416 | PLEKHA5 | Q9HAU0-6 | 416 | VIQRTNSMQQLEQ | 6.6763 | 6.0925 | 7.3589 | 6.0030 | 5.1432 | 6.1137 | 6.2746 | 5.9815 | 6.7491 | 6.0836 | 6.5158 | 6.7996 | 5.5915 | 6.5683 | 5.3378 | 6.7106 |
| PLEKHA5_S55 | PLEKHA5 | Q9HAU0-6 | 140 | YAVHPMSPVGRTS | 5.7779 | 5.9616 | 7.2261 | 7.9787 | 5.4628 | 7.3837 | 6.9417 | 6.8870 | 6.6267 | 7.3224 | 7.0205 | 6.6289 | 4.3624 | 5.5372 | 4.2246 | 4.6577 |
| PLEKHA5_S573 | PLEKHA5 | Q9HAU0-6 | 55 | TGHRRQSTDLPTG | 5.6238 | 5.0643 | 5.6760 | 5.4711 | 5.8596 | 6.2241 | 5.8134 | 5.8591 | 6.8170 | 6.4821 | 7.2045 | 7.0169 | 6.3841 | 6.9292 | 6.6604 | 6.9145 |
| PLEKHA5_S602;S613 | PLEKHA5 | Q9HAU0-6 | 1256 | MAVKSLSPSPESS | 6.5311 | 3.3710 | 6.0333 | 5.0709 | 6.0764 | 4.8314 | 5.3247 | 6.2775 | 6.7085 | 7.6997 | 6.9910 | 8.5664 | 7.7140 | 8.7052 | 7.7891 | 5.2097 |
| PLEKHA5_S975 | PLEKHA5 | Q9HAU0-6 | 573 | TYRSEVSSPIQRG | 7.3142 | 7.0467 | 6.9170 | 6.2118 | 5.0419 | 6.1470 | 7.7601 | 6.8711 | 7.1062 | 7.2602 | 7.5709 | 6.9414 | 4.4880 | 4.8392 | 5.0257 | 3.4585 |
| PLEKHA5_T444 | PLEKHA5 | Q9HAU0-6 | 602;613 | NA | 7.2532 | 7.6676 | 6.6482 | 6.1457 | 5.2274 | 6.6690 | 8.1010 | 5.9990 | 7.3178 | 6.4725 | 7.2182 | 7.6105 | 4.0609 | 5.3352 | 3.5531 | 4.7205 |
| PLEKHA5_T466 | PLEKHA5 | Q9HAU0-6 | 1099;1103 | NA | 6.0466 | 6.8473 | 6.5002 | 5.4459 | 5.4107 | 7.3921 | 7.2014 | 7.8232 | 7.3940 | 6.4380 | 6.5567 | 6.8555 | 4.8651 | 4.9811 | 5.4390 | 5.7032 |
| PLEKHF1_S227 | PLEKHF1 | Q96S99 | 227 | EEQGAGSPGQPAH | 6.4013 | 5.3958 | 6.5451 | 6.0813 | 5.8432 | 6.5517 | 6.6882 | 6.5856 | 6.5341 | 6.9420 | 6.9809 | 6.2144 | 5.5777 | 6.4194 | 5.5364 | 6.4029 |
| PLEKHF2_S226 | PLEKHF2 | Q9H8W4 | 226 | QPARSDSYSQSLK | 5.5457 | 5.1281 | 6.0264 | 6.2771 | 5.7747 | 5.2318 | 5.3112 | 5.9328 | 5.7373 | 5.8043 | 5.5309 | 5.4319 | 8.5630 | 8.4499 | 8.2831 | 6.9718 |
| PLEKHG2_S1049 | PLEKHG2 | Q9H7P9 | 469 | RRNTAPSPGPSVI | 6.3621 | 6.5594 | 6.9150 | 7.1738 | 6.4883 | 6.9614 | 6.7790 | 6.7574 | 6.7492 | 7.0793 | 7.0539 | 7.0591 | 4.2739 | 5.2695 | 3.9451 | 4.5737 |
| PLEKHG2_S1227 | PLEKHG2 | Q9H7P9 | 1227 | LDIQGLSPTPVQT | 6.5224 | 6.6641 | 6.9160 | 6.6522 | 6.2914 | 7.0310 | 6.9281 | 7.4047 | 7.3363 | 6.9087 | 7.1993 | 7.2835 | 3.9099 | 4.7009 | 3.8388 | 4.4125 |
| PLEKHG2_S1310 | PLEKHG2 | Q9H7P9 | 1318 | PTSRASSPPPQPQ | 6.1328 | 5.9890 | 6.5363 | 5.7904 | 5.9446 | 6.4269 | 5.9677 | 6.8083 | 6.3840 | 6.5347 | 6.6256 | 6.4190 | 6.3489 | 6.3242 | 6.0827 | 5.6849 |
| PLEKHG2_S1318 | PLEKHG2 | Q9H7P9 | 90 | PIRLHLSPVGIPG | 6.4846 | 6.5233 | 7.1008 | 6.5126 | 6.3408 | 7.2062 | 7.0059 | 7.3714 | 7.4104 | 7.2519 | 7.3734 | 6.7262 | 3.9743 | 4.5700 | 3.7058 | 4.4424 |
| PLEKHG2_S469 | PLEKHG2 | Q9H7P9 | 445 | PVLEPLTPPLGSP | 5.8882 | 5.9684 | 6.4359 | 6.5388 | 6.3657 | 5.8406 | 6.0793 | 6.3697 | 6.6441 | 6.2335 | 6.4510 | 6.4189 | 6.2272 | 6.5711 | 6.2133 | 5.9578 |
| PLEKHG2_S610 | PLEKHG2 | Q9H7P9 | 610 | MDERGPSPLHVLE | 5.9115 | 6.4964 | 6.1068 | 6.0724 | 5.9360 | 6.3305 | 6.1421 | 6.3482 | 6.3011 | 6.0037 | 7.0019 | 7.0244 | 4.9790 | 6.5887 | 6.4228 | 6.3345 |
| PLEKHG2_S90 | PLEKHG2 | Q9H7P9 | 1310 | SRGSWSSAPTSRA | 6.0446 | 6.0886 | 6.6810 | 6.5278 | 5.9889 | 6.7404 | 6.1363 | 7.4882 | 6.9992 | 7.0396 | 7.3728 | 7.4243 | 4.6278 | 5.7935 | 4.3826 | 4.6714 |
| PLEKHG2_T1257;S1261 | PLEKHG2 | Q9H7P9 | 1049 | GHLDSESPTNIPL | 5.7195 | 5.9390 | 6.3718 | 6.5155 | 5.8321 | 6.3307 | 5.2500 | 6.1067 | 6.2909 | 6.9337 | 6.3383 | 7.0426 | 6.4547 | 6.8053 | 5.6594 | 6.4098 |
| PLEKHG2_T445 | PLEKHG2 | Q9H7P9 | 445;450 | NA | 6.7776 | 6.9831 | 6.9370 | 6.6900 | 6.4295 | 7.0011 | 6.1554 | 7.4693 | 6.5348 | 7.2528 | 6.9848 | 7.1205 | 4.4246 | 4.9374 | 4.0108 | 4.2914 |
| PLEKHG2_T445;S450 | PLEKHG2 | Q9H7P9 | 1257;1261 | NA | 5.2885 | 5.5657 | 7.0067 | 6.3864 | 6.4746 | 5.9623 | 5.8754 | 6.5293 | 6.2853 | 6.2660 | 6.4402 | 7.0317 | 6.1436 | 6.4183 | 6.0631 | 6.2630 |
| PLEKHG3_S1028 | PLEKHG3 | A1L390 | 827 | MESSGGSPGKGPG | 6.2873 | 6.5857 | 6.5666 | 6.8969 | 6.4704 | 6.1952 | 6.1216 | 5.9973 | 6.0237 | 6.4230 | 6.6639 | 6.8620 | 5.7464 | 5.7985 | 5.3376 | 5.4239 |
| PLEKHG3_S1037;S1040 | PLEKHG3 | A1L390 | 1028 | VSQRTTSPGGRPS | 6.3806 | 6.6881 | 6.6950 | 6.4945 | 6.5473 | 6.6721 | 6.0773 | 5.8938 | 6.0025 | 6.2190 | 6.4668 | 6.0380 | 5.8966 | 6.4365 | 5.8959 | 5.5960 |
| PLEKHG3_S1040 | PLEKHG3 | A1L390 | 76 | NVKGPLSPFNSRA | 6.3694 | 6.7070 | 6.4096 | 6.7842 | 6.3888 | 6.6477 | 6.3031 | 6.4396 | 6.3227 | 6.7284 | 6.4217 | 6.7138 | 5.1536 | 5.5747 | 5.3839 | 5.6517 |
| PLEKHG3_S1154 | PLEKHG3 | A1L390 | 631 | KSSGFGSPRLVSR | 6.5175 | 6.6607 | 6.4696 | 6.7903 | 6.2531 | 6.3555 | 6.4201 | 5.8545 | 6.5132 | 6.8070 | 6.6046 | 6.3506 | 5.5544 | 5.6924 | 5.4529 | 5.7037 |
| PLEKHG3_S1154;S1169 | PLEKHG3 | A1L390 | 614 | RFVSSFSRRSSVA | 7.8652 | 6.5413 | 7.0214 | 6.8426 | 6.2349 | 5.2740 | 5.9078 | 5.5672 | 8.2499 | 6.9422 | 7.1149 | 7.2917 | 5.0857 | 5.5099 | 3.9940 | 4.5574 |
| PLEKHG3_S1169 | PLEKHG3 | A1L390 | 1154 | EKGPLPSPTAGLE | 5.8207 | 6.0446 | 6.3034 | 6.4596 | 6.5268 | 6.6722 | 6.0899 | 5.3928 | 6.7861 | 6.0890 | 6.3125 | 6.5562 | 6.2214 | 6.1585 | 6.0778 | 6.4885 |
| PLEKHG3_S488 | PLEKHG3 | A1L390 | 741 | SVRRRESLSYIPK | 6.1932 | 6.4253 | 6.9155 | 6.4163 | 7.0230 | 5.7528 | 6.9295 | 5.1670 | 7.4708 | 6.7438 | 6.9277 | 6.5975 | 6.3135 | 6.1737 | 5.6089 | 5.3417 |
| PLEKHG3_S614 | PLEKHG3 | A1L390 | 1040 | SARSPLSPTETFS | 6.9843 | 5.7785 | 6.8401 | 6.8157 | 6.2991 | 6.1943 | 6.5495 | 6.6196 | 6.7590 | 6.9583 | 6.5952 | 6.3676 | 4.9232 | 5.0043 | 5.7265 | 5.5847 |
| PLEKHG3_S631 | PLEKHG3 | A1L390 | 488 | RRPSGRSPTSTEK | 6.3706 | 6.5038 | 6.5508 | 6.7205 | 6.2292 | 7.1504 | 6.0360 | 6.1817 | 7.0282 | 6.7171 | 6.9611 | 6.9065 | 5.0309 | 5.4143 | 5.1186 | 5.0803 |
| PLEKHG3_S741 | PLEKHG3 | A1L390 | 962 | PERDGKSPTVPCL | 7.1465 | 7.3969 | 6.7691 | 7.5781 | 6.5395 | 6.0399 | 6.2871 | 7.0189 | 7.2034 | 7.2269 | 5.9261 | 6.5775 | 4.3303 | 5.1654 | 4.6030 | 5.0915 |
| PLEKHG3_S76 | PLEKHG3 | A1L390 | 1169 | SGQGPSSPVALLG | 6.0931 | 5.8861 | 6.9872 | 6.9872 | 5.4630 | 5.1731 | 6.6909 | 6.2859 | 5.7166 | 7.3987 | 6.4502 | 6.1410 | 5.8744 | 6.9639 | 6.1889 | 5.6998 |
| PLEKHG3_S827 | PLEKHG3 | A1L390 | 1037;1040 | NA | 6.1032 | 6.0411 | 5.9174 | 5.8496 | 6.3548 | 6.1486 | 6.8523 | 6.6823 | 6.5792 | 5.7778 | 5.6641 | 5.5162 | 6.4167 | 7.0157 | 6.2925 | 6.7886 |
| PLEKHG3_S962 | PLEKHG3 | A1L390 | 1154;1169 | NA | 5.5243 | 6.4862 | 5.7034 | 6.0604 | 6.9948 | 7.4713 | 6.6375 | 6.5172 | 5.4488 | 5.8195 | 5.2272 | 5.5212 | 6.4686 | 6.7383 | 6.2604 | 7.1210 |
| PLEKHG4_S64 | PLEKHG4 | Q58EX7 | 64 | DEELQGSPLSRKF | 6.5008 | 6.9114 | 7.3308 | 6.2905 | 6.0621 | 6.8986 | 6.9622 | 6.8489 | 6.9044 | 6.3471 | 7.2858 | 6.6912 | 4.5813 | 5.4236 | 4.2653 | 4.6958 |
| PLEKHG5_S237 | PLEKHG5 | O94827-7 | 262 | AQSRRESLDILAP | 6.5660 | 5.0227 | 5.5765 | 4.6377 | 5.8225 | 5.5544 | 4.9763 | 6.0265 | 8.0630 | 7.3139 | 7.2599 | 7.4353 | 7.1253 | 7.4066 | 6.1633 | 5.0502 |
| PLEKHG5_S262 | PLEKHG5 | O94827-7 | 237 | KDSKSLSLPILRP | 8.3008 | 6.1832 | 6.5861 | 5.8761 | 6.4436 | 5.6673 | 5.9277 | 6.4559 | 7.7161 | 6.8563 | 5.8835 | 3.9919 | 5.9302 | 7.3378 | 5.2350 | 5.6084 |
| PLEKHG5_S972 | PLEKHG5 | O94827-7 | 972 | APSRSLSELCLAV | 5.7384 | 6.4482 | 6.2485 | 6.9345 | 6.4601 | 6.1126 | 6.2418 | 6.1847 | 6.8187 | 6.3547 | 6.6220 | 5.9953 | 6.2752 | 6.4067 | 5.7281 | 5.4303 |
| PLEKHM1_S432 | PLEKHM1 | Q9Y4G2 | 435 | VVSSPTSPKNKSW | 6.7272 | 6.8013 | 7.2113 | 6.6762 | 5.9247 | 6.2285 | 6.9386 | 6.2563 | 7.0552 | 7.6869 | 7.6899 | 7.5856 | 4.1729 | 4.4840 | 3.8419 | 4.7194 |
| PLEKHM1_S432;S435 | PLEKHM1 | Q9Y4G2 | 432 | LKLVVSSPTSPKN | 7.1187 | 7.2111 | 6.6682 | 5.9775 | 6.7496 | 6.1077 | 6.4866 | 6.4000 | 7.2853 | 6.3238 | 6.8016 | 6.3518 | 4.7484 | 5.6982 | 4.1228 | 5.9488 |
| PLEKHM1_S435 | PLEKHM1 | Q9Y4G2 | 432;435 | NA | 7.3611 | 7.2634 | 6.8168 | 7.7669 | 6.6150 | 6.5520 | 7.0754 | 6.6556 | 7.9077 | 6.7674 | 7.6029 | 7.2031 | 3.3522 | 3.6817 | 3.0871 | 4.2918 |
| PLEKHM2_S264 | PLEKHM2 | Q8IWE5 | 441 | QSFRTGSPGDAPE | 6.0362 | 5.9810 | 7.5025 | 6.3224 | 6.1541 | 5.9969 | 6.4831 | 5.8506 | 6.8990 | 6.9170 | 7.0086 | 6.9509 | 5.4477 | 5.5690 | 5.3191 | 5.5617 |
| PLEKHM2_S441 | PLEKHM2 | Q8IWE5 | 264 | SKASTRSPTQRQN | 6.7018 | 7.0972 | 7.9611 | 7.3863 | 6.9300 | 6.8377 | 5.7056 | 5.3434 | 5.6551 | 6.9126 | 6.8342 | 6.9770 | 4.8992 | 5.3625 | 4.6589 | 4.7373 |
| PLEKHM3_S152 | PLEKHM3 | Q6ZWE6 | 192 | PSFLLPSPNKIED | 7.3892 | 5.5050 | 6.3611 | 7.6306 | 6.2962 | 6.5369 | 8.4352 | 7.4048 | 9.4611 | 6.0034 | 5.9565 | 6.8975 | 3.5448 | 4.0567 | 4.9954 | 4.4255 |
| PLEKHM3_S192 | PLEKHM3 | Q6ZWE6 | 152 | GHARSRSDITQVD | 6.6215 | 6.8006 | 6.3887 | 6.5156 | 6.1310 | 5.6444 | 6.4026 | 6.3315 | 6.5141 | 6.2182 | 6.4175 | 6.6636 | 5.5575 | 6.5310 | 5.7022 | 5.5599 |
| PLEKHO2_S439 | PLEKHO2 | Q8TD55 | 232 | VGERAPTPVSASS | 6.1498 | 6.1777 | 6.3703 | 6.5986 | 6.2248 | 5.8346 | 5.8082 | 6.6548 | 6.3411 | 6.7443 | 6.3950 | 6.4904 | 6.0171 | 6.3907 | 5.8314 | 5.9711 |
| PLEKHO2_T232 | PLEKHO2 | Q8TD55 | 364 | AEGTPGTPPKDAT | 5.7999 | 5.7430 | 6.7524 | 6.1280 | 6.0962 | 5.7066 | 5.6392 | 5.8328 | 6.6487 | 6.8387 | 6.1442 | 6.8109 | 6.3749 | 6.5710 | 6.1744 | 6.7393 |
| PLEKHO2_T364 | PLEKHO2 | Q8TD55 | 439 | SEPAPVSAETLLS | 5.7071 | 6.2980 | 6.1090 | 6.1923 | 6.1478 | 5.9520 | 6.6786 | 6.7185 | 6.6050 | 6.4000 | 6.2493 | 6.2964 | 5.8651 | 6.0660 | 6.5880 | 6.1271 |
| PLIN3_S31 | PLIN3 | O60664 | 31 | VVDRVASMPLISS | 3.7284 | 3.8403 | 3.9683 | 3.8253 | 4.2608 | 4.3732 | 4.1622 | 4.2240 | 4.0272 | 4.0059 | 4.0578 | 3.8090 | 13.5556 | 12.4616 | 13.7369 | 11.9637 |
| PLK1_S565 | PLK1 | P53350 | 210 | DGERKKTLCGTPN | 8.1992 | 7.8016 | 8.0927 | 8.7535 | 6.7907 | 6.9227 | 7.4154 | 6.4010 | 7.1117 | 6.6067 | 7.0410 | 6.2794 | 2.8500 | 3.7942 | 2.7617 | 3.1785 |
| PLK1_T210 | PLK1 | P53350 | 565 | FRTYRLSLLEEYG | 8.2922 | 7.8142 | 7.5680 | 7.2919 | 7.0648 | 5.8216 | 7.1089 | 5.3057 | 7.4686 | 6.4211 | 6.0551 | 6.7335 | 4.6651 | 4.0751 | 4.1019 | 4.2124 |
| PLPP6_S70 | PLPP6 | Q8IY26 | 70 | PVHRRGSFPLAAA | 5.9348 | 5.1852 | 5.4678 | 5.4614 | 6.2698 | 5.4876 | 5.8837 | 6.3301 | 7.2650 | 7.6983 | 6.9318 | 7.1491 | 6.5567 | 6.8806 | 5.6015 | 5.8966 |
| PLRG1_S201 | PLRG1 | O43660 | 201 | KLYRVISGHLGWV | 6.3470 | 6.6376 | 6.8536 | 5.8442 | 6.5826 | 5.2394 | 5.9974 | 7.0579 | 5.5300 | 4.6502 | 4.6895 | 6.6297 | 6.4452 | 7.0500 | 6.9872 | 7.4585 |
| PLXNA1_S1632 | PLXNA1 | Q9UIW2 | 1635 | TASSPDSLRSRTP | 7.2353 | 7.0402 | 6.3226 | 6.3753 | 6.5820 | 6.4598 | 6.4495 | 6.3068 | 7.0548 | 6.8822 | 6.3424 | 6.0262 | 5.6048 | 5.7625 | 4.7279 | 4.8277 |

| | | | | | | | | | | | | | | | | | | | | | |
|---|---|---|---|---|---|---|---|---|---|---|---|---|---|---|---|---|---|---|---|---|---|
| PLXNA1_S1635 | PLXNA1 | Q9UIW2 | | 1632 | MLRTASSPDSLRS | 7.0936 | 7.1664 | 6.4757 | 6.7631 | 6.4279 | 6.9010 | 6.5544 | 6.8299 | 6.9121 | 6.9183 | 6.6118 | 6.5955 | 4.5791 | 5.0989 | 4.4042 | 4.6680 |
| PMF1-BGLAP_S196 | PMF1-BGLAP | U3KQ54 | | 196 | RSPAVQSPAKVQP | 6.5056 | 5.5614 | 6.0837 | 5.5156 | 5.9355 | 5.1819 | 5.9097 | 6.7090 | 5.4582 | 6.6985 | 6.5552 | 7.7005 | 6.1456 | 7.0290 | 6.3742 | 6.6364 |
| PML_S36 | PML | P29590-2 | | 583 | QPRTPASPHFRSQ | 6.4631 | 6.3501 | 5.7497 | 6.4541 | 6.4778 | 5.8371 | 6.0278 | 6.1139 | 6.0630 | 5.8220 | 5.8833 | 6.1166 | 7.0142 | 6.4708 | 6.5864 | 6.5702 |
| PML_S38 | PML | P29590 | | 527 | HLDGPPSPRSPVI | 6.6496 | 6.8551 | 6.5997 | 6.8336 | 6.7464 | 6.0555 | 6.5892 | 6.9603 | 6.6599 | 6.4413 | 6.6760 | 6.6020 | 5.1416 | 5.2414 | 5.0299 | 4.9185 |
| PML_S403 | PML | P29590 | | 36 | SEGRQPSPSPSPT | 6.1938 | 6.1402 | 5.9451 | 6.6015 | 5.9514 | 5.9794 | 6.1273 | 6.5360 | 6.8550 | 6.2558 | 6.0362 | 6.3247 | 5.9241 | 6.3073 | 6.1531 | 6.6692 |
| PML_S504;S505 | PML | P29590 | | 530 | GPPSPRSPVIGSE | 5.9652 | 6.7959 | 5.4220 | 6.9622 | 6.7993 | 6.1648 | 5.6684 | 7.6617 | 6.4313 | 5.8252 | 6.0689 | 5.8651 | 6.1746 | 6.3923 | 6.2760 | 5.5271 |
| PML_S505 | PML | P29590 | | 8 | EPAPARSPRPQQD | 5.6348 | 6.0682 | 6.1968 | 6.5826 | 6.1213 | 5.5177 | 5.6723 | 6.1729 | 6.7748 | 6.8588 | 5.7648 | 5.9875 | 6.7757 | 6.5855 | 6.5976 | 6.6888 |
| PML_S518 | PML | P29590 | | 403 | AVSKKASPEAAST | 6.3338 | 6.1484 | 5.5656 | 6.0074 | 6.3220 | 5.7670 | 6.0473 | 6.3491 | 6.2350 | 6.1188 | 5.9755 | 5.8589 | 7.0843 | 6.9588 | 6.7662 | 6.4618 |
| PML_S518;S527 | PML | P29590-8 | | 720 | QLQRGISPPHRIR | 6.8190 | 6.7984 | 5.7955 | 6.4748 | 6.4731 | 6.3675 | 6.8337 | 6.4886 | 6.7999 | 6.5182 | 6.9686 | 6.6999 | 5.2265 | 5.6896 | 4.9971 | 5.0495 |
| PML_S518;S527;S530 | PML | P29590-8 | | 819 | PERPPISPVPGAR | 6.4594 | 6.2528 | 6.5168 | 6.4782 | 6.1224 | 6.2361 | 5.9554 | 6.6055 | 6.6660 | 6.0953 | 6.7181 | 6.1589 | 5.5892 | 5.5788 | 5.8657 | 6.7014 |
| PML_S518;S530;S535 | PML | P29590 | | 518 | STSKAVSPPHLDG | 6.8250 | 6.9684 | 5.9670 | 6.9105 | 6.5701 | 5.4777 | 6.0886 | 6.5961 | 6.7158 | 6.3945 | 5.9169 | 6.5417 | 6.0101 | 6.0727 | 5.6670 | 5.2776 |
| PML_S527 | PML | P29590 | | 867 | RAEGVSTPLAGRG | 6.7821 | 7.0596 | 6.3125 | 6.8665 | 5.8937 | 6.4243 | 6.7718 | 6.3006 | 6.7713 | 6.9395 | 6.7710 | 6.6958 | 4.9763 | 5.5377 | 4.8304 | 5.0669 |
| PML_S530 | PML | P29590 | | 38 | GRQPSPSPSPTER | 6.5927 | 6.7717 | 5.7938 | 6.7563 | 6.1500 | 6.1983 | 6.6936 | 6.7480 | 6.7619 | 6.6499 | 6.8677 | 6.4580 | 5.3237 | 5.6987 | 5.0860 | 5.4496 |
| PML_S583 | PML | P29590 | | 196 | SNPNHRTPTLTSI | 6.2474 | 5.9899 | 6.4684 | 6.7612 | 6.5025 | 5.9447 | 5.6612 | 6.0598 | 6.6696 | 6.0041 | 6.2053 | 6.6609 | 5.7447 | 6.6492 | 6.0066 | 6.6244 |
| PML_S720 | PML | P29590 | | 505 | ARLARSSPEQPRP | 6.5637 | 6.7551 | 5.7882 | 6.5606 | 6.6202 | 5.6824 | 6.0389 | 6.0218 | 6.0236 | 5.8723 | 5.9779 | 6.1463 | 7.3498 | 6.4103 | 6.4343 | 7.5745 |
| PML_S8 | PML | P29590 | 518;527 | | NA | 6.7255 | 6.4558 | 6.2974 | 6.4349 | 6.4380 | 5.6078 | 6.0910 | 6.1569 | 6.7632 | 5.9002 | 6.0304 | 6.3231 | 6.6633 | 6.2781 | 5.7448 | 6.0894 |
| PML_S819 | PML | P29590 | 504;505 | | NA | 6.6660 | 6.3784 | 6.2543 | 6.3520 | 7.0607 | 6.3193 | 6.1852 | 6.4998 | 5.8251 | 5.7141 | 6.0285 | 6.6850 | 6.4468 | 6.0640 | 5.6616 | 5.8594 |
| PML_T196 | PML | P29590 | 518;527;530 | | NA | 6.9463 | 5.6142 | 6.9101 | 6.9287 | 5.7602 | 6.4966 | 5.8579 | 5.7287 | 6.2214 | 5.3528 | 6.1865 | 7.1947 | 6.6171 | 6.5160 | 5.3094 | 6.3594 |
| PML_T867 | PML | P29590 | 518;530;535 | | NA | 5.4941 | 6.4263 | 6.1542 | 6.6836 | 5.3282 | 6.7701 | 6.1592 | 6.8250 | 6.4173 | 5.7239 | 4.7366 | 5.8601 | 7.8218 | 6.8713 | 6.5617 | 6.1664 |
| PMS2_S445 | PMS2 | P54278 | | 445 | TPEPRRSPLGQKR | 5.8891 | 6.2265 | 6.1731 | 5.4911 | 6.2108 | 6.4298 | 6.1722 | 6.4014 | 6.2793 | 6.1604 | 6.2722 | 6.0910 | 6.2682 | 6.7668 | 6.6899 | 6.4782 |
| PMS2_T576 | PMS2 | P54278 | | 576 | NLATPNTKRFKKE | 7.4052 | 7.0046 | 6.1063 | 6.9629 | 6.1037 | 6.7014 | 7.2723 | 6.3582 | 6.1480 | 6.3600 | 7.2140 | 6.9742 | 4.4954 | 5.2939 | 4.5832 | 5.0167 |
| PNISR_S211 | PNISR | Q8TF01 | | 211 | FRDRQRSPIALPV | 6.1772 | 6.4299 | 5.6924 | 6.0564 | 6.4642 | 5.7716 | 5.9239 | 6.1381 | 6.4038 | 6.1434 | 5.9828 | 5.8348 | 6.9566 | 6.6104 | 6.8752 | 6.5392 |
| PNISR_S561;S565 | PNISR | Q8TF01 | | 726 | TFSRSGSISVKII | 7.5427 | 6.4499 | 5.8004 | 6.2496 | 6.7020 | 5.4250 | 6.0400 | 5.1080 | 6.7385 | 5.6441 | 5.5145 | 5.4278 | 7.4212 | 7.4401 | 6.1179 | 6.3784 |
| PNISR_S601 | PNISR | Q8TF01 | | 601 | RRSNRRNSIERERR | 6.1729 | 6.5357 | 6.2128 | 6.3804 | 6.1474 | 6.0886 | 6.1162 | 5.8666 | 6.4774 | 6.1938 | 6.4122 | 6.2445 | 6.3393 | 6.3201 | 6.2972 | 6.1949 |
| PNISR_S706 | PNISR | Q8TF01 | | 706 | KDFKFSSQDDRLK | 6.1525 | 7.2938 | 7.2733 | 6.8738 | 7.4900 | 8.2337 | 6.9149 | 5.0068 | 4.3496 | 5.8858 | 6.5283 | 4.8689 | 7.0277 | 5.6794 | 5.1335 | 5.2889 |
| PNISR_S726 | PNISR | Q8TF01 | 561;565 | | NA | 6.1786 | 6.2944 | 5.9503 | 6.3778 | 6.1559 | 5.6202 | 6.0309 | 6.0167 | 6.3700 | 6.3253 | 6.1378 | 6.2343 | 6.5172 | 6.4293 | 6.5838 | 6.7775 |
| PNKP_T118 | PNKP | Q96T60 | | 118 | PESQPDTPPGTPL | 5.5972 | 6.0723 | 6.1195 | 5.7021 | 5.6223 | 6.0069 | 5.7197 | 6.3578 | 6.3197 | 6.2699 | 6.5962 | 6.5098 | 6.4709 | 6.6466 | 7.0752 | 6.9140 |
| PNMA8A_S428 | PNMA8A | Q86V59 | | 428 | KAKPEGSPRRATN | 6.5611 | 6.7613 | 6.3792 | 6.6623 | 6.0925 | 6.3196 | 6.2099 | 6.3132 | 6.7996 | 6.6913 | 6.7186 | 6.6773 | 4.8048 | 5.7735 | 5.5602 | 5.6755 |
| PNN_S100 | PNN | Q9H307 | | 347 | EIAIVHSDAEKEQ | 6.2681 | 6.3267 | 5.9517 | 6.4938 | 6.5021 | 5.9579 | 6.3128 | 6.0879 | 6.1268 | 5.9777 | 5.7729 | 5.7383 | 6.6945 | 6.5589 | 6.6315 | 6.5986 |
| PNN_S120 | PNN | Q9H307 | | 381 | SEKQQDSQPEEVM | 5.9436 | 6.2225 | 6.1481 | 6.5754 | 5.9441 | 6.0938 | 6.2092 | 5.9050 | 6.1641 | 5.8856 | 5.7835 | 6.1358 | 6.6860 | 6.6939 | 6.8475 | 7.7619 |
| PNN_S347 | PNN | Q9H307 | | 66 | LLRRGFSDSGGGP | 3.9019 | 3.8772 | 3.7530 | 4.0361 | 4.8896 | 4.4146 | 4.5885 | 4.6079 | 3.9064 | 3.9610 | 3.6069 | 3.7349 | 12.9807 | 11.3151 | 13.5291 | 12.8969 |
| PNN_S381 | PNN | Q9H307 | | 450 | PGKENVSALDMEK | 6.4750 | 6.6962 | 6.2107 | 7.0056 | 7.0458 | 6.6890 | 7.3784 | 6.8141 | 6.6281 | 6.1823 | 6.6660 | 6.4890 | 4.6334 | 5.0991 | 4.7516 | 5.2356 |
| PNN_S413 | PNN | Q9H307 | | 100 | RESRQESDPEDDD | 6.5087 | 6.8066 | 6.4753 | 6.9836 | 5.9036 | 6.5278 | 6.8720 | 5.7322 | 6.3457 | 6.2823 | 6.2733 | 6.1162 | 5.8659 | 6.0813 | 5.8662 | 6.0593 |
| PNN_S450 | PNN | Q9H307 | | 552 | LTVHPESKSKTKT | 6.4087 | 6.6440 | 6.1911 | 6.6601 | 6.6846 | 6.5377 | 7.1439 | 6.0815 | 6.4699 | 6.6972 | 5.9649 | 6.1748 | 5.6817 | 6.1154 | 5.6976 | 5.8471 |
| PNN_S552 | PNN | Q9H307 | | 413 | ETNRVESVEPSEN | 5.7753 | 5.7871 | 6.0604 | 6.3310 | 5.3151 | 5.0350 | 6.0739 | 5.9657 | 5.7852 | 5.9516 | 5.7146 | 6.0371 | 7.3788 | 7.2276 | 8.1178 | 7.4438 |
| PNN_S66 | PNN | Q9H307 | | 690 | RSDRKRSISESSR | 6.4348 | 6.4013 | 5.9260 | 6.0643 | 6.5270 | 5.8626 | 6.0407 | 6.3491 | 6.4429 | 6.2189 | 6.1634 | 6.1571 | 6.9536 | 6.2273 | 6.1817 | 6.0491 |
| PNN_S690 | PNN | Q9H307 | | 695 | RSISESSRSGKRS | 7.5720 | 7.6043 | 6.0833 | 7.3046 | 5.6631 | 5.8288 | 6.2868 | 5.7749 | 6.4677 | 5.1579 | 6.2561 | 6.7003 | 5.7374 | 6.0258 | 5.5407 | 5.9961 |
| PNN_S695 | PNN | Q9H307 | | 120 | SSVVATSKERTRR | 6.3010 | 6.4212 | 6.3432 | 6.4714 | 6.1631 | 6.7762 | 7.1453 | 7.1084 | 6.6097 | 6.1890 | 6.7402 | 6.6182 | 5.3371 | 5.1947 | 5.1820 | 5.3993 |
| PNN_T558;S560 | PNN | Q9H307 | 558;560 | | NA | 5.6795 | 6.3633 | 6.5000 | 6.7566 | 6.7671 | 5.4048 | 5.8986 | 6.0726 | 6.9585 | 6.5255 | 6.2024 | 5.5794 | 6.0794 | 6.5998 | 5.7718 | 6.8405 |
| PNPLA2_S404 | PNPLA2 | Q96AD5 | | 404 | ELRRVQSLPSVPL | 5.6691 | 5.7430 | 5.9089 | 5.6704 | 6.7301 | 6.7453 | 6.6031 | 6.2416 | 5.9613 | 6.0117 | 5.9082 | 6.1543 | 6.7364 | 6.8574 | 6.3804 | 6.6791 |
| PNPLA2_S428 | PNPLA2 | Q96AD5 | | 428 | WMRNNLSLGDALA | 2.8761 | 2.6539 | 2.8667 | 3.2083 | 5.2722 | 5.0964 | 4.9635 | 4.9891 | 2.8342 | 2.9386 | 2.9973 | 3.0638 | 14.0589 | 12.8525 | 15.0308 | 14.2977 |
| PNPLA6_S354 | PNPLA6 | Q8IY17 | | 420 | LLSRCVSMPGDIS | 3.9950 | 4.2028 | 4.2511 | 4.4350 | 4.9615 | 4.9013 | 4.7841 | 4.7651 | 4.6240 | 4.4262 | 4.6057 | 4.5995 | 11.1188 | 10.7683 | 11.7729 | 11.7887 |
| PNPLA6_S354;S362 | PNPLA6 | Q8IY17 | | 354 | PLRLFPSPGLPTR | 5.9467 | 6.2923 | 5.6096 | 6.3111 | 5.8032 | 5.8227 | 6.4353 | 6.1647 | 7.0874 | 6.9975 | 6.9929 | 7.0130 | 5.4473 | 5.9718 | 5.7837 | 6.3210 |
| PNPLA6_S362 | PNPLA6 | Q8IY17 | | 464 | LAAPARTPTQEPR | 5.6934 | 5.4918 | 6.3126 | 7.1486 | 5.7277 | 5.3971 | 7.0463 | 6.4424 | 5.5590 | 6.4729 | 5.8392 | 7.0447 | 5.4147 | 7.1325 | 6.8607 | 6.4164 |
| PNPLA6_S420 | PNPLA6 | Q8IY17 | | 362 | GLPTRTSPVRGSK | 6.0121 | 6.4502 | 6.6831 | 6.5904 | 6.4476 | 7.1087 | 6.4668 | 5.4643 | 5.6940 | 6.2725 | 6.5765 | 6.3997 | 6.1930 | 6.2944 | 6.1048 | 5.2420 |
| PNPLA6_T464 | PNPLA6 | Q8IY17 | 354;362 | | NA | 6.0883 | 7.0552 | 6.5591 | 7.1965 | 5.8233 | 6.4697 | 6.1750 | 6.0104 | 6.1847 | 7.4212 | 6.3807 | 6.8970 | 5.3612 | 6.1765 | 5.0506 | 5.1505 |
| PNPO_S241 | PNPO | Q9NVS9 | | 241 | GLPTGDRSPLGPMT | 5.9742 | 5.7899 | 6.0869 | 5.8193 | 7.0934 | 6.0575 | 5.7585 | 7.6677 | 6.1081 | 6.7989 | 6.3866 | 7.1882 | 5.7527 | 5.6388 | 6.4455 | 5.4339 |
| PNRC1_S105 | PNRC1 | Q12796 | | 105 | KKKVRASPAGQLP | 5.8403 | 7.8135 | 7.4192 | 5.7507 | 6.6102 | 6.4788 | 6.0584 | 6.0555 | 7.3513 | 8.5777 | 7.5565 | 7.1317 | 4.3957 | 5.2423 | 4.8031 | 4.7150 |
| PNRC2_S103 | PNRC2 | Q9NPJ4 | | 103 | KFSEPPSPSVLPK | 6.6342 | 6.1789 | 7.0008 | 6.8918 | 6.2684 | 6.7790 | 7.0411 | 7.2436 | 6.8129 | 6.4825 | 6.7600 | 6.7099 | 4.7979 | 4.6624 | 4.8813 | 4.8553 |
| POC1B_S321 | POC1B | Q8TC44 | | 321 | KRLHFDSPPHLLD | 5.9558 | 6.1104 | 6.2784 | 6.2929 | 6.1702 | 6.3728 | 6.5544 | 6.6378 | 6.4372 | 6.6167 | 6.5181 | 6.3932 | 5.5214 | 6.1509 | 5.9157 | 6.0741 |
| POC1B_S398 | POC1B | Q8TC44 | | 398 | LNPSLMSPECLPT | 6.1562 | 6.0797 | 6.0379 | 6.7485 | 6.2859 | 6.7259 | 5.9380 | 6.1909 | 6.7804 | 5.8827 | 6.7344 | 6.1102 | 5.6425 | 6.3043 | 5.9047 | 6.4779 |
| POC5_S109 | POC5 | Q8NA72 | | 78 | HSQGNNSEVRETA | 6.1469 | 5.9087 | 6.1446 | 6.2726 | 7.0386 | 6.3913 | 5.8757 | 6.2312 | 5.7041 | 6.4189 | 5.9186 | 6.5437 | 5.8995 | 6.2097 | 6.3645 | 6.9314 |
| POC5_S512 | POC5 | Q8NA72 | | 512 | LAIMGVSPPMSSV | 5.8064 | 6.4127 | 6.1972 | 6.9376 | 6.5970 | 6.1832 | 5.5868 | 6.6036 | 7.1161 | 6.7031 | 7.7298 | 7.1449 | 5.1195 | 5.2569 | 5.2157 | 5.3893 |
| POC5_S78 | POC5 | Q8NA72 | | 109 | ESSPVLSPRKPSH | 5.8682 | 5.9730 | 6.3657 | 5.8747 | 6.1776 | 6.4818 | 6.5096 | 6.6578 | 6.7400 | 6.4906 | 5.9138 | 6.0684 | 6.9758 | 6.0642 | 6.2790 | 6.5597 |
| POGZ_S1338 | POGZ | Q7Z3K3 | | 445 | TPIPALSPPTKVP | 6.2684 | 6.9532 | 6.7697 | 7.2751 | 6.5892 | 6.4877 | 6.8819 | 6.4819 | 6.9703 | 6.7721 | 6.6641 | 6.5306 | 4.6040 | 5.2186 | 4.7980 | 4.7350 |
| POGZ_S289 | POGZ | Q7Z3K3 | | 425 | SAAKPPSPEKTAP | 6.0758 | 6.0546 | 6.1637 | 6.8182 | 5.9485 | 5.9797 | 6.2753 | 5.4487 | 6.6221 | 6.4644 | 6.2249 | 6.2739 | 6.4826 | 6.3982 | 6.0518 | 6.7175 |
| POGZ_S425 | POGZ | Q7Z3K3 | | 1338 | PDGNINSPTRNAD | 6.2014 | 6.3586 | 5.9926 | 6.7118 | 6.6679 | 6.3914 | 6.0469 | 6.7939 | 5.9886 | 6.2872 | 6.5191 | 5.7315 | 5.8424 | 6.6485 | 6.2728 | 5.5455 |
| POGZ_S434;S445 | POGZ | Q7Z3K3 | | 289 | NPKLAPSFPSPPA | 6.4304 | 7.5921 | 7.7136 | 8.0069 | 5.8959 | 7.6452 | 7.2789 | 6.2882 | 6.9721 | 7.8386 | 8.0393 | 7.9313 | 3.2287 | 4.0574 | 2.1705 | 2.9110 |
| POGZ_S445 | POGZ | Q7Z3K3 | 434;445 | | NA | 5.9258 | 6.2776 | 5.7130 | 6.0938 | 6.2264 | 6.0068 | 6.0444 | 6.0120 | 6.1644 | 5.9034 | 6.2702 | 5.8505 | 7.3452 | 7.0540 | 6.4278 | 6.6844 |
| POLA1_S209 | POLA1 | P09884 | | 219 | RKEPPLTPVPLKR | 6.5312 | 6.9701 | 6.6781 | 6.8995 | 6.1791 | 6.7169 | 6.9378 | 6.9102 | 7.1076 | 6.4396 | 6.5292 | 6.6193 | 4.5852 | 5.6762 | 4.3609 | 4.8590 |
| POLA1_T219 | POLA1 | P09884 | | 209 | PSGKIASPVSRKE | 6.3631 | 6.7455 | 6.3556 | 5.9184 | 6.0711 | 6.7557 | 6.6456 | 6.5559 | 6.7494 | 6.7006 | 6.6711 | 6.8309 | 4.9681 | 5.9734 | 5.0996 | 5.5960 |
| POLA1_T404 | POLA1 | P09884 | | 404 | LNTGKETGTPISM | 6.0295 | 6.9035 | 6.2914 | 6.1119 | 6.2336 | 5.5137 | 6.0330 | 5.6267 | 5.6906 | 6.4219 | 5.8185 | 5.6451 | 7.2643 | 6.8755 | 5.9770 | 7.5638 |
| POLA2_S126 | POLA2 | Q14181 | | 130 | AISTPETPLTKRS | 7.1756 | 7.2678 | 6.6720 | 6.7881 | 6.4477 | 6.5525 | 6.5329 | 6.1628 | 7.0729 | 5.3568 | 5.9220 | 5.9763 | 5.4450 | 5.9377 | 5.3099 | 5.3800 |
| POLA2_S126;T130 | POLA2 | Q14181 | | 141 | RSVSTRSPHQLLS | 5.3750 | 6.1965 | 6.1044 | 6.3216 | 6.0986 | 6.5279 | 6.0535 | 7.3619 | 5.8441 | 5.7624 | 6.4393 | 6.3331 | 6.1311 | 6.7497 | 6.4900 | 6.2111 |
| POLA2_S141 | POLA2 | Q14181 | | 126 | SQKRAISTPETPL | 6.0504 | 6.2955 | 6.7406 | 5.8733 | 6.8310 | 7.0717 | 6.7854 | 7.1110 | 6.5804 | 6.9609 | 7.2786 | 6.8480 | 4.6338 | 5.0095 | 4.8053 | 5.1247 |
| POLA2_S141;S147;S149 | POLA2 | Q14181 | | 152 | LSPSSFSPSATPS | 4.8124 | 7.3618 | 8.1598 | 5.0456 | 6.2896 | 7.6237 | 5.5899 | 8.4380 | 7.6523 | 5.8600 | 6.0196 | 7.4354 | 4.1290 | 4.8083 | 5.7208 | 5.0538 |
| POLA2_S141;S149 | POLA2 | Q14181 | | 127 | QKRAISTPETPLT | 5.3162 | 5.9910 | 6.5381 | 5.8671 | 6.2304 | 6.6349 | 7.1583 | 6.3838 | 6.7661 | 7.4340 | 6.9818 | 7.0280 | 4.8298 | 5.8728 | 5.2068 | 5.7608 |
| POLA2_S141;S152 | POLA2 | Q14181 | 141;152 | | NA | 5.1809 | 6.1341 | 6.1669 | 5.4901 | 6.4673 | 6.9353 | 6.1040 | 7.1435 | 6.3432 | 6.8618 | 7.1209 | 7.2288 | 4.9814 | 5.8953 | 5.7030 | 6.2435 |
| POLA2_S141;S152;T156 | POLA2 | Q14181 | 126;130 | | NA | 6.7309 | 7.4229 | 6.9468 | 6.7446 | 6.7164 | 7.2664 | 6.8910 | 7.3272 | 7.3508 | 6.6025 | 7.2692 | 7.4672 | 3.5262 | 4.1481 | 3.2252 | 4.3646 |
| POLA2_S152 | POLA2 | Q14181 | 141;149 | | NA | 5.2727 | 5.9117 | 5.2868 | 5.4420 | 7.2054 | 7.4650 | 6.7143 | 7.6353 | 7.0732 | 5.8991 | 6.7424 | 6.5223 | 5.5050 | 5.7616 | 5.7021 | 5.8612 |
| POLA2_T127 | POLA2 | Q14181 | 141;147;149 | | NA | 5.4404 | 6.6421 | 6.7868 | 6.5361 | 5.7606 | 7.9812 | 7.2340 | 7.6187 | 6.7227 | 8.5162 | 7.6582 | 8.2137 | 3.0213 | 4.3661 | 3.2760 | 4.2260 |
| POLA2_T127;T130 | POLA2 | Q14181 | 141;152;156 | | NA | 6.8492 | 7.3052 | 7.9202 | 6.8653 | 8.2602 | 6.1373 | 5.9662 | 6.2405 | 5.6643 | 5.1442 | 5.9805 | 5.8876 | 4.9119 | 6.3848 | 5.3257 | 5.1569 |
| POLA2_T130 | POLA2 | Q14181 | 127;130 | | NA | 6.0565 | 6.6291 | 7.6045 | 6.3804 | 7.7883 | 8.7211 | 8.2104 | 6.0013 | 7.0125 | 7.0060 | 8.2680 | 6.4619 | 3.8052 | 4.1592 | 2.6560 | 3.2397 |
| POLD3_S307 | POLD3 | Q15054 | | 307 | GKRVALSDDETKE | 5.5907 | 5.8873 | 6.3234 | 5.3685 | 5.7421 | 5.8524 | 5.6491 | 6.0356 | 6.0994 | 5.9636 | 5.8008 | 5.7443 | 7.1192 | 7.4850 | 7.7777 | 7.5607 |
| POLD3_T277 | POLD3 | Q15054 | | 277 | TEPKLATPAGLKK | 7.1610 | 7.4335 | 7.0242 | 6.8613 | 5.6764 | 7.0299 | 7.4366 | 6.6264 | 7.4653 | 7.3760 | 7.3794 | 7.7973 | 3.1597 | 4.4247 | 3.0303 | 4.1180 |
| POLDIP2_T292 | POLDIP2 | Q9Y2S7 | | 292 | IFSLSGTLETVRG | 6.6396 | 6.8479 | 6.0746 | 6.8859 | 6.5289 | 5.8797 | 6.2182 | 6.3894 | 6.5618 | 6.4832 | 6.2834 | 6.3390 | 5.7973 | 5.6565 | 5.8923 | 5.5222 |
| POLDIP3_S127 | POLDIP3 | Q9BY77 | | 368 | LLRLSDSPSMKKE | 7.1727 | 7.5541 | 6.4748 | 7.2507 | 6.7396 | 6.7575 | 7.2978 | 6.7077 | 6.8503 | 7.0280 | 6.9631 | 6.9614 | 3.7278 | 4.4044 | 3.7170 | 4.3932 |
| POLDIP3_S275 | POLDIP3 | Q9BY77 | | 127 | ISLKRSSPAAFIN | 6.0148 | 6.4825 | 6.0769 | 6.9383 | 6.1980 | 6.2080 | 6.3969 | 6.3441 | 6.4421 | 6.5845 | 6.2032 | 6.0779 | 5.7261 | 6.2969 | 5.9438 | 6.0658 |
| POLDIP3_S368 | POLDIP3 | Q9BY77 | | 140 | PPIGTVTPALKLT | 9.2092 | 7.3386 | 6.8021 | 8.2332 | 6.8778 | 6.2355 | 7.5608 | 5.9520 | 6.9145 | 5.9562 | 6.5142 | 6.5114 | 4.0069 | 4.5735 | 3.0907 | 4.2234 |
| POLDIP3_S44 | POLDIP3 | Q9BY77 | | 275 | AAEPVLSPLEGTK | 7.4175 | 6.6474 | 7.1047 | 7.7152 | 6.6552 | 6.1159 | 6.7041 | 6.2963 | 6.8525 | 7.7758 | 7.3927 | 7.3273 | 3.4823 | 4.6936 | 3.8916 | 3.9279 |
| POLDIP3_T140 | POLDIP3 | Q9BY77 | | 44 | QGLLSQSTRTATF | 6.0744 | 6.5993 | 6.0083 | 6.6974 | 6.3174 | 5.7331 | 6.4637 | 5.4673 | 6.1365 | 7.3786 | 6.8637 | 6.9098 | 6.0477 | 6.3328 | 5.6416 | 5.3284 |
| POLE_S1184 | POLE | Q07864 | | 1184 | YKQKKISELFTLE | 7.0381 | 7.6874 | 6.8310 | 7.7226 | 6.6148 | 7.1085 | 8.0456 | 6.9632 | 7.0492 | 7.0622 | 7.2211 | 6.1563 | 2.8395 | 4.3003 | 3.2674 | 4.0925 |
| POLE_T2023 | POLE | Q07864 | | 2023 | RSAPGSTPVRRRG | 6.8970 | 6.8565 | 6.4211 | 6.8605 | 5.5845 | 6.5544 | 7.2446 | 6.3427 | 6.9179 | 6.8160 | 6.9176 | 7.2041 | 4.8431 | 5.2241 | 4.3556 | 4.9603 |
| POLH_S512 | POLH | Q9Y253 | | 512 | QAPMSNSPSKPSL | 6.9877 | 6.4668 | 6.1520 | 6.9564 | 6.0519 | 5.7211 | 6.2861 | 6.5665 | 6.9015 | 7.1356 | 7.0217 | 6.5404 | 5.0609 | 6.0000 | 4.9605 | 5.1909 |
| POLI_S290 | POLI | Q9UNA4 | | 290 | RDLQTFSPKILEK | 6.4411 | 7.7330 | 6.4672 | 6.7814 | 6.6419 | 6.1270 | 6.7332 | 6.3721 | 7.8360 | 7.4567 | 8.1762 | 7.9950 | 2.9592 | 4.7008 | 2.9175 | 4.6617 |
| POLQ_S1776 | POLQ | O75417 | | 1776 | ESYLFGSPSDIKN | 6.5572 | 6.8018 | 6.8365 | 6.9648 | 6.5158 | 6.4857 | 6.5975 | 6.2172 | 7.2081 | 7.3618 | 7.3619 | 7.1662 | 4.6101 | 4.8773 | 3.9887 | 4.4494 |
| POLQ_S2267 | POLQ | O75417 | | 952 | DSYIKHSPNIVQD | 7.1889 | 7.5012 | 7.0949 | 7.7006 | 6.6230 | 7.0234 | 7.6201 | 5.8901 | 7.3613 | 6.3295 | 6.9338 | 7.0873 | 3.8244 | 4.5453 | 3.3187 | 3.9576 |
| POLQ_S952 | POLQ | O75417 | | 2267 | PTLVGESPPSQAV | 6.3494 | 5.9897 | 6.5880 | 7.6038 | 5.9114 | 5.4933 | 6.6861 | 6.6388 | 6.3626 | 6.8972 | 6.3972 | 6.9145 | 5.3091 | 5.8387 | 5.0207 | 5.9994 |
| POLR1A_T357 | POLR1A | O95602 | | 357 | LPEEVATPTTDEE | 4.7581 | 5.4039 | 5.1607 | 5.4707 | 5.7485 | 4.9006 | 5.9490 | 6.0400 | 5.4699 | 5.0931 | 5.5996 | 5.0118 | 8.7343 | 8.7255 | 8.0990 | 9.8351 |
| POLR1D_S104 | POLR1D | P0DPB5 | | 104 | RGSASYSPPRKRS | 6.3699 | 6.6235 | 6.1014 | 7.2548 | 5.9843 | 6.2014 | 6.1789 | 6.2250 | 6.3473 | 6.6067 | 6.5978 | 6.9158 | 5.3153 | 5.7409 | 5.6490 | 5.8879 |
| POLR1E_S259 | POLR1E | Q9GZS1-1 | | 97 | SNGKLQSPGNMRF | 6.9318 | 7.6035 | 6.8339 | 8.2062 | 6.2609 | 6.3646 | 6.6302 | 6.8853 | 6.7115 | 7.1723 | 7.2063 | 7.1456 | 3.6624 | 4.1924 | 3.8370 | 4.3563 |
| POLR1E_S283 | POLR1E | Q9GZS1-1 | | 358 | RVVKRKSALGPGV | 6.1388 | 6.6291 | 6.9361 | 7.0167 | 5.8891 | 7.4374 | 7.6151 | 6.4097 | 7.1056 | 7.1433 | 7.3120 | 6.5237 | 4.2634 | 5.3135 | 3.8392 | 4.4274 |
| POLR1E_S358 | POLR1E | Q9GZS1-1 | | 283 | KFEDLLSPAEYEA | 6.0252 | 6.4856 | 6.5776 | 6.1478 | 6.0202 | 6.1478 | 5.7587 | 6.5140 | 6.5480 | 5.9310 | 5.7615 | 6.1038 | 6.5553 | 6.5731 | 6.6261 | 5.7849 |
| POLR1E_S97 | POLR1E | Q9GZS1-1 | | 259 | NDLQDDSLYLPPC | 6.2119 | 6.7688 | 6.8015 | 6.8150 | 6.6173 | 5.8314 | 6.9164 | 6.1091 | 6.3844 | 6.2746 | 6.1909 | 6.0943 | 5.3400 | 5.6920 | 5.5027 | 5.2496 |
| POLR1F_S304 | POLR1F | Q3B726 | | 304 | DSSGYQSDHKKKK | 6.1107 | 6.0288 | 6.2217 | 7.2525 | 6.5147 | 6.3630 | 6.3330 | 6.0150 | 5.6878 | 5.8464 | 5.8975 | 5.9163 | 6.2647 | 6.9479 | 6.2466 | 6.3534 |
| POLR1F_S316 | POLR1F | Q3B726 | | 60 | QRHIALSPRYLNR | 6.1423 | 7.0127 | 6.3773 | 7.1255 | 5.5340 | 5.8107 | 6.4432 | 5.8705 | 5.9647 | 6.1391 | 5.9265 | 6.8545 | 5.5910 | 6.2696 | 6.3052 | 6.6334 |
| POLR1F_S60 | POLR1F | Q3B726 | | 322 | SEEAEFTPPLKCS | 6.0195 | 6.5393 | 6.5409 | 7.0261 | 6.3378 | 6.0641 | 6.5358 | 5.8921 | 6.5400 | 6.2406 | 6.0096 | 6.7412 | 5.8405 | 5.9582 | 5.8010 | 5.9133 |
| POLR1F_T322 | POLR1F | Q3B726 | | 316 | KKKRKHSEEAEFT | 5.6695 | 5.9929 | 5.0394 | 6.7029 | 6.1685 | 7.1717 | 6.1932 | 6.8289 | 5.1109 | 5.3193 | 5.8629 | 5.0407 | 7.0364 | 7.3961 | 7.8275 | 6.6390 |
| POLR1G_S126;S138 | POLR1G | O15446-2 | | 130 | QQSLSGSPLQPIP | 6.0383 | 6.1164 | 6.3036 | 6.4955 | 6.0740 | 6.1161 | 6.4726 | 6.1387 | 6.3818 | 6.3216 | 6.3394 | 6.2402 | 5.8251 | 6.5427 | 6.1753 | 6.4186 |
| POLR1G_S130 | POLR1G | O15446-2 | | 361 | EMKPLESPGGTMA | 6.2425 | 5.8421 | 6.4983 | 7.3086 | 5.7565 | 5.6553 | 6.4367 | 5.8395 | 6.2515 | 6.5907 | 6.5446 | 6.8655 | 5.1553 | 6.3818 | 5.8137 | 6.8173 |
| POLR1G_S130;S138 | POLR1G | O15446-2 | | 138 | LQPIPASPPPQJP | 6.2738 | 6.4655 | 6.4757 | 6.8171 | 6.1804 | 6.2753 | 6.4340 | 6.5920 | 6.5215 | 6.4389 | 6.5427 | 6.5837 | 5.2035 | 5.9561 | 5.4381 | 5.8017 |
| POLR1G_S138 | POLR1G | O15446-2 | | 287 | LEDTVLSPTKKRK | 6.4118 | 6.7453 | 6.6123 | 7.3020 | 6.7497 | 7.0737 | 6.9127 | 6.5840 | 7.0249 | 6.6154 | 7.0872 | 6.9092 | 4.4559 | 4.9380 | 4.0326 | 4.5453 |
| POLR1G_S174 | POLR1G | O15446-2 | | 492 | PPLNSESGEEAPT | 6.0496 | 6.1953 | 6.3154 | 6.2938 | 5.9900 | 5.7181 | 6.3985 | 6.3012 | 6.1705 | 6.3463 | 5.5824 | 5.8228 | 6.3525 | 6.8907 | 6.4017 | 7.1711 |
| POLR1G_S287 | POLR1G | O15446-2 | | 174 | APNLLTSGKKKKE | 6.9179 | 7.6517 | 7.0210 | 7.2247 | 5.9204 | 7.3265 | 7.3600 | 6.7193 | 7.1599 | 7.2690 | 7.7481 | 7.7462 | 3.0381 | 4.2447 | 2.9298 | 3.7228 |
| POLR1G_S29 | POLR1G | O15446-2 | | 422 | PQAAPTSTKKKKK | 6.7205 | 6.5180 | 6.4185 | 6.6155 | 5.8392 | 6.3849 | 6.6030 | 6.3081 | 6.8427 | 6.6052 | 6.5695 | 6.5531 | 5.1805 | 5.7658 | 5.1191 | 5.9562 |
| POLR1G_S361 | POLR1G | O15446-2 | | 29 | KPPASESPRFSLE | 5.9546 | 6.7013 | 6.2551 | 7.1054 | 5.7100 | 5.9604 | 6.3633 | 5.8200 | 6.5528 | 5.9984 | 6.3542 | 6.6484 | 5.6206 | 6.4724 | 6.0630 | 6.4201 |
| POLR1G_S422 | POLR1G | O15446-2 | | 461 | ARATPGSTKKRKK | 6.8372 | 7.4369 | 6.8697 | 7.7506 | 6.2918 | 7.2758 | 7.1767 | 6.9800 | 6.9777 | 7.3450 | 7.2715 | 7.6397 | 2.7422 | 4.3227 | 2.9238 | 4.1585 |
| POLR1G_S461 | POLR1G | O15446-2 | 130;138 | | NA | 6.4586 | 6.7262 | 6.9573 | 7.4261 | 6.1176 | 6.5881 | 6.5168 | 6.7342 | 6.5666 | 6.6028 | 6.8801 | 6.8955 | 4.7294 | 5.0296 | 4.6149 | 5.1564 |
| POLR1G_S492 | POLR1G | O15446-2 | 126;138 | | NA | 6.1422 | 6.0378 | 6.2883 | 6.7431 | 6.2360 | 6.0549 | 6.4033 | 6.0083 | 6.4623 | 6.3071 | 6.7240 | 6.0367 | 5.6789 | 5.6436 | 5.3893 | 5.9443 |
| POLR2A_S1843 | POLR2A | P24928 | | 1864 | PKYSPTSPKYSPT | 7.0890 | 7.0760 | 6.5315 | 6.7094 | 6.1059 | 6.3565 | 6.8537 | 6.3744 | 6.8254 | 6.7384 | 6.5234 | 6.4480 | 5.0098 | 5.0756 | 4.8428 | 5.4401 |
| POLR2A_S1850 | POLR2A | P24928 | | 1920 | PTYSPTSPKYSPT | 6.3389 | 6.8020 | 6.4611 | 6.8868 | 5.6033 | 6.2203 | 6.5143 | 6.2759 | 6.2874 | 6.7956 | 6.5344 | 6.9277 | 5.4181 | 5.6237 | 5.3157 | 5.9948 |
| POLR2A_S1857 | POLR2A | P24928 | | 1966 | AISPDDSDEENxx | 6.0053 | 6.6300 | 6.0331 | 6.4931 | 6.6204 | 6.2312 | 6.1413 | 6.2777 | 6.1122 | 6.7990 | 6.2627 | 6.5997 | 5.5625 | 6.3216 | 6.2238 | 5.6864 |
| POLR2A_S1861 | POLR2A | P24928 | | 1875 | PTSPKYSPTSPTY | 6.1930 | 6.3716 | 6.5404 | 6.8442 | 5.7008 | 6.3546 | 6.6626 | 6.3765 | 6.8215 | 6.3761 | 6.8215 | 7.1123 | 5.3638 | 5.7600 | 5.2987 | 5.7585 |
| POLR2A_S1864 | POLR2A | P24928 | | 1861 | PTSPKYSPTSPKY | 6.3041 | 6.0765 | 5.8408 | 6.6087 | 5.8826 | 5.9203 | 6.2325 | 6.0476 | 6.7947 | 6.0835 | 5.7852 | 6.0322 | 6.6226 | 6.5884 | 6.5054 | 6.6750 |
| POLR2A_S1864;S1868 | POLR2A | P24928 | | 1882 | PTSPTYSPTTPKY | 6.5480 | 6.3572 | 6.5285 | 6.8972 | 5.8625 | 6.3435 | 6.4104 | 6.2870 | 6.7734 | 6.7387 | 6.4239 | 6.8620 | 5.4315 | 5.7823 | 5.2295 | 5.5244 |
| POLR2A_S1875 | POLR2A | P24928 | | 217 | EKKILLSPERVHE | 7.1350 | 7.2542 | 6.6195 | 7.0815 | 6.5572 | 6.7467 | 7.0553 | 6.7434 | 6.6408 | 6.6041 | 6.6629 | 6.6969 | 4.3428 | 4.6885 | 4.4458 | 4.7254 |
| POLR2A_S1875;S1882 | POLR2A | P24928 | | 1917 | PTSPTYSPTSPKY | 6.5179 | 6.7463 | 6.6106 | 6.6691 | 5.6210 | 5.7239 | 6.3569 | 6.7912 | 6.4721 | 6.0851 | 6.3058 | 6.3824 | 5.9910 | 6.3878 | 5.4657 | 5.8732 |
| POLR2A_S1875;T1885 | POLR2A | P24928 | | 1885 | PTYSPTTPKYSPT | 6.6969 | 6.6320 | 6.8839 | 6.6969 | 5.7223 | 6.2004 | 6.7133 | 6.0606 | 6.7160 | 6.6501 | 6.7519 | 6.6514 | 5.2731 | 5.6504 | 5.3736 | 5.3273 |
| POLR2A_S1882 | POLR2A | P24928 | | 1850 | PSYSPSSPEYTPT | 5.8871 | 6.1422 | 6.1911 | 6.1505 | 5.7219 | 6.6108 | 6.2304 | 6.0848 | 6.8309 | 6.4234 | 6.3868 | 7.0669 | 5.9101 | 6.1507 | 5.7705 | 6.4419 |
| POLR2A_S1885 | POLR2A | P24928 | | 1843 | PKYTPTSPSYSPS | 5.7875 | 5.9871 | 6.3385 | 6.2367 | 6.0734 | 6.5063 | 6.2042 | 6.4051 | 6.0809 | 6.5866 | 6.6112 | 7.5847 | 5.8718 | 6.2553 | 5.5233 | 5.9473 |
| POLR2A_S1892 | POLR2A | P24928 | | 1906 | PVYTPTSPKYSPT | 6.3330 | 6.1648 | 6.9744 | 6.0054 | 5.2325 | 6.9930 | 5.9868 | 6.1373 | 6.5441 | 6.8483 | 6.4463 | 6.4507 | 5.9706 | 6.1931 | 5.3980 | 6.3217 |
| POLR2A_S1892;S1899 | POLR2A | P24928 | | 1857 | PEYTPTSPKYSPT | 6.3859 | 6.7403 | 6.5365 | 7.7402 | 5.0677 | 6.0300 | 6.2869 | 5.3443 | 6.3573 | 7.1797 | 7.6404 | 6.6805 | 5.5914 | 6.0594 | 4.5636 | 5.7960 |
| POLR2A_S1892;S1906 | POLR2A | P24928 | | 1892 | PKYSPTSPTYSPT | 5.7753 | 6.8157 | 6.6550 | 6.5331 | 5.2773 | 6.1452 | 6.4683 | 6.5500 | 5.9830 | 6.8424 | 6.5630 | 6.3870 | 5.5380 | 6.2047 | 6.0828 | 6.1792 |
| POLR2A_S1892;T1899;S1 | POLR2A | P24928 | | 1913 | PKYSPTSPTYSPT | 6.4765 | 6.5973 | 6.6308 | 6.4050 | 5.4912 | 6.8133 | 6.4667 | 6.6070 | 5.9821 | 6.7218 | 6.5144 | 6.1787 | 5.4673 | 6.1898 | 5.3525 | 6.1055 |
| POLR2A_S1906 | POLR2A | P24928 | 1892;1906 | | NA | 6.3437 | 6.5146 | 6.5236 | 7.2227 | 5.5431 | 6.3030 | 6.2229 | 6.1828 | 6.9370 | 6.5848 | 6.9367 | 6.7471 | 5.3649 | 5.8027 | 5.1910 | 5.5793 |
| POLR2A_S1910;S1917 | POLR2A | P24928 | 1864;1868 | | NA | 6.9168 | 7.0665 | 6.5837 | 6.8061 | 6.1728 | 6.7537 | 6.8364 | 6.1150 | 6.5870 | 6.4630 | 7.0929 | 6.3295 | 5.0032 | 5.4029 | 5.1754 | 4.6948 |
| POLR2A_S1910;S1920 | POLR2A | P24928 | 1910;1917 | | NA | 6.1061 | 6.4568 | 6.6821 | 7.0544 | 5.9156 | 5.7334 | 5.9015 | 6.1913 | 6.4768 | 6.2784 | 6.3244 | 6.0532 | 6.1591 | 6.3789 | 5.9304 | 6.3577 |
| POLR2A_S1913 | POLR2A | P24928 | 1913;1917 | | NA | 6.9467 | 6.7377 | 7.0051 | 6.9529 | 6.1262 | 6.0310 | 6.7974 | 6.5318 | 6.6969 | 6.5826 | 6.5086 | 5.9872 | 5.2221 | 5.4108 | 4.9282 | 5.5348 |
| POLR2A_S1917 | POLR2A | P24928 | 1892;1899 | | NA | 6.1606 | 6.6563 | 6.9351 | 6.5106 | 5.8398 | 6.3178 | 6.7471 | 6.7800 | 6.0666 | 7.0216 | 6.8251 | 7.0996 | 5.3467 | 5.3540 | 4.9191 | 5.4200 |
| POLR2A_S1920 | POLR2A | P24928 | 1875;1885 | | NA | 6.7521 | 7.1578 | 6.5765 | 6.3734 | 6.2256 | 6.3621 | 6.7047 | 6.8416 | 7.2080 | 6.7159 | 7.4230 | 6.2839 | 4.5346 | 5.2571 | 4.2357 | 5.3480 |
| POLR2A_S1966 | POLR2A | P24928 | 1875;1882 | | NA | 7.6098 | 7.4829 | 6.2743 | 7.5333 | 6.7990 | 7.1164 | 6.1931 | 6.2069 | 7.2343 | 6.4951 | 6.9159 | 7.0200 | 4.3616 | 4.2281 | 4.1188 | 4.4105 |
| POLR2A_S217 | POLR2A | P24928 | 1913;1920 | | NA | 5.6160 | 7.1983 | 7.0153 | 6.3398 | 6.2831 | 6.0176 | 6.9245 | 5.8436 | 6.0501 | 7.1150 | 7.3696 | 6.9448 | 4.8605 | 4.8136 | 4.9725 | 5.7358 |
| POLR2A_T1843;S1857 | POLR2A | P24928 | 1892;1899;19 | | NA | 6.4666 | 6.8972 | 6.3006 | 7.6720 | 5.9629 | 6.4100 | 6.2504 | 6.3459 | 6.6820 | 7.2029 | 6.8779 | 6.6794 | 4.8110 | 5.7468 | 4.4374 | 5.2572 |
| POLR2A_T1913;S1917 | POLR2A | P24928 | 1843;1857 | | NA | 6.6415 | 6.3567 | 6.7418 | 6.9723 | 5.8147 | 6.8323 | 6.3504 | 6.3202 | 6.7651 | 7.3543 | 6.7585 | 6.4091 | 5.1350 | 5.1338 | 4.9939 | 5.4203 |
| POLR2A_T1913;S1920 | POLR2A | P24928 | 1910;1920 | | NA | 6.7338 | 6.7786 | 6.9692 | 6.9608 | 6.1373 | 6.0724 | 6.9173 | 6.9467 | 6.5470 | 6.3268 | 6.5359 | 6.5088 | 5.4545 | 4.8198 | 4.8625 | 5.4286 |
| POLR3C_S205 | POLR3C | Q9BUI4 | | 205 | GKRRRSSDEDAAG | 6.1083 | 6.4848 | 6.7376 | 5.8417 | 5.7280 | 6.3524 | 6.3080 | 5.7328 | 6.7350 | 6.4697 | 7.0828 | 7.2519 | 5.5744 | 6.6339 | 5.3744 | 5.5843 |
| POM121_S108 | POM121 | Q96HA1 | | 697 | QATSAPSPAPKQS | 6.2458 | 5.3463 | 5.7298 | 6.3609 | 6.3642 | 6.4059 | 6.6795 | 5.8180 | 6.2730 | 6.1190 | 5.9416 | 6.1860 | 6.6337 | 6.9157 | 5.9631 | 7.0175 |
| POM121_S179 | POM121 | Q96HA1 | | 485 | EKAADTTPRKKQN | 7.1999 | 7.0118 | 6.1293 | 6.6034 | 6.1866 | 6.1349 | 6.4441 | 5.6463 | 5.9982 | 5.9392 | 5.7945 | 6.1479 | 6.1682 | 6.3944 | 6.0552 | 6.1461 |
| POM121_S184;S188 | POM121 | Q96HA1 | | 435 | RNGPSSSPFSSPA | 6.4573 | 6.1336 | 5.8246 | 6.3305 | 6.2624 | 5.5565 | 6.3604 | 5.7406 | 6.4691 | 5.9770 | 5.6714 | 5.8241 | 7.3476 | 6.7688 | 6.6013 | 6.6750 |
| POM121_S188 | POM121 | Q96HA1 | | 447 | ASSRSQTPERPAK | 6.4464 | 6.6609 | 5.9155 | 6.6636 | 6.2897 | 6.0369 | 6.5106 | 6.2431 | 6.5408 | 6.4185 | 6.1449 | 6.0855 | 6.0447 | 6.0334 | 5.8879 | 6.0776 |
| POM121_S269 | POM121 | Q96HA1 | | 393 | KRSRSSSMSSLTG | 6.0606 | 5.6747 | 5.9121 | 6.4950 | 6.4340 | 6.4037 | 6.4655 | 6.2376 | 6.8582 | 6.5125 | 6.3161 | 6.0463 | 6.5884 | 6.4563 | 5.3262 | 6.2129 |
| POM121_S298 | POM121 | Q96HA1 | | 269 | NSRMVCSPVTVRI | 6.5597 | 6.8116 | 6.4688 | 7.0118 | 5.7375 | 6.3227 | 6.6066 | 6.0406 | 6.6633 | 6.5094 | 6.7020 | 6.4929 | 5.4528 | 5.6054 | 5.1942 | 5.8208 |
| POM121_S314 | POM121 | Q96HA1 | | 108 | RRTLFASPLAKST | 7.7555 | 7.8545 | 6.8326 | 7.5736 | 6.6473 | 6.7585 | 7.4977 | 6.7785 | 7.3724 | 7.1511 | 7.7718 | 6.9655 | 2.7852 | 3.8942 | 2.7739 | 3.5878 |
| POM121_S345;S348 | POM121 | Q96HA1 | | 958 | FGSSAKSPLPSYP | 6.0809 | 5.5367 | 5.2693 | 5.9001 | 6.0340 | 6.4988 | 6.7171 | 6.2888 | 5.9054 | 5.6717 | 5.5027 | 5.4425 | 7.2172 | 7.4125 | 6.7427 | 7.7797 |
| POM121_S345;S348;S35 | POM121 | Q96HA1 | | 314 | AKETVLSALKEKE | 7.8583 | 8.1637 | 6.7804 | 8.2571 | 6.5263 | 6.8834 | 7.9598 | 7.0604 | 7.4715 | 6.9954 | 7.3694 | 7.2627 | 2.4311 | 3.4717 | 2.3076 | 3.2012 |
| POM121_S345;S351 | POM121 | Q96HA1 | | 298 | ISSTLSSPSSNAP | 7.1763 | 7.3475 | 7.5468 | 7.6876 | 6.6717 | 6.6896 | 7.1804 | 7.1995 | 6.7435 | 6.7674 | 7.1966 | 6.7383 | 3.1563 | 4.4724 | 3.2915 | 4.1348 |
| POM121_S381 | POM121 | Q96HA1 | | 179 | RSPPPPSRPPPRSP | 6.1393 | 6.2874 | 6.3919 | 7.0582 | 6.3876 | 6.0968 | 6.6973 | 6.5105 | 6.2868 | 5.8948 | 6.2151 | 5.9483 | 6.1819 | 5.7581 | 6.1499 | 5.9962 |
| POM121_S393 | POM121 | Q96HA1 | | 188 | PRSPPPSPPTHRA | 7.1722 | 6.8183 | 6.7832 | 6.3383 | 6.6531 | 6.4096 | 6.7030 | 6.8548 | 6.0192 | 6.4591 | 6.4321 | 5.8993 | 5.6006 | 5.7602 | 5.0299 | 5.0672 |
| POM121_S393;S396 | POM121 | Q96HA1 | | 381 | GLNSQSSDDHLNK | 3.6655 | 3.3500 | 3.1034 | 3.7350 | 3.9692 | 4.0696 | 4.3267 | 4.6108 | 3.4304 | 3.3857 | 3.2257 | 3.6365 | 12.7482 | 12.4631 | 15.6827 | 14.5974 |

| | | | | | | | | | | | | | | | | | | | | |
|---|---|---|---|---|---|---|---|---|---|---|---|---|---|---|---|---|---|---|---|---|
| POM121_S435 | POM121 | Q96HA1 | 80 | RGSRPLSSFVRKA | 7.1251 | 7.1116 | 5.8739 | 7.1492 | 6.8168 | 6.1920 | 6.5131 | 6.5236 | 5.5221 | 6.1732 | 5.4205 | 6.0657 | 6.4116 | 6.1567 | 5.1183 | 5.8266 |
| POM121_S697 | POM121 | Q96HA1 | 712 | FGTQNTSPSSPAA | 5.9258 | 5.7484 | 5.7661 | 5.8420 | 5.0574 | 6.2560 | 5.7464 | 6.0894 | 7.0544 | 5.5374 | 6.0638 | 6.4344 | 6.9548 | 7.0593 | 7.7601 | 6.7045 |
| POM121_S712 | POM121 | Q96HA1 | 345;348 | NA | 5.5690 | 6.2684 | 6.2405 | 6.3882 | 6.0102 | 6.2980 | 6.1997 | 6.5749 | 6.4597 | 6.9540 | 6.9055 | 6.8029 | 5.7645 | 6.0185 | 5.6014 | 5.9445 |
| POM121_S80 | POM121 | Q96HA1 | 184;188 | NA | 6.3770 | 6.1826 | 6.1398 | 6.9717 | 5.8746 | 6.0897 | 6.1386 | 5.9672 | 6.0944 | 6.6224 | 5.8490 | 6.4990 | 6.3508 | 6.6511 | 6.0514 | 6.1409 |
| POM121_S958 | POM121 | Q96HA1 | 345;348;351 | NA | 6.6635 | 6.5745 | 6.4435 | 7.3571 | 6.5492 | 5.7180 | 6.2561 | 6.0167 | 6.2164 | 5.6385 | 5.9086 | 6.2301 | 6.1014 | 6.4594 | 5.8802 | 5.9869 |
| POM121_T447 | POM121 | Q96HA1 | 393;396 | NA | 5.9336 | 5.9899 | 7.4004 | 7.3097 | 7.6520 | 5.6846 | 6.5808 | 6.5218 | 6.3510 | 6.6780 | 6.2210 | 5.7748 | 5.4893 | 5.7708 | 5.2550 | 5.3872 |
| POM121_T485 | POM121 | Q96HA1 | 345;351 | NA | 5.6626 | 6.4667 | 6.5892 | 6.7693 | 6.0040 | 6.0136 | 5.9678 | 6.3866 | 7.0600 | 7.1074 | 6.8298 | 6.6175 | 5.6900 | 5.5588 | 5.8689 | 5.4078 |
| POM121C_S190 | POM121C | A8CG34 | 689 | FGTQNTSPSSPAA | 5.5378 | 5.9899 | 5.5883 | 6.0437 | 6.1573 | 5.7406 | 5.9733 | 6.3232 | 6.3105 | 6.2530 | 5.9464 | 6.1094 | 6.8402 | 6.8290 | 7.0265 | 7.3310 |
| POM121C_S322;S325 | POM121C | A8CG34 | 370 | KRSRSSSMSSLTG | 5.3899 | 5.6833 | 6.3363 | 6.5278 | 6.3677 | 6.5698 | 6.3272 | 5.8952 | 6.6037 | 6.6008 | 6.7460 | 6.6125 | 5.8161 | 6.1327 | 6.3440 | 6.0470 |
| POM121C_S322;S325;S3 | POM121C | A8CG34 | 190 | RPSGRPSPRDRGT | 5.9925 | 5.7229 | 5.7891 | 6.4968 | 5.8910 | 5.9296 | 6.4932 | 5.9458 | 7.2108 | 7.0196 | 6.9752 | 7.0855 | 5.6162 | 6.1049 | 5.4251 | 6.3018 |
| POM121C_S322;S328 | POM121C | A8CG34 | 80 | RGSRPLSSFVQKA | 6.7400 | 7.0996 | 6.5482 | 7.0694 | 6.5627 | 6.0450 | 7.3132 | 6.7014 | 6.8643 | 6.7473 | 6.3358 | 6.4046 | 4.1929 | 6.3141 | 4.0602 | 5.0013 |
| POM121C_S370 | POM121C | A8CG34 | 81 | GSRPLSSFVQKAR | 3.6900 | 3.5020 | 3.7296 | 4.0489 | 3.9199 | 3.1074 | 3.1477 | 3.1493 | 3.3870 | 3.1656 | 3.2355 | 3.3148 | 13.8432 | 13.4999 | 16.9296 | 14.3296 |
| POM121C_S370;S373 | POM121C | A8CG34 | 322;325 | NA | 5.9031 | 6.2532 | 6.1872 | 6.3205 | 5.9345 | 6.1470 | 6.1009 | 6.5837 | 6.3857 | 6.4366 | 6.7308 | 6.5070 | 5.8689 | 6.4364 | 6.0229 | 6.1814 |
| POM121C_S689 | POM121C | A8CG34 | 322;325;328 | NA | 6.5396 | 6.2001 | 6.0004 | 7.2489 | 6.4442 | 6.1677 | 6.3583 | 5.8647 | 6.5085 | 5.8959 | 5.9679 | 6.2103 | 6.3576 | 6.3077 | 5.7405 | 6.1877 |
| POM121C_S80 | POM121C | A8CG34 | 322;328 | NA | 5.4835 | 5.4838 | 5.8529 | 6.1214 | 5.9125 | 6.2584 | 6.3014 | 6.6358 | 6.6786 | 6.6151 | 6.4544 | 6.7649 | 6.0009 | 6.5803 | 6.5414 | 6.3146 |
| POM121C_S81 | POM121C | A8CG34 | 370;373 | NA | 6.2881 | 5.8460 | 7.4730 | 6.0316 | 5.1062 | 6.6350 | 6.5910 | 6.4293 | 7.3162 | 6.0448 | 6.1258 | 6.1907 | 6.2340 | 5.7970 | 5.2291 | 6.6622 |
| POP1_S367 | POP1 | Q99575 | 730 | EPSVASSPNGKES | 5.9425 | 6.1329 | 6.2612 | 6.5390 | 6.5852 | 6.1297 | 6.0130 | 6.1197 | 6.3861 | 6.2834 | 5.9784 | 6.1609 | 6.2410 | 6.6954 | 6.3816 | 6.1499 |
| POP1_S730 | POP1 | Q99575 | 367 | DPLPTPSQEKSQT | 5.8172 | 6.3838 | 6.5852 | 6.4191 | 5.9632 | 5.9000 | 5.9836 | 5.8195 | 6.6142 | 6.6610 | 6.4322 | 6.4523 | 5.8424 | 6.2965 | 6.4567 | 6.3733 |
| POP4_S10 | POP4 | O95707 | 10 | VIYHALSQKEAND | 7.2545 | 6.6541 | 6.4645 | 6.2805 | 7.4014 | 5.9162 | 5.9545 | 6.6768 | 7.0586 | 5.3812 | 5.3043 | 5.4875 | 6.7608 | 6.6876 | 5.4273 | 5.2901 |
| POP5_S154 | POP5 | Q969H6 | 154 | LEEEEESGEEAAE | 6.4434 | 6.3908 | 6.4062 | 6.1804 | 6.2319 | 6.2210 | 6.4639 | 6.7207 | 6.2520 | 6.2356 | 6.2743 | 5.9101 | 6.1289 | 6.2915 | 5.7114 | 6.1377 |
| POR_S62 | POR | P16435 | 62 | KIQTLTSSVRESS | 6.9299 | 7.0018 | 6.1252 | 7.2635 | 6.3858 | 6.3116 | 6.4646 | 6.0255 | 6.6625 | 5.8934 | 6.3416 | 6.4437 | 5.4806 | 5.4539 | 5.5688 | 5.6476 |
| POU2F1_S358 | POU2F1 | P14859-6 | 471 | SGGTSSSPIKAIF | 5.8239 | 5.9686 | 6.2171 | 5.5740 | 6.0789 | 6.1577 | 5.9563 | 6.5165 | 6.2915 | 6.4760 | 6.2065 | 6.0697 | 6.6624 | 6.6869 | 6.5390 | 6.7749 |
| POU2F1_S393 | POU2F1 | P14859-6 | 293 | LPQSQSTPKRIDT | 6.3872 | 6.3215 | 6.8975 | 6.6823 | 5.9670 | 6.1010 | 6.4529 | 6.6294 | 6.8445 | 6.7732 | 7.4971 | 6.9846 | 4.9336 | 5.6392 | 4.6637 | 5.2254 |
| POU2F1_S471 | POU2F1 | P14859-6 | 358 | FEALNLSFKNMCK | 6.8101 | 7.7386 | 6.4798 | 7.4376 | 6.4759 | 7.2226 | 7.2148 | 7.3653 | 6.2179 | 6.9704 | 6.8707 | 6.1729 | 4.7382 | 5.9325 | 3.3249 | 3.0278 |
| POU2F1_T293 | POU2F1 | P14859-6 | 393 | SPSALNSPGIEGL | 6.2387 | 5.9524 | 6.7196 | 5.9495 | 5.9067 | 6.8000 | 6.1693 | 7.3394 | 6.2221 | 6.2925 | 6.6911 | 6.6501 | 5.9072 | 6.3443 | 5.3378 | 5.4796 |
| POU4F1_S361 | POU4F1 | Q01851 | 361 | KKRKRTSIAAPEK | 6.4915 | 6.4569 | 6.1159 | 6.6246 | 5.1408 | 4.9671 | 5.8939 | 5.6523 | 6.1648 | 5.9277 | 5.5173 | 6.1024 | 6.1186 | 6.8041 | 7.5648 | 8.4574 |
| PPA2_S331 | PPA2 | Q9H2U2-2 | 332 | VESVSSSPNKESN | 5.9458 | 5.8367 | 6.5697 | 5.7009 | 6.0435 | 6.3871 | 5.8095 | 6.3333 | 6.1590 | 5.9260 | 6.7697 | 6.3777 | 6.4201 | 6.5029 | 6.8945 | 6.3235 |
| PPA2_S332 | PPA2 | Q9H2U2-2 | 331 | LVESVSSSPNKES | 5.1529 | 5.3614 | 6.2170 | 5.6649 | 6.2847 | 6.1052 | 5.5969 | 6.3670 | 6.6506 | 6.5455 | 6.4799 | 6.4097 | 6.3580 | 7.1867 | 6.7649 | 6.8547 |
| PPAN_S238 | PPAN | Q9NQ55-3 | 359 | KARVGGSDEEASG | 6.5575 | 6.5506 | 5.9798 | 6.7570 | 6.2649 | 5.7424 | 6.7802 | 6.0678 | 6.0946 | 5.6685 | 5.7705 | 5.6263 | 6.3409 | 6.5373 | 6.5701 | 6.6915 |
| PPAN_S359 | PPAN | Q9NQ55-3 | 238 | ATGAGLSESEAEP | 5.4260 | 6.1766 | 6.3331 | 6.9689 | 6.3186 | 6.2028 | 6.3955 | 6.3662 | 6.5765 | 6.4372 | 6.8141 | 6.3736 | 5.6924 | 6.3923 | 5.5568 | 5.9695 |
| PPAN_T233;S240 | PPAN | Q9NQ55-3 | 389 | EDDDIEYFCQAVG | 5.8460 | 6.4978 | 5.7461 | 6.9825 | 6.4819 | 5.9687 | 5.9326 | 6.4122 | 6.7918 | 6.0485 | 6.0384 | 6.0989 | 6.4357 | 6.5541 | 5.6369 | 6.5281 |
| PPAN_Y389 | PPAN | Q9NQ55-3 | 233;240 | NA | 5.9788 | 6.0264 | 6.7575 | 5.8250 | 5.6574 | 5.9274 | 7.1157 | 5.6862 | 5.5782 | 6.9327 | 7.5168 | 6.9917 | 5.6779 | 6.2600 | 5.9048 | 6.1634 |
| PPARGC1B_S638 | PPARGC1B | Q86YN6 | 638 | IALSLPSPEGLSL | 6.0978 | 5.9950 | 7.0758 | 6.4422 | 6.2333 | 6.5851 | 6.8561 | 6.4305 | 7.3126 | 6.4902 | 6.1599 | 7.1150 | 5.2259 | 5.4700 | 5.4891 | 5.0214 |
| PPFIA1_S1172 | PPFIA1 | Q13136 | 1172 | VTSSMSSPSMQPK | 5.3986 | 5.7966 | 5.9329 | 5.8963 | 5.5416 | 6.3924 | 6.4550 | 6.0277 | 6.2756 | 6.1801 | 6.1384 | 5.9417 | 6.9923 | 7.1715 | 6.9110 | 6.9482 |
| PPFIA1_S239;S242 | PPFIA1 | Q13136 | 763 | GALHTVSHEDIRD | 5.9512 | 6.0371 | 6.0260 | 5.8013 | 5.4952 | 5.9056 | 6.2229 | 6.0561 | 6.8285 | 6.1174 | 6.0936 | 6.4509 | 6.4015 | 7.0338 | 6.3043 | 7.2746 |
| PPFIA1_S708 | PPFIA1 | Q13136 | 708 | PRRIPHSPAREVD | 6.8169 | 6.7910 | 6.7884 | 6.6707 | 6.5947 | 6.6810 | 6.5684 | 6.5321 | 7.4202 | 6.7125 | 6.8074 | 6.7676 | 4.6126 | 4.9947 | 4.4325 | 4.8093 |
| PPFIA1_S763 | PPFIA1 | Q13136 | 230 | NHEQENTPSTSGK | 6.7366 | 7.4550 | 5.3141 | 6.2347 | 7.2195 | 7.9997 | 8.5040 | 6.7318 | 7.7428 | 7.1695 | 7.6405 | 7.7714 | 3.6109 | 3.4825 | 3.3564 | 3.0305 |
| PPFIA1_T230 | PPFIA1 | Q13136 | 239;242 | NA | 5.8118 | 5.6129 | 7.0031 | 6.1049 | 6.4495 | 6.1573 | 5.5350 | 5.9852 | 6.9513 | 5.8185 | 5.9527 | 6.5234 | 6.6855 | 6.5724 | 6.4543 | 6.3821 |
| PPFIA3_S1164 | PPFIA3 | O75145 | 1164 | AAGALGSPGLPLR | 6.0040 | 5.8799 | 6.6893 | 6.6023 | 5.9413 | 6.1324 | 6.0677 | 6.8310 | 6.5011 | 6.1543 | 5.9607 | 6.6610 | 5.8898 | 6.2126 | 6.0854 | 6.3872 |
| PPFIA3_S508 | PPFIA3 | O75145 | 683 | PRLAPPSPAREGT | 7.3325 | 6.8029 | 6.4297 | 6.8466 | 6.3787 | 6.6957 | 6.4277 | 7.1310 | 6.0783 | 6.2894 | 7.2800 | 6.0885 | 5.1255 | 5.2051 | 4.7673 | 5.1211 |
| PPFIA3_S683 | PPFIA3 | O75145 | 508 | PSSYSRSLPGSAL | 6.7479 | 6.5919 | 7.3575 | 6.2944 | 6.2991 | 6.7938 | 6.2875 | 6.3443 | 6.1842 | 6.6037 | 7.1459 | 6.7517 | 4.9947 | 5.6467 | 5.1233 | 4.8333 |
| PPFIBP1_S37 | PPFIBP1 | Q86W92 | 37 | GIFDCQSPTSPFM | 6.0014 | 6.1384 | 6.6483 | 6.4949 | 6.4394 | 6.7895 | 6.4636 | 6.3501 | 6.8431 | 6.5491 | 6.5675 | 6.1613 | 5.6149 | 5.9722 | 5.5691 | 5.3973 |
| PPFIBP1_S37;S40 | PPFIBP1 | Q86W92 | 466 | SLQKSSSLGNLKK | 6.1029 | 5.9199 | 6.4483 | 5.7785 | 5.5683 | 6.3892 | 5.9966 | 6.4035 | 6.3180 | 6.1505 | 5.9231 | 5.8322 | 6.7186 | 6.7789 | 6.4588 | 6.2127 |
| PPFIBP1_S40 | PPFIBP1 | Q86W92 | 601 | KLRRSQSTTFNPD | 6.6281 | 6.3859 | 7.3018 | 5.8702 | 6.0826 | 6.4096 | 6.0568 | 6.2964 | 6.2968 | 6.6862 | 6.6724 | 6.8144 | 5.5510 | 6.1536 | 5.2611 | 5.5333 |
| PPFIBP1_S466 | PPFIBP1 | Q86W92 | 579 | FQIPPPSPDSKKK | 6.2855 | 6.4749 | 6.4651 | 6.1250 | 6.1576 | 6.4581 | 6.3255 | 6.2834 | 6.7432 | 6.4553 | 6.7850 | 6.8270 | 5.9084 | 5.8423 | 5.3002 | 5.5635 |
| PPFIBP1_S534 | PPFIBP1 | Q86W92 | 636 | SRDLGQSNSDLDM | 5.9472 | 6.7957 | 7.1824 | 6.4085 | 6.0402 | 6.7348 | 6.3612 | 7.0367 | 6.1447 | 6.3119 | 6.9626 | 6.7013 | 4.9275 | 5.4093 | 5.4900 | 5.5461 |
| PPFIBP1_S540 | PPFIBP1 | Q86W92 | 40 | DCQSPTSPFMGSL | 5.1370 | 5.7833 | 5.9623 | 6.2413 | 7.0848 | 6.0109 | 6.8883 | 6.2887 | 6.0325 | 5.7658 | 5.9641 | 5.8389 | 6.4192 | 6.7666 | 7.4685 | 6.3478 |
| PPFIBP1_S571;S579 | PPFIBP1 | Q86W92-2 | 534 | DRKRSASAPTLAE | 6.6606 | 6.0085 | 7.3581 | 5.6828 | 6.5166 | 6.6404 | 6.4460 | 6.6379 | 6.3608 | 6.3192 | 6.3814 | 6.3806 | 5.7082 | 6.2396 | 5.1183 | 5.5407 |
| PPFIBP1_S579 | PPFIBP1 | Q86W92 | 794 | CLRRRPSDENTIA | 5.9912 | 5.0923 | 6.6548 | 4.7690 | 6.5254 | 6.8572 | 5.6294 | 4.8561 | 7.2490 | 6.9041 | 6.7616 | 6.7645 | 6.5761 | 7.6643 | 5.7944 | 5.9106 |
| PPFIBP1_S601 | PPFIBP1 | Q86W92 | 540 | SNKRTASAPNLAE | 6.8097 | 7.2072 | 7.7554 | 5.7092 | 5.9768 | 6.2398 | 5.8349 | 6.2199 | 7.3657 | 6.2592 | 6.1321 | 6.8684 | 5.2682 | 5.7887 | 5.3366 | 5.2280 |
| PPFIBP1_S636 | PPFIBP1 | Q86W92 | 37;40 | NA | 6.0721 | 6.3718 | 6.7636 | 6.8614 | 6.4529 | 6.8783 | 6.7464 | 6.7863 | 6.7196 | 6.6290 | 6.3003 | 6.2587 | 5.4198 | 5.2753 | 5.1497 | 5.3147 |
| PPFIBP1_S794 | PPFIBP1 | Q86W92 | 571;579 | NA | 5.7748 | 6.6166 | 7.1346 | 6.8291 | 5.3102 | 6.8406 | 7.6540 | 6.7916 | 7.0065 | 7.0585 | 7.2391 | 6.8558 | 3.8748 | 5.4604 | 3.9045 | 5.6488 |
| PPHLN1_S110 | PPHLN1 | Q8NEY8 | 133 | NTFFRESPVGRKD | 5.9704 | 6.1034 | 5.6975 | 6.4903 | 6.2701 | 6.0869 | 6.0960 | 6.2157 | 6.1369 | 6.0920 | 5.9062 | 5.9672 | 7.0298 | 6.5808 | 6.6875 | 6.6693 |
| PPHLN1_S133 | PPHLN1 | Q8NEY8 | 205 | KTSRDTSPSSGSA | 5.5363 | 5.6386 | 6.0631 | 6.7721 | 6.8607 | 6.3296 | 7.0886 | 6.6792 | 6.4451 | 6.0654 | 6.6780 | 6.2594 | 6.1456 | 5.9707 | 5.6827 | 5.7847 |
| PPHLN1_S140 | PPHLN1 | Q8NEY8 | 197 | KERPVQSLKTSRD | 5.1559 | 5.8094 | 5.2422 | 5.9745 | 4.6217 | 4.6413 | 4.7436 | 5.0030 | 5.6915 | 5.8045 | 4.9995 | 5.8350 | 8.7773 | 8.3336 | 9.1850 | 10.1818 |
| PPHLN1_S178 | PPHLN1 | Q8NEY8 | 178 | SVRPGASYKRQNE | 5.6987 | 6.2662 | 7.0102 | 7.0135 | 6.8864 | 6.6442 | 6.2477 | 7.2965 | 6.9848 | 8.0007 | 7.7338 | 7.9243 | 4.1295 | 4.5664 | 3.2874 | 4.3098 |
| PPHLN1_S197 | PPHLN1 | Q8NEY8 | 110 | DGFRRKSFYSSHY | 6.3634 | 6.2657 | 5.5268 | 6.9518 | 7.9041 | 6.4805 | 6.0844 | 6.5752 | 7.0668 | 7.1609 | 6.5515 | 7.1633 | 5.1871 | 5.6258 | 3.9698 | 5.1229 |
| PPHLN1_S205 | PPHLN1 | Q8NEY8 | 140 | PVGRKDSPHSRSG | 6.1042 | 6.3572 | 5.8624 | 6.3879 | 6.9877 | 7.3614 | 6.8679 | 6.6018 | 6.4415 | 5.7523 | 6.9823 | 6.0337 | 6.3388 | 5.6290 | 5.8290 | 4.4629 |
| PPIE_S97 | PPIE | Q9UNP9-3 | 97 | SSRPVWSDDDWLK | 6.1707 | 6.1300 | 7.6270 | 7.9844 | 6.4449 | 6.2282 | 6.8673 | 6.8627 | 6.1163 | 6.0750 | 6.0460 | 7.0733 | 5.0468 | 4.9390 | 4.8586 | 5.5299 |
| PPIG_S173 | PPIG | Q13427 | 687 | KADRDQSPFSKIK | 6.3149 | 6.2372 | 5.6758 | 6.0918 | 6.2089 | 5.5599 | 6.1764 | 5.8105 | 6.1043 | 5.7450 | 5.6959 | 5.5696 | 6.9872 | 7.2673 | 7.4122 | 7.1432 |
| PPIG_S356;T358 | PPIG | Q13427 | 717 | DEKIRSSVEKENQ | 6.4834 | 6.5900 | 5.7640 | 6.2557 | 6.4448 | 6.1488 | 6.2199 | 6.0029 | 5.8978 | 5.8696 | 5.4472 | 5.3959 | 6.7444 | 6.9894 | 6.9956 | 6.7506 |
| PPIG_S358 | PPIG | Q13427 | 358 | RFRRSETPPHWRQ | 6.5520 | 6.5619 | 6.1715 | 6.5284 | 6.1175 | 6.0450 | 6.2117 | 6.2283 | 6.0698 | 6.2535 | 6.4212 | 6.3161 | 5.8469 | 6.3560 | 6.2526 | 6.0676 |
| PPIG_S376 | PPIG | Q13427 | 611 | RSRERRTPPGRSR | 7.1140 | 7.0548 | 6.3822 | 6.3303 | 6.5712 | 6.5302 | 6.2061 | 5.1916 | 5.0905 | 5.2963 | 6.3981 | 6.7949 | 6.3261 | 7.0031 | 5.8860 | 5.8247 |
| PPIG_S397 | PPIG | Q13427 | 173 | AEVRILSCGELIP | 6.2634 | 5.8027 | 5.8442 | 6.6041 | 5.8564 | 5.8133 | 5.8553 | 5.7373 | 6.1226 | 6.0455 | 6.1108 | 5.8706 | 6.3288 | 7.1459 | 7.6150 | 6.9840 |
| PPIG_S415 | PPIG | Q13427 | 415 | HRNVSESPNRKNE | 6.1903 | 6.6109 | 6.0329 | 6.6220 | 5.7115 | 5.7544 | 6.0980 | 5.3711 | 6.1194 | 6.1643 | 6.0151 | 6.0381 | 6.4767 | 7.2274 | 6.8032 | 6.7646 |
| PPIG_S544;S546 | PPIG | Q13427 | 744 | KKFDHESSPGTDE | 5.8906 | 6.7537 | 4.6984 | 7.8114 | 6.2931 | 6.2752 | 5.2324 | 7.7038 | 6.7706 | 6.4571 | 6.8287 | 5.6458 | 4.0671 | 7.1313 | 6.9652 | 5.4755 |
| PPIG_S546 | PPIG | Q13427 | 397 | IKENQRSPVRVKE | 6.7268 | 6.6979 | 6.3789 | 6.4862 | 6.2035 | 5.1632 | 6.7376 | 4.9723 | 5.7380 | 6.0471 | 6.0589 | 5.9308 | 6.5689 | 6.8114 | 7.5151 | 5.9634 |
| PPIG_S687 | PPIG | Q13427 | 546 | RRAQSRSRECDIT | 5.1034 | 6.0384 | 5.9656 | 5.8967 | 5.9440 | 5.5999 | 6.4436 | 6.2923 | 5.9292 | 5.7671 | 5.4494 | 6.0643 | 7.0914 | 7.3139 | 7.3968 | 7.7040 |
| PPIG_S716;S717 | PPIG | Q13427 | 376 | QRMRVSSGERWIK | 5.7040 | 6.5396 | 5.5142 | 6.5696 | 5.5713 | 4.6956 | 5.2309 | 5.1288 | 10.7429 | 5.8039 | 5.2922 | 6.9464 | 5.2337 | 5.6505 | 8.9981 | 6.3784 |
| PPIG_S717 | PPIG | Q13427 | 356;358 | NA | 6.2553 | 6.1195 | 5.9808 | 6.3415 | 6.0932 | 6.0731 | 6.0800 | 6.0580 | 5.8408 | 6.0301 | 6.0384 | 6.1915 | 6.6921 | 6.8562 | 6.6919 | 6.6576 |
| PPIG_S744 | PPIG | Q13427 | 716;717 | NA | 6.9433 | 6.8972 | 6.1198 | 7.7204 | 6.1762 | 6.0094 | 7.1546 | 6.2868 | 4.8712 | 5.0573 | 5.0826 | 4.9725 | 6.4651 | 6.7679 | 6.7159 | 6.7598 |
| PPIG_S744;S745;T748 | PPIG | Q13427 | 544;546 | NA | 6.3748 | 6.6262 | 6.7478 | 6.0842 | 6.7687 | 6.5154 | 5.8911 | 4.6640 | 5.3939 | 5.4826 | 6.1406 | 6.2254 | 6.8495 | 6.7169 | 6.9939 | 6.5251 |
| PPIG_T611 | PPIG | Q13427 | 744;745;748 | NA | 5.9655 | 6.2186 | 5.7809 | 7.6156 | 5.6823 | 5.9268 | 5.7658 | 5.8727 | 5.5528 | 5.8424 | 5.5191 | 6.2040 | 6.4877 | 6.7430 | 7.2156 | 7.6071 |
| PPIL2_S519 | PPIL2 | Q13356 | 519 | GFGDFSSWxxxxx | 6.1730 | 7.2741 | 6.9434 | 6.5150 | 5.8314 | 6.6396 | 6.5208 | 7.1827 | 6.8703 | 7.1106 | 6.6406 | 7.2747 | 4.3808 | 5.2975 | 4.2809 | 5.0644 |
| PPIL4_S178 | PPIL4 | Q8WUA2 | 178 | LLIPDRSPEPTRE | 5.5521 | 5.9222 | 6.0356 | 6.2182 | 6.0093 | 5.9560 | 6.1452 | 6.0763 | 6.2685 | 6.1233 | 6.2151 | 6.2311 | 6.3289 | 7.0828 | 6.8883 | 6.9469 |
| PPIL4_S204 | PPIL4 | Q8WUA2 | 204 | DDFKGRSAEEVEE | 6.4821 | 6.8106 | 7.3681 | 6.6795 | 6.6906 | 6.4474 | 6.6819 | 6.0067 | 5.7477 | 5.7012 | 5.8717 | 7.3365 | 4.7465 | 6.0510 | 5.9199 | 5.4587 |
| PPIP5K1_S1152 | PPIP5K1 | Q6PFW1 | 944 | MKTNQGSMENLCP | 5.4635 | 5.7488 | 6.0719 | 5.3368 | 6.1694 | 6.7353 | 5.9777 | 6.5312 | 5.5037 | 6.1123 | 5.9552 | 5.7994 | 7.5311 | 7.4900 | 6.9326 | 6.6412 |
| PPIP5K1_S944 | PPIP5K1 | Q6PFW1 | 1152 | PPRTLHSPPLQLQ | 5.8594 | 5.7085 | 6.5484 | 6.1403 | 6.4218 | 6.9242 | 6.3705 | 6.5208 | 6.3633 | 7.1996 | 7.1247 | 6.8744 | 5.0879 | 5.8442 | 5.3862 | 5.6259 |
| PPIP5K1_S964 | PPIP5K1 | Q6PFW1 | 964 | DRALQTSPQPPEG | 6.9452 | 6.3939 | 6.6605 | 5.7794 | 6.3097 | 6.8406 | 7.1283 | 7.5996 | 6.7180 | 6.7621 | 6.3021 | 6.1882 | 5.5953 | 5.2020 | 4.9717 | 4.6035 |
| PPIP5K2_S1006 | PPIP5K2 | O43314 | 1006 | GRRRRRSGEQITS | 5.3603 | 5.4369 | 5.4694 | 4.7522 | 5.9295 | 5.9720 | 5.9161 | 5.9166 | 5.6839 | 5.8576 | 5.8951 | 5.3793 | 7.6533 | 8.4740 | 8.2270 | 8.0765 |
| PPIP5K2_S1180 | PPIP5K2 | O43314 | 1192 | PAKILPTPPATLK | 5.8731 | 6.8069 | 6.9701 | 7.8439 | 6.7676 | 7.6735 | 7.6425 | 7.4493 | 7.0620 | 7.4793 | 6.7160 | 7.7561 | 3.0378 | 4.2628 | 2.9913 | 3.6679 |
| PPIP5K2_S492 | PPIP5K2 | O43314 | 1185 | VSLNTYTPAKILP | 7.5389 | 7.7321 | 7.8845 | 7.2007 | 5.5319 | 8.1403 | 8.2471 | 6.8082 | 7.9614 | 7.3001 | 6.7987 | 7.6076 | 2.2805 | 4.0868 | 1.9417 | 2.9398 |
| PPIP5K2_T1185 | PPIP5K2 | O43314 | 1180 | IMRKKVSLNTYTP | 4.6718 | 3.9519 | 5.6489 | 4.4471 | 5.8669 | 5.5517 | 4.7137 | 6.0193 | 4.5195 | 4.5719 | 4.2767 | 4.8690 | 10.2350 | 12.1719 | 10.0874 | 8.3971 |
| PPIP5K2_T1192 | PPIP5K2 | O43314 | 492 | HGCPKTSSEEEDS | 5.5721 | 5.4009 | 6.5316 | 6.2406 | 5.5528 | 6.2124 | 7.1290 | 6.3483 | 6.4464 | 6.7528 | 6.4400 | 6.7130 | 6.2579 | 6.3015 | 6.2067 | 5.8939 |
| PPL_S14 | PPL | O60437 | 14 | RNKGKYSPTVQTR | 5.7242 | 5.5682 | 5.2769 | 5.2630 | 6.4891 | 6.1240 | 6.0720 | 6.1325 | 5.6604 | 5.4055 | 5.5615 | 5.3456 | 8.3698 | 7.7146 | 7.9384 | 7.3544 |
| PPL_S1657 | PPL | O60437 | 1657 | ENHLRRSIVVIHP | 6.7763 | 6.0856 | 6.3818 | 5.9621 | 6.7254 | 6.1486 | 6.1961 | 6.4108 | 6.7397 | 5.5533 | 6.4511 | 5.7239 | 6.8788 | 6.2209 | 5.9058 | 5.8398 |
| PPM1D_S40 | PPM1D | O15297 | 40 | TAEEKPSPRRSLS | 6.7256 | 6.6645 | 6.2242 | 6.6233 | 5.6254 | 5.3446 | 6.2603 | 6.6855 | 6.4819 | 6.0356 | 6.8370 | 6.0226 | 6.5451 | 6.1149 | 5.2774 | 6.5321 |
| PPM1D_T529 | PPM1D | O15297 | 529 | AQEIERTPPTNFK | 7.3014 | 7.3585 | 6.3215 | 7.6298 | 6.8411 | 7.6063 | 7.1447 | 6.4848 | 7.5860 | 6.8119 | 6.7074 | 7.2465 | 3.4354 | 4.7034 | 3.1743 | 3.6469 |
| PPM1F_S3 | PPM1F | P49593 | 3 | xxxxMSSGAPQKS | 7.4860 | 7.7037 | 6.6732 | 7.3851 | 6.2197 | 6.0285 | 6.2955 | 7.0635 | 7.3458 | 7.0189 | 6.7369 | 6.9577 | 3.9601 | 4.1534 | 4.2325 | 4.7396 |
| PPME1_S42 | PPME1 | Q9Y570-4 | 42 | GRKRDFSPVPWSQ | 6.2297 | 6.0186 | 7.1094 | 6.2991 | 6.1345 | 6.6274 | 6.1466 | 6.4938 | 6.7257 | 6.5266 | 6.7840 | 6.9209 | 5.2672 | 5.7954 | 5.2658 | 5.6554 |
| PPP1CA_T331 | PPP1CA | P62136-2 | 331 | PGGRPITPPRNSA | 6.8105 | 7.0707 | 6.9585 | 7.0546 | 6.1434 | 6.7275 | 6.7411 | 6.8588 | 6.6975 | 6.9123 | 7.3547 | 7.1125 | 4.2431 | 4.7160 | 4.2161 | 4.3827 |
| PPP1CB_T316 | PPP1CB | P62140 | 316 | NSGRPVTPPRTAN | 6.8814 | 6.7106 | 7.0039 | 6.9900 | 6.1969 | 6.6299 | 7.1942 | 6.9229 | 6.7483 | 7.1842 | 6.7603 | 7.0784 | 4.1653 | 5.2285 | 3.8691 | 4.4361 |
| PPP1R10_S313 | PPP1R10 | Q96QC0 | 545 | PGGSGGSPDGAGG | 6.1939 | 6.4532 | 6.6726 | 6.7266 | 6.3106 | 5.7589 | 6.1776 | 5.9538 | 6.5144 | 6.6938 | 6.3839 | 6.4727 | 5.7565 | 6.3381 | 5.8307 | 5.7628 |
| PPP1R10_S337 | PPP1R10 | Q96QC0 | 313 | KKKKVLSPTAAKP | 6.0555 | 6.0034 | 5.7698 | 6.4179 | 6.5272 | 6.0260 | 6.4245 | 6.0856 | 6.0133 | 5.7200 | 5.5623 | 5.5062 | 7.0760 | 6.8382 | 7.2065 | 6.7678 |
| PPP1R10_S337;S344 | PPP1R10 | Q96QC0 | 382 | DAKPVESPGDPNQ | 6.5291 | 6.0401 | 6.1457 | 6.3511 | 5.7810 | 5.3013 | 6.2086 | 5.3160 | 6.3606 | 6.2369 | 5.4423 | 5.7306 | 6.7769 | 6.7765 | 7.2244 | 7.7789 |
| PPP1R10_S382 | PPP1R10 | Q96QC0 | 337 | STAKPSSPEPAPP | 6.1452 | 5.8827 | 6.2790 | 6.7805 | 5.9759 | 5.7622 | 5.4100 | 5.7377 | 5.7507 | 6.2905 | 6.1537 | 6.8041 | 6.9622 | 6.5229 | 6.8918 | 6.6509 |
| PPP1R10_S398 | PPP1R10 | Q96QC0 | 398 | KGRKRKSVTWPEE | 5.8171 | 6.6698 | 6.4629 | 6.9540 | 6.9792 | 6.2309 | 7.4461 | 7.3809 | 6.9950 | 7.5806 | 7.6292 | 7.0443 | 4.4015 | 4.9353 | 3.4942 | 3.9791 |
| PPP1R10_S527 | PPP1R10 | Q96QC0 | 527 | PLDEECSMDETPY | 6.1813 | 7.3031 | 6.4864 | 5.4179 | 6.3646 | 5.1963 | 5.8007 | 6.1871 | 5.6244 | 7.1163 | 6.4574 | 7.5885 | 5.4237 | 6.6384 | 5.9295 | 6.2845 |
| PPP1R10_S545 | PPP1R10 | Q96QC0 | 337;344 | NA | 5.6250 | 6.3825 | 6.9756 | 6.6132 | 6.4546 | 5.1583 | 6.0052 | 5.8162 | 7.1155 | 6.7271 | 3.9895 | 6.2075 | 6.5605 | 6.5716 | 6.8890 | 6.9088 |
| PPP1R11_T109 | PPP1R11 | O60927 | 109 | LGPTPTTPPQPPD | 6.0333 | 6.7002 | 6.5299 | 6.6855 | 6.6247 | 6.5027 | 6.0610 | 7.1260 | 6.8254 | 6.6184 | 7.2002 | 6.6862 | 5.3524 | 5.1353 | 4.6017 | 5.3170 |
| PPP1R12A_S299 | PPP1R12A | O14974 | 299 | EKRDKKSPLIEST | 6.6896 | 6.6017 | 7.1925 | 6.2867 | 6.9616 | 6.7742 | 6.4844 | 7.0585 | 6.3457 | 5.7995 | 6.2870 | 5.8793 | 5.4456 | 5.5710 | 5.2957 | 5.3271 |
| PPP1R12A_S432 | PPP1R12A | O14974 | 696 | RQSRRSTQGVTLT | 9.5201 | 8.4169 | 9.9115 | 8.8228 | 7.7909 | 6.1182 | 6.8683 | 5.5378 | 4.4916 | 4.4163 | 5.2420 | 4.9772 | 4.2997 | 5.3301 | 4.1842 | 4.0724 |
| PPP1R12A_S445 | PPP1R12A | O14974 | 871 | NEQEQQSDTEEGS | 5.2881 | 5.5348 | 5.7392 | 5.5883 | 5.9925 | 5.8468 | 5.4194 | 5.9439 | 6.8406 | 6.8413 | 6.4968 | 6.5627 | 6.9204 | 7.1842 | 6.9798 | 6.8214 |
| PPP1R12A_S479 | PPP1R12A | O14974 | 445 | GLRKTGSYGALAE | 6.6340 | 5.5773 | 7.0297 | 5.4892 | 5.9820 | 6.4187 | 6.4581 | 6.0841 | 6.1974 | 6.0689 | 6.1281 | 6.3624 | 6.6772 | 6.8892 | 5.1973 | 6.8064 |
| PPP1R12A_S507 | PPP1R12A | O14974 | 507 | IPRRLASTSDIEE | 6.2048 | 5.5790 | 6.0208 | 5.3523 | 6.2295 | 6.8431 | 6.4802 | 7.0673 | 5.5408 | 4.9683 | 4.9620 | 5.6450 | 7.9445 | 7.9306 | 5.9431 | 7.2888 |
| PPP1R12A_S527 | PPP1R12A | O14974 | 432 | EERKDESPATWRL | 6.9005 | 7.7824 | 7.0932 | 7.4529 | 5.8648 | 7.5415 | 7.5814 | 7.0197 | 7.8523 | 7.5921 | 7.9025 | 7.3545 | 2.7098 | 3.8194 | 2.2659 | 3.2673 |
| PPP1R12A_S871 | PPP1R12A | O14974 | 910 | LLGRSGSYSYLEE | 3.9911 | 3.9099 | 4.0062 | 3.3818 | 4.4893 | 4.7175 | 4.2300 | 4.5532 | 3.7988 | 4.2731 | 3.7850 | 4.0016 | 12.9861 | 12.8217 | 12.4669 | 12.5878 |
| PPP1R12A_S903 | PPP1R12A | O14974 | 479 | SPRLSSSLDNKEK | 7.0665 | 7.3135 | 7.4671 | 6.9167 | 5.8265 | 6.2693 | 6.2487 | 6.3247 | 6.6662 | 7.0473 | 7.3854 | 7.1101 | 4.6856 | 4.5348 | 4.4716 | 4.6660 |
| PPP1R12A_S910 | PPP1R12A | O14974 | 508 | PRRLASTSDIEEK | 7.8420 | 7.5679 | 8.0318 | 6.0340 | 5.9702 | 7.5750 | 7.0775 | 7.2029 | 5.2430 | 6.2696 | 6.1289 | 5.6561 | 4.7295 | 5.6326 | 4.5229 | 4.5161 |
| PPP1R12A_S995 | PPP1R12A | O14974 | 995 | ALERRISEMEEEL | 5.3903 | 5.2146 | 6.1844 | 5.6204 | 5.7639 | 5.1244 | 5.9894 | 5.0913 | 5.2652 | 5.3876 | 5.2532 | 5.9415 | 8.2659 | 9.0047 | 8.7683 | 7.7349 |
| PPP1R12A_T508 | PPP1R12A | O14974 | 903 | AGDRYDSLLGRSG | 8.2897 | 7.1705 | 7.4670 | 7.3864 | 6.3027 | 7.5838 | 6.4614 | 7.0577 | 6.8063 | 6.1689 | 5.6573 | 5.9022 | 4.6106 | 4.7806 | 4.2496 | 4.1054 |
| PPP1R12A_T696 | PPP1R12A | O14974 | 527 | SLRTSSSYTRRKW | 5.6331 | 3.9540 | 4.9129 | 4.2073 | 5.4931 | 5.2138 | 5.2197 | 5.2778 | 5.2180 | 3.9223 | 3.7258 | 4.0424 | 12.2780 | 12.2088 | 9.2388 | 9.4542 |
| PPP1R12C_S407 | PPP1R12C | Q9BZL4 | 499 | PCLENSSPPSRIP | 5.9936 | 6.3289 | 6.4070 | 6.4187 | 6.3728 | 6.2565 | 6.0752 | 6.4995 | 6.8208 | 6.5251 | 6.5910 | 6.5557 | 5.8150 | 5.8605 | 5.6675 | 5.8124 |
| PPP1R12C_S407 | PPP1R12C | Q9BZL4 | 509 | RIPEPESPAKPNV | 6.2652 | 6.3098 | 6.5448 | 6.3425 | 6.3332 | 6.4480 | 6.3901 | 6.4391 | 6.6835 | 6.2791 | 6.5753 | 6.4619 | 5.7564 | 5.7934 | 5.6736 | 5.7042 |
| PPP1R12C_S499 | PPP1R12C | Q9BZL4 | 407 | PHPSPKSPVQLEE | 6.6295 | 6.4537 | 6.9059 | 6.7619 | 7.2396 | 6.8473 | 6.0869 | 6.4906 | 6.3696 | 6.2754 | 5.6054 | 5.8385 | 6.2155 | 5.8411 | 5.1597 | 5.2794 |
| PPP1R12C_S509 | PPP1R12C | Q9BZL4 | 560 | RQSRRSTQGVTLT | 10.4561 | 8.6216 | 9.8358 | 8.2776 | 7.3278 | 5.3691 | 6.0286 | 5.2923 | 4.7011 | 3.8666 | 5.0179 | 5.2266 | 4.8387 | 6.0094 | 5.0148 | 4.1161 |
| PPP1R12C_T560 | PPP1R12C | Q9BZL4-3 | 407 | PHPSPKSPVLEEA | 6.7371 | 5.8816 | 6.6077 | 6.6757 | 6.6303 | 5.9339 | 6.3662 | 6.9534 | 6.4214 | 5.9155 | 6.1107 | 6.3508 | 5.7805 | 5.9490 | 5.9796 | 5.7069 |
| PPP1R13B_S332 | PPP1R13B | Q96KQ4 | 671 | SLPRPLSPTKLTP | 6.2905 | 6.1314 | 6.3033 | 6.0648 | 6.3715 | 6.3612 | 5.7765 | 6.0462 | 6.6330 | 6.4573 | 6.4222 | 6.1166 | 6.5912 | 6.4335 | 6.1725 | 5.8285 |
| PPP1R13B_S478 | PPP1R13B | Q96KQ4 | 478 | GPGSTSSLERRKE | 6.0796 | 5.8946 | 6.7177 | 5.8148 | 5.9861 | 6.6287 | 6.5513 | 6.3857 | 6.4559 | 6.4990 | 6.1464 | 6.2275 | 6.5494 | 6.2181 | 6.0924 | 5.7528 |
| PPP1R13B_S486 | PPP1R13B | Q96KQ4 | 332 | RVNGTSSPQSPLS | 6.0664 | 5.8840 | 5.8505 | 5.9967 | 5.9837 | 5.9144 | 6.4954 | 6.4601 | 6.2751 | 5.9353 | 5.7553 | 5.8030 | 6.9709 | 7.4343 | 6.7964 | 6.3783 |
| PPP1R13B_S491 | PPP1R13B | Q96KQ4 | 681 | LTPIVHSPLRYQS | 7.0887 | 7.0396 | 6.8052 | 7.1003 | 6.6775 | 6.9865 | 6.9819 | 7.2367 | 7.2542 | 6.6713 | 7.2079 | 7.3449 | 3.6201 | 4.0680 | 3.6507 | 4.2664 |
| PPP1R13B_S527 | PPP1R13B | Q96KQ4 | 486 | ERRKEGSLPRPSA | 6.1590 | 5.8483 | 6.4822 | 6.1551 | 5.9831 | 5.7255 | 6.2330 | 6.0902 | 6.2914 | 6.4742 | 6.3081 | 6.4781 | 7.1081 | 6.7155 | 5.7823 | 6.1659 |
| PPP1R13B_S566 | PPP1R13B | Q96KQ4 | 710 | PLKKRSSITEPEG | 6.4110 | 6.0713 | 6.0866 | 6.2104 | 5.8254 | 6.1394 | 6.5418 | 5.9791 | 6.4768 | 6.3992 | 6.1361 | 6.2519 | 6.5003 | 6.9093 | 5.5929 | 6.4685 |
| PPP1R13B_S671 | PPP1R13B | Q96KQ4 | 566 | KGSRPQSPRKGPQ | 6.5245 | 6.0685 | 6.6369 | 6.2221 | 5.8859 | 6.7464 | 6.7895 | 6.2586 | 6.1341 | 5.9786 | 5.6621 | 6.4243 | 6.3878 | 6.8242 | 6.1181 | 5.3382 |
| PPP1R13B_S681 | PPP1R13B | Q96KQ4 | 491 | GSLPRPSAGLPSR | 6.1192 | 5.0477 | 6.2791 | 4.7378 | 6.5449 | 5.4610 | 5.3853 | 7.7486 | 4.8103 | 6.7937 | 6.0616 | 6.2424 | 5.5318 | 8.4991 | 6.9415 | 7.7959 |
| PPP1R13B_S710 | PPP1R13B | Q96KQ4 | 527 | RISVPPSPTYPPA | 5.1168 | 5.2060 | 6.0560 | 6.1183 | 6.1472 | 6.3953 | 5.8656 | 6.5587 | 6.4471 | 6.0553 | 5.2375 | 5.5841 | 7.0588 | 6.8337 | 8.2763 | 7.0431 |
| PPP1R13L_S102;S113 | PPP1R13L | Q8WUF5 | 526 | PLKRRGSMEQAPA | 5.9638 | 5.0941 | 6.2952 | 5.0206 | 5.8510 | 6.1390 | 5.6551 | 6.2927 | 7.9909 | 7.7920 | 7.6456 | 7.8344 | 5.9228 | 6.5643 | 4.5608 | 5.3776 |
| PPP1R13L_S110;S113 | PPP1R13L | Q8WUF5 | 358 | IPMPPSSPQPRGA | 6.7288 | 6.6445 | 6.6354 | 6.3671 | 6.0973 | 7.3407 | 6.6989 | 6.8844 | 7.1545 | 6.7583 | 7.0460 | 7.1118 | 4.6018 | 5.0323 | 3.9525 | 4.9458 |
| PPP1R13L_S113 | PPP1R13L | Q8WUF5 | 280 | VFARPASPSLQLL | 6.8962 | 6.5587 | 7.3453 | 6.8382 | 5.8400 | 7.4285 | 6.8525 | 6.5701 | 7.3890 | 7.2989 | 7.6874 | 7.5238 | 3.6337 | 4.5434 | 3.3660 | 4.2283 |
| PPP1R13L_S119;T123 | PPP1R13L | Q8WUF5 | 597 | PAPSQSSPPEQPQ | 6.2268 | 6.1974 | 6.7668 | 7.1241 | 6.1207 | 6.4566 | 6.3701 | 6.7806 | 6.4794 | 6.6389 | 7.0782 | 6.1448 | 5.5510 | 5.7584 | 4.8521 | 5.4539 |
| PPP1R13L_S120 | PPP1R13L | Q8WUF5 | 158 | SLGRAPSPRPGPG | 6.5980 | 6.6768 | 6.5915 | 6.4412 | 6.1908 | 8.3201 | 7.3487 | 6.6407 | 6.0376 | 6.7348 | 7.3987 | 7.0524 | 4.5061 | 4.9931 | 4.0596 | 4.4098 |
| PPP1R13L_S134 | PPP1R13L | Q8WUF5 | 316 | PRNYKVSPLASDR | 7.7704 | 7.5347 | 7.2230 | 7.3316 | 6.0732 | 6.8368 | 7.3260 | 7.2930 | 8.1646 | 7.5351 | 7.3082 | 7.4885 | 2.7284 | 3.4538 | 2.5224 | 3.4105 |
| PPP1R13L_S158 | PPP1R13L | Q8WUF5 | 113 | HPYSPLSPKGRPS | 6.4237 | 6.5427 | 7.1826 | 6.6531 | 6.5804 | 7.0641 | 6.2803 | 7.4595 | 7.4936 | 7.0754 | 7.1583 | 6.8461 | 4.3218 | 4.6864 | 3.7374 | 4.4946 |
| PPP1R13L_S183;S187 | PPP1R13L | Q8WUF5 | 395 | RAMLPGSPLFTRA | 8.0009 | 7.2628 | 7.0046 | 7.0681 | 5.5572 | 6.4846 | 7.0538 | 6.3679 | 7.2036 | 6.8913 | 7.2472 | 7.3232 | 4.2733 | 4.6485 | 3.5917 | 4.0212 |
| PPP1R13L_S187 | PPP1R13L | Q8WUF5 | 332 | AGSYRRSLGSAGP | 8.0584 | 7.1415 | 6.9816 | 6.7126 | 5.3047 | 6.7830 | 7.0898 | 6.0747 | 7.9331 | 7.1453 | 8.2165 | 8.1879 | 3.4392 | 4.5488 | 2.6989 | 3.6840 |
| PPP1R13L_S203 | PPP1R13L | Q8WUF5 | 187 | AGSPRGSPLAEGP | 6.3635 | 6.5534 | 6.7654 | 6.7749 | 6.1083 | 6.4897 | 6.4539 | 6.3811 | 7.0557 | 6.7042 | 7.1072 | 6.9648 | 4.9929 | 5.0639 | 4.9089 | 5.3122 |
| PPP1R13L_S280 | PPP1R13L | Q8WUF5 | 173 | RQQGPPTPFDFLG | 8.6506 | 7.5684 | 7.2535 | 6.6310 | 5.9903 | 6.6056 | 5.6826 | 6.4476 | 6.4801 | 7.0243 | 6.5899 | 7.4116 | 4.7042 | 4.2167 | 4.2457 | 4.4979 |
| PPP1R13L_S316 | PPP1R13L | Q8WUF5 | 134 | QPDAYGSLDRATS | 6.5683 | 6.0013 | 6.6675 | 5.5482 | 5.8799 | 6.5020 | 6.1906 | 6.5399 | 5.9967 | 6.3684 | 6.3414 | 6.0146 | 6.5679 | 6.8316 | 5.7001 | 6.2818 |
| PPP1R13L_S332 | PPP1R13L | Q8WUF5 | 567 | GPEPELSPITEGS | 6.3747 | 7.2187 | 7.4064 | 6.1786 | 6.0414 | 6.6000 | 5.7133 | 6.7993 | 6.6207 | 7.1675 | 7.1701 | 6.4421 | 4.6539 | 5.4506 | 4.7646 | 5.3981 |
| PPP1R13L_S358 | PPP1R13L | Q8WUF5 | 53 | ESLWSDSPAPPGP | 6.9450 | 7.4138 | 7.1794 | 6.3509 | 6.5120 | 7.6547 | 6.8607 | 6.3087 | 6.9738 | 6.5882 | 6.5753 | 6.8372 | 4.8021 | 4.4277 | 4.0110 | 4.5595 |
| PPP1R13L_S395 | PPP1R13L | Q8WUF5 | 71 | SRPPRYSSSSIPE | 7.9333 | 5.4882 | 6.6791 | 6.8906 | 6.0256 | 6.8481 | 7.1541 | 5.7101 | 6.8620 | 6.8256 | 7.0960 | 6.7675 | 4.0893 | 6.4035 | 4.2080 | 5.0193 |
| PPP1R13L_S526 | PPP1R13L | Q8WUF5 | 120 | PKGRPSSPRTPLY | 6.5609 | 5.9710 | 6.3514 | 5.9021 | 6.3209 | 6.9263 | 6.5298 | 7.0774 | 6.4084 | 6.3466 | 6.5586 | 6.5853 | 5.4884 | 5.9942 | 5.4863 | 5.4923 |
| PPP1R13L_S53 | PPP1R13L | Q8WUF5 | 203 | FPERGPSPRPPAT | 5.5085 | 7.0541 | 6.3884 | 6.1917 | 5.7151 | 7.1499 | 7.1058 | 7.2533 | 6.0801 | 6.1292 | 6.9650 | 6.2434 | 5.2860 | 5.8384 | 5.7843 | 5.3068 |
| PPP1R13L_S567 | PPP1R13L | Q8WUF5 | 341 | SAGPSGTLPRSWQ | 9.4929 | 7.6213 | 7.8503 | 7.2528 | 6.5283 | 6.3067 | 5.6802 | 5.5911 | 6.1513 | 6.1453 | 6.2250 | 6.2190 | 5.2510 | 4.8171 | 3.9793 | 4.8885 |
| PPP1R13L_S597 | PPP1R13L | Q8WUF5 | 102;113 | NA | 5.2748 | 5.4273 | 5.5324 | 5.2071 | 6.1473 | 6.8678 | 6.1453 | 6.3652 | 4.8754 | 4.9192 | 5.4485 | 5.7604 | 7.7770 | 7.7904 | 7.6661 | 8.7957 |
| PPP1R13L_S71 | PPP1R13L | Q8WUF5 | 110;113 | NA | 6.3944 | 7.6266 | 7.1206 | 7.8282 | 6.1237 | 6.9307 | 7.1434 | 7.0401 | 7.6771 | 7.3007 | 8.2287 | 7.7225 | 2.7294 | 4.0519 | 2.6883 | 3.3937 |
| PPP1R13L_T173 | PPP1R13L | Q8WUF5 | 119;123 | NA | 6.7135 | 7.4710 | 6.7773 | 6.7068 | 5.8959 | 7.7694 | 6.5400 | 6.5066 | 7.1580 | 7.0979 | 6.9434 | 6.7986 | 4.3561 | 4.6688 | 4.1266 | 4.4701 |
| PPP1R13L_T341 | PPP1R13L | Q8WUF5 | 183;187 | NA | 7.0587 | 7.2792 | 7.5531 | 7.0840 | 6.4431 | 8.1261 | 6.7437 | 6.5541 | 7.5122 | 7.0406 | 8.0531 | 7.6285 | 3.1826 | 3.8025 | 2.8312 | 3.1075 |
| PPP1R14A_S128 | PPP1R14A | Q96A00 | 26 | ARGPGGSPGGLQK | 6.2682 | 6.3475 | 7.1299 | 6.2725 | 6.2601 | 6.9378 | 6.4037 | 6.8789 | 7.0341 | 7.3308 | 7.3709 | 7.2544 | 4.6375 | 5.2095 | 4.0984 | 4.5658 |

| | | | | | | | | | | | | | | | | | | | | | |
|---|---|---|---|---|---|---|---|---|---|---|---|---|---|---|---|---|---|---|---|---|---|
| PPP1R14A_S128;S136 | PPP1R14A | Q96A00 | | 128 | PGLRQPSPSHDGS | 6.2147 | 5.7578 | 6.4717 | 6.6492 | 5.8546 | 5.8401 | 5.8807 | 6.4423 | 6.8967 | 6.4096 | 5.9121 | 6.1613 | 6.1439 | 7.1906 | 5.8107 | 6.3641 |
| PPP1R14A_S26 | PPP1R14A | Q96A00 | 128;136 | | NA | 6.5365 | 5.6064 | 7.1493 | 6.9214 | 6.2196 | 7.9352 | 6.5459 | 6.9190 | 5.4640 | 7.3922 | 7.8642 | 6.1124 | 5.0834 | 5.5082 | 4.7772 | 3.9649 |
| PPP1R14B_S32 | PPP1R14B | Q96C90 | | 32 | PRVYFQSPPGAAG | 7.2860 | 6.5757 | 7.1715 | 7.3404 | 6.2913 | 6.7919 | 6.7419 | 7.1187 | 6.5997 | 6.8060 | 7.3025 | 7.4997 | 4.0134 | 4.2073 | 3.7298 | 4.5242 |
| PPP1R15A_S143 | PPP1R15A | O75807 | | 143 | GQRAPLSPSLLIR | 5.2635 | 5.6980 | 6.4694 | 6.0060 | 6.1292 | 5.1551 | 5.6146 | 5.8257 | 6.3282 | 5.0242 | 4.9831 | 5.6852 | 8.8714 | 7.4307 | 7.8215 | 7.6944 |
| PPP1R15B_S653 | PPP1R15B | Q5SWA1 | | 653 | VTEYYISGDEDRK | 5.9299 | 6.4670 | 6.2546 | 6.5380 | 6.1350 | 6.1931 | 6.6696 | 6.1029 | 5.6075 | 5.4568 | 5.7480 | 5.6688 | 6.8167 | 6.7283 | 6.9910 | 6.6929 |
| PPP1R16A_S353 | PPP1R16A | Q96I34 | | 353 | KVVRRVSLTQRTD | 7.6321 | 6.5773 | 6.1512 | 5.8027 | 5.2470 | 5.9911 | 6.3485 | 6.2006 | 6.7585 | 7.4233 | 6.7083 | 7.1325 | 6.0085 | 6.4338 | 4.3377 | 5.2470 |
| PPP1R16A_S433 | PPP1R16A | Q96I34 | | 433 | RLDRSVSYQLSPL | 7.9023 | 6.6027 | 7.1949 | 6.2857 | 6.1705 | 6.5045 | 6.4863 | 6.3228 | 6.9379 | 6.5309 | 6.9732 | 6.3469 | 5.2866 | 5.6977 | 4.1617 | 4.5957 |
| PPP1R18_S125 | PPP1R18 | Q6NYC8 | | 125 | ARERRPSPGEMRD | 6.3782 | 6.4686 | 5.9008 | 6.0742 | 6.2554 | 6.2688 | 5.8052 | 6.2308 | 6.3157 | 6.3827 | 6.0725 | 6.3122 | 6.6178 | 6.4479 | 6.2957 | 6.1736 |
| PPP1R18_S145 | PPP1R18 | Q6NYC8 | | 224 | EVESRLSPGESAY | 6.4353 | 6.4865 | 6.2671 | 6.4702 | 6.7235 | 6.4335 | 6.3550 | 6.4201 | 6.3518 | 6.2486 | 6.2845 | 6.1128 | 6.1716 | 5.6984 | 5.8142 | 5.7269 |
| PPP1R18_S187 | PPP1R18 | Q6NYC8 | | 195 | AWKWRLSPGETPE | 6.5889 | 7.0046 | 6.4880 | 6.7926 | 6.5901 | 6.4615 | 6.2758 | 6.6573 | 6.7392 | 6.2301 | 6.5167 | 6.2806 | 5.3785 | 5.4735 | 5.1648 | 5.3579 |
| PPP1R18_S195 | PPP1R18 | Q6NYC8 | | 432 | PPPAPLSPPPPAP | 6.3227 | 6.2991 | 6.0157 | 6.4131 | 6.5672 | 6.2128 | 6.6422 | 6.5931 | 6.5917 | 6.1887 | 6.5230 | 6.3409 | 5.6979 | 5.9717 | 5.8278 | 5.7925 |
| PPP1R18_S195;T199 | PPP1R18 | Q6NYC8 | | 484 | RSVPPATPATPTS | 5.9171 | 5.3753 | 6.1528 | 6.4663 | 6.3342 | 6.0283 | 5.9574 | 6.1650 | 6.0140 | 5.8638 | 5.7871 | 6.0585 | 7.5541 | 6.7357 | 6.7715 | 6.8190 |
| PPP1R18_S199 | PPP1R18 | Q6NYC8 | | 368 | AEKLLESPGVEAG | 5.7119 | 6.3027 | 5.6347 | 5.7543 | 5.9843 | 6.3258 | 5.8673 | 6.5233 | 6.2797 | 5.7072 | 5.2768 | 5.4860 | 7.5706 | 7.2448 | 6.9704 | 7.3601 |
| PPP1R18_S224 | PPP1R18 | Q6NYC8 | | 401 | NCCSVPSPLPPED | 6.4249 | 5.7994 | 6.1704 | 6.2250 | 6.4575 | 6.6455 | 6.5800 | 6.2486 | 5.3317 | 6.1110 | 6.2269 | 5.7494 | 6.5550 | 6.5772 | 6.3632 | 6.5343 |
| PPP1R18_S338 | PPP1R18 | Q6NYC8 | | 199 | RLSPGETPERSLR | 8.1995 | 8.4598 | 6.8475 | 6.8959 | 6.4783 | 7.6238 | 7.3848 | 8.1386 | 7.1771 | 6.8149 | 7.7129 | 7.8907 | 2.6373 | 3.1208 | 1.7188 | 2.8992 |
| PPP1R18_S368 | PPP1R18 | Q6NYC8 | | 145 | SREERLSPRETRE | 6.7581 | 7.0272 | 5.7598 | 6.2689 | 6.7059 | 6.2470 | 5.9879 | 6.0885 | 6.3365 | 6.1108 | 6.1397 | 6.1875 | 6.5577 | 6.0555 | 6.0535 | 5.7157 |
| PPP1R18_S401 | PPP1R18 | Q6NYC8 | | 338 | WKWTLNSGKAREW | 9.2459 | 5.9635 | 6.5150 | 4.4314 | 6.2277 | 6.2660 | 6.6069 | 8.0470 | 9.9353 | 5.8332 | 5.9788 | 6.7946 | 3.8952 | 6.8291 | 2.7079 | 4.7225 |
| PPP1R18_S432 | PPP1R18 | Q6NYC8 | | 187 | DRSSRLSEAWKWR | 9.3664 | 7.2261 | 7.1249 | 6.7299 | 5.5017 | 7.2067 | 6.9433 | 5.6625 | 7.7792 | 6.3847 | 8.7953 | 6.7465 | 4.1460 | 5.2370 | 2.4729 | 2.6767 |
| PPP1R18_T484 | PPP1R18 | Q6NYC8 | | 490 | TPATPTSPATVDA | 5.6487 | 6.2225 | 5.4369 | 6.2317 | 6.0489 | 6.2551 | 6.4007 | 5.8172 | 6.2378 | 6.1380 | 6.0321 | 6.5545 | 6.7124 | 6.8208 | 6.0902 | 7.3524 |
| PPP1R18_T487 | PPP1R18 | Q6NYC8 | | 489 | ATPATPTSPATVD | 5.7813 | 5.7902 | 5.8095 | 6.1882 | 6.3841 | 6.0261 | 5.6466 | 6.3552 | 6.4323 | 5.9003 | 5.8280 | 6.3821 | 7.4050 | 6.4703 | 6.7882 | 6.8128 |
| PPP1R18_T487;S490 | PPP1R18 | Q6NYC8 | | 487 | PPATPATPTSPAT | 5.9310 | 5.7719 | 5.7760 | 6.0243 | 5.9999 | 6.1146 | 6.2437 | 6.3307 | 5.8019 | 6.4441 | 6.1241 | 6.1423 | 7.0510 | 6.7933 | 6.9811 | 6.4702 |
| PPP1R18_T487;T489 | PPP1R18 | Q6NYC8 | 487;489 | | NA | 6.7981 | 8.1230 | 6.9719 | 6.6858 | 6.0584 | 6.4657 | 6.8734 | 6.0908 | 7.1810 | 7.0344 | 7.3183 | 7.9515 | 4.1547 | 4.5750 | 3.5340 | 4.1840 |
| PPP1R18_T489 | PPP1R18 | Q6NYC8 | 195;199 | | NA | 8.1907 | 8.4412 | 6.9574 | 7.2368 | 5.9221 | 7.1044 | 7.0175 | 6.6119 | 6.9285 | 7.1131 | 7.7975 | 7.2858 | 3.0845 | 3.8990 | 2.9464 | 3.4634 |
| PPP1R18_T490 | PPP1R18 | Q6NYC8 | 487;490 | | NA | 6.9351 | 6.9258 | 6.0346 | 7.6510 | 5.8952 | 7.4922 | 6.5812 | 6.4011 | 6.5493 | 7.9777 | 7.8411 | 7.2350 | 3.9802 | 4.3503 | 4.0302 | 4.1197 |
| PPP1R2_S87 | PPP1R2 | P41236 | | 87 | DDEDACSDTEATE | 6.0763 | 6.2678 | 6.3981 | 5.7451 | 6.3727 | 6.3432 | 5.8481 | 6.3884 | 6.0071 | 6.1050 | 6.1080 | 6.4054 | 6.3892 | 6.9655 | 6.2010 | 6.3792 |
| PPP1R35_S45;S52 | PPP1R35 | Q8TAP8 | | 87 | FRLTPPSPVRSEP | 6.6032 | 6.5587 | 7.2248 | 6.1066 | 6.2711 | 6.6855 | 6.1471 | 7.1935 | 7.0529 | 6.6786 | 6.9527 | 7.2040 | 4.7217 | 5.1389 | 4.4964 | 4.9643 |
| PPP1R35_S52 | PPP1R35 | Q8TAP8 | | 52 | LSPRPDSPQPRHG | 6.2359 | 6.3177 | 6.5514 | 6.5884 | 6.2575 | 6.2286 | 6.2357 | 6.6637 | 6.5016 | 6.7410 | 6.5108 | 6.7877 | 5.5278 | 5.5491 | 5.4179 | 5.8853 |
| PPP1R35_T84 | PPP1R35 | Q8TAP8 | | 84 | QVRFRLTPPSPVR | 6.1664 | 6.8955 | 6.8427 | 5.4546 | 6.1715 | 7.3511 | 6.5112 | 6.3393 | 6.2640 | 6.4857 | 7.5443 | 6.6383 | 5.5349 | 5.9895 | 4.8597 | 4.9512 |
| PPP1R35_T84;S87 | PPP1R35 | Q8TAP8 | 84;87 | | NA | 5.9981 | 5.8062 | 6.5079 | 5.6513 | 6.2159 | 6.6998 | 6.1973 | 6.8551 | 6.9551 | 6.6643 | 6.5580 | 7.1185 | 5.9346 | 5.3578 | 5.5773 | 5.9028 |
| PPP1R35_T87 | PPP1R35 | Q8TAP8 | 45;52 | | NA | 5.4387 | 5.8418 | 6.3404 | 6.1621 | 5.7713 | 6.3494 | 6.1566 | 6.6975 | 6.7484 | 6.5143 | 6.6826 | 6.7359 | 5.3532 | 6.4611 | 6.3759 | 6.3709 |
| PPP1R37_S50 | PPP1R37 | O75864 | | 561 | QRISVSSPGRGHK | 6.4844 | 6.6719 | 6.9000 | 6.0377 | 5.9823 | 6.6791 | 6.2434 | 6.5980 | 6.6697 | 6.9827 | 6.5572 | 6.6104 | 5.1775 | 6.0128 | 4.9998 | 5.3931 |
| PPP1R37_S561 | PPP1R37 | O75864 | | 50 | KRVTFPSDEDIVS | 4.8905 | 5.3123 | 5.8359 | 5.4255 | 5.1568 | 5.5469 | 5.3509 | 5.9544 | 5.8034 | 5.7781 | 5.8863 | 5.9009 | 8.0941 | 8.1438 | 8.8350 | 8.0851 |
| PPP1R37_T588;S597;S60 | PPP1R37 | O75864 | 588;597;601 | | NA | 5.5149 | 6.4599 | 6.6102 | 6.3943 | 5.6008 | 5.9803 | 6.1287 | 6.8954 | 6.3300 | 6.0868 | 5.8811 | 6.2755 | 6.2541 | 6.5323 | 6.0614 | 6.9943 |
| PPP1R3D_S74;S78 | PPP1R3D | O95685 | 74;78 | | NA | 6.5169 | 6.4899 | 7.2942 | 6.4096 | 6.0810 | 6.4908 | 6.0544 | 6.9733 | 6.8863 | 6.9224 | 6.7710 | 6.7207 | 4.9872 | 5.4137 | 4.8085 | 5.1801 |
| PPP1R7_S37 | PPP1R7 | Q15435 | | 37 | EGKKHSSGIVADL | 6.0993 | 5.7544 | 6.8417 | 6.3088 | 6.5244 | 7.2858 | 6.6415 | 7.2042 | 6.2919 | 5.7889 | 6.3961 | 6.5549 | 5.3714 | 5.9549 | 4.9578 | 6.0239 |
| PPP1R9A_S1149;S1153 | PPP1R9A | Q9ULJ8-3 | | 199 | SRTEAVSPTVSQL | 4.9903 | 5.7353 | 5.8256 | 5.9166 | 5.8288 | 5.4941 | 6.2549 | 6.2644 | 5.7812 | 5.4310 | 5.7342 | 6.1015 | 7.7863 | 7.4224 | 7.7257 | 7.7078 |
| PPP1R9A_S1153 | PPP1R9A | Q9ULJ8-3 | | 338 | PKSEIPSPQSQLL | 6.0371 | 6.8253 | 5.8922 | 6.6297 | 6.9881 | 6.5728 | 6.2562 | 5.7322 | 6.3337 | 6.2445 | 6.2328 | 6.5728 | 5.3185 | 6.4719 | 5.7115 | 6.1809 |
| PPP1R9A_S199 | PPP1R9A | Q9ULJ8-3 | | 862 | IFERRTSLGEVSK | 5.9789 | 6.1471 | 5.6675 | 6.2200 | 6.2936 | 5.7881 | 6.2127 | 6.1929 | 6.5878 | 6.2683 | 6.1422 | 6.1967 | 6.7487 | 6.8345 | 6.5582 | 6.1628 |
| PPP1R9A_S284 | PPP1R9A | Q9ULJ8-3 | | 1153 | TSSLQPSPETLIS | 6.3379 | 6.3504 | 6.6030 | 8.0735 | 6.8711 | 6.1610 | 6.4025 | 6.8552 | 6.3805 | 6.9708 | 6.9457 | 6.7837 | 4.8982 | 5.1143 | 4.3155 | 4.9367 |
| PPP1R9A_S338 | PPP1R9A | Q9ULJ8-3 | | 284 | PVPEVASKSTSLA | 6.9190 | 6.9771 | 7.1440 | 6.9027 | 6.0522 | 6.2354 | 6.4196 | 6.1054 | 6.2418 | 6.5196 | 6.4684 | 6.7796 | 5.0536 | 5.5150 | 5.0920 | 5.5746 |
| PPP1R9A_S371 | PPP1R9A | Q9ULJ8-3 | | 371 | AGGDFTSPDASAS | 7.0210 | 8.5719 | 6.2061 | 6.8515 | 5.5503 | 6.8632 | 7.2836 | 6.5735 | 7.5914 | 6.5644 | 6.6032 | 6.3574 | 3.9981 | 5.1752 | 4.4250 | 4.3642 |
| PPP1R9A_S862 | PPP1R9A | Q9ULJ8-3 | 1149;1153 | | NA | 7.2723 | 7.8294 | 6.1174 | 6.8906 | 6.0775 | 6.7706 | 7.2558 | 6.3741 | 6.8581 | 6.2078 | 5.7197 | 6.6848 | 4.9154 | 5.5833 | 5.1241 | 4.3192 |
| PPP1R9B_S100 | PPP1R9B | Q96SB3 | | 192 | VVRFNGSTEALDK | 5.9922 | 6.8508 | 6.3360 | 6.7905 | 5.7892 | 6.3391 | 6.6032 | 6.2382 | 5.8900 | 6.6071 | 6.8829 | 6.3751 | 5.6157 | 5.6794 | 5.9567 | 6.0541 |
| PPP1R9B_S192 | PPP1R9B | Q96SB3 | | 205 | LDADAVSPTVSQL | 6.3238 | 6.4997 | 6.3211 | 6.7067 | 6.3353 | 6.2323 | 7.0328 | 5.8378 | 5.4656 | 6.2660 | 6.9928 | 6.4659 | 6.1488 | 5.9818 | 5.8485 | 5.5411 |
| PPP1R9B_S205 | PPP1R9B | Q96SB3 | | 100 | SLPRASSLNENVD | 4.5036 | 5.6320 | 6.0717 | 5.5735 | 5.0200 | 5.8142 | 4.7459 | 5.8294 | 5.2554 | 5.4444 | 5.5386 | 5.9722 | 8.4339 | 8.2122 | 9.1690 | 8.7840 |
| PPP2R3A_S562 | PPP2R3A | Q06190 | | 562 | PKSKVSSPIEKVS | 6.3950 | 6.4710 | 6.8660 | 6.3511 | 6.2452 | 6.6362 | 6.4569 | 6.8095 | 6.3525 | 6.0479 | 6.5937 | 6.6235 | 5.4884 | 5.7194 | 5.4698 | 5.4738 |
| PPP2R3A_S568 | PPP2R3A | Q06190 | | 568 | SPIEKVSPSCLTR | 6.9721 | 7.2882 | 6.8833 | 6.5409 | 5.9923 | 6.5491 | 6.5362 | 6.6468 | 6.6152 | 6.9040 | 7.0340 | 6.7575 | 4.7195 | 5.2214 | 4.5843 | 4.7551 |
| PPP2R5C_S39 | PPP2R5C | Q13362-4 | | 513 | PLARRKSELPQDP | 7.7868 | 5.6571 | 7.1408 | 5.5569 | 5.8387 | 6.1319 | 6.0472 | 5.4418 | 7.8854 | 6.6519 | 7.4634 | 7.8468 | 5.3583 | 6.7873 | 3.6092 | 4.7965 |
| PPP2R5C_S513 | PPP2R5C | Q13362-4 | | 39 | ISVRKNSLVAVPS | 6.2846 | 4.8555 | 6.1658 | 5.1920 | 5.9088 | 5.8883 | 5.9127 | 5.1986 | 9.4695 | 7.7035 | 7.8052 | 7.4461 | 6.6901 | 7.1536 | 3.3094 | 5.0164 |
| PPP2R5D_S573 | PPP2R5D | Q14738 | | 573 | VLLRRKSELPQDV | 7.6446 | 5.9230 | 6.9306 | 5.7507 | 6.3054 | 5.7497 | 5.8743 | 5.6919 | 7.4437 | 7.0066 | 7.3315 | 7.3197 | 5.1047 | 6.5983 | 4.4279 | 4.8973 |
| PPP2R5D_S89;S90 | PPP2R5D | Q14738 | | 461 | KSHWNKTIHGLIY | 4.9887 | 6.3378 | 6.4490 | 5.8506 | 7.1704 | 6.3764 | 7.2008 | 6.2394 | 6.1879 | 5.5262 | 5.5157 | 5.5332 | 7.5744 | 6.1469 | 6.6141 | 6.2886 |
| PPP2R5D_T461 | PPP2R5D | Q14738 | 89;90 | | NA | 8.0770 | 7.8121 | 8.7827 | 7.8724 | 5.9769 | 7.1002 | 7.4346 | 6.5927 | 6.9450 | 7.2157 | 8.2084 | 7.9608 | 1.9495 | 3.2086 | 2.0510 | 2.8123 |
| PPP2R5E_S33 | PPP2R5E | Q16537 | | 46 | GKPIELTPLPLLK | 5.6577 | 6.6635 | 7.3428 | 5.5569 | 6.8147 | 6.5441 | 6.1487 | 6.9554 | 6.6608 | 5.5410 | 5.8700 | 6.6396 | 7.2155 | 6.1327 | 4.6971 | 5.5595 |
| PPP2R5E_T46 | PPP2R5E | Q16537 | | 33 | QKRSQSSSQFRSQ | 5.8353 | 6.3506 | 6.3066 | 5.8105 | 6.1056 | 6.5736 | 5.9515 | 6.8191 | 6.5241 | 6.4930 | 6.5721 | 6.6270 | 5.9056 | 6.4708 | 5.4764 | 6.1781 |
| PPP3CC_S492 | PPP3CC | P48454-3 | | 492 | MPPRKDSIHAGGP | 6.5208 | 6.3184 | 6.1091 | 5.6152 | 6.1177 | 7.4126 | 7.1125 | 5.3494 | 8.3113 | 7.3835 | 7.6373 | 6.5053 | 4.9858 | 6.8432 | 3.4319 | 4.3460 |
| PPP4C_S47 | PPP4C | P60510 | | 47 | NVQRVDSPVTVCG | 5.4988 | 5.2657 | 6.2019 | 5.2908 | 6.1153 | 6.0316 | 5.8254 | 6.1207 | 6.3051 | 5.8120 | 6.0131 | 5.9622 | 7.3239 | 7.2944 | 7.2795 | 7.6596 |
| PPP4R2_S159 | PPP4R2 | Q9NY27 | | 159 | VMFPGNSPSYTER | 6.2346 | 6.5286 | 6.5230 | 6.4137 | 6.3459 | 6.6111 | 6.1300 | 6.6609 | 6.4897 | 6.3008 | 6.4775 | 6.3864 | 5.9461 | 5.6923 | 5.6983 | 5.5612 |
| PPP4R3A_S117 | PPP4R3A | Q6IN85 | | 117 | QDLVDESEEERFD | 5.9363 | 6.1639 | 6.6429 | 5.8354 | 5.9025 | 5.6645 | 6.4059 | 5.8225 | 5.9159 | 5.7743 | 6.1207 | 6.0173 | 6.8671 | 7.2760 | 7.2045 | 6.4501 |
| PPP4R3A_S127 | PPP4R3A | Q6IN85 | | 768 | TTNLPGSPGSPGS | 5.9001 | 6.1830 | 6.6968 | 6.4595 | 5.7012 | 5.4670 | 5.6896 | 6.0322 | 6.3677 | 6.9444 | 6.6487 | 6.1919 | 6.5717 | 5.9206 | 7.1074 | 6.1182 |
| PPP4R3A_S698 | PPP4R3A | Q6IN85 | | 127 | RFDDMSSPGLELP | 6.1069 | 5.4109 | 5.5616 | 5.2670 | 5.6071 | 6.0161 | 5.7578 | 6.5758 | 6.1251 | 5.3041 | 6.3167 | 5.7267 | 7.9126 | 7.4506 | 7.2953 | 7.5657 |
| PPP4R3A_S758;S780 | PPP4R3A | Q6IN85 | | 777 | SPGSPGSPGSPGS | 6.8027 | 5.3156 | 6.3685 | 6.7454 | 5.5645 | 5.5598 | 6.3308 | 6.2138 | 6.6496 | 6.1903 | 5.7694 | 6.5122 | 6.4188 | 6.2578 | 6.6669 | 6.6339 |
| PPP4R3A_S761 | PPP4R3A | Q6IN85 | | 780 | SPGSPGSPGSVPK | 5.8481 | 6.3563 | 6.5446 | 6.3517 | 5.5113 | 5.6736 | 5.8772 | 6.2132 | 6.1160 | 6.6012 | 6.7910 | 6.6073 | 5.8741 | 6.6540 | 6.7627 | 6.2178 |
| PPP4R3A_S761;S780 | PPP4R3A | Q6IN85 | | 698 | DGEAVVSPSDKTK | 6.3289 | 5.8233 | 5.9211 | 5.5543 | 6.1844 | 6.0088 | 6.0111 | 6.5812 | 5.6621 | 5.5432 | 6.1355 | 6.5801 | 7.0368 | 7.1413 | 7.1102 | 6.3778 |
| PPP4R3A_S768 | PPP4R3A | Q6IN85 | | 761 | NLTSQSSTTNLPG | 6.4087 | 7.1025 | 6.2181 | 6.0408 | 5.5122 | 4.9263 | 5.7005 | 6.0012 | 6.3261 | 6.5463 | 6.6598 | 5.8481 | 5.6730 | 6.9329 | 7.1135 | 6.9902 |
| PPP4R3A_S768;S774;S78 | PPP4R3A | Q6IN85 | 774;780 | | NA | 5.6463 | 5.7985 | 6.4723 | 6.5266 | 6.1371 | 5.7420 | 5.6709 | 5.8741 | 5.6525 | 6.5476 | 6.6816 | 6.5898 | 6.5939 | 6.4075 | 6.8302 | 6.8292 |
| PPP4R3A_S768;S777 | PPP4R3A | Q6IN85 | 758;774;780 | | NA | 5.8841 | 5.5646 | 6.0645 | 6.3844 | 6.0054 | 6.5889 | 6.5130 | 5.7431 | 5.6111 | 6.5252 | 5.7935 | 7.0104 | 6.7640 | 7.3759 | 6.1115 | 6.0602 |
| PPP4R3A_S774;S777;S78 | PPP4R3A | Q6IN85 | 768;774;780 | | NA | 4.4536 | 6.2137 | 6.2258 | 5.4161 | 6.3424 | 6.4719 | 5.5073 | 6.1467 | 6.4216 | 6.8560 | 6.5901 | 6.0080 | 7.2428 | 6.4678 | 6.7053 | 6.9310 |
| PPP4R3A_S774;S780 | PPP4R3A | Q6IN85 | 777;780 | | NA | 5.4987 | 5.6027 | 7.0927 | 5.9469 | 6.1938 | 6.8239 | 6.3225 | 5.8305 | 5.5195 | 5.5693 | 5.5719 | 6.9512 | 6.3640 | 6.7465 | 6.8617 | 7.1042 |
| PPP4R3A_S777 | PPP4R3A | Q6IN85 | 774;777;780 | | NA | 5.0909 | 5.3615 | 6.3895 | 5.9553 | 5.7673 | 5.8292 | 6.1118 | 5.7420 | 6.5139 | 5.9872 | 6.6636 | 6.4653 | 7.1041 | 7.3594 | 6.7078 | 6.9513 |
| PPP4R3A_S777;S780 | PPP4R3A | Q6IN85 | 768;777 | | NA | 5.9062 | 5.5829 | 6.2404 | 6.4901 | 6.4599 | 5.5088 | 6.2680 | 6.1530 | 6.1154 | 5.8944 | 6.2379 | 6.3537 | 6.9901 | 6.2118 | 6.6885 | 6.8991 |
| PPP4R3A_S780 | PPP4R3A | Q6IN85 | 758;780 | | NA | 6.5127 | 6.4766 | 6.3286 | 5.9528 | 5.9268 | 6.3109 | 6.6681 | 5.7319 | 5.8712 | 5.8129 | 5.6542 | 6.8565 | 6.3549 | 6.4505 | 7.0734 | 6.0178 |
| PPP4R3A_T758;S774;S78 | PPP4R3A | Q6IN85 | 761;780 | | NA | 5.3775 | 6.1703 | 5.9999 | 6.3410 | 5.6979 | 5.7695 | 6.0120 | 5.8890 | 6.0103 | 6.2294 | 6.8232 | 6.2531 | 6.9748 | 6.8850 | 6.1627 | 7.4044 |
| PPP4R3B_S129 | PPP4R3B | Q5MIZ7 | | 840 | DEEEESSPRKRPR | 5.6596 | 5.9606 | 6.5054 | 6.1991 | 5.9284 | 6.1824 | 6.6123 | 6.6935 | 6.3849 | 6.5380 | 6.4331 | 6.7789 | 5.7681 | 6.1688 | 6.2030 | 5.9840 |
| PPP4R3B_S840 | PPP4R3B | Q5MIZ7 | | 129 | EEMPETSHLIDLP | 5.9064 | 5.8008 | 6.2185 | 6.6490 | 6.1239 | 7.1769 | 6.2158 | 6.1439 | 5.3713 | 6.5425 | 7.0504 | 7.0140 | 5.9027 | 5.9291 | 5.6397 | 6.3150 |
| PPP4R4_S775 | PPP4R4 | Q6NUP7 | | 775 | KLIRSQSFNNQAF | 8.1217 | 6.7048 | 7.6632 | 6.3708 | 6.5496 | 6.6340 | 6.1145 | 5.9999 | 6.6535 | 5.9085 | 6.1844 | 6.4295 | 5.0523 | 5.8267 | 4.5362 | 5.2505 |
| PPP6R1_S529;S530;S531 | PPP6R1 | Q9UPN7 | | 726 | PTDAPTSPRVSGE | 6.4697 | 6.0681 | 6.7177 | 6.8186 | 6.2414 | 6.3290 | 6.3748 | 6.7705 | 6.4720 | 6.1497 | 6.4285 | 6.6856 | 5.7068 | 5.6028 | 5.6633 | 5.5015 |
| PPP6R1_S635;S638 | PPP6R1 | Q9UPN7 | | 759 | PTQSLASPPARDA | 6.2779 | 6.3986 | 6.9168 | 6.7352 | 6.6848 | 6.3435 | 6.9586 | 7.1216 | 7.0674 | 6.8032 | 7.3232 | 6.8236 | 4.5186 | 5.0249 | 4.3285 | 4.6736 |
| PPP6R1_S726 | PPP6R1 | Q9UPN7 | | 877 | APEGPASPGSQxx | 5.8513 | 6.3417 | 6.2008 | 7.3535 | 6.5420 | 7.5324 | 6.4341 | 6.2636 | 7.2623 | 7.1627 | 6.0704 | 6.5622 | 5.7707 | 5.5527 | 4.3123 | 5.6973 |
| PPP6R1_S759 | PPP6R1 | Q9UPN7 | 529;530;531 | | NA | 5.8960 | 6.7965 | 6.9185 | 6.2953 | 6.4310 | 6.3489 | 6.0062 | 6.4135 | 6.8082 | 6.5684 | 6.2592 | 6.6519 | 5.6013 | 5.7250 | 5.7299 | 5.5502 |
| PPP6R1_S877 | PPP6R1 | Q9UPN7 | 635;638 | | NA | 6.8037 | 5.9097 | 6.4998 | 7.2508 | 6.0924 | 5.7426 | 5.8782 | 5.8141 | 6.9309 | 5.4201 | 6.0538 | 6.5560 | 6.6890 | 6.4977 | 5.4592 | 6.4021 |
| PPP6R3_S315 | PPP6R3 | Q5H9R7-5 | | 859 | PRPPSSSPEQRTG | 6.6606 | 6.7078 | 7.4817 | 6.9614 | 6.0587 | 7.1721 | 6.7953 | 6.5904 | 6.6842 | 6.9250 | 7.4080 | 7.2555 | 4.0605 | 4.5480 | 4.2115 | 4.4794 |
| PPP6R3_S617 | PPP6R3 | Q5H9R7-5 | | 617 | QFDDGGSDEEDIW | 5.6796 | 6.5109 | 6.9708 | 5.9618 | 6.7296 | 6.8146 | 6.0575 | 6.5148 | 6.1695 | 6.1168 | 6.4898 | 6.5835 | 5.8447 | 6.0243 | 6.6383 | 5.8935 |
| PPP6R3_S859 | PPP6R3 | Q5H9R7-5 | | 315 | IRGRLGSFHELLL | 5.2743 | 5.2231 | 6.2485 | 6.1611 | 5.5258 | 6.3658 | 6.3253 | 6.0524 | 6.1888 | 6.5449 | 6.1537 | 6.8487 | 7.0533 | 7.4413 | 6.1430 | 6.4500 |
| PPP6R3_T631 | PPP6R3 | Q5H9R7-5 | | 631 | EKHIAFTPESQRR | 6.5197 | 5.7295 | 7.1238 | 5.7742 | 5.4033 | 6.9003 | 6.3106 | 6.9667 | 6.3988 | 7.7847 | 7.7581 | 8.2209 | 4.2095 | 5.7621 | 4.4028 | 4.7351 |
| PPRC1_S1076 | PPRC1 | Q5VV67 | | 548 | RSAGQSSPAKEGP | 6.8838 | 7.2482 | 6.9078 | 7.7981 | 6.3781 | 6.1104 | 6.9691 | 6.1719 | 6.8681 | 6.5434 | 6.9946 | 7.1883 | 4.2534 | 4.4348 | 4.3921 | 4.8578 |
| PPRC1_S1413 | PPRC1 | Q5VV67 | | 828 | VGPSPASPSPEPP | 6.2994 | 6.9209 | 6.5913 | 6.5377 | 6.5428 | 6.2533 | 6.7537 | 6.5309 | 6.5683 | 6.2338 | 6.4400 | 6.4551 | 5.5297 | 4.8779 | 5.3529 | 6.1123 |
| PPRC1_S206 | PPRC1 | Q5VV67 | | 237 | RPRWGQSPPPQQR | 6.8809 | 6.8002 | 7.5854 | 7.0809 | 6.1136 | 6.6366 | 6.6606 | 6.7105 | 6.8014 | 6.5950 | 6.8926 | 6.2560 | 4.5184 | 4.7239 | 4.7171 | 5.0268 |
| PPRC1_S237 | PPRC1 | Q5VV67 | | 1391 | NDMNTRTPPEPSA | 6.4099 | 7.2721 | 6.8956 | 8.0265 | 5.9160 | 6.4976 | 6.9720 | 6.4822 | 7.2288 | 6.1197 | 6.7727 | 6.9119 | 4.2651 | 4.6863 | 4.8330 | 4.7105 |
| PPRC1_S536 | PPRC1 | Q5VV67 | | 1076 | VSALVQSPQMKAL | 6.9271 | 6.8125 | 6.6446 | 7.1006 | 6.6812 | 6.3246 | 6.7649 | 6.8943 | 6.9860 | 6.3475 | 6.7973 | 7.5437 | 4.0914 | 5.7232 | 4.2816 | 4.0795 |
| PPRC1_S548 | PPRC1 | Q5VV67 | | 1229 | ANVAGLTPPATPP | 5.5145 | 5.9188 | 6.2438 | 7.1494 | 6.2728 | 6.2110 | 6.1896 | 5.9037 | 5.3344 | 5.5359 | 5.2424 | 5.6354 | 7.0751 | 6.6746 | 7.5991 | 7.4996 |
| PPRC1_S828 | PPRC1 | Q5VV67 | | 1233 | GLTPPATPPHQLW | 6.2392 | 6.4436 | 7.2408 | 8.0586 | 6.1113 | 6.7216 | 7.1634 | 7.6382 | 6.8012 | 6.4122 | 6.9444 | 6.8472 | 3.8409 | 4.7935 | 4.2238 | 4.5201 |
| PPRC1_T1229 | PPRC1 | Q5VV67 | | 1413 | KACRSASPSSQGW | 7.2762 | 6.9164 | 6.5835 | 6.9368 | 5.7684 | 6.1493 | 6.1508 | 6.2142 | 6.5478 | 6.2790 | 5.7509 | 6.6170 | 5.5222 | 5.5193 | 6.5500 | 5.2184 |
| PPRC1_T1233 | PPRC1 | Q5VV67 | | 536 | AALENSSPKNLER | 6.3761 | 5.5073 | 7.3905 | 6.5862 | 5.9826 | 6.2518 | 7.5677 | 6.0283 | 6.9675 | 7.2213 | 6.9393 | 7.6100 | 5.0118 | 5.4569 | 4.4277 | 4.6748 |
| PPRC1_T1391 | PPRC1 | Q5VV67 | | 206 | MSLPDPSWDFSPP | 6.9272 | 6.2976 | 5.8378 | 6.8125 | 6.1605 | 5.8350 | 6.5858 | 6.5075 | 6.8750 | 7.2443 | 7.2313 | 6.6791 | 5.7585 | 5.4535 | 4.0442 | 5.7501 |
| PQBP1_S247 | PQBP1 | O60828 | | 247 | QQRPYPSPGAVLR | 6.2619 | 6.1966 | 6.8607 | 6.6395 | 6.1908 | 6.9056 | 6.9046 | 6.4008 | 7.1169 | 7.0258 | 7.2660 | 7.3487 | 4.5239 | 5.1110 | 4.2131 | 5.0341 |
| PQBP1_S95 | PQBP1 | O60828 | | 95 | AKKLRSSNADAEE | 6.0325 | 6.6077 | 6.8213 | 7.0346 | 6.6564 | 6.9876 | 5.9155 | 6.4410 | 6.5683 | 6.2875 | 7.0548 | 6.8899 | 4.7888 | 5.5376 | 4.8521 | 5.5244 |
| PRAG1_S696 | PRAG1 | Q86YV5 | | 696 | GMSKSASFAFEFP | 4.9634 | 5.5979 | 5.5103 | 5.1218 | 5.9247 | 5.3071 | 5.9292 | 4.5064 | 5.6496 | 5.7967 | 5.5182 | 4.9994 | 8.9286 | 9.5574 | 7.7787 | 8.9106 |
| PRC1_S513 | PRC1 | O43663 | | 481 | RGLAPNTPGKARK | 7.0568 | 7.5890 | 7.4536 | 6.0975 | 4.4784 | 7.4053 | 7.9736 | 6.6621 | 7.4274 | 7.3356 | 8.1848 | 8.0937 | 2.9361 | 4.1587 | 2.7763 | 4.3711 |
| PRC1_S521 | PRC1 | O43663 | | 532 | VAASTCSGKKTPR | 7.0329 | 6.3672 | 7.2115 | 7.1483 | 4.7881 | 7.3075 | 7.4209 | 6.2964 | 7.3721 | 7.5648 | 8.0517 | 8.2886 | 3.2686 | 4.4901 | 3.0122 | 4.3792 |
| PRC1_S532 | PRC1 | O43663 | | 513 | GGTVYHSPVSRLP | 6.6796 | 6.5161 | 6.5823 | 6.6984 | 5.3403 | 6.9275 | 7.3052 | 6.8245 | 6.9430 | 6.8295 | 6.6108 | 6.9896 | 4.7304 | 5.4554 | 4.4430 | 5.1244 |
| PRC1_S554 | PRC1 | O43663 | | 571 | PLQRNFSINSVAS | 6.7312 | 6.1032 | 6.6484 | 6.2853 | 5.2172 | 6.3079 | 6.1886 | 5.8811 | 7.4261 | 7.2990 | 7.0497 | 7.3974 | 5.1086 | 6.2523 | 4.6723 | 5.4317 |
| PRC1_S571 | PRC1 | O43663 | | 554 | NLELNGSILSGGY | 6.6579 | 6.9145 | 6.9626 | 6.5114 | 5.7065 | 6.3497 | 6.3263 | 6.9719 | 6.9544 | 6.4904 | 5.9560 | 6.7446 | 5.1475 | 5.6467 | 5.2858 | 5.3737 |
| PRC1_S571 | PRC1 | O43663 | | 521 | VSRLPPSGSKPVA | 7.1183 | 7.4091 | 7.1728 | 7.1764 | 5.1847 | 7.5763 | 8.4116 | 7.2598 | 7.2089 | 7.3570 | 7.7571 | 7.0556 | 2.9286 | 4.1413 | 2.5351 | 3.7073 |
| PRC1_T481 | PRC1 | O43663-4 | | 571 | PLQRNFSINSVAS | 5.9038 | 5.3885 | 6.9884 | 5.9835 | 6.5663 | 6.1099 | 7.0791 | 5.3087 | 7.1307 | 6.1688 | 7.6202 | 7.5000 | 6.1712 | 6.0056 | 5.1443 | 4.9309 |
| PRCC_S157;S159 | PRCC | Q92733 | | 267 | TQEEDDSDEEVAP | 5.8172 | 6.0957 | 6.2122 | 6.2638 | 6.1780 | 5.9180 | 6.0685 | 6.1124 | 6.1621 | 6.2532 | 6.3946 | 6.3435 | 6.6658 | 6.5218 | 6.5343 | 6.4588 |
| PRCC_S209 | PRCC | Q92733 | | 212 | RKPSDGSPDTKPS | 6.6578 | 6.7175 | 6.8032 | 6.4035 | 6.5123 | 6.2377 | 6.7667 | 6.0271 | 7.1690 | 7.1384 | 6.8997 | 7.4274 | 4.7830 | 4.8854 | 4.3932 | 5.1782 |
| PRCC_S212 | PRCC | Q92733 | | 209 | AFSRKPSDGSPDT | 4.7315 | 4.5345 | 4.5492 | 5.5577 | 7.4661 | 4.7974 | 4.4977 | 4.9041 | 8.0658 | 8.7135 | 7.8626 | 8.7105 | 7.5594 | 7.5975 | 4.6192 | 5.8335 |
| PRCC_S267 | PRCC | Q92733 | 157;159 | | NA | 6.3773 | 6.0061 | 5.9095 | 5.8388 | 5.6526 | 6.3184 | 6.1407 | 6.0528 | 6.0890 | 5.9877 | 7.0088 | 6.4139 | 7.1983 | 6.0390 | 6.2760 | 6.6911 |
| PRDM10_S424 | PRDM10 | Q9NQV6-6 | | 424 | SENGEKSDDGTQD | 6.4525 | 6.9229 | 7.1417 | 7.0296 | 5.8737 | 5.7211 | 6.7505 | 6.0513 | 6.2197 | 6.1804 | 5.2897 | 6.1184 | 5.7654 | 5.6465 | 6.1796 | 6.6571 |
| PRDM2_S447 | PRDM2 | Q13029 | | 643 | KKRRTASPPALPK | 6.4492 | 6.5275 | 6.2127 | 6.7148 | 6.0853 | 5.7914 | 6.2690 | 5.8982 | 6.3314 | 6.1981 | 5.9371 | 6.0253 | 6.2788 | 6.3986 | 6.4573 | 6.4253 |
| PRDM2_S643 | PRDM2 | Q13029 | | 796 | DERETVSPPCFDE | 6.1163 | 6.1963 | 6.1817 | 6.6490 | 5.9982 | 5.8639 | 6.2624 | 6.3449 | 6.5362 | 6.4269 | 6.0614 | 6.3688 | 5.8900 | 6.3428 | 6.2647 | 6.4965 |
| PRDM2_S796 | PRDM2 | Q13029 | | 447 | ASKDDSSPPSLGP | 6.0454 | 6.4121 | 6.0717 | 6.8138 | 5.7858 | 5.8684 | 5.7054 | 5.6575 | 6.4170 | 6.3297 | 6.1101 | 6.4882 | 6.0348 | 7.2989 | 6.6627 | 6.2985 |
| PRDM2_S912;S914 | PRDM2 | Q13029 | 912;914 | | NA | 6.4987 | 6.7689 | 6.2970 | 6.9376 | 6.2266 | 6.0105 | 6.1721 | 5.6677 | 6.2360 | 6.1676 | 6.2589 | 6.1142 | 6.0515 | 6.3517 | 6.1575 | 6.0836 |
| PRDM8_S375 | PRDM8 | Q9NQV8 | | 375 | RLFAPPSPETGEA | 5.9816 | 6.0272 | 7.2404 | 5.6083 | 5.6392 | 6.3079 | 6.1153 | 6.1837 | 6.4335 | 6.2410 | 6.3439 | 6.6290 | 6.1800 | 6.9604 | 6.1800 | 5.9287 |
| PRDX6_T44 | PRDX6 | P30041 | | 44 | SHPRDFTPVCTTE | 6.0829 | 5.6111 | 5.9969 | 5.6319 | 5.2995 | 5.1987 | 5.5755 | 5.8010 | 6.2075 | 7.4063 | 5.3708 | 6.8574 | 5.9375 | 7.7921 | 7.5665 | 7.6645 |
| PRICKLE1_S353 | PRICKLE1 | Q96MT3 | | 353 | RQSLLLSPALNYK | 6.3265 | 6.1278 | 7.7293 | 6.7977 | 5.6988 | 6.5973 | 7.4957 | 6.6436 | 5.6897 | 6.5095 | 6.6080 | 7.4668 | 4.3482 | 5.1150 | 5.5887 | 5.2574 |
| PRICKLE2_S806 | PRICKLE2 | Q7Z3G6 | | 806 | RLRYVTSDELLHK | 6.2378 | 7.5204 | 5.7212 | 5.0684 | 5.4785 | 6.4360 | 5.6821 | 6.1043 | 5.9411 | 6.0638 | 6.0220 | 5.0819 | 7.0983 | 7.2884 | 7.1967 | 7.0591 |
| PRICKLE3_S437 | PRICKLE3 | O43900 | | 475 | QSTPRVSFRDPLV | 6.8891 | 6.7703 | 8.2327 | 6.3010 | 5.8887 | 6.3461 | 6.5334 | 6.2536 | 6.6874 | 6.8273 | 6.9743 | 7.0723 | 4.7525 | 5.5980 | 4.1458 | 4.7275 |
| PRICKLE3_S475 | PRICKLE3 | O43900 | | 437 | GAPHRHSMPELGL | 5.8518 | 5.9892 | 7.2703 | 6.9018 | 6.0706 | 6.3824 | 6.4476 | 6.3295 | 6.1640 | 6.7567 | 6.1235 | 7.2695 | 5.7785 | 5.3313 | 5.5782 | 5.7553 |
| PRICKLE3_S80 | PRICKLE3 | O43900 | | 80 | SDFQRHSISDDDS | 5.8476 | 6.3424 | 6.5365 | 6.0240 | 6.3650 | 5.9562 | 6.2525 | 5.6300 | 6.3960 | 6.3891 | 6.2579 | 6.3950 | 6.3012 | 6.8505 | 6.2162 | 6.2397 |
| PRIM2_T470 | PRIM2 | P49643 | | 470 | EPIQPETPQPKPS | 6.4447 | 6.8738 | 6.8906 | 6.6863 | 6.4853 | 6.8098 | 7.0544 | 6.7155 | 7.0728 | 6.6075 | 6.5036 | 6.8697 | 4.5831 | 5.3069 | 4.1434 | 4.9525 |
| PRIMPOL_S499 | PRIMPOL | Q96LW4 | | 499 | QNPHKPSPSRLST | 5.9124 | 6.5116 | 6.9511 | 6.5747 | 5.7749 | 6.4199 | 5.8451 | 6.8850 | 6.7113 | 7.2318 | 7.0122 | 7.1005 | 4.9166 | 5.6395 | 5.1707 | 5.3427 |
| PRKAA1_S371 | PRKAA1 | Q13131-2 | | 371 | DFYLATSPPDSFL | 6.4993 | 6.6384 | 7.4138 | 6.2663 | 6.3262 | 6.7641 | 6.2568 | 6.7620 | 6.8507 | 6.4272 | 6.8919 | 7.1637 | 4.6013 | 5.3324 | 4.8147 | 4.9910 |
| PRKAA1_S501 | PRKAA1 | Q13131-2 | | 198 | DGEFLRTSCGSPN | 6.0470 | 5.7368 | 6.2583 | 7.3918 | 6.5792 | 4.5530 | 5.1574 | 4.8996 | 4.9642 | 4.8642 | 4.9652 | 5.1203 | 7.8775 | 8.5166 | 8.9648 | 8.1041 |
| PRKAA1_S511 | PRKAA1 | Q13131-2 | | 397 | PFLVAETPRARHT | 9.0112 | 8.0659 | 7.0799 | 7.4649 | 6.5716 | 6.8053 | 8.3879 | 5.9876 | 7.4600 | 5.4606 | 6.7315 | 5.8618 | 3.2078 | 4.8638 | 3.1107 | 3.9295 |
| PRKAA1_T198 | PRKAA1 | Q13131-2 | | 505 | AKSGTATPQRSGS | 4.8873 | 4.6966 | 5.1542 | 4.4715 | 6.0251 | 4.3300 | 5.0853 | 5.2120 | 4.5613 | 4.9303 | 4.6056 | 5.8394 | 9.5260 | 9.9197 | 10.8103 | 9.9455 |
| PRKAA1_T397 | PRKAA1 | Q13131-2 | | 511 | TPQRSGSVSNYRS | 5.4289 | 4.9824 | 5.7049 | 5.2346 | 5.7333 | 5.6127 | 5.0762 | 5.5631 | 5.2894 | 5.3317 | 5.4601 | 5.4963 | 8.9337 | 8.8653 | 9.0843 | 8.2032 |
| PRKAA1_T505 | PRKAA1 | Q13131-2 | | 501 | EITEAKSGTATPQ | 5.0336 | 5.5372 | 5.2565 | 5.4122 | 5.5891 | 6.4121 | 5.9852 | 6.4625 | 5.6256 | 5.5694 | 5.6154 | 5.3187 | 7.7313 | 8.0222 | 7.9360 | 8.4929 |
| PRKAA2_S377 | PRKAA2 | P54646 | | 377 | PPLIADSPKARCP | 6.7421 | 6.2759 | 6.9966 | 6.3489 | 6.1177 | 5.9685 | 6.1582 | 6.2837 | 7.1908 | 6.3270 | 6.5933 | 6.7523 | 5.4059 | 5.9209 | 5.5500 | 5.3682 |
| PRKAB1_S108 | PRKAB1 | Q9Y478 | | 40 | PKILMDSPEDADL | 6.9587 | 7.4979 | 7.4922 | 7.6656 | 6.1285 | 6.7938 | 6.4851 | 6.7489 | 7.2198 | 7.6668 | 7.9592 | 8.2713 | 2.8529 | 3.7196 | 2.9790 | 3.5609 |
| PRKAB1_S182 | PRKAB1 | Q9Y478 | | 182 | VSELSSSPPGPYH | 5.5525 | 5.6126 | 6.2394 | 5.7252 | 5.6148 | 5.9345 | 5.9040 | 6.2048 | 7.7332 | 5.9493 | 6.3547 | 6.4239 | 7.3832 | 6.2365 | 6.3354 | 6.7961 |
| PRKAB1_S40 | PRKAB1 | Q9Y478 | | 108 | KLPLTRSHNNFVA | 3.3431 | 3.4855 | 3.5791 | 3.2354 | 5.8372 | 5.9604 | 5.6172 | 6.3211 | 4.1568 | 3.9431 | 4.3788 | 4.3422 | 12.0047 | 11.3719 | 11.4465 | 10.9770 |
| PRKAB1_S96 | PRKAB1 | Q9Y478 | | 96 | EVYLSGSFNNWSK | 2.6945 | 2.7402 | 2.8991 | 3.4818 | 3.4634 | 3.6460 | 3.4984 | 3.3652 | 3.0664 | 3.4814 | 2.5550 | 3.1368 | 15.0965 | 15.1974 | 16.2885 | 15.3894 |
| PRKAB2_S108 | PRKAB2 | O43741 | | 108 | KIPLIKSHNDFVA | 3.6001 | 3.8741 | 3.7415 | 3.5360 | 6.0963 | 5.8495 | 5.4432 | 6.3319 | 3.9648 | 3.7685 | 4.3957 | 4.0603 | 11.4069 | 11.2659 | 11.9034 | 10.7617 |
| PRKAB2_S158 | PRKAB2 | O43741 | | 39 | HKIMVGSTDDPSV | 5.6450 | 6.0653 | 5.8169 | 5.6070 | 6.5989 | 6.0947 | 5.9662 | 6.8716 | 5.9790 | 6.4183 | 6.4602 | 6.5821 | 6.1554 | 7.0596 | 6.1474 | 6.5324 |
| PRKAB2_S183 | PRKAB2 | O43741 | | 184 | CRDLSSSPPGPYG | 5.6849 | 5.5454 | 6.2634 | 6.0784 | 5.8865 | 6.4595 | 5.8732 | 6.7205 | 6.5284 | 6.3865 | 6.4724 | 6.3110 | 6.4254 | 6.8221 | 6.2886 | 6.2537 |
| PRKAB2_S184 | PRKAB2 | O43741 | | 183 | SCRDLSSSPPGPY | 5.7192 | 5.7926 | 6.1565 | 6.0586 | 6.4906 | 6.4369 | 5.9283 | 6.2901 | 6.2257 | 5.7203 | 6.1374 | 6.1241 | 6.7072 | 6.0629 | 7.0732 | 7.0764 |
| PRKAB2_S39 | PRKAB2 | O43741 | | 158 | LIHVKKSDFEVFD | 1.2030 | 1.1399 | 1.1877 | 1.3224 | 1.9601 | 1.7461 | 2.3591 | 1.8015 | 1.7649 | 1.3607 | 0.5329 | 1.2789 | 20.0888 | 19.6395 | 22.6483 | 19.9661 |
| PRKACA_S339 | PRKACA | P17612 | | 339 | EEEIRVSINEKCG | 5.8691 | 6.1550 | 6.3355 | 6.1841 | 6.2024 | 6.2057 | 6.0292 | 6.3449 | 6.2572 | 6.2946 | 6.2358 | 6.1417 | 6.3865 | 6.7384 | 6.3700 | 6.2498 |
| PRKACB_S386 | PRKACB | P22694-2 | | 245 | VKGRTWTLCGTPE | 6.9878 | 6.7945 | 7.2878 | 6.8269 | 6.6784 | 5.9193 | 6.4325 | 5.4571 | 5.4206 | 5.3342 | 5.5094 | 4.5062 | 7.1020 | 6.8569 | 7.5677 | 5.3188 |
| PRKACB_T245 | PRKACB | P22694-2 | | 386 | EEDIRVSITEKCA | 6.5193 | 5.9906 | 5.9267 | 6.0249 | 6.2874 | 6.0387 | 5.9436 | 5.9646 | 6.3588 | 6.2588 | 6.2404 | 6.3659 | 6.7390 | 6.7589 | 6.4362 | 6.1464 |
| PRKAG2_S196 | PRKAG2 | Q9UGJ0 | | 196 | RIYASSSPPDTGQ | 7.2145 | 7.0574 | 6.7818 | 7.3893 | 6.4068 | 5.8543 | 6.2116 | 6.5924 | 7.6985 | 6.7862 | 7.1081 | 7.5452 | 4.2623 | 4.4720 | 4.0659 | 4.5537 |
| PRKAR1A_S83 | PRKAR1A | P10644 | | 83 | SREDEISPPPPNP | 5.8048 | 5.8879 | 6.4948 | 5.8776 | 5.9816 | 6.0561 | 5.9590 | 6.2511 | 6.4412 | 6.4684 | 6.5029 | 6.3725 | 6.4362 | 6.6493 | 6.5227 | 6.2938 |
| PRKAR2A_S78;S80 | PRKAR2A | P13861 | | 99 | RFNRRVSVCAETY | 6.7932 | 6.7596 | 6.3563 | 6.1121 | 6.0948 | 6.4550 | 5.9880 | 6.2358 | 6.1046 | 5.9523 | 6.2288 | 6.2059 | 6.0269 | 6.8381 | 6.0276 | 5.8211 |
| PRKAR2A_S99 | PRKAR2A | P13861 | | 54 | SVLPAATPRQSLG | 8.1779 | 8.4970 | 7.0108 | 7.4883 | 6.5159 | 6.9666 | 7.7906 | 7.0742 | 6.5504 | 5.4055 | 5.9373 | 5.6312 | 3.9062 | 4.6867 | 4.0811 | 4.2804 |
| PRKAR2A_T54 | PRKAR2A | P13861 | 78;80 | | NA | 5.8025 | 5.0473 | 6.5038 | 5.5369 | 6.0388 | 5.7661 | 6.4338 | 6.7843 | 6.6696 | 6.1370 | 7.1341 | 6.8708 | 6.6549 | 7.0065 | 5.8057 | 5.8080 |
| PRKAR2B_S364 | PRKAR2B | P31323 | | 69 | AAAGGGTPSKGVN | 6.7779 | 6.9205 | 6.5065 | 6.6879 | 6.7953 | 7.0647 | 7.5515 | 6.8787 | 7.1815 | 6.6951 | 6.7635 | 7.1592 | 4.2208 | 4.7542 | 3.7946 | 4.2479 |
| PRKAR2B_T69 | PRKAR2B | P31323 | | 364 | NKPRAASAHAIGT | 6.7220 | 4.5463 | 5.4291 | 6.1013 | 5.9250 | 5.9359 | 6.1340 | 5.4836 | 8.2754 | 6.7438 | 5.1436 | 6.4678 | 6.9481 | 6.8015 | 7.0027 | 6.3399 |
| PRKCA_S226 | PRKCA | P17252 | | 497 | DGVTTRTFCGTPD | 7.9979 | 7.8613 | 8.7433 | 8.2191 | 7.0411 | 5.4348 | 5.9448 | 5.6483 | 5.4061 | 4.9292 | 5.9849 | 4.8435 | 5.3886 | 6.0411 | 5.9210 | 4.5950 |

| | | | | | | | | | | | | | | | | | | | | |
|---|---|---|---|---|---|---|---|---|---|---|---|---|---|---|---|---|---|---|---|---|
| PRKCA_S319 | PRKCA | P17252 | 319 | AGNKVISPSEDRK | 7.0718 | 7.6135 | 7.5926 | 7.2254 | 6.1273 | 7.2342 | 7.4916 | 6.7634 | 6.3364 | 7.2293 | 7.1682 | 6.8356 | 3.3734 | 4.3328 | 3.5475 | 4.0567 |
| PRKCA_T497 | PRKCA | P17252 | 226 | NPQWNESFTFKLK | 5.1979 | 4.7961 | 6.6000 | 7.0474 | 8.2699 | 7.2640 | 5.1523 | 6.8166 | 6.7140 | 7.0274 | 7.2013 | 5.5085 | 5.2891 | 6.3207 | 5.3489 | 5.4458 |
| PRKCD_S304 | PRKCD | Q05655-2 | 676 | NEKARLSYSDKNL | 7.0414 | 7.2144 | 7.2284 | 7.0594 | 6.7295 | 6.7392 | 6.6395 | 6.7418 | 5.5206 | 5.9240 | 5.9699 | 6.0430 | 5.3625 | 5.6382 | 5.0957 | 5.0525 |
| PRKCD_S676 | PRKCD | Q05655-2 | 538 | GESRASTFCGTPD | 7.9012 | 7.5855 | 7.5763 | 8.0223 | 7.1621 | 6.0084 | 5.9632 | 5.7276 | 6.0656 | 5.2975 | 5.6712 | 5.1012 | 5.8938 | 5.3967 | 5.7463 | 4.8812 |
| PRKCD_S695 | PRKCD | Q05655-2 | 695 | SAFAGFSFVNPKF | 8.2413 | 7.4106 | 8.2548 | 8.3930 | 7.1410 | 5.2847 | 6.1999 | 5.6156 | 5.8054 | 4.9116 | 5.4035 | 4.7882 | 5.3441 | 5.9149 | 6.1071 | 5.1840 |
| PRKCD_T538 | PRKCD | Q05655-2 | 304 | ASRRSDSASSEPV | 6.3165 | 6.7018 | 7.3542 | 6.8742 | 6.0722 | 6.8471 | 6.0373 | 6.2481 | 7.0600 | 5.9554 | 6.2374 | 6.7246 | 5.2332 | 5.2724 | 5.3878 | 5.6777 |
| PRKCE_S368 | PRKCE | Q02156 | 729 | EEFKGFSYFGEDL | 6.8872 | 7.1672 | 7.2365 | 7.2168 | 6.9974 | 5.9467 | 6.3131 | 6.5299 | 6.2304 | 5.6339 | 5.8695 | 5.5699 | 5.8398 | 5.9605 | 5.5286 | 5.0727 |
| PRKCE_S729 | PRKCE | Q02156 | 309 | LADLGVTPDKITN | 8.1271 | 8.3563 | 7.6888 | 7.7197 | 6.0999 | 7.1483 | 7.1977 | 7.3758 | 7.9975 | 6.8397 | 7.1351 | 7.6598 | 2.0457 | 3.6108 | 2.4848 | 2.5130 |
| PRKCE_T309 | PRKCE | Q02156 | 368 | NIRKALSFDNRGE | 6.2002 | 5.1310 | 5.6972 | 5.7473 | 6.4059 | 5.7498 | 5.9554 | 6.5123 | 5.8098 | 5.8471 | 5.2393 | 5.3812 | 7.6588 | 7.8183 | 7.8779 | 6.9687 |
| PRKCH_S317 | PRKCH | P24723 | 675 | DEFRNFSYVSPEL | 6.8694 | 6.9038 | 6.6251 | 7.2075 | 6.2304 | 6.1527 | 6.5699 | 6.3194 | 6.8825 | 5.6462 | 5.8938 | 6.4332 | 5.5084 | 5.8927 | 5.6706 | 5.1943 |
| PRKCH_S675 | PRKCH | P24723 | 656 | KEEPVLTPIDEGH | 6.0546 | 6.7460 | 6.9029 | 6.9049 | 6.2584 | 6.2175 | 6.5512 | 5.7988 | 6.1775 | 6.6940 | 6.4123 | 6.9398 | 5.6929 | 5.1234 | 5.6210 | 5.9047 |
| PRKCH_T656 | PRKCH | P24723 | 317 | LQPGNISPTSKLV | 6.1481 | 6.0705 | 7.2806 | 6.4881 | 6.7044 | 7.1007 | 6.0308 | 6.7995 | 5.8914 | 6.8645 | 6.4350 | 6.2995 | 5.4745 | 5.7511 | 4.9927 | 5.6687 |
| PRKCI_T412 | PRKCI | P41743 | 564 | NEPVQLTPDDDDI | 5.5323 | 5.7734 | 6.5245 | 6.1894 | 5.7647 | 5.9748 | 6.1157 | 6.0735 | 6.2896 | 6.3107 | 6.0561 | 6.4942 | 6.1667 | 7.4983 | 6.4117 | 6.8243 |
| PRKCI_T564 | PRKCI | P41743 | 412 | PGDTTSTFCGTPN | 7.0451 | 6.9920 | 7.7695 | 8.3474 | 7.3455 | 5.1062 | 5.9533 | 5.3886 | 5.4881 | 4.8008 | 5.1505 | 4.1079 | 7.1090 | 6.2772 | 7.8630 | 5.2558 |
| PRKCZ_S591 | PRKCZ | Q05513 | 560 | SEPVQLTPDDEDA | 6.4926 | 6.2796 | 7.6624 | 5.8123 | 6.0032 | 6.6682 | 5.5657 | 5.9451 | 6.4086 | 6.2552 | 6.2009 | 7.6334 | 5.7428 | 5.2862 | 5.8352 | 6.2086 |
| PRKCZ_T408 | PRKCZ | Q05513 | 591 | LLSTEESVxxxxx | 6.0379 | 7.0102 | 6.3096 | 6.9235 | 6.5342 | 6.1641 | 6.2133 | 7.2369 | 5.7491 | 6.7278 | 6.5609 | 5.6411 | 5.0977 | 6.5834 | 5.8464 | 5.3640 |
| PRKCZ_T560 | PRKCZ | Q05513 | 408 | LGPGDTTSTFCGT | 6.6705 | 7.4891 | 7.0635 | 7.0659 | 7.5895 | 5.3560 | 6.1347 | 5.3838 | 4.9170 | 6.0876 | 6.8579 | 5.2157 | 6.5580 | 6.2532 | 6.1130 | 5.2447 |
| PRKD1_S205 | PRKD1 | Q15139 | 742 | EKSFRRSVVGTPA | 6.4133 | 6.4368 | 6.5661 | 6.5765 | 6.7215 | 5.7172 | 6.2567 | 6.5874 | 6.5003 | 6.4588 | 6.0632 | 6.7603 | 6.2317 | 6.3270 | 5.3108 | 5.0724 |
| PRKD1_S205;S208 | PRKD1 | Q15139 | 397 | DANRTISPSTSNN | 6.5480 | 6.1248 | 6.6412 | 6.4589 | 6.2989 | 6.5135 | 6.1972 | 6.1817 | 6.8577 | 6.6916 | 6.8589 | 6.6019 | 5.5361 | 5.4007 | 5.6934 | 5.3956 |
| PRKD1_S237 | PRKD1 | Q15139 | 249 | REKRSNSQSYIGR | 5.9128 | 5.0274 | 6.3243 | 5.4991 | 7.5968 | 5.8535 | 4.7893 | 6.0809 | 7.3855 | 7.4208 | 7.1234 | 6.9491 | 6.7721 | 7.3456 | 5.0600 | 4.8595 |
| PRKD1_S249 | PRKD1 | Q15139 | 205 | VRRRRLSNVSLTG | 6.1408 | 6.2287 | 6.8607 | 6.6401 | 5.7445 | 6.1575 | 5.7579 | 6.0351 | 5.6393 | 6.7415 | 6.4584 | 6.4929 | 6.2729 | 6.4007 | 6.0685 | 6.3603 |
| PRKD1_S397 | PRKD1 | Q15139 | 237 | LLQKSPSESFIGR | 4.4601 | 4.4134 | 4.0790 | 4.9040 | 5.8536 | 4.5174 | 4.2203 | 5.4015 | 5.3312 | 4.9171 | 4.3075 | 5.1276 | 10.0459 | 11.3149 | 11.2862 | 9.8205 |
| PRKD1_S397;S401 | PRKD1 | Q15139 | 205;208 | NA | 6.4146 | 5.9040 | 6.4587 | 6.2271 | 6.9617 | 5.8801 | 6.5580 | 6.5138 | 7.1234 | 6.2503 | 6.6002 | 6.3568 | 5.6336 | 6.1880 | 5.4857 | 5.4441 |
| PRKD1_S742 | PRKD1 | Q15139 | 397;401 | NA | 6.1809 | 6.1463 | 5.9743 | 6.2549 | 7.0252 | 5.9597 | 6.3254 | 6.9046 | 6.0982 | 6.1423 | 5.5791 | 6.0272 | 6.0723 | 6.4074 | 5.8451 | 6.0569 |
| PRKD2_S197 | PRKD2 | Q9BZL6-3 | 710 | EKSFRRSVVGTPA | 6.3302 | 5.7805 | 6.7583 | 6.8620 | 6.7109 | 6.5309 | 6.4057 | 6.1577 | 6.4031 | 6.7321 | 5.7945 | 6.7334 | 5.4877 | 5.9660 | 5.2550 | 5.8219 |
| PRKD2_S396 | PRKD2 | Q9BZL6-3 | 886 | GLAERISVLxxxx | 7.2354 | 6.4820 | 7.2600 | 7.6502 | 7.2128 | 6.3524 | 5.3682 | 6.2655 | 8.1131 | 7.4369 | 7.1861 | 7.4608 | 4.2044 | 5.0769 | 3.3987 | 3.2965 |
| PRKD2_S710 | PRKD2 | Q9BZL6-3 | 197 | ARKRRLSSTSLAS | 5.4154 | 6.5507 | 6.9934 | 6.6073 | 5.9522 | 6.9036 | 5.8315 | 6.5265 | 6.8766 | 6.5418 | 7.0259 | 6.7573 | 5.3400 | 6.0536 | 5.2129 | 5.4115 |
| PRKD2_S886 | PRKD2 | Q9BZL6-3 | 396 | HTTRKSSTTLREG | 6.7504 | 6.3025 | 7.0177 | 6.4435 | 6.2070 | 6.4825 | 6.1803 | 6.4669 | 6.2936 | 6.9930 | 7.1841 | 6.8732 | 5.3243 | 5.6696 | 5.3321 | 4.4794 |
| PRKD3_S213 | PRKD3 | O94806 | 213 | VRKRRLSNVSLPG | 5.4571 | 5.7649 | 6.5282 | 5.8826 | 5.9668 | 5.7330 | 6.1631 | 6.7098 | 6.7354 | 6.8934 | 6.8797 | 6.9156 | 6.0054 | 6.4693 | 5.6839 | 6.2119 |
| PRKD3_S213;S216 | PRKD3 | O94806 | 535 | QALMPVTPQASVC | 5.8336 | 5.3715 | 6.3943 | 6.2084 | 6.4922 | 5.8794 | 5.8706 | 6.7211 | 6.5214 | 6.2295 | 6.3373 | 6.4772 | 6.5642 | 6.7753 | 6.1336 | 6.1903 |
| PRKD3_S27 | PRKD3 | O94806 | 27 | AVLPAASPCSSPK | 6.5610 | 6.4351 | 6.1985 | 6.0149 | 6.4847 | 6.5745 | 6.3587 | 6.4953 | 6.4413 | 6.4111 | 6.5339 | 6.6268 | 6.0816 | 6.1735 | 5.2667 | 5.3424 |
| PRKD3_S41 | PRKD3 | O94806 | 41 | GLSARLSNGSFSA | 4.9014 | 5.7941 | 6.7019 | 6.6517 | 5.6083 | 5.9909 | 6.7531 | 6.1978 | 6.7481 | 7.4510 | 6.2550 | 6.6577 | 5.4808 | 6.4408 | 5.9026 | 6.4649 |
| PRKD3_T535 | PRKD3 | O94806 | 213;216 | NA | 5.7922 | 5.9099 | 6.2553 | 6.3636 | 6.2948 | 5.9071 | 5.8009 | 6.4023 | 7.0839 | 6.3512 | 6.1527 | 6.8422 | 6.6750 | 6.4115 | 5.7576 | 5.9998 |
| PRKDC_S1065 | PRKDC | P78527 | 893 | DREKRLSFAVPFR | 6.2004 | 6.2846 | 5.9398 | 6.2097 | 6.3408 | 5.9597 | 6.0647 | 5.8371 | 6.6441 | 6.5099 | 6.6780 | 6.2954 | 5.9120 | 6.9051 | 6.1537 | 6.0649 |
| PRKDC_S2672 | PRKDC | P78527 | 2671 | DPLVDHTSPSSDS | 6.0236 | 6.0384 | 6.2684 | 6.4129 | 6.0604 | 6.0616 | 6.5930 | 6.1488 | 6.1166 | 6.7711 | 6.2517 | 5.9041 | 6.1064 | 6.6186 | 6.4615 | 6.1628 |
| PRKDC_S3205 | PRKDC | P78527 | 687 | KYFEGVSPKSLKH | 6.5552 | 6.7815 | 6.3354 | 6.3997 | 6.3048 | 6.0261 | 6.4415 | 6.0371 | 6.6572 | 6.5327 | 6.4779 | 6.4185 | 5.5796 | 5.9809 | 6.5547 | 5.8173 |
| PRKDC_S4026 | PRKDC | P78527 | 2672 | PLVDHTSPSSDSL | 6.2258 | 6.1843 | 5.9948 | 6.5280 | 6.2791 | 6.6201 | 6.3777 | 6.1464 | 5.7673 | 6.1353 | 6.0639 | 6.5053 | 6.0186 | 6.5605 | 6.2845 | 6.3084 |
| PRKDC_S687 | PRKDC | P78527 | 4026 | MLKKGGSWIQEIN | 5.5080 | 5.7914 | 5.2879 | 5.8090 | 5.7124 | 5.7988 | 6.0995 | 5.6413 | 5.7247 | 5.7645 | 5.5599 | 5.3274 | 7.6466 | 8.2191 | 7.8893 | 8.2201 |
| PRKDC_S893 | PRKDC | P78527 | 1065 | LFKRLYSLALHPN | 6.2160 | 6.4380 | 6.1830 | 6.1950 | 6.4897 | 6.2286 | 5.8775 | 5.8885 | 6.0933 | 5.8878 | 5.5157 | 5.2313 | 7.1018 | 7.1519 | 7.2604 | 6.2416 |
| PRKDC_T2671 | PRKDC | P78527 | 3205 | PLPEDNSMNVDQD | 6.9049 | 6.1114 | 7.8311 | 6.3409 | 6.6630 | 7.4219 | 6.5594 | 5.9829 | 6.9892 | 7.1233 | 6.8786 | 6.7418 | 4.6945 | 5.2378 | 5.1451 | 4.2743 |
| PRKRA_S167 | PRKRA | O75569 | 18 | PLEREDSGTFSLG | 5.0494 | 5.0242 | 5.5682 | 5.0767 | 5.4722 | 5.4157 | 5.2561 | 5.6313 | 5.2803 | 5.4693 | 5.2758 | 5.3379 | 9.3784 | 8.9814 | 9.1166 | 8.6667 |
| PRKRA_S18 | PRKRA | O75569 | 167 | TICRLESFMETGK | 6.1030 | 5.0859 | 3.7665 | 6.7001 | 6.7332 | 4.7210 | 4.2161 | 5.9077 | 5.0527 | 2.2628 | 4.4888 | 3.8623 | 12.7294 | 12.1213 | 9.4494 | 6.7996 |
| PRMT3_S25;S27 | PRMT3 | O60678 | 25;27 | NA | 6.6549 | 5.9609 | 6.1396 | 7.0168 | 5.9222 | 6.0446 | 6.4881 | 6.6672 | 5.9226 | 6.2350 | 6.2897 | 6.2017 | 5.8556 | 6.8925 | 5.9153 | 5.7932 |
| PROSER1_S613 | PROSER1 | Q86XN7 | 613 | IKTEPTSPTPSAF | 5.8391 | 5.4901 | 6.3615 | 5.2047 | 6.7650 | 6.1704 | 6.2743 | 6.7086 | 6.0909 | 6.4561 | 6.2202 | 6.6262 | 6.8105 | 6.3971 | 6.3078 | 6.2775 |
| PROSER2_S179 | PROSER2 | Q86WR7 | 312 | APGGGSSPERVAR | 6.4305 | 5.9309 | 6.3424 | 6.2929 | 6.6390 | 6.2425 | 5.8683 | 6.5761 | 6.6984 | 6.7316 | 6.7941 | 6.6442 | 5.7889 | 5.7036 | 5.6715 | 5.6452 |
| PROSER2_S212 | PROSER2 | Q86WR7 | 179 | RELRAPSPPVEHP | 6.0721 | 6.4392 | 6.6717 | 6.5749 | 6.3360 | 6.2397 | 6.1693 | 6.4724 | 6.3178 | 6.8990 | 6.7900 | 6.5567 | 5.0703 | 5.8154 | 5.8105 | 5.7648 |
| PROSER2_S312 | PROSER2 | Q86WR7 | 212 | AGNEALSPTSPFR | 5.8838 | 6.1852 | 6.8735 | 6.3723 | 6.8171 | 6.2548 | 6.6527 | 7.2034 | 6.4772 | 5.8167 | 6.4961 | 6.6929 | 5.5726 | 5.6309 | 5.3590 | 5.7117 |
| PROSER2_S43 | PROSER2 | Q86WR7 | 43 | SSSRSRSFTLDDE | 5.5470 | 5.8466 | 7.7042 | 6.3097 | 6.4664 | 6.1978 | 5.7898 | 7.1722 | 6.4061 | 6.7677 | 7.3806 | 6.1582 | 4.8946 | 5.9620 | 4.5761 | 6.8210 |
| PROSER3_S210 | PROSER3 | Q2NL68 | 356 | QPEVPLSPAEQAT | 5.7687 | 5.8626 | 6.6578 | 6.3786 | 6.4976 | 6.1464 | 6.3717 | 6.5593 | 6.7636 | 6.5112 | 7.0615 | 7.0145 | 6.0026 | 5.6861 | 5.3330 | 5.3848 |
| PROSER3_S356 | PROSER3 | Q2NL68 | 459 | WSPPAGSPPRSPR | 5.8564 | 6.4525 | 6.5084 | 6.9969 | 6.5429 | 6.2803 | 4.9365 | 5.9629 | 7.6048 | 6.4148 | 7.1981 | 7.1949 | 5.1441 | 5.2797 | 5.3421 | 6.2846 |
| PROSER3_S459 | PROSER3 | Q2NL68-2 | 210 | LLKRSPSDASTSS | 6.5015 | 6.0033 | 6.2710 | 5.4573 | 6.3525 | 6.5401 | 5.6069 | 6.7748 | 6.5085 | 6.9273 | 7.4775 | 6.5282 | 5.9983 | 6.1023 | 5.5409 | 5.4095 |
| PRPF18_T116 | PRPF18 | Q99633 | 116 | RKIEILTPEVNKG | 5.6382 | 6.7471 | 6.7178 | 6.5802 | 6.6794 | 6.9156 | 6.6187 | 6.4091 | 6.8847 | 6.6837 | 6.4210 | 6.6236 | 4.4865 | 4.9676 | 5.5970 | 6.0299 |
| PRPF3_S133 | PRPF3 | O43395 | 619 | GDDDEESDEEAVK | 5.4127 | 5.9395 | 5.8717 | 6.5303 | 6.2389 | 6.6892 | 6.4139 | 6.0125 | 6.2901 | 5.9800 | 7.0363 | 7.0160 | 6.7296 | 6.7142 | 5.6184 | 5.5069 |
| PRPF3_S619 | PRPF3 | O43395 | 133 | PGPPSESPGMLTK | 6.4339 | 6.5385 | 6.2654 | 6.9324 | 6.2291 | 6.8740 | 6.5122 | 6.4429 | 6.5858 | 6.9271 | 6.0822 | 6.4810 | 5.2939 | 5.7607 | 4.8548 | 5.7862 |
| PRPF3_T167 | PRPF3 | O43395 | 167 | SFISPPTPQPKTP | 7.0763 | 7.0485 | 7.0340 | 7.0768 | 5.3984 | 6.2105 | 7.0738 | 6.0858 | 7.3493 | 6.3009 | 6.2790 | 6.3012 | 5.3357 | 5.7667 | 4.5662 | 5.0969 |
| PRPF31_S379 | PRPF31 | Q8WWY3 | 450 | RSSGTASSVAFTP | 7.0651 | 6.1861 | 5.7680 | 6.7740 | 6.1063 | 5.9270 | 6.5006 | 5.4726 | 6.6398 | 5.1762 | 4.9304 | 5.2300 | 7.7905 | 7.2891 | 6.2080 | 6.9363 |
| PRPF31_S439;T440 | PRPF31 | Q8WWY3 | 455 | ASSVAFTPLQGLE | 5.6213 | 5.9186 | 5.5080 | 6.0630 | 6.0133 | 6.2354 | 6.5309 | 5.6613 | 6.5458 | 5.5156 | 5.5288 | 6.2447 | 7.6163 | 7.0972 | 6.2553 | 7.6444 |
| PRPF31_S450 | PRPF31 | Q8WWY3 | 379 | KQANRMSFGEIEE | 5.6393 | 6.4758 | 6.5308 | 6.6229 | 5.8931 | 7.2079 | 6.1906 | 6.2355 | 6.5437 | 7.5660 | 7.1304 | 7.3465 | 5.1088 | 5.3142 | 4.3481 | 5.8465 |
| PRPF31_T450;T455 | PRPF31 | Q8WWY3 | 450;455 | NA | 9.3091 | 6.7535 | 5.7420 | 7.9597 | 7.3017 | 4.7213 | 5.5329 | 4.9671 | 5.6967 | 4.3095 | 4.3359 | 4.6361 | 7.7095 | 7.4971 | 6.7403 | 6.7877 |
| PRPF31_T455 | PRPF31 | Q8WWY3 | 439;440 | NA | 6.4527 | 6.4972 | 6.4711 | 7.9522 | 7.1237 | 5.6083 | 6.2509 | 4.7143 | 5.5160 | 6.4603 | 4.9731 | 4.8823 | 7.8126 | 7.8784 | 6.7417 | 6.4852 |
| PRPF38A_S193;S194 | PRPF38A | Q8NAV1 | 209 | KLERVPSPDHRRR | 6.4184 | 6.4350 | 5.8468 | 6.6680 | 6.0289 | 5.9133 | 6.2040 | 5.8890 | 6.1967 | 6.0537 | 5.9409 | 5.8556 | 6.4952 | 6.6000 | 6.7270 | 6.7274 |
| PRPF38A_S209 | PRPF38A | Q8NAV1 | 193;194 | NA | 5.8998 | 6.0962 | 6.0564 | 6.4495 | 5.9113 | 5.7291 | 6.4320 | 5.8146 | 6.2375 | 6.0548 | 6.0735 | 6.2512 | 6.7474 | 6.6776 | 6.6645 | 6.9047 |
| PRPF38B_S513 | PRPF38B | Q5VTL8 | 513 | SHKRDHSDSKDQS | 5.8783 | 6.3572 | 6.0587 | 6.8984 | 6.1439 | 5.2813 | 6.5879 | 6.0976 | 5.9002 | 5.3969 | 6.3936 | 5.9441 | 5.8957 | 7.1720 | 7.1296 | 6.8645 |
| PRPF38B_S527;S529 | PRPF38B | Q5VTL8 | 527;529 | NA | 5.4123 | 5.9025 | 5.1206 | 5.5155 | 5.7922 | 6.0226 | 6.4781 | 5.2095 | 5.9639 | 5.6781 | 5.9438 | 6.0250 | 7.7365 | 7.8938 | 7.6613 | 7.6442 |
| PRPF4_S285 | PRPF4 | O43172 | 483 | WTHPGWSPLKTLA | 5.7920 | 6.2799 | 5.9746 | 6.7796 | 6.3257 | 6.3092 | 6.3236 | 6.4172 | 6.1664 | 6.6493 | 6.5873 | 6.4388 | 6.5517 | 6.0267 | 5.9237 | 6.3544 |
| PRPF4_S483 | PRPF4 | O43172 | 285 | IVFHPKSTVSLDP | 7.1571 | 6.8005 | 6.7580 | 7.3359 | 5.8049 | 6.3491 | 6.9689 | 6.8525 | 7.2453 | 6.7175 | 6.4908 | 7.6392 | 3.7342 | 4.5640 | 4.1189 | 5.4630 |
| PRPF40A_S13 | PRPF40A | O75400 | 373 | ETVADFTPKKEEE | 5.6787 | 5.8816 | 5.7210 | 5.9594 | 6.3685 | 6.1306 | 6.5362 | 5.8731 | 6.2113 | 5.8365 | 5.7986 | 5.7337 | 7.3227 | 6.9084 | 7.0471 | 6.9926 |
| PRPF40A_S34 | PRPF40A | O75400 | 34 | DGERRLSGSSLCS | 5.3116 | 5.5678 | 6.0024 | 5.5656 | 6.3149 | 5.7277 | 5.5767 | 6.2320 | 7.0464 | 7.6032 | 5.7713 | 6.7451 | 7.1108 | 6.8376 | 5.2946 | 5.4923 |
| PRPF40A_S883;S885;S888 | PRPF40A | O75400-2 | 13 | RRRSSLSPTMRPG | 7.1212 | 6.8278 | 5.8200 | 7.3317 | 7.0851 | 5.5176 | 6.2803 | 6.5886 | 6.0336 | 6.0381 | 6.1193 | 5.4725 | 5.2754 | 6.2983 | 5.7628 | 6.4277 |
| PRPF40A_T373 | PRPF40A | O75400 | 883;885;888 | NA | 5.6162 | 5.7265 | 6.3306 | 6.8308 | 6.5177 | 6.0058 | 6.5787 | 5.3154 | 6.5115 | 5.4481 | 5.7300 | 6.0869 | 7.4692 | 6.6578 | 7.0897 | 6.9850 |
| PRPF4B_S832 | PRPF4B | Q6NWY9 | 832 | AELPNRSPGFGIK | 5.9741 | 6.5048 | 6.5288 | 6.0098 | 6.1879 | 5.9455 | 6.1789 | 6.5839 | 6.5879 | 6.0427 | 5.8010 | 5.8500 | 6.6007 | 6.6869 | 6.9851 | 6.4321 |
| PRPF4B_S142;S144 | PRPF4B | Q13523 | 93 | SDKEGMSPAKRTK | 6.1421 | 6.4051 | 5.7343 | 6.4025 | 5.4928 | 5.2289 | 5.7624 | 5.5519 | 5.9889 | 6.0647 | 5.5919 | 5.9542 | 6.5750 | 7.5061 | 7.6402 | 7.9589 |
| PRPF4B_S20 | PRPF4B | Q13523 | 87 | KEIIDASDKEGMS | 6.4014 | 6.3957 | 5.9035 | 6.4521 | 5.9104 | 5.1810 | 6.1979 | 5.8265 | 5.5538 | 5.6142 | 4.9707 | 5.2941 | 6.8092 | 7.6247 | 7.7861 | 8.0787 |
| PRPF4B_S23 | PRPF4B | Q13523 | 277 | SKDRKKSPIINES | 6.1985 | 6.3430 | 5.7064 | 6.3624 | 5.9128 | 5.5302 | 5.8837 | 5.9440 | 5.9541 | 5.8028 | 5.7969 | 5.8664 | 6.4496 | 7.5262 | 7.2010 | 7.3221 |
| PRPF4B_S23;S32 | PRPF4B | Q13523 | 849 | DNDITPYLVSRFY | 6.0537 | 6.4965 | 5.9593 | 6.8385 | 6.3820 | 6.1355 | 5.7686 | 6.2546 | 6.0092 | 5.5583 | 5.9019 | 5.7531 | 6.1855 | 7.0089 | 6.7860 | 6.9082 |
| PRPF4B_S232 | PRPF4B | Q13523 | 20 | EMEDANSEKSINE | 6.0593 | 5.9077 | 5.8735 | 6.3245 | 5.9912 | 5.4582 | 6.2315 | 5.5161 | 5.8775 | 6.0079 | 5.7584 | 5.4580 | 6.8719 | 7.4329 | 7.4711 | 7.7601 |
| PRPF4B_S241 | PRPF4B | Q13523 | 431 | ERSKDASPINRWS | 6.2666 | 6.1263 | 5.4914 | 6.4845 | 5.6147 | 5.0727 | 5.8827 | 6.1648 | 6.4404 | 6.0966 | 5.4361 | 5.8218 | 6.2350 | 7.6443 | 7.6249 | 7.5972 |
| PRPF4B_S257 | PRPF4B | Q13523 | 294 | RGKKSRSPVDLRG | 6.0332 | 6.1868 | 5.3156 | 6.5754 | 6.6198 | 5.1730 | 5.2292 | 4.7998 | 5.3841 | 5.4674 | 5.5323 | 5.1403 | 8.0618 | 9.0278 | 8.5982 | 8.4553 |
| PRPF4B_S277 | PRPF4B | Q13523 | 241 | QARKSKSPTLRRR | 6.0403 | 6.0694 | 5.3915 | 5.8798 | 6.2230 | 5.7286 | 5.8321 | 5.6990 | 5.9434 | 5.7627 | 5.9898 | 5.7226 | 7.2174 | 8.2643 | 7.5401 | 6.6960 |
| PRPF4B_S292;S294 | PRPF4B | Q13523 | 368 | KSRRSRSPLLNDR | 5.8585 | 6.5228 | 5.6773 | 6.6154 | 5.1923 | 5.0952 | 5.4129 | 5.1990 | 5.7830 | 6.3798 | 5.6884 | 5.8728 | 7.0379 | 7.7329 | 7.9167 | 8.0151 |
| PRPF4B_S294 | PRPF4B | Q13523 | 349 | SRRPGRSPKRRSL | 6.5438 | 6.5255 | 5.3947 | 6.7913 | 5.9281 | 5.6890 | 6.2896 | 6.0588 | 5.5447 | 5.7742 | 6.3354 | 5.9905 | 6.0141 | 7.7193 | 6.6337 | 7.6672 |
| PRPF4B_S349 | PRPF4B | Q13523 | 23 | DANSEKSINEENG | 6.3376 | 6.3159 | 5.8738 | 6.5861 | 5.7762 | 5.3654 | 5.8759 | 5.7783 | 6.6303 | 6.7252 | 6.6173 | 6.2614 | 6.1416 | 7.0871 | 6.1971 | 6.4308 |
| PRPF4B_S354;S356 | PRPF4B | Q13523 | 257 | KIGKARSPTDDKV | 6.4554 | 6.5299 | 5.6307 | 6.0584 | 6.1879 | 5.7895 | 5.8497 | 5.5050 | 5.7449 | 5.7631 | 5.4467 | 5.5680 | 6.8366 | 7.8018 | 7.3000 | 7.5325 |
| PRPF4B_S366;S368 | PRPF4B | Q13523 | 636 | PDMFTESDDMFAA | 6.4115 | 6.2389 | 6.0160 | 8.4726 | 7.3474 | 3.6041 | 6.0885 | 5.6985 | 7.5813 | 6.7123 | 3.5873 | 6.3113 | 4.0553 | 6.6566 | 6.5564 | 8.6620 |
| PRPF4B_S368 | PRPF4B | Q13523 | 507 | EDKFKGSLSEGMK | 5.7772 | 6.1344 | 5.7594 | 6.0089 | 5.9795 | 6.0528 | 5.8430 | 6.0487 | 5.5722 | 6.3826 | 4.8270 | 5.3241 | 7.2409 | 7.6749 | 7.7640 | 7.6104 |
| PRPF4B_S387 | PRPF4B | Q13523 | 852 | ITPYLVSRFYRAP | 5.8779 | 6.1237 | 5.9078 | 6.6324 | 6.1244 | 6.2113 | 5.8874 | 5.9164 | 5.9699 | 5.6831 | 5.5090 | 5.5501 | 6.2276 | 7.5002 | 7.5258 | 7.3529 |
| PRPF4B_S431 | PRPF4B | Q13523 | 232 | PSKRSKSQDQARK | 6.0482 | 9.2913 | 8.0209 | 6.6377 | 4.4962 | 10.7432 | 5.9523 | 4.3829 | 8.3353 | 5.4948 | 4.3465 | 3.7535 | 5.8985 | 5.8659 | 5.5055 | 5.2273 |
| PRPF4B_S431;S437 | PRPF4B | Q13523 | 387 | SPSRTLSPGRRAK | 6.2016 | 6.2850 | 6.1307 | 6.4650 | 6.0857 | 5.6697 | 5.8723 | 6.2033 | 6.0124 | 6.3883 | 5.8606 | 5.5088 | 6.6860 | 6.4450 | 6.8335 | 7.3520 |
| PRPF4B_S451;S453 | PRPF4B | Q13523 | 509 | KFKGSLSEGMKVE | 6.8718 | 6.4247 | 6.0849 | 6.6392 | 6.5885 | 6.3471 | 6.6545 | 6.0988 | 5.8477 | 5.4199 | 5.5331 | 5.4014 | 6.0081 | 6.9452 | 6.4584 | 6.6767 |
| PRPF4B_S507 | PRPF4B | Q13523 | 140;144 | NA | 5.9270 | 5.8178 | 5.7441 | 6.2377 | 6.0963 | 5.7622 | 6.1014 | 5.7814 | 6.0050 | 6.1371 | 5.8801 | 5.7734 | 6.7090 | 7.5476 | 7.1408 | 7.3392 |
| PRPF4B_S509 | PRPF4B | Q13523 | 142;144 | NA | 5.9595 | 6.0446 | 5.9065 | 6.6594 | 6.1641 | 5.9641 | 6.1114 | 5.7194 | 6.1188 | 5.9181 | 5.8306 | 5.8983 | 6.4651 | 7.3198 | 6.7824 | 7.1379 |
| PRPF4B_S636 | PRPF4B | Q13523 | 366;368 | NA | 5.9967 | 5.9483 | 5.6874 | 5.9514 | 5.6582 | 5.0946 | 5.8888 | 5.6337 | 6.1288 | 5.8842 | 5.6403 | 5.8266 | 7.0899 | 7.8195 | 7.7986 | 7.9531 |
| PRPF4B_S852 | PRPF4B | Q13523 | 354;356 | NA | 6.0837 | 6.0445 | 5.1620 | 6.3259 | 5.1703 | 6.0261 | 6.5784 | 5.0759 | 5.7665 | 5.5956 | 6.2048 | 6.0080 | 7.1492 | 7.5144 | 6.5029 | 6.7916 |
| PRPF4B_S87 | PRPF4B | Q13523 | 431;437 | NA | 6.5296 | 6.4041 | 6.0340 | 6.3896 | 6.1900 | 5.8225 | 6.1376 | 5.5716 | 5.7521 | 5.9090 | 6.0517 | 6.5762 | 6.8140 | 7.4077 | 6.7274 | 6.5828 |
| PRPF4B_S93 | PRPF4B | Q13523 | 292;294 | NA | 6.0761 | 6.1408 | 5.6151 | 6.1912 | 5.8535 | 5.3842 | 6.0635 | 5.6581 | 6.0894 | 5.8507 | 5.7759 | 6.0231 | 6.7729 | 7.6655 | 7.2848 | 7.5554 |
| PRPF4B_Y140;S142 | PRPF4B | Q13523 | 140;142 | NA | 5.3871 | 5.5854 | 6.1883 | 6.1276 | 5.7438 | 5.6305 | 5.9599 | 6.2435 | 5.7938 | 6.1042 | 5.8814 | 6.0806 | 7.1485 | 7.2378 | 7.4523 | 7.4354 |
| PRPF4B_Y140;S144 | PRPF4B | Q13523 | 23;32 | NA | 6.7700 | 6.9417 | 6.0270 | 6.6399 | 5.4116 | 6.7291 | 6.0317 | 4.9927 | 5.6784 | 5.5492 | 5.4998 | 5.8762 | 6.2468 | 7.0046 | 7.0995 | 7.5020 |
| PRPF4B_Y849 | PRPF4B | Q13523 | 451;453 | NA | 6.4371 | 6.4143 | 6.6014 | 6.1891 | 6.0992 | 5.3104 | 5.9693 | 5.8865 | 6.2286 | 6.1676 | 6.4711 | 6.1593 | 6.3200 | 6.5012 | 6.7172 | 6.5276 |
| PRPF6_S143 | PRPF6 | O94906 | 275 | DPKGYLTDLNSMI | 8.5750 | 5.9540 | 6.8798 | 8.2673 | 7.2549 | 4.0320 | 5.1378 | 3.8053 | 5.1596 | 3.5935 | 3.8231 | 3.7833 | 9.4954 | 8.3111 | 8.2734 | 7.4726 |
| PRPF6_S279 | PRPF6 | O94906 | 212 | QFGGLNTPYPGGL | 6.4274 | 6.2564 | 6.0805 | 5.7255 | 6.0090 | 6.3260 | 5.9934 | 6.2347 | 6.6301 | 6.2042 | 6.4280 | 6.5608 | 6.2527 | 6.6214 | 6.4591 | 6.6910 |
| PRPF6_T180 | PRPF6 | O94906 | 143 | KIQQQFSDLKRKL | 5.9703 | 6.4252 | 8.4695 | 6.4497 | 6.0526 | 5.3457 | 6.3170 | 6.7056 | 6.3833 | 7.7861 | 6.9640 | 5.6231 | 5.5642 | 5.7867 | 5.0135 | 5.1437 |
| PRPF6_T212 | PRPF6 | O94906 | 180 | PRYEKLTPVPDSF | 6.3434 | 7.1755 | 6.3654 | 6.9367 | 6.1950 | 6.4252 | 6.6841 | 6.4095 | 6.3257 | 5.8668 | 6.2423 | 5.9910 | 5.2291 | 5.8868 | 5.6716 | 6.2518 |
| PRPF6_T275 | PRPF6 | O94906 | 279 | YLTDLNSMIPTHG | 8.2902 | 6.1553 | 5.7457 | 8.8760 | 6.9882 | 5.6542 | 4.3293 | 4.9654 | 4.4734 | 3.9541 | 4.0250 | 3.8259 | 8.5488 | 8.5374 | 6.7571 | 6.8738 |
| PRPF6_T275;S279 | PRPF6 | O94906 | 222 | GGLNTPYPGGMTP | 6.0875 | 6.4611 | 6.6294 | 7.7175 | 5.8670 | 5.2334 | 6.7430 | 6.0589 | 5.8022 | 5.7391 | 5.7526 | 6.4704 | 6.4460 | 5.9427 | 6.3214 | 6.7279 |
| PRPF6_Y222 | PRPF6 | O94906 | 275;279 | NA | 7.6234 | 6.1588 | 6.4682 | 9.0287 | 6.5674 | 3.5969 | 4.8497 | 3.0663 | 4.2868 | 4.2839 | 4.1621 | 2.5820 | 11.0013 | 10.1544 | 8.4780 | 7.6922 |
| PRPF8_S1539 | PRPF8 | Q6P2Q9 | 823 | LESRRFSPIPFPP | 6.1187 | 7.3521 | 5.7031 | 6.5583 | 6.7995 | 5.8757 | 6.6214 | 6.0020 | 6.3592 | 6.3683 | 6.6828 | 6.3063 | 5.9413 | 5.1618 | 6.4290 | 5.7205 |
| PRPF8_S823 | PRPF8 | Q6P2Q9 | 1404 | AQNRRLTLEDLED | 4.1875 | 5.5397 | 7.8396 | 5.4742 | 7.1230 | 7.1847 | 7.1307 | 4.3763 | 6.7263 | 7.3465 | 5.5628 | 5.7516 | 5.1391 | 5.8209 | 4.1143 | 10.6826 |
| PRPF8_T1404 | PRPF8 | Q6P2Q9 | 1539 | RFTLWWSPTINRA | 6.8868 | 8.3918 | 7.6864 | 5.4120 | 4.2933 | 6.7827 | 5.8588 | 7.0548 | 7.7132 | 10.0951 | 7.5486 | 5.9563 | 4.3269 | 4.0414 | 4.1287 | 3.8230 |
| PRPSAP1_S244 | PRPSAP1 | Q14558-2 | 244 | MDDGRHSPPMVKN | 6.1050 | 6.3374 | 6.8654 | 6.1269 | 6.3904 | 6.5945 | 5.9278 | 6.4599 | 6.7210 | 6.5877 | 6.5497 | 6.4656 | 5.8705 | 5.8315 | 5.6066 | 5.5600 |
| PRPSAP2_S227 | PRPSAP2 | O60256 | 227 | LVDGRHSPPMVRS | 6.0867 | 6.1248 | 6.9413 | 6.1198 | 6.1324 | 6.5131 | 6.1456 | 6.6054 | 6.5907 | 6.5687 | 6.4631 | 6.3374 | 5.7871 | 6.1548 | 5.6484 | 5.7807 |
| PRR11_S311 | PRR11 | Q96HE9 | 287 | VTNVLITPGKSQM | 7.5899 | 8.0935 | 6.9969 | 8.0186 | 5.7852 | 6.9106 | 7.8343 | 7.1650 | 7.4781 | 7.5002 | 7.3756 | 7.5370 | 2.5523 | 3.4973 | 2.4668 | 3.1985 |
| PRR11_S344 | PRR11 | Q96HE9 | 307 | KVDVERSPGGTPL | 6.9979 | 7.3924 | 6.4117 | 7.7833 | 6.9861 | 6.7248 | 7.5235 | 7.4222 | 7.5686 | 7.1219 | 6.9961 | 7.1276 | 3.1353 | 3.8932 | 3.1673 | 3.7482 |
| PRR11_S344;T346 | PRR11 | Q96HE9 | 40 | PPPPPPSPERVGI | 6.8846 | 7.4468 | 6.9342 | 7.3277 | 5.8655 | 6.4007 | 7.6273 | 6.3209 | 7.2142 | 7.2586 | 7.1768 | 6.7860 | 4.0082 | 4.7683 | 3.8582 | 5.0220 |
| PRR11_S40 | PRR11 | Q96HE9 | 326 | ETGTGLTPVMTQA | 4.4861 | 7.7648 | 6.9438 | 7.5720 | 6.0199 | 6.6397 | 5.6677 | 7.3324 | 7.2058 | 7.1834 | 7.1084 | 6.4498 | 3.9871 | 4.5238 | 3.7888 | 4.4265 |
| PRR11_T287 | PRR11 | Q96HE9 | 344 | QLAHPRSPTPTLP | 6.3210 | 5.5076 | 7.0560 | 7.3942 | 6.4101 | 6.7027 | 6.0149 | 6.9771 | 6.5830 | 6.3897 | 6.6372 | 6.8418 | 5.1215 | 5.8077 | 5.0560 | 5.1795 |
| PRR11_T326 | PRR11 | Q96HE9 | 344;346 | NA | 6.7602 | 9.3537 | 6.6524 | 7.5028 | 6.0183 | 7.7823 | 8.0196 | 7.0124 | 8.8161 | 7.5651 | 7.0507 | 5.8621 | 2.5475 | 4.5598 | 1.1690 | 3.3279 |
| PRR12_S1135 | PRR12 | Q9ULL5 | 651 | HLLQAPSPPRTSG | 6.2330 | 6.4078 | 6.0825 | 6.7969 | 6.1058 | 5.6606 | 6.5986 | 5.9617 | 6.4724 | 6.3484 | 6.3631 | 6.6075 | 6.2471 | 6.1491 | 5.9576 | 6.0078 |
| PRR12_S1308 | PRR12 | Q9ULL5 | 1135 | RSRPALSPLGDID | 5.5717 | 5.9327 | 5.4501 | 6.3067 | 6.2125 | 6.0710 | 6.2073 | 6.0667 | 5.7754 | 5.4452 | 6.1474 | 5.3510 | 7.5001 | 7.2904 | 7.5621 | 7.1094 |
| PRR12_S1381;S1382 | PRR12 | Q9ULL5 | 917 | GAYRSPSPQGTKA | 6.7117 | 6.6617 | 5.9309 | 6.4589 | 6.4642 | 5.8226 | 6.0665 | 6.0417 | 6.1187 | 6.0952 | 5.7072 | 5.5886 | 6.4812 | 6.3502 | 6.8796 | 6.2211 |
| PRR12_S651 | PRR12 | Q9ULL5 | 865 | LEPAAPSPRLRPE | 6.0042 | 5.9240 | 5.2733 | 5.7260 | 6.6397 | 5.9992 | 6.0581 | 6.0923 | 6.8439 | 6.1282 | 4.8324 | 6.0715 | 7.5434 | 7.4907 | 6.8523 | 6.5208 |
| PRR12_S865 | PRR12 | Q9ULL5 | 1308 | GLTPPLSPPKSVP | 6.4856 | 7.0031 | 5.8721 | 6.5986 | 6.1760 | 5.9804 | 6.0817 | 6.3852 | 6.2566 | 6.2823 | 6.1041 | 6.5191 | 6.2399 | 6.0598 | 5.9060 | 6.0494 |
| PRR12_S917 | PRR12 | Q9ULL5 | 1060 | EPKGGLTSPIFCS | 7.4597 | 7.6236 | 6.6041 | 8.0259 | 5.9173 | 6.8296 | 7.6822 | 6.5584 | 7.1060 | 6.8839 | 7.0027 | 7.2862 | 3.4856 | 4.0419 | 3.2512 | 4.2419 |
| PRR12_T1060 | PRR12 | Q9ULL5 | 1381;1382 | NA | 6.0800 | 6.3345 | 6.7102 | 6.5842 | 6.0211 | 6.0215 | 6.6368 | 5.9001 | 6.4577 | 5.9420 | 5.7732 | 6.3298 | 6.3587 | 6.6513 | 6.1624 | 6.0365 |
| PRR14_S277 | PRR14 | Q9BWN1 | 194 | RIVRQPTPPPGDL | 6.4874 | 6.0080 | 6.2209 | 5.5457 | 6.2335 | 6.0714 | 6.1376 | 6.3598 | 6.3274 | 5.9147 | 6.6957 | 6.4047 | 6.6045 | 6.5800 | 5.9266 | 6.4821 |
| PRR14_S42 | PRR14 | Q9BWN1 | 42 | QLPGAPSPLEKAS | 7.8939 | 7.7214 | 6.8947 | 7.3019 | 5.9003 | 7.5090 | 7.4078 | 6.8016 | 7.9266 | 7.6472 | 8.1850 | 8.1312 | 2.4566 | 3.2149 | 1.9270 | 3.0810 |
| PRR14_S93 | PRR14 | Q9BWN1 | 277 | PQPPPPSPPMKLE | 6.7057 | 6.7469 | 6.3706 | 6.8732 | 6.4927 | 6.0853 | 6.4533 | 6.3672 | 6.6073 | 6.4142 | 6.0035 | 6.2020 | 5.8096 | 5.6032 | 5.4530 | 5.8122 |
| PRR14_T194 | PRR14 | Q9BWN1 | 93 | HRQPPASPPRQAG | 7.2322 | 7.0973 | 6.4061 | 6.6855 | 6.1532 | 6.5249 | 6.5267 | 6.3440 | 6.6810 | 6.3347 | 6.2036 | 6.3259 | 5.6140 | 5.7033 | 5.5197 | 5.5479 |
| PRR14L_S1029 | PRR14L | Q5THK1 | 1029 | NSLPCGSPKKCNL | 5.9299 | 6.1138 | 6.5806 | 6.9533 | 6.3891 | 6.6861 | 6.4909 | 6.6788 | 6.3046 | 6.1214 | 6.2683 | 6.1066 | 5.8498 | 5.8261 | 5.3109 | 6.3899 |
| PRR14L_S1994 | PRR14L | Q5THK1 | 600 | TALLLPSPEFDYR | 5.5572 | 6.1544 | 6.5277 | 6.8680 | 6.2565 | 6.5463 | 6.0216 | 5.9985 | 6.2565 | 5.8492 | 6.1039 | 6.5474 | 5.8953 | 6.8471 | 5.8196 | 6.7505 |
| PRR14L_S600 | PRR14L | Q5THK1 | 1994 | CPCPQSSPPEQKE | 5.8690 | 6.4484 | 6.1883 | 6.9843 | 6.4443 | 6.6540 | 6.7119 | 6.4110 | 6.5431 | 6.9461 | 6.5015 | 6.9570 | 5.2705 | 5.3965 | 5.1330 | 5.5408 |
| PRR3_S33 | PRR3 | P79522-2 | 33 | GDEEDGSPIALHR | 6.4428 | 7.3001 | 6.4566 | 6.5667 | 6.0588 | 6.0144 | 6.0190 | 5.9210 | 6.0097 | 6.4416 | 6.6446 | 5.9579 | 5.9452 | 6.1395 | 6.0916 | 5.9907 |
| PRRC1_S408 | PRRC1 | Q96M27-2 | 408 | KSLLNVSRTDWHM | 6.1188 | 6.1611 | 6.2499 | 6.2316 | 6.5735 | 7.1092 | 7.0163 | 6.8934 | 6.2341 | 5.8104 | 6.6830 | 6.4292 | 5.7599 | 5.8696 | 5.2070 | 5.6531 |
| PRRC2A_S1085;S1089 | PRRC2A | P48634 | 610 | EVPPPTTPPVPKV | 6.0610 | 6.3904 | 6.3805 | 6.2615 | 6.2615 | 6.5674 | 6.3394 | 6.5842 | 6.3569 | 6.3523 | 6.2423 | 6.3266 | 5.9769 | 6.1733 | 5.9490 | 5.7769 |
| PRRC2A_S1147 | PRRC2A | P48634 | 1219 | LIPGPLSPVARGG | 6.4084 | 6.6485 | 6.6610 | 6.5683 | 6.5658 | 6.2347 | 6.7733 | 6.7566 | 6.3058 | 5.9792 | 6.2274 | 6.0869 | 5.4738 | 5.8506 | 5.5856 | 5.8738 |
| PRRC2A_S1219 | PRRC2A | P48634 | 1691 | KRPGGSSPLNAVP | 6.1909 | 6.6438 | 6.3587 | 6.6512 | 6.2066 | 6.2201 | 6.5879 | 6.1922 | 6.5728 | 6.7332 | 6.7122 | 6.5587 | 5.5385 | 5.8768 | 5.4549 | 5.5015 |
| PRRC2A_S1306 | PRRC2A | P48634 | 825 | KGMRSETPPVPPP | 6.3155 | 6.5840 | 6.3107 | 6.4793 | 6.6475 | 6.1545 | 6.0942 | 6.2941 | 6.7810 | 6.4857 | 6.6780 | 6.4556 | 5.5389 | 5.7339 | 5.5534 | 5.8938 |
| PRRC2A_S1386 | PRRC2A | P48634 | 878 | EVAKIQTPPPKKE | 7.2195 | 7.2669 | 7.0688 | 6.9511 | 6.7145 | 6.8836 | 6.8703 | 6.4942 | 7.5289 | 7.3832 | 6.9702 | 7.0472 | 3.7177 | 4.3937 | 3.5651 | 3.9251 |
| PRRC2A_S1435 | PRRC2A | P48634 | 1147 | PPGAPPSPAPARF | 6.7888 | 5.9314 | 6.4236 | 6.5836 | 5.9320 | 6.7722 | 6.7574 | 5.9006 | 6.6334 | 6.3894 | 6.6622 | 6.5898 | 5.4350 | 6.2209 | 5.6339 | 5.3459 |
| PRRC2A_S1619 | PRRC2A | P48634 | 1386 | LSKRSFSSQRPGM | 6.9411 | 6.4319 | 6.0512 | 5.4203 | 5.0867 | 6.4083 | 6.3911 | 5.8794 | 6.8119 | 6.5090 | 6.4716 | 6.6195 | 6.4491 | 6.7329 | 5.5993 | 6.1969 |
| PRRC2A_S1691 | PRRC2A | P48634 | 1347 | PPPVLLTPKAVGT | 6.9397 | 6.7849 | 6.7222 | 6.4381 | 7.1000 | 6.8533 | 7.2879 | 6.9811 | 7.2223 | 6.3802 | 6.7214 | 6.3111 | 4.4897 | 4.8091 | 4.1812 | 4.7778 |
| PRRC2A_S2036 | PRRC2A | P48634 | 808 | EPRPLTSPLRQAA | 6.8554 | 6.6676 | 7.1280 | 6.7931 | 6.6803 | 6.7768 | 6.6726 | 6.5740 | 6.5842 | 6.3083 | 6.8894 | 6.4156 | 5.4997 | 5.3583 | 4.5727 | 4.8039 |
| PRRC2A_S2113 | PRRC2A | P48634 | 782 | MLRERGTPPVDPK | 6.1935 | 6.2365 | 6.7326 | 6.6209 | 6.0269 | 6.2000 | 6.3926 | 6.3986 | 6.7169 | 6.4755 | 6.4881 | 6.3795 | 5.7340 | 5.9605 | 5.7338 | 5.7101 |
| PRRC2A_S342;S350 | PRRC2A | P48634 | 456 | RQRRKQSSSEISL | 5.3462 | 5.1042 | 5.6260 | 5.6713 | 7.0210 | 5.2522 | 4.9516 | 5.8852 | 6.7890 | 7.1135 | 6.8891 | 6.6506 | 6.8604 | 7.7105 | 6.7207 | 6.4088 |
| PRRC2A_S363 | PRRC2A | P48634 | 2036 | ATRVLPSPARPFP | 7.2195 | 7.8753 | 6.8531 | 7.0951 | 4.8562 | 7.3137 | 7.6322 | 7.1872 | 7.0964 | 7.4535 | 7.7056 | 8.1289 | 3.0729 | 3.8692 | 2.8850 | 3.7563 |
| PRRC2A_S380 | PRRC2A | P48634 | 1004 | PPNGNLSPAPRLR | 6.9064 | 6.9772 | 6.4899 | 6.4092 | 6.5638 | 6.2727 | 6.3820 | 6.8544 | 7.0939 | 6.5733 | 6.2901 | 6.3560 | 5.3692 | 5.4676 | 4.9766 | 5.0178 |
| PRRC2A_S380;S383 | PRRC2A | P48634 | 2113 | DLYQQASPPDALR | 6.8337 | 7.0562 | 7.0950 | 7.1131 | 6.2069 | 6.9691 | 6.2117 | 7.0432 | 6.9752 | 7.1128 | 6.4399 | 7.4106 | 3.9795 | 5.2629 | 3.9486 | 4.3415 |
| PRRC2A_S383 | PRRC2A | P48634 | 1306 | RRAAAKSPDLSNQ | 6.1116 | 7.3448 | 5.7909 | 7.0246 | 6.1419 | 6.7998 | 6.2347 | 6.3454 | 7.2069 | 6.9882 | 6.8893 | 6.4091 | 5.0337 | 5.2823 | 5.0527 | 5.3441 |
| PRRC2A_S456 | PRRC2A | P48634 | 761 | PRERSDSGGSSSE | 5.9526 | 6.5447 | 6.7148 | 6.3148 | 6.0656 | 5.9165 | 6.0040 | 6.6730 | 6.3471 | 6.2512 | 6.6019 | 6.1708 | 5.8597 | 6.0650 | 6.3307 | 6.1876 |
| PRRC2A_S680 | PRRC2A | P48634 | 932 | WGPRPGSSRRGIP | 5.9145 | 6.8684 | 7.5182 | 6.4459 | 5.3080 | 7.0439 | 6.8349 | 6.5287 | 6.6459 | 8.0845 | 8.4431 | 8.0409 | 3.7571 | 5.1003 | 3.4068 | 4.0591 |

| | | | | | | | | | | | | | | | | | | | | |
|---|---|---|---|---|---|---|---|---|---|---|---|---|---|---|---|---|---|---|---|---|
| PRRC2A_S761 | PRRC2A | P48634 | 997 | LGGPKETPPNGNL | 5.2670 | 10.0218 | 7.2694 | 10.1672 | 9.8632 | 9.1231 | 6.4169 | 5.6205 | 7.0414 | 7.9897 | 6.9455 | 5.6569 | 3.3076 | 1.6257 | 1.9264 | 1.7579 |
| PRRC2A_S808 | PRRC2A | P48634 | 1546 | VRGVGGTPRDSAG | 8.4743 | 9.0764 | 7.5799 | 7.8405 | 5.0221 | 7.3680 | 7.8087 | 6.9013 | 8.0007 | 7.4409 | 7.6711 | 7.8080 | 1.6933 | 3.1381 | 1.6358 | 2.5409 |
| PRRC2A_S932 | PRRC2A | P48634 | 1435 | DKRSWPSPKNRSR | 7.4864 | 7.0250 | 7.7303 | 7.2015 | 7.2141 | 6.8515 | 6.6663 | 4.5949 | 4.8250 | 6.0086 | 6.6777 | 6.9247 | 5.2747 | 5.3590 | 4.2649 | 5.8955 |
| PRRC2A_T1004 | PRRC2A | P48634 | 380 | DGKKGNSPNSEPP | 5.9937 | 6.2265 | 6.9666 | 6.2824 | 6.4272 | 6.0137 | 6.1685 | 7.2522 | 7.4220 | 6.8897 | 6.1745 | 6.9086 | 4.9041 | 5.3256 | 4.8941 | 6.1506 |
| PRRC2A_T1347 | PRRC2A | P48634 | 680 | PTPVPPSPPQPVT | 5.9954 | 5.9186 | 6.7554 | 6.3256 | 6.0452 | 6.0321 | 6.1509 | 6.2549 | 6.8008 | 7.2812 | 6.3920 | 6.3972 | 5.5884 | 5.9797 | 5.7587 | 6.3239 |
| PRRC2A_T1546 | PRRC2A | P48634 | 383 | KGNSPNSEPPTPK | 5.6977 | 5.9716 | 6.2618 | 6.7868 | 6.4995 | 6.6253 | 6.7734 | 6.1331 | 6.3089 | 6.1128 | 6.5154 | 6.3569 | 5.4293 | 6.3099 | 5.8419 | 6.3757 |
| PRRC2A_T610 | PRRC2A | P48634 | 363 | GHRDSQSASGEER | 5.9919 | 6.3965 | 7.0228 | 6.6310 | 6.1929 | 6.0651 | 5.7871 | 6.7135 | 6.6068 | 6.4100 | 5.9522 | 6.0142 | 5.7511 | 5.7793 | 6.2077 | 6.4780 |
| PRRC2A_T782 | PRRC2A | P48634 | 1619 | RLHTATSRKSYRP | 6.5622 | 7.6978 | 9.0622 | 6.0242 | 5.0060 | 7.9901 | 6.7960 | 5.5377 | 7.0738 | 8.6719 | 9.0442 | 8.8796 | 2.7926 | 3.7082 | 1.9814 | 3.1720 |
| PRRC2A_T825 | PRRC2A | P48634 | 342;350 | NA | 6.3709 | 6.0298 | 6.5336 | 6.3309 | 6.1049 | 6.2137 | 6.3012 | 6.5124 | 6.6385 | 6.3363 | 6.5054 | 6.6040 | 5.9580 | 6.2132 | 5.7430 | 5.6042 |
| PRRC2A_T878 | PRRC2A | P48634 | 1085;1089 | NA | 5.3296 | 5.7218 | 6.3448 | 5.8811 | 5.9643 | 6.1392 | 5.4535 | 5.9582 | 6.5807 | 7.7427 | 7.8314 | 7.8659 | 5.4799 | 5.8052 | 5.8112 | 6.0905 |
| PRRC2A_T997 | PRRC2A | P48634 | 997;1004 | NA | 7.5701 | 7.6713 | 6.9663 | 7.2723 | 5.2067 | 7.7864 | 7.8885 | 6.1040 | 8.0122 | 8.1776 | 8.5921 | 7.8843 | 2.1017 | 3.4737 | 1.9741 | 3.3189 |
| PRRC2A_T997;S1004 | PRRC2A | P48634 | 380;383 | NA | 5.6824 | 6.3994 | 6.0887 | 6.4788 | 6.4674 | 6.6312 | 6.4640 | 6.2626 | 6.0824 | 6.3816 | 6.3984 | 6.2516 | 5.9344 | 6.1312 | 5.8620 | 6.4840 |
| PRRC2B_S1132;S1136;S1 | PRRC2B | Q5JSZ5 | 556 | EKEVPWSPSAEKA | 6.9036 | 6.8499 | 6.8195 | 6.5096 | 6.5897 | 6.2982 | 6.5163 | 6.7941 | 6.9292 | 6.7333 | 6.8423 | 6.6015 | 4.8298 | 5.1381 | 4.8766 | 4.7683 |
| PRRC2B_S1422 | PRRC2B | Q5JSZ5 | 388 | SEKLKFSDDEEEE | 6.6383 | 6.4478 | 6.8445 | 5.9265 | 6.5991 | 5.9105 | 6.3687 | 6.1537 | 6.9387 | 6.4258 | 6.4221 | 6.2746 | 5.8155 | 5.9609 | 5.5346 | 5.7388 |
| PRRC2B_S1453 | PRRC2B | Q5JSZ5 | 613 | EAREAGSPAQEFK | 6.4671 | 6.9504 | 6.6220 | 6.4863 | 6.1339 | 6.3360 | 6.0730 | 6.3193 | 7.0015 | 6.8066 | 6.7804 | 6.5889 | 5.3282 | 5.5653 | 5.1882 | 5.3530 |
| PRRC2B_S166 | PRRC2B | Q5JSZ5 | 480 | SMFRQQSIEDKED | 5.7290 | 5.7246 | 6.6130 | 5.5049 | 6.3486 | 6.1041 | 5.9063 | 6.3237 | 6.3234 | 6.4479 | 6.3094 | 5.7861 | 6.8145 | 6.7390 | 6.9432 | 6.3824 |
| PRRC2B_S1691 | PRRC2B | Q5JSZ5 | 226 | ATSLSTSPTELGS | 6.3832 | 6.3292 | 6.4374 | 6.4794 | 6.0610 | 6.1862 | 6.7848 | 7.1819 | 6.8490 | 6.4867 | 5.8414 | 6.6434 | 5.7023 | 6.0301 | 5.4025 | 5.2015 |
| PRRC2B_S1808 | PRRC2B | Q5JSZ5 | 1453 | PGGGDTSPRYESQ | 6.2900 | 5.3884 | 6.5704 | 6.4164 | 5.9452 | 6.4300 | 6.1905 | 5.5880 | 6.5112 | 6.7512 | 6.8898 | 7.1563 | 6.0628 | 6.1029 | 5.5937 | 5.8132 |
| PRRC2B_S1843 | PRRC2B | Q5JSZ5 | 853 | TQNLRCSPLEPDF | 5.8286 | 5.7339 | 6.4201 | 5.8143 | 6.2756 | 5.7843 | 6.0541 | 5.9325 | 6.7225 | 6.3544 | 6.6077 | 6.3471 | 7.1209 | 6.3727 | 6.3336 | 6.2976 |
| PRRC2B_S2153 | PRRC2B | Q5JSZ5 | 1843 | QRAIGLSPMSFPT | 7.0841 | 7.3223 | 6.7438 | 7.0363 | 5.4595 | 7.0885 | 7.2605 | 7.2210 | 7.4314 | 7.3259 | 7.4560 | 7.4471 | 3.3111 | 4.4772 | 3.4991 | 3.8360 |
| PRRC2B_S2161;S2163 | PRRC2B | Q5JSZ5 | 980 | AKEKEQSPTAEKD | 5.5638 | 6.2566 | 6.5437 | 6.6920 | 5.5549 | 6.0880 | 6.1218 | 5.7859 | 6.1933 | 6.6151 | 6.3674 | 6.5358 | 6.3846 | 6.6045 | 6.3713 | 6.3215 |
| PRRC2B_S2163 | PRRC2B | Q5JSZ5 | 1422 | LAKRSFSSQRPVV | 5.3604 | 5.4103 | 6.6019 | 5.8338 | 5.7342 | 6.5180 | 6.1126 | 6.3289 | 6.7327 | 6.6265 | 6.8782 | 6.8538 | 6.3844 | 6.6416 | 6.1278 | 5.8548 |
| PRRC2B_S226 | PRRC2B | Q5JSZ5 | 416 | RRQRQLSMSSADS | 6.3600 | 6.3701 | 7.1338 | 6.2740 | 5.9267 | 5.6217 | 5.9478 | 6.0056 | 7.0246 | 7.1781 | 6.9673 | 6.9939 | 5.4999 | 6.2570 | 5.2030 | 5.2365 |
| PRRC2B_S388 | PRRC2B | Q5JSZ5 | 166 | GSSRLLSFSPEEF | 6.4874 | 5.8506 | 6.4898 | 6.3986 | 6.4088 | 6.0554 | 6.0050 | 5.7470 | 7.7446 | 7.6540 | 7.6383 | 7.0982 | 4.9674 | 6.6503 | 4.3033 | 4.5011 |
| PRRC2B_S416 | PRRC2B | Q5JSZ5 | 745 | DSPPVWSPEGYMA | 6.2984 | 6.2870 | 6.8176 | 6.8811 | 6.3764 | 6.8115 | 6.4954 | 5.9155 | 6.5475 | 6.7562 | 7.0520 | 6.9173 | 5.2567 | 5.8210 | 4.8475 | 4.9189 |
| PRRC2B_S480 | PRRC2B | Q5JSZ5 | 740 | KVTPIDSPPVWSP | 5.5584 | 6.7518 | 6.6108 | 6.3161 | 5.9213 | 6.2015 | 6.0489 | 6.7289 | 6.4348 | 6.2613 | 6.5923 | 6.7369 | 5.5614 | 6.4089 | 5.9836 | 5.8829 |
| PRRC2B_S556 | PRRC2B | Q5JSZ5 | 563 | PSAEKASPQENGP | 6.8467 | 6.7956 | 6.6851 | 6.7797 | 6.2625 | 6.1632 | 6.3068 | 6.3761 | 6.4271 | 6.6540 | 6.6958 | 6.6071 | 5.4028 | 5.7008 | 5.0531 | 5.2436 |
| PRRC2B_S563 | PRRC2B | Q5JSZ5 | 1808 | ASGPTGSPVVKLQ | 6.3817 | 7.1016 | 6.9178 | 6.6559 | 6.0769 | 6.7167 | 6.9606 | 6.8794 | 7.1937 | 7.5807 | 6.7194 | 7.2168 | 3.9497 | 4.8704 | 4.2708 | 4.5076 |
| PRRC2B_S613 | PRRC2B | Q5JSZ5 | 2163 | YRPSSASPSGKPS | 7.1550 | 6.9966 | 5.7953 | 5.7841 | 6.7009 | 6.8631 | 4.9097 | 8.4834 | 6.3292 | 6.2968 | 6.3055 | 6.7658 | 5.3612 | 6.0348 | 5.7167 | 4.5018 |
| PRRC2B_S740 | PRRC2B | Q5JSZ5 | 2153 | SGGPVPSPQTYRP | 6.1048 | 6.7895 | 5.5774 | 6.8407 | 5.7681 | 6.4609 | 6.6033 | 6.7176 | 7.8681 | 7.3392 | 6.8830 | 6.6730 | 4.2905 | 5.8311 | 5.2095 | 5.0434 |
| PRRC2B_S740;S745 | PRRC2B | Q5JSZ5 | 1691 | TSSQRSSPYGTLK | 5.9667 | 5.9667 | 6.1869 | 6.1558 | 6.4528 | 6.6923 | 6.0841 | 5.9524 | 6.5143 | 6.3981 | 7.0998 | 6.7395 | 5.6328 | 6.3761 | 5.7614 | 6.0204 |
| PRRC2B_S745 | PRRC2B | Q5JSZ5 | 221;226 | NA | 4.9459 | 7.6485 | 7.0549 | 5.6129 | 6.2920 | 6.4939 | 6.4524 | 6.7031 | 6.8563 | 6.2035 | 6.9695 | 6.1724 | 4.4994 | 5.4677 | 6.1670 | 6.4606 |
| PRRC2B_S853 | PRRC2B | Q5JSZ5 | 740;745 | NA | 7.0536 | 7.8859 | 7.6466 | 7.1036 | 5.7063 | 7.8078 | 6.8659 | 6.0171 | 7.7577 | 7.2124 | 8.2857 | 7.2729 | 3.0188 | 2.8977 | 3.1562 | 4.3119 |
| PRRC2B_S980 | PRRC2B | Q5JSZ5 | 2161;2163 | NA | 5.4154 | 6.0019 | 6.4098 | 5.8527 | 6.1634 | 6.0343 | 6.5360 | 6.9784 | 5.7536 | 6.5938 | 6.1220 | 6.4458 | 5.6552 | 7.2871 | 5.9731 | 6.7776 |
| PRRC2B_T221;S226 | PRRC2B | Q5JSZ5 | 1132;1136;11 | NA | 6.0993 | 6.3591 | 8.2566 | 6.2845 | 6.1749 | 6.4196 | 6.0554 | 5.1751 | 7.5757 | 6.7825 | 7.2108 | 6.8383 | 4.6398 | 5.7218 | 5.0865 | 5.3202 |
| PRRC2C_S1478 | PRRC2C | Q9Y520-7 | 1546 | IAKRSFSSQRPVD | 6.8073 | 6.5241 | 6.6033 | 6.1642 | 5.5473 | 6.2362 | 6.5821 | 6.4478 | 6.8482 | 6.2175 | 6.2439 | 6.3602 | 6.0151 | 6.2912 | 5.2552 | 5.8562 |
| PRRC2C_S1546 | PRRC2C | Q9Y520-7 | 2015 | KKLGPISPPQPPS | 6.3097 | 6.8786 | 6.9331 | 7.3824 | 6.2579 | 7.0267 | 6.4044 | 6.8525 | 6.8132 | 7.2869 | 7.0939 | 7.3018 | 4.0023 | 4.6504 | 4.1059 | 4.7002 |
| PRRC2C_S189 | PRRC2C | Q9Y520-7 | 2684 | IAGRSTTPTSSPF | 6.3117 | 6.1488 | 6.1661 | 5.7864 | 6.9187 | 7.1155 | 6.8339 | 7.1068 | 6.5139 | 6.2966 | 6.3062 | 6.0722 | 5.6913 | 5.9363 | 5.3304 | 5.4650 |
| PRRC2C_S2015 | PRRC2C | Q9Y520-7 | 2107 | QKLPDLSPVENKE | 7.2169 | 7.3483 | 6.8938 | 7.0797 | 6.8228 | 6.9942 | 7.2578 | 6.7349 | 6.5591 | 6.5776 | 6.1796 | 6.2042 | 4.5637 | 4.8507 | 4.1863 | 4.5305 |
| PRRC2C_S2107 | PRRC2C | Q9Y520-7 | 2696 | FRATSTSPNSQSS | 6.2438 | 6.4145 | 6.4209 | 6.3868 | 6.7459 | 6.7629 | 6.4728 | 6.9786 | 6.8163 | 6.7257 | 6.6431 | 6.6489 | 5.0322 | 5.4473 | 5.0655 | 5.1945 |
| PRRC2C_S2145 | PRRC2C | Q9Y520-7 | 2675 | IDIKPGTPPIAGR | 6.3146 | 6.5313 | 6.6748 | 6.4487 | 6.6900 | 7.2801 | 6.9699 | 7.1062 | 6.7023 | 6.8857 | 6.7950 | 6.4586 | 4.6976 | 5.3409 | 4.5723 | 4.5322 |
| PRRC2C_S2694 | PRRC2C | Q9Y520-7 | 824 | EPQQATTPKATEE | 6.1035 | 6.6194 | 6.6366 | 6.7275 | 6.3515 | 6.7931 | 6.6793 | 6.9737 | 6.3363 | 6.7532 | 6.3770 | 6.2515 | 5.1724 | 5.5481 | 5.2068 | 5.4700 |
| PRRC2C_S2694;S2696 | PRRC2C | Q9Y520-7 | 2694 | SPFRATSTSPNSQ | 6.3243 | 6.6987 | 6.7964 | 6.7289 | 6.3959 | 6.9267 | 6.3703 | 7.2249 | 6.4745 | 6.7652 | 6.7786 | 6.5583 | 4.8530 | 5.2791 | 4.9602 | 4.8651 |
| PRRC2C_S2696 | PRRC2C | Q9Y520-7 | 880 | QKFLSRSVEDVRP | 6.0699 | 6.9321 | 6.2386 | 6.7632 | 6.5534 | 6.9194 | 6.0402 | 7.1885 | 6.1170 | 5.9739 | 6.8614 | 6.1190 | 5.4060 | 5.7782 | 5.5691 | 5.4700 |
| PRRC2C_S502 | PRRC2C | Q9Y520-7 | 1982 | SGKSIQTPQSHGT | 6.5782 | 7.4334 | 6.3832 | 8.7042 | 5.7786 | 7.6599 | 6.7239 | 7.5119 | 6.3432 | 6.1504 | 6.8775 | 6.7747 | 3.9142 | 5.4162 | 3.7057 | 4.0450 |
| PRRC2C_S67 | PRRC2C | Q9Y520-7 | 502 | ILEKQPSPEEIRE | 6.3941 | 6.7905 | 6.2023 | 6.8267 | 6.1292 | 5.9261 | 6.1330 | 6.8361 | 6.4929 | 6.6663 | 6.5086 | 6.2260 | 5.2569 | 5.9599 | 5.9556 | 5.6959 |
| PRRC2C_S781 | PRRC2C | Q9Y520-7 | 1478 | NTLGDISGNKTPD | 6.4614 | 7.1608 | 6.8885 | 7.6083 | 6.6039 | 6.4323 | 7.4087 | 6.3741 | 6.3380 | 7.5245 | 7.3063 | 7.0177 | 3.7300 | 4.6488 | 3.8842 | 4.6127 |
| PRRC2C_S880 | PRRC2C | Q9Y520-7 | 189 | GGKAAGSPSSSDQ | 7.6211 | 7.6399 | 7.5713 | 7.3864 | 6.3711 | 6.8529 | 7.2962 | 6.8166 | 7.1880 | 6.9629 | 7.0085 | 6.4490 | 3.4238 | 4.1630 | 3.5180 | 3.7313 |
| PRRC2C_T1967 | PRRC2C | Q9Y520-7 | 2145 | EGQEKPSPATVRS | 6.6889 | 7.6650 | 7.0236 | 7.3252 | 6.2419 | 7.1660 | 6.8557 | 6.6205 | 7.2008 | 7.2381 | 7.1382 | 7.0589 | 3.4102 | 4.5599 | 3.6544 | 4.1526 |
| PRRC2C_T1982 | PRRC2C | Q9Y520-7 | 781 | RDQMEGSPNSSES | 5.5980 | 6.9534 | 6.3933 | 6.6308 | 6.4051 | 5.8887 | 6.1688 | 6.0825 | 6.5800 | 7.1543 | 7.3718 | 6.6816 | 5.5365 | 5.2434 | 5.4952 | 5.8166 |
| PRRC2C_T2675 | PRRC2C | Q9Y520-7 | 1967 | RLPSAQTPNGTDY | 7.2112 | 7.9702 | 6.5588 | 7.5395 | 6.9839 | 6.1450 | 8.0432 | 7.3290 | 6.8352 | 7.4988 | 6.8506 | 7.2940 | 3.6014 | 3.9157 | 3.2240 | 2.9995 |
| PRRC2C_T2684 | PRRC2C | A0A2R8Y7F4 | 67 | SKKCPISPPQPPS | 6.0512 | 4.8558 | 6.6597 | 6.9066 | 6.2658 | 6.6962 | 6.7178 | 6.1228 | 8.8630 | 6.7056 | 7.6204 | 8.6026 | 4.3283 | 6.0445 | 3.7346 | 3.8250 |
| PRRC2C_T2684;S2688 | PRRC2C | Q9Y520-7 | 2694;2696 | NA | 5.8483 | 6.8336 | 6.1898 | 6.2758 | 6.7875 | 7.4548 | 6.4066 | 6.8329 | 6.0479 | 7.4575 | 6.6613 | 6.3982 | 4.8942 | 5.8270 | 5.5881 | 4.4964 |
| PRRC2C_T824 | PRRC2C | Q9Y520-7 | 2684;2688 | NA | 6.9169 | 7.4345 | 6.3961 | 6.7937 | 6.1635 | 7.2221 | 7.4397 | 7.3798 | 7.5118 | 7.0587 | 7.0560 | 7.1555 | 3.5716 | 4.5838 | 3.3838 | 3.9326 |
| PRRT3_S815 | PRRT3 | Q5FWE3 | 815 | PINLSRSIDAALF | 6.6011 | 6.8355 | 6.1439 | 5.8201 | 5.8183 | 6.4320 | 6.0980 | 5.9989 | 6.0437 | 6.3012 | 6.1689 | 5.8311 | 6.3324 | 6.8959 | 6.2652 | 6.4140 |
| PSAT1_S331 | PSAT1 | Q9Y617 | 331 | LELNMLSLKGHRS | 6.4981 | 5.1765 | 6.7981 | 5.1080 | 7.1990 | 6.8473 | 6.4723 | 6.6533 | 6.4399 | 5.6744 | 5.4920 | 6.7593 | 5.9938 | 6.6417 | 6.4994 | 5.7468 |
| PSEN1_S367 | PSEN1 | P49768 | 43 | EHNDRRSLGHPEP | 5.9258 | 6.7293 | 6.5935 | 6.4278 | 5.9572 | 6.5535 | 6.5068 | 6.3262 | 5.9985 | 6.6777 | 6.3634 | 6.2102 | 5.5332 | 6.2363 | 5.6375 | 6.3231 |
| PSEN1_S43 | PSEN1 | P49768 | 367 | VQELSSSILAGED | 6.0060 | 6.5565 | 6.3742 | 6.6373 | 6.9652 | 5.6246 | 5.5945 | 6.9877 | 6.3085 | 6.5180 | 5.9139 | 6.5697 | 6.8928 | 5.8500 | 5.7166 | 5.4846 |
| PSIP1_S106 | PSIP1 | O75475 | 141 | KAVDITTPKAARR | 7.8415 | 7.9290 | 6.9531 | 7.5268 | 7.0507 | 6.9340 | 7.7799 | 6.9800 | 7.6055 | 7.2078 | 6.9957 | 6.8165 | 2.6825 | 3.8877 | 2.6164 | 3.1927 |
| PSIP1_S115;T122 | PSIP1 | O75475 | 129 | DHEEKASNEDVTK | 5.4944 | 5.8560 | 5.6735 | 6.2688 | 5.9047 | 5.5715 | 5.9082 | 5.7553 | 6.1069 | 6.2668 | 6.0075 | 6.1475 | 6.8988 | 7.3281 | 7.5451 | 7.2670 |
| PSIP1_S129 | PSIP1 | O75475 | 514 | TKKKPSSEERETE | 5.8126 | 6.8907 | 5.8736 | 7.1080 | 5.3254 | 5.4997 | 6.2483 | 5.6174 | 6.0420 | 6.3837 | 6.1948 | 6.5742 | 5.7438 | 6.8455 | 6.5940 | 7.2461 |
| PSIP1_S177 | PSIP1 | O75475 | 106 | KQSNASSDVEVEE | 5.8875 | 6.0960 | 6.1005 | 6.7858 | 6.4265 | 6.1856 | 6.8014 | 6.2526 | 5.8905 | 6.0339 | 5.8543 | 5.7907 | 6.1830 | 6.5216 | 6.6862 | 6.5039 |
| PSIP1_S514 | PSIP1 | O75475 | 177 | SVNLKVSPKRGRP | 5.8999 | 5.6197 | 6.2235 | 7.3835 | 7.1813 | 7.4382 | 6.4667 | 6.6155 | 6.3115 | 6.1454 | 6.9331 | 6.1966 | 5.8543 | 6.1167 | 5.2282 | 4.3858 |
| PSIP1_S522 | PSIP1 | O75475 | 522 | ERETEISLKDSTL | 6.3917 | 6.8689 | 6.4263 | 7.1048 | 6.2123 | 7.0669 | 7.5675 | 6.9868 | 5.9934 | 6.1720 | 6.6585 | 6.0457 | 4.7321 | 5.7796 | 4.7404 | 5.2530 |
| PSIP1_T141 | PSIP1 | O75475 | 212 | SESDIITEEDKSK | 6.1595 | 6.8266 | 7.8688 | 6.6926 | 6.6583 | 5.2201 | 6.2534 | 6.1000 | 6.0643 | 5.9987 | 6.8072 | 6.7194 | 5.5417 | 5.4270 | 5.6876 | 5.9749 |
| PSIP1_T212 | PSIP1 | O75475 | 271;275 | NA | 5.6290 | 5.5087 | 6.4206 | 6.2070 | 5.5673 | 6.0274 | 6.0843 | 5.4057 | 6.4634 | 6.4350 | 6.1377 | 5.9965 | 6.9200 | 7.3595 | 7.0383 | 6.7995 |
| PSIP1_T271;S275 | PSIP1 | O75475 | 115;122 | NA | 6.2789 | 6.5786 | 6.2054 | 7.1027 | 6.0510 | 6.6302 | 6.6864 | 6.1570 | 6.0720 | 6.1031 | 6.4635 | 6.6766 | 5.5619 | 6.2991 | 5.4571 | 5.6763 |
| PSMA1_S116 | PSMA1 | P25786-2 | 116 | PVSRLVSLIGSKT | 6.0842 | 6.1009 | 6.3641 | 5.7024 | 6.3374 | 6.2676 | 5.9163 | 6.3018 | 6.1194 | 6.0303 | 6.0840 | 6.2258 | 6.7765 | 6.4290 | 6.9034 | 6.3567 |
| PSMA2_T222 | PSMA2 | P25787 | 222 | AGFRRLTPTEVKD | 5.9967 | 6.4095 | 6.4199 | 6.2766 | 6.0763 | 6.5749 | 5.9688 | 6.5271 | 6.3180 | 5.6424 | 6.2789 | 6.3483 | 6.0606 | 6.4484 | 6.4701 | 6.1835 |
| PSMA3_S250 | PSMA3 | P25788 | 250 | LKEEDESDDDNMx | 5.7090 | 5.8407 | 6.3162 | 5.7934 | 6.6429 | 6.5840 | 6.1453 | 6.4187 | 6.4394 | 6.1564 | 6.4951 | 6.5548 | 6.2574 | 6.6500 | 6.1626 | 5.8341 |
| PSMA4_S13 | PSMA4 | P25789 | 13 | SRTTIFSPEGRLY | 7.8642 | 7.9211 | 7.3484 | 7.6183 | 5.3751 | 7.4911 | 7.9126 | 7.1474 | 8.3114 | 7.6998 | 8.2722 | 8.4630 | 1.6605 | 2.7004 | 1.5907 | 2.6238 |
| PSMA5_S16 | PSMA5 | P28066 | 56 | VEKRITSPLMEPS | 5.0238 | 4.1698 | 5.1434 | 4.3169 | 5.6425 | 5.4885 | 5.5255 | 6.4928 | 4.9493 | 5.0714 | 5.1927 | 5.8382 | 8.1280 | 9.4597 | 9.4812 | 10.0763 |
| PSMA5_S56 | PSMA5 | P28066 | 16 | RGVNTFSPEGRLF | 7.1786 | 6.9390 | 6.7866 | 5.8525 | 6.3327 | 7.1196 | 7.6742 | 7.5570 | 7.3763 | 7.7575 | 8.1304 | 8.3698 | 2.4490 | 3.7200 | 2.7452 | 4.0115 |
| PSMA6_S17 | PSMA6 | P60900 | 17 | RHITIFSPEGRLY | 7.0193 | 8.7027 | 6.7870 | 8.1207 | 6.0549 | 6.1415 | 7.0906 | 8.5448 | 6.2401 | 6.2949 | 6.9360 | 7.7306 | 3.1979 | 4.5864 | 2.6686 | 3.8839 |
| PSMA6_S63 | PSMA6 | P60900 | 63 | PDKLLDSSTVTHL | 6.3893 | 5.1504 | 6.3347 | 5.7793 | 6.2540 | 6.7121 | 5.8410 | 7.4502 | 6.6931 | 4.4170 | 6.5910 | 6.6907 | 5.8885 | 6.8023 | 7.6448 | 5.3616 |
| PSMB10_S230 | PSMB10 | P40306 | 230 | LLRTLSSPTEPVK | 5.9826 | 6.8958 | 6.9154 | 6.8417 | 6.1836 | 5.5316 | 5.7505 | 7.1090 | 6.5002 | 6.3511 | 6.4003 | 6.1339 | 6.1853 | 5.8591 | 5.9337 | 5.4261 |
| PSMC1_T434 | PSMC1 | P62191 | 434 | YKKQEGTPEGLYL | 6.0847 | 6.8751 | 7.3970 | 6.7054 | 6.3294 | 6.4069 | 6.5442 | 7.1432 | 6.3831 | 7.0145 | 6.1495 | 6.8756 | 4.4328 | 5.3626 | 4.9364 | 5.3595 |
| PSMC1_Y25 | PSMC1 | P62191 | 25 | KDKKKKYEPPVPT | 6.4290 | 5.8713 | 7.5516 | 5.5924 | 5.5031 | 6.5061 | 6.1774 | 6.4734 | 6.3971 | 6.4311 | 6.8471 | 7.0135 | 6.4688 | 6.4785 | 5.1127 | 5.1470 |
| PSMC4_S28 | PSMC4 | P43686 | 28 | RPQTGLSFLGPEP | 6.1275 | 6.2812 | 6.6297 | 6.6307 | 6.5063 | 6.8072 | 6.8622 | 6.5613 | 6.5891 | 6.2168 | 6.5314 | 6.4137 | 5.5750 | 5.6867 | 5.1483 | 5.4328 |
| PSMD1_T273 | PSMD1 | Q99460 | 273 | NLRTVGTPIASVP | 5.5723 | 5.9354 | 6.0307 | 5.9165 | 6.0636 | 6.4941 | 6.0573 | 6.0305 | 5.8977 | 6.3970 | 6.1680 | 6.2084 | 6.5199 | 6.8300 | 6.9973 | 6.8813 |
| PSMD1_T311 | PSMD1 | Q99460 | 311 | SAFVGKTPEASPE | 5.7941 | 6.6185 | 6.6026 | 6.4912 | 6.1148 | 6.1552 | 6.5238 | 6.3868 | 6.0329 | 6.2651 | 6.5592 | 6.1732 | 5.9663 | 6.0960 | 6.2172 | 6.0031 |
| PSMD1_T311;S315 | PSMD1 | Q99460 | 830 | EKEKVSTAVLSIT | 6.3517 | 6.5888 | 6.9432 | 6.6010 | 6.7037 | 5.8165 | 6.6303 | 5.6797 | 6.8625 | 7.1216 | 5.5721 | 5.8434 | 6.3053 | 6.0340 | 6.2711 | 4.6752 |
| PSMD1_T830 | PSMD1 | Q99460 | 311;315 | NA | 5.8148 | 5.5998 | 6.5147 | 6.3519 | 6.2190 | 6.1414 | 6.3065 | 5.4829 | 5.8530 | 6.4405 | 6.4115 | 6.2016 | 7.2830 | 6.3372 | 6.3633 | 6.6788 |
| PSMD11_S14 | PSMD11 | O00231-2 | 23 | STDREASIDILHS | 4.8771 | 5.6911 | 8.2365 | 7.6406 | 7.5214 | 6.7682 | 7.0627 | 7.1212 | 6.9930 | 8.2050 | 7.3550 | 5.3268 | 2.9771 | 4.3532 | 4.8389 | 5.0323 |
| PSMD11_S23 | PSMD11 | O00231-2 | 14 | EFQRAQSLLSTDR | 5.5231 | 5.3151 | 6.6879 | 6.1922 | 6.3151 | 5.6541 | 5.3136 | 5.8321 | 5.9474 | 6.6405 | 6.2038 | 5.6519 | 7.4312 | 7.2099 | 7.2872 | 6.7951 |
| PSMD2_S16 | PSMD2 | Q13200 | 16 | PVQPQQSPAAAPG | 5.7867 | 6.1124 | 6.4337 | 6.3956 | 6.2605 | 6.7519 | 6.1578 | 6.5249 | 6.4438 | 6.4835 | 6.4123 | 6.2264 | 6.0323 | 6.2514 | 5.7328 | 5.9941 |
| PSMD8_T295 | PSMD8 | P48556 | 295 | RILFFNTPKKMTD | 5.0994 | 6.0106 | 7.4749 | 5.4332 | 6.6027 | 5.3430 | 6.5101 | 7.3165 | 5.4478 | 6.2080 | 6.6611 | 8.0085 | 5.3236 | 6.0179 | 7.2118 | 5.3309 |
| PSME2_S10 | PSME2 | Q9UL46 | 10 | PCGVRLSGEARKQ | 6.0105 | 5.6275 | 7.3393 | 6.4035 | 6.8394 | 7.7691 | 6.6096 | 5.5442 | 5.5573 | 6.8611 | 7.0971 | 6.6944 | 5.5883 | 6.1584 | 4.8305 | 5.0700 |
| PSME3_S24 | PSME3 | P61289-2 | 24 | FRERITSEAEDLV | 5.1312 | 5.5091 | 5.7499 | 5.2444 | 5.5639 | 6.0989 | 5.6096 | 5.8397 | 5.9514 | 5.4848 | 6.0275 | 5.6912 | 7.6425 | 8.4668 | 8.2963 | 7.6929 |
| PSME4_S1121 | PSME4 | Q14997 | 1121 | INQILLSPEKIKE | 5.7875 | 6.4264 | 6.4147 | 5.8925 | 5.6359 | 6.8239 | 5.9239 | 6.3914 | 5.9427 | 6.6803 | 5.9293 | 6.5861 | 6.8562 | 6.8014 | 6.6372 | 5.2707 |
| PSMF1_S153 | PSMF1 | Q92530 | 252 | FGPIGTSPPGPNP | 6.0576 | 6.4220 | 7.4328 | 6.4372 | 6.6037 | 6.6606 | 6.5749 | 7.5151 | 7.2332 | 7.7398 | 7.8405 | 7.9270 | 3.2468 | 4.5484 | 3.5885 | 4.1719 |
| PSMF1_S252 | PSMF1 | Q92530 | 153 | EKANVSSPHREFP | 5.6318 | 5.7962 | 6.1727 | 5.6183 | 6.1647 | 6.4341 | 6.3398 | 7.5774 | 6.5499 | 7.0580 | 7.1521 | 7.7020 | 4.2570 | 5.7105 | 5.4503 | 6.3851 |
| PSMF1_S252 | PSMF1 | Q92530 | 251 | PFGPIGTSPPGPN | 6.0219 | 6.3096 | 7.4663 | 6.4063 | 6.6267 | 6.9474 | 6.7546 | 7.3651 | 7.3362 | 7.3572 | 7.8929 | 8.5138 | 3.2841 | 4.3783 | 3.3317 | 4.0079 |
| PSMF1_T251 | PSMF1 | Q5QPM7 | 252 | FGPIGTSPPGPRP | 6.2872 | 6.4541 | 7.1575 | 6.5255 | 6.3300 | 6.6708 | 6.8629 | 6.7800 | 6.2837 | 6.7867 | 7.1278 | 7.4624 | 4.9283 | 5.1316 | 4.4861 | 4.7256 |
| PSPC1_S164 | PSPC1 | Q8WXF1 | 477 | PPSQMGSPMGSRT | 6.3225 | 6.3148 | 6.3356 | 6.4928 | 6.5753 | 6.1165 | 6.7765 | 6.9362 | 7.1793 | 6.6853 | 6.8158 | 6.6711 | 4.9743 | 5.3843 | 4.9875 | 5.4323 |
| PSPC1_S477 | PSPC1 | Q8WXF1 | 164 | LTVKNLSPVVSNE | 7.5104 | 7.0385 | 6.1535 | 5.3575 | 6.6595 | 7.3114 | 6.5306 | 6.0815 | 6.6424 | 5.2419 | 5.9810 | 7.7738 | 4.9937 | 6.7959 | 5.4011 | 4.5275 |
| PSRC1_S122 | PSRC1 | Q6PGN9-2 | 122 | TFVLKDSPVRDLL | 6.9044 | 6.6926 | 7.2346 | 6.9255 | 6.5580 | 7.4123 | 7.2806 | 6.5152 | 6.9580 | 6.9229 | 7.9378 | 7.5083 | 3.5626 | 4.1578 | 3.2651 | 4.1644 |
| PSRC1_S140 | PSRC1 | Q6PGN9-2 | 212 | SGRAAASPPTPIR | 6.0403 | 6.3028 | 7.1382 | 6.6701 | 6.0577 | 7.0083 | 6.7104 | 7.2222 | 6.5702 | 6.8957 | 6.7711 | 6.7766 | 4.4039 | 5.2842 | 4.9619 | 5.1865 |
| PSRC1_S150 | PSRC1 | Q6PGN9-2 | 65 | PQGVRLSLGPLSP | 6.4553 | 7.3209 | 6.5136 | 8.3554 | 7.2379 | 5.8164 | 6.1133 | 6.6050 | 6.9709 | 6.3763 | 6.4326 | 6.5152 | 4.6825 | 5.2524 | 4.8031 | 4.5492 |
| PSRC1_S175 | PSRC1 | Q6PGN9-2 | 175 | SNMKRESPTCNLF | 6.5291 | 6.9118 | 6.7814 | 6.8478 | 6.4995 | 6.9293 | 6.8543 | 6.7306 | 7.3645 | 7.3594 | 7.4550 | 7.1488 | 4.5343 | 5.1954 | 3.2096 | 3.6491 |
| PSRC1_S186 | PSRC1 | Q6PGN9-2 | 150 | LTPRLRSNDRKGS | 5.7947 | 6.2901 | 7.3935 | 6.3744 | 5.3984 | 7.0461 | 6.4049 | 6.5357 | 6.3675 | 7.1059 | 7.9416 | 8.4144 | 4.3855 | 5.1406 | 4.2810 | 5.1257 |
| PSRC1_S212 | PSRC1 | Q6PGN9-2 | 116 | PSPRRETFVLKDS | 11.8442 | 7.8861 | 7.2420 | 10.5557 | 7.0290 | 4.0143 | 5.5950 | 4.2478 | 7.3366 | 5.2454 | 5.8469 | 5.5122 | 4.8414 | 4.8909 | 3.5861 | 4.3264 |
| PSRC1_S65 | PSRC1 | Q6PGN9-2 | 70 | LSLGPLSPEKLEE | 6.6147 | 5.5141 | 7.8977 | 6.3326 | 6.2437 | 5.9201 | 6.2887 | 7.0149 | 7.6591 | 7.8161 | 6.9349 | 7.2312 | 3.8390 | 4.3208 | 4.6729 | 5.6996 |
| PSRC1_S70 | PSRC1 | Q6PGN9-2 | 186 | LFPASKSPASSPL | 6.6886 | 7.5650 | 6.4543 | 5.5062 | 6.9490 | 7.1247 | 6.3340 | 6.9751 | 6.5578 | 6.6865 | 7.2178 | 5.9611 | 5.3833 | 5.2238 | 4.9707 | 4.4022 |
| PSRC1_T116 | PSRC1 | Q6PGN9-2 | 140 | LTRSTPSPSSLTP | 6.1565 | 6.1966 | 6.5803 | 6.4672 | 6.2169 | 6.7566 | 6.9612 | 6.9967 | 6.8785 | 6.8988 | 6.4995 | 6.6255 | 4.8374 | 5.4773 | 5.1209 | 5.3301 |
| PTAR1_S45 | PTAR1 | Q7Z6K3 | 45 | EARYNRSPIVLVE | 7.0718 | 7.7353 | 6.8918 | 7.2623 | 6.1732 | 7.1914 | 6.5201 | 7.0810 | 7.3109 | 6.3125 | 7.2071 | 7.3044 | 4.1304 | 3.8900 | 3.6180 | 4.2998 |
| PTBP1_S141 | PTBP1 | P26599-3 | 16 | VGTKRGSDELFST | 5.6434 | 5.8479 | 6.1351 | 5.7257 | 6.2320 | 6.3369 | 5.8524 | 5.8355 | 6.3605 | 6.3669 | 6.4859 | 6.8045 | 6.4471 | 6.7124 | 6.4144 | 6.7994 |
| PTBP1_S16 | PTBP1 | P26599-3 | 141 | ELKTDSSPNQARA | 5.8773 | 6.0258 | 6.4915 | 6.7201 | 5.5752 | 5.8116 | 5.9522 | 6.1953 | 6.4107 | 6.0488 | 6.5132 | 7.0768 | 5.9667 | 6.4285 | 6.4896 | 6.4166 |
| PTBP1_S53 | PTBP1 | P26599-3 | 53 | FKGDSRSAGVPSR | 5.1720 | 6.7677 | 5.0787 | 5.3460 | 5.6649 | 6.2572 | 6.2539 | 5.3223 | 5.5971 | 5.8245 | 5.8941 | 6.3870 | 7.7836 | 7.2674 | 7.2147 | 8.1689 |
| PTBP3_S62 | PTBP3 | O95758-4 | 62 | RDRPPCSPSRVLH | 6.5144 | 6.4915 | 6.3186 | 6.7594 | 6.2982 | 6.4051 | 6.4021 | 5.9231 | 6.3483 | 6.4370 | 6.3521 | 6.4310 | 5.5666 | 5.9817 | 5.6316 | 6.1393 |
| PTDSS1_S442 | PTDSS1 | P48651 | 442 | TKGSEDSPPKHAG | 6.3631 | 6.6365 | 6.2623 | 7.1605 | 5.6201 | 5.8364 | 6.4191 | 6.0842 | 6.0960 | 6.2807 | 6.3014 | 6.5487 | 5.5669 | 6.0788 | 6.2239 | 6.5213 |
| PTDSS2_S16 | PTDSS2 | Q9BVG9 | 16 | GGPRPESPVPAGR | 6.5161 | 7.0057 | 6.0226 | 6.5227 | 6.2366 | 6.0448 | 6.6903 | 6.3089 | 6.4275 | 6.3232 | 6.1944 | 6.4177 | 5.7713 | 6.2857 | 5.5575 | 5.6750 |
| PTGES3_S113 | PTGES3 | Q15185 | 113 | KDWEDDSDEDMSN | 5.9737 | 6.1384 | 6.6566 | 5.9860 | 6.1956 | 6.2946 | 5.9043 | 6.4388 | 6.3425 | 6.3978 | 6.4203 | 6.5720 | 6.5969 | 6.2727 | 5.7818 | 6.0280 |
| PTGES3_S148;S151 | PTGES3 | Q15185 | 151 | DDDSQDSDDEKMP | 5.9641 | 6.1289 | 6.7512 | 5.9457 | 6.0190 | 6.0621 | 5.5150 | 6.0400 | 6.8471 | 6.9544 | 7.2198 | 7.0020 | 6.1620 | 6.3045 | 5.4049 | 5.6791 |
| PTGES3_S151 | PTGES3 | Q15185 | 148;151 | NA | 6.0772 | 5.9617 | 6.6398 | 5.9100 | 6.2345 | 6.1977 | 5.9674 | 6.2113 | 6.3816 | 6.5715 | 6.4942 | 6.5797 | 6.6253 | 6.3001 | 5.8212 | 6.0269 |
| PTK2_S29 | PTK2 | Q05397 | 722 | SRPGYPSPRSSEG | 6.3211 | 5.9394 | 7.1152 | 6.2355 | 6.4813 | 6.5245 | 6.4527 | 6.7058 | 7.0753 | 6.3476 | 7.0322 | 6.9495 | 5.1133 | 5.4433 | 4.9531 | 5.3102 |
| PTK2_S722 | PTK2 | Q05397 | 397 | VSETDDYAEIIDE | 6.0457 | 5.7755 | 6.6308 | 6.5627 | 6.3598 | 6.0859 | 5.7636 | 6.1001 | 6.4713 | 6.1718 | 6.2145 | 6.6779 | 6.4012 | 6.3152 | 6.0996 | 6.3243 |
| PTK2_S840;S843 | PTK2 | Q05397 | 575 | RYMEDSTYYKASK | 6.2388 | 5.5776 | 6.5413 | 5.9141 | 6.9169 | 6.2429 | 5.4974 | 6.3707 | 7.3831 | 6.2045 | 6.2706 | 6.6635 | 6.4054 | 6.3836 | 5.6946 | 5.6950 |
| PTK2_S843 | PTK2 | Q05397 | 843 | VRLSRGSIDREDG | 5.8186 | 6.3220 | 6.4671 | 5.7316 | 6.5308 | 5.7540 | 6.2120 | 6.6096 | 6.4205 | 7.2033 | 7.0214 | 6.1519 | 6.1125 | 6.9314 | 5.5996 | 5.1137 |
| PTK2_S910 | PTK2 | Q05397 | 910 | LQPQEISPPPTAN | 5.6504 | 5.9907 | 6.5150 | 6.0910 | 5.8978 | 6.6545 | 6.4858 | 6.9098 | 6.0030 | 6.2328 | 5.5993 | 6.5588 | 6.0372 | 7.0449 | 6.1783 | 6.1505 |
| PTK2_T575 | PTK2 | Q05397 | 29 | GTGMERSPGAMER | 6.6789 | 7.6515 | 6.9862 | 7.4185 | 6.0921 | 6.8099 | 6.3804 | 6.7418 | 6.5832 | 7.8716 | 6.8831 | 7.0403 | 3.8108 | 4.6081 | 3.9764 | 4.4671 |
| PTK2_Y397 | PTK2 | Q05397 | 840;843 | NA | 6.9015 | 6.1541 | 7.8791 | 6.2822 | 6.9657 | 6.3434 | 6.9445 | 7.3233 | 6.2342 | 5.7399 | 6.8541 | 7.1933 | 4.8575 | 4.8488 | 4.2273 | 5.2511 |
| PTMA_S84 | PTMA | P06454-2 | 84 | EDEEAESATGKRA | 5.5606 | 6.3300 | 6.4255 | 6.2836 | 7.0394 | 5.6642 | 7.0312 | 5.8552 | 6.5155 | 6.0189 | 6.0026 | 6.1363 | 7.5196 | 6.3218 | 5.9180 | 5.3778 |
| PTMS_S5 | PTMS | P20962 | 5 | xxMSEKSVEAAAE | 6.0401 | 6.1384 | 6.3458 | 6.5599 | 6.3808 | 6.5882 | 5.6694 | 6.2539 | 6.2648 | 6.0905 | 6.4053 | 6.2945 | 6.5845 | 6.9881 | 6.1707 | 6.1250 |
| PTMS_T52 | PTMS | P20962 | 52 | EEEEEETAEDGEE | 5.5119 | 5.2753 | 6.8767 | 6.0147 | 7.0707 | 5.3715 | 5.4106 | 8.0985 | 5.7731 | 5.2753 | 4.9639 | 5.8182 | 6.6912 | 7.8637 | 6.6469 | 7.3378 |
| PTOV1_S36 | PTOV1 | Q86YD1 | 36 | RAVRSRSWPASPR | 6.9139 | 7.1976 | 7.3592 | 7.0439 | 6.4921 | 7.3002 | 6.6472 | 6.5141 | 6.8392 | 6.7199 | 7.2703 | 7.3071 | 3.7718 | 4.8237 | 3.7050 | 4.0948 |
| PTOV1_S53 | PTOV1 | Q86YD1 | 53 | PRIRARSAPPMEG | 6.8641 | 7.0736 | 7.2540 | 6.1741 | 6.6995 | 6.5915 | 6.5305 | 6.7393 | 7.2432 | 7.0331 | 7.3554 | 7.6688 | 3.7211 | 4.6588 | 3.9395 | 4.4535 |
| PTPDC1_S446 | PTPDC1 | A2A3K4-2 | 446 | KRRLSYSDSDLKR | 6.8326 | 6.5264 | 7.2238 | 6.0935 | 5.6136 | 6.6470 | 6.5939 | 6.6572 | 6.3561 | 6.3866 | 6.5467 | 6.5236 | 5.5469 | 5.5439 | 5.2971 | 5.6111 |
| PTPN1_S352 | PTPN1 | P18031 | 386 | RGAQAASPAKGEP | 7.5200 | 8.0713 | 6.7917 | 8.2057 | 6.3085 | 6.8231 | 7.3740 | 6.9847 | 7.1187 | 7.4804 | 7.6268 | 7.3864 | 2.5374 | 3.5876 | 2.5960 | 3.5879 |
| PTPN1_S378 | PTPN1 | P18031 | 378 | SRVVGGSLRGAQA | 5.6440 | 5.9974 | 6.7248 | 6.4012 | 5.5860 | 5.9109 | 5.8373 | 6.0675 | 6.2638 | 7.3242 | 7.1204 | 6.7860 | 5.9092 | 6.3073 | 6.0902 | 6.0300 |
| PTPN1_S386 | PTPN1 | P18031 | 50 | NRYRDVSPFDHSR | 5.9401 | 6.0393 | 5.0298 | 5.4061 | 6.4033 | 6.1654 | 6.0963 | 5.2547 | 6.0428 | 6.0852 | 5.5474 | 5.6851 | 7.8155 | 7.9605 | 7.5324 | 6.9960 |
| PTPN1_S50 | PTPN1 | P18031 | 352 | IKEEKGSPLNAAP | 6.3874 | 6.2559 | 6.6430 | 7.2352 | 5.6246 | 6.0823 | 7.3262 | 6.4700 | 6.2686 | 6.5785 | 6.9252 | 6.0503 | 5.3924 | 5.8977 | 4.7135 | 6.1492 |
| PTPN11_T557 | PTPN11 | Q06124-1 | 557 | YSLADQTSGDQSP | 6.5044 | 6.7220 | 7.2995 | 6.4587 | 6.0186 | 6.0351 | 6.3296 | 6.5011 | 7.2995 | 6.7052 | 6.3094 | 6.6734 | 4.9202 | 5.7842 | 5.1397 | 5.2996 |
| PTPN11_Y584 | PTPN11 | Q06124-1 | 584 | EDSARVYENVGLM | 8.8592 | 6.8705 | 6.8380 | 7.0773 | 6.6040 | 6.7528 | 6.0606 | 7.2545 | 8.9957 | 6.6955 | 6.7889 | 6.7495 | 3.5451 | 4.3652 | 2.9695 | 3.5735 |
| PTPN12_S332 | PTPN12 | Q05209 | 435 | KLERNLSFEIKKV | 5.1883 | 4.7883 | 5.5112 | 4.4100 | 5.6833 | 5.5126 | 5.2269 | 5.8069 | 5.2097 | 5.3106 | 5.1096 | 4.9417 | 9.7629 | 9.1115 | 9.6556 | 8.7710 |
| PTPN12_S369 | PTPN12 | Q05209 | 332 | EPEKQDSPPPKPP | 6.4739 | 6.5130 | 7.0217 | 6.1616 | 6.0160 | 6.3867 | 6.0664 | 6.1910 | 6.6285 | 6.7861 | 6.8408 | 6.8114 | 5.5841 | 5.8435 | 5.1973 | 5.4780 |
| PTPN12_S435 | PTPN12 | Q05209 | 369 | PPILTPSPPSAFP | 6.1571 | 5.8364 | 6.4063 | 6.2108 | 6.0209 | 6.5761 | 5.9939 | 6.5692 | 6.9343 | 6.1670 | 6.5537 | 6.5644 | 5.8121 | 6.3119 | 5.8348 | 6.0512 |
| PTPN12_S468 | PTPN12 | Q05209 | 468 | IKIKSASPCIADK | 6.4729 | 7.1137 | 6.9731 | 7.4069 | 5.9684 | 7.2800 | 7.4686 | 6.8175 | 7.2998 | 7.1756 | 7.4780 | 7.6024 | 3.2476 | 4.3140 | 3.2473 | 4.1339 |
| PTPN12_S567 | PTPN12 | Q05209 | 571 | TVSLTPSPTTQVE | 5.6978 | 6.0002 | 6.5052 | 6.0470 | 6.0059 | 6.6587 | 5.8407 | 6.8706 | 6.2314 | 6.4241 | 6.4271 | 6.0504 | 6.2801 | 6.6346 | 6.0047 | 6.3215 |
| PTPN12_S571;T573 | PTPN12 | Q05209 | 567 | QTRKTVSLTPSPT | 5.6075 | 5.7358 | 6.8218 | 6.1242 | 6.3763 | 6.7089 | 5.8480 | 6.6861 | 6.3897 | 6.0658 | 6.1048 | 5.6888 | 6.4784 | 7.1565 | 5.9885 | 6.2190 |
| PTPN12_S603;S606 | PTPN12 | Q05209 | 573 | SLTPSPTTQVETP | 5.2941 | 5.8019 | 6.7045 | 6.3361 | 6.2496 | 5.9423 | 6.0104 | 6.5601 | 6.5913 | 6.5503 | 6.8733 | 6.5370 | 6.2982 | 6.4666 | 5.4509 | 6.3334 |
| PTPN12_S603;S606;S608 | PTPN12 | Q05209 | 693 | LASEHNTPVRSEW | 6.1580 | 6.9309 | 5.8534 | 6.4450 | 6.0607 | 7.1465 | 6.7636 | 6.5416 | 6.7272 | 6.8021 | 6.5234 | 7.4286 | 5.0854 | 5.6644 | 4.1760 | 5.6931 |
| PTPN12_T519 | PTPN12 | Q05209 | 519 | ESQNSDTPPRPDR | 6.5589 | 6.3093 | 6.8902 | 7.6233 | 6.2336 | 6.3542 | 6.8505 | 6.1751 | 6.9794 | 6.7260 | 6.9337 | 7.6665 | 4.0193 | 5.5155 | 5.0912 | 4.9735 |
| PTPN12_T569 | PTPN12 | Q05209 | 569 | RKTVSLTPSPTTQ | 6.3248 | 5.2472 | 6.7778 | 5.7073 | 7.4667 | 6.4047 | 6.1651 | 6.2294 | 5.8611 | 5.7812 | 6.0971 | 6.6526 | 6.9446 | 6.3737 | 6.1436 | 5.8230 |
| PTPN12_T569;S571 | PTPN12 | Q05209 | 588 | VDHDNTSPLFRTP | 6.3625 | 6.1063 | 6.1670 | 5.4208 | 7.3132 | 7.6076 | 6.9154 | 6.5288 | 6.9895 | 5.3984 | 6.8232 | 6.5378 | 7.0705 | 5.7714 | 4.2949 | 4.6927 |
| PTPN12_T569;T573 | PTPN12 | Q05209 | 603;606 | NA | 6.4715 | 6.5639 | 7.0383 | 6.3236 | 6.6703 | 6.9469 | 5.7722 | 6.2135 | 6.5145 | 6.4907 | 6.9643 | 6.7157 | 5.5434 | 5.7850 | 5.1415 | 4.8446 |
| PTPN12_T571 | PTPN12 | Q05209 | 569;571 | NA | 6.1610 | 6.9245 | 6.6650 | 5.9091 | 5.9384 | 6.9079 | 6.2140 | 6.1347 | 6.5838 | 5.9353 | 5.9767 | 6.5549 | 6.2868 | 5.9885 | 6.2453 | 5.5743 |
| PTPN12_T573 | PTPN12 | Q05209 | 569;573 | NA | 6.5485 | 6.7812 | 6.7185 | 5.6613 | 6.2697 | 6.5343 | 5.9253 | 6.4879 | 6.4214 | 5.8377 | 6.3917 | 6.4865 | 6.0153 | 6.0262 | 5.5807 | 6.3137 |
| PTPN12_T573;S588 | PTPN12 | Q05209 | 603;606;608 | NA | 5.6046 | 5.7919 | 6.3640 | 5.7641 | 6.4652 | 6.5177 | 5.7573 | 6.9970 | 6.5047 | 6.4335 | 6.8826 | 6.4424 | 6.2173 | 6.5648 | 6.0133 | 5.6797 |

| | | | | | | | | | | | | | | | | | | | | |
|---|---|---|---|---|---|---|---|---|---|---|---|---|---|---|---|---|---|---|---|---|
| PTPN12_T588 | PTPN12 | Q05209 | 573;588 | NA | 5.5828 | 5.1932 | 6.4619 | 5.4124 | 5.5234 | 6.0768 | 5.7373 | 6.5588 | 6.4008 | 6.2593 | 6.4919 | 5.9723 | 7.9768 | 7.2054 | 6.6652 | 6.4815 |
| PTPN12_T693 | PTPN12 | Q05209 | 571;573 | NA | 6.4275 | 5.1958 | 5.5422 | 5.9323 | 6.8128 | 6.2618 | 6.7339 | 7.1390 | 6.0624 | 5.4280 | 5.9565 | 6.0162 | 6.6352 | 6.6421 | 6.4881 | 6.7262 |
| PTPN13_S1082 | PTPN13 | Q12923-4 | 240 | NKGLSKSMGFLSI | 7.2160 | 6.7214 | 6.6555 | 7.0616 | 5.7911 | 6.0553 | 6.3669 | 5.5337 | 5.9620 | 5.6888 | 6.6252 | 6.5360 | 5.6759 | 6.2792 | 5.4453 | 6.3863 |
| PTPN13_S1082;S1085 | PTPN13 | Q12923-4 | 938 | LKRLSCSELSLYQ | 6.2555 | 5.9577 | 5.9331 | 6.5218 | 6.3647 | 6.3785 | 6.7314 | 5.8963 | 6.5723 | 6.3520 | 6.3724 | 6.7580 | 6.0755 | 6.2097 | 5.3984 | 6.2228 |
| PTPN13_S1358 | PTPN13 | Q12923-4 | 2037 | TYQIKGSPNLTLP | 6.5028 | 6.6598 | 6.3876 | 6.6001 | 6.6202 | 6.4551 | 6.6772 | 6.8403 | 6.5500 | 6.6855 | 6.6172 | 6.6338 | 5.0339 | 5.3857 | 5.1105 | 5.2404 |
| PTPN13_S1519 | PTPN13 | Q12923-4 | 908 | NMGRAISTGSLAS | 5.0211 | 5.9547 | 6.3363 | 6.0467 | 5.3332 | 6.8264 | 6.1473 | 6.3746 | 6.2716 | 6.6311 | 6.9468 | 7.6584 | 5.4977 | 6.2740 | 5.6995 | 6.9806 |
| PTPN13_S2037 | PTPN13 | Q12923-4 | 1082 | VLHKRWSIVSSPE | 7.1731 | 6.3167 | 6.1112 | 6.2330 | 6.8455 | 5.8915 | 5.6643 | 5.9311 | 7.0318 | 6.3828 | 6.1752 | 6.2720 | 6.4155 | 6.9504 | 5.4205 | 5.1855 |
| PTPN13_S240 | PTPN13 | Q12923-4 | 345 | KKEARYSDGSIAL | 7.1595 | 5.8145 | 7.0977 | 6.4675 | 6.4974 | 5.8869 | 5.7333 | 6.0419 | 6.6894 | 6.8608 | 6.8691 | 6.9346 | 5.6716 | 6.2962 | 4.7470 | 5.2326 |
| PTPN13_S345 | PTPN13 | Q12923-4 | 1358 | NFKTFSSSPPKPG | 6.1290 | 6.4433 | 7.2065 | 6.9970 | 6.3333 | 5.9232 | 6.3016 | 6.2744 | 7.0088 | 6.6326 | 6.5746 | 7.1691 | 5.3805 | 5.6255 | 4.8607 | 5.1397 |
| PTPN13_S908 | PTPN13 | Q12923-4 | 1519 | SSGLGFSFSREDN | 5.8615 | 5.6379 | 5.6391 | 5.9538 | 5.6496 | 5.4071 | 6.3393 | 6.0402 | 6.1003 | 5.1814 | 5.7264 | 5.3268 | 7.9446 | 7.3559 | 8.4171 | 7.4188 |
| PTPN13_S908;S911 | PTPN13 | Q12923-4 | 1082;1085 | NA | 6.8139 | 6.7066 | 6.5538 | 6.7676 | 6.6213 | 5.7565 | 7.1832 | 6.2891 | 6.3980 | 5.7297 | 5.6123 | 5.5141 | 6.1463 | 6.1349 | 6.2346 | 5.5380 |
| PTPN13_S938 | PTPN13 | Q12923-4 | 908;911 | NA | 7.6959 | 6.3689 | 6.9179 | 7.7650 | 5.2649 | 5.4091 | 6.5644 | 5.4334 | 5.8743 | 5.4981 | 6.6823 | 6.0914 | 5.8832 | 6.6895 | 5.2829 | 6.5787 |
| PTPN14_S314 | PTPN14 | Q15678 | 594 | KYVSGSSPDLVTR | 6.6063 | 6.7484 | 6.7463 | 6.4548 | 6.0270 | 6.4185 | 6.2683 | 6.3085 | 6.8951 | 7.0812 | 6.8739 | 6.7303 | 4.9311 | 5.5831 | 5.0265 | 5.3007 |
| PTPN14_S486 | PTPN14 | Q15678 | 578 | RPRPATSTPDLAS | 6.0084 | 5.4121 | 6.7409 | 5.9856 | 6.0284 | 6.6886 | 6.0316 | 5.6536 | 6.2007 | 6.4254 | 6.5549 | 6.6391 | 6.6562 | 6.4611 | 5.9570 | 6.5566 |
| PTPN14_S512 | PTPN14 | Q15678 | 831 | VKERPVSEMFSLE | 6.7911 | 5.8290 | 6.8296 | 6.2154 | 5.9706 | 6.4555 | 5.7471 | 6.0507 | 6.7941 | 5.9545 | 6.1270 | 6.5246 | 6.4993 | 6.6307 | 5.5296 | 6.0512 |
| PTPN14_S578 | PTPN14 | Q15678 | 642 | TVNKRHSLEVMNS | 6.3358 | 6.1841 | 6.6903 | 5.8892 | 6.3099 | 6.2495 | 5.7895 | 6.3738 | 7.0679 | 6.4297 | 6.3140 | 6.6101 | 6.1437 | 6.2516 | 5.7985 | 5.5625 |
| PTPN14_S594 | PTPN14 | Q15678 | 486 | PEDLVYSQPEMRE | 5.9984 | 6.1692 | 6.8703 | 6.3337 | 5.9747 | 6.2636 | 6.3347 | 6.5230 | 6.5003 | 6.1675 | 6.7986 | 6.7151 | 6.1266 | 5.9759 | 5.5220 | 5.7264 |
| PTPN14_S620 | PTPN14 | Q15678 | 809 | SLGPSISEPDLTS | 6.0611 | 6.0458 | 6.5467 | 5.6545 | 6.2220 | 6.2903 | 6.1023 | 6.2978 | 6.3695 | 6.5656 | 6.5799 | 6.5922 | 6.3339 | 6.4680 | 5.8894 | 5.9807 |
| PTPN14_S642 | PTPN14 | Q15678 | 620 | SPVVHQSLQEVSE | 6.5664 | 5.8054 | 6.1753 | 6.0498 | 6.1114 | 6.2537 | 5.8652 | 5.6305 | 5.9369 | 5.0542 | 5.9108 | 5.6523 | 7.1179 | 7.6486 | 7.2204 | 7.0011 |
| PTPN14_S760 | PTPN14 | Q15678 | 314 | CTEQSNSPPPIRR | 6.0798 | 6.3333 | 6.5423 | 6.5320 | 6.3074 | 6.4999 | 6.7659 | 5.7631 | 6.8849 | 6.7679 | 6.7338 | 6.2464 | 5.1308 | 5.4630 | 5.5996 | 6.3499 |
| PTPN14_S809 | PTPN14 | Q15678 | 760 | GPRKSVSNGALRQ | 6.3139 | 6.0624 | 6.5610 | 6.2255 | 5.6283 | 6.0545 | 5.7899 | 6.4210 | 6.5238 | 6.1227 | 6.5782 | 6.7237 | 6.3000 | 6.4345 | 6.0892 | 6.1713 |
| PTPN14_S831 | PTPN14 | Q15678 | 512 | YSNKLVSPSDQRN | 6.9041 | 7.4451 | 6.5330 | 7.3463 | 6.3865 | 6.5240 | 7.2558 | 6.9171 | 7.3463 | 7.0650 | 7.4150 | 6.9661 | 3.5669 | 4.4544 | 3.4518 | 4.4228 |
| PTPN2_S298 | PTPN2 | P17706 | 304 | SPAFDHSPNKIMT | 6.0863 | 6.4066 | 6.8530 | 6.5146 | 6.1401 | 6.4336 | 6.1308 | 6.5135 | 6.7943 | 6.7912 | 7.2747 | 7.2126 | 4.7201 | 5.4460 | 5.1130 | 5.5696 |
| PTPN2_S304 | PTPN2 | P17706 | 298 | LSKEDLSPAFDHS | 5.9320 | 6.6616 | 6.3074 | 6.0424 | 6.1401 | 7.3283 | 6.1749 | 6.3439 | 5.6415 | 7.6129 | 7.0200 | 6.5520 | 4.9076 | 5.5421 | 5.8657 | 5.9277 |
| PTPN21_S330 | PTPN21 | Q16825 | 637 | QLHKRNSIEVAGL | 6.0914 | 5.8905 | 7.0201 | 6.0841 | 6.4067 | 6.3548 | 5.9687 | 6.6163 | 7.2430 | 6.5178 | 6.5102 | 7.0341 | 5.5113 | 5.9719 | 5.3070 | 5.4721 |
| PTPN21_S492 | PTPN21 | Q16825 | 492 | PAALVYSQPEIRE | 6.3946 | 5.8620 | 6.7993 | 6.7187 | 6.2926 | 6.7978 | 6.2695 | 5.9716 | 6.5489 | 6.1578 | 6.4806 | 6.7816 | 5.8360 | 6.0245 | 4.9713 | 6.0933 |
| PTPN21_S577 | PTPN21 | Q16825 | 590 | HLYISSSNPDLIT | 6.3689 | 6.2737 | 7.3024 | 6.4430 | 6.3295 | 6.5753 | 6.3612 | 6.6370 | 6.5418 | 6.2522 | 6.5960 | 6.7480 | 5.1653 | 5.9791 | 5.0442 | 5.3824 |
| PTPN21_S590 | PTPN21 | Q16825 | 673 | RAVSVGSQPSVFT | 6.3209 | 7.0012 | 7.3696 | 6.2616 | 6.5377 | 6.6444 | 6.0139 | 6.3503 | 6.2601 | 6.4624 | 6.7032 | 6.1156 | 5.8499 | 5.8235 | 5.1945 | 5.0913 |
| PTPN21_S602;S616 | PTPN21 | Q16825 | 330 | RSSSRMSLPKPQP | 6.3532 | 7.6219 | 6.7470 | 6.9051 | 5.7491 | 4.8459 | 6.1254 | 5.9074 | 6.2108 | 6.3507 | 5.5879 | 6.5268 | 5.5738 | 5.9203 | 6.8773 | 6.6972 |
| PTPN21_S637 | PTPN21 | Q16825 | 577 | PPRPANSTPDLSR | 6.4121 | 6.1029 | 7.4325 | 6.8323 | 6.6131 | 6.3893 | 6.6287 | 5.9091 | 6.7116 | 5.9587 | 6.1975 | 6.9778 | 5.0779 | 6.3684 | 5.0218 | 5.3663 |
| PTPN21_S673 | PTPN21 | Q16825 | 602;616 | NA | 6.7169 | 6.1833 | 7.7455 | 6.3396 | 6.8660 | 6.7508 | 6.7852 | 7.9222 | 5.6825 | 5.7927 | 6.3098 | 6.3930 | 4.7700 | 5.7124 | 4.8293 | 5.2009 |
| PTPN23_S1122 | PTPN23 | Q9H3S7 | 1122 | AADLLSSSPESQH | 7.0545 | 7.4692 | 7.1325 | 7.2406 | 6.7763 | 6.6990 | 5.5489 | 7.6641 | 6.4515 | 6.3711 | 6.6547 | 6.4225 | 4.0167 | 5.1525 | 4.4798 | 4.8661 |
| PTPN23_S1533 | PTPN23 | Q9H3S7 | 1533 | PGLPPASLPESTP | 5.3777 | 6.1818 | 6.5988 | 6.2854 | 5.7986 | 5.8878 | 6.4393 | 6.6193 | 6.6597 | 6.1131 | 6.4199 | 6.4438 | 6.1602 | 6.3928 | 6.0450 | 6.5767 |
| PTPN3_S357 | PTPN3 | P26045 | 359 | AMRRSLSVEHLET | 6.6478 | 6.3542 | 6.3957 | 5.9792 | 6.0190 | 6.0738 | 6.0169 | 6.3428 | 6.7569 | 6.4515 | 6.3359 | 6.1994 | 5.8094 | 6.2469 | 6.0864 | 6.2841 |
| PTPN3_S359 | PTPN3 | P26045 | 357 | NPAMRRSLSVEHL | 6.4141 | 5.4014 | 6.4325 | 6.1896 | 7.1334 | 6.2910 | 5.5403 | 6.9110 | 7.9685 | 7.2088 | 6.6216 | 6.9412 | 5.4380 | 6.1662 | 4.5288 | 4.8138 |
| PTPN3_S395 | PTPN3 | P26045 | 395 | RKPRHSSADNLAN | 5.4773 | 6.1453 | 6.0912 | 7.1375 | 5.8557 | 6.1269 | 6.4535 | 5.7504 | 6.9525 | 6.9032 | 6.3999 | 6.3137 | 5.5499 | 6.2036 | 6.2578 | 6.3815 |
| PTPRA_S189 | PTPRA | P18433 | 213 | LLARSPSTNRKYP | 7.4773 | 7.6808 | 6.5351 | 6.8592 | 6.9093 | 6.5663 | 6.3233 | 5.5818 | 5.2532 | 5.9153 | 5.3872 | 5.3955 | 6.2572 | 6.8108 | 5.4995 | 5.5482 |
| PTPRA_S213 | PTPRA | P18433 | 189 | AGSHSNSFRLSNG | 7.5379 | 7.6024 | 7.0655 | 6.6160 | 6.8681 | 6.8597 | 6.0055 | 6.6919 | 6.5526 | 6.1882 | 5.7376 | 5.9383 | 4.8602 | 5.9906 | 4.6075 | 4.8779 |
| PTPRK_S873 | PTPRK | Q15262-4 | 873 | LCEGTESPYQTGQ | 5.7285 | 5.9778 | 5.8811 | 5.8720 | 5.8495 | 7.2316 | 6.6503 | 6.4278 | 6.0952 | 6.0398 | 5.1977 | 5.5771 | 6.4399 | 7.5490 | 6.8042 | 6.6783 |
| PTPRZ1_S2055 | PTPRZ1 | P23471 | 2055 | REKNRTSSIIPVE | 6.5219 | 6.7194 | 6.7910 | 6.9446 | 6.5540 | 6.0680 | 6.3306 | 5.8060 | 6.4160 | 6.2460 | 5.9778 | 6.2226 | 5.7248 | 5.6535 | 6.1158 | 5.9081 |
| PTS_S19 | PTS | Q03393 | 19 | QVSRRISFSASHR | 6.5784 | 6.0358 | 6.3701 | 5.8588 | 7.5901 | 7.0762 | 5.5103 | 7.7015 | 6.4567 | 6.2717 | 6.3800 | 6.6775 | 5.5066 | 5.9199 | 5.2101 | 4.8564 |
| PTTG1_S165 | PTTG1 | O95997 | 165 | FQLGPPSPVKMPS | 6.5907 | 6.1475 | 7.1390 | 5.7539 | 7.0003 | 6.8434 | 6.4049 | 7.6247 | 7.0638 | 6.5525 | 6.8061 | 6.9251 | 4.6761 | 4.8741 | 4.7398 | 4.8581 |
| PTTG1_S87 | PTTG1 | O95997 | 87 | LKQKQPSFSAKKM | 7.9880 | 5.8967 | 7.5647 | 6.0116 | 6.1801 | 5.8885 | 6.8003 | 6.0982 | 7.9662 | 6.7831 | 8.1462 | 8.2825 | 4.6754 | 4.6807 | 3.3416 | 3.6963 |
| PUF60_S244 | PUF60 | Q9UHX1 | 244 | SDDDIKSVFEAFG | 6.1803 | 7.3065 | 7.6026 | 8.0545 | 6.0577 | 5.6875 | 6.9623 | 5.8036 | 5.7415 | 5.6198 | 4.9921 | 5.0695 | 6.1832 | 6.0185 | 7.5466 | 5.1739 |
| PUF60_T314 | PUF60 | Q9UHX1 | 314 | PLLTPATPGGLPP | 5.9466 | 6.7001 | 6.7914 | 5.9075 | 5.8264 | 5.9017 | 5.4293 | 7.0899 | 7.7868 | 5.6539 | 5.8408 | 5.9640 | 6.1654 | 6.9073 | 6.1306 | 5.9582 |
| PUM1_S124 | PUM1 | Q14671-3 | 229 | EYVLSSSPGDSCL | 6.1917 | 6.7026 | 6.4456 | 6.3508 | 6.2341 | 6.3929 | 6.2273 | 6.8193 | 6.0371 | 6.5036 | 6.2463 | 6.2876 | 5.3170 | 6.0585 | 6.0545 | 6.1311 |
| PUM1_S19 | PUM1 | Q14671-3 | 709 | SGSRRDSLTGSSD | 5.8710 | 5.2014 | 6.0290 | 5.4625 | 6.7797 | 5.9025 | 5.4410 | 6.2171 | 7.3364 | 7.1274 | 7.1877 | 7.0484 | 6.5630 | 6.8065 | 5.6024 | 5.4241 |
| PUM1_S212 | PUM1 | Q14671-3 | 124 | AEHQVRSMDELNH | 5.6875 | 6.1254 | 6.1240 | 7.0348 | 6.8498 | 7.1308 | 6.0168 | 7.1617 | 6.2135 | 6.4485 | 6.6375 | 6.4167 | 5.1704 | 5.9889 | 5.9128 | 5.0810 |
| PUM1_S229 | PUM1 | Q14671-3 | 19 | LWQDSFSPHLKHH | 7.1328 | 7.4865 | 6.5767 | 7.4728 | 5.8091 | 7.5064 | 7.9592 | 7.1336 | 7.2700 | 7.4082 | 7.5579 | 7.7092 | 2.9381 | 4.1501 | 2.3885 | 3.5008 |
| PUM1_S709 | PUM1 | Q14671-3 | 212 | SVLSPRSESGGLG | 6.4012 | 6.7036 | 6.2754 | 6.3260 | 6.0525 | 6.5672 | 6.4146 | 6.5819 | 6.8294 | 6.0821 | 6.8941 | 6.3661 | 5.5484 | 5.8191 | 5.6673 | 5.4711 |
| PUM1_S806 | PUM1 | Q14671-3 | 806 | SASSLFSPSSTLF | 6.5973 | 6.9800 | 6.8229 | 6.7106 | 6.2394 | 5.8444 | 7.3112 | 6.4174 | 6.1052 | 7.1656 | 6.4383 | 6.3213 | 4.7260 | 6.2689 | 4.7526 | 5.2990 |
| PUM2_S102 | PUM2 | Q8TB72 | 136 | DQKGKASPFEEDQ | 6.3420 | 6.3804 | 6.3078 | 6.0240 | 6.3852 | 6.4361 | 6.2464 | 6.5185 | 6.5331 | 6.1511 | 6.3512 | 6.1989 | 6.2176 | 6.2240 | 5.8358 | 5.8478 |
| PUM2_S136 | PUM2 | Q8TB72 | 102 | EYVLSSSPADKLD | 6.3576 | 6.3298 | 6.6970 | 6.2131 | 5.8465 | 6.6022 | 6.2619 | 6.2054 | 7.2103 | 6.8150 | 6.6986 | 7.5816 | 5.3996 | 5.4234 | 5.1185 | 5.2396 |
| PUM2_S182 | PUM2 | Q8TB72 | 82 | EVNAILSPRSESG | 6.1485 | 6.7528 | 6.8120 | 6.8725 | 6.4512 | 6.7612 | 6.1050 | 7.6197 | 6.3686 | 6.2554 | 6.7955 | 6.3886 | 5.4780 | 5.5833 | 4.6109 | 4.9968 |
| PUM2_S700 | PUM2 | Q8TB72 | 182 | PGSRQASPTEVVE | 6.3515 | 6.5032 | 6.6139 | 6.0061 | 6.1831 | 6.1090 | 5.9225 | 6.5008 | 6.1518 | 6.1301 | 6.3172 | 6.3577 | 6.3249 | 6.0521 | 6.2663 | 6.2099 |
| PUM2_S82 | PUM2 | Q8TB72 | 700 | RLRYNRSDIMPSG | 3.9836 | 4.7686 | 4.5881 | 4.0647 | 4.7522 | 4.5338 | 4.5921 | 4.5268 | 4.4314 | 4.4801 | 4.1603 | 4.1553 | 11.6863 | 12.0067 | 11.9424 | 11.3277 |
| PURB_S100 | PURB | Q96QR8 | 101 | AQLGPSSPEQLAA | 6.2320 | 6.7163 | 6.3382 | 6.2142 | 6.2229 | 6.6598 | 6.1618 | 6.4718 | 6.5153 | 6.4803 | 6.7540 | 6.6056 | 5.8680 | 6.6850 | 5.5058 | 5.5689 |
| PURB_S101 | PURB | Q96QR8 | 100 | YAQLGPSSPEQLA | 6.0226 | 6.3799 | 6.4419 | 6.3661 | 6.1340 | 6.7751 | 6.1280 | 6.2517 | 6.2532 | 7.1176 | 6.7625 | 6.4513 | 5.7489 | 6.1333 | 5.5254 | 5.5084 |
| PUS1_T426 | PUS1 | Q9Y606 | 426 | SEGDGDTDxxxxx | 5.9483 | 6.0082 | 5.6662 | 6.0243 | 6.1193 | 5.5651 | 5.9145 | 5.9392 | 6.5016 | 6.1770 | 6.4114 | 6.3272 | 6.1905 | 6.9451 | 6.9594 | 7.3027 |
| PUS7_T616 | PUS7 | Q96PZ0-2 | 616 | DNLEGKTPPVFAS | 5.0074 | 6.2623 | 5.9423 | 5.3256 | 5.6600 | 6.0539 | 5.4830 | 6.2143 | 6.3687 | 6.7464 | 6.3078 | 6.0195 | 7.0756 | 7.4592 | 7.2061 | 6.8680 |
| PWP1_S485 | PWP1 | Q13610 | 50 | QEEGGGSDEEETG | 6.0894 | 6.0508 | 6.6139 | 6.6814 | 6.5276 | 5.8815 | 6.0662 | 6.6040 | 7.1582 | 6.2198 | 6.6916 | 6.9386 | 5.3717 | 5.7736 | 5.5962 | 5.7353 |
| PWP1_S50 | PWP1 | Q13610 | 86 | DPEDDRTLDDDEL | 6.7877 | 6.5458 | 6.8221 | 7.1572 | 6.5249 | 6.2941 | 7.1433 | 6.9356 | 6.9858 | 6.9818 | 7.1293 | 7.0889 | 4.0923 | 4.3785 | 4.1053 | 5.0275 |
| PWP1_T86 | PWP1 | Q13610 | 485 | GSARNSSISGPFG | 5.7419 | 4.5476 | 5.7668 | 5.1729 | 6.9639 | 5.3926 | 5.4564 | 5.8471 | 6.9277 | 7.1356 | 6.5462 | 7.0018 | 7.6295 | 7.1425 | 6.7907 | 5.9368 |
| PWP2_T262 | PWP2 | Q15269 | 262 | TIRGKATPAEEEK | 6.6348 | 8.1921 | 6.5512 | 7.0865 | 6.7346 | 7.1121 | 6.6429 | 6.0274 | 6.2295 | 5.5988 | 7.9018 | 6.5488 | 5.6662 | 4.7197 | 3.8127 | 4.5409 |
| PWWP2A_S81 | PWWP2A | Q96N64 | 81 | PGELARSPEAVGP | 6.2691 | 6.2829 | 5.8846 | 6.5429 | 6.1256 | 5.6670 | 6.2732 | 6.1074 | 6.3602 | 6.4109 | 6.3318 | 6.1788 | 6.2945 | 6.3293 | 6.3270 | 6.6150 |
| PWWP2B_S186 | PWWP2B | Q6NUJ5 | 186 | KCKSTLSPPEASP | 5.9932 | 5.8171 | 7.6843 | 6.8528 | 5.7671 | 6.3277 | 6.1683 | 5.6338 | 7.5791 | 6.8049 | 7.2477 | 7.0539 | 5.0982 | 5.5608 | 4.8148 | 5.5962 |
| PXDC1_S150 | PXDC1 | Q5TGL8 | 150 | IQPSFQSPVKISE | 7.4124 | 6.8449 | 7.2785 | 6.7621 | 7.0245 | 7.3535 | 7.1036 | 7.2749 | 7.5414 | 6.8866 | 7.4486 | 7.3710 | 3.1371 | 3.7757 | 3.1628 | 3.6223 |
| PXN_S106 | PXN | P49023-3 | 336 | GKTGSSSPPGGPP | 6.3261 | 6.1002 | 6.6741 | 6.2636 | 6.4713 | 6.4682 | 6.1918 | 6.5388 | 6.5413 | 6.4408 | 6.7951 | 6.6976 | 5.9762 | 5.8328 | 5.2379 | 5.4443 |
| PXN_S126;S130 | PXN | P49023-3 | 354 | LDSMLGSLQSDLN | 5.3168 | 5.6928 | 6.5182 | 6.0432 | 5.5956 | 6.3339 | 6.4745 | 6.1435 | 6.5372 | 5.9492 | 6.7015 | 6.6350 | 7.3920 | 6.8975 | 5.4842 | 6.2847 |
| PXN_S244 | PXN | P49023-3 | 84 | HQQPQSSSPVYGS | 7.0061 | 6.9463 | 6.7386 | 7.1749 | 7.3076 | 7.1082 | 6.7971 | 7.6734 | 6.9524 | 6.6023 | 6.9582 | 7.3326 | 4.1383 | 4.0993 | 3.3273 | 3.8375 |
| PXN_S258 | PXN | P49023-3 | 118 | GEEEHVYSFPNKQ | 6.3471 | 5.6331 | 6.7350 | 5.3476 | 6.4337 | 5.8658 | 5.6421 | 6.2136 | 7.4777 | 6.3273 | 6.0682 | 6.1742 | 7.4724 | 7.0201 | 5.7294 | 5.5127 |
| PXN_S303 | PXN | P49023 | 303 | RDGGRSSPGGQDE | 6.7760 | 6.2138 | 6.8549 | 5.7036 | 6.7076 | 7.0189 | 6.2438 | 7.1742 | 6.6997 | 6.3847 | 7.2695 | 6.3948 | 5.6579 | 5.0997 | 4.8936 | 4.9072 |
| PXN_S316 | PXN | P49023-3 | 316 | RTPSLRSPDQPPP | 6.0366 | 5.7905 | 6.7488 | 6.3581 | 5.9891 | 6.8092 | 6.0778 | 6.7567 | 5.9775 | 6.8996 | 6.3910 | 6.1965 | 6.6451 | 6.1012 | 5.7017 | 5.5206 |
| PXN_S334 | PXN | P49023-3 | 83 | IHQQPQSSSPVYG | 6.5218 | 6.6437 | 7.0564 | 6.9658 | 6.5592 | 7.2618 | 6.4031 | 7.4847 | 6.8349 | 6.9029 | 7.1716 | 7.1096 | 4.4724 | 4.3947 | 4.0349 | 4.1825 |
| PXN_S336 | PXN | P49023-3 | 106 | PQDSVGSPCSRVG | 7.6939 | 6.6519 | 6.3503 | 7.6025 | 5.9664 | 6.0405 | 5.9948 | 7.2267 | 7.1089 | 6.9210 | 6.2335 | 6.6387 | 4.3019 | 5.4007 | 5.0290 | 4.8391 |
| PXN_S354 | PXN | P49023-3 | 258 | QQQTRISASSATR | 7.0195 | 6.1045 | 7.1631 | 6.0024 | 7.2919 | 6.5940 | 6.2412 | 7.4718 | 6.3186 | 5.1851 | 6.3439 | 6.3324 | 6.3457 | 5.1927 | 5.0007 | 5.3924 |
| PXN_S83 | PXN | P49023-3 | 244 | NQGEMSSPQRVTS | 6.0678 | 5.8386 | 6.4697 | 7.6473 | 7.3194 | 7.4816 | 5.4049 | 7.3582 | 5.8386 | 7.3758 | 8.4864 | 6.5014 | 5.2604 | 4.1251 | 4.3155 | 4.5094 |
| PXN_S83;Y88 | PXN | P49023-3 | 334 | AQGKTGSSSPPGG | 6.5757 | 6.4565 | 7.1074 | 5.6836 | 6.8938 | 6.8787 | 6.2456 | 6.6248 | 6.4827 | 6.5592 | 6.7356 | 6.9072 | 6.0204 | 5.3554 | 4.7338 | 4.7396 |
| PXN_S84 | PXN | P49023-3 | 126;130 | NA | 5.7344 | 5.6490 | 7.0891 | 7.3714 | 6.3039 | 5.3482 | 7.3802 | 7.2484 | 6.3107 | 4.9539 | 4.6920 | 7.5728 | 5.2041 | 7.1473 | 5.8173 | 6.1772 |
| PXN_Y118 | PXN | P49023-3 | 83;88 | NA | 6.5051 | 4.7495 | 5.6141 | 5.8934 | 8.3781 | 8.3680 | 4.5600 | 5.1033 | 7.8815 | 8.9413 | 7.0486 | 5.4966 | 5.8368 | 5.7653 | 5.9736 | 3.8850 |
| PYCR2_S304 | PYCR2 | Q96C36 | 304 | STLTPSSPGKLLT | 6.2472 | 6.4865 | 6.1739 | 7.5073 | 5.8308 | 6.0532 | 6.3219 | 6.6746 | 6.3417 | 6.6537 | 6.6969 | 7.0173 | 5.4658 | 5.6513 | 5.0360 | 5.8418 |
| PYGB_S430 | PYGB | P11216 | 430 | DRLRRMSVIEEGD | 4.4850 | 5.0927 | 7.2111 | 5.4864 | 4.9900 | 6.8830 | 5.1641 | 6.8317 | 6.7846 | 12.6605 | 10.0271 | 7.4009 | 4.6947 | 4.3866 | 4.3067 | 3.5949 |
| PYGO1_S34 | PYGO1 | Q9Y3Y4-2 | 34 | PGVQLGSPDKKKR | 6.0523 | 5.9217 | 6.5367 | 6.0450 | 6.1866 | 6.1617 | 5.9385 | 6.1511 | 6.4241 | 6.7340 | 6.5222 | 6.7995 | 6.4033 | 6.3001 | 5.9750 | 5.8481 |
| PYGO2_S97 | PYGO2 | Q9BRQ0 | 97 | APPFLGSPVPFGG | 6.4437 | 6.8402 | 6.9169 | 6.0354 | 6.5282 | 6.5221 | 6.2718 | 7.6180 | 7.2713 | 7.3795 | 7.7874 | 7.4755 | 4.2026 | 4.5243 | 3.6155 | 4.5676 |
| PYGO2_T302 | PYGO2 | Q9BRQ0 | 302 | SGRGGGTPDANSL | 5.6373 | 5.2198 | 5.8266 | 4.9918 | 6.4167 | 5.7906 | 5.8079 | 6.3250 | 6.0517 | 6.4945 | 6.8002 | 6.4857 | 7.1660 | 7.3732 | 6.8961 | 6.7170 |
| PYGO2_T50 | PYGO2 | Q9BRQ0 | 50 | KRRKSNTQGPAYS | 5.2355 | 5.4597 | 5.8584 | 5.3603 | 6.5784 | 6.1462 | 5.5741 | 6.6351 | 6.3773 | 6.0445 | 6.4062 | 6.4190 | 6.0769 | 7.3979 | 7.6452 | 6.7853 |
| PYM1_S117 | PYM1 | Q9BRP8 | 72 | EATAPVTPSRPEG | 7.3006 | 7.6493 | 6.8723 | 7.3605 | 5.6034 | 7.3800 | 7.1607 | 6.7063 | 7.4633 | 7.8752 | 7.5019 | 7.6256 | 2.9406 | 4.1935 | 2.7489 | 3.6178 |
| PYM1_T72 | PYM1 | Q9BRP8 | 117 | RTLDKVSLEETAQ | 4.8527 | 5.3099 | 7.6073 | 5.2337 | 7.1883 | 6.7165 | 7.0476 | 8.4396 | 6.6110 | 6.8952 | 5.2073 | 6.8396 | 4.4835 | 6.5289 | 5.9897 | 5.0491 |
| Q9UFV3_T4 | Q9UFV3 | Q9UFV3 | 4 | xxxMAETYRRSRQ | 6.8633 | 7.4429 | 5.4554 | 7.2577 | 6.3941 | 6.4350 | 5.4586 | 7.8246 | 6.0096 | 6.4824 | 5.8826 | 6.2165 | 4.8075 | 6.1314 | 6.1335 | 5.2050 |
| QARS1_S14 | QARS1 | A0A1B0GVU9 | 708 | GKRRGLTLAQGGV | 7.6110 | 8.0699 | 6.5635 | 8.0031 | 6.4044 | 7.1939 | 7.9933 | 7.2566 | 6.3596 | 6.9921 | 7.1964 | 7.3127 | 2.7464 | 4.1305 | 2.5849 | 3.5817 |
| QARS1_S439 | QARS1 | A0A1B0GVU9 | 14 | RDTRRLSFLVSYI | 5.3845 | 5.8446 | 6.6402 | 6.1437 | 6.4583 | 6.0822 | 5.7405 | 5.2122 | 7.0274 | 6.9981 | 6.8888 | 7.2156 | 6.1012 | 6.4500 | 6.2629 | 5.5500 |
| QARS1_T708 | QARS1 | A0A1B0GVU9 | 439 | LHYAVVSKRKILQ | 6.3276 | 6.4114 | 5.8070 | 5.6191 | 6.6237 | 6.0558 | 6.2735 | 6.2843 | 5.6313 | 6.0261 | 6.3343 | 7.0131 | 6.4844 | 7.1483 | 6.6129 | 5.3473 |
| QKI_S211 | QKI | Q96PU8-3 | 211 | RDANIKSPTAQAA | 6.8828 | 6.2987 | 7.0015 | 6.4351 | 6.1168 | 5.8694 | 6.2959 | 6.1931 | 6.3999 | 6.5697 | 7.2398 | 6.8208 | 5.6351 | 5.6255 | 5.0320 | 5.5837 |
| QSER1_S1211 | QSER1 | Q2KHR3 | 931 | QVTVNLSPVPALQ | 6.0017 | 6.1277 | 7.1158 | 6.6908 | 6.5029 | 6.6057 | 6.5944 | 5.9477 | 6.3741 | 6.5052 | 6.5990 | 6.5271 | 5.3739 | 5.7788 | 5.3825 | 5.8726 |
| QSER1_S236 | QSER1 | Q2KHR3 | 236 | QSSLSCSPIGDST | 5.7642 | 6.3286 | 6.6539 | 6.7909 | 6.5068 | 6.7026 | 6.9752 | 6.4844 | 6.2619 | 6.3013 | 6.7807 | 6.4293 | 5.5434 | 5.7856 | 5.4178 | 5.2734 |
| QSER1_S615 | QSER1 | Q2KHR3 | 615 | SVIRSNSRLEDQV | 4.6779 | 4.9575 | 5.5164 | 5.0536 | 5.3542 | 5.4609 | 5.4977 | 6.1981 | 5.3755 | 5.3936 | 5.2897 | 5.1945 | 8.7305 | 8.4276 | 9.8037 | 9.0685 |
| QSER1_S931 | QSER1 | Q2KHR3 | 1211 | SETGGNSPSDKVD | 5.5249 | 5.1975 | 6.1596 | 6.0142 | 7.1738 | 6.0248 | 6.4481 | 6.0436 | 6.0248 | 5.1659 | 6.0692 | 5.5676 | 7.4778 | 6.8359 | 6.9868 | 7.2853 |
| QSER1_T1341;S1348 | QSER1 | Q2KHR3 | 1341;1348 | NA | 5.7422 | 6.5000 | 6.3598 | 6.4909 | 6.3309 | 6.3414 | 5.9726 | 6.4014 | 6.1170 | 6.3342 | 6.1107 | 6.5046 | 6.5374 | 6.3229 | 5.7402 | 6.1938 |
| QSOX2_S405 | QSOX2 | Q6ZRP7 | 405 | NNKMRISGIFLTN | 6.8276 | 7.0539 | 5.9642 | 7.9635 | 5.6163 | 5.4654 | 5.6792 | 7.4647 | 6.9282 | 5.1679 | 5.7630 | 7.5904 | 3.8350 | 5.0798 | 6.0313 | 7.5695 |
| R3HCC1_S236 | R3HCC1 | Q9Y3T6 | 236 | MATRFGSTLQLDL | 6.2878 | 6.2434 | 6.6123 | 5.8654 | 5.1455 | 6.1154 | 7.3389 | 5.9678 | 7.3057 | 6.2325 | 6.9024 | 7.1349 | 6.1426 | 6.0092 | 4.4332 | 6.2630 |
| R3HDM1_S142 | R3HDM1 | Q15032-3 | 381 | AVTKASSFSGISV | 6.5403 | 6.9698 | 6.8113 | 6.5490 | 6.4773 | 6.0315 | 5.3923 | 6.4413 | 7.0540 | 6.9680 | 6.3342 | 6.3299 | 5.7158 | 5.8778 | 5.4791 | 5.0284 |
| R3HDM1_S380 | R3HDM1 | Q15032-3 | 142 | MLSRDSSQEYTDS | 6.2408 | 6.6105 | 8.3521 | 6.9326 | 6.2669 | 6.0953 | 6.0192 | 6.2835 | 6.1650 | 7.3066 | 7.0569 | 6.7092 | 4.1242 | 6.0183 | 5.2801 | 4.5387 |
| R3HDM1_S381 | R3HDM1 | Q15032-3 | 380 | PAVTKASSFSGIS | 7.2032 | 6.1051 | 6.4668 | 6.6515 | 5.7293 | 6.3633 | 5.5381 | 5.9663 | 6.6415 | 5.9987 | 7.2126 | 7.1956 | 5.0282 | 5.9993 | 6.1416 | 5.7587 |
| R3HDM2_S349 | R3HDM2 | Q9Y2K5 | 349 | PVTKASSFSGISI | 6.4895 | 6.4243 | 7.5393 | 6.7814 | 6.6143 | 6.2855 | 5.7387 | 6.4925 | 6.3656 | 6.9134 | 6.6870 | 6.6940 | 4.9747 | 6.0425 | 5.1004 | 4.8569 |
| RAB10_T73 | RAB10 | P61026 | 73 | GQERFHTITTSYY | 8.0844 | 8.1450 | 7.3322 | 8.1998 | 6.4086 | 6.2436 | 6.8044 | 6.2349 | 6.1628 | 5.6723 | 6.0837 | 5.8882 | 4.5847 | 4.9811 | 4.6493 | 4.5250 |
| RAB10_Y6 | RAB10 | P61026 | 6 | xMAKKTYDLLFKL | 6.2933 | 6.7072 | 6.4633 | 6.7985 | 6.1088 | 5.9036 | 6.4727 | 5.7287 | 6.3680 | 6.3599 | 6.7460 | 6.7647 | 5.3992 | 6.0130 | 5.7202 | 6.1528 |
| RAB11A_S190 | RAB11A | P62491 | 190 | RRENDMSPSNNVV | 6.1326 | 6.8652 | 6.5270 | 7.7300 | 6.2764 | 7.2053 | 7.3321 | 6.7809 | 6.7322 | 7.2270 | 7.9700 | 7.2602 | 3.5836 | 4.6918 | 3.4898 | 4.1959 |
| RAB11A_S78 | RAB11A | P62491 | 78 | RYRAITSAYYRGA | 6.2512 | 6.0685 | 5.7045 | 5.6102 | 6.1641 | 5.8548 | 6.0461 | 6.0106 | 5.7089 | 6.2247 | 5.9124 | 5.9918 | 6.8571 | 6.7420 | 7.6109 | 7.2423 |
| RAB11FIP1_S156 | RAB11FIP1 | Q6WKZ4 | 435 | PESRRSSLLSLMT | 5.4031 | 4.5224 | 6.0169 | 5.4215 | 6.6759 | 5.3816 | 5.0290 | 5.9605 | 7.9898 | 7.9295 | 7.8670 | 8.0231 | 6.0649 | 6.9150 | 5.4894 | 5.3104 |
| RAB11FIP1_S202 | RAB11FIP1 | Q6WKZ4 | 156 | RNNMTASMFDLSM | 6.6574 | 6.6520 | 6.9000 | 7.1457 | 6.4064 | 6.0581 | 6.3006 | 5.6314 | 6.2526 | 5.5935 | 6.2763 | 6.2005 | 5.9758 | 6.1979 | 5.6501 | 6.1015 |
| RAB11FIP1_S234 | RAB11FIP1 | Q6WKZ4 | 500 | VVSRQGSSLNLFE | 6.1389 | 6.3269 | 6.4633 | 6.4036 | 5.8674 | 5.8392 | 5.8460 | 5.8544 | 6.4382 | 6.2878 | 6.0334 | 6.5053 | 6.2424 | 6.5077 | 6.2749 | 6.9705 |
| RAB11FIP1_S280 | RAB11FIP1 | Q6WKZ4 | 234 | KTPLSQSMSVLPT | 7.8937 | 7.1029 | 6.8755 | 7.4869 | 7.0871 | 5.3653 | 6.1106 | 4.9099 | 5.8177 | 5.0374 | 5.8185 | 5.5115 | 6.3703 | 6.3825 | 6.3712 | 5.8590 |
| RAB11FIP1_S315 | RAB11FIP1 | Q6WKZ4 | 528 | EPRPPISSPRAPQ | 6.1890 | 6.4786 | 6.3191 | 6.4658 | 6.1045 | 6.6638 | 6.7779 | 6.5880 | 6.4116 | 6.3734 | 6.6212 | 6.7569 | 5.1709 | 5.7627 | 5.6754 | 5.6412 |
| RAB11FIP1_S356 | RAB11FIP1 | Q6WKZ4 | 315 | NDVLSRSNVCING | 6.8688 | 6.3612 | 5.7026 | 6.6605 | 5.4722 | 6.2033 | 5.9498 | 4.9405 | 6.9417 | 5.4112 | 6.5122 | 6.9123 | 6.5544 | 6.5609 | 5.3505 | 7.5980 |
| RAB11FIP1_S434 | RAB11FIP1 | Q6WKZ4 | 496 | DTAAVVSRQGSSL | 6.4923 | 7.1113 | 6.8811 | 6.6943 | 5.6492 | 5.9888 | 6.3344 | 6.0506 | 6.6363 | 6.4032 | 6.7544 | 6.7106 | 5.3448 | 5.8521 | 5.2537 | 5.8430 |
| RAB11FIP1_S435 | RAB11FIP1 | Q6WKZ4 | 202 | TTPSVDSDDESVV | 5.5760 | 6.1272 | 6.8790 | 6.2849 | 6.3213 | 6.4275 | 6.0127 | 6.5879 | 6.0592 | 6.6556 | 6.6549 | 6.9460 | 5.4334 | 6.4848 | 5.5763 | 5.9733 |
| RAB11FIP1_S496 | RAB11FIP1 | Q6WKZ4 | 280 | SHKRTASTDLKQL | 6.5018 | 6.6926 | 7.1609 | 6.5533 | 4.9310 | 5.2830 | 5.8192 | 6.0326 | 5.8938 | 5.9519 | 6.5407 | 6.6601 | 6.3869 | 6.1773 | 6.8222 | 6.5928 |
| RAB11FIP1_S500 | RAB11FIP1 | Q6WKZ4 | 356 | RKKHLFSSTENLA | 5.8358 | 6.5744 | 6.2513 | 6.0381 | 5.9346 | 6.3063 | 6.3023 | 6.3812 | 6.0585 | 6.2581 | 6.4100 | 6.7762 | 5.8532 | 5.9879 | 6.2844 | 6.7477 |
| RAB11FIP1_S528 | RAB11FIP1 | Q6WKZ4 | 434 | KPESRRSSLLSLM | 5.1795 | 4.7768 | 6.2892 | 5.5720 | 6.1102 | 5.2235 | 4.5503 | 6.2213 | 7.4517 | 7.7236 | 8.5508 | 8.8829 | 6.1738 | 6.7075 | 5.2089 | 5.3779 |
| RAB11FIP2_S162 | RAB11FIP2 | Q7L804-2 | 277 | PHRRTLSFDTSKM | 6.4212 | 6.3061 | 6.3864 | 6.6635 | 5.3272 | 5.4188 | 5.8025 | 5.9749 | 6.0761 | 6.0813 | 6.1957 | 6.3713 | 6.6253 | 6.8244 | 6.7428 | 6.7826 |
| RAB11FIP2_S277 | RAB11FIP2 | Q7L804-2 | 388 | SPCDLKSPNAFSE | 6.0865 | 5.9948 | 6.4451 | 6.5228 | 6.2601 | 6.5425 | 6.8502 | 6.4063 | 6.0816 | 6.3170 | 6.4582 | 6.2439 | 5.6129 | 5.9308 | 5.8617 | 6.3855 |
| RAB11FIP2_S388 | RAB11FIP2 | Q7L804-2 | 162 | MKDKTRSPFAKLK | 6.9299 | 7.5127 | 6.8188 | 7.1391 | 6.1908 | 7.5920 | 7.0215 | 6.8193 | 5.8626 | 6.7455 | 7.5930 | 6.3622 | 3.7885 | 4.8596 | 4.0145 | 4.7499 |
| RAB11FIP3_S467 | RAB11FIP3 | O75154-3 | 467 | SDLEMNSPTKRLS | 6.9756 | 6.6499 | 6.6055 | 6.5929 | 6.9357 | 6.8879 | 6.6841 | 6.7444 | 6.8355 | 6.0577 | 7.0394 | 6.8503 | 4.9154 | 5.0543 | 4.6899 | 4.4815 |
| RAB11FIP5_S1194 | RAB11FIP5 | A0A1B0GTL5 | 1209 | AAPVEGSPDRKQS | 6.2246 | 5.6322 | 6.7159 | 6.7011 | 6.5354 | 6.7584 | 6.3933 | 5.9788 | 6.4586 | 6.9173 | 7.0821 | 7.0006 | 5.3286 | 5.6978 | 5.0115 | 5.5637 |
| RAB11FIP5_S1209 | RAB11FIP5 | A0A1B0GTL5 | 640 | SPTPLASPGKALP | 6.5447 | 6.4472 | 6.5287 | 6.2044 | 6.5130 | 6.3990 | 6.2910 | 6.4446 | 6.3276 | 6.1023 | 6.4410 | 6.4957 | 5.5981 | 5.6965 | 5.9553 | 6.0110 |
| RAB11FIP5_S1218 | RAB11FIP5 | A0A1B0GTL5 | 307 | THKRTYSDEANQM | 6.2266 | 6.4359 | 7.0553 | 6.4976 | 5.9078 | 6.2168 | 5.9778 | 5.8118 | 6.0899 | 6.2438 | 6.5171 | 6.5110 | 6.0253 | 6.1405 | 5.9660 | 6.3767 |
| RAB11FIP5_S174 | RAB11FIP5 | A0A1B0GTL5 | 209 | YDLESASAILPSS | 8.5130 | 6.8426 | 6.9480 | 7.3795 | 6.9701 | 4.9226 | 5.9439 | 4.5799 | 5.8968 | 4.9182 | 5.2907 | 5.4968 | 7.2538 | 7.1397 | 5.6714 | 6.2329 |
| RAB11FIP5_S209 | RAB11FIP5 | A0A1B0GTL5 | 482 | ELGRRSSLGEKGG | 5.9667 | 5.6336 | 5.8162 | 5.5820 | 6.4187 | 5.9950 | 5.5840 | 6.3494 | 6.7643 | 7.1117 | 7.0734 | 6.9852 | 6.2789 | 6.7616 | 5.8108 | 5.8685 |
| RAB11FIP5_S244 | RAB11FIP5 | A0A1B0GTL5 | 296 | GRDSAQSPKLFTH | 6.1726 | 6.5444 | 6.5947 | 6.5114 | 5.6866 | 6.1829 | 6.2664 | 5.9607 | 6.1993 | 6.6146 | 6.7952 | 6.7811 | 5.6401 | 6.0198 | 6.0152 | 6.0152 |
| RAB11FIP5_S283 | RAB11FIP5 | A0A1B0GTL5 | 1194 | EPQPSASPHPVKP | 6.3031 | 6.1241 | 6.5509 | 6.3884 | 6.9593 | 6.7005 | 6.6257 | 6.9705 | 6.3336 | 5.7283 | 6.2569 | 6.1477 | 5.5051 | 6.1020 | 5.5777 | 5.7262 |
| RAB11FIP5_S296 | RAB11FIP5 | A0A1B0GTL5 | 866 | PQVQPESPETVSP | 5.5152 | 5.5494 | 6.5889 | 6.5020 | 6.0739 | 6.1704 | 6.1951 | 6.2493 | 5.4267 | 6.2158 | 6.7945 | 6.6694 | 6.1808 | 6.3458 | 6.6152 | 6.9077 |
| RAB11FIP5_S307 | RAB11FIP5 | A0A1B0GTL5 | 899 | KGLQPSTPPPKPP | 6.4100 | 6.5135 | 6.7248 | 6.5741 | 6.3045 | 6.3226 | 6.0379 | 6.0521 | 6.3998 | 6.5397 | 6.5847 | 6.5139 | 5.4431 | 5.8965 | 5.7322 | 5.9507 |
| RAB11FIP5_S482 | RAB11FIP5 | A0A1B0GTL5 | 174 | FTRNNLSASMFDL | 7.4310 | 6.8421 | 6.3511 | 6.9767 | 6.4415 | 5.8446 | 6.0120 | 5.4459 | 6.4691 | 5.9377 | 6.3009 | 5.8849 | 6.3872 | 6.4806 | 5.4697 | 5.7250 |
| RAB11FIP5_S634 | RAB11FIP5 | A0A1B0GTL5 | 244 | NKLRKSSLTQSNT | 5.8547 | 5.3829 | 5.5965 | 6.1874 | 7.5009 | 6.0086 | 6.2942 | 6.7328 | 6.6492 | 8.0421 | 7.1111 | 8.8690 | 5.1702 | 5.4417 | 5.0469 | 4.1117 |
| RAB11FIP5_S634;S640 | RAB11FIP5 | A0A1B0GTL5 | 283 | RSPSRSSWLSTEG | 5.7477 | 5.7797 | 5.7162 | 5.3352 | 5.5469 | 5.4982 | 5.5336 | 5.3136 | 5.3453 | 5.5841 | 5.1841 | 5.2863 | 8.5872 | 8.6260 | 8.8665 | 8.0495 |
| RAB11FIP5_S640 | RAB11FIP5 | A0A1B0GTL5 | 634 | PSASRASPTPLAS | 6.4503 | 6.7343 | 6.7192 | 6.1998 | 6.4348 | 6.2476 | 6.2340 | 6.7479 | 6.0545 | 6.4744 | 6.5292 | 5.9814 | 5.4596 | 6.0403 | 5.8008 | 5.8917 |
| RAB11FIP5_S780 | RAB11FIP5 | A0A1B0GTL5 | 913 | LFTPSRSQEEEEE | 4.8067 | 4.9217 | 5.1249 | 5.0220 | 5.2684 | 5.3942 | 5.2889 | 5.1728 | 4.8412 | 4.7159 | 5.1206 | 5.0024 | 10.3980 | 9.3608 | 9.8900 | 9.6715 |
| RAB11FIP5_S866 | RAB11FIP5 | A0A1B0GTL5 | 938 | TEGEDASPSALVV | 5.9271 | 6.3887 | 6.5675 | 6.4293 | 6.2166 | 6.8184 | 6.3587 | 6.3010 | 7.3634 | 6.0602 | 6.5728 | 6.9109 | 5.4279 | 5.5023 | 5.6258 | 5.5295 |
| RAB11FIP5_S913 | RAB11FIP5 | A0A1B0GTL5 | 780 | EVEAGQSPADSGT | 5.8504 | 5.9465 | 6.5109 | 6.3373 | 6.1862 | 6.2662 | 6.0152 | 5.9178 | 6.1363 | 6.7318 | 7.0827 | 6.0439 | 5.7330 | 5.5658 | 6.8979 | 6.7780 |
| RAB11FIP5_S938 | RAB11FIP5 | A0A1B0GTL5 | 1218 | RKQSRSSLSIALS | 4.8288 | 4.4557 | 5.4761 | 4.8708 | 5.3683 | 4.9246 | 5.3893 | 4.9791 | 5.5801 | 5.9949 | 5.1382 | 5.4138 | 8.4980 | 9.7702 | 9.9179 | 9.3942 |
| RAB11FIP5_T899 | RAB11FIP5 | A0A1B0GTL5 | 634;640 | NA | 6.3222 | 6.6159 | 6.4462 | 6.3906 | 6.4314 | 6.4142 | 6.6145 | 6.5060 | 6.3866 | 6.5780 | 6.9216 | 6.3700 | 4.9280 | 5.7104 | 5.4315 | 5.9330 |
| RAB12_S106 | RAB12 | Q6IQ22 | 21 | GGLGAGSPALSGG | 6.2150 | 6.5529 | 6.2724 | 6.5244 | 5.6895 | 6.2973 | 6.7413 | 6.6137 | 6.5568 | 6.4554 | 6.2843 | 6.4347 | 5.5581 | 5.9598 | 5.8239 | 6.0205 |
| RAB12_S21 | RAB12 | Q6IQ22 | 106 | GQERFNSITSAYY | 6.5702 | 6.5334 | 6.8847 | 6.5566 | 6.0080 | 6.2457 | 6.0371 | 6.1479 | 6.8222 | 6.6287 | 6.6680 | 6.5054 | 5.3816 | 5.4717 | 5.9432 | 5.5956 |
| RAB12_S220 | RAB12 | Q6IQ22 | 220 | RNELSNSILSLQP | 5.5068 | 6.4778 | 6.6324 | 5.9834 | 6.3365 | 5.9933 | 6.5535 | 6.8607 | 7.0924 | 6.5076 | 6.7676 | 6.3899 | 6.1767 | 5.8628 | 5.4397 | 5.4189 |
| RAB12_S220;S223 | RAB12 | Q6IQ22 | 220;223 | NA | 5.9966 | 6.3656 | 6.1348 | 6.5786 | 5.8459 | 6.3389 | 6.2516 | 5.8841 | 6.6179 | 6.6337 | 6.6877 | 5.7767 | 5.7831 | 6.6940 | 5.9841 | 6.4266 |
| RAB13_S178 | RAB13 | P51153 | 5 | xxMAKAYDHLFKL | 6.9292 | 7.7350 | 7.2072 | 8.4148 | 5.9904 | 6.7939 | 6.7828 | 6.7744 | 6.9608 | 7.7197 | 7.5739 | 8.6251 | 2.7804 | 3.2894 | 2.9617 | 3.4613 |
| RAB13_S186 | RAB13 | P51153 | 178 | KSGGRRSGNGNKP | 6.0293 | 5.8360 | 6.8065 | 6.9073 | 6.5700 | 6.4384 | 6.8929 | 6.3828 | 6.5462 | 6.8584 | 6.9283 | 6.7658 | 4.8930 | 5.8517 | 5.0639 | 5.2294 |
| RAB13_Y5 | RAB13 | P51153 | 186 | NGNKPPSTDLKTC | 5.5491 | 5.5927 | 5.9754 | 6.6654 | 6.3459 | 5.8822 | 6.0599 | 5.8037 | 6.4979 | 6.1368 | 5.6279 | 6.0083 | 6.7661 | 7.2697 | 7.0363 | 6.7828 |
| RAB14_S180 | RAB14 | P61106 | 180 | QNIQDGSLDLNAA | 6.1039 | 5.6411 | 4.7916 | 5.6108 | 5.5862 | 5.7517 | 5.4248 | 5.5485 | 5.8418 | 5.8221 | 6.1564 | 6.2227 | 8.1331 | 7.4966 | 7.5441 | 8.3248 |

| | | | | | | | | | | | | | | | | | | | | | |
|---|---|---|---|---|---|---|---|---|---|---|---|---|---|---|---|---|---|---|---|---|---|
| RAB18_S173 | RAB18 | Q9NP72-2 | | 173 | KFARKHSMLFIEA | 4.8394 | 4.7729 | 5.3084 | 5.5828 | 4.9379 | 5.0068 | 5.5731 | 4.7083 | 5.4312 | 5.4728 | 5.1073 | 5.6749 | 9.3781 | 8.5590 | 10.1003 | 9.5470 |
| RAB1A_T195 | RAB1A | P62820 | | 195 | NVKIQSTPVKQSG | 6.8158 | 7.2644 | 6.6111 | 7.1200 | 6.7860 | 7.3260 | 7.5768 | 7.1696 | 7.1888 | 7.0547 | 7.1603 | 6.9663 | 3.3236 | 4.2991 | 3.2171 | 4.1205 |
| RAB1A_T75 | RAB1A | P62820 | | 75 | GQERFRTITSSYY | 8.5531 | 7.3681 | 6.3714 | 7.2580 | 6.1836 | 5.4633 | 6.2603 | 4.9826 | 6.5316 | 5.1761 | 5.2958 | 5.6755 | 6.4798 | 7.0979 | 5.7721 | 5.5308 |
| RAB23_S187 | RAB23 | Q9ULC3 | | 187 | PELTHSSSNKIGV | 5.0327 | 5.3482 | 5.7053 | 6.5942 | 5.5912 | 5.4409 | 6.1848 | 5.9644 | 5.8709 | 5.8740 | 5.5986 | 5.2126 | 7.6898 | 8.3461 | 8.3477 | 7.1987 |
| RAB31_S35 | RAB31 | Q13636 | | 35 | HFDHNISPTIGAS | 5.8656 | 6.8177 | 5.7647 | 6.1331 | 6.9838 | 6.6136 | 6.8455 | 6.4608 | 6.3776 | 6.9645 | 6.4777 | 6.4650 | 5.5066 | 5.9171 | 5.3847 | 5.4219 |
| RAB32_S71 | RAB32 | Q13637 | | 71 | KVLNWDSRTLVRL | 6.4274 | 6.6655 | 6.1814 | 6.3936 | 5.9056 | 6.0340 | 6.3922 | 6.1517 | 6.2425 | 6.5179 | 6.1509 | 6.1677 | 5.8894 | 5.9708 | 6.6487 | 6.2608 |
| RAB34_S123 | RAB34 | Q9BZG1 | | 244 | RINSDDSNLYLTA | 5.6999 | 5.5438 | 5.5338 | 5.4536 | 6.4679 | 5.9019 | 5.8659 | 5.5307 | 5.5863 | 5.4693 | 5.3138 | 4.8157 | 8.4144 | 8.2949 | 8.6509 | 7.4572 |
| RAB34_S244 | RAB34 | P0DI83 | | 123 | PRVIVGSPRARVA | 6.8131 | 6.8726 | 5.9188 | 6.2858 | 5.4869 | 7.0801 | 7.0608 | 7.3729 | 7.0301 | 7.1273 | 7.5378 | 7.3467 | 4.0438 | 4.9005 | 4.2772 | 4.8456 |
| RAB3B_S190 | RAB3B | P20337 | | 190 | CDKMSDSLDTDPS | 6.5168 | 6.7413 | 6.6786 | 6.8872 | 5.8926 | 6.1245 | 6.0927 | 6.1932 | 6.3879 | 6.8261 | 6.8377 | 7.0733 | 4.9370 | 5.6926 | 5.6315 | 5.4869 |
| RAB3GAP1_S537 | RAB3GAP1 | Q15042-3 | | 537 | DEGKKTSASDVTN | 1.8607 | 2.2641 | 2.2180 | 2.1304 | 2.5368 | 2.9689 | 2.9631 | 2.9053 | 2.1724 | 2.3913 | 2.1927 | 2.3805 | 17.7081 | 16.9051 | 20.3887 | 16.0142 |
| RAB3IL1_S179 | RAB3IL1 | Q8TBN0 | | 179 | LHPQLLSPTKAGP | 6.8111 | 6.8413 | 7.4369 | 6.8980 | 6.0539 | 7.1364 | 6.2504 | 7.0984 | 6.4678 | 7.0145 | 7.4781 | 7.7584 | 3.8167 | 4.6631 | 3.6560 | 4.6189 |
| RAB3IP_S163 | RAB3IP | Q96QF0 | | 262 | KTLVLSSSPTSPT | 6.0899 | 6.4829 | 6.8980 | 6.6916 | 6.6721 | 6.8842 | 6.7496 | 6.9132 | 6.6389 | 6.3951 | 6.9422 | 6.8842 | 4.7330 | 5.4084 | 4.5584 | 5.0986 |
| RAB3IP_S262 | RAB3IP | Q96QF0 | | 163 | SLSRLRSPSVLEV | 7.1660 | 7.6283 | 6.5476 | 7.4326 | 6.0451 | 7.7442 | 7.9859 | 7.1575 | 7.2152 | 7.3616 | 7.8512 | 7.8479 | 2.6396 | 3.4334 | 2.1695 | 3.7741 |
| RAB5B_S123 | RAB5B | P61020 | | 123 | ELQRQASPSIVIA | 5.6236 | 6.3832 | 5.8756 | 6.3632 | 6.0555 | 5.9329 | 6.7572 | 6.9586 | 6.2880 | 6.2275 | 6.8371 | 6.2175 | 6.2764 | 6.1218 | 5.8840 | 6.1980 |
| RAB7A_S72 | RAB7A | P51149 | | 72 | GQERFQSLGVAFY | 6.0454 | 5.9404 | 6.4693 | 6.8405 | 6.6125 | 5.7937 | 6.5711 | 6.4168 | 6.4437 | 5.6809 | 5.9945 | 5.8372 | 6.3638 | 6.6738 | 6.1707 | 6.1457 |
| RAB8A_S181 | RAB8A | P61006 | | 185 | GNSPQGSNQGVKI | 5.3003 | 5.9695 | 6.3277 | 6.3165 | 5.5861 | 5.9896 | 6.1480 | 6.1592 | 6.1451 | 6.4479 | 5.9265 | 6.1396 | 6.0218 | 6.8364 | 6.9403 | 7.4544 |
| RAB8A_S181;S185 | RAB8A | P61006 | | 72 | GQERFRTITTAYY | 7.5643 | 7.2737 | 7.2377 | 7.8997 | 6.1167 | 6.0883 | 6.4757 | 5.8600 | 6.2209 | 6.5064 | 6.3221 | 6.3096 | 5.1045 | 5.4208 | 4.6060 | 4.9938 |
| RAB8A_S185 | RAB8A | P61006 | | 181 | KKLEGNSPQGSNQ | 6.2572 | 7.4539 | 7.1150 | 7.8900 | 5.9655 | 6.6727 | 7.1718 | 7.0197 | 7.8588 | 7.6382 | 6.8669 | 6.9351 | 3.3305 | 4.5866 | 2.9040 | 4.3341 |
| RAB8A_T192 | RAB8A | P61006 | | 192 | NQGVKITPDQQKR | 7.2055 | 6.7608 | 5.3422 | 6.7415 | 6.3232 | 6.8449 | 6.8064 | 5.6452 | 5.9265 | 6.2559 | 7.1478 | 6.6982 | 5.7149 | 5.8327 | 4.9936 | 5.7606 |
| RAB8A_T72 | RAB8A | P61006 | 181;185 | | NA | 7.3267 | 8.3574 | 6.7291 | 7.1724 | 7.2549 | 7.0984 | 8.2242 | 6.9742 | 6.6370 | 6.8550 | 7.5650 | 6.2520 | 2.9770 | 4.3123 | 2.8957 | 3.3687 |
| RAB8B_S182 | RAB8B | Q92930 | | 182 | KMNDSNSAGAGGP | 6.0767 | 5.9285 | 5.6069 | 6.6672 | 5.9453 | 6.1488 | 5.1371 | 6.3004 | 5.8054 | 5.8975 | 6.8054 | 6.2744 | 6.5918 | 7.6194 | 6.4896 | 7.6057 |
| RAB9A_S179 | RAB9A | P51151 | | 179 | LATEDRSDHLIQT | 5.6099 | 5.8065 | 6.1776 | 6.3456 | 6.0161 | 6.3138 | 6.2268 | 6.0189 | 6.4684 | 6.3810 | 5.9884 | 6.5272 | 6.4732 | 6.6440 | 6.2103 | 6.7922 |
| RAB9A_Y107 | RAB9A | P51151 | | 107 | WKKEFIYYADVKE | 6.5275 | 6.3350 | 6.4428 | 7.2658 | 4.9292 | 5.8198 | 6.8300 | 6.5349 | 6.5507 | 5.7839 | 6.8088 | 6.6713 | 5.0995 | 6.5560 | 5.6210 | 6.2239 |
| RABEP1_S377 | RABEP1 | Q15276 | | 377 | TRGSVHSLDAGLL | 5.8796 | 5.8890 | 5.6957 | 7.9465 | 6.5218 | 6.5312 | 6.1758 | 6.6808 | 4.4611 | 6.7089 | 7.5693 | 7.5786 | 7.1266 | 6.2319 | 4.2959 | 4.7074 |
| RABEP1_S407 | RABEP1 | Q15276 | | 407 | GLRRAQSTDSLGT | 6.5640 | 6.1641 | 6.7952 | 6.6559 | 6.2744 | 6.8286 | 6.4911 | 6.8532 | 7.8135 | 6.9600 | 6.6620 | 7.4672 | 4.2637 | 5.0838 | 4.0087 | 5.1145 |
| RABEP1_S407;S410 | RABEP1 | Q15276 | 407;410 | | NA | 6.2690 | 5.9833 | 7.3031 | 6.2224 | 5.7986 | 5.4894 | 5.7340 | 7.1189 | 6.8631 | 5.5169 | 7.4205 | 6.1949 | 6.0825 | 6.2421 | 5.9662 | 5.6950 |
| RABEPK_S133 | RABEPK | Q7Z6M1 | | 133 | TTPEVTSPPPSPR | 5.3834 | 5.7295 | 6.7570 | 7.3084 | 6.3159 | 6.1098 | 6.2718 | 7.4499 | 6.6863 | 6.5737 | 6.4520 | 7.4309 | 5.6116 | 4.9362 | 5.5660 | 5.4176 |
| RABEPK_S61 | RABEPK | Q7Z6M1 | | 61 | GANPNRSFSDVHT | 6.3064 | 6.1644 | 7.4758 | 5.9751 | 6.1783 | 5.5407 | 6.2980 | 7.1834 | 6.7769 | 5.5073 | 5.0312 | 6.7992 | 5.4934 | 6.6071 | 6.3704 | 6.2925 |
| RABGAP1_S352 | RABGAP1 | Q9Y3P9 | | 491 | RRKTTASPSVRLP | 6.8395 | 7.5807 | 6.9663 | 7.1985 | 6.3183 | 7.1882 | 6.9815 | 7.0181 | 7.2457 | 7.5040 | 7.7738 | 7.8091 | 2.9434 | 3.7921 | 2.9599 | 3.8808 |
| RABGAP1_S491 | RABGAP1 | Q9Y3P9 | | 352 | CFGLLLSPGKDVR | 6.6213 | 9.2102 | 7.0985 | 6.3410 | 6.7705 | 7.4684 | 6.6094 | 7.3073 | 8.7569 | 6.2396 | 7.3431 | 6.4304 | 3.4359 | 3.6209 | 3.3882 | 3.3584 |
| RABGAP1L_S119 | RABGAP1L | Q5R372-3 | | 119 | STPRPSSPGGLPE | 5.6207 | 5.0278 | 7.0861 | 6.4635 | 5.8116 | 6.1458 | 6.1440 | 6.5808 | 6.1862 | 6.8865 | 6.5324 | 6.7143 | 5.9860 | 6.6318 | 6.0010 | 6.1813 |
| RABGEF1_S164 | RABGEF1 | Q9UJ41-3 | | 430 | YMSGQTSPRKQEA | 6.2992 | 6.5858 | 6.6889 | 6.5805 | 6.0649 | 6.5439 | 6.4390 | 6.7799 | 6.7340 | 6.3771 | 6.8496 | 6.9418 | 4.9561 | 5.3704 | 5.1455 | 5.6434 |
| RABGEF1_S430 | RABGEF1 | Q9UJ41-3 | | 164 | QEAKAPSPSINRQ | 7.8448 | 8.0254 | 7.0316 | 6.7214 | 6.4121 | 7.1529 | 7.2748 | 7.5338 | 7.8710 | 6.9450 | 7.4789 | 6.9807 | 2.7478 | 3.7381 | 2.7686 | 3.4731 |
| RABGEF1_S440 | RABGEF1 | Q9UJ41-3 | | 440 | QEAESWSPDACLG | 5.9919 | 6.6511 | 7.1622 | 7.4193 | 6.2333 | 6.7922 | 7.2514 | 6.1526 | 6.4524 | 6.5987 | 7.4541 | 7.1628 | 4.6103 | 4.7133 | 3.8126 | 5.5419 |
| RABL6_S403 | RABL6 | Q3YEC7-2 | | 403 | DDRLDRSFLEDTT | 5.6693 | 6.0571 | 7.0339 | 6.9916 | 5.9703 | 6.1600 | 6.6464 | 7.0454 | 7.6842 | 7.0517 | 6.9487 | 7.1253 | 4.1587 | 5.6349 | 4.4081 | 5.4144 |
| RABL6_S597;T600 | RABL6 | Q3YEC7-2 | | 641 | EAGPKSSEEGKE | 5.9666 | 6.1516 | 7.4364 | 5.9476 | 5.9489 | 5.9020 | 6.3457 | 6.1994 | 7.1923 | 6.7692 | 6.7340 | 6.7299 | 5.2365 | 6.4360 | 5.1639 | 5.8399 |
| RABL6_S641 | RABL6 | Q3YEC7-2 | 597;600 | | NA | 6.0776 | 5.8510 | 6.6149 | 6.2633 | 5.8747 | 6.3170 | 6.0253 | 6.2286 | 6.4290 | 6.2138 | 6.6518 | 6.8106 | 5.9153 | 6.7367 | 6.0212 | 5.9689 |
| RABL6_S641;S642 | RABL6 | Q3YEC7-2 | 641;642 | | NA | 6.0771 | 5.7950 | 6.7071 | 6.3732 | 5.9855 | 5.6732 | 6.6790 | 5.7084 | 6.0227 | 6.2523 | 6.1164 | 6.2709 | 6.4866 | 6.7049 | 6.3890 | 6.7588 |
| RAC1_S71 | RAC1 | P63000-2 | | 71 | DRLRPLSYPQTVG | 8.0024 | 8.6792 | 8.2191 | 7.9453 | 7.3407 | 6.4205 | 7.9510 | 6.1562 | 5.5859 | 5.9946 | 6.2817 | 6.6506 | 3.5439 | 4.2227 | 3.5896 | 3.4165 |
| RACGAP1_S123 | RACGAP1 | Q9H0H5 | | 628 | QGNFFASPMLKxx | 7.6936 | 7.8561 | 7.0672 | 7.0904 | 5.4247 | 7.4520 | 7.7249 | 6.9348 | 7.7867 | 7.5962 | 7.4704 | 7.7939 | 2.4830 | 3.7948 | 2.4936 | 3.3378 |
| RACGAP1_S149;S157 | RACGAP1 | Q9H0H5 | | 342 | IPTLIGTPVKIGE | 6.8184 | 6.6782 | 6.6977 | 7.0437 | 6.3447 | 6.7025 | 7.0453 | 6.3323 | 6.8708 | 7.2034 | 6.7808 | 6.6213 | 4.2743 | 5.0913 | 4.6004 | 4.8951 |
| RACGAP1_S154;S157 | RACGAP1 | Q9H0H5 | | 580 | LLGPVTTPEHQLL | 6.8104 | 7.3153 | 6.8473 | 7.4204 | 6.4057 | 6.7435 | 7.1247 | 6.9012 | 7.1033 | 7.1818 | 6.6450 | 7.2512 | 3.7232 | 4.5876 | 3.7092 | 4.2302 |
| RACGAP1_S157 | RACGAP1 | Q9H0H5 | | 203 | PVKKTRSIGSAVD | 6.0749 | 6.3997 | 6.1323 | 6.8693 | 5.6043 | 6.1554 | 6.4017 | 6.3461 | 6.4083 | 6.6105 | 6.2643 | 6.6472 | 5.8343 | 6.0491 | 5.8424 | 6.3602 |
| RACGAP1_S203 | RACGAP1 | Q9H0H5 | | 123 | LSEEQKSALAFLN | 7.8046 | 7.5300 | 6.1779 | 7.1451 | 6.4322 | 6.3232 | 7.4771 | 6.0754 | 6.0491 | 5.8794 | 6.4378 | 5.4361 | 5.3335 | 5.6348 | 4.9702 | 5.2935 |
| RACGAP1_S573 | RACGAP1 | Q9H0H5 | | 573 | TPDIKVSLLGPVT | 7.3386 | 6.4971 | 6.1923 | 8.0249 | 4.9583 | 6.2816 | 6.8238 | 5.7888 | 6.7703 | 5.9549 | 5.7311 | 5.7476 | 6.0098 | 6.1882 | 5.8231 | 5.8698 |
| RACGAP1_S573;T580 | RACGAP1 | Q9H0H5 | | 157 | SILSDISFDKTDE | 6.2889 | 6.8789 | 5.9458 | 6.1559 | 6.7847 | 5.4066 | 5.9319 | 5.8159 | 7.1253 | 7.5181 | 6.9969 | 7.3043 | 5.3284 | 5.5737 | 5.1416 | 5.8030 |
| RACGAP1_S628 | RACGAP1 | Q9H0H5 | | 243 | ETVPYWTRSRRKT | 6.2698 | 8.0596 | 6.1607 | 6.6953 | 6.1479 | 6.8279 | 7.0139 | 6.8749 | 6.2698 | 6.3168 | 5.3524 | 5.3866 | 5.1450 | 5.9234 | 5.2989 | 6.2569 |
| RACGAP1_T243 | RACGAP1 | Q9H0H5 | | 249 | TRSRRKTGTLQPW | 7.4627 | 7.1345 | 6.3350 | 7.0405 | 5.5516 | 6.5118 | 7.0913 | 6.7808 | 7.1252 | 6.4060 | 6.4644 | 6.8950 | 4.4813 | 5.4864 | 5.4668 | 4.6666 |
| RACGAP1_T249 | RACGAP1 | Q9H0H5 | 573;580 | | NA | 7.5611 | 6.8805 | 7.9095 | 7.9993 | 5.9945 | 6.6993 | 8.0729 | 5.7961 | 8.0581 | 6.6231 | 6.2032 | 6.0637 | 3.8350 | 3.9906 | 3.6466 | 4.6665 |
| RACGAP1_T342 | RACGAP1 | Q9H0H5 | 149;157 | | NA | 6.4996 | 6.8581 | 7.1480 | 7.3735 | 6.0295 | 6.7329 | 6.5628 | 7.4203 | 6.3290 | 6.4858 | 6.7116 | 6.9839 | 4.5660 | 4.9278 | 4.2406 | 5.1304 |
| RACGAP1_T580 | RACGAP1 | Q9H0H5 | 588;592 | | NA | 7.0859 | 7.3769 | 6.2572 | 6.9172 | 5.2297 | 5.9317 | 6.9076 | 6.9225 | 7.4913 | 7.6492 | 7.0868 | 7.7166 | 3.9724 | 4.7589 | 3.6567 | 5.0395 |
| RACGAP1_T588;S592 | RACGAP1 | Q9H0H5 | 154;157 | | NA | 5.9440 | 8.0074 | 8.1629 | 6.8570 | 8.0778 | 7.1047 | 6.8110 | 4.6869 | 6.8714 | 4.7790 | 5.7100 | 8.4460 | 4.9252 | 4.1463 | 4.5126 | 4.9577 |
| RACK1_S276 | RACK1 | P63244 | | 276 | LKQEVISTSSKAE | 6.1295 | 6.6165 | 6.6929 | 6.4018 | 6.5879 | 6.1965 | 6.1060 | 6.1306 | 6.7724 | 6.0821 | 6.7728 | 6.7724 | 5.1605 | 5.6777 | 5.7761 | 5.8744 |
| RAD18_S103 | RAD18 | Q9NS91 | | 471 | LQDTEISPRQNRR | 6.4729 | 6.5791 | 6.7353 | 6.7630 | 6.3309 | 6.0143 | 6.5484 | 6.1818 | 6.7029 | 6.3404 | 6.3772 | 6.7635 | 5.5033 | 5.4455 | 5.4331 | 5.8086 |
| RAD18_S403 | RAD18 | Q9NS91 | | 99 | LQFALESPAKSPA | 6.8337 | 6.3876 | 6.8662 | 6.6370 | 6.4240 | 6.3542 | 6.9264 | 6.0197 | 6.9004 | 6.6970 | 6.6535 | 6.8511 | 5.0680 | 5.1759 | 4.6080 | 5.5973 |
| RAD18_S471 | RAD18 | Q9NS91 | | 118 | LAVKVYTPVASRQ | 6.1486 | 6.2222 | 7.2287 | 6.9048 | 6.2937 | 6.6271 | 6.5415 | 6.7711 | 7.4278 | 7.2751 | 7.2296 | 7.2452 | 4.7589 | 4.9506 | 3.7235 | 4.6517 |
| RAD18_S99 | RAD18 | Q9NS91 | | 403 | SVTNHFSQSKLDS | 5.9481 | 6.1068 | 5.2603 | 6.0369 | 6.4730 | 6.1711 | 5.9701 | 5.4825 | 5.3586 | 5.4862 | 6.0468 | 5.8162 | 7.1670 | 7.1400 | 8.3077 | 7.2288 |
| RAD18_T118 | RAD18 | Q9NS91 | | 103 | LESPAKSPASSSS | 6.8461 | 7.2333 | 7.0176 | 7.9657 | 6.0730 | 6.3246 | 6.6157 | 6.7213 | 7.4034 | 7.0523 | 7.5411 | 7.8847 | 3.4068 | 3.9623 | 3.4610 | 4.4910 |
| RAD21_S153 | RAD21 | O60216 | | 153 | EEVGNISILQENO | 6.2592 | 5.5508 | 6.5859 | 7.6213 | 6.6281 | 5.7988 | 6.1791 | 6.7345 | 7.1093 | 6.2139 | 6.7623 | 6.8225 | 5.6815 | 5.5035 | 4.7642 | 5.7851 |
| RAD21_S175 | RAD21 | O60216 | | 175 | EIMREGSAFEDDD | 5.7322 | 6.2528 | 6.2824 | 6.3191 | 5.9491 | 5.9414 | 6.6830 | 5.9664 | 6.1617 | 6.1593 | 5.9235 | 6.0910 | 6.9459 | 6.5331 | 6.3894 | 6.6697 |
| RAD21_T394 | RAD21 | O60216 | | 394 | LFTRCLTPLVPED | 6.1229 | 7.2202 | 7.0851 | 6.3631 | 6.0386 | 6.0567 | 5.6940 | 5.4190 | 6.1222 | 6.1403 | 6.2802 | 6.0513 | 6.4621 | 6.2855 | 6.6241 | 6.0346 |
| RAD23A_S123 | RAD23A | P54725 | | 295 | GELADISDVEGEV | 6.6012 | 6.3753 | 7.1708 | 6.1248 | 6.3706 | 6.4010 | 6.5139 | 7.0104 | 6.1014 | 6.4238 | 6.5073 | 6.5948 | 5.3224 | 5.7680 | 5.1624 | 5.5518 |
| RAD23A_S295 | RAD23A | P54725 | | 123 | AAREDKSPSEESA | 5.7730 | 5.7528 | 6.3857 | 5.5540 | 5.6332 | 5.9810 | 6.0383 | 7.0435 | 6.3658 | 6.4698 | 6.4656 | 6.8722 | 5.8602 | 7.0698 | 6.5883 | 6.1468 |
| RAD23B_S160 | RAD23B | P54727 | | 160 | ETPVATSPTATDS | 5.1726 | 4.7277 | 5.7176 | 5.3157 | 5.9297 | 6.2583 | 5.7965 | 5.6830 | 5.6923 | 6.2109 | 6.1850 | 5.9768 | 8.1398 | 7.6070 | 7.6900 | 7.8972 |
| RAD50_S641 | RAD50 | Q92878-2 | | 641 | LFDVCGSQDFESD | 6.6586 | 7.7171 | 6.5471 | 6.8879 | 6.2181 | 7.5281 | 5.8428 | 5.0868 | 5.9787 | 6.1926 | 6.6851 | 5.8256 | 6.2864 | 6.0263 | 6.0484 | 5.4703 |
| RAD50_T696 | RAD50 | Q92878-2 | | 696 | CQRVFQTEAELQE | 7.1186 | 6.8780 | 7.2395 | 9.7626 | 7.8221 | 3.3513 | 5.2295 | 3.1016 | 4.7068 | 3.3721 | 3.9829 | 2.9183 | 9.2266 | 6.7957 | 12.0174 | 6.4770 |
| RAD51AP1_S19;S21 | RAD51AP1 | Q96B01 | | 294 | KPLEIRSPSAESK | 7.4542 | 7.0562 | 8.3676 | 6.8982 | 6.9470 | 8.6632 | 7.0540 | 5.2359 | 6.5621 | 7.7507 | 8.7011 | 7.9816 | 2.6763 | 3.5428 | 2.2021 | 2.9070 |
| RAD51AP1_S294 | RAD51AP1 | Q96B01 | | 327 | VVVSVKSPNQSLR | 7.2705 | 7.8674 | 7.2065 | 7.7611 | 6.2717 | 6.7726 | 7.1971 | 7.2282 | 7.5577 | 7.9950 | 8.6215 | 8.3493 | 1.9507 | 3.0908 | 1.8718 | 2.9881 |
| RAD51AP1_S318 | RAD51AP1 | Q96B01 | | 318 | GSRSSSSPLVVVS | 6.7260 | 7.8820 | 7.3743 | 7.0976 | 6.0242 | 7.5702 | 7.7283 | 6.7275 | 7.8869 | 7.8282 | 7.7975 | 8.6185 | 2.1692 | 3.2949 | 2.0743 | 3.2004 |
| RAD51AP1_S327 | RAD51AP1 | Q96B01 | 19;21 | | NA | 5.7377 | 6.2352 | 6.6206 | 6.1679 | 5.7880 | 6.4275 | 5.6858 | 6.6409 | 6.4397 | 6.4622 | 6.8046 | 6.7789 | 5.5196 | 6.6312 | 5.9003 | 6.1598 |
| RAD51C_S20 | RAD51C | O43502 | | 20 | LVSFPLSPAVRVK | 7.2496 | 6.7164 | 6.6380 | 7.1397 | 6.2640 | 6.6370 | 7.1986 | 6.9006 | 7.0229 | 7.0593 | 6.9405 | 7.0898 | 3.5851 | 4.9041 | 4.0159 | 4.6385 |
| RAD54B_S14 | RAD54B | Q9Y620 | | 14 | SQLQGNSFKKPKF | 6.0703 | 6.3599 | 7.3404 | 6.5626 | 5.5509 | 6.3838 | 5.5165 | 6.5452 | 7.5325 | 8.7747 | 8.9317 | 9.9972 | 3.0262 | 5.0535 | 2.9279 | 3.4268 |
| RAD54L_S38 | RAD54L | Q92698 | | 38 | PRKRSSSETQIQ | 3.0079 | 3.4090 | 4.2586 | 3.6849 | 5.7765 | 5.0110 | 4.9357 | 4.8678 | 4.2935 | 4.0831 | 5.0951 | 3.9204 | 12.7428 | 12.4338 | 11.7325 | 10.7975 |
| RAD54L2_S1155 | RAD54L2 | Q9Y4B4 | | 1260 | HKRKLATPPAAQE | 5.5676 | 5.9193 | 5.8915 | 5.4912 | 6.1229 | 6.4178 | 5.9286 | 5.8775 | 6.0385 | 5.7228 | 6.3165 | 5.7123 | 6.9277 | 7.3444 | 7.3929 | 7.3284 |
| RAD54L2_S1169 | RAD54L2 | Q9Y4B4 | | 1169 | GLARPVSPDSPEI | 5.3714 | 6.6518 | 7.1792 | 6.7375 | 6.2363 | 5.8259 | 6.2651 | 6.7458 | 6.4396 | 6.1168 | 7.3102 | 7.1862 | 5.2001 | 5.1324 | 5.6361 | 5.9659 |
| RAD54L2_S1169;S1172 | RAD54L2 | Q9Y4B4 | | 1155 | GRHSASSPKAPDP | 5.7046 | 6.2470 | 6.2979 | 6.2143 | 5.8549 | 5.8218 | 5.8382 | 5.9124 | 5.7532 | 6.2181 | 6.3442 | 6.6106 | 6.6742 | 6.7172 | 6.8450 | 6.9464 |
| RAD54L2_S1195 | RAD54L2 | Q9Y4B4 | | 1195 | RESRQSSPSTNAA | 4.0081 | 6.6982 | 5.6037 | 5.9560 | 5.3650 | 6.7019 | 6.0422 | 6.1996 | 5.9735 | 6.7494 | 4.9202 | 5.9722 | 7.3766 | 7.9726 | 8.3649 | 6.0959 |
| RAD54L2_S1450;S1451 | RAD54L2 | Q9Y4B4 | 1450;1451 | | NA | 5.6689 | 5.9764 | 6.2408 | 5.7869 | 6.3077 | 5.8601 | 6.3929 | 6.3762 | 5.9273 | 6.7413 | 6.2685 | 6.2173 | 6.4237 | 6.3482 | 6.6111 | 6.8526 |
| RAD54L2_T1260 | RAD54L2 | Q9Y4B4 | 1169;1172 | | NA | 6.0617 | 5.9664 | 6.2552 | 6.4074 | 6.1061 | 6.0491 | 6.1316 | 6.2817 | 6.0260 | 5.3281 | 6.4012 | 6.8386 | 6.1661 | 6.8857 | 6.6897 | 6.4052 |
| RAD9A_S272;S277 | RAD9A | Q99638 | | 277 | HSQDLSPERHQP | 5.6140 | 6.0644 | 6.1860 | 6.3063 | 6.1843 | 6.6203 | 6.1275 | 6.2647 | 6.2221 | 6.6344 | 6.3709 | 6.2832 | 6.0703 | 6.6178 | 6.1140 | 6.3196 |
| RAD9A_S277 | RAD9A | Q99638 | | 328 | LPSISLSPGPQPP | 5.7545 | 6.2316 | 6.3850 | 5.7859 | 5.8395 | 6.1008 | 5.8082 | 6.2228 | 6.1379 | 6.4281 | 6.0180 | 6.0097 | 6.1708 | 6.8769 | 7.1414 | 7.0889 |
| RAD9A_S328 | RAD9A | Q99638 | 272;277 | | NA | 5.8815 | 6.4024 | 6.7350 | 4.4812 | 6.5833 | 5.9131 | 4.9141 | 6.7398 | 5.7623 | 5.3736 | 6.7258 | 4.6631 | 6.4105 | 7.9743 | 7.0912 | 8.3490 |
| RAE1_S209 | RAE1 | P78406 | | 209 | EFRRIESPLKHQH | 7.1411 | 7.2561 | 6.8081 | 7.3275 | 6.7604 | 6.5204 | 6.8024 | 7.0408 | 6.9670 | 6.9866 | 7.2371 | 7.4179 | 3.7172 | 4.0033 | 3.7165 | 4.2976 |
| RAF1_S233 | RAF1 | P04049-2 | | 259 | QRQRSTSTPNVHM | 5.8610 | 6.5949 | 6.7958 | 6.3401 | 6.4868 | 6.6744 | 6.1991 | 6.5222 | 6.4636 | 6.5278 | 6.4121 | 6.5558 | 5.8167 | 5.9777 | 5.3985 | 5.3733 |
| RAF1_S259 | RAF1 | P04049-2 | | 29 | DGSSCISPTIVQQ | 6.1823 | 6.1605 | 7.3588 | 6.7992 | 6.9032 | 7.0300 | 6.2949 | 6.4070 | 7.0331 | 5.9885 | 6.8905 | 6.4476 | 5.4822 | 5.3077 | 4.6086 | 5.1058 |
| RAF1_S29 | RAF1 | P04049-2 | | 662 | ACTLTTSPRLPVF | 6.0920 | 6.5289 | 7.3330 | 6.6197 | 6.1819 | 6.2579 | 7.4022 | 7.7860 | 6.3183 | 5.6607 | 6.6224 | 6.9524 | 5.5685 | 5.5136 | 4.9463 | 4.9164 |
| RAF1_S315 | RAF1 | P04049-2 | | 233 | SSQHRYSTPHAFT | 5.8997 | 4.2223 | 6.1828 | 5.2782 | 6.6389 | 5.0916 | 5.2943 | 6.8787 | 6.3687 | 6.6631 | 5.9877 | 6.1773 | 6.9878 | 8.1781 | 7.4878 | 6.6631 |
| RAF1_S662 | RAF1 | P04049-2 | | 315 | SPSALSSSPNNLS | 6.4995 | 7.1791 | 6.0441 | 6.5951 | 7.8255 | 6.6079 | 6.2002 | 8.0202 | 6.8815 | 5.6584 | 6.5914 | 7.0836 | 5.1938 | 6.1818 | 3.8090 | 4.5289 |
| RAI1_S1108;S1122 | RAI1 | Q7Z5J4 | | 892 | GMQDPLSPKAPLI | 5.9947 | 5.9219 | 6.1098 | 6.4325 | 6.2759 | 6.5934 | 6.2790 | 5.8030 | 6.3219 | 6.2448 | 6.2661 | 6.1016 | 6.9703 | 6.6183 | 6.4570 | 6.3098 |
| RAI1_S1122 | RAI1 | Q7Z5J4 | | 1266 | EERPEGSPTLFKR | 6.6358 | 6.7637 | 6.2020 | 7.2615 | 6.1789 | 6.0832 | 6.6043 | 6.4595 | 6.4150 | 6.5391 | 6.3206 | 6.4718 | 5.0379 | 5.7602 | 5.6137 | 5.6527 |
| RAI1_S1192 | RAI1 | Q7Z5J4 | | 1122 | PVASDSSPMGSKT | 6.3216 | 6.3935 | 6.3759 | 6.7661 | 6.2791 | 6.3336 | 6.4963 | 6.2733 | 6.6174 | 6.4220 | 6.4379 | 6.4492 | 5.5259 | 6.1707 | 5.4434 | 5.6943 |
| RAI1_S1266 | RAI1 | Q7Z5J4 | | 805 | PGEKVASLPGDFK | 6.3394 | 6.3699 | 5.7856 | 6.5484 | 5.9484 | 5.7501 | 6.6080 | 6.1864 | 5.9405 | 6.1005 | 6.1260 | 6.1761 | 6.1399 | 6.7394 | 6.4918 | 6.7496 |
| RAI1_S1338 | RAI1 | Q7Z5J4 | | 683 | GNAKDSPGLFED | 5.5389 | 5.8615 | 5.3008 | 6.0598 | 5.8619 | 6.0366 | 6.1350 | 5.6475 | 5.7936 | 6.0589 | 5.6024 | 5.7120 | 7.6701 | 7.7711 | 7.8519 | 7.0979 |
| RAI1_S1352;S1358 | RAI1 | Q7Z5J4 | | 1374 | LKGAGGSPVGVEE | 6.1762 | 6.6994 | 6.0300 | 6.8864 | 6.5698 | 6.0056 | 6.9421 | 6.1933 | 6.3976 | 6.3998 | 5.9611 | 6.2846 | 5.8046 | 6.3205 | 5.6332 | 5.6957 |
| RAI1_S1358 | RAI1 | Q7Z5J4 | | 1476 | GPALLLTPRDRAS | 7.6053 | 8.4900 | 7.4641 | 7.1603 | 5.8559 | 6.6175 | 6.9517 | 6.8588 | 6.7486 | 6.7126 | 7.7689 | 7.0113 | 3.2258 | 4.1449 | 2.9204 | 4.4638 |
| RAI1_S1374 | RAI1 | Q7Z5J4 | | 1616 | ICTVVNSPGDAPK | 5.7963 | 6.1232 | 6.4852 | 7.4119 | 6.4931 | 6.0275 | 6.7926 | 5.6442 | 6.4571 | 6.8539 | 6.0572 | 6.5942 | 5.2379 | 6.0501 | 5.7564 | 6.2192 |
| RAI1_S1431 | RAI1 | Q7Z5J4 | | 528 | EDPLERSFLYCNQ | 8.4037 | 8.2012 | 6.5651 | 9.2751 | 7.4798 | 6.1208 | 7.3789 | 6.3236 | 7.0701 | 5.6460 | 6.5466 | 6.5066 | 3.1176 | 4.3919 | 3.0263 | 3.9467 |
| RAI1_S1550 | RAI1 | Q7Z5J4 | | 1358 | SPGNPLSPSLSDK | 6.5979 | 6.7727 | 6.8007 | 7.7756 | 6.1856 | 6.3391 | 6.2391 | 6.5734 | 6.6947 | 6.8810 | 6.6711 | 6.7597 | 4.3272 | 4.9410 | 5.1805 | 5.2607 |
| RAI1_S1590 | RAI1 | Q7Z5J4 | | 1176 | ETDSPSTPGKDQR | 7.1318 | 6.8343 | 5.9698 | 6.7950 | 6.5233 | 5.9752 | 6.4824 | 6.0996 | 6.7240 | 6.2064 | 5.9457 | 5.9983 | 5.8797 | 5.8773 | 5.7542 | 5.8031 |
| RAI1_S1596 | RAI1 | Q7Z5J4 | | 1192 | TFKVSSSPQKEGR | 6.4101 | 6.9791 | 6.3413 | 7.4007 | 5.8084 | 5.5115 | 5.8206 | 5.8897 | 6.0171 | 6.4223 | 6.5440 | 6.7170 | 5.5467 | 6.1807 | 6.2343 | 6.1768 |
| RAI1_S1616 | RAI1 | Q7Z5J4 | | 1431 | KTEAFTSPEALQP | 6.7987 | 7.3705 | 6.6005 | 7.6462 | 6.0915 | 6.2299 | 7.2045 | 6.6363 | 7.0815 | 6.8947 | 7.3469 | 7.6148 | 3.6983 | 4.7082 | 3.7753 | 4.3022 |
| RAI1_S528 | RAI1 | Q7Z5J4 | | 1297 | KLPPPETPDACLK | 6.9031 | 6.7228 | 6.7523 | 7.6209 | 6.4025 | 6.3031 | 6.9490 | 6.8472 | 6.8028 | 5.9696 | 5.9342 | 6.2778 | 5.2090 | 5.1270 | 4.6019 | 5.5767 |
| RAI1_S683 | RAI1 | Q7Z5J4 | | 1068 | ALSEPRTPGPPGL | 7.2863 | 6.4222 | 5.8740 | 7.2181 | 6.6310 | 5.8903 | 6.2793 | 6.2067 | 6.2266 | 6.4784 | 5.9134 | 5.4134 | 6.2505 | 6.7965 | 5.7306 | 6.3828 |
| RAI1_S805 | RAI1 | Q7Z5J4 | | 696 | PSVAFATPDPKKT | 6.0660 | 7.9214 | 7.8086 | 8.5108 | 8.2904 | 5.9455 | 9.5692 | 5.8943 | 6.0275 | 6.1300 | 6.6375 | 5.3279 | 3.8133 | 4.3746 | 2.6883 | 4.9947 |
| RAI1_S880 | RAI1 | Q7Z5J4 | | 1596 | SRTPAFSPFVRVE | 7.2563 | 7.1531 | 6.8792 | 9.0051 | 6.5484 | 5.8889 | 6.9824 | 7.4354 | 8.3182 | 5.7478 | 6.4557 | 7.1952 | 4.5068 | 3.9337 | 3.9547 | 2.7390 |
| RAI1_S880;S892 | RAI1 | Q7Z5J4 | | 880 | LCELLGSPEQRPG | 5.9491 | 4.7580 | 6.2981 | 7.4282 | 6.0059 | 7.2050 | 5.9754 | 4.6647 | 6.8033 | 7.1502 | 6.4807 | 6.5639 | 5.8537 | 7.7042 | 5.7746 | 5.3850 |
| RAI1_S892 | RAI1 | Q7Z5J4 | | 1590 | KRLRSDSRTPAFS | 5.2292 | 6.0477 | 6.3050 | 6.2120 | 6.7596 | 4.9257 | 6.0227 | 5.9917 | 6.7759 | 7.7223 | 7.1159 | 6.1462 | 6.7803 | 6.0956 | 5.6192 | 6.2511 |
| RAI1_T1068 | RAI1 | Q7Z5J4 | | 1338 | IVQKITSPSLKKF | 6.9834 | 7.5437 | 6.7443 | 7.5721 | 6.1766 | 6.9054 | 6.8334 | 6.8705 | 6.9198 | 6.8987 | 7.1825 | 7.3090 | 3.7111 | 4.6318 | 3.4854 | 4.2321 |
| RAI1_T1136 | RAI1 | Q7Z5J4 | | 1548 | RGRAKNTTSSPCK | 6.4459 | 6.3904 | 6.4145 | 6.8578 | 5.8308 | 5.9276 | 6.1819 | 5.4417 | 6.6796 | 6.7642 | 5.9673 | 6.5697 | 6.1679 | 6.0705 | 6.3140 | 5.9762 |
| RAI1_T1297 | RAI1 | Q7Z5J4 | | 1550 | RAKNTTSSPCKGR | 6.6115 | 6.5107 | 6.2207 | 7.3606 | 6.2282 | 5.6135 | 6.2903 | 5.5121 | 6.6339 | 6.2909 | 6.2731 | 6.4077 | 5.6361 | 5.9353 | 6.2153 | 6.2602 |
| RAI1_T1476 | RAI1 | Q7Z5J4 | 880;892 | | NA | 6.4641 | 7.0402 | 6.7844 | 5.9934 | 7.1753 | 6.9562 | 7.0007 | 6.5706 | 6.8264 | 5.8888 | 6.1574 | 5.8766 | 5.6086 | 5.6758 | 5.6816 | 4.2999 |
| RAI1_T1548 | RAI1 | Q7Z5J4 | 1352;1358 | | NA | 6.8107 | 7.5958 | 7.2558 | 6.9822 | 6.5077 | 6.4797 | 7.8266 | 7.1627 | 7.5209 | 7.2925 | 6.7877 | 7.0429 | 3.1393 | 4.3300 | 3.0049 | 4.2603 |
| RAI1_T696 | RAI1 | Q7Z5J4 | 1108;1122 | | NA | 5.6306 | 6.5930 | 6.8207 | 7.4858 | 6.3095 | 6.5206 | 6.8003 | 6.0899 | 6.8538 | 6.5202 | 6.7439 | 6.3965 | 6.0693 | 5.7942 | 5.5037 | 4.8679 |
| RAI14_S284 | RAI14 | Q9P0K7-2 | | 300 | PRSITSTPLSGKE | 6.1042 | 5.4827 | 5.2399 | 5.0694 | 6.3335 | 5.6361 | 5.7002 | 5.7654 | 6.3676 | 5.3166 | 4.5195 | 4.8015 | 9.5861 | 8.2570 | 8.1571 | 7.6631 |
| RAI14_S296 | RAI14 | Q9P0K7-2 | | 422 | PSVLIHSLGKSTT | 6.0258 | 6.2412 | 6.2909 | 7.1631 | 6.5532 | 6.6912 | 5.9944 | 6.4700 | 6.1542 | 6.5625 | 6.3130 | 6.2156 | 6.0025 | 6.4302 | 5.8686 | 5.9336 |
| RAI14_S296;S303 | RAI14 | Q9P0K7-2 | | 296 | DVSSPRSITSTPL | 6.0073 | 6.5431 | 6.6921 | 6.5918 | 6.4458 | 6.0649 | 5.7267 | 6.5227 | 6.6010 | 6.8348 | 6.8966 | 7.0621 | 5.5870 | 5.6491 | 5.4627 | 5.3123 |
| RAI14_S296;T300 | RAI14 | Q9P0K7-2 | | 321 | FKAEISSIRENKD | 6.0280 | 5.4619 | 5.3612 | 6.2292 | 6.4334 | 5.9181 | 5.9283 | 5.6667 | 5.8460 | 5.1961 | 5.4008 | 5.4258 | 7.8555 | 7.6013 | 7.3718 | 7.2759 |
| RAI14_S300;S303 | RAI14 | Q9P0K7-2 | | 284 | APPPPISPTQLSD | 5.6642 | 5.8396 | 6.1245 | 6.0147 | 6.2804 | 6.2383 | 6.0563 | 5.9078 | 6.5312 | 6.5272 | 6.8338 | 6.5409 | 6.0005 | 6.4516 | 5.9459 | 6.1433 |
| RAI14_S303 | RAI14 | Q9P0K7-2 | | 415 | PPDSKSSPSVLIH | 6.7673 | 7.1525 | 5.9943 | 6.5310 | 7.3319 | 7.0600 | 5.8214 | 7.5485 | 6.9450 | 6.3862 | 6.3123 | 6.2904 | 4.8014 | 5.4269 | 5.0305 | 4.6004 |
| RAI14_S321 | RAI14 | Q9P0K7-2 | | 670 | DYRKRKSLEDVTA | 6.5651 | 5.9732 | 6.6995 | 5.0174 | 6.7862 | 6.4560 | 5.4551 | 6.0131 | 6.6457 | 6.7754 | 6.1155 | 5.9458 | 7.1878 | 7.1057 | 5.9038 | 6.3547 |
| RAI14_S330 | RAI14 | Q9P0K7-2 | | 303 | ITSTPLSGKESVF | 5.7903 | 5.7847 | 6.2771 | 5.7129 | 6.9723 | 6.6441 | 6.5528 | 6.5696 | 6.7353 | 6.0370 | 6.0905 | 5.8471 | 6.3182 | 6.0553 | 6.5612 | 6.0516 |
| RAI14_S361 | RAI14 | Q9P0K7-2 | | 361 | LQAKVASLTLHNK | 6.0723 | 5.7165 | 5.7366 | 6.1629 | 6.7838 | 6.1559 | 6.7319 | 6.4545 | 5.7071 | 5.5928 | 5.9855 | 5.7465 | 6.5922 | 6.9388 | 6.6413 | 6.9813 |
| RAI14_S408 | RAI14 | Q9P0K7-2 | | 330 | ENKDRLSDSTTGA | 6.0243 | 5.7266 | 6.2442 | 6.0569 | 5.2622 | 5.6043 | 5.4022 | 6.3231 | 5.8587 | 6.7559 | 5.9879 | 6.4739 | 7.0507 | 7.2598 | 6.7845 | 7.1848 |
| RAI14_S415 | RAI14 | Q9P0K7-2 | | 408 | GKPGETSPPDSKS | 6.1238 | 6.2227 | 6.2358 | 6.1414 | 5.7686 | 5.4784 | 6.2760 | 5.3342 | 6.5589 | 6.8186 | 5.9076 | 7.2129 | 5.8761 | 6.3680 | 6.5554 | 7.1216 |
| RAI14_S415;S422 | RAI14 | Q9P0K7-2 | 296;300 | | NA | 6.5543 | 6.7712 | 7.0191 | 6.5531 | 6.4040 | 6.0854 | 5.5379 | 6.0880 | 7.2631 | 6.3126 | 5.3872 | 6.1631 | 6.2665 | 6.2095 | 5.9898 | 5.3952 |
| RAI14_S422 | RAI14 | Q9P0K7-2 | 296;303 | | NA | 7.4172 | 6.3649 | 5.5515 | 6.1915 | 7.5361 | 5.8777 | 5.4790 | 6.9814 | 6.6534 | 5.8287 | 7.1387 | 7.3871 | 5.3757 | 5.2186 | 5.4551 | 5.5436 |
| RAI14_S670 | RAI14 | Q9P0K7-2 | 300;303 | | NA | 6.3313 | 5.4548 | 6.6859 | 5.3767 | 8.2673 | 7.5060 | 6.8300 | 5.8470 | 7.4146 | 6.4807 | 5.9650 | 6.2564 | 5.5776 | 6.0759 | 5.2819 | 4.6490 |
| RAI14_T300 | RAI14 | Q9P0K7-2 | 415;422 | | NA | 6.2627 | 6.2235 | 6.6348 | 7.2733 | 6.7367 | 6.7651 | 7.0995 | 6.9755 | 6.2351 | 5.9863 | 6.4172 | 6.2971 | 5.1695 | 5.5846 | 4.9255 | 5.4136 |
| RALA_S11 | RALA | P11233 | | 11 | KPKGQNSLALHKV | 5.9312 | 6.4531 | 6.3569 | 7.1727 | 5.9895 | 5.7856 | 5.7306 | 6.1186 | 6.0548 | 6.5317 | 6.4455 | 6.2518 | 6.6034 | 5.5672 | 6.9461 | 6.9612 |
| RALBP1_S29 | RALBP1 | Q15311 | | 463 | QEIASLSKEDVSK | 6.0817 | 6.3718 | 6.0266 | 6.1520 | 5.9144 | 6.0099 | 6.3140 | 5.7430 | 6.5638 | 6.7173 | 6.7433 | 6.6012 | 5.9184 | 6.4483 | 6.1410 | 6.2533 |
| RALBP1_S29;S34 | RALBP1 | Q15311 | | 29 | GLTRTPSSEEISP | 5.3753 | 6.8483 | 6.1010 | 6.5516 | 5.5257 | 5.8824 | 5.1982 | 5.2589 | 6.1475 | 7.7744 | 6.4069 | 6.1168 | 6.3362 | 7.4070 | 6.2672 | 6.8026 |
| RALBP1_S463 | RALBP1 | Q15311 | | 645 | KEPAKPSPSRDRK | 6.2880 | 6.8169 | 6.5593 | 6.1775 | 5.8984 | 6.4189 | 5.7559 | 6.1517 | 6.9991 | 7.2440 | 6.7884 | 7.5964 | 4.9465 | 5.5772 | 5.0766 | 5.7049 |
| RALBP1_S645 | RALBP1 | Q15311 | 29;34 | | NA | 6.0143 | 5.8161 | 6.0258 | 6.4690 | 6.6667 | 6.3046 | 6.5423 | 5.7919 | 6.5654 | 7.2359 | 6.5808 | 7.3124 | 4.9732 | 5.9959 | 5.6902 | 6.0158 |
| RALGAPA1_S773 | RALGAPA1 | Q6GYQ0-2 | | 773 | TRVRHFSQSEETG | 6.1341 | 6.2777 | 6.6790 | 6.5172 | 6.1195 | 5.7247 | 6.5287 | 5.9370 | 6.2502 | 6.7485 | 6.3097 | 6.4339 | 5.7923 | 6.0929 | 6.0003 | 6.4544 |
| RALGAPA1_S797 | RALGAPA1 | Q6GYQ0-2 | | 754 | SIVRQKTVDIDDA | 6.0847 | 6.3052 | 6.9284 | 6.2296 | 5.7298 | 6.4663 | 6.3646 | 6.7699 | 6.5713 | 6.4639 | 6.5178 | 6.3797 | 5.5410 | 6.0195 | 5.4728 | 6.1554 |
| RALGAPA1_S860 | RALGAPA1 | Q6GYQ0-2 | | 797 | PLPRSSSTSDILE | 5.7872 | 5.7908 | 7.8571 | 5.8627 | 5.6558 | 7.1420 | 6.4079 | 6.6517 | 6.2334 | 6.0202 | 7.5027 | 6.7479 | 4.8129 | 5.8987 | 5.8681 | 5.7611 |
| RALGAPA1_S860;S861 | RALGAPA1 | Q6GYQ0-2 | | 860 | TMTRRGSSPGSLE | 6.6314 | 6.2077 | 6.7501 | 6.0517 | 6.4757 | 6.3021 | 5.9404 | 6.1741 | 6.0134 | 7.4831 | 6.7984 | 6.3654 | 5.5184 | 5.6501 | 5.8581 | 5.7801 |
| RALGAPA1_T754 | RALGAPA1 | Q6GYQ0-2 | 860;861 | | NA | 5.1447 | 5.3403 | 6.0306 | 6.2414 | 6.4431 | 6.0459 | 6.4886 | 6.0215 | 6.7486 | 6.4754 | 6.6720 | 6.1496 | 6.3264 | 6.9968 | 6.0948 | 6.7802 |
| RALGAPA2_S696 | RALGAPA2 | Q2PPJ7 | | 820 | ILVRRSSSPAELD | 5.9500 | 6.5213 | 6.7164 | 6.5544 | 6.2486 | 6.0920 | 5.7013 | 6.3185 | 6.6263 | 6.5845 | 6.5725 | 6.6071 | 6.0024 | 6.5850 | 6.0282 | 5.7914 |
| RALGAPA2_S820 | RALGAPA2 | Q2PPJ7 | | 696 | TVGRSFSLSWRSH | 7.4532 | 6.5084 | 6.9970 | 6.2794 | 7.1232 | 6.5127 | 5.8638 | 5.8841 | 6.5550 | 5.4015 | 5.7655 | 5.5940 | 6.5521 | 6.7228 | 6.6775 | 5.1099 |
| RALGAPB_S357 | RALGAPB | Q86X10 | | 379 | SAAVSTTPPHNRR | 6.3265 | 6.1094 | 6.7577 | 6.1594 | 6.2224 | 6.3183 | 6.1167 | 6.8534 | 6.7412 | 6.7493 | 6.6811 | 6.6386 | 5.5952 | 5.7621 | 5.1877 | 5.7807 |
| RALGAPB_T379 | RALGAPB | Q86X10 | | 357 | GISRPRSDSAPPT | 6.2663 | 6.9488 | 6.2825 | 6.1908 | 6.0298 | 5.8571 | 6.0713 | 6.2042 | 6.6379 | 6.3446 | 6.7063 | 6.2949 | 6.0180 | 6.1718 | 5.9266 | 6.0491 |
| RALGPS2_S343 | RALGPS2 | Q86X27 | | 343 | GHRKCHSLGYNFI | 6.4835 | 6.5107 | 7.0180 | 5.8385 | 6.2882 | 6.5988 | 6.2441 | 6.0732 | 6.1661 | 6.5690 | 6.5554 | 6.4083 | 5.6723 | 6.2414 | 6.0474 | 5.9850 |
| RALGPS2_T361 | RALGPS2 | Q86X27 | | 361 | AEFKSATFPNAGP | 6.3192 | 6.5763 | 6.7802 | 5.5392 | 6.2387 | 6.7457 | 6.3907 | 6.4966 | 6.2618 | 6.1728 | 6.4194 | 6.1693 | 5.9470 | 6.1352 | 5.9414 | 5.8665 |
| RALY_S106 | RALY | Q9UKM9 | | 135 | DYRGRLSPVPVPR | 6.9840 | 7.0029 | 5.9573 | 6.7343 | 6.4453 | 5.8927 | 6.2946 | 6.2698 | 6.5371 | 6.3159 | 6.1246 | 6.0890 | 5.7775 | 6.0214 | 5.7487 | 5.8048 |
| RALY_S106 | RALY | Q9UKM9 | | 106 | GLKRAASAIYSGY | 5.8180 | 6.2146 | 6.0096 | 6.6033 | 6.2000 | 6.1621 | 5.9681 | 6.0646 | 5.8053 | 5.9315 | 6.1133 | 6.0992 | 6.7951 | 6.8780 | 6.7281 | 6.6092 |
| RALY_S135 | RALY | Q9UKM9 | | 288 | EGLLTHSEEELEH | 6.1901 | 5.9288 | 5.9211 | 6.5730 | 6.0683 | 5.3995 | 5.8816 | 5.1855 | 5.9557 | 6.2515 | 6.4506 | 6.5792 | 6.8851 | 7.0567 | 6.4720 | 7.2013 |
| RALY_S177 | RALY | Q9UKM9-2 | | 106 | GLKRAASAIYRLF | 6.6440 | 6.6774 | 6.2914 | 6.7251 | 6.3927 | 5.7201 | 6.2889 | 5.8399 | 6.4509 | 6.0744 | 6.0735 | 5.8556 | 6.2709 | 6.2796 | 6.0613 | 6.3544 |
| RALY_S288 | RALY | Q9UKM9 | | 286 | DEEGLLTHSEEEL | 5.8974 | 6.3030 | 5.7349 | 7.3799 | 6.1579 | 5.0412 | 5.4601 | 6.4031 | 6.5543 | 5.9975 | 6.9171 | 6.2070 | 7.0979 | 5.7554 | 6.7945 | 6.2989 |

| | | | | | | | | | | | | | | | | | | | | | |
|---|---|---|---|---|---|---|---|---|---|---|---|---|---|---|---|---|---|---|---|---|---|
| RALY_T151 | RALY | Q9UKM9 | | 177 | TAVTTSSAKIKLK | 6.6864 | 6.3544 | 6.5458 | 6.5086 | 6.4461 | 6.1595 | 6.2654 | 6.3765 | 6.5982 | 6.0164 | 6.3076 | 6.3864 | 6.2602 | 5.7985 | 5.5937 | 5.6963 |
| RALY_T286 | RALY | Q9UKM9 | | 298 | LEHSQDTDADDGA | 5.8943 | 6.3793 | 6.6435 | 6.6025 | 5.8696 | 6.2900 | 6.2852 | 6.9053 | 6.0102 | 5.8775 | 5.9999 | 6.0132 | 6.5066 | 6.5283 | 6.0675 | 6.1272 |
| RALY_T286;T298 | RALY | Q9UKM9 | | 151 | VKRPRVTVPLVRR | 5.3443 | 5.0923 | 5.5467 | 5.9113 | 5.3950 | 4.8156 | 5.7504 | 4.9570 | 5.0119 | 4.7412 | 4.8540 | 4.6768 | 9.6590 | 8.5976 | 10.1851 | 9.4619 |
| RALY_T298 | RALY | Q9UKM9 | 286;298 | | NA | 5.8556 | 6.8657 | 6.1507 | 6.4362 | 5.5338 | 6.0725 | 6.0890 | 5.9711 | 6.0519 | 6.7716 | 6.4028 | 6.8736 | 6.3224 | 6.4787 | 6.2157 | 5.9085 |
| RAMAC_S36 | RAMAC | Q9BTL3 | | 36 | LKRPPESPPIVEE | 5.7256 | 5.7401 | 6.3710 | 5.7206 | 6.4732 | 5.5853 | 5.6935 | 6.3816 | 5.8006 | 6.6054 | 6.3529 | 6.5917 | 6.7440 | 6.9997 | 6.5978 | 6.6170 |
| RANBP1_S188 | RANBP1 | P43487 | | 188 | EKLEALSVKEETK | 6.5267 | 6.1222 | 6.5971 | 5.7070 | 6.4896 | 6.1936 | 5.9973 | 6.3857 | 6.0959 | 5.7936 | 6.1718 | 6.1310 | 6.7687 | 6.6065 | 6.1641 | 6.2492 |
| RANBP1_S60 | RANBP1 | P43487 | | 60 | KLFRFASENDLPE | 6.6995 | 4.6726 | 5.4709 | 3.8365 | 6.7948 | 5.7171 | 4.6726 | 4.6615 | 6.0253 | 6.3047 | 4.7701 | 6.1673 | 9.4826 | 8.0145 | 8.3627 | 8.3472 |
| RANBP1_T13;S14 | RANBP1 | P43487 | 13;14 | | NA | 5.8831 | 7.6782 | 6.5171 | 7.3186 | 6.7587 | 7.7408 | 6.7756 | 7.3523 | 6.3271 | 6.0092 | 7.3391 | 6.7315 | 4.0937 | 4.8033 | 4.1595 | 4.5123 |
| RANBP2_S1103 | RANBP2 | P49792 | | 2900 | GEDEDGSDEEVVH | 5.8667 | 5.9595 | 6.2577 | 6.3341 | 6.0419 | 5.8137 | 6.1058 | 5.9690 | 6.4434 | 6.1516 | 6.2561 | 6.4419 | 6.6019 | 6.6052 | 6.3551 | 6.7965 |
| RANBP2_S1107 | RANBP2 | P49792 | | 2743 | CGVCSDTDEDNGN | 6.0038 | 6.0502 | 5.9270 | 6.5097 | 6.2223 | 6.2811 | 6.1650 | 6.0467 | 6.2574 | 6.1367 | 6.2827 | 6.0055 | 6.4109 | 6.6361 | 6.4456 | 6.6194 |
| RANBP2_S1128 | RANBP2 | P49792 | | 955 | PATGILSPRGDDY | 6.0848 | 5.9618 | 5.8534 | 6.4035 | 6.4819 | 6.2374 | 6.6839 | 6.5540 | 6.4377 | 5.8681 | 5.9441 | 6.1336 | 6.3376 | 6.3493 | 6.2236 | 6.4455 |
| RANBP2_S1160 | RANBP2 | P49792 | | 2510 | TPKAVVSPPKFVF | 6.8231 | 7.1169 | 6.7067 | 7.4101 | 6.1649 | 6.2827 | 6.6955 | 6.2027 | 6.9262 | 6.7195 | 6.7719 | 7.1103 | 4.4276 | 4.8617 | 4.5844 | 5.1957 |
| RANBP2_S128 | RANBP2 | P49792 | | 2153 | LDIPLQTPHKLVD | 8.5374 | 7.7744 | 6.8281 | 8.3805 | 6.9236 | 6.6287 | 7.3165 | 5.6597 | 6.7073 | 5.2237 | 5.7293 | 5.3677 | 4.8738 | 5.2189 | 4.1413 | 4.6891 |
| RANBP2_S1374 | RANBP2 | P49792 | | 3207 | KIESFGSPKGSVC | 5.9916 | 6.5613 | 6.4960 | 6.8680 | 5.6088 | 5.6842 | 6.0259 | 5.8624 | 6.2245 | 6.4898 | 6.3045 | 6.3943 | 6.0656 | 6.3446 | 6.1986 | 6.8800 |
| RANBP2_S1447 | RANBP2 | P49792 | | 788 | PTRYSLSPSKSYK | 6.6257 | 6.3188 | 5.6339 | 6.0554 | 6.9944 | 6.3612 | 6.6465 | 6.5180 | 6.3405 | 5.9620 | 5.7762 | 5.4896 | 6.3395 | 6.2095 | 6.4912 | 6.2377 |
| RANBP2_S1486 | RANBP2 | P49792 | | 1396 | LAETVFTPKTSPE | 6.0656 | 6.3315 | 6.5045 | 7.2568 | 5.6610 | 5.8474 | 6.7196 | 6.3920 | 6.5749 | 6.9315 | 6.1288 | 6.4812 | 5.2043 | 5.8673 | 5.4980 | 6.5355 |
| RANBP2_S1509 | RANBP2 | P49792 | | 781 | IKHSTPSPTRYSL | 6.1978 | 6.8847 | 5.5513 | 6.8745 | 6.4716 | 6.2886 | 6.0358 | 6.7372 | 5.9335 | 5.6875 | 6.0329 | 5.8178 | 6.2258 | 6.4418 | 6.7200 | 6.0992 |
| RANBP2_S1573 | RANBP2 | P49792 | | 2626 | KKKPEDSPSDDDV | 6.0007 | 6.0719 | 5.9181 | 6.6380 | 6.0732 | 6.1180 | 6.1401 | 6.0318 | 5.9505 | 6.2406 | 6.2269 | 6.2207 | 6.5682 | 6.3618 | 6.6700 | 6.7696 |
| RANBP2_S1613 | RANBP2 | P49792 | | 1993 | ANKANTSGDFEKD | 7.1111 | 7.0778 | 6.1966 | 6.8934 | 6.9127 | 6.5203 | 7.0366 | 6.9064 | 6.9778 | 6.3454 | 6.4397 | 6.2918 | 4.5579 | 4.9384 | 4.7465 | 5.0474 |
| RANBP2_S1869 | RANBP2 | P49792 | | 2668 | NRPDYVSEEEEDD | 6.3417 | 6.3512 | 6.2500 | 6.3713 | 6.0024 | 6.0484 | 6.1085 | 5.8932 | 6.0713 | 6.1752 | 6.1429 | 5.8282 | 6.8580 | 6.3835 | 6.4994 | 6.6750 |
| RANBP2_S1955 | RANBP2 | P49792 | | 896 | RPAANVTPTKGPV | 6.6840 | 5.5933 | 6.1996 | 6.8308 | 6.9159 | 6.4601 | 6.8893 | 6.3653 | 7.2991 | 6.5274 | 6.4209 | 6.3792 | 5.2107 | 5.3250 | 5.0009 | 5.8984 |
| RANBP2_S1993 | RANBP2 | P49792 | | 1412 | DRFALVTPKKEGH | 6.7990 | 6.4459 | 6.2521 | 6.7612 | 6.8870 | 6.8421 | 7.1170 | 6.6013 | 7.0713 | 6.3751 | 6.1471 | 6.0387 | 5.0723 | 5.3674 | 4.6393 | 5.5834 |
| RANBP2_S21 | RANBP2 | P49792 | | 2613 | INYTFKTPEKAKE | 6.1054 | 6.2274 | 6.2962 | 6.8802 | 7.0974 | 7.3266 | 7.2881 | 6.4682 | 6.8194 | 6.3562 | 7.1860 | 6.7356 | 5.3236 | 5.2888 | 4.2452 | 4.3558 |
| RANBP2_S2207 | RANBP2 | P49792 | | 1869 | HVDQENSPSFMFQ | 6.0293 | 6.5348 | 5.9353 | 7.0820 | 6.2734 | 7.0302 | 6.6157 | 6.3232 | 6.1662 | 6.3458 | 6.2946 | 6.3466 | 5.6113 | 6.0854 | 5.3375 | 5.9887 |
| RANBP2_S2270 | RANBP2 | P49792 | | 1573 | QNPDKPSPSTSVP | 5.7665 | 6.5775 | 6.1696 | 6.4001 | 6.3093 | 5.9402 | 6.2680 | 6.2380 | 6.0461 | 6.0677 | 6.0074 | 6.0943 | 6.4761 | 6.4374 | 6.4775 | 6.7243 |
| RANBP2_S2276 | RANBP2 | P49792 | | 1447 | TKSANKSGSSFVH | 5.2158 | 5.5301 | 5.6910 | 6.3466 | 6.3116 | 6.5242 | 6.1247 | 6.4645 | 5.9539 | 5.0177 | 5.5945 | 5.6610 | 7.3497 | 7.5573 | 7.0895 | 7.5678 |
| RANBP2_S2278 | RANBP2 | P49792 | | 2276 | SFKSALSPSKSPA | 6.5129 | 6.5324 | 5.8283 | 6.5752 | 6.2993 | 6.2413 | 6.5458 | 6.2115 | 6.5670 | 6.2053 | 6.2131 | 6.2866 | 5.9171 | 5.9376 | 6.0146 | 6.1121 |
| RANBP2_S2457;T2458;S | RANBP2 | P49792 | | 2804 | SITKSISSPSVSS | 4.1481 | 4.7161 | 4.3400 | 5.0058 | 5.0998 | 5.6309 | 4.9661 | 5.1828 | 4.9174 | 4.4074 | 5.4519 | 4.3753 | 10.3806 | 9.5947 | 10.9147 | 10.8685 |
| RANBP2_S2510 | RANBP2 | P49792 | | 2526 | SVKSIFSSEKSKP | 7.3179 | 7.8230 | 6.5028 | 7.4752 | 6.3358 | 7.2194 | 7.6232 | 6.8464 | 7.3635 | 7.4493 | 7.8795 | 7.3751 | 2.9363 | 3.8131 | 2.6649 | 3.3745 |
| RANBP2_S2518 | RANBP2 | P49792 | | 128 | AKLFPGSPAIYKL | 7.4433 | 7.6762 | 6.4335 | 7.2806 | 6.7523 | 7.0612 | 7.4677 | 6.6401 | 7.6197 | 7.1827 | 6.7967 | 6.7685 | 3.1502 | 4.5093 | 3.2530 | 3.9651 |
| RANBP2_S2520 | RANBP2 | P49792 | | 2270 | TTGFNFSFKSALS | 7.2132 | 7.0558 | 7.1394 | 7.5766 | 6.5481 | 7.1476 | 7.5286 | 6.3261 | 6.4958 | 6.2121 | 6.6538 | 6.5886 | 4.1580 | 4.9881 | 3.6735 | 4.6948 |
| RANBP2_S2526 | RANBP2 | P49792 | | 2008 | AYKTEDSDDIHFE | 6.0369 | 5.9058 | 6.0151 | 7.0672 | 6.2931 | 6.0052 | 6.2762 | 6.3420 | 6.2191 | 6.0929 | 6.6821 | 6.3223 | 6.2785 | 6.1937 | 6.0769 | 6.1930 |
| RANBP2_S2599 | RANBP2 | P49792 | | 837 | NSSNSASPHRWPT | 6.1732 | 5.9139 | 6.0686 | 6.2849 | 5.6458 | 5.8599 | 6.2233 | 6.6984 | 6.2718 | 6.2413 | 5.9967 | 5.4437 | 6.7692 | 7.8768 | 6.2614 | 6.2712 |
| RANBP2_S2626 | RANBP2 | P49792 | | 21 | VQGSTPSPRQKSM | 6.4699 | 6.7374 | 6.2124 | 7.0289 | 6.3414 | 6.7899 | 6.6883 | 6.4644 | 6.6534 | 6.4510 | 7.0041 | 6.7164 | 4.6282 | 5.3678 | 4.9038 | 5.5426 |
| RANBP2_S2668 | RANBP2 | P49792 | | 2766 | VQEAQKSQTEEIT | 5.9221 | 6.5023 | 6.5603 | 7.2938 | 5.7656 | 6.2067 | 6.9444 | 6.6395 | 6.4725 | 6.3852 | 6.8545 | 7.1301 | 4.7229 | 5.7006 | 5.0304 | 5.8690 |
| RANBP2_S2766 | RANBP2 | P49792 | | 1144 | GKSVFGTPTLETA | 7.2938 | 7.4033 | 6.8026 | 7.2992 | 6.0908 | 7.4725 | 7.1466 | 6.9113 | 7.7913 | 7.4314 | 7.0388 | 7.1653 | 3.1075 | 3.9400 | 3.0625 | 4.0433 |
| RANBP2_S2804 | RANBP2 | P49792 | | 2207 | TGAAGASDTTIKP | 7.0060 | 6.7089 | 6.4603 | 6.8055 | 6.2456 | 6.7483 | 6.2810 | 6.4822 | 6.8358 | 6.9192 | 6.2261 | 6.4105 | 4.5728 | 5.3867 | 5.0909 | 5.8203 |
| RANBP2_S2900 | RANBP2 | P49792 | | 2293 | SGTSVGTDEESDV | 6.6002 | 7.3342 | 6.4920 | 6.2797 | 6.5634 | 6.1683 | 6.1659 | 6.6233 | 6.5615 | 6.0889 | 6.2286 | 6.1172 | 5.8226 | 6.3627 | 5.2674 | 5.3241 |
| RANBP2_S3207 | RANBP2 | P49792 | | 2278 | KSALSPSKSPAKL | 6.7535 | 6.6265 | 5.6190 | 6.1236 | 6.9837 | 6.5022 | 6.8193 | 6.5803 | 6.4459 | 5.9486 | 5.9820 | 5.7222 | 5.9579 | 5.7793 | 6.1511 | 6.0047 |
| RANBP2_S781 | RANBP2 | P49792 | | 2518 | PKFVFGSESVKSI | 9.2519 | 9.2750 | 8.1475 | 8.7204 | 6.0630 | 6.3871 | 7.7158 | 5.4485 | 6.6590 | 5.3272 | 6.8157 | 5.2197 | 3.0983 | 5.1659 | 2.9078 | 3.7972 |
| RANBP2_S788 | RANBP2 | P49792 | | 2520 | FVFGSESVKSIFS | 7.6739 | 6.7854 | 6.3998 | 8.7594 | 5.5574 | 6.2154 | 6.8650 | 7.1920 | 6.2322 | 6.0268 | 6.8064 | 5.0335 | 4.3755 | 5.9765 | 6.0520 | 4.0486 |
| RANBP2_S837 | RANBP2 | P49792 | | 1613 | EGQWDCSVCLVRN | 6.0479 | 5.6734 | 6.1264 | 6.1946 | 6.3347 | 6.5628 | 6.5904 | 6.4100 | 6.5327 | 6.1245 | 5.5978 | 6.5661 | 6.1628 | 5.9869 | 6.1478 | 6.9412 |
| RANBP2_S955 | RANBP2 | P49792 | | 1128 | NSGFRRSDDMFTF | 6.2731 | 7.2763 | 6.6176 | 8.1162 | 5.9712 | 6.2323 | 6.2488 | 6.4041 | 6.4515 | 6.5175 | 7.2328 | 7.2328 | 4.4306 | 5.0471 | 4.5724 | 5.3757 |
| RANBP2_T1144 | RANBP2 | P49792 | | 2454 | LITPHVSRSSTPR | 6.1351 | 7.2277 | 6.2646 | 5.9867 | 6.2308 | 6.6275 | 6.2864 | 6.7381 | 6.1725 | 5.9432 | 6.4143 | 6.5124 | 6.1858 | 5.7763 | 5.9806 | 5.5180 |
| RANBP2_T1255 | RANBP2 | P49792 | | 1374 | TAKKCVSCQNLNP | 5.0201 | 5.0105 | 5.0232 | 5.3854 | 4.8927 | 4.7502 | 6.0222 | 5.6879 | 5.6353 | 5.0670 | 5.7277 | 5.2740 | 8.9887 | 8.9998 | 9.6692 | 8.8462 |
| RANBP2_T1393;T1400 | RANBP2 | P49792 | | 1103 | NTFNFGSKNVSGI | 6.2508 | 6.4647 | 6.4096 | 6.2764 | 6.7189 | 6.0571 | 7.6565 | 5.9670 | 6.7687 | 4.5640 | 6.2280 | 6.5131 | 5.8379 | 5.5352 | 5.3657 | 7.3861 |
| RANBP2_T1396 | RANBP2 | P49792 | | 1509 | QNPRKQSLPATSI | 5.0949 | 5.4010 | 5.4063 | 4.7468 | 4.2405 | 5.7334 | 5.4834 | 5.4782 | 6.3784 | 6.2999 | 6.3653 | 7.1556 | 7.2629 | 7.4395 | 8.4797 | 9.0344 |
| RANBP2_T1396;S1400 | RANBP2 | P49792 | | 1486 | EGQWDCSACLVQN | 6.0049 | 5.7606 | 6.5081 | 6.1522 | 6.6963 | 6.9212 | 6.7928 | 6.5459 | 7.1052 | 6.4646 | 7.2394 | 6.5695 | 5.5121 | 6.2717 | 4.7767 | 4.6786 |
| RANBP2_T1412 | RANBP2 | P49792 | | 1255 | SPDMKLTPNAGSD | 5.8229 | 6.3121 | 5.6653 | 6.0950 | 6.3105 | 5.6853 | 5.8798 | 5.6720 | 6.4971 | 5.5957 | 5.4577 | 6.2517 | 7.1210 | 6.8979 | 7.3133 | 7.4227 |
| RANBP2_T2005;S2008 | RANBP2 | P49792 | | 1160 | HETDGGSAHGDDD | 5.8796 | 6.4716 | 6.1898 | 6.9329 | 6.3787 | 6.3105 | 6.2562 | 5.4695 | 5.8562 | 5.8562 | 6.2479 | 6.0660 | 6.9165 | 6.1323 | 6.2688 | 6.7673 |
| RANBP2_T2008 | RANBP2 | P49792 | | 2599 | VKNLFASFPTEES | 5.6937 | 6.2438 | 6.7677 | 6.6939 | 6.7926 | 6.1433 | 6.5178 | 5.6506 | 6.0066 | 5.8232 | 6.6717 | 6.5563 | 5.7711 | 6.4453 | 6.2773 | 5.9450 |
| RANBP2_T2153 | RANBP2 | P49792 | | 1955 | DLAKSTSGEGFQF | 1.4237 | 2.2244 | 1.3996 | 1.3374 | 1.8921 | 2.2672 | 1.8189 | 1.8009 | 1.4901 | 1.4486 | 0.9830 | 1.2759 | 19.0171 | 18.9335 | 23.4770 | 19.2105 |
| RANBP2_T2293 | RANBP2 | P49792 | | 1107 | FGSKNVSGISFTE | 5.9343 | 7.2468 | 6.9477 | 7.4014 | 5.8107 | 6.9544 | 6.7283 | 6.3695 | 8.2115 | 7.1863 | 7.2418 | 8.5274 | 3.5747 | 3.9747 | 3.5654 | 4.3251 |
| RANBP2_T2450;S2454 | RANBP2 | P49792 | 2450;2454 | | NA | 6.9252 | 6.7883 | 6.2821 | 7.0058 | 6.4903 | 6.2444 | 6.6371 | 6.1730 | 6.4476 | 5.9210 | 6.1022 | 6.0737 | 5.7150 | 5.7501 | 5.7311 | 5.7130 |
| RANBP2_T2454 | RANBP2 | P49792 | 2458;2462 | | NA | 6.7974 | 6.1809 | 6.2908 | 6.7828 | 6.5101 | 6.7442 | 7.0631 | 6.1569 | 6.6015 | 6.5254 | 6.6066 | 6.7539 | 5.1433 | 5.2653 | 5.2387 | 5.3393 |
| RANBP2_T2458;S2462 | RANBP2 | P49792 | 1396;1400 | | NA | 6.3102 | 6.5398 | 6.2014 | 6.8901 | 6.2563 | 5.8537 | 6.7648 | 6.2824 | 6.3434 | 6.2370 | 6.2973 | 6.8860 | 5.6846 | 6.6137 | 5.4467 | 5.3927 |
| RANBP2_T2613 | RANBP2 | P49792 | 1393;1400 | | NA | 6.2371 | 6.5801 | 6.3756 | 6.8210 | 6.0912 | 6.4714 | 6.7155 | 6.2369 | 6.1890 | 6.3195 | 6.4265 | 6.6147 | 5.7161 | 6.1953 | 5.0939 | 5.9163 |
| RANBP2_T2743 | RANBP2 | P49792 | 2457;2458;24 | | NA | 6.2221 | 7.3994 | 6.4451 | 7.4157 | 6.0511 | 6.0580 | 6.2413 | 6.2944 | 6.0014 | 6.4646 | 5.9913 | 6.4126 | 5.7885 | 5.6313 | 5.8667 | 5.7167 |
| RANBP2_T896 | RANBP2 | P49792 | 2005;2008 | | NA | 6.2943 | 6.5555 | 7.0734 | 7.0633 | 5.9182 | 6.7792 | 6.7530 | 5.7656 | 6.5580 | 6.3854 | 5.7756 | 6.3486 | 6.2596 | 5.5104 | 5.9290 | 5.9311 |
| RANBP3_S108 | RANBP3 | Q9H6Z4 | | 211 | GAVPAASPDTAAW | 6.3644 | 5.9394 | 7.2010 | 5.9882 | 5.7077 | 6.1725 | 5.7850 | 6.0158 | 6.3968 | 6.7332 | 6.1762 | 6.8602 | 6.0505 | 6.8979 | 5.6631 | 6.0480 |
| RANBP3_S125 | RANBP3 | Q9H6Z4 | | 125 | VKRERTSSLTQFP | 5.1447 | 5.2158 | 6.1613 | 5.8761 | 6.0411 | 5.9551 | 5.7170 | 6.0406 | 5.9998 | 6.2213 | 6.1510 | 6.2142 | 6.9909 | 7.4623 | 7.3715 | 7.4374 |
| RANBP3_S211 | RANBP3 | Q9H6Z4 | | 108 | PEGGEDSDREDGN | 5.6332 | 5.9579 | 6.2744 | 5.2070 | 6.0909 | 5.6677 | 5.8220 | 5.8699 | 5.7387 | 6.5426 | 6.3815 | 6.1182 | 7.0449 | 7.2678 | 7.0066 | 7.3766 |
| RANBP3_S219 | RANBP3 | Q9H6Z4 | | 219 | DTAAWRSPSEAAD | 5.0340 | 5.8169 | 6.3735 | 5.4757 | 6.0544 | 5.7960 | 5.8907 | 6.4957 | 6.3903 | 6.4223 | 5.9625 | 5.9326 | 7.0732 | 6.8185 | 6.9780 | 7.4858 |
| RANBP9_S467 | RANBP9 | Q96S59 | | 467 | RCLGGRSPKSQDS | 5.6255 | 5.9668 | 6.8647 | 5.8380 | 6.7459 | 6.2144 | 6.3398 | 6.2845 | 7.9296 | 5.1611 | 4.8664 | 6.4984 | 6.3394 | 6.7302 | 6.3477 | 6.2476 |
| RANBP9_S477 | RANBP9 | Q96S59 | | 477 | QDSYPVSPRPFSS | 6.9755 | 6.5390 | 6.3994 | 6.7418 | 5.7351 | 5.8280 | 6.4611 | 6.8768 | 7.0893 | 6.0274 | 6.2016 | 6.9827 | 4.9323 | 5.4551 | 5.6576 | 6.0971 |
| RANBP9_S483 | RANBP9 | Q96S59 | | 487 | FSSPSMSPSHGMN | 6.1935 | 5.7917 | 6.1603 | 6.7579 | 7.5979 | 7.7551 | 6.3821 | 7.2366 | 5.7238 | 5.9547 | 7.1542 | 6.1548 | 5.3054 | 5.3809 | 5.7328 | 4.7183 |
| RANBP9_S483;S485 | RANBP9 | Q96S59 | | 485 | RPFSSPSMSPSHG | 6.8885 | 5.6709 | 6.2126 | 7.7045 | 7.9242 | 6.3931 | 5.9914 | 7.3070 | 5.4708 | 5.9508 | 6.3105 | 6.7612 | 6.1762 | 5.3029 | 4.5429 | 5.3924 |
| RANBP9_S485 | RANBP9 | Q96S59 | | 483 | SPRPFSSPSMSPS | 5.1409 | 7.1090 | 7.2750 | 5.6056 | 6.8624 | 6.5778 | 7.5405 | 7.0948 | 5.4207 | 2.9071 | 6.3312 | 8.1239 | 6.5968 | 7.4267 | 6.1131 | 3.8746 |
| RANBP9_S487 | RANBP9 | Q96S59 | 483;485 | | NA | 6.3635 | 6.2283 | 6.7344 | 7.3546 | 6.6788 | 6.1028 | 5.7493 | 6.0197 | 6.9527 | 6.6185 | 7.1933 | 7.3480 | 5.0168 | 5.2420 | 5.3179 | 5.0792 |
| RANGAP1_S428 | RANGAP1 | P46060 | | 442 | TFLAFPSPEKLLR | 6.1852 | 5.9846 | 6.8313 | 6.5495 | 6.1962 | 6.4182 | 6.5505 | 6.7082 | 6.2355 | 6.3650 | 6.4451 | 6.5497 | 5.5303 | 5.8350 | 5.6376 | 5.9780 |
| RANGAP1_S428;S442 | RANGAP1 | P46060 | | 409 | QGEKSATPSRKIL | 7.3155 | 7.6895 | 6.8310 | 7.3255 | 6.6274 | 7.6509 | 7.4492 | 6.6557 | 7.1047 | 6.9488 | 7.7785 | 7.6204 | 2.9884 | 3.6359 | 2.8224 | 3.5563 |
| RANGAP1_S442 | RANGAP1 | P46060 | | 428 | PAPVLSSPPPADV | 5.8898 | 6.6402 | 6.1705 | 6.0847 | 5.4025 | 6.0097 | 6.3837 | 6.2004 | 6.2521 | 6.3162 | 6.6627 | 7.2066 | 6.5728 | 6.7693 | 5.5366 | 5.9022 |
| RANGAP1_T409 | RANGAP1 | P46060 | 428;442 | | NA | 6.5372 | 6.4759 | 6.0685 | 7.6206 | 6.4222 | 7.6639 | 6.5823 | 6.1532 | 6.8489 | 7.1511 | 7.1693 | 7.3070 | 4.5003 | 4.2533 | 4.4016 | 4.8445 |
| RAP1GAP_S505 | RAP1GAP | P47736-4 | | 505 | IRSRSQSMDAMGL | 6.2902 | 5.4508 | 6.6456 | 4.8724 | 6.7760 | 6.3901 | 5.9879 | 6.9622 | 6.6797 | 6.1184 | 6.7535 | 6.7715 | 6.0014 | 7.3428 | 5.8197 | 5.1378 |
| RAP1GAP2_S418 | RAP1GAP2 | Q684P5 | | 418 | FGPPLPSPPVFQK | 5.9659 | 6.0247 | 7.2905 | 6.8686 | 5.6561 | 5.0953 | 5.8858 | 8.9462 | 5.3731 | 7.0769 | 4.5666 | 6.8472 | 7.4240 | 5.2983 | 5.8324 | 5.8484 |
| RAPGEF1_S235 | RAPGEF1 | Q13905-3 | | 299 | IRVVDNSPPPALP | 6.5480 | 6.5023 | 6.9176 | 6.8041 | 6.2863 | 6.5365 | 6.5089 | 6.7852 | 7.2469 | 6.9255 | 6.8552 | 7.1520 | 4.3570 | 5.0928 | 4.3350 | 5.1467 |
| RAPGEF1_S299 | RAPGEF1 | Q13905-3 | | 367 | GESPRLSPCSSIG | 5.4626 | 5.6746 | 5.7436 | 5.3330 | 5.7908 | 6.0183 | 6.0115 | 6.8098 | 5.4542 | 5.6806 | 5.6997 | 5.6501 | 7.6694 | 7.4643 | 8.0244 | 7.5131 |
| RAPGEF1_S314 | RAPGEF1 | Q13905-3 | | 314 | KRQSAPSPTRVAV | 6.0441 | 6.4446 | 6.8756 | 5.5377 | 6.0640 | 6.4608 | 5.7956 | 6.7305 | 6.5134 | 6.5320 | 6.5620 | 6.5809 | 5.8683 | 6.3142 | 5.9218 | 5.7547 |
| RAPGEF1_S329 | RAPGEF1 | Q13905-3 | | 329 | PMSRATSGSSLPV | 5.0946 | 4.6767 | 5.2358 | 4.7296 | 4.7672 | 5.2994 | 5.1071 | 4.8730 | 5.1942 | 4.6181 | 4.8835 | 4.9024 | 10.5002 | 10.1927 | 10.2907 | 9.6348 |
| RAPGEF1_S367 | RAPGEF1 | Q13905-3 | | 235 | GRPSPTSPVKPSS | 6.7204 | 5.6029 | 7.1666 | 6.1277 | 5.5965 | 5.8998 | 6.1798 | 6.4880 | 7.5093 | 6.9219 | 6.6810 | 7.1968 | 5.2117 | 5.4298 | 5.4012 | 5.8664 |
| RAPGEF2_S1080 | RAPGEF2 | Q9Y4G8 | | 423 | DSGSKATEAGLKR | 5.9528 | 5.9534 | 5.3078 | 5.8323 | 7.3462 | 6.8676 | 6.5623 | 6.9422 | 6.3694 | 6.2845 | 7.5011 | 5.8541 | 6.0159 | 6.1198 | 5.5551 | 5.5356 |
| RAPGEF2_S1325 | RAPGEF2 | Q9Y4G8 | | 1080 | PPFGINSPQALKK | 5.8734 | 6.4123 | 6.3591 | 6.5488 | 6.5212 | 6.6477 | 7.1810 | 7.0495 | 6.9534 | 7.3172 | 6.0088 | 6.7637 | 4.7253 | 5.8120 | 4.7941 | 5.0327 |
| RAPGEF2_T423 | RAPGEF2 | Q9Y4G8 | | 1325 | QAQSRASWASSTG | 6.5026 | 6.4807 | 6.9357 | 5.8344 | 6.6357 | 6.5857 | 6.3771 | 6.6969 | 6.6867 | 6.6189 | 6.2674 | 6.5164 | 5.5615 | 6.3434 | 4.7683 | 5.1886 |
| RAPGEF3_S864 | RAPGEF3 | O95398 | | 864 | HNPVPLSPLRSRV | 5.7570 | 5.4925 | 6.3055 | 6.6046 | 5.5962 | 6.8570 | 6.5730 | 5.7720 | 7.6054 | 6.9021 | 5.7450 | 7.1050 | 5.5677 | 6.0260 | 5.7465 | 6.3446 |
| RAPGEF5_S225 | RAPGEF5 | Q92565 | | 225 | ALFHQFSLKENWL | 7.6913 | 6.3327 | 6.3827 | 6.4526 | 6.2595 | 6.5896 | 6.2165 | 5.6068 | 6.8132 | 6.4419 | 5.6541 | 6.0758 | 6.1628 | 6.4901 | 5.5124 | 5.3180 |
| RAPGEF6_S1070 | RAPGEF6 | Q8TEU7 | | 1507 | YREPPPTPPGYLG | 6.4491 | 6.3934 | 6.8426 | 7.3430 | 6.2755 | 6.7817 | 6.5372 | 6.9569 | 7.0047 | 6.4350 | 7.1530 | 6.6189 | 5.0497 | 5.2477 | 4.2498 | 4.6618 |
| RAPGEF6_S1094 | RAPGEF6 | Q8TEU7 | | 1494 | IVYCVTSPKKDDR | 6.5032 | 6.4071 | 7.6658 | 6.5665 | 6.5297 | 6.9582 | 6.3546 | 6.5979 | 6.7231 | 6.6490 | 7.1628 | 6.8116 | 4.6015 | 5.2064 | 4.3233 | 4.9393 |
| RAPGEF6_S1157 | RAPGEF6 | Q8TEU7 | | 1094 | KRARRSSLLNAKK | 7.0246 | 6.2260 | 6.8614 | 6.2484 | 6.2316 | 6.3007 | 5.8668 | 6.3109 | 7.0639 | 6.4420 | 6.4912 | 6.9162 | 5.4410 | 5.9374 | 5.1522 | 5.4857 |
| RAPGEF6_S1494 | RAPGEF6 | Q8TEU7 | | 230 | PEGPVDSEDDEEE | 6.2296 | 5.7341 | 6.8653 | 6.2738 | 6.7694 | 6.0834 | 6.0804 | 5.9260 | 5.8368 | 6.7686 | 6.5835 | 5.7206 | 6.6772 | 7.0452 | 5.5624 | 5.8435 |
| RAPGEF6_S230 | RAPGEF6 | Q8TEU7 | | 1157 | AKSSEMSPVPMRS | 6.7526 | 6.8082 | 7.0023 | 6.3419 | 5.9794 | 7.3864 | 6.4698 | 6.8214 | 6.3864 | 6.2975 | 6.4807 | 6.3414 | 5.1190 | 5.5786 | 5.0729 | 5.1615 |
| RAPGEF6_T1507 | RAPGEF6 | Q8TEU7 | | 1070 | FRQRSLSQGSTNS | 5.3149 | 6.4366 | 5.8662 | 5.3715 | 6.0151 | 6.7218 | 6.9588 | 6.6886 | 6.4824 | 6.0134 | 5.6292 | 5.9186 | 5.7864 | 7.1974 | 6.8773 | 6.7218 |
| RAPH1_S1012 | RAPH1 | Q70E73 | | 894 | ANVAPQSPPAVKA | 6.1151 | 5.7057 | 7.1717 | 6.1162 | 5.9212 | 6.3447 | 6.1217 | 6.3160 | 6.1095 | 6.8704 | 6.4614 | 7.0974 | 5.7000 | 6.0455 | 5.8229 | 6.0807 |
| RAPH1_S1154 | RAPH1 | Q70E73 | | 571 | GHVRSQSIVSSVF | 6.5155 | 5.7691 | 6.8734 | 6.2556 | 6.0395 | 5.7800 | 5.9587 | 5.6202 | 7.1970 | 6.5528 | 6.4758 | 6.4955 | 6.4478 | 7.1631 | 5.1444 | 5.7115 |
| RAPH1_S571 | RAPH1 | Q70E73 | | 1012 | PPAESGSPSKETL | 6.4865 | 6.5423 | 7.2101 | 6.5586 | 6.7824 | 6.8444 | 6.7254 | 7.0264 | 7.8464 | 6.6286 | 6.9393 | 7.0552 | 3.9362 | 4.8665 | 4.0523 | 4.4994 |
| RAPH1_S610 | RAPH1 | Q70E73 | | 1154 | VPQVPTSPKSSLS | 5.6781 | 6.1072 | 7.5105 | 5.8965 | 6.3896 | 6.3430 | 6.5528 | 6.2956 | 6.4586 | 6.7214 | 6.7359 | 6.4844 | 5.9958 | 5.5649 | 5.5155 | 5.7501 |
| RAPH1_S853;S856 | RAPH1 | Q70E73 | | 610 | SLVPPLSPQPKIV | 4.8283 | 4.7633 | 6.5449 | 6.4269 | 7.4073 | 6.0568 | 6.0771 | 7.9971 | 6.4859 | 6.0263 | 8.6926 | 6.6710 | 5.2636 | 6.6649 | 5.2453 | 4.8487 |
| RAPH1_S894 | RAPH1 | Q70E73 | 853;856 | | NA | 5.8177 | 5.7578 | 6.2875 | 5.5641 | 5.9217 | 7.5961 | 7.5139 | 7.3741 | 6.7664 | 6.8121 | 6.1757 | 6.5316 | 4.5700 | 6.4279 | 5.3274 | 5.5558 |
| RARS1_S200 | RARS1 | P54136 | | 38 | CGCLGASPNLEQL | 6.3458 | 6.5485 | 7.0870 | 7.0965 | 6.3032 | 6.4324 | 6.9301 | 6.2449 | 6.9540 | 6.6269 | 7.0086 | 7.3469 | 4.7503 | 5.1223 | 4.3992 | 4.8033 |
| RARS1_S38 | RARS1 | P54136 | | 200 | VIVDFSSPNIAKE | 7.9341 | 7.1667 | 7.2702 | 7.0808 | 6.6668 | 5.0916 | 7.1667 | 8.7545 | 6.9243 | 6.3513 | 8.1587 | 7.9290 | 3.1403 | 3.3650 | 3.8900 | 3.1100 |
| RASA3_S207 | RASA3 | Q14644 | | 207 | EVTRPCSYSKKSH | 6.1903 | 6.3894 | 6.0816 | 6.3783 | 6.4741 | 6.2143 | 6.1782 | 6.1562 | 6.2417 | 5.8296 | 5.7861 | 5.5551 | 7.3027 | 6.5469 | 6.5969 | 6.0786 |
| RASA3_S77 | RASA3 | Q14644 | | 90 | DVFRRDSIIGKVA | 6.3631 | 6.1548 | 6.1030 | 6.5240 | 6.5661 | 5.9559 | 6.2888 | 6.0123 | 6.2304 | 5.4695 | 5.6800 | 6.0479 | 7.1395 | 6.2525 | 6.8875 | 6.3247 |
| RASA3_S809 | RASA3 | Q14644 | | 809 | KKTKYGSQEHPIG | 5.6657 | 5.9584 | 6.3949 | 5.7829 | 5.7729 | 5.7756 | 6.0788 | 5.7192 | 6.3703 | 5.7209 | 6.0974 | 5.4676 | 7.7229 | 7.2297 | 6.9279 | 7.3150 |
| RASA3_S90 | RASA3 | Q14644 | | 77 | RSFRHLSFYIFDR | 6.2125 | 7.2993 | 8.7287 | 6.5804 | 5.8911 | 5.1214 | 6.0010 | 5.5231 | 6.3901 | 3.9584 | 4.0895 | 4.8803 | 7.4093 | 5.4089 | 10.0694 | 6.4366 |
| RASAL2_S134 | RASAL2 | Q9UJF2-2 | | 134 | CLRRTVSVPSEGQ | 5.8342 | 6.0709 | 6.6910 | 6.0119 | 6.6391 | 6.4303 | 5.3716 | 6.3504 | 6.6748 | 6.0684 | 6.1082 | 6.5502 | 6.0658 | 6.8852 | 5.9725 | 6.2755 |
| RASAL2_S237 | RASAL2 | Q9UJF2-2 | | 237 | KLKESRSHESLLS | 6.9975 | 6.1154 | 5.8998 | 7.5631 | 5.9809 | 6.1945 | 7.0865 | 7.6699 | 6.4219 | 6.3943 | 7.3574 | 6.0679 | 4.0684 | 5.5201 | 5.4548 | 5.2076 |
| RASAL2_S877 | RASAL2 | Q9UJF2-2 | | 877 | PIRLTGSQLSITQ | 6.0307 | 6.8938 | 7.0583 | 5.9109 | 5.8923 | 6.5011 | 6.5927 | 6.0159 | 5.9478 | 6.1587 | 6.5362 | 6.3297 | 6.1305 | 5.9435 | 6.0511 | 6.0068 |
| RASSF1_S182 | RASSF1 | Q9NS23-5 | | 182 | RPVSVPSSKKPPS | 7.4108 | 10.1640 | 7.2550 | 5.8466 | 3.8914 | 7.8591 | 6.7050 | 9.5505 | 6.0883 | 7.9386 | 6.1932 | 8.9814 | 3.5671 | 2.5021 | 2.7787 | 3.2683 |
| RASSF1_S207 | RASSF1 | Q9NS23-5 | | 207 | SVRRRTSFYLPKD | 6.3890 | 6.5580 | 6.4863 | 5.8657 | 6.1622 | 5.8607 | 5.6836 | 5.6924 | 6.8082 | 6.6080 | 6.8035 | 7.2703 | 6.2129 | 6.1062 | 5.2475 | 6.2454 |
| RASSF8_S129 | RASSF8 | Q8NHQ8 | | 387 | APFQSGSLKRPGS | 7.4889 | 7.1193 | 7.4419 | 6.2370 | 6.0294 | 6.3850 | 6.4280 | 6.3121 | 6.9371 | 6.9274 | 6.9112 | 6.7913 | 4.6069 | 5.5132 | 4.1422 | 4.7290 |
| RASSF8_S387 | RASSF8 | Q8NHQ8 | | 131 | PKRKSLTFTGGAK | 7.6243 | 6.2814 | 7.8185 | 7.6822 | 7.1876 | 6.0394 | 6.2168 | 5.5307 | 6.3308 | 5.8347 | 6.6664 | 6.2343 | 5.3577 | 6.0217 | 4.5192 | 4.6544 |
| RASSF8_T131 | RASSF8 | Q8NHQ8 | | 129 | REPKRKSLTFTGG | 9.5153 | 6.8233 | 7.1459 | 6.9019 | 6.7917 | 6.1177 | 6.7524 | 5.5984 | 6.4875 | 5.9921 | 7.0444 | 6.8382 | 4.6508 | 5.5351 | 3.4387 | 4.3667 |
| RAVER1_S567 | RAVER1 | Q8IY67-2 | | 617 | PSRHKMSPPPSGF | 5.9961 | 6.1786 | 5.9301 | 6.2280 | 6.2466 | 6.0612 | 6.0440 | 6.4437 | 6.3053 | 6.2474 | 6.2905 | 6.2984 | 6.5118 | 6.4584 | 6.4144 | 6.3454 |
| RAVER1_S617 | RAVER1 | Q8IY67-2 | | 463 | PPAAQLTPPPAPV | 5.3825 | 5.6158 | 5.7976 | 5.9298 | 6.2827 | 6.4948 | 6.2689 | 6.4422 | 5.9152 | 6.3006 | 6.0577 | 6.3906 | 6.6498 | 6.4223 | 6.9877 | 7.0619 |
| RAVER1_S667;S680 | RAVER1 | Q8IY67-2 | | 567 | LKSRLLSPLSSAR | 6.8949 | 7.0336 | 6.8496 | 6.5797 | 6.8504 | 6.8225 | 6.7906 | 7.0317 | 7.2610 | 7.0387 | 7.2243 | 7.1427 | 4.0576 | 4.4083 | 3.7310 | 4.2833 |
| RAVER1_S668 | RAVER1 | Q8IY67-2 | | 488 | DSGPLPTPPGVSL | 6.7146 | 7.1091 | 6.7529 | 7.1793 | 6.0503 | 6.8125 | 6.4336 | 7.0713 | 7.0325 | 6.9295 | 7.2227 | 7.2377 | 4.2875 | 4.8790 | 3.7236 | 4.5638 |
| RAVER1_T463 | RAVER1 | Q8IY67-2 | | 668 | SKAIGSSPLGSGE | 5.4246 | 5.9226 | 6.6080 | 6.6947 | 6.7345 | 6.6088 | 6.1887 | 6.3041 | 6.1805 | 6.1329 | 6.4836 | 6.9151 | 5.6868 | 6.3764 | 5.8553 | 5.8835 |
| RAVER1_T488 | RAVER1 | Q8IY67-2 | | 692 | HSHLLKTPLGGQK | 7.7103 | 7.8041 | 6.9443 | 7.3905 | 5.2237 | 7.6601 | 7.6689 | 6.8350 | 7.4229 | 7.5871 | 8.2721 | 8.5198 | 2.1778 | 3.3089 | 2.1097 | 3.3650 |
| RAVER1_T692 | RAVER1 | Q8IY67-2 | | 647 | SGSPTSYFTSGLQ | 6.5721 | 8.3185 | 6.3173 | 8.1120 | 6.5170 | 7.7623 | 6.1983 | 8.0225 | 7.5439 | 6.6778 | 6.9670 | 6.8982 | 3.3281 | 4.0457 | 3.5150 | 3.2042 |
| RAVER1_Y647 | RAVER1 | Q8IY67-2 | 667;680 | | NA | 6.9038 | 7.5319 | 6.8494 | 7.1030 | 5.6788 | 6.6555 | 6.8729 | 6.5369 | 6.8997 | 7.0441 | 7.7425 | 7.5778 | 4.1763 | 4.3808 | 3.6974 | 4.3494 |
| RB1_S249 | RB1 | P06400 | | 807 | GGNIYISPLKSPY | 5.1013 | 5.7586 | 6.2953 | 5.7395 | 6.0844 | 5.8368 | 5.4277 | 6.0571 | 6.1491 | 6.3115 | 6.1942 | 5.9219 | 6.5192 | 7.5134 | 8.0241 | 7.0658 |
| RB1_S249;T252 | RB1 | P06400 | | 821 | ISEGLPTPTKMTP | 5.7179 | 5.7565 | 6.6029 | 5.5110 | 5.3484 | 6.1802 | 5.8471 | 6.1703 | 6.4773 | 6.3019 | 6.4570 | 6.5633 | 6.0040 | 7.2340 | 6.6639 | 7.1644 |
| RB1_S608 | RB1 | P06400 | | 373 | NVIPPHTPVRTVM | 5.6242 | 5.8462 | 6.3739 | 6.2077 | 5.5628 | 6.0049 | 6.0809 | 6.0741 | 6.1276 | 6.1389 | 6.2112 | 6.3698 | 6.2066 | 7.4353 | 6.8407 | 6.8953 |
| RB1_S608;S612 | RB1 | P06400 | | 249 | VIPINGSPRTPRR | 6.3317 | 6.6391 | 7.2666 | 6.4124 | 6.0689 | 6.5881 | 6.4708 | 7.0159 | 6.8637 | 6.9709 | 7.3549 | 7.2978 | 3.9841 | 5.1770 | 4.6675 | 4.8906 |
| RB1_S612 | RB1 | P06400 | | 780 | TRPPTLSPIPHIP | 5.5524 | 6.2766 | 6.6164 | 6.1647 | 5.8598 | 6.2805 | 6.0148 | 6.5217 | 6.4592 | 6.9673 | 6.9511 | 6.5128 | 5.5437 | 6.2282 | 5.9709 | 6.0798 |
| RB1_S780 | RB1 | P06400 | | 795 | PYKFPSSPLRIPG | 5.9323 | 6.1160 | 6.3180 | 5.2882 | 5.2059 | 6.4780 | 5.9029 | 6.6091 | 6.0368 | 6.3255 | 6.6037 | 6.7627 | 5.8651 | 7.0373 | 6.9235 | 6.5949 |
| RB1_S795 | RB1 | P06400 | | 356 | SFETQRTPRKSNL | 5.6655 | 5.5358 | 6.5726 | 5.6091 | 5.6188 | 6.1049 | 6.1079 | 6.1111 | 6.2345 | 6.0023 | 6.5941 | 6.7596 | 6.1818 | 7.5496 | 6.7238 | 6.6286 |
| RB1_S807 | RB1 | P06400 | | 855 | NQMVCNSDRVLKR | 7.2102 | 6.6357 | 9.0583 | 6.9018 | 5.2379 | 7.7116 | 7.4566 | 7.4438 | 7.0470 | 6.4431 | 8.5198 | 8.2619 | 2.2355 | 3.5238 | 2.7775 | 3.5355 |
| RB1_S807;S811 | RB1 | P06400 | | 612 | YLSPVRSPKKKGS | 5.1870 | 5.8773 | 5.5852 | 4.5176 | 5.2577 | 4.8017 | 5.2384 | 4.8306 | 5.2705 | 4.5738 | 5.5659 | 4.6894 | 8.9918 | 10.0722 | 10.9536 | 8.5873 |
| RB1_S855 | RB1 | P06400 | | 608 | AADMYLSPVRSPK | 5.8343 | 5.7720 | 6.1865 | 6.4743 | 5.6724 | 6.6171 | 6.1873 | 6.5102 | 6.4743 | 5.8541 | 6.7196 | 6.4062 | 6.7028 | 6.4545 | 5.7222 | 6.4121 |
| RB1_T356 | RB1 | P06400 | 807;811 | | NA | 5.5714 | 5.6117 | 6.6318 | 5.2707 | 5.2573 | 6.6802 | 6.1653 | 6.2153 | 6.5933 | 6.3411 | 7.1083 | 7.2111 | 6.0255 | 6.8715 | 5.8251 | 6.6206 |
| RB1_T373 | RB1 | P06400 | 249;252 | | NA | 7.1983 | 7.3122 | 8.3080 | 7.0925 | 5.4501 | 7.6219 | 7.2871 | 6.4424 | 6.5019 | 7.0951 | 8.3566 | 8.2228 | 2.7578 | 3.8866 | 3.0096 | 3.4572 |
| RB1_T780;S788 | RB1 | P06400 | 821;826 | | NA | 6.2016 | 5.7906 | 6.7653 | 5.6375 | 5.9670 | 7.1434 | 6.2566 | 6.5847 | 7.0275 | 5.8202 | 7.1493 | 6.4175 | 5.6552 | 6.9719 | 5.8074 | 4.8043 |
| RB1_T821 | RB1 | P06400 | 608;612 | | NA | 5.4295 | 4.9586 | 6.5925 | 6.2769 | 5.8036 | 6.1357 | 5.5509 | 6.7304 | 6.9540 | 7.4502 | 6.4690 | 6.9557 | 5.1493 | 6.8329 | 6.2240 | 6.4869 |
| RB1_T821;T826 | RB1 | P06400 | 780;788 | | NA | 6.2554 | 6.1214 | 6.9148 | 6.6201 | 6.4749 | 6.2905 | 5.5635 | 5.0827 | 6.5493 | 7.0990 | 7.0941 | 7.1191 | 5.4381 | 6.3880 | 5.3673 | 5.6217 |
| RB1CC1_S1160 | RB1CC1 | Q8TDY2 | | 237 | KAEMKRSTELVLS | 6.8706 | 6.9472 | 7.5427 | 6.4743 | 5.8147 | 5.8792 | 5.5913 | 6.0563 | 7.1316 | 7.5616 | 6.6112 | 7.1578 | 4.6456 | 5.4302 | 4.7765 | 5.5091 |
| RB1CC1_S222 | RB1CC1 | Q8TDY2 | | 222 | CLGRLDSLPEHED | 4.8405 | 4.8504 | 5.6710 | 5.0471 | 6.1243 | 5.7781 | 5.8158 | 5.9430 | 5.2259 | 5.5350 | 5.5488 | 5.6496 | 8.8640 | 8.1074 | 8.6842 | 8.3148 |
| RB1CC1_S237 | RB1CC1 | Q8TDY2 | | 243 | STELVLSPDMPRT | 6.1585 | 6.1056 | 6.5593 | 6.7934 | 6.5559 | 6.7301 | 6.4580 | 6.5476 | 6.3905 | 6.5913 | 6.3465 | 5.8818 | 5.3904 | 5.9037 | 5.8454 | 5.7420 |
| RB1CC1_S243 | RB1CC1 | Q8TDY2 | | 1160 | ELNKVTSLHNQAF | 7.3295 | 7.3055 | 6.9191 | 7.4296 | 6.4706 | 5.4946 | 6.0602 | 6.6958 | 6.6808 | 5.5096 | 7.4146 | 5.9481 | 5.1342 | 5.9711 | 4.8309 | 4.8059 |
| RB1CC1_S257 | RB1CC1 | Q8TDY2 | | 647 | ASVSQTSPQSASS | 5.6859 | 7.6465 | 6.2516 | 7.4722 | 5.8172 | 5.8457 | 5.9634 | 5.7900 | 6.1926 | 6.2244 | 5.5773 | 5.6545 | 6.2533 | 6.5733 | 6.6100 | 6.4420 |
| RB1CC1_S624 | RB1CC1 | Q8TDY2 | | 624 | LVKAAQSLDEMSQ | 6.3140 | 7.6943 | 7.7366 | 6.6957 | 5.4522 | 5.1682 | 6.2543 | 5.7785 | 6.4605 | 7.7740 | 6.4798 | 7.4754 | 4.7738 | 5.5294 | 4.6447 | 5.7688 |
| RB1CC1_S629 | RB1CC1 | Q8TDY2 | | 629 | QSLDEMSQTITDL | 8.1246 | 7.1777 | 7.5464 | 6.7614 | 6.8177 | 6.9570 | 6.6723 | 5.8812 | 6.7779 | 7.1353 | 6.7095 | 6.2283 | 4.2791 | 4.5206 | 3.9346 | 4.4764 |
| RB1CC1_S647 | RB1CC1 | Q8TDY2 | | 257 | NESLLTSFPKSVE | 5.6035 | 6.3070 | 7.1208 | 6.6450 | 6.3606 | 6.3832 | 6.2138 | 6.6534 | 7.2418 | 7.2856 | 7.0992 | 6.7504 | 4.8868 | 5.2637 | 5.4550 | 4.7303 |
| RB1CC1_S667 | RB1CC1 | Q8TDY2 | | 667 | GITTTTSPRTPPP | 5.4918 | 6.3650 | 6.2292 | 6.1067 | 6.2753 | 5.8086 | 6.5300 | 7.1476 | 6.7545 | 6.3437 | 6.9799 | 5.9461 | 5.3294 | 6.6640 | 5.9390 | 6.0890 |
| RB1CC1_T631 | RB1CC1 | Q8TDY2 | | 631 | LDEMSQTITDLLS | 9.2914 | 7.7374 | 8.0207 | 9.2226 | 6.4141 | 4.6093 | 5.8395 | 5.7100 | 5.6291 | 4.8723 | 6.4222 | 4.6174 | 4.5567 | 4.9492 | 7.2680 | 4.8399 |
| RBAK_S78 | RBAK | Q9NYW8 | | 78 | EFPCQHSPEAWRV | 5.3321 | 5.6704 | 5.3056 | 6.4983 | 6.6617 | 5.5989 | 6.0307 | 6.4504 | 6.5191 | 6.2761 | 5.3287 | 6.3257 | 6.9250 | 7.1351 | 7.3805 | 6.5618 |
| RBBP4_S110 | RBBP4 | Q09028 | | 355 | KIGEEQSPEDAED | 5.9287 | 5.9268 | 6.9853 | 5.4119 | 6.6933 | 6.0036 | 5.6828 | 6.1112 | 5.7001 | 5.7250 | 7.3830 | 5.3927 | 7.1947 | 6.2553 | 7.1294 | 6.4762 |
| RBBP4_S355 | RBBP4 | Q09028 | | 110 | EFGGFGSVSGKIE | 6.5526 | 6.2812 | 5.8615 | 6.0411 | 6.7312 | 6.7058 | 6.4351 | 6.2054 | 5.7851 | 5.9864 | 6.0912 | 5.6038 | 6.9315 | 6.7072 | 6.5192 | 5.5618 |
| RBBP5_S350 | RBBP5 | Q15291 | | 350 | EYEERESEFDIED | 6.0597 | 6.3656 | 5.9195 | 6.4353 | 6.1621 | 5.7068 | 6.3322 | 5.7629 | 6.3972 | 5.7798 | 5.6592 | 6.0788 | 6.8169 | 6.7879 | 6.7665 | 6.9695 |
| RBBP5_S515 | RBBP5 | Q15291 | | 515 | GLPLEGSAKGKVQ | 5.7582 | 9.5809 | 6.8671 | 7.5180 | 6.9463 | 6.6880 | 7.8624 | 8.6751 | 7.2459 | 7.1426 | 6.1094 | 6.4297 | 3.3853 | 3.2786 | 3.0582 | 3.4542 |

| | | | | | | | | | | | | | | | | | | | | | |
|---|---|---|---|---|---|---|---|---|---|---|---|---|---|---|---|---|---|---|---|---|---|
| RBBP6_S1179 | RBBP6 | Q7Z6E9 | | 984 | PSRDDATPVRDEP | 5.8465 | 6.1939 | 5.8776 | 6.4712 | 6.0002 | 5.7086 | 6.0514 | 5.7673 | 5.9641 | 6.0350 | 5.8262 | 5.9746 | 6.6065 | 7.4454 | 7.0138 | 7.2176 |
| RBBP6_S1221 | RBBP6 | Q7Z6E9 | | 1179 | TEIVKPSPKRKME | 5.8985 | 5.8894 | 5.9586 | 6.7416 | 5.8817 | 5.9149 | 6.2962 | 5.8257 | 6.1905 | 5.9543 | 6.0848 | 6.0628 | 6.3123 | 7.0717 | 6.7243 | 7.1925 |
| RBBP6_S1273;S1277 | RBBP6 | Q7Z6E9 | | 861 | ANRENFSPERFLP | 6.2564 | 6.4555 | 5.8134 | 6.7635 | 5.9150 | 5.5919 | 6.1914 | 5.8172 | 6.0560 | 6.1214 | 5.7492 | 5.9290 | 6.2114 | 7.2539 | 6.7586 | 7.1163 |
| RBBP6_S1277 | RBBP6 | Q7Z6E9 | | 1194 | TEKMDRTPEKDKI | 6.2753 | 6.5855 | 5.9706 | 6.4370 | 6.0855 | 5.7080 | 6.0576 | 6.0797 | 6.1938 | 6.2339 | 5.9025 | 6.0119 | 6.1630 | 6.9611 | 6.7709 | 6.5638 |
| RBBP6_S1328 | RBBP6 | Q7Z6E9 | | 1328 | DKDDFESEEEDVK | 6.1722 | 6.2727 | 6.3813 | 6.6426 | 5.6628 | 5.5186 | 6.4532 | 5.5403 | 6.0084 | 6.1726 | 5.7984 | 6.1287 | 6.3277 | 6.9794 | 6.7254 | 7.2158 |
| RBBP6_S1625 | RBBP6 | Q7Z6E9 | | 516 | KLGYLVSPPQQIR | 6.3956 | 7.0482 | 6.8631 | 7.1599 | 6.6831 | 6.4118 | 6.8551 | 6.5872 | 6.8055 | 6.3641 | 6.7620 | 6.4827 | 4.5936 | 5.3639 | 4.6034 | 5.0210 |
| RBBP6_S516 | RBBP6 | Q7Z6E9 | | 1277 | TSSTGGSPVRKSE | 6.0896 | 5.8122 | 5.8176 | 6.9638 | 6.1213 | 5.8620 | 5.9365 | 5.5172 | 6.3164 | 6.3196 | 6.0646 | 6.4549 | 6.4177 | 7.1366 | 6.5902 | 6.5799 |
| RBBP6_S770;S772 | RBBP6 | Q7Z6E9 | | 873 | PLNIRNSPFTRGR | 6.6558 | 7.0757 | 6.1681 | 6.9371 | 6.3191 | 6.6264 | 6.1852 | 5.3068 | 5.7229 | 6.1712 | 6.1302 | 5.6673 | 6.3701 | 7.5131 | 5.6580 | 5.4930 |
| RBBP6_S815 | RBBP6 | Q7Z6E9 | | 1221 | TGKKIGSTENISN | 6.1381 | 6.3639 | 6.3944 | 6.6281 | 5.7819 | 5.7075 | 6.4830 | 5.7240 | 6.2197 | 6.5335 | 6.0029 | 6.6353 | 5.9547 | 6.6362 | 5.9077 | 6.8892 |
| RBBP6_S861 | RBBP6 | Q7Z6E9 | | 815 | KAYYGRSVDFRDP | 7.2520 | 7.5159 | 7.1685 | 7.2169 | 6.8856 | 5.8360 | 6.1767 | 5.2417 | 6.6515 | 6.3084 | 6.5710 | 7.1325 | 5.0877 | 5.1725 | 4.4754 | 5.3076 |
| RBBP6_S873 | RBBP6 | Q7Z6E9 | | 1625 | SHSSRLSSDLTRE | 3.7992 | 3.7885 | 3.2044 | 3.8737 | 4.7633 | 4.2604 | 5.0298 | 5.0333 | 3.6252 | 3.1627 | 2.9362 | 3.3532 | 12.1550 | 12.4211 | 14.4423 | 14.1517 |
| RBBP6_T1194 | RBBP6 | Q7Z6E9 | 770;772 | | NA | 6.3681 | 6.2374 | 5.8093 | 6.4241 | 5.6563 | 5.5223 | 6.0321 | 6.2716 | 6.2743 | 6.1509 | 5.7333 | 5.9902 | 6.3221 | 7.1020 | 6.6006 | 7.5053 |
| RBBP6_T984 | RBBP6 | Q7Z6E9 | 1273;1277 | | NA | 6.1013 | 5.5588 | 6.0393 | 6.1514 | 5.9567 | 5.4391 | 5.7272 | 5.6478 | 5.7126 | 5.7352 | 5.9383 | 5.7252 | 6.7497 | 7.8593 | 7.9859 | 7.6723 |
| RBBP7_S143 | RBBP7 | Q16576-2 | | 189 | IATKTPSSDVLVF | 5.7937 | 5.6317 | 5.5415 | 5.8453 | 6.2173 | 6.0443 | 5.7155 | 6.1336 | 6.1824 | 6.7037 | 6.1673 | 6.9637 | 7.1418 | 6.7103 | 6.3436 | 6.8642 |
| RBBP7_S189 | RBBP7 | Q16576-2 | | 143 | DASHCDSDKGEFG | 5.9597 | 6.0142 | 6.1889 | 6.4946 | 6.5655 | 5.7522 | 5.9863 | 5.9469 | 6.4818 | 5.5376 | 6.2865 | 6.2059 | 6.6013 | 6.2865 | 6.3217 | 7.3703 |
| RBBP8_S163 | RBBP8 | Q99708-2 | | 327 | LPTRVSSPVFGAT | 5.7532 | 6.0498 | 5.9014 | 6.2935 | 6.5108 | 6.1996 | 6.3162 | 6.4011 | 6.2531 | 5.7896 | 6.1057 | 6.1251 | 6.7127 | 6.3996 | 6.5511 | 6.6374 |
| RBBP8_S327 | RBBP8 | Q99708-2 | | 315 | SDSTSKTPPQEEL | 5.4539 | 6.3567 | 5.9989 | 6.1109 | 6.7326 | 6.7312 | 6.7471 | 6.2520 | 6.7340 | 6.4293 | 6.5328 | 6.2839 | 6.3341 | 6.3572 | 5.8475 | 5.0979 |
| RBBP8_S345 | RBBP8 | Q99708-2 | | 163 | EDVIPDSPITAFS | 5.7546 | 5.4824 | 7.5160 | 5.8152 | 5.8819 | 6.1741 | 7.0050 | 6.2922 | 6.1536 | 6.1613 | 5.9405 | 6.4576 | 6.4293 | 6.3492 | 5.8465 | 6.7405 |
| RBBP8_T315 | RBBP8 | Q99708-2 | | 345 | GLDLNTSLSPSLL | 6.3942 | 6.1811 | 5.5374 | 6.4687 | 6.8263 | 7.4791 | 6.1066 | 5.6865 | 7.2472 | 6.6411 | 6.8196 | 6.7902 | 5.5705 | 5.7505 | 5.0631 | 5.4380 |
| RBFOX2_T67 | RBFOX2 | O43251-8 | | 67 | MQNEPLTPGYHGF | 6.8766 | 4.9107 | 5.3660 | 6.3781 | 5.7998 | 5.9028 | 6.9597 | 7.3236 | 7.1059 | 6.2568 | 5.9626 | 5.7881 | 7.2555 | 5.5804 | 5.9277 | 6.6057 |
| RBL1_S1009 | RBL1 | P28749 | | 650 | VHERYSSPTAGSA | 6.4275 | 6.6811 | 7.3780 | 6.1310 | 6.3392 | 6.8083 | 6.4963 | 7.1186 | 7.0520 | 6.9460 | 7.5743 | 7.1276 | 3.9545 | 4.7322 | 4.5168 | 4.7168 |
| RBL1_S1041 | RBL1 | P28749 | | 964 | MDAPPLSPFPHIK | 6.3285 | 6.5603 | 7.4177 | 6.2478 | 5.8272 | 6.7070 | 6.3216 | 6.7883 | 7.0069 | 6.9361 | 7.5773 | 7.3306 | 4.5757 | 4.7258 | 4.6274 | 5.0217 |
| RBL1_S640 | RBL1 | P28749 | | 915 | LEDATKTPDCSSG | 6.3639 | 6.7716 | 7.5536 | 6.5507 | 5.8213 | 6.4482 | 6.2629 | 6.1897 | 6.9023 | 7.0740 | 7.1860 | 7.1925 | 4.5259 | 4.9975 | 4.6565 | 5.5035 |
| RBL1_S649 | RBL1 | P28749 | | 1009 | LYKFNGSPSKSLK | 5.9608 | 6.3079 | 6.7261 | 5.6534 | 5.1289 | 6.4821 | 6.0436 | 6.1708 | 6.6666 | 6.7466 | 6.9664 | 7.2104 | 5.5766 | 6.1748 | 5.8043 | 6.3807 |
| RBL1_S650 | RBL1 | P28749 | | 369 | RSFAPSTPLTGRR | 6.1454 | 6.2589 | 7.0648 | 6.1487 | 6.3126 | 6.6601 | 6.3463 | 6.5459 | 7.0885 | 6.6642 | 7.3255 | 7.0314 | 4.8598 | 5.3995 | 4.9277 | 5.2208 |
| RBL1_S964 | RBL1 | P28749 | | 1041 | IDSDAESPAKRVC | 5.8252 | 6.6963 | 6.6424 | 5.9864 | 5.0742 | 5.8542 | 5.7190 | 6.3056 | 6.9102 | 6.5776 | 7.0454 | 7.0375 | 5.8018 | 6.2542 | 5.8320 | 6.4379 |
| RBL1_S975 | RBL1 | P28749 | | 332 | AEEEIGTPRKFTR | 5.3634 | 6.1810 | 6.1852 | 5.7655 | 5.4799 | 6.0162 | 5.9650 | 5.8816 | 6.1467 | 6.3534 | 5.8645 | 6.3632 | 6.9770 | 7.2912 | 7.1181 | 7.0480 |
| RBL1_S988 | RBL1 | P28749 | | 997 | KNGSGLTPRSALL | 6.7113 | 6.6050 | 7.7093 | 5.9772 | 5.2248 | 6.6944 | 6.4421 | 6.4696 | 7.2719 | 6.8099 | 7.8830 | 7.9756 | 4.3736 | 4.5858 | 4.3343 | 4.9324 |
| RBL1_T332 | RBL1 | P28749 | | 649 | SVHERYSSPTAGS | 6.6781 | 6.7031 | 7.2998 | 6.1852 | 6.0620 | 6.7916 | 6.4296 | 7.2089 | 6.8246 | 6.9707 | 7.8216 | 7.5127 | 4.1101 | 4.5350 | 4.1290 | 4.7381 |
| RBL1_T340 | RBL1 | P28749 | | 975 | IKQQPGSPRRISQ | 6.0248 | 6.5853 | 6.0028 | 5.5117 | 5.6058 | 6.0536 | 5.8327 | 5.9895 | 6.2192 | 6.1440 | 6.2712 | 6.7368 | 7.0518 | 7.0244 | 6.5766 | 6.3698 |
| RBL1_T369 | RBL1 | P28749 | | 385 | EKEAVITPVASAT | 5.9084 | 6.5988 | 7.7751 | 6.7254 | 5.9645 | 6.4595 | 6.7731 | 7.0233 | 7.2087 | 6.7284 | 7.3536 | 7.0622 | 4.1906 | 4.8945 | 4.4867 | 4.8472 |
| RBL1_T385 | RBL1 | P28749 | | 640 | RDMQPLSPISVHE | 6.6292 | 6.9528 | 6.1997 | 6.4456 | 6.2228 | 7.1115 | 6.1020 | 7.0580 | 6.5588 | 5.2949 | 6.0566 | 6.5271 | 5.1405 | 6.8955 | 4.9656 | 5.8394 |
| RBL1_T915 | RBL1 | P28749 | | 988 | QHSIYISPHKNGS | 5.9257 | 6.4110 | 6.0039 | 5.8665 | 6.8020 | 7.7112 | 5.7143 | 7.7088 | 6.7482 | 6.2310 | 7.1832 | 6.1219 | 5.4701 | 5.9891 | 5.2760 | 4.8372 |
| RBL1_T997 | RBL1 | P28749 | | 340 | RKFTRDTPLGKLT | 6.0328 | 6.3569 | 6.2064 | 5.1006 | 5.1185 | 6.3924 | 5.8625 | 5.7010 | 6.3826 | 5.8958 | 5.7795 | 6.4059 | 7.4222 | 7.4070 | 6.8136 | 7.1224 |
| RBL2_S1080 | RBL2 | Q08999 | | 401 | KALRISTPLTGVR | 6.3161 | 6.6671 | 6.9876 | 5.9726 | 6.0166 | 6.5948 | 6.3697 | 6.2920 | 6.8926 | 6.6529 | 6.6773 | 6.5346 | 5.3228 | 5.8952 | 5.2187 | 5.5894 |
| RBL2_S1112 | RBL2 | Q08999 | | 662 | LGRSITSPTTLYD | 6.0284 | 6.5879 | 6.8080 | 6.3131 | 6.0367 | 6.7324 | 6.1885 | 6.6148 | 6.9442 | 6.7879 | 6.9472 | 6.8631 | 5.0357 | 5.7735 | 4.9986 | 5.3400 |
| RBL2_S662 | RBL2 | Q08999 | | 1080 | FYYFSNSPSKRLR | 5.4402 | 5.9879 | 6.6339 | 5.2276 | 6.2225 | 6.4284 | 6.3388 | 6.7078 | 6.5385 | 6.2754 | 5.9241 | 6.2844 | 6.3407 | 7.2255 | 6.1190 | 6.3054 |
| RBL2_S965 | RBL2 | Q08999 | | 1112 | LEDGSESPAKRIC | 6.5100 | 6.5567 | 7.0191 | 5.9040 | 5.8501 | 6.5337 | 6.2025 | 6.6085 | 6.9422 | 7.0662 | 6.8351 | 6.7868 | 4.9712 | 5.5548 | 4.8964 | 5.7628 |
| RBL2_S982;T986 | RBL2 | Q08999 | | 1097 | MIRTGETPTKKRG | 6.4103 | 6.5579 | 7.3027 | 6.2436 | 5.5968 | 6.6016 | 6.5685 | 5.8507 | 7.1503 | 7.2443 | 7.1815 | 6.9235 | 5.0813 | 5.1651 | 4.7535 | 5.3686 |
| RBL2_T1097 | RBL2 | Q08999 | | 965 | DRTSRDSSPVMRS | 5.1232 | 5.8171 | 5.6617 | 5.5233 | 5.5941 | 5.5442 | 6.5738 | 7.0270 | 7.3273 | 6.9329 | 6.2711 | 5.7374 | 6.1616 | 7.2533 | 7.0987 | 6.3532 |
| RBL2_T401 | RBL2 | Q08999 | 982;986 | | NA | 5.5127 | 7.2263 | 5.1952 | 5.4822 | 5.2722 | 7.2177 | 6.5859 | 7.7148 | 6.2323 | 7.4644 | 5.8932 | 5.9231 | 4.8173 | 7.0576 | 6.4777 | 5.9274 |
| RBM10_S115 | RBM10 | P98175-5 | | 154 | RRRHRHSPTGPPG | 6.6005 | 6.6176 | 6.2859 | 6.8736 | 6.2364 | 5.8778 | 6.3169 | 5.9630 | 6.3979 | 6.2204 | 6.2487 | 6.1759 | 6.0705 | 5.9766 | 5.9850 | 6.1533 |
| RBM10_S154 | RBM10 | P98175-5 | | 788 | ASDDRPSPPRGLV | 6.2115 | 6.6478 | 5.8157 | 6.4206 | 6.1009 | 6.1849 | 6.2282 | 6.1154 | 6.1058 | 6.0072 | 5.7905 | 5.9890 | 6.5189 | 6.4618 | 6.7949 | 6.6069 |
| RBM10_S30 | RBM10 | P98175-5 | | 115 | YPREYGSQEGKHD | 6.5323 | 6.8768 | 6.5656 | 7.0337 | 5.9200 | 5.8205 | 6.0725 | 6.1384 | 6.3674 | 6.3957 | 6.1155 | 6.9027 | 5.5900 | 5.6369 | 5.7092 | 6.3228 |
| RBM10_S788 | RBM10 | P98175-5 | | 862 | HRRAHLSENELEA | 7.2382 | 7.9119 | 7.2302 | 8.1273 | 6.5861 | 6.7011 | 7.2182 | 6.7366 | 6.5079 | 6.5452 | 6.3019 | 6.4788 | 3.9102 | 4.3357 | 4.1327 | 4.0379 |
| RBM10_S801;S803 | RBM10 | P98175-5 | | 30 | ESLQEASPRLADH | 6.8891 | 7.1836 | 5.9759 | 6.9530 | 6.4643 | 6.5052 | 6.3172 | 6.0859 | 6.6261 | 6.4116 | 6.4737 | 6.7147 | 5.2613 | 5.0942 | 5.2322 | 5.8120 |
| RBM10_S862 | RBM10 | P98175-5 | | 910 | GGISTASVDFEQP | 5.5641 | 6.0419 | 6.7152 | 5.8580 | 6.2512 | 6.0904 | 6.4043 | 6.8637 | 6.8806 | 7.4254 | 6.8175 | 7.1946 | 4.8215 | 6.0119 | 5.6341 | 5.4256 |
| RBM10_S910 | RBM10 | P98175-5 | 801;803 | | NA | 5.8883 | 5.6324 | 6.4291 | 7.0406 | 6.0042 | 5.5061 | 6.1478 | 5.8282 | 6.6166 | 6.0719 | 6.2282 | 6.9318 | 6.4464 | 6.3427 | 6.5116 | 6.3743 |
| RBM12_S352 | RBM12 | Q9NTZ6 | | 424 | KRSRSRSPHEAGF | 5.3408 | 5.6857 | 5.7596 | 5.3376 | 6.0015 | 6.0956 | 4.7918 | 6.1991 | 6.0344 | 5.8137 | 5.9731 | 5.6717 | 7.9547 | 7.5596 | 8.1250 | 7.6560 |
| RBM12_S420;S422;S424 | RBM12 | Q9NTZ6 | | 352 | GLVKFLSPQDTFE | 5.4676 | 5.8731 | 7.1178 | 6.4635 | 5.8658 | 6.4870 | 5.2285 | 5.9102 | 6.2107 | 6.9571 | 6.3577 | 6.3747 | 6.1558 | 6.6194 | 6.4555 | 6.4555 |
| RBM12_S422;S424 | RBM12 | Q9NTZ6 | 422;424 | | NA | 5.5603 | 6.4133 | 6.3014 | 6.0452 | 5.8594 | 6.1142 | 5.6495 | 6.0752 | 6.1422 | 6.2714 | 6.3980 | 6.4355 | 6.4769 | 6.6295 | 7.0248 | 6.6034 |
| RBM12_S424 | RBM12 | Q9NTZ6 | 420;422;424 | | NA | 5.7684 | 5.9369 | 6.4655 | 5.5663 | 6.3854 | 7.1907 | 5.9253 | 5.7337 | 6.0638 | 6.1066 | 6.7019 | 6.1974 | 6.8390 | 6.6923 | 6.1251 | 6.3016 |
| RBM12B_S105 | RBM12B | Q8IXT5 | | 575 | PEDFRHSPEDFRR | 6.1734 | 6.2139 | 5.9088 | 6.5720 | 6.0066 | 5.9876 | 6.2057 | 5.9616 | 6.1916 | 6.1788 | 6.0608 | 6.0992 | 6.5119 | 6.7021 | 6.6279 | 6.5984 |
| RBM12B_S250;S254 | RBM12B | Q8IXT5 | | 280 | RRTRSRSPLGFYV | 6.2080 | 6.4534 | 5.9191 | 6.6936 | 6.0570 | 5.7877 | 5.6849 | 5.9372 | 6.2122 | 5.9823 | 5.8304 | 5.8147 | 6.8781 | 6.9105 | 7.0344 | 6.5964 |
| RBM12B_S254 | RBM12B | Q8IXT5 | | 638 | EEDFRRSPTEDFR | 6.1671 | 6.3604 | 5.9605 | 6.5444 | 5.8729 | 5.6204 | 6.1594 | 6.0911 | 6.1392 | 6.1208 | 5.8254 | 6.2294 | 6.5596 | 6.5851 | 6.9568 | 6.8077 |
| RBM12B_S278;S280 | RBM12B | Q8IXT5 | | 98 | GRGRPGSGTSGVD | 5.7356 | 5.9689 | 5.7739 | 6.3971 | 6.1364 | 5.9881 | 6.1774 | 6.2528 | 5.8766 | 5.7898 | 5.8814 | 5.9784 | 7.0716 | 6.7688 | 7.1985 | 7.0048 |
| RBM12B_S280 | RBM12B | Q8IXT5 | | 874 | PGEDFRSPPDDFR | 6.5772 | 6.9447 | 6.1169 | 7.5345 | 6.4561 | 6.3074 | 6.8543 | 6.6080 | 7.0529 | 6.9832 | 6.7709 | 6.8721 | 4.4609 | 5.1288 | 4.4376 | 4.8945 |
| RBM12B_S280;Y285 | RBM12B | Q8IXT5 | | 710 | EEDFRHSPEEDFR | 6.0928 | 6.4752 | 5.9683 | 6.6165 | 6.2296 | 6.0309 | 6.4840 | 6.2187 | 6.3799 | 6.1357 | 6.1583 | 5.9860 | 6.5099 | 6.1958 | 6.2407 | 6.2777 |
| RBM12B_S562 | RBM12B | Q8IXT5 | | 839 | EDFRCPSDEDFRQ | 5.9491 | 6.0418 | 5.9213 | 6.3441 | 6.0847 | 5.8222 | 6.4265 | 5.9342 | 5.9191 | 6.1712 | 5.8500 | 6.2073 | 6.4625 | 6.9972 | 6.8707 | 6.9980 |
| RBM12B_S575 | RBM12B | Q8IXT5 | | 718 | EEDFRQSPQEHFR | 6.2257 | 6.7907 | 6.1860 | 6.7016 | 6.2646 | 5.9380 | 6.8345 | 5.9343 | 6.3930 | 6.2391 | 6.0333 | 6.1726 | 5.8260 | 6.2405 | 5.9096 | 6.3105 |
| RBM12B_S638 | RBM12B | Q8IXT5 | | 105 | GTSGVDSLSNFIE | 6.4280 | 5.9918 | 5.9840 | 6.1763 | 6.7005 | 6.5901 | 6.4108 | 6.1349 | 5.2219 | 4.9150 | 4.7150 | 5.9159 | 7.5833 | 7.3419 | 7.2910 | 6.5996 |
| RBM12B_S710 | RBM12B | Q8IXT5 | | 562 | EDFRHSSEDFRFP | 5.8395 | 5.8646 | 5.7461 | 6.8615 | 6.0056 | 5.6466 | 6.4608 | 6.2440 | 6.1443 | 6.1935 | 5.9357 | 6.5132 | 6.4619 | 6.5744 | 6.3919 | 7.1164 |
| RBM12B_S710;S718 | RBM12B | Q8IXT5 | | 254 | RRSEEHSPPRGIN | 7.4776 | 7.8442 | 5.5498 | 7.8034 | 6.4528 | 6.0046 | 5.9487 | 6.7092 | 6.5710 | 6.3189 | 6.3963 | 6.5116 | 4.6189 | 5.7181 | 5.1137 | 4.9612 |
| RBM12B_S718 | RBM12B | Q8IXT5 | 278;280 | | NA | 6.4675 | 6.5533 | 5.9764 | 6.5517 | 6.1317 | 5.7948 | 6.0686 | 5.8969 | 6.2785 | 6.2786 | 6.2102 | 6.0884 | 6.7961 | 6.1681 | 6.4737 | 6.2656 |
| RBM12B_S839 | RBM12B | Q8IXT5 | 710;718 | | NA | 6.1280 | 6.0676 | 6.0309 | 6.8175 | 6.2940 | 6.3711 | 6.1072 | 6.2887 | 5.7301 | 6.3758 | 6.0468 | 6.2161 | 6.3833 | 6.2117 | 6.7328 | 6.1983 |
| RBM12B_S874 | RBM12B | Q8IXT5 | 276;278;280 | | NA | 6.4280 | 6.5263 | 6.1817 | 6.7192 | 6.5514 | 6.3037 | 6.4070 | 6.0273 | 5.8222 | 5.7757 | 5.9140 | 5.9893 | 6.3379 | 6.4431 | 6.3893 | 6.1842 |
| RBM12B_S98 | RBM12B | Q8IXT5 | 98;105 | | NA | 5.8300 | 5.8354 | 5.0827 | 6.8369 | 6.4342 | 5.7836 | 5.9429 | 6.2495 | 5.6998 | 5.2697 | 5.1443 | 5.0422 | 7.8529 | 7.4354 | 7.5033 | 8.0572 |
| RBM12B_S98;S105 | RBM12B | Q8IXT5 | 280;285 | | NA | 6.5909 | 7.2828 | 5.6678 | 7.4253 | 6.3318 | 6.9343 | 5.9050 | 7.9370 | 5.6288 | 6.5121 | 5.8877 | 5.7158 | 4.7001 | 5.7298 | 6.3403 | 5.4105 |
| RBM12B_T276;S278;S28 | RBM12B | Q8IXT5 | 250;254 | | NA | 6.2051 | 8.6088 | 5.5289 | 7.0169 | 6.5654 | 5.7170 | 5.4281 | 6.7636 | 5.5479 | 5.6147 | 6.0856 | 5.8684 | 6.2090 | 6.0252 | 6.8696 | 5.9458 |
| RBM14_S215 | RBM14 | Q96PK6 | | 206 | GQARQPTPPFFGR | 6.2155 | 6.1918 | 5.6654 | 6.4678 | 6.1890 | 5.7964 | 6.1022 | 6.1708 | 6.1006 | 6.0628 | 5.9632 | 6.0117 | 6.8703 | 6.6816 | 6.8594 | 6.6517 |
| RBM14_S215;S220 | RBM14 | Q96PK6 | | 582 | YERTRLSPPRASY | 6.4546 | 6.4894 | 5.9203 | 6.7217 | 6.0053 | 5.8101 | 6.1154 | 6.1265 | 6.3103 | 6.1908 | 6.1233 | 6.3589 | 6.3467 | 6.2782 | 6.4200 | 6.3285 |
| RBM14_S256 | RBM14 | Q96PK6 | | 572 | VANANSTPPPYER | 6.0683 | 6.1055 | 6.0491 | 6.5929 | 6.5218 | 6.1740 | 6.2266 | 6.3147 | 5.9067 | 6.0993 | 6.3667 | 6.1417 | 6.8870 | 6.6546 | 5.7886 | 6.1026 |
| RBM14_S520 | RBM14 | Q96PK6 | | 618 | SDYRRLSESQLSF | 5.4229 | 5.0710 | 5.8069 | 6.4798 | 8.1882 | 5.6748 | 5.3513 | 6.0609 | 6.8685 | 7.8250 | 7.0824 | 7.2825 | 6.2129 | 6.3582 | 4.9390 | 5.3757 |
| RBM14_S582 | RBM14 | Q96PK6 | | 215 | FFGRDRSPLRRSP | 6.7226 | 6.8033 | 5.8695 | 6.5732 | 6.1968 | 6.0931 | 5.9234 | 5.9667 | 5.9632 | 6.1695 | 6.0375 | 6.3382 | 6.5597 | 6.3557 | 6.1938 | 6.2337 |
| RBM14_S618 | RBM14 | Q96PK6 | | 256 | RAQPSASLGVGYR | 6.2890 | 5.6425 | 5.3847 | 6.4023 | 5.7055 | 5.1925 | 5.8559 | 5.3250 | 5.8532 | 5.2394 | 5.5762 | 5.7931 | 8.8219 | 7.9726 | 7.3442 | 7.6020 |
| RBM14_S623 | RBM14 | Q96PK6 | | 623 | LSESQLSFRRSPT | 5.6420 | 5.6549 | 6.4803 | 7.0055 | 5.5703 | 6.5139 | 6.0631 | 5.9548 | 6.9728 | 7.3381 | 8.5031 | 8.6939 | 5.2732 | 5.4719 | 3.6533 | 5.2089 |
| RBM14_S649 | RBM14 | Q96PK6 | | 520 | APYRTQSSASLAA | 7.6303 | 6.2762 | 5.4065 | 7.2498 | 6.7374 | 5.5300 | 6.1589 | 5.0462 | 5.7024 | 5.3560 | 4.9833 | 5.5929 | 8.0954 | 7.3221 | 6.2288 | 6.6838 |
| RBM14_T206 | RBM14 | Q96PK6 | | 649 | SDYARYSGSYNDY | 7.3657 | 7.2912 | 6.0405 | 7.8786 | 6.5436 | 6.8034 | 7.4250 | 6.6635 | 7.1505 | 6.4900 | 7.3352 | 7.0326 | 3.9618 | 4.3898 | 3.3565 | 4.2722 |
| RBM14_T572 | RBM14 | Q96PK6 | 215;220 | | NA | 7.1701 | 7.1587 | 6.9300 | 6.9865 | 6.7996 | 7.4807 | 6.3916 | 4.4680 | 5.3596 | 6.0994 | 6.3431 | 6.3880 | 6.3668 | 6.1777 | 5.0753 | 4.8048 |
| RBM15_S109 | RBM15 | Q96T37 | | 700 | LLLERPSPIRDRR | 6.4108 | 6.5588 | 5.9441 | 6.7484 | 6.0028 | 5.8007 | 6.2443 | 6.2311 | 6.2285 | 6.1568 | 5.7361 | 5.9111 | 6.2609 | 6.5212 | 6.6654 | 6.5790 |
| RBM15_S127 | RBM15 | Q96T37 | | 294 | GGQRSLSPGGAAL | 6.5559 | 6.7152 | 5.8537 | 6.5662 | 6.2856 | 5.7843 | 6.2602 | 6.1846 | 6.2631 | 5.9195 | 5.8522 | 5.6893 | 6.4424 | 6.4476 | 6.7488 | 6.4314 |
| RBM15_S128 | RBM15 | Q96T37 | | 568 | RGARDRTPPLLYR | 6.2293 | 6.3447 | 6.0133 | 6.5562 | 5.9846 | 5.7992 | 6.3604 | 5.7430 | 6.2375 | 6.1071 | 6.1142 | 6.2035 | 6.4946 | 6.7086 | 6.5684 | 6.5352 |
| RBM15_S257;S259 | RBM15 | Q96T37 | | 259 | SRRRSRSPLDKDT | 6.6280 | 6.9061 | 6.1470 | 6.9774 | 5.6855 | 5.4886 | 5.8374 | 5.2902 | 6.4889 | 6.4589 | 6.2022 | 6.1933 | 6.2125 | 6.6643 | 6.6219 | 6.1980 |
| RBM15_S259 | RBM15 | Q96T37 | | 622 | RRRDGWSLDRDRG | 5.7386 | 6.0796 | 6.0216 | 6.8670 | 6.2996 | 5.9781 | 6.8771 | 6.9263 | 6.3335 | 6.5313 | 6.5631 | 6.2594 | 5.3357 | 5.9800 | 6.0884 | 6.1208 |
| RBM15_S292;S294 | RBM15 | Q96T37 | | 935 | SQQFLDSPAKALA | 5.9108 | 6.3040 | 5.8387 | 6.9612 | 6.1002 | 6.2380 | 6.1832 | 6.1526 | 6.1712 | 6.2654 | 6.6712 | 6.6411 | 6.0808 | 6.4464 | 6.0405 | 5.9948 |
| RBM15_S294 | RBM15 | Q96T37 | | 656 | GRHLDRSPESDRP | 6.1912 | 6.4281 | 6.0586 | 6.8093 | 6.0168 | 5.8714 | 5.9299 | 5.6682 | 6.2186 | 6.2494 | 6.0783 | 5.9993 | 6.5431 | 6.6876 | 6.7544 | 6.4958 |
| RBM15_S622 | RBM15 | Q96T37 | | 781 | APVASASPKLCLA | 6.2585 | 6.7622 | 6.4720 | 7.0548 | 6.0305 | 5.8194 | 6.2168 | 5.6583 | 6.1483 | 6.2404 | 6.1287 | 5.9678 | 5.9860 | 6.3731 | 6.6359 | 6.2474 |
| RBM15_S656 | RBM15 | Q96T37 | | 128 | RLHSYSSPSTKNS | 6.6854 | 6.7489 | 5.8718 | 6.9393 | 6.5843 | 6.1566 | 6.7116 | 6.4503 | 6.5445 | 6.0168 | 6.1464 | 6.0946 | 5.7434 | 6.0025 | 5.7539 | 5.5498 |
| RBM15_S656;S659 | RBM15 | Q96T37 | | 127 | SRLHSYSSPSTKN | 6.4901 | 7.0712 | 5.9177 | 6.7959 | 6.7422 | 6.3672 | 6.3533 | 6.5658 | 6.8486 | 5.8906 | 6.0103 | 6.0233 | 5.6005 | 6.2754 | 5.4768 | 5.5713 |
| RBM15_S670;S674 | RBM15 | Q96T37 | | 109 | SSSRGGSREYDTG | 6.3435 | 6.5545 | 5.9328 | 6.7155 | 6.7985 | 7.4869 | 6.4909 | 5.5810 | 5.9312 | 5.8998 | 6.2799 | 6.4063 | 6.6358 | 6.2002 | 5.5270 | 5.2162 |
| RBM15_S700 | RBM15 | Q96T37 | | 765 | ASSKLKSPSQKQD | 7.4366 | 7.4847 | 6.5002 | 7.2047 | 7.8387 | 7.0534 | 7.3341 | 7.1879 | 7.5415 | 6.2667 | 6.3802 | 5.7549 | 4.0106 | 4.8219 | 3.8072 | 3.3768 |
| RBM15_S765 | RBM15 | Q96T37 | | 84 | GGSNGSSSGKTDS | 6.0823 | 6.1968 | 6.0946 | 6.8531 | 5.9994 | 6.0758 | 6.6604 | 6.8478 | 5.8649 | 6.1416 | 6.5746 | 5.3378 | 6.3355 | 6.4783 | 6.3061 | 6.1510 |
| RBM15_S781 | RBM15 | Q96T37 | 257;259 | | NA | 6.0515 | 6.3678 | 5.9629 | 6.7462 | 6.1555 | 5.3862 | 6.3787 | 5.4541 | 6.0264 | 6.3610 | 6.1121 | 6.3527 | 6.6768 | 6.4398 | 6.6650 | 6.8633 |
| RBM15_S84 | RBM15 | Q96T37 | 656;659 | | NA | 6.6806 | 5.8960 | 6.5267 | 7.4520 | 5.7439 | 5.5342 | 6.5966 | 6.5534 | 5.6439 | 6.3259 | 6.2824 | 5.7807 | 5.7846 | 6.3032 | 6.7029 | 6.1929 |
| RBM15_S935 | RBM15 | Q96T37 | 670;674 | | NA | 5.6122 | 6.5592 | 6.0690 | 6.3142 | 6.1613 | 6.0096 | 6.7758 | 6.0774 | 5.6580 | 6.1249 | 6.3845 | 5.4218 | 6.2904 | 6.6928 | 6.6888 | 7.1599 |
| RBM15_T568 | RBM15 | Q96T37 | 292;294 | | NA | 6.0743 | 7.2492 | 6.0488 | 6.5023 | 6.5074 | 5.6455 | 6.7632 | 6.5922 | 5.2654 | 6.0877 | 6.3552 | 6.1585 | 5.9469 | 6.2775 | 6.3582 | 6.1678 |
| RBM15B_S109 | RBM15B | Q8NDT2 | | 532 | DLVRDRTPPHLLY | 6.1202 | 6.5496 | 6.2788 | 6.9281 | 5.7856 | 6.4183 | 6.1945 | 6.2585 | 6.8690 | 6.0396 | 5.8760 | 5.8979 | 6.4712 | 6.3792 | 5.9965 | 5.9369 |
| RBM15B_S113 | RBM15B | Q8NDT2 | | 267 | YKQRSLSPVAAPP | 6.2179 | 6.5643 | 5.9182 | 6.6944 | 5.9918 | 5.6139 | 5.9088 | 5.9234 | 6.5205 | 6.3251 | 6.1184 | 5.8543 | 6.4564 | 6.5953 | 6.6562 | 6.6410 |
| RBM15B_S147 | RBM15B | Q8NDT2 | | 109 | PGASGMSPRASPL | 6.1965 | 6.7051 | 5.8074 | 6.7099 | 5.8607 | 5.9354 | 6.2480 | 5.9897 | 6.2909 | 6.1461 | 5.8418 | 6.1139 | 6.5033 | 6.3024 | 6.7630 | 6.5859 |
| RBM15B_S265;S267 | RBM15B | Q8NDT2 | | 552 | LEGDWTSPSKSSD | 6.4324 | 6.4890 | 6.1530 | 6.7840 | 6.1888 | 5.9360 | 6.5101 | 5.9213 | 6.4335 | 6.8514 | 6.2451 | 6.3683 | 5.8091 | 5.8727 | 5.8962 | 6.1091 |
| RBM15B_S267 | RBM15B | Q8NDT2 | | 113 | GMSPRASPLPPPP | 5.7398 | 6.2361 | 5.6910 | 6.5373 | 5.8520 | 5.9156 | 6.4974 | 6.1351 | 6.0431 | 5.8668 | 5.8284 | 6.0782 | 6.7453 | 6.7572 | 7.1939 | 6.8828 |
| RBM15B_S504 | RBM15B | Q8NDT2 | | 562 | SSDRRNSLEGYSR | 5.1626 | 5.7114 | 5.2161 | 6.2198 | 6.6557 | 5.6772 | 5.9410 | 5.6428 | 5.7455 | 6.2299 | 5.6144 | 5.9354 | 7.6531 | 7.5281 | 7.5634 | 7.5036 |
| RBM15B_S552 | RBM15B | Q8NDT2 | | 238 | SSAAASTPPPGPP | 6.0570 | 6.3333 | 5.4399 | 6.6244 | 6.1432 | 5.7275 | 6.4299 | 6.5363 | 5.9656 | 5.9906 | 6.2667 | 5.7791 | 6.4637 | 6.7460 | 6.8003 | 6.6965 |
| RBM15B_S562 | RBM15B | Q8NDT2 | | 504 | PQQYQPSPLPVHY | 5.0510 | 6.6929 | 6.6502 | 7.3448 | 6.2673 | 7.0422 | 7.1309 | 6.2942 | 5.8006 | 6.4048 | 6.6620 | 6.6615 | 5.6142 | 6.0236 | 4.7478 | 5.6120 |
| RBM15B_S609 | RBM15B | Q8NDT2 | | 147 | LLISSLSPALPAE | 6.5592 | 7.1140 | 6.1818 | 6.5061 | 5.6812 | 6.8745 | 6.6423 | 6.1756 | 5.8738 | 6.6632 | 6.5219 | 6.7457 | 5.8524 | 5.8772 | 4.8896 | 5.8416 |
| RBM15B_T238 | RBM15B | Q8NDT2 | | 609 | RGRTTHSPYEERS | 6.2915 | 6.7786 | 6.4039 | 7.1591 | 6.2138 | 6.0851 | 5.8038 | 5.9711 | 6.3641 | 6.5346 | 5.9274 | 6.5298 | 5.7672 | 5.9595 | 6.0523 | 6.1580 |
| RBM15B_T532 | RBM15B | Q8NDT2 | 265;267 | | NA | 5.3445 | 5.9921 | 6.1488 | 5.9271 | 6.2381 | 6.9436 | 6.5273 | 6.0027 | 6.5302 | 6.2577 | 6.7468 | 6.3515 | 6.5145 | 6.1192 | 5.4478 | 6.9081 |
| RBM17_S169 | RBM17 | Q96I25 | | 222 | EQDRPRSPTGPSN | 6.1140 | 6.3387 | 5.8856 | 6.6827 | 6.3207 | 6.2183 | 5.9617 | 6.3898 | 6.4685 | 5.9469 | 6.1417 | 6.0235 | 6.5081 | 6.3715 | 6.4355 | 6.1929 |
| RBM17_S222 | RBM17 | Q96I25 | | 71 | DRQIVDTPPHVAA | 6.0844 | 6.1859 | 6.1524 | 6.7596 | 6.0818 | 5.8267 | 6.1430 | 6.1464 | 6.4929 | 6.3080 | 6.0719 | 6.0980 | 6.4999 | 6.5258 | 6.5600 | 6.0634 |
| RBM17_T71 | RBM17 | Q96I25 | | 169 | RERRKRSMGGAAI | 5.9512 | 5.6577 | 6.3767 | 6.3835 | 7.9532 | 6.3804 | 5.9171 | 6.5414 | 7.7841 | 7.8386 | 7.7013 | 7.5280 | 5.1385 | 5.2375 | 3.6816 | 3.9292 |
| RBM19_S677 | RBM19 | Q9Y4C8 | | 710 | EGADNSSAKMEEE | 5.8377 | 6.5897 | 5.8923 | 6.9225 | 5.6020 | 5.9380 | 6.1414 | 6.1010 | 6.0291 | 6.2396 | 5.5021 | 6.2048 | 6.0766 | 6.8020 | 6.9926 | 7.1287 |
| RBM19_S710 | RBM19 | Q9Y4C8 | | 677 | KLQDTPSEPMEKD | 5.4489 | 8.3351 | 6.7244 | 5.9563 | 5.8609 | 4.4899 | 5.5373 | 6.7593 | 5.1137 | 5.6700 | 6.1914 | 5.9540 | 6.2007 | 6.0750 | 7.4436 | 8.2396 |
| RBM22_T141 | RBM22 | Q9NW64 | | 141 | GMLGKATSTSDML | 6.0876 | 6.0434 | 5.9372 | 6.0405 | 5.1291 | 6.2149 | 6.0329 | 6.0356 | 6.1829 | 6.1477 | 6.5014 | 6.8797 | 6.1515 | 7.2663 | 6.0733 | 7.2761 |
| RBM23_S128 | RBM23 | Q86U06 | | 149 | PHFREKSPVREPV | 6.2142 | 6.3832 | 6.4006 | 6.2454 | 5.9020 | 6.5864 | 6.1358 | 6.0926 | 6.3108 | 6.6841 | 6.3243 | 6.7276 | 6.0151 | 6.0678 | 5.7627 | 6.1474 |
| RBM23_S149 | RBM23 | Q86U06 | | 128 | DRVHYRSPPLATG | 5.7127 | 6.4874 | 6.5927 | 6.6790 | 5.8458 | 6.4625 | 5.7902 | 7.0338 | 7.0329 | 7.0394 | 7.0739 | 7.0343 | 5.2418 | 5.6792 | 4.9604 | 5.3340 |
| RBM25_S583 | RBM25 | P49756 | | 683 | SPGQPNSVKRKKL | 6.3136 | 6.6157 | 6.2858 | 6.5449 | 6.6359 | 6.1036 | 6.5329 | 6.2427 | 6.4542 | 6.4685 | 5.9242 | 5.8008 | 6.1092 | 6.0849 | 5.9948 | 5.8885 |
| RBM25_S675;S683 | RBM25 | P49756 | | 583 | IKQEPESEEEEEE | 5.5587 | 5.8534 | 6.2605 | 6.6651 | 5.8526 | 5.7318 | 5.8527 | 5.9206 | 6.5137 | 6.3937 | 6.3226 | 6.5790 | 6.5038 | 6.5586 | 6.7100 | 6.7234 |
| RBM25_S683 | RBM25 | P49756 | 675;683 | | NA | 7.9162 | 8.3401 | 7.1400 | 7.6348 | 6.5340 | 6.5255 | 7.9950 | 6.7750 | 7.8089 | 6.9717 | 7.0517 | 7.3008 | 2.7179 | 3.6935 | 2.2814 | 3.3135 |
| RBM26_S127 | RBM26 | Q5T8P6 | | 127 | SRRLNHSPPQSSS | 5.9270 | 6.1379 | 5.5982 | 6.1153 | 6.1244 | 5.7872 | 6.3208 | 6.4223 | 6.2357 | 6.0954 | 5.7997 | 5.8686 | 6.6443 | 6.6970 | 7.1847 | 7.0416 |
| RBM26_S518 | RBM26 | Q5T8P6 | | 518 | NFNRTNSPGFQKK | 6.7668 | 6.5042 | 6.1368 | 5.7786 | 6.2377 | 6.4582 | 6.8626 | 6.0318 | 7.0404 | 6.4741 | 6.5042 | 6.4282 | 5.7641 | 5.8392 | 5.4561 | 5.7170 |
| RBM27_S1020 | RBM27 | Q9P2N5 | | 888 | TSSAVSTPSKVKT | 7.0414 | 6.5365 | 5.8524 | 6.2143 | 6.6418 | 6.7391 | 6.9285 | 6.4024 | 7.4873 | 6.4344 | 6.4551 | 6.2849 | 5.5232 | 5.6133 | 4.6390 | 5.2065 |
| RBM27_S120 | RBM27 | Q9P2N5 | | 1020 | HKPKVPSISTETE | 5.2315 | 5.1892 | 5.1199 | 5.7677 | 5.9038 | 5.8543 | 6.0742 | 5.8467 | 6.2252 | 4.9237 | 5.9241 | 5.9182 | 8.0434 | 7.9077 | 7.8603 | 8.2100 |
| RBM27_S566 | RBM27 | Q9P2N5 | | 447 | ERLQLGTPPPLLA | 6.1443 | 6.6521 | 6.7705 | 6.5345 | 5.8563 | 6.3915 | 6.1587 | 6.1046 | 6.7536 | 6.7554 | 6.7348 | 6.2482 | 5.8920 | 5.7866 | 5.2549 | 5.9620 |
| RBM27_S884 | RBM27 | Q9P2N5 | | 120 | RKKKYPSPQKTRS | 5.9612 | 6.3762 | 6.0399 | 6.5023 | 5.8952 | 5.9806 | 5.9384 | 5.8080 | 6.3735 | 6.1208 | 6.2278 | 6.6914 | 6.5856 | 6.5092 | 6.6000 | 6.3900 |
| RBM27_T447 | RBM27 | Q9P2N5 | | 566 | SRKRAMSGLEGPL | 6.5495 | 6.6229 | 5.8537 | 6.4895 | 5.0613 | 6.9266 | 7.4461 | 6.1251 | 5.5598 | 5.8830 | 5.3709 | 5.6910 | 6.6454 | 7.2722 | 5.7863 | 6.7166 |
| RBM27_T888 | RBM27 | Q9P2N5 | | 884 | DELKTSSAVSTPS | 6.3234 | 6.2938 | 5.5905 | 6.2700 | 6.0288 | 5.4310 | 5.4201 | 5.0922 | 6.1414 | 5.9482 | 5.6768 | 6.1590 | 7.7667 | 7.8152 | 6.6769 | 7.3659 |
| RBM3_S147 | RBM3 | P98179 | | 147 | GGYDRYSGGNYRD | 6.9322 | 7.1155 | 6.3880 | 7.4299 | 6.3765 | 6.2303 | 6.9338 | 7.8604 | 7.4591 | 6.8119 | 6.9938 | 7.1697 | 4.0356 | 4.1499 | 3.5717 | 4.5416 |
| RBM33_S205 | RBM33 | Q96EV2 | | 741 | KPIVSASPPSRAV | 6.1289 | 5.7083 | 6.3301 | 6.5627 | 6.3058 | 6.4465 | 6.3699 | 5.6988 | 6.3784 | 6.5648 | 6.3170 | 6.3837 | 6.3151 | 6.4204 | 5.7043 | 6.3654 |
| RBM33_S41 | RBM33 | Q96EV2 | | 765 | VKVKPASPVAQPK | 5.7147 | 6.1271 | 5.6270 | 6.1206 | 6.2215 | 6.1674 | 6.1001 | 6.0284 | 6.2031 | 6.1336 | 6.1505 | 6.0000 | 6.8104 | 7.0394 | 6.8012 | 6.7550 |
| RBM33_S741 | RBM33 | Q96EV2 | | 205 | KDIKEESDEEEED | 5.6129 | 6.0524 | 6.4210 | 5.9755 | 5.5767 | 5.7437 | 6.0090 | 5.5941 | 6.1941 | 6.3843 | 6.3908 | 6.4262 | 7.1297 | 7.2033 | 6.7245 | 6.5617 |
| RBM33_S765 | RBM33 | Q96EV2 | | 41 | ADEDWDSELEDDL | 5.8716 | 6.0327 | 5.9445 | 6.3044 | 5.7705 | 6.0355 | 5.7918 | 6.3852 | 6.6665 | 6.3319 | 6.3965 | 6.7192 | 6.4576 | 6.6491 | 5.9047 | 6.7383 |
| RBM33_S849 | RBM33 | Q96EV2 | | 851 | EQALSPSPTNGNP | 5.9753 | 5.2721 | 6.0501 | 6.1134 | 6.4531 | 6.5059 | 5.6919 | 6.4839 | 6.7558 | 6.3484 | 5.9154 | 5.8899 | 6.9362 | 7.6174 | 6.0826 | 5.9084 |
| RBM33_S849;S851 | RBM33 | Q96EV2 | | 849 | QEEQALSPSPTNG | 6.1733 | 6.6102 | 5.5316 | 5.6231 | 6.8027 | 5.7817 | 5.4099 | 6.6800 | 6.3290 | 6.3309 | 5.9619 | 7.1613 | 6.6630 | 6.3809 | 5.5495 | 7.0112 |
| RBM33_S851 | RBM33 | Q96EV2 | 849;851 | | NA | 6.7074 | 6.0402 | 6.2752 | 6.4575 | 5.4238 | 5.4466 | 5.7311 | 6.9263 | 5.6574 | 5.3658 | 5.7233 | 6.1257 | 6.8624 | 7.2780 | 6.6958 | 7.2836 |
| RBM34_S28 | RBM34 | P42696 | | 28 | DDGVRGSPPEDYR | 6.3459 | 7.0213 | 6.0938 | 7.3505 | 5.9403 | 5.7774 | 6.3620 | 6.0617 | 6.2017 | 6.0530 | 6.3501 | 6.0399 | 5.5166 | 6.6127 | 6.0111 | 6.2621 |
| RBM39_S100 | RBM39 | Q14498 | | 136 | PFRKDKSPVREPI | 5.8616 | 6.2079 | 6.0385 | 6.3416 | 6.1389 | 5.9556 | 6.1890 | 5.8107 | 6.2058 | 6.3278 | 5.8748 | 6.2100 | 6.5355 | 6.7664 | 6.8327 | 6.7032 |
| RBM39_S117 | RBM39 | Q14498 | | 97 | FRGRYRSPYSGPK | 5.9011 | 6.1775 | 5.6528 | 6.1690 | 6.3866 | 6.0363 | 6.1294 | 6.2448 | 6.2004 | 6.0718 | 5.6747 | 5.6457 | 6.8599 | 6.9432 | 7.1157 | 6.7910 |
| RBM39_S136 | RBM39 | Q14498 | | 117 | KIGLPHSIKLSRR | 6.5814 | 6.5412 | 7.2400 | 6.9866 | 4.8190 | 5.9579 | 6.2903 | 5.5473 | 7.9396 | 8.2905 | 9.2105 | 9.8281 | 3.2087 | 5.1574 | 2.4535 | 3.9480 |
| RBM39_S97 | RBM39 | Q14498 | | 100 | RYRSPYSGPKFNS | 6.2014 | 6.1140 | 6.6171 | 7.1514 | 5.4817 | 6.2947 | 6.7417 | 6.2853 | 6.2085 | 6.2414 | 6.4385 | 7.1060 | 5.2028 | 5.9895 | 5.3181 | 6.6078 |
| RBM4_S86 | RBM4 | Q9BWF3 | | 86 | LHVGNISPTCTNK | 6.2356 | 6.1509 | 6.4775 | 7.0685 | 5.8686 | 6.3310 | 6.0205 | 6.0950 | 7.0949 | 6.4675 | 6.5054 | 6.9627 | 5.7541 | 5.6848 | 5.3614 | 5.9216 |
| RBM41_S6 | RBM41 | Q96IZ5 | | 6 | xMKRVNSCVKSDE | 5.7738 | 6.3556 | 7.5559 | 6.2673 | 5.0802 | 6.7860 | 6.1273 | 6.3991 | 6.6755 | 8.3842 | 8.1914 | 8.4214 | 4.0465 | 5.3011 | 3.5575 | 5.0773 |
| RBM42_S135 | RBM42 | Q9BTD8 | | 135 | DRSHLDSPEAREA | 7.1794 | 8.0987 | 6.8332 | 7.7786 | 5.9110 | 7.0950 | 7.4985 | 7.1679 | 7.1265 | 7.6043 | 7.7382 | 7.7525 | 2.5702 | 3.7988 | 2.4868 | 3.3606 |
| RBM4B_S86 | RBM4B | Q9BQ04 | | 86 | LHVGNISPTCTNQ | 5.8261 | 6.6865 | 6.7327 | 5.6020 | 6.3124 | 6.1554 | 6.2297 | 6.7439 | 7.1278 | 5.9887 | 6.2437 | 6.0475 | 5.3287 | 5.5459 | 6.0531 | 7.3759 |
| RBM5_S576 | RBM5 | P52756 | | 59 | DYRDYDSPERERE | 6.4070 | 6.7808 | 6.3900 | 6.6024 | 5.8768 | 5.9608 | 6.4311 | 6.0879 | 6.4739 | 6.7632 | 6.5732 | 6.5040 | 5.6815 | 6.1189 | 5.7207 | 5.6279 |
| RBM5_S59 | RBM5 | P52756 | | 576 | EEERRESAAADAG | 5.2911 | 5.3311 | 5.8214 | 6.2324 | 7.1776 | 5.7554 | 5.5551 | 5.7586 | 8.9478 | 8.9706 | 8.0922 | 8.7852 | 5.1756 | 5.7248 | 2.8820 | 4.4991 |
| RBM5_S621;S624 | RBM5 | P52756 | | 624 | AAYSGDSDNEEEL | 6.8917 | 6.4292 | 8.7532 | 7.7880 | 7.5582 | 9.5921 | 7.6788 | 3.2117 | 4.0793 | 5.9437 | 7.3571 | 6.1994 | 4.7946 | 4.7630 | 4.5705 | 4.3895 |
| RBM5_S624 | RBM5 | P52756 | 621;624 | | NA | 6.9232 | 6.9948 | 6.9348 | 7.5845 | 5.9972 | 6.5551 | 6.6046 | 6.2810 | 7.0363 | 5.9307 | 6.2824 | 6.0540 | 5.1686 | 5.6284 | 4.8670 | 5.1574 |
| RBM6_S1025 | RBM6 | P78332 | | 1025 | KLQSFDSPERKRI | 6.6810 | 7.1494 | 6.3883 | 7.1036 | 6.0804 | 5.9616 | 6.3279 | 6.5936 | 7.0507 | 6.4125 | 5.9113 | 6.1136 | 5.2803 | 5.5607 | 5.6658 | 5.7191 |
| RBM6_S362 | RBM6 | P78332 | | 891 | TVKKEESPPPPKV | 5.5848 | 5.6974 | 5.6991 | 5.9000 | 5.7514 | 5.7410 | 5.4853 | 5.4979 | 5.4688 | 5.7627 | 5.6363 | 5.9167 | 8.2688 | 7.7370 | 8.2181 | 7.6345 |

| | | | | | | | | | | | | | | | | | | | | | |
|---|---|---|---|---|---|---|---|---|---|---|---|---|---|---|---|---|---|---|---|---|---|
| RBM6_S891 | RBM6 | P78332 | | 362 | DFQNSQSPVQDQD | 5.7276 | 6.4503 | 6.0643 | 8.0287 | 6.5031 | 7.2585 | 5.5936 | 6.8061 | 6.3353 | 6.0525 | 6.1017 | 5.6973 | 5.6950 | 5.9558 | 6.2149 | 5.5153 |
| RBM7_S137 | RBM7 | Q9Y580 | | 204 | RKVRMNSYPYLAD | 4.6181 | 4.6785 | 4.5925 | 4.8552 | 5.5913 | 5.3588 | 5.1301 | 4.9825 | 4.4640 | 4.3691 | 4.4006 | 4.8861 | 11.3169 | 9.2905 | 10.8796 | 10.5861 |
| RBM7_S196 | RBM7 | Q9Y580 | | 137 | IQRSFSSPENFQR | 5.8760 | 6.1368 | 5.9166 | 6.0498 | 5.8716 | 6.1431 | 6.0236 | 6.2219 | 6.1368 | 6.0674 | 6.1828 | 6.4654 | 6.9037 | 6.5614 | 6.6387 | 6.8043 |
| RBM7_S204 | RBM7 | Q9Y580 | | 196 | RQGTPSSQRKVRM | 7.9695 | 7.6663 | 6.7311 | 7.7583 | 5.1060 | 7.4634 | 7.3028 | 6.3884 | 7.0813 | 7.1049 | 7.5218 | 8.3303 | 3.2883 | 3.7980 | 2.8002 | 3.6896 |
| RBM8A_S42 | RBM8A | Q9Y5S9 | | 56 | MREDYDSVEQDGD | 6.7010 | 6.9208 | 5.9697 | 6.5223 | 6.1199 | 6.0479 | 5.9198 | 6.8876 | 5.5514 | 6.3104 | 6.2994 | 5.9610 | 5.7293 | 6.3231 | 6.1847 | 6.5516 |
| RBM8A_S42 | RBM8A | Q9Y5S9 | | 42 | KGRGFGSEEGSRA | 5.6897 | 5.7457 | 5.2430 | 5.6433 | 5.4942 | 5.4746 | 6.1099 | 5.9321 | 5.6519 | 5.5991 | 5.4979 | 5.6023 | 7.9312 | 7.7652 | 8.3163 | 8.3038 |
| RBM8A_S56 | RBM8A | Q9Y5S9-2 | | 42 | KGRGFGSEGSRAR | 5.9247 | 6.5843 | 5.9000 | 6.5349 | 6.6167 | 6.0406 | 6.1013 | 6.2666 | 5.9071 | 6.5313 | 6.1624 | 6.2725 | 5.8806 | 6.8306 | 6.2006 | 6.2460 |
| RBMS1_S273 | RBMS1 | P29558 | | 38 | HPMAPPSPSTTSS | 6.4625 | 6.4907 | 6.4937 | 6.2458 | 6.4502 | 6.4571 | 6.2507 | 6.5477 | 6.2513 | 6.3304 | 6.3691 | 6.2134 | 5.7104 | 6.1042 | 5.8680 | 5.7548 |
| RBMS1_S38 | RBMS1 | P29558 | | 273 | QNGFYPSPYSIAT | 6.7479 | 6.7727 | 6.8861 | 6.8248 | 6.3177 | 6.6174 | 6.6353 | 6.4155 | 7.1462 | 6.7790 | 6.5926 | 7.3955 | 5.0947 | 4.9409 | 4.0278 | 4.8058 |
| RBMS2_S106 | RBMS2 | Q15434 | | 106 | GFVDFDSPSAAQK | 6.2454 | 6.9110 | 6.8756 | 6.9663 | 6.4634 | 6.4855 | 6.1316 | 6.7954 | 6.4786 | 7.0682 | 7.1682 | 7.4158 | 4.4786 | 5.1713 | 4.3992 | 4.9461 |
| RBMS2_S37 | RBMS2 | Q15434 | | 37 | MAPPSPSNSTPNS | 5.4074 | 5.7129 | 6.4833 | 6.0790 | 5.7989 | 6.8629 | 6.7839 | 6.6936 | 6.4480 | 6.6336 | 7.1169 | 7.3025 | 5.8180 | 6.0035 | 5.2183 | 5.6374 |
| RBMS3_S111 | RBMS3 | Q6XE24 | | 111 | GFVDFDSPAAAQK | 6.4173 | 6.6082 | 6.9409 | 6.7668 | 6.2581 | 6.6048 | 6.2543 | 6.4886 | 6.4228 | 6.9841 | 7.0752 | 6.9416 | 4.8083 | 5.6805 | 4.6524 | 5.0959 |
| RBMS3_S39 | RBMS3 | Q6XE24 | | 39 | HPMAPPSPSTNSS | 6.3770 | 6.6432 | 6.1659 | 6.4597 | 7.2243 | 6.0697 | 6.1801 | 7.3297 | 5.9112 | 5.9593 | 6.2543 | 5.7674 | 6.3589 | 6.0847 | 6.0782 | 5.1362 |
| RBMX_S135 | RBMX | P38159 | | 352 | ERGLPPSMERGYP | 6.7890 | 7.0348 | 6.6417 | 7.5994 | 6.3586 | 6.1879 | 6.7372 | 6.9914 | 6.5616 | 6.6074 | 6.7690 | 7.1967 | 4.2519 | 4.7416 | 4.5264 | 5.0055 |
| RBMX_S141 | RBMX | P38159 | | 208 | RRDVYLSPRDDGY | 7.2318 | 7.7374 | 6.5093 | 7.5646 | 6.4875 | 6.6080 | 6.6919 | 6.3635 | 6.7308 | 6.6614 | 6.7088 | 6.5764 | 4.3097 | 4.7443 | 4.5282 | 4.5463 |
| RBMX_S165 | RBMX | P38159 | | 189 | VSRGRDSYGGPPR | 7.0424 | 7.2906 | 6.2155 | 7.6110 | 6.2236 | 6.4040 | 7.1815 | 6.8022 | 6.7583 | 6.4416 | 6.7446 | 6.7809 | 4.1959 | 5.0752 | 4.1125 | 5.1201 |
| RBMX_S168 | RBMX | P38159 | | 174 | SGPVRSSSGMGGR | 6.6705 | 6.7063 | 6.2021 | 6.6705 | 6.2941 | 6.0110 | 6.4978 | 6.1461 | 6.1001 | 5.9456 | 6.4752 | 6.3990 | 6.0301 | 5.5681 | 5.9319 | 6.3514 |
| RBMX_S174 | RBMX | P38159 | | 88 | EQATKPSFESGRR | 5.9563 | 5.9668 | 5.8979 | 6.5470 | 6.5756 | 6.1362 | 7.1358 | 6.6311 | 5.9629 | 5.8875 | 5.6363 | 5.7672 | 6.3893 | 6.5900 | 6.4656 | 6.4545 |
| RBMX_S189 | RBMX | P38159 | | 307 | PPSYGGSSRYDDY | 7.1077 | 5.9816 | 5.7067 | 7.5188 | 6.2713 | 5.7490 | 6.8973 | 5.4245 | 6.7180 | 5.7253 | 6.7150 | 6.7898 | 6.0194 | 5.4578 | 5.8386 | 6.0794 |
| RBMX_S208 | RBMX | P38159 | | 261 | YPSRGYSDRDGYG | 6.4010 | 6.7208 | 5.5959 | 6.8678 | 6.4699 | 6.5363 | 6.8993 | 6.4287 | 5.6096 | 5.7376 | 5.6231 | 5.6876 | 6.5957 | 6.2152 | 6.1257 | 6.4857 |
| RBMX_S215 | RBMX | P38159 | | 332 | SYSSSRSDLYSSG | 6.3503 | 6.5954 | 5.5932 | 6.7021 | 6.3583 | 5.8085 | 6.4155 | 6.6848 | 6.2620 | 5.6675 | 6.0585 | 6.5383 | 5.6815 | 6.0149 | 6.3665 | 6.9026 |
| RBMX_S261 | RBMX | P38159 | | 58 | AFVTFESPADAKD | 6.7362 | 6.7010 | 5.9973 | 7.2578 | 5.8487 | 6.3144 | 6.5936 | 6.1050 | 6.8365 | 7.1362 | 6.4417 | 5.8239 | 5.2309 | 5.6794 | 5.6987 | 5.5987 |
| RBMX_S307 | RBMX | P38159 | | 165 | GPPPKRSAPSGPV | 6.2217 | 6.5915 | 5.8430 | 7.3864 | 6.5542 | 6.2283 | 7.1140 | 6.5660 | 6.0778 | 5.5982 | 6.2545 | 6.5242 | 5.5000 | 5.7024 | 5.6321 | 6.2058 |
| RBMX_S308 | RBMX | P38159 | | 168 | PKRSAPSGPVRSS | 6.1823 | 6.6264 | 6.0967 | 6.7988 | 5.9599 | 6.0763 | 6.9928 | 5.9478 | 5.8076 | 6.2949 | 5.3469 | 6.2066 | 6.2286 | 6.2562 | 6.4684 | 6.7098 |
| RBMX_S332 | RBMX | P38159 | | 135 | MDDGGYSMNFNMS | 8.2903 | 6.9951 | 7.7053 | 8.5819 | 6.6633 | 5.6110 | 6.0524 | 5.3670 | 6.2891 | 6.2291 | 6.4927 | 5.8012 | 5.9139 | 5.7826 | 3.5332 | 4.6920 |
| RBMX_S352 | RBMX | P38159 | | 141 | SMNFNMSSSRGPL | 7.3067 | 8.8510 | 5.8834 | 7.3124 | 7.5660 | 8.0500 | 6.7938 | 6.9091 | 5.5664 | 5.6241 | 6.4193 | 7.0070 | 4.0567 | 4.3102 | 4.9729 | 3.3710 |
| RBMX_S58 | RBMX | P38159 | | 308 | PSYGGSSRYDDYS | 7.8151 | 6.4837 | 6.3068 | 7.9761 | 6.8488 | 7.0204 | 6.5003 | 5.9796 | 5.6703 | 5.4262 | 6.6998 | 6.4671 | 5.3990 | 5.9782 | 4.7632 | 4.6654 |
| RBMX_S88 | RBMX | P38159 | | 215 | PRDDGYSTKDSYS | 8.4173 | 8.1670 | 6.5508 | 9.3343 | 6.8090 | 5.2977 | 6.5420 | 5.2789 | 6.0284 | 5.3298 | 6.0200 | 6.5142 | 4.8465 | 5.0973 | 4.4559 | 5.3109 |
| RBMX2_S188 | RBMX2 | Q9Y388 | | 188 | EQPSSSSPRRKTV | 6.2116 | 6.4434 | 5.7657 | 6.6744 | 5.8101 | 5.5427 | 6.2500 | 5.8592 | 5.9287 | 6.1287 | 6.1277 | 6.1111 | 6.1631 | 7.0573 | 6.6851 | 7.2413 |
| RBMXL1_S161 | RBMXL1 | Q96E39 | | 161 | PRSGGPSPKRSAP | 6.5137 | 6.3684 | 5.7296 | 6.1270 | 6.3275 | 5.7484 | 6.5047 | 5.9198 | 6.2507 | 5.9681 | 5.7656 | 5.8099 | 6.9019 | 6.6188 | 6.8187 | 6.6272 |
| RBMXL1_S88 | RBMXL1 | Q96E39 | | 88 | EQATKPSFERGRH | 5.9006 | 6.2457 | 5.5688 | 6.5927 | 6.4103 | 6.2870 | 6.8398 | 6.6054 | 6.0288 | 6.0434 | 5.6424 | 6.1076 | 6.4350 | 6.3443 | 6.4165 | 6.5318 |
| RC3H1_S462 | RC3H1 | Q5TC82 | | 772 | EERKVISPPPFAP | 5.7339 | 6.0259 | 6.4270 | 6.3866 | 6.1661 | 6.0488 | 5.7352 | 6.0386 | 6.4917 | 6.4821 | 6.7542 | 6.3401 | 6.3536 | 6.4645 | 6.0740 | 6.4777 |
| RC3H1_S530;S535 | RC3H1 | Q5TC82 | | 462 | RRPLSASLGQLNE | 5.5946 | 6.3273 | 6.5498 | 5.7656 | 5.7701 | 6.3492 | 6.6559 | 6.1019 | 6.2327 | 5.6650 | 6.6422 | 5.8647 | 6.9877 | 6.9210 | 6.1655 | 6.4068 |
| RC3H1_S535 | RC3H1 | Q5TC82 | | 690 | EEIFRESPIPIEI | 6.1013 | 6.2104 | 6.0067 | 6.5377 | 4.9269 | 7.1737 | 6.1813 | 6.6645 | 7.0230 | 6.2644 | 6.6978 | 6.5440 | 6.2104 | 5.9464 | 5.8217 | 5.6897 |
| RC3H1_S690 | RC3H1 | Q5TC82 | | 535 | SSSAPGSPPDLLE | 6.3192 | 8.0502 | 6.5799 | 7.6443 | 6.4240 | 6.5074 | 4.6447 | 7.6927 | 6.3407 | 5.8354 | 7.1041 | 6.8245 | 4.0130 | 6.0074 | 4.9376 | 5.0747 |
| RC3H1_S772 | RC3H1 | Q5TC82 | 530;535 | | NA | 5.3268 | 6.1070 | 6.0519 | 5.8683 | 8.5872 | 5.6986 | 5.8239 | 7.8596 | 4.5628 | 3.4989 | 6.6796 | 6.4487 | 5.1585 | 8.0643 | 7.8526 | 6.4113 |
| RC3H2_S580 | RC3H2 | Q9HBD1 | | 808 | QSPTPPSPLFSVD | 4.8746 | 5.9767 | 6.6557 | 5.6641 | 5.7705 | 5.6278 | 6.4189 | 6.0163 | 6.3727 | 6.6870 | 6.0955 | 5.9949 | 6.8042 | 6.9436 | 6.4437 | 7.6539 |
| RC3H2_S808 | RC3H2 | Q9HBD1 | | 580 | SVPQKSSPFLTRV | 5.8887 | 5.9996 | 6.8055 | 6.7660 | 7.5347 | 5.9394 | 12.1760 | 7.4592 | 6.5188 | 6.1433 | 7.1003 | 7.1483 | 3.2495 | 3.8262 | 3.4383 | 4.0062 |
| RCAN1_S163;S167 | RCAN1 | P53805 | | 167 | LISPPASPPVGWK | 5.1723 | 6.3278 | 7.3334 | 5.6790 | 6.3382 | 6.7263 | 6.2431 | 7.6829 | 6.3011 | 7.4682 | 7.0354 | 6.2846 | 4.9880 | 5.3995 | 5.6819 | 5.3383 |
| RCAN1_S167 | RCAN1 | P53805 | 163;167 | | NA | 7.3085 | 6.5506 | 7.4305 | 6.9507 | 6.2155 | 6.6662 | 5.9877 | 6.2371 | 6.8284 | 6.2210 | 6.6036 | 7.3345 | 5.7174 | 4.6562 | 4.4261 | 4.8659 |
| RCC1_S11 | RCC1 | P18754-2 | | 11 | RIAKRRSPPADAI | 6.0114 | 6.1313 | 5.8992 | 6.3033 | 6.1745 | 5.8980 | 6.1922 | 5.8606 | 6.0752 | 6.1066 | 5.9786 | 6.1206 | 6.8489 | 6.8799 | 6.8347 | 6.6847 |
| RCC1L_T274 | RCC1L | Q96I51 | | 274 | NITSSPTKLGGDL | 5.8910 | 7.3822 | 7.0658 | 7.3201 | 5.3592 | 7.1668 | 6.1686 | 6.5836 | 6.0923 | 7.5678 | 7.4253 | 5.4221 | 4.7668 | 5.1405 | 6.0004 | 4.6475 |
| RCOR1_S127 | RCOR1 | Q9UKL0 | | 260 | KREREESEDELEE | 5.8712 | 6.3235 | 6.0887 | 6.5143 | 5.9188 | 6.3057 | 6.2810 | 5.7736 | 6.2130 | 6.1005 | 6.3605 | 6.6622 | 6.3550 | 6.6543 | 6.1891 | 6.3885 |
| RCOR1_S139 | RCOR1 | Q9UKL0 | | 460 | NQKPVKSPDNSIK | 6.3698 | 6.5775 | 5.7505 | 6.1314 | 6.2182 | 6.0137 | 6.3018 | 6.0639 | 6.1529 | 6.1461 | 5.9094 | 5.8439 | 6.6299 | 6.6094 | 6.7603 | 6.5213 |
| RCOR1_S235 | RCOR1 | Q9UKL0 | | 127 | AKLARRSQERDNL | 6.6685 | 6.6736 | 6.5741 | 6.8545 | 6.8404 | 7.5374 | 6.5419 | 5.5011 | 5.0432 | 6.8002 | 7.0428 | 6.1060 | 6.1518 | 5.8024 | 4.4542 | 5.4079 |
| RCOR1_S260 | RCOR1 | Q9UKL0 | | 139 | LGMLVWSPNQNLS | 5.5287 | 6.4435 | 6.4770 | 6.6633 | 5.9451 | 6.4692 | 5.9010 | 6.4419 | 6.4906 | 6.8039 | 6.8500 | 6.5376 | 5.8939 | 5.9914 | 5.6808 | 5.8818 |
| RCOR1_S460 | RCOR1 | Q9UKL0 | | 235 | LVKFYYSWKKTRT | 6.5898 | 6.9811 | 6.8497 | 6.4653 | 6.2107 | 7.4066 | 6.3613 | 6.1081 | 5.7236 | 6.4817 | 8.4671 | 5.9686 | 4.9067 | 5.8030 | 5.2201 | 4.4566 |
| RECQL_S634 | RECQL | P46063 | | 634 | ANMLQQSGSKNTG | 7.0015 | 7.3765 | 6.9508 | 6.8360 | 6.4958 | 7.4926 | 7.4737 | 6.9670 | 7.3771 | 7.4718 | 7.2641 | 7.0525 | 3.1650 | 4.1903 | 3.0737 | 3.8116 |
| RECQL_Y61 | RECQL | P46063 | | 61 | AGASNEYDSSPAA | 5.1792 | 5.3959 | 5.7793 | 5.8147 | 5.7488 | 5.6949 | 5.9027 | 5.9445 | 6.0502 | 5.9787 | 6.6427 | 6.4401 | 7.2013 | 7.7214 | 7.1222 | 7.3834 |
| RECQL4_S251 | RECQL4 | O94761 | | 251 | VSIRVGSPQPSSS | 6.4870 | 6.0324 | 6.0459 | 5.2881 | 5.9241 | 5.7117 | 6.4983 | 5.5109 | 6.8378 | 6.7409 | 6.1634 | 6.4843 | 6.7060 | 6.6426 | 6.0548 | 6.8720 |
| RECQL4_S323 | RECQL4 | O94761 | | 323 | PRYHGLSPSSQAR | 5.9685 | 6.6320 | 6.5701 | 7.4421 | 6.7594 | 6.0829 | 6.7019 | 6.6899 | 6.5648 | 7.3080 | 7.1967 | 6.9803 | 4.6943 | 5.0978 | 4.5002 | 4.8110 |
| RECQL5_S727 | RECQL5 | O94762 | | 815 | DGAGGHSPAPPQT | 5.8900 | 5.8952 | 5.8738 | 6.9971 | 6.1786 | 6.2190 | 6.1319 | 5.7515 | 6.7357 | 6.2456 | 6.3009 | 5.8897 | 6.3294 | 6.9748 | 6.0285 | 6.5581 |
| RECQL5_S815 | RECQL5 | O94762 | | 727 | AHYGGPSPEKKAK | 5.8112 | 6.2727 | 6.0947 | 6.8619 | 6.4650 | 5.9560 | 5.8410 | 6.6785 | 6.8124 | 6.3217 | 6.3051 | 6.1779 | 5.6361 | 6.3452 | 6.1303 | 6.2905 |
| RECQL5_T839 | RECQL5 | O94762 | | 839 | PPRDQGTPEVQPT | 5.7887 | 8.4026 | 7.1787 | 6.0801 | 5.9448 | 6.7245 | 6.2396 | 6.7960 | 6.2505 | 7.2216 | 6.5277 | 6.7982 | 4.6538 | 4.3701 | 6.0317 | 4.9914 |
| REEP2_S152 | REEP2 | Q9BRK0-2 | | 152 | EKLRSFSMQDLTL | 5.5833 | 6.0322 | 6.2021 | 5.6325 | 6.0807 | 5.7660 | 5.7896 | 5.6595 | 5.6063 | 5.3380 | 6.0443 | 5.5584 | 7.4132 | 8.0515 | 7.6559 | 7.5864 |
| REEP3_S152 | REEP3 | Q6NUK4 | | 152 | ERLRSFSMHDLTT | 5.6479 | 5.9523 | 6.0295 | 6.4615 | 6.6306 | 5.9879 | 6.0182 | 6.2820 | 5.7169 | 5.8934 | 5.8635 | 5.8100 | 6.9181 | 7.2193 | 6.6512 | 6.9176 |
| REEP4_S152 | REEP4 | Q9H6H4 | | 152 | GRLRSFSMQDLRS | 5.9514 | 6.1768 | 5.6583 | 6.2068 | 6.2874 | 5.6526 | 5.8570 | 5.8716 | 5.8777 | 5.7313 | 5.9659 | 5.8417 | 7.0980 | 7.3228 | 7.1644 | 7.3365 |
| RELA_S338 | RELA | Q2TAM5 | | 359 | WNDRFSSVQLRSS | 6.1647 | 5.9679 | 5.3153 | 6.3401 | 6.5613 | 5.5979 | 6.8138 | 6.8962 | 6.5047 | 5.2806 | 5.5242 | 5.8284 | 6.8170 | 6.6200 | 6.7304 | 7.0374 |
| RELA_S359 | RELA | Q2TAM5 | | 338 | AVPSRSSASVPKP | 6.3652 | 6.5037 | 6.9989 | 6.6410 | 6.3048 | 6.7391 | 6.9880 | 6.3393 | 7.8830 | 6.9503 | 7.1630 | 6.7405 | 4.7215 | 5.4418 | 3.7309 | 4.4892 |
| RELB_S116 | RELB | Q01201 | | 573 | FGGGLLSPGPEAT | 6.1681 | 6.3065 | 6.4594 | 6.3403 | 6.3328 | 6.8389 | 6.3754 | 7.0347 | 6.7680 | 6.3725 | 6.5645 | 6.7782 | 5.3163 | 5.6811 | 5.1520 | 5.5114 |
| RELB_S37 | RELB | Q01201 | | 116 | PLGRLVSPAPGPG | 6.9760 | 7.2614 | 7.1386 | 7.4696 | 6.0137 | 7.4831 | 7.3198 | 6.8591 | 7.5397 | 7.7665 | 7.8920 | 7.8981 | 2.8829 | 3.5765 | 2.6341 | 3.2888 |
| RELB_S573 | RELB | Q01201 | | 37 | ELGALGSPDLSSL | 6.1680 | 6.4026 | 6.6513 | 6.6159 | 6.3691 | 6.5198 | 6.6079 | 6.5638 | 6.6163 | 7.1280 | 7.0597 | 6.8807 | 4.9109 | 5.4081 | 4.8286 | 5.2694 |
| RELCH_S170 | RELCH | Q9P260-2 | | 453 | TIPKENSPNSFPR | 6.4379 | 6.8300 | 6.8437 | 6.2080 | 6.3666 | 6.7921 | 6.3358 | 6.7068 | 6.3486 | 6.4585 | 6.4339 | 6.6719 | 5.4807 | 5.6659 | 5.1631 | 5.2565 |
| RELCH_S180 | RELCH | Q9P260-2 | | 180 | QLNRAGSISTLDS | 6.2824 | 5.7720 | 6.6057 | 6.1550 | 5.8917 | 5.8303 | 6.0206 | 6.4008 | 6.6109 | 6.3604 | 6.2841 | 6.2420 | 6.4591 | 6.8291 | 5.8647 | 6.3913 |
| RELCH_S193 | RELCH | Q9P260-2 | | 170 | REPSTASGGGQLN | 5.7238 | 5.9523 | 6.4723 | 6.5079 | 6.5574 | 6.2640 | 5.6912 | 6.6060 | 6.4513 | 6.3582 | 6.0678 | 6.1050 | 6.0614 | 6.5771 | 6.1726 | 6.4317 |
| RELCH_S453 | RELCH | Q9P260-2 | | 143 | FERQSGTPPGMGA | 6.5514 | 6.4524 | 6.8212 | 6.2204 | 6.9473 | 6.5949 | 6.0268 | 6.4984 | 6.7891 | 6.1397 | 6.1970 | 6.1970 | 5.7752 | 5.9846 | 5.6990 | 5.1057 |
| RELCH_S54;S56 | RELCH | Q9P260-2 | | 193 | LDFARYSDDGNRE | 6.0606 | 6.1537 | 6.6783 | 5.6036 | 6.2744 | 6.1548 | 5.3046 | 6.3260 | 6.9185 | 6.3761 | 6.3688 | 6.5054 | 6.3412 | 6.3604 | 6.5401 | 6.0334 |
| RELCH_S56 | RELCH | Q9P260-2 | | 56 | GSAGSLSPQDPVA | 6.3985 | 6.9153 | 6.9287 | 6.7414 | 6.5152 | 6.6536 | 6.4916 | 6.4841 | 6.9511 | 6.9342 | 7.5861 | 6.6161 | 4.5725 | 5.2575 | 4.0577 | 4.8965 |
| RELCH_T143 | RELCH | Q9P260-2 | 54;56 | | NA | 6.9697 | 6.4726 | 7.1848 | 6.3013 | 5.9430 | 6.4829 | 6.4727 | 6.1317 | 6.2025 | 6.5300 | 7.6158 | 7.2580 | 5.1650 | 5.2906 | 4.5595 | 5.4201 |
| RELL1_S109 | RELL1 | Q8IUW5 | | 244 | NQKERRSLMSVSG | 5.9921 | 6.2453 | 7.0794 | 6.4729 | 5.6249 | 6.3497 | 5.8569 | 6.8940 | 6.6972 | 7.7557 | 7.0916 | 7.6270 | 4.9647 | 5.5267 | 4.7982 | 5.0236 |
| RELL1_S148 | RELL1 | Q8IUW5 | | 261 | NGEVPATPVKRER | 6.7604 | 6.6788 | 6.1382 | 7.2135 | 6.5758 | 6.3735 | 6.5860 | 6.2566 | 6.2673 | 5.8097 | 5.7108 | 6.0377 | 5.8875 | 6.0604 | 5.6269 | 6.0168 |
| RELL1_S157 | RELL1 | Q8IUW5 | | 109 | KIELNDSVNENSD | 6.4051 | 6.9524 | 6.0484 | 6.6647 | 6.3560 | 7.0811 | 6.2681 | 6.6390 | 6.9173 | 5.7729 | 6.2616 | 6.6889 | 5.6497 | 5.7165 | 5.1642 | 5.4140 |
| RELL1_S244 | RELL1 | Q8IUW5 | | 148 | SLYDPESPVTPST | 5.8552 | 5.5576 | 5.8456 | 5.8369 | 5.9008 | 5.5055 | 6.1122 | 6.1098 | 5.5935 | 5.8656 | 5.9339 | 5.7304 | 7.9194 | 6.8792 | 7.7270 | 7.6276 |
| RELL1_S244;S247 | RELL1 | Q8IUW5 | | 157 | TPSTPGSPPVSPG | 6.1852 | 5.7947 | 5.8015 | 6.0469 | 5.6914 | 6.7672 | 6.4926 | 5.9577 | 5.1802 | 5.8835 | 6.0843 | 5.8195 | 6.7468 | 7.2405 | 7.5019 | 6.8060 |
| RELL1_S244;T261 | RELL1 | Q8IUW5 | | 247 | ERRSLMSVSGAET | 6.7887 | 6.4999 | 6.9760 | 6.5574 | 6.4909 | 5.9905 | 6.2544 | 6.6123 | 6.3484 | 6.6513 | 6.8999 | 7.1364 | 4.9321 | 6.6066 | 4.3115 | 4.9436 |
| RELL1_S247 | RELL1 | Q8IUW5 | 148;161 | | NA | 5.3033 | 6.3805 | 6.3763 | 7.0720 | 6.2009 | 6.8215 | 6.6515 | 6.9448 | 6.2319 | 6.3034 | 7.2711 | 7.0092 | 5.2480 | 5.0773 | 5.5500 | 5.5583 |
| RELL1_T148;S161 | RELL1 | Q8IUW5 | 244;261 | | NA | 6.7671 | 7.0820 | 6.7383 | 6.3796 | 5.9183 | 6.2884 | 6.5550 | 6.9895 | 6.5769 | 6.5780 | 6.8795 | 7.7937 | 4.3523 | 5.7371 | 4.5808 | 4.7834 |
| RELL1_T261 | RELL1 | Q8IUW5 | 244;247 | | NA | 7.8991 | 7.2153 | 7.0909 | 6.9567 | 7.2186 | 5.2207 | 6.1342 | 6.1012 | 7.0007 | 6.1486 | 6.3458 | 6.3552 | 5.5414 | 5.4800 | 4.7671 | 4.5245 |
| REPIN1_S386 | REPIN1 | Q9BWE0-4 | | 386 | AAPGPGSPQLPAG | 5.4196 | 5.8698 | 6.2715 | 5.9225 | 5.3304 | 5.7408 | 5.3318 | 6.1460 | 6.6224 | 6.4967 | 6.1833 | 6.0613 | 6.4143 | 6.8537 | 8.1614 | 7.1744 |
| REPS1_S118 | REPS1 | Q96D71 | | 401 | EAPPSKSPSMPSL | 6.0345 | 6.3192 | 6.2516 | 5.7736 | 6.5340 | 6.1349 | 5.8811 | 6.4744 | 6.2148 | 6.4846 | 6.5812 | 6.8387 | 6.2204 | 6.5133 | 5.7827 | 5.9609 |
| REPS1_S143 | REPS1 | Q96D71 | | 392 | EVGYSGSPAEAPP | 5.8734 | 6.0574 | 6.1661 | 6.3117 | 6.0537 | 6.0090 | 6.0782 | 6.2468 | 6.3684 | 6.4797 | 6.4582 | 6.5079 | 6.3624 | 6.4601 | 6.3184 | 6.2486 |
| REPS1_S166;S174 | REPS1 | Q96D71 | | 562 | SHSRSSSLDMNRT | 6.3772 | 6.1434 | 6.3703 | 5.9613 | 5.8790 | 5.8485 | 5.8577 | 6.4943 | 6.7468 | 6.3951 | 6.5501 | 6.6898 | 6.1416 | 6.2545 | 6.2304 | 6.0598 |
| REPS1_S170;S174 | REPS1 | Q96D71 | | 272 | IEIRRQSSSYDDP | 5.8615 | 5.6712 | 6.1226 | 5.8596 | 6.3724 | 5.8373 | 5.7255 | 6.1486 | 7.3063 | 7.0569 | 7.1275 | 7.1320 | 6.0976 | 6.2608 | 5.5998 | 5.8202 |
| REPS1_S272 | REPS1 | Q96D71 | | 694 | NVSKGTTPLAPPP | 7.3594 | 7.7739 | 6.7895 | 7.0269 | 6.4036 | 7.0648 | 7.4265 | 6.9156 | 7.5060 | 7.4364 | 7.3118 | 7.2732 | 2.9462 | 4.1089 | 2.8031 | 3.8542 |
| REPS1_S272;S274 | REPS1 | Q96D71 | | 430 | FSERSSSSQTLTQ | 4.8507 | 5.0771 | 5.4881 | 4.9963 | 4.9198 | 5.0993 | 5.6939 | 5.4113 | 5.0610 | 5.2176 | 5.9124 | 5.3315 | 8.9511 | 9.1394 | 9.5315 | 9.3191 |
| REPS1_S392 | REPS1 | Q96D71-4 | | 401 | EAPPSKSPSMPSL | 6.7574 | 6.1195 | 6.3967 | 6.5009 | 6.2469 | 6.0377 | 5.9590 | 6.4036 | 6.1060 | 6.2876 | 6.3660 | 6.1323 | 6.1047 | 6.4253 | 5.9376 | 6.2189 |
| REPS1_S401 | REPS1 | Q96D71 | | 709 | VRRRLKSEDELRP | 5.7379 | 5.9429 | 7.5553 | 5.9383 | 5.9306 | 7.2179 | 6.1130 | 6.3197 | 6.7205 | 6.3905 | 7.4425 | 7.1315 | 5.3176 | 5.3814 | 5.5615 | 5.2987 |
| REPS1_S401 | REPS1 | Q96D71 | | 143 | GQVKKGSVSHDTV | 5.4254 | 5.0783 | 5.3441 | 5.7192 | 6.9311 | 6.1972 | 5.1569 | 6.7461 | 6.8025 | 7.3062 | 7.3516 | 6.9401 | 6.0980 | 7.6026 | 6.1548 | 5.1460 |
| REPS1_S403 | REPS1 | Q96D71 | | 482 | DHTNPTSPLLVKP | 6.4625 | 6.7311 | 5.6675 | 6.7516 | 6.8323 | 6.8300 | 5.9336 | 7.3847 | 6.8897 | 5.9856 | 6.3459 | 6.7944 | 5.2357 | 5.8217 | 5.2152 | 5.1184 |
| REPS1_S430 | REPS1 | Q96D71 | | 475 | ELKRTGSDHTNPT | 5.2479 | 5.1114 | 5.3454 | 5.3563 | 5.4805 | 4.6083 | 5.1199 | 5.1438 | 6.0481 | 6.0055 | 5.7073 | 5.5126 | 9.0443 | 8.8673 | 9.0532 | 8.3481 |
| REPS1_S475 | REPS1 | Q96D71 | | 118 | HAASYSSDSENQG | 6.0002 | 6.5118 | 6.8273 | 7.0177 | 7.3973 | 6.9125 | 6.1477 | 7.1670 | 5.6418 | 5.4514 | 5.8590 | 6.6499 | 5.6804 | 5.5298 | 5.2099 | 5.9965 |
| REPS1_S482 | REPS1 | Q96D71-4 | | 403 | PPSKSPSMPSLNQ | 5.4065 | 5.4868 | 6.2805 | 4.7683 | 6.0464 | 6.1750 | 5.3285 | 6.0661 | 5.3611 | 5.7916 | 5.8921 | 5.7596 | 7.1309 | 8.2540 | 7.9560 | 8.2967 |
| REPS1_S537;T539 | REPS1 | Q96D71 | | 520 | SSSDSFTSDPEQI | 5.8401 | 6.7654 | 5.8736 | 5.5923 | 6.0532 | 5.4966 | 6.1178 | 5.6929 | 6.5044 | 6.7989 | 6.2231 | 6.8647 | 6.8815 | 6.7845 | 6.4889 | 6.0220 |
| REPS1_S562 | REPS1 | Q96D71 | 170;174 | | NA | 5.5595 | 5.2435 | 6.0662 | 5.5236 | 5.5226 | 5.8750 | 5.2280 | 6.5117 | 6.1946 | 5.9437 | 4.7545 | 5.4937 | 7.7782 | 8.6827 | 7.8337 | 7.7889 |
| REPS1_S709 | REPS1 | Q96D71 | 272;274 | | NA | 3.4194 | 3.5117 | 4.1358 | 3.6518 | 5.0384 | 4.9748 | 4.2566 | 4.4537 | 4.8542 | 4.0278 | 4.1664 | 5.0117 | 13.1076 | 10.9046 | 12.1418 | 12.3436 |
| REPS1_T520 | REPS1 | Q96D71 | 166;174 | | NA | 6.1984 | 5.4955 | 6.1430 | 5.9374 | 6.3094 | 6.1153 | 5.0432 | 5.4024 | 5.7714 | 5.8304 | 6.2472 | 6.0736 | 6.8960 | 7.8389 | 7.5556 | 7.1420 |
| REPS1_T539;S558 | REPS1 | Q96D71 | 539;558 | | NA | 5.6061 | 5.8252 | 8.6567 | 7.1620 | 5.4046 | 6.1693 | 6.1893 | 6.7575 | 6.1696 | 6.2513 | 5.6293 | 6.4632 | 6.4636 | 7.0855 | 4.7345 | 5.4323 |
| REPS1_T694 | REPS1 | Q96D71 | 537;539 | | NA | 8.0890 | 6.6553 | 6.7861 | 6.2933 | 6.3171 | 6.5628 | 5.9515 | 6.0212 | 7.0704 | 7.3986 | 5.9267 | 6.3631 | 5.6207 | 5.4213 | 4.3258 | 5.1970 |
| RER1_S95 | RER1 | O15258 | | 95 | PSLMEDSDDGPSL | 6.1591 | 6.7758 | 6.3352 | 6.1653 | 5.7521 | 6.1360 | 5.8700 | 6.2980 | 5.9423 | 6.3998 | 5.7482 | 5.9443 | 6.3662 | 6.6689 | 6.6310 | 6.8076 |
| RERE_S1266 | RERE | Q9P2R6 | | 1266 | ARPHVMSPTNRNH | 6.5225 | 6.0929 | 6.6024 | 6.7318 | 6.2067 | 6.5526 | 6.2769 | 5.7811 | 6.5574 | 6.9416 | 7.1742 | 7.1784 | 5.6710 | 5.6056 | 4.6466 | 5.4583 |
| RERE_S304 | RERE | Q9P2R6 | | 642 | VKEEASSPLKSNK | 6.3683 | 6.6753 | 6.0573 | 6.1003 | 6.1197 | 5.8567 | 6.0265 | 6.0399 | 6.4584 | 6.3190 | 6.2420 | 6.1646 | 6.4774 | 6.3411 | 6.3399 | 6.4135 |
| RERE_S594 | RERE | Q9P2R6 | | 594 | RKKQPASPDGRTS | 5.6912 | 6.5307 | 6.1998 | 6.7827 | 5.8415 | 5.8131 | 5.8930 | 5.8705 | 6.2699 | 6.4236 | 6.2314 | 6.4701 | 6.3761 | 6.3416 | 6.5512 | 6.7137 |
| RERE_S594;T599 | RERE | Q9P2R6 | | 600 | SPDGRTSPINEDI | 5.9664 | 6.1934 | 6.0239 | 6.0852 | 6.0268 | 5.6558 | 6.1728 | 6.0659 | 6.5124 | 6.3071 | 6.1856 | 6.4433 | 6.6948 | 6.4390 | 6.5187 | 6.7090 |
| RERE_S600 | RERE | Q9P2R6 | | 120 | ESRRPNTPYFICS | 6.0750 | 6.2729 | 6.4006 | 4.8077 | 5.8707 | 7.5978 | 5.6377 | 5.6856 | 6.0942 | 5.2450 | 4.0607 | 3.8851 | 7.1189 | 9.1109 | 8.7821 | 7.3551 |
| RERE_S642 | RERE | Q9P2R6 | | 304 | DLQPFPSPDGDTV | 6.2153 | 5.2928 | 6.5319 | 6.5641 | 5.6892 | 6.1500 | 6.5705 | 6.5568 | 6.6028 | 6.7333 | 6.4022 | 6.2628 | 5.7899 | 6.4038 | 5.8713 | 6.3635 |
| RERE_T120 | RERE | Q9P2R6 | 594;599 | | NA | 6.6724 | 6.4771 | 5.9736 | 5.9082 | 6.1972 | 6.8346 | 6.0548 | 5.3863 | 5.8899 | 6.5435 | 6.9020 | 5.6248 | 5.8556 | 6.1637 | 7.6569 | 5.8594 |
| RESF1_S1708 | RESF1 | Q9HCM1 | | 1708 | LSKRSFSADGFEM | 5.9695 | 6.1691 | 6.1741 | 6.9671 | 6.1854 | 6.0414 | 6.1389 | 6.3496 | 6.3356 | 6.4723 | 6.2152 | 6.1711 | 5.8771 | 6.3114 | 6.1094 | 6.5129 |
| RESF1_S1744 | RESF1 | Q9HCM1 | | 764 | LQIAVVSPLVLSE | 6.5605 | 7.2841 | 5.8530 | 7.8389 | 5.7212 | 6.0283 | 7.4707 | 7.7922 | 6.4641 | 6.6136 | 6.8130 | 5.7646 | 4.7789 | 5.4752 | 5.1793 | 4.3624 |
| RESF1_S764 | RESF1 | Q9HCM1 | | 1744 | SRSLGSSPVKxxx | 8.0683 | 8.0060 | 7.8466 | 9.0447 | 5.8788 | 6.3925 | 6.6883 | 6.4636 | 7.6160 | 7.3888 | 7.8220 | 8.0375 | 2.3409 | 3.0490 | 2.3016 | 3.0555 |
| REST_S766 | REST | Q13127 | | 864 | EDLSPPSPPLPKE | 5.6685 | 6.2756 | 5.9443 | 6.6578 | 6.3521 | 5.8582 | 5.9358 | 5.9844 | 6.2021 | 6.0420 | 5.9626 | 6.0754 | 6.0940 | 6.9166 | 7.0097 | 7.0208 |
| REST_S864 | REST | Q13127 | | 971 | TVEEPVSPMLPPS | 5.8556 | 6.3214 | 5.8315 | 6.1796 | 5.8399 | 5.5217 | 6.5697 | 5.8568 | 6.1344 | 6.2782 | 6.4674 | 6.0378 | 6.7612 | 6.7449 | 7.0852 | 6.5146 |
| REST_S971 | REST | Q13127 | | 766 | PVKIELSPPIEVV | 5.8185 | 6.2727 | 5.8806 | 6.6283 | 6.2408 | 5.7657 | 5.8182 | 6.0384 | 6.0115 | 6.3313 | 6.0587 | 5.8124 | 6.4069 | 6.8340 | 6.8389 | 7.2432 |
| RETREG2_S138 | RETREG2 | Q8NC44 | | 330 | SVSRATTPQLTDV | 5.8367 | 5.7842 | 6.1620 | 6.0832 | 6.7285 | 6.2352 | 6.1951 | 5.9502 | 6.4411 | 6.2323 | 6.0672 | 6.1503 | 6.3509 | 6.4599 | 6.4010 | 6.9221 |
| RETREG2_S379 | RETREG2 | Q8NC44 | | 385 | SRQALDSEEEEED | 6.1264 | 6.5603 | 6.0563 | 6.6764 | 6.6930 | 5.9533 | 5.9441 | 6.6207 | 5.6299 | 6.5370 | 5.9240 | 7.0533 | 5.7995 | 5.9542 | 6.2886 | 6.1830 |
| RETREG2_S385 | RETREG2 | Q8NC44 | | 379 | GRPQALSRQALDS | 6.3527 | 6.1012 | 5.9218 | 6.6417 | 6.2942 | 6.0099 | 7.2175 | 6.5769 | 6.9717 | 7.2928 | 6.5120 | 6.9148 | 5.5102 | 5.4333 | 5.5374 | 4.7118 |
| RETREG2_T330 | RETREG2 | Q8NC44 | | 138 | SPEEPHSDSEGAG | 5.6341 | 6.3103 | 6.6296 | 5.8575 | 6.6467 | 6.6339 | 7.2827 | 6.2606 | 6.5089 | 6.2820 | 5.5981 | 7.0011 | 5.5031 | 5.3122 | 6.7931 | 5.7462 |
| RETREG2_T330;S344 | RETREG2 | Q8NC44 | 330;344 | | NA | 5.9329 | 5.8624 | 6.0940 | 5.9317 | 6.1253 | 5.8947 | 7.1715 | 6.1665 | 6.0621 | 6.2985 | 5.7925 | 5.8081 | 6.4292 | 7.1265 | 6.5545 | 6.7497 |
| RETREG3_S258;S260 | RETREG3 | Q86VR2 | | 436 | FLRSPSSDLDTDA | 4.7080 | 5.9817 | 5.3157 | 5.9069 | 5.5318 | 6.1937 | 5.6290 | 6.8785 | 7.3831 | 6.6414 | 5.9158 | 6.7190 | 6.7983 | 7.2944 | 6.5885 | 6.5141 |
| RETREG3_S320 | RETREG3 | Q86VR2 | | 320 | EDLDGHSDPEESF | 5.5585 | 5.7704 | 5.8589 | 6.2519 | 6.4797 | 6.7918 | 6.3262 | 5.9282 | 6.0675 | 6.2987 | 5.5526 | 5.5785 | 7.6105 | 7.0263 | 6.8519 | 6.0483 |
| RETREG3_S436 | RETREG3 | Q86VR2 | 258;260 | | NA | 6.1315 | 5.9355 | 6.4196 | 6.0799 | 6.0261 | 6.4206 | 5.7542 | 5.7304 | 6.3903 | 6.1420 | 6.0585 | 5.8046 | 7.1611 | 6.5359 | 6.8650 | 6.5448 |
| RETREG3_S436;T440 | RETREG3 | Q86VR2 | 436;440 | | NA | 5.9818 | 6.7095 | 5.2673 | 6.2268 | 6.8204 | 6.3686 | 6.6463 | 7.4410 | 6.6247 | 5.4833 | 6.2011 | 5.5773 | 6.1818 | 6.4875 | 6.3479 | 5.6346 |
| REV1_S1097 | REV1 | Q9UBZ9 | | 1108 | LPGACGSPQKLID | 6.7252 | 7.0139 | 6.7826 | 7.1375 | 6.2452 | 5.9532 | 6.3722 | 6.4333 | 6.5965 | 6.6239 | 6.5992 | 6.7372 | 5.0935 | 5.4890 | 5.0164 | 5.1811 |
| REV1_S1108 | REV1 | Q9UBZ9 | | 1097 | NNKLLNSPAKTLP | 6.9296 | 7.0255 | 6.3114 | 6.7108 | 6.3761 | 6.1702 | 6.7724 | 6.4647 | 6.8220 | 6.4069 | 6.2774 | 6.3058 | 5.1532 | 5.4136 | 5.4419 | 5.4186 |
| REV1_S901 | REV1 | Q9UBZ9 | | 901 | PAHLPTSPDTNKA | 6.1301 | 6.0191 | 7.0532 | 6.2159 | 6.3934 | 6.6256 | 6.7292 | 6.1478 | 7.5842 | 5.9674 | 6.5428 | 6.3786 | 5.1301 | 4.6316 | 6.3786 | 6.0724 |
| REV3L_S1030 | REV3L | O60673 | | 1041 | YPIYPLTPKKSHR | 6.7959 | 6.6323 | 6.8129 | 6.9799 | 6.4749 | 5.9645 | 6.4949 | 5.9700 | 6.5547 | 6.0866 | 6.4540 | 6.3332 | 5.3915 | 5.9859 | 5.4336 | 5.6353 |
| REV3L_S1086 | REV3L | O60673 | | 1030 | WPKVPDSPATKYP | 6.8987 | 6.9946 | 5.9763 | 7.0937 | 6.2884 | 5.6801 | 6.3858 | 6.1565 | 6.4201 | 5.9069 | 6.5137 | 6.1615 | 5.4620 | 5.9192 | 5.9659 | 6.1767 |
| REV3L_S1724 | REV3L | O60673 | | 2147 | GDDRPSSPVEELP | 6.3382 | 6.8651 | 6.1071 | 6.4079 | 6.3635 | 6.1735 | 5.7212 | 6.2789 | 6.3801 | 6.1921 | 6.3527 | 6.0760 | 6.1469 | 6.2511 | 6.2274 | 6.1183 |
| REV3L_S203 | REV3L | O60673 | | 2281 | TYGFKVSIQNLQE | 6.4072 | 6.6555 | 8.9425 | 7.9596 | 5.3763 | 6.2851 | 7.5444 | 6.5897 | 7.2992 | 7.1323 | 7.1990 | 8.2873 | 2.6438 | 4.7190 | 2.9631 | 3.9960 |
| REV3L_S2147 | REV3L | O60673 | | 203 | SCKNHLSGNSLAD | 6.7259 | 6.1720 | 6.0065 | 5.7869 | 6.0065 | 5.9365 | 7.1492 | 5.9747 | 6.1561 | 6.4298 | 7.2193 | 7.5344 | 4.1698 | 7.6872 | 5.5227 | 5.5227 |
| REV3L_S2171 | REV3L | O60673 | | 2171 | KDGIQKSPCSEPQ | 6.2781 | 6.4299 | 5.8155 | 6.8593 | 5.9709 | 6.0245 | 6.1281 | 6.3540 | 6.3985 | 5.7107 | 6.2889 | 6.7442 | 6.1889 | 6.7334 | 5.9023 | 6.1727 |
| REV3L_S2281 | REV3L | O60673 | | 1724 | IRSGTLSPEIFEK | 6.8048 | 5.5864 | 5.7710 | 7.6098 | 6.1030 | 5.4033 | 6.1107 | 6.2616 | 6.0875 | 6.7002 | 6.6006 | 6.4342 | 5.8889 | 6.3984 | 5.7829 | 6.4567 |
| REV3L_T1041 | REV3L | O60673 | | 1086 | RSHAILSPPSPSY | 6.4038 | 6.1854 | 7.0478 | 6.6472 | 5.7791 | 6.6390 | 6.0504 | 6.2774 | 6.3246 | 6.7527 | 6.4498 | 5.9925 | 5.6658 | 6.3266 | 5.7992 | 5.6588 |
| REXO1_S358 | REXO1 | Q8N1G1 | | 459 | RPARRPSPTSGDS | 6.3854 | 6.7457 | 6.0888 | 6.7883 | 5.9987 | 5.8601 | 5.9841 | 6.0140 | 6.2406 | 6.1966 | 6.1302 | 6.3626 | 5.8768 | 6.2381 | 6.4860 | 6.6040 |
| REXO1_S422 | REXO1 | Q8N1G1 | | 358 | PPAKPASPAQVQS | 5.5886 | 5.8537 | 6.4904 | 6.8159 | 6.0776 | 6.2817 | 6.4925 | 5.7154 | 6.2302 | 6.0063 | 6.9856 | 6.5995 | 6.2241 | 6.5152 | 5.9017 | 6.2216 |
| REXO1_S459 | REXO1 | Q8N1G1 | | 422 | KGPQASSPRRKAE | 6.2931 | 6.4575 | 5.8302 | 6.8389 | 5.8791 | 5.6832 | 6.0684 | 5.9752 | 6.2374 | 6.0212 | 6.5294 | 6.5696 | 6.2612 | 6.7948 | 6.2829 | 6.2780 |
| REXO4_S111 | REXO4 | Q9GZR2 | | 15 | SKRAPSSPVAKPG | 6.7129 | 6.9204 | 6.1121 | 6.9458 | 6.3364 | 6.0448 | 6.4764 | 6.4729 | 6.7666 | 6.2039 | 6.2118 | 6.1491 | 5.1512 | 5.8819 | 5.7480 | 5.8659 |
| REXO4_S15 | REXO4 | Q9GZR2 | | 111 | QNKKETSPQVKGE | 6.2879 | 7.0119 | 6.6990 | 7.2339 | 5.6343 | 5.7949 | 6.0179 | 6.2283 | 6.6138 | 6.2577 | 6.3405 | 6.3014 | 5.5789 | 6.4346 | 5.7465 | 5.8186 |
| RFC1_S108 | RFC1 | P35251 | | 312 | KIGEVSSPKASSK | 7.2290 | 7.3821 | 7.2045 | 6.9035 | 6.9288 | 6.7244 | 7.5716 | 6.6437 | 7.7096 | 7.6376 | 7.2844 | 7.2418 | 2.7247 | 4.0444 | 2.9287 | 3.8410 |
| RFC1_S108;T110 | RFC1 | P35251 | | 506 | ESKLERTPQKNVQ | 6.3525 | 6.7086 | 6.1818 | 6.2694 | 6.1740 | 6.0391 | 6.1139 | 5.8444 | 6.2040 | 6.5451 | 6.3243 | 5.8992 | 5.7467 | 6.9341 | 6.3523 | 6.3104 |
| RFC1_S110 | RFC1 | P35251 | | 156 | TKNKPLSPIKLTP | 6.0228 | 6.1401 | 5.9310 | 6.0969 | 6.2847 | 6.0886 | 6.3883 | 5.8973 | 6.2703 | 6.4938 | 6.2286 | 6.3317 | 5.9468 | 6.8612 | 6.3124 | 6.7055 |
| RFC1_S156 | RFC1 | P35251 | | 368 | AKKESVSPEDSEK | 5.6905 | 6.2520 | 5.8484 | 6.1684 | 6.1219 | 5.9912 | 6.2438 | 6.1701 | 6.3099 | 6.4592 | 6.2694 | 6.1096 | 5.9871 | 6.7434 | 6.6694 | 6.9656 |
| RFC1_S312 | RFC1 | P35251 | | 161 | LSPIKLTPTSVLD | 5.7820 | 6.6046 | 6.9562 | 6.4154 | 5.8354 | 6.1685 | 6.5637 | 6.2231 | 6.4130 | 6.8306 | 6.2553 | 6.4974 | 5.2914 | 6.0661 | 5.9276 | 6.1697 |
| RFC1_S366;S372 | RFC1 | P35251 | | 110 | VTYISETDEEDDF | 5.3546 | 5.5469 | 5.4988 | 5.2833 | 5.6206 | 5.7974 | 5.3715 | 5.4690 | 6.2976 | 6.8610 | 6.3689 | 5.9651 | 6.9676 | 8.5623 | 7.6323 | 7.4030 |
| RFC1_S368 | RFC1 | P35251 | | 108 | DPVTYISETDEED | 5.2009 | 4.9241 | 5.1612 | 6.0905 | 5.7062 | 4.9732 | 5.8053 | 5.6585 | 7.2810 | 6.7138 | 7.0096 | 6.2635 | 7.3866 | 7.4999 | 7.3193 | 7.0063 |
| RFC1_S69;S71 | RFC1 | P35251 | 366;372 | | NA | 6.0125 | 6.6413 | 6.4635 | 6.8201 | 6.1368 | 6.1213 | 6.0498 | 6.3276 | 6.2773 | 7.1133 | 6.7202 | 7.0768 | 4.7874 | 6.1133 | 5.3005 | 6.0382 |
| RFC1_T161 | RFC1 | P35251 | 108;110 | | NA | 5.7347 | 7.4861 | 5.8187 | 6.1665 | 6.3373 | 5.7896 | 6.0411 | 7.2265 | 7.0767 | 6.0290 | 5.5929 | 5.6003 | 5.4774 | 6.8757 | 6.0580 | 6.6894 |
| RFC1_T506 | RFC1 | P35251 | 69;71 | | NA | 5.7714 | 5.7488 | 6.8288 | 6.1404 | 4.9748 | 5.8327 | 5.4321 | 5.8487 | 6.6510 | 6.9405 | 6.9110 | 6.0028 | 5.9311 | 6.5680 | 6.8256 | 7.5923 |
| RFC2_S30 | RFC2 | P35250 | | 30 | FSKAPGSAGHYEL | 5.7356 | 6.6399 | 5.8514 | 6.8918 | 5.9570 | 5.7422 | 5.7541 | 6.7235 | 6.1325 | 6.1677 | 6.4077 | 5.7147 | 6.6470 | 6.9097 | 6.3253 | 6.3999 |
| RFC3_S12 | RFC3 | P40938 | | 12 | DKYRPCSLGRLDY | 6.3821 | 6.4972 | 6.6852 | 6.3224 | 5.6092 | 6.0228 | 6.5976 | 5.9525 | 6.5430 | 6.3213 | 6.3866 | 6.6057 | 6.1796 | 6.3134 | 5.7297 | 5.8516 |

| | | | | | | | | | | | | | | | | | | | | | |
|---|---|---|---|---|---|---|---|---|---|---|---|---|---|---|---|---|---|---|---|---|---|
| RFC3_T75 | RFC3 | P40938 | | 75 | IEHQTITTPSKKK | 6.5421 | 6.7962 | 5.6793 | 7.1374 | 6.5863 | 6.4685 | 5.6743 | 5.6707 | 7.2516 | 7.3375 | 5.9984 | 7.1927 | 4.9158 | 5.4006 | 5.7738 | 5.5749 |
| RFFL_S240 | RFFL | Q8WZ73 | | 240 | VPGRRASLSDLTD | 6.0525 | 6.3514 | 6.3906 | 6.5933 | 6.2655 | 5.9623 | 5.9219 | 6.3080 | 6.4775 | 6.4557 | 6.4643 | 6.3008 | 5.8300 | 6.3187 | 6.2922 | 6.0154 |
| RFFL_S39 | RFFL | Q8WZ73 | | 39 | GYSSFPSPTGLEP | 6.4765 | 7.4511 | 6.9661 | 7.1908 | 7.0129 | 6.7104 | 6.8868 | 6.2284 | 6.5701 | 7.2895 | 6.8771 | 6.5467 | 4.2355 | 5.0103 | 3.9594 | 4.5883 |
| RFLNA_S25 | RFLNA | Q6ZTI6 | | 33 | DSGLPPSPSPSPP | 6.0053 | 6.1656 | 5.8485 | 5.8134 | 6.6309 | 6.0968 | 5.7118 | 5.5617 | 6.9833 | 6.3488 | 6.6327 | 7.0022 | 6.3971 | 6.4703 | 6.1823 | 6.1494 |
| RFLNA_S33 | RFLNA | Q6ZTI6 | | 25 | REGLLDSPDSGLP | 6.6917 | 6.3969 | 7.0296 | 6.5117 | 6.0890 | 5.5222 | 5.6453 | 6.2351 | 6.9884 | 6.5721 | 7.0219 | 6.6098 | 5.8729 | 5.5481 | 5.3988 | 5.8665 |
| RFTN1_S220 | RFTN1 | Q14699 | | 486 | NLEDQSSKAGDMG | 6.8074 | 6.5640 | 6.5207 | 6.6948 | 6.3068 | 5.6488 | 6.2371 | 5.7263 | 6.3539 | 5.6748 | 5.8239 | 6.0079 | 6.2171 | 6.4666 | 6.6248 | 6.3249 |
| RFTN1_S239 | RFTN1 | Q14699 | | 555 | GICTGHSNPGEDA | 4.7399 | 4.9857 | 4.8065 | 6.2123 | 5.1525 | 5.7082 | 5.8724 | 5.9540 | 4.7342 | 5.6085 | 5.7356 | 5.6251 | 8.5295 | 8.8384 | 8.3317 | 9.1653 |
| RFTN1_S486 | RFTN1 | Q14699 | | 220 | PAGRNQSPEPSSG | 7.2384 | 6.8131 | 6.8187 | 7.4922 | 6.3724 | 6.2083 | 6.8192 | 6.3297 | 7.2463 | 6.9478 | 7.2111 | 7.2391 | 4.0894 | 4.4985 | 4.0704 | 4.6051 |
| RFTN1_S555 | RFTN1 | Q14699 | | 239 | LAKQPSSPSGEGD | 6.1422 | 6.9860 | 6.8090 | 7.1375 | 6.8924 | 6.4974 | 7.3390 | 6.2812 | 6.5242 | 6.7582 | 7.3970 | 6.8237 | 4.1968 | 5.0928 | 4.2994 | 4.8231 |
| RFX1_S159 | RFX1 | P22670 | | 159 | AKPGHVSPLQLTN | 5.8001 | 6.2785 | 5.9588 | 6.2367 | 6.4522 | 5.9888 | 5.9633 | 5.8106 | 6.2665 | 6.1214 | 5.8841 | 6.1441 | 6.8693 | 7.0108 | 6.5945 | 6.6201 |
| RFX1_S204 | RFX1 | P22670 | | 204 | GTQQVHSPPEQSP | 6.6525 | 7.0349 | 5.0674 | 7.2096 | 6.7146 | 6.9011 | 5.6603 | 7.7411 | 5.8227 | 5.0162 | 6.8104 | 5.7671 | 5.7532 | 7.4224 | 5.3540 | 5.0725 |
| RFX1_S979 | RFX1 | P22670 | | 979 | VQALPSSxxxxxx | 5.9906 | 6.2380 | 6.1694 | 6.7901 | 6.5276 | 6.6692 | 7.2528 | 6.0406 | 5.6817 | 6.7472 | 5.7853 | 5.6001 | 6.3389 | 6.2309 | 5.9255 | 6.0120 |
| RFX7_S1028 | RFX7 | Q2KHR2 | | 225 | QEQKLQSPLPGES | 5.8219 | 6.3103 | 6.2698 | 6.6978 | 5.9120 | 6.0337 | 5.7165 | 6.1321 | 6.1047 | 6.3077 | 6.4345 | 6.2235 | 6.8914 | 6.5187 | 6.3211 | 6.3044 |
| RFX7_S1081 | RFX7 | Q2KHR2 | | 202 | QVKTLPSPIDAKQ | 7.1195 | 7.1433 | 6.9782 | 7.5355 | 6.0567 | 6.1041 | 6.4885 | 6.6719 | 6.6989 | 6.2943 | 6.9657 | 6.8790 | 4.7037 | 4.8393 | 4.6676 | 4.8539 |
| RFX7_S1128 | RFX7 | Q2KHR2 | | 349 | SALTIRSPTTVLF | 4.6349 | 5.1647 | 5.5811 | 5.4437 | 5.3930 | 6.0033 | 5.8680 | 5.6783 | 5.7243 | 5.0926 | 5.8622 | 5.0916 | 8.9893 | 8.0896 | 8.8827 | 8.5007 |
| RFX7_S1296 | RFX7 | Q2KHR2 | | 1128 | IAQRSQSVPLTVM | 6.3495 | 6.5158 | 6.4317 | 6.8491 | 6.1090 | 6.0646 | 6.3445 | 5.8520 | 6.5651 | 6.2325 | 6.7890 | 6.3662 | 6.2981 | 5.9927 | 5.4490 | 5.7912 |
| RFX7_S202 | RFX7 | Q2KHR2 | | 1296 | SSELSGSINDLNT | 5.6225 | 6.4648 | 6.3241 | 7.9046 | 5.5049 | 8.5021 | 6.1294 | 7.0874 | 6.4147 | 6.1699 | 6.4128 | 6.4995 | 4.9864 | 5.8557 | 4.6896 | 5.4317 |
| RFX7_S225 | RFX7 | Q2KHR2 | | 467 | EAKAPQTPSALLG | 7.0250 | 6.9654 | 7.1235 | 7.7215 | 6.3603 | 6.0612 | 6.3333 | 6.9072 | 6.5585 | 6.2481 | 6.7951 | 6.1174 | 4.7939 | 5.3403 | 4.7543 | 4.8951 |
| RFX7_S349 | RFX7 | Q2KHR2 | | 1081 | HRQRNLSGSTLYP | 6.4850 | 6.1395 | 6.3025 | 6.6547 | 6.6144 | 6.7448 | 5.9704 | 6.2112 | 6.1681 | 6.4940 | 6.7919 | 6.8311 | 5.8181 | 5.9435 | 5.5706 | 5.2604 |
| RFX7_S349;S358 | RFX7 | Q2KHR2 | | 424 | SVVSLQSPGSRSS | 7.4430 | 7.5090 | 7.9984 | 7.9548 | 5.9179 | 6.4007 | 6.8180 | 6.3262 | 7.2470 | 6.8518 | 7.6370 | 7.5481 | 3.8839 | 4.0350 | 3.0303 | 3.3990 |
| RFX7_S358 | RFX7 | Q2KHR2 | | 358 | TVLFTSSPIKTAV | 6.8123 | 7.1316 | 6.9724 | 7.6920 | 6.6107 | 6.4097 | 6.2606 | 6.2719 | 7.4426 | 6.8894 | 6.6765 | 7.0501 | 4.2662 | 5.1920 | 4.0747 | 4.2473 |
| RFX7_S424 | RFX7 | Q2KHR2 | | 1028 | GRLTPVSPVQHQG | 8.0760 | 7.9422 | 7.3691 | 8.0364 | 6.5340 | 7.7009 | 6.1965 | 6.2474 | 7.9497 | 7.4068 | 8.1495 | 7.1881 | 2.8032 | 3.9419 | 2.3791 | 2.0793 |
| RFX7_T467 | RFX7 | Q2KHR2 | 349;358 | | NA | 5.5984 | 5.7303 | 5.3845 | 6.9972 | 5.6056 | 5.8251 | 5.9478 | 5.7505 | 6.4399 | 5.8915 | 6.3817 | 5.4923 | 5.6605 | 8.7629 | 5.9037 | 8.6280 |
| RGL1_S555 | RGL1 | Q9NZL6-2 | | 555 | SMVKRLSLLFLGS | 6.8698 | 6.0804 | 6.5325 | 5.6989 | 5.8349 | 7.2689 | 5.8084 | 6.5430 | 6.3417 | 6.6260 | 7.1011 | 5.4269 | 6.1951 | 6.4848 | 5.4040 | 5.7837 |
| RGL2_S13 | RGL2 | O15211 | | 13 | RLLLDTSPPGGVV | 6.4706 | 6.4983 | 7.1254 | 6.4875 | 5.9091 | 6.4674 | 6.3528 | 7.0398 | 7.5020 | 6.6852 | 6.6296 | 7.3532 | 4.2881 | 5.0368 | 4.7929 | 5.3612 |
| RGL2_S409 | RGL2 | O15211 | | 409 | QEVKLQSPLEPHS | 6.0922 | 5.8426 | 6.8876 | 6.5776 | 7.1993 | 6.9652 | 7.1784 | 6.2991 | 7.0731 | 7.3784 | 7.6820 | 7.7050 | 4.1524 | 4.4635 | 3.9156 | 4.5880 |
| RGL2_S619 | RGL2 | O15211 | | 752 | GPSASGTPPSEGG | 5.3298 | 5.7901 | 6.8661 | 6.2273 | 6.4811 | 6.7242 | 7.2801 | 6.3903 | 6.9366 | 5.5614 | 6.5756 | 6.5515 | 6.1269 | 5.7390 | 5.2458 | 6.1742 |
| RGL2_T752 | RGL2 | O15211 | | 619 | GHRRSASCGSPLS | 6.4880 | 6.0497 | 5.4298 | 4.8294 | 6.2235 | 5.9369 | 6.5010 | 6.7332 | 6.4816 | 5.6334 | 5.7022 | 7.5566 | 7.1455 | 6.3921 | 5.4052 | 7.4918 |
| RGPD8_S1302 | RGPD8 | O14715 | | 21 | VLGLTPSPRQKSM | 5.8558 | 6.6521 | 5.9195 | 8.1066 | 7.8160 | 5.9290 | 6.1394 | 6.0515 | 6.4049 | 6.1520 | 6.0459 | 6.1425 | 5.3578 | 5.5008 | 6.0752 | 5.8511 |
| RGPD8_S1621 | RGPD8 | O14715 | | 781 | IKHSTPSPTKYSL | 6.3784 | 6.9545 | 5.6482 | 6.9857 | 8.8625 | 6.4939 | 6.2763 | 6.8929 | 5.9274 | 5.5862 | 5.7776 | 5.6801 | 5.7314 | 6.5266 | 6.5949 | 5.6835 |
| RGPD8_S1621;S1629 | RGPD8 | O14715 | | 1621 | SKVKNLSASFPTE | 6.2231 | 6.2551 | 7.0509 | 7.0552 | 6.2901 | 6.6680 | 6.7339 | 6.2888 | 6.4301 | 6.9726 | 7.1561 | 6.7761 | 4.5664 | 5.1020 | 4.9378 | 5.4938 |
| RGPD8_S1742 | RGPD8 | O14715 | | 1742 | NTMLQLSPEEKGK | 6.1123 | 6.5901 | 6.4342 | 7.3147 | 6.1931 | 5.8100 | 6.8249 | 5.9741 | 6.3414 | 6.3042 | 6.6885 | 6.7264 | 5.0917 | 5.8680 | 5.7191 | 6.0075 |
| RGPD8_S21 | RGPD8 | O14715 | | 1302 | KSALSLSKSPAKL | 7.4084 | 6.1056 | 6.5133 | 7.4248 | 6.7457 | 6.9591 | 7.1077 | 6.8814 | 7.0931 | 6.3354 | 5.9647 | 4.5859 | 4.7517 | 4.2792 | 5.5500 | 6.2939 |
| RGPD8_S781 | RGPD8 | O14715 | 1621;1629 | | NA | 6.7998 | 5.8611 | 7.1774 | 6.6859 | 6.8793 | 6.1213 | 7.8556 | 4.8758 | 7.3947 | 7.1088 | 7.2182 | 6.2147 | 5.0596 | 4.8984 | 4.4075 | 5.4419 |
| RGS10_S10 | RGS10 | O43665 | | 10 | RAVSRLSRKRPPS | 6.7322 | 6.1589 | 7.2660 | 5.5843 | 6.1699 | 6.8514 | 5.7525 | 6.3434 | 6.3556 | 7.7706 | 8.3582 | 9.8811 | 3.9276 | 5.5300 | 3.2399 | 4.0784 |
| RGS12_S1294 | RGS12 | O14924 | | 506 | KHLGPASPVEVPP | 5.8858 | 6.7403 | 7.3043 | 6.5549 | 7.4029 | 6.5129 | 7.1478 | 7.0545 | 7.4081 | 6.0185 | 6.8757 | 6.4984 | 3.8401 | 6.1145 | 4.0728 | 4.5685 |
| RGS12_S1294;S1304 | RGS12 | O14924 | | 16 | RPLPGPSPPPRVRS | 6.6346 | 6.8093 | 6.9159 | 6.9361 | 6.4605 | 6.9786 | 6.1786 | 6.2474 | 6.5220 | 7.2472 | 7.5206 | 7.0015 | 4.7319 | 5.0722 | 4.1373 | 4.6063 |
| RGS12_S16 | RGS12 | O14924 | | 1294 | TPPGQKSPSGPFC | 6.3170 | 6.4206 | 6.5140 | 5.9631 | 6.3929 | 5.9682 | 5.8215 | 5.9412 | 6.2944 | 6.5447 | 6.6644 | 6.3425 | 6.0557 | 6.2871 | 6.2353 | 6.2375 |
| RGS12_S506 | RGS12 | O14924 | | 850 | SQQVPSSPASKHS | 6.4482 | 6.1083 | 6.6231 | 6.3249 | 6.5512 | 6.2622 | 6.2493 | 6.9519 | 6.6287 | 6.4176 | 6.2545 | 6.0024 | 5.8747 | 6.3559 | 5.3809 | 5.5662 |
| RGS12_S671 | RGS12 | O14924 | | 671 | LESATVSDGELTG | 6.0720 | 7.8092 | 7.0145 | 5.9364 | 6.3231 | 6.1702 | 7.2241 | 6.6582 | 7.0236 | 6.0508 | 6.1672 | 5.3006 | 5.9880 | 6.0538 | 4.9858 | 5.2227 |
| RGS12_S850 | RGS12 | O14924 | 1294;1304 | | NA | 6.0732 | 6.4907 | 7.2820 | 6.4305 | 6.0062 | 6.5616 | 6.4789 | 6.7294 | 7.2195 | 7.1470 | 6.9950 | 7.0085 | 4.7900 | 5.5549 | 4.3573 | 4.8754 |
| RGS19_S65 | RGS19 | P49795 | | 65 | WQASRESKLQPLP | 7.3132 | 6.4148 | 7.2641 | 7.1336 | 7.3448 | 5.7865 | 5.4485 | 6.9752 | 6.6474 | 6.4881 | 6.0168 | 6.2405 | 5.3055 | 5.6315 | 4.5813 | 5.4081 |
| RHBDF1_S261 | RHBDF1 | Q96CC6 | | 390 | YRKRIDSFVKRQI | 5.5472 | 5.5240 | 5.1574 | 5.6856 | 6.1631 | 6.0677 | 6.3371 | 6.4450 | 5.4834 | 5.1562 | 5.3136 | 5.5594 | 8.1380 | 7.6081 | 8.1849 | 7.6293 |
| RHBDF1_S390 | RHBDF1 | Q96CC6 | | 92 | ADWFGVSKDSDST | 3.9075 | 6.3179 | 8.2133 | 6.8178 | 5.6449 | 6.3248 | 5.3977 | 6.6942 | 5.8921 | 6.3852 | 6.2149 | 7.8919 | 4.7741 | 7.0760 | 6.1778 | 6.2698 |
| RHBDF1_S92 | RHBDF1 | Q96CC6 | | 261 | PDELDTSFFAREG | 6.4306 | 6.9727 | 6.9332 | 8.3041 | 7.1783 | 5.8137 | 6.7214 | 6.5926 | 6.5926 | 6.4783 | 5.6330 | 5.0909 | 4.8063 | 5.2653 | 4.7523 | 6.4347 |
| RHBDF2_S102 | RHBDF2 | Q6PJF5 | | 102 | SRWQESSEKRPGF | 8.1455 | 7.3856 | 8.0505 | 7.7969 | 6.5629 | 6.0163 | 6.0596 | 6.3157 | 6.9166 | 7.1544 | 7.4313 | 7.3000 | 3.5185 | 4.8962 | 3.0382 | 3.4118 |
| RHBDF2_S113 | RHBDF2 | Q6PJF5 | | 177 | SFQGTESPKPCKM | 6.3709 | 6.4309 | 6.3987 | 6.5747 | 6.2567 | 6.3040 | 6.8408 | 5.9502 | 6.1600 | 5.9328 | 5.9452 | 6.1635 | 6.2313 | 6.2243 | 5.6938 | 6.5222 |
| RHBDF2_S177 | RHBDF2 | Q6PJF5 | | 313 | FESPPLSASYFRG | 6.2538 | 6.6427 | 6.0921 | 6.7840 | 6.5554 | 6.2515 | 5.7538 | 6.1378 | 6.5145 | 5.8081 | 5.6998 | 6.3237 | 6.1685 | 6.6129 | 5.8534 | 6.5480 |
| RHBDF2_S309 | RHBDF2 | Q6PJF5 | | 325 | GIPHSASPVSPDG | 5.7582 | 5.9835 | 5.8613 | 6.4677 | 5.9569 | 6.1841 | 6.2036 | 6.3721 | 6.2590 | 6.1316 | 5.3706 | 5.9277 | 6.5591 | 7.0735 | 7.0855 | 6.8058 |
| RHBDF2_S313 | RHBDF2 | Q6PJF5 | | 113 | GFRRQASLSQSIR | 5.9974 | 7.6789 | 8.0423 | 6.2496 | 5.0740 | 6.2935 | 6.1057 | 6.1771 | 5.9905 | 7.6333 | 6.8378 | 6.3398 | 4.5231 | 5.9989 | 5.4019 | 5.6563 |
| RHBDF2_S325 | RHBDF2 | Q6PJF5 | | 309 | PDDVFESPPLSAS | 7.0977 | 7.7828 | 6.6705 | 7.6723 | 7.4974 | 6.4366 | 5.1051 | 6.8141 | 6.5950 | 5.4458 | 7.0185 | 6.8417 | 5.5636 | 4.5673 | 4.1713 | 4.7202 |
| RHBDF2_S325;S328 | RHBDF2 | Q6PJF5 | | 90 | AYLKSVSLQEPRS | 5.7678 | 7.0029 | 8.1683 | 6.7563 | 5.3802 | 5.6567 | 6.1837 | 5.5774 | 5.5837 | 7.1091 | 6.7363 | 6.5015 | 5.1707 | 6.5279 | 6.1738 | 5.7035 |
| RHBDF2_S90 | RHBDF2 | Q6PJF5 | 325;328 | | NA | 5.3122 | 5.7978 | 6.0470 | 5.9888 | 5.8293 | 6.3366 | 6.5568 | 6.5704 | 5.6792 | 5.5403 | 5.8025 | 5.6260 | 7.2616 | 6.9398 | 7.2014 | 7.5104 |
| RHPN2_S616 | RHPN2 | Q8IUC4 | | 654 | NRQKSASTLCLPS | 7.4664 | 6.6407 | 7.0185 | 6.5139 | 6.5336 | 6.2384 | 6.3436 | 6.1181 | 6.9039 | 5.8986 | 6.2338 | 6.3496 | 5.5665 | 5.9257 | 4.9158 | 5.3327 |
| RHPN2_S639 | RHPN2 | Q8IUC4 | | 606 | MHNKSATYSVGMQ | 7.5325 | 7.3905 | 8.1428 | 7.5276 | 7.3918 | 5.3383 | 6.0975 | 5.7742 | 5.9738 | 5.6010 | 5.7374 | 5.0544 | 6.0734 | 5.9007 | 5.9786 | 4.4856 |
| RHPN2_S642 | RHPN2 | Q8IUC4 | | 31 | CNPLAQTGRSKLQ | 10.7088 | 6.5945 | 6.5643 | 8.6038 | 7.4359 | 3.9417 | 5.3741 | 3.8534 | 5.2465 | 3.4098 | 3.8901 | 3.8534 | 8.9199 | 7.1262 | 7.8037 | 6.6740 |
| RHPN2_S654 | RHPN2 | Q8IUC4 | | 642 | KKLSFLSWGTNKN | 6.6762 | 6.3591 | 5.7767 | 5.8984 | 6.0225 | 6.9221 | 6.8635 | 6.5694 | 5.3849 | 5.3860 | 6.2270 | 6.7176 | 6.6165 | 5.7238 | 6.1271 | 6.7291 |
| RHPN2_T31 | RHPN2 | Q8IUC4 | | 639 | KISKKLSFLSWGT | 7.7067 | 5.8625 | 6.2836 | 4.9746 | 7.5200 | 5.7131 | 5.8189 | 5.0534 | 7.2337 | 5.0430 | 6.5201 | 7.9432 | 6.2048 | 7.5386 | 5.4413 | 5.1426 |
| RHPN2_T606 | RHPN2 | Q8IUC4 | | 616 | GMQKTYSMICLAI | 6.7278 | 5.7766 | 7.3427 | 6.7227 | 6.3030 | 5.4271 | 5.9250 | 5.5112 | 6.7673 | 6.1334 | 6.1385 | 7.5725 | 6.8046 | 6.0581 | 6.2372 | 6.3725 |
| RIC1_S1017 | RIC1 | Q4ADV7 | | 992 | GSGESETPPSTPT | 6.1312 | 7.3004 | 6.6501 | 6.5514 | 6.3084 | 6.3969 | 6.5149 | 5.8227 | 7.0044 | 7.0515 | 6.5717 | 6.8244 | 5.1171 | 5.6874 | 4.8182 | 5.2494 |
| RIC1_S1037 | RIC1 | Q4ADV7 | | 1017 | FRNRSISLSQSAE | 5.1236 | 4.5052 | 5.5095 | 6.2218 | 6.7008 | 5.9491 | 5.6597 | 5.7237 | 6.2903 | 6.5269 | 6.0690 | 6.5313 | 7.5207 | 7.4284 | 7.4395 | 6.8003 |
| RIC1_S992;T996 | RIC1 | Q4ADV7 | | 1037 | SLQKTLSMPSGPS | 4.9251 | 4.8244 | 5.5058 | 5.1561 | 5.6610 | 5.5359 | 5.1490 | 5.0356 | 5.1664 | 4.8309 | 5.1750 | 5.8295 | 9.1011 | 8.8264 | 9.9765 | 9.3012 |
| RIC1_T992 | RIC1 | Q4ADV7 | 992;996 | | NA | 5.6835 | 5.3864 | 6.0565 | 5.8126 | 6.2564 | 5.8222 | 5.7793 | 6.7495 | 6.5255 | 6.1172 | 6.1248 | 5.9872 | 6.7455 | 6.5240 | 7.1202 | 7.3092 |
| RIC8A_S442;T447 | RIC8A | Q9NPQ8-3 | | 508 | IQPMGMSPRGHLT | 6.1095 | 6.5221 | 6.8956 | 6.7243 | 6.2061 | 6.6286 | 6.4981 | 6.5358 | 6.8184 | 6.6352 | 7.0501 | 7.4274 | 4.5094 | 5.4319 | 4.7623 | 5.2452 |
| RIC8A_S508 | RIC8A | Q9NPQ8-3 | 442;447 | | NA | 5.9504 | 6.1882 | 7.2618 | 5.8135 | 5.8403 | 6.8831 | 5.8672 | 6.5742 | 6.3907 | 6.8015 | 7.2056 | 6.5179 | 5.7907 | 5.8112 | 5.6558 | 5.4478 |
| RICTOR_S1023 | RICTOR | Q6R327-3 | | 1615 | SASSTKSTELLLG | 5.9364 | 7.0028 | 7.3538 | 6.5317 | 5.4543 | 6.7616 | 6.1831 | 6.5321 | 6.2789 | 7.3622 | 7.1846 | 7.2797 | 4.4931 | 5.1571 | 4.9251 | 5.5634 |
| RICTOR_S1144 | RICTOR | Q6R327-3 | | 1412 | KALSYASLDKEDL | 6.7497 | 6.0233 | 6.9363 | 6.5960 | 6.6383 | 6.1060 | 5.9365 | 6.6155 | 7.0353 | 6.7854 | 6.2418 | 6.7148 | 5.6113 | 5.9218 | 4.9433 | 5.1448 |
| RICTOR_S1307 | RICTOR | Q6R327-3 | | 1409 | RFMKALSYASLDK | 6.3309 | 5.7031 | 5.0330 | 5.0228 | 5.2469 | 5.6093 | 5.3753 | 4.6941 | 5.4903 | 4.6185 | 5.2173 | 5.3580 | 9.0432 | 9.4985 | 8.2775 | 9.4813 |
| RICTOR_S1409 | RICTOR | Q6R327-3 | | 291 | KMGIIATFRSWAG | 2.1316 | 2.5197 | 3.8936 | 3.9967 | 2.6723 | 2.3986 | 7.5726 | 6.2792 | 14.9593 | 1.7829 | 17.5561 | 13.6068 | 14.6438 | 2.3479 | 2.0752 | 1.5637 |
| RICTOR_S1412 | RICTOR | Q6R327-3 | | 21 | VRGRNDSGEENVP | 6.6713 | 6.8719 | 7.0312 | 6.8478 | 6.6542 | 6.5973 | 7.7602 | 6.0837 | 6.2089 | 6.5728 | 7.2125 | 7.1959 | 4.4163 | 4.8712 | 4.9882 | 5.0166 |
| RICTOR_S1420 | RICTOR | Q6R327-3 | | 1332 | DAFGYATLKRLQQ | 9.8834 | 6.8459 | 8.7819 | 8.1340 | 5.3690 | 4.8612 | 6.9354 | 4.6192 | 6.3695 | 5.4233 | 5.3805 | 6.0341 | 5.6225 | 6.2799 | 5.0937 | 4.3667 |
| RICTOR_S1494 | RICTOR | Q6R327-3 | | 1420 | DKEDLLSPINQNT | 6.3243 | 6.3359 | 6.2403 | 6.6463 | 6.0790 | 6.2735 | 6.8184 | 6.9452 | 6.0180 | 6.5256 | 6.2450 | 6.7805 | 5.1321 | 6.0742 | 6.1197 | 5.4419 |
| RICTOR_S1615 | RICTOR | Q6R327-3 | | 1307 | QSLKAPSIATIKS | 6.5139 | 5.8677 | 5.8790 | 5.8863 | 6.0631 | 6.2906 | 6.1358 | 5.5156 | 6.8594 | 6.0624 | 5.8118 | 5.7575 | 7.3728 | 7.5807 | 6.3152 | 6.0881 |
| RICTOR_S21 | RICTOR | Q6R327-3 | | 1023 | SIPSTLSLNSEST | 5.3130 | 4.4991 | 8.4113 | 6.9443 | 4.7122 | 7.0316 | 6.2910 | 6.7207 | 6.6508 | 6.4343 | 6.2456 | 5.7811 | 5.9173 | 7.1364 | 5.4876 | 6.4238 |
| RICTOR_T1103 | RICTOR | Q6R327-3 | | 1144 | SVDFNHSDDFTPI | 6.0853 | 6.3206 | 6.8031 | 5.9972 | 6.8330 | 7.3100 | 6.9521 | 7.7080 | 7.3482 | 9.0162 | 7.3582 | 7.7189 | 3.0227 | 4.4454 | 3.3579 | 3.7232 |
| RICTOR_T1332 | RICTOR | Q6R327-3 | | 1103 | RILNSLTLPNKKH | 8.3363 | 7.5021 | 9.3034 | 8.7014 | 6.1207 | 5.0293 | 6.7295 | 5.2181 | 6.7748 | 6.3307 | 6.5359 | 6.4704 | 4.0169 | 5.1093 | 4.0131 | 3.8079 |
| RICTOR_T291 | RICTOR | Q6R327-3 | | 1494 | DAGGLPSGTGGLV | 7.1344 | 7.0275 | 6.6225 | 7.6688 | 7.2323 | 7.8340 | 7.0976 | 4.7140 | 4.9125 | 6.1097 | 7.0868 | 7.2484 | 4.9224 | 5.1631 | 4.3000 | 4.9260 |
| RIF1_S1046 | RIF1 | Q5UIP0 | | 2205 | NKVRRVSFADPIY | 5.3312 | 4.8009 | 5.1052 | 5.4407 | 6.6192 | 6.0127 | 5.4536 | 5.9947 | 8.4455 | 8.4911 | 8.1049 | 8.3727 | 6.2127 | 6.8674 | 3.8992 | 4.8481 |
| RIF1_S1236 | RIF1 | Q5UIP0 | | 2144 | IEDNNASPQKLRE | 6.1082 | 6.3286 | 6.2726 | 6.5914 | 6.3297 | 5.5762 | 6.4517 | 6.2642 | 6.7722 | 5.9614 | 6.3133 | 6.3357 | 6.0007 | 6.2444 | 6.0931 | 6.3567 |
| RIF1_S1238 | RIF1 | Q5UIP0 | | 2243 | GKSVKTSPTTQSK | 7.4720 | 7.2581 | 6.6159 | 7.3801 | 6.8450 | 6.5118 | 7.1625 | 6.5425 | 7.2293 | 6.7727 | 7.0424 | 7.1318 | 3.8486 | 4.2965 | 3.6237 | 4.2670 |
| RIF1_S1384 | RIF1 | Q5UIP0 | | 1236 | SENRPFSPSPLNN | 6.2977 | 6.3090 | 6.7292 | 6.8645 | 5.3359 | 6.4005 | 6.8679 | 6.5803 | 6.9991 | 6.7798 | 6.7221 | 6.9423 | 5.0349 | 5.4310 | 5.0467 | 5.6590 |
| RIF1_S1422 | RIF1 | Q5UIP0 | | 1046 | LLEEEKSTDFVFI | 3.9047 | 4.8690 | 7.0217 | 7.3848 | 4.2735 | 5.6139 | 6.3404 | 5.7336 | 5.1828 | 5.8141 | 6.2724 | 6.1650 | 6.9107 | 8.5134 | 9.1391 | 6.8609 |
| RIF1_S1542 | RIF1 | Q5UIP0 | | 2161 | LVSANDSPSGMQT | 6.3916 | 6.5897 | 6.7117 | 6.9149 | 6.4278 | 6.0796 | 6.7610 | 6.6135 | 6.7997 | 7.0220 | 6.4703 | 6.6401 | 5.1524 | 5.2546 | 4.8831 | 5.2880 |
| RIF1_S1542;S1554 | RIF1 | Q5UIP0 | | 1384 | VEINLESKENTPP | 5.5757 | 5.8660 | 6.1092 | 6.1429 | 5.7878 | 5.6316 | 5.5575 | 5.6749 | 5.8726 | 6.1925 | 6.2333 | 6.7872 | 6.9827 | 7.1342 | 7.0868 | 7.3652 |
| RIF1_S1554 | RIF1 | Q5UIP0 | | 409 | PYGAPGTPRMNLS | 6.9002 | 6.8461 | 6.4191 | 6.9127 | 6.9175 | 6.7268 | 6.7165 | 6.7657 | 6.8861 | 6.6216 | 6.2184 | 6.4228 | 4.9022 | 5.3804 | 4.4406 | 4.9234 |
| RIF1_S1579 | RIF1 | Q5UIP0 | | 2348 | KTLPIRSPKVSNV | 7.7999 | 7.9713 | 6.7242 | 7.9410 | 6.9536 | 6.9947 | 7.3149 | 6.7973 | 7.7087 | 7.4598 | 7.1912 | 7.2667 | 2.5373 | 3.4340 | 2.4941 | 3.4111 |
| RIF1_S1608 | RIF1 | Q5UIP0 | | 2172 | QTRCVWSPLASPS | 6.5925 | 6.8090 | 6.2009 | 7.4766 | 6.8637 | 6.1861 | 6.3741 | 6.4705 | 7.2832 | 6.8572 | 6.5082 | 6.5761 | 4.8812 | 5.2724 | 4.7425 | 4.9057 |
| RIF1_S1616 | RIF1 | Q5UIP0 | | 2260 | SAKGFLSPGSRSP | 8.2709 | 8.4512 | 6.9041 | 7.7180 | 7.0637 | 7.0104 | 7.5495 | 6.8790 | 7.8209 | 7.2723 | 7.1417 | 7.5054 | 2.2053 | 3.1537 | 1.9989 | 3.0549 |
| RIF1_S1663 | RIF1 | Q5UIP0 | | 1097 | DTLFTQYSQEEPM | 5.0800 | 5.8901 | 6.4720 | 6.5950 | 5.4501 | 6.4038 | 6.5091 | 6.3584 | 5.8007 | 6.7971 | 7.0165 | 7.1055 | 5.6888 | 6.3646 | 6.5835 | 5.8848 |
| RIF1_S1688 | RIF1 | Q5UIP0 | | 1688 | RLHKRDSFDNCSL | 4.7248 | 5.5510 | 5.5945 | 6.0058 | 7.1151 | 5.3755 | 5.4972 | 5.8399 | 6.7984 | 7.5834 | 7.8620 | 7.2586 | 5.2831 | 7.0974 | 6.5373 | 5.8760 |
| RIF1_S1706 | RIF1 | Q5UIP0 | | 1408 | DSQVQITPNQKTL | 6.7722 | 6.7769 | 6.2804 | 6.9852 | 6.3971 | 6.2030 | 6.5351 | 6.5422 | 7.1961 | 7.1001 | 6.4854 | 7.0028 | 5.0471 | 5.0468 | 4.6727 | 4.9567 |
| RIF1_S1873 | RIF1 | Q5UIP0 | | 782 | YQPKVKSPQRPSD | 7.0859 | 6.9897 | 6.7404 | 8.2150 | 6.0059 | 5.9874 | 6.4046 | 6.3327 | 6.9534 | 6.5807 | 7.2891 | 7.6187 | 3.8702 | 5.0300 | 3.9943 | 4.9019 |
| RIF1_S2144 | RIF1 | Q5UIP0 | | 1663 | SKYAEYSFTSLPV | 6.7393 | 7.0046 | 6.3489 | 7.6348 | 5.8751 | 6.3515 | 6.3184 | 6.4303 | 6.7842 | 6.4396 | 7.3792 | 6.6880 | 5.3639 | 5.3686 | 4.5642 | 4.7096 |
| RIF1_S2161 | RIF1 | Q5UIP0 | | 2465 | LQSRWRSPSHENS | 6.5516 | 7.1404 | 6.2980 | 8.2132 | 6.2671 | 6.4877 | 6.4327 | 5.9938 | 6.6566 | 6.3267 | 6.5046 | 6.7650 | 4.8418 | 5.6220 | 4.8108 | 5.0878 |
| RIF1_S2172 | RIF1 | Q5UIP0 | | 2176 | VWSPLASPSTSIL | 8.0723 | 7.1279 | 6.4249 | 7.2687 | 6.4138 | 6.2448 | 7.1191 | 6.4200 | 7.3531 | 6.8657 | 6.4457 | 6.6099 | 4.4052 | 4.5363 | 3.8926 | 4.8001 |
| RIF1_S2172;S2176 | RIF1 | Q5UIP0 | | 1238 | NRPFSPSPLNNIS | 6.4072 | 6.3399 | 6.2315 | 6.6286 | 6.0794 | 6.4133 | 6.7271 | 6.1464 | 6.4673 | 7.0734 | 6.5247 | 6.6157 | 5.2461 | 5.9113 | 5.4121 | 5.7760 |
| RIF1_S2176 | RIF1 | Q5UIP0 | | 2234 | SHSSNSSPIGKSV | 7.6629 | 8.5420 | 6.2801 | 7.7537 | 6.8086 | 6.9733 | 6.5382 | 7.3475 | 7.4635 | 6.4104 | 6.5364 | 6.9645 | 3.1235 | 4.2109 | 3.5317 | 3.8528 |
| RIF1_S2205 | RIF1 | Q5UIP0 | | 1542 | YQTRRASQGLLSS | 5.4045 | 4.4458 | 5.1057 | 5.4484 | 6.2726 | 5.3500 | 5.4638 | 4.9527 | 8.0591 | 7.6885 | 8.2156 | 8.0442 | 7.0665 | 7.7335 | 5.0470 | 5.7022 |
| RIF1_S2234 | RIF1 | Q5UIP0 | | 1422 | RSSRRRSEVVEST | 4.1885 | 4.0104 | 4.7371 | 5.5003 | 7.5647 | 5.5361 | 5.2274 | 5.8054 | 7.4097 | 7.8987 | 7.5402 | 8.5054 | 6.6507 | 8.2711 | 5.7276 | 5.4266 |
| RIF1_S2243 | RIF1 | Q5UIP0 | | 1873 | GDSKNVSQESLET | 5.8113 | 6.8705 | 6.5853 | 8.0218 | 6.0829 | 6.3113 | 6.4199 | 6.1211 | 6.2335 | 6.0656 | 6.6921 | 7.7243 | 4.5828 | 5.3446 | 4.9816 | 6.1514 |
| RIF1_S2260 | RIF1 | Q5UIP0 | | 1616 | EDVSIKSPICEKQ | 6.4850 | 6.9534 | 6.3595 | 7.1432 | 6.9361 | 6.8673 | 7.2830 | 6.5947 | 7.2672 | 6.7543 | 7.3691 | 6.9777 | 4.3801 | 4.5380 | 4.0280 | 4.0633 |
| RIF1_S2348 | RIF1 | Q5UIP0 | | 1980 | SLSDNTTPVKLNA | 6.5744 | 6.7762 | 6.3842 | 7.3157 | 6.1398 | 6.4730 | 6.3681 | 7.1416 | 8.1180 | 6.3047 | 6.2332 | 6.7174 | 4.9478 | 5.4666 | 4.4337 | 4.6055 |
| RIF1_S2393 | RIF1 | Q5UIP0 | | 1608 | HDYKATSEEDVSI | 7.2711 | 7.7249 | 7.0082 | 8.0950 | 6.0306 | 6.2842 | 6.9148 | 6.3573 | 6.1987 | 6.4616 | 6.1107 | 6.2693 | 4.6947 | 4.7110 | 4.8899 | 4.9779 |
| RIF1_S2465 | RIF1 | Q5UIP0 | | 2393 | IENKSLSPDEERL | 6.6578 | 6.5658 | 6.5799 | 6.8955 | 6.1245 | 5.3823 | 6.8089 | 5.9583 | 6.2347 | 5.9916 | 5.8954 | 5.2381 | 6.3649 | 6.1190 | 6.8034 | 6.3800 |
| RIF1_S782 | RIF1 | Q5UIP0 | | 1706 | IGISDISSLSEKT | 9.1295 | 7.5856 | 6.9653 | 7.7820 | 5.5478 | 6.1410 | 7.5262 | 5.7797 | 7.3865 | 6.4177 | 7.6756 | 7.4561 | 3.4408 | 4.3009 | 2.7535 | 4.1116 |
| RIF1_T1408 | RIF1 | Q5UIP0 | | 1554 | SIENSESDSSEAK | 6.5460 | 7.5590 | 7.4774 | 8.1283 | 5.3151 | 6.6676 | 6.9578 | 7.6497 | 4.6246 | 4.7020 | 5.3050 | 5.4012 | 5.0516 | 5.4865 | 6.2746 | 6.8537 |
| RIF1_T1980 | RIF1 | Q5UIP0 | | 1579 | KNKSNESVDIQDQ | 4.8892 | 5.6824 | 6.6613 | 6.8690 | 5.4773 | 6.2517 | 5.9136 | 5.5641 | 6.1487 | 7.9405 | 7.5740 | 7.8502 | 4.5869 | 6.5309 | 6.2203 | 5.8399 |
| RIF1_T409 | RIF1 | Q5UIP0 | 2172;2176 | | NA | 6.9368 | 7.1774 | 6.7297 | 7.1878 | 6.6245 | 7.0554 | 6.9192 | 6.8264 | 7.4777 | 7.8585 | 6.8880 | 7.0336 | 3.4488 | 4.2195 | 3.5160 | 4.1007 |
| RIF1_Y1097 | RIF1 | Q5UIP0 | 1542;1554 | | NA | 5.0814 | 4.7927 | 5.8478 | 5.6006 | 7.3913 | 6.4511 | 6.7359 | 6.5056 | 7.5499 | 7.8321 | 7.6844 | 7.8482 | 5.4935 | 6.1454 | 3.7241 | 5.3161 |
| RIMS2_S1044 | RIMS2 | D6RA03 | | 1044 | DLQRRFSQDDTSV | 5.8848 | 5.5050 | 6.1809 | 6.2219 | 6.6614 | 6.0507 | 5.3977 | 6.4280 | 6.7831 | 6.4924 | 6.8992 | 6.9420 | 6.0185 | 7.0968 | 5.9285 | 5.5089 |
| RIMS2_S231 | RIMS2 | D6RA03 | | 231 | DLRRTDSLRKQHH | 7.2053 | 6.8235 | 7.3440 | 6.2798 | 5.4696 | 6.7460 | 6.5923 | 5.9534 | 7.1992 | 7.0679 | 7.4542 | 7.1118 | 4.3037 | 6.0037 | 3.9953 | 4.4503 |
| RIMS2_S826 | RIMS2 | D6RA03 | | 826 | TGSVQTSPSSTPV | 5.0403 | 10.6818 | 6.5356 | 7.3942 | 5.3984 | 5.0281 | 5.1512 | 8.2415 | 7.4318 | 6.7889 | 6.4960 | 6.2630 | 3.5827 | 5.5388 | 5.9131 | 4.5144 |
| RIN1_S254 | RIN1 | Q13671 | | 351 | PLLRSMSAAFCSL | 6.9419 | 6.4373 | 6.6773 | 6.7479 | 6.5644 | 6.5277 | 6.0114 | 6.1323 | 6.5227 | 5.7912 | 6.2514 | 6.4513 | 6.0117 | 5.9413 | 5.1217 | 5.8685 |
| RIN1_S333 | RIN1 | Q13671 | | 337 | RGSPATSPHLGRR | 5.9531 | 6.0344 | 5.7393 | 5.6390 | 6.2537 | 6.5465 | 6.0415 | 6.8277 | 5.7011 | 5.6276 | 5.6859 | 5.6919 | 7.0916 | 7.1361 | 7.2710 | 6.7596 |
| RIN1_S333;S337 | RIN1 | Q13671 | | 254 | RVSTETSSPLSPP | 5.8843 | 6.0973 | 6.0533 | 6.1563 | 6.8173 | 6.4746 | 6.1563 | 4.6625 | 5.9040 | 6.0834 | 6.0921 | 6.4891 | 6.6205 | 6.4874 | 5.9341 | 6.2877 |
| RIN1_S337 | RIN1 | Q13671 | | 333 | VPGSRGSPATSPH | 6.9710 | 6.7762 | 7.0224 | 6.8372 | 6.3590 | 7.1570 | 5.1914 | 7.5101 | 7.3074 | 6.9298 | 6.6899 | 5.9237 | 4.8566 | 5.6577 | 4.2526 | 4.5581 |
| RIN1_S351 | RIN1 | Q13671 | | 36 | PAQDPLYDVPNAS | 5.8137 | 6.1772 | 6.5504 | 6.2951 | 5.2278 | 5.9010 | 6.8437 | 6.4354 | 6.7564 | 6.1451 | 6.1294 | 5.9294 | 7.0094 | 6.0063 | 6.4839 | 6.2959 |
| RIN1_Y36 | RIN1 | Q13671 | 333;337 | | NA | 6.0194 | 6.1207 | 6.4757 | 5.7011 | 6.4481 | 6.4970 | 6.2614 | 7.2959 | 6.7499 | 6.5775 | 6.3137 | 6.6922 | 5.4465 | 6.0426 | 5.4000 | 5.9584 |
| RIN2_S533 | RIN2 | Q8WYP3-2 | | 533 | KSKKKRSSSFVLP | 6.9754 | 7.1278 | 6.9825 | 7.2352 | 6.1876 | 7.0735 | 6.7563 | 6.8782 | 7.1476 | 7.2383 | 7.2255 | 6.6380 | 3.7192 | 4.6880 | 3.8836 | 4.2432 |
| RING1_S140 | RING1 | Q06587 | | 248 | GTLGPPSPPGAPS | 5.8541 | 6.1472 | 6.5051 | 6.5729 | 6.3276 | 6.1651 | 6.2311 | 6.0848 | 6.4173 | 6.5617 | 6.6773 | 6.3179 | 6.3409 | 6.0363 | 5.9699 | 5.7908 |
| RING1_S248 | RING1 | Q06587 | | 243 | GTLGGGTLGPPSP | 6.1703 | 6.3248 | 5.9966 | 6.4880 | 6.2925 | 5.8156 | 5.9006 | 6.2964 | 6.4583 | 6.2894 | 6.1098 | 6.4422 | 6.4399 | 6.3051 | 6.0259 | 6.6445 |
| RING1_S248;S254 | RING1 | Q06587 | | 41 | IAVSPRSLHSELM | 6.1244 | 7.6571 | 5.4412 | 5.8800 | 6.8829 | 6.7795 | 4.9334 | 6.1088 | 7.6070 | 6.7011 | 5.6950 | 6.3814 | 6.1307 | 7.1713 | 5.6888 | 4.8174 |
| RING1_S41 | RING1 | Q06587 | | 140 | QQALSSSIEEGLR | 6.5856 | 6.4714 | 6.2354 | 5.5176 | 7.2516 | 5.7816 | 6.5331 | 7.1605 | 6.4427 | 7.0415 | 6.5267 | 7.0926 | 5.5379 | 5.5968 | 4.8215 | 5.4035 |
| RING1_T243 | RING1 | Q06587 | 243;254 | | NA | 5.7676 | 6.0817 | 5.9375 | 5.8940 | 6.0434 | 5.9294 | 6.1987 | 6.0956 | 6.9417 | 6.4311 | 6.1118 | 6.6436 | 5.9679 | 6.4499 | 6.3855 | 7.1208 |
| RING1_T243;S254 | RING1 | Q06587 | 248;254 | | NA | 6.0180 | 6.3180 | 6.3323 | 6.5084 | 5.7665 | 5.8211 | 6.4290 | 6.7975 | 6.3860 | 6.2117 | 6.4729 | 6.0442 | 6.0981 | 6.8018 | 5.8986 | 6.0958 |
| RIOK1_S22 | RIOK1 | Q9BRS2 | | 22 | FDDADSSDSENRD | 5.9152 | 6.3746 | 6.0587 | 6.5402 | 6.6399 | 6.8834 | 5.6197 | 6.5143 | 6.3014 | 6.1464 | 6.6067 | 6.3274 | 6.0779 | 6.0340 | 6.0881 | 5.8722 |
| RIOK2_S350 | RIOK2 | Q9BVS4 | | 390 | ESADARSFEMTEF | 5.8468 | 6.6143 | 7.2938 | 6.2723 | 6.4055 | 6.6038 | 6.6081 | 6.7991 | 6.4101 | 6.6958 | 6.8212 | 6.5433 | 5.2917 | 5.3210 | 5.2946 | 5.1786 |
| RIOK2_S350;S354 | RIOK2 | Q9BVS4 | | 442 | GGVPAGSDEYEDE | 5.9714 | 6.3598 | 6.7118 | 6.4136 | 6.5716 | 6.6026 | 6.4993 | 6.2846 | 6.4745 | 6.1874 | 6.1368 | 6.3521 | 5.7955 | 6.3855 | 5.4964 | 5.7572 |
| RIOK2_S380 | RIOK2 | Q9BVS4 | | 483 | YRTRTLSITSSGS | 6.5430 | 5.5306 | 6.1096 | 5.9722 | 6.3584 | 6.7554 | 6.3165 | 6.3311 | 6.8366 | 6.8380 | 7.4103 | 6.9892 | 6.5534 | 6.0408 | 5.0129 | 5.3021 |
| RIOK2_S390 | RIOK2 | Q9BVS4 | | 350 | KAEVYGSENESER | 6.7536 | 6.5587 | 7.0828 | 6.2784 | 6.3870 | 6.9183 | 6.1927 | 7.2882 | 6.0850 | 5.9575 | 6.3798 | 6.0592 | 5.2904 | 5.8333 | 5.4157 | 5.5195 |
| RIOK2_S442 | RIOK2 | Q9BVS4 | | 380 | EQIKEDSLSEESA | 6.6166 | 6.1619 | 7.2330 | 7.3278 | 6.5636 | 6.3729 | 7.3556 | 5.8583 | 7.2839 | 7.2999 | 8.0898 | 7.0477 | 4.4476 | 4.4252 | 3.9030 | 4.0133 |
| RIOK2_S483 | RIOK2 | Q9BVS4 | 350;354 | | NA | 5.8762 | 5.6932 | 6.0784 | 7.3099 | 5.8990 | 6.1286 | 7.3896 | 7.4525 | 5.5681 | 5.5249 | 6.1847 | 6.1010 | 5.9009 | 6.2488 | 6.2651 | 6.3790 |
| RIOX1_S109 | RIOX1 | Q9H6W3 | | 109 | AELLEASPAARSL | 6.6125 | 7.4193 | 6.3512 | 7.1762 | 6.5490 | 6.5721 | 6.7681 | 6.6214 | 7.2277 | 6.6765 | 6.7748 | 7.0375 | 4.0299 | 5.2723 | 3.9175 | 4.9941 |
| RIOX2_S47 | RIOX2 | Q8IUF8 | | 47 | LFESLISPIKTET | 5.7597 | 6.1989 | 6.2284 | 6.4803 | 6.1334 | 5.9649 | 6.4328 | 6.4328 | 5.9828 | 6.3109 | 6.1047 | 6.3037 | 5.8950 | 6.6363 | 6.1863 | 6.9491 |
| RIPK1_S320 | RIPK1 | Q13546 | | 320 | VVKRMQSLQLDCV | 6.0652 | 5.6906 | 6.1124 | 6.3270 | 6.3545 | 5.8827 | 5.6701 | 5.8942 | 6.3871 | 6.2398 | 5.9954 | 6.6298 | 7.1413 | 6.5670 | 6.4675 | 6.5755 |
| RIPK1_S416 | RIPK1 | Q13546 | | 416 | ERRRRVSHDPFAQ | 5.7850 | 4.6747 | 6.6470 | 4.8195 | 6.1385 | 6.2064 | 5.6190 | 6.3674 | 8.1713 | 7.5623 | 7.9207 | 6.7726 | 6.7174 | 6.8996 | 4.3646 | 5.3340 |
| RIPK2_S531 | RIPK2 | O43353 | | 531 | VVSRSPSLNLLQN | 3.9438 | 3.7740 | 4.5756 | 3.9102 | 4.9013 | 4.6169 | 4.0771 | 4.6263 | 4.6508 | 4.0051 | 4.2129 | 4.1804 | 11.7974 | 11.2182 | 13.3926 | 12.1173 |
| RIPK4_S415 | RIPK4 | P57078 | | 415 | KLPSSGSGKRLSG | 5.7849 | 5.6126 | 5.8423 | 5.5999 | 6.1706 | 6.1252 | 6.2675 | 6.6134 | 5.6028 | 5.7884 | 5.7083 | 5.5151 | 7.3830 | 7.6686 | 7.3249 | 6.9923 |
| RIPK4_S434 | RIPK4 | P57078 | | 434 | AFSSRGSLSLSFE | 7.4216 | 6.9086 | 5.8794 | 5.8263 | 6.7749 | 5.6870 | 5.6450 | 5.8108 | 8.1369 | 6.9429 | 5.8307 | 5.6958 | 6.4830 | 6.8799 | 5.3299 | 4.7473 |
| RIPOR1_S1141 | RIPOR1 | Q6ZS17-4 | | 42 | RVNRSQSFAGVLG | 6.8209 | 6.5751 | 6.8907 | 6.6506 | 6.1665 | 6.6692 | 6.0463 | 6.9207 | 6.6272 | 6.3920 | 6.6720 | 6.5796 | 5.2879 | 5.6447 | 5.1175 | 4.9391 |
| RIPOR1_S367 | RIPOR1 | Q6ZS17-4 | | 61 | RSFPVFSPPGPPR | 6.5634 | 6.7151 | 6.9795 | 6.8296 | 6.3808 | 6.8400 | 6.9904 | 6.9703 | 7.0758 | 6.8291 | 7.0716 | 7.2816 | 4.1160 | 4.8453 | 3.9687 | 4.5429 |
| RIPOR1_S367;T371 | RIPOR1 | Q6ZS17-4 | | 367 | FSTYSQSPPDTPS | 6.2084 | 6.3535 | 6.9262 | 6.7519 | 6.2075 | 6.7732 | 6.2206 | 6.8496 | 6.5984 | 7.0941 | 6.8439 | 6.7984 | 5.3659 | 5.2740 | 4.7882 | 4.9461 |
| RIPOR1_S42 | RIPOR1 | Q6ZS17-4 | | 80 | RVSRMFSVAHPAA | 9.6668 | 6.9866 | 9.3075 | 7.0073 | 6.0358 | 5.1682 | 5.3106 | 4.8407 | 6.5361 | 5.9221 | 6.8062 | 5.9645 | 5.6598 | 5.9595 | 4.4640 | 4.3642 |
| RIPOR1_S61 | RIPOR1 | Q6ZS17-4 | | 1141 | TQLRSLSLGPTFR | 6.0151 | 6.7610 | 6.8492 | 6.1715 | 6.6198 | 6.6019 | 6.6725 | 5.7168 | 5.4551 | 6.5486 | 6.4543 | 6.5873 | 5.7768 | 6.4028 | 5.6788 | 5.6883 |
| RIPOR1_S723;S727 | RIPOR1 | Q6ZS17-4 | 367;371 | | NA | 6.7210 | 6.0456 | 5.8746 | 7.3671 | 5.5000 | 7.5139 | 7.7311 | 6.0653 | 7.1035 | 5.9491 | 6.0016 | 7.1618 | 5.4019 | 5.1356 | 4.5276 | 5.9003 |
| RIPOR1_S80 | RIPOR1 | Q6ZS17-4 | 723;727 | | NA | 6.6296 | 5.7191 | 6.6955 | 7.2158 | 6.8994 | 5.8282 | 5.6268 | 5.8832 | 7.3949 | 6.7900 | 6.7328 | 6.2564 | 5.6065 | 5.6630 | 5.2749 | 5.7843 |
| RLF_S41 | RLF | Q13129 | | 41 | RGHRPVSPAPGAS | 6.7767 | 7.4883 | 6.3115 | 6.6456 | 7.0246 | 5.9478 | 5.7975 | 5.9836 | 6.6814 | 6.1338 | 5.9493 | 6.0561 | 6.0217 | 6.2185 | 5.7129 | 5.2507 |
| RLF_S632 | RLF | Q13129 | | 632 | KGICPKSPSAIPE | 5.8263 | 7.1809 | 6.2792 | 6.7077 | 6.3295 | 5.4789 | 5.8625 | 6.5331 | 6.9930 | 6.9215 | 5.7745 | 7.1526 | 5.4679 | 6.1495 | 5.4883 | 5.8547 |
| RLIM_S230 | RLIM | Q9NVW2 | | 230 | RAERSRSPLHPMS | 6.2758 | 6.9798 | 6.7731 | 6.7266 | 6.5334 | 6.4220 | 6.5170 | 6.6429 | 6.7161 | 6.8989 | 6.5135 | 6.5612 | 4.8793 | 5.2579 | 5.0742 | 5.2284 |

| | | | | | | | | | | | | | | | | | | | | | |
|---|---|---|---|---|---|---|---|---|---|---|---|---|---|---|---|---|---|---|---|---|---|
| RMDN3_S212 | RMDN3 | Q96TC7 | | 212 | KMGRKDSLDLEEE | 5.8717 | 6.2372 | 5.8057 | 6.5576 | 6.1686 | 6.1415 | 5.7217 | 5.5788 | 6.4676 | 6.3774 | 6.0161 | 6.1601 | 6.8731 | 7.1000 | 6.4454 | 6.4776 |
| RMDN3_S46 | RMDN3 | Q96TC7 | | 46 | RHGRSQSLPNSLD | 5.3354 | 5.0519 | 4.6982 | 5.3105 | 5.7868 | 6.1742 | 5.6348 | 5.1870 | 5.5513 | 6.6648 | 6.1200 | 5.8135 | 8.6304 | 7.6609 | 7.9382 | 8.4420 |
| RMDN3_S89 | RMDN3 | H0YMB1 | | 89 | EGQGPHSPQLRKK | 6.5484 | 7.4387 | 6.8912 | 7.0151 | 6.5922 | 6.8505 | 6.6965 | 6.4063 | 7.2502 | 7.2706 | 6.8820 | 7.1572 | 3.9931 | 4.4753 | 4.1033 | 4.4294 |
| RMI1_S225 | RMI1 | Q9H9A7 | | 292 | EPEFVISRPKEE | 6.8329 | 7.1691 | 6.9594 | 7.3951 | 6.7639 | 6.9169 | 6.2719 | 6.4248 | 7.3242 | 7.0325 | 7.2729 | 7.2839 | 4.0589 | 4.5323 | 3.8195 | 3.9419 |
| RMI1_S284 | RMI1 | Q9H9A7 | | 225 | SVIPNSNSNENIPR | 6.4451 | 6.3896 | 6.5260 | 6.8261 | 6.2769 | 6.4192 | 6.2522 | 6.8812 | 6.7207 | 6.6359 | 6.8474 | 6.7260 | 4.9687 | 5.7785 | 4.9399 | 5.3666 |
| RMI1_S292 | RMI1 | Q9H9A7 | | 284 | IPTRQSSFEPEFV | 4.4973 | 4.7664 | 4.7323 | 4.8583 | 5.8406 | 5.0246 | 5.3622 | 5.2248 | 4.6972 | 4.6116 | 4.7151 | 4.7699 | 10.3722 | 9.5692 | 10.5826 | 10.3758 |
| RMI1_S388 | RMI1 | Q9H9A7 | | 388 | MTNEDKSFGCPSV | 7.4783 | 8.0680 | 7.0596 | 8.2561 | 6.0282 | 6.9244 | 7.6297 | 6.7140 | 7.3756 | 6.7837 | 7.2621 | 6.8222 | 2.7268 | 3.9704 | 2.9188 | 3.9820 |
| RNF10_S128 | RNF10 | Q8N5U6-2 | | 128 | AQRAEFSPAQFSG | 7.3936 | 8.1253 | 7.4863 | 7.4288 | 5.9016 | 7.5283 | 7.1654 | 7.2061 | 9.0358 | 6.5968 | 8.2763 | 7.3144 | 2.4857 | 3.2250 | 2.4031 | 2.4275 |
| RNF113A_S253 | RNF113A | O15541 | | 253 | ENYEVGSDDEEIP | 5.4364 | 5.8470 | 6.8781 | 6.6740 | 5.8786 | 6.0340 | 5.6701 | 6.6363 | 6.0395 | 5.5304 | 6.3898 | 6.7360 | 5.7748 | 6.8036 | 6.6429 | 6.0282 |
| RNF123_S675 | RNF123 | Q5XPI4-2 | | 675 | RPGWLSSPTLGRA | 6.3061 | 5.8372 | 7.0642 | 6.0483 | 6.2156 | 6.5835 | 6.3926 | 6.1139 | 6.6031 | 6.6552 | 6.6058 | 6.7745 | 5.8227 | 6.0532 | 5.2230 | 5.7013 |
| RNF13_T380 | RNF13 | O43567 | | 380 | DYNIANTVxxxxx | 6.9497 | 7.0284 | 7.3746 | 7.5462 | 6.0806 | 6.6776 | 6.6616 | 6.0048 | 6.3117 | 6.8128 | 6.6863 | 6.9049 | 4.0394 | 5.3375 | 4.6221 | 4.9620 |
| RNF14_S348 | RNF14 | Q9UBS8 | | 348 | LTYHGVSPCKVTA | 5.9530 | 6.3525 | 7.3090 | 6.6614 | 6.5944 | 7.0692 | 6.6377 | 6.9986 | 6.1344 | 6.4638 | 6.6002 | 6.7055 | 4.9137 | 5.5318 | 4.8448 | 5.2300 |
| RNF168_S197 | RNF168 | Q8IYW5 | | 204 | PLNSRKSDPVTPK | 5.9737 | 6.6728 | 6.0363 | 7.3485 | 6.3762 | 6.7628 | 6.8730 | 6.8944 | 6.6967 | 7.2209 | 6.9112 | 6.6876 | 4.8045 | 5.3411 | 4.5973 | 4.8029 |
| RNF168_S204 | RNF168 | Q8IYW5 | | 208 | RKSDPVTPKSEKK | 6.5565 | 6.3769 | 5.8770 | 6.6785 | 6.1300 | 6.2125 | 7.0945 | 7.3753 | 7.4524 | 6.0574 | 6.3220 | 6.7739 | 5.1783 | 5.9889 | 4.0655 | 5.8603 |
| RNF168_S481 | RNF168 | Q8IYW5 | | 197 | EGSISASPLNSRK | 5.3984 | 6.4770 | 7.7738 | 7.4555 | 5.8574 | 7.7949 | 7.4513 | 6.3629 | 7.2049 | 6.6417 | 5.5082 | 5.8335 | 4.4635 | 5.5153 | 4.7817 | 5.4801 |
| RNF168_T208 | RNF168 | Q8IYW5 | | 481 | HLRATSSPPDKVL | 6.9721 | 7.3256 | 6.7699 | 7.1597 | 6.2413 | 6.6453 | 6.8135 | 6.5761 | 6.8366 | 6.7921 | 6.6678 | 7.1053 | 4.2302 | 5.0250 | 4.0980 | 4.7416 |
| RNF169_S12 | RNF169 | Q8NCN4 | | 403 | PDGRVLSPLIIKS | 7.2379 | 7.2848 | 6.4604 | 6.6168 | 6.1699 | 6.6228 | 6.9148 | 6.6752 | 6.6999 | 6.4315 | 6.5513 | 6.6424 | 4.7230 | 5.0089 | 4.7026 | 5.2577 |
| RNF169_S368 | RNF169 | Q8NCN4 | | 693 | VSRRKGSVDQYLL | 4.1020 | 3.9017 | 4.5455 | 5.0786 | 7.4207 | 5.0073 | 4.6894 | 5.7870 | 7.3711 | 8.5693 | 7.5426 | 7.9500 | 8.0378 | 7.9172 | 5.7433 | 6.3365 |
| RNF169_S368;S388;T391 | RNF169 | Q8NCN4 | | 368 | KPERSVSPESNDS | 5.9713 | 5.7448 | 5.4961 | 6.3868 | 5.7497 | 5.2843 | 5.9552 | 5.6081 | 5.8286 | 6.0925 | 5.3403 | 5.7892 | 6.9076 | 7.6694 | 8.1123 | 8.0635 |
| RNF169_S388;T391 | RNF169 | Q8NCN4 | | 391 | IVCSPCTPPKRLP | 5.0653 | 4.3184 | 4.7335 | 5.0748 | 5.5662 | 4.9494 | 4.7846 | 4.6142 | 5.4976 | 4.7387 | 4.3088 | 4.7955 | 11.0854 | 9.7451 | 10.2423 | 10.4801 |
| RNF169_S391 | RNF169 | Q8NCN4 | | 485 | AVNTRLSGGQVLS | 6.3426 | 7.2369 | 5.8775 | 7.0221 | 6.6685 | 6.2965 | 7.3902 | 7.1404 | 6.9930 | 6.9372 | 7.6433 | 6.6666 | 3.9861 | 5.2815 | 3.8425 | 4.6752 |
| RNF169_S403 | RNF169 | Q8NCN4 | | 554 | FEGLGSTPDAKLD | 6.9579 | 7.3261 | 7.2416 | 7.8098 | 6.2500 | 6.4984 | 7.7208 | 6.4888 | 6.7876 | 6.6778 | 6.7125 | 7.0784 | 3.7284 | 4.4561 | 3.8010 | 4.4648 |
| RNF169_S485 | RNF169 | Q8NCN4 | | 12 | PSTRASSAAAAAA | 4.4697 | 4.9482 | 4.2350 | 4.8088 | 4.3453 | 5.4939 | 4.9943 | 5.3655 | 4.1979 | 4.7526 | 3.9702 | 4.6192 | 10.8082 | 11.0870 | 10.4561 | 11.4481 |
| RNF169_S693 | RNF169 | Q8NCN4 | 368;388;391 | | NA | 6.3216 | 6.3334 | 6.5812 | 7.9788 | 6.1234 | 6.7609 | 6.8165 | 6.4825 | 6.7524 | 6.6831 | 6.9289 | 7.2892 | 4.5951 | 4.7967 | 4.6718 | 4.8845 |
| RNF169_T554 | RNF169 | Q8NCN4 | 388;391 | | NA | 7.0266 | 5.8652 | 7.2103 | 6.8578 | 5.7113 | 6.5229 | 6.9418 | 6.1950 | 7.5219 | 7.4018 | 6.7830 | 7.2261 | 4.0158 | 4.6285 | 4.4437 | 5.6482 |
| RNF185_T106 | RNF185 | Q96GF1 | | 106 | QDPREKTPPRPQG | 6.3608 | 6.2539 | 6.3052 | 5.6142 | 5.9836 | 5.4620 | 7.1233 | 7.3684 | 6.2782 | 6.9814 | 6.0205 | 6.6661 | 4.3825 | 5.7351 | 6.1780 | 7.2869 |
| RNF19A_S599 | RNF19A | Q9NV58 | | 66 | PKKRRISIGSLFR | 6.6601 | 5.3279 | 5.5288 | 5.4315 | 5.8007 | 6.2067 | 6.1762 | 6.6900 | 8.9362 | 7.4612 | 7.4752 | 7.7291 | 5.5974 | 5.8826 | 3.8669 | 5.2296 |
| RNF19A_S66 | RNF19A | Q9NV58 | | 599 | TKAMAGSILNSYI | 6.8270 | 6.0044 | 6.2570 | 7.2373 | 6.4401 | 5.6900 | 6.1250 | 5.8908 | 6.7152 | 6.4357 | 5.7724 | 6.4729 | 5.7562 | 6.5505 | 6.2221 | 5.6033 |
| RNF19A_S69 | RNF19A | Q9NV58 | | 69 | RRISIGSLFRRKK | 13.2800 | 12.7952 | 12.2659 | 10.7578 | 4.7036 | 5.8861 | 7.6807 | 5.9541 | 4.0476 | 3.9116 | 4.3224 | 3.5175 | 2.2406 | 2.7891 | 2.2814 | 3.5663 |
| RNF19B_S14 | RNF19B | Q6ZMZ0 | | 42 | HSVFSASARGRRA | 5.5479 | 8.1578 | 9.3588 | 6.2962 | 8.2271 | 7.0445 | 5.0721 | 5.3585 | 5.1737 | 7.5065 | 7.9222 | 5.8712 | 4.4669 | 6.3285 | 3.7324 | 3.9357 |
| RNF19B_S42 | RNF19B | Q6ZMZ0 | | 14 | ESPRSTSLHAAAP | 7.7989 | 7.2570 | 7.6470 | 7.1780 | 5.9557 | 6.8328 | 6.2320 | 6.5117 | 6.9358 | 8.2900 | 8.9382 | 8.2070 | 3.0789 | 3.6496 | 2.4941 | 2.9932 |
| RNF20_S524 | RNF20 | Q5VTR2 | | 525 | LLQSQSSTEDPKD | 3.9942 | 4.4606 | 4.7910 | 4.5234 | 5.5939 | 5.4483 | 5.7151 | 4.9919 | 4.6783 | 5.2221 | 5.1281 | 4.9415 | 11.1158 | 9.4226 | 10.5512 | 9.4220 |
| RNF20_S524;S525 | RNF20 | Q5VTR2 | | 524 | ALLQSQSSTEDPK | 4.2515 | 4.1027 | 4.6218 | 4.0162 | 5.9054 | 5.4417 | 5.5997 | 5.0055 | 4.8446 | 5.2194 | 5.3517 | 4.6491 | 10.9837 | 9.2847 | 10.8186 | 9.8538 |
| RNF20_S524;T526 | RNF20 | Q5VTR2 | | 526 | LQSQSSTEDPKDE | 4.4066 | 4.1741 | 5.2790 | 4.7450 | 5.0101 | 6.0663 | 6.1539 | 5.3413 | 4.7574 | 5.0107 | 5.1627 | 5.5440 | 9.1120 | 9.6707 | 10.2813 | 9.2848 |
| RNF20_S525 | RNF20 | Q5VTR2 | 524;526 | | NA | 4.7469 | 5.1171 | 5.5490 | 5.3071 | 6.6280 | 6.0949 | 6.5261 | 6.4745 | 5.0516 | 5.1526 | 5.1264 | 4.8055 | 9.3004 | 7.4877 | 8.4293 | 8.2028 |
| RNF20_S526 | RNF20 | Q5VTR2 | 524;525 | | NA | 5.2423 | 5.0660 | 6.1005 | 5.3515 | 6.0327 | 5.4314 | 6.7717 | 6.4168 | 5.0129 | 5.3293 | 5.1776 | 5.3735 | 8.1728 | 7.7480 | 8.7302 | 8.0428 |
| RNF213_S88 | RNF213 | Q63HN8 | | 303 | KQPPATTPPFKTH | 6.7405 | 7.1846 | 6.7454 | 6.9664 | 6.1621 | 6.9110 | 6.5166 | 6.4754 | 6.4941 | 6.7721 | 7.1213 | 6.9435 | 4.2116 | 5.4602 | 4.4242 | 4.8710 |
| RNF213_T303 | RNF213 | Q63HN8 | | 88 | SWTVQESKKKKRK | 5.3527 | 8.1465 | 7.9610 | 6.4891 | 5.3126 | 5.8025 | 6.0470 | 5.9743 | 7.7592 | 8.0906 | 8.5251 | 8.0667 | 3.8006 | 5.0717 | 3.2298 | 3.4708 |
| RNF214_S176 | RNF214 | Q8ND24 | | 176 | DFRPVVSPANGVE | 6.2809 | 6.2011 | 6.7381 | 6.6075 | 6.3428 | 6.8576 | 5.9886 | 6.7627 | 6.7154 | 6.8252 | 6.7824 | 6.9937 | 5.5704 | 5.4047 | 4.9254 | 5.0034 |
| RNF214_S200 | RNF214 | Q8ND24-2 | | 47 | AQKQKNSPLLSTD | 6.2376 | 6.5369 | 6.7218 | 5.2089 | 6.2005 | 7.1473 | 5.7768 | 6.2657 | 6.2305 | 7.7472 | 6.6703 | 6.2395 | 6.7827 | 6.5285 | 5.1832 | 5.4226 |
| RNF214_S47 | RNF214 | Q8ND24 | | 200 | SSSLKLSQNIAVQ | 5.9226 | 5.7962 | 7.2180 | 5.6502 | 6.3297 | 6.0231 | 6.2556 | 6.1574 | 6.4598 | 6.0316 | 6.0823 | 6.7670 | 6.7582 | 6.3755 | 5.9376 | 6.2354 |
| RNF216_S476 | RNF216 | Q9NWF9-1 | | 476 | KKWQELSPETSGK | 6.4513 | 7.0876 | 6.8877 | 6.6603 | 5.6433 | 6.2922 | 5.9019 | 6.8718 | 7.0153 | 6.9826 | 6.7967 | 7.0097 | 4.7282 | 5.3887 | 4.9127 | 5.3700 |
| RNF220_S390 | RNF220 | Q5VTB9 | | 390 | GKENPDSDADLDV | 5.7354 | 6.0610 | 6.5427 | 6.3529 | 6.4751 | 6.6338 | 5.5440 | 6.5357 | 6.4588 | 6.5067 | 6.8712 | 6.2188 | 6.1591 | 6.6281 | 5.5039 | 5.7729 |
| RNF26_T348 | RNF26 | Q9BY78 | | 348 | ARTIRVTPVRGRE | 6.8012 | 7.3377 | 7.2318 | 7.7214 | 5.5323 | 6.0155 | 5.5335 | 6.2368 | 6.8922 | 7.4002 | 7.5788 | 7.3583 | 4.0938 | 4.9957 | 4.2183 | 4.0526 |
| RNF34_S255 | RNF34 | Q969K3-2 | | 255 | KERVRASLSDLSS | 6.3555 | 5.9014 | 6.6308 | 6.1013 | 5.7455 | 5.7817 | 6.1242 | 7.0062 | 6.5669 | 6.3431 | 6.5777 | 6.3874 | 6.2409 | 6.2481 | 5.8619 | 6.1275 |
| RNF4_S90;S95 | RNF4 | P78317 | | 94 | ADSCVVSSDDEEL | 6.4149 | 6.7476 | 6.2626 | 7.5699 | 5.2083 | 4.9154 | 7.0334 | 5.3207 | 7.5910 | 5.8549 | 6.7335 | 5.9650 | 5.7097 | 7.6824 | 5.5925 | 5.3981 |
| RNF4_S94 | RNF4 | P78317 | 94;95 | | NA | 7.3781 | 6.7101 | 6.9138 | 6.3376 | 6.2014 | 6.4758 | 6.2267 | 7.9877 | 6.6577 | 5.7936 | 6.6291 | 6.7405 | 5.0212 | 5.1346 | 5.0944 | 4.6976 |
| RNF4_S94;S95 | RNF4 | P78317 | 90;95 | | NA | 6.0233 | 6.1359 | 6.5643 | 6.3867 | 5.2784 | 7.4942 | 6.7586 | 6.2920 | 6.4498 | 6.8732 | 7.1754 | 6.3979 | 4.8023 | 5.7368 | 5.3878 | 6.2435 |
| RNF6_S191 | RNF6 | Q9Y252 | | 191 | RQQRSTSPVARRT | 7.0529 | 7.3096 | 6.3819 | 7.4675 | 6.1079 | 6.5926 | 6.6862 | 6.0278 | 6.4797 | 6.9089 | 6.5245 | 6.7215 | 4.9449 | 4.9859 | 4.9745 | 5.5336 |
| RNF8_S157 | RNF8 | O76064 | | 157 | RTKRKFSLDELAG | 5.3017 | 5.7313 | 6.8061 | 6.1061 | 5.9537 | 5.9260 | 6.2930 | 5.6530 | 6.1024 | 6.2366 | 6.5728 | 6.7377 | 6.8949 | 6.6187 | 6.2449 | 6.8213 |
| RNH1_T82 | RNH1 | P13489 | | 82 | VLQGLQTPSCKIQ | 7.4641 | 6.4671 | 7.8500 | 7.5331 | 6.7660 | 6.4608 | 7.3425 | 5.1109 | 6.1095 | 6.1416 | 7.1534 | 7.2844 | 4.3846 | 4.5188 | 4.6592 | 4.7541 |
| RNMT_T77 | RNMT | O43148-2 | | 77 | SSCGKDTPSKKRK | 7.5189 | 7.5480 | 6.4832 | 6.9742 | 6.5778 | 6.4898 | 6.5869 | 6.1087 | 6.6921 | 6.4087 | 6.9975 | 6.9669 | 4.4975 | 5.2911 | 4.1721 | 4.6865 |
| RNPC3_S108 | RNPC3 | Q96LT9 | | 108 | EQDRVHSPCPTSG | 6.4754 | 6.8812 | 5.9988 | 6.6119 | 5.6196 | 6.3717 | 6.3593 | 5.8114 | 6.7039 | 6.4256 | 6.6309 | 6.2922 | 5.8695 | 6.2373 | 5.6041 | 6.1073 |
| RNPS1_S132;S134 | RNPS1 | Q15287-2 | | 4 | xxxMAPSPTKRKD | 6.4003 | 6.9274 | 5.9996 | 6.9731 | 6.1827 | 6.4214 | 6.8020 | 6.4632 | 6.0043 | 6.3790 | 6.3197 | 6.4093 | 5.5305 | 5.6456 | 5.6817 | 5.8602 |
| RNPS1_S4 | RNPS1 | Q15287-2 | 132;134 | | NA | 6.1364 | 6.2993 | 5.7422 | 6.3190 | 6.2264 | 5.5702 | 6.3703 | 6.0754 | 6.3645 | 5.9647 | 5.9055 | 5.8412 | 7.0517 | 6.5343 | 6.9376 | 6.6613 |
| RO60_S19 | RO60 | P10155 | | 19 | EKQIANSQDGYVW | 5.5861 | 5.7541 | 6.4155 | 6.8112 | 6.6260 | 6.2637 | 5.6643 | 6.7353 | 6.7583 | 6.3177 | 6.2004 | 6.3005 | 5.1996 | 6.1233 | 6.5432 | 6.7008 |
| ROBO1_S1059 | ROBO1 | Q9Y6N7-2 | | 1446 | QCPRPTSPVSTDS | 6.2130 | 5.7035 | 6.3286 | 6.1746 | 5.6116 | 5.6430 | 6.2743 | 6.1179 | 7.2013 | 6.4128 | 6.2530 | 6.5291 | 6.3804 | 6.3103 | 6.2955 | 6.5510 |
| ROBO1_S1446 | ROBO1 | Q9Y6N7-2 | | 1496 | PPPAIKSPTAQSK | 6.1769 | 6.5175 | 6.1471 | 6.6199 | 6.1807 | 6.7828 | 7.0361 | 6.7286 | 6.2494 | 6.3900 | 6.7843 | 6.8080 | 4.8992 | 5.5511 | 5.2754 | 5.8528 |
| ROBO1_S1496 | ROBO1 | Q9Y6N7-2 | | 1059 | EMKTFNSPNLKDG | 6.0208 | 6.6100 | 5.6191 | 5.6171 | 5.6759 | 6.8442 | 7.0697 | 6.4097 | 5.8013 | 6.0065 | 5.9806 | 6.0861 | 5.8682 | 6.3310 | 6.9568 | 7.1030 |
| ROBO1_S944 | ROBO1 | Q9Y6N7-2 | | 944 | GIRKVPSFTFTPT | 5.9787 | 6.6322 | 6.4358 | 5.5406 | 6.3592 | 6.4023 | 5.5987 | 5.7898 | 6.0036 | 6.4768 | 6.2584 | 5.9179 | 6.7491 | 7.0036 | 6.5765 | 6.2767 |
| ROBO1_T1244 | ROBO1 | Q9Y6N7-2 | | 1244 | EDERGPTPPVRGA | 5.4971 | 6.2363 | 7.2142 | 6.7413 | 6.3194 | 6.0818 | 5.8857 | 7.2297 | 7.2297 | 5.3153 | 5.4033 | 6.7140 | 5.7324 | 5.5114 | 6.1780 | 6.7104 |
| ROCK1_S1098 | ROCK1 | Q13464 | | 1098 | AKLLDLSDSTSVA | 6.5235 | 6.1033 | 7.1455 | 5.7160 | 5.6875 | 5.6989 | 6.1439 | 6.5070 | 6.2755 | 6.7835 | 5.5141 | 7.2585 | 5.9870 | 6.0166 | 5.8959 | 5.4735 |
| ROCK1_S1147 | ROCK1 | Q13464 | | 1105 | DSTSVASFPSADE | 7.2365 | 5.6470 | 6.2559 | 6.8060 | 5.9147 | 6.1991 | 5.7931 | 7.3140 | 5.7666 | 5.5745 | 6.8149 | 6.7737 | 5.4921 | 6.8109 | 6.1540 | 5.4470 |
| ROCK1_T1101 | ROCK1 | Q13464 | | 1101 | LDLSDSTSVASFP | 5.9971 | 6.5094 | 6.6512 | 6.0979 | 5.7422 | 5.7143 | 6.6473 | 6.4140 | 6.5094 | 6.7597 | 6.3273 | 6.0072 | 6.5814 | 6.2792 | 5.8724 | 5.8902 |
| ROCK1_T1105 | ROCK1 | Q13464 | | 1147 | QYVVVSSKKILFY | 6.3587 | 5.0548 | 6.1583 | 6.8065 | 7.6181 | 5.8342 | 6.3018 | 6.2226 | 5.7253 | 7.3484 | 5.2998 | 4.9336 | 5.1736 | 7.3979 | 6.3018 | 7.4647 |
| ROCK2_S1133;S1134 | ROCK2 | O75116 | | 1374 | KIQQNQSIRRPSR | 11.2213 | 7.8723 | 7.9011 | 6.9410 | 5.9407 | 5.9807 | 6.5185 | 5.8419 | 7.0790 | 4.8716 | 5.8635 | 6.0618 | 5.3491 | 5.4129 | 3.2929 | 3.8518 |
| ROCK2_S1374 | ROCK2 | O75116 | 1133;1134 | | NA | 6.2645 | 6.3143 | 7.9447 | 6.8391 | 6.9271 | 5.0806 | 7.0926 | 7.5330 | 5.6263 | 6.3043 | 5.3968 | 6.2801 | 5.8490 | 5.2307 | 5.4669 | 5.8499 |
| RPA1_S135 | RPA1 | P27694 | | 135 | PPAPAASPAASSR | 6.3736 | 6.3824 | 6.2873 | 6.7273 | 6.0717 | 6.7439 | 6.8982 | 6.8469 | 7.6757 | 7.4828 | 6.8624 | 7.3465 | 4.2772 | 4.6881 | 4.2761 | 5.0599 |
| RPA1_S38 | RPA1 | P27694 | | 384 | IKGARVSDFGGRS | 6.7234 | 7.3338 | 6.7235 | 7.1598 | 6.5075 | 7.0708 | 7.6457 | 6.9342 | 6.8861 | 7.0222 | 7.1293 | 6.7615 | 3.5434 | 4.4603 | 3.6619 | 4.4367 |
| RPA1_S384 | RPA1 | P27694 | | 38 | PITTGNSPPRYRL | 5.4914 | 4.9863 | 5.8652 | 5.6309 | 5.0565 | 5.8482 | 5.8228 | 5.5579 | 6.0291 | 6.3017 | 5.9295 | 6.1226 | 7.2875 | 7.8604 | 7.9577 | 8.2524 |
| RPA1_S395 | RPA1 | P27694 | | 395 | RSLSVLSSSTIIA | 5.5693 | 5.0597 | 5.5512 | 5.7128 | 6.2432 | 6.0171 | 6.4394 | 6.0587 | 6.3874 | 5.7017 | 5.9811 | 5.2864 | 6.4089 | 8.3536 | 7.7511 | 7.4786 |
| RPA2_S326 | RPA2 | P15927-3 | | 326 | NQLKHMSVSSIKQ | 4.9265 | 4.9628 | 5.7523 | 5.0331 | 5.5283 | 6.3330 | 5.7187 | 6.1802 | 5.2668 | 5.5739 | 5.8905 | 5.8603 | 7.9917 | 8.3472 | 8.7687 | 7.8659 |
| RPA2_S328 | RPA2 | P15927-3 | | 328 | LKHMSVSSIKQAV | 5.4533 | 5.7483 | 6.7330 | 6.0544 | 6.2191 | 6.8416 | 5.6724 | 6.4992 | 6.1455 | 6.1181 | 6.6273 | 6.6751 | 5.9482 | 6.5959 | 6.5336 | 6.1347 |
| RPAIN_S18 | RPAIN | Q86UA6-8 | | 18 | LYKLVGSPPWKEA | 6.8727 | 6.9188 | 7.1327 | 6.5478 | 6.4134 | 6.9147 | 6.8637 | 7.2014 | 7.2470 | 7.5126 | 7.8289 | 7.8096 | 3.4098 | 4.1477 | 3.2490 | 3.9301 |
| RPAP1_S601 | RPAP1 | Q9BWH6 | | 72 | PPALVPSPPKRAR | 5.8498 | 6.2583 | 6.8755 | 6.1105 | 6.4852 | 6.9300 | 6.7894 | 6.8971 | 6.6893 | 6.7573 | 6.9075 | 7.3837 | 4.8301 | 5.3370 | 4.5269 | 5.3723 |
| RPAP1_S72 | RPAP1 | Q9BWH6 | | 601 | REFLPTSWSPVGA | 5.9189 | 6.5835 | 7.3053 | 6.1120 | 5.6948 | 6.6736 | 5.7745 | 6.6681 | 6.4385 | 5.6872 | 5.7161 | 6.2667 | 6.0852 | 6.3258 | 6.8269 | 5.9230 |
| RPAP1_T321 | RPAP1 | Q9BWH6 | | 321 | ALALPVTPQKEWL | 6.5562 | 6.3829 | 7.1497 | 6.6603 | 6.4306 | 7.3422 | 7.0667 | 6.9510 | 7.0353 | 6.9634 | 6.7677 | 7.1662 | 4.2902 | 4.5898 | 3.8902 | 4.7575 |
| RPGR_S518 | RPGR | Q92834 | | 961 | LFKRVPSINQKIV | 7.1745 | 7.0384 | 6.2963 | 6.3493 | 6.7767 | 6.6926 | 5.8638 | 6.4952 | 6.7866 | 6.6074 | 5.7402 | 6.3322 | 5.9043 | 6.1130 | 4.8579 | 4.9716 |
| RPGR_S961 | RPGR | Q92834 | | 518 | EKSLKLSPVQKQK | 6.5125 | 6.8679 | 6.5388 | 6.7574 | 7.3939 | 8.3045 | 6.8003 | 7.1787 | 6.9535 | 6.2083 | 7.1167 | 6.8514 | 4.4118 | 4.8526 | 3.4506 | 3.2011 |
| RPL12_S126 | RPL12 | P30050 | | 38 | IGPLGLSPKKVGD | 7.5782 | 7.6893 | 7.7459 | 8.0308 | 5.4918 | 5.4975 | 5.7277 | 5.1106 | 7.1215 | 6.7618 | 7.2920 | 7.0072 | 4.7438 | 4.7968 | 4.7486 | 4.6566 |
| RPL12_S38 | RPL12 | P30050 | | 126 | SLARELSGTIKEI | 6.2775 | 6.2765 | 6.3421 | 5.7172 | 6.4197 | 6.3096 | 5.9950 | 6.4108 | 6.3603 | 6.1938 | 6.0687 | 6.0715 | 6.4383 | 6.4798 | 6.5808 | 6.0583 |
| RPL13_S77 | RPL13 | P26373 | | 77 | RAGRGFSLEELRV | 6.4519 | 6.0384 | 6.5562 | 6.0806 | 6.7558 | 6.1829 | 5.7802 | 7.3774 | 8.1471 | 8.2971 | 7.5011 | 7.1832 | 4.3299 | 5.0078 | 4.2811 | 4.0292 |
| RPL15_S187 | RPL15 | P61313 | | 34 | WQYRQLSALHRAP | 6.2827 | 6.4555 | 5.8706 | 6.1335 | 6.0626 | 5.9834 | 6.3230 | 6.6977 | 6.6870 | 6.1881 | 6.1610 | 6.3738 | 6.0188 | 6.3503 | 6.0242 | 6.3879 |
| RPL15_S34 | RPL15 | P61313 | | 187 | HHTIGGSRRAAWR | 6.0794 | 7.1063 | 7.5319 | 7.0985 | 4.7548 | 5.9497 | 6.9146 | 8.4600 | 8.6570 | 8.9408 | 8.2201 | 8.1116 | 2.8877 | 3.6681 | 2.1997 | 3.4198 |
| RPL15_T183 | RPL15 | P61313 | | 197 | AWRRRNTLQLHRY | 7.1817 | 5.9515 | 7.1441 | 5.2337 | 6.6340 | 6.3459 | 6.2617 | 6.2795 | 9.7606 | 8.5025 | 7.6683 | 6.3742 | 5.6811 | 4.6426 | 2.7409 | 3.5976 |
| RPL15_T197 | RPL15 | P61313 | | 183 | GHKFHHTIGGSRR | 7.2178 | 7.6689 | 8.8250 | 5.9481 | 5.1056 | 5.8525 | 8.4337 | 4.8815 | 5.2460 | 6.0347 | 5.4671 | 5.1624 | 5.7479 | 5.7270 | 5.9003 | 6.7816 |
| RPL17_S141 | RPL17 | P18621-3 | | 142 | INPYMSSPCHIEM | 6.3021 | 6.6531 | 6.6824 | 6.6552 | 6.4072 | 6.6186 | 6.2464 | 6.4174 | 6.3368 | 6.5516 | 6.6188 | 6.2506 | 5.2988 | 6.0147 | 5.5405 | 5.4057 |
| RPL17_S142 | RPL17 | P18621-3 | | 141 | RINPYMSSPCHIE | 6.8338 | 6.2282 | 6.7610 | 5.8177 | 7.0044 | 7.2177 | 6.1897 | 6.7780 | 6.8961 | 6.1800 | 5.6655 | 5.9853 | 4.6144 | 5.8968 | 5.7555 | 5.2759 |
| RPL17_S87 | RPL17 | P18621-3 | | 87 | GRWPKKSAEFLLH | 4.0373 | 5.5072 | 8.9051 | 4.9786 | 5.7589 | 5.1800 | 6.3529 | 7.5862 | 7.7876 | 8.8145 | 8.9152 | 6.9922 | 3.5792 | 5.4820 | 4.6363 | 5.4870 |
| RPL18_S130 | RPL18 | Q07020 | | 130 | DQLALDSPKGCGT | 6.5011 | 6.6236 | 6.0242 | 6.5051 | 6.3262 | 6.7265 | 6.9006 | 6.7567 | 6.6939 | 6.8118 | 6.5361 | 6.9511 | 4.9328 | 5.9314 | 5.4583 | 5.2307 |
| RPL18A_S123 | RPL18A | Q02543 | | 123 | HRARAHSIQIMKV | 6.5663 | 6.4428 | 6.7933 | 6.3894 | 6.5917 | 6.4592 | 6.4610 | 6.3528 | 6.7466 | 6.5087 | 6.3966 | 6.1550 | 5.2443 | 5.7439 | 5.5240 | 5.6243 |
| RPL18A_S49 | RPL18A | Q02543 | | 49 | RFWYFVSQLKKMK | 5.9659 | 6.5941 | 6.6279 | 5.8816 | 4.1783 | 7.4205 | 7.3868 | 7.3573 | 5.7256 | 7.1338 | 8.1246 | 7.4838 | 4.6294 | 5.3293 | 4.5535 | 5.6076 |
| RPL19_S13 | RPL19 | P84098 | | 13 | QKRLASVLRCGK | 6.6779 | 8.0300 | 7.8331 | 7.4028 | 5.9678 | 6.1257 | 7.6035 | 6.1930 | 7.1830 | 6.4062 | 6.7297 | 6.0136 | 4.2324 | 4.9965 | 3.7117 | 4.8930 |
| RPL21_S104 | RPL21 | P46778 | | 124 | EAKEKGTWVQLKR | 7.4127 | 6.4596 | 7.2309 | 8.7563 | 9.0511 | 5.0701 | 6.6950 | 4.2796 | 6.4041 | 4.9935 | 4.8767 | 3.4470 | 7.9161 | 7.3955 | 5.0854 | 4.9265 |
| RPL21_T124 | RPL21 | P46778 | | 104 | HSKSRDSFLKRVK | 5.7399 | 6.8922 | 7.0680 | 6.4108 | 5.6487 | 6.9768 | 5.8153 | 7.0627 | 6.6633 | 7.1132 | 7.2332 | 7.8661 | 5.1332 | 4.6748 | 4.2970 | 5.4047 |
| RPL23_S41 | RPL23 | P62829 | | 41 | KNLYIISVKGIKG | 6.9250 | 7.3349 | 7.7355 | 9.1725 | 4.2722 | 5.4053 | 6.8938 | 5.4583 | 4.9814 | 5.0094 | 7.7604 | 6.9624 | 5.4349 | 6.5416 | 5.3866 | 4.7258 |
| RPL23_S9 | RPL23 | P62829 | | 9 | KRGRGGSSGAKFR | 5.9635 | 6.7853 | 7.4268 | 7.5939 | 5.4561 | 5.7040 | 6.1457 | 6.2866 | 6.5294 | 5.8266 | 6.8075 | 7.4992 | 6.0713 | 6.0053 | 4.4990 | 5.4000 |
| RPL23A_S43 | RPL23A | P62750 | | 85 | FPLTTESAMKKIE | 5.9756 | 5.9568 | 5.7595 | 6.3029 | 6.1255 | 6.5131 | 6.5071 | 6.1244 | 6.0237 | 6.4141 | 5.7017 | 5.4124 | 6.7814 | 6.8658 | 7.3283 | 6.2077 |
| RPL23A_S85 | RPL23A | P62750 | | 43 | KKKIRTSPTFRRP | 6.1982 | 6.6652 | 6.6885 | 6.4238 | 6.4423 | 6.7783 | 6.5946 | 5.6683 | 5.3162 | 6.2628 | 6.3377 | 6.5660 | 5.9414 | 6.7348 | 5.7829 | 5.5989 |
| RPL24_S42 | RPL24 | P83731 | | 42 | CESAFLSKRNPRQ | 6.4475 | 6.4189 | 7.4472 | 7.9860 | 5.4483 | 5.3490 | 5.9806 | 4.9200 | 5.5947 | 5.5768 | 6.4977 | 6.7650 | 4.9954 | 7.6436 | 6.4632 | 6.4660 |
| RPL24_S86 | RPL24 | P83731 | | 86 | RAITGASLADIMA | 6.3719 | 6.1703 | 6.3455 | 6.1475 | 6.2302 | 5.9062 | 6.0769 | 5.6446 | 6.2285 | 5.4670 | 5.5880 | 5.6870 | 6.8885 | 7.9471 | 6.4535 | 6.8472 |
| RPL24_T83 | RPL24 | P83731 | | 83 | KFQRAITGASLAD | 7.0689 | 6.6077 | 7.4787 | 7.6379 | 5.9887 | 5.0360 | 6.6989 | 4.7991 | 6.5532 | 5.3949 | 5.8188 | 5.9413 | 6.3113 | 6.8068 | 6.1112 | 5.7467 |
| RPL26_S32 | RPL26 | P61254 | | 32 | RRKIMSSPLSKEL | 8.1986 | 7.9703 | 7.0464 | 7.5541 | 7.0824 | 7.0926 | 7.3555 | 6.6585 | 7.7180 | 7.2895 | 7.0581 | 6.7243 | 2.7571 | 3.5945 | 2.7108 | 3.1893 |
| RPL26_T139 | RPL26 | P61254 | | 139 | GKYKEETIEKMQE | 6.9477 | 7.3044 | 6.4625 | 6.8966 | 6.6837 | 6.9538 | 7.3944 | 6.6284 | 6.7934 | 6.7354 | 7.0910 | 6.7000 | 4.0141 | 4.8770 | 3.9017 | 4.6158 |
| RPL27A_S68 | RPL27A | P46776 | | 68 | HLKRNQSFCPTVN | 7.0909 | 6.7083 | 6.6399 | 7.4324 | 7.6063 | 5.7433 | 6.5725 | 5.7188 | 8.1736 | 6.6544 | 6.0663 | 5.4528 | 6.4614 | 5.3040 | 4.0006 | 4.3743 |
| RPL28_S115 | RPL28 | P46779 | | 115 | AAIRRASAILRSQ | 5.7256 | 5.7603 | 6.8890 | 5.5342 | 5.4581 | 6.5264 | 6.0043 | 7.4356 | 7.2434 | 8.4784 | 8.1809 | 7.0999 | 4.9750 | 5.3150 | 4.3633 | 5.0109 |
| RPL28_S15 | RPL28 | P46779 | | 15 | VVRNCSSFLIKRN | 5.2501 | 6.9036 | 8.2738 | 7.7768 | 5.5372 | 6.7252 | 7.3804 | 6.4525 | 7.1034 | 7.6166 | 7.2774 | 6.8778 | 4.8537 | 4.0244 | 4.1626 | 3.7844 |
| RPL29_S142 | RPL29 | P47914 | | 142 | QAAAPASVPAQAP | 6.3766 | 7.4938 | 6.6412 | 7.0569 | 6.8960 | 6.6005 | 6.7752 | 6.9312 | 7.4667 | 6.3679 | 6.8858 | 6.5084 | 4.0358 | 4.9137 | 4.2893 | 4.7609 |
| RPL3_S13 | RPL3 | P39023 | | 13 | SAPRHGSLGFLPR | 6.3988 | 5.7574 | 6.4015 | 6.4236 | 8.5596 | 5.5030 | 5.7165 | 5.9790 | 8.1904 | 7.5587 | 6.1159 | 5.3112 | 7.4687 | 5.3295 | 4.7862 | 4.5000 |
| RPL30_S10 | RPL30 | P62888 | | 10 | AKKTKKSLESINS | 6.6490 | 7.2678 | 6.8376 | 6.9529 | 6.2851 | 6.8592 | 6.6309 | 6.7006 | 6.5835 | 7.0591 | 6.6809 | 6.5542 | 4.4673 | 4.5145 | 4.9206 | 5.0367 |
| RPL31_S15 | RPL31 | P62899-2 | | 15 | EKKGRSAINEVV | 5.4828 | 5.3072 | 5.1168 | 5.3782 | 5.5633 | 5.5525 | 5.9922 | 6.9815 | 5.2375 | 5.7710 | 5.5512 | 5.2871 | 9.7708 | 6.5579 | 7.8783 | 9.3717 |
| RPL34_S12 | RPL34 | P49207 | | 12 | TYRRRLSYNTASN | 4.2285 | 4.0318 | 5.1138 | 5.1132 | 8.7668 | 5.5566 | 5.1174 | 6.0342 | 8.8212 | 8.4614 | 7.0585 | 5.9879 | 8.4513 | 7.6210 | 4.8404 | 4.7960 |
| RPL35_S42 | RPL35 | P42766 | | 42 | VTGGAAKLSKIR | 3.9882 | 5.0610 | 5.3609 | 5.0679 | 4.7820 | 5.6789 | 5.8115 | 7.3069 | 6.6959 | 6.5982 | 6.8898 | 6.8465 | 8.1173 | 7.1213 | 7.2120 | 7.4617 |
| RPL36A_S32 | RPL36A | P83881 | | 32 | YKKGKDSLYAQGK | 6.2928 | 8.2304 | 8.6149 | 8.3327 | 6.5520 | 5.9466 | 7.0438 | 7.3117 | 7.2149 | 6.8586 | 6.8889 | 6.3659 | 3.5173 | 4.2810 | 2.8928 | 3.6557 |
| RPL36A_S46 | RPL36A | P83881 | | 46 | RYDRKQSGYGGQT | 5.9081 | 7.4819 | 8.4708 | 7.5892 | 6.2189 | 6.5310 | 6.8928 | 5.4521 | 8.8267 | 8.2717 | 8.4090 | 6.0752 | 4.5539 | 4.1558 | 2.0458 | 3.1168 |
| RPL37_S6 | RPL37 | P61927 | | 6 | xMTKGTSSFGKRR | 6.6347 | 6.8429 | 6.1586 | 7.1309 | 5.8971 | 6.0571 | 6.8878 | 5.4320 | 6.2885 | 5.9071 | 6.1227 | 6.1528 | 5.5393 | 6.6919 | 6.0591 | 6.1974 |
| RPL37_S96 | RPL37 | P61927 | | 96 | AVAASSSSxxxxx | 5.3432 | 7.7213 | 7.9484 | 6.2678 | 7.5396 | 6.5566 | 6.4073 | 5.4795 | 7.1503 | 8.0295 | 5.6514 | 6.4593 | 4.9864 | 4.8436 | 3.9288 | 5.6871 |
| RPL37A_S21 | RPL37A | P61513 | | 21 | GTRYGASLRKMVK | 5.7052 | 7.9467 | 9.0185 | 8.7393 | 4.8017 | 6.0912 | 7.0481 | 5.9747 | 6.8513 | 7.2568 | 9.0449 | 8.0404 | 3.3498 | 3.9824 | 2.6018 | 3.5471 |
| RPL38_Y43 | RPL38 | P63173 | | 43 | RCSRYLYTLVITD | 6.5465 | 8.1674 | 9.1909 | 8.9604 | 9.0896 | 7.9333 | 4.5169 | 5.4356 | 6.5151 | 6.0959 | 6.9273 | 6.6583 | 2.7667 | 4.1990 | 3.9370 | 3.0602 |
| RPL4_S12 | RPL4 | P36578 | | 12 | PLISVYSEKGESS | 7.5075 | 6.5922 | 7.8177 | 5.7502 | 5.3797 | 6.4534 | 6.9434 | 6.2806 | 6.6334 | 8.6250 | 8.0348 | 8.9293 | 4.0073 | 4.1551 | 3.7070 | 4.9833 |
| RPL4_S275 | RPL4 | P36578 | | 420 | AEKKPTTEEKKPA | 7.6189 | 8.2115 | 6.7183 | 7.6008 | 6.6746 | 6.8091 | 7.9136 | 7.4820 | 7.9827 | 6.9421 | 6.7380 | 7.0612 | 2.6620 | 3.7950 | 2.3200 | 3.4703 |
| RPL4_S295 | RPL4 | P36578 | | 295 | LSRILKSPEIQRA | 3.8062 | 7.5773 | 7.8546 | 6.9157 | 4.3268 | 10.5370 | 5.7156 | 7.6695 | 11.0235 | 5.6216 | 4.7884 | 6.3020 | 4.5989 | 5.3970 | 3.9329 | 3.9332 |
| RPL4_T23 | RPL4 | P36578 | | 23 | SSGKNVTLPAVFK | 5.2930 | 6.9565 | 5.0142 | 6.4583 | 6.5740 | 7.5466 | 6.0254 | 6.1914 | 6.0402 | 7.7215 | 6.9921 | 5.7911 | 5.4472 | 5.7466 | 6.2596 | 5.9424 |
| RPL4_T420 | RPL4 | P36578 | | 275 | KAASLKSNYNLPM | 5.8482 | 6.9614 | 6.4626 | 6.2903 | 5.8553 | 5.9595 | 6.5072 | 6.0008 | 6.3462 | 6.8517 | 7.0615 | 6.5310 | 5.3769 | 6.0859 | 5.3960 | 6.4655 |
| RPL5_S185 | RPL5 | P46777 | | 185 | RFPGYDSESKEFN | 6.8109 | 6.7077 | 6.6929 | 6.3984 | 6.4391 | 6.1315 | 6.4105 | 6.1424 | 6.6038 | 6.1655 | 6.2352 | 6.1665 | 5.4400 | 6.1703 | 5.8072 | 5.6781 |
| RPL5_Y79 | RPL5 | P46777 | | 79 | MIVCAAYAHELPK | 5.8662 | 5.7880 | 6.9350 | 6.4539 | 6.2233 | 6.4576 | 6.0088 | 6.7044 | 5.4297 | 6.3956 | 6.3646 | 5.0528 | 6.8023 | 7.2586 | 6.1848 | 6.0745 |
| RPL7_S102 | RPL7 | P18124 | | 17 | VPAVPETLKKKRR | 9.7241 | 9.4524 | 7.9930 | 9.7869 | 7.0532 | 5.6758 | 7.7657 | 5.4407 | 7.6457 | 5.3912 | 6.6273 | 5.5365 | 2.9324 | 3.6787 | 2.2976 | 2.9990 |
| RPL7_T17 | RPL7 | P18124 | | 102 | RGINGVSPKVRKV | 6.4568 | 6.8443 | 6.0583 | 6.3556 | 6.0780 | 6.1814 | 6.6008 | 6.5077 | 7.5673 | 6.3395 | 6.8648 | 6.3531 | 5.5198 | 5.7211 | 5.0870 | 5.4644 |
| RPL8_S14 | RPL8 | P62917 | | 14 | QRKGAGSVFRAHV | 8.3112 | 8.3269 | 8.2269 | 7.9571 | 5.2422 | 5.7998 | 7.5675 | 6.1451 | 8.4936 | 7.1981 | 7.4009 | 6.4151 | 3.3136 | 3.6518 | 2.7568 | 3.1934 |
| RPLP0_S304;S307 | RPLP0 | P05388 | | 307 | KEESEESDEDMGF | 6.0675 | 6.4313 | 7.4014 | 6.3532 | 6.1874 | 6.4730 | 5.8122 | 6.0119 | 7.2797 | 6.9874 | 7.2815 | 7.4231 | 4.7942 | 5.7250 | 4.8548 | 4.9165 |
| RPLP0_S307 | RPLP0 | P05388 | | 24 | IQLLDDYPKCFIV | 5.8647 | 5.8792 | 6.8431 | 6.0547 | 6.1508 | 6.1142 | 6.9303 | 6.6075 | 6.8859 | 6.7581 | 6.1086 | 5.9792 | 5.9192 | 5.9403 | 6.6475 | 6.3169 |
| RPLP0_Y24 | RPLP0 | P05388 | 304;307 | | NA | 5.8852 | 6.2066 | 6.6564 | 6.1440 | 6.2452 | 6.2846 | 6.2003 | 6.1145 | 6.6228 | 6.5259 | 6.5860 | 6.6851 | 6.0244 | 6.2733 | 5.8063 | 5.7495 |
| RPLP2_S102 | RPLP2 | P05387 | | 105 | KEESEESDDDMGF | 6.5693 | 5.4422 | 6.5453 | 5.7354 | 5.4719 | 5.3249 | 6.0459 | 6.3627 | 7.3683 | 7.9695 | 7.6919 | 8.2977 | 4.5552 | 6.5863 | 4.3249 | 5.7085 |
| RPLP2_S102;S105 | RPLP2 | P05387 | | 102 | DEKKEESEESDDD | 4.9900 | 4.4593 | 4.1923 | 4.9879 | 4.2675 | 4.0093 | 5.2474 | 5.6053 | 4.6437 | 5.8347 | 4.7550 | 6.6617 | 10.6164 | 12.1748 | 9.4568 | 12.6979 |
| RPLP2_S105 | RPLP2 | P05387 | 102;105 | | NA | 6.0763 | 5.2526 | 6.0120 | 5.4794 | 5.6972 | 4.7811 | 5.5361 | 6.2460 | 5.6763 | 6.2219 | 5.9643 | 6.9243 | 5.8358 | 8.8578 | 7.0554 | 8.3834 |
| RPP25_S162 | RPP25 | Q9BUL9 | | 162 | PHPGPSSPPAAPA | 5.9821 | 6.3599 | 5.9989 | 6.4868 | 6.2745 | 6.1200 | 6.3120 | 6.4309 | 6.1427 | 6.2801 | 6.4194 | 6.0547 | 6.2070 | 6.4481 | 6.2988 | 6.1843 |
| RPP25L_S100 | RPP25L | Q8N5L8 | | 16 | VELPAPSPMPQLP | 7.6761 | 6.1978 | 5.2730 | 6.1543 | 5.7223 | 6.6667 | 6.1435 | 6.2542 | 6.7687 | 5.6247 | 7.9822 | 5.3077 | 6.1587 | 6.7296 | 6.1088 | 5.2317 |
| RPP25L_S16 | RPP25L | Q8N5L8 | | 100 | DSWVPASPDTGLD | 6.0458 | 5.7607 | 6.6747 | 6.7643 | 6.4405 | 6.7353 | 6.5143 | 6.1990 | 6.1415 | 7.2510 | 6.9338 | 6.8400 | 5.0319 | 5.9556 | 5.3358 | 5.3757 |
| RPP30_S251 | RPP30 | P78346-2 | | 251 | VKKPRPSEGDEDC | 5.2960 | 5.5806 | 6.9235 | 8.1083 | 5.2679 | 6.3908 | 6.0747 | 5.5156 | 5.6857 | 4.9012 | 5.6919 | 5.6355 | 6.9620 | 7.7244 | 7.3588 | 6.8834 |
| RPP38_S235 | RPP38 | P78345 | | 235 | RELLDTSFEDLSK | 5.9996 | 6.3485 | 6.2030 | 6.6662 | 6.3759 | 6.3622 | 6.7431 | 6.5066 | 6.6085 | 6.5962 | 6.4033 | 6.5168 | 5.3587 | 5.8564 | 5.5723 | 5.8828 |
| RPP38_S253 | RPP38 | P78345 | | 253 | ADGRQASVTLQPL | 5.0365 | 5.1142 | 5.5973 | 5.9850 | 8.1192 | 5.7483 | 5.1295 | 5.7011 | 7.3615 | 7.6154 | 7.3760 | 7.0281 | 6.0965 | 7.0505 | 5.2841 | 5.7569 |
| RPRD1A_S156 | RPRD1A | Q96P16 | | 156 | NCSSLGSPSEPPQ | 6.4069 | 6.1052 | 6.6574 | 6.2462 | 5.8831 | 6.6102 | 6.1448 | 5.9625 | 6.3975 | 6.1509 | 6.5241 | 6.6668 | 6.0434 | 6.4430 | 6.0040 | 5.7540 |
| RPRD1B_S134 | RPRD1B | Q9NQG5 | | 161 | EEEDDDYPGSYSP | 5.4666 | 5.9911 | 6.0475 | 5.9596 | 6.0884 | 5.8697 | 5.7585 | 6.0775 | 6.3487 | 6.3348 | 6.2169 | 6.5719 | 6.8504 | 7.0985 | 6.5992 | 6.7209 |
| RPRD1B_Y161 | RPRD1B | Q9NQG5 | | 134 | SMEDSKSPPPKAT | 6.1784 | 6.3687 | 6.2262 | 6.3103 | 6.7654 | 6.9613 | 6.2296 | 6.3623 | 6.3816 | 6.2963 | 7.2230 | 6.6941 | 5.9761 | 6.0532 | 5.1691 | 4.8043 |
| RPRD2_S1099 | RPRD2 | Q5VT52 | | 1099 | ASNRRMSGEPIQT | 5.0716 | 5.1228 | 5.2847 | 5.5185 | 6.8638 | 5.6324 | 5.3634 | 5.7348 | 5.9714 | 6.2748 | 6.0157 | 6.0010 | 8.0172 | 7.6533 | 7.9203 | 7.5542 |
| RPRD2_S436 | RPRD2 | Q5VT52 | | 798 | FGLGSESPYKQPS | 6.7059 | 6.3769 | 6.2676 | 6.6374 | 6.1251 | 6.3467 | 6.1490 | 5.9324 | 6.3673 | 6.4376 | 6.2320 | 6.1158 | 6.2334 | 6.2115 | 5.7780 | 6.0834 |
| RPRD2_S485 | RPRD2 | Q5VT52 | | 763 | ISPGSSTPSSTRS | 6.1903 | 6.5189 | 5.5276 | 6.8529 | 5.9964 | 6.0121 | 6.0922 | 5.8372 | 6.3786 | 6.0704 | 5.7345 | 5.8244 | 7.1408 | 7.0869 | 6.7138 | 7.0231 |
| RPRD2_S593 | RPRD2 | Q5VT52 | | 732 | RDERSGTPTQDEM | 6.0329 | 6.2722 | 6.0411 | 6.3809 | 5.7553 | 5.9453 | 6.1472 | 5.5117 | 6.4845 | 6.3592 | 6.3204 | 6.5846 | 6.7242 | 6.5025 | 6.2376 | 6.7002 |
| RPRD2_S614 | RPRD2 | Q5VT52 | | 826 | KFYPDTSFQEDED | 6.3278 | 6.9385 | 6.8190 | 7.0235 | 6.1768 | 6.6918 | 6.6334 | 6.9723 | 6.7102 | 6.8880 | 7.2585 | 7.3132 | 4.3362 | 5.0259 | 4.4058 | 4.4790 |

| | | | | | | | | | | | | | | | | | | | | | |
|---|---|---|---|---|---|---|---|---|---|---|---|---|---|---|---|---|---|---|---|---|---|
| RPRD2_S758 | RPRD2 | Q5VT52 | | 485 | SPGTPTSPSNLTS | 5.9738 | 5.8954 | 6.4789 | 6.3769 | 6.3944 | 6.0084 | 6.0765 | 5.5409 | 6.1704 | 6.4239 | 6.3176 | 6.3423 | 6.7749 | 6.6156 | 6.1567 | 6.4533 |
| RPRD2_S758;T763 | RPRD2 | Q5VT52 | | 723 | IDNIDGTPVRDER | 6.5706 | 6.4306 | 6.0316 | 7.0164 | 5.9462 | 6.1654 | 6.1607 | 6.2620 | 6.5392 | 6.0906 | 5.9827 | 6.3995 | 5.6904 | 6.6205 | 6.1999 | 5.8935 |
| RPRD2_S769 | RPRD2 | Q5VT52 | | 976 | PHRSLFSPQNTLA | 6.6302 | 7.1114 | 6.3968 | 6.8886 | 6.4423 | 6.5372 | 6.6779 | 6.4805 | 6.8213 | 6.6548 | 6.7482 | 7.4423 | 4.5222 | 4.9247 | 4.5095 | 5.2121 |
| RPRD2_S798 | RPRD2 | Q5VT52 | | 965 | QKQYPDSPHPVPH | 7.4635 | 7.7498 | 6.2048 | 7.0217 | 5.8652 | 6.5750 | 7.0392 | 6.2494 | 7.1519 | 6.7232 | 6.7782 | 6.8783 | 4.6904 | 4.9885 | 3.6870 | 4.9339 |
| RPRD2_S826 | RPRD2 | Q5VT52 | | 769 | TPSSTRSPPPGRD | 5.7970 | 5.9287 | 5.9326 | 6.4350 | 6.5433 | 6.1106 | 6.0815 | 6.1088 | 6.0216 | 5.9208 | 6.0751 | 6.6033 | 6.5225 | 7.0636 | 6.2624 | 6.5932 |
| RPRD2_S909 | RPRD2 | Q5VT52 | | 426 | PVTMTATPPLPKP | 5.9784 | 6.7136 | 6.3687 | 6.5996 | 6.6050 | 6.7253 | 6.6372 | 5.8840 | 6.6576 | 6.3900 | 5.9258 | 6.5654 | 5.7538 | 5.9972 | 5.5471 | 5.6515 |
| RPRD2_S914 | RPRD2 | Q5VT52 | | 593 | PKSFNYSPNSSTS | 5.6328 | 6.2537 | 5.6754 | 5.9297 | 6.5016 | 5.6852 | 5.7508 | 6.0319 | 7.0860 | 5.9044 | 6.7782 | 5.7195 | 6.6759 | 6.6363 | 7.2267 | 6.5118 |
| RPRD2_S928;S930 | RPRD2 | Q5VT52 | | 614 | KASIGQSPGLPST | 6.0428 | 6.5116 | 5.8818 | 6.6535 | 6.1288 | 6.4342 | 6.2983 | 6.3829 | 6.5463 | 6.6824 | 6.3433 | 6.1317 | 6.1566 | 6.2616 | 5.7050 | 5.8394 |
| RPRD2_S965 | RPRD2 | Q5VT52 | | 909 | CDRLSSSPGLFGA | 6.8914 | 7.0267 | 6.9418 | 7.5683 | 5.2982 | 6.3103 | 6.5233 | 6.3432 | 7.3387 | 7.0361 | 7.4501 | 7.2898 | 4.2269 | 5.1772 | 4.0550 | 4.5230 |
| RPRD2_S976 | RPRD2 | Q5VT52 | | 436 | PKPVNTSLSPSPA | 5.8449 | 5.4424 | 5.7853 | 6.7852 | 6.1285 | 6.3892 | 6.3473 | 5.7232 | 6.6257 | 6.5444 | 6.0258 | 6.5633 | 6.5589 | 6.7147 | 6.0714 | 6.4500 |
| RPRD2_T426 | RPRD2 | Q5VT52 | | 758 | LLSKIISPGSSTP | 6.3246 | 6.7056 | 6.2190 | 7.1821 | 6.5138 | 5.3922 | 7.0526 | 6.1882 | 7.1350 | 6.1372 | 6.7816 | 5.9323 | 5.6054 | 5.5236 | 5.4828 | 5.8241 |
| RPRD2_T723 | RPRD2 | Q5VT52-5 | | 914 | PCRSHGSPTHVRR | 6.7607 | 6.7117 | 6.4956 | 6.8040 | 5.8658 | 5.7903 | 6.0991 | 6.6516 | 5.9692 | 6.0390 | 6.5485 | 6.0991 | 5.4625 | 6.3098 | 6.3095 | 6.0837 |
| RPRD2_T732 | RPRD2 | Q5VT52 | 928;930 | | NA | 6.1255 | 6.3060 | 5.8041 | 6.9161 | 6.1212 | 6.1181 | 5.8898 | 5.9369 | 6.2085 | 6.2524 | 6.3630 | 6.4061 | 6.3592 | 6.4990 | 6.3441 | 6.3501 |
| RPRD2_T763 | RPRD2 | Q5VT52 | 758;763 | | NA | 6.4400 | 7.1689 | 7.4026 | 6.4954 | 6.1956 | 6.1483 | 6.4721 | 6.7184 | 6.6678 | 6.5120 | 6.1535 | 5.5628 | 5.5270 | 5.7407 | 5.4984 | 5.2963 |
| RPS10_S130 | RPS10 | P46783 | | 146 | AEAGAGSATEFQF | 7.1952 | 7.1467 | 7.6095 | 7.1655 | 6.5808 | 6.8212 | 6.7350 | 7.5111 | 6.5990 | 6.2738 | 7.0573 | 6.7507 | 3.9932 | 4.4355 | 3.9137 | 4.2117 |
| RPS10_S146 | RPS10 | P46783 | | 12 | KNRIAIYELLFKE | 6.4581 | 6.6304 | 6.7422 | 6.6390 | 6.2786 | 6.5113 | 5.9822 | 6.2974 | 6.6763 | 6.7595 | 6.8968 | 7.2016 | 5.1245 | 5.3566 | 5.2138 | 5.2314 |
| RPS10_Y12 | RPS10 | P46783 | | 130 | RDTYRRSAVPPGA | 8.0338 | 8.0717 | 6.9000 | 7.3958 | 6.8571 | 6.9943 | 7.2179 | 6.0895 | 6.7869 | 6.4672 | 6.9893 | 7.2370 | 3.7214 | 4.3623 | 3.1392 | 3.7365 |
| RPS11_S67 | RPS11 | P62280 | | 67 | PFTGNVSIRGRIL | 4.1086 | 6.7811 | 7.6367 | 10.2893 | 7.5479 | 5.2151 | 6.7080 | 5.9153 | 7.3141 | 6.4965 | 6.8477 | 6.3051 | 4.5730 | 4.7859 | 3.6703 | 5.8053 |
| RPS11_T28 | RPS11 | P62280 | | 28 | RVLLGETGKEKLP | 6.1371 | 6.4602 | 5.5017 | 6.7249 | 6.0895 | 6.8785 | 6.7302 | 7.0109 | 6.2907 | 4.5909 | 4.5539 | 6.0577 | 7.5351 | 5.6818 | 7.1803 | 6.5766 |
| RPS19BP1_S84 | RPS19BP1 | Q86WX3 | | 84 | FLTRTRSTVAESV | 6.5628 | 6.8865 | 6.8381 | 7.3074 | 6.0917 | 5.7101 | 7.1883 | 6.0931 | 7.2848 | 6.7671 | 7.0223 | 6.2191 | 4.5516 | 5.1851 | 4.9845 | 5.3074 |
| RPS2_S264 | RPS2 | P15880 | | 264 | ETVFTKSPYQEFT | 6.6169 | 6.7966 | 6.9447 | 6.8374 | 6.5381 | 6.0882 | 6.3655 | 6.5999 | 7.2575 | 7.2631 | 7.0959 | 6.8505 | 4.4313 | 4.9476 | 4.5266 | 4.8401 |
| RPS2_S281 | RPS2 | P15880 | | 281 | KTHTRVSVQRTQA | 7.5489 | 7.0908 | 7.0810 | 6.6986 | 6.9972 | 7.0943 | 6.8827 | 6.0746 | 6.1043 | 6.6014 | 7.2230 | 7.1750 | 4.5895 | 4.8028 | 3.8369 | 4.1991 |
| RPS2_S77 | RPS2 | P15880 | | 77 | KDMKIKSLEEIYL | 5.8987 | 6.2960 | 6.5758 | 5.9401 | 5.6933 | 6.1170 | 5.4292 | 6.4100 | 6.7977 | 7.3077 | 6.5620 | 6.8930 | 5.9683 | 6.3230 | 5.4870 | 6.3011 |
| RPS2_T252 | RPS2 | P15880 | | 252 | KTYSYLTPDLWKE | 6.9166 | 6.8052 | 6.0483 | 6.6127 | 5.7553 | 7.0888 | 6.5400 | 6.5155 | 6.9892 | 6.4598 | 6.7481 | 5.9290 | 5.1938 | 5.4420 | 5.3670 | 5.5887 |
| RPS2_Y266 | RPS2 | P15880 | | 266 | VFTKSPYQEFTDH | 10.2365 | 8.3997 | 10.7523 | 9.6117 | 6.6762 | 5.1623 | 5.9507 | 5.8081 | 8.0097 | 6.0891 | 6.7433 | 8.8610 | 2.6336 | 1.2874 | 1.3545 | 2.4239 |
| RPS20_S93 | RPS20 | P60866-2 | | 93 | RLIDLHSPSEIVK | 6.1104 | 6.3589 | 7.0092 | 6.8274 | 6.0145 | 6.7595 | 5.8831 | 6.6776 | 6.3686 | 6.7731 | 6.9484 | 7.0214 | 4.9618 | 5.7052 | 5.2784 | 5.3026 |
| RPS20_S95 | RPS20 | P60866-2 | | 95 | IDLHSPSEIVKQI | 3.8216 | 5.6266 | 6.5541 | 5.5743 | 5.9548 | 5.7265 | 6.3329 | 8.3520 | 6.2045 | 7.4721 | 6.9441 | 7.2771 | 5.7170 | 5.3841 | 6.5541 | 6.5042 |
| RPS25_S93 | RPS25 | P62851 | | 93 | ALQELLSKGLIKL | 5.3604 | 5.6701 | 6.3745 | 6.3255 | 4.7410 | 6.6932 | 6.2282 | 6.3606 | 6.1898 | 7.6901 | 6.6033 | 6.4791 | 6.0215 | 6.3010 | 6.3173 | 6.6442 |
| RPS26_S54 | RPS26 | P62854 | | 96 | EARKDRTPPPRFR | 6.1446 | 6.4459 | 6.7853 | 6.2674 | 6.2741 | 6.8606 | 6.2391 | 6.2284 | 6.4925 | 6.2659 | 6.5237 | 6.4198 | 5.5221 | 6.5667 | 5.2639 | 5.7001 |
| RPS26_T96 | RPS26 | P62854 | | 54 | AAVRDISEASVFD | 6.5276 | 6.3354 | 6.5842 | 6.2225 | 6.5906 | 5.8600 | 6.2223 | 6.0658 | 6.5946 | 6.2018 | 6.3963 | 6.2312 | 5.8493 | 6.1957 | 6.2737 | 5.8491 |
| RPS27_S11 | RPS27 | P42677 | | 11 | KDLLHPSPEEEKR | 6.1635 | 6.3057 | 7.0639 | 6.9342 | 6.0255 | 6.2153 | 6.0809 | 5.8229 | 6.8768 | 6.6462 | 6.9552 | 6.8310 | 5.4563 | 5.9763 | 5.1881 | 5.4581 |
| RPS27_S27 | RPS27 | P42677 | | 27 | KKRLVQSPNSYFM | 5.9582 | 6.5870 | 6.7975 | 6.5613 | 6.1630 | 6.4419 | 6.3586 | 6.8476 | 6.5247 | 6.9963 | 6.7290 | 6.4994 | 4.9847 | 5.7450 | 5.2029 | 5.6029 |
| RPS27_S78 | RPS27 | P42677 | | 78 | RLTEGCSFRRKQH | 4.3168 | 7.1445 | 7.3199 | 8.2322 | 6.1756 | 5.7149 | 6.1562 | 6.3909 | 6.7303 | 6.0601 | 6.5881 | 5.7765 | 5.1715 | 6.3615 | 5.5564 | 6.3045 |
| RPS27_Y31 | RPS27 | P42677 | | 31 | VQSPNSYFMDVKC | 6.4240 | 6.2414 | 7.3328 | 5.7326 | 5.9570 | 6.2976 | 5.6620 | 6.3933 | 6.4087 | 6.2154 | 6.9272 | 6.8645 | 5.4483 | 6.3329 | 5.6365 | 6.1257 |
| RPS3_S240 | RPS3 | P23396-2 | | 237 | DEILPTTPISEQK | 6.0755 | 6.1729 | 6.5691 | 6.2711 | 6.2802 | 6.2754 | 6.1205 | 6.4076 | 6.5562 | 6.5688 | 6.4652 | 6.7318 | 5.6056 | 6.1100 | 5.8649 | 5.9253 |
| RPS3_T237 | RPS3 | P23396-2 | | 240 | LPTTPISEQKGGK | 6.0899 | 6.1133 | 6.2582 | 6.5742 | 5.9549 | 6.0597 | 6.3432 | 6.9752 | 6.9704 | 6.8071 | 6.7886 | 6.8590 | 5.5246 | 5.6614 | 5.2371 | 5.7833 |
| RPS3A_S236 | RPS3A | P61247 | | 263 | EPPVQESVxxxxx | 6.4436 | 6.9549 | 6.8729 | 6.3701 | 6.3697 | 6.2931 | 6.3919 | 6.9165 | 6.3959 | 6.5108 | 6.4561 | 6.5391 | 5.1693 | 5.5324 | 5.3248 | 5.4589 |
| RPS3A_S237 | RPS3A | P61247 | | 238 | HGEGSSSGKATGD | 5.9942 | 6.7938 | 6.5132 | 6.3102 | 6.3105 | 6.7239 | 6.1365 | 6.6395 | 6.3361 | 6.5468 | 6.8612 | 6.4999 | 5.2586 | 5.9766 | 5.5370 | 5.5620 |
| RPS3A_S238 | RPS3A | P61247 | | 236 | ELHGEGSSSGKAT | 6.1489 | 6.0178 | 6.2599 | 6.5214 | 6.5094 | 6.5962 | 5.8657 | 7.1377 | 6.3140 | 6.7746 | 6.3465 | 6.2893 | 5.4827 | 6.2654 | 5.8839 | 5.5867 |
| RPS3A_S263 | RPS3A | P61247 | | 242 | SSSGKATGDETGA | 6.5519 | 6.0695 | 7.4051 | 7.2451 | 6.0337 | 6.5235 | 6.0148 | 6.7461 | 6.7459 | 6.3646 | 7.1611 | 6.9789 | 4.7742 | 5.2470 | 4.9022 | 5.2364 |
| RPS3A_T242 | RPS3A | P61247 | | 237 | LHGEGSSSGKATG | 5.8210 | 5.9268 | 6.3951 | 6.1862 | 6.4716 | 6.7205 | 6.4923 | 6.7425 | 6.5064 | 6.2446 | 6.4307 | 5.8694 | 5.6403 | 6.4936 | 6.1103 | 5.9487 |
| RPS4X_S223 | RPS4X | P62701 | | 223 | SFATRLSNIFVIG | 5.8541 | 6.0736 | 6.0756 | 6.3914 | 6.4811 | 5.8066 | 6.0419 | 6.6304 | 7.1276 | 5.6384 | 5.9549 | 6.0965 | 6.3779 | 6.3820 | 6.8027 | 6.2654 |
| RPS5_S142 | RPS5 | P46782 | | 142 | RQAVDVSPLRRVN | 6.5826 | 6.8830 | 6.4150 | 7.0875 | 6.1391 | 5.8873 | 6.4296 | 6.2957 | 6.8041 | 6.3235 | 6.3279 | 6.5546 | 5.5989 | 5.4423 | 5.4750 | 5.7538 |
| RPS6_S235;S236;S240 | RPS6 | P62753 | | 82 | LLSKGHSCYRPRR | 8.1547 | 8.9061 | 9.4799 | 10.1881 | 4.6964 | 5.9780 | 8.1158 | 5.9650 | 8.8625 | 6.0209 | 6.2788 | 4.1305 | 3.8618 | 3.4101 | 1.9999 | 3.9515 |
| RPS6_S236;S240 | RPS6 | P62753 | | 244 | RASTSKSESSQKx | 4.9887 | 6.4739 | 5.0723 | 5.4129 | 5.9797 | 5.4420 | 5.8834 | 5.9406 | 6.5701 | 5.1358 | 4.7516 | 6.1805 | 8.4504 | 8.3360 | 7.1597 | 8.2224 |
| RPS6_S244 | RPS6 | P62753 | 236;240 | | NA | 7.8195 | 7.4663 | 8.6957 | 7.9384 | 6.3382 | 6.8766 | 7.7436 | 7.9110 | 8.3606 | 7.4947 | 9.0901 | 8.9001 | 1.0585 | 1.5653 | 1.1283 | 1.6130 |
| RPS6_S82 | RPS6 | P62753 | 235;236;240 | | NA | 9.8770 | 9.0322 | 11.2142 | 9.7628 | 5.9246 | 5.6842 | 7.0107 | 7.1321 | 8.5597 | 6.5377 | 8.9923 | 8.3484 | 0.2523 | 0.4906 | 0.5345 | 0.6467 |
| RPS6KA1_S230 | RPS6KA1 | Q15418-2 | | 230 | HEKKAYSFCGTVE | 5.8372 | 6.0580 | 6.6324 | 6.3781 | 6.5041 | 6.4998 | 6.5487 | 6.4474 | 6.1549 | 6.0629 | 6.2599 | 5.9314 | 6.0475 | 6.4480 | 6.1694 | 6.0203 |
| RPS6KA1_S389 | RPS6KA1 | Q15418-2 | | 582 | ENGLLMTPCYTAN | 5.9457 | 5.9737 | 6.0448 | 6.4776 | 6.9471 | 6.7841 | 6.0697 | 5.9576 | 5.8072 | 6.0408 | 6.3714 | 6.0963 | 6.6332 | 6.6526 | 5.8364 | 6.3618 |
| RPS6KA1_T582 | RPS6KA1 | Q15418-2 | | 389 | QLFRGFSFVATGL | 7.1372 | 6.0920 | 6.6634 | 6.9535 | 7.0009 | 6.1649 | 5.9053 | 5.9555 | 6.5086 | 5.9664 | 6.3968 | 6.1800 | 5.8164 | 6.4631 | 5.6553 | 5.1409 |
| RPS6KA3_S369 | RPS6KA3 | P51812 | | 715 | ALNRNQSPVLEPV | 6.4540 | 6.0096 | 6.8570 | 6.1585 | 6.3352 | 6.3565 | 6.6211 | 7.1332 | 6.2425 | 5.8955 | 6.2986 | 6.5265 | 5.6214 | 5.9773 | 5.8227 | 5.6905 |
| RPS6KA3_S715 | RPS6KA3 | P51812 | | 369 | AKTPKDSPGIPPS | 6.4168 | 6.5182 | 6.1169 | 6.7009 | 6.2352 | 6.5188 | 5.6934 | 6.3162 | 6.3996 | 6.2210 | 6.7538 | 6.6254 | 5.9655 | 6.1556 | 5.7084 | 5.6544 |
| RPS6KA3_T365;S369 | RPS6KA3 | P51812 | 365;369 | | NA | 6.1906 | 5.5019 | 6.0301 | 6.3020 | 6.6027 | 6.7141 | 6.2215 | 6.6686 | 6.1163 | 5.5821 | 6.7583 | 6.3962 | 6.5387 | 6.2429 | 6.3518 | 5.7822 |
| RPS6KA4_S347 | RPS6KA4 | O75676 | | 634 | IREGRFSLDGEAW | 4.1409 | 4.0658 | 4.5644 | 3.9566 | 5.4359 | 5.1452 | 4.6978 | 5.6089 | 5.0045 | 4.6513 | 4.7628 | 4.8871 | 10.2979 | 10.3754 | 11.5869 | 10.8187 |
| RPS6KA4_S634 | RPS6KA4 | O75676 | | 745 | TASRRGSPAPANP | 5.9232 | 5.6058 | 6.1909 | 5.8754 | 5.3906 | 5.7203 | 5.7825 | 6.2469 | 6.1796 | 6.3005 | 6.3098 | 6.9441 | 6.3406 | 7.0522 | 7.0571 | 7.0804 |
| RPS6KA4_S678 | RPS6KA4 | O75676 | | 347 | VYSPPGSPPPGDP | 6.2388 | 6.1344 | 6.7578 | 6.1849 | 6.4963 | 6.4158 | 6.4870 | 6.2726 | 6.5414 | 6.7895 | 6.7187 | 6.7263 | 4.8190 | 6.2025 | 5.6078 | 5.6069 |
| RPS6KA4_S745 | RPS6KA4 | O75676 | | 678 | SWLQDGSARSSPP | 3.4881 | 3.6814 | 4.3360 | 4.0169 | 4.7985 | 4.6707 | 4.3848 | 5.1597 | 4.0986 | 3.8313 | 3.7392 | 3.9746 | 11.7694 | 11.9441 | 13.7177 | 12.3891 |
| RPS6KB1_S427 | RPS6KB1 | P23443 | | 447 | SPRTPVSPVKFSP | 6.4056 | 5.9795 | 6.7400 | 5.7133 | 6.2733 | 6.3993 | 6.1747 | 6.8058 | 6.5341 | 6.1995 | 6.2086 | 6.3038 | 6.0667 | 6.2853 | 6.0182 | 5.8922 |
| RPS6KB1_S441;T444;S44 | RPS6KB1 | P23443 | | 503 | PIRQPNSGPYKKQ | 7.1743 | 7.7585 | 7.2911 | 7.6871 | 5.8934 | 7.3734 | 7.4322 | 7.0335 | 7.6748 | 7.6603 | 8.0020 | 7.7160 | 2.3611 | 3.4342 | 2.2602 | 3.2478 |
| RPS6KB1_S447 | RPS6KB1 | P23443 | | 452 | VSPVKFSPGDFWG | 7.8254 | 7.4921 | 8.8977 | 7.8118 | 6.0638 | 6.2192 | 6.4917 | 7.0225 | 8.1934 | 7.7193 | 8.3557 | 7.9933 | 1.6479 | 2.7504 | 2.4000 | 3.1159 |
| RPS6KB1_S452 | RPS6KB1 | P23443 | | 427 | SVKEKFSFEPKIR | 7.7328 | 7.2249 | 9.4704 | 7.1749 | 7.0093 | 7.2804 | 6.9836 | 7.7423 | 8.7276 | 7.2889 | 8.1930 | 7.5133 | 1.7334 | 2.3076 | 1.6143 | 2.0033 |
| RPS6KB1_S503 | RPS6KB1 | P23443 | 441;444;447 | | NA | 6.6596 | 6.5301 | 8.3925 | 6.5362 | 5.6957 | 6.9581 | 7.0100 | 6.9555 | 7.2337 | 7.3621 | 8.3827 | 9.0858 | 2.9792 | 3.8964 | 2.9889 | 3.3335 |
| RPS6KB2_S423 | RPS6KB2 | Q9UBS0 | | 423 | SPRAPVSPLKFSP | 6.2144 | 6.1779 | 6.5961 | 5.8859 | 6.1711 | 6.2321 | 6.0949 | 6.6334 | 6.3990 | 6.1490 | 6.2754 | 6.2697 | 5.9558 | 6.5362 | 6.2581 | 6.1512 |
| RPS6KC1_S209 | RPS6KC1 | Q96S38 | | 664 | DTISRGSDDSVPV | 6.4309 | 5.5480 | 6.1792 | 6.9267 | 5.5613 | 5.5896 | 5.8470 | 6.1776 | 5.4891 | 5.8246 | 6.3038 | 7.1136 | 6.6726 | 6.6917 | 6.8171 | 6.8271 |
| RPS6KC1_S282 | RPS6KC1 | Q96S38 | | 528 | EKIEPGSLNEEPF | 6.5650 | 7.6234 | 6.3835 | 5.9525 | 6.1758 | 7.1461 | 5.9141 | 6.3154 | 6.1252 | 6.2483 | 7.8005 | 7.3712 | 5.5572 | 5.7212 | 4.6629 | 4.4377 |
| RPS6KC1_S423;S427 | RPS6KC1 | Q96S38 | | 823 | SLFRICSPLSGAN | 6.0226 | 6.6996 | 6.5782 | 6.7848 | 6.3981 | 6.9929 | 6.2401 | 6.4179 | 7.1376 | 6.6878 | 6.9773 | 7.4250 | 4.7797 | 4.9495 | 4.8590 | 5.0497 |
| RPS6KC1_S528 | RPS6KC1 | Q96S38 | | 209 | ALGAVASDSEQSK | 6.0407 | 6.0648 | 7.0060 | 5.9911 | 6.2542 | 5.5336 | 6.7273 | 6.7046 | 6.1003 | 7.1975 | 6.4854 | 5.8436 | 5.7946 | 5.8436 | 6.2450 | 6.1677 |
| RPS6KC1_S661;S664 | RPS6KC1 | Q96S38 | | 282 | GVQGESSPTRREA | 6.6547 | 6.6176 | 6.3109 | 7.1722 | 6.0922 | 5.8656 | 6.5454 | 6.1941 | 6.6305 | 6.6488 | 7.0411 | 6.4301 | 5.7691 | 5.4342 | 5.1878 | 5.4055 |
| RPS6KC1_S664 | RPS6KC1 | Q96S38 | 423;427 | | NA | 6.1303 | 6.2050 | 6.3906 | 5.7062 | 6.3167 | 6.3307 | 6.7809 | 5.0924 | 6.7849 | 6.6932 | 6.6710 | 6.3079 | 6.3965 | 5.9610 | 5.9017 | 6.3309 |
| RPS6KC1_S823 | RPS6KC1 | Q96S38 | 661;664 | | NA | 5.2733 | 7.8731 | 6.1391 | 6.0006 | 6.5266 | 5.5213 | 6.1100 | 5.8665 | 5.8705 | 6.2447 | 6.1305 | 6.6180 | 5.9899 | 7.3141 | 6.3956 | 6.1262 |
| RPS7_T121 | RPS7 | P62081 | | 121 | KRPRSRTLTAVHD | 4.3865 | 6.3164 | 7.0086 | 7.2147 | 6.0970 | 6.2788 | 6.0044 | 5.7503 | 6.1454 | 7.6218 | 8.6756 | 8.6654 | 4.7157 | 5.8337 | 4.2491 | 5.0367 |
| RPS7_T123 | RPS7 | P62081 | | 123 | PRSRTLTAVHDAI | 9.0199 | 8.0332 | 8.7921 | 11.2179 | 9.5628 | 4.5241 | 5.1724 | 4.2829 | 6.6331 | 5.0115 | 5.3390 | 5.6244 | 4.1888 | 4.7462 | 4.0675 | 3.7842 |
| RPS8_S159 | RPS8 | P62241 | | 130 | KKGAKLTPEEEEI | 5.4075 | 6.0664 | 7.0066 | 6.0430 | 6.5024 | 5.8495 | 5.9673 | 6.2981 | 6.5778 | 6.3917 | 6.6075 | 6.3663 | 5.8407 | 6.7134 | 6.0859 | 6.2759 |
| RPS8_T130 | RPS8 | P62241 | | 159 | KKNAKISSLLEEQ | 5.8043 | 5.4933 | 6.6982 | 6.5657 | 6.6001 | 5.6179 | 6.3327 | 6.4826 | 6.3509 | 7.2370 | 6.1061 | 5.9724 | 5.6309 | 6.2769 | 6.7021 | 6.1288 |
| RPS9_S163 | RPS9 | P46781 | | 163 | IDFSLRSPYGGGR | 7.1794 | 7.6699 | 7.2059 | 7.3355 | 5.8194 | 6.2920 | 7.0199 | 6.7005 | 7.3136 | 6.8620 | 7.5818 | 7.6147 | 3.7211 | 4.3075 | 3.3286 | 4.0483 |
| RPSA_S43 | RPSA | P08865 | | 43 | YIYKRKSDGIYII | 5.6428 | 5.1957 | 9.2699 | 4.6761 | 7.1716 | 5.2756 | 5.2557 | 5.8102 | 6.8818 | 5.3955 | 6.3997 | 5.8077 | 6.0400 | 6.4397 | 7.3714 | 7.3664 |
| RPTOR_S719;S722 | RPTOR | Q8N122 | | 722 | RLRSVSSYGNIRA | 5.3007 | 5.4404 | 5.0200 | 4.7517 | 5.8366 | 5.8382 | 5.5403 | 5.9313 | 5.0616 | 5.2266 | 5.7087 | 6.1306 | 8.4073 | 8.7818 | 8.2453 | 8.7790 |
| RPTOR_S722 | RPTOR | Q8N122 | | 877 | IHQAGGSPPASST | 6.8668 | 6.8571 | 6.7246 | 6.6452 | 6.3852 | 6.7211 | 6.5875 | 7.3252 | 6.8034 | 6.3101 | 6.5683 | 6.4939 | 4.7091 | 5.5581 | 4.7117 | 4.7326 |
| RPTOR_S863 | RPTOR | Q8N122 | | 863 | TQSAPASPTNKGV | 6.3135 | 5.9681 | 6.0115 | 6.2682 | 6.5060 | 6.3173 | 6.2168 | 6.7881 | 6.5804 | 6.7214 | 6.1674 | 6.5730 | 5.8245 | 6.3104 | 5.7705 | 5.6629 |
| RPTOR_S877 | RPTOR | Q8N122 | 719;722 | | NA | 5.1446 | 5.2726 | 6.0175 | 6.3052 | 6.5158 | 7.5350 | 5.8129 | 5.4564 | 6.6901 | 5.0097 | 5.7156 | 5.3607 | 6.5275 | 9.0662 | 6.2773 | 7.2929 |
| RPUSD2_S68 | RPUSD2 | Q8IZ73 | | 68 | DAKVELSPGPPKP | 6.0941 | 6.5702 | 6.5605 | 5.9553 | 6.0671 | 6.6855 | 6.3891 | 6.4625 | 6.7434 | 6.6585 | 7.0987 | 6.8036 | 4.9784 | 5.9264 | 5.4310 | 5.5757 |
| RRAD_S301 | RRAD | P55042 | | 301 | FRAKSKSCHDLSV | 6.6635 | 6.3753 | 7.0747 | 6.3446 | 6.1606 | 6.3873 | 5.9925 | 6.4396 | 6.7909 | 6.5210 | 6.5135 | 6.4200 | 5.8309 | 5.8708 | 5.0159 | 5.5989 |
| RRAD_S39 | RRAD | P55042 | | 39 | PPLHRRSMPVDER | 6.6995 | 6.8441 | 6.8885 | 6.0017 | 6.3960 | 6.3587 | 5.8944 | 6.6346 | 6.8672 | 6.3670 | 6.7226 | 6.6922 | 5.4371 | 5.7178 | 5.1225 | 5.3560 |
| RRAGC_T394 | RRAGC | Q9HB90 | | 394 | ALTHNGTPRNAIx | 8.1996 | 8.4746 | 8.6363 | 8.7814 | 5.3009 | 5.9590 | 6.3551 | 6.3552 | 8.1263 | 7.6174 | 8.1790 | 8.0604 | 1.9475 | 2.8406 | 2.3979 | 2.7688 |
| RRAGD_S96 | RRAGD | Q9NQL2 | | 96 | ETLFLESTNKICR | 5.8775 | 6.3236 | 6.4330 | 6.4702 | 6.0048 | 5.4586 | 5.5493 | 5.3524 | 6.4115 | 6.7546 | 6.4976 | 6.3199 | 6.8731 | 6.6202 | 6.9410 | 6.1126 |
| RRAGD_T391 | RRAGD | Q9NQL2 | | 391 | QKKKRATPNGTPR | 6.7045 | 6.5141 | 7.0304 | 7.1616 | 6.4896 | 5.9839 | 6.7821 | 6.1709 | 6.7815 | 6.1832 | 6.8033 | 6.7776 | 5.6094 | 5.4414 | 4.5389 | 5.0278 |
| RRAS_S201 | RRAS | P10301 | | 201 | EQELPPSPPSAPR | 6.0834 | 6.2912 | 6.8336 | 7.2176 | 6.2217 | 5.8975 | 6.7465 | 6.6065 | 7.3254 | 7.1925 | 7.2586 | 7.1105 | 4.5474 | 5.5677 | 4.0794 | 5.0205 |
| RRAS2_S192 | RRAS2 | P62070-4 | | 192 | EQECPPSPEPTRK | 6.9596 | 7.0431 | 6.9798 | 6.8191 | 6.7299 | 6.9001 | 6.7049 | 5.8996 | 6.3444 | 6.2302 | 6.4123 | 6.2025 | 5.2158 | 5.1742 | 5.0329 | 5.3514 |
| RRBP1_S1277 | RRBP1 | Q9P2E9 | | 225 | GRKAEGTPNQGKK | 6.9439 | 7.4643 | 6.2605 | 7.4855 | 6.5903 | 6.9960 | 7.5050 | 6.8477 | 6.6007 | 6.9245 | 7.1094 | 6.7961 | 3.4178 | 4.9734 | 3.6246 | 4.4603 |
| RRBP1_S552 | RRBP1 | Q9P2E9 | | 275 | GKKVDTTPNQGKK | 6.5548 | 7.4007 | 6.5220 | 7.8161 | 6.0126 | 6.0444 | 6.6657 | 6.5189 | 6.6875 | 6.8872 | 6.4264 | 6.4831 | 4.1745 | 5.5044 | 4.6777 | 5.6240 |
| RRBP1_S573 | RRBP1 | Q9P2E9 | | 583 | GKKAEGSPNQGKK | 6.8349 | 6.7895 | 5.7227 | 6.4103 | 7.0794 | 6.8837 | 6.5738 | 6.2905 | 6.2534 | 5.8489 | 6.0963 | 5.9570 | 5.9497 | 6.4036 | 5.6047 | 5.3015 |
| RRBP1_S583 | RRBP1 | Q9P2E9 | | 959 | LTERIRSIEALLE | 6.5548 | 6.6279 | 5.5652 | 6.1699 | 6.4579 | 5.8057 | 5.4910 | 5.6417 | 5.9765 | 6.1219 | 5.8087 | 5.7031 | 6.7580 | 7.6016 | 6.9751 | 6.7409 |
| RRBP1_S959 | RRBP1 | Q9P2E9 | | 1277 | AGAPASSPEAPPA | 5.8891 | 6.2366 | 6.5548 | 6.4954 | 5.8164 | 5.7952 | 6.0948 | 5.7104 | 6.2709 | 6.6083 | 6.7391 | 6.3005 | 5.7589 | 7.0279 | 6.3691 | 6.3327 |
| RRBP1_T225 | RRBP1 | Q9P2E9 | | 573 | GKKAEGSPSEGKK | 5.7946 | 6.3114 | 6.0727 | 6.8835 | 6.2976 | 6.5298 | 5.9025 | 6.1075 | 6.0994 | 6.3272 | 6.2538 | 6.2858 | 5.8830 | 6.8957 | 6.1395 | 6.2162 |
| RRBP1_T235 | RRBP1 | Q9P2E9 | | 235 | GKKTEGTPNQGKK | 7.0556 | 7.4171 | 6.6556 | 7.2457 | 6.1697 | 6.3125 | 7.4375 | 6.6524 | 6.4493 | 6.9684 | 6.6498 | 6.5958 | 3.7492 | 5.2404 | 4.3925 | 5.0084 |
| RRBP1_T275 | RRBP1 | Q9P2E9 | | 552 | QGKKADSVANQGT | 5.4802 | 5.7650 | 6.0549 | 7.2171 | 6.5236 | 6.5492 | 6.2157 | 5.8727 | 5.3630 | 6.3645 | 6.1242 | 5.5127 | 6.2225 | 7.1530 | 6.8896 | 6.6920 |
| RREB1_S1122 | RREB1 | Q92766-2 | | 36 | VTENGGSPQGIKS | 6.6019 | 7.0758 | 6.3276 | 6.8307 | 6.5497 | 6.1123 | 6.8602 | 6.6276 | 7.0137 | 6.6719 | 6.0974 | 6.1776 | 5.2712 | 5.3193 | 5.0575 | 5.4056 |
| RREB1_S1167 | RREB1 | Q92766-2 | | 161 | TATAPPSPLKRRR | 6.0034 | 5.8008 | 6.0421 | 6.6375 | 6.5039 | 6.1977 | 6.8740 | 6.4889 | 6.3176 | 6.0841 | 6.2201 | 5.9799 | 6.3534 | 6.2741 | 6.2781 | 5.9444 |
| RREB1_S1219;S1225 | RREB1 | Q92766-2 | | 1708 | GDLNPESPAALGQ | 5.5713 | 6.2818 | 6.2637 | 6.2752 | 6.4607 | 5.9772 | 6.1479 | 6.4012 | 6.7054 | 6.4158 | 6.3287 | 6.0571 | 6.1129 | 6.6882 | 5.9231 | 6.3898 |
| RREB1_S1238 | RREB1 | Q92766-2 | | 1122 | TPAATTSPKESSE | 6.1154 | 6.2451 | 6.3296 | 7.7497 | 6.0070 | 6.0253 | 6.0996 | 6.0712 | 5.9521 | 6.6504 | 6.6447 | 6.7613 | 5.8345 | 6.0774 | 5.7161 | 5.7206 |
| RREB1_S1375 | RREB1 | Q92766-2 | | 1167 | SRPRANSGGVDLD | 5.5638 | 5.5908 | 5.7997 | 6.1411 | 6.5597 | 5.7489 | 6.0817 | 6.2785 | 6.0716 | 6.1923 | 6.3553 | 5.8897 | 6.6870 | 7.0381 | 6.7811 | 7.2207 |
| RREB1_S161 | RREB1 | Q92766-2 | | 1238 | LRAKRNSYTNCLQ | 5.6686 | 5.6666 | 5.3511 | 6.3828 | 10.3189 | 6.0199 | 5.8066 | 6.0225 | 7.3952 | 7.2400 | 7.1136 | 6.4087 | 5.6339 | 5.6906 | 4.6395 | 4.6414 |
| RREB1_S1708 | RREB1 | Q92766-2 | | 1375 | ATAETASPVHREE | 8.0099 | 8.8364 | 7.0944 | 7.3986 | 5.5769 | 5.2620 | 5.5025 | 4.9962 | 4.3650 | 5.4334 | 7.3548 | 7.9900 | 4.8500 | 5.6380 | 5.9396 | 5.7523 |
| RREB1_S36 | RREB1 | Q92766-2 | 1219;1225 | | NA | 5.7960 | 6.0489 | 6.9927 | 6.8081 | 6.3359 | 6.0129 | 6.9290 | 5.9496 | 6.1846 | 6.2780 | 5.7348 | 6.8103 | 5.8392 | 6.1813 | 5.5406 | 6.5582 |
| RRM1_S119 | RRM1 | P23921 | | 119 | SPMVAKSTLDIVL | 6.1093 | 6.7314 | 7.1998 | 5.7873 | 6.4136 | 6.7706 | 5.8811 | 6.9127 | 6.3845 | 7.1596 | 6.3648 | 6.1409 | 5.8811 | 5.8763 | 5.1806 | 5.2064 |
| RRM1_T721 | RRM1 | P23921 | | 721 | PNYGKLTSMHFYG | 6.3683 | 6.6380 | 6.6898 | 6.2646 | 7.0062 | 8.2871 | 5.0926 | 7.7841 | 7.1514 | 6.7624 | 6.3683 | 4.1176 | 7.0321 | 4.7762 | 5.3259 | 4.3354 |
| RRM2_S437 | RRM2 | P31350-2 | | 80 | PQQLQLSPLKGLS | 5.6911 | 5.0848 | 5.9058 | 5.0667 | 5.8765 | 7.3087 | 6.6955 | 6.9011 | 6.3135 | 5.7152 | 5.8709 | 6.1658 | 6.4240 | 7.0318 | 6.6692 | 7.2794 |
| RRM2_S80 | RRM2 | P31350-2 | | 437 | RMGVMSSPTENSF | 6.9289 | 7.5014 | 6.6218 | 6.2627 | 6.6159 | 5.7281 | 6.8144 | 6.5304 | 6.7378 | 6.2002 | 5.9660 | 6.8523 | 5.1139 | 5.3482 | 5.0626 | 5.7155 |
| RRM2_T93 | RRM2 | P31350-2 | | 93 | LVDKENTPPALSG | 7.2776 | 7.1709 | 8.1746 | 6.6047 | 6.0632 | 6.2721 | 6.8538 | 6.7614 | 8.0938 | 7.1940 | 7.8755 | 8.0566 | 3.3652 | 3.8080 | 2.7437 | 3.6848 |
| RRN3_S44 | RRN3 | Q9NYV6 | | 44 | ENDFFNSPPRKTV | 7.5079 | 7.5778 | 8.3436 | 7.8015 | 6.1972 | 7.1803 | 7.1026 | 6.9201 | 7.1871 | 7.2843 | 8.3504 | 7.7027 | 2.2914 | 2.8703 | 2.5807 | 3.1022 |
| RRP1_S383 | RRP1 | P56182 | | 383 | GEKEPPSPGMERK | 7.1475 | 6.8554 | 6.3580 | 7.2114 | 6.7800 | 6.2961 | 6.8762 | 6.8491 | 7.3148 | 6.7704 | 6.8321 | 6.8088 | 4.2513 | 4.7450 | 4.1266 | 4.7773 |
| RRP12_S1080 | RRP12 | Q5JTH9 | | 1080 | EEILADSEDEEDN | 5.9937 | 6.0320 | 5.8317 | 6.4998 | 6.2273 | 5.9407 | 6.0608 | 6.1763 | 6.3371 | 6.2211 | 5.6500 | 6.2642 | 6.4344 | 7.0172 | 6.9584 | 6.3553 |
| RRP12_S1284 | RRP12 | Q5JTH9 | | 1284 | KAARRGSQVGHKN | 3.9901 | 3.4979 | 4.0330 | 4.4870 | 6.4464 | 5.2590 | 5.2780 | 5.4565 | 9.8617 | 10.3132 | 8.1017 | 8.5263 | 8.3841 | 8.0299 | 3.3353 | 4.9998 |
| RRP15_S206 | RRP15 | Q9Y3B9 | | 206 | KLISTVSKKDFIS | 5.9578 | 5.6789 | 7.1149 | 6.7098 | 4.6017 | 7.0021 | 6.1737 | 6.1674 | 8.5369 | 8.3351 | 8.6350 | 7.9938 | 4.0859 | 5.2528 | 2.7641 | 4.9900 |
| RRP15_S58;S67 | RRP15 | Q9Y3B9 | 58;67 | | NA | 5.7188 | 6.1097 | 6.4748 | 6.5908 | 6.4993 | 6.1257 | 6.1227 | 5.9301 | 6.4461 | 5.8608 | 6.2614 | 6.3156 | 6.5201 | 6.3585 | 6.1792 | 6.4865 |
| RRP1B_S245 | RRP1B | Q14684 | | 350 | SFAEDISADEDDQ | 6.0560 | 6.5931 | 6.3464 | 7.0729 | 6.5623 | 6.3124 | 6.6907 | 6.1923 | 6.0918 | 6.0762 | 6.4860 | 6.4015 | 5.8213 | 5.9838 | 5.7478 | 5.5656 |
| RRP1B_S338 | RRP1B | Q14684 | | 513 | PQGPRGSPTGGAQ | 6.4226 | 7.1295 | 6.5483 | 7.3963 | 6.5217 | 6.6223 | 6.9443 | 6.8684 | 7.0892 | 6.8352 | 7.0945 | 6.9938 | 3.9403 | 4.8576 | 4.2332 | 4.5027 |
| RRP1B_S339 | RRP1B | Q14684 | | 339 | DLSEGSSISQLSF | 6.2130 | 6.5666 | 6.2901 | 7.2646 | 6.1791 | 5.8012 | 6.2758 | 6.5364 | 6.6381 | 6.3132 | 5.6970 | 6.1088 | 5.6975 | 6.1369 | 6.0905 | 6.1914 |
| RRP1B_S344 | RRP1B | Q14684 | | 706 | DKSILVSPTGPSR | 7.7033 | 8.0744 | 7.2456 | 7.8850 | 6.8301 | 6.6738 | 7.5440 | 6.7976 | 7.6879 | 7.1421 | 7.4333 | 7.2903 | 2.5526 | 3.3238 | 2.5838 | 3.2324 |
| RRP1B_S350 | RRP1B | Q14684 | | 344 | SSISQLSFAEDIS | 6.6137 | 6.7560 | 6.4737 | 6.8850 | 6.4835 | 6.3187 | 6.2892 | 6.2637 | 5.4480 | 6.0687 | 5.6372 | 7.0436 | 5.5909 | 6.0167 | 6.3048 | 5.8067 |
| RRP1B_S513 | RRP1B | Q14684 | | 245 | VGDGDLSAEEIPE | 5.6179 | 6.8839 | 6.8615 | 7.1349 | 6.7021 | 6.2715 | 5.8646 | 6.5378 | 6.9279 | 5.5535 | 7.1628 | 6.1736 | 4.5601 | 6.6567 | 5.2641 | 5.8269 |
| RRP1B_S630 | RRP1B | Q14684 | | 747 | KKPLTTTPRRRPR | 7.8004 | 8.5152 | 7.4355 | 6.9100 | 5.2432 | 7.6128 | 7.5325 | 6.9721 | 8.3085 | 8.1330 | 7.9306 | 8.1786 | 1.9259 | 2.9073 | 1.7153 | 2.8790 |
| RRP1B_S706 | RRP1B | Q14684 | | 630 | SSGTCSSLKKQKL | 6.8253 | 6.8550 | 7.3324 | 7.5948 | 5.2149 | 6.8653 | 8.0424 | 6.9159 | 6.3517 | 7.6598 | 7.4629 | 7.7011 | 3.4257 | 4.5532 | 3.0806 | 4.1190 |
| RRP1B_S731;S736 | RRP1B | Q14684 | | 338 | QDLSEGSSISQLS | 5.7313 | 5.9867 | 6.8541 | 7.1910 | 6.4644 | 6.1429 | 6.3135 | 6.1871 | 6.2058 | 5.2824 | 6.4185 | 6.5113 | 5.8992 | 5.9323 | 6.0586 | 6.8210 |
| RRP1B_S736 | RRP1B | Q14684 | | 736 | TSSPASSPLVAKK | 8.0025 | 7.3570 | 6.2638 | 7.5324 | 8.8651 | 6.6824 | 8.3260 | 7.4360 | 6.9783 | 5.6728 | 6.1102 | 5.7263 | 3.8608 | 3.9172 | 3.6355 | 3.6337 |
| RRP1B_T728;S731 | RRP1B | Q14684 | | 731 | VLKTPTSSPASSP | 5.8271 | 6.0855 | 5.9663 | 7.2641 | 6.2563 | 5.9714 | 6.8021 | 6.2614 | 6.3124 | 5.5390 | 6.2130 | 6.3547 | 5.9186 | 6.3875 | 6.2851 | 6.5554 |
| RRP1B_T731 | RRP1B | Q14684 | 731;736 | | NA | 6.5023 | 6.3342 | 6.0522 | 7.0312 | 6.8616 | 6.2817 | 6.3877 | 6.5970 | 6.6703 | 6.0135 | 6.2018 | 6.6141 | 5.7779 | 5.8804 | 5.6333 | 5.1610 |
| RRP1B_T747 | RRP1B | Q14684 | 728;731 | | NA | 5.7276 | 6.7779 | 6.0823 | 6.5641 | 5.2308 | 6.1134 | 6.0536 | 6.3531 | 6.7138 | 6.8105 | 6.7257 | 7.3361 | 5.4845 | 5.9588 | 5.6319 | 6.4359 |
| RRP36_S73 | RRP36 | Q96EU6 | | 73 | QLVAGNSPKKQAS | 6.0633 | 5.8538 | 6.2159 | 6.4945 | 6.5026 | 5.8090 | 6.6109 | 6.2473 | 6.8382 | 6.1385 | 5.9215 | 6.0452 | 6.2586 | 6.0612 | 6.2922 | 6.6475 |
| RRP7A_S19 | RRP7A | Q9Y3A4 | | 99 | KPDLAESPKESRS | 6.0199 | 6.1683 | 6.7172 | 7.0217 | 6.1958 | 6.4375 | 6.5163 | 6.0711 | 6.3870 | 6.4235 | 6.5729 | 6.1507 | 5.7656 | 5.8485 | 5.6134 | 6.0906 |
| RRP7A_S99 | RRP7A | Q9Y3A4 | | 19 | PEDRIPSPLGYAA | 5.8511 | 6.2261 | 6.4360 | 6.4602 | 6.0826 | 5.8782 | 6.7019 | 6.6245 | 6.6563 | 6.3145 | 6.3298 | 6.2335 | 5.7641 | 6.1197 | 6.1178 | 6.2036 |
| RRP8_S104;S106 | RRP8 | O43159 | | 223 | TEKTEVSPVPRTD | 6.2749 | 6.9869 | 6.3851 | 7.2816 | 5.9598 | 6.0408 | 6.4606 | 6.3385 | 6.5523 | 6.6124 | 6.6795 | 6.8954 | 4.8693 | 5.5316 | 5.3566 | 5.7747 |
| RRP8_S176 | RRP8 | O43159 | | 176 | QSPGSTSPKPPHT | 6.9031 | 6.6615 | 6.0880 | 7.4438 | 6.7321 | 6.2336 | 6.1701 | 6.0968 | 6.9034 | 6.4384 | 6.3083 | 6.6761 | 5.0980 | 5.5616 | 5.0972 | 5.5880 |
| RRP8_S223 | RRP8 | O43159 | 104;106 | | NA | 6.4491 | 6.6583 | 6.2730 | 7.0251 | 6.1379 | 6.0554 | 6.2779 | 5.1220 | 5.8199 | 6.1661 | 6.2645 | 5.9492 | 6.6447 | 6.4189 | 6.4286 | 6.3093 |
| RRP8_S62;S64 | RRP8 | O43159 | 62;64 | | NA | 5.9922 | 6.5757 | 5.6254 | 7.4303 | 6.9193 | 6.8264 | 6.0191 | 7.5469 | 5.8484 | 5.3396 | 6.1386 | 5.8323 | 6.1114 | 6.0796 | 6.0050 | 5.7096 |
| RSBN1_S81 | RSBN1 | Q5VWQ0 | | 81 | KPHAGVSPRGVKR | 6.3577 | 5.8185 | 6.0433 | 6.7636 | 5.9125 | 5.7093 | 6.1814 | 5.5654 | 6.4073 | 6.6743 | 5.9572 | 6.4495 | 5.8826 | 6.6012 | 6.4891 | 7.1871 |
| RSBN1L_S41 | RSBN1L | Q6PCB5 | | 41 | DPPGSLSAKKVRT | 7.2394 | 7.1108 | 6.6730 | 7.7087 | 6.3949 | 7.1479 | 7.5179 | 7.3371 | 7.5789 | 7.2149 | 6.6207 | 7.0568 | 3.0222 | 4.2283 | 2.9614 | 4.1871 |
| RSBN1L_S76 | RSBN1L | Q6PCB5 | | 342 | NLKRLDTLEFKQL | 4.4859 | 4.4328 | 4.7328 | 4.9894 | 5.6262 | 4.8933 | 5.5197 | 5.0189 | 4.7988 | 4.3923 | 4.1941 | 3.9590 | 10.1972 | 10.1527 | 11.6187 | 10.9882 |
| RSBN1L_T285 | RSBN1L | Q6PCB5 | | 285 | ICSGLLTDVEDQA | 8.0013 | 7.9466 | 7.6985 | 9.7404 | 7.2920 | 7.1820 | 7.8809 | 6.2369 | 4.6064 | 4.4566 | 3.9966 | 4.5972 | 4.5106 | 4.8318 | 4.8644 | 6.1577 |
| RSBN1L_T342 | RSBN1L | Q6PCB5 | | 76 | QPPAAPSPQSYGS | 6.1728 | 6.8125 | 5.8397 | 8.1538 | 4.4572 | 6.5472 | 8.0182 | 5.5626 | 5.9104 | 5.4948 | 4.6163 | 5.7395 | 6.4293 | 6.2612 | 7.3637 | 6.6209 |
| RSF1_S1345 | RSF1 | Q96T23 | | 397 | LSDDFDSPVKGPL | 7.2748 | 7.1145 | 6.7293 | 7.0859 | 6.6161 | 5.9707 | 7.0889 | 6.6921 | 7.6870 | 6.5228 | 6.5110 | 6.8084 | 4.4597 | 4.7326 | 4.0913 | 4.6149 |
| RSF1_S1359 | RSF1 | Q96T23 | | 604 | KDAQRLSPIPEEV | 5.8696 | 6.1134 | 5.6427 | 6.2713 | 6.1716 | 5.7690 | 6.2066 | 6.0756 | 6.2397 | 6.0781 | 5.6891 | 5.9096 | 7.1960 | 6.8191 | 7.0246 | 6.9241 |
| RSF1_S1359;S1375 | RSF1 | Q96T23 | | 1345 | KPYRIESDEEEDF | 5.9641 | 6.1015 | 6.3216 | 6.5644 | 5.9900 | 6.0845 | 6.2671 | 5.5989 | 6.4649 | 6.2801 | 6.1828 | 6.5791 | 6.6470 | 6.2925 | 6.1735 | 6.4882 |
| RSF1_S1375 | RSF1 | Q96T23 | | 1375 | PSTNGQSPGKAIE | 6.3694 | 6.5704 | 6.4450 | 6.6868 | 6.0603 | 6.0029 | 6.3111 | 6.1890 | 6.8229 | 6.4670 | 6.5058 | 6.5757 | 6.0354 | 5.7639 | 5.5213 | 5.6731 |
| RSF1_S227 | RSF1 | Q96T23 | | 748 | RKKKPDSPPKVLE | 5.9048 | 6.1649 | 6.1202 | 6.8150 | 6.2647 | 5.9792 | 6.2184 | 5.7403 | 6.3049 | 6.2824 | 6.0259 | 6.3871 | 6.3367 | 6.2630 | 6.4947 | 6.6978 |
| RSF1_S397 | RSF1 | Q96T23 | | 622 | ESEKPGSPEAAET | 6.0204 | 6.4251 | 6.0849 | 6.9559 | 6.3557 | 6.2721 | 5.8766 | 5.9499 | 6.2289 | 6.4497 | 6.4167 | 6.3267 | 6.4965 | 5.8912 | 6.0083 | 6.2416 |
| RSF1_S429 | RSF1 | Q96T23 | | 473 | TKEESYSPSKDRN | 6.7509 | 6.8800 | 5.9373 | 6.6473 | 6.2935 | 5.8633 | 6.2660 | 6.0196 | 6.2771 | 6.0572 | 6.0430 | 6.1709 | 6.3675 | 6.1726 | 5.9522 | 6.3016 |
| RSF1_S473 | RSF1 | Q96T23 | | 1359 | NVGKVGSPLDYSL | 5.1662 | 5.5920 | 6.6514 | 5.7901 | 5.7844 | 5.5584 | 6.3010 | 5.3300 | 6.2224 | 6.2994 | 5.8237 | 6.5442 | 7.5054 | 6.9781 | 7.0453 | 7.4080 |
| RSF1_S604 | RSF1 | Q96T23 | | 429 | ETCKRISTITALG | 7.4404 | 6.6523 | 6.4574 | 8.1691 | 5.1693 | 6.0125 | 6.5082 | 6.8472 | 8.4869 | 5.2244 | 6.4997 | 12.3088 | 1.7245 | 4.7498 | 3.3049 | 4.4447 |

| | | | | | | | | | | | | | | | | | | | | |
|---|---|---|---|---|---|---|---|---|---|---|---|---|---|---|---|---|---|---|---|---|
| RSF1_S622 | RSF1 | Q96T23 | 227 | DNSSRESPSLEDE | 5.4811 | 5.5673 | 5.6941 | 6.3419 | 5.9602 | 5.4479 | 5.9277 | 5.5766 | 6.4295 | 6.5595 | 5.8826 | 6.3216 | 7.1648 | 6.8682 | 7.6122 | 7.1648 |
| RSF1_S622;S629 | RSF1 | Q96T23 | 1359;1375 | NA | 5.8983 | 5.8164 | 6.1930 | 6.4133 | 5.7571 | 5.8580 | 6.5222 | 5.5793 | 6.3397 | 6.1018 | 6.4022 | 6.6034 | 6.7488 | 6.3640 | 6.2237 | 7.1787 |
| RSF1_S748 | RSF1 | Q96T23 | 622;629 | NA | 6.1758 | 6.5088 | 6.0216 | 6.9224 | 6.0572 | 5.8905 | 6.0801 | 5.6825 | 6.2033 | 5.8140 | 6.0150 | 5.9487 | 6.8140 | 6.4785 | 6.5064 | 6.8812 |
| RSL1D1_S314 | RSL1D1 | O76021 | 361 | VKATNESEDEIPQ | 6.1904 | 6.3887 | 6.1994 | 7.1038 | 6.1575 | 6.2016 | 6.4685 | 6.3375 | 5.6340 | 5.5728 | 5.6163 | 5.6375 | 6.0804 | 6.8353 | 6.6880 | 6.8884 |
| RSL1D1_S317 | RSL1D1 | O76021 | 415 | ALPASETPKAAES | 7.9646 | 7.9693 | 7.0898 | 7.4421 | 6.5131 | 6.9526 | 8.0943 | 6.8050 | 7.9749 | 7.4815 | 7.4132 | 7.2573 | 2.2094 | 3.4279 | 2.2255 | 3.1796 |
| RSL1D1_S361 | RSL1D1 | O76021 | 427 | SETPGKSPEKKPK | 6.3225 | 6.3750 | 6.2441 | 6.9508 | 6.3981 | 6.1689 | 6.8924 | 5.9960 | 6.6156 | 6.4761 | 6.6168 | 6.7329 | 5.0565 | 6.0605 | 5.2350 | 5.8587 |
| RSL1D1_S396;T401 | RSL1D1 | O76021 | 401 | KSPNPSTPRGKKR | 7.4602 | 8.0563 | 6.6875 | 7.2790 | 6.1412 | 6.7141 | 7.3394 | 6.8248 | 7.6651 | 6.7298 | 7.0988 | 7.2847 | 3.9302 | 3.8557 | 2.8809 | 4.0522 |
| RSL1D1_S427 | RSL1D1 | O76021 | 464 | PEAKFFTTPSKSV | 7.9606 | 8.1575 | 6.9317 | 7.4012 | 5.6256 | 6.9169 | 8.2833 | 7.0409 | 7.6695 | 7.7761 | 7.2898 | 7.6995 | 2.0455 | 3.4852 | 2.3421 | 3.3746 |
| RSL1D1_S443 | RSL1D1 | O76021 | 465 | EAKFFTTPSKSVR | 8.1286 | 8.3500 | 6.9989 | 7.3360 | 5.8099 | 7.1594 | 8.3420 | 7.1743 | 7.8649 | 7.7867 | 7.3265 | 7.6722 | 1.8381 | 3.2866 | 1.9316 | 2.9943 |
| RSL1D1_S92 | RSL1D1 | O76021 | 92 | HSIRSDSEDICLF | 6.0393 | 6.2862 | 5.8280 | 6.8316 | 5.7675 | 5.8704 | 6.2270 | 5.7638 | 6.0657 | 5.9904 | 6.2366 | 6.1712 | 5.9777 | 6.9531 | 6.7913 | 7.2002 |
| RSL1D1_T340 | RSL1D1 | O76021 | 340 | ESKKEQTPEHGKK | 6.9150 | 7.5114 | 6.7035 | 7.6808 | 6.6105 | 7.1072 | 7.4984 | 6.9181 | 7.0599 | 7.3717 | 7.4777 | 7.0544 | 3.0725 | 4.2567 | 3.0065 | 3.7556 |
| RSL1D1_T401 | RSL1D1 | O76021 | 454 | KKDARQTPKKPEA | 8.1656 | 8.5615 | 7.5538 | 7.9484 | 5.6710 | 7.7182 | 7.7869 | 6.3593 | 6.9087 | 7.5003 | 8.2898 | 8.3285 | 1.8657 | 3.1782 | 1.6500 | 2.5141 |
| RSL1D1_T415 | RSL1D1 | O76021 | 317 | KTASVLSKDDVAP | 5.7633 | 5.7238 | 5.8739 | 6.4916 | 6.8163 | 5.7297 | 6.8826 | 6.4006 | 5.8855 | 5.2470 | 5.6255 | 5.5860 | 6.2680 | 7.1469 | 7.0239 | 7.5354 |
| RSL1D1_T423 | RSL1D1 | O76021 | 314 | QARKTASVLSKDD | 5.7696 | 6.6849 | 6.7583 | 6.6150 | 5.2676 | 6.2935 | 6.5206 | 6.5534 | 6.5361 | 7.5951 | 7.5700 | 8.1855 | 4.2297 | 5.4824 | 4.5224 | 5.4159 |
| RSL1D1_T423;S427 | RSL1D1 | O76021 | 443 | EAVKEKSPSLGKK | 5.1230 | 5.6564 | 6.3868 | 7.7133 | 6.7935 | 7.2368 | 7.3928 | 6.3444 | 6.9763 | 7.0364 | 6.6013 | 7.5256 | 4.4791 | 5.8013 | 4.3821 | 4.5509 |
| RSL1D1_T454 | RSL1D1 | O76021 | 423 | KAAESETPGKSPE | 7.3146 | 7.2497 | 6.7763 | 7.7273 | 5.6992 | 6.0281 | 6.6880 | 6.2660 | 6.9353 | 6.8488 | 6.6052 | 6.7034 | 4.3124 | 5.0918 | 4.4243 | 5.3297 |
| RSL1D1_T464 | RSL1D1 | O76021 | 423;427 | NA | 6.6518 | 7.0675 | 7.8414 | 8.4606 | 5.2823 | 7.4271 | 7.2272 | 5.7172 | 8.1419 | 7.7240 | 8.1907 | 8.6360 | 2.3656 | 3.8959 | 2.1691 | 3.2018 |
| RSL1D1_T465 | RSL1D1 | O76021 | 396;401 | NA | 6.9386 | 4.4821 | 7.4718 | 8.1703 | 6.2082 | 7.2883 | 6.9923 | 6.0855 | 6.5262 | 8.8328 | 7.8360 | 6.5150 | 4.2899 | 5.1633 | 3.2666 | 3.9331 |
| RSRC1_S331 | RSRC1 | Q96IZ7 | 331 | QERLMGSPVAxxx | 7.1835 | 7.4467 | 7.0291 | 7.4072 | 6.5360 | 6.4032 | 6.5862 | 6.7524 | 7.1621 | 6.7773 | 6.7643 | 6.9996 | 4.0852 | 4.4730 | 4.0193 | 4.3750 |
| RSRC2_S17 | RSRC2 | Q7L4I2 | 32 | QSEVSVSPRASKH | 5.9969 | 6.3569 | 6.0447 | 5.7117 | 6.0874 | 6.0028 | 5.8985 | 5.8675 | 6.3492 | 6.1600 | 6.1323 | 6.1084 | 6.7365 | 6.7461 | 6.9581 | 6.8431 |
| RSRC2_S32 | RSRC2 | Q7L4I2 | 54 | RERKRKSDNEGRK | 6.3122 | 6.9653 | 6.1664 | 6.3952 | 5.7536 | 6.6075 | 6.5533 | 6.5911 | 6.2293 | 6.2449 | 6.6938 | 6.4515 | 4.9921 | 5.9799 | 5.8280 | 6.2358 |
| RSRC2_S54 | RSRC2 | Q7L4I2 | 17 | LAPEKTSPDRDKK | 5.9354 | 5.6762 | 6.2452 | 5.3635 | 6.1901 | 5.9962 | 6.1249 | 5.3559 | 6.1706 | 6.3034 | 5.7492 | 5.6815 | 7.6670 | 7.2848 | 7.1232 | 7.1329 |
| RSRP1_S107 | RSRP1 | Q9BUV0 | 282 | KIDQKKSPYGLWI | 7.5815 | 8.3899 | 7.8159 | 9.2379 | 5.0766 | 6.2547 | 5.3342 | 6.2558 | 7.4324 | 6.4087 | 7.2393 | 6.6081 | 4.2471 | 4.0741 | 3.8487 | 4.1952 |
| RSRP1_S282 | RSRP1 | Q9BUV0 | 107 | TRRYYRSPSRYRS | 6.6998 | 7.6717 | 7.4462 | 9.0052 | 5.5601 | 6.0812 | 5.7279 | 6.0578 | 6.6331 | 6.9091 | 6.6951 | 6.4512 | 4.9245 | 4.8177 | 4.4279 | 4.8917 |
| RTEL1_S303 | RTEL1 | Q9NZ71-2 | 779 | AVSEAKSPGPFFS | 6.7625 | 6.8016 | 6.7793 | 6.4475 | 6.2420 | 6.5319 | 6.8236 | 6.6621 | 7.2927 | 7.2593 | 6.8414 | 7.0735 | 4.2891 | 5.0415 | 4.0850 | 5.0671 |
| RTEL1_S779 | RTEL1 | Q9NZ71-2 | 791 | STRKAKSLDLHVP | 6.0225 | 6.9702 | 7.2457 | 6.3967 | 6.5238 | 6.2215 | 6.3414 | 6.7465 | 6.5331 | 7.3785 | 7.1082 | 6.7274 | 4.1716 | 5.4934 | 5.1653 | 4.9544 |
| RTEL1_S791 | RTEL1 | Q9NZ71-2 | 303 | PEFSADSPSPGLN | 5.6393 | 5.9947 | 5.9081 | 6.6306 | 6.6127 | 6.2840 | 5.3557 | 5.9332 | 6.5456 | 6.3597 | 6.6055 | 6.0208 | 6.1452 | 6.2036 | 6.9818 | 6.7796 |
| RTF1_S675 | RTF1 | Q92541 | 697 | DGAPRRSLNLEDY | 6.4310 | 7.1275 | 6.5894 | 7.3159 | 6.5825 | 6.8029 | 6.9941 | 6.7608 | 7.6131 | 7.5559 | 7.7409 | 7.6734 | 3.1834 | 4.3604 | 3.1873 | 4.0812 |
| RTF1_S697 | RTF1 | Q92541 | 675 | DLQVPSSESKALA | 7.8280 | 7.8695 | 7.1735 | 7.5291 | 6.3282 | 6.9178 | 7.7590 | 6.4509 | 7.1690 | 6.2749 | 6.6216 | 6.7385 | 3.5554 | 4.4109 | 3.1714 | 4.2022 |
| RTF1_S77;S79 | RTF1 | Q92541 | 77;79 | NA | 4.2713 | 4.5383 | 5.3692 | 4.3730 | 5.1401 | 5.5699 | 5.0780 | 5.5380 | 4.6779 | 4.9350 | 5.1954 | 5.0165 | 9.7022 | 9.8206 | 10.8738 | 9.9008 |
| RTKN_S106 | RTKN | Q9BST9 | 561 | PRTWLQSPVxxxx | 6.8217 | 7.3921 | 7.7083 | 6.9668 | 5.8783 | 6.5690 | 6.6900 | 7.1313 | 7.7611 | 7.2824 | 7.2924 | 6.9274 | 3.8953 | 4.5906 | 3.2982 | 3.7950 |
| RTKN_S220 | RTKN | Q9BST9 | 106 | KTSRRPSDSGPPA | 5.6510 | 5.5806 | 5.8872 | 5.3656 | 6.9858 | 6.2010 | 5.8845 | 6.6484 | 6.4856 | 7.4211 | 6.5956 | 6.6005 | 6.4331 | 6.6840 | 5.8269 | 5.7493 |
| RTKN_S520 | RTKN | Q9BST9 | 220 | ATKLSSSLGRSSG | 6.9327 | 6.5578 | 6.7718 | 5.5621 | 6.4055 | 6.5492 | 6.3928 | 6.7582 | 6.6780 | 6.6727 | 6.2159 | 6.5720 | 5.4837 | 5.7417 | 5.3053 | 5.4006 |
| RTKN_S561 | RTKN | Q9BST9 | 520 | GRPRTFSLDAVPP | 5.8289 | 5.4845 | 6.8446 | 6.2938 | 6.5798 | 6.0704 | 6.0486 | 6.7538 | 6.7415 | 6.6902 | 6.5649 | 6.7896 | 6.2328 | 6.0068 | 5.2261 | 5.8438 |
| RTN1_S352 | RTN1 | Q16799 | 279 | RAPQITTPVKITL | 6.4152 | 6.8165 | 6.2507 | 6.9675 | 6.2134 | 6.5018 | 6.9950 | 6.4148 | 6.7568 | 6.6177 | 6.6919 | 6.7570 | 4.9508 | 5.3748 | 4.9130 | 5.3631 |
| RTN1_S48 | RTN1 | Q16799 | 352 | QGKGSISEDELIT | 6.4203 | 6.7111 | 6.0490 | 6.8034 | 5.8322 | 6.2390 | 6.3544 | 6.5754 | 5.9361 | 6.5000 | 5.9314 | 5.7501 | 5.9258 | 6.5530 | 6.3222 | 6.0967 |
| RTN1_S563 | RTN1 | Q16799 | 48 | PQAGEPSPGLGAR | 6.4037 | 6.8223 | 6.6166 | 7.1931 | 6.4085 | 6.3938 | 7.1083 | 6.8683 | 6.3532 | 6.5329 | 6.5902 | 6.6891 | 4.5474 | 5.4686 | 4.8073 | 5.1968 |
| RTN1_T279 | RTN1 | Q16799 | 563 | DSSSNQSPAATKG | 6.4252 | 7.3091 | 6.7216 | 7.8243 | 6.2731 | 7.0500 | 7.2550 | 6.6631 | 6.4636 | 7.3917 | 7.1094 | 6.6448 | 3.4503 | 4.8034 | 3.8575 | 4.7578 |
| RTN3_S229 | RTN3 | O95197 | 229 | IYTYSLSPSKVSG | 6.4229 | 6.2918 | 6.6273 | 6.4177 | 6.0466 | 6.6716 | 6.6333 | 6.6417 | 6.3041 | 6.4872 | 6.6831 | 6.7671 | 5.4350 | 5.6365 | 5.2165 | 5.7176 |
| RTN4_S107 | RTN4 | Q9NQC3-2 | 181 | APKRRGSSGSVVV | 5.4929 | 5.4857 | 5.1937 | 5.7149 | 5.5477 | 5.3653 | 5.4059 | 5.7897 | 6.9328 | 6.9833 | 6.7533 | 7.0015 | 6.6254 | 7.4084 | 6.8043 | 7.4951 |
| RTN4_S107;S121 | RTN4 | Q9NQC3 | 107 | APERQPSWDPSPV | 4.3450 | 4.3620 | 4.6781 | 4.8681 | 5.2423 | 5.0801 | 4.9888 | 5.2606 | 4.7728 | 4.4640 | 4.5517 | 4.4849 | 10.4308 | 10.0207 | 11.6464 | 10.8038 |
| RTN4_S107;S129 | RTN4 | Q9NQC3 | 181 | APKRRGSSGSVDE | 5.9268 | 6.2576 | 5.8536 | 6.4228 | 6.2179 | 5.8929 | 6.0505 | 6.2328 | 6.3832 | 6.2289 | 6.2918 | 6.4148 | 6.5502 | 6.4675 | 6.4404 | 6.3683 |
| RTN4_S111 | RTN4 | Q9NQC3 | 121 | STVPAPSPLSAAA | 6.1022 | 6.3954 | 6.2877 | 6.6491 | 6.3971 | 6.4117 | 6.6226 | 6.5371 | 6.3705 | 6.4872 | 6.4016 | 6.4776 | 5.5120 | 6.0067 | 5.6682 | 5.6734 |
| RTN4_S115;S129 | RTN4 | Q9NQC3 | 129 | LSAAAVSPSKLPE | 6.8413 | 7.3037 | 6.7635 | 7.4380 | 6.0765 | 6.7223 | 6.8894 | 6.6198 | 7.2714 | 7.6875 | 6.9868 | 7.4913 | 3.6754 | 4.3585 | 3.5296 | 4.3450 |
| RTN4_S121 | RTN4 | Q9NQC3 | 111 | QPSWDPSPVSSTV | 5.3509 | 5.7757 | 5.7711 | 5.8381 | 5.5687 | 6.1547 | 5.9302 | 5.7207 | 5.2591 | 5.3831 | 5.5323 | 5.5279 | 8.3901 | 7.9431 | 8.2857 | 7.5687 |
| RTN4_S121;S129 | RTN4 | Q9NQC3 | 152 | PPPASVSPQAEPV | 6.9770 | 7.2045 | 6.7588 | 7.7369 | 5.7192 | 6.5105 | 6.9040 | 6.9065 | 6.6173 | 7.3017 | 6.9875 | 7.1418 | 3.8070 | 4.6361 | 4.4852 | 4.3058 |
| RTN4_S129 | RTN4 | Q9NQC3 | 172 | PAAPPSTPAAPKR | 6.9247 | 6.2376 | 6.3243 | 7.0426 | 6.4681 | 6.3554 | 6.3794 | 6.5174 | 6.6004 | 6.0944 | 5.8952 | 5.9529 | 5.6528 | 5.5382 | 5.9225 | 6.0941 |
| RTN4_S152 | RTN4 | Q9NQC3 | 307 | GSSFSVSPKAESA | 5.9338 | 6.3479 | 6.7591 | 6.3980 | 6.2761 | 6.9459 | 5.7077 | 5.8859 | 6.9283 | 6.6013 | 6.1946 | 6.2174 | 6.2960 | 6.2157 | 5.1244 | 6.1679 |
| RTN4_S152;T172 | RTN4 | Q9NQC3 | 991 | TEKEDRSPSAIFS | 5.0710 | 5.7283 | 5.3486 | 5.7907 | 6.4347 | 6.1123 | 6.2981 | 6.1873 | 6.1821 | 5.7459 | 6.6051 | 6.9745 | 6.4928 | 6.9415 | 7.1251 | 6.9620 |
| RTN4_S172 | RTN4 | Q9NQC3 | 107;121 | NA | 4.5281 | 4.4983 | 4.6640 | 4.7700 | 5.2721 | 5.2019 | 5.7763 | 5.2463 | 4.6860 | 4.8310 | 4.5485 | 4.7627 | 9.8576 | 9.2698 | 11.2425 | 10.8449 |
| RTN4_S181 | RTN4 | Q9NQC3 | 121;129 | NA | 6.9659 | 7.7019 | 7.1255 | 7.8561 | 5.9249 | 6.4265 | 7.5973 | 6.8527 | 7.2950 | 7.3399 | 7.2980 | 7.2175 | 2.7437 | 4.2226 | 3.1287 | 4.3038 |
| RTN4_S181 | RTN4 | Q9NQC3 | 181;184 | NA | 6.1990 | 5.9513 | 6.2092 | 6.1234 | 5.8851 | 6.0846 | 6.5524 | 6.2223 | 6.2806 | 6.0809 | 6.1426 | 6.6375 | 6.4587 | 6.3070 | 6.2867 | 6.5790 |
| RTN4_S181;S184 | RTN4 | Q9NQC3 | 152;172 | NA | 7.4465 | 7.8329 | 7.1139 | 8.1294 | 6.0018 | 6.4885 | 7.3844 | 6.6079 | 6.9735 | 7.2610 | 6.8707 | 6.9099 | 3.1544 | 4.4786 | 2.9833 | 4.3632 |
| RTN4_S307 | RTN4 | Q9NQC3 | 115;129 | NA | 6.8276 | 7.7958 | 6.7847 | 8.0495 | 6.6042 | 7.2648 | 7.3475 | 6.7153 | 7.7044 | 7.9148 | 6.8302 | 6.4494 | 2.7466 | 4.0918 | 2.9961 | 3.8771 |
| RTN4_S991 | RTN4 | Q9NQC3 | 107;129 | NA | 6.0590 | 5.8896 | 5.8047 | 7.2311 | 6.1253 | 5.1211 | 5.2576 | 5.6924 | 5.7396 | 6.4443 | 6.1031 | 6.0282 | 7.5705 | 6.5454 | 7.1164 | 7.2717 |
| RUBCN_S197 | RUBCN | Q92622 | 266 | ESRGHVSPAEDQT | 6.0258 | 6.0667 | 6.6767 | 6.1723 | 5.8457 | 6.1200 | 6.3130 | 5.8615 | 6.6980 | 6.5525 | 6.6799 | 6.8720 | 5.9577 | 5.9899 | 5.8828 | 6.2856 |
| RUBCN_S266 | RUBCN | Q92622 | 388 | SVLRRSSFSEGQT | 5.9314 | 6.2463 | 6.2597 | 6.7370 | 5.9368 | 5.8955 | 5.9249 | 6.5490 | 6.7439 | 6.8913 | 6.9250 | 6.7166 | 5.7331 | 5.7317 | 5.3704 | 6.4073 |
| RUBCN_S388 | RUBCN | Q92622 | 197 | FARKHESPLLVTK | 6.8673 | 7.0253 | 6.9180 | 7.0261 | 6.6232 | 6.4690 | 6.4871 | 6.2252 | 7.3983 | 6.7502 | 6.4769 | 7.0948 | 4.5450 | 5.3216 | 3.9603 | 4.8118 |
| RUBCN_S410 | RUBCN | Q92622 | 469 | GMFRRPSEGQSLI | 5.2576 | 5.5790 | 5.5217 | 5.8934 | 6.1298 | 5.0645 | 4.6131 | 6.0558 | 7.9130 | 7.1454 | 7.6026 | 7.0448 | 6.5570 | 6.6466 | 6.7385 | 6.2373 |
| RUBCN_S469 | RUBCN | Q92622 | 410 | SHIRSHSDTSIAS | 6.5952 | 6.5822 | 5.4517 | 5.4171 | 5.5100 | 7.0989 | 6.3250 | 7.1075 | 5.5792 | 8.4456 | 7.2631 | 7.6522 | 4.7210 | 4.6648 | 5.7305 | 5.8559 |
| RUBCN_S671 | RUBCN | Q92622 | 671 | PDSLPISPDDGQH | 5.3399 | 7.9444 | 7.8042 | 7.3833 | 6.7848 | 8.2110 | 7.8135 | 4.7087 | 8.3700 | 7.9819 | 9.1789 | 6.2892 | 2.1650 | 3.1282 | 3.4742 | 3.4228 |
| RUNX1_S239 | RUNX1 | Q01196-8 | 48 | PPSTALSPGKMSE | 6.7691 | 6.7479 | 6.4482 | 7.3978 | 5.9868 | 6.3691 | 6.2411 | 6.3613 | 7.1049 | 6.8870 | 7.0622 | 7.1254 | 4.7604 | 4.9051 | 4.7869 | 5.0468 |
| RUNX1_T196 | RUNX1 | Q01196-8 | 239 | RTAMRVSPHHPAP | 5.3031 | 6.0830 | 5.4567 | 5.6949 | 6.1614 | 6.3459 | 6.2941 | 6.1412 | 5.9969 | 6.5292 | 5.8033 | 5.9512 | 7.4391 | 7.0794 | 6.9690 | 6.7516 |
| RUNX1_T41;S48 | RUNX1 | Q01196-8 | 196 | HRAIKITVDGPRE | 5.9966 | 5.6023 | 5.7944 | 5.5352 | 6.9966 | 5.4423 | 6.0131 | 5.9856 | 5.9074 | 6.3689 | 6.2291 | 5.4111 | 7.4498 | 6.9213 | 6.7963 | 7.5500 |
| RUNX1_T48 | RUNX1 | Q01196-8 | 41;48 | NA | 5.9440 | 5.9785 | 5.6056 | 6.0681 | 6.5054 | 6.5492 | 6.4730 | 6.2765 | 6.6081 | 6.2266 | 6.4083 | 6.4085 | 6.1456 | 6.4233 | 6.0599 | 6.3194 |
| RUNX2_S340 | RUNX2 | Q13950 | 340 | DVPRRISDDDTAT | 3.1879 | 3.8491 | 4.3279 | 5.5872 | 4.7919 | 6.0365 | 4.8764 | 5.9534 | 3.9618 | 3.8022 | 2.7359 | 4.0892 | 13.4517 | 10.9760 | 12.3037 | 10.0693 |
| RUNX2_S340 | RUNX2 | Q13950-3 | 340 | DVPRRISGASELG | 5.6598 | 5.8353 | 6.5085 | 6.9132 | 6.3761 | 5.6787 | 6.4344 | 6.2405 | 6.8139 | 6.7650 | 7.0872 | 6.8291 | 5.8306 | 6.5669 | 5.0848 | 5.3760 |
| RUSC2_S1347 | RUSC2 | Q8N2Y8 | 536 | AMAGPGSPPRRVT | 6.3410 | 6.0739 | 6.6333 | 6.2379 | 6.4039 | 6.5581 | 6.2251 | 7.0071 | 6.6794 | 6.1885 | 6.5924 | 6.4704 | 5.7948 | 5.8712 | 5.4438 | 5.4793 |
| RUSC2_S519 | RUSC2 | Q8N2Y8 | 542 | SPPRRVTSFAELA | 4.1864 | 3.9570 | 4.5259 | 3.8503 | 5.4511 | 5.1185 | 5.2271 | 5.6236 | 4.3114 | 3.5910 | 3.8293 | 3.8036 | 12.4590 | 10.8433 | 12.2940 | 10.9284 |
| RUSC2_S536 | RUSC2 | Q8N2Y8 | 519 | SLERMLSCPVRLS | 4.9565 | 4.5415 | 5.3955 | 4.5264 | 5.7871 | 5.8915 | 5.2873 | 5.8216 | 4.5362 | 4.0646 | 4.7810 | 4.6046 | 10.8496 | 9.0815 | 11.0137 | 8.8613 |
| RUSC2_S559 | RUSC2 | Q8N2Y8 | 890 | SNANHLSPQALKW | 5.9657 | 6.0258 | 6.2680 | 6.7406 | 7.4821 | 6.7456 | 5.8270 | 7.8111 | 8.2838 | 6.2496 | 6.6821 | 7.1364 | 4.8701 | 4.9786 | 3.6926 | 5.2408 |
| RUSC2_S781 | RUSC2 | Q8N2Y8 | 1347 | WPFWMGSPPDSVL | 6.2406 | 6.3942 | 7.2520 | 6.4046 | 6.4710 | 6.5534 | 5.6833 | 6.7762 | 6.9146 | 6.6700 | 6.4710 | 6.9345 | 5.4473 | 5.6653 | 4.6189 | 5.5032 |
| RUSC2_S890 | RUSC2 | Q8N2Y8 | 559 | KTGGSGSPPLRVS | 6.4006 | 5.7034 | 6.0317 | 5.7000 | 8.2120 | 7.6932 | 6.6666 | 6.7364 | 5.8842 | 5.4020 | 7.3506 | 6.8044 | 6.2291 | 6.0718 | 4.7128 | 4.4011 |
| RUSC2_T542 | RUSC2 | Q8N2Y8 | 781 | SPLGSYSPIRSVG | 5.6646 | 5.4595 | 6.7987 | 6.5996 | 6.5246 | 6.8518 | 6.3864 | 6.3357 | 6.1189 | 5.8389 | 6.4664 | 6.3190 | 6.3854 | 6.0268 | 6.0850 | 6.1387 |
| RWDD1_T144 | RWDD1 | Q9H446 | 144 | KQLFHGTPVTIEN | 7.1099 | 7.2482 | 7.6753 | 7.8017 | 6.7334 | 6.6529 | 6.5451 | 7.9800 | 7.7053 | 6.2990 | 7.4279 | 7.1026 | 3.1771 | 3.3747 | 3.1837 | 3.9833 |
| RYBP_S99 | RYBP | Q8N488 | 99 | EKDKEISPSVTKK | 6.3137 | 7.1590 | 6.5745 | 6.9836 | 6.0284 | 6.1006 | 6.5040 | 6.2525 | 6.5810 | 6.6967 | 6.8292 | 6.8949 | 4.8659 | 5.5275 | 5.1806 | 5.5080 |
| S1PR1_S351 | S1PR1 | P21453 | 351 | IAGMEFSRSKSDN | 6.4669 | 6.5160 | 7.4212 | 6.9376 | 6.6070 | 6.2616 | 6.2018 | 6.3927 | 6.3797 | 6.2007 | 6.3939 | 6.7937 | 5.1652 | 5.8835 | 5.0686 | 5.3099 |
| SAAL1_S56 | SAAL1 | Q96ER3 | 6 | xMDRNPSPPPPGR | 5.3584 | 5.5483 | 5.8569 | 5.2329 | 5.3209 | 5.6148 | 5.7227 | 5.9536 | 6.1520 | 6.1496 | 5.8394 | 6.2666 | 7.5138 | 7.8497 | 7.8537 | 7.7667 |
| SAAL1_S6 | SAAL1 | Q96ER3 | 56 | ENTKSSSDDEEQL | 5.5260 | 5.6337 | 6.2829 | 5.8788 | 5.6063 | 6.7301 | 5.9028 | 5.3347 | 5.4440 | 6.1297 | 6.9727 | 7.2270 | 6.5189 | 6.9371 | 7.3040 | 6.5711 |
| SACS_S1779 | SACS | Q9NZJ4 | 1779 | RIADLQSPLFRGP | 6.0146 | 6.1019 | 6.4677 | 5.9064 | 6.3865 | 6.4925 | 6.5179 | 5.9404 | 5.9919 | 5.6689 | 5.9967 | 5.9692 | 7.1507 | 6.6178 | 6.6833 | 6.0936 |
| SACS_S2511 | SACS | Q9NZJ4 | 2511 | ALERYASNVCFTT | 9.2303 | 8.1533 | 8.6380 | 8.0244 | 7.5589 | 5.1259 | 5.4583 | 5.0588 | 7.7549 | 5.8279 | 6.2902 | 5.9216 | 5.1291 | 5.2505 | 3.2980 | 3.2799 |
| SACS_T4266 | SACS | Q9NZJ4 | 4266 | STPTSPTEFLTPG | 5.5993 | 5.8763 | 6.1373 | 5.6652 | 5.7488 | 6.3813 | 6.7580 | 5.9616 | 6.3138 | 6.4480 | 5.9396 | 5.1871 | 7.0232 | 6.7766 | 7.4750 | 6.7090 |
| SAFB_S195 | SAFB | Q15424-3 | 604 | EKRSVVSFDKVKE | 5.8936 | 5.8441 | 5.8956 | 6.7495 | 6.5574 | 5.9713 | 7.0520 | 7.1308 | 6.2351 | 6.1043 | 6.0608 | 6.4501 | 5.8311 | 5.7856 | 5.9317 | 6.5069 |
| SAFB_S383;S384 | SAFB | Q15424-3 | 601 | ASREKRSVVSFDK | 6.9867 | 6.6473 | 6.0277 | 7.5327 | 6.9936 | 5.9045 | 6.2242 | 5.2809 | 6.6678 | 5.6049 | 5.7649 | 5.3815 | 6.8085 | 6.6768 | 5.6705 | 5.8274 |
| SAFB_S384 | SAFB | Q15424-3 | 195 | INNLDTSSSDFTI | 6.0035 | 6.3027 | 6.1976 | 6.5246 | 5.8484 | 6.0611 | 6.0239 | 6.3055 | 6.2274 | 6.2128 | 5.9945 | 6.1435 | 6.2299 | 6.7186 | 6.5527 | 6.6533 |
| SAFB_S576 | SAFB | Q15424-3 | 384 | ADQKMSSPEDDSD | 5.4610 | 6.1426 | 5.5569 | 6.1967 | 5.5992 | 5.8649 | 6.1355 | 5.9407 | 6.4155 | 6.5504 | 6.3095 | 6.6337 | 6.7798 | 6.8692 | 7.0920 | 6.4523 |
| SAFB_S601 | SAFB | Q15424-3 | 576 | KGVPVISVKTSGS | 6.4070 | 6.6509 | 5.4287 | 6.4540 | 7.7641 | 5.4733 | 6.3054 | 6.2063 | 5.2194 | 4.3142 | 5.1105 | 5.9005 | 7.3320 | 6.5655 | 6.9233 | 7.9449 |
| SAFB_S604 | SAFB | Q15424-3 | 383;384 | NA | 6.2999 | 6.2507 | 6.0671 | 6.4552 | 5.8338 | 5.7171 | 6.3199 | 5.5027 | 6.4241 | 6.3463 | 6.2598 | 6.1933 | 6.8674 | 6.5831 | 6.3902 | 6.4893 |
| SAFB2_S194 | SAFB2 | Q14151 | 246 | FAQDTSSVGPDRK | 5.9995 | 5.9061 | 5.9480 | 6.4619 | 6.1814 | 5.8618 | 6.4503 | 6.0321 | 5.9518 | 6.0010 | 5.9362 | 6.1083 | 7.0467 | 6.6411 | 6.8221 | 6.6516 |
| SAFB2_S195 | SAFB2 | Q14151 | 233 | EPVKEESSELEQP | 5.7865 | 6.0089 | 6.1416 | 6.3136 | 6.1188 | 5.5793 | 6.2032 | 6.1720 | 6.5206 | 6.2868 | 6.0927 | 6.1439 | 6.6382 | 6.7912 | 6.3926 | 6.8100 |
| SAFB2_S207 | SAFB2 | Q14151 | 287 | QSSKADSLLAVVK | 6.0305 | 6.0654 | 6.0765 | 6.5154 | 6.7855 | 6.3388 | 6.9727 | 6.5152 | 6.2626 | 5.8203 | 6.0106 | 5.9906 | 6.0591 | 6.2891 | 5.8244 | 6.4432 |
| SAFB2_S233 | SAFB2 | Q14151 | 195 | NTLETSSLNFKVT | 6.7613 | 6.4915 | 6.1299 | 6.8699 | 6.1735 | 5.7227 | 6.5418 | 5.5917 | 5.8654 | 5.3994 | 5.2099 | 5.3681 | 7.1486 | 7.0828 | 6.8683 | 6.7753 |
| SAFB2_S245 | SAFB2 | Q14151 | 194 | KNTLETSSLNFKV | 7.2520 | 7.4081 | 6.7445 | 8.3956 | 7.2448 | 6.2310 | 7.3414 | 5.6277 | 6.9662 | 6.0497 | 6.1180 | 6.2617 | 4.5850 | 5.0633 | 4.0090 | 4.7019 |
| SAFB2_S246 | SAFB2 | Q14151 | 245 | PFAQDTSSVGPDR | 7.3255 | 6.6993 | 6.4112 | 7.3858 | 7.0582 | 7.3334 | 7.5897 | 5.5066 | 6.1954 | 6.5691 | 7.0278 | 6.5470 | 4.4487 | 5.5705 | 3.8864 | 4.4455 |
| SAFB2_S287 | SAFB2 | Q14151 | 613 | EKRDILSFDKIKE | 6.0753 | 6.2647 | 5.6246 | 6.8850 | 6.2789 | 5.8840 | 6.3435 | 5.9095 | 5.7344 | 5.7208 | 5.7640 | 6.1522 | 6.5469 | 6.6755 | 6.5985 | 7.5422 |
| SAFB2_S613 | SAFB2 | Q14151 | 207 | TPDIEESLLEPEN | 7.8545 | 6.5852 | 7.8116 | 8.2920 | 6.0891 | 6.0110 | 7.8467 | 6.5266 | 5.4954 | 5.3236 | 6.6906 | 4.6987 | 4.5776 | 7.3585 | 4.8822 | 3.9566 |
| SALL2_S243 | SALL2 | Q9Y467 | 243 | PLLPLFSPIKPVQ | 7.0008 | 5.7310 | 6.9008 | 7.3192 | 7.1480 | 6.6224 | 6.9553 | 6.2467 | 7.0123 | 6.6261 | 7.1164 | 7.3803 | 4.7350 | 4.6397 | 3.9546 | 4.6115 |
| SALL3_S1177 | SALL3 | Q9BXA9 | 1177 | RRGRRLSVENPMA | 5.3873 | 5.2396 | 6.0077 | 6.3214 | 6.8798 | 6.2171 | 6.3536 | 6.6569 | 7.4143 | 6.4799 | 6.9407 | 7.2864 | 6.0441 | 5.6141 | 5.2064 | 5.9508 |
| SALL3_S45 | SALL3 | Q9BXA9 | 919 | ESFRSKSPGLGAP | 6.3963 | 6.5166 | 6.5350 | 7.4044 | 6.2960 | 6.1990 | 6.4514 | 6.4565 | 6.5887 | 6.0948 | 6.2630 | 6.5898 | 5.6792 | 5.1475 | 5.5111 | 5.8706 |
| SALL3_S552 | SALL3 | Q9BXA9 | 1265 | GMDKARTGSSPPI | 6.6604 | 6.2964 | 6.4678 | 6.8540 | 6.6305 | 6.1461 | 6.6484 | 6.7013 | 6.3612 | 5.7472 | 6.1901 | 6.4458 | 6.0292 | 5.1260 | 5.7257 | 5.9699 |
| SALL3_S919 | SALL3 | Q9BXA9 | 45 | SGPESRSGGEETS | 7.4664 | 7.0547 | 6.7692 | 7.7296 | 6.2848 | 6.2374 | 6.3933 | 6.5457 | 6.3076 | 5.9605 | 5.8731 | 6.4165 | 5.3901 | 4.8026 | 5.2119 | 5.5567 |
| SALL3_T1265 | SALL3 | Q9BXA9 | 702 | MHYRTHTGERPFK | 7.0690 | 7.4936 | 7.8391 | 8.2896 | 5.8387 | 7.0956 | 6.9947 | 6.3373 | 7.3254 | 7.6600 | 8.5809 | 8.5245 | 2.1739 | 3.0959 | 2.2910 | 3.3907 |
| SALL3_T702 | SALL3 | Q9BXA9 | 552 | ATPASRSPQRPSP | 7.3637 | 6.7383 | 6.9988 | 7.8323 | 7.0804 | 6.2367 | 6.4045 | 7.2684 | 5.7433 | 5.4884 | 6.7860 | 6.3412 | 4.8777 | 5.4811 | 5.2472 | 4.1120 |
| SAMD1_S161 | SAMD1 | Q6SPF0 | 161 | AATAPPSPGPAQP | 6.7965 | 6.1484 | 6.0163 | 6.8983 | 6.1930 | 6.2769 | 6.3532 | 6.5297 | 6.6962 | 6.4786 | 6.0027 | 6.2439 | 5.7189 | 5.8959 | 5.8804 | 5.8710 |
| SAMD1_S425 | SAMD1 | Q6SPF0 | 425 | VATGPDSPSPVPL | 5.9371 | 6.7101 | 6.0439 | 6.5152 | 6.2664 | 5.8767 | 6.0045 | 6.2426 | 6.4754 | 6.6191 | 6.3636 | 6.1237 | 6.0432 | 6.2728 | 6.3935 | 6.1122 |
| SAMD4A_S419 | SAMD4A | Q9UPU9 | 419 | TPIKAYSSPSTTP | 6.3995 | 6.4228 | 6.4987 | 6.4497 | 6.2795 | 6.2803 | 6.0912 | 6.6469 | 6.9862 | 6.3058 | 6.1620 | 6.7030 | 5.5421 | 5.5888 | 5.7897 | 5.8539 |
| SAMD4B_S172 | SAMD4B | Q5PRF9 | 238 | PLKRSMSLIPTSP | 6.3935 | 5.8312 | 6.5286 | 6.0774 | 6.2062 | 6.3408 | 5.8716 | 5.8469 | 6.5065 | 6.8381 | 6.7623 | 6.7659 | 6.3614 | 6.7070 | 5.2257 | 5.7368 |
| SAMD4B_S238 | SAMD4B | Q5PRF9 | 271 | PDHAPLSPQSSVA | 5.8548 | 6.0508 | 6.3527 | 6.2651 | 6.9620 | 6.2685 | 6.1038 | 6.5565 | 6.4275 | 6.4460 | 6.5949 | 6.7286 | 6.1157 | 5.8101 | 5.6182 | 5.8447 |
| SAMD4B_S238;S252 | SAMD4B | Q5PRF9 | 557 | WAFGSNSLPIAGS | 6.3100 | 6.2035 | 6.7089 | 6.4856 | 6.2693 | 6.4282 | 5.9952 | 6.8910 | 6.6960 | 6.6105 | 6.6045 | 6.5218 | 5.4465 | 6.2061 | 5.3696 | 5.2533 |
| SAMD4B_S238;T242 | SAMD4B | Q5PRF9 | 172 | YHSRQGSDEWGGP | 5.1177 | 5.5330 | 5.8177 | 5.5167 | 6.3221 | 6.1490 | 6.0542 | 6.2699 | 6.2996 | 6.8407 | 5.8924 | 6.3375 | 6.9677 | 7.2942 | 6.2634 | 7.3243 |
| SAMD4B_S243 | SAMD4B | Q5PRF9 | 592 | GRMGLLSPSGIGG | 6.1614 | 5.9586 | 6.2041 | 6.2571 | 6.5051 | 6.5660 | 6.0720 | 6.6567 | 6.5792 | 6.1874 | 6.5555 | 6.4649 | 5.6404 | 6.2903 | 5.8624 | 6.0390 |
| SAMD4B_S243;S252 | SAMD4B | Q5PRF9 | 243 | MSLIPTSPQVPGE | 6.0093 | 6.3177 | 6.2644 | 6.1600 | 6.3689 | 6.4428 | 6.8014 | 6.4102 | 6.4723 | 6.6076 | 6.4386 | 6.7138 | 5.6448 | 5.9553 | 5.6090 | 5.7839 |
| SAMD4B_S252 | SAMD4B | Q5PRF9 | 628 | NPGGSNSMPSQSR | 6.3153 | 6.8747 | 6.6811 | 6.2127 | 6.0573 | 6.4278 | 5.7844 | 6.5003 | 6.4008 | 6.5713 | 6.6858 | 6.7744 | 5.5118 | 6.0099 | 5.4343 | 5.7581 |
| SAMD4B_S271 | SAMD4B | Q5PRF9 | 407 | VAAATTTPTAKDG | 5.9695 | 6.5951 | 6.4463 | 6.1301 | 6.2805 | 6.8405 | 6.7899 | 6.1410 | 6.1353 | 6.0940 | 6.5822 | 6.9630 | 5.7251 | 6.4153 | 5.2751 | 5.6171 |
| SAMD4B_S557 | SAMD4B | Q5PRF9 | 252 | VPGEWPSPEELGA | 6.3673 | 6.9861 | 6.3805 | 6.6751 | 4.8483 | 8.3132 | 6.0661 | 6.9729 | 7.5189 | 5.3943 | 5.9371 | 7.2972 | 4.7688 | 5.0634 | 4.9509 | 6.4599 |
| SAMD4B_S592 | SAMD4B | Q5PRF9 | 606 | SPRHALTSPSLGG | 4.7383 | 4.1818 | 5.6931 | 4.6166 | 5.3179 | 5.3701 | 4.2704 | 5.6078 | 5.4090 | 4.1876 | 4.7681 | 4.5238 | 9.7523 | 11.3083 | 11.3116 | 8.9433 |
| SAMD4B_S592;S600 | SAMD4B | Q5PRF9 | 607 | PRHALTSPSLGGQ | 6.2108 | 6.2486 | 6.6532 | 7.4272 | 5.8370 | 7.5606 | 7.9029 | 7.4404 | 6.5505 | 7.6703 | 7.4386 | 6.6962 | 3.1541 | 4.9383 | 3.7211 | 4.5504 |
| SAMD4B_S607 | SAMD4B | Q5PRF9 | 238;252 | NA | 6.2760 | 6.2116 | 6.6085 | 6.4938 | 6.3034 | 6.1420 | 6.1968 | 5.7584 | 7.0169 | 6.5696 | 7.0375 | 6.4786 | 5.9284 | 5.9545 | 5.5332 | 5.4910 |
| SAMD4B_S628 | SAMD4B | Q5PRF9 | 243;252 | NA | 6.5104 | 7.2158 | 7.2581 | 7.9020 | 5.9643 | 5.6564 | 6.8493 | 6.6465 | 6.4383 | 7.0978 | 8.8799 | 6.3984 | 4.1415 | 4.2761 | 4.7122 | 4.0530 |
| SAMD4B_T407 | SAMD4B | Q5PRF9 | 238;242 | NA | 6.4909 | 5.0585 | 8.3965 | 5.1233 | 7.0044 | 7.7140 | 5.9417 | 6.0940 | 7.7931 | 5.7754 | 6.5865 | 6.6645 | 6.1030 | 6.8082 | 4.4989 | 3.9470 |
| SAMD4B_T606 | SAMD4B | Q5PRF9 | 592;600 | NA | 6.3973 | 7.0889 | 6.2581 | 6.7589 | 6.0054 | 7.3107 | 6.4516 | 5.7725 | 6.1046 | 7.1867 | 6.9142 | 6.0116 | 5.5550 | 5.6873 | 5.5126 | 4.9847 |
| SAMD8_S15 | SAMD8 | A0A5F9ZGS6 | 15 | PRLHSGSPPRAPP | 6.2711 | 6.3692 | 5.9937 | 6.4168 | 6.3156 | 5.8351 | 6.3775 | 6.5578 | 6.4922 | 5.8120 | 5.8071 | 6.0230 | 6.4894 | 6.7536 | 6.1661 | 6.3198 |
| SAMHD1_S278 | SAMHD1 | Q9Y3Z3 | 592 | VIAPLITPQKKEW | 5.2038 | 5.1925 | 6.2912 | 5.3977 | 5.8732 | 6.0007 | 5.6880 | 5.8598 | 6.3781 | 6.1551 | 6.7092 | 6.6710 | 7.1166 | 7.7228 | 6.7313 | 7.0090 |
| SAMHD1_S33 | SAMHD1 | Q9Y3Z3 | 33 | SAEADWSPGLELH | 5.2902 | 5.7121 | 6.0248 | 5.6202 | 5.7331 | 6.2164 | 5.6421 | 6.0083 | 5.8276 | 6.0725 | 6.0425 | 5.7731 | 7.1636 | 7.8104 | 7.6104 | 7.4526 |
| SAMHD1_T592 | SAMHD1 | Q9Y3Z3 | 278 | IVGPLESPVEDSL | 5.7309 | 6.2605 | 6.6486 | 5.7520 | 6.3623 | 6.1724 | 5.6653 | 5.4867 | 5.8348 | 6.2545 | 6.0096 | 6.6923 | 6.5062 | 7.3817 | 6.8392 | 6.4030 |
| SAP130_S442 | SAP130 | Q9H0E3-3 | 442 | QQITHTSPRIQPD | 6.8581 | 6.7571 | 6.2115 | 6.5525 | 6.4406 | 6.5239 | 6.7083 | 6.7393 | 7.0505 | 6.1045 | 6.7323 | 6.3554 | 5.2356 | 5.2916 | 5.2460 | 5.1929 |
| SAP130_S465 | SAP130 | Q9H0E3-3 | 709 | MRSTSGSPRPAGA | 6.2203 | 6.8357 | 6.0360 | 6.1062 | 7.6884 | 7.0368 | 6.4464 | 5.3751 | 5.5544 | 5.9552 | 5.7895 | 6.4740 | 6.5409 | 6.5144 | 6.0471 | 5.3794 |
| SAP130_S709 | SAP130 | Q9H0E3-3 | 866 | DLPPGASPRKKPR | 6.1428 | 5.3476 | 5.2746 | 5.9891 | 7.4048 | 6.7709 | 6.5174 | 5.7932 | 6.8343 | 5.5896 | 5.1555 | 5.3380 | 7.8331 | 6.0391 | 5.9680 | 8.0022 |
| SAP130_S866 | SAP130 | Q9H0E3-3 | 465 | ISGHRASPNPVAM | 6.3552 | 6.1547 | 4.9756 | 6.1127 | 5.6922 | 6.4248 | 5.9160 | 5.8841 | 5.5274 | 5.8496 | 6.6125 | 5.5577 | 6.5864 | 8.2260 | 7.1117 | 7.0133 |
| SAP30_S131 | SAP30 | O75446 | 131 | RRKRKGSDDDGGD | 5.9383 | 5.9457 | 5.8349 | 6.5229 | 6.2142 | 5.9078 | 6.4146 | 6.3197 | 5.9872 | 6.0991 | 6.3266 | 6.1689 | 6.7915 | 6.6960 | 6.4117 | 6.4209 |
| SAP30_S131;S138 | SAP30 | O75446 | 138 | DDDGGDSPVQDID | 5.9360 | 5.7801 | 5.2722 | 7.2773 | 5.8819 | 5.1365 | 6.0189 | 5.8882 | 6.8725 | 6.7757 | 6.3005 | 6.8738 | 6.1912 | 6.1170 | 6.7594 | 6.9190 |
| SAP30_S138 | SAP30 | O75446 | 131;138 | NA | 6.2373 | 6.3528 | 6.2488 | 6.9738 | 6.0377 | 5.8778 | 6.3092 | 5.2572 | 6.0659 | 6.2073 | 5.8271 | 6.8845 | 6.4817 | 6.5651 | 6.2508 | 6.4231 |
| SAP30BP_S104 | SAP30BP | Q9UHR5 | 113 | ERVRNMSPDEIKI | 6.4022 | 6.3877 | 6.4376 | 6.0991 | 6.1304 | 6.1004 | 6.1762 | 6.1622 | 6.6857 | 6.5012 | 6.7842 | 6.2719 | 6.1328 | 5.9795 | 5.7535 | 5.9956 |
| SAP30BP_S113 | SAP30BP | Q9UHR5 | 104 | PQELVASFSERVR | 7.5077 | 6.6267 | 6.3204 | 6.8145 | 5.3432 | 7.7984 | 6.4344 | 6.7117 | 6.2812 | 5.9637 | 6.5675 | 6.4668 | 5.4908 | 4.9977 | 5.1319 | 5.5433 |
| SAP30BP_S69;S72;S77 | SAP30BP | Q9UHR5 | 72;77 | NA | 5.3611 | 5.6135 | 6.7006 | 5.4781 | 5.5392 | 5.6888 | 5.8553 | 5.2675 | 6.2559 | 6.5089 | 6.9110 | 7.0377 | 7.2481 | 6.8941 | 6.7275 | 6.9126 |
| SAP30BP_S72;S77 | SAP30BP | Q9UHR5 | 69;72;77 | NA | 5.4533 | 5.2701 | 6.6394 | 5.6064 | 5.6331 | 5.5147 | 5.8991 | 5.1709 | 6.4025 | 6.7193 | 5.9977 | 6.3151 | 7.7646 | 7.2712 | 7.2508 | 7.0919 |
| SAP30L_S93 | SAP30L | Q9HAJ7 | 99 | SDDGGDSPEHDTD | 6.2636 | 6.6042 | 6.2469 | 6.7355 | 5.8911 | 5.7752 | 5.7809 | 5.8123 | 6.4274 | 6.6538 | 6.7463 | 6.6622 | 6.0346 | 6.1150 | 6.1802 | 6.0708 |
| SAP30L_S93;S99 | SAP30L | Q9HAJ7 | 93 | KRKRKTSDDGGDS | 5.9743 | 6.7244 | 6.4598 | 7.1623 | 6.8271 | 6.7672 | 6.1789 | 6.0509 | 6.3506 | 5.8903 | 6.5762 | 5.9477 | 5.3406 | 6.0873 | 6.1121 | 5.5504 |
| SAP30L_S99 | SAP30L | Q9HAJ7 | 93;99 | NA | 5.5608 | 6.1879 | 5.7277 | 6.8564 | 6.4321 | 6.0824 | 6.2475 | 6.5010 | 5.9512 | 5.7131 | 5.3598 | 5.8544 | 6.9750 | 6.6930 | 6.9844 | 6.8731 |
| SARM1_S548 | SARM1 | Q6SZW1 | 548 | AREMLHSPLPCTG | 5.8763 | 5.6652 | 6.0690 | 5.6334 | 6.1545 | 6.8985 | 5.3820 | 5.5714 | 6.1025 | 5.9970 | 7.8251 | 6.6103 | 6.9471 | 6.2265 | 7.2453 | 5.7959 |
| SARNP_S115 | SARNP | P82979 | 60 | DVLGDETEEEETK | 6.8595 | 7.3339 | 6.8478 | 7.3411 | 6.9466 | 6.8196 | 6.6792 | 7.2110 | 6.0192 | 6.2416 | 6.0692 | 6.2114 | 4.8714 | 5.0309 | 4.5520 | 4.9658 |
| SARNP_S163 | SARNP | P82979 | 163 | FGLNVSSISRKSE | 6.9733 | 6.9534 | 7.2287 | 6.5385 | 4.8083 | 6.0058 | 6.5106 | 5.8628 | 7.7908 | 7.9068 | 8.2129 | 8.3707 | 3.9503 | 5.5069 | 3.2719 | 4.1081 |
| SARNP_T60 | SARNP | P82979 | 115 | RFNVPVSLESKKA | 7.7147 | 7.7811 | 7.1548 | 7.7347 | 5.9598 | 7.1636 | 8.3242 | 7.6315 | 7.2444 | 6.4269 | 7.1812 | 6.6301 | 3.1875 | 3.6994 | 2.7331 | 3.4330 |
| SART1_S448 | SART1 | O43290 | 448 | RGRRRVSEVEEEK | 5.0765 | 4.9909 | 5.3143 | 5.8933 | 6.1141 | 5.3610 | 5.8213 | 5.5703 | 5.4920 | 5.3559 | 5.2521 | 5.3919 | 8.1576 | 8.5399 | 8.8659 | 8.8029 |
| SART1_S474;S486 | SART1 | O43290 | 392 | LASEYLTPEEMVT | 6.1770 | 6.8472 | 6.6253 | 7.3356 | 5.9116 | 6.8097 | 7.0476 | 6.4576 | 6.3916 | 7.3313 | 7.0537 | 6.7199 | 4.2342 | 5.3801 | 4.5555 | 5.1220 |
| SART1_S598 | SART1 | O43290 | 762 | LKKMSSSDTPLGT | 6.6337 | 6.2818 | 6.9883 | 6.5658 | 5.6459 | 6.8401 | 6.7886 | 6.1133 | 7.3360 | 7.4115 | 6.7768 | 7.4668 | 4.5112 | 5.6596 | 4.3202 | 4.6606 |
| SART1_S762 | SART1 | O43290 | 787 | TPYIVLSGSGKSM | 6.2903 | 6.0286 | 6.9048 | 6.1344 | 6.4185 | 6.0168 | 6.4330 | 8.3148 | 6.1700 | 4.8552 | 5.3930 | 5.7829 | 5.5450 | 7.1123 | 6.1212 | 6.4793 |
| SART1_S787 | SART1 | O43290 | 598 | ANGGSESDGEENI | 5.8510 | 5.8618 | 6.6833 | 6.3347 | 5.6222 | 5.7986 | 6.5198 | 5.8412 | 6.4226 | 6.8709 | 6.6683 | 5.5702 | 5.8063 | 6.5897 | 6.3866 | 6.1728 |
| SART1_T392 | SART1 | O43290 | 474;486 | NA | 5.7803 | 6.1652 | 6.0949 | 6.5116 | 6.0631 | 5.6809 | 6.1166 | 6.2616 | 6.1320 | 6.2236 | 6.0758 | 6.2430 | 6.1807 | 6.9895 | 6.6946 | 6.7864 |

| | | | | | | | | | | | | | | | | | | | | | |
|---|---|---|---|---|---|---|---|---|---|---|---|---|---|---|---|---|---|---|---|---|---|
| SART3_S250 | SART3 | A0A499FI31 | | 250 | PVAGFLSPFDREQ | 5.8015 | 6.0019 | 6.1536 | 6.7987 | 6.1199 | 6.2649 | 6.5334 | 6.1344 | 6.5960 | 6.9002 | 7.0350 | 6.6210 | 5.7502 | 5.9446 | 5.3234 | 6.0214 |
| SART3_S796 | SART3 | A0A499FI31 | | 796 | GRPMFVSPCVDKS | 7.8370 | 6.2529 | 8.1831 | 8.1146 | 6.8861 | 7.0554 | 8.1964 | 5.8877 | 6.7302 | 6.9964 | 7.4547 | 6.1445 | 3.0884 | 4.2199 | 2.5369 | 4.4158 |
| SASH1_S101 | SASH1 | O94885 | | 407 | SHGRTCSFGGFDL | 6.8627 | 6.2001 | 6.4037 | 5.5903 | 5.9251 | 6.0699 | 5.8123 | 6.0792 | 6.6528 | 5.8182 | 6.0848 | 6.0815 | 6.9683 | 6.9345 | 6.2853 | 6.2312 |
| SASH1_S1028 | SASH1 | O94885 | | 923 | ASGRGLSPPQCLP | 6.8939 | 6.8895 | 7.2284 | 6.8768 | 6.1660 | 6.7019 | 6.6532 | 6.5617 | 7.0281 | 7.3625 | 7.1504 | 7.0645 | 4.0496 | 4.7163 | 4.0725 | 4.5847 |
| SASH1_S1051 | SASH1 | O94885 | | 743 | GKTRKASLLSAKS | 6.3440 | 4.5584 | 6.0068 | 5.4692 | 7.5381 | 5.5565 | 5.2443 | 6.5475 | 7.6149 | 8.0701 | 7.2907 | 7.1745 | 6.5212 | 6.8915 | 4.8649 | 4.3072 |
| SASH1_S320 | SASH1 | O94885 | | 486 | LDGMPGSPPPSQP | 5.2836 | 6.0628 | 5.9927 | 6.4786 | 5.8743 | 6.5934 | 6.2003 | 6.4003 | 6.1672 | 6.5360 | 6.2886 | 6.7825 | 6.9081 | 6.2941 | 5.9429 | 6.1948 |
| SASH1_S320;S335 | SASH1 | O94885 | | 1028 | KRGSPASPTSPSD | 6.2934 | 6.3671 | 6.9627 | 6.0107 | 6.2152 | 6.0952 | 6.4528 | 6.8759 | 7.3087 | 6.5984 | 7.3616 | 6.6759 | 5.3570 | 5.4032 | 5.0386 | 4.9838 |
| SASH1_S407 | SASH1 | O94885 | | 320 | PLFFDGSPEKPPE | 6.2236 | 7.1206 | 8.0191 | 7.1543 | 6.2211 | 6.8858 | 6.6982 | 6.6148 | 7.4519 | 7.6549 | 7.2879 | 7.0793 | 3.9720 | 4.4260 | 3.5535 | 3.6369 |
| SASH1_S486 | SASH1 | O94885 | | 721 | QGLSGCSPRDSGC | 7.0647 | 6.5781 | 7.5514 | 6.7810 | 6.8436 | 6.3784 | 6.6485 | 6.7785 | 7.3899 | 5.9787 | 6.3680 | 7.3174 | 4.6280 | 4.5754 | 4.2784 | 4.8400 |
| SASH1_S614 | SASH1 | O94885 | | 839 | TWPRSHSLDDLQV | 6.4066 | 6.0994 | 5.7752 | 5.8227 | 6.3176 | 6.6452 | 6.1279 | 6.9294 | 6.9742 | 6.7723 | 6.5793 | 7.4208 | 5.0247 | 6.1164 | 5.2334 | 5.7548 |
| SASH1_S721 | SASH1 | O94885 | | 1051 | SGQAPGSPPSTRP | 6.0111 | 5.3838 | 6.4803 | 5.8059 | 6.5871 | 6.7386 | 5.8027 | 6.2481 | 6.3165 | 6.4494 | 6.3593 | 6.4462 | 6.5792 | 6.4588 | 5.9480 | 6.3849 |
| SASH1_S743 | SASH1 | O94885 | | 90 | LRKRRVSQDLEVE | 6.5073 | 6.3351 | 6.3022 | 5.9472 | 6.1023 | 5.3949 | 5.8814 | 6.0984 | 7.1687 | 6.2601 | 7.0582 | 5.8801 | 6.8439 | 6.2509 | 5.8709 | 6.0984 |
| SASH1_S813 | SASH1 | O94885 | | 614 | IYVDVLSEDEEKP | 5.5166 | 5.7664 | 7.3534 | 6.5079 | 6.8450 | 7.3280 | 6.5460 | 6.6983 | 6.7316 | 5.7744 | 6.5873 | 6.2550 | 5.9917 | 6.0773 | 5.1899 | 4.8314 |
| SASH1_S839 | SASH1 | O94885 | | 813 | LNKNRRSLPVSIC | 6.6047 | 6.7548 | 6.9167 | 6.2504 | 6.3651 | 6.2827 | 6.5869 | 7.0889 | 6.9061 | 6.5925 | 6.9552 | 7.0992 | 4.9877 | 5.0136 | 4.8193 | 4.7761 |
| SASH1_S90 | SASH1 | O94885 | | 101 | VEKPDASPTSLQL | 6.1779 | 7.3767 | 7.3086 | 6.5327 | 5.8441 | 7.9991 | 7.1814 | 6.6754 | 6.3151 | 6.7530 | 6.2043 | 7.2187 | 4.3341 | 5.4298 | 3.6700 | 4.9791 |
| SASH1_S923 | SASH1 | O94885 | 320;335 | | NA | 5.5100 | 6.1868 | 7.4831 | 7.0731 | 6.5155 | 6.2038 | 7.0375 | 6.1984 | 6.9772 | 7.4717 | 7.6005 | 8.4350 | 4.8452 | 4.6921 | 3.4513 | 4.3187 |
| SASH1_Y233;S244 | SASH1 | O94885 | 233;244 | | NA | 5.6997 | 6.1849 | 6.7716 | 5.7880 | 6.0952 | 6.6931 | 5.2894 | 7.5421 | 5.7589 | 5.7481 | 6.9903 | 6.8987 | 5.5980 | 6.7600 | 5.8791 | 6.3029 |
| SASS6_S510 | SASS6 | Q6UVJ0 | | 510 | TIRSGISPNLNVV | 6.3447 | 6.8005 | 6.7962 | 6.6195 | 6.1377 | 6.9602 | 7.5276 | 7.5152 | 7.0814 | 6.7781 | 7.3105 | 7.1207 | 3.8668 | 4.7019 | 4.1029 | 4.3361 |
| SATB1_S47 | SATB1 | Q01826-2 | | 47 | GRGRLGSTGAKMQ | 4.9099 | 5.0835 | 5.4060 | 6.0198 | 6.0102 | 5.9358 | 6.1939 | 6.3132 | 5.6542 | 5.5030 | 5.2025 | 5.9018 | 8.5615 | 7.5750 | 7.7526 | 7.9772 |
| SATB2_S20 | SATB2 | Q9UPW6 | | 20 | PDRRSGSPDVKGP | 6.2723 | 6.4378 | 5.8776 | 6.1909 | 6.5360 | 5.9678 | 6.5882 | 6.1052 | 6.4010 | 6.3169 | 6.0676 | 6.0612 | 6.3854 | 6.2648 | 6.1659 | 6.3614 |
| SATB2_S39 | SATB2 | Q9UPW6 | | 587 | VLHRQQSQPAKES | 4.8707 | 5.3156 | 5.1298 | 5.3352 | 5.3012 | 5.4674 | 5.3826 | 5.9311 | 5.3082 | 5.1087 | 5.5401 | 5.7404 | 8.5991 | 8.5716 | 9.6010 | 8.7971 |
| SATB2_S587 | SATB2 | Q9UPW6 | | 39 | RLEQNGSPMGARG | 6.8880 | 6.7430 | 7.2249 | 7.2468 | 6.0181 | 6.9986 | 7.1988 | 6.9255 | 7.1414 | 6.2601 | 6.6460 | 6.8431 | 4.1991 | 5.2088 | 3.9613 | 4.4964 |
| SATB2_S594 | SATB2 | Q9UPW6 | | 594 | QPAKESSPPREEA | 6.0963 | 5.8418 | 6.1709 | 6.7292 | 6.1009 | 5.8647 | 6.1768 | 5.7713 | 5.8798 | 6.1919 | 6.7860 | 6.3183 | 6.2459 | 6.3228 | 6.7425 | 6.7608 |
| SATB2_S7 | SATB2 | Q9UPW6 | | 7 | MERRSESPCLRDS | 5.5865 | 5.9997 | 6.2577 | 6.4152 | 5.3241 | 6.2025 | 6.0468 | 6.1658 | 5.5364 | 6.2440 | 5.8926 | 5.4330 | 6.5716 | 7.0653 | 7.3625 | 7.8962 |
| SAV1_S27 | SAV1 | Q9H4B6 | | 94 | RESNRLSAPSYLA | 6.4450 | 6.4584 | 7.3725 | 6.5694 | 6.7567 | 6.9086 | 6.4480 | 7.2667 | 7.5797 | 6.4160 | 6.5636 | 6.5467 | 4.9787 | 5.1049 | 4.0358 | 4.5496 |
| SAV1_S94 | SAV1 | Q9H4B6 | | 27 | YVKKETSPLLRNL | 7.5389 | 7.4290 | 6.3371 | 6.9629 | 6.4122 | 6.4374 | 6.9097 | 6.9963 | 6.9376 | 6.0266 | 6.5418 | 7.0475 | 4.6180 | 4.8446 | 4.2468 | 4.7136 |
| SBDS_S233 | SBDS | Q9Y3A5 | | 233 | ETKGKGSLEVLNL | 5.4309 | 7.3904 | 4.5600 | 5.0529 | 6.5709 | 6.3592 | 6.7221 | 6.4409 | 7.3148 | 5.0438 | 5.9510 | 5.7998 | 5.7696 | 6.9489 | 7.3904 | 7.2543 |
| SBDS_S96 | SBDS | Q9Y3A5 | | 96 | KGEVQVSDKERHT | 6.5362 | 7.0530 | 7.5671 | 6.0141 | 6.4326 | 6.8511 | 6.1124 | 7.7704 | 5.6528 | 5.9344 | 5.8892 | 5.9849 | 5.0111 | 6.5123 | 5.4880 | 5.1904 |
| SBF1_T1138 | SBF1 | O95248-4 | | 1138 | QRLGLGTLSSSLS | 6.6880 | 7.7090 | 7.4402 | 6.3225 | 7.5791 | 6.2349 | 6.2983 | 7.6879 | 5.9207 | 6.0053 | 5.1686 | 4.8091 | 5.3800 | 5.8120 | 6.8783 | 4.0660 |
| SBF2_T1105 | SBF2 | Q86WG5 | | 1105 | KASEKSTMEQLVE | 6.4935 | 6.0459 | 6.4647 | 6.4995 | 6.0765 | 5.7009 | 6.1749 | 6.0027 | 6.0399 | 6.6957 | 6.5055 | 5.8725 | 6.1797 | 6.5859 | 6.5835 | 6.0783 |
| SBNO1_S1062 | SBNO1 | A3KN83 | | 214 | MKMRSFSPTMKVP | 5.9885 | 6.4337 | 5.9462 | 5.5828 | 5.9072 | 6.4725 | 6.0538 | 6.3570 | 6.3181 | 6.6981 | 6.6409 | 6.2560 | 5.7862 | 6.8532 | 6.4988 | 6.2070 |
| SBNO1_S214 | SBNO1 | A3KN83 | | 815 | LGSKRPSFSSTPV | 4.9221 | 4.9657 | 5.5186 | 5.0300 | 8.2667 | 4.3610 | 4.5015 | 5.0721 | 7.7989 | 7.8972 | 6.3989 | 6.7271 | 7.1181 | 8.7162 | 6.0131 | 6.6931 |
| SBNO1_S692 | SBNO1 | A3KN83 | | 794 | EKKKKKSIDPDSI | 5.4815 | 6.9690 | 5.7326 | 7.0622 | 7.3403 | 5.9949 | 5.6739 | 6.9457 | 6.7021 | 6.6165 | 6.5751 | 7.8220 | 5.4672 | 6.1226 | 4.6390 | 4.8554 |
| SBNO1_S794 | SBNO1 | A3KN83 | | 1062 | LDSPMVSPPPDYP | 6.1924 | 5.6099 | 7.2237 | 6.4646 | 5.8176 | 4.8794 | 5.8606 | 5.5001 | 5.7985 | 6.2115 | 7.6558 | 6.5027 | 7.1258 | 6.3189 | 5.9680 | 6.8704 |
| SBNO1_S815 | SBNO1 | A3KN83 | | 692 | TAPSNNSSPRDSP | 5.3503 | 5.5339 | 7.0543 | 5.4994 | 6.4818 | 8.1044 | 7.9695 | 6.7226 | 6.1275 | 6.1933 | 7.4893 | 7.3687 | 4.8734 | 5.2794 | 4.8795 | 5.0727 |
| SCAF1_S239 | SCAF1 | Q9H7N4 | | 239 | PTDEAYSPPPAPE | 5.8806 | 6.4674 | 5.7915 | 6.6996 | 6.2536 | 6.1657 | 6.1819 | 6.1612 | 6.0497 | 5.9614 | 5.8573 | 5.8412 | 5.9520 | 7.1273 | 6.7190 | 6.8907 |
| SCAF1_S498;S500 | SCAF1 | Q9H7N4 | | 725 | SPAPASSPKREVL | 6.0920 | 6.2196 | 5.5742 | 6.2294 | 6.0405 | 5.6624 | 6.1246 | 5.6987 | 5.8409 | 5.6755 | 5.7087 | 5.5770 | 7.0698 | 7.5946 | 7.3846 | 7.5074 |
| SCAF1_S548 | SCAF1 | Q9H7N4 | | 874 | RDRESRSPFLKPD | 6.3741 | 6.4190 | 5.9281 | 6.8055 | 6.1392 | 5.8921 | 6.2288 | 6.2375 | 6.2865 | 6.0930 | 6.0370 | 5.9597 | 5.7408 | 6.8070 | 6.3524 | 6.6991 |
| SCAF1_S689 | SCAF1 | Q9H7N4 | | 548 | APPAPASPWDSKK | 6.2207 | 6.1564 | 6.1111 | 6.5622 | 6.4839 | 6.4068 | 6.1989 | 6.1706 | 6.5876 | 6.2246 | 6.4161 | 5.8788 | 5.7718 | 6.8197 | 6.0441 | 5.9465 |
| SCAF1_S719;S725 | SCAF1 | Q9H7N4 | | 989 | PPPPALTPDSQTV | 6.0158 | 5.9595 | 5.9958 | 6.8962 | 5.5559 | 5.5911 | 6.1472 | 5.7014 | 5.8735 | 6.1945 | 5.6489 | 5.8505 | 6.4336 | 7.3241 | 7.2869 | 7.5252 |
| SCAF1_S725 | SCAF1 | Q9H7N4 | | 976 | AAKVPSTPPPKAA | 6.2281 | 6.3415 | 5.9015 | 6.8490 | 6.0777 | 5.9955 | 6.4145 | 6.0677 | 5.9369 | 5.9650 | 5.9426 | 6.0277 | 6.0296 | 6.7650 | 6.6248 | 6.8329 |
| SCAF1_S734;S738 | SCAF1 | Q9H7N4 | | 1001 | VDSSCKTPEVSFL | 5.7743 | 5.8110 | 6.2878 | 6.6684 | 6.0040 | 6.1396 | 6.0271 | 6.3539 | 6.2553 | 6.1705 | 6.1532 | 6.2784 | 5.9029 | 6.7548 | 6.5423 | 6.8765 |
| SCAF1_S874 | SCAF1 | Q9H7N4 | | 335 | PTPAPGTPPQVDS | 6.1358 | 7.5223 | 6.0736 | 6.7942 | 6.2976 | 6.6187 | 6.7593 | 6.2018 | 7.0344 | 6.2267 | 7.2074 | 6.6448 | 4.6373 | 5.7226 | 5.3480 | 4.7755 |
| SCAF1_S965 | SCAF1 | Q9H7N4 | | 965 | AEETSWSGEERAA | 6.3599 | 6.9204 | 6.0141 | 6.7976 | 5.9582 | 5.8748 | 6.2270 | 6.1293 | 5.7517 | 5.5928 | 5.4729 | 5.7622 | 6.1383 | 7.0807 | 6.8148 | 7.1052 |
| SCAF1_T1001 | SCAF1 | Q9H7N4 | | 689 | RGAVPPSIQDLTD | 6.2771 | 6.4391 | 6.3522 | 7.3143 | 7.0249 | 6.3566 | 6.3117 | 6.1507 | 6.1052 | 5.9111 | 5.5649 | 6.3705 | 5.6760 | 6.4411 | 5.4231 | 6.2815 |
| SCAF1_T335 | SCAF1 | Q9H7N4 | | 849 | EGVSSTTPAKDAA | 7.4853 | 7.8492 | 6.5026 | 7.6463 | 6.1151 | 6.2133 | 7.0123 | 6.6332 | 7.4857 | 7.2320 | 7.1548 | 7.5743 | 3.1106 | 4.5280 | 3.4258 | 4.0315 |
| SCAF1_T849 | SCAF1 | Q9H7N4 | 719;725 | | NA | 5.9490 | 5.9999 | 5.7159 | 6.4906 | 6.0170 | 5.7488 | 6.4513 | 5.8510 | 6.9997 | 6.1538 | 5.9215 | 5.9905 | 6.2028 | 6.7893 | 6.8597 | 6.8593 |
| SCAF1_T976 | SCAF1 | Q9H7N4 | 734;738 | | NA | 5.8400 | 6.3675 | 6.1621 | 6.7442 | 6.1110 | 6.0354 | 6.3077 | 6.0039 | 5.9188 | 6.2879 | 5.8747 | 6.0266 | 6.2803 | 6.8314 | 6.3451 | 6.8634 |
| SCAF1_T989 | SCAF1 | Q9H7N4 | 498;500 | | NA | 5.5582 | 6.0160 | 5.9197 | 5.9760 | 6.7003 | 5.7230 | 6.3402 | 5.5634 | 5.8276 | 5.5203 | 5.8602 | 5.9794 | 6.6918 | 7.4094 | 7.5086 | 7.4060 |
| SCAF11_S1122 | SCAF11 | Q99590 | | 608 | TEELIESPKLESS | 5.9550 | 6.0867 | 6.2260 | 6.3634 | 5.8444 | 5.6064 | 6.3996 | 5.3912 | 6.5029 | 6.2272 | 6.3192 | 6.4637 | 6.7615 | 6.7986 | 6.2826 | 6.7717 |
| SCAF11_S1169 | SCAF11 | Q99590 | | 816 | EKKRPQSPSPRRE | 6.3365 | 6.4205 | 5.8691 | 6.6817 | 5.3617 | 5.0432 | 5.4551 | 5.5366 | 6.0154 | 5.8968 | 5.6440 | 5.9981 | 6.9674 | 7.1980 | 7.7454 | 7.8305 |
| SCAF11_S338 | SCAF11 | Q99590 | | 963 | IDRDSYSPRWKGR | 7.2758 | 7.1658 | 6.4063 | 7.2382 | 6.4591 | 6.4678 | 6.7740 | 6.2666 | 6.6055 | 6.5485 | 6.5017 | 6.7276 | 4.7994 | 5.2562 | 4.6251 | 4.8824 |
| SCAF11_S405 | SCAF11 | Q99590 | | 318 | RKPAMTTPTRRST | 6.8071 | 6.2777 | 6.5380 | 7.2317 | 6.1848 | 6.1497 | 6.6492 | 6.2437 | 6.8424 | 6.6920 | 6.3583 | 6.8114 | 4.9816 | 5.2837 | 5.0324 | 5.9163 |
| SCAF11_S533 | SCAF11 | Q99590 | | 1169 | SRRGRNSSGPQSG | 6.4070 | 6.8074 | 6.8844 | 7.2382 | 6.0553 | 6.3534 | 7.3126 | 6.4371 | 7.1661 | 6.7262 | 6.8290 | 7.0096 | 4.2873 | 5.2626 | 4.2411 | 4.9827 |
| SCAF11_S588 | SCAF11 | Q99590 | | 338 | ASQSQRSPISDNS | 5.8784 | 6.1021 | 5.8554 | 6.7831 | 5.9740 | 6.0539 | 6.0902 | 5.8871 | 6.6954 | 6.2450 | 6.5122 | 6.4134 | 5.5372 | 6.6787 | 6.7147 | 6.5792 |
| SCAF11_S608 | SCAF11 | Q99590 | | 902 | PRVKDSSPGEKSR | 6.3538 | 6.5842 | 5.9410 | 6.5275 | 5.8165 | 5.6263 | 6.5830 | 6.0206 | 7.3099 | 6.1345 | 5.8501 | 5.9791 | 6.1071 | 6.2834 | 6.4509 | 6.4321 |
| SCAF11_S614 | SCAF11 | Q99590 | | 405 | KSSSNDSVDEETA | 6.0855 | 6.1981 | 6.6086 | 6.8862 | 5.9563 | 5.7131 | 6.4833 | 5.8423 | 6.8095 | 6.4627 | 6.5815 | 6.7727 | 5.8880 | 5.9824 | 5.6951 | 6.0347 |
| SCAF11_S771;S776 | SCAF11 | Q99590 | | 818 | KRPQSPSPRRETG | 6.3743 | 6.8748 | 5.9289 | 6.8131 | 5.5766 | 5.1821 | 5.3821 | 5.5366 | 6.3110 | 5.8933 | 5.4081 | 5.5231 | 6.6963 | 7.5203 | 7.6006 | 7.3789 |
| SCAF11_S776 | SCAF11 | Q99590 | | 832 | ESRKSQSPSPKNE | 6.0642 | 5.9260 | 5.1938 | 5.9114 | 5.7035 | 5.3194 | 5.7599 | 5.1936 | 5.5402 | 5.3922 | 5.2906 | 5.4222 | 7.9240 | 8.5702 | 8.6528 | 8.1360 |
| SCAF11_S796 | SCAF11 | Q99590 | | 796 | RRSRFHSPSTTWS | 5.8217 | 6.2712 | 5.4574 | 6.3434 | 5.6751 | 5.5710 | 5.9046 | 6.1925 | 5.6167 | 6.0003 | 5.3938 | 5.5768 | 7.2807 | 7.7212 | 7.8216 | 7.3521 |
| SCAF11_S796;S802 | SCAF11 | Q99590 | | 588 | EKITESSLVEITE | 6.8637 | 6.9454 | 6.6586 | 7.8865 | 6.7507 | 6.7842 | 7.1544 | 6.6804 | 6.7214 | 6.7983 | 6.9392 | 7.1833 | 3.8257 | 4.6830 | 3.9106 | 4.2147 |
| SCAF11_S816 | SCAF11 | Q99590 | | 776 | VSQPSESPKDTID | 6.6644 | 6.2995 | 6.1702 | 6.7994 | 5.4362 | 5.9134 | 6.1735 | 5.2003 | 6.5376 | 6.3118 | 6.2012 | 6.6491 | 6.3840 | 6.1112 | 6.3530 | 6.7950 |
| SCAF11_S816;S818 | SCAF11 | Q99590 | | 1122 | KFVEQQSYKRKSE | 7.2140 | 7.0899 | 7.9811 | 6.6215 | 4.8964 | 5.7689 | 6.1999 | 4.9913 | 6.7667 | 7.6040 | 8.0467 | 9.2155 | 4.3846 | 4.7571 | 3.5965 | 4.8660 |
| SCAF11_S818 | SCAF11 | Q99590 | | 533 | DQISGLSQSEVKT | 6.1603 | 6.5013 | 6.6110 | 6.6067 | 6.1741 | 6.0164 | 6.0755 | 5.6963 | 6.5875 | 6.4412 | 5.8174 | 5.6903 | 5.9619 | 7.2209 | 6.0812 | 6.3579 |
| SCAF11_S830;S834 | SCAF11 | Q99590 | | 614 | SPKLESSEGEIIQ | 5.5819 | 6.4324 | 5.9009 | 6.2732 | 6.3677 | 6.0111 | 6.8453 | 6.2012 | 6.3059 | 6.0057 | 5.7962 | 6.5602 | 6.0553 | 6.6837 | 6.5838 | 6.3955 |
| SCAF11_S832 | SCAF11 | Q99590 | 816;818 | | NA | 6.3072 | 6.5718 | 6.3368 | 6.9511 | 5.7330 | 5.7508 | 6.1619 | 6.1037 | 6.3528 | 6.3218 | 5.7842 | 6.0698 | 6.0442 | 6.3255 | 6.3285 | 6.8570 |
| SCAF11_S832;S834 | SCAF11 | Q99590 | 961;963 | | NA | 6.2782 | 7.6142 | 6.8855 | 7.3109 | 6.4659 | 6.2398 | 7.2933 | 6.6330 | 6.3352 | 7.1889 | 7.1237 | 7.0259 | 3.9523 | 4.7255 | 4.1539 | 4.7736 |
| SCAF11_S902 | SCAF11 | Q99590 | 832;834 | | NA | 6.2075 | 6.7415 | 5.8346 | 6.6960 | 5.5953 | 6.1824 | 6.8429 | 5.5445 | 6.9419 | 6.1135 | 6.0856 | 5.8297 | 6.1560 | 6.2344 | 6.4902 | 6.5041 |
| SCAF11_S961;S963 | SCAF11 | Q99590 | 796;802 | | NA | 5.9980 | 6.1480 | 6.4429 | 7.2771 | 6.7470 | 6.2639 | 6.6516 | 6.3760 | 6.3890 | 6.3986 | 6.2169 | 6.3966 | 5.3683 | 5.5953 | 5.7524 | 5.9784 |
| SCAF11_S963 | SCAF11 | Q99590 | 830;834 | | NA | 6.3523 | 6.2961 | 6.1684 | 6.8357 | 5.9306 | 5.2302 | 5.7438 | 6.0102 | 6.2469 | 5.8745 | 6.3452 | 6.3338 | 6.9824 | 6.4151 | 6.4558 | 6.7788 |
| SCAF11_T318 | SCAF11 | Q99590 | 771;776 | | NA | 6.2133 | 6.1320 | 6.4653 | 7.0232 | 5.3597 | 5.6508 | 6.3485 | 5.7743 | 6.7111 | 6.2413 | 5.7720 | 6.8025 | 5.7139 | 6.2737 | 6.1610 | 7.3574 |
| SCAF4_S154 | SCAF4 | O95104 | | 656 | SHTEPVSPIPKPL | 5.9809 | 6.2679 | 5.8601 | 6.3624 | 6.3446 | 6.3441 | 6.3085 | 6.3338 | 6.1892 | 5.9770 | 5.7790 | 5.9914 | 6.5482 | 6.4086 | 6.6203 | 6.6841 |
| SCAF4_S656 | SCAF4 | O95104 | | 154 | VTNNEGSPPPPVK | 6.0562 | 6.1074 | 6.3051 | 6.5257 | 5.9234 | 6.1028 | 5.8506 | 5.9896 | 6.3915 | 6.8507 | 6.1382 | 6.0793 | 6.2255 | 6.4281 | 6.7359 | 6.2901 |
| SCAF4_T237 | SCAF4 | O95104 | | 237 | TAQLKTTPTQPSE | 5.8229 | 6.7918 | 6.1366 | 6.6668 | 5.6873 | 6.0913 | 7.2321 | 5.9008 | 6.1242 | 6.3433 | 6.7798 | 7.0366 | 5.4451 | 5.7840 | 5.4823 | 6.6750 |
| SCAF8_S683 | SCAF8 | Q9UPN6-2 | | 683 | TVQTTQSPTPVEK | 6.1290 | 6.2195 | 6.0276 | 6.2529 | 6.2710 | 6.0455 | 5.9457 | 6.2224 | 6.0780 | 6.1037 | 5.8155 | 6.2017 | 6.6554 | 6.5955 | 6.8208 | 6.6159 |
| SCAMP1_T45 | SCAMP1 | O15126 | | 45 | PFSDSRTPPPGGV | 6.6769 | 6.9393 | 6.6239 | 7.1216 | 6.2045 | 6.7197 | 6.9467 | 6.8671 | 6.7117 | 6.9686 | 7.2045 | 6.7453 | 4.2937 | 4.9882 | 4.2903 | 4.6979 |
| SCAMP2_S319 | SCAMP2 | O15127 | | 319 | TFHRAASSAAQGA | 2.3103 | 2.2690 | 2.2857 | 2.2214 | 2.5639 | 2.8110 | 2.8064 | 2.6317 | 2.3400 | 2.6362 | 2.1323 | 2.0762 | 17.3322 | 17.0054 | 19.9004 | 16.6779 |
| SCAMP3_S32 | SCAMP3 | O14828 | | 32 | VIQHRPSRQYATL | 6.0034 | 5.0717 | 5.7050 | 5.7155 | 7.0860 | 5.1939 | 5.5298 | 5.5640 | 7.6239 | 7.1305 | 6.6857 | 6.6563 | 7.4301 | 7.5183 | 5.8082 | 5.2779 |
| SCAMP3_S76 | SCAMP3 | O14828 | | 76 | QPSRKLSPTEPKN | 5.2257 | 6.0108 | 6.9679 | 7.5329 | 5.7373 | 7.5213 | 6.7220 | 5.7471 | 7.1676 | 6.1133 | 6.7550 | 6.7238 | 5.5555 | 5.5056 | 5.0029 | 5.7114 |
| SCAMP4_S212 | SCAMP4 | Q969E2 | | 221 | EYPTVPSYPGSGQ | 4.9897 | 4.6028 | 4.8211 | 5.0830 | 4.7926 | 4.1447 | 4.7393 | 4.8307 | 4.8532 | 5.0232 | 4.7172 | 4.5353 | 10.5087 | 11.4533 | 10.6217 | 10.2835 |
| SCAMP4_S221 | SCAMP4 | Q969E2 | | 212 | NNFSGNSLPEYPT | 5.2461 | 6.3270 | 5.8568 | 4.9169 | 5.7956 | 5.8265 | 5.8291 | 5.2944 | 5.5682 | 6.5119 | 5.6275 | 5.8542 | 7.5870 | 7.4221 | 8.8567 | 7.4801 |
| SCAMP4_S225 | SCAMP4 | Q969E2 | | 194 | TEWNTGTWRNPPS | 7.7367 | 8.3829 | 7.5755 | 7.0650 | 7.4076 | 5.8183 | 6.3938 | 5.7035 | 6.4380 | 6.3044 | 5.8516 | 5.8677 | 5.6286 | 5.7456 | 4.0622 | 4.0186 |
| SCAMP4_T194 | SCAMP4 | Q969E2 | | 225 | VPSYPGSGQWPxx | 6.0272 | 5.7772 | 6.3645 | 6.1024 | 5.4869 | 5.8229 | 6.8940 | 7.0176 | 4.8191 | 6.4397 | 6.3497 | 6.8993 | 7.2353 | 7.6868 | 5.0959 | 5.9815 |
| SCAND1_S52 | SCAND1 | P57086 | | 52 | PAPEPSSPNAAVP | 5.6671 | 5.6917 | 6.7217 | 6.0786 | 6.3231 | 6.2972 | 6.3074 | 6.3241 | 6.7959 | 6.9652 | 6.7872 | 7.0051 | 5.7960 | 6.0836 | 5.4370 | 5.7189 |
| SCAP_S822 | SCAP | Q12770 | | 851 | PEEPGDSPPLRHR | 6.3513 | 6.6249 | 6.1343 | 6.4781 | 6.4561 | 6.2501 | 6.3182 | 6.3920 | 5.9654 | 6.0254 | 6.1737 | 5.9936 | 5.9044 | 6.5291 | 6.0815 | 6.3218 |
| SCAP_S851 | SCAP | Q12770 | | 822 | GRQRRDSGVGSGL | 5.3252 | 5.4148 | 5.4011 | 6.4772 | 6.0303 | 5.5063 | 5.5082 | 6.4069 | 7.2446 | 6.4150 | 6.6272 | 6.9067 | 5.9164 | 6.8513 | 6.5556 | 7.4133 |
| SCAP_S943 | SCAP | Q12770 | | 943 | SPGPVLSQAPEDE | 6.0442 | 6.7111 | 6.0411 | 5.8354 | 5.1716 | 5.8354 | 5.7747 | 6.6488 | 6.0084 | 5.6936 | 6.7953 | 6.3746 | 6.9558 | 7.5245 | 6.2857 | 6.2998 |
| SCAPER_S533 | SCAPER | Q9BY12 | | 533 | MHEKLSSPSRKRT | 7.4441 | 7.2600 | 6.9462 | 5.5305 | 7.3089 | 7.3032 | 6.4945 | 5.8309 | 5.5291 | 6.2186 | 6.4252 | 6.2932 | 5.5366 | 5.7420 | 4.9074 | 5.2296 |
| SCARB1_S502 | SCARB1 | Q8WTV0-5 | | 489 | DRPPSPYTPLLPD | 6.4812 | 6.4013 | 6.3234 | 6.0558 | 6.5550 | 5.1661 | 6.3059 | 5.1671 | 6.1060 | 6.1224 | 6.8574 | 6.4648 | 6.6462 | 6.6134 | 6.3603 | 6.3736 |
| SCARB1_Y489 | SCARB1 | Q8WTV0-2 | | 502 | TSAPKGSVLQEAK | 8.0440 | 6.3205 | 7.1138 | 5.9751 | 7.6397 | 5.6411 | 6.0838 | 6.3896 | 6.1350 | 5.9904 | 6.8988 | 7.7612 | 5.5784 | 5.1677 | 4.6061 | 4.6547 |
| SCARF2_S543 | SCARF2 | Q96GP6 | | 841 | DLPAPETPRKKTP | 7.0383 | 6.8792 | 6.4111 | 6.8596 | 5.9663 | 5.8782 | 6.5711 | 6.0616 | 6.5528 | 6.2859 | 6.2480 | 6.4121 | 5.7531 | 5.9054 | 5.3243 | 5.8529 |
| SCARF2_S653 | SCARF2 | Q96GP6 | | 742 | TALAAPSPPRARA | 5.8350 | 6.5919 | 5.8191 | 6.7790 | 6.2956 | 5.3891 | 6.0245 | 6.5688 | 6.9616 | 6.7940 | 6.4469 | 6.4257 | 6.1420 | 6.0001 | 5.9831 | 5.9434 |
| SCARF2_S687 | SCARF2 | Q96GP6 | | 687 | AAGRAPSPPPPGS | 5.9765 | 6.0615 | 6.6264 | 6.8717 | 5.7303 | 6.2315 | 5.2838 | 5.9038 | 7.0298 | 6.1249 | 5.7092 | 5.7378 | 6.2288 | 6.6533 | 7.0738 | 6.7568 |
| SCARF2_S687;S698 | SCARF2 | Q96GP6 | | 543 | SGLEQPSPSWSSR | 5.4642 | 6.4230 | 6.5140 | 6.4808 | 6.3957 | 6.4158 | 5.9049 | 5.5513 | 6.6349 | 6.2774 | 6.3359 | 6.2689 | 6.4496 | 6.5796 | 6.0070 | 6.2969 |
| SCARF2_S718 | SCARF2 | Q96GP6 | | 846 | ETPRKKTPIQKPP | 6.7978 | 7.0679 | 5.8921 | 7.0995 | 5.7262 | 6.0051 | 6.9896 | 7.1917 | 14.9457 | 6.6298 | 6.7101 | 6.9923 | 2.6113 | 3.4854 | 2.5091 | 3.3463 |
| SCARF2_S742 | SCARF2 | Q96GP6 | | 718 | HTVEHGSPRTRDP | 4.7067 | 6.6572 | 6.5245 | 7.5661 | 9.1284 | 5.0801 | 5.6500 | 7.2074 | 7.2123 | 4.6477 | 5.8858 | 6.9225 | 6.1855 | 5.5223 | 5.0408 | 6.0627 |
| SCARF2_T841 | SCARF2 | Q96GP6 | | 653 | GLSLSPSPERRKP | 6.7066 | 6.5897 | 6.9705 | 7.1441 | 6.7774 | 5.0752 | 5.9196 | 5.5549 | 6.8709 | 6.7567 | 6.2550 | 6.3378 | 5.1827 | 6.1962 | 6.1427 | 5.5201 |
| SCARF2_T846 | SCARF2 | Q96GP6 | 687;698 | | NA | 6.5750 | 6.6541 | 5.7943 | 7.0390 | 5.9887 | 6.1422 | 6.1113 | 6.1215 | 6.1591 | 6.0293 | 6.1046 | 6.6979 | 5.9476 | 6.3941 | 6.1014 | 6.1400 |
| SCFD1_S303 | SCFD1 | Q8WVM8 | | 303 | SSGVENSPAGARP | 6.5382 | 6.6522 | 6.5779 | 6.8830 | 6.9454 | 6.8443 | 7.1539 | 6.8535 | 6.5451 | 6.6501 | 6.7712 | 6.3756 | 4.6968 | 5.0545 | 4.7044 | 4.7537 |
| SCFD1_S316 | SCFD1 | Q8WVM8 | | 316 | KRKNKKSYDLTPV | 6.6400 | 6.0836 | 6.5144 | 6.2222 | 5.4122 | 6.2722 | 5.9035 | 6.0291 | 6.0341 | 6.5034 | 6.6520 | 6.3403 | 6.2967 | 6.7850 | 6.2557 | 6.0556 |
| SCFD1_S388 | SCFD1 | Q8WVM8 | | 388 | LTSAVSSLPELLE | 5.5567 | 5.5546 | 5.5183 | 5.0009 | 6.3970 | 5.3579 | 5.4306 | 5.4926 | 5.2638 | 5.3280 | 5.2574 | 5.7556 | 8.2314 | 7.7824 | 9.6703 | 8.4025 |
| SCLT1_S681 | SCLT1 | Q96NL6 | | 681 | QRRKAASLMNLEN | 6.4711 | 6.1655 | 6.6113 | 6.0458 | 6.3185 | 6.4082 | 6.2088 | 6.4116 | 6.1464 | 6.3437 | 6.7833 | 7.1792 | 5.7288 | 5.9388 | 5.5623 | 5.6767 |
| SCLY_S129 | SCLY | Q96I15 | | 129 | HTGGHHSPVKGAK | 6.1114 | 5.9068 | 6.5128 | 5.6068 | 5.6975 | 6.5077 | 4.8596 | 5.9573 | 5.8594 | 6.6049 | 6.9250 | 6.9360 | 6.5390 | 7.0067 | 6.8408 | 6.1282 |
| SCML1_S138 | SCML1 | Q9UN30 | | 138 | SYPESYSPTLPVS | 5.7181 | 6.3122 | 6.5016 | 6.5221 | 6.1802 | 5.8603 | 6.0885 | 6.1959 | 6.3219 | 6.3309 | 6.1148 | 6.3917 | 6.5513 | 6.6396 | 6.1695 | 6.1013 |
| SCML2_S499 | SCML2 | Q9UQR0 | | 511 | PYVVPLSPKLPKT | 6.3691 | 6.3883 | 6.5531 | 6.8851 | 6.5186 | 6.1111 | 6.4772 | 6.1298 | 6.5710 | 6.3862 | 6.7048 | 6.6533 | 5.6536 | 5.5889 | 5.2751 | 5.7349 |
| SCML2_S499;S511 | SCML2 | Q9UQR0 | | 590 | PLVGDISPKSSPH | 6.5348 | 6.8350 | 6.9158 | 7.9978 | 6.4826 | 6.2484 | 6.8690 | 5.9543 | 6.7418 | 6.8605 | 6.9123 | 6.8371 | 4.7203 | 4.8282 | 4.2835 | 4.9785 |
| SCML2_S511 | SCML2 | Q9UQR0 | | 499 | RQSTKRSPQQTVP | 5.8733 | 6.2986 | 5.6063 | 5.7064 | 5.2688 | 6.3728 | 5.2082 | 5.7151 | 6.3839 | 6.2059 | 4.5827 | 5.8832 | 8.0256 | 6.9105 | 8.1047 | 7.8538 |
| SCML2_S590 | SCML2 | Q9UQR0 | 499;511 | | NA | 6.3652 | 6.8950 | 6.6751 | 7.0647 | 6.2408 | 6.2452 | 6.7318 | 5.9042 | 6.7643 | 6.6547 | 6.5892 | 6.3736 | 4.9685 | 5.7656 | 5.3530 | 5.4091 |
| SCNM1_S183 | SCNM1 | Q9BWG6 | | 183 | SAPAPMSPTRRRA | 5.9649 | 6.5162 | 6.9522 | 6.5609 | 6.6076 | 5.9745 | 6.1091 | 6.5331 | 6.3961 | 6.3273 | 7.1642 | 6.6585 | 5.5241 | 6.1153 | 5.3208 | 5.2751 |
| SCRIB_S1140 | SCRIB | A0A0G2JNZ2 | | 1309 | QPPSPPSPDELPA | 6.2219 | 6.0544 | 5.9149 | 6.1079 | 6.4570 | 5.9876 | 6.1214 | 5.9341 | 6.3250 | 6.0399 | 6.0064 | 5.8843 | 7.2215 | 6.5845 | 6.5779 | 6.5611 |
| SCRIB_S1220 | SCRIB | A0A0G2JNZ2 | | 835 | RPEDDYSPRERRG | 6.3898 | 6.3308 | 6.2126 | 6.5857 | 6.3005 | 6.3460 | 6.3209 | 6.6280 | 6.4158 | 6.7454 | 6.5171 | 6.3733 | 6.0377 | 5.8238 | 5.3678 | 5.6046 |
| SCRIB_S1223;S1226 | SCRIB | A0A0G2JNZ2 | | 853 | PLLPPESPGPLRQ | 6.8589 | 6.7457 | 6.2582 | 6.5814 | 6.5064 | 6.3603 | 6.4595 | 6.2962 | 6.6973 | 6.3143 | 6.0503 | 6.1021 | 6.0143 | 5.7964 | 5.2090 | 5.7498 |
| SCRIB_S1276 | SCRIB | A0A0G2JNZ2 | | 1566 | TSPGRLSPDFAEE | 5.9581 | 6.1928 | 5.8864 | 6.2338 | 5.9051 | 5.6465 | 5.9064 | 6.6105 | 6.3281 | 5.9842 | 5.9081 | 6.1972 | 6.8966 | 6.5574 | 6.9696 | 6.8192 |
| SCRIB_S1279 | SCRIB | A0A0G2JNZ2 | | 1448 | SRQSPASPPPLGG | 6.5668 | 6.3932 | 6.4571 | 6.6063 | 5.9791 | 6.3397 | 6.5748 | 6.4524 | 6.7342 | 6.3539 | 6.7483 | 6.7825 | 5.4631 | 5.6025 | 5.3889 | 5.5572 |
| SCRIB_S1285 | SCRIB | A0A0G2JNZ2 | | 1220 | GIGHRNSLESISS | 6.1726 | 5.5487 | 5.5324 | 5.6039 | 6.0257 | 5.5458 | 5.7686 | 5.7277 | 6.5243 | 6.0865 | 5.7202 | 6.0738 | 7.9055 | 7.0550 | 7.4167 | 7.2927 |
| SCRIB_S1306 | SCRIB | A0A0G2JNZ2 | | 1348 | TPGPAASPEQLSF | 6.4658 | 6.5256 | 6.1086 | 6.5323 | 6.8270 | 6.4230 | 5.9665 | 6.7377 | 6.9486 | 6.6741 | 7.0609 | 6.9248 | 5.3841 | 5.4026 | 4.8332 | 5.1851 |
| SCRIB_S1306;S1309 | SCRIB | A0A0G2JNZ2 | | 1285 | SVQRVPSGAAGGK | 6.2758 | 5.2781 | 5.1345 | 5.3890 | 6.6535 | 5.7900 | 5.7645 | 5.7941 | 6.1887 | 5.1125 | 4.7855 | 5.1753 | 8.8621 | 7.8929 | 8.0228 | 7.8807 |
| SCRIB_S1309 | SCRIB | A0A0G2JNZ2 | | 1378 | GPPKRVSLVGADD | 6.7303 | 5.8599 | 6.0918 | 5.9646 | 6.6648 | 5.4809 | 6.0821 | 6.5110 | 7.6682 | 6.3512 | 5.9999 | 6.3601 | 6.5023 | 6.3756 | 5.7258 | 5.6316 |
| SCRIB_S1378 | SCRIB | A0A0G2JNZ2 | | 703 | AVVSAPSVKGVSF | 7.0873 | 7.3294 | 7.2004 | 7.1942 | 7.0586 | 6.0697 | 6.4840 | 6.1963 | 6.8293 | 6.1508 | 6.7362 | 5.8829 | 5.0606 | 5.7481 | 4.4611 | 4.5108 |
| SCRIB_S1437 | SCRIB | A0A0G2JNZ2 | | 504 | CQPDSGSPLPAEE | 6.5047 | 6.7423 | 6.5358 | 7.1653 | 6.4935 | 6.4751 | 6.4134 | 6.2936 | 7.0681 | 6.6728 | 6.7865 | 7.1823 | 5.2255 | 5.0888 | 4.5091 | 4.8432 |
| SCRIB_S1439 | SCRIB | A0A0G2JNZ2 | | 1486 | APERALSPAELRA | 7.1449 | 7.3606 | 6.0275 | 6.8076 | 6.9460 | 6.8607 | 7.1679 | 7.1574 | 7.0077 | 6.6373 | 6.8378 | 6.7024 | 4.0990 | 4.6688 | 4.0556 | 4.5187 |
| SCRIB_S1448 | SCRIB | A0A0G2JNZ2 | | 1279 | AVPSAGSVQRVPS | 7.4316 | 7.5731 | 6.8897 | 7.1754 | 5.5984 | 6.3725 | 6.7860 | 6.2430 | 7.2542 | 6.8658 | 6.8106 | 6.7825 | 4.7694 | 5.1192 | 4.0002 | 4.3284 |
| SCRIB_S1475 | SCRIB | A0A0G2JNZ2 | | 939 | SLLTAASPTIALL | 6.2637 | 6.3877 | 6.1348 | 6.5134 | 6.1235 | 6.2243 | 6.7473 | 6.2911 | 6.6953 | 6.3099 | 6.3781 | 6.5919 | 5.8269 | 6.0133 | 5.4696 | 6.0292 |
| SCRIB_S1486 | SCRIB | A0A0G2JNZ2 | | 1579 | LRSLEPSPSPGPQ | 6.1502 | 5.8861 | 5.7855 | 5.8419 | 6.3877 | 6.1222 | 5.8982 | 6.1137 | 6.1062 | 6.3033 | 5.8798 | 5.7999 | 6.9138 | 7.1296 | 6.8116 | 6.8702 |
| SCRIB_S1508 | SCRIB | A0A0G2JNZ2 | | 1439 | PPWASPSPTSRQS | 5.6408 | 6.5492 | 6.4004 | 6.6254 | 6.1017 | 6.2656 | 6.0535 | 5.7897 | 6.0382 | 6.3539 | 6.1672 | 6.4123 | 6.4521 | 6.8126 | 6.3786 | 5.9588 |
| SCRIB_S1541 | SCRIB | A0A0G2JNZ2 | | 1581 | SLEPSPSPGPQEE | 6.0490 | 6.7336 | 6.4618 | 6.4595 | 6.4238 | 6.2770 | 6.5901 | 6.0688 | 6.5253 | 6.1610 | 6.2619 | 6.1579 | 5.6883 | 6.2273 | 6.1121 | 5.8026 |
| SCRIB_S1541;S1566 | SCRIB | A0A0G2JNZ2 | | 1276 | ALAAVPSAGSVQR | 3.7810 | 3.3543 | 3.3718 | 3.6592 | 4.2963 | 3.8160 | 3.7757 | 3.9123 | 3.7630 | 3.3794 | 3.4462 | 3.3201 | 13.5480 | 12.5572 | 15.0562 | 14.9633 |
| SCRIB_S1547 | SCRIB | A0A0G2JNZ2 | | 1475 | ERLRVQSPEPPAP | 6.4989 | 6.6688 | 6.1148 | 6.5954 | 6.6596 | 6.6550 | 6.7174 | 6.8432 | 7.1784 | 6.7924 | 6.8359 | 7.0440 | 4.7301 | 5.0257 | 4.6606 | 4.9800 |
| SCRIB_S1547;S1566 | SCRIB | A0A0G2JNZ2 | | 1508 | RAARMKSLEQDAL | 9.9332 | 8.0463 | 7.8998 | 5.6238 | 6.2603 | 7.5150 | 6.9290 | 6.7875 | 6.0070 | 5.5908 | 5.5851 | 5.9719 | 4.8572 | 5.2884 | 3.5952 | 4.1095 |
| SCRIB_S1566 | SCRIB | A0A0G2JNZ2 | | 1306 | SGQQPPSPPSPDE | 6.9133 | 7.6557 | 7.2711 | 7.2711 | 5.6720 | 6.3416 | 6.0298 | 5.8549 | 7.1288 | 6.9493 | 7.2550 | 7.2435 | 4.9105 | 5.1826 | 4.1632 | 4.1575 |
| SCRIB_S1579 | SCRIB | A0A0G2JNZ2 | | 1140 | IFISKVSPTGAAG | 6.6535 | 6.9248 | 5.8343 | 6.6233 | 5.9885 | 5.9072 | 5.9977 | 6.0142 | 6.4779 | 6.5210 | 6.0930 | 6.3270 | 6.2655 | 6.0769 | 6.2281 | 6.0671 |
| SCRIB_S1581 | SCRIB | A0A0G2JNZ2 | | 1437 | EQPPWASPSPTSR | 5.7322 | 6.5252 | 6.7447 | 6.3248 | 5.9462 | 6.5893 | 6.8357 | 6.0181 | 6.0468 | 6.4584 | 6.6087 | 6.6990 | 5.8222 | 5.9772 | 5.4463 | 6.2250 |
| SCRIB_S23 | SCRIB | A0A0G2JNZ2 | | 875 | ERGLGFSIAGGKG | 10.9515 | 5.1928 | 8.5959 | 7.1853 | 7.9979 | 4.2056 | 5.1467 | 3.5716 | 4.8497 | 4.5588 | 6.8061 | 4.0892 | 8.0782 | 7.8916 | 5.7486 | 5.1306 |
| SCRIB_S504 | SCRIB | A0A0G2JNZ2 | | 23 | VDKRHCSLQAVPE | 5.0999 | 5.7589 | 6.6675 | 5.3061 | 6.3117 | 6.5520 | 5.6263 | 6.8103 | 6.2117 | 7.6248 | 6.4944 | 6.3507 | 6.5992 | 6.3131 | 6.3419 | 5.9315 |
| SCRIB_S668 | SCRIB | A0A0G2JNZ2 | | 607 | RLIRKDTPHYKKH | 9.4690 | 7.7844 | 8.3858 | 8.2880 | 6.0779 | 5.4768 | 6.5252 | 4.1256 | 6.2022 | 5.4512 | 7.7634 | 7.0369 | 4.4079 | 5.2438 | 3.4090 | 4.3530 |
| SCRIB_S703 | SCRIB | A0A0G2JNZ2 | | 1342 | APAQPPTPGPAAS | 6.2767 | 7.0314 | 5.5797 | 5.7527 | 5.6286 | 6.0077 | 5.7859 | 7.0384 | 6.9581 | 6.2226 | 5.6636 | 6.0409 | 7.4227 | 6.5301 | 6.1545 | 5.9064 |
| SCRIB_S835 | SCRIB | A0A0G2JNZ2 | | 1541 | RLAEAPSPAPTPS | 5.5669 | 5.7014 | 6.4298 | 6.2868 | 5.9073 | 5.6094 | 5.8545 | 6.3702 | 6.1745 | 6.2085 | 6.9863 | 6.1030 | 8.0993 | 6.5761 | 5.6673 | 6.4587 |
| SCRIB_S853 | SCRIB | A0A0G2JNZ2 | | 475 | GLQRRATPHPSEL | 8.2104 | 8.0436 | 6.5675 | 7.0731 | 5.7687 | 7.9336 | 6.1696 | 8.3314 | 7.0071 | 7.0153 | 7.5728 | 7.3050 | 2.9712 | 4.4216 | 2.3493 | 3.2598 |
| SCRIB_S875 | SCRIB | A0A0G2JNZ2 | | 668 | EEEEEGSPQEEEE | 6.5901 | 5.4731 | 6.0118 | 6.2945 | 5.2976 | 5.2412 | 5.6468 | 6.2353 | 6.5495 | 5.7937 | 7.5417 | 6.7887 | 6.6511 | 6.6843 | 6.8636 | 6.3370 |
| SCRIB_S939 | SCRIB | A0A0G2JNZ2 | | 1547 | SPAPTPSPTPVED | 4.3663 | 5.3409 | 4.7152 | 5.1842 | 7.1308 | 6.2502 | 5.1242 | 5.7867 | 6.8895 | 6.2215 | 5.3682 | 5.0956 | 8.9970 | 8.2241 | 8.4790 | 6.8268 |
| SCRIB_T1342 | SCRIB | A0A0G2JNZ2 | | 749 | AGGKGSTPYKGDD | 6.7831 | 7.2695 | 7.2573 | 7.9300 | 7.0431 | 7.0997 | 7.3915 | 6.6991 | 7.3420 | 6.8384 | 7.4454 | 6.5198 | 3.3802 | 4.8999 | 2.7414 | 3.3599 |
| SCRIB_T1342;S1348 | SCRIB | A0A0G2JNZ2 | 1306;1309 | | NA | 5.7275 | 6.0567 | 6.2678 | 6.5657 | 6.1969 | 6.1536 | 5.9275 | 6.4991 | 6.1584 | 6.1627 | 6.2169 | 6.3877 | 6.6232 | 6.3039 | 6.5105 | 6.2419 |
| SCRIB_T1348 | SCRIB | A0A0G2JNZ2 | 1342;1348 | | NA | 7.2253 | 6.4201 | 6.3895 | 7.2127 | 6.7499 | 6.4299 | 6.5175 | 6.6895 | 6.9265 | 6.4722 | 6.4445 | 6.4774 | 5.4336 | 5.4158 | 4.6709 | 4.5246 |
| SCRIB_T475 | SCRIB | A0A0G2JNZ2 | 1547;1566 | | NA | 5.7527 | 5.3368 | 5.7158 | 6.0380 | 6.1582 | 6.3841 | 6.6717 | 6.1821 | 6.2565 | 6.7124 | 5.6635 | 5.7446 | 7.0842 | 6.9854 | 6.6287 | 6.6852 |
| SCRIB_T607 | SCRIB | A0A0G2JNZ2 | 1541;1566 | | NA | 6.7277 | 5.9833 | 6.1900 | 5.7786 | 6.0166 | 5.3082 | 6.8219 | 6.5796 | 5.8715 | 6.1217 | 5.2130 | 5.8753 | 7.2674 | 6.6298 | 6.9466 | 6.6687 |
| SCRIB_T749 | SCRIB | A0A0G2JNZ2 | 1223;1226 | | NA | 6.6269 | 6.3313 | 7.0073 | 6.8338 | 7.2968 | 5.3159 | 6.3657 | 7.0842 | 7.3718 | 5.8351 | 5.4039 | 5.8318 | 5.8445 | 6.0227 | 5.2041 | 5.6244 |
| SCYL1_S559 | SCYL1 | Q96KG9 | | 559 | DVHAASSPGMGGA | 5.7849 | 6.8165 | 6.6710 | 7.0243 | 6.2129 | 7.2676 | 6.5093 | 7.1755 | 6.4628 | 7.2759 | 6.4354 | 6.8026 | 4.6785 | 4.9940 | 4.6789 | 5.2100 |
| SCYL1_S754 | SCYL1 | Q96KG9 | | 754 | TQPRPDSWGEDNW | 5.4057 | 5.2341 | 8.2087 | 4.9971 | 6.6683 | 7.4814 | 8.5192 | 4.9726 | 4.2453 | 8.0534 | 6.6234 | 8.2659 | 7.4242 | 3.9103 | 4.4374 | 5.5528 |
| SCYL2_S677 | SCYL2 | Q6P3W7 | | 708 | LKPQVHTPVATVK | 7.4334 | 6.6659 | 7.4496 | 7.3126 | 5.7452 | 6.5796 | 7.4933 | 6.2566 | 7.8858 | 7.2799 | 7.6454 | 7.8867 | 3.1778 | 4.2251 | 2.8357 | 4.1274 |

| | | | | | | | | | | | | | | | | | | | | |
|---|---|---|---|---|---|---|---|---|---|---|---|---|---|---|---|---|---|---|---|---|
| SCYL2_T708 | SCYL2 | Q6P3W7 | 677 | NKHKRASLTLEEK | 6.4904 | 6.7490 | 6.7405 | 6.9924 | 5.9434 | 6.3884 | 6.5378 | 6.1642 | 6.8867 | 7.1798 | 7.2255 | 7.3390 | 4.5761 | 5.3453 | 4.4893 | 4.9522 |
| SCYL2_T810 | SCYL2 | Q6P3W7 | 810 | TKMTLGTPPTLPN | 7.1385 | 7.8070 | 7.9672 | 7.0257 | 6.2779 | 6.8774 | 7.1277 | 7.4041 | 6.4278 | 7.1415 | 7.7320 | 7.0703 | 3.1182 | 3.6345 | 3.9958 | 3.6694 |
| SDAD1_S585 | SDAD1 | Q9NVU7 | 585 | KYIEIDSDEEPRG | 6.1855 | 6.4557 | 6.1287 | 6.8419 | 6.3684 | 6.1145 | 6.2557 | 6.3297 | 5.8303 | 6.1708 | 6.3347 | 6.3290 | 5.9501 | 6.5409 | 6.0160 | 6.1484 |
| SDC3_S428 | SDC3 | O75056 | 428 | EEPKQASVTYQKP | 6.3123 | 6.2296 | 6.3256 | 6.3148 | 6.2916 | 6.2385 | 5.6427 | 6.9086 | 6.3635 | 7.2727 | 7.6220 | 7.6456 | 5.0651 | 6.1229 | 4.8688 | 4.7758 |
| SDC4_Y197 | SDC4 | P31431 | 197 | APTNEFYAxxxxx | 7.2853 | 6.7607 | 7.2433 | 6.6591 | 5.7028 | 6.6563 | 6.4081 | 6.5666 | 7.9076 | 6.2535 | 7.1787 | 7.4650 | 4.6538 | 4.4434 | 3.9997 | 4.8161 |
| SDCBP_S205 | SDCBP | O00560 | 205 | ITMHKDSTGHVGF | 5.9704 | 5.3657 | 5.1526 | 4.8625 | 6.0592 | 5.8556 | 5.8682 | 5.9790 | 6.6600 | 5.1471 | 5.3067 | 6.7111 | 7.7796 | 7.8276 | 7.3692 | 8.0855 |
| SDCCAG8_S4 | SDCCAG8 | Q86SQ7 | 4 | xxxMAKSPENSTL | 5.7700 | 6.6606 | 6.1653 | 6.1862 | 5.8195 | 5.8725 | 7.1211 | 6.4277 | 6.5190 | 6.2366 | 6.2566 | 6.0845 | 6.2922 | 6.6545 | 5.9420 | 5.9916 |
| SDE2_S278 | SDE2 | Q6IQ49 | 278 | VNTDHGSPEQLQI | 6.2045 | 6.5954 | 6.5924 | 7.0270 | 6.4242 | 6.7504 | 6.3124 | 6.4052 | 6.9516 | 6.8511 | 6.9079 | 6.7531 | 5.0660 | 5.0111 | 4.8062 | 5.3415 |
| SEC14L1_T234 | SEC14L1 | Q92503-2 | 234 | PEPVVGTPDDKLD | 6.0346 | 7.4283 | 8.9580 | 9.0096 | 5.4269 | 6.1519 | 6.8605 | 6.8135 | 7.2359 | 8.1650 | 6.5085 | 8.0242 | 2.9000 | 3.3575 | 3.4631 | 3.6625 |
| SEC16A_S1069 | SEC16A | O15027-5 | 1069 | PPQQQASPPQLPK | 6.1022 | 6.5238 | 6.6373 | 6.5095 | 6.1060 | 6.4920 | 5.9431 | 6.6027 | 6.6694 | 6.8060 | 6.8726 | 6.7199 | 5.3307 | 5.8200 | 5.2451 | 5.6196 |
| SEC16A_S1327 | SEC16A | O15027-5 | 2054 | GKFANLTPSRTVP | 7.1871 | 7.3732 | 6.9748 | 7.0712 | 6.6168 | 7.2121 | 7.1839 | 7.0592 | 7.2012 | 6.9433 | 7.3095 | 7.2094 | 3.3264 | 4.0929 | 3.2823 | 3.9568 |
| SEC16A_S1398 | SEC16A | O15027-5 | 314 | IVNHWASPELRQN | 6.8637 | 7.4478 | 6.9696 | 6.9858 | 6.4318 | 7.0470 | 6.8121 | 7.2535 | 7.2522 | 7.4617 | 7.3473 | 7.4856 | 3.2080 | 4.1617 | 3.2296 | 4.0425 |
| SEC16A_S1437;S1441 | SEC16A | O15027-5 | 536 | HGRLSGSARPQEL | 6.5024 | 7.1032 | 6.8837 | 7.1693 | 6.3258 | 6.5740 | 6.7860 | 6.7021 | 6.5739 | 7.3100 | 7.2795 | 7.0191 | 4.1176 | 5.0586 | 3.9414 | 4.6534 |
| SEC16A_S1441 | SEC16A | O15027-5 | 1964 | PDGPLASPARVPM | 6.0236 | 5.8165 | 6.5646 | 6.2493 | 6.4423 | 6.6955 | 6.9637 | 7.3213 | 6.5972 | 6.3066 | 6.3456 | 6.5598 | 5.3258 | 5.9932 | 5.0358 | 5.7592 |
| SEC16A_S1964 | SEC16A | O15027-5 | 1398 | APLPPGSFHGDFA | 7.7070 | 7.0162 | 7.2989 | 6.3153 | 5.9156 | 7.2830 | 7.5816 | 6.3178 | 6.6449 | 6.1957 | 6.1414 | 6.9995 | 4.6460 | 5.0240 | 4.2102 | 4.7028 |
| SEC16A_S2022 | SEC16A | O15027-5 | 1441 | VSSRPTSPEKFSV | 5.6447 | 5.9938 | 6.1281 | 5.6860 | 6.2971 | 6.4044 | 5.8889 | 6.0071 | 5.8386 | 6.3637 | 6.1050 | 6.8146 | 6.5879 | 7.2068 | 6.7463 | 6.2871 |
| SEC16A_S2083 | SEC16A | O15027-5 | 2313 | SQGGELSRCSSMS | 5.9342 | 6.0246 | 6.4403 | 6.3266 | 5.1801 | 6.4452 | 6.8095 | 5.7539 | 6.3061 | 5.8757 | 6.8631 | 5.8676 | 6.3764 | 7.1240 | 5.5855 | 7.0872 |
| SEC16A_S2313 | SEC16A | O15027-5 | 957 | LPGFANSPAGSTS | 7.1463 | 7.7795 | 6.4948 | 7.4268 | 6.2353 | 7.2318 | 6.5273 | 7.2113 | 7.1750 | 6.5995 | 6.9706 | 6.6025 | 4.1935 | 4.4491 | 3.6758 | 4.2807 |
| SEC16A_S2317 | SEC16A | O15027-5 | 2317 | ELSRCSSMSSLSR | 5.7960 | 5.7348 | 6.5582 | 5.5115 | 5.3721 | 6.0526 | 5.8264 | 5.8304 | 5.9642 | 5.9325 | 6.2331 | 6.4317 | 7.1594 | 7.3901 | 6.9508 | 7.2561 |
| SEC16A_S29 | SEC16A | O15027-5 | 589 | NYRGSVSQPSTPS | 5.7758 | 6.2122 | 7.1571 | 6.3844 | 5.9766 | 6.2674 | 5.4554 | 6.6547 | 6.6403 | 6.6846 | 7.2274 | 7.2518 | 5.3872 | 5.9867 | 5.6151 | 5.3233 |
| SEC16A_S314 | SEC16A | O15027-5 | 569 | FKQIDSSPVGGET | 6.5487 | 6.1407 | 6.3418 | 6.5213 | 6.4893 | 7.0351 | 6.0452 | 6.1875 | 5.9823 | 6.2304 | 6.3190 | 6.7568 | 5.4612 | 6.1635 | 5.7709 | 5.8223 |
| SEC16A_S536 | SEC16A | O15027-5 | 2022 | LLQEARSPDPGIV | 5.1989 | 6.0415 | 6.3768 | 5.7327 | 5.7304 | 7.3698 | 6.2749 | 7.7088 | 6.2084 | 5.9079 | 6.4372 | 7.1372 | 5.4216 | 6.3323 | 5.9819 | 6.1397 |
| SEC16A_S569 | SEC16A | O15027-5 | 2083 | QPPLSLSPAPETK | 5.7651 | 5.7623 | 5.9014 | 5.6200 | 6.3865 | 6.0139 | 6.0632 | 6.7502 | 6.4567 | 6.4767 | 6.6274 | 6.0902 | 6.7697 | 6.5009 | 6.4079 | 6.4079 |
| SEC16A_S589 | SEC16A | O15027-5 | 65 | RQALQSTPLGSSS | 6.7246 | 6.7047 | 7.2349 | 7.3846 | 5.8514 | 7.2510 | 7.2548 | 7.3268 | 7.0856 | 7.5263 | 7.7035 | 7.5867 | 3.4952 | 4.5510 | 2.7196 | 3.5993 |
| SEC16A_S957 | SEC16A | O15027-5 | 29 | SVFWASSPYRRRA | 6.9911 | 7.6052 | 6.5205 | 6.3091 | 5.3143 | 7.1515 | 7.3195 | 7.1368 | 7.0905 | 7.2712 | 7.8516 | 7.8560 | 3.2936 | 4.6204 | 3.4391 | 4.2294 |
| SEC16A_T2054 | SEC16A | O15027-5 | 2076 | RADSGPTQPPLSL | 5.5329 | 5.6179 | 6.5229 | 6.7007 | 6.1503 | 6.3229 | 6.2911 | 6.8192 | 5.7888 | 7.2347 | 6.1743 | 6.6208 | 5.8850 | 6.4560 | 5.9880 | 5.8944 |
| SEC16A_T2076 | SEC16A | O15027-5 | 1327 | DPRFTGSFDDDPD | 6.3430 | 6.3347 | 8.5016 | 7.7179 | 6.5361 | 6.2805 | 6.7922 | 7.1450 | 6.9848 | 7.0172 | 7.8999 | 7.4353 | 3.4824 | 4.2879 | 3.5501 | 3.6914 |
| SEC16A_T65 | SEC16A | O15027-5 | 1437;1441 | NA | 5.9218 | 6.4337 | 6.5042 | 6.2423 | 6.2071 | 6.7247 | 6.0859 | 6.2145 | 6.7717 | 6.5796 | 6.6478 | 6.7080 | 5.9198 | 6.0411 | 5.2601 | 5.7376 |
| SEC16A_Y1423;S1441 | SEC16A | O15027-5 | 1423;1441 | NA | 5.1588 | 6.3048 | 6.8452 | 6.2617 | 6.0774 | 6.0593 | 5.8650 | 6.4599 | 6.3451 | 6.7171 | 6.4545 | 7.3016 | 6.1474 | 6.2968 | 5.7732 | 5.9321 |
| SEC22B_S137 | SEC22B | O75396 | 137 | ARRNLGSINTELQ | 5.1872 | 4.9447 | 5.7582 | 5.8257 | 6.4523 | 5.0862 | 5.4423 | 6.3014 | 7.0230 | 7.1605 | 6.8328 | 6.6889 | 6.7154 | 7.7009 | 6.9396 | 5.9409 |
| SEC22B_S164 | SEC22B | O75396 | 168 | ALSALDSKANNLS | 2.1557 | 2.2069 | 2.5107 | 2.3696 | 2.5950 | 2.3522 | 2.6654 | 2.4883 | 2.2313 | 2.4324 | 2.0080 | 2.1517 | 17.6931 | 17.7897 | 18.8655 | 17.4846 |
| SEC22B_S168 | SEC22B | O75396 | 164 | QRGEALSALDSKA | 6.5723 | 7.1095 | 6.8188 | 6.9324 | 6.0159 | 5.6366 | 7.1538 | 6.2122 | 5.9908 | 5.8211 | 5.5834 | 5.7074 | 5.6454 | 7.0077 | 6.2196 | 5.5731 |
| SEC22B_S174 | SEC22B | O75396 | 175 | KANNLSSLSKKYR | 6.2356 | 6.1922 | 5.7285 | 6.0824 | 5.5026 | 6.0008 | 6.1603 | 6.2797 | 6.1816 | 6.4847 | 5.8671 | 6.7492 | 6.5398 | 6.5549 | 6.4164 | 7.0244 |
| SEC22B_S175 | SEC22B | O75396 | 174 | SKANNLSSLSKKY | 5.8791 | 5.9946 | 6.3098 | 5.3447 | 6.7587 | 7.3736 | 6.6082 | 6.4889 | 6.5356 | 6.4577 | 6.4655 | 6.4279 | 5.2915 | 6.4538 | 5.8260 | 5.7844 |
| SEC23A_S588 | SEC23A | Q15436 | 588 | FHLRRSSFLQVFN | 5.5591 | 6.0223 | 6.4253 | 6.1523 | 5.5516 | 7.1464 | 6.3963 | 6.3049 | 6.7359 | 6.2082 | 7.1690 | 6.6252 | 5.9095 | 6.4167 | 5.2443 | 6.1330 |
| SEC23IP_S134 | SEC23IP | Q9Y6Y8 | 926 | AEKVVESPDFSKD | 6.0348 | 6.2522 | 6.7417 | 6.3598 | 6.3589 | 6.7847 | 6.8431 | 6.9624 | 6.6859 | 6.7279 | 6.9125 | 6.5714 | 4.9039 | 5.6428 | 4.4929 | 5.2251 |
| SEC23IP_S926 | SEC23IP | Q9Y6Y8 | 134 | DVSNAFSPSISKA | 6.9298 | 6.0426 | 6.4020 | 6.6626 | 6.3760 | 8.0941 | 7.5265 | 6.7441 | 7.7057 | 7.3883 | 7.4407 | 7.7196 | 3.6857 | 4.0311 | 3.1909 | 4.0603 |
| SEC24A_T225 | SEC24A | O95486 | 225 | PPVRALTPLTSSY | 5.9152 | 5.7883 | 7.3058 | 5.9037 | 6.8201 | 8.0415 | 6.7211 | 6.9673 | 6.9490 | 6.5759 | 7.7559 | 6.1864 | 5.1440 | 5.5142 | 3.9082 | 4.5035 |
| SEC24B_S1254 | SEC24B | O95487-3 | 1254 | RDSRPLSPILHIV | 5.4051 | 5.6384 | 6.1496 | 6.0380 | 5.7992 | 6.1633 | 5.7502 | 6.2242 | 6.1869 | 6.2913 | 5.6337 | 6.5583 | 6.6942 | 7.2788 | 7.7223 | 7.3665 |
| SEC24B_T360 | SEC24B | O95487-3 | 360 | GSSSTRTPPTANH | 6.0950 | 6.6214 | 6.5456 | 6.8594 | 6.2011 | 6.8999 | 6.6330 | 7.0892 | 6.2177 | 6.9024 | 7.0900 | 6.7459 | 4.8098 | 5.7061 | 4.5193 | 5.0643 |
| SEC24C_S862 | SEC24C | P53992 | 862 | YRGVLNSPVKAVR | 6.2784 | 6.3588 | 6.6104 | 6.3488 | 5.6828 | 6.6307 | 5.9545 | 6.3669 | 6.2096 | 7.1495 | 6.3669 | 6.8948 | 5.6482 | 6.1535 | 5.6210 | 5.7253 |
| SEC31A_S1176 | SEC31A | O94979-8 | 527 | SDQVAQSDGEESP | 6.0881 | 6.1211 | 6.3209 | 6.3503 | 6.1999 | 6.0874 | 6.1081 | 6.4236 | 6.2194 | 6.1656 | 6.2798 | 6.2428 | 6.2890 | 6.5029 | 6.3145 | 6.2865 |
| SEC31A_S527 | SEC31A | O94979-8 | 1176 | LREQTLSPTITSG | 5.7643 | 6.1296 | 6.1588 | 6.4720 | 6.4569 | 6.5218 | 6.0367 | 6.5820 | 6.5395 | 6.5498 | 6.5513 | 6.7509 | 5.3565 | 6.2785 | 6.0515 | 5.7997 |
| SEC31A_S527;S532 | SEC31A | O94979-8 | 799 | PVAGHESPKIPYE | 5.5854 | 6.7444 | 6.7993 | 6.0798 | 5.5081 | 6.6775 | 6.1792 | 6.2881 | 6.4606 | 7.0112 | 7.3111 | 6.8557 | 5.3040 | 6.2035 | 5.6989 | 5.2931 |
| SEC31A_S532 | SEC31A | O94979-8 | 532 | QSDGEESPAAEEQ | 6.2013 | 5.5308 | 7.3309 | 6.9844 | 5.0763 | 5.3373 | 6.6694 | 5.4048 | 6.5074 | 5.8166 | 6.1226 | 6.9641 | 6.5006 | 7.5897 | 6.4534 | 5.5106 |
| SEC31A_S799 | SEC31A | O94979-8 | 527;532 | NA | 5.7606 | 5.8626 | 6.6667 | 5.7677 | 5.9185 | 6.2968 | 6.2398 | 6.4331 | 5.7711 | 5.7917 | 6.4884 | 6.4123 | 6.7466 | 6.9489 | 6.3910 | 6.5042 |
| SEC61A1_S408 | SEC61A1 | P61619 | 408 | RGHRETSMVHELN | 6.6833 | 6.4276 | 5.9309 | 6.7813 | 6.2684 | 5.9703 | 6.0729 | 6.0031 | 5.8753 | 5.9131 | 6.1418 | 6.1554 | 6.3312 | 6.5155 | 6.8943 | 6.0357 |
| SEC61B_S49 | SEC61B | P60468 | 49 | SAGRTTSAGTGGM | 5.9108 | 6.3705 | 6.1913 | 6.6107 | 5.1250 | 5.0345 | 5.3993 | 5.5159 | 6.1302 | 6.3945 | 5.9400 | 6.9973 | 7.1737 | 6.8001 | 7.3903 | 7.0159 |
| SEC61B_T30 | SEC61B | P60468 | 30 | ARAAGSTVRQRKN | 7.5332 | 8.0050 | 7.8307 | 7.6995 | 5.3697 | 6.8366 | 7.7175 | 6.5868 | 7.3262 | 7.7634 | 8.2433 | 7.8824 | 2.2739 | 3.7427 | 2.1265 | 3.1204 |
| SEC62_S307;S309 | SEC62 | Q99442 | 158 | KEETPGTPKKKET | 6.1938 | 6.3496 | 6.0759 | 6.4621 | 6.0262 | 5.8182 | 6.2545 | 5.7998 | 6.2264 | 6.3113 | 6.1544 | 6.1687 | 6.4281 | 6.7372 | 6.5180 | 6.4760 |
| SEC62_S341 | SEC62 | Q99442 | 375 | EELEQQTDGDCEE | 5.9397 | 6.2350 | 6.2909 | 6.6967 | 5.9237 | 5.9604 | 6.4049 | 6.5962 | 6.0250 | 5.8239 | 5.9518 | 5.9279 | 6.7215 | 6.7685 | 6.9352 | 6.6988 |
| SEC62_T155;T158 | SEC62 | Q99442 | 341 | SGGERHSDTDSDR | 6.1996 | 6.8504 | 6.1470 | 7.2893 | 5.3437 | 7.0622 | 5.8182 | 5.6217 | 6.7453 | 6.6114 | 6.6029 | 6.8758 | 4.9556 | 5.2742 | 7.2334 | 5.3691 |
| SEC62_T158 | SEC62 | Q99442 | 155;158 | NA | 6.3103 | 6.6264 | 5.9262 | 6.4171 | 6.7847 | 6.2351 | 6.9289 | 6.0183 | 6.0887 | 6.5628 | 6.3885 | 6.1984 | 5.5612 | 6.2338 | 5.8323 | 5.8874 |
| SEC62_T375 | SEC62 | Q99442 | 307;309 | NA | 6.2181 | 6.5667 | 6.7322 | 7.5785 | 5.7615 | 5.8824 | 6.4781 | 5.1475 | 6.1833 | 6.3234 | 5.7191 | 6.4445 | 6.3207 | 6.7013 | 5.8783 | 6.4692 |
| SEC63_S742;S748 | SEC63 | Q9UGP8 | 537 | PLKKKPTPVLLPQ | 1.7291 | 1.7469 | 1.6080 | 2.3575 | 2.2206 | 1.4430 | 1.7668 | 1.6785 | 1.4382 | 1.8462 | 1.7428 | 1.2623 | 16.9051 | 18.6685 | 22.8112 | 20.7754 |
| SEC63_T537 | SEC63 | Q9UGP8 | 742;748 | NA | 6.2129 | 5.9204 | 6.3311 | 6.2020 | 6.2661 | 6.5399 | 5.8139 | 6.3393 | 6.8040 | 5.9418 | 6.3613 | 6.1901 | 6.4673 | 6.3526 | 6.0775 | 6.1798 |
| SEC63_T731;S742 | SEC63 | Q9UGP8 | 731;742 | NA | 6.0719 | 6.5792 | 5.6036 | 6.5315 | 6.5095 | 6.5327 | 5.9712 | 6.7427 | 5.7348 | 6.6129 | 5.7264 | 6.2788 | 6.7829 | 6.3868 | 6.1945 | 5.7405 |
| SECISBP2L_S851 | SECISBP2L | Q93073 | 851 | GHSRNPSAASAIS | 6.2472 | 6.6194 | 8.1908 | 6.4933 | 5.8985 | 5.1444 | 5.1216 | 6.5432 | 6.5029 | 6.5756 | 5.8258 | 6.6694 | 6.3023 | 5.8766 | 5.6541 | 6.3347 |
| SEH1L_T258 | SEH1L | Q96EE3-1 | 258 | PVRKELTSSGGPT | 6.2076 | 6.5757 | 6.2302 | 6.9016 | 6.0964 | 6.2041 | 6.3692 | 5.9885 | 6.1132 | 6.4367 | 6.3515 | 6.2911 | 6.1128 | 6.0294 | 6.0779 | 6.0141 |
| SELENON_S13 | SELENON | Q9NZV5 | 13 | GQRGPPSPGPAAQ | 6.3834 | 6.6919 | 6.6617 | 6.4087 | 6.1030 | 6.3950 | 7.1173 | 6.9168 | 6.1624 | 6.3941 | 7.2321 | 7.0413 | 4.5914 | 6.1147 | 4.4425 | 5.3438 |
| SELENOS_S140 | SELENOS | Q9BQE4 | 140 | KPQEEDSPGPSTS | 7.2140 | 7.3392 | 6.7389 | 7.4522 | 7.0196 | 7.1711 | 7.7741 | 7.3803 | 7.4075 | 6.8385 | 6.9557 | 6.9965 | 3.1735 | 3.9256 | 3.1921 | 3.4211 |
| SEMA3F_T522 | SEMA3F | Q13275 | 522 | DPAPVKTMTISSK | 6.4842 | 6.8991 | 5.7210 | 6.7421 | 6.3061 | 5.9809 | 5.8721 | 6.2157 | 6.0978 | 5.8515 | 5.7502 | 5.7459 | 6.0406 | 6.6772 | 7.1623 | 6.4531 |
| SEMA4B_S793 | SEMA4B | Q9NPR2 | 793 | YQSLSDSPPGSRV | 5.9233 | 6.0148 | 5.9943 | 6.0883 | 6.0386 | 6.0949 | 6.0420 | 6.0908 | 5.9126 | 6.2927 | 6.0928 | 6.2397 | 6.3884 | 7.1592 | 6.5934 | 7.0342 |
| SEMA4B_S818 | SEMA4B | Q9NPR2 | 818 | DSFVEVSPVCPRP | 6.2821 | 6.1851 | 5.9643 | 6.3745 | 5.7840 | 6.3157 | 6.7858 | 6.3546 | 6.1620 | 6.7321 | 6.5327 | 6.6286 | 5.2451 | 6.7165 | 5.5288 | 6.4081 |
| SEMA4B_T780 | SEMA4B | Q9NPR2 | 780 | GLGPPSTPLDHRG | 5.8899 | 5.7921 | 6.1170 | 7.0783 | 5.9291 | 6.6559 | 6.2871 | 6.8112 | 6.9426 | 6.8469 | 6.6185 | 7.0987 | 4.8416 | 6.3153 | 4.5611 | 6.2148 |
| SEMA4C_S717 | SEMA4C | Q9C0C4 | 772 | PGQPLPSPTRLHL | 6.4928 | 6.3434 | 6.5762 | 6.3998 | 5.9137 | 6.2639 | 6.4752 | 5.5360 | 6.4627 | 6.6797 | 6.0859 | 6.6695 | 6.0539 | 6.2068 | 5.6335 | 6.2071 |
| SEMA4C_S760;S772 | SEMA4C | Q9C0C4 | 717 | LPKEPTSPPFRPC | 6.5594 | 6.9589 | 7.1430 | 6.9970 | 6.2435 | 7.0556 | 6.9151 | 6.8108 | 6.6356 | 6.5004 | 7.6449 | 7.0101 | 4.1556 | 5.1791 | 4.1348 | 4.9561 |
| SEMA4C_S772 | SEMA4C | Q9C0C4 | 760;772 | NA | 5.6722 | 6.4614 | 6.7709 | 6.4523 | 6.3207 | 6.1772 | 6.5509 | 7.1543 | 6.8063 | 6.5269 | 6.1064 | 6.5725 | 5.5973 | 5.7638 | 5.1406 | 5.9263 |
| SEMA6B_S794 | SEMA6B | Q9H3T3 | 794 | PLTPHASPDRRRV | 5.9153 | 6.0585 | 7.2195 | 6.4138 | 5.8203 | 6.3456 | 5.7005 | 6.3094 | 5.5519 | 5.9635 | 6.8300 | 5.9601 | 6.0096 | 6.6052 | 5.8651 | 7.4316 |
| SEMA6B_T709 | SEMA6B | Q9H3T3 | 709 | DSGLLPTPEQTPL | 6.2014 | 7.4144 | 6.7446 | 7.5285 | 5.4419 | 5.8413 | 6.0517 | 6.1495 | 6.1888 | 5.9739 | 6.3867 | 6.1481 | 6.3437 | 5.8487 | 6.3711 | 5.3656 |
| SENP1_S132 | SENP1 | Q9P0U3 | 157 | PIKPVPSPSWSGS | 5.9454 | 5.6381 | 6.1390 | 6.1478 | 6.0575 | 5.8797 | 6.2252 | 5.6116 | 6.4453 | 5.9407 | 5.6240 | 5.7667 | 7.4109 | 6.7573 | 6.9021 | 7.5088 |
| SENP1_S157 | SENP1 | Q9P0U3 | 57 | QGHLDRSFTCSTR | 7.9696 | 7.4277 | 6.1829 | 8.2073 | 6.4979 | 6.0451 | 6.9559 | 6.2636 | 7.5397 | 6.1229 | 6.6971 | 7.2070 | 3.7575 | 4.5655 | 3.5519 | 5.0085 |
| SENP1_S40 | SENP1 | Q9P0U3 | 132 | SSGLSNSFAGKSN | 7.3028 | 7.0436 | 6.9889 | 7.4182 | 6.0033 | 6.5407 | 6.9013 | 6.4875 | 7.3105 | 6.8464 | 7.3322 | 7.4077 | 3.9196 | 4.3418 | 3.5836 | 4.5720 |
| SENP1_S57 | SENP1 | Q9P0U3 | 49 | FPEQDLSLSDQQI | 5.5584 | 5.9625 | 6.3275 | 7.2245 | 6.8680 | 5.8997 | 6.1493 | 6.4321 | 6.1407 | 5.9095 | 6.8231 | 6.7818 | 5.8206 | 6.4950 | 5.9266 | 5.6806 |
| SENP1_S80 | SENP1 | Q9P0U3 | 80 | DNPSSDSFLGSGD | 7.2322 | 8.6884 | 6.3645 | 9.0571 | 7.5302 | 5.5471 | 6.8300 | 6.3831 | 6.8412 | 6.4651 | 7.1391 | 6.8878 | 4.1058 | 4.3609 | 3.3107 | 3.2567 |
| SENP2_S344 | SENP2 | Q9HC62 | 344 | LLRRKVSIIETKE | 6.1784 | 5.1216 | 5.6263 | 5.1251 | 5.5491 | 5.1242 | 5.2246 | 5.0485 | 7.4117 | 7.5257 | 6.7155 | 7.4892 | 7.2269 | 7.5530 | 7.5816 | 7.2985 |
| SENP3_S169 | SENP3 | Q9H4L4 | 26 | IPPAYSSPRRERL | 6.3262 | 6.3600 | 6.3267 | 6.5403 | 6.2456 | 6.0246 | 6.5223 | 6.3544 | 6.1110 | 6.3070 | 6.1456 | 6.1329 | 6.2652 | 6.3821 | 5.9693 | 5.9869 |
| SENP3_S169;S181 | SENP3 | Q9H4L4 | 232 | LRWTPKSPLDPDS | 6.0925 | 6.1177 | 6.3168 | 6.3276 | 6.3910 | 6.1316 | 6.2447 | 6.4016 | 6.3754 | 6.0832 | 6.1465 | 5.8831 | 6.3242 | 6.6190 | 6.3146 | 6.2306 |
| SENP3_S169;T176;S181 | SENP3 | Q9H4L4 | 169 | HPKNHLSPQQGGA | 5.8435 | 6.3520 | 6.0655 | 6.7032 | 6.2502 | 6.1684 | 6.2753 | 6.4393 | 6.3998 | 5.8539 | 6.0932 | 6.0108 | 6.1486 | 6.6173 | 6.1568 | 6.6225 |
| SENP3_S212 | SENP3 | Q9H4L4 | 212 | EDGVRGSPPVPSG | 6.6653 | 7.0592 | 6.0564 | 6.7154 | 6.6518 | 6.5775 | 6.5565 | 5.9567 | 6.5922 | 6.5068 | 6.6163 | 6.2805 | 5.4774 | 5.7444 | 5.4895 | 5.0543 |
| SENP3_S232 | SENP3 | Q9H4L4 | 353 | FQQEFSTPSRKGL | 6.6004 | 6.7972 | 6.0879 | 7.0293 | 6.2171 | 6.2973 | 6.7253 | 5.9189 | 6.1231 | 6.5067 | 6.3799 | 6.7038 | 5.6954 | 5.4198 | 5.5383 | 5.9599 |
| SENP3_S26 | SENP3 | Q9H4L4 | 169;181 | NA | 6.3526 | 6.0750 | 6.3503 | 6.1909 | 6.3619 | 6.2523 | 6.3819 | 6.4170 | 6.3880 | 6.1622 | 6.2219 | 6.0049 | 6.1014 | 6.2452 | 5.9814 | 6.5131 |
| SENP3_S69;S73;S75 | SENP3 | Q9H4L4 | 169;176;181 | NA | 6.3705 | 6.7906 | 6.5872 | 6.9780 | 4.9154 | 6.1502 | 6.1179 | 5.0504 | 6.4443 | 7.6032 | 6.1211 | 5.8665 | 6.1698 | 6.6234 | 6.1861 | 6.0255 |
| SENP3_S73;S75 | SENP3 | Q9H4L4 | 73;75 | NA | 6.3479 | 5.8057 | 6.6010 | 6.8913 | 6.2415 | 5.8891 | 6.3541 | 5.9305 | 6.5687 | 6.6574 | 6.2668 | 7.5071 | 5.6995 | 5.9980 | 5.6872 | 5.5543 |
| SENP3_T353 | SENP3 | Q9H4L4 | 69;73;75 | NA | 5.6503 | 5.2193 | 5.5407 | 5.9608 | 6.3454 | 6.2693 | 5.4847 | 6.1979 | 5.1576 | 4.8568 | 6.0425 | 5.3903 | 8.7228 | 7.8980 | 7.1084 | 8.1553 |
| SENP5_S228 | SENP5 | Q96HI0 | 363 | DQSSCSSPKWECT | 5.8977 | 5.6683 | 6.1374 | 6.4817 | 5.7137 | 5.7516 | 6.0337 | 5.5540 | 6.2667 | 6.2533 | 5.9986 | 6.1992 | 6.9871 | 6.9429 | 7.0041 | 7.1099 |
| SENP5_S363 | SENP5 | Q96HI0 | 465 | DHPYCKSPLEAPL | 5.9830 | 6.3712 | 5.6606 | 6.2606 | 6.1857 | 6.3768 | 6.7065 | 5.5873 | 5.7834 | 6.0712 | 5.7867 | 5.5167 | 7.0876 | 6.9434 | 6.4055 | 7.2738 |
| SENP5_S465 | SENP5 | Q96HI0 | 228 | HRRIKLSPLMMYE | 6.3643 | 7.5551 | 5.8378 | 6.9850 | 7.2293 | 6.1979 | 6.5743 | 6.7625 | 7.0140 | 6.9018 | 8.0256 | 6.8240 | 3.7676 | 4.7737 | 4.6868 | 4.5005 |
| SENP6_S352 | SENP6 | Q9GZR1 | 919 | LNYSDESPEAGKM | 5.5343 | 6.0866 | 6.8053 | 6.0409 | 6.4177 | 6.6395 | 6.3762 | 5.7140 | 6.4855 | 6.3457 | 6.0519 | 6.5804 | 6.3769 | 6.4025 | 6.4702 | 6.4023 |
| SENP6_S919 | SENP6 | Q9GZR1 | 352 | NRRESISPQPADS | 4.9337 | 6.0545 | 7.0832 | 6.0099 | 6.1723 | 6.1096 | 6.2131 | 6.8552 | 5.8588 | 5.4048 | 6.1561 | 6.2473 | 6.2245 | 7.4803 | 6.4857 | 6.7109 |
| SENP7_S11 | SENP7 | Q9BQF6 | 11 | KLGRRPSSSEIIT | 4.6518 | 4.4013 | 4.7199 | 5.5670 | 6.8862 | 5.3558 | 5.0463 | 5.5270 | 7.4415 | 7.7718 | 7.0391 | 7.0954 | 7.7739 | 8.1138 | 6.1917 | 6.4175 |
| SEPTIN10_S434 | SEPTIN10 | Q9P0V9-2 | 434 | EIFHSQSFLATGS | 8.5518 | 6.0021 | 6.8448 | 6.6818 | 6.1751 | 5.5837 | 6.2675 | 5.5229 | 7.3456 | 4.6544 | 6.0146 | 5.7550 | 7.3223 | 6.9729 | 5.5096 | 4.7959 |
| SEPTIN2_S253 | SEPTIN2 | Q15019-2 | 253 | HLPDAESDEDEDF | 5.9676 | 5.9295 | 6.5516 | 6.3572 | 6.2499 | 6.2444 | 6.1362 | 6.3615 | 6.2763 | 6.4067 | 6.4849 | 6.3363 | 6.1695 | 6.6664 | 6.0131 | 6.2417 |
| SEPTIN2_T49 | SEPTIN2 | Q15019-2 | 49 | QFINPETPGYVGF | 7.9593 | 6.8684 | 7.6140 | 6.9307 | 6.5151 | 6.2402 | 7.2296 | 6.1619 | 7.5052 | 6.7095 | 7.0113 | 6.5137 | 3.3555 | 5.0796 | 3.7167 | 4.5893 |
| SEPTIN4_S195 | SEPTIN4 | O43236-7 | 195 | VPRRILSYPKDEA | 6.0486 | 6.6498 | 6.6345 | 6.6664 | 5.6831 | 6.7482 | 6.1383 | 5.9152 | 6.1386 | 6.6448 | 7.1605 | 7.3731 | 4.6208 | 6.4042 | 5.4149 | 5.7590 |
| SEPTIN5_S225 | SEPTIN5 | Q99719 | 225 | QFPECDSDEDEDF | 5.8326 | 5.6439 | 6.9601 | 6.4420 | 6.1703 | 6.6142 | 5.9132 | 5.9597 | 6.4242 | 6.1327 | 6.3087 | 6.3562 | 6.4201 | 6.6645 | 6.3599 | 5.7977 |
| SEPTIN5_S327 | SEPTIN5 | Q99719 | 327 | QDSRMESPIPILP | 6.2667 | 7.7466 | 6.4964 | 6.2507 | 8.7242 | 6.2614 | 5.8179 | 4.7067 | 5.0326 | 7.9549 | 7.9335 | 6.8757 | 3.9107 | 5.0379 | 6.2186 | 4.7655 |
| SEPTIN6_T418 | SEPTIN6 | Q14141 | 418 | QAGGSQTLKRDKE | 6.5297 | 6.3387 | 6.6763 | 6.4813 | 5.7557 | 6.3099 | 6.3733 | 5.7269 | 6.2275 | 5.1530 | 6.5919 | 5.6423 | 7.1496 | 6.8287 | 6.4007 | 6.7146 |
| SEPTIN7_S334 | SEPTIN7 | Q16181 | 334 | KGQLTKSPLAQME | 6.5656 | 7.1888 | 6.9361 | 6.9814 | 6.2205 | 6.8464 | 6.6938 | 6.6378 | 6.5624 | 7.0309 | 7.0093 | 6.7809 | 4.3937 | 5.0429 | 4.4439 | 4.6658 |
| SEPTIN7_T426 | SEPTIN7 | Q16181 | 426 | QQNSSRTLEKNKK | 8.1237 | 7.3011 | 7.5454 | 5.9355 | 6.2642 | 8.3050 | 7.1611 | 5.4110 | 5.9725 | 5.3871 | 6.2163 | 6.4604 | 5.4153 | 5.8984 | 4.2213 | 4.3817 |
| SEPTIN8_S20 | SEPTIN8 | Q92599 | 20 | PEPRSLSLGGHVG | 5.9226 | 5.5899 | 6.7303 | 5.9722 | 6.8039 | 6.3497 | 5.0845 | 5.9962 | 6.9814 | 7.0598 | 6.7639 | 6.6504 | 6.9462 | 6.3481 | 5.5564 | 5.2445 |
| SEPTIN9_S111 | SEPTIN9 | Q9UHD8 | 49 | PPRRVQTPLLRAT | 7.1014 | 7.8221 | 7.2495 | 7.1982 | 5.4701 | 7.3434 | 7.7487 | 7.0803 | 7.2369 | 7.4885 | 7.7219 | 7.4895 | 2.6654 | 3.9569 | 2.8179 | 3.6092 |
| SEPTIN9_S203 | SEPTIN9 | Q9UHD8 | 30 | GPALKRSFEVEEV | 6.7397 | 6.8734 | 7.5713 | 7.1213 | 6.6453 | 6.4781 | 6.0658 | 6.9543 | 6.4743 | 6.9719 | 6.1188 | 6.4357 | 4.6748 | 5.6015 | 4.6897 | 4.5842 |
| SEPTIN9_S30 | SEPTIN9 | Q9UHD8 | 111 | SRRTELSIDISSK | 5.4252 | 6.6311 | 8.1657 | 6.8628 | 7.4289 | 6.6661 | 5.4884 | 6.7985 | 6.4163 | 7.8727 | 6.9598 | 6.6161 | 3.6936 | 5.7732 | 4.6151 | 4.5865 |
| SEPTIN9_T42 | SEPTIN9 | Q9UHD8 | 42 | VETPNSTPPRRVQ | 5.8897 | 6.9820 | 6.8392 | 6.8008 | 7.0284 | 6.5788 | 7.3742 | 6.7161 | 7.1478 | 7.0624 | 7.1985 | 7.1194 | 3.8689 | 5.3009 | 3.6890 | 4.4040 |
| SEPTIN9_T49 | SEPTIN9 | Q9UHD8 | 203 | EAPTAPSPAQTLE | 5.6474 | 6.3176 | 6.0379 | 6.1910 | 5.6918 | 6.1973 | 6.7176 | 7.2664 | 6.4696 | 6.8833 | 6.0559 | 6.2395 | 5.7298 | 6.7250 | 5.7266 | 6.1034 |
| SERBP1_S197 | SERBP1 | Q8NC51 | 234 | KDELTESPKYIQK | 6.7314 | 6.9453 | 7.2815 | 6.7530 | 6.2917 | 6.5057 | 7.6939 | 6.1617 | 7.1932 | 7.3909 | 7.4898 | 7.0472 | 3.8878 | 4.3672 | 3.7811 | 4.4786 |
| SERBP1_S202 | SERBP1 | Q8NC51-3 | 237 | LTDLDQSNVTEET | 6.6019 | 7.2559 | 6.5031 | 7.5033 | 6.7640 | 7.3727 | 6.7016 | 7.0044 | 7.1758 | 6.3292 | 7.1114 | 6.6243 | 3.8230 | 4.5183 | 4.0949 | 4.6162 |
| SERBP1_S203 | SERBP1 | Q8NC51 | 25 | QLFDDESDPFEVL | 6.2853 | 6.6296 | 7.2929 | 6.5182 | 6.3168 | 6.5992 | 6.4878 | 6.4962 | 6.1731 | 6.6152 | 6.4533 | 6.3588 | 5.3029 | 5.7115 | 5.3190 | 5.4852 |
| SERBP1_S219 | SERBP1 | Q8NC51 | 243 | YIQKQISYNYSDL | 6.0119 | 5.2347 | 5.8335 | 6.0768 | 5.6701 | 6.4591 | 6.1785 | 5.9999 | 5.8856 | 5.3855 | 5.4357 | 5.5708 | 7.7841 | 7.7427 | 7.7457 | 6.9853 |
| SERBP1_S234 | SERBP1 | Q8NC51 | 330 | VLHKSKSEEAHAE | 4.7193 | 6.2106 | 7.2524 | 5.3892 | 6.0247 | 6.3854 | 5.5891 | 6.3268 | 5.5949 | 7.8667 | 6.8502 | 5.8139 | 5.8707 | 6.8245 | 7.1207 | 6.1609 |
| SERBP1_S237 | SERBP1 | Q8NC51 | 197 | REFDRHSGSDRSS | 7.2948 | 7.8236 | 8.2571 | 8.0983 | 6.8696 | 7.1910 | 7.0083 | 7.0257 | 6.5033 | 7.2163 | 8.1573 | 7.5017 | 2.6309 | 3.1139 | 2.5171 | 2.7912 |
| SERBP1_S237;T240 | SERBP1 | Q8NC51 | 252 | YSDLDQSNVTEET | 7.3549 | 7.5468 | 6.7724 | 7.7625 | 6.9535 | 7.2279 | 7.4776 | 6.5098 | 7.3559 | 6.1483 | 7.2829 | 6.9301 | 3.8147 | 4.3437 | 3.1555 | 3.9034 |
| SERBP1_S243 | SERBP1 | Q8NC51 | 219 | EDKRGGSGSHNWG | 7.7326 | 8.0380 | 7.7724 | 7.7004 | 6.6882 | 6.8831 | 6.9005 | 7.3962 | 7.0602 | 7.0224 | 7.7742 | 7.1649 | 2.5302 | 3.5263 | 2.5016 | 3.3088 |
| SERBP1_S25 | SERBP1 | Q8NC51 | 203 | SGSDRSSFSHYSG | 7.4552 | 6.5508 | 5.4759 | 6.5788 | 8.2175 | 6.8499 | 5.1182 | 8.6741 | 8.1953 | 7.0349 | 7.3013 | 6.7874 | 3.7314 | 5.9555 | 3.2768 | 2.7971 |
| SERBP1_S252 | SERBP1 | Q8NC51 | 202 | HSGSDRSSFSHYS | 6.4007 | 6.3814 | 5.3648 | 7.1590 | 7.3687 | 7.3338 | 5.1737 | 8.4263 | 8.0708 | 7.8451 | 7.8749 | 7.5442 | 3.4286 | 5.3070 | 3.2086 | 2.9146 |
| SERBP1_S330 | SERBP1 | Q8NC51-3 | 237;240 | NA | 5.9705 | 6.5563 | 7.0664 | 6.4801 | 5.7140 | 6.7755 | 7.0712 | 6.0052 | 7.2304 | 7.2685 | 6.8254 | 6.6206 | 4.8292 | 5.2267 | 5.1407 | 5.5793 |
| SERF2_S21 | SERF2 | P84101-2 | 21 | MKKQSDSVKGKRR | 7.6116 | 7.0561 | 7.9989 | 6.8892 | 6.2576 | 6.5054 | 6.3093 | 5.8062 | 7.2281 | 7.2229 | 8.1991 | 7.9453 | 4.0791 | 4.4334 | 2.9860 | 3.4718 |
| SERPINB6_S151 | SERPINB6 | P35237 | 151 | KIAELLSPGSVDP | 6.9200 | 6.8097 | 6.2730 | 5.9057 | 6.6144 | 6.8357 | 6.0517 | 5.7330 | 7.1600 | 6.6732 | 6.6800 | 6.3947 | 5.8525 | 5.5626 | 5.6407 | 4.8930 |
| SERPINB7_S183 | SERPINB7 | O75635 | 183 | FKGKWQSAFTKSE | 6.1345 | 6.8630 | 6.0007 | 6.2110 | 5.2433 | 5.7236 | 5.3213 | 6.2143 | 6.5465 | 6.1656 | 6.3680 | 5.8664 | 6.5855 | 6.6660 | 6.8242 | 7.2662 |
| SET_S15 | SET | Q01105 | 7 | MAPKRQSPLPPQK | 5.3593 | 5.8285 | 5.3373 | 4.8073 | 5.7795 | 6.4571 | 5.5463 | 5.8158 | 5.6942 | 6.0016 | 6.3471 | 5.8907 | 7.7030 | 8.0838 | 7.7553 | 7.5931 |
| SET_S161 | SET | Q01105 | 63 | RLNEQASEEILKV | 5.2160 | 5.5710 | 6.0795 | 5.9261 | 6.2850 | 6.0724 | 5.7625 | 6.2200 | 6.0258 | 6.2348 | 5.5998 | 6.1732 | 6.9544 | 7.4481 | 6.6170 | 6.8145 |
| SET_S24 | SET | Q01105-2 | 24 | DGADETSEKEQQE | 5.2619 | 6.0313 | 5.6572 | 5.8959 | 5.9658 | 6.7634 | 5.2012 | 6.9712 | 5.7705 | 5.5534 | 6.9629 | 6.2211 | 6.3108 | 7.4504 | 7.2914 | 6.6916 |
| SET_S63 | SET | Q01105 | 161 | EFHLNESGDPSSK | 5.3636 | 5.1779 | 5.8328 | 5.2560 | 5.9020 | 6.3003 | 5.7823 | 6.0218 | 6.1256 | 6.0300 | 6.7854 | 6.5476 | 6.7124 | 7.6903 | 7.3414 | 7.1305 |
| SET_S7 | SET | Q01105-2 | 15 | SKKELNSNHDGAD | 5.4167 | 6.5143 | 6.4262 | 6.1362 | 5.0698 | 6.5109 | 6.3581 | 5.9566 | 6.5266 | 6.3157 | 6.7597 | 6.3615 | 6.4429 | 6.7050 | 5.9945 | 6.5054 |
| SETD1A_S1103 | SETD1A | O15047 | 1153 | PDERPSSPIPLLP | 5.7388 | 5.4140 | 5.7856 | 6.2590 | 5.9987 | 5.3944 | 6.2228 | 5.8161 | 6.4155 | 6.0705 | 5.8307 | 6.2775 | 7.0975 | 6.8082 | 7.1242 | 7.4465 |
| SETD1A_S1153 | SETD1A | O15047 | 1185 | PEPPPATPPQAKF | 5.8232 | 6.7150 | 5.3151 | 5.9831 | 5.7699 | 7.1627 | 6.6901 | 6.1465 | 5.4466 | 7.6778 | 7.0880 | 6.4734 | 6.2957 | 5.5461 | 6.3881 | 5.4786 |
| SETD1A_T1185 | SETD1A | O15047 | 1103 | PERVAGSPVTPLP | 5.9931 | 5.8410 | 5.9365 | 6.1533 | 6.5507 | 5.5650 | 6.1324 | 5.4153 | 6.2451 | 7.2302 | 6.4922 | 6.8045 | 6.4817 | 7.0454 | 6.4256 | 5.6882 |
| SETD1A_T279 | SETD1A | O15047 | 279 | YTSRGSTPYSQDS | 6.3718 | 6.5381 | 6.0066 | 6.4045 | 5.3175 | 6.6842 | 7.1926 | 5.8874 | 6.8295 | 7.2118 | 6.6130 | 6.9773 | 5.6510 | 4.9975 | 4.9927 | 4.9245 |
| SETD1A_T916 | SETD1A | O15047 | 916 | KRPRPSTPAEEDE | 5.9943 | 5.1734 | 6.3831 | 5.8254 | 5.9604 | 5.4711 | 5.8450 | 5.5640 | 6.5066 | 6.1298 | 5.7918 | 6.4419 | 7.4472 | 7.2998 | 6.3601 | 7.8062 |
| SETD1B_S1659;S1663 | SETD1B | Q9UPS6 | 909 | MAKASLTPVKSGE | 6.6691 | 6.9405 | 6.4970 | 6.8106 | 6.6363 | 6.1828 | 6.2566 | 6.4230 | 6.3287 | 6.3609 | 6.2745 | 5.8698 | 5.8018 | 5.9057 | 5.5194 | 5.5233 |
| SETD1B_S398 | SETD1B | Q9UPS6 | 398 | GFKSAFSPYQTPV | 5.4873 | 6.0098 | 5.5504 | 6.4352 | 6.6152 | 5.8715 | 6.0159 | 5.9552 | 6.5618 | 6.4093 | 6.0470 | 6.4259 | 7.0705 | 7.1207 | 6.2407 | 6.1837 |
| SETD1B_S986 | SETD1B | Q9UPS6 | 450 | AKEKPGTPPGPPP | 6.1381 | 6.2862 | 6.5311 | 6.0862 | 6.3597 | 6.4175 | 5.9270 | 6.1403 | 6.9307 | 6.6547 | 5.9851 | 5.7143 | 5.9854 | 6.5244 | 7.1304 | 7.0889 |
| SETD1B_T381 | SETD1B | Q9UPS6 | 381 | SATFAHTPPPAQA | 5.8656 | 6.5012 | 6.3456 | 6.5482 | 6.2301 | 6.1231 | 6.3332 | 5.7659 | 6.3449 | 6.2557 | 6.1867 | 6.2606 | 6.0782 | 6.6474 | 6.1701 | 6.3434 |
| SETD1B_T450 | SETD1B | Q9UPS6 | 986 | RLRPSTSVDEEDE | 5.3118 | 5.3201 | 5.9611 | 5.8905 | 5.6224 | 5.5145 | 5.9520 | 5.8183 | 6.1107 | 7.4922 | 7.0836 | 5.5715 | 7.1808 | 6.5505 | 7.4504 | 7.1694 |
| SETD1B_T909 | SETD1B | Q9UPS6 | 1659;1663 | NA | 5.6487 | 5.9209 | 6.4831 | 6.9979 | 6.2022 | 6.5620 | 6.7703 | 6.2940 | 6.2239 | 6.0300 | 6.8262 | 5.8785 | 5.8293 | 6.4567 | 6.4428 | 5.4333 |
| SETD2_S1228 | SETD2 | Q9BYW2 | 624 | SNRLNDSPTLKKL | 6.7238 | 6.5544 | 5.7236 | 6.6724 | 6.4550 | 5.9017 | 6.1806 | 6.0846 | 6.3896 | 6.3573 | 5.9627 | 5.9379 | 6.3496 | 6.3738 | 6.1526 | 6.1804 |
| SETD2_S131 | SETD2 | Q9BYW2 | 676 | ELNINGSPGAESD | 6.3999 | 6.8383 | 6.1438 | 6.7767 | 6.1631 | 6.0032 | 6.2839 | 6.0084 | 6.5657 | 6.4738 | 6.2666 | 6.4946 | 5.5650 | 6.1888 | 5.9022 | 5.9261 |
| SETD2_S1885 | SETD2 | Q9BYW2 | 532 | AIKRCCSPPNELG | 5.9814 | 6.1105 | 5.7060 | 6.2550 | 5.4791 | 5.2700 | 5.6729 | 5.3890 | 5.6534 | 5.8586 | 5.9651 | 5.9451 | 7.7724 | 7.7611 | 7.6906 | 7.4898 |
| SETD2_S2080;S2082 | SETD2 | Q9BYW2 | 744 | LHKKSESPFRETE | 5.6673 | 6.5012 | 6.0480 | 7.0372 | 6.2148 | 5.7813 | 6.4253 | 6.4787 | 6.6386 | 6.1732 | 6.2359 | 6.3021 | 5.8937 | 6.1866 | 5.8858 | 5.6304 |
| SETD2_S532 | SETD2 | Q9BYW2 | 614 | EREKAGSPAPSNR | 6.6004 | 6.6264 | 6.1308 | 6.7827 | 5.8694 | 5.8378 | 6.3280 | 5.7286 | 6.2637 | 6.0748 | 6.2507 | 6.2417 | 6.1158 | 6.4772 | 6.3973 | 6.2748 |
| SETD2_S543 | SETD2 | Q9BYW2 | 131 | STAEESSPPKSRV | 6.5710 | 6.5666 | 6.5957 | 6.5956 | 6.1451 | 6.1147 | 6.6319 | 6.3206 | 6.5512 | 6.2035 | 6.0205 | 6.5507 | 5.4306 | 5.9692 | 5.7361 | 5.9970 |
| SETD2_S614 | SETD2 | Q9BYW2 | 1228 | FQNRPDSRLGKTE | 5.5018 | 5.9079 | 5.7882 | 6.0280 | 6.0486 | 5.9347 | 6.0976 | 5.5030 | 5.1877 | 5.7562 | 5.2768 | 5.5421 | 7.8515 | 7.2953 | 8.3164 | 7.9640 |
| SETD2_S624 | SETD2 | Q9BYW2 | 543 | LGFRRGSSYSKHD | 5.5359 | 4.6844 | 4.5837 | 5.8741 | 7.4265 | 5.2668 | 4.8635 | 4.5180 | 8.8373 | 8.3895 | 7.8084 | 9.1238 | 6.6569 | 7.2655 | 4.0822 | 5.0835 |
| SETD2_S676 | SETD2 | Q9BYW2 | 754 | ETEPLVSPHQDKL | 6.2222 | 6.6656 | 6.0438 | 6.6077 | 6.1976 | 6.1067 | 6.5866 | 6.2790 | 6.5629 | 6.0446 | 6.2567 | 6.3313 | 5.7379 | 6.2351 | 6.4674 | 6.5549 |
| SETD2_S744 | SETD2 | Q9BYW2 | 1872 | TEADTDTPKKLMF | 6.8944 | 6.7850 | 6.3993 | 7.1156 | 5.9481 | 5.5307 | 6.4183 | 5.5457 | 6.4173 | 6.6190 | 6.2921 | 6.5558 | 5.8514 | 6.0838 | 5.4121 | 6.1313 |
| SETD2_S744;S754 | SETD2 | Q9BYW2 | 1885 | RRLKIISENSMDS | 4.6954 | 5.8833 | 6.4612 | 6.6619 | 5.4750 | 5.3478 | 6.0495 | 6.7778 | 6.3552 | 6.7349 | 6.6515 | 6.3927 | 5.8093 | 7.3458 | 7.1387 | 6.2201 |
| SETD2_S754 | SETD2 | Q9BYW2 | 905 | KCGENTSPVLDAV | 6.5890 | 7.4902 | 7.1209 | 7.6153 | 6.1543 | 6.2399 | 7.0690 | 6.1173 | 6.8195 | 6.6647 | 7.0569 | 7.3577 | 3.8369 | 4.6533 | 4.2288 | 4.9862 |
| SETD2_S905 | SETD2 | Q9BYW2 | 2080;2082 | NA | 5.6211 | 6.1779 | 6.2046 | 7.3490 | 6.1273 | 6.8095 | 5.7204 | 6.6355 | 6.3556 | 5.9719 | 6.1592 | 5.8369 | 6.0001 | 6.2858 | 5.9789 | 6.7663 |

| | | | | | | | | | | | | | | | | | | | | |
|---|---|---|---|---|---|---|---|---|---|---|---|---|---|---|---|---|---|---|---|---|
| SETD2_T1872 | SETD2 | Q9BYW2 | 744;754 | NA | 6.6347 | 6.6514 | 6.3056 | 7.3324 | 6.6240 | 6.4362 | 6.3783 | 5.4333 | 6.2290 | 6.2156 | 6.9248 | 6.4809 | 5.7174 | 5.5073 | 5.6599 | 5.4691 |
| SETD5_S1043 | SETD5 | Q9C0A6 | 1125 | SARTPSSPHKKFS | 7.0182 | 6.9471 | 6.3600 | 7.0335 | 6.2598 | 5.9000 | 6.1491 | 6.2002 | 6.1963 | 6.1233 | 6.0130 | 6.1971 | 5.6927 | 5.8610 | 6.1715 | 5.8773 |
| SETD5_S1125 | SETD5 | Q9C0A6 | 1233 | KGATVYSPSRYSY | 6.8685 | 6.5410 | 6.5447 | 6.8780 | 5.7759 | 5.9761 | 6.3515 | 6.9403 | 6.5762 | 6.7146 | 6.1481 | 5.9079 | 5.1476 | 5.9181 | 5.9457 | 5.7658 |
| SETD5_S1233 | SETD5 | Q9C0A6 | 829 | DNADLLSPLKKWK | 8.3660 | 10.1957 | 6.8619 | 8.5523 | 7.1501 | 6.3559 | 6.4461 | 5.9970 | 8.0875 | 6.1088 | 7.1639 | 6.3990 | 2.3729 | 3.3750 | 3.2280 | 3.3397 |
| SETD5_S72 | SETD5 | Q9C0A6 | 72 | DLNGLPSPVEERC | 5.4641 | 5.8070 | 6.9653 | 6.4399 | 5.4520 | 5.7119 | 5.8768 | 5.7079 | 6.1509 | 6.5483 | 6.7900 | 6.7776 | 6.2939 | 6.7549 | 6.3562 | 6.9033 |
| SETD5_S829 | SETD5 | Q9C0A6 | 1043 | LSRGSLSPGGERA | 6.7691 | 6.8040 | 5.9485 | 6.6822 | 5.8025 | 5.8181 | 6.1450 | 6.2083 | 6.2894 | 5.9244 | 5.9489 | 6.5468 | 6.0966 | 6.4048 | 6.4509 | 6.1606 |
| SETD5_S852;T855 | SETD5 | Q9C0A6 | 852;855 | NA | 6.3107 | 6.3417 | 5.6338 | 6.5446 | 6.2167 | 5.9200 | 5.8154 | 6.0623 | 6.2335 | 6.1853 | 6.1077 | 6.3231 | 6.7013 | 6.8193 | 6.5717 | 6.2130 |
| SETDB1_S1066 | SETDB1 | E9PQM8 | 468 | PPAPPLSPQAGDS | 6.2809 | 5.6787 | 6.5031 | 7.1779 | 5.8579 | 5.6652 | 6.3336 | 6.2411 | 6.0004 | 6.7282 | 5.8691 | 6.1081 | 6.1354 | 6.8248 | 6.0784 | 6.5172 |
| SETDB1_S468 | SETDB1 | Q15047 | 1066 | NYGYNPSPVKPEG | 5.9242 | 5.9722 | 6.3383 | 6.6049 | 6.2445 | 5.9591 | 6.3239 | 5.6375 | 6.5199 | 6.1965 | 6.2610 | 6.2094 | 6.3141 | 6.6948 | 6.3924 | 6.4073 |
| SETDB1_S468 | SETDB1 | Q15047 | 528 | INQTYRSPLGSTA | 6.1056 | 6.4664 | 5.9930 | 6.6738 | 6.5082 | 6.2710 | 6.1673 | 6.2029 | 5.9546 | 6.3362 | 6.4207 | 7.4780 | 5.8154 | 6.3393 | 5.6351 | 5.6326 |
| SETDB1_S528 | SETDB1 | Q15047 | 468 | PPAPPLSPQAGDS | 6.2677 | 6.6764 | 6.2759 | 6.2184 | 6.9364 | 5.8590 | 6.6992 | 5.5250 | 5.5196 | 5.7413 | 6.5432 | 5.8206 | 5.9785 | 6.7722 | 6.3407 | 6.8260 |
| SETX_S1430 | SETX | Q7Z333-4 | 947 | EQAPDISPKSDTL | 6.0087 | 6.4481 | 6.2038 | 6.7303 | 5.8700 | 5.9869 | 6.0846 | 5.8573 | 6.4246 | 6.7116 | 6.5240 | 6.4979 | 5.8279 | 6.3536 | 5.9108 | 6.5600 |
| SETX_S1686 | SETX | Q7Z333-4 | 1686 | KYFPSSSPVNILL | 6.5990 | 6.6160 | 6.5422 | 6.7371 | 6.1789 | 6.1254 | 6.2127 | 6.3265 | 6.5890 | 6.6943 | 6.8004 | 6.6796 | 5.2524 | 5.9210 | 5.5617 | 5.1639 |
| SETX_S2668 | SETX | Q7Z333-4 | 2668 | REARAFSEGEQEK | 6.1214 | 5.8871 | 5.9513 | 6.9122 | 6.1926 | 7.3320 | 7.1173 | 7.3418 | 5.0935 | 5.0321 | 5.0754 | 5.1633 | 6.6542 | 7.2609 | 5.9680 | 6.8969 |
| SETX_S947 | SETX | Q7Z333-4 | 1430 | IKAKHGSPATDDA | 5.8595 | 6.0330 | 7.3255 | 6.4444 | 6.4720 | 5.3345 | 6.7652 | 5.8042 | 4.4871 | 7.1229 | 6.3676 | 6.2249 | 5.9332 | 7.3133 | 5.8319 | 6.6808 |
| SETX_S956 | SETX | Q7Z333-4 | 956 | SDTLTDSQIDRDL | 6.1933 | 6.6860 | 6.4548 | 7.5936 | 6.7913 | 5.6714 | 6.3636 | 6.0288 | 5.8847 | 6.1224 | 6.2514 | 5.8922 | 6.0278 | 6.5174 | 5.9363 | 5.5850 |
| SEZ6_S956 | SEZ6 | Q53EL9 | 956 | RLQGKSSLQLPRP | 5.5381 | 4.3996 | 6.4047 | 4.6648 | 6.7358 | 5.3273 | 4.2676 | 6.6761 | 6.7018 | 8.8543 | 8.1865 | 9.1976 | 6.2486 | 6.9918 | 4.8634 | 4.9421 |
| SF1_S112 | SF1 | Q15637-6 | 463 | CGLWGLSPASRKR | 6.9562 | 7.1173 | 6.7393 | 7.2619 | 6.3799 | 6.9944 | 6.8217 | 7.1851 | 7.3945 | 6.9148 | 7.4212 | 7.4645 | 3.5490 | 4.3724 | 3.4859 | 3.9418 |
| SF1_S205;S207 | SF1 | Q15637-5 | 112 | PLYQRVSPPQPPP | 5.6443 | 6.3260 | 6.3277 | 5.9084 | 6.4429 | 6.6896 | 6.1459 | 5.5553 | 5.7710 | 5.9467 | 6.8001 | 6.2135 | 7.2493 | 6.5122 | 6.0092 | 6.4577 |
| SF1_S207 | SF1 | Q15637-5 | 207 | PEDRSPSPEPIYN | 6.2329 | 6.1806 | 6.6346 | 6.5549 | 6.1078 | 6.3183 | 5.9705 | 6.4382 | 6.9545 | 6.1337 | 6.1657 | 6.6786 | 6.0464 | 5.6804 | 6.0967 | 5.8061 |
| SF1_S456 | SF1 | Q15637-5 | 456 | VPASVGSTSGPAT | 7.1058 | 7.1430 | 6.9160 | 6.7084 | 6.1088 | 6.4930 | 7.1941 | 6.1646 | 6.3459 | 6.3342 | 6.8742 | 6.0639 | 4.8183 | 5.7238 | 4.9887 | 5.0174 |
| SF1_S463 | SF1 | Q15637-5 | 205;207 | NA | 5.7428 | 5.9343 | 6.2501 | 6.1318 | 6.1235 | 6.0732 | 5.9886 | 5.9139 | 6.2053 | 6.3469 | 6.4334 | 6.4520 | 6.8582 | 6.5801 | 6.4639 | 6.5021 |
| SF3A1_S329 | SF3A1 | Q15459 | 329 | VEMEVESDEEDDK | 5.8421 | 5.7901 | 6.2564 | 6.2936 | 6.3896 | 5.9810 | 6.3183 | 5.7780 | 6.3873 | 6.2039 | 6.7386 | 6.5828 | 7.1131 | 6.3417 | 5.8865 | 6.0970 |
| SF3A1_S359 | SF3A1 | Q15459 | 359 | QDMDEGSDDEEEG | 5.9786 | 5.9970 | 6.2191 | 6.3979 | 6.2695 | 5.9484 | 6.2093 | 6.1305 | 6.2934 | 6.1493 | 6.2036 | 6.2648 | 6.6299 | 6.4909 | 6.4405 | 6.3774 |
| SF3A1_S413 | SF3A1 | Q15459 | 383 | PPPLPPTPDQVIV | 6.1444 | 6.4695 | 6.5414 | 6.6270 | 6.2855 | 6.2615 | 6.4383 | 5.9738 | 6.4423 | 6.5240 | 6.9618 | 6.5863 | 5.5911 | 5.7337 | 5.5446 | 5.8749 |
| SF3A1_T383 | SF3A1 | Q15459 | 413 | PDEYLVSPITGEK | 7.3473 | 6.6652 | 6.7336 | 7.3850 | 6.7282 | 7.2905 | 6.9717 | 6.9794 | 7.4964 | 7.6331 | 7.7598 | 8.4097 | 2.8567 | 3.8745 | 2.5702 | 3.2984 |
| SF3A3_S299 | SF3A3 | Q12874 | 299 | LESLDTSLFAKNP | 6.6662 | 7.0648 | 6.5740 | 7.4806 | 6.3565 | 6.1871 | 6.8697 | 6.5686 | 6.7446 | 6.3372 | 6.6124 | 6.6861 | 4.8140 | 5.2584 | 4.7919 | 4.9878 |
| SF3A3_S365;S367;S369 | SF3A3 | Q12874 | 365;367;369 | NA | 6.6867 | 5.5377 | 6.6827 | 6.3684 | 6.1058 | 5.6441 | 6.0807 | 6.0247 | 5.8693 | 6.7114 | 6.5702 | 6.1357 | 6.7600 | 6.0374 | 6.2712 | 6.5138 |
| SF3B1_S129 | SF3B1 | O75533 | 129 | RRTMIISPERLDP | 6.5064 | 6.4580 | 5.9879 | 6.6329 | 6.1984 | 5.9442 | 6.5175 | 6.4540 | 6.5627 | 6.1775 | 5.8469 | 5.9792 | 6.2887 | 6.1538 | 6.0705 | 6.2214 |
| SF3B1_S129;T142 | SF3B1 | O75533 | 244 | RAKGSETPGATPG | 5.1693 | 5.5585 | 5.7030 | 5.9149 | 5.5186 | 6.4925 | 6.1806 | 6.5966 | 6.7234 | 7.0750 | 6.2408 | 6.9162 | 6.2630 | 6.4519 | 6.3351 | 6.8605 |
| SF3B1_S194 | SF3B1 | O75533 | 350 | SQMGGSTPVLTPG | 5.9420 | 5.8047 | 5.8475 | 6.3798 | 6.0466 | 5.7376 | 6.1019 | 5.5702 | 6.3358 | 6.1255 | 5.8460 | 6.2396 | 7.0449 | 6.9540 | 6.7961 | 7.2277 |
| SF3B1_S400 | SF3B1 | O75533 | 235 | SLRWDETPGRAKG | 7.1138 | 6.5963 | 6.4249 | 6.7957 | 6.3000 | 6.0723 | 6.5142 | 6.1218 | 6.4197 | 5.8067 | 5.8619 | 5.8152 | 6.3426 | 6.2244 | 5.7199 | 5.8706 |
| SF3B1_S488 | SF3B1 | O75533 | 211 | DQTPGATPKKLSS | 6.7223 | 6.2681 | 5.9628 | 6.6254 | 6.1756 | 6.0881 | 6.6804 | 6.0198 | 6.9207 | 6.5317 | 6.3691 | 6.3565 | 5.9625 | 5.9079 | 5.4460 | 5.9630 |
| SF3B1_T142 | SF3B1 | O75533 | 227 | AETPGHTPSLRWD | 5.9316 | 6.0956 | 6.3970 | 6.6424 | 6.2026 | 6.3548 | 6.2039 | 6.6583 | 6.5398 | 6.1015 | 5.9563 | 6.4204 | 6.1248 | 6.4052 | 5.9174 | 6.0484 |
| SF3B1_T207 | SF3B1 | O75533 | 296 | KNRWDETPKTERD | 6.2339 | 6.0218 | 6.0599 | 5.8571 | 6.2446 | 5.7721 | 6.4375 | 5.2657 | 6.2340 | 6.1078 | 5.8432 | 6.0889 | 7.4640 | 6.8688 | 6.4738 | 7.0268 |
| SF3B1_T207;T211 | SF3B1 | O75533 | 436 | KLTATPTPLGGMT | 6.8371 | 6.9016 | 6.8272 | 7.0467 | 6.3174 | 6.7677 | 6.4858 | 6.1929 | 6.7214 | 6.3150 | 6.3839 | 6.7327 | 5.1544 | 5.4015 | 4.7972 | 5.1174 |
| SF3B1_T211 | SF3B1 | O75533 | 194 | AASQPPSKRKRRW | 7.2068 | 7.7089 | 7.1882 | 7.6432 | 6.6222 | 6.9371 | 7.6154 | 6.5397 | 6.5392 | 5.9368 | 6.0873 | 6.2125 | 4.3846 | 4.6841 | 4.0006 | 4.6933 |
| SF3B1_T223 | SF3B1 | O75533 | 326 | GDSIGETPTPGAS | 6.8280 | 6.2784 | 5.7319 | 6.7105 | 6.1133 | 6.0922 | 6.1975 | 6.3034 | 6.5537 | 5.8917 | 5.8332 | 6.2501 | 6.1514 | 6.5306 | 5.9062 | 6.6280 |
| SF3B1_T223;T227 | SF3B1 | O75533 | 434 | ARKLTATPTPLGG | 6.1597 | 6.5668 | 6.2014 | 6.7598 | 6.4817 | 6.3595 | 6.6390 | 6.2075 | 6.9347 | 6.6740 | 6.4360 | 6.6526 | 4.8801 | 5.8507 | 5.1067 | 6.0900 |
| SF3B1_T227 | SF3B1 | O75533 | 142 | FADGGKTPDPKMN | 6.9936 | 6.2330 | 6.3437 | 6.9154 | 6.1829 | 6.0983 | 6.8925 | 6.3158 | 6.9578 | 6.2489 | 6.0621 | 6.3801 | 5.7526 | 6.0932 | 5.0094 | 5.5208 |
| SF3B1_T235 | SF3B1 | O75533 | 426 | GYVPIRTPARKLT | 7.3348 | 6.9379 | 6.5927 | 7.8775 | 6.5004 | 6.7954 | 6.9416 | 5.7439 | 6.3147 | 5.8861 | 6.6016 | 6.4855 | 4.9217 | 5.3170 | 4.6811 | 5.0682 |
| SF3B1_T244 | SF3B1 | O75533 | 223 | SWDQAETPGHTPS | 6.2370 | 6.2324 | 6.3572 | 6.9037 | 5.5810 | 6.1310 | 6.5425 | 6.5356 | 6.6501 | 6.5665 | 6.1792 | 6.4392 | 5.7734 | 6.0548 | 5.8285 | 5.9879 |
| SF3B1_T244;T248 | SF3B1 | O75533 | 261 | DPTPSHTPAGAAT | 5.4353 | 5.9291 | 6.0561 | 6.2091 | 5.8670 | 5.9842 | 5.8617 | 6.0386 | 6.3411 | 6.6373 | 6.1699 | 6.1097 | 6.5228 | 6.9589 | 6.9699 | 6.9094 |
| SF3B1_T261 | SF3B1 | O75533 | 488 | VDESTLSPEEQKE | 5.9392 | 6.1943 | 6.3170 | 6.5334 | 5.8507 | 5.7896 | 6.3074 | 5.9530 | 6.6186 | 6.3999 | 6.4827 | 6.5250 | 6.1236 | 6.6857 | 6.1112 | 6.1688 |
| SF3B1_T267 | SF3B1 | O75533 | 400 | ERNRPLSDEELDA | 5.4798 | 6.3965 | 5.4240 | 6.8478 | 5.4753 | 6.1106 | 6.3798 | 5.4785 | 6.2151 | 5.8118 | 5.6170 | 5.6413 | 7.3202 | 7.2061 | 7.5203 | 7.0760 |
| SF3B1_T278 | SF3B1 | O75533 | 267 | TPAGAATPGRGDT | 4.9365 | 5.8519 | 5.0170 | 6.9900 | 5.1328 | 6.3107 | 5.5916 | 6.5379 | 7.4179 | 6.9338 | 6.4529 | 6.8886 | 6.3614 | 5.7405 | 6.8147 | 7.0220 |
| SF3B1_T296 | SF3B1 | O75533 | 207 | DQTADQTPGATPK | 6.9332 | 5.9710 | 6.0824 | 5.8931 | 6.0985 | 5.4257 | 6.6156 | 6.9534 | 7.1407 | 6.7257 | 6.2456 | 6.5941 | 6.1261 | 5.9401 | 6.1549 | 5.1000 |
| SF3B1_T313 | SF3B1 | O75533 | 313 | GSGWAETPRTDRG | 6.5353 | 6.8311 | 5.7906 | 6.4497 | 6.2571 | 5.5881 | 5.6166 | 5.8669 | 7.2581 | 5.7000 | 5.3581 | 6.5754 | 7.0748 | 6.0875 | 6.1319 | 6.8789 |
| SF3B1_T326 | SF3B1 | O75533 | 354 | GSTPVLTPGKTPI | 6.1108 | 7.5214 | 6.7025 | 8.0411 | 4.8887 | 6.9575 | 6.8591 | 6.6138 | 7.8360 | 7.7182 | 8.2392 | 7.3690 | 3.2882 | 5.0563 | 3.4642 | 3.3339 |
| SF3B1_T341;T350 | SF3B1 | O75533 | 362 | GKTPIGTPAMNMA | 5.8314 | 5.9463 | 6.0653 | 6.4937 | 6.1353 | 6.3152 | 6.0882 | 6.3747 | 6.5162 | 6.7675 | 6.2429 | 7.0426 | 6.2301 | 6.5441 | 5.4954 | 5.9115 |
| SF3B1_T350 | SF3B1 | O75533 | 278 | DTPGHATPGHGGA | 6.1470 | 6.3016 | 5.0762 | 6.7145 | 6.5829 | 6.2621 | 5.6930 | 5.9299 | 7.7442 | 6.5730 | 5.2637 | 5.7917 | 6.7112 | 6.4924 | 6.1075 | 6.6092 |
| SF3B1_T350;T354 | SF3B1 | O75533 | 434;436 | NA | 6.4300 | 6.7072 | 6.6662 | 7.7497 | 6.2489 | 6.6988 | 7.3408 | 6.8955 | 6.5695 | 5.6825 | 6.2216 | 5.9646 | 5.1356 | 5.8740 | 4.9222 | 4.8926 |
| SF3B1_T354 | SF3B1 | O75533 | 223;227 | NA | 6.3548 | 6.2399 | 5.9875 | 6.8606 | 6.5609 | 6.0418 | 6.4964 | 5.7326 | 6.4049 | 5.6804 | 5.6604 | 6.1595 | 6.4509 | 6.5967 | 6.1366 | 6.6364 |
| SF3B1_T362 | SF3B1 | O75533 | 207;211 | NA | 6.5511 | 6.3285 | 6.4091 | 7.0773 | 6.1203 | 6.2590 | 6.6306 | 6.6716 | 6.8033 | 5.8744 | 5.6649 | 6.2777 | 6.0647 | 6.2733 | 5.5317 | 5.4624 |
| SF3B1_T426 | SF3B1 | O75533 | 129;142 | NA | 8.1369 | 6.9997 | 6.7331 | 7.4659 | 6.4340 | 6.4044 | 6.4882 | 5.1143 | 6.6133 | 6.2135 | 6.2916 | 6.1618 | 5.4319 | 5.6768 | 4.2672 | 5.5673 |
| SF3B1_T434 | SF3B1 | O75533 | 341;350 | NA | 6.4603 | 5.7366 | 6.7228 | 6.1573 | 6.1199 | 6.0877 | 6.3305 | 6.0120 | 6.7493 | 6.4288 | 5.8597 | 5.9748 | 6.5833 | 6.7681 | 5.7939 | 6.2150 |
| SF3B1_T434;T436 | SF3B1 | O75533 | 244;248 | NA | 6.1180 | 6.6211 | 6.7759 | 7.1895 | 6.3623 | 6.4649 | 6.0433 | 6.9305 | 5.8455 | 6.1028 | 6.0546 | 5.9898 | 5.7013 | 6.1784 | 5.9089 | 5.7132 |
| SF3B1_T436 | SF3B1 | O75533 | 350;354 | NA | 4.8669 | 5.6739 | 6.2397 | 5.0197 | 6.3784 | 6.5238 | 5.7159 | 6.9864 | 6.9496 | 6.6316 | 5.8612 | 7.4337 | 6.0506 | 6.4873 | 5.5601 | 7.6211 |
| SF3B2_S307 | SF3B2 | Q13435 | 780 | AMDGSETPQLFTV | 6.4659 | 6.1874 | 5.7621 | 6.6190 | 6.1267 | 5.6900 | 6.0889 | 5.6677 | 5.9912 | 5.2745 | 5.2584 | 5.4843 | 7.5436 | 7.5250 | 6.9729 | 7.3424 |
| SF3B2_S307;S309 | SF3B2 | Q13435 | 362 | RSRGSDSPAADVE | 5.7757 | 5.9665 | 6.2303 | 6.7083 | 6.1740 | 5.8121 | 6.0439 | 6.2191 | 6.0879 | 6.3351 | 6.1734 | 6.2165 | 6.3204 | 6.7331 | 6.7461 | 6.4577 |
| SF3B2_S346 | SF3B2 | Q13435 | 307 | RSSLGQSASETEE | 6.2045 | 6.3356 | 6.3345 | 7.0515 | 6.3148 | 5.7741 | 6.5049 | 5.9950 | 6.5262 | 6.4040 | 6.2247 | 6.3153 | 5.9795 | 6.3189 | 5.7834 | 5.9332 |
| SF3B2_S362 | SF3B2 | Q13435 | 861 | VEKEDFSDMVAEH | 5.6455 | 5.4054 | 5.7688 | 5.7423 | 6.1753 | 5.2092 | 6.1467 | 5.7852 | 6.2287 | 5.5123 | 5.2118 | 5.2427 | 8.5958 | 7.6091 | 7.3562 | 8.3649 |
| SF3B2_S431;S435;S436 | SF3B2 | Q13435 | 346 | RGVSSESSGDREK | 6.3264 | 7.3147 | 7.0442 | 7.4498 | 6.7371 | 7.4243 | 7.3618 | 6.3937 | 5.3352 | 6.0301 | 6.5197 | 5.4753 | 5.1050 | 5.2837 | 4.8284 | 5.3708 |
| SF3B2_S861 | SF3B2 | Q13435 | 307;309 | NA | 6.6168 | 6.2376 | 7.0660 | 7.0977 | 6.3825 | 6.2244 | 6.5406 | 6.2787 | 6.4743 | 6.2864 | 6.3736 | 6.3711 | 5.4311 | 5.7983 | 5.2593 | 5.5617 |
| SF3B2_T780 | SF3B2 | Q13435 | 431;435;436 | NA | 5.6283 | 5.7739 | 6.8321 | 6.5235 | 5.3734 | 5.6117 | 6.1323 | 5.0364 | 6.5159 | 5.8543 | 6.3730 | 6.7324 | 7.3841 | 6.2197 | 6.5768 | 7.4322 |
| SF3B3_S156 | SF3B3 | Q15393 | 1200 | SEELDRTPPEVSK | 5.9367 | 6.2622 | 5.8075 | 6.7794 | 5.9840 | 6.2693 | 6.3458 | 6.5386 | 6.4374 | 6.5972 | 6.5813 | 6.4257 | 5.8268 | 6.0599 | 6.0542 | 6.0939 |
| SF3B3_T1200 | SF3B3 | Q15393 | 156 | ARLTISSPLEAHK | 6.6417 | 6.6773 | 5.7458 | 6.6297 | 6.7552 | 6.4358 | 6.1240 | 5.8058 | 6.2819 | 6.2087 | 6.4542 | 6.7484 | 5.6759 | 5.6311 | 6.3551 | 5.8294 |
| SFMBT1_S775 | SFMBT1 | Q9UHJ3 | 775 | DENKPPSPKEIRI | 4.9745 | 5.2928 | 4.3208 | 4.8263 | 5.0724 | 5.0452 | 4.9513 | 4.1633 | 6.3531 | 7.0200 | 5.6266 | 5.5696 | 9.3974 | 10.1378 | 8.4986 | 8.7504 |
| SFPQ_S273 | SFPQ | P23246 | 33 | GLHDFRSPPPGMG | 7.1072 | 7.2562 | 7.0213 | 7.6638 | 5.9697 | 6.4464 | 6.9553 | 6.9999 | 7.6740 | 7.3408 | 7.4416 | 7.5795 | 3.2586 | 4.1057 | 3.2093 | 3.9707 |
| SFPQ_S283 | SFPQ | P23246 | 687 | RGMGPGTPAGYGR | 8.1212 | 7.9597 | 6.8406 | 7.2856 | 5.5473 | 6.9741 | 7.0618 | 6.5305 | 7.4141 | 7.0283 | 7.2437 | 7.3942 | 3.5587 | 4.1598 | 3.0488 | 3.8315 |
| SFPQ_S33 | SFPQ | P23246 | 420 | GIVEFASKPAARK | 5.4516 | 5.6633 | 5.5451 | 5.7327 | 5.4632 | 5.8661 | 5.4773 | 6.4377 | 6.6641 | 6.1740 | 6.4970 | 6.6174 | 7.2819 | 6.2076 | 7.5499 | 7.3712 |
| SFPQ_S374 | SFPQ | P23246 | 496 | YSQRWKSLDEMEK | 6.3261 | 6.4803 | 6.4531 | 6.2819 | 6.0819 | 6.1836 | 6.2584 | 6.3509 | 5.8054 | 6.6078 | 6.3543 | 6.3302 | 6.0868 | 6.1501 | 5.9634 | 6.2857 |
| SFPQ_S420 | SFPQ | P23246 | 273 | RSEEKISDSEGFK | 6.2259 | 6.6249 | 6.8131 | 7.2329 | 7.0593 | 6.7809 | 6.9320 | 6.7291 | 5.2972 | 5.4349 | 5.5366 | 5.5158 | 6.3001 | 5.5157 | 5.6865 | 6.3152 |
| SFPQ_S496 | SFPQ | P23246 | 226 | GGPGLSTPGGHPK | 7.0917 | 6.5883 | 6.3233 | 8.0450 | 6.1759 | 7.6513 | 7.0791 | 7.1047 | 7.2768 | 6.4237 | 7.7517 | 7.5992 | 3.4556 | 4.3384 | 3.0685 | 4.0267 |
| SFPQ_T226 | SFPQ | P23246 | 283 | GFKANLSLLRRPG | 7.5171 | 7.1358 | 6.3080 | 7.9837 | 6.7333 | 5.9701 | 7.0428 | 6.0776 | 6.8322 | 6.4277 | 6.9516 | 6.5505 | 4.6306 | 4.9012 | 4.4581 | 4.4799 |
| SFPQ_T687 | SFPQ | P23246 | 374 | THAAALSVRNLSP | 4.7428 | 5.5136 | 6.5750 | 4.8139 | 4.5577 | 7.7849 | 7.9937 | 9.0982 | 6.3360 | 6.4974 | 5.6126 | 5.8817 | 7.4103 | 5.3349 | 6.6417 | 5.2057 |
| SFSWAP_S283 | SFSWAP | Q12872-2 | 743 | PPDRPSSKSKDPP | 6.1184 | 6.1441 | 6.1491 | 6.8365 | 6.2981 | 5.8132 | 6.5513 | 5.8637 | 5.9725 | 6.2795 | 6.0331 | 6.2132 | 6.4116 | 6.6646 | 6.1778 | 6.4731 |
| SFSWAP_S743 | SFSWAP | Q12872-2 | 807 | RSRVTASPGTLRA | 5.9915 | 6.5089 | 5.9058 | 6.7486 | 5.9056 | 5.8984 | 6.1509 | 5.9076 | 6.0286 | 6.3579 | 6.2076 | 6.0348 | 5.9082 | 6.7915 | 6.7993 | 6.8546 |
| SFSWAP_S807 | SFSWAP | Q12872-2 | 283 | KKSGVSSDNEDDD | 6.4243 | 7.4860 | 5.6389 | 7.3891 | 6.5421 | 6.7727 | 5.5843 | 7.2251 | 5.5368 | 5.7245 | 6.0910 | 5.7968 | 5.1425 | 6.5604 | 6.4816 | 5.6040 |
| SFSWAP_S884;S886 | SFSWAP | Q12872-2 | 886 | AYRVSRSPGASRK | 6.2028 | 7.0325 | 6.9956 | 5.8256 | 5.9513 | 5.8939 | 6.9587 | 4.2209 | 5.5631 | 5.8557 | 5.7709 | 6.4967 | 6.9970 | 7.7939 | 5.6834 | 6.7578 |
| SFSWAP_S886 | SFSWAP | Q12872-2 | 918;924 | NA | 6.5509 | 6.3033 | 6.3917 | 6.6995 | 5.8548 | 5.9913 | 6.0051 | 5.6288 | 6.1311 | 6.3903 | 5.8663 | 6.2051 | 5.9742 | 6.9914 | 6.3985 | 6.6175 |
| SFSWAP_S918;S924 | SFSWAP | Q12872-2 | 884;886 | NA | 6.5762 | 5.9323 | 6.8015 | 6.2302 | 8.1532 | 6.8008 | 6.5837 | 5.1264 | 5.1082 | 6.2288 | 5.9319 | 6.1762 | 5.7707 | 7.1772 | 5.9525 | 5.4502 |
| SGK1_S496 | SGK1 | O00141-2 | 496 | PNSIGKSPDSVLV | 5.7688 | 5.6742 | 6.6028 | 5.5484 | 6.1750 | 6.6032 | 5.7968 | 6.4159 | 6.1333 | 5.4154 | 6.1495 | 5.8745 | 7.2316 | 6.3692 | 7.5778 | 6.6636 |
| SGK3_S121 | SGK3 | Q96BR1 | 121 | AFLQMDSPKHQSD | 5.9531 | 6.1526 | 7.3865 | 6.0186 | 5.1580 | 6.5572 | 7.0987 | 7.2241 | 6.3578 | 5.9702 | 6.0899 | 7.3409 | 5.7109 | 4.8559 | 6.7909 | 5.3347 |
| SGMS1_S8 | SGMS1 | Q86VZ5 | 8 | KEVVYWSPKKVAD | 6.1170 | 7.2296 | 6.3480 | 6.7008 | 5.8337 | 5.6077 | 5.9549 | 6.0736 | 6.7686 | 6.4887 | 6.0497 | 5.8826 | 5.6840 | 6.4393 | 6.6304 | 6.1914 |
| SGO1_S193 | SGO1 | Q5FBB7 | 507 | DLCFLNSPIFKQK | 8.1248 | 7.4936 | 6.9069 | 7.6930 | 5.8184 | 7.0283 | 7.1928 | 6.0147 | 7.4619 | 6.8886 | 6.6297 | 7.2636 | 3.5514 | 4.5375 | 2.9511 | 4.4436 |
| SGO1_S256 | SGO1 | Q5FBB7 | 468 | PKKNKASPAVALP | 7.4632 | 8.0303 | 6.6858 | 7.9379 | 5.7769 | 6.7714 | 7.1648 | 6.6813 | 7.1139 | 7.3179 | 7.3992 | 7.9419 | 2.6906 | 4.1959 | 2.9245 | 3.9044 |
| SGO1_S468 | SGO1 | Q5FBB7 | 256 | KDQVNLSPKLIQP | 6.7450 | 6.6951 | 5.9833 | 6.6027 | 6.4091 | 5.9051 | 6.7697 | 5.9454 | 6.3693 | 5.9391 | 5.9475 | 5.8728 | 5.9207 | 6.2569 | 6.1897 | 6.4484 |
| SGO1_S507 | SGO1 | Q5FBB7 | 346 | EEGVHLTPFRQKV | 6.7310 | 7.5497 | 6.4490 | 7.4356 | 6.3036 | 6.1266 | 6.5253 | 6.7010 | 6.4941 | 5.6615 | 5.4394 | 6.3214 | 4.7356 | 6.5826 | 5.2289 | 5.7149 |
| SGO1_T346 | SGO1 | Q5FBB7 | 193 | RTVSVRSSLKKHC | 6.3603 | 5.5580 | 6.2012 | 5.7125 | 5.8063 | 6.3557 | 6.3248 | 6.4685 | 6.4965 | 6.6877 | 6.9623 | 6.3477 | 5.4366 | 6.5108 | 5.7646 | 7.0064 |
| SGO2_S1166 | SGO2 | Q562F6 | 845 | DNFFSLTPKDKET | 6.6212 | 7.2615 | 6.4266 | 6.5950 | 6.2759 | 6.1437 | 6.7145 | 6.0965 | 6.2769 | 6.4048 | 6.5568 | 6.8235 | 5.2276 | 5.7819 | 5.0809 | 5.7127 |
| SGO2_S1187 | SGO2 | Q562F6 | 1219 | RPLQDLSNTSFVS | 10.9243 | 8.1854 | 8.3521 | 8.4628 | 5.6932 | 5.9692 | 8.1697 | 6.1553 | 7.3776 | 5.5443 | 5.9264 | 5.2829 | 3.6235 | 3.9077 | 2.7143 | 3.7113 |
| SGO2_S1219 | SGO2 | Q562F6 | 1187 | SPAFQVSDDEHEK | 6.9316 | 6.4878 | 7.1635 | 7.3219 | 6.4926 | 6.4190 | 6.3544 | 5.8448 | 5.3406 | 5.7546 | 6.6605 | 6.1153 | 5.4017 | 5.7434 | 5.2937 | 6.6747 |
| SGO2_S47 | SGO2 | Q562F6 | 47 | TKILNNSSIFKIS | 7.2951 | 6.3157 | 4.8761 | 6.4001 | 6.3706 | 5.6909 | 6.2777 | 7.4725 | 5.1210 | 4.5510 | 6.5774 | 6.2946 | 5.0112 | 8.0846 | 4.8845 | 8.7770 |
| SGO2_T845 | SGO2 | Q562F6 | 1166 | VPLSVSSGKNVII | 6.8954 | 7.7390 | 7.0189 | 7.7774 | 5.7809 | 6.4212 | 7.6395 | 7.0974 | 6.8166 | 7.5154 | 7.5600 | 7.9328 | 3.1719 | 3.9794 | 2.8431 | 3.8112 |
| SGPL1_S564 | SGPL1 | O95470 | 564 | GSQMNGSPKPHxx | 5.9151 | 5.7943 | 6.8190 | 6.4634 | 6.2856 | 6.0454 | 6.3168 | 6.0495 | 7.1068 | 6.1309 | 6.6955 | 6.5978 | 6.0182 | 5.8309 | 5.7644 | 6.1662 |
| SGPP1_S101 | SGPP1 | Q9BX95 | 112 | GALRRNSLTGEEG | 6.1782 | 6.1939 | 6.1638 | 6.3277 | 6.2703 | 5.9101 | 6.2051 | 5.9323 | 6.4538 | 6.0887 | 6.0137 | 6.1438 | 6.5308 | 6.5252 | 6.5289 | 6.5338 |
| SGPP1_S112 | SGPP1 | Q9BX95 | 101 | AELGPASPRRAGA | 6.5910 | 6.6566 | 6.6454 | 7.3563 | 6.0745 | 6.5605 | 7.4603 | 6.5306 | 6.5403 | 6.6912 | 6.7440 | 6.9286 | 4.5040 | 5.1733 | 4.4830 | 5.0603 |
| SGTA_S77 | SGTA | O43765 | 77 | MPQDLRSPARTPP | 6.5126 | 6.1471 | 6.3071 | 5.5553 | 5.9088 | 6.2922 | 6.0625 | 6.5435 | 6.1598 | 7.2165 | 6.8638 | 7.2383 | 5.5884 | 6.2793 | 5.6889 | 5.6356 |
| SGTA_S77;T81 | SGTA | O43765 | 77;81 | NA | 6.2806 | 5.9457 | 6.5358 | 5.7499 | 6.6587 | 6.2264 | 5.8225 | 6.6279 | 6.2884 | 6.4420 | 6.2265 | 6.6380 | 6.4646 | 6.4380 | 5.9047 | 5.7502 |
| SGTB_S297 | SGTB | Q96EQ0 | 297 | IRSRSFSSSAEEH | 4.9595 | 4.8374 | 5.5864 | 4.8997 | 6.4824 | 6.2020 | 5.8042 | 7.3839 | 5.3700 | 4.6227 | 5.3098 | 5.7478 | 8.4712 | 8.1839 | 8.5324 | 7.6065 |
| SH2B3_S103 | SH2B3 | Q9UQQ2 | 103 | PAKAEASPEPGPG | 6.5112 | 6.9356 | 6.0704 | 7.0537 | 7.1441 | 7.2219 | 7.1183 | 6.9997 | 6.9595 | 7.2766 | 6.7504 | 6.3548 | 4.5113 | 4.8517 | 3.8885 | 4.3525 |
| SH2B3_S520 | SH2B3 | Q9UQQ2 | 520 | GLPGRSSPPEQIF | 5.1194 | 6.9603 | 8.2749 | 7.0834 | 6.0831 | 6.4601 | 6.0647 | 7.3505 | 7.4107 | 5.5646 | 6.5151 | 6.4366 | 4.8942 | 6.6461 | 4.9073 | 4.2291 |
| SH2D4A_S315 | SH2D4A | Q9H788 | 315 | GVVRTLSSSAQED | 5.0614 | 4.8719 | 5.5187 | 5.2163 | 5.9771 | 5.4951 | 5.2780 | 5.9220 | 5.2501 | 4.9129 | 4.7695 | 5.1378 | 9.0482 | 9.1233 | 9.7945 | 8.6232 |
| SH2D5_S12 | SH2D5 | Q6ZV89 | 231 | LGNPYCSPTLVRK | 6.3479 | 6.3445 | 6.8381 | 6.3768 | 6.4009 | 6.5437 | 6.0216 | 6.3416 | 6.6104 | 6.4907 | 6.3869 | 6.6419 | 5.6884 | 5.4692 | 5.8160 | 5.6813 |
| SH2D5_S197 | SH2D5 | Q6ZV89 | 12 | AGGRRASDCGLAP | 5.8027 | 5.8132 | 6.4623 | 6.0624 | 6.5758 | 6.0802 | 5.5991 | 6.0087 | 7.1123 | 6.5513 | 7.1487 | 6.9959 | 6.0037 | 6.7962 | 5.3688 | 5.6186 |
| SH2D5_S231 | SH2D5 | Q6ZV89 | 415 | PLSHAKSEAELQG | 5.8686 | 6.1614 | 6.0421 | 6.7879 | 6.2297 | 5.5209 | 6.2902 | 6.4401 | 5.8961 | 5.9095 | 6.4378 | 6.2313 | 5.6999 | 6.8562 | 6.6003 | 7.0281 |
| SH2D5_S415 | SH2D5 | Q6ZV89 | 197 | NVHALVSFRRLPA | 7.5068 | 6.8317 | 7.5894 | 7.3049 | 7.4965 | 5.8869 | 5.8683 | 6.0206 | 5.7961 | 4.9377 | 5.7086 | 5.1937 | 6.0422 | 6.6853 | 6.1483 | 4.9829 |
| SH3BP1_S262 | SH3BP1 | Q9Y3L3 | 544 | VPPRPASPKVTRS | 6.1280 | 6.2918 | 7.1125 | 6.0076 | 6.0167 | 5.9834 | 6.1487 | 6.0895 | 7.0335 | 7.0097 | 6.5062 | 6.8174 | 6.2184 | 5.7447 | 5.3372 | 5.5545 |
| SH3BP1_S544 | SH3BP1 | Q9Y3L3 | 626 | PPPLPPTPPQPAR | 6.2862 | 6.8595 | 7.3671 | 6.0925 | 6.1890 | 7.0526 | 6.5559 | 6.9573 | 7.0941 | 5.9665 | 6.7969 | 6.7534 | 4.8389 | 5.3676 | 4.8006 | 5.0218 |
| SH3BP1_S586 | SH3BP1 | Q9Y3L3 | 262 | HGQADHSPSMTAT | 6.3096 | 6.0324 | 5.8578 | 6.5840 | 6.5678 | 6.3123 | 5.8219 | 6.5273 | 6.7946 | 7.2103 | 6.4022 | 6.7001 | 5.7022 | 5.9883 | 5.5474 | 5.6419 |
| SH3BP1_T626 | SH3BP1 | Q9Y3L3 | 586 | VSGSRSSPPAPPL | 6.2186 | 6.4436 | 7.1912 | 6.4840 | 5.2639 | 5.0584 | 6.6426 | 5.0309 | 6.3740 | 6.7251 | 6.0908 | 7.5116 | 6.2316 | 6.6135 | 6.0390 | 6.0811 |
| SH3BP2_S282 | SH3BP2 | P78314-4 | 282 | DMPRAHSFTSKGP | 7.5016 | 6.9372 | 6.6876 | 6.1940 | 6.1504 | 6.9051 | 6.5088 | 6.6339 | 6.4106 | 6.3094 | 6.8324 | 6.3988 | 5.1770 | 5.8132 | 4.2815 | 5.2585 |
| SH3BP4_S246 | SH3BP4 | Q9P0V3 | 637 | VGKIILSPFATTT | 7.6894 | 8.1802 | 7.0430 | 8.2855 | 6.9118 | 6.9043 | 7.9097 | 6.4517 | 7.9198 | 7.3196 | 7.4273 | 7.4861 | 2.1750 | 3.3377 | 2.0787 | 2.8800 |
| SH3BP4_S296 | SH3BP4 | Q9P0V3 | 296 | HRKLARSCHDLDL | 5.8069 | 5.5320 | 5.6757 | 6.9522 | 6.7873 | 5.6903 | 5.8786 | 5.3900 | 5.7083 | 5.5709 | 6.2950 | 5.8892 | 6.9288 | 6.8917 | 7.3557 | 7.6473 |
| SH3BP4_S42 | SH3BP4 | Q9P0V3 | 42 | NDIKVPSPSALLV | 6.6633 | 6.7740 | 6.5615 | 6.6095 | 6.3142 | 6.6767 | 6.2837 | 6.3041 | 6.9413 | 6.6039 | 7.0374 | 6.6997 | 4.9088 | 5.5802 | 4.9215 | 5.1201 |
| SH3BP4_S637 | SH3BP4 | Q9P0V3 | 246 | RSKRSYSLSELSV | 5.0344 | 5.1239 | 6.0015 | 6.1596 | 6.1945 | 5.8796 | 5.9170 | 6.0745 | 5.8503 | 5.7176 | 7.0118 | 6.0370 | 6.5533 | 7.6690 | 7.4277 | 7.3483 |
| SH3BP5L_S343 | SH3BP5L | Q7L8J4 | 343 | SLRTVASDLQKCD | 4.5799 | 5.4230 | 6.3524 | 4.8367 | 5.1733 | 7.3246 | 5.8371 | 7.7636 | 6.4680 | 7.8445 | 7.7209 | 8.3235 | 5.5940 | 5.4541 | 5.2317 | 6.0726 |
| SH3BP5L_S350 | SH3BP5L | Q7L8J4 | 362 | GLSDHVSLDGQEL | 6.2361 | 7.0174 | 6.6580 | 6.2094 | 6.5724 | 6.8822 | 6.0419 | 7.2764 | 6.5888 | 6.2517 | 7.0199 | 6.1442 | 5.5367 | 5.9367 | 4.9665 | 4.6617 |
| SH3BP5L_S362 | SH3BP5L | Q7L8J4 | 350 | DLQKCDSVEHLRG | 6.3316 | 6.3669 | 6.7438 | 6.5843 | 5.9746 | 6.2342 | 6.8068 | 5.9414 | 6.6186 | 6.4149 | 6.3178 | 6.4839 | 6.0171 | 6.0195 | 5.3387 | 5.8056 |
| SH3BP5L_S49 | SH3BP5L | Q7L8J4 | 49 | SSEAKLSPREEEE | 6.2466 | 6.8360 | 7.0927 | 6.3572 | 6.0551 | 6.6457 | 6.0543 | 6.4147 | 6.5956 | 6.4642 | 6.5105 | 6.7192 | 5.6538 | 5.9466 | 5.0695 | 5.3383 |
| SH3D19_S51 | SH3D19 | A0A0U1RQE4 | 51 | PEHRSSSQGPLSS | 6.2043 | 5.9755 | 6.2853 | 5.5416 | 5.6247 | 6.5758 | 6.1739 | 5.9776 | 6.3871 | 6.3255 | 6.3352 | 6.8996 | 6.5939 | 6.8471 | 5.8105 | 6.4421 |
| SH3D19_S555 | SH3D19 | A0A0U1RQE4 | 555 | LHEDPQSPPPLPA | 4.7835 | 5.1167 | 5.3210 | 5.0874 | 6.2380 | 6.6441 | 6.1554 | 6.4882 | 6.8048 | 5.6133 | 6.0971 | 6.3658 | 7.2035 | 7.0519 | 7.7764 | 7.2529 |
| SH3GL1_S288 | SH3GL1 | Q99961 | 288 | KIAASSSFRSSDK | 7.8427 | 6.4865 | 6.7375 | 5.8656 | 6.9589 | 6.5624 | 6.0016 | 6.7043 | 7.4787 | 6.2613 | 5.9331 | 5.9773 | 6.5193 | 5.8232 | 4.4422 | 4.4054 |
| SH3GLB2_S404 | SH3GLB2 | Q9NR46-2 | 404 | TYLELLSxxxxxx | 6.4715 | 7.1360 | 7.3712 | 5.4759 | 6.2112 | 6.5765 | 6.5994 | 6.0194 | 6.0970 | 7.0469 | 6.7638 | 5.5855 | 5.4668 | 5.9463 | 5.4713 | 5.7613 |
| SH3KBP1_S108 | SH3KBP1 | Q96B97 | 230 | IKLRPRSIEVEND | 6.2115 | 6.2388 | 7.2817 | 6.3154 | 5.8407 | 6.3245 | 6.0397 | 6.7456 | 6.6887 | 6.4411 | 6.6362 | 7.2151 | 5.4669 | 5.8787 | 5.4431 | 5.2323 |
| SH3KBP1_S230 | SH3KBP1 | Q96B97 | 511 | DIFDSPSPEEDKE | 6.1637 | 5.8270 | 6.4650 | 5.6364 | 6.0560 | 5.8393 | 6.1922 | 6.0121 | 6.4155 | 6.1682 | 6.2424 | 6.6967 | 6.6960 | 6.6910 | 6.6429 | 6.2556 |
| SH3KBP1_S410 | SH3KBP1 | Q96B97 | 493 | TGRRPPSQSLTSS | 6.7164 | 6.5315 | 6.9965 | 6.3385 | 6.5332 | 6.3656 | 6.5380 | 6.4618 | 7.0470 | 6.2711 | 6.3154 | 6.5241 | 5.2456 | 5.9957 | 4.6827 | 5.4369 |
| SH3KBP1_S493 | SH3KBP1 | Q96B97 | 503 | TSSSLSSPDIFDS | 5.7606 | 5.7331 | 6.3077 | 6.7584 | 5.9671 | 6.6765 | 5.8889 | 6.1935 | 6.0706 | 6.6878 | 6.1918 | 6.7708 | 6.0846 | 6.7061 | 6.0409 | 6.1617 |
| SH3KBP1_S503 | SH3KBP1 | Q96B97 | 108 | RCQVAFSYLPQND | 6.4373 | 6.0965 | 7.5150 | 7.2086 | 6.5755 | 5.3852 | 5.9118 | 6.6951 | 6.2917 | 5.5794 | 6.4213 | 5.7476 | 6.7151 | 5.8202 | 6.0990 | 5.5008 |
| SH3KBP1_S509;S511 | SH3KBP1 | Q96B97 | 410 | RPPKTNSLSRPGA | 6.9613 | 7.2145 | 6.8221 | 5.8573 | 7.3209 | 6.5824 | 6.4659 | 5.7424 | 6.9060 | 5.9880 | 6.1046 | 6.8632 | 6.3074 | 5.4213 | 4.6962 | 4.7465 |
| SH3KBP1_S511 | SH3KBP1 | Q96B97 | 79 | PLHEVPSGNSLLS | 4.8179 | 4.5991 | 4.7735 | 4.4433 | 4.5671 | 5.2482 | 4.7766 | 4.4031 | 4.2957 | 4.4877 | 4.8241 | 4.4082 | 10.2704 | 10.4798 | 13.6302 | 9.9752 |
| SH3KBP1_S587 | SH3KBP1 | Q96B97 | 587 | AGHRANSPSLFGT | 6.9134 | 6.4506 | 6.8633 | 6.6371 | 5.9405 | 6.7794 | 6.5005 | 6.3914 | 6.6095 | 7.6634 | 6.6853 | 6.4847 | 4.8571 | 5.3699 | 4.8571 | 4.9969 |
| SH3KBP1_S79 | SH3KBP1 | Q96B97 | 509;511 | NA | 6.2868 | 6.2527 | 7.2988 | 5.7580 | 5.9595 | 6.3751 | 6.1852 | 6.1054 | 6.1178 | 6.5265 | 6.3350 | 6.6945 | 6.1254 | 6.2132 | 5.8787 | 5.8873 |
| SH3PXD2A_S1056 | SH3PXD2A | Q5TCZ1 | 421 | QRAQISSPNLRTR | 6.9836 | 7.2280 | 6.5785 | 7.2486 | 6.5292 | 6.2485 | 7.0214 | 6.5470 | 7.1760 | 6.8119 | 6.7827 | 7.0860 | 4.2563 | 4.6518 | 4.1604 | 4.6903 |
| SH3PXD2A_S421 | SH3PXD2A | Q5TCZ1 | 256 | RLDRRWTLGGMVN | 8.6604 | 7.6616 | 7.0311 | 7.1323 | 6.8536 | 5.6255 | 6.1802 | 5.7828 | 6.3962 | 5.3089 | 6.5331 | 5.3150 | 5.7828 | 6.4820 | 4.5806 | 4.6740 |
| SH3PXD2A_T256 | SH3PXD2A | Q5TCZ1 | 1056 | RNSIPVSPVRPKP | 5.9460 | 5.9651 | 6.3880 | 6.7729 | 5.4487 | 6.8356 | 6.4783 | 6.4583 | 6.7923 | 6.2138 | 6.4988 | 6.8278 | 5.4934 | 5.8550 | 5.6334 | 6.3927 |
| SH3PXD2B_S291 | SH3PXD2B | A1X283 | 291 | PKPGPGSPSHPGA | 5.9848 | 5.9955 | 6.8332 | 6.3816 | 5.9864 | 6.2170 | 6.6416 | 6.4333 | 6.8699 | 6.6027 | 7.0605 | 7.0168 | 5.4967 | 5.7474 | 5.2187 | 5.5140 |
| SH3RF1_S727 | SH3RF1 | Q7Z6J0 | 727 | KLLSGASTKRKPR | 8.9376 | 6.7170 | 8.2803 | 6.3047 | 5.7986 | 6.3577 | 6.6822 | 6.3674 | 8.1779 | 6.9852 | 7.5401 | 6.9099 | 4.5834 | 4.8600 | 2.7376 | 2.7603 |
| SH3TC2_T21 | SH3TC2 | Q8TF17 | 21 | RGPGKETPSKDPT | 6.2666 | 6.6318 | 6.5146 | 7.8094 | 5.2450 | 5.6879 | 5.7096 | 5.8257 | 6.0667 | 6.6269 | 6.5017 | 6.9095 | 4.4863 | 7.0710 | 5.8153 | 6.8320 |
| SHANK3_S1121 | SHANK3 | Q9BYB0 | 1121 | RERALASQAPSRS | 6.6992 | 7.9583 | 5.5848 | 7.4086 | 6.3515 | 7.3550 | 6.0220 | 7.7950 | 6.6201 | 6.3957 | 6.5122 | 6.2560 | 3.9727 | 5.6215 | 5.0924 | 4.3549 |
| SHARPIN_S131 | SHARPIN | Q9H0F6 | 165 | EADLPRSPGNLTE | 5.9807 | 4.8148 | 5.6767 | 5.1262 | 5.0935 | 5.0501 | 5.6171 | 7.5979 | 5.2172 | 5.6893 | 4.8731 | 6.3052 | 4.5273 | 9.8601 | 8.0132 | 10.5577 |
| SHARPIN_S165 | SHARPIN | Q9H0F6 | 131 | NGSKSNSPPALGP | 5.1532 | 5.2752 | 5.8161 | 5.2576 | 5.6805 | 5.2881 | 5.3986 | 6.8804 | 5.1328 | 5.5118 | 5.3931 | 6.5780 | 5.6697 | 9.7735 | 8.5717 | 8.6198 |
| SHB_S102 | SHB | Q15464 | 388 | KPIKHGSPEFCGI | 6.3323 | 6.5512 | 7.0012 | 6.2513 | 6.3585 | 6.5643 | 6.8884 | 6.0105 | 6.8331 | 7.0219 | 6.7232 | 7.0065 | 5.2139 | 5.3971 | 4.7396 | 5.1071 |
| SHB_S18 | SHB | Q15464 | 211 | AGGRTWSPTACGG | 6.7676 | 6.5739 | 6.2609 | 5.9143 | 6.8900 | 6.7207 | 6.6710 | 6.5588 | 6.9941 | 6.5467 | 6.5835 | 6.3230 | 5.5199 | 5.5041 | 5.0444 | 5.1273 |
| SHB_S211 | SHB | Q15464 | 102 | YNGPGSSLRKLRA | 7.6617 | 6.8959 | 7.5353 | 7.1171 | 5.9534 | 6.5153 | 6.1622 | 6.8523 | 7.4872 | 6.7307 | 6.7508 | 6.9329 | 4.4837 | 5.4147 | 3.7044 | 3.8024 |
| SHB_S310 | SHB | Q15464 | 18 | GNSKTKSPPQPPR | 6.2583 | 5.8866 | 6.2053 | 6.9821 | 6.6434 | 6.8114 | 6.7505 | 6.6127 | 7.0709 | 6.5373 | 7.3505 | 6.6891 | 5.1226 | 5.1790 | 5.2041 | 4.6962 |
| SHB_S388 | SHB | Q15464 | 310 | EGQSVDSDSESTV | 5.9427 | 5.8652 | 6.4958 | 6.0569 | 6.3850 | 5.7032 | 6.3416 | 6.8180 | 6.6313 | 5.8617 | 6.3407 | 6.1481 | 5.9342 | 7.5739 | 5.9154 | 5.9862 |
| SHB_Y246 | SHB | Q15464 | 246 | VTIADDYSDPFDA | 7.3162 | 5.5723 | 6.4005 | 5.9888 | 6.3780 | 6.3165 | 7.2476 | 6.5022 | 6.5294 | 5.8113 | 6.8666 | 6.2940 | 5.2044 | 6.1532 | 5.8373 | 5.5818 |

| | | | | | | | | | | | | | | | | | | | | | |
|---|---|---|---|---|---|---|---|---|---|---|---|---|---|---|---|---|---|---|---|---|---|
| SHC1_S115 | SHC1 | P29353-6 | | 139 | EWTRHGSFVNKPT | 7.1342 | 6.5422 | 7.4639 | 7.0465 | 7.1536 | 6.2498 | 6.2702 | 6.3167 | 6.8278 | 6.6199 | 6.3019 | 6.2171 | 5.5096 | 6.0200 | 3.9382 | 4.3884 |
| SHC1_S139 | SHC1 | P29353-6 | | 428 | LFDDPSYVNVQNL | 6.6904 | 6.7974 | 7.1192 | 6.9182 | 7.7376 | 6.2503 | 5.5246 | 6.2194 | 7.2162 | 6.1743 | 6.3869 | 6.6069 | 5.3689 | 5.5417 | 4.7737 | 4.6741 |
| SHC1_Y428 | SHC1 | P29353-6 | | 115 | QDMNKLSGGGGRR | 7.0333 | 6.6355 | 7.1556 | 5.9753 | 6.0373 | 7.5844 | 6.5079 | 6.6208 | 5.7550 | 6.9876 | 7.2971 | 6.7980 | 4.0009 | 5.7030 | 4.7217 | 5.1867 |
| SHCBP1_S273 | SHCBP1 | Q8NEM2 | | 273 | SLSNCNSDSEQEN | 6.2161 | 6.2250 | 7.3096 | 6.6636 | 6.1447 | 6.3582 | 6.0285 | 6.4625 | 6.3773 | 5.9730 | 5.9657 | 6.1987 | 5.9249 | 5.6776 | 6.2147 | 6.2598 |
| SHCBP1_S42;S44;S47 | SHCBP1 | Q8NEM2 | | 634 | IKKKRLSELGITQ | 5.5541 | 5.9836 | 6.7727 | 6.9144 | 6.0528 | 6.4769 | 6.3978 | 7.0248 | 6.4188 | 6.6208 | 6.0962 | 7.0020 | 5.4460 | 6.2459 | 5.2780 | 5.7154 |
| SHCBP1_S47 | SHCBP1 | Q8NEM2 | | 666 | KWNGKGSFGTFLF | 4.2172 | 4.3265 | 6.0909 | 7.6041 | 5.8997 | 5.4791 | 5.2933 | 4.3319 | 3.9222 | 5.6211 | 7.7570 | 6.4023 | 8.5491 | 7.3091 | 8.7403 | 8.4562 |
| SHCBP1_S55 | SHCBP1 | Q8NEM2 | | 47 | DSCSDCSYRDKPG | 8.4006 | 6.7032 | 7.7456 | 7.7641 | 6.5672 | 5.5022 | 6.4333 | 5.9204 | 7.0204 | 5.9904 | 7.7044 | 6.6105 | 3.9511 | 4.9048 | 4.3692 | 4.4125 |
| SHCBP1_S634 | SHCBP1 | Q8NEM2 | | 55 | RDKPGSSLQSFMP | 7.2493 | 6.0099 | 5.9152 | 7.1366 | 5.7564 | 5.4338 | 7.1494 | 5.4184 | 7.3210 | 6.2122 | 6.4734 | 5.9485 | 6.2634 | 6.5349 | 4.6041 | 6.5733 |
| SHCBP1_S666 | SHCBP1 | Q8NEM2 | 42;44;47 | | NA | 6.5432 | 6.8471 | 7.7113 | 6.3259 | 5.0747 | 5.6393 | 7.7715 | 5.7236 | 6.6685 | 6.1648 | 4.9878 | 5.1661 | 6.5577 | 6.0397 | 5.7394 | 7.0394 |
| SHKBP1_S144 | SHKBP1 | Q8TBC3 | | 144 | KRRNRHSLVGPQQ | 6.7532 | 6.6234 | 6.6347 | 6.2577 | 6.8283 | 6.5374 | 6.4800 | 6.5742 | 6.8734 | 6.4523 | 6.6227 | 6.5807 | 5.3104 | 5.7413 | 4.7582 | 4.9723 |
| SHKBP1_S649 | SHKBP1 | Q8TBC3 | | 163 | PVRRSNTMPPNLG | 7.2220 | 6.4959 | 6.8207 | 5.7381 | 6.0491 | 6.2959 | 6.4292 | 5.6756 | 6.7400 | 6.3620 | 6.7817 | 6.9633 | 5.8227 | 6.0419 | 4.8975 | 5.6646 |
| SHKBP1_S662 | SHKBP1 | Q8TBC3 | | 662 | RRRGGGSFVERCQ | 7.0945 | 6.8629 | 7.4735 | 6.3873 | 5.7430 | 6.6775 | 6.6764 | 7.0545 | 6.8959 | 7.1885 | 7.0688 | 6.9515 | 4.3405 | 5.6282 | 3.7824 | 4.1747 |
| SHKBP1_T163 | SHKBP1 | Q8TBC3 | | 696 | GLGTPLTPPKMKL | 5.9141 | 6.8015 | 6.4379 | 6.3868 | 6.4423 | 7.0790 | 6.8641 | 6.7832 | 7.0118 | 6.7355 | 7.0043 | 6.6495 | 4.6107 | 5.1810 | 4.8346 | 5.2637 |
| SHKBP1_T696 | SHKBP1 | Q8TBC3 | | 649 | GHRGSPSPPQAEA | 6.1410 | 6.8390 | 6.2930 | 6.0359 | 6.3409 | 6.7382 | 6.2353 | 5.8815 | 6.5937 | 7.2880 | 6.2973 | 5.9535 | 5.4601 | 6.4479 | 5.3724 | 6.0823 |
| SHMT2_S266 | SHMT2 | P34897 | | 266 | AAKVIPSPFKHAD | 6.1811 | 6.1835 | 6.2923 | 6.8463 | 6.6660 | 6.1117 | 6.3916 | 6.1871 | 6.3515 | 6.5930 | 6.4081 | 6.6045 | 5.8260 | 5.6727 | 5.9083 | 5.7766 |
| SHMT2_T420 | SHMT2 | P34897 | | 420 | GDRSAITPGGLRL | 5.7337 | 5.6592 | 7.0711 | 6.7642 | 6.3404 | 6.4680 | 6.2765 | 5.9209 | 6.8638 | 6.4168 | 6.0082 | 6.3722 | 5.8240 | 6.2977 | 5.8042 | 6.1791 |
| SHPRH_S530 | SHPRH | Q149N8 | | 530 | KKQAVGSPRKIQK | 6.4181 | 6.7510 | 6.4519 | 7.0587 | 6.1496 | 5.9009 | 6.2880 | 6.1956 | 6.1937 | 5.9692 | 6.2453 | 6.4467 | 5.9391 | 5.6850 | 5.9997 | 6.3074 |
| SHROOM1_S224 | SHROOM1 | Q2M3G4 | | 224 | QRKWCFSEPGKLD | 6.7000 | 6.4368 | 7.3881 | 6.5307 | 6.2815 | 7.1739 | 6.3171 | 6.4852 | 6.3087 | 6.0155 | 5.9623 | 6.3766 | 5.7140 | 5.5847 | 5.2610 | 5.4639 |
| SHROOM1_S49 | SHROOM1 | Q2M3G4 | | 82 | DAALCTSPRPRPA | 5.8176 | 7.7328 | 6.2050 | 5.0141 | 6.0752 | 7.6049 | 7.1797 | 6.4680 | 6.9545 | 5.2050 | 6.0788 | 6.1815 | 6.6229 | 5.7870 | 5.3077 | 5.7654 |
| SHROOM1_S82 | SHROOM1 | Q2M3G4 | | 49 | PEPRTQSPGTDLL | 6.4144 | 6.5735 | 6.7263 | 6.3182 | 6.3812 | 6.9595 | 6.3570 | 6.6025 | 5.9571 | 7.0515 | 6.2885 | 7.2797 | 5.7067 | 5.5151 | 5.0939 | 4.7750 |
| SHTN1_S101 | SHTN1 | A0MZ66-3 | | 494 | AEADSSSPTGILA | 5.8824 | 6.6335 | 6.9390 | 6.1700 | 6.0232 | 6.2180 | 6.4255 | 6.4890 | 6.4467 | 6.8602 | 6.9954 | 6.7411 | 5.4124 | 6.1801 | 5.1005 | 5.4831 |
| SHTN1_S493 | SHTN1 | A0MZ66-3 | | 506 | ATSESKSMPVLGS | 5.7882 | 6.1732 | 6.8589 | 5.9131 | 6.0135 | 6.7635 | 5.9280 | 6.6572 | 6.5771 | 6.2767 | 7.2634 | 7.0192 | 5.7288 | 6.2178 | 4.8533 | 5.9681 |
| SHTN1_S494 | SHTN1 | A0MZ66-3 | | 101 | KTLKRISMLYMAK | 7.9204 | 5.9631 | 7.4718 | 6.6442 | 6.3031 | 6.6708 | 6.6698 | 6.0184 | 8.3639 | 6.9802 | 7.0498 | 7.8702 | 4.0795 | 5.5626 | 3.0502 | 3.3820 |
| SHTN1_S506 | SHTN1 | A0MZ66-3 | | 493 | TAEADSSSPTGIL | 5.4471 | 6.4978 | 6.9548 | 6.3624 | 6.3811 | 6.5438 | 6.7755 | 7.5945 | 5.8811 | 6.4373 | 7.3870 | 6.2342 | 5.5992 | 5.3671 | 5.3947 | 5.1426 |
| SIK2_S587 | SIK2 | Q9H0K1 | | 587 | REGRRASDTSLTQ | 6.3145 | 6.2714 | 6.6621 | 6.3946 | 6.5014 | 6.5170 | 6.0912 | 6.4894 | 6.6912 | 6.8173 | 6.5931 | 6.6505 | 5.8771 | 5.8609 | 4.9540 | 5.3144 |
| SIK2_T484 | SIK2 | Q9H0K1 | | 484 | SGQRRHTLSEVTN | 6.2366 | 5.6628 | 6.9619 | 6.4854 | 6.3962 | 6.3713 | 5.9951 | 5.9449 | 7.0907 | 7.1114 | 7.8295 | 7.4262 | 5.5617 | 6.4927 | 4.4204 | 4.0130 |
| SIK3_S551 | SIK3 | Q9Y2K2 | | 626 | SPVRRFSDGAASI | 5.8701 | 4.8117 | 6.3106 | 5.5731 | 6.3041 | 5.7254 | 5.4809 | 6.1127 | 7.4110 | 7.0263 | 6.9955 | 7.7072 | 6.2838 | 7.1491 | 5.3963 | 5.8422 |
| SIK3_S626 | SIK3 | Q9Y2K2 | | 731 | LRIQPSSPPPNHP | 6.2992 | 6.0843 | 6.7452 | 6.4118 | 6.3479 | 6.9438 | 6.4379 | 6.8540 | 6.8850 | 6.4065 | 6.5502 | 6.9221 | 5.1701 | 5.8822 | 4.9842 | 5.0755 |
| SIK3_S731 | SIK3 | Q9Y2K2 | | 551 | PLGRRASDGGANI | 6.6120 | 6.7902 | 6.1944 | 6.0130 | 6.4159 | 6.6194 | 6.3273 | 6.4062 | 6.4856 | 6.9964 | 7.1321 | 6.8428 | 5.4477 | 5.7855 | 4.7560 | 5.1755 |
| SIK3_T221 | SIK3 | Q9Y2K2 | | 221 | PGQLLKTWCGSPP | 6.5497 | 6.6544 | 7.5135 | 6.7280 | 7.1802 | 6.3219 | 5.7964 | 6.0462 | 6.1565 | 6.0308 | 6.3911 | 5.4490 | 5.9184 | 5.9657 | 6.0336 | 5.2648 |
| SIK3_T469 | SIK3 | Q9Y2K2 | | 469 | LSMRRHTVGVADP | 7.0984 | 6.9798 | 6.8270 | 6.6539 | 6.3814 | 5.5394 | 6.3761 | 5.9049 | 6.0929 | 6.3045 | 6.7031 | 6.5642 | 5.5618 | 6.0491 | 5.7691 | 5.1943 |
| SIN3A_S10 | SIN3A | Q96ST3 | | 277 | PYASPRSPPVQPH | 6.5124 | 6.9289 | 6.2494 | 7.0239 | 6.3263 | 6.4098 | 6.0418 | 6.1452 | 6.2281 | 6.1324 | 6.3517 | 6.6748 | 5.6682 | 5.8487 | 5.5861 | 5.8723 |
| SIN3A_S1112 | SIN3A | Q96ST3 | | 1112 | MNSDTTSPELREH | 6.2480 | 6.3655 | 6.1243 | 6.5769 | 6.1184 | 5.8187 | 6.2721 | 6.3736 | 5.9669 | 6.1267 | 6.0767 | 5.9856 | 6.3658 | 6.5748 | 6.7123 | 6.2936 |
| SIN3A_S274 | SIN3A | Q96ST3 | | 860 | HNGVGGSPPKSKL | 6.3256 | 6.4213 | 5.9663 | 6.8131 | 6.1718 | 5.8996 | 6.4180 | 6.1470 | 6.1989 | 6.5429 | 6.3420 | 6.2875 | 6.0104 | 6.1780 | 6.0429 | 6.2346 |
| SIN3A_S277 | SIN3A | Q96ST3 | | 832 | AQRGDLSDVEEEE | 6.1807 | 6.4119 | 6.0996 | 5.7433 | 5.9579 | 6.0761 | 6.3805 | 6.2806 | 6.0481 | 6.1851 | 6.1825 | 5.6552 | 6.3491 | 6.8528 | 6.2305 | 7.3662 |
| SIN3A_S277;T284 | SIN3A | Q96ST3 | | 266 | TPASQQTPPLPPY | 5.8868 | 5.5791 | 6.0807 | 6.3063 | 5.8165 | 6.3433 | 6.4233 | 5.4722 | 6.4271 | 6.4695 | 6.5587 | 6.3004 | 6.2429 | 6.4357 | 6.2219 | 7.4357 |
| SIN3A_S277;T287 | SIN3A | Q96ST3 | | 434 | RRHPTGTTPPVKK | 6.3055 | 6.5704 | 5.9437 | 7.1256 | 6.7209 | 6.9694 | 7.1247 | 6.7182 | 6.5098 | 6.5481 | 6.2868 | 6.4239 | 5.0457 | 5.5433 | 4.8021 | 5.3619 |
| SIN3A_S832 | SIN3A | Q96ST3 | | 10 | RLDDQESPVYAAQ | 6.9301 | 6.0235 | 6.3516 | 7.5450 | 7.3219 | 7.1270 | 6.4919 | 7.0896 | 6.2012 | 6.0618 | 7.0190 | 6.4879 | 4.6342 | 5.5510 | 4.3757 | 4.7887 |
| SIN3A_S860 | SIN3A | Q96ST3 | | 274 | PLPPYASPRSPPV | 5.8322 | 5.9006 | 6.8360 | 6.6103 | 6.0155 | 6.9002 | 5.8664 | 5.1923 | 6.3245 | 7.0548 | 6.9577 | 5.6326 | 5.3413 | 7.4253 | 6.3259 | 5.7844 |
| SIN3A_S940 | SIN3A | Q96ST3 | | 940 | KRDKSDSPAIQLR | 6.3675 | 7.0055 | 6.1446 | 6.5618 | 6.1861 | 5.8396 | 6.5852 | 5.9467 | 6.2536 | 6.1788 | 5.9944 | 6.0829 | 6.1518 | 6.3458 | 6.0867 | 6.2692 |
| SIN3A_T266 | SIN3A | Q96ST3 | 277;284 | | NA | 5.5517 | 5.6225 | 5.2945 | 6.1830 | 5.9238 | 6.0634 | 6.0863 | 6.4166 | 5.3263 | 5.4867 | 5.6875 | 5.4162 | 7.0855 | 7.6614 | 8.2236 | 7.9711 |
| SIN3A_T434 | SIN3A | Q96ST3 | 277;287 | | NA | 4.9497 | 5.4371 | 4.9908 | 6.0999 | 5.9871 | 6.3942 | 5.8000 | 6.5899 | 5.1289 | 5.9197 | 4.9710 | 5.0778 | 8.3056 | 8.1461 | 8.1667 | 8.0353 |
| SIN3B_S1003 | SIN3B | O75182 | | 1003 | TEGASSSPTEGFL | 5.7591 | 6.1927 | 6.0978 | 6.2688 | 6.2871 | 5.8992 | 6.3138 | 5.7802 | 5.9703 | 5.9782 | 6.3185 | 5.9548 | 6.6324 | 6.7734 | 6.8156 | 6.9580 |
| SIN3B_S279 | SIN3B | O75182 | | 740 | APGPHSSPPEEKG | 5.8632 | 6.3426 | 6.4203 | 7.1793 | 6.9146 | 6.3041 | 6.3913 | 6.0346 | 5.8895 | 6.5410 | 5.9196 | 6.0532 | 5.7415 | 6.2005 | 6.1276 | 6.0771 |
| SIN3B_S740 | SIN3B | O75182 | | 279 | SLLRPVSAPAKKK | 6.1521 | 5.4643 | 6.6243 | 7.1192 | 5.7031 | 6.7515 | 7.2112 | 6.0906 | 6.2172 | 6.6124 | 7.6918 | 7.2804 | 5.5879 | 5.2291 | 4.7629 | 5.5020 |
| SIN3B_T262 | SIN3B | O75182 | | 262 | KNEHDKTPEHSRK | 6.4739 | 6.6175 | 6.7888 | 6.8584 | 7.1228 | 6.7367 | 6.9208 | 6.4487 | 7.1733 | 6.5422 | 7.0173 | 6.8323 | 4.6245 | 5.0036 | 4.1031 | 4.7362 |
| SIN3B_T997 | SIN3B | O75182 | | 997 | VEQYVGTEGASSS | 6.3536 | 6.0417 | 6.3682 | 6.5936 | 6.0240 | 5.7403 | 6.6300 | 5.3646 | 6.1767 | 6.5702 | 6.1928 | 6.0918 | 6.0821 | 6.1993 | 7.2344 | 6.3367 |
| SIPA1_S314 | SIPA1 | Q96FS4 | | 817 | CLQDGGSPPGPGD | 6.3795 | 6.3073 | 6.2953 | 6.5356 | 6.2682 | 6.1806 | 6.3190 | 6.3164 | 6.5600 | 6.6037 | 6.4702 | 7.0743 | 5.1740 | 6.0985 | 5.5078 | 5.9096 |
| SIPA1_S67 | SIPA1 | Q96FS4 | | 839 | HSQNSLSPRSSLS | 5.8403 | 6.2627 | 6.5213 | 6.5322 | 6.3898 | 6.5656 | 6.0823 | 6.2671 | 6.7190 | 6.8344 | 6.6559 | 6.7213 | 5.3467 | 6.0512 | 5.6380 | 5.5721 |
| SIPA1_S817 | SIPA1 | Q96FS4 | | 884 | TQDRPGSPSGSED | 5.6650 | 5.3762 | 6.3376 | 6.2901 | 5.7312 | 6.3416 | 5.8802 | 5.3611 | 6.1146 | 6.7277 | 7.0662 | 6.8477 | 6.3449 | 6.7464 | 6.3304 | 6.8390 |
| SIPA1_S839 | SIPA1 | Q96FS4 | | 314 | LRLGSASPKVPRT | 6.5812 | 6.7854 | 7.1611 | 6.6686 | 6.0303 | 6.7517 | 6.6150 | 6.4446 | 6.7032 | 6.7730 | 6.7560 | 6.6175 | 4.7659 | 5.4104 | 4.8366 | 5.0995 |
| SIPA1_S843 | SIPA1 | Q96FS4 | | 843 | SLSPRSSLSDEAP | 5.7367 | 5.8517 | 5.6802 | 6.0641 | 5.8693 | 6.1005 | 5.8451 | 5.8938 | 6.3537 | 5.9231 | 6.1551 | 6.2646 | 6.8704 | 7.6710 | 6.8910 | 6.8296 |
| SIPA1_S884 | SIPA1 | Q96FS4 | | 67 | RPPTPASPRARAH | 6.3688 | 7.3757 | 6.2169 | 6.5471 | 5.9386 | 6.2366 | 6.9501 | 6.6529 | 6.1653 | 6.4323 | 6.3793 | 6.5494 | 5.1422 | 5.6689 | 5.3817 | 5.9944 |
| SIPA1L1_S1255 | SIPA1L1 | O43166 | | 162 | PEAYPSSPRKALR | 6.2997 | 6.5419 | 6.2596 | 6.7631 | 6.3222 | 6.3449 | 6.4679 | 6.3889 | 6.6727 | 6.3409 | 6.3031 | 6.2805 | 5.7400 | 6.0101 | 5.4358 | 5.8288 |
| SIPA1L1_S1270 | SIPA1L1 | O43166 | | 1270 | DPVVHLSPNKQGH | 6.5496 | 6.4286 | 5.7459 | 6.3401 | 6.5883 | 6.1218 | 6.6967 | 6.3187 | 6.3079 | 5.8902 | 6.0471 | 5.8420 | 6.2592 | 6.2382 | 6.2492 | 6.3765 |
| SIPA1L1_S1366 | SIPA1L1 | O43166 | | 1549 | KLIDLESPTPESQ | 6.3310 | 6.4307 | 6.0099 | 6.6306 | 6.1219 | 5.9774 | 6.2105 | 6.2010 | 6.0607 | 5.9603 | 5.7812 | 5.9989 | 6.3809 | 6.7219 | 6.5403 | 6.6429 |
| SIPA1L1_S1390;S1393 | SIPA1L1 | O43166 | | 1632 | LPLRRPSYTLGMK | 5.6387 | 5.0114 | 5.2999 | 5.7549 | 7.0508 | 5.4424 | 5.7299 | 5.8140 | 7.1287 | 6.6318 | 6.4674 | 6.5572 | 7.2305 | 7.5566 | 6.2535 | 6.4323 |
| SIPA1L1_S1393 | SIPA1L1 | O43166 | | 1433 | SSRHSASPVVFTS | 6.0444 | 6.2328 | 6.0255 | 5.9033 | 6.2989 | 6.6342 | 6.6893 | 6.7026 | 6.1076 | 6.5042 | 6.7037 | 6.0873 | 5.9431 | 6.2921 | 6.0984 | 5.7327 |
| SIPA1L1_S1431;S1433 | SIPA1L1 | O43166 | | 176 | IRQRSNSDITISE | 6.0400 | 6.4282 | 6.6239 | 6.4378 | 5.7708 | 5.7925 | 6.2013 | 5.6748 | 6.7541 | 6.6951 | 5.9589 | 6.4335 | 6.0715 | 6.6434 | 5.8528 | 6.6215 |
| SIPA1L1_S1433 | SIPA1L1 | O43166 | | 1585 | ALHRTLSDESIYN | 5.9190 | 6.2827 | 6.0488 | 6.2715 | 6.0441 | 5.7545 | 6.1807 | 5.8931 | 6.1212 | 6.2493 | 6.2927 | 6.1904 | 6.6207 | 6.6871 | 6.5398 | 6.9045 |
| SIPA1L1_S1433;S1439 | SIPA1L1 | O43166 | | 1366 | APLWHSSSEVISM | 6.1606 | 6.5370 | 6.3715 | 6.5704 | 6.3922 | 5.9682 | 6.0986 | 6.3719 | 6.3418 | 5.8004 | 6.1961 | 5.9596 | 6.4420 | 6.3881 | 6.1830 | 6.2185 |
| SIPA1L1_S1528 | SIPA1L1 | O43166 | | 255 | TGGGKGSGFSLDV | 6.5333 | 6.1798 | 6.2620 | 6.5161 | 6.5661 | 6.6802 | 6.5613 | 6.5671 | 6.2716 | 6.1938 | 6.6855 | 6.2969 | 5.6684 | 6.2152 | 5.3915 | 5.4112 |
| SIPA1L1_S1549 | SIPA1L1 | O43166 | | 1528 | TRDLRASPKPTSK | 6.4542 | 6.4716 | 5.9966 | 6.9422 | 6.1234 | 6.2203 | 6.6528 | 5.5944 | 5.8541 | 5.9646 | 6.1031 | 6.2995 | 6.2136 | 6.5512 | 6.2264 | 6.3320 |
| SIPA1L1_S1565 | SIPA1L1 | O43166 | | 318 | DLEDNRSEDSVRP | 6.1255 | 6.0361 | 6.3169 | 7.1760 | 5.7378 | 6.0281 | 6.2832 | 6.0198 | 6.4020 | 6.3673 | 6.9234 | 6.6775 | 5.8824 | 5.8617 | 6.0569 | 6.1055 |
| SIPA1L1_S1565;S1568 | SIPA1L1 | O43166 | | 239 | KSDRGPTPTKLSD | 6.5508 | 6.5332 | 6.0633 | 6.5545 | 6.1618 | 6.3510 | 6.6337 | 5.8288 | 5.9745 | 5.7982 | 5.8675 | 6.1168 | 6.3883 | 6.4806 | 6.2193 | 6.4778 |
| SIPA1L1_S1565;S1568;T1 | SIPA1L1 | O43166 | | 1734 | SSSKDSSPTLASK | 6.3151 | 6.6493 | 5.8584 | 6.7214 | 6.3984 | 6.1040 | 6.4024 | 6.2201 | 6.4978 | 6.3963 | 6.1412 | 6.4305 | 5.7995 | 6.0357 | 5.8733 | 6.1566 |
| SIPA1L1_S1565;T1573 | SIPA1L1 | O43166 | | 1565 | KFHALSSPQSPFP | 5.9198 | 6.6830 | 5.6758 | 6.1361 | 6.5260 | 5.7189 | 6.1236 | 6.0760 | 6.5748 | 6.3672 | 6.1660 | 6.9167 | 6.3157 | 6.0635 | 6.0181 | 6.7189 |
| SIPA1L1_S1568;T1573 | SIPA1L1 | O43166 | | 1621 | FSSTYPSLPKSLP | 6.1596 | 7.9450 | 6.2605 | 7.1126 | 6.0126 | 6.1313 | 7.0640 | 6.3054 | 6.6635 | 6.2076 | 5.7271 | 5.9486 | 5.3801 | 5.9806 | 5.5579 | 5.5438 |
| SIPA1L1_S1585 | SIPA1L1 | O43166 | | 1255 | LPAFRESPSGRLM | 7.0745 | 7.4628 | 6.0321 | 6.6268 | 6.7484 | 6.7929 | 7.4040 | 6.7152 | 6.4885 | 6.5514 | 6.2983 | 5.9431 | 4.7497 | 5.6656 | 4.5974 | 4.8492 |
| SIPA1L1_S1603 | SIPA1L1 | O43166 | | 1603 | FFTSRASLLDQAL | 6.2240 | 6.1469 | 5.7300 | 6.8568 | 6.8061 | 5.4113 | 5.5007 | 5.6340 | 7.2027 | 6.8208 | 7.0196 | 6.7396 | 5.9615 | 7.3434 | 5.4076 | 5.1949 |
| SIPA1L1_S162 | SIPA1L1 | O43166 | | 26 | VVGTDGTPKVHTD | 6.0102 | 6.5381 | 6.8726 | 7.7001 | 6.0962 | 6.8003 | 7.3240 | 6.2036 | 7.1235 | 7.3667 | 7.4225 | 7.5281 | 4.0754 | 4.6834 | 3.5291 | 4.7261 |
| SIPA1L1_S1621 | SIPA1L1 | O43166 | | 1393 | KTESSLSLDIHSK | 4.9237 | 5.1902 | 5.3230 | 5.0750 | 4.9881 | 5.5553 | 5.5590 | 5.2339 | 5.0634 | 4.9897 | 5.5159 | 5.0508 | 9.5990 | 8.9827 | 9.5146 | 9.4357 |
| SIPA1L1_S1632 | SIPA1L1 | O43166 | | 208 | LHREYGSTSSIDK | 6.2744 | 6.5495 | 5.6849 | 6.9303 | 6.1459 | 6.8788 | 5.7515 | 5.6321 | 6.2053 | 5.9660 | 5.5265 | 6.0527 | 6.9277 | 7.0790 | 6.1616 | 6.2339 |
| SIPA1L1_S1734 | SIPA1L1 | O43166 | | 288 | LKRRSKSETGDSS | 6.1938 | 6.0668 | 7.4788 | 6.3501 | 6.1399 | 5.7467 | 6.1863 | 6.3890 | 6.8067 | 6.3912 | 6.6601 | 6.4622 | 5.4223 | 6.4149 | 5.0886 | 6.2025 |
| SIPA1L1_S176 | SIPA1L1 | O43166 | | 1201 | GGSGKSTPSWQRS | 6.7517 | 7.0568 | 6.0800 | 7.3375 | 5.8971 | 6.2391 | 6.9291 | 6.4703 | 6.7895 | 6.5644 | 6.5996 | 6.5308 | 5.0536 | 5.2780 | 5.2159 | 5.2068 |
| SIPA1L1_S208 | SIPA1L1 | O43166 | 1565;1568;15 | | NA | 6.5315 | 6.6774 | 6.6253 | 7.4630 | 6.6826 | 6.0029 | 6.9412 | 6.6939 | 6.8470 | 6.3831 | 6.0744 | 6.3131 | 5.0498 | 5.4458 | 4.9460 | 5.3230 |
| SIPA1L1_S255 | SIPA1L1 | O43166 | 1565;1568 | | NA | 6.2466 | 5.9740 | 6.0864 | 6.1393 | 6.5088 | 6.1721 | 7.5057 | 6.8698 | 6.0066 | 7.0655 | 6.2176 | 6.7331 | 5.2934 | 5.9059 | 5.7669 | 5.5082 |
| SIPA1L1_S288 | SIPA1L1 | O43166 | 1390;1393 | | NA | 5.3048 | 4.3176 | 5.6214 | 4.9643 | 4.8315 | 4.6073 | 5.1131 | 4.6183 | 5.0428 | 4.9528 | 4.6323 | 5.1124 | 10.3194 | 9.3431 | 9.4781 | 11.7408 |
| SIPA1L1_S318 | SIPA1L1 | O43166 | 1431;1433 | | NA | 5.3141 | 6.6961 | 2.9620 | 5.1202 | 6.8721 | 5.2738 | 5.6790 | 6.3610 | 5.6337 | 9.4864 | 4.0589 | 6.8901 | 7.5364 | 6.2505 | 7.6271 | 8.2388 |
| SIPA1L1_T1201 | SIPA1L1 | O43166 | 1433;1439 | | NA | 5.7629 | 7.0819 | 6.2801 | 6.2283 | 6.1465 | 5.6136 | 6.5673 | 6.5328 | 7.0300 | 6.0314 | 6.0647 | 5.6392 | 6.0017 | 5.6900 | 6.2586 | 7.0709 |
| SIPA1L1_T239 | SIPA1L1 | O43166 | 1565;1573 | | NA | 6.7589 | 6.4587 | 6.6799 | 7.0387 | 6.5187 | 5.3352 | 5.8875 | 6.4707 | 6.8900 | 6.1613 | 6.3576 | 6.4138 | 5.7458 | 6.0443 | 5.3864 | 5.8525 |
| SIPA1L1_T26 | SIPA1L1 | O43166 | 1568;1573 | | NA | 6.6248 | 6.6530 | 6.2050 | 7.3508 | 6.7105 | 5.8241 | 6.4017 | 5.7319 | 6.5734 | 6.2139 | 7.1138 | 6.3612 | 5.6264 | 5.5915 | 5.6442 | 5.3739 |
| SIPA1L3_S100 | SIPA1L3 | O60292 | | 1702 | GPPTPRTTPTMSE | 5.6145 | 5.6293 | 5.4643 | 5.8408 | 5.8516 | 5.9395 | 6.1728 | 6.0317 | 5.7901 | 5.8266 | 5.8598 | 5.8305 | 7.5879 | 7.5825 | 7.3998 | 7.5784 |
| SIPA1L3_S1109 | SIPA1L3 | O60292 | | 1364 | DRRREVSPAPAVA | 5.6228 | 5.7180 | 5.3479 | 5.6552 | 6.1128 | 5.8145 | 6.0971 | 5.9796 | 6.0050 | 5.3416 | 5.2350 | 5.3308 | 7.9375 | 7.7282 | 8.1619 | 7.9120 |
| SIPA1L3_S1135 | SIPA1L3 | O60292 | | 1544 | GLQRTLSDESLCS | 6.1956 | 6.4463 | 5.9106 | 6.1501 | 5.9641 | 5.8044 | 6.2654 | 6.2483 | 6.4373 | 6.1088 | 6.0626 | 6.2424 | 6.3605 | 6.2661 | 6.7946 | 6.7430 |
| SIPA1L3_S1144 | SIPA1L3 | O60292 | | 1319 | TTLYTSSPSCMSL | 6.0234 | 6.4628 | 6.1305 | 6.6085 | 6.4533 | 6.3384 | 6.2304 | 6.6681 | 5.6982 | 5.7070 | 6.2299 | 6.1767 | 6.3194 | 6.7826 | 5.9032 | 6.2676 |
| SIPA1L3_S1160 | SIPA1L3 | O60292 | | 204 | LFREYGSTSSIDV | 6.2213 | 6.5893 | 6.1608 | 6.8465 | 6.3773 | 5.8681 | 6.3839 | 5.9836 | 6.2869 | 5.8868 | 6.1180 | 5.6853 | 6.2138 | 6.3909 | 6.5647 | 6.4228 |
| SIPA1L3_S1175 | SIPA1L3 | O60292 | | 1175 | YVRYKPSPERYTA | 6.5945 | 7.0237 | 6.0977 | 7.1102 | 6.4297 | 6.8571 | 7.0704 | 6.7248 | 6.5292 | 6.3772 | 6.6643 | 6.4420 | 4.6610 | 5.2288 | 4.8440 | 5.3453 |
| SIPA1L3_S1233 | SIPA1L3 | O60292 | | 1160 | YRQPSGSFSTPGS | 5.9841 | 5.7766 | 5.6869 | 6.1069 | 6.3967 | 5.6227 | 5.9515 | 6.0173 | 6.4851 | 5.5137 | 5.1420 | 5.2945 | 7.7635 | 7.8526 | 7.4263 | 6.9797 |
| SIPA1L3_S1239 | SIPA1L3 | O60292 | | 400 | GLDPAFTSTEDLN | 5.4908 | 6.1505 | 5.8122 | 6.3269 | 5.8055 | 6.2445 | 6.3028 | 6.3110 | 6.0593 | 6.0297 | 6.3576 | 6.3123 | 6.5537 | 6.6175 | 6.8323 | 6.7934 |
| SIPA1L3_S1319 | SIPA1L3 | O60292 | | 1135 | HSKRPVSFPETPY | 6.2120 | 6.5372 | 6.9018 | 6.9918 | 5.8996 | 6.0721 | 6.4106 | 6.6109 | 6.6672 | 6.2666 | 6.2585 | 6.5708 | 5.4752 | 5.8381 | 5.6976 | 5.5902 |
| SIPA1L3_S1364 | SIPA1L3 | O60292 | | 1144 | ETPYTVSPAGADR | 6.7118 | 6.7756 | 6.6546 | 7.3092 | 6.1551 | 6.8512 | 6.7486 | 6.7071 | 6.7612 | 7.1699 | 6.8843 | 6.9578 | 4.4799 | 4.6758 | 4.2326 | 4.9254 |
| SIPA1L3_S1501 | SIPA1L3 | O60292 | | 1387 | YSSGSSTPTGLAG | 6.9893 | 6.6308 | 6.2529 | 6.5561 | 5.7507 | 6.1170 | 6.4231 | 7.0989 | 6.9774 | 6.6393 | 6.6051 | 6.6520 | 4.7457 | 5.8261 | 5.1418 | 5.5939 |
| SIPA1L3_S1534 | SIPA1L3 | O60292 | | 172 | LRHRSSSEITLSE | 6.5806 | 6.6067 | 5.5344 | 6.0564 | 6.0532 | 6.4138 | 6.3105 | 6.2221 | 6.3504 | 6.2807 | 6.1784 | 5.9334 | 5.6255 | 6.8421 | 6.6024 | 6.4096 |
| SIPA1L3_S1544 | SIPA1L3 | O60292 | | 1117 | SRPLKQTPIVPFR | 7.7893 | 7.2178 | 6.7750 | 6.8649 | 5.6815 | 6.8602 | 7.4152 | 6.9300 | 6.7815 | 7.6812 | 7.3482 | 7.8227 | 3.7170 | 4.0858 | 3.2395 | 3.7904 |
| SIPA1L3_S1544;S1547 | SIPA1L3 | O60292 | | 1501 | SLRDLRSPRKNYK | 7.0312 | 7.1188 | 6.4351 | 6.3101 | 7.5573 | 8.2781 | 7.3166 | 5.6831 | 5.4338 | 5.8284 | 6.7535 | 6.1685 | 5.4657 | 5.5266 | 4.6713 | 4.4219 |
| SIPA1L3_S172 | SIPA1L3 | O60292 | | 391 | TASRAHSLGGLDP | 6.5468 | 6.3589 | 6.1616 | 7.2529 | 6.6150 | 6.0247 | 6.5801 | 6.2536 | 7.9374 | 7.2363 | 6.8256 | 6.5679 | 4.7859 | 5.7531 | 4.6114 | 4.4889 |
| SIPA1L3_S204 | SIPA1L3 | O60292 | | 1233 | PAQARLSAIAGSS | 6.9250 | 5.3216 | 5.0237 | 5.4390 | 8.6001 | 4.9501 | 5.6020 | 7.4682 | 9.4307 | 5.6160 | 4.1441 | 5.2025 | 7.8169 | 6.8514 | 5.8088 | 5.8000 |
| SIPA1L3_S204;S207 | SIPA1L3 | O60292 | | 493 | SRPRQYSIEHVDL | 5.6410 | 6.2540 | 6.5266 | 6.3016 | 6.5006 | 6.6910 | 5.5256 | 7.5997 | 6.0463 | 6.5064 | 7.3833 | 7.6963 | 4.9530 | 5.8501 | 5.8703 | 4.6544 |
| SIPA1L3_S391 | SIPA1L3 | O60292 | | 277 | LLPLQPTKEKEKA | 7.5569 | 6.2930 | 7.6558 | 8.9438 | 6.4883 | 6.3894 | 6.9611 | 6.0180 | 5.1786 | 6.9008 | 5.8998 | 6.3629 | 4.1993 | 5.2872 | 5.2269 | 4.6383 |
| SIPA1L3_S451 | SIPA1L3 | O60292 | | 1534 | ERDTGQSPQKGLQ | 6.5335 | 6.1539 | 5.6072 | 6.2566 | 6.5762 | 6.4084 | 7.0822 | 6.6401 | 6.3154 | 5.7288 | 5.9875 | 5.7476 | 5.7843 | 5.7978 | 6.7013 | 6.6792 |
| SIPA1L3_S493 | SIPA1L3 | O60292 | | 451 | SRASVGSPSSGEG | 6.1282 | 7.6795 | 6.6537 | 8.0002 | 6.7451 | 6.3274 | 6.2735 | 6.6343 | 5.7593 | 5.5013 | 6.8128 | 6.2416 | 4.9408 | 5.6948 | 5.0141 | 5.5934 |
| SIPA1L3_T1100;T1103 | SIPA1L3 | O60292 | | 1109 | TPGHAQSLSRPLK | 9.6232 | 7.1478 | 6.7318 | 7.3013 | 5.4789 | 5.9067 | 6.5843 | 5.2911 | 6.3532 | 5.6008 | 6.5007 | 6.2450 | 5.7464 | 5.8713 | 4.9164 | 4.7010 |
| SIPA1L3_T1103 | SIPA1L3 | O60292 | | 1103 | KWATPTTPGHAQS | 6.7071 | 7.2914 | 6.9092 | 7.3767 | 6.3993 | 6.8575 | 7.4284 | 7.4269 | 7.6697 | 6.4965 | 6.6272 | 7.1653 | 3.6151 | 4.4712 | 3.3888 | 4.1696 |
| SIPA1L3_T1117 | SIPA1L3 | O60292 | | 129 | GQPPTSTPASSGS | 8.2070 | 8.5020 | 7.0224 | 7.8350 | 5.8386 | 6.3049 | 7.6161 | 7.2498 | 7.2098 | 7.5371 | 6.6750 | 7.6818 | 2.9231 | 3.5645 | 2.3313 | 3.5017 |
| SIPA1L3_T129 | SIPA1L3 | O60292 | | 1239 | SAIAGSSGNKHPS | 6.6375 | 6.8049 | 7.0056 | 7.2723 | 5.8633 | 6.5533 | 7.1506 | 6.7576 | 7.3454 | 7.2631 | 7.9463 | 7.1687 | 3.6362 | 4.9824 | 3.4816 | 4.1312 |
| SIPA1L3_T1387 | SIPA1L3 | O60292 | | 100 | REHSNPSPSQDTD | 5.9366 | 5.7599 | 5.2787 | 5.8096 | 5.5738 | 5.3952 | 6.2376 | 6.2478 | 5.6376 | 8.9821 | 5.8529 | 5.2502 | 6.7700 | 7.1652 | 7.3642 | 6.7387 |
| SIPA1L3_T1702 | SIPA1L3 | O60292 | 1100;1103 | | NA | 7.4038 | 7.1166 | 6.8356 | 7.3416 | 5.7540 | 7.0417 | 6.7849 | 6.7185 | 7.3129 | 6.3345 | 6.6870 | 6.8194 | 4.3335 | 4.7329 | 4.1528 | 4.6305 |
| SIPA1L3_T277 | SIPA1L3 | O60292 | 1544;1547 | | NA | 6.4849 | 6.2931 | 5.3406 | 7.7897 | 6.6149 | 5.0623 | 6.4485 | 5.0330 | 5.9721 | 5.4117 | 5.7688 | 5.4560 | 7.6858 | 7.1269 | 6.4859 | 7.0257 |
| SIPA1L3_T400 | SIPA1L3 | O60292 | 204;207 | | NA | 5.8206 | 6.5857 | 6.9850 | 6.9152 | 6.3481 | 6.0267 | 6.8948 | 5.8107 | 5.9715 | 6.1229 | 6.3599 | 6.3716 | 5.8763 | 5.4286 | 5.8216 | 6.6610 |
| SIRT1_S27 | SIRT1 | Q96EB6 | | 615 | EKNERTSVAGTVR | 5.7019 | 6.3929 | 6.2457 | 5.5654 | 5.9409 | 6.1347 | 6.4050 | 6.3266 | 6.0486 | 6.0459 | 6.0451 | 5.9878 | 6.7879 | 6.8232 | 6.6178 | 6.9306 |
| SIRT1_S47 | SIRT1 | Q96EB6 | | 47 | GPGLERSPGEPGG | 5.2917 | 6.0649 | 6.1682 | 5.5485 | 5.4210 | 6.1747 | 5.6308 | 6.3051 | 6.1511 | 6.6100 | 6.2049 | 6.9006 | 6.8477 | 6.7887 | 6.9222 | 6.9698 |
| SIRT1_S615 | SIRT1 | Q96EB6 | | 719 | GGAGFGTDGDDQE | 5.3243 | 5.5174 | 6.7494 | 6.5650 | 5.8424 | 6.2811 | 6.1124 | 7.5017 | 5.7865 | 5.7616 | 6.0058 | 6.2615 | 6.6782 | 7.1810 | 6.3580 | 6.0737 |
| SIRT1_T719 | SIRT1 | Q96EB6 | | 27 | DREAASSPAGEPL | 5.3233 | 5.8248 | 5.9033 | 5.9263 | 5.4923 | 6.1663 | 5.7586 | 5.9876 | 6.6408 | 6.7410 | 6.1702 | 6.4485 | 6.1608 | 6.6509 | 7.6346 | 7.1708 |
| SIRT6_S338 | SIRT6 | Q8N6T7 | | 338 | KRERPTSPAPHRP | 5.0856 | 5.1141 | 5.8657 | 5.8852 | 5.6683 | 5.8873 | 5.5761 | 5.8650 | 6.6719 | 6.7077 | 6.5173 | 6.7731 | 6.8289 | 7.1688 | 7.0450 | 7.3401 |
| SIRT6_T294 | SIRT6 | Q8N6T7 | | 294 | PLPRPPTPKLEPK | 5.1585 | 5.8922 | 5.8896 | 5.5681 | 5.8719 | 5.8512 | 5.5898 | 5.6775 | 6.1901 | 6.1730 | 5.9740 | 6.1154 | 7.5472 | 7.7176 | 7.4852 | 7.2986 |
| SIX4_S44 | SIX4 | Q9UIU6 | | 44 | GAAVGLSPPAPAP | 5.8888 | 6.1337 | 6.2917 | 6.3617 | 6.2687 | 6.9550 | 6.4148 | 6.1717 | 6.1629 | 6.8156 | 6.5242 | 7.2553 | 6.0039 | 5.9893 | 5.3495 | 5.4132 |
| SKA2_S101 | SKA2 | Q8WVK7 | | 101 | QTDLELSPLTKEE | 6.2900 | 6.5478 | 6.8417 | 6.4814 | 5.9501 | 6.8193 | 6.7264 | 7.0250 | 6.4883 | 6.8691 | 6.9666 | 6.8150 | 4.7204 | 5.5150 | 5.0758 | 4.8680 |
| SKA3_S155 | SKA3 | Q8IX90 | | 155 | SEKSPRSPQLSDF | 6.6093 | 6.4312 | 6.7611 | 6.3073 | 6.5556 | 6.6835 | 6.1120 | 7.0785 | 6.5423 | 6.4072 | 7.1297 | 7.4913 | 5.0117 | 5.0459 | 4.5658 | 5.2675 |
| SKA3_T190 | SKA3 | Q8IX90 | | 217 | DDFECVTPKLEHF | 7.9978 | 7.4203 | 7.2548 | 6.7975 | 6.0756 | 7.0785 | 7.0325 | 6.5998 | 7.3916 | 8.0540 | 7.4249 | 8.0510 | 3.1339 | 3.4331 | 2.8075 | 3.4474 |
| SKA3_T217 | SKA3 | Q8IX90 | | 360 | NLLRTPTPPEVTK | 6.8370 | 6.3177 | 7.6377 | 6.2891 | 7.0180 | 6.1473 | 6.4428 | 6.9359 | 6.5612 | 6.3873 | 6.7807 | 7.3148 | 4.8093 | 5.1179 | 4.4632 | 4.9403 |
| SKA3_T265;S267 | SKA3 | Q8IX90 | | 190 | EEPVIVTPPTKQS | 7.7193 | 7.0188 | 7.0992 | 6.9628 | 7.1923 | 6.5901 | 6.6829 | 7.2542 | 7.1489 | 7.1336 | 7.7165 | 7.5991 | 3.4362 | 3.7789 | 2.8826 | 3.7845 |
| SKA3_T360 | SKA3 | Q8IX90 | 265;267 | | NA | 7.8522 | 7.9833 | 7.2385 | 7.2457 | 6.7412 | 7.2111 | 6.9718 | 6.9619 | 7.4660 | 8.0338 | 8.2716 | 7.8067 | 2.3777 | 3.4003 | 1.8437 | 2.5945 |
| SKI_S394 | SKI | P12755 | | 394 | ASEKELSPHLPAL | 7.7026 | 8.3057 | 7.1434 | 8.3819 | 6.3440 | 7.1603 | 6.4937 | 6.5234 | 7.9568 | 8.4031 | 8.2195 | 7.2097 | 2.0860 | 2.2441 | 1.9603 | 3.8654 |
| SKI_S480 | SKI | P12755 | | 515 | SAKDLGSPGARAL | 7.8729 | 8.2391 | 7.9158 | 8.2905 | 6.2390 | 6.2136 | 7.3489 | 6.9271 | 7.2625 | 7.8111 | 8.1508 | 8.0231 | 1.9840 | 3.1458 | 1.9487 | 2.6272 |
| SKI_S515 | SKI | P12755 | | 480 | PEEDKDSEAEVEV | 6.9154 | 6.8968 | 7.1805 | 7.3359 | 6.3024 | 6.5947 | 6.6350 | 6.3391 | 6.7333 | 6.4954 | 7.0310 | 6.8452 | 4.6782 | 4.7485 | 4.0730 | 5.1956 |
| SKIV2L_S245 | SKIV2L | Q15477 | | 256 | PLARASSLEDLVL | 6.4730 | 6.2424 | 6.8284 | 5.9423 | 6.4580 | 6.4102 | 6.1046 | 6.7481 | 6.4093 | 6.3981 | 6.5455 | 6.5724 | 5.5598 | 6.0911 | 5.6765 | 5.5400 |
| SKIV2L_S256 | SKIV2L | Q15477 | | 245 | GDTVSASPCSAPL | 6.7059 | 7.4311 | 7.0086 | 6.7015 | 6.3312 | 6.8351 | 7.0003 | 6.9303 | 7.0211 | 6.6609 | 6.9893 | 7.1867 | 3.9478 | 4.6314 | 3.7968 | 4.8221 |
| SKP1_T131 | SKP1 | P63208 | | 131 | NMIKGKTPEEIRK | 5.7661 | 4.9867 | 4.9174 | 4.5967 | 5.7073 | 5.3598 | 5.5334 | 7.1003 | 4.8742 | 5.9681 | 4.9866 | 5.7836 | 7.1579 | 9.6421 | 8.3411 | 9.2789 |
| SKP2_S64 | SKP2 | Q13309 | | 64 | NLGHPESPPRKRL | 5.8178 | 6.3334 | 6.1347 | 6.8608 | 6.2650 | 5.9662 | 5.9123 | 6.0843 | 6.1452 | 6.1845 | 6.3360 | 6.1834 | 6.1537 | 6.5134 | 6.5948 | 6.5145 |
| SLAIN2_S147 | SLAIN2 | Q9P270 | | 63 | PSSGAASPRGFPL | 6.6164 | 6.6154 | 6.9471 | 6.2365 | 6.6449 | 6.8748 | 6.3147 | 6.9589 | 7.2094 | 6.8260 | 7.2591 | 7.2769 | 4.3747 | 4.8814 | 4.2395 | 4.7241 |
| SLAIN2_S160 | SLAIN2 | Q9P270 | | 48 | GSLGPGSPVRAGA | 7.2091 | 6.5871 | 7.1948 | 6.3280 | 6.6974 | 6.8347 | 6.5885 | 7.0271 | 7.3713 | 6.9839 | 7.3529 | 7.1773 | 4.1535 | 4.4906 | 3.7188 | 4.2849 |
| SLAIN2_S179 | SLAIN2 | Q9P270 | | 377 | IEYSRVSPQPMIS | 6.7028 | 6.5964 | 6.6665 | 6.0680 | 6.7288 | 6.9254 | 6.3860 | 6.8660 | 6.7664 | 6.5742 | 7.1205 | 6.6556 | 5.3372 | 5.2832 | 4.5135 | 4.8096 |
| SLAIN2_S234 | SLAIN2 | Q9P270 | | 456 | QASAIPSPGKFRS | 6.7187 | 6.4598 | 6.7804 | 6.2943 | 6.3745 | 6.8863 | 6.6745 | 6.7267 | 7.4673 | 6.7861 | 7.1810 | 7.0920 | 4.6672 | 4.9790 | 4.0965 | 4.8156 |
| SLAIN2_S282 | SLAIN2 | Q9P270 | | 147 | PMQKSVSPLVWCR | 7.4063 | 7.5067 | 7.4163 | 7.1216 | 6.8163 | 7.2477 | 6.8303 | 7.2105 | 7.5783 | 7.4070 | 7.8891 | 7.5387 | 2.6626 | 3.3697 | 2.7002 | 3.2987 |
| SLAIN2_S289 | SLAIN2 | D6RIF6 | | 289 | STKRVPSPGKFRS | 5.9243 | 5.8039 | 5.6944 | 5.4279 | 5.6610 | 6.4526 | 6.0812 | 6.5090 | 6.0675 | 5.9313 | 6.0679 | 5.8825 | 7.4933 | 7.3097 | 6.9970 | 6.6965 |
| SLAIN2_S349;S353 | SLAIN2 | Q9P270 | | 353 | RNSPRPSPKQSPR | 6.2521 | 5.7159 | 5.5559 | 6.3962 | 8.2833 | 5.2397 | 5.1101 | 5.9674 | 6.0451 | 4.6588 | 5.0514 | 5.8574 | 8.3052 | 7.3969 | 7.4430 | 6.7215 |
| SLAIN2_S353 | SLAIN2 | Q9P270 | | 72 | GFPLGLSAKSGGG | 7.4066 | 7.7303 | 7.9442 | 7.5047 | 6.7562 | 6.7915 | 8.4278 | 6.4688 | 6.8847 | 7.5920 | 7.8314 | 7.9314 | 2.4869 | 3.3021 | 1.9630 | 2.9783 |
| SLAIN2_S377 | SLAIN2 | Q9P270 | | 413 | EEKLRRSLPNLSR | 7.1181 | 7.2421 | 6.5588 | 6.5310 | 6.4054 | 6.4480 | 6.2008 | 6.5912 | 7.5683 | 7.9235 | 7.8273 | 7.6804 | 3.8157 | 4.5751 | 3.3902 | 4.1241 |
| SLAIN2_S391 | SLAIN2 | Q9P270 | | 179 | KLDQTMSALKRQN | 7.9835 | 8.4441 | 7.2220 | 6.9180 | 6.4708 | 6.5737 | 7.0909 | 5.8462 | 8.0379 | 6.2432 | 7.2804 | 7.5061 | 3.6442 | 4.2864 | 2.6974 | 3.7552 |
| SLAIN2_S413 | SLAIN2 | Q9P270 | | 87 | SGPRRTSSEELRD | 6.9241 | 6.3853 | 6.3599 | 5.6541 | 6.9403 | 6.4531 | 6.0859 | 6.4577 | 6.3353 | 5.9728 | 6.4846 | 6.4538 | 6.8810 | 5.9843 | 5.3155 | 5.3124 |
| SLAIN2_S433 | SLAIN2 | Q9P270 | | 462 | SPGKFRSPAAPSP | 8.0357 | 7.5577 | 6.7821 | 7.4743 | 7.8756 | 5.9208 | 7.8035 | 7.8373 | 7.3661 | 7.2308 | 6.3785 | 5.9343 | 3.0123 | 3.8938 | 3.0619 | 3.8352 |
| SLAIN2_S456 | SLAIN2 | Q9P270 | | 234 | LILPGNSGNLKSS | 7.8162 | 8.3922 | 6.9449 | 7.7733 | 6.4757 | 7.1388 | 7.2983 | 7.1510 | 7.3606 | 8.0796 | 8.0089 | 7.9155 | 2.1660 | 2.9465 | 1.7869 | 2.7454 |
| SLAIN2_S462 | SLAIN2 | Q9P270 | | 160 | QVLDYPSPDVECA | 7.5927 | 7.3033 | 7.4727 | 7.3411 | 6.7964 | 8.0514 | 7.0335 | 6.9934 | 8.3167 | 7.7580 | 8.3053 | 7.7598 | 2.0078 | 2.6568 | 1.7684 | 2.8427 |
| SLAIN2_S462;S467 | SLAIN2 | Q9P270 | | 391 | LQQPRLSLQGHPT | 5.9722 | 5.6115 | 6.4537 | 5.7856 | 6.6987 | 7.3675 | 6.5351 | 6.8709 | 7.8736 | 6.6819 | 8.0298 | 7.4579 | 5.3769 | 5.3259 | 3.7689 | 4.1900 |
| SLAIN2_S48 | SLAIN2 | Q9P270 | | 433 | SVKSSRSDSNFQV | 5.5765 | 5.8410 | 7.2093 | 6.4583 | 6.1042 | 6.7298 | 5.2099 | 7.0935 | 5.7680 | 6.1621 | 6.4735 | 7.3347 | 5.4414 | 6.2778 | 6.3054 | 6.0146 |
| SLAIN2_S63 | SLAIN2 | Q9P270 | | 132 | EEESWLYSSPKKK | 6.4265 | 6.3143 | 6.5380 | 6.4800 | 7.4550 | 7.1629 | 6.4883 | 7.1419 | 7.3729 | 6.9498 | 8.5409 | 7.3841 | 4.0584 | 4.1415 | 3.4701 | 4.0754 |
| SLAIN2_S72 | SLAIN2 | D6RIF6 | | 282 | KQLLQTSSTKRVP | 8.2154 | 8.8256 | 7.0007 | 7.4265 | 5.6689 | 7.3001 | 7.8680 | 7.0337 | 7.7871 | 6.9511 | 7.3876 | 7.6506 | 2.2701 | 3.4830 | 2.1510 | 2.9806 |
| SLAIN2_S87 | SLAIN2 | Q9P270 | 462;467 | | NA | 7.0137 | 7.0274 | 7.3189 | 6.9101 | 5.6729 | 6.6987 | 7.4522 | 6.7769 | 7.5777 | 7.2262 | 7.3560 | 7.2209 | 3.8518 | 4.4722 | 3.3741 | 4.0504 |

| | | | | | | | | | | | | | | | | | | | | | |
|---|---|---|---|---|---|---|---|---|---|---|---|---|---|---|---|---|---|---|---|---|---|
| SLAIN2_Y132 | SLAIN2 | Q9P270 | 349;353 | | NA | 6.0625 | 6.0105 | 7.3983 | 5.7037 | 5.7831 | 5.8570 | 6.1867 | 6.4932 | 7.7209 | 7.5221 | 7.3075 | 7.4522 | 5.4862 | 4.9857 | 4.6365 | 5.3939 |
| SLBP_S182 | SLBP | Q14493 | | 62 | RPESFTTPEGPKP | 6.7503 | 6.8552 | 7.3804 | 5.7197 | 6.0361 | 6.7116 | 5.9447 | 6.6053 | 7.6629 | 7.6497 | 8.0335 | 7.9022 | 3.9116 | 4.5233 | 3.7602 | 4.5534 |
| SLBP_S20 | SLBP | Q14493 | | 182 | KKYSRRSWDQQIK | 5.8779 | 6.2544 | 7.0717 | 6.1974 | 6.0697 | 6.5046 | 5.7824 | 6.4462 | 6.5116 | 6.1408 | 6.7926 | 6.5923 | 5.8444 | 6.0782 | 5.9304 | 5.9054 |
| SLBP_S23 | SLBP | Q14493 | | 23 | GDASPPSPARWSL | 5.7335 | 5.5641 | 6.3779 | 4.6545 | 5.2672 | 4.6183 | 5.3441 | 5.2249 | 6.6592 | 6.3479 | 6.3791 | 5.8694 | 8.1816 | 7.7493 | 8.3214 | 7.7076 |
| SLBP_T238 | SLBP | Q14493 | | 20 | RCDGDASPPSPAR | 5.1218 | 4.9382 | 6.1846 | 5.5954 | 5.8275 | 5.5704 | 5.7798 | 6.4058 | 6.1753 | 5.7603 | 5.8596 | 5.3563 | 7.9602 | 8.5775 | 7.8140 | 7.0732 |
| SLBP_T62 | SLBP | Q14493 | | 238 | FDVYSGTPTKVRH | 5.0102 | 5.8586 | 6.7323 | 5.2421 | 6.5268 | 6.2888 | 5.3544 | 5.7695 | 6.3142 | 6.6067 | 7.3245 | 6.8750 | 6.0671 | 6.7677 | 6.5409 | 6.7211 |
| SLC12A1_S145 | SLC12A1 | A0A2R8Y6V7 | | 145 | AKGLTLSPRLECS | 6.7829 | 7.2499 | 6.2057 | 7.7073 | 6.3417 | 6.4121 | 5.9798 | 7.0362 | 6.9804 | 6.1539 | 7.5527 | 6.5133 | 4.5689 | 5.1703 | 4.5948 | 4.7502 |
| SLC12A2_S242 | SLC12A2 | P55011 | | 217 | RTFGHNTMDAVPR | 7.4064 | 7.4843 | 7.6162 | 8.5464 | 4.9416 | 6.2222 | 6.9852 | 6.0155 | 6.2906 | 5.5262 | 6.1311 | 5.8422 | 4.7097 | 6.1892 | 4.6629 | 5.4303 |
| SLC12A2_S77 | SLC12A2 | P55011 | | 940 | QEELLSSQEKSPG | 5.7443 | 6.0376 | 5.9418 | 5.8917 | 5.9098 | 6.6757 | 5.9976 | 6.0192 | 6.6746 | 5.8704 | 6.3124 | 6.6137 | 6.6321 | 6.6729 | 6.7973 | 6.2090 |
| SLC12A2_S940 | SLC12A2 | P55011 | | 77 | PLGPTPSQSRFQV | 3.4869 | 3.3846 | 3.6675 | 4.1093 | 4.9837 | 4.4616 | 4.4649 | 4.7798 | 4.0025 | 3.6095 | 3.6091 | 3.9836 | 12.6221 | 11.7624 | 13.1005 | 13.9719 |
| SLC12A2_T212;T217 | SLC12A2 | P55011 | | 242 | EKLLRPSLAELHD | 8.0044 | 5.1122 | 5.4197 | 5.4389 | 6.0755 | 4.3675 | 5.1912 | 4.9204 | 6.3012 | 5.8984 | 5.6910 | 6.3232 | 7.6917 | 8.4790 | 6.6399 | 8.4459 |
| SLC12A2_T217 | SLC12A2 | P55011 | 212;217 | | NA | 7.5956 | 6.3184 | 7.0708 | 7.4835 | 5.6607 | 5.6021 | 6.0753 | 4.9947 | 6.4213 | 7.2097 | 5.3057 | 6.4470 | 6.4560 | 6.6990 | 5.0582 | 5.6021 |
| SLC12A4_S47 | SLC12A4 | Q9UP95 | | 47 | GNHRESSPFLSPL | 6.8092 | 6.8064 | 7.1602 | 6.9695 | 6.0862 | 6.0252 | 7.2823 | 6.0815 | 6.6814 | 6.8312 | 7.0334 | 6.4344 | 5.6656 | 4.5535 | 4.6355 | 4.9445 |
| SLC12A4_S88 | SLC12A4 | Q9UP95 | | 88 | LLGKLVSYTNLTQ | 4.4176 | 4.5879 | 4.8880 | 4.7683 | 4.3474 | 4.7815 | 4.9746 | 4.9810 | 4.5337 | 4.4654 | 4.5965 | 5.0598 | 10.8701 | 10.2607 | 11.4886 | 10.9790 |
| SLC12A4_S967 | SLC12A4 | Q9UP95 | | 967 | RLESLYSDEEDES | 6.4270 | 6.3780 | 6.5839 | 6.4795 | 6.3151 | 6.6676 | 6.3561 | 6.1119 | 6.7077 | 5.9627 | 6.3086 | 6.3000 | 5.2189 | 6.0407 | 6.3350 | 5.8072 |
| SLC12A4_T983 | SLC12A4 | Q9UP95 | | 983 | ADKIQMTWTRDKY | 7.6181 | 7.9719 | 8.1907 | 8.3553 | 5.3655 | 5.0558 | 6.1060 | 5.1428 | 5.2794 | 4.4582 | 4.9031 | 4.0828 | 6.8186 | 7.5901 | 7.6949 | 5.3667 |
| SLC12A6_S45 | SLC12A6 | Q9UHW9-2 | | 940 | AYTYERTLMMEQR | 7.0114 | 6.7209 | 7.1407 | 8.1951 | 6.5546 | 4.8746 | 7.1936 | 6.3867 | 6.0377 | 5.1361 | 5.0129 | 4.5658 | 6.1843 | 6.4185 | 7.4982 | 5.0689 |
| SLC12A6_S972 | SLC12A6 | Q9UHW9-2 | | 972 | LVKDRNSMLRLTS | 6.0126 | 7.3144 | 5.9622 | 6.5615 | 5.9442 | 6.1398 | 6.1101 | 6.1928 | 5.4334 | 6.1695 | 6.9044 | 6.1863 | 5.7685 | 6.6700 | 6.4562 | 6.1740 |
| SLC12A6_T940 | SLC12A6 | Q9UHW9-2 | | 45 | PDLSQNSITGEHS | 6.0756 | 6.3407 | 6.3478 | 6.3966 | 5.7687 | 6.4580 | 6.8506 | 5.6059 | 5.8096 | 5.8647 | 5.9599 | 6.0992 | 6.4658 | 7.3714 | 6.4052 | 6.1802 |
| SLC15A4_S298 | SLC15A4 | Q8N697 | | 298 | KQSLFDSCKMSHG | 6.2522 | 6.2891 | 5.8362 | 6.2331 | 5.6887 | 5.5021 | 5.9197 | 5.4380 | 6.1019 | 6.3190 | 6.2937 | 6.2023 | 6.8643 | 6.9134 | 7.2623 | 6.8840 |
| SLC16A1_S467 | SLC16A1 | P53985 | | 467 | SKEEETSIDVAGK | 5.7469 | 6.1664 | 5.9670 | 5.9898 | 6.5278 | 6.4313 | 6.0607 | 5.9099 | 6.2318 | 6.1420 | 6.3099 | 6.0257 | 6.6667 | 7.0434 | 6.4218 | 6.3587 |
| SLC16A1_S483 | SLC16A1 | P53985 | | 466 | ESKEEETSIDVAG | 6.0874 | 6.0877 | 5.5588 | 5.9886 | 5.7761 | 5.8416 | 5.9725 | 5.8970 | 6.1797 | 6.2371 | 5.5295 | 5.9224 | 6.7977 | 7.4737 | 7.4647 | 7.1855 |
| SLC16A1_S498 | SLC16A1 | P53985 | | 483 | VTKAAESPDQKDT | 7.0267 | 7.5168 | 6.5358 | 7.0458 | 6.4020 | 6.1297 | 6.7637 | 6.4949 | 6.4248 | 6.6144 | 6.3917 | 6.5576 | 4.7408 | 5.2035 | 4.9550 | 5.1970 |
| SLC16A1_T466 | SLC16A1 | P53985 | | 498 | GPKEEESPVxxxx | 5.7330 | 6.1841 | 6.1970 | 6.8464 | 6.1750 | 6.2325 | 6.1869 | 6.1856 | 6.1608 | 5.9616 | 6.1876 | 6.3224 | 6.5338 | 6.9529 | 6.0704 | 6.0702 |
| SLC16A3_S436 | SLC16A3 | O15427 | | 464 | VHTPETSVxxxxx | 6.0285 | 6.4628 | 6.1429 | 6.3386 | 6.2331 | 6.1711 | 6.6765 | 6.3443 | 6.0336 | 5.9451 | 6.0357 | 6.2168 | 6.3603 | 6.4678 | 6.4203 | 6.1225 |
| SLC16A3_S464 | SLC16A3 | O15427 | | 436 | HKPPADSGVDLRE | 3.8774 | 4.2683 | 3.6376 | 4.0152 | 4.6668 | 4.4152 | 4.4487 | 3.9928 | 4.3036 | 4.0561 | 3.5204 | 3.5567 | 12.6658 | 12.9541 | 13.6063 | 12.0148 |
| SLC19A1_S499 | SLC19A1 | P41440 | | 499 | GLQPAQSPPLSPE | 6.7295 | 6.7837 | 6.3066 | 7.1353 | 6.8465 | 6.0406 | 6.9128 | 6.9406 | 7.0654 | 6.0292 | 7.0547 | 7.4020 | 4.4516 | 5.0821 | 4.1507 | 5.0685 |
| SLC1A4_S507 | SLC1A4 | P43007 | | 507 | KSEEETSPLVTHQ | 5.6899 | 6.1494 | 5.8398 | 6.0716 | 6.2544 | 6.7013 | 6.6553 | 6.3643 | 6.6946 | 6.6282 | 6.3570 | 6.4719 | 5.7628 | 6.2147 | 5.6781 | 6.4668 |
| SLC1A5_S493 | SLC1A5 | Q15758 | | 493 | DRTESRSTEPELI | 5.8853 | 6.5581 | 6.0468 | 6.4699 | 6.1917 | 5.8363 | 5.9864 | 6.2544 | 6.5156 | 6.1179 | 6.0445 | 6.0784 | 6.4344 | 6.2940 | 6.7010 | 6.5853 |
| SLC1A5_S535 | SLC1A5 | Q15758 | | 535 | GDATVASEKESVM | 4.9218 | 6.1350 | 6.1885 | 7.0793 | 6.4235 | 5.9341 | 6.3863 | 5.9645 | 6.9655 | 5.8330 | 6.8199 | 6.7736 | 5.8357 | 6.3346 | 6.0643 | 6.3405 |
| SLC20A1_S265 | SLC20A1 | Q8WUM9 | | 417 | PLRRNNSYTSYTM | 6.0768 | 5.9339 | 6.2809 | 6.3186 | 6.0192 | 5.8589 | 6.3056 | 6.2273 | 6.0604 | 6.1166 | 6.2603 | 6.4510 | 6.5395 | 6.7694 | 6.2220 | 6.5595 |
| SLC20A1_S267 | SLC20A1 | Q8WUM9 | | 335 | ERERLPSVDLKEE | 5.5043 | 5.7485 | 5.8644 | 5.9192 | 5.9692 | 6.4514 | 7.0623 | 6.4365 | 6.2222 | 6.2419 | 6.2937 | 6.5711 | 6.7344 | 6.8064 | 5.9823 | 6.1922 |
| SLC20A1_S269 | SLC20A1 | Q8WUM9 | | 267 | EIKCSPSESPLME | 2.0505 | 2.3314 | 2.4756 | 2.6420 | 3.2993 | 2.9971 | 3.1106 | 2.8871 | 2.7003 | 2.6948 | 2.2548 | 2.6024 | 16.3068 | 15.1847 | 20.2171 | 16.2455 |
| SLC20A1_S335 | SLC20A1 | Q8WUM9 | | 449 | EEMEKLTWPNADS | 7.3521 | 7.2144 | 7.1707 | 6.7771 | 6.1847 | 6.5388 | 7.3171 | 7.0613 | 6.3508 | 6.9083 | 5.1769 | 5.2949 | 5.5900 | 5.1681 | 4.4423 | 5.4523 |
| SLC20A1_S417 | SLC20A1 | Q8WUM9 | | 265 | EREIKCSPSESPL | 5.4684 | 5.7747 | 5.2976 | 6.6444 | 5.5711 | 5.7876 | 5.9084 | 4.6513 | 6.2725 | 6.8860 | 6.4684 | 6.1276 | 7.6082 | 7.0516 | 6.9447 | 7.5374 |
| SLC20A1_T449 | SLC20A1 | Q8WUM9 | | 269 | KCSPSESPLMEKK | 5.6847 | 5.5926 | 5.8522 | 6.4412 | 6.1035 | 6.7637 | 7.4845 | 6.7056 | 6.7443 | 6.0205 | 6.5569 | 6.1401 | 5.3963 | 6.7252 | 5.5566 | 6.2320 |
| SLC20A2_S268 | SLC20A2 | Q08357 | | 268 | KVQEAESPVFKEL | 6.5278 | 6.4042 | 6.3971 | 6.2766 | 6.0277 | 5.7638 | 6.2419 | 6.4412 | 6.9484 | 7.6746 | 7.3501 | 7.9637 | 5.1794 | 5.4196 | 4.7966 | 4.5871 |
| SLC20A2_S316;S321 | SLC20A2 | Q08357 | | 324 | THGSVKSPISNGT | 5.5230 | 6.7950 | 6.0980 | 6.6298 | 5.4910 | 7.8154 | 6.9982 | 7.0838 | 6.7751 | 6.4854 | 6.8080 | 6.2485 | 4.7638 | 5.7495 | 4.9332 | 5.8023 |
| SLC20A2_S324 | SLC20A2 | Q08357 | 316;321 | | NA | 6.2757 | 5.6321 | 8.3012 | 6.4402 | 6.3827 | 6.3252 | 5.7809 | 5.8049 | 7.0357 | 7.1201 | 9.0339 | 8.5839 | 3.7116 | 6.0902 | 3.6179 | 3.8637 |
| SLC23A2_S81 | SLC23A2 | Q9UGH3 | | 81 | TLDSTGSLDPQRS | 5.7384 | 6.3985 | 5.8220 | 6.5227 | 6.3471 | 6.2618 | 6.1760 | 5.7579 | 6.5936 | 6.5021 | 6.2458 | 6.2338 | 7.0867 | 6.4243 | 6.1011 | 5.7882 |
| SLC25A1_S156 | SLC25A1 | P53007 | | 156 | FIHDQTSPNPKYR | 5.1430 | 6.9388 | 7.1815 | 7.1052 | 5.3123 | 5.7936 | 6.0684 | 5.6393 | 6.2607 | 6.4460 | 6.7357 | 6.5957 | 6.0857 | 6.5223 | 6.1023 | 6.0696 |
| SLC25A3_S297 | SLC25A3 | Q00325 | | 297 | LNKEKGSSASLVL | 3.5975 | 2.9834 | 4.0637 | 4.1593 | 4.5960 | 5.5528 | 6.4660 | 5.1318 | 3.3348 | 3.7958 | 3.6688 | 2.5902 | 11.0655 | 11.9179 | 13.1843 | 13.8923 |
| SLC25A46_S34 | SLC25A46 | Q96AG3 | | 34 | FPARSFSTGSDLG | 2.8717 | 3.1844 | 3.0639 | 2.9886 | 5.7201 | 4.8953 | 5.0853 | 5.4081 | 3.4993 | 2.5934 | 4.0881 | 3.1537 | 13.7107 | 12.8428 | 13.2424 | 13.6522 |
| SLC25A46_S34;S37 | SLC25A46 | Q96AG3 | 34;37 | | NA | 2.8929 | 3.5199 | 3.4917 | 3.4533 | 5.7751 | 5.5825 | 7.0855 | 5.9978 | 4.3081 | 3.1553 | 4.2559 | 3.7114 | 11.6395 | 10.7769 | 12.6693 | 11.6849 |
| SLC25A5_S42 | SLC25A5 | P05141 | | 42 | LQVQHASKQITAD | 3.1297 | 3.0041 | 2.7322 | 3.2123 | 5.2955 | 5.4044 | 5.9804 | 5.3680 | 3.4875 | 3.1646 | 3.1297 | 3.3415 | 13.5311 | 12.1062 | 14.1741 | 12.9386 |
| SLC26A2_S35 | SLC26A2 | P50443 | | 35 | LELQRESSTDFKQ | 6.1298 | 5.4791 | 4.8665 | 5.3028 | 6.8477 | 5.7540 | 5.7324 | 5.8737 | 6.7924 | 5.7700 | 5.0233 | 5.4129 | 7.9705 | 7.8485 | 7.9793 | 7.2171 |
| SLC26A6_S752 | SLC26A6 | Q9BXS9 | | 752 | PRPVPDSPVSVTR | 6.1329 | 5.6538 | 5.9824 | 6.3832 | 6.1769 | 5.9599 | 6.1688 | 5.4831 | 6.3876 | 6.2845 | 6.2901 | 6.1301 | 7.0259 | 6.7193 | 6.3541 | 6.8675 |
| SLC29A1_S16 | SLC29A1 | Q99808-2 | | 16 | QAWEFPSPTKSGC | 6.4298 | 7.1427 | 6.6722 | 6.2111 | 6.6980 | 6.8198 | 6.3225 | 7.0292 | 6.9869 | 7.2892 | 8.3179 | 7.2830 | 4.0733 | 4.4519 | 3.6946 | 4.5778 |
| SLC29A2_S252 | SLC29A2 | Q14542 | | 252 | ENGIPSSPQKVAL | 6.2123 | 6.4464 | 6.4696 | 6.0238 | 5.7889 | 6.6040 | 6.6171 | 6.3537 | 7.9205 | 6.4719 | 7.2699 | 7.5156 | 4.2538 | 5.6437 | 4.6702 | 5.7387 |
| SLC2A1_S490 | SLC2A1 | P11166 | | 490 | HPLGADSQVxxxx | 6.2372 | 6.9024 | 5.9562 | 6.6493 | 6.9629 | 6.0798 | 6.6499 | 5.9714 | 5.6797 | 6.6061 | 5.7369 | 5.2737 | 6.4486 | 6.7562 | 6.4452 | 5.6445 |
| SLC2A13_S6 | SLC2A13 | Q96QE2 | | 6 | xMSRKASENVEYT | 5.8496 | 4.2802 | 5.5443 | 4.9082 | 6.9704 | 5.3664 | 4.6992 | 5.4873 | 7.7349 | 8.4972 | 7.8609 | 7.8590 | 6.9685 | 7.9433 | 5.5808 | 4.4498 |
| SLC2A6_S496 | SLC2A6 | Q9UGQ3 | | 496 | SLEQIESFFRTGR | 3.7017 | 3.0006 | 3.7046 | 3.1748 | 4.1238 | 4.0441 | 3.2530 | 3.5556 | 3.8699 | 3.9585 | 3.6515 | 3.5762 | 14.5883 | 12.2415 | 16.2118 | 13.3441 |
| SLC30A1_S167 | SLC30A1 | Q9Y6M5 | | 172 | KSTRPGSSDINVA | 6.0291 | 6.3677 | 6.5013 | 6.6152 | 6.2214 | 5.8718 | 6.2624 | 6.1942 | 6.4380 | 6.5048 | 6.3287 | 6.7323 | 6.3905 | 5.7575 | 5.8281 | 5.9571 |
| SLC30A1_S172 | SLC30A1 | Q9Y6M5 | | 167 | KGPRVKSTRPGSS | 7.2178 | 6.0257 | 7.4129 | 6.7692 | 5.8305 | 5.7368 | 6.4688 | 6.6317 | 7.1765 | 6.8016 | 6.8268 | 6.8723 | 5.1262 | 5.3901 | 4.7603 | 4.9528 |
| SLC30A1_S468 | SLC30A1 | Q9Y6M5 | | 449 | LKQCCGTLPQAPS | 7.2285 | 7.4547 | 6.1923 | 6.9595 | 6.9315 | 5.7577 | 6.1855 | 5.9257 | 6.8539 | 5.2601 | 5.6814 | 5.5336 | 6.4819 | 6.5225 | 5.6621 | 5.3691 |
| SLC30A1_T449 | SLC30A1 | Q9Y6M5 | | 468 | TPAVSISCLELSN | 6.2889 | 6.5118 | 6.4571 | 7.0133 | 5.5217 | 5.9001 | 5.7103 | 5.6012 | 5.9668 | 5.1917 | 5.8965 | 6.0100 | 6.5870 | 6.9254 | 7.5687 | 6.8495 |
| SLC30A6_S422 | SLC30A6 | Q6NXT4-2 | | 422 | TPAKPSSPPPEFS | 5.4585 | 5.5109 | 5.1560 | 5.8791 | 5.9102 | 5.8637 | 5.8914 | 5.9501 | 5.4810 | 6.1221 | 5.3198 | 5.9379 | 7.4887 | 7.2739 | 7.8751 | 8.8814 |
| SLC30A6_S428 | SLC30A6 | Q6NXT4-2 | | 431 | PEFSFNTPGKNVN | 6.1798 | 4.9735 | 6.9648 | 6.3062 | 6.1022 | 6.9560 | 7.2917 | 6.0551 | 6.6622 | 6.4449 | 6.8858 | 6.6653 | 5.2533 | 6.6123 | 5.3779 | 5.2691 |
| SLC30A6_T431 | SLC30A6 | Q6NXT4-2 | | 428 | SPPPEFSFNTPGK | 4.9344 | 5.5594 | 5.8547 | 6.5261 | 6.0735 | 6.1424 | 5.3310 | 5.8806 | 6.1015 | 5.7728 | 5.5077 | 5.8655 | 8.0283 | 8.0250 | 7.4658 | 6.9313 |
| SLC35B1_S29 | SLC35B1 | P78383-2 | | 29 | VPVVSGSPVGSSG | 6.1791 | 6.4068 | 6.0994 | 6.9029 | 6.3386 | 6.0087 | 6.3552 | 6.5046 | 6.1568 | 6.5030 | 6.2026 | 6.6321 | 5.4939 | 6.0578 | 6.0011 | 6.1575 |
| SLC35B2_S427 | SLC35B2 | Q8TB61 | | 427 | KAVPVESPVQKVx | 5.9858 | 6.4649 | 6.3249 | 7.0272 | 6.1551 | 6.3059 | 6.4789 | 6.4796 | 6.3237 | 6.5782 | 6.5636 | 6.4813 | 5.4244 | 5.9350 | 5.5952 | 5.8763 |
| SLC35B3_Y43 | SLC35B3 | Q9H1N7 | | 43 | FNNSRKYISITVP | 6.1270 | 6.1888 | 6.5038 | 7.3271 | 6.6735 | 7.1240 | 7.2661 | 7.1577 | 6.4560 | 6.5821 | 6.5472 | 6.7864 | 4.5772 | 5.0059 | 4.4271 | 5.2500 |
| SLC35C2_S364 | SLC35C2 | Q9NQQ7-3 | | 364 | ALKGLGSSPDLEL | 5.5068 | 5.9802 | 5.5181 | 5.7939 | 6.4413 | 6.2475 | 6.4590 | 6.4462 | 5.8126 | 5.9214 | 5.9505 | 5.9524 | 7.0171 | 7.0631 | 7.0494 | 6.8404 |
| SLC35E1_S398 | SLC35E1 | Q96K37 | | 398 | FQYSRQSYPNSYS | 5.2776 | 5.4690 | 5.3790 | 4.7909 | 5.9288 | 5.3183 | 4.6589 | 5.5313 | 6.7356 | 6.9645 | 6.2734 | 6.7874 | 7.5009 | 8.6042 | 7.7705 | 7.0097 |
| SLC35F2_S371 | SLC35F2 | Q8IXU6 | | 371 | NLQETHSAVLxxx | 5.4929 | 6.1334 | 6.3151 | 5.9026 | 6.0179 | 6.1206 | 5.9735 | 6.1765 | 5.6666 | 6.1465 | 5.8205 | 5.5120 | 6.8626 | 7.7429 | 7.1446 | 6.9719 |
| SLC35F6_T365 | SLC35F6 | Q8N357 | | 365 | LLGGTRTPINDAS | 6.9180 | 6.9697 | 6.9966 | 7.2245 | 6.6575 | 6.6117 | 6.5051 | 5.9229 | 6.1423 | 6.4554 | 6.4085 | 6.2734 | 4.8835 | 5.6905 | 5.1658 | 5.1747 |
| SLC35G2_S409 | SLC35G2 | Q8TBE7 | | 409 | YQEILDSPIKxxx | 5.8020 | 6.2416 | 5.9676 | 6.5345 | 6.1071 | 6.1825 | 6.2882 | 6.2200 | 6.1578 | 5.9300 | 5.7812 | 5.9671 | 6.6008 | 6.8363 | 6.5531 | 6.8301 |
| SLC38A1_S49 | SLC38A1 | Q9H2H9 | | 52 | DRESRRSLTNSHL | 6.3560 | 5.7819 | 6.1100 | 5.7445 | 6.7770 | 5.9204 | 5.9009 | 6.6173 | 7.6274 | 7.3076 | 7.2821 | 7.1480 | 5.5418 | 6.0982 | 4.8380 | 4.9489 |
| SLC38A1_S52 | SLC38A1 | Q9H2H9 | | 49 | FISDRESRRSLTN | 5.9571 | 6.5077 | 7.2000 | 6.7304 | 5.9063 | 6.4702 | 6.4695 | 6.2206 | 6.3072 | 6.7146 | 7.4397 | 7.2010 | 5.2981 | 5.7063 | 5.0030 | 4.8681 |
| SLC38A1_S52;S56 | SLC38A1 | Q9H2H9 | 17;25 | | NA | 6.0567 | 6.6207 | 6.0988 | 6.5352 | 6.4874 | 5.9951 | 6.6204 | 6.2161 | 6.3259 | 6.6892 | 6.3426 | 6.3150 | 5.4193 | 6.2224 | 6.0979 | 5.9571 |
| SLC38A1_T17;S25 | SLC38A1 | Q9H2H9 | 52;56 | | NA | 5.8085 | 6.0963 | 6.6467 | 6.7296 | 5.5168 | 5.6421 | 6.1686 | 5.8917 | 6.7432 | 6.7484 | 7.4644 | 7.7088 | 5.4428 | 5.8173 | 5.5666 | 6.0081 |
| SLC38A2_S10 | SLC38A2 | Q96QD8 | | 10 | AEMGRFSISPDED | 5.8887 | 6.0219 | 6.2848 | 6.2214 | 6.2343 | 5.9871 | 5.6856 | 5.8336 | 6.9940 | 7.2535 | 6.7660 | 6.6089 | 5.9277 | 6.8167 | 5.6313 | 5.8444 |
| SLC38A2_Y10;S28 | SLC38A2 | Q96QD8 | | 41 | AALKSHYADVDPE | 6.1974 | 6.6315 | 7.0965 | 7.5747 | 6.8456 | 6.5592 | 6.2608 | 6.5560 | 7.1496 | 5.9677 | 5.8413 | 6.2503 | 5.3413 | 5.3491 | 5.0979 | 5.2811 |
| SLC38A2_Y28 | SLC38A2 | Q96QD8 | | 28 | SNSDFNYSYPTKQ | 5.8194 | 5.8990 | 6.0340 | 6.4639 | 5.5034 | 6.0365 | 5.7373 | 5.9932 | 6.2086 | 5.7968 | 6.3296 | 5.9655 | 6.8003 | 7.1979 | 6.9456 | 7.2691 |
| SLC38A2_Y41 | SLC38A2 | Q96QD8 | 10;28 | | NA | 5.6481 | 6.2958 | 6.0286 | 6.5891 | 5.8804 | 5.8403 | 5.9610 | 5.9984 | 6.2084 | 5.6105 | 5.6873 | 5.5896 | 7.3462 | 6.8767 | 7.2782 | 7.1613 |
| SLC39A10_S591 | SLC39A10 | Q9ULF5 | | 556 | LNNTPDSDWLQLK | 8.7115 | 7.1525 | 4.2838 | 6.2506 | 6.9126 | 7.1477 | 6.2794 | 6.1547 | 6.5528 | 5.3727 | 6.1882 | 5.8908 | 5.5742 | 5.2096 | 5.7901 | 6.5288 |
| SLC39A10_T553;S556 | SLC39A10 | Q9ULF5 | | 591 | LEGQQESPPKNYL | 6.0203 | 6.6815 | 6.8019 | 6.6457 | 5.8533 | 6.2089 | 6.1931 | 6.7457 | 6.2485 | 6.4954 | 6.0923 | 6.6653 | 5.5776 | 6.0403 | 5.8071 | 5.9232 |
| SLC39A10_T556 | SLC39A10 | Q9ULF5 | 553;556 | | NA | 5.8049 | 8.8863 | 7.6666 | 8.7234 | 5.3173 | 6.8523 | 6.8115 | 6.0761 | 8.4743 | 7.5016 | 7.6771 | 7.6414 | 2.7345 | 4.5624 | 2.8427 | 2.4276 |
| SLC39A14_S309 | SLC39A14 | Q15043 | | 309 | EKVIVGSLSVQDL | 5.7867 | 6.3955 | 6.2004 | 6.5265 | 6.1675 | 6.8414 | 6.5936 | 5.8559 | 5.9884 | 5.9990 | 6.1250 | 6.5800 | 6.3538 | 6.3547 | 6.0333 | 6.1983 |
| SLC39A3_S125;S129 | SLC39A3 | Q9BRY0 | | 129 | AGSDVGSDSEYES | 6.1722 | 6.6691 | 6.5578 | 6.9022 | 5.9994 | 6.0407 | 6.1504 | 5.9407 | 7.0001 | 6.9470 | 7.3440 | 7.5443 | 4.9409 | 5.4129 | 4.9929 | 5.3855 |
| SLC39A3_S129 | SLC39A3 | Q9BRY0 | 125;129 | | NA | 5.7279 | 5.9762 | 5.6606 | 6.8339 | 6.0530 | 5.9886 | 6.7414 | 5.8260 | 7.5011 | 6.8623 | 6.7352 | 7.3119 | 5.4454 | 5.8837 | 5.7106 | 5.7421 |
| SLC39A6_S478 | SLC39A6 | Q13433 | | 478 | KYESQLSTNEEKV | 5.1205 | 5.2971 | 5.4680 | 5.4055 | 6.1594 | 5.7914 | 6.0709 | 5.9838 | 6.1431 | 5.5631 | 5.5642 | 5.3417 | 8.1854 | 7.0260 | 8.5558 | 8.3242 |
| SLC39A7_S275;S276 | SLC39A7 | Q92504 | | 293 | QKRRGGSTVPKDG | 5.3039 | 5.0357 | 4.8795 | 5.3329 | 5.8381 | 5.3908 | 5.8153 | 5.7482 | 6.2913 | 6.3037 | 5.8860 | 5.9023 | 8.1778 | 8.3159 | 8.3054 | 7.4732 |
| SLC39A7_S293 | SLC39A7 | Q92504 | 275;276 | | NA | 5.5621 | 6.3621 | 7.0392 | 7.0696 | 4.7441 | 5.0840 | 6.6442 | 5.6465 | 6.7035 | 6.0046 | 5.9528 | 6.1062 | 7.5464 | 6.5807 | 5.9153 | 7.0387 |
| SLC39A8_S275;S278 | SLC39A8 | Q9C0K1 | 275;278 | | NA | 5.8552 | 6.3742 | 6.9324 | 6.5383 | 5.8598 | 6.1694 | 5.9984 | 6.4756 | 6.2287 | 5.7621 | 6.3416 | 6.1415 | 6.2100 | 6.4085 | 6.1745 | 6.5299 |
| SLC3A2_S135 | SLC3A2 | J3KPF3 | | 135 | ASGAAMSLAGAEK | 6.8591 | 6.3002 | 6.3408 | 7.5093 | 6.2020 | 6.5134 | 6.0970 | 6.5023 | 6.3060 | 6.7477 | 6.0453 | 5.8036 | 5.4569 | 6.5345 | 6.1187 | 4.6631 |
| SLC3A2_S37 | SLC3A2 | J3KPF3 | | 37 | LSAGDDSETGSDC | 6.4081 | 5.9437 | 6.3022 | 6.3893 | 5.7367 | 6.2705 | 5.7149 | 6.0358 | 6.3269 | 5.8219 | 6.6280 | 6.2200 | 6.8904 | 5.7278 | 6.9587 | 6.6250 |
| SLC3A2_S37;S41 | SLC3A2 | J3KPF3 | | 41 | DDSETGSDCVTQA | 5.9193 | 6.1130 | 5.6023 | 6.5939 | 6.4191 | 6.4223 | 6.2724 | 6.1796 | 6.5128 | 5.7481 | 6.1284 | 6.4336 | 6.3779 | 5.9175 | 6.8614 | 6.4983 |
| SLC3A2_S41 | SLC3A2 | J3KPF3 | 37;41 | | NA | 6.1955 | 5.5520 | 6.2187 | 6.6370 | 5.7415 | 6.0257 | 6.6891 | 5.8655 | 6.3619 | 5.9965 | 5.8493 | 6.0614 | 6.4532 | 7.2060 | 6.7542 | 6.3923 |
| SLC41A3_S13 | SLC41A3 | Q96GZ6 | | 13 | RQRRLDSCGKPGE | 5.3032 | 4.4193 | 4.4489 | 4.7138 | 5.7477 | 4.8051 | 4.3938 | 5.1004 | 7.1102 | 6.7266 | 6.0783 | 6.7661 | 7.8977 | 8.8979 | 9.2978 | 8.2932 |
| SLC43A2_S274 | SLC43A2 | Q8N370-3 | | 274 | TVGRRLSVGSSMR | 6.1892 | 6.3862 | 6.0489 | 6.4355 | 5.7281 | 6.0294 | 5.9919 | 6.1296 | 7.1749 | 7.4041 | 6.8550 | 7.0834 | 5.5443 | 5.5508 | 5.5045 | 5.9442 |
| SLC44A2_T12 | SLC44A2 | Q8IWA5-3 | | 12 | KNGAYGTPQKYDP | 6.4077 | 6.8306 | 6.4077 | 7.1433 | 6.7322 | 6.5154 | 7.0285 | 5.8380 | 6.1068 | 6.6683 | 6.6093 | 6.2483 | 5.4177 | 5.4019 | 4.9879 | 5.6564 |
| SLC45A4_S238 | SLC45A4 | Q5BKX6-3 | | 238 | FPDEVQSEHELAL | 5.9871 | 7.0782 | 6.4093 | 6.0685 | 6.1515 | 5.7389 | 6.3122 | 5.8264 | 6.2472 | 6.1978 | 6.8929 | 6.0297 | 6.2860 | 6.8675 | 5.8301 | 6.0768 |
| SLC45A4_S338 | SLC45A4 | Q5BKX6-3 | | 338 | VPNGSGSPTKDAL | 6.0950 | 6.0872 | 6.6710 | 7.7487 | 7.3350 | 5.9758 | 6.3472 | 6.0285 | 6.5668 | 6.6717 | 7.4321 | 6.1204 | 4.8713 | 5.3913 | 4.7247 | 5.9334 |
| SLC45A4_S360 | SLC45A4 | Q5BKX6-3 | | 403 | LIKPSRSMSDLYD | 4.9483 | 7.8635 | 6.4037 | 4.7871 | 5.7722 | 5.0468 | 8.8666 | 4.3706 | 6.3051 | 7.8725 | 6.1887 | 5.6469 | 6.8515 | 6.1439 | 5.6200 | 7.3127 |
| SLC45A4_S373 | SLC45A4 | Q5BKX6-3 | | 373 | AFRRQASSTFSYY | 6.1045 | 6.6490 | 7.0467 | 6.0471 | 5.1986 | 6.7923 | 6.2714 | 6.4241 | 6.2283 | 7.7318 | 7.4184 | 7.7110 | 4.7890 | 5.4567 | 4.4484 | 5.6826 |
| SLC45A4_S403 | SLC45A4 | Q5BKX6-3 | | 296 | PATPRSTSQELAK | 5.9114 | 6.4293 | 6.3564 | 6.4601 | 6.1280 | 6.0503 | 6.1428 | 5.9213 | 6.3825 | 6.0136 | 6.5588 | 6.8527 | 6.0927 | 6.1759 | 6.1205 | 6.4039 |
| SLC45A4_T296 | SLC45A4 | Q5BKX6-3 | | 360 | PSATSSSMRRRRH | 5.7504 | 6.4164 | 7.1738 | 7.5335 | 5.6958 | 6.6018 | 6.1931 | 6.4799 | 6.8452 | 7.5697 | 7.7758 | 8.4596 | 4.3866 | 5.1707 | 3.8620 | 4.0858 |
| SLC49A4_S26 | SLC49A4 | Q96SL1 | | 26 | GPGLGASWRSREA | 6.2454 | 6.6064 | 5.9097 | 6.7726 | 6.2129 | 5.8771 | 5.9536 | 6.4695 | 6.2095 | 6.5333 | 6.3724 | 6.2075 | 5.9037 | 6.4037 | 6.5067 | 5.8159 |
| SLC4A1AP_S312 | SLC4A1AP | Q9BWU0 | | 82 | KPALPVSPAARSK | 5.8564 | 5.9686 | 6.3121 | 6.1322 | 6.6265 | 6.8563 | 6.4753 | 6.4710 | 6.7995 | 6.7363 | 6.9270 | 6.8923 | 5.3251 | 5.7958 | 5.1728 | 5.6527 |
| SLC4A1AP_S466 | SLC4A1AP | Q9BWU0 | | 530 | SQVLSESPSQDSL | 5.9516 | 6.8459 | 6.5671 | 6.5515 | 6.5463 | 6.8882 | 6.7324 | 6.8053 | 6.8911 | 6.9342 | 7.1074 | 6.7734 | 4.5827 | 5.3112 | 4.6477 | 4.8640 |
| SLC4A1AP_S530 | SLC4A1AP | Q9BWU0 | | 466 | DEDFYDSDDDTFL | 5.8029 | 5.7746 | 6.1699 | 6.1407 | 5.8010 | 6.4104 | 6.3351 | 6.2346 | 6.1651 | 6.2413 | 6.4429 | 6.3649 | 6.3179 | 6.8073 | 6.4940 | 6.4974 |
| SLC4A1AP_S712 | SLC4A1AP | Q9BWU0 | | 312 | KMLGEDSDEEEEM | 5.2809 | 5.4941 | 6.5233 | 6.3551 | 6.0453 | 6.0287 | 6.0561 | 5.3459 | 7.1819 | 6.7273 | 6.6449 | 6.8997 | 6.1244 | 6.6815 | 6.1910 | 6.4200 |
| SLC4A1AP_S82 | SLC4A1AP | Q9BWU0 | | 712 | ENMSQLSEEEQNK | 5.7977 | 6.4470 | 6.6722 | 6.6663 | 5.9063 | 6.5076 | 6.8397 | 6.4658 | 6.7666 | 6.8807 | 6.6718 | 6.5999 | 5.4053 | 5.7806 | 5.3209 | 5.2716 |
| SLC4A2_S144 | SLC4A2 | P04920 | | 173 | ERTSPSSPAPLPH | 6.3878 | 6.9868 | 6.9949 | 6.4929 | 6.3303 | 6.0828 | 6.7775 | 6.1053 | 6.8117 | 6.9963 | 6.8293 | 6.6716 | 4.9732 | 5.1281 | 5.1598 | 5.2717 |
| SLC4A2_S173 | SLC4A2 | P04920 | | 144 | RALTQPSPVSTPS | 5.9351 | 5.9698 | 5.9947 | 6.4845 | 6.0896 | 6.0200 | 5.9101 | 6.1254 | 6.1811 | 5.7224 | 5.9964 | 6.0530 | 6.9000 | 6.8317 | 7.1528 | 6.6333 |
| SLC4A2_S243 | SLC4A2 | P04920 | | 243 | ERRRIGSMTGAEQ | 6.1656 | 5.5196 | 6.1067 | 6.0508 | 5.6286 | 6.0052 | 6.2248 | 5.5230 | 7.2103 | 7.4055 | 6.8561 | 7.2706 | 5.9924 | 6.5335 | 5.3023 | 6.2048 |
| SLC4A2_S443;S449 | SLC4A2 | P04920 | | 141 | EGARALTQPSPVS | 6.1565 | 6.0685 | 6.6880 | 6.7083 | 6.1347 | 5.9276 | 6.5560 | 6.5169 | 6.6055 | 5.3871 | 5.9241 | 6.2939 | 6.5970 | 6.0596 | 5.8568 | 6.5194 |
| SLC4A2_T141 | SLC4A2 | P04920 | | 257 | LLPRVPTDEIEAQ | 2.9384 | 2.3822 | 2.8628 | 2.9411 | 3.5135 | 3.2705 | 3.1886 | 3.6188 | 2.7089 | 2.8970 | 3.3479 | 3.6647 | 14.3266 | 13.1548 | 18.7382 | 16.4460 |
| SLC4A2_T257 | SLC4A2 | P04920 | 443;449 | | NA | 4.6489 | 5.6431 | 7.4336 | 7.8730 | 5.7486 | 5.8991 | 6.6877 | 5.8836 | 6.6073 | 6.0322 | 6.1750 | 6.0842 | 5.0495 | 6.9679 | 6.3252 | 6.9411 |
| SLC4A4_S1069 | SLC4A4 | Q9Y6R1 | | 1069 | PSDRERSPTFLER | 6.7605 | 7.2518 | 6.9148 | 8.0135 | 5.8570 | 6.0582 | 6.7730 | 6.5486 | 6.9414 | 6.5832 | 6.5965 | 6.8909 | 3.9301 | 5.4278 | 4.3858 | 5.0669 |
| SLC4A7_S1061 | SLC4A7 | Q9Y6M7-13 | | 1134 | YVDAETSLxxxxx | 6.3463 | 6.2339 | 6.0939 | 6.4381 | 6.3916 | 6.5576 | 6.8583 | 6.3058 | 6.3302 | 6.0793 | 6.3998 | 5.9813 | 6.0220 | 6.4905 | 5.9408 | 5.5306 |
| SLC4A7_S1082 | SLC4A7 | Q9Y6M7-13 | | 1061 | LPFEGGSLLQIPV | 4.6237 | 4.7488 | 4.4755 | 5.1626 | 6.0687 | 5.5647 | 6.0370 | 5.7990 | 4.7250 | 4.3176 | 4.5440 | 4.3449 | 10.2798 | 9.8126 | 10.4868 | 9.0094 |
| SLC4A7_S1095 | SLC4A7 | Q9Y6M7-13 | | 242 | RIPLVRSFADIGK | 4.9041 | 5.1274 | 4.6976 | 4.7237 | 6.8739 | 6.4261 | 6.8646 | 6.3695 | 4.6333 | 4.6120 | 5.0952 | 5.1806 | 8.6158 | 8.6551 | 8.6307 | 8.5904 |
| SLC4A7_S1109 | SLC4A7 | Q9Y6M7-13 | | 267 | ILASPQSAPGNLD | 3.1535 | 3.2605 | 3.5722 | 3.4774 | 6.3735 | 6.2265 | 6.9538 | 6.0530 | 4.4105 | 3.9681 | 4.3882 | 4.4851 | 11.3320 | 10.7433 | 10.8086 | 10.7936 |
| SLC4A7_S1115 | SLC4A7 | Q9Y6M7-13 | | 251 | DIGKKHSDPHLLE | 4.6056 | 5.6693 | 5.0956 | 5.5819 | 5.6701 | 5.9685 | 6.1315 | 6.2108 | 4.8800 | 5.6576 | 5.8743 | 6.0812 | 7.3422 | 8.0577 | 8.3318 | 8.8418 |
| SLC4A7_S1119 | SLC4A7 | Q9Y6M7-13 | | 441 | PVFHNGSTPTLGE | 3.8780 | 4.2986 | 4.6066 | 4.4921 | 5.3207 | 5.5772 | 5.8740 | 5.6806 | 4.3812 | 4.6074 | 4.6897 | 4.6172 | 10.5403 | 10.2052 | 10.9146 | 10.3167 |
| SLC4A7_S1134 | SLC4A7 | Q9Y6M7-13 | | 159 | DGGDRWSKPYVAT | 5.6547 | 6.0000 | 6.6851 | 6.7381 | 6.3763 | 5.9667 | 6.8699 | 6.1141 | 6.3270 | 6.2269 | 6.1311 | 6.4867 | 6.2101 | 5.8363 | 6.0774 | 6.2994 |
| SLC4A7_S159 | SLC4A7 | Q9Y6M7-13 | | 1115 | SPDKPVSVKISFE | 6.5137 | 5.6696 | 7.0590 | 6.9516 | 6.3623 | 6.0418 | 6.1954 | 6.2120 | 6.6014 | 6.4299 | 6.2370 | 6.2981 | 5.9830 | 6.6435 | 5.1230 | 5.6786 |
| SLC4A7_S242 | SLC4A7 | Q9Y6M7-13 | | 1052 | LQDDDDTVHLPFE | 5.3968 | 5.6221 | 5.6059 | 5.7462 | 6.3217 | 5.3390 | 5.4425 | 6.1783 | 5.0583 | 4.7694 | 5.1711 | 4.6222 | 8.7129 | 8.5121 | 9.3118 | 8.1898 |
| SLC4A7_S251 | SLC4A7 | Q9Y6M7-13 | | 288 | GNGSGGSRENSTV | 7.9312 | 8.7988 | 7.7103 | 7.6261 | 6.2156 | 6.5107 | 6.3168 | 6.2619 | 7.3472 | 7.7961 | 7.3048 | 7.4038 | 2.6922 | 3.5294 | 3.2035 | 3.3518 |
| SLC4A7_S264;S267 | SLC4A7 | Q9Y6M7-13 | | 1109 | VTRSNMSPDKPVS | 6.7827 | 6.8532 | 6.6056 | 7.5334 | 6.6826 | 7.0541 | 7.1779 | 6.6185 | 7.0414 | 6.7353 | 7.1403 | 7.2539 | 3.6695 | 4.2983 | 3.8947 | 4.6588 |
| SLC4A7_S267 | SLC4A7 | Q9Y6M7-13 | | 442 | VFHNGSTPTLGET | 6.3206 | 6.7680 | 6.2678 | 7.3420 | 6.7019 | 7.2741 | 7.0628 | 7.6163 | 6.6077 | 6.6786 | 6.8885 | 6.6607 | 4.4472 | 4.5505 | 4.2381 | 4.5752 |
| SLC4A7_S288 | SLC4A7 | Q9Y6M7-13 | | 18 | PLLTRVTSRGPDE | 4.6111 | 4.7197 | 4.5811 | 5.1639 | 6.2268 | 5.2178 | 5.4905 | 5.5167 | 5.8013 | 4.3100 | 4.4328 | 5.5789 | 9.3668 | 9.4454 | 9.4866 | 10.0506 |
| SLC4A7_S288;S292 | SLC4A7 | Q9Y6M7-13 | | 1119 | PVSVKISFEDEPR | 2.1132 | 2.0569 | 2.1313 | 2.1346 | 2.6949 | 2.5157 | 2.7344 | 2.7068 | 2.2103 | 2.0157 | 1.9202 | 2.1983 | 18.1798 | 16.0804 | 20.4014 | 17.9061 |
| SLC4A7_S288;S292;T293 | SLC4A7 | Q9Y6M7-13 | | 53 | ESHRAVYIGVHVP | 6.3301 | 7.0214 | 6.7619 | 4.6145 | 6.6767 | 6.7484 | 6.2527 | 6.3960 | 7.3274 | 5.9777 | 6.5276 | 7.0621 | 5.5807 | 6.1268 | 5.3639 | 5.2322 |
| SLC4A7_S292 | SLC4A7 | Q9Y6M7-13 | | 1095 | AQWKALSMNTENA | 7.7971 | 7.0975 | 8.4162 | 6.8680 | 5.9931 | 6.1136 | 5.5040 | 6.4617 | 7.2061 | 6.6894 | 8.8087 | 7.3137 | 3.8757 | 4.8472 | 3.5547 | 3.4533 |
| SLC4A7_S441 | SLC4A7 | Q9Y6M7-13 | | 292 | GGSRENSTVDFSK | 5.4435 | 5.4394 | 5.1731 | 6.0324 | 6.1053 | 5.6134 | 6.1395 | 6.1780 | 5.0567 | 4.8604 | 5.3458 | 5.2245 | 8.3396 | 8.5359 | 8.1066 | 8.4059 |
| SLC4A7_S441;T442 | SLC4A7 | Q9Y6M7-13 | | 1082 | PSIVNISDEMAKT | 5.3329 | 5.8923 | 5.5678 | 5.6056 | 5.7463 | 5.5933 | 6.4262 | 5.9917 | 6.1553 | 5.0690 | 5.7208 | 6.1641 | 7.7500 | 7.8090 | 7.9330 | 7.2425 |
| SLC4A7_S441;T444 | SLC4A7 | Q9Y6M7-13 | | 93 | KEDGRESPSYDTP | 5.6425 | 5.8794 | 6.0927 | 5.7016 | 6.0240 | 6.9164 | 6.6742 | 6.6963 | 6.5214 | 6.3450 | 6.8092 | 5.6929 | 7.2306 | 6.5881 | 6.2796 | 4.9063 |
| SLC4A7_S93 | SLC4A7 | Q9Y6M7-13 | 441;442 | | NA | 4.9422 | 5.4501 | 5.2995 | 5.8007 | 5.9373 | 6.3686 | 6.4407 | 6.4039 | 6.0969 | 5.6591 | 6.0501 | 5.9404 | 7.1941 | 7.4710 | 7.0228 | 7.9227 |
| SLC4A7_T1052 | SLC4A7 | Q9Y6M7-13 | 288;292;293 | | NA | 3.7881 | 3.3868 | 1.5707 | 3.3656 | 6.0431 | 5.3689 | 7.0169 | 7.5453 | 2.8888 | 4.8247 | 2.4419 | 3.1101 | 13.1902 | 12.2615 | 13.1480 | 10.0494 |
| SLC4A7_T18 | SLC4A7 | Q9Y6M7-13 | 441;444 | | NA | 3.5141 | 4.1436 | 3.8777 | 4.5931 | 4.9224 | 5.4362 | 5.3087 | 5.2898 | 4.2576 | 4.0056 | 4.3042 | 4.3363 | 10.2741 | 11.4262 | 12.1545 | 12.1560 |
| SLC4A7_T442 | SLC4A7 | Q9Y6M7-13 | 264;267 | | NA | 6.1137 | 6.4858 | 7.3233 | 7.0252 | 6.4951 | 6.7747 | 6.6930 | 6.8620 | 7.2290 | 7.9092 | 7.8129 | 7.1387 | 3.6906 | 4.5414 | 3.5389 | 4.3666 |
| SLC4A7_Y53 | SLC4A7 | Q9Y6M7-13 | 288;292 | | NA | 4.5357 | 4.2751 | 3.7150 | 4.8210 | 6.7578 | 6.3608 | 6.3318 | 6.0156 | 4.2922 | 4.2402 | 4.7060 | 5.0059 | 9.8574 | 9.8942 | 9.1809 | 10.0102 |
| SLC4A8_S1050 | SLC4A8 | Q2Y0W8 | | 237 | LIPIVRSFAEVGK | 2.3989 | 2.7805 | 2.5411 | 2.7219 | 4.9957 | 5.6191 | 5.6612 | 5.0916 | 2.6984 | 2.6481 | 3.3499 | 3.5330 | 13.0178 | 13.4605 | 14.3858 | 15.0963 |
| SLC4A8_S1080 | SLC4A8 | Q2Y0W8 | | 1080 | TVWKALSMNSGNA | 6.6326 | 6.5976 | 7.0598 | 6.7431 | 6.6842 | 5.9927 | 5.6270 | 6.3016 | 6.6752 | 7.0711 | 6.1922 | 6.5838 | 5.4804 | 6.4867 | 4.9445 | 4.9275 |
| SLC4A8_S237 | SLC4A8 | Q2Y0W8 | | 1050 | SRKLLSSPGKNIS | 6.5151 | 6.8039 | 6.7263 | 7.2777 | 6.1555 | 6.3718 | 6.9321 | 6.6676 | 6.9344 | 6.9030 | 6.8832 | 7.3400 | 3.8836 | 5.0234 | 4.3196 | 5.2628 |
| SLC66A2_S110 | SLC66A2 | Q8N2U9 | | 110 | LNARRRSFTAADS | 4.8584 | 4.7930 | 4.7740 | 5.1031 | 5.3989 | 4.8274 | 5.3047 | 4.7303 | 6.2398 | 6.0655 | 6.1674 | 6.0548 | 8.7056 | 9.0516 | 9.3333 | 8.5924 |
| SLC6A15_S675 | SLC6A15 | Q9H2J7 | | 687 | IPSEMPSPNFGKN | 6.1377 | 6.6374 | 6.5868 | 6.9379 | 6.1779 | 6.6287 | 6.5295 | 6.6166 | 6.6184 | 6.6089 | 6.7259 | 6.7425 | 4.9880 | 5.5436 | 5.1358 | 5.3844 |
| SLC6A15_S683;S687 | SLC6A15 | Q9H2J7 | | 675 | LEGDDTSLIHGKI | 5.8614 | 6.2067 | 6.3808 | 6.5032 | 6.0332 | 6.2469 | 6.6318 | 6.2536 | 6.1501 | 6.6069 | 6.4215 | 6.5198 | 6.2392 | 6.1303 | 5.6014 | 6.2132 |
| SLC6A15_S687 | SLC6A15 | Q9H2J7 | 683;687 | | NA | 6.3611 | 6.8359 | 8.5707 | 7.4495 | 6.1887 | 5.6952 | 7.6688 | 5.9818 | 7.2526 | 7.3296 | 5.9290 | 6.5774 | 5.6574 | 4.9912 | 3.4670 | 4.0440 |
| SLC7A11_S26 | SLC7A11 | Q9UPY5 | | 26 | VNGRLPSLGNKEP | 7.1849 | 6.8933 | 6.9082 | 7.2577 | 6.6649 | 5.4815 | 5.8596 | 6.0743 | 7.4341 | 6.7948 | 5.6927 | 6.4617 | 5.4824 | 5.7889 | 4.8463 | 5.1750 |
| SLC7A2_S686 | SLC7A2 | P52569-3 | | 687 | HPRNLSSPFIFHE | 7.3015 | 7.2070 | 7.2736 | 7.5210 | 6.3868 | 7.4784 | 8.1637 | 7.2087 | 7.2116 | 7.0748 | 7.4204 | 7.5417 | 2.8394 | 3.5698 | 2.5151 | 3.2866 |
| SLC7A2_S687 | SLC7A2 | P52569-3 | | 686 | HHPRNLSSPFIFH | 5.5926 | 4.2119 | 6.1355 | 4.8507 | 5.3732 | 4.9966 | 4.6280 | 5.4295 | 7.4237 | 7.0726 | 7.0260 | 6.7853 | 8.4845 | 7.9344 | 7.2010 | 6.8548 |
| SLC7A5_S35 | SLC7A5 | Q01650 | | 35 | AKSADGSAPAGEG | 5.1950 | 6.3277 | 5.7824 | 5.8999 | 6.6460 | 7.0317 | 6.7779 | 5.9546 | 6.3196 | 5.4775 | 5.6488 | 6.0873 | 6.1214 | 7.0092 | 7.1527 | 6.5680 |
| SLC7A6_S20 | SLC7A6 | Q92536 | | 20 | HLVPNTSQSQVEE | 5.2193 | 4.2412 | 4.3510 | 5.0221 | 4.8686 | 4.8680 | 4.9938 | 5.8576 | 5.3849 | 4.8230 | 4.4319 | 4.6740 | 9.8387 | 9.9934 | 11.3498 | 10.0827 |
| SLC7A6_S36 | SLC7A6 | Q92536 | | 36 | SPPQRSSETMQLK | 5.9392 | 6.2598 | 6.3890 | 6.9406 | 5.4710 | 5.1210 | 6.4033 | 6.6385 | 6.2830 | 5.6747 | 6.6104 | 7.2283 | 5.7054 | 6.6282 | 5.8579 | 6.8497 |
| SLC7A6OS_S16 | SLC7A6OS | Q96CW6 | | 16 | RVKRKRSAEPAEA | 5.3649 | 5.1785 | 5.7020 | 5.5176 | 5.9639 | 5.8878 | 5.7304 | 5.5100 | 5.8256 | 5.9660 | 6.0884 | 6.1041 | 8.1636 | 7.3249 | 7.8914 | 7.7811 |

| | | | | | | | | | | | | | | | | | | | | | |
|---|---|---|---|---|---|---|---|---|---|---|---|---|---|---|---|---|---|---|---|---|---|
| SLC8A1_S285 | SLC8A1 | P32418 | | 392 | QARKAVSMHEVNT | 6.0440 | 6.3317 | 6.2045 | 6.3245 | 5.8219 | 6.6045 | 6.3776 | 5.6536 | 5.9429 | 5.8197 | 6.1590 | 5.8426 | 6.6322 | 7.0648 | 6.5231 | 6.6533 |
| SLC8A1_S392 | SLC8A1 | P32418 | | 285 | EGDRPSSKTEIEM | 6.0528 | 6.3656 | 6.6990 | 6.2270 | 6.1502 | 6.2643 | 6.0529 | 6.2682 | 6.4365 | 6.4215 | 6.6576 | 6.7030 | 5.8700 | 6.0699 | 5.8549 | 5.9068 |
| SLC9A1_S599 | SLC9A1 | P19634 | | 599 | GMGKIPSAVSTVS | 3.3457 | 3.5118 | 3.3857 | 3.5549 | 5.5984 | 6.3331 | 5.9321 | 6.3241 | 4.2039 | 4.5117 | 4.4416 | 4.5892 | 10.7991 | 10.3457 | 11.2362 | 11.8870 |
| SLC9A1_S599;S602 | SLC9A1 | P19634 | | 796 | RIQRCLSDPGPHP | 4.3204 | 4.4490 | 4.6454 | 4.7232 | 5.3686 | 5.3942 | 5.6469 | 5.4024 | 4.6714 | 4.4880 | 4.7886 | 4.5879 | 10.2895 | 9.4929 | 10.9625 | 10.7691 |
| SLC9A1_S599;S602;S605 | SLC9A1 | P19634 | | 693 | PAHKLDSPTMSRA | 6.4363 | 6.7257 | 6.1755 | 6.4393 | 5.9309 | 6.0489 | 6.2616 | 6.0109 | 6.4313 | 6.4835 | 5.9571 | 6.0587 | 6.0521 | 6.5133 | 6.1733 | 6.3015 |
| SLC9A1_S599;S605 | SLC9A1 | P19634 | | 602 | KIPSAVSTVSMQN | 6.2969 | 6.5705 | 6.4803 | 6.2524 | 6.8202 | 6.7947 | 6.7717 | 6.6449 | 6.3932 | 6.5010 | 7.1942 | 7.2184 | 4.8186 | 5.2590 | 4.7722 | 5.2117 |
| SLC9A1_S602 | SLC9A1 | P19634 | | 624 | RILPALSKDKEEE | 5.4899 | 5.1640 | 5.5686 | 6.1662 | 5.7449 | 5.9566 | 6.1883 | 7.0597 | 5.9910 | 5.1351 | 5.3003 | 5.3824 | 7.2825 | 8.5564 | 7.9743 | 7.0398 |
| SLC9A1_S602;S605 | SLC9A1 | P19634 | | 703 | SRARIGSDPLAYE | 5.6244 | 5.7877 | 6.0447 | 6.3243 | 5.7204 | 5.8885 | 6.8219 | 6.4020 | 5.9597 | 5.8685 | 6.0136 | 6.6600 | 6.9210 | 6.4197 | 6.4924 | 7.0512 |
| SLC9A1_S605 | SLC9A1 | P19634 | | 785 | TPAPSDSPSSQRI | 5.8032 | 6.0398 | 6.4421 | 6.3380 | 7.0871 | 6.0028 | 5.8510 | 6.0884 | 6.4228 | 6.9523 | 6.1970 | 6.2679 | 5.9280 | 6.1469 | 5.9781 | 6.4544 |
| SLC9A1_S624 | SLC9A1 | P19634 | | 718 | EDLPVITIDPASP | 5.8898 | 6.3573 | 6.3871 | 7.1510 | 6.1337 | 6.6518 | 6.4463 | 6.1730 | 6.5556 | 6.4653 | 7.0986 | 6.7955 | 5.0437 | 5.5961 | 5.3242 | 5.9309 |
| SLC9A1_S693 | SLC9A1 | P19634 | | 605 | SAVSTVSMQNIHP | 5.9465 | 6.3123 | 5.8784 | 6.4828 | 7.3798 | 8.0390 | 7.5923 | 7.0882 | 6.6489 | 6.2614 | 7.4337 | 6.8406 | 4.3240 | 5.1713 | 3.9377 | 4.6630 |
| SLC9A1_S703 | SLC9A1 | P19634 | 602;605 | | NA | 6.9678 | 6.7539 | 6.7492 | 7.1101 | 7.0279 | 6.3462 | 6.4522 | 5.9930 | 6.2737 | 5.8981 | 7.2861 | 7.2928 | 4.9497 | 5.2468 | 4.8372 | 4.8153 |
| SLC9A1_S785 | SLC9A1 | P19634 | 599;602 | | NA | 3.7426 | 3.8832 | 4.1431 | 3.3488 | 5.4785 | 6.1919 | 6.1940 | 6.4295 | 4.1142 | 4.6156 | 3.9941 | 4.4498 | 10.5687 | 10.2182 | 11.9085 | 10.7192 |
| SLC9A1_S796 | SLC9A1 | P19634 | 599;602;605 | | NA | 3.8501 | 3.4983 | 3.5667 | 4.2534 | 6.4284 | 6.1126 | 6.3275 | 5.7512 | 3.8744 | 3.4480 | 3.3601 | 3.2902 | 11.7928 | 11.5251 | 12.3425 | 10.5787 |
| SLC9A1_T718 | SLC9A1 | P19634 | 599;605 | | NA | 3.4132 | 3.8661 | 4.0335 | 3.2943 | 5.8311 | 5.8754 | 5.6959 | 6.3837 | 4.5283 | 3.2919 | 3.8194 | 3.5911 | 11.8710 | 11.5365 | 11.8264 | 11.1423 |
| SLC9A3R1_S269 | SLC9A3R1 | O14745 | | 280 | AEAALESPRPALV | 6.7319 | 6.7291 | 7.1573 | 6.7446 | 6.7246 | 7.1570 | 6.8031 | 7.3113 | 7.1536 | 7.5567 | 7.5527 | 7.5256 | 3.8654 | 4.3813 | 3.1405 | 3.4654 |
| SLC9A3R1_S280 | SLC9A3R1 | O14745 | | 269 | EIQKENSREALAE | 5.2748 | 5.1793 | 5.7219 | 5.0014 | 5.5626 | 6.5710 | 5.3414 | 4.6586 | 6.4185 | 5.9679 | 5.4149 | 5.8724 | 9.4271 | 8.2896 | 7.8891 | 7.4097 |
| SLC9A3R1_S290;S302 | SLC9A3R1 | O14745 | 290;302 | | NA | 5.8853 | 5.7401 | 5.7430 | 6.5113 | 5.5687 | 6.4643 | 7.1215 | 6.4219 | 5.7897 | 7.9446 | 6.5524 | 7.7583 | 5.1582 | 5.6413 | 6.1194 | 5.5800 |
| SLC9A3R2_S261 | SLC9A3R2 | Q15599 | | 280 | RSDLPGSDKDTED | 5.3081 | 5.9685 | 6.0006 | 6.2621 | 6.6201 | 6.8583 | 6.8510 | 6.3308 | 5.5725 | 5.0217 | 5.6383 | 6.1657 | 7.3433 | 6.1409 | 7.7144 | 6.2037 |
| SLC9A3R2_S280 | SLC9A3R2 | Q15599 | | 303 | ESGLHLSPTAAEA | 7.2230 | 6.7505 | 6.9998 | 7.3552 | 6.8777 | 7.0932 | 6.4243 | 7.4643 | 7.6997 | 6.7450 | 7.2411 | 7.0763 | 3.3888 | 4.5784 | 3.4298 | 3.6529 |
| SLC9A3R2_S303 | SLC9A3R2 | Q15599 | | 261 | PVTNGTSPAQLNG | 6.4701 | 6.6349 | 6.5004 | 6.5937 | 6.3465 | 7.5499 | 7.4675 | 6.4592 | 6.7454 | 7.0208 | 7.0468 | 5.3122 | 5.4575 | 4.2151 | 4.9241 | 5.2559 |
| SLC9A7_S545 | SLC9A7 | Q96T83 | | 545 | VGVEEPSEEDQNE | 6.5713 | 5.9811 | 6.3527 | 5.6266 | 5.9412 | 6.2353 | 6.1740 | 6.7657 | 5.9007 | 6.0680 | 7.3701 | 6.8575 | 6.0801 | 6.1997 | 5.8530 | 6.0231 |
| SLCO4A1_S40 | SLCO4A1 | Q96BD0 | | 40 | SPGTPLSPGSLRS | 5.7441 | 6.5552 | 6.3642 | 6.7033 | 5.1806 | 6.6682 | 6.4977 | 6.6744 | 6.4051 | 6.3135 | 7.5446 | 7.2019 | 5.0396 | 5.9728 | 5.2747 | 5.8601 |
| SLCO4C1_S16 | SLCO4C1 | Q6ZQN7 | | 16 | LAFVPSSPDILRR | 6.3127 | 6.4985 | 6.6389 | 6.8913 | 6.2619 | 6.6732 | 6.2828 | 6.4170 | 6.6165 | 6.7789 | 6.2802 | 6.8659 | 4.9774 | 5.8463 | 5.3034 | 5.3551 |
| SLF1_S159 | SLF1 | Q9BQI6 | | 159 | VILPKSSPSGITH | 5.5225 | 6.2118 | 7.2960 | 6.9476 | 7.5103 | 7.0026 | 5.7376 | 6.2990 | 6.1388 | 7.0303 | 5.8942 | 5.5663 | 5.3417 | 6.4081 | 5.8141 | 5.2791 |
| SLF2_S109 | SLF2 | Q8IX21-2 | | 109 | RVPPEKSPIIEAF | 6.8221 | 6.8225 | 6.6951 | 7.4631 | 6.8882 | 6.7834 | 6.9859 | 6.3642 | 6.8893 | 6.7317 | 6.7420 | 6.4451 | 4.2022 | 5.0321 | 4.1168 | 5.0163 |
| SLF2_S176 | SLF2 | Q8IX21-2 | | 444 | KPHLPLSQEKSAI | 5.4694 | 5.0076 | 5.9936 | 6.5695 | 6.0809 | 5.6728 | 5.8692 | 4.9905 | 5.9729 | 5.5486 | 5.6733 | 5.6680 | 8.4635 | 7.6368 | 7.5439 | 7.8395 |
| SLF2_S21 | SLF2 | Q8IX21-2 | | 21 | SPAPGSSPPRCHL | 6.1007 | 6.1963 | 6.0589 | 6.6155 | 6.0547 | 6.0896 | 6.4680 | 5.9706 | 7.1200 | 6.6246 | 6.3178 | 6.4400 | 6.1391 | 5.9346 | 5.7989 | 6.0708 |
| SLF2_S340 | SLF2 | Q8IX21-2 | | 68 | DRHMLDSPQKSNI | 6.2136 | 6.5184 | 6.3378 | 6.7583 | 5.9143 | 6.1871 | 5.9931 | 6.0606 | 6.5803 | 6.6080 | 6.2092 | 6.4373 | 5.8583 | 6.2164 | 5.9292 | 6.1781 |
| SLF2_S444 | SLF2 | Q8IX21-2 | | 555 | YGTPTKSPPAALE | 5.8415 | 6.0475 | 5.5481 | 6.8545 | 6.3992 | 5.9988 | 6.3433 | 5.7801 | 5.9554 | 6.5534 | 6.3628 | 6.4985 | 6.3676 | 6.4551 | 6.3600 | 6.6340 |
| SLF2_S555 | SLF2 | Q8IX21-2 | | 176 | SPERRKSLFIHEN | 5.9407 | 4.4516 | 4.8256 | 4.3150 | 6.3153 | 5.5936 | 5.5605 | 4.7262 | 10.3098 | 7.8567 | 7.5223 | 7.5279 | 7.7032 | 8.6668 | 3.6484 | 5.0364 |
| SLF2_S555;S568 | SLF2 | Q8IX21-2 | | 623 | EEETLKSLEEIMA | 5.1906 | 5.8808 | 6.9314 | 7.0565 | 6.3673 | 7.0733 | 5.9910 | 6.4570 | 6.4542 | 5.8014 | 6.1628 | 7.0211 | 5.9938 | 5.9807 | 6.0881 | 5.5502 |
| SLF2_S623 | SLF2 | Q8IX21-2 | | 340 | EKRKRNSVDSDLK | 4.0596 | 4.0559 | 4.3287 | 4.9937 | 5.7110 | 4.8884 | 5.6735 | 5.2926 | 4.9262 | 4.9303 | 4.7119 | 5.3301 | 9.8330 | 9.7023 | 10.8925 | 10.6704 |
| SLF2_S68 | SLF2 | Q8IX21-2 | | 551 | LRSEYGTPTKSPP | 7.0349 | 6.6933 | 6.0246 | 7.4563 | 6.4041 | 5.5929 | 6.1428 | 6.0651 | 6.6029 | 6.3591 | 6.0031 | 6.4129 | 5.9837 | 5.6006 | 5.6324 | 5.9913 |
| SLF2_T551 | SLF2 | Q8IX21-2 | 555;568 | | NA | 5.5877 | 6.3107 | 6.1732 | 5.8791 | 5.5102 | 7.0997 | 6.3580 | 5.5498 | 6.8489 | 6.6510 | 5.3694 | 6.3213 | 6.9037 | 6.9628 | 6.1297 | 6.3447 |
| SLIRP_S102 | SLIRP | Q9GZT3 | | 102 | PKLPQTSDDEKKD | 6.5152 | 6.2821 | 6.3899 | 7.0368 | 6.3226 | 5.7038 | 6.0945 | 6.0365 | 6.6614 | 6.4922 | 6.5620 | 7.1099 | 5.8303 | 5.5706 | 5.5523 | 5.8399 |
| SLK_S189 | SLK | Q9H2G2 | | 189 | TIQRRDSFIGTPY | 6.2076 | 6.2498 | 6.7410 | 6.4079 | 6.4682 | 6.3930 | 6.1855 | 6.1545 | 6.3247 | 6.3460 | 6.3681 | 6.2732 | 6.1268 | 6.4721 | 5.6052 | 5.6765 |
| SLK_S655 | SLK | Q9H2G2 | | 779 | KTKDSGSISLQET | 6.3576 | 5.9306 | 6.0081 | 6.3491 | 5.1979 | 6.3543 | 6.3118 | 6.0288 | 6.4091 | 4.8403 | 5.7584 | 5.8535 | 7.3227 | 7.3853 | 6.9197 | 6.9728 |
| SLK_S728 | SLK | Q9H2G2 | | 655 | EEMEVRSVVADTD | 6.4461 | 7.1261 | 6.1545 | 6.0548 | 5.9921 | 6.4810 | 6.6833 | 6.3555 | 6.7030 | 5.2260 | 5.6323 | 5.7000 | 6.4068 | 7.1779 | 6.0098 | 5.8507 |
| SLK_S779 | SLK | Q9H2G2 | | 1097 | RINSTATPDQDRD | 6.7331 | 8.0524 | 6.8400 | 7.3958 | 6.1782 | 7.3313 | 7.6480 | 6.7319 | 6.8697 | 7.4614 | 8.0239 | 7.2144 | 2.8307 | 4.0618 | 2.7818 | 3.8456 |
| SLK_T1097 | SLK | Q9H2G2 | | 728 | NKEEIGSLSKTET | 5.6582 | 6.1712 | 6.5079 | 5.9791 | 5.8721 | 6.3546 | 5.8990 | 6.2772 | 5.7025 | 5.6051 | 6.3746 | 6.0696 | 7.1179 | 6.7337 | 7.0764 | 6.6008 |
| SLMAP_S148 | SLMAP | Q14BN4 | | 452 | DFSDTLSPSKEKS | 6.2805 | 6.3874 | 6.0668 | 7.3064 | 6.7667 | 6.7024 | 7.2867 | 6.6317 | 6.5342 | 6.2053 | 7.1008 | 6.4157 | 4.7953 | 5.4906 | 4.7292 | 5.3003 |
| SLMAP_S402 | SLMAP | Q14BN4 | | 148 | IHAPLPSPVDKVA | 5.3729 | 5.6564 | 5.5845 | 6.7112 | 6.5219 | 6.1737 | 6.7533 | 8.8837 | 5.8027 | 6.2005 | 5.4762 | 5.7890 | 5.6690 | 6.1003 | 6.6723 | 6.6323 |
| SLMAP_S452 | SLMAP | Q14BN4 | | 402 | EHLLSKSGGDCTF | 7.6799 | 9.1862 | 8.1187 | 8.6721 | 6.0066 | 5.9277 | 5.3017 | 5.9052 | 6.9029 | 7.8919 | 7.8217 | 8.3380 | 2.5265 | 3.8638 | 2.8528 | 3.0042 |
| SLTM_S1002 | SLTM | Q9NWH9 | | 553 | KRISSKSPGHMVI | 6.3104 | 6.4033 | 5.9708 | 6.5878 | 6.3700 | 6.1542 | 6.4085 | 6.2181 | 6.2462 | 6.1275 | 6.0900 | 6.1490 | 6.0976 | 6.4385 | 6.3114 | 6.1167 |
| SLTM_S1014 | SLTM | Q9NWH9 | | 289 | QDAIAQSPEKESK | 6.1851 | 6.1900 | 6.0389 | 6.5447 | 6.0781 | 5.8108 | 6.2454 | 5.7811 | 6.1651 | 6.0593 | 6.0960 | 6.0781 | 6.4811 | 6.9631 | 6.7658 | 6.5175 |
| SLTM_S126 | SLTM | Q9NWH9 | | 590 | KEKERASLDKKRD | 5.9670 | 6.3994 | 5.7343 | 6.9489 | 5.8904 | 5.8445 | 6.3105 | 6.0813 | 5.7683 | 6.0966 | 6.1288 | 6.5131 | 5.9357 | 6.7766 | 6.7085 | 6.8961 |
| SLTM_S199 | SLTM | Q9NWH9 | | 1014 | VQISGNSMPRGSG | 6.6877 | 6.8103 | 6.5462 | 7.6247 | 6.0305 | 6.0682 | 6.7996 | 6.2965 | 6.6830 | 6.5993 | 6.6482 | 6.8689 | 4.7090 | 5.6665 | 4.6038 | 5.3575 |
| SLTM_S289 | SLTM | Q9NWH9 | | 344 | LKKGPSSTGASGQ | 6.7026 | 6.7102 | 5.7451 | 6.0896 | 6.5326 | 5.6830 | 6.5199 | 6.3382 | 6.1713 | 6.1341 | 5.6710 | 5.6035 | 6.0262 | 6.7175 | 6.5517 | 6.8035 |
| SLTM_S344 | SLTM | Q9NWH9 | | 126 | NDELKDSEEFGEN | 5.1557 | 5.9788 | 5.9859 | 6.1503 | 5.8302 | 5.3677 | 6.3024 | 5.2430 | 5.8277 | 6.3316 | 5.8115 | 5.8499 | 7.7495 | 7.1744 | 7.4105 | 7.8309 |
| SLTM_S371;T375 | SLTM | Q9NWH9 | | 93 | HEADELSGDASVE | 5.4004 | 6.5874 | 5.8607 | 6.8907 | 6.3088 | 6.1615 | 6.5341 | 6.5845 | 6.0138 | 6.3447 | 6.8239 | 5.8311 | 6.0898 | 6.5966 | 6.2769 | 5.6950 |
| SLTM_S551;S553 | SLTM | Q9NWH9 | | 1002 | QHSSNASPINRIV | 5.7909 | 8.0912 | 6.0862 | 6.6181 | 6.1791 | 7.3551 | 6.8013 | 7.3113 | 6.3167 | 6.2904 | 6.0161 | 6.2562 | 4.4930 | 5.9460 | 5.2887 | 5.1599 |
| SLTM_S553 | SLTM | Q9NWH9 | | 890 | KSEGSMSTDKRET | 6.7869 | 6.4317 | 6.1803 | 6.9326 | 6.0358 | 6.3944 | 6.6664 | 5.6721 | 6.7093 | 6.7392 | 6.5444 | 6.2594 | 5.0744 | 6.4055 | 5.4921 | 5.6754 |
| SLTM_S590 | SLTM | H7BXE3 | | 199 | ENEKEGSLAEADH | 6.5125 | 6.8200 | 6.0030 | 6.6800 | 6.4955 | 6.0445 | 6.3737 | 6.5365 | 6.0404 | 5.2879 | 5.9011 | 5.4542 | 6.3901 | 6.5734 | 6.4615 | 6.4258 |
| SLTM_S890 | SLTM | Q9NWH9 | 551;553 | | NA | 5.9863 | 6.1634 | 8.0883 | 5.3055 | 7.3075 | 9.6292 | 7.0315 | 6.3552 | 9.4746 | 4.7410 | 7.3808 | 6.2178 | 4.9906 | 4.4493 | 3.9144 | 2.9647 |
| SLTM_S93 | SLTM | Q9NWH9 | 371;375 | | NA | 6.3775 | 6.0957 | 6.3156 | 6.2256 | 5.9338 | 5.5291 | 5.9308 | 6.1808 | 5.7941 | 5.1093 | 5.3384 | 5.3186 | 6.6582 | 8.3128 | 7.8465 | 7.0332 |
| SLU7_S215 | SLU7 | O95391 | | 215 | LVEQANSPKHQWG | 6.1845 | 6.3652 | 6.1507 | 6.4737 | 6.2199 | 5.8494 | 6.3327 | 5.8393 | 6.5570 | 6.2573 | 5.9604 | 6.0138 | 6.2991 | 6.5682 | 6.5916 | 6.3373 |
| SLX4_S1028 | SLX4 | Q8IY92 | | 1028 | LAPWQASPPHPCR | 6.5866 | 6.3625 | 6.2155 | 7.1271 | 6.2517 | 6.0163 | 6.2925 | 6.2976 | 6.1744 | 6.4814 | 6.6101 | 6.4755 | 5.5455 | 5.9468 | 5.7341 | 5.8824 |
| SLX4_S1044 | SLX4 | Q8IY92 | | 1185 | KKALEISPRSCEL | 6.4721 | 6.9449 | 5.9589 | 7.1448 | 6.0168 | 5.8101 | 6.1696 | 6.4630 | 6.3894 | 6.5088 | 6.1184 | 6.3770 | 5.6272 | 6.1112 | 5.8742 | 6.0136 |
| SLX4_S1070 | SLX4 | Q8IY92 | | 1044 | GPPQGGSPRGSHH | 6.8820 | 6.9613 | 5.9249 | 7.2683 | 6.0662 | 6.5395 | 6.4531 | 6.6680 | 6.8203 | 7.0474 | 6.6540 | 6.7936 | 4.6091 | 5.2200 | 4.4270 | 5.6653 |
| SLX4_S1170 | SLX4 | Q8IY92 | | 467 | KKKPPVSPPLLLV | 5.6904 | 6.3291 | 6.2776 | 6.9168 | 5.4886 | 5.7579 | 5.7438 | 6.3575 | 6.6706 | 6.3485 | 6.4336 | 6.1410 | 6.0960 | 6.5148 | 6.6538 | 6.5799 |
| SLX4_S1185 | SLX4 | Q8IY92 | | 199 | LTTAVPSPSKPRT | 6.4498 | 6.1788 | 5.7640 | 6.7004 | 6.2001 | 6.2262 | 6.3956 | 6.5094 | 7.3132 | 6.0398 | 6.1569 | 6.4233 | 5.8598 | 6.3101 | 6.0159 | 5.4567 |
| SLX4_S1244 | SLX4 | Q8IY92 | | 1453 | GPLSTSSPSRRMN | 6.1628 | 6.6879 | 6.1833 | 6.7163 | 5.9960 | 6.0443 | 6.4240 | 5.8163 | 7.3759 | 6.8794 | 6.4587 | 6.7817 | 5.6440 | 5.6899 | 5.2667 | 5.8727 |
| SLX4_S1412;S1434 | SLX4 | Q8IY92 | | 1469 | DSRDCRSPGLLDT | 6.2678 | 6.4857 | 6.3458 | 6.5435 | 5.7809 | 6.0100 | 5.9522 | 5.9602 | 6.4496 | 5.9927 | 6.0333 | 6.1365 | 5.9288 | 6.6403 | 6.8336 | 6.6390 |
| SLX4_S1453 | SLX4 | Q8IY92 | | 1291 | QAVTQHTPRASVG | 7.2325 | 7.1912 | 7.1779 | 7.7210 | 5.9053 | 6.8947 | 7.5665 | 7.1358 | 7.6992 | 7.3964 | 6.9165 | 7.1561 | 3.4848 | 3.6830 | 3.1477 | 3.6916 |
| SLX4_S1469 | SLX4 | Q8IY92 | | 884 | DWLEGGSPVSGQL | 5.5018 | 6.6095 | 6.4870 | 6.8116 | 5.9084 | 6.7372 | 6.4959 | 5.6761 | 6.4208 | 6.7360 | 6.4759 | 6.1215 | 6.0264 | 6.1730 | 5.5748 | 6.2440 |
| SLX4_S199 | SLX4 | Q8IY92 | | 1070 | GTSQVGSPTLLSP | 5.3600 | 6.2381 | 5.2844 | 5.9358 | 7.3377 | 6.5247 | 7.4394 | 6.3606 | 6.9521 | 6.2746 | 6.9078 | 6.1730 | 5.2297 | 5.5971 | 6.0687 | 6.3163 |
| SLX4_S467 | SLX4 | Q8IY92 | | 1244 | FCDRESSPSEAST | 5.4741 | 6.4564 | 6.8919 | 6.1651 | 6.4607 | 5.9053 | 6.3108 | 6.5663 | 7.0347 | 6.8319 | 6.4792 | 6.7862 | 6.1623 | 5.9210 | 4.7631 | 5.7911 |
| SLX4_S884 | SLX4 | Q8IY92 | | 1170 | EQTKMKSISSDPL | 6.8186 | 6.0215 | 6.4463 | 7.0789 | 7.0211 | 6.6445 | 7.4326 | 6.1083 | 6.7433 | 6.5240 | 6.0968 | 6.8783 | 5.3497 | 4.7219 | 4.8472 | 5.2672 |
| SLX4_T1291 | SLX4 | Q8IY92 | 1315;1329 | | NA | 5.7731 | 5.7586 | 5.8777 | 6.7742 | 5.9684 | 6.9149 | 6.1465 | 6.0459 | 6.3422 | 6.2805 | 6.4640 | 6.9194 | 6.3949 | 6.2474 | 6.0467 | 6.0455 |
| SLX4_T1315;S1329 | SLX4 | Q8IY92 | 1412;1434 | | NA | 5.3390 | 6.0213 | 6.1121 | 6.4308 | 6.0242 | 5.9569 | 6.3637 | 6.2247 | 7.8046 | 6.2337 | 5.9373 | 6.2676 | 6.0446 | 6.0514 | 6.3146 | 6.8736 |
| SLX4IP_S396 | SLX4IP | Q5VYV7 | | 396 | LSTIQNSPTKKRK | 6.4261 | 6.3930 | 5.8764 | 6.4112 | 6.2408 | 6.1809 | 6.3585 | 6.1624 | 6.4075 | 6.2149 | 6.1115 | 6.1500 | 6.2564 | 6.4583 | 6.3214 | 6.0309 |
| SMAD2_T172 | SMAD2 | Q15796 | | 172 | HYQRVETPVLPPV | 6.0773 | 6.5402 | 6.0960 | 6.0730 | 5.9608 | 5.7495 | 7.5021 | 6.3246 | 6.2311 | 6.7847 | 5.5712 | 6.1492 | 6.7875 | 7.1110 | 5.6201 | 5.4217 |
| SMAD3_S416 | SMAD3 | P84022 | | 416 | VLTQMGSPSIRCS | 6.5158 | 7.4790 | 6.9635 | 7.0606 | 6.2639 | 6.6832 | 6.8966 | 6.7734 | 6.6917 | 7.2291 | 7.1946 | 6.9918 | 3.8738 | 5.0569 | 3.8681 | 4.4580 |
| SMAD5_S133 | SMAD5 | Q99717 | | 133 | HYKRVESPVLPPV | 6.0185 | 6.8133 | 6.8620 | 6.4049 | 6.5130 | 6.7345 | 6.9307 | 6.9180 | 6.7832 | 5.9591 | 7.5586 | 6.1012 | 4.5230 | 5.5160 | 5.0502 | 5.3138 |
| SMAD5_S463;S465 | SMAD5 | Q99717 | 463;465 | | NA | 5.4565 | 5.8677 | 5.9702 | 6.2541 | 5.3902 | 6.7683 | 5.7647 | 6.2786 | 6.1798 | 5.6889 | 6.4484 | 6.7658 | 6.4967 | 6.3358 | 7.6285 | 6.7058 |
| SMAGP_S77 | SMAGP | Q0VAQ4 | | 77 | PTEGEPSAIVQME | 4.2271 | 4.4897 | 4.5580 | 4.5949 | 4.8967 | 4.3548 | 6.1932 | 5.6521 | 4.1291 | 4.1447 | 4.1497 | 4.6421 | 6.8107 | 13.0698 | 10.0472 | 14.0404 |
| SMAP_S17 | SMAP | O00193 | | 17 | GVKRSASPDDDLG | 5.9169 | 5.5782 | 6.1825 | 6.1525 | 6.2853 | 5.9358 | 5.7298 | 6.0034 | 6.3236 | 5.6686 | 6.2764 | 5.9843 | 6.9615 | 7.3998 | 6.5158 | 7.0854 |
| SMAP1_S152 | SMAP1 | Q8IYB5 | | 152 | LQPLVSSPSLQAA | 7.0200 | 6.7717 | 6.7121 | 6.9337 | 5.9107 | 6.7093 | 6.4513 | 7.1651 | 6.8818 | 6.8313 | 6.6462 | 7.7562 | 4.9445 | 5.2425 | 3.6411 | 4.3822 |
| SMAP2_S219 | SMAP2 | Q8WU79 | | 240 | TAGSAGSVPENLN | 5.0671 | 6.0879 | 6.3847 | 5.2694 | 5.3207 | 5.8852 | 6.5163 | 6.3114 | 5.5000 | 6.0155 | 5.9770 | 6.2734 | 6.6537 | 7.0053 | 7.5557 | 8.1767 |
| SMAP2_S240 | SMAP2 | Q8WU79 | | 219 | LLASVPSPSSSGS | 6.6353 | 6.4505 | 6.8537 | 6.8971 | 6.7034 | 6.5265 | 6.4936 | 6.4844 | 6.5579 | 7.0182 | 6.9015 | 6.8207 | 4.9218 | 5.0666 | 4.6172 | 5.0517 |
| SMARCA1_S116 | SMARCA1 | P28370 | | 119 | AQKSPTSPLNMKL | 7.0795 | 7.0948 | 6.7560 | 7.1740 | 6.6174 | 6.2532 | 6.8291 | 6.5285 | 7.1425 | 6.8452 | 6.9419 | 6.7944 | 4.3373 | 4.7333 | 4.2614 | 4.6115 |
| SMARCA1_S116;S119 | SMARCA1 | P28370 | | 62 | KKEKNVSSFQLKL | 6.3745 | 5.3096 | 5.5532 | 6.4009 | 6.5942 | 6.1886 | 6.4247 | 6.6696 | 6.3632 | 5.4778 | 6.7148 | 5.3246 | 6.5302 | 6.7449 | 6.6746 | 6.6545 |
| SMARCA1_S119 | SMARCA1 | P28370 | | 116 | QPSAQKSPTSPLN | 5.9741 | 5.9064 | 6.0023 | 7.1336 | 5.5939 | 5.3954 | 5.4039 | 5.5854 | 5.4958 | 5.6407 | 6.0488 | 5.8446 | 7.1107 | 7.4220 | 8.2607 | 7.1817 |
| SMARCA1_S62 | SMARCA1 | P28370 | 116;119 | | NA | 6.9269 | 7.0458 | 6.2915 | 7.0091 | 6.6319 | 6.2442 | 6.6131 | 6.2944 | 7.0122 | 6.4969 | 6.4959 | 6.4487 | 5.1353 | 5.1903 | 5.0403 | 5.1236 |
| SMARCA2_S1377 | SMARCA2 | P51531 | | 1377 | PPAEKLSPNPPKL | 6.5809 | 6.6282 | 5.9786 | 6.2916 | 5.9642 | 5.4466 | 6.1716 | 5.8008 | 6.1277 | 6.2711 | 5.6010 | 5.7080 | 6.5494 | 6.7669 | 7.0890 | 7.0244 |
| SMARCA2_S329 | SMARCA2 | P51531 | | 329 | QKQSRISPIQKPQ | 5.5374 | 7.2278 | 5.9765 | 6.3476 | 6.5827 | 8.0263 | 6.3049 | 5.9318 | 6.5283 | 5.8347 | 5.8522 | 4.7525 | 5.5433 | 9.2038 | 5.4442 | 4.9060 |
| SMARCA2_T274 | SMARCA2 | P51531 | | 274 | ELSGPSTPQKLPV | 6.0993 | 5.8970 | 6.8642 | 7.2567 | 5.9538 | 6.6355 | 6.4600 | 5.9149 | 7.3615 | 6.3114 | 5.6866 | 6.5120 | 5.6060 | 6.1212 | 5.3285 | 5.9914 |
| SMARCA4_S1452 | SMARCA4 | P51532 | | 1452 | PPAEKLSPNPPNL | 6.8708 | 6.4087 | 5.5075 | 7.1602 | 6.0400 | 5.7578 | 6.3624 | 6.1606 | 5.9755 | 6.3171 | 5.4576 | 6.7711 | 6.0964 | 6.2623 | 6.3591 | 6.4928 |
| SMARCA5_S116 | SMARCA5 | O60264 | | 116 | AQKTPTSPLKMKP | 6.7733 | 6.6011 | 5.9073 | 6.5917 | 6.5348 | 5.7757 | 6.4884 | 6.2268 | 6.3945 | 6.0578 | 5.9312 | 5.9804 | 6.1056 | 6.0236 | 6.3714 | 6.2363 |
| SMARCA5_S66 | SMARCA5 | O60264 | | 66 | EIFDDASPGKQKE | 5.9652 | 6.3907 | 6.4085 | 6.8414 | 6.1603 | 6.1589 | 6.5760 | 6.3741 | 6.3194 | 6.4219 | 6.4071 | 6.6808 | 5.6879 | 5.9057 | 5.7724 | 5.9298 |
| SMARCAD1_S124;S127 | SMARCAD1 | Q9H4L7-2 | | 79 | DNERKASISYFKN | 4.5997 | 4.1978 | 4.9357 | 5.5900 | 9.8914 | 4.8159 | 4.4370 | 5.3461 | 8.3774 | 8.4643 | 7.2432 | 7.2167 | 7.3506 | 7.6236 | 4.8582 | 5.0524 |
| SMARCAD1_S146 | SMARCAD1 | Q9H4L7-2 | | 146 | RRNDDISELEDLS | 6.1747 | 6.1200 | 6.5099 | 6.3001 | 6.0087 | 6.4286 | 6.3145 | 6.2203 | 6.2382 | 5.8103 | 6.1399 | 6.3837 | 6.1732 | 6.5626 | 6.1551 | 6.4601 |
| SMARCAD1_S146;S152 | SMARCAD1 | Q9H4L7-2 | | 152 | SELEDLSELEDLK | 5.8123 | 6.3289 | 6.3068 | 6.0209 | 6.3243 | 6.1321 | 6.0615 | 5.7115 | 5.9992 | 5.8615 | 5.8075 | 6.0903 | 6.8333 | 7.2472 | 6.6565 | 6.8064 |
| SMARCAD1_S152 | SMARCAD1 | Q9H4L7-2 | | 211 | PRKRKLSSSSEPY | 5.5426 | 6.0211 | 6.1098 | 6.1139 | 6.3265 | 5.7834 | 5.8276 | 6.0854 | 7.0425 | 7.3911 | 6.8767 | 6.9819 | 5.7565 | 6.2162 | 5.8811 | 6.0437 |
| SMARCAD1_S211 | SMARCAD1 | Q9H4L7-2 | | 96 | QYIDLSSDSEDVV | 5.9762 | 6.2548 | 6.4908 | 6.0678 | 6.2408 | 6.4734 | 6.0702 | 5.7739 | 6.5958 | 6.4897 | 6.5385 | 6.4459 | 6.1606 | 6.2648 | 5.9544 | 6.2024 |
| SMARCAD1_S211;Y217 | SMARCAD1 | Q9H4L7-2 | | 24 | EEAPEATPQPSQP | 5.4890 | 6.3890 | 5.9049 | 6.0878 | 6.2039 | 6.0152 | 6.5122 | 5.9283 | 6.8766 | 6.8802 | 6.4508 | 6.3141 | 5.5231 | 7.1061 | 6.3729 | 5.9457 |
| SMARCAD1_S79 | SMARCAD1 | Q9H4L7-2 | | 54 | EVSRANTPDSDIT | 5.7770 | 7.1960 | 6.0898 | 6.6995 | 5.9284 | 5.5230 | 6.4266 | 5.7953 | 6.5772 | 6.8730 | 6.4794 | 6.0238 | 6.0612 | 6.1854 | 5.9578 | 6.4066 |
| SMARCAD1_S95;S96 | SMARCAD1 | Q9H4L7-2 | 124;127 | | NA | 5.7301 | 6.4554 | 6.5155 | 6.2465 | 6.2845 | 6.3065 | 5.9585 | 6.3411 | 6.3338 | 6.4347 | 6.4136 | 6.4136 | 5.9173 | 6.3714 | 6.0353 | 6.2421 |
| SMARCAD1_S96 | SMARCAD1 | Q9H4L7-2 | 211;217 | | NA | 5.5961 | 6.2282 | 6.1667 | 5.9817 | 5.9739 | 5.9638 | 5.8649 | 5.8857 | 7.0982 | 6.9168 | 7.0308 | 7.0865 | 5.7977 | 6.0438 | 5.9870 | 6.3784 |
| SMARCAD1_T24 | SMARCAD1 | Q9H4L7-2 | 95;96 | | NA | 5.3905 | 5.0374 | 5.9985 | 5.6997 | 6.6787 | 5.6380 | 6.0125 | 5.9817 | 5.9671 | 6.2315 | 6.4104 | 5.9810 | 7.6879 | 7.0854 | 7.0215 | 7.1783 |
| SMARCAD1_T54 | SMARCAD1 | Q9H4L7-2 | 146;152 | | NA | 5.3722 | 7.1874 | 6.6866 | 5.9823 | 6.7102 | 7.1700 | 5.9692 | 7.2704 | 4.9836 | 7.4896 | 5.3613 | 4.7008 | 7.1809 | 6.1871 | 6.1812 | 5.5672 |
| SMARCAL1_S112 | SMARCAL1 | Q9NZC9 | | 112 | TACPGHSPRSQMA | 5.7765 | 6.2774 | 6.6624 | 6.5850 | 5.8304 | 6.4800 | 6.0533 | 6.4968 | 6.8460 | 6.7105 | 7.6031 | 7.7253 | 4.8616 | 5.6019 | 5.3258 | 5.1641 |
| SMARCAL1_S129 | SMARCAL1 | Q9NZC9 | | 934 | ESSLTASPQKKRR | 4.9391 | 5.4055 | 6.7317 | 5.4202 | 5.5343 | 6.1018 | 5.5631 | 6.7024 | 5.4272 | 6.0490 | 6.5317 | 6.5138 | 7.3187 | 7.3753 | 7.1986 | 7.1877 |
| SMARCAL1_S934 | SMARCAL1 | Q9NZC9 | | 129 | SPPLAQSPPEVPK | 6.3935 | 6.6340 | 6.3763 | 6.6383 | 6.4618 | 6.5490 | 6.0137 | 6.3367 | 5.9579 | 7.1495 | 7.8012 | 6.8256 | 5.1253 | 5.4660 | 4.8173 | 5.4540 |
| SMARCC1_S310 | SMARCC1 | Q92922 | | 310 | NEEPVRSPERRDR | 6.7262 | 6.7527 | 6.8080 | 6.6158 | 6.2387 | 6.3116 | 6.3851 | 5.7767 | 6.6588 | 6.7338 | 6.9329 | 6.9920 | 5.6571 | 5.4109 | 4.8543 | 5.1453 |
| SMARCC1_S319 | SMARCC1 | Q92922 | | 573 | VPLHLRSPQVPAA | 6.4446 | 6.9042 | 6.6062 | 7.0052 | 6.4252 | 6.5413 | 6.7016 | 6.9308 | 6.9302 | 6.7768 | 6.7959 | 7.0188 | 4.5965 | 5.0689 | 4.2933 | 4.9607 |
| SMARCC1_S328;S330 | SMARCC1 | Q92922 | | 319 | RRDRKASANARKR | 3.3868 | 3.6954 | 3.5262 | 4.1066 | 6.0806 | 5.8237 | 5.4072 | 5.9910 | 3.6209 | 3.5828 | 3.8609 | 2.1817 | 16.1471 | 14.9780 | 4.7315 | 12.8794 |
| SMARCC1_S330 | SMARCC1 | Q92922 | | 398 | KKDSENTPVKGGT | 7.2624 | 7.0426 | 6.5878 | 6.3724 | 7.2013 | 6.5077 | 6.9433 | 6.6157 | 7.2789 | 6.5748 | 6.4483 | 6.4682 | 4.8368 | 4.7816 | 4.5069 | 4.5712 |
| SMARCC1_S573 | SMARCC1 | Q92922 | | 330 | KRKHSPSPPPPTP | 5.3532 | 5.7150 | 5.9731 | 6.7708 | 5.5172 | 6.0652 | 5.0125 | 7.3379 | 6.2577 | 6.1170 | 5.7721 | 5.8578 | 6.3762 | 6.9052 | 7.9346 | 7.0343 |
| SMARCC1_T398 | SMARCC1 | Q92922 | 328;330 | | NA | 5.8919 | 6.4347 | 6.3682 | 6.6056 | 6.2818 | 6.2338 | 6.2460 | 5.7919 | 6.1874 | 6.2404 | 6.2045 | 6.2645 | 6.3203 | 6.4012 | 6.3498 | 6.1779 |
| SMARCC2_S283 | SMARCC2 | Q8TAQ2 | | 347 | KDMDEPSPVPNVE | 5.7850 | 6.0855 | 5.9610 | 6.2859 | 6.1447 | 6.0424 | 6.0965 | 6.0880 | 6.0529 | 5.9339 | 5.8551 | 6.0008 | 6.8138 | 7.0797 | 6.9819 | 6.7927 |
| SMARCC2_S302 | SMARCC2 | Q8TAQ2 | | 283 | LTDEVNSPDSDRR | 6.2992 | 6.8812 | 6.7148 | 7.2368 | 6.2711 | 6.3358 | 6.5614 | 6.3298 | 6.4226 | 6.7702 | 6.6973 | 6.8509 | 5.0650 | 5.2982 | 5.0307 | 5.2351 |
| SMARCC2_S302;S304 | SMARCC2 | Q8TAQ2 | | 321 | NAKKGPSTPYTKS | 7.2841 | 7.5106 | 6.4012 | 6.7276 | 6.5270 | 6.6070 | 6.6167 | 6.4893 | 7.0108 | 6.4904 | 6.3752 | 6.3838 | 5.0069 | 5.1722 | 4.3104 | 5.0868 |
| SMARCC2_S302;S304;T3 | SMARCC2 | Q8TAQ2 | | 548 | VPLQPKTPQQTSA | 5.7908 | 6.2115 | 5.9738 | 6.3734 | 5.9917 | 6.3190 | 6.4913 | 6.1501 | 5.4709 | 6.0411 | 6.1180 | 6.2668 | 6.9467 | 6.5431 | 6.6955 | 6.6163 |
| SMARCC2_S304 | SMARCC2 | Q8TAQ2 | | 322 | AKKGPSTPYTKSK | 6.9066 | 7.1192 | 6.6503 | 6.7133 | 6.7223 | 6.6257 | 6.7547 | 6.4220 | 6.7960 | 6.3529 | 6.2736 | 6.3681 | 5.1517 | 5.4629 | 4.5930 | 5.0877 |
| SMARCC2_S321 | SMARCC2 | Q8TAQ2 | | 399 | KDEDENSTGNKGE | 5.4574 | 6.5798 | 5.9202 | 6.2295 | 5.8275 | 5.8630 | 6.0179 | 5.9715 | 5.6284 | 5.9701 | 6.4360 | 6.1711 | 7.4693 | 6.7854 | 6.7104 | 6.9624 |
| SMARCC2_S347 | SMARCC2 | Q8TAQ2 | | 813 | EKESEKSDGDPIV | 6.3406 | 7.1174 | 6.2133 | 6.7282 | 6.6371 | 5.8973 | 6.3955 | 5.7739 | 5.5485 | 6.3694 | 5.4561 | 5.9888 | 6.9392 | 6.6113 | 5.5141 | 6.4692 |
| SMARCC2_S399 | SMARCC2 | Q8TAQ2 | | 841 | LKEVVESEGERKT | 6.4739 | 6.7283 | 6.6184 | 6.7649 | 6.3832 | 6.4192 | 6.8506 | 6.1286 | 5.5363 | 5.5854 | 5.7603 | 5.8729 | 6.2775 | 6.1443 | 6.3227 | 6.1336 |
| SMARCC2_S806;S810;S8 | SMARCC2 | Q8TAQ2 | | 302 | YKKRKRSPSPSPT | 6.0445 | 6.2969 | 6.2651 | 6.8192 | 6.1936 | 6.0777 | 5.9642 | 6.0244 | 6.6153 | 6.0700 | 6.5858 | 6.0244 | 6.2436 | 6.4603 | 6.4727 | 5.8424 |
| SMARCC2_S813 | SMARCC2 | Q8TAQ2 | | 304 | KRKRSPSPSPTPE | 5.1269 | 5.6729 | 5.8354 | 5.7851 | 5.1863 | 5.5973 | 6.5333 | 6.1608 | 5.8013 | 6.4630 | 5.7048 | 5.9909 | 7.2068 | 7.2278 | 7.7498 | 7.9577 |
| SMARCC2_S841 | SMARCC2 | Q8TAQ2 | 302;304 | | NA | 5.5554 | 6.0655 | 5.8851 | 6.1886 | 5.9533 | 5.7634 | 5.9006 | 5.9680 | 5.9896 | 6.1439 | 6.0496 | 6.1491 | 6.9468 | 7.0368 | 7.2911 | 7.1133 |
| SMARCC2_T322 | SMARCC2 | Q8TAQ2 | 302;304;308 | | NA | 6.8882 | 6.4579 | 6.3387 | 6.8346 | 6.4551 | 6.8514 | 6.1039 | 6.5843 | 6.9645 | 6.4511 | 7.3538 | 6.7773 | 5.1114 | 5.6429 | 4.2579 | 4.9270 |
| SMARCC2_T548 | SMARCC2 | Q8TAQ2 | 806;810;813 | | NA | 5.6121 | 5.5258 | 6.5085 | 6.2377 | 5.5798 | 5.4626 | 6.5978 | 4.7129 | 6.3562 | 5.9149 | 6.6943 | 6.5804 | 8.1903 | 6.6602 | 6.3329 | 7.0337 |
| SMARCD2_S15 | SMARCD2 | Q92925 | | 203 | YISNTFSPSKAEG | 6.2252 | 6.6214 | 6.0738 | 6.7233 | 6.3623 | 6.2330 | 6.3507 | 6.0487 | 6.3750 | 6.3965 | 6.3927 | 6.0742 | 5.8508 | 5.9624 | 6.1638 | 6.1461 |
| SMARCD2_S203 | SMARCD2 | Q92925 | | 15 | FPLPPLSPGGGAV | 5.0603 | 6.1672 | 7.4279 | 8.2025 | 6.7984 | 6.2843 | 5.6916 | 7.9367 | 7.1429 | 6.1916 | 5.1600 | 6.8194 | 5.6566 | 5.1740 | 5.0778 | 5.2090 |
| SMARCD2_T217 | SMARCD2 | Q92925 | | 217 | SAGTAGTPGGTPA | 6.4858 | 7.7457 | 6.6611 | 6.5398 | 5.7674 | 7.3144 | 7.1878 | 6.3200 | 7.0011 | 6.6794 | 7.2076 | 6.7117 | 4.5825 | 4.7004 | 4.2086 | 4.8867 |
| SMARCE1_T363 | SMARCE1 | Q969G3 | | 363 | GEEGTSTPEDKES | 6.0681 | 5.8262 | 5.8564 | 5.9338 | 6.3471 | 6.2463 | 5.8951 | 6.0155 | 6.0298 | 6.3332 | 6.0198 | 5.9159 | 7.0172 | 6.5243 | 6.8179 | 7.1534 |
| SMC1A_S970 | SMC1A | Q14683 | | 970 | SGSQRISSIYARE | 7.0855 | 7.5506 | 6.3738 | 7.5949 | 6.5977 | 6.9317 | 7.5752 | 7.0411 | 6.9665 | 7.0325 | 6.9711 | 6.8515 | 3.1895 | 4.5964 | 3.3486 | 4.2935 |
| SMC3_S1067 | SMC3 | Q9UQE7 | | 1083 | SERGSGSQSSVPS | 6.3459 | 5.7104 | 6.7734 | 5.9992 | 6.9276 | 5.8627 | 6.0474 | 5.9273 | 6.1723 | 6.0831 | 6.3328 | 5.9833 | 6.5157 | 6.4750 | 6.2093 | 6.6347 |
| SMC3_S1083 | SMC3 | Q9UQE7 | | 1067 | DVEGSQSQDEGEG | 6.9497 | 7.0457 | 6.5207 | 6.7143 | 6.5555 | 6.6079 | 7.0419 | 7.0648 | 5.4482 | 5.7976 | 6.2221 | 5.9202 | 5.5409 | 5.4056 | 5.6376 | 5.5272 |
| SMC4_S1056 | SMC4 | Q9NTJ3 | | 28 | SPDGASSDAEPEP | 6.2661 | 6.1075 | 7.3577 | 6.2954 | 6.2717 | 6.1006 | 6.2261 | 6.5913 | 6.6311 | 6.4580 | 6.5518 | 6.6211 | 5.4086 | 6.1939 | 5.4218 | 5.4974 |
| SMC4_S22;S28 | SMC4 | Q9NTJ3 | | 1056 | NPIEEISVLSPED | 7.0401 | 7.8718 | 7.2180 | 8.5453 | 7.3197 | 6.4848 | 6.5165 | 7.0415 | 5.8413 | 5.5589 | 6.0638 | 5.2294 | 4.8528 | 5.3985 | 4.8360 | 4.1817 |
| SMC4_S28 | SMC4 | Q9NTJ3 | | 41 | PSGRTESPATAAE | 5.7795 | 6.0362 | 6.4108 | 6.5797 | 6.1317 | 6.1916 | 6.3464 | 6.7554 | 6.3191 | 6.1523 | 6.8413 | 6.3460 | 5.7643 | 6.5383 | 5.9118 | 5.8956 |
| SMC4_S41 | SMC4 | Q9NTJ3 | 22;28 | | NA | 6.3404 | 6.4240 | 6.2626 | 6.4443 | 6.4718 | 5.2631 | 5.8500 | 5.2359 | 7.2630 | 7.3579 | 7.1336 | 5.4022 | 5.6804 | 6.2457 | 5.7942 | 6.8308 |
| SMC5_S12 | SMC5 | Q8IY18 | | 12 | KKTSTPSPQPSKR | 6.1447 | 6.3336 | 5.7911 | 6.2120 | 6.2457 | 6.2007 | 6.3446 | 5.8479 | 6.3002 | 5.9546 | 5.9148 | 6.1795 | 6.4376 | 6.8073 | 6.5908 | 6.6950 |
| SMC5_S35 | SMC5 | Q8IY18 | | 35 | PSKRKNSAPQLPL | 4.8495 | 4.7650 | 5.3096 | 5.9264 | 8.8465 | 5.2691 | 4.9923 | 5.7885 | 7.6802 | 7.9320 | 7.2546 | 7.2536 | 6.6601 | 6.5025 | 5.3683 | 5.6018 |
| SMC5_T10;S12 | SMC5 | Q8IY18 | 10;12 | | NA | 6.4906 | 6.7982 | 6.7768 | 7.0258 | 7.2683 | 7.0884 | 7.2079 | 6.7735 | 6.3441 | 6.3525 | 6.7745 | 7.0593 | 4.3532 | 5.0396 | 4.1799 | 4.4673 |
| SMC6_S11 | SMC6 | Q96SB8-2 | | 11 | KEENFSSPKNAKR | 7.3948 | 7.5377 | 7.2883 | 7.3897 | 6.2082 | 6.1873 | 7.0969 | 6.2687 | 7.4493 | 7.3779 | 7.0676 | 7.4828 | 3.4849 | 3.9679 | 3.4932 | 4.3047 |
| SMCHD1_S1697 | SMCHD1 | A6NHR9 | | 1962 | EEKLGMTPIRKCN | 7.7884 | 7.5064 | 6.1596 | 7.6879 | 6.4245 | 6.4984 | 6.9469 | 7.3287 | 7.7368 | 6.3238 | 6.3247 | 6.7500 | 3.9891 | 4.4152 | 3.5827 | 4.5368 |
| SMCHD1_S1709 | SMCHD1 | A6NHR9 | | 1709 | KKKPRRSCTLPNY | 6.1776 | 6.3374 | 6.3595 | 6.9907 | 6.3755 | 6.7678 | 6.7370 | 6.1761 | 6.4654 | 6.4602 | 6.2837 | 6.0965 | 5.5211 | 5.5965 | 5.6125 | 6.0424 |
| SMCHD1_T1962 | SMCHD1 | A6NHR9 | | 1697 | LLKRKLSEQEELK | 5.9030 | 6.1501 | 5.3886 | 5.2913 | 6.3632 | 5.8420 | 5.8612 | 5.7440 | 6.6755 | 6.4712 | 6.1111 | 5.6506 | 7.3458 | 6.9834 | 7.3157 | 6.9032 |
| SMCHD1_T493 | SMCHD1 | A6NHR9 | | 493 | EDFDWCTPPKKRG | 7.5342 | 5.3248 | 4.7901 | 9.1692 | 7.0305 | 6.2086 | 6.2749 | 4.5868 | 6.9730 | 6.3588 | 6.8891 | 6.4825 | 5.5281 | 4.9669 | 5.1701 | 6.7123 |
| SMCR8_S417 | SMCR8 | Q8TEV9 | | 417 | VLISVGSYKSSVE | 6.4880 | 6.4438 | 6.3004 | 5.6051 | 6.3884 | 6.4911 | 6.6030 | 6.4661 | 6.2499 | 5.2954 | 5.4080 | 5.6678 | 6.8874 | 6.7558 | 7.1373 | 5.8124 |
| SMCR8_S421 | SMCR8 | Q8TEV9 | | 485 | KSTSVLSKSDSQA | 6.0902 | 6.3579 | 6.2482 | 8.0979 | 5.4266 | 6.9687 | 7.5397 | 7.6778 | 6.6540 | 6.7779 | 8.5835 | 6.5387 | 3.6011 | 5.0037 | 3.4616 | 4.9724 |
| SMCR8_S485 | SMCR8 | Q8TEV9 | | 421 | VGSYKSSVESVLI | 6.2090 | 6.6865 | 7.3242 | 5.9987 | 6.3717 | 6.8751 | 6.1886 | 6.7205 | 6.3797 | 6.7081 | 7.2320 | 7.1566 | 5.0772 | 5.0852 | 4.8168 | 5.1700 |
| SMG1_S3515 | SMG1 | Q96Q15 | | 3515 | NLVSFASPLVTDA | 5.9412 | 6.2019 | 6.7276 | 6.5369 | 5.5319 | 6.5096 | 5.8019 | 6.6669 | 6.3969 | 6.0830 | 6.4155 | 6.1716 | 5.8551 | 6.6644 | 6.4706 | 6.0248 |
| SMG1_S3556 | SMG1 | Q96Q15 | | 3588 | GKSVACSPKKAVR | 6.5318 | 6.6135 | 6.8528 | 6.2009 | 6.4938 | 6.5021 | 6.8955 | 6.4889 | 7.1189 | 6.5278 | 7.0627 | 6.8602 | 4.6326 | 5.0337 | 4.6921 | 5.4928 |
| SMG1_S3570 | SMG1 | Q96Q15 | | 3570 | QKNLATSADTPPS | 5.8666 | 5.8377 | 7.1459 | 5.7829 | 5.6772 | 6.7002 | 6.4317 | 6.3584 | 6.8589 | 6.3527 | 6.8884 | 6.5432 | 5.7857 | 6.4289 | 6.0519 | 5.2898 |

| | | | | | | | | | | | | | | | | | | | | |
|---|---|---|---|---|---|---|---|---|---|---|---|---|---|---|---|---|---|---|---|---|
| SMG1_S3570;T3573 | SMG1 | Q96Q15 | 3556 | TQPDVMSQNARKL | 6.0336 | 6.2650 | 6.3450 | 6.2126 | 6.7611 | 5.9055 | 6.1055 | 6.1817 | 6.6031 | 6.0350 | 6.1079 | 5.9203 | 6.8564 | 6.2583 | 6.2331 | 6.1760 |
| SMG1_S3588 | SMG1 | Q96Q15 | 3570;3573 | NA | 5.7006 | 5.5156 | 5.9884 | 5.9524 | 5.9424 | 6.3496 | 6.5226 | 6.2300 | 6.1222 | 5.5399 | 6.3182 | 6.4437 | 6.9557 | 7.0055 | 6.8496 | 6.5636 |
| SMG6_S240 | SMG6 | Q86US8 | 240 | ARGRPGSAKRYSR | 6.5158 | 6.2391 | 6.9874 | 6.5378 | 5.7718 | 6.2077 | 5.8005 | 6.5330 | 6.7718 | 7.2600 | 6.9531 | 7.1800 | 5.2346 | 6.1608 | 4.9069 | 4.9397 |
| SMG6_S831 | SMG6 | Q86US8 | 479 | QLHFLDTDDEVSP | 6.9762 | 7.1171 | 8.0274 | 7.1432 | 6.5104 | 7.7082 | 6.8957 | 7.8338 | 5.3285 | 5.2134 | 5.6760 | 5.8572 | 4.5437 | 5.1422 | 4.6668 | 5.3605 |
| SMG6_T479 | SMG6 | Q86US8 | 831 | HEEFDLSPDQWRK | 6.1316 | 6.6421 | 6.6682 | 6.3215 | 5.7950 | 6.2351 | 6.4773 | 6.2502 | 6.8732 | 6.7265 | 6.9094 | 6.7426 | 5.2473 | 5.6523 | 5.2936 | 6.0341 |
| SMG6_T479;S484 | SMG6 | Q86US8 | 479;484 | NA | 5.9913 | 6.4377 | 6.9254 | 6.2083 | 6.3783 | 6.9030 | 7.0526 | 7.0407 | 6.2497 | 6.1249 | 6.2657 | 6.3483 | 5.4728 | 5.6763 | 5.3053 | 5.6198 |
| SMG7_S520 | SMG7 | Q92540-4 | 520 | SLAADGSPGLKSV | 6.3101 | 6.8004 | 7.2447 | 7.3800 | 6.0754 | 6.7632 | 6.8698 | 6.8461 | 6.7945 | 7.3084 | 7.2327 | 7.3111 | 4.0558 | 4.4261 | 3.9170 | 4.6649 |
| SMG8_S469 | SMG8 | Q8ND04-2 | 469 | KEEDLGSPTGELT | 5.8648 | 6.4859 | 6.5673 | 6.5421 | 6.3220 | 6.6467 | 6.7591 | 6.4475 | 6.6856 | 5.6451 | 6.8084 | 7.0355 | 5.6608 | 5.9529 | 5.3824 | 5.1939 |
| SMG8_S742 | SMG8 | Q8ND04-2 | 742 | FVDRQASTVEYLP | 4.4657 | 4.3398 | 4.9392 | 4.7859 | 5.8746 | 5.2062 | 4.9528 | 5.4864 | 4.4363 | 4.0521 | 4.1813 | 4.3785 | 10.9413 | 10.4857 | 11.8446 | 9.6294 |
| SMG8_T657 | SMG8 | Q8ND04-2 | 657 | PVFEPSTPDPAPA | 5.1383 | 5.4818 | 6.1559 | 4.8324 | 6.1807 | 6.5099 | 6.0360 | 6.1072 | 5.7217 | 5.7457 | 6.5421 | 5.0843 | 8.5181 | 7.4653 | 7.2710 | 7.2095 |
| SMG9_S32 | SMG9 | Q9H0W8 | 118 | AVTGASTPEGTAP | 6.1041 | 6.1514 | 6.7699 | 6.3400 | 6.5230 | 6.2567 | 6.1991 | 6.7332 | 6.3682 | 6.2456 | 6.5399 | 6.9884 | 5.3741 | 5.8741 | 5.6138 | 5.9184 |
| SMG9_S450 | SMG9 | Q9H0W8 | 450 | PRAGPGSSPLFSL | 6.8472 | 6.2502 | 6.2335 | 7.1344 | 6.6894 | 6.7227 | 5.9711 | 6.7726 | 6.9064 | 6.2607 | 7.1636 | 7.3989 | 4.8041 | 5.0755 | 4.5210 | 5.2487 |
| SMG9_S53 | SMG9 | Q9H0W8 | 32 | GGPQNLSGPGGRE | 5.1337 | 5.4142 | 6.8895 | 5.8434 | 5.6520 | 6.1791 | 5.9157 | 6.4525 | 6.9053 | 6.0519 | 6.1233 | 6.2488 | 6.4880 | 7.1386 | 6.7633 | 6.8007 |
| SMG9_S69 | SMG9 | Q9H0W8 | 53 | RERRDASEETSTS | 6.0559 | 5.0163 | 5.5507 | 5.7621 | 6.0893 | 6.3313 | 6.0430 | 6.2925 | 5.7505 | 6.0866 | 6.4241 | 6.4800 | 7.1685 | 7.2183 | 6.6074 | 7.1235 |
| SMG9_T118 | SMG9 | Q9H0W8 | 69 | KTPIILSKPPAER | 6.8339 | 6.8314 | 8.0096 | 6.1674 | 5.2869 | 6.6190 | 7.0416 | 7.0629 | 8.0510 | 7.2352 | 8.4586 | 8.3407 | 3.2782 | 4.0531 | 3.0785 | 3.6521 |
| SMIM8_S22 | SMIM8 | Q96KF7 | 22 | KEKEFQSPGLRGV | 7.5240 | 7.1823 | 6.2716 | 7.5772 | 6.6092 | 5.6269 | 7.4053 | 7.5097 | 6.7197 | 6.4322 | 6.0014 | 6.1273 | 4.1166 | 5.6842 | 3.9262 | 5.2862 |
| SMN1_S175 | SMN1 | Q16637 | 175 | ESENSRSPGNKSD | 7.0891 | 6.2078 | 6.1514 | 6.6879 | 6.3713 | 6.3739 | 6.1522 | 5.8877 | 6.9499 | 7.4969 | 7.2947 | 7.7302 | 4.9521 | 5.2648 | 4.5626 | 4.8276 |
| SMN1_S28 | SMN1 | Q16637 | 28 | RRGTGQSDDSDIW | 6.3071 | 5.6972 | 6.9430 | 6.3932 | 6.1922 | 5.7888 | 6.6148 | 7.9522 | 5.7573 | 6.0773 | 7.0688 | 7.1632 | 5.2719 | 5.9734 | 5.1037 | 5.6958 |
| SMN1_S28;S31 | SMN1 | Q16637 | 167 | NESQVSTDESENS | 6.0249 | 6.0705 | 7.3273 | 7.1124 | 6.4206 | 4.6457 | 6.2626 | 6.8135 | 5.5866 | 5.5837 | 6.5900 | 6.3907 | 6.1758 | 6.3509 | 7.0554 | 5.5894 |
| SMN1_T167 | SMN1 | Q16637 | 28;31 | NA | 6.5449 | 5.9350 | 6.6179 | 6.1495 | 6.5418 | 6.4762 | 6.0035 | 6.6502 | 5.9590 | 6.3603 | 6.5151 | 6.5047 | 6.0020 | 5.9258 | 5.9285 | 5.8855 |
| SMNDC1_S201 | SMNDC1 | O75940 | 201 | KRSIFASPESVTG | 7.2599 | 7.4461 | 7.2673 | 7.2991 | 6.0827 | 7.0554 | 7.5048 | 7.2967 | 7.3635 | 7.2768 | 7.5664 | 7.5062 | 2.9301 | 3.7301 | 2.8284 | 3.5867 |
| SMPD4_S169 | SMPD4 | Q9NXE4-4 | 838 | ASRHLLSPVGRRQ | 7.5446 | 7.8226 | 6.6327 | 7.8090 | 6.4234 | 6.7050 | 7.2657 | 6.6818 | 7.1364 | 7.1919 | 7.2947 | 7.4057 | 3.0393 | 4.0657 | 3.1244 | 3.8571 |
| SMPD4_S248 | SMPD4 | Q9NXE4-4 | 754 | EDGLILTPLGRYQ | 6.3975 | 7.0577 | 6.4855 | 7.1010 | 6.1905 | 6.3936 | 6.7648 | 6.4810 | 6.7597 | 6.8892 | 7.1648 | 6.7318 | 4.8254 | 5.2904 | 4.7845 | 4.6825 |
| SMPD4_S838 | SMPD4 | Q9NXE4-4 | 248 | PPPLSSSPGGTSP | 6.1498 | 6.5361 | 5.9539 | 7.1490 | 6.1050 | 6.3643 | 6.3524 | 6.0539 | 6.4802 | 6.5131 | 6.9635 | 6.7568 | 5.7191 | 5.8661 | 5.7617 | 5.2750 |
| SMPD4_T595 | SMPD4 | Q9NXE4-4 | 169 | ECILPDSPLYHNK | 6.5839 | 6.8949 | 5.9452 | 7.4396 | 6.6286 | 6.5232 | 6.7147 | 6.9258 | 6.6664 | 6.5638 | 6.6554 | 6.2446 | 4.7280 | 5.0820 | 5.0023 | 5.4016 |
| SMPD4_T754 | SMPD4 | Q9NXE4-4 | 595 | FTAPTFTGSFLSP | 7.8510 | 6.6146 | 6.1560 | 7.7560 | 6.3984 | 4.7948 | 7.2074 | 5.5531 | 6.6792 | 5.6227 | 5.9870 | 5.7751 | 5.8360 | 5.5414 | 5.4638 | 6.7634 |
| SMTN_S245 | SMTN | P53814-6 | 304 | RSLSVLSPRQPAQ | 6.8520 | 7.6770 | 7.1107 | 7.0602 | 6.1361 | 7.1634 | 7.3354 | 7.1377 | 7.4997 | 7.3657 | 7.6834 | 7.5564 | 2.8450 | 3.8221 | 2.9545 | 3.8005 |
| SMTN_S277 | SMTN | P53814-6 | 301 | PCQRSLSVLSPRQ | 9.4585 | 7.2052 | 7.0967 | 6.3016 | 5.1684 | 6.0781 | 6.4161 | 5.4240 | 7.4111 | 6.3743 | 7.3240 | 7.0644 | 5.0149 | 5.8710 | 3.5102 | 4.2817 |
| SMTN_S301 | SMTN | P53814-6 | 245 | SPPKTTSPEPQES | 4.7721 | 5.5334 | 5.6772 | 5.4915 | 5.1510 | 5.2363 | 5.6467 | 5.4345 | 5.7854 | 5.5668 | 6.0371 | 5.7112 | 7.9824 | 8.2846 | 8.9782 | 8.7118 |
| SMTN_S304 | SMTN | P53814-6 | 792 | AVQRSTSFGVPNA | 7.4245 | 7.0785 | 7.3400 | 5.8295 | 6.4075 | 6.8365 | 6.2393 | 6.0466 | 8.0301 | 7.1075 | 7.5264 | 7.0039 | 4.5842 | 5.1913 | 3.4274 | 3.9268 |
| SMTN_S341 | SMTN | P53814-6 | 341 | HKFTSDSPMAARL | 7.2895 | 8.0438 | 6.8682 | 7.3150 | 5.9228 | 6.9087 | 7.6504 | 6.7146 | 7.4270 | 7.3758 | 7.7626 | 7.7062 | 2.9317 | 3.5907 | 2.5865 | 3.9064 |
| SMTN_S792 | SMTN | P53814-6 | 277 | ETPAAQSPTRGPS | 7.2290 | 8.1549 | 7.7270 | 6.4578 | 5.3470 | 6.9684 | 6.8231 | 6.4429 | 8.1105 | 7.5345 | 7.9410 | 7.7654 | 2.7179 | 4.1276 | 2.8194 | 3.8336 |
| SNAP23_S110 | SNAP23 | O00161 | 110 | GDGGENSPCNVVS | 6.6988 | 6.8720 | 6.6010 | 6.8170 | 6.5379 | 6.4961 | 7.0329 | 6.5516 | 6.7647 | 6.7406 | 6.8024 | 6.6048 | 4.5892 | 5.2183 | 4.4896 | 5.1831 |
| SNAP23_S136 | SNAP23 | O00161 | 161 | NLTQVGSILGNLK | 3.9832 | 4.1349 | 4.0101 | 4.1002 | 4.0653 | 3.5436 | 3.8014 | 3.4829 | 3.5451 | 3.1402 | 3.4439 | 3.5363 | 13.5877 | 13.3490 | 15.3531 | 12.9232 |
| SNAP23_S161 | SNAP23 | O00161 | 34 | LGLAIESQDAGIK | 6.2282 | 6.5929 | 6.1268 | 6.3810 | 6.0691 | 5.8277 | 5.9433 | 5.7340 | 5.6999 | 6.0927 | 6.5056 | 6.4816 | 6.2947 | 6.8121 | 7.0481 | 6.1622 |
| SNAP23_S34 | SNAP23 | O00161 | 136 | PTTGAASGGYIKR | 6.1811 | 6.2399 | 6.0685 | 6.3970 | 5.6362 | 5.5259 | 6.4188 | 5.9534 | 6.5651 | 5.4682 | 5.7664 | 6.0207 | 7.0859 | 6.7014 | 6.8492 | 7.1223 |
| SNAP23_Y139 | SNAP23 | O00161 | 139 | GAASGGYIKRITN | 5.9256 | 6.0420 | 6.4971 | 6.1832 | 6.7632 | 6.7401 | 6.0855 | 7.2075 | 7.5595 | 7.2000 | 7.0240 | 6.6843 | 4.6716 | 5.8526 | 3.8329 | 5.7308 |
| SNAP91_S298 | SNAP91 | O60641-4 | 298 | NKSGAPSPLSKSS | 5.9636 | 6.6947 | 6.7286 | 6.5957 | 5.7841 | 6.5045 | 6.5921 | 6.5550 | 6.1088 | 7.2172 | 7.0113 | 7.1056 | 4.7927 | 5.8910 | 5.0068 | 5.4484 |
| SNAPC1_S355 | SNAPC1 | Q16533 | 355 | EENESLSGTEFTA | 6.1127 | 6.4180 | 6.7777 | 6.9404 | 6.2944 | 6.3125 | 7.5458 | 6.6934 | 5.5670 | 6.0665 | 6.1477 | 6.1713 | 5.7710 | 5.9513 | 5.6849 | 5.5455 |
| SNAPC4_S599 | SNAPC4 | Q5SXM2 | 599 | GPAASLSPPKGSS | 5.8129 | 6.2086 | 6.3640 | 6.1304 | 6.0364 | 6.2724 | 6.4767 | 5.7590 | 6.4823 | 6.0257 | 6.3279 | 6.4889 | 6.0443 | 6.5988 | 6.3736 | 6.5981 |
| SNAPIN_S126 | SNAPIN | O95295 | 133 | GIYPPGSPGKxxx | 5.9207 | 4.9745 | 6.6706 | 5.4804 | 6.1113 | 5.4122 | 5.3626 | 7.0858 | 5.7084 | 6.9784 | 6.3663 | 7.4669 | 4.8144 | 7.8337 | 6.5672 | 7.2468 |
| SNAPIN_S133 | SNAPIN | O95295 | 126 | RRAMLDSGIYPPG | 7.0109 | 5.3374 | 5.8539 | 5.9579 | 5.3078 | 4.4840 | 5.2932 | 5.1070 | 5.3428 | 6.3353 | 6.5325 | 6.6728 | 6.7629 | 8.7152 | 7.4996 | 7.7868 |
| SND1_S426 | SND1 | Q7KZF4 | 426 | DYIRPASPATETV | 6.0676 | 5.9541 | 6.4028 | 6.2503 | 6.0554 | 6.1830 | 5.9241 | 6.2054 | 6.2859 | 6.1665 | 6.4304 | 6.2148 | 6.5586 | 6.4817 | 6.5421 | 6.2773 |
| SND1_S781 | SND1 | Q7KZF4 | 781 | TRLGTLSPAFSTR | 5.9111 | 6.3610 | 5.7904 | 6.0383 | 5.7447 | 6.1071 | 6.3344 | 6.1841 | 6.1368 | 6.5899 | 6.1323 | 6.1375 | 6.2892 | 6.6057 | 6.7932 | 6.8443 |
| SNIP1_S153 | SNIP1 | Q8TAD8 | 35 | VKQERLSPEVAPP | 6.0462 | 6.0024 | 5.9499 | 6.6792 | 5.9380 | 5.8410 | 6.2701 | 6.0541 | 6.2869 | 6.0629 | 5.7975 | 6.2543 | 6.1884 | 7.0458 | 6.7785 | 6.8048 |
| SNIP1_S35 | SNIP1 | Q8TAD8 | 52 | PDHSGGSPSPPTS | 6.1876 | 6.6647 | 6.3814 | 7.1121 | 6.0996 | 5.6902 | 5.7713 | 5.7054 | 5.9470 | 6.2920 | 6.0383 | 6.0810 | 6.6906 | 6.4081 | 6.4057 | 6.5251 |
| SNIP1_S396 | SNIP1 | Q8TAD8 | 153 | SHQRRTSNERPGS | 6.5369 | 6.4247 | 6.0107 | 6.1456 | 5.6938 | 5.6757 | 6.5084 | 5.5164 | 6.2729 | 5.6437 | 5.5362 | 5.7903 | 6.4473 | 7.3149 | 7.0637 | 7.4190 |
| SNIP1_S52 | SNIP1 | Q8TAD8 | 396 | EEEVSDSxxxxxx | 6.3148 | 7.5640 | 4.9871 | 7.4940 | 6.9359 | 6.6162 | 5.5059 | 8.1469 | 5.6603 | 5.9672 | 5.1822 | 5.4419 | 4.8421 | 6.8575 | 7.0874 | 5.3967 |
| SNIP1_S52;S54 | SNIP1 | Q8TAD8 | 52;54 | NA | 6.4689 | 6.2130 | 5.9100 | 6.4335 | 5.8651 | 5.5026 | 5.7876 | 5.7956 | 6.4047 | 5.9706 | 6.1819 | 5.7739 | 6.5903 | 7.0419 | 7.1745 | 6.8859 |
| SNIP1_S52;S54;T57 | SNIP1 | Q8TAD8 | 74;76 | NA | 6.0795 | 6.5123 | 5.9266 | 6.8413 | 5.8248 | 5.4854 | 6.0907 | 5.7966 | 5.8414 | 6.1429 | 5.9944 | 6.2237 | 6.1073 | 7.0990 | 6.7529 | 7.2812 |
| SNIP1_S74;S76 | SNIP1 | Q8TAD8 | 52;54;57 | NA | 6.4391 | 7.0007 | 5.8303 | 7.6880 | 5.7793 | 6.6326 | 5.4563 | 4.4993 | 7.3331 | 6.7172 | 5.8695 | 5.5465 | 5.9011 | 5.8462 | 6.0074 | 7.4533 |
| SNRK_S162 | SNRK | Q9NRH2 | 518 | LKMNIASPGTVHK | 5.8197 | 5.8772 | 6.9476 | 6.0390 | 6.6667 | 6.7331 | 6.4515 | 6.8944 | 5.9848 | 6.3070 | 6.3715 | 6.3534 | 6.0675 | 5.9798 | 5.4856 | 6.0211 |
| SNRK_S351 | SNRK | Q9NRH2 | 495 | IFEEGESDDEFDM | 6.1012 | 6.9021 | 6.5874 | 7.1021 | 6.1499 | 6.6473 | 5.7004 | 7.3516 | 5.8585 | 7.4647 | 6.3522 | 6.6863 | 4.9685 | 5.3625 | 5.5326 | 5.2329 |
| SNRK_S390 | SNRK | Q9NRH2 | 351 | IQTRSASPSNIKA | 6.4136 | 6.3900 | 6.9401 | 6.0936 | 5.8703 | 6.8202 | 6.2834 | 6.6717 | 6.5667 | 6.4837 | 6.8037 | 7.0232 | 5.2097 | 5.8045 | 5.3475 | 5.2780 |
| SNRK_S495 | SNRK | Q9NRH2 | 390 | HATVPQSPARAAD | 5.9996 | 6.7561 | 6.4213 | 6.3611 | 6.9037 | 6.4821 | 5.9894 | 6.5702 | 6.6577 | 6.1185 | 5.6785 | 5.9962 | 5.8104 | 6.3017 | 6.1739 | 5.7796 |
| SNRK_S518 | SNRK | Q9NRH2 | 162 | LTDFGFSNKFQPG | 6.5136 | 4.5928 | 5.2782 | 7.2881 | 5.9250 | 4.9878 | 5.6306 | 5.9482 | 3.3962 | 4.4185 | 6.0566 | 5.0265 | 9.2282 | 9.0927 | 7.5669 | 9.0501 |
| SNRNP200_S207 | SNRNP200 | O75643 | 457 | KPKPFGSEEQLLP | 5.8680 | 5.3164 | 6.5013 | 6.5999 | 5.9904 | 6.0339 | 6.2371 | 5.4203 | 6.2794 | 5.9560 | 6.6325 | 6.5061 | 7.1450 | 6.3357 | 6.4902 | 6.6876 |
| SNRNP200_S457 | SNRNP200 | O75643 | 207 | VNVQFESDEEEGD | 6.2846 | 6.3545 | 6.0782 | 6.6712 | 6.1563 | 5.7276 | 6.4259 | 6.2361 | 6.2246 | 6.3121 | 6.2038 | 6.3105 | 5.9265 | 6.4062 | 6.4222 | 6.2598 |
| SNRNP200_T1428 | SNRNP200 | O75643 | 1428 | GNIIISTPEKWDI | 6.3994 | 6.6902 | 6.1501 | 6.6943 | 6.2036 | 5.7233 | 6.4792 | 6.0573 | 6.1795 | 6.4889 | 5.8931 | 6.8544 | 5.5383 | 5.7549 | 6.4090 | 6.4844 |
| SNRNP27_S132 | SNRNP27 | Q8WVK2 | 132 | AYAINVSQKRKYR | 4.9968 | 5.1424 | 7.0608 | 6.0206 | 7.7817 | 5.5533 | 4.5601 | 7.0561 | 6.6710 | 10.1158 | 10.5267 | 9.1225 | 3.9590 | 5.1753 | 2.1579 | 4.0998 |
| SNRNP70_S117 | SNRNP70 | P08621 | 226 | DERPGPSPLPHRD | 6.0552 | 6.1765 | 5.8729 | 6.5089 | 6.2042 | 5.9590 | 6.3666 | 6.2261 | 6.3585 | 6.0139 | 5.9310 | 6.0722 | 6.1634 | 6.6032 | 6.6573 | 6.8310 |
| SNRNP70_S226 | SNRNP70 | P08621 | 320 | GDMAEPSEAGDAP | 6.6514 | 6.2381 | 5.9832 | 7.0951 | 6.4502 | 5.7032 | 6.2965 | 5.7847 | 6.0519 | 5.9400 | 5.7786 | 5.9059 | 6.3620 | 6.8229 | 6.5044 | 6.4320 |
| SNRNP70_S320 | SNRNP70 | P08621 | 410 | EGLGNDSRDMYME | 5.5462 | 6.6394 | 5.7390 | 6.6069 | 5.8251 | 5.9638 | 6.3749 | 6.6754 | 5.9694 | 6.2857 | 6.1016 | 5.9807 | 5.9704 | 6.5724 | 6.6325 | 7.1166 |
| SNRNP70_S410 | SNRNP70 | P08621 | 126 | RREFEVYGPIKRI | 6.1818 | 6.4291 | 6.1772 | 6.7345 | 5.9106 | 5.3947 | 6.2322 | 6.1437 | 6.1234 | 5.9522 | 5.7030 | 6.0338 | 6.2891 | 6.7987 | 7.0731 | 6.8230 |
| SNRNP70_Y126 | SNRNP70 | P08621 | 117 | NYDTTESKLRREF | 6.3050 | 5.9816 | 6.2481 | 7.0497 | 5.6737 | 5.5500 | 5.9304 | 5.2968 | 6.7787 | 6.3655 | 5.6179 | 5.5183 | 7.3012 | 6.8586 | 7.3208 | 6.2037 |
| SNRPA_T118 | SNRPA | P09012 | 131 | VQGGGATPVVGAV | 6.2177 | 6.7038 | 5.9711 | 6.6613 | 6.1391 | 5.9768 | 6.0179 | 7.1233 | 5.7380 | 6.2524 | 6.2765 | 6.3722 | 5.6912 | 7.0829 | 5.4475 | 6.3283 |
| SNRPA_T131 | SNRPA | P09012 | 118 | KPKSQETPATKKA | 8.2347 | 7.9902 | 6.7220 | 7.3944 | 6.5232 | 6.7644 | 7.6061 | 6.6495 | 7.3878 | 6.2260 | 6.8664 | 6.9597 | 3.4961 | 4.4337 | 2.9902 | 3.7555 |
| SNRPB_T30 | SNRPB | P14678-3 | 30 | GRIFIGTFKAFDK | 6.9183 | 6.6235 | 5.1823 | 6.6999 | 7.6570 | 5.5572 | 4.9931 | 6.6198 | 5.9138 | 5.8665 | 4.6110 | 5.5208 | 8.9053 | 7.0420 | 6.0485 | 5.8410 |
| SNRPC_S17 | SNRPC | P09234 | 17 | TYLTHDSPSVRKT | 6.9122 | 5.8467 | 6.6865 | 6.8351 | 7.2949 | 6.2144 | 6.2576 | 7.4445 | 7.3307 | 7.4746 | 6.9799 | 8.2918 | 3.9103 | 5.0464 | 3.3545 | 4.1200 |
| SNRPD2_S105 | SNRPD2 | P62316 | 12 | KPKSEMTPEELQK | 5.9654 | 6.1652 | 6.4243 | 6.3035 | 5.8397 | 5.8886 | 6.4011 | 5.9114 | 6.2671 | 6.6558 | 6.2831 | 6.4333 | 6.3176 | 6.3995 | 6.3406 | 6.4039 |
| SNRPD2_T12 | SNRPD2 | P62316 | 105 | MFLRGDSVIVVLR | 9.3186 | 7.7628 | 9.8840 | 7.6584 | 5.0224 | 5.1976 | 6.2648 | 5.4845 | 7.4365 | 7.1643 | 8.3760 | 7.1438 | 3.3158 | 3.5222 | 3.2093 | 3.2389 |
| SNTA1_S189 | SNTA1 | Q13424 | 189 | TSVGWDSPPASPL | 7.0003 | 6.2598 | 6.6352 | 6.9448 | 6.1528 | 5.9203 | 5.9070 | 6.3421 | 6.9180 | 5.6540 | 5.9913 | 6.3030 | 6.6660 | 6.5971 | 5.5820 | 5.1263 |
| SNTA1_S201 | SNTA1 | Q13424 | 201 | LQRQPSSPGPTPR | 6.9083 | 7.5601 | 6.6052 | 6.9691 | 6.3824 | 6.2756 | 5.6250 | 6.4754 | 7.3003 | 6.6986 | 6.6232 | 7.5867 | 4.5013 | 4.9969 | 4.7176 | 4.7744 |
| SNTB1_S219 | SNTB1 | Q13884 | 219 | ETPPPESPRLGGS | 5.9072 | 6.6622 | 6.0405 | 6.5404 | 6.4182 | 6.3976 | 6.2256 | 5.9174 | 6.3819 | 6.3062 | 6.2335 | 6.9521 | 6.1423 | 6.1708 | 5.8524 | 5.8516 |
| SNTB1_S269 | SNTB1 | Q13884 | 389 | SGPGKGSPQAGVD | 5.6743 | 6.0238 | 6.6703 | 6.3571 | 6.5204 | 5.8502 | 5.7643 | 6.2162 | 6.4144 | 6.1379 | 6.4376 | 5.7763 | 6.4649 | 6.9978 | 5.8784 | 6.8162 |
| SNTB1_S389 | SNTB1 | Q13884 | 269 | RQLEIHSPDAKHT | 7.3633 | 5.5924 | 6.8233 | 6.9247 | 6.6506 | 5.4636 | 6.7575 | 6.0694 | 7.3113 | 4.4794 | 6.2750 | 6.2284 | 5.6993 | 6.3024 | 5.3649 | 6.6944 |
| SNTB2_S110 | SNTB2 | Q13425 | 393 | SGSGCRSPSLGSD | 6.3227 | 6.3244 | 6.3334 | 6.5880 | 6.2927 | 6.4081 | 6.7631 | 6.5881 | 6.5326 | 6.4181 | 6.2996 | 6.5072 | 5.4478 | 5.7202 | 5.6234 | 5.8305 |
| SNTB2_S129 | SNTB2 | Q13425 | 110 | AGEAGASPPVRRV | 6.4467 | 6.7496 | 6.0856 | 6.5196 | 6.1800 | 6.2048 | 6.1307 | 5.8138 | 6.1060 | 6.0572 | 5.9098 | 5.9840 | 6.4719 | 6.4526 | 6.5957 | 6.2920 |
| SNTB2_S208 | SNTB2 | Q13425 | 233 | GSEDSGSPKHQNS | 6.0063 | 6.4024 | 6.7016 | 6.2813 | 6.1284 | 6.1483 | 6.3432 | 6.2828 | 6.4219 | 6.5508 | 6.4533 | 6.6891 | 6.0344 | 5.8744 | 5.6177 | 6.0641 |
| SNTB2_S211 | SNTB2 | Q13425 | 222 | EGAAPQSPSFSGS | 6.1066 | 6.1511 | 5.9133 | 6.2822 | 6.4563 | 6.0833 | 6.6010 | 6.2383 | 5.8014 | 6.1193 | 6.2443 | 6.6697 | 6.2979 | 6.4703 | 6.1973 | 6.3677 |
| SNTB2_S222 | SNTB2 | Q13425 | 95 | RGLGPPSPPAPPR | 6.9380 | 6.7139 | 6.4797 | 6.5540 | 6.5254 | 6.4248 | 6.9796 | 6.4703 | 6.7761 | 6.2620 | 6.5166 | 6.3848 | 4.9293 | 5.4760 | 5.1616 | 5.4077 |
| SNTB2_S222;S233 | SNTB2 | Q13425 | 129 | AGGLGISIKGGRE | 5.9643 | 5.8222 | 6.8916 | 6.7389 | 6.8836 | 5.4917 | 5.6363 | 5.3234 | 5.4786 | 4.9206 | 4.5137 | 4.5438 | 8.4643 | 7.8133 | 8.0896 | 7.4240 |
| SNTB2_S233 | SNTB2 | Q13425 | 208 | PYIKKPSLVSDLP | 5.4109 | 5.3432 | 4.7922 | 5.8171 | 5.2082 | 5.5770 | 6.1241 | 5.6990 | 5.5307 | 5.3768 | 5.8167 | 5.3625 | 8.5482 | 8.5541 | 8.4809 | 8.3581 |
| SNTB2_S393 | SNTB2 | Q13425 | 201 | KFIREVTPYIKKP | 6.2762 | 6.5272 | 6.2043 | 6.6323 | 6.7246 | 7.0068 | 6.6211 | 6.3191 | 6.5181 | 6.0707 | 5.7151 | 6.0948 | 5.6566 | 5.8899 | 5.6485 | 6.0948 |
| SNTB2_S95 | SNTB2 | Q13425 | 211 | KKPSLVSDLPWEG | 6.4490 | 6.1142 | 6.0000 | 5.7520 | 6.1869 | 5.9573 | 6.1711 | 5.7044 | 6.3323 | 5.8937 | 6.8918 | 5.8437 | 6.2602 | 6.7513 | 6.7922 | 6.8998 |
| SNTB2_T201 | SNTB2 | Q13425 | 222;233 | NA | 7.0063 | 6.2826 | 7.1610 | 7.3914 | 7.5551 | 7.0223 | 7.2877 | 6.7397 | 6.9218 | 6.9387 | 6.8866 | 7.3072 | 4.5250 | 4.1587 | 3.9269 | 4.6890 |
| SNU13_S122 | SNU13 | P55769 | 122 | IQSIQQSIERLLV | 6.4910 | 6.5824 | 5.5675 | 6.5554 | 5.8646 | 5.8961 | 6.1363 | 5.8586 | 5.7847 | 5.9795 | 6.3771 | 6.7512 | 6.7477 | 6.7052 | 6.5094 | 6.1933 |
| SNW1_S190 | SNW1 | Q13573 | 190 | QGVAFNSGAKQRV | 6.6328 | 7.3866 | 6.3930 | 7.1983 | 5.6405 | 7.4133 | 7.1508 | 6.9905 | 6.7621 | 7.1487 | 7.7611 | 7.2595 | 3.8136 | 4.2102 | 3.8622 | 4.3768 |
| SNW1_S224 | SNW1 | Q13573 | 224 | IPRGPPSPPAPVM | 5.1092 | 5.2883 | 6.1491 | 5.3346 | 5.5752 | 5.7299 | 5.1508 | 5.6461 | 6.2158 | 6.1241 | 5.9861 | 5.6606 | 8.1176 | 8.1936 | 7.9926 | 7.7265 |
| SNW1_S224;S232 | SNW1 | Q13573 | 420 | QSKGMDSGFAGGE | 6.9524 | 7.1728 | 7.0922 | 9.6838 | 6.3609 | 4.3069 | 7.0008 | 6.4362 | 5.6673 | 4.4306 | 5.5221 | 5.5974 | 5.1403 | 5.3124 | 6.9201 | 6.4039 |
| SNW1_S420 | SNW1 | Q13573 | 446 | GKDMAQSIYRPSK | 12.4793 | 8.5127 | 11.7089 | 10.5915 | 5.2751 | 5.5361 | 7.3475 | 4.9822 | 4.5556 | 3.5751 | 5.8672 | 4.7880 | 3.6610 | 5.0713 | 2.0916 | 3.9571 |
| SNW1_S446 | SNW1 | Q13573 | 224;232 | NA | 6.3019 | 6.5879 | 6.3844 | 6.5990 | 6.2125 | 6.5212 | 6.3057 | 6.1755 | 6.0873 | 6.0929 | 6.2888 | 6.4433 | 6.1026 | 6.1856 | 5.8466 | 5.8648 |
| SNX1_S188 | SNX1 | Q13596-3 | 72 | LPINNGSKENGIH | 6.3291 | 6.2114 | 6.4540 | 6.4140 | 5.8677 | 6.3844 | 6.2262 | 6.4068 | 6.3581 | 6.7726 | 6.7909 | 7.2940 | 5.4416 | 6.2431 | 5.3606 | 5.4454 |
| SNX1_S72 | SNX1 | Q13596-3 | 188 | AVKRRFSDFLGLY | 5.3462 | 5.5202 | 6.5015 | 5.0297 | 5.6529 | 6.0886 | 5.3678 | 6.5083 | 7.3571 | 9.4783 | 8.6493 | 8.4697 | 5.1780 | 5.3359 | 4.3770 | 5.1394 |
| SNX11_S228 | SNX11 | Q9Y5W9 | 228 | TLPPLSSPLCCDF | 6.4452 | 6.3192 | 6.6572 | 6.7984 | 6.6635 | 6.4078 | 6.8398 | 6.1909 | 6.2288 | 6.6835 | 6.2207 | 6.6857 | 4.9505 | 5.6631 | 5.8555 | 5.3901 |
| SNX12_S73 | SNX12 | Q9UMY4-1 | 73 | CVRRRYSDFEWLK | 6.1273 | 5.7039 | 6.8079 | 5.4344 | 6.3334 | 6.5554 | 5.5222 | 6.9577 | 6.4960 | 6.5817 | 7.0499 | 7.3900 | 5.8913 | 6.1099 | 5.4440 | 5.5950 |
| SNX14_S485 | SNX14 | Q9Y5W7 | 485 | SKLNRGSLSLDDF | 7.6809 | 6.5339 | 6.8347 | 6.2621 | 7.7062 | 6.4071 | 5.2094 | 6.3165 | 6.7278 | 6.4651 | 7.6881 | 6.2567 | 5.8309 | 5.7910 | 3.8993 | 4.3904 |
| SNX15_S179 | SNX15 | Q9NRS6 | 227 | EEGAAPSPTHVAE | 6.0415 | 6.1755 | 6.4956 | 6.2187 | 6.1397 | 6.6558 | 5.9324 | 6.7146 | 6.5123 | 6.4987 | 6.6201 | 6.7475 | 5.6791 | 6.2802 | 5.4675 | 5.8209 |
| SNX15_S227 | SNX15 | Q9NRS6 | 148 | PPPLIPTPPPDDP | 5.7684 | 6.2825 | 6.4786 | 6.8213 | 6.5539 | 6.7746 | 6.2192 | 6.2199 | 6.1145 | 6.6686 | 5.9258 | 6.6353 | 5.1523 | 6.4512 | 5.8071 | 6.1265 |
| SNX15_T148 | SNX15 | Q9NRS6 | 179 | PVDPPPSSPAQEA | 6.5008 | 5.8281 | 7.8665 | 5.9751 | 6.5819 | 4.7547 | 6.1205 | 6.3943 | 5.6540 | 5.8855 | 7.8192 | 6.6411 | 4.6381 | 6.6986 | 6.1306 | 6.5109 |
| SNX16_S108 | SNX16 | P57768 | 70 | TSSVCSSPLIRTK | 6.3907 | 6.2878 | 7.1231 | 6.5779 | 6.6575 | 7.1860 | 6.8520 | 6.5434 | 6.7500 | 6.9325 | 6.6515 | 6.3971 | 5.0290 | 5.0721 | 4.5320 | 5.0174 |
| SNX16_S286 | SNX16 | P57768 | 286 | EESLDVSETEGEQ | 6.0841 | 6.2519 | 6.8046 | 6.4346 | 6.1514 | 6.3639 | 6.4203 | 6.7591 | 5.9451 | 6.1446 | 6.4479 | 6.3239 | 5.6968 | 6.3366 | 5.6981 | 6.1371 |
| SNX16_S321 | SNX16 | P57768 | 321 | DNKPCLSFSEPEN | 6.1141 | 5.9275 | 7.3721 | 6.8165 | 5.6663 | 6.0653 | 6.3275 | 6.5964 | 6.8898 | 7.7788 | 7.2965 | 6.6764 | 5.1007 | 5.6430 | 4.3728 | 5.3563 |
| SNX16_S70 | SNX16 | P57768 | 108 | NWEDRPSTPTILG | 6.2010 | 5.6519 | 6.8887 | 6.1755 | 6.1524 | 5.8581 | 6.7675 | 6.5012 | 5.6692 | 5.8844 | 6.3404 | 6.6644 | 6.4731 | 6.8648 | 6.0848 | 5.8226 |
| SNX17_S336 | SNX17 | Q15036 | 421 | SPPLLESPDATRE | 6.1490 | 6.6288 | 6.9102 | 6.5362 | 5.7770 | 6.3943 | 6.4667 | 6.8202 | 6.9246 | 6.8561 | 7.5543 | 6.9558 | 4.9627 | 5.5434 | 4.6645 | 4.8561 |
| SNX17_S409 | SNX17 | Q15036 | 437 | KLSSKLSAVSLRG | 3.7060 | 3.7545 | 3.6037 | 3.6047 | 4.2746 | 3.9330 | 3.7841 | 4.3970 | 3.8799 | 3.9534 | 3.8378 | 4.0056 | 11.6574 | 11.6953 | 15.0001 | 14.9129 |
| SNX17_S421 | SNX17 | Q15036 | 336 | PSGSTSSPGRGRG | 6.3481 | 6.0569 | 7.2019 | 6.3109 | 6.2711 | 6.7168 | 6.9471 | 7.0148 | 7.1926 | 6.7838 | 7.5200 | 7.2678 | 4.0773 | 5.0796 | 4.6346 | 4.5766 |
| SNX17_S437 | SNX17 | Q15036 | 409 | TLRRSDSQQAVKS | 6.8989 | 6.2489 | 7.0520 | 5.7307 | 6.0962 | 6.6921 | 6.4337 | 6.5975 | 6.9827 | 6.7857 | 6.9305 | 6.6939 | 5.4640 | 5.5769 | 4.8356 | 4.9807 |
| SNX2_S119 | SNX2 | O60749 | 185 | SVKRRFSDFLGLH | 5.2978 | 5.4431 | 5.8089 | 5.3104 | 5.6925 | 6.6081 | 5.4970 | 6.3474 | 6.7349 | 7.7385 | 6.7809 | 6.9790 | 6.7567 | 6.3857 | 6.5222 | 6.0968 |
| SNX2_S185 | SNX2 | O60749 | 119 | IESKSMSAPVIFD | 3.2098 | 3.2078 | 3.1974 | 3.1435 | 3.6116 | 3.6840 | 3.5193 | 3.7659 | 3.3381 | 3.1918 | 3.1503 | 3.2877 | 14.9955 | 14.7684 | 16.0225 | 13.9064 |
| SNX2_S277 | SNX2 | O60749 | 277 | VNTQALSGAGILR | 6.9235 | 6.9041 | 7.8378 | 7.9823 | 6.3251 | 6.5493 | 7.0426 | 7.1484 | 6.9891 | 6.8699 | 7.1417 | 6.8237 | 3.8734 | 4.5420 | 3.1257 | 3.9216 |
| SNX24_S60 | SNX24 | Q9Y343 | 60 | KTPEIPSKHVRNW | 6.7116 | 6.9632 | 6.3459 | 7.2259 | 5.8867 | 5.4266 | 6.3149 | 6.0869 | 6.2329 | 6.4012 | 6.1473 | 6.5718 | 5.5820 | 5.8047 | 6.0371 | 6.2614 |
| SNX3_S72 | SNX3 | O60493 | 72 | TVRRRYSDFEWLR | 6.5073 | 6.4669 | 6.7473 | 6.0538 | 6.3048 | 6.5565 | 6.4032 | 7.5394 | 6.6997 | 7.8262 | 7.5622 | 7.7193 | 3.8701 | 4.9905 | 3.9989 | 4.7539 |
| SNX30_S67 | SNX30 | Q5VWJ9 | 67 | TPAGTSSPASSSS | 6.2125 | 5.9760 | 6.5630 | 6.3125 | 6.3847 | 6.2342 | 6.6671 | 6.3095 | 6.6674 | 6.5964 | 5.9717 | 6.2755 | 5.9708 | 6.0088 | 5.6793 | 6.1706 |
| SNX33_S169 | SNX33 | Q8WV41 | 169 | PLERQDSLASAKR | 7.2945 | 6.8334 | 8.1795 | 6.3009 | 6.1233 | 6.3911 | 6.7014 | 6.2820 | 6.5299 | 6.4019 | 6.8646 | 5.9260 | 4.6630 | 6.0940 | 4.4666 | 4.9477 |
| SNX4_S22 | SNX4 | O95219 | 22 | PLEPLGSPDAGLG | 6.8041 | 7.5044 | 6.9645 | 7.4935 | 6.1777 | 7.0422 | 7.4073 | 6.6212 | 7.1798 | 7.5306 | 7.7293 | 7.3903 | 2.8015 | 4.1840 | 3.0300 | 4.1397 |
| SNX8_T452 | SNX8 | Q9Y5X2 | 452 | GPHSTLTPPCSPP | 5.7420 | 7.3179 | 7.0982 | 6.8247 | 5.9749 | 6.9108 | 6.0057 | 7.1626 | 6.7814 | 6.7380 | 7.1430 | 6.9297 | 4.8536 | 5.1215 | 4.9096 | 4.4864 |
| SNX8_T452;S456 | SNX8 | Q9Y5X2 | 452;456 | NA | 6.2033 | 6.6463 | 6.7395 | 6.5757 | 6.1778 | 7.2875 | 6.6503 | 7.5081 | 6.5286 | 6.8546 | 7.2701 | 6.5261 | 4.3267 | 4.9089 | 4.7389 | 5.0575 |
| SNX9_S198 | SNX9 | Q9Y5X1 | 200 | SRASSSSMKIPLN | 8.5646 | 6.7949 | 7.8787 | 6.5464 | 5.4494 | 6.1946 | 5.4744 | 6.5671 | 7.9622 | 7.2582 | 7.0848 | 7.7054 | 4.6467 | 5.2829 | 3.1131 | 3.4768 |
| SNX9_S200 | SNX9 | Q9Y5X1 | 198 | GNSRASSSSMKIP | 7.3146 | 6.0960 | 6.6900 | 6.0980 | 6.0907 | 6.7313 | 5.8150 | 6.7512 | 7.0742 | 7.3006 | 7.2486 | 7.2759 | 5.4721 | 6.3524 | 3.6030 | 4.0864 |
| SNX9_T140 | SNX9 | Q9Y5X1 | 140 | AQRNTNTPNNWDT | 6.3357 | 6.9225 | 6.6990 | 6.3342 | 6.2569 | 6.9822 | 6.8852 | 5.8760 | 5.8015 | 7.7082 | 8.3224 | 7.1558 | 4.8900 | 5.0531 | 4.1964 | 4.5808 |
| SOCS4_S125 | SOCS4 | Q8WXH5 | 125 | KRKIHISELMLDK | 5.7270 | 6.1520 | 6.6836 | 5.9953 | 7.3392 | 6.8954 | 5.8373 | 6.9343 | 4.6616 | 5.3822 | 6.1595 | 5.4850 | 6.2899 | 6.5118 | 5.9025 | 6.2435 |
| SOCS5_S211 | SOCS5 | O75159 | 211 | KRKIHLSELMLEK | 6.1056 | 7.2798 | 6.8930 | 7.0562 | 6.1896 | 5.9886 | 6.7648 | 6.4009 | 6.7297 | 6.1799 | 5.7395 | 6.2836 | 5.2683 | 5.3340 | 5.5202 | 6.2664 |
| SOCS5_S271 | SOCS5 | O75159 | 271 | RERRRLSIEEGVD | 6.4353 | 6.8383 | 7.0691 | 6.9121 | 6.1959 | 5.8667 | 6.4513 | 6.3681 | 7.1755 | 6.6748 | 6.3514 | 6.8875 | 5.1526 | 5.6764 | 4.9450 | 5.0000 |
| SOCS6_S311 | SOCS6 | O14544 | 311 | DDVPPLSPLLPPM | 5.3464 | 4.9615 | 6.2818 | 6.7851 | 5.3345 | 5.3108 | 6.5719 | 8.3037 | 5.9562 | 7.3890 | 7.2114 | 4.9378 | 6.5749 | 6.4032 | 6.9923 | 5.6394 |
| SOCS6_S70 | SOCS6 | O14544 | 76 | SESLMGTLKRRLS | 9.4611 | 7.5850 | 9.2812 | 7.6687 | 5.7802 | 4.4344 | 7.1012 | 5.3709 | 6.9152 | 6.5741 | 7.6842 | 4.6639 | 4.3134 | 4.7445 | 4.4468 | 3.9754 |
| SOCS6_T76 | SOCS6 | O14544 | 70 | GKNRSKSESLMGT | 5.2740 | 5.9559 | 5.3124 | 5.8170 | 7.4257 | 5.7274 | 4.5574 | 6.8188 | 6.1826 | 5.0236 | 5.1241 | 5.4879 | 7.4878 | 8.4659 | 7.2886 | 8.0509 |
| SOGA1_S1164 | SOGA1 | O94964-2 | 1164 | GLKRTKSVSSMSE | 6.2254 | 6.0957 | 6.2377 | 6.5648 | 6.3930 | 6.0405 | 6.2881 | 6.3393 | 6.1975 | 6.4817 | 6.4115 | 6.6265 | 5.8087 | 6.7108 | 5.6722 | 5.9066 |
| SOGA1_S1255 | SOGA1 | O94964-2 | 1255 | GIRVYYSPPVARR | 6.4740 | 6.9155 | 6.9108 | 6.5846 | 5.9983 | 6.7707 | 6.4184 | 6.8018 | 6.8571 | 7.0484 | 6.7330 | 6.9115 | 4.5541 | 5.3436 | 4.6429 | 5.0353 |
| SOGA1_S142 | SOGA1 | O94964-2 | 175 | PPRPPASPDEPSV | 6.5434 | 6.2644 | 7.0117 | 6.1028 | 5.7456 | 6.1970 | 7.0069 | 5.8923 | 6.3173 | 6.6745 | 6.8052 | 6.6048 | 5.5287 | 6.3277 | 5.3126 | 5.6650 |
| SOGA1_S175 | SOGA1 | O94964-2 | 35 | KSHRAPSPARPKD | 6.5830 | 6.7676 | 6.6485 | 5.8440 | 5.1279 | 6.3638 | 6.3340 | 5.8395 | 6.9815 | 7.0351 | 6.9781 | 6.9010 | 5.7816 | 6.3659 | 4.8973 | 5.5513 |
| SOGA1_S35 | SOGA1 | O94964-2 | 142 | LELALSSDAESAA | 6.2681 | 4.8958 | 6.8524 | 6.5625 | 7.0335 | 4.9139 | 7.1060 | 6.5082 | 4.1803 | 7.5045 | 7.3596 | 6.4402 | 5.7156 | 6.1685 | 6.0281 | 6.4629 |
| SOGA3_S569 | SOGA3 | Q5TF21 | 569 | FYGDLDSPLPKGE | 5.6066 | 5.4849 | 5.5296 | 6.4754 | 5.8254 | 6.7142 | 5.7885 | 7.0683 | 6.0106 | 6.3129 | 6.2410 | 5.9161 | 6.2578 | 7.3557 | 6.3627 | 7.0502 |
| SON_S1066 | SON | P18583-9 | 1737 | TLAALLSPKESSG | 6.1020 | 6.0355 | 6.0339 | 6.4449 | 5.8515 | 5.5174 | 6.1248 | 5.7283 | 6.3022 | 5.9275 | 5.9272 | 6.1023 | 6.6953 | 7.1906 | 6.8635 | 7.1529 |
| SON_S152 | SON | P18583-9 | 1809 | RDRSAASPVVSSM | 5.8013 | 5.9767 | 5.4367 | 6.0824 | 5.9571 | 5.3811 | 5.7806 | 5.6253 | 6.6952 | 6.4835 | 6.4313 | 6.3858 | 6.7045 | 7.2550 | 6.9890 | 7.0144 |
| SON_S154 | SON | P18583-9 | 1610 | EKILPTSETKQRT | 6.8636 | 7.2714 | 6.3998 | 7.6484 | 6.3880 | 6.5754 | 7.2512 | 6.5609 | 7.0318 | 6.8848 | 6.8222 | 7.0156 | 3.6877 | 5.0410 | 3.7631 | 4.7949 |
| SON_S1596 | SON | P18583-9 | 152 | GNIDLESDSFLKF | 6.1687 | 8.0691 | 6.2242 | 6.3565 | 5.7661 | 5.5459 | 6.3401 | 7.8503 | 6.0981 | 5.1259 | 5.1748 | 5.4296 | 5.9756 | 6.8754 | 5.9428 | 7.0570 |
| SON_S1610 | SON | P18583-9 | 2169 | VNKPHVSDEEEEE | 6.3655 | 5.9189 | 6.1592 | 6.5538 | 5.5585 | 5.0886 | 6.0612 | 5.3958 | 6.0393 | 6.1058 | 5.4400 | 5.7441 | 7.0164 | 7.4672 | 7.0915 | 7.9942 |
| SON_S1737 | SON | P18583-9 | 94 | RKSRCVSVQTDPT | 4.2336 | 3.9451 | 4.4397 | 4.6563 | 4.9360 | 4.8911 | 5.0698 | 4.8893 | 4.3459 | 4.0216 | 4.0272 | 4.3487 | 11.0476 | 11.1618 | 12.1134 | 11.8731 |
| SON_S1809 | SON | P18583-9 | 154 | IDLESDSFLKFDS | 6.1189 | 6.7048 | 6.6272 | 7.3020 | 6.3907 | 6.1822 | 6.9296 | 7.0037 | 6.3713 | 5.8356 | 6.1404 | 6.3931 | 4.7396 | 5.8544 | 5.3848 | 6.0216 |
| SON_S1822 | SON | P18583-9 | 1596 | VCAAGTSPVGEIG | 5.9398 | 6.6524 | 5.4058 | 6.7059 | 6.5595 | 6.3106 | 6.0228 | 5.8692 | 5.8676 | 5.3652 | 5.8273 | 5.2547 | 6.0318 | 7.7981 | 7.6569 | 6.7324 |
| SON_S2049 | SON | P18583-9 | 1595 | TVCAAGTSPVGEI | 5.8525 | 5.9762 | 5.6447 | 6.9945 | 6.0126 | 6.2438 | 5.4922 | 5.8874 | 6.5890 | 5.4836 | 6.4386 | 5.4638 | 6.4241 | 7.9536 | 6.8438 | 6.6997 |
| SON_S2049;S2051;S205[illegible] | SON | P18583-9 | 1685 | HDMVISTSPSGGS | 6.0797 | 6.1716 | 6.0795 | 6.5926 | 5.7034 | 5.8651 | 6.2831 | 6.2925 | 6.1241 | 6.4870 | 6.0200 | 5.9578 | 6.4928 | 6.9927 | 6.2788 | 6.5793 |
| SON_S2069;S2071 | SON | P18583-9 | 2203 | IAAAKPTPPKSQV | 7.0656 | 6.0741 | 6.6223 | 7.1539 | 6.3855 | 6.9610 | 7.1298 | 5.9559 | 6.7779 | 6.9903 | 7.2020 | 7.4848 | 4.0905 | 5.4251 | 4.0923 | 4.5889 |
| SON_S2071 | SON | P18583-9 | 2049 | SVVRRRSFSISPV | 5.7583 | 6.5625 | 5.5025 | 5.8522 | 5.5950 | 5.6958 | 5.5861 | 5.7044 | 5.9331 | 6.2141 | 6.0584 | 5.6937 | 6.6462 | 7.2592 | 7.6384 | 8.3000 |
| SON_S2169 | SON | P18583-9 | 281 | VLKSVESTSPEPS | 6.5168 | 6.0651 | 5.9028 | 6.5146 | 6.4545 | 6.0869 | 6.7680 | 6.1633 | 6.0497 | 5.7477 | 5.8150 | 5.3918 | 5.9838 | 7.1501 | 6.8939 | 6.4962 |
| SON_S2278 | SON | P18583-9 | 2071 | RRRFSRSPIRRKR | 7.3541 | 7.2361 | 6.1157 | 7.2392 | 5.5383 | 5.6301 | 5.5359 | 5.3543 | 6.0130 | 6.1684 | 6.3942 | 7.1786 | 5.2999 | 6.4733 | 5.7580 | 6.7110 |
| SON_S281 | SON | P18583-9 | 2278 | LAQKRLSENAFDL | 5.1962 | 5.3058 | 5.8782 | 6.0947 | 6.3376 | 6.0186 | 6.0025 | 6.0934 | 6.3785 | 6.6032 | 5.6991 | 5.8417 | 7.2787 | 7.0944 | 7.0761 | 7.1013 |
| SON_S354 | SON | P18583-9 | 999 | HDPYRLTPDPYRM | 7.0427 | 6.8532 | 7.0401 | 7.4546 | 6.2294 | 6.9523 | 7.2810 | 6.5139 | 7.5695 | 6.5456 | 6.1878 | 7.1477 | 3.4765 | 5.1256 | 3.6864 | 4.8938 |

| | | | | | | | | | | | | | | | | | | | | |
|---|---|---|---|---|---|---|---|---|---|---|---|---|---|---|---|---|---|---|---|---|
| SON_S94 | SON | P18583-9 | 1066 | YERSMMSPMAERS | 7.1747 | 7.7706 | 6.8165 | 6.9930 | 6.7677 | 6.2278 | 7.0689 | 5.8035 | 6.6581 | 6.6134 | 6.9904 | 7.3460 | 4.1097 | 4.9650 | 3.9659 | 4.7288 |
| SON_T1595 | SON | P18583-9 | 354 | SEIADSSMTRPQE | 6.2882 | 7.5270 | 6.5156 | 8.0784 | 6.9816 | 6.3262 | 7.5938 | 6.7017 | 6.3152 | 6.2425 | 6.9900 | 6.1452 | 3.8671 | 5.5381 | 4.1030 | 4.7863 |
| SON_T1685 | SON | P18583-9 | 2062 | RLRRSRTPLRRRF | 6.0350 | 6.9175 | 6.4665 | 6.4318 | 6.1470 | 5.9511 | 6.0328 | 5.3621 | 5.2349 | 6.4665 | 5.8991 | 6.9196 | 6.7462 | 7.1459 | 6.2944 | 5.9497 |
| SON_T1685;S1686 | SON | P18583-9 | 1822 | PERASESSSEEKD | 5.5714 | 5.7650 | 6.3729 | 6.9953 | 6.1460 | 5.1090 | 6.0046 | 5.5173 | 6.6233 | 6.6389 | 6.2240 | 6.4099 | 6.0562 | 6.9102 | 6.3120 | 7.3440 |
| SON_T2062 | SON | P18583-9 | 2069;2071 | NA | 6.2003 | 6.2672 | 5.6257 | 6.6110 | 5.9212 | 5.4307 | 6.0315 | 6.0359 | 6.1144 | 5.8758 | 5.8495 | 5.8481 | 6.5382 | 7.2598 | 7.1350 | 7.2559 |
| SON_T2203 | SON | P18583-9 | 2049;2051;20 | NA | 6.1978 | 6.1988 | 6.2857 | 6.5453 | 6.3096 | 6.1237 | 6.3904 | 5.2394 | 6.1914 | 5.5705 | 5.6278 | 6.5881 | 6.1800 | 7.0655 | 6.8054 | 6.6803 |
| SON_T999 | SON | P18583-9 | 1685;1686 | NA | 6.5671 | 5.9960 | 5.6918 | 6.5028 | 5.8403 | 5.5854 | 6.2524 | 5.9564 | 5.8051 | 5.9179 | 5.7391 | 6.1491 | 6.3564 | 7.3678 | 7.4525 | 6.8200 |
| SORBS1_S325 | SORBS1 | Q9BX66 | 325 | EQQKRLSSLSDPV | 6.0138 | 7.1902 | 6.1421 | 6.9091 | 7.1291 | 6.9152 | 4.7273 | 6.5974 | 7.9756 | 6.1238 | 5.2834 | 5.9374 | 5.4912 | 6.2215 | 5.3629 | 5.9801 |
| SORBS3_S373 | SORBS3 | O60504 | 395 | FDFQAQSPKELTL | 7.8064 | 8.0400 | 7.3760 | 7.3113 | 6.6179 | 6.9921 | 7.4320 | 6.9216 | 7.6821 | 7.5380 | 7.7700 | 7.1558 | 2.4658 | 3.3851 | 2.2717 | 3.2342 |
| SORBS3_S395 | SORBS3 | O60504 | 530 | GPQLPTSPRLTAA | 6.2264 | 6.3362 | 6.9966 | 6.1561 | 6.2603 | 6.6004 | 6.2325 | 6.2709 | 6.6534 | 6.9240 | 6.6714 | 6.5852 | 5.4859 | 5.7502 | 5.4472 | 5.4034 |
| SORBS3_S530 | SORBS3 | O60504 | 373 | ASNGGGSPARREE | 6.5882 | 7.3882 | 6.6557 | 6.3679 | 6.2757 | 7.0334 | 6.0502 | 6.4860 | 7.2460 | 6.6810 | 6.6557 | 7.4956 | 5.1128 | 4.8256 | 4.6118 | 4.5260 |
| SORBS3_S545 | SORBS3 | O60504 | 545 | SARHPSSPSALRS | 5.5325 | 6.0686 | 6.8980 | 6.6914 | 6.2087 | 6.6156 | 6.2923 | 6.8907 | 7.1004 | 5.6430 | 6.4490 | 6.4418 | 5.8439 | 6.4127 | 5.2351 | 5.6763 |
| SORT1_S793 | SORT1 | Q99523 | 793 | FLVHRYSVLQQHA | 5.0180 | 6.0605 | 7.1547 | 7.0756 | 6.2093 | 5.4365 | 5.3516 | 7.0303 | 5.7902 | 6.6730 | 7.0720 | 6.5788 | 5.9764 | 6.7715 | 6.0037 | 5.7981 |
| SORT1_S814 | SORT1 | Q99523 | 814 | DALDTASHTNKSG | 6.3100 | 6.5269 | 7.2374 | 6.8316 | 5.7219 | 6.4825 | 6.6213 | 6.0266 | 6.0710 | 5.8369 | 6.9737 | 6.7971 | 4.7411 | 6.3971 | 5.7613 | 5.6636 |
| SOS1_S1134 | SOS1 | Q07889 | 1275 | TRRHLPSPPLTQE | 5.7606 | 5.7494 | 6.3941 | 6.1431 | 6.4144 | 6.0235 | 6.0892 | 6.3156 | 6.1408 | 6.1510 | 6.1786 | 6.0958 | 6.6956 | 7.0535 | 6.2593 | 6.5353 |
| SOS1_S1167 | SOS1 | Q07889 | 1210 | ISDPPESPPLLPP | 5.4416 | 5.7634 | 6.3902 | 5.9388 | 6.0138 | 6.1475 | 6.0593 | 6.3569 | 5.9027 | 5.7168 | 5.8090 | 6.2749 | 7.0388 | 6.9934 | 6.8384 | 7.3145 |
| SOS1_S1178 | SOS1 | Q07889 | 1167 | SAPAESSPSKIMS | 6.3787 | 6.5983 | 6.1017 | 6.6457 | 6.1390 | 6.8949 | 6.0555 | 7.0683 | 7.4448 | 6.1668 | 6.8529 | 7.2856 | 3.9760 | 5.8809 | 5.0202 | 5.4908 |
| SOS1_S1210 | SOS1 | Q07889 | 1134 | HGPRSASVSSISL | 5.5537 | 4.9302 | 5.9951 | 5.5491 | 6.2287 | 6.4829 | 6.3455 | 6.9960 | 6.7042 | 6.0799 | 6.1805 | 6.3575 | 7.4926 | 7.4511 | 6.0704 | 5.5827 |
| SOS1_S1275 | SOS1 | Q07889 | 1178 | MSKHLDSPPAIPP | 6.6135 | 5.9166 | 6.0565 | 6.2115 | 5.9263 | 7.2048 | 6.3385 | 6.7631 | 7.3458 | 6.5708 | 6.0802 | 6.5471 | 5.2648 | 5.6808 | 5.7842 | 5.6956 |
| SOWAHC_S213 | SOWAHC | Q53LP3 | 213 | DLVMGSSPQLKRS | 6.4499 | 6.4872 | 6.6829 | 5.9768 | 6.1728 | 6.7311 | 6.3538 | 6.8127 | 6.7821 | 6.4786 | 6.5431 | 6.5427 | 5.7318 | 5.8547 | 5.1401 | 5.2598 |
| SOWAHC_T485 | SOWAHC | Q53LP3 | 485 | TQIVHTTPSFRDP | 6.9294 | 6.6138 | 5.4692 | 5.4102 | 8.6050 | 6.5725 | 8.6315 | 7.1507 | 6.4191 | 6.2539 | 6.3660 | 5.9117 | 5.0090 | 5.5016 | 4.6314 | 4.5252 |
| SOX13_S310 | SOX13 | Q9UN79 | 386 | LISLDSSPAKERL | 6.1782 | 7.2050 | 7.2351 | 6.6521 | 5.9792 | 6.3269 | 6.0359 | 6.2174 | 6.8228 | 6.9273 | 6.9108 | 7.1557 | 4.6873 | 5.4376 | 4.7672 | 5.4618 |
| SOX13_S386 | SOX13 | Q9UN79 | 310 | LPNTSSSPSLKMS | 6.6200 | 6.5971 | 7.0150 | 5.7505 | 6.5847 | 6.4094 | 6.2167 | 6.5295 | 6.9002 | 6.9598 | 7.2446 | 7.0080 | 5.2058 | 5.3798 | 4.5699 | 5.0091 |
| SOX9_S199 | SOX9 | P48436 | 199 | TEQTHISPNAIFK | 5.7534 | 5.6345 | 6.6034 | 6.1395 | 6.4273 | 6.2790 | 6.1014 | 6.5976 | 6.6255 | 6.9279 | 6.8824 | 7.0174 | 6.0324 | 6.1160 | 5.2572 | 5.6051 |
| SP1_S42 | SP1 | P08047 | 59 | GQESQPSPLALLA | 5.5087 | 5.6091 | 6.2405 | 5.7478 | 5.9292 | 6.4022 | 5.7618 | 5.9682 | 6.3301 | 5.7335 | 5.7875 | 6.3767 | 7.4341 | 7.4080 | 6.9538 | 6.8089 |
| SP1_S59 | SP1 | P08047 | 73 | TCSRIESPNENSN | 5.4613 | 5.9263 | 5.5176 | 5.4446 | 6.5446 | 6.2073 | 5.9892 | 6.1302 | 6.1200 | 6.2579 | 6.2709 | 6.0811 | 7.3231 | 7.0546 | 6.9333 | 6.7380 |
| SP1_S73 | SP1 | P08047 | 42 | SQARSSSTGSSSS | 5.1871 | 6.5314 | 6.0230 | 6.5060 | 6.1753 | 6.0484 | 6.6454 | 6.4352 | 5.5513 | 6.1607 | 5.9581 | 6.7140 | 7.2160 | 6.6713 | 5.8257 | 6.3509 |
| SP1_T117 | SP1 | P08047 | 651 | AHLRWHTGERPFM | 6.7301 | 7.0751 | 7.1733 | 7.3903 | 5.7022 | 7.3722 | 6.7840 | 6.0175 | 7.1848 | 7.5137 | 8.2036 | 8.2983 | 3.0453 | 4.9468 | 3.0563 | 3.5066 |
| SP1_T651 | SP1 | P08047 | 117 | SSSSGATPTSKEQ | 5.9747 | 6.9244 | 5.5508 | 5.3254 | 5.7440 | 5.5240 | 8.2014 | 6.6695 | 8.8373 | 5.2181 | 5.8191 | 6.8466 | 4.2684 | 5.8647 | 6.2859 | 6.9459 |
| SP100_S157 | SP100 | P23497-4 | 18 | RLNECISPVANEM | 5.5918 | 6.1057 | 6.0796 | 5.9133 | 6.1397 | 6.4392 | 5.7494 | 6.2882 | 6.3912 | 6.2965 | 6.2735 | 6.4002 | 6.7956 | 6.4990 | 6.5990 | 6.4380 |
| SP100_S18 | SP100 | P23497-4 | 451 | SRKRRFSSSDFSD | 6.2013 | 6.4664 | 5.9807 | 5.7440 | 5.7937 | 5.4610 | 5.3644 | 6.0158 | 7.7563 | 8.3683 | 7.6366 | 7.7488 | 5.3948 | 6.0795 | 4.8043 | 5.1841 |
| SP100_S189 | SP100 | P23497-4 | 157 | KLPLQESEEEERE | 6.2227 | 6.1782 | 6.3163 | 6.3722 | 5.9615 | 6.3923 | 6.0754 | 6.4257 | 5.9386 | 6.0754 | 6.4051 | 6.2166 | 6.3312 | 6.5590 | 6.2074 | 6.3224 |
| SP100_S274 | SP100 | P23497-4 | 362 | NDNPLESNDEKEG | 6.3015 | 6.2612 | 6.7689 | 6.1372 | 6.2292 | 6.3371 | 7.0128 | 5.9246 | 6.3001 | 6.4240 | 6.2237 | 6.8034 | 5.6680 | 6.3709 | 5.7051 | 5.5321 |
| SP100_S362 | SP100 | P23497-4 | 394 | LSTFRESFKKRVI | 5.7754 | 6.6656 | 7.2700 | 6.3447 | 6.2181 | 6.4323 | 5.9906 | 6.3308 | 7.2523 | 8.2210 | 8.5080 | 8.2523 | 4.4294 | 4.8450 | 3.4300 | 4.0346 |
| SP100_S394 | SP100 | P23497-4 | 274 | LPCDEESPEAELH | 7.3416 | 5.4488 | 5.0215 | 5.8797 | 6.4129 | 7.5641 | 6.8720 | 5.6678 | 6.6777 | 6.6919 | 6.1798 | 6.6248 | 8.4293 | 5.9927 | 4.4883 | 4.7073 |
| SP100_S451 | SP100 | P23497-4 | 189 | SLTWPPSGSPSHA | 6.2529 | 7.2507 | 6.3275 | 5.2925 | 7.3603 | 7.3209 | 6.5271 | 6.9240 | 5.7585 | 5.8374 | 6.0402 | 6.3078 | 5.8604 | 5.9273 | 6.3560 | 4.6565 |
| SP100_S451;S453 | SP100 | P23497-4 | 451;453 | NA | 4.5798 | 3.7351 | 5.9847 | 5.3825 | 6.0835 | 5.2726 | 4.6548 | 6.9619 | 5.1283 | 7.5871 | 4.9756 | 6.0338 | 7.7674 | 9.5476 | 8.5369 | 7.7684 |
| SP110_S171 | SP110 | Q9HB58-6 | 244 | PQEMPHSPLGSMP | 5.7951 | 6.6342 | 6.2779 | 6.5051 | 6.1794 | 6.5544 | 6.5138 | 6.1932 | 6.2370 | 6.3000 | 6.1341 | 6.0510 | 5.8163 | 7.0381 | 6.0154 | 5.7550 |
| SP110_S244 | SP110 | Q9HB58-6 | 256 | PEIRDNSPEPNDP | 5.5142 | 5.9992 | 5.9449 | 6.4616 | 5.4159 | 4.6360 | 6.1195 | 5.1016 | 6.2669 | 6.5243 | 5.9022 | 6.6271 | 7.3055 | 7.5060 | 6.6032 | 8.0718 |
| SP110_S256 | SP110 | Q9HB58-6 | 287 | KRCIWSTPKRRHK | 6.8883 | 7.5173 | 6.5584 | 7.1487 | 5.9901 | 6.7686 | 7.0877 | 6.1433 | 6.8597 | 7.0964 | 7.5211 | 7.3707 | 3.8096 | 4.7466 | 3.8017 | 4.6920 |
| SP110_T287 | SP110 | Q9HB58-6 | 171 | QSDEILSESPSPS | 6.1670 | 5.7010 | 5.8693 | 6.4114 | 6.4141 | 6.5914 | 6.4982 | 6.1116 | 6.2442 | 6.3039 | 6.1404 | 7.1543 | 5.3214 | 6.7544 | 6.5440 | 5.7735 |
| SP2_S78 | SP2 | Q02086 | 78 | PAPLPLSPGKNSF | 6.7030 | 5.8253 | 6.5468 | 6.5096 | 6.5653 | 6.8922 | 6.8682 | 6.1206 | 7.0334 | 6.7599 | 6.7814 | 6.7056 | 5.2616 | 5.5460 | 4.4205 | 5.4608 |
| SP2_T238 | SP2 | Q02086 | 238 | LTESPPTPLSKTN | 6.5284 | 6.2581 | 6.3196 | 6.3273 | 6.4669 | 6.3294 | 7.0671 | 5.8832 | 6.6146 | 6.7707 | 5.5847 | 6.8573 | 5.7119 | 6.1192 | 5.2112 | 5.9504 |
| SP3_S73 | SP3 | Q02447 | 115 | WEVLSATPTTIKD | 6.8441 | 6.1750 | 6.0808 | 6.2437 | 6.1712 | 6.5253 | 6.5581 | 6.4792 | 6.5557 | 6.2987 | 6.3003 | 5.9972 | 6.0287 | 5.8510 | 5.6019 | 6.2890 |
| SP3_T115 | SP3 | Q02447 | 73 | SKIGPPSPGDDEE | 5.5932 | 5.5001 | 6.1646 | 6.8048 | 6.3561 | 5.8122 | 6.4782 | 6.4446 | 6.4210 | 6.2219 | 6.4324 | 5.6992 | 5.7374 | 6.8406 | 7.2717 | 6.2219 |
| SP4_S770 | SP4 | Q02446 | 770 | AAISQDSNPATPN | 6.5041 | 6.1142 | 7.0976 | 6.1314 | 5.5703 | 6.1577 | 5.9643 | 5.9795 | 6.5193 | 5.3313 | 5.6301 | 5.5885 | 6.6773 | 6.9284 | 6.6854 | 7.1209 |
| SPAG1_S423 | SPAG1 | Q07617 | 423 | RSPRRASAAAAAG | 5.6777 | 4.7519 | 6.3469 | 5.4248 | 6.0083 | 5.7571 | 5.4269 | 5.9032 | 7.3153 | 7.4528 | 7.4085 | 7.3486 | 6.3796 | 7.6470 | 5.7134 | 5.4382 |
| SPAG1_S565 | SPAG1 | Q07617 | 565 | DSVNRLSRILMEL | 6.3848 | 5.9741 | 6.2164 | 7.1413 | 6.3080 | 5.7564 | 6.1228 | 6.3661 | 6.3799 | 5.5958 | 5.7426 | 6.5395 | 6.3789 | 6.5808 | 6.1839 | 6.3287 |
| SPAG5_S12 | SPAG5 | Q96R06 | 135 | TIVLVPSPLGQQQ | 6.0885 | 6.3527 | 6.0926 | 6.8580 | 6.1566 | 6.3671 | 6.0997 | 6.9207 | 6.6336 | 6.3607 | 6.6430 | 6.8233 | 5.2934 | 6.0752 | 5.6668 | 5.5681 |
| SPAG5_S12;S14 | SPAG5 | Q96R06 | 12 | KLSLSLSPSPQTG | 6.7104 | 6.5266 | 7.0616 | 6.4970 | 6.3438 | 7.1152 | 7.3767 | 7.6559 | 8.0280 | 6.6645 | 7.1563 | 7.3587 | 3.8686 | 4.3714 | 3.0697 | 4.1956 |
| SPAG5_S135 | SPAG5 | Q96R06 | 43 | TNSGKRSPACSSL | 5.9448 | 6.1804 | 6.4996 | 6.7779 | 6.6219 | 6.7305 | 6.9054 | 7.4891 | 7.0549 | 6.5274 | 7.1383 | 6.9326 | 4.9039 | 5.1502 | 4.4090 | 4.7341 |
| SPAG5_S362 | SPAG5 | Q96R06 | 39 | PGALTNSGKRSPA | 7.0783 | 7.1797 | 6.7859 | 8.5819 | 6.7319 | 6.8902 | 7.3125 | 7.2090 | 7.5459 | 7.2744 | 7.8844 | 8.2995 | 2.1089 | 3.5689 | 2.1822 | 3.3661 |
| SPAG5_S39 | SPAG5 | Q96R06 | 362 | LENLRQSLSLPSM | 7.0439 | 7.0379 | 6.7477 | 7.6441 | 6.1259 | 6.8432 | 6.9009 | 6.6669 | 7.4741 | 7.1518 | 7.5383 | 7.3740 | 3.5110 | 4.4303 | 3.3917 | 4.1183 |
| SPAG5_S43 | SPAG5 | Q96R06 | 398 | APQEKSTNTSQTG | 6.5939 | 6.7918 | 6.4570 | 7.7521 | 6.3245 | 7.7274 | 8.3035 | 6.8242 | 6.0964 | 7.1282 | 7.5482 | 7.3382 | 3.0911 | 4.5401 | 3.0801 | 4.4033 |
| SPAG5_S43;T50 | SPAG5 | Q96R06 | 43;50 | NA | 7.6218 | 7.5610 | 7.3967 | 7.5654 | 5.8851 | 6.9827 | 7.8464 | 6.8657 | 7.4391 | 7.0461 | 7.8449 | 7.9077 | 2.6220 | 3.8245 | 2.1955 | 3.3955 |
| SPAG5_T398 | SPAG5 | Q96R06 | 12;14 | NA | 9.1458 | 6.5499 | 6.7437 | 7.3131 | 5.0052 | 8.8096 | 6.9942 | 7.3920 | 9.2121 | 5.5752 | 6.7328 | 9.2696 | 2.6608 | 3.2818 | 2.4859 | 2.8284 |
| SPAG7_S158 | SPAG7 | O75391 | 158 | QGPVVVSPASDYK | 6.2217 | 6.6535 | 7.3614 | 6.6207 | 5.8807 | 6.8957 | 6.3785 | 6.7776 | 6.8157 | 6.7746 | 7.0296 | 7.2937 | 4.4305 | 5.7288 | 4.4460 | 4.6914 |
| SPAG9_S1188 | SPAG9 | O60271 | 217 | AGDGLLTPDAQKG | 5.8625 | 5.9750 | 6.5783 | 6.3068 | 6.0749 | 6.5781 | 6.2657 | 6.7389 | 6.0132 | 5.8923 | 6.3598 | 5.9352 | 6.2745 | 6.4640 | 6.4636 | 6.2171 |
| SPAG9_S1244 | SPAG9 | O60271 | 365 | GYKGSSTPTKGIE | 6.5328 | 6.6342 | 6.8622 | 6.3058 | 6.1012 | 6.7627 | 6.0804 | 6.6227 | 6.5086 | 6.6447 | 6.8447 | 7.1236 | 5.0858 | 5.5413 | 5.0637 | 5.2856 |
| SPAG9_S183 | SPAG9 | O60271 | 226 | AQKGGETPGSEQW | 7.0107 | 6.8300 | 7.4147 | 6.7968 | 6.2277 | 6.6803 | 6.5805 | 6.8795 | 6.4637 | 5.9178 | 6.3562 | 5.9979 | 4.7568 | 5.6820 | 5.0855 | 5.3200 |
| SPAG9_S238 | SPAG9 | O60271 | 251 | SLKVSNSPEPQKA | 6.3753 | 6.4870 | 6.8463 | 6.5779 | 6.1485 | 6.7891 | 6.0112 | 6.6509 | 6.4341 | 6.6969 | 6.8434 | 7.1176 | 4.9915 | 5.5661 | 5.0683 | 5.3961 |
| SPAG9_S251 | SPAG9 | O60271 | 733 | RSASQSSLDKLDQ | 6.2850 | 5.9918 | 7.0006 | 6.2185 | 6.3076 | 6.4112 | 6.3111 | 6.5310 | 6.6124 | 6.1855 | 6.5537 | 6.8082 | 5.7172 | 6.1263 | 5.4523 | 5.4875 |
| SPAG9_S339 | SPAG9 | O60271 | 586 | APTSHVTPSVKKR | 7.7226 | 8.0182 | 7.1321 | 7.6640 | 6.0530 | 7.6713 | 7.4654 | 6.7968 | 7.4311 | 6.5606 | 7.2020 | 7.1557 | 2.9184 | 4.2021 | 2.5863 | 3.4206 |
| SPAG9_S730 | SPAG9 | O60271 | 238 | WKFQELSQPRSHT | 6.7371 | 7.3191 | 6.6693 | 7.4348 | 6.5368 | 6.8704 | 7.1749 | 7.0850 | 7.1654 | 6.7743 | 6.8663 | 6.7901 | 3.5885 | 4.4088 | 3.8697 | 4.7096 |
| SPAG9_S733 | SPAG9 | O60271 | 183 | TKLHQLSGSDQLE | 6.0333 | 7.0408 | 7.1917 | 7.6464 | 6.5270 | 7.4438 | 6.5142 | 7.4639 | 7.0551 | 6.6490 | 7.2327 | 7.2561 | 3.7278 | 4.5358 | 3.5405 | 4.1418 |
| SPAG9_T217 | SPAG9 | O60271 | 292 | ATIPTDTPLKEEN | 6.3638 | 6.7357 | 7.5235 | 6.6639 | 7.0965 | 5.8875 | 6.8252 | 6.3331 | 7.4703 | 7.0080 | 6.1824 | 7.0937 | 4.8465 | 4.7130 | 4.4206 | 4.8365 |
| SPAG9_T226 | SPAG9 | O60271 | 348 | QAIIESTPELDMD | 5.6883 | 6.8138 | 6.9550 | 7.2118 | 5.9317 | 6.8826 | 6.3757 | 6.9099 | 6.5404 | 6.9807 | 7.4581 | 7.4946 | 4.3612 | 4.8146 | 4.5182 | 5.0636 |
| SPAG9_T292 | SPAG9 | O60271 | 595 | VKKRSSTLSQLPG | 6.8166 | 5.6655 | 6.6464 | 5.4194 | 5.8694 | 5.7887 | 5.9526 | 6.1801 | 6.7243 | 6.4454 | 6.7946 | 6.8985 | 6.7609 | 7.2026 | 5.3214 | 5.5137 |
| SPAG9_T330 | SPAG9 | O60271 | 339 | AENEEKSEVQAII | 6.7579 | 7.8889 | 6.9156 | 7.2271 | 6.6756 | 6.4212 | 6.4144 | 6.6098 | 6.6427 | 6.8227 | 6.2528 | 7.0636 | 4.4156 | 4.8684 | 4.0160 | 5.0077 |
| SPAG9_T348 | SPAG9 | O60271 | 1188 | PGNRPGSVIRVYG | 6.1965 | 6.3014 | 8.3106 | 6.3769 | 7.0530 | 6.2800 | 5.5651 | 6.0351 | 7.2108 | 6.3123 | 6.6322 | 7.0143 | 5.2682 | 5.7919 | 4.7381 | 4.9136 |
| SPAG9_T365 | SPAG9 | O60271 | 730 | SKQRSASQSSLDK | 6.1285 | 5.6136 | 6.1596 | 5.8520 | 6.0335 | 6.1658 | 6.1673 | 6.5181 | 5.7347 | 5.5740 | 6.2390 | 6.1944 | 6.7782 | 7.1897 | 6.8264 | 6.8252 |
| SPAG9_T586 | SPAG9 | O60271 | 1244 | SPQSSSSGTDLTG | 5.3078 | 5.7143 | 6.0386 | 5.1145 | 5.6720 | 5.3700 | 4.9501 | 5.2400 | 5.8709 | 6.3673 | 4.8335 | 4.5703 | 9.2039 | 8.8496 | 9.2361 | 7.6612 |
| SPAG9_T595 | SPAG9 | O60271 | 330 | ETRNVSTGSAENE | 6.3405 | 6.3865 | 7.0552 | 5.6252 | 5.6600 | 6.1492 | 5.7627 | 6.3181 | 6.6210 | 6.8678 | 6.4140 | 6.5441 | 5.8754 | 5.8036 | 5.8356 | 6.7411 |
| SPART_S470 | SPART | Q8N0X7 | 470 | EKPVEVSPAVTKG | 6.9971 | 6.5236 | 7.6556 | 7.2644 | 6.0271 | 7.0946 | 7.4406 | 6.4649 | 7.8175 | 7.6656 | 7.6623 | 8.4083 | 2.8083 | 3.7395 | 2.4600 | 3.9708 |
| SPAST_S268 | SPAST | Q9UBP0 | 268 | GHHRAPSYSGLSM | 6.6034 | 6.5598 | 7.0029 | 6.7386 | 4.6950 | 6.0415 | 5.5125 | 5.7903 | 7.0430 | 6.5609 | 6.8031 | 7.5938 | 5.6908 | 6.4193 | 5.4815 | 5.4638 |
| SPAST_T303 | SPAST | Q9UBP0 | 303 | RTNKPSTPTTATR | 6.2579 | 6.6304 | 7.1740 | 6.2366 | 6.0931 | 6.9201 | 6.8741 | 7.1546 | 6.8926 | 7.4177 | 7.2278 | 6.9357 | 4.3390 | 5.3323 | 4.1888 | 4.3251 |
| SPATA13_S254 | SPATA13 | Q96N96-6 | 254 | GYRRSKSTDNLAF | 6.1185 | 5.9940 | 7.4440 | 6.3307 | 5.1094 | 6.4344 | 6.3425 | 5.9514 | 5.8529 | 6.2279 | 6.7166 | 7.6504 | 5.5915 | 6.2720 | 5.6763 | 6.2876 |
| SPATA2L_S252 | SPATA2L | Q8IUW3 | 252 | PSPGPDSPPAELA | 6.1539 | 5.8951 | 6.6423 | 5.9659 | 6.3725 | 7.1552 | 5.9671 | 6.2785 | 7.2516 | 6.3053 | 6.8072 | 6.6594 | 5.5068 | 6.2858 | 5.4836 | 5.2699 |
| SPATA5L1_T368 | SPATA5L1 | Q9BVQ7 | 368 | REVVIGTPTLKQR | 5.8744 | 7.2093 | 6.6643 | 6.6675 | 6.5602 | 7.0833 | 6.6444 | 6.9817 | 6.4773 | 6.2404 | 6.5970 | 6.6163 | 4.9778 | 4.9953 | 4.9516 | 5.4592 |
| SPATA6_S217 | SPATA6 | Q9NWH7 | 217 | ISSKSHSPSPYTK | 6.0640 | 6.5547 | 6.3052 | 6.4720 | 5.6534 | 7.2811 | 6.4452 | 6.7132 | 6.6215 | 6.6124 | 6.2173 | 6.4185 | 5.3152 | 6.4104 | 4.9933 | 5.9226 |
| SPATA7_S194 | SPATA7 | Q9P0W8 | 194 | SGPRKLSSGALYG | 6.4479 | 6.1778 | 5.8110 | 5.2203 | 6.0522 | 5.3531 | 5.5825 | 5.7415 | 8.0946 | 8.2365 | 7.3548 | 7.8923 | 5.4723 | 5.7695 | 5.6801 | 5.1137 |
| SPATS2_S408 | SPATS2 | Q86XZ4 | 408 | SSSTCASPPSLTS | 5.8226 | 6.0795 | 7.1185 | 6.9472 | 6.4114 | 7.0237 | 6.6220 | 6.0240 | 7.4035 | 6.8468 | 6.7994 | 7.1624 | 4.3768 | 5.4545 | 4.5608 | 5.3469 |
| SPATS2_S47 | SPATS2 | Q86XZ4 | 47 | AIVPNKSNNEIIL | 2.5938 | 3.4668 | 3.1695 | 3.3886 | 3.4417 | 2.7143 | 3.5466 | 3.5231 | 3.9893 | 3.1085 | 2.6674 | 4.2443 | 16.1152 | 13.8890 | 16.0996 | 14.0423 |
| SPATS2L_S189 | SPATS2L | Q9NUQ6-3 | 561 | GSVGRVSQCNLCP | 6.2604 | 5.5261 | 6.4507 | 5.3966 | 5.6586 | 5.8471 | 5.8210 | 6.2682 | 7.5182 | 8.1378 | 7.5140 | 6.8751 | 5.5820 | 6.7847 | 5.0494 | 5.3101 |
| SPATS2L_S368 | SPATS2L | Q9NUQ6-3 | 368 | GKAARFSCDIEQL | 6.1792 | 6.1678 | 6.6306 | 5.9956 | 6.6385 | 6.4111 | 5.7682 | 6.3643 | 6.1593 | 6.4091 | 6.9491 | 6.2802 | 5.8627 | 6.0991 | 5.8164 | 6.2688 |
| SPATS2L_S556 | SPATS2L | Q9NUQ6-3 | 189 | LLQQKLSLDGNPK | 5.7720 | 5.5896 | 5.8701 | 5.3902 | 6.8463 | 6.3093 | 6.0676 | 7.0483 | 7.1590 | 7.0442 | 7.0332 | 6.2500 | 5.5371 | 6.6261 | 5.6700 | 5.7871 |
| SPATS2L_S561 | SPATS2L | Q9NUQ6-3 | 556 | ARPFRGSVGRVSQ | 6.3463 | 5.2839 | 6.0712 | 6.5118 | 5.6992 | 7.1265 | 6.2334 | 5.1603 | 6.7230 | 6.8119 | 6.8151 | 6.7531 | 6.5899 | 6.8476 | 5.9875 | 5.0394 |
| SPATS2L_T528 | SPATS2L | Q9NUQ6-3 | 528 | ASLGMKTPEAPAH | 7.1331 | 7.8795 | 7.1524 | 6.9282 | 5.4355 | 7.0631 | 7.3805 | 7.1634 | 7.4014 | 7.5689 | 7.4228 | 7.7429 | 3.0346 | 4.1185 | 2.8973 | 3.6778 |
| SPDL1_S555 | SPDL1 | Q96EA4 | 555 | SGNTPNSPRLAAE | 5.7373 | 6.3334 | 6.4473 | 5.6928 | 6.0583 | 6.4219 | 6.3380 | 6.1213 | 6.1538 | 6.3444 | 6.9843 | 6.2684 | 6.6649 | 6.1999 | 6.1822 | 6.0519 |
| SPECC1_S134 | SPECC1 | Q5M775 | 810 | VCVSRTSPTPPES | 6.6178 | 6.2743 | 6.3915 | 7.0208 | 6.4320 | 6.0157 | 6.5470 | 6.3863 | 6.4556 | 6.1316 | 6.3170 | 6.4704 | 6.1650 | 5.5101 | 5.4113 | 5.8537 |
| SPECC1_S151 | SPECC1 | Q5M775 | 847 | VHKTPRSPLSGIP | 6.7242 | 6.6822 | 5.9535 | 6.8993 | 6.8578 | 6.9263 | 6.8641 | 7.1005 | 6.7433 | 6.2073 | 6.4938 | 6.5320 | 5.1092 | 4.8490 | 4.8727 | 5.1848 |
| SPECC1_S810 | SPECC1 | Q5M775 | 914 | HLRKSPSLESLSR | 5.5477 | 6.2529 | 6.0842 | 6.0788 | 5.7103 | 6.6330 | 6.2807 | 6.4453 | 6.0422 | 5.7929 | 5.5226 | 6.3600 | 6.5354 | 6.8626 | 6.3728 | 7.4784 |
| SPECC1_S847 | SPECC1 | Q5M775 | 134 | PRKSVSSPTSSNT | 6.5891 | 7.1089 | 6.9907 | 7.6292 | 6.7408 | 6.3936 | 6.6332 | 7.0582 | 6.9101 | 7.7081 | 6.8975 | 7.4701 | 3.8111 | 4.1682 | 3.7277 | 4.1637 |
| SPECC1_S914 | SPECC1 | Q5M775 | 151 | KHLRTPSTKPKQE | 7.9269 | 7.3759 | 7.0743 | 6.8157 | 5.0449 | 6.7306 | 6.0388 | 6.2927 | 6.8187 | 6.9473 | 7.7914 | 8.3981 | 3.9905 | 4.8687 | 3.4946 | 4.3911 |
| SPECC1L_S107 | SPECC1L | Q69YQ0 | 881 | PPAAAVSPMQRHS | 6.5295 | 6.4827 | 6.6828 | 6.9654 | 6.5643 | 6.5246 | 6.8756 | 6.7244 | 7.3981 | 6.6478 | 7.2441 | 6.9779 | 4.2557 | 4.9252 | 4.3351 | 4.8668 |
| SPECC1L_S171 | SPECC1L | Q69YQ0 | 870 | RTPLSPSPMKTPP | 6.2390 | 6.1309 | 6.3673 | 6.0074 | 6.6115 | 6.4573 | 6.5183 | 6.3728 | 7.0053 | 6.3757 | 6.6409 | 6.6296 | 5.6473 | 5.9213 | 5.4297 | 5.6458 |
| SPECC1L_S220 | SPECC1L | Q69YQ0 | 868 | IPRTPLSPSPMKT | 6.1276 | 6.3616 | 6.2785 | 6.4929 | 6.6892 | 6.0572 | 6.4911 | 6.4445 | 6.7239 | 5.8455 | 6.0611 | 5.9242 | 6.0838 | 6.1999 | 5.8719 | 6.3471 |
| SPECC1L_S384 | SPECC1L | Q69YQ0 | 220 | GINEDHSEGDEKS | 6.3598 | 6.3480 | 7.2144 | 6.5874 | 6.3604 | 6.5180 | 6.6724 | 6.4838 | 6.3314 | 6.0435 | 5.7662 | 6.0843 | 6.0112 | 5.8415 | 5.6059 | 5.7717 |
| SPECC1L_S384;S385 | SPECC1L | Q69YQ0 | 887 | SPMQRHSISGPIS | 6.8649 | 5.1109 | 5.9657 | 5.0279 | 7.0375 | 5.6743 | 5.5408 | 6.2730 | 8.6879 | 7.1225 | 7.6121 | 6.8670 | 6.4072 | 7.2149 | 4.4358 | 4.1577 |
| SPECC1L_S54 | SPECC1L | Q69YQ0 | 384 | ERSRKGSSGNASE | 5.1638 | 6.1585 | 6.1557 | 6.0133 | 5.7412 | 6.2380 | 6.2359 | 5.7779 | 6.7621 | 6.4960 | 7.1858 | 6.5451 | 7.1978 | 6.7009 | 5.7864 | 5.8415 |
| SPECC1L_S832 | SPECC1L | Q69YQ0 | 54 | SLSKTKSSDDLLA | 6.5586 | 6.5614 | 7.3296 | 6.8754 | 5.3691 | 6.3383 | 6.4428 | 6.4093 | 6.5996 | 6.8714 | 6.9480 | 7.0004 | 4.7663 | 5.0305 | 5.0463 | 5.8531 |
| SPECC1L_S868 | SPECC1L | Q69YQ0 | 902 | KPLTALSDKRPNY | 9.1830 | 6.6749 | 7.0189 | 6.8915 | 5.5514 | 7.7155 | 7.8939 | 5.2138 | 7.9704 | 8.8305 | 8.1572 | 6.3606 | 3.0242 | 4.1604 | 2.0919 | 3.2620 |
| SPECC1L_S868;S870 | SPECC1L | Q69YQ0 | 171 | RMSKSKSDNQISD | 1.9679 | 2.7181 | 2.1744 | 2.2729 | 2.2898 | 2.3197 | 2.1614 | 2.4186 | 2.2500 | 1.8352 | 1.4758 | 2.0605 | 18.4090 | 19.3195 | 18.5308 | 17.7964 |
| SPECC1L_S870 | SPECC1L | Q69YQ0 | 107 | STGTASSTKRSTS | 6.8630 | 6.7466 | 9.2014 | 6.3292 | 4.4509 | 6.9250 | 6.6288 | 6.1756 | 8.0780 | 8.3198 | 8.2537 | 8.3453 | 2.9080 | 3.8930 | 3.3006 | 3.5810 |
| SPECC1L_S881 | SPECC1L | Q69YQ0 | 832 | GLSRRSSTSSEPT | 5.3825 | 5.0416 | 5.8314 | 5.4421 | 6.7447 | 5.5435 | 5.5712 | 6.8817 | 7.5530 | 7.4235 | 7.7308 | 7.3059 | 6.3076 | 6.8376 | 5.1193 | 5.2836 |
| SPECC1L_S887 | SPECC1L | Q69YQ0 | 868;870 | NA | 6.1691 | 6.4337 | 6.6560 | 7.0063 | 6.7098 | 6.7347 | 6.6320 | 6.3734 | 6.7800 | 7.0472 | 6.8422 | 6.7805 | 4.6356 | 5.2148 | 4.6836 | 5.3011 |
| SPECC1L_S902 | SPECC1L | Q69YQ0 | 384;385 | NA | 5.6549 | 5.2533 | 6.5985 | 6.0198 | 5.8867 | 5.8693 | 6.5980 | 5.7392 | 6.8975 | 6.8148 | 6.8848 | 6.6258 | 6.3721 | 6.0998 | 5.9660 | 6.7194 |
| SPEG_S2047 | SPEG | Q15772 | 853 | TPTMKPSPSQNRR | 6.2029 | 6.2905 | 6.1092 | 7.3291 | 5.9061 | 6.1946 | 6.1829 | 5.7704 | 6.1524 | 6.8164 | 6.7683 | 6.6544 | 5.8943 | 6.0479 | 5.7677 | 5.9130 |
| SPEG_S853 | SPEG | Q15772 | 2047 | PQRRSPSPGATRL | 7.4800 | 7.2547 | 6.8077 | 6.2916 | 5.5930 | 5.9671 | 6.8383 | 6.9222 | 6.8873 | 6.7422 | 6.2653 | 6.4955 | 4.5558 | 5.9965 | 4.3526 | 5.5501 |
| SPEN_S1062 | SPEN | Q96T58 | 727 | PIERSQSPVHLRR | 6.4348 | 6.7467 | 6.0120 | 7.0310 | 5.8214 | 5.9385 | 6.0882 | 6.0434 | 6.2290 | 6.1506 | 5.9674 | 6.0702 | 5.8626 | 6.7888 | 6.4737 | 6.3417 |
| SPEN_S1222 | SPEN | Q96T58 | 1918 | GDHENRSPVKEPV | 6.3954 | 6.7072 | 6.1195 | 7.2399 | 6.3998 | 6.1697 | 6.3896 | 5.9738 | 6.3681 | 5.9328 | 5.9686 | 6.3179 | 5.2502 | 6.5113 | 5.9852 | 6.2711 |
| SPEN_S1261 | SPEN | Q96T58 | 2393 | SEKPHSTPPQSCT | 5.8531 | 5.4765 | 5.9660 | 7.0064 | 5.9507 | 5.9865 | 6.0773 | 5.2457 | 6.0758 | 6.1780 | 6.1228 | 6.3343 | 6.5373 | 7.3611 | 6.6781 | 7.1505 |
| SPEN_S1380;S1382 | SPEN | Q96T58 | 1983 | PPEGWRSPRSQKT | 7.2820 | 7.4053 | 6.9198 | 7.8183 | 6.2860 | 6.7561 | 7.0394 | 6.4944 | 6.8500 | 6.9880 | 6.7313 | 7.4402 | 3.5366 | 4.7115 | 3.6577 | 4.0833 |
| SPEN_S1918 | SPEN | Q96T58 | 2120 | SGVVAVSPEKSES | 6.8057 | 6.3711 | 6.0864 | 6.6868 | 6.1264 | 5.6679 | 6.1885 | 5.8172 | 6.3007 | 5.9384 | 5.7886 | 6.3510 | 6.1361 | 6.9950 | 6.1629 | 6.5772 |
| SPEN_S1983 | SPEN | Q96T58 | 1633 | KDSELKTPPSVGP | 6.4898 | 6.6342 | 6.6008 | 7.2834 | 6.0729 | 5.8208 | 6.3049 | 6.2631 | 7.0389 | 6.7827 | 6.4551 | 6.6105 | 4.8510 | 5.8815 | 5.3509 | 5.5594 |
| SPEN_S2101 | SPEN | Q96T58 | 1619 | TENHPKTPESAPE | 6.2288 | 6.3328 | 5.7886 | 6.6066 | 5.5773 | 5.0659 | 5.8138 | 5.7377 | 5.7669 | 6.0018 | 5.5055 | 5.5425 | 6.6133 | 7.7314 | 7.7622 | 7.9249 |
| SPEN_S2120 | SPEN | Q96T58 | 309 | DDSPARSVQSAAV | 5.8607 | 6.3082 | 6.3020 | 7.0116 | 6.2965 | 5.8259 | 6.2452 | 6.2166 | 6.1455 | 6.0558 | 5.6308 | 5.9384 | 6.2257 | 6.9164 | 6.3545 | 6.6662 |
| SPEN_S2481;S2493 | SPEN | Q96T58 | 1730 | SGDQPPYLDAKPP | 5.9740 | 5.9346 | 6.1257 | 6.6674 | 5.7301 | 5.8680 | 6.0047 | 5.9016 | 6.0167 | 6.3173 | 6.2711 | 6.2390 | 6.4594 | 6.9296 | 6.9227 | 6.6379 |
| SPEN_S2493 | SPEN | Q96T58 | 2421 | ISVEERTPTKASV | 6.9755 | 7.0952 | 6.2518 | 7.2334 | 6.2035 | 6.2001 | 6.1928 | 5.9210 | 6.3607 | 6.1342 | 6.4345 | 6.4260 | 5.1380 | 5.8971 | 5.6877 | 5.8486 |
| SPEN_S309 | SPEN | Q96T58 | 1947 | AAAVPTTPRRGRP | 7.6254 | 6.9421 | 6.8410 | 7.4725 | 6.1839 | 6.0077 | 6.5165 | 5.9370 | 7.0178 | 6.1885 | 6.8351 | 7.5698 | 4.3383 | 5.5959 | 4.1039 | 4.8245 |
| SPEN_S725;S727 | SPEN | Q96T58 | 3467 | TTSGPSTPPGLVL | 5.6117 | 5.9827 | 6.1578 | 6.6054 | 6.2582 | 5.7606 | 6.1633 | 6.5311 | 5.9255 | 5.8216 | 5.7589 | 5.8969 | 6.3703 | 7.2642 | 6.9218 | 6.9699 |
| SPEN_S727 | SPEN | Q96T58 | 2101 | NVDAAVSPRGAAA | 6.4331 | 6.8166 | 6.0947 | 6.9303 | 5.9812 | 5.4723 | 6.2018 | 6.0966 | 6.2128 | 6.3306 | 6.1831 | 5.8895 | 5.7396 | 6.8211 | 6.4156 | 6.3811 |
| SPEN_S736 | SPEN | Q96T58 | 1826 | KSKRSKTPVQAAA | 6.1308 | 6.5267 | 5.7747 | 6.6427 | 5.7279 | 6.2233 | 6.2327 | 5.9907 | 5.7726 | 5.7798 | 6.0494 | 5.5367 | 6.6383 | 7.0666 | 6.9846 | 6.9226 |
| SPEN_S736;S740 | SPEN | Q96T58 | 1062 | RLNTVASPKDCQE | 6.7691 | 6.8079 | 5.8141 | 6.5790 | 5.8587 | 5.8858 | 6.1863 | 5.8220 | 6.0121 | 6.1972 | 5.4378 | 5.8562 | 6.2279 | 7.0749 | 6.7414 | 6.7294 |
| SPEN_S749 | SPEN | Q96T58 | 1222 | QDVTDDSPPSKKK | 6.1594 | 7.0766 | 6.0054 | 7.8184 | 6.2001 | 6.6272 | 5.5636 | 7.0921 | 5.7274 | 6.0339 | 6.1461 | 5.8398 | 5.3449 | 6.6654 | 5.7929 | 5.9068 |
| SPEN_T1140 | SPEN | Q96T58 | 1140 | CSLRDETPERKSG | 6.9050 | 6.8952 | 7.1415 | 7.3906 | 6.3892 | 6.4443 | 6.4385 | 6.1730 | 6.3143 | 6.5700 | 6.7138 | 6.6497 | 4.5345 | 5.3718 | 4.8535 | 5.2151 |
| SPEN_T1619 | SPEN | Q96T58 | 2460 | IESDPVTPPSDPS | 5.6301 | 6.0209 | 5.4595 | 6.3642 | 5.4929 | 5.7297 | 5.8925 | 6.1143 | 6.1336 | 5.1302 | 5.4042 | 6.4498 | 7.6264 | 7.6938 | 6.9799 | 7.8779 |
| SPEN_T1633 | SPEN | Q96T58 | 3236 | GKEAAKTPDAKAA | 6.8037 | 7.9140 | 8.5491 | 7.3304 | 6.7314 | 7.7306 | 8.1541 | 6.2150 | 5.9826 | 6.5661 | 7.3562 | 5.6624 | 4.1829 | 4.1261 | 4.0254 | 2.6698 |
| SPEN_T1633;S1636 | SPEN | Q96T58 | 736 | HLRRPQSPGASPS | 6.3767 | 7.1912 | 7.1712 | 7.4608 | 6.0633 | 6.0676 | 6.3100 | 5.9928 | 6.4891 | 6.4829 | 6.8592 | 7.4180 | 4.0060 | 5.2278 | 5.1278 | 5.7556 |
| SPEN_T1826 | SPEN | Q96T58 | 1261 | SERTGGSPSVRHG | 5.8810 | 7.1450 | 5.5717 | 6.8257 | 5.9447 | 5.8757 | 6.2152 | 5.4697 | 5.9494 | 6.7284 | 6.0295 | 6.5010 | 5.8451 | 6.1662 | 6.9747 | 6.8769 |
| SPEN_T1947 | SPEN | Q96T58 | 2493 | GGIPHQSPPTKVT | 6.6487 | 6.8257 | 6.6344 | 7.0963 | 6.5230 | 5.9889 | 6.0845 | 6.0825 | 7.0953 | 6.5932 | 6.2676 | 5.8291 | 5.0437 | 5.7717 | 5.7188 | 5.7966 |
| SPEN_T2393 | SPEN | Q96T58 | 749 | QAERLPSDSERRL | 6.3768 | 6.4964 | 5.8407 | 7.0190 | 6.3823 | 5.7882 | 6.2202 | 6.1230 | 5.7971 | 5.7247 | 5.4221 | 5.3660 | 6.3749 | 7.2229 | 6.9150 | 6.9309 |
| SPEN_T2421 | SPEN | Q96T58 | 1633;1636 | NA | 6.3662 | 6.5816 | 6.3568 | 6.8065 | 6.1971 | 5.7508 | 6.1643 | 5.9819 | 6.3861 | 6.2781 | 6.3566 | 6.8473 | 5.9332 | 6.5637 | 5.5661 | 5.8638 |
| SPEN_T2460 | SPEN | Q96T58 | 736;740 | NA | 6.2170 | 6.5059 | 5.9787 | 7.3694 | 6.0013 | 4.8993 | 6.4101 | 6.4512 | 5.9292 | 6.0464 | 6.3184 | 6.2701 | 5.9316 | 6.7563 | 6.9475 | 5.9677 |
| SPEN_T3236 | SPEN | Q96T58 | 725;727 | NA | 6.3255 | 7.1423 | 6.1175 | 7.2031 | 5.8457 | 5.8878 | 6.4811 | 5.6659 | 6.6024 | 6.4209 | 6.2060 | 6.0954 | 5.5982 | 6.0792 | 6.0712 | 6.2576 |
| SPEN_T3467 | SPEN | Q96T58 | 1380;1382 | NA | 5.6452 | 6.3360 | 6.3238 | 7.3810 | 6.1852 | 6.3854 | 6.1744 | 5.9467 | 6.5837 | 6.4240 | 6.0152 | 5.4027 | 5.7432 | 6.2748 | 6.6905 | 6.4880 |
| SPEN_Y1730 | SPEN | Q96T58 | 2481;2493 | NA | 5.9363 | 7.3103 | 6.6244 | 7.8640 | 5.9413 | 6.1059 | 5.9825 | 6.2231 | 6.0184 | 6.2904 | 5.9886 | 6.3971 | 5.2750 | 6.0465 | 5.8523 | 6.1441 |
| SPG11_S1957 | SPG11 | Q96JI7 | 832 | HKSVLDSFLKYDC | 8.9302 | 7.3725 | 7.3988 | 8.1145 | 6.7358 | 6.0201 | 6.7358 | 6.5095 | 6.8568 | 5.7044 | 7.3199 | 6.1832 | 3.1890 | 5.0992 | 3.5468 | 4.2835 |
| SPG11_S1958 | SPG11 | Q96JI7 | 1957 | RRVHSTSSLDSQK | 7.1254 | 6.6290 | 6.9824 | 6.7272 | 7.1399 | 7.1495 | 6.3329 | 7.1505 | 8.1389 | 7.1355 | 7.2761 | 6.8048 | 3.9096 | 4.5355 | 3.4281 | 3.5345 |
| SPG11_S832 | SPG11 | Q96JI7 | 1958 | RVHSTSSLDSQKF | 7.2638 | 7.6703 | 7.3171 | 7.1874 | 6.8771 | 7.2511 | 6.3923 | 6.4814 | 6.7361 | 7.3180 | 6.9700 | 7.6505 | 3.2167 | 4.2617 | 3.6015 | 3.8052 |
| SPG11_Y1854;S1859;S18 | SPG11 | Q96JI7 | 1854;1859;18 | NA | 6.4306 | 5.1562 | 7.9856 | 8.2803 | 4.9297 | 7.5048 | 6.0511 | 7.5224 | 8.4157 | 7.3275 | 7.5664 | 8.6254 | 3.9824 | 4.2559 | 2.9112 | 3.0548 |
| SPHK2_S449 | SPHK2 | Q9NRA0-5 | 546 | PKAALHSPVSEGA | 4.2757 | 6.0554 | 6.0273 | 5.7228 | 5.8118 | 8.3267 | 4.3085 | 6.5892 | 6.5330 | 7.3713 | 6.9124 | 5.6105 | 6.9639 | 6.1631 | 5.2592 | 8.0691 |
| SPHK2_S546 | SPHK2 | Q9NRA0-5 | 449 | ATVEPASPTPAHS | 6.0176 | 7.1371 | 7.6615 | 6.7507 | 6.0296 | 6.2867 | 7.0396 | 6.5868 | 6.7791 | 7.0888 | 6.5903 | 6.8292 | 4.6384 | 5.3077 | 4.4918 | 4.7652 |
| SPICE1_S477 | SPICE1 | Q8N0Z3 | 477 | GRRVMDSPERPVV | 6.4200 | 6.4225 | 6.8272 | 6.4642 | 6.0241 | 6.6043 | 6.7973 | 7.3903 | 6.7334 | 6.3633 | 6.9431 | 6.8521 | 4.5385 | 5.6326 | 4.4647 | 5.5224 |
| SPICE1_S640 | SPICE1 | Q8N0Z3 | 760 | LVGLNLSPPMSPV | 5.7860 | 6.0889 | 6.2376 | 6.4911 | 6.0686 | 6.0517 | 6.1638 | 6.3244 | 6.3843 | 6.4388 | 6.4569 | 6.9160 | 6.2061 | 6.2166 | 5.6371 | 6.5322 |
| SPICE1_S760 | SPICE1 | Q8N0Z3 | 235 | ATQSQITPPGTPS | 5.9759 | 5.9417 | 5.7117 | 5.7821 | 6.7447 | 6.4147 | 6.3824 | 6.5570 | 6.4759 | 6.2659 | 6.8093 | 6.3343 | 6.1163 | 6.2474 | 5.9359 | 6.3047 |
| SPICE1_S800 | SPICE1 | Q8N0Z3 | 640 | NTQQSRSPTFSEE | 6.3176 | 6.4764 | 6.9447 | 6.4667 | 6.0651 | 6.1367 | 6.5914 | 6.2348 | 6.1011 | 5.8059 | 5.9438 | 6.9675 | 5.8718 | 6.0082 | 5.5384 | 6.5296 |
| SPICE1_T235 | SPICE1 | Q8N0Z3 | 537 | EPAVLLTPPRQKS | 6.2443 | 6.7900 | 7.0364 | 6.7359 | 6.6049 | 5.7286 | 5.4366 | 6.0375 | 6.9450 | 5.8849 | 7.4992 | 6.0388 | 5.7286 | 5.8008 | 5.5688 | 5.9198 |
| SPICE1_T45 | SPICE1 | Q8N0Z3 | 45 | LTVHRATPEDLVR | 6.4624 | 6.6862 | 6.0303 | 7.6625 | 6.1718 | 6.4601 | 6.6661 | 7.3904 | 6.6360 | 6.7867 | 6.5942 | 6.7713 | 4.6018 | 5.1212 | 4.6899 | 5.2689 |

| | | | | | | | | | | | | | | | | | | | | | |
|---|---|---|---|---|---|---|---|---|---|---|---|---|---|---|---|---|---|---|---|---|---|
| SPICE1_T537 | SPICE1 | Q8N0Z3 | | 800 | SKCSTVSPVSGIN | 6.6465 | 6.8989 | 6.5485 | 7.0647 | 5.9848 | 6.7006 | 6.6110 | 6.4650 | 7.6392 | 6.9542 | 6.3568 | 6.9602 | 4.4777 | 4.8863 | 4.7090 | 5.0966 |
| SPIDR_S857 | SPIDR | Q14159 | | 248 | TAKFPRTPENSAK | 6.4933 | 7.1935 | 7.8388 | 6.7845 | 5.8118 | 6.8597 | 6.8382 | 6.0922 | 6.7336 | 7.1101 | 6.5863 | 6.8311 | 4.7648 | 5.2953 | 3.8792 | 4.8875 |
| SPIDR_T248 | SPIDR | Q14159 | | 857 | LLQRSISSLLRFA | 6.2805 | 5.9195 | 5.6526 | 5.0001 | 5.4206 | 5.6925 | 4.6165 | 5.4525 | 5.8455 | 5.2169 | 4.4954 | 5.0342 | 8.8669 | 8.3470 | 9.8551 | 8.3042 |
| SPIN1_S124 | SPIN1 | Q9Y657 | | 199 | MPDSNDSPPAERE | 5.6935 | 6.0309 | 5.8030 | 6.1148 | 6.0049 | 6.1393 | 5.8848 | 5.9761 | 5.9097 | 6.3712 | 6.3255 | 6.2197 | 6.7157 | 7.1719 | 6.9905 | 6.6485 |
| SPIN1_S199 | SPIN1 | Q9Y657 | | 124 | VATSRISDAHLAD | 6.4137 | 7.2447 | 6.4795 | 7.3542 | 6.5199 | 6.9495 | 7.5542 | 7.0147 | 7.1370 | 7.2544 | 7.4041 | 7.2459 | 3.4286 | 4.3240 | 3.4302 | 4.2454 |
| SPIN3_S120 | SPIN3 | Q5JUX0 | | 120 | VASSRISDTHLAE | 6.1395 | 7.3999 | 6.2958 | 7.2568 | 6.2917 | 6.9838 | 7.0256 | 6.7454 | 6.7304 | 7.2387 | 6.4432 | 6.6451 | 4.2664 | 5.3006 | 4.5473 | 4.6898 |
| SPINDOC_S248;S251 | SPINDOC | Q9BUA3 | | 308 | GLPRAESPSPAPP | 6.5838 | 6.5763 | 6.2426 | 6.4549 | 6.2866 | 5.9635 | 5.9581 | 6.3379 | 6.5345 | 6.2306 | 6.5233 | 6.3798 | 6.3342 | 5.8743 | 5.8728 | 5.8470 |
| SPINDOC_S251 | SPINDOC | Q9BUA3 | | 49 | VTQQEKTPPPRPS | 6.2510 | 6.6275 | 6.8781 | 6.7170 | 6.3508 | 5.9813 | 6.3035 | 6.0708 | 6.4795 | 6.3800 | 6.9995 | 6.8963 | 5.7193 | 5.5443 | 5.3888 | 5.4123 |
| SPINDOC_S292 | SPINDOC | Q9BUA3 | | 292 | GPDSKDSPKDREV | 7.0789 | 7.0571 | 6.5677 | 6.5574 | 6.6061 | 6.1437 | 6.4631 | 5.8860 | 7.1144 | 7.0598 | 7.0025 | 6.8188 | 5.5472 | 4.8758 | 4.4682 | 4.7533 |
| SPINDOC_S308 | SPINDOC | Q9BUA3 | | 251 | EPPSPDSPTETFA | 5.7576 | 5.2917 | 5.6543 | 7.0575 | 6.8584 | 6.6048 | 6.8377 | 7.6962 | 5.8590 | 6.8809 | 6.1164 | 6.3644 | 6.4771 | 5.6712 | 5.3199 | 5.5528 |
| SPINDOC_T49 | SPINDOC | Q9BUA3 | 248;251 | | NA | 5.5405 | 6.2753 | 5.8230 | 5.3293 | 5.9412 | 6.0980 | 6.4675 | 6.2844 | 6.4777 | 6.1732 | 5.5062 | 6.2371 | 7.5332 | 6.0801 | 7.1257 | 7.1075 |
| SPIRE1_S387 | SPIRE1 | Q08AE8 | | 387 | RKLRPVSPEEIRR | 6.4393 | 6.6973 | 6.6960 | 6.4388 | 6.0659 | 6.3926 | 6.1838 | 6.4072 | 6.5671 | 6.5509 | 6.5418 | 6.6375 | 5.4387 | 6.0546 | 5.3945 | 5.4940 |
| SPIRE1_S477 | SPIRE1 | Q08AE8 | | 509 | FLPISSTPQPERR | 6.7245 | 6.1833 | 6.8840 | 6.3779 | 6.2456 | 6.3716 | 6.3997 | 6.4498 | 6.9989 | 6.1504 | 6.2819 | 6.4214 | 5.5596 | 5.6348 | 5.6508 | 5.6657 |
| SPIRE1_S735 | SPIRE1 | Q08AE8 | | 477 | TLHKSTSSSSVSP | 4.7461 | 5.9207 | 5.6302 | 5.1323 | 6.0372 | 5.6430 | 5.2855 | 6.2638 | 5.6797 | 5.0046 | 6.3515 | 6.7409 | 6.7026 | 8.1692 | 8.4389 | 8.2538 |
| SPIRE1_T509 | SPIRE1 | Q08AE8 | | 735 | LKRKTQSFYMSSP | 6.8750 | 5.7407 | 8.2577 | 6.7434 | 5.7230 | 6.9164 | 5.5618 | 5.4198 | 6.5201 | 6.4313 | 6.7419 | 7.3719 | 4.5118 | 6.5585 | 4.6612 | 5.9655 |
| SPIRE2_S442 | SPIRE2 | Q8WWL2 | | 442 | KRDRSFSEHDLAQ | 6.8417 | 8.5906 | 7.5106 | 6.6054 | 5.6377 | 5.9160 | 6.0988 | 7.1073 | 6.8110 | 7.4846 | 7.6213 | 7.1434 | 3.3081 | 4.6335 | 4.2797 | 4.4103 |
| SPOP_S6 | SPOP | O43791 | | 6 | xMSRVPSPPPPAE | 5.6637 | 5.8908 | 5.8897 | 6.2365 | 5.9412 | 6.1819 | 5.9719 | 5.6944 | 6.1496 | 6.2031 | 5.6016 | 5.7196 | 7.5359 | 6.5357 | 7.5350 | 7.2492 |
| SPRED1_S176 | SPRED1 | Q7Z699 | | 238 | KSIRHVSFQDEDE | 6.1383 | 6.9876 | 7.7828 | 6.8600 | 6.2422 | 6.2594 | 6.0481 | 6.6254 | 6.0191 | 6.8017 | 6.6508 | 6.8292 | 5.1880 | 5.8189 | 5.5322 | 5.2162 |
| SPRED1_S238 | SPRED1 | Q7Z699 | | 176 | SSNIRPSPFEDLN | 8.1071 | 7.1885 | 7.4162 | 6.9106 | 6.1867 | 7.6910 | 6.9812 | 6.2526 | 7.1053 | 6.7316 | 6.2417 | 6.8038 | 3.6194 | 5.2069 | 3.2127 | 4.3448 |
| SPRED1_S317 | SPRED1 | Q7Z699 | | 317 | FKTQPSSLKIKKS | 6.8834 | 7.4582 | 8.3121 | 5.8958 | 4.6901 | 6.6196 | 6.5764 | 6.5881 | 6.5501 | 7.7318 | 7.9698 | 7.3882 | 3.7349 | 5.7645 | 3.1815 | 4.6556 |
| SPRY2_S112 | SPRY2 | O43597 | | 167 | GELKPLSKEDLGL | 5.1183 | 4.7648 | 4.8495 | 5.8216 | 5.8570 | 5.4652 | 5.9387 | 5.7679 | 5.4703 | 5.8746 | 4.6654 | 4.9769 | 9.3205 | 8.7365 | 8.4375 | 8.9353 |
| SPRY2_S139 | SPRY2 | O43597 | | 112 | PLSRSISTVSSGS | 6.5208 | 7.6197 | 6.0303 | 5.9800 | 5.6183 | 5.8107 | 7.0913 | 5.3756 | 5.5473 | 5.3021 | 5.2064 | 5.2022 | 6.8758 | 8.2034 | 6.1583 | 7.4578 |
| SPRY2_S167 | SPRY2 | O43597 | | 139 | QRLLGSSFSSGPV | 6.5592 | 5.9369 | 6.0482 | 6.7662 | 6.7513 | 5.7163 | 6.4554 | 6.4567 | 7.6675 | 5.7767 | 5.3275 | 5.7462 | 7.1442 | 6.5898 | 5.9091 | 5.1490 |
| SPRY4_S125 | SPRY4 | Q9C004 | | 125 | PVADQASPRAVRI | 6.1301 | 6.4106 | 6.2475 | 6.5989 | 6.4239 | 6.4595 | 6.6045 | 6.8040 | 6.3301 | 6.5012 | 6.1918 | 6.3883 | 5.4589 | 5.9334 | 5.7364 | 5.7809 |
| SPRY4_S280 | SPRY4 | Q9C004 | | 280 | RCKHTNSVICKAA | 6.3252 | 6.8259 | 9.4396 | 8.3379 | 6.7870 | 6.4377 | 5.6449 | 6.4299 | 6.0465 | 7.3936 | 7.0384 | 6.9905 | 3.5930 | 5.4605 | 4.1030 | 3.1464 |
| SPTAN1_S1031 | SPTAN1 | Q13813-2 | | 1217 | LNERWRSLQQLAE | 6.3605 | 6.5131 | 6.0132 | 5.8176 | 6.8269 | 6.2131 | 5.8333 | 5.7856 | 5.9846 | 5.9634 | 5.7926 | 5.7828 | 7.0377 | 6.7898 | 6.8530 | 6.4327 |
| SPTAN1_S1217 | SPTAN1 | Q13813-2 | | 1031 | DPAQSASRENLLE | 6.1793 | 7.1435 | 6.8408 | 5.6931 | 5.1879 | 6.3946 | 5.6689 | 5.7302 | 5.7994 | 7.1420 | 7.6168 | 7.4220 | 4.0128 | 6.8643 | 5.6896 | 6.6149 |
| SPTBN1_S14 | SPTBN1 | Q01082 | | 2138 | LPAEQGSPRMAET | 6.3670 | 6.3939 | 6.8982 | 6.9595 | 6.2715 | 6.3379 | 6.4158 | 6.4654 | 6.7021 | 6.7769 | 7.0271 | 6.9395 | 4.9294 | 5.5358 | 4.8250 | 5.1547 |
| SPTBN1_S2102 | SPTBN1 | Q01082 | | 2102 | RKRRPPSPEPSTK | 5.8554 | 5.9359 | 6.1385 | 6.0465 | 6.1154 | 6.1315 | 6.0859 | 5.6524 | 6.1779 | 6.4120 | 6.2796 | 6.3555 | 6.6961 | 6.8976 | 6.5988 | 6.6212 |
| SPTBN1_S2138 | SPTBN1 | Q01082-3 | | 7 | MELQRTSSISGPL | 6.1968 | 5.9288 | 6.4164 | 6.0837 | 6.0654 | 6.0339 | 6.1537 | 5.9639 | 6.4747 | 6.3025 | 6.2874 | 6.2196 | 6.7391 | 6.7811 | 6.4267 | 5.9262 |
| SPTBN1_S2164 | SPTBN1 | Q01082 | | 2165 | TSSKESSPIPSPT | 5.7521 | 6.1546 | 5.7897 | 5.4464 | 5.5157 | 5.4082 | 5.5049 | 5.4014 | 5.7330 | 5.6176 | 5.3880 | 5.5647 | 8.4243 | 8.3884 | 8.2022 | 7.7090 |
| SPTBN1_S2164;S2169 | SPTBN1 | Q01082 | | 2187 | PAQSAATLPARTQ | 8.5914 | 7.4494 | 7.6277 | 7.4438 | 7.0471 | 5.2160 | 5.4988 | 4.9138 | 6.8645 | 5.3196 | 5.8615 | 5.3272 | 6.1642 | 6.4991 | 5.3423 | 4.8336 |
| SPTBN1_S2165 | SPTBN1 | Q01082 | | 2169 | ESSPIPSPTSDRK | 5.1862 | 5.0913 | 5.2764 | 5.4455 | 5.4133 | 5.2237 | 6.0332 | 5.5120 | 5.1071 | 5.6549 | 5.4727 | 5.3223 | 8.4372 | 8.8692 | 9.0873 | 8.8677 |
| SPTBN1_S2165;S2169 | SPTBN1 | Q01082 | | 2358 | DKEKRFSLFGKKK | 10.9106 | 6.9266 | 7.6697 | 5.9672 | 5.7029 | 5.5872 | 5.9787 | 5.5060 | 7.7957 | 6.2788 | 6.8277 | 7.2645 | 4.8072 | 5.9333 | 3.2820 | 3.5620 |
| SPTBN1_S2169 | SPTBN1 | Q01082 | | 2328 | ASSRAQTLPTSVV | 6.8480 | 6.6191 | 6.5461 | 6.6919 | 5.8994 | 6.4206 | 5.8384 | 6.1651 | 7.1270 | 6.2317 | 6.4382 | 6.5964 | 5.3120 | 6.1645 | 5.2477 | 5.8537 |
| SPTBN1_S2341 | SPTBN1 | Q01082 | | 2341 | TITSESSPGKREK | 6.2711 | 6.5388 | 6.6608 | 6.3572 | 5.9577 | 6.5286 | 6.2760 | 6.8915 | 6.7705 | 6.4929 | 6.8155 | 6.4795 | 5.4408 | 5.4356 | 4.9181 | 6.1655 |
| SPTBN1_S2358 | SPTBN1 | Q01082 | | 2164 | RTSSKESSPIPSP | 5.9745 | 5.7663 | 5.3513 | 5.5667 | 5.1080 | 4.9580 | 5.3787 | 5.5436 | 6.2116 | 5.2927 | 4.8343 | 4.6926 | 8.5194 | 8.9735 | 9.4160 | 8.4128 |
| SPTBN1_S7 | SPTBN1 | Q01082-3 | | 14 | SISGPLSPAYTGQ | 5.6089 | 6.2816 | 6.1582 | 5.7856 | 7.1914 | 6.7438 | 6.5066 | 7.2447 | 5.4371 | 6.6639 | 5.9235 | 6.3953 | 6.3978 | 5.5557 | 6.6494 | 5.4565 |
| SPTBN1_S7;S14 | SPTBN1 | Q01082 | 2165;2169 | | NA | 6.1531 | 6.1059 | 6.0504 | 6.1653 | 6.3400 | 6.0897 | 5.8473 | 6.8154 | 6.3277 | 6.1690 | 6.7155 | 6.0617 | 6.5040 | 6.5975 | 6.2770 | 5.7805 |
| SPTBN1_T2187 | SPTBN1 | Q01082 | 2164;2169 | | NA | 6.2624 | 6.6155 | 6.9207 | 6.0027 | 6.4424 | 6.1616 | 6.0772 | 5.9039 | 5.9796 | 6.4110 | 6.8898 | 6.3900 | 5.8610 | 5.9534 | 5.9918 | 6.1369 |
| SPTBN1_T2328 | SPTBN1 | Q01082-3 | 7;14 | | NA | 5.3295 | 6.6764 | 5.0037 | 5.2081 | 6.6044 | 6.0231 | 6.0268 | 5.3537 | 6.3490 | 7.6371 | 5.6542 | 5.6639 | 6.5897 | 7.3199 | 7.4491 | 7.1113 |
| SPTY2D1_S471 | SPTY2D1 | Q68D10 | | 471 | PGRPVSSPHELRR | 7.0003 | 6.7281 | 5.8827 | 6.6538 | 6.1982 | 6.1908 | 6.3179 | 6.3433 | 6.2260 | 6.7194 | 6.2897 | 6.0828 | 5.3911 | 6.5644 | 5.6522 | 5.7592 |
| SQSTM1_S152 | SQSTM1 | Q13501 | | 361 | SLQMPESEGPSSL | 5.4578 | 5.8089 | 6.3676 | 6.3552 | 6.4215 | 6.5273 | 6.6908 | 7.2136 | 6.2531 | 6.3874 | 6.4532 | 6.8217 | 5.8636 | 5.9993 | 5.5597 | 5.8191 |
| SQSTM1_S207 | SQSTM1 | Q13501 | | 24 | REIRRFSFCCSPE | 5.4287 | 5.5963 | 6.0903 | 6.0980 | 6.0763 | 5.8036 | 5.7924 | 5.7884 | 6.6788 | 7.0766 | 6.7417 | 7.0589 | 6.7419 | 6.5740 | 6.3958 | 6.0584 |
| SQSTM1_S24 | SQSTM1 | Q13501 | | 269 | GKRSRLTPVSPES | 5.8291 | 5.6078 | 6.0008 | 5.6073 | 6.8640 | 6.8557 | 6.5606 | 6.8467 | 6.1509 | 5.8507 | 5.9466 | 5.8755 | 6.9847 | 6.5644 | 6.3300 | 6.1252 |
| SQSTM1_S284 | SQSTM1 | Q13501 | | 366 | ESEGPSSLDPSQE | 5.6335 | 5.7576 | 6.2524 | 6.3010 | 6.2693 | 6.6824 | 6.8061 | 6.6878 | 6.2238 | 6.1681 | 6.6927 | 6.7081 | 5.8330 | 5.8745 | 6.0879 | 6.0217 |
| SQSTM1_S332 | SQSTM1 | Q13501 | | 272 | SRLTPVSPESSST | 5.0828 | 5.3095 | 6.1074 | 6.0974 | 6.0457 | 5.8807 | 6.0495 | 6.2825 | 5.8402 | 5.7265 | 6.1622 | 6.8079 | 7.1852 | 6.6396 | 7.6152 | 7.1678 |
| SQSTM1_S361 | SQSTM1 | Q13501 | | 284 | TEEKSSSQPSSCC | 5.3950 | 5.5094 | 5.7422 | 5.3311 | 6.3191 | 6.3280 | 6.3584 | 6.0800 | 5.2135 | 5.5429 | 5.6751 | 6.5081 | 7.6650 | 7.3294 | 7.2047 | 7.7982 |
| SQSTM1_S366 | SQSTM1 | Q13501 | | 207 | GPPGNWSPRPPRA | 6.5073 | 6.5173 | 6.1919 | 6.2962 | 5.8180 | 6.6663 | 5.8565 | 6.8527 | 6.5496 | 7.0663 | 7.1843 | 6.4129 | 6.0329 | 5.4155 | 5.2826 | 5.3497 |
| SQSTM1_T269 | SQSTM1 | Q13501 | | 332 | MESDNCSGGDDDW | 5.5032 | 6.5451 | 4.9407 | 6.2501 | 7.1023 | 7.4123 | 5.9551 | 8.0356 | 5.0054 | 5.2553 | 6.6294 | 5.3887 | 6.5281 | 7.4117 | 6.8956 | 5.1415 |
| SQSTM1_T269;S272 | SQSTM1 | Q13501 | | 152 | PDYDLCSVCEGKG | 6.6951 | 5.5680 | 6.1159 | 5.8105 | 6.5559 | 6.4930 | 5.2402 | 5.2717 | 7.3013 | 4.9888 | 5.1145 | 5.4468 | 9.9686 | 7.0274 | 5.0741 | 7.3282 |
| SQSTM1_T272 | SQSTM1 | Q13501 | 269;272 | | NA | 5.4952 | 5.5392 | 5.7689 | 6.1716 | 6.7059 | 6.6813 | 6.6326 | 6.8667 | 6.4703 | 5.8206 | 6.3431 | 6.1213 | 6.6925 | 6.3354 | 6.2212 | 6.1342 |
| SRA1_S60 | SRA1 | Q9HD15 | | 69 | VPASETSPGPPPM | 6.5117 | 6.3795 | 6.7736 | 6.5870 | 6.6223 | 6.2744 | 6.7973 | 6.8766 | 6.8117 | 6.7467 | 6.9179 | 6.8015 | 4.8122 | 5.2488 | 4.5480 | 5.2908 |
| SRA1_S69 | SRA1 | Q9HD15 | | 60 | AAPQDGSPRVPAS | 7.2373 | 7.1646 | 7.2564 | 6.7221 | 6.3487 | 7.2743 | 7.2848 | 8.0417 | 7.4751 | 7.7809 | 8.1217 | 8.1124 | 2.3524 | 3.2648 | 2.1681 | 3.3946 |
| SRC_S17 | SRC | P12931-2 | | 17 | ASQRRRSLEPAEN | 5.4138 | 5.7810 | 6.2166 | 6.5132 | 6.1384 | 6.0753 | 5.7600 | 6.1472 | 6.7121 | 6.6032 | 6.3267 | 5.8585 | 6.5972 | 6.7811 | 6.3993 | 6.6765 |
| SRCAP_S1859 | SRCAP | Q6ZRS2 | | 3148 | GAPVGGSPGLAKR | 6.6153 | 6.9151 | 6.2204 | 7.0491 | 6.2809 | 6.1812 | 6.8128 | 6.4590 | 6.6565 | 6.3352 | 6.3797 | 6.3683 | 5.0415 | 5.6149 | 5.3616 | 5.7085 |
| SRCAP_S2725 | SRCAP | Q6ZRS2 | | 1859 | LRSGPPSPPSTAT | 6.0725 | 6.1538 | 6.1329 | 6.7581 | 6.2352 | 5.9831 | 6.1613 | 6.1857 | 6.4418 | 6.1646 | 5.9918 | 6.1228 | 6.1037 | 6.6170 | 6.3851 | 6.4906 |
| SRCAP_S274 | SRCAP | Q6ZRS2 | | 274 | SSSAASSPPPPAS | 5.5692 | 5.8283 | 6.0594 | 5.9139 | 6.2011 | 6.0464 | 6.2301 | 5.9265 | 6.0403 | 5.9478 | 5.9674 | 5.9088 | 7.0384 | 6.8290 | 7.2786 | 7.2147 |
| SRCAP_S2790 | SRCAP | Q6ZRS2 | | 2869 | PPKKNRSPADAGR | 6.8970 | 6.9158 | 5.9422 | 7.1104 | 5.8960 | 5.6780 | 6.5848 | 5.8721 | 6.1932 | 6.4736 | 6.1390 | 6.2847 | 5.9610 | 5.7306 | 6.1001 | 6.2213 |
| SRCAP_S2840 | SRCAP | Q6ZRS2 | | 3016 | PPKNPPSPRPSQL | 6.1815 | 6.5493 | 6.1134 | 7.0859 | 6.2907 | 6.1302 | 6.1666 | 5.4480 | 5.9805 | 6.1992 | 5.9720 | 6.0821 | 6.6701 | 6.6245 | 6.3058 | 6.2001 |
| SRCAP_S2840;T2852 | SRCAP | Q6ZRS2 | | 2840 | GVTGGGSPENGDG | 6.5148 | 7.4466 | 6.9161 | 7.8150 | 5.8555 | 5.3476 | 6.5882 | 6.7692 | 6.1150 | 6.8574 | 6.1164 | 6.4405 | 5.1685 | 5.7327 | 5.1417 | 5.1749 |
| SRCAP_S2869 | SRCAP | Q6ZRS2 | | 2790 | TSASPGSPSVRSM | 7.0384 | 7.0179 | 5.2396 | 8.2389 | 6.3798 | 5.4669 | 7.1501 | 6.0100 | 5.7802 | 6.6609 | 6.6494 | 6.1820 | 6.4799 | 6.3541 | 4.9058 | 4.4462 |
| SRCAP_S3016 | SRCAP | Q6ZRS2 | | 2725 | PPRRRTSADVEIR | 5.6281 | 5.4913 | 5.4229 | 5.8706 | 6.1779 | 5.1733 | 5.8590 | 5.6325 | 5.4780 | 6.6549 | 6.1735 | 5.4504 | 7.5388 | 7.9420 | 7.5405 | 7.9660 |
| SRCAP_S3148 | SRCAP | Q6ZRS2 | 2840;2852 | | NA | 6.4858 | 7.2669 | 6.9255 | 7.0635 | 6.3811 | 6.0338 | 5.9989 | 6.5024 | 6.7442 | 7.1687 | 7.4639 | 6.9421 | 4.6521 | 4.8569 | 5.0349 | 4.4793 |
| SRCIN1_S844 | SRCIN1 | Q9C0H9 | | 844 | NNLLSQSPKKVTA | 5.8514 | 5.9562 | 6.5205 | 6.1807 | 5.7262 | 6.6514 | 5.8845 | 6.2801 | 6.5679 | 6.2895 | 6.1452 | 6.5596 | 5.8702 | 6.6089 | 6.0407 | 6.8672 |
| SREBF1_S147 | SREBF1 | P36956-4 | | 279 | HFIKADSLLLTAM | 8.6379 | 9.0663 | 8.2406 | 6.8250 | 6.4277 | 4.7970 | 6.5415 | 6.4919 | 8.3958 | 6.9182 | 7.6404 | 6.5250 | 3.0421 | 3.5160 | 2.9593 | 3.9754 |
| SREBF1_S279 | SREBF1 | P36956-4 | | 147 | PSMPAFSPGPGIK | 7.0495 | 6.7458 | 6.5262 | 7.8334 | 6.7102 | 7.0338 | 6.1411 | 6.9236 | 7.5967 | 8.2239 | 8.0858 | 7.6890 | 2.6978 | 4.4386 | 2.8571 | 3.4475 |
| SRF_S103 | SRF | P11831 | | 224 | CLNSPDSPPRSDP | 5.0933 | 5.6498 | 6.1590 | 5.0288 | 5.5893 | 6.4226 | 5.7122 | 6.3519 | 6.2061 | 6.1931 | 6.1179 | 6.2365 | 6.6577 | 7.6076 | 7.7438 | 7.2303 |
| SRF_S224 | SRF | P11831 | | 103 | GLKRSLSEMEIGM | 5.6894 | 5.8027 | 6.1619 | 5.6901 | 6.0874 | 6.2330 | 5.3682 | 6.1295 | 6.3352 | 6.0896 | 6.4736 | 6.2201 | 6.6260 | 7.2284 | 7.0165 | 6.8483 |
| SRF_T25 | SRF | P11831 | | 25 | GGSLNRTPTGRPG | 6.6210 | 7.0477 | 7.1544 | 7.3701 | 6.0816 | 6.5180 | 8.2623 | 8.0659 | 6.3616 | 7.4017 | 8.1023 | 7.6611 | 2.7420 | 3.7772 | 3.2875 | 3.5457 |
| SRFBP1_S203 | SRFBP1 | Q8NEF9 | | 203 | AVTIANSPSKPSE | 5.5593 | 5.6858 | 5.9114 | 6.7146 | 6.1473 | 5.8388 | 6.3843 | 6.1593 | 6.5766 | 6.0531 | 6.0070 | 6.3151 | 6.4459 | 6.5160 | 6.6884 | 6.9972 |
| SRFBP1_S264 | SRFBP1 | Q8NEF9 | | 264 | KEYFDDSTEERFY | 6.5904 | 6.5395 | 5.6904 | 6.6612 | 5.9689 | 5.8613 | 6.1156 | 6.1974 | 6.3336 | 5.5495 | 6.1368 | 5.8488 | 6.6558 | 6.5227 | 6.8931 | 6.4349 |
| SRFBP1_S349 | SRFBP1 | Q8NEF9 | | 351 | FHSLSGSKSSRRN | 5.6509 | 6.3727 | 7.3982 | 7.5944 | 6.3500 | 6.1252 | 6.1296 | 7.0683 | 6.1070 | 9.1163 | 9.0202 | 7.8917 | 3.8278 | 4.8966 | 2.7165 | 3.7346 |
| SRFBP1_S351 | SRFBP1 | Q8NEF9 | | 349 | VFFHSLSGSKSSR | 6.4762 | 8.4322 | 6.3522 | 6.9205 | 6.2701 | 6.5343 | 7.9590 | 6.8114 | 8.4362 | 7.4117 | 7.3647 | 7.0295 | 2.2681 | 4.4202 | 3.6258 | 3.6878 |
| SRGAP1_S1029 | SRGAP1 | Q7Z6B7 | | 932 | SLKKIDSPPIRRS | 6.2676 | 6.5039 | 7.0094 | 6.2840 | 6.1079 | 6.4519 | 6.7613 | 6.6800 | 7.0256 | 6.9544 | 6.9016 | 6.8774 | 4.9706 | 5.5351 | 4.7038 | 4.9655 |
| SRGAP1_S917 | SRGAP1 | Q7Z6B7 | | 940 | PIRRSTSSGQYTG | 4.8850 | 4.4697 | 5.3355 | 4.7751 | 5.5089 | 4.9391 | 4.9116 | 5.7748 | 5.2151 | 4.4813 | 4.6671 | 4.4280 | 10.0742 | 10.0654 | 10.9954 | 9.4739 |
| SRGAP1_S932 | SRGAP1 | Q7Z6B7 | | 1029 | QIRRSTSSSSDTM | 6.5822 | 6.4535 | 7.2033 | 6.3639 | 6.2486 | 6.5664 | 6.0171 | 6.4128 | 6.6283 | 6.9461 | 6.0594 | 6.4830 | 5.5224 | 6.4041 | 4.9708 | 5.1380 |
| SRGAP1_S940 | SRGAP1 | Q7Z6B7 | | 917 | RRPGHGSLTNISR | 5.5883 | 5.7422 | 6.1883 | 5.4060 | 5.4047 | 6.1612 | 5.8097 | 6.3458 | 6.2009 | 5.8349 | 6.0649 | 5.8696 | 7.2473 | 7.7258 | 7.7294 | 6.6811 |
| SRGAP2_S206 | SRGAP2 | O75044 | | 206 | DRQTPRSPDSTAN | 6.6786 | 7.1043 | 6.6350 | 6.8757 | 6.0311 | 6.5277 | 6.9233 | 6.3622 | 6.7478 | 7.0880 | 7.0066 | 7.1496 | 4.3530 | 4.8960 | 4.3370 | 5.2842 |
| SRGAP2_S424 | SRGAP2 | O75044 | | 424 | SMESVKSTVSETF | 5.6259 | 5.9017 | 6.4617 | 6.0902 | 5.8491 | 6.0351 | 6.0186 | 6.4182 | 5.8940 | 6.8505 | 6.0241 | 6.4556 | 6.7471 | 6.4345 | 6.8548 | 6.3390 |
| SRGAP2_S877 | SRGAP2 | O75044 | | 930 | TAGRSKSFNNHRP | 6.3664 | 6.8929 | 5.8148 | 5.6499 | 5.0198 | 7.7424 | 5.9774 | 6.7564 | 5.7591 | 6.8372 | 6.7371 | 7.1181 | 5.7056 | 6.5608 | 5.8910 | 5.1711 |
| SRGAP2_S930 | SRGAP2 | O75044 | | 983 | LEPLKTSPVVAPT | 6.0505 | 5.9930 | 7.6451 | 6.8766 | 5.1890 | 7.0294 | 6.0926 | 6.9807 | 6.8589 | 6.0107 | 5.6895 | 7.4569 | 5.0584 | 5.7206 | 4.6841 | 6.6640 |
| SRGAP2_S983 | SRGAP2 | O75044 | | 877 | ASCRPSSQPIMSQ | 6.4505 | 4.7509 | 6.6691 | 7.1094 | 6.2258 | 6.9308 | 5.9764 | 5.4529 | 6.8262 | 6.5244 | 6.7923 | 5.5207 | 6.0041 | 6.4013 | 5.4868 | 6.8785 |
| SRGAP2_S983;S994 | SRGAP2 | O75044 | 983;994 | | NA | 5.5505 | 6.0866 | 6.2946 | 6.1299 | 6.2541 | 6.5457 | 6.2104 | 6.8585 | 6.2365 | 5.8717 | 6.3687 | 6.3364 | 5.9747 | 6.4306 | 6.4032 | 6.4479 |
| SRGAP2_S983;T989;S994 | SRGAP2 | O75044 | 983;989;994 | | NA | 5.6351 | 6.5059 | 6.8159 | 6.1403 | 5.9228 | 5.6871 | 8.2061 | 7.2779 | 5.2082 | 7.7301 | 6.2341 | 7.2175 | 5.7727 | 3.8186 | 5.4645 | 6.3630 |
| SRP68_S627 | SRP68 | Q9UHB9 | | 627 | GIFGFRSxxxxxx | 6.2449 | 7.1250 | 7.1139 | 7.2527 | 6.1764 | 5.7624 | 6.9348 | 7.3847 | 6.6323 | 6.7968 | 6.1832 | 7.0419 | 4.4512 | 5.4248 | 4.3947 | 5.0803 |
| SRP72_S621 | SRP72 | O76094 | | 621 | ASKTVSSPPTSPR | 5.8338 | 5.5588 | 5.2709 | 5.5577 | 5.2783 | 4.4818 | 4.6326 | 5.1734 | 6.2147 | 5.4279 | 4.8688 | 5.0272 | 8.3798 | 9.4803 | 9.6607 | 9.1532 |
| SRP72_S621;S625;S630 | SRP72 | O76094 | 621;625 | | NA | 6.0896 | 6.1993 | 6.1841 | 6.6219 | 5.9825 | 6.0840 | 6.2590 | 6.2401 | 6.3525 | 6.3828 | 6.4689 | 6.6374 | 5.8929 | 6.3854 | 5.9754 | 6.2442 |
| SRP72_S621;S630 | SRP72 | O76094 | 621;630 | | NA | 5.9183 | 5.8072 | 6.7508 | 6.4242 | 6.0285 | 6.5383 | 6.6580 | 6.2616 | 6.3659 | 6.2908 | 6.5875 | 6.3028 | 5.5053 | 6.5931 | 5.8877 | 6.0800 |
| SRP72_S621;T625 | SRP72 | O76094 | 621;625;630 | | NA | 6.1774 | 5.0060 | 6.8042 | 6.4597 | 6.2534 | 6.0196 | 6.9026 | 6.5275 | 6.4095 | 7.1824 | 5.8544 | 6.9460 | 6.0828 | 6.5894 | 5.0630 | 5.7220 |
| SRPK1_S222 | SRPK1 | Q96SB4-3 | | 222 | EEEILGSDDDEQE | 5.7337 | 5.7005 | 6.1003 | 6.0341 | 6.9624 | 6.1276 | 6.2100 | 5.5393 | 6.8041 | 6.4574 | 6.5310 | 6.9352 | 5.8787 | 6.2564 | 6.3079 | 6.4213 |
| SRPK1_S480;S482 | SRPK1 | Q96SB4-3 | 480;482 | | NA | 5.6440 | 5.9573 | 6.0224 | 5.9534 | 6.2572 | 6.3583 | 5.8724 | 6.7687 | 6.0205 | 6.5672 | 6.1695 | 6.2450 | 6.0561 | 7.0419 | 6.7124 | 6.3537 |
| SRPK2_S391 | SRPK2 | P78362-2 | | 391 | DPTWIESPKTNGH | 6.3578 | 6.4447 | 6.9008 | 6.4669 | 5.5882 | 6.1018 | 6.5914 | 5.5716 | 6.9224 | 6.9087 | 6.9766 | 7.3473 | 5.2514 | 5.6778 | 5.0033 | 5.8892 |
| SRPRA_S296;S297;S298 | SRPRA | P08240 | | 269 | TPTTNGTPEAALS | 6.1076 | 6.1924 | 6.6580 | 6.5001 | 5.6990 | 6.5264 | 6.3069 | 6.4476 | 6.6857 | 6.1753 | 5.6687 | 6.4736 | 5.7502 | 6.3594 | 5.5189 | 6.0302 |
| SRPRA_T269 | SRPRA | P08240 | 296;297;298 | | NA | 5.6349 | 5.8603 | 6.2670 | 6.2872 | 6.8347 | 6.9936 | 6.2434 | 5.9101 | 6.1041 | 5.9502 | 6.6323 | 6.3057 | 6.8678 | 6.7252 | 5.9525 | 5.4311 |
| SRRM1_S187 | SRRM1 | A9Z1X7 | | 220 | PEKKEKTPELPEP | 5.7815 | 6.2436 | 5.8343 | 6.4211 | 5.9025 | 6.0019 | 6.0437 | 5.8297 | 6.2204 | 6.0246 | 5.9225 | 5.9464 | 6.5098 | 7.2124 | 6.9248 | 7.1807 |
| SRRM1_S207;S209;S211 | SRRM1 | A9Z1X7 | | 705 | APQTSSSPPPVRR | 6.1361 | 6.4929 | 5.9838 | 6.4763 | 5.9494 | 5.9103 | 6.1432 | 5.8843 | 6.1728 | 6.1144 | 5.9735 | 6.1283 | 6.2573 | 7.0143 | 6.5761 | 6.7868 |
| SRRM1_S234 | SRRM1 | A9Z1X7 | | 722 | SPQRRQSPSPSTR | 6.3626 | 6.7606 | 6.2977 | 6.8050 | 5.8715 | 5.7980 | 5.9954 | 5.9231 | 6.3012 | 6.4878 | 6.2881 | 6.3185 | 5.6488 | 6.5998 | 6.1441 | 6.3977 |
| SRRM1_S260 | SRRM1 | A9Z1X7 | | 606 | PSPRRYSPPIQRR | 6.0412 | 6.3202 | 5.8043 | 6.3303 | 6.0523 | 5.8215 | 5.9029 | 5.9973 | 6.0100 | 5.9638 | 5.8252 | 6.0203 | 6.4150 | 7.1982 | 7.0836 | 7.2139 |
| SRRM1_S375 | SRRM1 | A9Z1X7 | | 662 | PVTKRRSPSLSSK | 6.0907 | 6.4888 | 6.0965 | 6.4804 | 5.9781 | 5.8811 | 6.0800 | 6.1351 | 6.3658 | 6.5512 | 6.2297 | 6.3256 | 5.8197 | 6.5883 | 6.3027 | 6.5862 |
| SRRM1_S384;S386;S388 | SRRM1 | A9Z1X7 | | 409 | PPKTRHSPTPQQS | 6.4849 | 7.0210 | 6.5313 | 7.1716 | 5.8666 | 5.9257 | 6.1713 | 6.4116 | 6.4621 | 6.6789 | 6.8085 | 6.9543 | 4.7817 | 5.7000 | 5.3003 | 5.7302 |
| SRRM1_S384;S388 | SRRM1 | A9Z1X7 | | 625 | PKRRTASPPPPPK | 6.2281 | 6.4530 | 6.1049 | 6.5539 | 5.9079 | 5.6104 | 6.0749 | 5.6611 | 6.3529 | 6.2505 | 6.0604 | 6.0176 | 6.2657 | 6.7715 | 6.6773 | 7.0098 |
| SRRM1_S397 | SRRM1 | A9Z1X7 | | 547 | MGKRWQSPVTKSG | 6.2515 | 6.4086 | 5.6452 | 6.1072 | 6.1762 | 5.7689 | 5.9809 | 6.1479 | 6.1775 | 5.8192 | 5.6912 | 5.5962 | 6.7751 | 7.2746 | 7.0686 | 7.1110 |
| SRRM1_S397;T401 | SRRM1 | A9Z1X7 | | 883 | PKKETESEAEDNL | 5.9413 | 6.1764 | 5.8049 | 6.1436 | 5.8889 | 5.5329 | 5.8620 | 5.6435 | 6.4264 | 6.3079 | 6.1300 | 6.3360 | 6.6933 | 7.1864 | 6.8094 | 7.1170 |
| SRRM1_S409 | SRRM1 | A9Z1X7 | | 445 | GTEKRESPSPAPK | 6.4356 | 6.7543 | 6.4617 | 6.8870 | 5.8071 | 6.4900 | 6.1030 | 5.9838 | 6.9175 | 6.3561 | 6.0154 | 6.3980 | 5.4309 | 6.1948 | 5.7678 | 5.9969 |
| SRRM1_S409;T411 | SRRM1 | A9Z1X7 | | 647 | KRRVSHSPPPKQR | 6.6426 | 7.0224 | 6.6675 | 7.1627 | 5.9744 | 5.7635 | 5.9606 | 5.6930 | 6.7956 | 6.7798 | 6.7321 | 6.5842 | 5.0995 | 6.0845 | 5.3944 | 5.6432 |
| SRRM1_S424;S426 | SRRM1 | A9Z1X7 | | 375 | PPKRTSSPPRKTR | 6.2539 | 6.3154 | 5.8240 | 6.2032 | 5.9444 | 5.6576 | 5.9457 | 5.8131 | 6.1074 | 6.0653 | 5.8646 | 5.9742 | 6.7120 | 7.2086 | 7.0300 | 7.0806 |
| SRRM1_S426 | SRRM1 | A9Z1X7 | | 747 | KIKKAASPSPQSV | 6.4653 | 6.5768 | 5.8600 | 6.1084 | 5.7816 | 5.5985 | 5.6045 | 5.5833 | 6.3296 | 6.0126 | 5.8420 | 6.0977 | 6.6084 | 7.3880 | 7.1017 | 7.0414 |
| SRRM1_S445 | SRRM1 | A9Z1X7 | | 426 | KSRVSVSPGRTSG | 6.8431 | 7.4062 | 6.3216 | 7.2379 | 5.9771 | 5.4632 | 5.9363 | 6.1073 | 6.2488 | 6.3721 | 6.4418 | 6.4502 | 5.0155 | 6.0137 | 5.8726 | 6.2926 |
| SRRM1_S458 | SRRM1 | A9Z1X7 | | 458 | PRKVELSESEEDK | 5.3978 | 5.6373 | 5.8033 | 6.0702 | 6.4056 | 5.9353 | 6.2190 | 5.6259 | 6.6403 | 6.3815 | 6.5259 | 6.2401 | 6.9724 | 7.2507 | 6.4352 | 6.4593 |
| SRRM1_S458;S460 | SRRM1 | A9Z1X7 | | 910 | MRKAQVSPQSxxx | 6.2015 | 6.4988 | 5.8048 | 6.7359 | 6.0781 | 5.4685 | 6.1888 | 5.9377 | 5.9880 | 5.8549 | 5.8882 | 5.6117 | 6.2311 | 7.3245 | 7.0734 | 7.1141 |
| SRRM1_S473 | SRRM1 | A9Z1X7 | | 260 | PEPKEPSPEKNSK | 6.3415 | 6.0702 | 6.4036 | 6.2196 | 6.2024 | 6.5162 | 6.8644 | 4.8196 | 5.6726 | 6.1700 | 6.8780 | 6.0917 | 7.2527 | 6.9277 | 5.7192 | 5.8507 |
| SRRM1_S547 | SRRM1 | A9Z1X7 | | 778 | KPPAPPSPVQSQS | 5.8554 | 5.9803 | 6.2674 | 6.7294 | 5.8038 | 5.7286 | 5.9335 | 5.7677 | 5.8127 | 6.1145 | 5.6708 | 6.0583 | 6.6968 | 7.3603 | 6.9308 | 7.2897 |
| SRRM1_S558;S560 | SRRM1 | A9Z1X7 | | 397 | RRRHRPSPPATPP | 6.1395 | 6.7500 | 6.4274 | 6.9194 | 5.9443 | 5.6958 | 6.0230 | 6.1856 | 6.0163 | 6.6404 | 6.4506 | 6.4328 | 5.5613 | 6.3359 | 5.9845 | 6.4932 |
| SRRM1_S558;T564 | SRRM1 | A9Z1X7 | | 623 | PPPKRRTASPPPP | 6.3959 | 6.4190 | 5.9771 | 6.8019 | 5.9999 | 5.3648 | 6.0525 | 5.8462 | 6.2106 | 6.2214 | 5.9291 | 6.1083 | 6.2576 | 6.7589 | 6.6255 | 7.0313 |
| SRRM1_S569;S571 | SRRM1 | A9Z1X7 | | 736 | IRRVSRTPEPKKI | 6.2719 | 6.3577 | 5.8039 | 6.3240 | 5.5418 | 5.7147 | 5.5775 | 5.0672 | 5.7051 | 5.9965 | 5.6131 | 5.7006 | 7.1543 | 8.1156 | 7.4015 | 7.6546 |
| SRRM1_S606 | SRRM1 | A9Z1X7 | | 187 | SPRRRSSPVRRER | 5.7080 | 5.8707 | 5.7323 | 6.5023 | 6.1341 | 6.2250 | 6.3021 | 5.8028 | 5.7678 | 6.1241 | 6.2690 | 6.0800 | 6.7406 | 6.8777 | 7.0311 | 6.8323 |
| SRRM1_S614 | SRRM1 | A9Z1X7 | | 765 | SRSVSGSPEPAAK | 6.9727 | 7.7239 | 6.7548 | 7.2933 | 5.5860 | 5.6990 | 5.9172 | 5.9955 | 7.0483 | 7.8977 | 7.3034 | 7.1226 | 4.0081 | 4.9963 | 4.8129 | 4.8683 |
| SRRM1_S614;S616 | SRRM1 | A9Z1X7 | | 635 | PPKRRASPSPPPK | 6.5640 | 6.8753 | 6.5437 | 7.3092 | 5.7900 | 5.6784 | 5.9677 | 5.9169 | 6.4461 | 6.7628 | 6.8812 | 6.9505 | 5.0154 | 5.9339 | 5.6358 | 5.7289 |
| SRRM1_S635 | SRRM1 | A9Z1X7 | | 692 | QPNKRHSPSPRPR | 6.9618 | 7.5956 | 6.7277 | 8.0467 | 5.3645 | 5.3486 | 6.0543 | 6.7288 | 6.9099 | 7.4597 | 7.1778 | 8.1460 | 3.6933 | 4.3703 | 4.5251 | 4.8896 |
| SRRM1_S635;S637 | SRRM1 | A9Z1X7 | | 881 | PEPKKETESEAED | 5.6198 | 5.9560 | 5.9107 | 5.4451 | 5.8574 | 5.7036 | 6.4299 | 5.9057 | 6.2238 | 6.6691 | 6.0857 | 6.1703 | 6.3111 | 7.0455 | 7.0430 | 7.6232 |
| SRRM1_S645;S647 | SRRM1 | A9Z1X7 | | 614 | PIQRRYSPSPPPK | 6.1087 | 6.8823 | 6.9743 | 7.1963 | 5.5881 | 5.5486 | 6.0623 | 5.9611 | 6.2137 | 6.1671 | 6.1739 | 6.4646 | 5.7726 | 6.3560 | 6.1219 | 6.4084 |
| SRRM1_S647 | SRRM1 | A9Z1X7 | | 674 | KHRKGSSPSRSTR | 7.8756 | 11.4023 | 10.5433 | 7.3776 | 3.3822 | 9.8662 | 5.0992 | 4.2873 | 7.3631 | 4.8768 | 3.9345 | 3.0592 | 4.8645 | 5.1628 | 5.5231 | 5.3823 |
| SRRM1_S655 | SRRM1 | A9Z1X7 | | 684 | STREARSPQPNKR | 5.9521 | 5.6300 | 6.7747 | 6.9670 | 6.3580 | 7.7069 | 6.6959 | 4.4739 | 5.2137 | 5.9194 | 6.4219 | 7.1241 | 5.5819 | 7.1444 | 5.7469 | 6.2891 |
| SRRM1_S662 | SRRM1 | A9Z1X7 | | 473 | KMAAADSVQQRRQ | 6.8436 | 7.0810 | 6.0008 | 7.4690 | 6.3416 | 6.9423 | 6.8033 | 6.1586 | 6.8327 | 6.6247 | 6.1941 | 6.5349 | 4.6265 | 5.3417 | 4.6928 | 5.5124 |
| SRRM1_S662;S664 | SRRM1 | A9Z1X7 | | 234 | VKVKEPSVQEATS | 5.3241 | 6.2141 | 5.4905 | 5.8384 | 5.5956 | 6.5626 | 5.5082 | 5.3560 | 5.2729 | 5.6816 | 5.8320 | 5.4675 | 9.2819 | 7.6764 | 7.3424 | 7.5558 |
| SRRM1_S664 | SRRM1 | A9Z1X7 | | 855 | AAAAAVTPAAIAA | 5.8482 | 6.8238 | 6.5746 | 5.7263 | 5.4718 | 6.2525 | 5.6321 | 6.3970 | 7.8060 | 6.6740 | 5.8058 | 6.7920 | 6.4208 | 5.8310 | 5.8774 | 6.0669 |
| SRRM1_S674 | SRRM1 | A9Z1X7 | | 761 | RVSSSRSVSGSPE | 7.1503 | 6.6204 | 6.3811 | 7.7931 | 5.9163 | 5.9814 | 6.7659 | 6.3149 | 6.4255 | 5.8832 | 5.8595 | 6.1107 | 4.6335 | 6.1625 | 5.4905 | 6.5113 |
| SRRM1_S684 | SRRM1 | A9Z1X7 | | 716 | RRGASSSPQRRQS | 4.3522 | 6.2546 | 6.3252 | 7.8138 | 6.5624 | 5.6011 | 6.2874 | 6.0830 | 5.9468 | 6.0855 | 6.0401 | 5.7676 | 6.7844 | 6.6154 | 7.1074 | 6.3732 |
| SRRM1_S692 | SRRM1 | A9Z1X7 | | 655 | PPKQRSSPVTKRR | 5.7115 | 7.0365 | 7.7212 | 7.7334 | 7.1375 | 7.3469 | 7.7522 | 4.2778 | 3.1320 | 4.7974 | 5.8626 | 6.4411 | 6.5331 | 7.0896 | 5.3176 | 6.1096 |
| SRRM1_S692;S694 | SRRM1 | A9Z1X7 | | 664 | TKRRSPSLSSKHR | 5.9999 | 6.1994 | 5.6271 | 6.0006 | 6.3248 | 5.2920 | 6.1557 | 5.8569 | 5.8725 | 5.8662 | 5.0812 | 4.8885 | 7.3284 | 7.7514 | 7.8803 | 7.8752 |
| SRRM1_S703;S705 | SRRM1 | A9Z1X7 | 569;571 | | NA | 6.0433 | 6.2215 | 5.9524 | 6.3129 | 5.9418 | 5.8451 | 5.9721 | 5.6400 | 6.1544 | 5.9566 | 5.8735 | 6.0807 | 6.7271 | 7.2661 | 6.9070 | 7.1054 |
| SRRM1_S705 | SRRM1 | A9Z1X7 | 458;460 | | NA | 6.1987 | 6.7269 | 6.7368 | 6.6381 | 5.8597 | 6.0443 | 6.2586 | 5.5630 | 5.9698 | 5.9488 | 5.5275 | 5.6708 | 6.4284 | 6.8238 | 6.6288 | 6.9759 |
| SRRM1_S716 | SRRM1 | A9Z1X7 | 645;647 | | NA | 5.8751 | 6.1622 | 5.8927 | 6.3652 | 6.0782 | 5.7762 | 6.1786 | 5.3994 | 5.8491 | 5.8835 | 5.8004 | 5.9932 | 6.7593 | 7.4760 | 7.1733 | 7.3376 |
| SRRM1_S722 | SRRM1 | A9Z1X7 | 558;560 | | NA | 6.1924 | 6.3188 | 6.0985 | 6.3461 | 5.8016 | 5.9417 | 6.0744 | 5.6632 | 6.2418 | 6.0981 | 5.8745 | 5.9856 | 6.6121 | 7.1396 | 6.6774 | 6.9342 |
| SRRM1_S722;S724 | SRRM1 | A9Z1X7 | 590;592 | | NA | 6.1161 | 6.3809 | 5.9348 | 6.3303 | 6.0233 | 5.8171 | 5.9805 | 5.6576 | 5.9585 | 5.9949 | 5.8614 | 6.0568 | 6.5866 | 7.2134 | 6.8871 | 7.2007 |
| SRRM1_S722;S726 | SRRM1 | A9Z1X7 | 581;583 | | NA | 6.0843 | 6.4751 | 5.8247 | 6.5928 | 5.7869 | 5.7101 | 6.1662 | 5.7756 | 6.1836 | 6.1657 | 5.7937 | 5.9505 | 6.4339 | 7.0865 | 6.8679 | 7.1025 |
| SRRM1_S722;T727 | SRRM1 | A9Z1X7 | 397;401 | | NA | 5.9765 | 6.3335 | 5.9461 | 6.6254 | 6.0250 | 5.8975 | 5.9893 | 6.0337 | 6.2173 | 6.0160 | 5.8360 | 5.8701 | 6.2842 | 7.0522 | 6.7932 | 7.1041 |
| SRRM1_S734;T736 | SRRM1 | A9Z1X7 | 623;625 | | NA | 5.6769 | 6.0964 | 5.6042 | 6.3540 | 6.0292 | 5.6349 | 6.1104 | 5.8160 | 5.9977 | 5.8789 | 5.6878 | 5.7367 | 6.7297 | 7.4987 | 7.2998 | 7.8486 |
| SRRM1_S736 | SRRM1 | A9Z1X7 | 635;637 | | NA | 6.0272 | 6.4048 | 5.7898 | 6.4428 | 5.8881 | 5.8235 | 6.0204 | 5.7507 | 6.0115 | 5.9253 | 5.9516 | 6.2086 | 6.5056 | 7.1834 | 6.9321 | 7.1347 |
| SRRM1_S747 | SRRM1 | A9Z1X7 | 614;616 | | NA | 5.8841 | 6.1405 | 5.7277 | 6.5495 | 5.8493 | 5.8434 | 6.0771 | 5.9790 | 6.0499 | 6.0631 | 5.9312 | 5.8358 | 6.3071 | 7.4759 | 6.9547 | 7.3315 |
| SRRM1_S747;S749 | SRRM1 | A9Z1X7 | 424;426 | | NA | 6.1093 | 6.3799 | 5.8361 | 6.2843 | 6.1086 | 5.9259 | 5.9993 | 5.8585 | 5.8140 | 5.6122 | 5.8676 | 5.8739 | 6.8842 | 7.2295 | 7.1023 | 7.1142 |
| SRRM1_S747;S752 | SRRM1 | A9Z1X7 | 778;784 | | NA | 5.9305 | 6.4291 | 5.9603 | 6.3756 | 6.0056 | 5.9351 | 5.7692 | 5.7282 | 6.1065 | 6.1635 | 6.1991 | 6.2699 | 6.4424 | 6.9447 | 6.6962 | 7.0441 |
| SRRM1_S761 | SRRM1 | A9Z1X7 | 692;694 | | NA | 6.3142 | 6.5113 | 5.7131 | 6.4213 | 6.2021 | 5.7523 | 5.8435 | 5.7087 | 5.9813 | 5.9742 | 5.7915 | 5.8247 | 6.6968 | 6.9718 | 6.8273 | 7.4659 |
| SRRM1_S761;S765 | SRRM1 | A9Z1X7 | 778;782;784;7 | | NA | 5.7648 | 6.1777 | 5.9521 | 6.0023 | 6.2129 | 5.5785 | 6.0049 | 5.8827 | 6.2691 | 5.9904 | 5.8412 | 5.8696 | 6.7256 | 7.3261 | 6.9036 | 7.4985 |
| SRRM1_S765 | SRRM1 | A9Z1X7 | 384;388 | | NA | 6.6045 | 6.9930 | 6.2397 | 7.1583 | 6.0556 | 5.7813 | 6.2419 | 6.2458 | 6.3897 | 6.4425 | 6.1558 | 6.4783 | 5.1906 | 6.1612 | 5.7034 | 6.1584 |
| SRRM1_S778 | SRRM1 | A9Z1X7 | 409;411 | | NA | 5.4715 | 5.2669 | 4.7216 | 5.4375 | 5.9125 | 5.5512 | 5.3427 | 4.9144 | 5.2808 | 4.9147 | 4.3752 | 4.3054 | 9.6787 | 10.0379 | 9.4014 | 9.3874 |
| SRRM1_S778;S782 | SRRM1 | A9Z1X7 | 747;749 | | NA | 6.0580 | 6.4444 | 5.9586 | 6.5352 | 5.6524 | 5.8213 | 5.8972 | 5.7784 | 6.3152 | 6.3114 | 5.8916 | 6.2006 | 6.0845 | 6.9334 | 6.9064 | 7.2113 |
| SRRM1_S778;S782;S784 | SRRM1 | A9Z1X7 | 778;782 | | NA | 5.9371 | 6.3051 | 6.1123 | 6.3147 | 5.5787 | 5.7710 | 5.7952 | 5.7315 | 6.2477 | 6.1343 | 6.2853 | 6.2170 | 6.6532 | 7.1495 | 6.7506 | 7.0166 |

| | | | | | | | | | | | | | | | | | | | | |
|---|---|---|---|---|---|---|---|---|---|---|---|---|---|---|---|---|---|---|---|---|
| SRRM1_S778;S782;S784; | SRRM1 | A9Z1X7 | 722;724 | NA | 5.8352 | 6.1235 | 5.8484 | 6.3436 | 5.9660 | 5.7672 | 6.0934 | 5.8036 | 6.0794 | 6.0102 | 6.1755 | 6.0260 | 6.4786 | 7.3428 | 6.9449 | 7.1615 |
| SRRM1_S778;S782;S790 | SRRM1 | A9Z1X7 | 384;386;388 | NA | 5.8417 | 6.4505 | 6.0587 | 6.1326 | 6.1094 | 5.9841 | 6.3295 | 5.6327 | 6.1850 | 6.1092 | 5.8030 | 5.6657 | 6.7520 | 7.1025 | 6.5449 | 7.2984 |
| SRRM1_S778;S784 | SRRM1 | A9Z1X7 | 703;705 | NA | 5.8194 | 6.4988 | 5.7023 | 6.4816 | 6.4660 | 6.0662 | 6.7390 | 5.9363 | 5.7812 | 5.9463 | 5.8568 | 5.8873 | 5.8351 | 7.2376 | 6.9606 | 6.7856 |
| SRRM1_S778;S784;S790 | SRRM1 | A9Z1X7 | 722;727 | NA | 5.7492 | 5.8597 | 5.5879 | 6.2922 | 6.1818 | 5.4898 | 6.0159 | 5.7805 | 6.2700 | 5.9855 | 5.7967 | 6.0453 | 6.5925 | 7.4538 | 7.1071 | 7.7920 |
| SRRM1_S778;S790 | SRRM1 | A9Z1X7 | 778;782;784 | NA | 5.7735 | 5.8827 | 5.7350 | 6.2298 | 5.8751 | 5.5801 | 6.1388 | 5.9681 | 6.0175 | 6.3763 | 6.0033 | 5.8612 | 6.6497 | 7.1576 | 7.0629 | 7.6885 |
| SRRM1_S910 | SRRM1 | A9Z1X7 | 881;883 | NA | 6.1525 | 5.8928 | 6.5765 | 6.7646 | 5.7875 | 5.9614 | 6.3852 | 5.6703 | 5.5566 | 5.9637 | 6.2201 | 6.2694 | 6.3859 | 6.7484 | 6.7302 | 6.9349 |
| SRRM1_T220 | SRRM1 | A9Z1X7 | 778;790 | NA | 5.7944 | 6.2876 | 6.4430 | 6.7404 | 6.0982 | 5.6867 | 5.8715 | 5.7797 | 6.4361 | 5.7209 | 5.7001 | 6.2010 | 6.3794 | 7.0729 | 6.5945 | 7.1937 |
| SRRM1_T581;T583 | SRRM1 | A9Z1X7 | 558;564 | NA | 5.8577 | 6.6731 | 6.2766 | 6.1929 | 6.4052 | 5.9126 | 6.7267 | 5.8427 | 5.9650 | 6.4858 | 6.0971 | 6.1397 | 6.2016 | 6.5257 | 6.2509 | 6.4468 |
| SRRM1_T590;S592 | SRRM1 | A9Z1X7 | 722;726 | NA | 5.7908 | 6.3772 | 5.9664 | 6.1587 | 5.3845 | 5.5264 | 6.2979 | 6.2477 | 6.1641 | 5.8794 | 5.6938 | 6.2014 | 6.4530 | 7.2010 | 7.6377 | 7.0201 |
| SRRM1_T623 | SRRM1 | A9Z1X7 | 747;752 | NA | 5.9974 | 5.9210 | 6.2561 | 7.1026 | 6.0692 | 6.0336 | 6.5628 | 6.3448 | 6.3687 | 5.7374 | 5.4312 | 5.4636 | 6.4671 | 6.9555 | 6.5205 | 6.7684 |
| SRRM1_T623;S625 | SRRM1 | A9Z1X7 | 778;782;790 | NA | 5.8483 | 6.0728 | 5.7259 | 6.8853 | 5.7856 | 6.4874 | 6.1163 | 5.9914 | 6.3996 | 6.0291 | 5.8682 | 5.9705 | 6.2547 | 6.9047 | 6.8115 | 6.8488 |
| SRRM1_T625 | SRRM1 | A9Z1X7 | 761;765 | NA | 7.3064 | 7.2858 | 7.0587 | 8.2538 | 5.6792 | 5.8256 | 5.7869 | 5.3199 | 7.6358 | 7.7005 | 7.6798 | 7.0334 | 3.2817 | 5.2648 | 4.0269 | 4.8608 |
| SRRM1_T855 | SRRM1 | A9Z1X7 | 734;736 | NA | 5.9245 | 6.3413 | 5.8957 | 6.5381 | 5.8908 | 5.5407 | 5.8476 | 5.7046 | 5.9523 | 6.0751 | 6.2468 | 6.6252 | 6.1945 | 7.3897 | 6.4940 | 7.3392 |
| SRRM1_T881 | SRRM1 | A9Z1X7 | 662;664 | NA | 6.0540 | 5.9318 | 5.8227 | 6.3429 | 6.0105 | 6.1040 | 6.1829 | 5.2730 | 5.5072 | 5.6678 | 5.7547 | 5.6687 | 7.3078 | 7.8227 | 6.9494 | 7.5997 |
| SRRM1_T881;S883 | SRRM1 | A9Z1X7 | 778;784;790 | NA | 6.7273 | 5.7550 | 5.9121 | 6.7429 | 6.0747 | 6.3110 | 5.6925 | 6.3567 | 5.8891 | 7.1415 | 6.1552 | 6.9122 | 5.3850 | 6.0779 | 6.2056 | 6.6614 |
| SRRM1_T883 | SRRM1 | A9Z1X7 | 207;209;211 | NA | 6.5877 | 6.6034 | 5.5090 | 6.2794 | 6.1059 | 5.6341 | 6.4922 | 5.7453 | 6.3360 | 5.5023 | 6.3227 | 5.5547 | 7.0311 | 6.6623 | 6.9982 | 6.6359 |
| SRRM2_S1014 | SRRM2 | Q9UQ35 | 1124 | RGEFSASPMLKSG | 6.3638 | 6.5280 | 6.0901 | 6.5762 | 6.3313 | 6.1227 | 6.2207 | 6.1372 | 6.3958 | 6.3233 | 6.2420 | 6.1958 | 6.1907 | 6.2112 | 6.0592 | 6.0119 |
| SRRM2_S1032 | SRRM2 | Q9UQ35 | 2449 | AQDQPRSPVPSAF | 5.9137 | 5.9804 | 5.7966 | 6.1701 | 6.2121 | 5.8860 | 6.2576 | 6.1275 | 6.3485 | 5.8952 | 5.8598 | 5.8637 | 7.1017 | 6.9066 | 6.8049 | 6.8754 |
| SRRM2_S1064 | SRRM2 | Q9UQ35 | 2132 | PLDRCRSPGMLEP | 6.1813 | 6.2469 | 5.9141 | 6.1498 | 5.8625 | 5.8439 | 6.2366 | 5.9870 | 6.3419 | 6.1095 | 5.9152 | 6.0320 | 6.9703 | 6.7428 | 6.5930 | 6.8732 |
| SRRM2_S1083 | SRRM2 | Q9UQ35 | 1179 | QEDATASPPRQKD | 6.2726 | 6.3713 | 6.3676 | 6.5444 | 6.1827 | 6.0160 | 6.2115 | 6.1030 | 6.2367 | 6.3202 | 6.3368 | 6.2748 | 6.2412 | 6.2340 | 6.0727 | 6.2147 |
| SRRM2_S1099;S1101;S1 | SRRM2 | Q9UQ35 | 2123 | FSRPSMSPTPLDR | 6.2084 | 5.6454 | 5.9919 | 6.3547 | 6.1603 | 5.7977 | 5.9361 | 5.2364 | 6.1356 | 5.9856 | 5.9422 | 5.8453 | 7.7229 | 7.0602 | 6.9012 | 7.0762 |
| SRRM2_S1101 | SRRM2 | Q9UQ35 | 1208 | FKDTLRTPPRERS | 6.3627 | 6.6081 | 5.9351 | 6.6495 | 6.1384 | 6.0176 | 6.1220 | 5.9286 | 6.2274 | 6.2662 | 6.2148 | 6.2836 | 6.3288 | 6.3411 | 6.1875 | 6.3887 |
| SRRM2_S1101;S1103 | SRRM2 | Q9UQ35 | 2702 | VERRRPSPQPSPR | 6.0911 | 6.4456 | 5.8911 | 6.6627 | 5.8887 | 5.5391 | 5.7294 | 5.9258 | 6.1673 | 6.0233 | 5.8865 | 6.0924 | 7.0146 | 6.8384 | 6.8911 | 6.9128 |
| SRRM2_S1101;S1112 | SRRM2 | Q9UQ35 | 1132 | MLKSGMSPEQSRF | 6.4024 | 6.6562 | 5.9495 | 6.5237 | 6.0837 | 6.0121 | 5.7722 | 6.0832 | 6.1294 | 6.1836 | 6.0789 | 6.1249 | 6.5657 | 6.4196 | 6.5264 | 6.4886 |
| SRRM2_S1103 | SRRM2 | Q9UQ35 | 2581 | ALKRVPSPTPAPK | 6.2958 | 6.4820 | 5.9270 | 6.4344 | 6.2153 | 5.9131 | 6.0401 | 6.2317 | 6.3872 | 6.1138 | 5.9359 | 5.8522 | 6.6594 | 6.4241 | 6.5854 | 6.5028 |
| SRRM2_S1103;S1112 | SRRM2 | Q9UQ35 | 1404 | SNQSISSPVLDAV | 6.2773 | 6.2711 | 6.2460 | 6.4923 | 6.1867 | 5.9844 | 5.9782 | 6.1561 | 6.6823 | 6.4072 | 6.5818 | 6.7119 | 5.9119 | 6.3726 | 5.6261 | 6.1140 |
| SRRM2_S1112 | SRRM2 | Q9UQ35 | 2104 | NHSGSRTPPVALN | 7.0686 | 7.3213 | 6.5973 | 7.2571 | 5.9360 | 5.7947 | 6.2890 | 6.5402 | 6.9715 | 6.5932 | 6.5634 | 6.4620 | 5.0021 | 5.1762 | 5.1937 | 5.2336 |
| SRRM2_S1122;S1124 | SRRM2 | Q9UQ35 | 2409 | DRARSRTPPSAPS | 5.9656 | 6.2581 | 5.7896 | 6.3550 | 6.0825 | 6.0351 | 5.9573 | 5.8951 | 5.9196 | 5.8603 | 5.8410 | 5.8991 | 7.1584 | 7.0240 | 6.9833 | 6.9760 |
| SRRM2_S1124 | SRRM2 | Q9UQ35 | 866 | ANEQSVTPQRRSC | 6.3463 | 6.3712 | 6.2880 | 6.8296 | 6.2577 | 6.0602 | 6.1981 | 5.7666 | 5.8689 | 6.1439 | 6.0760 | 6.2846 | 6.4288 | 6.5729 | 6.1502 | 6.3570 |
| SRRM2_S1132 | SRRM2 | Q9UQ35 | 876 | RSCFESSPDPELK | 5.9570 | 6.0784 | 5.9492 | 6.4164 | 6.1583 | 5.8495 | 6.0940 | 5.8247 | 6.3937 | 6.2349 | 6.1505 | 6.0826 | 6.7905 | 6.5892 | 6.6967 | 6.7345 |
| SRRM2_S1152 | SRRM2 | Q9UQ35 | 323 | STQRPSSPETATK | 5.8090 | 5.8820 | 5.7938 | 6.3669 | 6.0680 | 5.7916 | 5.9099 | 5.9720 | 6.5043 | 6.2573 | 6.0710 | 6.1597 | 6.9180 | 6.7051 | 6.8397 | 6.9516 |
| SRRM2_S1179 | SRRM2 | Q9UQ35 | 1014 | SLSGSKSPCPQEK | 6.5364 | 6.4355 | 5.8224 | 6.3113 | 6.2829 | 5.9803 | 6.1818 | 6.0020 | 6.2505 | 6.0727 | 5.8035 | 5.8048 | 6.6769 | 6.5107 | 6.6953 | 6.6328 |
| SRRM2_S1188 | SRRM2 | Q9UQ35 | 333 | ATKQPSSPYEDKD | 5.7275 | 5.8754 | 5.9401 | 6.5171 | 6.2108 | 5.9564 | 6.0916 | 5.8874 | 6.2607 | 6.1771 | 6.0979 | 6.1985 | 6.8227 | 6.6037 | 6.8710 | 6.7620 |
| SRRM2_S1198 | SRRM2 | Q9UQ35 | 1329 | RENSFGSPLEFRN | 6.4764 | 6.5734 | 6.2619 | 6.7477 | 5.9636 | 5.6719 | 6.0895 | 6.3016 | 6.7416 | 6.2926 | 6.2021 | 6.3882 | 5.9604 | 6.1722 | 5.9503 | 6.2066 |
| SRRM2_S1199 | SRRM2 | Q9UQ35 | 1492 | PKALPQTPRPRSR | 6.6372 | 6.8703 | 5.9198 | 6.5513 | 6.2661 | 6.0285 | 6.1280 | 6.0375 | 6.2133 | 6.0830 | 5.9958 | 6.0205 | 6.3721 | 6.4006 | 6.2634 | 6.2126 |
| SRRM2_S1218;S1219 | SRRM2 | Q9UQ35 | 1003 | PKVKAQTPPGPSL | 5.7817 | 6.1898 | 6.0218 | 6.5870 | 6.1374 | 6.1504 | 6.1911 | 6.0234 | 6.3057 | 6.2080 | 6.0754 | 6.1603 | 6.4089 | 6.5893 | 6.5432 | 6.6266 |
| SRRM2_S1219 | SRRM2 | Q9UQ35 | 1434 | KDGLPRTPSRRSR | 7.0313 | 6.8858 | 6.2796 | 6.4910 | 6.1650 | 6.3657 | 6.4707 | 5.2372 | 5.9092 | 5.8418 | 6.2231 | 6.3863 | 6.3795 | 6.2329 | 6.0211 | 6.0797 |
| SRRM2_S1233 | SRRM2 | Q9UQ35 | 1472 | MKDIPRTPSRGRS | 6.4867 | 6.7905 | 5.8742 | 6.5585 | 6.1118 | 6.0457 | 6.0456 | 5.7201 | 6.1274 | 6.0291 | 6.0791 | 6.1908 | 6.5920 | 6.4588 | 6.4127 | 6.4769 |
| SRRM2_S1257 | SRRM2 | Q9UQ35 | 2398 | RVSGRTSPPLLDR | 6.3968 | 6.5919 | 6.1506 | 6.3563 | 6.1491 | 5.9583 | 6.1083 | 5.8067 | 6.2803 | 6.1908 | 6.1213 | 6.2431 | 6.7078 | 6.3550 | 6.2451 | 6.3385 |
| SRRM2_S1311 | SRRM2 | Q9UQ35 | 1188 | RQKDKFSPFPVQD | 5.9869 | 6.4996 | 6.2471 | 6.7133 | 6.0684 | 6.3146 | 6.3843 | 6.2854 | 6.4829 | 6.6730 | 6.6773 | 6.5132 | 5.5756 | 5.9727 | 5.7215 | 5.8843 |
| SRRM2_S1318;S1320 | SRRM2 | Q9UQ35 | 1531 | QKTVARTPLGQRS | 7.0992 | 6.8279 | 6.3436 | 6.4206 | 6.7325 | 6.8308 | 6.3596 | 5.1897 | 5.5324 | 5.8476 | 6.1836 | 6.2047 | 6.6031 | 6.3749 | 5.7851 | 5.6647 |
| SRRM2_S1318;S1320;S1 | SRRM2 | Q9UQ35 | 2382 | SGANLTSPRVPLS | 6.5320 | 6.5178 | 6.1107 | 6.5040 | 6.4240 | 6.4975 | 7.1103 | 6.5824 | 6.8970 | 6.6714 | 6.3237 | 6.3287 | 5.2261 | 5.6812 | 5.0650 | 5.5282 |
| SRRM2_S1318;S1326 | SRRM2 | Q9UQ35 | 983 | TKVKPETPPRQSH | 6.1197 | 6.4222 | 6.0171 | 6.6348 | 6.1229 | 5.8570 | 6.0629 | 6.1044 | 6.2652 | 6.1940 | 6.0245 | 6.2315 | 6.5361 | 6.4056 | 6.4883 | 6.5139 |
| SRRM2_S1320 | SRRM2 | Q9UQ35 | 2289 | NLADPRTPTAPAV | 6.5906 | 7.3466 | 6.4675 | 7.0530 | 6.0978 | 6.4651 | 7.2415 | 7.4569 | 8.1059 | 7.0805 | 7.1481 | 7.1766 | 3.5889 | 4.3700 | 3.5696 | 4.2412 |
| SRRM2_S1320;S1326 | SRRM2 | Q9UQ35 | 2694 | SRSLSYSPVERRR | 7.1120 | 7.1865 | 5.9834 | 6.5743 | 6.2932 | 6.0731 | 6.1384 | 5.8335 | 6.3559 | 5.9694 | 5.8810 | 5.8470 | 6.3243 | 6.4063 | 5.9328 | 6.0890 |
| SRRM2_S1320;S1326;S1 | SRRM2 | Q9UQ35 | 1103 | GRSRSSSPVTELA | 6.1982 | 6.4552 | 5.8053 | 6.3634 | 6.0457 | 5.5083 | 6.1010 | 6.0445 | 6.3230 | 6.0823 | 5.9642 | 6.0994 | 6.7394 | 6.5405 | 6.9415 | 6.7881 |
| SRRM2_S1320;S1329 | SRRM2 | Q9UQ35 | 895 | HSCSGSSPPRVKS | 6.3524 | 6.5892 | 5.9253 | 6.7019 | 5.9811 | 5.8205 | 5.8773 | 5.8471 | 6.2956 | 6.1775 | 6.2526 | 6.1642 | 6.4780 | 6.4410 | 6.5502 | 6.5461 |
| SRRM2_S1326 | SRRM2 | Q9UQ35 | 792 | PKQKSQTPPRRSR | 6.4551 | 6.7248 | 6.1110 | 6.6970 | 6.1513 | 6.0130 | 6.1283 | 6.3501 | 6.3670 | 6.2364 | 6.2474 | 6.2625 | 5.9527 | 6.1020 | 6.1008 | 6.1007 |
| SRRM2_S1329 | SRRM2 | Q9UQ35 | 1320 | HKELSNSPLRENS | 6.2810 | 6.4170 | 6.1400 | 6.5586 | 6.2824 | 6.1393 | 6.0065 | 5.6226 | 5.6997 | 5.7155 | 5.9266 | 5.9984 | 6.9925 | 6.7015 | 6.8857 | 6.6328 |
| SRRM2_S1329;S1336 | SRRM2 | Q9UQ35 | 1869 | RSRSRASPATHRR | 6.4829 | 6.5425 | 6.0692 | 6.4515 | 5.9442 | 5.6661 | 5.8248 | 6.3246 | 6.5803 | 6.0647 | 6.1541 | 6.2192 | 6.3557 | 6.1716 | 6.5587 | 6.5900 |
| SRRM2_S1368 | SRRM2 | Q9UQ35 | 820 | RSRSSSSPPPKQK | 6.9867 | 7.3360 | 6.6592 | 7.1735 | 5.8463 | 5.6383 | 5.8789 | 5.8835 | 6.7125 | 6.7993 | 6.6485 | 6.7488 | 5.3952 | 5.4895 | 5.4172 | 5.3864 |
| SRRM2_S1382;S1387 | SRRM2 | Q9UQ35 | 398 | NPPSEASPTRDRS | 6.2280 | 6.7332 | 6.6418 | 7.0558 | 5.8724 | 5.9743 | 5.9955 | 6.3535 | 6.8042 | 6.3468 | 6.5962 | 6.2878 | 5.3689 | 5.5783 | 6.0360 | 6.1272 |
| SRRM2_S1398 | SRRM2 | Q9UQ35 | 783 | RSLSGSSPCPKQK | 7.6757 | 7.8433 | 7.4181 | 7.8717 | 6.2550 | 6.1922 | 6.1123 | 5.9580 | 7.7346 | 7.6216 | 7.5251 | 7.2304 | 3.4698 | 4.2653 | 3.2150 | 3.6119 |
| SRRM2_S1398;S1404 | SRRM2 | Q9UQ35 | 1219 | RSGAGSSPETKEQ | 6.2212 | 6.4898 | 5.9264 | 6.6313 | 6.0401 | 5.7931 | 6.1183 | 5.9986 | 6.3421 | 6.2699 | 6.3308 | 6.5118 | 6.3547 | 6.3052 | 6.3112 | 6.3555 |
| SRRM2_S1401;S1404 | SRRM2 | Q9UQ35 | 1043 | AGVKSSTPPGESY | 6.1959 | 6.5331 | 6.0865 | 6.9110 | 6.2653 | 6.1759 | 6.2481 | 6.2656 | 6.4447 | 6.1270 | 6.4081 | 6.2690 | 6.1262 | 6.2994 | 5.7375 | 5.9067 |
| SRRM2_S1401;S1404;T1 | SRRM2 | Q9UQ35 | 994 | SHSGSISPYPKVK | 6.4256 | 6.6588 | 5.8264 | 6.6974 | 6.5486 | 6.4034 | 6.1172 | 6.4645 | 6.3039 | 5.8112 | 6.0021 | 6.0908 | 6.1321 | 6.3006 | 6.2698 | 5.9476 |
| SRRM2_S1401;T1413 | SRRM2 | Q9UQ35 | 1916 | RSRSRASPVSRRR | 6.3297 | 6.8287 | 6.4109 | 6.9777 | 6.3966 | 6.0534 | 6.3986 | 5.7708 | 5.3576 | 6.1881 | 6.3324 | 6.9053 | 6.0766 | 5.9571 | 5.9512 | 6.0656 |
| SRRM2_S1404 | SRRM2 | Q9UQ35 | 1987 | RSRSRTSPVTRRR | 6.0296 | 7.3725 | 7.1383 | 7.4364 | 6.7716 | 6.8951 | 7.1635 | 5.4128 | 4.3340 | 5.6412 | 6.4298 | 7.3457 | 5.9314 | 5.6190 | 5.1398 | 5.3393 |
| SRRM2_S1404;S1415 | SRRM2 | Q9UQ35 | 1424 | RSSSASSPEMKDG | 7.1175 | 7.1757 | 6.5203 | 7.2062 | 6.4199 | 6.4011 | 6.5238 | 6.1408 | 7.3732 | 7.4362 | 7.1686 | 7.2648 | 4.2239 | 4.8215 | 3.9496 | 4.2568 |
| SRRM2_S1404;T1413 | SRRM2 | Q9UQ35 | 1112 | TELASRSPIRQDR | 6.4307 | 6.7060 | 5.8198 | 6.6672 | 6.1614 | 5.8948 | 6.0180 | 5.9777 | 6.0789 | 6.1707 | 5.9934 | 6.1597 | 6.5280 | 6.4186 | 6.3487 | 6.6265 |
| SRRM2_S1419;S1424 | SRRM2 | Q9UQ35 | 2272 | AAMNLASPRTAVA | 5.9370 | 6.2277 | 6.8410 | 6.8208 | 6.2260 | 6.6044 | 6.8623 | 6.4808 | 6.7008 | 6.2745 | 6.4397 | 6.2331 | 5.6185 | 5.9779 | 5.1253 | 5.6304 |
| SRRM2_S1421 | SRRM2 | Q9UQ35 | 1463 | HSLSGSSPGMKDI | 6.5182 | 6.9032 | 5.7679 | 6.7209 | 6.1604 | 6.2166 | 5.9578 | 6.1664 | 6.6679 | 6.6820 | 6.5623 | 6.4667 | 5.7412 | 6.0350 | 5.7410 | 5.6924 |
| SRRM2_S1421;S1424 | SRRM2 | Q9UQ35 | 250 | KRKRSRSTTPAPK | 5.4829 | 6.7001 | 6.2186 | 6.1437 | 5.7366 | 6.1323 | 5.8361 | 6.5517 | 6.0279 | 6.9893 | 6.8413 | 6.8038 | 5.9154 | 5.8335 | 6.7324 | 6.0543 |
| SRRM2_S1424 | SRRM2 | Q9UQ35 | 1233 | SALPTSSQDEELM | 5.8748 | 6.3424 | 6.1537 | 6.6850 | 6.1597 | 5.8771 | 6.1414 | 6.3029 | 6.2590 | 6.5092 | 6.2329 | 6.0695 | 6.3105 | 6.3065 | 6.2959 | 6.4795 |
| SRRM2_S1443;S1444 | SRRM2 | Q9UQ35 | 1326 | SPLRENSFGSPLE | 5.8862 | 5.9603 | 6.0837 | 6.2152 | 5.8168 | 5.8270 | 5.9796 | 6.0870 | 6.4610 | 5.9550 | 6.2084 | 5.7527 | 6.8648 | 6.6961 | 7.0453 | 7.1608 |
| SRRM2_S1444 | SRRM2 | Q9UQ35 | 1601 | RSRSGSSPEVKDK | 6.2389 | 6.2559 | 5.8346 | 6.1811 | 6.2275 | 5.7971 | 5.9409 | 5.6097 | 6.8926 | 6.3622 | 6.5624 | 6.3826 | 6.6018 | 6.4257 | 6.3211 | 6.3659 |
| SRRM2_S1458;S1463 | SRRM2 | Q9UQ35 | 914 | SPSRSSSPQPKVK | 6.1712 | 6.2658 | 5.6747 | 6.3339 | 6.2013 | 5.9185 | 6.0912 | 5.9902 | 6.1299 | 6.0537 | 5.9610 | 5.9270 | 6.9469 | 6.7666 | 6.8222 | 6.7460 |
| SRRM2_S1463 | SRRM2 | Q9UQ35 | 2599 | GRPPEPTPAKRKR | 8.1842 | 8.7164 | 7.7016 | 7.9820 | 6.2270 | 7.1746 | 6.4608 | 5.4073 | 7.0414 | 5.0610 | 5.3765 | 4.8019 | 5.1890 | 5.4273 | 4.4228 | 4.8261 |
| SRRM2_S1483 | SRRM2 | Q9UQ35 | 2329 | YPSSSRTPQAPAS | 6.8297 | 7.4131 | 6.4113 | 7.2524 | 5.7643 | 6.8844 | 7.4530 | 7.3848 | 8.1177 | 7.6602 | 7.4846 | 7.0387 | 3.2643 | 4.1505 | 3.1487 | 3.7422 |
| SRRM2_S1497;S1499;S1 | SRRM2 | Q9UQ35 | 778 | RLSLRRSLSGSSP | 6.2470 | 6.7441 | 6.2192 | 7.0032 | 6.0769 | 5.9039 | 6.1689 | 6.3723 | 6.2851 | 6.5984 | 6.7268 | 6.9136 | 5.4728 | 5.4340 | 5.9093 | 5.9246 |
| SRRM2_S1497;S1502 | SRRM2 | Q9UQ35 | 2688 | IDSLRDSRSLSYS | 5.6268 | 6.2593 | 5.4791 | 6.4555 | 7.6746 | 6.9793 | 6.3180 | 7.1385 | 6.0570 | 6.0584 | 6.6498 | 5.9326 | 6.3573 | 6.5033 | 5.5801 | 4.9303 |
| SRRM2_S1499;S1502 | SRRM2 | Q9UQ35 | 1368 | QLETDPSLDMKEQ | 5.3953 | 5.8729 | 5.6209 | 5.6708 | 6.1998 | 5.3980 | 5.6950 | 6.5298 | 6.1393 | 6.1222 | 5.2807 | 5.9679 | 7.3642 | 7.3127 | 7.9942 | 7.4364 |
| SRRM2_S1519 | SRRM2 | Q9UQ35 | 486 | RMGRSRSPATAKR | 6.2009 | 6.0734 | 5.1359 | 5.7149 | 5.8025 | 5.6420 | 5.3575 | 4.9329 | 5.6404 | 5.4024 | 5.2061 | 5.4419 | 8.9246 | 8.3639 | 8.3358 | 7.8253 |
| SRRM2_S1539 | SRRM2 | Q9UQ35 | 1413 | LDAVPRTPSRERS | 7.4699 | 6.7684 | 6.9486 | 6.8620 | 6.3751 | 6.4012 | 6.3325 | 4.8002 | 5.9455 | 5.5909 | 6.0061 | 6.1162 | 6.7647 | 6.2881 | 5.7844 | 5.5460 |
| SRRM2_S1541 | SRRM2 | Q9UQ35 | 1444 | RSRSGSSPGLRDG | 7.2119 | 8.0492 | 6.7542 | 7.6152 | 5.9821 | 5.8964 | 6.0516 | 6.1089 | 7.3863 | 7.5881 | 7.1923 | 7.3048 | 3.9668 | 4.7224 | 3.8311 | 4.3388 |
| SRRM2_S1541;S1552 | SRRM2 | Q9UQ35 | 780 | SLRRSLSGSSPCP | 6.9737 | 7.1650 | 6.6150 | 6.7117 | 5.8727 | 5.6909 | 5.9216 | 6.1172 | 6.2906 | 6.2902 | 6.4479 | 6.2779 | 6.0352 | 5.7241 | 6.1268 | 5.7394 |
| SRRM2_S1552 | SRRM2 | Q9UQ35 | 1083 | RQSHSESPSLQSK | 6.0312 | 6.2607 | 6.2063 | 6.6963 | 6.1986 | 5.9894 | 6.2602 | 6.2922 | 6.1676 | 6.1182 | 6.4067 | 6.2959 | 6.4828 | 6.3684 | 5.7998 | 6.4256 |
| SRRM2_S1579 | SRRM2 | Q9UQ35 | 2365 | AALAPASLTSARM | 7.6250 | 6.9640 | 6.7968 | 7.4366 | 6.1205 | 6.2000 | 7.3421 | 6.0204 | 7.4047 | 6.1575 | 6.4195 | 6.6898 | 4.6314 | 5.1582 | 4.1359 | 4.8976 |
| SRRM2_S1581;S1582 | SRRM2 | Q9UQ35 | 1857 | RSRSRTSPAPWKR | 6.1407 | 6.6413 | 6.4023 | 6.8527 | 5.6978 | 5.5885 | 5.9556 | 6.1983 | 6.5948 | 6.5320 | 6.4397 | 6.9020 | 5.9945 | 6.0904 | 5.7919 | 6.1776 |
| SRRM2_S1582 | SRRM2 | Q9UQ35 | 817 | PPRRSRSSSSPPP | 6.9990 | 7.1639 | 6.6618 | 7.3084 | 5.7830 | 5.5024 | 5.8524 | 5.9552 | 6.7280 | 6.6418 | 6.4295 | 6.7012 | 5.6081 | 5.6086 | 5.4220 | 5.6345 |
| SRRM2_S1598 | SRRM2 | Q9UQ35 | 326 | RPSSPETATKQPS | 5.9231 | 6.3450 | 6.1036 | 6.2462 | 6.3721 | 6.1348 | 5.9891 | 6.2466 | 6.1611 | 6.3386 | 6.4891 | 6.4267 | 6.3514 | 6.4714 | 6.0578 | 6.3436 |
| SRRM2_S1598;S1601 | SRRM2 | Q9UQ35 | 2032 | IRRRSRSRTPLLP | 5.5474 | 6.4605 | 6.1001 | 6.6390 | 7.4292 | 5.3619 | 5.9094 | 6.5472 | 5.7865 | 5.9999 | 6.2619 | 6.0547 | 6.4650 | 5.9739 | 7.2254 | 6.2378 |
| SRRM2_S1601 | SRRM2 | Q9UQ35 | 1311 | WGGPHFSPEHKEL | 5.9060 | 6.1429 | 5.9062 | 6.1872 | 6.5412 | 6.5617 | 5.9016 | 6.2113 | 6.3942 | 6.0521 | 6.5715 | 6.3028 | 6.3637 | 6.8055 | 6.2473 | 5.9049 |
| SRRM2_S1691;S1694 | SRRM2 | Q9UQ35 | 536 | RRGRSRSPQRPGW | 6.1734 | 6.3670 | 6.0320 | 6.9362 | 6.2834 | 5.7696 | 5.7628 | 6.6410 | 6.5876 | 6.0437 | 5.9972 | 5.8844 | 6.3703 | 6.0568 | 6.6716 | 6.4231 |
| SRRM2_S1691;T1698 | SRRM2 | Q9UQ35 | 424 | SESSPPSPQPTKV | 6.1465 | 6.4700 | 6.8290 | 7.1311 | 6.6114 | 6.0047 | 6.9245 | 6.3353 | 7.4779 | 7.0814 | 7.4902 | 7.3777 | 4.3930 | 4.8001 | 4.2007 | 4.7267 |
| SRRM2_S1694 | SRRM2 | Q9UQ35 | 1694 | RRSSRSSPELTRK | 7.8226 | 8.1452 | 6.7056 | 7.1145 | 5.4700 | 5.6458 | 5.8025 | 6.1602 | 7.5887 | 7.5716 | 7.0454 | 7.0133 | 4.1835 | 4.8763 | 4.3448 | 4.5100 |
| SRRM2_S1762;S1764 | SRRM2 | Q9UQ35 | 1975 | RSRSRTSPITRRR | 6.0167 | 7.1423 | 6.9596 | 7.5584 | 6.6005 | 6.4781 | 7.0015 | 5.6667 | 4.5246 | 6.0649 | 6.7756 | 7.7343 | 5.3894 | 5.3913 | 5.2159 | 5.4802 |
| SRRM2_S1822;S1824 | SRRM2 | Q9UQ35 | 449 | PAPAPGSHREISS | 6.2262 | 5.9383 | 6.4418 | 6.9736 | 6.0723 | 5.6532 | 6.1060 | 6.1956 | 6.3365 | 6.3187 | 5.9988 | 6.3368 | 6.4239 | 6.4930 | 6.1299 | 6.3554 |
| SRRM2_S1842;T1844 | SRRM2 | Q9UQ35 | 367 | PEPPAPTPLLAER | 6.1799 | 6.1985 | 5.8597 | 6.4610 | 6.3695 | 5.4790 | 6.6594 | 5.8356 | 6.4183 | 5.0382 | 5.3026 | 5.2481 | 7.9368 | 7.4013 | 6.5868 | 7.0253 |
| SRRM2_S1854;S1857 | SRRM2 | Q9UQ35 | 2144 | PLGSSRTPMSVLQ | 6.0850 | 6.2047 | 6.6979 | 6.9905 | 5.6521 | 6.8936 | 6.3144 | 6.5397 | 6.9984 | 6.2531 | 6.7414 | 6.7691 | 4.8050 | 5.6986 | 5.7865 | 5.5701 |
| SRRM2_S1857 | SRRM2 | Q9UQ35 | 1152 | PTVDSNSLLGQSR | 5.7066 | 6.1106 | 5.4241 | 6.7644 | 6.0977 | 5.8153 | 6.0256 | 5.9288 | 6.0711 | 6.7555 | 6.2093 | 5.8913 | 6.9382 | 6.5994 | 7.2019 | 6.4602 |
| SRRM2_S1866;S1869 | SRRM2 | Q9UQ35 | 839 | QSHSSSSPHPKVK | 6.6262 | 7.0345 | 5.5516 | 6.8583 | 6.6273 | 6.6923 | 5.4705 | 6.9279 | 6.0024 | 5.7472 | 6.3477 | 5.9176 | 5.7205 | 6.3982 | 6.5020 | 5.5759 |
| SRRM2_S1869 | SRRM2 | Q9UQ35 | 1064 | KGQSQTSPDHRSD | 5.8069 | 6.1450 | 5.9497 | 6.6640 | 5.8331 | 5.2997 | 6.0284 | 6.0284 | 6.1161 | 6.6188 | 6.1423 | 6.1830 | 6.9560 | 6.2951 | 7.6105 | 6.3231 |
| SRRM2_S1876;S1878;S1 | SRRM2 | Q9UQ35 | 954 | RRSRSVSPCSNVE | 5.2480 | 6.2208 | 5.6818 | 6.3446 | 6.0132 | 5.7722 | 5.6962 | 5.6534 | 5.9701 | 6.2551 | 5.8651 | 6.1746 | 7.3755 | 7.3917 | 7.1834 | 7.1543 |
| SRRM2_S1878;S1884 | SRRM2 | Q9UQ35 | 454 | GSHREISSSPTSK | 5.9759 | 6.5181 | 6.6940 | 7.1524 | 5.8590 | 6.1399 | 6.4163 | 6.0198 | 6.8993 | 6.3543 | 6.7713 | 6.6817 | 5.4447 | 5.4742 | 5.3795 | 6.2194 |
| SRRM2_S1878;T1880 | SRRM2 | Q9UQ35 | 456 | HREISSSPTSKNR | 6.5361 | 7.0064 | 6.7434 | 7.6568 | 6.0180 | 5.9065 | 6.0245 | 6.3473 | 6.8103 | 7.1464 | 7.0897 | 7.0919 | 4.5601 | 5.0861 | 4.8174 | 5.1590 |
| SRRM2_S1890;S1893 | SRRM2 | Q9UQ35 | 1421 | SRERSSSASSPEM | 5.5246 | 5.4663 | 5.4499 | 5.9232 | 5.8223 | 5.7495 | 5.8689 | 6.2021 | 5.4943 | 5.2567 | 5.8306 | 5.5159 | 7.9898 | 7.7855 | 8.4854 | 7.6350 |
| SRRM2_S1893 | SRRM2 | Q9UQ35 | 322 | TSTQRPSSPETAT | 5.7698 | 5.4146 | 5.8831 | 6.1573 | 6.0323 | 5.0032 | 7.0825 | 5.6747 | 6.5130 | 6.2865 | 5.6849 | 5.4610 | 7.0138 | 6.3824 | 7.5958 | 8.0453 |
| SRRM2_S1902;S1905 | SRRM2 | Q9UQ35 | 1539 | LGQRSRSGSSQEL | 6.5795 | 6.4871 | 6.4986 | 6.6569 | 6.1743 | 5.7093 | 6.1285 | 6.1951 | 6.4911 | 6.5311 | 6.4848 | 6.9218 | 5.6737 | 5.8913 | 5.5347 | 6.0422 |
| SRRM2_S1913;S1916 | SRRM2 | Q9UQ35 | 358 | PSPERSSTGPEPP | 5.8187 | 5.6709 | 6.1854 | 6.1414 | 6.1683 | 6.2820 | 6.4919 | 5.9812 | 6.2329 | 5.7393 | 6.2205 | 5.9397 | 7.0902 | 6.4102 | 6.3983 | 7.2291 |
| SRRM2_S1916 | SRRM2 | Q9UQ35 | 1198 | VQDRPESSLVFKD | 6.4507 | 6.3140 | 5.6360 | 7.2746 | 6.5552 | 5.2803 | 6.2317 | 5.6428 | 5.6542 | 5.0436 | 5.4539 | 5.6921 | 7.2647 | 7.5606 | 6.8182 | 7.1274 |
| SRRM2_S1925;T1927 | SRRM2 | Q9UQ35 | 359 | SPERSSTGPEPPA | 6.0088 | 7.2188 | 6.7599 | 7.0932 | 5.7923 | 6.4958 | 5.5867 | 5.6798 | 6.5521 | 6.4027 | 6.5954 | 5.4677 | 6.6343 | 6.1300 | 5.2707 | 6.3118 |
| SRRM2_S1925;T1931 | SRRM2 | Q9UQ35 | 1978 | SRTSPITRRRSRS | 5.5767 | 6.4194 | 6.5305 | 6.9511 | 5.8544 | 6.1739 | 6.7464 | 6.5028 | 9.1776 | 5.8622 | 6.4110 | 7.0336 | 5.1184 | 5.4811 | 5.0093 | 5.1516 |
| SRRM2_S1972;S1975 | SRRM2 | Q9UQ35 | 400 | PSEASPTRDRSPP | 6.5289 | 6.5443 | 5.8004 | 7.1966 | 6.5730 | 5.9759 | 6.4716 | 6.3933 | 6.2667 | 6.8100 | 6.4793 | 7.5274 | 5.3389 | 5.1725 | 5.4151 | 5.5060 |
| SRRM2_S1972;T1978 | SRRM2 | Q9UQ35 | 510 | GHSRSRSPQWRRS | 6.9740 | 7.4079 | 6.7477 | 6.9728 | 6.0291 | 6.0066 | 5.8044 | 6.3720 | 6.6119 | 6.8011 | 6.5675 | 6.7428 | 5.1875 | 5.4937 | 5.1345 | 5.1464 |
| SRRM2_S1975 | SRRM2 | Q9UQ35 | 2738 | HKRRRETPSPRPM | 6.6935 | 6.7855 | 6.8635 | 6.6795 | 5.7419 | 5.8502 | 6.0307 | 5.4228 | 6.5898 | 5.8712 | 6.1647 | 6.4151 | 6.6713 | 6.7284 | 5.6883 | 5.8036 |
| SRRM2_S1978 | SRRM2 | Q9UQ35 | 353 | ATRPSPSPERSST | 7.2290 | 7.2386 | 6.9171 | 7.6969 | 6.0065 | 6.7439 | 6.8044 | 6.6402 | 6.7469 | 6.6010 | 6.9367 | 7.2332 | 3.5806 | 4.8986 | 4.0407 | 4.6855 |
| SRRM2_S1984;S1987 | SRRM2 | Q9UQ35 | 220 | KKKKHRSESESKK | 5.9501 | 7.8150 | 6.4010 | 7.8199 | 6.4843 | 5.8276 | 6.9107 | 7.1656 | 4.4062 | 6.0628 | 7.4695 | 4.9355 | 4.9404 | 5.2272 | 6.8911 | 5.6928 |
| SRRM2_S1987 | SRRM2 | Q9UQ35 | 351 | KSATRPSPSPERS | 6.1928 | 6.3324 | 6.5963 | 7.2442 | 5.4401 | 5.4871 | 6.4855 | 6.3398 | 6.5788 | 6.1834 | 6.0659 | 5.9296 | 6.0027 | 6.7635 | 5.7476 | 6.6104 |
| SRRM2_S2020;T2022 | SRRM2 | Q9UQ35 | 1032 | VQSCPGSLSLCAG | 6.4667 | 6.6062 | 6.3729 | 6.4006 | 6.4120 | 6.6482 | 7.0540 | 7.0576 | 7.0888 | 6.3038 | 6.5448 | 6.8893 | 4.6508 | 5.2675 | 4.9441 | 5.2927 |
| SRRM2_S2030;S2032;T2 | SRRM2 | Q9UQ35 | 1101 | KGGRSRSSSPVTE | 6.9621 | 6.3136 | 6.9342 | 7.2504 | 6.4860 | 6.0270 | 7.1510 | 6.2145 | 6.6469 | 5.9026 | 5.9011 | 5.5677 | 6.0886 | 5.9812 | 5.1571 | 5.4161 |
| SRRM2_S2032 | SRRM2 | Q9UQ35 | 2692 | RDSRSLSYSPVER | 6.6793 | 6.4209 | 5.8444 | 5.8565 | 6.4683 | 6.3485 | 6.2332 | 5.0600 | 6.2754 | 5.0909 | 6.1609 | 6.1217 | 7.0810 | 6.7336 | 6.1903 | 7.4351 |
| SRRM2_S2032;T2034 | SRRM2 | Q9UQ35 | 111 | PGGKEETPGQRPA | 6.3034 | 6.2060 | 7.1896 | 6.6175 | 6.4403 | 6.2631 | 6.0867 | 5.6813 | 7.7319 | 6.5807 | 6.3499 | 5.5313 | 6.2639 | 5.8173 | 5.0952 | 5.8419 |
| SRRM2_S2044;S2046 | SRRM2 | Q9UQ35 | 1519 | QRERSGSESSVDQ | 4.8558 | 4.8629 | 5.6479 | 5.8959 | 6.7254 | 5.9306 | 6.3350 | 5.5708 | 6.9633 | 7.4522 | 6.8973 | 6.6911 | 6.9973 | 6.9609 | 6.0658 | 6.1477 |
| SRRM2_S2067;T2069;S2 | SRRM2 | Q9UQ35 | 1552 | DVKPSASPQERSE | 6.6129 | 7.0685 | 6.7366 | 7.6022 | 5.6862 | 5.9388 | 6.0577 | 5.9874 | 7.7317 | 8.1156 | 7.6657 | 7.7379 | 3.8934 | 4.8293 | 3.9034 | 4.4328 |
| SRRM2_S2100;S2102;T2 | SRRM2 | Q9UQ35 | 875 | RRSCFESSPDPEL | 6.3328 | 6.4093 | 6.9065 | 6.6432 | 6.1053 | 6.6278 | 6.0099 | 6.4679 | 6.8141 | 6.0268 | 6.4013 | 5.8689 | 6.1043 | 6.4550 | 5.5291 | 5.2977 |
| SRRM2_S2102;T2104 | SRRM2 | Q9UQ35 | 1582 | RSRSGSSPEVDSK | 6.3798 | 7.0091 | 6.8520 | 6.9334 | 6.0492 | 5.8142 | 5.8810 | 5.5952 | 8.1934 | 7.8258 | 8.1954 | 7.9917 | 3.6436 | 4.9761 | 4.1578 | 4.5022 |
| SRRM2_S2121;S2123 | SRRM2 | Q9UQ35 | 810 | PKAKSRTPPRRSR | 6.6416 | 6.7462 | 5.3009 | 6.6844 | 5.5909 | 6.4705 | 7.4926 | 4.9491 | 6.1615 | 4.5450 | 5.5054 | 6.6131 | 7.3120 | 5.8049 | 7.5021 | 6.6797 |
| SRRM2_S2123 | SRRM2 | Q9UQ35 | 871 | VTPQRRSCFESSP | 5.9274 | 6.4419 | 6.0284 | 6.6070 | 6.6419 | 6.1060 | 6.0160 | 5.7349 | 6.1184 | 5.9777 | 5.8227 | 5.9338 | 6.5436 | 6.5151 | 6.7999 | 6.7853 |
| SRRM2_S2132 | SRRM2 | Q9UQ35 | 1598 | SPRRSRSGSSPEV | 6.5949 | 6.1311 | 6.2779 | 6.0278 | 6.4530 | 6.0699 | 6.1353 | 5.4833 | 6.9694 | 6.2327 | 6.3734 | 6.5268 | 6.1492 | 6.1915 | 6.1060 | 6.2778 |
| SRRM2_S220 | SRRM2 | Q9UQ35 | 1541 | QRSRSGSSQELDV | 5.8492 | 6.4064 | 5.5133 | 6.6712 | 5.7399 | 5.4500 | 5.6489 | 5.1061 | 6.0448 | 5.5871 | 5.3487 | 5.9216 | 7.4603 | 7.4169 | 7.5023 | 8.3331 |
| SRRM2_S220;S222 | SRRM2 | Q9UQ35 | 1199 | QDRPESSLVFKDT | 6.3794 | 6.7414 | 6.6868 | 7.0188 | 6.4094 | 5.7827 | 5.9969 | 6.5146 | 6.3265 | 6.2459 | 6.8196 | 6.0736 | 5.4579 | 5.9740 | 5.4587 | 6.1139 |
| SRRM2_S2272 | SRRM2 | Q9UQ35 | 857 | RQGSITSPQANEQ | 5.9598 | 7.0151 | 6.3401 | 5.9392 | 5.4778 | 6.4863 | 5.8912 | 6.4257 | 5.8627 | 6.3549 | 6.2841 | 5.7724 | 6.3641 | 6.2236 | 7.1544 | 6.4486 |
| SRRM2_S2365 | SRRM2 | Q9UQ35 | 2092 | DRSRSATPPATRN | 6.4872 | 6.5252 | 5.9294 | 6.3764 | 5.6954 | 5.8599 | 6.1990 | 5.9446 | 6.5258 | 6.0356 | 5.7806 | 6.5660 | 6.1926 | 6.4633 | 6.8479 | 6.5713 |
| SRRM2_S2382 | SRRM2 | Q9UQ35 | 1257 | QILSHLSSELKEM | 5.4820 | 5.8170 | 5.4375 | 6.3796 | 6.0817 | 5.8578 | 5.8434 | 5.8929 | 5.2747 | 6.0772 | 5.0848 | 5.4311 | 7.8391 | 7.5762 | 7.7042 | 8.2208 |
| SRRM2_S2398 | SRRM2 | Q9UQ35 | 1579 | LRQRSRSGSSPEV | 5.6269 | 5.7790 | 5.6893 | 6.2212 | 5.7694 | 6.4437 | 5.8174 | 6.0908 | 7.9729 | 7.2968 | 7.9297 | 6.8661 | 5.7155 | 6.1293 | 5.5909 | 5.0612 |
| SRRM2_S2407;S2412 | SRRM2 | Q9UQ35 | 1893 | RSRSRTSPVSRRR | 5.5216 | 7.5067 | 7.2220 | 7.4785 | 7.0378 | 6.8253 | 7.7324 | 5.1584 | 3.5868 | 6.3476 | 7.0193 | 7.7271 | 5.0130 | 5.6556 | 4.4832 | 5.6847 |
| SRRM2_S2407;T2409 | SRRM2 | Q9UQ35 | 1398 | EKAGMSSNQSISS | 8.0248 | 8.2454 | 6.8210 | 8.4605 | 6.1390 | 6.6196 | 7.0758 | 6.4076 | 6.4978 | 6.3504 | 6.6568 | 7.1344 | 3.6408 | 4.3417 | 3.4974 | 4.0872 |
| SRRM2_S2449 | SRRM2 | Q9UQ35 | 1483 | RSECDSSPEPKAL | 5.7636 | 7.0324 | 6.3703 | 6.5722 | 5.6436 | 6.5416 | 5.4314 | 5.8910 | 6.2821 | 6.4857 | 6.9436 | 6.4574 | 6.0404 | 6.1627 | 5.9303 | 6.4517 |
| SRRM2_S250 | SRRM2 | Q9UQ35 | 2121;2123 | NA | 6.3060 | 6.6568 | 6.6211 | 6.9653 | 6.6164 | 6.4165 | 6.3489 | 5.8894 | 6.0493 | 6.2390 | 6.3718 | 6.0526 | 6.1307 | 6.0454 | 5.6907 | 5.6002 |
| SRRM2_S2581 | SRRM2 | Q9UQ35 | 2032;2034 | NA | 6.1416 | 5.8842 | 6.2571 | 5.9824 | 6.0882 | 5.5450 | 6.5674 | 5.4508 | 6.2704 | 6.1160 | 6.2375 | 6.1792 | 7.2533 | 6.5762 | 6.4525 | 6.9980 |
| SRRM2_S2688 | SRRM2 | Q9UQ35 | 2702;2706 | NA | 6.2565 | 6.5044 | 6.0630 | 6.7250 | 6.3985 | 6.1642 | 6.3903 | 6.1952 | 6.3043 | 6.1448 | 6.1615 | 6.1995 | 6.0709 | 6.1504 | 6.0575 | 6.2139 |
| SRRM2_S2690;S2694 | SRRM2 | Q9UQ35 | 377;398 | NA | 6.0583 | 6.1854 | 6.4755 | 6.9830 | 6.3578 | 6.1419 | 6.2649 | 5.8528 | 6.5775 | 6.3580 | 6.3089 | 6.3155 | 6.1846 | 6.3176 | 5.7085 | 5.9099 |
| SRRM2_S2692 | SRRM2 | Q9UQ35 | 1842;1844 | NA | 6.1786 | 6.2607 | 6.0713 | 6.4933 | 5.9641 | 5.7515 | 6.1838 | 5.4308 | 6.3027 | 6.1885 | 6.1210 | 6.2295 | 7.0924 | 6.6031 | 6.4418 | 6.6866 |
| SRRM2_S2692;S2694 | SRRM2 | Q9UQ35 | 508;510 | NA | 6.6550 | 6.5733 | 5.9111 | 6.4464 | 6.1934 | 5.9264 | 6.1175 | 6.1923 | 6.4306 | 6.0402 | 6.3235 | 6.1194 | 6.5701 | 6.3636 | 5.9578 | 6.1794 |
| SRRM2_S2694 | SRRM2 | Q9UQ35 | 2738;2740 | NA | 6.4177 | 6.6516 | 6.1532 | 6.7792 | 6.4569 | 6.4684 | 5.9427 | 5.6900 | 5.9307 | 6.1501 | 6.4123 | 6.1378 | 6.7705 | 6.3419 | 6.0045 | 5.6924 |
| SRRM2_S2702 | SRRM2 | Q9UQ35 | 351;353 | NA | 6.4381 | 6.3720 | 5.9824 | 6.4856 | 5.9954 | 5.8846 | 6.1841 | 5.8684 | 6.1242 | 6.1826 | 6.2254 | 6.4407 | 6.4796 | 6.2346 | 6.5234 | 6.5787 |
| SRRM2_S2702;S2706 | SRRM2 | Q9UQ35 | 2044;2046 | NA | 6.2347 | 6.4797 | 6.0062 | 6.2851 | 6.1117 | 5.5967 | 5.9386 | 5.8646 | 6.1962 | 6.2521 | 6.2554 | 6.1801 | 6.7481 | 6.4369 | 6.6594 | 6.7545 |
| SRRM2_S2727;S2729 | SRRM2 | Q9UQ35 | 2067;2069;20 | NA | 6.3969 | 6.6069 | 6.1951 | 6.4915 | 6.0055 | 5.8984 | 6.1772 | 6.0548 | 6.2249 | 6.2269 | 6.1067 | 6.0883 | 6.3664 | 6.4895 | 6.2772 | 6.3937 |
| SRRM2_S295;S297 | SRRM2 | Q9UQ35 | 454;456 | NA | 5.9272 | 6.3244 | 5.9280 | 6.3294 | 6.0155 | 5.7586 | 5.9815 | 5.6849 | 6.4909 | 6.3327 | 6.2419 | 6.2289 | 6.6857 | 6.6885 | 6.6735 | 6.7085 |
| SRRM2_S295;S300 | SRRM2 | Q9UQ35 | 903;908 | NA | 6.1396 | 6.3766 | 6.0374 | 6.3304 | 6.3771 | 5.6517 | 5.9801 | 5.7627 | 6.3112 | 5.8965 | 5.9656 | 6.1090 | 7.0171 | 6.6105 | 6.8268 | 6.6075 |
| SRRM2_S311;S323 | SRRM2 | Q9UQ35 | 1404;1413 | NA | 6.6015 | 6.4287 | 6.6243 | 6.8290 | 6.7508 | 6.5577 | 6.0646 | 5.3113 | 6.2622 | 6.3408 | 6.5289 | 6.4780 | 6.1782 | 5.8173 | 5.7441 | 5.4825 |
| SRRM2_S322 | SRRM2 | Q9UQ35 | 2020;2022 | NA | 6.1169 | 5.9773 | 6.2004 | 6.3970 | 5.8039 | 5.7309 | 6.3566 | 5.6831 | 6.3856 | 6.2932 | 6.2476 | 6.3922 | 6.8058 | 6.4731 | 6.3052 | 6.8312 |
| SRRM2_S322;S323 | SRRM2 | Q9UQ35 | 778;780 | NA | 5.5494 | 5.7518 | 5.5770 | 6.0183 | 6.1319 | 5.8958 | 5.8822 | 5.7901 | 5.9132 | 5.5497 | 5.7283 | 5.6823 | 7.9163 | 7.3526 | 7.8535 | 7.4075 |
| SRRM2_S322;T326 | SRRM2 | Q9UQ35 | 952;954 | NA | 5.9558 | 6.4163 | 5.9865 | 6.7445 | 6.1860 | 5.8900 | 5.9443 | 6.2520 | 6.3892 | 6.2369 | 6.0831 | 6.0237 | 6.4573 | 6.3750 | 6.5778 | 6.4815 |
| SRRM2_S323 | SRRM2 | Q9UQ35 | 857;866 | NA | 6.4167 | 6.3180 | 6.4965 | 6.8344 | 6.3994 | 6.1929 | 6.2559 | 6.2195 | 6.6494 | 6.3497 | 6.0279 | 6.2039 | 5.7945 | 5.9804 | 5.9267 | 5.9341 |
| SRRM2_S333 | SRRM2 | Q9UQ35 | 1925;1927 | NA | 6.1114 | 6.2359 | 6.3104 | 6.1907 | 5.7275 | 5.5839 | 6.3386 | 5.6153 | 6.1812 | 6.4986 | 6.3995 | 6.4724 | 6.8839 | 6.5073 | 6.1518 | 6.7917 |
| SRRM2_S351 | SRRM2 | Q9UQ35 | 534;536 | NA | 6.5452 | 6.6109 | 6.3700 | 6.2976 | 6.2399 | 5.9647 | 6.0357 | 5.5457 | 6.1429 | 6.3092 | 6.1147 | 6.2314 | 6.8346 | 6.5717 | 6.1828 | 6.0032 |
| SRRM2_S351;S353 | SRRM2 | Q9UQ35 | 2692;2694 | NA | 6.6555 | 6.9468 | 6.3354 | 6.7910 | 6.1053 | 6.0220 | 6.1738 | 6.2460 | 6.3428 | 6.0793 | 6.4227 | 6.1936 | 5.8864 | 5.8803 | 5.8312 | 6.0878 |

| | | | | | | | | | | | | | | | | | | | | |
|---|---|---|---|---|---|---|---|---|---|---|---|---|---|---|---|---|---|---|---|---|
| SRRM2_S351;S353;S358 | SRRM2 | Q9UQ35 | 1984;1987 | NA | 6.4833 | 6.4694 | 5.9096 | 6.7132 | 6.2275 | 5.7291 | 6.1590 | 5.9400 | 6.3995 | 5.9970 | 6.3850 | 6.2753 | 6.3325 | 6.3235 | 6.3370 | 6.3189 |
| SRRM2_S353 | SRRM2 | Q9UQ35 | 1101;1103 | NA | 6.2048 | 6.3484 | 6.2875 | 6.6944 | 6.0204 | 5.6461 | 6.0630 | 6.0295 | 6.2628 | 6.3350 | 6.0878 | 5.9835 | 6.5116 | 6.4772 | 6.3435 | 6.7046 |
| SRRM2_S358 | SRRM2 | Q9UQ35 | 972;974 | NA | 6.1989 | 6.2653 | 6.0162 | 6.7618 | 5.9356 | 6.1634 | 5.9813 | 5.8768 | 6.6402 | 6.4763 | 6.6382 | 6.7567 | 5.8839 | 6.1764 | 6.0999 | 6.1289 |
| SRRM2_S377;S398 | SRRM2 | Q9UQ35 | 295;297 | NA | 6.2222 | 6.8426 | 5.9641 | 7.0657 | 6.3352 | 6.3302 | 6.0341 | 6.7522 | 6.4712 | 5.8493 | 6.0839 | 6.1521 | 5.2410 | 6.5084 | 6.1807 | 5.9671 |
| SRRM2_S377;T395;S398 | SRRM2 | Q9UQ35 | 1320;1329 | NA | 6.3335 | 6.9316 | 6.8557 | 6.6938 | 5.9495 | 6.7622 | 6.2537 | 5.7390 | 6.2301 | 6.2714 | 6.6348 | 6.3504 | 5.9419 | 6.0810 | 5.4995 | 5.4719 |
| SRRM2_S395;S398 | SRRM2 | Q9UQ35 | 2100;2102;21 | NA | 5.9805 | 5.9693 | 5.7532 | 6.3035 | 5.8841 | 5.4975 | 6.2173 | 5.7254 | 6.0472 | 5.7077 | 5.7518 | 6.1097 | 7.3088 | 7.1717 | 7.1098 | 7.4626 |
| SRRM2_S398 | SRRM2 | Q9UQ35 | 2407;2409 | NA | 6.1717 | 6.4225 | 6.4953 | 6.5218 | 5.8762 | 5.6021 | 6.3517 | 5.4221 | 6.6683 | 6.2052 | 6.4143 | 6.2563 | 6.6474 | 6.2693 | 6.2217 | 6.4541 |
| SRRM2_S398;T400;S408 | SRRM2 | Q9UQ35 | 1925;1931 | NA | 6.2554 | 6.2177 | 6.1041 | 6.0149 | 5.7938 | 5.4333 | 6.2043 | 5.3558 | 6.4841 | 6.0865 | 6.2106 | 6.4133 | 7.3786 | 6.7004 | 6.4871 | 6.8601 |
| SRRM2_S400 | SRRM2 | Q9UQ35 | 1598;1601 | NA | 6.6509 | 6.5749 | 6.4877 | 6.6754 | 6.0257 | 5.5928 | 6.4366 | 5.8454 | 6.4218 | 6.0962 | 6.1038 | 6.2112 | 6.4394 | 6.1494 | 6.0265 | 6.2624 |
| SRRM2_S424 | SRRM2 | Q9UQ35 | 322;323 | NA | 6.2587 | 6.3387 | 6.1789 | 6.9504 | 6.5067 | 6.3858 | 6.2760 | 6.1087 | 5.8325 | 6.3949 | 6.0185 | 6.0212 | 6.3465 | 6.1611 | 5.9635 | 6.2576 |
| SRRM2_S435;S436;S437; | SRRM2 | Q9UQ35 | 817;820 | NA | 6.1876 | 6.4982 | 6.1686 | 6.6677 | 5.7846 | 5.6833 | 5.9598 | 5.7055 | 6.6986 | 6.3935 | 6.2160 | 6.3407 | 6.4440 | 6.2731 | 6.2989 | 6.6798 |
| SRRM2_S435;S440 | SRRM2 | Q9UQ35 | 1421;1424 | NA | 6.6758 | 6.9663 | 6.5649 | 7.2652 | 6.4019 | 6.1098 | 6.1866 | 5.8383 | 7.0178 | 6.9800 | 6.8875 | 7.1370 | 4.5937 | 5.4198 | 4.7622 | 5.1932 |
| SRRM2_S449 | SRRM2 | Q9UQ35 | 534;543 | NA | 6.4172 | 6.4428 | 6.0292 | 6.4925 | 5.7904 | 5.9368 | 5.9224 | 5.7346 | 6.0273 | 6.4614 | 6.4618 | 6.2428 | 6.6831 | 6.7530 | 6.3400 | 6.2647 |
| SRRM2_S454 | SRRM2 | Q9UQ35 | 1854;1857 | NA | 6.2975 | 6.6398 | 5.8804 | 6.4548 | 6.1790 | 6.0892 | 6.0179 | 5.7201 | 6.0932 | 6.2387 | 6.3419 | 6.3353 | 6.6311 | 6.5341 | 6.4323 | 6.1148 |
| SRRM2_S454;S456 | SRRM2 | Q9UQ35 | 1878;1880 | NA | 6.1540 | 6.1907 | 6.4720 | 6.2313 | 5.8708 | 5.3352 | 6.6339 | 5.2232 | 6.2093 | 6.4200 | 6.3034 | 6.4969 | 6.9902 | 6.2000 | 6.4723 | 6.7968 |
| SRRM2_S456 | SRRM2 | Q9UQ35 | 1099;1101;11 | NA | 5.6562 | 6.3298 | 6.1316 | 6.9000 | 6.0663 | 5.4040 | 6.0968 | 6.2906 | 6.2505 | 6.4969 | 5.9409 | 6.6009 | 6.2589 | 6.3790 | 6.4901 | 6.7076 |
| SRRM2_S484;T486 | SRRM2 | Q9UQ35 | 1218;1219 | NA | 6.6304 | 6.2234 | 6.5461 | 7.1291 | 5.8062 | 5.8749 | 5.9321 | 5.8710 | 6.1538 | 5.7428 | 6.3350 | 6.0767 | 6.2801 | 6.5670 | 6.1758 | 6.6555 |
| SRRM2_S486 | SRRM2 | Q9UQ35 | 2030;2032;20 | NA | 6.2569 | 6.2178 | 6.1896 | 6.5093 | 6.0435 | 5.9486 | 6.0165 | 5.8229 | 6.3767 | 6.1810 | 6.4297 | 6.4205 | 6.4677 | 6.6458 | 6.1713 | 6.3021 |
| SRRM2_S508;S510 | SRRM2 | Q9UQ35 | 778;783 | NA | 6.4884 | 6.5427 | 6.1587 | 6.6997 | 6.3675 | 5.9982 | 6.3623 | 6.2074 | 6.4555 | 6.2605 | 6.2533 | 6.2684 | 5.8853 | 5.9084 | 5.9662 | 6.1775 |
| SRRM2_S510 | SRRM2 | Q9UQ35 | 1972;1978 | NA | 6.3265 | 6.6620 | 5.9731 | 6.6837 | 6.0150 | 5.7692 | 5.9936 | 5.9445 | 6.1264 | 6.2219 | 6.2368 | 6.0596 | 6.5390 | 6.5192 | 6.5510 | 6.3786 |
| SRRM2_S534;S536 | SRRM2 | Q9UQ35 | 1398;1404 | NA | 6.3657 | 6.1632 | 6.0475 | 6.6564 | 6.7102 | 6.0061 | 5.7140 | 6.3957 | 6.7537 | 5.4699 | 6.6031 | 6.0303 | 6.3008 | 6.6247 | 5.9810 | 6.1776 |
| SRRM2_S534;S543 | SRRM2 | Q9UQ35 | 1122;1124 | NA | 5.6511 | 5.4434 | 7.0745 | 6.3275 | 7.1782 | 5.8590 | 6.3712 | 6.4658 | 6.3732 | 5.9441 | 5.7225 | 6.8583 | 6.9682 | 5.8262 | 5.9704 | 5.9666 |
| SRRM2_S536 | SRRM2 | Q9UQ35 | 1413;1415 | NA | 6.3629 | 6.0390 | 6.6785 | 6.8075 | 6.4988 | 6.2485 | 6.6846 | 6.0349 | 6.7765 | 6.4196 | 6.4442 | 6.9816 | 5.6186 | 5.7372 | 5.2627 | 5.4049 |
| SRRM2_S724;S726;S727 | SRRM2 | Q9UQ35 | 1320;1326;13 | NA | 6.1101 | 6.5142 | 6.5433 | 7.3059 | 6.4657 | 6.5381 | 6.6063 | 6.3043 | 6.4910 | 6.7924 | 7.1993 | 6.1213 | 4.9271 | 5.3889 | 5.2857 | 5.4063 |
| SRRM2_S743;S746 | SRRM2 | Q9UQ35 | 1101;1112 | NA | 6.9631 | 6.9495 | 6.0283 | 6.1784 | 6.3783 | 6.2158 | 6.0609 | 5.5358 | 5.8494 | 6.2570 | 6.2796 | 5.9566 | 6.5683 | 6.2296 | 6.3165 | 6.2330 |
| SRRM2_S778 | SRRM2 | Q9UQ35 | 1762;1764 | NA | 5.9749 | 6.3982 | 5.8850 | 6.5494 | 6.9228 | 6.2354 | 6.3418 | 6.4760 | 6.0006 | 6.4504 | 6.7477 | 5.8879 | 6.8073 | 6.1313 | 5.9265 | 5.2649 |
| SRRM2_S778;S780 | SRRM2 | Q9UQ35 | 1822;1824 | NA | 7.1637 | 7.5892 | 6.1116 | 7.3491 | 6.0816 | 6.2334 | 6.2098 | 5.7724 | 6.4731 | 6.3574 | 6.3313 | 5.9581 | 5.3952 | 5.9082 | 5.6423 | 5.4237 |
| SRRM2_S778;S780;S783 | SRRM2 | Q9UQ35 | 1878;1884 | NA | 6.3173 | 6.3127 | 5.9354 | 6.5441 | 5.8037 | 5.2386 | 6.1085 | 5.7782 | 6.4132 | 6.0130 | 6.1426 | 6.1960 | 7.0201 | 6.5516 | 6.5295 | 7.0955 |
| SRRM2_S778;S783 | SRRM2 | Q9UQ35 | 2690;2694 | NA | 6.9865 | 6.6004 | 6.1108 | 6.9546 | 6.3680 | 5.7623 | 5.9006 | 6.4492 | 6.4170 | 6.0900 | 5.7703 | 6.3035 | 5.9586 | 6.2614 | 5.9229 | 6.1440 |
| SRRM2_S780 | SRRM2 | Q9UQ35 | 1443;1444 | NA | 5.8560 | 5.7794 | 5.9127 | 6.2604 | 6.1548 | 5.6777 | 6.3437 | 5.8702 | 5.6602 | 5.7342 | 5.7258 | 5.5759 | 7.5006 | 7.0050 | 7.5469 | 7.3963 |
| SRRM2_S780;S783 | SRRM2 | Q9UQ35 | 780;783 | NA | 6.3470 | 7.0438 | 6.6533 | 7.3607 | 6.1398 | 6.0494 | 6.5587 | 6.1097 | 6.6344 | 7.1027 | 6.8735 | 6.9767 | 4.5181 | 5.2854 | 5.0786 | 5.2684 |
| SRRM2_S783 | SRRM2 | Q9UQ35 | 970;973;974 | NA | 6.7327 | 6.6980 | 6.1667 | 6.9896 | 6.1133 | 5.9384 | 6.0202 | 5.9275 | 5.6387 | 5.6920 | 6.5828 | 5.6046 | 7.1745 | 6.1640 | 5.9877 | 6.5693 |
| SRRM2_S817 | SRRM2 | Q9UQ35 | 435;436;437; | NA | 5.5754 | 5.7447 | 5.7800 | 6.0756 | 5.6739 | 5.4764 | 5.5420 | 4.9561 | 6.6076 | 5.8420 | 6.1632 | 6.0496 | 8.1425 | 7.1562 | 7.4163 | 7.7985 |
| SRRM2_S817;S820 | SRRM2 | Q9UQ35 | 778;780;783 | NA | 6.3242 | 6.2537 | 5.7707 | 5.8522 | 6.1140 | 5.8004 | 6.0058 | 6.2538 | 6.1490 | 5.9985 | 5.9407 | 5.9337 | 6.7801 | 6.9084 | 6.9867 | 6.9281 |
| SRRM2_S820 | SRRM2 | Q9UQ35 | 992;994 | NA | 5.7390 | 5.8630 | 6.1384 | 6.1928 | 6.3613 | 6.2971 | 6.2435 | 6.4525 | 6.3890 | 5.9945 | 5.5148 | 5.7351 | 7.1209 | 6.6156 | 6.3622 | 6.9804 |
| SRRM2_S837;S839 | SRRM2 | Q9UQ35 | 1972;1975 | NA | 6.4196 | 6.5022 | 5.7885 | 6.0529 | 6.1320 | 5.4798 | 6.1271 | 6.0124 | 6.3130 | 6.0853 | 5.9852 | 6.4543 | 6.9132 | 6.6308 | 6.7624 | 6.3413 |
| SRRM2_S839 | SRRM2 | Q9UQ35 | 983;994 | NA | 6.5457 | 7.1175 | 6.4592 | 6.2627 | 6.9557 | 7.0874 | 5.6176 | 4.5867 | 5.3325 | 5.7733 | 6.3177 | 6.2798 | 6.9289 | 6.8033 | 5.8983 | 6.0337 |
| SRRM2_S854;T866 | SRRM2 | Q9UQ35 | 837;839 | NA | 5.8743 | 6.3733 | 5.8200 | 6.4685 | 6.2878 | 5.9154 | 6.3201 | 5.9935 | 6.0889 | 6.0656 | 6.1799 | 6.3082 | 6.4654 | 6.5885 | 6.5684 | 6.6821 |
| SRRM2_S857 | SRRM2 | Q9UQ35 | 983;990;992; | NA | 5.3353 | 5.6564 | 5.5378 | 5.7900 | 6.2853 | 5.8695 | 5.5596 | 5.1079 | 5.4559 | 5.3734 | 5.1611 | 5.3669 | 8.9306 | 8.6769 | 7.9169 | 7.9765 |
| SRRM2_S857;T866 | SRRM2 | Q9UQ35 | 1401;1413 | NA | 6.5601 | 5.9155 | 6.5461 | 6.7961 | 6.2155 | 6.0382 | 6.6086 | 5.7718 | 6.7543 | 6.5856 | 6.5499 | 5.9774 | 6.6517 | 6.4714 | 4.9226 | 5.6350 |
| SRRM2_S866 | SRRM2 | Q9UQ35 | 395;398 | NA | 6.0908 | 5.7247 | 6.4009 | 6.8471 | 6.1851 | 5.4387 | 5.7491 | 6.4444 | 6.7550 | 6.1727 | 6.3848 | 6.1605 | 6.2609 | 6.2333 | 6.7645 | 6.3874 |
| SRRM2_S871 | SRRM2 | Q9UQ35 | 1318;1320;13 | NA | 6.6707 | 6.6880 | 6.1036 | 6.8880 | 6.3646 | 5.4265 | 6.1427 | 6.4502 | 6.1657 | 6.3795 | 6.6925 | 6.5668 | 5.7259 | 6.0344 | 5.7645 | 5.9362 |
| SRRM2_S871;S876 | SRRM2 | Q9UQ35 | 377;395;398 | NA | 6.6176 | 5.2685 | 6.0740 | 6.7883 | 6.3929 | 5.6771 | 5.9012 | 6.3863 | 6.5032 | 5.9106 | 7.1618 | 6.0985 | 6.7449 | 5.7633 | 6.6712 | 6.0405 |
| SRRM2_S875 | SRRM2 | Q9UQ35 | 1866;1869 | NA | 6.0926 | 6.1272 | 6.3172 | 6.3664 | 6.3298 | 5.4112 | 6.3494 | 6.1792 | 6.8307 | 6.0336 | 5.8184 | 6.5599 | 6.6086 | 6.3341 | 6.2391 | 6.4025 |
| SRRM2_S875;S876 | SRRM2 | Q9UQ35 | 854;866 | NA | 6.1957 | 6.5502 | 5.4686 | 6.6375 | 6.2736 | 6.4991 | 5.0916 | 5.5170 | 5.9241 | 4.9654 | 6.0051 | 5.6199 | 7.2914 | 7.7092 | 7.1368 | 7.1149 |
| SRRM2_S876 | SRRM2 | Q9UQ35 | 1497;1499;15 | NA | 6.3882 | 6.3501 | 5.6525 | 6.2194 | 5.6469 | 5.9169 | 6.3155 | 5.5800 | 6.2993 | 6.3030 | 6.7139 | 6.3383 | 6.7083 | 6.2114 | 6.6359 | 6.7202 |
| SRRM2_S895 | SRRM2 | Q9UQ35 | 311;323 | NA | 6.1248 | 7.0396 | 6.4656 | 6.8324 | 6.3942 | 6.4878 | 6.6414 | 6.5032 | 6.0338 | 5.3098 | 5.4639 | 5.9400 | 6.2020 | 5.4285 | 6.3196 | 6.8134 |
| SRRM2_S913;S914 | SRRM2 | Q9UQ35 | 1691;1694 | NA | 5.7008 | 5.2715 | 5.2483 | 6.0275 | 5.9194 | 5.3656 | 5.9842 | 5.8051 | 6.0915 | 6.1115 | 6.0938 | 5.9012 | 7.8095 | 7.7980 | 7.4916 | 7.3806 |
| SRRM2_S914 | SRRM2 | Q9UQ35 | 743;746 | NA | 6.2301 | 6.6829 | 6.5750 | 7.8619 | 5.2078 | 6.1606 | 6.2675 | 5.3762 | 7.3081 | 7.3357 | 7.3521 | 6.9975 | 4.6521 | 5.1263 | 5.2405 | 5.6256 |
| SRRM2_S952;S954 | SRRM2 | Q9UQ35 | 1581;1582 | NA | 5.9558 | 6.5100 | 6.7722 | 7.2880 | 5.5329 | 5.6158 | 6.5989 | 6.1870 | 6.4471 | 6.9627 | 6.4459 | 6.4842 | 6.0480 | 6.2470 | 5.2388 | 5.6658 |
| SRRM2_S952;S957 | SRRM2 | Q9UQ35 | 1419;1424 | NA | 5.9696 | 6.1489 | 5.8809 | 6.5230 | 5.4424 | 5.8918 | 6.1188 | 5.9959 | 6.9498 | 5.4842 | 6.5769 | 6.3978 | 6.6904 | 6.7011 | 6.5611 | 6.6675 |
| SRRM2_S954 | SRRM2 | Q9UQ35 | 322;326 | NA | 7.1626 | 6.4495 | 6.3153 | 6.8599 | 6.8096 | 5.2689 | 6.3393 | 6.6810 | 6.4550 | 5.7333 | 5.9267 | 5.9308 | 5.7319 | 6.2975 | 6.1617 | 5.8770 |
| SRRM2_S970;S973;S974 | SRRM2 | Q9UQ35 | 1382;1387 | NA | 6.7265 | 7.7639 | 6.5108 | 8.1093 | 7.5392 | 7.1699 | 5.6931 | 8.1114 | 6.1196 | 5.9761 | 6.4049 | 5.8290 | 3.8085 | 4.5676 | 4.7356 | 4.9348 |
| SRRM2_S970;S974 | SRRM2 | Q9UQ35 | 2407;2412 | NA | 6.8609 | 6.5477 | 6.2489 | 6.0490 | 5.7025 | 4.9498 | 5.8347 | 5.5990 | 6.7851 | 6.6442 | 6.8036 | 6.3850 | 6.5514 | 6.0075 | 6.1891 | 6.8417 |
| SRRM2_S972;S974 | SRRM2 | Q9UQ35 | 1876;1878;18 | NA | 6.3638 | 6.4459 | 6.7886 | 6.2318 | 6.1547 | 5.4713 | 5.8117 | 5.5791 | 6.4837 | 6.0584 | 6.0975 | 6.1995 | 6.9512 | 6.4812 | 6.2925 | 6.5890 |
| SRRM2_S990;S994 | SRRM2 | Q9UQ35 | 484;486 | NA | 6.3924 | 6.1802 | 5.8572 | 6.4418 | 5.6592 | 6.0487 | 5.7924 | 5.9403 | 6.2503 | 6.4313 | 6.4263 | 7.1046 | 6.3877 | 6.2247 | 6.2910 | 6.5719 |
| SRRM2_S992;S994 | SRRM2 | Q9UQ35 | 1497;1502 | NA | 7.1147 | 7.2683 | 6.7061 | 8.1447 | 5.9940 | 5.6248 | 6.4723 | 6.1042 | 6.4552 | 7.1652 | 6.9295 | 6.9640 | 4.6830 | 5.2648 | 4.2045 | 4.9047 |
| SRRM2_S994 | SRRM2 | Q9UQ35 | 871;876 | NA | 6.3005 | 7.1598 | 6.5498 | 7.4803 | 6.3315 | 5.7288 | 5.2139 | 6.2384 | 6.7385 | 5.9804 | 6.3104 | 5.6464 | 6.0868 | 6.6038 | 6.0687 | 5.5620 |
| SRRM2_T1003 | SRRM2 | Q9UQ35 | 1541;1552 | NA | 5.0700 | 5.5647 | 4.9941 | 4.9986 | 5.4713 | 4.9376 | 5.9245 | 5.5096 | 5.6524 | 5.6074 | 5.9483 | 5.7461 | 8.8769 | 8.4627 | 8.7939 | 8.4418 |
| SRRM2_T1003;S1012 | SRRM2 | Q9UQ35 | 1691;1698 | NA | 6.2893 | 6.1131 | 5.8366 | 6.8890 | 6.3556 | 5.6859 | 6.2742 | 5.7406 | 5.7495 | 6.1559 | 5.6524 | 5.8572 | 7.2100 | 6.3489 | 6.8507 | 6.9911 |
| SRRM2_T1003;S1014 | SRRM2 | Q9UQ35 | 913;914 | NA | 5.9021 | 6.4854 | 6.3258 | 6.8863 | 6.5135 | 5.2727 | 6.2035 | 6.6286 | 5.3335 | 6.6531 | 5.9652 | 5.6433 | 6.6906 | 5.5712 | 7.1903 | 6.7348 |
| SRRM2_T1043 | SRRM2 | Q9UQ35 | 1320;1326 | NA | 6.0790 | 5.8158 | 5.2355 | 6.4759 | 6.4413 | 5.7334 | 6.6011 | 5.6239 | 5.1460 | 6.2329 | 6.3240 | 6.4077 | 6.5001 | 6.4539 | 6.8211 | 8.1083 |
| SRRM2_T111 | SRRM2 | Q9UQ35 | 220;222 | NA | 7.9216 | 6.8690 | 4.9691 | 7.7728 | 5.2184 | 7.6456 | 6.5968 | 5.6500 | 6.1240 | 4.2125 | 3.9340 | 6.1418 | 5.8367 | 7.9742 | 7.0857 | 6.0480 |
| SRRM2_T1208 | SRRM2 | Q9UQ35 | 1902;1905 | NA | 6.7388 | 6.1967 | 4.5417 | 8.0443 | 5.6902 | 5.9355 | 4.9403 | 6.3168 | 7.1864 | 6.3459 | 6.7039 | 6.3911 | 7.7280 | 4.2873 | 6.9630 | 5.9903 |
| SRRM2_T1413 | SRRM2 | Q9UQ35 | 983;990 | NA | 6.0752 | 6.1382 | 5.2103 | 6.3127 | 5.8574 | 5.9951 | 5.4563 | 5.4221 | 5.7470 | 5.9659 | 6.1485 | 5.7532 | 7.7105 | 7.5167 | 7.6273 | 7.0638 |
| SRRM2_T1413;S1415 | SRRM2 | Q9UQ35 | 1913;1916 | NA | 5.7927 | 6.9021 | 5.7833 | 6.2426 | 6.6256 | 5.4051 | 6.0354 | 6.1714 | 6.8132 | 6.5749 | 5.7990 | 5.6130 | 6.6343 | 6.1862 | 6.5199 | 6.9012 |
| SRRM2_T1434 | SRRM2 | Q9UQ35 | 295;300 | NA | 5.3651 | 6.4865 | 5.2346 | 5.7662 | 7.1369 | 6.8208 | 5.6273 | 6.0308 | 7.5187 | 6.1913 | 5.8485 | 6.0564 | 7.1263 | 6.7106 | 5.5929 | 6.4871 |
| SRRM2_T1472 | SRRM2 | Q9UQ35 | 1890;1893 | NA | 6.3040 | 6.2391 | 6.6617 | 7.5411 | 6.0958 | 5.3080 | 6.6214 | 6.4463 | 6.2461 | 5.2621 | 6.6212 | 5.3830 | 6.7047 | 6.9017 | 5.8870 | 5.7766 |
| SRRM2_T1492 | SRRM2 | Q9UQ35 | 1103;1112 | NA | 6.1866 | 5.9138 | 5.9578 | 6.3342 | 5.8109 | 6.1986 | 6.3788 | 5.5834 | 6.0897 | 6.2946 | 5.8609 | 6.1969 | 7.2423 | 6.6232 | 6.1500 | 7.1781 |
| SRRM2_T1531 | SRRM2 | Q9UQ35 | 435;440 | NA | 5.7224 | 6.0673 | 7.0304 | 4.7690 | 5.7619 | 6.8017 | 5.5719 | 6.8449 | 6.0770 | 6.1284 | 5.8991 | 5.6467 | 7.4710 | 6.4242 | 6.6589 | 7.1252 |
| SRRM2_T2092 | SRRM2 | Q9UQ35 | 1404;1415 | NA | 7.1075 | 7.6234 | 7.0565 | 7.2136 | 6.3628 | 6.2093 | 6.6975 | 6.5612 | 6.7764 | 6.1305 | 6.7347 | 6.2542 | 4.5571 | 4.8663 | 5.1422 | 4.7069 |
| SRRM2_T2104 | SRRM2 | Q9UQ35 | 2727;2729 | NA | 6.4585 | 8.1032 | 7.1436 | 7.0701 | 5.8865 | 7.6904 | 6.0622 | 4.7815 | 6.3010 | 6.1080 | 5.2036 | 6.3177 | 6.3280 | 4.8969 | 5.5156 | 6.1331 |
| SRRM2_T2144 | SRRM2 | Q9UQ35 | 724;726;727 | NA | 6.6779 | 6.8132 | 5.7176 | 6.2996 | 7.1821 | 6.1598 | 5.9678 | 5.0010 | 5.8663 | 5.6998 | 7.0089 | 5.9924 | 6.7222 | 6.1267 | 6.4001 | 6.3648 |
| SRRM2_T2289 | SRRM2 | Q9UQ35 | 970;974 | NA | 6.5156 | 6.9890 | 5.6324 | 6.9197 | 5.8356 | 5.4801 | 6.1732 | 6.4492 | 6.0761 | 6.4053 | 6.3514 | 6.6922 | 5.8236 | 6.3787 | 6.1295 | 6.1484 |
| SRRM2_T2329 | SRRM2 | Q9UQ35 | 1003;1014 | NA | 6.0646 | 6.6991 | 7.6524 | 6.3352 | 5.8101 | 12.1485 | 5.6754 | 6.7715 | 12.9783 | 4.2213 | 6.3254 | 3.9953 | 5.0112 | 4.1466 | 3.6093 | 2.5558 |
| SRRM2_T2409 | SRRM2 | Q9UQ35 | 983;992;994 | NA | 6.1666 | 6.3722 | 6.1268 | 6.9097 | 7.0717 | 7.6676 | 6.5845 | 5.2769 | 5.3023 | 5.8290 | 6.6807 | 6.0722 | 6.3129 | 6.9500 | 5.2944 | 5.3827 |
| SRRM2_T2599 | SRRM2 | Q9UQ35 | 1329;1336 | NA | 5.6710 | 6.1131 | 6.0465 | 7.0235 | 6.3648 | 6.2409 | 6.3393 | 5.7359 | 6.2020 | 6.8972 | 5.7020 | 6.3011 | 6.1457 | 6.2128 | 6.5316 | 6.4724 |
| SRRM2_T2738 | SRRM2 | Q9UQ35 | 990;994 | NA | 5.6957 | 5.9750 | 5.4032 | 6.6972 | 5.5132 | 5.2514 | 5.7387 | 5.6846 | 6.0412 | 6.2955 | 5.5437 | 5.6663 | 7.4614 | 7.6926 | 7.6435 | 7.6968 |
| SRRM2_T2738;S2740 | SRRM2 | Q9UQ35 | 398;400;408 | NA | 7.5955 | 6.8011 | 6.7035 | 6.1309 | 6.8822 | 6.2206 | 6.2637 | 4.7455 | 6.0030 | 6.6001 | 6.5868 | 7.5083 | 6.6166 | 4.7744 | 4.7248 | 5.8431 |
| SRRM2_T326 | SRRM2 | Q9UQ35 | 1003;1012 | NA | 5.4575 | 6.4588 | 6.3321 | 6.3871 | 6.0115 | 6.4077 | 5.8931 | 6.4874 | 6.5295 | 6.6843 | 6.7332 | 7.0767 | 5.4188 | 6.1551 | 5.8348 | 6.1324 |
| SRRM2_T359 | SRRM2 | Q9UQ35 | 2102;2104 | NA | 6.5484 | 6.3989 | 5.4050 | 6.8101 | 6.7498 | 6.7821 | 6.4322 | 5.6187 | 6.1490 | 7.2508 | 5.7279 | 5.9445 | 6.0704 | 6.1575 | 6.0050 | 5.9499 |
| SRRM2_T367 | SRRM2 | Q9UQ35 | 351;353;358 | NA | 5.7982 | 5.1587 | 6.7643 | 6.6801 | 5.4052 | 6.0173 | 5.8861 | 5.2099 | 6.1230 | 5.9470 | 5.8220 | 6.1055 | 6.9462 | 7.4540 | 7.5555 | 7.1270 |
| SRRM2_T792 | SRRM2 | Q9UQ35 | 1318;1326 | NA | 5.5007 | 5.7758 | 5.9505 | 6.2213 | 5.9013 | 5.3672 | 6.2108 | 7.3431 | 5.6516 | 5.4373 | 5.9088 | 5.5899 | 7.3239 | 6.8026 | 7.3251 | 7.6901 |
| SRRM2_T810 | SRRM2 | Q9UQ35 | 952;957 | NA | 5.9335 | 7.0170 | 6.1117 | 6.4830 | 5.9211 | 5.6103 | 6.3372 | 5.6560 | 6.5143 | 6.3532 | 5.9845 | 6.0201 | 6.4884 | 6.4547 | 6.4765 | 6.6387 |
| SRRM2_T903;S908 | SRRM2 | Q9UQ35 | 1318;1320 | NA | 5.4615 | 6.5505 | 5.4087 | 6.2319 | 6.5678 | 6.0664 | 6.0361 | 5.9532 | 5.9023 | 5.0504 | 6.0600 | 5.4077 | 7.1541 | 7.6372 | 7.7057 | 6.8065 |
| SRRM2_T983 | SRRM2 | Q9UQ35 | 1401;1404 | NA | 5.8842 | 5.2214 | 5.1164 | 6.6901 | 6.6303 | 5.5758 | 5.5326 | 6.8056 | 6.4150 | 5.8781 | 6.8202 | 6.2106 | 7.0370 | 6.5309 | 5.7376 | 7.9143 |
| SRRM2_T983;S990 | SRRM2 | Q9UQ35 | 1401;1404;14 | NA | 6.0703 | 7.4307 | 5.2465 | 5.2314 | 6.4373 | 7.9507 | 7.5583 | 4.7954 | 5.1498 | 6.3762 | 6.7842 | 5.8654 | 6.3194 | 7.4724 | 5.1422 | 6.1699 |
| SRRM2_T983;S990;S992; | SRRM2 | Q9UQ35 | 1458;1463 | NA | 5.7532 | 5.9474 | 5.8962 | 6.5306 | 6.1842 | 5.8678 | 6.2619 | 6.4855 | 6.4654 | 6.2774 | 6.2783 | 6.1407 | 6.0568 | 5.8774 | 6.9974 | 6.9799 |
| SRRM2_T983;S992;S994 | SRRM2 | Q9UQ35 | 875;876 | NA | 6.1944 | 6.8595 | 6.7259 | 7.1527 | 6.6720 | 6.6814 | 6.6022 | 5.4046 | 6.1518 | 6.4672 | 5.6758 | 5.6283 | 5.7588 | 5.8044 | 5.8045 | 6.4163 |
| SRRM2_T983;S994 | SRRM2 | Q9UQ35 | 1499;1502 | NA | 5.1725 | 5.6726 | 6.6888 | 6.3580 | 6.4819 | 6.0581 | 6.0288 | 5.6183 | 6.3199 | 6.8224 | 6.4024 | 6.7814 | 5.9657 | 6.6532 | 6.4434 | 6.5324 |
| SRRT_S136 | SRRT | Q9BXP5 | 74 | PPRHELSPPQKRM | 5.6460 | 5.9311 | 5.8744 | 6.5587 | 5.9447 | 5.6724 | 5.8167 | 6.2332 | 6.5078 | 6.3128 | 5.9867 | 6.1260 | 6.7947 | 6.5576 | 6.9349 | 7.1025 |
| SRRT_S67 | SRRT | Q9BXP5 | 67 | NRRERFSPPRHEL | 6.0334 | 6.2649 | 5.8123 | 6.4045 | 6.0321 | 5.7499 | 6.0561 | 6.2344 | 6.1576 | 5.9909 | 5.7899 | 6.0696 | 6.8273 | 6.8741 | 6.8988 | 6.8040 |
| SRRT_S67;S74 | SRRT | Q9BXP5 | 718 | KWLCPLSGKKFKG | 6.6977 | 7.8364 | 6.2310 | 6.9543 | 6.4611 | 6.5043 | 7.3690 | 6.6636 | 6.7487 | 6.2589 | 6.6812 | 6.3356 | 4.4941 | 5.3477 | 4.5146 | 4.9017 |
| SRRT_S718 | SRRT | Q9BXP5 | 544 | WASEPGTPPLPTS | 5.9977 | 5.8543 | 5.9616 | 6.0449 | 6.5036 | 6.3775 | 6.0409 | 6.2587 | 5.8471 | 6.5196 | 6.1379 | 6.1840 | 6.7014 | 6.3729 | 6.7309 | 6.4669 |
| SRRT_S74 | SRRT | Q9BXP5 | 136 | IQARLGSIAEIDL | 4.8375 | 5.2624 | 5.7123 | 5.8355 | 7.0275 | 6.0555 | 5.5344 | 5.7764 | 6.8262 | 7.5586 | 7.1198 | 7.6810 | 6.8433 | 6.6410 | 5.4404 | 5.8480 |
| SRRT_T544 | SRRT | Q9BXP5 | 67;74 | NA | 6.1824 | 6.7629 | 6.4851 | 6.4297 | 6.1659 | 6.8195 | 6.1264 | 5.6494 | 6.1553 | 5.9060 | 6.0918 | 6.0062 | 6.5640 | 6.3841 | 6.3866 | 5.8846 |
| SRSF1_S199 | SRSF1 | Q07955 | 199 | KVDGPRSPSYGRS | 8.5261 | 7.6433 | 7.4167 | 6.9204 | 6.5319 | 7.3355 | 6.2243 | 4.1733 | 5.9541 | 5.9923 | 6.6833 | 7.1808 | 5.4374 | 5.1970 | 4.5518 | 4.2315 |
| SRSF1_S199;S201 | SRSF1 | Q07955 | 199;201 | NA | 6.1159 | 6.2861 | 5.7841 | 6.4116 | 6.4700 | 6.0647 | 6.0577 | 5.9564 | 6.0269 | 5.8751 | 5.8048 | 5.6412 | 7.3705 | 6.6868 | 6.7698 | 6.6784 |
| SRSF1_S199;S201;S205 | SRSF1 | Q07955 | 238;242 | NA | 6.3280 | 6.4165 | 6.0121 | 6.5224 | 6.0717 | 5.6831 | 5.8224 | 5.6722 | 6.0985 | 6.2161 | 6.2723 | 6.3059 | 7.3096 | 6.3406 | 6.2589 | 6.6698 |
| SRSF1_S234;S238 | SRSF1 | Q07955 | 199;201;205 | NA | 6.2423 | 6.1041 | 5.9900 | 6.3404 | 6.1727 | 5.8047 | 6.4438 | 5.7145 | 5.8333 | 5.8654 | 5.5063 | 6.2530 | 7.4508 | 7.0279 | 6.6753 | 6.5756 |
| SRSF1_S238;S242 | SRSF1 | Q07955 | 234;238 | NA | 7.5858 | 6.6783 | 6.0596 | 6.2071 | 6.4082 | 7.0097 | 6.3647 | 4.9865 | 5.3735 | 5.7856 | 6.5784 | 6.6697 | 6.8090 | 6.2636 | 5.6877 | 5.5325 |
| SRSF10_S106 | SRSF10 | O75494 | 133 | RRSRSRSFDYNYR | 6.6660 | 6.5452 | 5.8821 | 6.3496 | 6.5751 | 5.8058 | 5.9863 | 6.4918 | 6.3818 | 5.9880 | 5.9235 | 5.8835 | 6.7225 | 6.2544 | 6.3948 | 6.1494 |
| SRSF10_S129;S131;S133 | SRSF10 | O75494-3 | 158 | RPRRSRSHSDNDR | 5.9752 | 6.5253 | 6.1541 | 6.9416 | 6.1037 | 6.1019 | 5.7584 | 6.7644 | 6.3032 | 6.4848 | 6.9374 | 6.3447 | 6.0624 | 5.9455 | 5.9042 | 5.6933 |
| SRSF10_S131;S133 | SRSF10 | O75494 | 235 | KKEPPRSKSQSRS | 6.7810 | 6.2796 | 5.6427 | 6.3585 | 6.7096 | 6.0626 | 6.1362 | 5.7858 | 5.7473 | 5.9951 | 5.6230 | 5.6138 | 7.3411 | 6.5510 | 6.6442 | 6.7284 |
| SRSF10_S133 | SRSF10 | O75494 | 106 | EGRNVYSSSRYDD | 6.6094 | 6.7824 | 6.3172 | 7.6468 | 6.1220 | 6.1613 | 6.4176 | 6.5833 | 6.1928 | 6.2536 | 6.6872 | 6.8389 | 5.0050 | 5.6138 | 4.9056 | 5.8634 |
| SRSF10_S156;S158 | SRSF10 | O75494 | 131;133 | NA | 6.5477 | 6.2413 | 5.9153 | 6.3313 | 6.1766 | 5.7046 | 6.0016 | 5.8099 | 6.2161 | 6.0256 | 6.1090 | 6.2643 | 7.1112 | 6.3990 | 6.5061 | 6.6405 |
| SRSF10_S156;S158;S160 | SRSF10 | O75494 | 129;131;133 | NA | 6.4130 | 6.4075 | 5.7409 | 6.3618 | 6.3357 | 5.4122 | 6.1388 | 5.8310 | 6.2911 | 6.2685 | 5.7912 | 5.6153 | 6.8760 | 7.1209 | 6.4323 | 6.9637 |
| SRSF10_S158 | SRSF10 | O75494-3 | 156;158;160 | NA | 7.1047 | 5.9696 | 5.4969 | 5.6649 | 6.6773 | 6.1277 | 5.5714 | 6.1359 | 6.3227 | 6.1560 | 6.3315 | 4.8840 | 5.5271 | 7.0130 | 7.6116 | 7.4056 |
| SRSF10_S235 | SRSF10 | O75494-3 | 156;158 | NA | 6.3158 | 6.2395 | 6.0351 | 7.2015 | 6.4651 | 6.1061 | 6.5991 | 6.8847 | 5.1907 | 6.5783 | 7.0619 | 6.5737 | 5.2770 | 5.5585 | 6.3243 | 5.5888 |
| SRSF11_S207 | SRSF11 | Q05519 | 449 | KPIETGSPKTKEC | 6.7741 | 6.8906 | 6.5137 | 6.2887 | 5.8196 | 5.7357 | 5.9632 | 5.4310 | 6.1117 | 6.1420 | 6.1964 | 6.4612 | 6.3073 | 6.5303 | 6.2434 | 6.5912 |
| SRSF11_S366;S368;S370 | SRSF11 | Q05519 | 434 | EEQGYDSEKEKKE | 5.3874 | 5.8485 | 6.1405 | 6.2415 | 5.8575 | 5.6074 | 6.1164 | 5.5750 | 6.0951 | 6.0306 | 5.9103 | 6.1265 | 7.1229 | 7.1130 | 7.0726 | 7.7547 |
| SRSF11_S434 | SRSF11 | Q05519 | 207 | VAAGLVSPSLKSD | 6.2455 | 6.1656 | 5.9863 | 6.9165 | 6.2542 | 6.8242 | 6.8343 | 6.5035 | 6.0257 | 6.2019 | 6.0123 | 5.8793 | 5.8480 | 6.1875 | 5.6568 | 6.4585 |
| SRSF11_S449 | SRSF11 | Q05519 | 468 | GDSLRESKVNGDD | 6.6216 | 7.6280 | 6.6462 | 6.9703 | 6.5271 | 6.8881 | 5.9715 | 5.9582 | 6.0017 | 5.6313 | 6.2323 | 5.9261 | 5.4857 | 5.8845 | 5.4432 | 6.1841 |
| SRSF11_S468 | SRSF11 | Q05519 | 366;368;370 | NA | 6.9825 | 6.4723 | 6.5084 | 6.4952 | 6.1378 | 6.2721 | 5.6629 | 5.6937 | 5.6770 | 6.4450 | 6.5269 | 6.0825 | 6.1731 | 6.2735 | 6.3573 | 6.2398 |
| SRSF12_S150 | SRSF12 | Q8WXF0 | 150 | RFSYSQSKSRSKS | 7.0941 | 8.0423 | 7.3217 | 7.1836 | 6.6390 | 5.9092 | 5.9350 | 5.9441 | 6.1626 | 6.1550 | 6.4205 | 6.8377 | 6.7527 | 5.2857 | 3.9885 | 4.3284 |
| SRSF12_S219;S223 | SRSF12 | Q8WXF0 | 219;223 | NA | 6.5005 | 6.3049 | 6.3343 | 7.1544 | 6.4644 | 6.2952 | 6.6032 | 5.6396 | 5.9348 | 5.7743 | 5.8584 | 5.9089 | 6.8755 | 6.6052 | 6.1101 | 5.6363 |
| SRSF2_S101 | SRSF2 | Q01130-2 | 179 | SRSRSRSPPPVSK | 5.5525 | 6.4290 | 5.8352 | 5.9915 | 6.7790 | 5.8345 | 5.6934 | 6.3400 | 6.3683 | 6.2417 | 5.5865 | 5.4593 | 6.9850 | 6.6409 | 7.1649 | 7.0984 |
| SRSF2_S109 | SRSF2 | Q01130-2 | 26 | NLTYRTSPDTLRR | 5.9369 | 6.6558 | 6.4554 | 5.9588 | 6.0383 | 6.6536 | 5.7988 | 5.2584 | 5.9321 | 6.3679 | 6.4700 | 6.6948 | 6.8988 | 6.6590 | 6.0518 | 6.1694 |
| SRSF2_S177;S179 | SRSF2 | Q01130-2 | 109 | RGPPPRSPRRRRR | 5.0517 | 5.8329 | 5.5502 | 5.0974 | 5.9072 | 5.9410 | 5.1603 | 6.3460 | 7.2854 | 5.7198 | 5.8736 | 5.7341 | 7.4471 | 7.8591 | 8.1738 | 7.0205 |
| SRSF2_S179 | SRSF2 | Q01130-2 | 101 | PPDSHHSRRGPPP | 6.9124 | 7.2416 | 6.9674 | 7.3646 | 6.8806 | 6.4993 | 5.5217 | 6.5844 | 7.4176 | 8.1792 | 8.2878 | 6.9169 | 4.0683 | 4.9311 | 2.7256 | 3.5014 |
| SRSF2_S196;S200;S208 | SRSF2 | Q01130-2 | 177;179 | NA | 5.2414 | 6.1515 | 6.1090 | 6.0666 | 5.9018 | 5.4111 | 5.5539 | 6.3265 | 6.4771 | 6.2495 | 6.4752 | 6.1978 | 7.1359 | 6.5306 | 6.8855 | 7.2867 |
| SRSF2_S26 | SRSF2 | Q01130-2 | 196;200;208 | NA | 6.9557 | 6.6929 | 6.9471 | 6.5427 | 5.8902 | 5.7683 | 5.5068 | 5.0898 | 6.1533 | 6.1240 | 5.9990 | 5.9818 | 6.9413 | 6.7805 | 5.9733 | 6.6534 |
| SRSF3_S108 | SRSF3 | P84103 | 108 | DDYRRRSPPPRRR | 6.8933 | 6.5176 | 6.6806 | 6.4285 | 6.6103 | 6.6837 | 6.3530 | 4.4580 | 5.1434 | 5.6737 | 6.1669 | 6.9274 | 7.0128 | 6.5780 | 5.9076 | 5.9653 |
| SRSF3_S138;S140 | SRSF3 | P84103 | 5 | xxMHRDSCPLDCK | 5.4597 | 5.5670 | 5.5949 | 6.0496 | 5.9550 | 6.3757 | 6.1845 | 6.0340 | 5.5494 | 5.8952 | 5.8954 | 5.8372 | 7.3382 | 6.9015 | 7.7787 | 7.5840 |
| SRSF3_S5 | SRSF3 | P84103 | 138;140 | NA | 6.6993 | 6.6301 | 6.3696 | 6.4128 | 6.3530 | 6.4357 | 5.8154 | 5.6977 | 6.5203 | 6.3113 | 6.7275 | 6.2628 | 5.8208 | 6.1098 | 6.1941 | 5.6398 |
| SRSF4_S113 | SRSF4 | Q08170 | 113 | IVENLSSRCSWQD | 5.8309 | 6.0664 | 5.7121 | 6.2587 | 6.2553 | 5.2395 | 5.9427 | 5.7973 | 5.2100 | 4.9539 | 4.8212 | 4.4603 | 8.8347 | 7.7172 | 9.6176 | 7.2823 |
| SRSF4_S116 | SRSF4 | Q08170 | 116 | NLSSRCSWQDLKD | 3.8474 | 3.3177 | 3.6913 | 3.8074 | 4.8261 | 4.1323 | 4.3044 | 3.2850 | 3.8988 | 4.6527 | 3.9655 | 4.3818 | 13.1688 | 12.0800 | 13.1161 | 13.5244 |
| SRSF4_S456 | SRSF4 | Q08170 | 456 | SKPNLPSESRSRS | 4.6184 | 5.6563 | 5.7700 | 5.9661 | 6.4610 | 6.4917 | 6.7869 | 6.2286 | 6.5992 | 5.3429 | 5.7965 | 6.0471 | 7.1576 | 7.0571 | 6.5385 | 7.4821 |
| SRSF4_S458 | SRSF4 | Q08170 | 458 | PNLPSESRSRSKS | 5.8819 | 6.2440 | 5.4612 | 6.0105 | 7.1310 | 5.8912 | 6.0527 | 6.4227 | 6.0781 | 5.7139 | 5.2993 | 5.3153 | 7.5372 | 6.6170 | 7.2578 | 7.0862 |
| SRSF4_S458;S460 | SRSF4 | Q08170 | 458;460;462 | NA | 6.2910 | 6.2745 | 7.0212 | 6.6277 | 6.0773 | 6.2957 | 6.2477 | 6.0695 | 5.8363 | 5.7939 | 6.0459 | 6.1052 | 6.3083 | 6.6677 | 6.0620 | 6.2763 |
| SRSF4_S458;S460;S462 | SRSF4 | Q08170 | 458;460 | NA | 5.2822 | 5.7329 | 5.7240 | 5.9616 | 6.6903 | 6.1036 | 6.3860 | 6.5080 | 6.3710 | 5.8887 | 5.6426 | 5.4304 | 6.5554 | 6.7750 | 7.4843 | 7.4640 |
| SRSF5_S120 | SRSF5 | Q13243 | 120 | NLSSRVSWQDLKD | 3.7465 | 3.9774 | 3.9960 | 3.1143 | 3.9052 | 3.8921 | 3.6082 | 3.7928 | 3.8429 | 3.8089 | 3.5386 | 3.6358 | 13.6352 | 12.6682 | 13.8291 | 15.0087 |
| SRSF5_S153 | SRSF5 | Q13243 | 153 | GVVEFASYGDLKN | 6.2795 | 5.5873 | 6.7249 | 6.6131 | 6.6466 | 5.2129 | 6.2091 | 6.4723 | 6.0426 | 6.0657 | 6.5150 | 6.0788 | 5.8380 | 6.7657 | 6.3223 | 6.6262 |
| SRSF6_S265 | SRSF6 | Q13247 | 303 | SQSRSNSPLPVPP | 6.1237 | 6.6456 | 6.2185 | 6.9150 | 6.2903 | 5.9302 | 6.3299 | 6.3320 | 6.5202 | 6.3970 | 6.3730 | 6.4671 | 5.6925 | 5.7574 | 5.9302 | 6.0773 |
| SRSF6_S303 | SRSF6 | Q13247 | 45 | FVEFEDSRDADDA | 5.7803 | 5.9939 | 5.6460 | 5.8527 | 6.3562 | 5.9286 | 6.0806 | 5.6934 | 5.9244 | 6.1007 | 6.1692 | 5.8094 | 7.4691 | 7.1137 | 7.0936 | 6.9881 |
| SRSF6_S314 | SRSF6 | Q13247 | 265 | SHSRSRSKDEYEK | 5.9479 | 6.4719 | 5.7273 | 6.5308 | 6.2030 | 5.4469 | 6.1412 | 6.1613 | 6.4239 | 6.3751 | 5.9922 | 6.0093 | 6.9064 | 6.3314 | 6.6427 | 6.6889 |
| SRSF6_S314;S316 | SRSF6 | Q13247 | 314 | PPSKARSVSPPPK | 6.0297 | 6.6486 | 5.5346 | 6.8112 | 6.3042 | 5.8958 | 5.9945 | 5.8771 | 5.9258 | 6.2284 | 6.3927 | 6.0206 | 6.2983 | 6.5240 | 6.5733 | 6.9411 |
| SRSF6_S316 | SRSF6 | Q13247 | 316 | SKARSVSPPPKRA | 6.3501 | 6.8465 | 6.1191 | 6.6254 | 5.7687 | 5.6655 | 6.2200 | 6.5566 | 6.3110 | 5.2341 | 5.4253 | 6.7738 | 6.6269 | 6.8105 | 6.3317 | 6.3348 |
| SRSF6_S45 | SRSF6 | Q13247 | 314;316 | NA | 6.2064 | 6.4337 | 6.5748 | 6.5029 | 6.4496 | 6.0340 | 6.3691 | 5.5514 | 6.1572 | 5.9269 | 6.1523 | 6.2227 | 6.4952 | 6.2933 | 6.2351 | 6.3953 |
| SRSF7_S134;S136 | SRSF7 | Q16629 | 192 | GSRYFQSPSRSRS | 6.2283 | 6.6283 | 5.3963 | 6.2235 | 6.1898 | 5.7807 | 5.4323 | 5.9508 | 6.2656 | 5.9466 | 5.6546 | 5.4669 | 7.4734 | 6.7577 | 7.4586 | 7.1467 |
| SRSF7_S179;S181 | SRSF7 | Q16629 | 192;194 | NA | 6.2389 | 7.3339 | 6.5516 | 6.8932 | 7.2152 | 5.8030 | 6.5161 | 6.0535 | 6.2863 | 5.9603 | 5.4236 | 5.6126 | 6.2938 | 5.5892 | 5.8968 | 6.3321 |
| SRSF7_S192 | SRSF7 | Q16629 | 179;181 | NA | 4.8761 | 5.5899 | 5.6666 | 5.7011 | 4.9001 | 5.9009 | 5.5227 | 6.3406 | 6.1209 | 7.5065 | 6.4885 | 7.7450 | 6.9077 | 5.9209 | 7.0239 | 7.7887 |
| SRSF7_S192;S194 | SRSF7 | Q16629 | 213;215;217 | NA | 6.3757 | 5.9757 | 5.9555 | 6.1794 | 6.5447 | 5.5718 | 5.9160 | 5.4283 | 6.8819 | 6.0620 | 5.9352 | 6.0934 | 7.3496 | 6.1793 | 6.5605 | 6.9910 |
| SRSF7_S192;S196 | SRSF7 | Q16629 | 192;196 | NA | 6.4906 | 6.6338 | 5.8228 | 6.6076 | 5.9402 | 5.5345 | 5.9255 | 6.0804 | 6.3831 | 6.1409 | 5.5056 | 5.4104 | 6.9463 | 6.9573 | 6.5638 | 7.0572 |
| SRSF7_S213;S215;S217 | SRSF7 | A0A0B4J1Z1 | 134;136 | NA | 7.6901 | 7.8318 | 7.2648 | 7.4524 | 6.5988 | 5.8242 | 6.4958 | 6.1300 | 6.9932 | 6.8152 | 7.2444 | 7.1972 | 4.2383 | 3.9458 | 4.2230 | 4.0549 |
| SRSF8_S156;S158 | SRSF8 | Q9BRL6 | 26 | NLTYRTSPDSLRR | 6.4278 | 6.7274 | 6.3392 | 6.9480 | 5.9146 | 5.8344 | 6.0998 | 6.2984 | 6.9086 | 6.1882 | 6.0962 | 6.3747 | 6.0709 | 5.8534 | 5.7590 | 6.1595 |
| SRSF8_S158 | SRSF8 | Q9BRL6 | 158 | GSRYSRSPYSRSP | 6.4262 | 7.2815 | 6.2335 | 6.4012 | 5.8615 | 6.8322 | 5.6952 | 5.2788 | 5.8064 | 5.9396 | 6.8228 | 7.0794 | 5.9128 | 6.2496 | 6.4106 | 5.7687 |
| SRSF8_S26 | SRSF8 | Q9BRL6 | 156;158 | NA | 6.1312 | 6.2612 | 6.0701 | 6.6290 | 5.9186 | 5.8969 | 5.5503 | 6.0865 | 6.6627 | 6.0707 | 6.1302 | 6.2993 | 6.5077 | 6.5569 | 6.6370 | 6.5917 |
| SRSF8_S273;S281 | SRSF8 | Q9BRL6 | 273;281 | NA | 6.5636 | 6.2833 | 5.9055 | 6.8180 | 6.2402 | 6.0191 | 6.2931 | 5.8256 | 6.6498 | 5.8195 | 6.1011 | 6.1521 | 6.8591 | 6.4556 | 6.0940 | 5.9204 |

| | | | | | | | | | | | | | | | | | | | | | |
|---|---|---|---|---|---|---|---|---|---|---|---|---|---|---|---|---|---|---|---|---|---|
| SRSF9_S187;S193 | SRSF9 | Q13242 | | 211 | PYQSRGSPHYFSP | 6.6188 | 6.4050 | 5.5563 | 6.2602 | 6.6953 | 5.9035 | 5.4906 | 6.8167 | 5.9476 | 5.7927 | 6.0220 | 5.9953 | 7.0245 | 6.5720 | 6.5604 | 6.3392 |
| SRSF9_S189 | SRSF9 | Q13242 | | 204 | GSRGRDSPYQSRG | 6.5153 | 6.4386 | 5.9374 | 6.4284 | 6.2802 | 5.7336 | 5.7644 | 6.4231 | 6.4091 | 6.2458 | 6.3179 | 6.2794 | 6.6552 | 6.1020 | 6.0522 | 6.4174 |
| SRSF9_S193 | SRSF9 | Q13242 | | 193 | STSYGYSRSRSGS | 6.0179 | 5.9822 | 5.7814 | 6.2729 | 6.8398 | 5.7104 | 6.0636 | 7.5445 | 5.9849 | 6.3707 | 6.3131 | 6.7768 | 6.2028 | 5.8182 | 5.9954 | 6.3254 |
| SRSF9_S204 | SRSF9 | Q13242 | | 189 | YPERSTSYGYSRS | 6.3531 | 6.0071 | 5.2678 | 5.7314 | 6.4028 | 5.4422 | 5.4088 | 5.9947 | 5.8566 | 5.7841 | 5.4678 | 5.5936 | 8.5199 | 7.2645 | 7.4797 | 7.4258 |
| SRSF9_S211 | SRSF9 | Q13242 | 211;216 | | NA | 6.8487 | 6.7272 | 6.0311 | 6.6158 | 7.1132 | 6.2992 | 6.1757 | 6.9771 | 6.5463 | 6.2719 | 6.5537 | 6.4641 | 5.5158 | 5.4988 | 5.0557 | 5.3056 |
| SRSF9_S211;S216 | SRSF9 | Q13242 | 187;193 | | NA | 6.0471 | 5.9888 | 5.3146 | 5.9261 | 6.7182 | 5.3138 | 5.6763 | 6.7178 | 6.0955 | 5.8963 | 5.9411 | 5.6859 | 7.6826 | 6.6510 | 7.1951 | 7.1496 |
| SSB_S350 | SSB | P05455 | | 366 | KKTKFASDDEHDE | 5.2209 | 5.5304 | 5.8834 | 5.1347 | 5.6063 | 5.8249 | 5.7656 | 5.9567 | 5.7337 | 6.0936 | 6.0098 | 5.8874 | 7.5675 | 8.0672 | 8.1036 | 7.6143 |
| SSB_S366 | SSB | P05455 | | 350 | KAAQPGSGKGKVQ | 5.9796 | 6.5666 | 6.6632 | 6.8587 | 6.9899 | 6.7411 | 6.5192 | 6.5601 | 7.1719 | 7.0037 | 6.2928 | 6.3585 | 5.1823 | 5.2457 | 4.9597 | 4.9071 |
| SSB_T389 | SSB | P05455 | | 389 | KRAREETDKEEPA | 4.4785 | 5.6648 | 5.3244 | 5.9633 | 5.3733 | 5.9695 | 5.9207 | 5.5199 | 3.6690 | 4.4065 | 4.5434 | 3.9915 | 9.7052 | 9.2671 | 9.8793 | 10.3236 |
| SSBP4_S342;T355 | SSBP4 | Q9BWG4 | | 355 | LSNAPGTPRDDGE | 5.8534 | 5.6612 | 6.2661 | 5.1100 | 6.0021 | 5.5045 | 5.6155 | 6.7800 | 5.7512 | 6.1868 | 5.9051 | 6.7740 | 6.6807 | 7.6232 | 7.4688 | 6.8173 |
| SSBP4_S355 | SSBP4 | Q9BWG4 | 342;355 | | NA | 5.5931 | 6.5431 | 6.3419 | 7.1496 | 5.3197 | 6.2796 | 6.3073 | 6.3750 | 6.2158 | 6.9658 | 5.6829 | 5.9478 | 5.4256 | 7.1684 | 6.2798 | 6.4046 |
| SSH1_S1042 | SSH1 | Q8WYL5 | | 971 | AGLTVSSPLKRSH | 6.7937 | 6.9578 | 6.4011 | 6.2799 | 6.7697 | 6.8719 | 6.8422 | 7.2129 | 7.4915 | 6.6183 | 6.4251 | 6.3689 | 4.4739 | 5.0352 | 4.7397 | 4.7183 |
| SSH1_S515 | SSH1 | Q8WYL5 | | 1042 | APENLKSPSWMSK | 6.0874 | 6.4848 | 6.2194 | 6.4008 | 6.3003 | 6.6435 | 6.6495 | 6.6074 | 6.8429 | 6.4786 | 6.6547 | 6.7577 | 5.2920 | 5.6990 | 5.2139 | 5.6682 |
| SSH1_S57 | SSH1 | Q8WYL5 | | 515 | CCFRRLSDPLLPS | 6.6717 | 6.7958 | 5.5153 | 6.6686 | 6.6477 | 6.0506 | 5.7965 | 7.1171 | 6.3780 | 6.0030 | 6.1889 | 5.7200 | 5.6239 | 6.4381 | 6.2299 | 6.1551 |
| SSH1_S897 | SSH1 | Q8WYL5 | | 897 | EGGSLKSPPPFFY | 5.9421 | 5.8392 | 5.8181 | 6.1205 | 6.0144 | 6.6224 | 6.2427 | 6.4710 | 6.1092 | 5.8199 | 6.5236 | 6.2031 | 6.3941 | 6.5475 | 6.4825 | 6.8496 |
| SSH1_S971 | SSH1 | Q8WYL5 | | 978 | PLKRSHSLAKLGS | 6.9296 | 6.7125 | 7.3044 | 6.6137 | 6.5532 | 6.6819 | 6.0525 | 4.4867 | 5.4965 | 5.4330 | 6.1473 | 6.1863 | 6.0070 | 6.9811 | 6.0049 | 6.4094 |
| SSH1_S978 | SSH1 | Q8WYL5 | | 57 | FLQQGSSPQGQRS | 7.2422 | 7.2605 | 6.0193 | 6.8772 | 6.3494 | 7.3041 | 7.2581 | 6.5851 | 6.4510 | 6.7875 | 6.7208 | 6.7851 | 4.7710 | 4.5544 | 4.4195 | 4.6147 |
| SSH2_S36 | SSH2 | Q76I76 | | 36 | CRSQPRSISESFL | 5.6943 | 5.4071 | 6.0870 | 5.8238 | 6.0263 | 6.0214 | 5.8429 | 6.0118 | 5.5301 | 5.4163 | 6.1101 | 6.1799 | 7.3197 | 7.3760 | 7.4240 | 7.7291 |
| SSH2_S784 | SSH2 | Q76I76 | | 784 | HTPKKNSIHELLL | 4.8039 | 4.2457 | 5.2282 | 4.8640 | 5.6039 | 5.0809 | 4.8431 | 6.0162 | 7.0811 | 5.6046 | 6.1817 | 5.7803 | 8.6495 | 8.7881 | 8.7820 | 8.4468 |
| SSH3_S37 | SSH3 | Q8TE77 | | 37 | RLQRRQSFAVLRG | 6.7042 | 6.2132 | 7.5071 | 5.5003 | 6.0677 | 6.6365 | 5.9841 | 6.6078 | 6.9806 | 6.8848 | 7.0250 | 7.0801 | 5.4747 | 6.5212 | 3.8163 | 4.9965 |
| SSH3_S87 | SSH3 | Q8TE77 | | 9 | LVTVSRSPPGSGA | 6.5033 | 5.8262 | 6.8097 | 6.2255 | 6.4845 | 6.1148 | 5.2195 | 6.5836 | 6.4672 | 6.1233 | 6.5118 | 6.7426 | 6.2753 | 6.5479 | 6.2345 | 5.3303 |
| SSH3_S9 | SSH3 | Q8TE77 | | 87 | FGQGSQSPQKQEE | 6.6253 | 6.1398 | 7.1119 | 6.4614 | 6.1255 | 6.2178 | 5.9599 | 6.2166 | 6.4683 | 6.7136 | 7.5080 | 6.8460 | 5.1844 | 5.8154 | 5.2170 | 5.3890 |
| SSR1_S246 | SSR1 | P43307 | | 246 | KVEMGTSSQNDVD | 5.0502 | 4.8081 | 4.7577 | 5.9741 | 5.7322 | 4.7519 | 4.8931 | 5.1078 | 4.6760 | 4.7150 | 4.4795 | 4.8218 | 10.1225 | 9.4818 | 10.5416 | 10.0867 |
| SSRP1_S444 | SSRP1 | Q08945 | | 444 | YDEYADSDEDQHD | 5.5949 | 6.3406 | 5.8972 | 6.1860 | 5.8322 | 6.0323 | 6.2029 | 6.2031 | 6.9870 | 6.0352 | 5.9429 | 5.9851 | 6.6000 | 6.8740 | 6.7857 | 6.5007 |
| SSRP1_S657 | SSRP1 | Q08945 | | 659 | SRQLSESFKSKEF | 6.7704 | 7.1404 | 6.7460 | 6.9600 | 6.6408 | 6.9500 | 7.4846 | 6.6620 | 7.2557 | 6.9391 | 7.1830 | 6.9270 | 3.5495 | 4.5659 | 3.7315 | 4.4939 |
| SSRP1_S659 | SSRP1 | Q08945 | | 657 | SSSRQLSESFKSK | 6.4746 | 6.9409 | 6.6774 | 6.4566 | 6.2260 | 6.5418 | 6.9853 | 6.0739 | 6.7401 | 7.2679 | 7.2373 | 7.3468 | 4.3208 | 5.1545 | 4.7373 | 4.8189 |
| SSX2IP_S29 | SSX2IP | Q9Y2D8 | | 293 | EMISLLSPQKKKP | 6.3068 | 6.5092 | 6.6137 | 6.8028 | 6.6886 | 6.7316 | 7.0187 | 6.5858 | 7.5268 | 6.8940 | 6.6457 | 6.6963 | 4.4835 | 4.8522 | 4.5648 | 5.0795 |
| SSX2IP_S293 | SSX2IP | Q9Y2D8 | | 341 | REQLTNSIRKQWR | 7.9764 | 8.2057 | 6.9134 | 8.6291 | 6.9451 | 6.9182 | 7.5097 | 6.4057 | 7.4192 | 6.2911 | 7.1474 | 6.9474 | 2.9201 | 3.5835 | 2.9221 | 3.2660 |
| SSX2IP_S341 | SSX2IP | Q9Y2D8 | | 29 | TSETKMSPSSLYS | 6.6011 | 6.6578 | 7.0220 | 6.6384 | 6.3441 | 7.1967 | 6.4047 | 5.9527 | 6.7821 | 6.5274 | 7.0663 | 6.6850 | 4.8897 | 5.4962 | 4.8074 | 4.9285 |
| ST13_S75;S76;S79 | ST13 | P50502 | | 79 | EPSSEESDLEIDK | 6.1102 | 5.7364 | 6.7671 | 6.0259 | 6.1954 | 5.5933 | 5.5267 | 6.9249 | 7.0459 | 6.6687 | 6.8215 | 6.6985 | 5.8128 | 5.8882 | 6.2194 | 5.9651 |
| ST13_S79 | ST13 | P50502 | 75;76;79 | | NA | 6.0167 | 6.3758 | 7.3295 | 6.0404 | 6.3722 | 6.7222 | 5.9623 | 6.0833 | 6.0075 | 6.5693 | 6.7139 | 6.7805 | 5.7760 | 6.0438 | 5.4610 | 5.7456 |
| STAG1_S1062 | STAG1 | Q8WVM7 | | 1062 | GEDDRMSVNSGSS | 6.2169 | 6.6563 | 6.6951 | 5.9778 | 5.9002 | 6.7355 | 6.5073 | 6.1300 | 5.2341 | 4.7822 | 6.7153 | 5.0571 | 6.1657 | 7.1330 | 7.4932 | 6.6004 |
| STAG1_S1062;S1065 | STAG1 | Q8WVM7 | | 1108 | TDTMIQTPGPLPA | 6.7177 | 6.0692 | 5.8878 | 6.3420 | 6.0757 | 5.8016 | 6.9261 | 6.2751 | 6.4179 | 7.6493 | 6.9429 | 6.2442 | 5.3834 | 6.4745 | 5.0412 | 5.7514 |
| STAG1_S1138;S1142 | STAG1 | Q8WVM7 | 1062;1065 | | NA | 5.1763 | 5.6113 | 5.2146 | 6.1737 | 6.5617 | 5.9484 | 5.3143 | 5.3323 | 5.8898 | 6.5806 | 5.4456 | 5.4799 | 8.8491 | 7.8870 | 7.4231 | 7.1123 |
| STAG1_S1142;S1155 | STAG1 | Q8WVM7 | 1142;1155 | | NA | 6.7447 | 6.4424 | 6.6860 | 6.6058 | 5.9975 | 6.2120 | 6.6387 | 6.0691 | 6.4296 | 5.7542 | 5.7598 | 5.4566 | 6.3064 | 6.6166 | 6.2087 | 6.0719 |
| STAG1_T1108 | STAG1 | Q8WVM7 | 1138;1142 | | NA | 5.4407 | 5.6311 | 6.5641 | 6.5706 | 6.1131 | 6.2346 | 6.7081 | 6.1259 | 6.1399 | 6.2289 | 5.4988 | 5.4556 | 6.8433 | 7.1963 | 7.2762 | 5.9726 |
| STAG2_S1198 | STAG2 | Q8N3U4-2 | | 1112 | REQTLHTPVMMQT | 5.3880 | 5.3746 | 5.4867 | 5.5224 | 5.5669 | 5.8216 | 5.7819 | 5.7594 | 5.5610 | 5.7283 | 5.3801 | 5.2950 | 8.8418 | 8.3867 | 8.0381 | 8.0675 |
| STAG2_T1112 | STAG2 | Q8N3U4-2 | | 1198 | AIRRGTSLMEDDE | 6.2003 | 6.6803 | 6.4131 | 6.4043 | 5.8907 | 6.0051 | 6.4243 | 6.2363 | 5.8563 | 6.2091 | 6.6155 | 6.3747 | 5.5564 | 5.8387 | 5.9923 | 7.3026 |
| STARD13_S314 | STARD13 | Q9Y3M8 | | 470 | GSHLYASTGDLLD | 6.2430 | 6.3648 | 7.2216 | 6.5778 | 6.2429 | 6.5389 | 6.4063 | 6.5320 | 6.3616 | 5.9452 | 6.7011 | 6.5943 | 5.5188 | 5.8196 | 5.2625 | 5.6696 |
| STARD13_S389 | STARD13 | Q9Y3M8 | | 389 | RMHEFHSQENLVV | 6.2856 | 6.0858 | 6.9879 | 5.9484 | 6.3563 | 6.5376 | 6.1113 | 6.7923 | 6.2260 | 6.4730 | 6.5395 | 6.7323 | 5.8669 | 5.9905 | 5.5113 | 5.5552 |
| STARD13_S411 | STARD13 | Q9Y3M8 | | 314 | NGDLQNSPPPACR | 6.3611 | 5.9628 | 6.4168 | 5.8989 | 6.4892 | 5.7703 | 6.6802 | 6.5205 | 6.2996 | 6.6085 | 6.5432 | 6.9092 | 5.9310 | 5.7213 | 6.0508 | 5.8366 |
| STARD13_S430 | STARD13 | Q9Y3M8 | | 430 | VNWRTGSISLGRE | 7.7046 | 6.1071 | 7.8068 | 7.1086 | 6.6017 | 5.8812 | 6.5047 | 5.8535 | 7.6750 | 6.6074 | 7.5327 | 6.9694 | 4.8340 | 5.4031 | 3.5243 | 3.8859 |
| STARD13_S470 | STARD13 | Q9Y3M8 | | 589 | RRLRWNSFQLSHQ | 6.6374 | 6.2969 | 6.0391 | 7.1391 | 6.8646 | 5.9105 | 6.3029 | 5.8205 | 7.2757 | 6.4928 | 5.2662 | 6.0091 | 6.6634 | 6.6167 | 5.5487 | 5.1163 |
| STARD13_S589 | STARD13 | Q9Y3M8 | | 411 | TFPKALSIESLSP | 4.8690 | 6.5419 | 6.2131 | 7.1221 | 4.6272 | 6.0439 | 6.4984 | 6.1696 | 7.8136 | 5.5362 | 7.5718 | 6.9577 | 5.1784 | 6.0052 | 6.6628 | 6.1890 |
| STARD3NL_S193 | STARD3NL | O95772 | | 193 | LIVQDASERAALI | 6.4524 | 6.8967 | 6.3217 | 6.6320 | 6.1663 | 5.8948 | 6.3278 | 6.2257 | 5.9402 | 6.1850 | 5.9783 | 5.8456 | 6.0986 | 6.4541 | 6.4575 | 6.1232 |
| STARD3NL_S218 | STARD3NL | O95772 | | 218 | PESEAGSEEAEEK | 6.7058 | 6.0805 | 6.5394 | 6.4213 | 6.0156 | 5.8227 | 5.6261 | 6.2770 | 6.9408 | 6.1317 | 6.6476 | 6.8005 | 5.7522 | 5.8488 | 6.1724 | 6.2174 |
| STARD3NL_Y209;S210;S2 | STARD3NL | O95772 | 209;210;218 | | NA | 5.9074 | 5.8571 | 5.8332 | 6.2232 | 6.4427 | 5.6361 | 6.8235 | 5.6850 | 6.0031 | 7.0921 | 5.8933 | 6.0224 | 6.6493 | 6.8799 | 6.7232 | 6.3286 |
| STARD8_S315 | STARD8 | Q92502-2 | | 826 | RIKSKRSLIGRPG | 7.2644 | 5.7702 | 8.2977 | 6.1496 | 6.2185 | 6.7558 | 6.5777 | 6.3637 | 8.1800 | 8.1705 | 9.4676 | 10.7933 | 2.0029 | 3.8383 | 1.5567 | 2.5931 |
| STARD8_S578 | STARD8 | Q92502-2 | | 97 | LEVHFQSKQNEDS | 7.5525 | 7.4839 | 7.0980 | 7.6945 | 5.9452 | 6.3971 | 7.1837 | 6.5701 | 6.8961 | 6.0864 | 7.4144 | 7.2920 | 3.5510 | 4.3415 | 4.0758 | 4.4180 |
| STARD8_S826 | STARD8 | Q92502-2 | | 578 | RKLRWHSFQNSHR | 6.7853 | 5.7621 | 7.5950 | 6.8464 | 8.0803 | 6.5219 | 7.1625 | 8.2384 | 6.9656 | 7.2262 | 6.9684 | 7.0294 | 3.8017 | 4.1927 | 3.2277 | 3.5965 |
| STARD8_S97 | STARD8 | Q92502-2 | | 315 | TFPRSLSIESLCP | 5.6997 | 5.5015 | 6.5541 | 6.5734 | 6.2198 | 6.7312 | 6.6171 | 6.7403 | 6.7581 | 7.0732 | 6.7129 | 7.5948 | 4.9010 | 5.5332 | 5.0617 | 5.7281 |
| STAT1_S727 | STAT1 | P42224 | | 727 | DNLLPMSPEEFDE | 5.7649 | 6.3210 | 6.4904 | 5.9793 | 5.7682 | 6.2378 | 6.2977 | 6.4857 | 6.1845 | 6.0156 | 6.6460 | 6.3458 | 6.2286 | 6.6652 | 6.2825 | 6.2864 |
| STAT3_S727 | STAT3 | P40763 | | 714 | TKFICVTPTTCSN | 5.6009 | 6.3923 | 6.0532 | 6.0146 | 5.8580 | 5.8374 | 6.2132 | 6.3261 | 6.1853 | 6.0299 | 6.0651 | 6.5787 | 7.2419 | 6.4447 | 6.3965 | 6.7624 |
| STAT3_T714 | STAT3 | P40763 | | 727 | TIDLPMSPRTLDS | 6.2020 | 6.6376 | 6.6185 | 7.2311 | 6.4182 | 5.8828 | 6.6925 | 6.4739 | 6.4223 | 6.8496 | 5.9626 | 5.9759 | 5.4722 | 5.2403 | 5.8828 | 6.0375 |
| STAT3_T714;S727 | STAT3 | P40763 | 714;727 | | NA | 5.6832 | 6.5040 | 6.1970 | 6.9986 | 5.9435 | 6.4665 | 6.4500 | 7.0741 | 6.8858 | 6.2455 | 6.6231 | 6.7605 | 5.3860 | 6.2055 | 5.3034 | 5.2734 |
| STAU1_S176 | STAU1 | O95793 | | 176 | EVNGRESEEENLN | 6.7976 | 7.0139 | 7.5793 | 6.6355 | 6.6449 | 6.8806 | 6.4779 | 7.2191 | 6.3378 | 6.0698 | 5.6099 | 5.9219 | 5.0636 | 5.2145 | 5.2404 | 5.2932 |
| STAU1_S278 | STAU1 | O95793 | | 390 | TFFEPGSGDENGT | 7.2675 | 7.5501 | 7.4509 | 7.0763 | 6.9443 | 7.3859 | 7.3775 | 7.6896 | 5.7562 | 5.8727 | 5.8898 | 5.7153 | 4.5297 | 4.6893 | 4.3041 | 4.5008 |
| STAU1_S390 | STAU1 | O95793 | | 278 | IVKPQTSPEYGQG | 6.1394 | 6.2305 | 6.9814 | 6.5105 | 6.0430 | 6.7577 | 6.4429 | 6.0303 | 6.2205 | 6.3578 | 5.9961 | 6.1682 | 5.3196 | 6.5748 | 5.6136 | 6.6137 |
| STAU2_S188 | STAU2 | Q9NUL3 | | 455 | SSFFSISPTSNSS | 6.5547 | 6.8595 | 7.0067 | 7.1517 | 6.3172 | 6.7612 | 7.0445 | 6.6186 | 7.1510 | 6.9728 | 6.8912 | 6.8705 | 4.1696 | 4.7166 | 4.1803 | 4.7338 |
| STAU2_S416 | STAU2 | Q9NUL3 | | 416 | KGILHLSPDVYQE | 6.5121 | 7.5226 | 7.1670 | 7.5488 | 6.3702 | 7.1602 | 7.2414 | 7.1513 | 7.0902 | 7.1947 | 7.5102 | 7.1388 | 3.3500 | 4.1852 | 3.0818 | 3.7752 |
| STAU2_S440 | STAU2 | Q9NUL3 | | 440 | TTLGYLSPKDMNQ | 6.4818 | 7.1549 | 7.0210 | 7.1193 | 6.6564 | 7.0899 | 7.5215 | 7.0161 | 6.9638 | 7.1181 | 7.0678 | 7.0238 | 3.5410 | 4.4785 | 3.5458 | 4.2003 |
| STAU2_S455 | STAU2 | Q9NUL3 | | 486 | IGLKGSSPTPPCS | 6.7135 | 6.5861 | 6.6829 | 6.6464 | 6.4370 | 6.7338 | 6.4765 | 6.4424 | 6.6153 | 6.5916 | 6.7472 | 6.6624 | 4.8116 | 5.4103 | 5.2750 | 5.1680 |
| STAU2_S486 | STAU2 | Q9NUL3 | | 408 | PEPTNNTPKGILH | 6.6091 | 6.3450 | 6.9593 | 7.8022 | 6.3840 | 7.3110 | 7.5001 | 6.2180 | 7.8528 | 7.8280 | 7.9955 | 7.8665 | 3.0332 | 3.8382 | 2.7281 | 3.7290 |
| STAU2_S492 | STAU2 | Q9NUL3 | | 188 | EPIPERSPQNGES | 6.2369 | 6.8108 | 6.7948 | 6.7788 | 6.1268 | 6.5700 | 6.5614 | 7.3507 | 6.5229 | 6.5676 | 6.5085 | 6.6460 | 4.5124 | 5.5485 | 5.2647 | 5.1991 |
| STAU2_T408 | STAU2 | Q9NUL3 | | 492 | SPTPPCSPVQPSK | 5.9202 | 5.3575 | 4.4911 | 6.1278 | 8.6881 | 5.8840 | 5.5052 | 5.8332 | 5.4090 | 5.0141 | 4.5255 | 4.4519 | 8.3806 | 7.9496 | 9.2113 | 7.2509 |
| STAU2_T486;S492 | STAU2 | Q9NUL3 | 486;492 | | NA | 5.6068 | 6.3075 | 6.4681 | 6.8083 | 5.5290 | 6.3088 | 5.9716 | 6.3098 | 6.5092 | 6.6214 | 6.7268 | 6.4259 | 5.5347 | 6.3225 | 6.1614 | 6.3882 |
| STBD1_S211 | STBD1 | O95210 | | 211 | MVPRHSSWGDVGV | 3.0956 | 3.3269 | 3.4258 | 3.6277 | 4.2443 | 4.3161 | 4.1031 | 4.0225 | 4.3215 | 4.7480 | 4.2074 | 4.3331 | 11.9674 | 12.8641 | 13.8072 | 13.5891 |
| STEAP3_S27;S30 | STEAP3 | Q658P3-2 | 27;30 | | NA | 4.7190 | 4.1394 | 4.5506 | 5.3052 | 7.2893 | 7.1264 | 5.9339 | 6.5957 | 4.4918 | 4.0416 | 4.6991 | 4.3440 | 8.7642 | 10.0729 | 9.4378 | 8.4891 |
| STIL_S409 | STIL | Q15468-2 | | 409 | SPRPIPSPHPVSQ | 6.5603 | 6.1514 | 6.9883 | 6.5001 | 5.5754 | 6.5670 | 6.7982 | 7.1488 | 7.5443 | 5.8744 | 7.1947 | 7.3141 | 4.0191 | 5.5973 | 4.4041 | 5.7626 |
| STIM1_S257 | STIM1 | Q13586 | | 575 | VEKLPDSPALAKK | 6.5173 | 6.8010 | 6.3237 | 6.7205 | 6.4143 | 6.4588 | 6.6768 | 6.5379 | 6.7950 | 6.9462 | 6.5642 | 6.5967 | 5.2145 | 5.4037 | 4.8677 | 5.1618 |
| STIM1_S512 | STIM1 | Q13586 | | 257 | LHRAEQSLHDLQE | 2.5799 | 2.6754 | 2.4677 | 2.5965 | 5.0805 | 4.9998 | 5.2801 | 4.7624 | 2.6099 | 2.7380 | 2.2565 | 2.5446 | 15.6169 | 13.5601 | 15.8205 | 14.4112 |
| STIM1_S575 | STIM1 | Q13586 | | 512 | PQHGLGSQRDLTH | 4.4424 | 4.5277 | 4.6936 | 4.8967 | 5.0105 | 5.2466 | 5.7971 | 4.7630 | 4.8620 | 4.7200 | 4.4734 | 4.7854 | 10.4339 | 10.2006 | 10.8957 | 10.2514 |
| STIM1_S618 | STIM1 | Q13586 | | 668 | GEETDSSPGRKKF | 6.8556 | 6.5288 | 6.3400 | 6.8780 | 7.0994 | 7.1239 | 7.1262 | 6.5699 | 6.3311 | 6.2904 | 6.8530 | 5.7435 | 5.2190 | 5.3183 | 4.7208 | 5.0021 |
| STIM1_S668 | STIM1 | Q13586 | | 618 | DSSRSHSPSSPDP | 5.9099 | 6.0796 | 6.5417 | 6.1779 | 5.8571 | 5.9886 | 6.2997 | 6.5685 | 6.1576 | 5.8766 | 6.3452 | 5.5850 | 6.4191 | 6.7196 | 6.1949 | 7.2791 |
| STIM2_S431 | STIM2 | Q9P246-2 | | 756 | GLTETKSMIFSPA | 4.5832 | 4.2772 | 3.7799 | 5.1837 | 7.1336 | 4.7000 | 5.2047 | 4.2672 | 3.7048 | 3.3209 | 4.1690 | 4.1779 | 11.5032 | 12.0308 | 10.9813 | 10.9827 |
| STIM2_S618 | STIM2 | Q9P246-2 | | 618 | RSIVPSSPQPQRA | 6.2436 | 6.7379 | 6.0914 | 6.7507 | 5.8957 | 6.6433 | 6.9554 | 6.6265 | 5.9013 | 5.9342 | 6.0653 | 6.1702 | 5.7323 | 6.0014 | 6.1586 | 6.0923 |
| STIM2_S704;S708 | STIM2 | Q9P246-2 | | 775 | ILEKSCSMNQLSS | 4.8344 | 5.5269 | 5.2002 | 5.3535 | 6.3403 | 6.3258 | 6.1045 | 5.5946 | 5.4539 | 5.2824 | 5.8917 | 6.0826 | 8.0672 | 8.1071 | 7.8924 | 7.9425 |
| STIM2_S716 | STIM2 | Q9P246-2 | | 760 | TKSMIFSPASKVY | 7.2366 | 8.5601 | 6.9689 | 8.2228 | 6.0926 | 6.7965 | 7.8926 | 7.7020 | 7.3111 | 6.9481 | 6.9042 | 7.3316 | 2.8715 | 3.3142 | 2.7749 | 3.0725 |
| STIM2_S735 | STIM2 | Q9P246-2 | | 735 | DSSRGDSPVTVDV | 4.9800 | 5.1622 | 5.0496 | 5.6404 | 5.8229 | 5.8351 | 6.1463 | 6.2050 | 5.7451 | 5.5737 | 5.3303 | 5.4091 | 8.0555 | 7.8632 | 8.7713 | 8.4104 |
| STIM2_S756 | STIM2 | Q9P246-2 | | 716 | EIYQTLSPRKISR | 6.3584 | 6.3806 | 6.6792 | 6.7788 | 6.0213 | 6.4945 | 6.2463 | 6.5968 | 6.6903 | 6.1908 | 6.7653 | 6.7880 | 5.4407 | 5.9588 | 5.2249 | 5.3851 |
| STIM2_S760 | STIM2 | Q9P246-2 | | 431 | EFELRSSWSVPDA | 4.1012 | 3.7549 | 4.1301 | 3.6819 | 3.4359 | 3.4171 | 3.6923 | 3.5027 | 3.6770 | 4.0053 | 3.7186 | 4.2033 | 14.0848 | 13.1735 | 14.5980 | 12.8233 |
| STIM2_S775 | STIM2 | Q9P246-2 | | 813 | PSVARISSIPHDL | 5.0671 | 5.3361 | 5.2937 | 5.9687 | 6.4583 | 6.3656 | 6.0655 | 6.3839 | 5.3000 | 5.7658 | 6.2905 | 6.6975 | 7.0858 | 7.7963 | 7.0868 | 7.0384 |
| STIM2_S813 | STIM2 | Q9P246-2 | 704;708 | | NA | 3.9347 | 4.1194 | 4.3215 | 4.8833 | 4.9742 | 5.1653 | 5.5641 | 5.9283 | 4.4893 | 3.9695 | 4.8667 | 5.2783 | 11.2197 | 9.4142 | 10.8677 | 11.0035 |
| STIP1_S528 | STIP1 | P31948-2 | | 528 | QYNRHDSPEDVKR | 5.9120 | 6.2616 | 7.0478 | 6.0483 | 6.1009 | 6.5242 | 6.3182 | 6.1328 | 6.5037 | 6.5174 | 6.9085 | 6.7728 | 6.0352 | 5.7688 | 5.4522 | 5.6956 |
| STK10_S13 | STK10 | O94804 | | 191 | TLQKRDSFIGTPY | 6.0365 | 6.2736 | 6.8424 | 6.1763 | 6.3773 | 6.4723 | 6.1557 | 6.2369 | 6.3870 | 6.3789 | 6.4599 | 6.3969 | 5.9591 | 6.3431 | 5.7012 | 5.8030 |
| STK10_S191 | STK10 | O94804 | | 514 | LNKEMGSLSIKDP | 7.7299 | 6.7060 | 7.8940 | 8.4027 | 5.5305 | 4.6642 | 5.4556 | 4.8456 | 6.4185 | 4.1424 | 5.1479 | 5.1343 | 7.2588 | 7.6753 | 7.2642 | 5.7302 |
| STK10_S417 | STK10 | O94804 | | 13 | RRILRLSTFEKRK | 10.0881 | 7.2635 | 7.6041 | 7.4850 | 8.0054 | 6.5067 | 6.4783 | 5.9435 | 7.6500 | 5.4483 | 5.4278 | 5.9873 | 5.0417 | 5.5693 | 2.7266 | 2.7746 |
| STK10_S438 | STK10 | O94804 | | 438 | EQGGDLSPAANRS | 6.4599 | 6.4789 | 6.1404 | 6.5129 | 6.6222 | 6.2074 | 6.1754 | 7.8280 | 6.4289 | 6.4305 | 5.8219 | 6.9097 | 5.8461 | 5.6915 | 4.9830 | 5.4633 |
| STK10_S450 | STK10 | O94804 | | 485 | GPSKRDSDCSSLC | 5.8284 | 5.0050 | 7.1046 | 5.4784 | 5.7755 | 6.6679 | 6.5106 | 7.4163 | 5.8754 | 6.3077 | 5.4976 | 6.2460 | 6.1033 | 6.5709 | 7.0826 | 6.5297 |
| STK10_S450;S454 | STK10 | O94804 | | 516 | KEMGSLSIKDPKL | 10.7039 | 5.4100 | 5.5769 | 6.5385 | 6.9993 | 3.6538 | 6.0196 | 6.3570 | 10.4608 | 6.7816 | 4.9093 | 6.7525 | 5.1560 | 5.4898 | 4.7896 | 4.4013 |
| STK10_S454 | STK10 | O94804 | | 417 | RKSRPVSMDARIQ | 6.4797 | 5.9502 | 6.4928 | 5.8952 | 5.3651 | 5.8004 | 6.7002 | 5.8647 | 6.5188 | 5.5004 | 5.8907 | 6.6046 | 6.8571 | 6.5346 | 6.6939 | 6.8517 |
| STK10_S485 | STK10 | O94804 | | 454 | SQSRPNSSALETL | 6.3720 | 5.8388 | 6.8609 | 5.9120 | 6.8347 | 5.9068 | 5.7151 | 6.7502 | 6.5420 | 5.7560 | 5.6131 | 5.6880 | 7.0787 | 5.6758 | 7.4943 | 5.9616 |
| STK10_S514 | STK10 | O94804 | | 259 | DPPTLLTPSKWSV | 7.2652 | 6.7803 | 7.8273 | 6.9906 | 5.4492 | 6.2474 | 5.9926 | 7.4943 | 6.6178 | 6.4449 | 5.7168 | 7.0444 | 4.9175 | 5.0706 | 5.0180 | 5.1232 |
| STK10_S516 | STK10 | O94804 | | 450 | SQKASQSRPNSSA | 5.4342 | 5.3995 | 6.6135 | 5.9776 | 6.4868 | 5.9161 | 5.1936 | 6.4333 | 6.2584 | 6.5611 | 6.6162 | 6.6381 | 7.2740 | 6.4119 | 5.7230 | 7.0627 |
| STK10_T259 | STK10 | O94804 | | 952 | ECPNPSTPSKAAK | 6.3862 | 6.5280 | 6.9896 | 6.3779 | 6.1864 | 6.5868 | 6.7310 | 7.7908 | 6.4657 | 6.3615 | 6.4074 | 6.9279 | 5.2415 | 5.2984 | 4.9091 | 4.8117 |
| STK10_T390 | STK10 | O94804 | | 390 | GDRSLQTTSPPVV | 5.8730 | 7.0793 | 6.9452 | 6.4404 | 6.0550 | 6.3030 | 7.0223 | 5.3335 | 6.7196 | 7.2312 | 7.3507 | 6.8838 | 5.0230 | 5.8517 | 4.3461 | 5.5423 |
| STK10_T952 | STK10 | O94804 | 450;454 | | NA | 6.2632 | 5.7925 | 7.6361 | 6.7763 | 6.6077 | 6.2016 | 5.9139 | 6.8883 | 6.6172 | 5.7604 | 6.1798 | 6.9959 | 5.8369 | 5.7263 | 5.3296 | 5.4744 |
| STK11_S31 | STK11 | Q15831 | | 31 | FIHRIDSTEVIYQ | 6.2673 | 6.1920 | 6.6960 | 6.1259 | 6.0553 | 6.1302 | 6.2453 | 6.2772 | 6.7581 | 6.3404 | 6.0828 | 6.3226 | 6.3437 | 6.3064 | 5.9523 | 5.9045 |
| STK11IP_S387 | STK11IP | Q8N1F8 | | 404 | PRTLNPSPAGWFV | 6.1725 | 6.4280 | 6.5959 | 6.6306 | 6.5352 | 6.6456 | 6.6623 | 6.9703 | 6.8151 | 6.3819 | 6.7755 | 6.7170 | 5.0628 | 5.4254 | 5.0065 | 5.1751 |
| STK11IP_S404 | STK11IP | Q8N1F8 | | 599 | EAQAQRSPRPTGS | 6.9744 | 6.4605 | 6.7887 | 7.3773 | 6.4003 | 6.4289 | 6.4820 | 6.7051 | 6.2322 | 6.8712 | 7.1158 | 6.7211 | 4.5300 | 4.7803 | 5.1210 | 5.0112 |
| STK11IP_S404;S421 | STK11IP | Q8N1F8 | | 482 | ARGPQESPQKMSE | 6.8297 | 6.6821 | 6.4140 | 6.8961 | 6.3448 | 6.2611 | 6.4113 | 6.3829 | 6.3855 | 6.0788 | 6.6234 | 6.5407 | 5.3755 | 5.4841 | 5.3732 | 5.9169 |
| STK11IP_S482 | STK11IP | Q8N1F8 | | 387 | VRVRRASISEPSD | 5.4445 | 5.6929 | 5.3441 | 5.5192 | 5.7350 | 5.7042 | 5.5858 | 6.0188 | 5.3119 | 5.5293 | 6.0723 | 5.6787 | 8.0860 | 8.1123 | 7.9917 | 8.1733 |
| STK11IP_S563 | STK11IP | Q8N1F8 | | 773 | STRDHGSWSLSPP | 6.7162 | 7.3733 | 6.7863 | 7.2673 | 6.3125 | 6.6349 | 6.3937 | 6.8833 | 6.0645 | 6.5833 | 7.1362 | 6.4373 | 4.6669 | 5.1407 | 4.6098 | 4.9938 |
| STK11IP_S599 | STK11IP | Q8N1F8 | | 761 | LDRAKNSPPQAPS | 6.4206 | 6.5226 | 6.3840 | 7.2248 | 6.3664 | 5.4266 | 6.3539 | 5.7213 | 6.3310 | 5.8709 | 6.2785 | 6.9203 | 5.9350 | 5.8491 | 5.9888 | 6.4062 |
| STK11IP_S761 | STK11IP | Q8N1F8 | | 563 | CFLRVTSAHLFEV | 2.0323 | 3.4622 | 3.0448 | 2.4750 | 2.7872 | 2.8343 | 4.2388 | 2.6347 | 2.9033 | 3.0920 | 2.5150 | 2.8416 | 15.9535 | 14.4475 | 19.2597 | 15.4781 |
| STK11IP_S773 | STK11IP | Q8N1F8 | 404;421 | | NA | 6.8764 | 7.1751 | 7.0470 | 7.9195 | 6.6089 | 6.9595 | 6.7357 | 6.5235 | 6.2377 | 5.8001 | 5.8233 | 5.8379 | 4.5212 | 5.3971 | 5.3409 | 5.1962 |
| STK25_T174 | STK25 | O00506 | | 174 | TQIKRNTFVGTPF | 11.2463 | 8.1014 | 8.0582 | 7.4166 | 8.3948 | 5.7403 | 5.5901 | 4.6972 | 6.6535 | 5.1947 | 5.1591 | 4.4835 | 5.3236 | 5.1450 | 4.3826 | 4.4131 |
| STK25_T277 | STK25 | O00506 | | 277 | TRYTKKTSFLTEL | 6.4422 | 5.6141 | 6.6721 | 6.1301 | 6.2145 | 6.4560 | 4.8313 | 6.4628 | 6.2570 | 5.3143 | 6.5355 | 6.5493 | 7.5825 | 6.4402 | 6.4189 | 6.0793 |
| STK3_S344 | STK3 | Q13188-2 | | 344 | EEEEENSDEDELD | 5.7211 | 6.1830 | 6.5963 | 6.5108 | 6.2562 | 6.0931 | 6.5015 | 5.7140 | 6.5670 | 5.8605 | 6.5104 | 7.1194 | 5.6504 | 6.5250 | 5.9445 | 6.2469 |
| STK3_S413 | STK3 | Q13188-2 | | 413 | MKRNATSPQVQRP | 6.1140 | 6.1480 | 7.2476 | 6.3772 | 6.5016 | 7.2313 | 7.1606 | 5.9434 | 6.6655 | 6.1283 | 6.0780 | 7.3360 | 5.8529 | 5.0865 | 4.9519 | 5.1770 |
| STK38_S281 | STK38 | Q15208 | | 281 | RRQLAFSTVGTPD | 7.2830 | 7.5367 | 7.7932 | 7.8546 | 6.2513 | 6.0067 | 5.9571 | 6.0384 | 5.9355 | 5.3694 | 6.3874 | 5.3429 | 5.5927 | 5.7159 | 5.9138 | 5.0214 |
| STK38L_S282 | STK38L | Q9Y2H1 | | 282 | RRQLAYSTVGTPD | 6.9760 | 7.2795 | 7.4503 | 7.8720 | 6.6543 | 5.6053 | 5.8220 | 5.6892 | 6.0853 | 5.5309 | 5.9728 | 5.5294 | 5.9020 | 6.0642 | 6.3454 | 5.2211 |
| STK39_S371 | STK39 | Q9UEW8 | | 371 | RRVPGSSGHLHKT | 6.4260 | 6.5572 | 6.7422 | 6.3263 | 6.1144 | 7.5128 | 6.2133 | 7.3809 | 6.5452 | 6.1901 | 6.2993 | 5.9045 | 5.2474 | 6.2904 | 5.2095 | 5.0407 |
| STK4_S410 | STK4 | Q13043 | | 414 | INSFGKSVPGPLK | 8.1131 | 7.4223 | 7.1575 | 7.5935 | 6.2646 | 6.3860 | 6.0792 | 5.5295 | 6.6282 | 6.1801 | 6.7017 | 7.5746 | 4.3204 | 4.7894 | 3.9467 | 5.3132 |
| STK4_S414 | STK4 | Q13043 | | 387 | MKRRDETMQPAKP | 6.3707 | 6.0805 | 6.3678 | 6.4104 | 6.4302 | 6.1985 | 6.4382 | 6.2888 | 6.6248 | 6.6863 | 5.9057 | 5.8980 | 6.0600 | 6.5624 | 6.0641 | 5.6136 |
| STK4_S438 | STK4 | Q13043 | | 410 | KENQINSFGKSVP | 7.8651 | 7.3421 | 6.8179 | 6.7485 | 6.6495 | 6.8593 | 5.7483 | 6.6129 | 7.2133 | 6.5274 | 6.9515 | 7.3103 | 3.9051 | 4.6768 | 4.2344 | 4.5375 |
| STK4_T387 | STK4 | Q13043 | | 438 | DYEFLKSWTVEDL | 6.5135 | 6.2731 | 6.6833 | 7.2234 | 7.1577 | 5.7899 | 7.3364 | 5.9301 | 6.6785 | 6.1112 | 6.1441 | 6.7971 | 6.0816 | 5.8140 | 4.7226 | 4.7435 |
| STMN1_S16 | STMN1 | P16949-2 | | 38 | VPEFPLSPPKKKD | 6.6298 | 5.7732 | 7.4958 | 6.1371 | 6.9108 | 6.3503 | 6.1251 | 7.5471 | 6.8042 | 7.7706 | 8.0137 | 8.8188 | 3.7270 | 4.4712 | 3.2715 | 4.1537 |
| STMN1_S16;S25 | STMN1 | P16949-2 | | 16 | ELEKRASGQAFEL | 7.1380 | 5.9313 | 7.6151 | 6.0602 | 7.0191 | 6.0125 | 5.7420 | 7.4328 | 6.9218 | 7.1027 | 7.3204 | 7.5879 | 4.6548 | 4.8210 | 4.1175 | 4.5229 |
| STMN1_S25 | STMN1 | P16949-2 | | 25 | AFELILSPRSKES | 6.8511 | 6.2515 | 6.9004 | 5.8650 | 6.8035 | 6.5344 | 6.2181 | 8.0529 | 6.7169 | 7.3829 | 7.6225 | 8.3045 | 3.8499 | 4.4860 | 3.6641 | 4.4963 |
| STMN1_S38 | STMN1 | P16949-2 | | 63 | AEERRKSHEAEVL | 7.4644 | 7.0013 | 7.1144 | 6.3330 | 6.9265 | 6.4834 | 6.6213 | 7.6752 | 7.0167 | 7.9809 | 8.3198 | 8.4093 | 2.8199 | 3.6442 | 2.5440 | 3.6458 |
| STMN1_S46 | STMN1 | P16949-2 | | 46 | PKKKDLSLEEIQK | 6.3507 | 5.4869 | 5.8840 | 5.0125 | 5.3040 | 5.3990 | 5.4391 | 6.1963 | 5.2678 | 6.2267 | 5.6022 | 6.0886 | 8.5334 | 7.9953 | 7.7868 | 7.4268 |
| STMN1_S63 | STMN1 | P16949-2 | 16;25 | | NA | 6.9432 | 6.6113 | 7.0807 | 6.4606 | 6.5377 | 6.6731 | 6.8070 | 7.4301 | 6.7404 | 7.4703 | 7.9064 | 7.7615 | 3.6785 | 4.5369 | 3.2552 | 4.1072 |
| STMN3_S50 | STMN3 | Q9NZ72 | | 50 | QLDKRASGQSFEV | 6.7521 | 6.3836 | 6.2983 | 6.1924 | 6.6085 | 6.3148 | 7.0311 | 6.6314 | 6.8474 | 6.9222 | 6.7821 | 7.3218 | 4.8704 | 5.7297 | 4.2295 | 5.0847 |
| STOML2_S30 | STOML2 | Q9UJZ1 | | 327 | KAPVPGTPDSLSS | 6.2178 | 5.8458 | 6.1961 | 5.8582 | 6.1643 | 5.8333 | 6.0143 | 6.4571 | 5.9614 | 6.5732 | 6.3215 | 6.7710 | 6.1875 | 6.6617 | 6.5442 | 6.3926 |
| STOML2_T327 | STOML2 | Q9UJZ1 | | 30 | APRRASSGLPRNT | 6.6792 | 6.8147 | 6.0249 | 6.3522 | 6.0880 | 5.7325 | 6.0026 | 6.1290 | 6.4761 | 6.0023 | 6.0179 | 6.3718 | 6.4040 | 6.2529 | 6.4583 | 6.1936 |
| STRADA_S313 | STRADA | Q7RTN6 | | 313 | AEELTMSPSRSVA | 6.0463 | 6.5824 | 6.8883 | 6.1234 | 6.2448 | 6.1781 | 6.2617 | 6.3560 | 6.5574 | 6.9961 | 6.5520 | 6.9654 | 5.5447 | 5.6920 | 5.5346 | 5.4766 |
| STRAP_T339 | STRAP | Q9Y3F4-2 | | 339 | IGFPETTEEELEE | 6.1283 | 6.8236 | 7.2527 | 6.4419 | 6.2684 | 6.0606 | 5.6268 | 6.7959 | 6.7782 | 6.6193 | 6.9448 | 6.6002 | 5.0844 | 6.0404 | 5.1909 | 5.3435 |
| STRBP_S465 | STRBP | Q96SI9 | | 465 | ADIECMSSDEKSD | 6.3982 | 6.5137 | 6.7552 | 7.2155 | 6.2538 | 6.0637 | 6.3347 | 5.8851 | 6.1815 | 6.2358 | 5.9091 | 6.0690 | 6.0877 | 5.8111 | 6.0403 | 6.2457 |
| STRBP_S465;S470 | STRBP | Q96SI9 | 465;470 | | NA | 5.9582 | 6.5016 | 6.4513 | 6.8283 | 6.0703 | 6.1354 | 6.6216 | 5.6972 | 6.2582 | 6.0836 | 6.6292 | 6.1428 | 6.2616 | 6.3932 | 6.3522 | 5.6153 |
| STRIP1_S335 | STRIP1 | Q5VSL9 | | 335 | RNMRAASPPASAS | 5.8796 | 6.1729 | 6.2154 | 6.0210 | 6.2823 | 6.2441 | 6.0406 | 6.4449 | 6.4671 | 6.1047 | 6.5318 | 6.2364 | 6.3720 | 6.4367 | 6.1670 | 6.3837 |
| STRIP1_S59 | STRIP1 | Q5VSL9 | | 59 | RNQRKDSEGYSES | 5.7466 | 5.9419 | 6.8408 | 6.5212 | 6.1505 | 6.2730 | 6.3885 | 6.1912 | 6.2566 | 6.4139 | 7.3183 | 7.1285 | 5.8049 | 5.6421 | 5.8260 | 5.5560 |
| STRN_S137 | STRN | O43815 | | 137 | KPPSYDSDEGNET | 5.8466 | 5.9538 | 6.3066 | 6.1486 | 6.3554 | 6.3183 | 6.0935 | 6.3746 | 6.4053 | 6.3354 | 6.2516 | 6.2498 | 6.4145 | 6.7062 | 6.3000 | 5.9397 |
| STRN_S227 | STRN | O43815 | | 376 | LQPSVGSPSRPSS | 6.5501 | 6.3711 | 6.7140 | 6.5095 | 6.1732 | 6.7912 | 6.8360 | 6.9704 | 7.0829 | 6.6210 | 6.6246 | 6.9811 | 4.6268 | 5.0947 | 4.7405 | 5.3129 |
| STRN_S245 | STRN | O43815 | | 245 | SAAADFSDEDEDD | 5.9565 | 6.5774 | 6.2196 | 6.4351 | 6.6163 | 6.4904 | 5.8649 | 6.5616 | 6.6972 | 6.3650 | 5.7257 | 6.0722 | 6.5836 | 6.3727 | 5.3771 | 6.0845 |
| STRN_S259 | STRN | O43815 | | 259 | VDGREKSVIDTST | 6.7675 | 6.9806 | 7.0501 | 7.3773 | 6.6813 | 6.6500 | 6.5677 | 6.3855 | 7.2142 | 6.3238 | 7.1383 | 6.9840 | 4.4622 | 4.9099 | 3.8069 | 4.7007 |
| STRN_S376 | STRN | O43815 | | 227 | KSELTDSASVLDN | 8.1976 | 7.2607 | 6.1677 | 7.0600 | 5.3876 | 5.0593 | 6.0483 | 5.3267 | 6.7268 | 6.1645 | 5.6834 | 6.3262 | 6.2051 | 5.7907 | 6.7544 | 5.8411 |
| STRN3_S229 | STRN3 | Q13033 | | 229 | ILNGGESPKQKGQ | 5.9778 | 6.0705 | 6.6606 | 6.4121 | 6.2086 | 6.1689 | 6.4091 | 5.9601 | 6.5857 | 6.5451 | 6.8688 | 7.1239 | 5.8119 | 5.6330 | 5.3802 | 6.1838 |
| STRN3_S257 | STRN3 | Q13033 | | 257 | LENADDSDEDEEN | 6.3684 | 6.3778 | 6.1733 | 5.8366 | 6.4663 | 6.4945 | 6.2297 | 6.8655 | 6.1418 | 6.7724 | 6.1126 | 6.4963 | 5.7482 | 6.1615 | 5.9076 | 5.8474 |
| STRN3_S335 | STRN3 | Q13033 | | 335 | WDKDDLSPTAEVW | 5.2254 | 6.2546 | 6.7916 | 6.8336 | 6.6194 | 5.2776 | 5.0937 | 5.9622 | 5.5396 | 6.3009 | 6.5542 | 6.5325 | 6.4399 | 6.8669 | 7.1665 | 6.5412 |
| STRN4_S167 | STRN4 | Q9NRL3-3 | | 377 | TGPPPGTPQPRPH | 6.4851 | 6.9037 | 6.5004 | 6.2470 | 5.3605 | 6.0893 | 6.2815 | 6.2502 | 7.1095 | 6.3558 | 7.1308 | 6.9248 | 5.8336 | 5.4126 | 5.2667 | 5.8485 |
| STRN4_S276 | STRN4 | Q9NRL3-3 | | 167 | SVTLENSPLVWKE | 6.3061 | 6.5038 | 6.6749 | 6.6357 | 6.1389 | 6.6799 | 6.6615 | 6.5911 | 6.7338 | 7.2083 | 7.1433 | 7.4054 | 4.6107 | 5.1735 | 4.5281 | 5.0050 |
| STRN4_S342 | STRN4 | Q9NRL3-3 | | 342 | RCTVDGSPHELES | 7.1581 | 7.0815 | 6.9161 | 7.1872 | 6.2699 | 6.3348 | 6.6735 | 7.2487 | 8.1508 | 6.7406 | 6.4510 | 7.2174 | 4.0290 | 4.4376 | 3.8306 | 4.2733 |
| STRN4_T377 | STRN4 | Q9NRL3-3 | | 276 | NCEDEDSDEDDEL | 6.2826 | 6.1655 | 6.6378 | 6.5011 | 5.5096 | 6.4657 | 6.0694 | 7.0583 | 6.4095 | 6.7431 | 6.9131 | 5.9720 | 6.2885 | 5.7254 | 5.4219 | 5.8366 |
| STT3B_S29 | STT3B | Q8TCJ2 | | 29 | LMALGNSRHGHHG | 5.7659 | 6.0845 | 7.7966 | 6.0592 | 7.8567 | 6.4673 | 5.5816 | 5.8759 | 6.2578 | 6.5526 | 6.2266 | 6.4213 | 5.6521 | 6.1214 | 5.6172 | 5.6632 |
| STT3B_S498;S499 | STT3B | Q8TCJ2 | 498;499 | | NA | 5.1998 | 6.3265 | 7.5899 | 9.1669 | 4.6906 | 6.3196 | 8.0540 | 5.4636 | 4.6580 | 6.4191 | 5.9283 | 4.8909 | 6.2991 | 6.0996 | 6.1767 | 6.7175 |
| STUB1_S19 | STUB1 | Q9UNE7 | | 19 | LGAGGGSPEKSPS | 6.6205 | 6.3101 | 6.6119 | 5.9273 | 6.6707 | 6.4561 | 6.2297 | 6.7588 | 6.7697 | 6.5237 | 6.4296 | 6.3570 | 5.9580 | 5.5056 | 5.4487 | 5.4225 |
| STUB1_S19;S23 | STUB1 | Q9UNE7 | | 23 | GGSPEKSPSAQEL | 6.3466 | 6.7440 | 6.4651 | 6.0241 | 5.9680 | 5.8577 | 5.6286 | 6.4703 | 6.3460 | 6.3433 | 6.4771 | 6.3659 | 6.3210 | 6.1536 | 6.1177 | 6.3711 |
| STUB1_S23 | STUB1 | Q9UNE7 | | 273 | FDPVTRSPLTQEQ | 6.2440 | 6.0888 | 6.4085 | 6.7271 | 6.5391 | 6.1853 | 6.4113 | 6.3818 | 6.2333 | 6.5975 | 6.5555 | 6.3529 | 5.8179 | 5.7791 | 5.6709 | 6.0069 |

| | | | | | | | | | | | | | | | | | | | | | |
|---|---|---|---|---|---|---|---|---|---|---|---|---|---|---|---|---|---|---|---|---|---|
| STUB1_S273 | STUB1 | Q9UNE7 | 19;23 | | NA | 6.0344 | 6.2373 | 6.7454 | 6.2513 | 5.7404 | 5.9274 | 6.1394 | 6.2087 | 6.3167 | 7.3458 | 7.9427 | 7.4574 | 5.4356 | 5.5343 | 5.4269 | 5.2562 |
| STX10_S108 | STX10 | O60499 | | 108 | MKDHMVSPTAVAF | 6.6759 | 6.4521 | 6.6288 | 6.9878 | 6.4737 | 6.5604 | 6.7612 | 6.1189 | 5.8221 | 6.6603 | 6.7795 | 6.9086 | 4.6036 | 6.1722 | 4.9546 | 5.4404 |
| STX10_S132 | STX10 | O60499 | | 132 | KPAAQKSPSDLLD | 5.6041 | 6.5054 | 6.0373 | 6.1967 | 6.7868 | 6.4006 | 6.3479 | 7.0199 | 6.7398 | 6.7106 | 6.9545 | 7.1539 | 5.3202 | 5.6943 | 4.9829 | 5.5450 |
| STX17_S289 | STX17 | P56962 | | 289 | MEKLTSSCPDLPS | 5.8522 | 6.1175 | 5.6749 | 6.0028 | 5.9603 | 5.3626 | 6.0351 | 5.5243 | 6.0233 | 5.5694 | 5.6715 | 5.6476 | 8.2891 | 7.5900 | 7.0575 | 7.6219 |
| STX18_S189 | STX18 | Q9P2W9 | | 189 | SEKVSQSPSKDSE | 7.1391 | 7.6879 | 6.7296 | 7.4374 | 6.3248 | 6.6726 | 6.9476 | 6.7324 | 7.0392 | 7.0417 | 7.0354 | 7.0926 | 3.4853 | 4.3453 | 3.9192 | 4.3696 |
| STX4_S117 | STX4 | Q12846 | | 36 | GTARLGSPDEEFF | 6.6520 | 6.7983 | 6.1505 | 6.8007 | 5.6306 | 5.5888 | 7.7459 | 6.4148 | 6.9519 | 7.5775 | 7.2321 | 7.6390 | 3.8373 | 4.7481 | 4.6940 | 5.5384 |
| STX4_S36 | STX4 | Q12846 | | 31 | LVVHPGTARLGSP | 7.3413 | 8.1305 | 8.5023 | 8.5584 | 8.6046 | 3.7741 | 6.3697 | 3.1533 | 6.3031 | 4.9211 | 5.9172 | 5.1947 | 5.5700 | 6.6047 | 6.0645 | 6.7906 |
| STX4_T31 | STX4 | Q12846 | | 117 | ADENYNSVNTRMR | 6.9867 | 6.9247 | 4.7474 | 7.3648 | 8.6913 | 8.2482 | 6.0709 | 5.1167 | 5.8405 | 6.2659 | 6.6499 | 5.2467 | 5.8641 | 6.8804 | 5.1492 | 3.9527 |
| STX6_S129 | STX6 | O43752 | | 129 | LLGDSGSQNWSTG | 5.3463 | 6.4082 | 6.3869 | 5.9370 | 5.9714 | 6.3507 | 5.9766 | 6.2419 | 5.7069 | 6.7979 | 6.0331 | 6.5779 | 6.0009 | 7.0423 | 6.7150 | 6.5069 |
| STX7_S126;S129 | STX7 | O15400 | | 205 | QGDVIDSIEANVE | 6.3968 | 6.0170 | 6.3299 | 6.6032 | 5.8236 | 5.7132 | 6.1042 | 6.0159 | 6.4157 | 6.2817 | 6.5839 | 6.4225 | 6.1552 | 6.8412 | 6.4562 | 5.8397 |
| STX7_S129 | STX7 | O15400 | | 131 | SSRVSGSFPEDSS | 7.0737 | 7.0381 | 7.6067 | 7.8041 | 6.3529 | 6.1092 | 6.5909 | 6.3587 | 7.4778 | 7.2302 | 7.6808 | 7.1366 | 3.7139 | 4.1378 | 3.7991 | 3.8894 |
| STX7_S131 | STX7 | O15400 | | 129 | RASSRVSGSFPED | 5.6500 | 5.7702 | 5.6535 | 5.9392 | 5.7118 | 5.8238 | 5.8605 | 5.8737 | 5.1796 | 5.5249 | 5.8199 | 6.0262 | 8.0778 | 7.8133 | 8.0972 | 7.1784 |
| STX7_S205 | STX7 | O15400 | | 79 | FGSLPTTPSEQRQ | 5.8951 | 5.9996 | 6.0079 | 6.3058 | 6.0933 | 6.6417 | 6.1485 | 6.3440 | 5.8377 | 6.4605 | 6.1571 | 6.2563 | 6.2971 | 6.9031 | 6.3560 | 6.2964 |
| STX7_T79 | STX7 | O15400 | 126;129 | | NA | 4.5915 | 4.5194 | 5.1659 | 6.0090 | 4.9609 | 5.2863 | 5.4710 | 5.9801 | 4.5503 | 4.5295 | 5.3558 | 5.5522 | 9.6977 | 9.0133 | 10.2457 | 9.0711 |
| STXBP3_S323 | STXBP3 | O00186 | | 587 | KPKDKVSLIKDEx | 5.8708 | 6.1932 | 5.0643 | 5.0889 | 6.7479 | 6.2828 | 6.1255 | 6.6043 | 5.5183 | 5.3479 | 5.4287 | 5.5035 | 7.7816 | 7.2417 | 8.0243 | 7.1763 |
| STXBP3_S508 | STXBP3 | O00186 | | 508 | PAVWNGSGAVSAR | 5.5313 | 5.6688 | 5.8124 | 7.6870 | 7.2385 | 6.7392 | 6.8020 | 6.2848 | 8.5003 | 5.8871 | 5.8751 | 5.9080 | 5.4506 | 5.0619 | 5.8991 | 5.6539 |
| STXBP3_S587 | STXBP3 | O00186 | | 323 | ATEGKTSLSALTQ | 5.0825 | 6.2198 | 4.4008 | 5.8065 | 7.4736 | 5.1602 | 4.9694 | 6.0290 | 6.3045 | 5.8348 | 7.9575 | 5.5558 | 7.5760 | 6.6895 | 7.3359 | 7.6043 |
| STXBP4_S10 | STXBP4 | Q6ZWJ1 | | 10 | NTSTVVSPSLLEK | 6.4084 | 6.7927 | 6.6412 | 6.7168 | 6.2715 | 6.5968 | 6.6715 | 6.8732 | 6.7223 | 7.0575 | 7.3215 | 7.1967 | 4.6729 | 4.8160 | 4.3165 | 4.9246 |
| STXBP4_S463 | STXBP4 | Q6ZWJ1 | | 465 | ASQTSLTPLGRNG | 6.1750 | 6.5659 | 6.5227 | 5.9511 | 6.1185 | 6.7498 | 6.7988 | 6.4203 | 7.3348 | 6.7284 | 7.3099 | 7.6870 | 4.8652 | 4.6449 | 4.5865 | 5.5411 |
| STXBP4_T465 | STXBP4 | Q6ZWJ1 | | 463 | VLASQTSLTPLGR | 6.2249 | 6.2473 | 6.3623 | 6.7531 | 5.8261 | 5.5807 | 5.8038 | 5.9337 | 6.0790 | 5.7063 | 6.1259 | 5.3219 | 7.6498 | 7.5348 | 6.2409 | 6.6094 |
| STXBP5_S759 | STXBP5 | Q5T5C0 | | 759 | KMSRKLSLPTDLK | 5.7400 | 5.3523 | 6.3550 | 5.5659 | 6.1222 | 5.7907 | 5.5235 | 5.9441 | 6.7935 | 7.3110 | 6.9717 | 7.0822 | 6.5336 | 6.7983 | 6.1353 | 5.9807 |
| STXBP5_S782 | STXBP5 | Q5T5C0 | | 782 | SRSRSSSVTSIDK | 6.1136 | 5.5734 | 6.1958 | 5.5046 | 6.7897 | 6.2669 | 5.9551 | 6.2840 | 7.5618 | 7.6009 | 6.7129 | 6.6701 | 6.0219 | 6.5363 | 5.0131 | 5.1997 |
| SUB1_S17 | SUB1 | P53999 | | 57 | SSKQSSSRDDNM | 5.7479 | 5.5505 | 5.8859 | 5.3847 | 5.8391 | 6.1302 | 5.6157 | 5.9665 | 5.7793 | 5.8230 | 6.1211 | 5.9228 | 7.4980 | 7.8782 | 7.8025 | 7.0547 |
| SUB1_S57 | SUB1 | P53999 | | 17 | SSSGSDSDSEVDK | 6.0428 | 5.9761 | 6.2704 | 5.4503 | 5.4246 | 5.1822 | 5.7669 | 5.6436 | 9.0892 | 8.8768 | 9.6519 | 8.8965 | 3.8185 | 5.6949 | 3.8399 | 4.3753 |
| SUCLA2_S124 | SUCLA2 | Q9P2R7 | | 124 | GVKIVFSPEEAKA | 5.2998 | 5.3622 | 6.2604 | 6.5118 | 7.2191 | 6.4075 | 7.1869 | 7.2298 | 6.3081 | 5.4041 | 5.5775 | 6.1961 | 5.8132 | 5.8785 | 6.4416 | 6.9034 |
| SUCO_S1224 | SUCO | Q9UBS9-2 | | 1377 | IQTKSGSLPSLHD | 3.7833 | 4.2844 | 3.9192 | 4.3932 | 5.3531 | 5.3807 | 4.9441 | 5.3582 | 3.8708 | 4.0863 | 4.0090 | 3.7600 | 11.6309 | 11.2174 | 12.7628 | 11.2466 |
| SUCO_S1232 | SUCO | Q9UBS9-2 | | 1232 | PLMRSKSLQLTGK | 5.9405 | 6.1556 | 5.5921 | 5.7450 | 5.8765 | 5.8168 | 5.4284 | 5.9054 | 5.5361 | 5.6014 | 5.4451 | 5.2084 | 7.8681 | 7.7982 | 8.3515 | 7.7309 |
| SUCO_S1254 | SUCO | Q9UBS9-2 | | 1254 | VEPLKFSPEKKKK | 5.9336 | 6.1326 | 5.8452 | 6.4016 | 6.8358 | 6.2013 | 6.9768 | 6.4172 | 6.3704 | 5.9055 | 6.8581 | 6.4462 | 6.2870 | 6.7537 | 5.8885 | 4.7467 |
| SUCO_S1377 | SUCO | Q9UBS9-2 | | 1224 | NLKRRTSFPLMRS | 4.1899 | 3.4954 | 3.9441 | 4.0524 | 4.7922 | 3.9052 | 4.0989 | 4.0080 | 5.7288 | 6.0803 | 5.3558 | 5.7974 | 11.4294 | 10.9610 | 11.3493 | 10.8119 |
| SUDS3_S234;S237 | SUDS3 | Q9H7L9 | 234;237 | | NA | 5.6810 | 6.0387 | 6.1922 | 6.8960 | 6.0214 | 6.0237 | 5.8842 | 6.4234 | 6.2896 | 6.1093 | 6.1962 | 6.0369 | 6.0246 | 6.9712 | 6.8153 | 6.3962 |
| SUFU_S346 | SUFU | Q9UMX1 | | 346 | APSRKDSLESDSS | 5.5297 | 6.5278 | 6.1563 | 5.2791 | 7.2439 | 7.1454 | 5.4917 | 4.6860 | 5.3059 | 6.6285 | 7.2349 | 6.2413 | 6.7650 | 6.5143 | 6.3197 | 6.9306 |
| SUGP1_S338 | SUGP1 | Q8IWZ8 | | 409 | QRDVDASPSPLSV | 6.1982 | 5.9203 | 6.4877 | 7.0238 | 6.6929 | 6.7442 | 6.0731 | 6.5466 | 6.6472 | 6.1134 | 6.2969 | 7.1845 | 5.8093 | 5.6653 | 5.0982 | 5.4984 |
| SUGP1_S409 | SUGP1 | Q8IWZ8 | | 338 | PGLKRKSPPEALS | 6.2496 | 6.8004 | 5.6427 | 6.5053 | 6.1559 | 6.0075 | 6.1738 | 6.2896 | 6.3993 | 6.1763 | 5.9791 | 6.9852 | 6.3093 | 5.9619 | 6.0376 | 6.3266 |
| SUGP1_S411 | SUGP1 | Q8IWZ8 | | 411 | DVDASPSPLSVQD | 6.1128 | 6.1491 | 6.5936 | 6.2708 | 5.2911 | 6.1454 | 6.3252 | 6.6076 | 6.3060 | 6.5050 | 7.3769 | 6.8070 | 5.2621 | 5.7584 | 5.6988 | 5.8900 |
| SUGP2_S277 | SUGP2 | Q8IX01 | | 757 | LEASGPSPKPAGV | 5.9280 | 5.7802 | 6.2251 | 6.6384 | 6.0284 | 5.7503 | 6.1430 | 5.7232 | 6.2289 | 6.0756 | 5.9860 | 5.9136 | 6.7678 | 6.6988 | 6.9781 | 7.1345 |
| SUGP2_S315 | SUGP2 | Q8IX01 | | 315 | LPRRKMSFDIIDK | 4.0293 | 4.1174 | 5.0745 | 5.5046 | 9.8596 | 4.7046 | 4.4068 | 5.5382 | 8.0149 | 9.1753 | 7.7768 | 9.6839 | 6.9914 | 6.8277 | 5.8516 | 5.1432 |
| SUGP2_S603 | SUGP2 | Q8IX01 | | 773 | EAPQTSSPCPSAD | 6.0792 | 6.2696 | 7.0385 | 7.0291 | 5.6468 | 5.8586 | 6.0853 | 5.7726 | 6.4054 | 6.8928 | 6.5309 | 6.4079 | 5.8525 | 5.8720 | 5.8778 | 6.3811 |
| SUGP2_S757 | SUGP2 | Q8IX01 | | 277 | IQKNTPSPDVTLG | 6.1445 | 5.7783 | 5.8850 | 6.3714 | 6.0767 | 5.8136 | 6.2057 | 6.0205 | 6.0256 | 6.4558 | 5.9683 | 6.2777 | 6.7881 | 6.8619 | 6.6495 | 6.6774 |
| SUGP2_S773 | SUGP2 | Q8IX01 | | 603 | VIEGSLSPKERTL | 6.1137 | 5.8547 | 5.9533 | 6.6409 | 5.7991 | 5.6956 | 6.4736 | 6.0782 | 6.8782 | 6.3956 | 5.7115 | 6.1550 | 6.6863 | 6.2152 | 6.6707 | 6.6785 |
| SUGP2_S846 | SUGP2 | Q8IX01 | | 262 | PTVNRITPKTQGT | 7.1556 | 7.6572 | 7.3423 | 7.0344 | 5.4879 | 6.0689 | 8.2757 | 5.6941 | 6.8686 | 7.1765 | 7.0456 | 6.7920 | 3.7143 | 4.5921 | 3.3716 | 6.0633 |
| SUGP2_T262 | SUGP2 | Q8IX01 | | 275 | NQIQKNTPSPDVT | 5.9830 | 6.2773 | 6.3016 | 5.9703 | 5.4272 | 6.7420 | 6.9463 | 5.3690 | 5.9735 | 5.9745 | 5.1128 | 5.9639 | 7.4152 | 7.5444 | 6.3831 | 6.6160 |
| SUGP2_T275 | SUGP2 | Q8IX01 | | 846 | SICFTSSPHNLHT | 5.7924 | 7.0566 | 6.9076 | 6.9936 | 7.2086 | 7.7561 | 6.7212 | 5.2606 | 5.5172 | 7.0882 | 7.5512 | 6.6897 | 5.0943 | 4.7473 | 4.5597 | 5.0685 |
| SUGT1_S281 | SUGT1 | Q9Y2Z0 | | 281 | NLYPSSSPYTRNW | 7.3712 | 8.0266 | 6.9918 | 7.2201 | 6.3678 | 7.2819 | 7.3864 | 6.8427 | 7.6395 | 7.3610 | 7.9204 | 7.1029 | 2.7653 | 3.7025 | 2.5881 | 3.4317 |
| SUGT1_T284 | SUGT1 | Q9Y2Z0 | | 284 | PSSSPYTRNWDKL | 7.7793 | 8.2621 | 6.8310 | 7.5541 | 5.8398 | 7.0863 | 7.5223 | 6.9978 | 7.9619 | 7.6401 | 7.6979 | 7.7131 | 2.5428 | 3.3844 | 2.1475 | 3.0398 |
| SUMO3_T37 | SUMO3 | P55854-2 | | 37 | FKIKRHTPLSKLM | 6.7747 | 5.9861 | 6.9322 | 6.0887 | 5.6893 | 6.1317 | 5.7069 | 7.0498 | 5.8797 | 7.3853 | 6.9514 | 8.1387 | 3.8638 | 6.5470 | 4.9749 | 5.8996 |
| SUN1_S114 | SUN1 | O94901-9 | | 333 | TAAHSQSPRLPGR | 6.7557 | 7.3792 | 6.9025 | 8.1833 | 6.3074 | 6.7105 | 6.8157 | 6.8703 | 7.2689 | 7.3682 | 6.8453 | 7.0311 | 3.5389 | 4.3516 | 3.6143 | 4.0571 |
| SUN1_S138 | SUN1 | O94901-9 | | 48 | LDPVFDSPRMSRR | 7.2719 | 7.5036 | 6.6859 | 7.2264 | 6.3213 | 6.5897 | 6.9351 | 6.6842 | 6.8269 | 6.8171 | 6.7250 | 6.7646 | 4.1476 | 4.7263 | 4.0886 | 4.6857 |
| SUN1_S333 | SUN1 | O94901-9 | | 138 | PPVLDESWIREQT | 7.9292 | 8.1386 | 6.9799 | 8.9740 | 7.3187 | 6.7180 | 7.6569 | 6.5188 | 6.8109 | 6.4443 | 6.9753 | 6.3075 | 2.9169 | 4.1068 | 2.7045 | 3.4999 |
| SUN1_S48 | SUN1 | O94901-9 | | 114 | SRQVTSSGVSHGG | 7.1694 | 8.5841 | 7.3462 | 8.2312 | 5.9712 | 6.9023 | 7.4501 | 7.3177 | 6.9147 | 7.4982 | 7.5976 | 6.8847 | 2.7067 | 3.6743 | 2.7175 | 3.0339 |
| SUN1_S79 | SUN1 | O94901-9 | | 79 | ADSGTSSAVSLKN | 6.8267 | 7.3684 | 6.6249 | 7.7237 | 6.2709 | 7.3735 | 7.3522 | 6.5333 | 6.7945 | 7.0879 | 6.9092 | 7.3194 | 3.9766 | 4.7704 | 3.0990 | 3.9695 |
| SUN2_S116 | SUN2 | Q9UH99-2 | | 116 | SESSRASGLVGRK | 5.6639 | 6.3264 | 5.2132 | 5.9034 | 5.5786 | 5.4706 | 5.9155 | 6.8594 | 6.4335 | 7.0054 | 5.8972 | 6.4526 | 6.3471 | 6.5213 | 7.0654 | 7.3465 |
| SUPT16H_S1039 | SUPT16H | Q9Y5B9 | | 1039 | RGSRHSSAPPKKK | 5.8150 | 6.6160 | 6.5268 | 6.6784 | 5.8368 | 6.6385 | 6.5840 | 6.8552 | 6.4239 | 6.5963 | 6.8090 | 7.1360 | 4.6778 | 5.5130 | 5.4365 | 5.8570 |
| SUPT16H_S188 | SUPT16H | Q9Y5B9 | | 650 | GIVKQDSLVINLN | 5.9773 | 5.5194 | 6.1460 | 6.4195 | 5.9539 | 5.7486 | 6.8226 | 6.2540 | 6.2456 | 6.2626 | 6.2402 | 6.0706 | 6.4455 | 6.4008 | 6.4872 | 7.0060 |
| SUPT16H_S650 | SUPT16H | Q9Y5B9 | | 188 | LMKKAASITSEVF | 5.3730 | 5.7477 | 5.5883 | 5.9415 | 5.9483 | 5.5659 | 5.9484 | 5.8525 | 5.4948 | 5.5785 | 5.8028 | 5.2957 | 7.8033 | 8.3171 | 8.1080 | 7.8342 |
| SUPT16H_T903 | SUPT16H | Q9Y5B9 | | 903 | VQSLNWTKIMKTI | 5.7335 | 6.6449 | 7.3725 | 7.4974 | 6.1125 | 5.4653 | 7.4555 | 6.3152 | 6.2463 | 5.7754 | 6.0085 | 5.3018 | 5.6011 | 5.5809 | 6.9414 | 5.9477 |
| SUPT20H_S492 | SUPT20H | Q8NEM7-3 | | 492 | TSSFLKSPTPPPS | 5.6989 | 5.1050 | 6.0351 | 5.7423 | 6.3620 | 6.1025 | 6.2060 | 6.2189 | 6.0360 | 6.2364 | 5.9242 | 5.5917 | 7.2986 | 6.6096 | 7.7272 | 7.1056 |
| SUPT20H_S509 | SUPT20H | Q8NEM7-3 | | 509 | SIPRKSSVDLNQV | 5.9770 | 5.8816 | 5.3018 | 6.2268 | 7.0057 | 5.3139 | 5.3419 | 5.9687 | 6.5516 | 6.4721 | 6.2200 | 7.0580 | 6.7196 | 6.8808 | 6.2594 | 6.8210 |
| SUPT5H_S104 | SUPT5H | O00267-2 | | 662 | GGFAPMSPRISSP | 7.6146 | 6.6973 | 6.3566 | 6.8548 | 5.9727 | 6.0767 | 6.7666 | 5.8774 | 6.7454 | 6.3658 | 6.4120 | 6.5501 | 5.4933 | 5.4542 | 4.9623 | 5.8002 |
| SUPT5H_S32 | SUPT5H | O00267-2 | | 104 | DILEKASNIDNVV | 6.7326 | 7.1354 | 7.2047 | 7.3806 | 6.1701 | 6.1740 | 6.6298 | 5.4947 | 7.1343 | 6.5980 | 6.2282 | 5.7461 | 5.4003 | 5.7042 | 4.8579 | 5.4092 |
| SUPT5H_S32;S36 | SUPT5H | O00267-2 | | 32 | VDEERRSAAGSEK | 4.3375 | 5.2599 | 4.9885 | 4.3101 | 5.2143 | 4.9261 | 4.7882 | 4.9905 | 6.4074 | 7.1606 | 6.3524 | 6.3083 | 8.1986 | 8.5945 | 9.5581 | 8.6051 |
| SUPT5H_S32;Y54 | SUPT5H | O00267-2 | | 824 | DPNNPNTPSRAEE | 5.8325 | 6.6949 | 6.7461 | 6.5422 | 5.8812 | 6.7499 | 6.8004 | 6.6600 | 6.8878 | 7.1689 | 6.8949 | 6.9120 | 4.5527 | 5.5038 | 4.6090 | 5.5637 |
| SUPT5H_S36 | SUPT5H | O00267-2 | | 36 | RRSAAGSEKEEEP | 5.6027 | 6.3328 | 6.0403 | 6.4674 | 6.3197 | 5.7469 | 6.5087 | 5.9470 | 5.7439 | 6.3367 | 6.0243 | 6.0969 | 6.1132 | 6.8216 | 6.8233 | 7.0747 |
| SUPT5H_S662 | SUPT5H | O00267-2 | | 787 | PMYGSQTPLQDGS | 7.1826 | 7.4584 | 8.3142 | 6.4337 | 5.7529 | 6.4572 | 6.4807 | 5.6085 | 7.2284 | 6.8997 | 6.7201 | 6.2823 | 5.2376 | 4.4969 | 4.0720 | 5.3749 |
| SUPT5H_T787 | SUPT5H | O00267-2 | | 802 | PHYGSQTPLHDGS | 7.8594 | 7.8053 | 7.0909 | 6.5415 | 5.6012 | 6.7075 | 7.4177 | 6.3072 | 7.8384 | 7.0636 | 7.9135 | 7.3725 | 3.7858 | 3.9864 | 2.9159 | 3.7931 |
| SUPT5H_T802 | SUPT5H | O00267-2 | 32;36 | | NA | 5.0618 | 5.2667 | 6.1270 | 5.1292 | 5.4865 | 5.7307 | 6.1038 | 5.7491 | 6.5425 | 6.3006 | 6.3043 | 6.1917 | 7.8495 | 7.5209 | 7.2006 | 7.4348 |
| SUPT5H_T824 | SUPT5H | O00267-2 | 32;54 | | NA | 4.8363 | 5.0809 | 5.4404 | 5.3719 | 5.5251 | 5.5831 | 6.1349 | 5.6836 | 6.0788 | 6.6350 | 6.4895 | 6.2619 | 7.5738 | 7.9224 | 7.5249 | 7.8574 |
| SUPT6H_S125 | SUPT6H | Q7KZ85 | | 1718 | SIAGDATPLLDEM | 6.1563 | 6.6326 | 6.6726 | 6.8407 | 6.3434 | 6.1597 | 6.4278 | 5.9942 | 6.3501 | 6.1910 | 6.1672 | 6.4209 | 5.9077 | 6.0389 | 5.9048 | 5.7920 |
| SUPT6H_S1701 | SUPT6H | Q7KZ85 | | 125 | RRVKKMSDDEDDD | 6.2207 | 6.4356 | 6.2619 | 6.3800 | 6.2511 | 5.8383 | 6.6237 | 5.4735 | 6.5816 | 5.9782 | 5.9823 | 5.9393 | 6.5198 | 6.3350 | 6.6677 | 6.5112 |
| SUPT6H_S1701;T1718 | SUPT6H | Q7KZ85 | | 619 | RAKLNITPTKKGR | 6.6765 | 7.1436 | 6.3182 | 7.2988 | 6.1689 | 6.0038 | 6.3900 | 6.1245 | 6.7489 | 6.5450 | 6.2352 | 6.5796 | 5.2272 | 5.4218 | 5.4835 | 5.6346 |
| SUPT6H_S91 | SUPT6H | Q7KZ85 | | 1535 | RTRTPASINATPA | 5.3977 | 6.0322 | 5.8285 | 7.4310 | 6.5769 | 6.6624 | 5.8654 | 6.3574 | 6.6181 | 6.7290 | 5.5044 | 6.9982 | 6.0280 | 6.0829 | 5.9847 | 5.9034 |
| SUPT6H_T1523 | SUPT6H | Q7KZ85 | | 1532 | SSSRTRTPASINA | 6.7120 | 6.5625 | 6.5767 | 7.0451 | 6.3520 | 6.3707 | 6.2469 | 6.1520 | 7.1248 | 6.7412 | 7.3016 | 6.4488 | 4.7896 | 5.5624 | 4.9907 | 5.0233 |
| SUPT6H_T1523;S1527 | SUPT6H | Q7KZ85 | | 91 | KKRKRTSFDDRLE | 5.1313 | 5.2705 | 5.0594 | 5.9068 | 6.3009 | 5.5822 | 6.0254 | 6.1810 | 5.3681 | 5.4789 | 5.1386 | 5.6323 | 8.2098 | 7.6320 | 8.9065 | 8.1763 |
| SUPT6H_T1532 | SUPT6H | Q7KZ85 | | 1701 | RLTPRPSPSPMIE | 5.2338 | 6.1392 | 6.1758 | 6.2924 | 6.6320 | 6.0400 | 6.4787 | 6.6111 | 6.5675 | 6.7730 | 5.4811 | 6.5397 | 6.9854 | 6.5780 | 5.5559 | 5.9164 |
| SUPT6H_T1532;S1535 | SUPT6H | Q7KZ85 | | 1523 | DPVPGITPSSSSR | 6.5884 | 6.0792 | 5.8881 | 6.3620 | 5.6912 | 6.1129 | 7.1144 | 6.7635 | 5.7796 | 6.0060 | 5.9184 | 5.5557 | 6.3822 | 6.7601 | 6.6641 | 6.3342 |
| SUPT6H_T1532;S1535;T1 | SUPT6H | Q7KZ85 | 1532;1535 | | NA | 6.5344 | 6.4070 | 6.1718 | 6.0554 | 6.1885 | 6.3278 | 6.4376 | 6.1123 | 6.5120 | 6.7312 | 6.3880 | 6.7182 | 5.9149 | 6.0502 | 5.6772 | 5.7735 |
| SUPT6H_T1535 | SUPT6H | Q7KZ85 | 1532;1535;15 | | NA | 7.1919 | 6.5960 | 6.5468 | 7.0467 | 6.6919 | 5.9187 | 6.3287 | 5.4834 | 5.4534 | 5.6412 | 5.8876 | 6.6587 | 7.0772 | 6.6481 | 5.9260 | 5.9036 |
| SUPT6H_T1718 | SUPT6H | Q7KZ85 | 1523;1527 | | NA | 6.5513 | 5.8981 | 6.7661 | 6.1581 | 6.6921 | 5.1268 | 6.3391 | 6.4437 | 7.1667 | 6.3374 | 6.0160 | 6.5807 | 5.3572 | 6.4724 | 6.4085 | 6.5855 |
| SUPT6H_T619 | SUPT6H | Q7KZ85 | 1701;1718 | | NA | 5.7639 | 6.1511 | 6.0653 | 6.8735 | 5.2384 | 5.9297 | 5.8259 | 7.1475 | 6.0967 | 6.5894 | 6.3656 | 5.7912 | 6.0869 | 7.2469 | 6.4450 | 6.3830 |
| SUPT7L_S108 | SUPT7L | O94864 | | 108 | LPSCPGSPPLPDD | 5.9372 | 5.8219 | 6.2439 | 6.1168 | 6.0185 | 5.5359 | 6.0117 | 6.0416 | 6.2514 | 6.3592 | 6.3085 | 6.3172 | 6.9266 | 6.8504 | 6.5793 | 6.6800 |
| SUPT7L_S323;S334 | SUPT7L | O94864 | 323;334 | | NA | 5.5539 | 5.7409 | 6.2355 | 5.8778 | 5.8506 | 6.5438 | 5.8303 | 5.9611 | 6.0117 | 5.9791 | 5.8468 | 6.2217 | 7.3894 | 6.5555 | 7.9092 | 6.4927 |
| SURF2_S163 | SURF2 | Q15527 | | 163 | DEGGAASDDSMTD | 5.6907 | 6.0276 | 8.0271 | 7.3631 | 5.9283 | 6.6917 | 6.5075 | 6.7523 | 6.0470 | 5.7925 | 7.1013 | 6.8250 | 4.9103 | 5.8846 | 4.7601 | 5.6907 |
| SURF2_T190;T195 | SURF2 | Q15527 | 190;195 | | NA | 5.8522 | 6.3017 | 8.1843 | 6.2016 | 5.9282 | 6.4723 | 6.9225 | 6.2665 | 5.7830 | 5.7009 | 6.2442 | 6.1031 | 5.1557 | 6.3677 | 6.1654 | 6.3507 |
| SURF6_S138 | SURF6 | O75683 | | 138 | GSAKELSPAALEK | 6.2513 | 6.7785 | 6.1003 | 7.2339 | 5.9005 | 5.7384 | 6.4703 | 6.2320 | 6.7820 | 6.3964 | 6.1427 | 6.5524 | 5.1544 | 6.2561 | 5.9065 | 6.1043 |
| SURF6_S206 | SURF6 | O75683 | | 206 | FNKVEVSEDEPAS | 6.5445 | 6.9222 | 7.5001 | 7.7775 | 6.9077 | 7.0660 | 7.2570 | 7.1124 | 5.5511 | 5.2055 | 5.4067 | 5.4891 | 4.9111 | 5.7471 | 5.1148 | 5.4871 |
| SURF6_T229 | SURF6 | O75683 | | 229 | RVKGNLTPLTGRN | 7.8848 | 8.6325 | 6.5907 | 7.4363 | 5.9408 | 7.1176 | 7.8694 | 7.2845 | 7.9098 | 7.4166 | 7.5696 | 7.4635 | 2.2458 | 3.3411 | 2.2074 | 3.0894 |
| SUV39H1_S402 | SUV39H1 | O43463-2 | | 402 | LAGLPGSPKKRVR | 6.6386 | 6.6747 | 6.4183 | 7.1049 | 6.5457 | 5.9041 | 6.9714 | 6.2050 | 6.7883 | 6.5703 | 6.7540 | 6.7606 | 4.9981 | 5.4062 | 4.8562 | 5.4037 |
| SUZ12_S546 | SUZ12 | Q15022 | | 583 | QEMEVDSEDEKDP | 6.3652 | 6.3488 | 6.4442 | 7.1456 | 6.0219 | 5.7739 | 6.3081 | 5.8234 | 6.3514 | 6.2276 | 6.2598 | 6.3582 | 6.1956 | 6.0385 | 5.9960 | 6.3417 |
| SUZ12_S583 | SUZ12 | Q15022 | | 546 | MSEFLESEDGEVE | 5.7808 | 5.9039 | 6.8984 | 6.4439 | 6.2223 | 5.8021 | 6.3292 | 6.0218 | 6.3480 | 6.4342 | 6.3549 | 6.5179 | 6.1735 | 6.4549 | 6.0437 | 6.2706 |
| SUZ12_S695 | SUZ12 | Q15022 | | 695 | EKGESASPANEEI | 6.0043 | 7.4143 | 6.1557 | 5.0965 | 6.4375 | 5.5230 | 6.5537 | 6.2404 | 6.0747 | 6.6407 | 6.0043 | 6.5785 | 6.5237 | 6.1774 | 6.6354 | 5.9398 |
| SVIL_S1093 | SVIL | O95425-3 | | 245 | FTEVPRSPKHAHS | 5.9889 | 5.8986 | 6.0037 | 6.1242 | 6.1325 | 6.0084 | 6.0997 | 5.8100 | 6.1266 | 6.1065 | 6.0198 | 5.9349 | 6.9376 | 7.0039 | 6.9942 | 6.8103 |
| SVIL_S1295 | SVIL | O95425-3 | | 1295 | DYNMPRSPVEMDE | 5.7974 | 5.9359 | 5.9219 | 5.8715 | 6.2797 | 6.0353 | 6.3188 | 6.0108 | 5.7389 | 5.6838 | 5.7131 | 5.5991 | 7.0983 | 7.2879 | 7.6289 | 7.0789 |
| SVIL_S193 | SVIL | O95425-3 | | 1093 | GEGLLDSPSKTMS | 7.0381 | 6.7214 | 6.4780 | 6.5980 | 6.9254 | 6.5678 | 6.8752 | 6.3804 | 7.3849 | 6.7354 | 6.5924 | 6.2376 | 5.1002 | 5.5198 | 4.2167 | 4.6290 |
| SVIL_S221 | SVIL | O95425-3 | | 221 | RQAHDLSPAAESS | 5.8783 | 6.1406 | 6.0411 | 6.2840 | 6.0994 | 6.5703 | 6.3578 | 6.3578 | 5.8201 | 5.9347 | 6.0785 | 6.0274 | 6.5116 | 7.2161 | 6.7348 | 6.5273 |
| SVIL_S238;S245 | SVIL | O95425-3 | | 270 | FGDPQLSPEARPR | 5.7520 | 5.9387 | 5.6939 | 6.2057 | 6.1701 | 5.9162 | 5.7819 | 5.9582 | 5.8303 | 5.6277 | 5.4540 | 5.4697 | 7.6570 | 7.4532 | 7.5499 | 7.5415 |
| SVIL_S245 | SVIL | O95425-3 | | 285 | MNARYQTQPVTLG | 9.8758 | 7.9504 | 8.6093 | 7.6191 | 7.6725 | 5.3163 | 6.1797 | 5.2157 | 5.6909 | 4.6504 | 5.0697 | 4.1005 | 5.8098 | 6.2860 | 5.2034 | 4.7504 |
| SVIL_S261;S270 | SVIL | O95425-3 | | 680 | FREMEKSFDEQNV | 7.4167 | 7.7251 | 8.4723 | 6.3838 | 6.1555 | 6.1703 | 6.3082 | 6.5154 | 6.2764 | 7.8223 | 6.4633 | 6.5492 | 3.6359 | 5.6405 | 4.4257 | 4.0395 |
| SVIL_S270 | SVIL | O95425-3 | | 630 | TSERFRTQPITSA | 9.1826 | 7.6387 | 7.1890 | 6.6506 | 6.0463 | 6.1395 | 6.4909 | 5.5261 | 6.0665 | 5.7403 | 5.9182 | 5.8135 | 5.3112 | 6.0349 | 4.7444 | 5.5073 |
| SVIL_S50 | SVIL | O95425-3 | | 893 | SIENSDSPVRSIL | 6.9274 | 6.1278 | 6.6002 | 6.6568 | 6.8074 | 6.7625 | 7.0419 | 7.0818 | 6.4194 | 7.3025 | 6.1484 | 6.8699 | 4.1821 | 5.4214 | 4.7887 | 4.8718 |
| SVIL_S680 | SVIL | O95425-3 | | 742 | VMARLPSPTVARS | 5.9652 | 6.5187 | 6.4098 | 6.5584 | 6.2560 | 6.1365 | 6.0335 | 6.2799 | 6.5555 | 6.5107 | 6.1657 | 6.2649 | 5.6792 | 6.3055 | 6.1174 | 6.2431 |
| SVIL_S742 | SVIL | O95425-4 | | 736 | LQDRSLTQPITTE | 8.4200 | 7.1390 | 7.0384 | 7.4114 | 6.6810 | 6.0805 | 6.3855 | 5.8030 | 5.9992 | 5.3349 | 5.8812 | 5.8239 | 5.6282 | 5.7032 | 5.0646 | 5.6059 |
| SVIL_S878 | SVIL | O95425-3 | | 941 | GMEKYGSFEEAEA | 6.0234 | 6.2963 | 8.2563 | 6.5172 | 6.4436 | 6.6000 | 5.8209 | 6.4929 | 6.2904 | 6.9281 | 7.1740 | 6.3739 | 5.1351 | 5.6749 | 4.5677 | 5.4054 |
| SVIL_S893 | SVIL | O95425-3 | | 50 | RASDPASPHIGRS | 6.6221 | 7.5702 | 6.4939 | 7.5758 | 6.3223 | 6.9124 | 6.9402 | 7.6060 | 6.7620 | 6.4401 | 6.6985 | 6.4109 | 3.9461 | 4.9978 | 3.8804 | 4.8214 |
| SVIL_S941 | SVIL | O95425-3 | | 193 | LHVGDGSSDPEVL | 5.6289 | 8.0603 | 5.6171 | 6.1827 | 5.7104 | 5.9785 | 6.1248 | 7.0390 | 6.7130 | 6.2030 | 4.3719 | 6.8623 | 5.6485 | 5.4600 | 6.2849 | 5.3146 |
| SVIL_T630 | SVIL | O95425-3 | | 878 | PLDHNASATDYKF | 7.0067 | 7.1626 | 7.3266 | 7.9140 | 6.6284 | 6.8294 | 6.6328 | 7.6176 | 7.6640 | 6.6140 | 7.3437 | 6.2195 | 3.8651 | 5.0689 | 3.4119 | 4.4950 |
| SVIL_T736 | SVIL | O95425-3 | 261;270 | | NA | 6.1441 | 6.2587 | 6.5870 | 6.4807 | 6.2661 | 6.3295 | 6.2161 | 6.0401 | 6.3397 | 6.1611 | 6.2468 | 6.2115 | 5.9393 | 6.4196 | 6.0519 | 6.3078 |
| SVIL_T825 | SVIL | O95425-3 | 238;245 | | NA | 6.0814 | 5.8930 | 6.9388 | 6.0164 | 5.6518 | 5.7431 | 7.2520 | 5.5546 | 5.9567 | 6.0186 | 6.2273 | 5.8564 | 7.0139 | 6.7964 | 6.2891 | 6.7105 |
| SWI5_S141 | SWI5 | Q1ZZU3 | | 141 | CRGAFRSPRPLPK | 7.0174 | 6.9479 | 6.4289 | 7.2725 | 5.8896 | 5.8569 | 6.5893 | 6.4537 | 6.7564 | 6.3565 | 6.7614 | 7.2250 | 4.9640 | 5.3639 | 4.9363 | 5.1803 |
| SYAP1_S269 | SYAP1 | Q96A49 | | 269 | EEEISTSPGVSEF | 6.3680 | 6.5640 | 6.8688 | 6.3513 | 6.6273 | 6.4003 | 6.3832 | 6.0147 | 6.3831 | 6.6159 | 6.7678 | 7.0385 | 5.4034 | 5.6191 | 5.0062 | 5.0884 |
| SYAP1_T248 | SYAP1 | Q96A49 | | 248 | EAVRPKTPPVVIK | 6.6467 | 6.2429 | 6.4844 | 5.5565 | 6.6331 | 6.4487 | 6.0198 | 7.0337 | 6.5012 | 6.3604 | 6.5035 | 6.5424 | 6.1943 | 5.8773 | 5.5233 | 5.4319 |
| SYAP1_T268 | SYAP1 | Q96A49 | | 268 | DEEEISTSPGVSE | 6.3470 | 6.4444 | 7.4274 | 6.4286 | 6.6245 | 6.8785 | 6.1624 | 6.2438 | 6.2336 | 6.7427 | 7.4177 | 7.1443 | 4.8767 | 5.6266 | 4.6403 | 4.7616 |
| SYCE1_S340 | SYCE1 | Q8N0S2 | | 340 | TLHPDLSPRGFQE | 6.1406 | 5.8531 | 6.5677 | 5.8250 | 6.1955 | 6.0903 | 5.8250 | 7.2489 | 6.4653 | 6.5887 | 7.4114 | 6.8128 | 6.0408 | 5.7677 | 5.0324 | 6.1349 |
| SYDE1_S645 | SYDE1 | Q6ZW31 | | 645 | GRGGPESPPSNRY | 6.6765 | 7.0822 | 7.2628 | 6.8732 | 5.9768 | 6.3589 | 6.3943 | 6.6212 | 7.1117 | 6.8774 | 6.9686 | 7.3073 | 4.4801 | 5.0066 | 4.2094 | 4.7930 |
| SYDE2_S104 | SYDE2 | Q5VT97 | | 212 | KGRARGTAPKVTG | 4.1713 | 4.4707 | 7.7990 | 6.5268 | 7.1327 | 6.8747 | 8.2291 | 5.6962 | 5.8669 | 6.3346 | 6.5635 | 6.3045 | 5.4163 | 6.3253 | 6.1405 | 6.1880 |
| SYDE2_S317 | SYDE2 | Q5VT97 | | 104 | PFQRWPSDSWIRC | 6.5544 | 6.6279 | 7.0113 | 6.4930 | 5.8536 | 6.1596 | 6.2378 | 6.6737 | 6.4240 | 6.1043 | 6.9294 | 6.7320 | 5.3366 | 5.8587 | 5.3195 | 5.6842 |
| SYDE2_T212 | SYDE2 | Q5VT97 | | 317 | RSHLSISPVSLPK | 8.0689 | 7.6247 | 6.6319 | 5.9886 | 6.1391 | 7.9160 | 6.9207 | 8.1830 | 7.7631 | 5.3744 | 6.7921 | 5.6366 | 3.5635 | 4.3500 | 4.0151 | 5.0322 |
| SYMPK_S1243 | SYMPK | Q92797 | | 494 | LIKRRLSAQGQAI | 5.1995 | 4.9784 | 5.1072 | 5.3914 | 6.9871 | 5.5021 | 5.1884 | 5.7518 | 7.4280 | 7.2543 | 6.7642 | 7.0379 | 7.5045 | 7.4383 | 6.0330 | 6.4339 |
| SYMPK_S1243;T1257;S12 | SYMPK | Q92797 | | 1243 | TLKEERSPQTLAP | 6.1807 | 6.2043 | 5.9907 | 6.2532 | 6.0312 | 6.0987 | 6.3268 | 6.1714 | 6.1994 | 6.3130 | 6.0222 | 6.2615 | 6.3101 | 6.7381 | 6.2052 | 6.6934 |
| SYMPK_S494 | SYMPK | Q92797 | | 547 | KKIFRLSDVLKPL | 7.3949 | 7.2103 | 6.4366 | 6.8468 | 6.2472 | 6.1298 | 6.4345 | 6.3122 | 6.2480 | 6.1385 | 5.5862 | 6.4817 | 5.0087 | 5.4526 | 5.7071 | 6.3650 |
| SYMPK_S547 | SYMPK | Q92797 | | 870 | TDKVPPSPELVKR | 6.4548 | 7.0360 | 6.3519 | 7.2274 | 6.4462 | 6.2102 | 6.2612 | 6.6051 | 6.0428 | 6.6583 | 7.0857 | 7.1525 | 4.9796 | 5.3077 | 4.6263 | 5.5541 |
| SYMPK_S870 | SYMPK | Q92797 | | 1083 | AHVRSFTPHQQAH | 6.1462 | 6.0387 | 5.9130 | 6.0864 | 7.3144 | 6.6796 | 7.2777 | 7.4462 | 6.6003 | 6.6259 | 6.6686 | 6.1413 | 5.1110 | 4.8815 | 5.7311 | 5.3381 |
| SYMPK_T1083 | SYMPK | Q92797 | 1243;1257;12 | | NA | 6.8091 | 6.4173 | 5.7078 | 6.1708 | 5.6489 | 6.1059 | 6.0532 | 5.6303 | 5.8806 | 5.9060 | 5.6285 | 5.8196 | 6.8537 | 6.7906 | 7.0623 | 7.5154 |
| SYNC_S325 | SYNC | Q9H7C4 | | 325 | QESRRLSAQFENL | 6.2271 | 5.6991 | 6.4729 | 5.8612 | 7.1293 | 5.7020 | 5.3226 | 6.7771 | 6.3338 | 7.6117 | 7.4871 | 7.0171 | 5.8767 | 6.4930 | 5.3536 | 4.7257 |
| SYNCRIP_S151 | SYNCRIP | O60506 | | 159 | YSGQQPSVGTEIF | 5.9789 | 5.9401 | 6.2914 | 6.8025 | 6.4678 | 6.0503 | 6.4575 | 6.1257 | 6.7080 | 6.2969 | 5.9774 | 6.2982 | 5.9378 | 6.3705 | 6.1305 | 6.1664 |
| SYNCRIP_S159 | SYNCRIP | O60506 | | 249 | NRLFVGSIPKSKT | 5.9151 | 6.8321 | 6.5545 | 5.9155 | 6.0248 | 6.4230 | 6.3423 | 6.6199 | 6.0067 | 6.1569 | 6.4240 | 6.2442 | 5.8999 | 6.4455 | 6.1865 | 6.0089 |
| SYNCRIP_S249 | SYNCRIP | O60506 | | 151 | GGPPPDSVYSGQQ | 5.8625 | 6.8177 | 5.9035 | 7.1451 | 6.5529 | 6.1043 | 6.1413 | 6.3105 | 6.0978 | 4.7195 | 5.2232 | 4.9989 | 7.1832 | 6.9168 | 7.6908 | 6.3321 |
| SYNE1_S5812 | SYNE1 | Q8NF91-4 | | 8312 | NIIRSKTPTGPEL | 6.3986 | 6.2803 | 5.7790 | 6.3506 | 6.3063 | 5.9339 | 6.3014 | 6.2082 | 6.1428 | 6.0839 | 5.9199 | 5.9311 | 6.5780 | 6.6857 | 6.6211 | 6.4792 |
| SYNE1_S6104 | SYNE1 | Q8NF91-4 | | 5812 | PACRSPSPVANTD | 5.9393 | 6.0563 | 5.9761 | 6.6671 | 6.0952 | 5.8541 | 5.9788 | 6.0946 | 6.3981 | 6.0058 | 5.6673 | 6.1688 | 6.8061 | 7.0264 | 6.1409 | 7.1250 |
| SYNE1_S6138 | SYNE1 | Q8NF91-4 | | 8234 | MSRALPSEDEEGQ | 7.2606 | 7.1876 | 6.3318 | 6.2515 | 6.4164 | 6.0672 | 6.6779 | 6.1826 | 6.5176 | 5.9339 | 5.6693 | 5.4443 | 6.3765 | 6.0384 | 5.6480 | 5.9962 |
| SYNE1_S8152 | SYNE1 | Q8NF91-4 | | 6104 | LRRLKGSLLELQR | 4.2819 | 4.7172 | 4.2345 | 4.0892 | 4.0405 | 4.3960 | 4.3686 | 4.5445 | 3.9761 | 4.7501 | 4.2970 | 4.5848 | 12.4657 | 11.9592 | 12.5099 | 10.7849 |
| SYNE1_S8187 | SYNE1 | Q8NF91-4 | | 8152 | DDEHDLSDRELEL | 6.0299 | 6.3978 | 6.2665 | 5.5939 | 7.0877 | 6.7062 | 5.7401 | 6.0572 | 6.3024 | 6.4052 | 6.4226 | 6.3396 | 6.0237 | 6.1042 | 6.5229 | 6.0002 |
| SYNE1_S8234 | SYNE1 | Q8NF91-4 | | 6138 | DLTATQSPGVQEW | 5.9647 | 7.1357 | 6.7612 | 6.6956 | 6.0923 | 6.5248 | 6.6845 | 5.8154 | 6.4697 | 6.4271 | 6.2946 | 6.8184 | 5.4548 | 6.0728 | 5.1110 | 5.6773 |
| SYNE1_T8312 | SYNE1 | Q8NF91-4 | | 8187 | QPSSNLSLSLAQP | 5.8166 | 6.5776 | 6.4790 | 6.2167 | 6.6413 | 7.0368 | 6.0026 | 6.0571 | 6.3871 | 6.4276 | 6.1203 | 6.5331 | 6.0003 | 6.1817 | 5.6221 | 5.9003 |
| SYNE2_S2276 | SYNE2 | Q8WXH0-2 | | 4108 | KLNRRGSMSYLAA | 6.1429 | 6.1663 | 6.3016 | 6.4425 | 6.2885 | 5.9824 | 6.0292 | 6.2412 | 6.5015 | 6.2674 | 5.8357 | 5.9936 | 6.8931 | 6.4075 | 6.3569 | 6.1496 |
| SYNE2_S4108 | SYNE2 | Q8WXH0-2 | | 2276 | KEFVSFSDKPVDQ | 7.2285 | 5.9722 | 5.7517 | 6.4782 | 6.2450 | 5.6090 | 7.1587 | 4.7873 | 6.0055 | 5.6803 | 5.5931 | 4.9808 | 6.9382 | 6.1435 | 8.0423 | 7.3856 |
| SYNE3_S763 | SYNE3 | Q6ZMZ3 | | 763 | QRMEVDSGKKMVF | 6.8841 | 7.7850 | 6.4530 | 7.6498 | 5.8366 | 6.9424 | 7.5421 | 6.6155 | 6.4941 | 7.6614 | 8.0292 | 7.1934 | 3.2236 | 4.5280 | 3.3889 | 3.7731 |
| SYNJ1_S1049;S1053 | SYNJ1 | O43426 | | 1392 | QENMRSSPNPFIT | 3.5655 | 5.5464 | 5.9321 | 5.4956 | 5.8222 | 6.2039 | 6.0843 | 6.1160 | 5.9492 | 6.5081 | 6.3470 | 6.1487 | 6.6736 | 7.4000 | 7.1834 | 7.2242 |
| SYNJ1_S1178 | SYNJ1 | O43426 | | 1565 | TLASKASPTLDFT | 5.4573 | 5.7841 | 6.5544 | 6.0835 | 5.8094 | 6.0516 | 6.2860 | 6.4752 | 6.2034 | 6.4522 | 6.3671 | 6.0469 | 6.5277 | 6.9423 | 6.5474 | 6.4117 |
| SYNJ1_S1345 | SYNJ1 | O43426 | | 1345 | QLSVQTSPVPTPD | 5.4032 | 5.2193 | 5.8187 | 6.1150 | 6.5196 | 6.2436 | 5.9614 | 6.3962 | 6.2313 | 6.3067 | 6.4198 | 6.3617 | 6.3550 | 7.2014 | 6.7320 | 7.6154 |
| SYNJ1_S1392 | SYNJ1 | O43426 | | 1178 | IGRSQPSPQAGLA | 5.9375 | 6.3538 | 7.3288 | 6.4030 | 5.9495 | 6.7435 | 5.4574 | 6.5595 | 6.3498 | 7.1186 | 7.0707 | 6.5374 | 5.5199 | 5.5983 | 5.7047 | 6.2674 |
| SYNJ1_S1439 | SYNJ1 | O43426 | | 1439 | EEPVTISPPPSLQ | 7.4635 | 7.0001 | 5.6793 | 6.3305 | 6.1570 | 7.3104 | 8.1147 | 6.9797 | 7.7963 | 8.1719 | 7.3716 | 6.1549 | 2.9458 | 4.5116 | 2.9458 | 5.0669 |
| SYNJ1_S1565 | SYNJ1 | O43426 | 1049;1053 | | NA | 5.8584 | 5.6945 | 6.3065 | 5.7081 | 5.5818 | 6.9037 | 5.8179 | 6.7060 | 6.3212 | 5.9533 | 6.2085 | 6.5232 | 5.9634 | 6.6168 | 6.8734 | 6.9634 |
| SYNJ2_S1191 | SYNJ2 | O15056 | | 1251 | RTGKPLSPEEQFE | 5.6157 | 5.6743 | 5.8101 | 6.4239 | 6.7740 | 6.4178 | 5.9180 | 5.9649 | 6.3177 | 5.9259 | 6.0075 | 6.3277 | 7.1154 | 7.3485 | 6.1279 | 6.2307 |
| SYNJ2_S1251 | SYNJ2 | O15056 | | 1191 | EARGGASEEALSA | 4.2568 | 3.9887 | 5.1173 | 5.2449 | 5.4293 | 4.9019 | 5.5212 | 5.3304 | 4.4418 | 5.2174 | 5.0464 | 5.0482 | 8.9339 | 9.9659 | 10.9709 | 10.5852 |
| SYNJ2BP_S21 | SYNJ2BP | P57105 | | 21 | NLTRGPSGLGFNI | 2.8617 | 3.2153 | 3.6285 | 3.5247 | 3.0809 | 4.0371 | 4.3289 | 4.4354 | 3.3028 | 3.2275 | 3.3394 | 3.6129 | 13.8158 | 14.1504 | 15.8863 | 13.5525 |
| SYNM_S1044 | SYNM | O15061 | | 598 | SPKGLQTPVKDAG | 7.2516 | 7.0309 | 6.2618 | 6.8373 | 7.1595 | 6.8705 | 6.7041 | 6.7142 | 7.1019 | 6.6048 | 6.3134 | 6.1360 | 4.9503 | 5.0375 | 4.3725 | 4.6538 |
| SYNM_S1044;S1049 | SYNM | O15061 | | 777 | EEVEDVSPGPWGL | 5.1804 | 5.2997 | 5.2086 | 5.5541 | 6.0096 | 5.8593 | 6.2683 | 5.8828 | 5.5888 | 5.2243 | 5.2793 | 5.5363 | 8.6794 | 8.0309 | 8.1493 | 8.2489 |
| SYNM_S1163 | SYNM | O15061 | | 1202 | CPEAWGSPEPGPA | 5.7124 | 6.1918 | 6.6212 | 6.3576 | 6.3672 | 6.2323 | 6.2547 | 6.1758 | 6.4417 | 6.8235 | 6.9705 | 6.4458 | 5.8567 | 6.1861 | 5.4586 | 5.9042 |
| SYNM_S1181 | SYNM | O15061 | | 429 | TNVRTFSPTYGLL | 6.8825 | 6.7576 | 6.4352 | 6.9465 | 6.4029 | 6.4263 | 6.3996 | 6.4365 | 6.7674 | 6.8836 | 6.7210 | 6.3268 | 5.2822 | 5.5349 | 5.4989 | 4.8481 |
| SYNM_S1202 | SYNM | O15061 | | 653 | LKQFTQSPETEAS | 6.3224 | 6.6890 | 6.0773 | 6.8643 | 6.1039 | 6.0982 | 6.2717 | 6.1335 | 6.2386 | 6.4660 | 6.4992 | 6.2165 | 5.8891 | 6.1508 | 5.8975 | 6.0822 |

| | | | | | | | | | | | | | | | | | | | | |
|---|---|---|---|---|---|---|---|---|---|---|---|---|---|---|---|---|---|---|---|---|
| SYNM_S429 | SYNM | O15061 | 1094 | HSSGQRTPQGPVS | 5.9246 | 6.4163 | 6.6303 | 6.1253 | 6.2900 | 6.2792 | 5.9925 | 5.3314 | 6.2671 | 7.0445 | 6.6502 | 6.7047 | 5.9549 | 6.7233 | 5.6483 | 6.0176 |
| SYNM_S653 | SYNM | O15061 | 1044 | SLSRQRSPAPGSP | 5.8327 | 5.9162 | 6.5023 | 6.4928 | 6.2639 | 5.9645 | 6.3313 | 6.0011 | 6.5431 | 6.5659 | 6.1302 | 5.9371 | 6.1108 | 6.8242 | 6.0681 | 6.5157 |
| SYNM_S754 | SYNM | O15061 | 1181 | TLGPGQSPLSREV | 6.7887 | 6.6111 | 6.7621 | 7.0857 | 6.7747 | 6.5243 | 6.2821 | 6.8493 | 7.1129 | 6.6724 | 6.8859 | 6.3962 | 4.7329 | 5.4880 | 4.5704 | 4.4635 |
| SYNM_S757 | SYNM | O15061 | 1163 | DLSQAASPTGASR | 5.9399 | 5.9901 | 5.9871 | 7.1548 | 6.6158 | 6.3908 | 5.8710 | 6.6158 | 6.1492 | 6.4516 | 6.2109 | 6.5337 | 6.2470 | 6.0611 | 5.5137 | 6.2674 |
| SYNM_S777 | SYNM | O15061 | 936 | PHEFHTSMKGISS | 5.9500 | 6.1249 | 7.0310 | 6.3303 | 6.6472 | 6.6409 | 5.8710 | 6.5019 | 7.3358 | 7.0253 | 7.1228 | 6.7521 | 5.4062 | 6.1761 | 4.4397 | 4.6450 |
| SYNM_S913 | SYNM | O15061 | 913 | WKEQARSGEFHAE | 4.8093 | 5.2566 | 4.3766 | 4.8719 | 4.7073 | 5.3760 | 4.3824 | 5.3017 | 3.7166 | 3.9468 | 4.4684 | 4.7976 | 12.3620 | 10.0702 | 11.8957 | 9.6608 |
| SYNM_S936 | SYNM | O15061 | 754 | IVEEPVSYVSGEK | 6.3925 | 6.9036 | 7.0720 | 7.4486 | 6.2150 | 6.4506 | 6.3951 | 6.3253 | 5.7733 | 6.5537 | 7.6699 | 7.0570 | 4.8665 | 4.0520 | 4.5395 | 4.4853 |
| SYNM_T1094 | SYNM | O15061 | 757 | EPVSYVSGEKPEE | 6.2809 | 6.8862 | 7.0748 | 6.1708 | 6.5166 | 6.2005 | 6.6457 | 6.0321 | 6.5951 | 6.3606 | 7.4784 | 6.9445 | 5.5310 | 5.2292 | 5.0268 | 5.0268 |
| SYNM_T1094;S1106 | SYNM | O15061 | 1044;1049 | NA | 6.2453 | 4.9654 | 5.8827 | 6.6270 | 5.8214 | 6.6649 | 6.6452 | 5.7507 | 5.1143 | 5.7885 | 6.4284 | 5.5584 | 7.5724 | 7.3827 | 6.7603 | 6.7925 |
| SYNM_T598 | SYNM | O15061 | 1094;1106 | NA | 6.3488 | 5.3759 | 6.7989 | 5.4184 | 6.2791 | 7.0787 | 6.6718 | 6.0379 | 6.6414 | 6.2101 | 5.9415 | 5.8090 | 6.7627 | 6.5899 | 5.8077 | 6.2283 |
| SYNPO_S281 | SYNPO | Q8N3V7-2 | 575 | PSLFVLSPIKEPA | 6.7370 | 6.1562 | 5.8109 | 6.4220 | 6.8409 | 6.5718 | 6.4718 | 6.4395 | 6.5999 | 6.5586 | 6.5651 | 6.1792 | 5.9214 | 5.6422 | 5.2036 | 5.8799 |
| SYNPO_S441 | SYNPO | Q8N3V7-2 | 784 | ENPRPFSPPRAPP | 6.3786 | 6.4034 | 5.9161 | 6.3248 | 6.1516 | 5.9334 | 6.0716 | 6.0615 | 6.5007 | 6.2961 | 6.1718 | 5.9857 | 6.8546 | 6.4724 | 6.3043 | 6.1735 |
| SYNPO_S510 | SYNPO | Q8N3V7-2 | 510 | ELARCPSPTMSLP | 5.3113 | 6.0866 | 5.8276 | 5.7150 | 6.0593 | 6.2180 | 5.4861 | 6.5143 | 6.0364 | 6.5500 | 6.2439 | 5.9626 | 7.6552 | 6.8003 | 7.2497 | 6.2837 |
| SYNPO_S510;S514 | SYNPO | Q8N3V7-2 | 871 | EDSGAKSPGILGY | 6.4399 | 6.6195 | 6.5035 | 6.3768 | 6.2741 | 6.6487 | 6.1902 | 6.6805 | 6.0471 | 5.4182 | 5.9379 | 5.9506 | 6.7640 | 5.7980 | 5.9999 | 6.3511 |
| SYNPO_S575 | SYNPO | Q8N3V7-2 | 441 | APRDRASPAAAEE | 5.8429 | 6.5461 | 5.8321 | 6.1549 | 6.3650 | 6.3633 | 6.1936 | 6.1160 | 6.4960 | 6.2749 | 6.3253 | 6.0621 | 7.0207 | 6.5423 | 6.0910 | 5.7737 |
| SYNPO_S595 | SYNPO | Q8N3V7-2 | 217 | GRAAATTPTKVYS | 6.0077 | 6.0661 | 5.6278 | 6.6088 | 6.3553 | 6.5945 | 6.2732 | 6.5254 | 6.1589 | 5.5653 | 5.8356 | 6.4736 | 6.5971 | 6.3182 | 6.3241 | 6.6683 |
| SYNPO_S638;S649 | SYNPO | Q8N3V7-2 | 649 | GDISPVSPSRAWS | 6.3074 | 6.4170 | 6.3907 | 6.4815 | 6.5717 | 6.9581 | 5.5876 | 5.8348 | 6.3645 | 6.5140 | 6.1390 | 6.1088 | 6.3759 | 6.6853 | 6.1327 | 6.1310 |
| SYNPO_S649 | SYNPO | Q8N3V7-2 | 894 | LRLKRGSLPAEAS | 4.9085 | 4.9789 | 5.9887 | 5.0882 | 6.4567 | 6.0031 | 5.3642 | 6.0106 | 7.1775 | 7.2403 | 6.7287 | 7.1996 | 6.8896 | 7.1916 | 6.5751 | 6.1987 |
| SYNPO_S718;S729 | SYNPO | Q8N3V7-2 | 281 | PSLPDRSPRPQRH | 6.1123 | 7.4487 | 6.3669 | 5.8083 | 6.6538 | 5.9280 | 7.6257 | 6.7927 | 7.4763 | 5.8734 | 5.5385 | 5.7892 | 6.0596 | 5.6095 | 5.4200 | 5.4970 |
| SYNPO_S784 | SYNPO | Q8N3V7-2 | 840 | SCTSPRSPLPAPP | 5.9613 | 5.6677 | 6.1235 | 5.5015 | 5.9531 | 6.5697 | 6.3213 | 6.8188 | 5.6027 | 5.8881 | 6.4315 | 6.3897 | 6.9379 | 6.0715 | 6.3603 | 6.5013 |
| SYNPO_S837 | SYNPO | Q8N3V7-2 | 837 | GPSSCTSPRSPLP | 5.6163 | 6.4831 | 5.7514 | 6.4456 | 6.0955 | 6.0354 | 6.0936 | 5.7770 | 6.8529 | 6.6340 | 7.5768 | 7.1221 | 5.6784 | 5.9170 | 6.2178 | 5.7030 |
| SYNPO_S840 | SYNPO | Q8N3V7-2 | 595 | SPAKPSSLDLVPN | 4.6512 | 5.5485 | 8.4461 | 6.9057 | 6.6253 | 5.4363 | 6.5356 | 4.7484 | 5.5111 | 6.5019 | 6.1018 | 6.8235 | 7.8217 | 5.8513 | 5.9149 | 6.5767 |
| SYNPO_S871 | SYNPO | Q8N3V7-2 | 510;514 | NA | 8.5949 | 6.5329 | 6.4352 | 7.5770 | 6.9717 | 6.5853 | 6.1762 | 5.3076 | 7.0395 | 5.2672 | 5.8549 | 6.7509 | 5.9276 | 5.7110 | 4.4777 | 5.7904 |
| SYNPO_S894 | SYNPO | Q8N3V7-2 | 718;729 | NA | 5.0019 | 5.5852 | 3.8165 | 6.5247 | 4.4804 | 6.0206 | 6.4032 | 7.5468 | 6.6716 | 5.8204 | 5.0358 | 5.6697 | 6.9227 | 6.6975 | 9.7436 | 8.0594 |
| SYNPO_T217 | SYNPO | Q8N3V7-2 | 638;649 | NA | 5.5855 | 6.8111 | 6.4020 | 5.6841 | 7.5897 | 5.3365 | 6.3226 | 7.3966 | 6.9302 | 5.5851 | 5.1873 | 5.7939 | 6.6792 | 6.1698 | 6.2145 | 6.3120 |
| SYNRG_S1075 | SYNRG | Q9UMZ2 | 752 | DVFRQLSLEGSGL | 5.8185 | 5.4229 | 6.5805 | 5.8051 | 6.2112 | 6.3857 | 5.8107 | 6.6361 | 6.0340 | 5.7246 | 5.9890 | 6.1041 | 7.2666 | 6.8126 | 6.8385 | 6.5600 |
| SYNRG_S1087 | SYNRG | Q9UMZ2 | 1075 | SHKRSLSLGDKEI | 6.1710 | 7.8429 | 6.7022 | 6.0289 | 5.4848 | 5.8254 | 5.5497 | 6.4416 | 5.9471 | 5.5626 | 5.9996 | 6.3077 | 6.6851 | 6.5897 | 6.6421 | 6.2196 |
| SYNRG_S578 | SYNRG | Q9UMZ2 | 789 | KFSSINSDKSLGE | 1.6586 | 1.8383 | 1.9417 | 2.0425 | 2.0841 | 2.1313 | 2.5806 | 2.5865 | 1.8997 | 1.6570 | 2.0503 | 1.4560 | 17.9231 | 18.8206 | 20.6895 | 18.6402 |
| SYNRG_S752 | SYNRG | Q9UMZ2 | 935 | KETSFGSSENITM | 6.3202 | 6.3206 | 6.6927 | 6.3889 | 6.2396 | 6.3375 | 6.1078 | 6.3983 | 6.4439 | 6.0112 | 6.5806 | 6.3716 | 6.0721 | 6.0592 | 5.4194 | 5.9364 |
| SYNRG_S789 | SYNRG | Q9UMZ2 | 812 | DSASVKSLDLPSI | 6.1442 | 6.0530 | 6.8980 | 6.3055 | 6.3355 | 6.3510 | 6.1676 | 7.0048 | 6.0993 | 6.3092 | 6.2919 | 6.4273 | 6.1642 | 5.9281 | 5.6010 | 5.9195 |
| SYNRG_S812 | SYNRG | Q9UMZ2 | 1087 | ISRSSPSPALEQP | 6.5629 | 6.4946 | 6.2394 | 6.3694 | 6.0023 | 6.8100 | 6.4359 | 7.0592 | 6.7773 | 6.0770 | 6.7308 | 6.3515 | 5.6318 | 5.5683 | 5.3993 | 5.4903 |
| SYNRG_S909;S919 | SYNRG | Q9UMZ2 | 919 | LSAGSGSPSATSI | 7.0004 | 6.8504 | 7.4719 | 7.3318 | 5.9242 | 7.1245 | 6.5933 | 6.4811 | 7.4176 | 7.3363 | 7.5216 | 7.1126 | 3.5155 | 5.0003 | 3.4716 | 3.8468 |
| SYNRG_S919 | SYNRG | Q9UMZ2 | 578 | DFKTADSVSPLEP | 5.7136 | 5.7009 | 6.3810 | 5.9124 | 6.1347 | 6.2698 | 6.5821 | 6.9349 | 5.6877 | 5.8376 | 6.3944 | 6.3692 | 6.5450 | 6.4281 | 6.8216 | 6.2869 |
| SYNRG_S935 | SYNRG | Q9UMZ2 | 909;919 | NA | 5.7465 | 6.0924 | 5.8014 | 6.6998 | 6.3487 | 6.8695 | 7.1201 | 6.9978 | 6.3519 | 6.2983 | 6.6698 | 6.1810 | 5.6260 | 6.1549 | 5.4913 | 5.5505 |
| SYTL4_S269 | SYTL4 | Q96C24 | 74 | ESLGRLSPKTNTC | 6.3921 | 6.7536 | 6.6577 | 6.5670 | 5.9076 | 6.4865 | 6.6090 | 5.2724 | 5.6812 | 6.4990 | 6.3453 | 6.4882 | 6.1258 | 6.5260 | 5.8632 | 5.8253 |
| SYTL4_S289 | SYTL4 | Q96C24 | 296 | LGDRSKSVPGLNV | 5.5190 | 6.3683 | 6.0988 | 6.1024 | 6.6954 | 6.5544 | 6.6376 | 5.7763 | 5.9004 | 6.5517 | 6.3692 | 6.2376 | 6.2707 | 7.0058 | 6.5466 | 6.3659 |
| SYTL4_S296 | SYTL4 | Q96C24 | 289 | VVHESGSLGDRSK | 6.1550 | 7.8748 | 5.8245 | 7.0967 | 5.9935 | 6.9126 | 5.8169 | 7.5299 | 6.5115 | 5.3256 | 7.0091 | 5.9257 | 4.8123 | 6.2885 | 5.7608 | 5.1627 |
| SYTL4_S74 | SYTL4 | Q96C24 | 269 | RKILRPSEYTKSV | 9.4303 | 7.3928 | 8.4791 | 7.4490 | 5.4657 | 5.6333 | 6.8260 | 5.1405 | 6.6949 | 6.4919 | 7.1355 | 7.0981 | 4.8950 | 5.4480 | 3.0320 | 3.3879 |
| SZRD1_S107 | SZRD1 | Q7Z422 | 107 | RILGSASPEEEQE | 5.8782 | 5.6009 | 6.7666 | 5.7203 | 6.3626 | 6.4516 | 6.1298 | 6.4684 | 6.2893 | 6.5759 | 6.3631 | 6.1832 | 6.8586 | 6.4629 | 5.8900 | 5.9989 |
| SZRD1_S39 | SZRD1 | Q7Z422 | 39 | ESRKSKSPPKVPI | 5.8367 | 5.3972 | 7.6099 | 5.6190 | 6.9964 | 7.3020 | 5.7466 | 6.5174 | 6.7984 | 6.1861 | 7.2545 | 6.9989 | 5.9285 | 6.0718 | 5.0458 | 4.6907 |
| SZT2_S1209 | SZT2 | Q5T011 | 1209 | QNQGELSPPFRRD | 5.6980 | 6.1259 | 6.3278 | 6.4821 | 6.3412 | 6.5759 | 5.8731 | 6.3560 | 6.6162 | 6.3003 | 6.7081 | 6.4297 | 5.6833 | 6.3640 | 6.0374 | 6.0810 |
| SZT2_S719 | SZT2 | Q5T011 | 723 | GSSPSKSPPVLGP | 5.4309 | 7.5753 | 6.7256 | 6.8958 | 5.1594 | 6.1601 | 7.1175 | 5.6638 | 6.1409 | 6.6645 | 6.4461 | 7.3424 | 5.4984 | 6.6156 | 5.9369 | 5.6269 |
| SZT2_S723 | SZT2 | Q5T011 | 719 | GAGGGSSPSKSPP | 6.1122 | 6.9216 | 7.0010 | 6.1303 | 6.4119 | 6.9990 | 6.1661 | 6.7061 | 6.3547 | 6.3501 | 6.4594 | 6.4891 | 5.0330 | 6.0736 | 5.6304 | 5.1617 |
| TAB2_S372 | TAB2 | Q9NYJ8 | 524 | SETRKLSMGSDDA | 5.6468 | 5.1497 | 5.8660 | 5.9559 | 6.3360 | 5.7383 | 5.4917 | 5.9537 | 7.0080 | 6.8123 | 6.6448 | 7.7767 | 6.3762 | 6.8702 | 6.3398 | 6.0338 |
| TAB2_S450 | TAB2 | Q9NYJ8 | 372 | TVYIAASPNTDE | 6.0451 | 6.7105 | 6.9182 | 5.9471 | 6.2856 | 6.5645 | 6.6604 | 6.5844 | 6.5007 | 6.4779 | 6.8576 | 6.7883 | 6.5908 | 5.4377 | 4.9245 | 5.6065 |
| TAB2_S524 | TAB2 | Q9NYJ8 | 450 | GNNSATSPRVVVT | 5.7451 | 5.0089 | 6.8367 | 5.5180 | 5.8943 | 6.1592 | 6.1888 | 7.3436 | 5.8051 | 5.6903 | 6.1404 | 5.9125 | 7.3125 | 7.3182 | 6.7251 | 6.4012 |
| TAB2_S582 | TAB2 | Q9NYJ8 | 582 | RLKRSNSISQIPS | 5.1632 | 5.9648 | 5.0939 | 5.9393 | 7.1552 | 6.2376 | 6.2532 | 5.5166 | 6.8583 | 8.0784 | 7.3644 | 5.4940 | 6.8145 | 6.7862 | 4.8945 | 6.3861 |
| TACC1_S147 | TACC1 | O75410 | 316 | GEESRSSPLKLEF | 6.2197 | 6.1533 | 6.3950 | 6.4066 | 6.8031 | 6.7908 | 6.2720 | 5.7820 | 6.4588 | 6.1193 | 6.5852 | 6.5410 | 5.9253 | 6.0105 | 5.8131 | 5.7243 |
| TACC1_S153 | TACC1 | O75410 | 350 | KLGSTLTPKIQKD | 8.5632 | 8.1864 | 6.9933 | 7.2706 | 5.5623 | 7.3934 | 7.8000 | 7.1154 | 8.2786 | 7.5430 | 7.8158 | 7.9465 | 2.1706 | 2.8601 | 1.8234 | 2.6774 |
| TACC1_S316 | TACC1 | O75410 | 465 | KSRLITSGCKVKK | 7.6313 | 8.2745 | 7.2603 | 7.9414 | 6.4248 | 6.8887 | 6.9598 | 7.0761 | 7.7419 | 7.3953 | 7.9761 | 7.6567 | 2.5123 | 3.1212 | 2.1651 | 2.9746 |
| TACC1_S350 | TACC1 | O75410 | 147 | HDFSKISIVRPFS | 7.3956 | 6.9635 | 9.3645 | 7.7143 | 6.6961 | 5.8399 | 6.3455 | 5.5737 | 6.7215 | 6.2444 | 7.7483 | 6.7768 | 4.4248 | 4.9595 | 3.2759 | 3.9555 |
| TACC1_S465 | TACC1 | O75410 | 153 | SIVRPFSIETKDS | 4.9903 | 4.9926 | 5.3730 | 5.3797 | 6.1089 | 5.1074 | 5.0580 | 5.8536 | 5.5902 | 4.8052 | 5.4838 | 5.0487 | 9.3836 | 8.4922 | 9.3866 | 8.9462 |
| TACC1_T350 | TACC1 | O75410 | 44 | DPEEEDSQAETKS | 5.8088 | 5.9642 | 6.0440 | 6.9503 | 6.7922 | 6.1342 | 5.8751 | 6.6632 | 6.2238 | 6.0627 | 6.0135 | 5.7642 | 6.6197 | 6.7430 | 6.3746 | 5.9663 |
| TACC2_S2072 | TACC2 | O95359 | 2569 | PVRMSESPTPCSG | 6.4707 | 6.3715 | 6.8902 | 5.9782 | 6.5189 | 6.5024 | 5.8775 | 6.2910 | 6.6175 | 6.4673 | 6.6421 | 6.9082 | 6.2186 | 5.8935 | 5.1266 | 5.2260 |
| TACC2_S2317 | TACC2 | O95359 | 2317 | LDNTPASPPRSPA | 6.0870 | 6.2040 | 6.5419 | 5.7713 | 6.0215 | 6.1870 | 5.5147 | 6.0739 | 6.9461 | 6.9151 | 6.6158 | 6.3565 | 6.2531 | 6.5292 | 6.0995 | 5.8833 |
| TACC2_S2359 | TACC2 | O95359 | 2390 | SSKTPSSPSKSPA | 5.7234 | 6.1926 | 6.6526 | 5.9452 | 6.0056 | 6.4630 | 5.7717 | 6.6424 | 6.4801 | 6.9306 | 6.4363 | 6.8372 | 6.1976 | 6.3489 | 5.7528 | 6.6198 |
| TACC2_S2390 | TACC2 | O95359 | 2394 | PSSPSKSPASFEI | 5.6424 | 7.0701 | 7.0942 | 6.5324 | 6.5336 | 6.1136 | 6.9356 | 7.6663 | 6.6593 | 7.1653 | 6.3796 | 6.6291 | 4.2302 | 5.9477 | 5.1341 | 5.1665 |
| TACC2_S2394 | TACC2 | O95359 | 2072 | VNTRRKSTDSVPI | 5.6174 | 5.3396 | 6.3189 | 5.6070 | 6.3760 | 5.8901 | 5.4026 | 6.2100 | 6.8975 | 7.1833 | 7.1359 | 6.7001 | 6.2489 | 7.1318 | 6.2360 | 7.7050 |
| TACC2_S2557 | TACC2 | O95359 | 2359 | TSKMQESPKLPQQ | 6.4026 | 5.5598 | 6.3268 | 5.9347 | 6.8445 | 6.8196 | 6.8541 | 6.3383 | 7.0785 | 5.8331 | 6.8417 | 5.9711 | 6.8762 | 6.5387 | 4.7343 | 5.0459 |
| TACC2_S2569 | TACC2 | O95359 | 2557 | FDTSQESPVKSSP | 5.9411 | 5.9442 | 6.6668 | 6.6369 | 6.0158 | 6.4896 | 6.5334 | 6.4813 | 6.3403 | 6.9193 | 6.4452 | 6.9478 | 5.0338 | 6.6352 | 5.4343 | 5.5350 |
| TACC2_T2317;S2321 | TACC2 | O95359 | 2317;2321 | NA | 5.9386 | 5.9418 | 6.6901 | 5.8645 | 6.6225 | 6.8704 | 6.0715 | 6.8059 | 6.4830 | 6.3629 | 6.3504 | 6.3730 | 6.2383 | 6.0781 | 5.8039 | 5.5050 |
| TACC3_S170 | TACC3 | Q9Y6A5 | 71 | QTHRILSPSMASK | 7.3567 | 7.0192 | 6.8550 | 6.5872 | 6.8773 | 7.1596 | 6.9836 | 7.2875 | 7.3003 | 6.9722 | 7.3654 | 7.1801 | 3.7662 | 4.0380 | 3.4055 | 3.8463 |
| TACC3_S177 | TACC3 | Q9Y6A5 | 317 | GRAMTLSPQEEVA | 6.3832 | 6.4369 | 7.2396 | 6.8757 | 6.6117 | 6.4622 | 6.6122 | 6.7518 | 6.9741 | 6.7739 | 7.3777 | 6.9960 | 4.5481 | 5.0554 | 4.2167 | 4.6847 |
| TACC3_S25 | TACC3 | Q9Y6A5 | 59 | MKVTFQTPLRDPQ | 7.9363 | 7.5250 | 7.6559 | 6.9094 | 5.1767 | 7.2839 | 7.5688 | 7.3826 | 8.0108 | 7.7706 | 8.1346 | 8.2527 | 2.2658 | 3.4189 | 1.8208 | 2.8872 |
| TACC3_S250 | TACC3 | Q9Y6A5 | 250 | PAAVATSPPGAIP | 7.0668 | 7.1960 | 7.3232 | 6.8783 | 6.5084 | 7.1737 | 6.7913 | 6.9142 | 7.3132 | 7.6195 | 7.9781 | 7.5430 | 3.4110 | 3.6397 | 3.0361 | 3.6077 |
| TACC3_S317 | TACC3 | Q9Y6A5 | 402 | MPASRGSYHLDWD | 4.9971 | 4.4565 | 4.8009 | 4.1793 | 5.1173 | 4.8663 | 4.6244 | 5.5104 | 5.1311 | 4.3679 | 5.1985 | 4.8326 | 10.8445 | 10.4195 | 11.0562 | 9.5974 |
| TACC3_S402 | TACC3 | Q9Y6A5 | 177 | PGKVSGSPEQAVE | 6.3779 | 6.4423 | 6.1001 | 6.4013 | 6.1744 | 6.3826 | 6.5277 | 6.5702 | 6.5707 | 6.6110 | 6.8832 | 6.7470 | 5.4805 | 5.9090 | 5.0978 | 5.7242 |
| TACC3_S558 | TACC3 | Q9Y6A5 | 25 | NCDFLFSPPEVTG | 6.6230 | 6.9725 | 6.9146 | 6.7477 | 6.3439 | 6.4241 | 6.8722 | 6.4130 | 7.1879 | 7.3196 | 7.2863 | 6.9297 | 4.0924 | 4.8049 | 4.1997 | 4.8685 |
| TACC3_S71 | TACC3 | Q9Y6A5 | 558 | SALRKQSLYLKFD | 8.0323 | 6.6819 | 7.5945 | 7.4696 | 6.2290 | 6.5724 | 6.3374 | 5.1660 | 7.5012 | 6.1598 | 8.0361 | 7.5779 | 4.6966 | 5.5994 | 3.0634 | 3.9827 |
| TACC3_T216 | TACC3 | Q9Y6A5 | 216 | SQHKAETPHGAEE | 6.9941 | 6.8026 | 6.6683 | 6.6473 | 6.0808 | 6.8089 | 6.9941 | 7.0846 | 6.8938 | 7.1306 | 7.5069 | 7.0716 | 4.2820 | 4.4826 | 3.9407 | 4.6110 |
| TACC3_T59 | TACC3 | Q9Y6A5 | 170 | SENQMVSPGKVSG | 6.8896 | 6.7861 | 6.9793 | 6.5853 | 6.5853 | 5.4726 | 6.4114 | 6.1323 | 5.7224 | 7.1120 | 6.2767 | 8.1016 | 6.4855 | 5.8981 | 5.6307 | 5.0432 | 5.3731 |
| TADA3_S280 | TADA3 | O75528 | 338 | DRPAEDSEDEVLA | 5.9397 | 6.3811 | 6.7297 | 6.8321 | 6.3419 | 6.5738 | 6.5053 | 6.5134 | 6.2547 | 6.1209 | 6.3455 | 6.0956 | 5.9330 | 6.2982 | 5.6857 | 5.4496 |
| TADA3_S338 | TADA3 | O75528 | 280 | ISPMEDSPIPDMS | 6.2567 | 5.5187 | 6.4119 | 6.0160 | 6.7575 | 6.6279 | 6.0248 | 6.6544 | 5.6139 | 5.8802 | 6.2779 | 5.9543 | 6.4075 | 6.2170 | 6.6262 | 6.7549 |
| TAF1_S307 | TAF1 | P21675-4 | 1680 | LDIPSATPKEQVT | 5.9185 | 6.2992 | 6.0972 | 6.6054 | 6.3165 | 6.1246 | 6.3845 | 6.3908 | 6.3763 | 6.3248 | 6.1736 | 6.5091 | 6.1076 | 6.0280 | 6.0527 | 6.2914 |
| TAF1_S524 | TAF1 | P21675-4 | 307 | PPEQCLSDDEITM | 5.7516 | 6.3945 | 6.2605 | 6.7188 | 6.2157 | 5.9906 | 6.1680 | 5.9712 | 6.3435 | 6.2888 | 6.3353 | 6.4704 | 6.0099 | 6.2241 | 6.2639 | 6.5932 |
| TAF1_T1680 | TAF1 | P21675-4 | 524 | EEATSNSPSKESK | 6.6221 | 6.9121 | 6.1776 | 6.7845 | 5.9740 | 6.7409 | 6.3938 | 6.2660 | 6.2019 | 6.0284 | 5.7953 | 6.2479 | 6.0259 | 6.1173 | 6.4579 | 6.2547 |
| TAF10_S44 | TAF10 | Q12962 | 44 | AAENKASPAGTAG | 5.6309 | 6.5320 | 6.0720 | 5.8415 | 5.7404 | 7.3026 | 6.6805 | 5.9278 | 6.7858 | 5.5162 | 5.3151 | 5.4593 | 7.7711 | 7.0805 | 6.0868 | 7.1574 |
| TAF12_S51 | TAF12 | Q16514 | 51 | GAGGRLSPENNQV | 6.1355 | 6.0738 | 6.2649 | 6.3771 | 6.2302 | 6.4810 | 6.1372 | 5.9959 | 6.0120 | 6.4017 | 6.3543 | 6.4791 | 6.4050 | 6.5156 | 6.1590 | 5.9776 |
| TAF12_T43 | TAF12 | Q16514 | 43 | VVKIPGTPGAGGR | 5.1552 | 6.3327 | 5.9803 | 7.7329 | 6.6591 | 7.1759 | 7.3961 | 6.3909 | 6.4492 | 7.0075 | 7.7173 | 7.5049 | 4.5128 | 5.0982 | 3.8962 | 4.9907 |
| TAF12_T43;S51 | TAF12 | Q16514 | 43;51 | NA | 6.5714 | 6.2175 | 6.4697 | 6.5205 | 6.8007 | 6.6710 | 6.3195 | 6.8585 | 7.2440 | 6.5735 | 6.8335 | 6.4245 | 4.9422 | 5.5349 | 4.6826 | 5.3359 |
| TAF15_S226 | TAF15 | Q92804 | 226 | PRTDADSESDNSD | 6.1322 | 6.3686 | 6.2384 | 6.7247 | 6.8698 | 5.9290 | 5.7382 | 6.2971 | 6.9459 | 6.1396 | 6.4578 | 6.1595 | 6.1462 | 5.9069 | 5.9830 | 5.9632 |
| TAF1C_T834 | TAF1C | Q15572 | 834 | TPGCATTPPHSQA | 6.2260 | 6.4017 | 6.0719 | 5.5187 | 6.2676 | 6.5728 | 5.8484 | 6.3185 | 8.3373 | 6.4264 | 5.7652 | 7.4543 | 5.6312 | 5.9609 | 5.4416 | 5.7575 |
| TAF1D_S138 | TAF1D | Q9H5J8 | 40 | TQCIPYSPKGEKR | 6.3713 | 6.3295 | 6.5907 | 6.6321 | 6.1826 | 5.9302 | 6.6479 | 5.7896 | 6.6868 | 6.2234 | 6.0165 | 6.1378 | 6.0428 | 5.9287 | 6.1863 | 6.3037 |
| TAF1D_S234 | TAF1D | Q9H5J8 | 138 | GFPFLESENEKNA | 6.1860 | 7.1751 | 7.1145 | 7.8212 | 7.1322 | 7.0775 | 7.7843 | 7.3846 | 6.1398 | 5.6056 | 5.7934 | 6.1651 | 4.4809 | 5.0497 | 4.0525 | 5.0376 |
| TAF1D_S40 | TAF1D | Q9H5J8 | 234 | IKLAGDSFIVSSE | 6.3890 | 6.1522 | 6.8326 | 6.3965 | 6.5043 | 6.0144 | 6.4704 | 5.8214 | 6.1422 | 6.3464 | 6.3251 | 5.3465 | 6.5557 | 6.0043 | 6.4379 | 6.2612 |
| TAF2_S1185 | TAF2 | Q6P1X5 | 1185 | EPFTFSSPASGRS | 6.6357 | 6.9258 | 6.3218 | 6.5484 | 5.8460 | 5.7920 | 6.3029 | 6.1665 | 6.6294 | 6.5988 | 6.0420 | 6.2378 | 5.4431 | 6.4265 | 5.8641 | 6.2194 |
| TAF2_T1070 | TAF2 | Q6P1X5 | 1070 | MLERPSTPGLSKY | 6.8013 | 6.0830 | 6.7353 | 6.0733 | 4.9947 | 6.0755 | 6.4170 | 5.7033 | 7.2754 | 6.8906 | 6.6374 | 6.9168 | 5.4115 | 6.2513 | 5.0522 | 6.6813 |
| TAF3_S183 | TAF3 | Q5VWG9 | 297 | SPAMVGSPIRSPK | 6.1667 | 6.4480 | 5.8438 | 6.4183 | 5.9462 | 5.5689 | 6.1074 | 5.8282 | 6.1774 | 6.0469 | 5.9388 | 5.9727 | 6.3033 | 7.1818 | 7.0394 | 7.0121 |
| TAF3_S229;S243 | TAF3 | Q5VWG9 | 183 | GKRPLDSPEAEEL | 6.0330 | 6.2108 | 6.1583 | 6.3843 | 6.0943 | 6.1082 | 6.3036 | 5.8270 | 5.9737 | 6.2192 | 6.0862 | 6.1566 | 6.3248 | 6.8939 | 6.4513 | 6.7749 |
| TAF3_S278 | TAF3 | Q5VWG9 | 278 | TKTKTSSPGQKTK | 6.2174 | 6.6758 | 6.0276 | 6.9455 | 5.7434 | 5.7763 | 6.1713 | 5.9737 | 6.2060 | 6.1111 | 6.2523 | 6.6196 | 5.5990 | 6.6627 | 6.3079 | 6.7104 |
| TAF3_S297 | TAF3 | Q5VWG9 | 755 | PVALAPSPVIPRL | 5.6690 | 6.7552 | 6.5876 | 6.7795 | 6.0255 | 6.3401 | 6.6706 | 5.8831 | 6.7829 | 6.2615 | 6.0348 | 6.6007 | 5.0548 | 5.8548 | 5.7217 | 6.0783 |
| TAF3_S297;S301 | TAF3 | Q5VWG9 | 433 | SGPECTTPKASTS | 6.7201 | 6.2802 | 6.2671 | 7.1191 | 5.9748 | 6.3233 | 6.9738 | 5.4941 | 6.4246 | 5.8187 | 6.0369 | 6.3432 | 5.7405 | 6.4689 | 5.8564 | 6.1584 |
| TAF3_S518 | TAF3 | Q5VWG9 | 667 | LALPLFSPATASR | 7.9201 | 7.4980 | 5.5373 | 7.5706 | 6.4859 | 6.8240 | 6.7877 | 7.2575 | 8.1448 | 7.1145 | 5.8777 | 5.9662 | 3.8988 | 5.8255 | 2.9207 | 4.3708 |
| TAF3_S667 | TAF3 | Q5VWG9 | 518 | KTKLPSSVEVKKK | 5.9236 | 8.0160 | 7.6859 | 6.9292 | 6.5415 | 8.3360 | 7.5048 | 7.5996 | 5.3192 | 7.4134 | 6.9901 | 8.3919 | 2.9372 | 4.7402 | 2.7663 | 2.9051 |
| TAF3_S755 | TAF3 | Q5VWG9 | 229;243 | NA | 6.1755 | 6.2802 | 6.1942 | 6.4543 | 5.9833 | 5.9560 | 6.0916 | 6.1352 | 6.2968 | 6.1660 | 6.2529 | 6.3554 | 5.9648 | 6.7760 | 6.3187 | 6.5991 |
| TAF3_T433 | TAF3 | Q5VWG9 | 297;301 | NA | 6.7685 | 6.5659 | 6.7828 | 7.1429 | 5.6028 | 6.1560 | 6.5703 | 6.4333 | 6.5143 | 6.5879 | 6.8599 | 7.1143 | 4.8642 | 5.8103 | 5.0260 | 5.2005 |
| TAF4_S109 | TAF4 | O00268 | 109 | QRPGPPSPRRPLV | 6.1095 | 5.4903 | 5.3777 | 7.2746 | 5.7191 | 5.6460 | 6.4366 | 5.4815 | 5.6653 | 6.4238 | 6.3833 | 7.0302 | 6.3714 | 6.6604 | 6.1789 | 6.8515 |
| TAF4B_S314 | TAF4B | Q92750 | 595 | ILSLQASPTQKNR | 6.3685 | 6.2819 | 6.3736 | 6.0110 | 7.6616 | 6.7626 | 6.5670 | 6.6423 | 6.7261 | 6.4332 | 5.7743 | 5.8384 | 5.9851 | 5.4370 | 5.7164 | 5.4212 |
| TAF4B_S595 | TAF4B | Q92750 | 314 | YVELKSSPQPHLV | 6.0213 | 5.8341 | 6.1907 | 6.3275 | 8.2831 | 6.3007 | 6.3869 | 7.0051 | 4.7850 | 6.1581 | 5.9738 | 7.4628 | 5.5726 | 6.8327 | 5.4477 | 5.4180 |
| TAF5L_S230 | TAF5L | O75529 | 230 | EPPDMPSPILQNE | 5.7905 | 5.9371 | 5.9703 | 6.9854 | 6.0528 | 6.1775 | 6.1969 | 6.0544 | 6.4058 | 6.0804 | 6.0893 | 6.1872 | 6.5782 | 6.3985 | 6.2382 | 6.8575 |
| TAF6_S569 | TAF6 | P49848-3 | 709 | SQPNSGSPQPAPx | 6.5263 | 6.1283 | 6.0185 | 6.7192 | 5.7406 | 5.9778 | 6.1123 | 6.0884 | 6.6940 | 6.5778 | 6.6673 | 6.3167 | 6.1586 | 6.5495 | 5.7063 | 6.0185 |
| TAF6_S690 | TAF6 | P49848-3 | 690 | KQEAGDSPPPAPG | 6.5308 | 7.0720 | 6.4038 | 6.4528 | 5.7542 | 6.0393 | 6.1454 | 6.2604 | 6.4365 | 6.7749 | 6.4575 | 6.8089 | 6.0837 | 5.6935 | 5.2887 | 5.7976 |
| TAF6_S690;T697 | TAF6 | P49848-3 | 697 | PPPAPGTPKANGS | 6.3129 | 6.1332 | 6.3327 | 6.0191 | 6.5296 | 6.1611 | 6.2674 | 6.5731 | 6.0766 | 6.1699 | 6.4627 | 6.6895 | 6.4159 | 5.9870 | 6.1845 | 5.6848 |
| TAF6_S697 | TAF6 | P49848-3 | 569 | AAPPQPSPPPTKF | 5.7187 | 6.0739 | 6.2487 | 6.6939 | 5.8734 | 6.0234 | 6.3092 | 6.6081 | 6.5776 | 6.2971 | 5.9376 | 6.7244 | 6.1908 | 6.2329 | 5.9887 | 6.5018 |
| TAF6_S709 | TAF6 | P49848-3 | 690;697 | NA | 6.0228 | 6.3163 | 6.5497 | 5.7985 | 6.2549 | 5.5638 | 5.6086 | 6.8785 | 6.4635 | 6.1244 | 5.9309 | 6.1754 | 6.9302 | 7.0651 | 6.0906 | 6.2268 |
| TAF7_S213 | TAF7 | Q15545 | 264 | QDKLNESDEQHQE | 5.9929 | 6.3885 | 6.9821 | 7.0439 | 6.3981 | 6.4296 | 6.3583 | 6.7723 | 6.5671 | 6.5092 | 6.8496 | 6.6419 | 5.4153 | 5.2119 | 5.0783 | 5.3609 |
| TAF7_S264 | TAF7 | Q15545 | 213 | HRQGHDSLEHDEL | 5.9455 | 6.5972 | 7.0248 | 6.2871 | 6.1964 | 7.1230 | 6.3056 | 6.9110 | 7.1609 | 6.2454 | 6.3643 | 7.0572 | 4.1571 | 5.0106 | 4.5829 | 5.0310 |
| TAF9_S149 | TAF9 | Q16594 | 149 | ITVPRLSVGSVTS | 5.0110 | 4.6299 | 5.3604 | 5.3678 | 6.0576 | 5.5212 | 5.8881 | 6.6480 | 5.3060 | 5.2382 | 5.3869 | 5.3699 | 8.3537 | 7.4253 | 8.7120 | 8.7241 |
| TAF9_T102 | TAF9 | Q16594 | 102 | ARQRNQTPLPLIK | 6.9471 | 7.2088 | 7.7294 | 7.0930 | 4.7553 | 7.5576 | 7.7532 | 6.1776 | 6.8982 | 8.0501 | 8.4730 | 6.8844 | 3.1115 | 4.3296 | 2.3435 | 2.8878 |
| TAF9_T178 | TAF9 | Q16594 | 178 | VSTKVGTPMSLTG | 4.9620 | 5.3280 | 6.6817 | 5.2143 | 5.1894 | 6.5023 | 7.9982 | 6.6284 | 5.0295 | 5.4115 | 6.9819 | 7.2822 | 6.1132 | 6.5982 | 6.1541 | 7.9253 |
| TAF9B_S147 | TAF9B | Q9HBM6 | 147 | RLVPRLSVGAVSS | 6.8495 | 6.2800 | 6.4679 | 6.2553 | 6.8598 | 6.7923 | 6.2807 | 6.3717 | 6.7089 | 6.6338 | 7.3400 | 7.1436 | 5.2063 | 5.4320 | 4.5138 | 5.4742 |
| TAGLN2_S184 | TAGLN2 | P37802-2 | 184 | ENPRNFSDNQLQE | 2.2448 | 2.3359 | 2.7859 | 2.5892 | 2.8215 | 2.6110 | 2.8907 | 3.1936 | 2.4876 | 2.6378 | 2.5096 | 2.6944 | 14.4378 | 17.6877 | 18.0886 | 17.9839 |
| TALDO1_S237 | TALDO1 | P37837 | 237 | TVMGASFRNTGE | 6.2385 | 6.3076 | 6.2756 | 6.6440 | 5.9260 | 6.3419 | 6.5739 | 6.3894 | 6.0333 | 6.0227 | 6.3684 | 6.1131 | 6.4845 | 6.6722 | 7.2725 | 5.8744 |
| TALDO1_S275 | TALDO1 | P37837 | 275 | KLVPVLSAKAAQA | 4.1498 | 8.7269 | 6.2247 | 5.9604 | 5.6005 | 4.9033 | 6.1347 | 5.9998 | 8.2883 | 5.5837 | 7.8104 | 6.3540 | 5.3138 | 6.7870 | 5.8142 | 6.3484 |
| TANC1_S1503 | TANC1 | Q9C0D5 | 313 | IMPRPNSVAATSS | 6.8211 | 5.5778 | 7.0381 | 6.3343 | 5.9938 | 5.6464 | 6.3445 | 5.8458 | 7.2259 | 6.0202 | 6.3674 | 6.5352 | 6.3036 | 6.2478 | 5.3953 | 6.1227 |
| TANC1_S214 | TANC1 | Q9C0D5 | 85 | SRVNKKSPGPVRK | 6.4869 | 6.7853 | 6.9510 | 6.1426 | 5.6313 | 6.7514 | 6.5413 | 6.3653 | 6.3542 | 6.7569 | 6.8134 | 6.8507 | 5.0774 | 5.8278 | 5.2011 | 5.4634 |
| TANC1_S242 | TANC1 | Q9C0D5 | 270 | RRADNCSPVAEEE | 6.4101 | 6.1963 | 6.4824 | 5.6112 | 6.0558 | 5.8703 | 7.0090 | 6.5162 | 6.7045 | 6.4256 | 6.2747 | 6.1302 | 6.4682 | 6.1621 | 5.9494 | 5.7341 |
| TANC1_S270 | TANC1 | Q9C0D5 | 347 | RLPWHNTAGGRAQ | 8.1172 | 6.6057 | 7.0695 | 8.9168 | 6.2330 | 2.8938 | 4.4653 | 2.9180 | 5.7311 | 4.0015 | 4.6580 | 4.1042 | 8.6503 | 7.1618 | 7.9614 | 10.5125 |
| TANC1_S313 | TANC1 | Q9C0D5 | 444 | RQIASNSPGSSPK | 6.2938 | 7.4616 | 7.2045 | 7.5961 | 6.5375 | 6.4117 | 6.3286 | 6.4011 | 6.7941 | 7.6316 | 7.3620 | 7.2949 | 3.9619 | 5.0132 | 4.1106 | 4.4970 |
| TANC1_S444 | TANC1 | Q9C0D5 | 67 | SMSLPSSPLLPRQ | 5.6255 | 6.0385 | 6.6885 | 6.1807 | 5.6061 | 6.4852 | 6.1468 | 6.6778 | 6.0953 | 6.6845 | 6.6750 | 6.3275 | 5.8330 | 6.2340 | 6.4144 | 6.2873 |
| TANC1_S63 | TANC1 | Q9C0D5 | 1503 | SKGRPVSPQSRAG | 6.2257 | 6.4077 | 6.8172 | 6.6201 | 6.3860 | 6.7952 | 6.8256 | 5.7639 | 6.5920 | 6.4130 | 6.7249 | 6.7242 | 5.4483 | 5.7097 | 4.9310 | 5.6154 |
| TANC1_S63;S67 | TANC1 | Q9C0D5 | 687 | NGKADATLIGKVS | 4.9396 | 4.9000 | 5.1110 | 5.1469 | 5.7182 | 5.6164 | 5.6740 | 5.4599 | 5.1059 | 4.6786 | 4.3152 | 4.1831 | 10.1528 | 9.7641 | 10.3497 | 8.8846 |
| TANC1_S67 | TANC1 | Q9C0D5 | 63 | AKGVSMSLPSSPL | 5.9349 | 6.1125 | 6.7138 | 6.5578 | 6.5163 | 6.6133 | 6.7398 | 7.1895 | 5.8405 | 5.8864 | 6.4339 | 7.1496 | 5.6074 | 5.9323 | 5.5970 | 5.1751 |
| TANC1_S85 | TANC1 | Q9C0D5 | 214 | PCETISSPSSTLE | 5.8279 | 6.3482 | 6.3870 | 6.8611 | 6.4322 | 5.9309 | 6.2204 | 6.4123 | 6.7630 | 6.4380 | 6.0854 | 6.4443 | 5.6803 | 5.8906 | 6.1131 | 6.1655 |
| TANC1_T347 | TANC1 | Q9C0D5 | 242 | NDDRSGSSLEWNK | 6.4913 | 5.4543 | 6.1956 | 6.1010 | 6.3079 | 6.1068 | 5.9978 | 6.0693 | 7.0526 | 6.5749 | 6.4717 | 6.2556 | 6.1510 | 6.1044 | 6.6248 | 6.0410 |
| TANC1_T687 | TANC1 | Q9C0D5 | 63;67 | NA | 7.0764 | 6.3737 | 7.2576 | 6.4749 | 5.6648 | 6.8649 | 6.7383 | 6.1451 | 6.8636 | 6.7161 | 6.6637 | 7.0492 | 4.6913 | 5.6605 | 4.6338 | 5.1263 |
| TANC2_S1468 | TANC2 | Q9HCD6-2 | 1987 | SRQGQTSPIKPKR | 6.4916 | 6.2576 | 6.2818 | 5.9162 | 5.9904 | 6.4265 | 6.2400 | 6.2930 | 6.6203 | 6.3290 | 6.5936 | 6.7139 | 5.9592 | 6.2252 | 5.6616 | 6.0002 |
| TANC2_S1486 | TANC2 | Q9HCD6-2 | 1527 | GGSYRFSPPPVGG | 6.2709 | 6.3686 | 6.9510 | 6.9468 | 6.3464 | 6.3115 | 6.4657 | 6.3032 | 6.8550 | 7.1429 | 6.3831 | 7.0038 | 5.0319 | 5.4539 | 5.1799 | 4.9852 |
| TANC2_S1527 | TANC2 | Q9HCD6-2 | 1486 | SSSSVGSPTRQTY | 6.3342 | 6.3571 | 7.1626 | 6.6048 | 6.2359 | 6.4731 | 6.6078 | 6.6766 | 6.9556 | 6.7790 | 7.2381 | 7.5270 | 4.6731 | 5.1499 | 4.4990 | 4.7261 |
| TANC2_S1544 | TANC2 | Q9HCD6-2 | 1468 | SAYISSSPLGSHQ | 6.4303 | 6.1955 | 6.5256 | 6.7699 | 6.1590 | 7.0608 | 6.1955 | 6.7731 | 6.8237 | 6.7576 | 6.8479 | 6.6300 | 5.4172 | 5.5057 | 4.7593 | 5.1487 |
| TANC2_S1555 | TANC2 | Q9HCD6-2 | 1555 | GPQYRASPPAESM | 5.7810 | 6.0052 | 6.9162 | 6.8149 | 5.9150 | 5.9790 | 6.4726 | 6.6765 | 6.6929 | 7.1841 | 6.8978 | 6.7718 | 5.3861 | 5.8480 | 5.2209 | 5.4382 |
| TANC2_S1570 | TANC2 | Q9HCD6-2 | 1570 | YRSQSGSPVRYQQ | 5.7909 | 6.9159 | 7.7427 | 6.1405 | 6.3271 | 5.8285 | 6.4998 | 6.7046 | 6.7808 | 6.0515 | 6.8794 | 6.4708 | 4.9963 | 6.2907 | 5.3062 | 5.2741 |
| TANC2_S1732 | TANC2 | Q9HCD6-2 | 400 | RQIASDSPHASPK | 5.3765 | 5.9876 | 6.5018 | 6.0689 | 6.6976 | 6.4294 | 6.5777 | 7.1328 | 7.3504 | 6.5755 | 6.0821 | 6.5129 | 6.5789 | 6.7284 | 5.8733 | 6.3261 |
| TANC2_S1987 | TANC2 | Q9HCD6-2 | 1544 | YPSPPPSPLRRGP | 6.1857 | 6.5191 | 6.2163 | 5.1317 | 5.9263 | 5.4484 | 5.2523 | 6.1021 | 5.6238 | 6.4860 | 6.3035 | 6.1273 | 7.1867 | 7.2336 | 7.5916 | 6.6657 |
| TANC2_S400 | TANC2 | Q9HCD6-2 | 1732 | QIGRSQSASYYPV | 6.0937 | 6.1910 | 8.3401 | 5.6098 | 5.7877 | 6.5312 | 6.4393 | 5.5825 | 7.7057 | 8.4025 | 8.8568 | 7.2346 | 3.4228 | 5.8993 | 3.6878 | 4.2151 |
| TANC2_S400;S404 | TANC2 | Q9HCD6-2 | 400;404 | NA | 6.0361 | 6.7936 | 5.8806 | 6.9139 | 6.6067 | 7.2033 | 5.8966 | 7.4950 | 6.3193 | 6.1566 | 5.7740 | 6.3158 | 6.4978 | 6.0802 | 5.5852 | 4.4453 |
| TANK_S117 | TANK | Q92844 | 129 | SARSLGSPLLHER | 7.0671 | 6.2198 | 6.6819 | 5.6515 | 7.4310 | 6.2671 | 5.9565 | 7.7894 | 6.2325 | 7.5709 | 7.4454 | 7.6116 | 4.4193 | 5.2203 | 3.9722 | 4.4635 |
| TANK_S129 | TANK | Q92844 | 225 | RDEEDTSFESLSK | 6.8307 | 7.0302 | 6.9306 | 6.0549 | 6.2359 | 6.2963 | 6.5697 | 6.9440 | 6.6511 | 6.7812 | 7.4835 | 7.5161 | 4.2768 | 5.4107 | 4.8746 | 4.1140 |
| TANK_S225 | TANK | Q92844 | 213 | PSITSVTPRGLCR | 7.7176 | 6.7543 | 6.8805 | 5.8860 | 6.9512 | 6.4199 | 6.5878 | 8.1763 | 6.8699 | 7.3451 | 7.4698 | 8.1621 | 3.6125 | 4.1233 | 3.0803 | 3.9633 |
| TANK_T213 | TANK | Q92844 | 117 | RRQEVSSPRKETS | 6.8943 | 5.5311 | 7.6425 | 5.1614 | 7.9008 | 6.7317 | 5.6523 | 7.1373 | 6.5807 | 7.3443 | 7.5163 | 8.1528 | 5.2143 | 4.9556 | 3.6443 | 3.9402 |
| TAOK1_S445 | TAOK1 | Q7L7X3 | 965 | SMGVRNSPQALRR | 5.9181 | 6.1550 | 6.3364 | 5.4448 | 5.6011 | 5.8281 | 6.2411 | 5.7788 | 6.5631 | 6.6521 | 6.4231 | 6.6622 | 6.5095 | 6.7078 | 6.5668 | 6.6120 |
| TAOK1_S9 | TAOK1 | Q7L7X3 | 445 | ATIRTASLVTRQM | 8.8383 | 7.4296 | 7.7612 | 7.2705 | 6.3645 | 5.8352 | 6.0014 | 6.2568 | 7.0538 | 6.3690 | 5.7864 | 5.9426 | 4.8713 | 5.9290 | 4.2204 | 4.0700 |
| TAOK1_S965 | TAOK1 | Q7L7X3 | 9 | STNRAGSLKDPEI | 5.9943 | 6.5240 | 8.7193 | 8.2750 | 5.8825 | 5.4249 | 6.7359 | 6.6682 | 7.1317 | 6.9846 | 6.8036 | 8.6090 | 5.1483 | 4.6289 | 2.9310 | 3.5386 |
| TAOK2_S1011 | TAOK2 | Q9UL54-2 | 453 | PTSTTSSARRRAY | 6.4277 | 6.5230 | 6.9217 | 6.7973 | 6.6384 | 6.7654 | 6.3427 | 6.9530 | 6.5190 | 6.3760 | 6.4636 | 6.1629 | 5.0227 | 5.9707 | 4.8867 | 5.2291 |
| TAOK2_S453 | TAOK2 | Q9UL54-2 | 9 | AGGRAGSLKDPDV | 7.1301 | 6.0480 | 7.8227 | 6.9255 | 6.3789 | 5.9824 | 5.9501 | 6.3161 | 7.5976 | 7.6332 | 7.1839 | 6.8038 | 4.9195 | 5.6136 | 4.0789 | 3.6156 |
| TAOK2_S473 | TAOK2 | Q9UL54-2 | 1011 | PLRRAASGGSGSE | 6.5703 | 5.2311 | 5.7143 | 6.6053 | 7.0329 | 5.8652 | 4.7516 | 5.6533 | 7.5070 | 6.0565 | 5.4750 | 5.2806 | 6.3698 | 7.8538 | 7.3096 | 6.7235 |
| TAOK2_S9 | TAOK2 | Q9UL54-2 | 473 | ATIRTASLVSRQI | 8.6809 | 7.5668 | 7.3666 | 7.2492 | 6.1972 | 6.1087 | 6.0347 | 6.2397 | 7.3627 | 6.6965 | 6.1047 | 6.1396 | 4.5665 | 5.9319 | 3.8702 | 3.8841 |
| TAOK3_S442 | TAOK3 | Q9H2K8 | 442 | ATIKSASLVTRQI | 7.1013 | 6.9671 | 6.8809 | 6.1691 | 6.1924 | 5.9401 | 5.9191 | 6.3058 | 7.0685 | 6.1790 | 6.1412 | 5.9705 | 6.2092 | 6.3535 | 5.6294 | 5.3329 |
| TAP1_S47 | TAP1 | Q03518 | 47 | PHRGAESPRPGRD | 6.6522 | 7.1214 | 6.3350 | 7.1457 | 6.3384 | 6.4059 | 6.8462 | 6.3907 | 6.5093 | 6.4161 | 6.7781 | 6.4848 | 4.9190 | 5.2534 | 4.9708 | 5.4332 |

| | | | | | | | | | | | | | | | | | | | | | |
|---|---|---|---|---|---|---|---|---|---|---|---|---|---|---|---|---|---|---|---|---|---|
| TAP1_T545 | TAP1 | Q03518 | | 545 | FEYLDRTPRCPPS | 6.0072 | 6.3109 | 6.2064 | 6.7620 | 6.6343 | 6.5131 | 6.4994 | 6.3090 | 6.2527 | 6.7772 | 6.0578 | 6.2421 | 5.4717 | 5.9107 | 5.7682 | 6.2774 |
| TARBP2_S142 | TARBP2 | Q15633 | | 142 | SVVLTRSPPMELQ | 5.8896 | 5.9346 | 6.2279 | 6.1339 | 6.5921 | 6.3846 | 5.6237 | 6.5613 | 6.7612 | 6.9102 | 6.7694 | 6.6824 | 5.5934 | 5.9315 | 6.1299 | 5.8742 |
| TARBP2_S142;S152 | TARBP2 | Q15633 | | 152 | ELQPPVSPQQSEC | 5.8653 | 5.4001 | 6.5154 | 6.1917 | 6.0724 | 6.1451 | 5.6003 | 6.5941 | 6.4484 | 5.7834 | 6.4798 | 6.5284 | 6.7892 | 6.3544 | 6.5200 | 6.7119 |
| TARBP2_S152 | TARBP2 | Q15633 | 142;152 | | NA | 5.9371 | 6.9445 | 7.3691 | 7.1905 | 5.9778 | 6.5627 | 6.6372 | 6.2869 | 6.9292 | 7.1930 | 7.2713 | 7.4694 | 4.1675 | 4.8430 | 4.3894 | 4.8314 |
| TARS1_S8 | TARS1 | P26639-2 | | 8 | FEEKASSPSGKMG | 6.3919 | 6.8759 | 7.0278 | 6.6050 | 6.2289 | 6.8902 | 6.5509 | 6.8475 | 6.6158 | 7.0032 | 6.9821 | 7.0869 | 4.3027 | 5.1467 | 4.5846 | 4.8599 |
| TASOR_S1103 | TASOR | Q9UK61 | | 979 | DYQFPSSPFTDTL | 6.7100 | 6.9815 | 6.6976 | 6.8670 | 6.1975 | 6.2801 | 6.3685 | 6.0105 | 6.7314 | 6.6913 | 6.6387 | 6.4813 | 5.4151 | 5.4914 | 5.3384 | 5.0996 |
| TASOR_S1139 | TASOR | Q9UK61 | | 927 | TGGNARSPEDQLG | 6.4200 | 6.4931 | 5.7910 | 6.5252 | 6.0078 | 5.9312 | 6.1491 | 6.1462 | 6.0363 | 6.0324 | 5.8610 | 5.9891 | 6.6365 | 6.5471 | 6.8391 | 6.5949 |
| TASOR_S1552 | TASOR | Q9UK61 | | 1103 | GNLPPVSPNDSGA | 5.9653 | 5.9141 | 6.1074 | 6.9715 | 5.9806 | 6.0050 | 6.2354 | 5.7682 | 6.2861 | 6.0190 | 6.0782 | 5.9848 | 6.6109 | 6.4948 | 6.8031 | 6.7755 |
| TASOR_S1651 | TASOR | Q9UK61 | | 1552 | QIELQSSPDVQNS | 5.9689 | 6.0979 | 6.0665 | 6.7347 | 6.1986 | 5.7989 | 6.1635 | 5.6912 | 6.4097 | 6.7653 | 6.4848 | 5.9972 | 6.5846 | 6.5806 | 6.1368 | 6.3206 |
| TASOR_S927 | TASOR | Q9UK61 | | 1651 | SLDRDKSPPPLSW | 6.8450 | 7.0141 | 5.9868 | 6.7102 | 5.7987 | 5.5738 | 6.3970 | 5.7478 | 6.7114 | 6.9388 | 5.5691 | 6.0590 | 6.3558 | 5.6001 | 6.4387 | 6.2537 |
| TASOR_S945 | TASOR | Q9UK61 | | 1139 | LLEPLCSDPLKDT | 5.9513 | 5.9460 | 6.1705 | 6.6072 | 5.9710 | 5.9315 | 6.3969 | 5.8637 | 6.7830 | 5.8023 | 5.7295 | 6.5656 | 6.4348 | 6.5033 | 6.3568 | 6.9866 |
| TASOR_S971 | TASOR | Q9UK61 | | 945 | QTPGMKSPEEQLV | 5.8826 | 5.8760 | 5.8062 | 6.0468 | 6.4561 | 6.4727 | 6.1823 | 6.1916 | 6.4667 | 6.6375 | 6.6886 | 6.4747 | 6.6621 | 5.9252 | 6.3418 | 5.8893 |
| TASOR_S971;S979 | TASOR | Q9UK61 | | 971 | VINRQRSSDYQFP | 5.1613 | 5.1556 | 5.1910 | 6.0069 | 5.9684 | 5.7306 | 6.0980 | 6.4222 | 5.4694 | 5.3294 | 5.0447 | 4.8007 | 8.6911 | 7.6488 | 8.6610 | 8.6209 |
| TASOR_S979 | TASOR | Q9UK61 | | 819 | EYELNSTPDKKDY | 6.7364 | 7.4224 | 6.6299 | 7.7797 | 6.1732 | 6.4802 | 6.4543 | 6.2401 | 7.1483 | 5.7551 | 7.0881 | 7.0871 | 4.3906 | 5.0856 | 4.2743 | 5.2546 |
| TASOR_T819 | TASOR | Q9UK61 | 971;979 | | NA | 5.2471 | 5.5649 | 4.7505 | 5.3049 | 5.9143 | 6.3604 | 5.9538 | 5.9182 | 4.9747 | 5.1120 | 5.2791 | 5.3556 | 8.5549 | 8.3984 | 9.0212 | 8.2900 |
| TASOR2_S1025 | TASOR2 | Q5VWN6 | | 340 | KRVFPLSPASNLR | 7.7778 | 7.5117 | 6.9195 | 7.8962 | 6.5698 | 6.6950 | 6.9394 | 6.7807 | 7.3100 | 6.9416 | 6.9880 | 7.0310 | 3.3248 | 3.9223 | 3.4460 | 3.9461 |
| TASOR2_S1087 | TASOR2 | Q5VWN6 | | 2037 | PAPRSRSPLLVTV | 5.8961 | 6.2934 | 5.8676 | 6.8623 | 6.3051 | 5.9606 | 6.2305 | 6.0570 | 6.1186 | 5.9049 | 6.0306 | 6.1681 | 6.4783 | 6.5617 | 6.5641 | 6.7009 |
| TASOR2_S1268 | TASOR2 | Q5VWN6 | | 19 | SENVLMSPWKGKL | 6.6575 | 6.7426 | 6.5572 | 7.0867 | 6.5787 | 5.3513 | 6.4554 | 5.8879 | 6.3288 | 5.7838 | 5.6261 | 5.7441 | 6.5077 | 6.2770 | 6.8040 | 5.6112 |
| TASOR2_S19 | TASOR2 | Q5VWN6 | | 1087 | ELIKQVSPAASLR | 6.7716 | 7.0899 | 6.3690 | 7.4415 | 6.2896 | 6.0939 | 6.9208 | 6.7898 | 6.5920 | 6.2615 | 6.5943 | 6.4928 | 4.7258 | 5.2013 | 4.9866 | 5.3797 |
| TASOR2_S2009 | TASOR2 | Q5VWN6 | | 685 | PRNGTKSPEAATP | 5.9781 | 6.3313 | 6.0256 | 7.2789 | 5.8354 | 5.9325 | 6.4350 | 5.9396 | 6.0005 | 6.4248 | 6.3782 | 6.8266 | 5.9193 | 6.0263 | 6.0863 | 6.5815 |
| TASOR2_S2037 | TASOR2 | Q5VWN6 | | 675 | QEREILSPLFPRN | 7.3080 | 7.5443 | 7.1264 | 8.2013 | 6.6642 | 6.9769 | 7.2371 | 6.7294 | 7.4960 | 6.8737 | 7.3542 | 7.2023 | 2.7911 | 3.8992 | 3.0049 | 3.5912 |
| TASOR2_S340 | TASOR2 | Q5VWN6 | | 1268 | QTSLSVSREVSLE | 5.6935 | 6.1318 | 5.3846 | 6.3378 | 5.7847 | 5.3492 | 5.9374 | 6.1358 | 5.5413 | 5.3034 | 5.1928 | 5.3900 | 7.9082 | 7.4628 | 8.2372 | 8.2095 |
| TASOR2_S675 | TASOR2 | Q5VWN6 | | 1025 | DLIQHGSPINNEC | 6.6545 | 7.1214 | 6.1331 | 6.9532 | 6.4023 | 5.7968 | 6.1001 | 5.4981 | 7.1691 | 6.0422 | 5.7945 | 7.3281 | 5.3583 | 6.3614 | 5.5231 | 5.7639 |
| TASOR2_S685 | TASOR2 | Q5VWN6 | | 2009 | NVFPKESPTQISI | 6.6250 | 6.3293 | 5.9955 | 5.8055 | 6.1845 | 6.5797 | 6.6458 | 6.6906 | 6.2465 | 5.8992 | 6.3256 | 5.7727 | 6.3725 | 6.2793 | 6.1887 | 6.0596 |
| TATDN2_S115 | TATDN2 | Q93075 | | 115 | FLSSGGSPLRPAN | 5.8258 | 5.7049 | 5.5881 | 6.0988 | 6.2184 | 6.3250 | 6.7760 | 6.1581 | 6.3663 | 5.8474 | 5.6395 | 5.8912 | 7.0723 | 6.9475 | 6.7365 | 6.8042 |
| TATDN2_S299 | TATDN2 | Q93075 | | 51 | RSGGPSSPKRLKA | 6.0503 | 6.2815 | 6.0885 | 6.4915 | 5.8491 | 5.4958 | 6.0818 | 5.8822 | 6.1394 | 5.9967 | 6.2110 | 6.1235 | 6.5325 | 6.8025 | 6.9171 | 7.0566 |
| TATDN2_S51 | TATDN2 | Q93075 | | 299 | TVIDKCSPPLEFL | 6.1438 | 5.4485 | 6.0225 | 5.8176 | 5.9745 | 6.2380 | 6.0366 | 5.8439 | 6.2122 | 6.1095 | 6.0548 | 6.5476 | 7.0487 | 6.8335 | 6.9403 | 6.7280 |
| TAX1BP1_S124 | TAX1BP1 | Q86VP1 | | 344 | TFLLTTSSKEDTC | 8.2556 | 7.5083 | 8.0845 | 8.2312 | 4.5434 | 7.1765 | 6.5794 | 5.9787 | 6.3698 | 6.4955 | 6.9286 | 7.5362 | 3.6633 | 3.9148 | 4.4805 | 4.2535 |
| TAX1BP1_S344 | TAX1BP1 | Q86VP1 | | 124 | FQFRASSPVEELL | 5.5597 | 6.1663 | 5.8756 | 5.7047 | 7.3319 | 6.2168 | 5.6729 | 7.0895 | 6.3899 | 6.8811 | 6.2471 | 6.4094 | 5.9946 | 5.7227 | 6.4808 | 6.2572 |
| TBC1D1_S235 | TBC1D1 | Q86TI0 | | 235 | RRPMRKSFSQPGL | 7.2073 | 5.9583 | 6.9852 | 6.3549 | 5.7880 | 5.9875 | 6.3737 | 6.3808 | 9.0878 | 8.0811 | 8.1456 | 7.7710 | 3.7526 | 5.0182 | 3.2276 | 3.8804 |
| TBC1D1_S237 | TBC1D1 | Q86TI0 | | 237 | PMRKSFSQPGLRS | 7.0101 | 6.5847 | 6.6777 | 5.8004 | 6.2585 | 6.6542 | 6.0374 | 6.3988 | 6.2711 | 6.2549 | 6.3795 | 6.5111 | 5.6212 | 6.1747 | 5.2317 | 6.1338 |
| TBC1D1_S559 | TBC1D1 | Q86TI0 | | 652 | LGDSGGTPVKTRR | 6.6470 | 6.2665 | 7.2310 | 6.4707 | 6.5528 | 6.9758 | 5.8114 | 7.2492 | 6.9326 | 6.0400 | 6.3418 | 6.1014 | 5.3661 | 5.4455 | 5.0397 | 5.5285 |
| TBC1D1_S614 | TBC1D1 | Q86TI0 | | 614 | PQPARGSPGVSQR | 7.1487 | 7.2398 | 7.7065 | 6.5243 | 6.0947 | 6.3309 | 6.0560 | 7.1287 | 7.2173 | 6.8627 | 7.0316 | 7.0222 | 4.3497 | 4.7951 | 3.7840 | 4.7077 |
| TBC1D1_T596 | TBC1D1 | Q86TI0 | | 596 | FRRRANTLSHFPI | 6.8056 | 5.8403 | 6.7305 | 5.4863 | 6.3613 | 5.2810 | 6.2431 | 5.6829 | 6.5811 | 5.8499 | 5.2602 | 5.4272 | 6.6298 | 8.3997 | 6.8408 | 6.5803 |
| TBC1D1_T652 | TBC1D1 | Q86TI0 | | 559 | LPISESSFKLLGS | 6.4629 | 7.0709 | 6.8048 | 7.3146 | 6.4776 | 7.3362 | 6.7554 | 6.4908 | 6.8790 | 6.9688 | 7.4135 | 7.5409 | 3.8565 | 4.9000 | 3.6897 | 4.0386 |
| TBC1D10A_S414 | TBC1D10A | Q9BXI6-2 | | 416 | ALQPSPSIRLPLD | 9.5573 | 6.2743 | 6.9145 | 7.4941 | 6.0823 | 5.1722 | 5.9372 | 5.0448 | 6.8016 | 5.3820 | 5.9727 | 6.2810 | 5.9924 | 7.1187 | 4.7322 | 5.2427 |
| TBC1D10A_S416 | TBC1D10A | Q9BXI6-2 | | 414 | RPALQPSPSIRLP | 5.9554 | 6.4619 | 7.3030 | 6.3395 | 6.5225 | 6.7023 | 6.5662 | 6.2140 | 6.6599 | 6.6568 | 6.5348 | 6.2272 | 5.5692 | 5.9996 | 5.3689 | 4.9186 |
| TBC1D10B_S111 | TBC1D10B | Q4KMP7 | | 678 | AAGGAPSPPPPVR | 6.1089 | 6.3161 | 6.0721 | 5.9704 | 6.1679 | 6.2087 | 6.2180 | 6.2510 | 6.8228 | 6.4492 | 6.2454 | 6.3166 | 5.9462 | 6.5531 | 6.1747 | 6.1788 |
| TBC1D10B_S132 | TBC1D10B | Q4KMP7 | | 22 | GAPAAPSPPPRGS | 5.8542 | 6.3578 | 6.3923 | 6.0085 | 6.0232 | 6.2752 | 5.9482 | 6.3284 | 6.3643 | 6.5779 | 6.1994 | 6.4300 | 6.0929 | 6.6986 | 6.1816 | 6.2676 |
| TBC1D10B_S22 | TBC1D10B | Q4KMP7 | | 132 | LAAGADSPKTEEA | 6.5210 | 6.6190 | 6.4603 | 6.0974 | 6.5877 | 6.2492 | 6.8828 | 6.2264 | 6.9830 | 6.7938 | 6.2653 | 6.1100 | 5.4753 | 5.6580 | 5.2963 | 5.7744 |
| TBC1D10B_S657 | TBC1D10B | Q4KMP7 | | 657 | PPLGPSSSLLSLP | 4.7454 | 5.9808 | 5.9917 | 4.9471 | 6.0731 | 6.8697 | 6.4552 | 6.7644 | 6.9554 | 6.9247 | 6.6615 | 6.7995 | 5.8080 | 6.5129 | 5.9231 | 6.5877 |
| TBC1D10B_S658 | TBC1D10B | Q4KMP7 | | 687 | PPVRRASAGPAPG | 5.7570 | 5.8672 | 5.9125 | 6.0495 | 5.8887 | 6.1580 | 5.4885 | 7.0328 | 6.6177 | 6.8534 | 6.9278 | 6.7435 | 5.9356 | 6.9921 | 5.8771 | 5.8987 |
| TBC1D10B_S658;S661 | TBC1D10B | Q4KMP7 | | 111 | LPSGPESPEPAAV | 5.7391 | 6.3558 | 6.3773 | 6.3958 | 5.8050 | 6.0692 | 6.9311 | 5.9840 | 6.3003 | 6.4139 | 6.9079 | 6.2138 | 5.6664 | 6.4160 | 5.8975 | 6.5270 |
| TBC1D10B_S678 | TBC1D10B | Q4KMP7 | | 658 | PLGPSSSLLSLPG | 5.1743 | 5.8910 | 5.8386 | 5.2899 | 5.6744 | 6.3118 | 6.1834 | 6.8117 | 7.1899 | 7.1237 | 6.7103 | 6.7608 | 6.1425 | 6.6829 | 5.7540 | 4.6608 |
| TBC1D10B_S687 | TBC1D10B | Q4KMP7 | | 150 | PGPGTPTGTPTRT | 6.3701 | 6.2845 | 6.5299 | 6.7411 | 6.0399 | 6.1664 | 6.8401 | 6.2522 | 6.6122 | 6.6230 | 7.0123 | 6.8358 | 5.5214 | 5.8882 | 4.9446 | 5.3382 |
| TBC1D10B_S687;S707 | TBC1D10B | Q4KMP7 | | 148 | PAPGPGTPTGTPT | 6.2699 | 5.6054 | 5.9574 | 6.4955 | 6.5770 | 6.2648 | 6.1457 | 6.2988 | 7.7931 | 7.0657 | 6.3694 | 6.8006 | 5.4578 | 5.6596 | 5.2781 | 5.9615 |
| TBC1D10B_T148 | TBC1D10B | Q4KMP7 | 687;707 | | NA | 6.2435 | 6.9600 | 6.4299 | 6.8653 | 6.7824 | 6.8610 | 5.7176 | 7.3383 | 6.9729 | 7.0144 | 7.6633 | 7.5355 | 4.3208 | 5.1102 | 4.0084 | 4.1764 |
| TBC1D10B_T150 | TBC1D10B | Q4KMP7 | 658;661 | | NA | 6.6110 | 7.1008 | 6.4129 | 6.5861 | 6.1697 | 6.1444 | 6.0596 | 5.3608 | 6.3958 | 5.3558 | 5.9327 | 6.2031 | 6.3195 | 6.9493 | 5.9327 | 6.4659 |
| TBC1D12_S770 | TBC1D12 | O60347 | | 770 | EGDKNSSPALKSx | 6.6789 | 7.2203 | 7.6318 | 6.7242 | 5.7906 | 6.6919 | 6.7601 | 6.7287 | 7.0083 | 7.1034 | 7.3255 | 7.3708 | 3.9125 | 5.0926 | 3.7401 | 4.2203 |
| TBC1D13_S184 | TBC1D13 | Q9NVG8 | | 184 | GVTNMSSPHKNSV | 6.3972 | 5.8197 | 6.8809 | 5.7940 | 6.3726 | 6.9543 | 6.5015 | 7.3734 | 6.5858 | 6.8128 | 6.8274 | 7.5846 | 4.2949 | 5.6871 | 4.8584 | 5.2555 |
| TBC1D15_S213 | TBC1D15 | Q8TC07-3 | | 78 | LYARKDSSSVVEW | 5.5778 | 4.6553 | 6.4066 | 5.2791 | 6.5682 | 5.3462 | 5.2863 | 5.8533 | 7.3357 | 7.5709 | 6.8006 | 6.7683 | 6.8762 | 6.9883 | 6.6372 | 6.0500 |
| TBC1D15_S666 | TBC1D15 | Q8TC07-3 | | 213 | NKSLSQSFENLLD | 5.9841 | 6.7683 | 7.0220 | 6.4570 | 5.7241 | 6.5456 | 6.0485 | 6.2559 | 6.7519 | 7.2258 | 7.6946 | 7.3801 | 4.6929 | 5.5121 | 4.8657 | 5.0714 |
| TBC1D15_S78 | TBC1D15 | Q8TC07-3 | | 666 | SGARNDSPTQIPV | 6.4893 | 6.3222 | 6.9163 | 6.4308 | 6.2909 | 5.9682 | 6.3318 | 6.0908 | 6.3132 | 6.3584 | 6.6922 | 6.7515 | 5.5649 | 6.1231 | 5.6010 | 5.7554 |
| TBC1D17_S20 | TBC1D17 | Q9HA65 | | 20 | GVYLHTSAKKYQD | 6.4795 | 7.2446 | 6.9375 | 6.6520 | 6.0281 | 6.5654 | 6.9255 | 6.7869 | 6.5607 | 6.7262 | 5.9406 | 5.8635 | 5.3922 | 5.3900 | 4.9639 | 5.5435 |
| TBC1D2_S436 | TBC1D2 | Q9BYX2 | | 436 | THPPDQSPLRPDA | 6.9802 | 7.0885 | 6.8714 | 7.5603 | 6.4152 | 6.1409 | 6.7781 | 6.6881 | 6.6787 | 7.4566 | 8.0601 | 7.2583 | 3.9707 | 4.4833 | 3.2997 | 4.2699 |
| TBC1D2_T186 | TBC1D2 | Q9BYX2 | | 186 | FLCPVKTPPGLVG | 5.5603 | 6.5594 | 7.0914 | 6.7454 | 5.6181 | 6.1050 | 5.8012 | 6.9918 | 7.0078 | 6.1269 | 5.9628 | 6.9399 | 5.5911 | 5.7189 | 5.9560 | 6.2239 |
| TBC1D22A_S132 | TBC1D22A | Q8WUA7 | | 51 | TAKMPTTPVKAKR | 6.4111 | 6.4577 | 7.2350 | 6.7863 | 5.7233 | 7.1236 | 7.2014 | 7.1585 | 7.4222 | 7.5693 | 7.4096 | 7.6052 | 3.5691 | 4.7532 | 3.4884 | 4.0860 |
| TBC1D22A_S167 | TBC1D22A | Q8WUA7 | | 167 | PLQRSQSLPHSAT | 5.1654 | 4.9271 | 5.6894 | 5.3264 | 5.9905 | 5.6308 | 6.3867 | 5.8409 | 5.8790 | 5.8990 | 5.3184 | 5.5092 | 7.8615 | 8.4157 | 8.6264 | 7.5336 |
| TBC1D22A_S288 | TBC1D22A | Q8WUA7 | | 288 | IDIPRMSPEALIL | 6.1217 | 6.4096 | 7.6142 | 5.0251 | 6.3310 | 6.8000 | 6.8240 | 5.0530 | 6.4589 | 7.0545 | 6.0858 | 5.3235 | 6.6254 | 6.9119 | 6.0644 | 5.2969 |
| TBC1D22A_T51 | TBC1D22A | Q8WUA7 | | 132 | PEAEPPSPPSGDL | 5.2890 | 5.1050 | 6.1076 | 6.0590 | 6.3107 | 6.3849 | 5.8969 | 6.1647 | 6.1812 | 6.6004 | 6.5316 | 6.4529 | 6.5727 | 7.0156 | 6.5132 | 6.8146 |
| TBC1D22B_S116 | TBC1D22B | Q9NU19 | | 154 | PLHKQQSLPLRPI | 6.7161 | 6.6668 | 7.4370 | 6.2424 | 6.3517 | 5.4593 | 5.0863 | 6.4853 | 6.4624 | 6.7861 | 6.5739 | 6.0359 | 5.7515 | 6.4588 | 5.4915 | 5.9952 |
| TBC1D22B_S154 | TBC1D22B | Q9NU19 | | 116 | KPERSQSTTSDVP | 5.8048 | 5.3266 | 6.4707 | 6.1487 | 5.9521 | 5.8327 | 6.1099 | 6.3757 | 5.7501 | 6.6374 | 5.8989 | 5.5993 | 7.3377 | 7.6656 | 7.0873 | 6.0024 |
| TBC1D23_S300 | TBC1D23 | Q9NUY8 | | 520 | TPVDRMSFNLPWP | 4.4120 | 4.3148 | 5.1942 | 5.2091 | 5.6412 | 5.2960 | 5.2395 | 5.6031 | 5.7036 | 5.3570 | 5.1229 | 6.1444 | 8.0312 | 9.4041 | 9.3011 | 10.0258 |
| TBC1D23_S507 | TBC1D23 | Q9NUY8 | | 507 | VREKVISFIENTS | 3.1241 | 3.1213 | 3.5726 | 3.1269 | 3.9485 | 3.9824 | 3.6809 | 4.1369 | 3.7798 | 3.2673 | 3.2946 | 3.9796 | 11.1389 | 13.5837 | 15.0497 | 17.2128 |
| TBC1D23_S520 | TBC1D23 | Q9NUY8 | | 300 | NHHLFGSTLLGIK | 4.1772 | 4.7063 | 3.8894 | 4.0386 | 4.8515 | 4.5886 | 4.7038 | 4.9697 | 4.3308 | 4.3617 | 3.6680 | 4.1857 | 11.3985 | 11.0629 | 13.8298 | 11.2375 |
| TBC1D23_Y560 | TBC1D23 | Q9NUY8 | | 560 | IGDEEEYDTDEID | 6.3841 | 5.4171 | 7.5949 | 6.6878 | 6.6145 | 6.8001 | 6.4960 | 6.0019 | 6.4755 | 6.4740 | 6.1577 | 6.8566 | 5.7056 | 6.1960 | 4.7844 | 5.3539 |
| TBC1D25_S506 | TBC1D25 | Q3MII6 | | 506 | RLLRQASLDGLQQ | 6.1191 | 6.1623 | 6.6160 | 5.6866 | 6.2916 | 6.7562 | 5.9701 | 7.0537 | 6.3867 | 6.5963 | 6.4690 | 6.4750 | 5.8083 | 6.1708 | 5.7747 | 5.6636 |
| TBC1D2B_S155 | TBC1D2B | Q9UPU7 | | 277 | EEKKKLTPEGNKG | 6.2082 | 6.5776 | 6.9320 | 6.9184 | 6.1488 | 6.6351 | 6.5422 | 6.3571 | 6.4179 | 6.5572 | 6.9938 | 6.2764 | 5.0124 | 5.6594 | 5.3211 | 5.4425 |
| TBC1D2B_S473 | TBC1D2B | Q9UPU7 | | 473 | PTVAPSSPSVVPV | 6.3548 | 5.2896 | 6.8597 | 6.5093 | 6.5129 | 7.0965 | 6.8692 | 6.8765 | 7.0317 | 6.9384 | 7.0317 | 6.6266 | 4.7301 | 5.2379 | 4.7446 | 5.2903 |
| TBC1D2B_T277 | TBC1D2B | Q9UPU7 | | 155 | KWDSRTSPTPGDF | 7.1447 | 6.7070 | 6.7373 | 6.5264 | 6.7456 | 6.4350 | 5.9770 | 6.8680 | 6.5008 | 6.6728 | 6.7693 | 7.6941 | 4.8801 | 5.1856 | 4.5815 | 4.5749 |
| TBC1D30_S713 | TBC1D30 | Q9Y2I9 | | 713 | HTGGKISPVPYED | 5.9701 | 6.4924 | 6.3839 | 6.2894 | 6.7390 | 6.3476 | 6.2052 | 6.0022 | 6.7414 | 5.9010 | 5.7489 | 6.2143 | 6.3130 | 6.2203 | 5.9458 | 6.4857 |
| TBC1D30_S80 | TBC1D30 | Q9Y2I9 | | 80 | RTSRTASLVSGLL | 5.9852 | 6.0457 | 6.7554 | 6.5692 | 5.3103 | 5.7199 | 5.0683 | 6.3365 | 7.4046 | 7.6374 | 5.1148 | 6.1248 | 5.6175 | 6.0433 | 6.7321 | 7.5350 |
| TBC1D4_S262 | TBC1D4 | O60343 | | 782 | IFLRVASPMNKSP | 6.9969 | 7.0177 | 7.1067 | 6.8451 | 6.3050 | 7.0396 | 7.0453 | 7.2644 | 7.2827 | 7.0174 | 7.3458 | 7.3357 | 3.5325 | 4.1251 | 3.4883 | 4.2520 |
| TBC1D4_S341 | TBC1D4 | O60343 | | 666 | RAQGVRSPLLRQS | 6.7353 | 6.4991 | 7.3479 | 5.9566 | 6.6991 | 7.1702 | 6.2899 | 6.2280 | 6.2116 | 6.3032 | 6.8011 | 6.7116 | 5.4062 | 5.5304 | 5.1157 | 4.9942 |
| TBC1D4_S370 | TBC1D4 | O60343 | | 105 | AGASGGTSPSATQ | 6.2062 | 6.3975 | 6.8502 | 6.3823 | 6.3209 | 6.6890 | 6.5362 | 6.4792 | 6.4206 | 6.6383 | 6.9200 | 7.0385 | 5.2814 | 5.3008 | 5.3644 | 5.1745 |
| TBC1D4_S566 | TBC1D4 | O60343 | | 570 | KRSLTSSLENIFS | 6.0505 | 6.0820 | 6.5973 | 6.1180 | 5.8811 | 6.6715 | 6.1629 | 6.9653 | 6.3165 | 6.2563 | 6.6303 | 6.3742 | 5.8417 | 6.1365 | 6.0611 | 5.8547 |
| TBC1D4_S569 | TBC1D4 | O60343 | | 642 | FRRRAHTFSHPPS | 6.7678 | 5.5155 | 6.8028 | 5.9098 | 6.4877 | 5.3777 | 6.6750 | 5.3224 | 6.0896 | 3.8094 | 4.2206 | 4.6100 | 9.1400 | 9.3339 | 7.2834 | 6.6543 |
| TBC1D4_S570 | TBC1D4 | O60343 | | 588 | MRGRLGSVDSFER | 5.6392 | 5.1369 | 6.4306 | 4.9061 | 5.9516 | 6.2967 | 6.1013 | 6.4697 | 6.8935 | 6.4284 | 6.6455 | 6.9304 | 7.0666 | 6.7969 | 5.9332 | 6.3734 |
| TBC1D4_S588 | TBC1D4 | O60343 | | 341 | PRRRHASAPSHVQ | 6.2550 | 6.9557 | 7.3828 | 6.9920 | 6.0876 | 6.9320 | 6.2003 | 6.6755 | 6.7362 | 6.7889 | 7.1937 | 7.4738 | 4.2425 | 4.7896 | 4.6275 | 4.6670 |
| TBC1D4_S591 | TBC1D4 | O60343 | | 370 | FEINLISPDTKSV | 6.4057 | 6.0309 | 7.8197 | 7.6868 | 7.0633 | 6.7396 | 6.2626 | 6.0615 | 6.6374 | 5.5436 | 7.7175 | 6.2898 | 4.1603 | 6.1501 | 5.0939 | 4.3375 |
| TBC1D4_S649 | TBC1D4 | O60343 | | 591 | RLGSVDSFERSNS | 8.2362 | 7.7503 | 8.5606 | 7.5979 | 5.9135 | 5.8628 | 5.7171 | 6.4197 | 7.0616 | 6.7171 | 6.1538 | 5.9128 | 4.0690 | 5.0357 | 4.6763 | 4.3154 |
| TBC1D4_S666 | TBC1D4 | O60343 | | 566 | KNKAKRSLTSSLE | 6.6385 | 5.7912 | 6.7699 | 6.5316 | 6.8724 | 6.1375 | 6.1575 | 6.0417 | 8.1039 | 7.3712 | 6.2566 | 6.6786 | 5.7355 | 5.9059 | 4.5998 | 4.4082 |
| TBC1D4_S782 | TBC1D4 | O60343 | | 809 | LPLSPLSPTMEEE | 4.8785 | 5.0437 | 6.4569 | 5.1474 | 6.2952 | 6.2697 | 6.3787 | 7.7356 | 5.8206 | 5.8127 | 5.3838 | 6.4877 | 7.9747 | 6.2926 | 6.4727 | 7.5493 |
| TBC1D4_S806 | TBC1D4 | O60343 | | 806 | NELLPLSPLSPTM | 6.2524 | 6.4508 | 5.1878 | 7.0170 | 6.7678 | 5.8523 | 6.2507 | 5.8133 | 6.0812 | 6.3609 | 6.5678 | 7.2696 | 4.7910 | 7.6053 | 5.8913 | 5.8405 |
| TBC1D4_S809 | TBC1D4 | O60343 | | 649 | FSHPPSSTKRKLN | 6.1913 | 6.6968 | 7.3918 | 6.3630 | 6.1874 | 7.1797 | 6.2795 | 6.9141 | 7.1597 | 7.7908 | 8.1846 | 7.4270 | 3.7337 | 4.7795 | 3.4190 | 4.3022 |
| TBC1D4_T105 | TBC1D4 | O60343 | | 262 | EVVVPGSPGDCLP | 5.9669 | 6.3220 | 5.9116 | 7.3446 | 6.5597 | 5.1530 | 5.3699 | 5.1875 | 7.2645 | 6.9356 | 6.5777 | 7.3750 | 6.1645 | 5.7472 | 6.5611 | 5.5592 |
| TBC1D4_T642 | TBC1D4 | O60343 | | 569 | AKRSLTSSLENIF | 6.0111 | 5.8939 | 6.8702 | 5.8861 | 5.9909 | 7.0509 | 6.4765 | 6.9192 | 5.8953 | 5.8897 | 6.3281 | 6.6364 | 5.8459 | 6.6008 | 5.9980 | 5.7069 |
| TBC1D4_T752 | TBC1D4 | O60343 | | 752 | RKRTSSTCSNESL | 4.8706 | 6.8000 | 5.8885 | 6.3244 | 5.9678 | 7.6260 | 5.6111 | 6.6728 | 5.6069 | 6.7082 | 6.6936 | 6.0074 | 5.5172 | 6.4663 | 7.4175 | 5.8217 |
| TBC1D5_S43 | TBC1D5 | Q92609-2 | | 576 | GREFTGSPPSSAT | 6.3950 | 6.3299 | 7.0821 | 6.4461 | 6.2900 | 6.4877 | 6.3145 | 6.8802 | 7.0267 | 7.0871 | 7.1033 | 7.1366 | 4.5677 | 5.3334 | 4.7232 | 4.7964 |
| TBC1D5_S522 | TBC1D5 | Q92609-2 | | 752 | GSSARGSFSGQAQ | 5.6432 | 4.9489 | 5.9991 | 5.6805 | 6.4520 | 6.1489 | 5.9477 | 6.2936 | 7.6561 | 7.7178 | 7.5081 | 7.2444 | 5.7535 | 7.0327 | 5.1734 | 4.8001 |
| TBC1D5_S561 | TBC1D5 | Q92609-2 | | 522 | RLMKSESMPVQLN | 3.6159 | 3.6257 | 3.5077 | 3.5136 | 3.2969 | 3.1814 | 2.9744 | 3.0132 | 3.5699 | 3.7802 | 3.4237 | 3.7156 | 15.1011 | 14.6379 | 15.5580 | 13.4848 |
| TBC1D5_S563 | TBC1D5 | Q92609-2 | | 580 | TGSPPSSATKKDS | 6.5647 | 6.7007 | 7.4448 | 6.7097 | 6.1315 | 6.1014 | 6.5698 | 6.5217 | 6.8499 | 7.3637 | 7.5438 | 7.3851 | 4.2458 | 4.8844 | 4.3531 | 4.6298 |
| TBC1D5_S576 | TBC1D5 | Q92609-2 | | 43 | KNGRRTSSTLDSE | 5.0233 | 5.0652 | 6.3289 | 5.6484 | 6.5747 | 5.5069 | 5.5602 | 6.4078 | 7.9606 | 7.5126 | 7.1062 | 7.5016 | 6.0225 | 6.9241 | 5.3780 | 5.4792 |
| TBC1D5_S580 | TBC1D5 | Q92609-2 | | 563 | NISSSPSVESLPG | 4.0156 | 4.1662 | 5.0775 | 4.5899 | 4.8648 | 5.1325 | 4.9765 | 4.7841 | 4.2869 | 4.7166 | 5.0207 | 4.7763 | 11.1476 | 10.0158 | 12.0893 | 10.3397 |
| TBC1D5_S752 | TBC1D5 | Q92609-2 | | 561 | SKNISSSPSVESL | 7.0734 | 7.1037 | 7.5775 | 6.8522 | 5.0513 | 6.0594 | 7.0804 | 6.7987 | 7.1409 | 7.2760 | 6.0641 | 6.8220 | 5.0327 | 4.5076 | 4.8930 | 4.6671 |
| TBC1D5_S798 | TBC1D5 | Q92609-2 | | 798 | SSNPSSSPDDDSS | 5.8635 | 5.7099 | 5.4516 | 6.0217 | 6.6988 | 5.8309 | 5.6924 | 6.7476 | 5.9973 | 5.1619 | 7.6330 | 5.9333 | 6.6266 | 7.5480 | 6.2183 | 6.8651 |
| TBC1D9B_S1138 | TBC1D9B | Q66K14 | | 435 | QPASPASPLSSRQ | 6.1790 | 6.3929 | 7.2118 | 7.2103 | 6.2097 | 6.7359 | 6.5717 | 7.1884 | 6.5585 | 6.7775 | 7.6577 | 6.6418 | 4.8696 | 4.4960 | 4.6281 | 4.6711 |
| TBC1D9B_S411 | TBC1D9B | Q66K14 | | 442 | PLSSRQSFCAQEA | 5.0241 | 4.1058 | 4.9837 | 4.5313 | 5.4236 | 4.7715 | 4.8856 | 4.5619 | 5.7053 | 5.3707 | 5.1813 | 4.4918 | 10.3642 | 10.1805 | 10.8934 | 9.5253 |
| TBC1D9B_S435 | TBC1D9B | Q66K14 | | 411 | IGSRKASVVDPST | 5.0618 | 5.5884 | 5.5528 | 5.6382 | 6.7580 | 5.4129 | 5.6098 | 5.9680 | 6.5951 | 6.6093 | 5.5078 | 6.6789 | 8.4123 | 8.0430 | 6.6995 | 5.8644 |
| TBC1D9B_S442 | TBC1D9B | Q66K14 | | 995 | EEKGTSSPDYRHY | 6.5437 | 7.1803 | 6.7778 | 6.8268 | 6.2915 | 6.6704 | 6.5622 | 6.6266 | 6.7390 | 6.5857 | 6.5275 | 7.2275 | 4.7970 | 5.0185 | 4.5452 | 5.0803 |
| TBC1D9B_S995 | TBC1D9B | Q66K14 | | 1138 | SGEGQGSPSQLLS | 6.4240 | 6.2158 | 6.8664 | 6.4325 | 6.5448 | 5.9460 | 6.5158 | 6.2200 | 5.9249 | 6.5635 | 6.1241 | 6.7940 | 6.0287 | 5.6262 | 5.9545 | 5.8187 |
| TBCB_S173 | TBCB | Q99426 | | 173 | VRAAGQSPRRGTV | 6.3987 | 6.6909 | 7.3827 | 5.9998 | 5.9143 | 6.3246 | 5.9534 | 6.6650 | 6.3849 | 6.8721 | 6.7717 | 6.5980 | 5.2906 | 5.7034 | 5.2935 | 5.7566 |
| TBCC_S168 | TBCC | Q15814 | | 168 | VESIQDSPLPKKA | 6.3546 | 6.7452 | 6.9942 | 6.9915 | 6.0751 | 7.3153 | 6.8559 | 6.8078 | 7.2837 | 7.1201 | 7.2774 | 7.4865 | 3.7939 | 4.7388 | 3.7616 | 4.3985 |
| TBCEL_S41 | TBCEL | Q5QJ74 | | 41 | PATPQGSPMKDRL | 5.9631 | 6.1963 | 6.7986 | 6.0700 | 6.7016 | 6.9494 | 6.3477 | 7.3025 | 6.8633 | 6.4740 | 7.0812 | 7.0882 | 4.8949 | 5.5071 | 4.8903 | 4.8718 |
| TBCK_S890 | TBCK | Q8TEA7 | | 890 | GLLTIPSPQIxxx | 6.0426 | 6.7260 | 6.0036 | 6.3107 | 6.2568 | 6.6855 | 6.5791 | 6.2447 | 6.4970 | 7.0893 | 7.0170 | 7.0160 | 5.6149 | 5.7013 | 4.8117 | 5.4037 |
| TBKBP1_S365 | TBKBP1 | A7MCY6 | | 504 | GPGRPLSPRRAFE | 6.0654 | 6.0813 | 6.5642 | 5.9246 | 6.5770 | 6.5199 | 6.1287 | 7.0177 | 6.6245 | 6.3798 | 6.6154 | 6.6659 | 5.8266 | 6.0023 | 5.5305 | 5.4762 |
| TBKBP1_S475 | TBKBP1 | A7MCY6 | | 365 | PCPPCQSPVPQRR | 6.5579 | 6.2488 | 7.2637 | 6.2467 | 6.5419 | 6.4792 | 6.4055 | 6.5790 | 7.1311 | 6.6973 | 6.6817 | 6.6401 | 5.6197 | 5.5393 | 4.5450 | 4.8230 |
| TBKBP1_S504 | TBKBP1 | A7MCY6 | | 64 | LERENATLRRRLK | 6.5714 | 5.6369 | 5.8823 | 7.1668 | 5.8674 | 6.7515 | 6.1427 | 6.5612 | 6.4872 | 6.5210 | 5.8563 | 5.8650 | 5.7966 | 6.5643 | 6.3763 | 5.9531 |
| TBKBP1_T64 | TBKBP1 | A7MCY6 | | 475 | AAYAGASPPWLQA | 6.5104 | 6.5129 | 6.5578 | 6.4055 | 6.4902 | 6.9482 | 6.8761 | 7.1276 | 7.4946 | 6.7315 | 7.4213 | 7.5243 | 3.9892 | 4.6127 | 4.3196 | 4.4782 |
| TBL3_S257 | TBL3 | Q12788 | | 257 | SQLGVKSPGLYFL | 5.6983 | 5.9090 | 6.6184 | 6.5131 | 6.2732 | 6.2614 | 6.2806 | 6.0012 | 6.4643 | 6.5020 | 6.1098 | 6.3226 | 5.8994 | 6.0286 | 6.1528 | 6.9652 |
| TBRG4_S553 | TBRG4 | Q969Z0 | | 553 | QPTGSQSPPPGSK | 6.3667 | 6.8126 | 6.6808 | 7.5318 | 6.0919 | 6.4551 | 8.4432 | 5.6979 | 5.2829 | 5.3264 | 5.5774 | 6.2419 | 5.5704 | 6.3106 | 6.0148 | 5.5956 |
| TBRG4_T65 | TBRG4 | Q969Z0 | | 65 | EKERASTPYIEKQ | 6.4561 | 6.3723 | 6.4251 | 6.0361 | 5.5633 | 6.0521 | 6.0206 | 5.8845 | 7.2578 | 6.5916 | 6.8616 | 7.0033 | 6.1830 | 6.5864 | 5.3575 | 5.3487 |
| TBX18_S117 | TBX18 | O95935 | | 543 | SPKLAASPEKIVS | 6.4061 | 6.8613 | 7.0180 | 6.3056 | 6.1563 | 6.4815 | 6.4946 | 6.5355 | 6.7795 | 6.9172 | 6.9753 | 7.0584 | 4.8918 | 5.0475 | 4.8243 | 5.2470 |
| TBX18_S543 | TBX18 | O95935 | | 8 | AEKRRGSPCSMLS | 5.3629 | 5.8597 | 5.5911 | 5.8952 | 5.8976 | 6.3308 | 5.7599 | 5.3904 | 6.0177 | 6.3525 | 6.6395 | 6.5105 | 7.1716 | 7.0822 | 7.0692 | 7.0693 |
| TBX18_S8 | TBX18 | O95935 | | 117 | LASPGGSPKGSPA | 6.0198 | 5.3820 | 5.4003 | 5.8875 | 6.2647 | 6.1276 | 6.2457 | 6.0781 | 6.4451 | 6.2090 | 6.4241 | 6.9207 | 7.4892 | 6.4092 | 6.2932 | 6.4038 |
| TBX18_T131 | TBX18 | O95935 | | 131 | SLARPGTPLPSPQ | 5.9417 | 6.0267 | 5.9502 | 6.5169 | 6.0918 | 6.4027 | 6.5039 | 6.2893 | 5.8367 | 6.5451 | 6.3989 | 6.3596 | 6.0879 | 6.4271 | 6.3826 | 6.2390 |
| TBX2_S336 | TBX2 | Q13207 | | 676 | PSRGALSPSGSAK | 6.9233 | 6.7759 | 6.0514 | 6.4958 | 6.5960 | 6.2613 | 6.1501 | 6.2043 | 6.6652 | 6.0594 | 6.4879 | 6.2188 | 5.7654 | 5.5974 | 5.7053 | 6.0426 |
| TBX2_S676 | TBX2 | Q13207 | | 336 | AREPPTSPGAAPS | 6.3130 | 6.9414 | 6.0711 | 6.4665 | 5.4584 | 6.0767 | 6.0151 | 5.7978 | 7.0691 | 7.0691 | 7.6829 | 5.5289 | 6.2559 | 5.0719 | 5.1537 | 7.0288 |
| TBX2_S705;S710 | TBX2 | Q13207 | 705;710 | | NA | 6.1689 | 6.4730 | 6.9959 | 6.6604 | 5.8269 | 6.0296 | 6.2761 | 5.7949 | 6.9995 | 6.8740 | 7.1662 | 7.0447 | 5.5324 | 5.0353 | 5.4807 | 5.6415 |
| TBXA2R_S331 | TBXA2R | P21731 | | 331 | TRPRSLSLQPQLT | 5.8913 | 6.3685 | 5.8310 | 5.8665 | 6.2605 | 5.5094 | 5.5026 | 6.1659 | 6.7964 | 6.9165 | 6.4061 | 5.6685 | 6.4531 | 6.9731 | 5.5606 | 5.9300 |
| TCEA1_S100 | TCEA1 | P23193 | | 100 | AITSQNSPEAREE | 5.7898 | 5.8559 | 6.2928 | 5.6798 | 5.7727 | 6.0727 | 5.8191 | 6.3113 | 6.1396 | 6.4882 | 6.1422 | 6.2794 | 6.5092 | 7.2952 | 6.8682 | 6.6838 |
| TCEA1_S81 | TCEA1 | P23193 | | 81 | KLLDGPSTEKDLD | 5.0474 | 5.7071 | 5.7478 | 4.6727 | 5.9124 | 6.5562 | 6.3818 | 5.9574 | 5.2818 | 5.3878 | 5.4492 | 6.6438 | 7.2754 | 7.5839 | 8.2914 | 8.1039 |
| TCEA3_S115 | TCEA3 | O75764 | | 115 | KPEAGLSPPRKKR | 5.6142 | 6.2315 | 6.5937 | 6.1615 | 6.0502 | 6.1447 | 6.2950 | 6.9350 | 6.1119 | 6.6585 | 6.1458 | 6.2273 | 5.8099 | 6.4034 | 6.2549 | 6.3624 |
| TCEA3_S140 | TCEA3 | O75764 | | 140 | KSSASSSPKRPSV | 6.7674 | 6.4090 | 6.4576 | 6.5866 | 5.7600 | 6.1074 | 5.5340 | 5.1067 | 6.9443 | 6.8111 | 6.4299 | 6.3773 | 6.2264 | 6.5805 | 5.8775 | 6.0243 |
| TCEA3_S164 | TCEA3 | O75764 | | 164 | SPKTPSSPLTPTF | 5.6930 | 5.9519 | 7.0975 | 6.9596 | 7.3588 | 5.3277 | 6.1297 | 5.7220 | 6.5434 | 6.3208 | 6.3825 | 6.8374 | 6.7140 | 5.7244 | 5.1377 | 6.0995 |
| TCEAL1_S28 | TCEAL1 | Q15170-2 | | 28 | RPPVEHSPEKQSP | 5.3976 | 5.3806 | 6.3925 | 5.1397 | 6.2114 | 5.7217 | 5.1117 | 5.7478 | 6.2543 | 7.2354 | 7.1573 | 7.2133 | 6.4796 | 7.3302 | 6.6441 | 6.5829 |
| TCEAL3_S30 | TCEAL3 | Q969E4 | | 30 | PDDEGKSDEEEKP | 5.5783 | 5.1676 | 6.9740 | 5.0965 | 5.4786 | 5.7080 | 5.8125 | 4.7469 | 6.1101 | 6.4771 | 6.7766 | 6.7609 | 7.7507 | 7.7936 | 6.8291 | 6.9395 |
| TCEAL4_S210 | TCEAL4 | Q96EI5-2 | | 210 | DKGKPESEGEAKE | 6.5469 | 6.5740 | 7.0656 | 5.7516 | 6.5914 | 6.4267 | 6.4024 | 6.7247 | 5.8163 | 6.5553 | 6.4770 | 6.5686 | 5.4437 | 5.9172 | 5.7720 | 5.3667 |
| TCEANC2_S14 | TCEANC2 | Q96MN5 | | 14 | TPRIQNSPQKKDS | 5.9029 | 6.1561 | 5.9049 | 5.5387 | 5.9669 | 6.0903 | 6.0037 | 6.1381 | 6.7527 | 6.5685 | 6.5006 | 6.4434 | 6.1306 | 6.6521 | 6.5037 | 6.7470 |
| TCEANC2_T55 | TCEANC2 | Q96MN5 | | 55 | LELPDQTKENLVE | 6.1159 | 6.3527 | 7.4734 | 6.5266 | 5.6522 | 4.3961 | 7.2271 | 8.1014 | 7.3092 | 6.6377 | 5.0821 | 6.8599 | 6.2560 | 6.1739 | 5.7295 | 4.1063 |
| TCF12_S386 | TCF12 | Q99081 | | 386 | GGQAPSSPSYENS | 5.5799 | 5.5907 | 6.3298 | 6.5833 | 5.6940 | 6.7192 | 6.2962 | 6.3922 | 6.6848 | 6.3279 | 7.7879 | 6.7192 | 5.5862 | 5.6197 | 6.0617 | 6.0273 |
| TCF19_T263 | TCF19 | Q9Y242 | | 263 | VDKAPLTPTGNRR | 6.9873 | 6.4137 | 6.6336 | 5.6967 | 5.6461 | 6.6899 | 6.0718 | 6.8285 | 7.3960 | 7.1637 | 7.9531 | 8.0674 | 4.4429 | 4.9989 | 4.3929 | 4.6174 |
| TCF20_S1005 | TCF20 | Q9UGU0 | | 1671 | KGQRSLTPPPSST | 6.2920 | 6.1311 | 5.8551 | 6.5584 | 5.9073 | 5.7825 | 6.2461 | 5.9728 | 6.0090 | 5.9157 | 5.8341 | 5.9264 | 6.5448 | 6.9960 | 7.0901 | 6.9386 |
| TCF20_S1019 | TCF20 | Q9UGU0 | | 574 | PPRLNASPAAREE | 6.2427 | 6.6305 | 5.9079 | 6.6028 | 5.9405 | 5.7714 | 6.0083 | 6.1974 | 6.3140 | 6.2220 | 5.8059 | 5.9213 | 6.3121 | 6.6527 | 6.7928 | 6.6777 |
| TCF20_S1322 | TCF20 | Q9UGU0 | | 1335 | PAVTLTSPAKTKI | 6.5460 | 6.5774 | 6.4922 | 6.9404 | 6.4129 | 6.0150 | 6.7730 | 6.0352 | 6.9490 | 6.5711 | 6.4154 | 6.9728 | 5.1979 | 5.5524 | 4.9673 | 5.5820 |
| TCF20_S1335 | TCF20 | Q9UGU0 | | 871 | SVICDISPLRQIV | 6.8029 | 6.7219 | 6.7658 | 7.1074 | 6.4223 | 5.9643 | 6.6092 | 6.9461 | 6.4851 | 7.1617 | 6.8783 | 7.3326 | 3.9135 | 5.4202 | 4.6178 | 4.8509 |
| TCF20_S1522 | TCF20 | Q9UGU0 | | 1322 | SSKDLPSPDSRNC | 5.8594 | 6.6418 | 6.1154 | 6.1948 | 6.4016 | 5.7604 | 6.5642 | 6.3334 | 6.1554 | 6.0164 | 6.1682 | 6.2489 | 6.3009 | 6.2848 | 6.5674 | 6.3870 |
| TCF20_S31 | TCF20 | Q9UGU0 | | 31 | SRLEEFSPRQAQM | 7.2030 | 7.9386 | 6.6104 | 7.4503 | 6.2876 | 6.5619 | 7.1245 | 6.7430 | 7.3321 | 7.3015 | 7.3458 | 7.3698 | 2.9882 | 4.3070 | 3.4121 | 4.0243 |
| TCF20_S559 | TCF20 | Q9UGU0 | | 1019 | AEKLKMSPGRSRG | 5.8791 | 6.3216 | 6.0809 | 8.3807 | 7.6073 | 6.9704 | 7.2894 | 5.8808 | 6.4597 | 5.0234 | 6.0262 | 5.8148 | 5.4253 | 6.9890 | 5.1255 | 4.7258 |
| TCF20_S574 | TCF20 | Q9UGU0 | | 1522 | NDTVTISPKQEGF | 6.1440 | 5.4578 | 5.5820 | 6.6420 | 6.5917 | 5.6505 | 6.7014 | 6.2488 | 6.9836 | 6.1826 | 5.6823 | 5.8936 | 6.6243 | 6.2846 | 6.7123 | 6.6184 |
| TCF20_S871 | TCF20 | Q9UGU0 | | 559 | SEKAGSSPAQGAQ | 5.8818 | 6.3623 | 6.2553 | 7.9478 | 5.3465 | 6.2451 | 6.4757 | 6.2502 | 5.1006 | 6.3623 | 5.8665 | 5.7085 | 6.9983 | 6.2591 | 6.8339 | 6.1061 |
| TCF20_T1360 | TCF20 | Q9UGU0 | | 1005 | EGMRGRSPSQYHD | 6.2142 | 6.9792 | 6.0647 | 6.7775 | 5.6153 | 6.7700 | 6.2263 | 6.5990 | 6.0217 | 5.5967 | 6.1917 | 6.0964 | 5.7826 | 7.1483 | 5.8657 | 6.0507 |
| TCF20_T1671 | TCF20 | Q9UGU0 | | 1360 | AIVQKITSPNIRR | 7.4996 | 7.5743 | 6.5662 | 7.8257 | 6.7204 | 6.8622 | 7.3116 | 6.8131 | 7.5071 | 7.1638 | 7.1482 | 6.8639 | 3.1468 | 3.9888 | 2.9666 | 4.0416 |
| TCF25_S602 | TCF25 | Q9BQ70 | | 602 | VRPERLSPISHGN | 6.1420 | 6.2518 | 5.9925 | 6.7941 | 6.6340 | 6.3505 | 6.3084 | 7.3495 | 6.1696 | 6.3742 | 6.0656 | 5.8585 | 5.4347 | 5.7922 | 6.0481 | 6.4342 |
| TCF25_T221 | TCF25 | Q9BQ70 | | 221 | KCTWLTTPKSTWP | 7.2121 | 9.2402 | 8.4174 | 8.8500 | 5.0836 | 6.2696 | 7.5289 | 5.9258 | 7.7877 | 8.1508 | 6.8838 | 5.9412 | 3.5269 | 3.6930 | 2.0010 | 3.4882 |
| TCF3_S328 | TCF3 | P15923-3 | | 328 | GAPGALSPSYDGG | 6.4906 | 6.5452 | 7.0495 | 6.8111 | 6.6819 | 6.4450 | 5.7374 | 6.7941 | 7.0278 | 7.3146 | 7.9546 | 7.3009 | 4.5508 | 4.8399 | 4.3052 | 4.1515 |
| TCF3_S88 | TCF3 | P15923-3 | | 88 | NSPGPLSPSGMKG | 6.3708 | 6.2456 | 6.3071 | 6.2359 | 6.4455 | 6.5273 | 5.9489 | 6.6002 | 6.7958 | 6.8033 | 7.6932 | 6.5040 | 4.9184 | 5.5754 | 5.8158 | 5.2129 |
| TCF4_S140 | TCF4 | P15884-3 | | 140 | GNPGTLSPTKPGS | 5.8637 | 5.8395 | 7.1312 | 5.8727 | 6.5569 | 6.8081 | 6.2664 | 5.5093 | 6.5473 | 6.1899 | 7.2314 | 7.3779 | 5.6860 | 6.1687 | 5.1816 | 5.7695 |
| TCF4_S515 | TCF4 | P15884-3 | | 515 | GSSEIKSDDEGDE | 6.9808 | 5.5140 | 6.9339 | 6.3269 | 6.1277 | 6.1670 | 6.1938 | 5.2785 | 7.4902 | 7.1494 | 6.7491 | 7.2135 | 5.8003 | 5.3273 | 5.2928 | 5.4546 |

| | | | | | | | | | | | | | | | | | | | | |
|---|---|---|---|---|---|---|---|---|---|---|---|---|---|---|---|---|---|---|---|---|
| TCF4_T483 | TCF4 | P15884-3 | 483 | LPVQSATSPDLNP | 5.0814 | 6.1891 | 4.6462 | 6.1125 | 8.7434 | 4.3420 | 5.7490 | 5.3805 | 6.9037 | 5.8330 | 8.8176 | 7.3686 | 6.7356 | 6.0234 | 6.5847 | 5.4893 |
| TCF7L2_S179 | TCF7L2 | Q9NQB0 | 179 | KDARSPSPAHIVS | 6.4197 | 6.8071 | 5.8591 | 5.8717 | 5.5697 | 6.6118 | 6.1774 | 6.2704 | 6.1275 | 6.2289 | 6.5304 | 6.3996 | 5.8964 | 6.9536 | 5.8973 | 6.3794 |
| TCOF1_S1073 | TCOF1 | Q13428-4 | 1293 | QAAGMLSPKTGGK | 6.6258 | 5.9833 | 7.0314 | 7.4857 | 6.5406 | 6.5385 | 7.1261 | 5.8261 | 7.0560 | 6.7351 | 7.0633 | 6.9557 | 4.3657 | 5.1641 | 4.2601 | 5.2425 |
| TCOF1_S1077 | TCOF1 | Q13428-4 | 1386 | SRKRKLSGDQPAA | 5.7418 | 5.9348 | 5.6879 | 6.3032 | 6.0763 | 5.8163 | 6.2495 | 6.0764 | 5.6848 | 5.7416 | 5.5011 | 5.7288 | 6.6886 | 7.7159 | 7.4613 | 7.5915 |
| TCOF1_S1078 | TCOF1 | Q13428-4 | 914 | RKGAAPTPPGKTG | 6.3259 | 6.7916 | 6.0666 | 7.0723 | 6.1389 | 6.3752 | 6.5350 | 6.3503 | 6.3410 | 6.4216 | 6.4423 | 6.4291 | 4.8522 | 6.2175 | 5.4806 | 6.1598 |
| TCOF1_S1147 | TCOF1 | Q13428-4 | 533 | GKVGPATPSAQVG | 6.3830 | 6.6884 | 5.9800 | 6.7526 | 5.9816 | 6.3693 | 6.9014 | 6.8914 | 6.8303 | 6.5070 | 6.2567 | 6.3992 | 4.7514 | 6.0073 | 5.2337 | 6.0667 |
| TCOF1_S1263 | TCOF1 | Q13428-4 | 249 | GKVGDVTPQVKGG | 7.2084 | 7.2079 | 6.0301 | 6.7289 | 7.0907 | 6.8055 | 7.3664 | 6.6740 | 6.9896 | 6.3279 | 6.0583 | 6.0316 | 4.3952 | 5.5557 | 4.3861 | 5.1437 |
| TCOF1_S1263;S1264 | TCOF1 | Q13428-4 | 1414 | GGEASVSPEKTST | 6.3423 | 6.3953 | 6.1442 | 6.4109 | 5.6639 | 5.6228 | 6.3086 | 5.8820 | 6.5267 | 5.9179 | 5.7923 | 6.1879 | 6.0522 | 7.1359 | 6.5652 | 7.0519 |
| TCOF1_S1264 | TCOF1 | Q13428-4 | 381 | SAPAKESPRKGAA | 6.0464 | 6.2784 | 5.6796 | 6.6127 | 6.2561 | 5.8482 | 6.2117 | 6.3882 | 6.2847 | 5.9241 | 5.9446 | 5.8006 | 6.3468 | 6.9689 | 6.6856 | 6.7235 |
| TCOF1_S1264;S1268 | TCOF1 | Q13428-4 | 1446 | GKGSLGSQGAKDE | 4.4187 | 4.5041 | 3.8324 | 4.4421 | 5.0003 | 4.3833 | 4.7890 | 4.5877 | 3.8883 | 3.7550 | 3.1858 | 3.6904 | 10.6190 | 11.7367 | 13.5852 | 13.5819 |
| TCOF1_S1293 | TCOF1 | Q13428-4 | 1019 | QAQAASTPRKARA | 6.2913 | 6.3794 | 6.0910 | 6.8908 | 6.3724 | 5.9811 | 6.3218 | 5.5587 | 5.7451 | 5.5275 | 5.6019 | 5.7877 | 7.0861 | 7.1606 | 6.4578 | 6.7468 |
| TCOF1_S1386 | TCOF1 | Q13428-4 | 583 | QAKPTSSPAKGPP | 5.4550 | 5.1198 | 5.9748 | 6.6968 | 6.0179 | 5.9522 | 6.1889 | 5.4045 | 6.3218 | 6.2041 | 6.0792 | 6.4289 | 6.5066 | 7.2939 | 6.8018 | 7.5540 |
| TCOF1_S1412;S1414 | TCOF1 | Q13428-7 | 1073 | GTLPATSPQSTSV | 6.2408 | 6.5865 | 6.2122 | 7.0953 | 5.8503 | 5.8799 | 6.1082 | 6.0163 | 6.3064 | 6.4305 | 6.1912 | 6.3563 | 5.4949 | 6.5757 | 5.8335 | 6.8220 |
| TCOF1_S1414 | TCOF1 | Q13428-4 | 1147 | GTLPATSPQSTSV | 5.8525 | 6.4262 | 6.1320 | 7.2399 | 6.0989 | 6.2607 | 6.3588 | 6.3084 | 6.4916 | 6.3733 | 6.1113 | 6.4429 | 5.4165 | 6.3725 | 5.7984 | 6.3161 |
| TCOF1_S1443 | TCOF1 | Q13428-4 | 233 | SVSTKESPARKAA | 6.3542 | 6.5441 | 5.8157 | 6.6707 | 7.2717 | 7.0404 | 6.8423 | 6.3193 | 6.1748 | 5.8222 | 6.5142 | 5.9414 | 5.7799 | 6.5261 | 5.3281 | 5.0549 |
| TCOF1_S1446 | TCOF1 | Q13428-4 | 328 | KAGAVASQTKAGK | 5.2246 | 5.4459 | 4.5430 | 5.1879 | 5.3429 | 5.2996 | 5.3328 | 4.8970 | 4.6805 | 4.4539 | 4.3061 | 4.2851 | 9.5231 | 10.5656 | 9.9514 | 10.9608 |
| TCOF1_S153 | TCOF1 | Q13428-4 | 846 | LARGPASVPSVGK | 6.2289 | 6.4081 | 5.7855 | 6.8491 | 6.0319 | 6.1981 | 6.6793 | 6.3884 | 6.1287 | 5.9081 | 6.4994 | 6.1622 | 5.3365 | 6.6070 | 6.1405 | 6.6481 |
| TCOF1_S156 | TCOF1 | Q13428-4 | 1264 | TPKLDSSPSVSST | 6.3152 | 6.6963 | 6.3091 | 7.1237 | 6.1248 | 5.9879 | 6.5569 | 6.0159 | 6.2851 | 6.3981 | 6.1098 | 6.1349 | 5.8840 | 6.1821 | 5.6823 | 6.1941 |
| TCOF1_S227 | TCOF1 | Q13428-4 | 316 | TPGKGATPAPPGK | 7.1846 | 7.9397 | 6.9600 | 6.9166 | 5.9062 | 7.3591 | 7.6675 | 6.6720 | 7.4627 | 6.7361 | 6.9625 | 7.1820 | 2.9648 | 4.6195 | 3.1494 | 4.3173 |
| TCOF1_S233 | TCOF1 | Q13428-4 | 153 | TVANLLSGKSPRK | 3.7711 | 3.5689 | 3.6494 | 4.0576 | 4.7531 | 4.0412 | 3.6223 | 4.5615 | 3.5689 | 3.5410 | 2.8162 | 3.0997 | 12.9911 | 12.3990 | 14.4308 | 15.1282 |
| TCOF1_S328 | TCOF1 | Q13428-4 | 1305 | KEAASGTTPQKSR | 6.3525 | 6.7719 | 5.9880 | 7.0398 | 5.7840 | 5.8519 | 6.2627 | 5.8438 | 6.4380 | 6.0386 | 6.1207 | 6.2750 | 5.5300 | 6.7064 | 6.3276 | 6.6693 |
| TCOF1_S381 | TCOF1 | Q13428-4 | 168 | EPSANTTLVSETE | 5.6246 | 6.0991 | 5.9742 | 7.0289 | 5.9606 | 5.9392 | 6.2980 | 6.3540 | 5.9544 | 5.6976 | 5.3709 | 6.1406 | 6.4258 | 7.0326 | 6.7865 | 7.3129 |
| TCOF1_S583 | TCOF1 | Q13428-4 | 906 | LAPAKESPRKGAA | 6.2313 | 6.6366 | 6.0459 | 6.7253 | 6.2707 | 5.7905 | 6.1944 | 6.4868 | 6.8814 | 6.1170 | 6.0908 | 5.8405 | 5.8899 | 6.2947 | 6.2347 | 6.2694 |
| TCOF1_S846 | TCOF1 | Q13428-4 | 1443 | EKKGKGSLGSQGA | 6.4997 | 7.0215 | 6.3526 | 6.7349 | 6.1279 | 6.5977 | 7.8944 | 7.0181 | 6.1703 | 6.8157 | 6.2821 | 6.2896 | 3.8467 | 5.8353 | 4.6185 | 5.8950 |
| TCOF1_S906 | TCOF1 | Q13428-4 | 227 | AQVKASSVSTKES | 6.3397 | 7.1486 | 5.8860 | 6.6767 | 5.3106 | 5.7281 | 6.2491 | 5.2675 | 5.9894 | 6.0140 | 5.7631 | 5.8835 | 6.2295 | 7.2729 | 6.7324 | 7.5090 |
| TCOF1_T1019 | TCOF1 | Q13428-4 | 1077 | ASMAGASSSKESS | 7.0435 | 7.3702 | 6.7096 | 6.8001 | 6.1455 | 7.1855 | 7.5560 | 6.5849 | 6.7852 | 6.6484 | 6.6378 | 6.5681 | 3.9663 | 5.2093 | 4.0273 | 4.7623 |
| TCOF1_T1222;Y1242 | TCOF1 | Q13428-4 | 1078 | SMAGASSSKESSR | 6.1284 | 7.2068 | 6.8352 | 7.8114 | 6.4370 | 7.7950 | 7.0443 | 7.4159 | 3.9134 | 4.7148 | 4.5032 | 4.8058 | 5.5820 | 6.5514 | 6.3025 | 6.9529 |
| TCOF1_T1305 | TCOF1 | Q13428-4 | 1306 | EAASGTTPQKSRK | 5.9897 | 5.8023 | 6.0790 | 6.7908 | 6.0806 | 6.5466 | 6.2425 | 6.0635 | 6.4778 | 6.4658 | 5.8619 | 5.9729 | 6.0033 | 6.7323 | 6.4978 | 6.3931 |
| TCOF1_T1306 | TCOF1 | Q13428-4 | 156 | NLLSGKSPRKSAE | 5.8492 | 8.6934 | 8.1287 | 7.0596 | 4.8859 | 10.8760 | 6.4607 | 5.4113 | 9.7552 | 4.9166 | 5.7688 | 4.5648 | 4.6843 | 5.0341 | 4.3273 | 3.5841 |
| TCOF1_T168 | TCOF1 | Q13428-4 | 1263 | ATPKLDSSPSVSS | 5.9237 | 6.5955 | 5.9944 | 7.2725 | 5.7448 | 5.5920 | 6.0709 | 5.8632 | 5.9201 | 6.4475 | 6.5446 | 6.8487 | 5.4291 | 6.9401 | 6.0839 | 6.7290 |
| TCOF1_T249 | TCOF1 | Q13428-4 | 1222;1242 | NA | 5.1557 | 6.4480 | 6.0964 | 6.0525 | 6.2994 | 6.1074 | 6.3747 | 6.2277 | 6.3334 | 6.5012 | 6.2432 | 6.2051 | 5.9668 | 6.7919 | 6.5577 | 6.6389 |
| TCOF1_T316 | TCOF1 | Q13428-4 | 1264;1268 | NA | 5.9207 | 6.3508 | 6.4032 | 6.5242 | 5.3077 | 6.1340 | 6.4213 | 6.0290 | 6.4185 | 6.2714 | 5.7330 | 5.8456 | 6.3170 | 7.0728 | 6.4276 | 6.8233 |
| TCOF1_T533 | TCOF1 | Q13428-4 | 1412;1414 | NA | 6.4045 | 5.9739 | 5.7762 | 6.6586 | 6.4585 | 5.8462 | 6.7496 | 6.5367 | 6.4864 | 5.8504 | 5.8310 | 5.9513 | 6.6310 | 6.7728 | 5.8637 | 6.2091 |
| TCOF1_T914 | TCOF1 | Q13428-4 | 1263;1264 | NA | 6.4569 | 6.3962 | 5.8285 | 5.7081 | 5.8028 | 6.0897 | 6.2256 | 6.2634 | 5.9072 | 5.3789 | 5.7953 | 6.0543 | 6.4533 | 7.6085 | 6.9105 | 7.1207 |
| TCP11L1_S56 | TCP11L1 | Q9NUJ3 | 56 | VQRPHSSPPRFVT | 6.4536 | 5.4429 | 6.6431 | 5.1606 | 5.7663 | 6.1386 | 6.0436 | 6.7362 | 6.2122 | 6.8353 | 6.6125 | 7.6762 | 5.4123 | 6.5518 | 5.9384 | 6.3765 |
| TDRD12_S211 | TDRD12 | Q587J7 | 211 | NYACYMSPTKNKN | 6.8598 | 7.0760 | 7.7258 | 7.1784 | 6.5320 | 7.1215 | 7.0958 | 6.9026 | 7.4225 | 7.0225 | 7.5717 | 7.4253 | 3.2507 | 3.7524 | 3.1283 | 3.9346 |
| TDRD12_S233 | TDRD12 | Q587J7 | 302 | MDSLRDSPKDKSE | 7.1406 | 6.7933 | 7.9408 | 6.6758 | 6.5737 | 6.9586 | 7.1252 | 6.6453 | 7.7643 | 7.6800 | 7.8564 | 7.9287 | 3.1007 | 3.5655 | 2.6984 | 3.5526 |
| TDRD12_S302 | TDRD12 | Q587J7 | 233 | NIKSAPSFNKLNP | 7.7546 | 6.1278 | 6.5064 | 5.2746 | 5.2056 | 5.9052 | 5.8270 | 5.8803 | 7.2726 | 6.1630 | 6.5767 | 6.9207 | 6.2044 | 6.5464 | 5.3893 | 6.4455 |
| TDRD12_S315 | TDRD12 | Q587J7 | 315 | KKHHCISLKDTNK | 7.4301 | 6.9776 | 7.7434 | 7.7000 | 5.4778 | 6.6147 | 6.5688 | 6.2313 | 7.7058 | 7.1083 | 7.9552 | 7.7621 | 3.4167 | 4.1307 | 3.3051 | 3.8724 |
| TDRD12_T1104 | TDRD12 | Q587J7 | 1104 | EESLSQTPPRVTG | 6.4346 | 7.1705 | 7.4372 | 7.7903 | 5.8726 | 6.8180 | 6.8256 | 6.8030 | 6.9598 | 7.6405 | 7.3948 | 8.0941 | 3.4956 | 3.9651 | 3.2465 | 4.0517 |
| TDRD12_T1129 | TDRD12 | Q587J7 | 1129 | EQGGQGTPPAEDA | 5.2849 | 6.6428 | 7.0689 | 7.3137 | 7.2340 | 6.8333 | 5.4801 | 7.3252 | 6.0793 | 5.6856 | 7.3864 | 6.5966 | 4.7618 | 6.0712 | 5.1718 | 5.0644 |
| TDRD3_S582 | TDRD3 | Q9H7E2-3 | 582 | NPVRSNSFIGVPN | 6.4547 | 5.2702 | 5.3827 | 5.4513 | 6.2074 | 5.3496 | 5.4852 | 5.8236 | 7.2365 | 8.0099 | 5.5572 | 5.1726 | 7.6261 | 8.5410 | 6.1503 | 6.2818 |
| TDRD7_S319 | TDRD7 | Q8NHU6 | 319 | TEKVPLSPLPGPK | 7.1352 | 7.0469 | 6.6900 | 6.7989 | 5.6855 | 7.5700 | 7.6319 | 6.7596 | 7.1538 | 6.8431 | 7.3368 | 7.3453 | 3.6191 | 4.4675 | 3.4113 | 4.5051 |
| TDRKH_S278 | TDRKH | Q9Y2W6 | 278 | EKPSDDSFQKSEA | 6.5802 | 6.9144 | 6.2308 | 8.2463 | 7.4828 | 6.9270 | 7.3609 | 6.9076 | 7.0218 | 7.7505 | 7.8945 | 7.3947 | 3.2297 | 3.9246 | 2.7448 | 3.3893 |
| TDRKH_T252 | TDRKH | Q9Y2W6 | 252 | PTAPLVTPPPKGG | 5.5797 | 6.2259 | 6.4331 | 6.0617 | 6.7191 | 6.9264 | 6.0486 | 6.6638 | 6.1958 | 6.7482 | 6.4987 | 6.6244 | 5.5975 | 5.9830 | 5.7420 | 5.9520 |
| TEAD1_S102 | TEAD1 | P28347 | 20 | ENMERMSDSADKP | 5.6237 | 6.5990 | 6.2675 | 6.7227 | 6.2483 | 6.2235 | 6.3368 | 6.1350 | 5.7503 | 6.1266 | 5.8548 | 6.5626 | 6.3792 | 6.4849 | 6.4953 | 6.1898 |
| TEAD1_S20 | TEAD1 | P28347 | 102 | VLARRKSRDFHSK | 5.1506 | 4.9363 | 5.6177 | 6.0167 | 7.0665 | 5.8462 | 5.7483 | 5.8143 | 7.8118 | 8.1545 | 7.8972 | 7.9313 | 6.1136 | 6.3616 | 4.3031 | 5.2303 |
| TEAD3_S148 | TEAD3 | Q99594 | 148 | NKFSPPSPLPQAV | 5.6324 | 5.8229 | 5.9686 | 7.2380 | 6.5349 | 6.4990 | 6.4663 | 6.0953 | 5.5959 | 6.1713 | 6.5465 | 6.3850 | 6.0319 | 6.4716 | 6.2552 | 6.2853 |
| TEAD3_S164 | TEAD3 | Q99594 | 164 | SSRFWSSPPLLGQ | 6.9354 | 7.8128 | 7.1540 | 7.7607 | 7.1258 | 7.4425 | 7.4618 | 6.6231 | 6.9161 | 8.0507 | 8.9459 | 7.8425 | 1.9658 | 3.2774 | 2.0625 | 2.6231 |
| TEAD4_S205 | TEAD4 | Q15561 | 205 | PAGPAPSPSAPPA | 5.8386 | 5.1626 | 6.1866 | 6.2752 | 7.4876 | 5.3634 | 6.7067 | 5.9322 | 6.8090 | 5.7788 | 6.1367 | 5.7738 | 7.5000 | 6.2328 | 6.6007 | 6.2153 |
| TECPR2_S412 | TECPR2 | O15040 | 412 | PRSRSSSLNSTDS | 5.4113 | 4.6393 | 6.7816 | 5.6426 | 6.6050 | 6.2650 | 5.0815 | 6.6153 | 7.1355 | 6.4917 | 6.6580 | 6.7360 | 6.1071 | 6.9776 | 6.4833 | 6.3691 |
| TELO2_S637 | TELO2 | Q9Y4R8 | 836 | RLLPPASPxxxxx | 6.6043 | 6.3242 | 6.7268 | 6.3857 | 6.1867 | 6.7813 | 6.5473 | 6.7925 | 6.3140 | 6.4806 | 6.5732 | 6.4345 | 5.4417 | 5.7880 | 5.2652 | 5.3539 |
| TELO2_S688 | TELO2 | Q9Y4R8 | 637 | RTPQPGSPSPNTP | 6.3271 | 5.9744 | 7.0690 | 6.0035 | 6.3008 | 7.0050 | 6.0634 | 6.3857 | 6.9414 | 6.4649 | 6.2750 | 6.2307 | 5.8013 | 5.2706 | 6.0183 | 5.8690 |
| TELO2_S836 | TELO2 | Q9Y4R8 | 688 | RQGPAGSPSRFNS | 6.7419 | 7.4496 | 5.4627 | 7.0316 | 5.9090 | 5.9497 | 5.9622 | 7.0926 | 7.4449 | 6.5947 | 7.9413 | 7.6203 | 3.5526 | 5.5598 | 4.7832 | 4.9038 |
| TENT4A_S771 | TENT4A | Q5XG87 | 771 | SLPVSLSRxxxxx | 6.8968 | 7.9195 | 6.8634 | 5.8622 | 5.2066 | 7.0875 | 6.0243 | 6.6226 | 7.1114 | 5.8145 | 6.9326 | 8.2938 | 4.7822 | 4.9539 | 3.7142 | 5.9146 |
| TEP1_S397 | TEP1 | Q99973 | 397 | PRRPPRSPGMEPP | 5.8119 | 6.1767 | 6.4806 | 6.4036 | 5.7984 | 6.0256 | 6.3994 | 6.1673 | 6.4944 | 7.0577 | 7.6656 | 6.9003 | 5.5901 | 6.3154 | 5.7056 | 5.9072 |
| TEPSIN_S249 | TEPSIN | Q96N21-2 | 249 | QELSMGSPGPVTN | 6.9614 | 6.6850 | 7.8569 | 7.3499 | 6.7370 | 6.9644 | 6.2938 | 6.9903 | 7.1760 | 7.2958 | 7.4580 | 7.8182 | 3.4308 | 3.9960 | 3.1943 | 3.7921 |
| TEPSIN_S272 | TEPSIN | Q96N21-2 | 272 | ASCGQLSPARGTS | 5.8284 | 5.9340 | 7.2674 | 7.1693 | 6.4753 | 6.8996 | 6.8097 | 6.9876 | 7.5346 | 6.8499 | 8.0331 | 6.7657 | 4.1479 | 4.4547 | 4.1316 | 4.7112 |
| TEPSIN_S371 | TEPSIN | Q96N21-2 | 371 | IPGGTDSPKRGPS | 5.9726 | 6.3056 | 8.0119 | 7.3157 | 5.9792 | 6.7434 | 7.6365 | 6.7493 | 7.2003 | 7.0744 | 7.7309 | 7.7881 | 3.7157 | 4.5943 | 3.3186 | 3.8633 |
| TEPSIN_S51 | TEPSIN | Q96N21-2 | 51 | MRPGPESPSTRRL | 6.5238 | 6.9224 | 7.3804 | 6.3989 | 6.3242 | 7.0649 | 7.0282 | 7.4111 | 5.9493 | 6.7312 | 7.3577 | 7.3289 | 4.1524 | 4.8838 | 3.9088 | 4.6341 |
| TERF1_T371 | TERF1 | P54274 | 371 | SKSQPVTPEKHRA | 6.7475 | 6.8623 | 6.2646 | 6.8610 | 6.3193 | 6.0940 | 6.5611 | 6.2295 | 6.6353 | 6.2931 | 6.2846 | 6.2592 | 5.6038 | 5.6533 | 5.6736 | 5.6579 |
| TERF2_S365 | TERF2 | Q15554 | 365 | TQALPASPALKNK | 6.3426 | 6.5104 | 6.4965 | 6.8235 | 6.2935 | 6.2045 | 6.3302 | 6.3905 | 6.6611 | 6.5411 | 6.6735 | 6.7740 | 5.2491 | 5.6461 | 5.3814 | 5.6819 |
| TERF2IP_S154 | TERF2IP | Q9NYB0 | 36 | SFYVRPSPAKRRL | 6.6677 | 6.8822 | 6.8269 | 7.4478 | 6.8902 | 7.0702 | 6.5871 | 6.8387 | 6.7747 | 6.8514 | 7.4461 | 7.1511 | 4.0986 | 4.6284 | 3.5434 | 4.2954 |
| TERF2IP_S203 | TERF2IP | Q9NYB0 | 203 | LGDAPVSPSSQKL | 6.6295 | 6.6526 | 6.5888 | 6.4976 | 6.6499 | 6.2548 | 6.2473 | 6.6496 | 6.8079 | 6.4876 | 6.7912 | 6.7601 | 5.5207 | 5.3579 | 5.1657 | 4.9386 |
| TERF2IP_S36 | TERF2IP | Q9NYB0 | 154 | VKENARSPSSVTG | 6.3532 | 7.0719 | 6.9099 | 7.4552 | 6.7553 | 7.1989 | 6.7125 | 6.4265 | 6.2040 | 7.1030 | 7.0515 | 6.9821 | 4.5218 | 4.8839 | 4.0767 | 4.2935 |
| TERF2IP_T230 | TERF2IP | Q9NYB0 | 230 | EPQNKRTPDLPEE | 5.9431 | 6.4840 | 6.5336 | 6.2021 | 4.9911 | 5.7505 | 5.6275 | 6.2341 | 5.8965 | 7.0012 | 6.0516 | 6.7147 | 6.1600 | 6.9920 | 6.5588 | 6.8591 |
| TESK2_S456 | TESK2 | Q96S53 | 456 | PIRRWRSLPGSPE | 5.4586 | 6.1808 | 6.5846 | 5.7003 | 6.1051 | 6.1759 | 5.7602 | 7.0519 | 6.2643 | 6.6016 | 6.0982 | 6.1637 | 6.9052 | 6.5954 | 5.9168 | 6.4372 |
| TET2_S1107 | TET2 | Q6N021 | 1122 | IKNLLDTPVKTQY | 6.7498 | 7.0286 | 6.3515 | 6.8533 | 6.7046 | 6.1454 | 6.1746 | 6.4230 | 6.5622 | 6.3484 | 6.6089 | 6.6789 | 5.5139 | 5.2085 | 5.0953 | 5.5531 |
| TET2_S15 | TET2 | Q6N021 | 1114 | PSKLLDTPIKNLL | 7.0805 | 7.6826 | 7.4195 | 7.2793 | 6.3885 | 6.4895 | 6.8934 | 6.5700 | 7.5457 | 7.2080 | 6.8990 | 6.6131 | 3.8363 | 4.2440 | 3.6355 | 4.2150 |
| TET2_T1114 | TET2 | Q6N021 | 15 | VEGNRLSPFLIPS | 5.8956 | 5.0600 | 6.3741 | 5.2243 | 6.5848 | 6.5883 | 4.7529 | 5.1814 | 5.1314 | 6.6955 | 6.3741 | 5.6349 | 6.9490 | 8.6916 | 6.7562 | 8.1060 |
| TET2_T1122 | TET2 | Q6N021 | 1107 | LNNFIESPSKLLD | 6.8120 | 6.5404 | 6.3884 | 6.6616 | 6.3074 | 6.0092 | 6.5984 | 6.2750 | 6.8360 | 7.3124 | 6.1784 | 6.2750 | 4.8777 | 6.1089 | 5.3635 | 5.4555 |
| TET3_T792;S801 | TET3 | O43151 | 792;801 | NA | 6.1223 | 5.9639 | 6.2240 | 7.3076 | 6.0021 | 6.0420 | 6.0791 | 6.2101 | 6.0682 | 6.8979 | 6.4797 | 6.7990 | 5.3279 | 5.9665 | 5.8978 | 6.6120 |
| TEX10_T29 | TEX10 | Q9NXF1 | 29 | PKLQNATPTNFKT | 7.1150 | 7.2936 | 6.0391 | 7.1484 | 6.5179 | 6.7214 | 7.1838 | 6.6248 | 7.3373 | 6.6320 | 6.4656 | 6.7745 | 4.2183 | 5.0714 | 4.0025 | 4.8544 |
| TEX2_S162 | TEX2 | Q8IWB9 | 732 | GLLPAHSRHNSPS | 7.0701 | 6.4716 | 6.7630 | 7.3305 | 6.1687 | 6.0684 | 6.2758 | 5.8365 | 6.6830 | 6.4646 | 6.2859 | 6.4417 | 5.3953 | 5.4262 | 5.5109 | 5.8078 |
| TEX2_S162;S166 | TEX2 | Q8IWB9 | 166 | SSSPLSSPSKSPI | 6.4480 | 6.7696 | 6.0295 | 7.0042 | 6.4088 | 7.3409 | 5.5321 | 5.5875 | 5.8548 | 5.6439 | 5.6895 | 5.4328 | 7.2924 | 6.0536 | 6.4183 | 6.4940 |
| TEX2_S166 | TEX2 | Q8IWB9 | 458 | DGVVLDSEDEVDS | 5.6794 | 6.0161 | 6.2786 | 6.5442 | 5.8475 | 6.6606 | 6.1891 | 6.3484 | 5.7922 | 6.2822 | 6.0881 | 6.6913 | 5.9525 | 6.6272 | 6.4477 | 6.5548 |
| TEX2_S196 | TEX2 | Q8IWB9 | 162 | QKTSSSSPLSSPS | 5.9593 | 6.3832 | 6.4343 | 7.4386 | 6.0114 | 6.2127 | 6.1336 | 5.8858 | 6.3331 | 6.2442 | 6.5119 | 6.9049 | 5.9213 | 5.8449 | 5.5192 | 6.2616 |
| TEX2_S458 | TEX2 | Q8IWB9 | 196 | SLVKSLSTEVEPK | 6.8104 | 6.4932 | 8.5421 | 7.1249 | 6.9632 | 5.5123 | 6.0185 | 6.9244 | 6.8023 | 6.8219 | 5.9117 | 6.2946 | 4.7519 | 6.0354 | 4.4896 | 4.5038 |
| TEX2_S718;S721 | TEX2 | Q8IWB9 | 718;721 | NA | 6.8826 | 6.5097 | 7.5399 | 7.7889 | 5.5623 | 5.0297 | 5.8658 | 6.3897 | 6.4752 | 6.3106 | 6.2890 | 6.1565 | 5.0396 | 5.1394 | 6.6000 | 6.4210 |
| TEX2_S732 | TEX2 | Q8IWB9 | 162;166 | NA | 6.9453 | 6.0340 | 6.0160 | 6.6355 | 6.1265 | 5.7159 | 6.3274 | 6.4996 | 6.1904 | 6.8784 | 5.0819 | 6.1031 | 6.2778 | 6.7819 | 6.0838 | 6.3024 |
| TEX264_S106 | TEX264 | Q9Y6I9 | 244 | TSAATLSPGASSR | 6.1735 | 6.8544 | 6.5549 | 6.8822 | 6.6472 | 6.8659 | 7.4359 | 6.8629 | 6.8730 | 6.2941 | 7.0663 | 6.4415 | 4.5142 | 5.2729 | 4.3523 | 4.9088 |
| TEX264_S244 | TEX264 | Q9Y6I9 | 106 | LSEGEESPSPELI | 5.7456 | 5.9427 | 6.2835 | 6.4401 | 6.3287 | 5.9339 | 6.5721 | 5.7456 | 6.0899 | 6.4994 | 6.3046 | 6.3739 | 6.2630 | 6.5023 | 6.5364 | 6.4384 |
| TFAM_S124 | TFAM | Q00059 | 122 | RFKEQLTPSQIMS | 6.2606 | 6.6642 | 6.3077 | 6.2861 | 7.0316 | 6.2992 | 7.8474 | 6.4389 | 6.2058 | 5.5004 | 6.7661 | 6.0075 | 5.4441 | 5.6648 | 5.4333 | 5.8423 |
| TFAM_S195 | TFAM | Q00059 | 124 | KEQLTPSQIMSLE | 4.6876 | 5.0874 | 5.4303 | 5.2201 | 5.9913 | 5.4707 | 6.1267 | 5.8109 | 4.9965 | 4.8198 | 5.6277 | 5.5961 | 9.2057 | 8.2981 | 8.9716 | 8.6594 |
| TFAM_T122 | TFAM | Q00059 | 195 | WKNLSDSEKELYI | 6.1189 | 6.0529 | 5.6562 | 6.7731 | 5.3564 | 5.3239 | 6.3337 | 5.3626 | 6.0947 | 6.4018 | 5.8820 | 6.5973 | 7.0658 | 6.9024 | 7.0128 | 7.0653 |
| TFAP2B_S251 | TFAP2B | Q92481-2 | 251 | RLSLLSSTSKYKV | 7.0552 | 7.0166 | 7.6474 | 6.3332 | 7.9559 | 5.7586 | 6.2853 | 6.0358 | 7.1225 | 5.9501 | 6.1196 | 5.6260 | 6.2614 | 5.5550 | 5.3055 | 3.9720 |
| TFAP4_S124 | TFAP4 | Q01664 | 124 | QELSGSSPKRRRA | 6.5125 | 5.8454 | 6.1462 | 6.1279 | 6.5985 | 5.8006 | 6.0676 | 6.5675 | 6.5913 | 6.2548 | 6.3485 | 6.5857 | 6.6621 | 5.9087 | 5.7321 | 6.2505 |
| TFAP4_T37 | TFAP4 | Q01664 | 37 | LANIPLTPETQRD | 6.9958 | 6.0283 | 6.6080 | 7.1956 | 5.2670 | 6.0724 | 6.6872 | 6.1126 | 6.1943 | 7.3681 | 6.6768 | 7.5963 | 4.9194 | 5.6548 | 5.6651 | 4.9583 |
| TFDP1_S23 | TFDP1 | Q14186 | 23 | FIDQNLSPGKGVV | 6.4312 | 6.4340 | 6.4005 | 6.0348 | 6.7240 | 6.3534 | 6.2336 | 6.4952 | 6.9160 | 6.4717 | 6.5559 | 6.2444 | 5.8796 | 5.7083 | 5.3615 | 5.7561 |
| TFDP2_S24 | TFDP2 | Q14188 | 24 | FIDQNLSPTKGNI | 6.5075 | 6.4470 | 6.6005 | 5.6785 | 6.4837 | 6.2770 | 6.0838 | 6.3902 | 6.5825 | 5.8159 | 6.0085 | 6.7243 | 5.9653 | 6.3311 | 5.9482 | 6.1558 |
| TFE3_S556 | TFE3 | P19532 | 568 | ASSRRSSFSMEEE | 5.6466 | 5.8164 | 5.7987 | 6.0582 | 6.2856 | 5.9936 | 5.6188 | 6.2999 | 6.3296 | 6.4094 | 5.9529 | 6.2205 | 7.3527 | 6.9342 | 6.9860 | 6.2968 |
| TFE3_S568 | TFE3 | P19532 | 556 | PLLSSVSPAVSKA | 5.5879 | 5.8690 | 6.1018 | 5.3160 | 6.2845 | 6.6170 | 5.9045 | 6.9519 | 6.3123 | 6.5660 | 6.4043 | 6.6717 | 6.2881 | 6.3068 | 6.4148 | 6.4034 |
| TFEB_T331 | TFEB | P19484 | 331 | VHGLPTTSPSGMN | 7.4364 | 7.0509 | 6.7119 | 6.1736 | 6.9792 | 5.8048 | 6.1453 | 6.7068 | 7.9254 | 6.8633 | 7.4422 | 7.3690 | 4.7427 | 4.6651 | 3.8776 | 4.1058 |
| TFG_S197 | TFG | Q92734 | 197 | QVSGPPSAPAEDR | 5.9957 | 5.8602 | 6.9307 | 6.4427 | 6.2322 | 6.2682 | 6.1412 | 6.7477 | 6.4347 | 5.7557 | 6.1347 | 6.5627 | 6.3542 | 6.2772 | 6.0177 | 5.8442 |
| TFIP11_S210 | TFIP11 | Q9UBB9 | 210 | DFPVVDSEEEAEE | 6.1315 | 5.7918 | 6.1791 | 5.8196 | 6.1393 | 5.9216 | 6.1746 | 6.2465 | 6.1455 | 6.1885 | 6.4050 | 6.2531 | 6.8679 | 6.7425 | 6.6493 | 6.3444 |
| TFIP11_S59 | TFIP11 | Q9UBB9 | 59 | VWAERDSDDERPS | 5.7539 | 6.2290 | 6.0608 | 6.0941 | 6.1407 | 6.0989 | 6.1746 | 6.1687 | 6.3944 | 6.0480 | 6.2012 | 5.8353 | 6.7875 | 6.8516 | 6.6993 | 6.4620 |
| TFIP11_S98 | TFIP11 | Q9UBB9 | 98 | EAELEDSDDEEKP | 6.0490 | 6.2478 | 6.3677 | 6.3475 | 6.3716 | 6.1859 | 6.2642 | 5.4799 | 6.5194 | 5.8852 | 6.8305 | 6.3420 | 6.2949 | 6.8842 | 6.0243 | 5.9060 |
| TFPT_S167 | TFPT | P0C1Z6 | 207 | RAGNALTPELAPV | 6.2170 | 6.2804 | 6.2151 | 6.4673 | 6.3504 | 6.5276 | 6.6703 | 6.4854 | 6.4664 | 6.6387 | 6.4992 | 6.6566 | 5.4318 | 5.7126 | 5.5898 | 5.7913 |
| TFPT_S180 | TFPT | P0C1Z6 | 252 | TLASPASDxxxxx | 6.0296 | 6.2753 | 5.6997 | 6.9622 | 6.3708 | 5.5878 | 6.2642 | 6.0816 | 6.1265 | 6.3118 | 5.8506 | 6.1302 | 6.4560 | 6.6494 | 6.3088 | 6.8955 |
| TFPT_S249 | TFPT | P0C1Z6 | 167 | PEKETLSPPRRTP | 6.4783 | 6.8841 | 6.1370 | 6.6979 | 6.0349 | 5.9523 | 6.2507 | 6.4576 | 6.3937 | 6.3530 | 6.1399 | 6.5600 | 5.9992 | 6.0025 | 5.7127 | 5.9461 |
| TFPT_S249;S252 | TFPT | P0C1Z6 | 180 | APPEPGSPAPGEG | 6.0357 | 6.7193 | 6.2341 | 6.3414 | 6.5414 | 6.6022 | 6.6384 | 6.4161 | 6.4884 | 6.6651 | 6.1676 | 5.5325 | 5.8677 | 5.4704 | 6.5127 | 5.7670 |
| TFPT_S252 | TFPT | P0C1Z6 | 249 | PYPTLASPASDxx | 7.0394 | 5.0975 | 5.5717 | 7.0207 | 5.9414 | 6.3560 | 5.2805 | 6.7555 | 6.2663 | 6.4680 | 7.7190 | 7.5584 | 5.0377 | 5.3775 | 7.2149 | 5.2954 |
| TFPT_T207 | TFPT | P0C1Z6 | 249;252 | NA | 5.6822 | 6.3562 | 6.6925 | 6.7819 | 6.0098 | 6.4693 | 5.7530 | 6.4137 | 6.5049 | 5.9134 | 6.4585 | 6.4666 | 6.0116 | 6.0723 | 6.4454 | 5.9686 |
| TGFB1I1_S137;S141 | TGFB1I1 | O43294 | 68 | TVCKPRSPKPAAP | 5.7262 | 5.4935 | 5.3237 | 6.6513 | 5.5901 | 6.7919 | 6.8724 | 5.7892 | 6.2078 | 6.7465 | 7.1812 | 6.6704 | 6.2781 | 6.8870 | 6.2034 | 5.5872 |
| TGFB1I1_S68 | TGFB1I1 | O43294 | 137;141 | NA | 6.5561 | 5.3313 | 6.6282 | 5.3820 | 7.2813 | 6.9211 | 6.3091 | 7.3744 | 6.3963 | 7.6359 | 7.6196 | 7.6664 | 4.8854 | 5.0177 | 4.1696 | 4.8256 |
| TGFBRAP1_S855 | TGFBRAP1 | Q8WUH2 | 855 | TNPSSSSPGTRTx | 4.9625 | 6.5095 | 7.4096 | 6.8209 | 4.8599 | 6.8004 | 6.8449 | 5.9790 | 7.4849 | 6.3247 | 6.1843 | 5.8284 | 5.0994 | 6.2117 | 6.2767 | 6.4034 |
| TGIF1_S286 | TGIF1 | Q15583 | 286 | TLGRPLSPKPSSP | 4.1105 | 4.1425 | 4.3612 | 4.6944 | 5.4841 | 5.2415 | 3.9423 | 4.8755 | 5.2017 | 5.4863 | 4.9171 | 5.3449 | 10.5297 | 9.6775 | 11.7447 | 10.2459 |
| TGIF1_S286;S291 | TGIF1 | Q15583 | 286;291 | NA | 5.7341 | 5.8518 | 5.7046 | 6.3443 | 6.2272 | 6.2555 | 6.5288 | 5.7498 | 6.1304 | 6.4733 | 5.9993 | 6.1997 | 6.5085 | 7.1852 | 6.3411 | 6.7666 |
| TGIF2_S174 | TGIF2 | Q9GZN2 | 174 | TRAEAGSPTGGLF | 5.7743 | 5.6839 | 7.4520 | 5.0611 | 7.1908 | 5.9029 | 5.6658 | 9.1055 | 5.8969 | 7.3555 | 6.5579 | 6.3630 | 5.5895 | 7.1968 | 4.1308 | 5.0731 |
| TGS1_S154 | TGS1 | Q96RS0 | 438 | IDENPASDFDDSG | 6.8146 | 6.9222 | 7.2984 | 7.1454 | 6.0731 | 7.0257 | 6.8095 | 7.1007 | 5.7894 | 5.8279 | 5.6710 | 6.0229 | 5.1853 | 5.3539 | 5.3648 | 5.5952 |
| TGS1_S294 | TGS1 | Q96RS0 | 298 | LVSFPSSPIMVDN | 5.5954 | 5.7299 | 5.9136 | 5.7807 | 6.0453 | 6.0204 | 6.2771 | 6.3782 | 6.0549 | 5.9542 | 6.5402 | 6.1109 | 6.7605 | 7.2702 | 7.0961 | 6.4724 |
| TGS1_S294;S298 | TGS1 | Q96RS0 | 294 | MKVDLVSFPSSPI | 5.2627 | 6.1287 | 6.0561 | 5.4544 | 5.8887 | 5.7024 | 6.7734 | 5.9952 | 5.9523 | 5.6675 | 5.9585 | 5.4248 | 7.7557 | 7.9510 | 7.1280 | 6.9006 |
| TGS1_S298 | TGS1 | Q96RS0 | 154 | EDDILASDDPSSI | 6.7494 | 6.1380 | 7.0311 | 6.7440 | 5.9679 | 6.1363 | 6.7954 | 7.9383 | 4.2420 | 7.0010 | 6.2231 | 5.6507 | 5.2432 | 6.1363 | 6.4924 | 5.5108 |
| TGS1_S438 | TGS1 | Q96RS0 | 445 | DFDDSGSLLGFKY | 6.1037 | 5.9340 | 5.8937 | 6.6066 | 6.8166 | 6.6617 | 6.8208 | 6.1144 | 6.7020 | 5.9001 | 7.0882 | 6.8145 | 5.7452 | 6.0847 | 5.6858 | 5.0281 |
| TGS1_S445 | TGS1 | Q96RS0 | 309 | DNDSSGTSDKDHS | 5.5469 | 5.2933 | 5.5749 | 5.5172 | 6.5416 | 5.7742 | 5.9455 | 5.9938 | 5.9336 | 6.0659 | 7.2531 | 5.8564 | 7.0123 | 7.2879 | 6.5179 | 7.8857 |
| TGS1_T309 | TGS1 | Q96RS0 | 294;298 | NA | 5.9720 | 6.4620 | 6.4451 | 7.0420 | 6.1000 | 5.8961 | 6.5793 | 6.3255 | 6.7237 | 5.8364 | 6.1174 | 6.0810 | 5.6164 | 6.2772 | 6.1571 | 6.3687 |
| THAP4_S128 | THAP4 | Q8WY91 | 128 | RGAAGWSPSSSGN | 7.0034 | 7.5817 | 5.5537 | 7.9227 | 6.9537 | 7.5034 | 7.0699 | 7.0732 | 7.1296 | 7.4327 | 7.8612 | 7.2770 | 3.8893 | 3.8599 | 3.0694 | 2.8194 |
| THAP7_S210 | THAP7 | Q9BT49 | 210 | LEPRPVSPSAYML | 5.6340 | 5.3276 | 6.6157 | 6.7738 | 6.1173 | 6.0852 | 6.5386 | 5.7499 | 6.1973 | 6.6776 | 6.7940 | 6.6187 | 6.3821 | 6.4277 | 6.0995 | 5.9610 |
| THOC2_S1219 | THOC2 | Q8NI27 | 1385 | VKGGEKTPVSGSL | 6.3065 | 6.8031 | 5.9798 | 7.1389 | 6.0068 | 6.2855 | 6.7862 | 6.2366 | 6.2781 | 6.1380 | 6.3231 | 6.2780 | 5.1177 | 6.4135 | 5.7448 | 6.1631 |
| THOC2_S1417 | THOC2 | Q8NI27 | 1417 | KIDTHPSPSHSST | 6.1945 | 7.2481 | 5.0356 | 7.4625 | 6.5546 | 7.2333 | 5.7499 | 7.6559 | 5.5622 | 5.4810 | 5.5700 | 5.4782 | 5.0818 | 7.1904 | 6.8727 | 5.6292 |
| THOC2_S1576 | THOC2 | Q8NI27 | 1443 | KLYINHTPPPLSK | 6.5148 | 7.4875 | 6.4173 | 7.7931 | 6.4903 | 6.6270 | 7.1720 | 6.8990 | 7.2973 | 7.0919 | 6.5182 | 6.5424 | 3.8991 | 4.8206 | 3.7260 | 4.7035 |
| THOC2_T1248 | THOC2 | Q8NI27 | 1285 | KEKKEKTPATTPE | 5.9083 | 6.1993 | 5.7621 | 6.5453 | 5.8481 | 5.9557 | 5.9597 | 5.6819 | 5.9430 | 5.7425 | 5.7472 | 5.6942 | 7.1936 | 7.6716 | 7.0109 | 7.1367 |
| THOC2_T1285 | THOC2 | Q8NI27 | 1219 | IGSASKSDESSTE | 6.5164 | 6.0173 | 6.6342 | 6.0605 | 5.9163 | 6.3449 | 6.3044 | 5.7361 | 6.0789 | 6.1200 | 6.0224 | 6.0938 | 6.7229 | 6.6480 | 6.2495 | 6.5342 |
| THOC2_T1385 | THOC2 | Q8NI27 | 1248 | NKASSTTPKGNSS | 6.2181 | 7.5782 | 6.1914 | 6.9005 | 6.2678 | 6.8774 | 7.3861 | 6.4297 | 6.7171 | 6.6568 | 6.3072 | 6.8017 | 4.7458 | 5.1230 | 4.3612 | 5.4379 |
| THOC2_T1443 | THOC2 | Q8NI27 | 1576 | KKEKRDSSGGKEE | 6.3473 | 7.1991 | 6.2893 | 7.0561 | 6.3819 | 6.7225 | 7.2165 | 6.8583 | 6.3380 | 6.5932 | 6.5673 | 6.5163 | 4.2851 | 5.4365 | 4.8163 | 5.3761 |
| THOC5_S307;S312;S314 | THOC5 | Q13769 | 328 | TKRRRPTLGVQLD | 5.4211 | 5.0936 | 4.8281 | 5.6685 | 6.7378 | 4.9419 | 5.6060 | 5.2642 | 4.9216 | 3.7184 | 3.7936 | 3.9351 | 10.6543 | 9.3909 | 10.4553 | 9.5696 |
| THOC5_S312;S314 | THOC5 | Q13769 | 312;314 | NA | 5.8414 | 6.1433 | 5.9361 | 6.5996 | 5.9833 | 5.8258 | 6.0890 | 6.0614 | 6.3146 | 6.1931 | 6.2330 | 6.1312 | 6.5532 | 6.6711 | 6.7434 | 6.6805 |
| THOC5_T328 | THOC5 | Q13769 | 307;312;314 | NA | 6.1587 | 6.4952 | 6.0155 | 6.0715 | 6.2143 | 6.1784 | 6.1466 | 6.3396 | 5.9481 | 5.9405 | 5.8165 | 6.0699 | 6.5953 | 6.8195 | 6.7982 | 6.3922 |
| THOP1_S16 | THOP1 | P52888 | 16 | DMADAASPCSVVN | 5.6923 | 5.8435 | 6.1250 | 5.6192 | 6.3054 | 6.3966 | 5.3267 | 6.4197 | 6.0311 | 6.4155 | 6.6218 | 6.5277 | 6.7101 | 6.8080 | 6.9028 | 6.2545 |
| THRAP3_S232 | THRAP3 | Q9Y2W1 | 682 | HRRIDISPSTFRK | 6.3327 | 6.4938 | 5.8542 | 6.5777 | 6.1984 | 5.8537 | 6.2923 | 6.1610 | 6.2406 | 6.0795 | 5.8891 | 5.9947 | 6.4646 | 6.4982 | 6.5327 | 6.5367 |
| THRAP3_S243 | THRAP3 | Q9Y2W1 | 248 | LSPRERSPALKSP | 6.4441 | 6.5075 | 5.6111 | 6.2463 | 6.2078 | 5.8318 | 6.1420 | 5.9093 | 6.0948 | 5.9068 | 5.7822 | 5.8452 | 6.9564 | 6.8778 | 6.8814 | 6.7555 |
| THRAP3_S248 | THRAP3 | Q9Y2W1 | 243 | SAVSELSPRERSP | 6.4220 | 6.6791 | 5.7894 | 6.5058 | 6.2429 | 5.9183 | 6.1592 | 6.1650 | 6.1670 | 6.0669 | 5.8966 | 5.8878 | 6.5313 | 6.4678 | 6.6101 | 6.4910 |
| THRAP3_S253 | THRAP3 | Q9Y2W1 | 535 | VQEKSSSPPPRKT | 6.3236 | 6.5946 | 5.8802 | 6.5773 | 6.1042 | 6.0201 | 6.0994 | 5.8920 | 6.2397 | 6.1853 | 6.0741 | 6.1482 | 6.5250 | 6.4405 | 6.4799 | 6.4159 |
| THRAP3_S264 | THRAP3 | Q9Y2W1 | 253 | RSPALKSPLQSVV | 7.2116 | 7.3649 | 6.4101 | 7.0725 | 6.8040 | 6.4344 | 6.9265 | 6.7513 | 6.8467 | 6.5787 | 6.5718 | 6.4617 | 4.4739 | 4.8002 | 4.4503 | 4.8413 |
| THRAP3_S264;S268 | THRAP3 | Q9Y2W1 | 320 | KSPVGKSPPSTGS | 5.9179 | 6.1502 | 5.8416 | 6.7808 | 6.1566 | 5.9724 | 6.0171 | 6.0381 | 6.1003 | 6.2206 | 6.0220 | 6.2495 | 6.4042 | 6.6887 | 6.7399 | 6.7000 |
| THRAP3_S315;S320 | THRAP3 | Q9Y2W1 | 379 | KEKGSFSDTGLGD | 6.3752 | 6.7026 | 6.3662 | 7.0437 | 6.9361 | 6.5403 | 7.0699 | 6.9963 | 5.0704 | 5.0933 | 5.0685 | 5.2884 | 6.4029 | 6.1299 | 6.4287 | 6.4876 |
| THRAP3_S315;S320;S323 | THRAP3 | Q9Y2W1 | 928 | WAHDKFSGEEGEI | 5.4845 | 5.8824 | 5.6775 | 5.8649 | 5.6508 | 6.0426 | 5.8031 | 5.5088 | 6.7363 | 7.3659 | 7.4774 | 7.2193 | 6.5010 | 6.6237 | 6.2344 | 5.9274 |
| THRAP3_S315;T323 | THRAP3 | Q9Y2W1 | 672 | AAKNKKSPEIHRR | 6.5639 | 6.5393 | 6.0200 | 6.7298 | 6.2311 | 6.0199 | 6.1544 | 5.6988 | 6.0946 | 6.1698 | 6.1886 | 6.3578 | 6.3009 | 6.2968 | 6.3206 | 6.3136 |
| THRAP3_S320 | THRAP3 | Q9Y2W1 | 939 | EIEDDESGTENRE | 6.3649 | 6.4065 | 6.3654 | 6.3659 | 5.8937 | 5.5798 | 5.9295 | 5.5365 | 6.8442 | 7.1029 | 6.7667 | 6.7663 | 6.1629 | 6.1290 | 5.7990 | 5.9869 |
| THRAP3_S320;S323 | THRAP3 | Q9Y2W1 | 874 | EWDPEYTPKSKKY | 7.4598 | 6.8327 | 6.3145 | 6.7112 | 6.3218 | 6.1381 | 6.8675 | 6.1914 | 7.2265 | 6.5535 | 6.2785 | 6.3189 | 5.5855 | 5.5923 | 4.5969 | 5.0109 |
| THRAP3_S320;S330 | THRAP3 | Q9Y2W1 | 622 | SAQSQRSPSELFA | 6.3989 | 6.7523 | 5.8591 | 6.7501 | 6.1088 | 6.1775 | 5.9474 | 6.2480 | 6.2929 | 6.3381 | 5.8664 | 6.0443 | 6.4247 | 6.3321 | 6.2950 | 6.1645 |
| THRAP3_S377 | THRAP3 | Q9Y2W1 | 624 | QSQRSPSELFAQH | 6.0144 | 6.3061 | 5.8547 | 6.5123 | 6.1812 | 5.9348 | 6.2930 | 5.8986 | 6.2450 | 6.2296 | 6.3333 | 6.1335 | 6.5999 | 6.3637 | 6.5647 | 6.5351 |
| THRAP3_S379 | THRAP3 | Q9Y2W1 | 264 | VVVRRRSPRPSPV | 6.5178 | 7.0757 | 5.8706 | 6.4309 | 6.9520 | 6.4257 | 5.9545 | 5.8974 | 5.5176 | 6.0356 | 5.9728 | 5.4339 | 7.1790 | 6.6882 | 6.3674 | 5.6808 |
| THRAP3_S406;S408 | THRAP3 | Q9Y2W1 | 232 | ATYGTGSASRASA | 6.2226 | 6.3864 | 6.6832 | 7.6033 | 6.5907 | 7.0224 | 6.9796 | 6.4198 | 6.8412 | 7.0317 | 7.0681 | 6.8633 | 4.3223 | 5.2113 | 3.8800 | 4.8744 |
| THRAP3_S535 | THRAP3 | Q9Y2W1 | 575 | VNVRMDSFDEDLA | 5.5445 | 6.3123 | 5.8878 | 6.2837 | 5.7727 | 5.2883 | 6.3729 | 5.6297 | 5.9717 | 6.8360 | 4.8086 | 5.8160 | 7.1274 | 6.7774 | 7.8034 | 7.7675 |
| THRAP3_S562 | THRAP3 | Q9Y2W1 | 377 | KIKEKGSFSDTGL | 4.9326 | 5.0875 | 4.7403 | 5.2010 | 5.8487 | 5.4409 | 5.5409 | 5.1655 | 5.5384 | 5.1951 | 4.5920 | 4.6798 | 8.5677 | 9.0601 | 10.0799 | 10.3296 |
| THRAP3_S575 | THRAP3 | Q9Y2W1 | 562 | TGKSSFSITREAQ | 6.2750 | 6.3874 | 6.0311 | 6.6552 | 6.2663 | 5.9713 | 6.5853 | 5.9230 | 6.1769 | 6.3237 | 6.1702 | 6.3216 | 6.2691 | 6.3661 | 5.8436 | 6.4343 |
| THRAP3_S622 | THRAP3 | Q9Y2W1 | 928;939 | NA | 6.3051 | 6.3076 | 5.9631 | 6.7563 | 6.5409 | 6.1647 | 6.5913 | 6.2394 | 5.5327 | 5.3637 | 5.3333 | 5.3815 | 7.0968 | 6.8159 | 6.7579 | 6.8498 |
| THRAP3_S624 | THRAP3 | Q9Y2W1 | 315;320 | NA | 5.9994 | 6.1806 | 5.7685 | 6.6336 | 7.1624 | 6.8563 | 6.4613 | 6.2380 | 6.1764 | 5.9733 | 6.9782 | 6.2984 | 6.4121 | 6.3004 | 5.6149 | 4.9460 |

| | | | | | | | | | | | | | | | | | | | | |
|---|---|---|---|---|---|---|---|---|---|---|---|---|---|---|---|---|---|---|---|---|
| THRAP3_S672 | THRAP3 | Q9Y2W1 | 264;268 | NA | 6.6272 | 6.3253 | 5.4315 | 5.9803 | 6.7315 | 5.7229 | 6.3691 | 6.2899 | 6.5958 | 5.7775 | 5.7888 | 5.5129 | 7.0939 | 6.5716 | 6.6248 | 6.5570 |
| THRAP3_S682 | THRAP3 | Q9Y2W1 | 315;320;323 | NA | 3.9857 | 5.2421 | 5.3520 | 4.7102 | 5.2117 | 6.7572 | 6.0310 | 7.0594 | 6.3401 | 7.8428 | 7.4356 | 7.0816 | 6.6859 | 6.6546 | 6.8461 | 7.7642 |
| THRAP3_S928 | THRAP3 | Q9Y2W1 | 320;330 | NA | 6.5274 | 7.3887 | 6.7542 | 6.6049 | 6.7796 | 5.9262 | 6.8064 | 7.3506 | 6.4306 | 6.8278 | 6.7294 | 6.1797 | 4.5053 | 4.8536 | 4.7178 | 5.6180 |
| THRAP3_S928;S939 | THRAP3 | Q9Y2W1 | 315;323 | NA | 5.9615 | 6.5128 | 5.7865 | 6.7608 | 7.3326 | 6.8853 | 6.3041 | 6.4340 | 6.0716 | 6.1275 | 6.7999 | 6.2475 | 6.3161 | 6.1161 | 5.4327 | 4.9108 |
| THRAP3_S939 | THRAP3 | Q9Y2W1 | 320;323 | NA | 4.2250 | 5.7291 | 5.5641 | 5.6766 | 4.5714 | 5.5715 | 6.1670 | 6.0006 | 5.4884 | 8.0976 | 7.3106 | 7.0546 | 6.4934 | 6.7676 | 7.0936 | 8.1889 |
| THRAP3_T874 | THRAP3 | Q9Y2W1 | 406;408 | NA | 6.9063 | 5.9205 | 5.9158 | 6.3468 | 6.6065 | 6.1845 | 6.9832 | 5.2832 | 6.3731 | 6.4511 | 5.7405 | 6.0830 | 6.8343 | 6.0612 | 5.8702 | 6.4397 |
| THSD1_S619 | THSD1 | Q9NS62 | 791 | QRVSSLSPSQCRK | 6.7087 | 6.5607 | 6.3566 | 6.6567 | 5.7723 | 6.0654 | 6.5584 | 6.3235 | 6.1795 | 6.1172 | 7.1613 | 6.7217 | 5.6717 | 5.6738 | 5.6210 | 5.8513 |
| THSD1_S791 | THSD1 | Q9NS62 | 619 | QASCAISPSQTLI | 6.3275 | 6.3327 | 5.7364 | 7.2709 | 5.5268 | 6.2576 | 6.3609 | 5.8450 | 5.9974 | 6.3698 | 6.5661 | 6.3438 | 6.1765 | 5.5847 | 6.5375 | 6.6665 |
| THUMPD1_S86;S88 | THUMPD1 | Q9NXG2 | 86;88 | NA | 5.2131 | 5.5309 | 6.1192 | 5.8004 | 5.6951 | 6.2253 | 5.5707 | 5.9174 | 5.6632 | 6.2403 | 5.8365 | 5.9330 | 6.8717 | 7.5810 | 8.4383 | 7.3639 |
| THUMPD3_S154 | THUMPD3 | Q9BV44 | 154 | KINQNSSKEKINN | 5.2761 | 6.5531 | 5.4203 | 4.9846 | 5.5504 | 6.0473 | 5.6096 | 5.1127 | 5.5829 | 5.4521 | 5.2898 | 5.7087 | 8.4485 | 8.8548 | 8.7290 | 8.2800 |
| TIAL1_S104 | TIAL1 | Q01085-2 | 102 | KVNWATTPSSQKK | 6.0734 | 6.7177 | 6.2095 | 6.6511 | 6.4409 | 6.5952 | 6.0332 | 6.5090 | 6.3750 | 6.0505 | 6.2059 | 6.6865 | 5.9005 | 5.9793 | 5.8823 | 5.6900 |
| TIAL1_T102 | TIAL1 | Q01085-2 | 104 | NWATTPSSQKKDT | 5.2410 | 6.0223 | 5.9448 | 5.2695 | 7.1704 | 6.5932 | 6.4034 | 6.5395 | 7.7018 | 6.1108 | 5.6902 | 4.6669 | 7.1056 | 7.0170 | 6.4967 | 6.9269 |
| TIAM1_S1321 | TIAM1 | Q13009 | 1321 | VGSHRLSIYEDWD | 6.4891 | 4.7922 | 5.5916 | 5.5193 | 6.5249 | 5.4489 | 5.5680 | 5.9268 | 8.2731 | 6.8101 | 6.8156 | 6.8904 | 6.7593 | 7.8526 | 5.0667 | 5.6714 |
| TIAM1_S1407 | TIAM1 | Q13009 | 231 | TCQRANSLGDLYA | 5.9399 | 6.0913 | 5.9002 | 5.8898 | 6.1274 | 6.1488 | 5.8842 | 6.3984 | 6.5044 | 6.3692 | 6.1986 | 6.1590 | 6.6248 | 7.0667 | 6.3376 | 6.3497 |
| TIAM1_S1466 | TIAM1 | Q13009 | 1466 | DAVSASSPEKESQ | 6.0341 | 6.3208 | 6.3384 | 6.3294 | 6.4193 | 6.4562 | 5.9449 | 6.0473 | 6.8879 | 6.4032 | 6.3664 | 6.4468 | 5.7711 | 6.0621 | 5.9910 | 6.1812 |
| TIAM1_S1506 | TIAM1 | Q13009 | 1407 | QLLKTESLPSSQQ | 6.5380 | 5.7526 | 7.8572 | 6.5804 | 6.9210 | 6.5228 | 6.8207 | 5.9418 | 6.8330 | 6.7722 | 6.2263 | 6.9498 | 5.8082 | 6.8764 | 4.4727 | 4.1269 |
| TIAM1_S231 | TIAM1 | Q13009 | 1506 | KETDILSDDDEFC | 6.2771 | 6.4411 | 6.7174 | 6.5137 | 6.1012 | 6.7433 | 5.7971 | 6.2523 | 6.0615 | 5.9114 | 5.9532 | 5.7762 | 6.1678 | 6.1907 | 6.6558 | 6.4401 |
| TIAM1_S358 | TIAM1 | Q13009 | 358 | ATNSSYSPTTGRA | 6.0613 | 6.9058 | 6.4121 | 6.1632 | 5.2409 | 5.9849 | 6.8011 | 6.7361 | 6.9355 | 6.6625 | 6.1886 | 6.2523 | 5.7615 | 6.4673 | 5.3838 | 6.0429 |
| TIAM1_S695 | TIAM1 | Q13009 | 695 | RRSRFSSLWGLDT | 7.7800 | 5.8416 | 7.4126 | 7.0359 | 6.2274 | 5.6211 | 5.5122 | 5.6093 | 6.8495 | 6.2498 | 5.8980 | 6.1303 | 5.5227 | 7.3338 | 5.8167 | 5.1591 |
| TICRR_S1009 | TICRR | Q7Z2Z1 | 1334 | SKIECPSPGELDQ | 6.6413 | 6.9201 | 6.5314 | 6.9055 | 5.9871 | 5.9295 | 6.3754 | 6.2505 | 6.5770 | 6.5331 | 6.6882 | 6.7740 | 4.9845 | 5.9249 | 5.3010 | 5.6766 |
| TICRR_S1045 | TICRR | Q7Z2Z1 | 820 | MTQENKSPLLSVP | 6.5393 | 6.5689 | 6.4188 | 7.3545 | 6.2088 | 6.1824 | 6.1257 | 6.1035 | 6.0383 | 6.8001 | 6.4819 | 6.6668 | 5.5718 | 5.8040 | 5.5239 | 5.6113 |
| TICRR_S1064 | TICRR | Q7Z2Z1 | 441 | QTAVADSPRDTAS | 6.5775 | 6.2910 | 6.0793 | 6.3993 | 5.9442 | 6.1734 | 6.2840 | 6.4905 | 6.4644 | 6.7342 | 6.1316 | 6.6085 | 5.5912 | 5.9449 | 6.0350 | 6.2509 |
| TICRR_S1334 | TICRR | Q7Z2Z1 | 599 | KLHPDGSPDVAGE | 6.5730 | 6.7559 | 6.4363 | 7.7228 | 5.3644 | 5.8962 | 6.6846 | 6.0357 | 6.9513 | 7.0284 | 7.1976 | 7.2579 | 4.6991 | 5.5011 | 6.4497 | 5.2462 |
| TICRR_S1346;T1360 | TICRR | Q7Z2Z1 | 1105 | VPAAYQTPKKSHQ | 7.3253 | 6.5944 | 6.8874 | 7.0722 | 5.7405 | 6.1258 | 6.8351 | 6.0351 | 7.9131 | 6.6390 | 6.9603 | 7.1984 | 4.4114 | 4.7951 | 4.0507 | 5.4162 |
| TICRR_S1623 | TICRR | Q7Z2Z1 | 923 | FNQELLSPSKRSL | 7.0324 | 7.5390 | 7.2390 | 6.8722 | 5.9103 | 6.0596 | 6.5886 | 6.6140 | 7.0769 | 6.6315 | 6.6246 | 7.3010 | 4.4648 | 4.5243 | 4.2317 | 5.2899 |
| TICRR_S441 | TICRR | Q7Z2Z1 | 1633 | PRLSHSTPGKSRG | 7.5422 | 7.5336 | 6.9085 | 7.2136 | 5.8418 | 6.5550 | 6.8535 | 6.3350 | 7.0890 | 6.6830 | 6.9882 | 6.9642 | 4.0402 | 4.7532 | 4.0151 | 4.6839 |
| TICRR_S599 | TICRR | Q7Z2Z1 | 1313 | TPKKLFTSPLCDV | 7.0744 | 7.4152 | 7.1043 | 7.5013 | 6.4770 | 6.4585 | 6.9677 | 6.5685 | 7.0955 | 7.4079 | 7.3424 | 7.0457 | 3.5324 | 4.2436 | 3.6782 | 4.0875 |
| TICRR_S820 | TICRR | Q7Z2Z1 | 1678 | KHSGKTTPDIIKD | 6.7074 | 6.8646 | 6.6801 | 6.8458 | 6.3155 | 6.0157 | 6.6557 | 6.1387 | 7.1152 | 6.5377 | 6.8912 | 6.9959 | 5.1254 | 5.0777 | 4.7641 | 5.2693 |
| TICRR_S838 | TICRR | Q7Z2Z1 | 1045 | SVQRVHSFQQDKS | 6.2244 | 6.7436 | 6.8615 | 6.7335 | 5.5956 | 5.8411 | 5.9130 | 6.0588 | 6.5384 | 7.5233 | 7.0364 | 7.6211 | 4.9913 | 5.5195 | 5.2368 | 5.5617 |
| TICRR_S865 | TICRR | Q7Z2Z1 | 1009 | EKGDEISLRRSPR | 7.7945 | 7.7864 | 7.6269 | 7.2689 | 6.0062 | 6.0445 | 6.5900 | 6.5115 | 7.6708 | 7.3166 | 6.9495 | 7.2596 | 4.3489 | 4.9013 | 3.0861 | 3.8083 |
| TICRR_S923 | TICRR | Q7Z2Z1 | 865 | ALIRHKSIAEVSQ | 7.0117 | 7.5752 | 7.2688 | 6.4670 | 4.7957 | 5.9410 | 6.7903 | 5.0161 | 6.4146 | 7.3842 | 7.6213 | 8.2013 | 3.9110 | 6.2849 | 3.7818 | 5.5349 |
| TICRR_T1105 | TICRR | Q7Z2Z1 | 1134 | PQTPLYTPERLQK | 6.0240 | 5.8597 | 6.7208 | 7.2491 | 6.2924 | 6.4752 | 6.5338 | 6.2887 | 6.4187 | 6.5565 | 6.7066 | 6.5925 | 5.4566 | 5.4471 | 5.7594 | 5.6188 |
| TICRR_T1134 | TICRR | Q7Z2Z1 | 1064 | PVQSIRSPKSLLF | 6.1707 | 6.5427 | 7.5017 | 7.9103 | 5.8653 | 6.2248 | 7.0902 | 5.8183 | 5.4823 | 6.1796 | 7.4075 | 7.2976 | 4.8672 | 5.0123 | 4.8114 | 5.8180 |
| TICRR_T1242;T1250 | TICRR | Q7Z2Z1 | 1623 | PEPTYVSPPCPRL | 7.1054 | 6.9318 | 6.7138 | 6.9821 | 6.8462 | 6.4831 | 7.4958 | 7.0197 | 7.5613 | 7.0821 | 7.1194 | 7.6876 | 3.5248 | 3.9801 | 3.3827 | 4.0840 |
| TICRR_T1313 | TICRR | Q7Z2Z1 | 838 | RRSVSGSPESDEL | 6.0631 | 6.6172 | 5.8388 | 7.1601 | 5.6239 | 6.0531 | 6.9290 | 6.2382 | 6.4532 | 6.4302 | 6.1848 | 7.1856 | 5.7667 | 5.4717 | 5.9214 | 6.0631 |
| TICRR_T1633 | TICRR | Q7Z2Z1 | 1346;1360 | NA | 5.6171 | 6.7827 | 6.5996 | 6.3982 | 6.5280 | 6.0156 | 6.2287 | 6.2878 | 7.0199 | 6.2283 | 5.6450 | 6.6732 | 5.7600 | 6.7177 | 5.5903 | 5.9078 |
| TICRR_T1678 | TICRR | Q7Z2Z1 | 1242;1250 | NA | 6.1285 | 7.4260 | 6.5575 | 7.3472 | 6.1321 | 6.7514 | 6.2504 | 5.9200 | 6.9275 | 6.8383 | 7.4977 | 7.0775 | 4.6474 | 5.0224 | 4.3537 | 5.1224 |
| TIMELESS_S1087 | TIMELESS | Q9UNS1 | 1149 | KKAGLASPEEEDA | 5.6998 | 6.6901 | 6.8159 | 6.2659 | 6.2547 | 6.2688 | 5.8886 | 6.3920 | 6.3195 | 6.4240 | 6.1623 | 6.1685 | 5.9290 | 6.3401 | 6.1233 | 6.2575 |
| TIMELESS_S1149 | TIMELESS | Q9UNS1 | 1173 | KRQLLDSDEEQEE | 5.3880 | 6.4193 | 6.6116 | 5.5373 | 5.3813 | 6.2027 | 6.2970 | 5.5855 | 5.8492 | 6.7909 | 6.5359 | 6.5250 | 6.3955 | 7.6351 | 6.1731 | 6.6727 |
| TIMELESS_S1173 | TIMELESS | Q9UNS1 | 962 | LPNGAESLKDFCQ | 7.3124 | 6.4878 | 6.1945 | 6.7458 | 5.8103 | 5.9753 | 6.5223 | 5.8941 | 6.9158 | 6.1917 | 6.1986 | 6.9529 | 5.4193 | 5.8928 | 5.0576 | 6.4286 |
| TIMELESS_S281 | TIMELESS | Q9UNS1 | 281 | HSRFGGSYIVQGL | 4.7781 | 6.3590 | 7.5441 | 5.4015 | 4.5176 | 6.8914 | 6.5334 | 6.0119 | 7.5186 | 7.9866 | 9.2204 | 8.1501 | 3.7317 | 6.8622 | 3.6678 | 4.8257 |
| TIMELESS_S962 | TIMELESS | Q9UNS1 | 1087 | RIPAKLSPTQLRR | 6.0008 | 7.2099 | 6.7638 | 6.5788 | 5.4207 | 6.2948 | 5.9984 | 6.4227 | 6.6360 | 6.9009 | 6.7181 | 6.6870 | 4.9646 | 5.9042 | 5.6046 | 5.8947 |
| TIMM44_S61 | TIMM44 | O43615 | 61 | NRKGFLSGLLDNV | 7.3770 | 5.3833 | 4.8942 | 5.8078 | 5.4886 | 6.7249 | 5.2882 | 5.8486 | 8.2193 | 6.2833 | 5.4954 | 5.4105 | 8.0698 | 6.9320 | 5.8010 | 6.9762 |
| TINF2_S295 | TINF2 | Q9BSI4 | 330 | STGKSKSPCQTLG | 6.4565 | 6.3725 | 5.8561 | 6.5116 | 7.1889 | 7.0354 | 6.9631 | 6.4055 | 7.9187 | 6.3253 | 6.9670 | 6.6209 | 5.4745 | 5.1628 | 4.1330 | 4.6083 |
| TINF2_S330 | TINF2 | Q9BSI4 | 295 | PFRNLGSPTQVIS | 6.2527 | 6.8248 | 6.9321 | 6.7762 | 6.4405 | 7.2280 | 6.3595 | 6.2829 | 6.9502 | 7.3503 | 7.0271 | 6.8917 | 4.3071 | 5.0432 | 4.1945 | 5.1391 |
| TIPIN_T233 | TIPIN | Q9BVW5 | 233 | NDMLMNTPRAHTV | 7.2293 | 7.0564 | 7.0433 | 7.7613 | 6.4689 | 5.8781 | 8.6426 | 6.9976 | 6.4330 | 8.5675 | 7.6243 | 5.4114 | 3.8644 | 3.7893 | 3.9329 | 3.2997 |
| TJAP1_S214 | TJAP1 | Q5JTD0 | 300 | LGTARGSPEEELP | 6.1025 | 5.8897 | 6.4680 | 6.4008 | 6.3076 | 6.2144 | 5.8046 | 6.4894 | 6.4291 | 6.4118 | 6.3736 | 6.5315 | 6.5664 | 6.1963 | 5.9375 | 5.8767 |
| TJAP1_S300 | TJAP1 | Q5JTD0 | 422 | RAFVDRTPPPAAV | 6.3320 | 6.1902 | 6.4762 | 6.6689 | 6.7334 | 6.8530 | 6.4969 | 6.9662 | 6.5402 | 6.6173 | 6.5878 | 6.8983 | 5.3607 | 5.5687 | 4.6732 | 5.0370 |
| TJAP1_S320 | TJAP1 | Q5JTD0 | 320 | NPYPTPSPPHPLY | 6.1999 | 6.5434 | 7.0169 | 6.8219 | 6.6060 | 6.7651 | 6.0906 | 6.8204 | 6.5715 | 6.6693 | 7.1760 | 6.7945 | 5.0668 | 5.2130 | 4.6583 | 4.9865 |
| TJAP1_S545 | TJAP1 | Q5JTD0 | 214 | SGQEAASPGPAPS | 5.4800 | 7.2938 | 6.1035 | 6.5501 | 6.1432 | 5.7328 | 6.3869 | 7.2632 | 6.4334 | 6.6771 | 6.7905 | 7.0081 | 4.3509 | 6.3517 | 5.4358 | 5.9992 |
| TJAP1_T422 | TJAP1 | Q5JTD0 | 545 | HLHRKDSLTQAQE | 6.0120 | 6.0736 | 5.5575 | 6.0894 | 7.1504 | 6.2170 | 5.2191 | 6.8379 | 5.8342 | 7.0859 | 6.9712 | 6.9741 | 6.0622 | 6.6830 | 5.4585 | 5.7740 |
| TJP1_S1111 | TJP1 | Q07157-2 | 912 | PIHRIDSPGFKPA | 6.7349 | 6.6492 | 6.4330 | 6.2940 | 6.2396 | 5.9867 | 6.8555 | 6.1791 | 6.7366 | 6.5777 | 6.5908 | 6.2617 | 5.6824 | 5.7423 | 5.6333 | 6.1031 |
| TJP1_S125 | TJP1 | Q07157 | 912 | PIHRIDSPGFKPA | 6.7432 | 6.6512 | 6.4388 | 6.2785 | 6.2169 | 5.9745 | 6.8683 | 6.1800 | 6.7455 | 6.5745 | 6.5549 | 6.2499 | 5.6900 | 5.7551 | 5.6553 | 6.1233 |
| TJP1_S125;S131 | TJP1 | Q07157 | 354 | KPGAVSTPVKHAD | 7.2353 | 6.4171 | 6.8711 | 6.4814 | 7.1185 | 6.7245 | 7.3531 | 6.1595 | 7.5167 | 6.8875 | 6.6061 | 6.5956 | 4.5086 | 4.8076 | 4.0581 | 4.6594 |
| TJP1_S1278 | TJP1 | Q07157 | 175 | PRSDRRSVASSQP | 6.6949 | 6.6274 | 6.8343 | 5.9412 | 5.0229 | 6.2965 | 6.1591 | 6.2528 | 6.9368 | 7.2925 | 6.8516 | 7.4008 | 5.0189 | 5.5133 | 4.9990 | 6.1581 |
| TJP1_S1594 | TJP1 | Q07157 | 1617 | PKAIPVSPSAVEE | 6.1545 | 6.2538 | 6.0547 | 6.4888 | 6.1331 | 6.2337 | 6.2398 | 6.3106 | 6.7092 | 6.5345 | 6.7693 | 6.4875 | 5.9983 | 6.0138 | 5.8602 | 5.7583 |
| TJP1_S1617 | TJP1 | Q07157 | 329 | SDHSRHSPQQPSN | 6.0032 | 6.5141 | 6.1501 | 6.3632 | 6.0061 | 5.8261 | 6.1779 | 6.2429 | 5.8848 | 6.2415 | 5.5814 | 6.3515 | 6.5306 | 6.6935 | 6.6866 | 6.7466 |
| TJP1_S175 | TJP1 | Q07157 | 617 | KRNLRKSREDLSA | 5.6172 | 6.2025 | 6.1019 | 5.9126 | 5.2221 | 6.1902 | 6.5903 | 6.1459 | 6.2366 | 6.0285 | 6.3328 | 6.4739 | 6.7960 | 6.3418 | 6.1269 | 7.6807 |
| TJP1_S175;S178 | TJP1 | Q07157 | 1111 | PFDNQHSQDLDSR | 5.3124 | 6.0961 | 7.8512 | 6.2047 | 5.6825 | 6.1189 | 5.6264 | 6.4406 | 5.8691 | 6.9761 | 6.9862 | 6.2016 | 5.6772 | 6.5944 | 6.2512 | 6.1115 |
| TJP1_S175;S178;S179 | TJP1 | Q07157 | 277 | VPDLSDSIHSANA | 5.7372 | 5.7138 | 5.9733 | 7.0705 | 6.4225 | 6.1895 | 6.3387 | 6.3055 | 6.8311 | 6.1242 | 6.3090 | 6.3276 | 6.1183 | 6.0940 | 6.1736 | 6.2714 |
| TJP1_S178;S179 | TJP1 | Q07157 | 402 | DVDLPVSPSDGVL | 6.1683 | 6.4215 | 6.6114 | 7.1042 | 6.6176 | 7.0026 | 6.7081 | 7.1661 | 6.4213 | 6.6598 | 7.3960 | 6.8214 | 4.6054 | 5.0984 | 4.6270 | 4.5710 |
| TJP1_S277 | TJP1 | Q07157 | 125 | PDPEPVSDNEEDS | 5.6157 | 6.8905 | 6.1474 | 7.0717 | 6.2260 | 6.4299 | 6.0682 | 6.2873 | 6.7543 | 6.6881 | 6.5731 | 6.9128 | 5.6883 | 5.6198 | 5.3184 | 5.7084 |
| TJP1_S277;S280 | TJP1 | Q07157 | 379 | ERNEKQTPSLPEP | 6.9344 | 7.3013 | 6.7552 | 6.9761 | 5.9407 | 6.5626 | 6.4380 | 6.4785 | 6.7378 | 6.4988 | 6.5983 | 6.5978 | 4.6738 | 5.4227 | 4.7952 | 5.2889 |
| TJP1_S329 | TJP1 | Q07157 | 337 | QQPSNGSLRSRDE | 6.0163 | 6.2378 | 7.4999 | 7.0479 | 5.5084 | 6.0583 | 6.0328 | 7.2698 | 6.3304 | 7.2583 | 6.7754 | 6.9685 | 5.3297 | 5.9196 | 4.5817 | 5.1650 |
| TJP1_S337 | TJP1 | Q07157 | 927 | QKAEASSPVPYLS | 5.9871 | 6.1192 | 5.3786 | 6.7397 | 7.3279 | 6.9414 | 5.1397 | 7.0278 | 5.9679 | 6.3868 | 6.7109 | 6.1276 | 6.4973 | 6.0844 | 6.5273 | 5.0365 |
| TJP1_S402 | TJP1 | Q07157 | 589 | LASVQYTLPKTAG | 7.2392 | 6.4370 | 6.5012 | 7.0356 | 6.2344 | 5.0707 | 5.3679 | 5.2980 | 5.6189 | 4.8792 | 4.8906 | 4.5987 | 8.1015 | 8.1612 | 7.8723 | 6.6937 |
| TJP1_S617 | TJP1 | Q07157 | 899 | QTYPPYSPQAQPQ | 6.5524 | 5.7958 | 6.2000 | 7.7516 | 6.9758 | 7.4664 | 6.6254 | 7.1290 | 6.1821 | 6.7362 | 7.1730 | 5.6735 | 5.5317 | 5.6898 | 4.0600 | 5.3573 |
| TJP1_S837 | TJP1 | Q07157 | 1594 | SGVETFSIHAEKP | 5.9506 | 5.9144 | 5.9631 | 5.9761 | 6.0843 | 6.1579 | 5.6421 | 5.7402 | 6.6847 | 5.8418 | 6.4265 | 5.9814 | 5.5648 | 7.0145 | 4.7580 | 6.5997 |
| TJP1_S899 | TJP1 | Q07157 | 1278 | ENKRSASLETKKD | 6.9047 | 6.4296 | 6.4897 | 6.6782 | 6.4135 | 5.5821 | 5.8757 | 6.4664 | 6.4436 | 6.6352 | 5.9818 | 5.9735 | 5.8661 | 6.1543 | 6.3619 | 5.7439 |
| TJP1_S912 | TJP1 | Q07157 | 837 | SMYSTDSRHTSDY | 5.9550 | 6.5744 | 7.4177 | 6.1316 | 5.0654 | 6.7861 | 6.7268 | 6.2084 | 6.1867 | 6.3310 | 6.3937 | 6.9286 | 5.7609 | 5.9785 | 5.9521 | 5.6032 |
| TJP1_S912 | TJP1 | Q07157 | 1521 | QTQKTVTPAYNRF | 7.3110 | 8.3839 | 7.1743 | 7.2710 | 5.5059 | 6.9195 | 7.3687 | 6.7394 | 7.2051 | 7.1348 | 7.4941 | 7.7133 | 2.9608 | 4.1058 | 2.5416 | 4.1709 |
| TJP1_S927 | TJP1 | Q07157 | 125;131 | NA | 6.2237 | 6.5118 | 6.3300 | 6.7133 | 6.5553 | 6.0798 | 5.8584 | 6.1944 | 6.0371 | 6.1538 | 6.0082 | 5.9704 | 6.2625 | 6.2262 | 6.5666 | 6.3086 |
| TJP1_T1521 | TJP1 | Q07157 | 277;280 | NA | 5.7985 | 6.4624 | 6.2308 | 6.7368 | 6.3692 | 6.1664 | 6.1117 | 6.6842 | 6.7936 | 6.2348 | 6.8043 | 5.8647 | 6.0825 | 6.3014 | 5.8668 | 5.4920 |
| TJP1_T354 | TJP1 | Q07157 | 175;178 | NA | 6.2845 | 6.9765 | 6.8483 | 6.8433 | 5.5115 | 6.6377 | 6.1445 | 6.5336 | 7.1245 | 7.8562 | 7.7657 | 8.0628 | 4.1691 | 4.5300 | 4.0367 | 4.6751 |
| TJP1_T379 | TJP1 | Q07157 | 175;178;179 | NA | 7.1381 | 7.4713 | 7.1084 | 7.0996 | 6.6435 | 6.0097 | 6.9346 | 6.2983 | 7.2509 | 6.9211 | 6.8682 | 6.7189 | 4.2148 | 4.8523 | 4.4031 | 4.0669 |
| TJP1_T589 | TJP1 | Q07157 | 178;179 | NA | 8.3372 | 8.5696 | 6.5660 | 7.6711 | 7.6394 | 6.6219 | 6.2785 | 5.5681 | 7.4222 | 6.2244 | 6.2349 | 6.3519 | 3.6797 | 4.8930 | 3.8087 | 4.1332 |
| TJP2_S1017 | TJP2 | Q9UDY2-7 | 297 | DYERAYSPEYRRG | 6.3893 | 6.7559 | 6.0791 | 5.9988 | 6.1917 | 6.1903 | 6.0878 | 6.2402 | 6.2345 | 6.0357 | 6.0089 | 6.0080 | 6.3864 | 6.6082 | 6.4961 | 6.2890 |
| TJP2_S1190 | TJP2 | Q9UDY2-7 | 1190 | TRGSYGSDAEEEE | 6.3104 | 6.6598 | 6.3505 | 6.6774 | 6.6199 | 6.8603 | 6.1003 | 6.4130 | 5.9601 | 5.8018 | 6.0518 | 6.0398 | 6.2806 | 6.4917 | 5.9338 | 5.4488 |
| TJP2_S205 | TJP2 | Q9UDY2-7 | 1017 | DQLRDNSPPPAFK | 5.8349 | 5.7498 | 6.2018 | 6.1738 | 6.1883 | 5.5328 | 6.0293 | 5.6491 | 6.7196 | 6.1951 | 6.1977 | 6.6194 | 6.8567 | 6.7417 | 6.6595 | 6.6504 |
| TJP2_S297 | TJP2 | Q9UDY2-7 | 997 | ESIRKPSPEPRAQ | 6.3213 | 6.5060 | 6.0861 | 6.2193 | 6.5922 | 6.1852 | 6.4846 | 6.4666 | 6.3896 | 6.1232 | 5.9291 | 6.0480 | 6.0112 | 6.3296 | 6.1696 | 6.1384 |
| TJP2_S997 | TJP2 | Q9UDY2-7 | 486 | LSRMGATPTPFKS | 7.7361 | 7.9848 | 7.1152 | 7.4483 | 5.9134 | 6.9846 | 7.1675 | 6.3268 | 7.7990 | 6.7491 | 7.5965 | 7.2584 | 3.3460 | 4.1105 | 3.4419 | 4.0219 |
| TJP2_T1085 | TJP2 | Q9UDY2-7 | 1085 | SLKPSTPIPPQE | 6.7036 | 6.5548 | 7.0034 | 6.4060 | 5.6964 | 6.1256 | 8.0603 | 5.7179 | 6.4211 | 6.2895 | 5.4224 | 5.8948 | 5.6532 | 6.2981 | 6.1622 | 5.5907 |
| TJP2_T486 | TJP2 | Q9UDY2-7 | 205 | SRSWEDSPERGRP | 6.7554 | 6.9443 | 6.4166 | 6.7068 | 6.3430 | 6.1459 | 6.1381 | 6.3869 | 6.2786 | 5.9195 | 6.0023 | 5.8774 | 6.1001 | 5.8854 | 5.9962 | 6.1033 |
| TJP3_S183 | TJP3 | O95049-3 | 183 | GSHGRRSPGGGSE | 6.1984 | 6.0000 | 5.7130 | 5.2653 | 6.5258 | 6.3534 | 5.3693 | 5.4213 | 6.9466 | 7.4165 | 5.7371 | 6.9090 | 6.0125 | 7.3876 | 6.2561 | 6.4882 |
| TJP3_S610 | TJP3 | O95049-3 | 610 | KKTTQRSREDLSA | 6.1915 | 5.9202 | 5.8771 | 6.3245 | 6.3640 | 5.9292 | 6.0154 | 5.7657 | 6.6029 | 5.8609 | 5.7193 | 5.8537 | 5.8645 | 7.2893 | 5.5087 | 7.1132 |
| TK1_S231 | TK1 | P04183 | 231 | QQILQCSPANxxx | 6.3867 | 6.5767 | 6.8850 | 7.2837 | 6.0657 | 7.1455 | 6.6554 | 6.8648 | 6.9087 | 6.8473 | 7.2142 | 6.9106 | 4.2393 | 5.1383 | 4.1868 | 4.6914 |
| TKT_T295 | TKT | P29401-2 | 295 | KKKILATPPQEDA | 5.0366 | 6.3701 | 6.5538 | 6.3551 | 6.4868 | 6.1106 | 6.1682 | 6.6705 | 6.4552 | 6.3874 | 6.1461 | 6.6003 | 5.6193 | 6.4205 | 6.6122 | 6.0073 |
| TLCD2_S235 | TLCD2 | A6NGC4 | 235 | SRPHPPSPGHEKT | 5.9050 | 5.3334 | 5.7672 | 6.3560 | 6.0104 | 5.7471 | 5.9590 | 5.7980 | 6.1363 | 6.3574 | 5.9379 | 6.4319 | 7.0095 | 6.7807 | 7.1318 | 7.3383 |
| TLE1_T312 | TLE1 | Q04724 | 319 | PVLKSSTPTPRSD | 5.7364 | 6.4256 | 5.5546 | 5.4291 | 5.6407 | 6.4364 | 6.1839 | 5.7897 | 6.0451 | 6.2648 | 6.2610 | 5.8457 | 6.7787 | 7.6156 | 7.0543 | 6.9385 |
| TLE1_T319 | TLE1 | Q04724 | 312 | LHEKASTPVLKSS | 6.6581 | 7.3773 | 6.5665 | 6.4039 | 6.1326 | 6.7397 | 6.8082 | 6.6423 | 6.6293 | 7.2360 | 7.3096 | 6.4729 | 4.4348 | 5.2628 | 4.0979 | 5.2282 |
| TLE3_S268;S272 | TLE3 | Q04726-4 | 317 | HNDKSSTPGLKSN | 6.5468 | 7.0111 | 6.7106 | 6.6624 | 6.1301 | 6.9090 | 6.8649 | 6.6367 | 6.9753 | 6.9033 | 7.4352 | 7.3448 | 4.2519 | 4.8820 | 4.2092 | 4.5270 |
| TLE3_S291 | TLE3 | Q04726-4 | 324 | PGLKSNTPTPRND | 5.9429 | 6.4526 | 5.8701 | 5.5912 | 5.8427 | 6.3692 | 6.3058 | 5.8294 | 6.3408 | 6.6568 | 6.0292 | 6.2765 | 6.7775 | 6.7803 | 6.2635 | 6.6716 |
| TLE3_T317 | TLE3 | Q04726-4 | 291 | KKDAPTSPASVAS | 5.0837 | 5.5225 | 5.9947 | 5.7052 | 6.0809 | 6.3449 | 5.4066 | 6.1532 | 6.2867 | 6.5028 | 5.4424 | 6.3158 | 7.1995 | 7.4702 | 7.3142 | 7.1765 |
| TLE3_T324 | TLE3 | Q04726-4 | 268;272 | NA | 5.6862 | 6.1402 | 6.5127 | 6.1449 | 6.2414 | 6.0032 | 6.6066 | 6.7291 | 6.0736 | 6.4228 | 6.6913 | 5.9418 | 6.1770 | 6.1797 | 6.1517 | 6.2980 |
| TLK1_S180 | TLK1 | Q9UKI8-2 | 764 | HMRRSNSSGNLHM | 5.5457 | 5.7445 | 5.4624 | 5.8175 | 5.6223 | 6.4858 | 6.1695 | 7.3955 | 6.1816 | 6.4210 | 6.5694 | 6.3453 | 6.2294 | 7.3165 | 6.3981 | 6.2955 |
| TLK1_S764 | TLK1 | Q9UKI8-2 | 180 | QGGNGSSPVRGIP | 5.9907 | 6.6359 | 5.8531 | 6.4249 | 6.5411 | 6.6481 | 5.9548 | 6.8759 | 6.4891 | 6.1497 | 6.1803 | 6.8181 | 5.2246 | 6.3405 | 5.6686 | 6.2046 |
| TLK2_S111 | TLK2 | Q86UE8 | 26 | FTGVGVSKGPLNS | 4.9428 | 6.5086 | 6.9082 | 6.4162 | 4.3759 | 6.6460 | 6.1704 | 5.9210 | 6.9975 | 6.4522 | 6.5685 | 6.2000 | 6.6704 | 6.9649 | 5.8275 | 6.4298 |
| TLK2_S223 | TLK2 | Q86UE8 | 46 | CSVGSLSDKEVET | 5.1561 | 5.5530 | 6.0285 | 5.3038 | 5.3718 | 6.0012 | 5.8426 | 5.4459 | 6.1673 | 6.3387 | 6.2478 | 6.2414 | 7.3133 | 7.9948 | 8.0027 | 6.9911 |
| TLK2_S26 | TLK2 | Q86UE8 | 99 | GSAPGTSPGRSVP | 6.2708 | 6.6809 | 6.0737 | 6.3696 | 6.3343 | 6.4526 | 6.2101 | 6.6282 | 6.4714 | 6.4738 | 6.4145 | 6.7826 | 5.6127 | 5.9557 | 5.6737 | 5.5954 |
| TLK2_S46 | TLK2 | Q86UE8 | 111 | PPVARSSPQHSLS | 6.3086 | 6.7647 | 6.1164 | 5.9595 | 6.4805 | 6.1048 | 6.7517 | 6.7115 | 7.1860 | 6.2782 | 6.3629 | 5.6246 | 6.1648 | 6.0269 | 5.8086 | 5.3504 |
| TLK2_S99 | TLK2 | Q86UE8 | 52 | SDKEVETPEKKQN | 5.6743 | 6.6928 | 6.5830 | 5.9605 | 5.8418 | 6.7299 | 6.2998 | 6.3394 | 6.8861 | 6.7904 | 6.4503 | 6.5643 | 5.6872 | 6.1150 | 5.5685 | 5.8168 |
| TLK2_T52 | TLK2 | Q86UE8 | 223 | ISALENSKNSDLE | 6.0205 | 7.1256 | 7.3840 | 6.3432 | 5.7895 | 5.3992 | 7.0651 | 6.0095 | 5.0858 | 6.8494 | 6.4854 | 6.0534 | 7.4500 | 5.1243 | 9.4678 | 8.7475 |
| TLN1_S1641 | TLN1 | Q9Y490 | 425 | MLEDSVSPKKSTV | 6.2735 | 6.3119 | 6.9085 | 6.5459 | 6.4010 | 6.7734 | 6.7782 | 6.4922 | 7.1293 | 6.8171 | 7.2797 | 7.3978 | 4.6464 | 4.9131 | 4.3755 | 4.9565 |
| TLN1_S2040 | TLN1 | Q9Y490 | 1641 | GHSRTVSDSIKKL | 5.5623 | 5.4437 | 6.1293 | 5.3348 | 5.5123 | 5.1678 | 5.5054 | 5.3669 | 5.3523 | 4.8156 | 5.0318 | 5.3871 | 8.8740 | 8.7933 | 9.0529 | 8.6703 |
| TLN1_S425 | TLN1 | Q9Y490 | 446 | GKVEHGSVALPAI | 6.7881 | 6.9970 | 7.7034 | 5.7161 | 4.6319 | 6.4349 | 5.8052 | 6.7574 | 8.5849 | 5.9987 | 5.9066 | 4.5367 | 6.1922 | 6.1031 | 6.4594 | 5.3844 |
| TLN1_S446 | TLN1 | Q9Y490 | 2040 | VQNAAGSQEKLAQ | 5.9123 | 6.1310 | 6.6406 | 6.1460 | 6.1923 | 6.3756 | 5.8119 | 6.6604 | 5.7735 | 6.0511 | 6.4363 | 7.0666 | 6.6472 | 6.0769 | 5.6473 | 6.4309 |
| TLN1_T2270 | TLN1 | Q9Y490 | 2270 | ELKQQLTGHSKRV | 6.2715 | 6.1302 | 7.5805 | 7.3170 | 7.0386 | 6.5242 | 6.5305 | 6.4665 | 6.2212 | 5.3886 | 6.5862 | 6.2960 | 5.1883 | 5.7055 | 5.3870 | 5.3683 |
| TLN2_T1843 | TLN2 | Q9Y4G6 | 1843 | SKLDEGTPPEPKG | 5.8771 | 5.9588 | 6.3111 | 6.0965 | 5.9527 | 5.8463 | 6.0682 | 6.4967 | 6.4760 | 6.6557 | 6.4916 | 6.2046 | 6.3657 | 6.7691 | 6.5064 | 5.9236 |
| TMBIM1_S83 | TMBIM1 | Q969X1 | 83 | ERAVSDSFGPGEW | 6.3553 | 6.8810 | 6.1277 | 6.9481 | 6.1762 | 6.3243 | 6.8958 | 6.5077 | 6.1478 | 6.3944 | 6.4194 | 6.7446 | 4.9899 | 5.8216 | 5.5925 | 5.6736 |
| TMC7_S72 | TMC7 | Q7Z402 | 72 | LLKPTDSYSSQLE | 1.6957 | 1.6299 | 1.8688 | 1.8540 | 2.4347 | 2.2735 | 2.2569 | 2.6366 | 2.0133 | 1.4540 | 1.2484 | 1.6688 | 19.2432 | 18.5486 | 21.6445 | 17.5290 |
| TMC8_S522 | TMC8 | Q8IU68 | 522 | KNSRASSRPFRAS | 7.4894 | 7.2164 | 6.3303 | 7.0693 | 6.2174 | 5.7397 | 6.2446 | 5.8963 | 5.9383 | 6.2085 | 6.1721 | 6.4771 | 5.8054 | 6.2927 | 5.4195 | 5.4830 |
| TMCC1_S134 | TMCC1 | O94876 | 382 | IRNKFGSADNIPN | 6.3623 | 6.0613 | 6.1087 | 5.7800 | 6.2660 | 5.7636 | 6.4704 | 5.7173 | 6.6980 | 6.1714 | 5.8419 | 5.8060 | 7.2221 | 6.3320 | 6.6503 | 6.7487 |
| TMCC1_S382 | TMCC1 | O94876 | 134 | PEAPKGSPQINRK | 6.0598 | 6.1514 | 5.9969 | 6.3850 | 5.8168 | 5.7646 | 6.2267 | 6.1687 | 6.5316 | 6.2870 | 5.9604 | 5.9365 | 6.5380 | 6.8511 | 6.8107 | 6.5747 |
| TMCC1_S414 | TMCC1 | O94876 | 414 | ISNFQSSPKYGSE | 6.2997 | 6.5353 | 6.3586 | 6.7135 | 6.1376 | 5.7729 | 6.6567 | 6.0079 | 6.0660 | 6.1010 | 6.0479 | 6.2347 | 6.1638 | 6.3601 | 6.3200 | 6.2244 |
| TMCC3_S216 | TMCC3 | Q9ULS5 | 216 | IRNKFGSADNIAH | 6.0391 | 6.1230 | 6.0273 | 6.5819 | 6.8391 | 5.9877 | 6.5114 | 6.5361 | 6.5864 | 6.1577 | 5.9386 | 5.6782 | 7.1248 | 6.6421 | 6.8867 | 7.1399 |
| TMCC3_S46 | TMCC3 | Q9ULS5 | 194 | MPGVSLTPPVFVF | 6.4969 | 7.4087 | 6.9921 | 6.8075 | 6.4458 | 6.7419 | 6.6154 | 6.9330 | 6.4559 | 6.9358 | 6.4500 | 6.7898 | 4.9953 | 5.2257 | 4.1989 | 4.5074 |
| TMCC3_T194 | TMCC3 | Q9ULS5 | 46 | NIRRGGSDTNLNF | 5.5200 | 6.0616 | 5.4135 | 5.6290 | 5.5687 | 6.4706 | 6.1504 | 5.6656 | 5.9056 | 5.1675 | 5.7148 | 5.7617 | 7.2329 | 7.7030 | 7.5401 | 8.4950 |
| TMCO1_S239 | TMCO1 | Q9UM00 | 239 | PPSGKFSxxxxxx | 6.0800 | 6.9362 | 6.5662 | 7.4503 | 6.2908 | 6.3087 | 6.5210 | 6.1918 | 6.4742 | 6.9964 | 6.9306 | 7.0864 | 4.6270 | 5.4741 | 5.0804 | 4.9859 |
| TMED8_S21 | TMED8 | Q6PL24 | 21 | PTARPGSAGGVGD | 5.9554 | 5.2315 | 7.0949 | 5.5247 | 5.7397 | 6.6307 | 6.1248 | 5.5303 | 7.3962 | 6.5236 | 6.6714 | 6.7146 | 6.5236 | 6.2543 | 6.0497 | 6.0346 |
| TMEM11_S17 | TMEM11 | P17152 | 17 | PGSSGGSARERVS | 4.8523 | 5.8192 | 6.1202 | 5.2200 | 4.8299 | 5.0103 | 5.2941 | 5.0828 | 5.1662 | 6.4608 | 6.3782 | 6.1914 | 7.5331 | 8.5702 | 9.1463 | 8.3248 |
| TMEM11_T152 | TMEM11 | P17152 | 152 | LPLHTLTSSTPVV | 7.0001 | 6.0199 | 7.1848 | 5.8510 | 5.1911 | 6.5126 | 6.4808 | 6.0410 | 6.4194 | 6.1695 | 6.5091 | 5.9618 | 6.4018 | 7.2024 | 6.9226 | 6.5320 |
| TMEM115_S320 | TMEM115 | Q12893 | 329 | PSDKAPTPPGKGA | 6.3281 | 6.6791 | 5.9445 | 6.8982 | 6.1110 | 6.1521 | 6.4139 | 6.3627 | 6.2805 | 6.3587 | 6.1963 | 6.4621 | 5.7310 | 6.0967 | 5.8770 | 6.1081 |
| TMEM115_T329 | TMEM115 | Q12893 | 320 | SGAKVDSPLPSDK | 5.7791 | 6.4452 | 6.5829 | 6.2389 | 6.1751 | 6.0150 | 6.9325 | 5.9913 | 5.9602 | 6.0304 | 5.7977 | 5.9154 | 6.7668 | 6.6035 | 6.3143 | 7.1815 |
| TMEM131_S1155 | TMEM131 | Q92545 | 1669 | TAGYDKSPGGNGF | 6.5508 | 7.0684 | 6.2250 | 6.8900 | 6.4383 | 6.7471 | 7.1337 | 6.7021 | 6.5580 | 6.2939 | 6.6460 | 6.2550 | 4.7264 | 5.2609 | 5.0058 | 5.4985 |
| TMEM131_S1342 | TMEM131 | Q92545 | 1649 | KLTKAASLPGKNG | 6.4529 | 6.2262 | 5.2543 | 6.0377 | 5.4386 | 4.3283 | 3.9114 | 4.7423 | 7.5570 | 7.1609 | 5.1922 | 6.3635 | 7.3314 | 8.0279 | 8.1846 | 7.7910 |
| TMEM131_S1649 | TMEM131 | Q92545 | 1863 | NPWRIWSPTIGRR | 6.8515 | 8.0006 | 7.3788 | 7.8526 | 6.2275 | 7.0413 | 7.5390 | 7.0151 | 6.7001 | 7.2076 | 7.0193 | 7.1971 | 2.9089 | 3.9567 | 3.2173 | 3.8865 |
| TMEM131_S1669 | TMEM131 | Q92545 | 1155 | PFRRRLSFEASNP | 5.2741 | 5.7194 | 6.1623 | 5.4064 | 5.3619 | 5.6523 | 5.5965 | 5.3719 | 6.9269 | 7.2914 | 6.6233 | 6.4145 | 7.2230 | 7.2901 | 7.0656 | 6.6202 |
| TMEM131_S1863 | TMEM131 | Q92545 | 1490 | SLELPYTPPLESK | 7.4971 | 6.6427 | 6.5455 | 7.1750 | 6.9089 | 6.9022 | 6.7298 | 7.3740 | 6.2600 | 6.4497 | 7.6568 | 6.8876 | 3.7978 | 4.9743 | 4.3521 | 3.8464 |
| TMEM131_T1490 | TMEM131 | Q92545 | 1342 | SSARHSSESDSIT | 5.7734 | 6.3047 | 6.1211 | 5.9652 | 5.9108 | 5.7022 | 6.3998 | 6.1184 | 6.6084 | 5.8630 | 7.0093 | 6.3042 | 6.2803 | 6.9088 | 6.5231 | 6.2075 |
| TMEM132A_S914 | TMEM132A | Q24JP5-2 | 914 | RQLDRQSPGPPKG | 7.2926 | 7.3860 | 7.2242 | 6.9651 | 6.7184 | 6.7498 | 6.5707 | 6.1919 | 6.8600 | 6.5417 | 7.1369 | 6.7712 | 4.6323 | 4.4875 | 3.9786 | 4.4930 |
| TMEM184B_S338 | TMEM184B | Q9Y519 | 388 | GLSRSHLSSGARD | 5.4833 | 6.3039 | 6.0021 | 5.4183 | 5.3159 | 6.0157 | 5.4826 | 5.8644 | 6.5068 | 7.2088 | 6.6094 | 6.9037 | 6.4117 | 7.3717 | 6.2464 | 6.8552 |
| TMEM184B_S388 | TMEM184B | Q9Y519 | 338 | PMKSISSSLKETM | 1.0937 | 1.5277 | 1.3455 | 1.3777 | 1.6422 | 1.3176 | 1.7097 | 1.6167 | 1.4674 | 1.1615 | 0.6277 | 1.1944 | 20.8466 | 19.5963 | 23.0439 | 20.4316 |
| TMEM184B_S402;S403 | TMEM184B | Q9Y519 | 402;403 | NA | 5.6518 | 5.7666 | 6.4377 | 7.2332 | 6.0701 | 6.1981 | 6.5808 | 6.3984 | 7.0208 | 6.2118 | 5.4398 | 6.8238 | 6.1969 | 6.1905 | 5.1866 | 6.5933 |
| TMEM185B_S333 | TMEM185B | Q9H7F4 | 333 | RVVITQSPGKYVP | 6.8081 | 7.3475 | 6.5977 | 7.0388 | 6.6829 | 6.5444 | 7.1357 | 7.0380 | 7.1543 | 7.0335 | 6.8909 | 6.9541 | 3.8398 | 4.6023 | 4.0291 | 4.3028 |
| TMEM192_S230 | TMEM192 | Q8IY95 | 230 | GFRTISSLEEIVE | 4.2769 | 4.6038 | 4.6255 | 5.0511 | 4.4018 | 4.3743 | 5.0891 | 4.5469 | 4.4068 | 4.6276 | 4.7037 | 4.7671 | 10.6509 | 11.1544 | 11.9568 | 10.7632 |
| TMEM201_S441 | TMEM201 | Q5SNT2 | 441 | RSPRRTSPSSLPG | 6.9703 | 7.1683 | 6.4144 | 7.4927 | 5.7520 | 6.8204 | 6.2824 | 5.9336 | 7.4872 | 7.2266 | 7.1804 | 7.4617 | 4.5151 | 4.9001 | 4.6818 | 4.7130 |
| TMEM201_S454 | TMEM201 | Q5SNT2 | 454 | RLSRALSLGTIPS | 5.7776 | 5.8426 | 5.8423 | 6.3173 | 6.1648 | 5.4153 | 5.7635 | 5.8772 | 6.0950 | 6.3341 | 6.2950 | 6.3465 | 6.9716 | 7.1191 | 6.7028 | 7.1353 |
| TMEM201_S477;S480 | TMEM201 | Q5SNT2 | 529 | HRRPLISPARLNL | 7.2293 | 7.8469 | 6.9333 | 7.8410 | 6.1809 | 6.4457 | 6.6739 | 6.6832 | 7.5179 | 7.0761 | 7.3544 | 7.3300 | 3.4485 | 4.1226 | 3.3373 | 3.9791 |
| TMEM201_S529 | TMEM201 | Q5SNT2 | 477;480 | NA | 7.4638 | 6.6958 | 6.8604 | 7.0194 | 6.5494 | 5.9987 | 6.5475 | 6.6232 | 7.0741 | 6.6440 | 7.3464 | 6.8272 | 4.6671 | 4.1760 | 5.2462 | 4.2608 |
| TMEM209_S147 | TMEM209 | Q96SK2 | 147 | SYSPSRSPSTSPK | 6.4230 | 6.7682 | 5.7318 | 6.6142 | 6.4884 | 5.8315 | 6.2174 | 6.7379 | 6.5412 | 6.1926 | 5.9502 | 5.9625 | 6.0361 | 5.9996 | 6.1512 | 6.3540 |
| TMEM209_S196 | TMEM209 | Q96SK2 | 221 | LRSYRSSPTVYN | 6.3847 | 6.4616 | 5.9650 | 6.6359 | 6.5905 | 6.3702 | 6.6078 | 5.9474 | 5.9593 | 5.9213 | 5.9120 | 6.2135 | 6.3641 | 6.2635 | 6.2713 | 6.1319 |
| TMEM209_S196;S201 | TMEM209 | Q96SK2 | 94 | YTVAPTSLVVSPG | 6.3457 | 6.5805 | 6.0042 | 6.7230 | 6.1350 | 6.3627 | 6.1389 | 6.5004 | 6.4077 | 6.4743 | 6.4119 | 6.2181 | 5.6803 | 6.5116 | 5.9928 | 5.5130 |
| TMEM209_S201 | TMEM209 | Q96SK2 | 201 | FSPSPPSPYPTTV | 5.8164 | 5.4380 | 6.1758 | 5.2822 | 5.6279 | 5.9135 | 6.4395 | 6.0364 | 5.8629 | 5.0417 | 5.5036 | 5.5309 | 7.7334 | 7.4861 | 7.8605 | 8.2512 |
| TMEM209_S221 | TMEM209 | Q96SK2 | 269 | STSPSSSPTFWNY | 5.9872 | 6.7800 | 6.2225 | 5.8770 | 5.9788 | 6.7286 | 6.3532 | 6.5773 | 6.5605 | 5.3392 | 7.2478 | 6.0292 | 6.5456 | 6.0442 | 5.4064 | 6.3224 |
| TMEM209_S260 | TMEM209 | Q96SK2 | 196 | NKLASFSPSPPSP | 6.3696 | 5.8136 | 6.0989 | 6.0998 | 6.0577 | 5.9324 | 5.9553 | 6.3802 | 6.4414 | 6.2263 | 6.0447 | 6.1043 | 6.4406 | 6.6120 | 6.5475 | 6.8757 |
| TMEM209_S260;S269 | TMEM209 | Q96SK2 | 115 | TAVVQTTPPHDLA | 5.5019 | 5.7541 | 5.9764 | 6.0109 | 6.0927 | 6.1531 | 5.9896 | 6.5314 | 5.7438 | 6.0547 | 5.8175 | 6.2683 | 6.7721 | 7.2476 | 7.5563 | 6.5297 |
| TMEM209_S269 | TMEM209 | Q96SK2 | 260 | HRVKLGSPDSTSP | 5.7687 | 5.3797 | 6.7447 | 7.0633 | 4.8095 | 5.2053 | 6.0337 | 3.6960 | 6.3858 | 5.6614 | 5.0141 | 7.9924 | 7.9924 | 6.3556 | 7.6201 | 8.2774 |
| TMEM209_S94 | TMEM209 | Q96SK2 | 196;201 | NA | 7.0496 | 6.8103 | 7.3502 | 8.4456 | 7.3414 | 8.3684 | 5.7447 | 6.9120 | 7.0202 | 7.7838 | 8.2765 | 7.7100 | 2.5210 | 3.2796 | 1.9312 | 3.4554 |
| TMEM209_T115 | TMEM209 | Q96SK2 | 260;269 | NA | 5.3431 | 5.6307 | 5.9078 | 7.0768 | 6.7868 | 6.5585 | 6.6014 | 6.3572 | 6.6158 | 6.0637 | 6.8326 | 6.4044 | 5.4292 | 6.7540 | 5.7149 | 5.9231 |

| | | | | | | | | | | | | | | | | | | | | | |
|---|---|---|---|---|---|---|---|---|---|---|---|---|---|---|---|---|---|---|---|---|---|
| TMEM214_T97 | TMEM214 | Q6NUQ4 | | 97 | QPKKVATPPNQNQ | 6.5498 | 6.8445 | 6.9644 | 7.7319 | 5.8375 | 6.0951 | 6.6294 | 6.3916 | 7.1334 | 7.2309 | 6.7166 | 7.4369 | 4.3623 | 4.8277 | 4.1247 | 5.1232 |
| TMEM230_S86 | TMEM230 | Q96A57-2 | | 87 | KYSRLSSTDDGYI | 4.9633 | 5.2618 | 5.5854 | 5.6250 | 5.5521 | 5.8744 | 5.6418 | 5.6609 | 5.6760 | 5.9880 | 5.4018 | 5.7486 | 8.1638 | 8.1700 | 8.7977 | 7.8895 |
| TMEM230_S87 | TMEM230 | Q96A57-2 | | 86 | VKYSRLSSTDDGY | 5.9806 | 6.0370 | 6.4948 | 5.7395 | 6.7974 | 6.4742 | 6.3486 | 5.7626 | 6.3332 | 6.5242 | 5.6279 | 6.1293 | 6.2011 | 7.9219 | 6.0139 | 5.6138 |
| TMEM237_S127 | TMEM237 | Q96Q45-3 | | 86 | TAGRRPSEGNEPS | 5.1633 | 4.9479 | 5.9126 | 5.7766 | 6.1372 | 5.2433 | 5.6576 | 5.9856 | 7.3383 | 7.5098 | 6.8694 | 7.6998 | 6.1846 | 7.1276 | 6.2103 | 6.2360 |
| TMEM237_S49 | TMEM237 | Q96Q45-3 | | 127 | STQKKSSSSSLLR | 6.1198 | 6.1093 | 6.4128 | 6.5437 | 5.5019 | 5.6791 | 5.7551 | 6.0007 | 6.0636 | 6.3656 | 5.8561 | 5.9567 | 6.7669 | 7.3274 | 6.8004 | 6.7407 |
| TMEM237_S73 | TMEM237 | Q96Q45-3 | | 49 | ALPPVPSQDDIPL | 4.0590 | 4.3706 | 4.3121 | 4.8020 | 4.2387 | 4.5645 | 4.3285 | 4.9924 | 3.3451 | 4.1709 | 3.6839 | 4.5945 | 10.3164 | 10.4676 | 13.7792 | 13.9746 |
| TMEM237_S86 | TMEM237 | Q96Q45-3 | | 73 | NTPASASLEGLAQ | 6.3383 | 6.1206 | 6.0849 | 6.1813 | 5.3095 | 6.0260 | 6.3339 | 5.9207 | 6.6301 | 7.7242 | 5.9520 | 6.8800 | 5.7039 | 6.8202 | 5.7199 | 6.2545 |
| TMEM238_S137 | TMEM238 | C9JI98 | | 137 | QRPAPGSRRARRA | 7.1027 | 6.5022 | 6.4333 | 7.0361 | 6.6351 | 6.4798 | 7.1362 | 6.1416 | 7.1020 | 6.9001 | 6.7471 | 7.1121 | 4.5435 | 5.1518 | 3.8232 | 5.1531 |
| TMEM245_S16 | TMEM245 | Q9H330 | | 332 | PSPTSPSPTLGRR | 5.8184 | 5.9882 | 5.6562 | 6.3240 | 5.8180 | 5.9289 | 6.4617 | 5.9937 | 6.4516 | 5.7333 | 5.6753 | 5.9075 | 6.6004 | 7.1601 | 7.4117 | 7.0711 |
| TMEM245_S327;S332 | TMEM245 | Q9H330 | | 16 | APSLRSSPGPAPR | 6.7547 | 6.8491 | 6.0512 | 6.7879 | 6.2897 | 6.0856 | 6.3872 | 6.3539 | 6.4494 | 6.4444 | 6.5564 | 6.6170 | 5.2492 | 5.8579 | 5.5982 | 5.6682 |
| TMEM245_S332 | TMEM245 | Q9H330 | | 36 | SGGGGETPRTAAL | 5.9121 | 8.1487 | 6.6658 | 6.4761 | 5.9628 | 6.9274 | 7.5650 | 6.3666 | 7.6844 | 7.0091 | 6.4647 | 6.8980 | 3.8520 | 4.8673 | 4.4455 | 4.7545 |
| TMEM245_T36 | TMEM245 | Q9H330 | 327;332 | | NA | 5.7233 | 5.9945 | 6.0438 | 7.2796 | 6.4819 | 6.3168 | 6.1997 | 5.9487 | 6.6414 | 5.4974 | 5.8568 | 5.9089 | 6.7456 | 6.8536 | 6.3634 | 6.1445 |
| TMEM248_S300 | TMEM248 | Q9NWD8 | | 300 | PSKLRQSNPEFCP | 5.8904 | 6.1682 | 5.9148 | 5.9353 | 6.9644 | 5.8847 | 5.6374 | 6.1118 | 6.0485 | 5.7651 | 6.0083 | 5.3740 | 6.8289 | 7.1402 | 7.3328 | 6.9950 |
| TMEM38B_S262 | TMEM38B | Q9NVV0 | | 262 | KKSEAKSPSNGVG | 6.4147 | 7.2866 | 6.8663 | 8.3078 | 6.6286 | 7.2598 | 7.4359 | 6.8402 | 6.9579 | 7.6847 | 8.1142 | 7.0343 | 2.9259 | 4.0687 | 2.8184 | 3.3559 |
| TMEM39B_S47 | TMEM39B | Q9GZU3 | | 47 | SRTRSSSGTGLSS | 4.2256 | 7.9798 | 7.1549 | 5.8698 | 5.1122 | 5.1740 | 5.6004 | 6.7284 | 6.3075 | 5.2245 | 7.4355 | 8.0191 | 6.2907 | 6.8575 | 5.1627 | 6.8575 |
| TMEM40_S137 | TMEM40 | Q8WWA1 | | 137 | GLRRRGSDPASGE | 6.1781 | 7.2269 | 6.4539 | 7.0748 | 6.2783 | 5.8241 | 6.1898 | 5.3754 | 5.6976 | 5.5738 | 5.9167 | 5.8185 | 6.4795 | 6.7996 | 6.9123 | 6.2009 |
| TMEM44_S333 | TMEM44 | Q2T9K0 | | 333 | AKEARESPDTQAL | 6.1932 | 6.1315 | 6.2355 | 7.0775 | 6.0069 | 5.6766 | 6.7797 | 5.9258 | 6.4055 | 6.1319 | 6.7634 | 6.6970 | 5.8943 | 6.1094 | 6.1051 | 5.8668 |
| TMEM51_S155 | TMEM51 | Q9NW97 | | 234 | PSIEPLTPPPQYD | 6.0234 | 6.1220 | 6.2506 | 6.7520 | 6.1776 | 5.7386 | 6.0622 | 5.9249 | 6.3925 | 6.0201 | 6.6455 | 6.9312 | 5.8305 | 6.5672 | 6.1955 | 6.3662 |
| TMEM51_S155;S163 | TMEM51 | Q9NW97 | | 155 | EQNPRLSISLPSY | 6.1686 | 6.1258 | 6.6821 | 6.9216 | 6.1994 | 5.9656 | 5.7810 | 5.7970 | 6.7501 | 6.9710 | 6.3919 | 6.8898 | 5.5076 | 6.2540 | 5.9575 | 5.6370 |
| TMEM51_T234 | TMEM51 | Q9NW97 | 155;163 | | NA | 5.7291 | 5.4317 | 6.6439 | 6.3934 | 6.1550 | 5.4258 | 5.9760 | 5.4448 | 7.5199 | 6.8359 | 6.3398 | 6.1144 | 6.5049 | 6.1421 | 6.7195 | 6.6238 |
| TMEM87A_S540 | TMEM87A | Q8NBN3 | | 540 | LPALLDSDEERMI | 6.4732 | 6.1143 | 6.6694 | 6.1685 | 6.0367 | 6.1459 | 6.3035 | 6.0622 | 6.0292 | 6.1029 | 6.3033 | 6.2869 | 6.4205 | 6.5512 | 6.0970 | 6.2352 |
| TMEM88_S145 | TMEM88 | Q6PEY1 | | 145 | PRQIRASPGSQAV | 6.4653 | 7.0297 | 6.3967 | 7.0196 | 6.6026 | 6.6787 | 6.5380 | 7.0260 | 6.7123 | 6.1103 | 6.1032 | 6.7955 | 5.2245 | 5.1426 | 4.9808 | 5.1742 |
| TMEM94_S1384 | TMEM94 | Q12767-2 | | 1384 | TKLGMNSPFxxxx | 6.3193 | 7.1598 | 7.2000 | 5.0641 | 4.1967 | 6.4620 | 6.7051 | 6.6997 | 5.9119 | 8.0192 | 7.5537 | 7.3398 | 4.4858 | 5.9433 | 5.1222 | 5.8176 |
| TMEM94_S221 | TMEM94 | Q12767-2 | | 798 | QNARRSSWSSDEG | 5.8495 | 6.5417 | 6.4501 | 6.1740 | 6.5591 | 5.8192 | 5.4055 | 6.6272 | 6.4033 | 5.9261 | 6.0899 | 6.2164 | 6.4221 | 7.0054 | 6.2988 | 6.2115 |
| TMEM94_S221;S225 | TMEM94 | Q12767-2 | | 221 | DLFPPFSPPPSPR | 6.1694 | 5.9014 | 6.2916 | 7.1240 | 5.9917 | 5.7892 | 6.2780 | 5.9581 | 6.3034 | 6.1493 | 6.7155 | 5.8170 | 6.5821 | 6.4345 | 6.4292 | 6.0655 |
| TMEM94_S225 | TMEM94 | Q12767-2 | | 225 | PFSPPPSPRGEVE | 4.7796 | 6.1734 | 5.9158 | 6.0834 | 6.9295 | 6.4269 | 5.8970 | 6.0886 | 5.8498 | 6.1127 | 6.8374 | 5.3021 | 6.8258 | 6.2331 | 6.4928 | 8.0522 |
| TMEM94_S513 | TMEM94 | Q12767-2 | | 513 | GRSKHPSGSNVSF | 6.8905 | 7.0305 | 6.2045 | 6.4846 | 7.0020 | 5.5484 | 4.9390 | 7.0331 | 6.9799 | 6.0217 | 6.3342 | 6.4457 | 5.1529 | 6.1099 | 6.0684 | 5.7546 |
| TMEM94_S798 | TMEM94 | Q12767-2 | 221;225 | | NA | 6.1817 | 6.5606 | 6.9310 | 6.5259 | 6.4282 | 4.9165 | 6.4382 | 6.1786 | 6.1431 | 6.1996 | 6.0833 | 7.1221 | 6.1611 | 5.8580 | 6.3512 | 5.9213 |
| TMF1_S199 | TMF1 | P82094-2 | | 401 | ESGRSATPVNCEQ | 6.1807 | 5.7088 | 5.8459 | 6.2493 | 6.5200 | 6.2939 | 5.8096 | 6.2797 | 6.2711 | 6.1250 | 6.1426 | 6.3617 | 6.9409 | 6.1666 | 6.6140 | 6.4901 |
| TMF1_S328 | TMF1 | P82094-2 | | 936 | SSTPTMSRSSSIS | 6.9464 | 5.6189 | 6.7558 | 6.8124 | 6.6145 | 6.2865 | 5.9544 | 6.1903 | 7.6459 | 7.4534 | 7.5213 | 7.9246 | 4.8543 | 4.7705 | 3.9508 | 4.7001 |
| TMF1_S333 | TMF1 | P82094-2 | | 77 | QSPPIASPKAITK | 5.4614 | 5.3344 | 5.2717 | 5.6890 | 6.5020 | 6.5288 | 6.4452 | 6.3188 | 6.6610 | 5.9631 | 6.0363 | 6.3471 | 7.3309 | 6.8456 | 6.0922 | 7.1723 |
| TMF1_S77 | TMF1 | P82094-2 | | 333 | DSFSVQSLDSRSV | 5.9937 | 6.1926 | 5.8148 | 5.9110 | 5.4518 | 5.8620 | 5.9789 | 6.3585 | 5.6832 | 5.1230 | 5.3845 | 6.0751 | 8.3087 | 6.8118 | 7.4546 | 7.5957 |
| TMF1_S936 | TMF1 | P82094-2 | | 364 | IIVNSSTPKSKTV | 6.6377 | 6.5793 | 6.4289 | 7.6872 | 6.9028 | 6.7336 | 5.9194 | 6.8970 | 7.3738 | 6.4145 | 7.1733 | 6.9578 | 4.3584 | 4.5166 | 4.2278 | 5.1919 |
| TMF1_S940 | TMF1 | P82094-2 | | 328 | AFERIDSFSVQSL | 5.6255 | 6.0654 | 6.3718 | 5.9593 | 6.0190 | 5.9862 | 5.7966 | 5.4179 | 6.9100 | 5.5341 | 6.5698 | 6.4383 | 7.3621 | 6.9448 | 5.9461 | 7.0531 |
| TMF1_T364 | TMF1 | P82094-2 | | 940 | TMSRSSSISGVDM | 5.7737 | 6.9355 | 7.0967 | 6.4652 | 6.5026 | 5.1796 | 4.6062 | 7.1585 | 6.3359 | 7.9456 | 7.1360 | 6.4783 | 4.5762 | 6.7331 | 4.6868 | 6.3902 |
| TMF1_T401 | TMF1 | P82094-2 | | 199 | SLKVSESVIDVKT | 1.7454 | 1.7032 | 1.9774 | 2.6700 | 3.3562 | 0.9862 | 2.3099 | 1.8330 | 1.0544 | 1.7308 | 0.9019 | 1.2117 | 22.7858 | 18.7662 | 19.7963 | 17.1717 |
| TMOD3_S25 | TMOD3 | Q9NYL9 | | 25 | ELLGNLSETELKQ | 6.9455 | 6.6222 | 7.0307 | 6.0780 | 6.8882 | 6.4414 | 6.8118 | 7.3028 | 5.6820 | 6.4388 | 6.5946 | 6.3309 | 5.3364 | 5.3922 | 4.9186 | 5.1858 |
| TMPO_S156 | TMPO | P42167 | | 306 | GNFKHASPILPIT | 6.4605 | 7.1664 | 6.9949 | 7.2975 | 6.1977 | 6.5325 | 6.6833 | 6.7579 | 6.9805 | 7.0626 | 6.5765 | 6.9196 | 4.4686 | 4.9130 | 4.1266 | 4.8620 |
| TMPO_S168 | TMPO | P42167 | | 385 | DFRMEESFSSKYV | 8.8580 | 8.6101 | 6.9857 | 9.2156 | 7.5190 | 5.7739 | 7.4110 | 5.9014 | 6.8186 | 5.7068 | 6.6212 | 5.8745 | 3.4450 | 4.5105 | 3.0401 | 3.7085 |
| TMPO_S180;S184 | TMPO | P42166 | | 74 | DEEREPTPVLGSG | 5.1229 | 5.1884 | 4.6881 | 5.4176 | 5.5660 | 5.0841 | 5.1844 | 5.0738 | 6.3284 | 6.3142 | 5.6973 | 5.4875 | 9.0252 | 8.7100 | 8.8671 | 8.2448 |
| TMPO_S184 | TMPO | P42166 | | 168 | LPTISSSAENTRQ | 6.0093 | 6.2897 | 5.9318 | 6.4138 | 5.9954 | 6.0016 | 6.2522 | 5.8013 | 6.3589 | 6.6034 | 6.3708 | 6.2783 | 6.2929 | 6.7976 | 6.3030 | 6.2999 |
| TMPO_S222 | TMPO | P42166 | | 424 | QETEFLSPPRKVP | 7.0559 | 7.9635 | 7.1763 | 7.7505 | 6.0327 | 7.1410 | 7.3320 | 6.9106 | 7.6158 | 7.6403 | 7.9485 | 8.0030 | 2.4164 | 3.4205 | 2.4545 | 3.1384 |
| TMPO_S306 | TMPO | P42166 | | 67 | GPPDFSSDEEREP | 6.2533 | 7.6112 | 7.1701 | 7.6480 | 6.5311 | 7.5129 | 6.8702 | 7.2396 | 6.2629 | 6.3262 | 6.5740 | 6.8137 | 3.4200 | 4.5066 | 4.3222 | 4.9380 |
| TMPO_S310 | TMPO | P42167 | | 355 | FPYEASTPTGISA | 6.3237 | 7.4227 | 6.6963 | 7.1937 | 6.5816 | 6.8354 | 6.9827 | 6.8514 | 7.2527 | 7.0218 | 7.2288 | 6.9651 | 3.8227 | 4.6716 | 3.5924 | 4.5576 |
| TMPO_S351 | TMPO | P42166 | | 66 | KGPPDFSSDEERE | 6.6989 | 7.3478 | 7.1930 | 7.5109 | 6.8111 | 7.1380 | 7.0686 | 6.7475 | 6.0864 | 6.2214 | 6.5703 | 6.5577 | 4.1612 | 4.6755 | 4.2642 | 4.9475 |
| TMPO_S378 | TMPO | P42166 | | 351 | SHISDQSPLSSKR | 7.5158 | 9.0634 | 6.2856 | 8.0723 | 6.9587 | 7.8253 | 6.5364 | 8.4143 | 7.0626 | 6.6098 | 7.1542 | 6.6125 | 2.5135 | 3.6665 | 2.7119 | 2.9973 |
| TMPO_S385 | TMPO | P42166 | | 442 | SVEERDSGSFVAF | 6.1132 | 6.1202 | 6.4389 | 6.8526 | 6.6048 | 6.3227 | 6.8150 | 6.5237 | 6.4577 | 6.7252 | 7.0228 | 6.9533 | 5.0999 | 5.7793 | 4.9469 | 5.2238 |
| TMPO_S424 | TMPO | P42166 | | 156 | REQGTESRSSTPL | 7.3102 | 7.2388 | 5.7811 | 6.8977 | 5.9556 | 5.1223 | 5.3629 | 4.1752 | 5.5644 | 5.3010 | 5.3052 | 5.5244 | 8.3480 | 8.1227 | 7.1799 | 6.8107 |
| TMPO_S442 | TMPO | P42167 | | 378 | GRPLELSDFRMEE | 6.9444 | 7.0112 | 6.6244 | 7.8195 | 6.3152 | 6.7903 | 7.7145 | 6.9158 | 7.0267 | 7.0019 | 7.3459 | 7.2129 | 3.3009 | 4.4143 | 3.1651 | 4.3971 |
| TMPO_S59 | TMPO | P42167 | | 184 | NDSDRYSDNEEDS | 5.7372 | 5.6842 | 5.4564 | 6.7872 | 5.9138 | 5.7518 | 6.1720 | 5.7680 | 6.1831 | 6.2163 | 6.3769 | 6.4098 | 7.0619 | 6.9893 | 6.6256 | 6.8663 |
| TMPO_S66 | TMPO | P42167 | | 222 | RVEHNQSYSQAGI | 7.7118 | 7.9319 | 6.8840 | 8.2709 | 6.4309 | 6.5852 | 7.7682 | 6.5584 | 7.3321 | 6.2082 | 6.7919 | 6.7449 | 3.1347 | 4.7851 | 2.7966 | 4.0650 |
| TMPO_S66;S67 | TMPO | P42167 | | 279 | PVISESTPIAETI | 6.5074 | 6.4240 | 6.1123 | 6.3635 | 6.8659 | 6.7126 | 7.3590 | 5.8705 | 7.0524 | 7.1989 | 6.6845 | 7.5514 | 4.8682 | 4.9202 | 5.0045 | 4.5046 |
| TMPO_S66;S67;T74 | TMPO | P42166 | | 310 | LVSTAASPSLIKE | 5.3615 | 6.0319 | 5.7550 | 6.8388 | 7.3248 | 6.9127 | 5.5030 | 6.9296 | 6.8937 | 7.1583 | 7.1875 | 7.0707 | 4.7967 | 5.8996 | 5.4340 | 4.9020 |
| TMPO_S67 | TMPO | P42166 | | 160 | TESRSSTPLPTIS | 6.3288 | 6.1401 | 6.5359 | 6.6488 | 7.0578 | 6.6411 | 6.2970 | 5.4337 | 6.0331 | 6.3346 | 6.8280 | 6.1815 | 6.4757 | 6.6370 | 5.2562 | 5.1707 |
| TMPO_S67;T74 | TMPO | P42166 | | 59 | LPAGTNSKGPPDF | 6.4779 | 6.1439 | 6.6252 | 6.9195 | 5.9583 | 6.0396 | 6.0984 | 6.5063 | 6.6104 | 6.5487 | 7.0978 | 6.7337 | 6.2337 | 5.7274 | 4.9188 | 5.3604 |
| TMPO_T160 | TMPO | P42166 | 66;67 | | NA | 7.3840 | 8.0068 | 7.7151 | 8.1124 | 6.3502 | 6.3830 | 6.7783 | 6.6946 | 6.7193 | 6.8481 | 6.9127 | 7.0434 | 3.5693 | 4.2972 | 3.3970 | 3.7887 |
| TMPO_T160;S168 | TMPO | P42166 | 66;67;74 | | NA | 7.2489 | 8.0798 | 7.2278 | 7.9649 | 6.2166 | 6.1885 | 6.3183 | 6.4166 | 7.0559 | 7.3364 | 7.3491 | 7.2304 | 3.4875 | 4.3030 | 3.4840 | 4.0923 |
| TMPO_T279 | TMPO | P42167 | 180;184 | | NA | 6.9533 | 6.5782 | 6.5316 | 6.9816 | 5.8077 | 6.2861 | 7.0443 | 6.0128 | 5.9487 | 5.9290 | 6.1401 | 6.3688 | 5.8449 | 5.3245 | 5.7445 | 6.5040 |
| TMPO_T355 | TMPO | P42166 | 67;74 | | NA | 6.3381 | 7.6154 | 6.5645 | 7.3021 | 6.5448 | 6.8851 | 6.6746 | 6.8931 | 6.0949 | 6.8585 | 7.0476 | 6.9742 | 3.8215 | 5.1448 | 3.9832 | 5.2575 |
| TMPO_T74 | TMPO | P42166 | 160;168 | | NA | 5.4605 | 6.7740 | 6.9025 | 6.5376 | 5.4975 | 6.6752 | 5.8434 | 5.6202 | 6.2446 | 6.5359 | 6.0875 | 6.1891 | 6.4718 | 6.6091 | 6.1935 | 6.3578 |
| TMUB1_S127 | TMUB1 | Q9BVT8 | | 127 | PHDTIGSLKRTQF | 6.3491 | 6.6775 | 5.5926 | 6.1440 | 6.2650 | 5.6276 | 6.5931 | 5.9537 | 5.3790 | 5.2969 | 6.0809 | 6.0624 | 5.0600 | 8.7520 | 6.2928 | 7.8735 |
| TMX1_S228 | TMX1 | Q9H3N1 | | 247 | ADEEDVSEEEAES | 6.1526 | 6.3803 | 6.0956 | 6.5131 | 6.3429 | 5.7904 | 6.3326 | 5.7667 | 6.0289 | 6.0989 | 5.8694 | 5.9476 | 6.7352 | 6.6495 | 6.6470 | 6.6495 |
| TMX1_S247 | TMX1 | Q9H3N1 | | 270 | NAIRQRSLGPSLA | 6.0155 | 6.2109 | 6.1054 | 6.4396 | 6.3991 | 5.7217 | 6.0441 | 6.0687 | 7.4706 | 7.7106 | 7.6729 | 7.3581 | 4.8464 | 5.9090 | 5.0673 | 4.9601 |
| TMX1_S270 | TMX1 | Q9H3N1 | | 280 | SLATDKSxxxxxx | 6.8072 | 7.7667 | 6.6499 | 6.6674 | 5.8835 | 6.3967 | 6.8780 | 6.1919 | 6.5674 | 6.9773 | 7.0536 | 7.3492 | 4.0831 | 5.6775 | 4.3504 | 4.7002 |
| TMX1_S280 | TMX1 | Q9H3N1 | | 228 | KKLLSESAQPLKK | 4.1146 | 4.7000 | 4.0617 | 4.8092 | 3.7942 | 4.1316 | 4.2219 | 4.2095 | 4.1285 | 4.3623 | 4.3317 | 4.1774 | 10.9900 | 12.2089 | 13.1463 | 12.6122 |
| TMX2_S211 | TMX2 | Q9Y320 | | 288 | STPTTVSDGENKK | 6.8994 | 6.8011 | 7.0545 | 7.3341 | 6.4429 | 6.2894 | 6.8648 | 6.5186 | 6.4952 | 6.4693 | 6.4298 | 6.6333 | 4.7398 | 5.1853 | 4.8757 | 4.9666 |
| TMX2_S243 | TMX2 | Q9Y320 | | 243 | KKGRAVSWTFSEE | 6.5263 | 6.3923 | 5.9685 | 6.8368 | 6.1109 | 6.1960 | 6.4547 | 5.6388 | 5.8146 | 6.1279 | 5.7345 | 6.3463 | 6.3044 | 6.3406 | 6.0989 | 7.1083 |
| TMX2_S288 | TMX2 | Q9Y320 | | 211 | RYKVSTSPLTKQL | 7.1570 | 7.1148 | 6.8811 | 7.3940 | 6.4969 | 6.3344 | 6.8214 | 6.3205 | 6.4744 | 6.4486 | 5.9956 | 5.7512 | 5.0610 | 5.2670 | 5.3862 | 5.0959 |
| TNFRSF10A_S424 | TNFRSF10A | O00220 | | 424 | KTGRNASIHTLLD | 4.8282 | 4.7015 | 4.9096 | 5.1445 | 5.5422 | 5.1944 | 5.8585 | 5.1863 | 5.1950 | 5.0515 | 4.6934 | 5.0949 | 9.2917 | 9.5836 | 9.7168 | 10.0082 |
| TNFRSF1A_S274 | TNFRSF1A | P19438 | | 274 | APNPSFSPTPGFT | 6.0631 | 5.5946 | 7.0177 | 6.6723 | 6.0703 | 6.4341 | 6.1998 | 5.4619 | 6.3501 | 6.3981 | 6.8274 | 6.1375 | 5.8049 | 6.5132 | 5.7010 | 6.7539 |
| TNIK_S548 | TNIK | Q9UKE5 | | 680 | QRTTSISPALARK | 6.0966 | 6.3168 | 6.0037 | 5.4300 | 6.2410 | 6.2790 | 5.8765 | 6.0202 | 6.3800 | 6.2821 | 6.2505 | 6.0903 | 6.6096 | 6.9363 | 6.6714 | 6.5159 |
| TNIK_S560 | TNIK | Q9UKE5 | | 581 | GVQPARTPPMLRP | 5.5348 | 6.0298 | 6.2843 | 6.0842 | 6.1922 | 6.5126 | 5.9478 | 6.5974 | 6.2826 | 6.6785 | 6.2311 | 6.1565 | 6.2192 | 6.5630 | 6.2846 | 6.4013 |
| TNIK_S570 | TNIK | Q9UKE5 | | 769 | NSKSEGSPVLPHE | 5.9214 | 5.9947 | 6.1630 | 5.6492 | 6.1481 | 6.0848 | 5.9402 | 6.0836 | 6.4408 | 6.5629 | 6.1903 | 6.2226 | 6.5621 | 6.9787 | 6.5615 | 6.4960 |
| TNIK_S680 | TNIK | Q9UKE5 | | 707 | SQPIRASNPDLRR | 5.9159 | 6.0686 | 6.1876 | 5.4921 | 5.7718 | 6.3698 | 5.9995 | 5.9078 | 6.1298 | 6.1918 | 6.2776 | 6.1673 | 6.6778 | 7.2884 | 6.5346 | 7.0198 |
| TNIK_S688 | TNIK | Q9UKE5 | | 548 | RLNRQSSPAMPHK | 5.5370 | 6.1539 | 6.5371 | 5.8895 | 5.4981 | 6.4416 | 5.9279 | 6.2851 | 6.2502 | 6.4177 | 6.5699 | 6.7116 | 5.9182 | 6.9822 | 6.4526 | 6.4273 |
| TNIK_S707 | TNIK | Q9UKE5 | | 560 | KVANRISDPNLPP | 6.2352 | 5.9343 | 6.6755 | 6.5133 | 5.5128 | 5.7682 | 6.3515 | 6.1392 | 6.5124 | 5.8282 | 6.0968 | 6.4592 | 6.7628 | 6.4358 | 6.2918 | 6.4829 |
| TNIK_S769 | TNIK | Q9UKE5 | | 187 | TVGRRNTFIGTPY | 6.6098 | 6.6989 | 6.7169 | 6.2010 | 6.3141 | 5.9049 | 6.4727 | 5.6475 | 6.0755 | 6.3337 | 6.3502 | 5.7167 | 6.6388 | 6.5591 | 6.3791 | 5.3811 |
| TNIK_T1036 | TNIK | Q9UKE5 | | 1036 | IRPHSDTPEIRKY | 5.3107 | 6.2418 | 6.5886 | 7.1176 | 6.2851 | 7.0284 | 6.1884 | 6.6822 | 5.9842 | 5.9622 | 7.4028 | 6.7400 | 5.2278 | 6.2826 | 5.6374 | 5.3201 |
| TNIK_T187 | TNIK | Q9UKE5 | | 688 | ALARKNSPGNGSA | 6.6648 | 7.2533 | 7.0235 | 6.2136 | 5.7986 | 6.6763 | 6.5804 | 5.8563 | 7.1699 | 7.2863 | 7.0476 | 7.1402 | 4.5596 | 4.9913 | 4.7435 | 4.9949 |
| TNIK_T581 | TNIK | Q9UKE5 | | 570 | LPPRSESFSISGV | 6.9927 | 4.8738 | 7.2657 | 5.8918 | 5.1700 | 5.7846 | 6.0333 | 6.1051 | 7.3162 | 5.9606 | 6.9422 | 5.6846 | 6.3397 | 7.3658 | 5.9353 | 6.3386 |
| TNIP1_S403 | TNIP1 | Q15025 | | 403 | YLQDQLSPLTRQR | 6.5991 | 6.5351 | 6.1525 | 6.2874 | 6.1469 | 6.5292 | 6.4144 | 6.5289 | 7.1603 | 6.8564 | 6.4541 | 6.5716 | 5.5897 | 6.0619 | 5.1001 | 5.0122 |
| TNIP1_S77 | TNIP1 | Q15025 | | 77 | ELLPPPSPSLGSF | 5.7519 | 6.4886 | 6.5399 | 6.6893 | 6.1565 | 7.0310 | 6.4569 | 6.2763 | 6.5953 | 6.3740 | 6.2626 | 6.6434 | 5.9785 | 5.7725 | 5.9067 | 5.0765 |
| TNIP1_T428;S435 | TNIP1 | Q15025 | 428;435 | | NA | 6.0852 | 6.1255 | 6.5066 | 6.2216 | 6.5080 | 5.9572 | 6.0546 | 6.3130 | 6.1212 | 6.1509 | 6.7360 | 6.8184 | 5.8076 | 6.6555 | 5.7964 | 6.1422 |
| TNK2_S102 | TNK2 | Q07912 | | 827 | FASDPKYATPQVI | 6.1894 | 6.9988 | 7.8799 | 8.0378 | 5.8588 | 6.4272 | 6.3049 | 6.6255 | 6.3188 | 6.8532 | 6.9221 | 6.8216 | 4.3341 | 5.0719 | 4.5932 | 4.7628 |
| TNK2_S810 | TNK2 | Q07912 | | 810 | PPRLSSSPGKTMP | 6.8582 | 6.9708 | 7.1670 | 6.7733 | 6.1954 | 6.5553 | 6.5848 | 6.8277 | 6.7552 | 6.6233 | 6.7318 | 6.5694 | 4.7330 | 5.0473 | 4.8172 | 4.7904 |
| TNK2_Y518 | TNK2 | Q07912 | | 102 | STFRKTSPAPGGP | 6.0496 | 6.2666 | 6.5236 | 6.1076 | 6.0485 | 6.6183 | 6.4359 | 6.0386 | 6.6829 | 6.5236 | 6.5823 | 6.1016 | 6.2860 | 6.1351 | 5.8557 | 5.7443 |
| TNK2_Y827 | TNK2 | Q07912 | | 518 | GVKKPTYDPVSED | 6.0041 | 6.6798 | 7.7096 | 7.9072 | 5.4968 | 6.7022 | 6.1635 | 6.6851 | 6.3848 | 7.0310 | 7.1003 | 7.3232 | 4.1921 | 5.0547 | 4.6122 | 4.9534 |
| TNKS1BP1_S1024 | TNKS1BP1 | Q9C0C2 | | 1545 | QTSRRPSQGPPAR | 5.6544 | 5.1168 | 5.7752 | 5.2089 | 6.5656 | 5.4238 | 4.9615 | 6.2471 | 7.0746 | 7.5317 | 7.1183 | 7.0647 | 6.9526 | 7.2085 | 6.1922 | 5.9040 |
| TNKS1BP1_S1029 | TNKS1BP1 | Q9C0C2 | | 872 | EFGKRDSLGTYSS | 6.3212 | 6.4956 | 6.8077 | 6.1991 | 6.1017 | 6.2502 | 5.8520 | 6.4652 | 6.5655 | 6.1515 | 6.7947 | 6.6408 | 6.0850 | 6.0037 | 5.7742 | 5.4918 |
| TNKS1BP1_S1103 | TNKS1BP1 | Q9C0C2 | | 983 | FGTRPLSSGFSPE | 5.5812 | 5.5973 | 6.5871 | 5.9167 | 5.9649 | 6.1583 | 5.8972 | 6.3466 | 6.3352 | 6.1048 | 6.2647 | 6.3320 | 7.1673 | 6.7033 | 6.6815 | 6.3618 |
| TNKS1BP1_S1138 | TNKS1BP1 | Q9C0C2 | | 1652 | NLFPGLSPSALKA | 6.9958 | 7.1140 | 6.9516 | 7.1145 | 6.6453 | 7.2596 | 6.9473 | 7.1298 | 7.3875 | 7.2356 | 7.2661 | 7.3096 | 3.5946 | 4.0884 | 3.3304 | 3.6300 |
| TNKS1BP1_S1158 | TNKS1BP1 | Q9C0C2 | | 178 | RKEVLASPDRLWG | 6.3213 | 6.6797 | 6.7466 | 6.2227 | 6.6417 | 7.0291 | 6.5335 | 7.0458 | 6.9196 | 6.7607 | 6.9699 | 7.0535 | 4.7942 | 5.0127 | 4.4575 | 4.8117 |
| TNKS1BP1_S1253 | TNKS1BP1 | Q9C0C2 | | 1138 | VSGAGFSPSSKME | 6.9273 | 6.6115 | 6.6944 | 6.5594 | 7.0445 | 6.6564 | 6.3329 | 7.0273 | 7.1521 | 7.0549 | 6.9880 | 6.6540 | 4.6436 | 4.9315 | 4.4178 | 4.3045 |
| TNKS1BP1_S1297 | TNKS1BP1 | Q9C0C2 | | 672 | QDLCRASPEPPGP | 5.8960 | 5.8561 | 6.1881 | 5.9171 | 5.9580 | 6.1682 | 5.8038 | 6.3038 | 6.7317 | 6.5018 | 6.6399 | 6.4266 | 6.5109 | 6.6532 | 6.4488 | 5.9958 |
| TNKS1BP1_S1385 | TNKS1BP1 | Q9C0C2 | | 1554 | PPARSPSQDFSFI | 5.9266 | 5.3715 | 6.5190 | 5.5508 | 6.2457 | 5.8318 | 5.3510 | 6.3153 | 6.3914 | 6.0902 | 6.1311 | 6.7032 | 7.0215 | 7.1550 | 6.9318 | 6.4640 |
| TNKS1BP1_S1473 | TNKS1BP1 | Q9C0C2 | | 221 | TLFRGWSQEGPVK | 5.0684 | 5.4736 | 6.3504 | 5.2388 | 5.2855 | 5.8002 | 5.0162 | 5.9551 | 5.5214 | 6.1961 | 5.8101 | 5.5787 | 8.5037 | 8.1537 | 8.4939 | 7.5542 |
| TNKS1BP1_S1473;S1503 | TNKS1BP1 | Q9C0C2 | | 836 | QLGTQRSQEADVQ | 4.8823 | 4.8304 | 4.9930 | 4.7116 | 6.0128 | 5.9246 | 5.3547 | 5.5426 | 4.9705 | 5.0498 | 5.0196 | 5.0448 | 10.3889 | 9.1343 | 9.5796 | 8.5604 |
| TNKS1BP1_S1533 | TNKS1BP1 | Q9C0C2 | | 691 | LDDLLASPPPSGG | 6.2966 | 6.6739 | 7.0088 | 6.9568 | 6.1801 | 6.3903 | 6.3687 | 6.7844 | 6.6304 | 6.5036 | 7.1139 | 6.9332 | 5.1717 | 5.2021 | 4.8954 | 4.8901 |
| TNKS1BP1_S1545 | TNKS1BP1 | Q9C0C2 | | 893 | EFGKRDSLGAYAS | 6.3782 | 6.3859 | 6.2045 | 5.8067 | 6.6801 | 6.3199 | 5.8247 | 6.1093 | 6.8837 | 6.6468 | 6.5412 | 6.3007 | 6.4341 | 6.1056 | 5.8937 | 5.4847 |
| TNKS1BP1_S1545;S1552 | TNKS1BP1 | Q9C0C2 | | 1385 | RHNGSLSPGLEAR | 6.1268 | 6.6998 | 6.1430 | 6.2710 | 5.7383 | 6.2676 | 6.0482 | 6.4970 | 6.1698 | 6.5307 | 6.5061 | 6.7720 | 6.1683 | 6.1488 | 6.1762 | 5.7364 |
| TNKS1BP1_S1552 | TNKS1BP1 | Q9C0C2 | | 1297 | APGAVCSPGESKE | 6.3871 | 6.5916 | 6.9231 | 7.0101 | 6.1251 | 6.9145 | 6.8596 | 6.3116 | 7.2921 | 7.3258 | 7.2253 | 7.3125 | 4.3356 | 4.7633 | 3.9313 | 4.6914 |
| TNKS1BP1_S1554 | TNKS1BP1 | Q9C0C2 | | 882 | YSSRDVSLGDWEF | 6.4981 | 7.0324 | 6.9673 | 7.4577 | 6.4509 | 7.1916 | 6.3801 | 6.8939 | 6.7312 | 6.6190 | 7.2184 | 7.0647 | 4.3816 | 4.9422 | 3.8042 | 4.3666 |
| TNKS1BP1_S1652 | TNKS1BP1 | Q9C0C2 | | 1282 | VGQADWTPDLGLR | 7.1396 | 7.4011 | 6.8053 | 6.7766 | 6.2401 | 7.0758 | 6.6951 | 6.6444 | 6.9704 | 7.3209 | 7.3304 | 7.0673 | 3.9732 | 4.6824 | 3.6929 | 4.1845 |
| TNKS1BP1_S1666 | TNKS1BP1 | Q9C0C2 | | 1103 | QREAAFSPGQQDW | 6.8364 | 6.8953 | 6.4618 | 6.8456 | 6.6443 | 6.9605 | 6.9492 | 7.0314 | 7.4385 | 6.5851 | 7.2070 | 7.0950 | 4.2283 | 4.6560 | 3.8311 | 4.3346 |
| TNKS1BP1_S1715 | TNKS1BP1 | Q9C0C2 | | 1024 | RPGERGSGGLFSP | 6.2158 | 6.1761 | 6.5221 | 6.2001 | 6.3848 | 6.4239 | 5.7532 | 7.0404 | 6.0897 | 5.9302 | 6.3432 | 6.1444 | 6.1642 | 6.4840 | 6.5138 | 5.6140 |
| TNKS1BP1_S178 | TNKS1BP1 | Q9C0C2 | | 601 | PLAGQESPLPLAT | 5.7244 | 6.2154 | 6.5194 | 6.8020 | 5.6598 | 6.0886 | 5.9469 | 6.5156 | 6.6433 | 6.6817 | 6.4304 | 6.5822 | 5.9456 | 6.2266 | 6.0083 | 6.0099 |
| TNKS1BP1_S221 | TNKS1BP1 | Q9C0C2 | | 1158 | GKTTAGSVDWTDQ | 6.0881 | 6.4053 | 6.8403 | 6.2540 | 5.6818 | 6.3460 | 6.2639 | 6.4036 | 6.0587 | 6.7933 | 6.5319 | 6.4316 | 5.8885 | 6.2730 | 5.8224 | 5.9176 |
| TNKS1BP1_S287;S297;S3 | TNKS1BP1 | Q9C0C2 | | 1552 | QGPPARSPSQDFS | 6.2855 | 6.6454 | 6.5639 | 6.7383 | 6.1602 | 6.3394 | 6.3540 | 6.4352 | 6.5517 | 6.5879 | 6.6442 | 7.3271 | 5.0395 | 5.5264 | 5.2808 | 5.5206 |
| TNKS1BP1_S297;S301 | TNKS1BP1 | Q9C0C2 | | 435 | SEGVLQSPSQDQE | 6.8312 | 7.9030 | 6.9526 | 7.1455 | 6.4222 | 7.6951 | 6.9237 | 6.9863 | 6.2962 | 6.2667 | 6.4112 | 6.5852 | 4.2886 | 4.5121 | 4.2731 | 4.5072 |
| TNKS1BP1_S32 | TNKS1BP1 | Q9C0C2 | | 131 | EEPPPLTPPARCA | 6.5018 | 6.7777 | 6.1481 | 6.0543 | 6.3672 | 6.6136 | 5.9318 | 6.9033 | 6.6480 | 6.1798 | 6.5064 | 6.5263 | 5.7398 | 5.5980 | 5.5846 | 5.9194 |
| TNKS1BP1_S353;S369 | TNKS1BP1 | Q9C0C2 | | 1533 | SSAARWSDQGPAQ | 5.5450 | 6.4008 | 7.9439 | 5.9109 | 6.3192 | 6.3134 | 6.3309 | 6.6052 | 6.0641 | 5.0895 | 7.0299 | 6.8417 | 5.9796 | 5.4419 | 6.3006 | 5.8835 |
| TNKS1BP1_S369 | TNKS1BP1 | Q9C0C2 | | 987 | PLSSGFSPEEAQQ | 6.2390 | 6.4372 | 6.5061 | 6.3256 | 6.4944 | 6.7073 | 6.3216 | 6.6168 | 6.0342 | 6.5817 | 7.2370 | 6.6407 | 5.7247 | 5.6309 | 5.1218 | 5.3809 |
| TNKS1BP1_S429 | TNKS1BP1 | Q9C0C2 | | 712 | ELKDTQSPSTCSE | 6.2031 | 6.4946 | 6.9730 | 6.9942 | 6.3014 | 6.8282 | 6.0006 | 7.0334 | 7.1814 | 7.0420 | 7.0375 | 6.4897 | 4.5981 | 5.4784 | 4.4296 | 4.9148 |
| TNKS1BP1_S429;S435 | TNKS1BP1 | Q9C0C2 | | 1715 | SGSEGSSPNWLQA | 6.7157 | 7.3632 | 6.8162 | 6.9008 | 6.6397 | 7.2191 | 6.8434 | 6.4598 | 7.4199 | 7.2161 | 7.1096 | 7.2669 | 3.2544 | 4.5086 | 3.5843 | 4.6822 |
| TNKS1BP1_S435 | TNKS1BP1 | Q9C0C2 | | 1666 | LRPRNRSAEEGEL | 5.1582 | 5.6769 | 6.5653 | 6.2212 | 5.8248 | 6.3135 | 5.5659 | 6.0682 | 6.3674 | 6.3714 | 6.5612 | 6.6622 | 6.6370 | 6.8872 | 6.8382 | 6.2813 |
| TNKS1BP1_S494 | TNKS1BP1 | Q9C0C2 | | 429 | LVQRRFSEGVLQS | 5.4438 | 5.2142 | 5.8102 | 5.5283 | 6.4099 | 5.6872 | 5.0618 | 5.8479 | 6.7010 | 7.7856 | 6.2823 | 7.0665 | 7.4963 | 7.8241 | 5.7083 | 6.1326 |
| TNKS1BP1_S601 | TNKS1BP1 | Q9C0C2 | | 1253 | HSQARESGVGQTD | 6.3700 | 6.2763 | 6.5517 | 6.5595 | 6.2833 | 5.8456 | 5.9300 | 6.5619 | 6.8984 | 6.1771 | 5.7281 | 6.5012 | 6.1926 | 6.3623 | 6.1537 | 5.6083 |
| TNKS1BP1_S672 | TNKS1BP1 | Q9C0C2 | | 1473 | AVARRESAASGLG | 5.4738 | 5.3356 | 6.0621 | 6.5966 | 5.9893 | 5.5831 | 5.9073 | 5.8162 | 6.2223 | 7.0992 | 6.9846 | 7.2427 | 6.2474 | 6.5959 | 6.7219 | 6.1221 |
| TNKS1BP1_S691 | TNKS1BP1 | Q9C0C2 | | 494 | GVWRLDSPPPSPI | 5.5008 | 5.8146 | 6.4948 | 6.2125 | 5.0503 | 5.5148 | 5.9023 | 5.2046 | 7.1784 | 5.9776 | 6.0618 | 8.4195 | 7.0960 | 6.9768 | 5.8917 | 6.7034 |
| TNKS1BP1_S691;S695 | TNKS1BP1 | Q9C0C2 | | 851 | EFRKRDSQGTYSS | 6.7872 | 6.1585 | 6.5729 | 5.8584 | 5.9534 | 6.3852 | 5.4493 | 6.4179 | 6.3717 | 6.0432 | 6.5470 | 6.8507 | 6.4760 | 6.1752 | 6.2774 | 5.6760 |
| TNKS1BP1_S712 | TNKS1BP1 | Q9C0C2 | | 1029 | GSGGLFSPSTAHV | 6.7078 | 7.6806 | 6.8428 | 6.3497 | 5.7820 | 7.1718 | 7.3296 | 7.0729 | 6.7541 | 7.2966 | 7.8109 | 6.9645 | 3.3252 | 4.4386 | 4.2651 | 4.2078 |
| TNKS1BP1_S836 | TNKS1BP1 | Q9C0C2 | | 369 | EAPRPSSPPPEVL | 5.5089 | 6.3163 | 7.0306 | 6.5759 | 6.2065 | 6.4117 | 6.4661 | 4.9277 | 6.4006 | 7.1803 | 6.2531 | 6.3496 | 6.0158 | 6.2520 | 5.9082 | 6.1966 |
| TNKS1BP1_S851 | TNKS1BP1 | A0A2R8Y5C4 | | 32 | EAGRLRSPEPRAL | 6.6346 | 6.9638 | 6.1783 | 6.5360 | 6.4293 | 6.7696 | 6.2451 | 6.3954 | 6.9217 | 7.0726 | 6.8272 | 6.9940 | 5.2925 | 4.9839 | 4.7763 | 4.9799 |
| TNKS1BP1_S872 | TNKS1BP1 | Q9C0C2 | 353;369 | | NA | 6.1987 | 6.2766 | 6.8469 | 6.2323 | 6.2058 | 6.0866 | 5.5325 | 6.2825 | 6.0794 | 6.7069 | 6.6450 | 6.7218 | 6.0917 | 6.4391 | 5.6251 | 6.0291 |
| TNKS1BP1_S882 | TNKS1BP1 | Q9C0C2 | 287;297;301 | | NA | 6.0899 | 5.7486 | 6.9846 | 6.6434 | 6.2470 | 6.5477 | 5.7829 | 5.9171 | 6.2200 | 6.4837 | 7.1327 | 7.1864 | 6.0217 | 5.5077 | 5.2881 | 6.1985 |
| TNKS1BP1_S893 | TNKS1BP1 | Q9C0C2 | 1473;1503 | | NA | 5.2748 | 5.5265 | 6.0866 | 6.1960 | 7.2816 | 5.4709 | 5.5445 | 6.4092 | 7.2511 | 6.5055 | 6.1764 | 6.0589 | 6.7013 | 7.4122 | 6.4782 | 5.6264 |
| TNKS1BP1_S983 | TNKS1BP1 | Q9C0C2 | 429;435 | | NA | 5.9413 | 5.0812 | 6.8926 | 5.1767 | 6.6224 | 5.9247 | 5.7302 | 5.6508 | 8.5283 | 8.2924 | 9.2629 | 8.6550 | 4.2229 | 5.5362 | 3.6670 | 4.8155 |
| TNKS1BP1_S987 | TNKS1BP1 | Q9C0C2 | 1545;1552 | | NA | 6.9650 | 6.2327 | 6.5288 | 6.6422 | 6.3780 | 6.5261 | 5.6806 | 6.1005 | 6.9805 | 6.1761 | 8.1373 | 6.6950 | 5.2187 | 6.2346 | 4.8112 | 4.6928 |
| TNKS1BP1_T1282 | TNKS1BP1 | Q9C0C2 | 691;695 | | NA | 6.5643 | 6.9161 | 6.3999 | 6.2161 | 6.6067 | 7.3866 | 6.3300 | 6.1905 | 6.6157 | 6.6168 | 6.2763 | 7.1782 | 6.1673 | 4.9427 | 4.6132 | 4.9797 |
| TNKS1BP1_T131 | TNKS1BP1 | Q9C0C2 | 297;301 | | NA | 5.2348 | 5.5851 | 7.2654 | 6.4143 | 6.7955 | 5.5351 | 5.8149 | 5.8116 | 6.8591 | 7.2643 | 7.6789 | 7.2070 | 5.7774 | 5.4848 | 5.3287 | 5.9431 |
| TNRC18_S1127 | TNRC18 | O15417 | | 1127 | PERLALSPEDKPI | 6.3568 | 7.1094 | 6.2890 | 6.9769 | 6.0220 | 5.8317 | 6.1586 | 6.1499 | 6.6114 | 6.7986 | 6.2513 | 6.5101 | 5.5239 | 5.6857 | 5.7949 | 5.9299 |
| TNRC18_S1136 | TNRC18 | O15417 | | 1857 | ARERALSPGLEES | 6.3796 | 6.7908 | 6.3815 | 6.7476 | 6.1280 | 5.9857 | 6.1128 | 6.4422 | 6.4701 | 6.2830 | 6.5403 | 6.0371 | 5.5198 | 6.3360 | 5.8108 | 6.0349 |
| TNRC18_S1232 | TNRC18 | O15417 | | 1957 | RGPDPSSPDKAKL | 6.8329 | 7.0675 | 6.3815 | 7.2050 | 6.0472 | 5.7421 | 6.5889 | 5.9755 | 6.6839 | 7.0381 | 6.6175 | 6.6361 | 4.9879 | 5.4856 | 5.0774 | 5.6330 |
| TNRC18_S1359 | TNRC18 | O15417 | | 999 | PPSPRASPVAALK | 5.0062 | 6.3182 | 6.8229 | 6.3142 | 5.5268 | 6.2599 | 6.1061 | 7.0439 | 6.9219 | 8.6353 | 8.8873 | 8.0372 | 4.2367 | 5.4140 | 3.8542 | 4.6152 |
| TNRC18_S1857 | TNRC18 | O15417 | | 2033 | AKGGPLSPRKDAG | 7.2792 | 7.2758 | 6.4442 | 7.3898 | 6.1254 | 6.0503 | 6.3973 | 6.2179 | 6.5661 | 6.4145 | 6.5080 | 6.5708 | 4.9758 | 5.0654 | 5.1334 | 5.5861 |
| TNRC18_S1957 | TNRC18 | O15417 | | 263 | RLAERLSPFLAES | 6.5658 | 6.4218 | 6.3856 | 6.3809 | 6.5408 | 6.4093 | 6.1714 | 6.2186 | 6.3266 | 7.0288 | 6.7601 | 6.4258 | 5.4831 | 6.0950 | 5.5845 | 5.2019 |
| TNRC18_S2033 | TNRC18 | O15417 | | 1232 | PEPRVDSPGRTEP | 6.8687 | 6.8825 | 6.8364 | 7.1817 | 6.3377 | 6.4273 | 6.8292 | 5.5372 | 6.3001 | 6.3334 | 6.7611 | 6.8017 | 5.2824 | 5.4041 | 4.6637 | 5.5529 |
| TNRC18_S2471 | TNRC18 | O15417 | | 2471 | DHEGVTSPKSKKA | 6.9139 | 7.3013 | 6.7604 | 7.1748 | 6.8420 | 6.4516 | 6.9881 | 6.0843 | 7.4852 | 6.9294 | 6.8467 | 6.8234 | 4.2266 | 4.6818 | 4.1175 | 4.3729 |
| TNRC18_S263 | TNRC18 | O15417 | | 2771 | KRQRLPSVENRPK | 4.5004 | 4.2863 | 4.6557 | 5.0829 | 5.7519 | 5.3272 | 5.8663 | 6.2620 | 4.2854 | 4.9828 | 4.1349 | 4.8881 | 9.7398 | 9.5426 | 10.8890 | 9.8045 |
| TNRC18_S2716 | TNRC18 | O15417 | | 1359 | QARPLPSPGAAGA | 6.5625 | 6.3104 | 6.2079 | 7.3364 | 6.2043 | 6.1715 | 6.1151 | 5.2941 | 6.9981 | 5.7441 | 5.6887 | 6.6773 | 6.2320 | 6.7532 | 6.0777 | 5.6267 |
| TNRC18_S2771 | TNRC18 | O15417 | | 2146 | AVERPLTPAPRSC | 6.8538 | 6.6698 | 5.9169 | 6.7529 | 6.0118 | 6.0209 | 6.0932 | 6.3741 | 6.2405 | 6.3195 | 6.2023 | 6.5983 | 6.1441 | 6.0799 | 5.8879 | 5.8340 |
| TNRC18_S416 | TNRC18 | O15417 | | 416 | LQAPPGSPRPLDR | 6.3060 | 6.5801 | 6.5115 | 7.0451 | 7.2251 | 6.2639 | 6.4001 | 7.4380 | 6.9336 | 6.5627 | 7.2342 | 5.9944 | 5.1338 | 5.3530 | 4.0895 | 4.9291 |
| TNRC18_S611 | TNRC18 | O15417 | | 611 | GGCGKKSPFGGLG | 6.0687 | 6.5756 | 6.9529 | 7.0015 | 4.8374 | 5.8141 | 7.0987 | 6.5131 | 6.5386 | 6.9645 | 7.7886 | 6.7099 | 5.1591 | 6.9483 | 6.5521 | 4.3768 |
| TNRC18_S999 | TNRC18 | O15417 | | 2716 | ARPSAHSPGKKTP | 6.7778 | 6.6067 | 5.6731 | 7.9604 | 7.2929 | 5.9644 | 5.9366 | 6.9328 | 5.9714 | 6.4978 | 6.8379 | 6.4338 | 5.0239 | 5.5242 | 5.3309 | 5.2355 |
| TNRC18_T2146 | TNRC18 | O15417 | | 1136 | DKPIRLSPSKITE | 7.3001 | 6.0384 | 6.6936 | 7.9402 | 6.3147 | 5.9576 | 6.2352 | 5.7274 | 6.5848 | 6.8868 | 7.1094 | 6.7674 | 5.1091 | 5.6883 | 4.3808 | 5.2662 |
| TNRC6A_S1217 | TNRC6A | Q8NDV7 | | 943 | SSKPVSSPDWNKQ | 6.4855 | 5.7224 | 6.9454 | 6.3799 | 6.3418 | 6.4550 | 6.3212 | 5.9978 | 6.4074 | 6.2794 | 6.5608 | 6.4792 | 6.1314 | 6.0388 | 5.5990 | 5.8549 |
| TNRC6A_S1585 | TNRC6A | Q8NDV7 | | 771 | ACIDKTSPNGNDT | 6.3476 | 6.4710 | 6.5678 | 6.8179 | 6.4957 | 5.9750 | 6.4706 | 6.9539 | 6.4712 | 6.2009 | 6.6394 | 6.8013 | 5.5357 | 5.5838 | 5.0019 | 5.6663 |
| TNRC6A_S1599 | TNRC6A | Q8NDV7 | | 1599 | GWPRAKSPNGSSS | 5.9010 | 6.9818 | 6.9238 | 5.5451 | 6.1224 | 5.7310 | 6.2072 | 6.8742 | 5.9290 | 6.8689 | 6.9749 | 6.6532 | 5.6364 | 6.0263 | 5.6466 | 5.9782 |
| TNRC6A_S1704 | TNRC6A | Q8NDV7 | | 991 | TGWEEPSPESIRR | 6.8931 | 7.0819 | 6.4956 | 5.9889 | 6.5132 | 6.2448 | 6.5120 | 6.5874 | 6.4775 | 6.8142 | 6.1675 | 6.4630 | 5.3425 | 5.4606 | 5.4873 | 5.4705 |

| | | | | | | | | | | | | | | | | | | | | |
|---|---|---|---|---|---|---|---|---|---|---|---|---|---|---|---|---|---|---|---|---|
| TNRC6A_S1942 | TNRC6A | Q8NDV7 | 1585 | SSTSPASPPGSIG | 6.0278 | 5.8430 | 6.7513 | 6.8567 | 6.3988 | 6.0333 | 6.0990 | 6.5103 | 6.4385 | 7.0771 | 6.6308 | 6.2961 | 5.6582 | 6.2961 | 5.2044 | 5.8786 |
| TNRC6A_S739 | TNRC6A | Q8NDV7 | 1495 | NVMPHTTPELQKG | 6.7997 | 6.9948 | 6.7383 | 7.9153 | 6.7548 | 6.6342 | 7.3750 | 7.4802 | 7.4013 | 8.2298 | 7.4484 | 6.6057 | 3.7313 | 3.8617 | 2.7878 | 3.2415 |
| TNRC6A_S771 | TNRC6A | Q8NDV7 | 1217 | ISFSRDSPEENVQ | 6.1596 | 7.0115 | 6.4371 | 7.3627 | 6.1809 | 6.9762 | 6.6987 | 6.3144 | 6.1457 | 7.1310 | 6.4542 | 7.5004 | 4.7515 | 5.3910 | 4.9533 | 4.5316 |
| TNRC6A_S943 | TNRC6A | Q8NDV7 | 1704 | DSKLTWSPGSVTN | 6.4037 | 7.6661 | 6.9846 | 7.0486 | 7.2575 | 6.9222 | 6.3723 | 7.9277 | 6.9377 | 6.6183 | 6.8624 | 6.6813 | 3.7247 | 4.3361 | 4.0802 | 4.1766 |
| TNRC6A_S991 | TNRC6A | Q8NDV7 | 739 | AWDTETSPRGERK | 6.1506 | 7.3886 | 6.2984 | 6.5492 | 6.1526 | 6.4616 | 6.0029 | 7.1100 | 6.5835 | 6.9985 | 7.2782 | 6.8139 | 5.0137 | 5.6501 | 4.7624 | 4.7858 |
| TNRC6A_T1495 | TNRC6A | Q8NDV7 | 1942 | DPRGISSPSPINA | 5.6060 | 6.1931 | 6.5484 | 5.3756 | 6.2214 | 7.4734 | 6.1720 | 6.4133 | 6.2570 | 6.4307 | 6.4692 | 6.2774 | 6.0318 | 6.3224 | 6.6327 | 5.5755 |
| TNRC6B_S1336 | TNRC6B | Q9UPQ9 | 1816 | DPHRMGSPAPLLP | 6.0940 | 6.0418 | 6.8369 | 5.7434 | 5.9492 | 6.5248 | 6.3379 | 6.0642 | 6.4569 | 6.6609 | 6.3477 | 6.5739 | 6.1810 | 5.9085 | 5.8985 | 6.3806 |
| TNRC6B_S1432 | TNRC6B | Q9UPQ9 | 1432 | GKTRGGSPYNQFD | 5.6505 | 6.0004 | 6.0553 | 5.9895 | 6.4496 | 6.5854 | 5.9647 | 7.2217 | 5.9255 | 6.1352 | 6.4479 | 6.1206 | 6.2617 | 6.7377 | 6.4639 | 5.9903 |
| TNRC6B_S1816 | TNRC6B | Q9UPQ9 | 879 | SGWEEPSPQSISR | 5.9685 | 5.9421 | 7.0868 | 5.5829 | 6.4702 | 7.0165 | 6.0225 | 5.7732 | 6.6187 | 6.7459 | 6.6289 | 6.5994 | 6.2921 | 5.5636 | 5.5331 | 6.1558 |
| TNRC6B_S803 | TNRC6B | Q9UPQ9 | 1336 | QPGMKHSPSHPVG | 6.4921 | 7.9702 | 6.3650 | 6.1263 | 6.1746 | 7.4723 | 6.6142 | 7.1301 | 6.9137 | 6.3749 | 6.6998 | 6.7413 | 5.0196 | 4.9352 | 4.2811 | 4.6897 |
| TNRC6B_S879 | TNRC6B | Q9UPQ9 | 803 | WEDCKRSPAWNET | 7.8068 | 6.3842 | 4.0742 | 8.7145 | 6.1394 | 8.1026 | 6.8431 | 7.4703 | 7.2102 | 6.1955 | 4.1099 | 5.6856 | 6.3485 | 3.7377 | 5.2878 | 5.8896 |
| TNRC6B_Y1499 | TNRC6B | Q9UPQ9 | 1499 | DPESDPYVTPGSV | 5.8739 | 7.2727 | 6.3346 | 6.8675 | 5.2502 | 6.4657 | 6.9331 | 6.3127 | 7.0575 | 7.0306 | 7.8443 | 6.3699 | 4.7694 | 4.9073 | 5.5730 | 5.1376 |
| TNRC6C_S1008 | TNRC6C | Q9HCJ0-2 | 1008 | PISKESSVDRPTF | 8.2696 | 7.1971 | 9.2894 | 7.8298 | 6.9785 | 5.7605 | 5.1407 | 6.8897 | 7.4789 | 8.1261 | 6.8479 | 6.3640 | 2.9245 | 4.9560 | 3.3565 | 2.5910 |
| TNRC6C_S568 | TNRC6C | Q9HCJ0-2 | 714 | TGWEEPSPPSIRR | 6.0198 | 6.6207 | 6.7090 | 7.0950 | 5.9909 | 6.7051 | 5.7924 | 6.1292 | 6.1831 | 6.3620 | 6.3870 | 6.9293 | 6.2104 | 6.0432 | 5.3267 | 5.4963 |
| TNRC6C_S714 | TNRC6C | Q9HCJ0-2 | 568 | ANQEDKSPTWGEP | 5.3873 | 6.3987 | 5.9747 | 6.0880 | 5.9387 | 6.6516 | 6.1420 | 6.4916 | 5.7846 | 5.5118 | 6.6190 | 6.3516 | 6.8281 | 7.1131 | 6.3370 | 6.3822 |
| TNS2_S1013 | TNS2 | Q63HR2-4 | 1013 | DGASPRSPVPTTL | 5.9421 | 6.3706 | 6.4386 | 6.2214 | 6.1913 | 6.3513 | 6.2822 | 6.5698 | 6.3451 | 6.4631 | 6.2362 | 6.2902 | 6.1445 | 6.2355 | 5.9387 | 5.9792 |
| TNS2_S1106 | TNS2 | Q63HR2-4 | 658 | RLCRSLSEGLYPY | 6.4344 | 6.1730 | 7.0474 | 5.9765 | 6.3273 | 6.1197 | 5.8641 | 6.4909 | 6.4769 | 6.8889 | 6.4521 | 6.5888 | 6.0881 | 6.7415 | 5.1773 | 5.1530 |
| TNS2_S112 | TNS2 | Q63HR2-4 | 130 | TLPRSFSLDPLME | 5.8437 | 5.9042 | 6.4288 | 6.2500 | 5.6611 | 6.1546 | 6.2816 | 6.6207 | 6.0932 | 6.4998 | 6.2272 | 6.4056 | 5.9492 | 6.9391 | 6.2612 | 6.4801 |
| TNS2_S130 | TNS2 | Q63HR2-4 | 112 | RIEHLGSTKSLNH | 8.7971 | 7.1550 | 7.5952 | 6.3438 | 5.6030 | 6.0448 | 5.9044 | 6.3398 | 7.0195 | 6.6863 | 6.3494 | 6.2341 | 4.8214 | 5.9877 | 4.3921 | 4.7262 |
| TNS2_S658 | TNS2 | Q63HR2-4 | 855 | PQSRKLSYEIPTE | 5.5100 | 6.3231 | 6.0492 | 5.3931 | 6.0903 | 5.6806 | 6.0145 | 5.8955 | 6.7886 | 5.9576 | 5.9976 | 6.8349 | 7.0834 | 6.1188 | 7.5763 | 6.6864 |
| TNS2_S842 | TNS2 | Q63HR2-4 | 941 | RRQDTRSPTSAPT | 6.9814 | 7.0619 | 6.0736 | 6.3747 | 6.0555 | 6.1322 | 6.1063 | 6.4829 | 6.8903 | 6.7677 | 6.2539 | 6.0804 | 5.3257 | 6.2070 | 5.4911 | 5.7155 |
| TNS2_S855 | TNS2 | Q63HR2-4 | 842 | PRAGSISPGSPPY | 6.7215 | 6.2696 | 6.2933 | 4.9029 | 5.8789 | 6.1013 | 5.9073 | 7.5464 | 6.1039 | 5.9400 | 5.7747 | 5.8681 | 6.6453 | 6.7456 | 6.6298 | 6.6715 |
| TNS2_S941 | TNS2 | Q63HR2-4 | 1106 | GPEQASSPARGIS | 7.0604 | 6.7048 | 6.6812 | 7.7059 | 6.4991 | 7.5424 | 7.0638 | 6.6980 | 6.0457 | 6.0913 | 7.2576 | 6.7621 | 4.1496 | 5.0345 | 4.1496 | 4.5541 |
| TNS3_S1115 | TNS3 | Q68CZ2 | 690 | TGPSPGSPTLDID | 6.1951 | 6.1303 | 6.5426 | 6.7709 | 6.2491 | 6.0871 | 6.6230 | 6.2829 | 6.1797 | 6.2076 | 6.1079 | 6.3969 | 6.1334 | 6.3218 | 5.5798 | 6.1920 |
| TNS3_S1115;S1122 | TNS3 | Q68CZ2 | 776 | GRLRKLSLGQYDN | 5.7413 | 5.8241 | 6.6288 | 6.3800 | 6.4101 | 6.0264 | 6.0546 | 6.0529 | 6.2813 | 7.0605 | 7.3877 | 6.9607 | 6.0616 | 6.5638 | 5.2945 | 5.2717 |
| TNS3_S1149 | TNS3 | Q68CZ2 | 332 | PLIRWDSYENLSA | 5.8361 | 6.0563 | 6.9139 | 5.8907 | 5.9924 | 6.2165 | 6.0690 | 6.1344 | 6.5685 | 6.6132 | 6.2985 | 6.1006 | 6.5524 | 6.7151 | 6.2622 | 5.7802 |
| TNS3_S1441 | TNS3 | Q68CZ2 | 660 | PDTQQPSPSKAFK | 5.9971 | 5.8941 | 6.8238 | 6.1292 | 6.0263 | 6.9914 | 6.3201 | 6.9905 | 6.5785 | 6.3658 | 6.7674 | 6.8259 | 5.1473 | 6.1888 | 5.3191 | 5.6349 |
| TNS3_S332 | TNS3 | Q68CZ2 | 1441 | SKVMIGSPKKVxx | 6.0562 | 6.1886 | 7.4481 | 6.4709 | 6.5484 | 6.2703 | 6.8491 | 6.1489 | 6.9439 | 6.5366 | 6.7096 | 6.1061 | 5.4312 | 5.9952 | 5.1606 | 5.1365 |
| TNS3_S660 | TNS3 | Q68CZ2 | 811 | NLPPFPSPADVKE | 5.8203 | 6.5837 | 7.0847 | 7.3684 | 6.4552 | 6.4534 | 7.6060 | 6.6440 | 7.0000 | 6.9238 | 7.0532 | 7.2629 | 4.4527 | 4.8535 | 4.2630 | 4.1752 |
| TNS3_S690 | TNS3 | Q68CZ2 | 1149 | KASEAASPLPDSP | 5.9406 | 5.0478 | 7.3386 | 5.9775 | 4.9232 | 6.0974 | 6.1413 | 7.0087 | 5.4653 | 5.4722 | 5.9337 | 6.4896 | 7.5947 | 5.9867 | 7.3778 | 7.2048 |
| TNS3_S776 | TNS3 | Q68CZ2 | 850 | TPAFPVSPETPYV | 5.7824 | 6.5677 | 6.3290 | 7.0166 | 6.7812 | 6.1669 | 6.1816 | 6.6581 | 6.5669 | 6.7494 | 6.9473 | 6.5058 | 5.3115 | 4.9482 | 5.6740 | 5.8133 |
| TNS3_S811 | TNS3 | Q68CZ2 | 1115 | RSLGSVSPSSSGF | 5.7303 | 6.7372 | 6.0147 | 6.8488 | 5.3054 | 6.9348 | 6.1271 | 7.7085 | 5.4576 | 6.5752 | 7.1927 | 6.5256 | 5.4683 | 6.5049 | 5.6766 | 5.1922 |
| TNS3_S850 | TNS3 | Q68CZ2 | 953 | RQWVESSPKPMVS | 6.3695 | 6.4277 | 7.4346 | 6.3626 | 6.6798 | 6.2740 | 7.0191 | 6.7656 | 6.3972 | 6.3294 | 7.8085 | 7.4235 | 4.8377 | 5.5386 | 3.9195 | 4.4126 |
| TNS3_S953 | TNS3 | Q68CZ2 | 632 | QPRVPLTPTRGTS | 6.7209 | 6.2735 | 6.9406 | 6.2553 | 6.0622 | 6.9959 | 6.7343 | 7.0168 | 7.0664 | 6.3669 | 7.0186 | 6.5614 | 4.8646 | 5.5692 | 4.9510 | 4.6024 |
| TNS3_T632 | TNS3 | Q68CZ2 | 1115;1122 | NA | 6.3506 | 7.3127 | 6.5982 | 6.7696 | 5.7695 | 7.6883 | 6.3853 | 6.9974 | 6.8950 | 6.0150 | 6.7500 | 6.1962 | 4.5781 | 5.4993 | 5.0662 | 5.1286 |
| TOB1_S268 | TOB1 | P50616 | 268 | QKTSALSPNAKEF | 7.4275 | 6.7431 | 8.0912 | 7.1964 | 6.7602 | 7.3394 | 6.0165 | 6.9032 | 8.2459 | 6.7953 | 6.8627 | 8.4113 | 2.8679 | 3.8689 | 2.8850 | 3.5856 |
| TOB2_S222 | TOB2 | Q14106 | 222 | QPRMARSPTNSLL | 6.4356 | 6.3384 | 6.4252 | 6.1114 | 6.7498 | 7.1235 | 6.6413 | 7.4191 | 6.3276 | 6.3812 | 6.6630 | 6.7493 | 5.1909 | 5.3273 | 4.9135 | 5.2029 |
| TOM1_S160 | TOM1 | O60784-2 | 160 | TDLDMLSPIHTPQ | 6.0055 | 6.3935 | 6.9970 | 5.9998 | 6.2683 | 6.7431 | 6.5771 | 7.1696 | 6.7152 | 6.5479 | 7.7058 | 6.7616 | 5.0627 | 4.9503 | 4.6787 | 5.4237 |
| TOM1_S160;T164 | TOM1 | O60784-2 | 463 | RLPNLSSPSAEGP | 6.0156 | 6.2216 | 6.6547 | 6.1607 | 6.6989 | 6.4205 | 6.4825 | 6.3927 | 6.6087 | 6.8091 | 6.7549 | 6.9557 | 5.6500 | 5.8965 | 4.9486 | 5.3294 |
| TOM1_S164 | TOM1 | O60784-2 | 164 | MLSPIHTPQRTVF | 5.7972 | 5.8830 | 7.0728 | 5.8061 | 6.7899 | 6.5595 | 6.5414 | 6.4458 | 7.1958 | 6.6830 | 7.2071 | 6.8169 | 5.2492 | 5.8355 | 4.7398 | 5.3771 |
| TOM1_S463 | TOM1 | O60784-2 | 160;164 | NA | 6.7517 | 6.6046 | 6.9736 | 6.0651 | 6.6837 | 6.7144 | 6.7889 | 7.1034 | 6.4015 | 6.5695 | 7.5535 | 7.0764 | 4.2205 | 5.5555 | 4.3638 | 4.5739 |
| TOM1L1_S321 | TOM1L1 | O75674 | 323 | LLDLSPSPRMPRA | 6.0951 | 5.8088 | 6.4295 | 6.1242 | 6.4617 | 6.3734 | 6.5790 | 6.1550 | 6.3021 | 6.5198 | 6.0390 | 6.7230 | 6.0937 | 6.4776 | 5.6274 | 6.1906 |
| TOM1L1_S323 | TOM1L1 | O75674 | 321 | QDLLDLSPSPRMP | 5.9352 | 9.4940 | 6.7986 | 6.5469 | 6.6405 | 5.7801 | 7.2727 | 6.4152 | 6.0172 | 5.9850 | 8.2209 | 8.0160 | 2.9003 | 4.9840 | 3.5968 | 5.3967 |
| TOM1L2_S160 | TOM1L2 | Q6ZVM7 | 423 | LDNRKQSSEGIPV | 5.3288 | 5.1764 | 5.5578 | 5.4106 | 5.5097 | 6.0746 | 5.3557 | 5.9619 | 6.5121 | 6.7457 | 6.9157 | 6.8696 | 7.2695 | 7.4466 | 6.8556 | 7.0097 |
| TOM1L2_S160;T164 | TOM1L2 | Q6ZVM7 | 160 | ADLDALSPIHTPQ | 6.1391 | 6.4742 | 6.8060 | 6.3159 | 6.5650 | 6.5526 | 6.4199 | 6.9144 | 7.1104 | 6.7162 | 7.0298 | 6.7300 | 4.8194 | 5.4171 | 4.7195 | 5.2705 |
| TOM1L2_S423 | TOM1L2 | Q6ZVM7 | 479 | MVPDLPSPPMEAP | 6.0947 | 6.3322 | 7.1047 | 6.3479 | 6.4266 | 6.1506 | 6.7268 | 6.8368 | 6.6158 | 6.9378 | 7.1023 | 6.7718 | 4.9306 | 5.4446 | 4.8485 | 5.3284 |
| TOM1L2_S479 | TOM1L2 | Q6ZVM7 | 160;164 | NA | 5.8508 | 6.4194 | 6.4787 | 6.9579 | 6.2906 | 6.3638 | 7.0254 | 5.9744 | 7.4719 | 6.4776 | 6.5386 | 6.5714 | 5.3271 | 6.8083 | 4.7525 | 4.6916 |
| TOMM20_S135 | TOMM20 | Q15388 | 135 | ISQRIVSAQSLAE | 4.4412 | 4.2259 | 6.2853 | 4.6118 | 5.1509 | 5.4041 | 5.3530 | 5.4024 | 4.9220 | 4.2864 | 5.1969 | 4.9948 | 9.4271 | 9.4156 | 10.7989 | 10.0836 |
| TOMM20_S135;S138 | TOMM20 | Q15388 | 138 | RIVSAQSLAEDDV | 4.4060 | 3.9218 | 5.9461 | 4.7862 | 5.5949 | 5.4147 | 6.0775 | 5.9904 | 4.6839 | 5.3757 | 4.6793 | 4.8626 | 10.7545 | 9.3976 | 9.5328 | 8.5760 |
| TOMM20_S138 | TOMM20 | Q15388 | 135;138 | NA | 4.7364 | 5.1362 | 4.1436 | 4.7151 | 4.2471 | 4.8084 | 4.7480 | 4.9719 | 4.7027 | 4.8921 | 3.9911 | 4.9002 | 11.8256 | 10.4720 | 11.4168 | 10.2930 |
| TOMM34_S186 | TOMM34 | Q15785 | 186 | TKNRVPSAGDVEK | 6.5528 | 6.3228 | 6.8270 | 5.6893 | 6.3172 | 6.2283 | 6.1059 | 5.9339 | 6.2839 | 5.8649 | 6.2366 | 6.4802 | 6.4215 | 6.3359 | 6.1008 | 6.2991 |
| TOMM34_S93 | TOMM34 | Q15785 | 93 | PLLRRASAYEALE | 4.6161 | 4.8290 | 6.0150 | 5.1730 | 4.1722 | 5.8839 | 5.1448 | 6.8838 | 6.5403 | 9.7585 | 8.2467 | 8.3989 | 5.6911 | 6.1619 | 6.2006 | 6.2843 |
| TOMM70_S279 | TOMM70 | O94826 | 91 | TPEGRASPAPGSG | 6.3539 | 6.6200 | 6.0215 | 6.9223 | 6.5234 | 6.2702 | 6.3073 | 6.3528 | 6.3194 | 6.1262 | 6.2832 | 6.0784 | 5.6828 | 6.1977 | 6.1119 | 5.8292 |
| TOMM70_S434 | TOMM70 | O94826 | 434 | IRLRPESALAQAQ | 5.8750 | 6.4298 | 6.5098 | 7.1572 | 6.2280 | 5.6666 | 6.1814 | 6.2157 | 6.0753 | 5.5772 | 6.6186 | 6.4224 | 5.8879 | 6.3972 | 6.4048 | 6.3529 |
| TOMM70_S91 | TOMM70 | O94826 | 279 | MLKGEKSDEDKDK | 6.0899 | 6.1785 | 5.8818 | 6.9268 | 5.7872 | 5.3805 | 6.3160 | 5.6942 | 6.2000 | 6.0748 | 5.7863 | 6.9486 | 6.7999 | 6.5109 | 6.5202 | 6.9044 |
| TONSL_S719 | TONSL | Q96HA7-2 | 719 | QAHVRVSPGQAAP | 6.3217 | 6.5240 | 6.2346 | 6.3522 | 6.3821 | 6.3801 | 6.4058 | 6.2739 | 6.4895 | 6.2897 | 6.3647 | 6.3095 | 5.9444 | 6.0173 | 5.7517 | 5.9588 |
| TONSL_S905 | TONSL | Q96HA7-2 | 905 | ASEPPGSPSTPRV | 6.1119 | 6.3995 | 6.4481 | 6.7072 | 6.5050 | 6.4579 | 6.0326 | 6.2410 | 6.4344 | 6.3717 | 6.5663 | 6.3160 | 6.0470 | 6.0657 | 5.7447 | 5.5510 |
| TONSL_T773 | TONSL | Q96HA7-2 | 773 | QRVAAWTPGPASN | 7.8906 | 8.0345 | 8.0228 | 7.0438 | 5.7686 | 6.6237 | 7.5190 | 6.2873 | 5.8472 | 6.8463 | 8.4311 | 8.6118 | 2.5705 | 4.0348 | 3.1228 | 3.3453 |
| TONSL_T862 | TONSL | Q96HA7-2 | 862 | NRRPSSTSGSDSE | 5.6645 | 4.1486 | 4.4784 | 8.8121 | 3.2112 | 4.2643 | 5.6645 | 4.9471 | 10.0503 | 10.5479 | 8.8873 | 7.2557 | 4.7156 | 5.9654 | 4.8834 | 6.5035 |
| TOP1_S112 | TOP1 | P11387 | 112 | KENGFSSPPQIKD | 6.6016 | 7.3695 | 6.6313 | 7.5660 | 5.7701 | 6.2286 | 6.9914 | 6.6114 | 6.8888 | 7.2532 | 7.2874 | 7.2288 | 3.6851 | 4.9243 | 4.1594 | 4.8032 |
| TOP1_S97 | TOP1 | P11387 | 125 | EPEDDGYFVPPKE | 6.2020 | 7.3592 | 6.7305 | 7.2389 | 6.4289 | 7.4436 | 7.2381 | 6.6026 | 6.9968 | 6.8276 | 7.9698 | 7.4413 | 3.7152 | 4.5591 | 3.5149 | 3.7316 |
| TOP1_Y125 | TOP1 | P11387 | 97 | EEKVRASGDAKIK | 6.5653 | 7.6693 | 6.4916 | 7.7317 | 5.8447 | 7.2112 | 7.5835 | 6.2935 | 6.2176 | 7.0345 | 7.7689 | 7.2193 | 3.2368 | 5.0142 | 3.4096 | 4.7082 |
| TOP2A_S1187 | TOP2A | P11388-4 | 1294 | MAEVLPSPRGQRV | 6.9894 | 7.1833 | 6.4161 | 7.4926 | 6.4719 | 6.4553 | 7.0344 | 6.5566 | 6.8354 | 6.8294 | 6.9418 | 6.9551 | 3.9822 | 4.9818 | 4.1871 | 4.6874 |
| TOP2A_S1294 | TOP2A | P11388-4 | 1606 | IKYLEESDEDDLF | 6.3773 | 6.4785 | 6.2891 | 6.6995 | 6.1833 | 6.0280 | 6.3005 | 6.2347 | 6.5073 | 6.3997 | 6.4312 | 6.3797 | 5.6923 | 6.1764 | 5.6602 | 6.1624 |
| TOP2A_S1328 | TOP2A | P11388-4 | 1458 | PQKSVVSDLEADD | 6.8252 | 6.9727 | 6.8055 | 7.2985 | 6.4171 | 6.2729 | 7.0176 | 6.6561 | 6.1237 | 6.0748 | 6.2130 | 6.2761 | 4.7538 | 5.5632 | 5.2470 | 5.4827 |
| TOP2A_S1455 | TOP2A | P11388-4 | 1474 | SVPLSSSPPATHF | 6.1206 | 6.3406 | 6.2102 | 7.1125 | 5.9349 | 6.0985 | 6.4922 | 6.3129 | 6.2090 | 6.5404 | 6.2810 | 6.5006 | 5.2029 | 6.2169 | 6.1821 | 6.2446 |
| TOP2A_S1455;S1458 | TOP2A | P11388-4 | 1328 | NENTEGSPQEDGV | 6.2307 | 6.5541 | 6.7054 | 6.3183 | 6.0372 | 5.9018 | 6.6374 | 5.7404 | 6.3669 | 6.2609 | 6.8433 | 6.0481 | 5.8467 | 6.1404 | 6.2092 | 6.1591 |
| TOP2A_S1458 | TOP2A | P11388-4 | 1455 | ELKPQKSVVSDLE | 5.9515 | 6.1651 | 6.9583 | 6.9313 | 5.7607 | 5.6084 | 6.2498 | 6.1905 | 6.2295 | 7.1789 | 7.1287 | 6.8212 | 4.9610 | 6.0917 | 5.7851 | 5.9883 |
| TOP2A_S1472;T1474 | TOP2A | P11388-4 | 1576 | KKSKGESDDFHMD | 6.1746 | 6.7603 | 6.1842 | 7.6683 | 6.3467 | 6.9119 | 6.8553 | 6.4389 | 6.6427 | 6.9411 | 7.3366 | 6.9643 | 4.1712 | 5.1294 | 4.4969 | 4.9777 |
| TOP2A_S1473 | TOP2A | P11388-4 | 1550 | RRKRKPSTSDDSD | 6.5734 | 6.5672 | 6.0378 | 7.3223 | 5.1810 | 4.9585 | 5.4479 | 5.2699 | 9.4448 | 9.5657 | 9.9415 | 10.1198 | 3.0312 | 4.0868 | 3.0252 | 3.4272 |
| TOP2A_S1473;S1474 | TOP2A | P11388-4 | 1187 | EEENEESDNEKET | 6.0331 | 6.4493 | 6.3046 | 7.2696 | 5.7489 | 6.0712 | 6.0876 | 5.7361 | 6.3235 | 6.2992 | 6.2229 | 6.4760 | 6.0871 | 6.5074 | 6.0214 | 6.3622 |
| TOP2A_S1474 | TOP2A | P11388-4 | 1473 | GSVPLSSSPPATH | 6.4376 | 6.3713 | 5.8280 | 7.2636 | 6.4287 | 5.8745 | 6.5344 | 6.0609 | 6.9889 | 6.0407 | 6.3612 | 5.9100 | 5.7068 | 6.2625 | 5.8757 | 6.0553 |
| TOP2A_S1550 | TOP2A | P11388-4 | 683 | EEWKSSTPNHKKW | 7.3561 | 7.8396 | 6.7212 | 8.7269 | 6.5295 | 6.7959 | 6.9950 | 7.1217 | 6.7005 | 7.0005 | 7.0411 | 7.0632 | 2.9339 | 4.0886 | 3.2730 | 3.8133 |
| TOP2A_S1550;S1552;S15 | TOP2A | P11388-4 | 1550;1552;15 | NA | 5.6290 | 5.7247 | 5.6336 | 6.0125 | 5.8872 | 5.8930 | 5.9424 | 5.2798 | 7.1892 | 6.5664 | 6.9128 | 6.8177 | 6.1286 | 6.8571 | 6.6155 | 6.9103 |
| TOP2A_S1550;S1555;S15 | TOP2A | P11388-4 | 1473;1474 | NA | 6.5804 | 6.9476 | 6.7915 | 7.3940 | 6.1118 | 6.1203 | 6.4992 | 6.1751 | 6.5572 | 6.6303 | 7.1043 | 6.4994 | 4.8087 | 5.5178 | 4.8405 | 5.4218 |
| TOP2A_S1550;T1551 | TOP2A | P11388-4 | 1550;1551 | NA | 5.9340 | 5.7241 | 4.3189 | 7.2529 | 5.2527 | 4.4405 | 4.0255 | 5.3396 | 8.0277 | 8.5467 | 7.7664 | 8.7120 | 5.1730 | 6.4957 | 6.5645 | 6.4258 |
| TOP2A_S1550;T1551;S15 | TOP2A | P11388-4 | 1472;1474 | NA | 6.5193 | 6.9215 | 6.4965 | 8.2292 | 6.0191 | 5.8317 | 6.3003 | 6.3065 | 6.6041 | 6.5518 | 7.1499 | 6.2799 | 4.2631 | 6.2598 | 5.1005 | 5.1668 |
| TOP2A_S1550;T1551;S15 | TOP2A | P11388-4 | 1455;1458 | NA | 5.9683 | 5.9571 | 7.1156 | 7.2871 | 6.0004 | 6.8185 | 6.4945 | 6.8681 | 6.4725 | 5.8857 | 6.5425 | 5.7724 | 5.0040 | 6.4697 | 5.5875 | 5.7560 |
| TOP2A_S1552;S1555 | TOP2A | P11388-4 | 1550;1551;15 | NA | 6.8985 | 5.8761 | 6.7033 | 6.9537 | 5.6059 | 5.9500 | 7.6155 | 6.0763 | 6.4759 | 5.9186 | 6.4622 | 6.1616 | 5.1167 | 6.3525 | 5.7111 | 6.1220 |
| TOP2A_S1576 | TOP2A | P11388-4 | 1550;1555;15 | NA | 6.2047 | 5.9556 | 5.7977 | 6.7166 | 5.4973 | 5.0774 | 5.5714 | 5.4323 | 6.8864 | 7.1087 | 6.6224 | 6.5709 | 6.7051 | 7.4261 | 6.0703 | 6.3573 |
| TOP2A_S1606 | TOP2A | P11388-4 | 1552;1555 | NA | 7.5646 | 8.3523 | 6.0161 | 7.6463 | 5.5196 | 6.3577 | 5.6830 | 5.6308 | 6.6557 | 7.9264 | 6.4906 | 7.0630 | 4.1001 | 5.9906 | 4.2375 | 4.7658 |
| TOP2A_T683 | TOP2A | P11388-4 | 1550;1551;15 | NA | 5.5304 | 6.1299 | 4.7439 | 7.4799 | 5.5060 | 4.9894 | 5.7812 | 5.8940 | 5.8612 | 6.6243 | 5.8623 | 6.5920 | 6.1574 | 6.6574 | 7.5229 | 8.6678 |
| TOP2B_S1212 | TOP2B | Q02880 | 1581 | TSFDQDSDVDIFP | 6.0028 | 6.0889 | 6.0129 | 6.6265 | 6.1373 | 5.8058 | 6.3715 | 6.0749 | 6.1813 | 6.2899 | 5.9123 | 6.1067 | 6.1828 | 6.8323 | 6.7579 | 6.6164 |
| TOP2B_S1236 | TOP2B | Q02880 | 1400 | ELKVKASPITNDG | 5.8050 | 5.9239 | 5.7055 | 6.3976 | 6.0864 | 5.8220 | 5.9584 | 5.6066 | 6.3607 | 6.4996 | 6.2276 | 6.3649 | 6.7717 | 7.1893 | 6.8052 | 6.4754 |
| TOP2B_S1279 | TOP2B | Q02880 | 1424 | KDEYTFSPGKSKA | 7.0574 | 6.9982 | 6.5104 | 7.0314 | 6.2048 | 5.6060 | 6.6766 | 6.0095 | 6.7213 | 6.6949 | 6.3000 | 6.6070 | 4.9540 | 5.4395 | 5.3144 | 5.8745 |
| TOP2B_S1375 | TOP2B | Q02880 | 1466 | DSAKFDSNEEDSA | 6.2575 | 6.1826 | 6.0895 | 6.7086 | 6.3122 | 6.2572 | 6.4358 | 6.2267 | 6.4650 | 6.2095 | 6.6078 | 6.2808 | 6.1592 | 6.5018 | 5.6223 | 5.6836 |
| TOP2B_S1400 | TOP2B | Q02880 | 1413 | EDEFVPSDGLDKD | 5.6069 | 6.3556 | 5.5379 | 6.1705 | 5.9915 | 5.3603 | 6.1308 | 6.5195 | 5.8439 | 5.9157 | 5.2373 | 5.4839 | 6.6883 | 7.6077 | 7.6098 | 7.9404 |
| TOP2B_S1400;S1413 | TOP2B | Q02880 | 1375 | KYTFDFSEEEDDD | 5.6579 | 6.0210 | 5.7221 | 6.5143 | 6.1550 | 5.9461 | 6.1331 | 5.3675 | 6.4330 | 6.3014 | 5.8407 | 6.3613 | 6.4768 | 6.9565 | 6.9253 | 7.1882 |
| TOP2B_S1400;S1413;S14 | TOP2B | Q02880 | 1613 | VKYFAESDEEEDD | 5.9814 | 6.0993 | 6.3318 | 6.4081 | 6.2148 | 5.8525 | 6.2436 | 5.9677 | 6.5928 | 6.5704 | 6.4342 | 6.6842 | 5.9949 | 6.2689 | 6.1702 | 6.1853 |
| TOP2B_S1400;S1424 | TOP2B | Q02880 | 1236 | LEETMPSPYGRRI | 7.3777 | 7.8374 | 6.8459 | 8.0291 | 6.3697 | 6.7181 | 7.3151 | 6.5568 | 7.3353 | 7.1849 | 6.8827 | 7.0049 | 3.3761 | 3.9806 | 3.4335 | 3.7522 |
| TOP2B_S1413 | TOP2B | Q02880 | 1317 | GTRVRKTPTSSGK | 7.4490 | 8.0314 | 6.4995 | 7.3815 | 6.3948 | 6.5629 | 7.3312 | 7.1441 | 7.8367 | 6.6649 | 7.1988 | 7.2351 | 2.7208 | 4.0400 | 3.3691 | 4.1404 |
| TOP2B_S1413;S1424 | TOP2B | Q02880 | 1212 | DVLAGMSGKAIKG | 6.1048 | 7.0276 | 7.3536 | 7.0636 | 7.2769 | 6.3589 | 7.1786 | 7.7491 | 6.7280 | 5.8316 | 6.7927 | 6.4835 | 5.3330 | 5.2132 | 3.2957 | 4.2089 |
| TOP2B_S1424 | TOP2B | Q02880 | 1441 | SLHDKKSQDFGNL | 5.4101 | 6.6357 | 6.9613 | 7.4137 | 6.2957 | 6.4066 | 6.9403 | 6.1086 | 6.7920 | 6.4335 | 7.1655 | 7.1104 | 4.7641 | 5.8092 | 4.7976 | 4.9558 |
| TOP2B_S1441 | TOP2B | Q02880 | 1279 | EFDEEFSGAPVEG | 5.4058 | 6.5743 | 6.3981 | 6.7267 | 5.8869 | 6.5702 | 5.9782 | 7.1100 | 5.9758 | 6.1690 | 5.9856 | 6.7740 | 5.9415 | 5.9195 | 6.2326 | 6.3517 |
| TOP2B_S1461 | TOP2B | Q02880 | 1552 | KRKASGSENEGDY | 7.0069 | 7.6416 | 6.3681 | 6.8159 | 6.2991 | 6.1416 | 6.5764 | 6.9860 | 5.8222 | 5.4446 | 5.1472 | 5.3924 | 6.1340 | 6.1657 | 6.3217 | 5.7366 |
| TOP2B_S1461;S1466 | TOP2B | Q02880 | 1431 | PGKSKATPEKSLH | 5.4108 | 6.5595 | 6.6493 | 7.2796 | 6.7455 | 6.8377 | 6.5514 | 6.6716 | 6.7584 | 7.0143 | 6.5015 | 6.6535 | 4.9223 | 5.7958 | 5.0622 | 5.4867 |
| TOP2B_S1466 | TOP2B | Q02880 | 1473 | NEEDSASVFSPSF | 6.6105 | 6.3199 | 6.6339 | 6.3732 | 5.9332 | 5.7031 | 5.7889 | 6.1769 | 7.0265 | 7.8383 | 7.5894 | 7.7070 | 4.5461 | 5.6127 | 4.5012 | 5.6394 |
| TOP2B_S1466;S1471 | TOP2B | Q02880 | 1461 | QKSEDDSAKFDSN | 4.9048 | 5.9405 | 5.7175 | 6.1176 | 5.8409 | 5.3317 | 6.0116 | 6.0195 | 6.8536 | 6.0773 | 5.6867 | 6.8449 | 7.9138 | 6.9603 | 7.0188 | 6.7603 |
| TOP2B_S1466;S1473 | TOP2B | Q02880 | 1292 | AGEEALTPSVPIN | 7.2020 | 6.0084 | 6.7318 | 7.1706 | 5.6226 | 5.8808 | 7.1853 | 5.7905 | 6.6985 | 7.2629 | 7.0999 | 6.8290 | 5.4214 | 5.1318 | 4.4408 | 5.5235 |
| TOP2B_S1473 | TOP2B | Q02880 | 1400;1413 | NA | 5.6552 | 5.8564 | 5.3386 | 6.3543 | 6.2779 | 5.9121 | 6.1331 | 6.0025 | 5.7662 | 6.0364 | 6.0710 | 5.8938 | 7.5533 | 7.9829 | 6.9705 | 6.1960 |
| TOP2B_S1522;S1524 | TOP2B | Q02880 | 1400;1413;14 | NA | 6.3601 | 6.4642 | 6.3798 | 7.3352 | 7.3512 | 7.1809 | 7.3293 | 6.6336 | 6.8896 | 6.5989 | 7.4460 | 6.8009 | 4.4924 | 5.2857 | 3.9789 | 3.4735 |
| TOP2B_S1550;S1552 | TOP2B | Q02880 | 1522;1524 | NA | 6.3618 | 5.8124 | 6.3401 | 6.4570 | 6.2594 | 6.4335 | 5.8159 | 5.9608 | 6.6855 | 6.8368 | 6.4244 | 6.2262 | 5.8965 | 6.4438 | 5.9968 | 6.0490 |
| TOP2B_S1552 | TOP2B | Q02880 | 1466;1473 | NA | 7.2987 | 6.5818 | 6.7293 | 7.7598 | 6.8570 | 4.9779 | 5.8872 | 5.1516 | 5.9070 | 4.9105 | 5.7592 | 5.3370 | 6.3879 | 6.6518 | 6.8032 | 6.9999 |
| TOP2B_S1581 | TOP2B | Q02880 | 1461;1466 | NA | 6.1787 | 5.8579 | 6.1835 | 6.7293 | 6.3771 | 6.3051 | 6.9273 | 5.9878 | 6.2765 | 5.8985 | 6.1037 | 6.4685 | 6.0086 | 6.3364 | 5.9531 | 6.4079 |
| TOP2B_S1613 | TOP2B | Q02880 | 1550;1552 | NA | 6.8757 | 6.5859 | 7.1038 | 7.1761 | 6.8983 | 6.2355 | 6.7865 | 6.8186 | 6.5767 | 6.1452 | 5.7408 | 5.9079 | 5.2024 | 5.1802 | 5.3311 | 5.4352 |
| TOP2B_T1292 | TOP2B | Q02880 | 1413;1424 | NA | 6.4035 | 6.3635 | 6.9101 | 6.9666 | 6.6365 | 6.7866 | 6.8517 | 6.0168 | 6.9559 | 6.1302 | 6.7191 | 6.4998 | 5.1300 | 5.5548 | 5.2872 | 4.7874 |
| TOP2B_T1317 | TOP2B | Q02880 | 1466;1471 | NA | 5.8635 | 5.7872 | 6.0874 | 6.0806 | 6.1389 | 5.4156 | 6.1778 | 5.4783 | 5.1833 | 5.3989 | 5.2533 | 5.3626 | 6.8816 | 7.3421 | 8.5427 | 9.0061 |
| TOP2B_T1317;S1321 | TOP2B | Q02880 | 1400;1424 | NA | 6.0444 | 6.9163 | 6.4761 | 5.9110 | 6.6401 | 6.4422 | 7.1169 | 6.1753 | 7.1501 | 6.8831 | 7.0796 | 7.3130 | 4.8500 | 5.2031 | 5.0572 | 4.7415 |
| TOP2B_T1431 | TOP2B | Q02880 | 1317;1321 | NA | 7.3529 | 7.0711 | 6.1664 | 7.4283 | 7.8916 | 7.1486 | 7.2771 | 7.2265 | 7.3716 | 8.1717 | 8.0064 | 8.3002 | 1.9987 | 3.3761 | 2.4231 | 2.7896 |
| TOP3A_S971 | TOP3A | Q13472 | 971 | KRPRASSSDMGST | 4.9458 | 5.0891 | 5.5972 | 5.7884 | 6.3062 | 5.4856 | 5.8298 | 6.1563 | 5.2052 | 6.0513 | 5.3915 | 6.0689 | 8.2372 | 7.5559 | 8.3370 | 7.9546 |
| TOP3A_T554 | TOP3A | Q13472 | 554 | RMYVGLTPDKRFL | 6.8455 | 6.9258 | 9.5236 | 6.3641 | 8.9268 | 7.1163 | 7.0612 | 7.9338 | 5.3310 | 7.8034 | 6.2387 | 6.8004 | 3.1344 | 3.0341 | 3.7462 | 3.2146 |
| TOPBP1_S1504 | TOPBP1 | Q92547 | 350 | LENLDVSAFQAPE | 6.4543 | 7.6278 | 7.0263 | 7.3595 | 6.5261 | 6.4908 | 6.4404 | 6.0956 | 6.4827 | 5.7919 | 6.3668 | 6.0602 | 4.9284 | 5.8122 | 5.3792 | 5.1579 |
| TOPBP1_S350 | TOPBP1 | Q92547 | 888 | AVALSASPQLKEA | 6.6835 | 6.5428 | 6.3931 | 6.8601 | 5.9692 | 5.7327 | 7.0145 | 5.9522 | 7.0778 | 6.6070 | 6.5761 | 6.1450 | 5.7124 | 5.8623 | 5.4058 | 5.4654 |
| TOPBP1_S805 | TOPBP1 | Q92547 | 1504 | ELGTGLSQKRKAP | 6.1661 | 5.9418 | 5.7503 | 5.8778 | 5.1710 | 5.3021 | 6.2295 | 4.8743 | 6.4343 | 5.6163 | 5.1660 | 5.1786 | 8.4067 | 8.8199 | 7.4559 | 7.6095 |
| TOPBP1_S836 | TOPBP1 | Q92547 | 860 | QQKRKPSTPLSEV | 4.8321 | 4.5046 | 5.0277 | 5.6347 | 8.6718 | 4.7702 | 4.7326 | 5.8039 | 8.2424 | 8.7818 | 7.6047 | 7.6049 | 6.6034 | 7.0291 | 5.0682 | 5.0878 |
| TOPBP1_S860 | TOPBP1 | Q92547 | 805 | ARQVAASPAVGQP | 7.8817 | 7.3294 | 6.9604 | 8.2524 | 5.4620 | 7.4305 | 7.3798 | 6.2584 | 8.2937 | 7.8153 | 7.5588 | 8.0325 | 2.6380 | 2.9744 | 2.4899 | 3.2427 |
| TOPBP1_S888 | TOPBP1 | Q92547 | 836 | DKLFKPSFDVKDA | 5.7085 | 5.9627 | 7.9454 | 7.3026 | 5.2418 | 6.5401 | 7.1955 | 4.9568 | 5.9536 | 9.2091 | 6.5927 | 6.9613 | 4.8715 | 5.2999 | 4.5955 | 5.6631 |
| TOPBP1_T298 | TOPBP1 | Q92547 | 298 | TMPNSSTPTSQIN | 5.2976 | 6.6466 | 5.4878 | 6.6767 | 6.9503 | 6.4209 | 5.4713 | 6.1363 | 6.3169 | 6.1267 | 7.4675 | 6.3757 | 6.5454 | 6.2088 | 5.2511 | 6.6206 |
| TOPBP1_T860;S864 | TOPBP1 | Q92547 | 860;864 | NA | 8.6376 | 8.3868 | 7.9203 | 9.0846 | 6.2045 | 7.0860 | 7.8698 | 6.5949 | 7.2201 | 6.9674 | 7.6086 | 7.7878 | 1.8014 | 2.7763 | 1.5976 | 2.4563 |
| TOPORS_S194 | TOPORS | Q9NS56 | 81 | FKMDNFSPKAGTS | 6.5473 | 6.7390 | 6.7272 | 7.2752 | 6.0771 | 6.0161 | 6.3739 | 5.7370 | 6.5026 | 6.8934 | 6.8045 | 6.9652 | 4.7975 | 5.6115 | 5.2128 | 5.7196 |
| TOPORS_S585 | TOPORS | Q9NS56 | 205 | VNRRTTTPPDSGV | 5.7116 | 6.6685 | 6.4149 | 6.5510 | 6.3119 | 6.0893 | 6.3419 | 6.3589 | 6.3286 | 6.4772 | 6.2708 | 6.2735 | 5.8097 | 6.3032 | 5.7315 | 6.3576 |
| TOPORS_S81 | TOPORS | Q9NS56 | 585 | RGDRVYSPYNHRH | 6.7348 | 6.7855 | 6.1806 | 7.2537 | 6.3135 | 5.8965 | 6.1588 | 6.1400 | 6.5543 | 6.1663 | 6.1900 | 6.1328 | 5.2923 | 6.3835 | 5.6399 | 6.1773 |
| TOPORS_T205 | TOPORS | Q9NS56 | 194 | RNASVYSPSGPVN | 6.6707 | 6.6203 | 6.6181 | 7.2417 | 6.0906 | 5.5646 | 6.3785 | 5.8309 | 7.1625 | 6.8611 | 6.2185 | 6.4995 | 5.3191 | 6.2585 | 5.2349 | 5.4307 |
| TOR1AIP1_S154;S156;S1 | TOR1AIP1 | Q5JTV8-3 | 79 | LVAKERSPVGKRT | 6.7182 | 7.0065 | 6.0081 | 6.9080 | 6.1259 | 5.8740 | 6.1088 | 6.0485 | 6.8904 | 6.5791 | 6.4959 | 6.5147 | 5.5678 | 5.8785 | 5.7068 | 5.5688 |
| TOR1AIP1_S156;S157 | TOR1AIP1 | Q5JTV8-3 | 306 | TRMQNDSILKSEL | 7.1777 | 6.5666 | 7.3230 | 8.0473 | 5.8816 | 7.1897 | 7.6007 | 7.0516 | 7.1809 | 7.4950 | 7.9109 | 7.3540 | 2.9989 | 3.8773 | 2.9464 | 3.3984 |
| TOR1AIP1_S254 | TOR1AIP1 | Q5JTV8-3 | 316 | SELGNQSPSTSSR | 7.2047 | 7.8340 | 7.0278 | 7.6988 | 6.5879 | 6.5036 | 6.7940 | 6.7873 | 7.3512 | 7.1147 | 7.3908 | 7.1507 | 3.1167 | 4.0231 | 3.1686 | 4.2462 |
| TOR1AIP1_S306 | TOR1AIP1 | Q5JTV8-3 | 254 | DKTTRSSSQYIES | 4.4865 | 4.5711 | 3.8346 | 4.9903 | 4.9783 | 5.0395 | 5.0349 | 4.7407 | 5.0387 | 5.0506 | 4.8137 | 4.8414 | 10.3797 | 10.3562 | 11.4138 | 10.4301 |
| TOR1AIP1_S316 | TOR1AIP1 | Q5JTV8-3 | 47 | DAPAYRTPPSRQG | 6.5225 | 6.9364 | 7.3842 | 8.2161 | 6.5984 | 7.0231 | 7.4852 | 5.8864 | 7.2975 | 7.0685 | 6.6465 | 6.2285 | 3.7748 | 4.6629 | 4.1372 | 4.1318 |
| TOR1AIP1_S79 | TOR1AIP1 | Q5JTV8-3 | 156;157 | NA | 6.4838 | 6.6022 | 6.3892 | 6.8632 | 6.3156 | 6.5005 | 6.0500 | 6.3525 | 6.3297 | 6.6179 | 6.5033 | 6.1896 | 5.5549 | 6.0414 | 5.6243 | 5.5819 |
| TOR1AIP1_T47 | TOR1AIP1 | Q5JTV8-3 | 154;156;157 | NA | 5.8814 | 5.9193 | 5.4819 | 6.1155 | 5.9058 | 5.7990 | 5.8211 | 6.5887 | 5.9751 | 6.1077 | 5.8888 | 5.7515 | 7.7643 | 7.0994 | 6.7768 | 7.1237 |
| TOR1AIP2_S72 | TOR1AIP2 | Q8NFQ8 | 72 | TGDKSESPDEANV | 5.7258 | 6.5370 | 6.3334 | 7.3805 | 6.4380 | 6.0890 | 6.5242 | 6.1998 | 6.1274 | 7.0684 | 6.2543 | 6.2368 | 5.4610 | 5.6945 | 5.7831 | 6.1468 |
| TOR4A_S63 | TOR4A | Q9NXH8 | 63 | APRPGCSPRAPRA | 7.8833 | 7.9043 | 8.1090 | 7.8652 | 6.8814 | 6.7381 | 6.7456 | 5.3910 | 6.4860 | 6.9220 | 7.3043 | 7.5484 | 3.7480 | 3.7500 | 3.0978 | 3.6255 |
| TOR4A_S81 | TOR4A | Q9NXH8 | 89 | AELPSRTPRKKRR | 7.8368 | 7.7997 | 6.9388 | 7.5865 | 6.3913 | 6.6429 | 6.9898 | 5.9593 | 7.0422 | 6.8000 | 6.6614 | 6.7788 | 3.8499 | 4.4346 | 3.8571 | 4.4309 |
| TOR4A_S97 | TOR4A | Q9NXH8 | 97 | RKKRRRSRLVLYP | 6.3703 | 5.8786 | 6.3886 | 6.6531 | 7.0246 | 5.5429 | 5.7284 | 5.6528 | 7.8933 | 8.8773 | 8.2977 | 8.1493 | 4.2806 | 5.1670 | 4.0984 | 3.9970 |
| TOR4A_T78;T89 | TOR4A | Q9NXH8 | 81 | KFFTFDSPAELPS | 7.6258 | 7.4608 | 6.8279 | 8.1170 | 5.6617 | 6.4169 | 7.1425 | 6.7404 | 8.0815 | 7.9766 | 7.1237 | 6.8596 | 2.9304 | 3.9019 | 2.7864 | 3.4470 |
| TOR4A_T89 | TOR4A | Q9NXH8 | 78;89 | NA | 8.3014 | 8.2411 | 7.7371 | 7.5016 | 5.7018 | 5.7590 | 6.4280 | 6.2434 | 9.1047 | 7.7524 | 7.1510 | 7.8722 | 2.8415 | 3.6427 | 2.6796 | 3.0426 |
| TOX4_S178 | TOX4 | O94842 | 178 | RLSTTPSPTSSLH | 7.3411 | 6.9309 | 5.9388 | 5.8645 | 7.1430 | 6.3954 | 6.8381 | 6.9449 | 7.7218 | 5.9280 | 6.3381 | 5.3646 | 4.7718 | 5.0210 | 5.6262 | 5.8320 |
| TOX4_T178;S182 | TOX4 | O94842 | 178;182 | NA | 6.6840 | 6.5535 | 6.0101 | 7.2339 | 6.3169 | 6.1635 | 6.5489 | 6.3704 | 6.3369 | 6.6422 | 6.3518 | 6.5627 | 5.4812 | 5.8439 | 5.7964 | 5.1037 |
| TP53BP1_S1004;S1007 | TP53BP1 | Q12888-2 | 1119 | VIQGPSSPQGEAM | 6.0892 | 6.1260 | 6.1224 | 6.4285 | 6.2381 | 6.0730 | 6.0312 | 6.0293 | 6.3017 | 6.2125 | 6.0082 | 6.0916 | 6.5494 | 6.7557 | 6.4652 | 6.4779 |
| TP53BP1_S1007 | TP53BP1 | Q12888-2 | 270 | SEDMPFSPKASVA | 6.4217 | 6.5120 | 5.9837 | 6.1094 | 6.2821 | 5.8922 | 6.3038 | 5.8855 | 6.4116 | 6.1801 | 6.1803 | 6.0695 | 6.6355 | 6.4775 | 6.3427 | 6.3122 |
| TP53BP1_S1033 | TP53BP1 | Q12888-2 | 1467 | ALRRSDSPEIPFQ | 6.0786 | 6.5852 | 6.3629 | 6.3902 | 6.1358 | 5.8774 | 6.2030 | 6.1985 | 6.2472 | 6.2200 | 6.2393 | 6.1353 | 6.5486 | 6.1219 | 6.5536 | 6.1024 |
| TP53BP1_S1072 | TP53BP1 | Q12888-2 | 557 | EDTEPMSPVLNSK | 6.0868 | 6.0926 | 6.1080 | 6.3911 | 5.9848 | 5.7706 | 6.1506 | 6.1809 | 6.3015 | 6.2067 | 5.9321 | 6.2304 | 6.6543 | 6.7239 | 6.4775 | 6.7083 |
| TP53BP1_S1072;S1073 | TP53BP1 | Q12888-2 | 385 | PFIVPSSPTEQEG | 6.1066 | 6.5116 | 6.1933 | 6.4405 | 6.1255 | 5.9644 | 6.0737 | 6.1313 | 6.2657 | 6.0667 | 6.1470 | 6.0977 | 6.4658 | 6.6312 | 6.3860 | 6.3930 |
| TP53BP1_S1073 | TP53BP1 | Q12888-2 | 1106 | PVKDPVSPASQKM | 6.0844 | 6.3060 | 6.1531 | 6.4860 | 6.0382 | 5.7735 | 5.9899 | 6.0235 | 6.2860 | 6.2378 | 5.7756 | 5.9953 | 6.4964 | 6.5140 | 6.8411 | 6.9994 |
| TP53BP1_S1099 | TP53BP1 | Q12888-2 | 732 | SVISIDSPQKLAI | 6.0596 | 6.1326 | 6.3913 | 6.4103 | 6.0453 | 5.9510 | 6.1973 | 5.9755 | 6.3490 | 6.4498 | 6.1755 | 6.2500 | 6.3663 | 6.4836 | 6.2774 | 6.4857 |
| TP53BP1_S1099;S1106 | TP53BP1 | Q12888-2 | 505 | EIEPKNSPEDLGL | 5.9579 | 6.1758 | 6.1930 | 6.4188 | 5.8200 | 5.6112 | 6.1600 | 5.8385 | 6.1698 | 6.3123 | 6.1702 | 6.2354 | 6.9461 | 6.7147 | 6.6314 | 6.6449 |
| TP53BP1_S1106 | TP53BP1 | Q12888-2 | 403 | MDTSVLSEEGGEP | 6.0021 | 5.9021 | 6.3594 | 6.8242 | 6.3790 | 6.1540 | 6.4065 | 6.1171 | 6.4050 | 6.4713 | 6.4722 | 6.4407 | 6.0384 | 6.0109 | 5.7542 | 6.2629 |
| TP53BP1_S1106;S1109 | TP53BP1 | Q12888-2 | 1670 | ASMGVLSGKRKLI | 7.8840 | 8.4381 | 7.1039 | 7.4698 | 6.5133 | 7.5517 | 7.6414 | 7.1795 | 7.7609 | 7.4264 | 7.6486 | 7.3179 | 2.0392 | 3.1974 | 1.9629 | 2.8649 |
| TP53BP1_S1119 | TP53BP1 | Q12888-2 | 585 | DGEVQLSQNDDKT | 6.1574 | 6.3766 | 6.3249 | 6.7870 | 6.5034 | 6.0765 | 6.5180 | 6.0101 | 6.6603 | 6.1924 | 6.2996 | 6.3070 | 6.0883 | 5.8811 | 5.9643 | 5.8532 |

| | | | | | | | | | | | | | | | | | | | | |
|---|---|---|---|---|---|---|---|---|---|---|---|---|---|---|---|---|---|---|---|---|
| TP53BP1_S1213;T1221 | TP53BP1 | Q12888-2 | 1435 | DISPNLSPDDKSF | 6.6367 | 6.7567 | 6.8928 | 7.0194 | 6.6448 | 6.6959 | 6.8029 | 6.7235 | 7.1443 | 6.9102 | 7.0341 | 6.8532 | 4.3154 | 4.6519 | 4.2694 | 4.6487 |
| TP53BP1_S1221;S1224 | TP53BP1 | Q12888-2 | 299 | GLQIQKSPEPEVL | 5.8352 | 6.0468 | 6.1355 | 6.3429 | 6.1237 | 5.8304 | 5.8761 | 5.8701 | 6.1118 | 5.9417 | 6.0151 | 6.1293 | 6.9942 | 6.9375 | 6.8527 | 6.9569 |
| TP53BP1_S124 | TP53BP1 | Q12888-2 | 1683 | TSEEERSPAKRGR | 7.0654 | 6.7098 | 6.3110 | 6.7180 | 6.7194 | 6.0850 | 6.4085 | 6.2388 | 6.9972 | 6.5595 | 6.2697 | 6.3889 | 5.4891 | 5.4349 | 5.2128 | 5.3920 |
| TP53BP1_S1295 | TP53BP1 | Q12888-2 | 1061 | SEDPPTTPIRGNL | 6.1482 | 6.3172 | 5.7421 | 6.4054 | 6.1240 | 5.8835 | 6.0282 | 5.8498 | 5.8938 | 5.8559 | 5.6138 | 5.8504 | 7.1978 | 6.8215 | 7.2232 | 7.0452 |
| TP53BP1_S1367 | TP53BP1 | Q12888-2 | 814 | DTKEEKSVEYEGD | 6.9161 | 7.7972 | 6.7094 | 7.7822 | 6.2858 | 6.5219 | 6.5611 | 6.4846 | 6.9061 | 7.0216 | 7.3569 | 7.2138 | 3.8001 | 4.2920 | 3.9840 | 4.3672 |
| TP53BP1_S1431 | TP53BP1 | Q12888-2 | 530 | TSEYSQSPKMESL | 6.2848 | 6.2816 | 6.0864 | 6.5172 | 6.0385 | 6.1591 | 6.5928 | 5.8536 | 6.6214 | 6.1895 | 5.9494 | 5.8241 | 6.1394 | 6.5061 | 6.3817 | 6.5743 |
| TP53BP1_S1431;S1435 | TP53BP1 | Q12888-2 | 1614 | GPYEAVTPLTKAA | 6.5477 | 7.0015 | 6.7538 | 6.8796 | 6.8248 | 6.9343 | 6.8137 | 6.4267 | 6.3101 | 6.4243 | 6.7415 | 6.2998 | 4.9897 | 5.3275 | 4.6436 | 5.0817 |
| TP53BP1_S1435 | TP53BP1 | Q12888-2 | 1099 | QPMKPISPVKDPV | 5.9911 | 5.8940 | 5.7352 | 6.4173 | 6.8016 | 5.9715 | 6.8650 | 6.0809 | 6.4741 | 5.5027 | 5.9614 | 5.6291 | 6.7541 | 7.1287 | 6.3959 | 6.3976 |
| TP53BP1_S1467 | TP53BP1 | Q12888-2 | 1431 | LGIEDISPNLSPD | 6.0324 | 6.2806 | 6.4371 | 6.3293 | 6.5314 | 5.8498 | 5.9684 | 5.9303 | 6.4958 | 6.3545 | 6.1126 | 6.3526 | 5.9830 | 6.5931 | 6.8953 | 6.7537 |
| TP53BP1_S1467;S1486 | TP53BP1 | Q12888-2 | 1764 | DGPTGSSEEEEEF | 5.7712 | 6.3042 | 6.0201 | 6.2457 | 6.0752 | 6.1362 | 6.0472 | 6.0266 | 6.2897 | 6.4333 | 6.0488 | 5.8977 | 6.8672 | 6.4528 | 6.7353 | 6.6488 |
| TP53BP1_S1486 | TP53BP1 | Q12888-2 | 697 | SVPLHLSLTETQS | 6.3962 | 7.2048 | 6.7948 | 7.4676 | 6.5719 | 7.0583 | 6.9373 | 6.4922 | 6.9598 | 6.4648 | 6.8243 | 6.8574 | 4.0430 | 4.7400 | 4.3897 | 4.7979 |
| TP53BP1_S162 | TP53BP1 | Q12888-2 | 836 | PVEQDSSQPSLPL | 6.4800 | 6.2446 | 6.5178 | 6.5532 | 6.2022 | 6.2453 | 6.3115 | 5.8474 | 6.4462 | 6.4369 | 6.1898 | 6.1382 | 6.1354 | 6.7294 | 5.9749 | 5.5472 |
| TP53BP1_S1623 | TP53BP1 | Q12888-2 | 635 | GSQAVPSPATRSE | 6.1374 | 6.0887 | 6.3901 | 6.2180 | 6.1530 | 6.1095 | 5.9363 | 6.3379 | 6.5300 | 6.5111 | 6.5682 | 6.3189 | 6.0779 | 6.2417 | 6.0759 | 6.3051 |
| TP53BP1_S1661 | TP53BP1 | Q12888-2 | 400 | DKPMDTSVLSEEG | 6.0981 | 6.3241 | 6.5255 | 6.6686 | 6.6170 | 6.3926 | 6.2892 | 6.1826 | 6.4702 | 6.5823 | 6.5604 | 6.2121 | 5.7004 | 5.8976 | 5.5781 | 5.9012 |
| TP53BP1_S1665 | TP53BP1 | Q12888-2 | 1367 | GGPGKLSPRKGVS | 5.7798 | 5.9200 | 6.0670 | 6.5574 | 6.5050 | 6.4817 | 6.3964 | 5.8770 | 5.9911 | 6.2000 | 6.1418 | 6.3828 | 6.6107 | 6.6678 | 6.2741 | 6.1474 |
| TP53BP1_S1670 | TP53BP1 | Q12888-2 | 1073 | LLHFPSSQGEEEK | 6.2349 | 6.3240 | 6.5008 | 7.1026 | 5.8917 | 5.9857 | 6.1953 | 5.8906 | 6.3648 | 6.0678 | 6.2761 | 6.3209 | 6.2264 | 6.4675 | 6.0730 | 6.0779 |
| TP53BP1_S1678 | TP53BP1 | Q12888-2 | 371 | SDLVAPSPDAFRS | 5.8868 | 5.8562 | 6.0395 | 6.3785 | 6.1987 | 5.8412 | 6.1809 | 6.2995 | 6.0759 | 6.1604 | 6.0679 | 6.1395 | 6.6533 | 6.7459 | 6.5377 | 6.9380 |
| TP53BP1_S1683 | TP53BP1 | Q12888-2 | 326 | SSDGCSTPSREEG | 5.9736 | 6.3648 | 5.7209 | 6.1647 | 5.9581 | 5.3340 | 5.5120 | 5.5676 | 6.0977 | 6.3207 | 5.8053 | 5.8669 | 7.6258 | 7.2166 | 7.3285 | 7.1427 |
| TP53BP1_S1706 | TP53BP1 | Q12888-2 | 124 | QPNRTSSVLGMSV | 5.5981 | 5.5761 | 5.3874 | 5.8285 | 6.0243 | 5.6396 | 6.0286 | 5.7609 | 5.5176 | 5.6152 | 5.5749 | 5.9742 | 8.3885 | 7.6564 | 8.1144 | 7.3154 |
| TP53BP1_S1763 | TP53BP1 | Q12888-2 | 1072 | NLLHFPSSQGEEE | 6.0536 | 6.6168 | 6.2941 | 6.9911 | 6.2718 | 6.2525 | 6.2504 | 6.2181 | 6.3711 | 6.5639 | 6.3701 | 6.1209 | 5.8661 | 6.2638 | 5.8172 | 5.6784 |
| TP53BP1_S1763;S1764 | TP53BP1 | Q12888-2 | 213 | SGYTRLSDVDANT | 5.7464 | 6.0518 | 6.0212 | 6.7319 | 5.6267 | 5.8034 | 6.1745 | 6.6395 | 6.1756 | 6.3119 | 6.5048 | 6.6334 | 6.1106 | 6.4021 | 6.3579 | 6.7084 |
| TP53BP1_S1764 | TP53BP1 | Q12888-2 | 1665 | TESPRASMGVLSG | 6.3422 | 6.9255 | 6.1029 | 6.8686 | 7.4803 | 5.5860 | 6.5336 | 6.6668 | 7.1070 | 6.5308 | 6.4413 | 6.5205 | 5.6757 | 6.0365 | 5.1131 | 5.8692 |
| TP53BP1_S181 | TP53BP1 | Q12888-2 | 596 | KTKGDDTDTRDDI | 6.1968 | 6.7255 | 6.8537 | 7.5022 | 6.2283 | 6.8115 | 7.1799 | 6.7103 | 6.5662 | 6.9734 | 6.9158 | 7.0022 | 4.2680 | 4.7825 | 4.3397 | 4.9439 |
| TP53BP1_S202 | TP53BP1 | Q12888-2 | 927 | PPLTGATPPLIGH | 6.5476 | 7.4435 | 6.6839 | 7.5475 | 6.5129 | 6.9018 | 7.0808 | 6.9084 | 6.8585 | 7.3357 | 7.2781 | 7.1592 | 3.7387 | 4.5531 | 3.4444 | 4.0059 |
| TP53BP1_S213 | TP53BP1 | Q12888-2 | 645 | RSEALSSVLDQEE | 6.5515 | 5.8660 | 6.1427 | 6.4550 | 6.1057 | 5.8904 | 5.8711 | 5.7538 | 6.3636 | 6.4291 | 5.9787 | 6.2966 | 6.4266 | 6.8851 | 6.5017 | 6.4824 |
| TP53BP1_S227 | TP53BP1 | Q12888-2 | 1653 | SSSSSTTPTRKIT | 6.9156 | 6.4072 | 6.6236 | 7.5815 | 6.6368 | 6.7514 | 6.4370 | 6.5906 | 6.4310 | 6.1440 | 6.4643 | 6.5508 | 5.5167 | 5.5202 | 4.9606 | 5.1688 |
| TP53BP1_S270 | TP53BP1 | Q12888-2 | 181 | FGVLELSQSQDVE | 6.2996 | 6.4413 | 6.4448 | 6.6158 | 6.6977 | 6.3797 | 6.0265 | 6.3819 | 6.9890 | 6.6971 | 6.6008 | 6.2891 | 5.9311 | 6.1104 | 6.4973 | 5.1480 |
| TP53BP1_S299 | TP53BP1 | Q12888-2 | 227 | IKHEEQSNEDIPI | 6.8888 | 7.3299 | 7.4071 | 7.2427 | 5.7534 | 6.4384 | 6.7637 | 5.5325 | 5.2830 | 5.4674 | 6.3990 | 6.4670 | 5.9400 | 6.0916 | 6.1688 | 4.8268 |
| TP53BP1_S321 | TP53BP1 | Q12888-2 | 571 | VPAENDSILMNPA | 6.8266 | 7.1493 | 6.4885 | 7.3639 | 7.2064 | 6.2096 | 6.5283 | 6.3806 | 6.5386 | 5.7034 | 6.5182 | 5.8496 | 5.8467 | 5.9443 | 5.6942 | 5.6518 |
| TP53BP1_S321;T326 | TP53BP1 | Q12888-2 | 1033 | VAESVASPQKTMS | 6.2307 | 6.1155 | 6.4898 | 6.2444 | 6.7791 | 5.8179 | 6.4129 | 5.9817 | 6.7099 | 6.5442 | 6.0358 | 5.9539 | 6.8039 | 6.0599 | 5.7585 | 6.0617 |
| TP53BP1_S371 | TP53BP1 | Q12888-2 | 1706 | GAGEFVSPCESGD | 6.0454 | 6.8693 | 6.3027 | 5.9007 | 5.8321 | 6.1515 | 6.0753 | 5.9241 | 6.3038 | 6.6097 | 6.6226 | 6.6941 | 5.7085 | 5.9763 | 5.9985 | 6.9853 |
| TP53BP1_S385 | TP53BP1 | Q12888-2 | 1763 | PDGPTGSSEEEEE | 6.2682 | 6.1723 | 5.5746 | 6.4086 | 6.1056 | 5.8610 | 5.5433 | 5.6905 | 6.2129 | 6.5057 | 6.1982 | 6.0827 | 6.8208 | 6.8238 | 7.0891 | 6.6427 |
| TP53BP1_S400 | TP53BP1 | Q12888-2 | 1678 | KRKLITSEEERSP | 6.1383 | 6.3856 | 6.0094 | 6.4801 | 5.9175 | 6.4602 | 6.3001 | 6.4958 | 5.2595 | 5.6645 | 5.6957 | 5.3268 | 6.6132 | 7.0674 | 7.1636 | 7.0222 |
| TP53BP1_S400;S403 | TP53BP1 | Q12888-2 | 162 | SGNTTHSLGAEDT | 5.2181 | 5.5418 | 5.5776 | 6.3443 | 5.9768 | 5.5497 | 5.4463 | 6.6719 | 5.7222 | 6.4265 | 5.9795 | 5.8614 | 6.8191 | 7.4531 | 7.4439 | 7.9678 |
| TP53BP1_S403 | TP53BP1 | Q12888-2 | 202 | DKEQLQSVTTNSG | 5.7506 | 6.7731 | 6.1253 | 6.4627 | 6.7228 | 6.7939 | 6.8393 | 6.3878 | 5.8638 | 6.1844 | 6.3338 | 6.3044 | 5.7469 | 6.2113 | 5.7920 | 5.7077 |
| TP53BP1_S505 | TP53BP1 | Q12888-2 | 1557 | ALSEDEYFSAGVV | 6.2236 | 6.5676 | 5.5321 | 6.6483 | 5.7846 | 6.5036 | 6.1316 | 5.8916 | 6.5889 | 5.8649 | 6.2068 | 6.2286 | 6.5822 | 6.4669 | 6.8748 | 6.8138 |
| TP53BP1_S523 | TP53BP1 | Q12888-2 | 1623 | TKAADISLDNLVE | 6.5490 | 6.8480 | 7.1471 | 7.2249 | 6.5376 | 7.1809 | 7.1233 | 6.9941 | 7.0829 | 7.0202 | 7.6809 | 7.4211 | 3.3452 | 4.7240 | 3.2954 | 3.8254 |
| TP53BP1_S528;S530 | TP53BP1 | Q12888-2 | 980 | GSGKGDSGAAPDV | 6.6882 | 7.2947 | 7.0226 | 8.0259 | 5.7721 | 6.5974 | 6.7174 | 7.1481 | 6.6910 | 7.4992 | 7.2214 | 7.8673 | 3.5260 | 4.2824 | 4.0692 | 4.4772 |
| TP53BP1_S530 | TP53BP1 | Q12888-2 | 776 | SPRVDVSCEPLEG | 6.3914 | 6.8835 | 7.4209 | 7.5680 | 5.8449 | 6.6782 | 6.7110 | 6.5578 | 6.7546 | 6.8903 | 7.1500 | 7.1907 | 4.3633 | 4.6320 | 4.1217 | 4.8538 |
| TP53BP1_S557 | TP53BP1 | Q12888-2 | 68 | PVLDVVSNPEQTA | 5.8742 | 6.5740 | 5.8977 | 6.4865 | 6.3024 | 6.1914 | 6.2205 | 6.3571 | 6.1786 | 6.2512 | 6.3336 | 6.5596 | 6.3878 | 6.4783 | 6.1438 | 6.3632 |
| TP53BP1_S571 | TP53BP1 | Q12888-2 | 1007 | TEASEESLQFNLE | 6.7202 | 6.9377 | 6.9890 | 7.8779 | 6.1247 | 5.9441 | 6.8819 | 7.1179 | 7.3009 | 7.8467 | 7.1934 | 6.8174 | 3.8581 | 4.5184 | 3.9480 | 3.9238 |
| TP53BP1_S585 | TP53BP1 | Q12888-2 | 1486 | DGLDASSPGNSFV | 6.5104 | 6.3069 | 6.0264 | 6.8420 | 6.1877 | 6.1214 | 6.4718 | 6.4306 | 5.8704 | 6.1115 | 6.6815 | 5.7154 | 5.9278 | 6.3284 | 6.8169 | 6.5509 |
| TP53BP1_S635 | TP53BP1 | Q12888-2 | 523 | SCKLMLSTSEYSQ | 7.6717 | 6.5231 | 6.4904 | 7.2797 | 10.2845 | 6.8442 | 6.5938 | 7.0275 | 7.8767 | 4.4854 | 4.7358 | 5.4743 | 4.8919 | 4.7957 | 4.5816 | 4.4437 |
| TP53BP1_S644;S645 | TP53BP1 | Q12888-2 | 1377 | PVSQTGTPVCEED | 6.4223 | 7.3147 | 6.6394 | 6.5463 | 6.0340 | 6.4576 | 6.1670 | 6.4509 | 6.4028 | 6.9091 | 6.8452 | 6.5628 | 4.4814 | 5.2551 | 5.3850 | 6.1264 |
| TP53BP1_S645 | TP53BP1 | Q12888-2 | 1661 | TRKITESPRASMG | 7.1438 | 9.4280 | 8.0383 | 7.4107 | 6.0502 | 7.9537 | 6.4307 | 6.4569 | 7.0532 | 6.2779 | 5.7585 | 5.0641 | 4.3860 | 4.3882 | 4.6331 | 3.5268 |
| TP53BP1_S664;S665;S66 | TP53BP1 | Q12888-2 | 321 | SNKTVSSDGCSTP | 5.9323 | 6.3183 | 5.5713 | 6.0783 | 6.1199 | 5.8393 | 5.6503 | 6.4413 | 6.0820 | 5.4973 | 5.7698 | 5.7024 | 7.2984 | 7.3301 | 7.4076 | 6.9612 |
| TP53BP1_S68 | TP53BP1 | Q12888-2 | 789 | VEKCSDSQSWEDI | 5.8345 | 6.2915 | 6.5420 | 6.4886 | 6.8590 | 6.3533 | 6.8299 | 5.7417 | 6.5915 | 7.0345 | 6.1925 | 6.4531 | 6.1152 | 5.7788 | 5.2183 | 6.6757 |
| TP53BP1_S697 | TP53BP1 | Q12888-2 | 867 | QEKTSNSLTEDSK | 6.5718 | 7.1234 | 6.9027 | 7.7298 | 6.4111 | 6.6277 | 7.4096 | 6.6035 | 6.8300 | 7.2353 | 6.9310 | 7.2056 | 3.6811 | 4.4064 | 3.9552 | 4.3757 |
| TP53BP1_S732 | TP53BP1 | Q12888-2 | 1761 | KLPDGPTGSSEEE | 6.2890 | 5.8654 | 6.6343 | 6.8824 | 5.8822 | 6.1310 | 5.7241 | 6.0195 | 6.4128 | 5.9340 | 5.8194 | 6.5015 | 6.5430 | 6.8902 | 6.1906 | 6.2806 |
| TP53BP1_S776 | TP53BP1 | Q12888-2 | 881 | ANAKQLSSDAEAQ | 6.8375 | 7.0573 | 6.4164 | 7.4629 | 6.0606 | 6.2888 | 6.6524 | 6.8210 | 6.5100 | 6.5700 | 6.5496 | 7.0622 | 4.6933 | 5.1150 | 4.9846 | 4.9184 |
| TP53BP1_S789 | TP53BP1 | Q12888-2 | 1295 | ETEVSPSQTGGSS | 6.1329 | 5.7491 | 5.9793 | 6.5682 | 5.8659 | 6.2088 | 6.3657 | 5.9219 | 6.0116 | 6.1916 | 6.1963 | 6.4033 | 6.1033 | 6.7713 | 6.8887 | 6.6420 |
| TP53BP1_S814 | TP53BP1 | Q12888-2 | 1099;1106 | NA | 5.8933 | 5.8399 | 6.0141 | 6.5147 | 6.9122 | 6.8850 | 6.3683 | 6.2685 | 6.4117 | 6.2193 | 6.9774 | 6.4747 | 6.6467 | 6.5375 | 5.2788 | 4.7578 |
| TP53BP1_S836 | TP53BP1 | Q12888-2 | 528;530 | NA | 5.8645 | 6.2088 | 6.8759 | 6.8461 | 5.9604 | 5.8386 | 6.1837 | 5.7755 | 6.2608 | 6.4666 | 6.5930 | 6.5643 | 5.8080 | 6.2456 | 5.9429 | 6.5654 |
| TP53BP1_S867 | TP53BP1 | Q12888-2 | 321;326 | NA | 5.8162 | 6.0306 | 6.3365 | 6.6794 | 6.1640 | 6.8020 | 6.0469 | 5.9155 | 6.9042 | 5.7413 | 6.0801 | 5.9112 | 6.8813 | 6.9533 | 6.3449 | 6.3926 |
| TP53BP1_S881 | TP53BP1 | Q12888-2 | 1431;1435 | NA | 6.2283 | 6.4394 | 5.8566 | 6.5709 | 6.2633 | 5.7337 | 5.7433 | 6.0933 | 6.5275 | 6.5156 | 6.2240 | 5.9953 | 6.3579 | 6.8577 | 6.2547 | 6.3385 |
| TP53BP1_S980 | TP53BP1 | Q12888-2 | 1221;1224 | NA | 5.9000 | 5.7670 | 6.1622 | 6.9954 | 5.9439 | 6.0267 | 5.7887 | 5.8324 | 6.1045 | 6.0420 | 6.0243 | 6.2687 | 7.2482 | 7.2900 | 6.4132 | 6.1928 |
| TP53BP1_T1061 | TP53BP1 | Q12888-2 | 1763;1764 | NA | 5.7394 | 6.2700 | 6.0930 | 6.9042 | 6.4922 | 6.2961 | 6.3309 | 6.3577 | 6.2134 | 6.3301 | 6.4498 | 6.3231 | 6.2098 | 6.6120 | 5.5034 | 5.8749 |
| TP53BP1_T1377 | TP53BP1 | Q12888-2 | 1106;1109 | NA | 6.6427 | 6.7455 | 6.6395 | 7.1979 | 6.5528 | 6.4659 | 6.1021 | 5.5114 | 6.4613 | 6.4400 | 6.1215 | 6.8793 | 5.7069 | 5.9095 | 6.0671 | 6.4565 |
| TP53BP1_T1614 | TP53BP1 | Q12888-2 | 664;665;667 | NA | 5.8852 | 6.4547 | 6.9191 | 6.3447 | 5.8716 | 6.1386 | 6.4342 | 6.0995 | 5.8751 | 6.4916 | 6.7707 | 6.2967 | 6.1213 | 5.9824 | 5.7043 | 6.6103 |
| TP53BP1_T1653 | TP53BP1 | Q12888-2 | 400;403 | NA | 5.8957 | 6.4559 | 6.5014 | 7.0954 | 6.5567 | 7.0968 | 7.8078 | 7.0145 | 6.4846 | 6.7266 | 7.8776 | 7.2757 | 4.1920 | 4.9370 | 3.8729 | 4.2093 |
| TP53BP1_T1761 | TP53BP1 | Q12888-2 | 1761;1764 | NA | 6.1544 | 6.0213 | 5.5973 | 6.1000 | 6.3857 | 5.9798 | 6.0677 | 6.3537 | 6.6949 | 6.7947 | 6.6638 | 6.5071 | 6.2372 | 6.6574 | 6.1203 | 5.6647 |
| TP53BP1_T1761;S1764 | TP53BP1 | Q12888-2 | 1004;1007 | NA | 6.9516 | 7.7243 | 6.4105 | 7.0924 | 5.7560 | 6.5019 | 8.5279 | 6.1298 | 6.2308 | 8.3577 | 8.1392 | 6.9243 | 4.0915 | 4.3067 | 3.2210 | 3.6343 |
| TP53BP1_T326 | TP53BP1 | Q12888-2 | 1467;1486 | NA | 5.7766 | 5.9465 | 6.6617 | 6.2426 | 6.5445 | 6.0423 | 6.5786 | 5.7885 | 6.4416 | 6.4843 | 6.7860 | 6.1791 | 6.0323 | 6.3891 | 6.5588 | 6.2476 |
| TP53BP1_T596 | TP53BP1 | Q12888-2 | 1213;1221 | NA | 5.2460 | 6.5672 | 5.6116 | 6.9660 | 7.3620 | 6.1718 | 5.4288 | 6.5555 | 5.8848 | 6.4382 | 7.6038 | 6.7180 | 5.7946 | 6.5413 | 5.4224 | 5.6879 |
| TP53BP1_T927 | TP53BP1 | Q12888-2 | 1072;1073 | NA | 5.4967 | 6.0888 | 7.5499 | 7.2254 | 6.0398 | 5.5858 | 7.5263 | 6.6834 | 7.1044 | 6.1379 | 5.9101 | 5.7986 | 5.9289 | 5.7534 | 4.8954 | 6.2751 |
| TP53BP1_Y1557 | TP53BP1 | Q12888-2 | 644;645 | NA | 5.7845 | 6.2268 | 6.1012 | 6.1221 | 5.4631 | 6.1283 | 5.5163 | 5.6247 | 5.3816 | 5.9200 | 6.9728 | 6.3384 | 7.3311 | 6.8669 | 6.5920 | 7.6303 |
| TP53BP2_S420 | TP53BP2 | Q13625-3 | 743 | PLKKRSSITEPEG | 6.8939 | 6.4057 | 6.6387 | 6.2167 | 6.3602 | 6.5490 | 6.5430 | 5.6660 | 6.7219 | 6.1954 | 6.3796 | 6.5728 | 6.2769 | 6.2297 | 4.8212 | 5.5292 |
| TP53BP2_S562 | TP53BP2 | Q13625-3 | 704 | RIPRPLSPTKLLP | 6.2906 | 6.7027 | 6.2790 | 6.4078 | 5.8198 | 6.1307 | 6.0541 | 6.4980 | 6.2935 | 6.7853 | 6.4935 | 6.7523 | 5.7906 | 6.0282 | 5.8091 | 5.8648 |
| TP53BP2_S562;S575 | TP53BP2 | Q13625-3 | 420 | PVGPDWSPSNADL | 6.1689 | 5.9678 | 6.6232 | 6.2626 | 7.1095 | 6.6590 | 6.6348 | 6.7615 | 6.4260 | 7.0824 | 6.5739 | 6.3612 | 4.5872 | 5.8711 | 4.9459 | 5.9649 |
| TP53BP2_S575 | TP53BP2 | Q13625-3 | 782 | YPSKSASVTASSE | 5.2107 | 5.7720 | 6.2034 | 6.4038 | 5.9593 | 6.4238 | 6.2039 | 6.5853 | 5.9162 | 5.3012 | 6.3554 | 6.3202 | 6.2776 | 7.5002 | 7.1414 | 6.4254 |
| TP53BP2_S582 | TP53BP2 | Q13625-3 | 582 | PGSKQESPPAAAV | 6.6269 | 6.6950 | 6.8139 | 7.3493 | 6.4099 | 6.4848 | 7.0349 | 6.6985 | 6.2927 | 6.1775 | 6.4281 | 6.8503 | 5.1074 | 5.0583 | 4.8061 | 5.1665 |
| TP53BP2_S679 | TP53BP2 | Q13625-3 | 679 | SQGKPGSPEPETE | 5.9618 | 5.4684 | 6.5804 | 6.1332 | 6.7062 | 7.1924 | 6.3594 | 6.0629 | 6.5406 | 5.6953 | 7.2628 | 7.0177 | 6.2134 | 6.0296 | 5.4385 | 5.3374 |
| TP53BP2_S704 | TP53BP2 | Q13625-3 | 562 | QPRVLLSPSIPSV | 6.2470 | 6.0975 | 6.1955 | 6.1157 | 6.0347 | 5.9727 | 6.0381 | 6.3298 | 6.1365 | 6.0797 | 7.1387 | 6.1044 | 6.5149 | 6.3259 | 6.8777 | 6.7910 |
| TP53BP2_S743 | TP53BP2 | Q13625-3 | 575 | GQDQTLSPGSKQE | 6.1672 | 6.7472 | 7.1009 | 6.2984 | 5.6358 | 6.3870 | 6.8684 | 6.6134 | 7.2121 | 5.9495 | 6.5335 | 6.5741 | 5.3035 | 5.4413 | 5.7850 | 5.3827 |
| TP53BP2_S782 | TP53BP2 | Q13625-3 | 562;575 | NA | 5.9520 | 6.7029 | 6.5417 | 6.6607 | 6.8983 | 7.0252 | 7.2264 | 6.7254 | 6.8986 | 7.7266 | 7.0310 | 7.2185 | 4.4379 | 4.6930 | 4.0474 | 4.2144 |
| TP53RK_T8 | TP53RK | Q96S44 | 8 | AAARATTPADGEE | 5.7000 | 6.0360 | 5.6220 | 6.2151 | 5.6804 | 6.6605 | 6.3225 | 5.9002 | 6.2259 | 6.2156 | 5.8089 | 6.6585 | 6.9584 | 7.1540 | 6.3699 | 6.4721 |
| TPCN1_S59 | TPCN1 | Q9ULQ1-3 | 59 | KPETISSPGYHIL | 5.8366 | 7.6787 | 6.8639 | 7.8447 | 6.0443 | 6.7321 | 7.0859 | 6.8383 | 6.8373 | 6.5338 | 6.8990 | 6.9929 | 3.6321 | 4.5940 | 4.5551 | 5.0313 |
| TPD52L2_S161 | TPD52L2 | O43399 | 21 | KGLLSDSMTDVPV | 6.0334 | 6.2965 | 6.6409 | 6.6607 | 6.4636 | 6.1340 | 6.0161 | 6.3506 | 6.9070 | 6.6577 | 6.5571 | 6.3597 | 5.6911 | 5.9271 | 5.4326 | 5.8718 |
| TPD52L2_S21 | TPD52L2 | O43399 | 161 | LGDMRNSATFKSF | 7.3919 | 7.1206 | 7.9501 | 6.2824 | 6.7184 | 7.3584 | 5.8263 | 5.2033 | 7.1962 | 7.8336 | 9.6427 | 8.4940 | 3.5268 | 3.7759 | 2.5803 | 3.0990 |
| TPI1_S117 | TPI1 | P60174-3 | 58 | MNGRKQSLGELIG | 5.7187 | 5.7047 | 6.3086 | 5.3915 | 6.1362 | 6.1610 | 5.5969 | 6.3091 | 6.8793 | 7.2774 | 7.0311 | 7.1567 | 5.9338 | 6.5248 | 5.9450 | 5.9252 |
| TPI1_S58 | TPI1 | P60174-3 | 117 | AFTGEISPGMIKD | 5.8090 | 6.2104 | 7.2517 | 5.7229 | 6.5785 | 6.4803 | 6.3138 | 7.4947 | 6.6771 | 6.4914 | 6.5207 | 6.4327 | 5.8576 | 5.9962 | 4.9246 | 5.2383 |
| TPM1_S252 | TPM1 | P09493-9 | 269 | AKEENLSMHQMLD | 7.4120 | 7.3731 | 6.7049 | 6.5656 | 7.0594 | 6.4655 | 7.0512 | 6.5615 | 7.3843 | 6.8506 | 7.1210 | 6.2855 | 3.7573 | 3.5551 | 4.0087 | 4.0442 |
| TPM1_S269 | TPM1 | P09493-9 | 252 | VTKLEKSIDDLEE | 7.4074 | 8.3393 | 7.7915 | 7.9920 | 5.9584 | 6.6086 | 6.4707 | 6.0111 | 7.7413 | 7.8108 | 8.5190 | 7.6741 | 2.7006 | 3.2631 | 2.5449 | 3.1673 |
| TPM1_S87 | TPM1 | P09493-9 | 87 | AEADVASLNRRIQ | 7.4601 | 6.6580 | 6.9631 | 6.9895 | 6.4494 | 6.3637 | 6.1288 | 6.3876 | 6.5929 | 6.3599 | 7.3629 | 6.4494 | 5.8365 | 5.2140 | 4.4009 | 4.9835 |
| TPM2_T252 | TPM2 | P07951-2 | 252 | VAKLEKTIDDLEE | 7.4300 | 7.7927 | 6.9519 | 7.3741 | 6.5760 | 7.2577 | 7.1264 | 6.5568 | 7.7215 | 7.4080 | 7.7422 | 7.0441 | 3.1667 | 3.5483 | 2.9087 | 3.3949 |
| TPR_S1185 | TPR | P12270 | 379 | EELAAMSPTAAAV | 6.5985 | 6.5497 | 6.2368 | 6.6376 | 6.3771 | 6.2591 | 6.4784 | 6.2482 | 6.7923 | 6.5101 | 6.5091 | 6.4417 | 5.7546 | 5.6980 | 5.4239 | 5.4848 |
| TPR_S2155 | TPR | P12270 | 2116 | GRGLQLTPGIGGM | 5.9019 | 5.0771 | 4.9893 | 5.4545 | 5.1443 | 6.2673 | 5.9295 | 6.3504 | 6.0828 | 4.7869 | 4.2787 | 4.3186 | 8.4746 | 8.8343 | 8.6822 | 9.4276 |
| TPR_S379 | TPR | P12270 | 1677 | PTPVVSTPSKVTA | 7.3871 | 6.4402 | 6.4677 | 6.8560 | 6.9789 | 6.3927 | 6.6342 | 6.0589 | 7.0318 | 6.5274 | 6.3689 | 6.5161 | 5.2102 | 5.3365 | 4.4304 | 5.3632 |
| TPR_S646 | TPR | P12270 | 2137 | DRTVPSTPTLVVP | 7.1297 | 5.6930 | 5.6853 | 6.2240 | 5.5877 | 5.7765 | 6.0939 | 5.9952 | 7.0781 | 5.6136 | 4.9510 | 5.2904 | 7.9540 | 7.5483 | 6.1210 | 7.2585 |
| TPR_T1677 | TPR | P12270 | 1185 | QGPLNVSLSEEGK | 6.4923 | 6.6114 | 6.0358 | 7.1806 | 6.5578 | 6.8427 | 6.7579 | 6.9667 | 6.7470 | 6.5910 | 7.0981 | 6.6217 | 4.6245 | 4.9199 | 4.7320 | 5.2206 |
| TPR_T2116 | TPR | P12270 | 2155 | FAEAIHSPQVAGV | 5.3300 | 4.9967 | 5.2160 | 5.9236 | 5.1641 | 5.9086 | 6.3925 | 6.0973 | 6.2278 | 5.9050 | 5.1881 | 5.1923 | 8.5688 | 7.9787 | 7.3383 | 8.5724 |
| TPR_T2137 | TPR | P12270 | 646 | STPKRPSTSQTVS | 5.3024 | 5.1675 | 5.0194 | 5.5741 | 6.9351 | 6.0432 | 6.0390 | 6.1072 | 5.7538 | 5.8135 | 5.4989 | 6.0326 | 7.9429 | 7.5699 | 7.3799 | 7.8206 |
| TPR_T653 | TPR | P12270 | 653 | TSQTVSTPAPVPV | 7.2391 | 6.9093 | 7.2439 | 7.5451 | 6.3133 | 6.8443 | 6.6580 | 5.9957 | 7.3939 | 6.7650 | 6.8664 | 7.1070 | 4.3309 | 4.6992 | 3.6791 | 4.4099 |
| TPRN_S241 | TPRN | Q4KMQ1 | 418 | RWQRPSSPPPFLP | 6.2935 | 6.3757 | 6.5534 | 7.0136 | 6.4383 | 5.9746 | 6.0919 | 6.4676 | 6.5939 | 6.2003 | 6.5348 | 6.8010 | 5.6274 | 5.9655 | 5.4778 | 5.5907 |
| TPRN_S274 | TPRN | Q4KMQ1 | 362 | GDLGPASPSQELG | 5.5992 | 6.2889 | 6.6043 | 6.6106 | 6.1770 | 6.0958 | 6.3580 | 6.4332 | 6.0507 | 5.7766 | 6.5413 | 6.4384 | 6.2674 | 6.1084 | 6.2061 | 6.4442 |
| TPRN_S362 | TPRN | Q4KMQ1 | 241 | ANRLAGSPPGSGQ | 6.8711 | 7.1618 | 6.7699 | 7.3990 | 6.1663 | 6.1025 | 6.4035 | 6.8413 | 6.9338 | 6.3750 | 6.8095 | 6.9632 | 4.6298 | 4.6283 | 4.9485 | 4.9964 |
| TPRN_S418 | TPRN | Q4KMQ1 | 274 | PSATPASPPASAT | 5.7164 | 6.5131 | 5.5527 | 5.8162 | 6.8951 | 7.6754 | 6.9393 | 6.3083 | 6.6017 | 5.7936 | 6.0468 | 6.1889 | 5.7051 | 5.2573 | 6.2558 | 6.7345 |
| TPT1_S46 | TPT1 | P13693 | 46 | EGNIDDSLIGGNA | 7.1452 | 6.4881 | 7.1365 | 7.1350 | 6.5203 | 6.2063 | 6.7773 | 6.5412 | 6.9900 | 6.5330 | 7.2286 | 7.4508 | 4.3201 | 4.8023 | 4.0256 | 4.6997 |
| TPX2_S121;S125 | TPX2 | Q9ULW0-2 | 405 | ICRDPQTPVLQTK | 7.9615 | 8.0634 | 6.9317 | 6.9045 | 4.8233 | 7.2201 | 8.2788 | 6.8894 | 7.8465 | 7.6061 | 7.7333 | 7.6804 | 2.4779 | 3.6389 | 2.2679 | 3.6762 |
| TPX2_S186 | TPX2 | Q9ULW0-2 | 522 | PITVPKSPAFALK | 7.7013 | 8.0584 | 7.0841 | 6.5935 | 5.0984 | 7.7695 | 8.4525 | 7.2272 | 6.9889 | 7.3558 | 7.6245 | 7.5649 | 2.5358 | 3.7470 | 2.3863 | 3.8118 |
| TPX2_S285 | TPX2 | Q9ULW0-2 | 186 | AEKNASSPEKAKG | 5.9941 | 6.0351 | 5.8901 | 6.4078 | 5.7556 | 5.6821 | 6.1965 | 5.4982 | 6.3667 | 6.2213 | 6.1675 | 6.3995 | 6.7731 | 6.6515 | 6.7040 | 7.2570 |
| TPX2_S293 | TPX2 | Q9ULW0-2 | 774 | PLTVPVSPKFSTR | 6.8618 | 7.0937 | 6.8505 | 7.2197 | 6.6626 | 6.8616 | 7.5748 | 6.8094 | 7.4410 | 7.2697 | 7.2054 | 7.3172 | 3.4815 | 4.1633 | 3.2306 | 3.9571 |
| TPX2_S310 | TPX2 | Q9ULW0-2 | 293 | LRKHPSSPARVTK | 6.8631 | 7.0762 | 6.8633 | 6.5940 | 6.2031 | 6.5022 | 6.4168 | 6.6942 | 6.1307 | 6.0024 | 6.4634 | 6.1646 | 5.4901 | 5.9568 | 5.6743 | 5.9049 |
| TPX2_S522 | TPX2 | Q9ULW0-2 | 147 | KAKRCATPVIIDE | 5.5949 | 5.8969 | 5.5970 | 5.4436 | 5.0798 | 5.9902 | 6.4090 | 5.6037 | 6.2152 | 6.1366 | 6.0465 | 6.2498 | 7.5588 | 7.2524 | 7.2110 | 7.7145 |
| TPX2_S690 | TPX2 | Q9ULW0-2 | 310 | VKPFNLSQGKKRT | 6.9346 | 6.3128 | 6.5210 | 6.7082 | 5.8552 | 6.2743 | 7.4333 | 6.1325 | 7.1703 | 6.9922 | 7.1083 | 6.9733 | 5.0345 | 5.3549 | 4.0499 | 5.1448 |
| TPX2_S774 | TPX2 | Q9ULW0-2 | 72 | LQQAIVTPLKPVD | 7.1552 | 7.9943 | 6.4950 | 7.1346 | 5.7876 | 6.9556 | 7.4669 | 6.8648 | 6.8022 | 7.3884 | 7.4318 | 7.8280 | 3.1899 | 4.4843 | 3.0095 | 4.0118 |
| TPX2_S9 | TPX2 | Q9ULW0-2 | 9 | QVKSSYSYDAPSD | 6.4514 | 6.8880 | 6.7534 | 6.9969 | 6.5649 | 6.1872 | 6.7777 | 5.8734 | 6.8830 | 6.2383 | 6.0426 | 5.7490 | 5.2709 | 6.2275 | 5.6426 | 5.4532 |
| TPX2_T113 | TPX2 | Q9ULW0-2 | 690 | AEGLSGSLVQEPF | 5.8701 | 6.6592 | 6.9388 | 7.7695 | 4.7076 | 7.0675 | 7.7504 | 6.8373 | 6.3650 | 6.5488 | 6.9371 | 6.8594 | 4.4324 | 5.6496 | 4.4699 | 5.1373 |
| TPX2_T147 | TPX2 | Q9ULW0-2 | 113 | AAISRKTPAQPQR | 7.4628 | 8.2077 | 7.3540 | 7.1563 | 4.2421 | 7.6356 | 8.9445 | 6.6197 | 6.8350 | 7.1060 | 7.8995 | 8.2765 | 2.2305 | 3.8266 | 2.3608 | 3.8425 |
| TPX2_T405 | TPX2 | Q9ULW0-2 | 285 | KEVNFTSELRKHP | 6.7031 | 6.9932 | 7.5902 | 9.6273 | 7.3817 | 4.6777 | 6.6591 | 6.1605 | 6.5451 | 4.5831 | 6.9608 | 4.2620 | 5.4197 | 5.8704 | 5.1154 | 5.4508 |
| TPX2_T72 | TPX2 | Q9ULW0-2 | 121;125 | NA | 7.4451 | 7.7746 | 6.7992 | 6.7808 | 6.1356 | 7.4615 | 8.4092 | 7.4275 | 7.5786 | 7.4682 | 7.8836 | 7.0367 | 2.3576 | 3.6721 | 2.4601 | 3.3095 |
| TRA2A_S20 | TRA2A | Q13595 | 202 | ITKRAHTPTPGIY | 6.2778 | 6.2006 | 5.7477 | 6.3187 | 6.2158 | 5.8737 | 5.9320 | 6.0831 | 6.1649 | 6.0994 | 6.0633 | 5.9262 | 7.0352 | 6.7668 | 6.7159 | 6.5789 |
| TRA2A_S20;T24 | TRA2A | Q13595 | 20 | SRSQSKSPTGTPA | 6.3008 | 7.2171 | 6.1682 | 6.6682 | 6.0863 | 5.2424 | 5.6507 | 5.9164 | 6.0050 | 6.6823 | 6.5719 | 5.7904 | 6.2690 | 6.1142 | 6.9152 | 7.1020 |
| TRA2A_S215 | TRA2A | Q13595 | 236 | GGRRRDSYYDRGY | 5.2419 | 5.5973 | 5.6012 | 5.7729 | 7.2466 | 5.7808 | 5.5361 | 5.6684 | 5.7690 | 6.5521 | 5.7083 | 5.6558 | 8.1097 | 6.9225 | 6.9926 | 7.8447 |
| TRA2A_S236 | TRA2A | Q13595 | 215 | MGRPTHSGGGGGG | 5.3203 | 6.6777 | 6.2378 | 8.7457 | 5.7941 | 6.9146 | 6.1776 | 6.3619 | 7.6440 | 6.8770 | 6.0047 | 8.3734 | 3.8088 | 4.8692 | 4.1510 | 6.0423 |
| TRA2A_S84;S86;T88 | TRA2A | Q13595 | 98;100 | NA | 5.5230 | 6.8275 | 6.0334 | 7.1194 | 5.7426 | 6.2885 | 5.3141 | 6.6466 | 6.4043 | 5.7516 | 6.3821 | 6.0004 | 6.2926 | 6.7807 | 6.7050 | 6.1883 |
| TRA2A_S98;S100 | TRA2A | Q13595 | 20;24 | NA | 5.7563 | 8.1637 | 7.3689 | 7.9603 | 7.5802 | 4.8914 | 6.0416 | 5.5805 | 6.0768 | 5.3604 | 4.6158 | 6.1216 | 6.0257 | 5.9059 | 6.0550 | 6.4960 |
| TRA2A_T202 | TRA2A | Q13595 | 84;86;88 | NA | 5.8802 | 7.3828 | 5.2537 | 6.2781 | 6.8012 | 5.9566 | 5.9731 | 4.7141 | 8.0542 | 5.7039 | 6.1006 | 5.5264 | 7.9340 | 6.1439 | 6.8596 | 6.4376 |
| TRA2B_S20;S26 | TRA2B | P62995 | 201 | ITKRPHTPTPGIY | 5.8801 | 6.0272 | 5.5782 | 6.3880 | 6.1303 | 5.9622 | 5.9530 | 6.2264 | 6.0205 | 5.7967 | 5.7088 | 5.6869 | 7.2382 | 7.0883 | 7.1996 | 7.1158 |
| TRA2B_S215 | TRA2B | P62995 | 264 | QIYRRRSPSPYYS | 5.7848 | 6.1591 | 6.0404 | 5.8503 | 5.8194 | 6.0791 | 5.6914 | 7.0194 | 5.9833 | 6.8410 | 7.0998 | 6.3949 | 6.2252 | 5.9936 | 7.2329 | 5.7855 |
| TRA2B_S22 | TRA2B | P62995 | 22 | SASRSGSAHGSGK | 6.5255 | 7.1306 | 5.8767 | 7.6336 | 6.2740 | 6.2717 | 5.7570 | 6.7036 | 6.0542 | 5.9306 | 6.8112 | 6.4313 | 5.3340 | 5.6857 | 5.8441 | 5.7361 |
| TRA2B_S22;S26 | TRA2B | P62995 | 99 | RHSHSHSPMSTRR | 6.6028 | 11.0769 | 6.5906 | 5.8878 | 5.6124 | 5.6974 | 4.5733 | 4.8772 | 6.6190 | 6.5805 | 6.9532 | 6.0883 | 4.9237 | 5.8878 | 6.3820 | 5.6468 |
| TRA2B_S264 | TRA2B | P62995 | 266 | YRRRSPSPYYSRG | 5.7579 | 6.2879 | 5.9965 | 5.5442 | 5.5989 | 5.8433 | 5.4361 | 7.0033 | 5.6854 | 7.4288 | 7.2223 | 6.5574 | 6.1852 | 5.8317 | 7.6747 | 5.5465 |
| TRA2B_S264;S266 | TRA2B | P62995 | 215 | GRPTYGSSRRRDY | 8.6154 | 7.9688 | 7.3216 | 8.1151 | 6.5231 | 6.4954 | 7.6620 | 6.4915 | 7.5596 | 7.4558 | 7.7513 | 8.1290 | 2.0264 | 3.2816 | 1.8560 | 2.7473 |
| TRA2B_S264;S270 | TRA2B | P62995 | 264;266 | NA | 6.4938 | 6.2728 | 5.9716 | 6.4368 | 6.2618 | 6.0304 | 6.1791 | 5.8242 | 6.0877 | 5.9936 | 5.9978 | 6.2767 | 6.7948 | 6.5959 | 6.3061 | 6.4770 |
| TRA2B_S266 | TRA2B | P62995 | 95;97;99 | NA | 6.3520 | 6.6613 | 5.9116 | 6.9542 | 6.3725 | 5.7873 | 5.8106 | 6.3126 | 6.0715 | 5.9775 | 6.1885 | 6.2195 | 6.5604 | 6.1324 | 6.3060 | 6.3820 |
| TRA2B_S266;S270 | TRA2B | P62995 | 264;270 | NA | 6.5512 | 6.2817 | 6.1126 | 6.4096 | 6.2035 | 6.1244 | 6.0264 | 5.8701 | 6.1750 | 5.8991 | 6.0664 | 6.1615 | 6.7247 | 6.4752 | 6.2920 | 6.6266 |
| TRA2B_S95;S97;S99 | TRA2B | P62995 | 95;99 | NA | 7.1689 | 7.0571 | 5.7445 | 7.5480 | 6.0583 | 6.3245 | 5.3704 | 6.3692 | 6.6734 | 6.6272 | 6.9123 | 6.6762 | 5.4105 | 5.7388 | 5.2390 | 5.0818 |
| TRA2B_S95;S99 | TRA2B | P62995 | 22;26 | NA | 5.4114 | 5.6379 | 5.8740 | 6.4770 | 6.2646 | 5.4313 | 6.0168 | 6.1168 | 6.2206 | 6.0751 | 6.1859 | 6.4976 | 6.8727 | 6.5965 | 7.0613 | 7.2605 |
| TRA2B_S97;S99 | TRA2B | P62995 | 97;99 | NA | 6.6856 | 8.0315 | 6.8694 | 6.5928 | 5.9514 | 4.4144 | 6.7746 | 5.1262 | 6.4170 | 6.6660 | 5.9886 | 6.0424 | 7.2488 | 5.2568 | 4.7990 | 7.1357 |
| TRA2B_S99 | TRA2B | P62995 | 266;270 | NA | 6.3886 | 6.6980 | 5.8801 | 6.2993 | 6.3590 | 5.8804 | 5.7960 | 6.4268 | 6.3093 | 6.0624 | 5.6473 | 5.7034 | 6.8283 | 6.3248 | 6.5996 | 6.7968 |
| TRA2B_T201 | TRA2B | P62995 | 20;26 | NA | 5.8022 | 6.5140 | 5.6965 | 6.4258 | 6.0163 | 5.3582 | 6.0838 | 5.3281 | 6.3717 | 6.0090 | 5.9465 | 6.3405 | 7.3367 | 6.5477 | 6.8562 | 7.3670 |
| TRAF1_S172 | TRAF1 | Q13077 | 66 | EDLQSISPGSRLR | 6.4275 | 6.4295 | 6.7914 | 6.4798 | 6.1694 | 6.5521 | 6.4378 | 6.5075 | 6.9101 | 6.6430 | 6.6209 | 6.9526 | 5.0690 | 5.4466 | 5.0272 | 5.5356 |
| TRAF1_S66 | TRAF1 | Q13077 | 172 | RAPCSESQEELAL | 6.1481 | 6.4807 | 6.3356 | 6.3522 | 6.2435 | 6.8392 | 5.5827 | 5.6015 | 7.0742 | 6.6556 | 7.2392 | 6.6947 | 5.5617 | 6.3252 | 5.2396 | 5.6263 |
| TRAF3_S9 | TRAF3 | Q13114 | 9 | SSKKMDSPGALQT | 6.6740 | 7.0219 | 7.4138 | 6.6541 | 6.0151 | 5.8847 | 6.8721 | 6.7187 | 6.3219 | 6.8999 | 6.5478 | 6.8153 | 4.5941 | 5.7065 | 4.3477 | 5.5126 |
| TRAF3IP1_S476 | TRAF3IP1 | Q8TDR0 | 476 | RVKRQDSMEALQM | 7.5224 | 5.9640 | 5.8887 | 6.7062 | 5.3473 | 5.7435 | 6.5873 | 6.3060 | 6.7485 | 7.0337 | 7.0416 | 6.3351 | 5.3354 | 5.7508 | 5.8861 | 5.7963 |
| TRAF7_S108 | TRAF7 | Q6Q0C0 | 13 | ARYNRFSGGPSNL | 5.1137 | 3.4810 | 8.5539 | 6.5656 | 5.5860 | 6.4198 | 6.5948 | 5.6793 | 4.5831 | 5.6910 | 8.4723 | 7.5044 | 11.9184 | 4.5656 | 4.0875 | 5.1837 |
| TRAF7_S13 | TRAF7 | Q6Q0C0 | 108 | SLRSTFSLPEEEE | 5.6161 | 4.6820 | 5.3025 | 5.7493 | 6.8918 | 5.7906 | 6.1355 | 6.6814 | 5.8186 | 5.9324 | 5.8432 | 5.7959 | 7.5204 | 7.1868 | 7.2920 | 7.7614 |
| TRAFD1_S327 | TRAFD1 | O14545 | 327 | LNTGSSSPRGVEE | 6.5612 | 6.6988 | 6.8312 | 6.0033 | 6.4273 | 6.7575 | 6.1447 | 6.7752 | 6.7256 | 6.2816 | 6.6946 | 6.4857 | 5.7185 | 5.4749 | 5.1879 | 5.2320 |
| TRAFD1_S415 | TRAFD1 | O14545 | 415 | SQPQETSPELPRR | 6.1735 | 6.3803 | 6.8810 | 6.2554 | 6.4353 | 6.6063 | 5.8451 | 6.6044 | 6.7324 | 6.7270 | 6.7523 | 6.6087 | 5.5743 | 5.8775 | 5.2005 | 5.3459 |
| TRAFD1_S480 | TRAFD1 | O14545 | 480 | PGCQPSSPCVPKL | 6.7005 | 6.3055 | 6.8575 | 6.7826 | 6.7813 | 7.1839 | 7.4038 | 7.2889 | 7.6203 | 7.5884 | 7.2221 | 7.5982 | 3.6396 | 3.7829 | 3.0899 | 4.1548 |
| TRAIP_S347 | TRAIP | Q9BWF2 | 469 | LDTFLWSxxxxxx | 7.2460 | 8.5167 | 6.4705 | 7.2974 | 5.4520 | 8.1430 | 7.3815 | 5.7977 | 5.8818 | 6.7414 | 6.8816 | 7.9421 | 5.3072 | 3.1114 | 2.7564 | 5.0736 |
| TRAIP_S469 | TRAIP | Q9BWF2 | 347 | CLEKSHSPIQDVP | 6.2790 | 7.6038 | 7.2260 | 7.9428 | 6.6429 | 6.3702 | 6.5222 | 7.0423 | 7.0446 | 6.9190 | 6.1970 | 6.4481 | 4.1271 | 4.7139 | 4.3037 | 4.6174 |
| TRAK1_S719 | TRAK1 | Q9UPV9 | 719 | CTPRRLSLAESFT | 7.9127 | 6.3341 | 6.4302 | 7.3162 | 6.4262 | 5.1533 | 4.5239 | 4.6296 | 8.2276 | 8.5487 | 7.0600 | 8.5088 | 5.6542 | 5.0156 | 3.5897 | 4.6691 |
| TRAK2_S420 | TRAK2 | O60296 | 420 | TRGRSISFPALLP | 4.2043 | 6.0853 | 5.5312 | 3.8924 | 7.4402 | 5.2286 | 3.3011 | 5.3171 | 5.6570 | 4.5535 | 3.5571 | 7.5240 | 10.8204 | 8.6321 | 7.1608 | 11.0951 |
| TRAM1_S365 | TRAM1 | Q15629 | 365 | TSNVADSPRNKKE | 6.1405 | 6.5763 | 6.5333 | 6.8850 | 6.7468 | 6.6791 | 6.8226 | 6.4363 | 6.4078 | 6.5575 | 6.7547 | 6.4636 | 4.9340 | 5.5301 | 5.1586 | 5.3738 |
| TRAP1_S195 | TRAP1 | Q12931 | 195 | NQAEASSKIIGQF | 6.6797 | 5.7512 | 6.7838 | 4.5546 | 5.8313 | 4.8547 | 5.4671 | 6.0274 | 5.0068 | 5.9073 | 7.1400 | 5.8433 | 8.3967 | 7.1240 | 7.5122 | 7.1200 |
| TRAPPC12_S182 | TRAPPC12 | Q8WVT3 | 184 | QMVKSPSFGGASE | 2.4308 | 3.2611 | 3.1271 | 3.2194 | 4.9130 | 5.2632 | 4.6511 | 4.9041 | 2.8387 | 3.1409 | 3.5095 | 3.1533 | 14.3665 | 12.8752 | 14.9302 | 13.4160 |
| TRAPPC12_S184 | TRAPPC12 | Q8WVT3 | 182 | EPQMVKSPSFGGA | 6.2080 | 6.6810 | 6.9316 | 6.9975 | 6.5352 | 6.8979 | 6.7499 | 6.9499 | 7.3718 | 6.7845 | 7.2109 | 6.8510 | 3.9483 | 4.8816 | 4.2435 | 4.7576 |

| | | | | | | | | | | | | | | | | | | | | |
|---|---|---|---|---|---|---|---|---|---|---|---|---|---|---|---|---|---|---|---|---|
| TRAPPC12_S536 | TRAPPC12 | Q8WVT3 | 536 | QEGRQASIRLWRS | 5.9077 | 6.1217 | 6.4162 | 5.9850 | 6.4213 | 6.6056 | 6.4402 | 6.6390 | 7.0803 | 6.7639 | 6.5113 | 6.8520 | 5.3398 | 6.1584 | 5.2439 | 5.5138 |
| TRAPPC14_S494 | TRAPPC14 | Q8WVR3 | 541 | MERRAITPPVASP | 6.3745 | 6.5667 | 6.6813 | 6.0761 | 6.1080 | 6.2249 | 6.0630 | 6.4620 | 7.1152 | 6.4286 | 6.0796 | 6.1216 | 5.7925 | 6.0787 | 5.7142 | 6.1130 |
| TRAPPC14_S533 | TRAPPC14 | Q8WVR3 | 533 | HLMRSGSVMERRA | 7.9782 | 7.7287 | 6.9316 | 7.1973 | 6.7720 | 5.9105 | 5.6719 | 5.2509 | 6.1982 | 5.6957 | 6.5216 | 6.1558 | 6.0743 | 5.4385 | 5.6303 | 4.8443 |
| TRAPPC14_T541 | TRAPPC14 | Q8WVR3 | 494 | RPLSRKSSPSSPA | 6.4159 | 6.9278 | 6.4092 | 5.7038 | 6.1506 | 6.0738 | 5.9691 | 5.9517 | 6.0785 | 6.1710 | 6.0757 | 6.0609 | 6.2729 | 6.8568 | 6.4022 | 6.4801 |
| TRAPPC8_S273 | TRAPPC8 | Q9Y2L5 | 273 | ITSNKNSDNNLLS | 2.5094 | 2.5456 | 2.4460 | 2.2176 | 5.0193 | 4.8084 | 5.0973 | 5.2485 | 2.6883 | 2.9348 | 3.0256 | 2.9999 | 15.6869 | 14.2536 | 14.9995 | 13.5192 |
| TRAPPC9_S1051 | TRAPPC9 | Q96Q05-2 | 1051 | FESFPESPGEKGQ | 5.6162 | 6.1620 | 6.2759 | 6.3531 | 6.1325 | 6.5250 | 6.3718 | 6.6344 | 6.1972 | 6.4293 | 6.6486 | 6.6873 | 5.8050 | 6.0890 | 5.9533 | 6.1192 |
| TRERF1_T634;S638 | TRERF1 | Q05GC8 | 735 | TPPPMLSPVRQGS | 6.2391 | 6.3111 | 6.1914 | 6.3244 | 6.2649 | 6.2518 | 6.2602 | 6.2775 | 6.5438 | 6.6610 | 6.6183 | 6.3649 | 6.0640 | 6.0597 | 5.8752 | 5.6927 |
| TRERF1_T646 | TRERF1 | Q05GC8 | 646 | QGAAPRTPAEIPR | 6.3237 | 6.8450 | 6.9896 | 6.3677 | 5.3956 | 7.0660 | 7.2920 | 7.4301 | 6.8257 | 6.6216 | 6.7337 | 6.7355 | 4.1931 | 5.0994 | 4.5144 | 5.5671 |
| TRERF1_T729;S735 | TRERF1 | Q05GC8 | 787 | GSNVTVTPGPGEQ | 8.0969 | 7.3553 | 7.6347 | 8.8233 | 7.2893 | 6.1743 | 5.8593 | 5.2092 | 6.6289 | 6.1870 | 7.9090 | 7.1800 | 5.0263 | 4.4294 | 2.6414 | 3.5557 |
| TRERF1_T735 | TRERF1 | Q05GC8 | 729;735 | NA | 5.5432 | 5.5670 | 6.3927 | 5.8262 | 6.2167 | 5.8851 | 6.0596 | 6.1976 | 5.9392 | 5.9839 | 6.1363 | 5.5260 | 7.4072 | 7.1662 | 7.0906 | 7.0624 |
| TRERF1_T787 | TRERF1 | Q05GC8 | 634;638 | NA | 6.1696 | 6.4343 | 7.4403 | 6.4933 | 6.6587 | 6.5650 | 7.0449 | 5.8920 | 6.0351 | 5.9508 | 6.6547 | 6.1106 | 5.9140 | 5.5851 | 5.4699 | 5.5816 |
| TRIM11_S85 | TRIM11 | Q96F44 | 85 | RRLHPPSPVPQGV | 5.3496 | 5.8285 | 5.8462 | 5.9641 | 5.9848 | 6.0617 | 6.1943 | 6.4507 | 6.2049 | 6.2363 | 6.1765 | 6.0188 | 7.1841 | 6.9089 | 6.8137 | 6.7768 |
| TRIM13_S278 | TRIM13 | O60858-3 | 278 | PSNLPASPLMKNF | 6.1045 | 6.9895 | 6.7871 | 7.8640 | 6.5038 | 7.0436 | 7.8232 | 7.1414 | 6.8496 | 6.9593 | 6.8995 | 6.4222 | 3.7657 | 4.6156 | 3.6807 | 4.5502 |
| TRIM24_S744 | TRIM24 | O15164 | 811 | LDPSQKSPLHVGE | 6.5402 | 6.3240 | 5.9374 | 6.5417 | 6.1283 | 6.0769 | 5.7287 | 5.8227 | 6.2076 | 5.9064 | 6.0414 | 6.2772 | 6.9089 | 6.4015 | 6.5061 | 6.6509 |
| TRIM24_S811 | TRIM24 | O15164 | 744 | VIVKQESDEESRP | 6.5875 | 5.5971 | 5.7129 | 7.1856 | 6.7354 | 6.3388 | 6.2874 | 6.3303 | 5.4964 | 6.7461 | 6.7011 | 5.7836 | 5.3506 | 6.9519 | 5.6786 | 6.5168 |
| TRIM25_S100 | TRIM25 | Q14258 | 100 | ARASAPSPNAQVA | 5.9719 | 6.3471 | 6.7570 | 6.3858 | 6.5778 | 6.9757 | 6.6993 | 6.7339 | 6.5827 | 6.7901 | 6.5234 | 6.4598 | 5.1038 | 5.6967 | 5.1451 | 5.2498 |
| TRIM25_S46 | TRIM25 | Q14258 | 91 | PPADVWTPPARAS | 6.6658 | 7.3204 | 6.5967 | 6.9469 | 6.3589 | 6.8909 | 7.2869 | 6.8744 | 6.8666 | 7.1593 | 7.0000 | 6.8699 | 3.7041 | 4.8211 | 3.8139 | 4.8243 |
| TRIM25_T91 | TRIM25 | Q14258 | 46 | TWAVQGSPYLCPQ | 6.3913 | 6.5634 | 6.7265 | 5.8756 | 5.9212 | 6.8251 | 5.8263 | 6.3866 | 6.0157 | 6.6803 | 6.3357 | 6.0545 | 5.8048 | 6.8734 | 6.1243 | 5.5955 |
| TRIM28_S258 | TRIM28 | Q13263 | 473 | GVKRSRSGEGEVS | 4.0706 | 4.1818 | 4.2484 | 4.3118 | 5.2193 | 4.9536 | 5.4620 | 5.0194 | 4.3022 | 4.1441 | 4.2423 | 4.4675 | 11.8824 | 10.2846 | 11.7905 | 11.4196 |
| TRIM28_S41 | TRIM28 | Q13263 | 594 | EGPRLASPSGSTS | 6.0974 | 6.2582 | 6.2426 | 6.6807 | 6.2966 | 6.0197 | 6.0407 | 6.1488 | 6.6487 | 6.2225 | 6.3635 | 6.3469 | 6.0936 | 6.2955 | 6.2748 | 5.9699 |
| TRIM28_S473 | TRIM28 | Q13263 | 498 | RLDLDLTADSQPP | 5.7035 | 6.0962 | 7.3776 | 7.3660 | 4.9656 | 6.5507 | 6.6409 | 6.1277 | 5.8639 | 6.3664 | 7.1959 | 6.8327 | 5.3096 | 7.1317 | 5.3211 | 5.1504 |
| TRIM28_S489 | TRIM28 | Q13263 | 41 | APSAAAASASASAA | 6.0498 | 6.4418 | 6.1133 | 6.5990 | 6.3172 | 6.6794 | 6.1859 | 6.2411 | 5.9863 | 6.1592 | 7.1012 | 6.7445 | 5.5449 | 6.3172 | 5.7860 | 5.7331 |
| TRIM28_S50 | TRIM28 | Q13263 | 501 | LDLTADSQPPVFK | 5.5026 | 6.2770 | 7.2988 | 7.0955 | 5.0819 | 5.8869 | 6.1557 | 5.9807 | 5.8545 | 6.0688 | 6.8647 | 6.2270 | 6.3904 | 7.7508 | 6.0893 | 5.4754 |
| TRIM28_S501 | TRIM28 | Q13263 | 531 | GAAAAATGQPGTA | 6.3911 | 5.7177 | 6.7323 | 6.7595 | 5.9115 | 6.6530 | 6.8759 | 6.1494 | 6.1830 | 5.8691 | 6.3483 | 6.2021 | 5.5282 | 6.3718 | 6.0853 | 6.2220 |
| TRIM28_S594 | TRIM28 | Q13263 | 697 | GVVAKLSPANQRK | 6.2662 | 6.7644 | 6.1612 | 6.4088 | 6.1908 | 6.1988 | 6.2060 | 6.2068 | 6.0509 | 6.4321 | 6.0435 | 5.8906 | 6.1713 | 6.4774 | 6.3361 | 6.1950 |
| TRIM28_S594;S600 | TRIM28 | Q13263 | 50 | ASAAASSPAGGGA | 6.7654 | 6.5888 | 5.9183 | 6.7628 | 6.2239 | 5.7328 | 6.3847 | 6.6400 | 6.9589 | 5.9881 | 6.3273 | 6.1126 | 5.4148 | 6.5843 | 5.5835 | 6.0137 |
| TRIM28_S697 | TRIM28 | Q13263 | 816 | VEPPPMSLPGAGL | 5.6194 | 6.1285 | 6.2357 | 6.4932 | 7.2312 | 5.5626 | 6.7326 | 6.7755 | 6.3059 | 6.1980 | 6.7054 | 6.5384 | 5.6513 | 5.7585 | 6.2096 | 5.8542 |
| TRIM28_S752 | TRIM28 | Q13263 | 258 | QRKLLASLVKRLG | 6.8277 | 5.6985 | 6.5804 | 6.6142 | 6.6031 | 6.6620 | 6.4137 | 5.6209 | 5.6758 | 6.5830 | 6.1503 | 5.4578 | 6.3870 | 6.4379 | 6.3775 | 5.9101 |
| TRIM28_S757 | TRIM28 | Q13263 | 752 | RLQEKLSPPYSSP | 6.2736 | 7.4042 | 7.0220 | 5.9279 | 5.6262 | 5.6058 | 5.8828 | 5.8044 | 5.4608 | 6.8652 | 5.7658 | 6.3649 | 6.2199 | 7.2496 | 6.9909 | 5.5360 |
| TRIM28_S816 | TRIM28 | Q13263 | 489 | RKVPRVSLERLDL | 6.1736 | 6.1315 | 6.4878 | 6.8709 | 6.9357 | 7.0024 | 6.8397 | 5.9683 | 5.2686 | 6.4638 | 7.0181 | 6.3918 | 5.6451 | 5.9635 | 5.5654 | 5.2737 |
| TRIM28_T498 | TRIM28 | Q13263 | 757 | LSPPYSSPQEFAQ | 5.0219 | 6.3798 | 6.4721 | 6.7023 | 4.8215 | 5.8812 | 6.5691 | 6.4218 | 5.7349 | 6.9605 | 6.1114 | 6.2214 | 6.1422 | 6.5271 | 6.8226 | 7.2103 |
| TRIM28_T531 | TRIM28 | Q13263 | 536 | ATGQPGTAPAGTP | 5.5463 | 6.2319 | 6.4774 | 6.9395 | 5.3756 | 6.1509 | 7.0627 | 5.9608 | 6.4519 | 6.3569 | 5.4539 | 5.9635 | 5.9503 | 6.3622 | 6.6526 | 7.0636 |
| TRIM28_T536 | TRIM28 | Q13263 | 594;600 | NA | 6.5007 | 5.9992 | 6.5346 | 6.2602 | 5.3986 | 6.6175 | 5.6549 | 6.1680 | 7.1317 | 6.9730 | 6.5090 | 6.4159 | 6.8897 | 6.1962 | 5.3922 | 5.3586 |
| TRIM32_S335 | TRIM32 | Q13049 | 335 | PEEVVASPRASPA | 5.8562 | 6.4320 | 6.3417 | 5.9328 | 6.9179 | 5.8217 | 6.1880 | 6.8811 | 5.7309 | 6.6835 | 6.1762 | 5.6420 | 6.3062 | 6.1460 | 6.6069 | 6.3370 |
| TRIM33_S1119 | TRIM33 | Q9UPN9 | 1119 | RRKRLKSDERPVH | 4.3766 | 4.8110 | 4.8824 | 4.9002 | 5.8283 | 5.7665 | 5.9948 | 5.7888 | 4.6217 | 4.5203 | 4.7339 | 4.9224 | 10.0896 | 8.7372 | 10.0079 | 10.0182 |
| TRIM33_T1102;S1105 | TRIM33 | Q9UPN9 | 1102;1105 | NA | 5.8148 | 6.1016 | 5.9288 | 5.7867 | 6.1400 | 5.3341 | 6.2428 | 6.0689 | 6.3598 | 6.0665 | 6.2983 | 6.9791 | 6.3507 | 6.6007 | 6.2715 | 7.6556 |
| TRIM36_S620 | TRIM36 | Q9NQ86 | 87 | PRLRLPSPSMDKI | 6.6077 | 6.9246 | 6.7491 | 7.0079 | 5.6107 | 6.5569 | 6.8927 | 6.2749 | 7.0427 | 6.9637 | 7.6170 | 8.6437 | 3.9372 | 4.9937 | 4.0829 | 4.0946 |
| TRIM36_S87 | TRIM36 | Q9NQ86 | 620 | QDSGHDSGSEDAC | 5.8994 | 6.1995 | 6.8225 | 6.7680 | 6.5265 | 6.1515 | 5.7059 | 7.0193 | 6.4191 | 6.1841 | 6.4606 | 6.4907 | 5.9889 | 5.9995 | 5.2879 | 6.0767 |
| TRIM37_S461 | TRIM37 | O94972 | 817 | RALIHGSIGDILP | 5.1155 | 4.7056 | 5.4874 | 4.6032 | 5.8896 | 5.3185 | 5.2008 | 4.5842 | 5.3621 | 5.2597 | 4.4419 | 5.7302 | 9.7642 | 9.0830 | 11.2271 | 8.2272 |
| TRIM37_S817 | TRIM37 | O94972 | 461 | PPDNHLSPQNDDA | 5.9546 | 6.6414 | 7.5811 | 7.1765 | 6.1422 | 6.3462 | 6.0753 | 6.9498 | 6.8943 | 6.1749 | 6.6708 | 8.1179 | 5.3901 | 4.8306 | 4.8404 | 4.2139 |
| TRIM38_S240 | TRIM38 | O00635 | 240 | EEKCQGSAQKLLQ | 6.6032 | 7.1909 | 6.3331 | 6.7312 | 5.7261 | 6.4810 | 6.4987 | 6.1521 | 7.0693 | 6.8250 | 6.8048 | 7.1184 | 4.8063 | 5.3644 | 4.7593 | 5.5360 |
| TRIM41_S447 | TRIM41 | Q8WV44 | 447 | HPALMLSPDRRGV | 6.0893 | 6.9818 | 6.3246 | 6.7365 | 6.3050 | 6.6176 | 6.3894 | 5.9524 | 6.7709 | 6.6747 | 6.5975 | 6.5962 | 5.1104 | 5.8038 | 5.4228 | 5.6270 |
| TRIM44_S320 | TRIM44 | Q96DX7 | 320 | KEQLDTSNESAEP | 6.7385 | 7.7133 | 6.7453 | 7.7157 | 6.3151 | 6.7530 | 7.3308 | 6.7130 | 6.9705 | 7.2271 | 7.8816 | 7.9217 | 3.0531 | 3.6748 | 3.1144 | 4.1321 |
| TRIM47_S588 | TRIM47 | Q96LD4 | 588 | RGGIPASPIDPFQ | 5.3719 | 5.5049 | 6.1450 | 5.5710 | 6.5201 | 6.4140 | 6.2374 | 6.7182 | 5.5689 | 5.1721 | 5.9472 | 5.6823 | 7.1969 | 7.4868 | 7.4190 | 7.0441 |
| TRIM5_S86 | TRIM5 | Q9C035 | 86 | LREVKLSPEGQKV | 6.2545 | 6.7485 | 7.2871 | 6.4565 | 6.2387 | 6.2956 | 6.4732 | 6.6034 | 6.8381 | 6.2705 | 6.7925 | 6.9073 | 5.1332 | 5.3489 | 5.1262 | 5.2259 |
| TRIM56_S469;S471 | TRIM56 | Q9BRZ2 | 442 | EEDRAQTPHEDGG | 6.2159 | 6.0275 | 6.5428 | 6.3287 | 6.1430 | 6.1872 | 6.0264 | 6.2335 | 6.5642 | 6.5159 | 5.9517 | 6.5367 | 5.8064 | 6.4040 | 6.2935 | 6.2226 |
| TRIM56_S469;S471;S475 | TRIM56 | Q9BRZ2 | 475 | SISREPSPALGPN | 5.5156 | 5.9351 | 6.2133 | 6.5181 | 6.6620 | 6.1743 | 6.6284 | 6.4064 | 6.7821 | 6.0910 | 6.1473 | 6.1048 | 5.7840 | 6.6412 | 6.3639 | 6.0326 |
| TRIM56_S471;S475 | TRIM56 | Q9BRZ2 | 418 | AGDGAQTPKEEKA | 6.3922 | 5.6559 | 6.6621 | 6.5811 | 6.0282 | 6.8184 | 6.5679 | 6.0503 | 6.3818 | 7.1665 | 6.3043 | 6.2828 | 5.3977 | 6.1257 | 5.7085 | 5.8766 |
| TRIM56_S475 | TRIM56 | Q9BRZ2 | 471;475 | NA | 5.9419 | 6.8145 | 6.4393 | 5.9311 | 6.3218 | 6.3802 | 5.8068 | 6.7113 | 6.5424 | 7.1133 | 6.3626 | 6.5189 | 5.6927 | 5.6033 | 5.5768 | 6.2431 |
| TRIM56_T418 | TRIM56 | Q9BRZ2 | 469;471 | NA | 5.5162 | 6.2011 | 5.0948 | 6.4366 | 6.7868 | 5.1335 | 6.2232 | 5.6949 | 5.1023 | 5.9758 | 6.2295 | 6.3831 | 7.2838 | 7.2568 | 7.5065 | 7.1750 |
| TRIM56_T442 | TRIM56 | Q9BRZ2 | 469;471;475 | NA | 6.1435 | 6.2617 | 6.3798 | 5.1101 | 6.2162 | 6.0731 | 5.8388 | 5.9452 | 5.2670 | 7.4066 | 7.0996 | 5.5217 | 5.6996 | 7.4166 | 6.1300 | 7.4906 |
| TRIM8_S321 | TRIM8 | Q9BZR9 | 321 | CTSSSLSPTKIGH | 5.8053 | 7.2365 | 6.9424 | 6.0886 | 6.1929 | 5.9401 | 6.2253 | 6.4103 | 7.3004 | 7.1736 | 7.1618 | 6.7536 | 4.4224 | 6.5558 | 5.0529 | 5.6381 |
| TRIO_S1724 | TRIO | O75962 | 1724 | IAHSRSSMEMEGI | 6.2787 | 6.1221 | 6.9708 | 6.2054 | 6.3346 | 6.3308 | 6.0035 | 6.6508 | 6.7887 | 6.2542 | 6.5922 | 6.6488 | 5.8896 | 5.8379 | 5.4820 | 5.6098 |
| TRIO_S1779 | TRIO | O75962 | 2455 | PRAGAASPLNSPL | 5.5294 | 5.7281 | 6.4320 | 6.0579 | 6.1349 | 6.1702 | 6.1939 | 6.5011 | 5.8190 | 6.1699 | 6.4745 | 6.8309 | 6.0485 | 6.6394 | 6.4334 | 6.8370 |
| TRIO_S2282 | TRIO | O75962 | 2631 | TLKKSTSWHTALR | 2.5748 | 2.1461 | 2.3096 | 2.2594 | 2.8906 | 2.5842 | 2.7203 | 3.3602 | 2.3503 | 2.4572 | 1.7750 | 2.3857 | 16.2326 | 16.1110 | 20.1616 | 17.6813 |
| TRIO_S2417 | TRIO | O75962 | 2426 | KGAANASGSSPDA | 6.2089 | 6.4706 | 6.7550 | 6.7891 | 6.4875 | 6.7449 | 6.6350 | 6.6026 | 6.8189 | 7.0246 | 6.8415 | 7.0976 | 4.6795 | 5.2307 | 4.6531 | 4.9605 |
| TRIO_S2426 | TRIO | O75962 | 2417 | KMKVLESPRKGAA | 6.6853 | 6.5636 | 7.3410 | 6.0320 | 6.7303 | 6.7370 | 6.8636 | 6.4578 | 6.5994 | 6.1200 | 6.5236 | 6.5817 | 5.0828 | 5.4498 | 5.1360 | 5.0961 |
| TRIO_S2440 | TRIO | O75962 | 2440 | AKDARASLGTLPL | 6.9353 | 7.2621 | 6.9619 | 7.4374 | 6.6533 | 6.9117 | 7.4230 | 7.1672 | 7.4912 | 6.9198 | 7.1198 | 6.9502 | 3.3812 | 4.2052 | 3.2250 | 3.9556 |
| TRIO_S2455 | TRIO | O75962 | 2477 | EPFPPSSPLQKGG | 6.6957 | 6.4305 | 6.4588 | 6.5277 | 6.1177 | 6.5703 | 6.2548 | 6.5114 | 6.4308 | 6.1782 | 6.1920 | 6.6836 | 5.6843 | 6.0304 | 5.5069 | 5.7270 |
| TRIO_S2455;S2459 | TRIO | O75962 | 2282 | FLNALTSPIEYQR | 6.1718 | 6.1834 | 6.5303 | 6.8798 | 6.4155 | 7.2151 | 6.8069 | 6.3053 | 6.5941 | 6.7695 | 6.5793 | 6.5245 | 5.2684 | 5.4218 | 5.0904 | 5.2438 |
| TRIO_S2477 | TRIO | O75962 | 1779 | LRKWLTSPVRRLS | 6.6421 | 6.5138 | 6.5997 | 5.8291 | 6.6446 | 6.4582 | 6.5394 | 7.1113 | 7.1381 | 6.5333 | 6.0902 | 6.7521 | 5.0893 | 5.5109 | 5.2741 | 5.2737 |
| TRIO_S2492 | TRIO | O75962 | 2492 | WSSIPASPASRPG | 6.2293 | 7.1167 | 6.3114 | 6.5519 | 5.6189 | 5.5936 | 6.2546 | 6.9084 | 5.7230 | 6.2664 | 6.4794 | 6.3114 | 6.1915 | 6.0795 | 6.1962 | 6.1678 |
| TRIO_S2631 | TRIO | O75962 | 2455;2459 | NA | 6.0340 | 6.3595 | 7.2417 | 7.0900 | 6.3126 | 6.7721 | 6.7747 | 7.0963 | 6.8984 | 6.4439 | 6.7356 | 7.0588 | 4.5691 | 5.2103 | 4.3679 | 5.0352 |
| TRIOBP_S1796 | TRIOBP | Q9H2D6 | 1955 | SPLTQASPQRART | 5.9936 | 6.1195 | 6.5990 | 6.5497 | 6.3546 | 6.0392 | 6.6502 | 7.0386 | 6.9104 | 7.1272 | 6.5949 | 6.7377 | 4.9844 | 5.6927 | 5.2012 | 5.4071 |
| TRIOBP_S1955 | TRIOBP | Q9H2D6 | 1796 | EPGEPPSPSLTTT | 5.8408 | 5.7324 | 5.9007 | 6.1255 | 6.6734 | 6.0637 | 6.3555 | 6.4299 | 6.5605 | 6.2057 | 6.2860 | 6.4415 | 5.9604 | 6.7744 | 6.6017 | 6.0478 |
| TRIP10_S296 | TRIP10 | Q15642 | 302 | SDSSLGTPSDGRP | 7.1389 | 7.1759 | 6.7820 | 7.1850 | 5.5717 | 7.4950 | 7.6563 | 7.1700 | 7.8097 | 7.8012 | 7.8447 | 7.9850 | 2.7699 | 3.3907 | 2.5412 | 3.6828 |
| TRIP10_S296;S299 | TRIP10 | Q15642 | 296 | PMNRAPSDSSLGT | 5.3030 | 5.8013 | 6.6522 | 6.0961 | 6.1638 | 5.9535 | 5.7864 | 6.8075 | 6.2162 | 5.8867 | 6.0898 | 6.4554 | 6.9429 | 6.4875 | 6.6315 | 6.7263 |
| TRIP10_T302 | TRIP10 | Q15642 | 296;299 | NA | 6.4353 | 6.7208 | 7.0061 | 6.8040 | 7.0175 | 6.5459 | 6.4828 | 6.5077 | 6.8889 | 6.4098 | 6.4887 | 6.4014 | 4.8307 | 5.4813 | 4.9776 | 5.0016 |
| TRIP11_S1341 | TRIP11 | Q15643 | 1891 | DMKPLDSPGRRKR | 6.3936 | 5.6826 | 6.3677 | 6.8470 | 6.3439 | 6.0362 | 6.0324 | 5.8683 | 6.5831 | 6.5507 | 6.6049 | 6.9902 | 6.4990 | 5.8760 | 5.4218 | 5.9027 |
| TRIP11_S1842 | TRIP11 | Q15643 | 1842 | LGGGSKSVPNTPL | 5.7327 | 5.9445 | 6.5717 | 6.2685 | 6.5776 | 6.6089 | 6.2764 | 6.8942 | 6.3680 | 6.7088 | 6.5213 | 6.4244 | 6.1547 | 5.7101 | 5.9323 | 5.3058 |
| TRIP11_S1842;T1846 | TRIP11 | Q15643 | 486 | EQELNQSISEKET | 7.0309 | 7.0140 | 6.7802 | 7.3741 | 7.0519 | 6.7015 | 6.9992 | 6.8631 | 7.3869 | 6.6614 | 7.1137 | 6.6514 | 4.1692 | 4.4127 | 3.7191 | 4.0707 |
| TRIP11_S1846 | TRIP11 | Q15643 | 464 | TATRDISLDSELH | 6.5756 | 6.0284 | 6.8146 | 6.9425 | 6.2956 | 7.5371 | 7.1937 | 6.6572 | 6.8914 | 6.9810 | 7.1072 | 7.1646 | 4.1486 | 4.9103 | 4.1600 | 4.5922 |
| TRIP11_S1891 | TRIP11 | Q15643 | 444 | KEELQMSLLKLNN | 7.9416 | 8.2955 | 7.4083 | 7.6852 | 9.0238 | 6.2030 | 6.2671 | 6.5697 | 5.8876 | 4.9414 | 6.2159 | 5.2132 | 5.1491 | 4.6696 | 3.5080 | 5.0209 |
| TRIP11_S444 | TRIP11 | Q15643 | 1846 | SKSVPNTPLRPNQ | 6.6847 | 6.3031 | 5.7721 | 6.6019 | 6.4158 | 5.6974 | 6.8375 | 6.2019 | 6.4579 | 4.9130 | 6.3038 | 6.4912 | 7.1811 | 6.1707 | 5.5208 | 6.4471 |
| TRIP11_S464 | TRIP11 | Q15643 | 1341 | SEVLSESSELLQQ | 5.8654 | 5.7748 | 6.5693 | 8.1138 | 6.1302 | 7.0012 | 6.3115 | 6.5408 | 5.8210 | 5.8156 | 6.6244 | 6.5088 | 6.0840 | 6.3258 | 3.9281 | 6.5853 |
| TRIP11_S486 | TRIP11 | Q15643 | 1842;1846 | NA | 7.2565 | 7.0480 | 6.2173 | 7.9193 | 6.8183 | 5.7295 | 6.7714 | 5.0260 | 5.6699 | 6.5066 | 7.2927 | 6.5828 | 5.5497 | 5.4214 | 4.3461 | 5.8448 |
| TRIP12_S1064 | TRIP12 | Q14669-3 | 354 | TKKRSESPPAELP | 6.2935 | 6.4245 | 6.2705 | 6.8106 | 6.0530 | 5.8938 | 6.2505 | 6.2982 | 6.5130 | 6.2905 | 6.2633 | 6.3237 | 5.7006 | 6.4334 | 6.0241 | 6.1568 |
| TRIP12_S1078 | TRIP12 | Q14669-3 | 127 | DYNRTNSPSSAKK | 6.5463 | 6.6922 | 6.5125 | 7.1275 | 5.9964 | 5.8986 | 6.4056 | 5.9885 | 6.4898 | 6.7180 | 6.6138 | 7.0081 | 5.0044 | 6.0243 | 5.1670 | 5.8072 |
| TRIP12_S1084 | TRIP12 | Q14669-3 | 119 | GVKRSASPDYNRT | 6.0569 | 6.7418 | 5.8855 | 6.5533 | 5.9378 | 5.4802 | 5.7013 | 5.7672 | 6.0797 | 5.9979 | 5.5999 | 5.8790 | 6.5437 | 7.0627 | 7.4538 | 7.2592 |
| TRIP12_S119 | TRIP12 | Q14669-3 | 1064 | PRRPKYSPPRDDD | 6.1295 | 6.4520 | 6.0571 | 6.7678 | 5.8499 | 5.8872 | 6.4256 | 5.7546 | 5.9917 | 6.2080 | 6.1447 | 6.5824 | 6.0049 | 6.7106 | 6.4451 | 6.5888 |
| TRIP12_S127 | TRIP12 | Q14669-3 | 1084 | SPTTTQSPKSSFL | 7.6328 | 7.4576 | 7.1839 | 7.6418 | 6.7293 | 6.3036 | 6.8906 | 6.4660 | 7.0864 | 6.8897 | 7.0025 | 7.0385 | 3.6011 | 4.3179 | 3.5938 | 4.1645 |
| TRIP12_S127;S130 | TRIP12 | Q14669-3 | 169 | QLKSAQSPSTSKA | 7.3675 | 7.8594 | 6.6994 | 7.8061 | 6.8402 | 6.4990 | 7.0842 | 6.7084 | 7.3781 | 7.2858 | 7.4597 | 7.2838 | 2.7965 | 3.9754 | 3.2435 | 3.7130 |
| TRIP12_S1365;S1370 | TRIP12 | Q14669-3 | 990 | SESLLTSPPKACT | 6.9869 | 7.3248 | 6.7446 | 7.8789 | 6.4133 | 6.3497 | 6.5338 | 6.6986 | 7.2243 | 6.9289 | 6.9670 | 6.9737 | 3.9615 | 4.6881 | 3.9695 | 4.3563 |
| TRIP12_S142 | TRIP12 | Q14669-3 | 382 | STSRRGSGLGKRG | 4.5408 | 4.2462 | 4.4473 | 4.7814 | 7.5272 | 5.1489 | 5.0036 | 5.4604 | 8.0455 | 9.1608 | 7.8164 | 8.1674 | 7.1683 | 8.1144 | 4.9220 | 5.4496 |
| TRIP12_S1424 | TRIP12 | Q14669-3 | 472 | PLGPRMSQLFHRT | 8.8048 | 7.5213 | 6.2999 | 8.0447 | 6.4191 | 5.8484 | 7.0501 | 5.8965 | 7.5787 | 6.5910 | 7.0686 | 7.1577 | 3.9491 | 4.7951 | 2.8178 | 4.1572 |
| TRIP12_S1424;T1425 | TRIP12 | Q14669-3 | 142 | ALQHTESPSETNK | 6.3924 | 6.7145 | 6.6274 | 6.9070 | 6.0292 | 6.0603 | 6.4443 | 6.3051 | 6.2112 | 6.3008 | 6.2235 | 6.0850 | 5.3973 | 6.1218 | 5.7165 | 6.4638 |
| TRIP12_S169 | TRIP12 | Q14669-3 | 1078 | VDNQAKSPTTTQS | 6.4580 | 6.9289 | 6.6961 | 7.1681 | 6.0963 | 6.3669 | 6.3648 | 6.7165 | 6.7456 | 6.9026 | 6.9851 | 7.0001 | 4.3984 | 5.2059 | 4.7301 | 5.2366 |
| TRIP12_S281 | TRIP12 | Q14669-3 | 45 | AKHKGYSPPESRK | 6.5715 | 6.9098 | 5.9008 | 7.0980 | 5.8802 | 6.0219 | 6.2222 | 6.0171 | 6.2308 | 6.2738 | 6.1515 | 6.4397 | 5.1977 | 6.6707 | 6.1179 | 6.2965 |
| TRIP12_S283 | TRIP12 | Q14669-3 | 1424 | AEDERESTDDESN | 5.9826 | 6.6283 | 6.7967 | 6.5161 | 5.1862 | 6.0447 | 6.1821 | 6.9899 | 5.3461 | 5.4674 | 5.6548 | 6.2764 | 6.4425 | 7.2763 | 7.0014 | 6.2085 |
| TRIP12_S352;S354 | TRIP12 | Q14669-3 | 283 | RSASSPSPRRSSR | 5.4351 | 5.3030 | 4.4735 | 5.1612 | 5.4469 | 5.0374 | 5.1645 | 5.2611 | 5.0728 | 4.8273 | 4.4264 | 4.9262 | 9.4696 | 9.9853 | 10.3365 | 9.6732 |
| TRIP12_S354 | TRIP12 | Q14669-3 | 281 | RSRSASSPSPRRS | 6.5309 | 6.5604 | 5.6525 | 6.6869 | 5.6024 | 5.4943 | 5.8825 | 5.6301 | 6.2633 | 6.2702 | 5.9806 | 5.8352 | 6.0833 | 7.2585 | 7.1829 | 7.0860 |
| TRIP12_S382 | TRIP12 | Q14669-3 | 1365;1370 | NA | 5.4997 | 5.5846 | 7.0303 | 6.0825 | 6.1744 | 6.4896 | 6.3926 | 6.3868 | 5.6690 | 6.2318 | 7.3770 | 6.2230 | 5.8668 | 6.2496 | 6.7942 | 5.9480 |
| TRIP12_S45 | TRIP12 | Q14669-3 | 127;130 | NA | 6.7632 | 6.8956 | 5.9823 | 6.8680 | 5.5830 | 7.2301 | 7.1974 | 6.9935 | 6.4291 | 6.7453 | 7.5093 | 7.2052 | 3.7079 | 5.9984 | 4.1737 | 4.7179 |
| TRIP12_S472 | TRIP12 | Q14669-3 | 352;354 | NA | 6.8380 | 6.6878 | 6.4652 | 6.6935 | 6.3270 | 6.1679 | 6.5079 | 6.6822 | 6.5650 | 6.8388 | 6.6415 | 6.9236 | 5.2272 | 5.9346 | 4.4584 | 5.0413 |
| TRIP12_S990 | TRIP12 | Q14669-3 | 1424;1425 | NA | 6.1375 | 6.9878 | 7.2657 | 6.5493 | 6.2685 | 6.6190 | 6.7864 | 5.9254 | 5.6520 | 6.1045 | 5.6246 | 6.5243 | 5.1260 | 6.1351 | 6.5030 | 5.7911 |
| TRIP13_S18 | TRIP13 | Q15645 | 18 | LPCVAESPTVHVE | 7.1304 | 6.4123 | 6.5548 | 6.8564 | 6.3587 | 6.6767 | 6.5989 | 6.8582 | 6.4364 | 6.0968 | 8.2158 | 6.2800 | 4.3235 | 5.0244 | 5.0278 | 5.1488 |
| TRIP6_S142 | TRIP6 | Q15654 | 92 | GGLRPGSLDAEID | 6.7068 | 6.0959 | 7.3311 | 6.3919 | 6.3762 | 6.5746 | 6.4386 | 7.5660 | 6.7701 | 5.9332 | 6.6482 | 6.9890 | 4.9418 | 5.2487 | 5.1650 | 4.8230 |
| TRIP6_S92 | TRIP6 | Q15654 | 142 | LKPNPASPLPASP | 6.2573 | 6.0696 | 6.6182 | 6.5149 | 6.3498 | 6.5516 | 6.8148 | 6.6619 | 6.9868 | 6.9350 | 6.8635 | 6.5387 | 5.2033 | 5.3294 | 5.0714 | 5.2338 |
| TRIR_S114 | TRIR | Q9BQ61 | 114 | GPGSTLSFVGKRR | 8.5219 | 6.3985 | 7.0740 | 5.5182 | 4.7560 | 5.9699 | 5.4745 | 5.5499 | 8.4696 | 7.6375 | 9.0700 | 9.8380 | 3.6720 | 5.4233 | 3.0417 | 3.5850 |
| TRIT1_S455 | TRIT1 | Q9H3H1 | 455 | EPKEKGSPGQNDQ | 4.4961 | 5.4094 | 7.1317 | 6.2064 | 7.4882 | 7.7145 | 8.5937 | 6.0783 | 7.1077 | 6.0222 | 9.2826 | 7.4201 | 5.3953 | 4.9607 | 3.4387 | 3.2544 |
| TRMT1_S625 | TRMT1 | Q9NXH9 | 625 | QCCYSHSPPTPRV | 5.4437 | 5.7497 | 6.1770 | 5.8725 | 5.6285 | 6.0129 | 5.8421 | 6.2322 | 5.6883 | 5.7750 | 6.2885 | 5.7905 | 6.8654 | 7.5677 | 7.5703 | 7.4955 |
| TRMT1_S643 | TRMT1 | Q9NXH9 | 643 | PDCPETSNQTPPG | 6.0327 | 6.3591 | 6.1968 | 6.7224 | 5.9718 | 6.5441 | 6.7617 | 6.6814 | 6.6091 | 6.2685 | 6.4490 | 6.4405 | 5.3163 | 5.9131 | 5.6347 | 6.0988 |
| TRMT1_T628 | TRMT1 | Q9NXH9 | 628 | YSHSPPTPRVSAD | 4.6574 | 5.6487 | 6.2573 | 5.5695 | 6.0119 | 5.8424 | 5.9102 | 5.2499 | 5.6616 | 5.8165 | 6.4042 | 5.9667 | 6.9159 | 8.1428 | 8.8515 | 7.0935 |
| TRMT1L_S612 | TRMT1L | Q7Z2T5 | 612 | AQGKRKSNEMITN | 6.0321 | 6.7993 | 7.1151 | 7.5814 | 6.0897 | 7.2812 | 6.9582 | 5.9388 | 7.2732 | 6.6041 | 7.0806 | 6.9919 | 4.4918 | 4.9054 | 4.0657 | 4.7913 |
| TRMT1L_S66 | TRMT1L | Q7Z2T5 | 66 | AQAPALSPSLASA | 6.0978 | 5.5978 | 5.9496 | 6.2990 | 5.6430 | 5.3256 | 5.6820 | 5.7476 | 5.5432 | 6.1625 | 6.0627 | 6.4308 | 7.0423 | 7.0540 | 7.6702 | 7.6920 |
| TRMT2A_S602 | TRMT2A | Q8IZ69 | 602 | GVLGPHSPPAQPT | 5.9811 | 5.9057 | 6.1576 | 6.2313 | 6.1489 | 6.0235 | 5.9765 | 6.2385 | 6.5981 | 6.5563 | 6.0132 | 6.0473 | 6.5245 | 7.1870 | 6.0596 | 6.3508 |
| TRMT5_T494 | TRMT5 | Q32P41 | 494 | PLKRQRTAEAFSD | 5.4011 | 5.8934 | 5.6641 | 5.3507 | 5.8979 | 5.6136 | 5.8509 | 5.4154 | 6.7289 | 6.6166 | 6.4936 | 6.7486 | 5.9359 | 7.3507 | 7.1812 | 7.8574 |
| TRMT61A_S263 | TRMT61A | Q96FX7 | 263 | PAGSDTSPFRSGT | 6.0103 | 6.0011 | 6.6410 | 6.4872 | 6.0744 | 6.7406 | 6.2298 | 6.0944 | 6.7760 | 6.9256 | 6.9374 | 6.7458 | 5.0331 | 5.8926 | 5.7365 | 5.6744 |
| TRO_S155 | TRO | Q12816 | 168 | GTIQLKSPLQVLK | 7.1786 | 7.2939 | 6.7850 | 6.9498 | 6.3784 | 6.8144 | 7.0597 | 7.0749 | 7.5905 | 7.3132 | 7.2422 | 7.4587 | 3.2573 | 4.1387 | 3.4373 | 4.0272 |
| TRO_S168 | TRO | Q12816 | 155 | AAQGSQSPTGHEG | 7.1776 | 7.3268 | 7.2942 | 5.8913 | 5.5738 | 6.8573 | 6.9089 | 6.1382 | 7.7908 | 7.0554 | 7.0256 | 6.9170 | 3.2889 | 5.9347 | 4.1817 | 4.6376 |
| TROAP_S376 | TROAP | Q12815 | 98 | LEELRPSPRGQNV | 7.1884 | 7.5800 | 7.0493 | 7.0656 | 5.8970 | 6.9788 | 7.4526 | 7.0429 | 7.3825 | 7.7727 | 7.4647 | 7.7208 | 2.9635 | 3.7832 | 2.9476 | 3.7105 |
| TROAP_S777 | TROAP | Q12815 | 376 | QWLRGVSPQSCSE | 6.1422 | 7.0319 | 7.9203 | 6.8832 | 6.1812 | 6.9709 | 5.7656 | 6.6626 | 7.4816 | 7.0709 | 7.9898 | 7.9483 | 3.2161 | 4.8358 | 3.4172 | 4.4824 |
| TROAP_S98 | TROAP | Q12815 | 777 | SAAPQGSPxxxxx | 5.8404 | 6.5293 | 6.0656 | 6.8951 | 6.2237 | 6.5775 | 6.5708 | 6.4328 | 6.2103 | 6.7758 | 5.7895 | 6.0749 | 5.1824 | 6.2023 | 6.8375 | 5.7922 |
| TRPM7_S1404;T1405 | TRPM7 | Q96QT4 | 1477 | TLKRVSSLAGFTD | 6.6374 | 4.8554 | 6.3478 | 6.7796 | 6.2671 | 6.0415 | 6.2729 | 6.4262 | 6.4581 | 5.4356 | 5.8166 | 6.1087 | 6.5628 | 6.9416 | 6.2813 | 6.7675 |
| TRPM7_S1476 | TRPM7 | Q96QT4 | 1851 | NLQPGNSTKESES | 5.8100 | 6.6978 | 6.7812 | 5.6822 | 5.9896 | 6.4765 | 6.5334 | 6.7699 | 5.9187 | 6.2766 | 5.7075 | 5.5975 | 5.9466 | 6.9115 | 6.7117 | 6.1894 |
| TRPM7_S1477 | TRPM7 | Q96QT4 | 1476 | NTLKRVSSLAGFT | 6.0496 | 5.0064 | 5.5771 | 6.6281 | 5.7994 | 6.0739 | 6.8572 | 6.9631 | 6.4984 | 5.8242 | 6.0078 | 6.1401 | 6.1468 | 7.1797 | 6.2730 | 6.9750 |
| TRPM7_S1488 | TRPM7 | Q96QT4 | 1488 | TDCHRTSIPVHSK | 5.1729 | 5.8343 | 6.2700 | 6.4492 | 5.9393 | 5.8709 | 5.9936 | 5.8693 | 6.0438 | 6.4896 | 6.2760 | 6.6726 | 6.7591 | 6.8474 | 6.2290 | 7.2829 |
| TRPM7_S1851 | TRPM7 | Q96QT4 | 1404;1405 | NA | 6.3689 | 7.4941 | 7.7841 | 8.6481 | 6.0804 | 4.9297 | 5.7683 | 6.1733 | 7.0982 | 6.5835 | 6.6683 | 7.7685 | 5.2994 | 6.4012 | 2.9663 | 3.9678 |
| TRPS1_S103 | TRPS1 | Q9UHF7-2 | 103 | AGFNYESPSKGGN | 6.8146 | 7.1974 | 6.5902 | 7.1396 | 6.2615 | 6.2157 | 6.7227 | 6.5503 | 6.8920 | 6.9170 | 6.8842 | 6.8900 | 4.7372 | 4.9688 | 4.3482 | 4.8706 |
| TRPS1_S1079 | TRPS1 | Q9UHF7-2 | 229 | PDPAPLSPELQDF | 6.1671 | 6.5044 | 6.2124 | 6.2124 | 5.8214 | 6.2087 | 6.3075 | 5.8857 | 6.4968 | 6.4084 | 6.4887 | 6.7458 | 6.0373 | 6.4373 | 5.8753 | 6.1908 |
| TRPS1_S191 | TRPS1 | Q9UHF7-2 | 191 | ANCQGLSPVSVAS | 6.2694 | 6.9464 | 6.6945 | 7.2740 | 6.5377 | 6.1376 | 6.5488 | 6.4650 | 6.6919 | 6.6465 | 6.5330 | 6.4668 | 5.3545 | 5.4860 | 4.8960 | 5.0520 |
| TRPS1_S229 | TRPS1 | Q9UHF7-2 | 1079 | GSSERGSPIEKYM | 6.0059 | 6.1167 | 6.0828 | 8.5167 | 6.3676 | 7.4527 | 5.2212 | 5.3411 | 6.0197 | 6.6249 | 5.8750 | 6.7522 | 6.2861 | 6.3777 | 5.4565 | 5.5032 |
| TRPS1_T764 | TRPS1 | Q9UHF7-2 | 764 | RVYNLLTPDSKMG | 5.9047 | 6.9751 | 7.4147 | 7.0475 | 7.0656 | 6.4390 | 7.5034 | 7.5017 | 6.9768 | 6.2882 | 7.5146 | 5.9891 | 3.8457 | 4.4533 | 4.4352 | 4.6455 |
| TRPT1_S242 | TRPT1 | Q86TN4-4 | 242 | ETECQSSPKHSSR | 5.3210 | 6.0478 | 5.8783 | 6.4171 | 6.3684 | 6.3701 | 5.4301 | 6.0663 | 6.6806 | 6.9106 | 6.7033 | 7.4737 | 6.5262 | 5.7692 | 6.2215 | 5.8161 |
| TRPV2_S741 | TRPV2 | Q9Y5S1 | 751 | EDEDGASEENYVP | 5.4877 | 5.6323 | 7.0301 | 6.3522 | 5.9504 | 6.2943 | 6.3262 | 6.3641 | 6.4219 | 6.2963 | 6.4434 | 6.2556 | 5.9953 | 6.4149 | 6.2805 | 6.4547 |
| TRPV2_S741;Y751 | TRPV2 | Q9Y5S1 | 741 | ENPVLASPPKEDE | 6.0844 | 5.9604 | 6.2150 | 6.0034 | 5.5678 | 5.9676 | 6.4177 | 6.0697 | 6.1864 | 6.4019 | 6.5820 | 6.7565 | 6.4174 | 6.5804 | 6.1503 | 6.6391 |
| TRPV2_S751 | TRPV2 | Q9Y5S1 | 741;751 | NA | 5.5552 | 6.9125 | 6.4654 | 6.5356 | 6.8435 | 6.3971 | 5.1774 | 5.1844 | 6.6376 | 7.1258 | 4.8789 | 5.8118 | 6.0622 | 6.1172 | 6.9528 | 7.3425 |
| TRRAP_S2077 | TRRAP | Q9Y4A5 | 2077 | VFGRSQSLPGADS | 4.3557 | 4.2115 | 4.4701 | 4.7055 | 5.2990 | 5.0433 | 5.3134 | 5.3383 | 4.7260 | 4.5882 | 5.0928 | 4.7441 | 10.5835 | 9.5679 | 11.2851 | 10.6755 |
| TSC1_S1080 | TSC1 | Q92574 | 505 | PRGGFDSPFYRDS | 6.6480 | 6.6099 | 6.8627 | 6.0377 | 5.8573 | 6.4523 | 6.0101 | 6.8776 | 6.7204 | 6.8732 | 6.9101 | 6.6611 | 5.4292 | 5.8179 | 4.9833 | 5.2491 |
| TSC1_S295 | TSC1 | Q92574 | 598 | TRVGFGSGQPPPY | 5.5977 | 8.1910 | 6.1182 | 7.2003 | 6.5565 | 9.5288 | 5.5155 | 6.3510 | 7.8486 | 6.2140 | 7.7984 | 7.0633 | 2.8399 | 4.6936 | 3.1778 | 5.3055 |
| TSC1_S505 | TSC1 | Q92574 | 1080 | IPTTVGSLPSSKS | 7.2258 | 6.9156 | 7.5889 | 7.3718 | 5.8196 | 5.7928 | 6.2414 | 6.8326 | 6.9314 | 6.6794 | 7.4042 | 7.3823 | 4.6357 | 4.3093 | 4.6825 | 4.1867 |
| TSC1_S598 | TSC1 | Q92574 | 295 | SADVTTSPYADTQ | 5.8419 | 6.2624 | 7.7961 | 7.0241 | 6.2021 | 6.8276 | 7.3239 | 6.5812 | 5.8333 | 7.4239 | 7.5531 | 6.6777 | 4.6925 | 5.0009 | 4.5339 | 4.4254 |
| TSC2_S1097 | TSC2 | P49815 | 1132 | DRVRSMSGGHGLR | 7.1469 | 7.3726 | 7.4691 | 7.2345 | 6.0230 | 6.8215 | 6.0195 | 7.1694 | 7.3101 | 7.5702 | 7.7398 | 8.0720 | 3.1846 | 4.0105 | 3.2327 | 3.6235 |
| TSC2_S1132 | TSC2 | P49815 | 1449 | GPLPSSSPRSPSG | 6.7123 | 6.5805 | 7.5933 | 6.9824 | 6.6775 | 6.4021 | 6.2260 | 6.7480 | 6.9088 | 7.1632 | 7.0406 | 7.1068 | 4.3740 | 4.8148 | 4.0192 | 4.6506 |
| TSC2_S1155 | TSC2 | P49815 | 1420 | VKARSQSGTLDGE | 6.2227 | 5.9432 | 6.0664 | 6.1043 | 6.1987 | 5.8206 | 6.6807 | 6.6967 | 6.1309 | 6.4029 | 6.1208 | 5.7379 | 6.6802 | 7.0700 | 5.8558 | 6.2680 |
| TSC2_S1387 | TSC2 | P49815 | 1388 | LSKSSSSPELQTL | 6.9721 | 7.0339 | 7.9563 | 7.9738 | 5.8047 | 6.5235 | 6.0606 | 6.7472 | 7.3971 | 7.4655 | 7.7434 | 7.3225 | 3.8303 | 3.8619 | 3.4273 | 3.8800 |
| TSC2_S1388 | TSC2 | P49815 | 1387 | PLSKSSSSPELQT | 5.7059 | 5.7737 | 5.8910 | 6.2378 | 7.1288 | 6.3218 | 6.1619 | 6.5064 | 6.5084 | 6.5366 | 6.3063 | 6.4782 | 6.2544 | 6.5935 | 5.8093 | 5.7861 |
| TSC2_S1420 | TSC2 | P49815 | 960 | KQGLNNSPPVKEF | 6.0192 | 6.2587 | 10.2612 | 6.0506 | 9.9710 | 7.5278 | 5.7216 | 5.6115 | 6.3391 | 6.1629 | 7.1738 | 6.4052 | 4.0054 | 4.8658 | 3.8971 | 3.7294 |
| TSC2_S1449 | TSC2 | P49815 | 927 | LLSFDDTPEKDSF | 6.6490 | 7.0033 | 6.7811 | 7.0646 | 6.3432 | 6.2563 | 6.3793 | 6.6657 | 7.5970 | 7.0499 | 7.4531 | 7.0484 | 4.0687 | 4.7916 | 4.2017 | 4.6472 |
| TSC2_S1452 | TSC2 | P49815 | 664 | KTSGPLSPPTGPP | 6.9792 | 7.2771 | 7.3586 | 6.6619 | 5.9178 | 7.2717 | 7.2345 | 6.8892 | 7.6721 | 7.2089 | 7.3997 | 6.3406 | 3.4015 | 4.5631 | 3.2611 | 4.5631 |
| TSC2_S664 | TSC2 | P49815 | 1452 | PSSSPRSPSGLRP | 6.5149 | 7.0427 | 6.6805 | 6.1940 | 6.5853 | 6.2094 | 6.3232 | 6.7800 | 7.1110 | 6.9437 | 6.8318 | 7.5004 | 4.5737 | 5.0187 | 4.6773 | 5.0133 |
| TSC2_S939 | TSC2 | P49815 | 1155 | FLGSATSPGPRTA | 6.2288 | 6.3825 | 7.0151 | 6.3685 | 5.8895 | 6.6150 | 6.7537 | 6.5511 | 7.0740 | 6.5820 | 5.7708 | 5.8217 | 5.8945 | 5.7648 | 5.3317 | 5.9564 |
| TSC2_S960 | TSC2 | P49815 | 1097 | GPESSSSPGVHVR | 6.9911 | 6.1504 | 6.2744 | 6.9333 | 6.0418 | 6.2709 | 6.9959 | 6.9179 | 5.6970 | 5.9946 | 7.0088 | 5.5872 | 5.6427 | 6.3075 | 5.8883 | 5.2980 |
| TSC2_T927 | TSC2 | P49815 | 939 | FRARSTSLNERPK | 5.8676 | 5.9812 | 6.3793 | 6.1096 | 6.0575 | 6.3601 | 6.3197 | 6.2195 | 6.4611 | 6.3984 | 6.4169 | 6.3244 | 6.5660 | 6.2206 | 6.3387 | 5.9792 |
| TSC22D1_S128 | TSC22D1 | H7C4J9 | 128 | SLFQCFSPTRGAR | 7.1698 | 6.3076 | 6.0988 | 6.8445 | 6.9471 | 6.6995 | 6.2446 | 7.3713 | 6.1625 | 5.8512 | 6.9669 | 6.4746 | 5.4695 | 5.1068 | 5.1530 | 5.1325 |
| TSC22D2_S552 | TSC22D2 | O75157 | 598 | DDTRRKSEPLPQP | 5.9510 | 6.0985 | 6.7050 | 6.0730 | 6.4285 | 5.2606 | 5.7653 | 6.2876 | 7.0630 | 6.9000 | 6.5626 | 7.3040 | 5.8607 | 6.1282 | 5.6193 | 5.9927 |
| TSC22D2_S598 | TSC22D2 | O75157 | 552 | PVSRSSSIIQHVG | 7.1130 | 6.0956 | 7.4359 | 5.9451 | 6.1693 | 5.2691 | 6.2245 | 7.2050 | 7.6133 | 6.5457 | 6.3115 | 7.0795 | 5.3979 | 5.3912 | 5.6890 | 4.5144 |
| TSC22D4_S165 | TSC22D4 | Q9Y3Q8 | 279 | ASLVHKSPDPFGA | 6.3490 | 5.7288 | 6.2110 | 5.7090 | 6.9306 | 6.8824 | 6.1328 | 6.9816 | 6.4627 | 6.1896 | 6.3807 | 6.4182 | 6.5262 | 5.5951 | 5.7124 | 5.7900 |
| TSC22D4_S203 | TSC22D4 | Q9Y3Q8 | 165 | PGPQARSFTGGLG | 5.5767 | 4.5987 | 5.9582 | 4.6317 | 4.7502 | 7.0047 | 7.4277 | 6.0401 | 5.7949 | 5.3568 | 5.7973 | 4.8676 | 7.5064 | 9.3234 | 7.9057 | 7.4600 |

| | | | | | | | | | | | | | | | | | | | | | |
|---|---|---|---|---|---|---|---|---|---|---|---|---|---|---|---|---|---|---|---|---|---|
| TSC22D4_S260;S264 | TSC22D4 | Q9Y3Q8 | | 49 | LPNGEPSPDPGGK | 6.6676 | 6.4637 | 6.7885 | 6.2094 | 6.2960 | 6.6160 | 5.8709 | 7.0660 | 6.8049 | 6.8167 | 6.7213 | 7.0098 | 5.0940 | 5.2152 | 5.1013 | 5.2585 |
| TSC22D4_S264 | TSC22D4 | Q9Y3Q8 | | 211 | GTSRAATPLPSLR | 6.4281 | 5.7495 | 6.0789 | 5.3412 | 6.5452 | 7.4169 | 7.4159 | 7.8989 | 6.9048 | 5.6964 | 6.4263 | 6.3732 | 5.7990 | 5.3750 | 4.9372 | 5.6135 |
| TSC22D4_S279 | TSC22D4 | Q9Y3Q8 | | 203 | EPETGESAGTSRA | 6.1740 | 5.6800 | 6.9750 | 5.4728 | 7.7808 | 6.4220 | 6.0582 | 7.2116 | 6.2241 | 5.8264 | 6.7903 | 6.5752 | 6.4824 | 6.1352 | 5.1060 | 5.0859 |
| TSC22D4_S49 | TSC22D4 | Q9Y3Q8 | | 264 | LDSRPSSPALYFT | 5.9441 | 6.8811 | 5.9842 | 4.8418 | 6.4703 | 7.2302 | 5.7771 | 7.2653 | 7.0849 | 5.8373 | 5.5650 | 5.8456 | 6.8460 | 6.2348 | 6.6490 | 5.5433 |
| TSC22D4_T157 | TSC22D4 | Q9Y3Q8 | | 19 | TSVTTDYEGPGSP | 6.7734 | 6.7474 | 5.6824 | 7.4942 | 5.4356 | 7.4001 | 5.7506 | 5.5168 | 6.3935 | 6.5916 | 6.2701 | 5.6759 | 6.8611 | 5.6239 | 6.2409 | 5.5427 |
| TSC22D4_T211 | TSC22D4 | Q9Y3Q8 | | 157 | WLRPPPTSPGPQA | 6.3325 | 5.6489 | 5.7982 | 6.1179 | 6.3800 | 7.6620 | 6.7860 | 6.9674 | 6.5559 | 5.5770 | 5.1489 | 5.8834 | 6.7827 | 5.7341 | 6.7008 | 5.9243 |
| TSC22D4_Y19 | TSC22D4 | Q9Y3Q8 | 260;264 | | NA | 6.2497 | 6.1734 | 6.2157 | 6.3365 | 6.4894 | 6.4368 | 6.3796 | 6.7294 | 6.4426 | 6.1200 | 6.5371 | 6.6571 | 5.7889 | 5.9998 | 5.7745 | 5.6697 |
| TSEN54_S230 | TSEN54 | Q7Z6J9 | | 249 | EKPQESSPMKGPG | 5.7307 | 5.7538 | 6.4581 | 5.7619 | 5.7984 | 5.6635 | 5.5636 | 5.4854 | 6.1597 | 6.3975 | 7.2916 | 6.6446 | 6.4970 | 6.8952 | 6.8006 | 7.0986 |
| TSEN54_S249 | TSEN54 | Q7Z6J9 | | 230 | KSLAASSPPPCSQ | 6.1273 | 6.2829 | 7.6281 | 6.7973 | 5.8227 | 6.6137 | 6.0467 | 6.3986 | 6.3817 | 6.9665 | 6.6094 | 7.0161 | 5.4202 | 5.2923 | 5.3237 | 5.2727 |
| TSEN54_S267 | TSEN54 | Q7Z6J9 | | 76 | RVERLGSLVAAEW | 4.8085 | 5.5133 | 7.3660 | 5.9748 | 5.5512 | 6.0793 | 5.4479 | 4.8315 | 6.4925 | 5.7831 | 5.6649 | 6.8552 | 7.5359 | 7.3913 | 7.3465 | 7.3580 |
| TSEN54_S76 | TSEN54 | Q7Z6J9 | | 267 | LGSLGPSPGPARE | 5.4455 | 6.2539 | 7.3508 | 5.5577 | 6.0661 | 6.8699 | 8.0125 | 6.4577 | 6.7577 | 6.4714 | 5.6264 | 6.8332 | 5.6768 | 5.5440 | 5.8508 | 5.2257 |
| TSHZ1_S671 | TSHZ1 | Q6ZSZ6 | | 671 | SLAKAASPIAKEN | 6.3308 | 7.0021 | 6.2499 | 6.6419 | 6.3338 | 6.2778 | 6.3245 | 6.2844 | 6.3982 | 6.7003 | 6.6993 | 6.4885 | 5.2108 | 5.9083 | 5.4802 | 5.6691 |
| TSHZ3_S463 | TSHZ3 | Q63HK5 | | 463 | NTPASISPKLNVE | 6.8801 | 8.4090 | 5.5434 | 8.7577 | 6.1469 | 6.9426 | 6.0217 | 5.1321 | 5.8384 | 6.9114 | 5.8071 | 7.1662 | 5.6551 | 4.2738 | 5.0740 | 5.4406 |
| TSHZ3_S584 | TSHZ3 | Q63HK5 | | 584 | GNSEIVSPTKNQT | 6.2889 | 6.3340 | 7.0753 | 7.2009 | 6.6516 | 6.5039 | 6.6912 | 5.6778 | 6.5637 | 7.5391 | 7.0326 | 7.1150 | 4.5675 | 5.4674 | 4.5430 | 4.7482 |
| TSHZ3_S600 | TSHZ3 | Q63HK5 | | 600 | PPSSQTSPMPKTN | 5.7591 | 6.0601 | 6.7353 | 6.2947 | 7.0558 | 7.4220 | 6.6781 | 5.3883 | 6.9162 | 6.6403 | 6.0967 | 7.7837 | 5.0667 | 6.0189 | 4.2964 | 5.7877 |
| TSHZ3_T340 | TSHZ3 | Q63HK5 | | 340 | PDSTGGTPKATIS | 6.3795 | 6.3508 | 6.7210 | 7.3232 | 6.1813 | 6.0131 | 6.9556 | 6.7108 | 6.5350 | 7.1760 | 6.3865 | 6.3769 | 4.5538 | 5.3420 | 5.4510 | 5.5434 |
| TSNAX_S32 | TSNAX | Q99598 | | 32 | GKDVNSSSPVMLA | 6.0064 | 6.1611 | 6.8681 | 5.7523 | 6.5052 | 6.8262 | 6.2896 | 6.6449 | 6.0191 | 6.5047 | 6.4444 | 6.5988 | 5.6505 | 5.8763 | 6.0179 | 5.8346 |
| TSPAN9_S14 | TSPAN9 | A0A1W2PQ04 | | 14 | DSSIHRSPSLDSK | 6.6797 | 6.8455 | 6.0402 | 6.8269 | 5.8096 | 5.9890 | 6.1696 | 6.0035 | 6.2750 | 6.6205 | 6.0316 | 6.1918 | 5.6375 | 6.3436 | 6.3571 | 6.1791 |
| TSPYL4_T328 | TSPYL4 | Q9UJ04 | | 328 | RVVSLSTPIRWHR | 6.1032 | 8.1476 | 6.7005 | 7.8439 | 6.0483 | 7.1689 | 7.0156 | 7.4141 | 7.7520 | 6.9386 | 7.1389 | 6.3976 | 2.8898 | 4.9712 | 3.3339 | 4.1358 |
| TSR3_S228 | TSR3 | Q9UJK0 | | 228 | LANAKESPQEEEI | 5.9903 | 5.7666 | 7.2275 | 6.4597 | 5.0362 | 6.9094 | 6.0693 | 6.1044 | 5.1568 | 6.6484 | 5.6218 | 7.0301 | 6.8787 | 6.4312 | 6.4510 | 6.2185 |
| TSSC4_S132 | TSSC4 | Q9Y5U2 | | 132 | LAPSGRSPVEGLG | 6.6169 | 7.0876 | 6.3888 | 6.6036 | 6.5784 | 6.6326 | 6.3622 | 6.8359 | 7.4096 | 6.4875 | 6.8998 | 7.2581 | 4.7668 | 4.8965 | 4.3701 | 4.8055 |
| TTC1_S67 | TTC1 | Q99614 | | 83 | DKVENKSNEDVNS | 6.5674 | 6.4569 | 6.7338 | 6.5689 | 5.9954 | 6.0801 | 6.3607 | 6.4514 | 6.7575 | 6.7622 | 6.5366 | 6.9488 | 5.4847 | 5.5844 | 5.2029 | 5.5084 |
| TTC1_S83 | TTC1 | Q99614 | | 67 | ECFHDCSASFEEE | 6.6337 | 6.2406 | 7.1370 | 6.3384 | 6.6590 | 6.7388 | 6.5748 | 6.3079 | 7.2297 | 6.6954 | 7.1694 | 7.5955 | 5.2284 | 4.3368 | 4.4890 | 4.6254 |
| TTC14_S454 | TTC14 | Q96N46 | | 670 | GSVRHSTSPASSE | 6.3086 | 6.9195 | 6.1750 | 7.1250 | 6.5778 | 6.2871 | 5.5533 | 6.7427 | 5.7749 | 6.2384 | 6.0030 | 6.2357 | 5.6868 | 6.0009 | 6.4515 | 5.9197 |
| TTC14_S629 | TTC14 | Q96N46 | | 454 | TKKIETSAEKLRK | 6.0122 | 6.6667 | 6.0306 | 6.9902 | 6.1950 | 6.3720 | 7.0626 | 6.1722 | 7.0847 | 7.4150 | 7.1948 | 7.5719 | 4.0878 | 4.9900 | 4.7435 | 5.4109 |
| TTC14_T670 | TTC14 | Q96N46 | | 629 | FSSRRNSSDSFCR | 5.0598 | 4.5778 | 5.0468 | 5.7042 | 7.3034 | 5.8301 | 5.3476 | 5.5622 | 5.9401 | 5.8883 | 5.7890 | 5.8098 | 8.4720 | 8.0660 | 7.8533 | 7.7496 |
| TTC28_S2079 | TTC28 | Q96AY4 | | 2398 | LSLLNLSPRHNKK | 6.9119 | 6.5359 | 7.1438 | 6.9837 | 6.9613 | 7.5103 | 6.3409 | 7.2994 | 6.8842 | 6.8300 | 6.8190 | 7.0208 | 3.9322 | 4.8049 | 3.9781 | 4.0437 |
| TTC28_S2117 | TTC28 | Q96AY4 | | 47 | DTIGQRSPDGPVL | 6.3566 | 6.8380 | 6.7900 | 6.7883 | 6.1565 | 6.7521 | 6.7238 | 6.7752 | 6.8562 | 6.9497 | 6.9918 | 7.0731 | 4.4855 | 4.9603 | 4.4829 | 5.0200 |
| TTC28_S2117;S2120 | TTC28 | Q96AY4 | | 28 | RRREPESPPASAP | 5.8575 | 6.6312 | 6.6960 | 5.9614 | 5.9843 | 6.4555 | 6.0583 | 6.5629 | 6.5963 | 6.2632 | 6.0440 | 6.7748 | 6.0173 | 6.5302 | 5.8635 | 5.7037 |
| TTC28_S2120 | TTC28 | Q96AY4 | | 2120 | LIPSPNSPFQKVG | 6.6438 | 6.7791 | 6.8279 | 6.4046 | 6.4528 | 7.1621 | 6.6386 | 7.3003 | 6.7351 | 6.5816 | 7.1133 | 7.2587 | 4.3714 | 4.9709 | 4.3104 | 4.4494 |
| TTC28_S2224 | TTC28 | Q96AY4 | | 2224 | VLSHQKSQPSPVT | 5.1422 | 4.4579 | 5.7222 | 4.6808 | 5.2654 | 5.3232 | 5.0232 | 5.0532 | 5.2628 | 5.7326 | 5.4327 | 5.4127 | 9.6111 | 9.6881 | 9.3474 | 8.8447 |
| TTC28_S2398 | TTC28 | Q96AY4 | | 2117 | KMTLIPSPNSPFQ | 6.5348 | 6.1994 | 7.1870 | 6.5761 | 7.0168 | 6.9337 | 5.6649 | 6.6068 | 6.3874 | 6.3227 | 6.1717 | 6.4993 | 5.5977 | 6.2319 | 5.0588 | 5.0111 |
| TTC28_S28 | TTC28 | Q96AY4 | | 2079 | NRNTCFSPDHKQP | 6.6993 | 6.1857 | 7.2884 | 6.7837 | 6.4943 | 7.1030 | 6.9670 | 6.4136 | 7.5970 | 7.3345 | 7.5349 | 7.7736 | 3.4756 | 4.1861 | 3.7358 | 4.4277 |
| TTC28_S47 | TTC28 | Q96AY4 | 2117;2120 | | NA | 5.8857 | 6.8196 | 6.5049 | 7.2493 | 6.4736 | 6.7006 | 6.8502 | 7.3387 | 7.1347 | 6.6363 | 7.0038 | 6.8872 | 4.5885 | 4.9835 | 4.1294 | 4.8141 |
| TTC3_S1244 | TTC3 | P53804 | | 1244 | KPVSANSPKPACE | 5.8878 | 6.1894 | 8.0551 | 8.0133 | 5.5354 | 7.2830 | 7.3738 | 6.3565 | 6.9724 | 9.3667 | 8.5238 | 7.2121 | 2.9993 | 4.5744 | 2.2708 | 3.3863 |
| TTC33_S197 | TTC33 | Q6PID6 | | 197 | AEVTHFSPKSIPD | 5.8596 | 6.1830 | 6.3225 | 6.1206 | 6.1035 | 6.1232 | 5.9993 | 6.2128 | 6.1073 | 6.1412 | 6.3306 | 6.3393 | 6.5076 | 6.7048 | 6.5019 | 6.4427 |
| TTC4_S267 | TTC4 | O95801 | | 47 | KVPLFMSRAPSEI | 8.1811 | 6.6688 | 7.6397 | 7.0482 | 5.6886 | 5.7679 | 6.2849 | 6.3635 | 6.9840 | 5.6017 | 6.3645 | 6.7307 | 5.2046 | 5.5127 | 4.7937 | 5.1653 |
| TTC4_S47 | TTC4 | O95801 | | 51 | FMSRAPSEIDPRE | 6.2492 | 6.2579 | 6.8021 | 6.2893 | 6.0955 | 6.7894 | 6.1709 | 6.7808 | 6.6732 | 6.2931 | 6.7131 | 6.8895 | 5.1733 | 5.7895 | 5.2575 | 5.7758 |
| TTC4_S51 | TTC4 | O95801 | | 267 | PHGARLSLDGQGR | 6.6903 | 6.9854 | 6.5799 | 6.7308 | 6.0014 | 6.5520 | 7.0680 | 7.3811 | 6.7467 | 6.4031 | 7.2430 | 7.1284 | 4.3272 | 4.9345 | 4.5888 | 4.6394 |
| TTC7A_S182 | TTC7A | Q9ULT0-4 | | 671 | GLEKDGSFGEGLT | 5.9880 | 6.2131 | 6.3196 | 6.0550 | 5.9481 | 5.8771 | 5.8821 | 5.4147 | 6.2967 | 6.2191 | 5.9922 | 6.2027 | 7.0202 | 6.7871 | 6.5965 | 7.1875 |
| TTC7A_S51 | TTC7A | Q9ULT0-4 | | 182 | LERLPNSIASRFR | 7.6303 | 7.2711 | 6.5564 | 6.5340 | 6.7764 | 6.0081 | 6.3378 | 5.9154 | 6.5620 | 5.8062 | 5.8864 | 5.7608 | 5.7235 | 6.0071 | 5.8020 | 5.4226 |
| TTC7A_S671 | TTC7A | Q9ULT0-4 | | 51 | GGNRRGSPSAAFT | 6.4023 | 6.1827 | 6.2020 | 6.7550 | 5.9640 | 6.4128 | 6.4010 | 6.3260 | 6.3576 | 6.2162 | 6.7392 | 6.4242 | 5.7018 | 6.1949 | 5.7499 | 5.9705 |
| TTC7B_S160 | TTC7B | Q86TV6-2 | | 160 | KLPISSSTSNLHV | 6.2588 | 6.4638 | 6.3694 | 6.9111 | 6.1175 | 6.2856 | 6.5065 | 6.4326 | 6.2988 | 6.2313 | 6.0231 | 6.0307 | 6.1621 | 6.0777 | 6.0916 | 5.7393 |
| TTC7B_S630 | TTC7B | Q86TV6-2 | | 695 | LSEVASSLQSSAP | 5.7149 | 5.9635 | 5.7548 | 5.8879 | 5.5768 | 6.7115 | 6.2994 | 6.5702 | 6.4692 | 7.5967 | 7.0144 | 7.4534 | 4.8535 | 6.3244 | 5.1705 | 6.6392 |
| TTC7B_S695 | TTC7B | Q86TV6-2 | | 630 | DSGRGSSLLDRTI | 7.0883 | 6.8483 | 6.6898 | 7.3403 | 6.0922 | 6.3856 | 6.7808 | 5.9464 | 6.7586 | 7.3307 | 6.5638 | 6.1126 | 4.6595 | 5.4621 | 4.9542 | 4.9867 |
| TTF1_S65 | TTF1 | Q15361 | | 65 | FQHLISSPLKKSR | 6.4417 | 6.6833 | 6.2809 | 7.3116 | 6.3922 | 5.7149 | 6.4216 | 6.2115 | 6.4716 | 6.1257 | 6.3135 | 6.1489 | 5.4008 | 6.0320 | 5.8270 | 6.2228 |
| TTF1_S872 | TTF1 | Q15361 | | 12 | SRFEIHTPVSDKK | 7.0886 | 6.7915 | 7.0146 | 7.0495 | 6.2761 | 6.0878 | 6.5641 | 6.6897 | 6.3806 | 6.2099 | 6.9390 | 7.2433 | 4.1048 | 5.1572 | 4.7862 | 5.6171 |
| TTF1_T12 | TTF1 | Q15361 | | 15 | EIHTPVSDKKKKK | 6.8740 | 7.3339 | 6.9071 | 7.5809 | 6.3794 | 7.0226 | 6.8372 | 6.5906 | 7.1335 | 6.8981 | 7.0796 | 6.8073 | 3.5007 | 4.5449 | 3.7497 | 4.7606 |
| TTF1_T12;S15 | TTF1 | Q15361 | | 872 | FYYEDDSEGEDIE | 5.9589 | 5.9990 | 6.3301 | 6.7996 | 6.2094 | 5.6963 | 6.5139 | 5.8418 | 6.6491 | 6.0369 | 5.9598 | 6.1797 | 6.5419 | 6.4121 | 6.4729 | 6.3986 |
| TTF1_T15 | TTF1 | Q15361 | 12;15 | | NA | 6.9957 | 7.0094 | 6.8467 | 7.2722 | 6.6334 | 6.7379 | 7.3226 | 6.6929 | 7.2472 | 7.3764 | 7.3543 | 7.2029 | 3.5747 | 4.5516 | 3.1333 | 4.0488 |
| TTF2_S368 | TTF2 | Q9UNY4 | | 883 | GNQSGRSPNNPFS | 6.8992 | 7.1111 | 6.3914 | 6.8826 | 5.9474 | 6.0143 | 6.4420 | 6.7043 | 7.2944 | 6.8976 | 6.6913 | 6.9681 | 4.4732 | 5.3895 | 4.7126 | 5.1810 |
| TTF2_S396 | TTF2 | Q9UNY4 | | 368 | VSSKPGSPLLFDS | 6.2499 | 6.0723 | 7.4773 | 6.6368 | 6.1775 | 5.4699 | 6.5311 | 5.7614 | 6.0764 | 6.4798 | 5.5659 | 6.5819 | 6.1000 | 6.1319 | 6.1965 | 6.4916 |
| TTF2_S396;S399 | TTF2 | Q9UNY4 | | 396 | SVQRKVSPASGVS | 6.6728 | 7.2064 | 7.1875 | 7.8589 | 5.9131 | 6.7538 | 6.9229 | 6.4053 | 7.3944 | 7.4065 | 7.2287 | 7.8896 | 3.3443 | 4.2638 | 3.2092 | 4.3427 |
| TTF2_S883 | TTF2 | Q9UNY4 | 396;399 | | NA | 6.0648 | 6.1896 | 6.2967 | 6.3804 | 6.0387 | 6.4157 | 6.2866 | 6.5619 | 7.1057 | 6.8036 | 6.1117 | 6.8404 | 5.7763 | 6.1882 | 5.2287 | 5.7110 |
| TTI1_S459 | TTI1 | O43156 | | 459 | SDDLNASPKTSAT | 5.5522 | 6.4094 | 7.5611 | 7.2088 | 6.0224 | 6.6712 | 5.7181 | 6.2970 | 7.4044 | 7.5441 | 7.0679 | 7.2689 | 4.5481 | 4.6854 | 4.3428 | 5.6980 |
| TTK_S258 | TTK | P33981 | | 436 | FSVSKQSPPISTS | 7.1818 | 7.3645 | 7.4421 | 7.1902 | 6.4129 | 6.6978 | 6.7814 | 6.8475 | 7.5169 | 7.0605 | 7.6924 | 7.3013 | 3.4000 | 4.0001 | 3.2802 | 3.8306 |
| TTK_S281 | TTK | P33981 | | 281 | PVNLLNSPDCDVK | 6.5574 | 6.5872 | 7.2682 | 7.5108 | 6.3551 | 6.7941 | 7.2332 | 6.7027 | 7.2624 | 7.1554 | 7.8123 | 7.2261 | 2.9567 | 4.5969 | 3.6578 | 4.3238 |
| TTK_S321 | TTK | P33981 | | 258 | EIGYRNSLRQTNK | 8.6246 | 5.7774 | 7.9905 | 7.2015 | 5.2454 | 6.6432 | 6.7868 | 5.5407 | 8.9716 | 7.4230 | 7.8529 | 8.5983 | 3.5178 | 4.1126 | 2.3830 | 3.3307 |
| TTK_S345 | TTK | P33981 | | 12 | LSGRELTIDSIMN | 10.6597 | 7.2251 | 5.1156 | 5.9471 | 5.5658 | 6.5752 | 6.4771 | 4.5856 | 9.6976 | 4.1862 | 5.0975 | 6.3247 | 6.3610 | 5.8672 | 4.1027 | 6.2121 |
| TTK_S436 | TTK | P33981 | | 321 | KPSGNDSCELRNL | 6.7688 | 7.0591 | 6.8824 | 7.6052 | 5.9736 | 8.3213 | 6.8767 | 7.1572 | 7.5476 | 6.9070 | 9.8024 | 8.2715 | 2.7798 | 3.4255 | 2.0492 | 2.5727 |
| TTK_S455 | TTK | P33981 | | 821 | QLVGLNSPNSILK | 7.4244 | 7.2675 | 6.8266 | 7.2681 | 6.6709 | 7.5962 | 7.1656 | 7.1078 | 7.8913 | 7.8166 | 7.3464 | 7.6228 | 2.7165 | 4.0874 | 2.2264 | 2.9657 |
| TTK_S821 | TTK | P33981 | | 345 | LVSDEKSSELIIT | 6.8324 | 6.9027 | 6.6455 | 7.4732 | 5.7093 | 6.4597 | 6.9662 | 6.2279 | 6.9621 | 6.7972 | 7.2304 | 7.5701 | 3.9466 | 4.9508 | 4.1929 | 5.1331 |
| TTK_T12 | TTK | P33981 | | 455 | SICKTPSSNTLDD | 5.9731 | 5.3616 | 4.8544 | 5.0617 | 5.2667 | 5.6428 | 5.0137 | 6.2777 | 6.3152 | 5.6884 | 5.6346 | 5.9567 | 7.5768 | 7.7689 | 8.2105 | 9.3972 |
| TTLL12_S16 | TTLL12 | Q14166 | | 16 | RPAERSSPGQTPE | 5.7522 | 5.9192 | 6.4858 | 5.8780 | 6.0362 | 6.4404 | 5.9304 | 6.6199 | 6.2256 | 6.3228 | 6.3009 | 6.4117 | 6.3262 | 6.6998 | 6.1801 | 6.4708 |
| TTLL4_S1143 | TTLL4 | Q14679 | | 207 | ISGKIPSPLSSSY | 6.6535 | 6.8875 | 7.0575 | 6.8851 | 6.2618 | 7.0876 | 7.2863 | 7.3495 | 7.0036 | 6.7200 | 7.1985 | 6.8512 | 3.8358 | 4.6013 | 3.7894 | 4.5316 |
| TTLL4_S207 | TTLL4 | Q14679 | | 1143 | KTQAGLSPYPQKP | 7.1992 | 6.8140 | 7.1645 | 6.3692 | 5.9298 | 7.0115 | 6.8006 | 7.9571 | 6.6654 | 6.7046 | 7.5656 | 7.2169 | 3.8162 | 4.7057 | 3.5217 | 4.5581 |
| TTYH2_S444 | TTYH2 | Q9BSA4 | | 444 | WRMAAHSPPRGQL | 7.0810 | 7.0511 | 6.6722 | 6.8904 | 5.3148 | 5.4227 | 6.2932 | 7.0121 | 7.5128 | 6.4517 | 7.1683 | 7.5610 | 4.9473 | 5.3584 | 4.8347 | 4.4282 |
| TTYH3_S496 | TTYH3 | Q9C0H2 | | 496 | PLIGRESPPPSYT | 6.5365 | 7.3063 | 6.5717 | 6.8022 | 6.2835 | 6.3296 | 6.3536 | 5.8895 | 6.8843 | 7.3653 | 7.6139 | 7.2019 | 4.8230 | 5.5756 | 4.1546 | 4.3085 |
| TTYH3_S522 | TTYH3 | Q9C0H2 | | 522 | RPDSSGSHxxxxx | 7.4455 | 7.9917 | 6.3497 | 7.2490 | 5.9594 | 6.6932 | 5.7814 | 7.4942 | 5.9469 | 7.0779 | 6.8686 | 6.6445 | 4.0459 | 5.0610 | 5.1334 | 4.2578 |
| TUBA1B_S158 | TUBA1B | P68363 | | 340 | TIKTKRSIQFVDW | 6.1216 | 5.8187 | 6.5682 | 5.8315 | 6.9222 | 6.6972 | 5.7639 | 6.2128 | 6.4204 | 6.9117 | 6.4426 | 6.7603 | 5.9816 | 6.1907 | 5.5298 | 5.8267 |
| TUBA1B_S340 | TUBA1B | P68363 | | 158 | LLMERLSVDYGKK | 4.5327 | 4.3698 | 4.8959 | 4.7508 | 6.8111 | 5.9035 | 4.3888 | 6.8148 | 7.4029 | 11.7834 | 7.5206 | 11.5137 | 5.5033 | 5.2964 | 3.6983 | 4.8141 |
| TUBA1B_S48 | TUBA1B | P68363 | | 48 | IGGGDDSFNTFFS | 6.0925 | 6.4693 | 6.1712 | 6.1182 | 7.0344 | 6.8917 | 6.4642 | 6.7539 | 6.4741 | 6.9024 | 6.7642 | 7.2744 | 4.7109 | 5.6793 | 4.2242 | 5.9752 |
| TUBB_S172 | TUBB | P07437 | | 172 | TFSVVPSPKVSDT | 5.4787 | 6.2045 | 6.9020 | 5.7435 | 7.3449 | 7.3700 | 6.1562 | 7.4932 | 6.2111 | 7.7004 | 6.8463 | 7.7282 | 5.2080 | 4.9910 | 3.9683 | 4.6536 |
| TUBB_S275 | TUBB | P07437 | | 275 | GFAPLTSRGSQQY | 5.0767 | 4.9074 | 5.3815 | 6.2248 | 6.3315 | 6.9530 | 5.4306 | 7.6286 | 4.9683 | 7.0681 | 8.0757 | 7.0342 | 6.3383 | 5.3358 | 7.1731 | 6.0724 |
| TUBG1_S80 | TUBG1 | P23258 | | 80 | IHSILNSPYAKLY | 5.4079 | 6.1378 | 7.1566 | 7.1643 | 6.8793 | 6.5615 | 6.5480 | 5.9760 | 7.2914 | 5.6486 | 6.3863 | 7.7151 | 5.7507 | 5.5119 | 5.1248 | 4.7396 |
| TULP3_S9 | TULP3 | O75386-2 | | 9 | ASRCRLSPSGDSV | 6.9874 | 7.3900 | 6.4263 | 7.6438 | 7.4271 | 7.3888 | 6.9439 | 7.1540 | 7.0001 | 6.9191 | 7.0870 | 7.1312 | 3.3467 | 4.4121 | 3.3311 | 3.4115 |
| TULP4_S1374 | TULP4 | Q9NRJ4 | | 1374 | FNSLISSPHLGRE | 6.7057 | 5.9803 | 7.2327 | 5.8556 | 5.1946 | 7.2493 | 7.7638 | 5.6311 | 6.6517 | 6.6330 | 6.6652 | 6.5000 | 5.5500 | 6.2006 | 4.9202 | 5.2663 |
| TUSC2_S50 | TUSC2 | O75896 | | 50 | VFTRRGSMFYDED | 5.9268 | 6.0916 | 6.6076 | 6.1697 | 6.2427 | 6.2649 | 6.0422 | 6.5217 | 6.6888 | 6.4539 | 6.6617 | 6.6074 | 6.2761 | 5.9993 | 5.8298 | 5.6156 |
| TUT1_S133 | TUT1 | Q9H6E5 | | 133 | EQKEFQSPASKSP | 6.1672 | 6.5386 | 6.6659 | 6.7493 | 6.2916 | 7.2295 | 7.1077 | 6.8871 | 6.9595 | 6.9101 | 7.5011 | 7.4527 | 3.9150 | 4.9368 | 3.9639 | 4.7240 |
| TUT4_S104 | TUT4 | Q5TAX3 | | 104 | AKKFPNSPVKAEK | 6.5155 | 6.5617 | 7.1071 | 6.5812 | 6.2481 | 6.8109 | 6.4318 | 6.7682 | 6.8615 | 7.0615 | 6.8409 | 6.9026 | 4.8357 | 5.3113 | 4.3910 | 4.7711 |
| TUT4_S156 | TUT4 | Q5TAX3 | | 209 | KCALQNSPRSQKQ | 6.5231 | 6.9599 | 7.0839 | 6.1004 | 6.1934 | 6.5405 | 6.4617 | 6.4715 | 6.7465 | 6.7606 | 6.9683 | 6.8591 | 5.1259 | 5.3865 | 4.6614 | 5.1574 |
| TUT4_S209 | TUT4 | Q5TAX3 | | 831 | CLKKELSQCNCID | 1.6920 | 3.5473 | 4.4961 | 2.9306 | 2.9728 | 2.1453 | 3.1731 | 3.9058 | 2.7409 | 2.7251 | 2.5511 | 3.0835 | 16.1132 | 14.6532 | 17.7630 | 15.5071 |
| TUT4_S831 | TUT4 | Q5TAX3 | | 156 | SEKVPSSPAEAEK | 6.4858 | 6.6696 | 6.6052 | 6.1080 | 5.5807 | 6.6919 | 6.6426 | 6.6008 | 7.1168 | 7.1020 | 6.6281 | 7.1215 | 5.3858 | 5.4660 | 4.8236 | 4.9716 |
| TUT7_S600 | TUT7 | Q5VYS8 | | 1414 | DKPIQCTPQKAKP | 6.7814 | 5.8438 | 7.1514 | 7.1090 | 5.4585 | 6.3513 | 6.7622 | 5.8721 | 7.9719 | 7.1960 | 7.1097 | 7.5330 | 4.3837 | 5.1747 | 4.0557 | 5.2455 |
| TUT7_S706 | TUT7 | Q5VYS8 | | 64 | PGNYGNTPRKGPC | 6.6403 | 7.9619 | 6.4521 | 7.0144 | 5.5952 | 6.1537 | 6.7687 | 6.4417 | 8.9254 | 7.7203 | 7.3645 | 8.0439 | 3.6526 | 4.0813 | 3.1099 | 4.0741 |
| TUT7_S939 | TUT7 | Q5VYS8 | | 939 | VCEEKNSPVDQSD | 5.2408 | 5.3785 | 7.3289 | 5.9071 | 5.9554 | 7.3524 | 7.3172 | 5.7987 | 6.9629 | 5.3785 | 6.7007 | 6.3185 | 5.9832 | 6.0198 | 5.8734 | 6.4840 |
| TUT7_T1414 | TUT7 | Q5VYS8 | | 600 | AIEDPYSVKRNVA | 5.9208 | 6.2621 | 6.9563 | 8.3165 | 6.2385 | 5.5726 | 6.8880 | 5.9844 | 6.7327 | 6.4762 | 5.6784 | 6.0032 | 4.5348 | 7.2951 | 5.9608 | 5.1796 |
| TUT7_T64 | TUT7 | Q5VYS8 | | 706 | TAESFGSPPKEEM | 6.7001 | 6.7771 | 6.7190 | 6.4656 | 5.9955 | 5.7473 | 5.7504 | 5.8369 | 6.4798 | 6.7676 | 6.3583 | 6.9752 | 5.7137 | 5.8329 | 5.6289 | 6.2517 |
| TWF1_S143 | TWF1 | Q12792-3 | | 143 | YLLSQSSPAPLTA | 6.0121 | 6.0794 | 6.2908 | 6.6981 | 6.4517 | 6.3784 | 7.0700 | 7.1218 | 6.4763 | 6.8550 | 6.4783 | 6.7204 | 4.6894 | 5.9934 | 5.0052 | 5.6796 |
| TXLNA_S515 | TXLNA | P40222 | | 515 | GAQAPSSPRVTEA | 6.3669 | 6.3932 | 6.7364 | 6.1949 | 6.3807 | 6.6878 | 6.1167 | 6.8274 | 6.5747 | 6.6027 | 6.7801 | 6.6711 | 5.5231 | 5.5484 | 5.2022 | 5.3937 |
| TXLNG_S517 | TXLNG | Q9NUQ3 | | 97 | ENRNLVSPAYCTQ | 6.6230 | 6.7808 | 7.2075 | 6.5295 | 6.3620 | 6.7584 | 6.9162 | 7.0910 | 6.5854 | 6.1836 | 7.2403 | 7.0248 | 4.4058 | 5.4291 | 4.7093 | 4.1533 |
| TXLNG_S97 | TXLNG | Q9NUQ3 | | 517 | AVQKPPSTGSAPA | 6.5705 | 6.9578 | 6.4286 | 5.9518 | 6.4084 | 6.7540 | 5.2541 | 6.7839 | 6.4585 | 6.7529 | 6.5801 | 7.2885 | 4.5660 | 5.5058 | 5.8216 | 5.9176 |
| TXNDC9_S188 | TXNDC9 | O14530 | | 188 | SDILNYSGNLMEP | 6.0611 | 6.4128 | 6.6072 | 6.5196 | 6.3524 | 6.9926 | 6.3843 | 6.9490 | 6.4765 | 6.8143 | 7.0701 | 6.4563 | 4.9762 | 5.9464 | 4.8678 | 5.1131 |
| TXNL4A_S132 | TXNL4A | P83876 | | 132 | GRGLVVSPKDYST | 5.9937 | 6.5331 | 7.0992 | 7.0542 | 5.9733 | 7.1968 | 7.0298 | 6.6901 | 7.3709 | 8.1738 | 7.8027 | 8.6900 | 3.3309 | 3.9812 | 2.9560 | 4.1241 |
| TXNL4B_S132 | TXNL4B | Q9NX01 | | 132 | GKLIVQSPIDPKN | 6.2741 | 5.9613 | 6.1286 | 5.6388 | 6.0128 | 6.0407 | 8.0866 | 9.5184 | 7.3677 | 7.4998 | 7.0312 | 7.9387 | 2.7539 | 5.1211 | 3.2443 | 5.3818 |
| TXNRD3_S32 | TXNRD3 | Q86VQ6 | | 32 | RGARVLSPPGRRA | 6.7847 | 5.9554 | 6.8686 | 5.9030 | 6.6202 | 5.7109 | 6.3257 | 7.8596 | 7.1194 | 5.7380 | 6.6714 | 6.1670 | 5.7155 | 6.3022 | 5.0915 | 5.1669 |
| TXNRD3_S4 | TXNRD3 | Q86VQ6 | | 4 | xxxMERSPPQSPG | 5.8428 | 6.1750 | 6.3182 | 5.5404 | 5.8205 | 5.8813 | 6.4453 | 5.8775 | 6.7446 | 6.2507 | 6.8203 | 6.4453 | 6.3597 | 6.8543 | 6.3610 | 6.2631 |
| TXNRD3_S42 | TXNRD3 | Q86VQ6 | | 42 | RRARLSSPGPSRS | 6.7659 | 6.5838 | 7.1612 | 5.6664 | 5.8812 | 6.8122 | 6.0725 | 6.7841 | 6.7113 | 6.3757 | 6.4120 | 6.1436 | 5.6491 | 5.9658 | 5.2648 | 5.7504 |
| TYK2_Y292 | TYK2 | P29597 | | 292 | AEGEPCYIRDSGV | 6.1648 | 6.1804 | 6.7614 | 6.2006 | 5.6946 | 6.2512 | 6.2622 | 6.8001 | 6.5428 | 6.3810 | 6.5635 | 6.3656 | 5.9189 | 6.2893 | 5.5411 | 6.0825 |
| TYMS_S114 | TYMS | P04818 | | 114 | IWDANGSRDFLDS | 5.9086 | 7.0111 | 7.9186 | 7.1964 | 6.0356 | 6.4250 | 7.1439 | 7.4333 | 6.8734 | 7.7218 | 7.8841 | 7.9890 | 3.0658 | 3.7724 | 3.5683 | 4.0527 |
| TYW1_S251 | TYW1 | Q9NV66 | | 251 | KGERKKSCGGHCK | 4.5860 | 5.0163 | 4.7680 | 4.9922 | 6.0296 | 5.2894 | 5.8506 | 5.0554 | 7.3980 | 7.2814 | 5.9807 | 8.2880 | 6.5126 | 8.0894 | 7.3078 | 7.5545 |
| U2AF1_S19 | U2AF1 | Q01081-2 | | 19 | KDKVNCSFYFKIG | 7.1993 | 8.0580 | 7.1196 | 7.9563 | 7.1196 | 6.6398 | 7.2961 | 6.7459 | 6.6517 | 6.8204 | 6.6657 | 5.3622 | 3.6384 | 4.3805 | 4.1914 | 4.1550 |
| U2AF2_S79 | U2AF2 | P26368 | | 79 | HGGLIRSPRHEKK | 6.4181 | 6.8001 | 6.2797 | 6.5512 | 6.4950 | 6.5804 | 6.3422 | 5.8481 | 6.1171 | 6.3835 | 6.4675 | 6.5250 | 5.7520 | 5.9933 | 5.6587 | 5.7880 |
| U2SURP_S485 | U2SURP | O15042 | | 485 | SILQGDSPTKWRT | 6.3037 | 6.5599 | 6.3391 | 6.5111 | 6.1625 | 5.9332 | 6.3427 | 6.1228 | 6.5451 | 6.2706 | 6.2101 | 6.2972 | 6.0111 | 6.3637 | 5.9460 | 6.0812 |
| U2SURP_S67 | U2SURP | O15042 | | 67 | RESLCDSPHQNLS | 7.3473 | 7.1231 | 6.6569 | 7.3533 | 6.5052 | 6.7063 | 6.6953 | 5.6129 | 6.8838 | 6.8367 | 6.6988 | 6.4856 | 4.9029 | 5.2489 | 4.3777 | 4.5652 |
| U2SURP_S788;S800 | U2SURP | O15042 | | 5 | xxMADKTPGGSQK | 6.8407 | 7.5482 | 6.4501 | 7.5098 | 6.0681 | 6.4019 | 7.1724 | 6.7107 | 7.1134 | 7.0598 | 6.6331 | 6.7573 | 3.8670 | 4.7948 | 4.1557 | 4.9170 |
| U2SURP_S788;S800;T80 | U2SURP | O15042 | | 974 | RSSHKDSPRDVSK | 6.4650 | 7.6598 | 5.4823 | 7.3980 | 7.4078 | 7.0243 | 5.7374 | 7.0409 | 5.7182 | 5.8268 | 6.6745 | 5.9065 | 5.3647 | 6.2075 | 5.6163 | 4.4700 |
| U2SURP_S811 | U2SURP | O15042 | | 811 | DTQSSKSEEHHLY | 6.1935 | 7.2968 | 6.4766 | 7.4489 | 6.3798 | 5.5512 | 6.2574 | 6.4405 | 5.4067 | 6.3663 | 5.0913 | 5.7391 | 5.5308 | 6.2375 | 6.7693 | 6.8143 |
| U2SURP_S974 | U2SURP | O15042 | 788;800;806;8 | | NA | 5.7549 | 5.8437 | 6.2290 | 6.6808 | 6.1611 | 5.9704 | 6.0677 | 6.0824 | 6.2151 | 5.9426 | 6.1490 | 6.3899 | 6.0964 | 7.2105 | 6.7534 | 6.4532 |
| U2SURP_T5 | U2SURP | O15042 | 788;800 | | NA | 6.3195 | 6.4139 | 6.3733 | 7.0908 | 6.3804 | 5.8764 | 6.3930 | 6.6339 | 6.3348 | 6.5417 | 6.6183 | 6.5544 | 5.3056 | 5.9350 | 5.7223 | 5.5068 |
| UACA_S1353 | UACA | Q9BZF9 | | 1353 | NPTKRQSQLIDTL | 6.8446 | 6.6051 | 7.7562 | 6.7284 | 7.2994 | 6.7274 | 6.4516 | 7.3501 | 6.9781 | 5.8551 | 6.6404 | 6.7190 | 4.7366 | 4.6948 | 4.1957 | 4.4175 |
| UAP1_S404 | UAP1 | Q16222 | | 404 | LREDEFSPLKNAD | 6.2677 | 6.1309 | 7.1905 | 6.8798 | 6.0814 | 6.6752 | 6.4689 | 6.5402 | 6.2693 | 6.3860 | 6.2640 | 6.8032 | 5.7253 | 5.2974 | 5.2264 | 5.7938 |
| UBA1_S13 | UBA1 | P22314 | | 13 | SKKRRVSGPDPKP | 5.0527 | 5.1427 | 5.5121 | 4.6937 | 5.2693 | 5.5102 | 5.4261 | 5.9253 | 5.6196 | 5.4371 | 5.2511 | 5.4371 | 8.0814 | 8.8032 | 9.7136 | 9.1246 |
| UBA1_S13;S24 | UBA1 | P22314 | | 24 | KPGSNCSPAQSVL | 5.6000 | 5.8574 | 6.8121 | 5.9499 | 5.5377 | 5.8304 | 6.1103 | 5.9542 | 6.4180 | 6.6847 | 6.5352 | 6.7071 | 6.1920 | 6.9494 | 6.2734 | 6.5882 |
| UBA1_S24 | UBA1 | P22314 | | 835 | LKATLPSPDKLPG | 5.8146 | 6.2864 | 6.7823 | 5.7146 | 5.4938 | 6.0812 | 5.9019 | 6.1873 | 6.0595 | 6.1990 | 6.1602 | 6.3569 | 6.5199 | 6.8363 | 6.7444 | 6.8615 |
| UBA1_S810 | UBA1 | P22314 | | 810 | GVKIHVSDQELQS | 4.5962 | 5.8533 | 7.3035 | 6.8681 | 4.5013 | 7.1234 | 5.5586 | 6.8616 | 6.2625 | 6.1184 | 7.3395 | 5.5161 | 7.0940 | 3.8956 | 6.6520 | 8.4558 |
| UBA1_S835 | UBA1 | P22314 | 13;24 | | NA | 5.0929 | 6.5667 | 6.2494 | 5.8159 | 6.4472 | 6.8588 | 6.3367 | 7.0126 | 5.5955 | 6.8510 | 7.0080 | 7.3538 | 5.9601 | 6.1601 | 5.3732 | 5.3182 |
| UBA2_T578 | UBA2 | Q9UBT2 | | 578 | DGAQPSTSTAQEQ | 5.1444 | 5.5702 | 6.0623 | 5.6208 | 6.1883 | 6.3362 | 5.5322 | 6.2071 | 6.2267 | 6.3966 | 6.3930 | 6.3129 | 6.6601 | 7.4869 | 7.2602 | 6.6022 |
| UBA3_S377 | UBA3 | Q8TBC4 | | 377 | PQNIQFSPSAKLQ | 5.5466 | 5.0571 | 7.2475 | 5.2507 | 5.9586 | 5.6522 | 5.0218 | 6.1664 | 4.3598 | 7.5433 | 7.3074 | 6.3917 | 7.0010 | 9.8077 | 6.8143 | 4.8739 |
| UBA6_S737 | UBA6 | A0AVT1 | | 737 | GSLFWQSPKRPPS | 6.3224 | 6.2273 | 6.9817 | 6.5613 | 6.5038 | 5.7902 | 5.5433 | 6.5741 | 6.0148 | 6.5102 | 6.9217 | 6.8003 | 5.3611 | 6.7100 | 5.4738 | 5.7039 |
| UBAC1_S98 | UBAC1 | Q9BSL1 | | 98 | IKKRAPSPLPKMA | 6.5529 | 6.2625 | 6.6484 | 5.7999 | 6.2949 | 6.6577 | 5.9657 | 6.7889 | 6.3940 | 6.4636 | 6.7078 | 6.7183 | 5.8978 | 5.7994 | 5.6162 | 5.4320 |
| UBAP1_S269 | UBAP1 | Q9NZ09-4 | | 317 | SLSSKVSLPPIPA | 7.3824 | 5.9022 | 7.3789 | 5.5384 | 5.4182 | 6.0741 | 5.0351 | 7.1049 | 8.9777 | 8.9554 | 7.3267 | 8.6837 | 4.2301 | 5.6200 | 3.1089 | 3.2632 |
| UBAP1_S317 | UBAP1 | Q9NZ09-4 | | 269 | NLPRGGSGSVLQD | 3.1826 | 4.1835 | 5.6137 | 3.2261 | 3.1042 | 4.2966 | 4.5519 | 4.6535 | 4.3982 | 5.3816 | 4.6883 | 5.6892 | 9.9162 | 11.9557 | 14.0300 | 11.1288 |
| UBAP2_S630 | UBAP2 | Q5T6F2 | | 630 | NRIPYQSPVSSSE | 6.2069 | 6.4523 | 7.0665 | 7.2892 | 6.8172 | 6.8776 | 6.6206 | 6.9204 | 6.9986 | 6.8101 | 6.9512 | 6.8614 | 4.4552 | 4.5599 | 4.3580 | 4.7551 |
| UBAP2_S987;S988 | UBAP2 | Q5T6F2 | | 657 | SQQTLDTPKTTGP | 6.8079 | 7.5358 | 7.3633 | 6.9212 | 5.9773 | 6.5812 | 7.8235 | 6.9227 | 7.1423 | 7.4485 | 7.2111 | 6.6072 | 3.6319 | 4.4073 | 3.4654 | 4.1535 |
| UBAP2_T657 | UBAP2 | Q5T6F2 | 987;988 | | NA | 5.5772 | 6.7818 | 9.0729 | 6.1566 | 6.6511 | 6.2647 | 7.3048 | 6.2321 | 6.1951 | 5.2017 | 9.0151 | 7.6929 | 3.8147 | 3.8573 | 4.9659 | 5.2160 |
| UBAP2L_S356 | UBAP2L | Q14157-5 | | 609 | PSSISSSPQKDLT | 5.7016 | 6.0492 | 6.2727 | 6.1305 | 6.2506 | 6.3517 | 5.8713 | 6.9155 | 6.7786 | 6.8596 | 6.7563 | 6.7929 | 5.6192 | 5.9307 | 5.8000 | 5.9197 |
| UBAP2L_S416 | UBAP2L | Q14157-5 | | 467 | TSAPQMSPGSSDN | 5.7081 | 5.8453 | 6.1892 | 5.5533 | 6.1610 | 6.2719 | 5.8989 | 6.6708 | 6.2549 | 6.3383 | 6.3645 | 6.2293 | 6.6434 | 6.8030 | 6.7003 | 6.3680 |
| UBAP2L_S439 | UBAP2L | Q14157-5 | | 425 | TKRQAFTPSSTMM | 6.6466 | 7.1493 | 6.7049 | 7.1086 | 6.4385 | 7.1359 | 7.0408 | 6.9651 | 6.9881 | 6.8720 | 7.2662 | 6.7228 | 3.9600 | 4.7087 | 3.7950 | 4.4976 |
| UBAP2L_S439;S454 | UBAP2L | Q14157-5 | | 416 | SDSAVHSPFTKRQ | 7.2854 | 7.3129 | 6.9107 | 7.1820 | 6.9407 | 7.4770 | 7.1827 | 7.7255 | 7.7361 | 7.1057 | 7.5078 | 7.2071 | 2.8284 | 3.7924 | 2.6108 | 3.1949 |
| UBAP2L_S454 | UBAP2L | Q14157-5 | | 454 | APPPPSSPLPSKS | 5.9348 | 5.9796 | 6.4089 | 5.8772 | 6.4522 | 6.4001 | 6.4201 | 6.8956 | 6.3897 | 6.5111 | 6.4574 | 6.7387 | 5.6866 | 6.1236 | 5.5056 | 6.2190 |
| UBAP2L_S467 | UBAP2L | Q14157-5 | | 844 | PFPTPTTPLTGRD | 6.0650 | 6.7434 | 6.3997 | 6.5517 | 4.9196 | 7.1453 | 7.1718 | 7.6191 | 6.9418 | 6.5376 | 7.3569 | 7.7669 | 4.0657 | 5.3868 | 4.0874 | 5.2414 |
| UBAP2L_S467;S477 | UBAP2L | Q14157-5 | | 477 | SDNQSSSPQPAQQ | 6.4410 | 5.9985 | 6.4575 | 5.8952 | 6.1713 | 6.4628 | 5.7855 | 6.3605 | 7.1204 | 6.7548 | 7.0696 | 7.5328 | 5.1231 | 6.2534 | 4.9837 | 5.5899 |
| UBAP2L_S477 | UBAP2L | Q14157-5 | | 439 | VFLQEKSPAVATS | 6.9490 | 7.1808 | 6.0318 | 7.7180 | 6.8361 | 6.9490 | 6.4403 | 7.6984 | 7.7877 | 6.8400 | 8.0381 | 6.6750 | 3.3989 | 3.7603 | 3.6817 | 4.0147 |
| UBAP2L_S605 | UBAP2L | Q14157-5 | | 605 | TRRYPSSISSSPQ | 7.0890 | 6.3571 | 7.2763 | 5.5292 | 7.0562 | 7.0609 | 6.4749 | 5.8101 | 5.9303 | 6.1675 | 5.7477 | 6.2962 | 6.4663 | 6.2322 | 4.5211 | 5.9849 |
| UBAP2L_S605;S609 | UBAP2L | Q14157-5 | | 356 | GEAKGGSTTGSQF | 5.8203 | 6.7020 | 7.5900 | 5.6620 | 5.9558 | 6.8772 | 6.8889 | 6.9632 | 7.0180 | 6.8549 | 7.3657 | 6.9010 | 4.5623 | 5.3945 | 4.7405 | 4.7036 |
| UBAP2L_S609 | UBAP2L | Q14157-5 | | 779 | TASTRSSVATTSG | 8.0575 | 5.4683 | 5.1061 | 5.4455 | 6.0437 | 7.5360 | 7.2110 | 5.6173 | 7.3311 | 6.1658 | 6.1948 | 6.6129 | 6.3935 | 6.2465 | 4.6735 | 5.8967 |
| UBAP2L_S779 | UBAP2L | Q14157-5 | 467;477 | | NA | 6.4309 | 6.8216 | 7.1632 | 6.6812 | 6.1348 | 6.6219 | 6.2376 | 6.7257 | 7.4225 | 7.1693 | 7.4866 | 7.3847 | 4.2937 | 4.8888 | 4.0415 | 4.4962 |
| UBAP2L_T425 | UBAP2L | Q14157-5 | 439;454 | | NA | 7.2044 | 7.1938 | 6.8644 | 7.0294 | 6.6666 | 7.1650 | 7.4311 | 7.0037 | 7.6151 | 7.1077 | 7.3060 | 7.2282 | 3.2696 | 3.9765 | 3.0259 | 3.9125 |
| UBAP2L_T844 | UBAP2L | Q14157-5 | 605;609 | | NA | 7.4545 | 7.3230 | 7.1162 | 6.5996 | 6.3494 | 6.9255 | 6.7196 | 6.0656 | 6.6076 | 6.5595 | 7.3829 | 6.8131 | 4.4476 | 4.8592 | 4.3092 | 4.4675 |
| UBC_S65 | UBC | P0CG48 | | 65 | YNIQKESTLHLVL | 6.7777 | 7.0391 | 6.0862 | 6.9697 | 7.4492 | 6.3807 | 7.3481 | 6.7553 | 5.6281 | 5.3398 | 4.9756 | 4.9006 | 5.9819 | 5.8875 | 6.9012 | 5.5792 |
| UBC_T66 | UBC | P0CG48 | | 66 | NIQKESTLHLVLR | 5.6470 | 6.1553 | 5.8093 | 6.6966 | 6.1976 | 5.9212 | 6.1804 | 7.2256 | 6.5507 | 5.4861 | 5.5390 | 5.5276 | 6.5917 | 6.7588 | 7.0744 | 6.6385 |
| UBE2J1_S251 | UBE2J1 | Q9Y385 | | 266 | QSQRRLSTSPDVI | 5.6913 | 5.4738 | 5.9464 | 6.0191 | 6.5446 | 5.7901 | 5.8961 | 6.2048 | 6.9928 | 6.7257 | 6.5631 | 6.7931 | 6.3111 | 6.5528 | 6.1448 | 6.3502 |
| UBE2J1_S266 | UBE2J1 | Q9Y385 | | 251 | AKNTSMSPRQRRA | 6.7642 | 6.5811 | 6.3651 | 6.7869 | 6.3375 | 5.9813 | 6.6650 | 6.0855 | 6.2516 | 6.5654 | 6.6272 | 6.5002 | 5.7219 | 5.3016 | 5.7418 | 5.7236 |
| UBE2K_S159 | UBE2K | P61086 | | 159 | AGAPVSSPEYTKK | 5.7283 | 6.2771 | 6.1337 | 5.2393 | 6.0644 | 7.6721 | 6.7202 | 6.8142 | 6.0068 | 6.0115 | 6.7084 | 7.0657 | 6.5968 | 5.8916 | 5.5461 | 5.5238 |
| UBE2M_S28 | UBE2M | P61081 | | 28 | GSSKKASAAQLRI | 1.6230 | 1.8541 | 1.6028 | 2.1551 | 2.5972 | 2.4392 | 2.2657 | 2.8128 | 2.0084 | 1.5536 | 1.2620 | 1.7144 | 21.3583 | 18.3341 | 20.9570 | 15.4621 |
| UBE2M_S50 | UBE2M | P61081 | | 50 | PKTCDISFSDPDD | 5.8169 | 6.7499 | 7.9691 | 4.7814 | 5.7621 | 5.9649 | 6.1701 | 6.4826 | 6.3872 | 6.2154 | 8.0741 | 5.6595 | 4.9365 | 6.1104 | 5.9744 | 6.9455 |
| UBE2O_S401 | UBE2O | Q9C0C9 | | 839 | LTGSPTSPTVEPE | 5.4864 | 5.5165 | 6.5787 | 5.9078 | 5.9735 | 6.0558 | 6.1115 | 6.3044 | 6.1025 | 6.2234 | 5.9492 | 6.3223 | 6.7897 | 6.7249 | 6.8473 | 7.1063 |
| UBE2O_S515 | UBE2O | Q9C0C9 | | 515 | GTSRKKSIPLSIK | 5.8819 | 4.6711 | 5.5870 | 5.6484 | 6.9704 | 5.2742 | 4.8433 | 6.1354 | 7.6364 | 7.7558 | 7.5451 | 7.9745 | 6.7507 | 7.4387 | 5.2492 | 4.6380 |

| | | | | | | | | | | | | | | | | | | | | | |
|---|---|---|---|---|---|---|---|---|---|---|---|---|---|---|---|---|---|---|---|---|---|
| UBE2O_S836;S839 | UBE2O | Q9C0C9 | | 896 | DKPEGQSPVKAEW | 6.6527 | 8.6501 | 8.3782 | 7.1513 | 5.9167 | 7.6502 | 5.6983 | 5.9902 | 6.3769 | 6.2845 | 7.2290 | 6.2786 | 4.3315 | 4.8155 | 5.0856 | 3.5109 |
| UBE2O_S839 | UBE2O | Q9C0C9 | | 401 | VRIMSCSPDTQCS | 6.2415 | 6.7529 | 6.8154 | 6.9267 | 6.1922 | 7.0163 | 6.7728 | 6.4037 | 7.3933 | 7.3794 | 6.9444 | 7.3296 | 4.2634 | 4.7538 | 4.6829 | 4.1317 |
| UBE2O_S896 | UBE2O | Q9C0C9 | 834;839 | | NA | 6.5105 | 6.5737 | 7.3775 | 6.8253 | 5.4860 | 7.4991 | 7.0152 | 7.1684 | 7.3942 | 7.0288 | 6.9778 | 7.3240 | 4.1710 | 4.3961 | 3.7060 | 4.5465 |
| UBE2O_T834;S839 | UBE2O | Q9C0C9 | 836;839 | | NA | 5.1347 | 6.7650 | 7.4586 | 6.7356 | 5.3782 | 5.2904 | 7.5480 | 5.8680 | 5.6861 | 8.8383 | 7.4235 | 6.5023 | 4.4801 | 6.7429 | 4.5889 | 5.5595 |
| UBE2T_S184 | UBE2T | Q9NPD8 | | 184 | TQKRKASQLVGIE | 7.4091 | 4.9106 | 6.2369 | 4.9845 | 8.7058 | 5.5348 | 5.3955 | 5.6921 | 7.6885 | 7.7259 | 7.4146 | 8.0380 | 6.2898 | 6.0338 | 3.9128 | 4.0272 |
| UBE3B_S419 | UBE3B | Q7Z3V4 | | 419 | AHAQPASPQNVLP | 6.5892 | 6.7769 | 6.6988 | 6.8847 | 5.9590 | 7.3084 | 7.3805 | 7.4612 | 6.7405 | 7.1763 | 6.7431 | 7.0468 | 3.6921 | 4.9035 | 4.0123 | 4.6264 |
| UBE3C_S15 | UBE3C | Q15386 | | 20 | VSLGGASRKEEKA | 6.9072 | 6.6507 | 7.6333 | 6.2524 | 4.8556 | 5.9274 | 6.6291 | 6.6488 | 8.0142 | 7.9992 | 8.7726 | 7.9860 | 4.1530 | 4.5995 | 3.1506 | 3.8204 |
| UBE3C_S20 | UBE3C | Q15386 | | 15 | KTRPKVSLGGASR | 6.7238 | 7.6756 | 7.0003 | 6.7209 | 5.9356 | 6.8290 | 6.9008 | 6.8281 | 7.2145 | 7.4221 | 7.1457 | 6.9515 | 4.4156 | 4.8432 | 2.8640 | 4.5294 |
| UBE3D_S381 | UBE3D | Q7Z6J8 | | 381 | SLRRVNSFQVAFL | 6.0771 | 5.7925 | 5.7625 | 6.6615 | 6.8622 | 5.4748 | 6.9881 | 7.5455 | 6.5716 | 5.8973 | 5.1542 | 6.9342 | 6.3049 | 6.0022 | 6.1820 | 5.7895 |
| UBE4B_S101 | UBE4B | O95155-4 | | 88 | VSSLSSSPSNSLE | 6.2872 | 6.8337 | 7.9489 | 6.4395 | 6.5387 | 7.2014 | 6.3101 | 6.4528 | 6.7880 | 7.2955 | 7.0565 | 6.9868 | 4.3320 | 4.9338 | 4.4902 | 4.1048 |
| UBE4B_S105 | UBE4B | O95155-4 | | 1316 | LRHLLNSPTDPFN | 6.2166 | 6.7213 | 6.7571 | 6.7832 | 5.6679 | 6.3030 | 6.4232 | 6.9826 | 6.7327 | 6.8773 | 6.9207 | 6.8836 | 4.7779 | 5.5324 | 4.8460 | 5.5745 |
| UBE4B_S1316 | UBE4B | O95155-4 | | 78 | VAHRSQSSEGVSS | 4.7971 | 5.1577 | 5.7584 | 5.2723 | 5.9932 | 5.7664 | 5.4278 | 5.4785 | 5.9904 | 5.3221 | 5.1352 | 5.4757 | 9.1602 | 8.4342 | 8.4572 | 8.3736 |
| UBE4B_S78 | UBE4B | O95155-4 | | 854 | ESQWKDSPLATRH | 6.6765 | 7.4351 | 7.0135 | 7.1035 | 5.5907 | 6.2446 | 6.6638 | 6.1650 | 6.5965 | 6.8890 | 7.0859 | 7.3520 | 4.3668 | 4.9697 | 4.5213 | 5.3262 |
| UBE4B_S84 | UBE4B | O95155-4 | | 101 | TQSQSLSRSQSMD | 5.3218 | 5.2786 | 4.9361 | 5.2542 | 5.8748 | 5.8050 | 5.4326 | 5.4858 | 5.5257 | 5.6720 | 4.7366 | 4.6402 | 9.5553 | 9.1064 | 8.9624 | 8.4127 |
| UBE4B_S84;S88 | UBE4B | O95155-4 | | 84 | SSEGVSSLSSSPS | 5.9706 | 6.0200 | 7.1477 | 6.6503 | 7.1256 | 7.0898 | 6.0745 | 7.3521 | 7.4407 | 7.2772 | 8.4100 | 6.5157 | 4.0900 | 4.2893 | 3.8328 | 4.7135 |
| UBE4B_S854 | UBE4B | O95155-4 | | 105 | SLSRSQSMDIDGV | 5.5587 | 5.8099 | 6.3363 | 5.6892 | 6.1942 | 6.1003 | 5.7822 | 6.0719 | 6.2398 | 7.0551 | 6.2781 | 5.7024 | 6.6994 | 7.1769 | 6.7277 | 6.5776 |
| UBE4B_S88 | UBE4B | O95155-4 | 84;88 | | NA | 6.3661 | 7.0479 | 7.4466 | 6.9481 | 5.1233 | 6.2832 | 6.1111 | 6.7549 | 6.7159 | 7.3411 | 6.1849 | 6.8792 | 4.7721 | 5.3811 | 5.1457 | 5.4988 |
| UBIAD1_S15 | UBIAD1 | Q9Y5Z9 | | 15 | EKINILSGETVKA | 5.4716 | 6.6594 | 5.6946 | 6.8174 | 5.9408 | 6.8329 | 7.3548 | 5.3616 | 6.1422 | 6.2490 | 6.2862 | 6.6068 | 6.0431 | 6.2769 | 5.6915 | 6.5712 |
| UBL4A_S90 | UBL4A | P11441 | | 90 | AQRLADSPPPQVW | 5.8557 | 6.5966 | 6.9454 | 6.0380 | 6.0293 | 5.5014 | 6.0538 | 6.6543 | 5.8170 | 7.0243 | 6.5343 | 7.0109 | 5.8415 | 6.2818 | 5.6624 | 6.1532 |
| UBL7_S230 | UBL7 | Q96S82 | | 230 | FLFEGLSDDEDDF | 5.6117 | 5.5977 | 5.7884 | 5.6088 | 6.0754 | 5.4966 | 5.9157 | 6.5378 | 4.9754 | 6.2339 | 5.6659 | 6.1727 | 6.2646 | 8.3483 | 7.4525 | 8.2547 |
| UBN1_S173 | UBN1 | Q9NPG3 | | 660 | PVNLEDSLDEDLI | 6.3417 | 6.3373 | 6.2429 | 6.5102 | 6.1027 | 6.1275 | 6.1965 | 5.8334 | 5.8449 | 6.3160 | 6.6223 | 5.9773 | 5.9505 | 6.7000 | 6.4132 | 6.4836 |
| UBN1_S231 | UBN1 | Q9NPG3 | | 764 | SGYKELSCQAPLN | 6.1206 | 7.5148 | 6.9750 | 7.0820 | 5.0542 | 5.7382 | 6.7975 | 5.3226 | 6.7823 | 5.8552 | 7.5450 | 6.4757 | 5.7120 | 7.0194 | 4.9281 | 5.0774 |
| UBN1_S660 | UBN1 | Q9NPG3 | | 231 | FTALNASKEKKKK | 7.1498 | 6.5105 | 6.1188 | 6.9547 | 6.1057 | 7.6481 | 8.0611 | 7.1940 | 6.5269 | 6.0909 | 7.9103 | 7.4497 | 2.6963 | 4.8140 | 4.1814 | 4.5878 |
| UBN1_S764 | UBN1 | Q9NPG3 | | 821 | KNFTPPSPFANKL | 6.9423 | 6.8198 | 6.5262 | 6.4872 | 6.0821 | 6.7071 | 7.5609 | 6.2626 | 6.9020 | 6.0897 | 6.4291 | 6.2016 | 5.5079 | 5.3138 | 4.5633 | 5.6045 |
| UBN1_S821 | UBN1 | Q9NPG3 | | 173 | LQFRQASESEDDF | 6.2155 | 5.8705 | 5.6999 | 6.5319 | 6.0478 | 6.5522 | 5.6462 | 5.5382 | 6.1970 | 6.0271 | 6.1589 | 6.2129 | 7.0509 | 7.9058 | 6.6793 | 6.5660 |
| UBN2_S1036 | UBN2 | Q6ZU65 | | 13 | VAFISLSPVRRRE | 6.4367 | 6.4295 | 6.1435 | 6.4743 | 6.1569 | 6.2360 | 6.1689 | 6.6121 | 6.3931 | 6.3157 | 6.0900 | 6.3928 | 6.1164 | 6.0804 | 5.9626 | 5.9910 |
| UBN2_S1123 | UBN2 | Q6ZU65 | | 1036 | PFSMAASPKLAAS | 6.5924 | 7.9014 | 5.1743 | 6.5106 | 5.5395 | 6.2699 | 6.7204 | 7.0429 | 6.1703 | 7.8112 | 7.8302 | 5.9142 | 6.0090 | 5.8051 | 4.9514 | 4.6573 |
| UBN2_S13 | UBN2 | Q6ZU65 | | 1123 | MNISRQSPTLNLL | 7.4538 | 7.2195 | 7.2550 | 7.0021 | 5.7811 | 5.2971 | 6.5473 | 6.7015 | 6.3577 | 5.8829 | 5.4343 | 6.7478 | 5.0535 | 5.8459 | 5.5237 | 5.8968 |
| UBR1_S1593 | UBR1 | Q8IWV7 | | 21 | SAELPQTPQRLAS | 5.6647 | 6.0207 | 6.2768 | 5.9062 | 6.1656 | 6.3685 | 6.1627 | 6.3444 | 6.2414 | 6.2312 | 6.3562 | 6.3627 | 6.3768 | 6.6762 | 6.5330 | 6.3129 |
| UBR1_T21 | UBR1 | Q8IWV7 | | 1593 | YPRKRNSLIELPD | 6.0689 | 5.7073 | 6.7389 | 5.8916 | 5.8516 | 5.9977 | 6.0036 | 6.0831 | 6.0750 | 6.3646 | 6.7142 | 6.6981 | 6.5357 | 6.7328 | 6.1063 | 6.4304 |
| UBR2_S1009 | UBR2 | Q8IWV8 | | 1009 | RESSPTSPVAETE | 5.6956 | 5.3386 | 5.4818 | 6.7881 | 6.2023 | 5.4626 | 7.1174 | 8.1115 | 6.7389 | 5.5331 | 6.9827 | 6.4054 | 5.4754 | 6.6855 | 7.0960 | 4.8853 |
| UBR4_S178 | UBR4 | Q5T4S7-2 | | 2719 | HVTLPSSPRSNTP | 6.4858 | 6.5615 | 6.5715 | 6.0775 | 6.4781 | 6.8507 | 6.0240 | 7.1191 | 6.4190 | 6.2317 | 6.5753 | 6.4423 | 5.1765 | 5.9314 | 5.5781 | 5.4774 |
| UBR4_S181 | UBR4 | Q5T4S7-2 | | 181 | ELASPVSPELRQK | 6.1407 | 6.3885 | 6.6054 | 6.2565 | 5.9101 | 6.5014 | 6.1700 | 6.3463 | 6.5955 | 6.4899 | 6.4315 | 6.5401 | 5.8684 | 6.1161 | 5.7271 | 5.9124 |
| UBR4_S2719;S2722 | UBR4 | Q5T4S7-2 | | 620 | PPPLESSPRVKSP | 5.8765 | 6.4257 | 7.0414 | 6.7603 | 5.5011 | 7.2717 | 6.4162 | 6.4043 | 6.8219 | 6.5353 | 6.9301 | 6.8945 | 5.1329 | 5.5037 | 5.0991 | 5.3851 |
| UBR4_S457 | UBR4 | Q5T4S7-2 | | 457 | TKEGVGSPKLGPG | 6.1958 | 6.3296 | 6.6693 | 5.9973 | 6.1885 | 6.8121 | 6.4915 | 6.5018 | 6.2886 | 6.3991 | 6.4697 | 6.3047 | 5.5680 | 6.1659 | 5.6397 | 5.9784 |
| UBR4_S620 | UBR4 | Q5T4S7-2 | | 178 | DQKELASPVSPEL | 6.0989 | 6.5733 | 6.8132 | 6.3630 | 6.1444 | 6.0131 | 5.8282 | 6.2467 | 6.4362 | 6.5786 | 6.2582 | 6.3999 | 6.1779 | 6.0099 | 6.0844 | 5.9740 |
| UBR4_T2715;S2719 | UBR4 | Q5T4S7-2 | | 905 | NSRRATTPLYHGF | 6.4095 | 6.6315 | 6.5118 | 6.1722 | 5.9961 | 6.1856 | 5.9507 | 6.9289 | 6.7127 | 6.3907 | 6.4708 | 6.4078 | 5.6059 | 6.0821 | 5.6540 | 5.8897 |
| UBR4_T2719 | UBR4 | Q5T4S7-2 | 2715;2719 | | NA | 6.2948 | 7.0158 | 7.6533 | 7.0500 | 6.1655 | 5.6775 | 7.0502 | 6.2625 | 6.3180 | 6.6951 | 7.6005 | 7.3386 | 4.2881 | 5.0502 | 4.7545 | 4.7853 |
| UBR4_T905 | UBR4 | Q5T4S7-2 | 2719;2722 | | NA | 6.7764 | 5.9631 | 8.8910 | 6.0663 | 6.1110 | 6.8023 | 5.2968 | 5.2184 | 2.9527 | 6.1883 | 4.5968 | 6.4401 | 5.9070 | 8.6661 | 7.1602 | 6.9636 |
| UBR5_S1227 | UBR5 | O95071 | | 1549 | PQQRRISQSQPVR | 5.4752 | 5.5345 | 5.4465 | 5.7839 | 6.5854 | 5.7041 | 6.1739 | 6.1250 | 5.7717 | 5.2970 | 5.3129 | 5.5549 | 7.8552 | 7.4561 | 8.0306 | 7.8931 |
| UBR5_S143 | UBR5 | O95071 | | 2028 | FFRRSDSMTFLGC | 6.3932 | 6.2298 | 6.4219 | 6.8908 | 6.1701 | 5.9810 | 6.4949 | 5.9904 | 6.0941 | 6.7237 | 6.0721 | 6.1844 | 5.6955 | 6.0487 | 6.4068 | 6.2026 |
| UBR5_S1549 | UBR5 | O95071 | | 2486 | RHGSSRSVVDMDL | 5.4906 | 5.9636 | 5.8436 | 5.9321 | 5.8107 | 6.4333 | 5.9924 | 6.3221 | 6.0935 | 5.8241 | 5.3949 | 5.6213 | 7.4340 | 7.3361 | 6.9725 | 7.5351 |
| UBR5_S1990 | UBR5 | O95071 | | 612 | EMGPPPSPASTCS | 5.9701 | 6.2877 | 6.6062 | 6.7737 | 6.3788 | 6.4986 | 6.3393 | 5.9655 | 6.9086 | 6.4636 | 6.4259 | 6.2180 | 5.9237 | 5.8125 | 5.4517 | 5.9761 |
| UBR5_S1990;S2011 | UBR5 | O95071 | | 2241 | QQSRDLSLEVDRD | 6.1761 | 6.0358 | 6.1637 | 6.4089 | 6.4096 | 6.0300 | 6.3088 | 5.8912 | 6.0621 | 6.0730 | 6.0363 | 6.3515 | 6.6876 | 6.5079 | 6.4603 | 6.3021 |
| UBR5_S2028 | UBR5 | O95071 | | 1990 | GIDNEDSEHENDD | 5.5874 | 6.4773 | 6.2159 | 6.2479 | 6.7559 | 6.2942 | 6.6090 | 6.0753 | 5.9907 | 6.1917 | 6.1170 | 6.1662 | 6.7479 | 6.5945 | 6.0803 | 5.8490 |
| UBR5_S2241 | UBR5 | O95071 | | 636 | PYKRRRSTPAPKE | 6.1825 | 6.3834 | 5.9559 | 6.0104 | 6.1563 | 6.0381 | 6.2082 | 6.2719 | 5.6265 | 5.6156 | 5.4999 | 5.5335 | 7.0791 | 7.1993 | 7.1365 | 7.1028 |
| UBR5_S2486 | UBR5 | O95071 | | 578 | CRFQLRSPESLKN | 6.8946 | 7.4553 | 6.2559 | 6.6775 | 7.0214 | 6.4921 | 6.4640 | 5.7951 | 5.8104 | 6.4871 | 7.0280 | 5.7700 | 5.8785 | 5.6531 | 5.2869 | 5.0301 |
| UBR5_S578 | UBR5 | O95071 | | 1227 | CKLKRTSPTAYCD | 6.6600 | 6.9067 | 7.1640 | 6.4658 | 6.0185 | 5.9031 | 6.2374 | 6.2117 | 5.7852 | 6.4454 | 6.3159 | 6.2532 | 6.1501 | 5.8411 | 5.7282 | 5.9140 |
| UBR5_S612 | UBR5 | O95071 | | 143 | GGSGGGSSGRSSA | 6.9592 | 7.2370 | 6.6585 | 6.5528 | 5.8425 | 6.5872 | 7.4574 | 6.1071 | 5.5377 | 7.0993 | 6.8945 | 6.8421 | 5.0396 | 5.6213 | 4.7897 | 4.7741 |
| UBR5_S636 | UBR5 | O95071 | 1990;2011 | | NA | 6.4736 | 5.7119 | 5.0714 | 5.3659 | 5.9458 | 5.5584 | 6.1648 | 5.7393 | 6.2692 | 5.3301 | 5.5786 | 6.0759 | 8.2083 | 7.2311 | 7.7328 | 7.5429 |
| UBTD1_S164 | UBTD1 | Q9HAC8 | | 164 | GKDVRLSASLPDT | 5.5678 | 6.2117 | 6.5078 | 7.1521 | 6.0449 | 6.1022 | 6.3032 | 6.6062 | 6.4862 | 6.3131 | 6.6362 | 6.9766 | 5.7090 | 5.3382 | 5.1273 | 5.9174 |
| UBTF_S389 | UBTF | P17480 | | 389 | NKKQATSPASKKP | 6.4416 | 6.6430 | 6.1507 | 6.6135 | 6.6423 | 6.4963 | 6.6130 | 6.3796 | 6.7909 | 6.4633 | 6.3788 | 6.3432 | 5.4037 | 5.6469 | 5.4051 | 5.5882 |
| UBTF_S449 | UBTF | P17480 | | 449 | RMWNDLSEKKKAK | 6.4474 | 7.2532 | 6.4328 | 7.1754 | 6.4907 | 6.6562 | 6.9973 | 6.4111 | 6.1383 | 7.1488 | 6.8350 | 6.1004 | 4.6981 | 5.3375 | 4.7247 | 5.1530 |
| UBTF_S638 | UBTF | P17480 | | 117 | FPKKPLTPYFRFF | 8.4667 | 7.9949 | 6.9850 | 8.1381 | 6.3501 | 7.7055 | 7.0534 | 7.2482 | 7.7992 | 7.4747 | 7.3660 | 7.1399 | 2.1420 | 3.3147 | 1.9738 | 2.6478 |
| UBTF_T117 | UBTF | P17480 | | 638 | LWVKSLSPQDRAA | 6.6098 | 7.8825 | 6.3708 | 7.5515 | 6.0866 | 5.9784 | 6.2950 | 6.5219 | 6.3451 | 7.2664 | 7.3840 | 6.5785 | 4.1171 | 5.1426 | 4.7658 | 5.1042 |
| UBXN1_S200 | UBXN1 | Q04323-2 | | 200 | PGPVPSSPSQEPP | 6.2070 | 6.3952 | 6.5694 | 6.1754 | 6.8440 | 7.0971 | 6.2284 | 7.1776 | 7.1654 | 7.0439 | 7.1451 | 7.2531 | 4.7215 | 4.8567 | 4.4049 | 4.7154 |
| UBXN2A_S53 | UBXN2A | P68543 | | 53 | VSSKCVSPAEQKK | 6.5449 | 6.7858 | 6.5894 | 7.4396 | 5.7499 | 5.7838 | 6.0135 | 6.4513 | 6.7625 | 7.2939 | 7.4882 | 7.6046 | 4.8572 | 5.4103 | 4.2029 | 5.0222 |
| UBXN2B_S17 | UBXN2B | Q14CS0 | | 235 | IVSTPSSPEEEDK | 5.3018 | 4.6102 | 5.7018 | 6.0198 | 6.5924 | 5.7143 | 5.2105 | 6.0178 | 6.2591 | 5.7216 | 5.8710 | 6.0562 | 7.9914 | 7.2892 | 8.3471 | 7.2958 |
| UBXN2B_S235 | UBXN2B | Q14CS0 | | 17 | EQERRSSGPRPPS | 6.8332 | 6.6923 | 5.9788 | 5.9398 | 6.1046 | 6.9590 | 6.3952 | 7.0132 | 8.7546 | 8.2124 | 8.4185 | 7.1477 | 4.2287 | 4.2613 | 3.4632 | 3.5977 |
| UBXN4_S331 | UBXN4 | Q92575 | | 331 | RLPDGSSFTNQFP | 4.8017 | 4.9419 | 5.0848 | 5.2506 | 4.9620 | 4.2984 | 4.6748 | 5.6518 | 4.6446 | 4.8594 | 4.5212 | 5.8880 | 10.1159 | 10.1347 | 11.2801 | 8.8900 |
| UBXN7_S280 | UBXN7 | O94888 | | 280 | KCARSESLIDASE | 6.0825 | 5.5817 | 6.0411 | 6.1920 | 6.5298 | 5.9718 | 5.6533 | 6.2045 | 6.4848 | 6.6248 | 6.1753 | 6.4585 | 7.1969 | 6.6341 | 6.2771 | 5.8918 |
| UCK1_S258 | UCK1 | Q9HA47-4 | | 258 | SYKRTFSEPGDHP | 4.2421 | 4.4287 | 5.7367 | 4.7308 | 5.3776 | 5.6645 | 5.4772 | 5.8537 | 5.1607 | 4.8534 | 5.1913 | 5.2733 | 9.5637 | 9.5470 | 9.9292 | 8.9701 |
| UCK2_T246 | UCK2 | Q9BZX2 | | 246 | GCLNGYTPSRKRQ | 6.7109 | 7.2616 | 8.3680 | 7.0610 | 6.4832 | 7.7647 | 7.1151 | 6.9607 | 8.2257 | 6.5786 | 6.2136 | 7.2631 | 3.4480 | 3.8428 | 3.0346 | 3.6685 |
| UCKL1_S16 | UCKL1 | Q9NWZ5 | | 16 | ADPSPTSPPTARD | 6.3478 | 6.8855 | 6.7137 | 6.3957 | 6.0027 | 6.4098 | 6.5035 | 5.9618 | 6.1950 | 6.9021 | 6.8322 | 6.4037 | 5.5985 | 5.6074 | 5.4562 | 5.7843 |
| UCKL1_S56 | UCKL1 | Q9NWZ5 | | 56 | PVGTGRSPRKRTT | 6.3878 | 6.5248 | 6.4246 | 6.0451 | 6.2537 | 6.6877 | 6.1531 | 6.0677 | 6.6112 | 6.4841 | 6.9274 | 7.2893 | 5.3066 | 5.8388 | 5.1527 | 5.8454 |
| UCKL1_T547 | UCKL1 | Q9NWZ5 | | 547 | EEEVAYTGxxxxx | 5.8225 | 6.4426 | 6.6640 | 6.3847 | 6.0740 | 6.9260 | 6.5261 | 6.5991 | 6.7336 | 6.0926 | 6.1783 | 6.2398 | 6.0219 | 6.4913 | 5.2626 | 5.5408 |
| UFD1_S165 | UFD1 | Q92890-1 | | 335 | FSGEGQSLRKKGR | 6.3049 | 6.8777 | 7.8750 | 7.3832 | 7.0241 | 6.2883 | 6.3312 | 6.5224 | 7.6274 | 7.3973 | 7.6840 | 6.8480 | 4.0242 | 4.8685 | 3.1490 | 3.7947 |
| UFD1_S253 | UFD1 | Q92890-1 | | 283 | GVEPSPSPIKPGD | 7.0566 | 7.0922 | 6.8223 | 6.5995 | 6.4085 | 6.4475 | 6.8533 | 6.7466 | 7.1020 | 6.9330 | 6.5904 | 6.7933 | 4.2088 | 4.7445 | 4.3904 | 5.2111 |
| UFD1_S283 | UFD1 | Q92890-1 | | 165 | SKFQPQSPDFLDI | 5.5690 | 6.4793 | 4.8941 | 6.5943 | 6.7620 | 5.5980 | 6.3241 | 6.0983 | 6.6317 | 4.9967 | 5.8059 | 6.6304 | 6.9407 | 6.6470 | 6.4890 | 7.5394 |
| UFD1_S335 | UFD1 | Q92890-1 | | 253 | EGEADHSGYAGEL | 6.4646 | 7.4687 | 6.3526 | 6.6574 | 6.3127 | 6.5094 | 7.4502 | 7.1356 | 6.9146 | 6.5737 | 6.8600 | 6.7470 | 4.2938 | 5.3232 | 4.2179 | 4.7185 |
| UFL1_S458 | UFL1 | O94874 | | 458 | GRKDDDSDDESQS | 5.9010 | 6.7603 | 5.9614 | 6.4270 | 6.7546 | 6.9756 | 5.9819 | 6.3956 | 6.6480 | 6.0173 | 6.5438 | 6.0712 | 6.0378 | 6.3229 | 6.1779 | 5.0237 |
| UFL1_S790 | UFL1 | O94874 | | 790 | LKSRKSSVTEExx | 6.1540 | 7.3713 | 6.3443 | 7.3114 | 5.7606 | 6.4192 | 5.9960 | 6.1195 | 6.3733 | 6.0193 | 6.6664 | 6.1961 | 5.7918 | 6.0810 | 6.5130 | 4.8828 |
| UGDH_S476 | UGDH | O60701 | | 91 | VFISVNTPTKTYG | 5.4742 | 5.8363 | 6.9188 | 5.5162 | 5.8672 | 6.6616 | 5.7022 | 6.5318 | 6.3534 | 6.3579 | 6.5390 | 6.3272 | 6.0721 | 6.8760 | 6.6817 | 6.2844 |
| UGDH_T91 | UGDH | O60701 | | 476 | RIPYAPSGEIPKF | 2.5236 | 2.8571 | 3.0704 | 2.9073 | 3.2030 | 2.9862 | 3.1576 | 3.1946 | 3.1490 | 2.9573 | 2.7691 | 2.7346 | 15.0707 | 17.1643 | 16.6708 | 15.5844 |
| UHRF1_S104 | UHRF1 | Q96T88-2 | | 300 | ERPGEGSPMVDNP | 6.0424 | 6.4695 | 6.1774 | 6.9292 | 6.1576 | 6.2628 | 6.2985 | 6.5321 | 6.6728 | 6.5258 | 6.4364 | 6.6383 | 5.4276 | 6.0434 | 5.7700 | 5.6163 |
| UHRF1_S300 | UHRF1 | Q96T88-2 | | 104 | ERDSELSDTDSGC | 6.0047 | 6.4374 | 6.6472 | 7.0440 | 6.8670 | 6.4986 | 6.1054 | 5.6879 | 5.8673 | 5.9312 | 5.8125 | 6.3144 | 5.6271 | 6.3264 | 6.4446 | 6.3843 |
| UHRF1_S664 | UHRF1 | Q96T88-2 | | 89 | QLLVRQSLVLPHS | 7.3739 | 8.0942 | 6.6532 | 7.6133 | 7.5602 | 7.4205 | 7.5130 | 7.3188 | 6.9251 | 6.2082 | 6.8510 | 8.5381 | 3.5572 | 4.2671 | 3.5303 | 3.2758 |
| UHRF1_S720 | UHRF1 | Q96T88-2 | | 664 | GKWKRKSAGGGPS | 6.4980 | 7.3608 | 6.8431 | 6.9130 | 6.0461 | 6.4338 | 6.7653 | 6.7019 | 7.7726 | 8.1265 | 8.3561 | 7.6450 | 3.3558 | 4.1431 | 3.0548 | 3.9842 |
| UHRF1_S89 | UHRF1 | Q96T88-2 | | 720 | LKDRPASGSPFQL | 5.7053 | 6.3428 | 7.0119 | 5.2677 | 6.6726 | 6.2127 | 5.6387 | 4.8237 | 5.4833 | 6.4284 | 7.7699 | 7.6969 | 5.9685 | 6.8216 | 6.2445 | 5.9115 |
| UHRF1BP1_S1106 | UHRF1BP1 | Q6BDS2 | | 1106 | RTVSQQSFDGVSL | 5.9939 | 5.6116 | 5.7716 | 6.8666 | 6.4372 | 6.1553 | 6.6699 | 6.2730 | 6.3852 | 6.0909 | 6.1227 | 7.0585 | 6.0888 | 6.8340 | 6.1899 | 5.4509 |
| UHRF1BP1_S950 | UHRF1BP1 | Q6BDS2 | | 950 | SQERPHSNGELQD | 6.0687 | 5.6958 | 6.2080 | 6.0001 | 6.8968 | 6.2266 | 6.9874 | 5.7829 | 5.5205 | 6.8805 | 6.4392 | 6.0315 | 5.8108 | 6.5147 | 6.4380 | 6.4984 |
| UHRF1BP1L_S1377 | UHRF1BP1L | A0JNW5 | | 1399 | RSVTQATQTSPGV | 5.8308 | 6.1313 | 6.6156 | 6.0912 | 6.2086 | 5.5924 | 6.0965 | 6.1209 | 6.0427 | 6.4501 | 6.7590 | 6.2458 | 6.1626 | 6.6214 | 6.1865 | 6.4446 |
| UHRF1BP1L_S887;S891 | UHRF1BP1L | A0JNW5 | | 935 | SDNRSMSVDLSHI | 6.0619 | 4.9570 | 6.6150 | 4.4183 | 6.5796 | 5.5053 | 5.4271 | 5.8809 | 7.4406 | 7.1870 | 6.9618 | 7.1227 | 7.2236 | 8.2567 | 5.4363 | 5.9263 |
| UHRF1BP1L_S891 | UHRF1BP1L | A0JNW5 | | 891 | PVSESVSPVVPDY | 6.1982 | 5.4741 | 6.6454 | 5.7427 | 5.7488 | 6.5082 | 6.1884 | 6.5082 | 6.6958 | 6.7409 | 7.3139 | 6.3809 | 5.9970 | 5.9262 | 6.2729 | 5.6582 |
| UHRF1BP1L_S935 | UHRF1BP1L | A0JNW5 | | 953 | LLFKSASDTNLQK | 5.0727 | 4.6755 | 5.2609 | 4.5964 | 6.3501 | 5.8733 | 5.0388 | 5.8836 | 5.4386 | 5.5658 | 6.1727 | 6.5509 | 7.9791 | 8.8839 | 7.8520 | 8.0457 |
| UHRF1BP1L_S953 | UHRF1BP1L | A0JNW5 | | 1377 | LKENVKSDSVLLT | 3.2539 | 3.7495 | 2.6462 | 3.1677 | 4.3701 | 2.6074 | 1.0257 | 4.6244 | 2.6936 | 4.7192 | 3.6547 | 2.4480 | 12.2786 | 16.3600 | 17.4331 | 14.9679 |
| UHRF1BP1L_T1399 | UHRF1BP1L | A0JNW5 | 887;891 | | NA | 5.3447 | 6.0313 | 6.2033 | 5.6656 | 6.0766 | 6.2413 | 5.7155 | 6.5351 | 6.2769 | 5.3279 | 6.7143 | 5.6903 | 7.0871 | 7.4614 | 6.6919 | 6.9368 |
| UHRF2_S643 | UHRF2 | Q96PU4 | | 95 | AKPCSNSPPKVKK | 5.7182 | 5.5139 | 6.2222 | 6.0801 | 6.0636 | 6.0713 | 6.1371 | 5.3096 | 6.4503 | 6.3988 | 6.4224 | 6.4921 | 6.9985 | 6.6695 | 6.4512 | 7.0011 |
| UHRF2_S667 | UHRF2 | Q96PU4 | | 643 | YPAGYPSDKEGKK | 6.4105 | 6.9303 | 6.6080 | 6.6748 | 6.2583 | 6.3039 | 6.8744 | 6.6159 | 6.0365 | 6.1384 | 5.9692 | 5.9703 | 5.3708 | 5.9407 | 5.5015 | 6.3965 |
| UHRF2_S95 | UHRF2 | Q96PU4 | | 667 | TTKRPISDDDCPS | 6.2818 | 6.6273 | 6.2104 | 6.7194 | 6.3816 | 6.1199 | 6.4616 | 6.4948 | 6.3491 | 6.1630 | 6.3624 | 6.2430 | 5.9432 | 6.0717 | 5.6863 | 5.8844 |
| UIMC1_S140 | UIMC1 | Q96RL1 | | 653 | GAFRVPSPGMEEA | 6.2779 | 6.5709 | 5.8503 | 6.6240 | 6.4096 | 6.2430 | 6.4315 | 5.8019 | 6.2588 | 6.0241 | 6.0375 | 6.0675 | 6.2112 | 7.0781 | 6.6823 | 6.2612 |
| UIMC1_S198 | UIMC1 | Q96RL1 | | 677 | RRDLNESPVKSFV | 6.2598 | 6.2993 | 5.7109 | 6.2443 | 6.2326 | 5.7332 | 6.0586 | 5.7809 | 6.2849 | 5.8754 | 5.8584 | 5.7207 | 6.6016 | 7.1958 | 7.2016 | 6.9422 |
| UIMC1_S343 | UIMC1 | Q96RL1 | | 343 | GQGEQASEKNECI | 5.7825 | 5.8024 | 5.7322 | 6.4028 | 6.0040 | 5.7901 | 6.3379 | 5.6668 | 5.8367 | 5.8604 | 5.6488 | 5.5782 | 7.0308 | 7.7548 | 7.2045 | 7.5542 |
| UIMC1_S44;S46 | UIMC1 | Q96RL1 | | 463 | DLEREVSPGSRDI | 5.3177 | 5.9145 | 6.5994 | 7.0970 | 6.1477 | 6.4252 | 5.7021 | 6.3878 | 7.0227 | 6.8671 | 6.1360 | 6.3688 | 5.3873 | 6.2853 | 5.9946 | 6.3468 |
| UIMC1_S463 | UIMC1 | Q96RL1 | | 140 | LATGPSSQSHQEK | 5.5172 | 5.7499 | 5.6523 | 6.5828 | 5.8775 | 5.8401 | 5.7701 | 5.6642 | 5.8698 | 5.2457 | 5.8945 | 5.8137 | 6.9710 | 7.9008 | 8.2467 | 7.4039 |
| UIMC1_S653 | UIMC1 | Q96RL1 | | 198 | EPVSGSGSWDQS | 5.6450 | 6.4134 | 6.2287 | 6.4689 | 6.4673 | 5.5499 | 6.2662 | 6.1385 | 6.3096 | 5.8732 | 5.7428 | 5.6912 | 6.6933 | 7.2966 | 6.9710 | 6.2444 |
| UIMC1_S677 | UIMC1 | Q96RL1 | 44;46 | | NA | 6.1747 | 5.4329 | 5.3004 | 6.5199 | 7.3718 | 6.2346 | 6.5277 | 6.6671 | 6.1220 | 6.2251 | 7.0658 | 6.2396 | 6.3128 | 6.1866 | 5.8908 | 5.7282 |
| ULK1_S450 | ULK1 | O75385 | | 638 | DFPKTPSSQNLLA | 5.9616 | 5.3106 | 6.5561 | 6.0083 | 6.1898 | 6.4442 | 6.3472 | 6.1161 | 6.1700 | 5.7277 | 6.6083 | 6.0579 | 6.5816 | 7.1888 | 6.1130 | 6.6189 |
| ULK1_S469 | ULK1 | O75385 | | 495 | VLARKMSLGGGRP | 5.6233 | 4.9454 | 5.7898 | 5.0507 | 6.0672 | 5.3656 | 5.3231 | 6.3656 | 7.8284 | 7.6499 | 6.9890 | 7.0322 | 6.8571 | 7.6436 | 5.4489 | 6.0202 |
| ULK1_S495 | ULK1 | O75385 | | 758 | VVFTVGSPPSGST | 5.1286 | 5.9529 | 6.3598 | 5.7507 | 5.9250 | 6.3440 | 6.3747 | 6.0238 | 6.4687 | 6.2691 | 6.0381 | 7.1630 | 6.5095 | 6.8426 | 6.5254 | 6.3239 |
| ULK1_S623 | ULK1 | O75385 | | 623 | LPPILGSPTKAVP | 6.0994 | 6.0376 | 5.8489 | 5.6050 | 6.2915 | 6.3821 | 6.6334 | 6.6295 | 6.1520 | 6.1366 | 6.0774 | 6.2198 | 6.8750 | 6.6024 | 6.0867 | 6.3229 |
| ULK1_S638 | ULK1 | O75385 | | 660 | TPPRNRTLPDLSE | 5.3523 | 5.4091 | 5.9336 | 5.1432 | 6.6307 | 6.5185 | 6.6266 | 6.4880 | 5.8130 | 5.9089 | 5.5817 | 6.2698 | 7.1293 | 7.3256 | 5.9253 | 6.9445 |
| ULK1_S758 | ULK1 | O75385 | | 450 | IERNLQSPTQFQT | 6.6630 | 6.0317 | 6.9894 | 5.9754 | 6.5156 | 6.4373 | 6.3421 | 6.5532 | 6.8516 | 6.5180 | 6.8997 | 6.6232 | 5.3634 | 5.9094 | 5.1213 | 5.2056 |
| ULK1_S758;T764 | ULK1 | O75385 | | 469 | RRSGSTSPLGFAR | 6.6902 | 6.6127 | 6.5326 | 6.4445 | 6.5087 | 6.2902 | 6.2783 | 6.8345 | 6.6952 | 6.5774 | 6.3326 | 6.6476 | 5.3338 | 5.7154 | 5.1585 | 5.3477 |
| ULK1_T660 | ULK1 | O75385 | 758;764 | | NA | 6.8861 | 6.9149 | 7.3099 | 6.0825 | 6.0725 | 5.9015 | 6.6233 | 5.4213 | 7.0878 | 6.3639 | 6.2791 | 7.2565 | 5.1156 | 5.5787 | 4.9651 | 6.1412 |
| ULK2_S600 | ULK2 | Q8IYT8 | | 600 | LPTIIGSPTKTTA | 6.3600 | 6.2497 | 6.2434 | 6.3820 | 6.3437 | 6.4332 | 6.5042 | 6.7622 | 6.6513 | 6.2591 | 6.8517 | 6.3628 | 5.4372 | 5.8714 | 5.3439 | 5.9443 |
| UMPS_S214 | UMPS | P11172 | | 214 | AANHNGSPLSIKE | 5.9372 | 6.6062 | 7.0694 | 6.5118 | 6.0570 | 6.6727 | 6.9199 | 6.9675 | 6.8630 | 6.8063 | 6.6370 | 7.5939 | 4.3026 | 5.2539 | 4.7251 | 5.0764 |
| UMPS_S335 | UMPS | P11172 | | 335 | GIFKIASWADLVN | 4.4824 | 4.8870 | 6.4873 | 4.7581 | 6.4390 | 6.5500 | 6.4425 | 5.9198 | 5.6048 | 5.6370 | 4.9944 | 6.0434 | 8.3096 | 7.3967 | 8.4833 | 7.5649 |
| UNC13B_S367 | UNC13B | O14795-2 | | 367 | PEENASSPFTQAR | 6.6328 | 6.0194 | 7.0616 | 7.2298 | 6.0464 | 7.1857 | 6.6679 | 5.9345 | 7.4430 | 6.9097 | 6.8337 | 6.3837 | 4.5795 | 5.2264 | 4.4660 | 5.3799 |
| UNC45A_T15 | UNC45A | Q9H3U1 | | 15 | PEPRPATPGASSV | 6.3451 | 6.0935 | 7.0622 | 6.2089 | 6.8398 | 6.5124 | 6.6017 | 5.8216 | 5.9937 | 6.4317 | 7.2267 | 6.8690 | 5.5587 | 5.8714 | 5.0270 | 5.5365 |
| UNC93B1_S547;S550 | UNC93B1 | Q9H1C4 | 547;550 | | NA | 6.1750 | 6.5097 | 5.9658 | 7.2389 | 6.0261 | 7.1528 | 7.4338 | 5.8871 | 6.4882 | 6.1286 | 5.2532 | 5.5943 | 6.2205 | 6.2526 | 6.0200 | 5.6534 |
| UNG_S12 | UNG | P13051 | | 23 | RKRHAPSPEPAVQ | 6.1322 | 5.7381 | 7.1324 | 5.7619 | 6.0221 | 5.9270 | 5.5402 | 6.0631 | 6.1700 | 6.1863 | 6.4679 | 6.2996 | 6.2986 | 7.0418 | 6.7387 | 6.4800 |
| UNG_S12;S14 | UNG | P13051 | | 60 | QEEPGTPPSSPL | 6.0852 | 5.5606 | 6.6247 | 5.9733 | 6.1851 | 6.4331 | 5.5881 | 6.1329 | 7.0588 | 6.7420 | 7.5488 | 7.1939 | 5.5948 | 6.3163 | 5.7531 | 5.2093 |
| UNG_S14 | UNG | P13051 | | 12 | TLYSFFSPSPARK | 6.6313 | 6.2433 | 6.6013 | 6.2912 | 5.7158 | 6.1836 | 6.9077 | 6.1940 | 6.5308 | 7.1724 | 7.3578 | 7.0558 | 5.0715 | 6.0044 | 4.9355 | 5.1036 |
| UNG_S23 | UNG | P13051 | | 14 | YSFFSPSPARKRH | 6.5413 | 7.0131 | 6.6615 | 6.0899 | 6.5809 | 6.6474 | 5.5146 | 6.9555 | 7.3660 | 6.2741 | 8.5857 | 7.8991 | 3.4829 | 4.8983 | 4.2206 | 5.2691 |
| UNG_T60 | UNG | P13051 | 60;64 | | NA | 6.2804 | 5.9830 | 6.6453 | 6.4226 | 5.9734 | 6.0975 | 5.6254 | 6.2148 | 8.0270 | 6.4344 | 6.8641 | 6.8896 | 5.4594 | 6.3144 | 5.2808 | 5.4878 |
| UNG_T60;S64 | UNG | P13051 | 12;14 | | NA | 9.8423 | 6.6285 | 8.5382 | 8.4848 | 5.6796 | 4.4781 | 7.5277 | 4.8831 | 9.8347 | 7.9918 | 8.9468 | 6.3841 | 2.0834 | 3.1533 | 2.4428 | 3.1006 |
| UNK_S240 | UNK | Q9C0B0 | | 385 | PSNLCGSPPGSIR | 6.4603 | 6.5434 | 6.7125 | 6.2997 | 6.5048 | 6.5426 | 5.9996 | 6.0213 | 6.3524 | 6.7745 | 6.5488 | 6.3239 | 6.4045 | 6.1943 | 4.9826 | 5.3351 |
| UNK_S375;S385 | UNK | Q9C0B0 | | 240 | CPYYHNSKDRRRS | 6.0517 | 6.7790 | 6.9235 | 6.3105 | 5.9557 | 6.9809 | 5.8057 | 6.5006 | 6.7803 | 6.4255 | 6.9376 | 6.3525 | 5.4945 | 5.8856 | 5.5302 | 5.2861 |
| UNK_S385 | UNK | Q9C0B0 | | 411 | SGLAPGSYKKAPG | 7.6068 | 7.5703 | 9.3524 | 7.0416 | 7.2604 | 6.1073 | 5.8429 | 6.2303 | 6.1848 | 7.0188 | 4.8585 | 7.5976 | 4.0290 | 5.4008 | 2.8258 | 5.0727 |
| UNK_S411 | UNK | Q9C0B0 | 375;385 | | NA | 5.9357 | 6.4313 | 6.5658 | 5.9382 | 6.4514 | 6.1335 | 6.2082 | 6.6754 | 6.8310 | 6.3697 | 6.8194 | 6.6352 | 5.7238 | 5.9450 | 5.6450 | 5.6912 |
| UNKL_S15 | UNKL | Q9H9P5-6 | | 15 | AALSGSPPQTEK | 5.8387 | 6.4036 | 6.5461 | 6.4701 | 5.9104 | 6.4989 | 6.2121 | 6.5903 | 6.4874 | 6.8206 | 6.7180 | 6.9340 | 5.4664 | 5.5638 | 5.7079 | 5.8317 |
| UPF1_S1107 | UPF1 | Q92900 | | 1107 | QIDVALSQDSTYQ | 6.1273 | 6.3580 | 6.4096 | 6.0783 | 6.2170 | 6.3877 | 6.3133 | 6.7907 | 6.6493 | 6.6371 | 6.8614 | 6.7145 | 5.4122 | 6.0804 | 5.3487 | 5.6144 |
| UPF1_S1127 | UPF1 | Q92900 | | 1127 | GGVTGLSQYxxxx | 6.7968 | 6.3943 | 6.9369 | 6.7951 | 6.3697 | 6.4467 | 6.3258 | 6.0894 | 6.4698 | 5.9137 | 6.3069 | 5.7902 | 6.0708 | 6.2684 | 5.4156 | 5.6099 |
| UPF3B_T177 | UPF3B | Q9BZI7 | | 177 | NEKMTSTPETLLE | 6.4255 | 6.8790 | 5.6881 | 7.5263 | 7.0943 | 6.9557 | 7.5761 | 6.2156 | 5.7177 | 6.7996 | 7.9569 | 6.9409 | 3.9603 | 5.3315 | 3.9616 | 4.9709 |
| UQCRC1_S107 | UQCRC1 | P31930 | | 107 | TKNRPGSALEKEV | 6.0196 | 5.5803 | 5.4492 | 5.7962 | 5.7924 | 5.1903 | 6.0665 | 5.8303 | 6.2817 | 6.4967 | 6.5197 | 6.9378 | 6.6609 | 6.8709 | 7.1883 | 7.3191 |
| UQCRC1_T266 | UQCRC1 | P31930 | | 266 | DAVPTLTPCRFTG | 6.1416 | 6.4441 | 5.9595 | 6.3672 | 6.4311 | 5.8730 | 6.5701 | 6.7969 | 6.5651 | 6.0511 | 6.4622 | 6.9846 | 6.5921 | 6.5989 | 6.0574 | 5.9053 |
| UQCRFS1P1_S108 | UQCRFS1P1 | P0C7P4 | | 108 | RLEVLDSTKSSRE | 6.2726 | 5.9316 | 5.6029 | 6.2416 | 5.7987 | 5.9842 | 6.1801 | 5.3643 | 5.9985 | 5.2944 | 5.5266 | 5.9144 | 7.4958 | 6.9442 | 7.7649 | 7.6852 |
| UQCRQ_S16 | UQCRQ | O14949 | | 16 | RMRHVISYSLSPF | 5.2081 | 5.7465 | 5.9199 | 4.9747 | 5.8432 | 4.1294 | 5.6398 | 5.8816 | 5.7282 | 6.1883 | 4.9597 | 5.9182 | 8.2755 | 8.5973 | 8.9107 | 8.0788 |
| URB1_S9 | URB1 | O60287 | | 9 | VPKRKASGGQDGA | 5.7506 | 5.6954 | 6.5515 | 8.0842 | 7.3710 | 6.5255 | 5.8343 | 6.4500 | 6.7870 | 7.7707 | 8.7640 | 6.9243 | 4.5225 | 5.0505 | 3.8110 | 4.1075 |
| URB2_S1414 | URB2 | Q14146 | | 1414 | RQKDKGSIDDLPT | 6.0529 | 6.1410 | 5.7874 | 6.6423 | 5.7054 | 6.5556 | 6.3328 | 5.8802 | 6.2584 | 5.9821 | 5.9192 | 6.2750 | 6.8904 | 6.4325 | 7.0002 | 7.0446 |
| URI1_S372 | URI1 | O94763 | | 492 | FVSPSLTPPPAIA | 6.3643 | 6.5530 | 6.6911 | 6.8576 | 6.4953 | 6.4727 | 6.6893 | 6.9464 | 7.1554 | 6.3219 | 6.9388 | 6.4757 | 5.0685 | 5.2028 | 4.6669 | 5.1003 |
| URI1_S442 | URI1 | O94763 | | 372 | KRKRKNSTGSGHS | 6.4744 | 5.7711 | 6.4854 | 6.5118 | 7.6180 | 7.0897 | 5.8051 | 7.0508 | 7.0907 | 6.9607 | 7.5747 | 7.6811 | 4.7221 | 5.1235 | 4.1613 | 3.8797 |
| URI1_T492 | URI1 | O94763 | | 442 | GVLRSISCEEATC | 5.8425 | 5.0827 | 6.7359 | 5.5070 | 6.2237 | 6.1822 | 5.2750 | 6.2354 | 6.5123 | 6.2270 | 5.9334 | 6.4535 | 7.3954 | 7.2823 | 6.8836 | 6.2281 |
| USF2_S155 | USF2 | Q15853-3 | | 155 | PRTHPYSPKIDGT | 6.9515 | 6.6410 | 6.1189 | 6.5602 | 6.6027 | 6.2087 | 6.2072 | 6.2355 | 6.8075 | 6.4336 | 6.3635 | 6.4092 | 6.1245 | 5.5622 | 5.1792 | 5.5947 |
| USF2_S88 | USF2 | Q15853-3 | | 88 | PFSNGGSPAAEAV | 5.6876 | 6.3684 | 4.9190 | 6.5711 | 6.7113 | 5.9663 | 6.0727 | 5.7450 | 7.0187 | 5.4984 | 5.0964 | 7.2501 | 5.7180 | 6.7163 | 7.7636 | 6.8971 |
| USO1_S862 | USO1 | O60763-2 | | 953 | EEDELESGDQEDE | 6.2254 | 6.2049 | 6.2170 | 6.0704 | 5.8051 | 5.9715 | 5.9690 | 6.3372 | 5.8256 | 5.9844 | 6.0765 | 6.3256 | 6.3394 | 7.3998 | 6.5417 | 6.7063 |
| USO1_S953 | USO1 | O60763-2 | | 862 | DVEGRLSALLQET | 4.8296 | 5.6401 | 6.5544 | 6.3549 | 5.9767 | 5.8022 | 5.7689 | 4.7922 | 6.1638 | 7.4564 | 7.3857 | 6.6542 | 5.0582 | 6.8911 | 7.4314 | 7.2402 |
| USP1_S327 | USP1 | O94782 | | 67 | VPAAQSSPINCEK | 5.6020 | 5.8316 | 6.8263 | 6.6620 | 6.4385 | 6.3422 | 6.2341 | 6.3661 | 6.5430 | 6.5828 | 6.9001 | 6.5774 | 5.5980 | 5.8278 | 5.5809 | 5.8172 |
| USP1_S475 | USP1 | O94782 | | 558 | INTPLLTPLKLSL | 7.7590 | 7.2647 | 7.9482 | 7.0171 | 6.0189 | 7.7354 | 7.9120 | 5.7414 | 7.2528 | 8.1666 | 7.5359 | 7.3640 | 2.7170 | 3.7609 | 2.4718 | 3.3343 |
| USP1_S67 | USP1 | O94782 | | 475 | EESSEISPEPKTE | 6.1166 | 6.9599 | 6.9925 | 6.3628 | 5.9328 | 5.9554 | 6.3443 | 6.1394 | 6.8274 | 7.0585 | 7.1751 | 6.9534 | 4.9929 | 5.6600 | 5.0874 | 5.4416 |
| USP1_T558 | USP1 | O94782 | | 327 | YISENESPRPSQK | 7.6226 | 8.5744 | 7.6107 | 7.1618 | 4.8171 | 6.1763 | 6.7636 | 6.5887 | 7.9545 | 7.9443 | 7.7266 | 8.1512 | 2.1670 | 4.2601 | 2.2698 | 4.2113 |
| USP10_S124 | USP10 | Q14694-2 | | 259 | TPRTCNSPQNSTD | 6.0850 | 6.1782 | 6.6319 | 6.3952 | 6.0315 | 5.9323 | 6.2917 | 6.0786 | 6.0194 | 6.3129 | 6.6494 | 6.2727 | 6.1070 | 6.7469 | 6.0250 | 6.2425 |
| USP10_S259 | USP10 | Q14694-2 | | 624 | EEQGEGSEDEWEQ | 6.0694 | 5.9776 | 6.7509 | 6.3835 | 6.1425 | 6.0122 | 6.5546 | 6.3009 | 6.1460 | 6.5492 | 6.4156 | 6.5399 | 5.8510 | 6.3003 | 5.9253 | 6.0812 |
| USP10_S413 | USP10 | Q14694-2 | | 413 | YVETKYSPPAISP | 5.9554 | 5.8265 | 6.7012 | 6.3001 | 6.4296 | 6.8087 | 6.2293 | 6.3539 | 6.2678 | 6.4546 | 6.9278 | 6.7429 | 5.9709 | 6.4488 | 5.1970 | 5.3854 |
| USP10_S413;S418 | USP10 | Q14694-2 | | 124 | TLPRTPSYSISST | 3.3159 | 3.4071 | 3.4312 | 3.4438 | 4.3102 | 4.1276 | 4.1562 | 4.4358 | 3.6052 | 3.5202 | 3.5512 | 3.5999 | 13.6987 | 13.5874 | 14.6725 | 13.1372 |
| USP10_S418 | USP10 | Q14694-2 | | 595 | NLKKLLSPSNEKL | 6.1573 | 6.6657 | 6.6211 | 6.4691 | 5.7925 | 6.3239 | 6.3763 | 6.4609 | 6.3887 | 6.5703 | 6.6186 | 6.6283 | 5.3657 | 6.0193 | 5.5972 | 5.9453 |
| USP10_S595 | USP10 | Q14694-2 | | 418 | YSPPAISPLVSEK | 6.9649 | 7.3896 | 7.1628 | 7.6006 | 6.4967 | 7.1781 | 7.5463 | 7.2514 | 6.8916 | 6.9396 | 7.0670 | 7.0230 | 3.1895 | 4.0968 | 3.2024 | 3.9997 |
| USP10_S624 | USP10 | Q14694-2 | | 148 | CTASKITPDGITK | 6.7819 | 7.3082 | 7.2438 | 7.4673 | 6.1663 | 6.8277 | 7.1149 | 6.9234 | 6.7745 | 7.3525 | 7.2562 | 7.4198 | 3.3889 | 4.2713 | 3.4805 | 4.2228 |

| | | | | | | | | | | | | | | | | | | | | |
|---|---|---|---|---|---|---|---|---|---|---|---|---|---|---|---|---|---|---|---|---|
| USP10_S766 | USP10 | Q14694-2 | | 122 | SDTLPRTPSYSIS | 6.3263 | 6.3948 | 6.2839 | 6.3597 | 6.8076 | 6.5325 | 6.7457 | 7.1169 | 6.7875 | 6.2650 | 5.8775 | 6.6205 | 5.3378 | 6.0503 | 5.1260 | 5.3680 |
| USP10_T122 | USP10 | Q14694-2 | | 766 | ISKELLSPGVKNK | 6.2572 | 7.1086 | 7.0655 | 6.8566 | 6.2580 | 6.7268 | 6.1050 | 6.6462 | 6.8368 | 7.0281 | 6.7758 | 6.8286 | 4.2233 | 5.1879 | 4.9900 | 5.1057 |
| USP10_T148 | USP10 | Q14694-2 | 413;418 | | NA | 5.9237 | 6.2914 | 5.9599 | 6.3891 | 6.2625 | 6.9598 | 6.9505 | 7.0544 | 6.4365 | 6.3621 | 6.2102 | 6.8572 | 5.0852 | 6.1007 | 5.0547 | 6.1021 |
| USP11_S648 | USP11 | P51784 | | 948 | SSGAPASPACSSP | 6.1531 | 6.1186 | 6.6853 | 6.4241 | 6.3431 | 6.5072 | 6.4322 | 6.2336 | 6.2607 | 6.8055 | 6.5460 | 6.6051 | 5.3243 | 6.0168 | 5.8532 | 5.6913 |
| USP11_S733 | USP11 | P51784 | | 648 | YVTKPNSDDEDDG | 5.0549 | 5.2924 | 5.8453 | 5.0878 | 6.1563 | 5.4082 | 5.8744 | 5.5407 | 6.8442 | 6.9622 | 7.4706 | 7.2009 | 7.2412 | 6.9407 | 6.1122 | 6.9678 |
| USP11_S943 | USP11 | P51784 | | 943 | LSPAGSSGAPASP | 6.0776 | 7.5083 | 6.4649 | 4.5385 | 6.9459 | 7.3011 | 6.1886 | 7.2419 | 6.0530 | 6.6425 | 6.6894 | 7.0544 | 3.6999 | 5.3204 | 6.6524 | 5.6213 |
| USP11_S943;S948 | USP11 | P51784 | | 733 | TSDRTTSPEEVHA | 4.6464 | 7.4166 | 5.9226 | 5.5061 | 7.4166 | 5.5367 | 5.9340 | 4.3063 | 6.5034 | 6.1633 | 5.7621 | 6.4155 | 6.6295 | 7.0498 | 8.2687 | 6.5225 |
| USP11_S948 | USP11 | P51784 | 948;952 | | NA | 5.5641 | 5.9214 | 6.4118 | 5.8788 | 5.8794 | 5.8567 | 6.0689 | 6.5523 | 6.1443 | 6.3168 | 6.2598 | 6.6642 | 6.5127 | 7.3128 | 6.2301 | 6.4258 |
| USP11_S948;S952 | USP11 | P51784 | 943;948 | | NA | 5.7728 | 4.2421 | 7.3060 | 6.7053 | 5.4302 | 6.2755 | 6.7020 | 7.0924 | 6.1403 | 6.4547 | 6.4071 | 6.1922 | 6.1974 | 6.2526 | 6.3911 | 6.4385 |
| USP13_T122 | USP13 | Q92995 | | 122 | IFLDLDTDDDLNS | 5.8934 | 5.7020 | 5.6665 | 6.0138 | 6.4854 | 6.3852 | 5.7451 | 7.5971 | 5.0238 | 6.2078 | 6.6172 | 6.1520 | 6.5234 | 6.5247 | 6.3434 | 7.1192 |
| USP14_S143 | USP14 | P54578 | | 143 | AGALRASGEMASA | 6.2172 | 6.0691 | 6.9803 | 6.4088 | 6.4666 | 6.7188 | 6.2757 | 6.8660 | 6.4217 | 6.1441 | 6.7917 | 6.5553 | 5.3802 | 5.9069 | 5.4338 | 5.3637 |
| USP15_S229 | USP15 | Q9Y4E8-3 | | 230 | WPRGPSTPNVKNS | 7.3113 | 6.9164 | 7.2019 | 6.4839 | 7.2989 | 6.6763 | 6.7749 | 6.2371 | 6.7179 | 6.4973 | 6.7873 | 6.6177 | 4.7611 | 4.7832 | 4.2852 | 4.6495 |
| USP15_T230 | USP15 | Q9Y4E8 | | 229 | GPSTPKSPGASNF | 6.3460 | 6.2490 | 6.2263 | 6.4634 | 6.2344 | 7.0120 | 7.0033 | 6.2015 | 6.5356 | 6.5539 | 6.1388 | 6.7654 | 5.5866 | 6.0403 | 5.1423 | 5.5012 |
| USP16_S189 | USP16 | Q9Y5T5 | | 552 | LEVLTSSPTRNLN | 6.1227 | 5.9492 | 6.4166 | 6.4696 | 6.0766 | 6.0481 | 6.5924 | 6.2581 | 6.5409 | 6.5439 | 6.6056 | 6.7170 | 5.5388 | 6.3259 | 5.7091 | 6.0854 |
| USP16_S415 | USP16 | Q9Y5T5 | | 415 | EDEDQDSEEEKDN | 5.8069 | 6.7352 | 5.8274 | 5.9180 | 6.1785 | 6.1826 | 6.4198 | 6.9674 | 6.3906 | 6.6552 | 6.7448 | 6.7169 | 5.7488 | 6.2338 | 5.6949 | 5.7790 |
| USP16_S552 | USP16 | Q9Y5T5 | | 189 | ENPPMNSPCQITV | 6.1555 | 6.1516 | 5.6461 | 8.8324 | 6.3891 | 7.0644 | 7.2456 | 6.8103 | 6.2218 | 6.5844 | 6.1963 | 6.6635 | 4.9772 | 5.1700 | 4.8215 | 5.0704 |
| USP19_S345 | USP19 | O94966-5 | | 345 | PGNDPVSPAMVRS | 6.1284 | 6.4337 | 6.8049 | 6.3604 | 6.3835 | 6.2690 | 6.4169 | 6.4922 | 6.8436 | 6.6434 | 6.4253 | 6.5222 | 5.4035 | 6.0456 | 5.4002 | 5.4273 |
| USP19_T508 | USP19 | O94966-5 | | 508 | PTPLDSTPPGGAP | 6.0957 | 7.3442 | 7.7026 | 9.4352 | 6.5246 | 6.0900 | 6.1205 | 7.7617 | 6.9973 | 5.6878 | 7.5730 | 6.6695 | 3.1718 | 4.7462 | 3.6349 | 4.4450 |
| USP20_S112 | USP20 | Q9Y2K6 | | 112 | KFSEQDSPPPSHP | 6.6069 | 7.1135 | 6.6043 | 7.5320 | 6.3212 | 7.0347 | 7.3456 | 6.9535 | 6.8482 | 7.1122 | 7.2151 | 7.0170 | 3.8382 | 4.5739 | 3.5362 | 4.3474 |
| USP20_S132 | USP20 | Q9Y2K6 | | 413 | SSPPRASPVRMAP | 6.2708 | 6.5385 | 6.3924 | 6.2489 | 5.9593 | 6.2468 | 6.1491 | 6.1035 | 5.9626 | 6.8053 | 6.8869 | 7.1834 | 5.5741 | 6.0911 | 5.6682 | 5.9191 |
| USP20_S265;S266 | USP20 | Q9Y2K6 | | 132 | VADEGESESEDDD | 6.2833 | 6.8445 | 6.1237 | 6.2900 | 5.9129 | 6.6615 | 6.4312 | 6.0708 | 7.0130 | 5.6481 | 6.3178 | 6.5058 | 6.2027 | 5.8372 | 6.1687 | 5.6887 |
| USP20_S413 | USP20 | Q9Y2K6 | 265;266 | | NA | 5.9643 | 6.3506 | 6.4613 | 6.8053 | 5.8147 | 5.8773 | 5.5486 | 6.2171 | 6.6778 | 6.4650 | 6.2845 | 6.5452 | 5.6828 | 5.9875 | 6.1242 | 7.1938 |
| USP22_S237 | USP22 | Q9UPT9 | | 237 | FYSGHRSPHIPYK | 6.1422 | 6.6503 | 7.5203 | 6.8488 | 6.0500 | 6.4876 | 6.0842 | 6.7145 | 6.5258 | 6.3257 | 6.5805 | 6.7029 | 4.6504 | 5.5220 | 5.4895 | 5.7053 |
| USP24_S1141 | USP24 | Q9UPU5 | | 1141 | SSQSSKSPSLSSK | 6.0858 | 6.4779 | 6.0929 | 6.4755 | 5.9821 | 5.8772 | 6.0877 | 6.1333 | 6.3555 | 6.5532 | 6.2315 | 6.3246 | 5.8218 | 6.6070 | 6.3077 | 6.5865 |
| USP24_S1145 | USP24 | Q9UPU5 | | 2561 | TFQRTISAQDTLA | 5.8613 | 5.9197 | 6.8502 | 6.4041 | 6.0395 | 6.2366 | 6.0372 | 6.4547 | 6.3229 | 6.2470 | 6.5023 | 6.5962 | 5.8429 | 6.4342 | 6.2790 | 5.9723 |
| USP24_S1305 | USP24 | Q9UPU5 | | 2047 | RVSDQNSPVLPKK | 6.1131 | 6.2545 | 6.6423 | 6.1102 | 6.2703 | 6.5031 | 6.2156 | 6.5210 | 6.7612 | 6.4060 | 6.3161 | 6.2220 | 5.8975 | 6.0955 | 6.0054 | 5.6663 |
| USP24_S1943 | USP24 | Q9UPU5 | | 1289 | QMSLCGTPEKSSY | 7.0488 | 7.2831 | 7.1973 | 7.0931 | 6.4181 | 7.4240 | 7.0045 | 6.9967 | 7.0734 | 7.2950 | 7.7310 | 6.8490 | 3.4717 | 4.1447 | 3.0486 | 3.9209 |
| USP24_S2047 | USP24 | Q9UPU5 | | 2604 | LQQGSESPMMIGE | 6.6069 | 6.9001 | 6.6047 | 6.4674 | 6.2800 | 6.7259 | 6.3096 | 6.3022 | 6.6229 | 6.8582 | 6.4938 | 6.3556 | 5.3991 | 5.8303 | 5.0105 | 5.2328 |
| USP24_S2561 | USP24 | Q9UPU5 | | 1305 | SVSDRSSIRVEEI | 8.5478 | 7.1500 | 8.2775 | 6.6282 | 5.8654 | 6.1563 | 7.4546 | 7.4270 | 7.4222 | 6.3733 | 6.9171 | 6.7583 | 3.8132 | 4.1191 | 3.2426 | 3.8472 |
| USP24_S2604 | USP24 | Q9UPU5 | | 1943 | DQGGGGSPRKKVA | 6.2533 | 6.0942 | 6.4068 | 6.0761 | 6.3188 | 6.4209 | 6.2354 | 6.2336 | 6.3359 | 6.1816 | 6.0252 | 6.5179 | 6.3433 | 6.3804 | 6.1350 | 6.0417 |
| USP24_T1289 | USP24 | Q9UPU5 | | 1145 | SKSPSLSSKQQHQ | 5.9999 | 6.1994 | 5.6271 | 6.0006 | 6.3248 | 5.2920 | 6.1557 | 5.8569 | 5.8725 | 5.8662 | 5.0812 | 4.8885 | 7.3284 | 7.7514 | 7.8803 | 7.8752 |
| USP28_S1053 | USP28 | Q96RU2 | | 1053 | PTIRPNSPYDLCS | 5.8567 | 6.1076 | 5.8892 | 5.7952 | 5.6986 | 6.2806 | 6.2687 | 5.4291 | 5.7154 | 6.8700 | 6.6377 | 6.7939 | 7.3379 | 6.8331 | 6.0979 | 6.3883 |
| USP28_S131 | USP28 | Q96RU2 | | 131 | RMHEATSAETKRS | 6.3413 | 7.4150 | 7.1110 | 6.9849 | 6.9606 | 6.4724 | 6.5106 | 6.8022 | 7.5992 | 7.6841 | 6.7637 | 7.3551 | 3.3526 | 4.6127 | 3.7696 | 4.2651 |
| USP28_S495 | USP28 | Q96RU2 | | 495 | TSTESSSQDVEST | 6.0550 | 6.4935 | 7.0474 | 6.9023 | 5.7580 | 6.0260 | 6.2051 | 6.3094 | 6.0964 | 6.5793 | 6.3774 | 6.3051 | 6.1896 | 6.2057 | 5.8401 | 5.6098 |
| USP31_S1174 | USP31 | Q70CQ4 | | 1174 | ASATSTSKPNSPR | 7.5074 | 6.7795 | 7.4771 | 6.6154 | 4.7885 | 6.2853 | 6.3621 | 6.9009 | 7.4486 | 7.1863 | 7.0043 | 7.8072 | 4.8581 | 4.7617 | 4.2229 | 3.9946 |
| USP32_S1350 | USP32 | Q8NFA0 | | 1416 | RLPQIGSKNKLSS | 2.8363 | 3.0043 | 3.4431 | 3.1230 | 4.1795 | 3.6243 | 4.0989 | 3.3806 | 3.0758 | 3.3415 | 2.8136 | 2.9376 | 15.7688 | 14.1347 | 15.6047 | 14.6335 |
| USP32_S1386 | USP32 | Q8NFA0 | | 1430 | KENLDASKENGAG | 5.6213 | 6.1078 | 6.8567 | 7.0646 | 6.1802 | 6.4341 | 7.0761 | 6.2942 | 6.5555 | 6.4431 | 6.7279 | 7.1419 | 5.1452 | 5.6851 | 4.9110 | 5.7553 |
| USP32_S1416 | USP32 | Q8NFA0 | | 1326 | CQHKPLTPQGDEL | 5.3369 | 5.2925 | 6.5746 | 5.9677 | 6.2786 | 6.8690 | 6.2919 | 6.4150 | 6.0802 | 6.4144 | 6.2503 | 5.8291 | 6.5547 | 6.7703 | 5.8751 | 7.1999 |
| USP32_S1430 | USP32 | Q8NFA0 | | 1386 | SRKSGTSCPSSKN | 5.4072 | 5.8970 | 6.4537 | 6.5371 | 6.7930 | 6.4682 | 6.7159 | 6.2679 | 6.3981 | 6.8423 | 6.7835 | 6.8088 | 5.5178 | 6.1365 | 5.3851 | 5.5880 |
| USP32_T1326 | USP32 | Q8NFA0 | | 1350 | KVDAQSSAGEEDV | 6.2960 | 6.7802 | 6.6251 | 5.4560 | 7.0366 | 6.7712 | 6.4991 | 6.6892 | 6.3739 | 5.6251 | 6.0340 | 5.6521 | 6.1030 | 6.0793 | 6.1176 | 5.8616 |
| USP33_S377 | USP33 | Q8TEY7 | | 439 | NPRLSASPPKSGN | 7.3984 | 7.5092 | 7.5644 | 8.0297 | 6.3272 | 6.2789 | 6.9050 | 6.5510 | 7.6107 | 7.5079 | 7.6131 | 7.7128 | 2.8911 | 3.4738 | 3.0355 | 3.5913 |
| USP33_S387 | USP33 | Q8TEY7 | | 461 | KKAQSASPKRKKQ | 6.8047 | 7.5494 | 7.0306 | 7.9098 | 6.1216 | 6.2222 | 6.5671 | 6.2689 | 6.8846 | 7.0134 | 6.9916 | 7.2418 | 4.0941 | 4.5410 | 4.0620 | 4.6973 |
| USP33_S439 | USP33 | Q8TEY7 | | 377 | KINKVNSEGEFDK | 2.3219 | 1.9858 | 2.6824 | 2.2333 | 3.0146 | 3.2560 | 3.3163 | 2.4288 | 2.1225 | 2.9925 | 2.1462 | 2.1317 | 17.1254 | 14.8479 | 18.5178 | 18.8768 |
| USP33_S461 | USP33 | Q8TEY7 | | 387 | FDKDRDSISETVD | 7.0029 | 7.0086 | 6.5399 | 7.4490 | 5.6234 | 6.5080 | 7.0443 | 5.7006 | 6.6115 | 7.1121 | 6.0356 | 6.0582 | 4.9741 | 5.5143 | 5.2188 | 5.5989 |
| USP34_S2488 | USP34 | Q70CQ2 | | 2488 | PQVEVLSEEEGEE | 5.9210 | 6.0042 | 6.1120 | 6.2759 | 6.1586 | 5.9115 | 6.3337 | 6.1712 | 5.9411 | 6.3390 | 6.4086 | 6.1115 | 6.2837 | 7.1121 | 6.6934 | 6.2224 |
| USP34_S3358 | USP34 | Q70CQ2 | | 3358 | IKRRRVSSDEEHT | 5.3189 | 6.2166 | 6.2708 | 6.1889 | 6.4511 | 6.4395 | 6.0786 | 6.0501 | 6.6948 | 7.0102 | 6.3380 | 6.5841 | 6.1842 | 6.2665 | 5.9188 | 5.9889 |
| USP34_S3359 | USP34 | Q70CQ2 | | 3359 | KRRRVSSDEEHTV | 7.8182 | 7.9973 | 7.8781 | 7.4694 | 6.1946 | 6.3998 | 6.9102 | 6.8375 | 7.3839 | 7.7353 | 7.6823 | 6.9530 | 3.0088 | 3.5167 | 2.8087 | 3.4064 |
| USP34_S3406 | USP34 | Q70CQ2 | | 3350 | KDDEGATPIKRRR | 5.9573 | 6.2093 | 6.5129 | 6.4189 | 6.4102 | 6.0244 | 6.2932 | 6.1649 | 6.7749 | 6.3652 | 5.8741 | 6.3122 | 5.7251 | 6.2581 | 5.8454 | 6.8539 |
| USP34_S487 | USP34 | Q70CQ2 | | 3406 | TEQDLPSPENSSV | 5.3656 | 6.0190 | 6.0448 | 5.5108 | 6.1104 | 6.7966 | 5.2110 | 5.9324 | 5.9652 | 8.1807 | 7.2369 | 6.8153 | 6.4546 | 5.9347 | 7.4477 | 4.9745 |
| USP34_T3350 | USP34 | Q70CQ2 | | 487 | QLSKQSSFASLLN | 4.8204 | 4.9426 | 5.1258 | 5.3508 | 5.2358 | 6.5195 | 5.2867 | 5.2032 | 6.0248 | 7.0092 | 6.1510 | 5.6634 | 7.7188 | 7.6536 | 7.9173 | 9.3772 |
| USP35_S613 | USP35 | Q9P2H5 | | 613 | RRRRLGSVMRPTE | 7.1187 | 4.5719 | 6.4854 | 5.6064 | 5.9136 | 5.9099 | 5.9964 | 6.3500 | 7.1701 | 6.2108 | 5.8259 | 6.0628 | 7.8723 | 7.7306 | 5.9613 | 5.2139 |
| USP36_S429 | USP36 | Q9P275 | | 513 | PKLPSGSPSPKLS | 6.9567 | 6.9527 | 6.1925 | 7.0896 | 6.6227 | 6.5184 | 7.0288 | 6.4684 | 6.6158 | 6.2634 | 6.1625 | 6.1302 | 4.8473 | 5.6857 | 5.0852 | 5.3801 |
| USP36_S464 | USP36 | Q9P275 | | 582 | DVVLSTSPKLLAT | 6.6130 | 7.2021 | 6.6194 | 7.7563 | 6.2210 | 6.2774 | 7.1907 | 6.3435 | 6.6945 | 6.5862 | 6.5751 | 6.6840 | 4.2769 | 5.2325 | 4.6793 | 5.0482 |
| USP36_S513 | USP36 | Q9P275 | | 742 | HRARAVSPAPQSS | 6.0631 | 6.4880 | 5.6083 | 6.6653 | 6.2319 | 6.0769 | 6.2938 | 6.5261 | 5.9745 | 5.7570 | 5.8026 | 5.6621 | 6.1214 | 6.9692 | 7.0013 | 6.7584 |
| USP36_S546 | USP36 | Q9P275 | | 682 | EPASTMSPPPAKK | 6.6301 | 6.5788 | 6.4513 | 7.0760 | 6.4346 | 6.0363 | 6.4916 | 6.1101 | 6.9658 | 6.2039 | 6.4771 | 6.3369 | 5.1244 | 6.0939 | 5.5018 | 5.4874 |
| USP36_S582 | USP36 | Q9P275 | | 807 | ASEPPQSPSEKRK | 6.3881 | 6.4399 | 6.0398 | 6.4234 | 6.1074 | 6.3414 | 6.5050 | 6.1807 | 6.3051 | 5.7734 | 6.2186 | 6.3913 | 5.9930 | 6.7532 | 6.2196 | 5.9202 |
| USP36_S667;S682 | USP36 | Q9P275 | | 546 | APPQHFSPRTAQG | 6.3469 | 6.3170 | 6.5339 | 7.6067 | 6.3225 | 6.5587 | 6.7634 | 5.8844 | 6.7091 | 6.7103 | 6.6362 | 7.2811 | 4.6370 | 5.5369 | 4.6270 | 5.5289 |
| USP36_S682 | USP36 | Q9P275 | | 871 | TQRQPGSPMYRRE | 6.7427 | 6.7969 | 6.1820 | 7.1540 | 6.1862 | 5.9497 | 6.3312 | 6.2224 | 6.4251 | 5.8070 | 6.0878 | 6.1442 | 5.5529 | 6.2936 | 6.0378 | 6.0863 |
| USP36_S713 | USP36 | Q9P275 | | 952 | PEAMEESPRKKKK | 5.9566 | 5.8322 | 5.8047 | 6.6360 | 6.0969 | 5.6348 | 5.7911 | 5.7562 | 5.7775 | 6.1129 | 6.2096 | 6.6461 | 6.5519 | 6.9527 | 6.8550 | 7.3857 |
| USP36_S742 | USP36 | Q9P275 | | 464 | GNGIISSPLTGKR | 7.0785 | 7.4306 | 6.2164 | 7.0853 | 6.2611 | 6.2256 | 6.8030 | 6.3794 | 6.4757 | 6.5740 | 6.5529 | 6.8229 | 4.6221 | 5.6473 | 4.5924 | 5.2329 |
| USP36_S807 | USP36 | Q9P275 | | 429 | IPGSKKSPEGLIS | 6.2794 | 7.1160 | 6.9682 | 7.6837 | 5.6933 | 5.9699 | 6.7093 | 6.0715 | 6.2149 | 6.0752 | 6.4262 | 6.8513 | 4.6934 | 5.9424 | 5.0484 | 6.2570 |
| USP36_S871 | USP36 | Q9P275 | | 713 | LRPPPPSPSSDLT | 5.4002 | 5.6144 | 5.8728 | 6.5366 | 5.8563 | 5.6464 | 6.2319 | 6.1853 | 5.7780 | 5.1453 | 5.4885 | 5.9440 | 6.5208 | 7.9761 | 7.9570 | 7.8464 |
| USP36_S952 | USP36 | Q9P275 | | 653 | TAGHSKTPPSGAD | 6.1839 | 6.7192 | 6.5406 | 6.9327 | 5.4253 | 6.0428 | 6.2758 | 6.3152 | 6.4698 | 6.7185 | 7.0515 | 6.8658 | 4.8706 | 5.8414 | 5.4037 | 6.3430 |
| USP36_T653 | USP36 | Q9P275 | 667;682 | | NA | 6.8372 | 7.1239 | 6.6830 | 8.2127 | 6.3739 | 6.9093 | 7.2100 | 6.9624 | 6.8252 | 6.9462 | 7.3568 | 7.3949 | 3.2640 | 4.1989 | 3.5096 | 4.1921 |
| USP37_S312 | USP37 | Q86T82 | | 312 | PQPVPLSVKKLRC | 7.8370 | 6.1580 | 6.4136 | 6.8854 | 6.0813 | 5.8296 | 6.6692 | 6.7753 | 7.1351 | 6.0518 | 6.3723 | 7.8351 | 3.1399 | 5.6586 | 5.9594 | 5.1985 |
| USP37_T631 | USP37 | Q86T82 | | 631 | CITSPSTPSKKFT | 6.4631 | 6.4132 | 5.9072 | 5.8546 | 5.8444 | 6.2125 | 6.2780 | 5.2826 | 7.8727 | 6.7276 | 6.3037 | 6.2715 | 5.4109 | 7.0345 | 6.0113 | 6.1122 |
| USP39_S82 | USP39 | Q53GS9 | | 82 | REVDEDSEPEREV | 7.4646 | 5.8421 | 5.8723 | 5.8502 | 6.2417 | 5.6725 | 6.0921 | 5.6925 | 6.1845 | 5.3723 | 6.2447 | 6.6774 | 6.2618 | 7.1493 | 6.9084 | 6.4736 |
| USP4_S244 | USP4 | Q13107 | | 244 | SRNFTTSPKSSAS | 6.6334 | 6.6421 | 7.3944 | 6.4443 | 5.9747 | 6.7542 | 6.7453 | 6.4194 | 6.8122 | 6.9006 | 7.3200 | 7.4527 | 4.3238 | 5.0289 | 4.2206 | 4.9331 |
| USP40_S1211 | USP40 | Q9NVE5-3 | | 1211 | ALGRRKSQEALHE | 6.2532 | 6.1204 | 6.4936 | 6.4738 | 6.4817 | 6.4422 | 5.6345 | 6.5695 | 6.4590 | 6.8385 | 6.7526 | 6.5774 | 5.5800 | 6.2090 | 5.6116 | 5.5030 |
| USP42_S1133 | USP42 | Q9H9J4 | | 936 | PKAPGPSPAKEKI | 6.3277 | 6.3829 | 5.8107 | 6.6104 | 5.9822 | 5.6910 | 6.2079 | 6.0491 | 6.2673 | 5.7949 | 5.8574 | 6.0207 | 6.1558 | 7.0293 | 6.7464 | 7.0662 |
| USP42_S754 | USP42 | Q9H9J4 | | 856 | EAPEGLSPAPPAR | 5.6351 | 5.5556 | 6.3767 | 6.5845 | 5.8484 | 5.7819 | 6.2973 | 5.4587 | 6.6545 | 6.2782 | 6.2651 | 6.6252 | 6.6352 | 6.5212 | 6.6489 | 6.8333 |
| USP42_S856 | USP42 | Q9H9J4 | | 754 | AEPQPGSPAAESL | 6.3503 | 6.0963 | 5.5934 | 6.5390 | 6.6248 | 6.1147 | 6.2316 | 5.7154 | 6.5788 | 5.5174 | 5.4497 | 6.0037 | 6.0610 | 7.2814 | 6.9633 | 6.8791 |
| USP42_S936 | USP42 | Q9H9J4 | | 1133 | PHPDRFSHDRTAL | 6.7505 | 7.1015 | 5.4666 | 7.0021 | 6.9420 | 5.9683 | 5.5157 | 7.2262 | 6.2156 | 6.1726 | 5.5284 | 5.7845 | 5.1652 | 6.6776 | 6.3735 | 6.1096 |
| USP42_T783 | USP42 | Q9H9J4 | | 783 | PPRDPGTPATKEG | 6.7188 | 6.9638 | 5.6888 | 6.4790 | 5.9927 | 5.7162 | 6.1798 | 5.4428 | 5.9498 | 5.3208 | 6.0273 | 5.9813 | 5.9643 | 7.3209 | 6.9798 | 7.2739 |
| USP43_S1041 | USP43 | Q70EL4 | | 1041 | DGQAPGSPPALRI | 6.0316 | 6.2899 | 6.1207 | 6.4293 | 5.8854 | 6.4621 | 5.9583 | 6.4699 | 6.2087 | 6.6689 | 6.1256 | 6.1427 | 5.9764 | 6.7648 | 6.0419 | 6.4239 |
| USP45_S523 | USP45 | Q70EL2 | | 523 | TGLFRSSSGSGVQ | 5.8132 | 4.8605 | 5.3134 | 4.6414 | 6.6341 | 5.8655 | 5.7097 | 6.1887 | 5.6889 | 5.0440 | 5.8841 | 5.3959 | 8.9901 | 8.3511 | 8.1462 | 7.4733 |
| USP47_S832 | USP47 | Q96K76 | | 832 | VLLPEQSPVSYSK | 6.0431 | 6.2047 | 6.7275 | 6.4923 | 5.8534 | 6.4941 | 6.2984 | 6.8342 | 6.4652 | 6.3843 | 6.9106 | 6.8900 | 5.3519 | 5.9874 | 5.6522 | 5.4109 |
| USP5_T623 | USP5 | P45974 | | 623 | IAPPLVTPDEPKG | 5.7733 | 5.8697 | 6.4632 | 6.3699 | 6.0248 | 5.8023 | 6.1913 | 6.5325 | 6.1204 | 5.9525 | 6.5533 | 6.0904 | 6.1365 | 6.8534 | 6.4488 | 6.8176 |
| USP54_S1036 | USP54 | Q70EL1 | | 632 | IKFGGPSPQYKRW | 6.6705 | 7.1835 | 6.6419 | 6.7184 | 6.3832 | 6.8310 | 7.5413 | 6.6780 | 7.1313 | 6.9078 | 6.2339 | 6.3683 | 4.4598 | 5.1474 | 4.1447 | 4.9589 |
| USP54_S1056 | USP54 | Q70EL1 | | 1036 | KDPANPSPVMPGI | 6.0072 | 5.5232 | 5.7648 | 6.1756 | 5.9993 | 6.3027 | 5.5619 | 5.7448 | 6.2176 | 6.0776 | 5.8456 | 5.9265 | 7.4794 | 7.1538 | 7.2318 | 6.9880 |
| USP54_S456 | USP54 | Q70EL1 | | 456 | HTSKKGSLIERKR | 8.7941 | 6.3801 | 6.8136 | 5.9887 | 6.1991 | 6.1149 | 5.7298 | 5.3118 | 9.0598 | 8.2139 | 6.3563 | 7.9249 | 4.9234 | 5.8056 | 3.1508 | 3.2332 |
| USP54_S632 | USP54 | Q70EL1 | | 1056 | EHSLGCSPSNSSA | 7.6172 | 7.2140 | 6.6072 | 7.8763 | 5.9631 | 7.0377 | 6.8969 | 6.1692 | 7.3067 | 6.2777 | 8.1602 | 7.5129 | 3.8901 | 3.9539 | 3.1715 | 4.3455 |
| USP54_S671 | USP54 | Q70EL1 | | 671 | SERNSSSPVSLDA | 5.9633 | 7.7122 | 6.6199 | 6.4937 | 5.0036 | 6.1117 | 7.0335 | 6.0707 | 5.2909 | 6.1306 | 6.3737 | 6.2601 | 5.4803 | 5.3856 | 6.2506 | 7.8196 |
| USP6NL_S408 | USP6NL | Q92738-2 | | 422 | ASGRRESGAPHRR | 5.6658 | 5.3634 | 6.5521 | 4.8408 | 6.6497 | 6.3166 | 5.8362 | 7.2622 | 8.3877 | 8.7572 | 7.9417 | 7.6221 | 5.2486 | 5.6735 | 3.1660 | 4.7164 |
| USP6NL_S408;S413 | USP6NL | Q92738-2 | | 602 | HALYPPSPRKHAE | 6.2084 | 6.2106 | 6.1374 | 6.3899 | 6.1414 | 6.5649 | 6.8601 | 6.1542 | 6.1070 | 5.9686 | 5.6663 | 5.3283 | 6.3053 | 6.5464 | 6.9747 | 6.4366 |
| USP6NL_S413 | USP6NL | Q92738-2 | | 408 | LSNGQRSVGRPSP | 7.5232 | 6.8558 | 7.3992 | 6.5613 | 5.0764 | 6.4862 | 5.7429 | 6.9088 | 7.2042 | 8.8318 | 7.9890 | 6.9880 | 3.3247 | 5.7054 | 3.6363 | 3.7669 |
| USP6NL_S422 | USP6NL | Q92738-2 | | 413 | RSVGRPSPLASGR | 4.6748 | 5.3851 | 8.9972 | 6.2777 | 13.2104 | 7.0598 | 5.7167 | 5.1515 | 6.1050 | 5.3588 | 5.4901 | 5.4970 | 5.6296 | 5.5523 | 4.4786 | 5.4155 |
| USP6NL_S452 | USP6NL | Q92738-2 | | 452 | QPPRRKSVEEESK | 5.9346 | 5.5934 | 6.2860 | 5.9091 | 6.0706 | 5.7806 | 5.3915 | 5.6864 | 7.1108 | 7.4018 | 7.3583 | 8.0395 | 5.6891 | 6.1934 | 5.8978 | 5.6572 |
| USP6NL_S602 | USP6NL | Q92738-2 | | 733 | YSGNSGSPKNGKL | 6.6250 | 5.0451 | 5.5551 | 6.4750 | 6.4528 | 6.1372 | 6.5228 | 5.7795 | 6.9105 | 6.7572 | 6.1983 | 6.4117 | 6.1050 | 7.0894 | 6.2350 | 5.7006 |
| USP6NL_S612 | USP6NL | Q92738-2 | | 612 | HAEPSSSPSKVSN | 6.0102 | 6.3690 | 6.3690 | 6.1097 | 6.3262 | 5.9932 | 5.9954 | 6.3410 | 6.3449 | 7.4566 | 6.3372 | 5.9585 | 6.1773 | 5.7470 | 5.7492 | 6.7158 |
| USP6NL_S733 | USP6NL | Q92738-2 | | 801 | PAVSVDSPVRYKA | 6.0067 | 6.5683 | 7.0180 | 6.6637 | 6.4487 | 7.1771 | 6.2721 | 7.3482 | 6.6264 | 6.5848 | 6.2656 | 6.7339 | 5.1171 | 4.4995 | 5.2356 | 5.4341 |
| USP6NL_S801 | USP6NL | Q92738-2 | 408;413 | | NA | 6.7528 | 6.8319 | 6.8561 | 7.0258 | 5.9349 | 6.5870 | 7.0437 | 7.6484 | 6.5646 | 7.5781 | 7.0183 | 7.1911 | 2.8659 | 5.4843 | 3.9806 | 4.6362 |
| USP7_S1026 | USP7 | Q93009 | | 18 | AGEQQLSEPEDME | 5.7186 | 6.2553 | 6.2926 | 6.1966 | 5.9487 | 6.2516 | 5.8855 | 6.4654 | 5.9912 | 6.2289 | 6.3160 | 6.1274 | 6.3935 | 6.5802 | 6.9084 | 6.4400 |
| USP7_S18 | USP7 | Q93009 | | 1026 | VMKRIQSLLDIQE | 5.3530 | 6.9102 | 4.9417 | 5.3908 | 5.4230 | 5.1767 | 5.4887 | 6.4709 | 5.4677 | 5.6160 | 5.7531 | 5.5027 | 8.5430 | 7.1453 | 8.4520 | 8.3653 |
| USP7_S963 | USP7 | Q93009 | | 963 | ELLECLSPATSRT | 5.6846 | 6.4502 | 7.7686 | 6.0263 | 6.2076 | 6.4126 | 6.4299 | 5.4984 | 5.7431 | 6.0134 | 6.0872 | 5.5796 | 6.6661 | 6.6171 | 7.1515 | 6.6637 |
| USP8_S452 | USP8 | P40818 | | 577 | RGKRCPTPEIQKK | 6.2973 | 6.7334 | 6.9697 | 6.2577 | 6.1566 | 6.5925 | 6.3443 | 6.5796 | 6.5847 | 6.8574 | 6.8190 | 6.9933 | 4.9542 | 5.6551 | 4.9499 | 5.2555 |
| USP8_S718 | USP8 | P40818 | | 718 | KLKRSYSSPDITQ | 6.1897 | 6.1732 | 6.9303 | 6.4958 | 6.1491 | 6.3183 | 6.1980 | 6.5287 | 6.5645 | 6.8755 | 6.5143 | 6.5330 | 5.4670 | 5.8978 | 5.5392 | 5.6256 |
| USP8_T577 | USP8 | P40818 | | 452 | TKPVVFSPTLMLT | 6.6535 | 6.3959 | 6.3358 | 7.3489 | 7.3063 | 7.8602 | 7.4496 | 6.3320 | 8.2147 | 7.3004 | 9.1522 | 7.3256 | 4.0405 | 2.9694 | 3.0469 | 2.2682 |
| USP9X_S1600 | USP9X | Q93008-3 | | 2443 | YTYNNWSPPVQSN | 6.2384 | 6.4252 | 6.9397 | 6.7431 | 6.1621 | 6.3128 | 5.9835 | 6.8188 | 6.4073 | 6.8943 | 7.0700 | 6.9234 | 4.9263 | 5.6566 | 5.2752 | 5.2233 |
| USP9X_S2443 | USP9X | Q93008-3 | | 1600 | DVDDDMSGDEKQD | 6.3123 | 5.9574 | 7.0933 | 6.6299 | 6.2930 | 6.6518 | 6.6252 | 6.1146 | 6.3602 | 6.1538 | 6.3602 | 6.3416 | 5.7797 | 6.2570 | 5.2617 | 5.8083 |
| USPL1_S1028 | USPL1 | Q5W0Q7 | | 909 | LAALMSSPQSRTV | 6.6902 | 6.7651 | 6.5442 | 6.9418 | 6.4403 | 6.1238 | 6.3194 | 6.7171 | 6.7661 | 6.4309 | 6.8531 | 6.5794 | 5.2949 | 5.4414 | 4.8856 | 5.2067 |
| USPL1_S1055 | USPL1 | Q5W0Q7 | | 1055 | RTLNLESPMKTDI | 6.6482 | 6.8772 | 7.0211 | 7.5268 | 6.5663 | 6.4904 | 6.4071 | 6.2928 | 6.9737 | 6.4498 | 6.6412 | 6.8119 | 4.3004 | 5.2380 | 4.5534 | 5.2017 |
| USPL1_S909 | USPL1 | Q5W0Q7 | | 1028 | QDHNYCSPTKKNP | 5.8817 | 6.4032 | 6.4857 | 6.9591 | 6.0989 | 6.2179 | 6.1569 | 5.8862 | 6.6078 | 6.4388 | 6.8086 | 6.2704 | 5.6701 | 6.1172 | 5.8923 | 6.1053 |
| UTP11_S241 | UTP11 | Q9Y3A2 | | 241 | KKETVNSPAIYKF | 7.2886 | 7.3532 | 6.4494 | 7.9010 | 6.6458 | 6.5789 | 7.0576 | 6.9438 | 7.1353 | 6.8384 | 6.8805 | 7.2389 | 3.4914 | 4.3168 | 3.6833 | 4.1972 |
| UTP11_T90 | UTP11 | Q9Y3A2 | | 90 | ETKEEVTPEQLKL | 7.4524 | 8.1976 | 6.9158 | 8.7156 | 7.4710 | 5.8390 | 6.8152 | 6.8078 | 6.0476 | 5.6713 | 6.2004 | 6.2228 | 3.9647 | 4.5571 | 4.3634 | 4.7584 |
| UTP14A_S405 | UTP14A | Q9BVJ6 | | 569 | NLLTTQSPSVKSL | 6.6073 | 7.1623 | 6.5568 | 7.8448 | 6.4819 | 6.6170 | 7.0178 | 6.9162 | 6.9732 | 6.9310 | 7.0454 | 7.3700 | 3.8735 | 4.4977 | 3.6930 | 4.4121 |
| UTP14A_S445 | UTP14A | Q9BVJ6 | | 405 | LEAHGVSESEGEE | 6.0408 | 6.5908 | 6.8533 | 6.0030 | 6.2853 | 5.9049 | 6.0425 | 5.9305 | 6.7084 | 6.4666 | 5.8796 | 6.2966 | 5.9450 | 6.6854 | 5.8548 | 6.5125 |
| UTP14A_S453 | UTP14A | Q9BVJ6 | | 52 | LLEAISSLDGKNR | 6.5770 | 7.1166 | 6.8310 | 7.3265 | 6.2579 | 6.7145 | 7.6853 | 7.1766 | 6.9507 | 6.9805 | 7.2316 | 7.5456 | 3.4135 | 4.1108 | 3.4765 | 4.6054 |
| UTP14A_S52 | UTP14A | Q9BVJ6 | | 541 | GQQSERTPNNRPD | 7.0773 | 7.8062 | 7.1072 | 8.1394 | 6.5660 | 7.5235 | 7.6524 | 6.5700 | 7.0440 | 7.1100 | 7.5929 | 7.3522 | 2.7351 | 3.9718 | 2.4064 | 3.3456 |
| UTP14A_S569 | UTP14A | Q9BVJ6 | | 453 | ETKDSGSQEVLSE | 5.7807 | 5.8719 | 6.6325 | 6.7693 | 6.2601 | 5.8429 | 6.3627 | 5.7405 | 5.7143 | 6.3295 | 5.8276 | 6.0836 | 6.3417 | 6.7820 | 6.7514 | 6.9095 |
| UTP14A_T541 | UTP14A | Q9BVJ6 | | 445 | DAEPAGSQETKDS | 6.9369 | 6.7910 | 5.4816 | 6.7010 | 6.2626 | 6.0229 | 5.4877 | 5.7258 | 6.6995 | 5.9655 | 6.7078 | 6.7003 | 5.7167 | 5.9957 | 6.6950 | 6.1099 |
| UTP18_S121 | UTP18 | Q9Y5J1 | | 121 | VQEHEDSGDSEVE | 4.4187 | 4.6182 | 5.4008 | 5.2448 | 5.3366 | 5.1728 | 5.4535 | 6.1284 | 6.7694 | 7.6988 | 7.5916 | 8.0843 | 6.8085 | 6.2023 | 7.0824 | 7.9887 |
| UTP18_S121;S124 | UTP18 | Q9Y5J1 | 121;124 | | NA | 5.9687 | 5.6335 | 6.1836 | 6.7608 | 6.1472 | 6.0187 | 6.3281 | 5.6620 | 6.3237 | 5.7159 | 5.8889 | 6.1170 | 6.8931 | 6.5561 | 6.8307 | 6.9719 |
| UTP20_S2601 | UTP20 | O75691 | | 2064 | CLLLPPTPVRGGQ | 6.2800 | 6.4714 | 5.8088 | 7.1291 | 6.2153 | 5.6494 | 6.7128 | 6.1179 | 6.9381 | 6.0077 | 6.3345 | 5.9844 | 5.5993 | 6.4962 | 5.9453 | 6.3097 |
| UTP20_S39 | UTP20 | O75691 | | 843 | PRSRELSPLFLRF | 8.3704 | 8.1782 | 6.2972 | 7.3670 | 6.6734 | 5.0959 | 7.1881 | 7.8437 | 6.7588 | 6.8252 | 6.7695 | 7.4682 | 3.3851 | 3.9124 | 3.8150 | 4.0521 |
| UTP20_S788 | UTP20 | O75691 | | 2601 | ADEKAESDGEEKE | 5.7185 | 5.9469 | 6.1530 | 7.2530 | 6.1412 | 5.0464 | 6.4158 | 5.7150 | 6.3831 | 6.3491 | 6.4258 | 6.1116 | 6.2524 | 6.7322 | 6.2593 | 7.0966 |
| UTP20_S843 | UTP20 | O75691 | | 788 | KSVGDESWEQTQE | 5.1110 | 5.5425 | 5.7418 | 6.1545 | 6.5055 | 5.9057 | 6.0100 | 6.5930 | 5.8316 | 5.8047 | 6.2690 | 5.9761 | 6.7962 | 6.9397 | 6.9090 | 7.9096 |
| UTP20_T2064 | UTP20 | O75691 | | 39 | RIDRTASYEEEVE | 4.7306 | 4.7114 | 5.2420 | 5.5314 | 5.2693 | 5.2792 | 5.7889 | 5.6692 | 5.4582 | 4.8381 | 4.9793 | 4.6440 | 10.2632 | 8.4994 | 9.5094 | 9.5862 |
| UTP3_S37 | UTP3 | Q9NQZ2 | | 37 | DLGLPPSPGDTSY | 5.8235 | 6.0482 | 5.8550 | 6.7946 | 6.2770 | 6.2698 | 6.1060 | 6.8866 | 6.4392 | 6.1086 | 5.9666 | 6.3964 | 5.7460 | 6.5901 | 6.3184 | 6.3741 |
| UTRN_S10 | UTRN | P46939 | | 881 | VQRGFDSFLGRYQ | 6.3198 | 6.0662 | 5.9088 | 6.4921 | 7.0517 | 5.7713 | 6.3485 | 5.9370 | 5.2849 | 4.8322 | 5.1471 | 4.9958 | 7.6854 | 7.4774 | 7.5081 | 7.1736 |
| UTRN_S1405 | UTRN | P46939 | | 295 | PEEEHESPRAETP | 5.5117 | 5.9883 | 6.4704 | 6.7346 | 5.6701 | 5.4121 | 6.0761 | 6.3121 | 6.5520 | 6.3873 | 6.4775 | 5.8433 | 6.5190 | 6.8843 | 6.4085 | 6.7526 |
| UTRN_S1866 | UTRN | P46939 | | 833 | SESSRQSLPSLKD | 6.2501 | 6.2099 | 5.8560 | 6.6184 | 6.3446 | 5.7852 | 6.5759 | 6.2147 | 6.4933 | 5.6627 | 5.8709 | 6.1523 | 6.5138 | 6.1105 | 6.8531 | 6.4887 |
| UTRN_S213 | UTRN | P46939 | | 2223 | VQSHRTSEISIPA | 5.5252 | 5.7499 | 6.5042 | 6.0977 | 6.1822 | 6.1116 | 6.8191 | 5.9324 | 5.8100 | 5.5389 | 5.6789 | 5.8728 | 7.6519 | 7.2439 | 6.6081 | 6.6732 |
| UTRN_S2223 | UTRN | H0Y337 | | 213 | LETPSQSPQLFHD | 5.7892 | 6.0087 | 6.1166 | 6.8879 | 5.5384 | 6.7700 | 5.9071 | 7.2704 | 6.2909 | 5.9899 | 6.0225 | 6.8653 | 6.3198 | 7.2642 | 5.4143 | 5.5447 |
| UTRN_S295 | UTRN | P46939 | | 10 | YGEHEASPDNGQN | 6.3806 | 7.0055 | 5.2242 | 7.3179 | 6.9649 | 5.9119 | 6.2792 | 6.5875 | 7.0826 | 6.8959 | 6.1615 | 6.3563 | 4.7981 | 5.9262 | 6.1169 | 4.9909 |
| UTRN_S3217;S3221 | UTRN | P46939 | | 1866 | ASGIRSSLLPTDY | 6.7744 | 5.9640 | 5.3730 | 5.1964 | 5.9292 | 5.9774 | 7.1595 | 5.6805 | 6.9081 | 6.4828 | 5.3462 | 7.0418 | 8.0741 | 5.5521 | 6.5337 | 6.0068 |
| UTRN_S3221 | UTRN | P46939 | | 3221 | SSSTTGSVEDEHA | 5.4942 | 5.3243 | 4.9908 | 6.6126 | 6.3539 | 5.9811 | 5.8264 | 5.3332 | 5.0631 | 6.2613 | 6.1738 | 5.3966 | 9.1092 | 6.4667 | 8.5260 | 7.0868 |
| UTRN_S833 | UTRN | P46939 | | 1405 | RSQPLTSPESRTA | 5.7560 | 6.3576 | 7.0520 | 7.5685 | 5.9553 | 6.4648 | 6.6911 | 6.5858 | 6.8356 | 6.8306 | 6.4127 | 7.1008 | 5.1371 | 5.2496 | 4.7041 | 5.2985 |
| UTRN_S881 | UTRN | H0Y337 | | 209 | EGDNLETPSQSPQ | 5.6956 | 5.3647 | 7.0214 | 5.8840 | 5.0184 | 6.7409 | 5.7043 | 5.8095 | 7.7248 | 6.6488 | 5.7876 | 5.9125 | 7.4947 | 6.3070 | 7.1397 | 5.7460 |
| UTRN_T209 | UTRN | H0Y337 | 209;213 | | NA | 6.3382 | 6.5828 | 6.5722 | 6.9246 | 6.7763 | 6.3210 | 5.6199 | 6.0734 | 7.1097 | 6.3060 | 6.4341 | 6.6595 | 5.8538 | 6.1429 | 4.7894 | 5.4963 |
| UTRN_T209;S213 | UTRN | P46939 | 3217;3221 | | NA | 6.2669 | 6.1258 | 6.0650 | 6.0993 | 5.8004 | 5.7136 | 5.1865 | 6.3651 | 6.9094 | 5.6697 | 5.5592 | 5.7030 | 7.9566 | 6.9085 | 6.4185 | 7.2525 |
| UVRAG_S549 | UVRAG | Q9P2Y5 | | 518 | ERLQYKTPPPSYN | 6.3308 | 6.8341 | 6.9032 | 6.7514 | 6.2639 | 6.1548 | 6.6321 | 6.1271 | 6.7000 | 6.1165 | 6.6925 | 6.3229 | 5.4525 | 6.0163 | 5.4589 | 5.2431 |
| UVRAG_S689 | UVRAG | Q9P2Y5 | | 549 | KITSLSSSLDTSL | 6.7311 | 6.6292 | 7.3017 | 6.8606 | 5.6179 | 6.1998 | 5.8209 | 5.7907 | 6.5844 | 7.0239 | 6.8839 | 6.3463 | 5.3683 | 5.5921 | 5.4057 | 5.8435 |
| UVRAG_T518 | UVRAG | Q9P2Y5 | | 689 | LNENVSSFRRPRR | 6.3028 | 5.7545 | 7.3053 | 6.6489 | 6.4861 | 6.3486 | 6.2323 | 6.6607 | 6.6001 | 6.4332 | 6.4732 | 6.5355 | 4.9835 | 5.7821 | 5.3443 | 6.1088 |
| UVSSA_S478 | UVSSA | Q2YD98 | | 478 | PPPSSASPSRALP | 6.1319 | 5.7710 | 6.7951 | 6.4477 | 7.0383 | 7.4220 | 6.1903 | 6.0167 | 6.7699 | 6.5746 | 5.6494 | 6.4371 | 5.3517 | 5.8055 | 5.6231 | 5.9757 |
| VAC14_S509 | VAC14 | Q08AM6 | | 509 | AGLLNTSGTKGLE | 6.9198 | 6.7706 | 6.3851 | 6.8391 | 6.2891 | 7.1284 | 7.6712 | 7.3921 | 7.1621 | 6.6399 | 6.6931 | 6.8204 | 3.7486 | 4.8308 | 3.7178 | 4.9920 |
| VAC14_S517 | VAC14 | Q08AM6 | | 499 | SELQVPTPGRAGL | 6.6329 | 6.4823 | 5.6427 | 7.7773 | 6.2473 | 6.6121 | 6.9548 | 6.5900 | 7.4799 | 5.9915 | 6.7173 | 6.1274 | 4.2032 | 5.8079 | 4.6488 | 6.0845 |
| VAC14_T11 | VAC14 | Q08AM6 | | 11 | KDFAPLTPNIVRA | 7.1652 | 7.4109 | 6.4486 | 6.7535 | 6.5367 | 7.8786 | 5.9216 | 6.4299 | 7.0331 | 6.6027 | 6.7908 | 7.0788 | 3.4157 | 4.2002 | 4.9542 | 5.3794 |
| VAC14_T499 | VAC14 | Q08AM6 | | 517 | TKGLECSPSTPTM | 5.6268 | 6.1476 | 6.1547 | 6.5858 | 5.9501 | 6.2414 | 6.5038 | 6.3156 | 5.8204 | 6.5475 | 7.2160 | 6.7037 | 6.3780 | 6.5147 | 5.5705 | 5.7236 |
| VAMP4_S30 | VAMP4 | O75379 | | 30 | NLLEDDSDEEEDF | 6.5084 | 5.9623 | 7.5332 | 5.9903 | 6.0968 | 5.8496 | 6.0408 | 5.6572 | 6.5245 | 7.0058 | 7.0482 | 5.4611 | 5.8440 | 6.0289 | 5.9424 | 6.5065 |
| VANGL1_S42 | VANGL1 | Q8TAA9 | | 523 | RLQSETSVxxxxx | 5.7649 | 6.1518 | 5.8097 | 6.4490 | 6.2380 | 5.8772 | 5.7637 | 6.0984 | 6.3501 | 6.1016 | 5.4167 | 5.9204 | 6.8281 | 7.1910 | 7.0837 | 6.9557 |
| VANGL1_S45 | VANGL1 | Q8TAA9 | | 42 | NKDGRGSEKSVTI | 6.4046 | 7.0342 | 6.8804 | 6.6894 | 4.9005 | 5.8861 | 6.5170 | 5.6606 | 7.0144 | 7.5251 | 8.0752 | 8.0857 | 4.7742 | 5.2527 | 4.4449 | 4.8550 |
| VANGL1_S523 | VANGL1 | Q8TAA9 | | 45 | GRGSEKSVTIQPP | 6.6654 | 6.5392 | 6.3056 | 6.3866 | 4.8708 | 6.3667 | 6.2963 | 6.6572 | 7.1671 | 7.0063 | 7.5885 | 7.7405 | 4.9805 | 5.9782 | 4.7699 | 4.6813 |
| VAPA_S198 | VAPA | Q9P0L0-2 | | 198 | AVPLNASKQDGPM | 7.1580 | 6.8143 | 6.6248 | 6.9824 | 7.4646 | 6.6299 | 7.7752 | 7.1955 | 6.8320 | 6.7047 | 6.9192 | 6.5347 | 3.8573 | 4.4880 | 3.7647 | 4.2548 |
| VAPA_S209 | VAPA | Q9P0L0-2 | | 261 | DKPGSTSTASFRD | 4.4964 | 4.2783 | 4.1562 | 4.6309 | 4.7415 | 4.6996 | 4.7104 | 4.8190 | 4.3883 | 4.3466 | 4.3342 | 4.5544 | 11.7010 | 11.2223 | 11.9479 | 10.9730 |
| VAPA_S261 | VAPA | Q9P0L0-2 | | 209 | PMPKPHSVSLNDT | 4.7653 | 4.6421 | 4.8498 | 4.7951 | 5.1682 | 4.9843 | 4.9062 | 5.0777 | 4.7642 | 4.7564 | 4.6833 | 5.4494 | 11.2063 | 9.7007 | 9.9573 | 10.2935 |
| VAPA_T215 | VAPA | Q9P0L0-2 | | 215 | SVSLNDTETRKLM | 5.9504 | 4.6432 | 5.4647 | 5.8234 | 6.4016 | 5.8740 | 6.6428 | 5.5906 | 5.7322 | 5.3886 | 5.5293 | 5.3023 | 8.7941 | 8.0594 | 7.9469 | 6.8566 |
| VAPA_T262 | VAPA | Q9P0L0-2 | | 262 | KPGSTSTASFRDN | 6.4106 | 6.5630 | 5.7925 | 6.7583 | 7.2878 | 6.3055 | 6.4836 | 6.5619 | 6.2461 | 6.2816 | 6.3796 | 6.1599 | 6.1619 | 6.0488 | 5.3722 | 5.1867 |
| VAPB_S146 | VAPB | O95292 | | 150 | TASKTETPIVSKS | 7.5941 | 6.9868 | 7.2508 | 6.9223 | 6.6638 | 5.8844 | 7.3814 | 7.0947 | 6.7852 | 7.9751 | 7.6988 | 8.4233 | 3.0221 | 4.0485 | 2.6085 | 3.6602 |

| | | | | | | | | | | | | | | | | | | | | | |
|---|---|---|---|---|---|---|---|---|---|---|---|---|---|---|---|---|---|---|---|---|---|
| VAPB_S159 | VAPB | O95292 | | 146 | IISTTASKTETPI | 5.4686 | 5.1045 | 5.7141 | 5.6478 | 6.5949 | 5.1866 | 6.0088 | 6.5038 | 5.6135 | 6.2605 | 5.9681 | 6.6482 | 6.5809 | 7.6734 | 7.7988 | 7.2276 |
| VAPB_S204 | VAPB | O95292 | | 159 | VSKSLSSSLDDTE | 6.3700 | 6.0022 | 6.4492 | 6.1799 | 6.2317 | 5.4869 | 5.7011 | 6.8297 | 5.3591 | 7.0809 | 6.5703 | 7.4487 | 6.1844 | 6.4619 | 5.4684 | 6.1758 |
| VAPB_T150 | VAPB | O95292 | | 204 | MRKTVQSNSPISA | 6.9339 | 6.9601 | 6.1046 | 6.3968 | 6.0526 | 6.5996 | 6.8988 | 7.2027 | 6.7550 | 7.6681 | 7.3914 | 7.6686 | 3.8200 | 5.0941 | 3.5376 | 4.9162 |
| VASH1_S313 | VASH1 | Q7L8A9 | | 313 | KGTGPPSPTKDRK | 6.0278 | 5.6471 | 6.9378 | 5.9270 | 6.1030 | 6.8969 | 5.6771 | 6.1104 | 6.5067 | 6.7119 | 7.1474 | 6.5625 | 5.9427 | 6.2891 | 5.5155 | 5.9970 |
| VASP_S305 | VASP | P50552 | | 322 | TLPRMKSSSSVTT | 4.7852 | 4.9993 | 4.4913 | 4.2721 | 4.7852 | 5.0099 | 4.3972 | 4.9637 | 5.2368 | 4.9264 | 4.5914 | 5.7020 | 11.2849 | 9.3668 | 11.1267 | 10.0610 |
| VASP_S322 | VASP | P50552 | | 305 | VPAQSESVRRPWE | 6.4498 | 6.3007 | 7.8802 | 6.6924 | 6.0157 | 6.1955 | 6.0917 | 6.7453 | 7.9420 | 7.6041 | 7.6135 | 6.4060 | 4.4769 | 5.3343 | 3.2462 | 5.0059 |
| VASP_T316 | VASP | P50552 | | 316 | WEKNSTTLPRMKS | 8.7789 | 7.7191 | 7.6847 | 6.8775 | 6.6038 | 6.3410 | 6.2723 | 5.7219 | 7.0136 | 6.2345 | 6.5029 | 6.3929 | 5.0926 | 5.3625 | 3.5274 | 3.8744 |
| VAV2_S566 | VAV2 | P52735-3 | | 771 | SRASSRSPVFTPR | 6.5565 | 7.5538 | 7.3458 | 6.6363 | 7.1571 | 8.1424 | 7.2946 | 5.4247 | 5.6440 | 6.1609 | 6.7343 | 6.9071 | 4.8020 | 5.0062 | 4.2863 | 4.3482 |
| VAV2_S769;S771 | VAV2 | P52735-3 | | 566 | PPCKFTSPADLDA | 7.1900 | 6.4157 | 8.0468 | 7.3569 | 6.2244 | 5.7652 | 6.3170 | 6.3658 | 6.3752 | 7.4129 | 6.0735 | 6.8539 | 4.7125 | 5.6520 | 4.2350 | 5.0031 |
| VAV2_S771 | VAV2 | P52735-3 | 769;771 | | NA | 6.0983 | 6.0519 | 6.8721 | 6.4238 | 6.1088 | 6.7320 | 6.0099 | 6.9823 | 6.4111 | 6.7887 | 6.2712 | 6.6286 | 5.5555 | 5.9028 | 5.2949 | 5.8683 |
| VCL_S272 | VCL | P18206 | | 721 | EAIDTKSLLDASE | 6.1419 | 6.2570 | 6.9154 | 6.1127 | 6.0183 | 6.0722 | 6.0831 | 6.7103 | 6.3953 | 6.3196 | 6.4151 | 6.7287 | 6.1582 | 5.9417 | 5.7074 | 6.0229 |
| VCL_S275 | VCL | P18206 | | 272 | MKRALASIDSKLN | 4.6039 | 3.7127 | 4.8527 | 4.6389 | 4.7985 | 5.3856 | 4.9864 | 5.4345 | 4.8241 | 4.6274 | 4.5428 | 5.3398 | 10.8105 | 10.8083 | 10.7792 | 9.8545 |
| VCL_S288 | VCL | P18206 | | 346 | ARGQGSSPVAMQK | 6.2783 | 6.1659 | 7.0255 | 5.7509 | 6.1527 | 6.7408 | 5.8765 | 7.1415 | 6.2487 | 6.4548 | 6.5287 | 6.5300 | 5.9172 | 5.8563 | 5.6251 | 5.7071 |
| VCL_S346 | VCL | P18206 | | 434 | RDDILRSLGEISA | 6.2995 | 6.6270 | 6.7655 | 6.1918 | 6.3996 | 7.1451 | 6.1673 | 6.7400 | 6.1459 | 6.5051 | 6.2449 | 6.3331 | 6.0422 | 5.9340 | 5.2580 | 5.2010 |
| VCL_S434 | VCL | P18206 | | 604 | VFSDTTTPIKLLA | 5.9915 | 6.0826 | 7.1157 | 6.4417 | 5.7816 | 6.5536 | 5.6854 | 6.7663 | 6.7647 | 7.2188 | 6.5336 | 6.5400 | 5.6343 | 5.9241 | 5.5296 | 5.4367 |
| VCL_S579 | VCL | P18206 | | 442 | GEISALTSKLADL | 5.9459 | 7.0670 | 7.1846 | 6.1714 | 6.2816 | 5.8271 | 7.2814 | 6.9971 | 6.3870 | 5.1287 | 5.9998 | 5.7058 | 5.5477 | 6.3331 | 6.3686 | 5.7732 |
| VCL_S721 | VCL | P18206 | | 579 | ASQLQDSLKDLKA | 5.6717 | 6.0281 | 6.8456 | 6.7894 | 5.6815 | 6.0025 | 6.7247 | 5.7185 | 6.6137 | 6.6735 | 6.7732 | 6.3406 | 5.8113 | 6.4920 | 5.6892 | 6.1445 |
| VCL_S795 | VCL | P18206 | | 288 | GWLRDPSASPGDA | 6.0113 | 6.6405 | 5.6496 | 5.7564 | 5.9441 | 6.0074 | 6.9713 | 5.9922 | 6.4817 | 6.5331 | 6.5832 | 6.9733 | 6.3263 | 6.1787 | 6.1517 | 5.7992 |
| VCL_T442 | VCL | P18206 | | 275 | ALASIDSKLNQAK | 5.9785 | 5.8854 | 6.2556 | 6.2812 | 6.2997 | 6.5515 | 6.4112 | 7.0475 | 6.4420 | 6.2776 | 6.4170 | 6.8997 | 5.8776 | 6.1833 | 5.7700 | 5.4222 |
| VCL_T604 | VCL | P18206 | | 795 | ELSKTISPMVMDA | 6.0240 | 5.4484 | 8.2900 | 5.9262 | 7.0550 | 5.7055 | 6.9070 | 7.5971 | 5.8480 | 5.3115 | 4.2470 | 6.8092 | 6.1861 | 6.1805 | 6.2559 | 6.2084 |
| VCL_T754 | VCL | P18206 | | 822 | SFLDSGYRILGAV | 6.7173 | 6.8459 | 6.7956 | 6.3495 | 6.2800 | 6.5919 | 5.9025 | 6.9110 | 6.3067 | 6.2464 | 6.5201 | 6.7050 | 5.3735 | 5.5693 | 5.6237 | 5.2615 |
| VCL_Y822 | VCL | P18206 | | 754 | MLVAGATSIARRA | 4.1465 | 6.8276 | 6.7822 | 8.2590 | 7.6796 | 8.3726 | 7.2309 | 6.2993 | 8.1852 | 5.4473 | 3.9591 | 4.7373 | 5.2712 | 6.2028 | 6.5947 | 4.0045 |
| VCP_S197 | VCP | P55072 | | 702 | KLAIRESIESEIR | 6.2884 | 6.7194 | 6.6276 | 6.6192 | 6.4865 | 6.6441 | 6.1805 | 5.9772 | 5.3586 | 6.3931 | 6.7819 | 6.1832 | 6.5400 | 6.2555 | 5.3408 | 5.6041 |
| VCP_S352 | VCP | P55072 | | 509 | FLKFGMTPSKGVL | 6.8037 | 6.8435 | 6.6672 | 6.8658 | 6.7049 | 6.1209 | 6.3598 | 6.3168 | 6.3239 | 6.2233 | 6.0734 | 5.6856 | 5.8665 | 5.7376 | 5.9915 | 5.4157 |
| VCP_S702 | VCP | P55072 | | 775 | GSFRFPSGNQGGA | 5.6314 | 6.3022 | 6.6416 | 6.3515 | 5.9960 | 6.4080 | 6.0148 | 6.1984 | 6.6682 | 6.5306 | 6.6009 | 6.5420 | 5.8453 | 6.2618 | 6.0492 | 5.9580 |
| VCP_S775 | VCP | P55072 | | 197 | REDEEESLNEVGY | 5.6699 | 5.7844 | 6.4776 | 7.0847 | 6.1940 | 6.2800 | 6.3325 | 5.9000 | 6.4898 | 6.4665 | 6.5219 | 6.4470 | 6.6087 | 5.7150 | 5.7032 | 6.3247 |
| VCP_T509 | VCP | P55072 | | 352 | ATNRPNSIDPALR | 5.6703 | 6.8815 | 6.6440 | 6.0151 | 6.7160 | 6.4922 | 5.8888 | 6.6302 | 6.5478 | 6.5733 | 6.6950 | 6.8992 | 5.8162 | 5.9124 | 5.1153 | 5.5027 |
| VCPIP1_S1198 | VCPIP1 | Q96JH7 | | 1198 | KAQRGNSVEELEE | 5.9235 | 5.6250 | 6.5168 | 6.0546 | 6.0710 | 5.8008 | 5.9725 | 6.1174 | 6.6206 | 6.2366 | 5.8952 | 6.2869 | 6.6369 | 7.0860 | 6.6287 | 6.5274 |
| VCPIP1_S710 | VCPIP1 | Q96JH7 | | 768 | PTKAPYSPTTSKE | 6.9133 | 6.9606 | 6.6914 | 6.5273 | 6.8319 | 6.8161 | 6.5710 | 6.8571 | 6.7803 | 6.5034 | 6.6751 | 6.6982 | 4.7484 | 5.0932 | 4.6040 | 4.7287 |
| VCPIP1_S747 | VCPIP1 | Q96JH7 | | 747 | GQPRTVSPSTIRD | 6.3231 | 6.5195 | 6.4911 | 6.2888 | 6.6106 | 6.4841 | 6.4734 | 6.4207 | 6.5602 | 6.3032 | 6.2961 | 6.4443 | 5.5540 | 5.8674 | 5.7051 | 5.6584 |
| VCPIP1_S757 | VCPIP1 | Q96JH7 | | 695 | PTKIILTGQKTKT | 6.6053 | 6.6512 | 7.3488 | 7.1213 | 6.0453 | 6.5734 | 6.7141 | 6.5841 | 7.8787 | 7.0733 | 8.0149 | 7.9705 | 3.5602 | 4.4111 | 3.3945 | 4.0533 |
| VCPIP1_S757;T761 | VCPIP1 | Q96JH7 | | 757 | IRDGPSSAPATPT | 5.9163 | 5.9352 | 6.2592 | 6.2151 | 6.7844 | 5.9021 | 5.7396 | 6.1009 | 6.5062 | 6.1115 | 6.2016 | 6.3626 | 6.7487 | 6.6018 | 6.0716 | 6.5434 |
| VCPIP1_S757;T763 | VCPIP1 | Q96JH7 | | 710 | KEELNMSKTERTI | 7.0067 | 7.5131 | 6.9255 | 7.6726 | 5.3631 | 7.1001 | 8.4591 | 7.1798 | 7.7295 | 6.7684 | 7.7591 | 7.1267 | 3.1667 | 4.0876 | 2.7711 | 3.3709 |
| VCPIP1_S768 | VCPIP1 | Q96JH7 | | 761 | PSSAPATPTKAPY | 7.4139 | 6.5687 | 6.7163 | 6.1698 | 7.4396 | 6.7658 | 7.3306 | 7.0550 | 6.9764 | 6.4277 | 6.5912 | 6.0335 | 4.6156 | 4.7564 | 4.6615 | 4.4779 |
| VCPIP1_T695 | VCPIP1 | Q96JH7 | 757;761 | | NA | 7.1533 | 6.9660 | 6.4060 | 6.5633 | 6.8252 | 5.8892 | 6.8786 | 6.1283 | 6.1579 | 6.7792 | 7.2767 | 6.8486 | 5.5886 | 5.0335 | 4.8313 | 4.6746 |
| VCPIP1_T761 | VCPIP1 | Q96JH7 | 757;763 | | NA | 6.8473 | 6.7157 | 6.8819 | 6.2907 | 6.2120 | 7.2029 | 6.6569 | 6.2633 | 7.2020 | 6.6897 | 6.9073 | 7.1502 | 4.5135 | 5.5497 | 4.0660 | 4.8508 |
| VDAC1_S104 | VDAC1 | P21796 | | 104 | TFDSSFSPNTGKK | 6.1075 | 6.2231 | 5.9031 | 6.4916 | 6.1123 | 5.8556 | 6.2177 | 6.2843 | 6.4276 | 6.4889 | 6.0732 | 6.4129 | 6.0122 | 6.5348 | 6.4473 | 6.4080 |
| VDAC1_S57 | VDAC1 | P21796 | | 57 | TTKVTGSLETKYR | 6.5632 | 5.9778 | 6.0882 | 6.6610 | 6.3428 | 5.7044 | 6.1750 | 6.2239 | 6.4828 | 6.2649 | 6.1092 | 6.3791 | 6.2424 | 6.1881 | 6.2137 | 6.3834 |
| VDAC1_T107 | VDAC1 | P21796 | | 107 | SSFSPNTGKKNAK | 5.7620 | 6.0948 | 5.7709 | 6.9370 | 5.9118 | 5.9027 | 6.3337 | 6.4845 | 6.5447 | 6.1649 | 6.0185 | 6.3463 | 6.1723 | 6.1376 | 6.6850 | 6.7333 |
| VDAC2_S130 | VDAC2 | P45880-1 | | 130 | TFDTTFSPNTGKK | 6.2102 | 6.2456 | 6.0157 | 6.6126 | 6.3600 | 5.9571 | 6.1783 | 6.3166 | 6.4356 | 6.2498 | 6.0349 | 6.2445 | 6.0644 | 6.1399 | 6.5778 | 6.3569 |
| VDAC2_T83 | VDAC2 | P45880-1 | | 83 | TGKVTGTLETKYK | 6.5140 | 6.7188 | 5.9350 | 6.7583 | 6.8162 | 6.1076 | 5.5606 | 6.1696 | 6.3565 | 5.5641 | 6.0809 | 6.3901 | 6.1279 | 5.9175 | 6.5781 | 6.4047 |
| VEPH1_S430 | VEPH1 | Q14D04 | | 449 | RFNRSKSLAFHTM | 9.2587 | 7.2429 | 7.5316 | 6.9907 | 7.0407 | 5.6551 | 6.1460 | 5.2166 | 6.1930 | 5.9661 | 7.0554 | 6.4317 | 4.7052 | 6.0978 | 4.2431 | 4.2255 |
| VEPH1_S449 | VEPH1 | Q14D04 | | 397 | ETKLIVTENEDHE | 6.5371 | 6.0820 | 6.8358 | 6.7143 | 5.9189 | 5.3167 | 5.6687 | 6.7230 | 6.6773 | 6.5334 | 6.4337 | 7.4325 | 5.5493 | 5.6179 | 5.8178 | 6.1416 |
| VEPH1_T397 | VEPH1 | Q14D04 | | 430 | GSIRRYSLGQVSK | 6.1152 | 5.6368 | 6.2289 | 5.9591 | 6.3836 | 5.9289 | 6.0695 | 6.0815 | 6.8202 | 6.9961 | 6.7876 | 7.1551 | 5.9938 | 6.8182 | 5.1204 | 5.9050 |
| VGLL4_S58 | VGLL4 | Q14135-4 | | 58 | TGPPPISPSKRKF | 5.9259 | 4.9363 | 6.0220 | 3.9646 | 6.5165 | 5.1553 | 5.0389 | 6.7963 | 5.5609 | 7.1908 | 6.6793 | 6.8986 | 7.2280 | 8.0271 | 7.0300 | 7.0296 |
| VIM_S10 | VIM | P08670 | | 56 | RSLYASSPGGVYA | 7.5509 | 7.3960 | 6.4088 | 7.8213 | 6.6457 | 6.3744 | 6.7952 | 6.4042 | 6.9406 | 6.5740 | 7.1473 | 7.2731 | 4.1612 | 4.7454 | 3.4087 | 4.3530 |
| VIM_S144 | VIM | P08670 | | 73 | AVRLRSSVPGVRL | 6.0797 | 5.6279 | 5.1355 | 6.3295 | 6.7513 | 5.5573 | 6.0150 | 5.7032 | 6.7366 | 6.5835 | 6.3277 | 6.6153 | 6.6874 | 7.2478 | 5.8564 | 6.7458 |
| VIM_S205 | VIM | P08670 | | 412 | GEESRISLPLPNF | 6.4690 | 7.0184 | 6.5849 | 7.5588 | 6.7579 | 7.1063 | 7.4188 | 6.8644 | 5.9765 | 6.2796 | 6.5234 | 6.4459 | 4.6414 | 5.3183 | 4.3549 | 4.6814 |
| VIM_S214 | VIM | P08670 | | 10 | RSVSSSSYRRMFG | 7.9525 | 7.1484 | 6.0416 | 7.9662 | 6.9524 | 6.0411 | 6.8580 | 5.9283 | 6.6553 | 5.8560 | 6.4728 | 6.8218 | 4.9002 | 5.1730 | 4.0164 | 5.2160 |
| VIM_S22 | VIM | P08670 | | 430 | RETNLDSLPLVDT | 6.2785 | 6.4609 | 6.5520 | 7.4283 | 6.6386 | 6.6374 | 7.2241 | 6.6786 | 6.2647 | 6.2184 | 6.7725 | 7.0513 | 5.1107 | 5.3154 | 4.4832 | 4.8856 |
| VIM_S22;S26 | VIM | P08670 | | 51 | RPSTSRSLYASSP | 7.6033 | 6.8183 | 6.3471 | 7.7350 | 6.9174 | 6.1753 | 6.9002 | 5.9605 | 6.2061 | 5.9344 | 6.7494 | 6.6233 | 5.2151 | 5.4727 | 4.4069 | 4.9350 |
| VIM_S25 | VIM | P08670 | | 20 | MFGGPGTASRPSS | 10.3722 | 8.1441 | 6.6135 | 9.7263 | 7.1032 | 4.8308 | 6.9279 | 5.3864 | 6.2191 | 5.4279 | 6.0488 | 5.6166 | 4.6188 | 4.8126 | 3.8462 | 4.3055 |
| VIM_S26 | VIM | P08670 | | 22 | GGPGTASRPSSSR | 6.5885 | 6.4215 | 5.3672 | 6.8970 | 6.4280 | 5.5464 | 6.1602 | 5.9774 | 6.1199 | 5.6646 | 5.9044 | 5.8984 | 7.0147 | 6.4243 | 6.6063 | 6.9813 |
| VIM_S29 | VIM | P08670 | | 83 | VRLLQDSVDFSLA | 7.2520 | 6.6564 | 6.8673 | 8.2454 | 6.9600 | 5.4089 | 5.7689 | 5.3002 | 5.5965 | 5.8246 | 5.7920 | 5.5681 | 6.5603 | 6.1226 | 6.2740 | 5.8028 |
| VIM_S325 | VIM | P08670 | | 42 | RTYSLGSALRPST | 9.9513 | 7.7966 | 6.0316 | 9.1208 | 7.0573 | 5.8852 | 7.7053 | 5.7746 | 6.9717 | 6.5385 | 6.7398 | 6.4543 | 3.7112 | 4.4831 | 2.8286 | 3.8500 |
| VIM_S34 | VIM | P08670 | | 459 | QVINETSQHHDDL | 6.4902 | 6.9819 | 6.1807 | 7.5233 | 6.8500 | 6.4552 | 6.9774 | 6.5747 | 6.9106 | 6.7426 | 7.0368 | 6.6860 | 4.6620 | 5.2152 | 4.1506 | 4.5628 |
| VIM_S39 | VIM | P08670 | | 39 | TSTRTYSLGSALR | 6.7678 | 5.7496 | 5.6791 | 6.6918 | 6.7828 | 5.2483 | 5.7370 | 5.3390 | 5.9096 | 5.4358 | 5.5520 | 5.4726 | 7.9817 | 7.8502 | 6.8910 | 6.9116 |
| VIM_S39;S42 | VIM | P08670 | | 34 | RSYVTTSTRTYSL | 8.7529 | 7.2164 | 5.8374 | 7.8477 | 7.1292 | 5.5624 | 6.5488 | 5.6551 | 6.5506 | 5.3224 | 6.1180 | 5.9641 | 5.7858 | 5.6164 | 4.7708 | 5.3221 |
| VIM_S412 | VIM | P08670 | | 205 | AENTLQSFRQDVD | 6.1850 | 6.1115 | 6.2864 | 6.7477 | 5.9953 | 5.9184 | 6.1766 | 6.5253 | 6.1279 | 5.9701 | 6.3691 | 6.0776 | 6.3726 | 6.6252 | 6.3522 | 6.1591 |
| VIM_S419 | VIM | P08670 | | 420 | PLPNFSSLNLRET | 6.1679 | 6.2360 | 5.8806 | 6.7401 | 6.4612 | 6.0410 | 6.6814 | 6.3906 | 5.9828 | 5.5211 | 5.8003 | 5.6623 | 6.7274 | 6.9072 | 6.2901 | 6.5101 |
| VIM_S42 | VIM | P08670 | | 436 | SLPLVDTHSKRTL | 7.9324 | 6.6750 | 6.1704 | 8.5079 | 7.2118 | 4.7867 | 5.9259 | 4.7311 | 5.6019 | 4.8086 | 5.3011 | 4.8760 | 7.5824 | 7.0951 | 6.5415 | 6.2523 |
| VIM_S420 | VIM | P08670 | | 25 | GTASRPSSSRSYV | 7.0789 | 6.8988 | 5.9846 | 7.5095 | 6.2235 | 6.0957 | 7.4687 | 5.8749 | 6.5913 | 5.7846 | 6.5651 | 6.3169 | 5.1783 | 5.9734 | 4.9610 | 5.4947 |
| VIM_S430 | VIM | P08670 | | 47 | GSALRPSTSRSLY | 7.4850 | 6.6061 | 5.6046 | 7.5398 | 6.7105 | 5.2906 | 6.0648 | 5.4117 | 5.7739 | 5.3625 | 5.6181 | 5.2177 | 7.3995 | 7.1220 | 6.3251 | 6.4682 |
| VIM_S459 | VIM | P08670 | | 5 | xxMSTRSVSSSSY | 9.2885 | 7.1327 | 6.2674 | 8.8684 | 7.3339 | 5.1194 | 6.3883 | 4.6577 | 6.1476 | 4.4850 | 5.5687 | 5.2790 | 6.7562 | 5.8609 | 5.1605 | 5.6858 |
| VIM_S47 | VIM | P08670 | | 214 | QDVDNASLARLDL | 6.7681 | 6.7338 | 5.9627 | 6.6005 | 6.3319 | 5.7211 | 6.0306 | 6.4147 | 6.1667 | 5.8931 | 5.9802 | 5.7375 | 6.9367 | 6.0870 | 6.9763 | 5.6591 |
| VIM_S49 | VIM | P08670 | | 325 | YRRQVQSLTCEVD | 5.7452 | 5.8962 | 6.5846 | 7.2746 | 6.1075 | 5.9425 | 6.5437 | 5.9360 | 6.1879 | 6.0672 | 7.5515 | 6.2572 | 6.3272 | 6.4961 | 6.4024 | 6.4802 |
| VIM_S5 | VIM | P08670 | | 63 | PGGVYATRSSAVR | 5.8945 | 6.2485 | 5.5851 | 6.1265 | 6.4013 | 6.0650 | 6.1811 | 6.4815 | 6.6172 | 7.1750 | 6.9447 | 7.0245 | 5.9646 | 5.9317 | 5.3295 | 6.0292 |
| VIM_S5;S8 | VIM | P08670 | | 419 | LPLPNFSSLNLRE | 3.4713 | 3.8208 | 3.2766 | 3.7079 | 3.9069 | 2.9302 | 3.7323 | 2.9057 | 3.1928 | 2.5023 | 2.3766 | 2.5826 | 16.3227 | 14.8929 | 15.5853 | 14.7932 |
| VIM_S51 | VIM | P08670 | | 26 | TASRPSSSRSYVT | 7.2786 | 6.9200 | 5.6544 | 7.4770 | 6.7582 | 5.8777 | 7.2498 | 6.1206 | 6.7261 | 5.7258 | 6.3253 | 6.0672 | 5.3994 | 5.7832 | 4.9811 | 5.6556 |
| VIM_S51;S56 | VIM | P08670 | | 144 | LKGQGKSRLGDLY | 6.6108 | 6.5364 | 6.1700 | 6.9732 | 6.3399 | 5.8977 | 6.4882 | 5.9457 | 5.7384 | 5.9813 | 6.0199 | 6.2812 | 6.5790 | 6.0624 | 6.0672 | 6.3087 |
| VIM_S56 | VIM | P08670 | | 29 | RPSSSRSYVTTST | 7.1345 | 6.8544 | 6.2688 | 6.8280 | 6.1797 | 5.5106 | 6.3684 | 5.0898 | 6.0851 | 5.6687 | 5.6582 | 5.8672 | 7.0823 | 7.2257 | 5.5631 | 6.6156 |
| VIM_S73 | VIM | P08670 | | 49 | ALRPSTSRSLYAS | 3.1446 | 3.1075 | 2.9183 | 3.9133 | 4.1106 | 3.2677 | 3.7379 | 3.4793 | 3.4222 | 3.2720 | 2.4525 | 2.2029 | 16.2502 | 15.4862 | 14.1195 | 15.1154 |
| VIM_S83 | VIM | P08670 | 22;26 | | NA | 8.5085 | 7.1491 | 5.9795 | 8.1005 | 6.7831 | 6.0564 | 6.7705 | 5.5500 | 6.6136 | 5.6798 | 6.2145 | 6.2217 | 5.6591 | 5.4160 | 4.1921 | 5.1055 |
| VIM_T20 | VIM | P08670 | 51;56 | | NA | 7.9780 | 7.0804 | 6.6092 | 8.1322 | 6.9280 | 6.1331 | 7.1336 | 5.9362 | 6.6249 | 6.5430 | 8.3372 | 7.4505 | 3.6132 | 4.4859 | 3.0683 | 3.9463 |
| VIM_T436 | VIM | P08670 | 5;8 | | NA | 7.0437 | 6.1125 | 5.6348 | 6.7325 | 6.8891 | 5.3966 | 6.2933 | 5.4353 | 5.8574 | 5.2846 | 4.8390 | 5.5064 | 7.5387 | 7.4424 | 6.5242 | 7.4694 |
| VIM_T63 | VIM | P08670 | 39;42 | | NA | 11.0486 | 7.9973 | 6.3538 | 8.6361 | 8.3251 | 5.3352 | 6.6764 | 5.3026 | 7.7852 | 4.8624 | 6.0202 | 5.4120 | 4.2700 | 5.0762 | 3.3104 | 3.5884 |
| VIPAS39_S121 | VIPAS39 | Q9H9C1 | | 121 | GRTRPGSFQSLSD | 4.0561 | 3.8512 | 4.5787 | 3.9052 | 5.4010 | 5.3689 | 5.0558 | 5.7227 | 4.7262 | 4.9271 | 4.7366 | 5.0445 | 10.7646 | 10.5262 | 10.7582 | 10.5769 |
| VIPAS39_S93 | VIPAS39 | Q9H9C1 | | 93 | QLKSRNSFSSYAQ | 5.4408 | 4.7465 | 5.8610 | 4.3866 | 5.3463 | 5.9850 | 5.7796 | 5.2962 | 7.0459 | 6.6384 | 6.1204 | 6.0052 | 8.3682 | 7.9153 | 7.2866 | 7.7779 |
| VIRMA_S138 | VIRMA | Q69YN4 | | 184 | QPRGPRTPPGPPP | 6.2133 | 6.5060 | 6.1011 | 6.6020 | 6.0105 | 5.9220 | 6.2850 | 5.8829 | 6.0598 | 6.2336 | 5.8882 | 6.3571 | 6.3272 | 6.3676 | 6.5573 | 6.6865 |
| VIRMA_S1579 | VIRMA | Q69YN4 | | 1579 | FLSEPSSPGRTKT | 6.5940 | 6.6464 | 5.8377 | 6.4371 | 6.4272 | 5.7142 | 6.2119 | 6.2680 | 6.2232 | 6.0667 | 5.9581 | 5.7374 | 6.5356 | 6.3449 | 6.6694 | 6.3283 |
| VIRMA_S173 | VIRMA | Q69YN4 | | 173 | EDQFNGSPPRPQP | 6.1478 | 6.2641 | 6.1972 | 6.7154 | 5.9731 | 5.6678 | 6.1879 | 6.2622 | 6.4221 | 6.2668 | 6.1535 | 6.0775 | 6.2414 | 6.4435 | 6.5211 | 6.4585 |
| VIRMA_S222 | VIRMA | Q69YN4 | | 1708 | SQNRFFTPPASKG | 7.0934 | 7.5201 | 7.0293 | 7.1687 | 6.4045 | 6.5224 | 7.3464 | 6.5755 | 7.6846 | 7.6767 | 7.2526 | 7.0571 | 3.3215 | 4.2585 | 3.1521 | 3.9365 |
| VIRMA_S63 | VIRMA | Q69YN4 | | 138 | ISHDRDSPPPPPP | 6.2269 | 6.5695 | 6.0760 | 6.6704 | 6.1562 | 5.9449 | 6.0616 | 5.7495 | 6.3208 | 6.0229 | 5.9783 | 6.3402 | 6.6222 | 6.4057 | 6.3679 | 6.4871 |
| VIRMA_T1708 | VIRMA | Q69YN4 | | 63 | RAYGETSPHTFQL | 5.4079 | 5.8364 | 5.9509 | 5.8049 | 5.8815 | 6.2881 | 6.0369 | 5.8483 | 5.8959 | 6.7360 | 5.7470 | 5.7958 | 6.6613 | 7.3006 | 7.1993 | 7.6091 |
| VIRMA_T184 | VIRMA | Q69YN4 | | 222 | DYFEPISPDRNSV | 6.2815 | 7.0442 | 6.1510 | 7.3306 | 6.5698 | 5.5918 | 5.9092 | 6.6347 | 6.6341 | 6.0273 | 5.7137 | 6.8209 | 5.2318 | 6.6453 | 5.3469 | 6.0669 |
| VPS13A_S1416 | VPS13A | Q96RL7 | | 1416 | PGPKQASFTDVRD | 5.4364 | 5.5760 | 5.6706 | 5.9794 | 5.5733 | 6.0684 | 5.8335 | 5.8663 | 5.8137 | 6.0296 | 6.1153 | 5.7818 | 7.3165 | 7.8941 | 7.4599 | 7.5851 |
| VPS13B_S1033 | VPS13B | Q7Z7G8 | | 1419 | LLVRPISKQDPFS | 6.2783 | 6.1207 | 6.4428 | 6.6141 | 6.0630 | 4.9635 | 6.2605 | 6.9659 | 6.0749 | 4.9940 | 6.3792 | 6.1580 | 6.0334 | 7.2007 | 6.8158 | 6.6352 |
| VPS13B_S1263 | VPS13B | Q7Z7G8 | | 1263 | AGPVPTSPVRSSI | 7.0273 | 6.0976 | 6.5634 | 6.7174 | 5.7232 | 7.1205 | 6.4326 | 7.7743 | 5.8791 | 6.3592 | 6.9180 | 5.4689 | 5.2665 | 6.1639 | 4.8831 | 5.6050 |
| VPS13B_S1419 | VPS13B | Q7Z7G8 | | 1033 | TESRPLSVPVKAM | 6.3433 | 6.9758 | 6.3497 | 6.3220 | 5.5288 | 6.3160 | 6.0861 | 5.8570 | 6.5002 | 7.2194 | 6.8540 | 7.6720 | 5.2672 | 5.6429 | 5.4006 | 5.6651 |
| VPS13B_S3052 | VPS13B | Q7Z7G8 | | 3052 | LVHNLTSPKWKDG | 6.0220 | 6.2752 | 6.9287 | 7.2390 | 5.9426 | 5.9993 | 6.4718 | 6.4905 | 5.6642 | 6.5021 | 6.2328 | 6.3601 | 5.2886 | 6.1574 | 6.3975 | 6.0281 |
| VPS13B_Y1011 | VPS13B | Q7Z7G8 | | 1011 | LAKQQSYQASEYA | 6.5296 | 6.8138 | 5.3032 | 6.7961 | 7.0344 | 7.3302 | 5.1327 | 8.4020 | 6.3688 | 5.1831 | 6.4151 | 5.4963 | 4.8064 | 6.6344 | 6.3311 | 5.4229 |
| VPS13C_S3641 | VPS13C | Q709C8 | | 3641 | HCAIPGSKKTILM | 5.9880 | 6.1219 | 6.8163 | 5.8769 | 6.8681 | 6.0629 | 6.4898 | 6.1939 | 6.5469 | 6.1228 | 6.5573 | 6.0328 | 6.1492 | 6.1667 | 5.9933 | 6.0132 |
| VPS13C_S737 | VPS13C | Q709C8 | | 737 | QKTTNSSLEEIMD | 4.0474 | 4.2689 | 4.1626 | 4.5639 | 5.3701 | 5.3309 | 5.2520 | 6.0175 | 4.7825 | 5.1060 | 4.9202 | 4.3277 | 10.8852 | 10.2011 | 11.2087 | 9.5552 |
| VPS13D_S1038 | VPS13D | Q5THJ4 | | 1724 | YPDMPRSLPSHME | 5.6832 | 6.2926 | 6.5287 | 6.4483 | 5.9362 | 5.7644 | 6.0419 | 6.0861 | 6.6188 | 6.4848 | 6.4530 | 6.3850 | 6.4979 | 6.3276 | 6.3013 | 6.1501 |
| VPS13D_S1138 | VPS13D | Q5THJ4 | | 1138 | KEKDDLSPQPLMT | 5.6819 | 6.3482 | 5.8268 | 4.8664 | 6.3415 | 4.8731 | 6.2256 | 6.6601 | 7.4088 | 6.5777 | 5.9560 | 7.5626 | 5.4435 | 5.9293 | 6.9543 | 7.3442 |
| VPS13D_S1573 | VPS13D | Q5THJ4 | | 2079 | ESVPSASPTGIPK | 7.0266 | 7.1283 | 7.0723 | 6.6521 | 7.1180 | 6.9229 | 7.2086 | 7.1318 | 5.7060 | 5.6104 | 5.6645 | 5.2916 | 5.6847 | 5.4406 | 5.2965 | 5.0451 |
| VPS13D_S1709 | VPS13D | Q5THJ4 | | 1709 | KEYLSQSCPSVSN | 5.9588 | 5.7525 | 6.7625 | 6.9071 | 6.5589 | 6.4340 | 6.0156 | 5.6616 | 6.4355 | 6.6355 | 6.5620 | 6.2492 | 5.8963 | 6.7331 | 5.7351 | 5.7021 |
| VPS13D_S1724 | VPS13D | Q5THJ4 | | 2692 | ASTSRDSPGAVAA | 5.1233 | 5.9170 | 6.1342 | 6.3819 | 5.9997 | 5.8943 | 6.8109 | 7.0116 | 6.8496 | 6.7775 | 6.1487 | 7.0024 | 5.7187 | 6.4684 | 5.5562 | 6.2053 |
| VPS13D_S2079 | VPS13D | Q5THJ4 | | 791 | SSNGEKTPPFSGV | 5.5696 | 6.0314 | 6.8082 | 7.1780 | 6.1184 | 5.0981 | 6.0777 | 5.1921 | 6.9106 | 6.1886 | 6.6384 | 6.5008 | 6.3429 | 6.3900 | 6.2931 | 6.6622 |
| VPS13D_S2692 | VPS13D | Q5THJ4 | | 1038 | TGSLRDSRAQSPV | 6.0281 | 6.1511 | 5.9979 | 5.9693 | 5.6601 | 6.2773 | 5.8470 | 5.7506 | 6.4797 | 6.2777 | 6.5919 | 6.7416 | 6.8403 | 6.6878 | 6.3957 | 6.3040 |
| VPS13D_T791 | VPS13D | Q5THJ4 | | 1573 | SSPCPDSPLPPLS | 5.9433 | 6.5486 | 6.7812 | 6.8981 | 6.2563 | 5.9913 | 5.9368 | 5.9705 | 7.4866 | 6.1238 | 5.6601 | 5.8523 | 6.6915 | 6.1992 | 5.5535 | 6.1069 |
| VPS26A_S315 | VPS26A | O75436 | | 315 | FHQRFESPESQAS | 6.0362 | 6.2920 | 6.5248 | 7.0493 | 6.7819 | 6.5909 | 6.4249 | 6.5335 | 6.5673 | 6.1289 | 6.4467 | 7.0296 | 5.0327 | 5.3581 | 5.1798 | 6.0235 |
| VPS26B_S304 | VPS26B | Q4G0F5 | | 304 | IVRKSMSHQAAIA | 2.7484 | 2.6096 | 3.0091 | 3.2762 | 3.0383 | 3.1021 | 3.1676 | 3.1297 | 3.2362 | 2.7913 | 3.1915 | 3.1044 | 15.8010 | 14.8635 | 17.8996 | 15.0314 |
| VPS41_S249 | VPS41 | P49754 | | 847 | KNRGPGSAILEMK | 6.6719 | 7.3180 | 6.7536 | 7.7384 | 4.9202 | 5.8739 | 5.1265 | 7.4192 | 6.8859 | 6.3410 | 6.6174 | 6.9755 | 4.9202 | 4.8501 | 5.5236 | 6.0646 |
| VPS41_S847 | VPS41 | P49754 | | 249 | VKERHASEMRDLP | 6.5189 | 6.5725 | 6.9410 | 6.3012 | 5.1933 | 7.2046 | 6.7371 | 5.3963 | 6.7687 | 6.3090 | 6.6119 | 6.9108 | 5.3560 | 5.6925 | 6.0514 | 5.4349 |
| VPS4B_S93 | VPS4B | O75351 | | 93 | VKEGQPSPADEKG | 6.3774 | 5.6971 | 6.8366 | 5.7864 | 5.9102 | 6.5973 | 6.3150 | 6.8558 | 6.6748 | 6.3063 | 5.9434 | 6.4056 | 6.0237 | 6.5704 | 5.8817 | 5.8182 |
| VPS50_S15 | VPS50 | Q96JG6 | | 546 | ASNGYESDEQEKS | 5.8891 | 5.6580 | 6.8782 | 5.9591 | 6.7322 | 6.8956 | 6.1413 | 6.4996 | 6.5748 | 6.5117 | 7.2400 | 6.3102 | 6.2304 | 6.2720 | 5.3415 | 4.8664 |
| VPS50_S494 | VPS50 | Q96JG6 | | 494 | FMEQSRSPSVSPS | 6.4190 | 6.4786 | 6.6821 | 5.9601 | 6.1947 | 6.0447 | 5.5356 | 6.5255 | 6.6731 | 6.4313 | 6.3118 | 6.1041 | 6.3180 | 6.2148 | 6.2842 | 5.8224 |
| VPS50_S494;S500 | VPS50 | Q96JG6 | | 15 | TRQGLKSPQESLS | 5.2150 | 5.5685 | 6.2803 | 5.7193 | 5.9623 | 5.8397 | 5.7457 | 6.3415 | 5.7617 | 6.0866 | 5.8370 | 6.0121 | 7.0765 | 7.8197 | 7.6135 | 7.1207 |
| VPS50_S546 | VPS50 | Q96JG6 | | 595 | LKSRKKSDYSLNK | 6.0019 | 4.5132 | 6.2755 | 5.5104 | 7.0869 | 5.9215 | 5.9789 | 5.4644 | 7.3609 | 7.2436 | 6.8884 | 6.5560 | 6.6403 | 7.5741 | 5.7768 | 5.2072 |
| VPS50_S595 | VPS50 | Q96JG6 | 494;500 | | NA | 6.1596 | 6.7789 | 6.5699 | 6.3000 | 6.5217 | 6.4935 | 6.7651 | 7.1638 | 6.7859 | 6.6292 | 6.4697 | 6.4166 | 4.8533 | 5.5653 | 5.2144 | 5.3132 |
| VPS51_S652 | VPS51 | Q9UID3 | | 652 | RTFSVYSSSRQQG | 6.5303 | 6.5196 | 6.9702 | 6.5014 | 5.5574 | 5.9144 | 6.1423 | 6.3524 | 5.7435 | 5.2067 | 5.8140 | 5.9120 | 6.5324 | 7.2781 | 6.2930 | 6.7321 |
| VPS53_S377 | VPS53 | Q5VIR6 | | 377 | TLKKLESPPPSTN | 6.0171 | 6.2500 | 6.4979 | 6.6671 | 6.0380 | 6.7360 | 6.3842 | 6.7024 | 6.5105 | 6.5448 | 6.5752 | 6.6180 | 5.3541 | 5.7970 | 5.7055 | 5.6023 |
| VPS72_T103 | VPS72 | Q15906-2 | | 103 | RPRKVNTPAGSSQ | 7.4026 | 7.5147 | 7.3345 | 7.5323 | 5.4918 | 6.6847 | 7.3624 | 6.0702 | 7.7140 | 7.4444 | 7.5456 | 7.7987 | 3.3190 | 4.1379 | 2.6871 | 3.9600 |
| VPS8_S26 | VPS8 | Q8N3P4 | | 26 | EEELNKSFNLEAS | 6.8430 | 7.1528 | 6.7903 | 6.9658 | 7.2420 | 7.4384 | 7.4724 | 6.2967 | 7.3000 | 7.0972 | 8.3570 | 7.6514 | 3.1292 | 3.9508 | 2.9368 | 3.3761 |
| VPS9D1_S147 | VPS9D1 | Q9Y2B5-2 | | 147 | RHRRVYSDEGGKL | 6.1033 | 5.6353 | 6.2903 | 5.9762 | 5.8850 | 5.3839 | 5.7310 | 6.2024 | 5.7920 | 5.5211 | 5.8464 | 6.7478 | 7.7197 | 7.5966 | 7.1223 | 6.4466 |
| VRK1_S342 | VRK1 | Q99986 | | 342 | DGKLDLSVVENGG | 6.4775 | 6.8821 | 6.7222 | 7.5388 | 6.5135 | 6.0446 | 6.7601 | 6.7803 | 6.2546 | 5.7549 | 6.1728 | 5.6891 | 5.1807 | 5.9749 | 5.8976 | 5.3563 |
| VRK1_S376 | VRK1 | Q99986 | | 376 | VEDTEWSNTQTEE | 5.1635 | 5.8466 | 6.1627 | 6.3454 | 5.7876 | 5.5849 | 5.1300 | 6.4811 | 5.5809 | 6.1307 | 6.1244 | 6.0717 | 6.3143 | 8.4659 | 7.4412 | 7.3693 |
| VRK3_S136 | VRK3 | Q8IV63 | | 136 | PQKTRQSPQTLKR | 5.8654 | 6.3577 | 6.8745 | 6.6937 | 6.0331 | 6.6000 | 6.0652 | 5.9302 | 6.4525 | 7.6086 | 8.0672 | 7.7660 | 4.4200 | 5.7483 | 4.5425 | 4.9749 |
| VRK3_S59 | VRK3 | Q8IV63 | | 71 | WSSTVTSPRLSLF | 6.7615 | 7.0194 | 7.1086 | 6.7379 | 6.0872 | 6.7784 | 6.4249 | 7.0277 | 7.0106 | 7.1870 | 7.4764 | 7.5516 | 3.8495 | 4.3528 | 3.7639 | 4.8626 |
| VRK3_S71 | VRK3 | Q8IV63 | | 59 | NSSFETSPKKVKW | 5.6401 | 5.9178 | 6.5723 | 6.6759 | 5.8410 | 6.5471 | 5.4392 | 5.8121 | 6.5406 | 6.3415 | 6.7866 | 7.6424 | 6.0202 | 5.9203 | 5.9600 | 6.3430 |
| VSIR_S305 | VSIR | Q9H7M9 | | 305 | LDPVPDSPNFEVI | 6.1398 | 5.5454 | 6.8089 | 6.0844 | 6.0481 | 7.0211 | 5.6237 | 6.5183 | 5.9716 | 6.3654 | 6.8681 | 5.9774 | 6.3081 | 5.3408 | 7.5047 | 5.8741 |
| VTI1B_T103 | VTI1B | Q9UEU0 | | 103 | STPLTATPGGRGD | 7.4765 | 8.0345 | 7.0593 | 8.2535 | 6.2170 | 6.6548 | 7.4925 | 6.7395 | 7.7835 | 7.2120 | 7.6970 | 7.1820 | 2.5377 | 3.4630 | 2.7137 | 3.4836 |
| VTI1B_T184 | VTI1B | Q9UEU0 | | 184 | KSRLVNTSENLSK | 6.2420 | 6.3305 | 6.0468 | 6.5924 | 5.6122 | 4.8364 | 6.1908 | 5.5772 | 5.8304 | 6.2557 | 7.2253 | 5.3397 | 6.8924 | 7.2746 | 7.4642 | 6.2894 |
| WAC_S456 | WAC | Q9BTA9 | | 511 | QGHEPVSPRSLQR | 6.0949 | 6.4597 | 5.9017 | 6.4761 | 6.1560 | 6.0822 | 6.0641 | 6.3558 | 6.1335 | 6.3043 | 6.0284 | 6.5027 | 6.6846 | 6.1762 | 6.2732 | 6.3065 |
| WAC_S511 | WAC | Q9BTA9 | | 53 | RDAGDPSPPNKML | 6.2465 | 6.3538 | 6.5336 | 6.2417 | 5.9251 | 6.1628 | 5.9570 | 6.0331 | 6.3722 | 6.3469 | 6.4710 | 6.5341 | 6.5372 | 5.7004 | 6.0901 | 6.4943 |
| WAC_S53 | WAC | Q9BTA9 | | 64 | MLRRSDSPENKYS | 6.4230 | 6.2627 | 5.8885 | 5.8433 | 6.7193 | 6.0986 | 6.0889 | 5.9702 | 6.2518 | 5.8325 | 6.3462 | 6.0028 | 7.2588 | 6.3021 | 6.5357 | 6.1757 |
| WAC_S64 | WAC | Q9BTA9 | | 482 | SQPKVSTPVVKQG | 7.1811 | 7.4093 | 6.6637 | 7.0025 | 6.3182 | 7.1292 | 6.9300 | 6.5700 | 7.3762 | 7.2722 | 7.2654 | 7.4778 | 3.8866 | 4.1548 | 3.3323 | 4.0307 |
| WAC_T293 | WAC | Q9BTA9 | | 456 | YVSPRISTPQTNT | 6.6927 | 6.2917 | 5.8227 | 5.8061 | 6.5338 | 6.2247 | 6.4934 | 6.3179 | 6.0840 | 5.8056 | 5.6707 | 5.6052 | 7.9457 | 6.3957 | 6.1739 | 6.1359 |
| WAC_T471 | WAC | Q9BTA9 | | 293 | TLSKLPTPTSSVP | 6.1748 | 5.8522 | 5.8125 | 6.4659 | 6.9370 | 5.9944 | 5.7554 | 6.4878 | 6.5312 | 5.4090 | 5.6041 | 6.5582 | 7.2783 | 6.5303 | 6.7373 | 5.8714 |
| WAC_T482 | WAC | Q9BTA9 | | 471 | IKPLISTPPVSSQ | 6.0976 | 5.4872 | 6.0981 | 6.0092 | 6.6626 | 7.5916 | 6.6369 | 6.4157 | 6.6905 | 6.7786 | 6.5051 | 6.7146 | 6.2625 | 5.9446 | 4.9415 | 5.1637 |
| WAPL_S1154 | WAPL | Q7Z5K2-3 | | 306 | FGKRPESPSEISP | 5.3568 | 5.7293 | 5.3195 | 5.6026 | 5.8653 | 5.6924 | 5.5286 | 5.8565 | 5.5186 | 5.9340 | 5.4580 | 5.1842 | 8.3352 | 8.3969 | 8.4766 | 7.7456 |
| WAPL_S162 | WAPL | Q7Z5K2-3 | | 1154 | TTQHDKSGEWQET | 6.5121 | 6.6478 | 6.3798 | 6.8005 | 6.1741 | 6.5503 | 6.5489 | 6.3186 | 6.4989 | 6.5046 | 6.8201 | 7.0268 | 5.2285 | 5.5585 | 5.0370 | 5.3935 |
| WAPL_S265 | WAPL | Q7Z5K2-3 | | 162 | DPFGFDSDDESLP | 5.3716 | 5.9374 | 6.3421 | 6.8224 | 6.0803 | 6.0488 | 6.2368 | 6.2699 | 6.2722 | 6.3428 | 6.4156 | 5.9674 | 6.1756 | 6.5680 | 6.6809 | 6.4681 |
| WAPL_S306 | WAPL | Q7Z5K2-3 | | 265 | EEHEKNSHHIHKN | 5.5182 | 4.3401 | 5.7258 | 4.4549 | 6.1647 | 5.2296 | 5.2454 | 5.5205 | 5.2566 | 4.1837 | 5.2358 | 4.7789 | 10.6066 | 9.1826 | 9.4577 | 9.0988 |
| WAPL_S306;S311 | WAPL | Q7Z5K2-3 | | 311 | ESPSEISPIKGSV | 6.6928 | 7.1922 | 7.8793 | 7.9772 | 5.9531 | 5.9126 | 6.1872 | 5.8751 | 7.2769 | 6.8735 | 7.7862 | 8.1101 | 3.8005 | 4.3445 | 3.5691 | 4.5697 |
| WAPL_S311 | WAPL | Q7Z5K2-3 | | 528 | GGDEGGSGSSNYK | 5.0380 | 5.4613 | 5.2404 | 5.8195 | 6.2266 | 6.4371 | 6.3929 | 6.3167 | 4.6891 | 5.2485 | 5.1624 | 4.7659 | 7.8989 | 7.7274 | 8.5990 | 8.9764 |
| WAPL_S432 | WAPL | Q7Z5K2-3 | | 570 | RTKTAPSPSLQPP | 6.2557 | 5.2951 | 5.0764 | 5.9675 | 6.3899 | 5.6793 | 6.6830 | 6.3037 | 6.1414 | 6.4478 | 6.3749 | 6.4362 | 6.9579 | 6.3269 | 6.4313 | 7.2329 |
| WAPL_S528 | WAPL | Q7Z5K2-3 | | 432 | GVSCGTSFRGTVG | 5.7222 | 6.1380 | 6.8022 | 5.6461 | 5.3087 | 7.0719 | 7.0446 | 5.9188 | 7.0072 | 7.5798 | 8.9884 | 8.3278 | 4.9950 | 5.7430 | 3.1860 | 4.5203 |
| WAPL_S544;S546 | WAPL | Q7Z5K2-3 | 306;311 | | NA | 5.7508 | 6.2870 | 6.0988 | 6.1966 | 6.2678 | 6.2265 | 6.3877 | 5.9965 | 6.2933 | 6.1969 | 6.3239 | 6.1898 | 6.2922 | 6.5060 | 6.3914 | 6.5949 |
| WAPL_S570 | WAPL | Q7Z5K2-3 | 544;546 | | NA | 5.7922 | 6.2604 | 5.7557 | 5.8951 | 6.9069 | 6.3456 | 6.3127 | 6.5024 | 6.1800 | 6.4431 | 6.2479 | 6.4391 | 6.0319 | 6.3272 | 6.3236 | 6.2360 |
| WARS1_S467 | WARS1 | P23381 | | 467 | MTPRKLSFDFQxx | 6.1355 | 5.9397 | 6.0431 | 5.1196 | 7.0721 | 6.4670 | 5.6826 | 6.9586 | 6.9233 | 6.7569 | 7.1564 | 7.0270 | 5.9556 | 5.7907 | 5.5041 | 5.4678 |
| WASF1_S310 | WASF1 | Q92558 | | 310 | IENRPQSPATGRT | 6.4537 | 6.0354 | 6.5777 | 6.4683 | 6.5133 | 6.4659 | 6.6130 | 6.9924 | 6.5133 | 6.8550 | 6.9182 | 6.6738 | 5.4419 | 5.9502 | 4.6369 | 4.8910 |
| WASF2_S284 | WASF2 | Q9Y6W5 | | 129 | TYNTCDTPPPLNN | 6.1057 | 6.5060 | 6.5543 | 6.3359 | 6.1112 | 6.3685 | 6.0544 | 6.1726 | 6.3843 | 6.1394 | 6.4888 | 6.4481 | 6.1186 | 6.4254 | 5.9469 | 5.8398 |
| WASF2_S293;S296;S308 | WASF2 | Q9Y6W5 | | 284 | VDNQRGSGLAGPK | 7.2347 | 6.7619 | 7.2639 | 6.4658 | 6.8969 | 6.4220 | 6.4826 | 6.5702 | 7.0552 | 5.8268 | 6.5744 | 6.3647 | 5.1430 | 5.2323 | 4.7545 | 4.9511 |
| WASF2_S296;S308 | WASF2 | Q9Y6W5 | | 308 | PAPPLGSPPGPKP | 7.5472 | 8.3617 | 6.7842 | 7.9218 | 7.2214 | 6.9397 | 6.2043 | 7.6300 | 7.3134 | 6.8385 | 7.7174 | 6.6360 | 3.1505 | 3.9172 | 2.5927 | 3.2241 |
| WASF2_S308 | WASF2 | Q9Y6W5 | 296;308 | | NA | 6.8037 | 7.5544 | 6.9073 | 7.3141 | 6.1371 | 7.3150 | 7.2654 | 7.0131 | 7.0212 | 7.4546 | 8.0634 | 7.6588 | 3.2343 | 3.8734 | 2.9068 | 3.4774 |
| WASF2_T129 | WASF2 | Q9Y6W5 | 293;296;308 | | NA | 6.1623 | 6.5213 | 6.4246 | 6.7620 | 5.7019 | 6.2684 | 6.3320 | 6.8072 | 6.9649 | 6.7922 | 6.5292 | 6.8080 | 5.5696 | 5.6546 | 4.6343 | 6.0675 |
| WASH4P_S230 | WASH4P | A8MWX3 | | 230 | LFDAPLSISKREQ | 7.5891 | 7.2867 | 6.6559 | 8.3762 | 7.2861 | 6.6665 | 7.4429 | 6.1017 | 6.4818 | 5.4035 | 6.7721 | 6.7726 | 3.7202 | 4.9138 | 3.9541 | 4.5768 |

| | | | | | | | | | | | | | | | | | | | | |
|---|---|---|---|---|---|---|---|---|---|---|---|---|---|---|---|---|---|---|---|---|
| WASHC2A_S1075 | WASHC2A | Q641Q2 | 539 | TQKGLFSDEEDSE | 5.9032 | 6.3806 | 6.7273 | 6.3217 | 6.3748 | 6.5090 | 6.2107 | 6.4508 | 6.5060 | 6.6656 | 6.8044 | 6.7046 | 5.5186 | 5.9716 | 5.2437 | 5.7074 |
| WASHC2A_S1114 | WASHC2A | Q641Q2 | 802 | RTKTVLSLFDEEE | 5.1211 | 5.4608 | 6.6075 | 6.5192 | 5.7213 | 5.9799 | 5.9663 | 5.8740 | 6.2986 | 6.5807 | 6.6663 | 6.7452 | 6.9778 | 6.9111 | 6.4134 | 6.1567 |
| WASHC2A_S1144 | WASHC2A | Q641Q2 | 1180 | ALGEASSDDDLFQ | 6.0488 | 6.3240 | 6.5556 | 6.0219 | 6.1909 | 6.2307 | 5.9257 | 6.4126 | 6.2753 | 6.7456 | 6.6691 | 6.2783 | 6.2310 | 6.3008 | 6.0435 | 5.7461 |
| WASHC2A_S1179 | WASHC2A | Q641Q2 | 498 | KEKAVASPEATVS | 6.7382 | 6.7498 | 6.8811 | 6.8129 | 6.3838 | 6.6785 | 6.7274 | 6.5433 | 7.1911 | 7.1231 | 7.3496 | 7.3866 | 4.1677 | 4.7619 | 4.0402 | 4.4648 |
| WASHC2A_S1179;S1180 | WASHC2A | Q641Q2 | 352 | LTDEDFSPFGSGG | 6.6753 | 7.2157 | 6.9974 | 7.0237 | 6.3299 | 6.8127 | 6.9253 | 6.5707 | 6.8395 | 7.0403 | 7.2820 | 7.1261 | 4.0490 | 4.5188 | 4.1582 | 4.4356 |
| WASHC2A_S1180 | WASHC2A | Q641Q2 | 6 | xMMNRTTPDQELA | 5.5071 | 5.9123 | 6.3530 | 5.6203 | 5.9005 | 6.3452 | 6.0961 | 5.7386 | 6.5219 | 6.5266 | 6.7631 | 6.5415 | 6.5368 | 6.5466 | 6.2918 | 6.7984 |
| WASHC2A_S284 | WASHC2A | Q641Q2 | 700 | DVDSGGSLFGSPP | 6.4866 | 5.7391 | 6.5577 | 6.0727 | 6.0085 | 6.0824 | 6.4932 | 6.4999 | 6.4566 | 6.3703 | 6.4832 | 6.7366 | 6.2168 | 6.3391 | 5.8532 | 5.6038 |
| WASHC2A_S284;S288 | WASHC2A | Q641Q2 | 704 | GGSLFGSPPTSVP | 5.8677 | 6.0419 | 6.9012 | 6.0871 | 6.2744 | 6.1094 | 5.7277 | 6.3092 | 7.6234 | 6.4436 | 7.0025 | 7.3585 | 5.3381 | 5.4321 | 5.5694 | 5.9138 |
| WASHC2A_S288 | WASHC2A | Q641Q2 | 1075 | WADGAISPNGHRP | 6.8127 | 6.8544 | 7.1755 | 7.0240 | 5.8584 | 6.6287 | 6.6639 | 6.4442 | 6.4306 | 6.6408 | 7.1331 | 7.6333 | 4.4815 | 5.2445 | 4.5459 | 4.4285 |
| WASHC2A_S352 | WASHC2A | Q641Q2 | 333 | KERRTPSDDEEDN | 5.8563 | 6.3457 | 6.8131 | 6.3169 | 5.8975 | 6.5228 | 5.9172 | 6.1501 | 6.7440 | 6.5319 | 6.8116 | 6.5178 | 5.7122 | 6.2995 | 5.5931 | 5.9704 |
| WASHC2A_S441 | WASHC2A | Q641Q2 | 284 | NTRPKRSRPTSFA | 6.0449 | 6.3791 | 5.8342 | 5.9260 | 5.4817 | 5.4039 | 6.2419 | 6.3183 | 6.0043 | 5.5488 | 6.2294 | 6.0539 | 6.4546 | 7.0153 | 7.3896 | 7.6744 |
| WASHC2A_S498 | WASHC2A | Q641Q2 | 1144 | TGTGSQSVERTKP | 5.7814 | 6.9082 | 7.3151 | 6.2385 | 6.1916 | 6.8379 | 6.7860 | 7.5193 | 7.1292 | 7.1242 | 6.5985 | 7.5378 | 4.0334 | 5.6927 | 4.4286 | 3.8777 |
| WASHC2A_S521 | WASHC2A | Q641Q2 | 288 | KRSRPTSFADELA | 5.3948 | 5.4500 | 6.1597 | 5.8513 | 5.7422 | 5.9737 | 5.9287 | 6.3450 | 6.0217 | 6.1696 | 5.4974 | 5.8972 | 7.4159 | 7.7479 | 7.2215 | 7.1834 |
| WASHC2A_S539 | WASHC2A | Q641Q2 | 1179 | PALGEASSDDDLF | 5.8780 | 6.3898 | 6.3191 | 6.6308 | 6.3971 | 6.0754 | 6.2442 | 6.7370 | 6.0201 | 6.1943 | 6.5242 | 6.5285 | 6.0811 | 6.0136 | 6.0035 | 5.9633 |
| WASHC2A_S619;S620 | WASHC2A | Q641Q2 | 1114 | VPGVDRSPFAKSL | 6.2107 | 5.6775 | 7.1014 | 5.9078 | 7.6151 | 6.2694 | 5.7724 | 6.1996 | 6.3727 | 7.4797 | 6.3461 | 5.6705 | 5.3089 | 5.8101 | 6.4173 | 5.8408 |
| WASHC2A_S620 | WASHC2A | Q641Q2 | 996 | SSEHRRSHGLESV | 5.9347 | 6.8728 | 5.8278 | 7.0081 | 6.5126 | 7.5012 | 6.0142 | 6.9454 | 6.9215 | 6.8972 | 7.5018 | 7.8126 | 4.6417 | 4.7532 | 4.3855 | 4.4697 |
| WASHC2A_S700 | WASHC2A | Q641Q2 | 441 | QPTPRKSPYGPPP | 6.0802 | 6.0549 | 7.0523 | 5.8057 | 6.8625 | 7.4019 | 7.0634 | 6.7810 | 6.3317 | 6.6734 | 6.6971 | 6.3697 | 4.8985 | 5.1279 | 4.9483 | 5.8516 |
| WASHC2A_S704 | WASHC2A | Q641Q2 | 620 | SALLFSSDEEDQW | 4.2294 | 5.3358 | 4.8631 | 5.7080 | 6.3291 | 5.5621 | 5.2579 | 5.9318 | 6.5906 | 7.8981 | 5.0567 | 7.2946 | 7.3273 | 9.0674 | 7.7221 | 5.8261 |
| WASHC2A_S787 | WASHC2A | Q641Q2 | 877 | LEPSVGSLFGDDE | 5.8681 | 6.2650 | 6.3229 | 6.9556 | 6.6350 | 6.3247 | 5.8356 | 5.9253 | 6.5252 | 6.2141 | 6.5722 | 6.6647 | 5.6838 | 6.2664 | 6.0567 | 5.8847 |
| WASHC2A_S802 | WASHC2A | Q641Q2 | 787 | DLFSSSSPWDKGT | 6.8474 | 7.7817 | 6.4450 | 7.2259 | 6.3369 | 7.5487 | 7.6459 | 7.1221 | 7.6997 | 7.7501 | 7.9054 | 7.4035 | 2.6386 | 3.6725 | 2.7179 | 3.2588 |
| WASHC2A_S874 | WASHC2A | Q641Q2 | 874 | TKLLEPSVGSLFG | 6.1582 | 6.6169 | 7.0596 | 6.4403 | 6.0865 | 6.0723 | 6.3893 | 5.5294 | 6.4053 | 6.8765 | 7.4277 | 6.0984 | 5.8364 | 6.2892 | 5.6041 | 5.1098 |
| WASHC2A_S877 | WASHC2A | Q641Q2 | 521 | KKVTLSSSKNLKP | 7.0673 | 7.8659 | 7.0350 | 7.5760 | 6.0373 | 6.4106 | 6.9013 | 6.6553 | 7.3805 | 7.7222 | 7.6560 | 8.5121 | 2.9269 | 3.7961 | 2.8774 | 3.5801 |
| WASHC2A_S996 | WASHC2A | Q641Q2 | 619;620 | NA | 6.6133 | 6.8122 | 6.3876 | 6.3647 | 6.6470 | 6.5054 | 6.3472 | 7.5301 | 6.8795 | 5.9896 | 6.6004 | 6.4743 | 4.8634 | 5.7058 | 5.1855 | 5.0940 |
| WASHC2A_T331;S333 | WASHC2A | Q641Q2 | 284;288 | NA | 5.3381 | 5.6522 | 6.0451 | 5.8848 | 5.6789 | 5.4228 | 5.7061 | 5.9679 | 5.5348 | 5.2853 | 6.1823 | 5.8428 | 7.8822 | 7.9376 | 7.6606 | 7.9786 |
| WASHC2A_T333 | WASHC2A | Q641Q2 | 331;333 | NA | 6.4672 | 6.5078 | 5.6445 | 7.4064 | 5.4163 | 6.1494 | 6.1735 | 5.4718 | 6.3444 | 6.1665 | 6.8273 | 6.5316 | 6.0233 | 7.1315 | 5.5482 | 6.1903 |
| WASHC2A_T6 | WASHC2A | Q641Q2 | 1179;1180 | NA | 6.8114 | 6.9366 | 6.9133 | 7.2761 | 6.0250 | 6.6876 | 6.4135 | 6.1868 | 6.3730 | 6.4547 | 6.0976 | 6.5801 | 5.2006 | 5.8799 | 4.8642 | 5.2998 |
| WASHC2C_S1093 | WASHC2C | Q9Y4E1 | 6 | xMMNRTTPDQELV | 5.6395 | 5.5875 | 6.2178 | 5.4705 | 5.9532 | 5.8756 | 5.8078 | 6.2150 | 6.8869 | 6.4498 | 6.4365 | 6.8347 | 6.7638 | 6.6130 | 6.5497 | 6.6986 |
| WASHC2C_S539 | WASHC2C | Q9Y4E1 | 1093 | VPGVDTSPFAKSL | 6.0764 | 6.5101 | 6.8286 | 5.8314 | 5.8575 | 6.8313 | 6.2783 | 6.6555 | 7.2199 | 7.0334 | 6.7935 | 7.1921 | 4.8754 | 5.9220 | 5.0845 | 5.0103 |
| WASHC2C_S663 | WASHC2C | Q9Y4E1 | 539 | TQKGLFSDEEDSE | 6.4055 | 6.4479 | 6.6802 | 6.4026 | 6.4018 | 6.6846 | 6.1299 | 6.4154 | 6.4151 | 6.2507 | 6.6158 | 6.8989 | 5.6158 | 5.9123 | 5.2515 | 5.4721 |
| WASHC2C_T6 | WASHC2C | Q9Y4E1 | 663 | KAVKKTSLFEEDK | 5.8641 | 6.6437 | 6.7956 | 5.2417 | 5.9997 | 5.8627 | 5.9170 | 6.5163 | 6.8417 | 7.0274 | 7.1196 | 7.9738 | 5.9007 | 6.6057 | 4.8797 | 4.8106 |
| WASHC4_S1157 | WASHC4 | Q2M389 | 1157 | EEETKTSNGDLSD | 5.7176 | 6.0683 | 6.1385 | 5.7219 | 5.6090 | 5.9302 | 6.0633 | 5.9696 | 6.4387 | 6.2884 | 5.8801 | 6.4218 | 7.0471 | 6.8099 | 6.6461 | 7.2494 |
| WASHC5_S917 | WASHC5 | Q12768 | 917 | TLMNAVSPLKSIV | 5.7297 | 5.9080 | 6.7112 | 5.7703 | 6.0205 | 6.3879 | 6.3439 | 6.2548 | 6.6914 | 6.4804 | 6.3597 | 6.4349 | 5.7817 | 6.6740 | 6.3526 | 6.0989 |
| WASL_Y256 | WASL | O00401 | 256 | ETSKVIYDFIEKT | 5.6957 | 6.3543 | 7.8585 | 7.3224 | 5.5378 | 6.1256 | 6.3114 | 6.8570 | 6.3664 | 6.6867 | 7.2977 | 6.9955 | 4.5924 | 5.5759 | 5.4016 | 5.0211 |
| WBP11_S171 | WBP11 | Q9Y2W2 | 237 | DEDMLYSPELAQR | 6.7002 | 7.0018 | 6.7560 | 7.0320 | 6.5080 | 6.5170 | 6.7472 | 6.6785 | 6.5828 | 6.5938 | 7.1731 | 6.9280 | 4.6916 | 4.9660 | 4.5938 | 4.5303 |
| WBP11_S237 | WBP11 | Q9Y2W2 | 328 | KNMKELTPLQAMM | 7.1230 | 5.8870 | 6.1867 | 6.9701 | 6.7040 | 7.2499 | 6.9713 | 6.4528 | 6.4864 | 7.6603 | 6.9179 | 6.3993 | 4.8262 | 5.8608 | 3.7667 | 4.5377 |
| WBP11_T328 | WBP11 | Q9Y2W2 | 171 | SILKKTSAYGPPT | 7.4994 | 6.9937 | 7.0060 | 6.3312 | 5.4350 | 5.8428 | 5.6179 | 5.9145 | 7.5023 | 7.0979 | 7.1809 | 6.8290 | 5.0704 | 5.6909 | 4.8075 | 5.1806 |
| WBP4_S262 | WBP4 | O75554 | 262 | KNSDGGSDPETQK | 5.9348 | 6.4401 | 6.0760 | 6.4045 | 6.2164 | 5.5013 | 5.9792 | 5.7630 | 6.2870 | 6.5457 | 6.0043 | 6.7117 | 6.4921 | 6.4142 | 6.2829 | 6.9469 |
| WDCP_S468 | WDCP | Q9H6R7 | 468 | KLIESLSPDFCHQ | 6.5996 | 6.7873 | 7.4147 | 7.2004 | 6.2612 | 7.2182 | 6.5374 | 6.9721 | 6.9519 | 7.0708 | 7.4904 | 6.9829 | 3.8846 | 4.5469 | 3.6960 | 4.3857 |
| WDCP_S690 | WDCP | Q9H6R7 | 690 | RDSFSHSPGAVSS | 5.1958 | 6.3591 | 6.4543 | 5.5093 | 6.5951 | 7.2292 | 6.4902 | 7.9279 | 5.8482 | 5.6768 | 6.4251 | 5.4797 | 5.8943 | 6.3766 | 6.4446 | 6.0939 |
| WDFY3_S2278 | WDFY3 | Q8IZQ1 | 3339 | RWSDQLSLDEKDG | 5.9870 | 5.6291 | 6.6001 | 5.7451 | 5.8279 | 6.3654 | 5.9814 | 5.9761 | 6.2404 | 6.7588 | 5.9725 | 6.3203 | 6.4104 | 7.3665 | 6.7426 | 6.0764 |
| WDFY3_S3335 | WDFY3 | Q8IZQ1 | 799 | SESSLPSPWGTPA | 5.5189 | 6.7111 | 6.0375 | 6.6541 | 7.2564 | 6.0999 | 6.3476 | 6.8216 | 4.9433 | 8.5787 | 6.9000 | 7.1922 | 5.5991 | 5.7043 | 4.3766 | 5.2587 |
| WDFY3_S3339 | WDFY3 | Q8IZQ1 | 2278 | KLSRVSSGFGLSK | 5.8749 | 5.3455 | 5.6263 | 5.1473 | 5.8444 | 5.9064 | 5.6916 | 5.5891 | 5.6116 | 5.3201 | 5.7551 | 6.5812 | 7.5070 | 8.6955 | 7.2924 | 8.2116 |
| WDFY3_S799 | WDFY3 | Q8IZQ1 | 817 | HAYHSVSTPPVYP | 6.6519 | 6.0046 | 6.6862 | 6.0461 | 6.3907 | 7.2646 | 6.5701 | 6.4796 | 7.4044 | 6.5558 | 6.2369 | 6.2787 | 5.3637 | 5.3471 | 5.4531 | 5.2664 |
| WDFY3_S817 | WDFY3 | Q8IZQ1 | 3335 | DDSRRWSDQLSLD | 5.5274 | 5.5819 | 6.4945 | 5.9098 | 6.1565 | 6.2527 | 6.0445 | 6.7348 | 6.9294 | 6.6605 | 6.8486 | 7.0070 | 5.9776 | 6.0604 | 5.7355 | 6.0789 |
| WDHD1_S333 | WDHD1 | O75717 | 333 | FDGDDMSNAGDFL | 5.9735 | 6.3892 | 6.8403 | 6.8695 | 5.8197 | 5.8700 | 6.2553 | 5.6588 | 6.2812 | 6.0795 | 6.3082 | 6.2801 | 6.2508 | 6.6604 | 6.4047 | 6.0590 |
| WDHD1_S383 | WDHD1 | O75717 | 407 | EDGQEGSIHNLPL | 5.8233 | 6.1499 | 6.5062 | 6.3680 | 6.1689 | 5.9515 | 6.0777 | 6.0680 | 5.9937 | 5.8915 | 6.1537 | 5.9436 | 7.0666 | 6.7864 | 6.4025 | 6.6484 |
| WDHD1_S407 | WDHD1 | O75717 | 383 | LEDDENSVDISML | 5.8705 | 6.1615 | 6.2669 | 6.2920 | 5.8713 | 5.8362 | 5.9906 | 5.7094 | 5.8951 | 5.8874 | 6.0859 | 5.5360 | 7.0364 | 7.4790 | 7.6849 | 6.3970 |
| WDHD1_S868 | WDHD1 | O75717 | 868 | EEDAEDSGEADDE | 5.8750 | 6.8083 | 6.5524 | 6.6510 | 6.3742 | 7.8052 | 6.1516 | 6.1828 | 5.7711 | 5.7338 | 6.6474 | 6.3064 | 6.1962 | 6.0309 | 5.8354 | 5.0783 |
| WDHD1_S984 | WDHD1 | O75717 | 984 | YFQKRNSQTNKTE | 6.9850 | 7.6294 | 6.8870 | 7.3541 | 6.0199 | 5.5271 | 5.2385 | 5.7077 | 7.0549 | 7.3999 | 7.3343 | 7.1043 | 4.5795 | 5.2791 | 4.7540 | 5.1454 |
| WDR11_S209 | WDR11 | Q9BZH6 | 402 | NSSSGVSPLYSPV | 6.0155 | 5.6648 | 6.4469 | 6.1707 | 6.1136 | 6.3281 | 6.0135 | 5.4840 | 6.2904 | 6.3874 | 6.4594 | 6.4798 | 6.5282 | 6.8579 | 6.5614 | 6.1984 |
| WDR11_S402 | WDR11 | Q9BZH6 | 209 | ISSPHSSPAHNKL | 6.2381 | 7.4255 | 5.9222 | 7.2145 | 7.0480 | 7.4741 | 5.6814 | 8.1859 | 6.4160 | 6.0600 | 6.5443 | 5.9324 | 4.7049 | 5.5660 | 5.0808 | 4.5058 |
| WDR13_S81 | WDR13 | Q9H1Z4 | 81 | ARAYSNSIVRSSR | 6.7166 | 6.5678 | 6.7680 | 6.6434 | 5.5836 | 6.0959 | 5.7732 | 6.1693 | 7.0323 | 6.3746 | 6.3376 | 6.5587 | 6.4941 | 6.5255 | 5.2720 | 5.0874 |
| WDR18_S368 | WDR18 | Q9BV38 | 368 | GLHQQGSEPSYLD | 6.1708 | 6.4728 | 6.0345 | 6.6546 | 7.1383 | 5.4786 | 6.2457 | 6.4167 | 5.9410 | 5.1045 | 5.4118 | 5.5936 | 7.9855 | 7.3173 | 6.9726 | 5.0617 |
| WDR20_S388 | WDR20 | Q8TBZ3-7 | 496 | SKFATLSLHDRKE | 8.1387 | 7.1815 | 8.5496 | 6.9275 | 5.4558 | 6.4487 | 6.0203 | 6.4883 | 7.3060 | 7.1550 | 7.5206 | 7.8653 | 3.6199 | 4.8908 | 3.1728 | 3.2591 |
| WDR20_S465 | WDR20 | Q8TBZ3-7 | 465 | PLPRSNSLPHSAV | 5.7954 | 5.7023 | 6.5137 | 6.3779 | 6.4077 | 6.1309 | 5.3042 | 6.5153 | 6.3281 | 6.0897 | 6.0528 | 6.1915 | 6.4434 | 6.9293 | 6.5927 | 6.6252 |
| WDR20_S479 | WDR20 | Q8TBZ3-7 | 479 | NAGSKSSVMDGAI | 5.6108 | 5.5385 | 6.4265 | 6.6412 | 5.8982 | 5.7802 | 6.1807 | 5.5940 | 6.5819 | 5.8919 | 6.0932 | 6.4983 | 7.3143 | 6.7298 | 6.3764 | 6.8442 |
| WDR20_S496 | WDR20 | Q8TBZ3-7 | 397 | SRPVSVTYRFGSV | 5.8462 | 5.6718 | 6.9728 | 6.8078 | 6.1579 | 5.6757 | 7.2382 | 6.4327 | 7.6575 | 6.3173 | 5.5907 | 6.1353 | 5.2150 | 5.7303 | 5.5626 | 6.9883 |
| WDR20_T397 | WDR20 | Q8TBZ3-7 | 388 | RLSKRNSTDSRPV | 6.1664 | 6.5500 | 7.2699 | 6.8777 | 5.8140 | 5.5937 | 6.2771 | 6.5027 | 6.9228 | 7.0013 | 6.5446 | 6.9003 | 5.8591 | 5.8935 | 4.4827 | 5.3444 |
| WDR24_T581 | WDR24 | Q96S15 | 581 | RHEIVDTPPGPEH | 6.0999 | 6.7899 | 6.7477 | 6.9297 | 5.9771 | 7.0497 | 6.8819 | 6.2033 | 6.3014 | 6.7799 | 6.8221 | 7.2403 | 4.9116 | 5.1354 | 4.6788 | 5.4512 |
| WDR26_S121 | WDR26 | Q9H7D7 | 121 | KKKKRLSQSDEDV | 6.1936 | 6.4272 | 6.4615 | 6.1856 | 6.4948 | 6.6200 | 5.5926 | 6.4578 | 6.1658 | 5.8510 | 6.8846 | 6.5220 | 5.8253 | 6.1715 | 5.9832 | 6.1636 |
| WDR26_S658 | WDR26 | Q9H7D7 | 658 | IEEECSSMDSxxx | 5.2667 | 6.1286 | 6.3356 | 8.4891 | 7.0155 | 7.6363 | 5.1189 | 6.0604 | 5.4191 | 6.4288 | 7.4112 | 6.1400 | 5.7284 | 5.6283 | 6.2128 | 4.9802 |
| WDR3_S241 | WDR3 | Q9UNX4 | 241 | KKIKGSSPGIQDT | 6.1752 | 6.4570 | 6.6470 | 6.5834 | 6.1082 | 6.0852 | 6.1281 | 6.2345 | 6.4071 | 6.3136 | 6.4365 | 6.0019 | 5.8376 | 6.3663 | 5.9468 | 6.2717 |
| WDR33_S1181 | WDR33 | Q9C0J8 | 1218 | ASRERSSSLQGMD | 5.1826 | 5.9221 | 6.1340 | 6.0343 | 6.2561 | 6.0333 | 5.7071 | 6.6019 | 5.9211 | 5.8817 | 5.8237 | 5.9407 | 7.3528 | 6.6619 | 7.7189 | 6.8278 |
| WDR33_S1210 | WDR33 | Q9C0J8 | 1181 | GGRKPDSWDGNRE | 7.8802 | 6.0243 | 6.3058 | 7.1455 | 6.1467 | 7.9661 | 6.6413 | 6.7670 | 6.3471 | 5.8271 | 5.0557 | 6.3248 | 5.6442 | 5.0955 | 4.8840 | 5.9448 |
| WDR33_S1218 | WDR33 | Q9C0J8 | 1210 | PPHDGHSPASRER | 6.6160 | 6.7036 | 7.3011 | 7.4041 | 6.3868 | 6.2271 | 6.4563 | 6.2117 | 5.8769 | 5.9593 | 4.3472 | 6.6907 | 5.9954 | 6.6727 | 5.8975 | 5.2537 |
| WDR33_S1279 | WDR33 | Q9C0J8 | 1279 | KSGRSSSLDGEHH | 4.5679 | 7.4808 | 4.6432 | 4.8625 | 5.7868 | 4.2088 | 5.9788 | 6.9065 | 4.9055 | 4.2261 | 4.8302 | 4.6912 | 7.9566 | 8.8379 | 10.0146 | 10.1024 |
| WDR36_T842 | WDR36 | Q8NI36 | 842 | VNNKYDTALNLLK | 7.4685 | 4.8845 | 6.0363 | 8.3545 | 6.9813 | 6.6771 | 5.1090 | 4.4327 | 3.4641 | 4.6512 | 4.7162 | 3.3607 | 9.1489 | 8.5169 | 8.6941 | 7.5040 |
| WDR4_S391 | WDR4 | P57081 | 391 | KKQRRRSPPPGPD | 5.0153 | 5.3003 | 5.9463 | 5.3431 | 5.1465 | 5.9309 | 5.3747 | 6.1282 | 6.0120 | 6.2297 | 6.2534 | 6.0999 | 7.0774 | 8.2165 | 8.1425 | 7.7833 |
| WDR43_S590 | WDR43 | Q15061 | 590 | KTKGATSPGQKAK | 5.6806 | 6.0906 | 6.1171 | 7.2801 | 6.3416 | 6.4151 | 6.4585 | 6.4505 | 6.5029 | 6.4471 | 6.3394 | 6.5717 | 5.5903 | 5.8267 | 5.7897 | 6.0981 |
| WDR43_T394 | WDR43 | Q15061 | 394 | AITKVRTPVMNSE | 6.7140 | 7.0208 | 6.5822 | 7.6271 | 6.4055 | 6.6741 | 7.0530 | 6.8557 | 6.7462 | 6.7073 | 6.9582 | 6.6353 | 4.1086 | 4.9383 | 4.2223 | 4.7514 |
| WDR43_T589 | WDR43 | Q15061 | 589 | EKTKGATSPGQKA | 6.4979 | 7.6343 | 6.7927 | 8.2564 | 6.2548 | 6.3718 | 6.3498 | 6.4137 | 6.2936 | 6.1212 | 6.2317 | 6.5231 | 4.9180 | 5.1105 | 5.1285 | 5.1019 |
| WDR44_S161 | WDR44 | Q5JSH3 | 50 | YKVGNESPVQELK | 6.3339 | 6.3096 | 6.5654 | 5.9599 | 6.1375 | 6.6802 | 6.4143 | 6.4263 | 6.8006 | 6.8187 | 6.5402 | 6.3892 | 5.6815 | 6.1027 | 5.4519 | 5.3881 |
| WDR44_S344 | WDR44 | Q5JSH3 | 344 | QRPRSNSGRELTD | 6.2436 | 6.2433 | 6.3140 | 6.1464 | 6.6199 | 6.1205 | 6.0570 | 6.1953 | 6.1521 | 7.1897 | 6.7016 | 6.4941 | 5.9601 | 6.6220 | 5.6804 | 5.2600 |
| WDR44_S50 | WDR44 | Q5JSH3 | 161 | TKLTQTSSTEQLN | 5.7708 | 6.2309 | 6.3861 | 5.9693 | 5.8559 | 6.3165 | 5.8184 | 6.3325 | 6.7259 | 7.0465 | 6.3417 | 6.4804 | 5.7086 | 7.0770 | 5.9392 | 6.0003 |
| WDR44_S561 | WDR44 | Q5JSH3 | 96 | SDQATASPIVART | 6.5964 | 6.7939 | 6.5696 | 6.8870 | 6.1852 | 6.8265 | 6.6679 | 6.8675 | 6.4076 | 6.6489 | 6.4275 | 6.8389 | 4.7144 | 5.4645 | 4.8732 | 5.2311 |
| WDR44_S96 | WDR44 | Q5JSH3 | 561 | NTEGRVSPSPSQE | 5.8861 | 6.2904 | 6.9454 | 6.5663 | 6.4471 | 6.5014 | 5.6615 | 5.8606 | 6.5878 | 7.3777 | 7.2999 | 7.0889 | 5.1099 | 5.7407 | 5.3605 | 5.2756 |
| WDR44_T219 | WDR44 | Q5JSH3 | 219 | KPPRHLTPEPDIV | 6.0894 | 6.9559 | 7.4343 | 6.7999 | 5.5853 | 6.4307 | 6.6972 | 7.1243 | 6.7076 | 6.5705 | 7.1708 | 7.1496 | 4.7156 | 5.3042 | 4.3402 | 4.9243 |
| WDR46_S41 | WDR46 | O15213 | 41 | PTTAGASPGPPRN | 6.1302 | 7.2395 | 6.5396 | 6.7428 | 6.3619 | 6.6626 | 8.1433 | 5.8491 | 6.2437 | 6.3875 | 6.2235 | 6.6656 | 4.9983 | 5.4926 | 4.7761 | 5.5437 |
| WDR47_S303 | WDR47 | O94967-4 | 292 | AYADLLTPLISKL | 7.3567 | 7.3576 | 7.3137 | 7.8066 | 6.4833 | 7.0050 | 6.9807 | 6.5068 | 6.7636 | 5.7624 | 6.5969 | 6.5166 | 4.3457 | 4.4727 | 4.5340 | 4.1977 |
| WDR47_S304 | WDR47 | O94967-4 | 311 | PMRRPQSADAYMT | 6.0220 | 6.8576 | 6.7365 | 6.5149 | 6.7099 | 6.8352 | 5.6815 | 5.7107 | 5.9443 | 5.7420 | 6.3496 | 7.1392 | 5.6648 | 5.9679 | 6.5519 | 6.4720 |
| WDR47_S311 | WDR47 | O94967-4 | 304 | LSPYPSSPMRRPQ | 6.5559 | 6.6902 | 6.9060 | 6.1657 | 6.8019 | 6.8764 | 6.5470 | 6.4669 | 6.7402 | 6.4545 | 6.8813 | 6.7542 | 4.7091 | 5.1241 | 5.1028 | 5.2237 |
| WDR47_T292 | WDR47 | O94967-4 | 303 | KLSPYPSSPMRRP | 6.2532 | 6.1957 | 6.4945 | 6.4409 | 6.1608 | 6.2282 | 7.5282 | 6.3931 | 6.7475 | 5.3825 | 5.9153 | 6.2228 | 5.3749 | 6.3518 | 6.1527 | 6.1581 |
| WDR48_S335 | WDR48 | Q8TAF3 | 335 | IHNFRASGDYDND | 6.0193 | 6.4459 | 6.7950 | 6.8589 | 6.4431 | 6.1493 | 6.3019 | 6.0557 | 7.0117 | 6.7339 | 6.6875 | 6.6834 | 5.4078 | 5.3683 | 5.5659 | 5.4725 |
| WDR48_S586 | WDR48 | Q8TAF3 | 586 | LKKDRLSASDMLQ | 5.9903 | 6.7687 | 6.3447 | 6.4784 | 5.1554 | 5.6250 | 6.6035 | 6.2773 | 5.3587 | 5.3456 | 6.7513 | 6.2697 | 6.7078 | 7.4308 | 7.0536 | 5.8392 |
| WDR55_S14 | WDR55 | Q9H6Y2 | 14 | RPAEDGSDEEDPD | 4.7797 | 8.0357 | 7.1592 | 5.7931 | 3.5539 | 6.3992 | 6.0739 | 5.3275 | 5.3480 | 5.9472 | 7.0565 | 8.1659 | 6.4882 | 7.2209 | 6.1458 | 6.5053 |
| WDR59_S564 | WDR59 | Q6PJI9 | 603 | LRLYSGSPTRSEK | 6.9166 | 6.9807 | 7.0456 | 6.4387 | 6.2761 | 6.5369 | 6.4067 | 6.7442 | 6.5951 | 6.6691 | 7.0719 | 6.6907 | 4.7134 | 5.3923 | 4.6836 | 4.8384 |
| WDR59_S603 | WDR59 | Q6PJI9 | 564 | TMHRAVSPTEPTP | 5.9716 | 6.8256 | 6.8798 | 6.2840 | 5.8687 | 6.2071 | 6.2873 | 6.3942 | 6.4271 | 6.3705 | 5.9831 | 6.8388 | 5.7762 | 6.1660 | 5.8983 | 5.8218 |
| WDR62_S1008 | WDR62 | O43379-4 | 1233 | RISRSISLGDSEG | 6.2538 | 5.6664 | 6.7281 | 6.0337 | 5.9628 | 6.3844 | 5.8716 | 6.6829 | 6.7319 | 6.2532 | 6.5265 | 6.6778 | 5.9789 | 6.2124 | 5.8159 | 6.2196 |
| WDR62_S1049;T1053 | WDR62 | O43379-4 | 33 | ARRGQSSPPPAPP | 6.1075 | 6.3466 | 6.8269 | 6.1163 | 6.1190 | 6.7067 | 5.9152 | 6.2371 | 6.6492 | 6.7068 | 6.5662 | 6.6595 | 5.4352 | 6.1560 | 5.7592 | 5.6926 |
| WDR62_S1053 | WDR62 | O43379-4 | 1149 | AGTGYASPDRTHV | 6.8839 | 6.7756 | 7.2400 | 6.6500 | 6.6359 | 7.1270 | 6.5532 | 6.7489 | 6.8360 | 7.0217 | 7.2735 | 7.0866 | 4.1129 | 4.9337 | 3.9174 | 4.2038 |
| WDR62_S1070 | WDR62 | O43379-4 | 1353 | SLPAPESPGLPAH | 5.8968 | 5.5308 | 6.5094 | 6.2581 | 6.3818 | 6.9217 | 5.9075 | 7.6072 | 6.5428 | 5.4315 | 6.8356 | 6.4981 | 5.8074 | 6.1257 | 5.6409 | 6.1047 |
| WDR62_S1098 | WDR62 | O43379-4 | 1410 | VEALPPSPLELSR | 6.1316 | 6.1511 | 6.5042 | 6.3155 | 6.4344 | 6.5264 | 6.3998 | 6.6425 | 6.4957 | 6.1846 | 6.5689 | 6.8312 | 5.7413 | 6.0477 | 5.2025 | 5.8228 |
| WDR62_S1149 | WDR62 | O43379-4 | 982 | LRVSSDSPKDQSP | 6.2735 | 6.3561 | 7.3639 | 6.3329 | 6.1932 | 6.2218 | 6.3119 | 6.2632 | 6.7614 | 6.8365 | 6.4864 | 6.8178 | 5.2735 | 5.5746 | 5.2013 | 5.7321 |
| WDR62_S1231 | WDR62 | O43379-4 | 1053 | SSSLPQTPEQEKF | 5.5061 | 5.5783 | 6.5600 | 6.3618 | 6.3380 | 6.2153 | 5.8349 | 6.2641 | 6.7891 | 6.6780 | 6.2286 | 5.9184 | 6.2160 | 6.6568 | 6.5812 | 6.2733 |
| WDR62_S1233 | WDR62 | O43379-4 | 1196 | TDRNLPTPTSAPT | 6.3789 | 6.5208 | 6.5417 | 6.2752 | 6.2512 | 6.6072 | 6.0310 | 6.7734 | 6.4140 | 6.2397 | 6.2120 | 6.2455 | 5.5706 | 5.8837 | 6.0212 | 6.0337 |
| WDR62_S1253 | WDR62 | O43379-4 | 1070 | FETLTESPCRELF | 6.7327 | 6.7905 | 7.5362 | 7.0513 | 6.1348 | 6.7542 | 6.3945 | 6.8933 | 6.9758 | 6.9578 | 7.3252 | 7.0759 | 4.2879 | 4.5711 | 3.9271 | 4.5919 |
| WDR62_S1254 | WDR62 | O43379-4 | 829 | DPRCLLTNGKLPL | 9.2782 | 8.4396 | 7.0335 | 8.0731 | 6.6292 | 6.1069 | 7.0996 | 5.5348 | 6.8932 | 5.7117 | 6.5865 | 6.1655 | 4.0177 | 4.5781 | 3.4304 | 4.4220 |
| WDR62_S1353 | WDR62 | O43379-4 | 49 | RRRTRLSTASEET | 6.1387 | 5.7765 | 6.5224 | 6.3060 | 6.4768 | 7.1387 | 6.2361 | 6.7811 | 7.0683 | 6.6921 | 6.8961 | 6.5106 | 5.5804 | 5.6358 | 5.2321 | 5.0083 |
| WDR62_S1410 | WDR62 | O43379-4 | 1254 | PLRRPSSVGELAS | 6.3917 | 5.9483 | 6.7907 | 5.5906 | 6.4050 | 7.2832 | 5.5696 | 5.9810 | 6.4011 | 6.9762 | 6.4619 | 6.1018 | 5.7184 | 7.0206 | 5.9951 | 5.3646 |
| WDR62_S33 | WDR62 | O43379-4 | 1008 | SFAAIHSPAPPPD | 6.6673 | 6.2790 | 6.3242 | 6.1991 | 5.6559 | 6.5732 | 6.5232 | 6.7626 | 7.9431 | 6.1515 | 6.0121 | 7.2474 | 5.3081 | 5.5952 | 6.1765 | 4.5815 |
| WDR62_S49 | WDR62 | O43379-4 | 1098 | FFNPRLSISTQFL | 7.0244 | 6.4665 | 5.7882 | 6.8560 | 6.3756 | 6.6177 | 7.0821 | 5.7332 | 7.3562 | 6.1162 | 7.7277 | 6.5620 | 4.7295 | 6.2123 | 4.7794 | 4.5732 |
| WDR62_S982 | WDR62 | O43379-4 | 1231 | RARISRSISLGDS | 6.6062 | 6.5923 | 7.4350 | 7.4228 | 6.1162 | 6.3161 | 6.9019 | 6.6395 | 7.0839 | 6.7874 | 6.9109 | 6.4900 | 4.6479 | 5.1240 | 4.2806 | 4.6454 |
| WDR62_T1196 | WDR62 | O43379-4 | 1253 | QPLRRPSSVGELA | 5.1253 | 5.8518 | 7.2076 | 7.1870 | 6.3368 | 5.9717 | 5.6084 | 6.1515 | 6.0279 | 6.9511 | 7.4978 | 7.8387 | 5.4455 | 5.5560 | 4.9174 | 6.3256 |
| WDR62_T829 | WDR62 | O43379-4 | 1049;1053 | NA | 6.2944 | 6.4789 | 7.0113 | 6.5011 | 6.1834 | 6.5910 | 6.1724 | 6.3808 | 6.3139 | 6.5406 | 6.9420 | 6.8245 | 5.2222 | 5.6699 | 5.4710 | 5.4026 |
| WDR7_S1456 | WDR7 | Q9Y4E6 | 935 | LSKARGSPPTSSN | 6.0182 | 6.8545 | 7.2160 | 7.1253 | 6.3438 | 6.9011 | 6.7627 | 6.3592 | 5.9927 | 7.1828 | 6.7647 | 6.8407 | 4.8589 | 5.2240 | 4.4857 | 5.0698 |
| WDR7_S935 | WDR7 | Q9Y4E6 | 1456 | PPVQPASPGSHNA | 6.2693 | 6.1967 | 6.7293 | 6.8257 | 6.4037 | 6.7937 | 6.7333 | 6.4682 | 6.9371 | 6.8289 | 6.6521 | 6.6062 | 4.9252 | 5.3811 | 5.3729 | 4.8765 |
| WDR7_T926 | WDR7 | Q9Y4E6 | 926 | RPPRPSTPDLSKA | 6.9186 | 7.4398 | 7.6742 | 7.2075 | 5.8347 | 6.3582 | 6.7972 | 6.4615 | 7.1680 | 7.7679 | 7.6634 | 7.1963 | 3.3452 | 4.5673 | 3.4955 | 4.1048 |
| WDR70_S638 | WDR70 | Q9NW82 | 638 | MFAQVESDDEEAK | 5.8982 | 6.1340 | 6.3581 | 5.6023 | 6.1890 | 6.7046 | 6.0824 | 5.8255 | 6.1223 | 6.4060 | 6.4196 | 6.7130 | 5.8209 | 6.5603 | 6.1998 | 6.9640 |
| WDR75_S194 | WDR75 | Q8IWA0 | 796 | EKVQDTSNTGLGE | 5.9684 | 7.0880 | 6.3146 | 7.1475 | 6.3470 | 6.8040 | 6.8276 | 6.6871 | 6.7021 | 6.4404 | 6.8677 | 6.4065 | 5.0855 | 5.0434 | 4.8415 | 5.4288 |
| WDR75_S779;S782 | WDR75 | Q8IWA0 | 194 | TSRFTLSSSRNKK | 6.2614 | 5.7695 | 5.6885 | 6.8823 | 6.7868 | 5.8447 | 6.7655 | 6.7753 | 6.5461 | 6.1197 | 5.9159 | 5.7319 | 5.7377 | 6.5584 | 5.9965 | 6.6199 |
| WDR75_S796 | WDR75 | Q8IWA0 | 779;782 | NA | 6.2188 | 5.9522 | 6.1641 | 6.5388 | 5.7867 | 5.3488 | 6.2421 | 5.3683 | 6.3949 | 5.9626 | 6.0688 | 6.2888 | 7.0937 | 6.5277 | 6.6489 | 7.3947 |
| WDR77_T5 | WDR77 | Q9BQA1 | 5 | xxMRKETPPPLVP | 6.0417 | 5.9704 | 6.3581 | 5.7162 | 6.2809 | 6.1833 | 5.9815 | 6.3314 | 6.2900 | 6.3116 | 6.2252 | 6.4701 | 6.4429 | 6.4840 | 6.2571 | 6.6557 |
| WDR81_S116 | WDR81 | Q562E7 | 116 | LRKRRLSYPLGGG | 9.2425 | 6.5379 | 7.2395 | 5.5719 | 4.1586 | 7.7275 | 6.0651 | 8.1546 | 6.7260 | 7.5394 | 8.4698 | 7.5241 | 2.6436 | 4.1179 | 4.8246 | 3.4570 |
| WDR81_S1269 | WDR81 | Q562E7 | 686 | GSSSQASPGLLSF | 5.8858 | 6.4264 | 6.9087 | 6.4033 | 6.9245 | 6.4136 | 6.8298 | 6.5283 | 6.6715 | 6.6976 | 6.9985 | 6.3669 | 5.4411 | 5.5042 | 4.6596 | 5.3403 |
| WDR81_S686 | WDR81 | Q562E7 | 1269 | QQFTVSSGESPPL | 5.9846 | 6.3547 | 6.8051 | 6.1131 | 6.3682 | 6.0554 | 5.7687 | 6.5088 | 6.6183 | 6.3742 | 6.3833 | 6.5239 | 6.0885 | 6.2979 | 5.7561 | 5.9991 |
| WDR91_S293 | WDR91 | A4D1P6 | 293 | SSAKKESFGGQGT | 6.6243 | 6.0796 | 6.0635 | 6.0694 | 6.5335 | 5.4080 | 6.0103 | 6.0843 | 7.3996 | 7.4459 | 7.2252 | 6.9268 | 5.5615 | 6.5830 | 5.3853 | 4.5998 |
| WDTC1_S353 | WDTC1 | Q8N5D0 | 353 | ESRGHVSPQVELP | 6.2909 | 5.9521 | 6.2886 | 6.3158 | 5.9676 | 6.3617 | 5.7932 | 6.2206 | 6.0932 | 6.0196 | 7.0454 | 6.4243 | 5.5994 | 7.1937 | 6.3927 | 6.0412 |
| WDTC1_S511 | WDTC1 | Q8N5D0 | 511 | STSRKDSISEDEM | 6.0997 | 5.6167 | 6.0991 | 6.0032 | 6.3487 | 6.3149 | 6.3341 | 6.2849 | 7.2601 | 5.8667 | 5.9902 | 6.2542 | 6.8130 | 6.2501 | 6.5297 | 5.9347 |
| WEE1_S127;S139 | WEE1 | P30291 | 190 | LFDTPHTPKSLLS | 6.1496 | 6.2160 | 6.3644 | 5.4413 | 4.8170 | 6.0989 | 5.8708 | 6.0038 | 7.3449 | 6.8780 | 6.9610 | 6.9690 | 6.4472 | 6.7960 | 5.3561 | 6.2860 |
| WEE1_S150 | WEE1 | P30291 | 642 | KMNRSVSLTIYxx | 6.7717 | 4.7041 | 6.1937 | 5.3207 | 5.2575 | 5.7060 | 5.3015 | 5.5877 | 6.0212 | 5.8907 | 5.6600 | 6.0110 | 7.7784 | 8.1243 | 8.5136 | 7.1579 |
| WEE1_S642 | WEE1 | P30291 | 150 | GGPGDASPRGCGA | 5.3403 | 5.8897 | 6.5110 | 5.3368 | 5.2661 | 5.9834 | 5.5795 | 5.4070 | 6.4615 | 6.0896 | 6.3540 | 6.6670 | 7.2922 | 7.4326 | 7.0643 | 7.3249 |
| WEE1_T187;T190 | WEE1 | P30291 | 187;190 | NA | 7.2650 | 6.3073 | 8.7101 | 8.0960 | 4.8993 | 5.5205 | 7.0934 | 6.2660 | 9.0702 | 7.4576 | 7.5324 | 7.2299 | 2.9011 | 4.1365 | 3.2441 | 4.2705 |
| WEE1_T190 | WEE1 | P30291 | 127;139 | NA | 5.1041 | 6.8882 | 6.5747 | 6.7117 | 6.3683 | 5.9060 | 5.5041 | 6.5528 | 6.7785 | 6.6589 | 7.0606 | 6.0661 | 5.3852 | 6.4757 | 5.2538 | 6.7114 |
| WFS1_S235 | WFS1 | O76024 | 235 | MLERLVSSESKNY | 6.4707 | 5.8941 | 6.7093 | 6.1132 | 6.0067 | 4.6787 | 4.5643 | 5.2564 | 6.3790 | 6.0882 | 4.7568 | 4.9031 | 7.9467 | 8.0142 | 8.5221 | 7.6963 |
| WFS1_S32 | WFS1 | O76024 | 32 | RLNATASLEQERS | 7.2378 | 6.0992 | 7.2128 | 7.9554 | 7.2102 | 5.3944 | 6.5503 | 5.8378 | 6.5669 | 7.0526 | 5.9794 | 6.2126 | 4.4871 | 6.2305 | 5.5200 | 4.4531 |
| WHRN_S685 | WHRN | Q9P202 | 685 | PFPRVQSPPHLKS | 5.9317 | 5.9893 | 8.0363 | 6.0575 | 7.7664 | 7.0522 | 5.9741 | 6.5139 | 6.0560 | 6.1606 | 6.2546 | 7.3524 | 4.8324 | 5.2387 | 5.7376 | 5.0462 |
| WIPF2_S235 | WIPF2 | Q8TF74 | 267 | VPNGPSSPTNESA | 6.1835 | 5.9833 | 6.4530 | 6.7700 | 6.3156 | 6.5853 | 6.4981 | 6.4150 | 6.3392 | 5.9488 | 6.5245 | 6.8664 | 5.8596 | 6.0897 | 5.5280 | 5.6400 |
| WIPF2_S267 | WIPF2 | Q8TF74 | 235 | PVKPPPSPVNIRT | 5.3910 | 5.1399 | 4.9085 | 4.8986 | 6.0991 | 6.8056 | 7.0479 | 7.0499 | 6.6300 | 5.7330 | 5.2948 | 5.2775 | 6.9576 | 7.4890 | 7.5719 | 7.7056 |
| WIPF2_S385 | WIPF2 | Q8TF74 | 385 | RNGHRDSITTVRS | 6.9124 | 5.4549 | 6.8187 | 5.7981 | 6.4597 | 6.5476 | 5.3670 | 5.5417 | 7.4862 | 7.4098 | 8.0141 | 7.3340 | 6.2534 | 7.2303 | 3.6331 | 3.7389 |
| WIPI2_S390 | WIPI2 | Q9Y4P8 | 413 | GTYVPSSPTRLAY | 6.4598 | 6.7577 | 7.0052 | 5.8676 | 6.2105 | 6.7410 | 6.5122 | 6.8501 | 6.6860 | 6.4633 | 6.8470 | 7.0399 | 4.9328 | 5.4292 | 4.9443 | 5.2535 |
| WIPI2_S413 | WIPI2 | Q9Y4P8 | 390 | EILDSASHDCPLV | 8.9272 | 9.6813 | 8.3186 | 7.8143 | 5.1886 | 5.4881 | 5.1675 | 5.1541 | 7.3183 | 7.5959 | 7.4592 | 7.2977 | 3.6029 | 4.1453 | 3.2799 | 3.5611 |
| WIZ_S1006 | WIZ | O95785 | 983 | LGHPPSSPLLKKT | 6.8934 | 6.8437 | 6.1524 | 6.8688 | 6.5877 | 6.4756 | 6.5395 | 6.7225 | 6.7911 | 6.6778 | 6.3841 | 6.4717 | 4.9829 | 5.4106 | 4.9537 | 5.2446 |
| WIZ_S1012 | WIZ | O95785 | 996 | PLALAGSPTPKNP | 7.9702 | 7.5275 | 7.1508 | 7.6678 | 6.9948 | 6.3477 | 7.0140 | 6.6832 | 7.3316 | 6.9003 | 6.9578 | 7.1863 | 3.4333 | 3.7446 | 3.2213 | 3.8687 |
| WIZ_S1012;S1017 | WIZ | A0A669KBH9 | 420 | KAAGMASPWGKQD | 7.9225 | 8.1687 | 7.2613 | 6.9967 | 5.5049 | 7.3454 | 7.7556 | 7.3287 | 7.5414 | 7.4413 | 7.1670 | 7.3858 | 2.6347 | 3.6794 | 2.4712 | 3.3954 |
| WIZ_S1025 | WIZ | O95785 | 1146 | LPPPPGSPLGHSP | 6.2648 | 6.8071 | 6.2952 | 7.1482 | 6.3625 | 6.3703 | 6.3986 | 6.5006 | 6.6162 | 6.4467 | 6.1983 | 6.3509 | 5.5059 | 5.6187 | 5.3315 | 5.7844 |
| WIZ_S106 | WIZ | O95785 | 1106 | PNPPGPSPKALAK | 6.9245 | 6.2835 | 6.9224 | 7.5329 | 6.5827 | 6.6480 | 6.7075 | 6.2306 | 7.5320 | 6.9296 | 6.8251 | 7.0356 | 4.6678 | 4.5851 | 3.7657 | 4.8269 |
| WIZ_S1079 | WIZ | O95785 | 1134 | PSDLHISPLAKKL | 7.3691 | 7.5417 | 7.0047 | 8.0119 | 7.2094 | 6.8764 | 6.8861 | 7.1170 | 7.0961 | 7.3367 | 7.2193 | 7.2423 | 2.9886 | 3.7881 | 2.8990 | 3.4135 |
| WIZ_S1106 | WIZ | O95785 | 1314 | QSPLPLSPLAGRP | 6.7353 | 6.6978 | 6.8162 | 7.4251 | 6.0286 | 5.8154 | 6.5850 | 6.8265 | 7.3778 | 7.0190 | 6.7021 | 7.0143 | 4.8292 | 4.9408 | 4.4533 | 4.7336 |
| WIZ_S1127;S1134 | WIZ | O95785 | 1006 | KNPEDKSPQLSLS | 6.4692 | 6.6229 | 6.4463 | 7.1932 | 6.2846 | 6.3607 | 6.4569 | 6.4677 | 6.4952 | 6.3850 | 6.6070 | 6.6317 | 5.3569 | 5.2172 | 5.2784 | 5.7271 |
| WIZ_S1134 | WIZ | O95785 | 1337 | PRELSLTPITGAK | 6.9817 | 7.1589 | 7.2004 | 7.3765 | 6.4117 | 7.1413 | 6.9799 | 7.1169 | 7.8453 | 7.2488 | 7.4930 | 7.5874 | 3.0875 | 3.8317 | 2.7880 | 3.7511 |
| WIZ_S1146 | WIZ | O95785 | 1335 | QVPRELSLTPITG | 7.1940 | 7.2613 | 6.2429 | 7.2929 | 6.1547 | 6.1982 | 6.1726 | 5.8935 | 6.5297 | 6.2378 | 6.8952 | 6.9756 | 4.7428 | 5.9982 | 4.8282 | 5.3825 |
| WIZ_S1146;S1151;S1155 | WIZ | O95785 | 1012 | SPQLSLSPRPASP | 7.7702 | 7.3908 | 7.0072 | 8.0646 | 5.9508 | 5.8998 | 6.6162 | 7.1767 | 7.9758 | 7.3690 | 7.5495 | 8.2088 | 2.8832 | 3.2668 | 3.1405 | 3.7301 |
| WIZ_S1263 | WIZ | O95785 | 1079 | EWYVNGSPIDTLR | 6.8258 | 7.0129 | 6.5337 | 6.1133 | 6.2552 | 5.9578 | 6.9047 | 6.4570 | 6.4612 | 6.6230 | 7.5657 | 7.0035 | 4.8595 | 4.7355 | 4.9247 | 5.7665 |
| WIZ_S1309 | WIZ | A0A669KBH9 | 361 | DPDAKGSPIDVLH | 8.1891 | 7.8111 | 5.9300 | 8.5783 | 6.4483 | 7.0694 | 7.5717 | 7.1476 | 7.2296 | 6.7678 | 7.6462 | 7.6867 | 2.6464 | 3.7190 | 2.1997 | 3.3589 |
| WIZ_S1314 | WIZ | A0A669KBH9 | 307 | KAQWPQSEDEGPL | 6.7683 | 7.0769 | 8.4460 | 8.1427 | 6.0268 | 7.6273 | 8.6703 | 6.7456 | 4.1196 | 5.2555 | 6.0233 | 5.9620 | 4.3457 | 4.4561 | 5.1608 | 5.1731 |
| WIZ_S1335 | WIZ | A0A669KBH9 | 106 | VAMDLGSPSLPKK | 6.3477 | 7.2657 | 6.9112 | 7.9910 | 6.7870 | 6.6175 | 6.8615 | 7.6698 | 6.5892 | 7.2408 | 7.1081 | 7.0593 | 3.8285 | 4.3697 | 3.3241 | 4.0289 |
| WIZ_S1480 | WIZ | O95785 | 1480 | DSDKRPSLGLAPG | 6.2617 | 6.9739 | 6.3947 | 6.9374 | 8.7286 | 6.0193 | 6.1104 | 6.3796 | 7.6388 | 7.3364 | 7.2999 | 7.4930 | 4.8684 | 4.6592 | 3.5577 | 3.3411 |
| WIZ_S1517 | WIZ | O95785 | 1025 | KAQWPQSEDEGPL | 6.6832 | 7.9357 | 7.8642 | 7.5711 | 7.5946 | 7.8429 | 8.4771 | 8.5741 | 4.3702 | 5.4041 | 5.5960 | 5.6290 | 3.5367 | 4.8275 | 3.7211 | 4.3723 |
| WIZ_S307 | WIZ | O95785 | 1309 | APGPVQSPLPLSP | 6.5429 | 6.7578 | 5.8690 | 6.4552 | 6.4105 | 6.2730 | 6.9366 | 6.8300 | 6.5102 | 7.6019 | 5.5114 | 5.9670 | 5.2914 | 5.4186 | 5.4066 | 6.2180 |

| | | | | | | | | | | | | | | | | | | | | |
|---|---|---|---|---|---|---|---|---|---|---|---|---|---|---|---|---|---|---|---|---|
| WIZ_S361 | WIZ | O95785 | 1263 | EWSVNGSPIDTLR | 6.4978 | 7.4383 | 6.8322 | 7.0443 | 5.9831 | 7.5606 | 6.9953 | 7.1014 | 6.7890 | 7.1185 | 7.6291 | 6.8257 | 3.8900 | 4.6135 | 3.6673 | 4.0140 |
| WIZ_S420 | WIZ | O95785 | 1517 | ERPLAASPPGTVK | 7.2997 | 7.4750 | 7.0603 | 7.8513 | 6.1698 | 6.4618 | 6.8965 | 6.9938 | 8.0786 | 7.6581 | 7.8068 | 7.6605 | 2.7254 | 3.7470 | 2.4002 | 3.7153 |
| WIZ_S983 | WIZ | O95785 | 1127;1134 | NA | 7.0902 | 7.6063 | 7.2442 | 8.0958 | 6.2416 | 7.0493 | 7.4081 | 7.0571 | 7.6255 | 7.3293 | 7.9182 | 7.3819 | 2.6141 | 3.4576 | 2.6214 | 3.2593 |
| WIZ_S996 | WIZ | O95785 | 1012;1017 | NA | 7.2253 | 7.4013 | 7.5374 | 8.3927 | 6.2167 | 6.4690 | 6.6237 | 6.4471 | 7.3985 | 7.3990 | 7.7988 | 8.0949 | 3.0772 | 3.4157 | 2.8661 | 3.6368 |
| WIZ_T1337 | WIZ | O95785 | 1146;1151;11 | NA | 7.0299 | 7.7853 | 6.5712 | 8.4254 | 6.5738 | 7.2432 | 7.5785 | 6.6285 | 7.9746 | 7.2384 | 7.7332 | 7.5806 | 2.4007 | 3.8365 | 2.5355 | 2.8647 |
| WNK1_S167 | WNK1 | Q9H4A3-7 | 2823 | NLQKSISNPPGSN | 4.6095 | 4.6894 | 5.6685 | 4.9324 | 5.7959 | 6.2509 | 5.8301 | 6.3371 | 5.6746 | 5.5813 | 5.6565 | 5.6769 | 8.4709 | 8.6480 | 8.3160 | 7.8620 |
| WNK1_S1674 | WNK1 | Q9H4A3-7 | 2425 | HLNGPSSDPEAAF | 6.2579 | 6.5132 | 6.4984 | 6.0606 | 6.6094 | 6.6936 | 5.9688 | 7.1216 | 6.2742 | 6.1974 | 6.3027 | 6.3380 | 5.7990 | 6.1826 | 5.6320 | 5.5508 |
| WNK1_S185 | WNK1 | Q9H4A3-7 | 2553 | KSSRSSSLGNKSP | 5.9146 | 5.3121 | 6.3800 | 6.0945 | 6.2662 | 6.4720 | 6.0400 | 6.5926 | 6.6863 | 6.9020 | 6.7799 | 6.8249 | 5.7462 | 6.5286 | 5.6260 | 5.8340 |
| WNK1_S19 | WNK1 | Q9H4A3-7 | 2391 | KEGPVASPPFMDL | 6.2971 | 5.6082 | 6.3932 | 6.4599 | 6.2338 | 5.9792 | 6.0449 | 6.7149 | 6.0937 | 6.4096 | 6.1734 | 6.8091 | 6.1304 | 6.5705 | 6.2193 | 5.8628 |
| WNK1_S2218 | WNK1 | Q9H4A3-7 | 19 | PGSLFLSPPAPAP | 6.6627 | 7.6850 | 7.0749 | 8.0604 | 6.2916 | 7.0060 | 7.5849 | 6.6526 | 6.8229 | 7.2486 | 7.6738 | 7.9357 | 2.9611 | 3.9736 | 2.8083 | 3.5578 |
| WNK1_S2262 | WNK1 | Q9H4A3-7 | 185 | PPARSGSGGGSAK | 6.2357 | 6.1826 | 6.6999 | 6.0826 | 5.8501 | 6.4144 | 5.9213 | 6.2166 | 6.3765 | 6.4784 | 6.4834 | 6.8036 | 5.8586 | 6.4816 | 5.9415 | 5.9732 |
| WNK1_S2391 | WNK1 | Q9H4A3-7 | 2534 | PPAAPLSGRRRRP | 6.5126 | 6.9254 | 6.0878 | 6.7297 | 5.8539 | 6.8745 | 6.6669 | 6.5277 | 6.4998 | 6.2365 | 6.7743 | 6.0806 | 5.1547 | 6.2135 | 5.3225 | 5.5397 |
| WNK1_S2418;S2425 | WNK1 | Q9H4A3-7 | 1674 | GRRFIVSPVPESR | 5.7388 | 5.3294 | 5.2718 | 5.2584 | 6.1742 | 6.9800 | 7.2521 | 6.5819 | 6.6339 | 5.4561 | 5.3589 | 5.3509 | 6.6999 | 7.3830 | 6.9973 | 7.5334 |
| WNK1_S2425 | WNK1 | Q9H4A3-7 | 167 | SKDRPVSQPSLVG | 5.7205 | 5.6734 | 6.6431 | 5.4655 | 6.5425 | 7.1105 | 6.5519 | 6.9297 | 6.6459 | 6.3702 | 6.8133 | 6.0859 | 5.4981 | 6.7839 | 5.7734 | 5.3922 |
| WNK1_S2534 | WNK1 | Q9H4A3-7 | 2262 | GPVLATSSGAGVF | 6.0165 | 6.7397 | 6.7784 | 5.8104 | 5.4483 | 6.8804 | 6.3331 | 7.9938 | 5.9476 | 5.5269 | 6.8987 | 6.7928 | 5.0977 | 7.3024 | 5.4309 | 5.0025 |
| WNK1_S2553 | WNK1 | Q9H4A3-7 | 2218 | QLRRTLSPEMITV | 5.5948 | 6.5344 | 6.4572 | 6.2897 | 5.4744 | 6.8162 | 6.2496 | 7.2003 | 6.6137 | 6.9325 | 5.2558 | 6.4692 | 6.0902 | 5.7312 | 6.1373 | 6.1534 |
| WNK1_S2823 | WNK1 | Q9H4A3-7 | 2418;2425 | NA | 6.2476 | 6.7729 | 7.8023 | 6.4676 | 6.5315 | 7.3222 | 6.5242 | 7.0962 | 5.6525 | 5.6648 | 6.3334 | 6.2425 | 5.0651 | 5.8638 | 5.3506 | 5.0629 |
| WNK3_S1070 | WNK3 | Q9BYP7 | 905 | GRHPLPSPKNTSN | 6.0845 | 5.9625 | 6.7694 | 6.0091 | 6.1001 | 6.5430 | 6.5173 | 6.7536 | 6.8993 | 6.2893 | 6.7904 | 6.9792 | 5.4949 | 5.7224 | 5.2254 | 5.8598 |
| WNK3_S1095 | WNK3 | Q9BYP7 | 1585 | IPLPPASPRRPRS | 5.7818 | 7.3762 | 6.7423 | 5.7393 | 6.0615 | 6.8756 | 6.1375 | 6.3043 | 6.2533 | 6.8442 | 6.4300 | 6.3926 | 5.4703 | 6.2032 | 5.1898 | 6.1980 |
| WNK3_S1395 | WNK3 | Q9BYP7 | 734 | IKQRRASCPRPEK | 5.8151 | 5.0046 | 6.4889 | 5.8045 | 6.6258 | 5.9183 | 5.2289 | 6.3245 | 7.5221 | 7.5178 | 7.9680 | 7.4765 | 5.8344 | 6.3849 | 5.2962 | 4.7897 |
| WNK3_S1585 | WNK3 | Q9BYP7 | 1095 | PTTVLESDGERPP | 6.9068 | 6.8276 | 7.4425 | 5.7161 | 6.5375 | 6.7130 | 6.7240 | 6.3230 | 5.7977 | 6.8495 | 6.6033 | 5.9610 | 5.3224 | 5.7928 | 5.0518 | 5.4309 |
| WNK3_S734 | WNK3 | Q9BYP7 | 1070 | VNTEASSPKTVIP | 5.4508 | 5.1005 | 6.3198 | 5.3821 | 7.3330 | 5.7324 | 6.8968 | 4.5681 | 6.2751 | 5.7977 | 6.4778 | 6.7457 | 6.8521 | 6.5361 | 7.9341 | 6.5980 |
| WNK3_S905 | WNK3 | Q9BYP7 | 1395 | NPKTSFSYEKLQA | 3.5374 | 3.6450 | 4.1825 | 3.5083 | 4.2235 | 4.4588 | 4.1874 | 4.4207 | 4.0074 | 4.1318 | 3.8438 | 3.9308 | 12.9925 | 12.9722 | 13.6539 | 12.3038 |
| WRAP53_S103 | WRAP53 | Q9BUR4 | 491 | FPEPTESGDEGEE | 5.9692 | 6.0919 | 6.2002 | 6.2551 | 6.0439 | 6.3654 | 6.1412 | 6.3466 | 6.3925 | 6.2162 | 6.2768 | 6.2459 | 6.5868 | 6.4302 | 6.1597 | 6.2783 |
| WRAP53_S16;S30 | WRAP53 | Q9BUR4 | 103 | ELSENTSLPAEEA | 6.5321 | 5.8092 | 6.7162 | 6.1485 | 5.8289 | 5.5255 | 6.2234 | 6.0253 | 6.5891 | 6.2344 | 6.2620 | 5.9876 | 6.5281 | 6.4970 | 6.6000 | 6.4926 |
| WRAP53_S26;S30 | WRAP53 | Q9BUR4 | 26;30 | NA | 5.9679 | 6.3645 | 6.6803 | 6.3389 | 6.2788 | 6.8038 | 6.3496 | 6.3484 | 6.4291 | 6.4173 | 6.4179 | 6.7584 | 5.9146 | 5.3447 | 5.7107 | 5.8751 |
| WRAP53_S491 | WRAP53 | Q9BUR4 | 16;30 | NA | 6.3021 | 5.8959 | 6.4801 | 6.4809 | 6.6532 | 6.7644 | 6.5950 | 6.3992 | 6.2483 | 6.6721 | 6.9252 | 6.4433 | 6.1425 | 5.8606 | 5.4052 | 4.7320 |
| WRN_S1133 | WRN | Q14191 | 1133 | KSIMVQSPEKAYS | 6.6083 | 6.6106 | 6.4416 | 6.5839 | 6.4435 | 6.0327 | 6.5017 | 6.4826 | 6.6852 | 6.4358 | 6.3691 | 6.2909 | 5.7035 | 5.6213 | 5.5598 | 5.6294 |
| WRNIP1_S153 | WRNIP1 | Q96S55 | 75 | AKRRRLSESSALK | 5.3751 | 5.4930 | 5.5189 | 5.5482 | 6.9404 | 6.1844 | 5.7197 | 6.5604 | 5.9778 | 6.1436 | 6.0123 | 4.0099 | 8.3924 | 7.6721 | 6.3083 | 8.1435 |
| WRNIP1_S65 | WRNIP1 | Q96S55 | 153 | AAAGSASPRSWDE | 5.7364 | 6.1642 | 5.9121 | 5.8680 | 6.0022 | 5.9698 | 5.9250 | 5.9825 | 6.2402 | 6.1936 | 6.0897 | 6.1115 | 6.8729 | 7.0131 | 7.0176 | 6.9012 |
| WRNIP1_S75 | WRNIP1 | Q96S55 | 65 | ERAKGPSPPGAKR | 6.0385 | 6.5506 | 6.1102 | 6.2150 | 6.2240 | 6.2952 | 6.0035 | 6.2274 | 6.3301 | 6.3194 | 6.2046 | 6.4185 | 6.0187 | 6.4787 | 6.2703 | 6.2952 |
| WRNIP1_S75;S78 | WRNIP1 | Q96S55 | 116 | SYDAPPTPSGARL | 6.3033 | 6.6256 | 5.8015 | 6.2165 | 6.1762 | 6.2236 | 6.1119 | 5.6735 | 5.8135 | 5.0682 | 5.7363 | 5.6069 | 7.2234 | 7.3518 | 6.9807 | 7.0869 |
| WRNIP1_T116 | WRNIP1 | Q96S55 | 75;78 | NA | 5.9651 | 6.2203 | 6.0230 | 5.8824 | 6.3219 | 6.3274 | 6.2446 | 6.0071 | 5.3119 | 5.1271 | 5.7375 | 5.2798 | 7.3894 | 7.1024 | 7.3948 | 7.6652 |
| WTAP_S14 | WTAP | Q15007 | 306 | EDDFPSSPGNGNK | 6.9112 | 6.4237 | 5.8424 | 6.9634 | 6.5180 | 6.2505 | 5.6103 | 5.9229 | 6.5467 | 6.3731 | 6.6621 | 6.4313 | 6.3302 | 5.8294 | 5.8713 | 5.5137 |
| WTAP_S306 | WTAP | Q15007 | 14 | PKKVRLSETDFKV | 7.1316 | 6.5955 | 6.4136 | 7.1150 | 6.8057 | 6.2728 | 6.2039 | 6.6336 | 6.3745 | 6.1813 | 7.0191 | 6.5645 | 5.6903 | 5.0867 | 4.7430 | 5.1689 |
| WTIP_S422 | WTIP | A6NIX2 | 88 | SAQPAGSPRASLA | 6.7405 | 6.4549 | 6.8954 | 6.9196 | 6.2376 | 6.1622 | 6.2216 | 6.7530 | 6.6358 | 5.9761 | 6.1012 | 6.6334 | 5.3878 | 5.6580 | 5.5324 | 5.6907 |
| WTIP_S88 | WTIP | A6NIX2 | 422 | QPGPLPSPTVHVT | 6.3563 | 7.6188 | 7.1156 | 7.2737 | 6.7270 | 6.9430 | 6.0790 | 6.7693 | 6.0946 | 5.8641 | 6.5277 | 6.5088 | 5.2418 | 4.7897 | 4.3043 | 5.7862 |
| WWC1_S876 | WWC1 | Q8IX03-2 | 876 | VFTEKASPDMDGY | 6.0558 | 6.6722 | 7.0759 | 7.2257 | 6.2625 | 5.8808 | 5.8403 | 6.5084 | 6.3233 | 6.2478 | 6.6998 | 6.8110 | 5.7974 | 5.4507 | 5.1500 | 5.9982 |
| WWC1_S974 | WWC1 | Q8IX03-2 | 974 | VRMKRPSPPPQPS | 6.1784 | 6.1198 | 6.8458 | 6.3992 | 6.7342 | 6.4535 | 6.5068 | 7.2164 | 6.8412 | 7.1499 | 7.3985 | 6.9481 | 4.6789 | 4.7344 | 4.8676 | 4.9274 |
| WWC1_T527 | WWC1 | Q8IX03-2 | 527 | LRSLSGTPKSMTS | 5.1594 | 6.3691 | 6.0983 | 6.4521 | 5.8511 | 6.5794 | 5.9704 | 7.1350 | 6.3182 | 7.0056 | 6.6932 | 7.3060 | 5.4826 | 5.5314 | 6.0083 | 6.0399 |
| WWC2_S1046 | WWC2 | Q6AWC2-6 | 269 | EPDLRCSPVNSHL | 6.6079 | 6.3034 | 6.7401 | 6.0166 | 5.9231 | 6.2760 | 6.4921 | 5.9642 | 6.6816 | 6.2051 | 6.6822 | 6.5136 | 5.7611 | 5.9203 | 5.7310 | 6.1818 |
| WWC2_S269 | WWC2 | Q6AWC2-6 | 533 | AAATGHTPPLAEA | 6.2728 | 5.8536 | 6.9698 | 6.3296 | 5.9250 | 6.5804 | 6.6909 | 6.3174 | 6.7446 | 6.1978 | 5.7168 | 7.2829 | 6.0898 | 5.8816 | 5.2286 | 5.9183 |
| WWC2_T1028 | WWC2 | Q6AWC2-6 | 1028 | VIVRSQTFSPGER | 6.2341 | 6.8352 | 8.1036 | 7.2235 | 6.4683 | 5.6702 | 5.8191 | 5.4720 | 6.1391 | 5.3586 | 5.2961 | 5.6838 | 6.8057 | 6.5591 | 6.4854 | 5.8462 |
| WWC2_T533 | WWC2 | Q6AWC2-6 | 1046 | RLNRSDSDSSTLA | 6.4810 | 6.2100 | 6.7241 | 5.9734 | 6.2874 | 6.5010 | 6.3837 | 6.2580 | 6.8491 | 6.6688 | 6.0085 | 6.7953 | 5.4409 | 6.0527 | 5.4038 | 5.9623 |
| WWP2_S211 | WWP2 | O00308 | 211 | PATGEQSPGARSR | 6.9173 | 6.9046 | 6.7430 | 6.5248 | 6.1403 | 6.5285 | 6.6228 | 6.5369 | 6.7597 | 6.4077 | 6.5487 | 6.6743 | 5.1703 | 5.3592 | 5.0166 | 5.1454 |
| WWTR1_S117 | WWTR1 | Q9GZV5 | 393 | ESALNKSEPFLTW | 8.1124 | 6.2377 | 7.2764 | 6.3003 | 5.9533 | 5.1371 | 6.7716 | 8.8453 | 5.8839 | 7.3165 | 7.3453 | 8.6392 | 3.0156 | 4.4191 | 3.8045 | 4.9418 |
| WWTR1_S296 | WWTR1 | Q9GZV5 | 117 | AHLRQQSYDVTDE | 6.4394 | 5.8228 | 6.4705 | 5.0062 | 6.2443 | 5.2325 | 6.4312 | 5.8671 | 6.0180 | 5.9885 | 6.2328 | 6.8805 | 4.9521 | 7.1379 | 8.2366 | 7.0396 |
| WWTR1_S393 | WWTR1 | Q9GZV5 | 296 | SITNNSSDPFLNG | 6.2048 | 4.5565 | 5.6183 | 5.9007 | 5.2996 | 6.0628 | 5.2780 | 4.6785 | 5.1546 | 5.6615 | 5.3845 | 6.2318 | 7.7704 | 9.2226 | 8.3684 | 8.6069 |
| XAGE1A_S20 | XAGE1A | Q9HD64 | 20 | GILHLGSRQKKIR | 8.1439 | 6.7779 | 8.1697 | 7.7787 | 5.9347 | 6.6391 | 6.4406 | 6.3806 | 8.2804 | 8.1810 | 8.3972 | 8.8372 | 2.1339 | 3.4389 | 2.0726 | 2.3936 |
| XAGE1A_S56 | XAGE1A | Q9HD64 | 56 | INLDLGSGVKVKI | 6.0745 | 5.7910 | 7.0236 | 7.4995 | 6.8963 | 6.6409 | 6.7870 | 5.8858 | 6.9831 | 6.9180 | 6.1664 | 6.3820 | 5.0409 | 5.6622 | 5.4185 | 4.8304 |
| XPC_S347 | XPC | Q01831 | 94 | IKDEALSDGDDLR | 6.0908 | 6.0720 | 5.6900 | 6.6877 | 6.9653 | 7.1401 | 6.6947 | 6.6182 | 5.7375 | 6.0667 | 6.7599 | 6.0955 | 6.0591 | 6.4436 | 5.6490 | 5.2299 |
| XPC_S351 | XPC | Q01831 | 351 | GSSETSSQVLENH | 5.8099 | 7.9205 | 6.2377 | 6.5201 | 5.3621 | 5.5646 | 5.8755 | 7.3044 | 5.7215 | 4.3923 | 5.3763 | 5.0655 | 7.3016 | 8.1601 | 6.1949 | 7.1932 |
| XPC_S398;S399 | XPC | Q01831 | 347 | ADPGGSSETSSQV | 6.1758 | 7.2234 | 5.7503 | 6.8130 | 5.7030 | 6.2571 | 5.9451 | 6.8395 | 6.8735 | 5.6804 | 4.6611 | 4.7179 | 6.7809 | 7.0021 | 6.1607 | 7.4162 |
| XPC_S94 | XPC | Q01831 | 398;399 | NA | 6.4374 | 6.1796 | 6.0380 | 7.0527 | 5.6142 | 5.1592 | 4.8158 | 5.4773 | 8.1800 | 8.1148 | 6.6758 | 7.6394 | 5.4206 | 6.4951 | 5.1468 | 5.5534 |
| XPNPEP1_T478 | XPNPEP1 | Q9NQW7-3 | 478 | RTMHFGTPTAYEK | 6.4838 | 6.1460 | 7.3654 | 5.9780 | 7.1116 | 7.0020 | 5.6995 | 7.1767 | 6.4608 | 6.5753 | 6.9567 | 6.2482 | 4.9218 | 5.1648 | 5.5174 | 5.1920 |
| XPO1_S1055 | XPO1 | O14980 | 1055 | KHKRQMSVPGIFN | 4.4464 | 5.6461 | 6.5346 | 4.8044 | 5.3368 | 6.2422 | 6.1747 | 5.8166 | 6.2871 | 5.6817 | 5.8373 | 6.2309 | 6.6339 | 9.1795 | 7.5318 | 7.6162 |
| XPO1_S391 | XPO1 | O14980 | 984 | VANLLKSAFPHLQ | 4.1321 | 4.0810 | 14.3308 | 11.8534 | 14.1142 | 8.2582 | 3.9922 | 3.3905 | 3.6072 | 8.2883 | 4.7338 | 3.8584 | 3.4462 | 4.3698 | 3.7877 | 3.7561 |
| XPO1_S984 | XPO1 | O14980 | 391 | PFSTSASPLLSGS | 6.4959 | 7.5287 | 6.1065 | 6.4557 | 5.8561 | 7.0278 | 5.8807 | 7.4912 | 6.0498 | 6.6851 | 7.2618 | 5.7546 | 4.6843 | 5.9365 | 5.3899 | 5.3954 |
| XPO4_S521 | XPO4 | Q9C0E2 | 521 | QQQLLASPGSSTV | 6.2058 | 6.5595 | 6.6493 | 6.4376 | 6.2249 | 6.8121 | 6.5291 | 6.4820 | 6.5390 | 6.7952 | 7.0050 | 6.8211 | 4.8469 | 5.5656 | 5.1381 | 5.3888 |
| XPO6_T204;S208 | XPO6 | Q96QU8 | 204;208 | NA | 5.4917 | 5.7246 | 6.1335 | 5.6542 | 6.0225 | 6.0271 | 5.7394 | 5.9549 | 6.5719 | 6.1519 | 6.0825 | 6.1674 | 6.8043 | 7.3612 | 7.1664 | 6.9466 |
| XPR1_S666 | XPR1 | Q9UBH6 | 467 | IQCLRRYRDTKRA | 5.9621 | 4.9898 | 5.4946 | 5.3544 | 5.7893 | 5.3548 | 4.8870 | 5.6827 | 5.8800 | 6.5331 | 5.6868 | 6.5817 | 7.8741 | 7.6239 | 8.0106 | 8.2951 |
| XPR1_Y467 | XPR1 | Q9UBH6 | 666 | SWKYNQSISLRRP | 8.7774 | 7.8927 | 7.6644 | 7.7946 | 6.4519 | 5.0003 | 6.4550 | 6.0358 | 6.1279 | 5.8176 | 5.5484 | 5.0769 | 5.7376 | 6.0404 | 4.9780 | 4.6008 |
| XRCC1_S199 | XRCC1 | P18887 | 447 | QAAGPSSPQKPPT | 5.7167 | 6.1486 | 6.4047 | 5.9336 | 5.7663 | 6.3228 | 5.8958 | 6.1414 | 6.7154 | 6.9323 | 7.1753 | 7.0262 | 5.5712 | 6.2870 | 5.8694 | 6.0932 |
| XRCC1_S226 | XRCC1 | P18887 | 257 | NQEEKKTPSKPPA | 6.6756 | 6.7273 | 7.2512 | 6.3003 | 5.8924 | 6.6713 | 6.0796 | 6.4989 | 8.1072 | 8.0774 | 8.0035 | 8.0430 | 3.6626 | 4.7314 | 3.3607 | 3.9176 |
| XRCC1_S241 | XRCC1 | P18887 | 241 | TSKPQESPKGKRK | 5.7528 | 5.8445 | 6.5107 | 5.3664 | 7.0345 | 7.0100 | 6.4297 | 6.3560 | 7.1315 | 7.0166 | 7.4956 | 6.8325 | 5.6491 | 6.1313 | 4.8407 | 4.5983 |
| XRCC1_S447 | XRCC1 | P18887 | 281 | LPAPTRTPATAPV | 6.2988 | 7.3302 | 7.3520 | 7.2248 | 5.2634 | 7.1818 | 6.8370 | 6.6797 | 7.7862 | 7.6876 | 7.5441 | 8.3172 | 3.1050 | 4.3601 | 2.8049 | 4.2272 |
| XRCC1_S447;T453 | XRCC1 | P18887 | 226 | SAASSASPVSRAI | 5.1027 | 6.5299 | 7.5446 | 6.1317 | 5.8422 | 5.9841 | 5.9070 | 6.1008 | 7.2551 | 7.1082 | 7.5006 | 7.1766 | 5.7393 | 5.8545 | 4.3588 | 5.8638 |
| XRCC1_T257 | XRCC1 | P18887 | 199 | SRINKTSPVTASD | 4.6151 | 5.4145 | 6.4216 | 6.1263 | 7.0457 | 5.7364 | 6.7642 | 6.8400 | 6.3675 | 5.4411 | 7.2564 | 6.0879 | 5.7876 | 6.5762 | 6.6933 | 6.8262 |
| XRCC1_T257;S266 | XRCC1 | P18887 | 447;453 | NA | 5.1359 | 5.2292 | 5.7717 | 5.1489 | 5.6207 | 5.7265 | 5.9234 | 5.3460 | 6.0321 | 6.3646 | 6.0933 | 6.2139 | 7.9609 | 7.8327 | 7.9064 | 7.6937 |
| XRCC1_T281 | XRCC1 | P18887 | 257;266 | NA | 6.3544 | 7.5026 | 7.1136 | 7.3374 | 6.1077 | 7.2050 | 7.3510 | 6.8796 | 7.6879 | 7.3509 | 7.8867 | 7.7802 | 2.9147 | 4.1504 | 2.6300 | 3.7479 |
| XRCC5_S145 | XRCC5 | P13010 | 145 | SSRFSKSQLDIII | 6.0108 | 5.8846 | 6.3511 | 5.9923 | 6.5063 | 6.2185 | 7.0181 | 5.9366 | 5.2470 | 6.1538 | 5.7018 | 6.2746 | 6.4113 | 6.7611 | 7.0181 | 6.5140 |
| XRCC5_S324 | XRCC5 | P13010 | 341 | SEGKCFSVLGFCK | 6.1531 | 5.8148 | 7.5720 | 6.9968 | 7.1593 | 5.5108 | 6.5023 | 5.2031 | 6.4010 | 5.6182 | 4.7256 | 5.2458 | 6.7893 | 7.0017 | 7.7344 | 5.5718 |
| XRCC5_S341 | XRCC5 | P13010 | 324 | SDIVPFSKVDEEQ | 4.0889 | 4.9826 | 5.4524 | 6.2647 | 5.6459 | 6.1388 | 6.3614 | 6.3000 | 5.8578 | 6.1173 | 5.2114 | 5.3250 | 7.1261 | 8.2761 | 8.8089 | 8.0427 |
| XRCC5_S436 | XRCC5 | P13010 | 437 | RQYMFSSLKNSKK | 5.7647 | 6.0613 | 6.5117 | 6.1166 | 6.6600 | 6.5914 | 5.3198 | 5.8001 | 6.3900 | 6.7231 | 5.4847 | 6.0193 | 6.9954 | 7.0197 | 7.1846 | 5.3575 |
| XRCC5_S437 | XRCC5 | P13010 | 436 | LRQYMFSSLKNSK | 4.3902 | 5.4151 | 5.2597 | 6.4498 | 5.2101 | 5.8746 | 6.4134 | 7.9606 | 6.9655 | 4.4464 | 6.4068 | 6.3308 | 7.2531 | 5.3159 | 8.3342 | 7.9738 |
| XRCC6_S237 | XRCC6 | P12956 | 455 | TEKIMATPEQVGK | 5.9908 | 6.5841 | 6.3738 | 6.4273 | 6.1869 | 5.9852 | 6.1651 | 6.3381 | 6.3811 | 6.3840 | 6.0995 | 5.9467 | 6.1369 | 6.3443 | 6.3286 | 6.3277 |
| XRCC6_S27 | XRCC6 | P12956 | 520 | MNKRLGSLVDEFK | 5.9560 | 5.9412 | 6.1916 | 5.8955 | 5.9439 | 5.9608 | 6.1113 | 5.3471 | 5.6610 | 5.7154 | 5.9987 | 5.7521 | 7.2819 | 7.6169 | 7.5474 | 7.0792 |
| XRCC6_S306 | XRCC6 | P12956 | 477 | FTYRSDSFENPVL | 3.6915 | 5.5951 | 7.9490 | 4.5152 | 4.3568 | 7.5352 | 5.5970 | 8.1529 | 5.2531 | 11.7443 | 8.5415 | 8.0331 | 3.8039 | 4.9892 | 4.8979 | 5.3442 |
| XRCC6_S477 | XRCC6 | P12956 | 237 | VHFEESSKLEDLL | 6.3683 | 6.4770 | 5.9859 | 6.9128 | 6.3531 | 5.7692 | 6.3538 | 5.7261 | 6.2657 | 6.1219 | 5.4081 | 5.6017 | 6.8964 | 5.7516 | 6.8399 | 7.1683 |
| XRCC6_S520 | XRCC6 | P12956 | 306 | TRTFNTSTGGLLL | 6.0589 | 6.0839 | 6.2439 | 6.9806 | 6.2679 | 6.3139 | 6.4085 | 5.8207 | 6.8200 | 6.2225 | 6.7381 | 6.2431 | 5.6393 | 6.1846 | 5.5980 | 6.3760 |
| XRCC6_T428 | XRCC6 | P12956 | 428 | DQKIQVTPPGFQL | 5.5613 | 5.7078 | 6.0751 | 5.8342 | 6.1186 | 6.8629 | 6.2221 | 5.7724 | 6.0757 | 6.1901 | 5.9166 | 5.8262 | 6.5614 | 6.9676 | 7.3068 | 7.0013 |
| XRCC6_T455 | XRCC6 | P12956 | 27 | EENLEASGDYKYS | 5.7596 | 6.1096 | 6.0730 | 5.6038 | 5.9302 | 6.1522 | 6.1696 | 6.3916 | 6.2445 | 6.5527 | 5.5089 | 6.3995 | 7.1534 | 5.9807 | 6.7625 | 7.2082 |
| XRN2_S448 | XRN2 | Q9H0D6 | 678 | EETRRNSLGGDVL | 5.8582 | 5.7993 | 5.8640 | 6.1829 | 6.1735 | 5.6800 | 6.0405 | 6.0084 | 7.2284 | 6.9134 | 6.9829 | 7.0544 | 6.1394 | 5.9431 | 5.6557 | 6.4760 |
| XRN2_S471 | XRN2 | Q9H0D6 | 471 | RMQNNSSPSISPN | 6.0498 | 6.1766 | 6.4789 | 6.6970 | 6.4884 | 6.6237 | 6.2841 | 6.3719 | 6.3365 | 6.3130 | 6.4177 | 6.1029 | 5.9140 | 6.2247 | 5.8673 | 5.6534 |
| XRN2_S471;S475 | XRN2 | Q9H0D6 | 448 | ALGSRNSPGSQVA | 6.5413 | 6.5880 | 6.2245 | 6.7151 | 6.3021 | 6.1318 | 6.7786 | 6.3736 | 6.8739 | 6.5444 | 6.8863 | 6.8665 | 5.4302 | 5.5851 | 4.8615 | 5.2972 |
| XRN2_S471;S475;S487 | XRN2 | Q9H0D6 | 485 | SFTSDGSPSPLGG | 6.4703 | 6.5047 | 6.3230 | 6.4477 | 6.9857 | 6.3314 | 6.0126 | 5.7543 | 6.5820 | 7.1700 | 6.4207 | 6.4124 | 5.9102 | 5.7667 | 5.3983 | 5.5100 |
| XRN2_S475 | XRN2 | Q9H0D6 | 439 | TPSGILTPHALGS | 6.1183 | 6.5887 | 7.0850 | 5.8887 | 5.4593 | 6.4035 | 6.7641 | 7.1161 | 7.1220 | 6.7932 | 6.6886 | 6.9354 | 4.5601 | 5.9926 | 4.4695 | 6.0151 |
| XRN2_S475;S482 | XRN2 | Q9H0D6 | 475 | NSSPSISPNTSFT | 5.7715 | 6.0518 | 5.8003 | 6.3876 | 6.0761 | 5.7567 | 6.1292 | 5.8807 | 6.2021 | 5.9559 | 5.9650 | 6.5499 | 7.1410 | 7.3644 | 6.6459 | 6.3219 |
| XRN2_S485 | XRN2 | Q9H0D6 | 487 | TSDGSPSPLGGIK | 6.3143 | 6.4387 | 5.9058 | 6.9396 | 6.0407 | 6.5645 | 6.3167 | 6.1019 | 6.1091 | 6.6329 | 6.5105 | 6.3507 | 5.9393 | 5.9302 | 5.8766 | 6.0287 |
| XRN2_S487 | XRN2 | Q9H0D6 | 499;501 | NA | 5.7614 | 6.1101 | 6.2698 | 6.2996 | 6.2361 | 6.3263 | 5.9820 | 6.1244 | 6.3861 | 6.3564 | 6.3152 | 6.2708 | 6.7540 | 6.6405 | 6.4563 | 5.7108 |
| XRN2_S499;S501 | XRN2 | Q9H0D6 | 471;475 | NA | 5.4554 | 6.0209 | 5.8060 | 6.3114 | 5.6420 | 5.9799 | 5.7537 | 5.6388 | 6.0112 | 6.5256 | 5.9606 | 6.4061 | 7.1233 | 6.8842 | 7.2269 | 7.2541 |
| XRN2_S678 | XRN2 | Q9H0D6 | 471;475;487 | NA | 5.3834 | 5.9272 | 7.0282 | 6.2022 | 6.2352 | 5.6409 | 5.2859 | 7.7447 | 6.7169 | 5.2539 | 6.4197 | 6.5576 | 6.9119 | 6.6026 | 6.2152 | 5.8746 |
| XRN2_T439 | XRN2 | Q9H0D6 | 475;482 | NA | 5.3883 | 6.4496 | 6.3826 | 6.8435 | 6.8882 | 5.9706 | 6.4259 | 6.6972 | 6.0331 | 6.6389 | 6.4542 | 7.2794 | 6.4061 | 5.4243 | 4.5981 | 6.1198 |
| YAP1_S109 | YAP1 | P46937-9 | 289 | PPLAPQSPQGGVM | 6.4946 | 5.3657 | 6.6110 | 5.9308 | 6.0411 | 5.3886 | 5.9858 | 7.5887 | 6.1089 | 6.8525 | 6.5661 | 7.4035 | 5.4346 | 6.2569 | 5.5040 | 6.4672 |
| YAP1_S164 | YAP1 | P46937-9 | 164 | QHLRQSSFEIPDD | 5.7254 | 4.8438 | 5.4159 | 4.5545 | 5.8319 | 4.9354 | 5.2707 | 7.1050 | 5.0347 | 6.7508 | 5.8128 | 7.1881 | 7.0832 | 9.0049 | 7.4256 | 8.0173 |
| YAP1_S276 | YAP1 | P46937-9 | 371 | TQNPVSSPGMSQE | 6.4885 | 6.1593 | 6.5424 | 6.2787 | 6.1975 | 5.8546 | 6.1145 | 6.5901 | 6.1621 | 6.8059 | 6.3184 | 7.5716 | 5.2672 | 6.2198 | 5.4352 | 5.9944 |
| YAP1_S289 | YAP1 | P46937-9 | 61 | VHVRGDSETDLEA | 5.6147 | 4.9109 | 5.2616 | 4.5765 | 5.5831 | 5.0261 | 5.1746 | 6.3068 | 4.9928 | 6.1511 | 5.5904 | 6.4786 | 7.7118 | 9.4153 | 8.6336 | 8.5720 |
| YAP1_S344 | YAP1 | P46937-9 | 276 | NQRISQSAPVKQP | 9.2815 | 6.9205 | 6.8500 | 6.4173 | 6.7053 | 5.2484 | 6.3616 | 7.6450 | 6.0408 | 7.0531 | 6.6968 | 7.9826 | 3.2669 | 5.4170 | 3.6074 | 4.5058 |
| YAP1_S371 | YAP1 | P46937-9 | 109 | SHSRQASTDAGTA | 6.3994 | 5.8717 | 6.3615 | 5.8825 | 5.6391 | 5.1818 | 5.4910 | 6.6444 | 5.9614 | 6.7909 | 6.1205 | 6.6029 | 6.1444 | 7.4353 | 6.4939 | 6.9793 |
| YAP1_S61 | YAP1 | P46937-9 | 344 | NPSTANSPKCQEL | 6.7436 | 6.6568 | 6.8068 | 7.0035 | 5.8099 | 5.9383 | 6.6464 | 6.7580 | 6.9357 | 8.3236 | 7.6473 | 9.1073 | 3.1428 | 4.7860 | 3.3738 | 4.3202 |
| YARS1_S205 | YARS1 | P54577 | 205 | LPALGYSKRVHLM | 5.7627 | 5.1405 | 4.9818 | 5.7284 | 5.4581 | 6.7368 | 6.4321 | 5.8099 | 6.9298 | 6.2605 | 7.1959 | 6.9470 | 6.9427 | 7.1058 | 6.3120 | 6.2562 |
| YARS1_S338 | YARS1 | P54577 | 338 | ALKKLASAAYPDP | 6.4381 | 6.6387 | 5.7125 | 5.9855 | 6.5084 | 6.5770 | 6.4961 | 6.4741 | 6.0338 | 6.4226 | 7.3729 | 6.5132 | 5.5109 | 5.6064 | 5.8525 | 5.8573 |
| YBX1_S165 | YBX1 | P67809 | 32 | KPGTTGSGAGSGG | 5.6190 | 5.3025 | 6.4300 | 5.6894 | 5.8829 | 5.9937 | 6.5722 | 6.0736 | 6.7723 | 5.8926 | 5.8602 | 6.5978 | 7.4110 | 6.4934 | 5.8703 | 7.5392 |
| YBX1_S165;S167 | YBX1 | P67809 | 167 | NYQNSESGEKNEG | 5.6260 | 5.3645 | 6.2279 | 5.3521 | 5.5301 | 6.0933 | 5.6306 | 5.9331 | 5.7614 | 6.4546 | 6.1335 | 6.3076 | 8.1272 | 6.4863 | 7.3605 | 7.6112 |
| YBX1_S167 | YBX1 | P67809 | 165 | QQNYQNSESGEKN | 5.3134 | 5.1939 | 5.9068 | 6.0419 | 5.8437 | 6.5775 | 6.3125 | 6.1325 | 5.5026 | 6.3515 | 6.4090 | 6.3036 | 6.9757 | 7.2997 | 6.8462 | 6.9895 |
| YBX1_S174 | YBX1 | P67809 | 174 | GEKNEGSESAPEG | 6.7722 | 5.9938 | 7.4104 | 5.8881 | 6.7377 | 7.5690 | 7.4409 | 6.7519 | 6.6198 | 5.9044 | 5.6035 | 6.3881 | 5.4613 | 5.1401 | 4.1727 | 6.1462 |
| YBX1_S176 | YBX1 | P67809 | 176 | KNEGSESAPEGQA | 5.8480 | 7.1893 | 6.9773 | 7.1272 | 7.4561 | 7.2697 | 5.5080 | 5.4240 | 7.2844 | 4.9890 | 5.0658 | 5.6944 | 5.1389 | 6.8713 | 4.9598 | 7.1966 |
| YBX1_S32 | YBX1 | P67809 | 165;167 | NA | 6.0685 | 6.2223 | 7.6818 | 6.6838 | 6.4505 | 6.8306 | 6.4908 | 6.1394 | 5.8296 | 6.0579 | 6.0760 | 6.2226 | 5.8131 | 5.9088 | 5.6564 | 5.8679 |
| YBX2_S189 | YBX2 | Q9Y2T7 | 189 | FIPRPPSVAPPPM | 6.4091 | 6.3226 | 6.6653 | 6.2813 | 6.6813 | 6.4229 | 6.5202 | 6.4101 | 6.6726 | 6.2025 | 6.6941 | 6.1417 | 5.3649 | 5.9226 | 5.7184 | 5.5705 |
| YBX2_T76 | YBX2 | Q9Y2T7 | 76 | ATAVSGTPAPPAR | 7.3543 | 6.9195 | 7.2608 | 6.9365 | 6.5623 | 5.9458 | 6.8611 | 7.1981 | 6.5188 | 6.0542 | 7.0854 | 6.8930 | 4.3320 | 4.6775 | 4.3714 | 5.0294 |
| YBX3_S203 | YBX3 | P16989 | 203 | EEEGSGSSEGFDP | 6.0157 | 6.0224 | 6.8046 | 6.9092 | 5.7854 | 6.1639 | 6.2073 | 6.5444 | 6.7545 | 6.5067 | 6.7120 | 6.6816 | 5.8310 | 6.1736 | 5.4947 | 5.3931 |
| YBX3_S324 | YBX3 | P16989 | 346 | RPPNAPSQDGKEA | 5.4626 | 6.7834 | 6.6023 | 6.0561 | 5.9173 | 6.0088 | 6.5108 | 6.5324 | 6.5802 | 6.3317 | 6.0536 | 6.7965 | 5.9384 | 6.3674 | 6.0707 | 5.9877 |
| YBX3_S346 | YBX3 | P16989 | 324 | SGPNQPSVRRGYR | 6.6332 | 6.9837 | 9.1089 | 6.7271 | 5.5081 | 6.5576 | 7.4687 | 6.4618 | 6.8539 | 8.6035 | 8.6354 | 6.8346 | 4.1275 | 4.3279 | 2.1107 | 3.0576 |
| YBX3_S370 | YBX3 | P16989 | 370 | PPTQQSSAExxxx | 6.4805 | 5.7635 | 6.2475 | 6.0633 | 6.4111 | 5.9391 | 7.0620 | 7.0047 | 5.6805 | 5.8244 | 6.1122 | 6.6115 | 6.2244 | 5.9948 | 6.3040 | 6.2766 |
| YEATS2_S118 | YEATS2 | Q9ULM3 | 627 | GHMIAVSPQKQVI | 6.6661 | 6.5342 | 6.0257 | 6.4162 | 6.5320 | 6.3459 | 6.4813 | 6.2335 | 6.4552 | 6.4103 | 6.3458 | 6.3306 | 5.8638 | 5.8372 | 5.8092 | 5.7129 |
| YEATS2_S447 | YEATS2 | Q9ULM3 | 447 | QVPNPESPGKSFQ | 6.3892 | 6.3411 | 6.2160 | 6.6303 | 6.4080 | 6.2863 | 6.5622 | 6.3859 | 6.4316 | 6.0697 | 6.3271 | 6.4115 | 5.9697 | 6.0436 | 5.8677 | 5.6601 |
| YEATS2_S465 | YEATS2 | Q9ULM3 | 118 | IKKFLESPSRSSS | 6.6883 | 6.8446 | 6.2657 | 6.7051 | 6.4942 | 6.0367 | 6.4413 | 6.2932 | 6.6089 | 6.3526 | 6.1818 | 6.1715 | 5.7113 | 5.7682 | 5.6611 | 5.7756 |
| YEATS2_S627 | YEATS2 | Q9ULM3 | 478 | PSPLPRTPTSTPV | 5.4908 | 6.1459 | 5.2974 | 6.1248 | 5.4382 | 6.0154 | 5.9719 | 6.2773 | 6.2395 | 5.9171 | 5.7193 | 5.7791 | 7.3192 | 7.2476 | 7.2721 | 7.7443 |
| YEATS2_T478 | YEATS2 | Q9ULM3 | 465 | CKIVSGSPISTPS | 5.3510 | 5.2341 | 5.7963 | 6.0310 | 6.5320 | 5.5452 | 5.5744 | 6.1083 | 6.2756 | 5.7294 | 6.3981 | 5.8369 | 7.1182 | 7.2803 | 7.5277 | 7.6615 |
| YES1_S195 | YES1 | P07947 | 427 | IEDNEYTARQGAK | 5.9500 | 5.8635 | 5.8135 | 5.5397 | 5.1314 | 6.6322 | 6.3842 | 6.5883 | 7.5132 | 7.8761 | 7.0665 | 7.2948 | 5.3943 | 6.6673 | 4.8659 | 5.4192 |
| YES1_T427 | YES1 | P07947 | 195 | TTKGAYSLSIRDW | 5.6833 | 5.9000 | 6.4385 | 6.3281 | 6.2328 | 5.9698 | 6.3354 | 5.8646 | 6.3781 | 5.9349 | 6.6219 | 5.9786 | 6.2547 | 6.9766 | 6.9703 | 6.1323 |
| YIPF1_S43 | YIPF1 | Q9Y548 | 43 | PKHQPGSPRGSGR | 6.0469 | 6.4891 | 5.9353 | 6.7639 | 6.1105 | 6.1553 | 6.5735 | 6.1113 | 6.2149 | 6.0747 | 6.4501 | 6.5423 | 5.8861 | 6.0709 | 6.2643 | 6.3107 |
| YKT6_S172 | YKT6 | O15498 | 172 | KLDDLVSKSEVLG | 6.0161 | 7.2465 | 7.9688 | 6.7357 | 6.0641 | 7.7939 | 5.8920 | 4.7632 | 5.4461 | 4.8818 | 6.9276 | 6.0952 | 7.4581 | 7.4807 | 5.2796 | 3.9506 |
| YLPM1_S1089 | YLPM1 | P49750 | 1089 | LVRPGSSREKVPG | 6.6482 | 6.8774 | 6.3834 | 7.2870 | 6.7489 | 6.3162 | 6.6838 | 6.3771 | 6.1320 | 6.2378 | 6.5031 | 6.7725 | 5.1962 | 5.3213 | 5.1895 | 5.3255 |
| YLPM1_S1249 | YLPM1 | P49750 | 756 | RSSYLESPRGPRF | 7.1497 | 7.2428 | 6.2647 | 7.0361 | 6.3468 | 6.5884 | 6.9672 | 6.8437 | 7.4491 | 6.6810 | 6.4104 | 6.9588 | 4.3020 | 4.8633 | 4.0401 | 4.8558 |
| YLPM1_S347 | YLPM1 | P49750 | 829 | TPSTSLSPRQSGP | 6.7613 | 7.3901 | 7.0816 | 6.7140 | 6.7442 | 6.6290 | 6.2737 | 6.1814 | 6.5495 | 7.0186 | 6.8258 | 6.9156 | 4.5744 | 4.9156 | 4.8839 | 4.5412 |
| YLPM1_S756 | YLPM1 | P49750 | 347 | QVPESPSSEEPPL | 5.4275 | 6.1596 | 5.7820 | 6.4805 | 6.0776 | 5.8468 | 5.6293 | 5.9324 | 6.2162 | 6.8164 | 6.7248 | 5.9324 | 6.9765 | 7.4077 | 6.1514 | 6.4388 |
| YLPM1_S829 | YLPM1 | P49750 | 1249 | NREDRFSAPPSRS | 6.2914 | 6.7908 | 6.1615 | 7.6164 | 6.1182 | 5.5247 | 10.8040 | 6.2042 | 6.5985 | 5.5859 | 5.2937 | 5.2637 | 5.7204 | 5.1188 | 4.9906 | 5.9173 |
| YOD1_S130 | YOD1 | Q5VVQ6 | 130 | QTRPRSSPAFTKR | 6.7070 | 7.0360 | 7.0449 | 6.6636 | 6.4201 | 6.9403 | 6.6913 | 7.0813 | 6.9056 | 6.8885 | 7.1732 | 7.2799 | 3.8760 | 4.7350 | 3.9804 | 4.5770 |
| YRDC_S37 | YRDC | Q86U90 | 37 | RLFRPPSPAPAAP | 6.1467 | 6.2327 | 6.3278 | 5.3521 | 6.4743 | 6.5499 | 6.1443 | 6.8476 | 6.2315 | 6.2204 | 6.3917 | 6.5869 | 6.2742 | 6.1718 | 6.0953 | 5.9529 |
| YRDC_S60 | YRDC | Q86U90 | 60 | GAVQAASPERAGW | 6.2543 | 5.8603 | 6.7399 | 6.3374 | 6.5911 | 6.7780 | 6.8984 | 7.0615 | 6.7249 | 6.7208 | 6.8971 | 7.3160 | 5.0812 | 5.0172 | 4.5294 | 5.1927 |
| YTHDC1_S118 | YTHDC1 | Q96MU7 | 308 | KRARGISPIVFDR | 6.2735 | 6.6654 | 5.9791 | 6.7063 | 6.3858 | 5.6436 | 6.3057 | 6.3103 | 6.5122 | 6.3070 | 5.9428 | 5.7910 | 6.1095 | 6.3072 | 6.4475 | 6.3128 |
| YTHDC1_S119 | YTHDC1 | Q96MU7 | 118 | DRKIRLSSSASRE | 6.2616 | 6.5655 | 6.1690 | 6.7249 | 6.0196 | 6.5647 | 7.0274 | 5.9963 | 5.7469 | 5.9477 | 6.0499 | 6.3555 | 5.4322 | 6.3585 | 6.2513 | 6.5290 |
| YTHDC1_S120 | YTHDC1 | Q96MU7 | 119 | RKIRLSSSASREP | 6.1585 | 6.7743 | 6.0802 | 6.1282 | 5.9728 | 6.5352 | 7.1836 | 6.0988 | 5.8903 | 5.5678 | 5.9634 | 6.3302 | 6.1659 | 6.3610 | 5.8814 | 6.9082 |
| YTHDC1_S146;S148 | YTHDC1 | Q96MU7 | 63 | TKRQKPSVHSRQL | 5.9944 | 6.0290 | 7.3748 | 7.3374 | 7.2134 | 6.6373 | 5.5372 | 5.7529 | 5.8966 | 6.4820 | 5.3781 | 6.0763 | 6.1568 | 6.3407 | 5.8629 | 5.9303 |
| YTHDC1_S308 | YTHDC1 | Q96MU7 | 120 | KIRLSSSASREPY | 5.5743 | 6.1599 | 5.8878 | 6.6850 | 5.3584 | 5.4570 | 6.2220 | 6.5913 | 5.6669 | 5.9798 | 6.1908 | 6.1267 | 8.3267 | 7.8575 | 5.9126 | 6.0036 |
| YTHDC1_S63 | YTHDC1 | Q96MU7 | 146;148 | NA | 6.4661 | 6.0533 | 6.0335 | 6.9474 | 6.6939 | 6.3000 | 6.2759 | 5.8726 | 6.1342 | 6.0870 | 6.4425 | 5.9270 | 6.8631 | 6.1754 | 6.2345 | 5.4936 |
| YTHDC2_S1183 | YTHDC2 | Q9H6S0 | 1221 | ERVLMKSPSPALH | 6.3364 | 6.3861 | 6.5319 | 6.3682 | 6.4264 | 6.7748 | 6.3505 | 7.0752 | 6.5213 | 6.1050 | 6.3461 | 6.4385 | 5.3036 | 6.0407 | 5.4853 | 5.5102 |
| YTHDC2_S1221 | YTHDC2 | Q9H6S0 | 1279 | SGKGSKSPSPRPN | 6.6916 | 6.2909 | 6.9365 | 6.4666 | 6.0446 | 6.3582 | 6.6727 | 6.3057 | 6.5903 | 5.5634 | 6.3678 | 6.1880 | 6.2214 | 5.8338 | 5.7137 | 5.7550 |
| YTHDC2_S1221;S1223 | YTHDC2 | Q9H6S0 | 1183 | QRPRPMSSEELPL | 5.5699 | 5.1727 | 6.7496 | 4.8666 | 6.3368 | 5.7075 | 6.1219 | 5.0162 | 5.5836 | 6.7547 | 6.4847 | 6.0273 | 6.7530 | 8.1304 | 6.0445 | 8.6807 |
| YTHDC2_S1263 | YTHDC2 | Q9H6S0 | 1281 | KGSKSPSPRPNMP | 6.6716 | 7.1920 | 7.3298 | 6.3306 | 6.2508 | 6.4769 | 5.7007 | 5.9140 | 6.3742 | 7.3941 | 5.9225 | 6.2636 | 6.0029 | 5.9497 | 5.3187 | 4.9080 |
| YTHDC2_S1263;S1267 | YTHDC2 | Q9H6S0 | 1263 | DSSSYPSPCASPS | 6.5372 | 6.1546 | 7.1074 | 6.6990 | 5.7168 | 6.8149 | 5.7003 | 6.8480 | 7.4201 | 6.0589 | 6.8719 | 7.2784 | 5.6120 | 5.7407 | 4.9001 | 4.5396 |
| YTHDC2_S1279 | YTHDC2 | Q9H6S0 | 1263;1267 | NA | 5.6497 | 7.2830 | 7.3774 | 6.1544 | 6.6971 | 5.8506 | 6.3037 | 6.7793 | 7.0118 | 7.0423 | 6.6992 | 5.6730 | 5.3098 | 5.6153 | 4.8217 | 5.7314 |
| YTHDC2_S1281 | YTHDC2 | Q9H6S0 | 1221;1223 | NA | 6.5281 | 6.8983 | 6.8412 | 6.6706 | 6.3148 | 6.2155 | 6.0279 | 6.1514 | 6.3067 | 6.5761 | 6.6468 | 6.4375 | 5.3961 | 5.3684 | 5.4923 | 6.1285 |
| YTHDF2_S238 | YTHDF2 | Q9Y5A9 | 238 | APPKPASWADIAS | 4.1707 | 3.5901 | 4.7601 | 4.1432 | 4.2849 | 3.9669 | 4.2738 | 4.1593 | 4.3742 | 4.3495 | 4.1039 | 4.5020 | 12.0762 | 11.4268 | 13.0790 | 12.7394 |
| YTHDF2_S39 | YTHDF2 | Q9Y5A9 | 39 | DFEPYLSPQARPN | 6.3420 | 6.6318 | 6.9590 | 8.2051 | 6.2859 | 5.9521 | 7.3740 | 6.3457 | 6.0970 | 6.5402 | 6.4158 | 6.5383 | 4.8247 | 5.1304 | 5.4623 | 4.8958 |
| YTHDF3_S385 | YTHDF3 | Q7Z739 | 385 | VVPVSASPSSVEV | 5.8019 | 6.0477 | 6.9453 | 6.8255 | 6.5736 | 5.9340 | 6.3616 | 6.6518 | 6.6441 | 6.7452 | 6.6612 | 6.7285 | 5.1587 | 6.2104 | 5.0881 | 5.6225 |
| YWHAE_S210 | YWHAE | P62258 | 210 | AELDTLSEESYKD | 6.0360 | 5.8981 | 6.6872 | 6.2982 | 5.9565 | 6.7598 | 6.1033 | 6.6998 | 6.5439 | 6.4798 | 6.9090 | 6.6738 | 5.5162 | 6.1871 | 5.5782 | 5.6732 |
| YWHAE_S46 | YWHAE | P62258 | 46 | EERNLLSVAYKNV | 5.7259 | 5.8620 | 6.7192 | 5.6723 | 5.9443 | 5.7663 | 6.0036 | 6.3661 | 5.9513 | 6.1732 | 6.0885 | 6.2916 | 6.9656 | 6.4342 | 7.2270 | 6.8090 |

| | | | | | | | | | | | | | | | | | | | | | |
|---|---|---|---|---|---|---|---|---|---|---|---|---|---|---|---|---|---|---|---|---|---|
| YWHAG_S215 | YWHAG | P61981 | | 215 | DTLNEDSYKDSTL | 4.9620 | 6.3394 | 7.2627 | 5.5669 | 6.3507 | 6.5851 | 6.3921 | 5.8093 | 6.2199 | 5.9168 | 5.7038 | 6.6111 | 6.6231 | 6.7260 | 6.6919 | 6.2392 |
| YWHAQ_S230 | YWHAQ | P27348 | | 230 | NLTLWTSDSAGEE | 6.5270 | 5.7674 | 6.4521 | 5.7128 | 6.5879 | 6.3252 | 6.3837 | 6.0029 | 6.3149 | 6.6362 | 6.2287 | 6.4979 | 5.9561 | 6.9390 | 5.8174 | 5.8508 |
| YWHAZ_S210 | YWHAZ | P63104 | | 210 | DTLSEESYKDSTL | 6.1318 | 5.7890 | 6.6523 | 6.2280 | 6.4119 | 6.1626 | 6.1407 | 6.2381 | 6.2619 | 6.3155 | 6.5136 | 6.3939 | 6.3755 | 6.4783 | 6.0768 | 5.8300 |
| YWHAZ_T232 | YWHAZ | P63104 | | 232 | TLWTSDTQGDEAE | 7.0844 | 6.5217 | 6.5888 | 6.7870 | 5.3952 | 6.8081 | 6.3124 | 6.6458 | 6.3885 | 6.2053 | 6.4716 | 6.1482 | 5.4012 | 6.4366 | 4.9145 | 5.8908 |
| YY1_S247 | YY1 | P25490 | | 247 | QIIGENSPPDYSE | 5.5563 | 6.0267 | 6.0172 | 5.9984 | 6.4792 | 6.4616 | 6.0495 | 6.1686 | 6.4250 | 6.4968 | 6.4872 | 6.2487 | 6.4570 | 6.9948 | 6.1617 | 5.9711 |
| YY1AP1_S816 | YY1AP1 | Q9H869-9 | | 816 | FPKVEHSPGPPPV | 6.1565 | 6.2192 | 6.0308 | 6.2020 | 6.2698 | 6.4109 | 6.2644 | 6.3071 | 6.4156 | 6.3709 | 6.4074 | 6.4153 | 5.7544 | 6.6109 | 6.0857 | 6.0789 |
| ZADH2_T288 | ZADH2 | Q8N4Q0 | | 288 | SPVKAGTLPAKLL | 6.8486 | 7.9720 | 7.6174 | 7.3985 | 5.9131 | 8.1913 | 6.8456 | 7.4523 | 7.6140 | 6.7631 | 8.4838 | 7.4286 | 2.5904 | 3.5057 | 2.2269 | 3.1488 |
| ZBED4_S1053 | ZBED4 | O75132 | | 1053 | HRCDAGSPSKDSA | 6.3661 | 6.4340 | 6.5413 | 6.8820 | 6.2496 | 5.7586 | 6.2307 | 5.1604 | 6.5951 | 6.3099 | 6.5228 | 7.0041 | 5.9041 | 5.8109 | 5.7450 | 6.4854 |
| ZBTB10_S210 | ZBTB10 | Q96DT7 | | 466 | KDALNISIKSEAP | 7.5325 | 6.7559 | 7.3039 | 8.1901 | 7.5131 | 6.4429 | 8.0019 | 6.2777 | 8.2581 | 7.0964 | 7.4991 | 7.3490 | 2.2149 | 3.7305 | 2.7230 | 3.1110 |
| ZBTB10_S466 | ZBTB10 | Q96DT7 | | 210 | CSSSRRSGGDGGD | 5.4815 | 6.1766 | 6.4000 | 6.5120 | 5.7744 | 6.6000 | 6.8923 | 6.1558 | 6.0287 | 6.1385 | 6.6364 | 6.0522 | 6.3575 | 5.9929 | 6.4857 | 6.3155 |
| ZBTB11_S1050 | ZBTB11 | O95625 | | 1050 | VEVAHISGGExxx | 6.0441 | 7.3510 | 6.9682 | 7.7758 | 6.0982 | 6.7304 | 6.4682 | 6.8819 | 5.7477 | 5.2154 | 5.5036 | 5.7914 | 5.6357 | 5.7463 | 6.4353 | 5.6069 |
| ZBTB14_S190 | ZBTB14 | O43829 | | 235 | KDLGSQTPQALTF | 5.8511 | 5.8278 | 5.6798 | 5.7332 | 5.6824 | 6.2442 | 6.2697 | 5.9680 | 6.0494 | 5.5296 | 6.2131 | 5.6454 | 7.1060 | 7.2090 | 7.4353 | 7.5560 |
| ZBTB14_T235 | ZBTB14 | O43829 | | 190 | SQEDGKSPTTTLR | 6.1920 | 6.8081 | 6.2044 | 6.4151 | 5.3934 | 5.4061 | 6.1562 | 6.0774 | 6.5171 | 6.0434 | 5.4394 | 5.7767 | 6.2598 | 6.6535 | 7.2119 | 7.4455 |
| ZBTB17_S120 | ZBTB17 | Q13105-2 | | 120 | LAEPATSPGGNAE | 5.9837 | 6.3868 | 5.6591 | 6.2807 | 6.1053 | 6.0001 | 6.1978 | 5.9782 | 6.5233 | 6.1612 | 6.2203 | 6.0948 | 6.7264 | 6.5916 | 6.6881 | 6.4024 |
| ZBTB17_S799 | ZBTB17 | Q13105-2 | | 609 | KHIIIHTGEKPYL | 8.8057 | 8.8972 | 7.1281 | 8.1268 | 5.9376 | 6.5089 | 6.9344 | 6.4240 | 7.8537 | 7.2027 | 6.8413 | 7.2361 | 3.2095 | 3.4900 | 2.1479 | 3.2562 |
| ZBTB17_T609 | ZBTB17 | Q13105-2 | | 798 | QPALAETSPTAPE | 6.3688 | 6.3515 | 6.4723 | 6.1808 | 5.8393 | 6.0964 | 6.8196 | 6.3361 | 6.6392 | 6.2796 | 6.4886 | 5.6504 | 6.1213 | 6.5328 | 5.8863 | 5.9371 |
| ZBTB17_T798 | ZBTB17 | Q13105-2 | | 799 | PALAETSPTAPEC | 5.6906 | 5.8183 | 6.1918 | 6.1421 | 6.1602 | 5.8428 | 6.3176 | 5.6487 | 6.4937 | 6.5285 | 5.9041 | 6.5008 | 6.2034 | 6.9904 | 6.3014 | 7.2658 |
| ZBTB2_S414 | ZBTB2 | Q8N680 | | 414 | HVCLNQSIDTYTM | 5.5070 | 6.1559 | 6.4452 | 6.6933 | 7.1707 | 6.7400 | 6.7732 | 6.8530 | 6.5274 | 5.6406 | 6.0903 | 5.8066 | 5.3126 | 5.6619 | 5.9631 | 6.6593 |
| ZBTB20_S432 | ZBTB20 | Q9HC78 | | 432 | LETGASSPERSNE | 5.7197 | 5.4392 | 6.5753 | 6.0811 | 5.7281 | 6.4522 | 5.9315 | 6.3032 | 5.9872 | 6.6400 | 6.7236 | 6.0927 | 6.4872 | 6.6964 | 6.3181 | 6.8246 |
| ZBTB21_S1003 | ZBTB21 | Q9ULJ3 | | 1003 | QPLEPDSPTGLSE | 6.1280 | 6.1375 | 6.2390 | 6.3835 | 6.2960 | 5.9994 | 6.1469 | 6.4690 | 6.2341 | 6.0040 | 6.0153 | 5.9425 | 6.6182 | 6.5630 | 6.4366 | 6.3869 |
| ZBTB21_S345 | ZBTB21 | Q9ULJ3 | | 345 | LLRRSLSMDSQVP | 5.5140 | 5.6432 | 5.9201 | 5.9684 | 5.7768 | 5.9928 | 6.1018 | 6.1445 | 6.2266 | 5.9120 | 5.8392 | 6.0478 | 6.9962 | 7.4180 | 7.1597 | 7.3388 |
| ZBTB21_S411 | ZBTB21 | Q9ULJ3 | | 983 | VPTNSPSPPPLPP | 5.3854 | 6.0544 | 5.7384 | 5.7702 | 5.8303 | 5.6886 | 5.7970 | 5.8976 | 5.9996 | 5.7464 | 5.7992 | 5.6740 | 7.7833 | 7.3760 | 7.8430 | 7.6166 |
| ZBTB21_S435 | ZBTB21 | Q9ULJ3 | | 411 | HRLRSFSASQSTD | 4.9436 | 5.3099 | 5.1348 | 5.7448 | 5.8719 | 5.9517 | 6.1442 | 6.2284 | 5.1907 | 5.0001 | 5.1135 | 5.1484 | 8.7190 | 8.2571 | 8.9759 | 8.2660 |
| ZBTB21_S714 | ZBTB21 | Q9ULJ3 | | 714 | EHAPLASPVENKE | 6.4906 | 6.7909 | 6.2387 | 6.9202 | 5.9584 | 6.1178 | 6.4841 | 6.3764 | 6.5140 | 5.9300 | 6.7195 | 6.6060 | 5.9329 | 5.6483 | 5.4613 | 5.8110 |
| ZBTB21_S981;S983 | ZBTB21 | Q9ULJ3 | | 435 | IKTEPSSPLSDPS | 5.8103 | 6.3429 | 5.9881 | 6.3619 | 6.6490 | 6.1765 | 6.5532 | 6.5207 | 5.9302 | 6.6486 | 6.5481 | 6.1490 | 5.9315 | 6.0865 | 6.0109 | 6.2926 |
| ZBTB21_S983 | ZBTB21 | Q9ULJ3 | 981;983 | | NA | 5.8672 | 5.7522 | 6.5551 | 6.7978 | 6.6482 | 6.2727 | 6.3985 | 5.7204 | 6.6252 | 6.0996 | 6.4172 | 6.4524 | 5.8994 | 6.0154 | 6.3445 | 6.1342 |
| ZBTB3_S549 | ZBTB3 | Q9H5J0 | | 549 | VQPLPGSPTADRQ | 5.8253 | 6.8184 | 6.1203 | 6.0082 | 5.8082 | 6.0763 | 5.7014 | 6.2914 | 6.0514 | 6.0319 | 5.9582 | 6.1656 | 7.2445 | 6.4559 | 6.4535 | 6.9897 |
| ZBTB33_S219 | ZBTB33 | Q86T24 | | 219 | TVAQVQSNPGPVA | 5.8710 | 6.2429 | 6.0806 | 6.9881 | 6.7013 | 5.6228 | 6.1932 | 6.0744 | 5.9842 | 6.0800 | 6.0924 | 6.1739 | 6.9396 | 6.5782 | 6.0980 | 6.2796 |
| ZBTB37_S310 | ZBTB37 | Q5TC79 | | 310 | SEVDRFSPSGSVV | 6.2683 | 5.4684 | 5.9264 | 6.3864 | 6.0963 | 6.9044 | 6.7801 | 5.7088 | 5.9429 | 7.7001 | 6.4071 | 6.3159 | 6.4506 | 6.7241 | 5.9056 | 5.0146 |
| ZBTB38_S130 | ZBTB38 | Q8NAP3 | | 130 | DRNFSNSPGPYVF | 6.0835 | 6.2812 | 5.9624 | 6.3367 | 6.1750 | 5.9097 | 5.8340 | 5.8884 | 6.2763 | 6.5483 | 6.1935 | 6.3325 | 6.2946 | 6.8624 | 6.6179 | 6.4034 |
| ZBTB40_S214 | ZBTB40 | Q9NUA8 | | 728 | EKEASASPDPAKK | 6.4258 | 6.6418 | 5.8917 | 6.8985 | 5.8610 | 6.1020 | 6.5472 | 6.1105 | 6.4217 | 6.5105 | 6.3168 | 6.4596 | 5.5533 | 5.9729 | 6.0097 | 6.2769 |
| ZBTB40_S728 | ZBTB40 | Q9NUA8 | | 214 | TTAEACSPSPAVQ | 6.2757 | 5.6104 | 5.8811 | 6.6215 | 6.1184 | 6.0040 | 5.8010 | 5.5582 | 6.8688 | 6.5180 | 6.1150 | 5.7652 | 6.8415 | 6.8365 | 6.3867 | 6.7982 |
| ZBTB44_S194 | ZBTB44 | Q8NCP5 | | 194 | IVMSPESPVKCGT | 6.2787 | 6.6231 | 6.4207 | 6.5522 | 6.3439 | 6.1354 | 6.1937 | 6.3826 | 6.5363 | 5.9704 | 6.0906 | 5.9579 | 5.9805 | 6.2577 | 6.1593 | 6.1170 |
| ZBTB46_S168 | ZBTB46 | Q86UZ6 | | 168 | MAGRSISPWLARR | 3.8105 | 4.1075 | 3.5722 | 4.0897 | 4.6621 | 4.9212 | 4.8145 | 4.8238 | 3.9852 | 4.6536 | 3.6265 | 3.7406 | 10.6788 | 12.7013 | 12.3496 | 13.4629 |
| ZBTB48_S171 | ZBTB48 | P10074 | | 171 | EPRGSHSPQRPQL | 6.1021 | 6.0448 | 5.7927 | 6.1091 | 5.5759 | 6.0892 | 7.1106 | 6.5201 | 6.3015 | 6.1445 | 5.9959 | 6.6919 | 5.9213 | 6.1246 | 6.6552 | 6.8205 |
| ZBTB49_S592 | ZBTB49 | Q6ZSB9 | | 592 | CKAGDESPDVLEE | 6.0580 | 6.4855 | 5.3762 | 6.4485 | 6.0738 | 5.8360 | 5.9533 | 5.4888 | 6.6240 | 5.7998 | 6.0769 | 6.3950 | 6.8760 | 8.0199 | 6.2509 | 6.2375 |
| ZBTB5_S234 | ZBTB5 | O15062 | | 234 | SREEFFSPDSLKI | 6.9118 | 6.8904 | 6.4966 | 7.1547 | 6.5841 | 6.2835 | 6.3747 | 5.9382 | 7.3120 | 7.2264 | 7.1277 | 6.0946 | 4.1567 | 4.9609 | 5.3341 | 5.1536 |
| ZBTB6_S202 | ZBTB6 | Q15916 | | 202 | ERKEMKSPELSTV | 6.6275 | 6.7870 | 6.4225 | 6.7145 | 6.2253 | 6.0467 | 6.4670 | 6.0959 | 6.3126 | 7.1261 | 6.5918 | 6.3399 | 4.9736 | 5.5804 | 5.6296 | 6.0595 |
| ZBTB7A_S511;S526 | ZBTB7A | O95365 | | 526 | APAQPSSPDARRN | 6.2058 | 6.4661 | 6.1743 | 6.7165 | 6.1780 | 5.8256 | 6.3363 | 6.2702 | 6.0876 | 6.2220 | 6.1915 | 6.5392 | 6.1656 | 6.4504 | 6.2419 | 5.9288 |
| ZBTB7A_S526 | ZBTB7A | O95365 | | 549 | EDEDVASPDGLGR | 5.6043 | 5.8954 | 6.4652 | 6.9524 | 5.4760 | 6.4842 | 6.2881 | 5.6957 | 6.5191 | 6.6130 | 6.5576 | 6.6479 | 6.2172 | 6.6623 | 6.0750 | 5.8466 |
| ZBTB7A_S549 | ZBTB7A | O95365 | 511;526 | | NA | 4.0431 | 7.6931 | 6.1498 | 7.4461 | 7.2247 | 5.1449 | 5.1085 | 7.4159 | 7.2220 | 5.6480 | 6.5466 | 5.0179 | 6.9955 | 6.6267 | 4.8301 | 6.8869 |
| ZBTB8A_S161 | ZBTB8A | Q96BR9 | | 161 | WQEGSSSPRSHLS | 6.1343 | 6.6304 | 6.1585 | 6.1343 | 5.9464 | 6.4563 | 6.2555 | 5.9104 | 6.3470 | 6.0703 | 6.4474 | 6.4878 | 5.7068 | 6.9375 | 6.1139 | 6.2631 |
| ZBTB9_S241 | ZBTB9 | Q96C00 | | 376 | PVKLGGTPPADGK | 7.0019 | 7.3935 | 6.7710 | 7.0964 | 6.3491 | 6.6204 | 6.5947 | 6.3381 | 6.6732 | 7.3192 | 7.4373 | 7.3711 | 3.6605 | 4.8259 | 4.0422 | 4.5054 |
| ZBTB9_T258 | ZBTB9 | Q96C00 | | 241 | PQPQRVSGVFPRP | 6.6844 | 7.1747 | 6.7289 | 6.6514 | 6.3109 | 6.2800 | 6.6880 | 6.8965 | 7.4732 | 6.2638 | 6.4324 | 7.4161 | 4.1423 | 4.9160 | 4.3707 | 5.5707 |
| ZBTB9_T376 | ZBTB9 | Q96C00 | | 258 | PLPMTATPRKLPE | 6.8757 | 6.0595 | 7.9855 | 7.9254 | 6.0080 | 7.2500 | 7.3083 | 5.9642 | 7.4731 | 6.7255 | 7.4705 | 6.7444 | 4.2528 | 4.9111 | 3.1465 | 3.8992 |
| ZC2HC1A_S179 | ZC2HC1A | Q96GY0 | | 223 | NKLQTLSPSHKGI | 6.9431 | 7.3431 | 7.1008 | 6.3161 | 7.6038 | 7.6119 | 7.4381 | 7.6618 | 7.0520 | 7.0095 | 6.4587 | 6.3761 | 3.4731 | 4.4595 | 3.3715 | 3.7812 |
| ZC2HC1A_S223 | ZC2HC1A | Q96GY0 | | 179 | ALKKSNSPGTASS | 7.3196 | 8.7667 | 8.2514 | 7.1478 | 5.6678 | 5.8030 | 7.8385 | 7.4073 | 8.3245 | 6.8116 | 7.0528 | 6.8262 | 1.4727 | 3.8809 | 3.4606 | 3.9686 |
| ZC2HC1A_T244 | ZC2HC1A | Q96GY0 | | 244 | VKPRNSTPPSLAR | 6.5146 | 6.5993 | 6.5290 | 6.3993 | 6.1053 | 6.5609 | 7.0214 | 6.1981 | 10.1403 | 7.0063 | 6.3869 | 6.4591 | 4.6078 | 4.9841 | 4.1235 | 4.3639 |
| ZC3H11A_S108 | ZC3H11A | O75152 | | 579 | REKSVLTPLRGDV | 7.1723 | 7.6104 | 6.7878 | 7.5478 | 5.8080 | 6.7103 | 7.3216 | 7.2533 | 7.6137 | 6.9615 | 7.2530 | 7.2380 | 3.2905 | 4.2417 | 3.2959 | 3.8943 |
| ZC3H11A_S132 | ZC3H11A | O75152 | | 132 | SVQSNPSPQLRSV | 6.8701 | 7.1956 | 6.1986 | 7.1627 | 6.6671 | 6.1769 | 7.0320 | 7.1849 | 6.3909 | 6.8520 | 6.7369 | 6.2566 | 4.5958 | 4.9571 | 4.3040 | 5.4188 |
| ZC3H11A_S149 | ZC3H11A | O75152 | | 758 | MKTRRLSSASTGK | 5.0486 | 4.9422 | 4.8721 | 5.7450 | 7.1341 | 5.3196 | 5.4364 | 5.9611 | 7.6893 | 7.4891 | 6.6746 | 6.9099 | 7.2639 | 7.3139 | 6.1217 | 6.0783 |
| ZC3H11A_S171 | ZC3H11A | O75152 | | 781 | KLIWEISGGKLEA | 6.0701 | 7.1000 | 7.0030 | 6.6113 | 6.3219 | 6.1845 | 7.5833 | 6.3192 | 6.4134 | 5.4589 | 7.1054 | 5.9045 | 5.0497 | 5.4428 | 5.0241 | 6.4080 |
| ZC3H11A_S290 | ZC3H11A | O75152 | | 108 | LPTVPESPEEEVK | 5.5293 | 5.6385 | 5.9875 | 6.2705 | 5.8607 | 6.3614 | 6.2189 | 6.1189 | 6.9829 | 6.0359 | 6.0574 | 5.7071 | 7.6208 | 6.3966 | 6.2378 | 6.9757 |
| ZC3H11A_S758 | ZC3H11A | O75152 | | 290 | LGKRKFSAGGDSD | 4.2942 | 4.8142 | 5.1360 | 5.8275 | 8.2132 | 5.3107 | 5.5708 | 5.9310 | 6.8815 | 7.3129 | 6.3748 | 6.7754 | 7.2303 | 7.3881 | 6.3226 | 6.6167 |
| ZC3H11A_S758;S761 | ZC3H11A | O75152 | | 171 | DDDDQFSEEGDET | 5.9290 | 5.8157 | 6.2594 | 6.3347 | 6.2316 | 6.4106 | 6.6678 | 6.1786 | 5.9215 | 5.9236 | 5.8361 | 6.0952 | 6.4914 | 6.9581 | 6.4113 | 6.5355 |
| ZC3H11A_S761 | ZC3H11A | O75152 | | 149 | SSENVPSPTHPPV | 6.1584 | 6.9247 | 5.5355 | 7.8808 | 7.0848 | 6.7611 | 5.3409 | 7.5595 | 6.1186 | 6.7129 | 6.3678 | 5.6295 | 4.3813 | 6.5041 | 6.0025 | 5.0375 |
| ZC3H11A_S781 | ZC3H11A | O75152 | | 761 | RRLSSASTGKPPL | 5.1551 | 6.3469 | 6.0806 | 6.3715 | 5.6945 | 5.9079 | 7.5052 | 6.1512 | 5.3543 | 4.2383 | 4.0161 | 5.5676 | 7.5651 | 6.6043 | 8.6370 | 8.8045 |
| ZC3H11A_T579 | ZC3H11A | O75152 | 758;761 | | NA | 4.3738 | 4.3775 | 4.6077 | 5.0218 | 6.0456 | 5.3560 | 5.7717 | 5.6116 | 4.9812 | 4.6318 | 4.5621 | 5.1767 | 9.6950 | 9.6513 | 10.3502 | 9.7860 |
| ZC3H12C_S124 | ZC3H12C | Q9C0D7-2 | | 124 | HRQLCRSPCLEPH | 6.6493 | 5.9868 | 7.2381 | 6.4868 | 6.6633 | 7.2472 | 6.1238 | 6.3341 | 5.9018 | 6.4188 | 6.9227 | 6.0335 | 5.9891 | 6.0332 | 5.2106 | 4.7611 |
| ZC3H12C_S613 | ZC3H12C | Q9C0D7-2 | | 613 | SPERRFSLDTDYR | 6.3220 | 5.4262 | 6.8885 | 5.6251 | 6.6846 | 5.9258 | 5.2361 | 6.7331 | 7.8542 | 7.6894 | 8.3834 | 7.4693 | 5.2684 | 5.9281 | 4.4274 | 4.1383 |
| ZC3H13_S1010;S1014;S1 | ZC3H13 | Q5T200 | | 77 | SSNYRRSPERPTG | 6.2286 | 6.5856 | 6.0015 | 6.6278 | 5.9356 | 5.7858 | 6.2154 | 6.0566 | 6.0487 | 5.9987 | 5.9134 | 5.8516 | 6.1261 | 7.1456 | 6.6668 | 6.8123 |
| ZC3H13_S1014;S1017 | ZC3H13 | Q5T200 | | 364 | SYQRTLTPPLRRS | 6.3539 | 6.3381 | 5.9373 | 6.6519 | 5.8816 | 5.6573 | 6.1487 | 5.6395 | 6.1244 | 5.9118 | 5.9608 | 5.9614 | 6.4777 | 7.1768 | 6.8033 | 6.9753 |
| ZC3H13_S110 | ZC3H13 | Q5T200 | | 198 | IIKKEVSPEVVRS | 6.3039 | 6.5491 | 5.6256 | 6.4228 | 6.2252 | 5.9027 | 6.0750 | 6.1005 | 6.0227 | 5.9939 | 5.7252 | 5.7433 | 6.5145 | 7.2124 | 6.8805 | 6.7027 |
| ZC3H13_S1191 | ZC3H13 | Q5T200 | | 1033 | KKRGPRTPPITTK | 6.1180 | 6.3315 | 5.3867 | 6.4092 | 5.9240 | 5.4953 | 6.0951 | 6.3368 | 6.2022 | 6.0141 | 5.5065 | 5.6197 | 6.3932 | 7.2360 | 7.4946 | 7.4371 |
| ZC3H13_S1208 | ZC3H13 | Q5T200 | | 1208 | TSGRLRSPSNDSA | 6.3057 | 6.8420 | 5.8627 | 6.9659 | 6.0984 | 6.0900 | 5.9311 | 6.3258 | 6.0348 | 5.8036 | 6.0667 | 6.0787 | 5.7444 | 6.8661 | 6.7240 | 6.2598 |
| ZC3H13_S1232 | ZC3H13 | Q5T200 | | 64 | RFVHGPSPRGKGY | 6.8705 | 7.0537 | 6.0772 | 6.8171 | 6.3004 | 6.1315 | 6.4923 | 6.3154 | 6.6618 | 6.1289 | 6.3432 | 6.3825 | 5.0918 | 6.0135 | 5.4744 | 5.8459 |
| ZC3H13_S1240 | ZC3H13 | Q5T200 | | 643 | RDQRPSSPIRHQG | 5.8998 | 6.4790 | 5.6740 | 6.8425 | 5.7062 | 6.6579 | 6.0310 | 5.7908 | 6.1637 | 6.0491 | 5.8580 | 6.1747 | 6.3311 | 7.1599 | 7.0529 | 7.1291 |
| ZC3H13_S1292 | ZC3H13 | Q5T200 | | 242 | KTSAVSSPLLDQQ | 6.4661 | 6.4669 | 5.8288 | 6.4399 | 6.3350 | 5.9034 | 6.3336 | 6.0526 | 6.6295 | 6.2311 | 5.8419 | 6.0698 | 6.2031 | 6.6192 | 6.2162 | 6.3629 |
| ZC3H13_S1366 | ZC3H13 | Q5T200 | | 25 | STSRRPSVFERLG | 5.9185 | 3.8116 | 4.0256 | 5.0160 | 7.6751 | 4.7142 | 4.5839 | 4.3524 | 10.0037 | 7.7531 | 7.3220 | 7.6197 | 8.0872 | 10.0098 | 3.8119 | 5.2952 |
| ZC3H13_S1465 | ZC3H13 | Q5T200 | | 207 | VVRSKLSPSPSLR | 6.4295 | 6.7800 | 6.4584 | 6.5507 | 6.0154 | 6.2482 | 6.1284 | 6.0765 | 6.6199 | 6.6597 | 6.5177 | 6.5876 | 5.4340 | 6.1377 | 5.9347 | 5.4215 |
| ZC3H13_S198 | ZC3H13 | Q5T200 | | 877 | QESRSLSPSHLTE | 6.2791 | 6.2608 | 6.3418 | 7.1432 | 6.0055 | 5.6575 | 5.9400 | 5.8941 | 6.3825 | 6.1188 | 6.2785 | 5.6687 | 5.9833 | 6.5966 | 6.0639 | 6.4856 |
| ZC3H13_S207 | ZC3H13 | Q5T200 | | 993 | EDGQVFSPKKGQK | 6.2999 | 6.1823 | 5.8053 | 6.5458 | 5.9172 | 5.7690 | 6.4281 | 5.8454 | 6.2785 | 6.4411 | 5.8150 | 6.3298 | 6.1836 | 6.8325 | 6.4640 | 6.8626 |
| ZC3H13_S207;S209 | ZC3H13 | Q5T200 | | 1465 | EGYEPISDDELDE | 5.6879 | 6.1333 | 6.3661 | 7.1773 | 6.5935 | 6.0275 | 6.5360 | 6.2510 | 6.3272 | 6.4865 | 6.7255 | 6.3881 | 5.1298 | 6.2160 | 6.0016 | 5.9528 |
| ZC3H13_S242 | ZC3H13 | Q5T200 | | 1232 | RVLHSGSRDREKT | 5.8379 | 6.1183 | 5.5194 | 6.1945 | 5.4033 | 5.6289 | 5.9437 | 5.8308 | 5.7177 | 5.5294 | 5.2251 | 5.9239 | 7.4182 | 7.7993 | 7.9964 | 7.9131 |
| ZC3H13_S25 | ZC3H13 | Q5T200 | | 110 | NTEESSSPVRKES | 5.6862 | 6.2789 | 5.7949 | 6.8883 | 5.9932 | 6.2895 | 6.0927 | 5.4765 | 6.0360 | 6.0977 | 6.2474 | 6.4581 | 6.3243 | 6.8157 | 6.5623 | 6.9585 |
| ZC3H13_S370;S372;S381 | ZC3H13 | Q5T200 | | 265 | KGPRTPSPPPPIP | 5.9395 | 6.1469 | 5.7344 | 6.2945 | 6.3729 | 6.1635 | 5.8679 | 5.7155 | 6.6267 | 6.0623 | 6.1642 | 5.7293 | 6.0391 | 7.0821 | 6.5834 | 7.4777 |
| ZC3H13_S64 | ZC3H13 | Q5T200 | | 1366 | ERLISDSVERDRD | 6.2768 | 6.5604 | 6.3800 | 6.9708 | 6.1764 | 5.8349 | 6.8003 | 6.9223 | 6.4029 | 6.5356 | 5.9469 | 6.1290 | 5.1350 | 5.9500 | 5.8941 | 6.0846 |
| ZC3H13_S642 | ZC3H13 | Q5T200 | | 1191 | DQKRSSSLGSNRS | 5.9992 | 6.3449 | 5.2922 | 6.3173 | 5.5906 | 5.5924 | 6.0818 | 6.0943 | 6.3015 | 6.0628 | 5.8147 | 5.9511 | 6.6026 | 7.2667 | 7.2181 | 7.4699 |
| ZC3H13_S643 | ZC3H13 | Q5T200 | | 642 | ERDQRPSSPIRHQ | 4.6145 | 5.6141 | 4.9237 | 5.6779 | 5.1977 | 5.2168 | 5.5289 | 5.9298 | 5.9761 | 6.4660 | 6.0678 | 6.5859 | 6.9898 | 8.4172 | 8.0724 | 8.7214 |
| ZC3H13_S77 | ZC3H13 | Q5T200 | | 873 | VVRPQESRSLSPS | 6.1808 | 6.1351 | 5.3243 | 6.3382 | 5.4751 | 4.8530 | 4.8054 | 4.4868 | 5.8302 | 5.5575 | 4.6958 | 5.9505 | 8.0792 | 9.2743 | 8.0037 | 9.0101 |
| ZC3H13_S873 | ZC3H13 | Q5T200 | | 1240 | DREKTKSLEITGE | 5.7847 | 6.4403 | 5.6968 | 6.3916 | 6.3505 | 6.1298 | 6.8416 | 6.5564 | 5.7177 | 5.9060 | 5.8219 | 5.9614 | 6.3114 | 6.5311 | 6.4737 | 7.0850 |
| ZC3H13_S877 | ZC3H13 | Q5T200 | | 943 | IIGHHQSEDRETS | 5.2372 | 6.0567 | 6.5316 | 5.8640 | 5.5218 | 5.9985 | 6.1469 | 6.1527 | 6.5327 | 6.1213 | 6.1114 | 6.2656 | 6.2179 | 7.2305 | 6.9902 | 7.0210 |
| ZC3H13_S943 | ZC3H13 | Q5T200 | | 1292 | VHSRSGSFDSRDR | 6.0791 | 6.2751 | 5.7515 | 6.4536 | 5.6023 | 5.4504 | 6.2705 | 6.0348 | 6.2581 | 6.4244 | 6.0441 | 6.3038 | 6.4398 | 7.2867 | 6.6148 | 6.7111 |
| ZC3H13_S986 | ZC3H13 | Q5T200 | | 986 | GNIETTSEDGQVF | 6.3833 | 6.3270 | 5.8393 | 6.8082 | 6.1709 | 5.9732 | 5.6897 | 5.9038 | 6.1563 | 6.0575 | 6.2826 | 6.3076 | 6.7625 | 6.3288 | 6.4777 | 6.5314 |
| ZC3H13_S986;S993 | ZC3H13 | Q5T200 | 263;265 | | NA | 5.8201 | 6.0130 | 5.9441 | 6.6454 | 5.9739 | 5.9842 | 6.2157 | 5.8601 | 5.9575 | 6.1347 | 6.0881 | 6.0305 | 6.3107 | 7.2037 | 6.9372 | 6.8811 |
| ZC3H13_S993 | ZC3H13 | Q5T200 | 207;209 | | NA | 6.2554 | 6.2289 | 5.6412 | 6.3046 | 5.8446 | 5.4670 | 6.0568 | 5.7185 | 5.9382 | 5.7811 | 5.4106 | 5.5027 | 6.9591 | 7.5755 | 7.5700 | 7.7458 |
| ZC3H13_T1033 | ZC3H13 | Q5T200 | 354;356 | | NA | 6.0307 | 6.6766 | 5.8946 | 6.8091 | 5.9958 | 5.8542 | 5.7215 | 5.8745 | 5.9628 | 6.2423 | 5.8829 | 6.2674 | 6.1352 | 6.9709 | 6.7834 | 6.8981 |
| ZC3H13_T263;S265 | ZC3H13 | Q5T200 | 986;993 | | NA | 5.6177 | 5.9829 | 5.9221 | 6.7450 | 6.7239 | 6.6520 | 6.4817 | 5.8258 | 5.9344 | 5.8692 | 6.7208 | 6.3381 | 6.3451 | 6.9110 | 5.9789 | 5.9513 |
| ZC3H13_T265 | ZC3H13 | Q5T200 | 1014;1017 | | NA | 6.8775 | 7.1129 | 6.6042 | 7.1448 | 6.0006 | 5.7999 | 6.0718 | 5.1191 | 7.0694 | 7.1568 | 6.8970 | 6.3485 | 5.0810 | 6.3243 | 4.7626 | 5.6297 |
| ZC3H13_T317;S325 | ZC3H13 | Q5T200 | 354;358 | | NA | 6.0106 | 6.9389 | 5.9019 | 6.7841 | 5.5717 | 5.6786 | 5.7503 | 5.6087 | 6.2831 | 6.9593 | 6.1311 | 5.7426 | 5.7549 | 6.4983 | 7.3375 | 7.0483 |
| ZC3H13_T354;S356 | ZC3H13 | Q5T200 | 317;325 | | NA | 5.3950 | 7.1350 | 6.7709 | 7.0915 | 5.8999 | 6.5767 | 5.1048 | 5.6870 | 5.9428 | 6.5317 | 5.8320 | 5.9459 | 6.3595 | 6.5565 | 6.2813 | 6.8894 |
| ZC3H13_T354;S358 | ZC3H13 | Q5T200 | 1010;1014;10: | | NA | 5.0208 | 5.9731 | 6.0684 | 6.1404 | 5.5542 | 5.4610 | 6.9611 | 5.8982 | 6.0608 | 5.8804 | 6.0315 | 6.3392 | 6.8439 | 7.2048 | 6.8559 | 7.7061 |
| ZC3H13_T364 | ZC3H13 | Q5T200 | 370;372;381 | | NA | 6.4420 | 6.8955 | 6.3747 | 6.7594 | 5.8234 | 6.2565 | 6.4953 | 5.4443 | 5.7154 | 6.1213 | 5.7253 | 6.1584 | 5.7627 | 6.8041 | 6.2147 | 7.0071 |
| ZC3H14_S195 | ZC3H14 | Q6PJT7 | | 533 | SPPGYMSDQEEDM | 5.7978 | 5.7271 | 5.9177 | 6.7300 | 6.1487 | 6.1112 | 5.9132 | 6.0756 | 6.2166 | 6.2886 | 6.4360 | 6.4778 | 6.7760 | 6.4588 | 6.5187 | 6.4061 |
| ZC3H14_S301 | ZC3H14 | Q6PJT7 | | 665 | RRIPVLSPKPAVA | 7.7040 | 7.4141 | 7.1399 | 7.5672 | 6.0465 | 6.3488 | 7.0893 | 6.8358 | 7.8087 | 7.3915 | 7.1109 | 7.4583 | 3.1702 | 3.5294 | 3.2631 | 4.1221 |
| ZC3H14_S343 | ZC3H14 | Q6PJT7 | | 527 | TLDGVPSPPGYMS | 5.8410 | 6.0443 | 6.2720 | 6.6227 | 6.0276 | 6.0880 | 6.4282 | 5.8704 | 6.8426 | 6.2648 | 6.3102 | 6.5192 | 6.0751 | 5.9949 | 6.4631 | 6.3359 |
| ZC3H14_S390 | ZC3H14 | Q6PJT7 | | 343 | KPERRPSLPPSKQ | 4.7684 | 4.4012 | 4.8774 | 5.0126 | 7.3760 | 5.6148 | 4.8940 | 5.5467 | 7.2012 | 7.4646 | 6.6919 | 6.9511 | 8.1962 | 7.3054 | 6.7470 | 6.9513 |
| ZC3H14_S409 | ZC3H14 | Q6PJT7 | | 515 | QPDKPASPKFIVT | 6.2590 | 6.2163 | 6.4366 | 5.9547 | 6.4783 | 5.7729 | 6.0522 | 6.0132 | 6.8797 | 6.4986 | 6.1861 | 6.3818 | 6.9302 | 6.1307 | 6.0030 | 5.8067 |
| ZC3H14_S515 | ZC3H14 | Q6PJT7 | | 195 | VQENPLSQKKPTV | 5.2483 | 5.5866 | 7.3452 | 6.6581 | 6.2237 | 8.2087 | 6.3619 | 6.2711 | 7.5005 | 7.4637 | 6.6278 | 7.5519 | 5.0745 | 5.3088 | 3.8687 | 4.7007 |
| ZC3H14_S527 | ZC3H14 | Q6PJT7 | | 409 | SRTPRISPPIKEE | 4.8921 | 5.5835 | 5.8283 | 6.1673 | 7.3708 | 8.1416 | 5.8442 | 6.3396 | 6.8417 | 5.9488 | 7.3024 | 6.1998 | 7.8843 | 6.4707 | 5.3089 | 3.8762 |
| ZC3H14_S527;Y533 | ZC3H14 | Q6PJT7 | | 579 | LEDPNGSFSNAEM | 6.1657 | 5.9004 | 6.0935 | 6.6632 | 6.9527 | 6.5414 | 6.9741 | 6.8285 | 7.1417 | 5.9031 | 6.7135 | 6.6897 | 5.0818 | 5.4643 | 4.9739 | 5.9126 |
| ZC3H14_S579 | ZC3H14 | Q6PJT7 | | 301 | KLPVVSSVVKVKK | 9.2567 | 8.2301 | 5.7993 | 7.5132 | 5.7314 | 7.0847 | 7.6913 | 6.3889 | 6.3889 | 7.0932 | 6.2023 | 7.3859 | 3.3939 | 3.8902 | 3.0460 | 4.9041 |
| ZC3H14_S665 | ZC3H14 | Q6PJT7 | | 390 | APRTRTSQEELLA | 4.3342 | 4.3621 | 4.6569 | 4.9662 | 4.7644 | 4.4563 | 5.9317 | 5.0814 | 4.9409 | 4.3981 | 4.3127 | 5.6331 | 11.2908 | 9.0183 | 11.3901 | 10.4627 |
| ZC3H14_Y533 | ZC3H14 | Q6PJT7 | 527;533 | | NA | 5.9112 | 6.0335 | 6.0404 | 7.0291 | 6.2601 | 6.0421 | 6.0942 | 6.2967 | 6.2240 | 6.2146 | 6.1884 | 6.2590 | 6.4889 | 6.2743 | 6.1864 | 6.4571 |
| ZC3H15_S328 | ZC3H15 | Q8WU90 | | 328 | EVDDSVSVNDIDL | 6.1498 | 6.4298 | 6.7817 | 6.8809 | 6.2974 | 6.0160 | 6.4939 | 5.9390 | 6.6555 | 6.3646 | 6.6399 | 6.6144 | 5.7550 | 5.8246 | 5.5569 | 5.6008 |
| ZC3H15_S360 | ZC3H15 | Q8WU90 | | 360 | RFSTYTSDKDENK | 6.0745 | 6.3553 | 6.7004 | 6.7285 | 5.7791 | 6.6117 | 6.4876 | 6.0722 | 5.7079 | 6.0293 | 6.0711 | 6.1962 | 6.2723 | 6.5978 | 6.1926 | 6.1233 |
| ZC3H18_S118 | ZC3H18 | Q86VM9 | | 852 | AKRPNTSPDRGSR | 6.3476 | 6.6426 | 5.8587 | 6.6320 | 5.9331 | 5.7048 | 6.0400 | 6.0572 | 6.1831 | 6.0359 | 6.0069 | 5.9848 | 6.5386 | 6.6215 | 6.6955 | 6.7176 |
| ZC3H18_S173 | ZC3H18 | Q86VM9 | | 534 | KLGVSVSPSRARR | 6.3987 | 6.5992 | 5.8086 | 6.5858 | 6.3325 | 6.0334 | 6.2514 | 6.1693 | 6.1400 | 6.0302 | 5.8055 | 5.8218 | 6.4518 | 6.4281 | 6.5952 | 6.5484 |
| ZC3H18_S179 | ZC3H18 | Q86VM9 | | 677 | KERPARTPPRRRT | 6.8937 | 6.9463 | 6.2766 | 6.7247 | 5.9190 | 6.0883 | 6.1138 | 5.3056 | 5.9280 | 5.8393 | 6.2997 | 6.4030 | 6.3893 | 6.3981 | 6.3178 | 6.1568 |
| ZC3H18_S46 | ZC3H18 | Q86VM9 | | 842 | SDKDRQSPPPAKR | 6.1143 | 6.3969 | 5.8482 | 6.5850 | 6.0495 | 5.8931 | 6.0270 | 6.0057 | 6.1270 | 6.1354 | 6.0294 | 5.9526 | 6.6629 | 6.7235 | 6.7522 | 6.6973 |
| ZC3H18_S487 | ZC3H18 | Q86VM9 | | 868 | SGGRLGSPKPERQ | 6.3069 | 6.2102 | 6.0112 | 6.0824 | 6.2207 | 5.6345 | 6.6469 | 5.4282 | 6.2643 | 6.0903 | 5.8303 | 5.9235 | 7.4254 | 6.4533 | 6.6093 | 6.8626 |
| ZC3H18_S532;S534 | ZC3H18 | Q86VM9 | | 893 | DRKRQLSPQSKSS | 6.5509 | 6.6250 | 5.9207 | 6.5893 | 6.4290 | 5.8918 | 6.2504 | 5.8613 | 6.0057 | 5.9470 | 5.8814 | 5.8222 | 6.6601 | 6.4310 | 6.5789 | 6.5556 |
| ZC3H18_S534 | ZC3H18 | Q86VM9 | | 487 | GKPKPRSPQPPSR | 6.1903 | 6.3252 | 6.0160 | 6.8484 | 6.4716 | 6.1699 | 5.9161 | 6.2487 | 6.2552 | 6.1998 | 6.0581 | 5.8723 | 6.2764 | 6.3375 | 6.2826 | 6.5318 |
| ZC3H18_S67;S74 | ZC3H18 | Q86VM9 | | 118 | LRDEASSVTRELD | 6.0277 | 6.4505 | 6.4793 | 7.1133 | 6.3079 | 6.2298 | 6.3584 | 5.8083 | 6.3827 | 5.9748 | 6.1276 | 6.4160 | 6.1561 | 5.9326 | 5.8267 | 6.4083 |
| ZC3H18_S746 | ZC3H18 | Q86VM9 | | 796 | SSQQPSTPQQAPP | 5.8621 | 6.6410 | 6.0877 | 7.1291 | 6.2700 | 6.3918 | 7.3329 | 6.8214 | 6.3244 | 6.2802 | 6.7899 | 6.4979 | 5.4958 | 5.2885 | 5.0791 | 5.7083 |
| ZC3H18_S775 | ZC3H18 | Q86VM9 | | 775 | KRKRDSSTQPPKS | 6.0368 | 6.3556 | 5.9482 | 6.6273 | 6.1496 | 5.9449 | 6.7711 | 6.2022 | 6.3555 | 6.1629 | 6.3050 | 6.1239 | 6.1897 | 6.7719 | 6.0899 | 5.9654 |
| ZC3H18_S842 | ZC3H18 | Q86VM9 | | 173 | GKAGVQSVGEKES | 6.0481 | 6.0643 | 5.8654 | 7.1751 | 5.8557 | 5.9163 | 6.1637 | 5.8403 | 5.6503 | 5.9664 | 4.8605 | 5.4062 | 7.5745 | 7.0199 | 6.9730 | 7.6205 |
| ZC3H18_S852 | ZC3H18 | Q86VM9 | | 746 | SSASSRSPAPAQT | 6.3012 | 7.0944 | 7.4770 | 8.0749 | 6.1233 | 5.7908 | 6.5447 | 6.3550 | 6.9394 | 6.4660 | 6.3466 | 6.5746 | 4.9451 | 4.8581 | 5.0041 | 5.1048 |
| ZC3H18_S868 | ZC3H18 | Q86VM9 | | 179 | SVGEKESLEAAKE | 5.7962 | 6.2161 | 5.9476 | 6.3814 | 6.5456 | 6.7660 | 6.4695 | 5.7745 | 5.7096 | 6.0310 | 6.2412 | 6.1649 | 6.5341 | 6.3955 | 6.2258 | 6.8010 |
| ZC3H18_S893 | ZC3H18 | Q86VM9 | | 46 | GAGVRASDLEDEE | 5.8055 | 6.3393 | 6.7522 | 8.0359 | 6.1617 | 5.7338 | 5.3017 | 6.5747 | 8.1836 | 6.7351 | 6.9127 | 6.1992 | 5.0995 | 4.9797 | 5.1230 | 6.0623 |
| ZC3H18_T677 | ZC3H18 | Q86VM9 | 532;534 | | NA | 5.5887 | 5.8544 | 6.0967 | 6.5966 | 6.1797 | 6.3253 | 5.9292 | 6.1119 | 6.3293 | 6.3635 | 6.6163 | 6.1820 | 6.6021 | 6.6154 | 6.2813 | 6.3273 |
| ZC3H18_T796 | ZC3H18 | Q86VM9 | 67;74 | | NA | 6.2294 | 5.9551 | 7.1380 | 6.4786 | 6.7573 | 6.5944 | 5.8428 | 6.0685 | 5.3503 | 4.7282 | 7.2022 | 5.5239 | 7.1375 | 6.9498 | 6.5655 | 5.4785 |
| ZC3H3_S382 | ZC3H3 | Q8IXZ2 | | 465 | DKKSGTSPAATAK | 6.6536 | 6.5239 | 6.6329 | 7.2853 | 6.2308 | 6.4608 | 6.9421 | 6.6340 | 7.0414 | 6.8573 | 7.1577 | 7.3564 | 4.3809 | 4.8019 | 4.3105 | 4.7304 |
| ZC3H3_S408 | ZC3H3 | Q8IXZ2 | | 408 | DHASQLSPVLSRS | 6.7849 | 6.6110 | 7.1691 | 6.9599 | 6.8895 | 5.6664 | 6.3335 | 6.0154 | 6.5955 | 7.0139 | 6.3776 | 6.0352 | 5.5431 | 5.1688 | 3.9776 | 6.8587 |
| ZC3H3_S453 | ZC3H3 | Q8IXZ2 | | 866 | CPGGSASPSSSKA | 6.5875 | 5.8564 | 6.5554 | 7.1956 | 6.1910 | 6.5119 | 6.3767 | 6.4233 | 6.5325 | 6.5638 | 6.8648 | 6.4195 | 4.9389 | 5.5669 | 5.6861 | 5.7296 |
| ZC3H3_S465 | ZC3H3 | Q8IXZ2 | | 549 | VKKTPASPLSAPP | 6.4819 | 5.7580 | 7.5337 | 6.7578 | 5.8532 | 6.9125 | 6.3173 | 6.9484 | 6.7257 | 6.2517 | 5.9250 | 7.2169 | 4.5539 | 5.9361 | 5.1801 | 5.6478 |
| ZC3H3_S549 | ZC3H3 | Q8IXZ2 | | 453 | IIRRRSSTSLPGD | 5.7109 | 4.9002 | 5.4151 | 5.2081 | 6.0739 | 6.1890 | 5.8051 | 6.0051 | 9.0577 | 8.4219 | 7.1733 | 7.7473 | 6.3415 | 6.1542 | 4.4465 | 5.3502 |
| ZC3H3_S866 | ZC3H3 | Q8IXZ2 | | 809 | HSRRAATSPAPGP | 6.1034 | 6.1231 | 6.2772 | 6.4682 | 5.8865 | 6.9514 | 6.4017 | 6.1324 | 5.9555 | 6.9994 | 6.7961 | 6.2932 | 5.7250 | 6.0073 | 5.2739 | 6.6057 |
| ZC3H3_T809 | ZC3H3 | Q8IXZ2 | | 382 | KYKWKASSPSASS | 7.1122 | 6.9564 | 6.8134 | 6.9628 | 5.9822 | 5.9703 | 6.9023 | 6.8563 | 7.0315 | 6.2441 | 6.9142 | 6.2228 | 4.2165 | 5.6304 | 5.1377 | 5.0469 |
| ZC3H4_S1104 | ZC3H4 | Q9UPT8 | | 1275 | SPFAGNSPAREGE | 6.5400 | 6.7137 | 6.4537 | 6.5810 | 6.6722 | 6.2930 | 6.8833 | 6.4747 | 6.4073 | 6.0635 | 6.5328 | 6.2785 | 5.6381 | 5.5682 | 5.4611 | 5.4389 |
| ZC3H4_S1114 | ZC3H4 | Q9UPT8 | | 159 | RKYREYSPPYAPS | 6.0263 | 7.0772 | 5.8759 | 6.7979 | 6.6718 | 6.5785 | 5.8722 | 6.9016 | 5.9480 | 6.0301 | 6.0352 | 5.6825 | 5.7778 | 6.6021 | 6.3442 | 5.7787 |
| ZC3H4_S1269 | ZC3H4 | Q9UPT8 | | 1114 | SPSGDASPPATAP | 6.2350 | 5.9072 | 6.2717 | 6.1181 | 6.6177 | 5.6906 | 5.8210 | 4.9904 | 6.0155 | 6.2407 | 5.8776 | 5.8669 | 7.2512 | 7.3403 | 6.7460 | 7.0100 |
| ZC3H4_S1269;S1275 | ZC3H4 | Q9UPT8 | | 1269 | PKTGSGSPFAGNS | 6.1609 | 5.8012 | 5.7817 | 5.5854 | 5.8074 | 5.5703 | 5.6216 | 5.6606 | 5.9753 | 6.1383 | 5.5737 | 5.6768 | 7.7243 | 7.7579 | 7.7349 | 7.4297 |
| ZC3H4_S1275 | ZC3H4 | Q9UPT8 | | 1104 | GPAEAPSPTASPS | 6.3757 | 6.5799 | 6.1873 | 6.6102 | 6.6783 | 5.8890 | 5.4431 | 5.5101 | 5.7634 | 5.9916 | 5.6828 | 5.8377 | 7.4016 | 6.4291 | 5.9539 | 7.6664 |
| ZC3H4_S1299 | ZC3H4 | Q9UPT8 | | 531 | GPQAPTSPNGRPM | 5.7727 | 6.8193 | 6.0171 | 5.8100 | 6.2273 | 6.3841 | 6.3639 | 6.7234 | 5.9912 | 6.1460 | 6.1186 | 5.7885 | 5.8800 | 6.9100 | 6.6194 | 6.4283 |
| ZC3H4_S159 | ZC3H4 | Q9UPT8 | | 1299 | GFDPTASPFCQxx | 7.0886 | 6.8817 | 6.7002 | 6.4474 | 5.1762 | 6.6061 | 7.4155 | 6.0398 | 6.8373 | 6.5614 | 6.4819 | 6.3324 | 5.3406 | 5.7478 | 4.8990 | 5.4442 |
| ZC3H4_S531 | ZC3H4 | Q9UPT8 | | 908 | EGSLHSSPVGPSS | 5.9591 | 7.0375 | 5.7816 | 7.5758 | 7.4425 | 7.1170 | 6.3801 | 7.8496 | 6.3271 | 6.0268 | 6.6114 | 6.0520 | 4.7847 | 5.7466 | 4.4491 | 4.8592 |
| ZC3H4_S908 | ZC3H4 | Q9UPT8 | | 1232 | AAAPAATTATPPP | 6.1430 | 6.4868 | 6.3510 | 8.0891 | 7.1918 | 6.0161 | 7.0244 | 5.6389 | 7.0120 | 6.1748 | 7.2306 | 6.4127 | 4.8492 | 4.9691 | 4.9532 | 5.4573 |
| ZC3H4_T1232 | ZC3H4 | Q9UPT8 | 1269;1275 | | NA | 5.8526 | 6.3806 | 5.6615 | 6.3065 | 5.2993 | 5.7162 | 5.5267 | 5.7219 | 6.0725 | 6.4718 | 5.8320 | 6.0462 | 7.1469 | 7.3886 | 7.3821 | 7.1945 |
| ZC3H4_T519;S531 | ZC3H4 | Q9UPT8 | 519;531 | | NA | 6.6016 | 6.9611 | 6.0315 | 6.1245 | 6.5509 | 5.4741 | 6.1738 | 7.5486 | 7.5953 | 7.3584 | 7.0793 | 6.4763 | 3.9670 | 4.7989 | 5.0724 | 6.1863 |
| ZC3H7B_S217 | ZC3H7B | Q9UGR2 | | 399 | AQKPAPSPEPCMP | 6.0463 | 5.6144 | 6.6470 | 6.2045 | 6.5121 | 6.4801 | 6.3970 | 6.0379 | 6.5385 | 6.7073 | 6.6418 | 6.3764 | 5.9405 | 6.1982 | 5.5733 | 6.0848 |
| ZC3H7B_S217;T224 | ZC3H7B | Q9UGR2 | | 224 | PALLPSTPTMPLF | 5.7515 | 5.5907 | 5.7834 | 6.1825 | 6.6274 | 6.5695 | 6.1092 | 6.5479 | 6.2807 | 5.9682 | 5.8447 | 6.3708 | 6.1729 | 6.4595 | 7.0041 | 6.7370 |
| ZC3H7B_S399 | ZC3H7B | Q9UGR2 | | 217 | YVDPRGSPALLPS | 6.0651 | 7.0550 | 6.8872 | 6.7065 | 6.5556 | 5.4019 | 5.9728 | 6.7006 | 5.7623 | 7.1831 | 6.9637 | 6.5298 | 5.8428 | 5.3334 | 5.3493 | 5.6909 |
| ZC3H7B_T224 | ZC3H7B | Q9UGR2 | 217;224 | | NA | 5.0751 | 7.0438 | 7.5361 | 5.2381 | 6.7241 | 5.4823 | 5.7367 | 7.9688 | 5.8709 | 6.4852 | 6.5226 | 5.7520 | 5.8851 | 7.1301 | 5.4914 | 6.0577 |
| ZC3H8_S59 | ZC3H8 | Q8N5P1 | | 59 | FRHSAISPKSSLH | 7.1105 | 7.2951 | 6.5400 | 7.0105 | 6.5756 | 6.3478 | 7.0633 | 6.3178 | 6.9806 | 6.9331 | 6.6998 | 6.6989 | 4.3064 | 4.7484 | 4.5266 | 4.8455 |
| ZC3H8_S77 | ZC3H8 | Q8N5P1 | | 77 | KDYDVYSDNDICS | 6.4210 | 6.8633 | 7.2440 | 7.4634 | 5.2543 | 6.1364 | 5.9205 | 6.1144 | 6.3954 | 6.4351 | 6.3373 | 6.5849 | 4.8666 | 5.4429 | 5.9989 | 6.5215 |
| ZC3HAV1_S257 | ZC3HAV1 | Q7Z2W4 | | 378 | RRKTVFSPTLPAA | 6.6378 | 7.1157 | 6.5957 | 6.9718 | 6.2316 | 7.0210 | 7.1863 | 6.9531 | 6.8731 | 6.9511 | 7.3389 | 7.4292 | 3.6829 | 4.6113 | 3.7416 | 4.6590 |
| ZC3HAV1_S275 | ZC3HAV1 | Q7Z2W4 | | 335 | RFLENGSQEDLLH | 5.6351 | 7.0667 | 6.6298 | 6.3122 | 5.2992 | 6.6924 | 6.3978 | 6.5070 | 5.1117 | 6.6498 | 7.7014 | 6.7836 | 5.4420 | 5.8342 | 6.0321 | 5.9049 |
| ZC3HAV1_S284 | ZC3HAV1 | Q7Z2W4 | | 387 | LPAARSSLGSLQT | 5.2306 | 5.2030 | 4.8480 | 5.3814 | 5.3597 | 5.1524 | 5.0916 | 5.0049 | 6.2151 | 6.0656 | 5.7687 | 5.4763 | 8.6730 | 8.7387 | 9.1034 | 8.6874 |

| | | | | | | | | | | | | | | | | | | | | | |
|---|---|---|---|---|---|---|---|---|---|---|---|---|---|---|---|---|---|---|---|---|---|
| ZC3HAV1_S302 | ZC3HAV1 | Q7Z2W4 | | 257 | DRFFQGSQEFLAS | 5.3996 | 5.8219 | 5.7264 | 5.9895 | 5.8107 | 6.3307 | 6.1119 | 7.0459 | 6.1587 | 6.2275 | 5.7863 | 6.2532 | 7.2433 | 6.4746 | 7.0030 | 6.6169 |
| ZC3HAV1_S310 | ZC3HAV1 | Q7Z2W4 | | 393 | SLGSLQTPEAVTT | 7.5927 | 7.8880 | 7.1427 | 7.4664 | 6.0616 | 6.7130 | 7.3926 | 6.8132 | 6.9330 | 7.1226 | 7.2579 | 7.2306 | 3.3799 | 4.0961 | 2.8989 | 4.0106 |
| ZC3HAV1_S335 | ZC3HAV1 | Q7Z2W4 | | 310 | DRARPPSGSSKAT | 6.7591 | 6.7852 | 6.3461 | 6.6736 | 6.2335 | 6.1656 | 6.4071 | 6.3553 | 6.7102 | 6.2672 | 6.3723 | 6.5255 | 5.7132 | 5.4927 | 5.6546 | 5.5389 |
| ZC3HAV1_S378 | ZC3HAV1 | Q7Z2W4 | | 284 | QISHRASLEDAPV | 5.5927 | 5.7965 | 5.9906 | 6.3373 | 6.3004 | 5.9475 | 5.7982 | 6.1036 | 6.5437 | 6.1908 | 5.8424 | 6.8111 | 6.6066 | 6.7953 | 6.5899 | 6.7534 |
| ZC3HAV1_S387 | ZC3HAV1 | Q7Z2W4 | | 302 | KFTYLGSQDRARP | 6.8389 | 8.3772 | 7.1764 | 7.6389 | 5.5467 | 6.2963 | 6.8026 | 6.5476 | 6.6369 | 7.4581 | 7.3459 | 7.6331 | 3.3815 | 4.3425 | 3.7292 | 4.2482 |
| ZC3HAV1_T273 | ZC3HAV1 | Q7Z2W4 | | 275 | ERSCTPSPDQISH | 6.1917 | 6.2143 | 5.9263 | 6.7346 | 6.1778 | 5.6690 | 6.5195 | 6.0529 | 6.6259 | 6.2894 | 6.9393 | 5.8338 | 5.4424 | 6.4970 | 5.9067 | 6.9792 |
| ZC3HAV1_T393 | ZC3HAV1 | Q7Z2W4 | | 273 | SAERSCTPSPDQI | 5.6813 | 6.3977 | 5.5370 | 6.3691 | 5.6024 | 6.1211 | 7.5606 | 7.1750 | 6.8405 | 5.0642 | 5.1334 | 5.7806 | 6.9758 | 7.3606 | 6.7082 | 5.6926 |
| ZC3HC1_S24 | ZC3HC1 | Q86WB0 | | 321 | LPLVPESPRRMMT | 6.3474 | 6.4413 | 5.9484 | 6.5009 | 6.3018 | 5.9730 | 6.3163 | 6.3842 | 6.3533 | 5.8752 | 6.0003 | 6.0265 | 6.3183 | 6.3405 | 6.4327 | 6.4398 |
| ZC3HC1_S24;T28 | ZC3HC1 | Q86WB0 | | 395 | GLEVPSSPLRKAK | 5.9484 | 6.0505 | 5.9177 | 6.3898 | 6.1806 | 6.0308 | 6.3324 | 6.1360 | 6.3028 | 5.8417 | 5.9461 | 5.9492 | 6.4810 | 6.9046 | 6.7427 | 6.8456 |
| ZC3HC1_S28 | ZC3HC1 | Q86WB0 | | 344 | SEQAEKSPGPIVS | 6.4375 | 6.3987 | 5.9766 | 6.2366 | 6.2677 | 6.0215 | 6.3295 | 6.3425 | 6.1996 | 5.8538 | 5.9304 | 6.0893 | 6.2888 | 6.5097 | 6.5821 | 6.5358 |
| ZC3HC1_S321 | ZC3HC1 | Q86WB0 | | 24 | WGAVVRSPEGTPQ | 6.5638 | 6.7660 | 6.3028 | 6.5182 | 6.0498 | 6.0797 | 6.0222 | 5.8721 | 6.4845 | 6.0670 | 6.2480 | 6.5688 | 5.9436 | 6.3413 | 6.0119 | 6.1602 |
| ZC3HC1_S335 | ZC3HC1 | Q86WB0 | | 62 | SQSVNGSPQAEQP | 6.3121 | 6.3775 | 5.7491 | 6.1628 | 6.0222 | 6.0618 | 5.8156 | 6.1870 | 6.4857 | 6.1370 | 6.4655 | 6.2774 | 6.5889 | 6.4365 | 6.6494 | 6.2715 |
| ZC3HC1_S344 | ZC3HC1 | Q86WB0 | | 28 | VRSPEGTPQKIRQ | 7.0722 | 6.8394 | 6.4284 | 6.6997 | 7.1928 | 6.9043 | 7.4577 | 7.2282 | 6.8983 | 6.3371 | 6.6926 | 6.4582 | 4.2077 | 4.9202 | 4.1483 | 4.5148 |
| ZC3HC1_S354 | ZC3HC1 | Q86WB0 | | 354 | IVSRTRSWDSSSP | 4.7408 | 4.1300 | 4.3820 | 4.2837 | 6.1313 | 5.5119 | 5.3238 | 5.4988 | 4.3274 | 3.8958 | 4.3204 | 4.4315 | 11.4233 | 10.3623 | 11.0703 | 10.1668 |
| ZC3HC1_S354;S358 | ZC3HC1 | Q86WB0 | | 407 | KRARLCSSSSSDT | 6.2496 | 6.2148 | 6.1749 | 6.3724 | 5.9077 | 5.9897 | 6.0942 | 6.3367 | 6.4092 | 5.5977 | 6.0421 | 5.9612 | 6.4214 | 6.4420 | 6.6501 | 7.1363 |
| ZC3HC1_S354;S370 | ZC3HC1 | Q86WB0 | | 87 | RVETFSSLKWAGK | 5.6797 | 6.3184 | 6.7416 | 6.0514 | 6.0311 | 5.2284 | 6.5714 | 5.5751 | 5.3924 | 5.9952 | 5.4736 | 6.0857 | 7.0836 | 7.5552 | 7.0243 | 7.1929 |
| ZC3HC1_S359 | ZC3HC1 | Q86WB0 | | 370 | PEPEAASPTTRTR | 7.9535 | 7.8167 | 7.1286 | 7.3281 | 6.3852 | 6.2997 | 6.6893 | 5.9274 | 7.5158 | 7.7662 | 8.0021 | 7.4699 | 2.9310 | 4.2305 | 3.1826 | 3.3734 |
| ZC3HC1_S370 | ZC3HC1 | Q86WB0 | | 359 | RSWDSSSPVDRPE | 7.5103 | 8.2468 | 8.4735 | 10.1405 | 6.1463 | 6.0490 | 4.8897 | 5.0027 | 8.0761 | 8.6284 | 8.0773 | 8.3152 | 2.3826 | 3.0186 | 2.5127 | 2.5302 |
| ZC3HC1_S395 | ZC3HC1 | Q86WB0 | | 335 | SQDATFSPGSEQA | 6.1986 | 6.8112 | 6.0080 | 7.2117 | 6.6942 | 6.4047 | 6.3940 | 6.6034 | 6.2563 | 6.8095 | 6.3716 | 7.1186 | 4.8934 | 5.5915 | 5.0880 | 5.5452 |
| ZC3HC1_S407 | ZC3HC1 | Q86WB0 | 24;28 | | NA | 6.7464 | 6.9903 | 6.7710 | 6.8247 | 6.3898 | 6.2386 | 6.1355 | 6.3913 | 6.8608 | 7.1368 | 6.7831 | 7.3593 | 4.4945 | 5.2321 | 4.8558 | 4.7899 |
| ZC3HC1_S62 | ZC3HC1 | Q86WB0 | 354;358 | | NA | 3.6792 | 5.2708 | 5.4046 | 5.4892 | 6.5307 | 5.5407 | 5.1393 | 6.1124 | 5.9858 | 4.3927 | 4.3442 | 4.5015 | 6.9827 | 9.2053 | 11.7753 | 9.6457 |
| ZC3HC1_S87 | ZC3HC1 | Q86WB0 | 354;370 | | NA | 4.5644 | 4.6162 | 4.8008 | 4.4588 | 6.0058 | 5.5373 | 6.0457 | 5.7515 | 5.4390 | 4.4450 | 4.3260 | 4.9027 | 10.7325 | 9.3501 | 9.1887 | 9.8356 |
| ZC3HC1_T387;S395 | ZC3HC1 | Q86WB0 | 387;395 | | NA | 7.2183 | 7.2249 | 7.2678 | 7.6856 | 6.0754 | 7.0351 | 7.5923 | 6.1317 | 7.4699 | 6.9547 | 7.0245 | 6.6465 | 3.7183 | 4.3574 | 3.5021 | 4.0953 |
| ZCCHC17_S114 | ZCCHC17 | Q9NP64 | | 170 | EKEEAKSAEFEKP | 6.0782 | 6.5566 | 6.2595 | 6.8471 | 5.8536 | 5.9061 | 6.0791 | 5.8685 | 6.4237 | 6.4244 | 6.4011 | 6.4412 | 5.5300 | 6.9254 | 6.0398 | 6.3657 |
| ZCCHC17_S170 | ZCCHC17 | Q9NP64 | | 114 | EERRRRSFQDYTG | 5.0735 | 5.5536 | 5.9872 | 6.3593 | 7.5969 | 6.3598 | 4.9649 | 5.6546 | 7.1696 | 8.0725 | 8.2033 | 8.1917 | 5.0905 | 6.1654 | 4.3962 | 5.1609 |
| ZCCHC3_S363 | ZCCHC3 | Q9NUD5 | | 363 | GEEGHLSPYCRKG | 6.5141 | 7.6130 | 6.7918 | 6.5568 | 5.6133 | 6.5403 | 6.8451 | 6.2642 | 7.2242 | 7.1260 | 7.1660 | 6.7202 | 4.4905 | 5.5349 | 4.4034 | 4.5962 |
| ZCCHC7_S481 | ZCCHC7 | Q8N3Z6 | | 482 | GPKTYSSPGSFKT | 7.0890 | 7.0487 | 6.1966 | 7.3202 | 6.8778 | 6.4110 | 7.1645 | 6.5493 | 6.9709 | 6.3311 | 6.3881 | 6.8018 | 4.4088 | 4.8184 | 4.6644 | 4.9595 |
| ZCCHC7_S482 | ZCCHC7 | Q8N3Z6 | | 481 | RGPKTYSSPGSFK | 7.7037 | 7.1721 | 6.2556 | 7.3763 | 6.9193 | 6.2113 | 6.7744 | 6.1845 | 7.3658 | 6.3243 | 6.7997 | 6.0537 | 3.8433 | 5.3333 | 4.7800 | 4.9027 |
| ZCCHC8_S597;S598 | ZCCHC8 | Q6NZY4 | | 649 | LFPADTSPSTATK | 6.2826 | 6.7230 | 6.7112 | 7.0444 | 6.4357 | 6.7214 | 6.6243 | 6.5476 | 6.6505 | 6.7491 | 6.7663 | 6.8697 | 5.1012 | 5.1045 | 4.5815 | 5.0870 |
| ZCCHC8_S649 | ZCCHC8 | Q6NZY4 | | 658 | TATKIHSPIPDMS | 6.4454 | 6.7384 | 6.4215 | 7.1080 | 6.3121 | 6.2705 | 6.6121 | 6.3330 | 6.9061 | 6.5093 | 6.6101 | 6.9087 | 5.2864 | 5.1879 | 4.9297 | 5.4208 |
| ZCCHC8_S658 | ZCCHC8 | Q6NZY4 | | 479 | PPLPRGTPPPVFT | 6.5097 | 6.6762 | 6.4176 | 6.7350 | 6.4422 | 5.9532 | 6.1384 | 6.3442 | 6.6442 | 6.7577 | 6.5760 | 6.3333 | 5.7011 | 5.7577 | 5.3954 | 5.6181 |
| ZCCHC8_T479 | ZCCHC8 | Q6NZY4 | | 492 | PPLPKGTPPLTPS | 6.1888 | 6.6351 | 6.7477 | 6.7182 | 5.9909 | 6.0120 | 5.7320 | 6.0951 | 6.3983 | 5.9941 | 6.1236 | 5.9425 | 6.2267 | 6.1783 | 6.9593 | 6.0573 |
| ZCCHC8_T479;T485 | ZCCHC8 | Q6NZY4 | 479;485 | | NA | 6.9844 | 7.1908 | 6.9493 | 7.7186 | 6.2278 | 6.9800 | 7.1037 | 6.7006 | 6.7978 | 7.0337 | 7.2831 | 7.5812 | 3.5217 | 4.1534 | 3.5446 | 4.2293 |
| ZCCHC8_T492 | ZCCHC8 | Q6NZY4 | 597;598 | | NA | 7.4128 | 5.8150 | 6.7477 | 6.8911 | 6.1470 | 6.4220 | 6.2085 | 6.9099 | 6.3143 | 5.9969 | 5.9182 | 6.5097 | 6.2367 | 5.6155 | 5.9145 | 4.9400 |
| ZCCHC9_S48 | ZCCHC9 | Q8N567 | | 48 | LEANRLSLKNDAP | 5.2760 | 6.5536 | 7.1175 | 7.2871 | 5.4048 | 6.7352 | 6.4767 | 7.0305 | 7.2483 | 9.3082 | 9.0707 | 9.0704 | 2.9357 | 4.2765 | 2.5970 | 3.6119 |
| ZCRB1_S155 | ZCRB1 | Q8TBF4 | | 155 | IEEVEESEDEGED | 6.1203 | 6.3273 | 5.9899 | 6.4575 | 6.1045 | 6.0037 | 6.0489 | 6.2767 | 6.4509 | 6.4380 | 6.4764 | 6.3565 | 6.1696 | 6.5958 | 6.0544 | 6.1296 |
| ZCRB1_S210;S216 | ZCRB1 | Q8TBF4 | 210;216 | | NA | 6.3005 | 6.6849 | 6.2561 | 6.5227 | 6.0532 | 5.9025 | 5.9018 | 5.9504 | 6.4832 | 6.3794 | 6.3068 | 6.3160 | 6.0519 | 6.5053 | 6.2033 | 6.1821 |
| ZDHHC14_S455 | ZDHHC14 | Q8IZN3 | | 455 | RLLAAGSPLAHSR | 5.8564 | 5.9864 | 6.4090 | 6.4066 | 6.0539 | 6.0672 | 6.4405 | 5.6896 | 6.2182 | 6.3491 | 6.4678 | 6.6644 | 6.5617 | 6.3733 | 5.8279 | 6.6281 |
| ZDHHC18_S19 | ZDHHC18 | Q9NUE0 | | 19 | AAPLPASPGARRP | 6.1853 | 6.6473 | 6.6570 | 6.6108 | 5.7420 | 5.9904 | 6.4781 | 6.3676 | 6.4864 | 6.7319 | 6.3313 | 6.5752 | 5.9254 | 5.6826 | 5.3929 | 6.1959 |
| ZDHHC18_S30 | ZDHHC18 | Q9NUE0 | | 366 | SDTVLPSPIRSDE | 7.0107 | 7.4620 | 6.9089 | 7.8295 | 6.1153 | 6.3737 | 6.8020 | 7.0954 | 7.5331 | 7.1493 | 7.0643 | 7.2226 | 3.4280 | 4.4190 | 3.4693 | 4.1170 |
| ZDHHC18_S366 | ZDHHC18 | Q9NUE0 | | 30 | RPGPAASPTPGPG | 5.6866 | 7.8422 | 7.6440 | 6.9557 | 6.3455 | 6.2981 | 5.8052 | 7.0855 | 7.4094 | 6.8538 | 6.6904 | 6.6890 | 4.9703 | 4.5152 | 4.2136 | 4.9955 |
| ZDHHC18_S61 | ZDHHC18 | Q9NUE0 | | 61 | SGSGSGSLGRRPR | 6.4583 | 6.9367 | 6.1588 | 6.2454 | 6.0891 | 6.0964 | 6.0978 | 5.9838 | 6.5870 | 6.1021 | 6.3558 | 6.2172 | 5.7966 | 6.0910 | 6.3090 | 6.4749 |
| ZDHHC20_S305 | ZDHHC20 | Q5W0Z9 | | 305 | MDPEQASVTNQNE | 5.3071 | 6.6716 | 6.7588 | 7.2523 | 6.1790 | 5.4610 | 6.4323 | 6.8080 | 6.5853 | 6.6586 | 5.5881 | 5.9555 | 5.8599 | 6.4879 | 6.3952 | 5.5993 |
| ZDHHC23_S206 | ZDHHC23 | Q8IYP9 | | 252 | SKMPAGSPTKAKE | 6.3994 | 6.8584 | 6.0151 | 6.7719 | 6.2809 | 6.3788 | 6.3689 | 6.4169 | 6.4034 | 6.4821 | 6.3287 | 6.4478 | 5.5966 | 5.7851 | 5.7366 | 5.7294 |
| ZDHHC23_S252 | ZDHHC23 | Q8IYP9 | | 206 | SGDRSLSSSQLEC | 3.1263 | 2.9879 | 3.0498 | 2.7110 | 3.7636 | 3.5878 | 3.5008 | 3.4616 | 3.5265 | 2.8419 | 2.8454 | 3.2023 | 15.6807 | 14.2694 | 16.9618 | 14.4832 |
| ZDHHC5_S274 | ZDHHC5 | Q9C0B5 | | 348 | AKDSPPTPTMYKY | 6.1119 | 6.4404 | 5.9695 | 6.3126 | 5.5386 | 6.3201 | 6.8418 | 5.8502 | 5.5089 | 5.4000 | 5.7913 | 5.8995 | 6.7216 | 6.6887 | 7.2697 | 7.3353 |
| ZDHHC5_S296;S299 | ZDHHC5 | Q9C0B5 | | 636 | YLGRSMSYSSQKA | 7.7730 | 7.1094 | 7.0319 | 6.8895 | 6.0628 | 6.3063 | 6.7228 | 5.8096 | 6.4407 | 5.8070 | 6.2728 | 6.3126 | 5.1662 | 6.3687 | 4.5474 | 5.3793 |
| ZDHHC5_S299 | ZDHHC5 | Q9C0B5 | | 694 | GQPPLSSPTRGGV | 6.1647 | 6.4944 | 6.3274 | 6.6182 | 6.5170 | 6.3597 | 6.7719 | 6.5424 | 6.8131 | 6.5794 | 6.7676 | 6.5591 | 5.3012 | 5.5263 | 5.2619 | 5.3957 |
| ZDHHC5_S345;T348 | ZDHHC5 | Q9C0B5 | | 425 | DPLSSGSRSSSLK | 6.3459 | 7.0489 | 5.9885 | 7.0460 | 6.6358 | 6.7049 | 5.9112 | 7.0634 | 6.2275 | 5.4395 | 6.0529 | 5.6888 | 5.8660 | 6.1510 | 6.3312 | 5.4986 |
| ZDHHC5_S371 | ZDHHC5 | Q9C0B5 | | 621 | RGRGVGSPEPGPT | 6.2534 | 6.7714 | 6.6576 | 6.9453 | 6.8128 | 7.0127 | 7.1093 | 7.1051 | 6.5914 | 6.9713 | 6.9717 | 6.7382 | 4.2925 | 4.9428 | 4.0153 | 4.8092 |
| ZDHHC5_S398 | ZDHHC5 | Q9C0B5 | | 529 | PTHREPSPVRYDN | 6.1074 | 6.2896 | 5.9342 | 6.4489 | 6.2462 | 7.0424 | 6.6470 | 5.7164 | 5.7783 | 5.8224 | 6.2829 | 5.8559 | 6.8376 | 6.8027 | 6.1025 | 6.0855 |
| ZDHHC5_S409 | ZDHHC5 | Q9C0B5 | | 409 | EPESFRSPTFGKS | 5.5997 | 6.2792 | 6.1266 | 6.5308 | 6.1177 | 7.1691 | 6.9110 | 6.6844 | 6.1825 | 6.5194 | 6.5281 | 6.6027 | 4.9845 | 6.2954 | 5.4069 | 6.0620 |
| ZDHHC5_S415;S425 | ZDHHC5 | Q9C0B5 | | 455 | EGTTSTSYKSLAN | 6.0421 | 6.4182 | 6.3289 | 6.4349 | 5.6346 | 6.0442 | 6.6377 | 5.9222 | 5.8146 | 6.2150 | 6.7031 | 6.4146 | 5.9837 | 6.4047 | 6.3490 | 6.6526 |
| ZDHHC5_S425 | ZDHHC5 | Q9C0B5 | | 554 | EKLLRQSPPLPGR | 6.0215 | 7.0117 | 6.2061 | 6.5442 | 6.0902 | 5.5802 | 6.2323 | 6.6556 | 6.5031 | 6.4017 | 6.0287 | 6.0636 | 6.0422 | 6.1481 | 6.0180 | 6.4528 |
| ZDHHC5_S432 | ZDHHC5 | Q9C0B5 | | 398 | RHPSYRSEPSLEP | 5.3332 | 6.3949 | 6.3215 | 6.1928 | 4.9001 | 5.3759 | 5.8851 | 6.1581 | 6.3917 | 6.3054 | 6.4552 | 6.6117 | 6.1908 | 6.9009 | 7.1011 | 7.4814 |
| ZDHHC5_S455 | ZDHHC5 | Q9C0B5 | | 659 | ALQPLLTPKDEVQ | 6.5779 | 6.6186 | 6.2679 | 7.5966 | 6.7105 | 6.7427 | 7.2700 | 6.2607 | 6.6038 | 6.5961 | 7.2268 | 7.0731 | 4.5363 | 5.4271 | 3.8506 | 4.6414 |
| ZDHHC5_S499 | ZDHHC5 | Q9C0B5 | | 432 | RSSSLKSAQGTGF | 5.8469 | 7.0530 | 6.4936 | 6.7462 | 6.2914 | 6.0638 | 6.2614 | 6.2512 | 6.6809 | 6.2986 | 6.2334 | 5.9774 | 6.2046 | 6.1090 | 5.7858 | 5.7028 |
| ZDHHC5_S529 | ZDHHC5 | Q9C0B5 | | 299 | RTKSKGSLEITES | 6.4274 | 6.7616 | 6.2477 | 6.7439 | 7.1135 | 7.0211 | 5.8794 | 5.7427 | 7.1628 | 6.4717 | 5.2212 | 5.9022 | 6.9768 | 6.4034 | 5.5933 | 4.3314 |
| ZDHHC5_S554 | ZDHHC5 | Q9C0B5 | | 274 | FLRPEVSDGQITV | 2.0929 | 2.8096 | 2.5596 | 2.9507 | 2.4599 | 2.9897 | 2.6938 | 3.0596 | 2.2477 | 2.7649 | 2.3750 | 2.6330 | 16.1296 | 16.2408 | 19.5619 | 16.4312 |
| ZDHHC5_S621 | ZDHHC5 | Q9C0B5 | | 499 | PPLGYTSPFLSAR | 6.4751 | 5.7956 | 5.8458 | 6.7580 | 6.8320 | 6.6681 | 5.2800 | 5.9727 | 4.6164 | 6.2847 | 5.8220 | 5.6713 | 6.7077 | 6.2239 | 6.8505 | 8.1963 |
| ZDHHC5_S636 | ZDHHC5 | Q9C0B5 | | 371 | AAMPHSSSAKLSR | 5.8209 | 5.5736 | 5.3990 | 6.5469 | 4.7836 | 6.4174 | 5.4586 | 5.3655 | 5.0062 | 3.5586 | 5.8253 | 5.3539 | 8.0221 | 9.1074 | 9.6021 | 8.1589 |
| ZDHHC5_S694 | ZDHHC5 | Q9C0B5 | 345;348 | | NA | 6.1312 | 6.2828 | 6.4718 | 6.4358 | 6.2741 | 6.3880 | 6.7616 | 6.2548 | 6.5667 | 6.4252 | 6.4195 | 6.4992 | 5.6505 | 6.3302 | 5.4612 | 5.6475 |
| ZDHHC5_T348 | ZDHHC5 | Q9C0B5 | 296;299 | | NA | 5.0062 | 5.9978 | 5.9226 | 5.8977 | 4.9498 | 5.7651 | 5.8226 | 5.5598 | 5.7941 | 6.6917 | 7.4436 | 7.0133 | 6.2694 | 6.8897 | 7.5487 | 7.4279 |
| ZDHHC5_T659 | ZDHHC5 | Q9C0B5 | 415;425 | | NA | 7.6811 | 7.3841 | 8.1213 | 7.3854 | 5.2905 | 6.1873 | 6.2168 | 5.9252 | 7.8316 | 7.5659 | 7.8578 | 7.1907 | 3.6539 | 5.0742 | 2.8298 | 3.8045 |
| ZDHHC7_S344 | ZDHHC7 | Q9NXF8-2 | | 344 | KGGPEFSVxxxxx | 7.1519 | 6.3492 | 6.5715 | 5.0799 | 6.8753 | 7.1765 | 5.4528 | 6.4036 | 6.9247 | 5.2108 | 6.6826 | 4.9194 | 5.8306 | 6.3221 | 7.0827 | 5.9665 |
| ZDHHC8_S337 | ZDHHC8 | Q9ULC8 | | 754 | TLVKKVSGVGGTT | 7.1148 | 6.6214 | 6.2023 | 6.9335 | 6.9984 | 6.2754 | 5.8363 | 6.2275 | 7.3112 | 6.8209 | 6.0388 | 6.3812 | 5.0413 | 6.0286 | 5.0101 | 5.1580 |
| ZDHHC8_S526 | ZDHHC8 | Q9ULC8 | | 351 | QRTSPPTPAMYKF | 7.3124 | 7.1119 | 6.7223 | 6.6272 | 6.1021 | 6.0694 | 6.6635 | 6.1715 | 6.2019 | 5.6837 | 6.0343 | 6.2398 | 5.9063 | 6.1681 | 5.3265 | 5.6592 |
| ZDHHC8_S627 | ZDHHC8 | Q9ULC8 | | 743 | PPGPSASPTRHTL | 6.6603 | 6.7395 | 6.4155 | 6.4704 | 7.3529 | 6.1744 | 6.7562 | 6.9005 | 6.6268 | 5.9538 | 6.8446 | 5.7574 | 4.9596 | 5.7136 | 5.2203 | 5.4539 |
| ZDHHC8_S675 | ZDHHC8 | Q9ULC8 | | 337 | QTPRPGSAESALS | 5.9892 | 4.7469 | 5.5929 | 7.6896 | 5.6001 | 5.6577 | 5.0366 | 5.7831 | 6.2183 | 8.6912 | 7.2761 | 7.0152 | 5.7903 | 6.0526 | 6.1880 | 6.6722 |
| ZDHHC8_S743 | ZDHHC8 | Q9ULC8 | | 526 | PLPRSFSPVLGPR | 6.7475 | 6.4589 | 5.9928 | 6.1352 | 5.9409 | 5.7833 | 6.5707 | 6.3499 | 6.0670 | 6.2478 | 6.3430 | 6.1600 | 6.0239 | 6.4378 | 6.3114 | 6.4301 |
| ZDHHC8_S754 | ZDHHC8 | Q9ULC8 | | 627 | NPALQTSLSSLSS | 6.1979 | 5.4127 | 6.6641 | 6.5252 | 5.9215 | 6.6105 | 7.2265 | 5.4457 | 6.6297 | 5.9421 | 7.1495 | 6.0123 | 6.5211 | 6.1525 | 5.5117 | 6.0769 |
| ZDHHC8_T351 | ZDHHC8 | Q9ULC8 | | 675 | ARQGLPSPPGTPH | 6.5875 | 6.0270 | 6.3549 | 5.9702 | 5.3089 | 6.4611 | 7.1060 | 6.0966 | 6.5069 | 6.9447 | 7.3624 | 6.5271 | 5.1037 | 6.9173 | 4.5724 | 6.1534 |
| ZEB1_S323 | ZEB1 | P37275-2 | | 522 | KSEKDKSFEGGVN | 6.1899 | 6.5523 | 6.7387 | 7.3603 | 6.3510 | 6.3539 | 6.6262 | 6.5516 | 6.8030 | 7.1731 | 6.8408 | 7.3526 | 4.4107 | 5.1080 | 4.4319 | 5.1560 |
| ZEB1_S46 | ZEB1 | P37275-2 | | 647 | QSSEPSSPEPGKV | 6.5833 | 6.5252 | 6.6197 | 7.0775 | 6.6020 | 6.6737 | 6.6939 | 6.6989 | 6.2426 | 6.8236 | 6.7962 | 6.5743 | 5.2213 | 5.3453 | 4.8474 | 4.6750 |
| ZEB1_S522 | ZEB1 | P37275-2 | | 323 | LSASPGSPTRPQI | 5.8526 | 5.9830 | 6.3341 | 6.6996 | 5.9735 | 6.0649 | 6.4177 | 6.3287 | 6.0222 | 6.0728 | 6.0603 | 6.3200 | 6.4762 | 6.3275 | 6.3557 | 6.7112 |
| ZEB1_S584 | ZEB1 | P37275-2 | | 584 | TGDGNLSPSQPPL | 6.0546 | 6.1815 | 6.7598 | 6.9188 | 6.6177 | 6.3311 | 5.9248 | 6.8275 | 6.4169 | 6.6889 | 6.3706 | 6.7954 | 5.0515 | 5.4391 | 5.5109 | 6.1109 |
| ZEB1_S639;S647 | ZEB1 | P37275-2 | | 680 | STVNLQSPLKMTN | 5.9902 | 5.7759 | 7.0022 | 6.4659 | 5.6708 | 6.2400 | 6.7272 | 6.5348 | 6.4065 | 5.8053 | 6.3826 | 7.2738 | 6.1786 | 5.7507 | 5.3351 | 6.4604 |
| ZEB1_S643;S647 | ZEB1 | P37275-2 | | 46 | HIVEEESVTDAAD | 5.9802 | 6.5398 | 6.4728 | 7.4706 | 5.6662 | 5.9740 | 5.9876 | 6.3140 | 6.0013 | 6.1527 | 6.9394 | 5.8859 | 6.2433 | 6.6739 | 6.6885 | 6.0100 |
| ZEB1_S647 | ZEB1 | P37275-2 | | 699 | GSTTNGSRSSTPS | 6.4705 | 6.7672 | 7.3576 | 7.6457 | 5.8927 | 6.0442 | 6.3747 | 6.0002 | 6.4469 | 6.6738 | 6.1839 | 5.8667 | 5.8353 | 5.4161 | 5.4271 | 5.5975 |
| ZEB1_S680 | ZEB1 | P37275-2 | | 687 | PLKMTNSPVLPVG | 5.0911 | 5.8828 | 6.1017 | 6.3356 | 6.1008 | 6.2924 | 6.4592 | 7.3630 | 7.4151 | 5.7513 | 6.2156 | 6.5642 | 6.4186 | 6.2871 | 5.5359 | 6.1856 |
| ZEB1_S687 | ZEB1 | P37275-2 | | 703 | NGSRSSTPSPSPL | 5.5520 | 7.0517 | 7.4688 | 7.5980 | 5.4228 | 6.3134 | 7.2446 | 6.5801 | 6.0550 | 6.2110 | 6.7093 | 6.0688 | 5.5908 | 5.8833 | 4.7445 | 5.5059 |
| ZEB1_S699 | ZEB1 | P37275-2 | 643;647 | | NA | 5.4997 | 5.8813 | 5.6447 | 6.1386 | 5.3272 | 5.7686 | 6.2068 | 5.9313 | 6.6891 | 6.3834 | 6.6073 | 6.5311 | 6.7308 | 7.0978 | 7.3885 | 6.1740 |
| ZEB1_T703 | ZEB1 | P37275-2 | 639;647 | | NA | 5.7247 | 6.5790 | 6.3195 | 5.9841 | 5.3852 | 5.6864 | 6.3171 | 5.3193 | 5.3824 | 6.3003 | 6.2383 | 7.2787 | 6.9297 | 7.4684 | 6.6699 | 6.4170 |
| ZEB2_S360 | ZEB2 | O60315 | | 360 | PNSVSSSPTNSAI | 5.8409 | 6.1654 | 7.4767 | 7.8395 | 5.5707 | 5.5234 | 5.1907 | 5.2119 | 7.4444 | 6.8961 | 7.1753 | 7.6480 | 5.7995 | 5.4276 | 5.4840 | 5.3056 |
| ZFAND2B_S173 | ZFAND2B | Q8WV99 | | 173 | TMPSCTSPSRATT | 6.0538 | 6.0932 | 7.4535 | 6.3844 | 6.5606 | 6.7453 | 6.4924 | 7.1880 | 6.6179 | 6.7948 | 6.8427 | 6.7129 | 5.4602 | 5.0431 | 4.6559 | 4.9010 |
| ZFAND2B_S183 | ZFAND2B | Q8WV99 | | 183 | ATTRSPSWTAPPV | 5.6943 | 5.5315 | 6.3130 | 5.9086 | 5.3177 | 5.7866 | 5.5283 | 5.8215 | 5.9581 | 5.8469 | 6.0362 | 5.9209 | 7.6791 | 7.3527 | 7.9779 | 7.3269 |
| ZFAND3_T112 | ZFAND3 | Q9H8U3 | | 112 | AHVSLITPTKRSC | 6.6125 | 8.0133 | 7.3833 | 8.5478 | 7.1395 | 7.0451 | 5.9900 | 8.5935 | 5.7010 | 5.7885 | 6.5130 | 5.9951 | 3.7738 | 4.5348 | 4.3437 | 4.0250 |
| ZFAND5_S48 | ZFAND5 | O76080 | | 48 | QNSGRMSPMGTAS | 5.9051 | 6.1874 | 6.3389 | 5.5097 | 6.2711 | 6.8269 | 6.1989 | 7.2078 | 6.6232 | 6.3151 | 6.3033 | 6.8879 | 6.4189 | 5.9220 | 5.5984 | 5.4854 |
| ZFC3H1_S1046 | ZFC3H1 | O60293 | | 352 | NLTRRISTSDILS | 4.5228 | 4.3313 | 4.7095 | 5.7601 | 6.3734 | 5.3432 | 5.4582 | 5.9216 | 6.4448 | 6.6845 | 6.3035 | 6.5717 | 7.8678 | 8.1886 | 7.8559 | 7.6632 |
| ZFC3H1_S1301 | ZFC3H1 | O60293 | | 1046 | SRERRRSFLESNY | 4.4995 | 4.5108 | 4.8389 | 5.5314 | 6.3563 | 5.5813 | 5.4108 | 6.0681 | 5.9860 | 6.6542 | 6.2703 | 6.2442 | 7.8229 | 7.9481 | 8.1335 | 8.1435 |
| ZFC3H1_S194 | ZFC3H1 | O60293 | | 655 | SNSDPPSPPVLNN | 6.0442 | 5.9507 | 6.0057 | 7.2473 | 6.1858 | 5.9876 | 6.5872 | 5.8200 | 6.3864 | 6.5598 | 5.9333 | 6.3830 | 6.0221 | 6.0583 | 6.3008 | 6.5278 |
| ZFC3H1_S352 | ZFC3H1 | O60293 | | 949 | MMRLDSSPVSSPR | 5.8359 | 6.0755 | 5.8109 | 6.4170 | 6.1071 | 5.7006 | 6.2422 | 6.0188 | 6.0336 | 5.8773 | 5.6626 | 5.7400 | 7.0022 | 6.8684 | 7.3581 | 7.2500 |
| ZFC3H1_S381 | ZFC3H1 | O60293 | | 1301 | KPISDNSFSSDEE | 5.7935 | 6.0610 | 6.4357 | 6.5487 | 6.1725 | 5.6508 | 6.3856 | 5.8832 | 6.3395 | 6.3907 | 6.0647 | 6.2329 | 6.6282 | 6.8549 | 6.1448 | 6.4132 |
| ZFC3H1_S655 | ZFC3H1 | O60293 | | 194 | QSWREPSPPRKSS | 6.1771 | 6.7004 | 5.9762 | 6.5690 | 6.0792 | 5.8887 | 6.0863 | 5.7444 | 6.2925 | 6.2186 | 5.9740 | 6.3817 | 6.2057 | 6.5218 | 6.5834 | 6.6011 |
| ZFC3H1_S949 | ZFC3H1 | O60293 | | 1333 | ALDTVVTPDDVRY | 5.6063 | 6.8116 | 6.2510 | 6.5003 | 5.9970 | 5.7095 | 6.6221 | 5.5853 | 7.0348 | 6.5591 | 5.9079 | 6.5624 | 6.3724 | 6.2178 | 6.0311 | 6.2314 |
| ZFC3H1_S949;S953 | ZFC3H1 | O60293 | | 766 | ENDPLRTPEALPE | 5.8258 | 6.1506 | 6.2420 | 6.6954 | 6.1073 | 5.8162 | 5.5528 | 5.8739 | 5.5576 | 6.6857 | 5.7428 | 6.2059 | 5.8980 | 7.0429 | 7.0538 | 7.5493 |
| ZFC3H1_S998 | ZFC3H1 | O60293 | | 998 | KEQQNISPVVEEE | 5.6173 | 6.4736 | 7.0992 | 6.7633 | 5.6889 | 6.1505 | 5.1478 | 6.4481 | 5.4582 | 6.3302 | 5.0791 | 5.7243 | 6.5433 | 7.2946 | 6.9911 | 7.1905 |
| ZFC3H1_T1333 | ZFC3H1 | O60293 | | 381 | RLLALQSASKKWQ | 9.0545 | 9.6249 | 7.4817 | 7.4498 | 6.9857 | 6.4721 | 7.4179 | 6.2595 | 7.7438 | 6.1320 | 5.8946 | 6.4048 | 2.5754 | 5.1862 | 2.5647 | 2.7525 |
| ZFC3H1_T766 | ZFC3H1 | O60293 | 949;953 | | NA | 7.1078 | 7.1080 | 6.4023 | 7.0142 | 6.1346 | 5.4395 | 6.8107 | 6.5685 | 6.8700 | 6.0232 | 6.1061 | 6.7959 | 5.3037 | 5.5847 | 5.1130 | 5.6176 |
| ZFHX3_S1189 | ZFHX3 | Q15911 | | 2230 | IDSRPPSPEPPKQ | 6.7217 | 6.4371 | 6.1883 | 6.7813 | 6.5509 | 6.1131 | 6.6682 | 6.1577 | 6.2447 | 6.5052 | 5.8886 | 5.9607 | 5.6498 | 5.6438 | 6.3967 | 6.0922 |
| ZFHX3_S1590 | ZFHX3 | Q15911 | | 2801 | SPRTLLSPSSIKV | 7.0018 | 7.3996 | 6.2259 | 7.3229 | 6.7217 | 6.5811 | 6.8754 | 6.3902 | 6.7549 | 6.8370 | 6.4186 | 6.5677 | 4.7243 | 4.9576 | 4.4735 | 4.7478 |
| ZFHX3_S2230 | ZFHX3 | Q15911 | | 1189 | EKELTDSPATSKR | 6.6156 | 7.4584 | 6.7268 | 7.2779 | 6.2934 | 6.1342 | 6.2546 | 6.4494 | 6.9189 | 6.8766 | 6.9847 | 7.2914 | 4.0671 | 5.3212 | 4.3010 | 5.0288 |
| ZFHX3_S2801 | ZFHX3 | Q15911 | | 426 | LGPLASSPTKSSE | 6.2084 | 6.3824 | 6.2093 | 6.7172 | 5.9823 | 6.1871 | 5.8977 | 6.4580 | 6.3941 | 6.1016 | 6.6201 | 6.2616 | 5.7574 | 6.4373 | 6.1416 | 6.2438 |
| ZFHX3_S3432 | ZFHX3 | Q15911 | | 3432 | NTPREVSPLLPKL | 5.9250 | 6.0754 | 6.2489 | 6.4049 | 5.4667 | 6.3484 | 7.1473 | 5.8622 | 6.0619 | 6.2481 | 6.2059 | 5.7452 | 6.0834 | 7.3271 | 5.9441 | 6.9054 |
| ZFHX3_S426 | ZFHX3 | Q15911 | | 533 | NQSISNSPLMPNV | 5.9013 | 6.5308 | 6.6001 | 6.3236 | 6.1342 | 6.3768 | 5.7941 | 7.3883 | 6.1939 | 6.1238 | 6.4728 | 6.7629 | 5.4991 | 6.2906 | 5.3718 | 6.2358 |
| ZFHX3_S533 | ZFHX3 | Q15911 | | 1590 | QPEPTSSPDNKPF | 6.3756 | 7.2541 | 6.1258 | 7.4559 | 6.2601 | 6.5288 | 6.1611 | 6.8322 | 6.8273 | 6.5167 | 7.6145 | 6.6462 | 5.2103 | 5.3136 | 3.9982 | 4.8796 |
| ZFHX4_S1095 | ZFHX4 | Q86UP3-5 | | 2769 | SPKNLLSPSSFKA | 6.9462 | 6.9376 | 6.7026 | 7.0842 | 6.3764 | 6.3791 | 6.5574 | 6.1290 | 6.6915 | 6.4686 | 6.5796 | 6.7109 | 4.9655 | 5.3462 | 5.0188 | 5.1062 |
| ZFHX4_S1500 | ZFHX4 | Q86UP3-5 | | 1174 | KDSGIITPEKELK | 6.3426 | 6.8170 | 6.3297 | 7.0616 | 6.2187 | 6.0476 | 6.5501 | 6.2575 | 6.3274 | 6.5811 | 6.3995 | 6.6982 | 5.3568 | 5.7467 | 5.5080 | 5.7576 |
| ZFHX4_S2191 | ZFHX4 | Q86UP3-5 | | 508 | LLNQSISPLSSSV | 6.2654 | 6.5722 | 6.7401 | 7.0655 | 6.0491 | 6.2579 | 6.5267 | 6.0476 | 6.2716 | 6.3346 | 6.6240 | 6.3774 | 5.5610 | 6.0098 | 5.8236 | 5.4734 |
| ZFHX4_S2394 | ZFHX4 | A0A494BZV7 | | 293 | STSGIITPEKELK | 6.4387 | 6.6504 | 6.5919 | 7.1118 | 6.0820 | 6.1489 | 6.8235 | 6.4539 | 6.5321 | 6.6502 | 6.8915 | 7.0890 | 4.7839 | 5.3522 | 5.1799 | 5.2201 |
| ZFHX4_S2403 | ZFHX4 | Q86UP3-5 | | 2191 | RQRNKDSPYNFSN | 6.2917 | 7.6937 | 6.7728 | 6.9655 | 6.4845 | 5.3282 | 8.3262 | 8.4115 | 7.8378 | 6.8743 | 7.6760 | 6.6110 | 4.4912 | 4.3176 | 3.3320 | 2.5861 |
| ZFHX4_S2763 | ZFHX4 | Q86UP3-5 | | 2772 | NLLSPSSFKAECS | 5.7266 | 5.9628 | 5.7650 | 6.1560 | 5.9684 | 5.0373 | 7.1956 | 5.4995 | 5.3639 | 4.0159 | 5.1244 | 5.7662 | 8.2894 | 7.8758 | 8.6329 | 7.6204 |
| ZFHX4_S2769 | ZFHX4 | Q86UP3-5 | | 2870 | FLFSLTSPSIHFN | 7.0319 | 7.3776 | 5.9795 | 7.7273 | 7.0522 | 7.1990 | 6.4109 | 7.2255 | 6.1937 | 6.0947 | 6.1140 | 6.2278 | 4.2959 | 5.2713 | 4.9101 | 4.8888 |
| ZFHX4_S2769;S2772 | ZFHX4 | Q86UP3-5 | | 2903 | DRSETSSIADPSS | 6.6058 | 6.5311 | 6.0098 | 6.4669 | 6.3141 | 5.9058 | 5.9347 | 6.1425 | 6.5051 | 6.4329 | 6.3288 | 6.2168 | 5.8134 | 6.4757 | 6.4493 | 5.8671 |
| ZFHX4_S2772 | ZFHX4 | Q86UP3-5 | | 2854 | LPKPATTPTTEVC | 5.5617 | 5.3539 | 6.2161 | 6.5737 | 5.8649 | 5.5516 | 6.0288 | 5.5935 | 5.5909 | 5.9246 | 5.9801 | 6.0585 | 7.4466 | 7.2588 | 7.2744 | 7.7219 |
| ZFHX4_S2870 | ZFHX4 | Q86UP3-5 | | 1095 | PEEDNLSEIFFVK | 6.1430 | 6.5418 | 6.7672 | 6.7602 | 6.4665 | 6.3396 | 6.4809 | 6.9929 | 6.1300 | 5.7851 | 6.1996 | 5.9495 | 5.5028 | 6.5479 | 6.1407 | 5.2524 |
| ZFHX4_S2903 | ZFHX4 | Q86UP3-5 | | 1500 | DHEGKASPVGSDS | 5.7317 | 6.7217 | 6.2228 | 6.2228 | 5.9297 | 6.2659 | 6.2993 | 6.1498 | 6.0505 | 6.7803 | 6.0128 | 5.5636 | 6.4088 | 6.1857 | 6.5572 | 6.8975 |
| ZFHX4_S2903;S2908 | ZFHX4 | Q86UP3-5 | | 2763 | SKTAELSPKNLLS | 5.8377 | 6.1465 | 6.2568 | 6.3686 | 5.5287 | 6.1626 | 6.4782 | 5.4879 | 5.8962 | 6.5324 | 6.0399 | 5.9564 | 6.6462 | 6.8668 | 6.5760 | 7.2192 |
| ZFHX4_S508 | ZFHX4 | Q86UP3-5 | | 2394 | STPLIPSPKPEPE | 6.3951 | 7.2639 | 7.5556 | 7.1735 | 6.8654 | 6.1969 | 7.3390 | 5.5393 | 6.2949 | 7.2475 | 6.0162 | 6.5116 | 4.9885 | 5.8409 | 3.9335 | 4.8382 |
| ZFHX4_S757 | ZFHX4 | Q86UP3-5 | | 757 | SGCGTPSPSKPKQ | 5.5536 | 6.4191 | 5.8295 | 7.0446 | 7.3268 | 6.5897 | 6.3349 | 6.3854 | 6.3328 | 7.1815 | 5.2756 | 7.1289 | 6.1369 | 6.3707 | 5.4146 | 4.6754 |
| ZFHX4_T1174 | ZFHX4 | Q86UP3-5 | | 2403 | PEPEKTSPKPEYP | 5.4793 | 6.0657 | 5.6546 | 6.6136 | 5.8460 | 5.8573 | 6.1919 | 5.2937 | 6.1627 | 6.8995 | 6.6296 | 6.6026 | 6.9313 | 6.6971 | 6.3547 | 6.7203 |
| ZFHX4_T2854 | ZFHX4 | Q86UP3-5 | | 478 | EDEEDAYSNELDD | 7.0830 | 6.3038 | 5.7402 | 6.7396 | 5.6522 | 6.2016 | 6.5847 | 5.9181 | 6.3170 | 5.8388 | 6.3628 | 6.6287 | 5.8961 | 6.0484 | 6.2641 | 6.4209 |
| ZFHX4_T293 | ZFHX4 | Q86UP3-5 | 2903;2908 | | NA | 6.3367 | 6.0985 | 5.7573 | 8.2548 | 5.5706 | 5.9306 | 5.6267 | 6.9600 | 6.1644 | 5.8760 | 6.3249 | 6.4248 | 6.2445 | 5.8051 | 6.3959 | 6.2291 |
| ZFHX4_Y478 | ZFHX4 | Q86UP3-5 | 2769;2772 | | NA | 5.6424 | 7.2947 | 6.4230 | 8.1279 | 6.5077 | 4.2696 | 7.6880 | 6.8279 | 6.7509 | 5.7536 | 7.0261 | 6.5838 | 5.3175 | 4.9068 | 4.5189 | 6.3612 |
| ZFP36_S186 | ZFP36 | P26651 | | 186 | VLRQSISFSGLPS | 5.1541 | 4.7813 | 5.0683 | 4.6404 | 6.0624 | 5.0310 | 4.9372 | 5.5489 | 4.9300 | 4.6182 | 4.9059 | 4.9268 | 10.9285 | 9.6846 | 9.1179 | 9.6644 |
| ZFP36_S60 | ZFP36 | P26651 | | 257 | PSCRRATPISVWG | 5.6308 | 6.7756 | 6.6940 | 7.1790 | 6.5891 | 5.3300 | 6.8736 | 7.6640 | 6.5052 | 5.9922 | 6.9388 | 7.0624 | 5.3370 | 5.0992 | 5.4373 | 4.8917 |
| ZFP36_S93 | ZFP36 | P26651 | | 93 | LSPSPTSPTATST | 6.1256 | 5.5027 | 5.9305 | 5.3812 | 6.7408 | 6.7110 | 5.3812 | 6.8261 | 6.6451 | 6.6464 | 5.7431 | 5.7108 | 6.7253 | 7.5084 | 6.6335 | 5.7883 |
| ZFP36_T257 | ZFP36 | P26651 | | 60 | LPGRSTSLVEGRS | 4.5668 | 4.8083 | 5.0130 | 4.7254 | 5.9235 | 5.5590 | 4.8614 | 6.1875 | 5.0832 | 4.8007 | 5.5941 | 5.3071 | 10.3459 | 9.3490 | 9.1892 | 8.6859 |
| ZFP36L1_S192 | ZFP36L1 | Q07352 | | 54 | GFPRRHSVTLPSS | 4.3778 | 3.8186 | 4.2799 | 3.9439 | 6.3386 | 5.8542 | 5.0445 | 6.2910 | 5.0230 | 5.1069 | 5.3134 | 5.4417 | 11.2375 | 10.0033 | 9.1416 | 8.7839 |
| ZFP36L1_S54 | ZFP36L1 | Q07352 | | 43 | DRKAVGTPAGGGF | 6.3295 | 6.8107 | 5.6832 | 6.6371 | 5.9395 | 6.4264 | 7.2122 | 7.5283 | 6.8392 | 6.4520 | 6.9873 | 6.9588 | 4.4190 | 5.2419 | 4.2311 | 6.3039 |
| ZFP36L1_T43 | ZFP36L1 | Q07352 | | 192 | AGARDLSADRPRL | 4.7718 | 4.6856 | 5.0599 | 5.1934 | 6.1911 | 5.8001 | 6.2031 | 6.3390 | 4.6472 | 5.1515 | 6.0007 | 5.4024 | 9.4903 | 7.8949 | 9.1957 | 7.9734 |
| ZFP36L2_S426 | ZFP36L2 | P47974 | | 438 | QLPRRLSDSPVFD | 6.3800 | 4.9535 | 6.5334 | 5.2815 | 6.7592 | 4.8211 | 6.6418 | 8.3602 | 8.7454 | 6.4432 | 6.8645 | 7.6680 | 4.3095 | 6.4582 | 4.7639 | 5.0167 |
| ZFP36L2_S438 | ZFP36L2 | P47974 | | 426 | GAAAPPSPPFSFQ | 5.8959 | 6.0489 | 6.9364 | 6.6197 | 5.9261 | 6.5185 | 5.7968 | 7.5525 | 5.5706 | 6.9794 | 7.7183 | 5.5771 | 6.9191 | 5.9692 | 5.0536 | 4.9179 |
| ZFP37_S289 | ZFP37 | Q9Y6Q3-2 | | 289 | GEKHEKSPSLSSS | 5.9958 | 6.2199 | 5.5916 | 6.5845 | 6.0162 | 6.2868 | 6.2856 | 6.5039 | 6.0817 | 6.2111 | 5.7181 | 6.3130 | 5.9513 | 7.0916 | 6.6432 | 6.5057 |
| ZFP64_S547 | ZFP64 | Q9NTW7-5 | | 547 | QILRQVSLIAPPQ | 5.9914 | 5.1012 | 6.7938 | 6.2889 | 5.0255 | 6.1353 | 7.0682 | 6.9889 | 6.5804 | 5.5633 | 6.1597 | 6.5621 | 6.9682 | 6.6914 | 6.5267 | 5.5548 |
| ZFP90_S406 | ZFP90 | Q8TF47 | | 406 | GRAFGQSPSLYKH | 6.6366 | 6.2506 | 6.9361 | 8.1222 | 5.8733 | 5.6663 | 7.4044 | 6.6043 | 6.8054 | 6.7144 | 8.7681 | 6.9111 | 4.0457 | 3.7609 | 4.2820 | 5.2186 |
| ZFP90_S511 | ZFP90 | Q8TF47 | | 511 | AFKRSTSFIEHHR | 5.7866 | 5.6495 | 6.0059 | 5.7191 | 4.6607 | 5.9224 | 5.5754 | 4.9236 | 5.7197 | 5.7575 | 5.6158 | 5.6694 | 8.7803 | 7.7491 | 8.5703 | 7.8948 |
| ZFP91_S101 | ZFP91 | Q96JP5 | | 83 | PRRRRSSPSARPP | 5.6017 | 6.0901 | 6.0873 | 6.1949 | 6.0451 | 5.7869 | 5.9984 | 5.8523 | 6.1898 | 6.3568 | 6.1722 | 6.0455 | 6.6973 | 7.0898 | 7.0026 | 6.7892 |
| ZFP91_S103 | ZFP91 | Q96JP5 | | 101 | QPQAAKSPSPVQG | 6.3878 | 6.9072 | 6.3764 | 6.6411 | 6.3815 | 6.5503 | 6.4227 | 6.4837 | 6.8833 | 6.8711 | 6.9604 | 6.8682 | 4.8025 | 5.5870 | 4.8850 | 4.9920 |
| ZFP91_S83 | ZFP91 | Q96JP5 | | 103 | QAAKSPSPVQGKK | 5.6056 | 5.9738 | 5.4183 | 5.5537 | 5.8531 | 5.8894 | 5.8181 | 5.6015 | 5.6475 | 5.7750 | 5.7203 | 5.8218 | 7.6500 | 7.7805 | 8.2166 | 7.6748 |
| ZFR_S1054 | ZFR | Q96KR1 | | 1054 | RKRRRDSDGVDGF | 5.5377 | 5.5780 | 5.4844 | 6.2809 | 6.0528 | 5.6291 | 6.1909 | 6.0743 | 5.8448 | 5.6615 | 5.5823 | 5.5489 | 7.6854 | 7.4175 | 7.8181 | 7.6133 |
| ZFR_T529 | ZFR | Q96KR1 | | 529 | TECVKSTPVTSAV | 6.8149 | 7.0111 | 6.7517 | 6.6727 | 6.0946 | 6.3501 | 6.7706 | 6.5675 | 6.8727 | 6.0067 | 6.8388 | 6.1690 | 4.7812 | 5.9913 | 4.8833 | 5.4236 |
| ZFYVE16_S142 | ZFYVE16 | Q7Z3T8 | | 815 | AFERMMSPTGSNL | 7.2542 | 7.8892 | 7.3378 | 7.5105 | 5.7384 | 7.1021 | 7.1658 | 6.8205 | 7.4652 | 8.0148 | 8.1926 | 7.9187 | 2.4785 | 3.4535 | 2.1549 | 3.5035 |
| ZFYVE16_S815 | ZFYVE16 | Q7Z3T8 | | 939 | RNEIIQSPISQVP | 6.2090 | 7.4387 | 7.3002 | 7.2021 | 5.6771 | 6.9965 | 7.5204 | 6.2861 | 7.5935 | 7.7103 | 8.2447 | 8.7321 | 3.0204 | 3.6194 | 2.6707 | 3.7789 |
| ZFYVE16_S939 | ZFYVE16 | Q7Z3T8 | | 946 | PISQVPSVEKLSM | 3.2578 | 3.7518 | 3.6459 | 3.8279 | 3.9015 | 3.8611 | 4.0046 | 3.8577 | 3.9948 | 3.8700 | 3.6097 | 3.8736 | 14.5121 | 13.1584 | 14.3594 | 12.5136 |
| ZFYVE16_S946 | ZFYVE16 | Q7Z3T8 | | 142 | LVHATNSEEDIKK | 4.9117 | 4.9608 | 5.4764 | 5.3494 | 5.1055 | 5.4413 | 5.2663 | 5.0556 | 5.2207 | 5.2819 | 5.6457 | 5.8498 | 9.1487 | 8.7566 | 8.9631 | 9.5666 |
| ZFYVE19_S144 | ZFYVE19 | Q96K21 | | 144 | ANASKWSPPQNYK | 6.2612 | 6.2173 | 6.8315 | 5.7241 | 6.0418 | 6.4890 | 6.1057 | 6.6948 | 6.5924 | 6.3245 | 6.3797 | 6.6997 | 5.9873 | 6.2836 | 5.6996 | 5.6679 |
| ZFYVE26_S1291 | ZFYVE26 | Q68DK2 | | 1742 | REKRSDSVIHLQE | 6.1448 | 5.0974 | 5.8669 | 5.8443 | 6.5760 | 5.9692 | 5.7046 | 6.7111 | 6.8146 | 7.0318 | 7.5833 | 6.7916 | 5.9829 | 6.6873 | 5.7458 | 5.4485 |
| ZFYVE26_S1742 | ZFYVE26 | Q68DK2 | | 297 | TASGKVSPDHLDP | 6.5484 | 6.6050 | 6.3081 | 6.3714 | 6.1037 | 6.1554 | 5.9830 | 6.0494 | 6.6800 | 6.9282 | 6.5177 | 6.1361 | 5.8633 | 6.1066 | 5.6852 | 5.9586 |

| | | | | | | | | | | | | | | | | | | | | |
|---|---|---|---|---|---|---|---|---|---|---|---|---|---|---|---|---|---|---|---|---|
| ZFYVE26_S297 | ZFYVE26 | Q68DK2 | 1291 | ERKPYSSPRDSSL | 6.9935 | 8.0579 | 6.7545 | 7.6660 | 6.0212 | 7.1170 | 7.3865 | 6.6669 | 7.7590 | 7.7548 | 7.7464 | 7.4943 | 2.7098 | 3.9826 | 2.7131 | 3.1765 |
| ZFYVE28_S569 | ZFYVE28 | Q9HCC9 | 569 | SCVCCGSCGDSRE | 7.2185 | 6.5276 | 8.5418 | 7.1315 | 6.6562 | 7.2466 | 6.3538 | 7.4652 | 7.8994 | 7.4574 | 7.9624 | 7.1742 | 3.0991 | 3.5004 | 2.8179 | 2.9481 |
| ZFYVE28_S586 | ZFYVE28 | Q9HCC9 | 586 | RLREKCSPGGVIG | 6.6639 | 6.6126 | 7.4682 | 6.3321 | 5.8544 | 6.3219 | 6.4228 | 6.7602 | 6.7941 | 6.9354 | 7.9518 | 6.9605 | 4.5829 | 5.3931 | 4.5361 | 4.4101 |
| ZGRF1_S877 | ZGRF1 | Q86YA3 | 877 | PFITVVSPKSPHL | 6.6223 | 6.1417 | 7.9615 | 7.0636 | 5.7475 | 6.8820 | 7.0970 | 6.1785 | 7.5109 | 7.1904 | 6.7637 | 7.6651 | 4.0013 | 4.3629 | 3.5335 | 5.2780 |
| ZHX1_S156 | ZHX1 | Q9UKY1 | 686 | VRTQWPSPEEYDK | 6.2283 | 6.1371 | 6.8514 | 6.8953 | 6.1089 | 6.0713 | 6.5661 | 6.3230 | 6.0409 | 6.4361 | 5.8847 | 6.4147 | 6.0136 | 6.5066 | 5.8883 | 5.6337 |
| ZHX1_S534 | ZHX1 | Q9UKY1 | 568 | NPFPDFTPQKFKE | 7.9263 | 7.2016 | 5.8053 | 7.6573 | 6.0802 | 6.0724 | 6.3847 | 5.5009 | 8.3839 | 7.1977 | 4.9706 | 6.9365 | 5.5637 | 4.9962 | 3.9200 | 5.4027 |
| ZHX1_S640 | ZHX1 | Q9UKY1 | 156 | DLTFDGSFVKEEN | 6.7182 | 6.1676 | 6.4825 | 6.7238 | 6.8794 | 6.0362 | 7.2912 | 6.0204 | 7.0825 | 7.1486 | 6.3893 | 5.6487 | 4.7403 | 5.3580 | 5.4046 | 5.9086 |
| ZHX1_S686 | ZHX1 | Q9UKY1 | 647 | KEEAGETSPADES | 5.6101 | 5.7435 | 6.6167 | 6.1209 | 5.9795 | 5.9742 | 6.1952 | 6.3557 | 6.0255 | 6.2514 | 5.5172 | 6.0077 | 6.8955 | 6.8810 | 6.7759 | 7.0499 |
| ZHX1_T568 | ZHX1 | Q9UKY1 | 640 | ESNAGSSKEEAGE | 6.1663 | 6.2187 | 6.8597 | 6.6433 | 6.9669 | 6.3652 | 6.7724 | 6.4703 | 6.0683 | 5.8786 | 5.1962 | 6.1509 | 5.8673 | 5.6544 | 5.8283 | 6.8932 |
| ZHX1_T647 | ZHX1 | Q9UKY1 | 534 | LHLNNDSSTTIII | 6.3987 | 6.3787 | 6.5974 | 7.0060 | 6.4776 | 6.5540 | 5.7245 | 6.2416 | 5.8755 | 6.9444 | 5.8382 | 6.6303 | 5.4998 | 5.5814 | 5.3836 | 6.8681 |
| ZHX3_S946 | ZHX3 | Q9H4I2 | 946 | EPFDTSSPQAGRQ | 5.8346 | 6.7074 | 6.0654 | 5.6771 | 5.6749 | 6.9742 | 6.1452 | 6.1966 | 7.2629 | 5.6662 | 5.8292 | 7.0923 | 5.5371 | 6.6242 | 6.3519 | 6.3607 |
| ZKSCAN1_S13 | ZKSCAN1 | P17029 | 208 | APPHEGSPRDQAM | 6.2666 | 6.5678 | 6.4023 | 7.1518 | 6.3587 | 6.4995 | 5.9918 | 6.6916 | 6.6320 | 6.7576 | 6.7302 | 6.5669 | 5.2025 | 5.5909 | 5.4341 | 5.1557 |
| ZKSCAN1_S208 | ZKSCAN1 | P17029 | 13 | REATGLSPQAAQE | 6.2135 | 6.4531 | 6.4574 | 6.2960 | 6.2062 | 6.3705 | 5.9954 | 6.3486 | 6.5999 | 6.7892 | 6.5452 | 6.3271 | 5.7432 | 6.1387 | 5.6594 | 5.8566 |
| ZKSCAN1_S548 | ZKSCAN1 | P17029 | 400 | RHKIIHTGEKPYE | 7.5077 | 7.9355 | 7.8242 | 8.4800 | 5.9302 | 7.3959 | 7.4690 | 6.3475 | 8.3243 | 7.4804 | 8.2967 | 8.0579 | 1.9358 | 2.8099 | 1.6637 | 2.5414 |
| ZKSCAN1_T400 | ZKSCAN1 | P17029 | 548 | ERASEYSPASLDA | 6.1947 | 6.9332 | 6.6232 | 7.3651 | 6.1959 | 7.4938 | 7.0825 | 6.1229 | 7.3799 | 7.5987 | 7.7389 | 7.6067 | 4.0124 | 4.3156 | 3.7162 | 3.6204 |
| ZKSCAN4_S44 | ZKSCAN4 | Q969J2 | 44 | EVRAPCSPARGPE | 6.2224 | 6.7569 | 6.0908 | 6.5330 | 6.1614 | 6.2707 | 6.1686 | 6.3654 | 6.1973 | 6.3684 | 6.2981 | 6.3485 | 6.0126 | 6.2331 | 5.9925 | 5.9802 |
| ZKSCAN4_S445 | ZKSCAN4 | Q969J2 | 445 | KAFRRNSHLLRHQ | 6.7074 | 7.0433 | 5.7695 | 6.9951 | 6.5924 | 6.0010 | 6.0812 | 8.9700 | 6.4144 | 6.3395 | 6.0451 | 6.0545 | 5.0496 | 6.2846 | 4.5693 | 5.0831 |
| ZKSCAN4_T214 | ZKSCAN4 | Q969J2 | 214 | MVASRLTPGSQGL | 6.4642 | 6.3988 | 5.8497 | 7.0002 | 6.5557 | 6.0944 | 5.3791 | 7.5064 | 6.9797 | 6.4250 | 6.5370 | 8.1433 | 5.1568 | 5.9805 | 4.6133 | 4.9159 |
| ZKSCAN8_S12 | ZKSCAN8 | Q15776 | 12 | RKPSAPSPPDQTP | 5.8602 | 6.1509 | 6.0765 | 6.3653 | 6.1937 | 6.2323 | 6.1556 | 6.1046 | 6.2215 | 6.5633 | 5.7941 | 5.8671 | 6.6239 | 6.3087 | 6.7591 | 6.7233 |
| ZKSCAN8_S193 | ZKSCAN8 | Q15776 | 193 | AMSTSQSPTRSQK | 6.2862 | 6.7102 | 6.3997 | 6.4245 | 5.8244 | 5.8541 | 6.0736 | 6.1388 | 5.8153 | 6.3016 | 6.5791 | 6.5817 | 6.5380 | 5.9708 | 6.0445 | 6.4575 |
| ZMIZ2_T580 | ZMIZ2 | Q8NF64 | 580 | SGTIPGTPGPNGE | 6.0666 | 5.6095 | 6.0349 | 6.3064 | 6.3382 | 6.8598 | 6.0194 | 6.1234 | 6.9715 | 6.3012 | 6.5195 | 6.3760 | 6.8710 | 6.0984 | 4.8971 | 6.6071 |
| ZMYM1_S127 | ZMYM1 | Q5SVZ6 | 127 | YISSASSPVPSKR | 6.2359 | 6.5733 | 7.7091 | 7.5286 | 6.0535 | 6.5897 | 7.0596 | 6.3375 | 6.5724 | 6.8646 | 7.1476 | 6.9135 | 3.9879 | 5.7660 | 4.3154 | 4.3453 |
| ZMYM2_S1056 | ZMYM2 | Q9UBW7 | 838 | GSAPPPSPTPNKE | 6.0243 | 6.2032 | 6.1207 | 6.4291 | 6.3607 | 6.3358 | 6.2605 | 6.3372 | 6.0635 | 6.1413 | 6.2568 | 6.1826 | 6.1983 | 6.2780 | 6.3182 | 6.4898 |
| ZMYM2_S305 | ZMYM2 | Q9UBW7 | 958 | TMTDMMSEDEGKT | 6.3422 | 6.5588 | 6.8059 | 6.5108 | 6.2259 | 6.1685 | 6.2257 | 6.3674 | 6.3734 | 6.3146 | 6.1285 | 6.0962 | 5.7503 | 5.7044 | 6.0928 | 6.3346 |
| ZMYM2_S838 | ZMYM2 | Q9UBW7 | 1056 | AVSGYQSHDDSSD | 6.0235 | 6.1779 | 6.9661 | 7.8274 | 7.1067 | 7.4179 | 6.7007 | 6.7133 | 4.7504 | 5.4482 | 6.1436 | 5.5774 | 4.4759 | 6.2672 | 6.1425 | 6.2614 |
| ZMYM2_S958 | ZMYM2 | Q9UBW7 | 305 | PGVDSLSPVASLP | 6.3528 | 6.4224 | 6.8483 | 7.2819 | 6.9426 | 6.5427 | 5.8231 | 6.2408 | 6.0522 | 6.7148 | 7.1990 | 7.1426 | 4.1061 | 5.8478 | 4.8833 | 5.5997 |
| ZMYM3_S172;S186;S192 | ZMYM3 | Q14202 | 774 | SLLNSQSPESKPQ | 5.9525 | 6.5171 | 6.3219 | 7.2948 | 6.5534 | 6.2864 | 6.4298 | 6.2254 | 6.4761 | 6.7660 | 6.4611 | 6.6624 | 5.3503 | 5.8101 | 5.3964 | 5.4963 |
| ZMYM3_S192;S194 | ZMYM3 | Q14202 | 464 | IYTKTGSPGPELL | 5.9894 | 6.0349 | 6.7075 | 6.3083 | 5.8968 | 6.0840 | 6.6063 | 6.1896 | 6.5235 | 6.4254 | 6.1564 | 6.5587 | 6.1145 | 5.9857 | 5.8199 | 6.5991 |
| ZMYM3_S220 | ZMYM3 | Q14202 | 51 | WAPPGPSPSSGAL | 5.5712 | 5.9366 | 5.7444 | 5.8644 | 5.9232 | 5.6329 | 5.9567 | 5.7737 | 5.7332 | 6.2179 | 5.9287 | 5.9420 | 7.5559 | 7.5036 | 7.4685 | 7.2471 |
| ZMYM3_S263 | ZMYM3 | Q14202 | 826 | PPPPPATPRKNKA | 6.3163 | 6.6330 | 6.7453 | 7.1943 | 6.2667 | 6.4072 | 6.5850 | 6.0487 | 6.7751 | 6.1571 | 6.1025 | 6.3379 | 5.4941 | 5.6526 | 5.4834 | 5.8008 |
| ZMYM3_S263;S267 | ZMYM3 | Q14202 | 960 | DLVSNQSAEGLLE | 6.3045 | 6.4899 | 6.4685 | 6.2622 | 6.1307 | 5.9437 | 6.7424 | 6.3499 | 6.2035 | 6.4512 | 6.8783 | 6.3608 | 5.6827 | 6.0849 | 5.8155 | 5.8311 |
| ZMYM3_S464 | ZMYM3 | Q14202 | 795 | NSNTVRTPEENGN | 6.8736 | 7.5529 | 6.7802 | 7.9663 | 6.1912 | 6.8286 | 7.0095 | 6.4351 | 6.9541 | 6.9925 | 7.4341 | 6.8551 | 3.6714 | 4.7453 | 3.7674 | 3.9426 |
| ZMYM3_S51 | ZMYM3 | Q14202 | 220 | SPPAHPSLPGDGL | 5.8429 | 6.6427 | 5.8807 | 7.1392 | 5.9239 | 6.4113 | 5.6417 | 6.6676 | 5.9979 | 6.2487 | 6.6090 | 6.2235 | 5.2770 | 6.5941 | 6.5407 | 6.3592 |
| ZMYM3_S774 | ZMYM3 | Q14202 | 950 | ELDKASSDLCDLV | 5.9062 | 5.7614 | 5.9527 | 6.4820 | 6.0589 | 6.1631 | 5.9429 | 5.9483 | 6.2369 | 5.5247 | 5.9777 | 6.5882 | 6.8279 | 6.5833 | 7.7643 | 7.2814 |
| ZMYM3_S950 | ZMYM3 | Q14202 | 817 | SAPTAPTPPPPPP | 5.8593 | 6.2741 | 5.6231 | 7.0297 | 6.2430 | 6.1035 | 6.4469 | 6.6602 | 5.9253 | 5.8000 | 6.2329 | 6.3608 | 6.2788 | 6.5078 | 6.4899 | 6.1647 |
| ZMYM3_S957;S960 | ZMYM3 | Q14202 | 263 | TESIPVSDEDSDA | 5.9457 | 5.6058 | 6.1846 | 5.0125 | 6.0389 | 5.9051 | 5.6124 | 6.0521 | 8.0878 | 6.6860 | 6.9853 | 7.1034 | 6.7319 | 6.9524 | 5.8972 | 5.1989 |
| ZMYM3_S960 | ZMYM3 | Q14202 | 263;267 | NA | 5.9583 | 6.2756 | 6.4148 | 6.3930 | 5.9561 | 5.8780 | 6.1943 | 6.3994 | 6.3628 | 5.9657 | 6.2143 | 6.4799 | 6.3057 | 6.4227 | 6.4660 | 6.3132 |
| ZMYM3_T795 | ZMYM3 | Q14202 | 957;960 | NA | 5.1055 | 7.1565 | 6.0019 | 7.5487 | 6.2384 | 5.8237 | 6.4739 | 6.9140 | 7.1697 | 6.3840 | 5.8520 | 5.7261 | 5.6260 | 5.5870 | 6.1922 | 6.2002 |
| ZMYM3_T817 | ZMYM3 | Q14202 | 192;194 | NA | 5.7013 | 5.9268 | 6.5300 | 6.8974 | 6.5567 | 5.5973 | 6.0665 | 6.7198 | 5.7941 | 7.7066 | 6.2507 | 6.0307 | 6.2563 | 5.9800 | 6.4589 | 6.5269 |
| ZMYM3_T826 | ZMYM3 | Q14202 | 172;186;192 | NA | 5.7439 | 6.3552 | 6.8408 | 5.7925 | 6.0090 | 5.5970 | 5.6557 | 6.5316 | 5.9828 | 6.6647 | 5.5553 | 6.3947 | 6.7441 | 7.2079 | 6.4402 | 7.3848 |
| ZMYM4_S1030 | ZMYM4 | Q5VZL5 | 122 | RVTQHESDNENEI | 6.3596 | 6.1548 | 6.1675 | 6.8222 | 6.0750 | 6.0739 | 5.9997 | 6.3268 | 6.1592 | 6.2278 | 6.0259 | 5.7820 | 6.0117 | 6.6030 | 6.6609 | 6.5501 |
| ZMYM4_S1071 | ZMYM4 | Q5VZL5 | 1181 | RRGRKKSIVAVEP | 5.3171 | 4.2673 | 4.3570 | 4.7140 | 6.5495 | 5.4403 | 5.1707 | 5.5062 | 8.9467 | 8.1213 | 7.3711 | 7.9411 | 7.5798 | 8.2891 | 4.4998 | 5.9290 |
| ZMYM4_S1088 | ZMYM4 | Q5VZL5 | 1030 | EDKVTESIEDIKE | 5.9401 | 5.7142 | 6.1508 | 6.7478 | 6.3017 | 5.6567 | 6.0233 | 5.9567 | 6.3757 | 6.3634 | 5.8657 | 6.0191 | 6.5696 | 6.7329 | 6.7424 | 6.8400 |
| ZMYM4_S1181 | ZMYM4 | Q5VZL5 | 1241 | VEQASSSPRSDPL | 6.2651 | 6.4890 | 6.1492 | 6.6567 | 5.9076 | 5.5486 | 6.2779 | 5.6850 | 6.3164 | 6.1288 | 6.0189 | 6.0301 | 6.9960 | 6.3719 | 6.6050 | 6.5538 |
| ZMYM4_S122 | ZMYM4 | Q5VZL5 | 1071 | QGESQTSEHELFL | 5.4842 | 6.1708 | 5.5779 | 6.4193 | 5.9580 | 6.0099 | 6.0665 | 6.3089 | 5.5225 | 5.6978 | 5.7198 | 5.7909 | 6.9096 | 7.2433 | 7.4303 | 7.6903 |
| ZMYM4_S1241 | ZMYM4 | Q5VZL5 | 1028 | DSEDKVTESIEDI | 6.6565 | 5.7978 | 6.0530 | 6.5333 | 6.1148 | 6.2810 | 6.1158 | 5.9367 | 5.8763 | 5.7882 | 6.1572 | 6.1427 | 6.3277 | 6.9151 | 6.5258 | 6.7781 |
| ZMYM4_T1028 | ZMYM4 | Q5VZL5 | 217 | HTHLPQTPETNFR | 5.2826 | 5.9354 | 6.2346 | 6.7434 | 5.8392 | 5.6595 | 6.4027 | 5.7315 | 6.3343 | 6.6658 | 5.8996 | 5.9671 | 6.6662 | 6.5770 | 7.0793 | 6.9817 |
| ZMYM4_T217 | ZMYM4 | Q5VZL5 | 1088 | FEKDQGSTYSGDL | 6.2933 | 6.5170 | 6.0597 | 6.5443 | 5.9380 | 5.6870 | 6.0730 | 6.4051 | 5.9108 | 6.0157 | 6.9205 | 6.4044 | 6.3003 | 5.6989 | 6.5254 | 6.7065 |
| ZMYM6_S397 | ZMYM6 | O95789 | 397 | GNTSAVSPSSIRG | 6.9711 | 7.2331 | 6.2470 | 7.1454 | 6.0032 | 5.4486 | 5.7058 | 6.5969 | 7.8475 | 6.2129 | 6.0910 | 7.7451 | 4.9720 | 5.1512 | 4.9659 | 5.6632 |
| ZMYND11_S403 | ZMYND11 | Q15326 | 403 | KGRRNQSVEPKKE | 5.8638 | 6.5105 | 6.4342 | 6.5953 | 6.2921 | 5.8174 | 7.0551 | 6.1068 | 6.2539 | 5.7781 | 6.0191 | 6.4324 | 6.1791 | 5.7705 | 6.4077 | 6.4838 |
| ZMYND11_S444 | ZMYND11 | Q15326 | 444 | TQTKKLSASSPRM | 5.6840 | 5.9415 | 5.4860 | 6.4977 | 5.4532 | 5.4894 | 5.4899 | 5.7062 | 5.7000 | 5.5618 | 5.6494 | 5.7752 | 7.9624 | 7.7271 | 8.2457 | 7.6305 |
| ZMYND8_S433 | ZMYND8 | Q9ULU4-19 | 433 | NFDMTASPKILMS | 6.5726 | 6.5520 | 6.2446 | 6.5454 | 6.5417 | 5.9665 | 6.8525 | 6.0696 | 6.5804 | 6.0841 | 5.9957 | 6.0516 | 5.8550 | 5.9925 | 5.7846 | 6.3113 |
| ZMYND8_S452 | ZMYND8 | Q9ULU4-19 | 452 | GTGRRISLSDMPR | 5.1191 | 5.0600 | 4.9495 | 5.8439 | 7.0827 | 5.3262 | 5.4893 | 6.1715 | 6.3258 | 6.5259 | 6.0266 | 6.0880 | 7.8157 | 7.6403 | 7.4915 | 7.0440 |
| ZMYND8_S487 | ZMYND8 | Q9ULU4-19 | 574 | TPVPLISPKRQIR | 6.6247 | 6.7402 | 6.5580 | 6.9750 | 6.3509 | 6.2947 | 6.5803 | 6.2443 | 6.7786 | 6.6100 | 6.8054 | 6.6514 | 5.1914 | 5.4270 | 5.0448 | 5.1232 |
| ZMYND8_S517 | ZMYND8 | Q9ULU4-19 | 517 | AGSLSGSPKPFSP | 6.1942 | 5.7704 | 6.0415 | 6.3793 | 6.3152 | 5.8695 | 6.4622 | 5.6021 | 6.4852 | 6.3305 | 6.0748 | 6.0991 | 6.8806 | 6.4465 | 6.4882 | 6.5607 |
| ZMYND8_S522 | ZMYND8 | Q9ULU4-19 | 783 | TTVGSHSPPETPV | 6.1658 | 6.4188 | 6.1507 | 6.7952 | 6.2676 | 6.4920 | 5.9784 | 6.2774 | 5.8539 | 6.1535 | 6.4139 | 6.6500 | 5.6803 | 6.3508 | 6.2648 | 6.0870 |
| ZMYND8_S574 | ZMYND8 | Q9ULU4-19 | 522 | GSPKPFSPQLSAP | 6.6736 | 6.8227 | 6.6208 | 6.9774 | 5.8775 | 5.9281 | 6.3318 | 5.8679 | 6.7599 | 6.9935 | 6.2402 | 6.9199 | 5.6077 | 5.1617 | 4.9119 | 6.3056 |
| ZMYND8_S679 | ZMYND8 | Q9ULU4-19 | 695 | AVKDKASPEPEKD | 6.2537 | 6.5090 | 6.1043 | 6.3904 | 5.8907 | 5.3031 | 5.9159 | 5.8557 | 6.1476 | 6.2715 | 5.5082 | 5.6709 | 6.9868 | 6.3920 | 7.3291 | 7.4711 |
| ZMYND8_S695 | ZMYND8 | Q9ULU4-19 | 709 | SEKAKPSPHPIKD | 6.4193 | 6.8047 | 5.8100 | 6.6560 | 6.5699 | 6.3830 | 6.7240 | 6.2894 | 6.1973 | 6.4728 | 5.7567 | 6.2197 | 5.7844 | 5.8175 | 6.1190 | 5.9761 |
| ZMYND8_S709 | ZMYND8 | Q9ULU4-19 | 487 | ATSSHFSASEESM | 6.3276 | 7.5978 | 6.0485 | 6.6301 | 7.4071 | 5.7554 | 6.1974 | 7.7421 | 4.4433 | 4.5084 | 5.6902 | 6.7324 | 5.8670 | 6.3416 | 7.4443 | 5.2668 |
| ZMYND8_S734;S736 | ZMYND8 | Q9ULU4-19 | 679 | EELKSTSPASEKA | 6.7316 | 6.8545 | 5.6327 | 6.5195 | 6.3566 | 5.8969 | 6.2134 | 6.0936 | 6.3218 | 5.9585 | 5.9383 | 5.8063 | 6.7412 | 6.3460 | 6.5207 | 6.0683 |
| ZMYND8_S783 | ZMYND8 | Q9ULU4-19 | 500 | DFLDKSTASPAST | 5.5151 | 6.9486 | 6.0153 | 7.1676 | 5.7369 | 6.1040 | 5.6663 | 6.6446 | 6.1418 | 6.0109 | 5.9300 | 6.5339 | 6.0207 | 6.4849 | 6.2046 | 6.8748 |
| ZMYND8_T500 | ZMYND8 | Q9ULU4-19 | 734;736 | NA | 6.3423 | 6.3823 | 5.6122 | 6.8527 | 6.5062 | 6.1156 | 6.3023 | 6.7363 | 5.9503 | 5.9459 | 6.2319 | 6.3354 | 6.5576 | 6.1549 | 6.2176 | 5.7568 |
| ZNF101_S152 | ZNF101 | Q8IZC7 | 152 | HGKASISPSSGAR | 6.2932 | 6.9440 | 6.6378 | 6.9073 | 6.2690 | 6.5738 | 6.6635 | 6.9055 | 6.9110 | 6.5649 | 6.5473 | 6.6524 | 4.6968 | 5.1792 | 4.9463 | 5.3081 |
| ZNF106_S1026 | ZNF106 | Q9H2Y7 | 452 | KMPSLKSPLLPCP | 6.6281 | 7.2676 | 7.0218 | 7.6228 | 6.6457 | 6.2968 | 6.9100 | 6.5973 | 7.1587 | 6.5832 | 7.0196 | 7.1597 | 3.8530 | 4.4077 | 4.1312 | 4.6967 |
| ZNF106_S1205 | ZNF106 | Q9H2Y7 | 1370 | AHVPENSDTEQDV | 5.9344 | 6.3996 | 5.8643 | 6.5948 | 5.8414 | 5.8908 | 6.2069 | 6.0574 | 6.4899 | 6.2866 | 6.7663 | 6.6415 | 5.9942 | 6.4166 | 6.3487 | 6.2665 |
| ZNF106_S1279 | ZNF106 | Q9H2Y7 | 1279 | TRNRENSPSSQSA | 6.1369 | 6.4235 | 6.5263 | 6.9631 | 6.1251 | 6.1418 | 6.7267 | 5.8240 | 6.4854 | 6.2564 | 6.2605 | 6.5220 | 5.6688 | 6.0674 | 5.7097 | 6.1625 |
| ZNF106_S1370 | ZNF106 | Q9H2Y7 | 861 | ARKRSLSESSVIM | 6.2479 | 6.4061 | 6.8999 | 7.3778 | 5.7451 | 5.9415 | 6.5918 | 6.3424 | 6.4869 | 6.6715 | 7.1304 | 7.1654 | 4.8165 | 5.3082 | 5.3138 | 5.5549 |
| ZNF106_S1370;T1372 | ZNF106 | Q9H2Y7 | 641 | SRELSTSPCNPIV | 6.2056 | 6.7823 | 5.9097 | 6.9864 | 6.2885 | 5.9801 | 6.2983 | 6.2904 | 6.4738 | 6.1287 | 6.2551 | 5.9849 | 6.0264 | 6.2722 | 5.9209 | 6.1967 |
| ZNF106_S400 | ZNF106 | Q9H2Y7 | 400 | TDETRNSPTQKTQ | 6.0234 | 5.6863 | 6.3754 | 6.9147 | 6.1835 | 6.0915 | 6.3775 | 5.7139 | 6.3865 | 6.2677 | 6.2992 | 6.6458 | 6.2192 | 6.4977 | 6.1149 | 6.2027 |
| ZNF106_S452 | ZNF106 | Q9H2Y7 | 761 | RRIRNISGHRKSE | 6.5275 | 7.2626 | 7.4397 | 7.8431 | 6.3922 | 7.4370 | 6.9459 | 5.5892 | 5.4335 | 6.3602 | 7.7832 | 7.1256 | 4.3644 | 4.8160 | 4.3917 | 4.2883 |
| ZNF106_S590;S595 | ZNF106 | Q9H2Y7 | 1205 | IKQEPMSPEQDEN | 5.5855 | 6.2690 | 6.0466 | 6.6891 | 5.8198 | 6.5361 | 6.1115 | 5.7071 | 6.7025 | 6.0929 | 6.0116 | 6.4988 | 6.2981 | 6.2265 | 6.7726 | 6.6323 |
| ZNF106_S641 | ZNF106 | Q9H2Y7 | 859 | GLARKRSLSESSV | 5.2694 | 5.7921 | 6.1202 | 6.0996 | 7.4288 | 4.8659 | 5.3743 | 5.4355 | 6.2098 | 7.4024 | 6.3955 | 6.5110 | 7.2294 | 7.2073 | 6.5158 | 6.1429 |
| ZNF106_S761 | ZNF106 | Q9H2Y7 | 1026 | ATGDGSSPELPSL | 5.3923 | 5.6411 | 5.8736 | 6.4918 | 6.4149 | 5.7223 | 6.0780 | 5.9681 | 7.0122 | 6.6462 | 7.0546 | 7.0825 | 5.9390 | 6.3816 | 6.0108 | 6.2908 |
| ZNF106_S859 | ZNF106 | Q9H2Y7 | 1370;1372 | NA | 7.0586 | 6.9716 | 7.5525 | 8.7976 | 7.7755 | 8.1066 | 6.9477 | 6.2344 | 6.0695 | 4.3985 | 4.2370 | 5.4516 | 5.0232 | 5.2133 | 5.0675 | 5.0947 |
| ZNF106_S861 | ZNF106 | Q9H2Y7 | 590;595 | NA | 5.9584 | 7.0099 | 7.6561 | 7.3547 | 6.3806 | 4.8957 | 6.8677 | 5.4629 | 7.7166 | 6.8256 | 6.4586 | 5.5940 | 4.8868 | 6.0676 | 5.4430 | 5.4218 |
| ZNF121_S91 | ZNF121 | P58317 | 91 | DKSFEYSDCEEAF | 6.2087 | 6.8009 | 6.7753 | 7.0499 | 6.7622 | 6.0209 | 6.5650 | 6.7402 | 5.7338 | 5.3289 | 5.5810 | 5.8426 | 6.1491 | 6.1528 | 6.7104 | 5.5784 |
| ZNF136_S292 | ZNF136 | P52737 | 405 | KPFHSLSPFRIHE | 7.4592 | 6.9238 | 5.7977 | 7.4619 | 6.2613 | 6.4914 | 6.6257 | 6.9769 | 6.0892 | 6.8109 | 6.2766 | 6.6186 | 4.7757 | 5.1237 | 4.8020 | 5.5056 |
| ZNF136_S405 | ZNF136 | P52737 | 292 | GKAFSCSPTLRIH | 6.1187 | 6.9150 | 6.5153 | 6.8611 | 6.2527 | 6.6242 | 6.4802 | 6.6438 | 6.3871 | 7.1375 | 6.5462 | 6.5644 | 4.8262 | 5.3981 | 5.0373 | 5.6921 |
| ZNF142_S1011 | ZNF142 | P52746 | 904 | GLPRPDSPIPLQP | 5.5621 | 6.6276 | 5.7023 | 6.4699 | 6.4902 | 6.4840 | 5.7705 | 6.4254 | 6.6417 | 6.4344 | 5.7007 | 6.6653 | 5.9563 | 6.4575 | 6.3674 | 6.2447 |
| ZNF142_S154 | ZNF142 | P52746 | 154 | LEGDVVSGTESLF | 6.7513 | 7.5848 | 8.2866 | 7.9390 | 7.2753 | 7.3612 | 7.4648 | 8.7096 | 4.9057 | 5.5760 | 5.9525 | 5.1497 | 4.3358 | 4.1436 | 3.7508 | 4.8132 |
| ZNF142_S904 | ZNF142 | P52746 | 1011 | PPAGNSSPTEALK | 5.8087 | 6.4572 | 6.4334 | 7.4359 | 6.0608 | 6.0205 | 6.6556 | 5.9664 | 6.8601 | 7.0360 | 6.7120 | 6.5010 | 5.1658 | 5.7965 | 5.5838 | 5.5063 |
| ZNF148_S306 | ZNF148 | Q9UQR1 | 665 | FRSGMNSPLRTTP | 6.3630 | 6.6483 | 5.8809 | 6.5111 | 5.8178 | 6.1758 | 5.9687 | 6.2780 | 6.1117 | 6.2483 | 6.1520 | 6.3662 | 6.3008 | 6.2125 | 6.2863 | 6.6785 |
| ZNF148_S412 | ZNF148 | Q9UQR1 | 783 | NRAGMTSSPDATT | 6.8305 | 6.3029 | 6.1371 | 6.5824 | 5.9875 | 6.2818 | 6.4352 | 6.1620 | 5.9857 | 6.1650 | 5.8567 | 5.7871 | 6.2680 | 6.3336 | 6.2968 | 6.5876 |
| ZNF148_S625 | ZNF148 | Q9UQR1 | 412 | EQNQTISPLSTYE | 5.9368 | 5.9562 | 6.7041 | 6.4226 | 6.1390 | 6.2127 | 6.3297 | 5.7108 | 6.3955 | 6.4938 | 6.3582 | 6.6052 | 6.4288 | 6.2087 | 6.0453 | 6.0527 |
| ZNF148_S665 | ZNF148 | Q9UQR1 | 306 | DSGFSTSPKDNSL | 5.9840 | 6.9422 | 6.3503 | 6.7930 | 5.9066 | 5.6916 | 6.1552 | 6.4158 | 7.0057 | 6.7504 | 6.5680 | 6.8937 | 5.3403 | 5.9777 | 5.3403 | 5.8853 |
| ZNF148_S665;T670 | ZNF148 | Q9UQR1 | 625 | NDAYLNSPSLNFV | 7.4814 | 6.8296 | 6.6825 | 7.1361 | 5.4348 | 6.8073 | 7.0679 | 6.1792 | 7.5686 | 6.7438 | 6.9588 | 7.1781 | 4.3348 | 5.3603 | 3.7808 | 4.4560 |
| ZNF148_S783 | ZNF148 | Q9UQR1 | 665;670 | NA | 6.8311 | 7.5076 | 6.4062 | 6.8099 | 6.1619 | 6.3085 | 7.6454 | 6.3514 | 7.2364 | 6.8604 | 6.5525 | 7.1172 | 4.2818 | 4.7493 | 4.1567 | 5.0235 |
| ZNF174_S188 | ZNF174 | Q15697 | 188 | GAYDRLSPHHWEK | 6.1089 | 6.6105 | 6.2136 | 7.2330 | 5.9915 | 5.3869 | 6.2898 | 6.4820 | 6.5830 | 6.4534 | 6.1032 | 6.8269 | 5.5972 | 5.9571 | 6.0354 | 6.1278 |
| ZNF174_S195 | ZNF174 | Q15697 | 195 | PHHWEKSPLLQEP | 6.1882 | 7.3964 | 6.7591 | 8.0546 | 6.6433 | 6.6628 | 6.9000 | 6.3489 | 7.0253 | 7.5799 | 7.3607 | 7.6528 | 3.2633 | 4.4061 | 3.6075 | 4.1511 |
| ZNF174_S287 | ZNF174 | Q15697 | 287 | RVEYISSPLKSHP | 8.1168 | 7.4727 | 7.3040 | 7.3592 | 6.0127 | 5.9263 | 6.6525 | 6.2561 | 7.8700 | 7.5050 | 6.9652 | 7.3670 | 3.7980 | 4.0094 | 3.4626 | 3.9225 |
| ZNF202_T387 | ZNF202 | O95125 | 387 | VNLRETTPVHPLL | 6.1350 | 5.6377 | 5.5615 | 6.8987 | 5.3028 | 6.8010 | 6.9151 | 5.9535 | 6.0614 | 5.1137 | 5.9776 | 5.9903 | 7.1752 | 6.6538 | 6.7807 | 7.0420 |
| ZNF217_S106 | ZNF217 | O75362 | 407 | RRAGAESPTMSVD | 6.0963 | 6.3912 | 6.0174 | 6.1365 | 5.9445 | 5.9299 | 6.0529 | 5.9233 | 6.0393 | 6.1676 | 5.9567 | 6.0099 | 6.8892 | 6.8118 | 7.0755 | 6.5580 |
| ZNF217_S407 | ZNF217 | O75362 | 593 | LGSAVLSPAHKDT | 6.3859 | 6.7693 | 6.8482 | 7.0997 | 6.4808 | 6.7375 | 6.5082 | 6.6805 | 7.1743 | 7.0041 | 7.4218 | 6.9831 | 4.4163 | 4.9684 | 4.0355 | 4.4865 |
| ZNF217_S421 | ZNF217 | O75362 | 795 | SAKGKQSPPGPGK | 6.0881 | 6.5680 | 7.3035 | 8.2338 | 5.6319 | 6.3453 | 6.1307 | 5.9599 | 6.6598 | 6.7070 | 7.1110 | 7.3455 | 4.5714 | 5.2280 | 4.8630 | 5.2530 |
| ZNF217_S441;S445 | ZNF217 | O75362 | 848 | FPKTSVSPAPDKT | 6.5358 | 7.0658 | 6.7274 | 6.6633 | 6.5778 | 6.6838 | 7.0589 | 6.6775 | 6.6953 | 6.6560 | 6.6326 | 6.5214 | 4.6652 | 5.1547 | 4.5512 | 5.1334 |
| ZNF217_S570 | ZNF217 | O75362 | 106 | VEAEYLSPLDKSQ | 7.6710 | 8.4092 | 7.2671 | 7.4338 | 7.1975 | 7.7707 | 8.2589 | 6.5839 | 8.2176 | 7.2516 | 7.6281 | 7.8867 | 1.8554 | 2.6220 | 1.6621 | 2.2843 |
| ZNF217_S593 | ZNF217 | O75362 | 662 | THNLEVSPKEKQT | 6.8978 | 6.6271 | 6.4659 | 7.2621 | 6.0973 | 6.8714 | 5.7037 | 6.3607 | 7.0546 | 6.3799 | 6.6646 | 6.8441 | 4.5574 | 6.3365 | 4.9473 | 4.9297 |
| ZNF217_S662 | ZNF217 | O75362 | 421 | RQPGTCSPDLAAP | 5.2607 | 6.7930 | 5.9642 | 6.7877 | 6.2860 | 5.4185 | 6.1679 | 6.9049 | 7.0010 | 5.9616 | 6.9164 | 6.3354 | 6.1908 | 6.3081 | 5.3348 | 6.3689 |
| ZNF217_S795 | ZNF217 | O75362 | 570 | AKDVTGSPPAKQL | 6.9368 | 7.0638 | 6.4507 | 7.2921 | 6.1244 | 6.1770 | 6.5508 | 6.5854 | 7.4525 | 6.8766 | 7.2642 | 7.6146 | 4.2838 | 4.4522 | 4.3453 | 4.5298 |
| ZNF217_S848 | ZNF217 | O75362 | 441;445 | NA | 5.2413 | 5.8270 | 5.9457 | 6.4357 | 6.0352 | 5.6153 | 5.8848 | 6.8914 | 6.0695 | 6.6107 | 6.0282 | 6.0016 | 6.1038 | 6.6081 | 7.1545 | 7.5471 |
| ZNF219_S114 | ZNF219 | Q9P2Y4 | 16 | SGHLAPSPPAFDG | 5.9474 | 6.1300 | 5.8710 | 6.4112 | 6.6533 | 6.2488 | 6.2159 | 6.2616 | 6.7175 | 6.5529 | 6.3757 | 6.2743 | 6.1862 | 6.4781 | 5.3864 | 6.2897 |
| ZNF219_S16 | ZNF219 | Q9P2Y4 | 114 | QPERPRSPAARLL | 6.0524 | 5.8071 | 5.6281 | 6.4994 | 6.4075 | 5.9075 | 5.6916 | 5.8525 | 5.5022 | 6.2939 | 5.3989 | 4.8544 | 8.4057 | 7.4880 | 6.9198 | 7.2911 |
| ZNF239_S191 | ZNF239 | Q16600 | 191 | GKILNTSPDGHPY | 6.0788 | 6.7401 | 6.5295 | 6.5431 | 6.1832 | 6.2184 | 6.3787 | 6.5353 | 6.5022 | 6.2318 | 6.4918 | 6.5170 | 5.2876 | 6.0980 | 5.6795 | 5.9851 |
| ZNF24_S39 | ZNF24 | P17028 | 39 | DGEEGSSIPWNHL | 4.3087 | 4.7165 | 6.0888 | 8.1426 | 4.9493 | 5.3217 | 7.3159 | 5.4557 | 5.6084 | 5.9529 | 6.3402 | 6.7387 | 7.8726 | 7.3308 | 6.3718 | 7.4853 |
| ZNF250_S134 | ZNF250 | P15622 | 134 | SQNTDLSPKPLIS | 5.8384 | 6.6050 | 6.1306 | 6.6172 | 6.0691 | 6.7600 | 6.6130 | 6.2910 | 6.1497 | 6.7820 | 6.3934 | 6.9060 | 5.6949 | 5.7535 | 5.5676 | 5.8284 |
| ZNF263_S153 | ZNF263 | O14978 | 153 | LETARESPSFKLE | 5.5168 | 6.8915 | 5.5707 | 7.5034 | 5.5273 | 5.6141 | 6.3485 | 5.9985 | 5.9267 | 6.9843 | 5.7008 | 5.5116 | 6.3605 | 6.7644 | 6.8527 | 5.9282 |
| ZNF267_S230 | ZNF267 | Q14586 | 230 | TLNQSSSPKNHQE | 7.0657 | 6.8646 | 7.4959 | 7.5439 | 6.7008 | 6.2246 | 6.9480 | 6.1889 | 7.2368 | 6.8605 | 6.7716 | 6.7646 | 4.0720 | 4.4534 | 4.1646 | 4.6441 |
| ZNF276_S599 | ZNF276 | Q8N554 | 599 | PPPGPPSPSVTTE | 5.6814 | 6.2235 | 6.1313 | 6.7308 | 5.7445 | 6.0889 | 6.4581 | 5.9628 | 6.0555 | 6.2890 | 5.9027 | 6.0699 | 6.8054 | 6.8047 | 6.3943 | 6.6572 |
| ZNF277_T49 | ZNF277 | Q9NRM2 | 49 | PESPGGTTTLEGS | 7.3600 | 6.5356 | 7.1176 | 7.8125 | 6.0370 | 5.4238 | 6.0796 | 6.5530 | 7.4321 | 7.1193 | 7.4034 | 7.3113 | 4.8505 | 4.9894 | 3.7030 | 4.2720 |
| ZNF280B_S114 | ZNF280B | Q86YH2 | 114 | ESRSTDSPIIIEP | 5.9471 | 6.2949 | 6.3029 | 6.6740 | 5.3457 | 5.8930 | 6.1147 | 5.9701 | 6.4198 | 6.6050 | 6.5688 | 6.9559 | 5.9938 | 6.0496 | 6.1570 | 6.7079 |
| ZNF280C_S209 | ZNF280C | Q8ND82 | 80 | IKSEPHSPGIPEI | 6.3721 | 6.5986 | 5.9769 | 7.1820 | 6.4091 | 6.1832 | 6.3662 | 6.9412 | 6.6624 | 6.4975 | 6.6675 | 6.5801 | 5.0456 | 5.7889 | 5.1020 | 5.6267 |
| ZNF280C_S226 | ZNF280C | Q8ND82 | 185 | SKVNSVTPKKPKT | 7.0354 | 6.8801 | 6.3635 | 7.3835 | 6.4474 | 6.5467 | 7.1610 | 6.5864 | 7.1410 | 6.7088 | 6.5765 | 6.7848 | 4.3838 | 4.8624 | 4.2060 | 4.9328 |
| ZNF280C_S76 | ZNF280C | Q8ND82 | 540 | TSFLQVTPPTSQN | 5.7050 | 6.4944 | 7.1160 | 7.0240 | 5.9806 | 5.9848 | 6.5616 | 5.9438 | 6.4224 | 6.4830 | 6.8372 | 6.3814 | 5.6369 | 5.8018 | 5.8947 | 5.7322 |
| ZNF280C_S80 | ZNF280C | Q8ND82 | 76 | SSKGIKSEPHSPG | 5.5318 | 7.9859 | 4.7473 | 6.6784 | 5.9427 | 5.1093 | 5.7358 | 5.6783 | 7.0807 | 4.7473 | 5.4025 | 6.3594 | 7.0835 | 6.7732 | 7.6238 | 7.5203 |
| ZNF280C_T185 | ZNF280C | Q8ND82 | 209 | SLPLIGSPPVTSS | 6.8671 | 7.0728 | 6.7606 | 7.2094 | 5.8700 | 6.9408 | 6.7398 | 6.7449 | 7.2515 | 7.5252 | 7.8253 | 7.9045 | 3.0112 | 4.5668 | 3.3715 | 4.3385 |
| ZNF280C_T540 | ZNF280C | Q8ND82 | 226 | SKGTNTSSPYDAG | 6.1474 | 6.9125 | 6.4715 | 6.7203 | 5.9082 | 6.2181 | 6.3878 | 6.4732 | 6.4262 | 6.6733 | 6.5693 | 6.6852 | 5.3682 | 5.5774 | 5.6532 | 5.8081 |
| ZNF280D_S545 | ZNF280D | Q6N043 | 545 | ASTLQLSPPRTKN | 6.0280 | 6.8957 | 6.6290 | 7.3136 | 5.9563 | 5.3016 | 6.9574 | 6.2966 | 5.8449 | 5.6777 | 6.0718 | 6.7006 | 5.5882 | 6.6509 | 5.8091 | 6.2787 |
| ZNF286A_S19 | ZNF286A | Q9HBT8 | 19 | ALSSQDSPHFQEK | 5.3863 | 6.8740 | 7.0624 | 6.0530 | 6.3261 | 5.9454 | 5.7278 | 6.1556 | 6.1063 | 6.6245 | 6.2532 | 6.6211 | 5.4659 | 5.8781 | 6.4764 | 7.0439 |
| ZNF286B_S19 | ZNF286B | P0CG31 | 19 | VLSSQDSPHFQEK | 6.3026 | 6.7391 | 7.0481 | 6.4718 | 6.9917 | 7.0539 | 6.5675 | 7.2913 | 5.8632 | 7.1597 | 7.8834 | 7.9545 | 3.3302 | 5.4220 | 3.6562 | 4.2648 |
| ZNF292_S1462 | ZNF292 | O60281 | 1810 | NKLPDSSPFSSFI | 6.4239 | 6.8037 | 6.3799 | 6.9521 | 6.1695 | 6.2891 | 6.7850 | 6.6175 | 6.3667 | 7.0380 | 6.9382 | 6.6317 | 4.7921 | 5.6829 | 4.8467 | 5.2830 |
| ZNF292_S1607 | ZNF292 | O60281 | 654 | FNDNDGSDDENDD | 5.5374 | 5.6866 | 6.3445 | 6.0566 | 6.6796 | 5.8355 | 6.1404 | 5.3396 | 6.4470 | 6.3016 | 6.3016 | 6.4423 | 6.6745 | 7.1853 | 6.6337 | 6.3938 |
| ZNF292_S1810 | ZNF292 | O60281 | 1872 | SVTLTPTPVKSTA | 6.1423 | 6.6690 | 6.1870 | 6.6705 | 6.7519 | 5.9007 | 6.0592 | 6.3048 | 6.5435 | 6.3870 | 6.5267 | 6.8056 | 5.9694 | 5.7515 | 5.3615 | 5.9694 |
| ZNF292_S1831 | ZNF292 | O60281 | 1870 | NTSVTLTPTPVKS | 5.7981 | 6.1732 | 5.9826 | 6.4662 | 6.0850 | 5.9310 | 6.2355 | 6.2944 | 6.5864 | 6.0208 | 6.3291 | 6.6195 | 5.9581 | 6.4627 | 6.7541 | 6.3033 |
| ZNF292_S654 | ZNF292 | O60281 | 1462 | LLAENRSPAFLPN | 5.7237 | 6.8850 | 7.4320 | 7.0898 | 5.7083 | 6.3787 | 6.2034 | 6.5778 | 6.1388 | 7.1192 | 6.8724 | 6.5302 | 4.6115 | 6.7013 | 4.2412 | 5.7868 |
| ZNF292_T1870 | ZNF292 | O60281 | 1607 | SNSSRVSVISGPQ | 6.1423 | 6.8236 | 6.0714 | 6.5161 | 5.8190 | 6.7585 | 6.8487 | 5.8368 | 5.9293 | 6.2135 | 6.6161 | 6.8671 | 5.1795 | 6.2109 | 5.5027 | 6.6646 |
| ZNF292_T1872 | ZNF292 | O60281 | 1831 | IPQSEVSHKEDQI | 6.7468 | 6.5248 | 6.0395 | 9.6333 | 7.4764 | 7.7365 | 7.1528 | 5.2243 | 5.1576 | 6.1822 | 7.9109 | 6.3059 | 3.6034 | 5.4971 | 4.1743 | 4.6343 |
| ZNF30_S299 | ZNF30 | P17039-2 | 302 | KAFSTSSPLAKHQ | 6.3845 | 6.3559 | 6.2944 | 6.2051 | 5.9863 | 6.8852 | 7.0237 | 5.8757 | 7.3532 | 6.5296 | 6.4739 | 7.2442 | 5.1283 | 5.4381 | 5.5093 | 5.3127 |
| ZNF30_S302 | ZNF30 | P17039-2 | 299 | ECGKAFSTSSPLA | 6.5452 | 6.1136 | 5.9723 | 7.1212 | 7.0383 | 6.8724 | 6.0905 | 5.7418 | 8.0966 | 6.2534 | 6.3640 | 5.0260 | 5.4761 | 5.1812 | 5.7495 | 6.3578 |
| ZNF316_S112 | ZNF316 | A6NFI3 | 112 | RGGDAKSPVLQEK | 6.0296 | 6.4064 | 6.0092 | 6.3663 | 6.1615 | 5.8686 | 6.0579 | 6.1398 | 6.2369 | 6.1423 | 5.9346 | 5.8973 | 6.5663 | 6.7987 | 6.9083 | 6.4764 |
| ZNF318_S1043 | ZNF318 | Q5VUA4 | 2101 | RSVRIPSPNILKT | 6.7328 | 6.6386 | 5.9228 | 6.4656 | 6.5074 | 5.9818 | 6.2362 | 5.9566 | 6.3978 | 6.0721 | 5.7303 | 5.7647 | 6.5515 | 6.4792 | 6.3089 | 6.2538 |
| ZNF318_S1238;S1243 | ZNF318 | Q5VUA4 | 173 | LSDRLGSPVDNLE | 6.0002 | 6.1961 | 6.5772 | 6.4247 | 5.8004 | 5.7299 | 6.4164 | 5.5129 | 6.4104 | 6.3076 | 6.2525 | 6.5909 | 6.8914 | 6.3161 | 6.2273 | 6.3460 |
| ZNF318_S1243 | ZNF318 | Q5VUA4 | 161 | DHFCVSTPERRRL | 7.0628 | 7.0057 | 6.4674 | 7.2097 | 5.8063 | 6.2849 | 6.5041 | 6.0464 | 7.0242 | 6.7067 | 6.4228 | 6.5366 | 5.3475 | 5.5028 | 4.8593 | 5.2127 |
| ZNF318_S1267 | ZNF318 | Q5VUA4 | 1847 | LMTGSETPSKVVI | 7.6906 | 7.6700 | 6.9592 | 7.8374 | 6.3680 | 6.4425 | 6.6787 | 6.4747 | 7.2588 | 7.1585 | 7.0756 | 7.1167 | 3.7230 | 4.1968 | 3.3096 | 4.0398 |
| ZNF318_S136 | ZNF318 | Q5VUA4 | 2035 | VSPAHRSPTVLCQ | 6.5102 | 6.8566 | 5.8151 | 6.5965 | 6.1554 | 5.7388 | 6.2400 | 6.3904 | 6.4378 | 6.2989 | 5.8242 | 5.5482 | 6.3622 | 6.3588 | 6.5931 | 6.2738 |
| ZNF318_S1558 | ZNF318 | Q5VUA4 | 1243 | LSEGRNSPEKAEN | 6.4618 | 6.2925 | 6.2905 | 6.5460 | 6.2847 | 6.0585 | 6.2591 | 5.8227 | 6.3708 | 6.5944 | 6.2783 | 6.2423 | 6.1356 | 6.2182 | 6.1141 | 6.0307 |
| ZNF318_S173 | ZNF318 | Q5VUA4 | 1896 | PELLLHSPARSAM | 7.1527 | 6.9989 | 6.6206 | 7.1623 | 6.7934 | 6.4866 | 6.5564 | 6.5834 | 7.1228 | 6.4891 | 6.2083 | 6.5460 | 4.2929 | 5.5874 | 4.5447 | 4.8547 |
| ZNF318_S1856 | ZNF318 | Q5VUA4 | 1856 | KVVIKLSPQACSF | 6.7994 | 6.9732 | 6.6389 | 7.3725 | 6.8031 | 6.5971 | 7.3549 | 6.3020 | 6.4026 | 6.5421 | 6.4355 | 6.0513 | 5.0593 | 4.9962 | 4.8715 | 4.8004 |
| ZNF318_S1896 | ZNF318 | Q5VUA4 | 136 | SGSRRRSPGLCSD | 6.6089 | 6.7437 | 6.3338 | 6.7420 | 6.1740 | 6.0911 | 6.3968 | 6.0517 | 6.4729 | 6.3646 | 6.3076 | 5.9627 | 5.9721 | 6.1578 | 5.7857 | 5.8346 |
| ZNF318_S1971 | ZNF318 | Q5VUA4 | 40 | GPARRSSPPPPPS | 6.2389 | 7.0254 | 6.2265 | 6.6380 | 6.1136 | 5.9777 | 6.1476 | 6.0547 | 5.9327 | 6.7750 | 6.1970 | 6.1393 | 6.2117 | 6.1982 | 5.9882 | 6.1355 |
| ZNF318_S2030;S2035 | ZNF318 | Q5VUA4 | 709 | YLLTKNSPPFLKS | 6.2783 | 6.4505 | 6.2114 | 6.4394 | 6.2316 | 5.8061 | 6.4638 | 5.8617 | 6.3082 | 6.4337 | 6.2305 | 6.3219 | 6.4661 | 6.1313 | 6.0816 | 6.2839 |
| ZNF318_S2035 | ZNF318 | Q5VUA4 | 527 | QEKRRRSFPDIED | 5.5644 | 5.8505 | 5.6229 | 6.0551 | 6.7967 | 5.7797 | 6.3039 | 5.8464 | 6.3568 | 6.2504 | 6.1631 | 6.1033 | 6.8560 | 6.8740 | 6.8139 | 6.7628 |
| ZNF318_S2050 | ZNF318 | Q5VUA4 | 1883 | LLNPQDTPVKISA | 8.6915 | 8.4459 | 7.1160 | 8.3993 | 5.4752 | 6.7903 | 7.2850 | 6.5917 | 7.8531 | 7.3168 | 7.4428 | 7.7503 | 2.3550 | 3.1347 | 2.1461 | 3.2063 |
| ZNF318_S2050;S2075 | ZNF318 | Q5VUA4 | 214 | QEEGPLSPFLGQL | 6.5658 | 6.3811 | 6.1003 | 6.6473 | 6.3325 | 6.1608 | 6.1192 | 6.0371 | 6.6716 | 6.7931 | 6.3159 | 6.6357 | 5.9675 | 6.0514 | 5.5639 | 5.6567 |
| ZNF318_S207 | ZNF318 | Q5VUA4 | 1971 | RPETWESPEKPKT | 6.9883 | 7.0414 | 6.9750 | 7.4087 | 5.9594 | 5.8056 | 6.3565 | 6.2557 | 6.7435 | 7.0648 | 6.3206 | 6.9677 | 4.9589 | 5.1357 | 4.7664 | 5.2518 |
| ZNF318_S2091 | ZNF318 | Q5VUA4 | 472 | LREKFGSFLCHKD | 7.0459 | 6.8560 | 6.9269 | 6.3251 | 6.4281 | 6.1473 | 7.0352 | 5.6572 | 8.1981 | 6.8273 | 6.6753 | 7.4491 | 4.2894 | 5.8336 | 3.2995 | 5.0059 |
| ZNF318_S2101 | ZNF318 | Q5VUA4 | 207 | GLERYISQEEGPL | 5.3498 | 6.0443 | 5.3700 | 5.2817 | 6.0114 | 5.5045 | 6.4952 | 5.9975 | 5.5969 | 5.7297 | 6.2954 | 5.4964 | 7.9690 | 7.5112 | 7.6146 | 7.7323 |
| ZNF318_S214 | ZNF318 | Q5VUA4 | 1043 | SPKPAESPQSATK | 6.5342 | 6.3544 | 6.6917 | 6.6854 | 6.3649 | 6.0381 | 6.6023 | 5.6528 | 6.6502 | 6.7196 | 6.1836 | 6.3013 | 5.8007 | 5.7872 | 5.1835 | 6.4502 |
| ZNF318_S2166 | ZNF318 | Q5VUA4 | 2050 | CEENSVSPIGCNS | 6.4282 | 6.4887 | 6.5929 | 7.7279 | 5.5718 | 5.8902 | 6.4045 | 6.3342 | 6.5312 | 6.1737 | 6.8643 | 6.3222 | 5.3702 | 5.9225 | 5.7590 | 5.6184 |

| | | | | | | | | | | | | | | | | | | | | | |
|---|---|---|---|---|---|---|---|---|---|---|---|---|---|---|---|---|---|---|---|---|---|
| ZNF318_S2189 | ZNF318 | Q5VUA4 | | 2189 | QKDKLCSPLSEPG | 6.9890 | 6.8954 | 6.3292 | 6.3737 | 6.3391 | 6.0374 | 6.7301 | 6.3493 | 6.3978 | 5.9833 | 5.5529 | 5.7813 | 6.6327 | 6.1377 | 5.6386 | 5.8325 |
| ZNF318_S305;S307 | ZNF318 | Q5VUA4 | | 1558 | KLEKRNSCLATAN | 5.3570 | 7.8346 | 9.5324 | 5.8030 | 6.0239 | 5.7349 | 5.2676 | 7.4918 | 6.4822 | 9.4372 | 9.3137 | 6.4156 | 3.6421 | 4.2042 | 3.9471 | 3.5128 |
| ZNF318_S40 | ZNF318 | Q5VUA4 | | 842 | RVIPTVTPDKPKQ | 7.2837 | 7.2377 | 6.6054 | 7.4205 | 6.9334 | 6.6222 | 8.0667 | 7.3535 | 7.0548 | 7.1428 | 7.7443 | 6.9809 | 3.1178 | 3.5728 | 2.4270 | 4.4367 |
| ZNF318_S472 | ZNF318 | Q5VUA4 | | 2166 | SAGLGPSPCLPDL | 6.0790 | 5.6674 | 5.1707 | 6.3044 | 6.1940 | 6.3803 | 6.7701 | 4.9913 | 6.6942 | 6.2596 | 5.0879 | 5.3949 | 7.7969 | 6.7367 | 6.3377 | 8.1349 |
| ZNF318_S501 | ZNF318 | Q5VUA4 | | 2091 | TEGERNSPNPRSV | 6.1246 | 6.2433 | 6.4515 | 6.5321 | 6.1618 | 6.3035 | 6.5037 | 6.6061 | 6.6868 | 6.8753 | 6.6651 | 6.1267 | 5.6697 | 5.9469 | 5.2613 | 5.8416 |
| ZNF318_S527 | ZNF318 | Q5VUA4 | | 1267 | EEVKKESPTSSSF | 5.3430 | 6.2212 | 7.2385 | 7.6578 | 5.7118 | 7.2946 | 5.9525 | 5.6585 | 7.0899 | 6.7915 | 6.7367 | 7.5040 | 4.9677 | 5.4495 | 5.1691 | 5.2136 |
| ZNF318_S69;S71 | ZNF318 | Q5VUA4 | | 501 | LPHERASQDGSGF | 4.9579 | 7.0020 | 7.0863 | 6.6060 | 4.5959 | 8.0879 | 7.7168 | 6.7699 | 6.2349 | 6.4921 | 7.4710 | 5.9117 | 5.5566 | 6.0892 | 4.9533 | 4.4685 |
| ZNF318_S709 | ZNF318 | Q5VUA4 | 69;71 | | NA | 6.6709 | 6.7120 | 6.2058 | 6.7120 | 5.9815 | 5.9635 | 5.7561 | 6.1683 | 6.3000 | 6.3980 | 6.1825 | 6.3515 | 5.8955 | 6.5335 | 6.0161 | 6.1527 |
| ZNF318_S79;S81 | ZNF318 | Q5VUA4 | 79;81 | | NA | 5.9388 | 5.7879 | 6.0710 | 6.3852 | 5.9901 | 6.0201 | 6.2574 | 6.3353 | 7.1786 | 6.3377 | 6.2553 | 6.1158 | 6.5457 | 6.3455 | 6.1647 | 6.2710 |
| ZNF318_S89;S91 | ZNF318 | Q5VUA4 | 2030;2035 | | NA | 6.3180 | 6.5914 | 6.5657 | 7.0926 | 5.6318 | 6.6359 | 6.2930 | 5.7148 | 5.5965 | 6.4208 | 6.6122 | 6.6869 | 6.0415 | 6.4055 | 5.4476 | 5.9459 |
| ZNF318_T161 | ZNF318 | Q5VUA4 | 305;307 | | NA | 6.9596 | 6.3832 | 6.2883 | 6.6337 | 6.0836 | 5.6432 | 6.1309 | 4.9483 | 5.8336 | 6.1247 | 5.8483 | 5.8047 | 6.9117 | 7.0626 | 7.0948 | 6.2488 |
| ZNF318_T1847 | ZNF318 | Q5VUA4 | 89;91 | | NA | 6.0598 | 6.9739 | 6.3738 | 6.9367 | 6.0997 | 5.9847 | 6.4377 | 5.7240 | 6.3082 | 5.9453 | 6.4179 | 6.3015 | 6.2055 | 6.0249 | 6.2056 | 6.0009 |
| ZNF318_T1883 | ZNF318 | Q5VUA4 | 1238;1243 | | NA | 6.7081 | 7.1134 | 6.4539 | 7.1088 | 6.2516 | 5.5361 | 6.8111 | 5.9535 | 6.5043 | 5.5934 | 6.2089 | 6.4161 | 5.9299 | 6.0198 | 5.5597 | 5.8313 |
| ZNF318_T842 | ZNF318 | Q5VUA4 | 2050;2075 | | NA | 6.9025 | 5.8155 | 6.4206 | 7.0903 | 5.1741 | 6.1957 | 6.2591 | 5.2719 | 6.3913 | 6.3650 | 6.1602 | 6.7813 | 5.8756 | 6.1126 | 6.2529 | 6.9313 |
| ZNF326_S155 | ZNF326 | Q5BKZ1 | | 162 | SYSSFSSPHMKPA | 6.5325 | 7.2478 | 6.2741 | 7.0936 | 6.1011 | 6.4554 | 6.5587 | 6.5366 | 6.5997 | 6.8562 | 6.9112 | 6.9452 | 4.9735 | 5.3724 | 4.4150 | 5.1272 |
| ZNF326_S162 | ZNF326 | Q5BKZ1 | | 155 | RGRENYSSYSSFS | 7.1061 | 7.7618 | 5.9986 | 7.4713 | 7.1468 | 7.7517 | 7.0689 | 7.4318 | 6.9233 | 6.8730 | 6.8493 | 7.0012 | 3.1060 | 4.6256 | 3.1385 | 3.7460 |
| ZNF335_S1007 | ZNF335 | Q9H4Z2 | | 1007 | GLAVPPSPPSAAT | 6.0656 | 6.3179 | 6.1757 | 6.5911 | 6.0277 | 5.8705 | 6.4205 | 6.2143 | 6.4693 | 5.9910 | 6.3695 | 6.2603 | 6.1811 | 6.4531 | 6.2068 | 6.3856 |
| ZNF335_S976 | ZNF335 | Q9H4Z2 | | 992 | SQSSASSPPATSK | 5.8842 | 6.9047 | 6.2269 | 8.2848 | 6.3667 | 4.5556 | 5.6425 | 7.8537 | 5.9069 | 5.9103 | 6.1149 | 6.6656 | 4.8695 | 6.1048 | 6.4197 | 5.3891 |
| ZNF335_S992 | ZNF335 | Q9H4Z2 | | 976 | DGPEPPSPAKTHC | 6.7277 | 6.5145 | 6.2032 | 6.2901 | 5.9958 | 5.7310 | 6.5322 | 5.9530 | 6.3519 | 6.3241 | 5.8824 | 6.4561 | 6.3648 | 6.2782 | 5.8328 | 6.5621 |
| ZNF358_S553 | ZNF358 | Q9NW07 | | 553 | ALPTGESPEWVQE | 6.1940 | 6.7842 | 7.5359 | 6.6537 | 5.4961 | 5.5499 | 5.9827 | 6.5460 | 6.6972 | 6.4507 | 7.1859 | 6.6454 | 5.6617 | 6.1877 | 4.8127 | 5.6162 |
| ZNF358_T485;S487 | ZNF358 | Q9NW07 | | 487 | GSRSTPSPTPVES | 6.8788 | 7.1890 | 6.5068 | 5.7514 | 6.7161 | 5.9128 | 5.0112 | 5.1272 | 6.3479 | 7.7388 | 6.4299 | 6.7993 | 5.6556 | 6.2987 | 6.3340 | 5.3025 |
| ZNF358_T487 | ZNF358 | Q9NW07 | 485;487 | | NA | 6.0104 | 5.2254 | 6.9166 | 6.0376 | 6.3930 | 4.4425 | 4.9673 | 6.5609 | 6.5065 | 8.3803 | 5.1506 | 5.9222 | 7.3163 | 6.5792 | 7.6947 | 5.8963 |
| ZNF367_S60 | ZNF367 | Q7RTV3 | | 60 | PPLIPTSPGFSDF | 5.0855 | 5.9078 | 5.7536 | 6.1274 | 6.5724 | 5.9393 | 5.2175 | 6.6903 | 6.4147 | 6.1916 | 6.8048 | 6.0281 | 7.8759 | 7.0466 | 6.4310 | 5.9136 |
| ZNF384_S214 | ZNF384 | Q8TF68 | | 234 | YRCRMCSLTFYSK | 7.1796 | 6.9399 | 6.7114 | 6.6177 | 7.0005 | 6.2685 | 6.4692 | 6.5312 | 6.4359 | 6.1940 | 6.5970 | 6.2937 | 5.6650 | 4.9882 | 5.2474 | 4.8608 |
| ZNF384_S234 | ZNF384 | Q8TF68 | | 214 | NDPYVLSPEDDDD | 6.9452 | 7.4059 | 7.3800 | 8.0547 | 5.7001 | 6.4112 | 6.7532 | 7.0944 | 6.7897 | 6.3331 | 7.0465 | 6.6629 | 4.1109 | 4.6620 | 4.2992 | 4.3510 |
| ZNF385A_S153 | ZNF385A | Q96PM9 | | 160 | PEKQPGSPSPPSI | 6.8108 | 6.6862 | 6.5074 | 6.7225 | 6.1283 | 6.1347 | 6.8023 | 6.1116 | 6.2873 | 6.3820 | 6.2166 | 6.3335 | 6.1040 | 5.7112 | 5.6086 | 5.4531 |
| ZNF385A_S160 | ZNF385A | Q96PM9 | | 153 | LGPAPGSPEKQPG | 5.6918 | 5.4241 | 6.2023 | 7.0858 | 6.2499 | 6.2118 | 6.1291 | 5.9810 | 6.5361 | 6.6782 | 6.6342 | 6.3933 | 6.3249 | 6.3147 | 5.4699 | 6.6729 |
| ZNF395_S248 | ZNF395 | Q9H8N7 | | 449 | VRSRSLSFSEPQQ | 6.2456 | 5.9182 | 6.0298 | 5.5256 | 6.2770 | 6.2245 | 5.7337 | 6.3753 | 6.9862 | 7.4330 | 7.2432 | 7.1830 | 6.0155 | 6.8967 | 4.7008 | 5.2117 |
| ZNF395_S449 | ZNF395 | Q9H8N7 | | 248 | LGDAFGSPQTDHG | 7.0785 | 5.5935 | 5.8602 | 7.0051 | 5.9639 | 7.0688 | 6.8301 | 6.7837 | 6.7631 | 6.8787 | 7.1714 | 6.7945 | 5.1571 | 5.7328 | 4.2478 | 5.0707 |
| ZNF397_S142 | ZNF397 | Q8NF99 | | 142 | GQQVPASPQGPAV | 5.3838 | 4.9867 | 8.5602 | 6.4139 | 7.8182 | 6.9414 | 7.7947 | 5.4637 | 6.0707 | 5.6282 | 5.7136 | 5.9376 | 5.6074 | 5.7348 | 6.2264 | 5.7187 |
| ZNF397_S31 | ZNF397 | Q8NF99 | | 31 | KVEDNFSWDEKFK | 6.0890 | 6.6797 | 6.5257 | 6.4538 | 6.5317 | 6.9955 | 6.9236 | 6.3837 | 5.4262 | 6.7380 | 6.6234 | 6.5152 | 5.9194 | 5.1200 | 4.9592 | 6.1160 |
| ZNF398_S284 | ZNF398 | Q8TD17 | | 284 | VPVPFSSPPAAAK | 6.6727 | 9.3183 | 6.9666 | 8.1082 | 6.4058 | 6.2164 | 6.3950 | 5.6754 | 7.8665 | 6.4094 | 8.2182 | 7.6537 | 3.0676 | 3.9026 | 3.3111 | 3.8124 |
| ZNF408_S338 | ZNF408 | Q9H9D4 | | 338 | FPTLSRSPPGPAG | 5.5350 | 6.6149 | 6.5160 | 6.4431 | 5.5278 | 6.0305 | 5.9673 | 5.3623 | 5.8332 | 5.5794 | 5.9181 | 7.1201 | 6.4088 | 6.6940 | 7.6373 | 6.8122 |
| ZNF414_S376 | ZNF414 | Q96IQ9-2 | | 376 | PLAPKVSPLLSEG | 5.5346 | 5.9969 | 6.2662 | 6.0920 | 5.6271 | 5.4717 | 4.4583 | 6.8128 | 6.6761 | 5.1863 | 6.0773 | 5.5655 | 7.8350 | 7.1156 | 6.9079 | 7.3768 |
| ZNF428_T108 | ZNF428 | Q96B54 | | 108 | GEAPPGTPPCRLC | 7.1038 | 7.0829 | 7.4119 | 7.1070 | 6.4617 | 6.7671 | 6.3521 | 6.7071 | 7.0698 | 7.2013 | 7.6675 | 7.5392 | 3.5915 | 4.2698 | 3.3109 | 4.3564 |
| ZNF444_S104 | ZNF444 | Q8N0Y2 | | 162 | YKQEPSSPPLAPG | 5.7137 | 6.2380 | 6.5179 | 6.5182 | 6.3398 | 6.2027 | 6.4083 | 5.9629 | 7.2584 | 6.4781 | 6.2112 | 6.5182 | 5.9396 | 5.5737 | 5.9493 | 6.1698 |
| ZNF444_S162 | ZNF444 | Q8N0Y2 | | 104 | LWGPAASPDGSSA | 8.3675 | 6.9229 | 6.4201 | 7.4025 | 6.6281 | 6.3161 | 5.4897 | 6.3739 | 8.7085 | 5.3337 | 5.4320 | 5.2875 | 4.1838 | 7.1078 | 4.7559 | 5.2702 |
| ZNF451_S149 | ZNF451 | Q9Y4E5-4 | | 149 | FQTLPSSPLLVPQ | 5.8670 | 5.9566 | 6.2658 | 6.8203 | 5.5169 | 6.2116 | 6.2729 | 6.3934 | 5.8083 | 6.2658 | 6.4505 | 6.0656 | 6.5595 | 6.4495 | 6.6443 | 6.4520 |
| ZNF451_S601 | ZNF451 | Q9Y4E5-4 | | 145 | SSQNFQTLPSSPL | 6.1050 | 5.6967 | 6.6620 | 6.8549 | 6.4307 | 6.4591 | 6.1401 | 6.6303 | 5.8762 | 5.8578 | 6.1175 | 5.9372 | 5.4888 | 6.6795 | 6.7731 | 6.2912 |
| ZNF451_T145 | ZNF451 | Q9Y4E5 | | 601 | HSPANSSPRGKWQ | 5.8185 | 5.9996 | 6.6537 | 7.1960 | 7.1893 | 6.1673 | 7.2988 | 6.2065 | 6.8930 | 6.3317 | 6.6123 | 6.8248 | 4.1481 | 5.1398 | 5.7078 | 5.8129 |
| ZNF462_S1090 | ZNF462 | Q96JM2-3 | | 688 | DFPLDLSPVKKRT | 6.4245 | 6.5403 | 6.3309 | 7.1374 | 6.3371 | 6.3101 | 6.3236 | 6.3029 | 6.5681 | 5.9935 | 6.0439 | 6.3517 | 5.8363 | 5.7419 | 5.9055 | 5.8522 |
| ZNF462_S1098 | ZNF462 | Q96JM2-3 | | 1747 | NKVIIPSPPKDDS | 6.3326 | 6.3915 | 6.4845 | 7.2024 | 5.8264 | 5.7257 | 6.6477 | 6.2768 | 6.7945 | 6.5501 | 6.4658 | 6.7810 | 5.4275 | 5.4129 | 5.4636 | 6.2168 |
| ZNF462_S1166 | ZNF462 | Q96JM2-3 | | 1451 | PEDARLSPEKSLQ | 5.8505 | 4.2350 | 5.9951 | 7.0563 | 6.6287 | 5.7100 | 7.1022 | 5.4757 | 5.9339 | 5.7323 | 6.1043 | 6.0362 | 7.4485 | 7.1412 | 6.6613 | 6.8887 |
| ZNF462_S1356 | ZNF462 | Q96JM2-3 | | 2462 | RELMRFSDHGAAL | 5.7658 | 5.8584 | 5.6463 | 7.4192 | 6.9799 | 6.1315 | 7.1381 | 7.1158 | 5.0842 | 4.9772 | 5.4243 | 5.3968 | 6.3600 | 6.1171 | 7.2182 | 7.3673 |
| ZNF462_S1451 | ZNF462 | Q96JM2-3 | | 1090 | EVGAPMSPKMSNM | 6.1245 | 6.0755 | 6.7317 | 6.9527 | 6.2307 | 6.3846 | 6.5689 | 6.0007 | 6.0232 | 6.5564 | 6.6915 | 6.7444 | 5.7086 | 6.1509 | 5.3210 | 5.7345 |
| ZNF462_S1747 | ZNF462 | Q96JM2-3 | | 1356 | WAGPDPSPPSLTM | 6.0202 | 6.5540 | 6.0532 | 6.4702 | 6.3216 | 5.7922 | 6.1364 | 6.1272 | 5.8989 | 5.6819 | 5.6126 | 6.0104 | 6.9122 | 6.4414 | 6.8403 | 7.1274 |
| ZNF462_S2232 | ZNF462 | Q96JM2-3 | | 1166 | VEGPQGSPRPPAP | 6.6972 | 6.9598 | 7.1528 | 7.6222 | 6.1568 | 6.1043 | 6.5105 | 6.2927 | 7.1546 | 6.7445 | 6.4215 | 6.5715 | 4.5464 | 5.3222 | 4.9128 | 4.8302 |
| ZNF462_S2462 | ZNF462 | Q96JM2-3 | | 1098 | KMSNMGSPPPPQP | 6.8622 | 6.7989 | 6.0030 | 8.4929 | 6.7563 | 6.5713 | 6.6371 | 6.0797 | 6.8123 | 7.1603 | 7.1117 | 6.6772 | 4.7143 | 4.7667 | 3.6812 | 4.8750 |
| ZNF462_S688 | ZNF462 | Q96JM2-3 | | 2232 | RNNSRVSPVPLSG | 6.9421 | 7.1952 | 6.6178 | 6.6489 | 6.2013 | 6.3233 | 6.7219 | 5.1521 | 5.4297 | 5.8412 | 6.1009 | 5.7270 | 7.5092 | 5.9039 | 6.1514 | 5.5341 |
| ZNF462_S934 | ZNF462 | Q96JM2-3 | | 792 | GAEIELTLSEDEE | 6.8806 | 5.9790 | 6.4934 | 6.2332 | 5.9574 | 6.1978 | 6.1456 | 5.8716 | 5.3908 | 6.5013 | 6.9466 | 6.8067 | 5.4598 | 6.4806 | 6.0096 | 6.6461 |
| ZNF462_T792 | ZNF462 | Q96JM2-3 | | 934 | RFCSYTSPNVRSL | 7.0033 | 7.7259 | 6.9723 | 7.1751 | 5.9619 | 6.6008 | 8.0708 | 6.7045 | 6.8354 | 7.1860 | 7.1720 | 7.3657 | 3.2712 | 4.5559 | 3.2450 | 4.1543 |
| ZNF48_S416 | ZNF48 | Q96MX3 | | 416 | PPLTPRSPSHSGE | 6.1396 | 6.5352 | 6.0790 | 7.0356 | 6.2743 | 6.1899 | 6.3091 | 6.5559 | 6.4067 | 6.5524 | 6.5420 | 6.4799 | 5.4184 | 5.7968 | 5.8068 | 5.8785 |
| ZNF496_S335 | ZNF496 | Q96IT1-2 | | 335 | QVSYLDSPSLQPF | 5.6735 | 6.3453 | 6.7452 | 5.9302 | 5.8815 | 6.9830 | 6.0654 | 6.7365 | 7.0892 | 6.1316 | 6.2348 | 7.1182 | 5.5884 | 6.3795 | 5.4234 | 5.6742 |
| ZNF507_S136 | ZNF507 | Q8TCN5 | | 195 | KAQQCVSPSSSLC | 6.4173 | 6.8162 | 6.6114 | 6.6894 | 6.4660 | 6.6467 | 6.0769 | 6.4659 | 6.9716 | 6.6335 | 7.0984 | 7.0982 | 4.6220 | 5.1896 | 4.7890 | 5.4080 |
| ZNF507_S195 | ZNF507 | Q8TCN5 | | 404 | IVERLPSAEETLS | 4.2204 | 3.9580 | 4.9834 | 3.6890 | 5.4022 | 5.1983 | 4.9044 | 5.3094 | 4.7105 | 4.0376 | 4.4619 | 4.3514 | 10.4149 | 10.8149 | 11.3735 | 12.1702 |
| ZNF507_S388 | ZNF507 | Q8TCN5 | | 884 | NENEKLSPTSNTS | 5.9902 | 6.8735 | 6.1479 | 6.1780 | 5.8318 | 5.4479 | 6.1293 | 6.5132 | 6.3497 | 7.0741 | 7.0547 | 7.0699 | 5.3707 | 5.8978 | 6.0274 | 6.0439 |
| ZNF507_S404 | ZNF507 | Q8TCN5 | | 388 | QKIISSSPNKKGH | 5.4071 | 5.1340 | 6.7735 | 6.2697 | 5.9950 | 6.4772 | 6.3458 | 5.7980 | 7.2204 | 6.2177 | 6.2393 | 7.0026 | 5.7475 | 6.2833 | 6.2497 | 6.8392 |
| ZNF507_S884 | ZNF507 | Q8TCN5 | | 136 | CKFLSSSFSVLKD | 5.6179 | 5.9717 | 7.0822 | 6.6032 | 6.9080 | 5.5906 | 4.4838 | 6.0352 | 6.3237 | 5.4310 | 6.8554 | 5.9554 | 6.1822 | 6.6884 | 8.4631 | 5.8084 |
| ZNF512B_S63 | ZNF512B | Q96KM6 | | 686 | GRVRRTSAQVAVF | 5.1627 | 5.2054 | 5.7464 | 5.7364 | 7.3702 | 5.8267 | 5.3291 | 6.2509 | 7.2521 | 7.5483 | 6.7832 | 7.2071 | 6.5754 | 7.3367 | 5.2846 | 5.3847 |
| ZNF512B_S665 | ZNF512B | Q96KM6 | | 685 | SGRVRRTSAQVAV | 5.2112 | 4.9892 | 6.1027 | 5.7977 | 6.8871 | 5.3102 | 5.5123 | 6.1892 | 6.6604 | 7.4583 | 7.1341 | 8.0103 | 6.6245 | 6.3508 | 5.7070 | 6.0550 |
| ZNF512B_S686 | ZNF512B | Q96KM6 | | 665 | EEPTDKSPEAEDP | 6.3514 | 6.6208 | 6.8482 | 6.4078 | 5.8446 | 5.4435 | 6.5082 | 6.1024 | 5.9633 | 6.5170 | 6.2666 | 5.7306 | 5.4646 | 6.4587 | 6.5448 | 6.9276 |
| ZNF512B_T685 | ZNF512B | Q96KM6 | | 63 | LCFDPGSPASDKT | 7.0371 | 8.1812 | 6.3590 | 6.6761 | 6.8297 | 6.6431 | 6.5993 | 7.4410 | 6.5318 | 7.1469 | 5.8583 | 7.3350 | 4.0184 | 4.3509 | 4.4899 | 4.5022 |
| ZNF516_S849 | ZNF516 | Q92618 | | 930 | GFSRSATPTPTVI | 5.7534 | 5.8636 | 6.3332 | 6.2248 | 5.0799 | 6.1283 | 6.3676 | 6.2253 | 6.1927 | 7.0348 | 6.4111 | 6.0902 | 6.3567 | 7.2311 | 6.4220 | 6.2851 |
| ZNF516_T930 | ZNF516 | Q92618 | | 849 | GMPGSKSGSSPLG | 6.6871 | 7.4615 | 6.5242 | 7.4667 | 6.1466 | 6.9344 | 7.1650 | 6.6246 | 6.8741 | 7.3012 | 7.5309 | 6.7465 | 3.6120 | 4.9705 | 3.7577 | 4.1969 |
| ZNF518A_S437 | ZNF518A | Q6AHZ1 | | 437 | NNKLAVSPNYNAT | 7.0694 | 6.4854 | 6.6407 | 7.0235 | 6.8944 | 5.3086 | 7.3603 | 7.0891 | 6.9382 | 6.9426 | 6.1508 | 6.2032 | 6.2426 | 4.8012 | 4.3921 | 4.4578 |
| ZNF518B_S317 | ZNF518B | Q9C0D4 | | 317 | VEVEVLSPAKEPV | 7.7637 | 7.4179 | 7.6756 | 7.8870 | 5.9546 | 6.4898 | 6.7842 | 7.1132 | 8.0104 | 6.9509 | 7.5236 | 7.8973 | 2.8767 | 3.3672 | 2.8011 | 3.4868 |
| ZNF518B_S836 | ZNF518B | Q9C0D4 | | 836 | RGPIDMSPNIETP | 6.1475 | 7.9311 | 6.2911 | 7.9683 | 6.0901 | 6.5878 | 7.5450 | 6.8580 | 7.5068 | 7.2682 | 8.0939 | 7.4163 | 3.5906 | 4.4575 | 2.9695 | 3.8183 |
| ZNF532_S1022 | ZNF532 | Q9HCE3 | | 314 | PRAADKSPESQNL | 6.3595 | 6.4351 | 6.1740 | 7.0558 | 5.8538 | 6.2524 | 6.7459 | 5.7628 | 6.0647 | 6.3085 | 6.4000 | 6.3051 | 5.8605 | 5.8485 | 6.0086 | 6.5646 |
| ZNF532_S314 | ZNF532 | Q9HCE3 | | 1022 | ETKKVASPGWTCW | 7.0880 | 7.4437 | 6.8111 | 7.3217 | 6.3320 | 4.7004 | 7.6774 | 6.6798 | 6.5393 | 6.5695 | 5.4236 | 6.3989 | 5.3816 | 5.0601 | 5.6022 | 4.9708 |
| ZNF532_S349 | ZNF532 | Q9HCE3 | | 418 | SAAVLSSPPRAPL | 6.2714 | 5.7501 | 6.2800 | 6.9651 | 5.9207 | 5.9884 | 6.3520 | 5.3385 | 5.5811 | 6.8184 | 6.3245 | 7.0860 | 5.6223 | 6.8982 | 6.2217 | 6.5818 |
| ZNF532_S418 | ZNF532 | Q9HCE3 | | 349 | SSKGSPSSPAGST | 5.9519 | 5.6319 | 6.0613 | 7.0382 | 6.6701 | 5.9983 | 6.4583 | 6.3968 | 6.3175 | 5.8027 | 6.3680 | 6.1301 | 6.3570 | 6.4347 | 6.3306 | 6.0528 |
| ZNF542P_S18;S21 | ZNF542P | Q5EBM4 | 18;21 | | NA | 6.5703 | 7.0893 | 5.6686 | 6.5061 | 5.4551 | 5.6936 | 6.3895 | 5.5843 | 6.8673 | 7.3566 | 8.1069 | 8.1039 | 4.8933 | 6.0595 | 4.5344 | 5.1214 |
| ZNF568_S35 | ZNF568 | Q3ZCX4 | | 35 | SQESALSEEEEDT | 6.0531 | 6.2175 | 6.6766 | 6.6538 | 6.0742 | 6.1738 | 6.0731 | 5.9618 | 5.7751 | 6.1137 | 5.7627 | 5.9764 | 6.6254 | 6.4932 | 6.7572 | 6.6122 |
| ZNF574_S202 | ZNF574 | Q6ZN55-2 | | 253 | APVVLGSPVVLGP | 6.2511 | 6.4049 | 6.2144 | 7.0183 | 7.0587 | 6.1599 | 6.2541 | 6.4233 | 6.4872 | 6.5284 | 6.8631 | 6.7285 | 5.1655 | 5.7767 | 5.1766 | 5.4892 |
| ZNF574_S253 | ZNF574 | Q6ZN55-2 | | 878 | DHRRLHTGERPFA | 6.7284 | 6.9329 | 7.1105 | 7.4513 | 5.7295 | 6.7456 | 7.3106 | 6.0181 | 8.3801 | 8.1702 | 9.1171 | 8.6304 | 2.5636 | 3.4680 | 2.4182 | 3.2255 |
| ZNF574_S623 | ZNF574 | Q6ZN55-2 | | 202 | AVPAEPSPKAPPL | 6.9721 | 6.4159 | 7.0578 | 7.2471 | 7.5715 | 6.7017 | 6.9644 | 6.1309 | 5.9046 | 6.4615 | 6.5333 | 6.3534 | 4.9753 | 5.7655 | 4.4748 | 4.4701 |
| ZNF574_T607 | ZNF574 | Q6ZN55-2 | | 623 | CSKPFNSPANLAR | 7.3657 | 7.1922 | 7.3638 | 7.3306 | 6.6207 | 6.6118 | 7.3851 | 6.2419 | 8.2145 | 7.4747 | 7.0727 | 7.8977 | 2.7207 | 3.9103 | 2.9627 | 3.6348 |
| ZNF574_T878 | ZNF574 | Q6ZN55-2 | | 607 | NHLRTHTGERPFP | 6.6305 | 7.2279 | 7.6715 | 7.8326 | 6.0541 | 7.3251 | 7.7275 | 6.1969 | 7.9659 | 8.0063 | 9.0785 | 8.1521 | 2.0883 | 3.3739 | 1.9452 | 2.7236 |
| ZNF576_S20 | ZNF576 | Q9H609 | | 23 | KERSPQSPGGNIC | 6.7376 | 6.0910 | 7.6223 | 6.4717 | 6.4057 | 6.7383 | 6.2914 | 6.6711 | 7.2139 | 6.5173 | 7.2458 | 7.2244 | 4.5058 | 5.0105 | 4.7102 | 4.5430 |
| ZNF576_S23 | ZNF576 | Q9H609 | | 20 | DSPKERSPQSPGG | 6.2678 | 6.5147 | 6.8900 | 5.5484 | 6.8475 | 7.2371 | 6.6772 | 6.5677 | 6.6294 | 6.3799 | 7.2085 | 7.4151 | 4.5992 | 5.4327 | 4.8608 | 4.9241 |
| ZNF579_S483 | ZNF579 | Q8NAF0 | | 483 | QAQAPTSPPPPPP | 6.0739 | 6.2202 | 6.2912 | 6.6021 | 6.0826 | 6.2040 | 6.1536 | 6.2237 | 6.3842 | 6.4061 | 6.2673 | 6.2043 | 5.9696 | 6.3951 | 6.2166 | 6.3056 |
| ZNF581_S30;S45 | ZNF581 | Q9P0T4 | 30;45 | | NA | 5.5598 | 5.1439 | 5.7632 | 6.4609 | 6.4639 | 6.3582 | 6.7189 | 6.8675 | 6.3335 | 5.5789 | 5.6959 | 6.9748 | 6.3173 | 7.1115 | 7.5681 | 5.0838 |
| ZNF589_S187 | ZNF589 | Q86UQ0-3 | | 187 | ILEIQLSPAQNAS | 6.2171 | 6.4813 | 6.2729 | 6.4220 | 6.1097 | 6.0160 | 6.3632 | 6.6197 | 7.4040 | 7.1490 | 6.8251 | 5.8315 | 5.4789 | 6.5012 | 5.5989 | 4.7097 |
| ZNF592_S142 | ZNF592 | Q92610 | | 348 | SIKPSDSPRSICS | 6.2617 | 6.6588 | 5.8576 | 7.0332 | 6.0616 | 6.0006 | 6.1707 | 5.9546 | 6.0256 | 6.1870 | 6.2472 | 6.1727 | 6.4139 | 6.2668 | 6.3662 | 6.3217 |
| ZNF592_S334 | ZNF592 | Q92610 | | 680 | APAPSSSPKHGLT | 6.1698 | 5.6375 | 6.4770 | 6.6109 | 6.5157 | 5.8657 | 6.2553 | 5.8077 | 6.8597 | 6.6504 | 6.2583 | 6.4532 | 5.8513 | 6.3169 | 5.9701 | 6.3006 |
| ZNF592_S348 | ZNF592 | Q92610 | | 687 | PKHGLTSGSASPP | 5.9870 | 6.6984 | 6.2194 | 7.2277 | 6.6463 | 6.3395 | 6.0381 | 6.4508 | 6.0409 | 6.1050 | 6.4423 | 6.0600 | 5.9768 | 5.9671 | 5.8201 | 5.9805 |
| ZNF592_S368 | ZNF592 | Q92610 | | 78 | NTSRQESFEAEKD | 5.2699 | 5.1476 | 5.0039 | 5.8082 | 6.1270 | 5.6897 | 6.1356 | 5.6488 | 5.1273 | 5.1149 | 5.1769 | 5.5847 | 8.6539 | 8.0111 | 8.8917 | 8.6087 |
| ZNF592_S680 | ZNF592 | Q92610 | | 142 | PTFNQFSPISSPE | 5.5244 | 5.6045 | 5.6025 | 5.8519 | 5.7350 | 5.9934 | 5.7644 | 5.4006 | 6.1178 | 5.6665 | 5.6736 | 5.5518 | 7.6938 | 8.1712 | 7.5665 | 8.0821 |
| ZNF592_S687 | ZNF592 | Q92610 | | 334 | SPKSPRSPLEATR | 6.6957 | 7.2493 | 7.0836 | 6.9604 | 6.2372 | 5.3885 | 6.5399 | 6.0068 | 6.9999 | 6.5765 | 6.6647 | 5.9997 | 5.2427 | 5.8854 | 5.1216 | 5.3480 |
| ZNF592_S78 | ZNF592 | Q92610 | | 368 | PSVAASSPPAIPK | 6.0381 | 6.2222 | 6.3094 | 7.2736 | 6.3257 | 5.5238 | 6.4053 | 6.6155 | 6.4728 | 7.2943 | 6.3257 | 6.1263 | 5.8725 | 5.9248 | 6.0065 | 5.2634 |
| ZNF593_S133 | ZNF593 | O00488 | | 133 | VPEMDTSTxxxxx | 6.0044 | 6.1322 | 6.6348 | 6.6870 | 5.9804 | 5.9199 | 6.2743 | 6.3183 | 6.3279 | 6.3070 | 6.3061 | 6.5547 | 5.5564 | 6.6163 | 6.1981 | 6.1821 |
| ZNF594_S50 | ZNF594 | Q96JF6 | | 50 | LLKHWVSPLKDAM | 6.4021 | 6.5228 | 7.2750 | 7.9864 | 6.2386 | 6.1425 | 6.8956 | 6.6348 | 6.6107 | 7.0036 | 7.8858 | 7.5183 | 4.2459 | 4.5570 | 3.8955 | 4.1854 |
| ZNF597_T109 | ZNF597 | Q96LX8 | | 109 | AKILSGTPTYKRR | 7.2116 | 7.5147 | 6.7442 | 7.4519 | 6.1044 | 6.7353 | 6.8626 | 6.5137 | 6.9363 | 7.5556 | 7.2282 | 7.2683 | 3.7342 | 4.8267 | 3.6935 | 4.4190 |
| ZNF598_S431 | ZNF598 | Q86UK7-3 | | 431 | GALPPPSPKLKDE | 6.2241 | 6.0178 | 6.5542 | 5.9799 | 6.4308 | 6.5245 | 6.6187 | 6.5755 | 6.5692 | 6.3473 | 6.6398 | 6.1120 | 5.5727 | 6.2707 | 5.7339 | 5.8291 |
| ZNF598_S437 | ZNF598 | Q86UK7 | | 437 | GALPPPSPKLKDE | 6.3347 | 6.1064 | 6.6008 | 6.3427 | 6.6982 | 6.3209 | 6.3977 | 5.9516 | 6.4826 | 6.4620 | 6.3370 | 6.2165 | 5.6717 | 5.9504 | 5.8116 | 6.3152 |
| ZNF608_S1430 | ZNF608 | Q9ULD9 | | 493 | AEDIKASPSSTNK | 6.5571 | 7.6516 | 6.6486 | 6.2543 | 5.8828 | 5.7171 | 6.1844 | 5.9586 | 6.1361 | 6.7896 | 5.9252 | 6.0697 | 5.5722 | 7.2757 | 5.7100 | 5.6672 |
| ZNF608_S421 | ZNF608 | Q9ULD9 | | 1430 | NQYRSKSPAPVEK | 6.1230 | 6.2890 | 5.3488 | 5.7253 | 5.1083 | 6.1900 | 6.4834 | 5.8799 | 6.3481 | 6.4259 | 5.5213 | 5.6241 | 6.5505 | 7.9775 | 6.5343 | 7.8706 |
| ZNF608_S493 | ZNF608 | Q9ULD9 | | 481 | NASGRRTPPNCAA | 5.9377 | 5.9017 | 5.3961 | 6.0899 | 6.2328 | 5.8479 | 6.6087 | 5.3794 | 6.3792 | 6.7013 | 6.2823 | 6.0710 | 6.0831 | 7.9745 | 6.4116 | 6.7028 |
| ZNF608_S596 | ZNF608 | Q9ULD9 | | 596 | DSEDKISDCEEGL | 6.6951 | 6.3636 | 6.3856 | 5.5341 | 5.8669 | 5.7903 | 5.5648 | 5.8512 | 5.3777 | 6.8094 | 6.4634 | 6.4446 | 5.9856 | 7.4436 | 6.3190 | 7.1052 |
| ZNF608_S964 | ZNF608 | Q9ULD9 | | 421 | PPRFCESPTSDLE | 6.2750 | 6.1841 | 6.2546 | 5.7549 | 7.1694 | 6.3611 | 5.4275 | 5.6938 | 6.5099 | 6.0510 | 7.1380 | 5.3445 | 6.4488 | 7.8445 | 5.9946 | 5.5482 |
| ZNF608_T481 | ZNF608 | Q9ULD9 | | 964 | MRSKASSPSDIIS | 6.5515 | 6.8404 | 5.7496 | 6.3876 | 6.2704 | 5.8650 | 5.5648 | 6.6644 | 6.5358 | 6.1919 | 6.2604 | 6.1619 | 5.7827 | 6.9978 | 6.3976 | 5.7783 |
| ZNF609_S1055 | ZNF609 | O15014 | | 722 | TVNPALTPAKDKK | 6.4779 | 6.0616 | 6.6147 | 6.9125 | 6.2469 | 6.3202 | 6.8529 | 5.9370 | 6.8410 | 6.4212 | 6.4521 | 6.6984 | 5.4834 | 5.9336 | 4.8939 | 5.8528 |
| ZNF609_S1285 | ZNF609 | O15014 | | 1055 | TLTKAPSLTDLVK | 6.4184 | 6.3598 | 5.9373 | 6.2877 | 6.6707 | 6.1894 | 6.3625 | 6.2542 | 5.0577 | 4.9492 | 4.8935 | 4.7544 | 7.1287 | 7.4767 | 7.6439 | 7.6160 |
| ZNF609_S1313 | ZNF609 | O15014 | | 491 | LDRNCPSPVLIDC | 5.8006 | 5.8996 | 6.0687 | 6.0943 | 6.2133 | 6.0197 | 6.0776 | 5.7267 | 5.9758 | 6.4396 | 6.2329 | 6.2624 | 6.5539 | 7.1886 | 6.5372 | 6.9088 |
| ZNF609_S252 | ZNF609 | O15014 | | 252 | KSEKMESPVSTPA | 5.6793 | 5.8693 | 6.1900 | 6.2555 | 5.6991 | 6.1261 | 6.0259 | 6.3608 | 5.3719 | 6.3226 | 5.8826 | 6.5765 | 6.8634 | 7.2467 | 6.4280 | 7.1024 |
| ZNF609_S358 | ZNF609 | O15014 | | 823 | TPTQPLTPLHVVT | 6.1561 | 6.0591 | 5.9795 | 6.6128 | 6.3918 | 6.3497 | 6.3473 | 6.3484 | 6.6175 | 6.3007 | 6.3044 | 6.2912 | 5.5307 | 6.7847 | 5.7801 | 6.1460 |
| ZNF609_S433 | ZNF609 | O15014 | | 758 | RAEEGKSPFRESS | 5.9228 | 6.0638 | 5.7399 | 6.0816 | 6.6084 | 6.6224 | 6.5618 | 6.3811 | 6.4450 | 6.0103 | 6.3807 | 5.8704 | 6.4087 | 6.7771 | 6.0629 | 6.0631 |
| ZNF609_S452 | ZNF609 | O15014 | | 746 | ELESPLTPGKVCR | 6.3627 | 6.5400 | 6.6592 | 6.8281 | 6.4718 | 5.8645 | 6.1248 | 6.1302 | 6.6536 | 6.3870 | 6.4436 | 6.1245 | 5.2865 | 6.0958 | 5.9601 | 6.0675 |
| ZNF609_S452;S453 | ZNF609 | O15014 | | 576 | RLVEPHSPSPSSK | 5.8211 | 5.9634 | 6.3398 | 6.5610 | 5.8757 | 6.2837 | 5.8628 | 6.1959 | 5.8897 | 5.8116 | 5.8702 | 6.1369 | 6.4936 | 7.0633 | 6.8176 | 7.0137 |
| ZNF609_S453 | ZNF609 | O15014 | | 1285 | ADKARASPSVTCK | 6.0747 | 6.6431 | 6.4248 | 6.3445 | 5.7790 | 6.6133 | 6.4184 | 6.2932 | 6.7276 | 6.9515 | 6.7256 | 6.9448 | 4.8672 | 6.0625 | 4.9722 | 6.1574 |
| ZNF609_S467 | ZNF609 | O15014 | | 452 | SDMELNSSSEDSK | 6.7280 | 7.5542 | 7.6899 | 7.8809 | 6.7302 | 7.3090 | 7.8422 | 6.3290 | 4.2288 | 4.9243 | 5.1532 | 5.4374 | 5.1091 | 5.6917 | 5.1991 | 6.1932 |
| ZNF609_S491 | ZNF609 | O15014 | | 467 | KRVRTNSMGSATG | 5.2956 | 5.5563 | 6.7964 | 6.2314 | 6.1433 | 6.1532 | 6.8214 | 5.8878 | 6.0295 | 5.6094 | 6.3968 | 6.2650 | 6.2086 | 7.2215 | 6.6858 | 6.6980 |
| ZNF609_S560 | ZNF609 | O15014 | | 433 | SEDVKASPSSANK | 6.0641 | 6.9927 | 6.2890 | 6.9576 | 5.8104 | 6.0894 | 6.0034 | 6.0466 | 5.9580 | 6.3345 | 6.1661 | 6.8738 | 5.8201 | 6.5395 | 5.5674 | 6.4874 |
| ZNF609_S576 | ZNF609 | O15014 | | 560 | SQKGSLSPARSAT | 6.1322 | 6.6005 | 5.6636 | 6.2701 | 5.9681 | 5.9511 | 6.2145 | 5.9568 | 6.0668 | 6.5817 | 6.1603 | 5.9872 | 6.0297 | 7.1325 | 6.6059 | 6.6791 |
| ZNF609_S576;S578 | ZNF609 | O15014 | | 1313 | SHYKSKSPTISDK | 5.9240 | 6.3651 | 5.8547 | 6.4683 | 5.5616 | 5.9414 | 5.8620 | 6.0158 | 6.0078 | 6.1275 | 5.9548 | 6.1668 | 5.9155 | 7.3088 | 6.9107 | 7.6152 |
| ZNF609_S578 | ZNF609 | O15014 | | 804 | FTDNAPSPSIGGS | 6.2035 | 7.2705 | 7.0998 | 6.7485 | 6.6777 | 6.1990 | 6.3264 | 5.6328 | 6.7251 | 6.7499 | 6.9662 | 6.6930 | 4.4990 | 5.6671 | 4.9984 | 5.5430 |
| ZNF609_S758 | ZNF609 | O15014 | | 899 | QGFESYYSPSYAQ | 5.9119 | 6.2593 | 6.3615 | 6.8569 | 6.7985 | 6.1395 | 6.1913 | 6.2633 | 5.8993 | 6.5483 | 6.1828 | 5.1442 | 5.9331 | 6.8820 | 6.6308 | 5.9975 |
| ZNF609_S804 | ZNF609 | O15014 | | 76 | DNIKFVTPVPGPQ | 6.7416 | 6.4981 | 6.6735 | 6.4875 | 6.0293 | 8.3385 | 7.9982 | 6.0233 | 7.1529 | 7.4569 | 7.6746 | 6.0626 | 3.1757 | 4.8861 | 3.9137 | 4.8876 |
| ZNF609_T722 | ZNF609 | O15014 | | 578 | VEPHSPSPSSKFS | 6.5615 | 7.0915 | 6.5413 | 7.0564 | 5.7602 | 5.4088 | 5.4143 | 5.1915 | 6.4492 | 6.2491 | 6.2814 | 6.6765 | 6.1194 | 6.5704 | 6.2900 | 6.3384 |
| ZNF609_T746 | ZNF609 | O15014 | | 358 | PPRFCDSPTSDLE | 5.2756 | 6.0553 | 6.4049 | 6.9452 | 6.8054 | 5.4536 | 6.8350 | 5.1019 | 5.9388 | 5.9664 | 6.4960 | 6.7863 | 5.7735 | 6.9749 | 5.7375 | 7.4495 |
| ZNF609_T76 | ZNF609 | O15014 | | 453 | DMELNSSSEDSKG | 6.5026 | 6.5157 | 6.7333 | 5.8512 | 5.3503 | 5.2047 | 6.4158 | 5.7268 | 7.2325 | 7.0099 | 6.7677 | 6.3340 | 6.7775 | 6.7055 | 5.7595 | 5.1130 |
| ZNF609_T823 | ZNF609 | O15014 | 576;578 | | NA | 6.2841 | 6.7072 | 5.9780 | 6.5035 | 6.4952 | 6.3281 | 6.2932 | 6.0395 | 6.4795 | 6.2775 | 6.2834 | 6.1083 | 5.5901 | 6.3795 | 6.0333 | 6.2197 |
| ZNF609_Y899 | ZNF609 | O15014 | 452;453 | | NA | 6.3812 | 6.7820 | 6.7540 | 7.7278 | 5.8982 | 7.0281 | 7.0247 | 6.0783 | 5.1048 | 6.2544 | 6.4544 | 6.2822 | 4.8990 | 6.3516 | 5.2488 | 5.7304 |
| ZNF622_S143 | ZNF622 | Q969S3 | | 143 | KAQPSMSPKKAPP | 5.5500 | 5.6681 | 5.5850 | 5.1038 | 6.0732 | 5.9163 | 5.9486 | 5.8325 | 6.3712 | 5.9706 | 5.8542 | 5.9867 | 7.5985 | 7.3118 | 7.6920 | 7.5376 |
| ZNF622_S78 | ZNF622 | Q969S3 | | 78 | CSKKFASFNAYEN | 6.2066 | 6.5290 | 7.8879 | 6.9289 | 4.9825 | 8.4816 | 6.4780 | 6.3005 | 5.9679 | 6.5127 | 5.8353 | 6.9391 | 5.7496 | 4.0990 | 5.2865 | 5.8149 |
| ZNF629_T561 | ZNF629 | Q9UEG4 | | 561 | GDPSLLTPPPGAK | 6.3288 | 7.1692 | 6.6933 | 7.1174 | 6.1801 | 6.4973 | 7.2272 | 6.6674 | 7.2712 | 7.4542 | 7.3987 | 7.7613 | 3.8565 | 4.7227 | 3.5340 | 4.1206 |
| ZNF638_S1100 | ZNF638 | Q14966 | | 1401 | KAVIVSSPKAKAT | 6.6019 | 6.4328 | 6.1006 | 6.7320 | 6.6715 | 5.9943 | 6.6907 | 6.0192 | 6.5643 | 6.3400 | 5.9264 | 5.8922 | 5.8449 | 6.0804 | 6.1477 | 5.9612 |
| ZNF638_S1119 | ZNF638 | Q14966 | | 605 | AADKGHSPAQKPK | 6.1776 | 6.3880 | 5.4969 | 6.4515 | 5.7184 | 5.9186 | 5.9329 | 5.9388 | 5.7994 | 6.1391 | 5.4640 | 5.8908 | 6.7319 | 7.4824 | 7.2623 | 7.2073 |
| ZNF638_S128 | ZNF638 | Q14966 | | 1656 | DQDDCISHSEPKD | 6.1179 | 6.1328 | 6.6756 | 6.6852 | 6.0694 | 5.9732 | 6.3033 | 5.3034 | 6.3975 | 5.8166 | 6.2767 | 5.8264 | 6.4048 | 6.8616 | 6.5499 | 6.6058 |
| ZNF638_S1401 | ZNF638 | Q14966 | | 552 | RNPFRGSPKCFRS | 6.5781 | 6.4122 | 5.9758 | 6.2926 | 6.1896 | 6.2269 | 6.4667 | 5.2111 | 6.2955 | 5.6490 | 5.5616 | 5.8088 | 7.2583 | 6.9397 | 6.6537 | 6.4805 |
| ZNF638_S1656 | ZNF638 | Q14966 | | 383 | ADIPIRSPFGIVK | 6.6844 | 6.6871 | 6.6405 | 6.7284 | 5.9453 | 6.3867 | 7.0275 | 6.3002 | 6.7710 | 6.8050 | 6.8341 | 6.6140 | 4.5363 | 5.8622 | 4.7646 | 5.4126 |
| ZNF638_S1910 | ZNF638 | Q14966 | | 283 | ESSNNRSFFSVES | 7.3937 | 7.3078 | 6.3985 | 7.5808 | 5.7416 | 6.6403 | 7.7684 | 6.0485 | 7.3690 | 6.3190 | 7.0860 | 7.3941 | 3.6687 | 4.9084 | 3.4010 | 4.9741 |
| ZNF638_S283 | ZNF638 | Q14966 | | 1100 | PELEKESPGLKNS | 5.8179 | 5.8993 | 5.3818 | 7.0495 | 6.6905 | 6.6277 | 6.6612 | 5.9831 | 6.5660 | 6.1707 | 7.0957 | 6.8823 | 6.2111 | 6.5840 | 5.5606 | 4.8186 |
| ZNF638_S383 | ZNF638 | Q14966 | | 650 | LEDDTLSECKQVS | 6.2629 | 6.1802 | 5.9052 | 7.7944 | 5.8466 | 5.0160 | 5.9569 | 4.9241 | 6.2480 | 6.6755 | 5.5724 | 4.7858 | 7.8262 | 7.4470 | 6.7558 | 7.8028 |
| ZNF638_S508;S510 | ZNF638 | Q14966 | | 1910 | DSSSGKSVASDVP | 5.6614 | 6.1901 | 6.6421 | 7.1098 | 5.6917 | 6.1981 | 7.2277 | 5.9648 | 6.1508 | 5.7536 | 6.3286 | 6.4168 | 5.4104 | 6.7867 | 6.1760 | 6.2914 |
| ZNF638_S552 | ZNF638 | Q14966 | | 128 | TQVTEQSPKVQSR | 6.0044 | 5.7282 | 6.1585 | 6.6572 | 6.1360 | 5.9805 | 7.0647 | 6.4196 | 6.1494 | 6.2455 | 7.1356 | 6.7042 | 5.7145 | 6.1157 | 5.8338 | 5.9521 |
| ZNF638_S605 | ZNF638 | Q14966 | | 624 | PSVKPTSATKSDS | 7.7678 | 7.0057 | 6.6813 | 7.9662 | 7.3565 | 6.7180 | 7.9175 | 6.2648 | 7.2409 | 6.8830 | 6.8592 | 6.8102 | 2.9717 | 4.1060 | 3.3087 | 4.1424 |
| ZNF638_S624 | ZNF638 | Q14966 | | 1119 | VQTATDSPSVKPN | 5.9590 | 5.9146 | 5.6480 | 6.6769 | 5.8764 | 5.9858 | 6.5263 | 4.8516 | 6.8842 | 6.0812 | 6.2885 | 6.2173 | 6.2798 | 6.4175 | 7.3231 | 7.0699 |
| ZNF638_S650 | ZNF638 | Q14966 | 508;510 | | NA | 6.0308 | 6.3607 | 5.9706 | 6.5878 | 6.1894 | 5.8557 | 6.1239 | 6.6256 | 5.9494 | 6.3519 | 6.5248 | 6.0484 | 6.0264 | 6.5982 | 6.4992 | 6.2574 |
| ZNF639_S100 | ZNF639 | Q9UID6 | | 88 | QNYLVPSPVLRIL | 5.6789 | 5.7346 | 6.0241 | 6.2166 | 5.7734 | 5.8556 | 6.2046 | 5.6159 | 6.2213 | 5.7804 | 5.9792 | 6.3344 | 7.3567 | 6.9060 | 6.8220 | 7.4964 |
| ZNF639_S182 | ZNF639 | Q9UID6 | | 182 | CKILDKSQALNVT | 8.0074 | 6.8113 | 6.0128 | 6.9826 | 6.6335 | 6.1971 | 6.8210 | 7.0214 | 5.7962 | 4.8296 | 5.3145 | 4.8038 | 6.5721 | 6.2294 | 5.7930 | 6.1744 |
| ZNF639_S88 | ZNF639 | Q9UID6 | | 100 | LDHTAFSTEKSAD | 5.6606 | 5.6922 | 7.1586 | 5.8005 | 6.5111 | 8.0273 | 8.4582 | 7.1970 | 5.6967 | 6.6465 | 5.9832 | 5.5433 | 4.7672 | 6.7841 | 4.6250 | 5.4485 |
| ZNF644_S1189 | ZNF644 | Q9H582 | | 394 | KKKCEESDSESPA | 6.4293 | 6.9625 | 6.5629 | 7.1211 | 6.6947 | 6.4342 | 6.7398 | 6.4250 | 5.8598 | 5.6781 | 5.8094 | 6.0477 | 5.7286 | 5.7814 | 5.6969 | 6.0287 |
| ZNF644_S1312 | ZNF644 | Q9H582 | | 165 | ADLQLSTPQKASQ | 7.5644 | 7.2564 | 6.5002 | 7.3552 | 6.2888 | 6.5428 | 6.9029 | 6.4367 | 7.1313 | 6.5838 | 6.4547 | 6.4341 | 4.6342 | 4.9902 | 4.1905 | 4.7337 |
| ZNF644_S394 | ZNF644 | Q9H582 | | 1189 | ERNSAISPQKIHN | 7.0542 | 6.8309 | 6.6707 | 6.7820 | 6.2203 | 5.7697 | 6.7278 | 5.8769 | 6.7875 | 6.2786 | 6.3794 | 6.2946 | 5.7988 | 5.4298 | 5.3439 | 5.7549 |
| ZNF644_T1020 | ZNF644 | Q9H582 | | 1020 | KRTGTGTPVKRVR | 7.0464 | 7.0056 | 6.6605 | 7.4374 | 6.1953 | 5.8393 | 6.4663 | 5.4651 | 6.5260 | 6.4989 | 6.2642 | 6.6779 | 5.6721 | 5.8856 | 4.9521 | 5.4073 |
| ZNF644_T165 | ZNF644 | Q9H582 | | 1312 | LLKKPASITETSF | 4.2950 | 4.3912 | 5.0061 | 6.1984 | 5.9496 | 4.9873 | 6.5762 | 5.7172 | 4.8465 | 4.1495 | 4.5344 | 5.7431 | 9.3198 | 8.5383 | 9.6109 | 10.1366 |
| ZNF646_S930 | ZNF646 | O15015 | | 930 | EAAPARSPPLQLS | 6.0684 | 6.4517 | 7.3439 | 6.4242 | 5.9664 | 6.9909 | 5.7416 | 6.2035 | 6.6379 | 6.3911 | 5.8381 | 6.5814 | 5.6492 | 6.1870 | 6.0077 | 5.5169 |
| ZNF652_S204 | ZNF652 | Q9Y2D9 | | 204 | VAAATTSPTPRTT | 6.7992 | 6.9162 | 6.2436 | 6.7395 | 6.1146 | 6.0752 | 6.5004 | 6.1057 | 6.8510 | 6.5267 | 6.4576 | 6.3492 | 5.5870 | 5.5735 | 5.4666 | 5.6940 |

| | | | | | | | | | | | | | | | | | | | | |
|---|---|---|---|---|---|---|---|---|---|---|---|---|---|---|---|---|---|---|---|---|
| ZNF654_S23 | ZNF654 | A0A1B0GWA0 | 23 | LVAIVESPLGPVG | 6.6311 | 7.1677 | 6.6894 | 6.4511 | 6.2867 | 6.3779 | 5.9843 | 5.8635 | 7.3271 | 6.8907 | 6.2284 | 5.8035 | 5.8635 | 6.4338 | 5.3104 | 4.6908 |
| ZNF655_S28 | ZNF655 | Q8N720 | 285 | ASDKSCSPSSGII | 6.3005 | 6.4188 | 6.3457 | 6.5009 | 6.9015 | 6.6347 | 6.6792 | 6.7151 | 6.2703 | 6.3944 | 6.3984 | 6.2640 | 5.3425 | 5.7582 | 5.6587 | 5.4171 |
| ZNF655_S285 | ZNF655 | Q8N720 | 28 | TQSECLSPEPQFV | 5.8284 | 7.0488 | 5.2489 | 6.0958 | 6.3721 | 6.4028 | 5.7657 | 6.1981 | 5.7798 | 6.3094 | 7.1345 | 6.9337 | 6.3030 | 7.4876 | 5.7632 | 5.3282 |
| ZNF668_S33 | ZNF668 | Q96K58-2 | 33 | EAAEARSPAPGYK | 6.0209 | 6.6762 | 6.5259 | 7.0887 | 6.1674 | 6.4876 | 6.7226 | 6.6573 | 6.9373 | 6.8944 | 6.9309 | 7.0794 | 4.3336 | 5.4151 | 4.8575 | 5.2052 |
| ZNF668_T623 | ZNF668 | Q96K58-2 | 623 | HPVPMGTPTPLEP | 5.5022 | 6.7632 | 6.2114 | 6.9347 | 5.7610 | 6.2644 | 6.8178 | 5.0190 | 5.6098 | 6.2738 | 6.4483 | 7.2246 | 5.9729 | 6.3533 | 6.3844 | 6.4593 |
| ZNF687_S1057 | ZNF687 | Q8N1G0 | 183 | LPPSAPSPTREGA | 5.7489 | 6.2751 | 6.1358 | 6.2111 | 5.9938 | 5.7819 | 5.9360 | 6.1335 | 6.6448 | 6.2477 | 6.1349 | 6.2025 | 6.5119 | 6.8516 | 6.5869 | 6.6034 |
| ZNF687_S1146 | ZNF687 | Q8N1G0 | 1057 | LQLGAQSPGRGTT | 6.3035 | 6.5774 | 5.7105 | 6.4709 | 6.2364 | 5.7551 | 6.2563 | 5.9615 | 6.3970 | 5.9723 | 5.7744 | 5.9571 | 6.8272 | 6.4951 | 6.7984 | 6.5069 |
| ZNF687_S1191 | ZNF687 | Q8N1G0 | 253 | DIPASASPPPVAG | 6.3030 | 6.4010 | 5.6113 | 6.6585 | 6.2485 | 6.1695 | 6.1989 | 5.9258 | 6.2708 | 6.1319 | 5.9386 | 6.2269 | 6.4228 | 6.5362 | 6.5340 | 6.4223 |
| ZNF687_S140 | ZNF687 | Q8N1G0 | 433 | PGGTATSPKMIAK | 6.5202 | 6.4159 | 6.1283 | 6.9362 | 6.7294 | 6.1138 | 6.4676 | 6.2826 | 6.6502 | 6.3851 | 6.1023 | 6.0735 | 5.7490 | 5.8276 | 5.7891 | 5.8292 |
| ZNF687_S174;S183 | ZNF687 | Q8N1G0 | 900 | AGGALLTPKTEPE | 5.8856 | 6.4849 | 5.3891 | 6.4639 | 6.4824 | 6.1464 | 6.2874 | 6.4901 | 6.0322 | 6.4708 | 5.7170 | 5.9452 | 6.5895 | 6.3837 | 6.3988 | 6.8332 |
| ZNF687_S183 | ZNF687 | Q8N1G0 | 191 | TREGALTPPPFPS | 6.1136 | 6.3822 | 5.6960 | 6.2842 | 5.9280 | 5.7970 | 6.2630 | 6.0540 | 6.0625 | 6.3682 | 5.8080 | 6.5103 | 6.5532 | 6.8259 | 6.7295 | 6.6246 |
| ZNF687_S227;S242 | ZNF687 | Q8N1G0 | 374 | ASLLKLSPATPTS | 6.0887 | 6.3577 | 6.2219 | 6.3741 | 6.3921 | 6.1854 | 6.3392 | 6.0295 | 6.6528 | 6.6347 | 6.0143 | 6.3188 | 6.0907 | 6.1524 | 6.2362 | 5.9117 |
| ZNF687_S253 | ZNF687 | Q8N1G0 | 140 | LPGTPHSPAPPSG | 5.9900 | 6.5506 | 5.9485 | 6.2215 | 6.0276 | 6.3113 | 5.8012 | 6.2784 | 6.1435 | 6.1701 | 5.6768 | 6.1165 | 6.8423 | 6.7978 | 6.7926 | 6.3315 |
| ZNF687_S266;S271 | ZNF687 | Q8N1G0 | 377 | LKLSPATPTSEGP | 5.5347 | 5.5414 | 5.2009 | 5.8713 | 5.9515 | 5.8010 | 5.8529 | 5.9668 | 5.6724 | 5.7209 | 5.4155 | 5.1501 | 7.8400 | 8.1288 | 8.4035 | 7.9486 |
| ZNF687_S266;S271;S275 | ZNF687 | Q8N1G0 | 1146 | CGLCFASPGSLSR | 6.2881 | 7.0959 | 6.5914 | 7.3183 | 6.2697 | 5.9058 | 6.3848 | 7.6901 | 7.2480 | 6.3980 | 7.0001 | 7.2409 | 4.2727 | 5.3327 | 4.5381 | 4.4256 |
| ZNF687_S374 | ZNF687 | Q8N1G0 | 1191 | DPDGGDSPLPASG | 4.9120 | 5.6424 | 6.0930 | 5.2533 | 6.0436 | 5.7501 | 6.5156 | 6.1123 | 6.1227 | 6.5261 | 5.7402 | 5.9870 | 7.2207 | 7.2471 | 7.2532 | 7.5807 |
| ZNF687_S433 | ZNF687 | Q8N1G0 | 191;215 | NA | 5.9230 | 6.2865 | 5.9528 | 6.7905 | 6.1579 | 5.8048 | 6.1497 | 6.0109 | 6.5567 | 6.5488 | 6.1577 | 6.0780 | 6.1161 | 6.3933 | 6.4649 | 6.6085 |
| ZNF687_T191 | ZNF687 | Q8N1G0 | 266;271;275 | NA | 6.9787 | 6.5357 | 7.3523 | 6.8510 | 6.0582 | 5.9075 | 5.9243 | 5.5199 | 5.6955 | 5.7920 | 6.0407 | 5.8548 | 6.2921 | 6.3938 | 6.6090 | 6.1946 |
| ZNF687_T191;S215 | ZNF687 | Q8N1G0 | 266;271 | NA | 5.0435 | 5.4179 | 5.2709 | 6.0927 | 5.6976 | 5.6381 | 5.3443 | 5.4817 | 5.4703 | 5.5437 | 5.3904 | 5.2437 | 8.5209 | 8.4678 | 8.9919 | 8.3847 |
| ZNF687_T377 | ZNF687 | Q8N1G0 | 227;242 | NA | 6.6055 | 6.6080 | 6.5755 | 6.7398 | 6.1818 | 6.5190 | 6.5813 | 6.5983 | 6.6258 | 6.1507 | 6.5185 | 6.3078 | 5.0491 | 5.9395 | 5.3795 | 5.6198 |
| ZNF687_T900 | ZNF687 | Q8N1G0 | 174;183 | NA | 6.1470 | 5.1585 | 6.8724 | 5.0424 | 6.7528 | 7.0177 | 6.4298 | 5.6969 | 6.7438 | 8.4886 | 6.5810 | 4.5232 | 7.5075 | 7.1328 | 4.3448 | 5.5610 |
| ZNF692_S236 | ZNF692 | Q9BU19-5 | 236 | APRLLPSPVTCTP | 5.9821 | 6.5173 | 6.2577 | 6.4232 | 6.1549 | 6.0578 | 6.0698 | 5.9829 | 6.1177 | 6.1639 | 6.1053 | 5.7773 | 6.6335 | 6.4804 | 6.8311 | 6.4452 |
| ZNF696_S86 | ZNF696 | Q9H7X3 | 86 | RERPGGSPRGPVT | 6.6478 | 6.6037 | 6.8425 | 6.6385 | 6.7551 | 5.6409 | 6.3870 | 5.8072 | 6.3628 | 6.6293 | 5.6523 | 6.2044 | 6.3678 | 6.1802 | 5.7717 | 5.5088 |
| ZNF7_S149 | ZNF7 | P17097-2 | 149 | LDSHLGSPGLKVT | 5.8223 | 5.3468 | 5.1694 | 5.5813 | 5.6143 | 6.4760 | 5.6865 | 6.4671 | 6.1523 | 5.7926 | 5.7998 | 5.8608 | 7.3144 | 8.5921 | 7.1168 | 7.2076 |
| ZNF708_S480 | ZNF708 | P17019 | 480 | GKAFNQSPNLTKH | 7.1603 | 6.7876 | 6.9771 | 6.9894 | 6.7370 | 6.0212 | 5.5688 | 6.0258 | 7.3840 | 7.0905 | 7.0107 | 6.7949 | 4.6504 | 5.0795 | 4.0686 | 4.6542 |
| ZNF710_S542 | ZNF710 | Q8N1W2 | 542 | THMIVHSPVKPFK | 6.9234 | 6.6068 | 6.3823 | 6.9354 | 6.4195 | 6.5598 | 6.7883 | 6.2666 | 7.0700 | 6.7967 | 6.6698 | 6.8995 | 5.0260 | 5.2607 | 4.8733 | 4.5218 |
| ZNF711_T221 | ZNF711 | Q9Y462-3 | 221 | LEHMGNTPLKIGS | 6.7551 | 6.9031 | 6.4143 | 7.4597 | 6.5891 | 6.3573 | 6.5368 | 6.8731 | 6.5947 | 6.3834 | 6.5329 | 6.8791 | 4.8172 | 4.9733 | 4.6449 | 5.2861 |
| ZNF740_S44 | ZNF740 | Q8NDX6 | 44 | NGERAGSPDVLRC | 6.3476 | 6.7514 | 5.8438 | 6.2746 | 6.1348 | 5.7866 | 6.3149 | 5.8585 | 6.3718 | 6.4691 | 6.1658 | 6.3095 | 6.2429 | 6.4515 | 6.2528 | 6.4244 |
| ZNF746_S323 | ZNF746 | Q6NUN9-2 | 323 | ATRFFPSPAQEGA | 5.6531 | 4.9735 | 5.3778 | 7.3218 | 6.2783 | 6.4089 | 5.9299 | 7.2829 | 6.1741 | 5.1399 | 5.8662 | 5.3327 | 7.7199 | 6.6126 | 7.1616 | 6.7666 |
| ZNF764_S131 | ZNF764 | Q96H86 | 131 | DPVAAGSPGLKSP | 6.2302 | 6.3072 | 5.9799 | 6.8156 | 6.3410 | 6.3539 | 6.9904 | 6.4683 | 7.4118 | 7.0803 | 6.8020 | 6.5423 | 4.9184 | 5.5621 | 4.9465 | 5.2500 |
| ZNF768_S125 | ZNF768 | Q9H5H4 | 83 | PRFEPESPGFESR | 5.9546 | 6.3679 | 5.9838 | 6.1230 | 6.5265 | 6.1029 | 6.3646 | 6.3771 | 6.5143 | 6.2890 | 5.9071 | 5.9798 | 6.1580 | 6.7853 | 6.4990 | 6.0672 |
| ZNF768_S139;S144 | ZNF768 | Q9H5H4 | 23 | SSDEMRSPEGYLR | 5.6887 | 6.5649 | 5.8431 | 6.5070 | 5.8497 | 6.0298 | 6.3269 | 6.2206 | 6.5360 | 6.6711 | 6.4108 | 6.7559 | 5.9163 | 6.5077 | 5.8138 | 6.3577 |
| ZNF768_S160 | ZNF768 | Q9H5H4 | 125 | PRYEPQSPGYEPR | 6.0873 | 6.4677 | 5.9766 | 6.1509 | 6.2912 | 5.9658 | 5.9376 | 6.0902 | 6.3660 | 6.3497 | 5.8926 | 5.9839 | 6.7750 | 6.8027 | 6.6355 | 6.2273 |
| ZNF768_S17 | ZNF768 | Q9H5H4 | 97 | PGLVPPSPEFAPR | 5.4484 | 5.6482 | 5.6711 | 5.9792 | 6.1674 | 5.7593 | 6.0949 | 5.8547 | 6.2774 | 6.4498 | 6.0700 | 5.9244 | 7.1462 | 7.2476 | 7.1266 | 7.1351 |
| ZNF768_S23 | ZNF768 | Q9H5H4 | 160 | TELKTQSPEFEAQ | 6.0827 | 6.4145 | 5.9596 | 6.4551 | 5.9157 | 6.2995 | 6.0610 | 6.0246 | 6.2091 | 6.7759 | 6.4449 | 6.3671 | 6.1979 | 6.5757 | 6.1927 | 6.0240 |
| ZNF768_S83 | ZNF768 | Q9H5H4 | 17 | EPQDVQSSDEMRS | 6.1620 | 6.9804 | 6.3418 | 5.9973 | 5.7772 | 6.7433 | 5.6168 | 5.8789 | 6.6641 | 7.0628 | 6.9274 | 6.8316 | 5.5598 | 6.1854 | 5.5396 | 5.7316 |
| ZNF768_S90;S97 | ZNF768 | Q9H5H4 | 90;97 | NA | 5.9998 | 6.8741 | 6.1389 | 6.6555 | 6.5795 | 6.2255 | 5.8238 | 6.4514 | 6.5709 | 6.2286 | 5.9612 | 6.4195 | 5.7544 | 6.2399 | 5.9207 | 6.1562 |
| ZNF768_S97 | ZNF768 | Q9H5H4 | 139;144 | NA | 6.3068 | 6.7731 | 6.6867 | 6.5822 | 6.9636 | 6.7898 | 7.3338 | 6.5187 | 5.4515 | 5.3444 | 5.8290 | 5.7532 | 6.1432 | 5.7688 | 5.5750 | 6.1801 |
| ZNF786_S578 | ZNF786 | Q8N393 | 578 | KGFTRQSKLTEHL | 8.5258 | 7.4983 | 9.2758 | 6.9126 | 5.6201 | 6.8121 | 4.4829 | 7.2101 | 7.3938 | 7.0728 | 8.6267 | 8.3447 | 3.7763 | 3.9963 | 2.2188 | 2.2327 |
| ZNF787_S216 | ZNF787 | Q6DD87 | 9 | LREEAWSPGPLDS | 5.6860 | 6.1995 | 6.0729 | 6.2494 | 5.8890 | 6.2225 | 5.9713 | 6.1261 | 6.1437 | 6.5693 | 6.3251 | 6.5150 | 6.3342 | 6.6478 | 6.4782 | 6.5700 |
| ZNF787_S9 | ZNF787 | Q6DD87 | 216 | AKVLAASVRRAKG | 5.5602 | 6.6727 | 7.8827 | 6.7890 | 4.2904 | 6.2135 | 5.9714 | 6.4358 | 8.0730 | 8.8610 | 9.7281 | 9.0305 | 3.2188 | 4.5951 | 2.5566 | 4.1213 |
| ZNF800_S317 | ZNF800 | Q2TB10 | 336 | LFLDSISPKKSFK | 6.4420 | 6.1910 | 6.7367 | 7.3176 | 6.4007 | 6.2236 | 6.6754 | 5.7714 | 6.7593 | 6.6640 | 6.7030 | 6.9134 | 5.4327 | 5.5688 | 4.8431 | 5.3571 |
| ZNF800_S317;S336 | ZNF800 | Q2TB10 | 462 | ESPKSTSPSAAGG | 6.8373 | 6.7957 | 6.1518 | 7.1330 | 6.1077 | 6.4197 | 6.8079 | 6.6835 | 6.4435 | 6.6475 | 6.3285 | 6.5466 | 4.8399 | 5.7743 | 5.0704 | 5.4125 |
| ZNF800_S336 | ZNF800 | Q2TB10 | 317 | RGLRRDSITPDIA | 5.0266 | 4.0025 | 4.6205 | 5.0473 | 7.9038 | 4.7844 | 5.9215 | 6.0660 | 9.2568 | 9.7003 | 7.6937 | 6.8192 | 6.7448 | 6.7073 | 4.3623 | 5.3432 |
| ZNF800_S426 | ZNF800 | Q2TB10 | 647 | KANASNSPEGNKT | 6.8303 | 5.8776 | 5.4993 | 8.3919 | 6.4310 | 5.4437 | 6.5337 | 6.1637 | 6.7433 | 6.8460 | 5.8441 | 5.2907 | 5.9206 | 6.1480 | 5.8692 | 6.1669 |
| ZNF800_S460 | ZNF800 | Q2TB10 | 460 | DSESPKSTSPSAA | 6.0717 | 6.7240 | 6.5331 | 7.3711 | 5.8677 | 5.9875 | 6.2934 | 6.3732 | 6.7502 | 6.5474 | 6.9386 | 7.2409 | 4.5475 | 5.5993 | 5.0213 | 6.1332 |
| ZNF800_S462 | ZNF800 | Q2TB10 | 426 | PPSITHSPQNELK | 6.3087 | 6.1825 | 6.0265 | 6.8477 | 5.7587 | 5.9947 | 6.4686 | 6.0781 | 5.8998 | 5.9942 | 6.4150 | 6.1462 | 6.2530 | 6.2535 | 6.8949 | 6.4781 |
| ZNF800_S647 | ZNF800 | Q2TB10 | 317;336 | NA | 5.2945 | 4.9578 | 5.4495 | 5.7058 | 8.2916 | 6.2266 | 6.4304 | 7.4251 | 8.0018 | 8.7257 | 8.3366 | 7.3162 | 5.3929 | 5.2413 | 2.8547 | 4.3495 |
| ZNF814_S87 | ZNF814 | B7Z6K7 | 87 | TPMAGVSPKKAHP | 6.7571 | 7.0629 | 6.9245 | 6.4869 | 7.1117 | 6.6828 | 6.7737 | 6.3612 | 6.5604 | 6.6398 | 6.9743 | 6.6450 | 4.7233 | 4.8928 | 4.4858 | 4.9177 |
| ZNF827_T328 | ZNF827 | Q17R98 | 328 | KKPEKVTPPPPPP | 6.1404 | 6.2652 | 6.2400 | 6.5662 | 6.2389 | 6.3385 | 6.4151 | 6.0430 | 6.1568 | 6.9208 | 5.2921 | 6.1623 | 5.8821 | 7.0872 | 5.9434 | 6.3079 |
| ZNF830_S351 | ZNF830 | Q96NB3 | 351 | EEENADSDDEGEL | 5.8800 | 6.3023 | 5.9713 | 6.5452 | 6.2756 | 6.2207 | 5.9306 | 5.9908 | 6.4189 | 6.3278 | 6.0617 | 6.1389 | 6.8272 | 6.5565 | 6.3019 | 6.2506 |
| ZNF830_S40 | ZNF830 | Q96NB3 | 40 | SRKRIESPFAKYN | 7.3698 | 7.9810 | 6.5951 | 7.1460 | 6.4640 | 7.4314 | 7.3831 | 6.5751 | 7.9335 | 7.6851 | 7.9072 | 7.6526 | 2.7688 | 3.3948 | 2.3561 | 3.3562 |
| ZNF839_S690 | ZNF839 | A8K0R7-5 | 690 | LEARRGSIGAALS | 4.9840 | 4.4222 | 5.8402 | 5.3223 | 7.3571 | 5.4009 | 5.3335 | 5.8331 | 8.2287 | 7.8141 | 7.6126 | 7.2418 | 7.1942 | 7.4425 | 5.0125 | 4.9601 |
| ZNF839_S786 | ZNF839 | A8K0R7-5 | 786 | LNHQQPSPTSVLP | 6.1296 | 6.7373 | 6.0616 | 6.6219 | 6.1323 | 6.3572 | 5.7551 | 6.1468 | 6.0886 | 6.3088 | 5.8938 | 6.3432 | 6.3408 | 6.4702 | 5.9301 | 6.6827 |
| ZNF841_S93 | ZNF841 | Q6ZN19-3 | 93 | GVITGISPKCVIK | 8.0952 | 7.2903 | 7.8124 | 8.4623 | 7.0034 | 6.1224 | 7.7933 | 7.1027 | 6.3766 | 7.5595 | 7.4657 | 6.7546 | 2.8688 | 3.3596 | 2.6567 | 3.2767 |
| ZNHIT2_T161 | ZNHIT2 | Q9UHR6 | 161 | ELAPARTPPDSVK | 6.0340 | 6.3868 | 7.0247 | 6.4221 | 6.4948 | 6.8461 | 6.3286 | 6.5159 | 6.7632 | 6.5702 | 6.7717 | 7.3808 | 4.9438 | 5.3602 | 4.7741 | 5.3830 |
| ZNHIT3_S80 | ZNHIT3 | Q15649 | 80 | IADFLNSDEEEDR | 5.8755 | 5.7891 | 6.4526 | 6.6696 | 7.0120 | 6.8856 | 6.3136 | 6.3469 | 6.6598 | 6.3184 | 7.2423 | 6.9794 | 6.1744 | 5.6991 | 4.8966 | 4.6851 |
| ZNHIT6_S25 | ZNHIT6 | Q9NWK9 | 25 | AEGVRLSPEPGRE | 6.4650 | 6.6186 | 6.3600 | 6.9658 | 6.3728 | 6.1170 | 6.1477 | 6.6923 | 6.5867 | 6.2574 | 6.5923 | 6.6341 | 5.4436 | 5.4234 | 5.6614 | 5.6618 |
| ZNRF1_S123 | ZNRF1 | Q8ND25-2 | 123 | YLGSRASLADALP | 6.2774 | 5.8911 | 7.0432 | 6.3074 | 6.2462 | 6.2469 | 6.6414 | 6.8551 | 6.7553 | 6.7330 | 6.8091 | 6.7486 | 4.8325 | 6.0310 | 5.0114 | 5.5705 |
| ZNRF1_S50 | ZNRF1 | Q8ND25-2 | 50 | MGLRSRSVSSVAG | 6.6852 | 5.8466 | 6.4247 | 6.1410 | 7.1233 | 5.5080 | 6.1062 | 6.1680 | 5.7193 | 6.7913 | 6.8734 | 6.8927 | 5.6768 | 5.9451 | 6.2413 | 5.8573 |
| ZNRF2_S135 | ZNRF2 | Q8NHG8 | 151 | SLPAHLSPHMFGG | 6.0991 | 5.7472 | 6.4136 | 5.5748 | 6.6812 | 6.3697 | 6.0286 | 7.8117 | 5.6121 | 8.5393 | 7.6274 | 8.7151 | 3.9096 | 5.2801 | 4.5217 | 5.0689 |
| ZNRF2_S145;S151 | ZNRF2 | Q8NHG8 | 135 | DRPVGGSPGGPRL | 5.6349 | 4.9995 | 6.5189 | 5.6840 | 5.9426 | 5.5908 | 6.1626 | 6.7303 | 5.7921 | 7.0665 | 6.0374 | 6.8679 | 6.5429 | 6.9759 | 6.4915 | 6.9622 |
| ZNRF2_S151 | ZNRF2 | Q8NHG8 | 82 | AAPRSRSLGGAVG | 6.8746 | 5.9530 | 6.9270 | 5.7428 | 5.8174 | 5.6293 | 5.9725 | 7.0821 | 5.7372 | 7.4328 | 6.6885 | 7.2106 | 5.1929 | 6.7746 | 5.6801 | 5.2846 |
| ZNRF2_S19 | ZNRF2 | Q8NHG8 | 19 | GRTRAYSGSDLPS | 6.0705 | 5.5029 | 6.5434 | 5.9474 | 5.5968 | 4.9698 | 5.2376 | 6.6149 | 5.7811 | 5.3040 | 6.5839 | 5.9077 | 6.9043 | 7.8000 | 7.4900 | 7.7458 |
| ZNRF2_S82 | ZNRF2 | Q8NHG8 | 145;151 | NA | 8.5840 | 6.3838 | 6.8873 | 6.9185 | 6.8759 | 5.9998 | 5.3808 | 6.1125 | 6.8061 | 6.2921 | 6.8817 | 7.8206 | 4.4207 | 6.2209 | 4.0207 | 4.3946 |
| ZRANB1_S216 | ZRANB1 | Q9UGI0 | 216 | GNSQRRSPPATKR | 8.0184 | 6.6783 | 7.0866 | 6.3732 | 6.4245 | 7.1710 | 6.6199 | 6.0516 | 6.9723 | 6.7752 | 7.3885 | 7.0448 | 4.4774 | 4.2780 | 4.2606 | 4.3797 |
| ZRANB2_S120 | ZRANB2 | O95218-2 | 120 | YIEREESDGEYDE | 5.530048 | 6.090451 | 5.979302 | 5.859051 | 5.945796 | 6.104048 | 5.731717 | 5.983897 | 6.164962 | 6.178837 | 6.335397 | 6.130825 | 6.996710 | 6.979856 | 6.929534 | 7.059569 |
| ZRANB2_S143 | ZRANB2 | O95218-2 | 188 | KYNLDASEEEDSN | 5.268876 | 5.734996 | 6.101871 | 5.912526 | 5.944951 | 6.195646 | 5.922941 | 6.071064 | 6.221945 | 6.163439 | 6.311695 | 6.298304 | 6.922831 | 7.101849 | 6.941385 | 6.885680 |
| ZRANB2_S153 | ZRANB2 | O95218-2 | 153 | EVEDKESEGEEED | 5.330276 | 5.746939 | 6.308154 | 5.523754 | 5.608232 | 5.897456 | 5.565270 | 5.396797 | 6.295319 | 6.306707 | 6.481206 | 6.310564 | 7.298263 | 7.307422 | 7.236683 | 7.386959 |
| ZRANB2_S181 | ZRANB2 | O95218-2 | 143 | KAVGPASILKEVE | 6.922122 | 6.567300 | 6.954954 | 5.764248 | 5.079419 | 7.065624 | 6.866325 | 5.989442 | 6.299719 | 6.269508 | 6.889291 | 6.747486 | 5.017610 | 6.270895 | 4.754345 | 6.541713 |
| ZRANB2_S188 | ZRANB2 | O95218-2 | 181 | EDDADLSKYNLDA | 6.747626 | 6.348652 | 5.707876 | 7.309144 | 6.695235 | 6.840316 | 7.076745 | 6.800016 | 7.537513 | 5.299499 | 7.041818 | 6.194167 | 4.343037 | 5.714593 | 5.390846 | 4.952916 |
| ZRANB2_S305;S307 | ZRANB2 | O95218-2 | 307 | RRTRSRSPESQVI | 5.691061 | 6.135316 | 5.473720 | 6.258826 | 5.728880 | 5.889252 | 5.169253 | 5.874412 | 5.375582 | 5.790635 | 6.041007 | 5.625476 | 7.438879 | 7.526964 | 7.939145 | 8.041591 |
| ZRANB2_S307 | ZRANB2 | O95218-2 | 305;307 | NA | 5.525735 | 5.872103 | 6.384024 | 5.599372 | 5.803140 | 5.565985 | 5.735154 | 5.978060 | 6.250682 | 6.408905 | 6.266418 | 6.121311 | 7.232236 | 7.274094 | 7.003938 | 6.978842 |
| ZRSR2_S349 | ZRSR2 | Q15696 | 384 | RGRRNPSPDHSYK | 5.867622 | 6.362509 | 5.932676 | 6.702836 | 5.776179 | 5.760971 | 5.998923 | 5.953300 | 7.271822 | 5.859223 | 5.811116 | 6.162725 | 6.076898 | 6.777134 | 6.796367 | 6.889699 |
| ZRSR2_S384 | ZRSR2 | Q15696 | 349 | NRDIYLSPDRTGS | 6.515445 | 6.874607 | 6.117618 | 6.750911 | 6.141893 | 5.873076 | 6.311184 | 6.272978 | 6.291307 | 6.109386 | 6.147593 | 6.178833 | 5.534072 | 6.536764 | 6.115613 | 6.228720 |
| ZRSR2P1_S475 | ZRSR2P1 | Q15695 | 475 | RDRTVQSPQSKxx | 5.863906 | 6.211456 | 5.924277 | 6.497797 | 5.907507 | 5.726815 | 6.244576 | 5.387650 | 6.111677 | 5.923019 | 6.647885 | 6.322135 | 6.304527 | 6.622730 | 6.846604 | 7.457437 |
| ZSCAN18_S70 | ZSCAN18 | Q8TBC5-3 | 70 | FASPRSSPAPPDL | 6.248110 | 6.471122 | 5.738459 | 6.275829 | 5.749168 | 5.781297 | 6.199602 | 6.262600 | 6.061637 | 6.985813 | 6.688464 | 6.548609 | 5.834215 | 6.512700 | 6.563098 | 6.079276 |
| ZSCAN21_S157 | ZSCAN21 | Q9Y5A6 | 157 | SGTAKESPSSMQP | 5.946257 | 6.738334 | 6.953186 | 6.872995 | 5.977274 | 6.416056 | 6.624856 | 5.691309 | 6.885100 | 6.800369 | 6.460691 | 7.005386 | 5.036162 | 5.328935 | 5.344066 | 5.919022 |
| ZSCAN29_S405 | ZSCAN29 | Q8IWY8 | 561 | APVLFQSPRGFEA | 7.221913 | 7.546826 | 6.614658 | 7.111582 | 5.517427 | 7.129826 | 7.549866 | 6.942176 | 7.258835 | 7.821785 | 7.702332 | 7.511641 | 2.938545 | 3.821628 | 3.244344 | 4.066616 |
| ZSCAN29_S561 | ZSCAN29 | Q8IWY8 | 405 | APVVFRSPGGVHW | 7.193741 | 7.724705 | 6.783716 | 7.151306 | 6.159926 | 5.878796 | 6.813420 | 7.451001 | 6.895107 | 6.920037 | 7.462140 | 6.963002 | 3.839809 | 4.652964 | 3.327941 | 4.782390 |
| ZSCAN5A_S296 | ZSCAN5A | Q9BUG6 | 296 | DALNLSSPKRSKP | 6.797462 | 6.788922 | 6.525955 | 6.954210 | 6.654236 | 6.376857 | 6.716152 | 6.122965 | 7.386202 | 6.531649 | 6.393404 | 6.545704 | 5.302572 | 5.337267 | 4.576655 | 4.989787 |
| ZSCAN5A_T209 | ZSCAN5A | Q9BUG6 | 209 | HKSIDVTGDPKSL | 6.831216 | 7.109282 | 6.516977 | 6.715625 | 5.405722 | 5.220614 | 6.012173 | 6.089975 | 6.820910 | 6.438973 | 5.970342 | 7.136766 | 5.552232 | 5.722386 | 5.559507 | 6.897297 |
| ZSWIM8_S1093 | ZSWIM8 | A7E2V4-4 | 1841 | LEMATFSPxxxxx | 6.880605 | 6.717672 | 6.693586 | 6.650232 | 6.586758 | 6.717105 | 6.983182 | 6.632663 | 7.146116 | 6.264859 | 6.895623 | 6.948046 | 4.514246 | 4.927955 | 4.499795 | 4.941557 |
| ZSWIM8_S1135 | ZSWIM8 | A7E2V4-4 | 593 | ALGGAGSGSKGSA | 6.530553 | 7.102826 | 7.083324 | 7.082132 | 6.361852 | 7.135783 | 7.358901 | 6.940938 | 6.662721 | 6.697892 | 6.981048 | 6.777942 | 3.792755 | 4.926401 | 3.958136 | 4.606797 |
| ZSWIM8_S1841 | ZSWIM8 | A7E2V4-4 | 1135 | NGRGWGSPGRPKK | 6.357364 | 6.039778 | 7.386860 | 6.025493 | 6.472638 | 6.291920 | 6.024164 | 6.449716 | 6.496557 | 6.382279 | 6.942373 | 7.122094 | 5.870022 | 5.611569 | 4.831557 | 5.695616 |
| ZSWIM8_S567 | ZSWIM8 | A7E2V4-4 | 567 | RGPRRLSAEGGDK | 6.278779 | 5.921046 | 6.631256 | 6.030531 | 6.307933 | 5.882132 | 6.103353 | 6.565315 | 6.800175 | 6.762888 | 6.932432 | 6.844845 | 5.857482 | 6.384134 | 5.198949 | 5.498749 |
| ZSWIM8_S593 | ZSWIM8 | A7E2V4-4 | 1093 | EKNVPESSPHSPC | 5.868002 | 6.430089 | 6.538405 | 6.549447 | 6.792369 | 6.856518 | 5.534640 | 7.206705 | 5.990514 | 5.730766 | 5.822256 | 7.043179 | 6.277605 | 6.730850 | 4.987275 | 5.641379 |
| ZXDB_T753 | ZXDB | P98169 | 753 | SVSELLTPTKAEW | 5.752548 | 6.868067 | 6.029399 | 5.849941 | 6.587609 | 6.354946 | 6.418072 | 6.144828 | 6.662458 | 6.106953 | 6.330598 | 5.815673 | 6.495626 | 6.105149 | 6.315267 | 6.162864 |
| ZXDC_T172 | ZXDC | Q2QGD7 | 172 | QASGPSTPGYRCP | 6.469333 | 6.522740 | 6.147136 | 6.072540 | 5.897608 | 6.098456 | 6.649343 | 6.532371 | 7.332085 | 6.872604 | 6.522390 | 6.562314 | 5.263020 | 5.703414 | 5.463692 | 5.890954 |
| ZYX_S143 | ZYX | Q15942 | 344 | NQNQVRSPGAPGP | 7.148754 | 6.682089 | 7.048914 | 6.403045 | 6.934447 | 6.841215 | 6.645806 | 7.533658 | 7.035108 | 7.151069 | 7.799695 | 7.801800 | 3.827146 | 3.968320 | 3.373423 | 3.805511 |
| ZYX_S169 | ZYX | Q15942 | 267 | APAPKFSPVTPKF | 7.389638 | 6.360390 | 7.180560 | 5.785751 | 7.622967 | 7.026007 | 6.941573 | 7.695564 | 7.325895 | 7.242671 | 7.223924 | 6.941179 | 4.096037 | 3.972997 | 3.389618 | 3.805228 |
| ZYX_S202 | ZYX | Q15942 | 308 | LSAGTGSPQPPSF | 6.923464 | 6.665217 | 7.236339 | 6.562833 | 6.837600 | 7.319259 | 7.126268 | 7.490825 | 7.144095 | 7.536456 | 7.731409 | 7.695428 | 3.188601 | 3.981008 | 2.980073 | 3.581125 |
| ZYX_S258 | ZYX | Q15942 | 259 | RGPPASSPAPAPK | 5.943871 | 5.339266 | 6.147743 | 4.985568 | 6.895694 | 7.330012 | 7.153938 | 8.065003 | 6.654773 | 6.540773 | 6.570730 | 6.623960 | 5.549089 | 5.760786 | 5.052125 | 5.386668 |
| ZYX_S259 | ZYX | Q15942 | 202 | PLPPWKSPSSSQP | 6.709626 | 7.164340 | 7.027869 | 7.239155 | 6.638946 | 6.814516 | 6.925986 | 7.726201 | 7.136708 | 7.078059 | 7.688042 | 7.139527 | 3.450869 | 4.249204 | 3.193153 | 3.817798 |
| ZYX_S267 | ZYX | Q15942 | 258 | PRGPPASSPAPAP | 6.260527 | 5.541770 | 6.160513 | 5.273743 | 6.894381 | 7.435039 | 6.956372 | 7.800269 | 6.569776 | 6.198185 | 6.666592 | 6.732799 | 5.436520 | 5.677031 | 4.919759 | 5.476723 |
| ZYX_S267;T270 | ZYX | Q15942 | 274 | PVTPKFTPVASKF | 8.163786 | 7.136479 | 7.283506 | 6.411599 | 6.789664 | 6.634213 | 7.612596 | 7.706655 | 8.264904 | 7.668347 | 7.636272 | 8.067923 | 2.440665 | 3.036681 | 1.945586 | 3.201123 |
| ZYX_S270 | ZYX | Q15942 | 169 | PFKARVSSGYVPP | 7.120928 | 6.474012 | 6.907047 | 6.718289 | 6.687060 | 7.152573 | 6.854584 | 6.844174 | 6.751321 | 7.073183 | 7.948830 | 7.808093 | 3.864005 | 4.413904 | 3.228748 | 4.153250 |
| ZYX_S281 | ZYX | Q15942 | 306 | EALSAGTGSPQPP | 6.525924 | 6.190495 | 6.854269 | 6.612168 | 6.537629 | 6.703649 | 6.965462 | 7.490013 | 6.522844 | 6.469865 | 7.632008 | 7.437342 | 4.697544 | 5.263061 | 3.728835 | 4.368892 |
| ZYX_S281;S288 | ZYX | Q15942 | 270 | PKFSPVTPKFTPV | 6.900278 | 6.556139 | 7.114537 | 6.351642 | 6.078362 | 6.810176 | 6.689329 | 7.439432 | 6.993634 | 7.045698 | 7.262426 | 7.739025 | 4.135611 | 4.399410 | 3.898406 | 4.585897 |
| ZYX_S308 | ZYX | Q15942 | 281 | PVASKFSPGAPGG | 7.066464 | 6.578844 | 7.259947 | 6.654179 | 6.999156 | 6.891505 | 6.661178 | 7.573637 | 7.017475 | 7.213223 | 7.369039 | 7.365539 | 3.983492 | 4.061915 | 3.559887 | 3.744520 |
| ZYX_S344 | ZYX | Q15942 | 143 | PREKVSSIDLEID | 5.547247 | 5.546597 | 6.945275 | 6.449138 | 6.464093 | 6.173434 | 6.460842 | 6.433532 | 5.981611 | 6.897157 | 6.795719 | 5.854163 | 5.951700 | 6.116862 | 5.806695 | 6.575936 |
| ZYX_T274 | ZYX | Q15942 | 267;270 | NA | 7.659464 | 6.915830 | 7.466231 | 6.378583 | 6.738449 | 6.797053 | 7.346940 | 7.480861 | 7.733752 | 7.941393 | 8.390460 | 8.285316 | 2.522089 | 3.217292 | 1.948985 | 3.177302 |
| ZYX_T306 | ZYX | Q15942 | 281;288 | NA | 6.729945 | 6.547330 | 7.626166 | 6.848064 | 5.832980 | 7.121876 | 7.184108 | 7.334617 | 7.019120 | 7.339311 | 7.578885 | 7.614911 | 3.713799 | 4.453524 | 2.911427 | 4.143937 |
| ZZEF1_S1463 | ZZEF1 | O43149 | 1463 | NKRQRTSSVVEEH | 5.720198 | 5.483562 | 6.310666 | 6.408236 | 6.349918 | 6.000011 | 5.796179 | 6.805807 | 6.826274 | 6.432629 | 6.397302 | 7.000387 | 6.269731 | 6.575900 | 5.997207 | 5.625992 |
| ZZEF1_S1518 | ZZEF1 | O43149 | 1518 | TAEEPLSPSTPTR | 5.141820 | 5.651416 | 6.502951 | 5.908762 | 6.162541 | 6.828582 | 6.096293 | 6.467279 | 6.091707 | 6.268537 | 6.009662 | 7.301487 | 6.863235 | 6.750614 | 6.150820 | 5.804295 |
| ZZZ3_S113 | ZZZ3 | Q8IYH5 | 113 | RQTEPVSPVLKRI | 6.040328 | 5.887848 | 6.310145 | 6.810569 | 6.338608 | 6.161247 | 6.266289 | 6.566117 | 6.176350 | 6.335994 | 6.558856 | 6.238988 | 5.919893 | 6.172671 | 5.965299 | 6.250799 |
| ZZZ3_S391 | ZZZ3 | Q8IYH5 | 391 | AAPTRGSPTKNSS | 6.691939 | 7.137373 | 6.710299 | 7.325770 | 6.337261 | 6.584493 | 6.261110 | 5.264047 | 5.506689 | 6.440535 | 6.975056 | 6.767673 | 5.638754 | 5.859072 | 5.218148 | 5.281781 |
| ZZZ3_S82 | ZZZ3 | Q8IYH5 | 82 | TRESWVSPRKRGL | 7.494865 | 7.650445 | 6.640741 | 6.890445 | 5.770971 | 5.677223 | 6.302170 | 6.235958 | 6.748633 | 6.361999 | 6.659140 | 6.685800 | 5.201221 | 5.043639 | 5.313494 | 5.323257 |

**IACS-10759 up-regulated phosphosites @ 6 hr treatment (AMPK consensus phosphorylation motif is derived from Gwinn et al. (2008) and Banko et al. (2011)**

| Phosphosite | Gene | Site | -6 | -5 | -4 | -3 | -2 | -1 | 0 | 1 | 2 | 3 | 4 | 5 | 6 | AMPK motif | PSP |
|---|---|---|---|---|---|---|---|---|---|---|---|---|---|---|---|---|---|
| BAD_S118 | BAD | S118 | R | E | L | R | R | M | S | D | E | F | V | D | S | FALSE | NO |
| BAD_S134 | BAD | S134 | G | L | P | R | P | K | S | A | G | T | A | T | Q | TRUE | NO |
| BAD_S146 | BAD | S146 | Q | M | R | Q | S | S | S | W | T | R | V | F | Q | TRUE | NO |
| DENND4C_S1038 | DENND4C | S1038 | S | I | V | K | V | P | S | G | I | F | D | V | N | FALSE | NO |
| DMXL1_S574 | DMXL1 | S574 | G | L | T | R | S | T | S | M | L | I | S | S | G | TRUE | NO |
| DMXL2_S1151 | DMXL2 | S1151 | L | F | V | Y | S | K | S | D | A | L | L | S | K | FALSE | NO |
| DOCK6_T1264 | DOCK6 | T1264 | L | W | V | L | K | N | T | E | P | A | L | L | Q | FALSE | NO |
| GPAT3_S68 | GPAT3 | S68 | I | L | K | N | S | A | S | V | G | I | I | Q | R | TRUE | NO |
| HMGCR_S524 | HMGCR | S524 | L | L | S | K | K | L | S | E | P | S | S | L | Q | TRUE | NO |
| PARP1_S257 | PARP1 | S257 | E | L | K | K | V | C | S | T | N | D | L | K | E | TRUE | NO |
| PHKA1_S758 | PHKA1 | S758 | H | L | P | R | D | Q | S | G | E | V | D | F | K | FALSE | NO |
| PNPLA2_S428 | PNPLA2 | S428 | W | M | R | N | N | L | S | L | G | D | A | L | A | TRUE | NO |
| PRKAB1_S108 | PRKAB1 | S108 | K | L | P | L | T | R | S | H | N | N | F | V | A | TRUE | YES |
| PRKAB2_S108 | PRKAB2 | S108 | K | I | P | L | I | K | S | H | N | D | F | V | A | TRUE | NO |
| SLC25A46_S34 | SLC25A46 | S34 | F | P | A | R | S | F | S | T | G | S | D | L | G | FALSE | NO |
| SLC25A5_S42 | SLC25A5 | S42 | L | Q | V | Q | H | A | S | K | Q | I | T | A | D | FALSE | NO |
| SLC4A7_S267 | SLC4A7 | S267 | I | L | A | S | P | Q | S | A | P | G | N | L | D | FALSE | NO |
| SLC4A8_S237 | SLC4A8 | S237 | L | I | P | I | V | R | S | F | A | E | V | G | K | TRUE | NO |
| SLC9A1_S599 | SLC9A1 | S599 | G | M | G | K | I | P | S | A | V | S | T | V | S | FALSE | NO |
| SMARCC1_S319 | SMARCC1 | S319 | R | R | D | R | K | A | S | A | N | A | R | K | R | FALSE | NO |
| STIM1_S257 | STIM1 | S257 | L | H | R | A | E | Q | S | L | H | D | L | Q | E | TRUE | YES |
| TRAPPC12_S184 | TRAPPC12 | S184 | Q | M | V | K | S | P | S | F | G | G | A | S | E | FALSE | NO |
| TRAPPC8_S273 | TRAPPC8 | S273 | I | T | S | N | K | N | S | D | N | N | L | L | S | FALSE | NO |

**KD025 down-regulated phosphosites @ 6 hr treatment**

| Phosphosite | Gene | Site | -6 | -5 | -4 | -3 | -2 | -1 | 0 | 1 | 2 | 3 | 4 | 5 | 6 | ROCK1/2 motif | PSP |
|---|---|---|---|---|---|---|---|---|---|---|---|---|---|---|---|---|---|
| RNF19A_S69 | RNF19A | S69 | R | R | I | S | I | G | S | L | F | R | R | K | K | FALSE | NO |
| CDC5L_T227 | CDC5L | T227 | A | L | G | F | Y | D | T | S | E | E | N | Y | Q | FALSE | NO |
| PHTF1_T222 | PHTF1 | T222 | S | N | K | G | T | E | T | D | N | D | P | S | C | FALSE | NO |
| SNW1_S446 | SNW1 | S446 | G | K | D | M | A | Q | S | I | Y | R | P | S | K | FALSE | NO |
| PPP1R12C_T560 | PPP1R12C | T560 | R | Q | S | R | R | S | T | Q | G | V | T | L | T | TRUE | YES |
| PPP1R12A_T696 | PPP1R12A | T696 | R | Q | S | R | R | S | T | Q | G | V | T | L | T | TRUE | NO |
| BAZ1B_T833 | BAZ1B | T833 | F | E | P | Q | V | D | T | E | A | E | D | M | I | FALSE | NO |
| CEP170_S1059 | CEP170 | S1059 | G | R | T | P | L | T | S | A | D | E | H | V | H | FALSE | NO |
| RSBN1L_T285 | RSBN1L | T285 | I | C | S | G | L | L | T | D | V | E | D | Q | A | FALSE | NO |
| IWS1_S527 | IWS1 | S527 | K | H | M | D | F | L | S | D | F | E | M | M | L | FALSE | NO |
| DTL_S488 | DTL | S488 | R | R | G | S | V | S | S | V | S | P | K | P | P | FALSE | NO |
| DTL_S490 | DTL | S490 | G | S | V | S | S | V | S | P | K | P | P | S | S | FALSE | NO |
| JPH1_S220 | JPH1 | S220 | R | G | S | L | L | G | S | M | K | L | R | K | S | FALSE | NO |
| GPATCH2_S284 | GPATCH2 | S284 | Q | G | D | D | E | Q | S | D | W | F | Y | E | K | FALSE | NO |
| NUMA1_T1812 | NUMA1 | T1812 | S | A | R | R | R | T | T | Q | I | I | N | I | T | TRUE | NO |
| SVIL_T825 | SVIL | T825 | M | N | A | R | Y | Q | T | Q | P | V | T | L | G | TRUE | NO |
| SLC12A4_T983 | SLC12A4 | T983 | A | D | K | I | Q | M | T | W | T | R | D | K | Y | FALSE | NO |
| H1-2_T165 | H1-2 | T165 | K | K | P | A | A | A | T | V | T | K | K | V | A | FALSE | NO |
| GPATCH2L_S248 | GPATCH2L | S248 | Q | G | D | D | E | Q | S | D | W | F | Y | E | G | FALSE | NO |
| ALS2_S492 | ALS2 | S492 | G | L | L | S | Q | V | S | P | R | L | L | R | K | FALSE | NO |
| CAMK1D_T180 | CAMK1D | T180 | K | G | D | V | M | S | T | A | C | G | T | P | G | FALSE | NO |
| TCOF1_S1078 | TCOF1 | S1078 | S | M | A | G | A | S | S | S | K | E | S | S | R | FALSE | NO |
| MKI67_S2638 | MKI67 | S2638 | T | H | K | E | P | A | S | G | D | E | G | I | K | FALSE | NO |
| PRKCA_T497 | PRKCA | T497 | D | G | V | T | T | R | T | F | C | G | T | P | D | FALSE | NO |
| PRKCD_S695 | PRKCD | S695 | S | A | F | A | G | F | S | F | V | N | P | K | F | FALSE | NO |
| PRKCI_T412 | PRKCI | T412 | P | G | D | T | T | S | T | F | C | G | T | P | N | FALSE | NO |
| PCM1_S1969 | PCM1 | S1969 | L | K | L | T | I | Y | S | E | A | D | L | R | K | FALSE | NO |
| ZNF609_S452 | ZNF609 | S452 | S | D | M | E | L | N | S | S | S | E | D | S | K | FALSE | NO |
| PLEKHA5_T444 | PLEKHA5 | T444 | G | V | I | S | Y | Q | T | L | P | R | N | M | P | FALSE | NO |
